# G protein-coupled receptor 151 regulates glucose metabolism and hepatic gluconeogenesis

**DOI:** 10.1101/2022.05.25.493489

**Authors:** Ewa Bielczyk-Maczynska, Meng Zhao, Peter-James H. Zushin, Theresia M. Schnurr, Hyun-Jung Kim, Jiehan Li, Pratima Nallagatla, Panjamaporn Sangwung, Chong Park, Cameron Cornn, Andreas Stahl, Katrin J. Svensson, Joshua W. Knowles

**Affiliations:** Division of Cardiovascular Medicine, Department of Medicine, Stanford University School of Medicine, Stanford, CA, USA; Stanford Diabetes Research Center, Stanford University School of Medicine, Stanford, CA, USA; Stanford Cardiovascular Institute, Stanford University School of Medicine, CA, USA; Department of Pathology, Stanford University School of Medicine, Stanford, CA, USA; Department of Nutritional Sciences and Toxicology, University of California at Berkeley, Berkeley, CA, USA; Genetics Bioinformatics Service Center, Stanford University School of Medicine, Stanford, CA, USA; Stanford Prevention Research Center, Stanford University School of Medicine, Stanford, CA, USA

## Abstract

Human genetics has been instrumental in identification of genetic variants linked to type 2 diabetes (T2D). Recently a rare, putative loss-of-function mutation in the orphan G-protein coupled receptor 151 (*GPR151*) was found to be associated with lower odds ratio for T2D, but the mechanism behind this association has remained elusive. Here for the first time we show that *Gpr151* is a fasting- and glucagon-responsive hepatic gene which regulates hepatic gluconeogenesis. *Gpr151* ablation in mice leads to suppression of hepatic gluconeogenesis genes and reduced hepatic glucose production in response to pyruvate. Importantly, the restoration of hepatic *Gpr151* levels in the *Gpr151* knockout mice reverses the reduced hepatic glucose production. Our findings establish a previously unknown role of *Gpr151* in the liver and provides an explanation to the lowered T2D risk in individuals with nonsynonymous mutations in *GPR151*. This study highlights the therapeutic potential of targeting GPR151 for treatment of metabolic disease.

---

Type 2 diabetes (T2D) is a major health problem worldwide. There is a great need for deeper understanding of molecular mechanisms and identification of novel drug targets to correct abnormal glucose metabolism that characterizes T2D^1,2^. Human genetics is being increasingly used to identify potential drug targets to combat T2D and cardiovascular disease, including identifying new functions for orphan G-protein coupled receptors (GPCRs)^3,4^. GPCRs are a superfamily of over 800 highly druggable receptors and are targeted by approximately 34% of the current FDA-approved drugs^5^.

Recently, a rare nonsynonymous, presumed inactivating, mutation (p.Arg95Ter) in the gene encoding an orphan G-protein coupled receptor 151 (*GPR151*), a Gα_o1_-linked GPCR, was associated with lower odds ratio for T2D, obesity and coronary artery disease^6^ and with reduced body-mass index (BMI)^6-8^. The function of Gpr151 was first described in the habenula, a brain structure with crucial role in processing reward-related aversive signals^9^. The nonsynonymous *GPR151* p.Arg95Ter variant is linked to 12% lower odds of clinical obesity^6^ and a 14% decrease in the odds of T2D^6,7^, which suggests the presence of a habenula-mediated regulation of appetite or other mechanisms which mediate the effect of *GPR151* loss-of-function variants on decreased odds of T2D. In addition to the brain, *Gpr151* is widely expressed in peripheral tissues in mice^10^. However, the peripheral functions of GPR151 and its mechanism of action in controlling glucose metabolism are entirely unknown.

Several mechanisms sustain glucose homeostasis in the body, including glucose production by the liver, kidney and gut, as well as glucose uptake by skeletal muscle, heart muscle and adipose tissue^2^. T2D is associated with increased rates of gluconeogenesis^11^ and impaired glucose uptake in peripheral tissues due to insulin resistance^12,13^. In diabetic patients^14^ and in mouse models^15^, excessive signaling through the glucagon receptor, a G^s^ alpha subunit-coupled GCPR, contributes to pathologically elevated hepatic gluconeogenesis. This effect is mediated through an increase in intercellular cyclic AMP (cAMP) levels^16^. Surprisingly, excessive signaling through the Gα_i_ class of G proteins, which inhibits cAMP production, can also trigger an increase in hepatic gluconeogenesis in mice^17^. Moreover, a lack of functional Gα_i_-type G proteins in mouse hepatocytes reduces blood glucose levels^17^. Therefore, the relationship between the dynamics of cAMP signaling in the liver and hepatic gluconeogenesis is complex. In addition, the identity and role of Gα_i_-linked GPCRs which regulate hepatic gluconeogenesis are currently unknown.

Here, we dissect the mechanism of action of *Gpr151 in vivo* and *in vitro*. In mice, *Gpr151* expression in liver is increased by fasting. Whole-body loss of *Gpr151* confers increased glucose tolerance in high-fat diet-induced obesity. Furthermore, we show that Gpr151 has a cell-autonomous role in hepatic gluconeogenesis and that loss of *Gpr151* is protective for metabolic health in diet-induced obesity through decreasing gluconeogenesis in hepatocytes. Liver-specific *Gpr151* overexpression abrogates these positive effects resulting in increased hepatic gluconeogenesis. Lastly, we query summary statistics from published genome-wide association studies (GWAS) in humans to identify associations between the *GPR151* p.Arg95Ter loss-of-function variant and selected metabolic traits. Our results demonstrate a new function for Gpr151 in regulating glucose metabolism, suggesting that inhibition of GPR151 may be a novel molecular target for the treatment of T2D.

## Results

### *Gpr151* loss improves glucose metabolism in diet-induced obesity (DIO)

Given the association between *GPR151* loss-of-function variant p.Arg96Ter and lower odds ratio for T2D, obesity and reduced BMI, we examined the function of Gpr151 with respect to metabolic health in diet-induced obesity. We employed the previously described whole-body *Gpr151* knockout (KO) mouse model, which was used to determine the conserved expression and role of Gpr151 in the habenula, a brain structure critical for processing of reward-related and aversive signals^9,18^. As the loss-of-function *GPR151* variant was associated with lower BMI in humans, we compared body weights of *Gpr151* wild-type (WT) and knockout littermates fed either standard diet (SD) or obesity-inducing high-fat diet (HFD). Body weights of *Gpr151* KO animals did not differ significantly from WT littermates, irrespective of sex or diet (**Fig. 1a, Extended Data Figs. 1-2**).

**Fig. 1.**
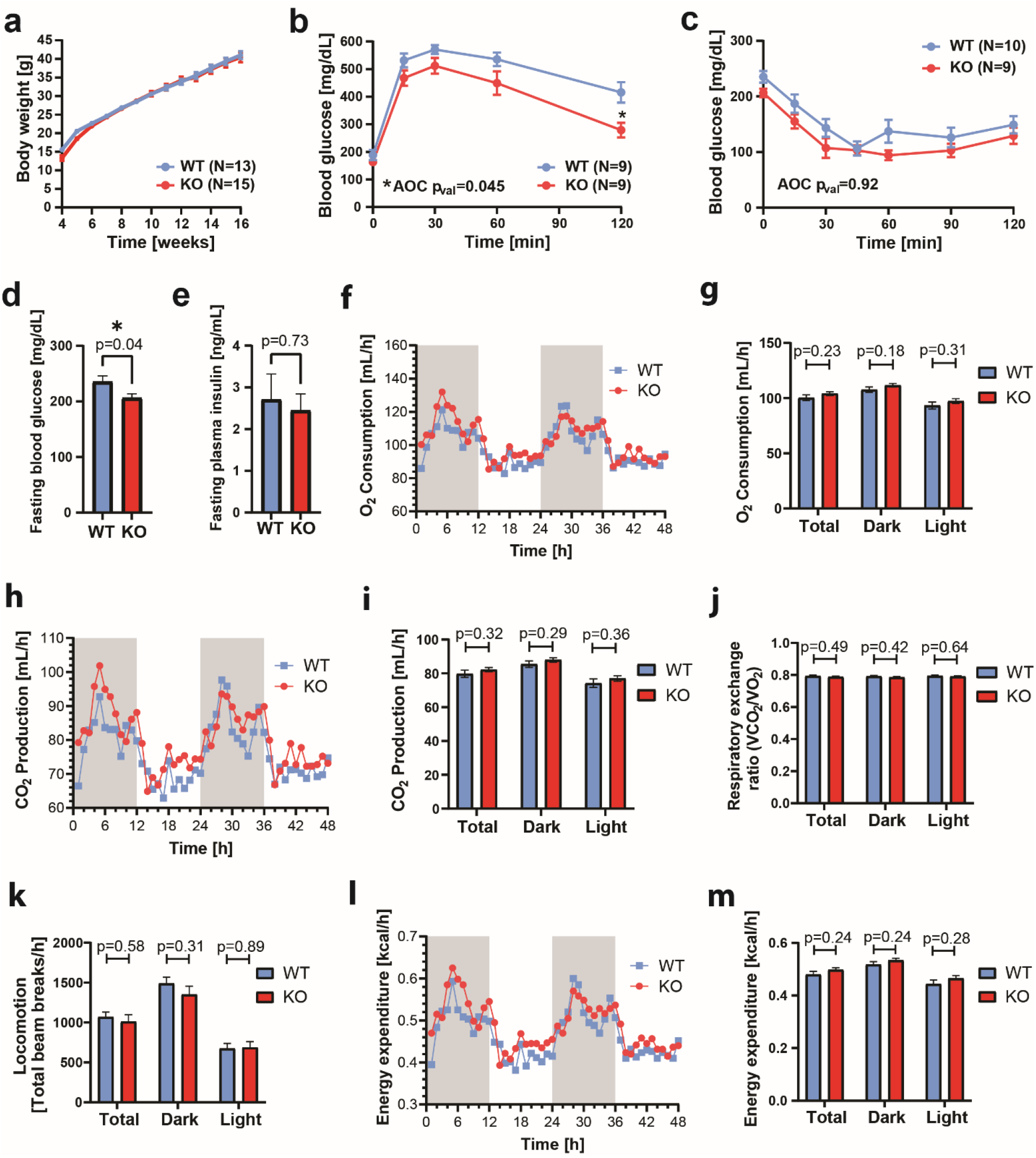
*Gpr151* loss improves glucose metabolism in DIO male mice which is not explained by behavioral differences. **a**, Body weight in male DIO KO and WT mice over 12 weeks of HFD (N=13, WT; N=15, KO). **b**, Blood glucose levels measured during glucose tolerance testing in *Gpr151* KO and WT DIO males. Area of the curve (AOC) compared using two-tailed Student *t* test. Student t test with Bonferroni correction used to test differences at every time point (N=9, WT; N=9, KO; * t=120 q_val_=0.02). **c**, Blood glucose levels measured during insulin tolerance testing in *Gpr151* KO and WT in DIO male mice (N=10, WT; N=9, KO). AOC compared using two-tailed Student *t* test. **d**, Fasting glucose levels in *Gpr151* WT and KO DIO male mice measured in whole blood (N=10, WT; N=9, KO). Two-tailed *t* Student test. **e**, Fasting insulin levels measured in blood plasma of DIO male mice (N=9, WT; N=8, KO). Two-tailed Student *t* tests. **f-m**, Metabolic phenotyping of DIO male mice using CLAMS, conducted at 23°C. **f**, Representative time course of oxygen consumption (N=5, WT; N=5, KO). **g**, Average oxygen consumption (N=10, WT, N=10, KO). Two-tailed Student *t* tests. **h**, Representative time course of CO_2_ production (N=5, WT; N=5, KO). **i**, Average CO_2_ production (N=10, WT; N=10, KO). Two-tailed Student *t* tests. **j**, Average respiratory exchange ratio (N=10, WT; N=10, KO). Two-tailed Student *t* tests. **k**, Average locomotion measured as average number of beam breaks per hour (N=10, WT; N=10, KO). Two-tailed Student *t* tests. **l**, Representative time course of energy expenditure (N=5, WT; N=5, KO). **m**, Average energy expenditure (N=10, WT; N=10, KO). Two-tailed Student *t* tests. *, p<0.05

Next, the effect of *Gpr151* KO on whole-body glucose metabolism was assessed using glucose and insulin tolerance testing. *Gpr151* KO mice showed dramatically improved glucose tolerance compared to WT littermates in diet-induced obesity (DIO) but not when fed standard diet (**Figure 1b, Extended Data Figs. 1-2**). There were no significant differences in insulin tolerance between *Gpr151* WT and KO mice in DIO conditions, suggesting that Gpr151 regulates blood glucose levels but not insulin action *per se* (**Figure 1c**). While *Gpr151* KO mice showed lower fasting blood glucose levels (**Figure 1d**), their plasma insulin levels did not differ from WT littermates in DIO (**Figure 1e**). Because of the known role of Gpr151 in the regulation of appetite^18^ we conducted an analysis of food intake, activity, and metabolic parameters in *Gpr151* WT and KO DIO mice using Comprehensive Lab Animal Monitoring System (CLAMS) to identify a mechanism which could explain the improved glucose metabolism in *Gpr151* KO mice. In spite of small increases in food intake in *Gpr151* KO mice compared to WT littermates in this cohort (**Extended Data Fig. 3**), there were no significant differences in oxygen consumption (**Figure 1f,g**), CO_2_ production (**Figure 1h,i**), or respiratory exchange ratio (**Figure 1j**), indicating no differences in the usage of carbohydrates and lipids for energy and no differences in sympathetic activity in adipose tissue. In addition, locomotion and energy expenditure in *Gpr151* KO and WT mice were comparable (**Figure 1k-m**), indicating that behavioral differences are not the cause of the differences in glucose metabolism.

In summary, whole-body *Gpr151* KO resulted in improved whole-body glucose metabolism in DIO mice, which could not be explained by differences in body composition or physical activity.

### *Gpr151* expression in liver is decreased by feeding

To determine which tissues may mediate the effect of Gpr151 on whole-body glucose metabolism, we quantified *Gpr151* expression in a tissue panel from WT mice. Compared to the brain, known to express *Gpr151*^9^, several tissues showed higher *Gpr151* expression, including skeletal muscle and liver (**Figure 2a**). To identify tissues in which *Gpr151* expression is regulated by changes in the metabolic state, we compared gene expression in SD-fed and DIO WT mice, focusing on the top ten *Gpr151*-expressing tissues, as well as adipose tissue due to its role in glucose metabolism^19^. In the DIO model, *Gpr151* was significantly upregulated in the hind brain and pituitary gland and downregulated in the liver and subcutaneous white adipose tissue (**Figure 2b**). Next, we focused on peripheral tissues involved in glucose metabolism (liver, skeletal muscle, brown and white adipose tissue) and compared *Gpr151* expression under fasting and feeding conditions. *Gpr151* expression was robustly downregulated in the liver following feeding (**Figure 2c**). This postprandial regulation of *Gpr151* gene expression in the liver resembled that of *Fgf21* and *Pck1* (**Figure 2d**), whose expression levels are tightly regulated by fasting and feeding^20^.

**Fig. 2.**
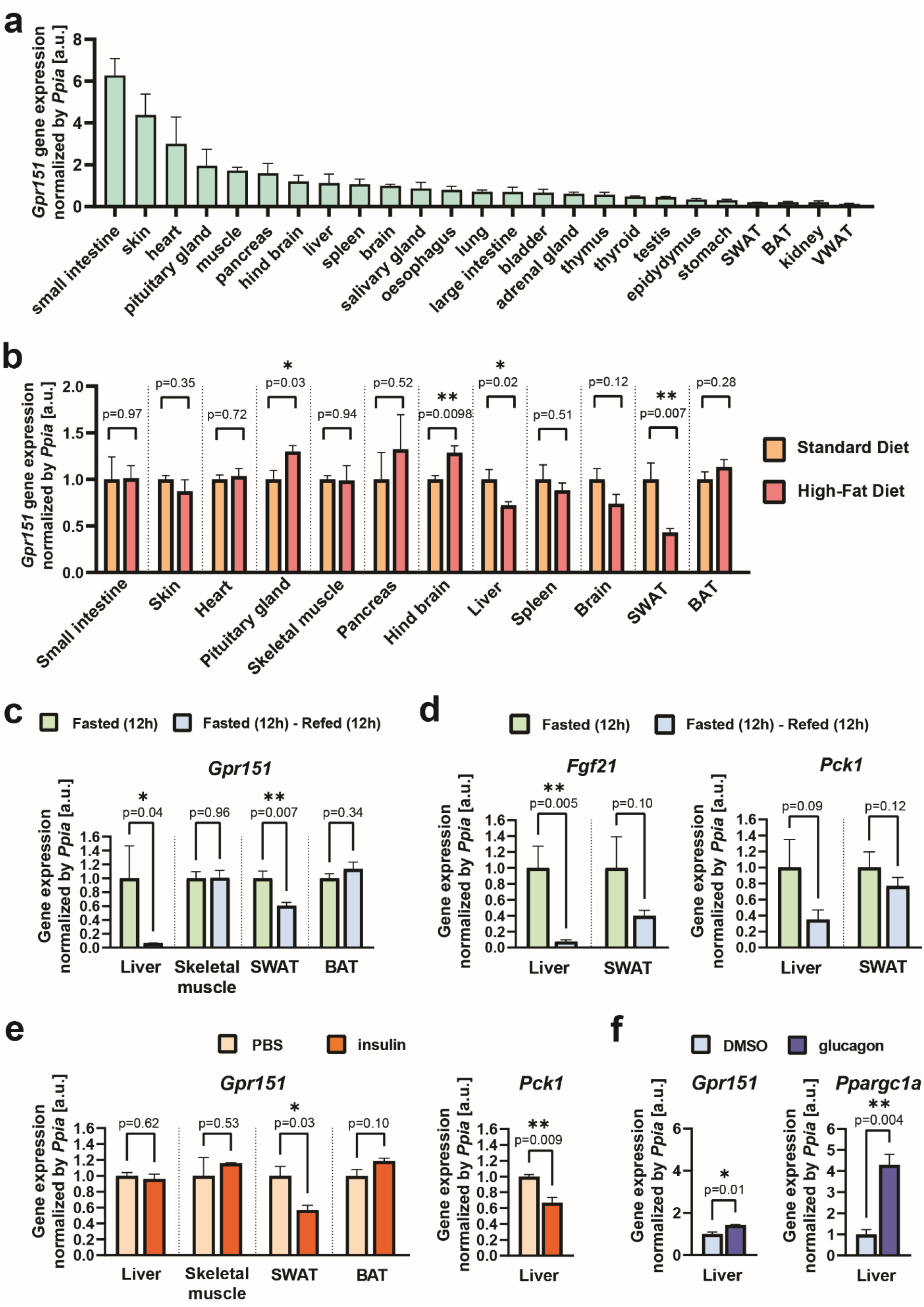
*Gpr151* expression is upregulated by fasting and glucagon in the liver. **a**, RT-qPCR quantification of *Gpr151* expression in the tissues of eight-week-old male c57BL/6J mice (N=5). **b**, RT-qPCR quantification of *Gpr151* expression in the 10 top-expressing tissues and the metabolically relevant tissues in HFD-fed mice compared to the same tissues from SD-fed mice. All mice were 16-week-old male c57BL/6J mice (N=4, SD; N=4, HFD). Two-tailed Student *t* tests. **c**, RT-qPCR quantification of *Gpr151* expression in the liver, SWAT, BAT and skeletal muscle of mice which were fasted for 12 h and then refed for 12 h, normalized by *Gpr151* expression in the same tissues of mice fasted for 12 h. All mice were eight-week-old male c57BL/6J. Two-tailed Student *t* tests (N=5, fasted; N=6, refed). **d**, Quantification of *Fgf21* expression in the liver and SWAT of fasted-refed and fasted mice. Two-tailed Student *t* tests (N=5, fasted; N=6, refed). **e**, *Gpr151* expression in the liver, skeletal muscle, SWAT and BAT in insulin-injected mice compared to the same tissues of PBS-injected mice, quantified by RT-qPCR. Quantification of *Pck1* expression in the liver shown as a positive control. Eight-week-old male c57BL/6J mice (N=3, PBS-injected; N=3, insulin-injected). Two-tailed Student *t* tests. **f**, *Gpr151* expression in the liver of glucagon-injected mice compared to the livers of control mice (vehicle-injected), quantified by RT-qPCR. Quantification of *Ppargc1a* expression in the liver shown as a positive control. Eight-week-old male c57BL/6J mice (N=3, vehicle-injected; N=3, glucagon-injected). Two-tailed Student *t* tests.

To further determine the physiological regulation of *Gpr151*, we assessed *Gpr151* expression following the injection of insulin and glucagon, the major hormones which regulate carbohydrate metabolism. Insulin injection led to a significant downregulation of *Gpr151* expression in the white adipose tissue but not the liver (**Figure 2e**). In contrast, glucagon induced a significant upregulation of *Gpr151* in the liver (**Figure 2f**). These data indicate that *Gpr151* expression is regulated by feeding hormones in peripheral tissues involved in glucose metabolism. However, the molecular mechanisms behind the regulation of *Gpr151* by feeding in adipose tissue and liver are distinct. Given the high hepatic expression of *Gpr151* compared with the adipose tissue, *Gpr151* in the liver is likely to be more physiologically important (**Figure 2a**).

To further characterize the regulation of *Gpr151* expression in the liver, we investigated and identified transcription factor binding sites upstream of the human *GPR151* transcript for the cAMP-responsive element-binding protein (CREB) and glucocorticoid receptor (GR) which are conserved in mice (**Extended Data Fig. 4a,b**). The activity of CREB is regulated by glucagon, insulin^21^ and glucocorticoids. Glucocorticoid blood levels increase during prolonged fasting and stimulate hepatic gluconeogenesis through GR^22^. We tested whether CREB and GR induce *Gpr151* expression in mouse hepatocyte AML12 cell line following cAMP induction by forskolin treatment, GR activation by dexamethasone treatment, or both. *Gpr151* expression increased by forskolin but not by dexamethasone (**Extended Data Fig. 4c**), and the induction of *Gpr151* expression by forskolin was abrogated by co-treatment with the CREB inhibitor 666-15^23^ (**Extended Data Fig. 4d**). Gene expression induction by forskolin was unique to *Gpr151*, compared to other genes encoding G_i_- interacting GPCRs that are known to be expressed in the liver (**Extended Data Fig. 4e**). We concluded that *Gpr151* expression in the liver may be at least partially upregulated by cAMP signaling through CREB. However, considering the strong upregulation of *Gpr151* expression observed in the livers of fasting mice (**Figure 2c**), additional regulators of *Gpr151* expression in addition to CREB might be present.

In conclusion, we found that *Gpr151* expression is downregulated in the liver and adipose tissue by feeding, which strongly suggests that the effects of *Gpr151* ablation on whole-body glucose metabolism is mediated by these peripheral metabolic tissues.

### *Gpr151* loss impairs hepatic glucose production by direct action on hepatocytes

To determine the mechanistic role of *Gpr151* in the liver, we verified the absence of *Gpr151* transcript in the livers of *Gpr151* KO mice using RT-qPCR (**Figure 3a, Extended Data Fig. 5**). We compared the liver transcriptomes from *Gpr151* WT and KO DIO mice using bulk RNA sequencing (RNA-Seq) (**Figure 3b**). The analysis revealed 79 significantly upregulated (p-val_adj_ <0.05, log2Fold change >1) and 338 significantly downregulated (p-val_adj_ <0.05, log2Fold change <(−1)) genes in *Gpr151* KO livers compared to WT littermates (**Figure 3c, Supplementary Table 1**). To gain biological insights from the transcriptome changes, we conducted Gene Set Enrichment Analysis (GSEA) using the Hallmark gene sets^24^, which revealed significant enrichment (FDR q-val < 0.25) in 32 gene sets in WT livers and no gene sets significantly enriched in KO livers (**Supplementary Table 1**). Surprisingly, the expression of genes in the glycolysis and gluconeogenesis pathway was decreased in KO livers (**Figure 3d**). Downregulation of the expression of several genes from this pathway in the livers of *Gpr151* KO mice was further validated using RT-qPCR in female littermate *Gpr151* WT and KO DIO mice (**Figure 3e**). Expression of *Ppargc1a* (*Pgc1a*), a gene encoding transcriptional coactivator PPARGC1A which regulates the expression of genes involved in energy metabolism^25^, was diminished in the livers but not in skeletal muscle or adipose tissue from the *Gpr151* KO mice compared to WT littermates in DIO, indicating liver-specific effects of *Gpr151* loss on energy metabolism (**Figure 3f**). Further, the expression of genes encoding hepatic gluconeogenesis enzymes which are directly regulated by PPARGC1A (PGC1-α), *Pck1* and *G6pc*^*26*^, was also significantly decreased in *Gpr151* KO DIO livers (**Figure 3g**). Altogether, these data revealed that *Gpr151* loss leads to transcriptional downregulation of genes involved in glycolysis and gluconeogenesis in the liver that is consistent with PGC1-α downregulation.

**Fig. 3.**
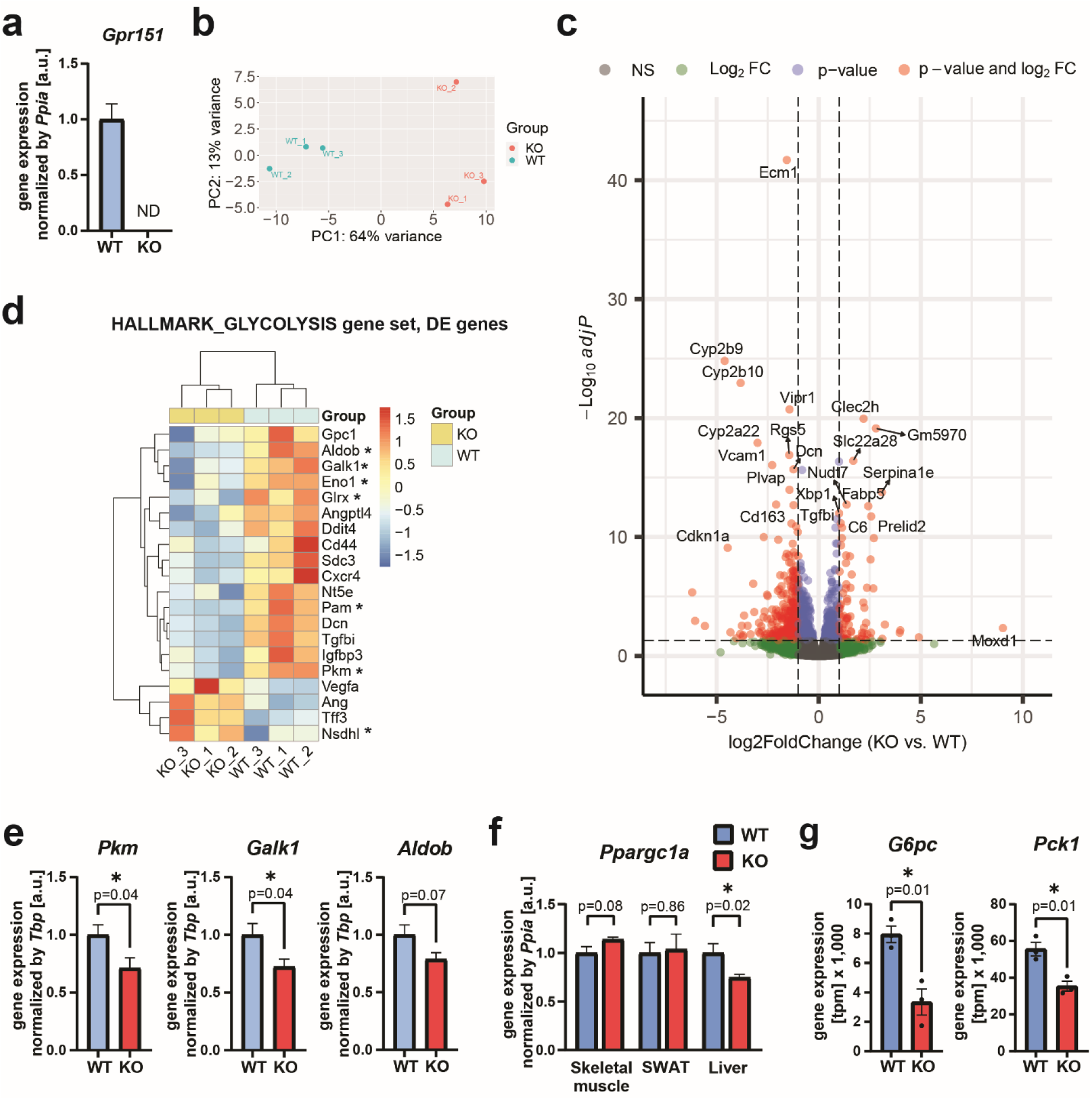
Gpr151 upregulates the expression of hepatic gluconeogenesis genes. **a**, RT-qPCR quantification of *Gpr151* gene expression in the livers of *Gpr151* WT and KO mice (N=3, WT; N=3, KO). **b**, Principal component analysis showing the clustering of samples analyzed by RNA-Seq. **c**, Volcano plot showing the statistical significance (-log10(p_adj_)) versus log2 of fold change of gene expression between *Gpr151* KO and WT livers. **d**, Heat map of transcriptional regulation patterns between DE genes (p-val_adj_<0.05) from the glycolysis/gluconeogenesis pathway within the HALLMARK gene sets. Genes which encode enzymes are marked with an asterisk. **e**, RT-qPCR quantification of selected genes from the glycolysis/gluconeogenesis pathway in 16-week-old DIO female mice (N=6, WT; N=6, KO). Two-tailed Student *t* tests. **f**, RT-qPCR quantification of *Ppargc1a* expression in the liver, skeletal muscle and adipose tissue of *Gpr151* WT and KO DIO male mice (N=4, WT; N=4, KO). Two-tailed Student *t* tests. **g**, Quantification of the expression of PPARGC1A target genes *G6pc* and *Pck1* in *Gpr151* WT and KO livers by RNA-Seq. N=3 mice / group. Two-tailed Student *t* tests (N=3, WT; N=3, KO).

To functionally test the direct effect of *Gpr151* loss on hepatic gluconeogenesis, we assessed glucose and lactate production in DIO *Gpr151* WT and KO mice following an injection of pyruvate. In line with the insights from the RNA-Seq in the liver, pyruvate tolerance testing resulted in lower blood glucose levels in KO compared to WT mice, consistent with a lower gluconeogenesis action in the liver of *Gpr151* KO mice (**Figure 4a**). Conversely, pyruvate administration resulted in elevated levels of blood lactate in *Gpr151* KO mice compared to WT littermates (**Figure 4b**), further supporting a decrease in hepatic gluconeogenesis in *Gpr151* KO mice. Plasma levels of glucagon, the pancreatic hormone which stimulates hepatic gluconeogenesis^27^, were not affected in *Gpr151* KO mice (**Figure 4c**), supporting that the effects of *Gpr151* loss on glucose metabolism are selective to the liver. In addition, there were no changes in the expression of glucagon receptor in the liver between *Gpr151* WT and KO mice (**Figure 4d**). To determine whether the impairment of hepatic gluconeogenesis by *Gpr151* loss is cell-autonomous, we assessed glucose secretion by glucagon-stimulated primary hepatocytes isolated from *Gpr151* WT and KO mice. *Gpr151* KO hepatocytes showed impaired glucose production compared to WT hepatocytes and did not increase glucose production in response to glucagon (**Figure 4e**), demonstrating that *Gpr151* KO hepatocytes have a cell-autonomous impairment in both basal and glucagon-induced hepatic gluconeogenesis. In addition, gross liver morphology and the levels of liver triglycerides, plasma triglycerides, total cholesterol, HDL-cholesterol and LDL-cholesterol were comparable in *Gpr151* KO mice and WT controls (**Extended Data Fig. 6**), indicating that in contrast to the differences in hepatic gluconeogenesis in *Gpr151* KO mice, there were no changes in liver lipid metabolism. In conclusion, we observe for the first time that loss of *Gpr151* directly impairs glucose production in primary hepatocytes *ex vivo* and in mice in a glucagon-independent way.

**Fig. 4.**
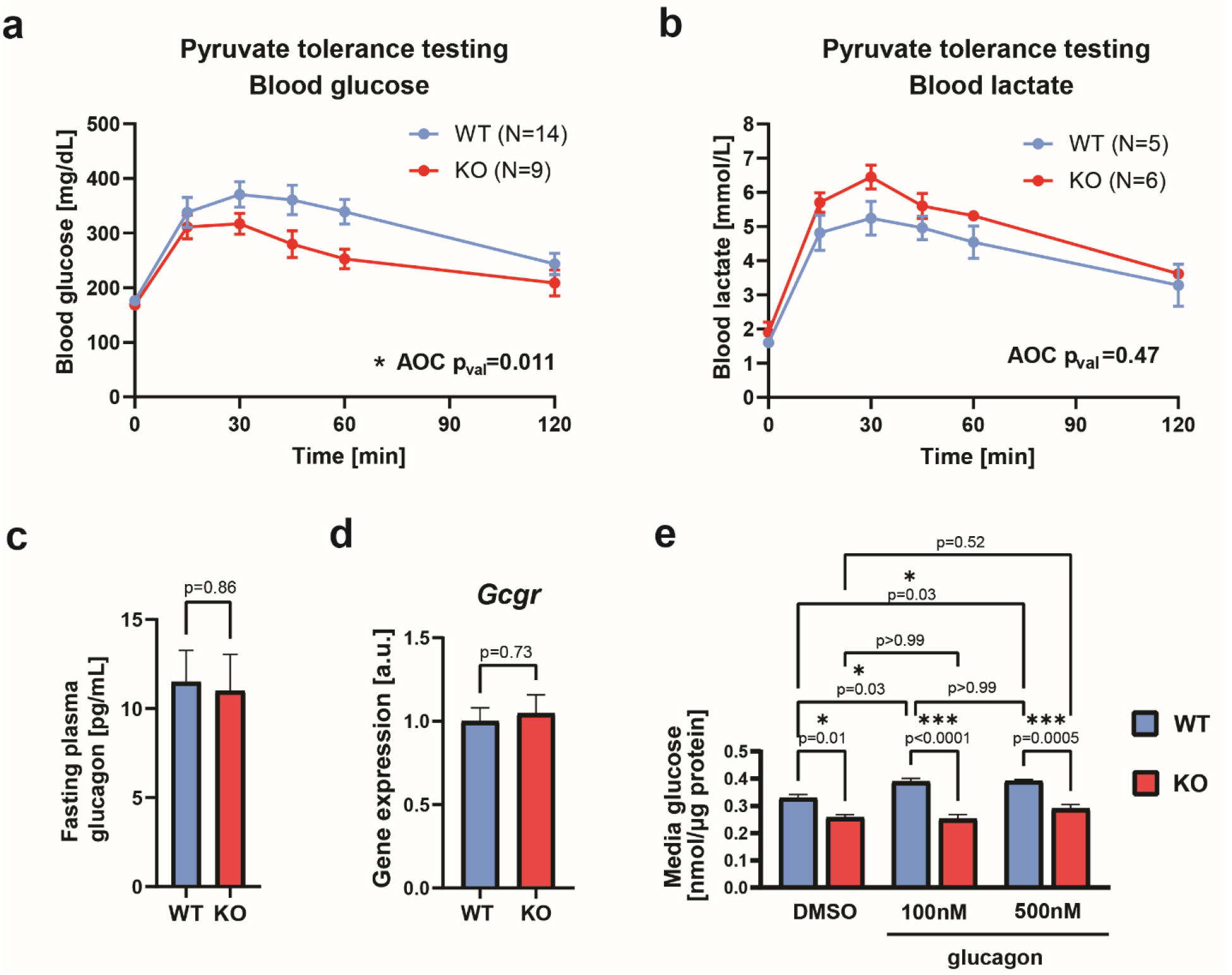
*Gpr151* regulates hepatic gluconeogenesis in a cell-autonomous manner. **a**, Blood glucose levels measured during pyruvate tolerance testing (PTT) in male DIO WT and KO mice. AOC compared using two-tailed Student *t* test (N=14, WT; N=9, KO). **b**, Blood lactate levels during PTT in male DIO WT and KO mice. AOC compared using two-tailed Student *t* test (N=5, WT; N=6, KO). **c**, Quantification of glucagon levels in blood plasma of fasted 16-week-old DIO mice by ELISA. Two-tailed Student *t* test (N=6, WT; N=7, KO). **d**, RT-qPCR quantification of glucagon receptor (*Gcgr*) gene expression in the livers from *Gpr151* WT and KO DIO male mice. Two-tailed Student *t* test (N=5, WT; N=5, KO). **e**, Quantification of media glucose produced by *Gpr151* WT and KO primary hepatocytes *ex vivo*, normalized by protein concentration. Results of one experiment representative for three independent experiments. Ordinary one-way ANOVA with Sidak’s multiple comparisons test. n=3 technical replicates.

### Gpr151 ablation results in a suppression of cAMP-dependent gene expression in the liver

To identify the molecular mechanism explaining the impairment in hepatic gluconeogenesis in *Gpr151* KO hepatocytes, we next focused on the regulation of hepatic gluconeogenesis gene expression in the liver. CREB regulates transcription of hepatic gluconeogenesis genes and is activated by cAMP through PKA^25^. In glucagon-injected mice, the levels of CREB phosphorylation were nominally but not significantly lower in *Gpr151* KO mice compared to WT controls (**Figure 5a, Extended Data Fig. 7a**). Additionally, there were no differences in insulin-dependent signaling between *Gpr151* WT and KO livers (**Extended Data Fig. 7b-d**), indicating specificity of the effect of *Gpr151* loss on glucagon signaling. To determine if there is orthogonal evidence for cAMP-dependent CREB activity dysregulation in the livers of *Gpr151* KO mice, we conducted a custom gene set enrichment analysis on liver RNA-Seq data, using a set of genes which were previously identified as regulated by cAMP signaling in mouse hepatocytes^28^ (**Supplementary Table 2**). The analysis revealed a significant (p-val = 9.5×10^−8^) enrichment of cAMP-regulated genes in WT livers, supporting an inhibition of cAMP-dependent transcription in *Gpr151* KO livers (**Figure 5b**). CREB-regulated genes, such as *Dusp1*^29^, *Ppp1r3c*^*30*^, and *Btg1*^31^ were significantly downregulated in *Gpr151* KO livers compared to WT (**Figure 5c**). Therefore, Gpr151 stimulates cAMP-dependent gene expression in the liver, including genes within hepatic gluconeogenesis pathway. In summary, *Gpr151* loss impairs hepatic glucogenesis regardless of the presence of glucagon, which likely explains its effects on whole-body glucose metabolism in diet-induced obesity (**Figure 5d**).

**Fig. 5.**
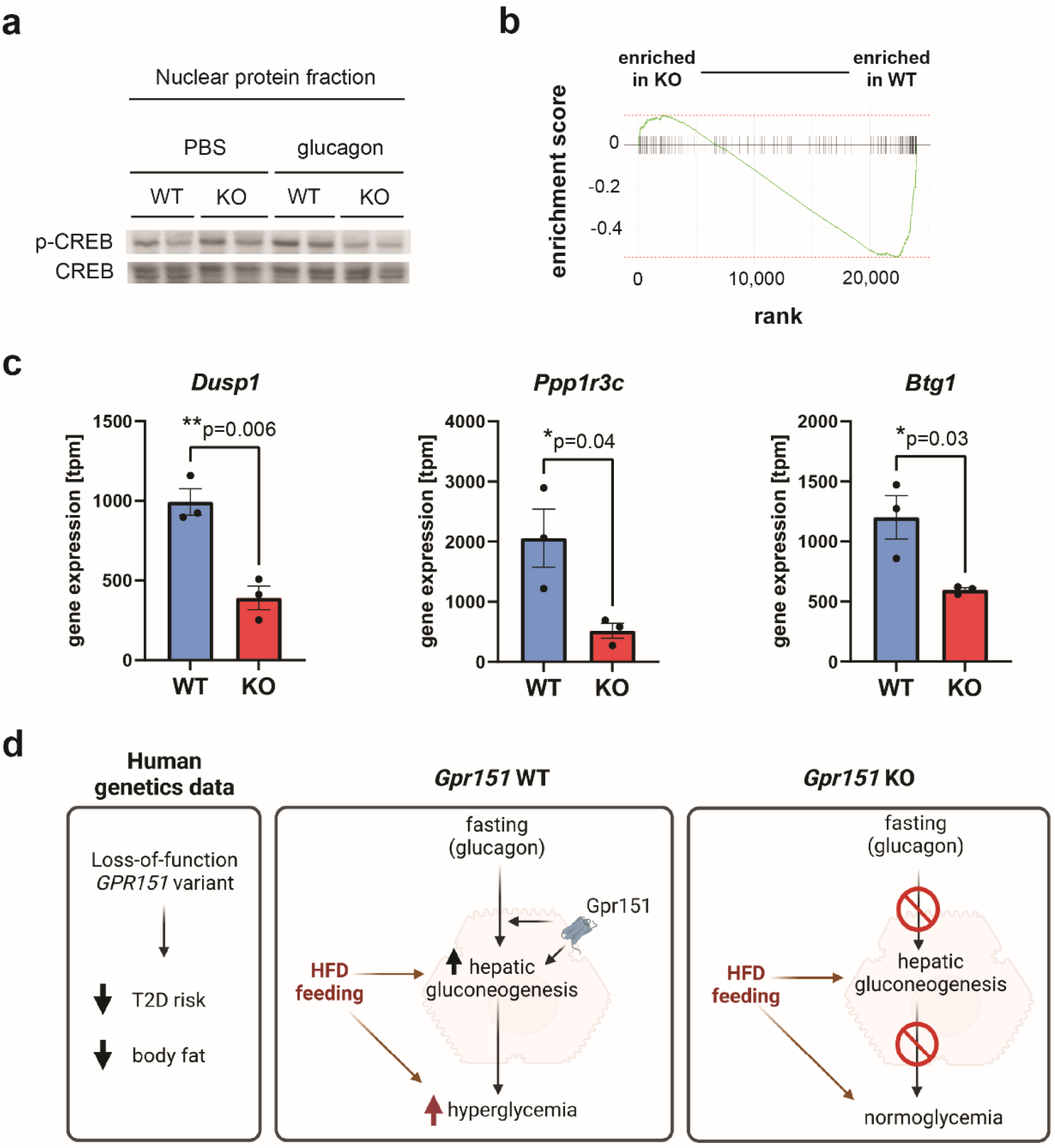
*Gpr151* loss results in a downregulation of cAMP-dependent gene expression. **a**, Representative Western blotting of phosphorylated CREB and total CREB in the nuclear protein fraction isolated from the livers of PBS- and glucagon-injected *Gpr151* WT and KO eight-week-old male mice. **b**, The results of custom gene set enrichment analysis of liver expression of 127 cAMP-responsive genes in the liver of *Gpr151* WT and KO mice. **c**, Quantification of the expression of CREB-regulated genes *Dusp1, Igfbp1, Btg1* in *Gpr151* WT and KO livers by RNA-Seq. N=3 mice / group. Two-tailed Student *t* tests (N=3, WT; N=3, KO). **d**, A model of the role of Gpr151 in metabolic health.

### Liver-specific overexpression of *Gpr151* reverses the metabolic phenotype associated with whole-body *Gpr151* loss

*Gpr151* loss leads to a decrease in basal and glucagon-dependent hepatic gluconeogenesis (**Figure 5e**). To determine whether improved glucose metabolism in *Gpr151* KO mice is attributable to Gpr151 function in the liver, recombinant adenovirus-associated virus serotype 8 (AAV8) was used to overexpress either *Gpr151* or green fluorescent protein (*GFP*) in the livers of DIO *Gpr151* KO mice (**Figure 6a**). As expected, viral transduction was restricted to the liver, and elevated *Gpr151* transcript levels about 100-fold compared to livers from wild-type mice (**Figure 6b**). Liver-specific overexpression of *Gpr151* did not lead to the expected decrease in glucose tolerance in *Gpr151* KO mice, although both groups of mice showed poor glucose tolerance (**Extended Data Fig. 8**).

**Fig. 6.**
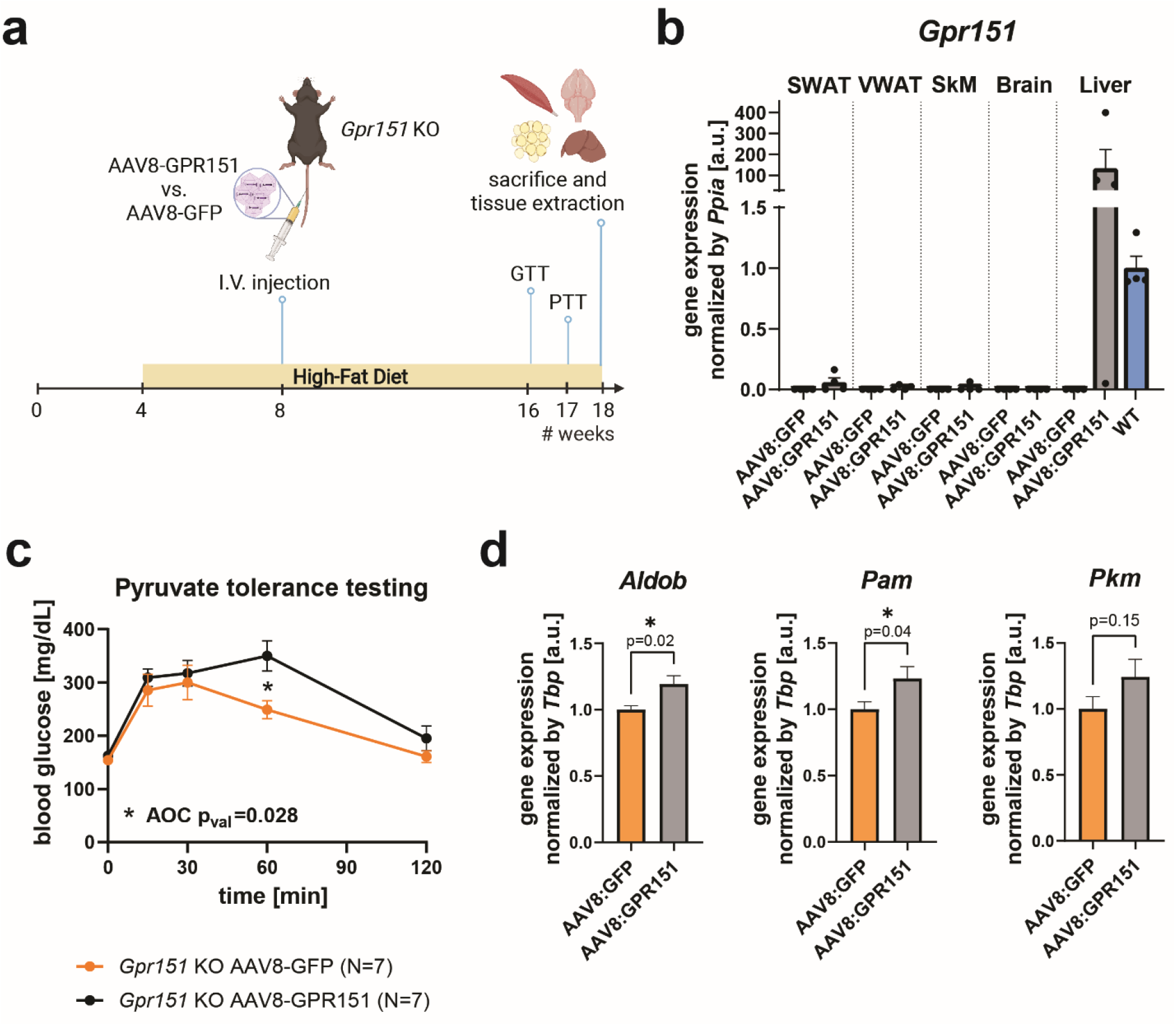
Overexpression of *Gpr151* in the liver of DIO *Gpr151* mice rescues the downregulation of gluconeogenesis. **a**, Schematic of the experiments involving liver overexpression of *Gpr151* using AAV8. **b**, Quantification of *Gpr151* overexpression by RT-qPCR in subcutaneous (SWAT) and visceral (VWAT) adipose tissue, skeletal muscle (SkM), brain and liver of *Gpr151* KO mice injected either with AAV8-GFP and AAV8-GPR151. Data normalized to *Gpr151* expression in the livers of age-, sex- and diet-matched wild-type mice (N=4, *Gpr151* KO AAV8:GFP; N=4, *Gpr151* KO AAV8:GPR151). **c**, Blood glucose levels in *Gpr151* KO DIO mice injected with AAV8-GPR151 or AAV8-GFP, subjected to pyruvate tolerance testing. AOC compared using two-tailed Student *t* test. Student *t* test with Bonferroni correction used to test differences at every time point (* t=60 q_val_=0.049; N=7 *Gpr151* KO AAV8:GFP; N=7, *Gpr151* KO AAV8:GPR151). **d**, RT-qPCR quantification of the expression of hepatic gluconeogenesis genes, identified as downregulated in *Gpr151* KO mice compared to WT, in the livers of *Gpr151* KO mice with liver-specific GPR151 and GFP overexpression by AAV8 (N=8, *Gpr151* KO AAV8:GFP; N=9, *Gpr151* KO AAV8:GPR151).

Remarkably, pyruvate tolerance testing revealed a significant increase in the amount of glucose present in the blood following an injection of pyruvate in the mice with liver-specific *Gpr151* over-expression compared to *GFP*-overexpressing controls (**Figure 6c**). This is strongly supporting a cell-autonomous role of liver Gpr151 in hepatic glucose production important to regulate physiological glucose metabolism. Furthermore, several hepatic gluconeogenesis genes were upregulated following *Gpr151* over-expression, consistent with the increased blood glucose levels following pyruvate injection (**Figure 6d**).

In summary, re-expression of *Gpr151* in the livers of whole-body *Gpr151* KO mice resulted in a reversal of the phenotype associated with the defect in hepatic gluconeogenesis, a pathway that can be successfully targeted in the treatment of T2D^11^. Therefore, due to its novel function in the liver that we have uncovered, GPR151 appears to be a promising target for antidiabetic treatments.

### Human *GPR151* loss-of-function carriers are characterized by favorable metabolic traits

Based on our observations *in vitro* and *in vivo*, we queried published GWAS studies for associations between the human *GPR151* p.Arg95Ter loss-of-function variant (*rs114285050*, allele frequency 0.8% in European ancestry) and metabolic traits as summarized in **Table 1**. Previous studies in UK Biobank found that being a carrier of the *GPR151* loss-of-function variant was associated with reduced BMI and lower risk of T2D^6,7^. Both associations were confirmed by larger independent GWAS studies^32,33^. In addition, there is indication of associations between the *GPR151* loss-of-function variant with improved lipid profile (reduced triglycerides and increased HDL cholesterol)^34^, as well as reduced waist-hip ratio adjusted by BMI (WHRadjBMI)^35^ (all p-value < 0.05). Furthermore, there is indication of directionality that *GPR151* loss-of-function is associated with improved glycemic traits (nominally reduced fasting glucose and reduced fasting insulin)^36^. These data strongly suggest that humans with a loss-of-function *GPR151* variant have improved metabolic health.

**Table 1.**
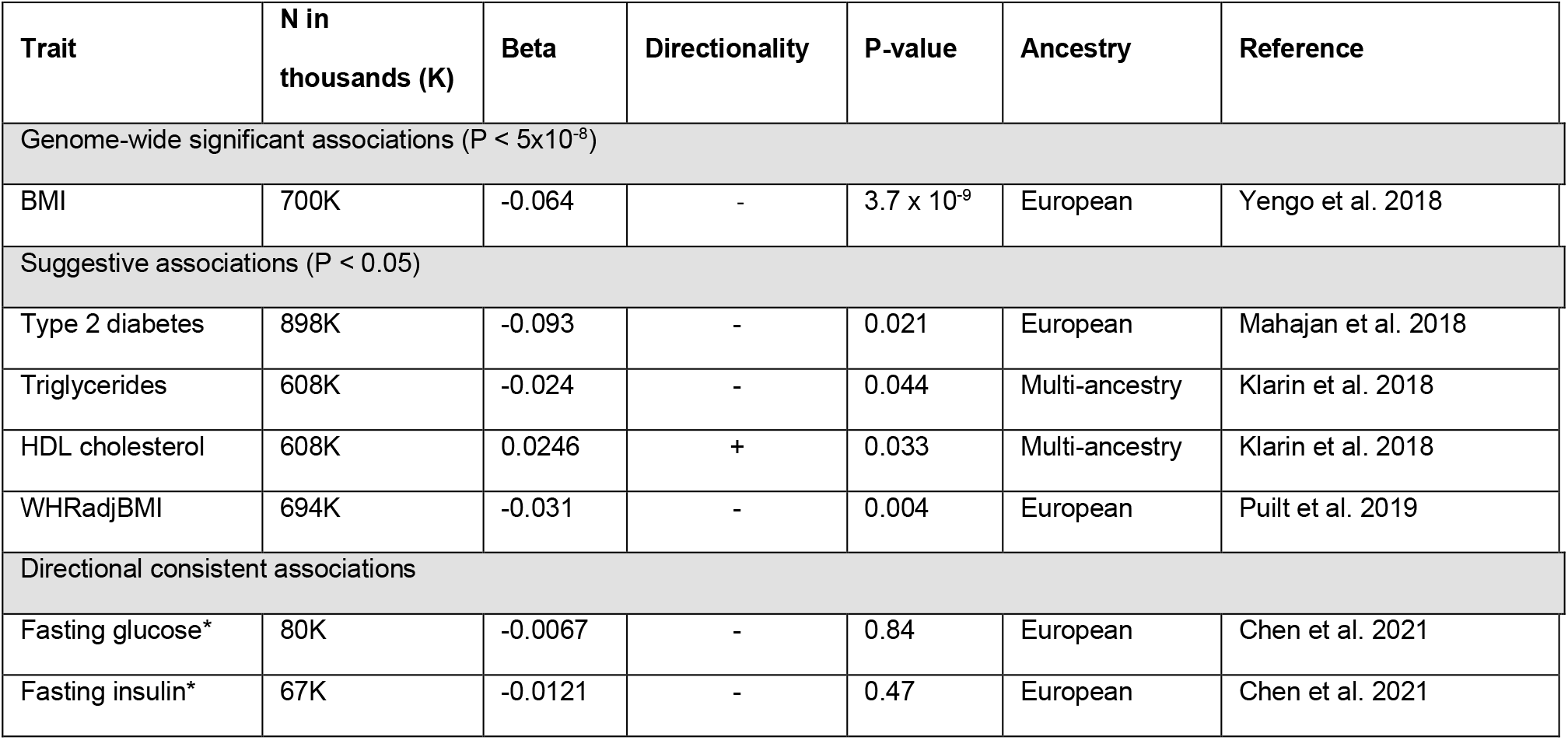
Results from query of GWAS studies with publicly available summary statistics. Directionality indicates the effect of the *GPR151* p.Arg95Ter loss-of-function variant (*rs114285050* A-allele is the effect allele, while the G-allele is the other allele) on the respective trait analyzed. *GPR151* rs114285050 chromosomal position is Chr5:146515831 (GRCh38). Abbreviations: N, Number of participants of respective GWAS study. *GWAS analyses were adjusted for BMI.

## Discussion

As the prevalence of IR and T2D keeps increasing worldwide, there is a need for the discovery of novel molecular targets and development of new therapies to target T2D, especially in those with insulin resistance. Here, we build on human genetics data to dissect the role of *Gpr151*in glucose metabolism and understand the mechanism behind the association between predicted loss-of-function variants in *GPR151* and lower risk of T2D.

We report that *Gpr151* knockout in mice leads to cell-autonomous lowering of hepatic glucose production. This pathway is also targeted by the first-line antidiabetic drug metformin.^37^ Although the molecular mechanism of metformin action in hepatocytes is not fully understood, it is known to affect mitochondrial respiration, leading to alterations in ATP:AMP ratios and inducing effects on AMPK signaling^38^. More recently AMPK-independent effects of metformin have also been identified^39^. Nevertheless, these pathways seem to be distinct from the Gpr151 function. The mechanism of hepatic gluconeogenesis regulation by GPR151 should be explored further to understand whether it can be utilized in an orthogonal way to augment currently available therapies for T2D. GPCRs are highly druggable molecular targets^40^. Therefore, GPR151 appears to be an exciting novel target candidate for T2D treatment.

In humans, the *GPR151* p.Arg95Ter loss-of-function variant is associated with reduced BMI and WHRadjBMI, lower risk of T2D, improved lipid profile and a directionally consistent reduction of fasting glucose and insulin. However, we are limited to searching for associations in available published GWAS studies that may not be sufficiently powered to detect genotype effects of *GPR151* p.Arg95Ter due to its low allele frequency (minor allele frequency < 1% in European populations), and therefore the reported effect sizes need careful interpretation. To quantify the clinical impact of the *GPR151* p.Arg95Ter loss-of-function variant on metabolic traits, future recall-by-genotype studies of carriers of this variant should be performed. Such a trial might include detailed oral glucose tolerance tests as a measure of glucose homeostasis and T2D prevalence as primary outcomes. If designed properly, studies of human *GPR151* p.Arg95Ter loss-of-function variant carriers may also provide information on molecular mechanisms of the role of GPR151 in its protective effect on metabolic traits.

Importantly, the role of *Gpr151* in the regulation of hepatic gluconeogenesis that we discovered is independent from the previously described role of *Gpr151* in the habenula, a brain structure that processes reward-related signals and affects appetite. Amongst peripheral metabolic tissues, *Gpr151* expression is regulated by feeding not only in the liver, but also in the white adipose tissue. Of note, the patterns of *Gpr151* expression in the liver and SWAT are distinct, indicating different mechanisms. In addition, liver-specific *Gpr151* overexpression reversed the hepatic gluconeogenesis phenotype in *Gpr151* KO mice but not the whole-body glucose tolerance. Therefore, while hepatic Gpr151 regulates hepatic gluconeogenesis, this receptor likely has additional functions in other tissues which contribute to its effects on whole-body glucose metabolism. Whether Gpr151 function in the fat contributes to whole-body glucose metabolism remains to be studied.

In mouse hepatocytes, Gpr151 is regulated at the level of gene expression. In particular, the expression of hepatic *Gpr151* is upregulated by fasting and the activity of the cAMP-inducing glucagon receptor and is at least partially mediated by CREB. Consequently, *Gpr151* loss would be predicted to primarily affect glucagon-induced hepatic gluconeogenesis. However, we instead observed that *Gpr151* KO hepatocytes show a general decrease in hepatic gluconeogenesis, independent of glucagon stimulation. This contradiction may be due to a chronic downregulation of hepatic gluconeogenesis in *Gpr151* KO hepatocytes and is supported by the decrease in hepatic gluconeogenesis when hepatic Gi signaling is inhibited in general^17^. Molecular mechanism underlying this paradoxical gluconeogenesis impairment by the loss of cAMP-inhibiting GPCRs remains to be understood and could be utilized to design antidiabetic treatments.

Taken together, we show here that GPR151 may be a promising target for the treatment of T2D by affecting hepatic gluconeogenesis. In the future, development of molecular tools, in particular of an inverse agonist to GPR151, would allow for assessment of the feasibility of such approach.

## Acknowledgements

We would like to thank Dr. Erik Ingelsson for his support of this project. We also thank the members of Knowles and Svensson labs for valuable discussions and feedback. We would like to thank Dr. Ines Ibañez-Tallon for sharing the *Gpr151* KO mice. We acknowledge the support of the Genetics Bioinformatics Service Center at Stanford University.

The study was supported by a grant from the National Institute of Diabetes and Digestive and Kidney Diseases of the National Institutes of Health (R01DK120565 to J.W.K.). K.J.S. was supported by NIH grants DK125260, DK120565, P30DK116074, Merck, the Jacob Churg Foundation, the McCormick and Gabilan Award, and the Stanford Cardiovascular Institute. E.B.M. and M.Z. were supported by the American Heart Association (AHA) postdoctoral fellowships (AHA Award Numbers: 18POST34030448 to E.B.M., 905674 to M.Z.). T.M.S was funded by a grant from the Novo Nordisk Foundation and the Stanford Bio-X Program (NNF19OC0054265). P.S. was funded by the Dean’s fellowship at Stanford University.

## Author contributions

E.B.M. designed the project, performed most experiments, analyzed data and wrote the manuscript with J.W.K and K.J.S. who both supervised the project; M.Z. performed major experiments; P.H.Z. conducted CLAMS experiments and analyzed the data; J.L., H.K., P.S., C.P., and C.C. contributed experimental work or data analysis; P.N. analyzed RNA-Seq data; T.M.S. queried GWAS summary statistics; A.S. supervised CLAMS experiments; E.B.M., J.W.K. and K.J.S. conceived the project. All authors reviewed the manuscript.

## Methods

### Animals

*Gpr151* KO mice on the C57BL/6J genetic background, described previously^9^, were crossed with C57BL/6J mice (#000664, Jax) to obtain heterozygous animals which were further in-crossed. For experiments conducted exclusively on wild-type mice, male C57BL/6J mice (#000664, Jax) were purchased. All animal studies were approved by the Administrative Panel on Laboratory Animal Care at Stanford University, and were performed according to the guidelines of the American Association for the Accreditation of Laboratory Animal Care.

Unless indicated otherwise, mice were housed with *ad libitum* access to chow and water in an air-conditioned room with a standard 12-hour light/dark cycle. For studies on diet-induced obesity, mice were fed a standard diet (18% protein / 6% fat, #2918, Envigo Teklad) for the first 4 weeks, followed by either high-fat diet (HFD, 60 kcal% Fat, D12492, Research Diets) or control sucrose-matched diet (SD, 10 kcal% Fat, D12450J, Research Diets) for the remainder of the experiment. Body weight and chow intake were monitored weekly.

To quantify the changes in gene expression in response to insulin *in vivo*, eight-week-old c57BL/6J male mice were fasted for 4 h, injected i.v. with either PBS or insulin (Humulin, Eli Lilly) at 1 U/kg body weight and sacrificed after 4 h. To assess protein phosphorylation in response to insulin, 16-week-old male *Gpr151* WT and KO mice fed HFD for 12 weeks were injected i.v. with 1 U/kg body weight (Humulin, Eli Lilly) diluted in PBS or vehicle control (PBS) and sacrificed after 10 min.

To assess gene expression changes in response to glucagon, eight-week-old male c57BL/6J mice were injected i.p. with 2 mg/kg body weight glucagon (Sigma-Aldrich, #G2044) in PBS or vehicle control and sacrificed after 60 min. To assess protein phosphorylation in response to glucagon, eight-week-old male *Gpr151* WT and KO mice fed HFD for 4 weeks were injected i.v. with either 2 mg/kg body weight glucagon (Sigma-Aldrich, #G2044) diluted in PBS or vehicle control (DMSO in PBS) and sacrificed after 10 min.

### Indirect Calorimetry

For measurements of metabolic rate and food intake, approximately 16-week-old male mice fed HFD for 12 weeks were placed within the CLAMS (Columbus Instruments) indirect calorimeter. Prior to the experiment, body composition of conscious mice was assessed with an EchoMRI 3-in-1 (Echo Medical Systems). Mice were acclimated to CLAMS for 13 h followed by 48 h measurement of VO_2_, VCO_2_, RER, locomotor and ambulatory activity, food intake at 23 ± 0.1°C while on HFD. Energy expenditure was calculated as previously described^41^. Calories consumed were calculated by multiplying hourly food intake by the 5.21 kcal/g caloric value of the 60% HFD. Energy balance was calculated by subtracting hourly food intake from hourly energy expenditure. Data from two separate experimental runs were combined. Analyses using ANOVA and ANCOVA were performed using CalR without the remove outliers feature^42^.

### Glucose, insulin and pyruvate tolerance testing

GTTs, ITTs and PTTs were performed on 16-week-old mice. GTTs were performed after an overnight fast (14-16h). 1.5 g glucose / kg body weight was injected i. p. ITTs were performed after 5-6h fasting. Mice were injected i.p. with insulin (Humulin, Eli Lilly) with various concentrations depending on the group (males receiving standard diet - 0.75 U/kg body weight, females receiving standard diet - 0.5 U/kg body weight, mice receiving high-fat diet - 1 U/kg body weight). PTTs were performed after an overnight fast (16-18h). 1 g sodium pyruvate / kg body weight was injected i.p. Blood glucose levels were measured from the tail vain using a glucometer (TRUEbalance, Nipro Diagnostics), and blood lactate levels were measured using Lactate Plus Meter (Nova Biomedical). For statistical comparisons, the area of the curve (AOC)^43^ was calculated after subtracting baseline level of the metabolite measured, followed by statistical testing using Student *t* test. If the difference in AOC was significantly significant, Student *t* test with Bonferroni correction was used for the follow-up comparison of metabolite levels at different time points.

### Systemic metabolic parameters

Blood was obtained by collection from vena cava from euthanized animals which had been fasted for 5-6h. Heparin (Sigma Alrich, #H3393) solution in PBS was used to wash syringe. Collected blood was immediately centrifuged at room temperature to obtain plasma. ELISA was used to measure plasma glucagon (Mouse glucagon ELISA kit, Crystal Chem, #81518) and insulin (Ultra Sensitive Mouse Insulin ELISA kit, Crystal Chem, #90080). Plasma triglycerides, HDL and LDL were measured by the Diagnostic Laboratory at the Department of Comparative Medicine at Stanford University.

#### Gpr151 overexpression in vivo

Adeno-associated viruses serotype 8 (AAV8) encoding *GFP* (AAV8-GFP) and *Gpr151* (AAV8-GPR151) were purchased from Vector Biolabs. HFD-fed *Gpr151* KO male mice were injected intravenously with 5 × 10^10^ vg / mouse at 8 weeks of age. Following the injection, GTT was performed at 16 weeks of age and PTT was performed at 17 weeks of age. Mice were sacrificed at 18 weeks of age.

### Cell culture

AML12 mouse hepatocyte cell line was obtained from ATCC and cultured in Dulbecco’s Modified Eagle Medium/Nutrient Mixture F-12 (Invitrogen, #11330057) with the addition of 10% Fetal Bovine Serum (BenchMark Fetal Bovine Serum, GeminiBio, #100-106), 1x Insulin-Transferrin-Selenium (ITS-G, Gibco, #41400045), 40 ng/ml dexamethasone (Sigma, #D4902), and 1x Pen/Strep (Thermo Fisher Scientific, #15140163). Cells were cultured in a humidified 5% CO_2_ incubator. Serum starvation was carried out overnight in DMEM/F-12 with the addition of 1x Pen/Strep.

Primary hepatocytes were isolated from eight-week-old male mice and cultured as previously described^44^. For each sample, hepatocytes isolated from two mice of the same genotype were pooled. All experiments were performed within 36 h of hepatocyte isolation.

Cell were stimulated with 666-15 (Tocris, #5661), forskolin (Sigma Aldrich, #F6886), glucagon (Sigma Aldrich, #G2044), dexamethasone (Sigma Aldrich, #D4902).

Media glucose was quantified using Glucose Assay Kit (Abcam, ab65333) according to manufacturer’s protocol. Glucose levels were normalized using Pierce BCA Protein Assay Kit (Thermo Fisher Scientific, #23209).

### Liver triglyceride measurement

Liver triglycerides were quantified using Triglyceride Assay Kit (Abcam, ab65336) according to manufacturer’s protocol, and normalized by tissue sample weight.

### Histology

For hematoxylin and Eosin (H&E) staining tissues were harvested, weighed and fixed in 10% Neutral Buffered Formalin (Sigma Aldrich, #HT501320) for 72h, following by dehydration in 70% ethanol. Dehydrated tissues underwent standard H&E staining at the Stanford Animal Histology facility, using the following steps: xylene (2 min x 3), 100% ethanol (2 min), 100% ethanol (1 min), 95% ethanol (1 min x 2), 80% ethanol (1 min), running tap water (1 min), Harris hematoxylin pH 2.5 (10 min), running tap water (1 min), 1% HCl/70% ethanol (20s), running tap water (5 min), 0.5% ammonium hydroxide/deionized water, running tap water (3 min), 95% ethanol (1 min), eosin (95% ethanol solution pH 4.6, 2 min), 95% ethanol (30s x 3), 100% ethanol (2 min x 3), xylene (2 min x 3), sealed with Cytoseal XYL. Histological images were collected using Zeiss Axioplan2 microscope at 20x magnification using Leica DC500 camera and NIS Elements software.

### RNA analyses

Total RNA was extracted from *in vitro* cell cultures using [RNAeasy Mini](https://www.zymoresearch.com/collections/direct-zol-rna-kits/products/direct-zol-rna-miniprep-kits) kit (QIAGEN, #74106) or from tissue using Ambion TRIzol Reagent (Thermo Fisher Scientific, #15-596-018) according to the manufacturer’s instructions. To remove any contaminating DNA, RNA was treated with Invitrogen TURBO DNA-free Kit (Invitrogen, #AM1907), according to the manufacturer’s instructions. Next, RNA was converted to cDNA using High-Capacity cDNA Reverse Transcription Kit (Applied Biosystems, #4374966). Quantitative PCR was conducted using TaqMan Fast Advanced Master Mix (Thermo Fisher Scientific, #4444557) and performed on ViiA 7 Real-Time PCR System (Thermo Fisher Scientific). All data were normalized to the expression of the housekeeping genes cyclophilin A (*Ppia*) or TATA-box binding protein (*Tbp*). The TaqMan assays (Integrated DNA Technologies) were: *Adra2a* (Mm.PT.58.33590743.g), *Adra2b* (Mm.PT.58.6098909.g), *Adra2c* (Mm.PT.58.30363072.g), *Aldob* (Mm.PT.58.42159810), *Cnr1* (Mm.PT.58.30057922), *Fgf21* (Mm.PT.58.29365871.g), *G6pc* (Mm.PT.58.11964858), *Galk1* (Mm.PT.58.7143435), *Gcgr* (Mm.PT.58.16192096), *Gpr151* (Mm00808987_s1), *Pam* (Mm.PT.58.8164623), *Pck1* (Mm.PT.58.11992693), *Pkm* (Mm.PT.58.6642152), *Ppargc1a* (Mm.PT.58.28716430), *Ppia* (Mm02342430_g1), *Tbp* (Mm.PT.39a.22214839). *LacZ* Taqman Gene Expression Assay was ordered from Thermo Fisher Scientific (Mr03987581_mr).

For RNA-Seq, RNA was extracted from snap-frozen liver tissue using TRIzol according to manufacturer’s protocol. Library preparation and sequencing was conducted by Novogene using the mRNA-Seq pipeline, with over 20 million raw reads per sample and Q30 of over 96%. Raw paired-end FASTQ files were filtered to remove reads with adaptor contamination, reads when uncertain nucleotides constitute more than 10% of either read (N > 10%), and reads when low quality nucleotides (Base Quality less than 5) constitute more than 50% of the read. STAR 2.6.1d^45^ was used to align clean reads to the mouse reference genome (Ensembl Mus Musculus GRCm38.p6 GCA_000001635) and to generate gene counts. STAR output for each sample was combined for differential expression testing using DESeq2_1.32.0 R package installed in R v4.1, using default settings. PCA was performed on rlog-transformed data using DESeq2^46^. Gene set enrichment analysis was performed using GSEA v4.1.0 on Hallmark gene sets^24^ with Permutation type set to Gene set. Additionally, gene set enrichment analysis was performed on a custom gene set of 127 cAMP-responsive genes^28^ using fgsea v1.18.0^47^; specifically, the fgseaMultilevel method was used with default settings. Both gene set enrichment analyses were performed using DESeq2 normalized counts. RNA-Seq data have been deposited with the Gene Expression Omnibus (#GSE196535).

### Western blotting

Tissue samples were snap-frozen in liquid nitrogen, ground using mortar and pestle and lysed in T-PER Tissue Protein Extraction Reagent (Thermo Fisher Scientific, #78510) with Halt Protease and Phosphatase Inhibitor Cocktail (Thermo Fisher Scientific, #78445). Nuclear and cytoplasmic protein fractions were isolated using Nuclear Extraction kit (Abcam, ab219177) according to manufacturer’s protocols. Protein concentrations were determined using Pierce BCA Protein Assay Kit (Thermo Fisher Scientific). Samples were subjected to SDS-PAGE in polyacrylamide gels (4-20% Mini-Protean TGX, Bio-Rad, #4561094) and transferred onto 0.45 µm PVDF membrane (Thermo Fisher Scientific, #88518) using Mini-PROTEAN system (Bio-Rad). Proteins were detected using primary antibodies against CREB (Cell Signaling, #9197, 1:1,000), p-CREB (Millipore Sigma, #06-519, 1:1,000), Akt (Cell Signaling, #58295, 1:1,000), phospho-Akt (Cell Signaling, #4060, 1:1,000), S6 (Cell Signaling, #2317, 1:1,000), phospho-S6 (Cell Signaling, #2211, 1:1,000). Equal protein loading was verified using antibody against β-actin (Santa Cruz, sc-47778, 1:15,000). Secondary antibodies were: anti-rabbit IRDye 680RD (goat anti-rabbit IgG, #925-68071, Li-cor, 1:10,000), anti-mouse IRDye 800CW (goat anti-mouse IgG, #925-32210, Li-cor, 1:10,000), anti-mouse HRP (Cell Signaling, #7076, 1:10,000), anti-rabbit HRP (Cell Signaling, #7074, 1:10,000). HRP was visualized using SuperSignal West Pico PLUS Chemiluminescent Substrate (Thermo Fisher Scientific, #34579) and Supersignal West Femto Maximum Sensitivity substrate (Thermo Fisher Scientific, #34096) according to manufacturer’s protocol. Blots were imaged using Odyssey Fc imager (Li-cor). Protein expression was quantified using ImageJ 1.53c.

### Analysis of promoter sequences

Conserved predicted transcription factor-binding sites were identified using ConTra v3^48^ by the analysis of 1,000 bp promoter sequence upstream of human *GPR151* (ENST00000311104) transcript. Initial search was conducted using position weight matrices (PWMs) for glucocorticoid receptor (GR TRANSFAC20113,V$GR_Q6,M00192), FOXO1 (FOXO1 TRANSFAC20113,V$FOXO1_Q5,M01216), and CREB (CREB TRANSFAC20113,V$CREB_Q4_01,M00917) using core = 0.95, similarity matrix = 0.85 settings.

### Query of published summary statistics from GWAS analysis in humans

We queried summary statistics for GWAS of BMI and T2D to confirm previous reports. We queried selected GWAS of selected traits (WHRadjBMI, triglycerides, HDL cholesterol, fasting glucose and fasting insulin) to explore if our findings in vivo translate to human variant carriers. Summary statistics for GWAS of BMI, WHRadjBMI, fasting glucose and fasting insulin were downloaded from GWAS catalogue (https://www.ebi.ac.uk/gwas/) with the following accession codes: GCST006900 (BMI)^32^, GCST008996 (WHRadjBMI)^35^, GCST90002232 (fasting glucose)^36^, GCST90002238 (fasting insulin)^36^. The summary statistics from the GWAS of T2D^33^ were downloaded from the DIAGRAM consortium website (http://diagram-consortium.org/), and the summary statistics from the GWAS of triglycerides and HDL cholesterol^34^ were downloaded through dbGaP, with accession number phs001672.v1.p1. Association results for the *GPR151* p.Arg64Ter loss-of-function variant was retrieved based on rs-number (*rs114285050*) or position (Chr5:145895394 for GRCh37, Chr5:146515831 for GRCh38). All GWAS studies were performed assuming additive genetic effects. We aligned the effect allele of the *rs114285050* across studies (A-allele is the effect allele; G-allele is the other allele) to indicate directionality of effect for *GPR151* p.Arg64Ter loss-of-function variant carriers.

### Statistical analysis

Unless indicated, all data are expressed as mean +/- standard error of the mean (SEM). Student’s *t* test was used for single variables and one-way ANOVA with Bonferroni post hoc correction (or equivalent) was used for multiple comparisons using GraphPad Prism 7 software.

## Extended Data Figures

**Extended Data Fig. 1.**
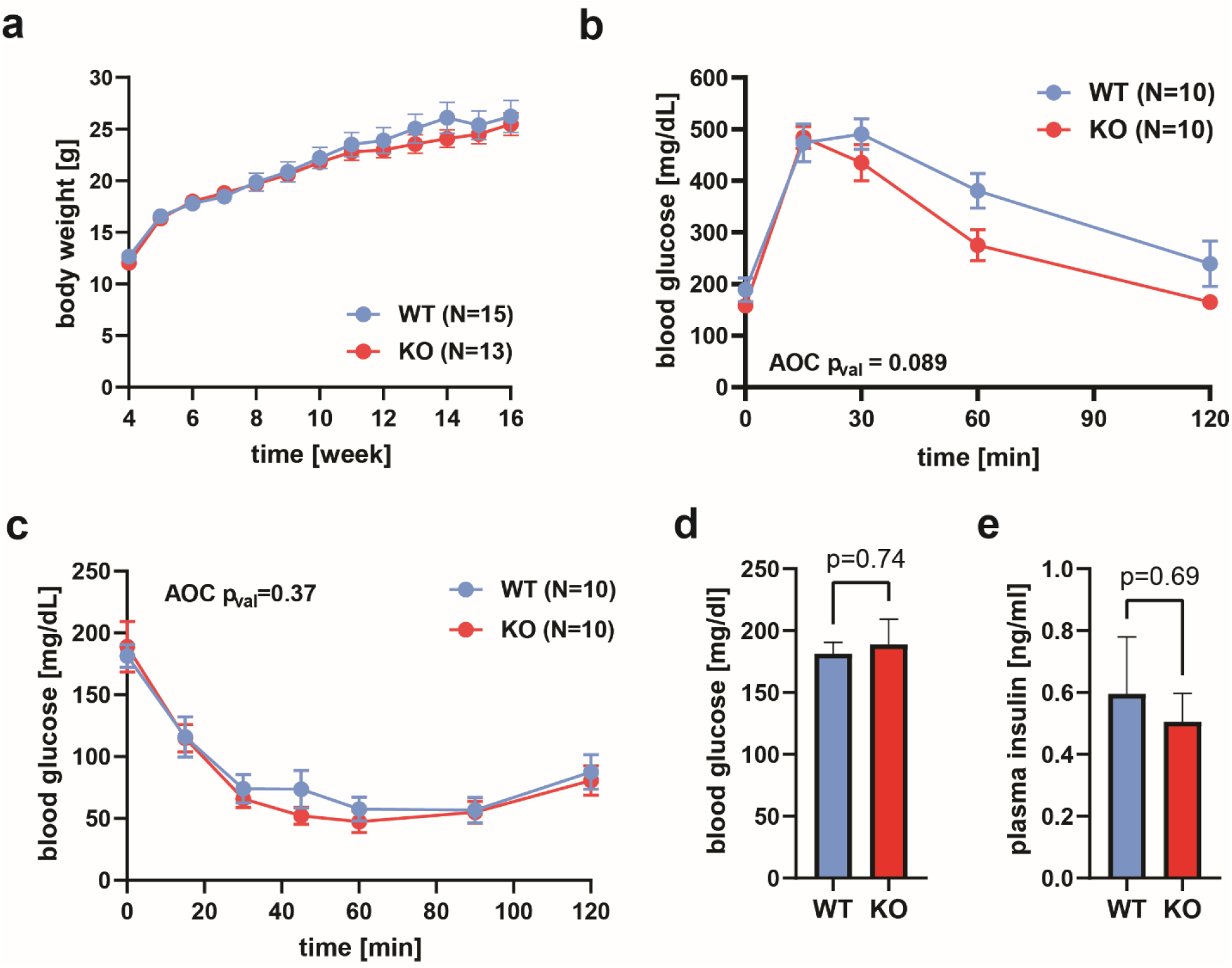
Supplementary data corresponding to Fig. 1 for DIO females. **a**, Body weight in female DIO KO and WT mice over 12 weeks of HFD (N=15, WT; N=13, KO). **b**, Blood glucose levels measured during glucose tolerance testing in *Gpr151* KO and WT DIO females. Area of the curve (AOC) compared using two-tailed Student *t* test (N=10, WT; N=10, KO). **c**, Blood glucose levels measured during insulin tolerance testing in *Gpr151* KO and WT in DIO female mice. AOC compared using two-tailed Student *t* test (N=10, WT; N=10, KO). **d**, Fasting glucose levels in *Gpr151* WT and KO DIO female mice measured in whole blood (N=10, WT; N=10, KO). Two-tailed Student *t* test. **e**, Fasting insulin levels measured in blood plasma of DIO female mice (N=7, WT; N=6, KO). Two-tailed Student *t* test.

**Extended Data Fig. 2.**
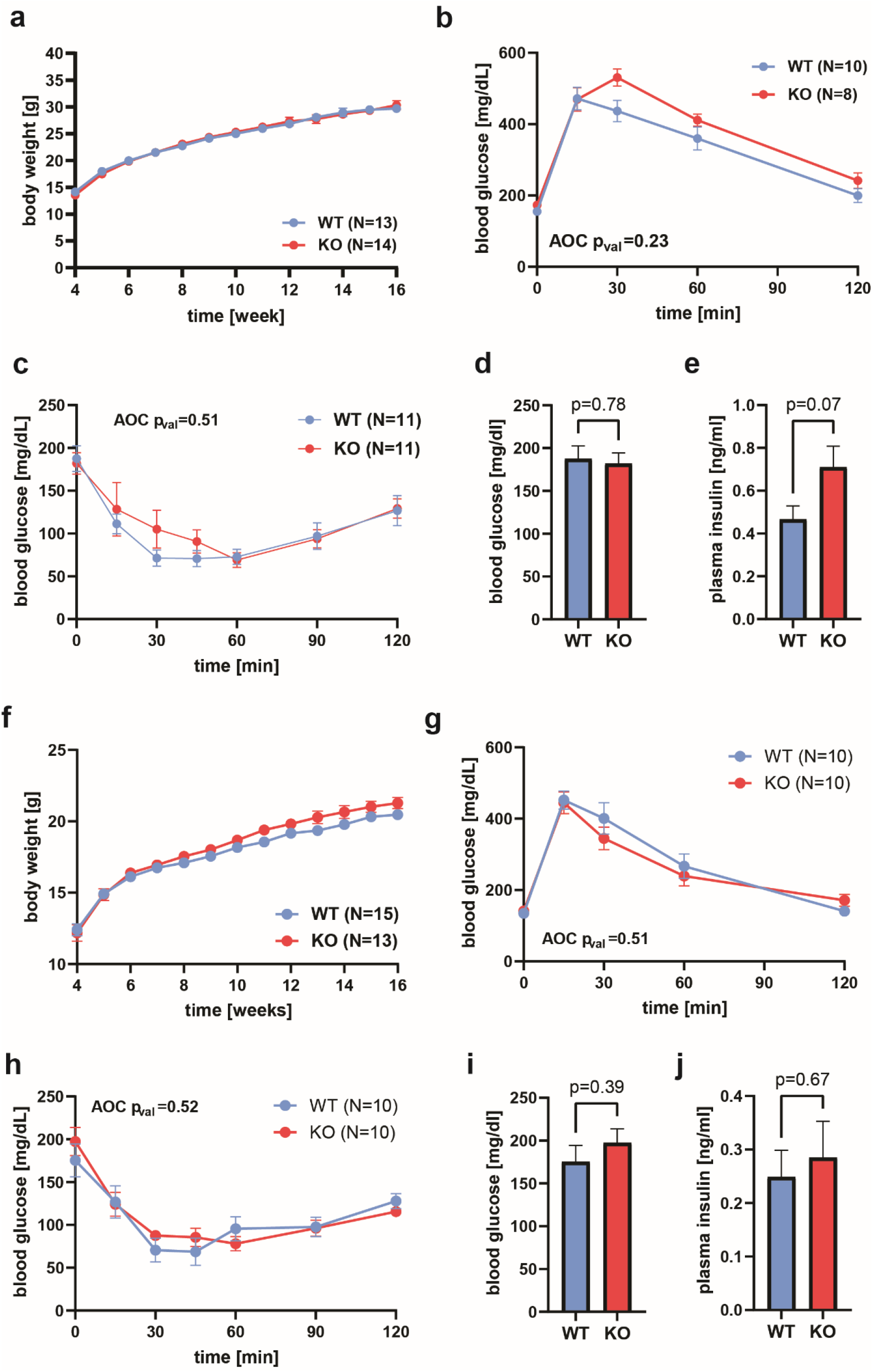
Supplementary data corresponding for Fig. 1 for SD-fed mice. **a-e**, SD-fed males. **a**, Body weight in male KO and WT mice over 12 weeks of SD. **b**, Blood glucose levels measured during glucose tolerance testing in *Gpr151* KO and WT SD males. Area of the curve (AOC) compared using two-tailed Student *t* test. **c**, Blood glucose levels measured during insulin tolerance testing in *Gpr151* KO and WT in SD male mice. AOC compared using two-tailed Student *t* test. **d**, Fasting glucose levels in *Gpr151* WT and KO DIO SD-fed male mice measured in whole blood (N=12, WT; N=12, KO). Two-tailed Student *t* test. **e**, Fasting insulin levels measured in blood plasma of SD male mice (N=6, WT; N=7, KO). Two-tailed Student *t* test. **f-j**, SD-fed females. **f**, Body weight in female KO and WT mice over 12 weeks of SD. **g**, Blood glucose levels measured during glucose tolerance testing in *Gpr151* KO and WT SD females. Area of the curve (AOC) compared using two-tailed Student *t* test. **h**, Blood glucose levels measured during insulin tolerance testing in *Gpr151* KO and WT in SD female mice. AOC compared using two-tailed Student *t* test. **i**, Fasting glucose levels in *Gpr151* WT and KO DIO SD-fed female mice measured in whole blood (N=10, WT; N=10, KO). Two-tailed Student *t* test. **j**, Fasting insulin levels measured in blood plasma of SD female mice (N=7, WT; N=5, KO). Two-tailed Student *t* test.

**Extended Data Fig. 3.**
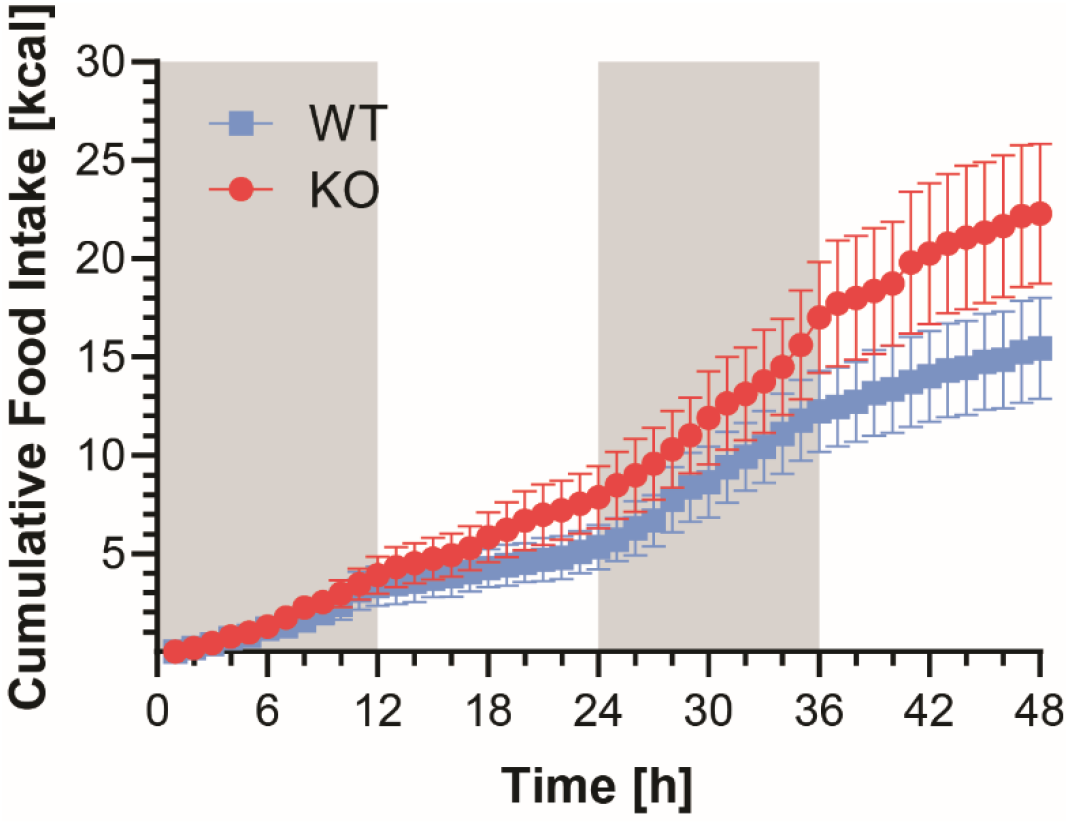
Increased food intake in *Gpr151* KO mice compared to WT littermates in the male DIO cohort of mice assessed using CLAMS. Representative graph of cumulative food intake over 48 h of measurement (N=5, WT; N=4, KO).

**Extended Data Fig. 4.**
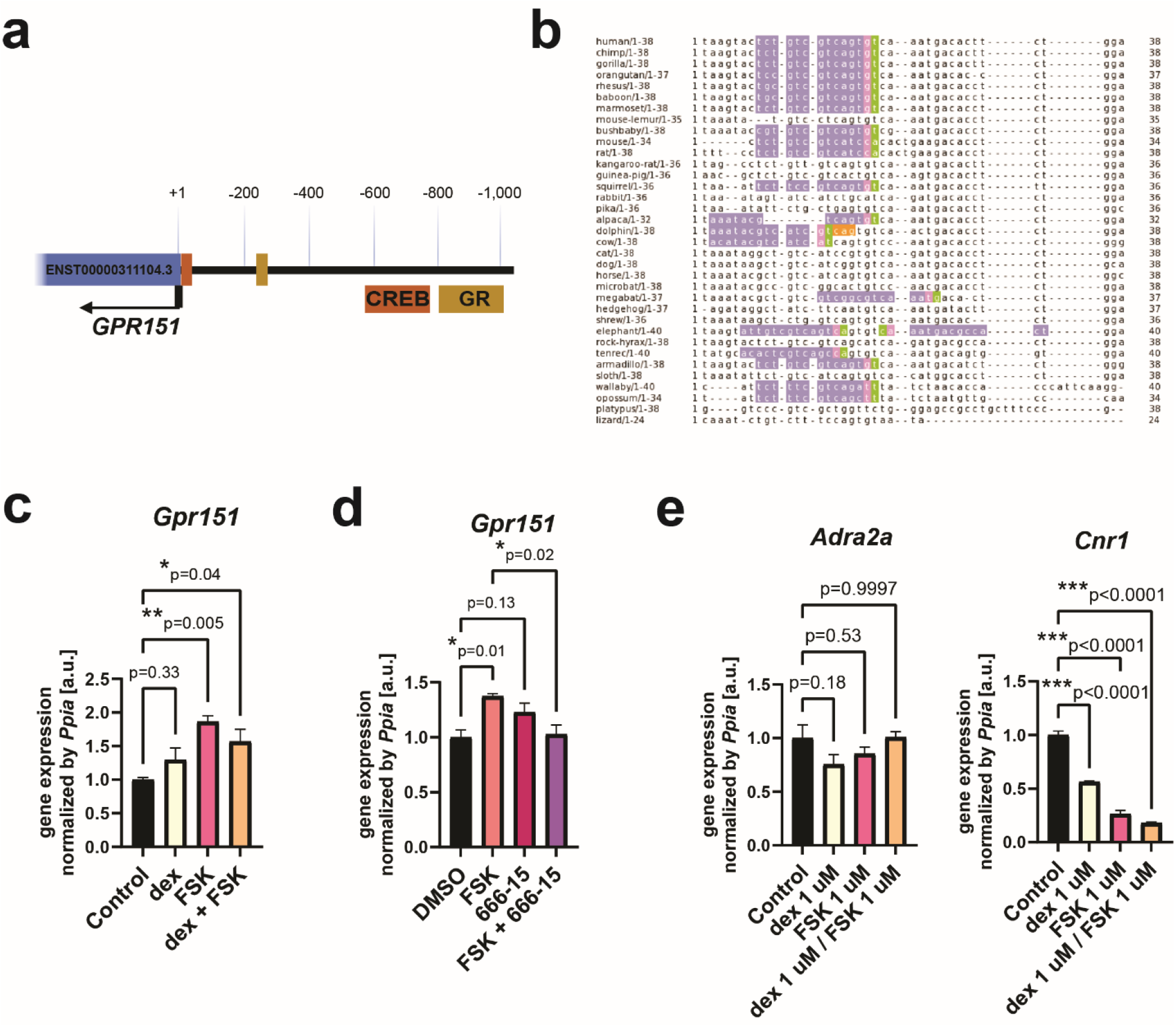
*Gpr151* expression is regulated by cAMP through CREB. **a**, Location of the conserved CREB and GR motifs in the proximal promoter of the human *GPR151* gene. The sites conserved between mouse and human genes are shown. **b**, Analysis of promoter sequence of *GPR151*, conducted using Contra v3 for 1kb upstream of the human transcript. The alignment of conserved sequences 1-38 bp downstream of TSS. CREB binding sites are highlighted. **c**, RT-qPCR quantification of *Gpr151* expression in AML12 cells following 2 h treatment with 1 μM dexamethasone (dex), 1 μM forskolin (FSK), or both. Data from one experiment representative for two independent experiments. n=3 technical replicates. Ordinary one-way ANOVA with Dunnett’s multiple comparisons test. **d**, RT-qPCR quantification of *Gpr151* expression following 3 h treatment with 10 μM FSK, 500 nM CREB inhibitor 666-15, or both. Data from one experiment representative for three independent experiments. n=3 technical replicates. Ordinary one-way ANOVA with Sidak’s multiple comparisons test. **e**, RT-qPCR quantification of the expression of *Adra2a* and *Cnr1* in response to 2 h treatment with dexamethasone, forskolin, or both. The expression of *Adra2b* and *Adra2c* was not detected. Data from one experiment representative for two independent experiments. n=3 technical replicates.

**Extended Data Fig. 5.**
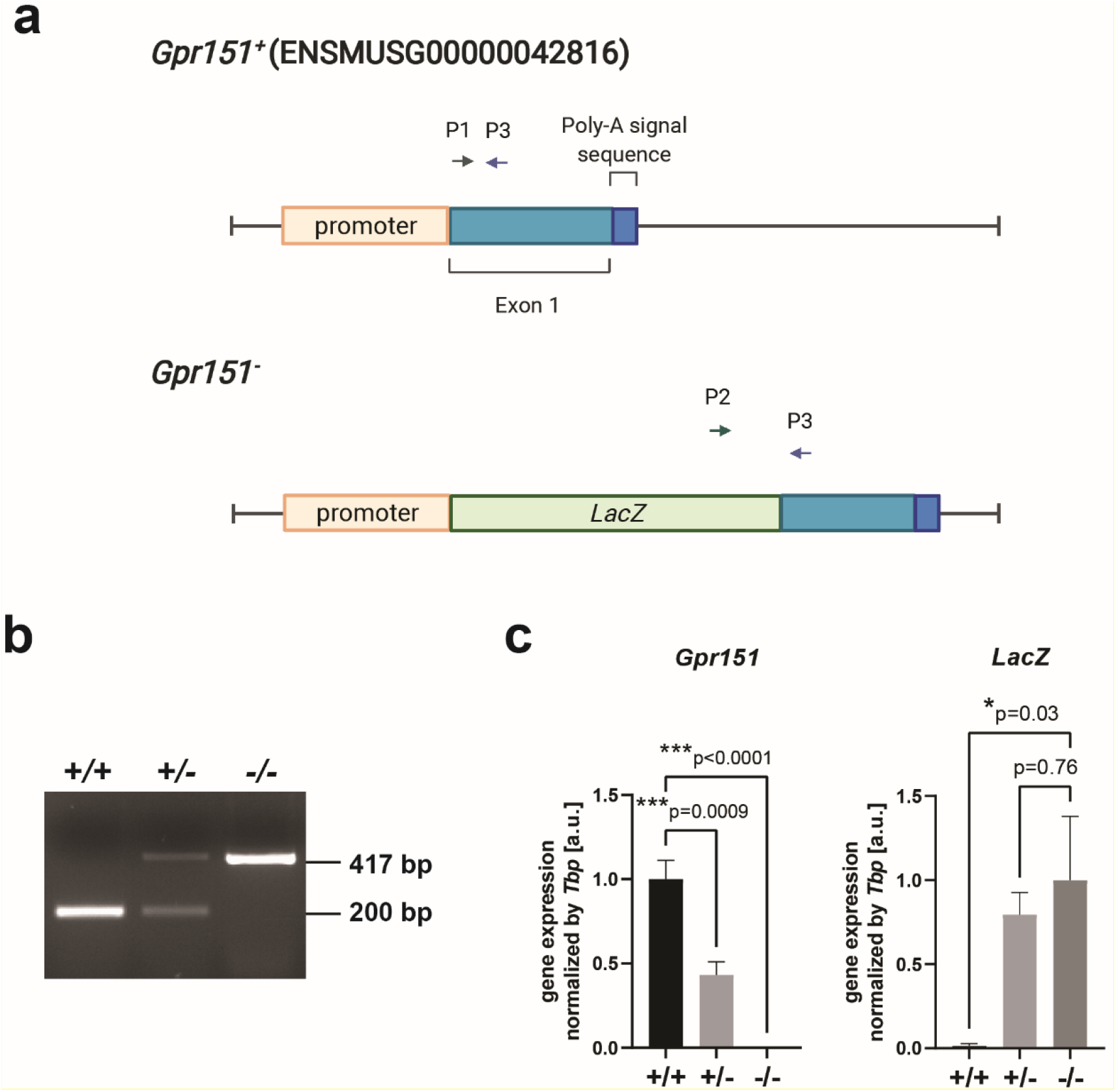
Analysis of the *Gpr151* KO allele in the liver. **a**, Schematic of the wild-type (*Gpr151*^*+*^) and knockout (*Gpr151*^*-*^) alleles of the *Gpr151* gene. Location of the primers used for genotyping (two forward primers P1 and P2 and a common reverse primer P3) is indicated. **b**, Representative image of agarose gel electrophoresis of the PCR genotyping product. Sanger sequencing of the PCR product for *Gpr151* KO (-/-) confirms the disruption of the gene. **c**, RT-qPCR quantification of *Gpr151* and *LacZ* expression in the livers of 16-week-old SD-fed male mice. Data normalized to the gene expression in WT and KO mice, respectively (N=4, WT; N=5, KO). One-way ANOVA with Dunnett’s multiple comparisons test.

**Extended Data Fig. 6.**
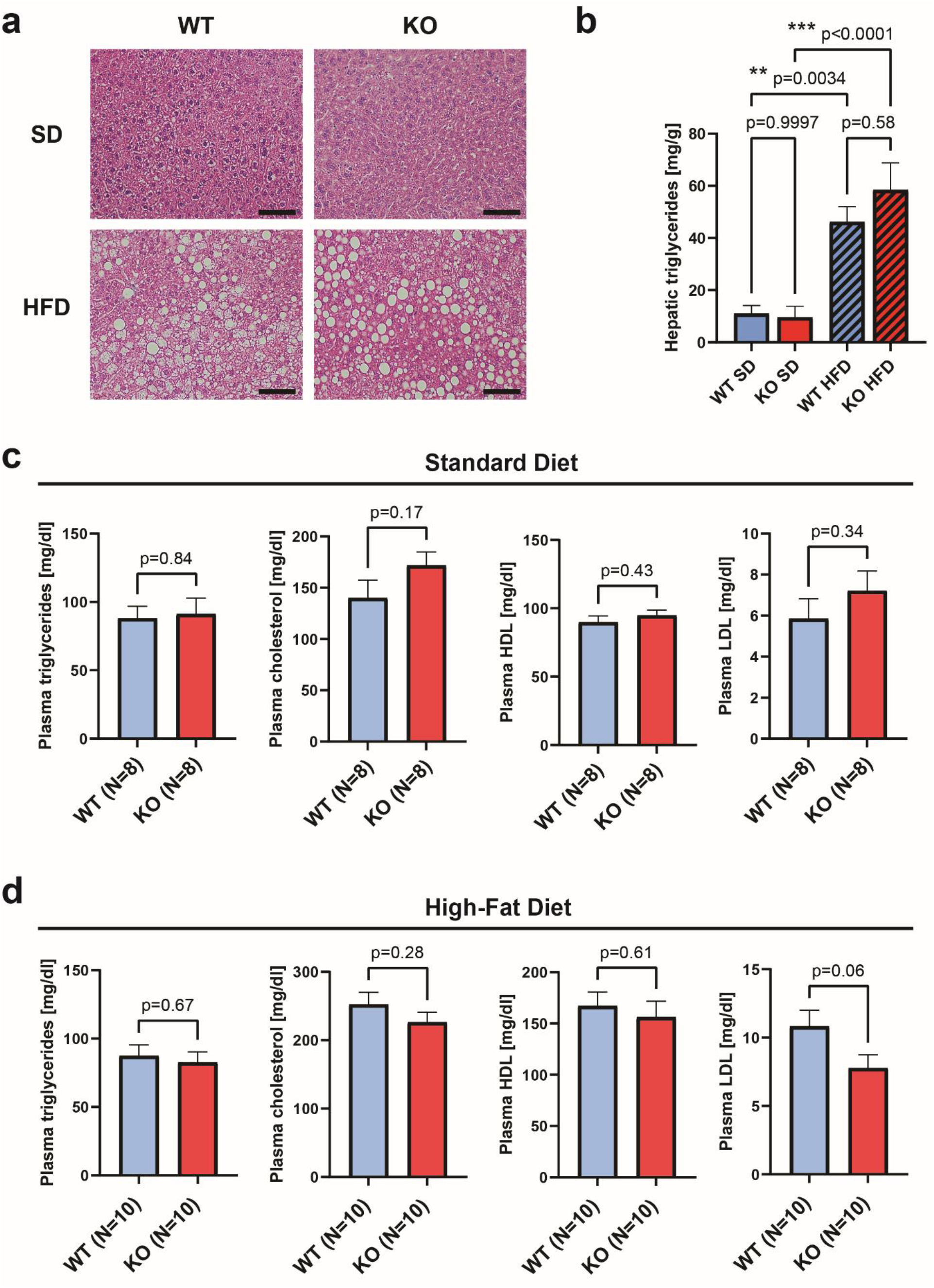
Analysis of liver morphology and lipid metabolism in *Gpr151* KO mice. **a**, Representative images of H&E staining of livers from SD- and HFD-fed 16-week-old male mice. Scale: 250 μm. Images representative for N=8, WT; N=8, KO. **b**, Quantification of liver triglycerides in SD- and HFD-fed 16-week-old male mice (N=7, WT; N=7, KO). Ordinary one-way ANOVA with Sidak’s multiple comparisons test. **c**, Quantification of total triglycerides, total cholesterol, HDL and LDL in the plasma of SD-fed male mice at 16 weeks of age. Two-tailed Student *t* tests. **d**, Quantification of total triglycerides, total cholesterol, HDL and LDL in the plasma of HFD-fed male mice at 16 weeks of age. Two-tailed Student *t* tests. The number of mice is indicated.

**Extended Data Fig. 7.**
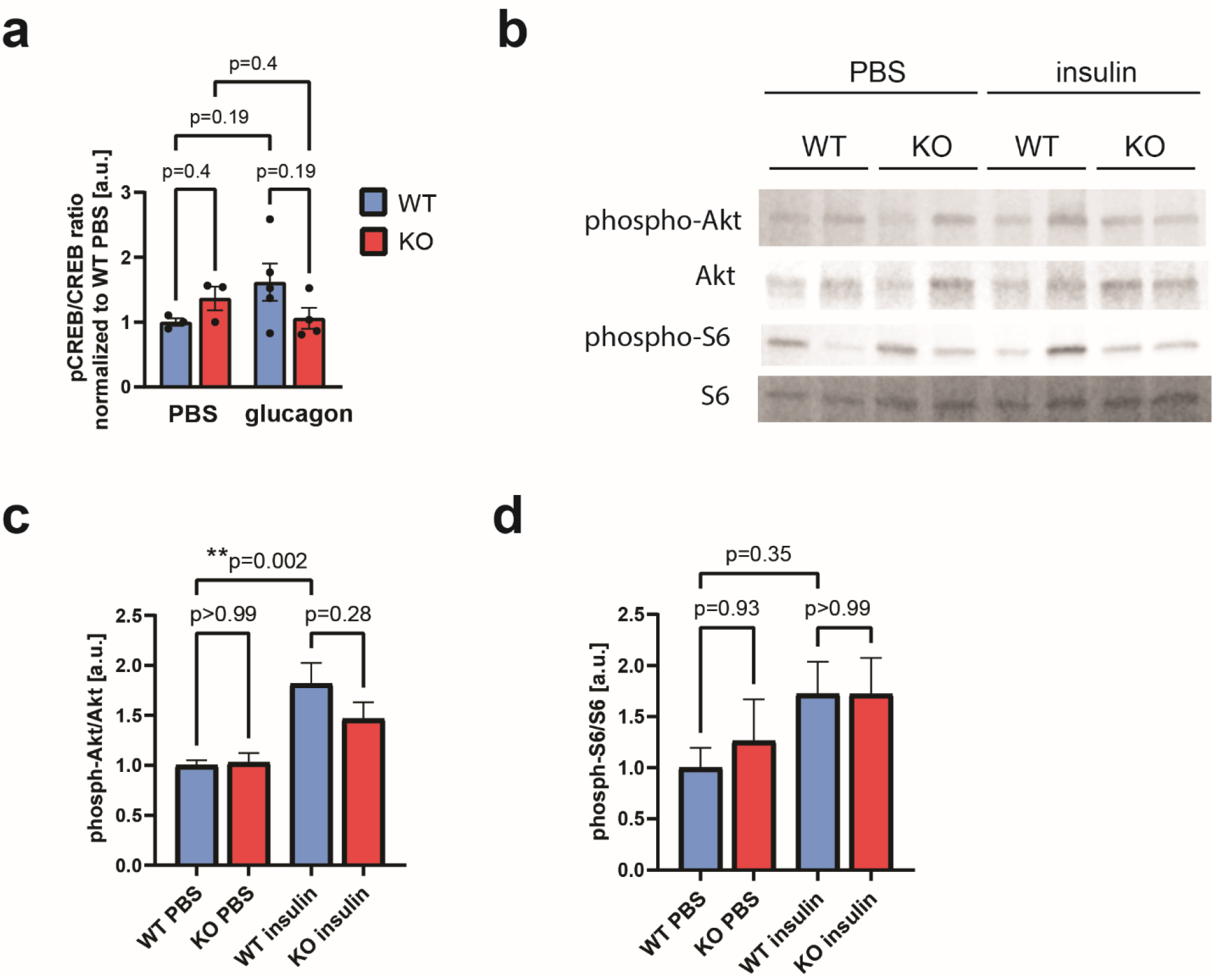
Downstream signaling from glucagon and insulin receptors in the livers of *Gpr151* WT and KO mice. **a**, Quantification of the ratio between phosphorylated CREB (Ser133) and total CREB, normalized to PBS-injected WT mice, in the livers of PBS- and glucagon-injected mice (N=3, WT PBS; N=3, KO PBS; N=5, WT glucagon; N=4, KO glucagon). Ordinary one-way ANOVA with Sidak’s multiple comparisons test. **b**, Representative Western blotting of phosphorylated S6, total S6, phosphorylated Akt and total Akt in the total protein fraction isolated from the livers of PBS- and insulin-injected *Gpr151* WT and KO eight-week-old male mice. **c**, Quantification of the phosphorylated to total Akt ratio in the livers of PBS- and insulin-injected mice (N=6, WT PBS; N=6, KO PBS; N=6, WT insulin; N=6, KO insulin). Ordinary one-way ANOVA with Sidak’s multiple comparisons test. **d**, Quantification of the phosphorylated to total S6 in the livers of PBS- and insulin-injected mice (N=6, WT PBS; N=6, KO PBS; N=6, WT insulin; N=6, KO insulin). Ordinary one-way ANOVA with Sidak’s multiple comparisons test.

**Extended Data Fig. 8.**
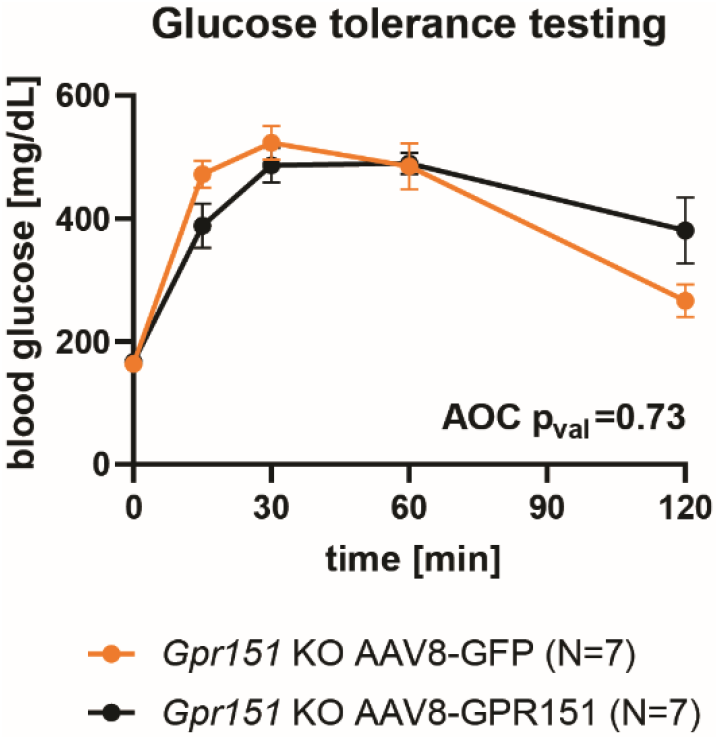
Supplementary data to Figure 6. Blood glucose levels of *Gpr151* KO mice injected with AAV8-GPR151 and AAV8-GFP subjected to glucose tolerance testing (N=7, *Gpr151* KO AAV8:GFP; N=7, *Gpr151* KO AAV8:GPR151). AOC compared using two-tailed Student *t* test.

**Supplementary Table S1** | **Results of RNA-Seq for WT and KO livers from male DIO mice**.

DESeq2 results for all genes and the results of GSEA on the Hallmark gene sets are shown.

**Supplementary Table S2** | **Gene list used for custom GSEA on cAMP-regulated genes**.

