## Supplementary Table 1 for "G protein-coupled receptor 151 regulates glucose metabolism and hepatic gluconeogenesis"

**Supplementary Table S1 |** Results of RNA-Seq for WT and KO livers from male DIO mice. DESeq2 results for all genes and the results of GSEA on the Hallmark gene sets are shown.

| | gene_ID | baseMean | log2FoldChange | lfcSE | stat | pvalue | padj | mgi_symbol | description | WT_1 | WT_2 | WT_3 | KO_1 | KO_2 | KO_3 |
|---|---|---|---|---|---|---|---|---|---|---|---|---|---|---|---|
| 1 | ENSMUSG00000000001 | 1661.798701 | 0.166296178 | 0.097134769 | 1.712014967 | 0.08689391 | 0.434457217 | Gnai3 | guanine nucleotide binding protein (G prc | 1584.5 | 1517.2 | 1595.9 | 1667.7 | 1779.9 | 1825.6 |
| 2 | ENSMUSG00000000003 | 0 | NA | NA | NA | NA | NA | Pbsn | probasin [Source:MGI Symbol;Acc:MGI:18 | 0 | 0 | 0 | 0 | 0 | 0 |
| 3 | ENSMUSG00000000028 | 23.40484971 | 0.258772847 | 0.497103909 | 0.520560878 | 0.602672709 | 0.915657359 | Cdc45 | cell division cycle 45 [Source:MGI Symbol; | 27.305 | 16.447 | 20.125 | 33.739 | 29.447 | 13.366 |
| 4 | ENSMUSG00000000031 | 2.182307175 | -1.085893717 | 1.481339325 | -0.733048599 | 0.463528789 | NA | H19 | H19, imprinted maternally expressed tran | 0.8031 | 5.1396 | 3.0187 | 1.928 | 1.0906 | 1.1138 |
| 5 | ENSMUSG00000000037 | 0.435408177 | -2.179616019 | 3.531869654 | -0.617128103 | 0.537150234 | NA | Scml2 | Scm polycomb group protein like 2 [Sourc | 1.6062 | 0 | 1.0062 | 0 | 0 | 0 |
| 6 | ENSMUSG00000000049 | 37093.42396 | 0.000179488 | 0.089741709 | 0.002000048 | 0.998404194 | 0.999764475 | Apoh | apolipoprotein H [Source:MGI Symbol;Acc | 35166 | 39538 | 36571 | 39117 | 36033 | 36135 |
| 7 | ENSMUSG00000000056 | 1004.872649 | 0.883000692 | 0.128658554 | 6.863132406 | 6.74E-12 | 2.50E-09 | Narf | nuclear prelamin A recognition factor [Sou | 801.5 | 629.09 | 687.27 | 1191.5 | 1312 | 1407.9 |
| 8 | ENSMUSG00000000058 | 46.51256249 | -0.151290405 | 0.346594799 | -0.436505123 | 0.662470283 | 0.9288455 | Cav2 | caveolin 2 [Source:MGI Symbol;Acc:MGI:1 | 52.202 | 39.061 | 55.343 | 38.559 | 35.99 | 57.92 |
| 9 | ENSMUSG00000000078 | 114.8152343 | -0.516760362 | 0.271824624 | -1.901080019 | 0.057291532 | 0.340239855 | Klf6 | Kruppel-like factor 6 [Source:MGI Symbol; | 176.68 | 137.74 | 90.562 | 97.362 | 79.615 | 106.93 |
| 10 | ENSMUSG00000000085 | 138.2254416 | 0.108996629 | 0.199677112 | 0.54586441 | 0.585159144 | 0.909199899 | Scmh1 | sex comb on midleg homolog 1 [Source:M | 122.87 | 130.55 | 145.91 | 143.63 | 150.51 | 135.89 |
| 11 | ENSMUSG00000000088 | 2043.915475 | -0.155574216 | 0.094491983 | -1.646427675 | 0.099675739 | 0.465067707 | Cox5a | cytochrome c oxidase subunit 5A [Source: | 2107.3 | 2083.6 | 2271.1 | 1888.4 | 1930.4 | 1982.6 |
| 12 | ENSMUSG00000000093 | 63.04950606 | -0.6796821 | 0.295858723 | -2.297319787 | 0.021600534 | 0.182976706 | Tbx2 | T-box 2 [Source:MGI Symbol;Acc:MGI:984 | 82.72 | 61.675 | 88.55 | 54.947 | 43.625 | 46.781 |
| 13 | ENSMUSG00000000094 | 0 | NA | NA | NA | NA | NA | Tbx4 | T-box 4 [Source:MGI Symbol;Acc:MGI:102 | 0 | 0 | 0 | 0 | 0 | 0 |
| 14 | ENSMUSG00000000103 | 0 | NA | NA | NA | NA | NA | Zfy2 | zinc finger protein 2, Y-linked [Source:MG | 0 | 0 | 0 | 0 | 0 | 0 |
| 15 | ENSMUSG00000000120 | 33.69250976 | -0.614628154 | 0.37227197 | -1.651019154 | 0.098734664 | 0.463365025 | Ngfr | nerve growth factor receptor (TNFR super | 40.958 | 40.089 | 41.256 | 26.991 | 28.356 | 24.505 |
| 16 | ENSMUSG00000000125 | 0 | NA | NA | NA | NA | NA | Wnt3 | wingless-type MMTV integration site fami | 0 | 0 | 0 | 0 | 0 | 0 |
| 17 | ENSMUSG00000000126 | 31.66921767 | 0.96836658 | 0.544681346 | 1.777858901 | 0.075427032 | 0.401870868 | Wnt9a | wingless-type MMTV integration site fami | 18.471 | 30.838 | 15.094 | 50.127 | 11.997 | 63.489 |
| 18 | ENSMUSG00000000127 | 103.8036697 | -0.186539726 | 0.230673599 | -0.808673932 | 0.418702727 | 0.837591673 | Fer | fer (fms/fps related) protein kinase [Sourc | 108.42 | 126.43 | 96.599 | 89.65 | 99.246 | 102.47 |
| 19 | ENSMUSG00000000131 | 629.2741486 | 0.038672038 | 0.120868913 | 0.319950241 | 0.749006051 | 0.951074124 | Xpo6 | exportin 6 [Source:MGI Symbol;Acc:MGI:2 | 660.15 | 633.2 | 568.53 | 636.23 | 629.29 | 648.26 |
| 20 | ENSMUSG00000000134 | 321.6769174 | 0.051146546 | 0.15990398 | 0.319857865 | 0.74907608 | 0.951074124 | Tfe3 | transcription factor E3 [Source:MGI Symb | 323.65 | 307.35 | 316.97 | 351.85 | 270.47 | 359.77 |
| 21 | ENSMUSG00000000142 | 126.1968356 | -0.173804374 | 0.216829127 | -0.801573001 | 0.422800001 | 0.84005258 | Axin2 | axin 2 [Source:MGI Symbol;Acc:MGI:1270 | 152.59 | 120.27 | 127.79 | 104.11 | 124.33 | 128.09 |
| 22 | ENSMUSG00000000148 | 247.9895627 | 0.141126994 | 0.162873361 | 0.866482971 | 0.386225361 | 0.817115886 | Brat1 | BRCA1-associated ATM activator 1 [Sourc | 256.99 | 211.75 | 238.48 | 272.81 | 261.75 | 246.16 |
| 23 | ENSMUSG00000000149 | 3240.803848 | 0.374083541 | 0.124907917 | 2.994874551 | 0.002745577 | 0.044003638 | Gna12 | guanine nucleotide binding protein, alpha | 2726.5 | 2587.3 | 3154.6 | 3125.2 | 3777.9 | 4073.3 |
| 24 | ENSMUSG00000000154 | 1818.386677 | 0.093919612 | 0.101392976 | 0.926293085 | 0.354293688 | 0.79449673 | Slc22a18 | solute carrier family 22 (organic cation tra | 1876.8 | 1677.6 | 1722.7 | 1965.6 | 1893.3 | 1774.4 |
| 25 | ENSMUSG00000000157 | 51.99224566 | 0.456188055 | 0.35991682 | 1.267481899 | 0.204983032 | 0.650619832 | Itgb2l | integrin beta 2-like [Source:MGI Symbol;A | 47.383 | 38.033 | 46.287 | 86.758 | 55.622 | 37.871 |
| 26 | ENSMUSG00000000159 | 185.6563473 | 0.317578114 | 0.179830208 | 1.765988694 | 0.077397746 | 0.40712138 | Igsf5 | immunoglobulin superfamily, member 5 [ | 151.79 | 172.69 | 172.07 | 222.68 | 189.77 | 204.95 |
| 27 | ENSMUSG00000000167 | 8.955506916 | 0.155986401 | 0.717099863 | 0.217523959 | 0.827800038 | 0.969794868 | Pih1d2 | PIH1 domain containing 2 [Source:MGI Sy | 7.2279 | 6.1675 | 12.075 | 10.604 | 7.6343 | 10.025 |
| 28 | ENSMUSG00000000168 | 1071.380247 | -0.103174992 | 0.127833575 | -0.807104016 | 0.419606558 | 0.837591673 | Dlat | dihydrolipoamide S-acetyltransferase (E2 | 1207.1 | 1178 | 942.85 | 958.19 | 1086.3 | 1055.9 |
| 29 | ENSMUSG00000000171 | 3955.075822 | 0.049105481 | 0.088400504 | 0.555488697 | 0.578560439 | 0.908137944 | Sdhd | succinate dehydrogenase complex, subun | 3764.1 | 3970.8 | 3928.4 | 3830.8 | 4109.4 | 4126.8 |
| 30 | ENSMUSG00000000182 | 0 | NA | NA | NA | NA | NA | Fgf23 | fibroblast growth factor 23 [Source:MGI S | 0 | 0 | 0 | 0 | 0 | 0 |
| 31 | ENSMUSG00000000183 | 0 | NA | NA | NA | NA | NA | Fgf6 | fibroblast growth factor 6 [Source:MGI Sy | 0 | 0 | 0 | 0 | 0 | 0 |
| 32 | ENSMUSG00000000184 | 68.40413655 | -0.872270979 | 0.277464468 | -3.14372138 | 0.001668142 | 0.030619294 | Ccnd2 | cyclin D2 [Source:MGI Symbol;Acc:MGI:88 | 99.585 | 76.066 | 89.556 | 48.199 | 46.897 | 50.123 |
| 33 | ENSMUSG00000000194 | 418.5362255 | 0.23166226 | 0.14149138 | 1.63728886 | 0.101570127 | 0.469189975 | Gpr107 | G protein-coupled receptor 107 [Source:N | 390.31 | 366.97 | 397.47 | 447.29 | 404.62 | 504.57 |
| 34 | ENSMUSG00000000197 | 0.856598198 | -3.132279849 | 3.123122476 | -1.002932121 | 0.315893613 | NA | Nalcn | sodium leak channel, non-selective [Sourc | 0 | 5.1396 | 0 | 0 | 0 | 0 |
| 35 | ENSMUSG00000000202 | 0 | NA | NA | NA | NA | NA | Btbd17 | BTB (POZ) domain containing 17 [Source:I | 0 | 0 | 0 | 0 | 0 | 0 |
| 36 | ENSMUSG00000000204 | 4.732335571 | -0.311567316 | 1.031244891 | -0.302127379 | 0.762554962 | NA | Slfn4 | schlafen 4 [Source:MGI Symbol;Acc:MGI:1 | 1.6062 | 6.1675 | 8.05 | 4.8199 | 2.1812 | 5.5692 |
| 37 | ENSMUSG00000000214 | 0 | NA | NA | NA | NA | NA | Th | tyrosine hydroxylase [Source:MGI Symbol | 0 | 0 | 0 | 0 | 0 | 0 |
| 38 | ENSMUSG00000000215 | 0 | NA | NA | NA | NA | NA | Ins2 | insulin II [Source:MGI Symbol;Acc:MGI:96 | 0 | 0 | 0 | 0 | 0 | 0 |
| 39 | ENSMUSG00000000216 | 0.315620088 | 0.165143421 | 4.063300985 | 0.040642675 | 0.967580763 | NA | Scnn1g | sodium channel, nonvoltage-gated 1 gam | 0.8031 | 0 | 0 | 0 | 1.0906 | 0 |
| 40 | ENSMUSG00000000223 | 0 | NA | NA | NA | NA | NA | Drp2 | dystrophin related protein 2 [Source:MGI | 0 | 0 | 0 | 0 | 0 | 0 |
| 41 | ENSMUSG00000000244 | 2.735507428 | 0.530488369 | 1.502478146 | 0.353075597 | 0.724031767 | NA | Tspan32 | tetraspanin 32 [Source:MGI Symbol;Acc:N | 1.6062 | 4.1117 | 1.0062 | 0.964 | 8.7249 | 0 |
| 42 | ENSMUSG00000000247 | 41.18511976 | -1.195484982 | 0.425385674 | -2.810355529 | 0.00494868 | 0.066398616 | Lhx2 | LIM homeobox protein 2 [Source:MGI Syn | 46.58 | 56.535 | 69.431 | 41.451 | 14.178 | 18.935 |
| 43 | ENSMUSG00000000248 | 0 | NA | NA | NA | NA | NA | Clec2g | C-type lectin domain family 2, member g [ | 0 | 0 | 0 | 0 | 0 | 0 |
| 44 | ENSMUSG00000000253 | 7.698001816 | -1.418764311 | 0.792451853 | -1.790347647 | 0.073398041 | 0.393487496 | Gmpr | guanosine monophosphate reductase [So | 11.243 | 12.335 | 10.062 | 4.8199 | 3.2719 | 4.4554 |
| 45 | ENSMUSG00000000263 | 0 | NA | NA | NA | NA | NA | Glra1 | glycine receptor, alpha 1 subunit [Source: | 0 | 0 | 0 | 0 | 0 | 0 |
| 46 | ENSMUSG00000000266 | 115.8785759 | -0.27598364 | 0.230946051 | -1.195013465 | 0.232081769 | 0.679520387 | Mid2 | midline 2 [Source:MGI Symbol;Acc:MGI:1 | 134.92 | 144.94 | 100.62 | 95.434 | 107.97 | 111.38 |
| 47 | ENSMUSG00000000275 | 1760.067113 | 0.102534867 | 0.127300638 | 0.805454463 | 0.420557474 | 0.837906635 | Trim25 | tripartite motif-containing 25 [Source:MG | 1911.4 | 1461.7 | 1717.7 | 1623.3 | 1924.9 | 1921.4 |
| 48 | ENSMUSG00000000276 | 42.91714558 | 0.014427611 | 0.362450509 | 0.03980574 | 0.968248 | 0.994231936 | Dgke | diacylglycerol kinase, epsilon [Source:MGI | 54.611 | 30.838 | 42.262 | 41.451 | 35.99 | 52.351 |
| 49 | ENSMUSG00000000278 | 495.1623273 | 0.21054025 | 0.146915263 | 1.433072688 | 0.151837066 | 0.567340036 | Scpep1 | serine carboxypeptidase 1 [Source:MGI Sy | 536.47 | 447.14 | 392.44 | 532.12 | 513.68 | 549.13 |
| 50 | ENSMUSG00000000282 | 237.6777893 | 0.054200649 | 0.177020072 | 0.306183634 | 0.759464832 | 0.95402891 | Mnt | max binding protein [Source:MGI Symbol; | 220.85 | 209.7 | 269.67 | 271.84 | 230.12 | 223.88 |
| 51 | ENSMUSG00000000290 | 57.18063815 | -1.52180769 | 0.369668649 | -4.116680417 | 3.84E-05 | 0.001694046 | Itgb2 | integrin beta 2 [Source:MGI Symbol;Acc:N | 82.72 | 111.02 | 60.375 | 18.316 | 29.447 | 41.212 |
| 52 | ENSMUSG00000000295 | 16.78015006 | -0.77646403 | 0.567894319 | -1.367268528 | 0.171541152 | 0.599034187 | Hddc2 | HD domain containing 2 [Source:MGI Sym | 24.896 | 12.335 | 26.162 | 8.6758 | 16.359 | 12.252 |

| | | | | | | | | | | | | | | | |
|---|---|---|---|---|---|---|---|---|---|---|---|---|---|---|---|
| 53 | ENSMUSG00000000296 | 169.4001676 | 0.06062593 | 0.197486624 | 0.306987527 | 0.758852866 | 0.95402891 | Tpd52l1 | tumor protein D52-like 1 [Source:MGI Syn | 146.97 | 165.49 | 185.15 | 147.49 | 190.86 | 180.44 |
| 54 | ENSMUSG00000000301 | 5048.728159 | 0.162726581 | 0.095216896 | 1.709009514 | 0.087449187 | 0.43616036 | Pemt | phosphatidylethanolamine N-methyltrans | 4821 | 4824 | 4646.8 | 4867.1 | 5677.8 | 5455.6 |
| 55 | ENSMUSG00000000303 | 387.0678243 | 0.983656938 | 0.250170156 | 3.931951576 | 8.43E-05 | 0.003149154 | Cdh1 | cadherin 1 [Source:MGI Symbol;Acc:MGI: | 179.89 | 234.37 | 366.27 | 347.03 | 590.02 | 604.82 |
| 56 | ENSMUSG00000000305 | 11.78409623 | 0.103813108 | 0.708769608 | 0.14646947 | 0.88355079 | 0.981769403 | Cdh4 | cadherin 4 [Source:MGI Symbol;Acc:MGI: | 13.653 | 12.335 | 8.05 | 14.46 | 3.2719 | 18.935 |
| 57 | ENSMUSG00000000308 | 1.254537109 | -3.692333964 | 2.19368904 | -1.683161969 | 0.092343741 | NA | Ckmt1 | creatine kinase, mitochondrial 1, ubiquito | 2.4093 | 4.1117 | 1.0062 | 0 | 0 | 0 |
| 58 | ENSMUSG00000000317 | 25.29213076 | -0.176566231 | 0.461143433 | -0.382887878 | 0.701802899 | 0.942189108 | Bcl6b | B cell CLL/lymphoma 6, member B [Source | 33.73 | 24.67 | 22.137 | 33.739 | 18.541 | 18.935 |
| 59 | ENSMUSG00000000318 | 5.617678995 | -2.788355873 | 1.113567709 | -2.503984131 | 0.012280354 | 0.124560339 | Clec10a | C-type lectin domain family 10, member A | 8.031 | 11.307 | 10.062 | 0.964 | 0 | 3.3415 |
| 60 | ENSMUSG00000000320 | 4.93262643 | 0.648013187 | 0.975126724 | 0.664542537 | 0.506343133 | NA | Alox12 | arachidonate 12-lipoxygenase [Source:MG | 2.4093 | 6.1675 | 3.0187 | 4.8199 | 8.7249 | 4.4554 |
| 61 | ENSMUSG00000000325 | 254.1744918 | -0.987858618 | 0.163616107 | -6.037661187 | 1.56E-09 | 3.06E-07 | NA | NA | 322.04 | 345.38 | 347.15 | 190.87 | 160.32 | 159.28 |
| 62 | ENSMUSG00000000326 | 15480.21523 | 0.476533267 | 0.099503661 | 4.789102873 | 1.68E-06 | 0.000124296 | Comt | catechol-O-methyltransferase [Source:MG | 14024 | 12083 | 12732 | 19542 | 17138 | 17363 |
| 63 | ENSMUSG00000000339 | 146.2555043 | 0.008767032 | 0.210973579 | 0.041555116 | 0.966853355 | 0.994231936 | Rtca | RNA 3'-terminal phosphate cyclase [Sourc | 175.08 | 139.8 | 121.76 | 137.85 | 152.69 | 150.37 |
| 64 | ENSMUSG00000000340 | 2275.922239 | -0.038332784 | 0.115123975 | -0.332969601 | 0.73915723 | 0.948310621 | Dbt | dihydrolipoamide branched chain transac | 2048.7 | 2265.5 | 2605.2 | 2161.2 | 2390.6 | 2184.2 |
| 65 | ENSMUSG00000000346 | 2246.95925 | 0.046116597 | 0.094114817 | 0.490003582 | 0.624131364 | 0.919503423 | Dazap2 | DAZ associated protein 2 [Source:MGI Syr | 2136.3 | 2315.9 | 2181.5 | 2276.9 | 2199.8 | 2371.4 |
| 66 | ENSMUSG00000000355 | 549.66196 | 0.106945992 | 0.121952808 | 0.876945717 | 0.380516123 | 0.812628154 | Mcts1 | malignant T cell amplified sequence 1 [Sou | 523.62 | 524.24 | 540.35 | 585.13 | 592.21 | 532.42 |
| 67 | ENSMUSG00000000359 | 3.998206513 | -0.929957917 | 1.146205103 | -0.811336396 | 0.417172514 | NA | Rem1 | rad and gem related GTP binding protein : | 5.6217 | 4.1117 | 6.0375 | 3.8559 | 4.3625 | 0 |
| 68 | ENSMUSG00000000365 | 0.486939728 | -0.698472448 | 3.137608162 | -0.222613026 | 0.823836697 | NA | Rnf17 | ring finger protein 17 [Source:MGI Symbo | 0.8031 | 1.0279 | 0 | 0 | 1.0906 | 0 |
| 69 | ENSMUSG00000000374 | 496.6435541 | -0.144237993 | 0.141146013 | -1.02190625 | 0.306825274 | 0.755905427 | Trappc10 | trafficking protein particle complex 10 [So | 473.03 | 566.38 | 526.27 | 527.3 | 458.06 | 428.83 |
| 70 | ENSMUSG00000000378 | 377.7799095 | 0.204413546 | 0.137422929 | 1.487477727 | 0.136888682 | 0.540676649 | Ccm2 | cerebral cavernous malformation 2 [Sourc | 358.18 | 360.8 | 334.07 | 409.69 | 378.44 | 425.49 |
| 71 | ENSMUSG00000000381 | 0 | NA | NA | NA | NA | NA | Wap | whey acidic protein [Source:MGI Symbol;/ | 0 | 0 | 0 | 0 | 0 | 0 |
| 72 | ENSMUSG00000000384 | 749.911905 | 0.004390069 | 0.128034299 | 0.034288228 | 0.972647312 | 0.99573532 | Tbrg4 | transforming growth factor beta regulate | 781.42 | 762.72 | 701.35 | 681.53 | 850.68 | 721.77 |
| 73 | ENSMUSG00000000385 | 84.58156782 | -0.441359408 | 0.279358321 | -1.579904285 | 0.114128788 | 0.496868018 | Tmprss2 | transmembrane protease, serine 2 [Sourc | 95.569 | 106.9 | 89.556 | 60.731 | 56.712 | 98.018 |
| 74 | ENSMUSG00000000386 | 22.99870148 | -0.607944634 | 0.494273536 | -1.229976095 | 0.218706057 | 0.663735712 | Mx1 | MX dynamin-like GTPase 1 [Source:MGI S | 26.502 | 23.642 | 33.206 | 17.352 | 27.265 | 10.025 |
| 75 | ENSMUSG00000000392 | 2.200446982 | -1.639531804 | 1.630534821 | -1.005517811 | 0.314647578 | NA | Fap | fibroblast activation protein [Source:MGI | 0.8031 | 7.1954 | 2.0125 | 0.964 | 0 | 2.2277 |
| 76 | ENSMUSG00000000394 | 0 | NA | NA | NA | NA | NA | Gcg | glucagon [Source:MGI Symbol;Acc:MGI:9 | 0 | 0 | 0 | 0 | 0 | 0 |
| 77 | ENSMUSG00000000399 | 2135.441243 | -0.083650875 | 0.093070884 | -0.898786726 | 0.368766272 | 0.806506847 | Ndufa9 | NADH:ubiquinone oxidoreductase subuni | 2124.2 | 2254.2 | 2213.7 | 1993.5 | 2147.4 | 2079.5 |
| 78 | ENSMUSG00000000402 | 3.179517392 | -3.98606895 | 1.62414482 | -2.454257096 | 0.014117601 | NA | Egfl6 | EGF-like-domain, multiple 6 [Source:MGI ! | 4.8186 | 4.1117 | 9.0562 | 0 | 1.0906 | 0 |
| 79 | ENSMUSG00000000409 | 8.489961198 | -1.337067152 | 0.862968369 | -1.549381414 | 0.121290059 | 0.510576081 | Lck | lymphocyte protein tyrosine kinase [Sourc | 8.031 | 21.586 | 7.0437 | 7.7118 | 5.4531 | 1.1138 |
| 80 | ENSMUSG00000000411 | 0.596071152 | -1.224343926 | 2.897583865 | -0.4225396 | 0.672631203 | NA | Tssk3 | testis-specific serine kinase 3 [Source:MG | 1.6062 | 0 | 1.0062 | 0.964 | 0 | 0 |
| 81 | ENSMUSG00000000416 | 2.39475491 | -0.222726924 | 1.490821106 | -0.149398827 | 0.881238937 | NA | Cttnbp2 | cortactin binding protein 2 [Source:MGI S | 1.6062 | 3.0838 | 3.0187 | 0 | 1.0906 | 5.5692 |
| 82 | ENSMUSG00000000420 | 1012.744579 | -0.092656978 | 0.110054893 | -0.841916017 | 0.399834964 | 0.82676382 | Galnt1 | polypeptide N-acetylgalactosaminyltransf | 978.18 | 1108.1 | 1050.5 | 998.68 | 1014.3 | 926.72 |
| 83 | ENSMUSG00000000435 | 0 | NA | NA | NA | NA | NA | Myf5 | myogenic factor 5 [Source:MGI Symbol;Ac | 0 | 0 | 0 | 0 | 0 | 0 |
| 84 | ENSMUSG00000000439 | 266.7528878 | 0.110093485 | 0.156484902 | 0.703540621 | 0.481718901 | 0.871274502 | Mkrn2 | makorin, ring finger protein, 2 [Source:MG | 248.96 | 259.04 | 261.62 | 252.56 | 302.1 | 276.23 |
| 85 | ENSMUSG00000000440 | 182.6502302 | -0.196356539 | 0.660050127 | -0.297487313 | 0.766094492 | 0.955091733 | Pparg | peroxisome proliferator activated recepto | 262.61 | 221 | 101.63 | 283.41 | 155.96 | 71.286 |
| 86 | ENSMUSG00000000441 | 490.5341182 | 0.210267658 | 0.124312788 | 1.691440288 | 0.090752743 | 0.444602826 | Raf1 | v-raf-leukemia viral oncogene 1 [Source:N | 433.68 | 464.62 | 466.9 | 539.83 | 519.13 | 519.05 |
| 87 | ENSMUSG00000000486 | 25.01241973 | -0.333869833 | 0.489393268 | -0.68221174 | 0.495105072 | 0.877919732 | Septin1 | septin 1 [Source:MGI Symbol;Acc:MGI:18 | 20.078 | 25.698 | 38.237 | 31.811 | 13.087 | 21.163 |
| 88 | ENSMUSG00000000489 | 20.73935001 | -0.982303067 | 0.508315841 | -1.932465975 | 0.053302018 | 0.324478276 | Pdgfb | platelet derived growth factor, B polypept | 25.699 | 32.893 | 24.15 | 17.352 | 7.6343 | 16.708 |
| 89 | ENSMUSG00000000530 | 47.79938838 | -0.523279458 | 0.351210202 | -1.489932396 | 0.136242012 | 0.538921153 | Acvrl1 | activin A receptor, type II-like 1 [Source:M | 58.626 | 46.256 | 64.4 | 50.127 | 26.175 | 41.212 |
| 90 | ENSMUSG00000000531 | 27.24412444 | -0.761883643 | 0.422550184 | -1.803060727 | 0.071378678 | 0.386964468 | Tamalin | trafficking regulator and scaffold protein t | 33.73 | 38.033 | 31.194 | 24.099 | 16.359 | 20.049 |
| 91 | ENSMUSG00000000532 | 351.4856218 | 0.330456669 | 0.150749582 | 2.192090123 | 0.028373 | 0.220401219 | Acvr1b | activin A receptor, type 1B [Source:MGI S | 324.45 | 302.21 | 306.9 | 364.38 | 362.09 | 448.88 |
| 92 | ENSMUSG00000000538 | 730.48091 | 0.063468067 | 0.142818815 | 0.444395698 | 0.656756523 | 0.9268505 | Tom1l2 | target of myb1-like 2 (chicken) [Source:M | 843.26 | 681.51 | 616.83 | 700.81 | 730.71 | 809.76 |
| 93 | ENSMUSG00000000544 | 0 | NA | NA | NA | NA | NA | Gpa33 | glycoprotein A33 (transmembrane) [Sourc | 0 | 0 | 0 | 0 | 0 | 0 |
| 94 | ENSMUSG00000000552 | 189.3511468 | -0.571533337 | 0.215225994 | -2.655503293 | 0.007919016 | 0.09134023 | Zfp385a | zinc finger protein 385A [Source:MGI Sym | 204.79 | 253.9 | 221.37 | 195.69 | 117.79 | 142.57 |
| 95 | ENSMUSG00000000555 | 274.0581895 | -0.51108617 | 0.194192471 | -2.63185369 | 0.008492042 | 0.096227639 | Itga5 | integrin alpha 5 (fibronectin receptor alph | 244.95 | 317.63 | 404.51 | 226.53 | 226.85 | 223.88 |
| 96 | ENSMUSG00000000560 | 0 | NA | NA | NA | NA | NA | Gabra2 | gamma-aminobutyric acid (GABA) A recep | 0 | 0 | 0 | 0 | 0 | 0 |
| 97 | ENSMUSG00000000561 | 225.7248857 | -0.016879336 | 0.173560942 | -0.097253081 | 0.922525415 | 0.986685804 | Wdr77 | WD repeat domain 77 [Source:MGI Symbo | 228.08 | 232.31 | 220.37 | 195.69 | 258.48 | 219.43 |
| 98 | ENSMUSG00000000562 | 0.181769715 | 1.126924175 | 4.080472857 | 0.276174898 | 0.782413736 | NA | Adora3 | adenosine A3 receptor [Source:MGI Symb | 0 | 0 | 0 | 0 | 1.0906 | 0 |
| 99 | ENSMUSG00000000563 | 6118.736176 | 0.014236263 | 0.094455597 | 0.150719103 | 0.880197303 | 0.981359269 | Atp5pb | ATP synthase peripheral stalk-membrane | 5936.5 | 6358.7 | 5970 | 5652.8 | 6502.3 | 6292.1 |
| 100 | ENSMUSG00000000567 | 85.50217132 | -0.317009309 | 0.383524322 | -0.826568982 | 0.408481404 | 0.832151413 | Sox9 | SRY (sex determining region Y)-box 9 [Sou | 54.611 | 76.066 | 153.96 | 44.343 | 92.703 | 91.335 |
| 101 | ENSMUSG00000000568 | 827.6612798 | 0.038240829 | 0.110287859 | 0.34673653 | 0.728789264 | 0.946248466 | Hnrnpd | heterogeneous nuclear ribonucleoprotein | 813.54 | 818.22 | 818.08 | 802.03 | 894.31 | 819.79 |
| 102 | ENSMUSG00000000579 | 9.00487932 | -1.038602584 | 0.803759257 | -1.292181179 | 0.196294408 | 0.637163895 | Dynlt1c | dynein light chain Tctex-type 1C [Source:N | 8.031 | 12.335 | 16.1 | 8.6758 | 1.0906 | 7.7969 |
| 103 | ENSMUSG00000000581 | 885.546832 | 0.006764812 | 0.128554086 | 0.052622305 | 0.958032845 | 0.991861398 | C1d | C1D nuclear receptor co-repressor [Sourc | 934.81 | 891.2 | 823.11 | 763.47 | 971.74 | 928.95 |
| 104 | ENSMUSG00000000594 | 1780.032057 | -0.036654459 | 0.112192666 | -0.326709934 | 0.743887286 | 0.949911274 | Gm2a | GM2 ganglioside activator protein [Source | 1707.4 | 1675.5 | 2025.6 | 1779.5 | 1658.8 | 1833.4 |
| 105 | ENSMUSG00000000600 | 321.7018227 | -0.034815689 | 0.157143665 | -0.221553246 | 0.824661682 | 0.96943211 | Krit1 | KRIT1, ankyrin repeat containing [Source:I | 298.75 | 323.79 | 355.2 | 358.6 | 305.37 | 288.49 |
| 106 | ENSMUSG00000000605 | 1097.987959 | -0.06956373 | 0.121794588 | -0.571156163 | 0.567893791 | 0.903352921 | Clcn4 | chloride channel, voltage-sensitive 4 [Sou | 1093.8 | 1206.8 | 1073.7 | 1147.1 | 1134.2 | 932.29 |
| 107 | ENSMUSG00000000606 | 0 | NA | NA | NA | NA | NA | Vmn2r88 | vomeronasal 2, receptor 88 [Source:MGI ! | 0 | 0 | 0 | 0 | 0 | 0 |
| 108 | ENSMUSG00000000617 | 0.167707431 | -0.796629176 | 4.080472857 | -0.195229623 | 0.84521319 | NA | Grm6 | glutamate receptor, metabotropic 6 [Sour | 0 | 0 | 1.0062 | 0 | 0 | 0 |

| | | | | | | | | | | | | | | | |
|---|---|---|---|---|---|---|---|---|---|---|---|---|---|---|---|
| 109 | ENSMUSG00000000627 | 1.199030887 | -1.291105934 | 2.361925445 | -0.546632806 | 0.584631026 | NA | Sema4f | sema domain, immunoglobulin domain (Ig | 0 | 5.1396 | 0 | 0.964 | 1.0906 | 0 |
| 110 | ENSMUSG00000000628 | 11.7979059 | -1.15625158 | 0.704698019 | -1.640775976 | 0.100843927 | 0.46735231 | Hk2 | hexokinase 2 [Source:MGI Symbol;Acc:MG | 15.259 | 22.614 | 11.069 | 9.6398 | 2.1812 | 10.025 |
| 111 | ENSMUSG00000000631 | 2130.999438 | 0.283404195 | 0.091805839 | 3.086995304 | 0.002021908 | 0.035277293 | Myo18a | myosin XVIIIA [Source:MGI Symbol;Acc:M | 1915.4 | 1996.2 | 1855.5 | 2324.2 | 2307.7 | 2387 |
| 112 | ENSMUSG00000000632 | 0 | NA | NA | NA | NA | NA | Sez6 | seizure related gene 6 [Source:MGI Symbo | 0 | 0 | 0 | 0 | 0 | 0 |
| 113 | ENSMUSG00000000673 | 9988.756526 | 0.17646261 | 0.081591411 | 2.162759617 | 0.030559673 | 0.231785586 | Haao | 3-hydroxyanthranilate 3,4-dioxygenase [S | 9600.3 | 9190.6 | 9344 | 10263 | 10763 | 10771 |
| 114 | ENSMUSG00000000682 | 44.94255023 | -1.632166378 | 0.439131181 | -3.716808209 | 0.000201755 | 0.006326357 | Cd52 | CD52 antigen [Source:MGI Symbol;Acc:M | 53.808 | 100.74 | 49.306 | 19.28 | 11.997 | 34.529 |
| 115 | ENSMUSG00000000686 | 518.1571821 | 0.074374005 | 0.124167475 | 0.598981374 | 0.549185305 | 0.895703112 | Abhd15 | abhydrolase domain containing 15 [Sourc | 494.71 | 516.01 | 504.13 | 546.58 | 551.85 | 495.66 |
| 116 | ENSMUSG00000000690 | 1.274125293 | -2.554690053 | 2.062976523 | -1.238351491 | 0.215585759 | NA | Hoxb6 | homeobox B6 [Source:MGI Symbol;Acc:M | 1.6062 | 2.0558 | 3.0187 | 0.964 | 0 | 0 |
| 117 | ENSMUSG00000000693 | 30.04079904 | -0.47002747 | 0.454052932 | -1.035182106 | 0.300583874 | 0.751875664 | Loxl3 | lysyl oxidase-like 3 [Source:MGI Symbol;A | 37.746 | 31.865 | 35.219 | 33.739 | 30.537 | 11.138 |
| 118 | ENSMUSG00000000701 | 0 | NA | NA | NA | NA | NA | Tcl1b5 | T cell leukemia/lymphoma 1B, 5 [Source:N | 0 | 0 | 0 | 0 | 0 | 0 |
| 119 | ENSMUSG00000000706 | 0 | NA | NA | NA | NA | NA | Btn1a1 | butyrophilin, subfamily 1, member A1 [So | 0 | 0 | 0 | 0 | 0 | 0 |
| 120 | ENSMUSG00000000708 | 1812.299315 | 0.127946646 | 0.119779686 | 1.068183185 | 0.285437888 | 0.738036041 | Kat2b | K(lysine) acetyltransferase 2B [Source:MG | 1475.3 | 1904.7 | 1817.3 | 1797.8 | 1975.1 | 1903.6 |
| 121 | ENSMUSG00000000711 | 2006.319416 | 0.128588668 | 0.094640408 | 1.35870788 | 0.174239174 | 0.604151393 | Rab5b | RAB5B, member RAS oncogene family [So | 2006.1 | 1830.7 | 1912.9 | 2033 | 2132.2 | 2123 |
| 122 | ENSMUSG00000000724 | 0.499690045 | -0.82080011 | 3.113553401 | -0.263621658 | 0.792071468 | NA | Cryba1 | crystallin, beta A1 [Source:MGI Symbol;Ac | 0 | 1.0279 | 1.0062 | 0.964 | 0 | 0 |
| 123 | ENSMUSG00000000730 | 0 | NA | NA | NA | NA | NA | Dnmt3l | DNA (cytosine-5-)-methyltransferase 3-lik | 0 | 0 | 0 | 0 | 0 | 0 |
| 124 | ENSMUSG00000000731 | 0 | NA | NA | NA | NA | NA | Aire | autoimmune regulator (autoimmune poly | 0 | 0 | 0 | 0 | 0 | 0 |
| 125 | ENSMUSG00000000732 | 10.81983943 | -0.304981689 | 0.709270017 | -0.42999377 | 0.667200173 | 0.929709374 | Icosl | icos ligand [Source:MGI Symbol;Acc:MGI: | 18.471 | 9.2513 | 8.05 | 12.532 | 4.3625 | 12.252 |
| 126 | ENSMUSG00000000738 | 966.7393212 | -0.013511916 | 0.103356329 | -0.130731389 | 0.895987801 | 0.982798755 | Spg7 | SPG7, paraplegin matrix AAA peptidase su | 958.9 | 988.86 | 965.99 | 935.06 | 985.92 | 965.7 |
| 127 | ENSMUSG00000000739 | 195.3462477 | 0.711324097 | 0.251740555 | 2.825623777 | 0.004718865 | 0.06420853 | Sult5a1 | sulfotransferase family 5A, member 1 [Sou | 207.2 | 111.02 | 124.77 | 186.05 | 285.74 | 257.3 |
| 128 | ENSMUSG00000000740 | 4147.476007 | 0.120807486 | 0.092955941 | 1.299620929 | 0.193730923 | 0.631851946 | Rpl13 | ribosomal protein L13 [Source:MGI Symbo | 3764.1 | 4172.3 | 3985.7 | 4097.9 | 4354.8 | 4510 |
| 129 | ENSMUSG00000000743 | 1561.014585 | 0.034048818 | 0.100080281 | 0.34021505 | 0.733694585 | 0.947937336 | Chmp1a | charged multivesicular body protein 1A [S | 1576.5 | 1550.1 | 1500.3 | 1478.7 | 1663.2 | 1597.3 |
| 130 | ENSMUSG00000000751 | 530.7679254 | 0.071172805 | 0.141036164 | 0.504642232 | 0.613810142 | 0.917353781 | Rpa1 | replication protein A1 [Source:MGI Symbo | 577.43 | 481.07 | 493.06 | 479.1 | 574.76 | 579.2 |
| 131 | ENSMUSG00000000753 | 9202.430075 | 0.115784391 | 0.079257032 | 1.460872148 | 0.144050531 | 0.554282225 | Serpinf1 | serine (or cysteine) peptidase inhibitor, cl | 8740.2 | 8779.4 | 8980.7 | 9589.7 | 9629.1 | 9495.5 |
| 132 | ENSMUSG00000000759 | 330.0554093 | -0.127068626 | 0.177675474 | -0.71517257 | 0.474502416 | 0.869199968 | Tubgcp3 | tubulin, gamma complex associated prote | 368.62 | 303.24 | 362.25 | 389.45 | 268.29 | 288.49 |
| 133 | ENSMUSG00000000766 | 0.17131964 | -0.796629176 | 4.080472857 | -0.195229623 | 0.84521319 | NA | Oprm1 | opioid receptor, mu 1 [Source:MGI Symbo | 0 | 1.0279 | 0 | 0 | 0 | 0 |
| 134 | ENSMUSG00000000776 | 113.2669875 | 0.065693149 | 0.216910077 | 0.302858909 | 0.761997388 | 0.954327666 | Polr3d | polymerase (RNA) III (DNA directed) polyp | 105.21 | 116.15 | 110.69 | 106.04 | 113.42 | 128.09 |
| 135 | ENSMUSG00000000782 | 84.16910306 | 0.124384229 | 0.254053783 | 0.489598018 | 0.624418379 | 0.919503423 | Tcf7 | transcription factor 7, T cell specific [Sour | 83.523 | 88.401 | 69.431 | 75.19 | 93.793 | 94.677 |
| 136 | ENSMUSG00000000787 | 3108.965546 | -0.090221764 | 0.120797054 | -0.74688712 | 0.455131701 | 0.857337687 | Ddx3x | DEAD box helicase 3, X-linked [Source:MG | 2769.1 | 3653.2 | 3197.8 | 3110.8 | 3163.9 | 2759 |
| 137 | ENSMUSG00000000791 | 29.09523611 | 0.130781812 | 0.469533102 | 0.278535873 | 0.780601032 | 0.959921066 | Il12rb1 | interleukin 12 receptor, beta 1 [Source:M | 32.124 | 26.726 | 24.15 | 16.388 | 26.175 | 49.009 |
| 138 | ENSMUSG00000000792 | 0.321325949 | 1.911575114 | 4.048267893 | 0.472195804 | 0.636787034 | NA | Slc5a5 | solute carrier family 5 (sodium iodide sym | 0 | 0 | 0 | 1.928 | 0 | 0 |
| 139 | ENSMUSG00000000794 | 0.301557804 | -1.64335916 | 4.042814367 | -0.406488899 | 0.684383412 | NA | Kcnn3 | potassium intermediate/small conductanc | 0.8031 | 0 | 1.0062 | 0 | 0 | 0 |
| 140 | ENSMUSG00000000804 | 556.4401598 | 0.080294432 | 0.129811299 | 0.618547329 | 0.53621461 | 0.890560262 | Usp32 | ubiquitin specific peptidase 32 [Source:M | 534.87 | 523.21 | 565.51 | 637.19 | 538.77 | 539.1 |
| 141 | ENSMUSG00000000805 | 0 | NA | NA | NA | NA | NA | Car4 | carbonic anhydrase 4 [Source:MGI Symbo | 0 | 0 | 0 | 0 | 0 | 0 |
| 142 | ENSMUSG00000000811 | 181.6861055 | 0.424741965 | 0.196185393 | 2.165003008 | 0.030387455 | 0.230721337 | Txnrd3 | thioredoxin reductase 3 [Source:MGI Sym | 160.62 | 181.94 | 122.76 | 208.22 | 209.4 | 207.17 |
| 143 | ENSMUSG00000000817 | 0.904673413 | -3.227537609 | 2.678727401 | -1.204877214 | 0.228250706 | NA | Fasl | Fas ligand (TNF superfamily, member 6) [S | 2.4093 | 0 | 3.0187 | 0 | 0 | 0 |
| 144 | ENSMUSG00000000823 | 427.9373076 | 0.304664318 | 0.162641766 | 1.873223135 | 0.061037579 | 0.352293512 | Zfp512b | zinc finger protein 512B [Source:MGI Sym | 384.69 | 429.67 | 335.08 | 562 | 435.16 | 421.03 |
| 145 | ENSMUSG00000000826 | 1275.843484 | 0.038703907 | 0.098996758 | 0.390961356 | 0.6958258 | 0.939992732 | Dnajc5 | DnaJ heat shock protein family (Hsp40) m | 1211.1 | 1290 | 1275.9 | 1325.5 | 1272.8 | 1279.8 |
| 146 | ENSMUSG00000000827 | 596.2141493 | 0.015046865 | 0.124661433 | 0.120701843 | 0.903927199 | 0.983821605 | Tpd52l2 | tumor protein D52-like 2 [Source:MGI Syn | 596.7 | 541.71 | 640.98 | 618.87 | 600.93 | 578.08 |
| 147 | ENSMUSG00000000838 | 472.5425629 | -0.143962893 | 0.139974522 | -1.028493544 | 0.303717726 | 0.75273701 | Fmr1 | FMRP translational regulator 1 [Source:M | 498.73 | 535.55 | 453.82 | 422.22 | 501.68 | 423.26 |
| 148 | ENSMUSG00000000861 | 2.857079428 | -1.545408277 | 1.550740269 | -0.996561647 | 0.31897733 | NA | Bcl11a | B cell CLL/lymphoma 11A (zinc finger prot | 5.6217 | 1.0279 | 6.0375 | 0 | 0 | 4.4554 |
| 149 | ENSMUSG00000000869 | 5.083537097 | -1.678524754 | 1.024738949 | -1.6380023 | 0.101421214 | NA | Il4 | interleukin 4 [Source:MGI Symbol;Acc:MG | 12.047 | 5.1396 | 6.0375 | 2.8919 | 3.2719 | 1.1138 |
| 150 | ENSMUSG00000000876 | 2241.316577 | -0.336898407 | 0.101916395 | -3.305635035 | 0.000947614 | 0.020109255 | Pxmp4 | peroxisomal membrane protein 4 [Source | 2283.2 | 2564.7 | 2658.5 | 1928.9 | 2066.7 | 1945.9 |
| 151 | ENSMUSG00000000881 | 364.3561838 | 0.335232363 | 0.151742821 | 2.20921399 | 0.02715976 | 0.21354341 | Dlg3 | discs large MAGUK scaffold protein 3 [Sou | 281.89 | 344.35 | 341.12 | 390.41 | 381.72 | 446.65 |
| 152 | ENSMUSG00000000884 | 25.71113425 | -0.067762371 | 0.448206181 | -0.151185713 | 0.87982922 | 0.98128671 | Gnb1l | guanine nucleotide binding protein (G pro | 24.896 | 31.865 | 22.137 | 21.208 | 19.631 | 34.529 |
| 153 | ENSMUSG00000000889 | 0 | NA | NA | NA | NA | NA | Dbh | dopamine beta hydroxylase [Source:MGI S | 0 | 0 | 0 | 0 | 0 | 0 |
| 154 | ENSMUSG00000000901 | 7.88286682 | -0.328695911 | 0.754577492 | -0.435602591 | 0.663125091 | 0.929111881 | Mmp11 | matrix metallopeptidase 11 [Source:MGI S | 8.031 | 11.307 | 7.0437 | 7.7118 | 7.6343 | 5.5692 |
| 155 | ENSMUSG00000000902 | 257.2275958 | 0.070483006 | 0.164401407 | 0.42872508 | 0.668123307 | 0.930136128 | Smarcb1 | SWI/SNF related, matrix associated, actin | 281.89 | 242.59 | 227.41 | 245.81 | 267.2 | 278.46 |
| 156 | ENSMUSG00000000903 | 11.36080005 | -0.354229452 | 0.88487406 | -0.400316235 | 0.688923612 | 0.936908509 | Vpreb3 | pre-B lymphocyte gene 3 [Source:MGI Syr | 4.8186 | 19.53 | 14.087 | 14.46 | 15.269 | 0 |
| 157 | ENSMUSG00000000915 | 416.2652129 | 0.208214779 | 0.150447026 | 1.383974047 | 0.166366402 | 0.5925379 | Hip1r | huntingtin interacting protein 1 related [S | 399.94 | 409.11 | 350.17 | 513.8 | 403.53 | 421.03 |
| 158 | ENSMUSG00000000916 | 177.1582511 | 0.193943221 | 0.195026372 | 0.994446134 | 0.320005718 | 0.767207475 | Nsun5 | NOL1/NOP2/Sun domain family, member | 155 | 160.36 | 181.12 | 223.64 | 167.96 | 174.87 |
| 159 | ENSMUSG00000000934 | 368.701923 | 0.007040791 | 0.144182038 | 0.048832653 | 0.96105266 | 0.992557454 | Top1mt | DNA topoisomerase 1, mitochondrial [Sou | 349.35 | 375.19 | 379.35 | 384.63 | 328.28 | 395.41 |
| 160 | ENSMUSG00000000938 | 0 | NA | NA | NA | NA | NA | Hoxa10 | homeobox A10 [Source:MGI Symbol;Acc:I | 0 | 0 | 0 | 0 | 0 | 0 |
| 161 | ENSMUSG00000000942 | 4.282089825 | -1.364155763 | 1.149953483 | -1.186270387 | 0.235515537 | NA | Hoxa4 | homeobox A4 [Source:MGI Symbol;Acc:M | 6.4248 | 3.0838 | 9.0562 | 3.8559 | 3.2719 | 0 |
| 162 | ENSMUSG00000000948 | 16.46657718 | 0.250003609 | 0.557214322 | 0.448666877 | 0.653671982 | 0.925797231 | Gm38393 | predicted gene, 38393 [Source:MGI Symb | 8.8341 | 14.391 | 22.137 | 19.28 | 17.45 | 16.708 |
| 163 | ENSMUSG00000000957 | 502.6225405 | -0.426349189 | 0.154414014 | -2.761078341 | 0.005761086 | 0.073396321 | Mmp14 | matrix metallopeptidase 14 (membrane-ir | 527.64 | 630.11 | 571.55 | 416.44 | 362.09 | 507.91 |
| 164 | ENSMUSG00000000958 | 30.71824816 | -0.802115858 | 0.469153656 | -1.709708211 | 0.087319844 | 0.435815811 | Slc7a7 | solute carrier family 7 (cationic amino acid | 28.912 | 52.424 | 36.225 | 34.703 | 11.997 | 20.049 |

| | | | | | | | | | | | | | | | |
|---|---|---|---|---|---|---|---|---|---|---|---|---|---|---|---|
| 165 | ENSMUSG00000000959 | 770.8790425 | 0.134674572 | 0.121092472 | 1.112163049 | 0.266068052 | 0.722063933 | Oxa1l | oxidase assembly 1-like [Source:MGI Syml | 710.75 | 796.64 | 697.33 | 741.3 | 856.14 | 823.13 |
| 166 | ENSMUSG00000000976 | 230.0879251 | -0.084783666 | 0.179172431 | -0.473195935 | 0.636073398 | 0.923478764 | Heatr6 | HEAT repeat containing 6 [Source:MGI Sy | 208 | 274.45 | 228.42 | 201.47 | 244.3 | 223.88 |
| 167 | ENSMUSG00000000982 | 0.439020386 | -2.189747911 | 3.52148864 | -0.621824499 | 0.534057275 | NA | Ccl3 | chemokine (C-C motif) ligand 3 [Source:M | 1.6062 | 1.0279 | 0 | 0 | 0 | 0 |
| 168 | ENSMUSG00000000983 | 0.633540418 | -1.320719359 | 2.623870149 | -0.503347835 | 0.61471974 | NA | Wfdc18 | WAP four-disulfide core domain 18 [Sourc | 0.8031 | 1.0279 | 1.0062 | 0.964 | 0 | 0 |
| 169 | ENSMUSG00000001014 | 0 | NA | NA | NA | NA | NA | Icam4 | intercellular adhesion molecule 4, Landste | 0 | 0 | 0 | 0 | 0 | 0 |
| 170 | ENSMUSG00000001016 | 277.07507 | 0.188564895 | 0.167236195 | 1.127536386 | 0.259515765 | 0.712728614 | Ilf2 | interleukin enhancer binding factor 2 [Sou | 237.72 | 282.68 | 256.59 | 251.6 | 309.74 | 324.13 |
| 171 | ENSMUSG00000001017 | 1233.798093 | 0.078482185 | 0.108959617 | 0.720286903 | 0.471348367 | 0.867794597 | Chtop | chromatin target of PRMT1 [Source:MGI S | 1194.2 | 1113.2 | 1293 | 1238.7 | 1224.8 | 1338.8 |
| 172 | ENSMUSG00000001018 | 374.8494208 | 0.380646576 | 0.157231862 | 2.420925198 | 0.015481063 | 0.145412036 | Snapin | SNAP-associated protein [Source:MGI Syn | 362.2 | 326.88 | 286.78 | 367.28 | 451.52 | 454.45 |
| 173 | ENSMUSG00000001020 | 0 | NA | NA | NA | NA | NA | S100a4 | S100 calcium binding protein A4 [Source:I | 0 | 0 | 0 | 0 | 0 | 0 |
| 174 | ENSMUSG00000001021 | 0 | NA | NA | NA | NA | NA | S100a3 | S100 calcium binding protein A3 [Source:I | 0 | 0 | 0 | 0 | 0 | 0 |
| 175 | ENSMUSG00000001023 | 0 | NA | NA | NA | NA | NA | S100a5 | S100 calcium binding protein A5 [Source:I | 0 | 0 | 0 | 0 | 0 | 0 |
| 176 | ENSMUSG00000001025 | 27.34277403 | -0.721884634 | 0.489765394 | -1.473939652 | 0.14049784 | 0.548157814 | S100a6 | S100 calcium binding protein A6 (calcyclin | 38.549 | 45.228 | 18.112 | 12.532 | 22.903 | 26.732 |
| 177 | ENSMUSG00000001027 | 0.315620088 | 0.165143421 | 4.063300985 | 0.040642675 | 0.967580763 | NA | Scn4a | sodium channel, voltage-gated, type IV, al | 0.8031 | 0 | 0 | 0 | 1.0906 | 0 |
| 178 | ENSMUSG00000001029 | 32.92566493 | -0.662573274 | 0.427498371 | -1.549884909 | 0.121169142 | 0.510364145 | Icam2 | intercellular adhesion molecule 2 [Source | 33.73 | 35.977 | 51.318 | 21.208 | 17.45 | 37.871 |
| 179 | ENSMUSG00000001034 | 20.8384259 | -0.502391362 | 0.516625074 | -0.972448661 | 0.330827394 | 0.774747658 | Mapk7 | mitogen-activated protein kinase 7 [Sourc | 17.668 | 29.81 | 26.162 | 26.027 | 11.997 | 13.366 |
| 180 | ENSMUSG00000001036 | 151.4853391 | -0.204430956 | 0.190865683 | -1.071072355 | 0.284136897 | 0.736651636 | Epn2 | epsin 2 [Source:MGI Symbol;Acc:MGI:133 | 155 | 166.52 | 165.02 | 129.17 | 145.05 | 148.14 |
| 181 | ENSMUSG00000001039 | 5.396314519 | -0.827506048 | 1.022301911 | -0.809453684 | 0.418254233 | 0.837575303 | B9d1 | B9 protein domain 1 [Source:MGI Symbol | 10.44 | 5.1396 | 5.0312 | 1.928 | 8.7249 | 1.1138 |
| 182 | ENSMUSG00000001052 | 338.89827 | 0.07443314 | 0.145106075 | 0.512956743 | 0.607981579 | 0.916376001 | Sec24b | Sec24 related gene family, member B (S. c | 338.91 | 354.63 | 296.84 | 349.92 | 353.36 | 339.72 |
| 183 | ENSMUSG00000001053 | 73.24163884 | 0.373729757 | 0.310006957 | 1.205552808 | 0.227989965 | 0.674239914 | N4bp3 | NEDD4 binding protein 3 [Source:MGI Syn | 67.461 | 41.117 | 82.512 | 67.478 | 83.978 | 96.904 |
| 184 | ENSMUSG00000001054 | 260.2724338 | -0.084254037 | 0.161442462 | -0.521882756 | 0.601751965 | 0.915461118 | Rmnd5b | required for meiotic nuclear division 5 hor | 236.92 | 284.73 | 282.75 | 262.2 | 238.85 | 256.18 |
| 185 | ENSMUSG00000001056 | 450.280465 | -0.076512975 | 0.13793869 | -0.55468828 | 0.579107894 | 0.908337213 | Nhp2 | NHP2 ribonucleoprotein [Source:MGI Sym | 503.55 | 457.42 | 424.64 | 404.87 | 466.78 | 444.42 |
| 186 | ENSMUSG00000001062 | 490.6614294 | -0.252189992 | 0.129087148 | -1.953641372 | 0.050743644 | 0.316199857 | Vps9d1 | VPS9 domain containing 1 [Source:MGI Sy | 538.88 | 561.24 | 500.1 | 440.54 | 476.6 | 426.6 |
| 187 | ENSMUSG00000001065 | 696.5745397 | -0.83989744 | 0.183434077 | -4.578742704 | 4.68E-06 | 0.000287913 | Zfp276 | zinc finger protein (C2H2 type) 276 [Sourc | 881.81 | 996.05 | 802.98 | 392.34 | 669.64 | 436.63 |
| 188 | ENSMUSG00000001076 | 1.5430864 | 2.003247258 | 1.813519248 | 1.104618691 | 0.269324849 | NA | C1ql4 | complement component 1, q subcompon | 0.8031 | 1.0279 | 0 | 1.928 | 3.2719 | 2.2277 |
| 189 | ENSMUSG00000001082 | 76.70406538 | 0.149134124 | 0.316364104 | 0.471400268 | 0.637354924 | 0.923478764 | Mfsd10 | major facilitator superfamily domain cont | 59.43 | 70.926 | 88.55 | 114.71 | 69.8 | 56.806 |
| 190 | ENSMUSG00000001089 | 277.2916424 | 0.084099166 | 0.148906045 | 0.564780063 | 0.57222337 | 0.905144228 | Luzp1 | leucine zipper protein 1 [Source:MGI Sym | 258.6 | 278.57 | 270.68 | 280.52 | 283.56 | 291.83 |
| 191 | ENSMUSG00000001095 | 6.858516215 | 0.315355463 | 0.995772986 | 0.316694134 | 0.7514757 | 0.952255112 | Slc13a2 | solute carrier family 13 (sodium-depender | 7.2279 | 1.0279 | 10.062 | 10.604 | 1.0906 | 11.138 |
| 192 | ENSMUSG00000001098 | 88.45882178 | -0.355530301 | 0.259360564 | -1.370795529 | 0.170438705 | 0.597556794 | Kctd10 | potassium channel tetramerisation domai | 115.65 | 97.652 | 84.525 | 90.614 | 64.346 | 77.969 |
| 193 | ENSMUSG00000001100 | 1786.096567 | 0.046668815 | 0.093266354 | 0.500382109 | 0.616806048 | 0.917509899 | Poldip2 | polymerase (DNA-directed), delta interact | 1750 | 1792.7 | 1728.7 | 1760.2 | 1812.6 | 1872.4 |
| 194 | ENSMUSG00000001103 | 5.449066968 | 0.798554386 | 0.918979206 | 0.868958058 | 0.384870072 | 0.816437825 | Sebox | SEBOX homeobox [Source:MGI Symbol;Ac | 4.8186 | 2.0558 | 5.0312 | 8.6758 | 6.5437 | 5.5692 |
| 195 | ENSMUSG00000001105 | 849.5592044 | 0.106801473 | 0.114576759 | 0.932139061 | 0.351264663 | 0.792511048 | Ift20 | intraflagellar transport 20 [Source:MGI Sy | 860.12 | 785.33 | 808.01 | 855.05 | 836.5 | 952.34 |
| 196 | ENSMUSG00000001119 | 121.7103722 | -0.61804891 | 0.220347557 | -2.80488206 | 0.005033499 | 0.066915933 | Col6a1 | collagen, type VI, alpha 1 [Source:MGI Syr | 152.59 | 150.08 | 139.87 | 114.71 | 87.249 | 85.766 |
| 197 | ENSMUSG00000001120 | 18.68686053 | -0.271190837 | 0.563926759 | -0.480897267 | 0.630589515 | 0.921020101 | Pcbp3 | poly(rC) binding protein 3 [Source:MGI Sy | 14.456 | 32.893 | 14.087 | 16.388 | 10.906 | 23.391 |
| 198 | ENSMUSG00000001123 | 6341.074164 | 0.232358229 | 0.140833159 | 1.649882958 | 0.098966877 | 0.463570426 | Lgals9 | lectin, galactose binding, soluble 9 [Sourc | 6964.5 | 5276.3 | 5251.6 | 6007.5 | 6497.9 | 8048.6 |
| 199 | ENSMUSG00000001127 | 1217.767444 | 0.181660612 | 0.110424202 | 1.645115915 | 0.099945909 | 0.465620067 | Araf | Araf proto-oncogene, serine/threonine kir | 1147.6 | 1203.7 | 1071.7 | 1185.7 | 1350.2 | 1347.8 |
| 200 | ENSMUSG00000001128 | 232.7632519 | -1.614972762 | 0.239683057 | -6.737951281 | 1.61E-11 | 5.05E-09 | Cfp | complement factor properdin [Source:MG | 314.01 | 484.15 | 254.58 | 102.18 | 106.88 | 134.78 |
| 201 | ENSMUSG00000001131 | 3.963598487 | -1.170861732 | 1.174379601 | -0.997004488 | 0.318762332 | NA | Timp1 | tissue inhibitor of metalloproteinase 1 [So | 7.2279 | 8.2233 | 1.0062 | 2.8919 | 1.0906 | 3.3415 |
| 202 | ENSMUSG00000001134 | 205.2311419 | 0.111068468 | 0.168485425 | 0.659217068 | 0.509756388 | 0.882793133 | Uxt | ubiquitously expressed prefoldin like chap | 205.59 | 185.03 | 201.25 | 221.71 | 203.95 | 213.86 |
| 203 | ENSMUSG00000001138 | 291.2060342 | 0.123867676 | 0.175317469 | 0.706533563 | 0.479856388 | 0.870278888 | Cnnm3 | cyclin M3 [Source:MGI Symbol;Acc:MGI:2 | 259.4 | 243.62 | 333.07 | 274.73 | 294.47 | 341.95 |
| 204 | ENSMUSG00000001143 | 238.5322625 | 0.197583895 | 0.171937309 | 1.149162429 | 0.25048901 | 0.702302228 | Lman2l | lectin, mannose-binding 2-like [Source:MC | 203.99 | 218.95 | 244.52 | 286.3 | 227.94 | 249.5 |
| 205 | ENSMUSG00000001150 | 395.558047 | 0.068522447 | 0.145283594 | 0.471646147 | 0.637179382 | 0.923478764 | Mcm3ap | minichromosome maintenance complex c | 363.81 | 367.99 | 426.65 | 365.35 | 432.98 | 416.58 |
| 206 | ENSMUSG00000001151 | 213.7235672 | 0.153065854 | 0.173242786 | 0.883533783 | 0.376947938 | 0.81030298 | Pcnt | pericentrin (kendrin) [Source:MGI Symbol | 208.81 | 181.94 | 216.34 | 230.39 | 241.03 | 203.83 |
| 207 | ENSMUSG00000001155 | 10171.26713 | 0.145088633 | 0.086576376 | 1.675845521 | 0.093768455 | 0.450886614 | Ftcd | formiminotransferase cyclodeaminase [Sc | 9422.8 | 9389 | 10169 | 10401 | 11138 | 10507 |
| 208 | ENSMUSG00000001156 | 105.6596598 | 0.393201675 | 0.221672664 | 1.773794155 | 0.076097205 | 0.403877794 | Mxd1 | MAX dimerization protein 1 [Source:MGI S | 85.932 | 97.652 | 90.562 | 111.82 | 123.24 | 124.75 |
| 209 | ENSMUSG00000001157 | 236.2587788 | 0.214657571 | 0.174854008 | 1.227638842 | 0.219582574 | 0.665321537 | Gmcl1 | germ cell-less, spermatogenesis associate | 206.4 | 202.5 | 247.54 | 252.56 | 283.56 | 225 |
| 210 | ENSMUSG00000001158 | 233.1013116 | 0.090469059 | 0.182474263 | 0.495790789 | 0.620042024 | 0.917977899 | Snrnp27 | small nuclear ribonucleoprotein 27 (U4/U | 206.4 | 236.42 | 234.45 | 194.72 | 272.65 | 253.96 |
| 211 | ENSMUSG00000001166 | 18.35180682 | 0.491288315 | 0.546114683 | 0.899606493 | 0.3683297 | 0.80627882 | Oas1c | 2'-5' oligoadenylate synthetase 1C [Source | 10.44 | 13.363 | 22.137 | 20.244 | 29.447 | 14.48 |
| 212 | ENSMUSG00000001168 | 0 | NA | NA | NA | NA | NA | Oas1h | 2'-5' oligoadenylate synthetase 1H [Sourc | 0 | 0 | 0 | 0 | 0 | 0 |
| 213 | ENSMUSG00000001173 | 111.5589366 | -0.00621793 | 0.251453804 | -0.024727923 | 0.980271983 | 0.99820894 | Ocrl | OCRL, inositol polyphosphate-5-phosphat | 93.16 | 133.63 | 108.67 | 86.758 | 111.24 | 135.89 |
| 214 | ENSMUSG00000001175 | 3563.234187 | -0.076293135 | 0.101592459 | -0.750972424 | 0.452669252 | 0.856769845 | Calm1 | calmodulin 1 [Source:MGI Symbol;Acc:MC | 3997.8 | 3601.8 | 3370.9 | 3300.7 | 3627.4 | 3480.8 |
| 215 | ENSMUSG00000001211 | 5995.506533 | -0.138796885 | 0.083488986 | -1.662457417 | 0.096421095 | 0.4584598 | Agpat3 | 1-acylglycerol-3-phosphate O-acyltransfer | 6154.2 | 6483.1 | 6214.6 | 5833 | 5573.1 | 5715.1 |
| 216 | ENSMUSG00000001225 | 2.033902574 | 1.019073505 | 1.885384065 | 0.54051242 | 0.588843698 | NA | Slc26a3 | solute carrier family 26, member 3 [Sourc | 0 | 4.1117 | 0 | 4.8199 | 3.2719 | 0 |
| 217 | ENSMUSG00000001227 | 186.2685457 | 0.078866734 | 0.219784113 | 0.358837282 | 0.71971682 | 0.9447719 | Sema6b | sema domain, transmembrane domain (TI | 195.15 | 160.36 | 188.17 | 237.14 | 137.42 | 199.38 |
| 218 | ENSMUSG00000001228 | 8.240049062 | -0.382659462 | 0.934126862 | -0.409643997 | 0.682067117 | 0.934933546 | Uhrf1 | ubiquitin-like, containing PHD and RING fi | 17.668 | 1.0279 | 9.0562 | 2.8919 | 6.5437 | 12.252 |
| 219 | ENSMUSG00000001229 | 847.4212856 | -0.008578973 | 0.10977825 | -0.078148207 | 0.93771016 | 0.988420305 | Dpp9 | dipeptidylpeptidase 9 [Source:MGI Symbo | 845.67 | 873.73 | 830.15 | 811.67 | 827.78 | 895.53 |
| 220 | ENSMUSG00000001240 | 87.22589142 | -0.811302018 | 0.261476579 | -3.102771273 | 0.001917178 | 0.033734861 | Ramp2 | receptor (calcitonin) activity modifying pro | 115.65 | 114.1 | 103.64 | 68.442 | 46.897 | 74.628 |

| | | | | | | | | | | | | | | | |
|---|---|---|---|---|---|---|---|---|---|---|---|---|---|---|---|
| 221 | ENSMUSG00000001247 | 1152.19475 | 0.28967497 | 0.126868201 | 2.283274837 | 0.022414184 | 0.187573567 | Lsr | lipolysis stimulated lipoprotein receptor [S | 900.28 | 1004.3 | 1207.5 | 1285.9 | 1249.8 | 1265.3 |
| 222 | ENSMUSG00000001248 | 233.3374746 | 0.052739515 | 0.171400707 | 0.307697182 | 0.758312764 | 0.953676708 | Gramd1a | GRAM domain containing 1A [Source:MGI | 215.23 | 228.2 | 244.52 | 266.06 | 238.85 | 207.17 |
| 223 | ENSMUSG00000001249 | 9933.347482 | 0.123518395 | 0.083128372 | 1.485875304 | 0.137312107 | 0.541266927 | Hpn | hepsin [Source:MGI Symbol;Acc:MGI:119 | 9487.8 | 9290.3 | 9746.5 | 10032 | 10277 | 10766 |
| 224 | ENSMUSG00000001260 | 0 | NA | NA | NA | NA | NA | Gabrg1 | gamma-aminobutyric acid (GABA) A recep | 0 | 0 | 0 | 0 | 0 | 0 |
| 225 | ENSMUSG00000001270 | 50.96981615 | -1.119220125 | 0.406221121 | -2.755199241 | 0.00586564 | 0.074161384 | Ckb | creatine kinase, brain [Source:MGI Symbo | 76.295 | 90.457 | 42.262 | 17.352 | 34.9 | 44.554 |
| 226 | ENSMUSG00000001280 | 692.7144749 | 0.095833827 | 0.13619973 | 0.703627144 | 0.481665003 | 0.871274502 | Sp1 | trans-acting transcription factor 1 [Source | 595.1 | 702.07 | 712.42 | 657.43 | 677.27 | 811.99 |
| 227 | ENSMUSG00000001281 | 18.61867902 | -0.931611643 | 0.520936449 | -1.788340293 | 0.073721123 | 0.39478057 | Itgb7 | integrin beta 7 [Source:MGI Symbol;Acc:M | 23.29 | 25.698 | 24.15 | 7.7118 | 15.269 | 15.594 |
| 228 | ENSMUSG00000001285 | 306.2810027 | 0.027370875 | 0.152848526 | 0.179071892 | 0.857881252 | 0.975277277 | Myg1 | melanocyte proliferating gene 1 [Source:M | 308.39 | 279.59 | 322 | 305.58 | 338.09 | 284.03 |
| 229 | ENSMUSG00000001288 | 69.38968132 | -0.391691347 | 0.364283821 | -1.075236735 | 0.282268753 | 0.735230549 | Rarg | retinoic acid receptor, gamma [Source:MG | 93.963 | 54.48 | 87.543 | 60.731 | 31.628 | 87.994 |
| 230 | ENSMUSG00000001289 | 860.6631345 | -0.085455571 | 0.113619479 | -0.752120777 | 0.45197843 | 0.856579667 | Pfdn5 | prefoldin 5 [Source:MGI Symbol;Acc:MGI: | 910.72 | 896.34 | 851.28 | 865.65 | 765.61 | 874.37 |
| 231 | ENSMUSG00000001300 | 36.38185435 | -0.801056524 | 0.436964306 | -1.833231028 | 0.066768221 | 0.37192178 | Efnb2 | ephrin B2 [Source:MGI Symbol;Acc:MGI:1 | 36.14 | 65.787 | 37.231 | 39.523 | 22.903 | 16.708 |
| 232 | ENSMUSG00000001305 | 128.9478728 | 0.032049451 | 0.211810454 | 0.151311943 | 0.879729648 | 0.98128671 | Rrp15 | ribosomal RNA processing 15 homolog [Sc | 115.65 | 124.38 | 142.89 | 132.06 | 142.87 | 115.84 |
| 233 | ENSMUSG00000001313 | 202.3267132 | 0.129757851 | 0.18203783 | 0.712807061 | 0.475965156 | 0.869483933 | Rnd2 | Rho family GTPase 2 [Source:MGI Symbol | 216.03 | 193.25 | 170.06 | 221.71 | 224.67 | 188.24 |
| 234 | ENSMUSG00000001323 | 1096.617534 | 0.088207443 | 0.101626761 | 0.86795488 | 0.385419033 | 0.816437825 | Srr | serine racemase [Source:MGI Symbol;Acc | 1053.7 | 1037.2 | 1098.8 | 1159.7 | 1128.8 | 1101.6 |
| 235 | ENSMUSG00000001333 | 0.48903338 | 0.95637552 | 3.456690196 | 0.276673773 | 0.782030612 | NA | Sync | syncoilin [Source:MGI Symbol;Acc:MGI:19 | 0 | 0 | 1.0062 | 1.928 | 0 | 0 |
| 236 | ENSMUSG00000001334 | 8.331716443 | -0.751649076 | 0.832909165 | -0.902438235 | 0.366824118 | 0.805239112 | Fndc5 | fibronectin type III domain containing 5 [S | 20.078 | 4.1117 | 7.0437 | 7.7118 | 4.3625 | 6.6831 |
| 237 | ENSMUSG00000001348 | 2134.678573 | -0.062861808 | 0.108854639 | -0.577483957 | 0.563612573 | 0.900821918 | Acp5 | acid phosphatase 5, tartrate resistant [Sou | 2119.4 | 2378.6 | 2046.7 | 2274 | 1955.5 | 2033.9 |
| 238 | ENSMUSG00000001349 | 1.331156114 | 0.754896071 | 2.025254231 | 0.372741387 | 0.70934093 | NA | Cnn1 | calponin 1 [Source:MGI Symbol;Acc:MGI: | 0 | 1.0279 | 2.0125 | 3.8559 | 1.0906 | 0 |
| 239 | ENSMUSG00000001366 | 1245.401415 | 0.078207853 | 0.109202947 | 0.716169804 | 0.473886506 | 0.868952133 | Fbxo9 | f-box protein 9 [Source:MGI Symbol;Acc:M | 1106.7 | 1241.7 | 1288 | 1337 | 1225.9 | 1273.1 |
| 240 | ENSMUSG00000001376 | 213.5790485 | -0.000479729 | 0.183046358 | -0.002620807 | 0.997908901 | 0.999705939 | Vps50 | VPS50 EARP/GARPII complex subunit [Sou | 203.99 | 201.47 | 235.46 | 201.47 | 250.84 | 188.24 |
| 241 | ENSMUSG00000001380 | 589.0227206 | 0.007454232 | 0.118378084 | 0.062969694 | 0.949790637 | 0.990766637 | Hars | histidyl-tRNA synthetase [Source:MGI Syn | 565.38 | 580.77 | 616.83 | 600.56 | 578.03 | 592.56 |
| 242 | ENSMUSG00000001383 | 660.0198633 | 0.048173151 | 0.137167863 | 0.351198525 | 0.725439416 | 0.945037892 | Zmat2 | zinc finger, matrin type 2 [Source:MGI Syr | 619.19 | 627.03 | 700.35 | 568.75 | 725.26 | 719.54 |
| 243 | ENSMUSG00000001403 | 17.25035419 | 0.306344742 | 0.600051896 | 0.510530413 | 0.609679913 | 0.916376001 | Ube2c | ubiquitin-conjugating enzyme E2C [Source | 22.487 | 18.503 | 5.0312 | 14.46 | 19.631 | 23.391 |
| 244 | ENSMUSG00000001415 | 676.160154 | 0.002612218 | 0.132557249 | 0.019706341 | 0.984277632 | 0.998580735 | Smg5 | Smg-5 homolog, nonsense mediated mRN | 603.13 | 758.6 | 666.13 | 712.38 | 620.56 | 696.15 |
| 245 | ENSMUSG00000001416 | 1441.366076 | 0.005358912 | 0.096732102 | 0.055399525 | 0.955820174 | 0.991678478 | Cct3 | chaperonin containing Tcp1, subunit 3 (ga | 1458.4 | 1427.8 | 1429.9 | 1484.5 | 1384 | 1463.6 |
| 246 | ENSMUSG00000001418 | 834.1380417 | 0.05704126 | 0.117881511 | 0.483886397 | 0.628466495 | 0.920328819 | Glmp | glycosylated lysosomal membrane proteir | 870.56 | 843.92 | 737.58 | 825.17 | 835.41 | 892.19 |
| 247 | ENSMUSG00000001419 | 412.031156 | -0.168483508 | 0.149383288 | -1.127860491 | 0.259378842 | 0.712728614 | Mef2d | myocyte enhancer factor 2D [Source:MGI | 379.87 | 467.7 | 461.87 | 425.11 | 353.36 | 384.28 |
| 248 | ENSMUSG00000001420 | 42.64142177 | 0.242253357 | 0.367447179 | 0.659287567 | 0.509711124 | 0.882793133 | Tmem79 | transmembrane protein 79 [Source:MGI S | 53.808 | 30.838 | 32.2 | 52.055 | 49.078 | 37.871 |
| 249 | ENSMUSG00000001424 | 2048.432283 | 0.285490332 | 0.116940476 | 2.441330337 | 0.014633262 | 0.140541507 | Snd1 | staphylococcal nuclease and tudor domai | 1939.5 | 1803 | 1797.2 | 2592.1 | 2103.8 | 2055 |
| 250 | ENSMUSG00000001435 | 18145.09422 | -0.016339674 | 0.095693091 | -0.170750822 | 0.864419699 | 0.977227145 | Col18a1 | collagen, type XVIII, alpha 1 [Source:MGI S | 17121 | 18410 | 19213 | 16961 | 17628 | 19538 |
| 251 | ENSMUSG00000001436 | 285.014311 | -0.357534939 | 0.168242679 | -2.125114401 | 0.033577068 | 0.245153552 | Slc19a1 | solute carrier family 19 (folate transporter | 285.1 | 343.32 | 333.07 | 279.55 | 256.3 | 212.74 |
| 252 | ENSMUSG00000001440 | 1645.386072 | 0.054615683 | 0.099247 | 0.550300592 | 0.582113218 | 0.908645447 | Kpnb1 | karyopherin (importin) beta 1 [Source:MG | 1579.7 | 1651.9 | 1611 | 1597.3 | 1641.4 | 1791.1 |
| 253 | ENSMUSG00000001441 | 539.3297236 | 0.177394642 | 0.122473361 | 1.448434506 | 0.14749557 | 0.55862161 | Npepps | aminopeptidase puromycin sensitive [Sou | 497.12 | 504.71 | 517.21 | 596.7 | 536.58 | 583.65 |
| 254 | ENSMUSG00000001444 | 2.196317325 | -1.654744586 | 1.676991002 | -0.986734326 | 0.323772898 | NA | Tbx21 | T-box 21 [Source:MGI Symbol;Acc:MGI:18 | 0.8031 | 8.2233 | 1.0062 | 0.964 | 2.1812 | 0 |
| 255 | ENSMUSG00000001445 | 755.2729558 | -0.035632973 | 0.113663167 | -0.313496215 | 0.753903696 | 0.953048681 | Mrpl10 | mitochondrial ribosomal protein L10 [Sou | 742.87 | 743.18 | 808.01 | 739.37 | 780.88 | 717.32 |
| 256 | ENSMUSG00000001467 | 1081.178941 | 1.341689374 | 0.197199198 | 6.803726328 | 1.02E-11 | 3.57E-09 | Cyp51 | cytochrome P450, family 51 [Source:MGI S | 742.87 | 692.82 | 398.47 | 1396.8 | 1396 | 1860.1 |
| 257 | ENSMUSG00000001472 | 2902.062179 | 0.141639642 | 0.098900158 | 1.432147782 | 0.152101532 | 0.567594572 | Tcf25 | transcription factor 25 (basic helix-loop-he | 2695.2 | 2791.8 | 2793.3 | 3297.8 | 3037.4 | 2796.9 |
| 258 | ENSMUSG00000001473 | 137.5455831 | -0.791476914 | 0.275956206 | -2.868125072 | 0.004129123 | 0.058667613 | Tubb6 | tubulin, beta 6 class V [Source:MGI Symbo | 196.76 | 180.91 | 144.9 | 74.226 | 150.51 | 77.969 |
| 259 | ENSMUSG00000001482 | 1111.052668 | -0.085526553 | 0.128627045 | -0.664918898 | 0.506102367 | 0.881592135 | Def8 | differentially expressed in FDCP 8 [Source | 1149.2 | 1074.2 | 1207.5 | 907.1 | 1185.5 | 1142.8 |
| 260 | ENSMUSG00000001493 | 0.863771898 | -1.920093103 | 2.569535405 | -0.747253025 | 0.454910842 | NA | Meox1 | mesenchyme homeobox 1 [Source:MGI Sy | 3.2124 | 0 | 1.0062 | 0.964 | 0 | 0 |
| 261 | ENSMUSG00000001494 | 0 | NA | NA | NA | NA | NA | Sost | sclerostin [Source:MGI Symbol;Acc:MGI:1 | 0 | 0 | 0 | 0 | 0 | 0 |
| 262 | ENSMUSG00000001496 | 0 | NA | NA | NA | NA | NA | Nkx2-1 | NK2 homeobox 1 [Source:MGI Symbol;Ac | 0 | 0 | 0 | 0 | 0 | 0 |
| 263 | ENSMUSG00000001497 | 6.361135157 | 0.183361263 | 0.969139283 | 0.189200114 | 0.849935978 | 0.973277966 | Pax9 | paired box 9 [Source:MGI Symbol;Acc:MG | 9.6372 | 1.0279 | 7.0437 | 2.8919 | 11.997 | 5.5692 |
| 264 | ENSMUSG00000001504 | 0 | NA | NA | NA | NA | NA | Irx2 | Iroquois homeobox 2 [Source:MGI Symbo | 0 | 0 | 0 | 0 | 0 | 0 |
| 265 | ENSMUSG00000001506 | 109.3101413 | -1.389846493 | 0.755438736 | -1.839787169 | 0.06579949 | 0.368337165 | Col1a1 | collagen, type I, alpha 1 [Source:MGI Sym | 291.53 | 140.82 | 42.262 | 63.623 | 73.071 | 44.554 |
| 266 | ENSMUSG00000001507 | 128.1866572 | 0.294750993 | 0.21703681 | 1.358069138 | 0.174441747 | 0.604196702 | Itga3 | integrin alpha 3 [Source:MGI Symbol;Acc: | 114.04 | 97.652 | 133.83 | 152.31 | 127.6 | 143.69 |
| 267 | ENSMUSG00000001508 | 0.17131964 | -0.796629176 | 4.080472857 | -0.195229623 | 0.84521319 | NA | Sgca | sarcoglycan, alpha (dystrophin-associated | 0 | 1.0279 | 0 | 0 | 0 | 0 |
| 268 | ENSMUSG00000001510 | 0 | NA | NA | NA | NA | NA | Dlx3 | distal-less homeobox 3 [Source:MGI Symb | 0 | 0 | 0 | 0 | 0 | 0 |
| 269 | ENSMUSG00000001517 | 7.235366441 | -1.275321495 | 0.875493634 | -1.456688484 | 0.14520239 | 0.555361204 | Foxm1 | forkhead box M1 [Source:MGI Symbol;Ac | 7.2279 | 18.503 | 5.0312 | 3.8559 | 5.4531 | 3.3415 |
| 270 | ENSMUSG00000001518 | 162.8543184 | 0.1453216 | 0.204488126 | 0.710660337 | 0.477294744 | 0.869688987 | Itfg2 | integrin alpha FG-GAP repeat containing 2 | 141.35 | 153.16 | 170.06 | 187.98 | 187.59 | 137 |
| 271 | ENSMUSG00000001520 | 3.97028741 | 0.049448988 | 1.112413286 | 0.044451993 | 0.964544118 | NA | Nrip2 | nuclear receptor interacting protein 2 [So | 5.6217 | 3.0838 | 3.0187 | 7.7118 | 3.2719 | 1.1138 |
| 272 | ENSMUSG00000001521 | 17.0395387 | -0.596379375 | 0.547018278 | -1.090236652 | 0.275608912 | 0.730038567 | Tulp3 | tubby-like protein 3 [Source:MGI Symbol; | 17.668 | 23.642 | 20.125 | 7.7118 | 15.269 | 17.822 |
| 273 | ENSMUSG00000001524 | 265.3262918 | 0.011703571 | 0.150790296 | 0.077614879 | 0.938134407 | 0.988420305 | Gtf2h4 | general transcription factor II H, polypepti | 265.83 | 262.12 | 264.64 | 260.27 | 261.75 | 277.35 |
| 274 | ENSMUSG00000001525 | 605.9140319 | 0.410212321 | 0.150418982 | 2.727131345 | 0.00638876 | 0.078845588 | Tubb5 | tubulin, beta 5 class I [Source:MGI Symbo | 531.65 | 565.35 | 463.88 | 697.92 | 576.94 | 799.74 |
| 275 | ENSMUSG00000001542 | 1698.325562 | -0.293597613 | 0.139599407 | -2.103143692 | 0.035453211 | 0.252635616 | Ell2 | elongation factor for RNA polymerase II 2 | 1664 | 2152.5 | 1795.1 | 1279.2 | 1729.7 | 1569.4 |
| 276 | ENSMUSG00000001552 | 840.6646889 | -0.20918114 | 0.12598658 | -1.66034461 | 0.096845146 | 0.458530879 | Jup | junction plakoglobin [Source:MGI Symbol | 936.42 | 807.94 | 959.96 | 817.45 | 699.09 | 823.13 |

| | | | | | | | | | | | | | | | |
|---|---|---|---|---|---|---|---|---|---|---|---|---|---|---|---|
| 277 | ENSMUSG00000001555 | 15.49875564 | -1.138919587 | 0.588939488 | -1.933848229 | 0.053131791 | 0.323714501 | Fkbp10 | FK506 binding protein 10 [Source:MGI Syr | 27.305 | 21.586 | 15.094 | 12.532 | 10.906 | 5.5692 |
| 278 | ENSMUSG00000001558 | 0.538988726 | 1.126003851 | 3.343747182 | 0.336749099 | 0.736306047 | NA | Klhl10 | kelch-like 10 [Source:MGI Symbol;Acc:MG | 0 | 0 | 1.0062 | 0 | 0 | 2.2277 |
| 279 | ENSMUSG00000001566 | 0 | NA | NA | NA | NA | NA | Mnx1 | motor neuron and pancreas homeobox 1 | 0 | 0 | 0 | 0 | 0 | 0 |
| 280 | ENSMUSG00000001569 | 440.8810358 | 0.014472926 | 0.146415744 | 0.098848157 | 0.921258832 | 0.986685804 | Nom1 | nucleolar protein with MIF4G domain 1 [S | 387.9 | 457.42 | 471.93 | 480.06 | 454.79 | 393.19 |
| 281 | ENSMUSG00000001576 | 1229.440311 | 0.007925731 | 0.11625997 | 0.068172483 | 0.945648331 | 0.990309554 | Ergic1 | endoplasmic reticulum-golgi intermediate | 1303.4 | 1174.9 | 1198.4 | 1127.9 | 1196.4 | 1375.6 |
| 282 | ENSMUSG00000001583 | 0.877498349 | -0.489540585 | 2.815048567 | -0.173901293 | 0.861943037 | NA | Tnk1 | tyrosine kinase, non-receptor, 1 [Source:N | 0 | 3.0838 | 0 | 0 | 2.1812 | 0 |
| 283 | ENSMUSG00000001588 | 35.16784057 | -0.113608503 | 0.37198502 | -0.3054115 | 0.760052764 | 0.95402891 | Acap1 | ArfGAP with coiled-coil, ankyrin repeat an | 34.533 | 41.117 | 34.212 | 40.487 | 28.356 | 32.301 |
| 284 | ENSMUSG00000001604 | 1445.573861 | 0.219504183 | 0.111941051 | 1.960890851 | 0.049891757 | 0.312937715 | Tcea3 | transcription elongation factor A (SII), 3 [S | 1292.2 | 1225.3 | 1490.2 | 1541.4 | 1621.7 | 1502.6 |
| 285 | ENSMUSG00000001622 | 0.133850373 | -0.796629176 | 4.080472857 | -0.195229623 | 0.84521319 | NA | Csn3 | casein kappa [Source:MGI Symbol;Acc:M( | 0.8031 | 0 | 0 | 0 | 0 | 0 |
| 286 | ENSMUSG00000001627 | 265.6404456 | 0.082468658 | 0.195334879 | 0.422191156 | 0.672885497 | 0.931454408 | Ifrd1 | interferon-related developmental regulato | 234.51 | 329.96 | 210.31 | 294.01 | 241.03 | 284.03 |
| 287 | ENSMUSG00000001630 | 351.5018815 | 0.030680769 | 0.162936042 | 0.188299464 | 0.850641906 | 0.973463808 | Stk38l | serine/threonine kinase 38 like [Source:M | 366.21 | 340.24 | 337.09 | 402.94 | 376.26 | 286.26 |
| 288 | ENSMUSG00000001632 | 297.9626594 | 0.021785238 | 0.150886928 | 0.144381218 | 0.885199447 | 0.981795936 | Brpf1 | bromodomain and PHD finger containing, | 276.27 | 319.68 | 291.81 | 312.33 | 281.38 | 306.31 |
| 289 | ENSMUSG00000001642 | 109.2529664 | 0.230427045 | 0.279491849 | 0.824449965 | 0.409683935 | 0.833241465 | Akr1b3 | aldo-keto reductase family 1, member B3 | 110.83 | 93.541 | 96.599 | 71.334 | 142.87 | 140.34 |
| 290 | ENSMUSG00000001655 | 0 | NA | NA | NA | NA | NA | Hoxc13 | homeobox C13 [Source:MGI Symbol;Acc:I | 0 | 0 | 0 | 0 | 0 | 0 |
| 291 | ENSMUSG00000001656 | 0 | NA | NA | NA | NA | NA | Hoxc11 | homeobox C11 [Source:MGI Symbol;Acc:I | 0 | 0 | 0 | 0 | 0 | 0 |
| 292 | ENSMUSG00000001657 | 0 | NA | NA | NA | NA | NA | Hoxc8 | homeobox C8 [Source:MGI Symbol;Acc:M | 0 | 0 | 0 | 0 | 0 | 0 |
| 293 | ENSMUSG00000001661 | 0.51034671 | -2.381743948 | 3.331081356 | -0.715006238 | 0.474605189 | NA | Hoxc6 | homeobox C6 [Source:MGI Symbol;Acc:M | 0 | 2.0558 | 1.0062 | 0 | 0 | 0 |
| 294 | ENSMUSG00000001663 | 5765.878387 | -0.177581289 | 0.095072503 | -1.867851212 | 0.061782813 | 0.355601514 | Gstt1 | glutathione S-transferase, theta 1 [Source | 6252.2 | 6062.7 | 6046.5 | 5735.7 | 5582.9 | 4915.4 |
| 295 | ENSMUSG00000001665 | 847.1316655 | -0.804831265 | 0.143905359 | -5.592781745 | 2.23E-08 | 3.11E-06 | Gstt3 | glutathione S-transferase, theta 3 [Source | 1057.7 | 965.21 | 1210.5 | 707.56 | 621.65 | 520.17 |
| 296 | ENSMUSG00000001666 | 934.4421741 | -0.006407538 | 0.110654269 | -0.057905928 | 0.953823561 | 0.9915542 | Ddt | D-dopachrome tautomerase [Source:MGI | 913.93 | 894.29 | 1001.2 | 895.54 | 924.84 | 976.84 |
| 297 | ENSMUSG00000001670 | 20990.83993 | -1.282315184 | 0.550049772 | -2.331271187 | 0.019739065 | 0.172018247 | Tat | tyrosine aminotransferase [Source:MGI Sy | 20783 | 45059 | 23409 | 6270.7 | 18145 | 12279 |
| 298 | ENSMUSG00000001672 | 13.51789412 | 0.182024843 | 0.680370064 | 0.267537995 | 0.789054964 | 0.960994657 | Marveld3 | MARVEL (membrane-associating) domain | 4.8186 | 9.2513 | 24.15 | 16.388 | 10.906 | 15.594 |
| 299 | ENSMUSG00000001674 | 305.6091656 | 0.01670967 | 0.164448692 | 0.101610233 | 0.919066057 | 0.986374671 | Ddx18 | DEAD box helicase 18 [Source:MGI Symbo | 268.24 | 327.91 | 315.96 | 293.05 | 352.27 | 276.23 |
| 300 | ENSMUSG00000001687 | 1337.938397 | -0.126116073 | 0.103254935 | -1.2214048 | 0.221932791 | 0.667949912 | Ubl3 | ubiquitin-like 3 [Source:MGI Symbol;Acc:I | 1334 | 1398 | 1458 | 1322.6 | 1201.9 | 1313.2 |
| 301 | ENSMUSG00000001700 | 726.7205975 | 0.239340672 | 0.123669483 | 1.935325246 | 0.052950396 | 0.323119557 | Gramd3 | GRAM domain containing 3 [Source:MGI ! | 742.07 | 613.67 | 642.99 | 819.38 | 753.62 | 788.6 |
| 302 | ENSMUSG00000001707 | 167.3659439 | 0.181382582 | 0.21209999 | 0.855174873 | 0.392454345 | 0.821365886 | Eef1e1 | eukaryotic translation elongation factor 1 | 195.96 | 134.66 | 138.86 | 159.06 | 195.22 | 180.44 |
| 303 | ENSMUSG00000001729 | 1100.914877 | 0.17449002 | 0.107241104 | 1.627081535 | 0.103719782 | 0.474047094 | Akt1 | thymoma viral proto-oncogene 1 [Source: | 1077 | 966.24 | 1059.6 | 1197.3 | 1108.1 | 1197.4 |
| 304 | ENSMUSG00000001739 | 14.13636672 | 0.409897692 | 0.648338195 | 0.632228202 | 0.527237771 | 0.888334265 | Cldn15 | claudin 15 [Source:MGI Symbol;Acc:MGI:1 | 7.2279 | 14.391 | 15.094 | 25.063 | 16.359 | 6.6831 |
| 305 | ENSMUSG00000001741 | 29.89452033 | -0.634646189 | 0.434761552 | -1.459756931 | 0.144356889 | 0.55464443 | Il16 | interleukin 16 [Source:MGI Symbol;Acc:M | 37.746 | 45.228 | 26.162 | 24.099 | 30.537 | 15.594 |
| 306 | ENSMUSG00000001750 | 728.7147014 | 0.039392066 | 0.125277361 | 0.314438822 | 0.753187775 | 0.952989051 | Tcirg1 | T cell, immune regulator 1, ATPase, H+ tra | 679.42 | 705.15 | 772.8 | 813.6 | 675.09 | 726.23 |
| 307 | ENSMUSG00000001751 | 499.1816838 | -0.033366545 | 0.131361196 | -0.254006101 | 0.799490845 | 0.962424862 | Naglu | alpha-N-acetylglucosaminidase (Sanfilippo | 546.11 | 463.59 | 504.13 | 471.39 | 488.6 | 521.28 |
| 308 | ENSMUSG00000001755 | 3454.105108 | 0.106588165 | 0.093201909 | 1.143626414 | 0.252778602 | 0.705781321 | Coasy | Coenzyme A synthase [Source:MGI Symbo | 3564.2 | 3273.9 | 3140.5 | 3610.1 | 3629.6 | 3506.4 |
| 309 | ENSMUSG00000001761 | 109.9135268 | -0.086846952 | 0.290921948 | -0.298523205 | 0.765303868 | 0.954980327 | Smo | smoothened, frizzled class receptor [Sour | 137.33 | 111.02 | 90.562 | 67.478 | 145.05 | 108.04 |
| 310 | ENSMUSG00000001763 | 232.4515134 | 0.245305033 | 0.231844054 | 1.058060488 | 0.290027866 | 0.740058153 | Tspan33 | tetraspanin 33 [Source:MGI Symbol;Acc:N | 159.01 | 190.16 | 289.8 | 269.91 | 200.67 | 285.14 |
| 311 | ENSMUSG00000001767 | 265.8571996 | 0.118066655 | 0.159339404 | 0.740975884 | 0.458708063 | 0.859322431 | Crnkl1 | crooked neck pre-mRNA splicing factor 1 [ | 246.55 | 256.98 | 261.62 | 271.84 | 308.64 | 249.5 |
| 312 | ENSMUSG00000001768 | 261.5265326 | 0.054773522 | 0.177859902 | 0.307958801 | 0.758113682 | 0.953675099 | Rin2 | Ras and Rab interactor 2 [Source:MGI Syn | 229.69 | 270.34 | 270.68 | 314.26 | 224.67 | 259.53 |
| 313 | ENSMUSG00000001773 | 0.342432689 | 1.990653086 | 3.900281263 | 0.51038706 | 0.609780321 | NA | Folh1 | folate hydrolase 1 [Source:MGI Symbol;A | 0 | 0 | 0 | 0.964 | 1.0906 | 0 |
| 314 | ENSMUSG00000001774 | 690.3227437 | -0.14882261 | 0.129383861 | -1.150240907 | 0.250044662 | 0.702261358 | Chordc1 | cysteine and histidine-rich domain (CHOR | 712.35 | 761.69 | 704.37 | 700.81 | 698 | 564.72 |
| 315 | ENSMUSG00000001783 | 1103.716087 | 0.106895754 | 0.113410657 | 0.942554757 | 0.345908691 | 0.786899236 | Rtcb | RNA 2',3'-cyclic phosphate and 5'-OH ligas | 1044.8 | 1090.6 | 1052.5 | 1019.9 | 1221.5 | 1192.9 |
| 316 | ENSMUSG00000001785 | 247.4902371 | -0.116631819 | 0.163884707 | -0.711669937 | 0.476669188 | 0.869535015 | Pwp1 | PWP1 homolog, endonuclein [Source:MG | 256.19 | 285.76 | 230.43 | 226.53 | 243.21 | 242.82 |
| 317 | ENSMUSG00000001786 | 342.7713992 | 0.268931866 | 0.150129458 | 1.791333091 | 0.073239859 | 0.392921239 | Fbxo7 | F-box protein 7 [Source:MGI Symbol;Acc:I | 332.48 | 321.74 | 277.72 | 344.14 | 378.44 | 402.1 |
| 318 | ENSMUSG00000001794 | 2044.405361 | -0.143432183 | 0.111523329 | -1.286118199 | 0.198401814 | 0.640554426 | Capns1 | calpain, small subunit 1 [Source:MGI Syml | 2259.1 | 2153.5 | 2025.6 | 2142.9 | 1953.3 | 1732 |
| 319 | ENSMUSG00000001802 | 621.8713859 | 0.1183951 | 0.117029421 | 1.011669536 | 0.311696086 | 0.759830222 | Lrp3 | low density lipoprotein receptor-related p | 568.6 | 603.39 | 617.83 | 663.22 | 651.1 | 627.09 |
| 320 | ENSMUSG00000001804 | 0 | NA | NA | NA | NA | NA | Dsg4 | desmoglein 4 [Source:MGI Symbol;Acc:M( | 0 | 0 | 0 | 0 | 0 | 0 |
| 321 | ENSMUSG00000001815 | 0 | NA | NA | NA | NA | NA | Evx2 | even-skipped homeobox 2 [Source:MGI S | 0 | 0 | 0 | 0 | 0 | 0 |
| 322 | ENSMUSG00000001819 | 0 | NA | NA | NA | NA | NA | Hoxd13 | homeobox D13 [Source:MGI Symbol;Acc:I | 0 | 0 | 0 | 0 | 0 | 0 |
| 323 | ENSMUSG00000001823 | 0 | NA | NA | NA | NA | NA | Hoxd12 | homeobox D12 [Source:MGI Symbol;Acc:I | 0 | 0 | 0 | 0 | 0 | 0 |
| 324 | ENSMUSG00000001827 | 3.077810336 | -0.50720119 | 1.348795429 | -0.376040116 | 0.706887073 | NA | Folr1 | folate receptor 1 (adult) [Source:MGI Sym | 5.6217 | 4.1117 | 1.0062 | 0 | 3.2719 | 4.4554 |
| 325 | ENSMUSG00000001829 | 908.4298471 | -0.160720629 | 0.106619048 | -1.507428852 | 0.131700768 | 0.532100533 | Clpb | ClpB caseinolytic peptidase B [Source:MG | 961.31 | 962.13 | 953.92 | 898.43 | 857.23 | 817.56 |
| 326 | ENSMUSG00000001833 | 750.975023 | -0.066446626 | 0.134139383 | -0.495355089 | 0.620349491 | 0.917977899 | Septin7 | septin 7 [Source:MGI Symbol;Acc:MGI:13 | 801.5 | 855.23 | 648.02 | 763.47 | 695.81 | 741.82 |
| 327 | ENSMUSG00000001844 | 307.1390823 | -0.234690339 | 0.154020059 | -1.523764772 | 0.127567483 | 0.523285632 | Zdhhc4 | zinc finger, DHHC domain containing 4 [Sc | 299.56 | 363.88 | 333.07 | 263.17 | 284.65 | 298.51 |
| 328 | ENSMUSG00000001847 | 2039.060276 | -0.030682474 | 0.094888694 | -0.323352263 | 0.746428478 | 0.950047651 | Rac1 | Rac family small GTPase 1 [Source:MGI Sy | 2034.3 | 2042.5 | 2105.1 | 1899 | 2091.8 | 2061.7 |
| 329 | ENSMUSG00000001855 | 194.2765144 | -0.111160432 | 0.169923176 | -0.654180519 | 0.512995511 | 0.884860213 | Nup214 | nucleoporin 214 [Source:MGI Symbol;Acc | 198.37 | 208.67 | 198.23 | 181.23 | 187.59 | 191.58 |
| 330 | ENSMUSG00000001864 | 1.625688923 | -0.479836157 | 1.909187649 | -0.251330013 | 0.801558972 | NA | Aif1l | allograft inflammatory factor 1-like [Sourc | 5.6217 | 0 | 0 | 1.928 | 1.0906 | 1.1138 |
| 331 | ENSMUSG00000001865 | 0.736965982 | -2.935629304 | 2.883220453 | -1.018177192 | 0.308593757 | NA | Cpa3 | carboxypeptidase A3, mast cell [Source:M | 2.4093 | 0 | 2.0125 | 0 | 0 | 0 |
| 332 | ENSMUSG00000001870 | 47.76900763 | -0.723862842 | 0.34619304 | -2.090922576 | 0.036535005 | 0.257902544 | Ltbp1 | latent transforming growth factor beta bir | 79.507 | 49.34 | 49.306 | 35.667 | 33.809 | 38.985 |

| | | | | | | | | | | | | | | | |
|---|---|---|---|---|---|---|---|---|---|---|---|---|---|---|---|
| 333 | ENSMUSG00000001891 | 10011.75564 | -0.046697586 | 0.119267941 | -0.391535112 | 0.69540174 | 0.939992732 | Ugp2 | UDP-glucose pyrophosphorylase 2 [Source | 11073 | 9091.9 | 10354 | 8525.4 | 10003 | 11023 |
| 334 | ENSMUSG00000001901 | 0.185640647 | 1.126924175 | 4.080472857 | 0.276174898 | 0.782413736 | NA | Kcnh6 | potassium voltage-gated channel, subfam | 0 | 0 | 0 | 0 | 0 | 1.1138 |
| 335 | ENSMUSG00000001909 | 594.2219452 | 0.104750372 | 0.11676629 | 0.897094289 | 0.369668606 | 0.80653433 | Trmt1 | tRNA methyltransferase 1 [Source:MGI Sy | 558.96 | 586.94 | 572.55 | 633.33 | 600.93 | 612.61 |
| 336 | ENSMUSG00000001910 | 614.6531173 | -0.016311328 | 0.115376112 | -0.14137526 | 0.887573497 | 0.982382091 | Nacc1 | nucleus accumbens associated 1, BEN and | 608.75 | 613.67 | 631.92 | 595.74 | 610.75 | 627.09 |
| 337 | ENSMUSG00000001911 | 2067.870595 | 0.393965573 | 0.110346478 | 3.570259605 | 0.000356628 | 0.009958034 | Nfix | nuclear factor I/X [Source:MGI Symbol;Ac | 1717.8 | 1678.6 | 1965.2 | 2144.9 | 2413.5 | 2487.2 |
| 338 | ENSMUSG00000001918 | 26.01627694 | -1.005585215 | 0.462692598 | -2.173333263 | 0.029755244 | 0.227593784 | Slc1a5 | solute carrier family 1 (neutral amino acid | 41.761 | 39.061 | 23.144 | 13.496 | 16.359 | 22.277 |
| 339 | ENSMUSG00000001924 | 2616.996868 | -0.002130867 | 0.106719391 | -0.019967006 | 0.984069693 | 0.998580735 | Uba1 | ubiquitin-like modifier activating enzyme 1 | 2445.4 | 2696.2 | 2716.9 | 2871.7 | 2583.7 | 2388.1 |
| 340 | ENSMUSG00000001930 | 70.21378372 | -0.763233055 | 0.293737323 | -2.598352322 | 0.009367233 | 0.103504702 | Vwf | Von Willebrand factor [Source:MGI Symbo | 88.341 | 84.289 | 92.575 | 58.803 | 34.9 | 62.375 |
| 341 | ENSMUSG00000001942 | 868.3515819 | 0.033324089 | 0.143245833 | 0.23263566 | 0.816044334 | 0.967767553 | Siae | sialic acid acetylesterase [Source:MGI Sym | 946.05 | 891.2 | 736.57 | 778.89 | 1017.5 | 839.84 |
| 342 | ENSMUSG00000001943 | 7.625505978 | -0.482757972 | 0.957019094 | -0.504439227 | 0.613952758 | 0.917378096 | Vsig2 | V-set and immunoglobulin domain contai | 5.6217 | 5.1396 | 16.1 | 14.46 | 1.0906 | 3.3415 |
| 343 | ENSMUSG00000001946 | 116.0620172 | -0.62409353 | 0.279976373 | -2.229093555 | 0.025807681 | 0.207494326 | Esam | endothelial cell-specific adhesion molecul | 163.83 | 127.46 | 130.81 | 98.326 | 54.531 | 121.41 |
| 344 | ENSMUSG00000001948 | 5.170579785 | -0.548377536 | 0.974141976 | -0.562933894 | 0.573479899 | 0.905547299 | Spa17 | sperm autoantigenic protein 17 [Source:M | 3.2124 | 10.279 | 5.0312 | 4.8199 | 5.4531 | 2.2277 |
| 345 | ENSMUSG00000001962 | 296.8729236 | -0.133397972 | 0.157970566 | -0.844448276 | 0.398418955 | 0.82588279 | Fam50a | family with sequence similarity 50, memb | 273.05 | 346.41 | 312.94 | 273.77 | 298.83 | 276.23 |
| 346 | ENSMUSG00000001964 | 262.4031887 | 0.027230395 | 0.160963887 | 0.169170834 | 0.865662271 | 0.977511053 | Emd | emerin [Source:MGI Symbol;Acc:MGI:108 | 253.78 | 245.67 | 280.74 | 289.19 | 239.94 | 265.09 |
| 347 | ENSMUSG00000001983 | 342.7281933 | -0.02885088 | 0.151890255 | -0.189945563 | 0.849351788 | 0.973071229 | Taco1 | translational activator of mitochondrially | 328.47 | 356.69 | 353.19 | 298.83 | 382.81 | 336.38 |
| 348 | ENSMUSG00000001985 | 0 | NA | NA | NA | NA | NA | Grik3 | glutamate receptor, ionotropic, kainate 3 | 0 | 0 | 0 | 0 | 0 | 0 |
| 349 | ENSMUSG00000001986 | 17.29397073 | 0.935489004 | 0.576545914 | 1.622575031 | 0.104680282 | 0.476887243 | Gria3 | glutamate receptor, ionotropic, AMPA3 (a | 7.2279 | 20.558 | 8.05 | 18.316 | 23.994 | 25.618 |
| 350 | ENSMUSG00000001988 | 1.87305964 | 0.316912849 | 1.734991202 | 0.182659629 | 0.855065094 | NA | Npas1 | neuronal PAS domain protein 1 [Source:M | 0 | 2.0558 | 3.0187 | 2.8919 | 3.2719 | 0 |
| 351 | ENSMUSG00000001995 | 172.4458309 | 0.776535793 | 0.219294792 | 3.541058977 | 0.000398524 | 0.010635163 | Sipa1l2 | signal-induced proliferation-associated 1 l | 122.87 | 153.16 | 105.66 | 240.03 | 170.14 | 242.82 |
| 352 | ENSMUSG00000001998 | 113.0787927 | 0.043182056 | 0.254745023 | 0.169510892 | 0.865394806 | 0.977361636 | Ap4e1 | adaptor-related protein complex AP-4, ep | 94.766 | 129.52 | 110.69 | 149.42 | 91.612 | 102.47 |
| 353 | ENSMUSG00000001999 | 250.4149294 | 0.089038035 | 0.166822943 | 0.533727754 | 0.593529904 | 0.913011006 | Blvra | biliverdin reductase A [Source:MGI Symbo | 233.7 | 240.53 | 253.57 | 226.53 | 255.2 | 292.94 |
| 354 | ENSMUSG00000002006 | 4.843341385 | -0.550648635 | 1.016035679 | -0.541957971 | 0.58784746 | NA | Pdzd4 | PDZ domain containing 4 [Source:MGI Syr | 4.0155 | 8.2233 | 5.0312 | 1.928 | 7.6343 | 2.2277 |
| 355 | ENSMUSG00000002007 | 3.436663198 | 0.284048622 | 1.134773408 | 0.250313076 | 0.802345245 | NA | Srpk3 | serine/arginine-rich protein specific kinase | 3.2124 | 4.1117 | 2.0125 | 5.7839 | 3.2719 | 2.2277 |
| 356 | ENSMUSG00000002010 | 1305.865975 | -0.073814295 | 0.10660084 | -0.692436332 | 0.488663356 | 0.874431125 | Idh3g | isocitrate dehydrogenase 3 (NAD+), gamm | 1399.8 | 1402.1 | 1215.5 | 1291.7 | 1262.9 | 1263.1 |
| 357 | ENSMUSG00000002012 | 0.633540418 | -1.320719359 | 2.623870149 | -0.503347835 | 0.61471974 | NA | Pnck | pregnancy upregulated non-ubiquitously | 0.8031 | 1.0279 | 1.0062 | 0.964 | 0 | 0 |
| 358 | ENSMUSG00000002014 | 1494.565746 | 0.200752613 | 0.106544785 | 1.884208726 | 0.059536748 | 0.347373747 | Ssr4 | signal sequence receptor, delta [Source:M | 1366.1 | 1404.1 | 1401.7 | 1499.9 | 1532.3 | 1763.2 |
| 359 | ENSMUSG00000002015 | 2328.634607 | -0.063963974 | 0.094394912 | -0.677620992 | 0.498012033 | 0.879195172 | Bcap31 | B cell receptor associated protein 31 [Sou | 2530.6 | 2327.2 | 2282.2 | 2319.3 | 2284.8 | 2227.7 |
| 360 | ENSMUSG00000002017 | 469.009619 | 0.002099995 | 0.125812241 | 0.016691502 | 0.986682726 | 0.99859176 | Fam98a | family with sequence similarity 98, memb | 482.66 | 455.37 | 467.9 | 485.84 | 470.06 | 452.22 |
| 361 | ENSMUSG00000002020 | 12.52880385 | -0.180211231 | 0.632228272 | -0.285041399 | 0.775612425 | 0.95893551 | Ltbp2 | latent transforming growth factor beta bir | 18.471 | 12.335 | 9.0562 | 15.424 | 7.6343 | 12.252 |
| 362 | ENSMUSG00000002028 | 342.7696854 | 0.070914536 | 0.151542344 | 0.467951953 | 0.639818949 | 0.923932929 | Kmt2a | lysine (K)-specific methyltransferase 2A [S | 337.3 | 321.74 | 344.14 | 375.95 | 301.01 | 376.48 |
| 363 | ENSMUSG00000002031 | 406.3626901 | 0.24922021 | 0.140183383 | 1.777815631 | 0.07543414 | 0.401870868 | Ift46 | intraflagellar transport 46 [Source:MGI Sy | 377.46 | 367.99 | 368.29 | 431.86 | 400.26 | 492.32 |
| 364 | ENSMUSG00000002032 | 801.3288214 | 0.451649632 | 0.177870973 | 2.539198077 | 0.01111069 | 0.11696144 | Tmem25 | transmembrane protein 25 [Source:MGI S | 636.86 | 564.33 | 829.15 | 705.63 | 1033.9 | 1038.1 |
| 365 | ENSMUSG00000002033 | 2.398392094 | -1.199048312 | 1.639421746 | -0.731384901 | 0.464544085 | NA | Cd3g | CD3 antigen, gamma polypeptide [Source | 4.8186 | 5.1396 | 0 | 0 | 1.0906 | 3.3415 |
| 366 | ENSMUSG00000002043 | 242.4570997 | 0.022785933 | 0.178896227 | 0.127369554 | 0.898647912 | 0.982798755 | Trappc6a | trafficking protein particle complex 6A [Sc | 285.1 | 222.03 | 213.32 | 221.71 | 251.93 | 260.64 |
| 367 | ENSMUSG00000002052 | 809.6868284 | -0.035293297 | 0.110151414 | -0.32040712 | 0.748659728 | 0.951074124 | Supt6 | SPT6, histone chaperone and transcriptior | 782.22 | 852.14 | 825.12 | 808.78 | 817.96 | 771.89 |
| 368 | ENSMUSG00000002055 | 7.984772701 | 0.597536836 | 0.770939015 | 0.775076659 | 0.438294363 | 0.847566655 | Spag5 | sperm associated antigen 5 [Source:MGI S | 8.8341 | 5.1396 | 5.0312 | 13.496 | 8.7249 | 6.6831 |
| 369 | ENSMUSG00000002057 | 0 | NA | NA | NA | NA | NA | Foxn1 | forkhead box N1 [Source:MGI Symbol;Acc | 0 | 0 | 0 | 0 | 0 | 0 |
| 370 | ENSMUSG00000002058 | 135.2459916 | -0.163782149 | 0.273992317 | -0.597761832 | 0.549998861 | 0.895703112 | Unc119 | unc-119 lipid binding chaperone [Source:I | 198.37 | 120.27 | 109.68 | 153.27 | 136.33 | 93.563 |
| 371 | ENSMUSG00000002059 | 111.5837794 | 0.03448607 | 0.25094297 | 0.137425928 | 0.890694138 | 0.982489666 | Rab34 | RAB34, member RAS oncogene family [So | 142.95 | 98.68 | 88.55 | 122.43 | 118.88 | 98.018 |
| 372 | ENSMUSG00000002064 | 553.339255 | 0.097483252 | 0.122068468 | 0.798594866 | 0.424525365 | 0.840450984 | Sdf2 | stromal cell derived factor 2 [Source:MGI | 571.01 | 516.01 | 516.2 | 572.6 | 561.67 | 582.54 |
| 373 | ENSMUSG00000002068 | 35.38025152 | 0.816089626 | 0.550123031 | 1.483467479 | 0.137950251 | 0.5424993 | Ccne1 | cyclin E1 [Source:MGI Symbol;Acc:MGI:88 | 44.171 | 4.1117 | 28.175 | 52.055 | 41.443 | 42.326 |
| 374 | ENSMUSG00000002076 | 4.677035701 | 0.485400626 | 1.114622852 | 0.435484186 | 0.663211015 | NA | Hsf2bp | heat shock transcription factor 2 binding p | 5.6217 | 0 | 6.0375 | 8.6758 | 3.2719 | 4.4554 |
| 375 | ENSMUSG00000002083 | 177.3515055 | 0.080376121 | 0.197632022 | 0.406695841 | 0.684231395 | 0.935528319 | Bbc3 | BCL2 binding component 3 [Source:MGI S | 181.5 | 144.94 | 190.18 | 156.16 | 191.95 | 199.38 |
| 376 | ENSMUSG00000002100 | 1.795510235 | 0.722507877 | 2.036434268 | 0.354790669 | 0.722746419 | NA | Mybpc3 | myosin binding protein C, cardiac [Source | 0 | 3.0838 | 1.0062 | 0 | 0 | 6.6831 |
| 377 | ENSMUSG00000002102 | 2064.084817 | 0.037268504 | 0.095974033 | 0.388318623 | 0.697780257 | 0.940502785 | Psmc3 | proteasome (prosome, macropain) 26S su | 2122.6 | 1994.2 | 1994.4 | 1990.6 | 2195.4 | 2087.3 |
| 378 | ENSMUSG00000002103 | 825.8583532 | -0.051383317 | 0.129291135 | -0.397423359 | 0.691055293 | 0.937657013 | Acp2 | acid phosphatase 2, lysosomal [Source:M | 791.06 | 838.78 | 891.53 | 695.99 | 880.13 | 857.66 |
| 379 | ENSMUSG00000002104 | 1.749833788 | 2.219837245 | 1.781498859 | 1.246050333 | 0.212745917 | NA | Rapsn | receptor-associated protein of the synaps | 0.8031 | 1.0279 | 0 | 0.964 | 4.3625 | 3.3415 |
| 380 | ENSMUSG00000002105 | 159.6546325 | -0.233395497 | 0.238487853 | -0.978647317 | 0.327754271 | 0.772937193 | Slc39a13 | solute carrier family 39 (metal ion transpo | 133.31 | 215.86 | 169.05 | 141.7 | 181.04 | 116.95 |
| 381 | ENSMUSG00000002107 | 372.791599 | -0.361686756 | 0.1681459 | -2.151029292 | 0.031473888 | 0.236119457 | Celf2 | CUGBP, Elav-like family member 2 [Source | 403.16 | 482.09 | 372.31 | 276.66 | 380.63 | 321.9 |
| 382 | ENSMUSG00000002108 | 2176.39631 | 0.019628272 | 0.101922583 | 0.19258021 | 0.847287745 | 0.972803041 | Nr1h3 | nuclear receptor subfamily 1, group H, me | 1980.5 | 2165.8 | 2339.5 | 2149.7 | 2256.5 | 2166.4 |
| 383 | ENSMUSG00000002109 | 83.87733083 | -0.117930406 | 0.269735001 | -0.43720839 | 0.661960226 | 0.928760569 | Ddb2 | damage specific DNA binding protein 2 [Sc | 77.098 | 96.624 | 88.55 | 89.65 | 58.893 | 92.449 |
| 384 | ENSMUSG00000002111 | 60.79428851 | -1.519763669 | 0.34655307 | -4.385370666 | 1.16E-05 | 0.000600274 | Spi1 | spleen focus forming virus (SFFV) proviral | 82.72 | 116.15 | 71.443 | 26.991 | 22.903 | 44.554 |
| 385 | ENSMUSG00000002129 | 375.0217585 | 0.069078987 | 0.141321515 | 0.488807294 | 0.624978134 | 0.919503423 | Sf3a1 | splicing factor 3a, subunit 1 [Source:MGI S | 403.96 | 336.13 | 357.22 | 384.63 | 382.81 | 385.39 |
| 386 | ENSMUSG00000002147 | 1105.898172 | 0.217347005 | 0.111836992 | 1.943426781 | 0.051964614 | 0.320245495 | Stat6 | signal transducer and activator of transcri | 987.82 | 1035.1 | 1046.5 | 1311 | 1125.5 | 1129.4 |
| 387 | ENSMUSG00000002190 | 0 | NA | NA | NA | NA | NA | Clgn | calmegin [Source:MGI Symbol;Acc:MGI:10 | 0 | 0 | 0 | 0 | 0 | 0 |
| 388 | ENSMUSG00000002204 | 7.221661407 | -0.954639345 | 0.875686463 | -1.090161131 | 0.275642172 | 0.730038567 | Napsa | napsin A aspartic peptidase [Source:MGI S | 11.243 | 9.2513 | 8.05 | 3.8559 | 9.8156 | 1.1138 |

| | | | | | | | | | | | | | | | |
|---|---|---|---|---|---|---|---|---|---|---|---|---|---|---|---|
| 389 | ENSMUSG00000002205 | 426.9913752 | -0.063385896 | 0.139295017 | -0.455047836 | 0.649074825 | 0.925643431 | Vrk3 | vaccinia related kinase 3 [Source:MGI Sym | 445.72 | 415.28 | 447.78 | 413.55 | 376.26 | 463.36 |
| 390 | ENSMUSG00000002210 | 162.5571701 | 0.00445327 | 0.18852948 | 0.023621081 | 0.981154857 | 0.99820894 | Smg9 | smg-9 homolog, nonsense mediated mRN | 150.98 | 155.22 | 181.12 | 170.62 | 159.23 | 158.17 |
| 391 | ENSMUSG00000002221 | 402.7550881 | -0.006672682 | 0.166520375 | -0.040071266 | 0.96803631 | 0.994231936 | Paxip1 | PAX interacting (with transcription-activat | 339.71 | 447.14 | 425.64 | 460.78 | 403.53 | 339.72 |
| 392 | ENSMUSG00000002222 | 3499.355148 | 0.036223315 | 0.096221115 | 0.376459103 | 0.706575614 | 0.94295036 | Rmnd5a | required for meiotic nuclear division 5 hor | 3462.2 | 3613.1 | 3290.4 | 3317 | 3772.4 | 3540.9 |
| 393 | ENSMUSG00000002227 | 348.0609261 | 0.318834356 | 0.158784415 | 2.007970085 | 0.044646471 | 0.291866583 | Mov10 | Mov10 RISC complex RNA helicase [Sourc | 278.68 | 329.96 | 320.99 | 345.1 | 446.06 | 367.57 |
| 394 | ENSMUSG00000002228 | 4.302298308 | -0.817545977 | 1.122443486 | -0.728362709 | 0.466391585 | NA | Ppm1j | protein phosphatase 1J [Source:MGI Symb | 2.4093 | 3.0838 | 11.069 | 4.8199 | 1.0906 | 3.3415 |
| 395 | ENSMUSG00000002233 | 104.8139229 | -0.165721642 | 0.234067823 | -0.708006939 | 0.478940949 | 0.870169438 | Rhoc | ras homolog family member C [Source:MC | 118.06 | 107.93 | 106.66 | 115.68 | 99.246 | 81.311 |
| 396 | ENSMUSG00000002240 | 0 | NA | NA | NA | NA | NA | Prr27 | proline rich 27 [Source:MGI Symbol;Acc:M | 0 | 0 | 0 | 0 | 0 | 0 |
| 397 | ENSMUSG00000002249 | 227.2517117 | 0.133615865 | 0.172923552 | 0.772687488 | 0.439707353 | 0.84844279 | Tead3 | TEA domain family member 3 [Source:MG | 219.25 | 192.22 | 238.48 | 230.39 | 220.3 | 262.87 |
| 398 | ENSMUSG00000002250 | 153.605193 | -0.165662853 | 0.196569416 | -0.842770235 | 0.399356958 | 0.8263661 | Ppard | peroxisome proliferator activator recepto | 159.82 | 165.49 | 162.01 | 148.45 | 123.24 | 162.62 |
| 399 | ENSMUSG00000002257 | 5.814291025 | -1.978385878 | 1.06002993 | -1.866349073 | 0.061992543 | 0.355884025 | Def6 | differentially expressed in FDCP 6 [Source | 9.6372 | 11.307 | 7.0437 | 5.7839 | 0 | 1.1138 |
| 400 | ENSMUSG00000002265 | 113.5313986 | 0.010964254 | 0.371222809 | 0.029535508 | 0.9764375 | 0.9970485 | Peg3 | paternally expressed 3 [Source:MGI Symb | 120.47 | 91.485 | 127.79 | 203.4 | 55.622 | 82.424 |
| 401 | ENSMUSG00000002266 | 0.829837731 | -1.7608533 | 2.399900965 | -0.733719152 | 0.463119924 | NA | Zim1 | zinc finger, imprinted 1 [Source:MGI Symb | 0.8031 | 2.0558 | 1.0062 | 0 | 0 | 1.1138 |
| 402 | ENSMUSG00000002274 | 7.932796952 | 0.404901925 | 0.842879721 | 0.480379246 | 0.630957749 | 0.921277295 | Metrn | meteorin, glial cell differentiation regulato | 11.243 | 1.0279 | 8.05 | 9.6398 | 8.7249 | 8.9108 |
| 403 | ENSMUSG00000002279 | 748.8129631 | 0.045670002 | 0.116366009 | 0.392468573 | 0.694712027 | 0.939895655 | Lmf1 | lipase maturation factor 1 [Source:MGI Sy | 794.27 | 701.04 | 714.43 | 731.66 | 789.61 | 761.87 |
| 404 | ENSMUSG00000002280 | 489.544011 | 0.074928022 | 0.140255983 | 0.534223357 | 0.593187011 | 0.913011006 | Ciao3 | cytosolic iron-sulfur assembly component | 427.25 | 495.46 | 508.15 | 459.82 | 490.78 | 555.81 |
| 405 | ENSMUSG00000002289 | 4357.990617 | -0.78368436 | 0.160663268 | -4.877806678 | 1.07E-06 | 8.67E-05 | Angptl4 | angiopoietin-like 4 [Source:MGI Symbol;A | 5278 | 5832.4 | 5429.7 | 2912.2 | 4219.6 | 2476.1 |
| 406 | ENSMUSG00000002297 | 38.75288822 | 0.403652379 | 0.380932243 | 1.05964351 | 0.289306811 | 0.739770202 | Dbf4 | DBF4 zinc finger [Source:MGI Symbol;Acc: | 45.777 | 30.838 | 23.144 | 50.127 | 42.534 | 40.098 |
| 407 | ENSMUSG00000002307 | 217.7775099 | 0.340629832 | 0.181297176 | 1.878847974 | 0.060265254 | 0.349789984 | Daxx | Fas death domain-associated protein [Sou | 208.81 | 180.91 | 186.16 | 227.5 | 217.03 | 286.26 |
| 408 | ENSMUSG00000002308 | 76.86191345 | 0.330537689 | 0.259051306 | 1.275954535 | 0.201971591 | 0.646452112 | Cd320 | CD320 antigen [Source:MGI Symbol;Acc:M | 61.036 | 70.926 | 72.45 | 78.082 | 82.887 | 95.791 |
| 409 | ENSMUSG00000002319 | 244.7596722 | -0.019193785 | 0.187019142 | -0.102630055 | 0.918256588 | 0.986374671 | Ipo4 | importin 4 [Source:MGI Symbol;Acc:MGI:1 | 295.54 | 197.36 | 245.52 | 255.45 | 253.02 | 221.65 |
| 410 | ENSMUSG00000002320 | 805.0175265 | 0.056146811 | 0.11488954 | 0.48870255 | 0.625052299 | 0.919503423 | Tm9sf1 | transmembrane 9 superfamily member 1 | 734.04 | 788.41 | 846.25 | 808.78 | 799.42 | 853.2 |
| 411 | ENSMUSG00000002324 | 26.48770887 | 0.542920438 | 0.481080273 | 1.128544378 | 0.259090086 | 0.712448479 | Rec8 | REC8 meiotic recombination protein [Sour | 19.274 | 15.419 | 30.187 | 42.415 | 33.809 | 17.822 |
| 412 | ENSMUSG00000002325 | 649.6790785 | 0.372644658 | 0.126352202 | 2.949253362 | 0.003185427 | 0.048652606 | Irf9 | interferon regulatory factor 9 [Source:MG | 602.33 | 506.76 | 588.65 | 709.49 | 707.81 | 783.03 |
| 413 | ENSMUSG00000002326 | 70.31133429 | 7.68E-05 | 0.272682939 | 0.00028147 | 0.999775419 | 0.999897338 | Gmpr2 | guanosine monophosphate reductase 2 [S | 63.445 | 74.01 | 73.456 | 58.803 | 73.071 | 79.083 |
| 414 | ENSMUSG00000002329 | 269.5314096 | -0.119949361 | 0.167606749 | -0.715659496 | 0.474201627 | 0.86899591 | Mdp1 | magnesium-dependent phosphatase 1 [Sc | 298.75 | 266.23 | 276.72 | 254.49 | 222.49 | 298.51 |
| 415 | ENSMUSG00000002332 | 2830.076827 | 0.225758939 | 0.093326994 | 2.419010078 | 0.015562809 | 0.14579046 | Dhrs1 | dehydrogenase/reductase (SDR family) m | 2692.8 | 2469.1 | 2664.5 | 3001.8 | 2958.8 | 3193.4 |
| 416 | ENSMUSG00000002341 | 0 | NA | NA | NA | NA | NA | Ncan | neurocan [Source:MGI Symbol;Acc:MGI:1 | 0 | 0 | 0 | 0 | 0 | 0 |
| 417 | ENSMUSG00000002342 | 291.1899262 | -0.012211775 | 0.145546616 | -0.083902846 | 0.933133676 | 0.988420305 | Tmem161a | transmembrane protein 161A [Source:MC | 287.51 | 298.1 | 291.81 | 290.16 | 296.65 | 282.92 |
| 418 | ENSMUSG00000002343 | 76.45389755 | 0.088612413 | 0.254102354 | 0.348727241 | 0.727294092 | 0.945907265 | Armc6 | armadillo repeat containing 6 [Source:MG | 77.901 | 66.815 | 77.481 | 80.01 | 77.434 | 79.083 |
| 419 | ENSMUSG00000002345 | 190.7762257 | -0.025547973 | 0.1797274 | -0.142148457 | 0.886962744 | 0.982382091 | Borcs8 | BLOC-1 related complex subunit 8 [Source | 176.68 | 208.67 | 192.19 | 171.59 | 203.95 | 191.58 |
| 420 | ENSMUSG00000002346 | 2712.244349 | -0.033059418 | 0.103151736 | -0.320493084 | 0.748594573 | 0.951074124 | Slc25a42 | solute carrier family 25, member 42 [Sour | 2751.4 | 2586.2 | 2893 | 2892.9 | 2689.5 | 2460.5 |
| 421 | ENSMUSG00000002365 | 1316.927686 | -0.002359635 | 0.108238676 | -0.021800296 | 0.982607258 | 0.998580735 | Snx9 | sorting nexin 9 [Source:MGI Symbol;Acc:M | 1413.5 | 1253 | 1286 | 1223.3 | 1349.1 | 1376.7 |
| 422 | ENSMUSG00000002372 | 595.668625 | -0.031864546 | 0.126553913 | -0.251786333 | 0.801206222 | 0.96255706 | Ranbp3 | RAN binding protein 3 [Source:MGI Symb | 590.28 | 605.44 | 611.8 | 650.69 | 535.49 | 580.31 |
| 423 | ENSMUSG00000002379 | 712.3839398 | -0.12215107 | 0.118947029 | -1.026936711 | 0.304450267 | 0.75326183 | Ndufa11 | NADH:ubiquinone oxidoreductase subuni | 723.6 | 741.13 | 763.74 | 740.33 | 676.18 | 629.32 |
| 424 | ENSMUSG00000002384 | 0.492645588 | 0.9565034 | 3.447648281 | 0.277436479 | 0.781444974 | NA | Bmp8b | bone morphogenetic protein 8b [Source:M | 0 | 1.0279 | 0 | 1.928 | 0 | 0 |
| 425 | ENSMUSG00000002393 | 1377.363732 | 0.246767597 | 0.121497907 | 2.031044015 | 0.04225053 | 0.283184274 | Nr2f6 | nuclear receptor subfamily 2, group F, me | 1099.4 | 1234.5 | 1447 | 1513.4 | 1504 | 1465.8 |
| 426 | ENSMUSG00000002395 | 781.6653093 | 0.069792272 | 0.117444747 | 0.594256224 | 0.552340753 | 0.896035963 | Use1 | unconventional SNARE in the ER 1 homolo | 751.7 | 781.22 | 755.69 | 805.89 | 867.04 | 728.45 |
| 427 | ENSMUSG00000002396 | 342.8095223 | -0.203387012 | 0.176414623 | -1.152892024 | 0.248954706 | 0.700833178 | Ocel1 | occludin/ELL domain containing 1 [Source | 342.12 | 313.51 | 445.77 | 349.92 | 332.64 | 272.89 |
| 428 | ENSMUSG00000002409 | 258.0115046 | 0.423376965 | 0.209254084 | 2.023267398 | 0.043045586 | 0.28638977 | Dyrk1b | dual-specificity tyrosine-(Y)-phosphorylati | 265.83 | 172.69 | 222.38 | 370.17 | 253.02 | 263.98 |
| 429 | ENSMUSG00000002413 | 353.7258477 | -0.091011537 | 0.160556705 | -0.566849803 | 0.570816231 | 0.904651508 | Braf | Braf transforming gene [Source:MGI Symb | 325.26 | 428.64 | 341.12 | 312.33 | 369.72 | 345.29 |
| 430 | ENSMUSG00000002416 | 152.1934966 | 0.141585591 | 0.191636214 | 0.738824818 | 0.460013379 | 0.859753653 | Ndufb2 | NADH:ubiquinone oxidoreductase subuni | 158.21 | 138.77 | 136.85 | 160.98 | 166.86 | 151.48 |
| 431 | ENSMUSG00000002428 | 247.6663778 | 0.102837921 | 0.171648718 | 0.599118492 | 0.549093871 | 0.895703112 | Hltf | helicase-like transcription factor [Source:M | 252.17 | 222.03 | 241.5 | 215.93 | 277.02 | 277.35 |
| 432 | ENSMUSG00000002455 | 1080.536821 | 0.004475581 | 0.137721557 | 0.032497315 | 0.974075457 | 0.996324688 | Prpf6 | pre-mRNA splicing factor 6 [Source:MGI S | 1147.6 | 1122.5 | 965.99 | 1049.8 | 1268.4 | 928.95 |
| 433 | ENSMUSG00000002458 | 14.96093718 | -1.07316214 | 0.679353464 | -1.57968156 | 0.114179811 | 0.496868018 | Rgs19 | regulator of G-protein signaling 19 [Sourc | 28.912 | 23.642 | 8.05 | 4.8199 | 7.6343 | 16.708 |
| 434 | ENSMUSG00000002459 | 0.335414862 | -1.779047242 | 4.030804179 | -0.441362855 | 0.658950329 | NA | Rgs20 | regulator of G-protein signaling 20 [Sourc | 0 | 0 | 2.0125 | 0 | 0 | 0 |
| 435 | ENSMUSG00000002475 | 1231.790486 | -0.085225006 | 0.106899855 | -0.797241545 | 0.425310762 | 0.840969906 | Abhd3 | abhydrolase domain containing 3 [Source | 1310.7 | 1222.2 | 1271.9 | 1289.8 | 1176.8 | 1119.4 |
| 436 | ENSMUSG00000002477 | 160.8067138 | 0.010806367 | 0.204896826 | 0.052740527 | 0.957938648 | 0.991861398 | Snrpd1 | small nuclear ribonucleoprotein D1 [Sourc | 139.74 | 193.25 | 147.92 | 145.56 | 167.96 | 170.42 |
| 437 | ENSMUSG00000002486 | 40.35550704 | -0.413769739 | 0.362916187 | -1.140124782 | 0.254234319 | 0.707434626 | Tchp | trichoplein, keratin filament binding [Sour | 53.005 | 41.117 | 44.275 | 45.307 | 28.356 | 30.074 |
| 438 | ENSMUSG00000002489 | 12.36305674 | -0.361705541 | 0.602009609 | -0.600830178 | 0.547953102 | 0.895600915 | Tiam1 | T cell lymphoma invasion and metastasis 1 | 15.259 | 13.363 | 13.081 | 11.568 | 11.997 | 8.9108 |
| 439 | ENSMUSG00000002496 | 723.4053858 | -0.094084255 | 0.114126333 | -0.824386914 | 0.409719748 | 0.833241465 | Tsc2 | TSC complex subunit 2 [Source:MGI Symb | 714.76 | 729.82 | 796.95 | 711.42 | 696.91 | 690.58 |
| 440 | ENSMUSG00000002500 | 5.390207261 | 1.282067874 | 0.977314781 | 1.311826956 | 0.189578528 | 0.626663447 | Rpl3l | ribosomal protein L3-like [Source:MGI Syr | 2.4093 | 1.0279 | 6.0375 | 9.6398 | 6.5437 | 6.6831 |
| 441 | ENSMUSG00000002504 | 348.3864909 | -0.397657052 | 0.178211688 | -2.231374695 | 0.02565632 | 0.206447904 | Slc9a3r2 | solute carrier family 9 (sodium/hydrogen e | 390.31 | 360.8 | 437.72 | 352.82 | 230.12 | 318.56 |
| 442 | ENSMUSG00000002524 | 994.2281308 | 0.017882607 | 0.103137567 | 0.173385965 | 0.862348056 | 0.976334255 | Puf60 | poly-U binding splicing factor 60 [Source:M | 961.31 | 1002.2 | 1001.2 | 1020.9 | 971.74 | 1008 |
| 443 | ENSMUSG00000002546 | 699.5919225 | 0.179822313 | 0.122378696 | 1.4693923 | 0.141726415 | 0.551204857 | Golga2 | golgi autoantigen, golgin subfamily a, 2 [S | 681.83 | 669.17 | 616.83 | 761.54 | 674 | 794.17 |
| 444 | ENSMUSG00000002550 | 1333.931258 | -0.403899477 | 0.125033348 | -3.230334006 | 0.001236457 | 0.024465268 | Uck1 | uridine-cytidine kinase 1 [Source:MGI Sym | 1750 | 1440.1 | 1367.5 | 1217.5 | 1163.7 | 1064.8 |

| | | | | | | | | | | | | | | | |
|---|---|---|---|---|---|---|---|---|---|---|---|---|---|---|---|
| 445 | ENSMUSG00000002565 | 0.167707431 | -0.796629176 | 4.080472857 | -0.195229623 | 0.84521319 | NA | Scin | scinderin [Source:MGI Symbol;Acc:MGI:1 | 0 | 0 | 1.0062 | 0 | 0 | 0 |
| 446 | ENSMUSG00000002578 | 16.8891166 | 1.53928982 | 0.76607259 | 2.009326324 | 0.044502538 | 0.291685757 | Ikzf4 | IKAROS family zinc finger 4 [Source:MGI S | 8.8341 | 3.0838 | 14.087 | 52.055 | 5.4531 | 17.822 |
| 447 | ENSMUSG00000002580 | 238.9922566 | 0.106822807 | 0.16405646 | 0.651134414 | 0.514959723 | 0.885627279 | Mien1 | migration and invasion enhancer 1 [Sourc | 245.75 | 207.64 | 236.47 | 239.07 | 239.94 | 265.09 |
| 448 | ENSMUSG00000002588 | 14095.36865 | -0.093735351 | 0.089871031 | -1.042998504 | 0.296948986 | 0.747770393 | Pon1 | paraoxonase 1 [Source:MGI Symbol;Acc:N | 13791 | 14542 | 15328 | 14438 | 13282 | 13191 |
| 449 | ENSMUSG00000002602 | 201.3635313 | -1.449024968 | 0.213053 | -6.801241809 | 1.04E-11 | 3.57E-09 | Axl | AXL receptor tyrosine kinase [Source:MGI | 266.63 | 376.22 | 241.5 | 102.18 | 104.7 | 116.95 |
| 450 | ENSMUSG00000002603 | 105.6145442 | -0.992376412 | 0.258745978 | -3.835330771 | 0.000125395 | 0.004328686 | Tgfb1 | transforming growth factor, beta 1 [Sourc | 153.39 | 146.99 | 121.76 | 91.578 | 67.618 | 52.351 |
| 451 | ENSMUSG00000002608 | 346.3650161 | 0.098061882 | 0.150520863 | 0.651483657 | 0.514734324 | 0.885488593 | Ccdc97 | coiled-coil domain containing 97 [Source:I | 317.23 | 309.4 | 377.34 | 351.85 | 338.09 | 384.28 |
| 452 | ENSMUSG00000002617 | 49.4004807 | 0.303557015 | 0.408394748 | 0.74329314 | 0.457304224 | 0.858923498 | Zfp40 | zinc finger protein 40 [Source:MGI Symbo | 54.611 | 30.838 | 47.293 | 88.686 | 35.99 | 38.985 |
| 453 | ENSMUSG00000002625 | 476.1786837 | 0.138275242 | 0.164161303 | 0.842313259 | 0.399612631 | 0.826649016 | Akap8l | A kinase (PRKA) anchor protein 8-like [Sou | 453.75 | 459.48 | 447.78 | 627.55 | 431.88 | 436.63 |
| 454 | ENSMUSG00000002633 | 11.14409738 | 1.001337245 | 0.688456335 | 1.454467326 | 0.145816786 | 0.556690797 | Shh | sonic hedgehog [Source:MGI Symbol;Acc: | 8.031 | 3.0838 | 11.069 | 11.568 | 14.178 | 18.935 |
| 455 | ENSMUSG00000002635 | 191.2485066 | 0.214180057 | 0.184119588 | 1.163266002 | 0.244721595 | 0.694594417 | Pdcd2l | programmed cell death 2-like [Source:MG | 163.03 | 187.08 | 181.12 | 176.41 | 214.85 | 225 |
| 456 | ENSMUSG00000002658 | 329.9563029 | -0.058922374 | 0.143448557 | -0.410756126 | 0.681251367 | 0.934452508 | Gtf2f1 | general transcription factor IIF, polypeptic | 333.29 | 350.52 | 326.02 | 299.8 | 333.73 | 336.38 |
| 457 | ENSMUSG00000002660 | 814.9734372 | 0.076855743 | 0.122026996 | 0.62982574 | 0.528808603 | 0.888334265 | Clpp | caseinolytic mitochondrial matrix peptida: | 828.8 | 788.41 | 761.73 | 794.32 | 777.61 | 938.97 |
| 458 | ENSMUSG00000002661 | 145.5043621 | 0.133274959 | 0.216812688 | 0.614700921 | 0.538752251 | 0.8916993 | Alkbh7 | alkB homolog 7 [Source:MGI Symbol;Acc:I | 110.83 | 150.08 | 155.97 | 141.7 | 140.69 | 173.76 |
| 459 | ENSMUSG00000002664 | 1.841445177 | -1.239977997 | 1.649098077 | -0.751912827 | 0.452103484 | NA | Pspn | persephin [Source:MGI Symbol;Acc:MGI:1 | 1.6062 | 4.1117 | 2.0125 | 0 | 1.0906 | 2.2277 |
| 460 | ENSMUSG00000002668 | 69.67709201 | 0.207531689 | 0.281746463 | 0.736590218 | 0.461371585 | 0.860071824 | Dennd1c | DENN/MADD domain containing 1C [Sour | 57.823 | 62.703 | 73.456 | 59.767 | 77.434 | 86.88 |
| 461 | ENSMUSG00000002679 | 158.1440118 | 0.187237729 | 0.202383449 | 0.925163245 | 0.354880999 | 0.794902164 | Med6 | mediator complex subunit 6 [Source:MGI | 163.03 | 119.24 | 161 | 169.66 | 177.77 | 158.17 |
| 462 | ENSMUSG00000002688 | 2.002546835 | -0.348143247 | 1.713699112 | -0.203153077 | 0.839015388 | NA | Prkd1 | protein kinase D1 [Source:MGI Symbol;Ac | 1.6062 | 5.1396 | 0 | 1.928 | 0 | 3.3415 |
| 463 | ENSMUSG00000002699 | 36.99028136 | -1.351171818 | 0.463284103 | -2.916508054 | 0.003539736 | 0.052782205 | Lcp2 | lymphocyte cytosolic protein 2 [Source:M | 48.989 | 82.233 | 28.175 | 18.316 | 15.269 | 28.96 |
| 464 | ENSMUSG00000002718 | 647.0497867 | 0.063315155 | 0.122569686 | 0.516564552 | 0.605460166 | 0.916376001 | Cse1l | chromosome segregation 1-like (S. cerevis | 595.9 | 685.62 | 617.83 | 669.96 | 691.45 | 621.52 |
| 465 | ENSMUSG00000002728 | 274.9316475 | -0.100524568 | 0.156487694 | -0.642380022 | 0.520626468 | 0.887802551 | Naa20 | N(alpha)-acetyltransferase 20, NatB cataly | 298.75 | 295.01 | 259.61 | 270.88 | 280.29 | 245.05 |
| 466 | ENSMUSG00000002731 | 313.0102963 | 0.356617278 | 0.162153324 | 2.19925975 | 0.027859458 | 0.217463141 | Prkra | protein kinase, interferon inducible doubl | 248.96 | 253.9 | 320.99 | 324.86 | 376.26 | 353.09 |
| 467 | ENSMUSG00000002732 | 51.80516552 | -0.464033764 | 0.347031537 | -1.337151571 | 0.181173166 | 0.614561135 | Fkbp7 | FK506 binding protein 7 [Source:MGI Sym | 73.885 | 63.731 | 42.262 | 41.451 | 33.809 | 55.692 |
| 468 | ENSMUSG00000002733 | 413.250561 | -0.097206022 | 0.137769039 | -0.705572331 | 0.480454136 | 0.870278888 | Plekha3 | pleckstrin homology domain-containing, f | 454.56 | 416.31 | 410.55 | 427.04 | 405.71 | 365.34 |
| 469 | ENSMUSG00000002741 | 1129.509898 | 0.129889738 | 0.106217173 | 1.222869464 | 0.221379004 | 0.667727967 | Ykt6 | YKT6 v-SNARE homolog (S. cerevisiae) [So | 1015.9 | 1138.9 | 1081.7 | 1129.8 | 1186.6 | 1224.1 |
| 470 | ENSMUSG00000002748 | 1212.587659 | 0.046714717 | 0.105930519 | 0.440993938 | 0.659217386 | 0.927763076 | Baz1b | bromodomain adjacent to zinc finger dom | 1215.1 | 1188.3 | 1175.3 | 1222.3 | 1327.3 | 1147.3 |
| 471 | ENSMUSG00000002763 | 2775.211496 | 0.243512534 | 0.105991458 | 2.297473197 | 0.02159179 | 0.182976706 | Pex6 | peroxisomal biogenesis factor 6 [Source:N | 2502.5 | 2302.5 | 2819.5 | 2908.3 | 3133.3 | 2985.1 |
| 472 | ENSMUSG00000002767 | 1025.951753 | -0.033207556 | 0.112875843 | -0.294195419 | 0.768608581 | 0.955903225 | Mrpl2 | mitochondrial ribosomal protein L2 [Sourc | 962.92 | 1133.8 | 1017.3 | 998.68 | 983.74 | 1059.3 |
| 473 | ENSMUSG00000002768 | 681.4421549 | 0.039243894 | 0.127109244 | 0.308741463 | 0.7575182 | 0.953466696 | Mea1 | male enhanced antigen 1 [Source:MGI Syı | 723.6 | 677.4 | 614.82 | 677.68 | 754.71 | 640.46 |
| 474 | ENSMUSG00000002769 | 25297.96388 | 0.125911197 | 0.104056398 | 1.210028398 | 0.226267996 | 0.672783802 | Gnmt | glycine N-methyltransferase [Source:MGI | 21356 | 24086 | 27143 | 26314 | 26113 | 26776 |
| 475 | ENSMUSG00000002771 | 3.241068807 | 0.625344275 | 1.445026609 | 0.432756235 | 0.665191871 | NA | Grin2d | glutamate receptor, ionotropic, NMDA2D | 1.6062 | 5.1396 | 1.0062 | 10.604 | 1.0906 | 0 |
| 476 | ENSMUSG00000002778 | 1474.391835 | 0.101847876 | 0.097560161 | 1.043949454 | 0.296508776 | 0.74713095 | Kdelr1 | KDEL (Lys-Asp-Glu-Leu) endoplasmic retic | 1425.5 | 1457.6 | 1383.6 | 1460.4 | 1546.5 | 1572.7 |
| 477 | ENSMUSG00000002781 | 534.7074751 | 0.226975118 | 0.136034143 | 1.6685158 | 0.095213378 | 0.455057116 | Tmem143 | transmembrane protein 143 [Source:MGI | 530.85 | 492.37 | 453.82 | 521.51 | 572.57 | 637.12 |
| 478 | ENSMUSG00000002791 | 3.346421123 | 1.057555632 | 1.309305499 | 0.807722593 | 0.419250295 | NA | Pdxk-ps | pyridoxal (pyridoxine, vitamin B6) kinase, | 2.4093 | 4.1117 | 0 | 5.7839 | 1.0906 | 6.6831 |
| 479 | ENSMUSG00000002797 | 145.8201781 | -0.099604394 | 0.237040987 | -0.420199035 | 0.67434006 | 0.932114677 | Ggct | gamma-glutamyl cyclotransferase [Source | 191.14 | 131.57 | 128.8 | 116.64 | 135.24 | 171.53 |
| 480 | ENSMUSG00000002799 | 1.427195874 | 0.88083156 | 1.960322273 | 0.449329976 | 0.653193637 | NA | Jag2 | jagged 2 [Source:MGI Symbol;Acc:MGI:10 | 0 | 1.0279 | 2.0125 | 0 | 2.1812 | 3.3415 |
| 481 | ENSMUSG00000002803 | 217.0053404 | -0.003990788 | 0.174969641 | -0.022808459 | 0.981803061 | 0.998368917 | Btbd6 | BTB (POZ) domain containing 6 [Source:M | 197.56 | 248.76 | 206.28 | 227.5 | 219.21 | 202.72 |
| 482 | ENSMUSG00000002804 | 224.3597234 | 0.220805291 | 0.183012304 | 1.206505171 | 0.227622766 | 0.673700979 | Nudt14 | nudix (nucleoside diphosphate linked moi | 171.86 | 216.89 | 233.45 | 215.93 | 256.3 | 251.73 |
| 483 | ENSMUSG00000002808 | 5.13365347 | 0.431632653 | 0.922887902 | 0.467697813 | 0.640000705 | 0.923932929 | Epdr1 | ependymin related protein 1 (zebrafish) [S | 4.0155 | 4.1117 | 5.0312 | 7.7118 | 4.3625 | 5.5692 |
| 484 | ENSMUSG00000002812 | 976.8031691 | -0.021639159 | 0.101989179 | -0.212171129 | 0.83197353 | 0.970533397 | Flii | flightless I actin binding protein [Source:N | 985.41 | 982.69 | 984.11 | 953.37 | 972.83 | 982.41 |
| 485 | ENSMUSG00000002814 | 140.0435575 | 0.172425181 | 0.200078623 | 0.861787121 | 0.388804673 | 0.818464676 | Top3a | topoisomerase (DNA) III alpha [Source:MC | 136.53 | 122.32 | 135.84 | 137.85 | 142.87 | 164.85 |
| 486 | ENSMUSG00000002820 | 622.9346875 | 0.285154522 | 0.127381489 | 2.238586816 | 0.025182813 | 0.204058378 | Atg4d | autophagy related 4D, cysteine peptidase | 503.55 | 559.19 | 622.87 | 669 | 698 | 685.01 |
| 487 | ENSMUSG00000002825 | 47.84826611 | -0.255218442 | 0.361836055 | -0.705342759 | 0.480596956 | 0.870278888 | Qtrt1 | queuine tRNA-ribosyltransferase catalytic | 53.005 | 35.977 | 67.418 | 56.875 | 38.172 | 35.643 |
| 488 | ENSMUSG00000002831 | 792.4927603 | -2.428342554 | 0.880645199 | -2.757458461 | 0.005825261 | 0.073974171 | Plin4 | perilipin 4 [Source:MGI Symbol;Acc:MGI:1 | 1651.2 | 1921.2 | 437.72 | 342.21 | 339.18 | 63.489 |
| 489 | ENSMUSG00000002833 | 580.1116109 | 0.066657477 | 0.116347519 | 0.572917043 | 0.566700865 | 0.902528221 | Hdgfl2 | HDGF like 2 [Source:MGI Symbol;Acc:MGI | 562.17 | 574.61 | 563.5 | 597.67 | 583.48 | 599.25 |
| 490 | ENSMUSG00000002835 | 34.59988001 | -0.301649061 | 0.402519531 | -0.749402296 | 0.453614768 | 0.857099982 | Chaf1a | chromatin assembly factor 1, subunit A (p | 52.202 | 37.005 | 25.156 | 33.739 | 30.537 | 28.96 |
| 491 | ENSMUSG00000002844 | 96.28022362 | -0.119471727 | 0.233114713 | -0.512501876 | 0.608299808 | 0.916376001 | Adprh | ADP-ribosylarginine hydrolase [Source:MC | 101.19 | 111.02 | 88.55 | 90.614 | 90.521 | 95.791 |
| 492 | ENSMUSG00000002845 | 150.82818 | 0.125800069 | 0.188414235 | 0.667678158 | 0.504339055 | 0.881510796 | Tmem39a | transmembrane protein 39a [Source:MGI | 151.79 | 142.88 | 137.86 | 156.16 | 159.23 | 157.05 |
| 493 | ENSMUSG00000002846 | 651.4606665 | 0.220158358 | 0.119542797 | 1.841669787 | 0.065523466 | 0.367765392 | Timmdc1 | translocase of inner mitochondrial membı | 618.39 | 577.69 | 608.78 | 657.43 | 751.44 | 695.04 |
| 494 | ENSMUSG00000002847 | 794.891453 | 0.349048035 | 0.135460139 | 2.576758281 | 0.009973167 | 0.108460061 | Pla1a | phospholipase A1 member A [Source:MGI | 625.62 | 644.5 | 828.14 | 858.9 | 891.04 | 921.15 |
| 495 | ENSMUSG00000002870 | 72.25352021 | -0.552598887 | 0.311973827 | -1.771298869 | 0.076511015 | 0.404923281 | Mcm2 | minichromosome maintenance complex c | 118.06 | 59.619 | 79.493 | 53.019 | 66.528 | 56.806 |
| 496 | ENSMUSG00000002871 | 373.5392815 | 0.21981116 | 0.136514079 | 1.610172087 | 0.107360293 | 0.48315751 | Tpra1 | transmembrane protein, adipocyte assoc | 364.61 | 332.02 | 338.1 | 387.52 | 403.53 | 415.46 |
| 497 | ENSMUSG00000002881 | 1295.265641 | 0.279971501 | 0.106672487 | 2.624589618 | 0.008675346 | 0.097864221 | Nab1 | Ngfi-A binding protein 1 [Source:MGI Sym | 1146.8 | 1211.9 | 1152.2 | 1542.4 | 1333.8 | 1384.5 |
| 498 | ENSMUSG00000002885 | 117.6314905 | -0.97585438 | 0.254595375 | -3.832961931 | 0.00012661 | 0.00434447 | Adgre5 | adhesion G protein-coupled receptor E5 [ | 126.89 | 174.75 | 167.04 | 100.25 | 58.893 | 77.969 |
| 499 | ENSMUSG00000002897 | 313.1903878 | 0.173616322 | 0.165548656 | 1.048732899 | 0.294301068 | 0.745164683 | Il17ra | interleukin 17 receptor A [Source:MGI Syr | 266.63 | 305.29 | 311.94 | 359.56 | 274.84 | 360.89 |
| 500 | ENSMUSG00000002900 | 108.9913111 | -1.310315173 | 0.235935416 | -5.553702767 | 2.80E-08 | 3.68E-06 | Lamb1 | laminin B1 [Source:MGI Symbol;Acc:MGI: | 172.67 | 163.44 | 129.81 | 64.587 | 61.075 | 62.375 |

| | | | | | | | | | | | | | | | |
|---|---|---|---|---|---|---|---|---|---|---|---|---|---|---|---|
| 501 | ENSMUSG00000002908 | 165.8156039 | 0.004373555 | 0.224739508 | 0.019460553 | 0.984473706 | 0.998580735 | Kcnn1 | potassium intermediate/small conductanc | 167.05 | 148.02 | 181.12 | 123.39 | 211.58 | 163.74 |
| 502 | ENSMUSG00000002910 | 374.2071796 | -1.13931018 | 0.205655249 | -5.53990323 | 3.03E-08 | 3.87E-06 | Arrdc2 | arrestin domain containing 2 [Source:MGI | 518 | 629.09 | 396.46 | 176.41 | 282.47 | 242.82 |
| 503 | ENSMUSG00000002930 | 0 | NA | NA | NA | NA | NA | Ppp1r17 | protein phosphatase 1, regulatory subunit | 0 | 0 | 0 | 0 | 0 | 0 |
| 504 | ENSMUSG00000002944 | 1494.331502 | -1.658641567 | 0.479659707 | -3.457954758 | 0.000544293 | 0.013168962 | Cd36 | CD36 molecule [Source:MGI Symbol;Acc:M | 2964.3 | 2635.6 | 1209.5 | 888.79 | 822.33 | 445.54 |
| 505 | ENSMUSG00000002948 | 384.7520197 | 0.016146105 | 0.135858796 | 0.118844755 | 0.905398351 | 0.983972587 | Map2k7 | mitogen-activated protein kinase kinase 7 | 390.31 | 382.39 | 375.33 | 415.47 | 372.99 | 372.02 |
| 506 | ENSMUSG00000002949 | 885.947685 | 0.144151745 | 0.1060527 | 1.359246341 | 0.174068541 | 0.604151393 | Timm44 | translocase of inner mitochondrial membr | 871.37 | 811.03 | 842.23 | 925.42 | 943.38 | 922.26 |
| 507 | ENSMUSG00000002957 | 1250.214458 | -0.128589239 | 0.110438072 | -1.164356068 | 0.244279744 | 0.694157605 | Ap2a2 | adaptor-related protein complex 2, alpha | 1230.4 | 1301.3 | 1386.6 | 1205 | 1100.4 | 1277.6 |
| 508 | ENSMUSG00000002963 | 379.5982734 | -0.013285095 | 0.148042687 | -0.089738273 | 0.928495201 | 0.988240056 | Pnkp | polynucleotide kinase 3'- phosphatase [Sc | 341.32 | 419.39 | 384.39 | 405.83 | 345.73 | 380.93 |
| 509 | ENSMUSG00000002968 | 748.6955155 | 0.09743052 | 0.117217316 | 0.83119562 | 0.405863129 | 0.829973745 | Med25 | mediator complex subunit 25 [Source:MG | 727.61 | 720.57 | 721.48 | 719.13 | 757.98 | 845.41 |
| 510 | ENSMUSG00000002980 | 81.42077198 | -0.127321338 | 0.271173435 | -0.469519951 | 0.638698027 | 0.923932929 | Bcam | basal cell adhesion molecule [Source:MGI | 97.978 | 91.485 | 65.406 | 85.794 | 65.437 | 82.424 |
| 511 | ENSMUSG00000002981 | 1634.301448 | -0.030549788 | 0.104453766 | -0.292471868 | 0.769925866 | 0.956472364 | Clptm1 | cleft lip and palate associated transmemb | 1604.6 | 1736.2 | 1615 | 1695.6 | 1674.1 | 1480.3 |
| 512 | ENSMUSG00000002983 | 24.5834164 | -0.184334094 | 0.434067261 | -0.424667121 | 0.671079355 | 0.93116767 | Relb | avian reticuloendotheliosis viral (v-rel) on | 25.699 | 24.67 | 28.175 | 26.027 | 23.994 | 18.935 |
| 513 | ENSMUSG00000002984 | 800.4728024 | 0.086321023 | 0.134129655 | 0.643564048 | 0.51985817 | 0.887802551 | Tomm40 | translocase of outer mitochondrial memb | 836.03 | 817.19 | 675.19 | 728.77 | 876.86 | 868.8 |
| 514 | ENSMUSG00000002985 | 487990.9782 | -0.130408574 | 0.090265623 | -1.444720251 | 0.148536479 | 0.560454158 | Apoe | apolipoprotein E [Source:MGI Symbol;Acc | 522265 | 493927 | 513902 | 505000 | 455055 | 437796 |
| 515 | ENSMUSG00000002992 | 38.2637251 | -0.683582691 | 0.364269923 | -1.876582853 | 0.060575289 | 0.351307532 | Apoc2 | apolipoprotein C-II [Source:MGI Symbol;A | 44.171 | 40.089 | 57.356 | 31.811 | 30.537 | 25.618 |
| 516 | ENSMUSG00000002996 | 708.4242605 | 0.239980796 | 0.123055725 | 1.950179848 | 0.051154687 | 0.317554244 | Hbp1 | high mobility group box transcription fact | 621.6 | 668.15 | 659.09 | 703.7 | 749.25 | 848.75 |
| 517 | ENSMUSG00000002997 | 8.49570651 | -0.768761109 | 0.807794826 | -0.951678674 | 0.341259968 | 0.783129922 | Prkar2b | protein kinase, cAMP dependent regulato | 13.653 | 15.419 | 3.0187 | 6.7478 | 5.4531 | 6.6831 |
| 518 | ENSMUSG00000003031 | 511.3877055 | 0.033896599 | 0.126815945 | 0.267289727 | 0.789246096 | 0.960994657 | Cdkn1b | cyclin-dependent kinase inhibitor 1B [Sou | 510.77 | 476.95 | 528.28 | 523.44 | 485.33 | 543.56 |
| 519 | ENSMUSG00000003032 | 19.28998551 | -0.352727857 | 0.587446067 | -0.600442963 | 0.548211062 | 0.895600915 | Klf4 | Kruppel-like factor 4 (gut) [Source:MGI Sy | 16.062 | 38.033 | 11.069 | 24.099 | 11.997 | 14.48 |
| 520 | ENSMUSG00000003033 | 519.936588 | -0.179980116 | 0.128935007 | -1.395897981 | 0.162745219 | 0.586288131 | Ap1m1 | adaptor-related protein complex AP-1, mu | 562.97 | 590.02 | 504.13 | 508.98 | 467.88 | 485.64 |
| 521 | ENSMUSG00000003037 | 1313.617729 | 0.053350739 | 0.104394726 | 0.511048217 | 0.609317295 | 0.916376001 | Rab8a | RAB8A, member RAS oncogene family [So | 1329.9 | 1251 | 1286 | 1237.7 | 1400.4 | 1376.7 |
| 522 | ENSMUSG00000003038 | 346.0511538 | -0.105295364 | 0.209904494 | -0.501634637 | 0.615924551 | 0.917509899 | Hmgn2 | high mobility group nucleosomal binding | 428.86 | 325.85 | 319.99 | 229.43 | 404.62 | 367.57 |
| 523 | ENSMUSG00000003039 | 911.9540247 | 0.07723366 | 0.125788002 | 0.613998626 | 0.539216235 | 0.891951467 | Fam32a | family with sequence similarity 32, memb | 867.35 | 838.78 | 955.93 | 828.06 | 937.93 | 1043.7 |
| 524 | ENSMUSG00000003051 | 4.047365687 | -0.982579207 | 1.107488472 | -0.887213936 | 0.374963731 | NA | Elf3 | E74-like factor 3 [Source:MGI Symbol;Acc | 8.031 | 3.0838 | 5.0312 | 4.8199 | 1.0906 | 2.2277 |
| 525 | ENSMUSG00000003053 | 10044.60064 | -0.338173993 | 0.174602398 | -1.9368233 | 0.052766947 | 0.32296796 | Cyp2c29 | cytochrome P450, family 2, subfamily c, p | 9880.6 | 9046.7 | 14723 | 10878 | 7590.7 | 8147.8 |
| 526 | ENSMUSG00000003062 | 174.1776292 | -0.141125388 | 0.198195875 | -0.712050075 | 0.476433768 | 0.86953366 | Stard3nl | STARD3 N-terminal like [Source:MGI Symb | 177.49 | 188.11 | 182.13 | 139.78 | 155.96 | 201.61 |
| 527 | ENSMUSG00000003068 | 2056.278499 | 0.206613098 | 0.109792525 | 1.881850327 | 0.05985634 | 0.348255071 | Stk11 | serine/threonine kinase 11 [Source:MGI S | 1864 | 1786.5 | 2076.9 | 2022.4 | 2200.9 | 2387 |
| 528 | ENSMUSG00000003070 | 3.111751671 | -0.957964007 | 1.20100429 | -0.797635792 | 0.425081874 | NA | Efna2 | ephrin A2 [Source:MGI Symbol;Acc:MGI:1 | 3.2124 | 3.0838 | 6.0375 | 1.928 | 2.1812 | 2.2277 |
| 529 | ENSMUSG00000003072 | 4927.174757 | -0.083930525 | 0.083425427 | -1.006054488 | 0.31438936 | 0.760895517 | Atp5d | ATP synthase, H+ transporting, mitochond | 5078 | 5003.9 | 5129.8 | 4930.7 | 4723.5 | 4697.1 |
| 530 | ENSMUSG00000003099 | 898.815218 | 0.051651558 | 0.112503114 | 0.459112248 | 0.646153561 | 0.925643431 | Ppp5c | protein phosphatase 5, catalytic subunit [ | 820.77 | 954.94 | 873.42 | 915.78 | 933.57 | 894.42 |
| 531 | ENSMUSG00000003119 | 310.0957669 | 0.166903552 | 0.160098907 | 1.042502754 | 0.297178649 | 0.748066826 | Cdk12 | cyclin-dependent kinase 12 [Source:MGI S | 265.83 | 328.93 | 282.75 | 363.42 | 297.74 | 321.9 |
| 532 | ENSMUSG00000003123 | 449.2463064 | 0.352926992 | 0.195847607 | 1.802049036 | 0.071537691 | 0.38710107 | Lipe | lipase, hormone sensitive [Source:MGI Sy | 522.02 | 311.46 | 349.17 | 589.95 | 441.7 | 481.18 |
| 533 | ENSMUSG00000003131 | 457.1627896 | -0.091186611 | 0.154505073 | -0.590185223 | 0.555066478 | 0.897628185 | Pafah1b2 | platelet-activating factor acetylhydrolase, | 458.57 | 510.88 | 445.77 | 473.31 | 359.9 | 494.55 |
| 534 | ENSMUSG00000003134 | 391.1820301 | -0.6303726 | 0.154254204 | -4.086582958 | 4.38E-05 | 0.001884509 | Tbc1d8 | TBC1 domain family, member 8 [Source:M | 401.55 | 530.41 | 495.07 | 322.93 | 308.64 | 288.49 |
| 535 | ENSMUSG00000003135 | 350.6686008 | -0.067726022 | 0.139538852 | -0.485356022 | 0.627423823 | 0.920328819 | Cnot11 | CCR4-NOT transcription complex, subunit | 365.41 | 351.55 | 359.23 | 319.08 | 350.09 | 358.66 |
| 536 | ENSMUSG00000003153 | 5.228379081 | -1.00280812 | 0.992303079 | -1.010586524 | 0.312214368 | 0.759830222 | Slc2a3 | solute carrier family 2 (facilitated glucose | 9.6372 | 9.2513 | 2.0125 | 3.8559 | 3.2719 | 3.3415 |
| 537 | ENSMUSG00000003154 | 180.3498698 | 0.054459404 | 0.191651266 | 0.284158854 | 0.776288646 | 0.959022803 | Foxj2 | forkhead box J2 [Source:MGI Symbol;Acc: | 153.39 | 169.61 | 208.29 | 177.37 | 195.22 | 178.22 |
| 538 | ENSMUSG00000003161 | 580.5025308 | 0.128473058 | 0.159010472 | 0.807953441 | 0.419117385 | 0.837591673 | Sri | sorcin [Source:MGI Symbol;Acc:MGI:9841 | 552.53 | 571.52 | 539.35 | 515.73 | 543.13 | 760.76 |
| 539 | ENSMUSG00000003166 | 895.7254177 | 0.206764725 | 0.125655379 | 1.645490446 | 0.099868711 | 0.465517577 | Dgcr2 | DiGeorge syndrome critical region gene 2 | 800.69 | 785.33 | 908.64 | 872.4 | 934.66 | 1072.6 |
| 540 | ENSMUSG00000003184 | 1147.896386 | 0.180279913 | 0.102491488 | 1.758974502 | 0.07858184 | 0.411069703 | Irf3 | interferon regulatory factor 3 [Source:MG | 1052.9 | 1044.4 | 1132 | 1240.6 | 1234.6 | 1182.9 |
| 541 | ENSMUSG00000003190 | 74.89523522 | 0.246714935 | 0.264255133 | 0.933624001 | 0.35049788 | 0.791877747 | Bcl2l12 | BCL2-like 12 (proline rich) [Source:MGI Sy | 71.476 | 58.591 | 75.468 | 90.614 | 75.253 | 77.969 |
| 542 | ENSMUSG00000003198 | 97.70084527 | 0.302184014 | 0.234598459 | 1.288090364 | 0.197714509 | 0.639763459 | Zfp959 | zinc finger protein 959 [Source:MGI Symb | 91.554 | 92.513 | 78.487 | 116.64 | 112.33 | 94.677 |
| 543 | ENSMUSG00000003199 | 1501.544606 | 0.235305376 | 0.100186366 | 2.348676623 | 0.018840261 | 0.166466427 | Mpnd | MPN domain containing [Source:MGI Sym | 1380.5 | 1381.5 | 1375.5 | 1547.2 | 1584.7 | 1739.8 |
| 544 | ENSMUSG00000003200 | 378.1832708 | 0.031678794 | 0.147756052 | 0.214399299 | 0.830235688 | 0.9702546 | Sh3gl1 | SH3-domain GRB2-like 1 [Source:MGI Sym | 367.02 | 390.61 | 365.27 | 437.65 | 358.81 | 349.75 |
| 545 | ENSMUSG00000003206 | 4.432883723 | -1.32308952 | 1.404567932 | -0.941990408 | 0.346197551 | NA | Ebi3 | Epstein-Barr virus induced gene 3 [Source | 5.6217 | 12.335 | 1.0062 | 0 | 7.6343 | 0 |
| 546 | ENSMUSG00000003208 | 77.12465755 | -0.136712761 | 0.277989701 | -0.491790742 | 0.622867282 | 0.919113687 | Yju2 | YJU2 splicing factor [Source:MGI Symbol;A | 104.4 | 71.954 | 65.406 | 73.262 | 71.981 | 75.741 |
| 547 | ENSMUSG00000003226 | 1698.739743 | -0.044838033 | 0.121575364 | -0.368808546 | 0.712270434 | 0.944239215 | Ranbp2 | RAN binding protein 2 [Source:MGI Symbo | 1574.1 | 1922.2 | 1680.4 | 1737.1 | 1802.8 | 1475.8 |
| 548 | ENSMUSG00000003227 | 1.472354105 | -3.941665183 | 2.621323271 | -1.503692897 | 0.132660459 | NA | Edar | ectodysplasin-A receptor [Source:MGI Syr | 8.8341 | 0 | 0 | 0 | 0 | 0 |
| 549 | ENSMUSG00000003228 | 36.27509476 | -0.698555095 | 0.396101233 | -1.763577176 | 0.077803193 | 0.408177507 | Grk5 | G protein-coupled receptor kinase 5 [Sour | 44.171 | 33.921 | 56.35 | 19.28 | 31.628 | 32.301 |
| 550 | ENSMUSG00000003233 | 345.5579111 | -0.04391924 | 0.144235998 | -0.304495691 | 0.760750275 | 0.95402891 | Dvl3 | dishevelled segment polarity protein 3 [Sc | 327.67 | 383.41 | 342.12 | 356.67 | 331.55 | 331.93 |
| 551 | ENSMUSG00000003234 | 844.7961662 | 0.118502899 | 0.111534664 | 1.062475959 | 0.288019666 | 0.73951397 | Abcf3 | ATP-binding cassette, sub-family F (GCN2( | 807.92 | 805.89 | 816.06 | 825.17 | 870.31 | 943.43 |
| 552 | ENSMUSG00000003235 | 715.3261988 | 0.143021872 | 0.113830884 | 1.256441722 | 0.208955863 | 0.65350014 | Eif2b5 | eukaryotic translation initiation factor 2B, | 664.97 | 670.2 | 704.37 | 710.45 | 765.61 | 776.35 |
| 553 | ENSMUSG00000003269 | 351.1781678 | 0.092000811 | 0.157926992 | 0.5825528 | 0.560194388 | 0.899475387 | Cyth2 | cytohesin 2 [Source:MGI Symbol;Acc:MGI | 401.55 | 313.51 | 303.89 | 373.06 | 367.54 | 347.52 |
| 554 | ENSMUSG00000003271 | 0.685278559 | -2.809870862 | 3.376578027 | -0.832165239 | 0.405315682 | NA | Sult2b1 | sulfotransferase family, cytosolic, 2B, mer | 0 | 4.1117 | 0 | 0 | 0 | 0 |
| 555 | ENSMUSG00000003273 | 11.86466418 | -0.294377744 | 0.641380391 | -0.458975279 | 0.646251919 | 0.925643431 | Car11 | carbonic anhydrase 11 [Source:MGI Symb | 16.865 | 10.279 | 12.075 | 15.424 | 7.6343 | 8.9108 |
| 556 | ENSMUSG00000003279 | 57.77797793 | 0.103684251 | 0.319461038 | 0.324559928 | 0.74551416 | 0.949997329 | Dlgap1 | DLG associated protein 1 [Source:MGI Syr | 55.414 | 49.34 | 62.387 | 59.767 | 77.434 | 42.326 |

| | | | | | | | | | | | | | | |
|---|---|---|---|---|---|---|---|---|---|---|---|---|---|---|
| 557 | ENSMUSG00000003282 | 8.371879073 | 0.636512735 | 0.832513001 | 0.764567922 | 0.444528877 | 0.851724159 | Plag1 | pleiomorphic adenoma gene 1 [Source:M | 10.44 | 4.1117 | 5.0312 | 17.352 | 3.2719 | 10.025 |
| 558 | ENSMUSG00000003283 | 41.20659152 | -1.102848358 | 0.422202549 | -2.612130982 | 0.008997977 | 0.100267137 | Hck | hemopoietic cell kinase [Source:MGI Syml | 48.186 | 82.233 | 38.237 | 22.171 | 18.541 | 37.871 |
| 559 | ENSMUSG00000003299 | 687.0967712 | -0.019720985 | 0.112392099 | -0.175465935 | 0.860713535 | 0.975943719 | Mrpl4 | mitochondrial ribosomal protein L4 [Sourc | 712.35 | 670.2 | 692.3 | 669.96 | 682.73 | 695.04 |
| 560 | ENSMUSG00000003308 | 810.819861 | 0.068694978 | 0.112824707 | 0.608864673 | 0.542614142 | 0.894095573 | Keap1 | kelch-like ECH-associated protein 1 [Sourc | 798.28 | 763.74 | 812.04 | 798.17 | 801.6 | 891.08 |
| 561 | ENSMUSG00000003309 | 2.166512577 | 0.265867089 | 1.560321348 | 0.170392522 | 0.864701453 | NA | Ap1m2 | adaptor protein complex AP-1, mu 2 subu | 4.8186 | 0 | 1.0062 | 3.8559 | 1.0906 | 2.2277 |
| 562 | ENSMUSG00000003316 | 628.4389434 | -0.107442725 | 0.121683427 | -0.882969252 | 0.377252885 | 0.81030298 | Glg1 | golgi apparatus protein 1 [Source:MGI Syr | 611.16 | 708.24 | 636.95 | 619.84 | 608.57 | 585.88 |
| 563 | ENSMUSG00000003341 | 0 | NA | NA | NA | NA | NA | Atp8b3 | ATPase, class I, type 8B, member 3 [Sourc | 0 | 0 | 0 | 0 | 0 | 0 |
| 564 | ENSMUSG00000003344 | 537.5325362 | 0.145319849 | 0.130817098 | 1.110862811 | 0.266627405 | 0.722732157 | Btbd2 | BTB (POZ) domain containing 2 [Source:M | 481.06 | 526.29 | 524.25 | 532.12 | 535.49 | 625.98 |
| 565 | ENSMUSG00000003345 | 1687.96153 | 0.110449122 | 0.09932008 | 1.112052285 | 0.26611567 | 0.722063933 | Csnk1g2 | casein kinase 1, gamma 2 [Source:MGI Sy | 1654.4 | 1566.5 | 1648.2 | 1636.8 | 1799.5 | 1822.2 |
| 566 | ENSMUSG00000003346 | 1353.990261 | 0.017647452 | 0.099116454 | 0.178047647 | 0.858685556 | 0.975439379 | Abhd17a | abhydrolase domain containing 17A [Sour | 1415.9 | 1305.5 | 1315.2 | 1386.2 | 1351.3 | 1350 |
| 567 | ENSMUSG00000003348 | 12.55909057 | 0.308381492 | 0.661101547 | 0.466466147 | 0.640881873 | 0.923932929 | Mob3a | MOB kinase activator 3A [Source:MGI Sym | 11.243 | 9.2513 | 13.081 | 9.6398 | 7.6343 | 24.505 |
| 568 | ENSMUSG00000003352 | 7.733120123 | -1.114206214 | 0.809182516 | -1.376952902 | 0.16852681 | 0.595692028 | Cacnb3 | calcium channel, voltage-dependent, beta | 14.456 | 5.1396 | 12.075 | 4.8199 | 5.4531 | 4.4554 |
| 569 | ENSMUSG00000003354 | 0.469265235 | -2.274672822 | 3.439794491 | -0.661281605 | 0.508431736 | NA | Ccdc65 | coiled-coil domain containing 65 [Source:I | 0.8031 | 0 | 2.0125 | 0 | 0 | 0 |
| 570 | ENSMUSG00000003355 | 162.9304966 | 0.701202032 | 0.276241529 | 2.538365733 | 0.011137153 | 0.117069607 | Fkbp11 | FK506 binding protein 11 [Source:MGI Syr | 175.88 | 90.457 | 104.65 | 147.49 | 199.58 | 259.53 |
| 571 | ENSMUSG00000003360 | 926.2627962 | -0.048657237 | 0.110340741 | -0.440972541 | 0.659232876 | 0.927763076 | Ddx23 | DEAD box helicase 23 [Source:MGI Symbc | 943.65 | 937.46 | 944.86 | 940.84 | 955.38 | 835.38 |
| 572 | ENSMUSG00000003363 | 333.4308792 | -0.008837699 | 0.143511568 | -0.061581788 | 0.95089588 | 0.990883058 | Pld3 | phospholipase D family, member 3 [Sourc | 322.85 | 322.77 | 358.22 | 350.89 | 321.73 | 324.13 |
| 573 | ENSMUSG00000003378 | 188.1131879 | 0.130559695 | 0.251196618 | 0.519751006 | 0.603237131 | 0.91593763 | Grik5 | glutamate receptor, ionotropic, kainate 5 | 256.99 | 141.85 | 138.86 | 186.05 | 233.39 | 171.53 |
| 574 | ENSMUSG00000003379 | 10.84801199 | -0.994745571 | 0.791084184 | -1.257445909 | 0.208592214 | 0.653455953 | Cd79a | CD79A antigen (immunoglobulin-associat | 13.653 | 16.447 | 13.081 | 1.928 | 3.2719 | 16.708 |
| 575 | ENSMUSG00000003380 | 1494.939435 | -0.022667158 | 0.112987922 | -0.200615759 | 0.840999035 | 0.971854408 | Rabac1 | Rab acceptor 1 (prenylated) [Source:MGI | 1497 | 1497.7 | 1526.5 | 1678.3 | 1408 | 1362.2 |
| 576 | ENSMUSG00000003382 | 345.1079249 | 0.120908879 | 0.167589606 | 0.721458105 | 0.470627708 | 0.867110762 | Etv3 | ets variant 3 [Source:MGI Symbol;Acc:MG | 327.67 | 287.82 | 376.34 | 354.74 | 309.74 | 414.35 |
| 577 | ENSMUSG00000003402 | 1895.276856 | 0.093884854 | 0.106118553 | 0.884716684 | 0.376309455 | 0.809731241 | Prkcsh | protein kinase C substrate 80K-H [Source: | 1947.5 | 1672.4 | 1879.7 | 1891.3 | 1890 | 2090.7 |
| 578 | ENSMUSG00000003410 | 0 | NA | NA | NA | NA | NA | Elavl3 | ELAV like RNA binding protein 3 [Source:N | 0 | 0 | 0 | 0 | 0 | 0 |
| 579 | ENSMUSG00000003411 | 0.877757073 | -0.466619568 | 2.366442592 | -0.197181867 | 0.84368522 | NA | Rab3b | RAB3B, member RAS oncogene family [So | 0 | 2.0558 | 1.0062 | 0 | 1.0906 | 1.1138 |
| 580 | ENSMUSG00000003418 | 0.829837731 | -1.7608533 | 2.399900965 | -0.733719152 | 0.463119924 | NA | St8sia6 | ST8 alpha-N-acetyl-neuraminide alpha-2,8 | 0.8031 | 2.0558 | 1.0062 | 0 | 0 | 1.1138 |
| 581 | ENSMUSG00000003420 | 3407.574165 | -0.112118941 | 0.086162621 | -1.301248034 | 0.193173569 | 0.63128389 | Fcgrt | Fc fragment of IgG receptor and transport | 3495.1 | 3586.4 | 3539 | 3383.6 | 3240.2 | 3201.2 |
| 582 | ENSMUSG00000003421 | 369.6238885 | 0.036623299 | 0.140096617 | 0.261414585 | 0.793772811 | 0.961915077 | Nosip | nitric oxide synthase interacting protein [S | 369.43 | 350.52 | 374.32 | 342.21 | 396.99 | 384.28 |
| 583 | ENSMUSG00000003423 | 679.0956094 | -0.008133552 | 0.120170669 | -0.067683334 | 0.946037716 | 0.990337231 | Pih1d1 | PIH1 domain containing 1 [Source:MGI Sy | 632.84 | 687.68 | 723.49 | 707.56 | 639.1 | 683.9 |
| 584 | ENSMUSG00000003429 | 2103.919796 | -0.017907726 | 0.098051387 | -0.182636132 | 0.855083532 | 0.974158771 | Rps11 | ribosomal protein S11 [Source:MGI Symbo | 2068 | 2179.2 | 2104.1 | 2095.7 | 1950 | 2226.6 |
| 585 | ENSMUSG00000003435 | 826.3071815 | -0.126229452 | 0.114394656 | -1.103455842 | 0.269829258 | 0.725011151 | Supt5 | suppressor of Ty 5, DSIF elongation factor | 856.11 | 868.59 | 863.36 | 861.8 | 739.44 | 768.55 |
| 586 | ENSMUSG00000003436 | 0.160662974 | 1.126924175 | 4.080472857 | 0.276174898 | 0.782413736 | NA | Dll3 | delta like canonical Notch ligand 3 [Source | 0 | 0 | 0 | 0.964 | 0 | 0 |
| 587 | ENSMUSG00000003437 | 492.1525016 | -0.056604263 | 0.128282256 | -0.441247796 | 0.659033615 | 0.927763076 | Paf1 | Paf1, RNA polymerase II complex compon | 476.24 | 519.1 | 510.17 | 454.03 | 478.78 | 514.6 |
| 588 | ENSMUSG00000003438 | 580.5193511 | 0.023979907 | 0.123348515 | 0.19440775 | 0.845856622 | 0.972803041 | Timm50 | translocase of inner mitochondrial membr | 529.24 | 582.83 | 615.82 | 588.99 | 573.67 | 592.56 |
| 589 | ENSMUSG00000003444 | 176.800911 | -0.117444127 | 0.194561278 | -0.603635669 | 0.546085893 | 0.89549457 | Med29 | mediator complex subunit 29 [Source:MG | 174.27 | 171.66 | 206.28 | 169.66 | 194.13 | 144.8 |
| 590 | ENSMUSG00000003452 | 4.385850334 | -0.619269065 | 1.005584604 | -0.615829899 | 0.538006792 | NA | Bicd1 | BICD cargo adaptor 1 [Source:MGI Symbo | 4.8186 | 5.1396 | 6.0375 | 4.8199 | 3.2719 | 2.2277 |
| 591 | ENSMUSG00000003458 | 700.1449416 | -0.012107254 | 0.112173418 | -0.107933363 | 0.914048552 | 0.98625062 | Ncstn | nicastrin [Source:MGI Symbol;Acc:MGI:18 | 681.83 | 720.57 | 707.39 | 714.31 | 677.27 | 699.49 |
| 592 | ENSMUSG00000003464 | 2671.675094 | 0.051241617 | 0.105643933 | 0.485040788 | 0.627647412 | 0.920328819 | Pex19 | peroxisomal biogenesis factor 19 [Source: | 2814.1 | 2620.2 | 2437.1 | 2561.3 | 2965.4 | 2632 |
| 593 | ENSMUSG00000003469 | 2.381338202 | 1.454875946 | 1.433034627 | 1.015241306 | 0.309990815 | NA | Phyhip | phytanoyl-CoA hydroxylase interacting pr | 0.8031 | 2.0558 | 1.0062 | 3.8559 | 5.4531 | 1.1138 |
| 594 | ENSMUSG00000003476 | 1.156610816 | 0.369776097 | 2.544046901 | 0.14534956 | 0.884434887 | NA | Crhr2 | corticotropin releasing hormone receptor | 0 | 3.0838 | 0 | 3.8559 | 0 | 0 |
| 595 | ENSMUSG00000003477 | 11940.70893 | -0.163852398 | 0.134584138 | -1.217471839 | 0.223424745 | 0.668933038 | Inmt | indolethylamine N-methyltransferase [Sou | 11064 | 12811 | 13981 | 12988 | 11582 | 9218.2 |
| 596 | ENSMUSG00000003484 | 7.8711578 | -0.906117783 | 0.856496175 | -1.057935587 | 0.290084809 | 0.740058153 | Cyp4f18 | cytochrome P450, family 4, subfamily f, pc | 13.653 | 3.0838 | 14.087 | 8.6758 | 3.2719 | 4.4554 |
| 597 | ENSMUSG00000003500 | 29.97806306 | -0.6607297 | 0.42212815 | -1.565234869 | 0.117527848 | 0.502806291 | Impdh1 | inosine monophosphate dehydrogenase 1 | 28.912 | 42.145 | 39.244 | 23.135 | 16.359 | 30.074 |
| 598 | ENSMUSG00000003505 | 0 | NA | NA | NA | NA | NA | Psg18 | pregnancy specific glycoprotein 18 [Sourc | 0 | 0 | 0 | 0 | 0 | 0 |
| 599 | ENSMUSG00000003518 | 722.2990211 | -0.455926377 | 0.123368909 | -3.695634345 | 0.000219338 | 0.006778871 | Dusp3 | dual specificity phosphatase 3 (vaccinia vi | 860.12 | 847 | 799.96 | 680.57 | 575.85 | 570.29 |
| 600 | ENSMUSG00000003526 | 6200.584537 | -0.070225831 | 0.105032315 | -0.668611666 | 0.503743229 | 0.881510796 | Prodh | proline dehydrogenase [Source:MGI Symb | 6220.8 | 6475.9 | 6357.5 | 5597.8 | 6869.8 | 5681.7 |
| 601 | ENSMUSG00000003527 | 118.8325613 | 0.182389816 | 0.22490047 | 0.810980146 | 0.417377073 | 0.837009933 | Ess2 | ess-2 splicing factor [Source:MGI Symbol;, | 110.83 | 99.708 | 123.77 | 148.45 | 119.97 | 110.27 |
| 602 | ENSMUSG00000003528 | 4119.409741 | 0.294651348 | 0.110426496 | 2.668302975 | 0.007623548 | 0.088790029 | Slc25a1 | solute carrier family 25 (mitochondrial car | 4202.6 | 3528.8 | 3366.9 | 4274.3 | 4613.3 | 4730.5 |
| 603 | ENSMUSG00000003531 | 942.1282328 | 0.05392826 | 0.10842419 | 0.497382177 | 0.618919572 | 0.917509899 | Dgcr6 | DiGeorge syndrome critical region gene 6 | 907.51 | 927.18 | 938.83 | 935.06 | 916.12 | 1028.1 |
| 604 | ENSMUSG00000003534 | 33.9214643 | -0.429893393 | 0.431098787 | -0.997203903 | 0.318665547 | 0.7652277 | Ddr1 | discoidin domain receptor family, membe | 51.399 | 41.117 | 24.15 | 30.847 | 37.081 | 18.935 |
| 605 | ENSMUSG00000003541 | 13.74632509 | -0.205206383 | 0.669843672 | -0.306349663 | 0.75933843 | 0.95402891 | Ier3 | immediate early response 3 [Source:MGI S | 18.471 | 18.503 | 7.0437 | 9.6398 | 6.5437 | 22.277 |
| 606 | ENSMUSG00000003545 | 3.424197706 | -4.103826214 | 1.584219396 | -2.590440582 | 0.009585317 | NA | Fosb | FBJ osteosarcoma oncogene B [Source:M | 7.2279 | 7.1954 | 5.0312 | 0 | 1.0906 | 0 |
| 607 | ENSMUSG00000003546 | 1824.670866 | 0.098968404 | 0.104384974 | 0.948109678 | 0.343073622 | 0.784828638 | Klc4 | kinesin light chain 4 [Source:MGI Symbol;, | 1825.5 | 1603.6 | 1856.5 | 1858.5 | 1822.4 | 1981.5 |
| 608 | ENSMUSG00000003549 | 49.20638801 | -0.192490543 | 0.324336084 | -0.593490989 | 0.552852612 | 0.896226672 | Ercc1 | excision repair cross-complementing rode | 61.839 | 43.173 | 52.325 | 53.019 | 41.443 | 43.44 |
| 609 | ENSMUSG00000003555 | 564.4047112 | -0.58589407 | 0.213476853 | -2.744532069 | 0.006059723 | 0.075885575 | Cyp17a1 | cytochrome P450, family 17, subfamily a, | 790.25 | 825.42 | 416.59 | 498.38 | 401.35 | 454.45 |
| 610 | ENSMUSG00000003559 | 866.9339465 | -0.250398678 | 0.135186629 | -1.852244409 | 0.063990734 | 0.362113452 | As3mt | arsenite methyltransferase [Source:MGI S | 1002.3 | 1040.3 | 782.86 | 775.04 | 756.89 | 844.29 |
| 611 | ENSMUSG00000003573 | 16.83716138 | -1.379955774 | 0.607025507 | -2.273307725 | 0.023007642 | 0.191034753 | Homer3 | homer scaffolding protein 3 [Source:MGI S | 24.093 | 34.949 | 14.087 | 12.532 | 10.906 | 4.4554 |
| 612 | ENSMUSG00000003575 | 165.337927 | -0.082489566 | 0.202363963 | -0.40762972 | 0.683545541 | 0.935452995 | Crtc1 | CREB regulated transcription coactivator 1 | 175.08 | 162.41 | 173.07 | 194.72 | 134.15 | 152.6 |

| | | | | | | | | | | | | | | | |
|---|---|---|---|---|---|---|---|---|---|---|---|---|---|---|---|
| 613 | ENSMUSG00000003581 | 249.1480027 | -0.047832346 | 0.182061853 | -0.262725798 | 0.792761931 | 0.961632721 | Rnf215 | ring finger protein 215 [Source:MGI Symb | 294.74 | 235.39 | 229.42 | 280.52 | 244.3 | 210.52 |
| 614 | ENSMUSG00000003585 | 12648.6212 | 0.015831768 | 0.088964824 | 0.177955371 | 0.858758025 | 0.975439379 | Sec14l2 | SEC14-like lipid binding 2 [Source:MGI Syr | 12160 | 12437 | 13142 | 12460 | 13537 | 12156 |
| 615 | ENSMUSG00000003604 | 294.3865631 | 0.14642452 | 0.158554151 | 0.923498497 | 0.355747483 | 0.795525652 | Aven | apoptosis, caspase activation inhibitor [Sc | 262.61 | 290.9 | 285.77 | 351.85 | 293.38 | 281.8 |
| 616 | ENSMUSG00000003617 | 19521.98065 | 0.045886506 | 0.085884463 | 0.534281802 | 0.593146581 | 0.913011006 | Cp | ceruloplasmin [Source:MGI Symbol;Acc:M | 20005 | 18596 | 19033 | 20827 | 19706 | 18964 |
| 617 | ENSMUSG00000003623 | 8811.740089 | 0.104622876 | 0.14037039 | 0.745334367 | 0.45606961 | 0.857702619 | Crot | carnitine O-octanoyltransferase [Source:M | 8347.4 | 7705.3 | 9423.5 | 7437.1 | 8872.2 | 11085 |
| 618 | ENSMUSG00000003644 | 407.3502203 | -0.243975535 | 0.159941394 | -1.525405829 | 0.127157906 | 0.522483099 | Rps6ka1 | ribosomal protein S6 kinase polypeptide 1 | 511.58 | 437.89 | 375.33 | 417.4 | 357.72 | 344.18 |
| 619 | ENSMUSG00000003657 | 0 | NA | NA | NA | NA | NA | Calb2 | calbindin 2 [Source:MGI Symbol;Acc:MGI: | 0 | 0 | 0 | 0 | 0 | 0 |
| 620 | ENSMUSG00000003660 | 1053.612861 | 0.053821022 | 0.126648906 | 0.424962391 | 0.670864091 | 0.93116767 | Snrnp200 | small nuclear ribonucleoprotein 200 (U5) | 924.37 | 1155.4 | 1023.4 | 1082.5 | 1165.9 | 970.16 |
| 621 | ENSMUSG00000003662 | 842.8216396 | 0.113546918 | 0.110911873 | 1.023758007 | 0.305949589 | 0.755069291 | Ciao1 | cytosolic iron-sulfur protein assembly 1 [S | 837.64 | 791.5 | 798.96 | 820.35 | 907.39 | 901.1 |
| 622 | ENSMUSG00000003665 | 0.133850373 | -0.796629176 | 4.080472857 | -0.195229623 | 0.84521319 | NA | Has1 | hyaluronan synthase 1 [Source:MGI Symb | 0.8031 | 0 | 0 | 0 | 0 | 0 |
| 623 | ENSMUSG00000003680 | 250.5480066 | 0.013570461 | 0.163300991 | 0.083100912 | 0.933771301 | 0.988420305 | Taf6l | TATA-box binding protein associated facto | 266.63 | 234.37 | 246.53 | 243.89 | 232.3 | 279.57 |
| 624 | ENSMUSG00000003721 | 3650.147074 | -1.022409234 | 0.136666511 | -7.48105169 | 7.37E-14 | 3.81E-11 | Insig2 | insulin induced gene 2 [Source:MGI Symb | 4728.7 | 4813.7 | 5133.9 | 2527.5 | 1883.5 | 2813.6 |
| 625 | ENSMUSG00000003731 | 1013.550363 | -0.049682515 | 0.10594963 | -0.468925796 | 0.639122677 | 0.923932929 | Kpna6 | karyopherin (importin) alpha 6 [Source:M | 1007.9 | 1006.3 | 1078.7 | 954.34 | 1038.3 | 995.78 |
| 626 | ENSMUSG00000003746 | 5905.132968 | -0.091274067 | 0.094388181 | -0.967007368 | 0.333540351 | 0.775366061 | Man1a | mannosidase 1, alpha [Source:MGI Symbo | 5624.1 | 6239.5 | 6413.8 | 5996.9 | 5637.4 | 5519.1 |
| 627 | ENSMUSG00000003752 | 51.43297993 | 0.245996286 | 0.339067952 | 0.725507334 | 0.468140847 | 0.865229581 | Itpkc | inositol 1,4,5-trisphosphate 3-kinase C [Sc | 32.927 | 48.312 | 60.375 | 58.803 | 45.806 | 62.375 |
| 628 | ENSMUSG00000003762 | 234.7549293 | 0.092951667 | 0.172767288 | 0.53801659 | 0.590565595 | 0.911722201 | Coq8b | coenzyme Q8B [Source:MGI Symbol;Acc:M | 226.47 | 238.48 | 216.34 | 213.04 | 280.29 | 233.91 |
| 629 | ENSMUSG00000003778 | 296.1764659 | 0.266563028 | 0.146245959 | 1.822703541 | 0.068348304 | 0.376605663 | Brd8 | bromodomain containing 8 [Source:MGI S | 275.46 | 264.17 | 266.65 | 310.4 | 320.64 | 339.72 |
| 630 | ENSMUSG00000003779 | 9.740624931 | -1.292771223 | 0.706625522 | -1.829499761 | 0.067324775 | 0.373457298 | Kif20a | kinesin family member 20A [Source:MGI S | 12.047 | 12.335 | 17.106 | 4.8199 | 5.4531 | 6.6831 |
| 631 | ENSMUSG00000003808 | 321.9310257 | 0.031519318 | 0.146585879 | 0.215022879 | 0.829749481 | 0.970053063 | Farsa | phenylalanyl-tRNA synthetase, alpha subu | 297.95 | 330.99 | 327.03 | 346.07 | 303.19 | 326.36 |
| 632 | ENSMUSG00000003809 | 14080.68815 | -0.067700977 | 0.087125176 | -0.77705412 | 0.437126841 | 0.846852749 | Gcdh | glutaryl-Coenzyme A dehydrogenase [Sou | 14147 | 14320 | 14767 | 13474 | 14673 | 13103 |
| 633 | ENSMUSG00000003810 | 268.3668442 | 0.016184002 | 0.197417874 | 0.081978401 | 0.934663889 | 0.988420305 | Mast2 | microtubule associated serine/threonine I | 319.63 | 195.3 | 284.77 | 269.91 | 244.3 | 296.28 |
| 634 | ENSMUSG00000003812 | 325.3076097 | 0.041085095 | 0.153882858 | 0.266989419 | 0.789477308 | 0.961009963 | Dnase2a | deoxyribonuclease II alpha [Source:MGI S | 307.59 | 344.35 | 310.93 | 374.02 | 308.64 | 306.31 |
| 635 | ENSMUSG00000003813 | 621.9557009 | -0.045864811 | 0.137878319 | -0.332647015 | 0.73940075 | 0.948310621 | Rad23a | RAD23 homolog A, nucleotide excision rep | 554.14 | 717.49 | 624.88 | 599.59 | 662.01 | 573.63 |
| 636 | ENSMUSG00000003814 | 9610.797237 | 0.217119257 | 0.119717543 | 1.813596006 | 0.069739946 | 0.381582516 | Calr | calreticulin [Source:MGI Symbol;Acc:MGI: | 9880.6 | 8648.9 | 8136.5 | 10982 | 8825.3 | 11192 |
| 637 | ENSMUSG00000003824 | 159.2294363 | 0.03020713 | 0.188990061 | 0.159834491 | 0.873011454 | 0.979691028 | Syce2 | synaptonemal complex central element pr | 155.8 | 156.24 | 161 | 179.3 | 153.78 | 149.26 |
| 638 | ENSMUSG00000003847 | 484.0844935 | 0.153432458 | 0.162063869 | 0.946740687 | 0.343770932 | 0.785513865 | Nfat5 | nuclear factor of activated T cells 5 [Sourc | 430.46 | 476.95 | 468.91 | 631.41 | 464.6 | 432.17 |
| 639 | ENSMUSG00000003848 | 176.6522497 | 0.045032548 | 0.178480823 | 0.252310288 | 0.800801237 | 0.962424862 | Nob1 | NIN1/RPN12 binding protein 1 homolog [S | 176.68 | 184 | 161 | 181.23 | 182.13 | 174.87 |
| 640 | ENSMUSG00000003849 | 127.6478717 | -0.57396881 | 0.211720989 | -2.710967927 | 0.006708712 | 0.08160729 | Nqo1 | NAD(P)H dehydrogenase, quinone 1 [Sour | 168.65 | 153.16 | 135.84 | 94.47 | 109.06 | 104.7 |
| 641 | ENSMUSG00000003863 | 0 | NA | NA | NA | NA | NA | Ppfia3 | protein tyrosine phosphatase, receptor ty | 0 | 0 | 0 | 0 | 0 | 0 |
| 642 | ENSMUSG00000003865 | 44.25127313 | 0.326719885 | 0.348352515 | 0.937900174 | 0.348295714 | 0.789711685 | Gys1 | glycogen synthase 1, muscle [Source:MGI | 30.518 | 49.34 | 38.237 | 49.163 | 41.443 | 56.806 |
| 643 | ENSMUSG00000003868 | 199.2852796 | -0.215152881 | 0.174323855 | -1.234213653 | 0.217123301 | 0.663070346 | Ruvbl2 | RuvB-like protein 2 [Source:MGI Symbol;A | 221.66 | 199.42 | 221.37 | 198.58 | 186.5 | 168.19 |
| 644 | ENSMUSG00000003872 | 0.17131964 | -0.796629176 | 4.080472857 | -0.195229623 | 0.84521319 | NA | Lin7b | lin-7 homolog B (C. elegans) [Source:MGI | 0 | 1.0279 | 0 | 0 | 0 | 0 |
| 645 | ENSMUSG00000003873 | 293.0379872 | -0.141210035 | 0.161446794 | -0.874653698 | 0.38176236 | 0.813226919 | Bax | BCL2-associated X protein [Source:MGI Sy | 349.35 | 265.2 | 306.9 | 290.16 | 274.84 | 271.78 |
| 646 | ENSMUSG00000003882 | 4.333149131 | -1.270924265 | 1.241504441 | -1.023696914 | 0.305978454 | NA | Il7r | interleukin 7 receptor [Source:MGI Symbo | 4.0155 | 11.307 | 3.0187 | 0 | 6.5437 | 1.1138 |
| 647 | ENSMUSG00000003923 | 1019.409166 | 0.130016038 | 0.112590363 | 1.154770569 | 0.248184392 | 0.699604734 | Tfam | transcription factor A, mitochondrial [Sou | 984.6 | 1019.7 | 915.68 | 990.97 | 1146.2 | 1059.3 |
| 648 | ENSMUSG00000003929 | 60.65745173 | 0.430923913 | 0.282521048 | 1.525280739 | 0.12718909 | 0.522483099 | Zfp81 | zinc finger protein 81 [Source:MGI Symbo | 56.217 | 47.284 | 51.318 | 71.334 | 67.618 | 70.172 |
| 649 | ENSMUSG00000003934 | 0.815516723 | -3.06336425 | 2.557025255 | -1.19801877 | 0.23090971 | NA | Efnb3 | ephrin B3 [Source:MGI Symbol;Acc:MGI:1 | 0.8031 | 3.0838 | 1.0062 | 0 | 0 | 0 |
| 650 | ENSMUSG00000003948 | 2552.026902 | 0.021676772 | 0.098567036 | 0.219919083 | 0.825934174 | 0.96943211 | Mmd | monocyte to macrophage differentiation- | 2494.4 | 2445.4 | 2658.5 | 2447.5 | 2771.3 | 2495 |
| 651 | ENSMUSG00000003949 | 1780.803121 | -0.169901481 | 0.124228004 | -1.367658461 | 0.171419008 | 0.599034187 | Hlf | hepatic leukemia factor [Source:MGI Sym | 1759.6 | 2199.7 | 1697.5 | 1553 | 1736.3 | 1738.7 |
| 652 | ENSMUSG00000003955 | 823.8703133 | 0.034951069 | 0.124664013 | 0.280362136 | 0.779199683 | 0.959870443 | Fam162a | family with sequence similarity 162, meml | 864.14 | 847 | 729.53 | 778.89 | 811.42 | 912.24 |
| 653 | ENSMUSG00000003970 | 4560.517426 | 0.052457696 | 0.09819768 | 0.53420504 | 0.593199683 | 0.913011006 | Rpl8 | ribosomal protein L8 [Source:MGI Symbol | 4113.5 | 4658.5 | 4661.9 | 4602 | 4402.8 | 4924.3 |
| 654 | ENSMUSG00000003974 | 0 | NA | NA | NA | NA | NA | Grm3 | glutamate receptor, metabotropic 3 [Sour | 0 | 0 | 0 | 0 | 0 | 0 |
| 655 | ENSMUSG00000003992 | 24.54127273 | 0.362892419 | 0.451554543 | 0.803651352 | 0.421598357 | 0.838959049 | Ssbp2 | single-stranded DNA binding protein 2 [Sc | 21.684 | 23.642 | 19.119 | 30.847 | 18.541 | 33.415 |
| 656 | ENSMUSG00000004018 | 150.4757073 | 0.218124953 | 0.200453197 | 1.088159014 | 0.276524915 | 0.730038567 | Fancl | Fanconi anemia, complementation group | 127.69 | 153.16 | 136.85 | 157.13 | 183.22 | 144.8 |
| 657 | ENSMUSG00000004031 | 0 | NA | NA | NA | NA | NA | Brinp2 | bone morphogenic protein/retinoic acid ir | 0 | 0 | 0 | 0 | 0 | 0 |
| 658 | ENSMUSG00000004032 | 112.0964664 | 0.182937012 | 0.27573143 | 0.663460861 | 0.507035439 | 0.88185929 | Gstm5 | glutathione S-transferase, mu 5 [Source:M | 145.36 | 77.094 | 91.568 | 107.97 | 105.79 | 144.8 |
| 659 | ENSMUSG00000004035 | 784.2681866 | 0.344651988 | 0.140245173 | 2.457496265 | 0.013990927 | 0.136282395 | Gstm7 | glutathione S-transferase, mu 7 [Source:M | 746.89 | 648.62 | 676.2 | 751.9 | 855.04 | 1027 |
| 660 | ENSMUSG00000004038 | 348.6158662 | -0.263891105 | 0.513958348 | -0.513448428 | 0.607637677 | 0.916376001 | Gstm3 | glutathione S-transferase, mu 3 [Source:M | 608.75 | 226.14 | 305.9 | 230.39 | 268.29 | 452.22 |
| 661 | ENSMUSG00000004040 | 1606.0978 | 0.082007824 | 0.113241931 | 0.7241825 | 0.468953701 | 0.865872289 | Stat3 | signal transducer and activator of transcri | 1382.1 | 1575.8 | 1724.7 | 1647.4 | 1592.3 | 1714.2 |
| 662 | ENSMUSG00000004043 | 376.148769 | -0.335956587 | 0.151621961 | -2.2157515 | 0.026708526 | 0.211364326 | Stat5a | signal transducer and activator of transcri | 479.45 | 373.13 | 405.52 | 315.22 | 329.37 | 354.2 |
| 663 | ENSMUSG00000004044 | 116.2854226 | -0.423670759 | 0.231945276 | -1.826597924 | 0.067760246 | 0.37493657 | Cavin1 | caveolae associated 1 [Source:MGI Symbo | 147.77 | 135.69 | 115.72 | 87.722 | 90.521 | 120.3 |
| 664 | ENSMUSG00000004054 | 798.4056605 | 0.027282777 | 0.126313312 | 0.215992886 | 0.828993293 | 0.969955221 | Map3k11 | mitogen-activated protein kinase kinase k | 766.96 | 780.19 | 826.13 | 878.18 | 696.91 | 842.07 |
| 665 | ENSMUSG00000004056 | 2117.114978 | 0.218055366 | 0.094264678 | 2.313224549 | 0.020710297 | 0.17746074 | Akt2 | thymoma viral proto-oncogene 2 [Source: | 1941.1 | 1894.5 | 2036.6 | 2253.8 | 2199.8 | 2376.9 |
| 666 | ENSMUSG00000004069 | 2576.835903 | -0.138850696 | 0.10758803 | -1.290577543 | 0.196850207 | 0.637822894 | Dnaja3 | DnaJ heat shock protein family (Hsp40) m | 2522.5 | 2915.2 | 2664.5 | 2222 | 2522.6 | 2614.2 |
| 667 | ENSMUSG00000004070 | 751.0738138 | -0.165571126 | 0.152043784 | -1.088970044 | 0.276167096 | 0.730038567 | Hmox2 | heme oxygenase 2 [Source:MGI Symbol;A | 708.34 | 791.5 | 882.48 | 612.13 | 655.46 | 856.55 |
| 668 | ENSMUSG00000004071 | 2595.29363 | -0.486977422 | 0.125304004 | -3.886367616 | 0.000101755 | 0.003694676 | Cdip1 | cell death inducing Trp53 target 1 [Source | 2685.6 | 3024.1 | 3378 | 1957.8 | 2059.1 | 2467.2 |

| | | | | | | | | | | | | | | | |
|---|---|---|---|---|---|---|---|---|---|---|---|---|---|---|---|
| 669 | ENSMUSG00000004085 | 236.2017078 | 0.225774901 | 0.182073986 | 1.240017346 | 0.214968978 | 0.659728602 | Map3k20 | mitogen-activated protein kinase kinase k | 242.54 | 204.56 | 205.27 | 228.46 | 232.3 | 304.08 |
| 670 | ENSMUSG00000004096 | 849.2504289 | 0.070095303 | 0.115732423 | 0.605666946 | 0.544735937 | 0.895347795 | Cwc15 | CWC15 spliceosome-associated protein [S | 880.2 | 843.92 | 760.72 | 823.24 | 903.03 | 884.39 |
| 671 | ENSMUSG00000004098 | 1078.377575 | 0.356689275 | 0.140105924 | 2.545854334 | 0.010901067 | 0.115391119 | Col5a3 | collagen, type V, alpha 3 [Source:MGI Sym | 1009.5 | 816.17 | 1011.3 | 1337 | 1015.4 | 1280.9 |
| 672 | ENSMUSG00000004099 | 173.9037992 | -0.155260222 | 0.228514148 | -0.679433741 | 0.496863077 | 0.878668241 | Dnmt1 | DNA methyltransferase (cytosine-5) 1 [Sou | 176.68 | 166.52 | 207.29 | 219.79 | 145.05 | 128.09 |
| 673 | ENSMUSG00000004100 | 228.5826479 | -0.049266339 | 0.17489064 | -0.281697973 | 0.778175109 | 0.959559604 | Ppan | peter pan homolog [Source:MGI Symbol;A | 252.98 | 227.17 | 217.35 | 255.45 | 221.4 | 197.15 |
| 674 | ENSMUSG00000004105 | 225.8044655 | -0.830286023 | 0.182228533 | -4.55628988 | 5.21E-06 | 0.000316416 | Angptl2 | angiopoietin-like 2 [Source:MGI Symbol;A | 318.83 | 305.29 | 242.5 | 152.31 | 179.95 | 155.94 |
| 675 | ENSMUSG00000004110 | 1.335900328 | 0.890498008 | 1.988601539 | 0.447801126 | 0.654296731 | NA | Cacna1e | calcium channel, voltage-dependent, R ty | 0.8031 | 0 | 2.0125 | 1.928 | 3.2719 | 0 |
| 676 | ENSMUSG00000004113 | 0 | NA | NA | NA | NA | NA | Cacna1b | calcium channel, voltage-dependent, N ty | 0 | 0 | 0 | 0 | 0 | 0 |
| 677 | ENSMUSG00000004127 | 109.8234593 | -0.084327203 | 0.217677026 | -0.387395971 | 0.698463086 | 0.940696964 | Trmt10a | tRNA methyltransferase 10A [Source:MGI | 112.43 | 106.9 | 119.74 | 107 | 99.246 | 113.61 |
| 678 | ENSMUSG00000004151 | 8.387048985 | -0.746192565 | 0.755797426 | -0.987291752 | 0.323499633 | 0.770475661 | Etv1 | ets variant 1 [Source:MGI Symbol;Acc:MG | 11.243 | 8.2233 | 12.075 | 7.7118 | 3.2719 | 7.7969 |
| 679 | ENSMUSG00000004187 | 255.2484838 | 0.143877143 | 0.202395786 | 0.710870248 | 0.477164644 | 0.869671045 | Kifc2 | kinesin family member C2 [Source:MGI Sy | 236.92 | 189.14 | 301.87 | 302.69 | 278.11 | 222.77 |
| 680 | ENSMUSG00000004207 | 7943.361999 | -0.268599657 | 0.085855541 | -3.128506956 | 0.001756969 | 0.031805748 | Psap | prosaposin [Source:MGI Symbol;Acc:MGI: | 8499.2 | 9184.4 | 8358.9 | 7246.2 | 7085.7 | 7285.7 |
| 681 | ENSMUSG00000004221 | 1263.002106 | -0.060907577 | 0.105789049 | -0.575745572 | 0.564787167 | 0.901542552 | Ikbkg | inhibitor of kappaB kinase gamma [Source | 1246.4 | 1341.4 | 1282 | 1296.6 | 1141.9 | 1269.8 |
| 682 | ENSMUSG00000004231 | 0 | NA | NA | NA | NA | NA | Pax2 | paired box 2 [Source:MGI Symbol;Acc:MG | 0 | 0 | 0 | 0 | 0 | 0 |
| 683 | ENSMUSG00000004233 | 84.83104858 | -0.147276434 | 0.252785072 | -0.582615234 | 0.560152348 | 0.899475387 | Wars2 | tryptophanyl tRNA synthetase 2 (mitocho | 89.144 | 102.79 | 75.468 | 76.154 | 77.434 | 87.994 |
| 684 | ENSMUSG00000004263 | 46.88551638 | 0.038031817 | 0.332552719 | 0.114363271 | 0.908949828 | 0.985098284 | Atn1 | atrophin 1 [Source:MGI Symbol;Acc:MGI: | 51.399 | 44.2 | 43.269 | 59.767 | 40.353 | 42.326 |
| 685 | ENSMUSG00000004264 | 3509.393803 | -0.07520964 | 0.091737263 | -0.819837411 | 0.412308801 | 0.834424863 | Phb2 | prohibitin 2 [Source:MGI Symbol;Acc:MGI | 3516.8 | 3701.5 | 3584.2 | 3271.7 | 3638.3 | 3343.8 |
| 686 | ENSMUSG00000004266 | 378.6058554 | -0.119899087 | 0.157556501 | -0.760991047 | 0.446662415 | 0.853099851 | Ptpn6 | protein tyrosine phosphatase, non-recept | 412.79 | 438.92 | 331.05 | 366.31 | 329.37 | 393.19 |
| 687 | ENSMUSG00000004267 | 1.939997044 | -0.932772183 | 1.761429371 | -0.529554122 | 0.596421111 | NA | Eno2 | enolase 2, gamma neuronal [Source:MGI | 5.6217 | 0 | 2.0125 | 2.8919 | 0 | 1.1138 |
| 688 | ENSMUSG00000004268 | 745.8735011 | 0.194259477 | 0.125421255 | 1.548856104 | 0.121416315 | 0.510744254 | Emg1 | EMG1 N1-specific pseudouridine methyltr | 632.04 | 758.6 | 697.33 | 776.97 | 747.07 | 863.23 |
| 689 | ENSMUSG00000004270 | 2037.874674 | 0.113560915 | 0.120512728 | 0.942314699 | 0.346031545 | 0.786949727 | Lpcat3 | lysophosphatidylcholine acyltransferase 3 | 2156.3 | 2008.6 | 1707.6 | 2205.6 | 2243.4 | 1905.8 |
| 690 | ENSMUSG00000004285 | 801.0195207 | 0.046330448 | 0.112694881 | 0.411114046 | 0.680988911 | 0.934384556 | Atp6v1f | ATPase, H+ transporting, lysosomal V1 sul | 778.21 | 826.45 | 759.71 | 785.64 | 792.88 | 863.23 |
| 691 | ENSMUSG00000004296 | 0 | NA | NA | NA | NA | NA | Il12b | interleukin 12b [Source:MGI Symbol;Acc:I | 0 | 0 | 0 | 0 | 0 | 0 |
| 692 | ENSMUSG00000004317 | 102.3571414 | -0.249791024 | 0.253652912 | -0.984774909 | 0.324734648 | 0.771309263 | Clcn5 | chloride channel, voltage-sensitive 5 [Sou | 95.569 | 142.88 | 95.593 | 97.362 | 101.43 | 81.311 |
| 693 | ENSMUSG00000004319 | 1019.353445 | -0.13430053 | 0.124157168 | -1.081697758 | 0.27938685 | 0.732074527 | Clcn3 | chloride channel, voltage-sensitive 3 [Sou | 974.97 | 1123.5 | 1102.8 | 986.15 | 1071 | 857.66 |
| 694 | ENSMUSG00000004328 | 3.663231462 | -4.201000545 | 1.561398787 | -2.690536575 | 0.007133721 | NA | Hif3a | hypoxia inducible factor 3, alpha subunit [ | 5.6217 | 8.2233 | 7.0437 | 0 | 1.0906 | 0 |
| 695 | ENSMUSG00000004341 | 35.80177025 | 0.838582761 | 0.426226428 | 1.967458387 | 0.049130389 | 0.309234962 | Gpx6 | glutathione peroxidase 6 [Source:MGI Syr | 20.078 | 34.949 | 22.137 | 29.883 | 65.437 | 42.326 |
| 696 | ENSMUSG00000004344 | 1.552869784 | 1.055416156 | 2.057642344 | 0.512924979 | 0.6080038 | NA | Gpx5 | glutathione peroxidase 5 [Source:MGI Syr | 0 | 3.0838 | 0 | 2.8919 | 0 | 3.3415 |
| 697 | ENSMUSG00000004347 | 3.502439535 | -0.325629626 | 1.248859945 | -0.260741509 | 0.794291854 | NA | Pde1c | phosphodiesterase 1C [Source:MGI Symb | 5.6217 | 0 | 6.0375 | 3.8559 | 3.2719 | 2.2277 |
| 698 | ENSMUSG00000004356 | 421.228023 | -0.270229519 | 0.16316482 | -1.656175145 | 0.097686355 | 0.460719048 | Utp20 | UTP20 small subunit processome compon | 445.72 | 510.88 | 424.64 | 311.36 | 447.15 | 387.62 |
| 699 | ENSMUSG00000004359 | 14.86026756 | -1.698763408 | 0.814527681 | -2.085580941 | 0.037016599 | 0.260159423 | Spic | Spi-C transcription factor (Spi-1/PU.1 rela | 8.8341 | 51.396 | 8.05 | 8.6758 | 2.1812 | 10.025 |
| 700 | ENSMUSG00000004360 | 0 | NA | NA | NA | NA | NA | 9330159F19 | RIKEN cDNA 9330159F19 gene [Source:M | 0 | 0 | 0 | 0 | 0 | 0 |
| 701 | ENSMUSG00000004364 | 1207.95721 | 0.144421982 | 0.100958804 | 1.43050409 | 0.152572393 | 0.568325287 | Cul3 | cullin 3 [Source:MGI Symbol;Acc:MGI:134 | 1129.2 | 1207.8 | 1105.9 | 1251.2 | 1272.8 | 1280.9 |
| 702 | ENSMUSG00000004366 | 0 | NA | NA | NA | NA | NA | Sst | somatostatin [Source:MGI Symbol;Acc:M | 0 | 0 | 0 | 0 | 0 | 0 |
| 703 | ENSMUSG00000004371 | 0.506734502 | -2.372847156 | 3.338060886 | -0.710845978 | 0.477179685 | NA | Il11 | interleukin 11 [Source:MGI Symbol;Acc:M | 0 | 1.0279 | 2.0125 | 0 | 0 | 0 |
| 704 | ENSMUSG00000004383 | 89.39735112 | -0.345226219 | 0.266876554 | -1.293580175 | 0.195810474 | 0.636484845 | Large1 | LARGE xylosyl- and glucuronyltransferase | 101.19 | 107.93 | 90.562 | 55.911 | 88.34 | 92.449 |
| 705 | ENSMUSG00000004393 | 417.8100534 | 0.010192993 | 0.132731834 | 0.076793885 | 0.938787515 | 0.988721002 | Ddx56 | DEAD box helicase 56 [Source:MGI Symbo | 405.57 | 437.89 | 405.52 | 396.19 | 438.43 | 423.26 |
| 706 | ENSMUSG00000004394 | 1114.56014 | -0.019510663 | 0.101054531 | -0.193070644 | 0.846903643 | 0.972803041 | Tmed4 | transmembrane p24 trafficking protein 4 [ | 1157.3 | 1097.8 | 1110.9 | 1122.1 | 1123.3 | 1076 |
| 707 | ENSMUSG00000004415 | 0 | NA | NA | NA | NA | NA | Col26a1 | collagen, type XXVI, alpha 1 [Source:MGI | 0 | 0 | 0 | 0 | 0 | 0 |
| 708 | ENSMUSG00000004446 | 324.9797626 | 0.392162369 | 0.191989754 | 2.042621346 | 0.041089936 | 0.277592171 | Bid | BH3 interacting domain death agonist [So | 340.52 | 232.31 | 269.67 | 400.05 | 416.62 | 290.71 |
| 709 | ENSMUSG00000004451 | 165.7729797 | -0.030489945 | 0.219439747 | -0.138944495 | 0.88949401 | 0.982382091 | Ralb | v-ral simian leukemia viral oncogene B [Sc | 177.49 | 160.36 | 164.02 | 126.28 | 154.87 | 211.63 |
| 710 | ENSMUSG00000004455 | 1002.409683 | -0.149718632 | 0.114609907 | -1.306332376 | 0.191439552 | 0.629099068 | Ppp1cc | protein phosphatase 1 catalytic subunit ga | 943.65 | 1137.9 | 1082.7 | 944.7 | 955.38 | 950.11 |
| 711 | ENSMUSG00000004460 | 1292.077337 | 0.4306738 | 0.13206718 | 3.261020648 | 0.00111012 | 0.022572786 | Dnajb11 | DnaJ heat shock protein family (Hsp40) m | 1044 | 1181.1 | 1077.7 | 1617.6 | 1221.5 | 1610.6 |
| 712 | ENSMUSG00000004462 | 509.8859805 | 0.232538861 | 0.1309196 | 1.776195936 | 0.075700628 | 0.402878781 | Tbccd1 | TBCC domain containing 1 [Source:MGI Sy | 433.68 | 491.34 | 481.99 | 517.66 | 539.86 | 594.79 |
| 713 | ENSMUSG00000004473 | 34.85369422 | -0.15133417 | 0.39345156 | -0.384632279 | 0.700509876 | 0.941739459 | Clec11a | C-type lectin domain family 11, member a | 38.549 | 47.284 | 24.15 | 30.847 | 35.99 | 32.301 |
| 714 | ENSMUSG00000004500 | 98.01314026 | 0.673845383 | 0.272151897 | 2.475990026 | 0.013286729 | 0.131539526 | Zfp324 | zinc finger protein 324 [Source:MGI Symb | 76.295 | 70.926 | 79.493 | 143.63 | 79.615 | 138.12 |
| 715 | ENSMUSG00000004508 | 36.86005461 | -1.098480691 | 0.394667699 | -2.783305284 | 0.005380815 | 0.069926422 | Gab2 | growth factor receptor bound protein 2-a | 53.005 | 53.452 | 44.275 | 25.063 | 14.178 | 31.188 |
| 716 | ENSMUSG00000004530 | 245.3723461 | -0.062533586 | 0.176902426 | -0.353491964 | 0.723719652 | 0.944786345 | Coro1c | coronin, actin binding protein 1C [Source: | 289.12 | 226.14 | 236.47 | 271.84 | 219.21 | 229.45 |
| 717 | ENSMUSG00000004535 | 3136.33459 | -0.014913518 | 0.101805493 | -0.146490311 | 0.883534338 | 0.981769403 | Tax1bp1 | Tax1 (human T cell leukemia virus type I) b | 3261.4 | 3283.2 | 2912.1 | 2905.4 | 3342.7 | 3113.2 |
| 718 | ENSMUSG00000004540 | 0 | NA | NA | NA | NA | NA | Psg17 | pregnancy specific glycoprotein 17 [Sourc | 0 | 0 | 0 | 0 | 0 | 0 |
| 719 | ENSMUSG00000004542 | 0 | NA | NA | NA | NA | NA | Psg19 | pregnancy specific glycoprotein 19 [Sourc | 0 | 0 | 0 | 0 | 0 | 0 |
| 720 | ENSMUSG00000004552 | 12.74263745 | -0.754755865 | 0.771370386 | -0.978461033 | 0.327846355 | 0.772937193 | Ctse | cathepsin E [Source:MGI Symbol;Acc:MGI | 10.44 | 28.782 | 9.0562 | 18.316 | 7.6343 | 2.2277 |
| 721 | ENSMUSG00000004558 | 26209.67168 | 0.14491833 | 0.101121718 | 1.433107867 | 0.151827014 | 0.567340036 | Ndrg2 | N-myc downstream regulated gene 2 [Sou | 22405 | 24620 | 27660 | 26399 | 27947 | 28228 |
| 722 | ENSMUSG00000004561 | 277.9733064 | 0.311763173 | 0.156981829 | 1.985982553 | 0.047035261 | 0.300361156 | Mettl17 | methyltransferase like 17 [Source:MGI Syr | 237.72 | 247.73 | 258.6 | 277.63 | 307.55 | 338.61 |
| 723 | ENSMUSG00000004562 | 239.4224817 | 0.158184574 | 0.196817246 | 0.803712973 | 0.42156276 | 0.838959049 | Arhgef40 | Rho guanine nucleotide exchange factor ( | 217.64 | 176.8 | 284.77 | 274.73 | 247.57 | 235.02 |
| 724 | ENSMUSG00000004565 | 554.2180427 | 0.127588125 | 0.183027301 | 0.69709887 | 0.485740922 | 0.872998689 | Pnpla6 | patatin-like phospholipase domain contai | 450.54 | 526.29 | 613.81 | 742.26 | 524.59 | 467.81 |

| | | | | | | | | | | | | | | |
|---|---|---|---|---|---|---|---|---|---|---|---|---|---|---|
| 725 | ENSMUSG00000004567 | 736.0053769 | 0.226072841 | 0.120637876 | 1.873978946 | 0.060933327 | 0.352269825 | Mcoln1 | mucolipin 1 [Source:MGI Symbol;Acc:MGI | 733.23 | 642.45 | 659.09 | 827.09 | 820.14 | 734.02 |
| 726 | ENSMUSG00000004568 | 522.5235957 | -0.171665354 | 0.130112892 | -1.319356996 | 0.187049791 | 0.623384374 | Arhgef18 | rho/rac guanine nucleotide exchange fact | 531.65 | 539.66 | 589.66 | 493.56 | 450.43 | 530.19 |
| 727 | ENSMUSG00000004591 | 801.8367483 | 0.038218269 | 0.121396831 | 0.314820974 | 0.752897586 | 0.952989051 | Pkn2 | protein kinase N2 [Source:MGI Symbol;Ac | 738.05 | 884.01 | 752.67 | 847.34 | 808.15 | 780.8 |
| 728 | ENSMUSG00000004609 | 12.63508129 | -0.760593126 | 0.701219282 | -1.084672292 | 0.278066824 | 0.730997543 | Cd33 | CD33 antigen [Source:MGI Symbol;Acc:M | 16.865 | 20.558 | 10.062 | 2.8919 | 8.7249 | 16.708 |
| 729 | ENSMUSG00000004610 | 821.6060407 | 0.068052753 | 0.119842702 | 0.567850622 | 0.570136403 | 0.90421633 | Etfb | electron transferring flavoprotein, beta po | 851.29 | 739.07 | 816.06 | 889.75 | 862.68 | 770.78 |
| 730 | ENSMUSG00000004612 | 4.93954609 | -2.213869895 | 1.090434228 | -2.030264492 | 0.042329661 | NA | Nkg7 | natural killer cell group 7 sequence [Sourc | 8.031 | 12.335 | 4.025 | 1.928 | 1.0906 | 2.2277 |
| 731 | ENSMUSG00000004626 | 611.191805 | -0.057007073 | 0.127550209 | -0.446938296 | 0.654919612 | 0.925797231 | Stxbp2 | syntaxin binding protein 2 [Source:MGI Sy | 680.23 | 612.64 | 576.58 | 643.94 | 583.48 | 570.29 |
| 732 | ENSMUSG00000004630 | 0.649696354 | 1.537929172 | 3.118369374 | 0.493183773 | 0.621882743 | NA | Pcp2 | Purkinje cell protein 2 (L7) [Source:MGI Sy | 0 | 0 | 1.0062 | 2.8919 | 0 | 0 |
| 733 | ENSMUSG00000004631 | 57.9631105 | -0.261684867 | 0.324690066 | -0.805952797 | 0.420270067 | 0.837906635 | Sgce | sarcoglycan, epsilon [Source:MGI Symbol; | 79.507 | 62.703 | 47.293 | 63.623 | 53.44 | 41.212 |
| 734 | ENSMUSG00000004633 | 801.3129315 | 0.091593458 | 0.13317299 | 0.687778043 | 0.491592565 | 0.876368691 | Chn2 | chimerin 2 [Source:MGI Symbol;Acc:MGI: | 861.73 | 769.91 | 695.32 | 848.3 | 901.94 | 730.68 |
| 735 | ENSMUSG00000004637 | 141.6110532 | 0.008346165 | 0.20628659 | 0.040459075 | 0.967727134 | 0.994231936 | Wwox | WW domain-containing oxidoreductase [S | 131.71 | 154.19 | 137.86 | 137.85 | 124.33 | 163.74 |
| 736 | ENSMUSG00000004642 | 194.2334141 | -0.010806446 | 0.171660411 | -0.062952467 | 0.949804355 | 0.990766637 | Slbp | stem-loop binding protein [Source:MGI Sy | 207.2 | 190.16 | 187.16 | 192.8 | 187.59 | 200.49 |
| 737 | ENSMUSG00000004651 | 0 | NA | NA | NA | NA | NA | Tyr | tyrosinase [Source:MGI Symbol;Acc:MGI:9 | 0 | 0 | 0 | 0 | 0 | 0 |
| 738 | ENSMUSG00000004654 | 1.110030071 | -2.325698304 | 2.174895496 | -1.069337956 | 0.284917414 | NA | Ghrhr | growth hormone releasing hormone recep | 1.6062 | 3.0838 | 1.0062 | 0.964 | 0 | 0 |
| 739 | ENSMUSG00000004655 | 874.3950414 | -0.446608847 | 0.175436087 | -2.545706837 | 0.010905673 | 0.115391119 | Aqp1 | aquaporin 1 [Source:MGI Symbol;Acc:MG | 917.14 | 1036.1 | 1072.7 | 686.35 | 560.58 | 973.5 |
| 740 | ENSMUSG00000004661 | 55.52798737 | -0.349953625 | 0.311943163 | -1.1218506 | 0.261925968 | 0.716026686 | Arid3b | AT rich interactive domain 3B (BRIGHT-like | 75.492 | 62.703 | 48.3 | 53.019 | 47.987 | 45.668 |
| 741 | ENSMUSG00000004665 | 53.97420146 | -0.506554083 | 0.315574555 | -1.605180376 | 0.108454115 | 0.48522218 | Cnn2 | calponin 2 [Source:MGI Symbol;Acc:MGI:1 | 57.823 | 72.982 | 59.368 | 45.307 | 34.9 | 53.465 |
| 742 | ENSMUSG00000004667 | 443.4784977 | 0.100680692 | 0.140230663 | 0.717964886 | 0.472778938 | 0.86857093 | Polr2e | polymerase (RNA) II (DNA directed) polyp | 437.69 | 450.23 | 395.45 | 411.62 | 462.42 | 503.46 |
| 743 | ENSMUSG00000004668 | 0 | NA | NA | NA | NA | NA | Abca13 | ATP-binding cassette, sub-family A (ABC1) | 0 | 0 | 0 | 0 | 0 | 0 |
| 744 | ENSMUSG00000004677 | 590.4021098 | -0.074061225 | 0.123279008 | -0.600761039 | 0.547999157 | 0.895600915 | Myo9b | myosin IXb [Source:MGI Symbol;Acc:MGI: | 646.5 | 602.36 | 567.52 | 605.38 | 569.3 | 551.35 |
| 745 | ENSMUSG00000004698 | 9.859278832 | -2.432173578 | 0.813348362 | -2.990322095 | 0.002786834 | 0.044426743 | Hdac9 | histone deacetylase 9 [Source:MGI Symbo | 18.471 | 19.53 | 12.075 | 5.7839 | 2.1812 | 1.1138 |
| 746 | ENSMUSG00000004707 | 18.41247264 | -1.293983309 | 0.546948515 | -2.365822877 | 0.017990049 | 0.160343553 | Ly9 | lymphocyte antigen 9 [Source:MGI Symbo | 23.29 | 34.949 | 20.125 | 6.7478 | 11.997 | 13.366 |
| 747 | ENSMUSG00000004709 | 3.779348569 | -1.347828303 | 1.151372966 | -1.170627019 | 0.241748732 | NA | Cd244a | CD244 molecule A [Source:MGI Symbol;Ac | 8.031 | 5.1396 | 3.0187 | 0.964 | 2.1812 | 3.3415 |
| 748 | ENSMUSG00000004730 | 121.9962877 | -1.746268651 | 0.708220091 | -2.465714648 | 0.013674022 | 0.134453469 | Adgre1 | adhesion G protein-coupled receptor E1 [S | 165.44 | 330.99 | 67.418 | 45.307 | 38.172 | 84.652 |
| 749 | ENSMUSG00000004748 | 523.1481927 | -0.050733079 | 0.141048712 | -0.359684807 | 0.719082855 | 0.9447719 | Mtfp1 | mitochondrial fission process 1 [Source:M | 465 | 610.58 | 522.24 | 491.63 | 513.68 | 535.76 |
| 750 | ENSMUSG00000004768 | 179.0813221 | 0.030615748 | 0.243349134 | 0.125809974 | 0.899882347 | 0.982798755 | Rab23 | RAB23, member RAS oncogene family [So | 146.97 | 150.08 | 234.45 | 133.03 | 206.13 | 203.83 |
| 751 | ENSMUSG00000004771 | 790.4570849 | 0.167245266 | 0.109767741 | 1.523628567 | 0.127601523 | 0.523285632 | Rab11a | RAB11A, member RAS oncogene family [S | 762.14 | 730.85 | 740.6 | 813.6 | 875.77 | 819.79 |
| 752 | ENSMUSG00000004788 | 310.2430486 | 0.091832091 | 0.153862299 | 0.596845957 | 0.550610232 | 0.895703112 | Eif2b2 | eukaryotic translation initiation factor 2B, | 315.62 | 317.63 | 267.66 | 329.68 | 293.38 | 337.49 |
| 753 | ENSMUSG00000004789 | 3766.148756 | -0.078886853 | 0.094228652 | -0.837185414 | 0.402488355 | 0.828953151 | Dlst | dihydrolipoamide S-succinyltransferase (E | 3807.5 | 3865 | 3934.4 | 3429.8 | 3951.3 | 3608.9 |
| 754 | ENSMUSG00000004791 | 0.535401493 | -2.48596965 | 3.684117979 | -0.674780141 | 0.499815459 | NA | Pgf | placental growth factor [Source:MGI Syml | 3.2124 | 0 | 0 | 0 | 0 | 0 |
| 755 | ENSMUSG00000004798 | 1831.111628 | 0.207402429 | 0.113568768 | 1.826227686 | 0.067815973 | 0.375101426 | Ulk2 | unc-51 like kinase 2 [Source:MGI Symbol;A | 1665.6 | 1586.1 | 1847.5 | 1896.1 | 2175.8 | 1815.6 |
| 756 | ENSMUSG00000004814 | 12.85212197 | -1.049883567 | 0.645490564 | -1.626489411 | 0.103845584 | 0.474113351 | Ccl24 | chemokine (C-C motif) ligand 24 [Source:M | 14.456 | 19.53 | 18.112 | 10.604 | 3.2719 | 11.138 |
| 757 | ENSMUSG00000004815 | 372.8208011 | 0.248758928 | 0.175464843 | 1.417713791 | 0.156274342 | 0.574894692 | Dgkq | diacylglycerol kinase, theta [Source:MGI S | 336.5 | 290.9 | 395.45 | 484.88 | 383.9 | 345.29 |
| 758 | ENSMUSG00000004821 | 0 | NA | NA | NA | NA | NA | Tmed11 | transmembrane p24 trafficking protein 11 | 0 | 0 | 0 | 0 | 0 | 0 |
| 759 | ENSMUSG00000004837 | 38.82829852 | -0.907631117 | 0.449244449 | -2.020350212 | 0.043347074 | 0.287075127 | Grap | GRB2-related adaptor protein [Source:MG | 64.248 | 47.284 | 40.25 | 27.955 | 10.906 | 42.326 |
| 760 | ENSMUSG00000004842 | 0 | NA | NA | NA | NA | NA | Pou1f1 | POU domain, class 1, transcription factor 1 | 0 | 0 | 0 | 0 | 0 | 0 |
| 761 | ENSMUSG00000004843 | 505.5554902 | -0.115900288 | 0.136818139 | -0.847112 | 0.396932697 | 0.824416217 | Chmp2b | charged multivesicular body protein 2B [S | 546.91 | 539.66 | 490.04 | 426.08 | 507.14 | 523.51 |
| 762 | ENSMUSG00000004846 | 398.0543434 | 0.178400627 | 0.178327554 | 1.000409765 | 0.317112246 | 0.763688233 | Plod3 | procollagen-lysine, 2-oxoglutarate 5-dioxy | 327.67 | 383.41 | 410.55 | 518.62 | 333.73 | 414.35 |
| 763 | ENSMUSG00000004849 | 618.227219 | -0.04371852 | 0.130735285 | -0.334404897 | 0.738074045 | 0.948137969 | Ap1s1 | adaptor protein complex AP-1, sigma 1 [So | 635.25 | 616.75 | 629.91 | 535.01 | 609.66 | 682.79 |
| 764 | ENSMUSG00000004864 | 3.056529148 | -0.961548002 | 1.208631184 | -0.795567759 | 0.426283313 | NA | Mapk13 | mitogen-activated protein kinase 13 [Sour | 4.0155 | 4.1117 | 4.025 | 2.8919 | 2.1812 | 1.1138 |
| 765 | ENSMUSG00000004865 | 391.5728518 | 0.058503753 | 0.150758733 | 0.388062115 | 0.697970067 | 0.940510439 | Srpk1 | serine/arginine-rich protein specific kinase | 420.02 | 342.3 | 388.41 | 449.21 | 371.9 | 377.59 |
| 766 | ENSMUSG00000004872 | 0 | NA | NA | NA | NA | NA | Pax3 | paired box 3 [Source:MGI Symbol;Acc:MG | 0 | 0 | 0 | 0 | 0 | 0 |
| 767 | ENSMUSG00000004880 | 115.9824707 | -0.071431207 | 0.229360773 | -0.311436022 | 0.755469173 | 0.953085143 | Lbr | lamin B receptor [Source:MGI Symbol;Acc | 114.84 | 128.49 | 113.71 | 133.03 | 115.61 | 90.221 |
| 768 | ENSMUSG00000004885 | 0 | NA | NA | NA | NA | NA | Crabp2 | cellular retinoic acid binding protein II [So | 0 | 0 | 0 | 0 | 0 | 0 |
| 769 | ENSMUSG00000004891 | 23.56868703 | -0.33421761 | 0.447337581 | -0.747126161 | 0.45498741 | 0.857337687 | Nes | nestin [Source:MGI Symbol;Acc:MGI:1017 | 28.109 | 23.642 | 27.169 | 25.063 | 20.722 | 16.708 |
| 770 | ENSMUSG00000004892 | 0 | NA | NA | NA | NA | NA | Bcan | brevican [Source:MGI Symbol;Acc:MGI:10 | 0 | 0 | 0 | 0 | 0 | 0 |
| 771 | ENSMUSG00000004894 | 0.401551119 | -2.072982726 | 4.008251979 | -0.517178744 | 0.605031388 | NA | Hapln2 | hyaluronan and proteoglycan link protein | 2.4093 | 0 | 0 | 0 | 0 | 0 |
| 772 | ENSMUSG00000004895 | 408.6581125 | 0.038673084 | 0.147803396 | 0.261652198 | 0.793589597 | 0.961915077 | Prcc | papillary renal cell carcinoma (translocatio | 393.52 | 406.03 | 409.54 | 361.49 | 403.53 | 477.84 |
| 773 | ENSMUSG00000004896 | 178.1268036 | -0.300161374 | 0.184349543 | -1.628218706 | 0.10347852 | 0.473607836 | Rrnad1 | ribosomal RNA adenine dimethylase doma | 201.58 | 196.33 | 192.19 | 182.19 | 148.32 | 148.14 |
| 774 | ENSMUSG00000004897 | 3256.517203 | 0.119688412 | 0.098807035 | 1.211334926 | 0.225767068 | 0.6725584 | Hdgf | heparin binding growth factor [Source:MG | 2899.2 | 3144.4 | 3322.6 | 3598.5 | 3326.4 | 3248 |
| 775 | ENSMUSG00000004902 | 5.121683184 | 0.532191611 | 1.008001765 | 0.527966943 | 0.59752228 | 0.914641631 | Slc25a18 | solute carrier family 25 (mitochondrial car | 6.4248 | 4.1117 | 2.0125 | 11.568 | 3.2719 | 3.3415 |
| 776 | ENSMUSG00000004929 | 170.3434357 | -0.037879431 | 0.187950106 | -0.201539824 | 0.840276493 | 0.971838962 | Thop1 | thimet oligopeptidase 1 [Source:MGI Sym | 172.67 | 170.63 | 174.08 | 145.56 | 187.59 | 171.53 |
| 777 | ENSMUSG00000004931 | 716.0405186 | 0.021981859 | 0.118017824 | 0.186258808 | 0.852241809 | 0.973781977 | Apba3 | amyloid beta (A4) precursor protein-bindi | 692.27 | 673.29 | 766.76 | 757.69 | 705.63 | 700.61 |
| 778 | ENSMUSG00000004933 | 6.76847446 | -0.822984285 | 0.857902503 | -0.959298151 | 0.337408567 | 0.77947253 | Matk | megakaryocyte-associated tyrosine kinase | 13.653 | 5.1396 | 7.0437 | 4.8199 | 3.2719 | 6.6831 |
| 779 | ENSMUSG00000004934 | 359.9863079 | 0.093950632 | 0.13677381 | 0.686905134 | 0.492142511 | 0.876368691 | Pias4 | protein inhibitor of activated STAT 4 [Sour | 367.02 | 337.16 | 340.11 | 367.28 | 372.99 | 375.37 |
| 780 | ENSMUSG00000004936 | 1002.138974 | 0.131942504 | 0.103695919 | 1.272398235 | 0.203231661 | 0.647739134 | Map2k1 | mitogen-activated protein kinase kinase 1 | 983.8 | 923.07 | 961.97 | 1078.7 | 1035 | 1030.3 |

| | | | | | | | | | | | | | | | |
|---|---|---|---|---|---|---|---|---|---|---|---|---|---|---|---|
| 781 | ENSMUSG00000004937 | 1224.601207 | -0.091862944 | 0.101361683 | -0.906288668 | 0.364783082 | 0.803322548 | Sgta | small glutamine-rich tetratricopeptide rep | 1334.8 | 1240.7 | 1214.5 | 1190.5 | 1194.2 | 1172.9 |
| 782 | ENSMUSG00000004939 | 0 | NA | NA | NA | NA | NA | Nmrk2 | nicotinamide riboside kinase 2 [Source:M | 0 | 0 | 0 | 0 | 0 | 0 |
| 783 | ENSMUSG00000004945 | 769.9058193 | 0.061882232 | 0.127326691 | 0.486011467 | 0.626959036 | 0.920273791 | Tmem242 | transmembrane protein 242 [Source:MGI | 748.49 | 741.13 | 769.78 | 674.78 | 836.5 | 848.75 |
| 784 | ENSMUSG00000004947 | 85.56547458 | 0.036086478 | 0.296946561 | 0.121525158 | 0.903275089 | 0.983821605 | Dtx2 | deltex 2, E3 ubiquitin ligase [Source:MGI S | 53.808 | 90.457 | 109.68 | 74.226 | 89.431 | 95.791 |
| 785 | ENSMUSG00000004948 | 0 | NA | NA | NA | NA | NA | Zp3 | zona pellucida glycoprotein 3 [Source:MG | 0 | 0 | 0 | 0 | 0 | 0 |
| 786 | ENSMUSG00000004951 | 516.5950382 | -0.151150951 | 0.422301679 | -0.357921739 | 0.720401882 | 0.9447719 | Hspb1 | heat shock protein 1 [Source:MGI Symbol | 714.76 | 403.97 | 512.18 | 732.62 | 323.91 | 412.12 |
| 787 | ENSMUSG00000004952 | 30.16211162 | -0.97715033 | 0.495815891 | -1.970792683 | 0.048747594 | 0.307494635 | Rasa4 | RAS p21 protein activator 4 [Source:MGI S | 38.549 | 58.591 | 23.144 | 30.847 | 10.906 | 18.935 |
| 788 | ENSMUSG00000004961 | 0.167707431 | -0.796629176 | 4.080472857 | -0.195229623 | 0.84521319 | NA | Syt5 | synaptotagmin V [Source:MGI Symbol;Acc | 0 | 0 | 1.0062 | 0 | 0 | 0 |
| 789 | ENSMUSG00000004980 | 5844.467778 | 0.12666611 | 0.092255513 | 1.372992313 | 0.169754736 | 0.596876778 | Hnrnpa2b1 | heterogeneous nuclear ribonucleoprotein | 5227.4 | 5623.7 | 5913.7 | 5820.5 | 6236.2 | 6245.3 |
| 790 | ENSMUSG00000004988 | 0.133850373 | -0.796629176 | 4.080472857 | -0.195229623 | 0.84521319 | NA | Fxyd4 | FXYD domain-containing ion transport reg | 0.8031 | 0 | 0 | 0 | 0 | 0 |
| 791 | ENSMUSG00000004994 | 76.89290827 | 0.27541434 | 0.2590136 | 1.063319998 | 0.287636859 | 0.73951397 | Ccdc130 | coiled-coil domain containing 130 [Source | 71.476 | 60.647 | 76.475 | 80.974 | 92.703 | 79.083 |
| 792 | ENSMUSG00000004996 | 414.639003 | -0.045377416 | 0.144921651 | -0.313116888 | 0.754191859 | 0.953048681 | Mri1 | methylthioribose-1-phosphate isomerase | 451.34 | 385.47 | 426.65 | 451.14 | 407.89 | 365.34 |
| 793 | ENSMUSG00000005034 | 1056.140754 | -0.067618801 | 0.109452463 | -0.617791495 | 0.536712791 | 0.890560262 | Prkacb | protein kinase, cAMP dependent, catalytic | 1079.4 | 1078.3 | 1084.7 | 983.26 | 1128.8 | 982.41 |
| 794 | ENSMUSG00000005043 | 110.2232133 | -0.019417619 | 0.261224754 | -0.074332999 | 0.940745421 | 0.989001277 | Sgsh | N-sulfoglucosamine sulfohydrolase (sulfar | 114.84 | 101.76 | 115.72 | 72.298 | 131.96 | 124.75 |
| 795 | ENSMUSG00000005045 | 0.185640647 | 1.126924175 | 4.080472857 | 0.276174898 | 0.782413736 | NA | Chd5 | chromodomain helicase DNA binding prot | 0 | 0 | 0 | 0 | 0 | 1.1138 |
| 796 | ENSMUSG00000005054 | 545.3716164 | -0.175431097 | 0.135644223 | -1.293317867 | 0.195901144 | 0.636484845 | Cstb | cystatin B [Source:MGI Symbol;Acc:MGI:1 | 646.5 | 581.8 | 506.14 | 496.45 | 527.86 | 513.48 |
| 797 | ENSMUSG00000005057 | 35.57609763 | 1.427387335 | 0.510324853 | 2.797017088 | 0.005157681 | 0.067944759 | Sh2b2 | SH2B adaptor protein 2 [Source:MGI Syml | 18.471 | 15.419 | 24.15 | 93.506 | 21.812 | 40.098 |
| 798 | ENSMUSG00000005069 | 2882.998637 | 0.049055969 | 0.095911578 | 0.51147078 | 0.609021444 | 0.916376001 | Pex5 | peroxisomal biogenesis factor 5 [Source:N | 2781.1 | 2893.6 | 2827.5 | 2893.9 | 3157.3 | 2744.5 |
| 799 | ENSMUSG00000005078 | 383.4569082 | 0.000630092 | 0.149317757 | 0.004219807 | 0.996633091 | 0.999154389 | Jkamp | JNK1/MAPK8-associated membrane prote | 433.68 | 381.36 | 334.07 | 368.24 | 401.35 | 382.05 |
| 800 | ENSMUSG00000005087 | 76.12461148 | -1.497078204 | 0.413952724 | -3.616543915 | 0.000298563 | 0.00865413 | Cd44 | CD44 antigen [Source:MGI Symbol;Acc:M | 102.8 | 181.94 | 52.325 | 27.955 | 33.809 | 57.92 |
| 801 | ENSMUSG00000005089 | 3745.134156 | 0.121456991 | 0.096213272 | 1.262372527 | 0.20681477 | 0.652194904 | Slc1a2 | solute carrier family 1 (glial high affinity gl | 3457.4 | 3524.7 | 3780.5 | 3611.1 | 4048.4 | 4048.8 |
| 802 | ENSMUSG00000005102 | 185.3015314 | -0.117795002 | 0.184171633 | -0.639593623 | 0.52243683 | 0.888334265 | Eif2ak4 | eukaryotic translation initiation factor 2 al | 196.76 | 179.89 | 202.26 | 203.4 | 165.77 | 163.74 |
| 803 | ENSMUSG00000005103 | 689.129434 | -0.020671525 | 0.123478977 | -0.167409269 | 0.867048033 | 0.977930657 | Wdr1 | WD repeat domain 1 [Source:MGI Symbol | 683.44 | 763.74 | 634.94 | 650.69 | 696.91 | 705.06 |
| 804 | ENSMUSG00000005107 | 929.4930007 | 0.218478099 | 0.108458572 | 2.014392174 | 0.043968375 | 0.2895349 | Slc2a9 | solute carrier family 2 (facilitated glucose | 835.23 | 885.04 | 858.33 | 1052.7 | 1002.3 | 943.43 |
| 805 | ENSMUSG00000005124 | 6.868777745 | 0.065795479 | 0.848350074 | 0.077556991 | 0.938180455 | 0.988420305 | Ccn4 | cellular communication network factor 4 [ | 8.8341 | 7.1954 | 4.025 | 6.7478 | 3.2719 | 11.138 |
| 806 | ENSMUSG00000005125 | 72.6866884 | -0.384263321 | 0.297475853 | -1.291746263 | 0.196445031 | 0.637356187 | Ndrg1 | N-myc downstream regulated gene 1 [Sou | 76.295 | 72.982 | 97.606 | 58.803 | 46.897 | 83.538 |
| 807 | ENSMUSG00000005131 | 2.152671899 | 3.575152246 | 1.972792005 | 1.812229691 | 0.069950713 | NA | 4930550C14 | RIKEN cDNA 4930550C14 gene [Source:M | 0.8031 | 0 | 0 | 0 | 6.5437 | 5.5692 |
| 808 | ENSMUSG00000005142 | 1850.998642 | -0.309928987 | 0.098081864 | -3.159901084 | 0.001578227 | 0.029454792 | Man2b1 | mannosidase 2, alpha B1 [Source:MGI Syr | 1978 | 2182.3 | 1987.3 | 1631.1 | 1714.5 | 1612.8 |
| 809 | ENSMUSG00000005148 | 1.920254642 | -1.299695502 | 1.964072455 | -0.661735008 | 0.508141065 | NA | Klf5 | Kruppel-like factor 5 [Source:MGI Symbol; | 0 | 6.1675 | 2.0125 | 0 | 0 | 3.3415 |
| 810 | ENSMUSG00000005150 | 310.8311789 | -0.031774907 | 0.156272668 | -0.203329909 | 0.83887718 | 0.971535186 | Wdr83 | WD repeat domain containing 83 [Source: | 328.47 | 318.65 | 294.83 | 266.06 | 322.82 | 334.15 |
| 811 | ENSMUSG00000005161 | 2247.408508 | 0.038260242 | 0.096163149 | 0.397868024 | 0.690727472 | 0.937606823 | Prdx2 | peroxiredoxin 2 [Source:MGI Symbol;Acc: | 2353.1 | 2129.8 | 2168.5 | 2181.5 | 2345.9 | 2305.7 |
| 812 | ENSMUSG00000005198 | 673.8064695 | -0.250234879 | 0.146898178 | -1.703458017 | 0.088482388 | 0.438432246 | Polr2a | polymerase (RNA) II (DNA directed) polyp | 634.45 | 798.69 | 764.75 | 700.81 | 617.29 | 526.85 |
| 813 | ENSMUSG00000005202 | 0.346303622 | 2.004339653 | 3.884003423 | 0.516049919 | 0.605819544 | NA | Shbg | sex hormone binding globulin [Source:MG | 0 | 0 | 0 | 0.964 | 0 | 1.1138 |
| 814 | ENSMUSG00000005204 | 684.4413718 | 0.148653141 | 0.118922468 | 1.250000469 | 0.211299376 | 0.655985013 | Senp3 | SUMO/sentrin specific peptidase 3 [Sourc | 642.48 | 695.9 | 609.78 | 756.72 | 706.72 | 695.04 |
| 815 | ENSMUSG00000005220 | 6.811396086 | -3.15551626 | 1.119960464 | -2.817524689 | 0.004839539 | 0.065358634 | Corin | corin, serine peptidase [Source:MGI Symb | 7.2279 | 22.614 | 7.0437 | 2.8919 | 1.0906 | 0 |
| 816 | ENSMUSG00000005225 | 634.3670437 | -0.019183565 | 0.131825451 | -0.145522464 | 0.884298381 | 0.981769403 | Plekha8 | pleckstrin homology domain containing, f | 697.09 | 648.62 | 569.53 | 648.76 | 667.46 | 574.74 |
| 817 | ENSMUSG00000005232 | 0.706696157 | 0.149852971 | 3.037999338 | 0.049326203 | 0.960659338 | NA | G6pc2 | glucose-6-phosphatase, catalytic, 2 [Sourc | 0 | 0 | 2.0125 | 0 | 0 | 2.2277 |
| 818 | ENSMUSG00000005233 | 123.9803562 | -0.324966203 | 0.248910835 | -1.305552661 | 0.191704728 | 0.629099068 | Spc25 | SPC25, NDC80 kinetochore complex comp | 155.8 | 109.99 | 147.92 | 131.1 | 118.88 | 80.197 |
| 819 | ENSMUSG00000005237 | 1.934478312 | -1.529593741 | 1.722984031 | -0.887758513 | 0.374670664 | NA | Dnah2 | dynein, axonemal, heavy chain 2 [Source: | 1.6062 | 3.0838 | 4.025 | 2.8919 | 0 | 0 |
| 820 | ENSMUSG00000005251 | 163.8985374 | -0.28679629 | 0.183781862 | -1.560525545 | 0.118635738 | 0.504308497 | Ripk4 | receptor-interacting serine-threonine kina | 177.49 | 187.08 | 176.09 | 155.2 | 149.41 | 138.12 |
| 821 | ENSMUSG00000005262 | 635.5148859 | 0.034168527 | 0.115695217 | 0.295332234 | 0.767740095 | 0.95578799 | Ufd1 | ubiquitin recognition factor in ER-associat | 598.31 | 632.17 | 654.06 | 646.83 | 642.37 | 639.35 |
| 822 | ENSMUSG00000005267 | 5.014226428 | -0.466450905 | 0.998828352 | -0.466998062 | 0.640501265 | NA | Zfp287 | zinc finger protein 287 [Source:MGI Symb | 3.2124 | 11.307 | 3.0187 | 4.8199 | 3.2719 | 4.4554 |
| 823 | ENSMUSG00000005268 | 1388.396627 | -0.435709404 | 0.146543905 | -2.973234568 | 0.002946792 | 0.046110081 | Prlr | prolactin receptor [Source:MGI Symbol;Ac | 1655.2 | 1518.2 | 1617 | 1480.7 | 1005.6 | 1053.7 |
| 824 | ENSMUSG00000005299 | 1717.451646 | 0.006466153 | 0.101349847 | 0.063800328 | 0.949129217 | 0.990731019 | Letm1 | leucine zipper-EF-hand containing transm | 1803.8 | 1637.5 | 1698.5 | 1695.6 | 1637 | 1832.3 |
| 825 | ENSMUSG00000005312 | 2572.237079 | -0.051530665 | 0.094827433 | -0.543415159 | 0.586843991 | 0.910448513 | Ubqln1 | ubiquilin 1 [Source:MGI Symbol;Acc:MGI: | 2525 | 2642.8 | 2686.7 | 2380.1 | 2532.4 | 2666.5 |
| 826 | ENSMUSG00000005320 | 571.809883 | 0.329929532 | 0.161183855 | 2.046914262 | 0.040666504 | 0.276020792 | Fgfr4 | fibroblast growth factor receptor 4 [Sourc | 414.4 | 485.18 | 621.86 | 685.39 | 579.12 | 644.92 |
| 827 | ENSMUSG00000005338 | 77.87951663 | -1.361795646 | 0.285060175 | -4.777221669 | 1.78E-06 | 0.000129183 | Cadm3 | cell adhesion molecule 3 [Source:MGI Syn | 134.92 | 99.708 | 101.63 | 48.199 | 33.809 | 49.009 |
| 828 | ENSMUSG00000005339 | 0 | NA | NA | NA | NA | NA | Fcer1a | Fc receptor, IgE, high affinity I, alpha poly | 0 | 0 | 0 | 0 | 0 | 0 |
| 829 | ENSMUSG00000005354 | 3181.509965 | -0.033541014 | 0.092947716 | -0.360858941 | 0.718204899 | 0.9447719 | Txn2 | thioredoxin 2 [Source:MGI Symbol;Acc:M | 3198 | 3192.7 | 3264.3 | 2990.3 | 3072.3 | 3371.6 |
| 830 | ENSMUSG00000005355 | 0 | NA | NA | NA | NA | NA | Casp14 | caspase 14 [Source:MGI Symbol;Acc:MGI: | 0 | 0 | 0 | 0 | 0 | 0 |
| 831 | ENSMUSG00000005357 | 0.133850373 | -0.796629176 | 4.080472857 | -0.195229623 | 0.84521319 | NA | Slc1a6 | solute carrier family 1 (high affinity aspart | 0.8031 | 0 | 0 | 0 | 0 | 0 |
| 832 | ENSMUSG00000005360 | 2.578797709 | -2.613028369 | 1.692766038 | -1.543644137 | 0.122674566 | NA | Slc1a3 | solute carrier family 1 (glial high affinity gl | 4.0155 | 8.2233 | 1.0062 | 0 | 0 | 2.2277 |
| 833 | ENSMUSG00000005362 | 533.11948 | 0.245934364 | 0.139371624 | 1.764594231 | 0.077631986 | 0.407722964 | Crbn | cereblon [Source:MGI Symbol;Acc:MGI:19 | 430.46 | 534.52 | 499.1 | 534.04 | 633.65 | 566.95 |
| 834 | ENSMUSG00000005364 | 0.267700746 | -1.497481649 | 4.057001436 | -0.369110456 | 0.712045396 | NA | Il5ra | interleukin 5 receptor, alpha [Source:MGI | 1.6062 | 0 | 0 | 0 | 0 | 0 |
| 835 | ENSMUSG00000005370 | 209.4659113 | 0.116607372 | 0.179432329 | 0.64986824 | 0.515777335 | 0.886238519 | Msh6 | mutS homolog 6 [Source:MGI Symbol;Acc | 183.91 | 182.97 | 236.47 | 216.9 | 212.67 | 223.88 |
| 836 | ENSMUSG00000005371 | 416.9347988 | 0.13906932 | 0.149633461 | 0.929399871 | 0.352681894 | 0.793259753 | Fbxo11 | F-box protein 11 [Source:MGI Symbol;Acc | 377.46 | 424.53 | 388.41 | 374.99 | 495.14 | 441.08 |

| | | | | | | | | | | | | | | | |
|---|---|---|---|---|---|---|---|---|---|---|---|---|---|---|---|
| 837 | ENSMUSG00000005373 | 4849.187189 | 0.260710249 | 0.165543275 | 1.574876713 | 0.115284905 | 0.498648584 | Mlxipl | MLX interacting protein-like [Source:MGI | 4578.5 | 3824.9 | 4834 | 6956.1 | 4191.2 | 4710.4 |
| 838 | ENSMUSG00000005374 | 367.9058715 | 0.079515047 | 0.148802606 | 0.534365957 | 0.593088367 | 0.913011006 | Tbl2 | transducin (beta)-like 2 [Source:MGI Symt | 372.64 | 356.69 | 344.14 | 410.65 | 398.08 | 325.24 |
| 839 | ENSMUSG00000005378 | 460.9968901 | 0.109681919 | 0.135526686 | 0.809301266 | 0.418341878 | 0.837575303 | Bud23 | BUD23, rRNA methyltransferase and ribos | 454.56 | 397.8 | 477.97 | 497.41 | 452.61 | 485.64 |
| 840 | ENSMUSG00000005397 | 130.4159229 | -0.863822064 | 0.250450449 | -3.449073723 | 0.000562513 | 0.013537753 | Nid1 | nidogen 1 [Source:MGI Symbol;Acc:MGI:9 | 216.84 | 141.85 | 145.91 | 101.22 | 71.981 | 104.7 |
| 841 | ENSMUSG00000005410 | 47.43703627 | -0.141300589 | 0.333748709 | -0.423374187 | 0.672022275 | 0.931191575 | Mcm5 | minichromosome maintenance complex c | 53.808 | 37.005 | 58.362 | 46.271 | 49.078 | 40.098 |
| 842 | ENSMUSG00000005413 | 186.6264475 | -0.899478195 | 0.256645814 | -3.504745238 | 0.000457044 | 0.011618085 | Hmox1 | heme oxygenase 1 [Source:MGI Symbol;A | 235.31 | 322.77 | 171.06 | 139.78 | 93.793 | 157.05 |
| 843 | ENSMUSG00000005417 | 1868.967991 | 0.087551121 | 0.093498619 | 0.936389453 | 0.349072706 | 0.79040715 | Mprip | myosin phosphatase Rho interacting prote | 1839.1 | 1787.5 | 1810.2 | 1993.5 | 1845.3 | 1938.1 |
| 844 | ENSMUSG00000005442 | 472.2852072 | 0.25562461 | 0.161366637 | 1.584123053 | 0.113165719 | 0.493951959 | Cic | capicua transcriptional repressor [Source: | 428.05 | 371.08 | 493.06 | 604.41 | 453.7 | 483.41 |
| 845 | ENSMUSG00000005447 | 39.36232902 | -0.113890697 | 0.378814876 | -0.300650011 | 0.763681391 | 0.954980327 | Pafah1b3 | platelet-activating factor acetylhydrolase, | 34.533 | 50.368 | 38.237 | 50.127 | 27.265 | 35.643 |
| 846 | ENSMUSG00000005465 | 11.03813058 | -1.216220282 | 0.661434167 | -1.838762408 | 0.06595014 | 0.368871933 | Il27ra | interleukin 27 receptor, alpha [Source:MG | 12.85 | 15.419 | 18.112 | 7.7118 | 5.4531 | 6.6831 |
| 847 | ENSMUSG00000005469 | 1069.412903 | -0.050808576 | 0.106125587 | -0.478758964 | 0.632110113 | 0.921791304 | Prkaca | protein kinase, cAMP dependent, catalyti | 1058.5 | 1126.6 | 1079.7 | 1003.5 | 1117.9 | 1030.3 |
| 848 | ENSMUSG00000005470 | 5.789930092 | -0.637926551 | 0.909236435 | -0.701606895 | 0.482924351 | 0.871710696 | Asf1b | anti-silencing function 1B histone chaperc | 4.0155 | 6.1675 | 11.069 | 5.7839 | 4.3625 | 3.3415 |
| 849 | ENSMUSG00000005474 | 0.17131964 | -0.796629176 | 4.080472857 | -0.195229623 | 0.84521319 | NA | Myl10 | myosin, light chain 10, regulatory [Source: | 0 | 1.0279 | 0 | 0 | 0 | 0 |
| 850 | ENSMUSG00000005481 | 224.7682573 | -0.219580504 | 0.181546702 | -1.209498725 | 0.2264713 | 0.67317733 | Ddx39a | DEAD box helicase 39a [Source:MGI Symb | 264.22 | 229.23 | 231.44 | 197.62 | 179.95 | 246.16 |
| 851 | ENSMUSG00000005483 | 381.0111567 | -0.108904675 | 0.14137518 | -0.770323865 | 0.441107804 | 0.849828766 | Dnajb1 | DnaJ heat shock protein family (Hsp40) m | 354.97 | 425.56 | 406.52 | 375.95 | 357.72 | 365.34 |
| 852 | ENSMUSG00000005493 | 0.469265235 | -2.274672822 | 3.439794491 | -0.661281605 | 0.508431736 | NA | Msh4 | mutS homolog 4 [Source:MGI Symbol;Acc | 0.8031 | 0 | 2.0125 | 0 | 0 | 0 |
| 853 | ENSMUSG00000005501 | 804.4799142 | 0.043722403 | 0.12048701 | 0.362880639 | 0.71669405 | 0.9447719 | Usp40 | ubiquitin specific peptidase 40 [Source:M | 823.98 | 788.41 | 764.75 | 905.18 | 748.16 | 796.4 |
| 854 | ENSMUSG00000005503 | 0 | NA | NA | NA | NA | NA | Evx1 | even-skipped homeobox 1 [Source:MGI S | 0 | 0 | 0 | 0 | 0 | 0 |
| 855 | ENSMUSG00000005505 | 99.36721117 | 0.222361769 | 0.243513673 | 0.913138739 | 0.361169578 | 0.800074801 | Kbtbd4 | kelch repeat and BTB (POZ) domain conta | 85.129 | 98.68 | 91.568 | 109.89 | 85.068 | 125.86 |
| 856 | ENSMUSG00000005506 | 1926.006709 | 0.059566729 | 0.101104366 | 0.589160803 | 0.555753409 | 0.897628185 | Celf1 | CUGBP, Elav-like family member 1 [Source | 1790.9 | 2026 | 1842.4 | 1908.7 | 2072.2 | 1915.8 |
| 857 | ENSMUSG00000005510 | 2295.203677 | -0.060456658 | 0.092926476 | -0.650584532 | 0.515314718 | 0.885846456 | Ndufs3 | NADH:ubiquinone oxidoreductase core su | 2293.7 | 2364.2 | 2371.7 | 2124.6 | 2294.7 | 2322.4 |
| 858 | ENSMUSG00000005514 | 12464.21696 | -0.359363801 | 0.146886181 | -2.446546016 | 0.014423238 | 0.139264166 | Por | P450 (cytochrome) oxidoreductase [Sourc | 14676 | 13749 | 13601 | 13256 | 11321 | 8182.3 |
| 859 | ENSMUSG00000005533 | 34.17313061 | -0.443807933 | 0.398644002 | -1.113293892 | 0.265582229 | 0.721565724 | Igf1r | insulin-like growth factor I receptor [Sour | 37.746 | 51.396 | 29.181 | 33.739 | 22.903 | 30.074 |
| 860 | ENSMUSG00000005534 | 2804.336157 | 0.025882811 | 0.086917762 | 0.297785062 | 0.765867216 | 0.954996635 | Insr | insulin receptor [Source:MGI Symbol;Acc: | 2753 | 2817.5 | 2767.2 | 2814.8 | 2897.8 | 2775.7 |
| 861 | ENSMUSG00000005540 | 3.20325932 | -1.834433378 | 1.403637079 | -1.306914306 | 0.191241817 | NA | Fcer2a | Fc receptor, IgE, low affinity II, alpha poly | 0.8031 | 8.2233 | 6.0375 | 1.928 | 0 | 2.2277 |
| 862 | ENSMUSG00000005547 | 12863.28373 | -0.259104421 | 0.134196464 | -1.930784264 | 0.053509737 | 0.325468814 | Cyp2a5 | cytochrome P450, family 2, subfamily a, p | 11584 | 13910 | 16553 | 10325 | 12684 | 12124 |
| 863 | ENSMUSG00000005553 | 8.015673981 | 1.51594732 | 0.861072887 | 1.760533101 | 0.07831746 | 0.410131696 | Atp4a | ATPase, H+/K+ exchanging, gastric, alpha | 2.4093 | 2.0558 | 8.05 | 5.7839 | 13.087 | 16.708 |
| 864 | ENSMUSG00000005566 | 1060.833474 | 0.113549422 | 0.111610449 | 1.017372682 | 0.308976173 | 0.757462943 | Trim28 | tripartite motif-containing 28 [Source:MG | 1002.3 | 1042.3 | 1013.3 | 1134.6 | 1177.9 | 994.66 |
| 865 | ENSMUSG00000005575 | 871.9349444 | -0.074317775 | 0.122409462 | -0.607124435 | 0.543768335 | 0.894702561 | Ube2m | ubiquitin-conjugating enzyme E2M [Sourc | 902.69 | 852.14 | 928.76 | 932.17 | 748.16 | 867.68 |
| 866 | ENSMUSG00000005580 | 411.919817 | -0.09888332 | 0.166188781 | -0.59500599 | 0.551839467 | 0.895935901 | Adcy9 | adenylate cyclase 9 [Source:MGI Symbol;A | 362.2 | 503.68 | 412.56 | 345.1 | 402.44 | 445.54 |
| 867 | ENSMUSG00000005583 | 33.33036656 | -0.504839698 | 0.392935702 | -1.284789585 | 0.198865823 | 0.640976045 | Mef2c | myocyte enhancer factor 2C [Source:MGI | 44.171 | 29.81 | 43.269 | 30.847 | 21.812 | 30.074 |
| 868 | ENSMUSG00000005609 | 482.1758889 | -0.231393211 | 0.140385066 | -1.648275119 | 0.099296228 | 0.464258353 | Ctr9 | CTR9 homolog, Paf1/RNA polymerase II cc | 512.38 | 561.24 | 489.03 | 453.07 | 488.6 | 388.73 |
| 869 | ENSMUSG00000005610 | 8793.763234 | -0.098679648 | 0.09794346 | -1.007516461 | 0.313686651 | 0.760476509 | Eif4g2 | eukaryotic translation initiation factor 4, g | 8324.2 | 9953.3 | 9006.9 | 8328.8 | 8859.1 | 8290.3 |
| 870 | ENSMUSG00000005611 | 16.43041739 | -0.417262267 | 0.532117417 | -0.7841545 | 0.43294946 | 0.845264432 | Irag1 | inositol 1,4,5-triphosphate receptor assoc | 17.668 | 22.614 | 16.1 | 13.496 | 11.997 | 16.708 |
| 871 | ENSMUSG00000005615 | 1622.299987 | -0.001973531 | 0.10439716 | -0.018904065 | 0.984917636 | 0.99859176 | Pcyt1a | phosphate cytidylyltransferase 1, choline, | 1595.8 | 1685.8 | 1588.9 | 1582.9 | 1765.7 | 1514.8 |
| 872 | ENSMUSG00000005621 | 527.9487257 | -0.030167725 | 0.125865348 | -0.239682527 | 0.810576382 | 0.966384503 | Zfp592 | zinc finger protein 592 [Source:MGI Symb | 501.14 | 536.57 | 563.5 | 545.61 | 495.14 | 525.73 |
| 873 | ENSMUSG00000005625 | 1569.739888 | -0.021614402 | 0.112537617 | -0.192063798 | 0.847692232 | 0.972803041 | Psmd4 | proteasome (prosome, macropain) 26S su | 1782.9 | 1480.2 | 1480.2 | 1615.6 | 1525.8 | 1533.8 |
| 874 | ENSMUSG00000005628 | 6.41992237 | 1.319050644 | 0.863272245 | 1.527966006 | 0.126520978 | 0.521187766 | Tmod4 | tropomodulin 4 [Source:MGI Symbol;Acc: | 4.8186 | 3.0838 | 3.0187 | 7.7118 | 7.6343 | 12.252 |
| 875 | ENSMUSG00000005640 | 0.867384107 | -1.926017826 | 2.566777421 | -0.750364177 | 0.4530354 | NA | Insrr | insulin receptor-related receptor [Source: | 3.2124 | 1.0279 | 0 | 0.964 | 0 | 0 |
| 876 | ENSMUSG00000005649 | 0 | NA | NA | NA | NA | NA | Cabp5 | calcium binding protein 5 [Source:MGI Sy | 0 | 0 | 0 | 0 | 0 | 0 |
| 877 | ENSMUSG00000005656 | 245.5568121 | -0.240398193 | 0.167725307 | -1.433285159 | 0.151776361 | 0.567340036 | Snx6 | sorting nexin 6 [Source:MGI Symbol;Acc:N | 264.22 | 264.17 | 269.67 | 228.46 | 254.11 | 192.69 |
| 878 | ENSMUSG00000005667 | 14.91642902 | -0.344415786 | 0.595788337 | -0.578084135 | 0.563207317 | 0.900821918 | Mthfd2 | methylenetetrahydrofolate dehydrogenas | 18.471 | 11.307 | 20.125 | 7.7118 | 19.631 | 12.252 |
| 879 | ENSMUSG00000005672 | 57.76897753 | -0.075058843 | 0.291718526 | -0.257298857 | 0.796948075 | 0.962236316 | Kit | KIT proto-oncogene receptor tyrosine kina | 65.854 | 53.452 | 58.362 | 59.767 | 51.259 | 57.92 |
| 880 | ENSMUSG00000005674 | 221.468832 | -0.083084261 | 0.230874929 | -0.359866968 | 0.71894662 | 0.9447719 | Tomm40l | translocase of outer mitochondrial memb | 193.55 | 227.17 | 263.64 | 255.45 | 248.66 | 140.34 |
| 881 | ENSMUSG00000005677 | 1432.485709 | -0.427682305 | 0.32590007 | -1.312311176 | 0.189415163 | 0.626663447 | Nr1i3 | nuclear receptor subfamily 1, group I, mer | 1513 | 1720.7 | 1696.5 | 1645.5 | 1269.5 | 749.62 |
| 882 | ENSMUSG00000005681 | 108155.5391 | -0.132983409 | 0.094364646 | -1.409250328 | 0.158761161 | 0.579879149 | Apoa2 | apolipoprotein A-II [Source:MGI Symbol;A | 109023 | 119711 | 110678 | 112449 | 99483 | 97589 |
| 883 | ENSMUSG00000005682 | 471.346095 | 0.496692144 | 0.141902279 | 3.500240787 | 0.000464838 | 0.011774812 | Pan2 | PAN2 poly(A) specific ribonuclease subun | 371.84 | 390.61 | 411.55 | 630.44 | 522.41 | 501.23 |
| 884 | ENSMUSG00000005683 | 2239.698453 | 0.057857413 | 0.129109188 | 0.448127775 | 0.654060984 | 0.925797231 | Cs | citrate synthase [Source:MGI Symbol;Acc: | 2476.8 | 2264.5 | 1841.4 | 2049.4 | 2361.2 | 2444.9 |
| 885 | ENSMUSG00000005686 | 11.61407 | -0.382655579 | 0.627327362 | -0.609977506 | 0.541876709 | 0.894000766 | Ampd3 | adenosine monophosphate deaminase 3 [ | 12.047 | 15.419 | 12.075 | 12.532 | 9.8156 | 7.7969 |
| 886 | ENSMUSG00000005687 | 517.7573885 | 0.037245283 | 0.127706397 | 0.291647746 | 0.770555966 | 0.956515747 | Bcas2 | breast carcinoma amplified sequence 2 [S | 508.36 | 527.32 | 497.08 | 478.13 | 540.95 | 554.69 |
| 887 | ENSMUSG00000005696 | 0.294513347 | 0.165143496 | 4.080472857 | 0.040471657 | 0.967717103 | NA | Sh2d1a | SH2 domain containing 1A [Source:MGI Sy | 0.8031 | 0 | 0 | 0.964 | 0 | 0 |
| 888 | ENSMUSG00000005698 | 579.6819211 | -0.121892413 | 0.142129431 | -0.857615574 | 0.391104772 | 0.820195366 | Ctcf | CCCTC-binding factor [Source:MGI Symbo | 534.87 | 663.01 | 614.82 | 482.95 | 596.57 | 585.88 |
| 889 | ENSMUSG00000005699 | 72.01130865 | 0.087579681 | 0.284093519 | 0.308277647 | 0.757871073 | 0.953466696 | Pard6a | par-6 family cell polarity regulator alpha [S | 78.704 | 78.122 | 52.325 | 68.442 | 68.709 | 85.766 |
| 890 | ENSMUSG00000005705 | 4.509915897 | 0.019808844 | 1.074130753 | 0.018441744 | 0.985286451 | NA | Agrp | agouti related neuropeptide [Source:MGI | 7.2279 | 4.1117 | 2.0125 | 4.8199 | 1.0906 | 7.7969 |
| 891 | ENSMUSG00000005716 | 0 | NA | NA | NA | NA | NA | Pvalb | parvalbumin [Source:MGI Symbol;Acc:MG | 0 | 0 | 0 | 0 | 0 | 0 |
| 892 | ENSMUSG00000005718 | 110.0443385 | 0.338648878 | 0.240903509 | 1.405744896 | 0.15979989 | 0.581763305 | Tfap4 | transcription factor AP4 [Source:MGI Sym | 102.8 | 77.094 | 111.69 | 140.74 | 123.24 | 104.7 |

| | | | | | | | | | | | | | | |
|---|---|---|---|---|---|---|---|---|---|---|---|---|---|---|
| 893 | ENSMUSG00000005732 | 230.0785565 | -0.096795152 | 0.163473159 | -0.592116484 | 0.55377259 | 0.896749523 | Ranbp1 | RAN binding protein 1 [Source:MGI Symbo | 232.1 | 245.67 | 235.46 | 203.4 | 238.85 | 225 |
| 894 | ENSMUSG00000005763 | 1.700809071 | -0.448317306 | 1.700036252 | -0.263710439 | 0.792003051 | NA | Cd247 | CD247 antigen [Source:MGI Symbol;Acc:N | 0.8031 | 3.0838 | 2.0125 | 0.964 | 0 | 3.3415 |
| 895 | ENSMUSG00000005774 | 161.9607203 | 0.080479101 | 0.216821238 | 0.371177204 | 0.71050555 | 0.944239215 | Rfx5 | regulatory factor X, 5 (influences HLA clas | 189.53 | 118.21 | 164.02 | 180.26 | 153.78 | 165.96 |
| 896 | ENSMUSG00000005779 | 2475.879284 | 0.082915765 | 0.097849671 | 0.847379089 | 0.396783856 | 0.824225434 | Psmb4 | proteasome (prosome, macropain) subun | 2549.8 | 2285.1 | 2377.8 | 2400.3 | 2589.1 | 2653.2 |
| 897 | ENSMUSG00000005800 | 0.506966596 | 2.558866737 | 3.339834942 | 0.766165628 | 0.443577757 | NA | Mmp8 | matrix metallopeptidase 8 [Source:MGI Sy | 0 | 0 | 0 | 1.928 | 0 | 1.1138 |
| 898 | ENSMUSG00000005802 | 249.000681 | 0.194034533 | 0.168846249 | 1.14917882 | 0.250482252 | 0.702302228 | Slc30a4 | solute carrier family 30 (zinc transporter), | 204.79 | 267.26 | 225.4 | 254.49 | 279.2 | 262.87 |
| 899 | ENSMUSG00000005803 | 3237.688236 | -0.015130007 | 0.103308989 | -0.146453922 | 0.883563063 | 0.981769403 | Sqor | sulfide quinone oxidoreductase [Source:N | 3000.4 | 3179.3 | 3585.2 | 3192.7 | 3383.1 | 3085.3 |
| 900 | ENSMUSG00000005804 | 308.7971826 | 0.061963818 | 0.154253242 | 0.401701882 | 0.68790344 | 0.936885029 | Bloc1s6 | biogenesis of lysosomal organelles comple | 292.33 | 312.49 | 301.87 | 306.54 | 356.63 | 282.92 |
| 901 | ENSMUSG00000005813 | 602.1659742 | 0.177596398 | 0.124177185 | 1.430185411 | 0.152663811 | 0.568519405 | Metap1 | methionyl aminopeptidase 1 [Source:MGI | 516.39 | 607.5 | 572.55 | 642.97 | 658.73 | 614.84 |
| 902 | ENSMUSG00000005823 | 480.0762008 | 0.282004089 | 0.145104492 | 1.943455271 | 0.051961175 | 0.320245495 | Gpr108 | G protein-coupled receptor 108 [Source:N | 401.55 | 472.84 | 426.65 | 595.74 | 463.51 | 520.17 |
| 903 | ENSMUSG00000005824 | 1.59492828 | -0.96611914 | 1.740592475 | -0.555051888 | 0.57885917 | NA | Tnfsf14 | tumor necrosis factor (ligand) superfamily | 3.2124 | 2.0558 | 1.0062 | 0 | 2.1812 | 1.1138 |
| 904 | ENSMUSG00000005836 | 190.8109755 | -1.038798978 | 0.181208731 | -5.732609965 | 9.89E-09 | 1.54E-06 | Gata6 | GATA binding protein 6 [Source:MGI Symb | 263.42 | 267.26 | 239.49 | 129.17 | 134.15 | 111.38 |
| 905 | ENSMUSG00000005846 | 716.3186308 | 0.012959685 | 0.114997204 | 0.112695649 | 0.910271851 | 0.985639471 | Rsl1d1 | ribosomal L1 domain containing 1 [Source | 727.61 | 733.93 | 677.2 | 686.35 | 717.63 | 755.19 |
| 906 | ENSMUSG00000005864 | 0 | NA | NA | NA | NA | NA | Cnga2 | cyclic nucleotide gated channel alpha 2 [S | 0 | 0 | 0 | 0 | 0 | 0 |
| 907 | ENSMUSG00000005871 | 590.4668826 | -0.067748743 | 0.125835824 | -0.538389947 | 0.590307865 | 0.911690353 | Apc | APC, WNT signaling pathway regulator [Sc | 602.33 | 651.7 | 559.47 | 614.05 | 561.67 | 553.58 |
| 908 | ENSMUSG00000005873 | 181.2491134 | -0.277029694 | 0.206379317 | -1.342332639 | 0.179488165 | 0.611676727 | Reep5 | receptor accessory protein 5 [Source:MGI | 230.49 | 198.39 | 166.03 | 141.7 | 155.96 | 194.92 |
| 909 | ENSMUSG00000005881 | 1361.352922 | 0.203577463 | 0.104764983 | 1.94318232 | 0.051994134 | 0.320290949 | Ergic3 | ERGIC and golgi 3 [Source:MGI Symbol;Ac | 1195.8 | 1273.6 | 1327.2 | 1364 | 1513.8 | 1493.7 |
| 910 | ENSMUSG00000005882 | 786.632251 | -0.063010112 | 0.11483129 | -0.548719015 | 0.583198295 | 0.908645447 | Uqcc1 | ubiquinol-cytochrome c reductase comple | 787.04 | 850.09 | 774.81 | 785.64 | 803.79 | 718.43 |
| 911 | ENSMUSG00000005883 | 2.007188199 | -0.837586909 | 1.725238184 | -0.485490593 | 0.627328384 | NA | Spo11 | SPO11 initiator of meiotic double strande | 5.6217 | 0 | 2.0125 | 0 | 2.1812 | 2.2277 |
| 912 | ENSMUSG00000005886 | 811.3958697 | 0.371296682 | 0.124220021 | 2.989024462 | 0.002798697 | 0.044483909 | Ncoa2 | nuclear receptor coactivator 2 [Source:MC | 699.5 | 702.07 | 720.47 | 794.32 | 967.38 | 984.64 |
| 913 | ENSMUSG00000005891 | 0 | NA | NA | NA | NA | NA | Prl4a1 | prolactin family 4, subfamily a, member 1 | 0 | 0 | 0 | 0 | 0 | 0 |
| 914 | ENSMUSG00000005892 | 0.267700746 | -1.497481649 | 4.057001436 | -0.369110456 | 0.712045396 | NA | Trh | thyrotropin releasing hormone [Source:M | 1.6062 | 0 | 0 | 0 | 0 | 0 |
| 915 | ENSMUSG00000005893 | 204.8135678 | 0.142272062 | 0.182493956 | 0.779598763 | 0.435627084 | 0.845940264 | Nr2c2 | nuclear receptor subfamily 2, group C, me | 166.24 | 200.44 | 218.36 | 227.5 | 220.3 | 196.04 |
| 916 | ENSMUSG00000005897 | 232.6896092 | 0.152345785 | 0.163413489 | 0.932271787 | 0.351196084 | 0.792511048 | Nr2c1 | nuclear receptor subfamily 2, group C, me | 220.85 | 217.92 | 222.38 | 236.17 | 270.47 | 228.34 |
| 917 | ENSMUSG00000005899 | 241.1850167 | 0.043521491 | 0.15949516 | 0.272870295 | 0.784952922 | 0.960670212 | Smpd4 | sphingomyelin phosphodiesterase 4 [Sour | 254.58 | 223.06 | 234.45 | 240.03 | 241.03 | 253.96 |
| 918 | ENSMUSG00000005907 | 655.2924079 | -0.270369679 | 0.124788127 | -2.166629836 | 0.030263091 | 0.230381759 | Pex1 | peroxisomal biogenesis factor 1 [Source:N | 689.86 | 746.27 | 714.43 | 647.79 | 598.75 | 534.65 |
| 919 | ENSMUSG00000005917 | 0 | NA | NA | NA | NA | NA | Otx1 | orthodenticle homeobox 1 [Source:MGI S | 0 | 0 | 0 | 0 | 0 | 0 |
| 920 | ENSMUSG00000005936 | 728.8776572 | 0.245392101 | 0.135238428 | 1.814514583 | 0.06959854 | 0.381135916 | Kctd20 | potassium channel tetramerisation domai | 665.77 | 638.34 | 696.32 | 696.96 | 753.62 | 922.26 |
| 921 | ENSMUSG00000005947 | 8.347382327 | -0.458280092 | 0.7314989 | -0.626494574 | 0.530990595 | 0.888794797 | Itgae | integrin alpha E, epithelial-associated [Sou | 9.6372 | 10.279 | 9.0562 | 6.7478 | 5.4531 | 8.9108 |
| 922 | ENSMUSG00000005949 | 325.8238092 | 0.046660776 | 0.154679467 | 0.301661085 | 0.762910437 | 0.95469451 | Ctns | cystinosis, nephropathic [Source:MGI Sym | 310.8 | 335.1 | 315.96 | 336.43 | 285.74 | 370.91 |
| 923 | ENSMUSG00000005950 | 3.681656025 | 0.414591798 | 1.150024794 | 0.360506834 | 0.718468147 | NA | P2rx5 | purinergic receptor P2X, ligand-gated ion | 2.4093 | 2.0558 | 5.0312 | 4.8199 | 1.0906 | 6.6831 |
| 924 | ENSMUSG00000005951 | 1632.112437 | 0.051599364 | 0.095827682 | 0.538459896 | 0.590259585 | 0.911690353 | Shpk | sedoheptulokinase [Source:MGI Symbol;A | 1639.9 | 1568.6 | 1599.9 | 1663.8 | 1725.4 | 1595 |
| 925 | ENSMUSG00000005952 | 1.468645325 | -1.76532237 | 1.978996813 | -0.89202891 | 0.372377437 | NA | Trpv1 | transient receptor potential cation channe | 0.8031 | 2.0558 | 4.025 | 1.928 | 0 | 0 |
| 926 | ENSMUSG00000005958 | 7.481593358 | -1.500049158 | 0.850443968 | -1.763842434 | 0.077758511 | 0.408091111 | Ephb3 | Eph receptor B3 [Source:MGI Symbol;Acc | 13.653 | 13.363 | 6.0375 | 1.928 | 5.4531 | 4.4554 |
| 927 | ENSMUSG00000005968 | 103.2327719 | 0.978308125 | 0.277839003 | 3.521133157 | 0.000429707 | 0.011198701 | Tuft1 | tuftelin 1 [Source:MGI Symbol;Acc:MGI:1( | 79.507 | 55.508 | 73.456 | 186.05 | 110.15 | 114.73 |
| 928 | ENSMUSG00000005973 | 85.75216451 | -0.820360709 | 0.293059207 | -2.799300239 | 0.005121349 | 0.067648582 | Rcn1 | reticulocalbin 1 [Source:MGI Symbol;Acc:I | 134.12 | 96.624 | 97.606 | 72.298 | 40.353 | 73.514 |
| 929 | ENSMUSG00000005980 | 15.41484455 | 0.248005063 | 0.60306259 | 0.411242658 | 0.680894611 | 0.934384556 | Dnase1 | deoxyribonuclease I [Source:MGI Symbol; | 13.653 | 18.503 | 10.062 | 9.6398 | 27.265 | 13.366 |
| 930 | ENSMUSG00000005981 | 3480.314374 | -0.08616963 | 0.090562056 | -0.951498169 | 0.341351548 | 0.783129922 | Trap1 | TNF receptor-associated protein 1 [Source | 3574.6 | 3755 | 3423.2 | 3371 | 3488.9 | 3269.1 |
| 931 | ENSMUSG00000005982 | 1645.440034 | -0.131163084 | 0.109294409 | -1.200089609 | 0.23010454 | 0.676880752 | Naa60 | N(alpha)-acetyltransferase 60, NatF cataly | 1663.2 | 1696.1 | 1802.2 | 1652.3 | 1659.9 | 1399 |
| 932 | ENSMUSG00000005983 | 10.64715072 | -0.321941409 | 0.650672121 | -0.494782854 | 0.620753408 | 0.918069106 | 1700037C18 | RIKEN cDNA 1700037C18 gene [Source:M | 12.047 | 14.391 | 9.0562 | 9.6398 | 8.7249 | 10.025 |
| 933 | ENSMUSG00000005986 | 4.725006886 | -1.226998155 | 1.072276047 | -1.144293168 | 0.252502076 | NA | Ankrd13d | ankyrin repeat domain 13 family, member | 1.6062 | 10.279 | 8.05 | 2.8919 | 2.1812 | 3.3415 |
| 934 | ENSMUSG00000005994 | 0 | NA | NA | NA | NA | NA | Tyrp1 | tyrosinase-related protein 1 [Source:MGI | 0 | 0 | 0 | 0 | 0 | 0 |
| 935 | ENSMUSG00000006005 | 1078.789856 | 0.078771487 | 0.112240386 | 0.701810549 | 0.482797319 | 0.871710696 | Tpr | translocated promoter region, nuclear ba: | 1114.7 | 1051.6 | 981.09 | 1101.8 | 1187.7 | 1035.9 |
| 936 | ENSMUSG00000006007 | 0 | NA | NA | NA | NA | NA | Pdc | phosducin [Source:MGI Symbol;Acc:MGI:! | 0 | 0 | 0 | 0 | 0 | 0 |
| 937 | ENSMUSG00000006010 | 724.123253 | 0.067655953 | 0.118716248 | 0.569896323 | 0.568748019 | 0.903527882 | Odr4 | odr4 GPCR localization factor homolog [Sc | 757.33 | 691.79 | 671.17 | 695.03 | 778.7 | 750.73 |
| 938 | ENSMUSG00000006014 | 879.4764979 | -0.1404223 | 0.114356353 | -1.227936155 | 0.219470936 | 0.665219534 | Prg4 | proteoglycan 4 (megakaryocyte stimulatin | 987.82 | 888.12 | 890.53 | 894.57 | 801.6 | 814.22 |
| 939 | ENSMUSG00000006019 | 198.9249948 | -0.055511834 | 0.18182554 | -0.305302732 | 0.760135595 | 0.95402891 | Dhx34 | DEAH (Asp-Glu-Ala-His) box polypeptide 3 | 213.63 | 174.75 | 219.36 | 182.19 | 189.77 | 213.86 |
| 940 | ENSMUSG00000006021 | 249.382817 | 0.085020673 | 0.171989211 | 0.494337247 | 0.621068024 | 0.918069106 | Kptn | kaptin [Source:MGI Symbol;Acc:MGI:189C | 247.36 | 244.64 | 233.45 | 212.08 | 279.2 | 279.57 |
| 941 | ENSMUSG00000006024 | 1171.409496 | -0.05638862 | 0.102832728 | -0.548352862 | 0.583449637 | 0.908645447 | Napa | N-ethylmaleimide sensitive fusion protein | 1247.2 | 1137.9 | 1197.4 | 1184.7 | 1147.3 | 1113.8 |
| 942 | ENSMUSG00000006050 | 1713.813971 | -0.013531227 | 0.103322035 | -0.130961679 | 0.895805622 | 0.982798755 | Sra1 | steroid receptor RNA activator 1 [Source:I | 1793.3 | 1732 | 1640.2 | 1792 | 1755.9 | 1569.4 |
| 943 | ENSMUSG00000006056 | 0 | NA | NA | NA | NA | NA | Calcoco2 | calcium binding and coiled-coil domain 2 [ | 0 | 0 | 0 | 0 | 0 | 0 |
| 944 | ENSMUSG00000006057 | 2965.588363 | -0.101859429 | 0.10522415 | -0.968023305 | 0.333032731 | 0.77523112 | Atp5g1 | ATP synthase, H+ transporting, mitochonc | 2936.1 | 3008.7 | 3266.3 | 2874.6 | 2592.4 | 3115.4 |
| 945 | ENSMUSG00000006058 | 614.6751967 | -0.087374624 | 0.121607523 | -0.718496867 | 0.472450979 | 0.868340109 | Snf8 | SNF8, ESCRT-II complex subunit, homolog | 658.54 | 581.8 | 659.09 | 603.45 | 573.67 | 611.5 |
| 946 | ENSMUSG00000006095 | 360.8234446 | 0.201156316 | 0.151813311 | 1.325024233 | 0.185163114 | 0.620815781 | Tbcb | tubulin folding cofactor B [Source:MGI Sy | 377.46 | 337.16 | 291.81 | 401.01 | 362.09 | 395.41 |
| 947 | ENSMUSG00000006127 | 280.4802295 | -0.21078545 | 0.164261116 | -1.283234005 | 0.199410106 | 0.641417519 | Inpp5k | inositol polyphosphate 5-phosphatase K [ | 341.32 | 267.26 | 293.82 | 283.41 | 247.57 | 249.5 |
| 948 | ENSMUSG00000006134 | 540.5943447 | -0.063296454 | 0.129409979 | -0.489115711 | 0.624759779 | 0.919503423 | Crkl | v-crk avian sarcoma virus CT10 oncogene | 605.54 | 529.38 | 521.23 | 497.41 | 544.22 | 545.78 |

| | | | | | | | | | | | | | | | |
|---|---|---|---|---|---|---|---|---|---|---|---|---|---|---|---|
| 949 | ENSMUSG00000006143 | 0 | NA | NA | NA | NA | NA | Upk3bl | uroplakin 3B-like [Source:MGI Symbol;Acc | 0 | 0 | 0 | 0 | 0 | 0 |
| 950 | ENSMUSG00000006154 | 34.16993665 | 0.254708816 | 0.608168897 | 0.418812632 | 0.675353072 | 0.932446243 | Eps8l1 | EPS8-like 1 [Source:MGI Symbol;Acc:MGI: | 59.43 | 15.419 | 18.112 | 14.46 | 71.981 | 25.618 |
| 951 | ENSMUSG00000006169 | 956.8375567 | 0.02290877 | 0.111154326 | 0.206098771 | 0.836713759 | 0.971073158 | Clint1 | clathrin interactor 1 [Source:MGI Symbol; | 909.91 | 988.86 | 948.89 | 890.72 | 995.73 | 1006.9 |
| 952 | ENSMUSG00000006179 | 0.319491021 | 0.16514334 | 4.044653141 | 0.040830038 | 0.967431393 | NA | Prss16 | protease, serine 16 (thymus) [Source:MGI | 0.8031 | 0 | 0 | 0 | 0 | 1.1138 |
| 953 | ENSMUSG00000006191 | 141.3816771 | 0.031882528 | 0.21330308 | 0.149470549 | 0.881182347 | 0.981549593 | Cdkal1 | CDK5 regulatory subunit associated protei | 154.2 | 125.41 | 139.87 | 166.77 | 142.87 | 119.18 |
| 954 | ENSMUSG00000006200 | 0 | NA | NA | NA | NA | NA | Rhox6 | reproductive homeobox 6 [Source:MGI Sy | 0 | 0 | 0 | 0 | 0 | 0 |
| 955 | ENSMUSG00000006204 | 0 | NA | NA | NA | NA | NA | Cdcp3 | CUB domain containing protein 3 [Source: | 0 | 0 | 0 | 0 | 0 | 0 |
| 956 | ENSMUSG00000006205 | 24.51188222 | -0.100054315 | 0.520857396 | -0.192095411 | 0.84766747 | 0.972803041 | Htra1 | HtrA serine peptidase 1 [Source:MGI Symb | 32.927 | 31.865 | 11.069 | 17.352 | 33.809 | 20.049 |
| 957 | ENSMUSG00000006215 | 201.5598315 | -0.101643862 | 0.191116707 | -0.531841846 | 0.594835537 | 0.913409341 | Zbtb17 | zinc finger and BTB domain containing 17 | 236.92 | 175.77 | 212.32 | 167.73 | 207.22 | 209.4 |
| 958 | ENSMUSG00000006216 | 0 | NA | NA | NA | NA | NA | Clcnkb | chloride channel, voltage-sensitive Kb [So | 0 | 0 | 0 | 0 | 0 | 0 |
| 959 | ENSMUSG00000006218 | 102.1340285 | -0.790643615 | 0.24057286 | -3.286503787 | 0.001014394 | 0.021080742 | Fam131c | family with sequence similarity 131, meml | 136.53 | 104.85 | 146.91 | 79.046 | 73.071 | 72.4 |
| 960 | ENSMUSG00000006219 | 16.86879695 | -1.022928026 | 0.556373315 | -1.83856414 | 0.06597932 | 0.368892493 | Fblim1 | filamin binding LIM protein 1 [Source:MGI | 28.109 | 21.586 | 18.112 | 13.496 | 6.5437 | 13.366 |
| 961 | ENSMUSG00000006221 | 0.825760208 | 0.783249908 | 2.404539989 | 0.325737942 | 0.744622635 | NA | Hspb7 | heat shock protein family, member 7 (carc | 0.8031 | 0 | 1.0062 | 0.964 | 2.1812 | 0 |
| 962 | ENSMUSG00000006235 | 19.0312999 | -0.877951087 | 0.560381213 | -1.566703284 | 0.117184063 | 0.502643331 | Epor | erythropoietin receptor [Source:MGI Sym | 22.487 | 17.475 | 34.212 | 20.244 | 13.087 | 6.6831 |
| 963 | ENSMUSG00000006241 | 87.7467302 | 0.025829194 | 0.291497495 | 0.088608632 | 0.929392948 | 0.988405116 | Ccdc159 | coiled-coil domain containing 159 [Source | 82.72 | 59.619 | 118.74 | 102.18 | 76.343 | 86.88 |
| 964 | ENSMUSG00000006262 | 956.5331883 | 0.137585823 | 0.127932726 | 1.075454473 | 0.282171305 | 0.735230549 | Mob1b | MOB kinase activator 1B [Source:MGI Syn | 807.92 | 1028.9 | 897.57 | 1066.2 | 1021.9 | 916.69 |
| 965 | ENSMUSG00000006269 | 0.449470461 | -0.545136533 | 3.562105917 | -0.15303771 | 0.878368537 | NA | Atp6v1b1 | ATPase, H+ transporting, lysosomal V1 sul | 1.6062 | 0 | 0 | 0 | 1.0906 | 0 |
| 966 | ENSMUSG00000006270 | 0 | NA | NA | NA | NA | NA | Vax1 | ventral anterior homeobox 1 [Source:MGI | 0 | 0 | 0 | 0 | 0 | 0 |
| 967 | ENSMUSG00000006273 | 743.1572922 | 0.099473221 | 0.125573626 | 0.792150587 | 0.428272877 | 0.842753542 | Atp6v1b2 | ATPase, H+ transporting, lysosomal V1 sul | 730.02 | 716.46 | 705.38 | 699.85 | 733.99 | 873.25 |
| 968 | ENSMUSG00000006276 | 240.8158099 | 0.0988716 | 0.159142808 | 0.621275952 | 0.534418073 | 0.889858671 | Eps15l1 | epidermal growth factor receptor pathwa | 236.11 | 230.25 | 231.44 | 264.13 | 230.12 | 252.84 |
| 969 | ENSMUSG00000006281 | 263.442754 | -0.089719106 | 0.154127821 | -0.582108445 | 0.560493637 | 0.899624792 | Tep1 | telomerase associated protein 1 [Source:I | 257.8 | 287.82 | 269.67 | 256.42 | 265.02 | 243.93 |
| 970 | ENSMUSG00000006288 | 165.5645186 | -0.037036959 | 0.19092139 | -0.19399062 | 0.846183227 | 0.972803041 | Ttc5 | tetratricopeptide repeat domain 5 [Source | 167.85 | 145.96 | 189.17 | 160.98 | 170.14 | 159.28 |
| 971 | ENSMUSG00000006289 | 279.7823768 | 0.119915841 | 0.191657415 | 0.625678066 | 0.531526123 | 0.888947136 | Osgep | O-sialoglycoprotein endopeptidase [Sourc | 255.39 | 304.26 | 244.52 | 233.28 | 277.02 | 364.23 |
| 972 | ENSMUSG00000006299 | 2426.845714 | 0.149292231 | 0.092224539 | 1.618790766 | 0.10549229 | 0.478619974 | Aamp | angio-associated migratory protein [Sourc | 2197.3 | 2345.7 | 2361.7 | 2464.9 | 2619.7 | 2571.9 |
| 973 | ENSMUSG00000006301 | 841.2672016 | 0.046948122 | 0.114723582 | 0.409228179 | 0.682372217 | 0.93502957 | Tmbim1 | transmembrane BAX inhibitor motif conta | 819.97 | 864.48 | 797.95 | 791.43 | 916.12 | 857.66 |
| 974 | ENSMUSG00000006304 | 1946.084387 | 0.065099689 | 0.091998404 | 0.707617585 | 0.479182769 | 0.870169438 | Arpc2 | actin related protein 2/3 complex, subuni | 1852.8 | 1884.2 | 1970.2 | 2019.5 | 1981.7 | 1968.2 |
| 975 | ENSMUSG00000006307 | 492.0811598 | 0.178036907 | 0.135646175 | 1.3125096 | 0.189348249 | 0.626663447 | Kmt2b | lysine (K)-specific methyltransferase 2B [S | 439.3 | 483.12 | 463.88 | 585.13 | 482.05 | 499 |
| 976 | ENSMUSG00000006310 | 33.65492612 | 0.094941084 | 0.410300619 | 0.231393957 | 0.81700876 | 0.967910123 | Zbtb32 | zinc finger and BTB domain containing 32 | 26.502 | 34.949 | 36.225 | 27.955 | 26.175 | 50.123 |
| 977 | ENSMUSG00000006311 | 0 | NA | NA | NA | NA | NA | Etv2 | ets variant 2 [Source:MGI Symbol;Acc:MG | 0 | 0 | 0 | 0 | 0 | 0 |
| 978 | ENSMUSG00000006313 | 0.465832987 | -0.698295221 | 3.18348759 | -0.219349126 | 0.826378096 | NA | Upk1a | uroplakin 1A [Source:MGI Symbol;Acc:MC | 0.8031 | 1.0279 | 0 | 0.964 | 0 | 0 |
| 979 | ENSMUSG00000006315 | 1089.443175 | 0.187032337 | 0.1133621 | 1.649866545 | 0.098970235 | 0.463570426 | Tmem147 | transmembrane protein 147 [Source:MGI | 1082.6 | 1035.1 | 937.82 | 1073.9 | 1191 | 1216.3 |
| 980 | ENSMUSG00000006333 | 4430.916624 | 0.085784416 | 0.100591434 | 0.852800409 | 0.393769997 | 0.822674956 | Rps9 | ribosomal protein S9 [Source:MGI Symbol | 3910.3 | 4656.5 | 4331.9 | 4364.9 | 4541.3 | 4780.6 |
| 981 | ENSMUSG00000006335 | 143.9381045 | 0.232099669 | 0.224168106 | 1.035382209 | 0.300490451 | 0.751875664 | Tfpt | TCF3 (E2A) fusion partner [Source:MGI Sy | 142.95 | 101.76 | 151.94 | 143.63 | 141.78 | 181.56 |
| 982 | ENSMUSG00000006342 | 27.2970415 | -0.48681398 | 0.415587327 | -1.171387931 | 0.241442881 | 0.691388521 | Susd2 | sushi domain containing 2 [Source:MGI Sy | 34.533 | 29.81 | 31.194 | 23.135 | 26.175 | 18.935 |
| 983 | ENSMUSG00000006344 | 42.5388027 | -0.852822739 | 0.36743459 | -2.321019202 | 0.020285808 | 0.174859309 | Ggt5 | gamma-glutamyltransferase 5 [Source:MC | 71.476 | 49.34 | 43.269 | 33.739 | 23.994 | 33.415 |
| 984 | ENSMUSG00000006345 | 0.520796785 | -0.820103173 | 3.072141206 | -0.266948398 | 0.789508892 | NA | Ggt1 | gamma-glutamyltransferase 1 [Source:MC | 0 | 1.0279 | 1.0062 | 0 | 1.0906 | 0 |
| 985 | ENSMUSG00000006356 | 724.9453478 | -0.702740483 | 0.190674978 | -3.685541173 | 0.000228217 | 0.006978717 | Crip2 | cysteine rich protein 2 [Source:MGI Symb | 654.53 | 867.56 | 1173.3 | 608.27 | 464.6 | 581.43 |
| 986 | ENSMUSG00000006360 | 43.98996756 | -0.201105689 | 0.341407269 | -0.58904923 | 0.55582825 | 0.897628185 | Crip1 | cysteine-rich protein 1 (intestinal) [Source | 51.399 | 54.48 | 35.219 | 42.415 | 41.443 | 38.985 |
| 987 | ENSMUSG00000006362 | 43.97276301 | -1.220359077 | 0.34394341 | -3.548139149 | 0.000387963 | 0.010477107 | Cbfa2t3 | CBFA2/RUNX1 translocation partner 3 [So | 67.461 | 55.508 | 61.381 | 22.171 | 28.356 | 28.96 |
| 988 | ENSMUSG00000006369 | 15.28589359 | -0.451960186 | 0.702555717 | -0.643308673 | 0.52002383 | 0.887802551 | Fbln1 | fibulin 1 [Source:MGI Symbol;Acc:MGI:95 | 26.502 | 16.447 | 10.062 | 21.208 | 15.269 | 2.2277 |
| 989 | ENSMUSG00000006373 | 21564.07696 | -0.098573748 | 0.084101342 | -1.172082933 | 0.241163761 | 0.691388521 | Pgrmc1 | progesterone receptor membrane compo | 22660 | 21572 | 22669 | 20409 | 21920 | 20154 |
| 990 | ENSMUSG00000006378 | 200.2239874 | 0.414322227 | 0.173044393 | 2.394311778 | 0.016651588 | 0.153190407 | Gcat | glycine C-acetyltransferase (2-amino-3-ke | 166.24 | 167.55 | 181.12 | 210.15 | 230.12 | 246.16 |
| 991 | ENSMUSG00000006386 | 100.1309637 | -0.488960714 | 0.247898979 | -1.972419234 | 0.048561768 | 0.306856582 | Tek | TEK receptor tyrosine kinase [Source:MGI | 119.66 | 121.29 | 109.68 | 80.01 | 65.437 | 104.7 |
| 992 | ENSMUSG00000006389 | 0.685278559 | -2.809870862 | 3.376578027 | -0.832165239 | 0.405315682 | NA | Mpl | myeloproliferative leukemia virus oncoger | 0 | 4.1117 | 0 | 0 | 0 | 0 |
| 993 | ENSMUSG00000006390 | 505.8514465 | 0.282970005 | 0.139349854 | 2.030644431 | 0.042291077 | 0.283193583 | Elovl1 | elongation of very long chain fatty acids (I | 502.74 | 431.73 | 433.69 | 542.72 | 506.05 | 618.18 |
| 994 | ENSMUSG00000006392 | 553.988478 | 0.006688464 | 0.136865622 | 0.048868835 | 0.961023825 | 0.992557454 | Med8 | mediator complex subunit 8 [Source:MGI | 570.2 | 538.63 | 548.4 | 498.38 | 527.86 | 640.46 |
| 995 | ENSMUSG00000006395 | 1347.723046 | 0.169426676 | 0.099066815 | 1.710226326 | 0.087224029 | 0.43548787 | Hyi | hydroxypyruvate isomerase (putative) [So | 1286.6 | 1306.5 | 1212.5 | 1383.3 | 1448.3 | 1449.1 |
| 996 | ENSMUSG00000006398 | 11.89853353 | -0.365158352 | 0.711638261 | -0.513123551 | 0.607864898 | 0.916376001 | Cdc20 | cell division cycle 20 [Source:MGI Symbol; | 25.699 | 8.2233 | 6.0375 | 11.568 | 8.7249 | 11.138 |
| 997 | ENSMUSG00000006403 | 2.187827562 | -0.978872854 | 1.716172621 | -0.570381349 | 0.568419077 | NA | Adamts4 | a disintegrin-like and metallopeptidase (re | 1.6062 | 1.0279 | 6.0375 | 0 | 0 | 4.4554 |
| 998 | ENSMUSG00000006411 | 2.342144914 | -0.752225548 | 1.703000749 | -0.441705941 | 0.65870201 | NA | Nectin4 | nectin cell adhesion molecule 4 [Source:N | 0.8031 | 0 | 8.05 | 1.928 | 3.2719 | 0 |
| 999 | ENSMUSG00000006412 | 731.3626709 | 0.108758522 | 0.11872138 | 0.916082022 | 0.359623876 | 0.799452417 | Pfdn2 | prefoldin 2 [Source:MGI Symbol;Acc:MGI: | 760.54 | 683.57 | 666.13 | 721.06 | 797.24 | 759.64 |
| 1000 | ENSMUSG00000006418 | 1233.838559 | 0.318083039 | 0.105142313 | 3.025261959 | 0.002484179 | 0.041111167 | Rnf114 | ring finger protein 114 [Source:MGI Symb | 1146.8 | 1009.4 | 1138.1 | 1364 | 1351.3 | 1393.4 |
| 1001 | ENSMUSG00000006423 | 776.6493956 | -0.071696244 | 0.109914335 | -0.652292025 | 0.514212804 | 0.885004045 | C330007P06 | RIKEN cDNA C330007P06 gene [Source:M | 791.86 | 781.22 | 815.06 | 784.68 | 727.44 | 759.64 |
| 1002 | ENSMUSG00000006435 | 4.948032523 | -0.783374173 | 1.085613822 | -0.721595614 | 0.470543136 | NA | Neurl1a | neuralized E3 ubiquitin protein ligase 1A [ | 2.4093 | 12.335 | 4.025 | 0.964 | 3.2719 | 6.6831 |
| 1003 | ENSMUSG00000006442 | 531.21686 | 0.14508285 | 0.135036954 | 1.074393677 | 0.282646275 | 0.735551588 | Srm | spermidine synthase [Source:MGI Symbol | 538.08 | 483.12 | 492.05 | 576.46 | 606.38 | 491.21 |
| 1004 | ENSMUSG00000006445 | 624.1272739 | -1.025984482 | 0.512180178 | -2.003171007 | 0.045158936 | 0.294092237 | Epha2 | Eph receptor A2 [Source:MGI Symbol;Acc | 748.49 | 765.8 | 997.19 | 364.38 | 675.09 | 193.81 |

| | | | | | | | | | | | | | | | |
|---|---|---|---|---|---|---|---|---|---|---|---|---|---|---|---|
| 1005 | ENSMUSG00000006456 | 182.4695307 | -0.195564464 | 0.183816787 | -1.063909706 | 0.287369606 | 0.73951397 | Rbm14 | RNA binding motif protein 14 [Source:MG | 214.43 | 194.28 | 175.09 | 157.13 | 171.23 | 182.67 |
| 1006 | ENSMUSG00000006457 | 0.722877068 | -0.331317305 | 2.989821669 | -0.110815073 | 0.911762993 | NA | Actn3 | actinin alpha 3 [Source:MGI Symbol;Acc:M | 2.4093 | 0 | 0 | 1.928 | 0 | 0 |
| 1007 | ENSMUSG00000006462 | 14.66743542 | 0.511192697 | 0.586818818 | 0.871125262 | 0.383685762 | 0.816002185 | A530013C23 | RIKEN cDNA A530013C23 gene [Source:M | 12.85 | 11.307 | 12.075 | 14.46 | 26.175 | 11.138 |
| 1008 | ENSMUSG00000006463 | 62.13707129 | 0.241869051 | 0.296129282 | 0.816768437 | 0.414060773 | 0.835748677 | Zdhhc24 | zinc finger, DHHC domain containing 24 [S | 62.642 | 55.508 | 52.325 | 58.803 | 58.893 | 84.652 |
| 1009 | ENSMUSG00000006464 | 21.4657731 | -0.652777894 | 0.483928429 | -1.348914127 | 0.17736455 | 0.608561375 | Bbs1 | Bardet-Biedl syndrome 1 (human) [Source | 21.684 | 30.838 | 26.162 | 11.568 | 20.722 | 17.822 |
| 1010 | ENSMUSG00000006469 | 0.17131964 | -0.796629176 | 4.080472857 | -0.195229623 | 0.84521319 | NA | Slc34a3 | solute carrier family 34 (sodium phosphat | 0 | 1.0279 | 0 | 0 | 0 | 0 |
| 1011 | ENSMUSG00000006471 | 72.88369016 | -0.137233413 | 0.290294771 | -0.472738151 | 0.636400004 | 0.923478764 | Ndor1 | NADPH dependent diflavin oxidoreductas | 69.067 | 68.87 | 91.568 | 88.686 | 55.622 | 63.489 |
| 1012 | ENSMUSG00000006476 | 1991.49098 | 0.227805328 | 0.099245567 | 2.295370308 | 0.021711911 | 0.183390105 | Nsmf | NMDA receptor synaptonuclear signaling | 1894.5 | 1684.8 | 1923.9 | 2176.7 | 2106 | 2163.1 |
| 1013 | ENSMUSG00000006488 | 0 | NA | NA | NA | NA | NA | Prl7a1 | prolactin family 7, subfamily a, member 1 | 0 | 0 | 0 | 0 | 0 | 0 |
| 1014 | ENSMUSG00000006490 | 0 | NA | NA | NA | NA | NA | Prl8a9 | prolactin family8, subfamily a, member 9 [ | 0 | 0 | 0 | 0 | 0 | 0 |
| 1015 | ENSMUSG00000006494 | 2416.519659 | -0.330061932 | 0.104436171 | -3.160417795 | 0.001575431 | 0.029454792 | Pdk1 | pyruvate dehydrogenase kinase, isoenzym | 2544.2 | 2853.5 | 2678.6 | 2200.8 | 2276.1 | 1945.9 |
| 1016 | ENSMUSG00000006498 | 1066.507262 | 0.189866049 | 0.113043938 | 1.679577455 | 0.093039557 | 0.449345765 | Ptbp1 | polypyrimidine tract binding protein 1 [So | 1068.9 | 1025.9 | 893.55 | 1121.1 | 1120.1 | 1169.5 |
| 1017 | ENSMUSG00000006517 | 97.74849391 | 1.355836393 | 0.311229534 | 4.356387312 | 1.32E-05 | 0.000675805 | Mvd | mevalonate (diphospho) decarboxylase [S | 72.279 | 53.452 | 38.237 | 110.86 | 114.51 | 197.15 |
| 1018 | ENSMUSG00000006519 | 96.05936125 | -1.346399325 | 0.388453233 | -3.466052564 | 0.00052816 | 0.012909222 | Cyba | cytochrome b-245, alpha polypeptide [Sou | 120.47 | 196.33 | 96.599 | 36.631 | 30.537 | 95.791 |
| 1019 | ENSMUSG00000006522 | 12141.02705 | 0.089365733 | 0.107864653 | 0.828498771 | 0.40738809 | 0.830977484 | Itih3 | inter-alpha trypsin inhibitor, heavy chain 3 | 10286 | 11982 | 13029 | 13234 | 12335 | 11979 |
| 1020 | ENSMUSG00000006526 | 319.280886 | -0.04163751 | 0.152709746 | -0.27265784 | 0.785116246 | 0.960670212 | Stimate | STIM activating enhancer [Source:MGI Sy | 354.17 | 334.07 | 282.75 | 305.58 | 325 | 314.1 |
| 1021 | ENSMUSG00000006527 | 560.5486323 | 0.112519837 | 0.119294859 | 0.943207767 | 0.345574642 | 0.786899236 | Sfmbt1 | Scm-like with four mbt domains 1 [Source | 526.03 | 538.63 | 551.42 | 560.07 | 585.66 | 601.48 |
| 1022 | ENSMUSG00000006529 | 9301.987546 | 0.158532008 | 0.095533937 | 1.659431333 | 0.097028907 | 0.458785913 | Itih1 | inter-alpha trypsin inhibitor, heavy chain 1 | 8003.7 | 9082.7 | 9289.7 | 10122 | 9359.7 | 9954.4 |
| 1023 | ENSMUSG00000006538 | 26.56588083 | 0.753700097 | 0.438349192 | 1.719405694 | 0.085540526 | 0.430973029 | Ihh | Indian hedgehog [Source:MGI Symbol;Acc | 16.865 | 15.419 | 27.169 | 33.739 | 29.447 | 36.757 |
| 1024 | ENSMUSG00000006542 | 6.184565708 | -2.595440119 | 1.013677998 | -2.560418718 | 0.010454612 | 0.112094516 | Prkag3 | protein kinase, AMP-activated, gamma 3 n | 6.4248 | 14.391 | 11.069 | 1.928 | 2.1812 | 1.1138 |
| 1025 | ENSMUSG00000006546 | 0 | NA | NA | NA | NA | NA | Cryba2 | crystallin, beta A2 [Source:MGI Symbol;Ac | 0 | 0 | 0 | 0 | 0 | 0 |
| 1026 | ENSMUSG00000006567 | 776.1214881 | -0.068519945 | 0.124602351 | -0.549908929 | 0.58238184 | 0.908645447 | Atp7b | ATPase, Cu++ transporting, beta polypept | 716.37 | 867.56 | 800.97 | 782.75 | 790.7 | 698.38 |
| 1027 | ENSMUSG00000006570 | 0 | NA | NA | NA | NA | NA | Defb2 | defensin beta 2 [Source:MGI Symbol;Acc: | 0 | 0 | 0 | 0 | 0 | 0 |
| 1028 | ENSMUSG00000006574 | 5.55438019 | -1.832535794 | 0.990648346 | -1.849834808 | 0.064337362 | 0.363364158 | Slc4a1 | solute carrier family 4 (anion exchanger), | 8.8341 | 5.1396 | 12.075 | 2.8919 | 3.2719 | 1.1138 |
| 1029 | ENSMUSG00000006575 | 314.7632574 | 0.162070371 | 0.157209825 | 1.030917571 | 0.302579476 | 0.752385931 | Rundc3a | RUN domain containing 3A [Source:MGI S | 323.65 | 260.06 | 306.9 | 322.93 | 364.27 | 310.76 |
| 1030 | ENSMUSG00000006576 | 7.901169667 | -0.738670736 | 0.782278806 | -0.94425508 | 0.345039315 | 0.786176536 | Slc4a3 | solute carrier family 4 (anion exchanger), | 6.4248 | 9.2513 | 14.087 | 7.7118 | 4.3625 | 5.5692 |
| 1031 | ENSMUSG00000006585 | 60.96845587 | 0.538808459 | 0.323651144 | 1.664781568 | 0.095956343 | 0.457453045 | Cdt1 | chromatin licensing and DNA replication f | 55.414 | 44.2 | 49.306 | 74.226 | 47.987 | 94.677 |
| 1032 | ENSMUSG00000006586 | 1.772108237 | -4.187305224 | 2.19492838 | -1.907718384 | 0.056427624 | NA | Runx1t1 | RUNX1 translocation partner 1 [Source:M | 2.4093 | 8.2233 | 0 | 0 | 0 | 0 |
| 1033 | ENSMUSG00000006587 | 1.711749964 | 2.202100797 | 1.919805976 | 1.147043412 | 0.251363679 | NA | Snai3 | snail family zinc finger 3 [Source:MGI Sym | 0.8031 | 0 | 1.0062 | 2.8919 | 0 | 5.5692 |
| 1034 | ENSMUSG00000006589 | 464.9971659 | 0.144536871 | 0.171309831 | 0.843716149 | 0.398828041 | 0.82594896 | Aprt | adenine phosphoribosyl transferase [Sour | 456.97 | 422.47 | 444.76 | 378.84 | 478.78 | 608.16 |
| 1035 | ENSMUSG00000006599 | 659.3408658 | -0.073853968 | 0.114810731 | -0.643267117 | 0.520050789 | 0.887802551 | Gtf2h1 | general transcription factor II H, polypepti | 668.98 | 693.84 | 666.13 | 653.58 | 662.01 | 611.5 |
| 1036 | ENSMUSG00000006611 | 770.2171488 | -0.088325934 | 0.129808126 | -0.680434549 | 0.496229351 | 0.878165817 | Hfe | homeostatic iron regulator [Source:MGI S | 873.78 | 805.89 | 700.35 | 681.53 | 766.7 | 793.06 |
| 1037 | ENSMUSG00000006638 | 6.011675668 | -0.458808039 | 1.017034346 | -0.451123446 | 0.65190058 | 0.925643431 | Abhd1 | abhydrolase domain containing 1 [Source | 8.8341 | 1.0279 | 11.069 | 9.6398 | 3.2719 | 2.2277 |
| 1038 | ENSMUSG00000006641 | 306.41082 | 0.745165581 | 0.222022903 | 3.356255469 | 0.000790056 | 0.017553553 | Slc5a6 | solute carrier family 5 (sodium-dependent | 302.77 | 222.03 | 161 | 367.28 | 464.6 | 320.79 |
| 1039 | ENSMUSG00000006642 | 3.021063743 | -0.109484735 | 1.45444761 | -0.075275819 | 0.93999526 | NA | Tcf23 | transcription factor 23 [Source:MGI Symb | 7.2279 | 1.0279 | 1.0062 | 0 | 2.1812 | 6.6831 |
| 1040 | ENSMUSG00000006649 | 0 | NA | NA | NA | NA | NA | Nphs1 | nephrosis 1, nephrin [Source:MGI Symbol | 0 | 0 | 0 | 0 | 0 | 0 |
| 1041 | ENSMUSG00000006651 | 3.880661547 | 0.542151456 | 1.0702638 | 0.506558715 | 0.612464482 | NA | Aplp1 | amyloid beta (A4) precursor-like protein 1 | 2.4093 | 2.0558 | 5.0312 | 3.8559 | 4.3625 | 5.5692 |
| 1042 | ENSMUSG00000006673 | 708.5797287 | 0.09378206 | 0.122271064 | 0.767001261 | 0.443080766 | 0.850769625 | Qrich1 | glutamine-rich 1 [Source:MGI Symbol;Acc | 697.9 | 724.68 | 633.93 | 718.16 | 791.79 | 685.01 |
| 1043 | ENSMUSG00000006675 | 2.746376187 | 1.137426797 | 1.393591899 | 0.81618356 | 0.414395158 | NA | P4htm | prolyl 4-hydroxylase, transmembrane (end | 4.0155 | 0 | 1.0062 | 4.8199 | 2.1812 | 4.4554 |
| 1044 | ENSMUSG00000006676 | 1312.442207 | 0.00015699 | 0.110585794 | 0.001419618 | 0.998867309 | 0.999764475 | Usp19 | ubiquitin specific peptidase 19 [Source:M | 1268.9 | 1377.4 | 1292 | 1445 | 1219.3 | 1272 |
| 1045 | ENSMUSG00000006678 | 59.76408905 | -0.18408485 | 0.304118059 | -0.605307198 | 0.544974899 | 0.8954353 | Pola1 | polymerase (DNA directed), alpha 1 [Sour | 66.657 | 60.647 | 63.393 | 56.875 | 69.8 | 41.212 |
| 1046 | ENSMUSG00000006699 | 2204.600902 | -0.015970726 | 0.103763104 | -0.153915268 | 0.877676545 | 0.98072613 | Cdc42 | cell division cycle 42 [Source:MGI Symbol; | 2374.8 | 2222.4 | 2051.7 | 2050.4 | 2221.6 | 2306.8 |
| 1047 | ENSMUSG00000006705 | 277.143567 | 0.296930771 | 0.163653653 | 1.81438523 | 0.069618438 | 0.381135916 | Pknox1 | Pbx/knotted 1 homeobox [Source:MGI Sy | 261.81 | 221 | 262.63 | 276.66 | 299.92 | 340.84 |
| 1048 | ENSMUSG00000006711 | 97.10152164 | 0.241884915 | 0.250097434 | 0.967162723 | 0.333462694 | 0.775366061 | D130043K22 | RIKEN cDNA D130043K22 gene [Source:M | 110.83 | 79.15 | 76.475 | 116.64 | 98.156 | 101.36 |
| 1049 | ENSMUSG00000006715 | 200.116666 | 0.135010393 | 0.193681285 | 0.697075054 | 0.485755825 | 0.872998689 | Gmnn | geminin [Source:MGI Symbol;Acc:MGI:19 | 209.61 | 203.53 | 158.99 | 231.35 | 176.68 | 220.54 |
| 1050 | ENSMUSG00000006717 | 867.172086 | -0.165696735 | 0.116891451 | -1.417526556 | 0.156329036 | 0.574915631 | Acot13 | acyl-CoA thioesterase 13 [Source:MGI Sym | 929.99 | 939.52 | 881.47 | 844.44 | 873.59 | 734.02 |
| 1051 | ENSMUSG00000006720 | 3.009458113 | -0.480375295 | 1.255759954 | -0.382537517 | 0.702062706 | NA | Zfp184 | zinc finger protein 184 (Kruppel-like) [Sou | 2.4093 | 2.0558 | 6.0375 | 0.964 | 4.3625 | 2.2277 |
| 1052 | ENSMUSG00000006724 | 0.36353943 | 2.065989254 | 4.034137301 | 0.512126658 | 0.608562368 | NA | Cyp27b1 | cytochrome P450, family 27, subfamily b, | 0 | 0 | 0 | 0 | 2.1812 | 0 |
| 1053 | ENSMUSG00000006728 | 593.5278162 | 0.056418642 | 0.119282015 | 0.472985317 | 0.636223655 | 0.923478764 | Cdk4 | cyclin-dependent kinase 4 [Source:MGI Sy | 607.95 | 585.91 | 551.42 | 605.38 | 586.75 | 623.75 |
| 1054 | ENSMUSG00000006731 | 1158.674737 | -0.017938623 | 0.138086767 | -0.129908343 | 0.896638943 | 0.982798755 | B4galnt1 | beta-1,4-N-acetyl-galactosaminyl transfer | 1039.2 | 1179 | 1279.9 | 1119.2 | 990.28 | 1344.4 |
| 1055 | ENSMUSG00000006732 | 176.5236183 | 0.086056954 | 0.184002894 | 0.467693482 | 0.640003803 | 0.923932929 | Mettl1 | methyltransferase like 1 [Source:MGI Sym | 176.68 | 174.75 | 162.01 | 170.62 | 170.14 | 204.95 |
| 1056 | ENSMUSG00000006736 | 3070.233636 | 0.07800139 | 0.105876978 | 0.736717195 | 0.461294348 | 0.860071824 | Tspan31 | tetraspanin 31 [Source:MGI Symbol;Acc:M | 3288.7 | 2923.4 | 2748.1 | 3025 | 3376.6 | 3059.7 |
| 1057 | ENSMUSG00000006740 | 945.3336585 | -0.07243721 | 0.105470919 | -0.686797941 | 0.492210068 | 0.876368691 | Kif5b | kinesin family member 5B [Source:MGI Sy | 950.07 | 1006.3 | 950.9 | 907.1 | 903.03 | 954.56 |
| 1058 | ENSMUSG00000006763 | 105.2228264 | 0.008272425 | 0.238908687 | 0.034625884 | 0.972378061 | 0.995715135 | Saal1 | serum amyloid A-like 1 [Source:MGI Symb | 88.341 | 113.07 | 113.71 | 98.326 | 124.33 | 93.563 |
| 1059 | ENSMUSG00000006764 | 0 | NA | NA | NA | NA | NA | Tph2 | tryptophan hydroxylase 2 [Source:MGI Sy | 0 | 0 | 0 | 0 | 0 | 0 |
| 1060 | ENSMUSG00000006777 | 116.0556733 | -0.138842634 | 0.262310872 | -0.529305678 | 0.596593418 | 0.914226274 | Krt23 | keratin 23 [Source:MGI Symbol;Acc:MGI:2 | 131.71 | 98.68 | 133.83 | 75.19 | 122.15 | 134.78 |

| | | | | | | | | | | | | | | | |
|---|---|---|---|---|---|---|---|---|---|---|---|---|---|---|---|
| 1061 | ENSMUSG00000006782 | 240.0748272 | 0.320063317 | 0.188585059 | 1.697182794 | 0.089662107 | 0.441714141 | Cnp | 2',3'-cyclic nucleotide 3' phosphodiesteras | 239.32 | 180.91 | 219.36 | 220.75 | 271.56 | 308.53 |
| 1062 | ENSMUSG00000006784 | 25.4760103 | -1.27422043 | 0.572662018 | -2.225082841 | 0.026075677 | 0.208952132 | Odad4 | outer dynein arm complex subunit 4 [Sour | 65.051 | 19.53 | 23.144 | 8.6758 | 14.178 | 22.277 |
| 1063 | ENSMUSG00000006800 | 420.784385 | -0.363808918 | 0.163796058 | -2.221109114 | 0.026343572 | 0.210051502 | Sulf2 | sulfatase 2 [Source:MGI Symbol;Acc:MGI: | 550.93 | 390.61 | 477.97 | 345.1 | 344.64 | 415.46 |
| 1064 | ENSMUSG00000006818 | 4405.770972 | -0.172268395 | 0.097120627 | -1.773757039 | 0.076103347 | 0.403877794 | Sod2 | superoxide dismutase 2, mitochondrial [S | 4416.3 | 4677 | 4913.5 | 4298.4 | 4311.2 | 3818.3 |
| 1065 | ENSMUSG00000006850 | 110.3416352 | 0.389144874 | 0.242615639 | 1.603956262 | 0.108723694 | 0.485684837 | Tmco6 | transmembrane and coiled-coil domains 6 | 94.766 | 85.317 | 106.66 | 137.85 | 94.884 | 142.57 |
| 1066 | ENSMUSG00000006906 | 215.1856047 | 0.191054248 | 0.169416593 | 1.127718627 | 0.259438768 | 0.712728614 | Stambp | STAM binding protein [Source:MGI Symbo | 185.52 | 212.78 | 205.27 | 247.74 | 217.03 | 222.77 |
| 1067 | ENSMUSG00000006920 | 286.8089347 | 0.278240545 | 0.169558122 | 1.640974444 | 0.10080272 | 0.46735231 | Ezh1 | enhancer of zeste 1 polycomb repressive | 248.16 | 251.84 | 277.72 | 301.73 | 269.38 | 372.02 |
| 1068 | ENSMUSG00000006930 | 27.10755768 | -0.93190237 | 0.429373366 | -2.170377681 | 0.029978246 | 0.22888973 | Hap1 | huntingtin-associated protein 1 [Source:M | 34.533 | 32.893 | 39.244 | 16.388 | 23.994 | 15.594 |
| 1069 | ENSMUSG00000006931 | 18.33300039 | -0.513797517 | 0.51240202 | -1.002723442 | 0.315994316 | 0.762646719 | P3h4 | prolyl 3-hydroxylase family member 4 (no | 20.078 | 16.447 | 28.175 | 14.46 | 16.359 | 14.48 |
| 1070 | ENSMUSG00000006932 | 2969.22807 | 0.113892976 | 0.092888203 | 1.226129607 | 0.220149906 | 0.666439564 | Ctnnb1 | catenin (cadherin associated protein), bet | 2713.7 | 2848.4 | 2994.6 | 2957.5 | 3126.8 | 3174.5 |
| 1071 | ENSMUSG00000006941 | 347.9511987 | -0.202499766 | 0.15191842 | -1.332950705 | 0.182547981 | 0.616333801 | Eif1b | eukaryotic translation initiation factor 1B | 335.7 | 416.31 | 365.27 | 295.94 | 337 | 337.49 |
| 1072 | ENSMUSG00000006948 | 0 | NA | NA | NA | NA | NA | Klk4 | kallikrein related-peptidase 4 (prostase, e | 0 | 0 | 0 | 0 | 0 | 0 |
| 1073 | ENSMUSG00000006958 | 136.9749289 | 0.579667217 | 0.201269768 | 2.880051097 | 0.003976107 | 0.056970643 | Chrd | chordin [Source:MGI Symbol;Acc:MGI:13 | 97.978 | 117.18 | 114.71 | 159.06 | 162.5 | 170.42 |
| 1074 | ENSMUSG00000006998 | 3015.710793 | 0.170380295 | 0.10520277 | 1.619541904 | 0.105330718 | 0.478186913 | Psmd2 | proteasome (prosome, macropain) 26S su | 3107.2 | 2813.4 | 2591.1 | 2986.4 | 3295.8 | 3300.3 |
| 1075 | ENSMUSG00000007021 | 2.398625843 | -0.217212459 | 1.622552918 | -0.133870801 | 0.89350474 | NA | Syngr3 | synaptogyrin 3 [Source:MGI Symbol;Acc:N | 1.6062 | 3.0838 | 3.0187 | 0 | 0 | 6.6831 |
| 1076 | ENSMUSG00000007029 | 596.1258026 | -0.175082203 | 0.122581882 | -1.428287773 | 0.153209046 | 0.570109503 | Vars | valyl-tRNA synthetase [Source:MGI Symbo | 651.32 | 599.28 | 646.01 | 567.78 | 592.21 | 520.17 |
| 1077 | ENSMUSG00000007030 | 4.681068258 | -0.5928078 | 1.225319908 | -0.483798391 | 0.628528957 | NA | Vwa7 | von Willebrand factor A domain containin | 0.8031 | 6.1675 | 10.062 | 7.7118 | 0 | 3.3415 |
| 1078 | ENSMUSG00000007033 | 8.466376199 | -0.065524968 | 0.772290461 | -0.084844979 | 0.932384635 | 0.988420305 | Hspa1l | heat shock protein 1-like [Source:MGI Syn | 8.8341 | 4.1117 | 13.081 | 11.568 | 7.6343 | 5.5692 |
| 1079 | ENSMUSG00000007034 | 3.255384302 | 1.068687155 | 1.248455668 | 0.856007291 | 0.391993746 | NA | Slc44a4 | solute carrier family 44, member 4 [Sourc | 3.2124 | 2.0558 | 1.0062 | 7.7118 | 1.0906 | 4.4554 |
| 1080 | ENSMUSG00000007035 | 11.61524915 | 0.442274668 | 0.665133904 | 0.664940796 | 0.506088361 | 0.881592135 | Msh5 | mutS homolog 5 [Source:MGI Symbol;Acc | 11.243 | 4.1117 | 14.087 | 11.568 | 13.087 | 15.594 |
| 1081 | ENSMUSG00000007036 | 823.4548436 | -0.044049745 | 0.121205937 | -0.363428941 | 0.716284486 | 0.9447719 | Abhd16a | abhydrolase domain containing 16A [Sour | 828 | 868.59 | 811.03 | 750.94 | 772.16 | 910.01 |
| 1082 | ENSMUSG00000007038 | 819.9633319 | 0.208276073 | 0.114390164 | 1.820751582 | 0.068644627 | 0.377662947 | Neu1 | neuraminidase 1 [Source:MGI Symbol;Acc | 800.69 | 761.69 | 719.46 | 864.69 | 836.5 | 936.74 |
| 1083 | ENSMUSG00000007039 | 40.70428434 | -0.495790138 | 0.360492598 | -1.375312949 | 0.169034444 | 0.596320535 | Ddah2 | dimethylarginine dimethylaminohydrolase | 50.595 | 37.005 | 55.343 | 38.559 | 35.99 | 26.732 |
| 1084 | ENSMUSG00000007041 | 186.6245127 | -0.220267768 | 0.194998083 | -1.129589403 | 0.258649279 | 0.712165697 | Clic1 | chloride intracellular channel 1 [Source:M | 207.2 | 216.89 | 178.11 | 160.98 | 150.51 | 206.06 |
| 1085 | ENSMUSG00000007050 | 208.6423425 | 0.248584617 | 0.212234473 | 1.17127351 | 0.241488856 | 0.691388521 | Lsm2 | LSM2 homolog, U6 small nuclear RNA and | 208.81 | 205.58 | 156.97 | 178.34 | 218.12 | 284.03 |
| 1086 | ENSMUSG00000007080 | 35.02734806 | -0.116098889 | 0.437506939 | -0.265364681 | 0.790728542 | 0.961019028 | Pole | polymerase (DNA directed), epsilon [Sour | 43.368 | 16.447 | 49.306 | 32.775 | 37.081 | 31.188 |
| 1087 | ENSMUSG00000007097 | 12.06268112 | -0.537761156 | 0.638504807 | -0.842219431 | 0.399665139 | 0.826649016 | Atp1a2 | ATPase, Na+/K+ transporting, alpha 2 poly | 19.274 | 13.363 | 10.062 | 8.6758 | 7.6343 | 13.366 |
| 1088 | ENSMUSG00000007107 | 0.995767881 | -1.008676959 | 2.44979061 | -0.411740071 | 0.680529951 | NA | Atp1a4 | ATPase, Na+/K+ transporting, alpha 4 poly | 0 | 1.0279 | 3.0187 | 1.928 | 0 | 0 |
| 1089 | ENSMUSG00000007122 | 2.735926112 | 0.7453132 | 1.335239042 | 0.558187093 | 0.576716633 | NA | Casq1 | calsequestrin 1 [Source:MGI Symbol;Acc:I | 4.0155 | 1.0279 | 1.0062 | 4.8199 | 1.0906 | 4.4554 |
| 1090 | ENSMUSG00000007207 | 9.750091705 | -1.188707418 | 0.737256709 | -1.612338556 | 0.106888288 | 0.482381339 | Stx1a | syntaxin 1A (brain) [Source:MGI Symbol;A | 19.274 | 7.1954 | 14.087 | 5.7839 | 4.3625 | 7.7969 |
| 1091 | ENSMUSG00000007209 | 0.181769715 | 1.126924175 | 4.080472857 | 0.276174898 | 0.782413736 | NA | Ceacam9 | carcinoembryonic antigen-related cell adh | 0 | 0 | 0 | 0 | 1.0906 | 0 |
| 1092 | ENSMUSG00000007216 | 55.28789645 | -0.140173867 | 0.315460521 | -0.44434678 | 0.656791885 | 0.9268505 | Zfp775 | zinc finger protein 775 [Source:MGI Symb | 44.974 | 64.759 | 64.4 | 45.307 | 62.165 | 50.123 |
| 1093 | ENSMUSG00000007279 | 0 | NA | NA | NA | NA | NA | Scube2 | signal peptide, CUB domain, EGF-like 2 [S | 0 | 0 | 0 | 0 | 0 | 0 |
| 1094 | ENSMUSG00000007338 | 640.819117 | 0.022913705 | 0.123914872 | 0.184914893 | 0.853295789 | 0.973825032 | Mrpl49 | mitochondrial ribosomal protein L49 [Sou | 665.77 | 618.81 | 621.86 | 609.23 | 712.17 | 617.07 |
| 1095 | ENSMUSG00000007379 | 5.643845195 | -0.420055471 | 0.890511788 | -0.471701191 | 0.637140087 | 0.923478764 | Dennd2c | DENN/MADD domain containing 2C [Sour | 7.2279 | 5.1396 | 7.0437 | 6.7478 | 4.3625 | 3.3415 |
| 1096 | ENSMUSG00000007411 | 785.8632377 | -0.127820729 | 0.109847763 | -1.16361704 | 0.244579243 | 0.694577264 | Mark3 | MAP/microtubule affinity regulating kinas | 797.48 | 850.09 | 815.06 | 777.93 | 737.26 | 737.36 |
| 1097 | ENSMUSG00000007415 | 1036.295253 | 0.065307187 | 0.116085019 | 0.56258066 | 0.573720464 | 0.905828271 | Gatad1 | GATA zinc finger domain containing 1 [So | 910.72 | 1059.8 | 1069.6 | 1127.9 | 1031.7 | 1018.1 |
| 1098 | ENSMUSG00000007440 | 0.342639279 | -1.805976596 | 4.028547714 | -0.448294702 | 0.653940524 | NA | Pcdha11 | protocadherin alpha 11 [Source:MGI Sym | 0 | 2.0558 | 0 | 0 | 0 | 0 |
| 1099 | ENSMUSG00000007457 | 0 | NA | NA | NA | NA | NA | 2310003L06 | RIKEN cDNA 2310003L06 gene [Source:M | 0 | 0 | 0 | 0 | 0 | 0 |
| 1100 | ENSMUSG00000007458 | 1899.814679 | 0.046377061 | 0.09807915 | 0.472853411 | 0.636317766 | 0.923478764 | M6pr | mannose-6-phosphate receptor, cation de | 1894.5 | 1879 | 1833.4 | 1789.1 | 2003.5 | 1999.3 |
| 1101 | ENSMUSG00000007476 | 159.9437029 | -0.217157346 | 0.189887979 | -1.143607655 | 0.252786385 | 0.705781321 | Lrrc8a | leucine rich repeat containing 8A VRAC su | 191.94 | 162.41 | 161 | 144.6 | 152.69 | 147.03 |
| 1102 | ENSMUSG00000007480 | 0 | NA | NA | NA | NA | NA | Mc5r | melanocortin 5 receptor [Source:MGI Sym | 0 | 0 | 0 | 0 | 0 | 0 |
| 1103 | ENSMUSG00000007564 | 1046.18731 | -0.020541902 | 0.105236485 | -0.19519753 | 0.845238313 | 0.972803041 | Ppp2r1a | protein phosphatase 2, regulatory subuni | 1113.9 | 1026.9 | 1019.3 | 1032.4 | 1071 | 1013.6 |
| 1104 | ENSMUSG00000007570 | 253.0268393 | 0.186680373 | 0.17394639 | 1.073206363 | 0.283178537 | 0.735802616 | Fance | Fanconi anemia, complementation group | 220.05 | 211.75 | 278.73 | 283.41 | 280.29 | 243.93 |
| 1105 | ENSMUSG00000007589 | 194.658815 | 0.068994759 | 0.193361496 | 0.356817465 | 0.721228461 | 0.9447719 | Tinf2 | Terf1 (TRF1)-interacting nuclear factor 2 [ | 204.79 | 167.55 | 197.22 | 197.62 | 166.86 | 233.91 |
| 1106 | ENSMUSG00000007591 | 1.188162118 | 1.685781406 | 2.287800744 | 0.736856744 | 0.461209472 | NA | Tssk4 | testis-specific serine kinase 4 [Source:MG | 1.6062 | 0 | 0 | 0 | 2.1812 | 3.3415 |
| 1107 | ENSMUSG00000007594 | 96.47203443 | 0.091184107 | 0.250440712 | 0.364094584 | 0.715787381 | 0.9447719 | Hapln4 | hyaluronan and proteoglycan link protein | 83.523 | 86.345 | 110.69 | 105.07 | 80.706 | 112.5 |
| 1108 | ENSMUSG00000007603 | 307.5515636 | 0.104298292 | 0.166170377 | 0.627658758 | 0.530227508 | 0.888565713 | Dus3l | dihydrouridine synthase 3-like (S. cerevisi | 281.89 | 315.57 | 292.82 | 379.81 | 291.2 | 284.03 |
| 1109 | ENSMUSG00000007610 | 181.0859088 | 0.052814603 | 0.181836248 | 0.290451455 | 0.771470884 | 0.956965056 | Gtpbp3 | GTP binding protein 3 [Source:MGI Symbo | 169.45 | 169.61 | 194.21 | 170.62 | 182.13 | 200.49 |
| 1110 | ENSMUSG00000007613 | 530.0214081 | 0.010507211 | 0.130579762 | 0.080465843 | 0.935866762 | 0.988420305 | Tgfbr1 | transforming growth factor, beta recepto | 504.35 | 575.63 | 505.13 | 563.93 | 538.77 | 492.32 |
| 1111 | ENSMUSG00000007617 | 200.7141326 | -0.112989436 | 0.186318362 | -0.606432099 | 0.544227859 | 0.89481775 | Homer1 | homer scaffolding protein 1 [Source:MGI | 199.17 | 221 | 206.28 | 226.53 | 188.68 | 162.62 |
| 1112 | ENSMUSG00000007646 | 5.084888903 | -0.778225268 | 1.111655577 | -0.700059699 | 0.483890023 | NA | Rad51c | RAD51 paralog C [Source:MGI Symbol;Acc | 3.2124 | 4.1117 | 12.075 | 6.7478 | 4.3625 | 0 |
| 1113 | ENSMUSG00000007653 | 0 | NA | NA | NA | NA | NA | Gabrb2 | gamma-aminobutyric acid (GABA) A recep | 0 | 0 | 0 | 0 | 0 | 0 |
| 1114 | ENSMUSG00000007655 | 57.56251444 | 0.454747618 | 0.365997203 | 1.242489328 | 0.214056076 | 0.658874792 | Cav1 | caveolin 1, caveolae protein [Source:MGI | 61.839 | 43.173 | 40.25 | 53.019 | 101.43 | 45.668 |
| 1115 | ENSMUSG00000007656 | 773.2406578 | -0.113384871 | 0.117431059 | -0.96554414 | 0.334272339 | 0.775945291 | Arpp19 | cAMP-regulated phosphoprotein 19 [Sour | 832.82 | 842.89 | 734.56 | 722.02 | 726.35 | 780.8 |
| 1116 | ENSMUSG00000007659 | 690.2648866 | -0.050242312 | 0.135073852 | -0.37196179 | 0.709921298 | 0.944239215 | Bcl2l1 | BCL2-like 1 [Source:MGI Symbol;Acc:MGI: | 657.74 | 737.02 | 712.42 | 668.04 | 590.02 | 776.35 |

| | | | | | | | | | | | | | | | |
|---|---|---|---|---|---|---|---|---|---|---|---|---|---|---|---|
| 1117 | ENSMUSG00000007670 | 704.7881751 | 0.068858841 | 0.12900339 | 0.533775434 | 0.593496912 | 0.913011006 | Khsrp | KH-type splicing regulatory protein [Sourc | 603.93 | 775.05 | 686.26 | 738.41 | 709.99 | 715.09 |
| 1118 | ENSMUSG00000007682 | 0.133850373 | -0.796629176 | 4.080472857 | -0.195229623 | 0.84521319 | NA | Dio2 | deiodinase, iodothyronine, type II [Source | 0.8031 | 0 | 0 | 0 | 0 | 0 |
| 1119 | ENSMUSG00000007721 | 569.4251977 | 0.024712179 | 0.125036437 | 0.197639819 | 0.843326878 | 0.972273952 | Ccdc124 | coiled-coil domain containing 124 [Source | 539.68 | 576.66 | 577.58 | 567.78 | 533.31 | 621.52 |
| 1120 | ENSMUSG00000007739 | 1734.798704 | 0.103123985 | 0.096643755 | 1.067052761 | 0.285948011 | 0.738431001 | Cct4 | chaperonin containing Tcp1, subunit 4 (de | 1676.9 | 1697.1 | 1644.2 | 1733.2 | 1905.3 | 1752.1 |
| 1121 | ENSMUSG00000007777 | 317.6977623 | 0.013250275 | 0.150617616 | 0.087972946 | 0.929898178 | 0.988405116 | 0610009B22 | RIKEN cDNA 0610009B22 gene [Source:M | 336.5 | 290.9 | 320.99 | 333.54 | 290.1 | 334.15 |
| 1122 | ENSMUSG00000007783 | 7.243590581 | 1.024188344 | 0.857330148 | 1.194625368 | 0.232233433 | 0.67964336 | Cpt1c | carnitine palmitoyltransferase 1c [Source: | 3.2124 | 3.0838 | 8.05 | 4.8199 | 9.8156 | 14.48 |
| 1123 | ENSMUSG00000007805 | 0.744190399 | -2.947713985 | 2.874421214 | -1.025498271 | 0.305128141 | NA | Twist2 | twist basic helix-loop-helix transcription fa | 2.4093 | 2.0558 | 0 | 0 | 0 | 0 |
| 1124 | ENSMUSG00000007812 | 831.6483132 | 0.6978344 | 0.155309997 | 4.493171152 | 7.02E-06 | 0.000404554 | Zfp655 | zinc finger protein 655 [Source:MGI Symb | 744.48 | 537.6 | 619.85 | 1134.6 | 851.77 | 1101.6 |
| 1125 | ENSMUSG00000007815 | 1956.307126 | -0.032394435 | 0.091790002 | -0.35291899 | 0.724149174 | 0.944786345 | Rhoa | ras homolog family member A [Source:MG | 1992.5 | 1941.7 | 2000.4 | 1942.4 | 1873.7 | 1987.1 |
| 1126 | ENSMUSG00000007817 | 738.7350028 | -0.259803314 | 0.172136817 | -1.509283827 | 0.131226266 | 0.530924954 | Zmiz1 | zinc finger, MIZ-type containing 1 [Source | 775.8 | 817.19 | 823.11 | 828.06 | 480.96 | 707.29 |
| 1127 | ENSMUSG00000007827 | 90.77237329 | -0.190854301 | 0.241270839 | -0.791037583 | 0.42892206 | 0.842753542 | Ankrd26 | ankyrin repeat domain 26 [Source:MGI Sy | 106.81 | 100.74 | 82.512 | 84.83 | 83.978 | 85.766 |
| 1128 | ENSMUSG00000007833 | 1664.05969 | 0.066488693 | 0.09372112 | 0.709431262 | 0.478056893 | 0.869794311 | Aldh16a1 | aldehyde dehydrogenase 16 family, meml | 1588.5 | 1644.7 | 1644.2 | 1684.1 | 1671.9 | 1751 |
| 1129 | ENSMUSG00000007836 | 1716.019534 | -0.002285052 | 0.095924001 | -0.023821485 | 0.980995003 | 0.99820894 | Hnrnpa0 | heterogeneous nuclear ribonucleoprotein | 1689.7 | 1666.3 | 1796.1 | 1687.9 | 1682.8 | 1773.2 |
| 1130 | ENSMUSG00000007837 | 589.5585772 | 0.129522289 | 0.126307345 | 1.025453348 | 0.305149327 | 0.75399118 | Prrg2 | proline-rich Gla (G-carboxyglutamic acid) | 564.58 | 532.46 | 591.67 | 560.07 | 634.74 | 653.83 |
| 1131 | ENSMUSG00000007850 | 1100.335438 | 0.059825603 | 0.111629704 | 0.535929062 | 0.592007578 | 0.912780899 | Hnrnph1 | heterogeneous nuclear ribonucleoprotein | 1061.7 | 1112.2 | 1059.6 | 1232 | 1036.1 | 1100.5 |
| 1132 | ENSMUSG00000007867 | 46.15186915 | -0.060020283 | 0.356962468 | -0.16814172 | 0.866471788 | 0.977925452 | Ift43 | intraflagellar transport 43 [Source:MGI Sy | 38.549 | 53.452 | 49.306 | 29.883 | 56.712 | 49.009 |
| 1133 | ENSMUSG00000007872 | 548.0837757 | -0.240829948 | 0.471238112 | -0.511057873 | 0.609310533 | 0.916376001 | Id3 | inhibitor of DNA binding 3 [Source:MGI Sy | 520.41 | 485.18 | 775.81 | 710.45 | 249.75 | 546.9 |
| 1134 | ENSMUSG00000007877 | 0 | NA | NA | NA | NA | NA | Tcap | titin-cap [Source:MGI Symbol;Acc:MGI:13 | 0 | 0 | 0 | 0 | 0 | 0 |
| 1135 | ENSMUSG00000007880 | 656.2202371 | 0.151529577 | 0.123920495 | 1.222796732 | 0.22140648 | 0.667727967 | Arid1a | AT rich interactive domain 1A (SWI-like) [S | 609.55 | 666.09 | 589.66 | 660.32 | 658.73 | 752.96 |
| 1136 | ENSMUSG00000007888 | 0.401551119 | -2.072982726 | 4.008251979 | -0.517178744 | 0.605031388 | NA | Crlf1 | cytokine receptor-like factor 1 [Source:MG | 2.4093 | 0 | 0 | 0 | 0 | 0 |
| 1137 | ENSMUSG00000007891 | 3176.942529 | -0.200105701 | 0.103381269 | -1.935608872 | 0.052915623 | 0.323119557 | Ctsd | cathepsin D [Source:MGI Symbol;Acc:MG | 3624.4 | 3485.7 | 3080.1 | 3001.8 | 3100.6 | 2769 |
| 1138 | ENSMUSG00000007892 | 5337.886691 | 0.169261536 | 0.104452388 | 1.620465933 | 0.105132227 | 0.477915593 | Rplp1 | ribosomal protein, large, P1 [Source:MGI S | 4968 | 5292.7 | 4813.9 | 4995.3 | 5950.4 | 6007 |
| 1139 | ENSMUSG00000007907 | 0 | NA | NA | NA | NA | NA | Cabs1 | calcium binding protein, spermatid specifi | 0 | 0 | 0 | 0 | 0 | 0 |
| 1140 | ENSMUSG00000007908 | 0.185640647 | 1.126924175 | 4.080472857 | 0.276174898 | 0.782413736 | NA | Hmgcll1 | 3-hydroxymethyl-3-methylglutaryl-Coenzy | 0 | 0 | 0 | 0 | 0 | 1.1138 |
| 1141 | ENSMUSG00000007944 | 0 | NA | NA | NA | NA | NA | Ttc9b | tetratricopeptide repeat domain 9B [Sour | 0 | 0 | 0 | 0 | 0 | 0 |
| 1142 | ENSMUSG00000007946 | 0.17131964 | -0.796629176 | 4.080472857 | -0.195229623 | 0.84521319 | NA | Phox2a | paired-like homeobox 2a [Source:MGI Syr | 0 | 1.0279 | 0 | 0 | 0 | 0 |
| 1143 | ENSMUSG00000007950 | 77.90434161 | -0.516838172 | 0.324398971 | -1.593217671 | 0.111111368 | 0.489450944 | Abhd8 | abhydrolase domain containing 8 [Source | 75.492 | 78.122 | 121.76 | 70.37 | 39.262 | 82.424 |
| 1144 | ENSMUSG00000007987 | 76.83209591 | -0.394966292 | 0.25970488 | -1.520827381 | 0.128303161 | 0.524378899 | Ift22 | intraflagellar transport 22 [Source:MGI Sy | 89.947 | 80.178 | 91.568 | 61.695 | 76.343 | 61.261 |
| 1145 | ENSMUSG00000007989 | 2.377673859 | 0.519131532 | 1.521740132 | 0.341143354 | 0.732995665 | NA | Fzd3 | frizzled class receptor 3 [Source:MGI Syml | 0.8031 | 4.1117 | 1.0062 | 2.8919 | 5.4531 | 0 |
| 1146 | ENSMUSG00000008028 | 2.036456027 | 0.611476597 | 1.568786419 | 0.38977683 | 0.696701578 | NA | 1700008O03 | RIKEN cDNA 1700008O03 gene [Source:M | 0.8031 | 2.0558 | 2.0125 | 2.8919 | 0 | 4.4554 |
| 1147 | ENSMUSG00000008035 | 695.4444784 | 0.153712961 | 0.132678954 | 1.15853311 | 0.246646548 | 0.696725814 | Mid1ip1 | Mid1 interacting protein 1 (gastrulation sp | 751.7 | 592.08 | 629.91 | 687.32 | 776.52 | 735.14 |
| 1148 | ENSMUSG00000008036 | 882.1881997 | 0.0030264 | 0.109983946 | 0.027516741 | 0.978047587 | 0.997776859 | Ap2s1 | adaptor-related protein complex 2, sigma | 892.25 | 876.81 | 874.43 | 876.26 | 829.96 | 943.43 |
| 1149 | ENSMUSG00000008090 | 848.1834457 | 0.426169039 | 0.138246289 | 3.082679778 | 0.002051458 | 0.035537681 | Fgfrl1 | fibroblast growth factor receptor-like 1 [S | 829.6 | 702.07 | 637.96 | 855.05 | 1024.1 | 1040.3 |
| 1150 | ENSMUSG00000008129 | 0.658052777 | 1.6393978 | 2.811180186 | 0.583170658 | 0.559778421 | NA | Brme1 | break repair meiotic recombinase recruitr | 0.8031 | 0 | 0 | 0.964 | 2.1812 | 0 |
| 1151 | ENSMUSG00000008136 | 5.508236475 | -2.117624957 | 1.062580994 | -1.992906866 | 0.046271646 | 0.297454708 | Fhl2 | four and a half LIM domains 2 [Source:MG | 13.653 | 7.1954 | 6.0375 | 2.8919 | 3.2719 | 0 |
| 1152 | ENSMUSG00000008140 | 1118.385485 | 0.034320331 | 0.105771856 | 0.324475075 | 0.74557839 | 0.949997329 | Emc10 | ER membrane protein complex subunit 10 | 1037.6 | 1112.2 | 1166.2 | 1137.5 | 1092.8 | 1164 |
| 1153 | ENSMUSG00000008153 | 579.9019816 | -0.24240353 | 0.640174418 | -0.378652322 | 0.704946063 | 0.94295036 | Clstn3 | calsyntenin 3 [Source:MGI Symbol;Acc:MG | 974.16 | 599.28 | 311.94 | 557.18 | 790.7 | 246.16 |
| 1154 | ENSMUSG00000008167 | 386.3472858 | -0.017751682 | 0.138456746 | -0.128211027 | 0.897981974 | 0.982798755 | Fbxw9 | F-box and WD-40 domain protein 9 [Sourc | 409.58 | 386.5 | 369.29 | 354.74 | 396.99 | 400.98 |
| 1155 | ENSMUSG00000008193 | 6.487744682 | -1.657925029 | 0.905339063 | -1.831275261 | 0.067059468 | 0.372627026 | Spib | Spi-B transcription factor (Spi-1/PU.1 relat | 11.243 | 11.307 | 7.0437 | 3.8559 | 4.3625 | 1.1138 |
| 1156 | ENSMUSG00000008200 | 376.6425076 | 0.121069729 | 0.20966324 | 0.577448526 | 0.563636502 | 0.900821918 | Fnbp4 | formin binding protein 4 [Source:MGI Sym | 345.33 | 412.2 | 326.02 | 540.79 | 337 | 298.51 |
| 1157 | ENSMUSG00000008206 | 37.39435594 | -0.62540328 | 0.379444047 | -1.648209494 | 0.099309689 | 0.464258353 | Cers4 | ceramide synthase 4 [Source:MGI Symbol | 49.792 | 46.256 | 40.25 | 39.523 | 21.812 | 26.732 |
| 1158 | ENSMUSG00000008226 | 1363.675248 | 0.14071983 | 0.107543493 | 1.308492284 | 0.19070639 | 0.628188847 | Scrn3 | secernin 3 [Source:MGI Symbol;Acc:MGI:1 | 1246.4 | 1320.9 | 1324.2 | 1316.8 | 1422.2 | 1551.6 |
| 1159 | ENSMUSG00000008301 | 420.6215654 | 0.077469505 | 0.143296289 | 0.540624641 | 0.58876633 | 0.911039598 | Phax | phosphorylated adaptor for RNA export [S | 468.21 | 377.25 | 381.37 | 424.15 | 441.7 | 431.06 |
| 1160 | ENSMUSG00000008305 | 943.2865004 | 0.221734559 | 0.121738014 | 1.82140772 | 0.068544902 | 0.37732204 | Tle1 | transducin-like enhancer of split 1 [Source | 913.93 | 941.57 | 756.7 | 987.11 | 1003.4 | 1057 |
| 1161 | ENSMUSG00000008307 | 15.24669059 | -0.359528808 | 0.582764184 | -0.616937036 | 0.537276257 | 0.890758111 | 1700109H08 | RIKEN cDNA 1700109H08 gene [Source:M | 12.85 | 20.558 | 18.112 | 12.532 | 19.631 | 7.7969 |
| 1162 | ENSMUSG00000008318 | 3.601606199 | -1.247711561 | 1.259267942 | -0.990822937 | 0.321772047 | NA | Relt | RELT tumor necrosis factor receptor [Sour | 4.0155 | 5.1396 | 6.0375 | 0.964 | 5.4531 | 0 |
| 1163 | ENSMUSG00000008333 | 226.219412 | 0.201031367 | 0.183049912 | 1.098232527 | 0.272102966 | 0.727618887 | Snrpb2 | U2 small nuclear ribonucleoprotein B [Sou | 168.65 | 237.45 | 226.41 | 247.74 | 245.39 | 231.68 |
| 1164 | ENSMUSG00000008348 | 1137.539439 | -0.187743407 | 0.160203135 | -1.171908446 | 0.241233816 | 0.691388521 | Ubc | ubiquitin C [Source:MGI Symbol;Acc:MGI: | 1272.9 | 1448.3 | 912.66 | 976.51 | 1180 | 1034.8 |
| 1165 | ENSMUSG00000008373 | 193.4888664 | -0.1989568 | 0.171366251 | -1.161003395 | 0.245640519 | 0.695774774 | Prpf31 | pre-mRNA processing factor 31 [Source:M | 206.4 | 206.61 | 207.29 | 171.59 | 191.95 | 177.1 |
| 1166 | ENSMUSG00000008384 | 93.59174001 | 0.024649689 | 0.247330671 | 0.099662888 | 0.920611965 | 0.986685804 | Sertad1 | SERTA domain containing 1 [Source:MGI S | 101.19 | 77.094 | 99.618 | 82.902 | 92.703 | 108.04 |
| 1167 | ENSMUSG00000008393 | 2187.326898 | 0.21586913 | 0.119837269 | 1.801352221 | 0.071647382 | 0.387549638 | Carhsp1 | calcium regulated heat stable protein 1 [S | 2140.3 | 1806.1 | 2124.2 | 2099.5 | 2323 | 2630.9 |
| 1168 | ENSMUSG00000008398 | 77.0226145 | -0.637698822 | 0.307440979 | -2.074215427 | 0.038059308 | 0.265040844 | Elk3 | ELK3, member of ETS oncogene family [So | 116.45 | 88.401 | 76.475 | 78.082 | 40.353 | 62.375 |
| 1169 | ENSMUSG00000008429 | 586.7462341 | 0.180933058 | 0.11879314 | 1.523093488 | 0.127735318 | 0.523389132 | Herpud2 | HERPUD family member 2 [Source:MGI Sy | 573.41 | 551.99 | 524.25 | 636.23 | 606.38 | 628.21 |
| 1170 | ENSMUSG00000008435 | 175.6173887 | 0.197043678 | 0.205810575 | 0.957403079 | 0.338363845 | 0.780780576 | Rdh13 | retinol dehydrogenase 13 (all-trans and 9- | 140.54 | 197.36 | 153.96 | 215.93 | 179.95 | 165.96 |
| 1171 | ENSMUSG00000008438 | 0.945548297 | -0.814460806 | 2.273503937 | -0.358240332 | 0.720163467 | NA | Adam21 | a disintegrin and metallopeptidase domai | 1.6062 | 0 | 2.0125 | 0.964 | 1.0906 | 0 |
| 1172 | ENSMUSG00000008450 | 196.9593275 | -0.023943772 | 0.184157754 | -0.130017724 | 0.896552404 | 0.982798755 | Nutf2 | nuclear transport factor 2 [Source:MGI Sy | 218.44 | 184 | 193.2 | 214 | 205.04 | 167.08 |

| | | | | | | | | | | | | | | | |
|---|---|---|---|---|---|---|---|---|---|---|---|---|---|---|---|
| 1173 | ENSMUSG00000008461 | 9.000459324 | 0.146909938 | 0.715093236 | 0.205441655 | 0.837227078 | 0.971073158 | Fut1 | fucosyltransferase 1 [Source:MGI Symbol; | 6.4248 | 8.2233 | 11.069 | 10.604 | 6.5437 | 11.138 |
| 1174 | ENSMUSG00000008475 | 1513.953911 | -0.030679078 | 0.104337383 | -0.294037257 | 0.768729434 | 0.95597133 | Arpc5 | actin related protein 2/3 complex, subunit | 1429.5 | 1548 | 1613 | 1432.5 | 1472.3 | 1588.3 |
| 1175 | ENSMUSG00000008482 | 0.562214094 | -1.122836648 | 3.276281221 | -0.34271681 | 0.731811515 | NA | Rnf151 | ring finger protein 151 [Source:MGI Symb | 2.4093 | 0 | 0 | 0.964 | 0 | 0 |
| 1176 | ENSMUSG00000008489 | 0 | NA | NA | NA | NA | NA | Elavl2 | ELAV like RNA binding protein 1 [Source:N | 0 | 0 | 0 | 0 | 0 | 0 |
| 1177 | ENSMUSG00000008496 | 27.63388927 | -1.477649817 | 0.550680466 | -2.683316203 | 0.007289603 | 0.086502219 | Pou2f2 | POU domain, class 2, transcription factor : | 24.093 | 75.038 | 23.144 | 19.28 | 11.997 | 12.252 |
| 1178 | ENSMUSG00000008540 | 60906.85218 | 0.09644576 | 0.106664608 | 0.904196452 | 0.365891241 | 0.804235439 | Mgst1 | microsomal glutathione S-transferase 1 [S | 66328 | 57927 | 52358 | 60761 | 66696 | 61371 |
| 1179 | ENSMUSG00000008575 | 1651.830066 | -0.035293426 | 0.105530443 | -0.334438343 | 0.73804881 | 0.948137969 | Nfib | nuclear factor I/B [Source:MGI Symbol;Ac | 1578.1 | 1810.2 | 1629.1 | 1704.3 | 1656.6 | 1532.6 |
| 1180 | ENSMUSG00000008590 | 0 | NA | NA | NA | NA | NA | Htr3b | 5-hydroxytryptamine (serotonin) receptor | 0 | 0 | 0 | 0 | 0 | 0 |
| 1181 | ENSMUSG00000008601 | 2.312306606 | 1.134532138 | 1.483988581 | 0.76451541 | 0.444560157 | NA | Rab25 | RAB25, member RAS oncogene family [So | 3.2124 | 0 | 1.0062 | 1.928 | 3.2719 | 4.4554 |
| 1182 | ENSMUSG00000008604 | 655.9610095 | -0.076045424 | 0.129324765 | -0.588019038 | 0.556519514 | 0.898080805 | Ubqln4 | ubiquilin 4 [Source:MGI Symbol;Acc:MGI: | 638.47 | 712.35 | 670.16 | 720.09 | 596.57 | 598.13 |
| 1183 | ENSMUSG00000008658 | 0.167707431 | -0.796629176 | 4.080472857 | -0.195229623 | 0.84521319 | NA | Rbfox1 | RNA binding protein, fox-1 homolog (C. el | 0 | 0 | 1.0062 | 0 | 0 | 0 |
| 1184 | ENSMUSG00000008668 | 4602.595196 | 0.071384114 | 0.08936979 | 0.798749938 | 0.424435424 | 0.840450984 | Rps18 | ribosomal protein S18 [Source:MGI Symbo | 4261.3 | 4652.4 | 4553.3 | 4589.5 | 4646 | 4913.2 |
| 1185 | ENSMUSG00000008682 | 8563.851822 | 0.069634008 | 0.083197226 | 0.836975114 | 0.402606557 | 0.828953151 | Rpl10 | ribosomal protein L10 [Source:MGI Symbo | 8068.8 | 8697.2 | 8306.5 | 8834.9 | 8600.6 | 8875.1 |
| 1186 | ENSMUSG00000008683 | 1366.132512 | 0.114351715 | 0.10774978 | 1.061270979 | 0.28856677 | 0.73951397 | Rps15a | ribosomal protein S15A [Source:MGI Syml | 1256.9 | 1427.8 | 1251.8 | 1357.3 | 1496.3 | 1406.8 |
| 1187 | ENSMUSG00000008690 | 899.1884636 | 0.106265451 | 0.111086094 | 0.956604447 | 0.338766943 | 0.780858357 | Ncaph2 | non-SMC condensin II complex, subunit H | 890.64 | 840.84 | 866.38 | 938.91 | 867.04 | 991.32 |
| 1188 | ENSMUSG00000008730 | 1427.712339 | -0.233845673 | 0.099687734 | -2.345781804 | 0.018987219 | 0.167458011 | Hipk1 | homeodomain interacting protein kinase : | 1476.1 | 1615.9 | 1538.5 | 1359.2 | 1297.8 | 1278.7 |
| 1189 | ENSMUSG00000008734 | 5.532468693 | -2.470759607 | 1.183952897 | -2.086873231 | 0.036899595 | 0.259514449 | Gprc5b | G protein-coupled receptor, family C, grou | 12.85 | 13.363 | 2.0125 | 3.8559 | 0 | 1.1138 |
| 1190 | ENSMUSG00000008763 | 718.2543702 | -0.083267522 | 0.117925339 | -0.706103731 | 0.480123632 | 0.870278888 | Man1a2 | mannosidase, alpha, class 1A, member 2 [ | 695.49 | 800.75 | 721.48 | 710.45 | 670.73 | 710.63 |
| 1191 | ENSMUSG00000008789 | 0 | NA | NA | NA | NA | NA | Ceacam5 | carcinoembryonic antigen-related cell adh | 0 | 0 | 0 | 0 | 0 | 0 |
| 1192 | ENSMUSG00000008813 | 0 | NA | NA | NA | NA | NA | Tppp2 | tubulin polymerization-promoting protein | 0 | 0 | 0 | 0 | 0 | 0 |
| 1193 | ENSMUSG00000008822 | 171.5593018 | -0.029609001 | 0.218161448 | -0.135720594 | 0.892042167 | 0.982489666 | Acyp1 | acylphosphatase 1, erythrocyte (common | 187.12 | 143.91 | 188.17 | 134.96 | 164.68 | 210.52 |
| 1194 | ENSMUSG00000008843 | 0.52105551 | -0.812028029 | 3.385364865 | -0.239864257 | 0.810435491 | NA | Cldn13 | claudin 13 [Source:MGI Symbol;Acc:MGI:: | 0 | 0 | 2.0125 | 0 | 0 | 1.1138 |
| 1195 | ENSMUSG00000008845 | 185.3077653 | -2.084846905 | 0.253804219 | -8.214390251 | 2.13E-16 | 1.87E-13 | Cd163 | CD163 antigen [Source:MGI Symbol;Acc:N | 256.99 | 420.42 | 222.38 | 66.514 | 69.8 | 75.741 |
| 1196 | ENSMUSG00000008855 | 1082.42049 | -0.096410082 | 0.142463883 | -0.676733499 | 0.498575058 | 0.879866965 | Hdac5 | histone deacetylase 5 [Source:MGI Symbo | 1102.7 | 934.38 | 1319.2 | 1151 | 1036.1 | 951.22 |
| 1197 | ENSMUSG00000008859 | 448.7911654 | -0.008452672 | 0.130821509 | -0.064612246 | 0.948482734 | 0.990731019 | Rala | v-ral simian leukemia viral oncogene A (ra | 480.26 | 445.09 | 424.64 | 465.6 | 443.88 | 433.29 |
| 1198 | ENSMUSG00000008874 | 0 | NA | NA | NA | NA | NA | Clec3a | C-type lectin domain family 3, member a [ | 0 | 0 | 0 | 0 | 0 | 0 |
| 1199 | ENSMUSG00000008892 | 1511.644286 | -0.004546244 | 0.09746379 | -0.046645467 | 0.962795794 | 0.992956886 | Vdac3 | voltage-dependent anion channel 3 [Sour | 1480.9 | 1527.5 | 1533.5 | 1433.4 | 1557.4 | 1537.1 |
| 1200 | ENSMUSG00000008932 | 0.658259367 | -1.329928433 | 2.812497761 | -0.472863819 | 0.636310339 | NA | Slc1a7 | solute carrier family 1 (glutamate transpo | 0.8031 | 2.0558 | 0 | 0 | 1.0906 | 0 |
| 1201 | ENSMUSG00000008958 | 279.4002817 | 0.018382173 | 0.173754198 | 0.10579412 | 0.915745702 | 0.98625062 | Vps72 | vacuolar protein sorting 72 [Source:MGI S | 312.41 | 232.31 | 287.79 | 306.54 | 291.2 | 246.16 |
| 1202 | ENSMUSG00000008976 | 1020.208427 | 0.003465241 | 0.127174919 | 0.027247832 | 0.978262066 | 0.997797092 | Gabpa | GA repeat binding protein, alpha [Source: | 913.13 | 1087.5 | 1056.6 | 895.54 | 1105.9 | 1062.6 |
| 1203 | ENSMUSG00000008999 | 79.57740644 | 0.400745232 | 0.289789951 | 1.382881738 | 0.166701131 | 0.592715132 | Bmp7 | bone morphogenetic protein 7 [Source:M | 85.932 | 64.759 | 54.337 | 68.442 | 97.065 | 106.93 |
| 1204 | ENSMUSG00000009013 | 141.6087043 | -0.140991371 | 0.231850159 | -0.608114188 | 0.543111743 | 0.894344786 | Dynll1 | dynein light chain LC8-type 1 [Source:MGI | 140.54 | 156.24 | 148.92 | 139.78 | 95.974 | 168.19 |
| 1205 | ENSMUSG00000009030 | 462.6692392 | -0.016883921 | 0.150069686 | -0.112507206 | 0.910421256 | 0.985650811 | Pdcl | phosducin-like [Source:MGI Symbol;Acc:N | 389.5 | 484.15 | 523.25 | 427.04 | 483.14 | 468.93 |
| 1206 | ENSMUSG00000009035 | 593.5364358 | 0.11070904 | 0.139969633 | 0.790950417 | 0.428972926 | 0.842753542 | Tmem184b | transmembrane protein 184b [Source:MG | 668.18 | 531.43 | 511.17 | 590.92 | 639.1 | 620.41 |
| 1207 | ENSMUSG00000009047 | 0 | NA | NA | NA | NA | NA | Gm5965 | predicted gene 5965 [Source:MGI Symbol | 0 | 0 | 0 | 0 | 0 | 0 |
| 1208 | ENSMUSG00000009070 | 0.356960287 | 0.16514264 | 3.880370363 | 0.042558474 | 0.966053499 | NA | Rsph14 | radial spoke head homolog 14 (Chlamydo | 0 | 1.0279 | 0 | 0 | 0 | 1.1138 |
| 1209 | ENSMUSG00000009073 | 692.0822141 | 0.107439067 | 0.129453147 | 0.829945575 | 0.406569556 | 0.830399297 | Nf2 | neurofibromin 2 [Source:MGI Symbol;Acc | 607.15 | 645.53 | 746.63 | 667.07 | 720.9 | 765.21 |
| 1210 | ENSMUSG00000009075 | 0.267700746 | -1.497481649 | 4.057001436 | -0.369110456 | 0.712045396 | NA | Cabp7 | calcium binding protein 7 [Source:MGI Sy | 1.6062 | 0 | 0 | 0 | 0 | 0 |
| 1211 | ENSMUSG00000009076 | 58.67134199 | 0.003403717 | 0.30214001 | 0.011265363 | 0.991011731 | 0.99859176 | Zmat5 | zinc finger, matrin type 5 [Source:MGI Syr | 54.611 | 72.982 | 48.3 | 55.911 | 55.622 | 64.603 |
| 1212 | ENSMUSG00000009079 | 1857.984942 | -0.081119918 | 0.101552708 | -0.798796205 | 0.424408591 | 0.840450984 | Ewsr1 | Ewing sarcoma breakpoint region 1 [Sourc | 1909.8 | 1897.5 | 1923.9 | 1907.7 | 1867.1 | 1641.8 |
| 1213 | ENSMUSG00000009090 | 930.4124688 | -0.149418114 | 0.112375225 | -1.329635728 | 0.183638321 | 0.617851749 | Ap1b1 | adaptor protein complex AP-1, beta 1 sub | 1047.2 | 913.82 | 974.04 | 910 | 899.76 | 837.61 |
| 1214 | ENSMUSG00000009092 | 73.58538739 | 0.612569529 | 0.335219225 | 1.827369921 | 0.067644169 | 0.374543844 | Derl3 | Der1-like domain family, member 3 [Sourc | 60.233 | 38.033 | 76.475 | 123.39 | 66.528 | 76.855 |
| 1215 | ENSMUSG00000009093 | 0 | NA | NA | NA | NA | NA | Gstt4 | glutathione S-transferase, theta 4 [Source | 0 | 0 | 0 | 0 | 0 | 0 |
| 1216 | ENSMUSG00000009097 | 2.626897899 | -4.77173913 | 1.778627662 | -2.682820712 | 0.007300412 | NA | Tbx1 | T-box 1 [Source:MGI Symbol;Acc:MGI:984 | 9.6372 | 4.1117 | 2.0125 | 0 | 0 | 0 |
| 1217 | ENSMUSG00000009108 | 12.74085133 | -0.117640432 | 0.652031452 | -0.180421407 | 0.85682175 | 0.974757731 | Gnat2 | guanine nucleotide binding protein, alpha | 6.4248 | 18.503 | 15.094 | 15.424 | 7.6343 | 13.366 |
| 1218 | ENSMUSG00000009112 | 893.951745 | -0.176000584 | 0.123393715 | -1.426333454 | 0.153772112 | 0.571029482 | Bcl2l13 | BCL2-like 13 (apoptosis facilitator) [Source | 925.17 | 992.97 | 927.76 | 874.33 | 915.03 | 728.45 |
| 1219 | ENSMUSG00000009114 | 1.493389778 | 1.968920358 | 1.855984966 | 1.060849303 | 0.288758392 | NA | 2610028H24 | RIKEN cDNA 2610028H24 gene [Source:M | 0.8031 | 0 | 1.0062 | 3.8559 | 2.1812 | 1.1138 |
| 1220 | ENSMUSG00000009115 | 6.303985223 | 0.680737568 | 0.874012321 | 0.778864956 | 0.436059269 | 0.84624805 | Spatc1l | spermatogenesis and centriole associated | 2.4093 | 8.2233 | 4.025 | 7.7118 | 6.5437 | 8.9108 |
| 1221 | ENSMUSG00000009145 | 259.1289736 | 0.118539887 | 0.169318197 | 0.700101284 | 0.483864054 | 0.872318294 | Dqx1 | DEAQ RNA-dependent ATPase [Source:M( | 243.34 | 226.14 | 275.71 | 239.07 | 303.19 | 267.32 |
| 1222 | ENSMUSG00000009185 | 0.17131964 | -0.796629176 | 4.080472857 | -0.195229623 | 0.84521319 | NA | Ccl8 | chemokine (C-C motif) ligand 8 [Source:M | 0 | 1.0279 | 0 | 0 | 0 | 0 |
| 1223 | ENSMUSG00000009207 | 67.30400122 | -0.072730104 | 0.284413442 | -0.255719642 | 0.798167328 | 0.962424862 | Lnpk | lunapark, ER junction formation factor [Sc | 83.523 | 67.843 | 55.343 | 70.37 | 63.256 | 63.489 |
| 1224 | ENSMUSG00000009210 | 0 | NA | NA | NA | NA | NA | Prr29 | proline rich 29 [Source:MGI Symbol;Acc:N | 0 | 0 | 0 | 0 | 0 | 0 |
| 1225 | ENSMUSG00000009214 | 0 | NA | NA | NA | NA | NA | Mymk | myomaker, myoblast fusion factor [Sourc | 0 | 0 | 0 | 0 | 0 | 0 |
| 1226 | ENSMUSG00000009216 | 1.34516075 | 0.243467075 | 1.955597237 | 0.124497555 | 0.900921336 | NA | Fam163b | family with sequence similarity 163, meml | 1.6062 | 2.0558 | 0 | 0 | 2.1812 | 2.2277 |
| 1227 | ENSMUSG00000009246 | 2.831868006 | -2.855231761 | 1.554412989 | -1.836855315 | 0.066231257 | NA | Trpm5 | transient receptor potential cation channe | 8.8341 | 3.0838 | 3.0187 | 0.964 | 1.0906 | 0 |
| 1228 | ENSMUSG00000009248 | 0 | NA | NA | NA | NA | NA | Ascl2 | achaete-scute family bHLH transcription fa | 0 | 0 | 0 | 0 | 0 | 0 |

| | | | | | | | | | | | | | | | |
|---|---|---|---|---|---|---|---|---|---|---|---|---|---|---|---|
| 1229 | ENSMUSG00000009281 | 6208.566084 | 0.068731371 | 0.095461078 | 0.719993664 | 0.471528896 | 0.867794597 | Rarres2 | retinoic acid receptor responder (tazarote | 6142.1 | 5559 | 6480.2 | 6101 | 6406.3 | 6562.8 |
| 1230 | ENSMUSG00000009291 | 4359.124929 | 0.030675623 | 0.09854447 | 0.311287106 | 0.755582369 | 0.953085143 | Pttg1ip | pituitary tumor-transforming 1 interacting | 4310.2 | 4180.5 | 4446.6 | 4019.8 | 4419.2 | 4778.4 |
| 1231 | ENSMUSG00000009292 | 66.68756751 | -1.925830907 | 0.341551937 | -5.638471632 | 1.72E-08 | 2.43E-06 | Trpm2 | transient receptor potential cation chann | 75.492 | 139.8 | 101.63 | 27.955 | 20.722 | 34.529 |
| 1232 | ENSMUSG00000009293 | 622.9066763 | -0.330154626 | 0.146431465 | -2.254669968 | 0.024154052 | 0.197940062 | Ube2g2 | ubiquitin-conjugating enzyme E2G 2 [Sour | 593.49 | 745.24 | 743.61 | 542.72 | 487.51 | 624.87 |
| 1233 | ENSMUSG00000009350 | 0.133850373 | -0.796629176 | 4.080472857 | -0.195229623 | 0.84521319 | NA | Mpo | myeloperoxidase [Source:MGI Symbol;Acc | 0.8031 | 0 | 0 | 0 | 0 | 0 |
| 1234 | ENSMUSG00000009356 | 0 | NA | NA | NA | NA | NA | Lpo | lactoperoxidase [Source:MGI Symbol;Acc: | 0 | 0 | 0 | 0 | 0 | 0 |
| 1235 | ENSMUSG00000009376 | 1796.32283 | 0.167184438 | 0.106246533 | 1.573551929 | 0.115591074 | 0.499823407 | Met | met proto-oncogene [Source:MGI Symbol | 1631.9 | 1699.1 | 1746.8 | 1953 | 2037.3 | 1709.8 |
| 1236 | ENSMUSG00000009378 | 1834.772561 | 0.065666193 | 0.134018689 | 0.489977879 | 0.624149553 | 0.919503423 | Slc16a12 | solute carrier family 16 (monocarboxylic a | 1813.4 | 1812.2 | 1753.9 | 2009.9 | 2127.8 | 1491.4 |
| 1237 | ENSMUSG00000009394 | 0.167707431 | -0.796629176 | 4.080472857 | -0.195229623 | 0.84521319 | NA | Syn2 | synapsin II [Source:MGI Symbol;Acc:MGI: | 0 | 0 | 1.0062 | 0 | 0 | 0 |
| 1238 | ENSMUSG00000009406 | 105.9423726 | 0.281747523 | 0.255227074 | 1.103909229 | 0.269632515 | 0.725011151 | Elk1 | ELK1, member of ETS oncogene family [So | 114.04 | 74.01 | 98.612 | 139.78 | 113.42 | 95.791 |
| 1239 | ENSMUSG00000009418 | 59.79935912 | -0.549351716 | 0.313331036 | -1.753263013 | 0.079556871 | 0.413924671 | Nav1 | neuron navigator 1 [Source:MGI Symbol;A | 81.916 | 71.954 | 59.368 | 61.695 | 37.081 | 46.781 |
| 1240 | ENSMUSG00000009470 | 1593.113371 | 0.212946792 | 0.096444589 | 2.207970335 | 0.027246341 | 0.213715335 | Tnpo1 | transportin 1 [Source:MGI Symbol;Acc:M | 1415.1 | 1523.4 | 1489.2 | 1658 | 1726.4 | 1746.5 |
| 1241 | ENSMUSG00000009471 | 0 | NA | NA | NA | NA | NA | Myod1 | myogenic differentiation 1 [Source:MGI S | 0 | 0 | 0 | 0 | 0 | 0 |
| 1242 | ENSMUSG00000009487 | 0 | NA | NA | NA | NA | NA | Otog | otogelin [Source:MGI Symbol;Acc:MGI:12 | 0 | 0 | 0 | 0 | 0 | 0 |
| 1243 | ENSMUSG00000009535 | 252.0718314 | -0.007618765 | 0.155767824 | -0.048911032 | 0.960990197 | 0.992557454 | Rnmt | RNA (guanine-7-) methyltransferase [Sour | 253.78 | 243.62 | 260.62 | 243.89 | 243.21 | 267.32 |
| 1244 | ENSMUSG00000009545 | 9.042056064 | -0.569986763 | 0.732259483 | -0.778394512 | 0.436336473 | 0.846530029 | Kcnq1 | potassium voltage-gated channel, subfam | 12.047 | 13.363 | 7.0437 | 9.6398 | 4.3625 | 7.7969 |
| 1245 | ENSMUSG00000009549 | 657.796893 | 0.026768043 | 0.1333218 | 0.200777693 | 0.840872406 | 0.971854408 | Srp14 | signal recognition particle 14 [Source:MGI | 702.71 | 633.2 | 617.83 | 588.03 | 658.73 | 746.28 |
| 1246 | ENSMUSG00000009551 | 0.976179698 | -2.098728468 | 2.289297049 | -0.916756726 | 0.359270133 | NA | 6330409D20 | RIKEN cDNA 6330409D20 gene [Source:M | 0.8031 | 3.0838 | 1.0062 | 0.964 | 0 | 0 |
| 1247 | ENSMUSG00000009555 | 604.9208551 | 0.022998888 | 0.122694969 | 0.187447689 | 0.851309635 | 0.973568625 | Cdk9 | cyclin-dependent kinase 9 (CDC2-related l | 570.2 | 586.94 | 644 | 645.87 | 593.3 | 589.22 |
| 1248 | ENSMUSG00000009563 | 643.3489739 | -0.031140667 | 0.12524118 | -0.24864559 | 0.80363494 | 0.963596811 | Tor2a | torsin family 2, member A [Source:MGI Sy | 650.51 | 701.04 | 599.72 | 664.18 | 657.64 | 587 |
| 1249 | ENSMUSG00000009566 | 3635.155674 | -0.618397404 | 0.126146162 | -4.902229251 | 9.48E-07 | 7.87E-05 | Fpgs | folylpolyglutamyl synthetase [Source:MGI | 3764.1 | 4466.3 | 4977.9 | 2708.8 | 3220.6 | 2673.2 |
| 1250 | ENSMUSG00000009569 | 430.8520993 | -0.490113514 | 0.175938078 | -2.785715968 | 0.005340965 | 0.069658895 | Mrtfb | myocardin related transcription factor B [ | 437.69 | 607.5 | 465.89 | 396.19 | 389.35 | 288.49 |
| 1251 | ENSMUSG00000009575 | 575.9241663 | 0.175571523 | 0.13305009 | 1.31958966 | 0.186972057 | 0.623323359 | Cbx5 | chromobox 5 [Source:MGI Symbol;Acc:M | 500.33 | 516.01 | 606.77 | 575.49 | 655.46 | 601.48 |
| 1252 | ENSMUSG00000009580 | 0 | NA | NA | NA | NA | NA | Odam | odontogenic, ameloblast asssociated [Sou | 0 | 0 | 0 | 0 | 0 | 0 |
| 1253 | ENSMUSG00000009585 | 27.76855923 | -1.419438221 | 0.46871216 | -3.028379337 | 0.002458692 | 0.040782715 | Apobec3 | apolipoprotein B mRNA editing enzyme, c | 37.746 | 48.312 | 35.219 | 15.424 | 7.6343 | 22.277 |
| 1254 | ENSMUSG00000009588 | 0 | NA | NA | NA | NA | NA | St6galnac1 | ST6 (alpha-N-acetyl-neuraminyl-2,3-beta-g | 0 | 0 | 0 | 0 | 0 | 0 |
| 1255 | ENSMUSG00000009596 | 0 | NA | NA | NA | NA | NA | Taf7l | TATA-box binding protein associated facto | 0 | 0 | 0 | 0 | 0 | 0 |
| 1256 | ENSMUSG00000009614 | 20585.28676 | -0.056633031 | 0.112723064 | -0.50240855 | 0.615380166 | 0.917509899 | Sardh | sarcosine dehydrogenase [Source:MGI Sy | 18588 | 20783 | 23597 | 19060 | 22160 | 19323 |
| 1257 | ENSMUSG00000009621 | 965.5678737 | 0.20883531 | 0.103670728 | 2.014409614 | 0.043966546 | 0.2895349 | Vav2 | vav 2 oncogene [Source:MGI Symbol;Acc: | 923.57 | 889.15 | 874.43 | 1057.5 | 1026.3 | 1022.5 |
| 1258 | ENSMUSG00000009628 | 0 | NA | NA | NA | NA | NA | Tex15 | testis expressed gene 15 [Source:MGI Sym | 0 | 0 | 0 | 0 | 0 | 0 |
| 1259 | ENSMUSG00000009630 | 544.3624354 | 0.162553559 | 0.138981616 | 1.169604751 | 0.242160065 | 0.692000002 | Ppp2cb | protein phosphatase 2 (formerly 2A), cata | 454.56 | 579.75 | 508.15 | 611.16 | 531.13 | 581.43 |
| 1260 | ENSMUSG00000009633 | 4116.229935 | -0.740111837 | 0.691400847 | -1.070452603 | 0.284415632 | 0.736709833 | G0s2 | G0/G1 switch gene 2 [Source:MGI Symbol | 5747.8 | 4366.6 | 5334.1 | 3431.8 | 4991.8 | 825.36 |
| 1261 | ENSMUSG00000009640 | 209.4194108 | 0.165005502 | 0.191442696 | 0.861905447 | 0.388739551 | 0.818464676 | Dmap1 | DNA methyltransferase 1-associated prot | 184.71 | 185.03 | 222.38 | 179.3 | 234.48 | 250.61 |
| 1262 | ENSMUSG00000009646 | 897.4313361 | 0.106206357 | 0.117227271 | 0.90598677 | 0.364942855 | 0.803552056 | Pla2g12b | phospholipase A2, group XIIB [Source:MG | 780.62 | 910.74 | 902.6 | 878.18 | 942.29 | 970.16 |
| 1263 | ENSMUSG00000009647 | 244.1955673 | -0.202179766 | 0.177999614 | -1.135843846 | 0.256021923 | 0.708998869 | Mcu | mitochondrial calcium uniporter [Source:I | 230.49 | 281.65 | 272.69 | 261.24 | 195.22 | 223.88 |
| 1264 | ENSMUSG00000009654 | 239.4223155 | -0.479574349 | 0.245521898 | -1.953285447 | 0.050785781 | 0.316214178 | Oit3 | oncoprotein induced transcript 3 [Source: | 288.31 | 304.26 | 243.51 | 166.77 | 138.51 | 295.17 |
| 1265 | ENSMUSG00000009670 | 0.181769715 | 1.126924175 | 4.080472857 | 0.276174898 | 0.782413736 | NA | Tex11 | testis expressed gene 11 [Source:MGI Sym | 0 | 0 | 0 | 0 | 1.0906 | 0 |
| 1266 | ENSMUSG00000009681 | 591.5301255 | 0.158195575 | 0.12474256 | 1.268176439 | 0.204734949 | 0.650460832 | Bcr | BCR activator of RhoGEF and GTPase [Sou | 535.67 | 563.3 | 578.59 | 605.38 | 584.57 | 681.67 |
| 1267 | ENSMUSG00000009687 | 35.49225203 | -1.263353879 | 0.40480857 | -3.120867424 | 0.001803192 | 0.03231892 | Fxyd5 | FXYD domain-containing ion transport reg | 51.399 | 52.424 | 46.287 | 16.388 | 15.269 | 31.188 |
| 1268 | ENSMUSG00000009731 | 2.340928631 | -0.272078483 | 1.337579267 | -0.203411109 | 0.838813717 | NA | Kcnd1 | potassium voltage-gated channel, Shal-rel | 1.6062 | 3.0838 | 3.0187 | 1.928 | 2.1812 | 2.2277 |
| 1269 | ENSMUSG00000009733 | 134.1019502 | 0.253177461 | 0.248470294 | 1.018944587 | 0.308229274 | 0.75704334 | Tfcp2 | transcription factor CP2 [Source:MGI Sym | 104.4 | 125.41 | 137.86 | 170.62 | 99.246 | 167.08 |
| 1270 | ENSMUSG00000009734 | 0 | NA | NA | NA | NA | NA | Pou6f2 | POU domain, class 6, transcription factor 2 | 0 | 0 | 0 | 0 | 0 | 0 |
| 1271 | ENSMUSG00000009739 | 87.61316009 | 0.066302013 | 0.258094929 | 0.256890028 | 0.797263669 | 0.962236316 | Pou6f1 | POU domain, class 6, transcription factor 1 | 73.885 | 106.9 | 76.475 | 96.398 | 81.796 | 90.221 |
| 1272 | ENSMUSG00000009741 | 1118.944611 | -0.023634884 | 0.120936031 | -0.195432938 | 0.845054033 | 0.972803041 | Ubp1 | upstream binding protein 1 [Source:MGI S | 1060.1 | 1159.5 | 1166.2 | 1259.9 | 1016.5 | 1051.5 |
| 1273 | ENSMUSG00000009772 | 235.9079142 | 0.249214313 | 0.198422809 | 1.255976137 | 0.209124622 | 0.65350014 | Nuak2 | NUAK family, SNF1-like kinase, 2 [Source:I | 188.73 | 180.91 | 277.72 | 275.7 | 258.48 | 233.91 |
| 1274 | ENSMUSG00000009828 | 310.1910943 | 0.143747238 | 0.166457616 | 0.863566605 | 0.387826016 | 0.817832847 | Cilk1 | ciliogenesis associated kinase 1 [Source:M | 352.56 | 260.06 | 270.68 | 344.14 | 317.37 | 316.33 |
| 1275 | ENSMUSG00000009863 | 4660.672992 | 0.060381297 | 0.093474395 | 0.645966171 | 0.518301264 | 0.887543815 | Sdhb | succinate dehydrogenase complex, subun | 4761.6 | 4436.5 | 4489.9 | 4416.9 | 4997.2 | 4861.9 |
| 1276 | ENSMUSG00000009876 | 1.213455634 | -3.651113421 | 2.181791776 | -1.673447238 | 0.094239285 | NA | Cox4i2 | cytochrome c oxidase subunit 4I2 [Source | 3.2124 | 2.0558 | 2.0125 | 0 | 0 | 0 |
| 1277 | ENSMUSG00000009894 | 1340.745259 | 0.175693348 | 0.120714154 | 1.455449444 | 0.145544877 | 0.556332213 | Snap47 | synaptosomal-associated protein, 47 [Sou | 1285.8 | 1208.8 | 1283 | 1437.3 | 1599.9 | 1229.7 |
| 1278 | ENSMUSG00000009900 | 0 | NA | NA | NA | NA | NA | Wnt3a | wingless-type MMTV integration site fami | 0 | 0 | 0 | 0 | 0 | 0 |
| 1279 | ENSMUSG00000009905 | 546.3782266 | -0.267236612 | 0.149989017 | -1.781707868 | 0.074796883 | 0.399358478 | Kdsr | 3-ketodihydrosphingosine reductase [Sou | 594.3 | 653.76 | 543.37 | 579.35 | 484.23 | 423.26 |
| 1280 | ENSMUSG00000009907 | 1118.35179 | -0.023624864 | 0.104324564 | -0.226455429 | 0.820847221 | 0.969160361 | Vps4b | vacuolar protein sorting 4B [Source:MGI S | 1118.7 | 1165.7 | 1097.8 | 1046.9 | 1156.1 | 1125 |
| 1281 | ENSMUSG00000009927 | 3062.188887 | -0.035546194 | 0.091376574 | -0.389007733 | 0.697270426 | 0.940268454 | Rps25 | ribosomal protein S25 [Source:MGI Symbo | 2979.5 | 3264.7 | 3056 | 2929.5 | 3109.4 | 3034.1 |
| 1282 | ENSMUSG00000009941 | 0.17131964 | -0.796629176 | 4.080472857 | -0.195229623 | 0.84521319 | NA | Nxf2 | nuclear RNA export factor 2 [Source:MGI | 0 | 1.0279 | 0 | 0 | 0 | 0 |
| 1283 | ENSMUSG00000009995 | 667.8995156 | 0.115705553 | 0.118367376 | 0.977512191 | 0.328315645 | 0.773161429 | Taz | tafazzin [Source:MGI Symbol;Acc:MGI:109 | 650.51 | 676.37 | 596.7 | 724.91 | 679.46 | 679.44 |
| 1284 | ENSMUSG00000010021 | 3.076994701 | -0.018767034 | 1.302716438 | -0.014406078 | 0.988506011 | NA | Kif19a | kinesin family member 19A [Source:MGI S | 3.2124 | 4.1117 | 2.0125 | 5.7839 | 0 | 3.3415 |

| | | | | | | | | | | | | | | | |
|---|---|---|---|---|---|---|---|---|---|---|---|---|---|---|---|
| 1285 | ENSMUSG00000010025 | 11531.32542 | -0.301118608 | 0.164023525 | -1.835825731 | 0.066383435 | 0.370435957 | Aldh3a2 | aldehyde dehydrogenase family 3, subfam | 14364 | 12999 | 10828 | 12637 | 10820 | 7539.6 |
| 1286 | ENSMUSG00000010044 | 5.850224607 | 0.276973574 | 0.876748651 | 0.315909895 | 0.752070901 | 0.952618751 | Zmynd10 | zinc finger, MYND domain containing 10 [S | 5.6217 | 6.1675 | 4.025 | 3.8559 | 7.6343 | 7.7969 |
| 1287 | ENSMUSG00000010045 | 474.3205247 | 0.088877392 | 0.134110663 | 0.662716824 | 0.507511933 | 0.882258527 | Tmem115 | transmembrane protein 115 [Source:MGI | 428.05 | 452.28 | 499.1 | 468.49 | 471.15 | 526.85 |
| 1288 | ENSMUSG00000010047 | 1100.804132 | 0.114502543 | 0.108860977 | 1.051823584 | 0.292880497 | 0.743850082 | Hyal2 | hyaluronoglucosaminidase 2 [Source:MGI | 1028.8 | 1003.2 | 1140.1 | 1201.1 | 1123.3 | 1108.3 |
| 1289 | ENSMUSG00000010048 | 864.5979007 | 0.112005536 | 0.117167063 | 0.955947285 | 0.339098868 | 0.780887145 | Ifrd2 | interferon-related developmental regulato | 918.75 | 788.41 | 784.87 | 912.89 | 864.86 | 917.81 |
| 1290 | ENSMUSG00000010051 | 544.3176828 | 0.180463615 | 0.171799999 | 1.0504285 | 0.293521148 | 0.744822787 | Hyal1 | hyaluronoglucosaminidase 1 [Source:MGI | 433.68 | 452.28 | 646.01 | 642.01 | 506.05 | 585.88 |
| 1291 | ENSMUSG00000010054 | 494.5198704 | 0.104076376 | 0.131111615 | 0.793799816 | 0.42731198 | 0.842374461 | Tusc2 | tumor suppressor 2, mitochondrial calciur | 493.91 | 462.56 | 472.93 | 488.74 | 483.14 | 565.83 |
| 1292 | ENSMUSG00000010057 | 251.7074149 | 0.167836965 | 0.174880144 | 0.959725676 | 0.337193297 | 0.77926821 | Nprl2 | NPR2 like, GATOR1 complex subunit [Sour | 244.95 | 194.28 | 271.69 | 261.24 | 256.3 | 281.8 |
| 1293 | ENSMUSG00000010064 | 26017.83998 | 0.669969953 | 0.120629278 | 5.553958055 | 2.79E-08 | 3.68E-06 | Slc38a3 | solute carrier family 38, member 3 [Sourc | 23112 | 19195 | 17939 | 28119 | 34116 | 33626 |
| 1294 | ENSMUSG00000010066 | 0 | NA | NA | NA | NA | NA | Cacna2d2 | calcium channel, voltage-dependent, alph | 0 | 0 | 0 | 0 | 0 | 0 |
| 1295 | ENSMUSG00000010067 | 59.12016363 | -0.343602632 | 0.327932821 | -1.047783602 | 0.294738319 | 0.745164683 | Rassf1 | Ras association (RalGDS/AF-6) domain far | 55.414 | 74.01 | 69.431 | 67.478 | 33.809 | 54.578 |
| 1296 | ENSMUSG00000010080 | 1.781965171 | -0.615311128 | 1.654787947 | -0.371836844 | 0.71001433 | NA | Epn3 | epsin 3 [Source:MGI Symbol;Acc:MGI:191 | 2.4093 | 1.0279 | 3.0187 | 0.964 | 3.2719 | 0 |
| 1297 | ENSMUSG00000010086 | 1.056559854 | -0.88642246 | 2.653260908 | -0.334087936 | 0.738313203 | NA | Rnf112 | ring finger protein 112 [Source:MGI Symb | 0 | 4.1117 | 0 | 0 | 0 | 2.2277 |
| 1298 | ENSMUSG00000010095 | 603.6077391 | -0.177462151 | 0.144335071 | -1.229515115 | 0.218878734 | 0.664120414 | Slc3a2 | solute carrier family 3 (activators of dibasi | 549.32 | 654.78 | 719.46 | 632.37 | 534.4 | 531.3 |
| 1299 | ENSMUSG00000010097 | 1260.361286 | 0.027067203 | 0.105720863 | 0.256025184 | 0.797931391 | 0.962377049 | Nxf1 | nuclear RNA export factor 1 [Source:MGI | 1206.3 | 1274.6 | 1265.9 | 1372.7 | 1199.7 | 1243 |
| 1300 | ENSMUSG00000010110 | 735.7998346 | 0.18521066 | 0.118092784 | 1.568348658 | 0.116799786 | 0.50225288 | Stx5a | syntaxin 5A [Source:MGI Symbol;Acc:MGI | 718.78 | 689.73 | 657.08 | 810.71 | 718.72 | 819.79 |
| 1301 | ENSMUSG00000010122 | 2654.347359 | -0.176068041 | 0.09957606 | -1.76817642 | 0.077031415 | 0.406191172 | Slc47a1 | solute carrier family 47, member 1 [Sourc | 2739.4 | 3022.1 | 2687.7 | 2547.8 | 2596.8 | 2332.4 |
| 1302 | ENSMUSG00000010136 | 0 | NA | NA | NA | NA | NA | Pifo | primary cilia formation [Source:MGI Symb | 0 | 0 | 0 | 0 | 0 | 0 |
| 1303 | ENSMUSG00000010142 | 2.913360706 | -1.191861201 | 1.274197297 | -0.935381989 | 0.349591476 | NA | Tnfrsf13b | tumor necrosis factor receptor superfamil | 4.0155 | 3.0838 | 5.0312 | 0.964 | 3.2719 | 1.1138 |
| 1304 | ENSMUSG00000010154 | 1.429645359 | 1.717828417 | 2.246887115 | 0.764537037 | 0.444547274 | NA | Spire2 | spire type actin nucleation factor 2 [Sourc | 0 | 1.0279 | 1.0062 | 0 | 6.5437 | 0 |
| 1305 | ENSMUSG00000010175 | 2447.30881 | -0.217282231 | 0.144515611 | -1.503520823 | 0.132704791 | 0.533623048 | Prox1 | prospero homeobox 1 [Source:MGI Symbo | 2294.5 | 2605.8 | 2993.6 | 1879.8 | 2698.2 | 2212.1 |
| 1306 | ENSMUSG00000010205 | 0 | NA | NA | NA | NA | NA | Raver1 | ribonucleoprotein, PTB-binding 1 [Source: | 0 | 0 | 0 | 0 | 0 | 0 |
| 1307 | ENSMUSG00000010277 | 3518.188558 | 0.084069735 | 0.090500573 | 0.928941467 | 0.352919421 | 0.793259753 | 2610507B11 | RIKEN cDNA 2610507B11 gene [Source:M | 3255 | 3454.8 | 3538 | 3529.1 | 3774.6 | 3557.6 |
| 1308 | ENSMUSG00000010290 | 214.2931829 | 0.222265893 | 0.174686536 | 1.272369911 | 0.203241719 | 0.647739134 | AI597479 | expressed sequence AI597479 [Source:M( | 181.5 | 224.09 | 188.17 | 217.86 | 225.76 | 248.39 |
| 1309 | ENSMUSG00000010307 | 215.5676488 | -0.235316806 | 0.20541476 | -1.145569122 | 0.251973479 | 0.704395904 | Tmem86a | transmembrane protein 86A [Source:MGI | 299.56 | 214.83 | 184.14 | 209.18 | 195.22 | 190.47 |
| 1310 | ENSMUSG00000010311 | 0.339027071 | -1.79174149 | 3.919585585 | -0.457125237 | 0.647581032 | NA | Optc | opticin [Source:MGI Symbol;Acc:MGI:215 | 0 | 1.0279 | 1.0062 | 0 | 0 | 0 |
| 1311 | ENSMUSG00000010342 | 0.503302254 | -0.837877584 | 3.426698643 | -0.244514523 | 0.806832356 | NA | Tex14 | testis expressed gene 14 [Source:MGI Sym | 0 | 2.0558 | 0 | 0.964 | 0 | 0 |
| 1312 | ENSMUSG00000010358 | 474.1365441 | 0.154000361 | 0.149947287 | 1.027029992 | 0.304406342 | 0.75326183 | Ifi35 | interferon-induced protein 35 [Source:MC | 532.46 | 406.03 | 406.52 | 470.42 | 513.68 | 515.71 |
| 1313 | ENSMUSG00000010362 | 176.4943374 | 0.241364899 | 0.206710254 | 1.16764841 | 0.242948614 | 0.692615513 | Rdm1 | RAD52 motif 1 [Source:MGI Symbol;Acc:M | 140.54 | 145.96 | 199.24 | 187.98 | 215.94 | 169.3 |
| 1314 | ENSMUSG00000010376 | 915.9075565 | 0.087390148 | 0.118441347 | 0.737834799 | 0.460614844 | 0.859817816 | Nedd8 | neural precursor cell expressed, developn | 890.64 | 861.4 | 911.66 | 840.59 | 959.74 | 1031.4 |
| 1315 | ENSMUSG00000010392 | 900.1043327 | 0.026418012 | 0.113526097 | 0.232704308 | 0.815991024 | 0.967767553 | Gosr1 | golgi SNAP receptor complex member 1 [S | 882.61 | 925.13 | 867.38 | 829.02 | 959.74 | 936.74 |
| 1316 | ENSMUSG00000010406 | 726.7979697 | 0.129677037 | 0.111181171 | 1.166357894 | 0.243469775 | 0.693217879 | Mrpl52 | mitochondrial ribosomal protein L52 [Sou | 703.52 | 681.51 | 697.33 | 762.51 | 785.25 | 730.68 |
| 1317 | ENSMUSG00000010435 | 0 | NA | NA | NA | NA | NA | Spaca7 | sperm acrosome associated 7 [Source:MG | 0 | 0 | 0 | 0 | 0 | 0 |
| 1318 | ENSMUSG00000010453 | 1152.968736 | 0.144886679 | 0.099701291 | 1.453207652 | 0.146166108 | 0.556855766 | Kansl3 | KAT8 regulatory NSL complex subunit 3 [S | 1068.1 | 1096.8 | 1121 | 1224.3 | 1223.7 | 1184 |
| 1319 | ENSMUSG00000010461 | 3.286993849 | -4.038794381 | 1.596612725 | -2.529601774 | 0.011419205 | NA | Eya4 | EYA transcriptional coactivator and phosp | 6.4248 | 5.1396 | 7.0437 | 0 | 0 | 1.1138 |
| 1320 | ENSMUSG00000010476 | 0.472877444 | -2.283655516 | 3.093270275 | -0.738265755 | 0.460352972 | NA | Ebf3 | early B cell factor 3 [Source:MGI Symbol;A | 0.8031 | 1.0279 | 1.0062 | 0 | 0 | 0 |
| 1321 | ENSMUSG00000010492 | 8.577700517 | -0.385413491 | 0.946727478 | -0.407100776 | 0.683933973 | 0.935528319 | Uckl1os | uridine-cytidine kinase 1-like 1, opposite s | 12.85 | 12.335 | 4.025 | 14.46 | 0 | 7.7969 |
| 1322 | ENSMUSG00000010505 | 0 | NA | NA | NA | NA | NA | Myt1 | myelin transcription factor 1 [Source:MGI | 0 | 0 | 0 | 0 | 0 | 0 |
| 1323 | ENSMUSG00000010517 | 580.5006946 | 0.146682605 | 0.124226771 | 1.180764856 | 0.237696149 | 0.686377492 | Faf1 | Fas-associated factor 1 [Source:MGI Symb | 541.29 | 512.93 | 598.72 | 588.03 | 622.74 | 619.3 |
| 1324 | ENSMUSG00000010529 | 0.513958919 | -2.394274821 | 3.756096013 | -0.637437066 | 0.523840191 | NA | Gm266 | predicted gene 266 [Source:MGI Symbol;A | 0 | 3.0838 | 0 | 0 | 0 | 0 |
| 1325 | ENSMUSG00000010538 | 15.43657398 | -0.178714656 | 0.668058324 | -0.267513554 | 0.78907378 | 0.960994657 | Tsacc | TSSK6 activating co-chaperone [Source:M | 4.8186 | 13.363 | 31.194 | 13.496 | 15.269 | 14.48 |
| 1326 | ENSMUSG00000010554 | 259.2719255 | -0.07532499 | 0.164104458 | -0.45900636 | 0.646229599 | 0.925643431 | Mettl16 | methyltransferase like 16 [Source:MGI Syr | 270.65 | 266.23 | 260.62 | 217.86 | 258.48 | 281.8 |
| 1327 | ENSMUSG00000010592 | 0 | NA | NA | NA | NA | NA | Dazl | deleted in azoospermia-like [Source:MGI S | 0 | 0 | 0 | 0 | 0 | 0 |
| 1328 | ENSMUSG00000010601 | 2152.6246 | 0.349524126 | 0.144448929 | 2.419707284 | 0.015533005 | 0.145605564 | Apol7a | apolipoprotein L 7a [Source:MGI Symbol;A | 1666.4 | 1936.6 | 2076.9 | 2107.3 | 2961 | 2167.5 |
| 1329 | ENSMUSG00000010607 | 378.1940993 | 0.006207218 | 0.134042576 | 0.046307809 | 0.963064915 | 0.992997643 | Pigyl | phosphatidylinositol glycan anchor biosyn | 378.26 | 379.3 | 374.32 | 363.42 | 378.44 | 395.41 |
| 1330 | ENSMUSG00000010608 | 1153.138342 | -0.153479776 | 0.158085096 | -0.970868096 | 0.331613967 | 0.774747658 | Rbm25 | RNA binding motif protein 25 [Source:MG | 1139.6 | 1352.7 | 1152.2 | 1290.8 | 1159.3 | 824.24 |
| 1331 | ENSMUSG00000010609 | 2592.839264 | -0.071813445 | 0.148145041 | -0.484750919 | 0.627853042 | 0.920328819 | Psen2 | presenilin 2 [Source:MGI Symbol;Acc:MGI | 2406.9 | 2406.4 | 3158.6 | 2158.3 | 2397.2 | 3029.7 |
| 1332 | ENSMUSG00000010651 | 32520.92068 | -0.446516008 | 0.159046166 | -2.80746163 | 0.004993363 | 0.066565898 | Acaa1b | acetyl-Coenzyme A acyltransferase 1B [So | 40921 | 36848 | 34773 | 32889 | 30153 | 19541 |
| 1333 | ENSMUSG00000010660 | 6.890256321 | -0.363537034 | 0.961660128 | -0.378030682 | 0.705407801 | 0.94295036 | Plcd1 | phospholipase C, delta 1 [Source:MGI Syn | 12.047 | 9.2513 | 2.0125 | 12.532 | 3.2719 | 2.2277 |
| 1334 | ENSMUSG00000010663 | 10480.20306 | 0.351758131 | 0.140879326 | 2.496875454 | 0.012529295 | 0.126529431 | Fads1 | fatty acid desaturase 1 [Source:MGI Symb | 10746 | 9926.6 | 6953.2 | 11683 | 11651 | 11921 |
| 1335 | ENSMUSG00000010721 | 116.8894059 | -0.234983036 | 0.221995489 | -1.05850365 | 0.289825887 | 0.740058153 | Lmbr1 | limb region 1 [Source:MGI Symbol;Acc:M( | 111.63 | 146.99 | 120.75 | 99.29 | 109.06 | 113.61 |
| 1336 | ENSMUSG00000010751 | 7.005008022 | -1.781671656 | 0.863949555 | -2.062240378 | 0.039184856 | 0.270922448 | Tnfrsf22 | tumor necrosis factor receptor superfamil | 11.243 | 8.2233 | 13.081 | 2.8919 | 4.3625 | 2.2277 |
| 1337 | ENSMUSG00000010755 | 345.5370966 | 0.217863703 | 0.184072356 | 1.183576437 | 0.236580772 | 0.685340334 | Cars | cysteinyl-tRNA synthetase [Source:MGI Sy | 391.91 | 298.1 | 267.66 | 412.58 | 307.55 | 395.41 |
| 1338 | ENSMUSG00000010760 | 1.028022105 | 1.080690112 | 2.389621529 | 0.45224321 | 0.651093782 | NA | Phlda2 | pleckstrin homology like domain, family A | 0 | 0 | 2.0125 | 1.928 | 0 | 2.2277 |
| 1339 | ENSMUSG00000010796 | 0.185640647 | 1.126924175 | 4.080472857 | 0.276174898 | 0.782413736 | NA | Asz1 | ankyrin repeat, SAM and basic leucine zip | 0 | 0 | 0 | 0 | 0 | 1.1138 |
| 1340 | ENSMUSG00000010797 | 36.29051969 | -0.351221233 | 0.456473561 | -0.769422948 | 0.44164227 | 0.849906689 | Wnt2 | wingless-type MMTV integration site fami | 33.73 | 43.173 | 45.281 | 35.667 | 11.997 | 47.895 |

| | | | | | | | | | | | | | | | |
|---|---|---|---|---|---|---|---|---|---|---|---|---|---|---|---|
| 1341 | ENSMUSG00000010803 | 0.160662974 | 1.126924175 | 4.080472857 | 0.276174898 | 0.782413736 | NA | Gabra1 | gamma-aminobutyric acid (GABA) A recep | 0 | 0 | 0 | 0.964 | 0 | 0 |
| 1342 | ENSMUSG00000010825 | 0.68303045 | 1.69421863 | 2.781340055 | 0.609137537 | 0.542433279 | NA | Grid2ip | glutamate receptor, ionotropic, delta 2 (G | 0.8031 | 0 | 0 | 0 | 2.1812 | 1.1138 |
| 1343 | ENSMUSG00000010830 | 2.849490893 | -0.076223454 | 1.347766679 | -0.056555378 | 0.954899381 | NA | Kdelr3 | KDEL (Lys-Asp-Glu-Leu) endoplasmic retic | 5.6217 | 3.0838 | 0 | 2.8919 | 3.2719 | 2.2277 |
| 1344 | ENSMUSG00000010841 | 0 | NA | NA | NA | NA | NA | Cfap97d1 | CFAP97 domain containing 1 [Source:MGI | 0 | 0 | 0 | 0 | 0 | 0 |
| 1345 | ENSMUSG00000010911 | 362.333004 | 0.018804338 | 0.1684027 | 0.111662923 | 0.911090679 | 0.98582838 | Apip | APAF1 interacting protein [Source:MGI Sy | 397.54 | 329.96 | 351.18 | 292.09 | 411.16 | 392.07 |
| 1346 | ENSMUSG00000010914 | 621.0721451 | -0.095469903 | 0.123212061 | -0.774842187 | 0.438432917 | 0.847566655 | Pdhx | pyruvate dehydrogenase complex, compo | 689.86 | 644.5 | 589.66 | 586.1 | 628.2 | 588.11 |
| 1347 | ENSMUSG00000010936 | 644.6313368 | -0.630512015 | 0.221966091 | -2.840578095 | 0.004503185 | 0.062269657 | Vac14 | Vac14 homolog (S. cerevisiae) [Source:MG | 697.09 | 875.79 | 777.83 | 593.81 | 632.56 | 290.71 |
| 1348 | ENSMUSG00000011008 | 1.745807871 | 0.0800449 | 2.024770098 | 0.039532834 | 0.968465576 | NA | Mcoln2 | mucolipin 2 [Source:MGI Symbol;Acc:MGI | 4.0155 | 0 | 1.0062 | 0 | 5.4531 | 0 |
| 1349 | ENSMUSG00000011034 | 10.70759379 | -0.091401283 | 0.713361229 | -0.128127629 | 0.898047972 | 0.982798755 | Slc5a1 | solute carrier family 5 (sodium/glucose co | 12.85 | 8.2233 | 12.075 | 14.46 | 3.2719 | 13.366 |
| 1350 | ENSMUSG00000011052 | 0 | NA | NA | NA | NA | NA | NA | NA | 0 | 0 | 0 | 0 | 0 | 0 |
| 1351 | ENSMUSG00000011096 | 1046.97795 | 0.154181284 | 0.113409579 | 1.359508475 | 0.173985519 | 0.604151393 | Akt1s1 | AKT1 substrate 1 (proline-rich) [Source:M | 1054.5 | 904.57 | 1013.3 | 1101.8 | 1042.6 | 1165.1 |
| 1352 | ENSMUSG00000011114 | 687.940995 | -0.034010496 | 0.113613204 | -0.299353372 | 0.764670435 | 0.954980327 | Tbrg1 | transforming growth factor beta regulatec | 724.4 | 665.06 | 698.33 | 692.14 | 681.64 | 666.08 |
| 1353 | ENSMUSG00000011118 | 0 | NA | NA | NA | NA | NA | Panx3 | pannexin 3 [Source:MGI Symbol;Acc:MGI: | 0 | 0 | 0 | 0 | 0 | 0 |
| 1354 | ENSMUSG00000011148 | 405.7627617 | -0.154676731 | 0.150704118 | -1.026360348 | 0.304721761 | 0.753556583 | Adssl1 | adenylosuccinate synthetase like 1 [Sourc | 397.54 | 398.83 | 486.02 | 345.1 | 387.17 | 419.92 |
| 1355 | ENSMUSG00000011154 | 0 | NA | NA | NA | NA | NA | Cfap161 | cilia and flagella associated protein 161 [S | 0 | 0 | 0 | 0 | 0 | 0 |
| 1356 | ENSMUSG00000011158 | 342.9785088 | 0.090013884 | 0.148908863 | 0.604489765 | 0.545518072 | 0.89549457 | Brf1 | BRF1, RNA polymerase III transcription ini | 337.3 | 306.32 | 353.19 | 358.6 | 383.9 | 318.56 |
| 1357 | ENSMUSG00000011171 | 0.328370405 | 0.16514316 | 4.003149536 | 0.041253308 | 0.967093956 | NA | Vipr2 | vasoactive intestinal peptide receptor 2 [S | 0 | 0 | 1.0062 | 0.964 | 0 | 0 |
| 1358 | ENSMUSG00000011179 | 1519.246792 | -0.121423949 | 0.112060881 | -1.083553404 | 0.27856286 | 0.731329747 | Odc1 | ornithine decarboxylase, structural 1 [Sou | 1403 | 1709.4 | 1638.2 | 1419.9 | 1520.3 | 1424.6 |
| 1359 | ENSMUSG00000011254 | 43.04053155 | -0.082479725 | 0.366700061 | -0.224924221 | 0.822038227 | 0.969160361 | Thg1l | tRNA-histidine guanylyltransferase 1-like ( | 61.036 | 39.061 | 32.2 | 36.631 | 42.534 | 46.781 |
| 1360 | ENSMUSG00000011256 | 7.779550509 | -0.617892623 | 0.847157681 | -0.729371447 | 0.465774479 | 0.863698893 | Adam19 | a disintegrin and metallopeptidase domai | 8.031 | 12.335 | 8.05 | 10.604 | 6.5437 | 1.1138 |
| 1361 | ENSMUSG00000011257 | 819.4032651 | 0.338177068 | 0.116413502 | 2.904964299 | 0.00367295 | 0.054127116 | Pabpc4 | poly(A) binding protein, cytoplasmic 4 [So | 790.25 | 700.01 | 680.22 | 937.95 | 931.39 | 876.6 |
| 1362 | ENSMUSG00000011263 | 85.03080384 | -0.213627849 | 0.254671591 | -0.83883659 | 0.401561015 | 0.828437957 | Exoc3l2 | exocyst complex component 3-like 2 [Sou | 93.963 | 88.401 | 91.568 | 83.866 | 62.165 | 90.221 |
| 1363 | ENSMUSG00000011267 | 9.467882463 | 0.014916592 | 0.707132414 | 0.021094482 | 0.983170287 | 0.998580735 | Zfp296 | zinc finger protein 296 [Source:MGI Symb | 12.85 | 7.1954 | 8.05 | 7.7118 | 7.6343 | 13.366 |
| 1364 | ENSMUSG00000011305 | 1790.712545 | -1.069737553 | 0.170887556 | -6.259891445 | 3.85E-10 | 8.71E-08 | Plin5 | perilipin 5 [Source:MGI Symbol;Acc:MGI:1 | 1935.5 | 3271.9 | 2070.9 | 1178.9 | 1183.3 | 1103.8 |
| 1365 | ENSMUSG00000011306 | 299.5858689 | -0.134468125 | 0.160522079 | -0.837692394 | 0.402203486 | 0.828953151 | Sugp1 | SURP and G patch domain containing 1 [S | 358.18 | 277.54 | 303.89 | 275.7 | 277.02 | 305.19 |
| 1366 | ENSMUSG00000011349 | 0 | NA | NA | NA | NA | NA | Dmrtc2 | doublesex and mab-3 related transcriptio | 0 | 0 | 0 | 0 | 0 | 0 |
| 1367 | ENSMUSG00000011350 | 1.035014429 | -2.151020407 | 2.418909279 | -0.889252204 | 0.373867552 | NA | Gm5893 | predicted gene 5893 [Source:MGI Symbol | 0 | 3.0838 | 2.0125 | 0 | 0 | 1.1138 |
| 1368 | ENSMUSG00000011382 | 3989.631563 | 0.054892294 | 0.104817041 | 0.523696273 | 0.600489809 | 0.91534406 | Dhdh | dihydrodiol dehydrogenase (dimeric) [Sou | 4067.7 | 4152.8 | 3519.8 | 3825.1 | 4359.2 | 4013.2 |
| 1369 | ENSMUSG00000011427 | 137.7272893 | 0.059750631 | 0.209966142 | 0.284572695 | 0.775971533 | 0.95893551 | Zfp790 | zinc finger protein 790 [Source:MGI Symb | 113.24 | 155.22 | 136.85 | 146.52 | 131.96 | 142.57 |
| 1370 | ENSMUSG00000011463 | 0 | NA | NA | NA | NA | NA | Cpb1 | carboxypeptidase B1 (tissue) [Source:MGI | 0 | 0 | 0 | 0 | 0 | 0 |
| 1371 | ENSMUSG00000011486 | 0.185640647 | 1.126924175 | 4.080472857 | 0.276174898 | 0.782413736 | NA | Slc25a41 | solute carrier family 25, member 41 [Sour | 0 | 0 | 0 | 0 | 0 | 1.1138 |
| 1372 | ENSMUSG00000011589 | 1.528765393 | 1.154518049 | 1.769806046 | 0.652341567 | 0.514180851 | NA | Fsd1 | fibronectin type 3 and SPRY domain-conta | 0.8031 | 2.0558 | 0 | 1.928 | 3.2719 | 1.1138 |
| 1373 | ENSMUSG00000011632 | 0.503122293 | -2.366675966 | 3.785596405 | -0.625179156 | 0.531853481 | NA | Pinlyp | phospholipase A2 inhibitor and LY6/PLAU | 0 | 0 | 3.0187 | 0 | 0 | 0 |
| 1374 | ENSMUSG00000011658 | 124.7826033 | -0.308451953 | 0.249371481 | -1.236917518 | 0.216117706 | 0.66165144 | Fuz | fuzzy planar cell polarity protein [Source:M | 124.48 | 127.46 | 163.01 | 146.52 | 100.34 | 86.88 |
| 1375 | ENSMUSG00000011751 | 1.266874246 | 0.147248781 | 2.008467279 | 0.073314005 | 0.941556247 | NA | Sptbn4 | spectrin beta, non-erythrocytic 4 [Source: | 1.6062 | 0 | 2.0125 | 2.8919 | 1.0906 | 0 |
| 1376 | ENSMUSG00000011752 | 1503.240594 | -0.156476911 | 0.108472172 | -1.442553494 | 0.149146292 | 0.561319344 | Pgam1 | phosphoglycerate mutase 1 [Source:MGI S | 1541.2 | 1701.2 | 1511.4 | 1309.1 | 1441.8 | 1514.8 |
| 1377 | ENSMUSG00000011831 | 1859.234311 | 0.103696616 | 0.118937929 | 0.871854901 | 0.383287539 | 0.815515 | Evi5 | ecotropic viral integration site 5 [Source:N | 1591.7 | 1723.8 | 2062.8 | 1913.5 | 1848.6 | 2014.9 |
| 1378 | ENSMUSG00000011832 | 91.34484152 | 0.345975077 | 0.257641032 | 1.342857053 | 0.179318265 | 0.611423712 | Evi5l | ecotropic viral integration site 5 like [Sour | 99.585 | 69.898 | 71.443 | 117.61 | 95.974 | 93.563 |
| 1379 | ENSMUSG00000011837 | 175.6066312 | 0.170346298 | 0.213629913 | 0.79738973 | 0.425224722 | 0.840969906 | Snapc2 | small nuclear RNA activating complex, pol | 180.7 | 144.94 | 170.06 | 217.86 | 139.6 | 200.49 |
| 1380 | ENSMUSG00000011877 | 196.7572426 | -0.045810222 | 0.196864589 | -0.232699147 | 0.815995032 | 0.967767553 | Git1 | GIT ArfGAP 1 [Source:MGI Symbol;Acc:MG | 214.43 | 192.22 | 193.2 | 234.25 | 154.87 | 191.58 |
| 1381 | ENSMUSG00000011884 | 47.42885765 | -1.069891195 | 0.375078547 | -2.852445721 | 0.004338422 | 0.060746309 | Gltp | glycolipid transfer protein [Source:MGI Sy | 71.476 | 78.122 | 43.269 | 35.667 | 35.99 | 20.049 |
| 1382 | ENSMUSG00000011958 | 819.1920952 | 0.035278517 | 0.108617086 | 0.324797124 | 0.745334622 | 0.949997329 | Bnip2 | BCL2/adenovirus E1B interacting protein 2 | 779.01 | 839.81 | 809.02 | 809.74 | 831.05 | 846.52 |
| 1383 | ENSMUSG00000011960 | 396.2521036 | 0.02637194 | 0.149689067 | 0.176178129 | 0.860154003 | 0.97570916 | Ccnt1 | cyclin T1 [Source:MGI Symbol;Acc:MGI:13 | 364.61 | 389.58 | 423.63 | 347.03 | 443.88 | 408.78 |
| 1384 | ENSMUSG00000012017 | 73.71015065 | -0.990631578 | 0.300243668 | -3.299425385 | 0.00096883 | 0.020487067 | Scarf2 | scavenger receptor class F, member 2 [Sou | 89.144 | 106.9 | 98.612 | 63.623 | 31.628 | 52.351 |
| 1385 | ENSMUSG00000012042 | 2.320695173 | -0.225566747 | 1.388427702 | -0.162462004 | 0.870942047 | NA | 4930579F01 | RIKEN cDNA 4930579F01 gene [Source:MG | 2.4093 | 2.0558 | 3.0187 | 0.964 | 4.3625 | 1.1138 |
| 1386 | ENSMUSG00000012076 | 71.83759632 | 0.082886606 | 0.263475063 | 0.314589947 | 0.753073013 | 0.952989051 | Brms1l | breast cancer metastasis-suppressor 1-like | 64.248 | 66.815 | 78.487 | 77.118 | 73.071 | 71.286 |
| 1387 | ENSMUSG00000012114 | 522.8579415 | 0.066464895 | 0.138262116 | 0.480716606 | 0.630717927 | 0.921020101 | Med15 | mediator complex subunit 15 [Source:MG | 578.23 | 502.65 | 450.8 | 562.96 | 525.68 | 516.82 |
| 1388 | ENSMUSG00000012117 | 511.2919992 | 0.086937109 | 0.138997158 | 0.62545962 | 0.531669443 | 0.888947136 | Dhdds | dehydrodolichyl diphosphate synthase [So | 554.94 | 457.42 | 473.94 | 489.7 | 514.77 | 576.97 |
| 1389 | ENSMUSG00000012123 | 82.71365885 | -0.518388477 | 0.345223203 | -1.501603807 | 0.133199459 | 0.534720228 | Crybg2 | crystallin beta-gamma domain containing | 146.97 | 91.485 | 53.331 | 74.226 | 54.531 | 75.741 |
| 1390 | ENSMUSG00000012126 | 1.049076712 | -0.889203617 | 2.241827177 | -0.396642358 | 0.691631212 | NA | Ubxn11 | UBX domain protein 11 [Source:MGI Symb | 0 | 3.0838 | 1.0062 | 0 | 1.0906 | 1.1138 |
| 1391 | ENSMUSG00000012187 | 31.37884341 | -0.962247562 | 0.441838194 | -2.177827938 | 0.029418849 | 0.225823158 | Mogat1 | monoacylglycerol O-acyltransferase 1 [Sou | 61.036 | 34.949 | 28.175 | 21.208 | 25.084 | 17.822 |
| 1392 | ENSMUSG00000012211 | 0 | NA | NA | NA | NA | NA | Tex22 | testis expressed gene 22 [Source:MGI Sym | 0 | 0 | 0 | 0 | 0 | 0 |
| 1393 | ENSMUSG00000012282 | 0 | NA | NA | NA | NA | NA | Wnt8a | wingless-type MMTV integration site fami | 0 | 0 | 0 | 0 | 0 | 0 |
| 1394 | ENSMUSG00000012296 | 105.807399 | -0.023039218 | 0.23795042 | -0.096823606 | 0.922866477 | 0.986938316 | Tjap1 | tight junction associated protein 1 [Source | 89.947 | 100.74 | 129.81 | 113.75 | 99.246 | 101.36 |
| 1395 | ENSMUSG00000012350 | 5.741234807 | -0.58235906 | 1.070344066 | -0.54408585 | 0.586382397 | 0.910220541 | Ehf | ets homologous factor [Source:MGI Symb | 2.4093 | 9.2513 | 9.0562 | 4.8199 | 0 | 8.9108 |
| 1396 | ENSMUSG00000012396 | 0 | NA | NA | NA | NA | NA | Nanog | Nanog homeobox [Source:MGI Symbol;Ac | 0 | 0 | 0 | 0 | 0 | 0 |

| | | | | | | | | | | | | | | | |
|---|---|---|---|---|---|---|---|---|---|---|---|---|---|---|---|
| 1397 | ENSMUSG00000012405 | 2838.836003 | 0.109652529 | 0.103226692 | 1.062249765 | 0.288122313 | 0.73951397 | Rpl15 | ribosomal protein L15 [Source:MGI Symbo | 2674.3 | 2756.9 | 2761.1 | 2627.8 | 3040.6 | 3172.2 |
| 1398 | ENSMUSG00000012422 | 778.6283291 | 0.141244656 | 0.115816813 | 1.219552257 | 0.222634655 | 0.668407528 | Tmem167 | transmembrane protein 167 [Source:MGI | 738.05 | 712.35 | 770.78 | 771.18 | 797.24 | 882.16 |
| 1399 | ENSMUSG00000012428 | 2537.887453 | -0.010004919 | 0.118199682 | -0.084644216 | 0.932544246 | 0.988420305 | Steap4 | STEAP family member 4 [Source:MGI Sym | 2354.7 | 2734.3 | 2551.8 | 2541 | 2207.4 | 2838.1 |
| 1400 | ENSMUSG00000012429 | 264.5990172 | 0.250965474 | 0.184696064 | 1.358802502 | 0.17420918 | 0.604151393 | Mplkip | M-phase specific PLK1 intereacting protei | 252.98 | 201.47 | 270.68 | 342.21 | 257.39 | 262.87 |
| 1401 | ENSMUSG00000012443 | 18.67469035 | -0.073193378 | 0.542112103 | -0.135015206 | 0.892599851 | 0.982512308 | Kif11 | kinesin family member 11 [Source:MGI Sy | 20.881 | 22.614 | 14.087 | 26.991 | 17.45 | 10.025 |
| 1402 | ENSMUSG00000012483 | 127.2700976 | -0.006764545 | 0.214928994 | -0.031473394 | 0.97489201 | 0.996336532 | Rpa3 | replication protein A3 [Source:MGI Symbo | 150.18 | 114.1 | 117.73 | 121.46 | 127.6 | 132.55 |
| 1403 | ENSMUSG00000012519 | 54.79704492 | -0.005287291 | 0.356111014 | -0.014847311 | 0.988153995 | 0.99859176 | Mlkl | mixed lineage kinase domain-like [Source: | 77.901 | 31.865 | 54.337 | 51.091 | 53.44 | 60.148 |
| 1404 | ENSMUSG00000012520 | 0 | NA | NA | NA | NA | NA | Phox2b | paired-like homeobox 2b [Source:MGI Syr | 0 | 0 | 0 | 0 | 0 | 0 |
| 1405 | ENSMUSG00000012535 | 747.0246874 | -0.115368096 | 0.125097245 | -0.922227313 | 0.356410019 | 0.795787977 | Tnpo3 | transportin 3 [Source:MGI Symbol;Acc:M( | 678.62 | 813.08 | 840.21 | 730.7 | 708.9 | 710.63 |
| 1406 | ENSMUSG00000012609 | 200.8291035 | 0.06295817 | 0.175818312 | 0.358086531 | 0.720278558 | 0.9447719 | Ttll5 | tubulin tyrosine ligase-like family, membe | 175.88 | 206.61 | 207.29 | 196.65 | 221.4 | 197.15 |
| 1407 | ENSMUSG00000012640 | 226.9647676 | 0.048090281 | 0.162863596 | 0.295279499 | 0.767780376 | 0.95578799 | Zfp715 | zinc finger protein 715 [Source:MGI Symb | 218.44 | 224.09 | 227.41 | 246.78 | 231.21 | 213.86 |
| 1408 | ENSMUSG00000012705 | 0 | NA | NA | NA | NA | NA | Retn | resistin [Source:MGI Symbol;Acc:MGI:188 | 0 | 0 | 0 | 0 | 0 | 0 |
| 1409 | ENSMUSG00000012777 | 0.185640647 | 1.126924175 | 4.080472857 | 0.276174898 | 0.782413736 | NA | Acp4 | acid phosphatase 4 [Source:MGI Symbol;/ | 0 | 0 | 0 | 0 | 0 | 1.1138 |
| 1410 | ENSMUSG00000012819 | 1.398785952 | 1.680204043 | 1.969343652 | 0.853179709 | 0.393559654 | NA | Cdh23 | cadherin 23 (otocadherin) [Source:MGI Sy | 0 | 2.0558 | 0 | 1.928 | 2.1812 | 2.2277 |
| 1411 | ENSMUSG00000012848 | 3129.144466 | 0.070077766 | 0.097645612 | 0.717674501 | 0.472958009 | 0.86857093 | Rps5 | ribosomal protein S5 [Source:MGI Symbol | 2871.9 | 3199.9 | 3088.2 | 3068.3 | 3114.8 | 3431.8 |
| 1412 | ENSMUSG00000012889 | 2.830290915 | -1.06137059 | 1.579789965 | -0.671842848 | 0.501683747 | NA | Podnl1 | podocan-like 1 [Source:MGI Symbol;Acc:N | 3.2124 | 8.2233 | 0 | 0 | 1.0906 | 4.4554 |
| 1413 | ENSMUSG00000013033 | 77.96361277 | -0.277817532 | 0.295055569 | -0.941576981 | 0.34640926 | 0.787313566 | Adgrl1 | adhesion G protein-coupled receptor L1 [! | 82.72 | 98.68 | 75.468 | 96.398 | 62.165 | 52.351 |
| 1414 | ENSMUSG00000013076 | 107.7140759 | -0.353925868 | 0.259442325 | -1.364179374 | 0.172511118 | 0.600818882 | Amotl1 | angiomotin-like 1 [Source:MGI Symbol;Ac | 135.72 | 96.624 | 129.81 | 89.65 | 73.071 | 121.41 |
| 1415 | ENSMUSG00000013083 | 0 | NA | NA | NA | NA | NA | 2200002J24I | RIKEN cDNA 2200002J24 gene [Source:M( | 0 | 0 | 0 | 0 | 0 | 0 |
| 1416 | ENSMUSG00000013089 | 76.67885318 | 0.411269476 | 0.297186981 | 1.38387447 | 0.166396895 | 0.5925379 | Etv5 | ets variant 5 [Source:MGI Symbol;Acc:MG | 61.839 | 51.396 | 84.525 | 108.93 | 67.618 | 85.766 |
| 1417 | ENSMUSG00000013091 | 1.786016064 | -0.076844098 | 1.625096695 | -0.047285862 | 0.962285396 | NA | Tmem190 | transmembrane protein 190 [Source:MGI | 2.4093 | 3.0838 | 0 | 1.928 | 2.1812 | 1.1138 |
| 1418 | ENSMUSG00000013150 | 183.0015559 | -0.039693445 | 0.205531617 | -0.193125736 | 0.846860498 | 0.972803041 | Gfod2 | glucose-fructose oxidoreductase domain ( | 195.15 | 166.52 | 195.21 | 224.61 | 148.32 | 168.19 |
| 1419 | ENSMUSG00000013155 | 12.79430072 | 0.817667337 | 0.667850176 | 1.224327501 | 0.220828706 | 0.667237604 | Enkd1 | enkurin domain containing 1 [Source:MGI | 9.6372 | 6.1675 | 12.075 | 27.955 | 10.906 | 10.025 |
| 1420 | ENSMUSG00000013158 | 1.027298067 | 3.575098506 | 2.566323775 | 1.393081629 | 0.163595108 | NA | 4933405L10I | RIKEN cDNA 4933405L10 gene [Source:M( | 0 | 0 | 0 | 2.8919 | 3.2719 | 0 |
| 1421 | ENSMUSG00000013160 | 1211.403586 | -0.043511141 | 0.09833542 | -0.442476793 | 0.65814422 | 0.927303282 | Atp6v0d1 | ATPase, H+ transporting, lysosomal V0 sul | 1259.3 | 1205.7 | 1223.6 | 1193.4 | 1176.8 | 1209.6 |
| 1422 | ENSMUSG00000013236 | 69.19885615 | -0.239512569 | 0.2732636 | -0.876489108 | 0.380764195 | 0.812628154 | Ptprs | protein tyrosine phosphatase, receptor ty | 79.507 | 67.843 | 77.481 | 70.37 | 66.528 | 53.465 |
| 1423 | ENSMUSG00000013275 | 113.8607834 | 0.744569653 | 0.263041269 | 2.830619153 | 0.0046458 | 0.063513096 | Slc41a1 | solute carrier family 41, member 1 [Sourc | 96.372 | 84.289 | 74.462 | 163.88 | 95.974 | 168.19 |
| 1424 | ENSMUSG00000013338 | 0 | NA | NA | NA | NA | NA | Fer1l4 | fer-1-like 4 (C. elegans) [Source:MGI Symb | 0 | 0 | 0 | 0 | 0 | 0 |
| 1425 | ENSMUSG00000013353 | 0 | NA | NA | NA | NA | NA | 4931406B18 | RIKEN cDNA 4931406B18 gene [Source:M | 0 | 0 | 0 | 0 | 0 | 0 |
| 1426 | ENSMUSG00000013367 | 6.13329341 | -1.126251684 | 0.948755174 | -1.187083575 | 0.235194656 | 0.683240713 | Iglon5 | IgLON family member 5 [Source:MGI Syml | 4.0155 | 8.2233 | 13.081 | 4.8199 | 1.0906 | 5.5692 |
| 1427 | ENSMUSG00000013415 | 0.51034671 | -2.381743948 | 3.331081356 | -0.715006238 | 0.474605189 | NA | Igf2bp1 | insulin-like growth factor 2 mRNA binding | 0 | 2.0558 | 1.0062 | 0 | 0 | 0 |
| 1428 | ENSMUSG00000013418 | 0.881705114 | -0.66605326 | 2.559736336 | -0.260203854 | 0.794706531 | NA | B4galnt2 | beta-1,4-N-acetyl-galactosaminyl transfer | 3.2124 | 0 | 0 | 0.964 | 0 | 1.1138 |
| 1429 | ENSMUSG00000013419 | 17.33208635 | -1.184390624 | 0.579676079 | -2.043193891 | 0.041033248 | 0.277468398 | Zfp651 | zinc finger protein 651 [Source:MGI Symb | 35.336 | 18.503 | 18.112 | 6.7478 | 15.269 | 10.025 |
| 1430 | ENSMUSG00000013465 | 450.9341855 | 0.106393953 | 0.147141471 | 0.723072513 | 0.469635336 | 0.8665398 | Nelfb | negative elongation factor complex meml | 421.63 | 389.58 | 492.05 | 494.52 | 416.62 | 491.21 |
| 1431 | ENSMUSG00000013483 | 3.052956931 | 0.913901323 | 1.422830852 | 0.642311995 | 0.520670628 | NA | Card14 | caspase recruitment domain family, meml | 3.2124 | 2.0558 | 1.0062 | 0 | 9.8156 | 2.2277 |
| 1432 | ENSMUSG00000013495 | 134.3570294 | 0.209052749 | 0.213721745 | 0.978153858 | 0.327998234 | 0.773161429 | Tmem175 | transmembrane protein 175 [Source:MGI | 141.35 | 113.07 | 118.74 | 123.39 | 159.23 | 150.37 |
| 1433 | ENSMUSG00000013523 | 0.845734943 | 0.605113486 | 2.575938408 | 0.23490992 | 0.81427865 | NA | Bcas1 | breast carcinoma amplified sequence 1 [S | 0 | 2.0558 | 0 | 1.928 | 1.0906 | 0 |
| 1434 | ENSMUSG00000013539 | 1149.157917 | -0.372184251 | 0.106366625 | -3.499069859 | 0.000466884 | 0.011805968 | Tango2 | transport and golgi organization 2 [Source | 1261.7 | 1232.5 | 1395.7 | 990.01 | 984.83 | 1030.3 |
| 1435 | ENSMUSG00000013562 | 0.87367955 | 2.066316271 | 2.569175551 | 0.804272121 | 0.421239835 | NA | 4833413E03 | RIKEN cDNA 4833413E03 gene [Source:M( | 0 | 0 | 1.0062 | 0.964 | 3.2719 | 0 |
| 1436 | ENSMUSG00000013584 | 4.405793454 | 1.926198033 | 1.084701693 | 1.775785957 | 0.075768203 | NA | Aldh1a2 | aldehyde dehydrogenase family 1, subfam | 2.4093 | 1.0279 | 2.0125 | 7.7118 | 4.3625 | 8.9108 |
| 1437 | ENSMUSG00000013593 | 4526.408971 | -0.003832202 | 0.085455789 | -0.04484426 | 0.964231446 | 0.993347771 | Ndufs2 | NADH:ubiquinone oxidoreductase core su | 4414.7 | 4477.6 | 4705.2 | 4433.3 | 4511.9 | 4615.8 |
| 1438 | ENSMUSG00000013611 | 0.877834182 | -0.676932822 | 2.563285057 | -0.264087999 | 0.791712111 | NA | Snx31 | sorting nexin 31 [Source:MGI Symbol;Acc: | 3.2124 | 0 | 0 | 0.964 | 1.0906 | 0 |
| 1439 | ENSMUSG00000013622 | 1170.736188 | -0.014750039 | 0.122428353 | -0.120478943 | 0.904103759 | 0.983821605 | Atraid | all-trans retinoic acid induced differentiat | 1034.4 | 1284.9 | 1212.5 | 1228.1 | 1200.8 | 1063.7 |
| 1440 | ENSMUSG00000013629 | 157.2380029 | 0.072905494 | 0.204489281 | 0.35652477 | 0.721447605 | 0.9447719 | Cad | carbamoyl-phosphate synthetase 2, aspar | 130.1 | 169.61 | 161 | 184.12 | 152.69 | 145.91 |
| 1441 | ENSMUSG00000013643 | 0 | NA | NA | NA | NA | NA | Lypd8 | LY6/PLAUR domain containing 8 [Source:N | 0 | 0 | 0 | 0 | 0 | 0 |
| 1442 | ENSMUSG00000013646 | 374.0264066 | 0.213150015 | 0.14975118 | 1.423361171 | 0.154631482 | 0.572750259 | Sh3bp5l | SH3 binding domain protein 5 like [Source | 358.99 | 350.52 | 329.04 | 345.1 | 441.7 | 418.81 |
| 1443 | ENSMUSG00000013653 | 0 | NA | NA | NA | NA | NA | 1810065E05 | RIKEN cDNA 1810065E05 gene [Source:M( | 0 | 0 | 0 | 0 | 0 | 0 |
| 1444 | ENSMUSG00000013662 | 1168.699812 | 0.106099611 | 0.112015639 | 0.947185701 | 0.34354416 | 0.785243795 | Atad1 | ATPase family, AAA domain containing 1 [ | 1040.8 | 1167.7 | 1169.3 | 1139.4 | 1308.7 | 1186.2 |
| 1445 | ENSMUSG00000013663 | 2147.275908 | 0.002491648 | 0.095639016 | 0.026052635 | 0.979215356 | 0.998055255 | Pten | phosphatase and tensin homolog [Source | 2035.9 | 2277.9 | 2123.2 | 2094.7 | 2185.6 | 2166.4 |
| 1446 | ENSMUSG00000013668 | 0 | NA | NA | NA | NA | NA | 4933402N03 | RIKEN cDNA 4933402N03 gene [Source:M | 0 | 0 | 0 | 0 | 0 | 0 |
| 1447 | ENSMUSG00000013698 | 175.685984 | -0.401305666 | 0.239524705 | -1.675424947 | 0.093850885 | 0.450983125 | Pea15a | phosphoprotein enriched in astrocytes 15 | 209.61 | 217.92 | 172.07 | 142.67 | 105.79 | 206.06 |
| 1448 | ENSMUSG00000013701 | 1389.004202 | -0.047897957 | 0.114744261 | -0.417432267 | 0.676362257 | 0.932771138 | Timm23 | translocase of inner mitochondrial membr | 1287.4 | 1467.9 | 1481.2 | 1234.9 | 1404.7 | 1458 |
| 1449 | ENSMUSG00000013707 | 15.94276139 | -1.383810473 | 0.57836219 | -2.392636479 | 0.016727805 | 0.153327616 | Tnfaip8l2 | tumor necrosis factor, alpha-induced prot | 15.259 | 29.81 | 24.15 | 7.7118 | 9.8156 | 8.9108 |
| 1450 | ENSMUSG00000013736 | 405.4750621 | 0.067580117 | 0.14091216 | 0.479590384 | 0.631518685 | 0.921549084 | Trnt1 | tRNA nucleotidyl transferase, CCA-adding | 422.43 | 379.3 | 385.39 | 385.59 | 459.15 | 400.98 |
| 1451 | ENSMUSG00000013766 | 1.188400852 | -3.603107576 | 2.463463935 | -1.462618358 | 0.143571837 | NA | Ly6g6e | lymphocyte antigen 6 complex, locus G6E | 0 | 4.1117 | 3.0187 | 0 | 0 | 0 |
| 1452 | ENSMUSG00000013787 | 1191.370635 | 0.313527967 | 0.109371386 | 2.866636124 | 0.004148597 | 0.058842478 | Ehmt2 | euchromatic histone lysine N-methyltrans | 1041.6 | 1142 | 1004.2 | 1374.6 | 1232.4 | 1353.3 |

| | | | | | | | | | | | | | | | |
|---|---|---|---|---|---|---|---|---|---|---|---|---|---|---|---|
| 1453 | ENSMUSG00000013822 | 594.8083969 | 0.11664658 | 0.130888266 | 0.891192033 | 0.372826158 | 0.808358199 | Elof1 | ELF1 homolog, elongation factor 1 [Source | 576.63 | 501.62 | 633.93 | 627.55 | 602.02 | 627.09 |
| 1454 | ENSMUSG00000013833 | 391.7216028 | 0.044413597 | 0.162909601 | 0.272627255 | 0.785139758 | 0.960670212 | Med16 | mediator complex subunit 16 [Source:MG | 423.23 | 390.61 | 342.12 | 329.68 | 454.79 | 409.89 |
| 1455 | ENSMUSG00000013846 | 1380.526008 | 0.028373946 | 0.120156511 | 0.236141561 | 0.813322825 | 0.966802988 | St3gal1 | ST3 beta-galactoside alpha-2,3-sialyltrans | 1479.3 | 1288 | 1333.3 | 1536.6 | 1225.9 | 1420.2 |
| 1456 | ENSMUSG00000013858 | 2433.374963 | 0.164459663 | 0.097003145 | 1.695405473 | 0.089998528 | 0.442644317 | Tmem259 | transmembrane protein 259 [Source:MGI | 2325.8 | 2183.3 | 2375.7 | 2743.5 | 2470.3 | 2501.7 |
| 1457 | ENSMUSG00000013878 | 326.9858985 | 0.059496676 | 0.145753981 | 0.408199318 | 0.683127349 | 0.935211432 | Rnf170 | ring finger protein 170 [Source:MGI Symb | 332.48 | 298.1 | 330.05 | 351.85 | 310.83 | 338.61 |
| 1458 | ENSMUSG00000013921 | 5.501630703 | -1.607251371 | 1.033642709 | -1.55493901 | 0.119960595 | 0.506897474 | Clip3 | CAP-GLY domain containing linker protein | 13.653 | 8.2233 | 3.0187 | 4.8199 | 2.1812 | 1.1138 |
| 1459 | ENSMUSG00000013928 | 0.346303622 | 2.004339653 | 3.884003423 | 0.516049919 | 0.605819544 | NA | 1700020L13I | RIKEN cDNA 1700020L13 gene [Source:MG | 0 | 0 | 0 | 0.964 | 0 | 1.1138 |
| 1460 | ENSMUSG00000013936 | 0 | NA | NA | NA | NA | NA | Myl2 | myosin, light polypeptide 2, regulatory, ca | 0 | 0 | 0 | 0 | 0 | 0 |
| 1461 | ENSMUSG00000013973 | 76.89847955 | 0.159366072 | 0.260426444 | 0.611942742 | 0.540575641 | 0.893324212 | Dedd | death effector domain-containing [Source | 76.295 | 77.094 | 64.4 | 78.082 | 73.071 | 92.449 |
| 1462 | ENSMUSG00000013974 | 1.750841526 | -0.484957103 | 1.850060474 | -0.262130406 | 0.793220906 | NA | Mcemp1 | mast cell expressed membrane protein 1 [ | 4.0155 | 1.0279 | 1.0062 | 0 | 0 | 4.4554 |
| 1463 | ENSMUSG00000013997 | 1766.788561 | 0.074723416 | 0.111473272 | 0.670325853 | 0.50265009 | 0.881220881 | Nit1 | nitrilase 1 [Source:MGI Symbol;Acc:MGI:1 | 1599.8 | 1732 | 1831.4 | 1623.3 | 1916.2 | 1898 |
| 1464 | ENSMUSG00000014030 | 4.165195119 | -2.385091058 | 1.30458515 | -1.828237166 | 0.067513965 | NA | Pax5 | paired box 5 [Source:MGI Symbol;Acc:MG | 4.8186 | 10.279 | 6.0375 | 3.8559 | 0 | 0 |
| 1465 | ENSMUSG00000014039 | 81.3265861 | -0.027328804 | 0.31133928 | -0.087778208 | 0.930052959 | 0.988411902 | Prdm15 | PR domain containing 15 [Source:MGI Syr | 92.357 | 97.652 | 56.35 | 95.434 | 92.703 | 53.465 |
| 1466 | ENSMUSG00000014074 | 298.4244806 | 0.046560645 | 0.144570575 | 0.3220617 | 0.747405953 | 0.950739825 | Rnf168 | ring finger protein 168 [Source:MGI Symb | 285.9 | 303.24 | 291.81 | 297.87 | 303.19 | 308.53 |
| 1467 | ENSMUSG00000014075 | 22.14576712 | -0.046707395 | 0.510961442 | -0.091410801 | 0.92716618 | 0.987712228 | Dynlt2b | dynein light chain Tctex-type 2B [Source:N | 26.502 | 22.614 | 18.112 | 10.604 | 30.537 | 24.505 |
| 1468 | ENSMUSG00000014077 | 7340.800177 | 0.127265643 | 0.105181276 | 1.209964814 | 0.226292395 | 0.672783802 | Chp1 | calcineurin-like EF hand protein 1 [Source | 6434.5 | 7034 | 7585.1 | 8319.1 | 7662.7 | 7009.4 |
| 1469 | ENSMUSG00000014104 | 0 | NA | NA | NA | NA | NA | Sval2 | seminal vesicle antigen-like 2 [Source:MG | 0 | 0 | 0 | 0 | 0 | 0 |
| 1470 | ENSMUSG00000014158 | 7.986194745 | -1.501272343 | 0.873845609 | -1.718006393 | 0.085795449 | 0.431795087 | Trpv4 | transient receptor potential cation chann | 12.047 | 12.335 | 11.069 | 5.7839 | 0 | 6.6831 |
| 1471 | ENSMUSG00000014164 | 181.6029849 | -0.316538602 | 0.193828418 | -1.633086653 | 0.102450773 | 0.470781813 | Klhl3 | kelch-like 3 [Source:MGI Symbol;Acc:MGI | 182.3 | 212.78 | 209.3 | 147.49 | 145.05 | 192.69 |
| 1472 | ENSMUSG00000014177 | 454.0129998 | 0.221629014 | 0.138792508 | 1.596837011 | 0.110302043 | 0.488191174 | Tvp23b | trans-golgi network vesicle protein 23B [S | 424.04 | 417.33 | 415.58 | 433.79 | 485.33 | 548.01 |
| 1473 | ENSMUSG00000014195 | 759.4848186 | 0.038104392 | 0.115797807 | 0.329059707 | 0.742110559 | 0.949043964 | Dnajc7 | DnaJ heat shock protein family (Hsp40) m | 720.38 | 799.72 | 728.52 | 738.41 | 814.69 | 755.19 |
| 1474 | ENSMUSG00000014198 | 0 | NA | NA | NA | NA | NA | Zfp385c | zinc finger protein 385C [Source:MGI Sym | 0 | 0 | 0 | 0 | 0 | 0 |
| 1475 | ENSMUSG00000014226 | 592.7655261 | 0.096271303 | 0.117331373 | 0.820507763 | 0.411926706 | 0.83423521 | Cacybp | calcyclin binding protein [Source:MGI Sym | 562.17 | 579.75 | 577.58 | 639.12 | 599.84 | 598.13 |
| 1476 | ENSMUSG00000014232 | 94.2745638 | 0.023027836 | 0.245900139 | 0.093647105 | 0.92538949 | 0.987546253 | Cluap1 | clusterin associated protein 1 [Source:MG | 90.751 | 80.178 | 109.68 | 93.506 | 106.88 | 84.652 |
| 1477 | ENSMUSG00000014243 | 186.9334673 | -0.011140782 | 0.180664324 | -0.061665646 | 0.950829098 | 0.990883058 | Zswim7 | zinc finger SWIM-type containing 7 [Sourc | 189.53 | 187.08 | 186.16 | 173.52 | 212.67 | 172.65 |
| 1478 | ENSMUSG00000014245 | 102.0056458 | -0.179147993 | 0.24191427 | -0.740543306 | 0.458970394 | 0.859322431 | Pigl | phosphatidylinositol glycan anchor biosyn | 110.83 | 99.708 | 114.71 | 111.82 | 100.34 | 74.628 |
| 1479 | ENSMUSG00000014294 | 1589.773441 | 0.137666889 | 0.102611188 | 1.341636248 | 0.179713967 | 0.612025777 | Ndufa2 | NADH:ubiquinone oxidoreductase subuni | 1551.6 | 1552.2 | 1437.9 | 1691.8 | 1549.8 | 1755.4 |
| 1480 | ENSMUSG00000014301 | 373.3712474 | 0.183107705 | 0.155942131 | 1.174202918 | 0.240313758 | 0.690347208 | Pam16 | presequence translocase-asssociated mot | 405.57 | 328.93 | 313.95 | 435.72 | 377.35 | 378.71 |
| 1481 | ENSMUSG00000014303 | 26.51172104 | -0.506105447 | 0.449348306 | -1.126309904 | 0.260034367 | 0.713552853 | Glis2 | GLIS family zinc finger 2 [Source:MGI Sym | 39.352 | 26.726 | 27.169 | 26.027 | 14.178 | 25.618 |
| 1482 | ENSMUSG00000014313 | 4067.087018 | 0.091798863 | 0.099042082 | 0.926867255 | 0.35399546 | 0.79449673 | Cox6c | cytochrome c oxidase subunit 6C [Source: | 3944.8 | 4152.8 | 3715.1 | 3979.3 | 4054.9 | 4555.6 |
| 1483 | ENSMUSG00000014329 | 48.00341755 | 0.027235854 | 0.339962105 | 0.080114381 | 0.936146285 | 0.988420305 | Bicc1 | BicC family RNA binding protein 1 [Source | 56.217 | 42.145 | 44.275 | 62.659 | 38.172 | 44.554 |
| 1484 | ENSMUSG00000014349 | 911.083416 | 0.076889021 | 0.108014007 | 0.711843057 | 0.476561966 | 0.86953366 | Ube2z | ubiquitin-conjugating enzyme E2Z [Source | 852.89 | 893.26 | 914.68 | 931.2 | 895.4 | 979.07 |
| 1485 | ENSMUSG00000014351 | 0 | NA | NA | NA | NA | NA | Gip | gastric inhibitory polypeptide [Source:MG | 0 | 0 | 0 | 0 | 0 | 0 |
| 1486 | ENSMUSG00000014353 | 698.7673023 | -0.211822977 | 0.140849742 | -1.503893259 | 0.132608853 | 0.533565316 | Tmem87b | transmembrane protein 87B [Source:MGI | 803.91 | 831.59 | 613.81 | 619.84 | 669.64 | 653.83 |
| 1487 | ENSMUSG00000014355 | 770.2266397 | 0.068153949 | 0.114294876 | 0.596299245 | 0.550975335 | 0.895729712 | Anapc1 | anaphase promoting complex subunit 1 [S | 753.31 | 714.4 | 787.89 | 743.23 | 800.51 | 822.02 |
| 1488 | ENSMUSG00000014361 | 348.5438571 | -0.245256463 | 0.161888493 | -1.514971566 | 0.129779605 | 0.527580721 | Mertk | MER proto-oncogene tyrosine kinase [Sou | 362.2 | 396.78 | 376.34 | 382.7 | 279.2 | 294.05 |
| 1489 | ENSMUSG00000014402 | 627.3436689 | 0.09671228 | 0.119789614 | 0.807351128 | 0.419464215 | 0.837591673 | Tsg101 | tumor susceptibility gene 101 [Source:MG | 574.22 | 636.28 | 608.78 | 615.98 | 680.55 | 648.26 |
| 1490 | ENSMUSG00000014418 | 333.4840657 | -0.000121833 | 0.149494249 | -0.000814968 | 0.999349749 | 0.999764475 | Hps5 | HPS5, biogenesis of lysosomal organelles | 335.7 | 355.66 | 308.92 | 312.33 | 368.63 | 319.67 |
| 1491 | ENSMUSG00000014426 | 364.5503895 | 0.1402664 | 0.140853486 | 0.995831939 | 0.319331809 | 0.766337562 | Map3k4 | mitogen-activated protein kinase kinase k | 322.04 | 364.91 | 354.2 | 381.74 | 403.53 | 360.89 |
| 1492 | ENSMUSG00000014444 | 68.76399441 | -0.360639579 | 0.289621089 | -1.245211738 | 0.213053929 | 0.657724029 | Piezo1 | piezo-type mechanosensitive ion channel | 59.43 | 83.261 | 89.556 | 57.839 | 53.44 | 69.058 |
| 1493 | ENSMUSG00000014453 | 6.856805026 | -0.210623055 | 0.839794242 | -0.250803166 | 0.801966293 | 0.962865482 | Blk | B lymphoid kinase [Source:MGI Symbol;A | 8.8341 | 3.0838 | 10.062 | 4.8199 | 6.5437 | 7.7969 |
| 1494 | ENSMUSG00000014470 | 269.5478498 | -0.153039877 | 0.17416401 | -0.878711259 | 0.379557852 | 0.812628154 | Rnf166 | ring finger protein 166 [Source:MGI Symb | 313.21 | 284.73 | 253.57 | 295.94 | 219.21 | 250.61 |
| 1495 | ENSMUSG00000014496 | 822.0988634 | -0.05273803 | 0.121798621 | -0.432993653 | 0.665019381 | 0.929426323 | Ankrd28 | ankyrin repeat domain 28 [Source:MGI Sy | 788.65 | 912.79 | 811.03 | 855.05 | 831.05 | 734.02 |
| 1496 | ENSMUSG00000014498 | 429.8594792 | 0.063257901 | 0.140135247 | 0.451406071 | 0.651696909 | 0.925643431 | Ankrd52 | ankyrin repeat domain 52 [Source:MGI Sy | 400.75 | 438.92 | 422.62 | 493.56 | 410.07 | 413.24 |
| 1497 | ENSMUSG00000014503 | 65.57474949 | -0.246350988 | 0.302898823 | -0.813311144 | 0.416039689 | 0.836590861 | Pkd2l2 | polycystic kidney disease 2-like 2 [Source: | 63.445 | 77.094 | 73.456 | 79.046 | 44.715 | 55.692 |
| 1498 | ENSMUSG00000014504 | 305.95353 | -0.113947759 | 0.158625957 | -0.718342456 | 0.472546159 | 0.868365854 | Srp19 | signal recognition particle 19 [Source:MGI | 298.75 | 362.85 | 292.82 | 284.37 | 318.46 | 278.46 |
| 1499 | ENSMUSG00000014529 | 0 | NA | NA | NA | NA | NA | Tmbim7 | transmembrane BAX inhibitor motif conta | 0 | 0 | 0 | 0 | 0 | 0 |
| 1500 | ENSMUSG00000014542 | 923.3439014 | -1.273013456 | 0.178093949 | -7.147988267 | 8.81E-13 | 3.64E-10 | Clec4f | C-type lectin domain family 4, member f [ | 1301 | 1566.5 | 1050.5 | 488.74 | 449.33 | 683.9 |
| 1501 | ENSMUSG00000014543 | 0.51034671 | -2.381743948 | 3.331081356 | -0.715006238 | 0.474605189 | NA | Klra17 | killer cell lectin-like receptor, subfamily A, | 0 | 2.0558 | 1.0062 | 0 | 0 | 0 |
| 1502 | ENSMUSG00000014547 | 348.7248369 | 0.12772423 | 0.143431465 | 0.890489614 | 0.373203043 | 0.808611351 | Wdfy2 | WD repeat and FYVE domain containing 2 | 319.63 | 361.83 | 318.98 | 380.77 | 342.45 | 368.68 |
| 1503 | ENSMUSG00000014550 | 265.0858199 | 0.192824802 | 0.1575152 | 1.224166318 | 0.220889492 | 0.667281875 | Rbsn | rabenosyn, RAB effector [Source:MGI Sym | 228.08 | 252.87 | 261.62 | 276.66 | 267.2 | 304.08 |
| 1504 | ENSMUSG00000014551 | 221.422316 | -0.027608895 | 0.16618438 | -0.166134116 | 0.868051407 | 0.978326885 | Mrps25 | mitochondrial ribosomal protein S25 [Sou | 232.1 | 206.61 | 231.44 | 204.36 | 229.03 | 225 |
| 1505 | ENSMUSG00000014554 | 218.1808265 | 0.136456357 | 0.169983868 | 0.802760629 | 0.422113103 | 0.839236277 | Dguok | deoxyguanosine kinase [Source:MGI Symb | 191.94 | 207.64 | 224.39 | 228.46 | 210.49 | 246.16 |
| 1506 | ENSMUSG00000014592 | 75.93804141 | 0.386945074 | 0.260433906 | 1.485770729 | 0.137339775 | 0.541266927 | Camta1 | calmodulin binding transcription activator | 67.461 | 61.675 | 68.425 | 98.326 | 82.887 | 76.855 |
| 1507 | ENSMUSG00000014599 | 156.6981671 | -0.418082614 | 0.228805265 | -1.827242106 | 0.067663376 | 0.374543844 | Csf1 | colony stimulating factor 1 (macrophage) | 227.28 | 153.16 | 156.97 | 159.06 | 114.51 | 129.21 |
| 1508 | ENSMUSG00000014601 | 276.7380533 | -0.185421025 | 0.160130853 | -1.157934412 | 0.246890803 | 0.696922048 | Strip1 | striatin interacting protein 1 [Source:MGI | 293.13 | 301.18 | 289.8 | 296.91 | 239.94 | 239.48 |

| | | | | | | | | | | | | | | | |
|---|---|---|---|---|---|---|---|---|---|---|---|---|---|---|---|
| 1509 | ENSMUSG00000014602 | 2.282733662 | -0.57059406 | 1.815268639 | -0.314330368 | 0.753270136 | NA | Kif1a | kinesin family member 1A [Source:MGI Sy | 4.0155 | 4.1117 | 0 | 0 | 0 | 5.5692 |
| 1510 | ENSMUSG00000014603 | 0 | NA | NA | NA | NA | NA | Alx3 | aristaless-like homeobox 3 [Source:MGI S | 0 | 0 | 0 | 0 | 0 | 0 |
| 1511 | ENSMUSG00000014606 | 1612.061354 | 0.088201596 | 0.114016495 | 0.773586278 | 0.43917549 | 0.848147242 | Slc25a11 | solute carrier family 25 (mitochondrial car | 1570.9 | 1585 | 1531.5 | 1441.1 | 1799.5 | 1744.3 |
| 1512 | ENSMUSG00000014609 | 4.323759456 | -0.274136267 | 1.024998834 | -0.267450321 | 0.789122459 | NA | Chrne | cholinergic receptor, nicotinic, epsilon pol | 4.0155 | 5.1396 | 5.0312 | 2.8919 | 2.1812 | 6.6831 |
| 1513 | ENSMUSG00000014633 | 159.2907533 | -0.303801121 | 0.208385528 | -1.45788013 | 0.144873586 | 0.555361204 | Cmc2 | COX assembly mitochondrial protein 2 [Sc | 177.49 | 174.75 | 176.09 | 159.06 | 160.32 | 108.04 |
| 1514 | ENSMUSG00000014668 | 346.7403266 | 0.123253421 | 0.140327792 | 0.87832509 | 0.379767324 | 0.812628154 | Chfr | checkpoint with forkhead and ring finger | 353.36 | 328.93 | 312.94 | 357.64 | 355.54 | 372.02 |
| 1515 | ENSMUSG00000014686 | 0.349477146 | 0.165142768 | 3.911021834 | 0.042224967 | 0.966319359 | NA | Ceacam16 | carcinoembryonic antigen-related cell adh | 0 | 0 | 1.0062 | 0 | 1.0906 | 0 |
| 1516 | ENSMUSG00000014704 | 3.246169443 | -3.119671659 | 1.561549881 | -1.99780468 | 0.04573784 | NA | Hoxa2 | homeobox A2 [Source:MGI Symbol;Acc:M | 10.44 | 3.0838 | 4.025 | 1.928 | 0 | 0 |
| 1517 | ENSMUSG00000014725 | 0 | NA | NA | NA | NA | NA | Adam28 | a disintegrin and metallopeptidase domai | 0 | 0 | 0 | 0 | 0 | 0 |
| 1518 | ENSMUSG00000014747 | 0.790591184 | -0.507346814 | 2.633409124 | -0.192657802 | 0.847226973 | NA | Ankrd53 | ankyrin repeat domain 53 [Source:MGI Sy | 0.8031 | 0 | 2.0125 | 1.928 | 0 | 0 |
| 1519 | ENSMUSG00000014748 | 1470.734267 | 0.06675141 | 0.117366764 | 0.568742015 | 0.56953123 | 0.903851186 | Tex261 | testis expressed gene 261 [Source:MGI Sy | 1272.1 | 1441.1 | 1597.9 | 1449.8 | 1605.4 | 1458 |
| 1520 | ENSMUSG00000014763 | 546.5320649 | -0.086850068 | 0.138194441 | -0.62846282 | 0.529700795 | 0.888334265 | Fam120b | family with sequence similarity 120, meml | 586.26 | 534.52 | 568.53 | 597.67 | 533.31 | 458.9 |
| 1521 | ENSMUSG00000014767 | 211.3072773 | -0.073335643 | 0.201914656 | -0.363201185 | 0.716454602 | 0.9447719 | Tbp | TATA box binding protein [Source:MGI Sy | 192.74 | 195.3 | 262.63 | 226.53 | 224.67 | 165.96 |
| 1522 | ENSMUSG00000014769 | 2560.529773 | 0.06027602 | 0.100637959 | 0.59893921 | 0.549213423 | 0.895703112 | Psmb1 | proteasome (prosome, macropain) subun | 2649.4 | 2380.7 | 2489.4 | 2427.3 | 2628.4 | 2788 |
| 1523 | ENSMUSG00000014771 | 0 | NA | NA | NA | NA | NA | Pdcd2 | programmed cell death 2 [Source:MGI Syr | 0 | 0 | 0 | 0 | 0 | 0 |
| 1524 | ENSMUSG00000014773 | 67.23679578 | -0.053972357 | 0.311136036 | -0.173468678 | 0.862283046 | 0.976334255 | Dll1 | delta like canonical Notch ligand 1 [Source | 51.399 | 65.787 | 88.55 | 60.731 | 54.531 | 82.424 |
| 1525 | ENSMUSG00000014776 | 9.065166438 | -0.44073959 | 0.809297966 | -0.544594956 | 0.586032125 | 0.909872118 | Nol3 | nucleolar protein 3 (apoptosis repressor v | 16.865 | 11.307 | 3.0187 | 6.7478 | 11.997 | 4.4554 |
| 1526 | ENSMUSG00000014778 | 104.5893214 | -0.554769607 | 0.268850631 | -2.063486353 | 0.039066443 | 0.270232919 | Fhod1 | formin homology 2 domain containing 1 [S | 114.04 | 114.1 | 145.91 | 115.68 | 66.528 | 71.286 |
| 1527 | ENSMUSG00000014782 | 0.771003 | -1.69715066 | 2.640552792 | -0.642725518 | 0.52040222 | NA | Plekhg4 | pleckstrin homology domain containing, fa | 1.6062 | 2.0558 | 0 | 0.964 | 0 | 0 |
| 1528 | ENSMUSG00000014786 | 3.261653712 | -1.885955193 | 1.316525235 | -1.432524909 | 0.151993655 | NA | Slc9a5 | solute carrier family 9 (sodium/hydrogen | 3.2124 | 8.2233 | 4.025 | 1.928 | 2.1812 | 0 |
| 1529 | ENSMUSG00000014791 | 480.8494136 | 0.082975673 | 0.141329888 | 0.58710634 | 0.55713229 | 0.898167794 | Elmo3 | engulfment and cell motility 3 [Source:MG | 496.32 | 445.09 | 459.85 | 563.93 | 476.6 | 443.31 |
| 1530 | ENSMUSG00000014813 | 1.023919079 | -2.141404304 | 2.462125713 | -0.869738004 | 0.384443598 | NA | Stc1 | stanniocalcin 1 [Source:MGI Symbol;Acc:N | 0 | 1.0279 | 4.025 | 0 | 1.0906 | 0 |
| 1531 | ENSMUSG00000014837 | 19.20886898 | 0.576155461 | 0.582268103 | 0.989502016 | 0.322417584 | 0.76988915 | 4931428F04 | RIKEN cDNA 4931428F04 gene [Source:M | 12.85 | 16.447 | 17.106 | 25.063 | 35.99 | 7.7969 |
| 1532 | ENSMUSG00000014846 | 2.821289217 | -0.726966404 | 1.264610135 | -0.574854165 | 0.56538993 | NA | Tppp3 | tubulin polymerization-promoting protein | 2.4093 | 5.1396 | 3.0187 | 1.928 | 1.0906 | 3.3415 |
| 1533 | ENSMUSG00000014850 | 248.1637287 | 0.079242186 | 0.159329585 | 0.497347598 | 0.618943952 | 0.917509899 | Msh3 | mutS homolog 3 [Source:MGI Symbol;Acc | 232.1 | 238.48 | 253.57 | 243.89 | 277.02 | 243.93 |
| 1534 | ENSMUSG00000014852 | 79.21382107 | -1.078745519 | 0.293736632 | -3.672492296 | 0.000240196 | 0.007268202 | Adamts13 | a disintegrin-like and metallopeptidase (re | 114.84 | 122.32 | 85.531 | 59.767 | 34.9 | 57.92 |
| 1535 | ENSMUSG00000014856 | 567.9016853 | 0.244161929 | 0.124987582 | 1.953489504 | 0.05076162 | 0.316199857 | Tmem208 | transmembrane protein 208 [Source:MGI | 479.45 | 536.57 | 544.38 | 588.03 | 611.84 | 647.14 |
| 1536 | ENSMUSG00000014859 | 310.9679298 | -0.016890302 | 0.154183319 | -0.10954688 | 0.91276874 | 0.98625062 | E2f4 | E2F transcription factor 4 [Source:MGI Sy | 309.19 | 320.71 | 308.92 | 341.25 | 268.29 | 317.45 |
| 1537 | ENSMUSG00000014867 | 4160.702301 | 0.040839813 | 0.088706694 | 0.460391564 | 0.645235189 | 0.925643431 | Surf4 | surfeit gene 4 [Source:MGI Symbol;Acc:M | 4300.6 | 4123 | 3881.1 | 4249.2 | 4251.2 | 4159.1 |
| 1538 | ENSMUSG00000014873 | 264.2836849 | -0.062052054 | 0.16207415 | -0.382862126 | 0.701821994 | 0.942189108 | Surf2 | surfeit gene 2 [Source:MGI Symbol;Acc:M | 301.16 | 244.64 | 263.64 | 272.81 | 261.75 | 241.7 |
| 1539 | ENSMUSG00000014905 | 810.6072132 | -0.041970344 | 0.159997392 | -0.262318926 | 0.793075572 | 0.961851842 | Dnajb9 | DnaJ heat shock protein family (Hsp40) m | 755.72 | 1033.1 | 679.22 | 830.95 | 848.5 | 716.2 |
| 1540 | ENSMUSG00000014907 | 110.8144153 | 0.143045705 | 0.23842079 | 0.599971611 | 0.548525155 | 0.895622715 | Naf1 | nuclear assembly factor 1 ribonucleoprote | 97.978 | 129.52 | 88.55 | 105.07 | 112.33 | 131.43 |
| 1541 | ENSMUSG00000014932 | 557.589707 | 0.089154404 | 0.131831833 | 0.67627372 | 0.498866875 | 0.880236544 | Yes1 | YES proto-oncogene 1, Src family tyrosine | 483.47 | 545.82 | 592.68 | 565.85 | 607.47 | 550.24 |
| 1542 | ENSMUSG00000014956 | 2071.924081 | 0.013112059 | 0.10028688 | 0.130745506 | 0.895976633 | 0.982798755 | Ppp1cb | protein phosphatase 1 catalytic subunit b | 2039.1 | 2157.6 | 1990.4 | 1930.8 | 2103.8 | 2209.9 |
| 1543 | ENSMUSG00000014959 | 1447.600814 | 0.084234298 | 0.10189408 | 0.826684909 | 0.408415677 | 0.832151413 | Gorasp2 | golgi reassembly stacking protein 2 [Sourc | 1464.9 | 1429.8 | 1321.2 | 1552 | 1490.9 | 1426.8 |
| 1544 | ENSMUSG00000014980 | 107.7155083 | 0.035214958 | 0.242547865 | 0.145187664 | 0.884562705 | 0.981795936 | Tsen15 | tRNA splicing endonuclease subunit 15 [S | 120.47 | 87.373 | 110.69 | 90.614 | 109.06 | 128.09 |
| 1545 | ENSMUSG00000015001 | 0 | NA | NA | NA | NA | NA | Oc90 | otoconin 90 [Source:MGI Symbol;Acc:MG | 0 | 0 | 0 | 0 | 0 | 0 |
| 1546 | ENSMUSG00000015002 | 828.1855581 | -0.00138482 | 0.128270467 | -0.010796091 | 0.991386133 | 0.99859176 | Efr3a | EFR3 homolog A [Source:MGI Symbol;Acc | 750.9 | 930.27 | 805 | 752.87 | 875.77 | 854.32 |
| 1547 | ENSMUSG00000015013 | 199.0073598 | 0.204740366 | 0.196158098 | 1.043751788 | 0.296600243 | 0.74713095 | Trappc2l | trafficking protein particle complex 2-like | 183.91 | 206.61 | 164.02 | 195.69 | 185.41 | 258.41 |
| 1548 | ENSMUSG00000015016 | 726.0027266 | -0.127493877 | 0.140582267 | -0.906898716 | 0.36446036 | 0.802897466 | Acsf3 | acyl-CoA synthetase family member 3 [So | 826.39 | 759.63 | 687.27 | 642.01 | 812.51 | 628.21 |
| 1549 | ENSMUSG00000015023 | 456.1473852 | 0.161139969 | 0.139022823 | 1.159090043 | 0.246419484 | 0.696725814 | Ddx19a | DEAD box helicase 19a [Source:MGI Symb | 436.89 | 388.55 | 466.9 | 524.4 | 465.69 | 454.45 |
| 1550 | ENSMUSG00000015027 | 85.95970402 | 0.351115926 | 0.285047246 | 1.231781507 | 0.218030713 | 0.663697152 | Galns | galactosamine (N-acetyl)-6-sulfate sulfata | 82.72 | 69.898 | 73.456 | 63.623 | 106.88 | 119.18 |
| 1551 | ENSMUSG00000015053 | 11.03577156 | -0.497847276 | 0.640558295 | -0.777208382 | 0.437035838 | 0.846852749 | Gata2 | GATA binding protein 2 [Source:MGI Syml | 15.259 | 12.335 | 11.069 | 7.7118 | 9.8156 | 10.025 |
| 1552 | ENSMUSG00000015083 | 8380.905501 | 0.120184169 | 0.09664593 | 1.243551262 | 0.213664763 | 0.658358849 | C8g | complement component 8, gamma polype | 8245.5 | 7268.4 | 8581.3 | 8391.4 | 8959.4 | 8839.5 |
| 1553 | ENSMUSG00000015085 | 59.47185723 | 0.049633207 | 0.291493462 | 0.17027211 | 0.864796144 | 0.977227145 | Entpd2 | ectonucleoside triphosphate diphosphohy | 58.626 | 49.34 | 67.418 | 64.587 | 56.712 | 60.148 |
| 1554 | ENSMUSG00000015087 | 522.2278701 | 0.029029363 | 0.150486178 | 0.192903847 | 0.847034272 | 0.972803041 | Rabl6 | RAB, member RAS oncogene family-like 6 | 521.21 | 520.13 | 510.17 | 594.77 | 564.94 | 422.15 |
| 1555 | ENSMUSG00000015090 | 28.84417841 | 1.243700044 | 0.562489743 | 2.211062618 | 0.027031501 | 0.212954048 | Ptgds | prostaglandin D2 synthase (brain) [Source | 12.85 | 14.391 | 24.15 | 11.568 | 59.984 | 50.123 |
| 1556 | ENSMUSG00000015092 | 3293.667104 | -0.10471355 | 0.092889567 | -1.127290764 | 0.259619566 | 0.712820312 | Edf1 | endothelial differentiation-related factor | 3658.9 | 3314 | 3265.3 | 3173.4 | 3121.3 | 3229 |
| 1557 | ENSMUSG00000015093 | 8.589389058 | 0.392431078 | 0.772155403 | 0.508228106 | 0.611293382 | 0.916376001 | Clic3 | chloride intracellular channel 3 [Source:M | 3.2124 | 7.1954 | 12.075 | 12.532 | 8.7249 | 7.7969 |
| 1558 | ENSMUSG00000015094 | 55.75773601 | -0.482861569 | 0.362607553 | -1.331636821 | 0.182979559 | 0.61680331 | Npdc1 | neural proliferation, differentiation and co | 90.751 | 58.591 | 45.281 | 44.343 | 62.165 | 33.415 |
| 1559 | ENSMUSG00000015095 | 627.9127903 | 0.03201781 | 0.118457249 | 0.270290004 | 0.786937155 | 0.960870081 | Fbxw5 | F-box and WD-40 domain protein 5 [Sourc | 656.94 | 604.42 | 600.73 | 627.55 | 615.11 | 662.74 |
| 1560 | ENSMUSG00000015112 | 5606.97463 | 0.029658866 | 0.096967477 | 0.305864058 | 0.759708153 | 0.95402891 | Slc25a13 | solute carrier family 25 (mitochondrial car | 5045.1 | 5665.9 | 5938.9 | 5872.6 | 5598.1 | 5521.3 |
| 1561 | ENSMUSG00000015120 | 1003.760069 | 0.132530875 | 0.117524068 | 1.12769135 | 0.259450292 | 0.712728614 | Ube2i | ubiquitin-conjugating enzyme E2I [Source | 958.1 | 950.82 | 964.99 | 1182.8 | 997.92 | 967.93 |
| 1562 | ENSMUSG00000015126 | 251.9579262 | 0.011199153 | 0.174197295 | 0.064290049 | 0.948739277 | 0.990731019 | Tsr3 | TSR3 20S rRNA accumulation [Source:MG | 273.05 | 236.42 | 242.5 | 211.11 | 283.56 | 265.09 |
| 1563 | ENSMUSG00000015127 | 160.0143659 | 0.237598618 | 0.221378459 | 1.073268915 | 0.283150478 | 0.735802616 | Unkl | unkempt family like zinc finger [Source:M | 148.57 | 113.07 | 179.11 | 199.54 | 151.6 | 168.19 |
| 1564 | ENSMUSG00000015133 | 304.6039623 | -0.114720083 | 0.156562217 | -0.732744371 | 0.463714356 | 0.861771097 | Lrrk1 | leucine-rich repeat kinase 1 [Source:MGI | 343.73 | 326.88 | 278.73 | 270.88 | 296.65 | 310.76 |

| | | | | | | | | | | | | | | | |
|---|---|---|---|---|---|---|---|---|---|---|---|---|---|---|---|
| 1565 | ENSMUSG00000015134 | 8.201118896 | -0.683705459 | 0.809484045 | -0.84461882 | 0.398323698 | 0.825882163 | Aldh1a3 | aldehyde dehydrogenase family 1, subfam | 12.047 | 6.1675 | 12.075 | 5.7839 | 10.906 | 2.2277 |
| 1566 | ENSMUSG00000015143 | 461.7358975 | 0.22288752 | 0.134254206 | 1.660190218 | 0.096876192 | 0.458530879 | Actn1 | actinin, alpha 1 [Source:MGI Symbol;Acc:I | 459.37 | 414.25 | 404.51 | 536.94 | 486.42 | 468.93 |
| 1567 | ENSMUSG00000015149 | 997.4801536 | -0.03635313 | 0.10946908 | -0.332085831 | 0.739824449 | 0.948393197 | Sirt2 | sirtuin 2 [Source:MGI Symbol;Acc:MGI:19 | 1022.3 | 1038.2 | 970.02 | 1043 | 996.83 | 914.47 |
| 1568 | ENSMUSG00000015165 | 1789.210134 | 0.04225471 | 0.100356174 | 0.421047436 | 0.673720444 | 0.931915649 | Hnrnpl | heterogeneous nuclear ribonucleoprotein | 1784.5 | 1776.2 | 1728.7 | 1914.5 | 1861.7 | 1669.7 |
| 1569 | ENSMUSG00000015176 | 535.3479887 | -0.426862745 | 0.146870311 | -2.906392328 | 0.003656227 | 0.054018047 | Nolc1 | nucleolar and coiled-body phosphoprotei | 598.31 | 657.87 | 585.63 | 418.37 | 543.13 | 408.78 |
| 1570 | ENSMUSG00000015189 | 511.5781401 | 0.285411201 | 0.143268267 | 1.992145276 | 0.04635512 | 0.297726671 | Casd1 | CAS1 domain containing 1 [Source:MGI Sy | 401.55 | 498.54 | 484 | 519.58 | 542.04 | 623.75 |
| 1571 | ENSMUSG00000015202 | 117.4956851 | -0.004854526 | 0.236968532 | -0.020485951 | 0.983655719 | 0.998580735 | Cnksr3 | Cnksr family member 3 [Source:MGI Symb | 122.07 | 106.9 | 123.77 | 107 | 95.974 | 149.26 |
| 1572 | ENSMUSG00000015214 | 629.4228468 | 0.033416065 | 0.125589169 | 0.266074421 | 0.790181894 | 0.961019028 | Mtmr1 | myotubularin related protein 1 [Source:M | 632.84 | 654.78 | 578.59 | 620.8 | 694.72 | 594.79 |
| 1573 | ENSMUSG00000015217 | 207.8405113 | 0.156365926 | 0.178052213 | 0.878202657 | 0.379833751 | 0.812628154 | Hmgb3 | high mobility group box 3 [Source:MGI Sy | 177.49 | 191.19 | 221.37 | 209.18 | 207.22 | 240.59 |
| 1574 | ENSMUSG00000015222 | 10.32171765 | -0.401090553 | 0.769321201 | -0.521356427 | 0.602118499 | 0.915461118 | Map2 | microtubule-associated protein 2 [Source | 16.865 | 3.0838 | 15.094 | 4.8199 | 9.8156 | 12.252 |
| 1575 | ENSMUSG00000015224 | 117.1709441 | -0.282566536 | 0.361338232 | -0.782000108 | 0.43421451 | 0.845264432 | Cyp2j9 | cytochrome P450, family 2, subfamily j, pc | 114.04 | 136.71 | 135.84 | 190.87 | 66.528 | 59.034 |
| 1576 | ENSMUSG00000015242 | 0 | NA | NA | NA | NA | NA | Nipsnap3a | nipsnap homolog 3A [Source:MGI Symbol | 0 | 0 | 0 | 0 | 0 | 0 |
| 1577 | ENSMUSG00000015243 | 2953.623334 | -0.024659122 | 0.117067794 | -0.210639676 | 0.83316845 | 0.970611245 | Abca1 | ATP-binding cassette, sub-family A (ABC1) | 3153 | 3100.2 | 2683.7 | 3221.6 | 2998.1 | 2565.2 |
| 1578 | ENSMUSG00000015247 | 1500.134754 | 0.074128707 | 0.117117186 | 0.632944738 | 0.526769731 | 0.888334265 | Nipsnap3b | nipsnap homolog 3B [Source:MGI Symbol | 1579.7 | 1488.4 | 1315.2 | 1392.9 | 1569.4 | 1655.2 |
| 1579 | ENSMUSG00000015289 | 292.6538983 | 0.099889543 | 0.146016381 | 0.684098196 | 0.493913161 | 0.877473722 | Lage3 | L antigen family, member 3 [Source:MGI S | 289.92 | 282.68 | 274.7 | 297.87 | 296.65 | 314.1 |
| 1580 | ENSMUSG00000015290 | 680.4640946 | 0.042280483 | 0.114786825 | 0.36833916 | 0.712620357 | 0.944239215 | Ubl4a | ubiquitin-like 4A [Source:MGI Symbol;Acc | 676.21 | 665.06 | 670.16 | 701.78 | 648.92 | 720.66 |
| 1581 | ENSMUSG00000015291 | 990.1985888 | 0.184266695 | 0.10889079 | 1.692215606 | 0.090604872 | 0.444541679 | Gdi1 | guanosine diphosphate (GDP) dissociation | 928.39 | 880.93 | 972.03 | 1084.5 | 1085.2 | 990.21 |
| 1582 | ENSMUSG00000015305 | 603.0312661 | 0.145432615 | 0.121321036 | 1.198741944 | 0.230628303 | 0.677871932 | Sash1 | SAM and SH3 domain containing 1 [Sourc | 611.16 | 550.96 | 555.45 | 663.22 | 631.47 | 605.93 |
| 1583 | ENSMUSG00000015312 | 20.5365641 | -2.268776559 | 0.669112098 | -3.390727154 | 0.000697075 | 0.015902977 | Gadd45b | growth arrest and DNA-damage-inducible | 30.518 | 50.368 | 21.131 | 4.8199 | 15.269 | 1.1138 |
| 1584 | ENSMUSG00000015314 | 2.189608702 | -1.178148657 | 1.676939801 | -0.702558706 | 0.482330804 | NA | Slamf6 | SLAM family member 6 [Source:MGI Symb | 4.0155 | 5.1396 | 0 | 2.8919 | 1.0906 | 0 |
| 1585 | ENSMUSG00000015316 | 0 | NA | NA | NA | NA | NA | Slamf1 | signaling lymphocytic activation molecule | 0 | 0 | 0 | 0 | 0 | 0 |
| 1586 | ENSMUSG00000015335 | 248.9291531 | 0.033769432 | 0.161955328 | 0.208510783 | 0.834830164 | 0.971073158 | Zdhhc12 | zinc finger, DHHC domain containing 12 [S | 247.36 | 248.76 | 241.5 | 225.57 | 251.93 | 278.46 |
| 1587 | ENSMUSG00000015337 | 684.6903953 | -0.034508235 | 0.139630487 | -0.247139691 | 0.804800117 | 0.963787787 | Endog | endonuclease G [Source:MGI Symbol;Acc | 662.56 | 629.09 | 787.89 | 760.58 | 657.64 | 610.39 |
| 1588 | ENSMUSG00000015340 | 120.0380792 | -1.363856482 | 0.277570536 | -4.913549196 | 8.94E-07 | 7.48E-05 | Cybb | cytochrome b-245, beta polypeptide [Sou | 165.44 | 229.23 | 123.77 | 52.055 | 81.796 | 67.944 |
| 1589 | ENSMUSG00000015341 | 1131.518802 | 0.089941586 | 0.109220834 | 0.823483785 | 0.410232935 | 0.833241465 | Golga7 | golgi autoantigen, golgin subfamily a, 7 [S | 1091.4 | 1111.2 | 1085.7 | 1079.7 | 1272.8 | 1148.4 |
| 1590 | ENSMUSG00000015342 | 114.1044755 | 0.246109686 | 0.243196953 | 1.01197685 | 0.311549122 | 0.759830222 | Xk | X-linked Kx blood group [Source:MGI Sym | 81.916 | 106.9 | 124.77 | 112.79 | 112.33 | 145.91 |
| 1591 | ENSMUSG00000015354 | 10.87017606 | -0.698807059 | 0.833888634 | -0.838010054 | 0.402025057 | 0.82893099 | Pcolce2 | procollagen C-endopeptidase enhancer 2 | 13.653 | 20.558 | 6.0375 | 3.8559 | 2.1812 | 18.935 |
| 1592 | ENSMUSG00000015355 | 20.01932263 | -1.786389729 | 0.620298542 | -2.879887035 | 0.003978177 | 0.056970643 | Cd48 | CD48 antigen [Source:MGI Symbol;Acc:M | 26.502 | 47.284 | 19.119 | 2.8919 | 8.7249 | 15.594 |
| 1593 | ENSMUSG00000015357 | 2684.166114 | -0.976431121 | 0.376062669 | -2.596458521 | 0.009419028 | 0.10399757 | Clpx | caseinolytic mitochondrial matrix peptida | 2067.2 | 3608 | 5003 | 1668.6 | 1568.3 | 2189.8 |
| 1594 | ENSMUSG00000015363 | 1941.1461 | 0.298157643 | 0.115529747 | 2.580786787 | 0.009857544 | 0.107526033 | Trabd | TraB domain containing [Source:MGI Sym | 1600.6 | 1841 | 1784.1 | 2412.8 | 2003.5 | 2004.9 |
| 1595 | ENSMUSG00000015365 | 0 | NA | NA | NA | NA | NA | Mov10l1 | Mov10 like RISC complex RNA helicase 1 [ | 0 | 0 | 0 | 0 | 0 | 0 |
| 1596 | ENSMUSG00000015377 | 147.3027148 | -0.070520208 | 0.207556254 | -0.339764312 | 0.734034025 | 0.948109711 | Dennd6b | DENN/MADD domain containing 6B [Sour | 141.35 | 139.8 | 172.07 | 164.84 | 125.42 | 140.34 |
| 1597 | ENSMUSG00000015396 | 14.0634518 | -1.044367519 | 0.609006408 | -1.714871148 | 0.086368854 | 0.433027696 | Cd83 | CD83 antigen [Source:MGI Symbol;Acc:M | 20.078 | 20.558 | 16.1 | 7.7118 | 5.4531 | 14.48 |
| 1598 | ENSMUSG00000015401 | 0.17131964 | -0.796629176 | 4.080472857 | -0.195229623 | 0.84521319 | NA | Cltrn | collectrin, amino acid transport regulator | 0 | 1.0279 | 0 | 0 | 0 | 0 |
| 1599 | ENSMUSG00000015405 | 2.475037729 | -0.417259668 | 1.327341701 | -0.314357387 | 0.753249617 | NA | Ace2 | angiotensin I converting enzyme (peptidyl | 2.4093 | 2.0558 | 4.025 | 1.928 | 1.0906 | 3.3415 |
| 1600 | ENSMUSG00000015437 | 1.614723054 | -1.79710927 | 1.773843138 | -1.013116228 | 0.311004648 | NA | Gzmb | granzyme B [Source:MGI Symbol;Acc:MGI | 2.4093 | 2.0558 | 3.0187 | 0 | 1.0906 | 1.1138 |
| 1601 | ENSMUSG00000015441 | 0 | NA | NA | NA | NA | NA | Gzmf | granzyme F [Source:MGI Symbol;Acc:MGI | 0 | 0 | 0 | 0 | 0 | 0 |
| 1602 | ENSMUSG00000015443 | 0 | NA | NA | NA | NA | NA | Gzmn | granzyme N [Source:MGI Symbol;Acc:MG | 0 | 0 | 0 | 0 | 0 | 0 |
| 1603 | ENSMUSG00000015451 | 2386.835675 | 0.97032664 | 0.371635304 | 2.610964649 | 0.009028723 | 0.100532289 | C4a | complement component 4A (Rodgers bloc | 1786.1 | 1296.2 | 1756.9 | 3151.2 | 4470.4 | 1860.1 |
| 1604 | ENSMUSG00000015452 | 1.513391143 | -0.296673934 | 1.904666913 | -0.155761583 | 0.876220953 | NA | Ager | advanced glycosylation end product-speci | 0 | 2.0558 | 3.0187 | 2.8919 | 0 | 1.1138 |
| 1605 | ENSMUSG00000015461 | 390.4244462 | -0.06153584 | 0.135487088 | -0.454182324 | 0.649697605 | 0.925643431 | Atf6b | activating transcription factor 6 beta [Sou | 371.03 | 418.36 | 407.53 | 387.52 | 386.08 | 372.02 |
| 1606 | ENSMUSG00000015467 | 8.177400925 | 0.792908561 | 0.874591433 | 0.906604537 | 0.364615962 | 0.803076788 | Egfl8 | EGF-like domain 8 [Source:MGI Symbol;Ac | 0.8031 | 11.307 | 6.0375 | 7.7118 | 8.7249 | 14.48 |
| 1607 | ENSMUSG00000015468 | 47.72781414 | -0.133559075 | 0.343316036 | -0.389026614 | 0.697256459 | 0.940268454 | Notch4 | notch 4 [Source:MGI Symbol;Acc:MGI:107 | 60.233 | 43.173 | 46.287 | 54.947 | 32.719 | 49.009 |
| 1608 | ENSMUSG00000015474 | 367.3060352 | -0.023066845 | 0.141466143 | -0.163055585 | 0.87047467 | 0.978802475 | Ppt2 | palmitoyl-protein thioesterase 2 [Source:I | 379.06 | 337.16 | 394.45 | 375.95 | 370.81 | 346.41 |
| 1609 | ENSMUSG00000015476 | 5.564414004 | -0.232915147 | 1.019871596 | -0.228376933 | 0.819353221 | 0.969160361 | Prrt1 | proline-rich transmembrane protein 1 [So | 12.85 | 2.0558 | 3.0187 | 7.7118 | 2.1812 | 5.5692 |
| 1610 | ENSMUSG00000015478 | 1053.711292 | 0.120153489 | 0.111374349 | 1.078825511 | 0.28066552 | 0.733530291 | Rnf5 | ring finger protein 5 [Source:MGI Symbol; | 970.95 | 1000.2 | 1059.6 | 1194.4 | 1053.5 | 1043.7 |
| 1611 | ENSMUSG00000015484 | 0 | NA | NA | NA | NA | NA | Fam163a | family with sequence similarity 163, meml | 0 | 0 | 0 | 0 | 0 | 0 |
| 1612 | ENSMUSG00000015488 | 609.454673 | 0.240470234 | 0.125642874 | 1.913918604 | 0.055630557 | 0.332908719 | Cacfd1 | calcium channel flower domain containing | 579.04 | 555.08 | 541.36 | 608.27 | 645.65 | 727.34 |
| 1613 | ENSMUSG00000015501 | 142.3271863 | 0.250229202 | 0.197081078 | 1.269676439 | 0.204199909 | 0.649416772 | Hivep2 | human immunodeficiency virus type I enh | 130.91 | 131.57 | 127.79 | 171.59 | 143.96 | 148.14 |
| 1614 | ENSMUSG00000015519 | 0.185640647 | 1.126924175 | 4.080472857 | 0.276174898 | 0.782413736 | NA | 2310057J18I | RIKEN cDNA 2310057J18 gene [Source:MC | 0 | 0 | 0 | 0 | 0 | 1.1138 |
| 1615 | ENSMUSG00000015522 | 695.2898855 | 0.089844135 | 0.150262083 | 0.597916208 | 0.549895844 | 0.895703112 | Arnt | aryl hydrocarbon receptor nuclear translo | 677.02 | 633.2 | 711.41 | 833.84 | 745.98 | 570.29 |
| 1616 | ENSMUSG00000015533 | 2.217579939 | -2.380130971 | 1.683639647 | -1.413681945 | 0.157455309 | NA | Itga2 | integrin alpha 2 [Source:MGI Symbol;Acc: | 4.0155 | 3.0838 | 4.025 | 0 | 2.1812 | 0 |
| 1617 | ENSMUSG00000015536 | 2153.628995 | 0.00996791 | 0.097050977 | 0.102707977 | 0.918194742 | 0.986374671 | Mocs2 | molybdenum cofactor synthesis 2 [Source | 2187.7 | 2076.4 | 2174.5 | 2207.5 | 2264.1 | 2011.6 |
| 1618 | ENSMUSG00000015542 | 579.2276377 | 0.027557448 | 0.129891587 | 0.212157296 | 0.831984322 | 0.970533397 | Nat9 | N-acetyltransferase 9 (GCN5-related, puta | 613.57 | 544.8 | 561.48 | 524.4 | 612.93 | 618.18 |
| 1619 | ENSMUSG00000015568 | 97.98439064 | -0.606058734 | 0.410382865 | -1.476812959 | 0.139725782 | 0.546014685 | Lpl | lipoprotein lipase [Source:MGI Symbol;Ac | 135.72 | 154.19 | 64.4 | 36.631 | 61.075 | 135.89 |
| 1620 | ENSMUSG00000015575 | 689.1731776 | -0.074887993 | 0.119105574 | -0.62875305 | 0.529510742 | 0.888334265 | Atp6v0e | ATPase, H+ transporting, lysosomal V0 sul | 726.81 | 713.37 | 680.22 | 619.84 | 719.81 | 674.99 |

| | | | | | | | | | | | | | | |
|---|---|---|---|---|---|---|---|---|---|---|---|---|---|---|
| 1621 | ENSMUSG00000015579 | 0 | NA | NA | NA | NA | NA | Nkx2-5 | NK2 homeobox 5 [Source:MGI Symbol;Ac | 0 | 0 | 0 | 0 | 0 | 0 |
| 1622 | ENSMUSG00000015597 | 458.0108678 | 0.213271474 | 0.13607542 | 1.567303443 | 0.117043781 | 0.502643331 | Zfp318 | zinc finger protein 318 [Source:MGI Symb | 426.45 | 399.86 | 446.77 | 541.76 | 478.78 | 454.45 |
| 1623 | ENSMUSG00000015599 | 132.2687648 | -0.262369703 | 0.219267445 | -1.196573907 | 0.231472676 | 0.678699825 | Ttbk1 | tau tubulin kinase 1 [Source:MGI Symbol; | 171.86 | 126.43 | 133.83 | 113.75 | 135.24 | 112.5 |
| 1624 | ENSMUSG00000015605 | 226.6614891 | 0.026169394 | 0.168595818 | 0.155219712 | 0.876648109 | 0.980632845 | Srf | serum response factor [Source:MGI Symb | 228.08 | 245.67 | 200.24 | 242.92 | 221.4 | 221.65 |
| 1625 | ENSMUSG00000015619 | 1.729321077 | -3.053230067 | 1.994397783 | -1.53090326 | 0.125793303 | NA | Gata3 | GATA binding protein 3 [Source:MGI Symb | 7.2279 | 1.0279 | 1.0062 | 0 | 0 | 1.1138 |
| 1626 | ENSMUSG00000015627 | 4.810291517 | 0.433808162 | 0.96413895 | 0.449943612 | 0.6527511 | NA | Gata5 | GATA binding protein 5 [Source:MGI Symb | 3.2124 | 4.1117 | 5.0312 | 7.7118 | 5.4531 | 3.3415 |
| 1627 | ENSMUSG00000015647 | 47.31359937 | -0.383045083 | 0.370524436 | -1.033791691 | 0.301233557 | 0.7521219 | Lama5 | laminin, alpha 5 [Source:MGI Symbol;Acc: | 49.792 | 68.87 | 42.262 | 51.091 | 25.084 | 46.781 |
| 1628 | ENSMUSG00000015652 | 1.644475959 | 1.440245623 | 1.855875835 | 0.776046326 | 0.437721634 | NA | Steap1 | six transmembrane epithelial antigen of th | 1.6062 | 0 | 1.0062 | 2.8919 | 4.3625 | 0 |
| 1629 | ENSMUSG00000015653 | 143.4778963 | -0.063508383 | 0.275615735 | -0.230423649 | 0.817762587 | 0.968565186 | Steap2 | six transmembrane epithelial antigen of p | 97.978 | 185.03 | 157.98 | 186.05 | 107.97 | 125.86 |
| 1630 | ENSMUSG00000015656 | 16774.04041 | -0.002208776 | 0.093199489 | -0.023699442 | 0.981092351 | 0.99820894 | Hspa8 | heat shock protein 8 [Source:MGI Symbol | 16860 | 17629 | 15871 | 15535 | 17033 | 17716 |
| 1631 | ENSMUSG00000015659 | 70.22703506 | 0.075527125 | 0.271239917 | 0.278451365 | 0.780665894 | 0.959921066 | Serac1 | serine active site containing 1 [Source:MG | 60.233 | 65.787 | 79.493 | 78.082 | 68.709 | 69.058 |
| 1632 | ENSMUSG00000015665 | 0 | NA | NA | NA | NA | NA | Awat1 | acyl-CoA wax alcohol acyltransferase 1 [So | 0 | 0 | 0 | 0 | 0 | 0 |
| 1633 | ENSMUSG00000015668 | 259.7284542 | -0.102129055 | 0.176047854 | -0.580120987 | 0.561833032 | 0.900565925 | Pdzd11 | PDZ domain containing 11 [Source:MGI Sy | 316.42 | 242.59 | 246.53 | 223.64 | 256.3 | 272.89 |
| 1634 | ENSMUSG00000015671 | 1355.062519 | 0.004375715 | 0.101311506 | 0.043190698 | 0.96554952 | 0.993576285 | Psma2 | proteasome subunit alpha 2 [Source:MGI | 1410.2 | 1302.4 | 1345.3 | 1305.2 | 1341.5 | 1425.7 |
| 1635 | ENSMUSG00000015672 | 420.8491578 | -0.087584613 | 0.139897563 | -0.626062467 | 0.53127397 | 0.888877582 | Mrpl32 | mitochondrial ribosomal protein L32 [Sou | 390.31 | 444.06 | 466.9 | 382.7 | 410.07 | 431.06 |
| 1636 | ENSMUSG00000015697 | 469.93944 | 0.194989223 | 0.135851517 | 1.435311342 | 0.151198397 | 0.566416373 | Setdb1 | SET domain, bifurcated 1 [Source:MGI Syr | 403.16 | 473.87 | 438.72 | 530.19 | 513.68 | 460.02 |
| 1637 | ENSMUSG00000015702 | 17.74360122 | -0.346001821 | 0.559270534 | -0.61866628 | 0.536136229 | 0.890560262 | Anxa9 | annexin A9 [Source:MGI Symbol;Acc:MGI | 17.668 | 29.81 | 12.075 | 11.568 | 14.178 | 21.163 |
| 1638 | ENSMUSG00000015709 | 0 | NA | NA | NA | NA | NA | Arnt2 | aryl hydrocarbon receptor nuclear translo | 0 | 0 | 0 | 0 | 0 | 0 |
| 1639 | ENSMUSG00000015711 | 284.5034954 | -0.091686282 | 0.177975445 | -0.515162537 | 0.606439445 | 0.916376001 | Prune1 | prune exopolyphosphatase [Source:MGI S | 350.15 | 285.76 | 243.51 | 246.78 | 290.1 | 290.71 |
| 1640 | ENSMUSG00000015714 | 5796.5912 | 0.088891655 | 0.090670554 | 0.980380631 | 0.326898267 | 0.772717198 | Cers2 | ceramide synthase 2 [Source:MGI Symbol | 5449 | 5537.4 | 5868.4 | 5996.9 | 6294 | 5633.8 |
| 1641 | ENSMUSG00000015721 | 0 | NA | NA | NA | NA | NA | Nlrp5 | NLR family, pyrin domain containing 5 [So | 0 | 0 | 0 | 0 | 0 | 0 |
| 1642 | ENSMUSG00000015733 | 1179.273559 | -0.00300716 | 0.104878074 | -0.02867291 | 0.977125462 | 0.997328536 | Capza2 | capping protein (actin filament) muscle Z- | 1182.2 | 1201.6 | 1157.2 | 1100.9 | 1253.1 | 1180.7 |
| 1643 | ENSMUSG00000015745 | 65.38590285 | -0.595191929 | 0.281947952 | -2.110999296 | 0.034772372 | 0.249973953 | Plekho1 | pleckstrin homology domain containing, f | 68.264 | 90.457 | 77.481 | 49.163 | 51.259 | 55.692 |
| 1644 | ENSMUSG00000015747 | 135.0628317 | -0.115714784 | 0.196270861 | -0.589566802 | 0.555481114 | 0.897628185 | Vps45 | vacuolar protein sorting 45 [Source:MGI S | 139.74 | 143.91 | 137.86 | 132.06 | 127.6 | 129.21 |
| 1645 | ENSMUSG00000015748 | 178.8973846 | 0.077914677 | 0.187647252 | 0.415218855 | 0.677981697 | 0.93325821 | Prpf3 | pre-mRNA processing factor 3 [Source:MG | 155 | 169.61 | 198.23 | 196.65 | 171.23 | 182.67 |
| 1646 | ENSMUSG00000015749 | 394.9387121 | -0.222268771 | 0.152501335 | -1.457487384 | 0.144981891 | 0.555361204 | Anp32e | acidic (leucine-rich) nuclear phosphoprote | 375.85 | 415.28 | 485.01 | 329.68 | 379.54 | 384.28 |
| 1647 | ENSMUSG00000015750 | 1256.412165 | -0.096078861 | 0.103642388 | -0.927022843 | 0.353914674 | 0.79449673 | Aph1a | aph1 homolog A, gamma secretase subun | 1232 | 1314.7 | 1348.4 | 1177 | 1185.5 | 1280.9 |
| 1648 | ENSMUSG00000015755 | 929.1160483 | -0.157888949 | 0.127615459 | -1.23722431 | 0.216003819 | 0.661643209 | Tab2 | TGF-beta activated kinase 1/MAP3K7 binc | 897.07 | 1026.9 | 1016.3 | 866.62 | 777.61 | 990.21 |
| 1649 | ENSMUSG00000015757 | 278.7493898 | -0.100545323 | 0.168899825 | -0.595295601 | 0.551645896 | 0.895935901 | Ppil4 | peptidylprolyl isomerase (cyclophilin)-like | 310.8 | 251.84 | 301.87 | 240.99 | 259.57 | 307.42 |
| 1650 | ENSMUSG00000015759 | 825.980936 | 0.054556449 | 0.125821406 | 0.43360228 | 0.664577279 | 0.929301301 | Cnih1 | cornichon family AMPA receptor auxiliary | 880.2 | 803.83 | 745.63 | 750.94 | 897.58 | 877.71 |
| 1651 | ENSMUSG00000015766 | 29.35883993 | -1.095593972 | 0.436689915 | -2.508860257 | 0.012112139 | 0.123460168 | Eps8 | epidermal growth factor receptor pathwa | 53.808 | 34.949 | 31.194 | 23.135 | 16.359 | 16.708 |
| 1652 | ENSMUSG00000015776 | 671.8614207 | 0.189343191 | 0.121910599 | 1.553131504 | 0.120391721 | 0.507977203 | Med22 | mediator complex subunit 22 [Source:MG | 647.3 | 609.56 | 625.88 | 660.32 | 784.15 | 703.95 |
| 1653 | ENSMUSG00000015787 | 0 | NA | NA | NA | NA | NA | Abo | ABO blood group (transferase A, alpha 1-3 | 0 | 0 | 0 | 0 | 0 | 0 |
| 1654 | ENSMUSG00000015790 | 704.709784 | -0.005643518 | 0.111330906 | -0.050691386 | 0.959571441 | 0.992148164 | Surf1 | surfeit gene 1 [Source:MGI Symbol;Acc:M | 686.65 | 725.71 | 706.38 | 715.27 | 693.63 | 700.61 |
| 1655 | ENSMUSG00000015804 | 339.7975169 | 0.121635242 | 0.155798138 | 0.780723334 | 0.434965233 | 0.845325456 | Med28 | mediator complex subunit 28 [Source:MG | 347.74 | 334.07 | 293.82 | 322.93 | 337 | 403.21 |
| 1656 | ENSMUSG00000015806 | 5246.380696 | 0.328261471 | 0.115355791 | 2.845643624 | 0.004432177 | 0.061582136 | Qdpr | quinoid dihydropteridine reductase [Sour | 5110.1 | 4449.9 | 4394.3 | 5235.4 | 5666.9 | 6621.8 |
| 1657 | ENSMUSG00000015812 | 5.227888791 | -0.179909217 | 0.924876602 | -0.1945224 | 0.845766858 | 0.972803041 | Gnrh1 | gonadotropin releasing hormone 1 [Sourc | 6.4248 | 7.1954 | 3.0187 | 4.8199 | 5.4531 | 4.4554 |
| 1658 | ENSMUSG00000015829 | 0.133850373 | -0.796629176 | 4.080472857 | -0.195229623 | 0.84521319 | NA | Tnr | tenascin R [Source:MGI Symbol;Acc:MGI:9 | 0.8031 | 0 | 0 | 0 | 0 | 0 |
| 1659 | ENSMUSG00000015837 | 6279.969938 | 0.001478154 | 0.104672446 | 0.014121711 | 0.988732879 | 0.99859176 | Sqstm1 | sequestosome 1 [Source:MGI Symbol;Acc | 6922.7 | 5911.6 | 5994.2 | 6105.8 | 5921 | 6824.5 |
| 1660 | ENSMUSG00000015839 | 1120.855144 | -0.008565532 | 0.125337964 | -0.068339483 | 0.945515395 | 0.990309554 | Nfe2l2 | nuclear factor, erythroid derived 2, like 2 [ | 1285.8 | 1020.7 | 1064.6 | 1094.1 | 1206.2 | 1053.7 |
| 1661 | ENSMUSG00000015843 | 153.8861213 | -0.530356349 | 0.203249424 | -2.609386727 | 0.009070467 | 0.100841841 | Rxrg | retinoid X receptor gamma [Source:MGI S | 199.17 | 196.33 | 149.93 | 131.1 | 127.6 | 119.18 |
| 1662 | ENSMUSG00000015846 | 8953.017933 | 0.142864425 | 0.099369963 | 1.437702299 | 0.150518541 | 0.564307976 | Rxra | retinoid X receptor alpha [Source:MGI Syr | 8059.9 | 8918.2 | 8552.1 | 8457 | 9924.6 | 9806.3 |
| 1663 | ENSMUSG00000015850 | 48.8268658 | -1.13546777 | 0.339263029 | -3.346865628 | 0.000817308 | 0.01799322 | Adamtsl4 | ADAMTS-like 4 [Source:MGI Symbol;Acc:M | 77.901 | 67.843 | 55.343 | 28.919 | 25.084 | 37.871 |
| 1664 | ENSMUSG00000015852 | 0.331982614 | 0.16514309 | 3.986751476 | 0.041422971 | 0.966958701 | NA | Fcrls | Fc receptor-like S, scavenger receptor [So | 0 | 1.0279 | 0 | 0.964 | 0 | 0 |
| 1665 | ENSMUSG00000015854 | 745.9697304 | -1.35593579 | 0.545414619 | -2.486064259 | 0.012916464 | 0.128577932 | Cd5l | CD5 antigen-like [Source:MGI Symbol;Acc | 823.98 | 1743.3 | 651.04 | 316.18 | 310.83 | 630.44 |
| 1666 | ENSMUSG00000015869 | 1282.41194 | 0.101083413 | 0.107797069 | 0.937719494 | 0.348388583 | 0.789711685 | Prpsap1 | phosphoribosyl pyrophosphate synthetas | 1257.7 | 1303.4 | 1151.1 | 1286.9 | 1418.9 | 1276.5 |
| 1667 | ENSMUSG00000015879 | 2.696111638 | 0.094088101 | 1.293450854 | 0.072741922 | 0.942011488 | NA | Fam184b | family with sequence similarity 184, meml | 0.8031 | 3.0838 | 4.025 | 3.8559 | 2.1812 | 2.2277 |
| 1668 | ENSMUSG00000015880 | 2.366008389 | -1.307384588 | 1.662275911 | -0.786502758 | 0.431573004 | NA | Ncapg | non-SMC condensin I complex, subunit G | 8.031 | 2.0558 | 0 | 1.928 | 2.1812 | 0 |
| 1669 | ENSMUSG00000015882 | 237.4023309 | 0.123152139 | 0.179372096 | 0.686573564 | 0.492351492 | 0.876368691 | Lcorl | ligand dependent nuclear receptor corepr | 245.75 | 215.86 | 220.37 | 293.05 | 237.75 | 211.63 |
| 1670 | ENSMUSG00000015889 | 643.8765691 | -0.032583491 | 0.118506724 | -0.274950563 | 0.78335422 | 0.960670212 | Lta4h | leukotriene A4 hydrolase [Source:MGI Syr | 617.59 | 654.78 | 681.23 | 602.49 | 650.01 | 657.17 |
| 1671 | ENSMUSG00000015890 | 3589.512271 | 0.135146704 | 0.130604123 | 1.034781298 | 0.300771059 | 0.751875664 | Amdhd1 | amidohydrolase domain containing 1 [Sou | 3294.3 | 3213.3 | 3756.3 | 3212 | 3594.7 | 4466.5 |
| 1672 | ENSMUSG00000015932 | 2452.269261 | 0.043963209 | 0.090771797 | 0.484326743 | 0.628153999 | 0.920328819 | Dstn | destrin [Source:MGI Symbol;Acc:MGI:192 | 2476 | 2410.5 | 2357.6 | 2431.2 | 2592.4 | 2446 |
| 1673 | ENSMUSG00000015937 | 1255.90472 | 0.075952893 | 0.109823208 | 0.691592367 | 0.489193356 | 0.874770056 | Macroh2a1 | macroH2A.1 histone [Source:MGI Symbol | 1198.2 | 1251 | 1219.6 | 1298.5 | 1167 | 1401.2 |
| 1674 | ENSMUSG00000015942 | 135.8926645 | -0.008785198 | 0.227439389 | -0.038626547 | 0.969188137 | 0.994602269 | Gtf2ird2 | GTF2I repeat domain containing 2 [Source | 135.72 | 121.29 | 151.94 | 144.6 | 102.52 | 159.28 |
| 1675 | ENSMUSG00000015943 | 392.2104239 | -0.036782256 | 0.150387731 | -0.244582822 | 0.806779466 | 0.964718338 | Bola1 | bolA-like 1 (E. coli) [Source:MGI Symbol;A | 446.52 | 401.92 | 342.12 | 362.46 | 394.8 | 405.44 |
| 1676 | ENSMUSG00000015944 | 81.65106088 | -0.208482058 | 0.307170201 | -0.678718369 | 0.497316325 | 0.878825085 | Castor2 | cytosolic arginine sensor for mTORC1 sub | 73.885 | 72.982 | 115.72 | 62.659 | 61.075 | 103.59 |

| | | | | | | | | | | | | | | |
|---|---|---|---|---|---|---|---|---|---|---|---|---|---|---|
| 1677 | ENSMUSG00000015947 | 26.46965811 | -1.293593724 | 0.543454585 | -2.380316146 | 0.01729779 | 0.156658967 | Fcgr1 | Fc receptor, IgG, high affinity I [Source:M( | 28.109 | 51.396 | 33.206 | 10.604 | 6.5437 | 28.96 |
| 1678 | ENSMUSG00000015950 | 40.98632017 | -1.122036659 | 0.419715204 | -2.673328599 | 0.007510264 | 0.08802955 | Ncf1 | neutrophil cytosolic factor 1 [Source:MGI | 56.217 | 68.87 | 43.269 | 22.171 | 14.178 | 41.212 |
| 1679 | ENSMUSG00000015957 | 8.476648886 | 0.618946975 | 0.973516242 | 0.63578495 | 0.524916598 | 0.888334265 | Wnt11 | wingless-type MMTV integration site fami | 4.8186 | 5.1396 | 10.062 | 0 | 16.359 | 14.48 |
| 1680 | ENSMUSG00000015961 | 1054.36636 | -0.08740803 | 0.118386949 | -0.738324876 | 0.460317054 | 0.859817816 | Adss | adenylosuccinate synthetase, non muscle | 1154.9 | 1050.5 | 1052.5 | 963.01 | 1146.2 | 959.02 |
| 1681 | ENSMUSG00000015962 | 0 | NA | NA | NA | NA | NA | 1700016C15 | RIKEN cDNA 1700016C15 gene [Source:M( | 0 | 0 | 0 | 0 | 0 | 0 |
| 1682 | ENSMUSG00000015966 | 136.3243222 | 0.259039913 | 0.209157847 | 1.23849005 | 0.215534409 | 0.660345201 | Il17rb | interleukin 17 receptor B [Source:MGI Syr | 113.24 | 121.29 | 137.86 | 128.21 | 161.41 | 155.94 |
| 1683 | ENSMUSG00000015968 | 65.51140792 | 0.173814287 | 0.292160868 | 0.594926651 | 0.551892501 | 0.895935901 | Cacna1d | calcium channel, voltage-dependent, L typ | 62.642 | 67.843 | 54.337 | 75.19 | 80.706 | 52.351 |
| 1684 | ENSMUSG00000015970 | 3549.689599 | -0.04683125 | 0.10752581 | -0.435534971 | 0.663174161 | 0.929111881 | Chdh | choline dehydrogenase [Source:MGI Symb | 3246.9 | 3833.1 | 3743.2 | 3487.7 | 3763.7 | 3223.5 |
| 1685 | ENSMUSG00000015971 | 517.758856 | 0.090125692 | 0.125314057 | 0.71919858 | 0.472018574 | 0.867870596 | Actr8 | ARP8 actin-related protein 8 [Source:MGI | 468.21 | 519.1 | 518.22 | 537.9 | 550.76 | 512.37 |
| 1686 | ENSMUSG00000015980 | 5.527081687 | 0.299478421 | 1.092672422 | 0.274078868 | 0.784024013 | 0.960670212 | Lrrc27 | leucine rich repeat containing 27 [Source: | 4.8186 | 2.0558 | 8.05 | 10.604 | 7.6343 | 0 |
| 1687 | ENSMUSG00000015981 | 0.924441557 | -0.875430577 | 2.469021705 | -0.354565768 | 0.722914925 | NA | Stk32c | serine/threonine kinase 32C [Source:MGI | 1.6062 | 0 | 2.0125 | 1.928 | 0 | 0 |
| 1688 | ENSMUSG00000015994 | 630.8302796 | 0.06752238 | 0.122757799 | 0.550045538 | 0.58228814 | 0.908645447 | Fnta | farnesyltransferase, CAAX box, alpha [Sou | 565.38 | 657.87 | 625.88 | 645.87 | 672.91 | 617.07 |
| 1689 | ENSMUSG00000016018 | 410.1407647 | 0.061644467 | 0.13413934 | 0.459555468 | 0.645835329 | 0.925643431 | Mtrex | Mtr4 exosome RNA helicase [Source:MGI | 386.29 | 392.66 | 425.64 | 428.97 | 432.98 | 394.3 |
| 1690 | ENSMUSG00000016024 | 1503.911447 | -0.534607849 | 0.150288445 | -3.557211931 | 0.000374812 | 0.010261584 | Lbp | lipopolysaccharide binding protein [Sourc | 1635.9 | 1876 | 1827.3 | 1393.9 | 928.12 | 1362.2 |
| 1691 | ENSMUSG00000016028 | 437.1255967 | -0.832015683 | 0.196006426 | -4.244838808 | 2.19E-05 | 0.001040792 | Celsr1 | cadherin, EGF LAG seven-pass G-type rece | 428.86 | 676.37 | 575.57 | 389.45 | 259.57 | 292.94 |
| 1692 | ENSMUSG00000016087 | 23.87387401 | -0.976495058 | 0.490243011 | -1.991859215 | 0.046386507 | 0.297796022 | Fli1 | Friend leukemia integration 1 [Source:MG | 25.699 | 32.893 | 36.225 | 8.6758 | 16.359 | 23.391 |
| 1693 | ENSMUSG00000016128 | 364.6840609 | -0.1147537 | 0.160154651 | -0.716518057 | 0.473671522 | 0.868952133 | Stard13 | StAR-related lipid transfer (START) domaiı | 323.65 | 362.85 | 451.8 | 350.89 | 341.36 | 357.54 |
| 1694 | ENSMUSG00000016150 | 0.36353943 | 2.065989254 | 4.034137301 | 0.512126658 | 0.608562368 | NA | Tenm1 | teneurin transmembrane protein 1 [Sourc | 0 | 0 | 0 | 0 | 2.1812 | 0 |
| 1695 | ENSMUSG00000016179 | 0 | NA | NA | NA | NA | NA | Camk1g | calcium/calmodulin-dependent protein ki | 0 | 0 | 0 | 0 | 0 | 0 |
| 1696 | ENSMUSG00000016181 | 149.3706424 | 0.276432071 | 0.205968881 | 1.342106003 | 0.179561628 | 0.611676727 | Utp25 | UTP25 small subunit processome compon | 128.5 | 138.77 | 137.86 | 133.03 | 184.31 | 173.76 |
| 1697 | ENSMUSG00000016194 | 13058.64048 | -0.290606135 | 0.127491389 | -2.279417752 | 0.022642244 | 0.188541979 | Hsd11b1 | hydroxysteroid 11-beta dehydrogenase 1 | 12687 | 13189 | 17234 | 12871 | 11322 | 11049 |
| 1698 | ENSMUSG00000016200 | 0.696064467 | -1.534444742 | 3.043667477 | -0.504143358 | 0.61416064 | NA | Syt14 | synaptotagmin XIV [Source:MGI Symbol;A | 3.2124 | 0 | 0 | 0.964 | 0 | 0 |
| 1699 | ENSMUSG00000016206 | 46.32782241 | -0.426925274 | 0.335276958 | -1.273351075 | 0.20289349 | 0.647682949 | H2-M3 | histocompatibility 2, M region locus 3 [Sou | 41.761 | 61.675 | 56.35 | 44.343 | 37.081 | 36.757 |
| 1700 | ENSMUSG00000016239 | 64.98440792 | -0.596176524 | 0.398262344 | -1.496944245 | 0.134407766 | 0.537569131 | Lonrf3 | LON peptidase N-terminal domain and rin | 66.657 | 70.926 | 97.606 | 79.046 | 55.622 | 20.049 |
| 1701 | ENSMUSG00000016252 | 3070.527225 | 0.006813097 | 0.088875477 | 0.076658906 | 0.938894896 | 0.988721002 | Atp5e | ATP synthase, H+ transporting, mitochonc | 3088.7 | 2990.2 | 3110.3 | 2995.1 | 3024.3 | 3214.6 |
| 1702 | ENSMUSG00000016253 | 696.4541145 | 0.005909793 | 0.134568357 | 0.043916665 | 0.964970832 | 0.993480943 | Nelfcd | negative elongation factor complex memt | 798.28 | 614.69 | 671.17 | 739.37 | 695.81 | 659.4 |
| 1703 | ENSMUSG00000016255 | 2.545199904 | 0.75250198 | 1.462385667 | 0.514571496 | 0.606852488 | NA | Tubb1 | tubulin, beta 1 class VI [Source:MGI Symb | 1.6062 | 3.0838 | 1.0062 | 2.8919 | 0 | 6.6831 |
| 1704 | ENSMUSG00000016256 | 3699.31637 | -0.262694742 | 0.109724548 | -2.394129173 | 0.01665988 | 0.153190407 | Ctsz | cathepsin Z [Source:MGI Symbol;Acc:MGI | 4062.9 | 4514.6 | 3527.9 | 3277.5 | 3419.1 | 3393.9 |
| 1705 | ENSMUSG00000016257 | 574.3519077 | -0.046792965 | 0.119624907 | -0.391164063 | 0.69567597 | 0.939992732 | Prelid3b | PRELI domain containing 3B [Source:MGI . | 595.9 | 605.44 | 549.41 | 559.11 | 569.3 | 566.95 |
| 1706 | ENSMUSG00000016262 | 9.713339119 | 2.191114305 | 0.806819359 | 2.715743346 | 0.006612716 | 0.080919055 | Sertad4 | SERTA domain containing 4 [Source:MGI S | 2.4093 | 5.1396 | 3.0187 | 14.46 | 7.6343 | 25.618 |
| 1707 | ENSMUSG00000016283 | 2.289830252 | -0.112501525 | 1.684533309 | -0.066784981 | 0.94675288 | NA | H2-M2 | histocompatibility 2, M region locus 2 [Sou | 4.0155 | 1.0279 | 2.0125 | 0 | 0 | 6.6831 |
| 1708 | ENSMUSG00000016308 | 464.7909107 | -0.074841481 | 0.128579599 | -0.582063418 | 0.560523965 | 0.899624792 | Ube2a | ubiquitin-conjugating enzyme E2A [Source | 484.27 | 470.79 | 474.95 | 427.04 | 446.06 | 485.64 |
| 1709 | ENSMUSG00000016319 | 8803.288265 | -0.269125418 | 0.098349676 | -2.736413875 | 0.006211285 | 0.077098412 | Slc25a5 | solute carrier family 25 (mitochondrial car | 9378.6 | 10486 | 9001.9 | 7750.4 | 8526.5 | 7676.6 |
| 1710 | ENSMUSG00000016327 | 0 | NA | NA | NA | NA | NA | Atp1b4 | ATPase, (Na+)/K+ transporting, beta 4 pol | 0 | 0 | 0 | 0 | 0 | 0 |
| 1711 | ENSMUSG00000016344 | 1273.567815 | 0.125895252 | 0.113360828 | 1.110571034 | 0.266753037 | 0.722778036 | Ppdpf | pancreatic progenitor cell differentiation a | 1244.8 | 1161.5 | 1246.7 | 1214.6 | 1296.7 | 1477 |
| 1712 | ENSMUSG00000016346 | 0 | NA | NA | NA | NA | NA | Kcnq2 | potassium voltage-gated channel, subfam | 0 | 0 | 0 | 0 | 0 | 0 |
| 1713 | ENSMUSG00000016349 | 0.185640647 | 1.126924175 | 4.080472857 | 0.276174898 | 0.782413736 | NA | Eef1a2 | eukaryotic translation elongation factor 1 | 0 | 0 | 0 | 0 | 0 | 1.1138 |
| 1714 | ENSMUSG00000016356 | 20.55945875 | 0.615334225 | 0.504445914 | 1.219822003 | 0.222532359 | 0.668407528 | Col20a1 | collagen, type XX, alpha 1 [Source:MGI Sy | 10.44 | 14.391 | 24.15 | 26.991 | 23.994 | 23.391 |
| 1715 | ENSMUSG00000016382 | 1679.052338 | -0.133383332 | 0.099465006 | -1.341007628 | 0.179917976 | 0.612025777 | Pls3 | plastin 3 (T-isoform) [Source:MGI Symbol; | 1656.8 | 1811.2 | 1802.2 | 1525 | 1622.8 | 1656.3 |
| 1716 | ENSMUSG00000016386 | 7.170408571 | -0.327613645 | 0.88244392 | -0.371257184 | 0.710445985 | 0.944239215 | Mpped2 | metallophosphoesterase domain containi | 13.653 | 2.0558 | 8.05 | 3.8559 | 8.7249 | 6.6831 |
| 1717 | ENSMUSG00000016409 | 190.75745 | 0.198923894 | 0.202352729 | 0.983055158 | 0.325580292 | 0.771619425 | Nkap | NFKB activating protein [Source:MGI Symb | 163.03 | 164.47 | 205.27 | 172.55 | 191.95 | 247.27 |
| 1718 | ENSMUSG00000016427 | 1100.318057 | 0.177839525 | 0.114705419 | 1.550402119 | 0.12104503 | 0.509989898 | Ndufa1 | NADH:ubiquinone oxidoreductase subuni | 1024 | 1015.6 | 1057.6 | 1055.6 | 1155 | 1294.3 |
| 1719 | ENSMUSG00000016458 | 11.59881119 | -0.044510459 | 0.700762274 | -0.063517203 | 0.949354661 | 0.990731019 | Wt1 | Wilms tumor 1 homolog [Source:MGI Sym | 16.865 | 13.363 | 5.0312 | 13.496 | 15.269 | 5.5692 |
| 1720 | ENSMUSG00000016477 | 140.7491321 | -0.136859514 | 0.201221476 | -0.680143674 | 0.496413493 | 0.878165817 | E2f3 | E2F transcription factor 3 [Source:MGI Syı | 141.35 | 141.85 | 158.99 | 121.46 | 149.41 | 131.43 |
| 1721 | ENSMUSG00000016481 | 738.2570968 | 0.060031372 | 0.113490312 | 0.528955916 | 0.596836031 | 0.91428215 | Cr1l | complement component (3b/4b) receptor | 692.27 | 746.27 | 730.53 | 733.59 | 793.97 | 732.91 |
| 1722 | ENSMUSG00000016487 | 586.8021838 | 0.03059452 | 0.167701267 | 0.18243464 | 0.855241643 | 0.974158771 | Ppfibp1 | PTPRF interacting protein, binding protein | 629.63 | 624.97 | 486.02 | 475.24 | 597.66 | 707.29 |
| 1723 | ENSMUSG00000016493 | 23.40200644 | -0.087445485 | 0.44598908 | -0.196070911 | 0.844554664 | 0.972803041 | Cd46 | CD46 antigen, complement regulatory prc | 26.502 | 24.67 | 21.131 | 25.063 | 18.541 | 24.505 |
| 1724 | ENSMUSG00000016494 | 26.26541285 | -0.449981932 | 0.500007781 | -0.899949859 | 0.368146935 | 0.80627882 | Cd34 | CD34 antigen [Source:MGI Symbol;Acc:M( | 41.761 | 30.838 | 18.112 | 16.388 | 34.9 | 15.594 |
| 1725 | ENSMUSG00000016495 | 101.9730712 | -0.033537337 | 0.227819832 | -0.147209912 | 0.882966337 | 0.981769403 | Plgrkt | plasminogen receptor, C-terminal lysine tı | 114.04 | 92.513 | 102.64 | 102.18 | 105.79 | 94.677 |
| 1726 | ENSMUSG00000016496 | 33.25923521 | -0.238053569 | 0.381780585 | -0.62353503 | 0.53293299 | 0.889143097 | Cd274 | CD274 antigen [Source:MGI Symbol;Acc:N | 41.761 | 28.782 | 37.231 | 28.919 | 29.447 | 33.415 |
| 1727 | ENSMUSG00000016498 | 0.133850373 | -0.796629176 | 4.080472857 | -0.195229623 | 0.84521319 | NA | Pdcd1lg2 | programmed cell death 1 ligand 2 [Source | 0.8031 | 0 | 0 | 0 | 0 | 0 |
| 1728 | ENSMUSG00000016503 | 99.02157091 | -0.210508713 | 0.253560284 | -0.830211693 | 0.406419105 | 0.830399297 | Gtf3a | general transcription factor III A [Source:N | 89.947 | 125.41 | 103.64 | 84.83 | 112.33 | 77.969 |
| 1729 | ENSMUSG00000016510 | 429.558845 | 0.157911358 | 0.142283884 | 1.109833063 | 0.267070969 | 0.722778036 | Mtif3 | mitochondrial translational initiation facto | 410.39 | 403.97 | 403.5 | 434.75 | 406.8 | 517.94 |
| 1730 | ENSMUSG00000016520 | 320.8418868 | -0.305635485 | 0.154879773 | -1.9733725 | 0.048453138 | 0.306305155 | Lnx2 | ligand of numb-protein X 2 [Source:MGI S | 314.82 | 352.58 | 397.47 | 297.87 | 269.38 | 292.94 |
| 1731 | ENSMUSG00000016524 | 0.133850373 | -0.796629176 | 4.080472857 | -0.195229623 | 0.84521319 | NA | Il19 | interleukin 19 [Source:MGI Symbol;Acc:M | 0.8031 | 0 | 0 | 0 | 0 | 0 |
| 1732 | ENSMUSG00000016526 | 25.40329713 | -0.787691213 | 0.437349218 | -1.801057784 | 0.071693773 | 0.387575723 | Dyrk3 | dual-specificity tyrosine-(Y)-phosphorylati | 26.502 | 31.865 | 38.237 | 18.316 | 17.45 | 20.049 |

| | | | | | | | | | | | | | | | |
|---|---|---|---|---|---|---|---|---|---|---|---|---|---|---|---|
| 1733 | ENSMUSG00000016528 | 1209.419257 | -0.081932496 | 0.112565181 | -0.727867133 | 0.466694924 | 0.863779479 | Mapkapk2 | MAP kinase-activated protein kinase 2 [Sc | 1237.6 | 1230.4 | 1262.8 | 1124 | 1087.3 | 1314.3 |
| 1734 | ENSMUSG00000016529 | 0.439020386 | -2.189747911 | 3.52148864 | -0.621824499 | 0.534057275 | NA | Il10 | interleukin 10 [Source:MGI Symbol;Acc:M | 1.6062 | 1.0279 | 0 | 0 | 0 | 0 |
| 1735 | ENSMUSG00000016534 | 12007.86346 | 0.050571045 | 0.082740407 | 0.611201306 | 0.541066319 | 0.893581096 | Lamp2 | lysosomal-associated membrane protein 2 | 11508 | 11900 | 11984 | 11724 | 12297 | 12634 |
| 1736 | ENSMUSG00000016541 | 1156.078465 | 0.078362686 | 0.126676844 | 0.618603079 | 0.536177873 | 0.890560262 | Atxn10 | ataxin 10 [Source:MGI Symbol;Acc:MGI:1 | 1262.5 | 1112.2 | 998.19 | 1157.7 | 1309.8 | 1096 |
| 1737 | ENSMUSG00000016552 | 3.393746795 | 0.067317763 | 1.125513072 | 0.059810734 | 0.952306376 | NA | Foxred2 | FAD-dependent oxidoreductase domain c | 4.8186 | 2.0558 | 3.0187 | 3.8559 | 3.2719 | 3.3415 |
| 1738 | ENSMUSG00000016554 | 648.6004932 | 0.172953182 | 0.11658108 | 1.483544173 | 0.137929889 | 0.5424993 | Eif3d | eukaryotic translation initiation factor 3, s | 634.45 | 612.64 | 581.61 | 668.04 | 716.54 | 678.33 |
| 1739 | ENSMUSG00000016559 | 3478.700859 | 0.015321229 | 0.105014533 | 0.145896275 | 0.884003272 | 0.981769403 | H3f3b | H3.3 histone B [Source:MGI Symbol;Acc:N | 3761.7 | 3107.4 | 3509.8 | 3299.7 | 3538 | 3655.6 |
| 1740 | ENSMUSG00000016619 | 738.5735638 | 0.223882465 | 0.133510192 | 1.676894186 | 0.093563175 | 0.450459278 | Nup50 | nucleoporin 50 [Source:MGI Symbol;Acc:I | 684.24 | 655.81 | 703.36 | 706.6 | 754.71 | 926.72 |
| 1741 | ENSMUSG00000016624 | 5.856007576 | 0.377468988 | 0.883930393 | 0.427034743 | 0.669354022 | 0.930837335 | Phf21b | PHD finger protein 21B [Source:MGI Symb | 8.031 | 4.1117 | 3.0187 | 6.7478 | 6.5437 | 6.6831 |
| 1742 | ENSMUSG00000016626 | 0 | NA | NA | NA | NA | NA | Nlrp14 | NLR family, pyrin domain containing 14 [S | 0 | 0 | 0 | 0 | 0 | 0 |
| 1743 | ENSMUSG00000016637 | 173.2456485 | -0.146982924 | 0.180416125 | -0.814688399 | 0.415250696 | 0.836001897 | Ift27 | intraflagellar transport 27 [Source:MGI Sy | 179.89 | 188.11 | 178.11 | 155.2 | 177.77 | 160.39 |
| 1744 | ENSMUSG00000016664 | 1176.400973 | 0.124862197 | 0.113623589 | 1.09891087 | 0.271806945 | 0.727526428 | Pacsin2 | protein kinase C and casein kinase substra | 1183 | 1134.8 | 1057.6 | 1111.5 | 1320.7 | 1250.8 |
| 1745 | ENSMUSG00000016756 | 4196.940256 | 0.149067547 | 0.085071974 | 1.752252128 | 0.079730464 | 0.414529629 | Cmah | cytidine monophospho-N-acetylneuramin | 3897.5 | 4001.7 | 4042.1 | 4346.6 | 4559.9 | 4334 |
| 1746 | ENSMUSG00000016757 | 122.6140089 | 0.243769186 | 0.242196036 | 1.00649536 | 0.314177343 | 0.760800449 | Ttll12 | tubulin tyrosine ligase-like family, membe | 126.09 | 86.345 | 123.77 | 114.71 | 122.15 | 162.62 |
| 1747 | ENSMUSG00000016758 | 116.0796846 | -0.067823339 | 0.272937873 | -0.248493689 | 0.803752452 | 0.963596811 | Bik | BCL2-interacting killer [Source:MGI Symbo | 110.83 | 87.373 | 157.98 | 88.686 | 110.15 | 141.46 |
| 1748 | ENSMUSG00000016763 | 3.090560653 | -0.550157467 | 1.227651577 | -0.448138117 | 0.65405352 | NA | Scube1 | signal peptide, CUB domain, EGF-like 1 [Sc | 4.8186 | 4.1117 | 2.0125 | 0.964 | 2.1812 | 4.4554 |
| 1749 | ENSMUSG00000016831 | 518.5264562 | 0.043433017 | 0.126421319 | 0.343557697 | 0.731178943 | 0.94705581 | Tox4 | TOX high mobility group box family memb | 489.89 | 536.57 | 506.14 | 533.08 | 490.78 | 554.69 |
| 1750 | ENSMUSG00000016833 | 270.8632876 | 0.046480663 | 0.152914838 | 0.303964375 | 0.761155033 | 0.95402891 | Mrps18c | mitochondrial ribosomal protein S18C [So | 277.87 | 246.7 | 274.7 | 280.52 | 280.29 | 265.09 |
| 1751 | ENSMUSG00000016918 | 53.82599677 | 0.31271491 | 0.386500317 | 0.809093542 | 0.418461343 | 0.837575303 | Sulf1 | sulfatase 1 [Source:MGI Symbol;Acc:MGI: | 35.336 | 69.898 | 39.244 | 66.514 | 77.434 | 34.529 |
| 1752 | ENSMUSG00000016921 | 1795.323264 | -0.087912986 | 0.112367841 | -0.782367844 | 0.433998426 | 0.845264432 | Srsf6 | serine and arginine-rich splicing factor 6 [S | 1808.6 | 1931.5 | 1811.2 | 1940.5 | 1730.8 | 1549.4 |
| 1753 | ENSMUSG00000016933 | 936.5460887 | 0.38647933 | 0.108824666 | 3.551394587 | 0.000383195 | 0.010379287 | Plcg1 | phospholipase C, gamma 1 [Source:MGI S | 753.31 | 845.98 | 837.2 | 1046.9 | 1066.6 | 1069.3 |
| 1754 | ENSMUSG00000016940 | 530.4795943 | 0.039711541 | 0.135428002 | 0.293229911 | 0.769346422 | 0.956153573 | Kctd2 | potassium channel tetramerisation domai | 515.59 | 538.63 | 516.2 | 613.09 | 511.5 | 487.86 |
| 1755 | ENSMUSG00000016942 | 10078.68565 | 0.145489111 | 0.090728667 | 1.603562759 | 0.108810465 | 0.485684837 | Tmprss6 | transmembrane serine protease 6 [Source | 9119.2 | 9555.5 | 10038 | 10121 | 10374 | 11264 |
| 1756 | ENSMUSG00000016946 | 91.32494076 | -0.123478544 | 0.284878086 | -0.433443463 | 0.664692631 | 0.929301301 | Kctd5 | potassium channel tetramerisation domai | 65.854 | 90.457 | 129.81 | 80.974 | 85.068 | 95.791 |
| 1757 | ENSMUSG00000016982 | 0 | NA | NA | NA | NA | NA | Pom121l2 | POM121 membrane glycoprotein-like 2 (ra | 0 | 0 | 0 | 0 | 0 | 0 |
| 1758 | ENSMUSG00000016984 | 120.1156209 | 0.23163105 | 0.213946402 | 1.082659242 | 0.278959701 | 0.731519729 | Etaa1 | Ewing tumor-associated antigen 1 [Source | 105.21 | 102.79 | 123.77 | 139.78 | 121.06 | 128.09 |
| 1759 | ENSMUSG00000016995 | 0.17131964 | -0.796629176 | 4.080472857 | -0.195229623 | 0.84521319 | NA | Matn4 | matrilin 4 [Source:MGI Symbol;Acc:MGI:1 | 0 | 1.0279 | 0 | 0 | 0 | 0 |
| 1760 | ENSMUSG00000016998 | 0 | NA | NA | NA | NA | NA | Svs4 | seminal vesicle secretory protein 4 [Sourc | 0 | 0 | 0 | 0 | 0 | 0 |
| 1761 | ENSMUSG00000017000 | 0 | NA | NA | NA | NA | NA | Svs6 | seminal vesicle secretory protein 6 [Sourc | 0 | 0 | 0 | 0 | 0 | 0 |
| 1762 | ENSMUSG00000017002 | 71.01250411 | 0.559047153 | 0.29342696 | 1.905234454 | 0.056749601 | 0.337927639 | Slpi | secretory leukocyte peptidase inhibitor [S | 61.036 | 62.703 | 48.3 | 70.37 | 110.15 | 73.514 |
| 1763 | ENSMUSG00000017003 | 0 | NA | NA | NA | NA | NA | Svs3a | seminal vesicle secretory protein 3A [Sour | 0 | 0 | 0 | 0 | 0 | 0 |
| 1764 | ENSMUSG00000017004 | 0 | NA | NA | NA | NA | NA | Svs5 | seminal vesicle secretory protein 5 [Sourc | 0 | 0 | 0 | 0 | 0 | 0 |
| 1765 | ENSMUSG00000017007 | 0 | NA | NA | NA | NA | NA | Rbpjl | recombination signal binding protein for i | 0 | 0 | 0 | 0 | 0 | 0 |
| 1766 | ENSMUSG00000017009 | 8128.338847 | -0.200411841 | 0.118456493 | -1.691860327 | 0.090672608 | 0.444602826 | Sdc4 | syndecan 4 [Source:MGI Symbol;Acc:MGI | 7611.8 | 9545.2 | 8919.4 | 6716 | 7607.1 | 8370.5 |
| 1767 | ENSMUSG00000017049 | 0 | NA | NA | NA | NA | NA | Ccdc70 | coiled-coil domain containing 70 [Source:I | 0 | 0 | 0 | 0 | 0 | 0 |
| 1768 | ENSMUSG00000017057 | 615.2183556 | 0.258630399 | 0.126767141 | 2.040200613 | 0.041330349 | 0.27856578 | Il13ra1 | interleukin 13 receptor, alpha 1 [Source:N | 544.5 | 582.83 | 553.43 | 635.26 | 747.07 | 628.21 |
| 1769 | ENSMUSG00000017064 | 0 | NA | NA | NA | NA | NA | Prl5a1 | prolactin family 5, subfamily a, member 1 | 0 | 0 | 0 | 0 | 0 | 0 |
| 1770 | ENSMUSG00000017119 | 4288.701977 | 0.20735785 | 0.091111771 | 2.275862353 | 0.022854249 | 0.19019785 | Nbr1 | NBR1, autophagy cargo receptor [Source: | 4206.6 | 3809.5 | 3925.4 | 4395.7 | 4667.8 | 4727.2 |
| 1771 | ENSMUSG00000017132 | 927.8508596 | 0.168983163 | 0.130326951 | 1.296609507 | 0.19476558 | 0.634479582 | Cyth1 | cytohesin 1 [Source:MGI Symbol;Acc:MGI | 929.99 | 863.45 | 826.13 | 898.43 | 906.3 | 1142.8 |
| 1772 | ENSMUSG00000017144 | 381.0161775 | 0.439792389 | 0.19363035 | 2.27129884 | 0.023128893 | 0.191931334 | Rnd3 | Rho family GTPase 3 [Source:MGI Symbol, | 356.58 | 301.18 | 311.94 | 459.82 | 311.92 | 544.67 |
| 1773 | ENSMUSG00000017146 | 54.99297589 | -0.409331634 | 0.335986344 | -1.218298426 | 0.223110588 | 0.668407528 | Brca1 | breast cancer 1, early onset [Source:MGI S | 70.673 | 74.01 | 43.269 | 38.559 | 58.893 | 44.554 |
| 1774 | ENSMUSG00000017165 | 0 | NA | NA | NA | NA | NA | Gast | gastrin [Source:MGI Symbol;Acc:MGI:104 | 0 | 0 | 0 | 0 | 0 | 0 |
| 1775 | ENSMUSG00000017167 | 131.7230091 | -0.978369097 | 0.367119301 | -2.664989539 | 0.007699072 | 0.089301824 | Cntnap1 | contactin associated protein-like 1 [Sourc | 314.01 | 108.96 | 100.62 | 81.938 | 109.06 | 75.741 |
| 1776 | ENSMUSG00000017176 | 20.42474135 | 0.246497886 | 0.483446788 | 0.50987594 | 0.610138379 | 0.916376001 | Nt5c3b | 5'-nucleotidase, cytosolic IIIB [Source:MG | 23.29 | 15.419 | 17.106 | 19.28 | 20.722 | 26.732 |
| 1777 | ENSMUSG00000017188 | 799.1941683 | 0.094515519 | 0.109663363 | 0.861869601 | 0.388759279 | 0.818464676 | Coa3 | cytochrome C oxidase assembly factor 3 [S | 788.65 | 737.02 | 792.92 | 808.78 | 819.05 | 848.75 |
| 1778 | ENSMUSG00000017195 | 3.676258839 | -1.027780784 | 1.187840679 | -0.865251377 | 0.386900832 | NA | Zpbp2 | zona pellucida binding protein 2 [Source:I | 9.6372 | 3.0838 | 2.0125 | 2.8919 | 1.0906 | 3.3415 |
| 1779 | ENSMUSG00000017204 | 0 | NA | NA | NA | NA | NA | Gsdma | gasdermin A [Source:MGI Symbol;Acc:MG | 0 | 0 | 0 | 0 | 0 | 0 |
| 1780 | ENSMUSG00000017210 | 261.4228915 | 0.035906654 | 0.178046899 | 0.201669638 | 0.840175 | 0.971838962 | Med24 | mediator complex subunit 24 [Source:MG | 264.22 | 256.98 | 253.57 | 303.65 | 208.31 | 281.8 |
| 1781 | ENSMUSG00000017211 | 0 | NA | NA | NA | NA | NA | Gsdma2 | gasdermin A2 [Source:MGI Symbol;Acc:M | 0 | 0 | 0 | 0 | 0 | 0 |
| 1782 | ENSMUSG00000017221 | 1805.238321 | 0.104821428 | 0.096411678 | 1.087227507 | 0.276936277 | 0.730041765 | Psmd3 | proteasome (prosome, macropain) 26S su | 1767.6 | 1682.7 | 1769 | 1974.2 | 1826.8 | 1811.1 |
| 1783 | ENSMUSG00000017264 | 424.2647417 | -0.108731354 | 0.14989183 | -0.725398804 | 0.468207407 | 0.865229581 | Exosc10 | exosome component 10 [Source:MGI Sym | 451.34 | 492.37 | 377.34 | 445.36 | 390.44 | 388.73 |
| 1784 | ENSMUSG00000017286 | 747.5016297 | 0.062624807 | 0.125378486 | 0.499486072 | 0.617436996 | 0.917509899 | Glod4 | glyoxalase domain containing 4 [Source:N | 726.81 | 725.71 | 740.6 | 663.22 | 825.6 | 803.08 |
| 1785 | ENSMUSG00000017288 | 515.8208567 | -0.047411178 | 0.129065568 | -0.3673418 | 0.713364078 | 0.944239215 | Vps53 | VPS53 GARP complex subunit [Source:MG | 501.94 | 531.43 | 539.35 | 459.82 | 533.31 | 529.08 |
| 1786 | ENSMUSG00000017291 | 927.817132 | 0.100557392 | 0.11425891 | 0.880083592 | 0.378814027 | 0.811969331 | Taok1 | TAO kinase 1 [Source:MGI Symbol;Acc:MG | 854.5 | 956.99 | 875.43 | 928.31 | 1036.1 | 915.58 |
| 1787 | ENSMUSG00000017299 | 182.568169 | 0.269871274 | 0.216103712 | 1.248804438 | 0.21173661 | 0.656595245 | Dnttip1 | deoxynucleotidyltransferase, terminal, int | 216.84 | 136.71 | 141.88 | 187.98 | 214.85 | 197.15 |
| 1788 | ENSMUSG00000017300 | 0 | NA | NA | NA | NA | NA | Tnnc2 | troponin C2, fast [Source:MGI Symbol;Acc | 0 | 0 | 0 | 0 | 0 | 0 |

| | | | | | | | | | | | | | | | |
|---|---|---|---|---|---|---|---|---|---|---|---|---|---|---|---|
| 1789 | ENSMUSG00000017307 | 382.2674278 | -0.238041844 | 0.1581716 | -1.504959449 | 0.132334503 | 0.533283479 | Acot8 | acyl-CoA thioesterase 8 [Source:MGI Syml | 449.74 | 367.99 | 423.63 | 408.73 | 327.19 | 316.33 |
| 1790 | ENSMUSG00000017309 | 171.1170432 | -0.655525112 | 0.209036946 | -3.135929439 | 0.001713104 | 0.031285781 | Cd300lg | CD300 molecule like family member G [So | 199.97 | 206.61 | 221.37 | 129.17 | 103.61 | 165.96 |
| 1791 | ENSMUSG00000017310 | 0 | NA | NA | NA | NA | NA | Spint4 | serine protease inhibitor, Kunitz type 4 [S | 0 | 0 | 0 | 0 | 0 | 0 |
| 1792 | ENSMUSG00000017311 | 0 | NA | NA | NA | NA | NA | Pyy | peptide YY [Source:MGI Symbol;Acc:MGI: | 0 | 0 | 0 | 0 | 0 | 0 |
| 1793 | ENSMUSG00000017314 | 5.670647697 | 0.936172413 | 1.255428347 | 0.745699598 | 0.455848902 | 0.857604173 | Mpp2 | membrane protein, palmitoylated 2 (MAG | 5.6217 | 2.0558 | 4.025 | 21.208 | 0 | 1.1138 |
| 1794 | ENSMUSG00000017316 | 0 | NA | NA | NA | NA | NA | Ppy | pancreatic polypeptide [Source:MGI Symb | 0 | 0 | 0 | 0 | 0 | 0 |
| 1795 | ENSMUSG00000017344 | 35663.41214 | 0.194584066 | 0.087554452 | 2.222434862 | 0.026253932 | 0.209626794 | Vtn | vitronectin [Source:MGI Symbol;Acc:MGI: | 30742 | 34756 | 34290 | 38591 | 37516 | 38086 |
| 1796 | ENSMUSG00000017376 | 227.9750228 | -0.022324499 | 0.184930773 | -0.120718139 | 0.903914291 | 0.983821605 | Nlk | nemo like kinase [Source:MGI Symbol;Acc | 212.02 | 270.34 | 207.29 | 223.64 | 256.3 | 198.26 |
| 1797 | ENSMUSG00000017386 | 776.3425493 | 0.329060383 | 0.114706132 | 2.868725309 | 0.004121296 | 0.058613981 | Traf4 | TNF receptor associated factor 4 [Source: | 713.15 | 670.2 | 681.23 | 930.24 | 824.51 | 838.72 |
| 1798 | ENSMUSG00000017390 | 139.3846718 | 0.869490455 | 0.27005281 | 3.21970527 | 0.001283225 | 0.025183934 | Aldoc | aldolase C, fructose-bisphosphate [Source | 111.63 | 120.27 | 63.393 | 133.03 | 194.13 | 213.86 |
| 1799 | ENSMUSG00000017400 | 1.559010242 | 1.09645564 | 1.761919995 | 0.62230728 | 0.533739835 | NA | Stac2 | SH3 and cysteine rich domain 2 [Source:N | 0 | 1.0279 | 2.0125 | 1.928 | 3.2719 | 1.1138 |
| 1800 | ENSMUSG00000017404 | 3577.119058 | 0.028478109 | 0.091226712 | 0.312168539 | 0.754912441 | 0.953085143 | Rpl19 | ribosomal protein L19 [Source:MGI Symbo | 3528.8 | 3625.5 | 3470.5 | 3382.6 | 3750.6 | 3704.6 |
| 1801 | ENSMUSG00000017405 | 124.7397178 | 0.449221511 | 0.232777856 | 1.929829233 | 0.053628 | 0.325777149 | Nek8 | NIMA (never in mitosis gene a)-related ex | 126.09 | 83.261 | 106.66 | 165.8 | 137.42 | 129.21 |
| 1802 | ENSMUSG00000017412 | 13.70408747 | 0.903744504 | 0.591524115 | 1.527823602 | 0.126556341 | 0.521187766 | Cacnb4 | calcium channel, voltage-dependent, beta | 11.243 | 9.2513 | 8.05 | 18.316 | 13.087 | 22.277 |
| 1803 | ENSMUSG00000017417 | 5.75713149 | -1.641980096 | 1.032468683 | -1.59034373 | 0.111757347 | 0.490460457 | Plxdc1 | plexin domain containing 1 [Source:MGI S | 4.8186 | 13.363 | 8.05 | 3.8559 | 0 | 4.4554 |
| 1804 | ENSMUSG00000017418 | 581.2704934 | 0.140157919 | 0.142433853 | 0.984021115 | 0.325105131 | 0.771501263 | Arl5b | ADP-ribosylation factor-like 5B [Source:M | 475.44 | 613.67 | 571.55 | 649.72 | 631.47 | 545.78 |
| 1805 | ENSMUSG00000017421 | 1111.229198 | -0.005267823 | 0.111100776 | -0.047414816 | 0.962182621 | 0.992956886 | Zfp207 | zinc finger protein 207 [Source:MGI Symb | 1068.1 | 1100.9 | 1171.3 | 1103.8 | 1198.6 | 1024.7 |
| 1806 | ENSMUSG00000017428 | 1615.158452 | -0.062722764 | 0.103943698 | -0.603430177 | 0.546222552 | 0.89549457 | Psmd11 | proteasome (prosome, macropain) 26S su | 1633.5 | 1690.9 | 1627.1 | 1693.7 | 1442.9 | 1602.8 |
| 1807 | ENSMUSG00000017446 | 105.634491 | -0.653366074 | 0.25473358 | -2.564899669 | 0.010320564 | 0.111317407 | C1qtnf1 | C1q and tumor necrosis factor related pro | 151.79 | 133.63 | 101.63 | 80.01 | 67.618 | 99.132 |
| 1808 | ENSMUSG00000017453 | 6606.162835 | -0.025913781 | 0.102737968 | -0.252231788 | 0.800861909 | 0.962424862 | Pipox | pipecolic acid oxidase [Source:MGI Symbo | 5861.8 | 6894.2 | 7241.9 | 6483.7 | 6555.7 | 6599.5 |
| 1809 | ENSMUSG00000017466 | 226.5538709 | -0.480568441 | 0.188695642 | -2.54679142 | 0.010871839 | 0.115286132 | Timp2 | tissue inhibitor of metalloproteinase 2 [So | 299.56 | 282.68 | 209.3 | 196.65 | 198.49 | 172.65 |
| 1810 | ENSMUSG00000017478 | 879.3848385 | 0.044832238 | 0.107963631 | 0.415253155 | 0.67795659 | 0.93325821 | Zc3h18 | zinc finger CCCH-type containing 18 [Sour | 836.03 | 896.34 | 865.37 | 909.03 | 853.95 | 915.58 |
| 1811 | ENSMUSG00000017485 | 1049.751815 | -0.038000168 | 0.117007431 | -0.324767136 | 0.745357319 | 0.949997329 | Top2b | topoisomerase (DNA) II beta [Source:MGI | 1134.8 | 939.52 | 1115.9 | 1082.5 | 1009.9 | 1015.8 |
| 1812 | ENSMUSG00000017491 | 15.56489228 | 0.483365056 | 0.730403172 | 0.661778418 | 0.508113239 | 0.882698762 | Rarb | retinoic acid receptor, beta [Source:MGI S | 18.471 | 10.279 | 10.062 | 21.208 | 2.1812 | 31.188 |
| 1813 | ENSMUSG00000017493 | 39322.23572 | 0.032154057 | 0.082041874 | 0.391922509 | 0.695115471 | 0.939992732 | Igfbp4 | insulin-like growth factor binding protein | 37067 | 38989 | 40597 | 39364 | 40314 | 39603 |
| 1814 | ENSMUSG00000017499 | 2.123112523 | -2.351405505 | 1.737542498 | -1.353293809 | 0.175961786 | NA | Cdc6 | cell division cycle 6 [Source:MGI Symbol;A | 1.6062 | 1.0279 | 8.05 | 0.964 | 1.0906 | 0 |
| 1815 | ENSMUSG00000017548 | 500.8230541 | 0.134276669 | 0.132774446 | 1.011314094 | 0.311866122 | 0.759830222 | Suz12 | SUZ12 polycomb repressive complex 2 sul | 440.9 | 530.41 | 461.87 | 508.98 | 514.77 | 548.01 |
| 1816 | ENSMUSG00000017550 | 26.20621567 | 0.454606601 | 0.421747568 | 1.077911611 | 0.281073203 | 0.733710968 | Atad5 | ATPase family, AAA domain containing 5 [ | 21.684 | 22.614 | 22.137 | 35.667 | 26.175 | 28.96 |
| 1817 | ENSMUSG00000017561 | 173.3001592 | -0.10476275 | 0.19041142 | -0.550191525 | 0.582188016 | 0.908645447 | Crlf3 | cytokine receptor-like factor 3 [Source:M | 200.78 | 171.66 | 166.03 | 175.44 | 178.86 | 147.03 |
| 1818 | ENSMUSG00000017588 | 0 | NA | NA | NA | NA | NA | Krt27 | keratin 27 [Source:MGI Symbol;Acc:MGI:1 | 0 | 0 | 0 | 0 | 0 | 0 |
| 1819 | ENSMUSG00000017607 | 5.198910942 | -0.542516646 | 1.015966704 | -0.533990576 | 0.593348054 | 0.913011006 | Tns4 | tensin 4 [Source:MGI Symbol;Acc:MGI:21 | 3.2124 | 12.335 | 3.0187 | 3.8559 | 6.5437 | 2.2277 |
| 1820 | ENSMUSG00000017615 | 582.2438494 | 0.002697033 | 0.128648281 | 0.02096439 | 0.983274062 | 0.998580735 | Tnfaip1 | tumor necrosis factor, alpha-induced prot | 571.01 | 604.42 | 569.53 | 558.14 | 538.77 | 651.6 |
| 1821 | ENSMUSG00000017631 | 66.34097462 | -0.641286955 | 0.308554887 | -2.078356173 | 0.037676566 | 0.263449005 | Abr | active BCR-related gene [Source:MGI Sym | 63.445 | 103.82 | 75.468 | 48.199 | 43.625 | 63.489 |
| 1822 | ENSMUSG00000017639 | 152.6616434 | -0.500204977 | 0.335166916 | -1.492405584 | 0.135592852 | 0.5381974 | Rab11fip4 | RAB11 family interacting protein 4 (class I | 116.45 | 222.03 | 198.23 | 101.22 | 71.981 | 206.06 |
| 1823 | ENSMUSG00000017652 | 3.291970157 | -1.839918954 | 1.307558805 | -1.407140502 | 0.15938573 | NA | Cd40 | CD40 antigen [Source:MGI Symbol;Acc:M | 7.2279 | 3.0838 | 5.0312 | 0 | 2.1812 | 2.2277 |
| 1824 | ENSMUSG00000017664 | 392.7802727 | 0.257845654 | 0.147587174 | 1.747073587 | 0.080624574 | 0.417261162 | Slc35c2 | solute carrier family 35, member C2 [Sour | 347.74 | 374.16 | 351.18 | 385.59 | 406.8 | 491.21 |
| 1825 | ENSMUSG00000017667 | 25.99636523 | -0.428778447 | 0.47552842 | -0.901688373 | 0.367222431 | 0.805261559 | Zfp334 | zinc finger protein 334 [Source:MGI Symb | 27.305 | 40.089 | 22.137 | 22.171 | 13.087 | 31.188 |
| 1826 | ENSMUSG00000017670 | 182.5138418 | -0.251134802 | 0.18481485 | -1.358845369 | 0.174195593 | 0.604151393 | Elmo2 | engulfment and cell motility 2 [Source:MG | 192.74 | 210.72 | 192.19 | 190.87 | 155.96 | 152.6 |
| 1827 | ENSMUSG00000017677 | 376.0838408 | 0.380766862 | 0.253802634 | 1.500247874 | 0.133550206 | 0.535830842 | Wsb1 | WD repeat and SOCS box-containing 1 [Sc | 286.71 | 384.44 | 309.92 | 604.41 | 238.85 | 432.17 |
| 1828 | ENSMUSG00000017679 | 274.3338151 | 0.232583924 | 0.153717569 | 1.513060124 | 0.130264387 | 0.528511668 | Ttpal | tocopherol (alpha) transfer protein-like [S | 233.7 | 249.78 | 273.7 | 288.23 | 303.19 | 297.4 |
| 1829 | ENSMUSG00000017686 | 758.5545857 | 0.10011809 | 0.114122721 | 0.877284464 | 0.380332149 | 0.812628154 | Rhot1 | ras homolog family member T1 [Source:M | 701.91 | 759.63 | 735.56 | 762.51 | 835.41 | 756.3 |
| 1830 | ENSMUSG00000017688 | 1.643861401 | 1.33770114 | 1.870945163 | 0.714986824 | 0.474617185 | NA | Hnf4g | hepatocyte nuclear factor 4, gamma [Sou | 0.8031 | 0 | 2.0125 | 4.8199 | 0 | 2.2277 |
| 1831 | ENSMUSG00000017692 | 3.795447386 | 0.661188431 | 1.178424879 | 0.561078133 | 0.574744274 | NA | Rhbdl3 | rhomboid like 3 [Source:MGI Symbol;Acc: | 5.6217 | 1.0279 | 2.0125 | 1.928 | 3.2719 | 8.9108 |
| 1832 | ENSMUSG00000017697 | 16.82343213 | -0.547476994 | 0.528008345 | -1.03687186 | 0.29979558 | 0.751124765 | Ada | adenosine deaminase [Source:MGI Symbo | 19.274 | 20.558 | 20.125 | 14.46 | 9.8156 | 16.708 |
| 1833 | ENSMUSG00000017707 | 5096.957784 | -0.46175588 | 0.0974246 | -4.739623046 | 2.14E-06 | 0.00015256 | Serinc3 | serine incorporator 3 [Source:MGI Symbo | 5399.3 | 6415.2 | 5903.6 | 4174 | 4370.1 | 4319.5 |
| 1834 | ENSMUSG00000017713 | 192.464674 | -0.001653353 | 0.191713389 | -0.008624086 | 0.99311906 | 0.99859176 | Tha1 | threonine aldolase 1 [Source:MGI Symbol | 216.03 | 169.61 | 191.19 | 168.7 | 186.5 | 222.77 |
| 1835 | ENSMUSG00000017715 | 183.2195617 | 0.207213609 | 0.190220782 | 1.089332127 | 0.27600745 | 0.730038567 | Pgs1 | phosphatidylglycerophosphate synthase 1 | 192.74 | 145.96 | 171.06 | 210.15 | 176.68 | 202.72 |
| 1836 | ENSMUSG00000017716 | 25.77313058 | -0.721598039 | 0.463149208 | -1.558024988 | 0.119227325 | 0.505144521 | Birc5 | baculoviral IAP repeat-containing 5 [Sourc | 20.078 | 41.117 | 35.219 | 16.388 | 22.903 | 18.935 |
| 1837 | ENSMUSG00000017718 | 851.2529243 | -0.274650171 | 0.203282406 | -1.351076938 | 0.176670788 | 0.60755261 | Afmid | arylformamidase [Source:MGI Symbol;Acc | 639.27 | 951.85 | 1206.5 | 910.96 | 786.34 | 612.61 |
| 1838 | ENSMUSG00000017720 | 0.96030799 | 1.298129309 | 2.443376799 | 0.531284945 | 0.595221336 | NA | Trp53tg5 | transformation related protein 53 target 5 | 1.6062 | 0 | 0 | 1.928 | 0 | 2.2277 |
| 1839 | ENSMUSG00000017721 | 636.158172 | 0.094050357 | 0.147472685 | 0.637747643 | 0.523637967 | 0.888334265 | Pigt | phosphatidylinositol glycan anchor biosyn | 575.82 | 583.86 | 686.26 | 548.5 | 674 | 748.5 |
| 1840 | ENSMUSG00000017723 | 171.3395499 | -1.194069076 | 0.308848426 | -3.866197703 | 0.000110545 | 0.003918941 | Wfdc2 | WAP four-disulfide core domain 2 [Source | 306.79 | 218.95 | 190.18 | 168.7 | 64.346 | 79.083 |
| 1841 | ENSMUSG00000017724 | 6.053546148 | -0.238617527 | 0.900117397 | -0.265096006 | 0.790935504 | 0.961019028 | Etv4 | ets variant 4 [Source:MGI Symbol;Acc:MG | 10.44 | 3.0838 | 6.0375 | 5.7839 | 7.6343 | 3.3415 |
| 1842 | ENSMUSG00000017733 | 0 | NA | NA | NA | NA | NA | Eppin | epididymal peptidase inhibitor [Source:M | 0 | 0 | 0 | 0 | 0 | 0 |
| 1843 | ENSMUSG00000017734 | 12.64774144 | -0.988811306 | 0.622117927 | -1.589427444 | 0.111963923 | 0.490954285 | Dbndd2 | dysbindin (dystrobrevin binding protein 1 | 12.85 | 19.53 | 18.112 | 7.7118 | 6.5437 | 11.138 |
| 1844 | ENSMUSG00000017737 | 2.699053068 | -1.485653934 | 1.488252019 | -0.998254271 | 0.318156076 | NA | Mmp9 | matrix metallopeptidase 9 [Source:MGI Sy | 0.8031 | 4.1117 | 7.0437 | 0.964 | 3.2719 | 0 |

| | | | | | | | | | | | | | | | |
|---|---|---|---|---|---|---|---|---|---|---|---|---|---|---|---|
| 1845 | ENSMUSG00000017740 | 22.28196255 | -0.103148245 | 0.457120142 | -0.225647997 | 0.821475207 | 0.969160361 | Slc12a5 | solute carrier family 12, member 5 [Sourc | 22.487 | 19.53 | 27.169 | 19.28 | 20.722 | 24.505 |
| 1846 | ENSMUSG00000017747 | 366.2717424 | 0.001236482 | 0.144284352 | 0.008569757 | 0.993162407 | 0.99859176 | Ghdc | GH3 domain containing [Source:MGI Syml | 375.05 | 340.24 | 382.37 | 331.61 | 375.17 | 393.19 |
| 1847 | ENSMUSG00000017754 | 385.781281 | -0.165387289 | 0.453234559 | -0.36490441 | 0.715182762 | 0.944595913 | Pltp | phospholipid transfer protein [Source:MG | 424.84 | 572.55 | 226.41 | 500.3 | 302.1 | 288.49 |
| 1848 | ENSMUSG00000017756 | 2283.219092 | 0.135146447 | 0.116102104 | 1.164030995 | 0.244411451 | 0.694303562 | Slc12a7 | solute carrier family 12, member 7 [Sourc | 2040.7 | 2255.3 | 2234.9 | 2734.8 | 2272.8 | 2160.9 |
| 1849 | ENSMUSG00000017760 | 2259.56392 | -0.166428922 | 0.111754043 | -1.489242961 | 0.136423402 | 0.5392807 | Ctsa | cathepsin A [Source:MGI Symbol;Acc:MGI | 2427 | 2653.1 | 2089 | 2071.6 | 2166 | 2150.8 |
| 1850 | ENSMUSG00000017764 | 60.01468279 | 0.378158774 | 0.342313413 | 1.104715033 | 0.269283088 | 0.724979259 | Zswim1 | zinc finger SWIM-type containing 1 [Sourc | 49.792 | 45.228 | 61.381 | 51.091 | 52.35 | 100.25 |
| 1851 | ENSMUSG00000017765 | 328.8842187 | 0.090191673 | 0.18505909 | 0.487366889 | 0.625998354 | 0.919638611 | Slc12a4 | solute carrier family 12, member 4 [Sourc | 313.21 | 349.49 | 293.82 | 402.94 | 360.99 | 252.84 |
| 1852 | ENSMUSG00000017767 | 0 | NA | NA | NA | NA | NA | Spata25 | spermatogenesis associated 25 [Source:M | 0 | 0 | 0 | 0 | 0 | 0 |
| 1853 | ENSMUSG00000017774 | 694.4514659 | -0.181869827 | 0.131028458 | -1.388017761 | 0.165131637 | 0.590574884 | Myo1c | myosin IC [Source:MGI Symbol;Acc:MGI:1 | 819.16 | 728.79 | 666.13 | 696.96 | 663.1 | 592.56 |
| 1854 | ENSMUSG00000017776 | 1582.261509 | 0.042752773 | 0.110858257 | 0.385652582 | 0.699753985 | 0.941362191 | Crk | v-crk avian sarcoma virus CT10 oncogene | 1410.2 | 1709.4 | 1557.7 | 1525 | 1682.8 | 1608.4 |
| 1855 | ENSMUSG00000017778 | 3935.98704 | -0.056543754 | 0.101923809 | -0.554764919 | 0.579055465 | 0.908337213 | Cox7c | cytochrome c oxidase subunit 7C [Source: | 4062.9 | 4208.3 | 3767.4 | 3633.2 | 3704.8 | 4239.3 |
| 1856 | ENSMUSG00000017781 | 1040.27715 | -0.145920475 | 0.110902488 | -1.31575474 | 0.188256375 | 0.625244593 | Pitpna | phosphatidylinositol transfer protein, alph | 1072.9 | 1133.8 | 1071.7 | 931.2 | 1078.6 | 953.45 |
| 1857 | ENSMUSG00000017801 | 535.7936769 | 0.184457596 | 0.13753399 | 1.341178251 | 0.179862586 | 0.612025777 | Mlx | MAX-like protein X [Source:MGI Symbol;A | 553.34 | 447.14 | 503.12 | 535.97 | 623.83 | 551.35 |
| 1858 | ENSMUSG00000017802 | 1906.81996 | 0.112922855 | 0.106075718 | 1.064549524 | 0.287079832 | 0.73951397 | Retreg3 | reticulophagy regulator family member 3 | 1766 | 1820.4 | 1909.9 | 1780.5 | 2064.5 | 2099.6 |
| 1859 | ENSMUSG00000017817 | 1.998831776 | -0.928431494 | 1.702414045 | -0.545361745 | 0.585504746 | NA | Jph2 | junctophilin 2 [Source:MGI Symbol;Acc:M | 4.8186 | 0 | 3.0187 | 1.928 | 0 | 2.2277 |
| 1860 | ENSMUSG00000017830 | 256.1859136 | -0.076275303 | 0.166140847 | -0.459100242 | 0.646162183 | 0.925643431 | Dhx58 | DEXH (Asp-Glu-X-His) box polypeptide 58 | 298.75 | 242.59 | 246.53 | 233.28 | 249.75 | 266.21 |
| 1861 | ENSMUSG00000017831 | 1069.351262 | 0.068647411 | 0.105761666 | 0.649076488 | 0.516288938 | 0.886716042 | Rab5a | RAB5A, member RAS oncogene family [So | 1038.4 | 1089.6 | 1004.2 | 1135.6 | 1112.4 | 1035.9 |
| 1862 | ENSMUSG00000017832 | 0.465832987 | -0.698295221 | 3.18348759 | -0.219349126 | 0.826378096 | NA | Hspb9 | heat shock protein, alpha-crystallin-relate | 0.8031 | 1.0279 | 0 | 0.964 | 0 | 0 |
| 1863 | ENSMUSG00000017837 | 300.907494 | 0.322193775 | 0.151572399 | 2.125675762 | 0.033530266 | 0.24494029 | Nkiras2 | NFKB inhibitor interacting Ras-like protein | 276.27 | 250.81 | 274.7 | 313.29 | 323.91 | 366.45 |
| 1864 | ENSMUSG00000017843 | 936.5222579 | -0.118476263 | 0.116484647 | -1.017097673 | 0.309106967 | 0.757655173 | Ppp2r5c | protein phosphatase 2, regulatory subunit | 949.27 | 1060.8 | 915.68 | 959.16 | 891.04 | 843.18 |
| 1865 | ENSMUSG00000017858 | 516.0150044 | 0.038120515 | 0.135115082 | 0.282133674 | 0.777841013 | 0.959559604 | Ift52 | intraflagellar transport 52 [Source:MGI Sy | 499.53 | 535.55 | 492.05 | 457.89 | 548.58 | 562.49 |
| 1866 | ENSMUSG00000017861 | 5.497922451 | 0.18114783 | 0.948334264 | 0.191016856 | 0.848512387 | 0.972803041 | Mybl2 | myeloblastosis oncogene-like 2 [Source:M | 2.4093 | 6.1675 | 7.0437 | 9.6398 | 3.2719 | 4.4554 |
| 1867 | ENSMUSG00000017868 | 849.3665408 | 0.553412344 | 0.201867148 | 2.741468091 | 0.00611653 | 0.076332607 | Sgk2 | serum/glucocorticoid regulated kinase 2 [ | 705.12 | 684.59 | 675.19 | 927.35 | 684.91 | 1419 |
| 1868 | ENSMUSG00000017897 | 0.79210974 | -1.697240781 | 2.618300099 | -0.648222403 | 0.516841114 | NA | Eya2 | EYA transcriptional coactivator and phosp | 1.6062 | 2.0558 | 0 | 0 | 1.0906 | 0 |
| 1869 | ENSMUSG00000017922 | 0 | NA | NA | NA | NA | NA | Prl3c1 | prolactin family 3, subfamily c, member 1 | 0 | 0 | 0 | 0 | 0 | 0 |
| 1870 | ENSMUSG00000017929 | 539.9684639 | 0.343268213 | 0.171939123 | 1.996452044 | 0.045884741 | 0.296229283 | B4galt5 | UDP-Gal:betaGlcNAc beta 1,4-galactosyltr | 388.7 | 438.92 | 601.73 | 674.78 | 544.22 | 591.45 |
| 1871 | ENSMUSG00000017943 | 0 | NA | NA | NA | NA | NA | Gdap1l1 | ganglioside-induced differentiation-associ | 0 | 0 | 0 | 0 | 0 | 0 |
| 1872 | ENSMUSG00000017950 | 10706.6944 | -0.127802588 | 0.105007149 | -1.217084634 | 0.223572016 | 0.66923544 | Hnf4a | hepatic nuclear factor 4, alpha [Source:M | 9677.4 | 12032 | 11834 | 10223 | 10355 | 10118 |
| 1873 | ENSMUSG00000017969 | 38.20105528 | -0.708549095 | 0.365302163 | -1.93962469 | 0.052425318 | 0.321850509 | Ptgis | prostaglandin I2 (prostacyclin) synthase [S | 58.626 | 40.089 | 43.269 | 29.883 | 27.265 | 30.074 |
| 1874 | ENSMUSG00000017978 | 451.8644001 | 0.054302395 | 0.138014016 | 0.393455653 | 0.693982972 | 0.939133502 | Cadps2 | Ca2+-dependent activator protein for secr | 425.64 | 498.54 | 406.52 | 480.06 | 450.43 | 449.99 |
| 1875 | ENSMUSG00000017999 | 422.9696723 | -0.068183321 | 0.129011967 | -0.528503845 | 0.597149679 | 0.914521894 | Ddx27 | DEAD box helicase 27 [Source:MGI Symbc | 421.63 | 433.78 | 443.75 | 414.51 | 423.16 | 400.98 |
| 1876 | ENSMUSG00000018001 | 73.5401202 | -0.536111238 | 0.270418138 | -1.982526917 | 0.047420298 | 0.302152946 | Cyth3 | cytohesin 3 [Source:MGI Symbol;Acc:MGI | 89.947 | 97.652 | 73.456 | 56.875 | 67.618 | 55.692 |
| 1877 | ENSMUSG00000018008 | 54.53769206 | -0.914226536 | 0.385401399 | -2.372141197 | 0.017685331 | 0.158586872 | Cyth4 | cytohesin 4 [Source:MGI Symbol;Acc:MGI | 73.885 | 88.401 | 51.318 | 32.775 | 21.812 | 59.034 |
| 1878 | ENSMUSG00000018012 | 23.30866476 | 0.480172532 | 0.571166964 | 0.840686808 | 0.400523413 | 0.827266295 | Rac3 | Rac family small GTPase 3 [Source:MGI Sy | 17.668 | 15.419 | 25.156 | 8.6758 | 27.265 | 45.668 |
| 1879 | ENSMUSG00000018040 | 420.4282904 | 0.112340194 | 0.136272196 | 0.824380887 | 0.409723171 | 0.833241465 | Rrp7a | ribosomal RNA processing 7 homolog A [S | 398.34 | 372.11 | 441.74 | 428.97 | 453.7 | 427.72 |
| 1880 | ENSMUSG00000018042 | 8993.049127 | 0.271292515 | 0.123833797 | 2.190779264 | 0.02846777 | 0.220942827 | Cyb5r3 | cytochrome b5 reductase 3 [Source:MGI S | 9463.8 | 8225.4 | 6758.9 | 9516.4 | 9557.1 | 10437 |
| 1881 | ENSMUSG00000018068 | 101.1603199 | -0.331728749 | 0.245374637 | -1.351927616 | 0.176398472 | 0.607400048 | Ints2 | integrator complex subunit 2 [Source:MG | 90.751 | 112.04 | 135.84 | 87.722 | 97.065 | 83.538 |
| 1882 | ENSMUSG00000018076 | 265.1078496 | -0.19595991 | 0.159760777 | -1.226583359 | 0.219979227 | 0.666201746 | Med13l | mediator complex subunit 13-like [Source | 283.5 | 308.38 | 257.6 | 255.45 | 257.39 | 228.34 |
| 1883 | ENSMUSG00000018102 | 2598.190998 | 0.223343104 | 0.112724576 | 1.981316863 | 0.04755575 | 0.302749285 | H2bc4 | H2B clustered histone 4 [Source:MGI Sym | 2629.4 | 2068.2 | 2493.5 | 2774.3 | 2718.9 | 2904.9 |
| 1884 | ENSMUSG00000018126 | 0.17131964 | -0.796629176 | 4.080472857 | -0.195229623 | 0.84521319 | NA | Baiap2l2 | BAI1-associated protein 2-like 2 [Source:N | 0 | 1.0279 | 0 | 0 | 0 | 0 |
| 1885 | ENSMUSG00000018143 | 293.672492 | 0.095895994 | 0.214245121 | 0.447599429 | 0.654442315 | 0.925797231 | Mafk | v-maf musculoaponeurotic fibrosarcoma | 386.29 | 249.78 | 214.33 | 288.23 | 278.11 | 345.29 |
| 1886 | ENSMUSG00000018160 | 501.2255834 | 0.19848426 | 0.123216438 | 1.610858617 | 0.107210542 | 0.48315751 | Med1 | mediator complex subunit 1 [Source:MGI | 487.48 | 462.56 | 449.79 | 526.33 | 539.86 | 541.33 |
| 1887 | ENSMUSG00000018166 | 2480.096391 | 0.325695484 | 0.133228901 | 2.44463087 | 0.014500046 | 0.139725963 | Erbb3 | erb-b2 receptor tyrosine kinase 3 [Source: | 2065.6 | 2000.3 | 2538.8 | 2933.4 | 2328.5 | 3014.1 |
| 1888 | ENSMUSG00000018167 | 327.4231542 | -0.036974252 | 0.140258958 | -0.26361419 | 0.792077223 | 0.961434599 | Stard3 | START domain containing 3 [Source:MGI S | 338.91 | 325.85 | 330.05 | 332.57 | 311.92 | 325.24 |
| 1889 | ENSMUSG00000018168 | 5.516102608 | -1.088678273 | 1.006214873 | -1.081954066 | 0.279272939 | 0.731908641 | Ikzf3 | IKAROS family zinc finger 3 [Source:MGI S | 11.243 | 5.1396 | 6.0375 | 1.928 | 7.6343 | 1.1138 |
| 1890 | ENSMUSG00000018169 | 18.25918527 | -0.598287684 | 0.519975027 | -1.150608497 | 0.249893336 | 0.702104867 | Mfng | MFNG O-fucosylpeptide 3-beta-N-acetylgl | 24.093 | 21.586 | 20.125 | 9.6398 | 19.631 | 14.48 |
| 1891 | ENSMUSG00000018171 | 2377.060799 | 0.223734479 | 0.153304607 | 1.459411325 | 0.144451931 | 0.55464443 | Vmp1 | vacuole membrane protein 1 [Source:MG | 2076 | 2443.4 | 2060.8 | 2918.9 | 1912.9 | 2850.3 |
| 1892 | ENSMUSG00000018189 | 331.5329739 | -0.095630566 | 0.153343997 | -0.623634233 | 0.532867823 | 0.889143097 | Uchl5 | ubiquitin carboxyl-terminal esterase L5 [S | 338.11 | 304.26 | 385.39 | 333.54 | 328.28 | 299.62 |
| 1893 | ENSMUSG00000018196 | 412.1656341 | 0.174583265 | 0.129954399 | 1.343419435 | 0.179136197 | 0.611235187 | Glrx2 | glutaredoxin 2 (thioltransferase) [Source:I | 383.08 | 387.53 | 391.43 | 448.25 | 442.79 | 419.92 |
| 1894 | ENSMUSG00000018199 | 82.42670397 | -0.537304928 | 0.264036069 | -2.034967912 | 0.041854102 | 0.281309353 | Ro60 | Ro60, Y RNA binding protein [Source:MGI | 79.507 | 117.18 | 96.599 | 72.298 | 63.256 | 65.717 |
| 1895 | ENSMUSG00000018209 | 400.6468143 | 0.040180269 | 0.15742992 | 0.255226385 | 0.798548253 | 0.962424862 | Stk4 | serine/threonine kinase 4 [Source:MGI Sy | 354.17 | 381.36 | 449.79 | 361.49 | 393.71 | 463.36 |
| 1896 | ENSMUSG00000018211 | 3.548394706 | -0.829318792 | 1.237957626 | -0.669908868 | 0.502915886 | NA | Wfdc15b | WAP four-disulfide core domain 15B [Sou | 2.4093 | 5.1396 | 6.0375 | 0 | 4.3625 | 3.3415 |
| 1897 | ENSMUSG00000018217 | 51.2631481 | -0.888342965 | 0.320378113 | -2.772795423 | 0.005557705 | 0.071518364 | Pmp22 | peripheral myelin protein 22 [Source:MGI | 79.507 | 61.675 | 58.362 | 38.559 | 32.719 | 36.757 |
| 1898 | ENSMUSG00000018238 | 4.447961441 | -0.584468292 | 1.100091888 | -0.53129043 | 0.595217536 | NA | Gdf9 | growth differentiation factor 9 [Source:M | 0.8031 | 9.2513 | 6.0375 | 2.8919 | 4.3625 | 3.3415 |
| 1899 | ENSMUSG00000018239 | 54.45925108 | 0.244874989 | 0.302262891 | 0.810139107 | 0.41786023 | 0.837344173 | Zcchc10 | zinc finger, CCHC domain containing 10 [S | 53.005 | 41.117 | 55.343 | 62.659 | 56.712 | 57.92 |
| 1900 | ENSMUSG00000018259 | 0 | NA | NA | NA | NA | NA | Prl8a2 | prolactin family 8, subfamily a, member 2 | 0 | 0 | 0 | 0 | 0 | 0 |

| | | | | | | | | | | | | | | | |
|---|---|---|---|---|---|---|---|---|---|---|---|---|---|---|---|
| 1901 | ENSMUSG00000018263 | 0 | NA | NA | NA | NA | NA | Tbx5 | T-box 5 [Source:MGI Symbol;Acc:MGI:102 | 0 | 0 | 0 | 0 | 0 | 0 |
| 1902 | ENSMUSG00000018286 | 2129.026378 | 0.076254376 | 0.107551309 | 0.709004623 | 0.478321608 | 0.869794311 | Psmb6 | proteasome (prosome, macropain) subun | 2278.4 | 1984.9 | 1953.1 | 2036.9 | 2208.5 | 2312.3 |
| 1903 | ENSMUSG00000018287 | 449.1524258 | 0.157766857 | 0.145159316 | 1.08685313 | 0.277101723 | 0.730041765 | Spag7 | sperm associated antigen 7 [Source:MGI S | 492.3 | 402.94 | 377.34 | 466.57 | 466.78 | 488.98 |
| 1904 | ENSMUSG00000018293 | 3233.072865 | -0.03159387 | 0.092584374 | -0.341244083 | 0.732919839 | 0.947867718 | Pfn1 | profilin 1 [Source:MGI Symbol;Acc:MGI:97 | 3409.2 | 3205 | 3189.8 | 3016.3 | 3253.3 | 3324.8 |
| 1905 | ENSMUSG00000018322 | 178.3238812 | -0.210473198 | 0.18828976 | -1.117815421 | 0.263645834 | 0.718555368 | Tomm34 | translocase of outer mitochondrial memb | 195.15 | 174.75 | 204.27 | 187.01 | 146.14 | 162.62 |
| 1906 | ENSMUSG00000018326 | 1950.904493 | -0.128185885 | 0.092037745 | -1.392753438 | 0.163694363 | 0.587804187 | Ywhab | tyrosine 3-monooxygenase/tryptophan 5- | 2103.3 | 1982.9 | 2025.6 | 1828.7 | 1872.6 | 1892.4 |
| 1907 | ENSMUSG00000018334 | 21.9008412 | -0.672891673 | 0.526949515 | -1.276956622 | 0.201617562 | 0.645999739 | Ksr1 | kinase suppressor of ras 1 [Source:MGI Sy | 16.062 | 21.586 | 43.269 | 17.352 | 13.087 | 20.049 |
| 1908 | ENSMUSG00000018339 | 93.56096401 | -0.745978915 | 0.312670704 | -2.385829265 | 0.01704066 | 0.155016422 | Gpx3 | glutathione peroxidase 3 [Source:MGI Syr | 147.77 | 121.29 | 82.512 | 84.83 | 42.534 | 82.424 |
| 1909 | ENSMUSG00000018340 | 2051.534178 | -0.068308327 | 0.101674669 | -0.671832303 | 0.501690461 | 0.880974509 | Anxa6 | annexin A6 [Source:MGI Symbol;Acc:MGI | 2051.9 | 2183.3 | 2064.8 | 1859.5 | 2169.2 | 1980.4 |
| 1910 | ENSMUSG00000018341 | 0.810301681 | -0.283528992 | 2.609307867 | -0.108660613 | 0.913471683 | NA | Il12rb2 | interleukin 12 receptor, beta 2 [Source:M | 1.6062 | 1.0279 | 0 | 0 | 0 | 2.2277 |
| 1911 | ENSMUSG00000018347 | 70.74769017 | 0.075959118 | 0.272605728 | 0.278640946 | 0.780520386 | 0.959905354 | Zkscan6 | zinc finger with KRAB and SCAN domains ( | 76.295 | 68.87 | 61.381 | 80.01 | 61.075 | 76.855 |
| 1912 | ENSMUSG00000018362 | 814.8104921 | 0.264226161 | 0.133824888 | 1.974417208 | 0.048334323 | 0.305821368 | Kpna2 | karyopherin (importin) alpha 2 [Source:M | 765.36 | 832.61 | 622.87 | 860.83 | 863.77 | 943.43 |
| 1913 | ENSMUSG00000018363 | 1042.825569 | 0.172237278 | 0.107424992 | 1.60332595 | 0.10886271 | 0.485684837 | Smurf2 | SMAD specific E3 ubiquitin protein ligase : | 954.89 | 979.61 | 1008.3 | 1160.6 | 1126.6 | 1027 |
| 1914 | ENSMUSG00000018372 | 109.8882553 | 0.231768145 | 0.21956385 | 1.055584266 | 0.291158192 | 0.740906421 | Cep95 | centrosomal protein 95 [Source:MGI Symb | 103.6 | 107.93 | 91.568 | 114.71 | 113.42 | 128.09 |
| 1915 | ENSMUSG00000018377 | 246.0703081 | 0.135845402 | 0.164875697 | 0.823926173 | 0.409981508 | 0.833241465 | Vezf1 | vascular endothelial zinc finger 1 [Source: | 216.84 | 244.64 | 242.5 | 266.06 | 229.03 | 277.35 |
| 1916 | ENSMUSG00000018378 | 43.16459891 | -0.054480307 | 0.391593532 | -0.139124633 | 0.889351663 | 0.982382091 | Cuedc1 | CUE domain containing 1 [Source:MGI Syr | 30.518 | 62.703 | 39.244 | 55.911 | 31.628 | 38.985 |
| 1917 | ENSMUSG00000018379 | 1702.29316 | 0.002912164 | 0.108545987 | 0.026828852 | 0.978596241 | 0.997797092 | Srsf1 | serine and arginine-rich splicing factor 1 [! | 1617.4 | 1686.8 | 1798.2 | 1702.4 | 1855.1 | 1553.8 |
| 1918 | ENSMUSG00000018381 | 12.7823142 | -1.11768546 | 0.631539076 | -1.76978037 | 0.076763735 | 0.405543261 | Abi3 | ABI family member 3 [Source:MGI Symbol | 16.865 | 20.558 | 15.094 | 8.6758 | 4.3625 | 11.138 |
| 1919 | ENSMUSG00000018387 | 708.9650285 | -0.152260088 | 0.112857242 | -1.349138834 | 0.177292377 | 0.608561375 | Shroom1 | shroom family member 1 [Source:MGI Syr | 731.63 | 730.85 | 776.82 | 674.78 | 691.45 | 648.26 |
| 1920 | ENSMUSG00000018395 | 49.83998368 | -0.110876476 | 0.327855689 | -0.33818683 | 0.7352224 | 0.948109711 | Kif3a | kinesin family member 3A [Source:MGI Sy | 47.383 | 59.619 | 48.3 | 41.451 | 61.075 | 41.212 |
| 1921 | ENSMUSG00000018398 | 50.01366435 | -0.974920122 | 0.336225547 | -2.899601563 | 0.003736373 | 0.054645736 | Septin8 | septin 8 [Source:MGI Symbol;Acc:MGI:89 | 75.492 | 66.815 | 56.35 | 31.811 | 26.175 | 43.44 |
| 1922 | ENSMUSG00000018401 | 885.5639288 | -0.061925634 | 0.140878305 | -0.439568275 | 0.660249822 | 0.927980329 | Mtmr4 | myotubularin related protein 4 [Source:M | 753.31 | 885.04 | 1076.7 | 895.54 | 848.5 | 854.32 |
| 1923 | ENSMUSG00000018405 | 71.04785152 | -0.114087266 | 0.3046467 | -0.374490405 | 0.708039494 | 0.943617731 | Mrm1 | mitochondrial rRNA methyltransferase 1 [ | 64.248 | 62.703 | 94.587 | 57.839 | 57.803 | 89.108 |
| 1924 | ENSMUSG00000018411 | 44.45463916 | -0.687973558 | 0.340385762 | -2.021158449 | 0.043263365 | 0.286918392 | Mapt | microtubule-associated protein tau [Sourc | 53.005 | 55.508 | 56.35 | 42.415 | 32.719 | 26.732 |
| 1925 | ENSMUSG00000018412 | 464.0431487 | -0.242863662 | 0.145537913 | -1.668731241 | 0.095170655 | 0.455057116 | Kansl1 | KAT8 regulatory NSL complex subunit 1 [S | 463.39 | 579.75 | 466.9 | 458.85 | 416.62 | 398.76 |
| 1926 | ENSMUSG00000018415 | 883.0544839 | -0.028475216 | 0.118644064 | -0.240005397 | 0.810326072 | 0.966245368 | Gid4 | GID complex subunit 4, VID24 homolog [S | 803.91 | 916.9 | 955.93 | 922.53 | 868.13 | 830.93 |
| 1927 | ENSMUSG00000018417 | 3964.40056 | -0.090343439 | 0.144553166 | -0.624984158 | 0.531981456 | 0.889135634 | Myo1b | myosin IB [Source:MGI Symbol;Acc:MGI:1 | 3282.3 | 4302.9 | 4681 | 3424 | 3626.3 | 4469.9 |
| 1928 | ENSMUSG00000018425 | 734.4945104 | -0.027414607 | 0.110058071 | -0.249092196 | 0.803289466 | 0.963596811 | Dhx40 | DEAH (Asp-Glu-Ala-His) box polypeptide 4 | 749.29 | 752.44 | 722.48 | 721.06 | 716.54 | 745.16 |
| 1929 | ENSMUSG00000018427 | 255.994191 | -0.478462544 | 0.187497106 | -2.551839624 | 0.010715583 | 0.114047236 | Ypel2 | yippee like 2 [Source:MGI Symbol;Acc:MG | 256.19 | 263.15 | 375.33 | 215.93 | 214.85 | 210.52 |
| 1930 | ENSMUSG00000018428 | 2758.870905 | -0.052243858 | 0.107392711 | -0.486474898 | 0.626630499 | 0.920067357 | Akap1 | A kinase (PRKA) anchor protein 1 [Source: | 2655.1 | 3000.5 | 2772.2 | 2846.6 | 2860.7 | 2418.2 |
| 1931 | ENSMUSG00000018433 | 202.6029474 | 0.03359975 | 0.168696604 | 0.199172651 | 0.842127692 | 0.971854408 | Nol11 | nucleolar protein 11 [Source:MGI Symbol; | 187.12 | 208.67 | 205.27 | 202.44 | 209.4 | 202.72 |
| 1932 | ENSMUSG00000018442 | 2055.175628 | 0.306404501 | 0.109269247 | 2.804123838 | 0.005045352 | 0.066950435 | Derl2 | Der1-like domain family, member 2 [Sourc | 2013.4 | 1749.5 | 1748.9 | 2244.1 | 2115.8 | 2459.4 |
| 1933 | ENSMUSG00000018446 | 1476.997476 | -0.196119842 | 0.108016577 | -1.815645777 | 0.069424727 | 0.380507483 | C1qbp | complement component 1, q subcompone | 1491.4 | 1688.9 | 1551.6 | 1281.1 | 1474.5 | 1374.5 |
| 1934 | ENSMUSG00000018449 | 397.7552707 | 0.204665231 | 0.148702928 | 1.376336253 | 0.168717554 | 0.595783862 | Rpain | RPA interacting protein [Source:MGI Symb | 407.17 | 312.49 | 388.41 | 436.68 | 431.88 | 409.89 |
| 1935 | ENSMUSG00000018451 | 0.160662974 | 1.126924175 | 4.080472857 | 0.276174898 | 0.782413736 | NA | 6330403K07 | RIKEN cDNA 6330403K07 gene [Source:M | 0 | 0 | 0 | 0.964 | 0 | 0 |
| 1936 | ENSMUSG00000018459 | 322.3874759 | -0.265267147 | 0.191385932 | -1.386032626 | 0.165736941 | 0.591759842 | Slc13a3 | solute carrier family 13 (sodium-depender | 432.07 | 312.49 | 309.92 | 249.67 | 273.75 | 356.43 |
| 1937 | ENSMUSG00000018470 | 8.424989608 | -0.170981748 | 0.814264629 | -0.209983022 | 0.833680924 | 0.970792812 | Kcnab3 | potassium voltage-gated channel, shaker- | 5.6217 | 8.2233 | 13.081 | 12.532 | 2.1812 | 8.9108 |
| 1938 | ENSMUSG00000018474 | 2896.21561 | 0.303471852 | 0.136253174 | 2.227264466 | 0.025929604 | 0.208275484 | Chd3 | chromodomain helicase DNA binding prot | 2753 | 2254.2 | 2771.2 | 3824.1 | 2954.5 | 2820.3 |
| 1939 | ENSMUSG00000018476 | 402.7234277 | 0.099643492 | 0.159431536 | 0.624992363 | 0.531976071 | 0.889135634 | Kdm6b | KDM1 lysine (K)-specific demethylase 6B [ | 351.76 | 355.66 | 459.85 | 455 | 370.81 | 423.26 |
| 1940 | ENSMUSG00000018479 | 0.321325949 | 1.911575114 | 4.048267893 | 0.472195804 | 0.636787034 | NA | 1700125H20 | RIKEN cDNA 1700125H20 gene [Source:M | 0 | 0 | 0 | 1.928 | 0 | 0 |
| 1941 | ENSMUSG00000018481 | 456.7055637 | -0.073429093 | 0.131489778 | -0.558439549 | 0.576544273 | 0.9077105 | Appbp2 | amyloid beta precursor protein (cytoplasr | 462.59 | 498.54 | 443.75 | 428.97 | 431.88 | 474.5 |
| 1942 | ENSMUSG00000018486 | 12.53068729 | -0.271350387 | 0.659564082 | -0.411408678 | 0.680772892 | 0.934384556 | Wnt9b | wingless-type MMTV integration site fami | 9.6372 | 15.419 | 16.1 | 9.6398 | 5.4531 | 18.935 |
| 1943 | ENSMUSG00000018500 | 0.17131964 | -0.796629176 | 4.080472857 | -0.195229623 | 0.84521319 | NA | Adora2b | adenosine A2b receptor [Source:MGI Sym | 0 | 1.0279 | 0 | 0 | 0 | 0 |
| 1944 | ENSMUSG00000018501 | 1877.986411 | -0.000337899 | 0.104741783 | -0.003226019 | 0.997426014 | 0.999498002 | Ncor1 | nuclear receptor co-repressor 1 [Source:N | 1831.1 | 1914 | 1890.7 | 2036.9 | 1894.4 | 1700.8 |
| 1945 | ENSMUSG00000018507 | 18.04555423 | -1.759894548 | 0.586331699 | -3.001534029 | 0.00268623 | 0.043509107 | Trpv2 | transient receptor potential cation channe | 23.29 | 35.977 | 24.15 | 3.8559 | 7.6343 | 13.366 |
| 1946 | ENSMUSG00000018509 | 400.2159375 | 0.064817745 | 0.143231536 | 0.452538224 | 0.650881291 | 0.925643431 | Cenpv | centromere protein V [Source:MGI Symbo | 439.3 | 383.41 | 350.17 | 417.4 | 412.25 | 398.76 |
| 1947 | ENSMUSG00000018537 | 72.25080879 | -0.106340486 | 0.322307174 | -0.329935214 | 0.741448915 | 0.948792385 | Pcgf2 | polycomb group ring finger 2 [Source:MGI | 62.642 | 84.289 | 78.487 | 100.25 | 62.165 | 45.668 |
| 1948 | ENSMUSG00000018541 | 202.5333028 | -0.122498569 | 0.195400531 | -0.626910115 | 0.530718157 | 0.888794797 | Cwc25 | CWC25 spliceosome-associated protein [S | 208.81 | 216.89 | 208.29 | 238.1 | 154.87 | 188.24 |
| 1949 | ENSMUSG00000018543 | 0.294513347 | 0.165143496 | 4.080472857 | 0.040471657 | 0.967717103 | NA | 1700001P01 | RIKEN cDNA 1700001P01 gene [Source:M | 0.8031 | 0 | 0 | 0.964 | 0 | 0 |
| 1950 | ENSMUSG00000018547 | 344.111084 | 0.13574144 | 0.149277385 | 0.909323539 | 0.363179369 | 0.802111222 | Pip4k2b | phosphatidylinositol-5-phosphate 4-kinas | 303.57 | 313.51 | 367.28 | 364.38 | 379.54 | 336.38 |
| 1951 | ENSMUSG00000018548 | 278.8034586 | 0.049258967 | 0.162399934 | 0.303318888 | 0.761646853 | 0.954216198 | Trim37 | tripartite motif-containing 37 [Source:MG | 238.52 | 283.71 | 300.87 | 297.87 | 290.1 | 261.75 |
| 1952 | ENSMUSG00000018554 | 2.224933599 | -1.182727417 | 1.683253524 | -0.70264366 | 0.482277846 | NA | Ybx2 | Y box protein 2 [Source:MGI Symbol;Acc:I | 7.2279 | 0 | 2.0125 | 1.928 | 2.1812 | 0 |
| 1953 | ENSMUSG00000018559 | 500.9988606 | 0.002057888 | 0.129290086 | 0.015916829 | 0.987300744 | 0.99859176 | Ctdnep1 | CTD nuclear envelope phosphatase 1 [Sou | 516.39 | 508.82 | 475.95 | 459.82 | 512.59 | 532.42 |
| 1954 | ENSMUSG00000018565 | 368.1035528 | 0.076419265 | 0.140091263 | 0.545496298 | 0.585412226 | 0.909202452 | Elp5 | elongator acetyltransferase complex subu | 375.85 | 341.27 | 357.22 | 350.89 | 401.35 | 382.05 |
| 1955 | ENSMUSG00000018566 | 28.40505602 | -0.992938396 | 0.509659191 | -1.948239949 | 0.051386259 | 0.318308714 | Slc2a4 | solute carrier family 2 (facilitated glucose | 46.58 | 34.949 | 32.2 | 34.703 | 13.087 | 8.9108 |
| 1956 | ENSMUSG00000018567 | 3589.531978 | 0.025132374 | 0.090433063 | 0.277911349 | 0.781080411 | 0.959993512 | Gabarap | gamma-aminobutyric acid receptor associ | 3528.8 | 3533 | 3612.4 | 3405.7 | 3657.9 | 3799.3 |

| | | | | | | | | | | | | | | | | |
|---|---|---|---|---|---|---|---|---|---|---|---|---|---|---|---|---|
| 1957 | ENSMUSG00000018569 | 6.425575569 | -0.040019067 | 0.882926179 | -0.045325496 | 0.963847866 | 0.993245949 | Cldn7 | claudin 7 [Source:MGI Symbol;Acc:MGI:1{ | 7.2279 | 10.279 | 2.0125 | 5.7839 | 5.4531 | 7.7969 |
| 1958 | ENSMUSG00000018570 | 1.224137803 | 0.310318343 | 2.061334224 | 0.150542469 | 0.880336647 | NA | 2810408A11 | RIKEN cDNA 2810408A11 gene [Source:M | 3.2124 | 0 | 0 | 1.928 | 1.0906 | 1.1138 |
| 1959 | ENSMUSG00000018572 | 304.1891808 | 0.167044652 | 0.162555974 | 1.027613125 | 0.304131848 | 0.752961647 | Phf23 | PHD finger protein 23 [Source:MGI Symbo | 273.05 | 303.24 | 283.76 | 324.86 | 272.65 | 367.57 |
| 1960 | ENSMUSG00000018574 | 4925.415315 | -0.046968724 | 0.091687907 | -0.512267381 | 0.608463891 | 0.916376001 | Acadvl | acyl-Coenzyme A dehydrogenase, very lor | 4658 | 5155 | 5205.3 | 4971.2 | 4888.2 | 4674.8 |
| 1961 | ENSMUSG00000018581 | 1.871018121 | 0.96264486 | 1.647809486 | 0.584196698 | 0.559087983 | NA | Dnah11 | dynein, axonemal, heavy chain 11 [Source | 0.8031 | 0 | 3.0187 | 1.928 | 4.3625 | 1.1138 |
| 1962 | ENSMUSG00000018583 | 1016.010187 | -0.154108553 | 0.12643294 | -1.218895597 | 0.222883821 | 0.668407528 | G3bp1 | GTPase activating protein (SH3 domain) b | 966.13 | 1132.8 | 1111.9 | 842.52 | 1048.1 | 994.66 |
| 1963 | ENSMUSG00000018585 | 1668.941409 | -0.256241564 | 0.116692245 | -2.19587483 | 0.028100904 | 0.21875752 | Atox1 | antioxidant 1 copper chaperone [Source:I | 2070.4 | 1749.5 | 1629.1 | 1532.7 | 1568.3 | 1463.6 |
| 1964 | ENSMUSG00000018589 | 0 | NA | NA | NA | NA | NA | Glra2 | glycine receptor, alpha 2 subunit [Source: | 0 | 0 | 0 | 0 | 0 | 0 |
| 1965 | ENSMUSG00000018593 | 577.4968246 | -0.966166157 | 0.216715981 | -4.458213708 | 8.26E-06 | 0.000455037 | Sparc | secreted acidic cysteine rich glycoprotein | 1046.4 | 708.24 | 536.33 | 394.27 | 313.01 | 466.7 |
| 1966 | ENSMUSG00000018595 | 2.684827743 | -0.965508093 | 1.614888463 | -0.59787912 | 0.549920593 | NA | Glra4 | glycine receptor, alpha 4 subunit [Source: | 2.4093 | 8.2233 | 0 | 0 | 4.3625 | 1.1138 |
| 1967 | ENSMUSG00000018599 | 596.141168 | -0.043796834 | 0.124421512 | -0.352003711 | 0.724835479 | 0.9448775 | Mief2 | mitochondrial elongation factor 2 [Source | 638.47 | 585.91 | 590.67 | 581.28 | 634.74 | 545.78 |
| 1968 | ENSMUSG00000018604 | 634.5501182 | 0.348625466 | 0.191214125 | 1.823220254 | 0.068270039 | 0.376317774 | Tbx3 | T-box 3 [Source:MGI Symbol;Acc:MGI:984 | 687.46 | 416.31 | 569.53 | 589.95 | 876.86 | 667.19 |
| 1969 | ENSMUSG00000018620 | 0 | NA | NA | NA | NA | NA | Mmp20 | matrix metallopeptidase 20 (enamelysin) | | 0 | 0 | 0 | 0 | 0 | 0 |
| 1970 | ENSMUSG00000018623 | 1.160062526 | -2.390610565 | 2.366072327 | -1.010370874 | 0.312317637 | NA | Mmp7 | matrix metallopeptidase 7 [Source:MGI Sy | 4.8186 | 1.0279 | 0 | 0 | 0 | 1.1138 |
| 1971 | ENSMUSG00000018634 | 0 | NA | NA | NA | NA | NA | Crhr1 | corticotropin releasing hormone receptor | 0 | 0 | 0 | 0 | 0 | 0 |
| 1972 | ENSMUSG00000018648 | 31.43526653 | -1.075104988 | 0.426956681 | -2.518065735 | 0.01180013 | 0.121391945 | Dusp14 | dual specificity phosphatase 14 [Source:M | 39.352 | 39.061 | 49.306 | 14.46 | 16.359 | 30.074 |
| 1973 | ENSMUSG00000018651 | 88.88832077 | -0.234460104 | 0.240568008 | -0.974610492 | 0.329753507 | 0.774655633 | Tada2a | transcriptional adaptor 2A [Source:MGI Sy | 97.175 | 100.74 | 90.562 | 90.614 | 79.615 | 74.628 |
| 1974 | ENSMUSG00000018654 | 17.91527086 | -1.695053581 | 0.541704628 | -3.129110393 | 0.001753364 | 0.031780276 | Ikzf1 | IKAROS family zinc finger 1 [Source:MGI S | 24.093 | 31.865 | 26.162 | 7.7118 | 7.6343 | 10.025 |
| 1975 | ENSMUSG00000018656 | 0 | NA | NA | NA | NA | NA | Tcaf3 | TRPM8 channel-associated factor 3 [Sourc | 0 | 0 | 0 | 0 | 0 | 0 |
| 1976 | ENSMUSG00000018659 | 2768.262642 | 0.111482019 | 0.097605791 | 1.142166041 | 0.253385009 | 0.706566564 | Pnpo | pyridoxine 5'-phosphate oxidase [Source:I | 2735.4 | 2628.4 | 2619.3 | 2671.2 | 3101.7 | 2853.7 |
| 1977 | ENSMUSG00000018661 | 329.2794113 | -0.000152752 | 0.151998981 | -0.001004953 | 0.999198163 | 0.999764475 | Cog1 | component of oligomeric golgi complex 1 | 336.5 | 314.54 | 337.09 | 356.67 | 345.73 | 285.14 |
| 1978 | ENSMUSG00000018666 | 539.0025269 | 0.176323463 | 0.125756036 | 1.40210736 | 0.16088319 | 0.583943402 | Cbx1 | chromobox 1 [Source:MGI Symbol;Acc:M( | 485.07 | 518.07 | 515.2 | 536.94 | 561.67 | 617.07 |
| 1979 | ENSMUSG00000018669 | 1162.508581 | 0.269975922 | 0.107254629 | 2.517149376 | 0.011830866 | 0.121535259 | Cdk5rap3 | CDK5 regulatory subunit associated protei | 1086.6 | 1043.3 | 1032.4 | 1375.6 | 1201.9 | 1235.3 |
| 1980 | ENSMUSG00000018672 | 292.5549659 | -0.178798654 | 0.172221988 | -1.038187145 | 0.299182936 | 0.750369688 | Copz2 | coatomer protein complex, subunit zeta 2 | 363 | 291.93 | 276.72 | 311.36 | 263.93 | 248.39 |
| 1981 | ENSMUSG00000018677 | 7181.249847 | -0.067100634 | 0.093301681 | -0.719179254 | 0.47203048 | 0.867870596 | Slc25a39 | solute carrier family 25, member 39 [Sour | 7011.9 | 7522.3 | 7510.6 | 6569.5 | 7536.2 | 6937 |
| 1982 | ENSMUSG00000018678 | 68.94381874 | -0.109251811 | 0.283457428 | -0.385425819 | 0.699921956 | 0.941372511 | Sp2 | Sp2 transcription factor [Source:MGI Sym | 70.673 | 66.815 | 77.481 | 82.902 | 54.531 | 61.261 |
| 1983 | ENSMUSG00000018697 | 232.2052176 | -0.023136342 | 0.158631585 | -0.145849531 | 0.884040174 | 0.981769403 | Aatf | apoptosis antagonizing transcription facto | 243.34 | 226.14 | 232.44 | 228.46 | 233.39 | 229.45 |
| 1984 | ENSMUSG00000018698 | 1.086881812 | -1.106124234 | 2.043719538 | -0.541230934 | 0.588348418 | NA | Lhx1 | LIM homeobox protein 1 [Source:MGI Syn | 2.4093 | 1.0279 | 1.0062 | 0.964 | 0 | 1.1138 |
| 1985 | ENSMUSG00000018707 | 1623.042975 | -0.068567715 | 0.116443973 | -0.588847264 | 0.555963737 | 0.897628185 | Dync1h1 | dynein cytoplasmic 1 heavy chain 1 [Sourc | 1635.9 | 1820.4 | 1529.5 | 1744.8 | 1558.5 | 1449.1 |
| 1986 | ENSMUSG00000018727 | 276.5676259 | 0.028173973 | 0.182029863 | 0.154776655 | 0.876997396 | 0.980632845 | Cpsf4l | cleavage and polyadenylation specific fact | 260.21 | 232.31 | 329.04 | 255.45 | 320.64 | 261.75 |
| 1987 | ENSMUSG00000018733 | 215.4933448 | 0.022138935 | 0.181250334 | 0.122145624 | 0.90278369 | 0.983821605 | Pex12 | peroxisomal biogenesis factor 12 [Source: | 216.03 | 210.72 | 214.33 | 185.08 | 257.39 | 209.4 |
| 1988 | ENSMUSG00000018736 | 590.171686 | -0.045217377 | 0.127443154 | -0.354804285 | 0.722736218 | 0.9447719 | Ndel1 | nudE neurodevelopment protein 1 like 1 [ | 620.8 | 597.22 | 580.6 | 643.94 | 564.94 | 533.53 |
| 1989 | ENSMUSG00000018740 | 25.02901683 | 0.251601756 | 0.457939478 | 0.549421414 | 0.582716282 | 0.908645447 | Slc25a35 | solute carrier family 25, member 35 [Sour | 30.518 | 20.558 | 17.106 | 20.244 | 29.447 | 32.301 |
| 1990 | ENSMUSG00000018750 | 268.2680157 | -0.044529839 | 0.161203702 | -0.27623335 | 0.782368844 | 0.960626736 | Zbtb4 | zinc finger and BTB domain containing 4 [! | 261.81 | 303.24 | 252.57 | 274.73 | 241.03 | 276.23 |
| 1991 | ENSMUSG00000018752 | 10.25135185 | -0.042716485 | 0.685627752 | -0.062302736 | 0.950321749 | 0.990815588 | Tnfsfm13 | tumor necrosis factor (ligand) superfamily | 12.85 | 9.2513 | 9.0562 | 10.604 | 14.178 | 5.5692 |
| 1992 | ENSMUSG00000018761 | 1291.888304 | -0.117838508 | 0.132877154 | -0.886822944 | 0.375174233 | 0.809332934 | Mpdu1 | mannose-P-dolichol utilization defect 1 [S | 1503.4 | 1291.1 | 1237.7 | 1061.3 | 1402.5 | 1255.3 |
| 1993 | ENSMUSG00000018765 | 809.758159 | 0.14473967 | 0.108737687 | 1.331090207 | 0.183159331 | 0.617101458 | Fxr2 | fragile X mental retardation, autosomal ho | 773.39 | 749.35 | 784.87 | 875.29 | 816.87 | 858.77 |
| 1994 | ENSMUSG00000018770 | 7543.168765 | -0.055236738 | 0.094644064 | -0.583626015 | 0.559471953 | 0.899423311 | Atp5g3 | ATP synthase, H+ transporting, mitochonc | 7619.8 | 7931.4 | 7510.6 | 6724.7 | 7739 | 7733.4 |
| 1995 | ENSMUSG00000018774 | 53.90567209 | -1.471145846 | 0.369173842 | -3.984967723 | 6.75E-05 | 0.002638284 | Cd68 | CD68 antigen [Source:MGI Symbol;Acc:M( | 78.704 | 104.85 | 54.337 | 34.703 | 18.541 | 32.301 |
| 1996 | ENSMUSG00000018776 | 0.356960287 | 0.16514264 | 3.880370363 | 0.042558474 | 0.966053499 | NA | Slc35g3 | solute carrier family 35, member G3 [Sour | 0 | 1.0279 | 0 | 0 | 0 | 1.1138 |
| 1997 | ENSMUSG00000018796 | 31290.50457 | -0.482512436 | 0.166412961 | -2.899488324 | 0.003737723 | 0.054645736 | Acsl1 | acyl-CoA synthetase long-chain family me | 29955 | 38229 | 41242 | 30413 | 29074 | 18831 |
| 1998 | ENSMUSG00000018800 | 137.8049715 | -0.544219931 | 0.235292848 | -2.312947185 | 0.020725543 | 0.177486238 | Abca5 | ATP-binding cassette, sub-family A (ABC1) | 148.57 | 188.11 | 153.96 | 103.15 | 145.05 | 87.994 |
| 1999 | ENSMUSG00000018809 | 59.87798372 | -0.103293152 | 0.283954985 | -0.363765939 | 0.7160328 | 0.9447719 | Smyd4 | SET and MYND domain containing 4 [Sour | 61.839 | 63.731 | 60.375 | 52.055 | 58.893 | 62.375 |
| 2000 | ENSMUSG00000018819 | 31.40914637 | -0.98903635 | 0.408588242 | -2.420618729 | 0.015494119 | 0.145412036 | Lsp1 | lymphocyte specific 1 [Source:MGI Symbo | 49.792 | 42.145 | 33.206 | 20.244 | 17.45 | 25.618 |
| 2001 | ENSMUSG00000018820 | 376.1436519 | 0.046608609 | 0.14545326 | 0.32043702 | 0.748637066 | 0.951074124 | Zfyve27 | zinc finger, FYVE domain containing 27 [Sc | 376.65 | 353.6 | 380.36 | 428.01 | 375.17 | 343.06 |
| 2002 | ENSMUSG00000018821 | 480.0219503 | 0.192692354 | 0.128922151 | 1.494641164 | 0.135008117 | 0.537616739 | Avpi1 | arginine vasopressin-induced 1 [Source:M | 474.63 | 433.78 | 434.7 | 480.06 | 526.77 | 530.19 |
| 2003 | ENSMUSG00000018822 | 2.814528459 | -1.194521597 | 1.398783124 | -0.853971982 | 0.393120514 | NA | Sfrp5 | secreted frizzled-related sequence proteir | 5.6217 | 5.1396 | 1.0062 | 2.8919 | 0 | 2.2277 |
| 2004 | ENSMUSG00000018830 | 16.03070618 | -0.457003911 | 0.583938265 | -0.782623675 | 0.433848135 | 0.845264432 | Myh11 | myosin, heavy polypeptide 11, smooth mu | 20.078 | 19.53 | 16.1 | 18.316 | 5.4531 | 16.708 |
| 2005 | ENSMUSG00000018841 | 314.058645 | 0.010828744 | 0.141402402 | 0.076581048 | 0.938956836 | 0.988721002 | Rad51d | RAD51 paralog D [Source:MGI Symbol;Acc | 309.19 | 316.6 | 312.94 | 316.18 | 308.64 | 320.79 |
| 2006 | ENSMUSG00000018844 | 0.838717116 | -1.825569927 | 2.582619186 | -0.706867639 | 0.479648736 | NA | Fndc8 | fibronectin type III domain containing 8 [S | 0 | 2.0558 | 2.0125 | 0.964 | 0 | 0 |
| 2007 | ENSMUSG00000018845 | 29.53385479 | -0.405939678 | 0.441976382 | -0.918464639 | 0.358375664 | 0.798191951 | Unc45b | unc-45 myosin chaperone B [Source:MGI | 24.896 | 34.949 | 41.256 | 24.099 | 16.359 | 35.643 |
| 2008 | ENSMUSG00000018846 | 1853.496221 | -0.110721215 | 0.133273085 | -0.830784509 | 0.406095376 | 0.830012328 | Pank3 | pantothenate kinase 3 [Source:MGI Symb | 1917.8 | 2176.1 | 1680.4 | 1889.4 | 1941.3 | 1515.9 |
| 2009 | ENSMUSG00000018848 | 520.2254101 | 0.086843643 | 0.123606675 | 0.702580531 | 0.482317199 | 0.871593699 | Rars | arginyl-tRNA synthetase [Source:MGI Sym | 513.18 | 500.6 | 500.1 | 562 | 543.13 | 502.34 |
| 2010 | ENSMUSG00000018849 | 1247.65763 | 0.103563695 | 0.123945303 | 0.835559656 | 0.403402673 | 0.828953151 | Wwc1 | WW, C2 and coiled-coil domain containing | 1090.6 | 1276.7 | 1242.7 | 1388.1 | 1122.2 | 1365.6 |
| 2011 | ENSMUSG00000018858 | 574.6853156 | -0.022661361 | 0.120236695 | -0.188472918 | 0.850505943 | 0.973432436 | Mrpl58 | mitochondrial ribosomal protein L58 [Sou | 595.1 | 561.24 | 580.6 | 543.68 | 567.12 | 600.36 |
| 2012 | ENSMUSG00000018861 | 414.7432156 | 0.40126679 | 0.143021005 | 2.805649357 | 0.00502153 | 0.066818229 | Fdxr | ferredoxin reductase [Source:MGI Symbol | 370.23 | 333.05 | 368.29 | 422.22 | 522.41 | 472.27 |

| | | | | | | | | | | | | | | | |
|---|---|---|---|---|---|---|---|---|---|---|---|---|---|---|---|
| 2013 | ENSMUSG00000018862 | 0 | NA | NA | NA | NA | NA | Otop3 | otopetrin 3 [Source:MGI Symbol;Acc:MGI | 0 | 0 | 0 | 0 | 0 | 0 |
| 2014 | ENSMUSG00000018865 | 0.160662974 | 1.126924175 | 4.080472857 | 0.276174898 | 0.782413736 | NA | Sult4a1 | sulfotransferase family 4A, member 1 [Sou | 0 | 0 | 0 | 0.964 | 0 | 0 |
| 2015 | ENSMUSG00000018868 | 2.300517935 | 1.548311773 | 1.527778561 | 1.013439914 | 0.310850083 | NA | Pnpla5 | patatin-like phospholipase domain contai | 2.4093 | 1.0279 | 0 | 4.8199 | 1.0906 | 4.4554 |
| 2016 | ENSMUSG00000018882 | 612.2069951 | -0.074346895 | 0.141906731 | -0.523913801 | 0.600338495 | 0.91534406 | Mrpl45 | mitochondrial ribosomal protein L45 [Sou | 708.34 | 638.34 | 536.33 | 582.24 | 652.19 | 555.81 |
| 2017 | ENSMUSG00000018893 | 0 | NA | NA | NA | NA | NA | Mb | myoglobin [Source:MGI Symbol;Acc:MGI: | 0 | 0 | 0 | 0 | 0 | 0 |
| 2018 | ENSMUSG00000018899 | 475.8269815 | 0.578132723 | 0.183750046 | 3.146299742 | 0.001653505 | 0.030466709 | Irf1 | interferon regulatory factor 1 [Source:MG | 387.9 | 295.01 | 461.87 | 523.44 | 499.5 | 687.24 |
| 2019 | ENSMUSG00000018900 | 878.6803512 | -0.276494742 | 0.141222305 | -1.957868777 | 0.050245412 | 0.314882862 | Slc22a5 | solute carrier family 22 (organic cation tra | 939.63 | 966.24 | 983.1 | 922.53 | 821.24 | 639.35 |
| 2020 | ENSMUSG00000018906 | 106.5995798 | -0.057136268 | 0.2753239 | -0.20752382 | 0.835600792 | 0.971073158 | P4ha2 | procollagen-proline, 2-oxoglutarate 4-dio | 100.39 | 89.429 | 136.85 | 139.78 | 80.706 | 92.449 |
| 2021 | ENSMUSG00000018907 | 1.193894609 | -0.207782015 | 2.285126421 | -0.090928018 | 0.927549788 | NA | Alox12e | arachidonate lipoxygenase, epidermal [So | 0.8031 | 0 | 3.0187 | 0 | 0 | 3.3415 |
| 2022 | ENSMUSG00000018909 | 134.5697171 | -0.678864174 | 0.209572425 | -3.239281947 | 0.001198311 | 0.023808192 | Arrb1 | arrestin, beta 1 [Source:MGI Symbol;Acc:I | 148.57 | 181.94 | 167.04 | 114.71 | 93.793 | 101.36 |
| 2023 | ENSMUSG00000018914 | 0 | NA | NA | NA | NA | NA | Il3 | interleukin 3 [Source:MGI Symbol;Acc:MG | 0 | 0 | 0 | 0 | 0 | 0 |
| 2024 | ENSMUSG00000018916 | 0 | NA | NA | NA | NA | NA | Csf2 | colony stimulating factor 2 (granulocyte-n | 0 | 0 | 0 | 0 | 0 | 0 |
| 2025 | ENSMUSG00000018919 | 0.524408994 | -0.837196744 | 3.377823722 | -0.247850928 | 0.804249749 | NA | Tm4sf5 | transmembrane 4 superfamily member 5 | 0 | 2.0558 | 0 | 0 | 1.0906 | 0 |
| 2026 | ENSMUSG00000018920 | 80.62616933 | -0.735813893 | 0.330946699 | -2.223360726 | 0.026191486 | 0.209416058 | Cxcl16 | chemokine (C-X-C motif) ligand 16 [Source | 106.01 | 139.8 | 56.35 | 56.875 | 53.44 | 71.286 |
| 2027 | ENSMUSG00000018921 | 192.0075298 | 0.030246603 | 0.173272408 | 0.174560991 | 0.861424603 | 0.976192255 | Pelp1 | proline, glutamic acid and leucine rich pro | 198.37 | 184 | 187.16 | 180.26 | 201.76 | 200.49 |
| 2028 | ENSMUSG00000018923 | 124.2820535 | 0.163056225 | 0.232403259 | 0.70160903 | 0.482923019 | 0.871710696 | Med11 | mediator complex subunit 11 [Source:MG | 148.57 | 102.79 | 99.618 | 131.1 | 122.15 | 141.46 |
| 2029 | ENSMUSG00000018924 | 7.966750592 | -0.474668189 | 0.895673446 | -0.52995675 | 0.596141918 | 0.913894721 | Alox15 | arachidonate 15-lipoxygenase [Source:MC | 14.456 | 7.1954 | 6.0375 | 4.8199 | 14.178 | 1.1138 |
| 2030 | ENSMUSG00000018925 | 0.167707431 | -0.796629176 | 4.080472857 | -0.195229623 | 0.84521319 | NA | Heatr9 | HEAT repeat containing 9 [Source:MGI Sy | 0 | 0 | 1.0062 | 0 | 0 | 0 |
| 2031 | ENSMUSG00000018927 | 71.77833082 | -1.198860813 | 0.27582854 | -4.346398721 | 1.38E-05 | 0.000702346 | Ccl6 | chemokine (C-C motif) ligand 6 [Source:M | 96.372 | 112.04 | 91.568 | 42.415 | 39.262 | 49.009 |
| 2032 | ENSMUSG00000018930 | 1.004383029 | -0.810797259 | 2.263328258 | -0.358232287 | 0.720169487 | NA | Ccl4 | chemokine (C-C motif) ligand 4 [Source:M | 0.8031 | 0 | 3.0187 | 0 | 1.0906 | 1.1138 |
| 2033 | ENSMUSG00000018931 | 123.2158995 | 0.014258888 | 0.261211979 | 0.054587421 | 0.956467161 | 0.991780128 | Natd1 | N-acetyltransferase domain containing 1 [ | 155 | 105.88 | 106.66 | 154.24 | 89.431 | 128.09 |
| 2034 | ENSMUSG00000018932 | 2252.167486 | 0.076064845 | 0.102854835 | 0.739535924 | 0.459581634 | 0.859724361 | Map2k3 | mitogen-activated protein kinase kinase 3 | 2291.3 | 2147.3 | 2139.3 | 2297.2 | 2512.8 | 2125.2 |
| 2035 | ENSMUSG00000018965 | 822.9882172 | -0.230182222 | 0.115460221 | -1.99360628 | 0.046195099 | 0.297429405 | Ywhah | tyrosine 3-monooxygenase/tryptophan 5- | 943.65 | 881.95 | 839.21 | 743.23 | 809.24 | 720.66 |
| 2036 | ENSMUSG00000018973 | 0 | NA | NA | NA | NA | NA | Hoxb1 | homeobox B1 [Source:MGI Symbol;Acc:M | 0 | 0 | 0 | 0 | 0 | 0 |
| 2037 | ENSMUSG00000018974 | 276.0343478 | -0.018986987 | 0.149820375 | -0.126731674 | 0.899152776 | 0.982798755 | Sart3 | squamous cell carcinoma antigen recogni | 267.43 | 287.82 | 278.73 | 283.41 | 274.84 | 263.98 |
| 2038 | ENSMUSG00000018983 | 20.68735373 | 0.921795968 | 0.574152254 | 1.605490464 | 0.10838591 | 0.48522218 | E2f2 | E2F transcription factor 2 [Source:MGI Sy | 14.456 | 6.1675 | 22.137 | 18.316 | 20.722 | 42.326 |
| 2039 | ENSMUSG00000018986 | 0.331982614 | 0.16514309 | 3.986751476 | 0.041422971 | 0.966958701 | NA | Slfn3 | schlafen 3 [Source:MGI Symbol;Acc:MGI: | 0 | 1.0279 | 0 | 0.964 | 0 | 0 |
| 2040 | ENSMUSG00000018995 | 378.8141445 | 0.06476406 | 0.145951138 | 0.443737958 | 0.657232049 | 0.927163206 | Nars2 | asparaginyl-tRNA synthetase 2 (mitochon | 350.96 | 389.58 | 370.3 | 341.25 | 423.16 | 397.64 |
| 2041 | ENSMUSG00000018999 | 419.1041022 | -0.063287709 | 0.138805686 | -0.455944642 | 0.648429787 | 0.925643431 | Slc35b4 | solute carrier family 35, member B4 [Sour | 432.87 | 431.73 | 420.61 | 446.32 | 363.18 | 419.92 |
| 2042 | ENSMUSG00000019027 | 0.997777256 | 1.176137242 | 2.241393702 | 0.524734785 | 0.599767572 | NA | Dnah1 | dynein, axonemal, heavy chain 1 [Source: | 0.8031 | 1.0279 | 0 | 1.928 | 0 | 2.2277 |
| 2043 | ENSMUSG00000019039 | 710.6116944 | 0.09718108 | 0.120510691 | 0.80641045 | 0.420006223 | 0.837906635 | Dalrd3 | DALR anticodon binding domain containir | 659.35 | 656.84 | 744.62 | 780.82 | 695.81 | 726.23 |
| 2044 | ENSMUSG00000019054 | 1190.789291 | 0.024834787 | 0.101173227 | 0.245467971 | 0.806094105 | 0.964428432 | Fis1 | fission, mitochondrial 1 [Source:MGI Sym | 1171.7 | 1191.4 | 1178.3 | 1164.5 | 1172.4 | 1266.4 |
| 2045 | ENSMUSG00000019055 | 1053.165291 | -0.182592891 | 0.111690725 | -1.634807998 | 0.102089304 | 0.47032223 | Plod1 | procollagen-lysine, 2-oxoglutarate 5-dioxy | 1105.9 | 1110.2 | 1144.1 | 1082.5 | 965.2 | 911.12 |
| 2046 | ENSMUSG00000019066 | 191.1399228 | -0.38047393 | 0.179053367 | -2.124919162 | 0.033593359 | 0.245153552 | Rab3d | RAB3D, member RAS oncogene family [So | 208 | 202.5 | 238.48 | 179.3 | 157.05 | 161.51 |
| 2047 | ENSMUSG00000019080 | 233.6300681 | 0.046321546 | 0.173586724 | 0.266849591 | 0.78958497 | 0.961009963 | Mfsd3 | major facilitator superfamily domain cont | 232.1 | 202.5 | 254.58 | 212.08 | 242.12 | 258.41 |
| 2048 | ENSMUSG00000019082 | 3704.744639 | 0.008179121 | 0.129300964 | 0.063256461 | 0.949562285 | 0.990731019 | Slc25a22 | solute carrier family 25 (mitochondrial car | 3720.8 | 3319.1 | 4042.1 | 3435.6 | 3308.9 | 4401.9 |
| 2049 | ENSMUSG00000019087 | 1139.99844 | -0.092919539 | 0.103921175 | -0.894134808 | 0.371249767 | 0.807214774 | Atp6ap1 | ATPase, H+ transporting, lysosomal acces | 1117.1 | 1240.7 | 1173.3 | 1134.6 | 1095 | 1079.3 |
| 2050 | ENSMUSG00000019088 | 73.70268996 | 0.010861836 | 0.281157939 | 0.038632508 | 0.969183384 | 0.994602269 | Dnase1l1 | deoxyribonuclease 1-like 1 [Source:MGI S | 61.036 | 85.317 | 74.462 | 83.866 | 79.615 | 57.92 |
| 2051 | ENSMUSG00000019102 | 0.832804665 | -0.381337792 | 2.590357773 | -0.147214333 | 0.882962848 | NA | Aldh3a1 | aldehyde dehydrogenase family 3, subfan | 0.8031 | 0 | 2.0125 | 0 | 2.1812 | 0 |
| 2052 | ENSMUSG00000019122 | 671.2970394 | 0.323187499 | 0.12362808 | 2.614191693 | 0.008943884 | 0.099758896 | Ccl9 | chemokine (C-C motif) ligand 9 [Source:M | 538.08 | 631.14 | 620.85 | 716.24 | 733.99 | 787.49 |
| 2053 | ENSMUSG00000019124 | 2.816487883 | 1.241482131 | 1.560896414 | 0.795364843 | 0.426401305 | NA | Scrn1 | secernin 1 [Source:MGI Symbol;Acc:MGI: | 0 | 3.0838 | 2.0125 | 2.8919 | 0 | 8.9108 |
| 2054 | ENSMUSG00000019132 | 3515.6348 | -0.158298836 | 0.1059833 | -1.493620568 | 0.13527482 | 0.538039097 | BC005537 | cDNA sequence BC005537 [Source:MGI S | 3669.4 | 3871.1 | 3585.2 | 3504.1 | 3540.1 | 2923.8 |
| 2055 | ENSMUSG00000019139 | 97.59325787 | -0.255375446 | 0.287822675 | -0.887266598 | 0.374935384 | 0.809332934 | Isyna1 | myo-inositol 1-phosphate synthase A1 [Sc | 94.766 | 78.122 | 145.91 | 94.47 | 68.709 | 103.59 |
| 2056 | ENSMUSG00000019143 | 528.4774817 | 0.22491303 | 0.128594959 | 1.749003478 | 0.080290418 | 0.41597095 | Hars2 | histidyl-tRNA synthetase 2 [Source:MGI S | 479.45 | 470.79 | 512.18 | 596.7 | 516.95 | 594.79 |
| 2057 | ENSMUSG00000019146 | 0 | NA | NA | NA | NA | NA | Cacng2 | calcium channel, voltage-dependent, gam | 0 | 0 | 0 | 0 | 0 | 0 |
| 2058 | ENSMUSG00000019158 | 417.4216969 | -0.007603509 | 0.147942721 | -0.051394954 | 0.959010805 | 0.992027194 | Tmem160 | transmembrane protein 160 [Source:MGI | 419.22 | 411.17 | 425.64 | 468.49 | 351.18 | 428.83 |
| 2059 | ENSMUSG00000019173 | 2064.418574 | -0.104717635 | 0.092494792 | -1.132146282 | 0.25757294 | 0.710872182 | Rab5c | RAB5C, member RAS oncogene family [So | 2109.7 | 2161.7 | 2146.3 | 1934.7 | 1950 | 2084 |
| 2060 | ENSMUSG00000019178 | 3.338834602 | -1.548284411 | 1.250682232 | -1.237951872 | 0.215733907 | NA | Styxl1 | serine/threonine/tyrosine interacting-like | 8.8341 | 3.0838 | 3.0187 | 2.8919 | 1.0906 | 1.1138 |
| 2061 | ENSMUSG00000019179 | 3700.132119 | -0.144540999 | 0.092615986 | -1.560648484 | 0.118606712 | 0.504308497 | Mdh2 | malate dehydrogenase 2, NAD (mitochon | 4066.1 | 3834.1 | 3754.3 | 3314.2 | 3560.9 | 3671.2 |
| 2062 | ENSMUSG00000019188 | 1884.83947 | 0.263595619 | 0.097748276 | 2.696677931 | 0.007003497 | 0.084204972 | H13 | histocompatibility 13 [Source:MGI Symbo | 1740.3 | 1627.2 | 1772 | 2155.5 | 2050.4 | 1963.7 |
| 2063 | ENSMUSG00000019189 | 337.6198556 | 0.205838051 | 0.162799909 | 1.264362196 | 0.206100055 | 0.651217828 | Rnf145 | ring finger protein 145 [Source:MGI Symb | 340.52 | 303.24 | 296.84 | 408.73 | 299.92 | 376.48 |
| 2064 | ENSMUSG00000019194 | 50.31178526 | -0.565074316 | 0.335180855 | -1.685878854 | 0.091819136 | 0.446623824 | Scn1b | sodium channel, voltage-gated, type I, bet | 44.171 | 74.01 | 62.387 | 45.307 | 40.353 | 35.643 |
| 2065 | ENSMUSG00000019210 | 1769.89539 | 0.00100853 | 0.133187193 | 0.007572273 | 0.993958258 | 0.998618081 | Atp6v1e1 | ATPase, H+ transporting, lysosomal V1 sul | 2034.3 | 1755.7 | 1516.4 | 1653.2 | 2001.3 | 1658.5 |
| 2066 | ENSMUSG00000019214 | 1.768368202 | -3.063643882 | 1.885230976 | -1.625076142 | 0.104146336 | NA | Chtf18 | CTF18, chromosome transmission fidelity | 2.4093 | 2.0558 | 5.0312 | 0 | 0 | 1.1138 |
| 2067 | ENSMUSG00000019230 | 1.444250066 | 0.711568442 | 2.365547075 | 0.300805023 | 0.763563178 | NA | Lhx9 | LIM homeobox protein 9 [Source:MGI Syn | 3.2124 | 0 | 0 | 0 | 5.4531 | 0 |
| 2068 | ENSMUSG00000019232 | 2267.399397 | -0.718553484 | 0.320665073 | -2.240822414 | 0.02503758 | 0.202995271 | Etnppl | ethanolamine phosphate phospholyase [S | 1779.7 | 3151.6 | 3530.9 | 1434.4 | 1734.1 | 1973.7 |

| | | | | | | | | | | | | | | | |
|---|---|---|---|---|---|---|---|---|---|---|---|---|---|---|---|
| 2069 | ENSMUSG00000019235 | 62.7525395 | -0.214331279 | 0.285919705 | -0.749620523 | 0.453483287 | 0.857073339 | Rps6kl1 | ribosomal protein S6 kinase-like 1 [Source | 68.264 | 63.731 | 70.437 | 69.406 | 53.44 | 51.237 |
| 2070 | ENSMUSG00000019254 | 493.6796375 | -0.157175219 | 0.146989795 | -1.069293405 | 0.284937482 | 0.737533238 | Ppp1r12c | protein phosphatase 1, regulatory subunit | 489.89 | 539.66 | 533.31 | 550.43 | 416.62 | 432.17 |
| 2071 | ENSMUSG00000019256 | 732.5142279 | -0.468132173 | 0.207483056 | -2.256242903 | 0.024055423 | 0.197338277 | Ahr | aryl-hydrocarbon receptor [Source:MGI Sy | 857.71 | 1058.8 | 633.93 | 449.21 | 792.88 | 602.59 |
| 2072 | ENSMUSG00000019261 | 85.2727724 | -0.228292481 | 0.242932529 | -0.939736155 | 0.347352915 | 0.788467131 | Map1s | microtubule-associated protein 1S [Source | 89.947 | 94.568 | 91.568 | 80.01 | 70.89 | 84.652 |
| 2073 | ENSMUSG00000019278 | 9.838895532 | -0.702643631 | 0.792636627 | -0.886463743 | 0.375367683 | 0.809332934 | Dpep1 | dipeptidase 1 [Source:MGI Symbol;Acc:M | 15.259 | 12.335 | 9.0562 | 13.496 | 1.0906 | 7.7969 |
| 2074 | ENSMUSG00000019295 | 445.8690003 | 0.240004158 | 0.133567636 | 1.796873597 | 0.072355689 | 0.389777537 | Tmem129 | transmembrane protein 129 [Source:MGI | 446.52 | 397.8 | 381.37 | 473.31 | 500.59 | 475.61 |
| 2075 | ENSMUSG00000019297 | 397.058842 | 0.253564768 | 0.154429347 | 1.641946772 | 0.100601036 | 0.46725495 | Nop9 | NOP9 nucleolar protein [Source:MGI Symb | 306.79 | 373.13 | 407.53 | 393.3 | 447.15 | 454.45 |
| 2076 | ENSMUSG00000019301 | 3.82747503 | 0.329943946 | 1.07252108 | 0.307633996 | 0.758360849 | NA | Hsd17b1 | hydroxysteroid (17-beta) dehydrogenase | 4.0155 | 3.0838 | 3.0187 | 2.8919 | 3.2719 | 6.6831 |
| 2077 | ENSMUSG00000019302 | 694.8915057 | -0.060808186 | 0.128319935 | -0.473879497 | 0.635585841 | 0.923478764 | Atp6v0a1 | ATPase, H+ transporting, lysosomal V0 sul | 721.19 | 764.77 | 641.98 | 614.05 | 707.81 | 719.54 |
| 2078 | ENSMUSG00000019303 | 10.85044464 | 0.694252879 | 0.804882176 | 0.862552184 | 0.38838373 | 0.818315288 | Psmc3ip | proteasome (prosome, macropain) 26S su | 10.44 | 8.2233 | 6.0375 | 1.928 | 23.994 | 14.48 |
| 2079 | ENSMUSG00000019312 | 1806.58897 | 0.40323136 | 0.12337211 | 3.268415852 | 0.001081513 | 0.022234986 | Grb7 | growth factor receptor bound protein 7 [S | 1460 | 1375.4 | 1832.4 | 2046.5 | 2112.5 | 2012.7 |
| 2080 | ENSMUSG00000019320 | 5.089372967 | -0.320119505 | 0.944704995 | -0.338856582 | 0.734717774 | 0.948109711 | Noxo1 | NADPH oxidase organizer 1 [Source:MGI S | 4.8186 | 5.1396 | 7.0437 | 5.7839 | 2.1812 | 5.5692 |
| 2081 | ENSMUSG00000019326 | 0.17131964 | -0.796629176 | 4.080472857 | -0.195229623 | 0.84521319 | NA | Aoc3 | amine oxidase, copper containing 3 [Sour | 0 | 1.0279 | 0 | 0 | 0 | 0 |
| 2082 | ENSMUSG00000019338 | 206.9546715 | 0.227718548 | 0.187466278 | 1.214717393 | 0.224473897 | 0.670547386 | Zfp687 | zinc finger protein 687 [Source:MGI Symb | 190.34 | 173.72 | 208.29 | 255.45 | 229.03 | 184.9 |
| 2083 | ENSMUSG00000019359 | 0 | NA | NA | NA | NA | NA | Gdpd2 | glycerophosphodiester phosphodiesterase | 0 | 0 | 0 | 0 | 0 | 0 |
| 2084 | ENSMUSG00000019362 | 1064.323794 | -0.006365208 | 0.108857391 | -0.058472902 | 0.953371947 | 0.9915542 | D8Ertd738e | DNA segment, Chr 8, ERATO Doi 738, expr | 1098.6 | 1018.7 | 1081.7 | 984.22 | 1121.2 | 1081.5 |
| 2085 | ENSMUSG00000019368 | 3980.929087 | -0.20477135 | 0.125051282 | -1.637499003 | 0.101526247 | 0.469189975 | Sec14l4 | SEC14-like lipid binding 4 [Source:MGI Syr | 3511.2 | 4793.2 | 4485.8 | 3484.8 | 3936 | 3674.6 |
| 2086 | ENSMUSG00000019370 | 875.3334059 | 0.025461468 | 0.115546283 | 0.220357312 | 0.825592889 | 0.96943211 | Calm3 | calmodulin 3 [Source:MGI Symbol;Acc:MG | 918.75 | 833.64 | 849.27 | 831.91 | 859.41 | 959.02 |
| 2087 | ENSMUSG00000019373 | 552.0782616 | 0.11049574 | 0.135156785 | 0.817537496 | 0.41362133 | 0.835385466 | Cops3 | COP9 signalosome subunit 3 [Source:MGI | 485.88 | 601.33 | 506.14 | 537.9 | 602.02 | 579.2 |
| 2088 | ENSMUSG00000019428 | 4986.862967 | -0.009624829 | 0.088272032 | -0.109035992 | 0.913173943 | 0.98625062 | Fkbp8 | FK506 binding protein 8 [Source:MGI Sym | 4911 | 5150.9 | 4949.7 | 5253.7 | 4811.8 | 4844.1 |
| 2089 | ENSMUSG00000019429 | 0 | NA | NA | NA | NA | NA | Ffar3 | free fatty acid receptor 3 [Source:MGI Syr | 0 | 0 | 0 | 0 | 0 | 0 |
| 2090 | ENSMUSG00000019432 | 912.1458675 | -0.161135677 | 0.111114745 | -1.45017367 | 0.147010095 | 0.558244689 | Ddx39b | DEAD box helicase 39b [Source:MGI Symb | 938.02 | 998.11 | 953.92 | 923.49 | 850.68 | 808.65 |
| 2091 | ENSMUSG00000019433 | 187.5577797 | -0.000660443 | 0.189621293 | -0.003482958 | 0.997221007 | 0.999498002 | Gipc1 | GIPC PDZ domain containing family, meml | 199.17 | 177.83 | 185.15 | 164.84 | 175.59 | 222.77 |
| 2092 | ENSMUSG00000019437 | 700.721168 | 0.224946271 | 0.137400391 | 1.637158882 | 0.101597276 | 0.469189975 | Tlcd1 | TLC domain containing 1 [Source:MGI Sym | 754.11 | 569.47 | 613.81 | 788.53 | 768.89 | 709.52 |
| 2093 | ENSMUSG00000019461 | 123.7793155 | 0.119842336 | 0.21233081 | 0.564413314 | 0.57247288 | 0.905173631 | Plscr3 | phospholipid scramblase 3 [Source:MGI S | 121.27 | 107.93 | 126.79 | 140.74 | 114.51 | 131.43 |
| 2094 | ENSMUSG00000019464 | 4.227480434 | -0.780790594 | 1.073200838 | -0.727534461 | 0.466898613 | NA | Ptger1 | prostaglandin E receptor 1 (subtype EP1) | 8.8341 | 5.1396 | 2.0125 | 3.8559 | 2.1812 | 3.3415 |
| 2095 | ENSMUSG00000019467 | 22.31659194 | -0.358125297 | 0.46077447 | -0.777224696 | 0.437026215 | 0.846852749 | Arhgef25 | Rho guanine nucleotide exchange factor ( | 31.321 | 22.614 | 21.131 | 20.244 | 18.541 | 20.049 |
| 2096 | ENSMUSG00000019470 | 363.519988 | 0.095607679 | 0.14881389 | 0.642464754 | 0.520571467 | 0.887802551 | Xab2 | XPA binding protein 2 [Source:MGI Symbo | 350.15 | 313.51 | 390.42 | 348 | 396.99 | 382.05 |
| 2097 | ENSMUSG00000019471 | 1680.100489 | 0.138056817 | 0.096974342 | 1.423642729 | 0.15454992 | 0.572594786 | Cdc37 | cell division cycle 37 [Source:MGI Symbol; | 1517.9 | 1632.3 | 1650.2 | 1802.6 | 1773.3 | 1704.2 |
| 2098 | ENSMUSG00000019478 | 594.4150549 | -0.346030055 | 0.136219341 | -2.540241734 | 0.011077587 | 0.116697906 | Rab4a | RAB4A, member RAS oncogene family [So | 660.15 | 740.1 | 595.7 | 506.09 | 488.6 | 575.86 |
| 2099 | ENSMUSG00000019487 | 202.0046595 | 0.356432365 | 0.202040172 | 1.764165811 | 0.077704067 | 0.407953403 | Trip10 | thyroid hormone receptor interactor 10 [S | 177.49 | 159.33 | 195.21 | 283.41 | 206.13 | 190.47 |
| 2100 | ENSMUSG00000019489 | 0 | NA | NA | NA | NA | NA | Cd70 | CD70 antigen [Source:MGI Symbol;Acc:M | 0 | 0 | 0 | 0 | 0 | 0 |
| 2101 | ENSMUSG00000019494 | 1805.404266 | 0.108568978 | 0.111575903 | 0.973050415 | 0.330528246 | 0.774747658 | Cops6 | COP9 signalosome subunit 6 [Source:MGI | 1886.5 | 1653.9 | 1670.4 | 1724.6 | 1842.1 | 2055 |
| 2102 | ENSMUSG00000019505 | 6145.575521 | -0.12867298 | 0.098587356 | -1.305167166 | 0.191835932 | 0.629099068 | Ubb | ubiquitin B [Source:MGI Symbol;Acc:MGI: | 6856.1 | 6537.6 | 5864.4 | 6224.4 | 5609 | 5782 |
| 2103 | ENSMUSG00000019518 | 166.0335973 | 0.478746975 | 0.192221144 | 2.490605166 | 0.012752575 | 0.127649303 | Ap4m1 | adaptor-related protein complex AP-4, mu | 149.38 | 141.85 | 124.77 | 201.47 | 208.31 | 170.42 |
| 2104 | ENSMUSG00000019528 | 103.5799252 | 0.039961372 | 0.279060645 | 0.143199598 | 0.886132547 | 0.982222482 | Gyg | glycogenin [Source:MGI Symbol;Acc:MGI: | 131.71 | 84.289 | 89.556 | 82.902 | 93.793 | 139.23 |
| 2105 | ENSMUSG00000019539 | 63.95329188 | -0.764610733 | 0.322485755 | -2.370990723 | 0.017740476 | 0.15881418 | Rcn3 | reticulocalbin 3, EF-hand calcium binding | 105.21 | 69.898 | 66.412 | 60.731 | 44.715 | 36.757 |
| 2106 | ENSMUSG00000019558 | 127.5768184 | -0.461511593 | 0.223488402 | -2.065035989 | 0.038919595 | 0.269932742 | Slc6a8 | solute carrier family 6 (neurotransmitter t | 163.03 | 126.43 | 153.96 | 119.53 | 114.51 | 87.994 |
| 2107 | ENSMUSG00000019564 | 32.95473695 | -0.41741562 | 0.423126257 | -0.986503704 | 0.323885999 | 0.77088057 | Arid3a | AT rich interactive domain 3A (BRIGHT-like | 32.124 | 30.838 | 50.312 | 34.703 | 17.45 | 32.301 |
| 2108 | ENSMUSG00000019577 | 94.74738901 | -1.279809615 | 0.323817607 | -3.952254565 | 7.74E-05 | 0.002946786 | Pdk4 | pyruvate dehydrogenase kinase, isoenzym | 149.38 | 156.24 | 97.606 | 84.83 | 41.443 | 38.985 |
| 2109 | ENSMUSG00000019578 | 293.3547953 | 0.047861234 | 0.153585724 | 0.311625539 | 0.755325123 | 0.953085143 | Ubxn6 | UBX domain protein 6 [Source:MGI Symbo | 264.22 | 281.65 | 319.99 | 290.16 | 294.47 | 309.65 |
| 2110 | ENSMUSG00000019579 | 1007.666192 | 0.155522416 | 0.103874854 | 1.497209476 | 0.134338761 | 0.537569131 | Mydgf | myeloid derived growth factor [Source:MG | 945.25 | 979.61 | 935.81 | 1100.9 | 1024.1 | 1060.4 |
| 2111 | ENSMUSG00000019590 | 115.7289326 | 1.396296313 | 0.271386142 | 5.145053845 | 2.67E-07 | 2.54E-05 | Cyb561 | cytochrome b-561 [Source:MGI Symbol;Ac | 48.989 | 87.373 | 55.343 | 130.14 | 185.41 | 187.13 |
| 2112 | ENSMUSG00000019647 | 52.55336643 | -0.667398483 | 0.336828807 | -1.981417473 | 0.047544476 | 0.302749285 | Sema6a | sema domain, transmembrane domain (TI | 55.414 | 68.87 | 69.431 | 45.307 | 26.175 | 50.123 |
| 2113 | ENSMUSG00000019659 | 255.7891418 | -0.144857152 | 0.165498577 | -0.875277324 | 0.381423029 | 0.8130053 | Ccdc12 | coiled-coil domain containing 12 [Source:I | 232.1 | 280.62 | 293.82 | 241.96 | 233.39 | 252.84 |
| 2114 | ENSMUSG00000019689 | 505.9195662 | 0.233363842 | 0.125811208 | 1.85487323 | 0.063614332 | 0.360558879 | Fmc1 | formation of mitochondrial complex V ass | 470.62 | 452.28 | 471.93 | 514.76 | 583.48 | 542.44 |
| 2115 | ENSMUSG00000019699 | 44.38590328 | -1.168266962 | 0.358805326 | -3.255991135 | 0.001129973 | 0.022858638 | Akt3 | thymoma viral proto-oncogene 3 [Source: | 61.036 | 74.01 | 49.306 | 26.991 | 33.809 | 21.163 |
| 2116 | ENSMUSG00000019710 | 810.7772615 | 0.087485128 | 0.114871025 | 0.761594383 | 0.446302128 | 0.852777619 | Mrpl24 | mitochondrial ribosomal protein L24 [Sou | 742.87 | 767.85 | 848.26 | 815.53 | 815.78 | 874.37 |
| 2117 | ENSMUSG00000019715 | 658.5533434 | 0.187938414 | 0.118522956 | 1.585670997 | 0.112813962 | 0.493122134 | Gle1 | GLE1 RNA export mediator (yeast) [Source | 653.73 | 582.83 | 609.78 | 696.96 | 674 | 734.02 |
| 2118 | ENSMUSG00000019718 | 729.1275502 | -0.29062764 | 0.131202744 | -2.21510336 | 0.026752971 | 0.211566414 | L3hypdh | L-3-hydroxyproline dehydratase (trans-) [S | 871.37 | 800.75 | 734.56 | 684.42 | 707.81 | 575.86 |
| 2119 | ENSMUSG00000019726 | 623.9645577 | 0.085237378 | 0.167668909 | 0.508367226 | 0.611195832 | 0.916376001 | Lyst | lysosomal trafficking regulator [Source:MG | 516.39 | 656.84 | 645 | 801.07 | 546.4 | 578.08 |
| 2120 | ENSMUSG00000019731 | 919.0680214 | 0.171859686 | 0.129690688 | 1.325150545 | 0.185121224 | 0.620815781 | Slc35e1 | solute carrier family 35, member E1 [Sour | 847.27 | 774.02 | 972.03 | 1001.6 | 872.49 | 1047 |
| 2121 | ENSMUSG00000019732 | 44.5516742 | 0.354238655 | 0.365761924 | 0.968495164 | 0.332797133 | 0.77523112 | Calr3 | calreticulin 3 [Source:MGI Symbol;Acc:MG | 40.958 | 29.81 | 46.287 | 37.595 | 44.715 | 67.944 |
| 2122 | ENSMUSG00000019734 | 41.3766934 | 0.163284691 | 0.355564428 | 0.459226734 | 0.646071354 | 0.925643431 | Tmc4 | transmembrane channel-like gene family | 36.14 | 35.977 | 45.281 | 55.911 | 37.081 | 37.871 |
| 2123 | ENSMUSG00000019737 | 22.00069971 | 0.023619863 | 0.465387968 | 0.050753058 | 0.959522297 | 0.992148164 | Syne4 | spectrin repeat containing, nuclear envelo | 20.881 | 22.614 | 22.137 | 28.919 | 19.631 | 17.822 |
| 2124 | ENSMUSG00000019738 | 351.7669641 | 0.156730193 | 0.158906655 | 0.98630352 | 0.323984193 | 0.77088057 | Polr2i | polymerase (RNA) II (DNA directed) polyp | 292.33 | 372.11 | 334.07 | 343.18 | 349 | 419.92 |

| | | | | | | | | | | | | | | | |
|---|---|---|---|---|---|---|---|---|---|---|---|---|---|---|---|
| 2125 | ENSMUSG00000019756 | 0 | NA | NA | NA | NA | NA | Prl8a1 | prolactin family 8, subfamily a, member 1 | 0 | 0 | 0 | 0 | 0 | 0 |
| 2126 | ENSMUSG00000019761 | 41.29720778 | 0.158146171 | 0.370394946 | 0.426966331 | 0.669403851 | 0.930837335 | Krt10 | keratin 10 [Source:MGI Symbol;Acc:MGI:9 | 32.927 | 31.865 | 52.325 | 33.739 | 51.259 | 45.668 |
| 2127 | ENSMUSG00000019762 | 1887.517766 | 0.050115288 | 0.097181426 | 0.515687922 | 0.606072392 | 0.916376001 | Iyd | iodotyrosine deiodinase [Source:MGI Sym | 1768.4 | 1864.6 | 1932 | 1930.8 | 1996.9 | 1832.3 |
| 2128 | ENSMUSG00000019763 | 667.365572 | -0.125511378 | 0.122333351 | -1.025978419 | 0.304901757 | 0.753627713 | Rmnd1 | required for meiotic nuclear division 1 hor | 718.78 | 666.09 | 704.37 | 673.82 | 665.28 | 575.86 |
| 2129 | ENSMUSG00000019767 | 0.17131964 | -0.796629176 | 4.080472857 | -0.195229623 | 0.84521319 | NA | Ccdc170 | coiled-coil domain containing 170 [Source | 0 | 1.0279 | 0 | 0 | 0 | 0 |
| 2130 | ENSMUSG00000019768 | 391.4763609 | -1.10740902 | 0.171503275 | -6.457072157 | 1.07E-10 | 2.91E-08 | Esr1 | estrogen receptor 1 (alpha) [Source:MGI S | 481.86 | 517.04 | 605.76 | 244.85 | 297.74 | 201.61 |
| 2131 | ENSMUSG00000019772 | 0 | NA | NA | NA | NA | NA | Vip | vasoactive intestinal polypeptide [Source: | 0 | 0 | 0 | 0 | 0 | 0 |
| 2132 | ENSMUSG00000019773 | 18.21119343 | 0.122494199 | 0.567829604 | 0.215723516 | 0.82920327 | 0.969955221 | Fbxo5 | F-box protein 5 [Source:MGI Symbol;Acc:I | 21.684 | 13.363 | 17.106 | 18.316 | 8.7249 | 30.074 |
| 2133 | ENSMUSG00000019774 | 110.8050118 | 0.075992209 | 0.245732838 | 0.309247268 | 0.75713344 | 0.953260973 | Mtrf1l | mitochondrial translational release factor | 85.932 | 129.52 | 108.67 | 104.11 | 135.24 | 101.36 |
| 2134 | ENSMUSG00000019775 | 0.785324008 | -0.358914107 | 2.435960446 | -0.147339875 | 0.88286376 | NA | Rgs17 | regulator of G-protein signaling 17 [Source | 1.6062 | 1.0279 | 0 | 0.964 | 0 | 1.1138 |
| 2135 | ENSMUSG00000019777 | 428.8050113 | -0.172355413 | 0.133921436 | -1.286988985 | 0.198098127 | 0.640288562 | Hdac2 | histone deacetylase 2 [Source:MGI Symbo | 425.64 | 486.21 | 451.8 | 396.19 | 424.25 | 388.73 |
| 2136 | ENSMUSG00000019779 | 111.6554751 | -0.118640511 | 0.233828294 | -0.507383041 | 0.611886083 | 0.916441371 | Frk | fyn-related kinase [Source:MGI Symbol;Ac | 126.09 | 90.457 | 131.82 | 101.22 | 113.42 | 106.93 |
| 2137 | ENSMUSG00000019782 | 284.2397689 | -0.076909353 | 0.147090366 | -0.522871453 | 0.601063712 | 0.915461118 | Rwdd1 | RWD domain containing 1 [Source:MGI Sy | 285.9 | 286.79 | 302.88 | 276.66 | 279.2 | 274.01 |
| 2138 | ENSMUSG00000019785 | 0 | NA | NA | NA | NA | NA | Clvs2 | clavesin 2 [Source:MGI Symbol;Acc:MGI:2 | 0 | 0 | 0 | 0 | 0 | 0 |
| 2139 | ENSMUSG00000019787 | 0 | NA | NA | NA | NA | NA | Trdn | triadin [Source:MGI Symbol;Acc:MGI:1924 | 0 | 0 | 0 | 0 | 0 | 0 |
| 2140 | ENSMUSG00000019789 | 3.079756171 | -1.776130267 | 1.299464536 | -1.366817038 | 0.171682661 | NA | Hey2 | hairy/enhancer-of-split related with YRPW | 3.2124 | 3.0838 | 8.05 | 1.928 | 1.0906 | 1.1138 |
| 2141 | ENSMUSG00000019790 | 245.8493667 | 0.091945517 | 0.16353272 | 0.562245381 | 0.573948846 | 0.9059841 | Stxbp5 | syntaxin binding protein 5 (tomosyn) [Sou | 225.67 | 259.04 | 229.42 | 237.14 | 246.48 | 277.35 |
| 2142 | ENSMUSG00000019791 | 468.105819 | 0.141863177 | 0.131610449 | 1.077902082 | 0.281077456 | 0.733710968 | Hint3 | histidine triad nucleotide binding protein | 444.12 | 443.03 | 447.78 | 449.21 | 534.4 | 490.09 |
| 2143 | ENSMUSG00000019792 | 108.5181226 | -0.055344387 | 0.232042977 | -0.238509211 | 0.811486172 | 0.966527993 | Trmt11 | tRNA methyltransferase 11 [Source:MGI S | 96.372 | 107.93 | 127.79 | 106.04 | 93.793 | 119.18 |
| 2144 | ENSMUSG00000019794 | 225.2218939 | 0.230801518 | 0.174771722 | 1.320588454 | 0.186638625 | 0.6230483 | Katna1 | katanin p60 (ATPase-containing) subunit A | 228.08 | 209.7 | 183.14 | 216.9 | 259.57 | 253.96 |
| 2145 | ENSMUSG00000019795 | 832.3931275 | 0.046543719 | 0.117473568 | 0.396205883 | 0.691953152 | 0.938171202 | Pcmt1 | protein-L-isoaspartate (D-aspartate) O-me | 769.37 | 864.48 | 824.11 | 881.08 | 882.31 | 773.01 |
| 2146 | ENSMUSG00000019796 | 67.85711042 | 0.305843491 | 0.349014167 | 0.876306809 | 0.380863265 | 0.812628154 | Lrp11 | low density lipoprotein receptor-related p | 61.036 | 44.2 | 76.475 | 42.415 | 88.34 | 94.677 |
| 2147 | ENSMUSG00000019797 | 260.0014233 | 0.032544435 | 0.159512478 | 0.204024381 | 0.838334446 | 0.971223091 | Mtres1 | mitochondrial transcription rescue factor | 244.14 | 277.54 | 249.55 | 240.03 | 279.2 | 269.55 |
| 2148 | ENSMUSG00000019802 | 2554.311021 | 0.190926791 | 0.09534288 | 2.002528043 | 0.04522797 | 0.294174243 | Sec63 | SEC63-like (S. cerevisiae) [Source:MGI Syn | 2293.7 | 2532.8 | 2330.5 | 2597 | 2798.5 | 2773.5 |
| 2149 | ENSMUSG00000019803 | 0 | NA | NA | NA | NA | NA | Nr2e1 | nuclear receptor subfamily 2, group E, me | 0 | 0 | 0 | 0 | 0 | 0 |
| 2150 | ENSMUSG00000019804 | 1179.579388 | -0.118164297 | 0.10400441 | -1.136146991 | 0.255895051 | 0.708918985 | Snx3 | sorting nexin 3 [Source:MGI Symbol;Acc:N | 1241.6 | 1162.6 | 1278.9 | 1106.6 | 1103.7 | 1184 |
| 2151 | ENSMUSG00000019806 | 161.4136183 | 0.200012303 | 0.204182529 | 0.979575988 | 0.327295463 | 0.772772049 | Aig1 | androgen-induced 1 [Source:MGI Symbol; | 153.39 | 168.58 | 128.8 | 182.19 | 145.05 | 190.47 |
| 2152 | ENSMUSG00000019808 | 44.73095425 | -0.142655405 | 0.374518121 | -0.380903878 | 0.703274575 | 0.942506832 | Adat2 | adenosine deaminase, tRNA-specific 2 [So | 32.124 | 58.591 | 50.312 | 32.775 | 56.712 | 37.871 |
| 2153 | ENSMUSG00000019809 | 556.4550785 | -0.137686563 | 0.126504369 | -1.088393742 | 0.276421323 | 0.730038567 | Pex3 | peroxisomal biogenesis factor 3 [Source:N | 627.22 | 573.58 | 547.4 | 517.66 | 563.85 | 509.03 |
| 2154 | ENSMUSG00000019810 | 157.5937546 | -0.356030052 | 0.191877425 | -1.855507764 | 0.063523752 | 0.36045804 | Fuca2 | fucosidase, alpha-L- 2, plasma [Source:MG | 181.5 | 192.22 | 156.97 | 135.92 | 134.15 | 144.8 |
| 2155 | ENSMUSG00000019813 | 92.30458272 | 0.413233421 | 0.24873252 | 1.661356628 | 0.096641843 | 0.458530879 | Cep57l1 | centrosomal protein 57-like 1 [Source:MG | 76.295 | 75.038 | 86.537 | 126.28 | 89.431 | 100.25 |
| 2156 | ENSMUSG00000019814 | 238.8430747 | 0.099205225 | 0.172539546 | 0.574970942 | 0.56531095 | 0.901542552 | Ltv1 | LTV1 ribosome biogenesis factor [Source:I | 238.52 | 233.34 | 219.36 | 208.22 | 257.39 | 276.23 |
| 2157 | ENSMUSG00000019815 | 0 | NA | NA | NA | NA | NA | Zc2hc1b | zinc finger, C2HC-type containing 1B [Sour | 0 | 0 | 0 | 0 | 0 | 0 |
| 2158 | ENSMUSG00000019817 | 12.34514903 | -0.047707535 | 0.611464248 | -0.07802179 | 0.93781072 | 0.988420305 | Plagl1 | pleiomorphic adenoma gene-like 1 [Sourc | 15.259 | 10.279 | 12.075 | 14.46 | 13.087 | 8.9108 |
| 2159 | ENSMUSG00000019818 | 4938.396594 | 0.050680188 | 0.097627882 | 0.519115925 | 0.603679904 | 0.916106578 | Cd164 | CD164 antigen [Source:MGI Symbol;Acc:N | 4690.1 | 4880.6 | 4983.9 | 4600.1 | 5022.3 | 5453.4 |
| 2160 | ENSMUSG00000019820 | 590.8127476 | 0.127708843 | 0.126242764 | 1.011613174 | 0.311723044 | 0.759830222 | Utrn | utrophin [Source:MGI Symbol;Acc:MGI:10 | 515.59 | 623.95 | 555.45 | 618.87 | 617.29 | 613.73 |
| 2161 | ENSMUSG00000019822 | 754.510282 | -0.120421471 | 0.138785087 | -0.867683074 | 0.385567854 | 0.816437825 | Smpd2 | sphingomyelin phosphodiesterase 2, neut | 808.72 | 785.33 | 762.73 | 589.95 | 795.06 | 785.26 |
| 2162 | ENSMUSG00000019823 | 10.14486119 | -1.545230354 | 0.729092656 | -2.119388176 | 0.034057675 | 0.247046246 | Mical1 | microtubule associated monooxygenase, | 16.865 | 18.503 | 10.062 | 7.7118 | 3.2719 | 4.4554 |
| 2163 | ENSMUSG00000019826 | 227.3388958 | 0.214862118 | 0.17595789 | 1.221099652 | 0.222048291 | 0.66813116 | Zbtb24 | zinc finger and BTB domain containing 24 | 204.79 | 213.81 | 213.32 | 284.37 | 210.49 | 237.25 |
| 2164 | ENSMUSG00000019828 | 0 | NA | NA | NA | NA | NA | Grm1 | glutamate receptor, metabotropic 1 [Sour | 0 | 0 | 0 | 0 | 0 | 0 |
| 2165 | ENSMUSG00000019831 | 20.26687273 | -0.777266967 | 0.564730249 | -1.376350865 | 0.168713032 | 0.595783862 | Wasf1 | WASP family, member 1 [Source:MGI Sym | 15.259 | 19.53 | 42.262 | 19.28 | 16.359 | 8.9108 |
| 2166 | ENSMUSG00000019832 | 475.404595 | 0.053391218 | 0.146331778 | 0.364864136 | 0.715212827 | 0.944595913 | Rab32 | RAB32, member RAS oncogene family [So | 407.17 | 472.84 | 520.23 | 435.72 | 492.96 | 523.51 |
| 2167 | ENSMUSG00000019834 | 0 | NA | NA | NA | NA | NA | Slc22a16 | solute carrier family 22 (organic cation tra | 0 | 0 | 0 | 0 | 0 | 0 |
| 2168 | ENSMUSG00000019836 | 2.777271825 | 0.591968711 | 1.442853407 | 0.410276406 | 0.681603197 | NA | Amd-ps4 | S-adenosylmethionine decarboxylase, pse | 1.6062 | 4.1117 | 1.0062 | 7.7118 | 0 | 2.2277 |
| 2169 | ENSMUSG00000019837 | 237.5595637 | -0.096244911 | 0.161133272 | -0.597300046 | 0.550307074 | 0.895703112 | Gtf3c6 | general transcription factor IIIC, polypepti | 239.32 | 255.95 | 241.5 | 241.96 | 210.49 | 236.13 |
| 2170 | ENSMUSG00000019838 | 1812.281158 | 0.262942668 | 0.132094638 | 1.990562772 | 0.046528977 | 0.298578135 | Slc16a10 | solute carrier family 16 (monocarboxylic a | 1510.6 | 1776.2 | 1656.3 | 1868.2 | 1718.8 | 2343.5 |
| 2171 | ENSMUSG00000019841 | 267.2160217 | 0.267190995 | 0.163629491 | 1.632902445 | 0.102489516 | 0.470781813 | Rev3l | REV3 like, DNA directed polymerase zeta c | 266.63 | 234.37 | 226.41 | 325.82 | 269.38 | 280.69 |
| 2172 | ENSMUSG00000019842 | 31.65942713 | -0.6699344 | 0.427747246 | -1.566192198 | 0.117303629 | 0.502643331 | Traf3ip2 | TRAF3 interacting protein 2 [Source:MGI S | 40.155 | 39.061 | 37.231 | 19.28 | 16.359 | 37.871 |
| 2173 | ENSMUSG00000019843 | 87.22553706 | -0.771312724 | 0.259493208 | -2.972381168 | 0.002954996 | 0.046110081 | Fyn | Fyn proto-oncogene [Source:MGI Symbol; | 102.8 | 132.6 | 94.587 | 58.803 | 62.165 | 72.4 |
| 2174 | ENSMUSG00000019845 | 17.80135922 | 0.120624551 | 0.52962783 | 0.227753424 | 0.819837938 | 0.969160361 | Tube1 | tubulin, epsilon 1 [Source:MGI Symbol;Ac | 17.668 | 13.363 | 20.125 | 18.316 | 25.084 | 12.252 |
| 2175 | ENSMUSG00000019846 | 52.11971608 | -0.727700195 | 0.340772822 | -2.13544082 | 0.032725019 | 0.241743964 | Lama4 | laminin, alpha 4 [Source:MGI Symbol;Acc: | 45.777 | 80.178 | 69.431 | 43.379 | 31.628 | 42.326 |
| 2176 | ENSMUSG00000019848 | 0 | NA | NA | NA | NA | NA | Popdc3 | popeye domain containing 3 [Source:MGI | 0 | 0 | 0 | 0 | 0 | 0 |
| 2177 | ENSMUSG00000019849 | 646.930636 | 0.034440014 | 0.13193196 | 0.261043751 | 0.794058768 | 0.961915077 | Prep | prolyl endopeptidase [Source:MGI Symbo | 578.23 | 638.34 | 701.35 | 591.88 | 702.36 | 669.42 |
| 2178 | ENSMUSG00000019850 | 63.20236792 | -0.676471615 | 0.32082702 | -2.108524449 | 0.03498565 | 0.250788285 | Tnfaip3 | tumor necrosis factor, alpha-induced prot | 72.279 | 93.541 | 67.418 | 40.487 | 67.618 | 37.871 |
| 2179 | ENSMUSG00000019851 | 2418.12411 | -0.075694733 | 0.090463309 | -0.83674513 | 0.402735846 | 0.828953151 | Perp | PERP, TP53 apoptosis effector [Source:MG | 2372.4 | 2512.2 | 2560.9 | 2370.4 | 2343.7 | 2349.1 |
| 2180 | ENSMUSG00000019852 | 1.310068835 | -0.687136541 | 2.222726139 | -0.309141342 | 0.757214012 | NA | Arfgef3 | ARFGEF family member 3 [Source:MGI Syı | 4.8186 | 0 | 0 | 1.928 | 0 | 1.1138 |

| | | | | | | | | | | | | | | | |
|---|---|---|---|---|---|---|---|---|---|---|---|---|---|---|---|
| 2181 | ENSMUSG00000019853 | 3.944679177 | 0.831910362 | 1.101184453 | 0.755468677 | 0.44996782 | NA | Hebp2 | heme binding protein 2 [Source:MGI Syml | 2.4093 | 2.0558 | 4.025 | 1.928 | 5.4531 | 7.7969 |
| 2182 | ENSMUSG00000019854 | 313.2375631 | 0.062479819 | 0.176063389 | 0.354871158 | 0.722686117 | 0.9447719 | Reps1 | RalBP1 associated Eps domain containing | 372.64 | 298.1 | 247.54 | 317.15 | 304.28 | 339.72 |
| 2183 | ENSMUSG00000019856 | 10.25727087 | 1.269767179 | 0.719131702 | 1.765694902 | 0.077447049 | 0.40712138 | Fam184a | family with sequence similarity 184, meml | 4.0155 | 5.1396 | 9.0562 | 20.244 | 14.178 | 8.9108 |
| 2184 | ENSMUSG00000019857 | 121.9318952 | -0.315411499 | 0.21090511 | -1.495513784 | 0.134780405 | 0.537616739 | Asf1a | anti-silencing function 1A histone chaperc | 146.16 | 121.29 | 137.86 | 107.97 | 104.7 | 113.61 |
| 2185 | ENSMUSG00000019861 | 587.5156425 | 0.106809439 | 0.139984136 | 0.763011021 | 0.445456823 | 0.852147532 | Gopc | golgi associated PDZ and coiled-coil motif | 617.59 | 558.16 | 520.23 | 524.4 | 660.91 | 643.8 |
| 2186 | ENSMUSG00000019863 | 109.444144 | 0.080324252 | 0.243462255 | 0.329924868 | 0.741456733 | 0.948792385 | Qrsl1 | glutaminyl-tRNA synthase (glutamine-hyd | 125.28 | 93.541 | 99.618 | 91.578 | 134.15 | 112.5 |
| 2187 | ENSMUSG00000019864 | 652.8094913 | 0.319693315 | 0.149401198 | 2.139830994 | 0.032368427 | 0.24029826 | Rtn4ip1 | reticulon 4 interacting protein 1 [Source:N | 597.51 | 516.01 | 627.9 | 601.52 | 834.32 | 739.59 |
| 2188 | ENSMUSG00000019865 | 0 | NA | NA | NA | NA | NA | Nmbr | neuromedin B receptor [Source:MGI Syml | 0 | 0 | 0 | 0 | 0 | 0 |
| 2189 | ENSMUSG00000019866 | 583.2170208 | -0.314241142 | 0.117808423 | -2.667391126 | 0.007644265 | 0.088879945 | Crybg1 | crystallin beta-gamma domain containing | 641.68 | 653.76 | 644 | 509.94 | 543.13 | 506.8 |
| 2190 | ENSMUSG00000019867 | 0 | NA | NA | NA | NA | NA | Gje1 | gap junction protein, epsilon 1 [Source:M | 0 | 0 | 0 | 0 | 0 | 0 |
| 2191 | ENSMUSG00000019868 | 192.1576132 | -0.043913081 | 0.187188344 | -0.234593032 | 0.814524618 | 0.967404825 | Vta1 | vesicle (multivesicular body) trafficking 1 [ | 223.26 | 182.97 | 178.11 | 168.7 | 207.22 | 192.69 |
| 2192 | ENSMUSG00000019872 | 698.6699997 | -0.369645207 | 0.135673099 | -2.724528367 | 0.00643934 | 0.079266903 | Smpdl3a | sphingomyelin phosphodiesterase, acid-lil | 770.98 | 889.15 | 703.36 | 637.19 | 545.31 | 646.03 |
| 2193 | ENSMUSG00000019873 | 2156.947983 | -0.270460203 | 0.096043528 | -2.816016963 | 0.00486231 | 0.065543757 | Reep3 | receptor accessory protein 3 [Source:MGI | 2228.6 | 2397.1 | 2451.2 | 2040.7 | 1926 | 1898 |
| 2194 | ENSMUSG00000019874 | 27.72845661 | -1.003056366 | 0.465480961 | -2.154881618 | 0.031171099 | 0.234334085 | Fabp7 | fatty acid binding protein 7, brain [Source | 22.487 | 49.34 | 39.244 | 14.46 | 17.45 | 23.391 |
| 2195 | ENSMUSG00000019876 | 10.48078082 | -1.432178655 | 0.779320814 | -1.837726684 | 0.06610269 | 0.369154172 | Pkib | protein kinase inhibitor beta, cAMP deper | 23.29 | 13.363 | 9.0562 | 3.8559 | 2.1812 | 11.138 |
| 2196 | ENSMUSG00000019877 | 4050.725054 | 0.036230938 | 0.092022001 | 0.393720387 | 0.693787488 | 0.939133502 | Serinc1 | serine incorporator 1 [Source:MGI Symbo | 3785.8 | 4117.8 | 4096.4 | 3912.8 | 4299.2 | 4092.3 |
| 2197 | ENSMUSG00000019878 | 408.8098939 | -0.154959724 | 0.140264727 | -1.104766159 | 0.269260928 | 0.724979259 | Hsf2 | heat shock factor 2 [Source:MGI Symbol;A | 470.62 | 405 | 415.58 | 359.56 | 412.25 | 389.85 |
| 2198 | ENSMUSG00000019880 | 59.21624113 | -0.631605637 | 0.366518163 | -1.723258764 | 0.084841741 | 0.429140871 | Rspo3 | R-spondin 3 [Source:MGI Symbol;Acc:MG | 97.978 | 50.368 | 67.418 | 58.803 | 27.265 | 53.465 |
| 2199 | ENSMUSG00000019883 | 325.4778565 | 0.273877942 | 0.156119466 | 1.754284385 | 0.07938179 | 0.413580628 | Echdc1 | enoyl Coenzyme A hydratase domain cont | 334.09 | 281.65 | 267.66 | 378.84 | 364.27 | 326.36 |
| 2200 | ENSMUSG00000019888 | 0 | NA | NA | NA | NA | NA | Mgat4c | MGAT4 family, member C [Source:MGI Sy | 0 | 0 | 0 | 0 | 0 | 0 |
| 2201 | ENSMUSG00000019889 | 747.9383139 | 0.14831174 | 0.115447143 | 1.284672236 | 0.198906844 | 0.640976045 | Ptprk | protein tyrosine phosphatase, receptor ty | 690.67 | 707.21 | 730.53 | 738.41 | 776.52 | 844.29 |
| 2202 | ENSMUSG00000019890 | 0.133850373 | -0.796629176 | 4.080472857 | -0.195229623 | 0.84521319 | NA | Nts | neurotensin [Source:MGI Symbol;Acc:MG | 0.8031 | 0 | 0 | 0 | 0 | 0 |
| 2203 | ENSMUSG00000019891 | 78.89957415 | -0.1434064 | 0.265458897 | -0.540220734 | 0.589044815 | 0.911039598 | Dcbld1 | discoidin, CUB and LCCL domain containir | 91.554 | 90.457 | 66.412 | 83.866 | 68.709 | 72.4 |
| 2204 | ENSMUSG00000019892 | 0 | NA | NA | NA | NA | NA | Lrriq1 | leucine-rich repeats and IQ motif containi | 0 | 0 | 0 | 0 | 0 | 0 |
| 2205 | ENSMUSG00000019893 | 0 | NA | NA | NA | NA | NA | Ros1 | Ros1 proto-oncogene [Source:MGI Symbo | 0 | 0 | 0 | 0 | 0 | 0 |
| 2206 | ENSMUSG00000019894 | 0 | NA | NA | NA | NA | NA | Slc6a15 | solute carrier family 6 (neurotransmitter t | 0 | 0 | 0 | 0 | 0 | 0 |
| 2207 | ENSMUSG00000019897 | 296.4350365 | 0.056712525 | 0.151732674 | 0.373766068 | 0.708578367 | 0.943888782 | Ccdc59 | coiled-coil domain containing 59 [Source:I | 298.75 | 267.26 | 305.9 | 324.86 | 294.47 | 287.37 |
| 2208 | ENSMUSG00000019899 | 17.92859968 | -0.429006761 | 0.584147537 | -0.734415081 | 0.462695798 | 0.860873685 | Lama2 | laminin, alpha 2 [Source:MGI Symbol;Acc: | 28.912 | 24.67 | 8.05 | 17.352 | 17.45 | 11.138 |
| 2209 | ENSMUSG00000019900 | 0 | NA | NA | NA | NA | NA | Rfx6 | regulatory factor X, 6 [Source:MGI Symbo | 0 | 0 | 0 | 0 | 0 | 0 |
| 2210 | ENSMUSG00000019905 | 0 | NA | NA | NA | NA | NA | Gprc6a | G protein-coupled receptor, family C, grou | 0 | 0 | 0 | 0 | 0 | 0 |
| 2211 | ENSMUSG00000019906 | 341.9591103 | 0.364414927 | 0.182135432 | 2.000790965 | 0.045414921 | 0.294697812 | Lin7a | lin-7 homolog A (C. elegans) [Source:MGI | 289.92 | 288.84 | 318.98 | 468.49 | 394.8 | 290.71 |
| 2212 | ENSMUSG00000019907 | 434.7460585 | 0.12436471 | 0.137389018 | 0.905201238 | 0.365358786 | 0.803870323 | Ppp1r12a | protein phosphatase 1, regulatory subunit | 375.05 | 457.42 | 416.59 | 466.57 | 439.52 | 453.33 |
| 2213 | ENSMUSG00000019909 | 0 | NA | NA | NA | NA | NA | Fam162b | family with sequence similarity 162, meml | 0 | 0 | 0 | 0 | 0 | 0 |
| 2214 | ENSMUSG00000019913 | 0 | NA | NA | NA | NA | NA | Sim1 | single-minded family bHLH transcription fa | 0 | 0 | 0 | 0 | 0 | 0 |
| 2215 | ENSMUSG00000019916 | 116.178766 | -0.518718721 | 0.236435043 | -2.193916407 | 0.02824142 | 0.219614996 | P4ha1 | procollagen-proline, 2-oxoglutarate 4-diox | 134.92 | 136.71 | 138.86 | 88.686 | 122.15 | 75.741 |
| 2216 | ENSMUSG00000019917 | 273.4245496 | 0.248039906 | 0.158289456 | 1.56700207 | 0.117114208 | 0.502643331 | Septin10 | septin 10 [Source:MGI Symbol;Acc:MGI:1! | 240.13 | 233.34 | 276.72 | 303.65 | 274.84 | 311.88 |
| 2217 | ENSMUSG00000019920 | 278.9763371 | -0.238529187 | 0.163964897 | -1.454757648 | 0.145736367 | 0.556690797 | Lims1 | LIM and senescent cell antigen-like domai | 293.94 | 329.96 | 281.75 | 221.71 | 280.29 | 266.21 |
| 2218 | ENSMUSG00000019923 | 1153.253927 | -0.021233088 | 0.100385098 | -0.211516331 | 0.83248439 | 0.97056812 | Zwint | ZW10 interactor [Source:MGI Symbol;Acc | 1179.8 | 1151.3 | 1154.2 | 1162.6 | 1172.4 | 1099.4 |
| 2219 | ENSMUSG00000019927 | 163.2942421 | -0.131311444 | 0.215595613 | -0.609063618 | 0.542482271 | 0.894095573 | Ube2d1 | ubiquitin-conjugating enzyme E2D 1 [Sour | 169.45 | 134.66 | 208.29 | 175.44 | 152.69 | 139.23 |
| 2220 | ENSMUSG00000019929 | 1823.054192 | -1.235443531 | 0.136800697 | -9.030973946 | 1.70E-19 | 2.05E-16 | Dcn | decorin [Source:MGI Symbol;Acc:MGI:948 | 2982.7 | 2427.9 | 2266.1 | 1101.8 | 940.11 | 1219.7 |
| 2221 | ENSMUSG00000019932 | 0 | NA | NA | NA | NA | NA | Kera | keratocan [Source:MGI Symbol;Acc:MGI:1 | 0 | 0 | 0 | 0 | 0 | 0 |
| 2222 | ENSMUSG00000019933 | 0.17131964 | -0.796629176 | 4.080472857 | -0.195229623 | 0.84521319 | NA | Mrln | myoregulin [Source:MGI Symbol;Acc:MGI | 0 | 1.0279 | 0 | 0 | 0 | 0 |
| 2223 | ENSMUSG00000019935 | 1271.738928 | -0.75701162 | 0.165753389 | -4.567095892 | 4.95E-06 | 0.000303087 | Slc17a8 | solute carrier family 17 (sodium-depender | 1281.8 | 1900.6 | 1613 | 1077.7 | 1008.8 | 748.5 |
| 2224 | ENSMUSG00000019936 | 0 | NA | NA | NA | NA | NA | Epyc | epiphycan [Source:MGI Symbol;Acc:MGI:1 | 0 | 0 | 0 | 0 | 0 | 0 |
| 2225 | ENSMUSG00000019942 | 24.65137005 | -0.314772846 | 0.529930214 | -0.593989242 | 0.552519308 | 0.896052884 | Cdk1 | cyclin-dependent kinase 1 [Source:MGI Sy | 44.171 | 13.363 | 24.15 | 21.208 | 30.537 | 14.48 |
| 2226 | ENSMUSG00000019943 | 606.9361793 | 0.146697976 | 0.125350605 | 1.170301294 | 0.241879741 | 0.691791181 | Atp2b1 | ATPase, Ca++ transporting, plasma memb | 530.85 | 623.95 | 574.57 | 664.18 | 600.93 | 647.14 |
| 2227 | ENSMUSG00000019944 | 1917.410352 | -0.560524089 | 0.142339306 | -3.937943119 | 8.22E-05 | 0.003087522 | Rhobtb1 | Rho-related BTB domain containing 1 [Sou | 2182 | 2657.2 | 2016.5 | 1384.3 | 1839.9 | 1424.6 |
| 2228 | ENSMUSG00000019945 | 22.63546663 | 0.022925853 | 0.506658443 | 0.045249129 | 0.963908735 | 0.993245949 | Cabcoco1 | ciliary associated calcium binding coiled-c | 18.471 | 34.949 | 14.087 | 24.099 | 16.359 | 27.846 |
| 2229 | ENSMUSG00000019947 | 1302.194802 | 0.971660508 | 0.510324129 | 1.904006595 | 0.056909325 | 0.338327228 | Arid5b | AT rich interactive domain 5B (MRF1-like) | 774.19 | 1100.9 | 763.74 | 2640.3 | 784.15 | 1749.8 |
| 2230 | ENSMUSG00000019948 | 257.015184 | -0.182560052 | 0.171646169 | -1.063583608 | 0.287517371 | 0.73951397 | Actr6 | ARP6 actin-related protein 6 [Source:MGI | 256.99 | 320.71 | 242.5 | 243.89 | 254.11 | 223.88 |
| 2231 | ENSMUSG00000019951 | 1008.243268 | -0.210080753 | 0.13678084 | -1.535893131 | 0.124564593 | 0.517879354 | Uhrf1bp1l | UHRF1 (ICBP90) binding protein 1-like [So | 1052.9 | 1240.7 | 951.91 | 1026.6 | 957.56 | 819.79 |
| 2232 | ENSMUSG00000019952 | 131.7258272 | -0.006399415 | 0.237557392 | -0.026938396 | 0.978508869 | 0.997797092 | Poc1b | POC1 centriolar protein B [Source:MGI Sy | 167.85 | 104.85 | 122.76 | 145.56 | 112.33 | 137 |
| 2233 | ENSMUSG00000019960 | 515.4867088 | -0.006230591 | 0.412686737 | -0.015097628 | 0.987954293 | 0.99859176 | Dusp6 | dual specificity phosphatase 6 [Source:M( | 705.12 | 391.64 | 452.81 | 570.67 | 666.37 | 306.31 |
| 2234 | ENSMUSG00000019961 | 730.5973205 | 0.069165778 | 0.119193665 | 0.580280654 | 0.561725371 | 0.900565925 | Tmpo | thymopoietin [Source:MGI Symbol;Acc:M | 721.99 | 660.95 | 755.69 | 698.88 | 781.97 | 764.1 |
| 2235 | ENSMUSG00000019966 | 50.45456716 | -0.124301571 | 0.330178362 | -0.376467948 | 0.706569039 | 0.94295036 | Kitl | kit ligand [Source:MGI Symbol;Acc:MGI:9( | 44.974 | 63.731 | 49.306 | 45.307 | 39.262 | 60.148 |
| 2236 | ENSMUSG00000019969 | 690.7992289 | 0.089614956 | 0.132664524 | 0.675500526 | 0.499357818 | 0.880660418 | Psen1 | presenilin 1 [Source:MGI Symbol;Acc:MGI | 638.47 | 703.1 | 666.13 | 608.27 | 754.71 | 774.12 |

| | | | | | | | | | | | | | | | |
|---|---|---|---|---|---|---|---|---|---|---|---|---|---|---|---|
| 2237 | ENSMUSG00000019970 | 230.0697829 | -1.166227299 | 0.213077688 | -5.473249252 | 4.42E-08 | 5.44E-06 | Sgk1 | serum/glucocorticoid regulated kinase 1 [ | 277.07 | 410.14 | 267.66 | 120.5 | 163.59 | 141.46 |
| 2238 | ENSMUSG00000019971 | 56.43341577 | -0.122369266 | 0.315256605 | -0.388157661 | 0.697899363 | 0.940502785 | Cep290 | centrosomal protein 290 [Source:MGI Syn | 46.58 | 68.87 | 61.381 | 53.983 | 64.346 | 43.44 |
| 2239 | ENSMUSG00000019975 | 39.51938722 | -0.371910747 | 0.365382695 | -1.017866339 | 0.30874148 | 0.75728287 | Ikbip | IKBKB interacting protein [Source:MGI Syr | 35.336 | 48.312 | 50.312 | 32.775 | 42.534 | 27.846 |
| 2240 | ENSMUSG00000019977 | 915.9613292 | 0.197020463 | 0.111628416 | 1.764966935 | 0.077569323 | 0.407541841 | Hbs1l | Hbs1-like (S. cerevisiae) [Source:MGI Sym | 852.09 | 877.84 | 830.15 | 916.74 | 963.02 | 1055.9 |
| 2241 | ENSMUSG00000019978 | 148.4551011 | -1.195820004 | 0.284133434 | -4.208656428 | 2.57E-05 | 0.001190933 | Epb41l2 | erythrocyte membrane protein band 4.1 l | 205.59 | 219.97 | 195.21 | 133.99 | 49.078 | 86.88 |
| 2242 | ENSMUSG00000019979 | 37.00770801 | -0.378173669 | 0.390429694 | -0.96860888 | 0.332740371 | 0.77523112 | Apaf1 | apoptotic peptidase activating factor 1 [So | 56.217 | 33.921 | 35.219 | 40.487 | 28.356 | 27.846 |
| 2243 | ENSMUSG00000019982 | 0.428363721 | -0.543730382 | 3.624677344 | -0.15000794 | 0.880758351 | NA | Myb | myeloblastosis oncogene [Source:MGI Syr | 1.6062 | 0 | 0 | 0.964 | 0 | 0 |
| 2244 | ENSMUSG00000019984 | 173.666694 | 0.211042656 | 0.197957965 | 1.066098326 | 0.286379196 | 0.738722073 | Med23 | mediator complex subunit 23 [Source:MG | 174.27 | 177.83 | 130.81 | 205.33 | 177.77 | 175.99 |
| 2245 | ENSMUSG00000019986 | 16.74908869 | -0.415623043 | 0.519904623 | -0.79942171 | 0.424045926 | 0.840450466 | Ahi1 | Abelson helper integration site 1 [Source: | 16.865 | 19.53 | 21.131 | 15.424 | 14.178 | 13.366 |
| 2246 | ENSMUSG00000019987 | 30065.82485 | -0.053484385 | 0.114590996 | -0.4667416 | 0.640684763 | 0.923932929 | Arg1 | arginase, liver [Source:MGI Symbol;Acc:M | 31154 | 30067 | 30647 | 24646 | 30858 | 33023 |
| 2247 | ENSMUSG00000019988 | 331.6927947 | 0.106153318 | 0.158925355 | 0.667944505 | 0.504169017 | 0.881510796 | Nedd1 | neural precursor cell expressed, developn | 310 | 279.59 | 369.29 | 370.17 | 338.09 | 323.01 |
| 2248 | ENSMUSG00000019989 | 981.8658259 | 0.244223207 | 0.135856677 | 1.797653318 | 0.072231963 | 0.389401088 | Enpp3 | ectonucleotide pyrophosphatase/phosph | 742.07 | 910.74 | 1045.5 | 1051.7 | 1079.7 | 1061.5 |
| 2249 | ENSMUSG00000019990 | 85.69543248 | -0.24052577 | 0.259464478 | -0.927008476 | 0.353922134 | 0.79449673 | Pde7b | phosphodiesterase 7B [Source:MGI Symb | 113.24 | 78.122 | 86.537 | 73.262 | 86.159 | 76.855 |
| 2250 | ENSMUSG00000019992 | 1.451991932 | 0.708226004 | 2.099467461 | 0.337336023 | 0.735863607 | NA | Mtfr2 | mitochondrial fission regulator 2 [Source: | 3.2124 | 0 | 0 | 0 | 3.2719 | 2.2277 |
| 2251 | ENSMUSG00000019996 | 475.7121361 | -0.143764569 | 0.138231027 | -1.04003112 | 0.298325443 | 0.74911959 | Map7 | microtubule-associated protein 7 [Source | 493.1 | 542.74 | 462.87 | 488.74 | 406.8 | 460.02 |
| 2252 | ENSMUSG00000019997 | 73.51239484 | -1.768126106 | 0.379503557 | -4.659050157 | 3.18E-06 | 0.00020791 | Ccn2 | cellular communication network factor 2 [ | 110.03 | 167.55 | 63.393 | 33.739 | 21.812 | 44.554 |
| 2253 | ENSMUSG00000019998 | 388.366149 | -0.301599289 | 0.140196125 | -2.151266948 | 0.031455135 | 0.236101235 | Stx7 | syntaxin 7 [Source:MGI Symbol;Acc:MGI:1 | 431.27 | 450.23 | 404.51 | 318.11 | 372.99 | 353.09 |
| 2254 | ENSMUSG00000020000 | 88.25137577 | 9.030732246 | 2.366274264 | 3.816435137 | 0.000135394 | 0.004586262 | Moxd1 | monooxygenase, DBH-like 1 [Source:MGI | 0 | 0 | 1.0062 | 0.964 | 333.73 | 193.81 |
| 2255 | ENSMUSG00000020003 | 1831.47903 | -0.05386364 | 0.095809345 | -0.562196102 | 0.573982417 | 0.9059841 | Pex7 | peroxisomal biogenesis factor 7 [Source:N | 1829.5 | 1915 | 1852.5 | 1726.5 | 1894.4 | 1771 |
| 2256 | ENSMUSG00000020007 | 0.321325949 | 1.911575114 | 4.048267893 | 0.472195804 | 0.636787034 | NA | Il20ra | interleukin 20 receptor, alpha [Source:MG | 0 | 0 | 0 | 1.928 | 0 | 0 |
| 2257 | ENSMUSG00000020009 | 266.4307465 | -0.28473876 | 0.168359982 | -1.691249465 | 0.090789133 | 0.444602826 | Ifngr1 | interferon gamma receptor 1 [Source:MG | 303.57 | 321.74 | 252.57 | 249.67 | 214.85 | 256.18 |
| 2258 | ENSMUSG00000020010 | 1466.071065 | -0.580141268 | 0.120333244 | -4.821122134 | 1.43E-06 | 0.000108104 | Vnn3 | vanin 3 [Source:MGI Symbol;Acc:MGI:134 | 1659.2 | 1848.2 | 1765 | 1322.6 | 1177.9 | 1023.6 |
| 2259 | ENSMUSG00000020014 | 16.00710545 | -0.489544349 | 0.639115787 | -0.765971297 | 0.443693381 | 0.851138072 | Cfap54 | cilia and flagella associated protein 54 [So | 19.274 | 28.782 | 8.05 | 12.532 | 20.722 | 6.6831 |
| 2260 | ENSMUSG00000020015 | 334.3126169 | 0.115770215 | 0.141382233 | 0.818845572 | 0.412874526 | 0.835103185 | Cdk17 | cyclin-dependent kinase 17 [Source:MGI S | 338.11 | 311.46 | 312.94 | 363.42 | 342.45 | 337.49 |
| 2261 | ENSMUSG00000020017 | 6034.687105 | -0.031013584 | 0.193311154 | -0.160433494 | 0.872539607 | 0.979419285 | Hal | histidine ammonia lyase [Source:MGI Sym | 4265.3 | 5393.5 | 8640.6 | 5215.1 | 6212.2 | 6481.5 |
| 2262 | ENSMUSG00000020018 | 96.6353353 | -0.16084153 | 0.236490459 | -0.680118475 | 0.496429447 | 0.878165817 | Snrpf | small nuclear ribonucleoprotein polypepti | 101.99 | 99.708 | 104.65 | 106.04 | 88.34 | 79.083 |
| 2263 | ENSMUSG00000020019 | 85.96180992 | -0.070305361 | 0.290629666 | -0.241907037 | 0.808852197 | 0.965682062 | Ntn4 | netrin 4 [Source:MGI Symbol;Acc:MGI:18 | 102.8 | 91.485 | 69.431 | 75.19 | 63.256 | 113.61 |
| 2264 | ENSMUSG00000020020 | 0 | NA | NA | NA | NA | NA | Usp44 | ubiquitin specific peptidase 44 [Source:M | 0 | 0 | 0 | 0 | 0 | 0 |
| 2265 | ENSMUSG00000020021 | 771.6411334 | -0.026967325 | 0.127313097 | -0.211818941 | 0.832248291 | 0.970533397 | Fgd6 | FYVE, RhoGEF and PH domain containing | 721.99 | 719.54 | 895.56 | 757.69 | 775.43 | 759.64 |
| 2266 | ENSMUSG00000020022 | 889.3611419 | -0.050328892 | 0.110644226 | -0.454871382 | 0.649201773 | 0.925643431 | Ndufa12 | NADH:ubiquinone oxidoreductase subuni | 913.93 | 855.23 | 944.86 | 831.91 | 875.77 | 914.47 |
| 2267 | ENSMUSG00000020023 | 290.461818 | -0.02502147 | 0.149327765 | -0.167560735 | 0.866928863 | 0.977930657 | Tmcc3 | transmembrane and coiled coil domains 3 | 277.87 | 308.38 | 292.82 | 271.84 | 294.47 | 297.4 |
| 2268 | ENSMUSG00000020024 | 135.5367436 | 0.004672526 | 0.199354484 | 0.023438278 | 0.981300672 | 0.99820894 | Cep83 | centrosomal protein 83 [Source:MGI Syml | 146.16 | 133.63 | 125.78 | 132.06 | 135.24 | 140.34 |
| 2269 | ENSMUSG00000020027 | 409.2647309 | 1.727147374 | 0.780990166 | 2.211484151 | 0.027002328 | 0.212875643 | Socs2 | suppressor of cytokine signaling 2 [Source | 296.34 | 165.49 | 107.67 | 1213.6 | 224.67 | 447.77 |
| 2270 | ENSMUSG00000020029 | 4039.187813 | -0.241053366 | 0.093059566 | -2.590312609 | 0.009588881 | 0.105150551 | Nudt4 | nudix (nucleoside diphosphate linked moi | 4360.8 | 4464.2 | 4301.7 | 3466.5 | 3671 | 3970.9 |
| 2271 | ENSMUSG00000020032 | 53.72619624 | 0.429142007 | 0.351793712 | 1.219868329 | 0.222514794 | 0.668407528 | Nuak1 | NUAK family, SNF1-like kinase, 1 [Source:I | 32.124 | 40.089 | 65.406 | 50.127 | 59.984 | 74.628 |
| 2272 | ENSMUSG00000020033 | 0 | NA | NA | NA | NA | NA | 4930463O16 | RIKEN cDNA 4930463O16 gene [Source:M | 0 | 0 | 0 | 0 | 0 | 0 |
| 2273 | ENSMUSG00000020034 | 409.1057646 | 0.197262261 | 0.138628919 | 1.422951738 | 0.154750146 | 0.572785168 | Tcp11l2 | t-complex 11 (mouse) like 2 [Source:MGI ! | 368.62 | 371.08 | 404.51 | 478.13 | 406.8 | 425.49 |
| 2274 | ENSMUSG00000020037 | 125.2532223 | -0.173509009 | 0.258259333 | -0.671840228 | 0.501685416 | 0.880974509 | Rfx4 | regulatory factor X, 4 (influences HLA clas | 139.74 | 86.345 | 172.07 | 117.61 | 122.15 | 113.61 |
| 2275 | ENSMUSG00000020038 | 375.6887203 | -1.154974245 | 0.246881433 | -4.678254785 | 2.89E-06 | 0.000191964 | Cry1 | cryptochrome 1 (photolyase-like) [Source | 315.62 | 543.77 | 697.33 | 293.05 | 207.22 | 197.15 |
| 2276 | ENSMUSG00000020042 | 6.882303717 | -1.941587197 | 0.910847406 | -2.131627301 | 0.033037498 | 0.243182883 | Btbd11 | BTB (POZ) domain containing 11 [Source:I | 7.2279 | 17.475 | 8.05 | 1.928 | 3.2719 | 3.3415 |
| 2277 | ENSMUSG00000020044 | 126.5073247 | -0.992757072 | 0.27599085 | -3.597065167 | 0.000321828 | 0.00921766 | Timp3 | tissue inhibitor of metalloproteinase 3 [So | 164.64 | 187.08 | 153.96 | 111.82 | 47.987 | 93.563 |
| 2278 | ENSMUSG00000020048 | 7684.977561 | 0.346498908 | 0.092274601 | 3.755084328 | 0.000173283 | 0.005582109 | Hsp90b1 | heat shock protein 90, beta (Grp94), mem | 7234.3 | 6682.5 | 6381.6 | 8822.3 | 8176.4 | 8812.7 |
| 2279 | ENSMUSG00000020051 | 24684.75187 | 0.198657845 | 0.100833675 | 1.970153779 | 0.048820749 | 0.307687718 | Pah | phenylalanine hydroxylase [Source:MGI S | 22458 | 22216 | 24289 | 23528 | 28555 | 27063 |
| 2280 | ENSMUSG00000020052 | 0.371281295 | 2.09253362 | 4.031852165 | 0.519000582 | 0.603760335 | NA | Ascl1 | achaete-scute family bHLH transcription fa | 0 | 0 | 0 | 0 | 0 | 2.2277 |
| 2281 | ENSMUSG00000020053 | 11626.26517 | 0.109941456 | 0.111686008 | 0.984379852 | 0.324928781 | 0.771501263 | Igf1 | insulin-like growth factor 1 [Source:MGI S | 11736 | 11184 | 10631 | 12768 | 13151 | 10289 |
| 2282 | ENSMUSG00000020056 | 357.3183676 | 0.05390044 | 0.152302874 | 0.353902971 | 0.7234116 | 0.9447719 | Washc3 | WASH complex subunit 3 [Source:MGI Syr | 400.75 | 335.1 | 314.95 | 349.92 | 354.45 | 388.73 |
| 2283 | ENSMUSG00000020057 | 9.602868401 | -0.563927626 | 0.78137546 | -0.721711463 | 0.470471893 | 0.867110762 | Dram1 | DNA-damage regulated autophagy modul | 16.865 | 12.335 | 5.0312 | 6.7478 | 3.2719 | 13.366 |
| 2284 | ENSMUSG00000020059 | 13.88186454 | -0.472214115 | 0.637761287 | -0.740424552 | 0.459042426 | 0.859322431 | Sycp3 | synaptonemal complex protein 3 [Source: | 8.8341 | 22.614 | 17.106 | 10.604 | 17.45 | 6.6831 |
| 2285 | ENSMUSG00000020061 | 1.540146097 | -0.833076984 | 1.905481017 | -0.437200359 | 0.66196605 | NA | Mybpc1 | myosin binding protein C, slow-type [Sour | 0.8031 | 3.0838 | 2.0125 | 0 | 0 | 3.3415 |
| 2286 | ENSMUSG00000020062 | 0.305170013 | -1.657836477 | 4.041480335 | -0.410205257 | 0.681655385 | NA | Slc5a8 | solute carrier family 5 (iodide transporter) | 0.8031 | 1.0279 | 0 | 0 | 0 | 0 |
| 2287 | ENSMUSG00000020063 | 240.2628 | -0.219344998 | 0.168479853 | -1.301906398 | 0.192948385 | 0.631239875 | Sirt1 | sirtuin 1 [Source:MGI Symbol;Acc:MGI:21 | 235.31 | 250.81 | 289.8 | 220.75 | 238.85 | 206.06 |
| 2288 | ENSMUSG00000020064 | 854.5782047 | 0.152769737 | 0.127325907 | 1.199832307 | 0.230204474 | 0.676901304 | Herc4 | hect domain and RLD 4 [Source:MGI Symb | 801.5 | 771.97 | 854.3 | 820.35 | 1031.7 | 847.64 |
| 2289 | ENSMUSG00000020067 | 2.002397892 | 1.612075924 | 1.689313071 | 0.954278962 | 0.339942454 | NA | Mypn | myopalladin [Source:MGI Symbol;Acc:MG | 0 | 2.0558 | 1.0062 | 6.7478 | 1.0906 | 1.1138 |
| 2290 | ENSMUSG00000020069 | 260.061849 | 0.599702179 | 0.173430942 | 3.457873037 | 0.000544458 | 0.013168962 | Hnrnph3 | heterogeneous nuclear ribonucleoprotein | 211.22 | 198.39 | 210.31 | 289.19 | 277.02 | 374.25 |
| 2291 | ENSMUSG00000020070 | 103.9622326 | 0.247555334 | 0.224754092 | 1.101449729 | 0.270700972 | 0.72615335 | Rufy2 | RUN and FYVE domain-containing 2 [Sour | 93.963 | 93.541 | 97.606 | 104.11 | 126.51 | 108.04 |
| 2292 | ENSMUSG00000020072 | 3978.462913 | -0.151639078 | 0.125122209 | -1.211927757 | 0.225540036 | 0.672301295 | Pbld2 | phenazine biosynthesis-like protein doma | 3745.7 | 4205.2 | 4612.6 | 4002.4 | 4120.4 | 3184.5 |

| | | | | | | | | | | | | | | | |
|---|---|---|---|---|---|---|---|---|---|---|---|---|---|---|---|
| 2293 | ENSMUSG00000020074 | 679.4203135 | -0.118216378 | 0.120664909 | -0.979708013 | 0.327230269 | 0.772772049 | Ccar1 | cell division cycle and apoptosis regulator | 672.2 | 741.13 | 708.4 | 606.34 | 646.74 | 701.72 |
| 2294 | ENSMUSG00000020075 | 668.8554902 | -0.154880965 | 0.141805645 | -1.092205917 | 0.274742602 | 0.729487777 | Ddx21 | DExD box helicase 21 [Source:MGI Symbo | 614.37 | 805.89 | 695.32 | 679.6 | 655.46 | 562.49 |
| 2295 | ENSMUSG00000020076 | 956.5762986 | -0.019295486 | 0.11624184 | -0.165994324 | 0.868161417 | 0.978326885 | Ddx50 | DExD box helicase 50 [Source:MGI Symbo | 888.23 | 985.77 | 1016.3 | 1016 | 954.29 | 878.82 |
| 2296 | ENSMUSG00000020077 | 73.59077766 | -1.040551202 | 0.298542126 | -3.48544179 | 0.000491325 | 0.012273793 | Srgn | serglycin [Source:MGI Symbol;Acc:MGI:97 | 114.04 | 89.429 | 93.581 | 54.947 | 31.628 | 57.92 |
| 2297 | ENSMUSG00000020078 | 998.3267458 | -0.077988698 | 0.106232239 | -0.734133991 | 0.462867079 | 0.860978434 | Vps26a | VPS26 retromer complex component A [S | 982.19 | 1064.9 | 1029.4 | 966.87 | 1013.2 | 933.4 |
| 2298 | ENSMUSG00000020079 | 361.8197196 | 0.046218508 | 0.140115915 | 0.329859087 | 0.741506438 | 0.948792385 | Supv3l1 | suppressor of var1, 3-like 1 (S. cerevisiae) | 367.82 | 362.85 | 337.09 | 358.6 | 394.8 | 349.75 |
| 2299 | ENSMUSG00000020080 | 2.648166792 | -2.709159643 | 1.58402035 | -1.710306084 | 0.087209287 | NA | Hkdc1 | hexokinase domain containing 1 [Source:I | 1.6062 | 6.1675 | 6.0375 | 0.964 | 0 | 1.1138 |
| 2300 | ENSMUSG00000020081 | 0 | NA | NA | NA | NA | NA | Tacr2 | tachykinin receptor 2 [Source:MGI Symbo | 0 | 0 | 0 | 0 | 0 | 0 |
| 2301 | ENSMUSG00000020083 | 55.2992324 | 0.051755967 | 0.333691977 | 0.155101024 | 0.876741675 | 0.980632845 | Fam241b | family with sequence similarity 241, meml | 73.885 | 47.284 | 41.256 | 54.947 | 66.528 | 47.895 |
| 2302 | ENSMUSG00000020085 | 924.6443965 | -0.43027929 | 0.118293212 | -3.637396282 | 0.000275408 | 0.008113038 | Aifm2 | apoptosis-inducing factor, mitochondrion | 1052.1 | 1108.1 | 1025.4 | 867.58 | 785.25 | 709.52 |
| 2303 | ENSMUSG00000020086 | 21.2042888 | -0.250622632 | 0.476453607 | -0.526016863 | 0.59887649 | 0.915055389 | Macroh2a2 | macroH2A.2 histone [Source:MGI Symbol | 23.29 | 23.642 | 22.137 | 16.388 | 26.175 | 15.594 |
| 2304 | ENSMUSG00000020087 | 1548.381211 | -0.140433292 | 0.110281101 | -1.273412137 | 0.202871833 | 0.647682949 | Tysnd1 | trypsin domain containing 1 [Source:MGI | 1607 | 1650.8 | 1614 | 1573.2 | 1547.6 | 1297.6 |
| 2305 | ENSMUSG00000020088 | 1656.619763 | 0.094942448 | 0.09481744 | 1.001318407 | 0.316672897 | 0.763392796 | Sar1a | secretion associated Ras related GTPase 1 | 1637.5 | 1549.1 | 1619 | 1659 | 1731.9 | 1743.2 |
| 2306 | ENSMUSG00000020089 | 3087.693612 | 0.055562147 | 0.099955061 | 0.555871278 | 0.578298855 | 0.908137944 | Ppa1 | pyrophosphatase (inorganic) 1 [Source:M | 3271.8 | 2817.5 | 2993.6 | 2971.9 | 3250 | 3221.2 |
| 2307 | ENSMUSG00000020090 | 0 | NA | NA | NA | NA | NA | Npffr1 | neuropeptide FF receptor 1 [Source:MGI S | 0 | 0 | 0 | 0 | 0 | 0 |
| 2308 | ENSMUSG00000020091 | 5410.101826 | 0.106061871 | 0.094401483 | 1.123519118 | 0.261217086 | 0.715305553 | Eif4ebp2 | eukaryotic translation initiation factor 4E I | 5176.8 | 5095.4 | 5361.3 | 5353 | 5364.8 | 6109.4 |
| 2309 | ENSMUSG00000020092 | 235.1458024 | -0.074040426 | 0.181866314 | -0.407114569 | 0.683923843 | 0.935528319 | Pald1 | phosphatase domain containing, paladin 1 | 219.25 | 213.81 | 290.8 | 230.39 | 211.58 | 245.05 |
| 2310 | ENSMUSG00000020096 | 0 | NA | NA | NA | NA | NA | Tbata | thymus, brain and testes associated [Sour | 0 | 0 | 0 | 0 | 0 | 0 |
| 2311 | ENSMUSG00000020097 | 940.5283555 | -0.050069284 | 0.116015577 | -0.431573801 | 0.666051203 | 0.929596279 | Sgpl1 | sphingosine phosphate lyase 1 [Source:M | 871.37 | 999.14 | 1001.2 | 955.3 | 861.59 | 954.56 |
| 2312 | ENSMUSG00000020098 | 4295.789994 | -0.078208528 | 0.103152611 | -0.758182722 | 0.448341604 | 0.854238022 | Pcbd1 | pterin 4 alpha carbinolamine dehydratase | 4911 | 4111.7 | 4212.1 | 4092.1 | 4057.1 | 4390.8 |
| 2313 | ENSMUSG00000020099 | 205.2098604 | -0.21224729 | 0.266945221 | -0.795096798 | 0.426557197 | 0.842053132 | Unc5b | unc-5 netrin receptor B [Source:MGI Syml | 326.86 | 154.19 | 179.11 | 231.35 | 155.96 | 183.78 |
| 2314 | ENSMUSG00000020100 | 102.7996851 | -0.354912521 | 0.231534998 | -1.5328677 | 0.125308456 | 0.51896773 | Slc29a3 | solute carrier family 29 (nucleoside transp | 110.03 | 121.29 | 114.71 | 80.974 | 83.978 | 105.82 |
| 2315 | ENSMUSG00000020101 | 83.79345098 | -0.56382294 | 0.256665372 | -2.196723838 | 0.028040175 | 0.21848271 | Vsir | V-set immunoregulatory receptor [Source | 97.978 | 92.513 | 109.68 | 79.046 | 56.712 | 66.831 |
| 2316 | ENSMUSG00000020102 | 2346.552715 | -0.771815193 | 0.148058608 | -5.21290322 | 1.86E-07 | 1.91E-05 | Slc16a7 | solute carrier family 16 (monocarboxylic a | 2983.5 | 3254.4 | 2642.4 | 2102.4 | 1722.1 | 1374.5 |
| 2317 | ENSMUSG00000020105 | 215.2484082 | 0.499374484 | 0.174903021 | 2.855150705 | 0.004301641 | 0.06046544 | Lrig3 | leucine-rich repeats and immunoglobulin- | 165.44 | 195.3 | 175.09 | 280.52 | 231.21 | 243.93 |
| 2318 | ENSMUSG00000020107 | 570.2066573 | -0.148049939 | 0.134761621 | -1.098606108 | 0.271939912 | 0.727618887 | Anapc16 | anaphase promoting complex subunit 16 | 674.61 | 551.99 | 570.54 | 531.15 | 510.41 | 582.54 |
| 2319 | ENSMUSG00000020108 | 444.4758316 | -2.70076592 | 0.612741344 | -4.407676983 | 1.04E-05 | 0.000555615 | Ddit4 | DNA-damage-inducible transcript 4 [Sour | 503.55 | 1070.1 | 737.58 | 55.911 | 202.86 | 96.904 |
| 2320 | ENSMUSG00000020109 | 999.7966048 | 0.093039107 | 0.103878223 | 0.895655558 | 0.370436753 | 0.807048832 | Dnajb12 | DnaJ heat shock protein family (Hsp40) m | 991.83 | 927.18 | 983.1 | 1008.3 | 1051.4 | 1037 |
| 2321 | ENSMUSG00000020111 | 481.7828546 | -0.115617805 | 0.155484582 | -0.743596592 | 0.457120565 | 0.858785798 | Micu1 | mitochondrial calcium uptake 1 [Source:M | 579.84 | 489.29 | 432.69 | 403.91 | 507.14 | 477.84 |
| 2322 | ENSMUSG00000020114 | 969.1214502 | 0.147524559 | 0.124857183 | 1.181546434 | 0.237385718 | 0.686297628 | Cand1 | cullin associated and neddylation disasso | 789.45 | 976.52 | 994.17 | 990.01 | 1068.8 | 995.78 |
| 2323 | ENSMUSG00000020115 | 522.0544485 | 0.173085277 | 0.132986341 | 1.301526726 | 0.193078223 | 0.631271165 | Tbk1 | TANK-binding kinase 1 [Source:MGI Symb | 534.06 | 492.37 | 444.76 | 514.76 | 564.94 | 581.43 |
| 2324 | ENSMUSG00000020116 | 604.2361779 | -0.060164459 | 0.120796376 | -0.498065097 | 0.61843816 | 0.917509899 | Pno1 | partner of NOB1 homolog [Source:MGI Sy | 655.33 | 618.81 | 575.57 | 574.53 | 606.38 | 594.79 |
| 2325 | ENSMUSG00000020120 | 33.46403 | -1.09487893 | 0.42714784 | -2.563231806 | 0.010370279 | 0.111734379 | Plek | pleckstrin [Source:MGI Symbol;Acc:MGI:1 | 48.989 | 58.591 | 29.181 | 23.135 | 15.269 | 25.618 |
| 2326 | ENSMUSG00000020121 | 7.420627706 | -1.164917619 | 0.896251698 | -1.299766151 | 0.19368113 | 0.631851946 | Srgap1 | SLIT-ROBO Rho GTPase activating protein | 2.4093 | 15.419 | 13.081 | 4.8199 | 5.4531 | 3.3415 |
| 2327 | ENSMUSG00000020122 | 6644.811579 | 1.117064708 | 0.413213681 | 2.703358478 | 0.006864267 | 0.082875426 | Egfr | epidermal growth factor receptor [Source | 4968 | 3446.6 | 4165.9 | 8325.9 | 13629 | 5333.1 |
| 2328 | ENSMUSG00000020123 | 284.9799206 | 0.44274424 | 0.276579721 | 1.600783452 | 0.10942489 | 0.486243197 | Avpr1a | arginine vasopressin receptor 1A [Source: | 153.39 | 208.67 | 363.25 | 279.55 | 263.93 | 441.08 |
| 2329 | ENSMUSG00000020124 | 772.1385162 | -0.009735472 | 0.120674781 | -0.080675283 | 0.935700194 | 0.988420305 | Usp15 | ubiquitin specific peptidase 15 [Source:M | 758.93 | 859.34 | 706.38 | 779.86 | 778.7 | 749.62 |
| 2330 | ENSMUSG00000020125 | 1.62202458 | -1.907363115 | 1.957675755 | -0.97429981 | 0.329907699 | NA | Elane | elastase, neutrophil expressed [Source:M | 5.6217 | 2.0558 | 0 | 0.964 | 1.0906 | 0 |
| 2331 | ENSMUSG00000020128 | 767.3498885 | 0.015415259 | 0.116513968 | 0.132303958 | 0.894743877 | 0.982798755 | Vps54 | VPS54 GARP complex subunit [Source:MG | 707.53 | 770.94 | 812.04 | 762.51 | 808.15 | 742.93 |
| 2332 | ENSMUSG00000020130 | 1069.861677 | 0.104521106 | 0.108025298 | 0.967561371 | 0.333263477 | 0.77523112 | Tbc1d15 | TBC1 domain family, member 15 [Source: | 1056.9 | 1045.4 | 991.15 | 1147.1 | 1153.9 | 1024.7 |
| 2333 | ENSMUSG00000020131 | 724.8532729 | 0.673132057 | 0.142207884 | 4.733436992 | 2.21E-06 | 0.000156516 | Pcsk4 | proprotein convertase subtilisin/kexin typ | 655.33 | 518.07 | 501.11 | 829.98 | 989.19 | 855.43 |
| 2334 | ENSMUSG00000020132 | 941.1468261 | -0.087436407 | 0.111657333 | -0.783078053 | 0.433581278 | 0.845264432 | Rab21 | RAB21, member RAS oncogene family [So | 917.14 | 1036.1 | 955.93 | 860.83 | 941.2 | 935.63 |
| 2335 | ENSMUSG00000020133 | 364.1462225 | 0.031431399 | 0.138593629 | 0.226788194 | 0.820588444 | 0.969160361 | 2310011J03I | RIKEN cDNA 2310011J03 gene [Source:MC | 367.02 | 334.07 | 379.35 | 374.99 | 371.9 | 357.54 |
| 2336 | ENSMUSG00000020134 | 391.2980424 | -0.018759253 | 0.169590384 | -0.110615077 | 0.911921591 | 0.98589875 | Peli1 | pellino 1 [Source:MGI Symbol;Acc:MGI:19 | 331.68 | 490.32 | 360.24 | 364.38 | 403.53 | 397.64 |
| 2337 | ENSMUSG00000020135 | 7.391824066 | -0.710641248 | 0.797013928 | -0.891629648 | 0.372591474 | 0.808133324 | Apc2 | APC regulator of WNT signaling pathway 2 | 7.2279 | 11.307 | 9.0562 | 5.7839 | 7.6343 | 3.3415 |
| 2338 | ENSMUSG00000020137 | 193.4845081 | 0.108840222 | 0.209342025 | 0.519915778 | 0.603122278 | 0.915859383 | Thap2 | THAP domain containing, apoptosis assoc | 161.42 | 236.42 | 161 | 173.52 | 222.49 | 206.06 |
| 2339 | ENSMUSG00000020140 | 75.95565432 | -0.02022909 | 0.267850298 | -0.075523866 | 0.939797909 | 0.988812319 | Lgr5 | leucine rich repeat containing G protein c | 79.507 | 85.317 | 64.4 | 64.587 | 85.068 | 76.855 |
| 2340 | ENSMUSG00000020142 | 37.5533785 | 0.213072451 | 0.375002318 | 0.56818969 | 0.569906171 | 0.904090127 | Slc1a4 | solute carrier family 1 (glutamate/neutral | 40.958 | 29.81 | 33.206 | 30.847 | 39.262 | 51.237 |
| 2341 | ENSMUSG00000020143 | 33.67347534 | -0.899994674 | 0.444691931 | -2.023861041 | 0.042984451 | 0.286246365 | Dock2 | dedicator of cyto-kinesis 2 [Source:MGI Sy | 42.564 | 58.591 | 30.187 | 15.424 | 19.631 | 35.643 |
| 2342 | ENSMUSG00000020149 | 2779.735574 | -0.095959086 | 0.094671639 | -1.013599072 | 0.310774101 | 0.759553327 | Rab1a | RAB1A, member RAS oncogene family [So | 3040.5 | 2732.2 | 2842.6 | 2653.8 | 2807.3 | 2601.9 |
| 2343 | ENSMUSG00000020150 | 4462.9821 | 0.02442281 | 0.101873247 | 0.23973723 | 0.810533971 | 0.966384503 | Gamt | guanidinoacetate methyltransferase [Sour | 4503.8 | 4394.3 | 4376.2 | 3982.2 | 4747.5 | 4773.9 |
| 2344 | ENSMUSG00000020151 | 0.603115608 | -2.642822926 | 3.106511347 | -0.850736608 | 0.394915683 | NA | Ptprr | protein tyrosine phosphatase, receptor ty | 1.6062 | 0 | 2.0125 | 0 | 0 | 0 |
| 2345 | ENSMUSG00000020152 | 2384.201629 | -0.0296385 | 0.097403473 | -0.304285867 | 0.760910111 | 0.95402891 | Actr2 | ARP2 actin-related protein 2 [Source:MGI | 2350.7 | 2374.5 | 2500.5 | 2206.5 | 2357.9 | 2515.1 |
| 2346 | ENSMUSG00000020153 | 1655.635401 | -0.063844536 | 0.092821234 | -0.687822536 | 0.491564542 | 0.876368691 | Ndufs7 | NADH:ubiquinone oxidoreductase core su | 1693.7 | 1708.4 | 1674.4 | 1583.8 | 1627.2 | 1646.3 |
| 2347 | ENSMUSG00000020154 | 587.6821393 | -0.532528321 | 0.164243569 | -3.242308501 | 0.001185656 | 0.02363684 | Ptprb | protein tyrosine phosphatase, receptor ty | 681.83 | 685.62 | 717.45 | 503.2 | 359.9 | 578.08 |
| 2348 | ENSMUSG00000020155 | 0.486939728 | -0.698472448 | 3.137608162 | -0.222613026 | 0.823836697 | NA | Kcnmb1 | potassium large conductance calcium-acti | 0.8031 | 1.0279 | 0 | 0 | 1.0906 | 0 |

| | | | | | | | | | | | | | | | |
|---|---|---|---|---|---|---|---|---|---|---|---|---|---|---|---|
| 2349 | ENSMUSG00000020156 | 203.8388616 | 0.234275387 | 0.172912065 | 1.354881669 | 0.175455261 | 0.605810563 | Pwwp3a | PWWP domain containing 3A, DNA repair | 179.89 | 191.19 | 191.19 | 229.43 | 196.31 | 235.02 |
| 2350 | ENSMUSG00000020159 | 0.167707431 | -0.796629176 | 4.080472857 | -0.195229623 | 0.84521319 | NA | Gabrp | gamma-aminobutyric acid (GABA) A recep | 0 | 0 | 1.0062 | 0 | 0 | 0 |
| 2351 | ENSMUSG00000020160 | 35.41583332 | -0.107618771 | 0.434669358 | -0.247587663 | 0.804453457 | 0.963771623 | Meis1 | Meis homeobox 1 [Source:MGI Symbol;Ac | 29.715 | 48.312 | 32.2 | 26.991 | 21.812 | 53.465 |
| 2352 | ENSMUSG00000020163 | 2455.666017 | 0.122225137 | 0.091806223 | 1.331338261 | 0.183077734 | 0.616970258 | Uqcr11 | ubiquinol-cytochrome c reductase, compl | 2399.7 | 2387.9 | 2267.1 | 2529.5 | 2481.2 | 2668.8 |
| 2353 | ENSMUSG00000020164 | 0 | NA | NA | NA | NA | NA | Kcnmb4os1 | potassium large conductance calcium-acti | 0 | 0 | 0 | 0 | 0 | 0 |
| 2354 | ENSMUSG00000020166 | 525.3521689 | 0.090702038 | 0.127424032 | 0.711812649 | 0.476580799 | 0.86953366 | Cnot2 | CCR4-NOT transcription complex, subunit | 525.23 | 522.18 | 478.97 | 547.54 | 575.85 | 502.34 |
| 2355 | ENSMUSG00000020167 | 394.5292821 | 0.323226794 | 0.139555165 | 2.316122034 | 0.020551607 | 0.176557791 | Tcf3 | transcription factor 3 [Source:MGI Symbo | 361.4 | 320.71 | 369.29 | 451.14 | 456.97 | 407.67 |
| 2356 | ENSMUSG00000020168 | 0 | NA | NA | NA | NA | NA | Olfr299 | olfactory receptor 299 [Source:MGI Symb | 0 | 0 | 0 | 0 | 0 | 0 |
| 2357 | ENSMUSG00000020169 | 0.167707431 | -0.796629176 | 4.080472857 | -0.195229623 | 0.84521319 | NA | Best3 | bestrophin 3 [Source:MGI Symbol;Acc:MG | 0 | 0 | 1.0062 | 0 | 0 | 0 |
| 2358 | ENSMUSG00000020170 | 307.0755536 | 0.016503253 | 0.157073432 | 0.105067118 | 0.916322551 | 0.986374671 | Frs2 | fibroblast growth factor receptor substrat | 276.27 | 345.38 | 294.83 | 295.94 | 332.64 | 297.4 |
| 2359 | ENSMUSG00000020171 | 379.2856796 | 0.31065579 | 0.148240787 | 2.095616171 | 0.036116254 | 0.255945856 | Yeats4 | YEATS domain containing 4 [Source:MGI S | 338.91 | 347.44 | 329.04 | 368.24 | 475.51 | 416.58 |
| 2360 | ENSMUSG00000020173 | 320.3990663 | 0.240652708 | 0.213781965 | 1.12569228 | 0.260295793 | 0.713728596 | Cobl | cordon-bleu WH2 repeat [Source:MGI Syr | 338.11 | 272.4 | 269.67 | 246.78 | 463.51 | 331.93 |
| 2361 | ENSMUSG00000020175 | 76.31564028 | 0.686029232 | 0.271657828 | 2.525343137 | 0.011558537 | 0.119930185 | Rab36 | RAB36, member RAS oncogene family [So | 50.595 | 65.787 | 59.368 | 86.758 | 82.887 | 112.5 |
| 2362 | ENSMUSG00000020176 | 32.94793346 | -0.042033356 | 0.405135841 | -0.103751265 | 0.917366742 | 0.986374671 | Grb10 | growth factor receptor bound protein 10 [ | 32.124 | 41.117 | 27.169 | 37.595 | 21.812 | 37.871 |
| 2363 | ENSMUSG00000020177 | 0 | NA | NA | NA | NA | NA | 9530003J23I | RIKEN cDNA 9530003J23 gene [Source:MC | 0 | 0 | 0 | 0 | 0 | 0 |
| 2364 | ENSMUSG00000020178 | 25.64163134 | -1.718341481 | 0.519569955 | -3.307237968 | 0.000942208 | 0.020070837 | Adora2a | adenosine A2a receptor [Source:MGI Sym | 45.777 | 30.838 | 41.256 | 11.568 | 4.3625 | 20.049 |
| 2365 | ENSMUSG00000020180 | 694.1474008 | 0.143342346 | 0.133281838 | 1.075482963 | 0.282158556 | 0.735230549 | Snrpd3 | small nuclear ribonucleoprotein D3 [Sourc | 613.57 | 669.17 | 696.32 | 653.58 | 699.09 | 833.16 |
| 2366 | ENSMUSG00000020181 | 0 | NA | NA | NA | NA | NA | Nav3 | neuron navigator 3 [Source:MGI Symbol;A | 0 | 0 | 0 | 0 | 0 | 0 |
| 2367 | ENSMUSG00000020182 | 734.3532511 | -0.592382437 | 0.170959683 | -3.465041737 | 0.000530149 | 0.012909222 | Ddc | dopa decarboxylase [Source:MGI Symbol; | 701.91 | 1062.9 | 884.49 | 481.02 | 605.29 | 670.53 |
| 2368 | ENSMUSG00000020183 | 60.48511814 | -0.423015132 | 0.329218628 | -1.284906431 | 0.198824984 | 0.640976045 | Cpm | carboxypeptidase M [Source:MGI Symbol | 56.217 | 75.038 | 76.475 | 32.775 | 57.803 | 64.603 |
| 2369 | ENSMUSG00000020184 | 483.2020556 | 0.096299042 | 0.133377727 | 0.722002426 | 0.470292986 | 0.867104619 | Mdm2 | transformed mouse 3T3 cell double minut | 483.47 | 488.26 | 429.67 | 539.83 | 466.78 | 491.21 |
| 2370 | ENSMUSG00000020185 | 7.098939045 | 0.343386733 | 0.990067113 | 0.346831774 | 0.728717705 | 0.946248466 | E2f7 | E2F transcription factor 7 [Source:MGI Syr | 9.6372 | 0 | 9.0562 | 10.604 | 3.2719 | 10.025 |
| 2371 | ENSMUSG00000020186 | 445.8322767 | -0.034149746 | 0.151693023 | -0.225124039 | 0.82188278 | 0.969160361 | Csrp2 | cysteine and glycine-rich protein 2 [Sourc | 392.72 | 467.7 | 494.07 | 481.02 | 382.81 | 456.68 |
| 2372 | ENSMUSG00000020189 | 744.2799279 | 0.151583466 | 0.110159656 | 1.376034306 | 0.168811013 | 0.595822959 | Osbpl8 | oxysterol binding protein-like 8 [Source:M | 698.7 | 722.63 | 694.31 | 764.43 | 808.15 | 777.46 |
| 2373 | ENSMUSG00000020190 | 3290.003964 | -0.100739699 | 0.13376711 | -0.753097672 | 0.451391223 | 0.85613987 | Mknk2 | MAP kinase-interacting serine/threonine k | 3822.8 | 3345.9 | 3043.9 | 2608.5 | 3462.7 | 3456.3 |
| 2374 | ENSMUSG00000020191 | 0 | NA | NA | NA | NA | NA | Spata48 | spermatogenesis associated 48 [Source:M | 0 | 0 | 0 | 0 | 0 | 0 |
| 2375 | ENSMUSG00000020193 | 1.331589285 | 1.899580413 | 2.208043411 | 0.860300302 | 0.389623527 | NA | Zpbp | zona pellucida binding protein [Source:MG | 1.6062 | 0 | 0 | 1.928 | 0 | 4.4554 |
| 2376 | ENSMUSG00000020196 | 454.8823076 | -0.009599864 | 0.133305417 | -0.072014055 | 0.942590723 | 0.989952964 | Cabin1 | calcineurin binding protein 1 [Source:MGI | 469.01 | 445.09 | 454.82 | 445.36 | 497.32 | 417.69 |
| 2377 | ENSMUSG00000020198 | 1104.126044 | 0.001549302 | 0.108960226 | 0.01421897 | 0.988655286 | 0.99859176 | Ap3d1 | adaptor-related protein complex 3, delta 1 | 1044 | 1131.7 | 1136.1 | 1170.3 | 1115.7 | 1027 |
| 2378 | ENSMUSG00000020205 | 1300.140178 | 1.457567546 | 0.788612572 | 1.848268208 | 0.064563553 | 0.364013257 | Phlda1 | pleckstrin homology like domain, family A | 385.49 | 970.35 | 726.51 | 2105.3 | 477.69 | 3135.5 |
| 2379 | ENSMUSG00000020211 | 287.6229571 | -0.181804907 | 0.167252962 | -1.087005602 | 0.277034334 | 0.730041765 | Sf3a2 | splicing factor 3a, subunit 2 [Source:MGI S | 352.56 | 282.68 | 280.74 | 240.03 | 287.92 | 281.8 |
| 2380 | ENSMUSG00000020212 | 18.5551145 | 0.310385401 | 0.500756728 | 0.619832713 | 0.535367929 | 0.889860001 | Mdm1 | transformed mouse 3T3 cell double minut | 16.062 | 17.475 | 16.1 | 15.424 | 23.994 | 22.277 |
| 2381 | ENSMUSG00000020213 | 0 | NA | NA | NA | NA | NA | Glipr1l1 | GLI pathogenesis-related 1 like 1 [Source:I | 0 | 0 | 0 | 0 | 0 | 0 |
| 2382 | ENSMUSG00000020214 | 0 | NA | NA | NA | NA | NA | Glipr1l2 | GLI pathogenesis-related 1 like 2 [Source:I | 0 | 0 | 0 | 0 | 0 | 0 |
| 2383 | ENSMUSG00000020216 | 0.181769715 | 1.126924175 | 4.080472857 | 0.276174898 | 0.782413736 | NA | Jsrp1 | junctional sarcoplasmic reticulum protein | 0 | 0 | 0 | 0 | 1.0906 | 0 |
| 2384 | ENSMUSG00000020218 | 9.870969338 | 1.333984002 | 0.828374505 | 1.610363421 | 0.107318541 | 0.48315751 | Wif1 | Wnt inhibitory factor 1 [Source:MGI Symb | 5.6217 | 2.0558 | 9.0562 | 3.8559 | 16.359 | 22.277 |
| 2385 | ENSMUSG00000020219 | 1298.139586 | 0.063541352 | 0.106189882 | 0.598374825 | 0.54958986 | 0.895703112 | Timm13 | translocase of inner mitochondrial membr | 1268.9 | 1279.8 | 1259.8 | 1311 | 1228 | 1441.3 |
| 2386 | ENSMUSG00000020220 | 1711.530349 | -0.120390296 | 0.110021995 | -1.094238438 | 0.273850417 | 0.728923893 | Vps13d | vacuolar protein sorting 13D [Source:MGI | 1829.5 | 1853.3 | 1666.3 | 1759.3 | 1688.3 | 1472.5 |
| 2387 | ENSMUSG00000020224 | 341.0171072 | -0.1739975 | 0.146135503 | -1.190658645 | 0.233787619 | 0.681342761 | Llph | LLP homolog, long-term synaptic facilitatio | 357.38 | 383.41 | 344.14 | 325.82 | 344.64 | 290.71 |
| 2388 | ENSMUSG00000020225 | 1064.097765 | -0.005241974 | 0.112790391 | -0.046475358 | 0.962931375 | 0.992956886 | Tmbim4 | transmembrane BAX inhibitor motif conta | 1081 | 1118.4 | 999.2 | 1131.7 | 1079.7 | 974.61 |
| 2389 | ENSMUSG00000020226 | 0.167707431 | -0.796629176 | 4.080472857 | -0.195229623 | 0.84521319 | NA | Slc5a4b | solute carrier family 5 (neutral amino acid | 0 | 0 | 1.0062 | 0 | 0 | 0 |
| 2390 | ENSMUSG00000020227 | 14.27386545 | -1.624228 | 0.694589255 | -2.338400699 | 0.019366472 | 0.1693571 | Irak3 | interleukin-1 receptor-associated kinase 3 | 21.684 | 35.977 | 7.0437 | 7.7118 | 6.5437 | 6.6831 |
| 2391 | ENSMUSG00000020228 | 252.8543181 | 0.207821484 | 0.162749796 | 1.276938526 | 0.201623951 | 0.645999739 | Helb | helicase (DNA) B [Source:MGI Symbol;Acc | 219.25 | 261.09 | 224.39 | 291.12 | 261.75 | 259.53 |
| 2392 | ENSMUSG00000020229 | 0 | NA | NA | NA | NA | NA | Slc5a4a | solute carrier family 5, member 4a [Sourc | 0 | 0 | 0 | 0 | 0 | 0 |
| 2393 | ENSMUSG00000020230 | 5.099977499 | -0.928810325 | 0.958435034 | -0.969090539 | 0.332500018 | 0.77523112 | Prmt2 | protein arginine N-methyltransferase 2 [S | 4.8186 | 9.2513 | 6.0375 | 3.8559 | 2.1812 | 4.4554 |
| 2394 | ENSMUSG00000020231 | 54.56223788 | -0.252682891 | 0.317561165 | -0.795698337 | 0.426207394 | 0.841707228 | Dip2a | disco interacting protein 2 homolog A [So | 70.673 | 54.48 | 52.325 | 38.559 | 54.531 | 56.806 |
| 2395 | ENSMUSG00000020232 | 684.0842468 | 0.048214745 | 0.132771006 | 0.363142121 | 0.716498721 | 0.9447719 | Hmg20b | high mobility group 20B [Source:MGI Sym | 722.79 | 630.11 | 665.13 | 786.61 | 679.46 | 620.41 |
| 2396 | ENSMUSG00000020234 | 0.6621303 | -1.329905799 | 2.807279375 | -0.473734752 | 0.635689068 | NA | 4930404N11 | RIKEN cDNA 4930404N11 gene [Source:M | 0.8031 | 2.0558 | 0 | 0 | 0 | 1.1138 |
| 2397 | ENSMUSG00000020235 | 258.2582407 | 0.218880287 | 0.16204772 | 1.350715008 | 0.176786743 | 0.607589534 | Fzr1 | fizzy and cell division cycle 20 related 1 [S | 262.61 | 244.64 | 208.29 | 278.59 | 280.29 | 275.12 |
| 2398 | ENSMUSG00000020238 | 1101.589616 | -0.067605134 | 0.113448676 | -0.595909414 | 0.551235744 | 0.89589637 | Ncln | nicalin [Source:MGI Symbol;Acc:MGI:192 | 1017.5 | 1164.6 | 1201.5 | 1125.9 | 1027.4 | 1072.6 |
| 2399 | ENSMUSG00000020241 | 60.74964467 | -0.775950814 | 0.343136025 | -2.261350477 | 0.023737563 | 0.19563539 | Col6a2 | collagen, type VI, alpha 2 [Source:MGI Syr | 108.42 | 71.954 | 49.306 | 44.343 | 40.353 | 50.123 |
| 2400 | ENSMUSG00000020246 | 206.2788155 | 0.22076992 | 0.180621257 | 1.222280937 | 0.221601405 | 0.667949912 | Hcfc2 | host cell factor C2 [Source:MGI Symbol;Ac | 172.67 | 202.5 | 197.22 | 250.63 | 195.22 | 219.43 |
| 2401 | ENSMUSG00000020248 | 547.1021243 | 0.259045858 | 0.129689993 | 1.997423644 | 0.045779181 | 0.296229283 | Nfyb | nuclear transcription factor-Y beta [Sourc | 513.18 | 493.4 | 488.03 | 661.29 | 546.4 | 580.31 |
| 2402 | ENSMUSG00000020250 | 2581.63277 | 0.098622648 | 0.088493367 | 1.114463731 | 0.265080296 | 0.720853222 | Txnrd1 | thioredoxin reductase 1 [Source:MGI Sym | 2435.8 | 2546.2 | 2498.5 | 2609.5 | 2670.9 | 2728.9 |
| 2403 | ENSMUSG00000020251 | 3.917019696 | -0.549126324 | 1.211401509 | -0.453298366 | 0.650333909 | NA | Glt8d2 | glycosyltransferase 8 domain containing 2 | 4.8186 | 3.0838 | 6.0375 | 1.928 | 7.6343 | 0 |
| 2404 | ENSMUSG00000020253 | 407.5193631 | -0.112447426 | 0.135700523 | -0.828644015 | 0.407305874 | 0.83094725 | Ppm1m | protein phosphatase 1M [Source:MGI Syn | 406.37 | 453.31 | 410.55 | 368.24 | 407.89 | 398.76 |

| | | | | | | | | | | | | | | | |
|---|---|---|---|---|---|---|---|---|---|---|---|---|---|---|---|
| 2405 | ENSMUSG00000020255 | 291.0779653 | 0.214945298 | 0.151265641 | 1.42097899 | 0.155322867 | 0.574114264 | D10Wsu102 | DNA segment, Chr 10, Wayne State Unive | 281.89 | 261.09 | 264.64 | 286.3 | 321.73 | 330.81 |
| 2406 | ENSMUSG00000020256 | 3.925020286 | -0.54222307 | 1.091317356 | -0.496851871 | 0.619293513 | NA | Aldh1l2 | aldehyde dehydrogenase 1 family, membe | 4.8186 | 2.0558 | 7.0437 | 1.928 | 4.3625 | 3.3415 |
| 2407 | ENSMUSG00000020257 | 972.2081501 | -0.045802301 | 0.112408151 | -0.407464233 | 0.683667059 | 0.935528319 | Wdr82 | WD repeat domain containing 82 [Source: | 924.37 | 1068 | 971.03 | 912.89 | 996.83 | 960.13 |
| 2408 | ENSMUSG00000020258 | 3449.985333 | -0.027300724 | 0.102915181 | -0.265274023 | 0.790798374 | 0.961019028 | Glyctk | glycerate kinase [Source:MGI Symbol;Acc: | 3557.7 | 3222.5 | 3666.8 | 3209.1 | 3720.1 | 3323.7 |
| 2409 | ENSMUSG00000020260 | 292.3874556 | 0.16604904 | 0.156370671 | 1.061893762 | 0.288283917 | 0.73951397 | Pofut2 | protein O-fucosyltransferase 2 [Source:M | 268.24 | 300.15 | 258.6 | 316.18 | 278.11 | 333.04 |
| 2410 | ENSMUSG00000020261 | 748.2294465 | 0.343922837 | 0.117230204 | 2.933739141 | 0.003349055 | 0.050564445 | Slc36a1 | solute carrier family 36 (proton/amino aci | 616.78 | 698.98 | 663.12 | 797.21 | 827.78 | 885.51 |
| 2411 | ENSMUSG00000020262 | 340.9332343 | 0.193083687 | 0.160797182 | 1.200790242 | 0.229832578 | 0.676539688 | Adarb1 | adenosine deaminase, RNA-specific, B1 [S | 344.53 | 261.09 | 348.16 | 359.56 | 344.64 | 387.62 |
| 2412 | ENSMUSG00000020263 | 404.7356551 | -0.251616217 | 0.135862929 | -1.851985804 | 0.064027861 | 0.362159575 | Appl2 | adaptor protein, phosphotyrosine interac | 428.86 | 469.76 | 421.62 | 377.88 | 383.9 | 346.41 |
| 2413 | ENSMUSG00000020264 | 0 | NA | NA | NA | NA | NA | Slc36a2 | solute carrier family 36 (proton/amino aci | 0 | 0 | 0 | 0 | 0 | 0 |
| 2414 | ENSMUSG00000020265 | 933.0431675 | -0.013362878 | 0.132254356 | -0.10103923 | 0.919519319 | 0.986685804 | Sumo3 | small ubiquitin-like modifier 3 [Source:MG | 819.16 | 955.96 | 1037.4 | 842.52 | 911.76 | 1031.4 |
| 2415 | ENSMUSG00000020267 | 2903.802701 | 0.077562224 | 0.08824517 | 0.878940156 | 0.379433724 | 0.81257468 | Hint1 | histidine triad nucleotide binding protein | 2920.9 | 2731.2 | 2824.5 | 2988.3 | 2931.6 | 3026.3 |
| 2416 | ENSMUSG00000020268 | 38.23918554 | 0.015293417 | 0.35738275 | 0.042792824 | 0.965866684 | 0.993761255 | Lyrm7 | LYR motif containing 7 [Source:MGI Symb | 33.73 | 45.228 | 35.219 | 33.739 | 42.534 | 38.985 |
| 2417 | ENSMUSG00000020270 | 0 | NA | NA | NA | NA | NA | Smim23 | small integral membrane protein 23 [Sour | 0 | 0 | 0 | 0 | 0 | 0 |
| 2418 | ENSMUSG00000020271 | 567.1579658 | 0.106077763 | 0.126757786 | 0.836854021 | 0.402674628 | 0.828953151 | Fbxw11 | F-box and WD-40 domain protein 11 [Sou | 566.19 | 504.71 | 567.52 | 582.24 | 551.85 | 630.44 |
| 2419 | ENSMUSG00000020272 | 46.15862209 | -0.946001859 | 0.379657108 | -2.491726977 | 0.012712371 | 0.127461989 | Stk10 | serine/threonine kinase 10 [Source:MGI S | 65.854 | 81.206 | 35.219 | 31.811 | 29.447 | 33.415 |
| 2420 | ENSMUSG00000020273 | 102.0246131 | -0.020186868 | 0.240252527 | -0.084023539 | 0.933037717 | 0.988420305 | Papolg | poly(A) polymerase gamma [Source:MGI S | 81.916 | 113.07 | 113.71 | 96.398 | 110.15 | 96.904 |
| 2421 | ENSMUSG00000020275 | 76.63703523 | -0.266406411 | 0.255181617 | -1.043987468 | 0.296491188 | 0.74713095 | Rel | reticuloendotheliosis oncogene [Source:M | 81.113 | 91.485 | 78.487 | 66.514 | 68.709 | 73.514 |
| 2422 | ENSMUSG00000020277 | 739.9218521 | -0.145695965 | 0.119050376 | -1.22381776 | 0.221020983 | 0.667400313 | Pfkl | phosphofructokinase, liver, B-type [Sourc | 724.4 | 789.44 | 819.08 | 747.08 | 702.36 | 657.17 |
| 2423 | ENSMUSG00000020279 | 0.965523032 | -0.944628791 | 2.267086424 | -0.416670834 | 0.67691919 | NA | Il9r | interleukin 9 receptor [Source:MGI Symbo | 0.8031 | 2.0558 | 1.0062 | 1.928 | 0 | 0 |
| 2424 | ENSMUSG00000020280 | 392.424834 | -0.06814918 | 0.154029771 | -0.442441611 | 0.658169674 | 0.927303282 | Pus10 | pseudouridylate synthase 10 [Source:MGI | 428.05 | 439.95 | 337.09 | 414.51 | 375.17 | 359.77 |
| 2425 | ENSMUSG00000020282 | 141.3885595 | -0.198057505 | 0.228402383 | -0.867142898 | 0.385863719 | 0.816790989 | Rhbdf1 | rhomboid 5 homolog 1 [Source:MGI Symb | 178.29 | 142.88 | 131.82 | 151.34 | 101.43 | 142.57 |
| 2426 | ENSMUSG00000020283 | 1151.898188 | -0.019208433 | 0.122618934 | -0.156651443 | 0.875519557 | 0.980527671 | Pex13 | peroxisomal biogenesis factor 13 [Source: | 1316.3 | 1138.9 | 1022.3 | 1164.5 | 1182.2 | 1087.1 |
| 2427 | ENSMUSG00000020284 | 129.9702017 | 0.0665018 | 0.229908324 | 0.289253554 | 0.772387353 | 0.957706548 | Cfap410 | cilia and flagella associated protein 410 [S | 134.92 | 146.99 | 98.612 | 112.79 | 145.05 | 141.46 |
| 2428 | ENSMUSG00000020286 | 0.589026695 | 0.241472922 | 3.228838524 | 0.074786311 | 0.940384733 | NA | 1700093K21 | RIKEN cDNA 1700093K21 gene [Source:M | 1.6062 | 0 | 0 | 1.928 | 0 | 0 |
| 2429 | ENSMUSG00000020287 | 92.60974493 | 0.187825674 | 0.250166518 | 0.750802608 | 0.452771459 | 0.856769845 | Mpg | N-methylpurine-DNA glycosylase [Source: | 84.326 | 77.094 | 98.612 | 111.82 | 103.61 | 80.197 |
| 2430 | ENSMUSG00000020288 | 352.0221562 | 0.181980331 | 0.152772848 | 1.191182424 | 0.233581978 | 0.681057972 | Ahsa2 | AHA1, activator of heat shock protein ATP | 307.59 | 309.4 | 373.32 | 409.69 | 347.91 | 364.23 |
| 2431 | ENSMUSG00000020289 | 253.3864661 | 0.109462646 | 0.181253658 | 0.603919653 | 0.545897062 | 0.89549457 | Nprl3 | nitrogen permease regulator-like 3 [Sourc | 260.21 | 206.61 | 264.64 | 308.47 | 246.48 | 233.91 |
| 2432 | ENSMUSG00000020290 | 583.1656598 | 0.19618638 | 0.13003206 | 1.508753916 | 0.131361682 | 0.531175668 | Xpo1 | exportin 1 [Source:MGI Symbol;Acc:MGI:2 | 575.02 | 487.23 | 567.52 | 581.28 | 664.19 | 623.75 |
| 2433 | ENSMUSG00000020295 | 0.6996517 | 1.656076801 | 2.758183935 | 0.600422901 | 0.548224429 | NA | Hbq1a | hemoglobin, theta 1A [Source:MGI Symbo | 0 | 0 | 1.0062 | 0.964 | 0 | 2.2277 |
| 2434 | ENSMUSG00000020297 | 0.167707431 | -0.796629176 | 4.080472857 | -0.195229623 | 0.84521319 | NA | Nsg2 | neuron specific gene family member 2 [So | 0 | 0 | 1.0062 | 0 | 0 | 0 |
| 2435 | ENSMUSG00000020299 | 0 | NA | NA | NA | NA | NA | 4930524B15 | RIKEN cDNA 4930524B15 gene [Source:M | 0 | 0 | 0 | 0 | 0 | 0 |
| 2436 | ENSMUSG00000020300 | 957.9509789 | -0.047519733 | 0.139902806 | -0.339662476 | 0.734110723 | 0.948109711 | Cpeb4 | cytoplasmic polyadenylation element binc | 847.27 | 990.91 | 1084.7 | 1091.2 | 869.22 | 864.34 |
| 2437 | ENSMUSG00000020303 | 4.100643801 | -0.645104964 | 1.283622604 | -0.502565912 | 0.615269501 | NA | Stc2 | stanniocalcin 2 [Source:MGI Symbol;Acc:M | 8.8341 | 2.0558 | 4.025 | 0.964 | 8.7249 | 0 |
| 2438 | ENSMUSG00000020305 | 127.1200239 | 0.237415022 | 0.23070882 | 1.02906782 | 0.303447807 | 0.752690022 | Asb3 | ankyrin repeat and SOCS box-containing 3 | 120.47 | 131.57 | 97.606 | 108.93 | 153.78 | 150.37 |
| 2439 | ENSMUSG00000020307 | 1208.838252 | -0.078317497 | 0.115293904 | -0.679285671 | 0.496956874 | 0.878676728 | Cdc34 | cell division cycle 34 [Source:MGI Symbol; | 1376.5 | 1157.4 | 1189.4 | 1125.9 | 1146.2 | 1257.5 |
| 2440 | ENSMUSG00000020308 | 476.8413971 | 0.097376508 | 0.132205433 | 0.736554508 | 0.461393309 | 0.860071824 | Tpgs1 | tubulin polyglutamylase complex subunit | 483.47 | 443.03 | 455.83 | 539.83 | 474.42 | 464.47 |
| 2441 | ENSMUSG00000020309 | 534.3247427 | 0.252532691 | 0.131010326 | 1.927578532 | 0.053907571 | 0.326515541 | Chac2 | ChaC, cation transport regulator 2 [Source | 497.12 | 443.03 | 522.24 | 537.9 | 605.29 | 600.36 |
| 2442 | ENSMUSG00000020310 | 0 | NA | NA | NA | NA | NA | Madcam1 | mucosal vascular addressin cell adhesion | 0 | 0 | 0 | 0 | 0 | 0 |
| 2443 | ENSMUSG00000020311 | 471.2861891 | 0.033020975 | 0.130127972 | 0.253757699 | 0.799682755 | 0.962424862 | Erlec1 | endoplasmic reticulum lectin 1 [Source:M | 432.87 | 496.48 | 468.91 | 466.57 | 498.41 | 464.47 |
| 2444 | ENSMUSG00000020312 | 2.063087013 | 0.905842508 | 1.582303304 | 0.572483484 | 0.566994472 | NA | Shc2 | SHC (Src homology 2 domain containing) t | 3.2124 | 1.0279 | 0 | 4.8199 | 1.0906 | 2.2277 |
| 2445 | ENSMUSG00000020315 | 1231.823932 | -0.234390406 | 0.114075918 | -2.054687882 | 0.039909161 | 0.273576354 | Sptbn1 | spectrin beta, non-erythrocytic 1 [Source: | 1356.4 | 1443.2 | 1195.4 | 1143.3 | 1183.3 | 1069.3 |
| 2446 | ENSMUSG00000020317 | 0 | NA | NA | NA | NA | NA | Theg | testicular haploid expressed gene [Source | 0 | 0 | 0 | 0 | 0 | 0 |
| 2447 | ENSMUSG00000020319 | 37.51994617 | 0.282146871 | 0.355373562 | 0.793944461 | 0.427227765 | 0.842342201 | Wdpcp | WD repeat containing planar cell polarity | 35.336 | 33.921 | 32.2 | 37.595 | 38.172 | 47.895 |
| 2448 | ENSMUSG00000020321 | 12896.61698 | 0.03385846 | 0.09077635 | 0.372987677 | 0.709157615 | 0.944237849 | Mdh1 | malate dehydrogenase 1, NAD (soluble) [S | 12907 | 12659 | 12669 | 12109 | 14065 | 12971 |
| 2449 | ENSMUSG00000020323 | 0 | NA | NA | NA | NA | NA | Prss57 | protease, serine 57 [Source:MGI Symbol;A | 0 | 0 | 0 | 0 | 0 | 0 |
| 2450 | ENSMUSG00000020325 | 1.002812338 | -2.141638232 | 2.477191155 | -0.864542983 | 0.387289679 | NA | Fstl3 | follistatin-like 3 [Source:MGI Symbol;Acc:I | 0 | 1.0279 | 4.025 | 0.964 | 0 | 0 |
| 2451 | ENSMUSG00000020326 | 767.7977668 | -0.04479896 | 0.121572993 | -0.368494346 | 0.71250466 | 0.944239215 | Ccng1 | cyclin G1 [Source:MGI Symbol;Acc:MGI:1( | 838.44 | 689.73 | 810.03 | 744.19 | 753.62 | 770.78 |
| 2452 | ENSMUSG00000020327 | 0 | NA | NA | NA | NA | NA | Fgf22 | fibroblast growth factor 22 [Source:MGI S | 0 | 0 | 0 | 0 | 0 | 0 |
| 2453 | ENSMUSG00000020328 | 570.562648 | -0.045009638 | 0.129455873 | -0.347683249 | 0.728078078 | 0.946158959 | Nudcd2 | NudC domain containing 2 [Source:MGI S | 586.26 | 591.05 | 560.48 | 507.05 | 623.83 | 554.69 |
| 2454 | ENSMUSG00000020329 | 477.9919821 | -0.108523337 | 0.136768725 | -0.793480654 | 0.427497837 | 0.842415355 | Polrmt | polymerase (RNA) mitochondrial (DNA dir | 480.26 | 513.96 | 494.07 | 462.71 | 510.41 | 406.55 |
| 2455 | ENSMUSG00000020330 | 5.35498392 | -0.539632682 | 1.150225465 | -0.46915383 | 0.638959685 | 0.923932929 | Hmmr | hyaluronan mediated motility receptor (R | 8.8341 | 9.2513 | 1.0062 | 8.6758 | 4.3625 | 0 |
| 2456 | ENSMUSG00000020331 | 9.020971866 | -1.142133666 | 0.729483324 | -1.565674811 | 0.117424766 | 0.502643331 | Hcn2 | hyperpolarization-activated, cyclic nucleo | 12.85 | 10.279 | 14.087 | 4.8199 | 7.6343 | 4.4554 |
| 2457 | ENSMUSG00000020332 | 0 | NA | NA | NA | NA | NA | Meikin | meiotic kinetochore factor [Source:MGI S | 0 | 0 | 0 | 0 | 0 | 0 |
| 2458 | ENSMUSG00000020333 | 1.758971897 | 2.130408155 | 1.826993604 | 1.166073133 | 0.243584878 | NA | Acsl6 | acyl-CoA synthetase long-chain family me | 0 | 0 | 2.0125 | 1.928 | 3.2719 | 3.3415 |
| 2459 | ENSMUSG00000020334 | 86.78458738 | 0.474994853 | 0.294816312 | 1.611155264 | 0.107145886 | 0.483091674 | Slc22a4 | solute carrier family 22 (organic cation tra | 65.854 | 50.368 | 101.63 | 87.722 | 97.065 | 118.07 |
| 2460 | ENSMUSG00000020335 | 74.51388049 | -0.161303333 | 0.276174609 | -0.584062866 | 0.559178017 | 0.899423311 | Zfp354b | zinc finger protein 354B [Source:MGI Sym | 73.885 | 69.898 | 92.575 | 81.938 | 71.981 | 56.806 |

| | | | | | | | | | | | | | | | |
|---|---|---|---|---|---|---|---|---|---|---|---|---|---|---|---|
| 2461 | ENSMUSG00000020340 | 18.42693587 | -0.558776916 | 0.509895752 | -1.095865015 | 0.273137851 | 0.728320076 | Cyfip2 | cytoplasmic FMR1 interacting protein 2 [S | 23.29 | 18.503 | 24.15 | 19.28 | 13.087 | 12.252 |
| 2462 | ENSMUSG00000020346 | 1510.308321 | 0.139766102 | 0.110857151 | 1.260776599 | 0.207389347 | 0.652364421 | Mgat1 | mannoside acetylglucosaminyltransferase | 1550.8 | 1307.5 | 1452 | 1550.1 | 1504 | 1697.5 |
| 2463 | ENSMUSG00000020349 | 2246.742129 | -0.114389404 | 0.100535257 | -1.137803869 | 0.255202389 | 0.708637501 | Ppp2ca | protein phosphatase 2 (formerly 2A), cata | 2373.2 | 2467 | 2166.4 | 2035 | 2239 | 2199.8 |
| 2464 | ENSMUSG00000020354 | 1.42753722 | 0.459331161 | 1.934778185 | 0.23740766 | 0.812340548 | NA | Sgcd | sarcoglycan, delta (dystrophin-associated | 1.6062 | 0 | 2.0125 | 3.8559 | 1.0906 | 0 |
| 2465 | ENSMUSG00000020357 | 203.5400389 | -0.647642815 | 0.241455199 | -2.682248378 | 0.007312914 | 0.086502219 | Flt4 | FMS-like tyrosine kinase 4 [Source:MGI Sy | 228.88 | 295.01 | 221.37 | 139.78 | 113.42 | 222.77 |
| 2466 | ENSMUSG00000020358 | 2379.114969 | 0.057524911 | 0.091384278 | 0.629483677 | 0.529032452 | 0.888334265 | Hnrnpab | heterogeneous nuclear ribonucleoprotein | 2258.3 | 2335.4 | 2401.9 | 2421.5 | 2508.4 | 2349.1 |
| 2467 | ENSMUSG00000020359 | 297.208076 | 0.007605775 | 0.173003969 | 0.04396301 | 0.964933889 | 0.993480943 | Phykpl | 5-phosphohydroxy-L-lysine phospholyase | 338.91 | 272.4 | 277.72 | 339.32 | 304.28 | 250.61 |
| 2468 | ENSMUSG00000020361 | 2322.732403 | 0.081038278 | 0.108888696 | 0.744230397 | 0.456737101 | 0.85828835 | Hspa4 | heat shock protein 4 [Source:MGI Symbol | 2349.1 | 2224.4 | 2197.6 | 2141 | 2366.6 | 2657.6 |
| 2469 | ENSMUSG00000020362 | 723.6231389 | 0.040654132 | 0.117476195 | 0.346062719 | 0.72929558 | 0.946534338 | Cnot6 | CCR4-NOT transcription complex, subunit | 660.15 | 761.69 | 719.46 | 746.12 | 749.25 | 705.06 |
| 2470 | ENSMUSG00000020363 | 3.731436396 | -2.11750837 | 1.248170974 | -1.696489034 | 0.089793305 | NA | Gfpt2 | glutamine fructose-6-phosphate transami | 4.0155 | 6.1675 | 8.05 | 1.928 | 0 | 2.2277 |
| 2471 | ENSMUSG00000020364 | 88.12833805 | 0.116641833 | 0.241831422 | 0.482327038 | 0.629573643 | 0.92057725 | Zfp354a | zinc finger protein 354A [Source:MGI Sym | 90.751 | 86.345 | 76.475 | 93.506 | 98.156 | 83.538 |
| 2472 | ENSMUSG00000020366 | 990.3492435 | -0.044029579 | 0.105315628 | -0.418072603 | 0.675894035 | 0.93265897 | Mapk9 | mitogen-activated protein kinase 9 [Sourc | 999.06 | 1042.3 | 975.05 | 959.16 | 1018.6 | 947.88 |
| 2473 | ENSMUSG00000020368 | 4863.0243 | 0.259993948 | 0.090172162 | 2.883306135 | 0.003935248 | 0.056579946 | Canx | calnexin [Source:MGI Symbol;Acc:MGI:88 | 4333.5 | 4671.9 | 4273.5 | 5514 | 5084.5 | 5300.8 |
| 2474 | ENSMUSG00000020372 | 4532.706492 | 0.085988679 | 0.094016539 | 0.91461225 | 0.360395225 | 0.799625179 | Rack1 | receptor for activated C kinase 1 [Source:I | 4062.1 | 4539.3 | 4592.5 | 4482.5 | 4714.7 | 4805.1 |
| 2475 | ENSMUSG00000020374 | 0 | NA | NA | NA | NA | NA | Rasgef1c | RasGEF domain family, member 1C [Sourc | 0 | 0 | 0 | 0 | 0 | 0 |
| 2476 | ENSMUSG00000020375 | 463.7034791 | -0.027076081 | 0.127558877 | -0.212263404 | 0.831901544 | 0.970533397 | Rufy1 | RUN and FYVE domain containing 1 [Sourc | 496.32 | 463.59 | 443.75 | 462.71 | 458.06 | 457.79 |
| 2477 | ENSMUSG00000020376 | 1351.88467 | 0.01009835 | 0.103303754 | 0.097753948 | 0.922127677 | 0.986685804 | Rnf130 | ring finger protein 130 [Source:MGI Symb | 1323.5 | 1340.4 | 1377.5 | 1338 | 1274.9 | 1456.9 |
| 2478 | ENSMUSG00000020377 | 6.53837299 | 0.087946071 | 0.859211251 | 0.10235675 | 0.918473512 | 0.986374671 | Ltc4s | leukotriene C4 synthase [Source:MGI Sym | 4.8186 | 6.1675 | 8.05 | 5.7839 | 3.2719 | 11.138 |
| 2479 | ENSMUSG00000020380 | 165.5847129 | 0.296119608 | 0.184119495 | 1.608301214 | 0.107769225 | 0.484241711 | Rad50 | RAD50 double strand break repair protein | 146.97 | 141.85 | 156.97 | 169.66 | 187.59 | 190.47 |
| 2480 | ENSMUSG00000020381 | 47.4642943 | 0.454884022 | 0.383193057 | 1.187088372 | 0.235192764 | 0.683240713 | Mrnip | MRN complex interacting protein [Source | 37.746 | 38.033 | 44.275 | 46.271 | 33.809 | 84.652 |
| 2481 | ENSMUSG00000020383 | 0 | NA | NA | NA | NA | NA | Il13 | interleukin 13 [Source:MGI Symbol;Acc:M | 0 | 0 | 0 | 0 | 0 | 0 |
| 2482 | ENSMUSG00000020385 | 615.4783511 | 0.247522671 | 0.165377552 | 1.496712627 | 0.134468049 | 0.537574866 | Clk4 | CDC like kinase 4 [Source:MGI Symbol;Acc | 624.01 | 571.52 | 493.06 | 818.42 | 645.65 | 540.21 |
| 2483 | ENSMUSG00000020386 | 4894.776606 | -0.116347958 | 0.094574595 | -1.230224221 | 0.218613152 | 0.663735712 | Sar1b | secretion associated Ras related GTPase 1 | 4995.3 | 4890.8 | 5390.5 | 4746.6 | 4954.7 | 4390.8 |
| 2484 | ENSMUSG00000020387 | 301.0059883 | -0.127066484 | 0.185658671 | -0.6844091 | 0.493716871 | 0.877333256 | Jade2 | jade family PHD finger 2 [Source:MGI Sym | 252.17 | 365.94 | 325.02 | 239.07 | 309.74 | 314.1 |
| 2485 | ENSMUSG00000020388 | 11.77250344 | 0.130368526 | 0.644819944 | 0.202178184 | 0.839777427 | 0.971838962 | Pdlim4 | PDZ and LIM domain 4 [Source:MGI Symb | 11.243 | 16.447 | 6.0375 | 11.568 | 13.087 | 12.252 |
| 2486 | ENSMUSG00000020389 | 5.199867075 | -0.345862254 | 0.983501877 | -0.351664051 | 0.725090224 | 0.9448775 | Cdkl3 | cyclin-dependent kinase-like 3 [Source:M( | 3.2124 | 11.307 | 3.0187 | 4.8199 | 3.2719 | 5.5692 |
| 2487 | ENSMUSG00000020390 | 1684.115313 | 0.157054878 | 0.114782789 | 1.368278987 | 0.171224766 | 0.598645156 | Ube2b | ubiquitin-conjugating enzyme E2B [Source | 1696.2 | 1513.1 | 1566.7 | 1572.2 | 1802.8 | 1953.7 |
| 2488 | ENSMUSG00000020392 | 253.65523 | -0.166692737 | 0.169492464 | -0.983481702 | 0.325370417 | 0.771501263 | Cdkn2aipnl | CDKN2A interacting protein N-terminal lik | 228.08 | 284.73 | 292.82 | 240.99 | 223.58 | 251.73 |
| 2489 | ENSMUSG00000020393 | 353.2981315 | -0.07262128 | 0.142724871 | -0.508820081 | 0.610878342 | 0.916376001 | Kremen1 | kringle containing transmembrane proteir | 387.1 | 366.97 | 332.06 | 348.96 | 328.28 | 356.43 |
| 2490 | ENSMUSG00000020395 | 3.08888071 | 0.211116302 | 1.392871285 | 0.15156914 | 0.879526775 | NA | Itk | IL2 inducible T cell kinase [Source:MGI Syr | 2.4093 | 5.1396 | 1.0062 | 0 | 2.1812 | 7.7969 |
| 2491 | ENSMUSG00000020396 | 1.925651061 | -0.324390327 | 1.705721857 | -0.19017774 | 0.849169854 | NA | Nefh | neurofilament, heavy polypeptide [Source | 2.4093 | 0 | 4.025 | 2.8919 | 0 | 2.2277 |
| 2492 | ENSMUSG00000020397 | 109.0908879 | -0.366546381 | 0.218943897 | -1.674156653 | 0.094099816 | 0.451490179 | Med7 | mediator complex subunit 7 [Source:MGI | 119.66 | 121.29 | 127.79 | 96.398 | 102.52 | 86.88 |
| 2493 | ENSMUSG00000020399 | 5.301039016 | 0.071260884 | 1.00787444 | 0.070704129 | 0.943633235 | 0.990077979 | Havcr2 | hepatitis A virus cellular receptor 2 [Sourc | 7.2279 | 4.1117 | 4.025 | 0.964 | 5.4531 | 10.025 |
| 2494 | ENSMUSG00000020400 | 140.5336485 | -0.217919576 | 0.263389558 | -0.82736604 | 0.408029623 | 0.832051384 | Tnip1 | TNFAIP3 interacting protein 1 [Source:MG | 187.12 | 125.41 | 139.87 | 91.578 | 123.24 | 175.99 |
| 2495 | ENSMUSG00000020401 | 0 | NA | NA | NA | NA | NA | Fam71b | family with sequence similarity 71, memb | 0 | 0 | 0 | 0 | 0 | 0 |
| 2496 | ENSMUSG00000020402 | 4176.664469 | 0.03138391 | 0.104623356 | 0.2999704 | 0.764199734 | 0.954980327 | Vdac1 | voltage-dependent anion channel 1 [Sour | 3930.4 | 4328.6 | 4134.7 | 3794.2 | 4636.2 | 4235.9 |
| 2497 | ENSMUSG00000020405 | 0 | NA | NA | NA | NA | NA | Fabp6 | fatty acid binding protein 6 [Source:MGI S | 0 | 0 | 0 | 0 | 0 | 0 |
| 2498 | ENSMUSG00000020407 | 41.90698585 | -0.280140976 | 0.384435864 | -0.728706665 | 0.466181115 | 0.863779479 | Upp1 | uridine phosphorylase 1 [Source:MGI Sym | 58.626 | 38.033 | 41.256 | 53.983 | 28.356 | 31.188 |
| 2499 | ENSMUSG00000020409 | 453.3376988 | -0.028586248 | 0.130508836 | -0.21903688 | 0.826621319 | 0.969648181 | Slu7 | SLU7 splicing factor homolog (S. cerevisia | 425.64 | 463.59 | 485.01 | 461.75 | 435.16 | 448.88 |
| 2500 | ENSMUSG00000020411 | 0 | NA | NA | NA | NA | NA | Nipal4 | NIPA-like domain containing 4 [Source:M( | 0 | 0 | 0 | 0 | 0 | 0 |
| 2501 | ENSMUSG00000020412 | 464.9041071 | 0.132781647 | 0.127841149 | 1.038645604 | 0.298969588 | 0.750094731 | Ascc2 | activating signal cointegrator 1 complex su | 448.93 | 420.42 | 460.86 | 467.53 | 506.05 | 485.64 |
| 2502 | ENSMUSG00000020413 | 127.6620789 | 0.211794427 | 0.244869633 | 0.864927285 | 0.387078701 | 0.817618864 | Hus1 | HUS1 checkpoint clamp component [Sour | 127.69 | 130.55 | 96.599 | 150.38 | 99.246 | 161.51 |
| 2503 | ENSMUSG00000020415 | 468.46613 | -0.023480973 | 0.137835841 | -0.170354626 | 0.864731254 | 0.977227145 | Pttg1 | pituitary tumor-transforming gene 1 [Sour | 519.61 | 449.2 | 446.77 | 425.11 | 473.33 | 496.77 |
| 2504 | ENSMUSG00000020419 | 7.958340847 | -0.97344588 | 0.886511904 | -1.098062954 | 0.272177001 | 0.727618887 | Hormad2 | HORMA domain containing 2 [Source:MG | 8.031 | 20.558 | 3.0187 | 2.8919 | 5.4531 | 7.7969 |
| 2505 | ENSMUSG00000020420 | 36.34838486 | 0.019757426 | 0.385936625 | 0.051193445 | 0.959171375 | 0.992094877 | Zfp607a | zinc finger protein 607A [Source:MGI Sym | 36.14 | 39.061 | 33.206 | 40.487 | 45.806 | 23.391 |
| 2506 | ENSMUSG00000020422 | 796.5060852 | 0.207548776 | 0.119719883 | 1.733619942 | 0.08298554 | 0.424735615 | Tns3 | tensin 3 [Source:MGI Symbol;Acc:MGI:24 | 693.08 | 795.61 | 730.53 | 924.45 | 825.6 | 809.76 |
| 2507 | ENSMUSG00000020423 | 179.5633239 | -0.422489498 | 0.225342147 | -1.874880061 | 0.060809226 | 0.352240545 | Btg2 | BTG anti-proliferation factor 2 [Source:M( | 247.36 | 208.67 | 159.99 | 119.53 | 162.5 | 179.33 |
| 2508 | ENSMUSG00000020424 | 34.87364592 | 0.190693024 | 0.364039067 | 0.523825713 | 0.600399767 | 0.91534406 | Castor1 | cytosolic arginine sensor for mTORC1 subu | 34.533 | 33.921 | 29.181 | 36.631 | 35.99 | 38.985 |
| 2509 | ENSMUSG00000020427 | 237.4060734 | -1.168635924 | 0.244177061 | -4.786018468 | 1.70E-06 | 0.000125543 | Igfbp3 | insulin-like growth factor binding protein : | 432.07 | 319.68 | 233.45 | 139.78 | 112.33 | 187.13 |
| 2510 | ENSMUSG00000020428 | 0 | NA | NA | NA | NA | NA | Gabra6 | gamma-aminobutyric acid (GABA) A recep | 0 | 0 | 0 | 0 | 0 | 0 |
| 2511 | ENSMUSG00000020429 | 516.6174259 | -1.710342075 | 0.548617893 | -3.117547015 | 0.001823629 | 0.032561965 | Igfbp1 | insulin-like growth factor binding protein | 689.86 | 1289 | 395.45 | 280.52 | 293.38 | 151.48 |
| 2512 | ENSMUSG00000020430 | 528.3823176 | -0.061607387 | 0.121448724 | -0.507270764 | 0.611964849 | 0.916441371 | Pes1 | pescadillo ribosomal biogenesis factor 1 [S | 530.85 | 547.88 | 540.35 | 514.76 | 496.23 | 540.21 |
| 2513 | ENSMUSG00000020431 | 5.59331551 | 0.783674487 | 1.004119036 | 0.780459745 | 0.435120313 | 0.845325456 | Adcy1 | adenylate cyclase 1 [Source:MGI Symbol;/ | 3.2124 | 4.1117 | 5.0312 | 6.7478 | 1.0906 | 13.366 |
| 2514 | ENSMUSG00000020432 | 1749.50362 | -0.50217455 | 0.096591669 | -5.198942686 | 2.00E-07 | 2.01E-05 | Tcn2 | transcobalamin 2 [Source:MGI Symbol;Ac | 2104.9 | 2110.3 | 1937 | 1413.2 | 1468 | 1463.6 |
| 2515 | ENSMUSG00000020434 | 0 | NA | NA | NA | NA | NA | 4921536K21 | RIKEN cDNA 4921536K21 gene [Source:M( | 0 | 0 | 0 | 0 | 0 | 0 |
| 2516 | ENSMUSG00000020435 | 14.53990018 | 0.738978724 | 0.588224229 | 1.256287461 | 0.209011766 | 0.65350014 | Osbp2 | oxysterol binding protein 2 [Source:MGI S | 11.243 | 9.2513 | 12.075 | 10.604 | 22.903 | 21.163 |

| | | | | | | | | | | | | | | | |
|---|---|---|---|---|---|---|---|---|---|---|---|---|---|---|---|
| 2517 | ENSMUSG00000020436 | 0 | NA | NA | NA | NA | NA | Gabrg2 | gamma-aminobutyric acid (GABA) A recep | 0 | 0 | 0 | 0 | 0 | 0 |
| 2518 | ENSMUSG00000020437 | 21.89890391 | -1.145123364 | 0.50543407 | -2.265623615 | 0.023474438 | 0.193908778 | Myo1g | myosin IG [Source:MGI Symbol;Acc:MGI:1 | 22.487 | 35.977 | 32.2 | 16.388 | 7.6343 | 16.708 |
| 2519 | ENSMUSG00000020439 | 59.43775877 | -0.640154708 | 0.318634167 | -2.009058586 | 0.044530921 | 0.291709801 | Smtn | smoothelin [Source:MGI Symbol;Acc:MGI | 84.326 | 54.48 | 78.487 | 57.839 | 43.625 | 37.871 |
| 2520 | ENSMUSG00000020440 | 1759.815715 | 0.13160645 | 0.113064704 | 1.163992352 | 0.244427111 | 0.694303562 | Arf5 | ADP-ribosylation factor 5 [Source:MGI Syr | 1748.4 | 1478.1 | 1811.2 | 1728.4 | 1878 | 1914.7 |
| 2521 | ENSMUSG00000020441 | 105.4988958 | -0.008999258 | 0.250813384 | -0.035880293 | 0.97137781 | 0.99516582 | 2310033P09 | RIKEN cDNA 2310033P09 gene [Source:M | 100.39 | 81.206 | 135.84 | 97.362 | 110.15 | 108.04 |
| 2522 | ENSMUSG00000020444 | 683.0456913 | 0.16466317 | 0.1224813 | 1.344394367 | 0.178820894 | 0.610879881 | Guk1 | guanylate kinase 1 [Source:MGI Symbol;A | 659.35 | 644.5 | 628.9 | 798.17 | 701.27 | 666.08 |
| 2523 | ENSMUSG00000020447 | 0 | NA | NA | NA | NA | NA | Npc1l1 | NPC1 like intracellular cholesterol transpo | 0 | 0 | 0 | 0 | 0 | 0 |
| 2524 | ENSMUSG00000020448 | 580.104199 | -0.035897153 | 0.127582582 | -0.281364056 | 0.778431184 | 0.959559604 | Rnf185 | ring finger protein 185 [Source:MGI Symb | 525.23 | 631.14 | 606.77 | 588.99 | 567.12 | 561.38 |
| 2525 | ENSMUSG00000020451 | 149.5852449 | 0.193841094 | 0.191911198 | 1.010056194 | 0.312468368 | 0.759830222 | Limk2 | LIM motif-containing protein kinase 2 [So | 151.79 | 130.55 | 135.84 | 154.24 | 163.59 | 161.51 |
| 2526 | ENSMUSG00000020453 | 98.73615318 | 0.184907508 | 0.232188006 | 0.796369764 | 0.425817147 | 0.841511028 | Patz1 | POZ (BTB) and AT hook containing zinc fin | 85.129 | 93.541 | 98.612 | 93.506 | 105.79 | 115.84 |
| 2527 | ENSMUSG00000020454 | 410.0885305 | 0.012524551 | 0.147514896 | 0.084903633 | 0.932338004 | 0.988420305 | Eif4enif1 | eukaryotic translation initiation factor 4E | 416.01 | 454.34 | 354.2 | 380.77 | 428.61 | 426.6 |
| 2528 | ENSMUSG00000020455 | 532.2915365 | 0.211567689 | 0.141302128 | 1.497271777 | 0.134322556 | 0.537569131 | Trim11 | tripartite motif-containing 11 [Source:MG | 538.88 | 418.36 | 522.24 | 612.13 | 549.67 | 552.47 |
| 2529 | ENSMUSG00000020456 | 3321.598923 | -0.051485964 | 0.096834679 | -0.531689317 | 0.594941192 | 0.913409341 | Ogdh | oxoglutarate (alpha-ketoglutarate) dehyd | 3144.9 | 3648.1 | 3350.8 | 3268.8 | 3355.8 | 3161.1 |
| 2530 | ENSMUSG00000020457 | 835.7608619 | 0.109411755 | 0.10990424 | 0.995518964 | 0.319483925 | 0.766337562 | Drg1 | developmentally regulated GTP binding p | 821.57 | 820.28 | 769.78 | 825.17 | 885.58 | 892.19 |
| 2531 | ENSMUSG00000020458 | 1326.978683 | -0.244377711 | 0.103780075 | -2.354765223 | 0.018534412 | 0.163964362 | Rtn4 | reticulon 4 [Source:MGI Symbol;Acc:MGI: | 1422.3 | 1512.1 | 1382.6 | 1149.1 | 1214.9 | 1280.9 |
| 2532 | ENSMUSG00000020459 | 634.7559292 | 0.255072731 | 0.136651116 | 1.866598228 | 0.061957715 | 0.355884025 | Mtif2 | mitochondrial translational initiation facto | 578.23 | 538.63 | 618.84 | 596.7 | 771.07 | 705.06 |
| 2533 | ENSMUSG00000020460 | 2891.459863 | 0.124609179 | 0.100939194 | 1.234497463 | 0.217017592 | 0.663063465 | Rps27a | ribosomal protein S27A [Source:MGI Syml | 2594 | 2853.5 | 2852.7 | 2790.7 | 3022.1 | 3235.7 |
| 2534 | ENSMUSG00000020461 | 0.428363721 | -0.543730382 | 3.624677344 | -0.15000794 | 0.880758351 | NA | Clhc1 | clathrin heavy chain linker domain contair | 1.6062 | 0 | 0 | 0.964 | 0 | 0 |
| 2535 | ENSMUSG00000020462 | 322.8016646 | -0.124240604 | 0.158910121 | -0.781829395 | 0.434314843 | 0.845264432 | Cfap36 | cilia and flagella associated protein 36 [So | 374.25 | 327.91 | 306.9 | 275.7 | 344.64 | 307.42 |
| 2536 | ENSMUSG00000020463 | 792.0636115 | 0.109500612 | 0.109184495 | 1.002895254 | 0.315911403 | 0.762646719 | Ppp4r3b | protein phosphatase 4 regulatory subunit | 756.52 | 763.74 | 765.75 | 813.6 | 792.88 | 859.89 |
| 2537 | ENSMUSG00000020464 | 467.0859087 | 0.03933513 | 0.127159182 | 0.309337708 | 0.75706465 | 0.953260973 | Pnpt1 | polyribonucleotide nucleotidyltransferase | 449.74 | 459.48 | 472.93 | 453.07 | 500.59 | 466.7 |
| 2538 | ENSMUSG00000020467 | 155.9537008 | -0.328307626 | 0.191302455 | -1.71617048 | 0.086130844 | 0.432482144 | Efemp1 | epidermal growth factor-containing fibuli | 176.68 | 177.83 | 166.03 | 124.35 | 151.6 | 139.23 |
| 2539 | ENSMUSG00000020469 | 6.770225649 | -0.178150408 | 0.885960844 | -0.201081582 | 0.840634784 | 0.971854408 | Myl7 | myosin, light polypeptide 7, regulatory [S | 7.2279 | 11.307 | 3.0187 | 4.8199 | 10.906 | 3.3415 |
| 2540 | ENSMUSG00000020471 | 195.4946071 | 0.193454089 | 0.182010774 | 1.062871632 | 0.287840169 | 0.73951397 | Pold2 | polymerase (DNA directed), delta 2, regul | 205.59 | 172.69 | 168.04 | 187.98 | 218.12 | 220.54 |
| 2541 | ENSMUSG00000020472 | 432.4239126 | 0.357322179 | 0.141941191 | 2.517395943 | 0.011822589 | 0.121535259 | Zkscan17 | zinc finger with KRAB and SCAN domains | 395.93 | 406.03 | 335.08 | 468.49 | 526.77 | 462.25 |
| 2542 | ENSMUSG00000020473 | 66.56177531 | -0.14902398 | 0.284774204 | -0.523305757 | 0.600761495 | 0.915429015 | Aebp1 | AE binding protein 1 [Source:MGI Symbol | 61.839 | 79.15 | 69.431 | 74.226 | 52.35 | 62.375 |
| 2543 | ENSMUSG00000020474 | 84.07230224 | 0.186171636 | 0.274171549 | 0.67903339 | 0.497116706 | 0.878687038 | Polm | polymerase (DNA directed), mu [Source:N | 65.854 | 64.759 | 105.66 | 89.65 | 90.521 | 87.994 |
| 2544 | ENSMUSG00000020475 | 10.44543077 | -0.33052499 | 0.843233365 | -0.39197333 | 0.69507792 | 0.939992732 | Pgam2 | phosphoglycerate mutase 2 [Source:MGI | 9.6372 | 4.1117 | 21.131 | 4.8199 | 19.631 | 3.3415 |
| 2545 | ENSMUSG00000020476 | 489.8684722 | -0.004958004 | 0.128909326 | -0.038461171 | 0.96931999 | 0.994602269 | Dbnl | drebrin-like [Source:MGI Symbol;Acc:MGI | 478.65 | 480.04 | 513.18 | 450.18 | 512.59 | 504.57 |
| 2546 | ENSMUSG00000020477 | 892.1149222 | 0.050823604 | 0.111758394 | 0.454763189 | 0.649279616 | 0.925643431 | Mrps24 | mitochondrial ribosomal protein S24 [Sou | 809.53 | 914.85 | 905.62 | 892.64 | 888.85 | 941.2 |
| 2547 | ENSMUSG00000020481 | 0.181769715 | 1.126924175 | 4.080472857 | 0.276174898 | 0.782413736 | NA | Ankrd36 | ankyrin repeat domain 36 [Source:MGI Sy | 0 | 0 | 0 | 0 | 1.0906 | 0 |
| 2548 | ENSMUSG00000020482 | 484.8332375 | 0.257295643 | 0.130018506 | 1.978915546 | 0.047825516 | 0.303663969 | Ccdc117 | coiled-coil domain containing 117 [Source | 409.58 | 456.4 | 459.85 | 531.15 | 497.32 | 554.69 |
| 2549 | ENSMUSG00000020483 | 4212.182533 | 0.089738125 | 0.113775992 | 0.788726369 | 0.430271952 | 0.843514979 | Dynll2 | dynein light chain LC8-type 2 [Source:MGI | 4066.1 | 4190.8 | 3985.7 | 3739.3 | 4365.7 | 4925.4 |
| 2550 | ENSMUSG00000020484 | 4525.209776 | 1.003174677 | 0.125664833 | 7.982938846 | 1.43E-15 | 1.03E-12 | Xbp1 | X-box binding protein 1 [Source:MGI Syml | 3121.7 | 2941.9 | 2973.5 | 6662.1 | 4880.5 | 6571.7 |
| 2551 | ENSMUSG00000020485 | 547.7998452 | 0.088107717 | 0.133208372 | 0.661427772 | 0.50833802 | 0.882793133 | Supt4a | SPT4A, DSIF elongation factor subunit [So | 545.31 | 501.62 | 545.38 | 499.34 | 575.85 | 619.3 |
| 2552 | ENSMUSG00000020486 | 116.4961148 | -0.94680117 | 0.245328727 | -3.859316365 | 0.000113705 | 0.004011279 | Septin4 | septin 4 [Source:MGI Symbol;Acc:MGI:12 | 173.47 | 126.43 | 159.99 | 80.01 | 62.165 | 96.904 |
| 2553 | ENSMUSG00000020490 | 0.810301681 | -0.283528992 | 2.609307867 | -0.108660613 | 0.913471683 | NA | Btnl10 | butyrophilin-like 10 [Source:MGI Symbol; | 1.6062 | 1.0279 | 0 | 0 | 0 | 2.2277 |
| 2554 | ENSMUSG00000020491 | 49.94051257 | 0.290283439 | 0.331655383 | 0.875256227 | 0.381434506 | 0.8130053 | 2810021J22I | RIKEN cDNA 2810021J22 gene [Source:M | 34.533 | 51.396 | 49.306 | 58.803 | 62.165 | 43.44 |
| 2555 | ENSMUSG00000020492 | 17.79817579 | -0.226137999 | 0.559550529 | -0.404142231 | 0.686108132 | 0.936390642 | Ska2 | spindle and kinetochore associated compl | 20.881 | 12.335 | 24.15 | 8.6758 | 21.812 | 18.935 |
| 2556 | ENSMUSG00000020493 | 1.732184271 | 0.039407798 | 1.707965082 | 0.023072953 | 0.98159208 | NA | Prr11 | proline rich 11 [Source:MGI Symbol;Acc:N | 4.0155 | 1.0279 | 0 | 0.964 | 3.2719 | 1.1138 |
| 2557 | ENSMUSG00000020495 | 192.3084228 | 0.018844465 | 0.181404575 | 0.103880869 | 0.91726389 | 0.986374671 | Smg8 | smg-8 homolog, nonsense mediated mRN | 174.27 | 181.94 | 217.35 | 197.62 | 179.95 | 202.72 |
| 2558 | ENSMUSG00000020496 | 2400.691707 | 0.010640791 | 0.095333929 | 0.111615995 | 0.911127889 | 0.98582838 | Rnf187 | ring finger protein 187 [Source:MGI Symb | 2262.3 | 2393 | 2520.6 | 2315.5 | 2480.1 | 2432.6 |
| 2559 | ENSMUSG00000020513 | 75.89099159 | -0.623088667 | 0.290301341 | -2.146351323 | 0.03184496 | 0.237793236 | Tubd1 | tubulin, delta 1 [Source:MGI Symbol;Acc:I | 105.21 | 80.178 | 90.562 | 60.731 | 76.343 | 42.326 |
| 2560 | ENSMUSG00000020514 | 374.2347648 | 0.174576607 | 0.157613667 | 1.107623531 | 0.268024437 | 0.723941261 | Mrpl22 | mitochondrial ribosomal protein L22 [Sou | 359.79 | 296.04 | 398.47 | 366.31 | 392.62 | 432.17 |
| 2561 | ENSMUSG00000020515 | 856.2347356 | 0.305528654 | 0.1505963 | 2.028792574 | 0.042479422 | 0.283892783 | Cnot8 | CCR4-NOT transcription complex, subunit | 758.13 | 682.54 | 856.31 | 779.86 | 943.38 | 1117.2 |
| 2562 | ENSMUSG00000020516 | 613.4669204 | -0.159512941 | 0.120367073 | -1.325220735 | 0.18509795 | 0.620815781 | Rps6kb1 | ribosomal protein S6 kinase, polypeptide | 627.22 | 669.17 | 646.01 | 576.46 | 618.38 | 543.56 |
| 2563 | ENSMUSG00000020519 | 541.5713154 | 0.05194687 | 0.137343077 | 0.378227072 | 0.705261916 | 0.94295036 | Sap30l | SAP30-like [Source:MGI Symbol;Acc:MGI: | 470.62 | 525.27 | 600.73 | 576.46 | 557.31 | 519.05 |
| 2564 | ENSMUSG00000020520 | 121.3371809 | -0.259737743 | 0.247111702 | -1.051094468 | 0.293215205 | 0.744176999 | Galnt10 | polypeptide N-acetylgalactosaminyltransf | 121.27 | 132.6 | 142.89 | 107.97 | 79.615 | 143.69 |
| 2565 | ENSMUSG00000020521 | 764.6474208 | 0.1477612 | 0.117700036 | 1.255404878 | 0.209331819 | 0.65350014 | Rnft1 | ring finger protein, transmembrane 1 [Sou | 677.02 | 742.16 | 757.7 | 747.08 | 844.14 | 819.79 |
| 2566 | ENSMUSG00000020522 | 194.6021936 | -0.084162848 | 0.195314943 | -0.430908391 | 0.666534982 | 0.929596279 | Mfap3 | microfibrillar-associated protein 3 [Source | 226.47 | 210.72 | 163.01 | 181.23 | 172.32 | 213.86 |
| 2567 | ENSMUSG00000020523 | 731.1767163 | 0.139572641 | 0.117174954 | 1.191147394 | 0.233595727 | 0.681057972 | Fam114a2 | family with sequence similarity 114, meml | 709.94 | 683.57 | 693.3 | 709.49 | 827.78 | 762.98 |
| 2568 | ENSMUSG00000020524 | 0 | NA | NA | NA | NA | NA | Gria1 | glutamate receptor, ionotropic, AMPA1 (a | 0 | 0 | 0 | 0 | 0 | 0 |
| 2569 | ENSMUSG00000020525 | 90.19412664 | 0.122208663 | 0.266553834 | 0.45847648 | 0.646610157 | 0.925643431 | Ppm1d | protein phosphatase 1D magnesium-depe | 63.445 | 94.568 | 101.63 | 90.614 | 83.978 | 106.93 |
| 2570 | ENSMUSG00000020526 | 105.6742268 | -0.187617634 | 0.237224134 | -0.790887636 | 0.429009564 | 0.842753542 | Znhit3 | zinc finger, HIT type 3 [Source:MGI Symbo | 130.91 | 92.513 | 113.71 | 95.434 | 110.15 | 91.335 |
| 2571 | ENSMUSG00000020527 | 188.8302607 | -0.017254843 | 0.171784773 | -0.100444541 | 0.91999141 | 0.986685804 | Myo19 | myosin XIX [Source:MGI Symbol;Acc:MGI: | 193.55 | 181.94 | 194.21 | 184.12 | 187.59 | 191.58 |
| 2572 | ENSMUSG00000020528 | 211.8937092 | -0.09383341 | 0.170321611 | -0.550918988 | 0.581689211 | 0.908645447 | Prpsap2 | phosphoribosyl pyrophosphate synthetas | 236.11 | 207.64 | 212.32 | 211.11 | 215.94 | 188.24 |

| | | | | | | | | | | | | | | | |
|---|---|---|---|---|---|---|---|---|---|---|---|---|---|---|---|
| 2573 | ENSMUSG00000020530 | 1509.885862 | 0.262152071 | 0.106621711 | 2.45871191 | 0.013943646 | 0.136178862 | Ggnbp2 | gametogenetin binding protein 2 [Source: | 1362.9 | 1284.9 | 1471.1 | 1583.8 | 1593.4 | 1763.2 |
| 2574 | ENSMUSG00000020532 | 1630.761808 | -0.004723857 | 0.172300318 | -0.027416417 | 0.978127604 | 0.997788114 | Acaca | acetyl-Coenzyme A carboxylase alpha [Sou | 1896.9 | 1840 | 1163.2 | 1882.6 | 1569.4 | 1432.4 |
| 2575 | ENSMUSG00000020534 | 4429.756087 | 0.067665144 | 0.109518245 | 0.617843569 | 0.53667846 | 0.890560262 | Shmt1 | serine hydroxymethyltransferase 1 (solub | 4858 | 3879.4 | 4238.3 | 4312.8 | 4469.4 | 4820.7 |
| 2576 | ENSMUSG00000020536 | 169.6936325 | 0.148430144 | 0.202098401 | 0.73444492 | 0.462677618 | 0.860873685 | Llgl1 | LLGL1 scribble cell polarity complex comp | 149.38 | 145.96 | 188.17 | 207.26 | 160.32 | 167.08 |
| 2577 | ENSMUSG00000020537 | 599.1138243 | -0.031259674 | 0.123449156 | -0.253219014 | 0.800098974 | 0.962424862 | Drg2 | developmentally regulated GTP binding pr | 644.09 | 554.05 | 617.83 | 575.49 | 615.11 | 588.11 |
| 2578 | ENSMUSG00000020538 | 5598.495304 | 0.047717149 | 0.137966403 | 0.345860642 | 0.729447449 | 0.946534338 | Srebf1 | sterol regulatory element binding transcri | 6611.1 | 4795.2 | 5109.7 | 5509.1 | 6504.4 | 5061.3 |
| 2579 | ENSMUSG00000020541 | 408.5029279 | 0.248587643 | 0.137567793 | 1.807019201 | 0.070759285 | 0.385049776 | Tom1l1 | target of myb1-like 1 (chicken) [Source:M( | 391.91 | 384.44 | 343.13 | 415.47 | 448.24 | 467.81 |
| 2580 | ENSMUSG00000020542 | 0.685278559 | -2.809870862 | 3.376578027 | -0.832165239 | 0.405315682 | NA | Myocd | myocardin [Source:MGI Symbol;Acc:MGI: | 0 | 4.1117 | 0 | 0 | 0 | 0 |
| 2581 | ENSMUSG00000020544 | 177.9860346 | 0.321940887 | 0.196941278 | 1.634704976 | 0.102110909 | 0.47032223 | Cox11 | cytochrome c oxidase assembly protein 1: | 160.62 | 129.52 | 184.14 | 180.26 | 202.86 | 210.52 |
| 2582 | ENSMUSG00000020545 | 0 | NA | NA | NA | NA | NA | Lrrc72 | leucine rich repeat containing 72 [Source: | 0 | 0 | 0 | 0 | 0 | 0 |
| 2583 | ENSMUSG00000020546 | 45.99087862 | -0.499485369 | 0.327133929 | -1.526852843 | 0.126797608 | 0.521635389 | Stxbp4 | syntaxin binding protein 4 [Source:MGI Sy | 53.005 | 49.34 | 59.368 | 40.487 | 41.443 | 32.301 |
| 2584 | ENSMUSG00000020547 | 159.0140276 | 0.003672774 | 0.201790661 | 0.018200914 | 0.985478574 | 0.99859176 | Bzw2 | basic leucine zipper and W2 domains 2 [So | 141.35 | 151.1 | 184.14 | 142.67 | 177.77 | 157.05 |
| 2585 | ENSMUSG00000020548 | 0 | NA | NA | NA | NA | NA | 1700086D15 | RIKEN cDNA 1700086D15 gene [Source:M | 0 | 0 | 0 | 0 | 0 | 0 |
| 2586 | ENSMUSG00000020549 | 474.3589644 | 0.074275712 | 0.126016009 | 0.589414888 | 0.555582992 | 0.897628185 | Elac2 | elaC ribonuclease Z 2 [Source:MGI Symbo | 469.01 | 448.17 | 468.91 | 468.49 | 511.5 | 480.07 |
| 2587 | ENSMUSG00000020553 | 2369.392929 | -0.183895269 | 0.096303052 | -1.909547668 | 0.056191479 | 0.335294368 | Pctp | phosphatidylcholine transfer protein [Sou | 2679.1 | 2505 | 2375.7 | 2277.9 | 2143.1 | 2235.5 |
| 2588 | ENSMUSG00000020561 | 309.7751907 | 0.04313585 | 0.147357285 | 0.292729678 | 0.769728783 | 0.956391815 | Polr1f | RNA polymerase I subunit F [Source:MGI S | 296.34 | 323.79 | 295.84 | 334.5 | 313.01 | 295.17 |
| 2589 | ENSMUSG00000020562 | 0 | NA | NA | NA | NA | NA | Efcab10 | EF-hand calcium binding domain 10 [Sour | 0 | 0 | 0 | 0 | 0 | 0 |
| 2590 | ENSMUSG00000020564 | 210.9300368 | 0.492436949 | 0.250731556 | 1.964000686 | 0.049530012 | 0.311073426 | Atxn7l1 | ataxin 7-like 1 [Source:MGI Symbol;Acc:M | 175.88 | 160.36 | 190.18 | 364.38 | 184.31 | 190.47 |
| 2591 | ENSMUSG00000020566 | 1.011684555 | -0.99386545 | 2.458038153 | -0.404332801 | 0.685968008 | NA | Atp6v1c2 | ATPase, H+ transporting, lysosomal V1 sul | 4.0155 | 0 | 0 | 0.964 | 1.0906 | 0 |
| 2592 | ENSMUSG00000020570 | 1170.025346 | -0.223028075 | 0.104884511 | -2.126415748 | 0.033468658 | 0.24471574 | Sypl | synaptophysin-like protein [Source:MGI S | 1214.3 | 1297.2 | 1269.9 | 1077.7 | 1143 | 1018.1 |
| 2593 | ENSMUSG00000020571 | 4450.885017 | 0.434856842 | 0.125247098 | 3.471991364 | 0.000516613 | 0.012751348 | Pdia6 | protein disulfide isomerase associated 6 [ | 4229.1 | 3577.2 | 3548 | 5429.1 | 4300.3 | 5621.6 |
| 2594 | ENSMUSG00000020572 | 2040.133289 | -0.305288059 | 0.171606651 | -1.778998993 | 0.075239927 | 0.401279611 | Nampt | nicotinamide phosphoribosyltransferase [ | 2013.4 | 2469.1 | 2284.2 | 2024.4 | 2184.5 | 1265.3 |
| 2595 | ENSMUSG00000020573 | 10.63911627 | -0.852040955 | 0.890777824 | -0.956513433 | 0.338812901 | 0.780858357 | Pik3cg | phosphatidylinositol-4,5-bisphosphate 3-k | 11.243 | 27.754 | 2.0125 | 2.8919 | 5.4531 | 14.48 |
| 2596 | ENSMUSG00000020576 | 336.90872 | 0.275956851 | 0.14961507 | 1.844445558 | 0.065118233 | 0.366343883 | Nbas | neuroblastoma amplified sequence [Sour | 296.34 | 311.46 | 306.9 | 388.48 | 321.73 | 396.53 |
| 2597 | ENSMUSG00000020577 | 11.90953485 | -0.61766041 | 0.689554421 | -0.895738453 | 0.370392468 | 0.807048832 | Tspan13 | tetraspanin 13 [Source:MGI Symbol;Acc:M | 20.881 | 6.1675 | 16.1 | 10.604 | 5.4531 | 12.252 |
| 2598 | ENSMUSG00000020580 | 448.0327837 | -0.059617151 | 0.165170352 | -0.360943415 | 0.718141747 | 0.9447719 | Rock2 | Rho-associated coiled-coil containing prot | 456.97 | 516.01 | 399.48 | 492.59 | 470.06 | 353.09 |
| 2599 | ENSMUSG00000020581 | 0 | NA | NA | NA | NA | NA | Agr2 | anterior gradient 2 [Source:MGI Symbol;A | 0 | 0 | 0 | 0 | 0 | 0 |
| 2600 | ENSMUSG00000020583 | 0 | NA | NA | NA | NA | NA | Matn3 | matrilin 3 [Source:MGI Symbol;Acc:MGI:1 | 0 | 0 | 0 | 0 | 0 | 0 |
| 2601 | ENSMUSG00000020585 | 3631.664989 | 0.056387019 | 0.085027984 | 0.663158361 | 0.507229137 | 0.882010367 | Laptm4a | lysosomal-associated protein transmembr | 3509.6 | 3645 | 3527.9 | 3713.2 | 3632.8 | 3761.5 |
| 2602 | ENSMUSG00000020589 | 42.479125 | -0.863997032 | 0.416967761 | -2.072095528 | 0.038256533 | 0.266030047 | Cyria | CYFIP related Rac1 interactor A [Source:M | 71.476 | 48.312 | 44.275 | 22.171 | 19.631 | 49.009 |
| 2603 | ENSMUSG00000020590 | 617.0041577 | -0.001664462 | 0.130592442 | -0.012745473 | 0.989830859 | 0.99859176 | Snx13 | sorting nexin 13 [Source:MGI Symbol;Acc: | 628.83 | 634.23 | 589.66 | 698.88 | 583.48 | 566.95 |
| 2604 | ENSMUSG00000020591 | 0.51014012 | 1.031855138 | 3.09002894 | 0.333930574 | 0.738431948 | NA | Ntsr2 | neurotensin receptor 2 [Source:MGI Symb | 0 | 0 | 1.0062 | 0.964 | 1.0906 | 0 |
| 2605 | ENSMUSG00000020592 | 3564.481923 | -0.150404754 | 0.112477962 | -1.337193093 | 0.181159616 | 0.614561135 | Sdc1 | syndecan 1 [Source:MGI Symbol;Acc:MGI | 3320.8 | 3726.2 | 4205.1 | 3606.2 | 3334 | 3194.5 |
| 2606 | ENSMUSG00000020593 | 5996.270142 | -1.405983362 | 0.483883972 | -2.905620859 | 0.003665253 | 0.054096138 | Lpin1 | lipin 1 [Source:MGI Symbol;Acc:MGI:1891 | 7309 | 12877 | 5934.8 | 2204.6 | 4745.3 | 2907.1 |
| 2607 | ENSMUSG00000020594 | 1070.689167 | 0.201930418 | 0.111292409 | 1.81441322 | 0.069614132 | 0.381135916 | Pum2 | pumilio RNA-binding family member 2 [So | 995.85 | 1083.4 | 908.64 | 1170.3 | 1132.1 | 1133.9 |
| 2608 | ENSMUSG00000020598 | 0.17131964 | -0.796629176 | 4.080472857 | -0.195229623 | 0.84521319 | NA | Nrcam | neuronal cell adhesion molecule [Source:I | 0 | 1.0279 | 0 | 0 | 0 | 0 |
| 2609 | ENSMUSG00000020599 | 2.508325205 | -1.342360737 | 1.501572951 | -0.893969711 | 0.371338097 | NA | Rgs9 | regulator of G-protein signaling 9 [Source: | 1.6062 | 7.1954 | 2.0125 | 0.964 | 3.2719 | 0 |
| 2610 | ENSMUSG00000020600 | 6.106679405 | 0.851752517 | 0.991238979 | 0.859280693 | 0.390185673 | 0.819725365 | Slc7a15 | solute carrier family 7 (cationic amino acid | 8.8341 | 2.0558 | 2.0125 | 10.604 | 10.906 | 2.2277 |
| 2611 | ENSMUSG00000020601 | 27.00513677 | -0.935202734 | 0.427850641 | -2.185815901 | 0.028829077 | 0.22274774 | Trib2 | tribbles pseudokinase 2 [Source:MGI Sym | 40.155 | 35.977 | 30.187 | 19.28 | 15.269 | 21.163 |
| 2612 | ENSMUSG00000020604 | 606.448189 | -0.289174439 | 0.187849289 | -1.539395977 | 0.123707656 | 0.515353551 | Arsg | arylsulfatase G [Source:MGI Symbol;Acc:M | 782.22 | 720.57 | 497.08 | 449.21 | 522.41 | 667.19 |
| 2613 | ENSMUSG00000020605 | 214.7923917 | 0.01111978 | 0.204433455 | 0.05439315 | 0.956621936 | 0.991798415 | Hs1bp3 | HCLS1 binding protein 3 [Source:MGI Sym | 240.13 | 203.53 | 197.22 | 166.77 | 212.67 | 268.44 |
| 2614 | ENSMUSG00000020607 | 18.8136175 | -0.651446247 | 0.551906678 | -1.180355796 | 0.237858735 | 0.686509684 | Lratd1 | LRAT domain containing 1 [Source:MGI Sy | 31.321 | 13.363 | 24.15 | 16.388 | 8.7249 | 18.935 |
| 2615 | ENSMUSG00000020608 | 428.960839 | 0.158195538 | 0.131379892 | 1.204107686 | 0.22854796 | 0.675041782 | Smc6 | structural maintenance of chromosomes ( | 399.94 | 420.42 | 396.46 | 472.35 | 461.33 | 423.26 |
| 2616 | ENSMUSG00000020609 | 94230.61911 | -0.043614237 | 0.105487971 | -0.413452235 | 0.679275311 | 0.934062537 | Apob | apolipoprotein B [Source:MGI Symbol;Acc | 83031 | 104027 | 99908 | 93868 | 96106 | 88443 |
| 2617 | ENSMUSG00000020610 | 251.1393364 | 0.012091021 | 0.170034957 | 0.071109034 | 0.943310979 | 0.990077979 | Amz2 | archaelysin family metallopeptidase 2 [So | 245.75 | 248.76 | 255.59 | 239.07 | 221.4 | 296.28 |
| 2618 | ENSMUSG00000020611 | 587.9684662 | 0.091265571 | 0.123115902 | 0.741297988 | 0.458512782 | 0.859322431 | Gna13 | guanine nucleotide binding protein, alpha | 583.86 | 522.18 | 601.73 | 607.31 | 586.75 | 625.98 |
| 2619 | ENSMUSG00000020612 | 2482.243136 | -0.001511417 | 0.097579445 | -0.015489095 | 0.987641984 | 0.99859176 | Prkar1a | protein kinase, cAMP dependent regulato | 2478.4 | 2622.2 | 2349.6 | 2349.2 | 2488.8 | 2605.3 |
| 2620 | ENSMUSG00000020614 | 503.0460699 | -0.092062397 | 0.17466026 | -0.527094125 | 0.598128227 | 0.915029425 | Fam20a | FAM20A, golgi associated secretory pathw | 583.05 | 506.76 | 466.9 | 507.05 | 368.63 | 585.88 |
| 2621 | ENSMUSG00000020617 | 0 | NA | NA | NA | NA | NA | 1700012B07 | RIKEN cDNA 1700012B07 gene [Source:M | 0 | 0 | 0 | 0 | 0 | 0 |
| 2622 | ENSMUSG00000020620 | 825.6926977 | 0.274004603 | 0.130481631 | 2.099947708 | 0.035733441 | 0.253977709 | Abca8b | ATP-binding cassette, sub-family A (ABC1) | 837.64 | 769.91 | 633.93 | 909.03 | 873.59 | 930.06 |
| 2623 | ENSMUSG00000020621 | 440.9215962 | 0.143374038 | 0.133489519 | 1.074047152 | 0.282801549 | 0.735558641 | Rdh14 | retinol dehydrogenase 14 (all-trans and 9- | 457.77 | 399.86 | 398.47 | 449.21 | 464.6 | 475.61 |
| 2624 | ENSMUSG00000020622 | 0.435408177 | -2.179616019 | 3.531869654 | -0.617128103 | 0.537150234 | NA | Nt5c1b | 5'-nucleotidase, cytosolic IB [Source:MGI : | 1.6062 | 0 | 1.0062 | 0 | 0 | 0 |
| 2625 | ENSMUSG00000020623 | 271.6715327 | -0.325845485 | 0.174748504 | -1.864653932 | 0.062229928 | 0.356472744 | Map2k6 | mitogen-activated protein kinase kinase 6 | 274.66 | 268.29 | 364.26 | 253.53 | 244.3 | 225 |
| 2626 | ENSMUSG00000020624 | 0 | NA | NA | NA | NA | NA | 4933434M1( | RIKEN cDNA 4933434M16 gene [Source:M | 0 | 0 | 0 | 0 | 0 | 0 |
| 2627 | ENSMUSG00000020627 | 5.115851749 | -3.730614053 | 1.391118983 | -2.681736141 | 0.00732412 | 0.086502219 | Klhl29 | kelch-like 29 [Source:MGI Symbol;Acc:MG | 7.2279 | 5.1396 | 16.1 | 0 | 0 | 2.2277 |
| 2628 | ENSMUSG00000020628 | 520.2827687 | -0.075609038 | 0.144773749 | -0.522256548 | 0.601491719 | 0.915461118 | Trappc12 | trafficking protein particle complex 12 [So | 568.6 | 519.1 | 513.18 | 453.07 | 595.48 | 472.27 |

| | | | | | | | | | | | | | | | |
|---|---|---|---|---|---|---|---|---|---|---|---|---|---|---|---|
| 2629 | ENSMUSG00000020629 | 3308.476098 | -0.042909665 | 0.087655435 | -0.489526575 | 0.624468945 | 0.919503423 | Adi1 | acireductone dioxygenase 1 [Source:MGI | 3508.8 | 3278 | 3285.4 | 3259.2 | 3288.2 | 3231.3 |
| 2630 | ENSMUSG00000020630 | 106.5649277 | 0.175407113 | 0.22567787 | 0.777245517 | 0.437013933 | 0.846852749 | Rnaseh1 | ribonuclease H1 [Source:MGI Symbol;Acc | 93.16 | 95.596 | 111.69 | 112.79 | 102.52 | 123.64 |
| 2631 | ENSMUSG00000020633 | 0 | NA | NA | NA | NA | NA | Dcdc2c | doublecortin domain containing 2C [Sourc | 0 | 0 | 0 | 0 | 0 | 0 |
| 2632 | ENSMUSG00000020634 | 522.1669122 | -0.035566701 | 0.133676934 | -0.266064606 | 0.790189453 | 0.961019028 | Ubxn2a | UBX domain protein 2A [Source:MGI Syml | 497.92 | 580.77 | 508.15 | 561.04 | 500.59 | 484.52 |
| 2633 | ENSMUSG00000020635 | 1.186126113 | 2.643610322 | 2.135807954 | 1.237756568 | 0.215806337 | NA | Fkbp1b | FK506 binding protein 1b [Source:MGI Syr | 0.8031 | 0 | 0 | 1.928 | 3.2719 | 1.1138 |
| 2634 | ENSMUSG00000020636 | 0 | NA | NA | NA | NA | NA | Allc | allantoicase [Source:MGI Symbol;Acc:MG | 0 | 0 | 0 | 0 | 0 | 0 |
| 2635 | ENSMUSG00000020638 | 105.4033448 | -0.367462846 | 0.269646868 | -1.362755847 | 0.172959471 | 0.601945568 | Cmpk2 | cytidine monophosphate (UMP-CMP) kina | 146.16 | 92.513 | 116.72 | 67.478 | 98.156 | 111.38 |
| 2636 | ENSMUSG00000020639 | 11.27105224 | 0.693401915 | 0.67735735 | 1.023687002 | 0.305983136 | 0.755069291 | Pfn4 | profilin family, member 4 [Source:MGI Syr | 10.44 | 6.1675 | 9.0562 | 7.7118 | 13.087 | 21.163 |
| 2637 | ENSMUSG00000020640 | 606.9779465 | 0.009915515 | 0.123166867 | 0.08050473 | 0.935835835 | 0.988420305 | Itsn2 | intersectin 2 [Source:MGI Symbol;Acc:MG | 605.54 | 631.14 | 577.58 | 571.64 | 596.57 | 659.4 |
| 2638 | ENSMUSG00000020641 | 160.3525091 | -0.166179185 | 0.261763771 | -0.634844097 | 0.5255301 | 0.888334265 | Rsad2 | radical S-adenosyl methionine domain coı | 195.96 | 103.82 | 208.29 | 122.43 | 171.23 | 160.39 |
| 2639 | ENSMUSG00000020642 | 35.09210913 | -0.460907001 | 0.424183581 | -1.086574356 | 0.277224963 | 0.730041765 | Rnf144a | ring finger protein 144A [Source:MGI Sym | 42.564 | 28.782 | 50.312 | 19.28 | 26.175 | 43.44 |
| 2640 | ENSMUSG00000020644 | 1733.206132 | 0.663524258 | 0.391311359 | 1.695642722 | 0.089953562 | 0.442644317 | Id2 | inhibitor of DNA binding 2 [Source:MGI Sy | 1378.1 | 911.76 | 1734.8 | 2916 | 1409.1 | 2049.5 |
| 2641 | ENSMUSG00000020646 | 1.107988552 | -1.056291367 | 2.031653537 | -0.519917076 | 0.603121373 | NA | Mboat2 | membrane bound O-acyltransferase domɛ | 2.4093 | 1.0279 | 1.0062 | 0 | 1.0906 | 1.1138 |
| 2642 | ENSMUSG00000020647 | 339.3030073 | 0.129565437 | 0.140686295 | 0.9209528 | 0.35707507 | 0.796166767 | Ncoa1 | nuclear receptor coactivator 1 [Source:M( | 313.21 | 346.41 | 312.94 | 354.74 | 360.99 | 347.52 |
| 2643 | ENSMUSG00000020648 | 93.74452604 | -0.02538375 | 0.25099779 | -0.101131368 | 0.919446178 | 0.986685804 | Dus4l | dihydrouridine synthase 4-like (S. cerevisi | 93.16 | 109.99 | 80.5 | 82.902 | 110.15 | 85.766 |
| 2644 | ENSMUSG00000020649 | 252.5944586 | -0.859671937 | 0.164607514 | -5.222555849 | 1.76E-07 | 1.82E-05 | Rrm2 | ribonucleotide reductase M2 [Source:MG | 334.89 | 317.63 | 324.01 | 159.06 | 201.76 | 178.22 |
| 2645 | ENSMUSG00000020650 | 221.7458699 | -0.041465489 | 0.171551241 | -0.241709061 | 0.809005608 | 0.965705794 | Bcap29 | B cell receptor associated protein 29 [Sou | 220.85 | 236.42 | 217.35 | 196.65 | 247.57 | 211.63 |
| 2646 | ENSMUSG00000020651 | 11.1518997 | -0.120348778 | 0.667795613 | -0.180217983 | 0.856981441 | 0.974809103 | Slc26a4 | solute carrier family 26, member 4 [Sourc | 14.456 | 13.363 | 7.0437 | 14.46 | 10.906 | 6.6831 |
| 2647 | ENSMUSG00000020652 | 57.15243011 | 0.27574114 | 0.30995676 | 0.889611634 | 0.373674457 | 0.80865857 | Cenpo | centromere protein O [Source:MGI Symbo | 45.777 | 43.173 | 66.412 | 68.442 | 55.622 | 63.489 |
| 2648 | ENSMUSG00000020653 | 179.0136543 | -0.089301798 | 0.182462866 | -0.489424504 | 0.624541192 | 0.919503423 | Klf11 | Kruppel-like factor 11 [Source:MGI Symbc | 190.34 | 197.36 | 166.03 | 182.19 | 177.77 | 160.39 |
| 2649 | ENSMUSG00000020654 | 12.44394144 | -0.273652259 | 0.68526785 | -0.3993362 | 0.689645497 | 0.937261276 | Adcy3 | adenylate cyclase 3 [Source:MGI Symbol;/ | 23.29 | 9.2513 | 8.05 | 7.7118 | 17.45 | 8.9108 |
| 2650 | ENSMUSG00000020656 | 30.64947777 | -0.253786644 | 0.476191436 | -0.532950878 | 0.594067584 | 0.913144152 | Grhl1 | grainyhead like transcription factor 1 [Sou | 46.58 | 34.949 | 18.112 | 21.208 | 20.722 | 42.326 |
| 2651 | ENSMUSG00000020657 | 16.02668781 | -0.462213656 | 0.616349366 | -0.749921524 | 0.45330197 | 0.857073339 | Dnajc27 | DnaJ heat shock protein family (Hsp40) m | 24.896 | 19.53 | 11.069 | 9.6398 | 7.6343 | 23.391 |
| 2652 | ENSMUSG00000020658 | 16.12919664 | 0.655759337 | 0.695778117 | 0.942483416 | 0.345945198 | 0.786899236 | Efr3b | EFR3 homolog B [Source:MGI Symbol;Acc | 19.274 | 7.1954 | 11.069 | 40.487 | 8.7249 | 10.025 |
| 2653 | ENSMUSG00000020659 | 102.4537661 | -0.024587141 | 0.227291613 | -0.108174433 | 0.913857325 | 0.98625062 | Cbll1 | Casitas B-lineage lymphoma-like 1 [Source | 103.6 | 92.513 | 113.71 | 95.434 | 101.43 | 108.04 |
| 2654 | ENSMUSG00000020660 | 5.643322234 | -0.371485639 | 0.89448042 | -0.415308855 | 0.67791582 | 0.93325821 | Pomc | pro-opiomelanocortin-alpha [Source:MGI | 8.8341 | 5.1396 | 5.0312 | 3.8559 | 6.5437 | 4.4554 |
| 2655 | ENSMUSG00000020661 | 352.2714412 | 0.516501852 | 0.158279569 | 3.263225029 | 0.00110152 | 0.022493298 | Dnmt3a | DNA methyltransferase 3A [Source:MGI S | 290.72 | 288.84 | 289.8 | 394.27 | 359.9 | 490.09 |
| 2656 | ENSMUSG00000020664 | 2870.830825 | -0.013197188 | 0.100124795 | -0.131807387 | 0.895136643 | 0.982798755 | Dld | dihydrolipoamide dehydrogenase [Source | 2743.4 | 3030.3 | 2878.9 | 2798.4 | 3094.1 | 2679.9 |
| 2657 | ENSMUSG00000020668 | 12.04830374 | -1.219470411 | 0.688818558 | -1.770379727 | 0.076663905 | 0.405435728 | Kif3c | kinesin family member 3C [Source:MGI Sy | 11.243 | 14.391 | 25.156 | 11.568 | 4.3625 | 5.5692 |
| 2658 | ENSMUSG00000020669 | 145.6270793 | 0.129274427 | 0.238306347 | 0.542471609 | 0.58749366 | 0.910765033 | Sh3yl1 | Sh3 domain YSC-like 1 [Source:MGI Symbo | 157.41 | 115.13 | 144.9 | 199.54 | 128.69 | 128.09 |
| 2659 | ENSMUSG00000020671 | 1614.302875 | -0.034493959 | 0.10590912 | -0.325693944 | 0.744655927 | 0.949997329 | Rab10 | RAB10, member RAS oncogene family [So | 1566.9 | 1731 | 1602.9 | 1477.8 | 1715.5 | 1591.7 |
| 2660 | ENSMUSG00000020672 | 261.9524345 | 0.257862015 | 0.167968742 | 1.53517858 | 0.124739968 | 0.51816166 | Sntg2 | syntrophin, gamma 2 [Source:MGI Symbo | 264.22 | 233.34 | 217.35 | 255.45 | 319.55 | 281.8 |
| 2661 | ENSMUSG00000020673 | 2.1660278 | 2.503113172 | 1.913696156 | 1.307999269 | 0.190873557 | NA | Tpo | thyroid peroxidase [Source:MGI Symbol;A | 0 | 0 | 2.0125 | 7.7118 | 3.2719 | 0 |
| 2662 | ENSMUSG00000020674 | 91.69534573 | -0.685468553 | 0.319463524 | -2.145686443 | 0.031898004 | 0.238066421 | Pxdn | peroxidasin [Source:MGI Symbol;Acc:MGI | 106.81 | 99.708 | 132.82 | 79.046 | 35.99 | 95.791 |
| 2663 | ENSMUSG00000020676 | 0 | NA | NA | NA | NA | NA | Ccl11 | chemokine (C-C motif) ligand 11 [Source:N | 0 | 0 | 0 | 0 | 0 | 0 |
| 2664 | ENSMUSG00000020677 | 335.0629195 | 0.366988959 | 0.151377722 | 2.424326076 | 0.015336828 | 0.14470442 | Ddx52 | DExD box helicase 52 [Source:MGI Symbo | 298.75 | 288.84 | 289.8 | 334.5 | 372.99 | 425.49 |
| 2665 | ENSMUSG00000020679 | 279.6530743 | 0.080380818 | 0.168275797 | 0.477673079 | 0.632882908 | 0.922107984 | Hnf1b | HNF1 homeobox B [Source:MGI Symbol;A | 225.67 | 285.76 | 304.89 | 271.84 | 289.01 | 300.74 |
| 2666 | ENSMUSG00000020680 | 1115.883349 | 0.093852433 | 0.104633521 | 0.89696334 | 0.369738479 | 0.80653433 | Taf15 | TATA-box binding protein associated facto | 1049.7 | 1036.1 | 1153.2 | 1129.8 | 1162.6 | 1164 |
| 2667 | ENSMUSG00000020681 | 13.04503946 | -0.361249687 | 0.600296901 | -0.601785028 | 0.547317242 | 0.895518681 | Ace | angiotensin I converting enzyme (peptidyl | 13.653 | 14.391 | 16.1 | 15.424 | 10.906 | 7.7969 |
| 2668 | ENSMUSG00000020682 | 1.727415082 | 1.314599518 | 1.700222262 | 0.773192745 | 0.439408319 | NA | Mmp28 | matrix metallopeptidase 28 (epilysin) [Sou | 0 | 1.0279 | 2.0125 | 2.8919 | 1.0906 | 3.3415 |
| 2669 | ENSMUSG00000020684 | 6.315567663 | -0.976511741 | 0.943627342 | -1.034848925 | 0.30073947 | 0.751875664 | Rasl10b | RAS-like, family 10, member B [Source:M( | 12.85 | 3.0838 | 9.0562 | 1.928 | 7.6343 | 3.3415 |
| 2670 | ENSMUSG00000020686 | 0 | NA | NA | NA | NA | NA | Gas2l2 | growth arrest-specific 2 like 2 [Source:MG | 0 | 0 | 0 | 0 | 0 | 0 |
| 2671 | ENSMUSG00000020687 | 495.4817393 | 0.064766713 | 0.140415981 | 0.461248877 | 0.644620058 | 0.925643431 | Cdc27 | cell division cycle 27 [Source:MGI Symbol; | 542.09 | 493.4 | 416.59 | 491.63 | 523.5 | 505.69 |
| 2672 | ENSMUSG00000020689 | 44.12854447 | -1.115817484 | 0.344996173 | -3.234289456 | 0.001219458 | 0.024195116 | Itgb3 | integrin beta 3 [Source:MGI Symbol;Acc:N | 61.839 | 66.815 | 52.325 | 23.135 | 28.356 | 32.301 |
| 2673 | ENSMUSG00000020690 | 0 | NA | NA | NA | NA | NA | Efcab3 | EF-hand calcium binding domain 3 [Sourc | 0 | 0 | 0 | 0 | 0 | 0 |
| 2674 | ENSMUSG00000020691 | 165.1021556 | 0.257801542 | 0.201205542 | 1.281284495 | 0.200093757 | 0.642574622 | Mettl2 | methyltransferase like 2 [Source:MGI Sym | 133.31 | 145.96 | 172.07 | 157.13 | 205.04 | 177.1 |
| 2675 | ENSMUSG00000020692 | 73.82174605 | -0.301620107 | 0.265826034 | -1.134652247 | 0.256521053 | 0.709880947 | Nle1 | notchless homolog 1 [Source:MGI Symbol | 69.87 | 85.317 | 89.556 | 63.623 | 62.165 | 72.4 |
| 2676 | ENSMUSG00000020694 | 407.5479087 | -0.103082303 | 0.141237811 | -0.729849195 | 0.46548237 | 0.863282087 | Tlk2 | tousled-like kinase 2 (Arabidopsis) [Source | 463.39 | 405 | 397.47 | 410.65 | 407.89 | 360.89 |
| 2677 | ENSMUSG00000020695 | 28.99174722 | -0.683285686 | 0.444196509 | -1.538250915 | 0.123987276 | 0.515924041 | Mrc2 | mannose receptor, C type 2 [Source:MGI ! | 24.093 | 33.921 | 49.306 | 20.244 | 18.541 | 27.846 |
| 2678 | ENSMUSG00000020696 | 469.7466995 | 0.116181577 | 0.148409748 | 0.782843301 | 0.433719136 | 0.845264432 | Rffl | ring finger and FYVE like domain containir | 465 | 421.45 | 465.89 | 508.02 | 407.89 | 550.24 |
| 2679 | ENSMUSG00000020697 | 270.9459104 | 0.152982372 | 0.151126636 | 1.012279347 | 0.311404507 | 0.759810187 | Lig3 | ligase III, DNA, ATP-dependent [Source:M | 268.24 | 249.78 | 251.56 | 294.98 | 273.75 | 287.37 |
| 2680 | ENSMUSG00000020698 | 19.63126253 | 0.294847614 | 0.489668062 | 0.602137728 | 0.547082463 | 0.895518681 | Cct6b | chaperonin containing Tcp1, subunit 6b (z | 15.259 | 19.53 | 18.112 | 16.388 | 23.994 | 24.505 |
| 2681 | ENSMUSG00000020700 | 233.4358833 | 0.056735929 | 0.174463574 | 0.325202151 | 0.745028081 | 0.949997329 | Map3k3 | mitogen-activated protein kinase kinase k | 212.02 | 256.98 | 218.36 | 266.06 | 235.57 | 211.63 |
| 2682 | ENSMUSG00000020701 | 27.90773011 | -0.475372903 | 0.49294369 | -0.964355386 | 0.334867782 | 0.776159726 | Tmem132e | transmembrane protein 132E [Source:MG | 44.171 | 26.726 | 26.162 | 18.316 | 13.087 | 38.985 |
| 2683 | ENSMUSG00000020702 | 0 | NA | NA | NA | NA | NA | Ccl1 | chemokine (C-C motif) ligand 1 [Source:M | 0 | 0 | 0 | 0 | 0 | 0 |
| 2684 | ENSMUSG00000020703 | 0 | NA | NA | NA | NA | NA | 5530401A14 | RIKEN cDNA 5530401A14 gene [Source:M | 0 | 0 | 0 | 0 | 0 | 0 |

| | | | | | | | | | | | | | | | |
|---|---|---|---|---|---|---|---|---|---|---|---|---|---|---|---|
| 2685 | ENSMUSG00000020704 | 0 | NA | NA | NA | NA | NA | Asic2 | acid-sensing (proton-gated) ion channel 2 | 0 | 0 | 0 | 0 | 0 | 0 |
| 2686 | ENSMUSG00000020705 | 748.4568746 | -0.01516292 | 0.118549142 | -0.12790409 | 0.898224874 | 0.982798755 | Ddx42 | DEAD box helicase 42 [Source:MGI Symbo | 780.62 | 774.02 | 702.36 | 755.76 | 788.52 | 689.47 |
| 2687 | ENSMUSG00000020706 | 672.6270838 | -0.073387627 | 0.115779038 | -0.633859362 | 0.526172608 | 0.888334265 | Ftsj3 | FtsJ RNA methyltransferase homolog 3 (E. | 652.92 | 715.43 | 701.35 | 643.94 | 680.55 | 641.57 |
| 2688 | ENSMUSG00000020707 | 222.2428135 | 0.311194752 | 0.181310532 | 1.716363344 | 0.086095561 | 0.432482144 | Rnf135 | ring finger protein 135 [Source:MGI Symb | 203.99 | 194.28 | 197.22 | 294.01 | 231.21 | 212.74 |
| 2689 | ENSMUSG00000020708 | 1267.413819 | 0.035272508 | 0.099325695 | 0.355119664 | 0.722499946 | 0.9447719 | Psmc5 | protease (prosome, macropain) 26S subu | 1256.9 | 1257.1 | 1241.7 | 1290.8 | 1225.9 | 1332.2 |
| 2690 | ENSMUSG00000020709 | 669.4074178 | -0.083919939 | 0.194668413 | -0.43109171 | 0.666401688 | 0.929596279 | Adap2 | ArfGAP with dual PH domains 2 [Source:N | 616.78 | 666.09 | 784.87 | 818.42 | 692.54 | 437.74 |
| 2691 | ENSMUSG00000020712 | 0 | NA | NA | NA | NA | NA | Tcam1 | testicular cell adhesion molecule 1 [Sourc | 0 | 0 | 0 | 0 | 0 | 0 |
| 2692 | ENSMUSG00000020713 | 0.17131964 | -0.796629176 | 4.080472857 | -0.195229623 | 0.84521319 | NA | Gh | growth hormone [Source:MGI Symbol;Acc | 0 | 1.0279 | 0 | 0 | 0 | 0 |
| 2693 | ENSMUSG00000020715 | 1218.958423 | 0.200303939 | 0.105580844 | 1.897161765 | 0.057806591 | 0.341829329 | Ern1 | endoplasmic reticulum (ER) to nucleus sig | 1130.8 | 1104 | 1169.3 | 1389.1 | 1306.6 | 1214.1 |
| 2694 | ENSMUSG00000020716 | 340.9461351 | -0.072884115 | 0.145835639 | -0.49976889 | 0.617237819 | 0.917509899 | Nf1 | neurofibromin 1 [Source:MGI Symbol;Acc | 361.4 | 345.38 | 342.12 | 367.28 | 305.37 | 324.13 |
| 2695 | ENSMUSG00000020717 | 179.760659 | -0.603459547 | 0.203681048 | -2.96276729 | 0.00304887 | 0.047316364 | Pecam1 | platelet/endothelial cell adhesion molecu | 204.79 | 235.39 | 210.31 | 139.78 | 113.42 | 174.87 |
| 2696 | ENSMUSG00000020718 | 238.9166401 | 0.391261363 | 0.170246898 | 2.298199665 | 0.021550428 | 0.182925697 | Polg2 | polymerase (DNA directed), gamma 2, acc | 231.29 | 192.22 | 196.22 | 294.98 | 273.75 | 245.05 |
| 2697 | ENSMUSG00000020719 | 8401.206237 | -0.115098782 | 0.116698545 | -0.986291494 | 0.323990093 | 0.77088057 | Ddx5 | DEAD box helicase 5 [Source:MGI Symbol; | 8524.1 | 8821.6 | 8864 | 9036.3 | 8442.5 | 6718.7 |
| 2698 | ENSMUSG00000020720 | 1286.623103 | 0.085202387 | 0.110004427 | 0.774535989 | 0.438613895 | 0.847803203 | Psmd12 | proteasome (prosome, macropain) 26S su | 1351.6 | 1270.5 | 1123 | 1349.6 | 1319.6 | 1305.4 |
| 2699 | ENSMUSG00000020721 | 224.2500511 | -0.148948951 | 0.168066577 | -0.886249688 | 0.375482994 | 0.809386889 | Helz | helicase with zinc finger domain [Source:N | 244.95 | 248.76 | 213.32 | 198.58 | 212.67 | 227.22 |
| 2700 | ENSMUSG00000020722 | 0.185640647 | 1.126924175 | 4.080472857 | 0.276174898 | 0.782413736 | NA | Cacng1 | calcium channel, voltage-dependent, gam | 0 | 0 | 0 | 0 | 0 | 1.1138 |
| 2701 | ENSMUSG00000020723 | 0 | NA | NA | NA | NA | NA | Cacng4 | calcium channel, voltage-dependent, gam | 0 | 0 | 0 | 0 | 0 | 0 |
| 2702 | ENSMUSG00000020728 | 3.588782092 | -1.724543393 | 1.360105968 | -1.267947817 | 0.204816586 | NA | Cep112 | centrosomal protein 112 [Source:MGI Syn | 10.44 | 5.1396 | 1.0062 | 3.8559 | 1.0906 | 0 |
| 2703 | ENSMUSG00000020732 | 0.824448242 | -0.598621533 | 2.872093747 | -0.208426878 | 0.834895672 | NA | Rab37 | RAB37, member RAS oncogene family [So | 0 | 0 | 3.0187 | 1.928 | 0 | 0 |
| 2704 | ENSMUSG00000020733 | 5929.872493 | 0.15272112 | 0.116981452 | 1.305515681 | 0.191717311 | 0.629099068 | Slc9a3r1 | solute carrier family 9 (sodium/hydrogen | 5166.4 | 5468.5 | 6215.6 | 6815.3 | 6495.7 | 5417.7 |
| 2705 | ENSMUSG00000020734 | 0.824421612 | 3.264491806 | 2.808832384 | 1.162223786 | 0.245144575 | NA | Grin2c | glutamate receptor, ionotropic, NMDA2C | 0 | 0 | 0 | 3.8559 | 1.0906 | 0 |
| 2706 | ENSMUSG00000020736 | 431.9544626 | 0.011531049 | 0.134516453 | 0.085722219 | 0.931687239 | 0.988420305 | Nt5c | 5',3'-nucleotidase, cytosolic [Source:MGI | 414.4 | 417.33 | 458.85 | 401.98 | 456.97 | 442.2 |
| 2707 | ENSMUSG00000020737 | 130.6970472 | 0.124508136 | 0.208586664 | 0.596913215 | 0.550565324 | 0.895703112 | Jpt1 | Jupiter microtubule associated homolog 1 | 118.86 | 135.69 | 120.75 | 133.03 | 122.15 | 153.71 |
| 2708 | ENSMUSG00000020738 | 402.3292417 | -0.011649677 | 0.14187821 | -0.082110401 | 0.934558922 | 0.988420305 | Sumo2 | small ubiquitin-like modifier 2 [Source:MG | 356.58 | 427.61 | 428.66 | 406.8 | 411.16 | 383.16 |
| 2709 | ENSMUSG00000020739 | 303.2742543 | 0.039906668 | 0.149091827 | 0.267665027 | 0.788957172 | 0.960994657 | Nup85 | nucleoporin 85 [Source:MGI Symbol;Acc:l | 310 | 289.87 | 296.84 | 292.09 | 294.47 | 336.38 |
| 2710 | ENSMUSG00000020740 | 121.5407194 | 0.123917179 | 0.215531803 | 0.574936863 | 0.565333998 | 0.901542552 | Gga3 | golgi associated, gamma adaptin ear cont | 116.45 | 99.708 | 132.82 | 130.14 | 127.6 | 122.52 |
| 2711 | ENSMUSG00000020741 | 4790.98854 | -0.065088571 | 0.118980469 | -0.547052571 | 0.584342615 | 0.908955096 | Cluh | clustered mitochondria (cluA/CLU1) homo | 4100.6 | 5297.9 | 5299.9 | 4398.6 | 4932.9 | 4716 |
| 2712 | ENSMUSG00000020743 | 928.4626715 | -0.379542999 | 0.148595034 | -2.554210529 | 0.010642888 | 0.113524143 | Mif4gd | MIF4G domain containing [Source:MGI Sy | 887.43 | 970.35 | 1293 | 852.16 | 807.06 | 760.76 |
| 2713 | ENSMUSG00000020744 | 256.7499997 | -0.134914335 | 0.15486821 | -0.871155767 | 0.383669108 | 0.816002185 | Slc25a19 | solute carrier family 25 (mitochondrial thi | 262.61 | 255.95 | 287.79 | 244.85 | 246.48 | 242.82 |
| 2714 | ENSMUSG00000020745 | 1563.998755 | 0.143599924 | 0.106240779 | 1.351646005 | 0.176488586 | 0.607400048 | Pafah1b1 | platelet-activating factor acetylhydrolase, | 1489.8 | 1512.1 | 1456 | 1486.5 | 1667.6 | 1772.1 |
| 2715 | ENSMUSG00000020747 | 637.8638997 | -0.01020416 | 0.135126042 | -0.075515866 | 0.939804275 | 0.988812319 | Tmem94 | transmembrane protein 94 [Source:MGI S | 577.43 | 622.92 | 720.47 | 609.23 | 598.75 | 698.38 |
| 2716 | ENSMUSG00000020752 | 312.8492229 | 0.172327541 | 0.161701151 | 1.06571623 | 0.286551939 | 0.73880343 | Recql5 | RecQ protein-like 5 [Source:MGI Symbol;/ | 318.83 | 263.15 | 299.86 | 309.44 | 380.63 | 305.19 |
| 2717 | ENSMUSG00000020755 | 331.3761023 | 0.096785457 | 0.150795917 | 0.641830755 | 0.520983078 | 0.887887809 | Sap30bp | SAP30 binding protein [Source:MGI Symb | 313.21 | 327.91 | 319.99 | 348 | 379.54 | 299.62 |
| 2718 | ENSMUSG00000020758 | 12.54564926 | -0.523118476 | 0.624577414 | -0.837555865 | 0.402280189 | 0.828953151 | Itgb4 | integrin beta 4 [Source:MGI Symbol;Acc:N | 12.85 | 14.391 | 17.106 | 6.7478 | 15.269 | 8.9108 |
| 2719 | ENSMUSG00000020766 | 1170.804518 | -0.483473158 | 0.123440189 | -3.91665926 | 8.98E-05 | 0.003321338 | Galk1 | galactokinase 1 [Source:MGI Symbol;Acc:l | 1349.2 | 1484.3 | 1262.8 | 1055.6 | 1020.8 | 852.09 |
| 2720 | ENSMUSG00000020770 | 172.3443901 | 0.086758953 | 0.202381993 | 0.428689091 | 0.6681495 | 0.930136128 | Unk | unkempt family zinc finger [Source:MGI S | 177.49 | 154.19 | 170.06 | 213.04 | 174.5 | 144.8 |
| 2721 | ENSMUSG00000020773 | 62.0457936 | -0.299706349 | 0.304253296 | -0.985055391 | 0.324596863 | 0.771309263 | Trim47 | tripartite motif-containing 47 [Source:MG | 85.129 | 54.48 | 65.406 | 56.875 | 46.897 | 63.489 |
| 2722 | ENSMUSG00000020774 | 302.3504056 | -0.046721158 | 0.151117449 | -0.309171168 | 0.757191325 | 0.953260973 | Aspa | aspartoacylase [Source:MGI Symbol;Acc:N | 282.69 | 328.93 | 310.93 | 314.26 | 298.83 | 278.46 |
| 2723 | ENSMUSG00000020775 | 1301.800159 | 0.020969922 | 0.102572763 | 0.204439478 | 0.838010083 | 0.971073158 | Mrpl38 | mitochondrial ribosomal protein L38 [Sou | 1343.6 | 1273.6 | 1259.8 | 1385.2 | 1303.3 | 1245.3 |
| 2724 | ENSMUSG00000020776 | 424.2784878 | -0.136304461 | 0.48865112 | -0.278940241 | 0.780290686 | 0.959892424 | Fbf1 | Fas (TNFRSF6) binding factor 1 [Source:M | 482.66 | 338.18 | 512.18 | 544.65 | 485.33 | 182.67 |
| 2725 | ENSMUSG00000020777 | 64089.38949 | -0.206176659 | 0.128442356 | -1.605207706 | 0.108448102 | 0.48522218 | Acox1 | acyl-Coenzyme A oxidase 1, palmitoyl [Sou | 71115 | 72598 | 62271 | 64580 | 66019 | 47953 |
| 2726 | ENSMUSG00000020778 | 68.03897756 | -0.372388766 | 0.309605973 | -1.202782885 | 0.22906035 | 0.67545951 | Ten1 | TEN1 telomerase capping complex subuni | 73.082 | 69.898 | 87.543 | 64.587 | 75.253 | 37.871 |
| 2727 | ENSMUSG00000020780 | 902.5873226 | 0.103577137 | 0.104419397 | 0.991933874 | 0.321229784 | 0.768486205 | Srp68 | signal recognition particle 68 [Source:MGI | 873.78 | 851.12 | 885.5 | 922.53 | 931.39 | 951.22 |
| 2728 | ENSMUSG00000020781 | 118.7111463 | -0.113501639 | 0.227011107 | -0.499982758 | 0.617087218 | 0.917509899 | Tsen54 | tRNA splicing endonuclease subunit 54 [S | 129.3 | 99.708 | 140.87 | 111.82 | 104.7 | 125.86 |
| 2729 | ENSMUSG00000020782 | 565.9129163 | 0.047417368 | 0.166134663 | 0.285415259 | 0.775326018 | 0.95893551 | Llgl2 | LLGL2 scribble cell polarity complex comp | 447.33 | 531.43 | 692.3 | 615.02 | 522.41 | 587 |
| 2730 | ENSMUSG00000020783 | 297.5148703 | 0.041854709 | 0.148821314 | 0.281241364 | 0.77852528 | 0.959559604 | Ncbp3 | nuclear cap binding subunit 3 [Source:MG | 287.51 | 303.24 | 288.79 | 290.16 | 287.92 | 327.47 |
| 2731 | ENSMUSG00000020785 | 1.422476634 | -0.840187361 | 1.937218141 | -0.433708183 | 0.664500363 | NA | Camkk1 | calcium/calmodulin-dependent protein ki | 2.4093 | 3.0838 | 0 | 1.928 | 0 | 1.1138 |
| 2732 | ENSMUSG00000020787 | 1.138206772 | 1.608612704 | 2.129587282 | 0.755363595 | 0.450030851 | NA | P2rx1 | purinergic receptor P2X, ligand-gated ion | 1.6062 | 0 | 0 | 1.928 | 2.1812 | 1.1138 |
| 2733 | ENSMUSG00000020788 | 18.06922165 | -1.210661895 | 0.522846615 | -2.315520193 | 0.020584481 | 0.176591892 | Atp2a3 | ATPase, Ca++ transporting, ubiquitous [Sc | 24.896 | 21.586 | 29.181 | 9.6398 | 13.087 | 10.025 |
| 2734 | ENSMUSG00000020790 | 570.2544259 | 0.094065027 | 0.127475154 | 0.737908714 | 0.460569924 | 0.859817816 | Ankfy1 | ankyrin repeat and FYVE domain containi | 542.9 | 548.91 | 563.5 | 594.77 | 642.37 | 529.08 |
| 2735 | ENSMUSG00000020792 | 515.7185693 | -0.02323476 | 0.129816998 | -0.17898088 | 0.857952715 | 0.975277277 | Exoc7 | exocyst complex component 7 [Source:M | 479.45 | 520.13 | 560.48 | 488.74 | 540.95 | 504.57 |
| 2736 | ENSMUSG00000020793 | 1.552456075 | -1.047537488 | 1.883233703 | -0.556244021 | 0.57804405 | NA | Galr2 | galanin receptor 2 [Source:MGI Symbol;A | 3.2124 | 3.0838 | 0 | 1.928 | 1.0906 | 0 |
| 2737 | ENSMUSG00000020794 | 1366.305932 | 0.052635316 | 0.103579891 | 0.508161532 | 0.611340066 | 0.916376001 | Ube2g1 | ubiquitin-conjugating enzyme E2G 1 [Sour | 1366.9 | 1283.9 | 1372.5 | 1297.5 | 1464.7 | 1412.4 |
| 2738 | ENSMUSG00000020798 | 2.25522418 | 2.005652901 | 1.593493516 | 1.258651435 | 0.20815626 | NA | Spns3 | spinster homolog 3 [Source:MGI Symbol;/ | 1.6062 | 0 | 1.0062 | 0.964 | 3.2719 | 6.6831 |
| 2739 | ENSMUSG00000020799 | 0 | NA | NA | NA | NA | NA | Tekt1 | tektin 1 [Source:MGI Symbol;Acc:MGI:133 | 0 | 0 | 0 | 0 | 0 | 0 |
| 2740 | ENSMUSG00000020801 | 70.17294349 | 0.172193948 | 0.291417506 | 0.590884023 | 0.554598132 | 0.897558119 | Med31 | mediator complex subunit 31 [Source:MG | 53.808 | 65.787 | 78.487 | 60.731 | 70.89 | 91.335 |

| | | | | | | | | | | | | | | | |
|---|---|---|---|---|---|---|---|---|---|---|---|---|---|---|---|
| 2741 | ENSMUSG00000020802 | 223.4325043 | -0.047973342 | 0.168318002 | -0.285016109 | 0.7756318 | 0.95893551 | Ube2o | ubiquitin-conjugating enzyme E2O [Source | 208 | 241.56 | 232.44 | 228.46 | 200.67 | 229.45 |
| 2742 | ENSMUSG00000020803 | 775.123338 | -0.03020456 | 0.10763313 | -0.280625123 | 0.778997945 | 0.959828458 | Txndc17 | thioredoxin domain containing 17 [Source | 775.8 | 778.13 | 795.94 | 771.18 | 766.7 | 762.98 |
| 2743 | ENSMUSG00000020804 | 0.845528352 | 3.296539104 | 2.749625613 | 1.198904712 | 0.230564999 | NA | Aanat | arylalkylamine N-acetyltransferase [Source | 0 | 0 | 0 | 2.8919 | 2.1812 | 0 |
| 2744 | ENSMUSG00000020805 | 26.39412675 | 0.917447044 | 0.44914349 | 2.042659115 | 0.041086195 | 0.277592171 | Slc13a5 | solute carrier family 13 (sodium-depender | 24.896 | 16.447 | 13.081 | 28.919 | 33.809 | 41.212 |
| 2745 | ENSMUSG00000020806 | 200.6438755 | -0.052582941 | 0.185992068 | -0.282716044 | 0.777394517 | 0.959559604 | Rhbdf2 | rhomboid 5 homolog 2 [Source:MGI Symb | 216.84 | 173.72 | 222.38 | 221.71 | 184.31 | 184.9 |
| 2746 | ENSMUSG00000020807 | 61.23355139 | -0.097031466 | 0.288230719 | -0.336645123 | 0.736384436 | 0.948109711 | 4933427D14 | RIKEN cDNA 4933427D14 gene [Source:M | 61.036 | 63.731 | 65.406 | 70.37 | 55.622 | 51.237 |
| 2747 | ENSMUSG00000020808 | 2.255564999 | -1.787055958 | 1.583110351 | -1.128825894 | 0.258971287 | NA | Pimreg | PICALM interacting mitotic regulator [Sou | 6.4248 | 3.0838 | 1.0062 | 1.928 | 1.0906 | 0 |
| 2748 | ENSMUSG00000020810 | 143.7734269 | -0.82587536 | 0.20184125 | -4.091707525 | 4.28E-05 | 0.001848838 | Cygb | cytoglobin [Source:MGI Symbol;Acc:MGI: | 195.15 | 191.19 | 165.02 | 103.15 | 112.33 | 95.791 |
| 2749 | ENSMUSG00000020811 | 4.410795864 | 0.1277934 | 1.016403942 | 0.125730917 | 0.899944928 | NA | Wscd1 | WSC domain containing 1 [Source:MGI Sy | 6.4248 | 3.0838 | 3.0187 | 2.8919 | 4.3625 | 6.6831 |
| 2750 | ENSMUSG00000020812 | 37.43498509 | 0.823805735 | 0.377124013 | 2.184442538 | 0.028929745 | 0.223286999 | Snhg16 | small nucleolar RNA host gene 16 [Source | 22.487 | 23.642 | 35.219 | 53.019 | 51.259 | 38.985 |
| 2751 | ENSMUSG00000020814 | 35.08307603 | -0.67266911 | 0.449210636 | -1.497446979 | 0.134276992 | 0.537569131 | Mxra7 | matrix-remodelling associated 7 [Source:N | 54.611 | 51.396 | 23.144 | 26.027 | 17.45 | 37.871 |
| 2752 | ENSMUSG00000020817 | 922.2630737 | 0.227259081 | 0.117055202 | 1.941469306 | 0.052201379 | 0.321125242 | Rabep1 | rabaptin, RAB GTPase binding effector pro | 799.09 | 892.23 | 858.33 | 945.66 | 944.48 | 1093.8 |
| 2753 | ENSMUSG00000020818 | 215.2984153 | 0.118571226 | 0.183733744 | 0.645342678 | 0.518705141 | 0.887592015 | Mfsd11 | major facilitator superfamily domain cont | 240.93 | 202.5 | 175.09 | 214 | 244.3 | 214.97 |
| 2754 | ENSMUSG00000020821 | 4725.785182 | 0.031975637 | 0.106140727 | 0.301256999 | 0.763218528 | 0.954864872 | Kif1c | kinesin family member 1C [Source:MGI Sy | 4270.9 | 4712 | 5038.3 | 4722.5 | 4408.3 | 5202.8 |
| 2755 | ENSMUSG00000020823 | 184.4497217 | -0.13446781 | 0.201639819 | -0.666871311 | 0.504854338 | 0.881592135 | Sec14l1 | SEC14-like lipid binding 1 [Source:MGI Syr | 150.98 | 203.53 | 225.4 | 175.44 | 186.5 | 164.85 |
| 2756 | ENSMUSG00000020826 | 0.305170013 | -1.657836477 | 4.041480335 | -0.410205257 | 0.681655385 | NA | Nos2 | nitric oxide synthase 2, inducible [Source: | 0.8031 | 1.0279 | 0 | 0 | 0 | 0 |
| 2757 | ENSMUSG00000020827 | 957.8581421 | 0.081346113 | 0.109589758 | 0.742278426 | 0.457918661 | 0.859322431 | Mink1 | misshapen-like kinase 1 (zebrafish) [Sourc | 896.26 | 919.99 | 977.06 | 1027.6 | 922.66 | 1003.6 |
| 2758 | ENSMUSG00000020828 | 123.2375259 | -0.000393629 | 0.249753453 | -0.00157607 | 0.998742478 | 0.999764475 | Pld2 | phospholipase D2 [Source:MGI Symbol;Ac | 93.963 | 137.74 | 138.86 | 143.63 | 93.793 | 131.43 |
| 2759 | ENSMUSG00000020829 | 763.0624477 | 0.107763813 | 0.114168357 | 0.943902642 | 0.345219402 | 0.786444837 | Slc46a1 | solute carrier family 46, member 1 [Source | 777.4 | 704.12 | 721.48 | 784.68 | 831.05 | 759.64 |
| 2760 | ENSMUSG00000020830 | 60.85275855 | 0.908680537 | 0.354561892 | 2.562826286 | 0.010382398 | 0.111734379 | Vmo1 | vitelline membrane outer layer 1 homolog | 32.927 | 27.754 | 66.412 | 68.442 | 91.612 | 77.969 |
| 2761 | ENSMUSG00000020831 | 7.081438243 | -0.810258092 | 0.943386867 | -0.858882098 | 0.390405567 | 0.819763149 | 0610010K14 | RIKEN cDNA 0610010K14 gene [Source:M | 16.865 | 3.0838 | 7.0437 | 6.7478 | 7.6343 | 1.1138 |
| 2762 | ENSMUSG00000020832 | 318.3424198 | -0.115621174 | 0.150428921 | -0.768610002 | 0.442124866 | 0.850193681 | Eral1 | Era (G-protein)-like 1 (E. coli) [Source:MGI | 299.56 | 341.27 | 353.19 | 311.36 | 320.64 | 284.03 |
| 2763 | ENSMUSG00000020834 | 73.34939995 | -0.138677652 | 0.27442563 | -0.505337828 | 0.613321578 | 0.916813074 | Dhrs13 | dehydrogenase/reductase (SDR family) m | 61.036 | 86.345 | 83.518 | 62.659 | 75.253 | 71.286 |
| 2764 | ENSMUSG00000020836 | 2.542207766 | 0.687013411 | 1.380109994 | 0.497796128 | 0.618627746 | NA | Coro6 | coronin 6 [Source:MGI Symbol;Acc:MGI:2 | 0.8031 | 4.1117 | 1.0062 | 3.8559 | 4.3625 | 1.1138 |
| 2765 | ENSMUSG00000020838 | 18.97654764 | 0.364737289 | 0.531415866 | 0.686350017 | 0.492492415 | 0.876429279 | Slc6a4 | solute carrier family 6 (neurotransmitter t | 11.243 | 25.698 | 13.081 | 23.135 | 23.994 | 16.708 |
| 2766 | ENSMUSG00000020839 | 21.2898975 | 0.467374132 | 0.507729327 | 0.92051829 | 0.357301979 | 0.796427158 | Tmigd1 | transmembrane and immunoglobulin don | 15.259 | 12.335 | 26.162 | 28.919 | 28.356 | 16.708 |
| 2767 | ENSMUSG00000020840 | 873.639297 | 0.252107886 | 0.12407972 | 2.031821846 | 0.042171695 | 0.282846725 | Blmh | bleomycin hydrolase [Source:MGI Symbol | 871.37 | 768.88 | 750.66 | 850.23 | 982.65 | 1018.1 |
| 2768 | ENSMUSG00000020841 | 650.6781674 | 0.054164642 | 0.134893341 | 0.401536809 | 0.688024943 | 0.936885029 | Cpd | carboxypeptidase D [Source:MGI Symbol; | 603.13 | 679.45 | 632.93 | 602.49 | 763.43 | 622.64 |
| 2769 | ENSMUSG00000020843 | 259.4967973 | 0.037346676 | 0.171017873 | 0.218378791 | 0.827133989 | 0.969748238 | Timm22 | translocase of inner mitochondrial memb | 250.57 | 272.4 | 245.52 | 264.13 | 221.4 | 302.97 |
| 2770 | ENSMUSG00000020844 | 22.05929246 | -0.670201254 | 0.490170818 | -1.367281015 | 0.17153724 | 0.599034187 | Nxn | nucleoredoxin [Source:MGI Symbol;Acc:N | 19.274 | 31.865 | 30.187 | 12.532 | 22.903 | 15.594 |
| 2771 | ENSMUSG00000020846 | 16.7900025 | -0.528649205 | 0.538521111 | -0.981668489 | 0.326263194 | 0.772018857 | Rflnb | refilin B [Source:MGI Symbol;Acc:MGI:192 | 25.699 | 17.475 | 16.1 | 10.604 | 15.269 | 15.594 |
| 2772 | ENSMUSG00000020847 | 38.4934861 | 0.317948123 | 0.358602075 | 0.886632132 | 0.375276989 | 0.809332934 | Rph3al | rabphilin 3A-like (without C2 domains) [Sc | 37.746 | 31.865 | 33.206 | 52.055 | 35.99 | 40.098 |
| 2773 | ENSMUSG00000020848 | 2.860846622 | -0.325476144 | 1.394432783 | -0.233411139 | 0.815442165 | NA | Doc2b | double C2, beta [Source:MGI Symbol;Acc: | 2.4093 | 3.0838 | 4.025 | 0.964 | 0 | 6.6831 |
| 2774 | ENSMUSG00000020849 | 3466.305808 | -0.035104235 | 0.094985888 | -0.369573168 | 0.711700547 | 0.944239215 | Ywhae | tyrosine 3-monooxygenase/tryptophan 5- | 3409.2 | 3430.2 | 3685.9 | 3210 | 3623 | 3439.5 |
| 2775 | ENSMUSG00000020850 | 3267.146849 | -0.044964081 | 0.100976018 | -0.445294652 | 0.656106832 | 0.92651636 | Prpf8 | pre-mRNA processing factor 8 [Source:MC | 3103.2 | 3419.9 | 3432.3 | 3266.9 | 3428.9 | 2951.7 |
| 2776 | ENSMUSG00000020857 | 10.52240774 | -0.187382413 | 0.665610144 | -0.281519768 | 0.778311768 | 0.959559604 | Nme2 | NME/NM23 nucleoside diphosphate kinas | 11.243 | 10.279 | 12.075 | 8.6758 | 14.178 | 6.6831 |
| 2777 | ENSMUSG00000020859 | 1413.814346 | 0.090345363 | 0.119842981 | 0.753864448 | 0.45093062 | 0.855818389 | Spag9 | sperm associated antigen 9 [Source:MGI S | 1403 | 1480.2 | 1224.6 | 1303.3 | 1526.9 | 1544.9 |
| 2778 | ENSMUSG00000020863 | 939.5827603 | 0.039107682 | 0.113156881 | 0.345605864 | 0.729638938 | 0.946534338 | Luc7l3 | LUC7-like 3 (S. cerevisiae) [Source:MGI Syı | 889.84 | 938.49 | 952.91 | 968.8 | 1017.5 | 869.91 |
| 2779 | ENSMUSG00000020864 | 900.4684106 | -0.275592493 | 0.120761384 | -2.282124337 | 0.022482001 | 0.187679993 | Ankrd40 | ankyrin repeat domain 40 [Source:MGI Sy | 952.48 | 1067 | 938.83 | 731.66 | 847.41 | 865.46 |
| 2780 | ENSMUSG00000020865 | 4474.542388 | -0.538055961 | 0.169120413 | -3.181496259 | 0.001465164 | 0.027884387 | Abcc3 | ATP-binding cassette, sub-family C (CFTR/ | 5189.6 | 5600.1 | 5109.7 | 4598.2 | 3836.8 | 2512.8 |
| 2781 | ENSMUSG00000020866 | 2.640992854 | -1.113835598 | 1.412869862 | -0.788349747 | 0.430492157 | NA | Cacna1g | calcium channel, voltage-dependent, T typ | 4.8186 | 2.0558 | 4.025 | 3.8559 | 1.0906 | 0 |
| 2782 | ENSMUSG00000020867 | 0 | NA | NA | NA | NA | NA | Spata20 | spermatogenesis associated 20 [Source:M | 0 | 0 | 0 | 0 | 0 | 0 |
| 2783 | ENSMUSG00000020868 | 415.4819199 | 0.035038332 | 0.161990988 | 0.216298033 | 0.828755442 | 0.969955221 | Xylt2 | xylosyltransferase II [Source:MGI Symbol; | 330.88 | 428.64 | 472.93 | 429.93 | 386.08 | 444.42 |
| 2784 | ENSMUSG00000020869 | 1152.961479 | -0.168652795 | 0.117121617 | -1.439980074 | 0.149873037 | 0.563095827 | Lrrc59 | leucine rich repeat containing 59 [Source: | 1312.3 | 1189.3 | 1159.2 | 1196.3 | 989.19 | 1071.5 |
| 2785 | ENSMUSG00000020870 | 0.892336804 | 0.740760821 | 2.797409738 | 0.264802403 | 0.791161686 | NA | Cdc34b | cell division cycle 34B [Source:MGI Symbo | 0 | 0 | 2.0125 | 0 | 0 | 3.3415 |
| 2786 | ENSMUSG00000020871 | 3.857817517 | 2.851239775 | 1.325496002 | 2.151073841 | 0.031470372 | NA | Dlx4 | distal-less homeobox 4 [Source:MGI Symb | 0.8031 | 0 | 2.0125 | 3.8559 | 10.906 | 5.5692 |
| 2787 | ENSMUSG00000020872 | 0.346303622 | 2.004339653 | 3.884003423 | 0.516049919 | 0.605819544 | NA | Tac4 | tachykinin 4 [Source:MGI Symbol;Acc:MG | 0 | 0 | 0 | 0.964 | 0 | 1.1138 |
| 2788 | ENSMUSG00000020873 | 624.7237525 | 0.126049654 | 0.165323993 | 0.762440176 | 0.445797335 | 0.852395554 | Slc35b1 | solute carrier family 35, member B1 [Sour | 667.38 | 571.52 | 553.43 | 789.5 | 508.23 | 658.28 |
| 2789 | ENSMUSG00000020875 | 0 | NA | NA | NA | NA | NA | Hoxb9 | homeobox B9 [Source:MGI Symbol;Acc:M | 0 | 0 | 0 | 0 | 0 | 0 |
| 2790 | ENSMUSG00000020876 | 139.4315888 | 0.024154968 | 0.201614267 | 0.11980783 | 0.904635379 | 0.983865503 | Snx11 | sorting nexin 11 [Source:MGI Symbol;Acc: | 150.98 | 124.38 | 138.86 | 129.17 | 145.05 | 148.14 |
| 2791 | ENSMUSG00000020877 | 909.4581138 | 0.082525435 | 0.117414499 | 0.702855579 | 0.482145757 | 0.871593699 | Scrn2 | secernin 2 [Source:MGI Symbol;Acc:MGI: | 791.06 | 899.43 | 960.96 | 924.45 | 961.93 | 918.92 |
| 2792 | ENSMUSG00000020878 | 23.19808595 | -0.058921519 | 0.504019799 | -0.116903184 | 0.906936775 | 0.984383096 | Lrrc46 | leucine rich repeat containing 46 [Source: | 24.093 | 23.642 | 23.144 | 17.352 | 13.087 | 37.871 |
| 2793 | ENSMUSG00000020882 | 5.704714277 | -0.438580913 | 0.914810971 | -0.479422446 | 0.631638128 | 0.921549084 | Cacnb1 | calcium channel, voltage-dependent, beta | 10.44 | 6.1675 | 3.0187 | 5.7839 | 4.3625 | 4.4554 |
| 2794 | ENSMUSG00000020883 | 882.2583686 | 0.19114301 | 0.124788248 | 1.531738869 | 0.125586887 | 0.519739264 | Fbxl20 | F-box and leucine-rich repeat protein 20 [ | 915.54 | 758.6 | 796.95 | 1030.5 | 898.67 | 893.3 |
| 2795 | ENSMUSG00000020884 | 11277.76896 | 0.066964512 | 0.08357653 | 0.801235845 | 0.422995123 | 0.84005258 | Asgr1 | asialoglycoprotein receptor 1 [Source:MG | 11040 | 10774 | 11234 | 11016 | 11961 | 11642 |
| 2796 | ENSMUSG00000020886 | 15.41767077 | -0.334505776 | 0.585753208 | -0.571069474 | 0.56795255 | 0.903352921 | Dlg4 | discs large MAGUK scaffold protein 4 [Sou | 20.881 | 21.586 | 9.0562 | 14.46 | 9.8156 | 16.708 |

| | | | | | | | | | | | | | | | |
|---|---|---|---|---|---|---|---|---|---|---|---|---|---|---|---|
| 2797 | ENSMUSG00000020887 | 0 | NA | NA | NA | NA | NA | A230052G05 | RIKEN cDNA A230052G05 gene [Source:M | 0 | 0 | 0 | 0 | 0 | 0 |
| 2798 | ENSMUSG00000020888 | 110.1772941 | -0.155112806 | 0.23863835 | -0.649991112 | 0.515697963 | 0.886238519 | Dvl2 | dishevelled segment polarity protein 2 [Sc | 104.4 | 140.82 | 103.64 | 120.5 | 99.246 | 92.449 |
| 2799 | ENSMUSG00000020889 | 531.7683017 | 1.849769656 | 0.65090955 | 2.841822887 | 0.00448564 | 0.062145883 | Nr1d1 | nuclear receptor subfamily 1, group D, me | 419.22 | 169.61 | 103.64 | 566.82 | 736.17 | 1195.2 |
| 2800 | ENSMUSG00000020890 | 0 | NA | NA | NA | NA | NA | Gucy2e | guanylate cyclase 2e [Source:MGI Symbol | 0 | 0 | 0 | 0 | 0 | 0 |
| 2801 | ENSMUSG00000020891 | 0 | NA | NA | NA | NA | NA | Alox8 | arachidonate 8-lipoxygenase [Source:MGI | 0 | 0 | 0 | 0 | 0 | 0 |
| 2802 | ENSMUSG00000020892 | 0.660353019 | -0.01500701 | 2.83315355 | -0.005296928 | 0.995773683 | NA | Aloxe3 | arachidonate lipoxygenase 3 [Source:MGI | 0 | 1.0279 | 1.0062 | 1.928 | 0 | 0 |
| 2803 | ENSMUSG00000020893 | 719.8260576 | -0.519165975 | 0.182237561 | -2.848841765 | 0.00438787 | 0.061260761 | Per1 | period circadian clock 1 [Source:MGI Syml | 885.02 | 886.07 | 772.8 | 585.13 | 767.8 | 422.15 |
| 2804 | ENSMUSG00000020894 | 371.2917485 | 0.000919995 | 0.135040304 | 0.006812743 | 0.994564259 | 0.998676967 | Vamp2 | vesicle-associated membrane protein 2 [S | 365.41 | 373.13 | 375.33 | 385.59 | 374.08 | 354.2 |
| 2805 | ENSMUSG00000020895 | 27.01859161 | -0.281863762 | 0.427050692 | -0.660024133 | 0.509238343 | 0.882793133 | Tmem107 | transmembrane protein 107 [Source:MGI | 24.093 | 34.949 | 30.187 | 29.883 | 20.722 | 22.277 |
| 2806 | ENSMUSG00000020897 | 5.005626116 | -1.855676792 | 1.206134698 | -1.538531969 | 0.123918598 | NA | Aurkb | aurora kinase B [Source:MGI Symbol;Acc: | 15.259 | 3.0838 | 5.0312 | 0 | 1.0906 | 5.5692 |
| 2807 | ENSMUSG00000020898 | 110.4083765 | 0.277072826 | 0.232227494 | 1.193109486 | 0.232826497 | 0.68004896 | Ctc1 | CTS telomere maintenance complex comp | 105.21 | 103.82 | 90.562 | 133.99 | 131.96 | 96.904 |
| 2808 | ENSMUSG00000020899 | 214.5269064 | 0.108504281 | 0.170305218 | 0.637116596 | 0.524048899 | 0.888334265 | Pfas | phosphoribosylformylglycinamidine synth | 209.61 | 191.19 | 218.36 | 212.08 | 244.3 | 211.63 |
| 2809 | ENSMUSG00000020900 | 289.9069409 | 0.658260971 | 0.184792336 | 3.562165973 | 0.000367808 | 0.010124897 | Myh10 | myosin, heavy polypeptide 10, non-muscl | 228.88 | 208.67 | 236.47 | 269.91 | 407.89 | 387.62 |
| 2810 | ENSMUSG00000020901 | 13.70180248 | -2.791511835 | 0.763698486 | -3.655253852 | 0.000256928 | 0.00766227 | Pik3r5 | phosphoinositide-3-kinase regulatory sub | 20.881 | 38.033 | 13.081 | 5.7839 | 1.0906 | 3.3415 |
| 2811 | ENSMUSG00000020902 | 144.3619754 | -0.98047455 | 0.218913537 | -4.478821008 | 7.51E-06 | 0.000427408 | Ntn1 | netrin 1 [Source:MGI Symbol;Acc:MGI:10 | 218.44 | 193.25 | 163.01 | 106.04 | 79.615 | 105.82 |
| 2812 | ENSMUSG00000020903 | 282.9570996 | 0.18030414 | 0.183638666 | 0.981841917 | 0.326177734 | 0.772018857 | Stx8 | syntaxin 8 [Source:MGI Symbol;Acc:MGI:1 | 300.36 | 250.81 | 243.51 | 236.17 | 328.28 | 338.61 |
| 2813 | ENSMUSG00000020904 | 0.133850373 | -0.796629176 | 4.080472857 | -0.195229623 | 0.84521319 | NA | Cfap52 | cilia and flagella associated protein 52 [So | 0.8031 | 0 | 0 | 0 | 0 | 0 |
| 2814 | ENSMUSG00000020905 | 1.040533161 | -0.903287288 | 2.649905161 | -0.340875327 | 0.73319744 | NA | Usp43 | ubiquitin specific peptidase 43 [Source:M | 4.0155 | 0 | 0 | 0 | 0 | 2.2277 |
| 2815 | ENSMUSG00000020907 | 0 | NA | NA | NA | NA | NA | Rcvrn | recoverin [Source:MGI Symbol;Acc:MGI:9 | 0 | 0 | 0 | 0 | 0 | 0 |
| 2816 | ENSMUSG00000020908 | 0 | NA | NA | NA | NA | NA | Myh3 | myosin, heavy polypeptide 3, skeletal mus | 0 | 0 | 0 | 0 | 0 | 0 |
| 2817 | ENSMUSG00000020910 | 328.4787711 | 0.068538832 | 0.152562137 | 0.449251915 | 0.653249941 | 0.925756399 | Adprm | ADP-ribose/CDP-alcohol diphosphatase, n | 326.06 | 308.38 | 327.03 | 305.58 | 319.55 | 384.28 |
| 2818 | ENSMUSG00000020911 | 16.97096286 | -0.209155625 | 0.542014292 | -0.385885813 | 0.699581238 | 0.941362191 | Krt19 | keratin 19 [Source:MGI Symbol;Acc:MGI:9 | 12.047 | 24.67 | 18.112 | 18.316 | 13.087 | 15.594 |
| 2819 | ENSMUSG00000020912 | 0 | NA | NA | NA | NA | NA | Krt12 | keratin 12 [Source:MGI Symbol;Acc:MGI:9 | 0 | 0 | 0 | 0 | 0 | 0 |
| 2820 | ENSMUSG00000020913 | 0 | NA | NA | NA | NA | NA | Krt24 | keratin 24 [Source:MGI Symbol;Acc:MGI:1 | 0 | 0 | 0 | 0 | 0 | 0 |
| 2821 | ENSMUSG00000020914 | 18.72102288 | -1.707766307 | 0.59576814 | -2.866494852 | 0.004150449 | 0.058842478 | Top2a | topoisomerase (DNA) II alpha [Source:MG | 44.171 | 17.475 | 24.15 | 7.7118 | 5.4531 | 13.366 |
| 2822 | ENSMUSG00000020916 | 0 | NA | NA | NA | NA | NA | Krt36 | keratin 36 [Source:MGI Symbol;Acc:MGI:1 | 0 | 0 | 0 | 0 | 0 | 0 |
| 2823 | ENSMUSG00000020917 | 3357.243047 | 0.099860078 | 0.110561002 | 0.903212488 | 0.366413131 | 0.804706883 | Acly | ATP citrate lyase [Source:MGI Symbol;Acc | 3544.1 | 3254.4 | 2923.1 | 3252.5 | 3367.8 | 3801.5 |
| 2824 | ENSMUSG00000020918 | 362.9991542 | -0.092266485 | 0.141038284 | -0.654194609 | 0.512986434 | 0.884860213 | Kat2a | K(lysine) acetyltransferase 2A [Source:MG | 379.87 | 352.58 | 391.43 | 362.46 | 367.54 | 324.13 |
| 2825 | ENSMUSG00000020919 | 1247.367769 | -0.160354708 | 0.14494193 | -1.106337608 | 0.26858042 | 0.7242258 | Stat5b | signal transducer and activator of transcri | 1145.2 | 1311.6 | 1493.3 | 1066.2 | 1062.3 | 1405.7 |
| 2826 | ENSMUSG00000020921 | 137.1352842 | 0.13308549 | 0.223451703 | 0.595589509 | 0.551449487 | 0.89589637 | Tmem101 | transmembrane protein 101 [Source:MGI | 163.03 | 104.85 | 123.77 | 145.56 | 135.24 | 150.37 |
| 2827 | ENSMUSG00000020922 | 403.4263716 | -0.162661686 | 0.13959759 | -1.165218439 | 0.243930584 | 0.693748143 | Lsm12 | LSM12 homolog [Source:MGI Symbol;Acc | 393.52 | 466.67 | 418.6 | 360.53 | 398.08 | 383.16 |
| 2828 | ENSMUSG00000020923 | 1234.433256 | 0.111383632 | 0.119527397 | 0.931866956 | 0.351405287 | 0.792511048 | Ubtf | upstream binding transcription factor, RN | 1271.3 | 1122.5 | 1165.2 | 1185.7 | 1212.8 | 1449.1 |
| 2829 | ENSMUSG00000020925 | 205.2215449 | 0.080423679 | 0.172719914 | 0.465630608 | 0.64147993 | 0.924019354 | Ccdc43 | coiled-coil domain containing 43 [Source:I | 191.14 | 210.72 | 197.22 | 233.28 | 199.58 | 199.38 |
| 2830 | ENSMUSG00000020926 | 64.88373564 | -1.233581157 | 0.356378809 | -3.461432399 | 0.000537309 | 0.013061576 | Adam11 | a disintegrin and metallopeptidase domai | 69.87 | 114.1 | 89.556 | 43.379 | 52.35 | 20.049 |
| 2831 | ENSMUSG00000020928 | 0.185640647 | 1.126924175 | 4.080472857 | 0.276174898 | 0.782413736 | NA | Higd1b | HIG1 domain family, member 1B [Source:I | 0 | 0 | 0 | 0 | 0 | 1.1138 |
| 2832 | ENSMUSG00000020929 | 382.1580932 | -0.028772005 | 0.143157449 | -0.20098154 | 0.840713009 | 0.971854408 | Eftud2 | elongation factor Tu GTP binding domain | 365.41 | 405 | 387.4 | 341.25 | 379.54 | 414.35 |
| 2833 | ENSMUSG00000020930 | 2.643798402 | -0.153833515 | 1.29510205 | -0.118780999 | 0.905448863 | NA | Ccdc103 | coiled-coil domain containing 103 [Source | 3.2124 | 3.0838 | 2.0125 | 0.964 | 4.3625 | 2.2277 |
| 2834 | ENSMUSG00000020932 | 7.133443931 | -1.26565862 | 0.8641384 | -1.464648046 | 0.143016966 | 0.552624717 | Gfap | glial fibrillary acidic protein [Source:MGI S | 8.031 | 5.1396 | 17.106 | 4.8199 | 4.3625 | 3.3415 |
| 2835 | ENSMUSG00000020935 | 463.107524 | -0.002389281 | 0.147504615 | -0.01619801 | 0.987076423 | 0.99859176 | Dcakd | dephospho-CoA kinase domain containing | 435.28 | 408.08 | 547.4 | 458.85 | 466.78 | 462.25 |
| 2836 | ENSMUSG00000020936 | 1269.949553 | 0.11959341 | 0.108954756 | 1.097642859 | 0.27236047 | 0.727618887 | Nmt1 | N-myristoyltransferase 1 [Source:MGI Syn | 1297.8 | 1159.5 | 1193.4 | 1262.8 | 1277.1 | 1429.1 |
| 2837 | ENSMUSG00000020937 | 13.16607689 | 0.054748192 | 0.596674208 | 0.091755586 | 0.926892232 | 0.987712228 | Plcd3 | phospholipase C, delta 3 [Source:MGI Syn | 15.259 | 9.2513 | 14.087 | 10.604 | 13.087 | 16.708 |
| 2838 | ENSMUSG00000020940 | 0 | NA | NA | NA | NA | NA | Efcab15 | EF-hand calcium binding domain 15 [Sour | 0 | 0 | 0 | 0 | 0 | 0 |
| 2839 | ENSMUSG00000020941 | 55.5476675 | -0.152882584 | 0.316848577 | -0.482509928 | 0.629443747 | 0.92057725 | Map3k14 | mitogen-activated protein kinase kinase k | 53.808 | 47.284 | 74.462 | 53.983 | 44.715 | 59.034 |
| 2840 | ENSMUSG00000020945 | 0 | NA | NA | NA | NA | NA | Lyzl6 | lysozyme-like 6 [Source:MGI Symbol;Acc:I | 0 | 0 | 0 | 0 | 0 | 0 |
| 2841 | ENSMUSG00000020946 | 1216.876387 | -0.134804685 | 0.122949582 | -1.096422477 | 0.272893932 | 0.728119873 | Gosr2 | golgi SNAP receptor complex member 2 [S | 1306.6 | 1106 | 1407.7 | 1123 | 1108.1 | 1249.7 |
| 2842 | ENSMUSG00000020948 | 121.2261329 | -0.209428562 | 0.21303066 | -0.983091175 | 0.325562567 | 0.771619425 | Klhl28 | kelch-like 28 [Source:MGI Symbol;Acc:MG | 140.54 | 119.24 | 129.81 | 103.15 | 123.24 | 111.38 |
| 2843 | ENSMUSG00000020949 | 244.1000169 | 0.138037948 | 0.160290692 | 0.861172576 | 0.389143002 | 0.818819375 | Fkbp3 | FK506 binding protein 3 [Source:MGI Sym | 229.69 | 237.45 | 230.43 | 271.84 | 231.21 | 263.98 |
| 2844 | ENSMUSG00000020950 | 0.133850373 | -0.796629176 | 4.080472857 | -0.195229623 | 0.84521319 | NA | Foxg1 | forkhead box G1 [Source:MGI Symbol;Acc | 0.8031 | 0 | 0 | 0 | 0 | 0 |
| 2845 | ENSMUSG00000020952 | 668.1174378 | 0.142962649 | 0.113370529 | 1.261021277 | 0.207301182 | 0.652364421 | Scfd1 | Sec1 family domain containing 1 [Source:I | 647.3 | 607.5 | 650.03 | 700.81 | 698 | 705.06 |
| 2846 | ENSMUSG00000020953 | 0.346303622 | 2.004339653 | 3.884003423 | 0.516049919 | 0.605819544 | NA | Coch | cochlin [Source:MGI Symbol;Acc:MGI:127 | 0 | 0 | 0 | 0.964 | 0 | 1.1138 |
| 2847 | ENSMUSG00000020954 | 432.6705704 | 0.064993074 | 0.143282977 | 0.45359941 | 0.650117179 | 0.925643431 | Strn3 | striatin, calmodulin binding protein 3 [Sou | 403.96 | 482.09 | 383.38 | 461.75 | 447.15 | 417.69 |
| 2848 | ENSMUSG00000020955 | 188.49749 | 0.100526952 | 0.190432658 | 0.527887142 | 0.597577669 | 0.914641631 | Ap4s1 | adaptor-related protein complex AP-4, sig | 183.11 | 188.11 | 174.08 | 158.09 | 214.85 | 212.74 |
| 2849 | ENSMUSG00000020956 | 265.1309179 | -0.155967563 | 0.155137296 | -1.005351822 | 0.31472747 | 0.761014653 | Dtd2 | D-tyrosyl-tRNA deacylase 2 [Source:MGI S | 293.94 | 275.48 | 268.67 | 256.42 | 232.3 | 263.98 |
| 2850 | ENSMUSG00000020961 | 196.4264976 | 0.186526276 | 0.193455246 | 0.964183083 | 0.334954144 | 0.776159726 | Ston2 | stonin 2 [Source:MGI Symbol;Acc:MGI:19 | 195.96 | 188.11 | 167.04 | 225.57 | 166.86 | 235.02 |
| 2851 | ENSMUSG00000020962 | 503.1659807 | 0.009419583 | 0.143750944 | 0.065527105 | 0.947754326 | 0.990677354 | Gtf2a1 | general transcription factor II A, 1 [Source | 512.38 | 482.09 | 509.16 | 428.01 | 511.5 | 575.86 |
| 2852 | ENSMUSG00000020963 | 0 | NA | NA | NA | NA | NA | Tshr | thyroid stimulating hormone receptor [So | 0 | 0 | 0 | 0 | 0 | 0 |

| | | | | | | | | | | | | | | | |
|---|---|---|---|---|---|---|---|---|---|---|---|---|---|---|---|
| 2853 | ENSMUSG00000020964 | 2266.258872 | 0.104725246 | 0.106963352 | 0.979075952 | 0.327542453 | 0.772937193 | Sel1l | sel-1 suppressor of lin-12-like (C. elegans) | 2101.7 | 2253.2 | 2198.6 | 2619.1 | 2164.9 | 2260 |
| 2854 | ENSMUSG00000020973 | 19.12463289 | 0.061807851 | 0.554477394 | 0.111470462 | 0.911243288 | 0.985839849 | Dnaaf2 | dynein, axonemal assembly factor 2 [Sour | 22.487 | 10.279 | 23.144 | 13.496 | 15.269 | 30.074 |
| 2855 | ENSMUSG00000020974 | 4.242220664 | 0.134236427 | 1.092526435 | 0.122867898 | 0.902211707 | NA | Pole2 | polymerase (DNA directed), epsilon 2 (p59 | 4.0155 | 6.1675 | 2.0125 | 7.7118 | 1.0906 | 4.4554 |
| 2856 | ENSMUSG00000020978 | 704.8739252 | 0.054425077 | 0.1349713 | 0.403234443 | 0.686775764 | 0.936582996 | Klhdc2 | kelch domain containing 2 [Source:MGI Sy | 694.68 | 614.69 | 764.75 | 644.9 | 791.79 | 718.43 |
| 2857 | ENSMUSG00000020982 | 483.6568851 | -0.004473901 | 0.145750725 | -0.030695566 | 0.975512327 | 0.996491676 | Nemf | nuclear export mediator factor [Source:M | 459.37 | 549.94 | 444.76 | 519.58 | 501.68 | 426.6 |
| 2858 | ENSMUSG00000020986 | 2105.92216 | -0.092844065 | 0.103128493 | -0.900275592 | 0.367973607 | 0.80627882 | Sec23a | SEC23 homolog A, COPII coat complex cor | 2261.5 | 2141.2 | 2118.1 | 2097.6 | 2170.3 | 1846.8 |
| 2859 | ENSMUSG00000020988 | 1531.861942 | 0.110483218 | 0.111741321 | 0.988740933 | 0.32278991 | 0.769926329 | L2hgdh | L-2-hydroxyglutarate dehydrogenase [Sou | 1329.1 | 1621 | 1471.1 | 1645.5 | 1619.6 | 1504.8 |
| 2860 | ENSMUSG00000020990 | 3.154693578 | 0.896775315 | 1.204699945 | 0.74439724 | 0.456636188 | NA | Cdkl1 | cyclin-dependent kinase-like 1 (CDC2-rela | 1.6062 | 2.0558 | 3.0187 | 6.7478 | 3.2719 | 2.2277 |
| 2861 | ENSMUSG00000020992 | 1.090494021 | -1.109926708 | 2.171201027 | -0.511204027 | 0.609208199 | NA | 4930512B01 | RIKEN cDNA 4930512B01 gene [Source:M | 2.4093 | 2.0558 | 0 | 0.964 | 0 | 1.1138 |
| 2862 | ENSMUSG00000020993 | 555.6280861 | -0.014611999 | 0.136005125 | -0.107437123 | 0.914442205 | 0.98625062 | Trappc6b | trafficking protein particle complex 6B [So | 547.72 | 574.61 | 552.43 | 479.1 | 558.4 | 621.52 |
| 2863 | ENSMUSG00000020994 | 657.6866587 | 0.243401537 | 0.125855851 | 1.933970766 | 0.053116722 | 0.323714501 | Pnn | pinin [Source:MGI Symbol;Acc:MGI:11005 | 588.67 | 589 | 629.91 | 749.01 | 757.98 | 631.55 |
| 2864 | ENSMUSG00000021000 | 7686.773893 | -0.131487234 | 0.09776646 | -1.344911474 | 0.178653824 | 0.610813333 | Mia2 | MIA SH3 domain ER export factor 2 [Sourc | 7513.8 | 8597.5 | 8000.7 | 7616.4 | 7622.3 | 6769.9 |
| 2865 | ENSMUSG00000021003 | 103.1647758 | -0.057573098 | 0.221205924 | -0.26026924 | 0.794656098 | 0.962189806 | Galc | galactosylceramidase [Source:MGI Symbo | 101.19 | 105.88 | 108.67 | 98.326 | 105.79 | 99.132 |
| 2866 | ENSMUSG00000021007 | 29.82548764 | 0.098130277 | 0.428684726 | 0.228910131 | 0.818938766 | 0.969160361 | Spata7 | spermatogenesis associated 7 [Source:MC | 34.533 | 17.475 | 34.212 | 28.919 | 37.081 | 26.732 |
| 2867 | ENSMUSG00000021009 | 300.0442885 | -0.144722172 | 0.154522471 | -0.936576869 | 0.348976255 | 0.79040715 | Ptpn21 | protein tyrosine phosphatase, non-recept | 299.56 | 350.52 | 295.84 | 305.58 | 278.11 | 270.66 |
| 2868 | ENSMUSG00000021010 | 1.216318829 | 0.500682538 | 2.087023804 | 0.239902648 | 0.810405728 | NA | Npas3 | neuronal PAS domain protein 3 [Source:M | 0 | 2.0558 | 1.0062 | 0.964 | 3.2719 | 0 |
| 2869 | ENSMUSG00000021012 | 694.9761279 | 0.096566171 | 0.112700613 | 0.856838034 | 0.391534403 | 0.820404559 | Zc3h14 | zinc finger CCCH type containing 14 [Sour | 644.89 | 689.73 | 681.23 | 735.52 | 705.63 | 712.86 |
| 2870 | ENSMUSG00000021013 | 92.45301696 | 0.034258393 | 0.232794511 | 0.147161515 | 0.883004536 | 0.981769403 | Ttc8 | tetratricopeptide repeat domain 8 [Source | 89.144 | 89.429 | 95.593 | 96.398 | 87.249 | 96.904 |
| 2871 | ENSMUSG00000021018 | 158.2040977 | 0.237089235 | 0.233237407 | 1.016514624 | 0.309384387 | 0.758141261 | Polr2h | polymerase (RNA) II (DNA directed) polyp | 156.6 | 103.82 | 175.09 | 187.01 | 140.69 | 186.01 |
| 2872 | ENSMUSG00000021022 | 171.5650321 | -0.062291739 | 0.186990428 | -0.333127954 | 0.739037699 | 0.948310621 | Ppp2r3c | protein phosphatase 2, regulatory subunit | 169.45 | 195.3 | 161 | 154.24 | 173.41 | 175.99 |
| 2873 | ENSMUSG00000021023 | 189.352724 | 0.214285495 | 0.181459465 | 1.18090007 | 0.237642423 | 0.686377492 | Prorp | protein only RNase P catalytic subunit [So | 158.21 | 197.36 | 171.06 | 214 | 206.13 | 189.35 |
| 2874 | ENSMUSG00000021024 | 1709.973739 | 0.014786459 | 0.109208484 | 0.13539661 | 0.892298304 | 0.982489666 | Psma6 | proteasome subunit alpha 6 [Source:MGI | 1851.2 | 1607.7 | 1643.2 | 1630.1 | 1664.3 | 1863.5 |
| 2875 | ENSMUSG00000021025 | 701.3619075 | -0.420694675 | 0.181527177 | -2.31752998 | 0.020474879 | 0.176278962 | Nfkbia | nuclear factor of kappa light polypeptide g | 762.95 | 904.57 | 741.6 | 603.45 | 763.43 | 432.17 |
| 2876 | ENSMUSG00000021027 | 164.0654861 | -0.325718281 | 0.201388053 | -1.617366451 | 0.105799203 | 0.479410925 | Ralgapa1 | Ral GTPase activating protein, alpha subu | 170.26 | 214.83 | 163.01 | 162.91 | 134.15 | 139.23 |
| 2877 | ENSMUSG00000021028 | 61.26287126 | 0.090392164 | 0.284702372 | 0.317497053 | 0.750866476 | 0.952074518 | Mbip | MAP3K12 binding inhibitory protein 1 [So | 56.217 | 53.452 | 68.425 | 61.695 | 66.528 | 61.261 |
| 2878 | ENSMUSG00000021032 | 4.473506513 | -1.697430064 | 1.066753662 | -1.591210908 | 0.111562119 | NA | Ngb | neuroglobin [Source:MGI Symbol;Acc:MG | 7.2279 | 9.2513 | 4.025 | 1.928 | 2.1812 | 2.2277 |
| 2879 | ENSMUSG00000021033 | 15871.18124 | 0.057978878 | 0.085371053 | 0.67913977 | 0.497049306 | 0.878676728 | Gstz1 | glutathione transferase zeta 1 (maleylacet | 15844 | 15341 | 15471 | 15494 | 17194 | 15883 |
| 2880 | ENSMUSG00000021036 | 311.4646267 | -0.188464557 | 0.169744821 | -1.11028163 | 0.266877687 | 0.722778036 | Sptlc2 | serine palmitoyltransferase, long chain ba | 356.58 | 326.88 | 310.93 | 248.71 | 274.84 | 350.86 |
| 2881 | ENSMUSG00000021037 | 1411.232627 | 0.04987395 | 0.113775473 | 0.438354145 | 0.661129582 | 0.928341045 | Ahsa1 | AHA1, activator of heat shock protein ATP | 1464.9 | 1290 | 1405.7 | 1593.5 | 1360 | 1353.3 |
| 2882 | ENSMUSG00000021038 | 273.3122297 | 0.184900001 | 0.155089299 | 1.192216368 | 0.233176417 | 0.680673483 | Vipas39 | VPS33B interacting protein, apical-basolat | 251.37 | 234.37 | 281.75 | 288.23 | 290.1 | 294.05 |
| 2883 | ENSMUSG00000021039 | 352.6310956 | 0.083369307 | 0.151551244 | 0.550106385 | 0.582246407 | 0.908645447 | Snw1 | SNW domain containing 1 [Source:MGI Sy | 307.59 | 339.21 | 381.37 | 374.02 | 383.9 | 329.7 |
| 2884 | ENSMUSG00000021040 | 48.62973772 | 0.463641561 | 0.310826541 | 1.491640836 | 0.135793327 | 0.538686971 | Slirp | SRA stem-loop interacting RNA binding pr | 39.352 | 42.145 | 41.256 | 58.803 | 54.531 | 55.692 |
| 2885 | ENSMUSG00000021044 | 261.296134 | 0.039823708 | 0.158479958 | 0.251285455 | 0.80159342 | 0.962796744 | Adck1 | aarF domain containing kinase 1 [Source:I | 272.25 | 244.64 | 255.59 | 246.78 | 290.1 | 258.41 |
| 2886 | ENSMUSG00000021047 | 0.17131964 | -0.796629176 | 4.080472857 | -0.195229623 | 0.84521319 | NA | Nova1 | NOVA alternative splicing regulator 1 [Sou | 0 | 1.0279 | 0 | 0 | 0 | 0 |
| 2887 | ENSMUSG00000021048 | 4818.170046 | 0.00929589 | 0.099253881 | 0.093657695 | 0.925381077 | 0.987546253 | Mthfd1 | methylenetetrahydrofolate dehydrogenas | 4643.5 | 4842.5 | 4921.5 | 4386.1 | 5259 | 4856.4 |
| 2888 | ENSMUSG00000021051 | 1067.858797 | 0.00302143 | 0.127975003 | 0.023609529 | 0.981164071 | 0.99820894 | Ppp2r5e | protein phosphatase 2, regulatory subunit | 913.13 | 1187.2 | 1100.8 | 1007.4 | 1052.4 | 1146.1 |
| 2889 | ENSMUSG00000021054 | 1423.112074 | -0.230413397 | 0.112051709 | -2.056313106 | 0.039752346 | 0.27301896 | Sgpp1 | sphingosine-1-phosphate phosphatase 1 [ | 1390.2 | 1585 | 1635.1 | 1262.8 | 1409.1 | 1256.4 |
| 2890 | ENSMUSG00000021055 | 1.350918216 | -3.807600627 | 2.14162527 | -1.777902362 | 0.075419892 | NA | Esr2 | estrogen receptor 2 (beta) [Source:MGI Sy | 4.0155 | 3.0838 | 1.0062 | 0 | 0 | 0 |
| 2891 | ENSMUSG00000021056 | 0 | NA | NA | NA | NA | NA | Tex21 | testis expressed gene 21 [Source:MGI Sym | 0 | 0 | 0 | 0 | 0 | 0 |
| 2892 | ENSMUSG00000021057 | 9.877414721 | -0.411195784 | 0.703392896 | -0.584589048 | 0.558824073 | 0.899349136 | Akap5 | A kinase (PRKA) anchor protein 5 [Source: | 6.4248 | 16.447 | 11.069 | 7.7118 | 9.8156 | 7.7969 |
| 2893 | ENSMUSG00000021061 | 28.86201773 | -0.263948629 | 0.485240212 | -0.543954565 | 0.586472739 | 0.910263086 | Sptb | spectrin beta, erythrocytic [Source:MGI Sy | 21.684 | 19.53 | 53.331 | 20.244 | 30.537 | 27.846 |
| 2894 | ENSMUSG00000021062 | 0.995947842 | -0.043351441 | 2.675688395 | -0.016201977 | 0.987073258 | NA | Rab15 | RAB15, member RAS oncogene family [So | 0 | 3.0838 | 0 | 2.8919 | 0 | 0 |
| 2895 | ENSMUSG00000021065 | 67.01458073 | -0.686504367 | 0.282107533 | -2.433484701 | 0.014954267 | 0.142208102 | Fut8 | fucosyltransferase 8 [Source:MGI Symbol; | 97.175 | 80.178 | 70.437 | 53.983 | 49.078 | 51.237 |
| 2896 | ENSMUSG00000021066 | 31.92627348 | 0.3945133 | 0.393914262 | 1.001520731 | 0.316575123 | 0.76328431 | Atl1 | atlastin GTPase 1 [Source:MGI Symbol;Acc | 33.73 | 25.698 | 23.144 | 38.559 | 40.353 | 30.074 |
| 2897 | ENSMUSG00000021067 | 175.6748577 | -0.282369984 | 0.263626716 | -1.071097754 | 0.284125478 | 0.736651636 | Sav1 | salvador family WW domain containing 1 | 203.18 | 169.61 | 206.28 | 238.1 | 122.15 | 114.73 |
| 2898 | ENSMUSG00000021068 | 40.86365974 | -0.466132528 | 0.389556301 | -1.19657294 | 0.231473053 | 0.678699825 | Nin | ninein [Source:MGI Symbol;Acc:MGI:1051 | 51.399 | 48.312 | 42.262 | 27.955 | 22.903 | 52.351 |
| 2899 | ENSMUSG00000021069 | 10432.02168 | -0.143957036 | 0.088420215 | -1.628100949 | 0.103503482 | 0.473607836 | Pygl | liver glycogen phosphorylase [Source:MG | 10589 | 10785 | 11483 | 10106 | 10278 | 9350.7 |
| 2900 | ENSMUSG00000021070 | 2.409695989 | 0.133853525 | 1.429505897 | 0.093636218 | 0.925398138 | NA | Bdkrb2 | bradykinin receptor, beta 2 [Source:MGI S | 0.8031 | 4.1117 | 2.0125 | 0.964 | 5.4531 | 1.1138 |
| 2901 | ENSMUSG00000021071 | 0.294513347 | 0.165143496 | 4.080472857 | 0.040471657 | 0.967717103 | NA | Trim9 | tripartite motif-containing 9 [Source:MGI | 0.8031 | 0 | 0 | 0.964 | 0 | 0 |
| 2902 | ENSMUSG00000021072 | 697.4196558 | 0.051249543 | 0.116407016 | 0.440261636 | 0.659747623 | 0.927787797 | Tmx1 | thioredoxin-related transmembrane prote | 636.86 | 721.6 | 697.33 | 697.92 | 702.36 | 728.45 |
| 2903 | ENSMUSG00000021076 | 646.0524515 | 0.047195575 | 0.114154364 | 0.413436454 | 0.679286871 | 0.934062537 | Actr10 | ARP10 actin-related protein 10 [Source:M | 632.84 | 645.53 | 627.9 | 636.23 | 654.37 | 679.44 |
| 2904 | ENSMUSG00000021078 | 0.17131964 | -0.796629176 | 4.080472857 | -0.195229623 | 0.84521319 | NA | Tomm20l | translocase of outer mitochondrial memb | 0 | 1.0279 | 0 | 0 | 0 | 0 |
| 2905 | ENSMUSG00000021079 | 566.1589316 | -0.070014556 | 0.143224965 | -0.488843243 | 0.624952681 | 0.919503423 | Timm9 | translocase of inner mitochondrial memb | 623.21 | 589 | 527.27 | 574.53 | 614.02 | 468.93 |
| 2906 | ENSMUSG00000021081 | 0 | NA | NA | NA | NA | NA | Serpina1f | serine (or cysteine) peptidase inhibitor, cl | 0 | 0 | 0 | 0 | 0 | 0 |
| 2907 | ENSMUSG00000021086 | 0.503095663 | 2.549494177 | 3.348715477 | 0.761334964 | 0.446457022 | NA | Ccdc175 | coiled-coil domain containing 175 [Source | 0 | 0 | 0 | 1.928 | 1.0906 | 0 |
| 2908 | ENSMUSG00000021087 | 15.11491446 | -1.379725104 | 0.580808539 | -2.375524826 | 0.017524014 | 0.157628937 | Rtn1 | reticulon 1 [Source:MGI Symbol;Acc:MGI: | 16.865 | 22.614 | 26.162 | 9.6398 | 8.7249 | 6.6831 |

| | | | | | | | | | | | | | | |
|---|---|---|---|---|---|---|---|---|---|---|---|---|---|---|
| 2909 | ENSMUSG00000021090 | 0.267700746 | -1.497481649 | 4.057001436 | -0.369110456 | 0.712045396 | NA | Lrrc9 | leucine rich repeat containing 9 [Source:M | 1.6062 | 0 | 0 | 0 | 0 | 0 |
| 2910 | ENSMUSG00000021091 | 9004.484977 | -0.193807559 | 0.092627088 | -2.09234213 | 0.036407922 | 0.257267344 | Serpina3n | serine (or cysteine) peptidase inhibitor, cla | 9561.7 | 9740.5 | 9522.1 | 7839.1 | 8223.3 | 9140.2 |
| 2911 | ENSMUSG00000021094 | 862.0820362 | -0.48959158 | 0.12109615 | -4.042998732 | 5.28E-05 | 0.00219337 | Dhrs7 | dehydrogenase/reductase (SDR family) m | 951.68 | 1053.6 | 1016.3 | 722.98 | 786.34 | 641.57 |
| 2912 | ENSMUSG00000021095 | 0.133850373 | -0.796629176 | 4.080472857 | -0.195229623 | 0.84521319 | NA | Gsc | goosecoid homeobox [Source:MGI Symbo | 0.8031 | 0 | 0 | 0 | 0 | 0 |
| 2913 | ENSMUSG00000021096 | 3066.321313 | 0.028043582 | 0.095551182 | 0.29349278 | 0.769145517 | 0.956045284 | Ppm1a | protein phosphatase 1A, magnesium depe | 3122.5 | 3154.7 | 2832.6 | 3252.5 | 3041.7 | 2994 |
| 2914 | ENSMUSG00000021097 | 3320.65401 | 0.433674474 | 0.102694776 | 4.2229458 | 2.41E-05 | 0.001126619 | Clmn | calmin [Source:MGI Symbol;Acc:MGI:2136 | 3034.1 | 2527.6 | 2913.1 | 3969.7 | 3715.7 | 3763.7 |
| 2915 | ENSMUSG00000021098 | 0 | NA | NA | NA | NA | NA | 4930447C04 | RIKEN cDNA 4930447C04 gene [Source:M | 0 | 0 | 0 | 0 | 0 | 0 |
| 2916 | ENSMUSG00000021099 | 0 | NA | NA | NA | NA | NA | Six6 | sine oculis-related homeobox 6 [Source:M | 0 | 0 | 0 | 0 | 0 | 0 |
| 2917 | ENSMUSG00000021101 | 0.713713984 | 3.041227437 | 2.670264816 | 1.138923532 | 0.254735049 | NA | 4930408O17 | RIKEN cDNA 4930408O17 gene [Source:M | 0 | 0 | 0 | 0.964 | 1.0906 | 2.2277 |
| 2918 | ENSMUSG00000021102 | 3099.393102 | -0.185839726 | 0.104210071 | -1.783318297 | 0.074534505 | 0.398693445 | Glrx5 | glutaredoxin 5 [Source:MGI Symbol;Acc:M | 3390.7 | 3208.1 | 3296.5 | 2688.5 | 3242.4 | 2770.1 |
| 2919 | ENSMUSG00000021103 | 146.4392975 | 0.007858972 | 0.230595474 | 0.034081208 | 0.972812393 | 0.995807393 | Mnat1 | menage a trois 1 [Source:MGI Symbol;Acc | 156.6 | 145.96 | 134.84 | 104.11 | 174.5 | 162.62 |
| 2920 | ENSMUSG00000021108 | 42.98805963 | -0.861794496 | 0.35971434 | -2.395774643 | 0.016585284 | 0.152893273 | Prkch | protein kinase C, eta [Source:MGI Symbol | 56.217 | 55.508 | 54.337 | 21.208 | 29.447 | 41.212 |
| 2921 | ENSMUSG00000021109 | 706.9607617 | 0.051908658 | 0.118334529 | 0.438660285 | 0.660907708 | 0.928184201 | Hif1a | hypoxia inducible factor 1, alpha subunit [ | 727.61 | 696.93 | 658.08 | 755.76 | 735.08 | 668.31 |
| 2922 | ENSMUSG00000021111 | 1639.185812 | 0.016367874 | 0.110363128 | 0.14830926 | 0.882098708 | 0.981738667 | Papola | poly (A) polymerase alpha [Source:MGI Sy | 1535.5 | 1796.8 | 1558.7 | 1765 | 1561.8 | 1617.3 |
| 2923 | ENSMUSG00000021112 | 381.8789928 | 0.266554066 | 0.156046701 | 1.708168547 | 0.087605073 | 0.43652818 | Mpp5 | membrane protein, palmitoylated 5 (MAG | 375.05 | 298.1 | 366.27 | 401.98 | 470.06 | 379.82 |
| 2924 | ENSMUSG00000021113 | 133.6972236 | -0.079922749 | 0.204340852 | -0.391124673 | 0.695705084 | 0.939992732 | Snapc1 | small nuclear RNA activating complex, pol | 138.94 | 128.49 | 144.9 | 141.7 | 115.61 | 132.55 |
| 2925 | ENSMUSG00000021114 | 736.9761045 | -0.069262699 | 0.126037569 | -0.549540103 | 0.582634851 | 0.908645447 | Atp6v1d | ATPase, H+ transporting, lysosomal V1 sul | 711.55 | 808.97 | 744.62 | 777.93 | 739.44 | 639.35 |
| 2926 | ENSMUSG00000021115 | 116.6898192 | 0.318610705 | 0.222669336 | 1.430869245 | 0.152467693 | 0.568249894 | Vrk1 | vaccinia related kinase 1 [Source:MGI Sym | 111.63 | 108.96 | 90.562 | 117.61 | 123.24 | 148.14 |
| 2927 | ENSMUSG00000021116 | 730.019766 | 0.027637406 | 0.11723677 | 0.235740086 | 0.813634362 | 0.966842541 | Eif2s1 | eukaryotic translation initiation factor 2, s | 721.19 | 731.88 | 715.44 | 670.93 | 773.25 | 767.44 |
| 2928 | ENSMUSG00000021118 | 8.645400586 | 0.557026868 | 0.739908857 | 0.752831734 | 0.451551035 | 0.856218428 | Plek2 | pleckstrin 2 [Source:MGI Symbol;Acc:MGI | 4.8186 | 6.1675 | 10.062 | 7.7118 | 13.087 | 10.025 |
| 2929 | ENSMUSG00000021120 | 30.29073116 | 0.311851546 | 0.41797954 | 0.746092849 | 0.455611329 | 0.857509728 | Pigh | phosphatidylinositol glycan anchor biosyn | 19.274 | 30.838 | 31.194 | 30.847 | 27.265 | 42.326 |
| 2930 | ENSMUSG00000021123 | 0.353089354 | 0.165142706 | 3.896102244 | 0.042386646 | 0.966190474 | NA | Rdh12 | retinol dehydrogenase 12 [Source:MGI Sy | 0 | 1.0279 | 0 | 0 | 1.0906 | 0 |
| 2931 | ENSMUSG00000021124 | 851.8990273 | 0.020965653 | 0.107170535 | 0.195628887 | 0.844900649 | 0.972803041 | Vti1b | vesicle transport through interaction with | 819.97 | 845.98 | 871.41 | 844.44 | 846.32 | 883.28 |
| 2932 | ENSMUSG00000021125 | 1.380956475 | -1.598416342 | 2.020048339 | -0.791276283 | 0.428782785 | NA | Arg2 | arginase type II [Source:MGI Symbol;Acc:I | 3.2124 | 0 | 3.0187 | 0.964 | 1.0906 | 0 |
| 2933 | ENSMUSG00000021127 | 1057.658106 | -0.074591955 | 0.183693511 | -0.406067444 | 0.684693046 | 0.935737593 | Zfp36l1 | zinc finger protein 36, C3H type-like 1 [Sou | 995.04 | 1014.6 | 1246.7 | 1389.1 | 904.12 | 796.4 |
| 2934 | ENSMUSG00000021130 | 0.535117793 | 1.111049431 | 3.044213722 | 0.364970903 | 0.715133126 | NA | Galnt16 | polypeptide N-acetylgalactosaminyltransf | 0 | 0 | 1.0062 | 0 | 1.0906 | 1.1138 |
| 2935 | ENSMUSG00000021131 | 440.5157146 | -0.131641114 | 0.138322813 | -0.951694887 | 0.341251743 | 0.783129922 | Erh | ERH mRNA splicing and mitosis factor [Sou | 491.5 | 480.04 | 409.54 | 398.12 | 439.52 | 424.37 |
| 2936 | ENSMUSG00000021133 | 1305.782493 | -0.597214386 | 0.144884206 | -4.122011665 | 3.76E-05 | 0.001672902 | Susd6 | sushi domain containing 6 [Source:MGI Sy | 1506.6 | 1925.3 | 1285 | 990.97 | 1074.3 | 1052.6 |
| 2937 | ENSMUSG00000021134 | 3770.891579 | 0.118359133 | 0.115141276 | 1.027947037 | 0.303974742 | 0.752961647 | Srsf5 | serine and arginine-rich splicing factor 5 [S | 3756.9 | 3465.1 | 3627.5 | 4557.7 | 3641.6 | 3576.6 |
| 2938 | ENSMUSG00000021135 | 14217.67852 | -0.097892525 | 0.100771909 | -0.971426719 | 0.331335829 | 0.774747658 | Slc10a1 | solute carrier family 10 (sodium/bile acid c | 13821 | 16296 | 13985 | 14230 | 14131 | 12844 |
| 2939 | ENSMUSG00000021136 | 2272.059087 | -0.182281549 | 0.121160583 | -1.504462463 | 0.132462332 | 0.533538058 | Smoc1 | SPARC related modular calcium binding 1 | 2337 | 2740.4 | 2168.5 | 1922.2 | 2145.2 | 2319 |
| 2940 | ENSMUSG00000021139 | 6.477418277 | 1.371335232 | 0.903287914 | 1.518159616 | 0.128974165 | 0.525636046 | Gm20498 | predicted gene 20498 [Source:MGI Symbo | 5.6217 | 3.0838 | 2.0125 | 10.604 | 13.087 | 4.4554 |
| 2941 | ENSMUSG00000021140 | 708.6880535 | 0.155867587 | 0.116477848 | 1.338173643 | 0.180839839 | 0.614186356 | Pcnx | pecanex homolog [Source:MGI Symbol;Ac | 661.76 | 636.28 | 713.43 | 757.69 | 711.08 | 771.89 |
| 2942 | ENSMUSG00000021143 | 405.9766771 | -0.074004763 | 0.136606287 | -0.541737608 | 0.587999278 | 0.910798643 | Pacs2 | phosphofurin acidic cluster sorting protei | 421.63 | 421.45 | 405.52 | 358.6 | 419.89 | 408.78 |
| 2943 | ENSMUSG00000021144 | 537.168066 | 0.195632834 | 0.127494876 | 1.534436834 | 0.124922222 | 0.518472028 | Mta1 | metastasis associated 1 [Source:MGI Syml | 485.07 | 533.49 | 484 | 559.11 | 542.04 | 619.3 |
| 2944 | ENSMUSG00000021147 | 210.717402 | 0.157963973 | 0.175162783 | 0.901812417 | 0.367156523 | 0.805239112 | Wdr37 | WD repeat domain 37 [Source:MGI Symbo | 198.37 | 190.16 | 209.3 | 237.14 | 237.75 | 191.58 |
| 2945 | ENSMUSG00000021148 | 0 | NA | NA | NA | NA | NA | Gm9745 | predicted gene 9745 [Source:MGI Symbol | 0 | 0 | 0 | 0 | 0 | 0 |
| 2946 | ENSMUSG00000021149 | 677.8645785 | 0.210176217 | 0.11670093 | 1.800981499 | 0.071705796 | 0.387575723 | Gtpbp4 | GTP binding protein 4 [Source:MGI Symbo | 636.06 | 623.95 | 625.88 | 753.83 | 754.71 | 672.76 |
| 2947 | ENSMUSG00000021156 | 869.9709548 | 0.083780956 | 0.112335262 | 0.745811733 | 0.455781151 | 0.857604173 | Zmynd11 | zinc finger, MYND domain containing 11 [S | 836.83 | 881.95 | 816.06 | 961.09 | 852.86 | 871.03 |
| 2948 | ENSMUSG00000021171 | 640.426537 | -0.118829271 | 0.122787592 | -0.967762862 | 0.333162815 | 0.77523112 | Esyt2 | extended synaptotagmin-like protein 2 [So | 638.47 | 691.79 | 671.17 | 671.89 | 588.93 | 580.31 |
| 2949 | ENSMUSG00000021175 | 2.326168939 | -1.190848361 | 1.502977849 | -0.792325956 | 0.428170641 | NA | Cdca7l | cell division cycle associated 7 like [Source | 1.6062 | 3.0838 | 5.0312 | 0.964 | 3.2719 | 0 |
| 2950 | ENSMUSG00000021176 | 1.836159426 | -3.15368553 | 2.040351209 | -1.545658177 | 0.122187143 | NA | Efcab11 | EF-hand calcium binding domain 11 [Sour | 4.0155 | 0 | 6.0375 | 0.964 | 0 | 0 |
| 2951 | ENSMUSG00000021177 | 54.53538867 | -0.317120487 | 0.314377176 | -1.008726178 | 0.313105974 | 0.760114938 | Tdp1 | tyrosyl-DNA phosphodiesterase 1 [Source | 67.461 | 49.34 | 64.4 | 41.451 | 58.893 | 45.668 |
| 2952 | ENSMUSG00000021178 | 1898.002359 | 0.141503875 | 0.097301894 | 1.454276671 | 0.145869616 | 0.556690797 | Psmc1 | protease (prosome, macropain) 26S subu | 1780.5 | 1838.9 | 1795.1 | 1853.7 | 2058 | 2061.7 |
| 2953 | ENSMUSG00000021179 | 198.6372371 | 0.18904162 | 0.177909272 | 1.062573176 | 0.287975557 | 0.73951397 | Nrde2 | nrde-2 necessary for RNA interference, do | 175.08 | 175.77 | 206.28 | 211.11 | 194.13 | 229.45 |
| 2954 | ENSMUSG00000021180 | 33.74481464 | 0.179570698 | 0.383559229 | 0.468169409 | 0.639663446 | 0.923932929 | Rps6ka5 | ribosomal protein S6 kinase, polypeptide 5 | 32.927 | 33.921 | 28.175 | 41.451 | 39.262 | 26.732 |
| 2955 | ENSMUSG00000021182 | 21.92456335 | -0.211692069 | 0.491391446 | -0.43080129 | 0.666612862 | 0.929596279 | Ccdc88c | coiled-coil domain containing 88C [Source | 24.896 | 17.475 | 28.175 | 19.28 | 28.356 | 13.366 |
| 2956 | ENSMUSG00000021185 | 1162.588201 | -0.066864759 | 0.137773624 | -0.485323367 | 0.627446982 | 0.920328819 | Dglucy | D-glutamate cyclase [Source:MGI Symbol; | 1433.5 | 1097.8 | 1035.4 | 1079.7 | 1200.8 | 1128.3 |
| 2957 | ENSMUSG00000021186 | 85.84303394 | -0.495068231 | 0.273466245 | -1.810344929 | 0.070242314 | 0.383245879 | Fbln5 | fibulin 5 [Source:MGI Symbol;Acc:MGI:13 | 120.47 | 108.96 | 71.443 | 66.514 | 74.162 | 73.514 |
| 2958 | ENSMUSG00000021187 | 47.14062727 | -0.241665092 | 0.321538964 | -0.751588825 | 0.452298365 | 0.856769845 | Tc2n | tandem C2 domains, nuclear [Source:MGI | 55.414 | 49.34 | 48.3 | 39.523 | 50.168 | 40.098 |
| 2959 | ENSMUSG00000021188 | 548.6794081 | -0.184236983 | 0.121396844 | -1.517642276 | 0.129104603 | 0.525983675 | Trip11 | thyroid hormone receptor interactor 11 [S | 575.82 | 606.47 | 568.53 | 489.7 | 519.13 | 532.42 |
| 2960 | ENSMUSG00000021189 | 376.4721593 | -0.07380996 | 0.142822619 | -0.516794617 | 0.605299538 | 0.916376001 | Atxn3 | ataxin 3 [Source:MGI Symbol;Acc:MGI:10 | 367.82 | 364.91 | 425.64 | 348 | 389.35 | 363.11 |
| 2961 | ENSMUSG00000021190 | 569.2579557 | -0.716471929 | 0.178322417 | -4.017845537 | 5.87E-05 | 0.00237374 | Lgmn | legumain [Source:MGI Symbol;Acc:MGI:1 | 701.91 | 867.56 | 553.43 | 377.88 | 404.62 | 510.14 |
| 2962 | ENSMUSG00000021192 | 537.3885191 | 0.0066564 | 0.124962506 | 0.053267178 | 0.957519031 | 0.991861398 | Golga5 | golgi autoantigen, golgin subfamily a, 5 [S | 531.65 | 558.16 | 519.22 | 578.39 | 526.77 | 510.14 |
| 2963 | ENSMUSG00000021193 | 610.014582 | -0.213097169 | 0.144706795 | -1.472613429 | 0.140855302 | 0.548849969 | Pitrm1 | pitrilysin metallepetidase 1 [Source:MGI S | 631.24 | 738.05 | 595.7 | 520.55 | 653.28 | 521.28 |
| 2964 | ENSMUSG00000021194 | 0 | NA | NA | NA | NA | NA | Chga | chromogranin A [Source:MGI Symbol;Acc: | 0 | 0 | 0 | 0 | 0 | 0 |

| | | | | | | | | | | | | | | | |
|---|---|---|---|---|---|---|---|---|---|---|---|---|---|---|---|
| 2965 | ENSMUSG00000021196 | 33.26151194 | -0.764928307 | 0.452303561 | -1.691183474 | 0.090801766 | 0.444602826 | Pfkp | phosphofructokinase, platelet [Source:MG | 31.321 | 50.368 | 44.275 | 33.739 | 10.906 | 28.96 |
| 2966 | ENSMUSG00000021198 | 17.79349571 | -0.779741015 | 0.662196648 | -1.177506738 | 0.23899331 | 0.688331189 | Unc79 | unc-79 homolog [Source:MGI Symbol;Acc | 8.031 | 17.475 | 42.262 | 20.244 | 8.7249 | 10.025 |
| 2967 | ENSMUSG00000021200 | 26.42297813 | -1.280263944 | 0.465119237 | -2.752549978 | 0.005913312 | 0.07450361 | Asb2 | ankyrin repeat and SOCS box-containing 2 | 40.958 | 41.117 | 30.187 | 16.388 | 8.7249 | 21.163 |
| 2968 | ENSMUSG00000021203 | 50.74869642 | -0.172757171 | 0.321894267 | -0.536689182 | 0.59148233 | 0.912621683 | Otub2 | OTU domain, ubiquitin aldehyde binding 2 | 60.233 | 52.424 | 48.3 | 36.631 | 53.44 | 53.465 |
| 2969 | ENSMUSG00000021207 | 4.221467553 | -0.106847712 | 1.411301955 | -0.075708612 | 0.939650925 | NA | Akr1c21 | aldo-keto reductase family 1, member C2 | 0 | 3.0838 | 10.062 | 0 | 3.2719 | 8.9108 |
| 2970 | ENSMUSG00000021208 | 173.1240016 | -1.072526856 | 0.274206029 | -3.911390503 | 9.18E-05 | 0.003377372 | Ifi27l2b | interferon, alpha-inducible protein 27 like | 347.74 | 199.42 | 155.97 | 96.398 | 116.7 | 122.52 |
| 2971 | ENSMUSG00000021209 | 221.4083844 | 0.488651384 | 0.22025808 | 2.218540105 | 0.026518028 | 0.210861331 | Ppp4r4 | protein phosphatase 4, regulatory subunit | 146.16 | 201.47 | 206.28 | 321.97 | 193.04 | 259.53 |
| 2972 | ENSMUSG00000021210 | 23535.52719 | 0.313701156 | 0.099413171 | 3.155529107 | 0.001602073 | 0.029784551 | Akr1c6 | aldo-keto reductase family 1, member C6 | 21465 | 20181 | 21313 | 23117 | 27634 | 27503 |
| 2973 | ENSMUSG00000021211 | 1630.440296 | 0.097585564 | 0.111859936 | 0.872390662 | 0.382995293 | 0.815133007 | Akr1c12 | aldo-keto reductase family 1, member C1 | 1438.4 | 1705.3 | 1583.8 | 1763.1 | 1749.4 | 1542.7 |
| 2974 | ENSMUSG00000021213 | 1753.543865 | 0.101685552 | 0.114717452 | 0.88640002 | 0.375402009 | 0.809332934 | Akr1c13 | aldo-keto reductase family 1, member C1 | 1504.2 | 1664.2 | 1907.8 | 1745.8 | 1822.4 | 1876.8 |
| 2975 | ENSMUSG00000021214 | 2.358680233 | 0.056617388 | 1.476847134 | 0.038336661 | 0.969419261 | NA | Akr1c18 | aldo-keto reductase family 1, member C1 | 4.8186 | 2.0558 | 0 | 2.8919 | 3.2719 | 1.1138 |
| 2976 | ENSMUSG00000021215 | 1273.913402 | -0.441250605 | 0.147528393 | -2.990953779 | 0.002781076 | 0.044398986 | Net1 | neuroepithelial cell transforming gene 1 [ | 1274.5 | 1543.9 | 1584.8 | 1241.6 | 1134.2 | 864.34 |
| 2977 | ENSMUSG00000021216 | 0 | NA | NA | NA | NA | NA | Tubal3 | tubulin, alpha-like 3 [Source:MGI Symbol; | 0 | 0 | 0 | 0 | 0 | 0 |
| 2978 | ENSMUSG00000021217 | 10.45582021 | -0.770249049 | 0.776012434 | -0.992573077 | 0.32091805 | 0.768121576 | Tshz3 | teashirt zinc finger family member 3 [Sour | 15.259 | 15.419 | 9.0562 | 16.388 | 3.2719 | 3.3415 |
| 2979 | ENSMUSG00000021218 | 4681.151789 | -0.020767611 | 0.092698832 | -0.224033148 | 0.82273151 | 0.969219124 | Gdi2 | guanosine diphosphate (GDP) dissociation | 4616.2 | 4772.6 | 4755.5 | 4389 | 5015.8 | 4537.8 |
| 2980 | ENSMUSG00000021219 | 0.356960287 | 0.16514264 | 3.880370363 | 0.042558474 | 0.966053499 | NA | Rgs6 | regulator of G-protein signaling 6 [Source: | 0 | 1.0279 | 0 | 0 | 0 | 1.1138 |
| 2981 | ENSMUSG00000021221 | 26.19656533 | 0.037091835 | 0.476464988 | 0.077847976 | 0.937948983 | 0.988420305 | Dpf3 | D4, zinc and double PHD fingers, family 3 | 17.668 | 41.117 | 19.119 | 30.847 | 27.265 | 21.163 |
| 2982 | ENSMUSG00000021222 | 55.60103567 | -0.812722874 | 0.380175356 | -2.137757906 | 0.032536397 | 0.241089369 | Dcaf4 | DDB1 and CUL4 associated factor 4 [Sourc | 79.507 | 85.317 | 47.293 | 22.171 | 43.625 | 55.692 |
| 2983 | ENSMUSG00000021223 | 61.79030239 | -0.329582228 | 0.313055874 | -1.05279043 | 0.292437052 | 0.743376015 | Papln | papilin, proteoglycan-like sulfated glycopr | 56.217 | 84.289 | 66.412 | 65.55 | 40.353 | 57.92 |
| 2984 | ENSMUSG00000021224 | 664.3761996 | -0.052665502 | 0.117077214 | -0.449835628 | 0.652828966 | 0.925756399 | Numb | NUMB endocytic adaptor protein [Source | 664.97 | 703.1 | 662.11 | 694.06 | 626.01 | 636 |
| 2985 | ENSMUSG00000021226 | 293.6073322 | -1.32250938 | 0.284613886 | -4.646679053 | 3.37E-06 | 0.000217655 | Acot2 | acyl-CoA thioesterase 2 [Source:MGI Sym | 507.56 | 471.81 | 279.74 | 249.67 | 155.96 | 96.904 |
| 2986 | ENSMUSG00000021228 | 819.6055988 | -0.994070989 | 0.924084842 | -1.075735629 | 0.282045508 | 0.735178632 | Acot3 | acyl-CoA thioesterase 3 [Source:MGI Sym | 1557.2 | 1107.1 | 609.78 | 1228.1 | 261.75 | 153.71 |
| 2987 | ENSMUSG00000021234 | 7.44992759 | -0.295138081 | 0.865893375 | -0.340848064 | 0.733217964 | 0.947915087 | Fam161b | family with sequence similarity 161, mem | 11.243 | 10.279 | 3.0187 | 4.8199 | 11.997 | 3.3415 |
| 2988 | ENSMUSG00000021235 | 270.6013115 | -0.09896355 | 0.163785936 | -0.604224959 | 0.545694089 | 0.89549457 | Coq6 | coenzyme Q6 monooxygenase [Source:M | 286.71 | 304.26 | 248.54 | 254.49 | 289.01 | 240.59 |
| 2989 | ENSMUSG00000021236 | 5422.331226 | -0.185526453 | 0.125219816 | -1.48160618 | 0.138445114 | 0.543828246 | Entpd5 | ectonucleoside triphosphate diphosphoh | 6205.6 | 6100.7 | 5005.1 | 5289.3 | 5593.8 | 4339.5 |
| 2990 | ENSMUSG00000021238 | 9266.250669 | -0.152054899 | 0.095783822 | -1.587479965 | 0.112403981 | 0.49222258 | Aldh6a1 | aldehyde dehydrogenase family 6, subfan | 9472.6 | 9954.4 | 9836 | 8805 | 9525.5 | 8004.1 |
| 2991 | ENSMUSG00000021239 | 0.160662974 | 1.126924175 | 4.080472857 | 0.276174898 | 0.782413736 | NA | Vsx2 | visual system homeobox 2 [Source:MGI Sy | 0 | 0 | 0 | 0.964 | 0 | 0 |
| 2992 | ENSMUSG00000021240 | 399.2007256 | -0.392952794 | 0.14074971 | -2.79185508 | 0.005240682 | 0.068745122 | Abcd4 | ATP-binding cassette, sub-family D (ALD), | 475.44 | 422.47 | 461.87 | 370.17 | 352.27 | 312.99 |
| 2993 | ENSMUSG00000021241 | 404.1781653 | -0.149936493 | 0.143032215 | -1.048270791 | 0.294513863 | 0.745164683 | Isca2 | iron-sulfur cluster assembly 2 [Source:MG | 376.65 | 447.14 | 452.81 | 405.83 | 380.63 | 362 |
| 2994 | ENSMUSG00000021242 | 1025.371622 | -0.128134418 | 0.114282184 | -1.121210786 | 0.26219815 | 0.716133499 | Npc2 | NPC intracellular cholesterol transporter 2 | 1061.7 | 1178 | 973.04 | 970.73 | 965.2 | 1003.6 |
| 2995 | ENSMUSG00000021243 | 367.008689 | -0.044796013 | 0.150141063 | -0.298359502 | 0.765428795 | 0.954980327 | Fcf1 | FCF1 rRNA processing protein [Source:MG | 400.75 | 370.05 | 347.15 | 381.74 | 388.26 | 314.1 |
| 2996 | ENSMUSG00000021244 | 482.042368 | -0.085077451 | 0.133243241 | -0.638512319 | 0.523140237 | 0.888334265 | Ylpm1 | YLP motif containing 1 [Source:MGI Symb | 505.15 | 487.23 | 496.08 | 449.21 | 522.41 | 432.17 |
| 2997 | ENSMUSG00000021245 | 245.8163531 | -0.128634507 | 0.15912687 | -0.808377032 | 0.41887357 | 0.837591673 | Mlh3 | mutL homolog 3 [Source:MGI Symbol;Acc | 258.6 | 255.95 | 255.59 | 224.61 | 257.39 | 222.77 |
| 2998 | ENSMUSG00000021248 | 2078.817601 | -0.082628651 | 0.095575812 | -0.864535163 | 0.387293973 | 0.817618864 | Tmed10 | transmembrane p24 trafficking protein 10 | 2182.8 | 2153.5 | 2078.9 | 2127.5 | 1909.7 | 2020.5 |
| 2999 | ENSMUSG00000021250 | 12.89074413 | -1.403521226 | 0.637183385 | -2.202695894 | 0.02761619 | 0.215913822 | Fos | FBJ osteosarcoma oncogene [Source:MGI | 18.471 | 16.447 | 21.131 | 4.8199 | 10.906 | 5.5692 |
| 3000 | ENSMUSG00000021252 | 976.9474584 | 0.258563878 | 0.118421871 | 2.18341322 | 0.029005393 | 0.223715404 | Erg28 | ergosterol biosynthesis 28 [Source:MGI Sy | 934.81 | 918.96 | 814.05 | 966.87 | 1131 | 1096 |
| 3001 | ENSMUSG00000021253 | 18.40443967 | -0.865534474 | 0.543076594 | -1.593761327 | 0.110989503 | 0.489438373 | Tgfb3 | transforming growth factor, beta 3 [Sourc | 31.321 | 24.67 | 15.094 | 9.6398 | 17.45 | 12.252 |
| 3002 | ENSMUSG00000021254 | 181.6035702 | 0.001434959 | 0.200256676 | 0.007165597 | 0.99428273 | 0.998676967 | Gpatch2l | G patch domain containing 2 like [Source: | 158.21 | 185.03 | 202.26 | 216.9 | 167.96 | 159.28 |
| 3003 | ENSMUSG00000021255 | 14.32618281 | -0.289938751 | 0.609451587 | -0.475737133 | 0.634261663 | 0.923398157 | Esrrb | estrogen related receptor, beta [Source:M | 12.85 | 16.447 | 18.112 | 16.388 | 5.4531 | 16.708 |
| 3004 | ENSMUSG00000021256 | 4.983252914 | -0.868511596 | 1.029095687 | -0.843956113 | 0.39869393 | NA | Vash1 | vasohibin 1 [Source:MGI Symbol;Acc:MGI | 12.047 | 4.1117 | 3.0187 | 1.928 | 5.4531 | 3.3415 |
| 3005 | ENSMUSG00000021257 | 92.95365814 | -0.102187329 | 0.237378524 | -0.430482621 | 0.666844606 | 0.929596279 | Angel1 | angel homolog 1 [Source:MGI Symbol;Acc | 105.21 | 94.568 | 88.55 | 80.974 | 95.974 | 92.449 |
| 3006 | ENSMUSG00000021258 | 321.9800409 | 0.300578773 | 0.156610933 | 1.919270689 | 0.054950085 | 0.330417929 | Ccnk | cyclin K [Source:MGI Symbol;Acc:MGI:127 | 263.42 | 325.85 | 276.72 | 323.9 | 357.72 | 384.28 |
| 3007 | ENSMUSG00000021259 | 113.6443112 | -0.710112767 | 0.341188945 | -2.081288911 | 0.03740747 | 0.262396527 | Cyp46a1 | cytochrome P450, family 46, subfamily a, | 153.39 | 164.47 | 105.66 | 114.71 | 106.88 | 36.757 |
| 3008 | ENSMUSG00000021260 | 61.3153914 | -0.353904321 | 0.285292325 | -1.240497167 | 0.214791561 | 0.659728602 | Hhipl1 | hedgehog interacting protein-like 1 [Sourc | 69.067 | 76.066 | 61.381 | 57.839 | 54.531 | 49.009 |
| 3009 | ENSMUSG00000021262 | 43.43526628 | -1.35209547 | 0.422078708 | -3.203420227 | 0.001358057 | 0.026260602 | Evl | Ena-vasodilator stimulated phosphoprote | 55.414 | 91.485 | 40.25 | 19.28 | 18.541 | 35.643 |
| 3010 | ENSMUSG00000021263 | 0.538988726 | 1.126003851 | 3.343747182 | 0.336749099 | 0.736306047 | NA | Degs2 | delta(4)-desaturase, sphingolipid 2 [Sourc | 0 | 0 | 1.0062 | 0 | 0 | 2.2277 |
| 3011 | ENSMUSG00000021264 | 1473.937684 | 0.137274091 | 0.097346599 | 1.410158061 | 0.158493016 | 0.579338637 | Yy1 | YY1 transcription factor [Source:MGI Sym | 1333.1 | 1450.4 | 1428.9 | 1537.5 | 1544.3 | 1549.4 |
| 3012 | ENSMUSG00000021265 | 75.24308941 | -0.539504375 | 0.276570513 | -1.950693764 | 0.051093486 | 0.317405278 | Slc25a29 | solute carrier family 25 (mitochondrial car | 96.372 | 78.122 | 92.575 | 50.127 | 76.343 | 57.92 |
| 3013 | ENSMUSG00000021266 | 292.0683295 | 0.018504519 | 0.161496157 | 0.114581792 | 0.908776613 | 0.985098284 | Wars | tryptophanyl-tRNA synthetase [Source:M | 267.43 | 303.24 | 299.86 | 254.49 | 299.92 | 327.47 |
| 3014 | ENSMUSG00000021268 | 8.848463631 | 0.339820227 | 0.747722979 | 0.454473431 | 0.649488111 | 0.925643431 | Meg3 | maternally expressed 3 [Source:MGI Symb | 7.2279 | 4.1117 | 12.075 | 8.6758 | 7.6343 | 13.366 |
| 3015 | ENSMUSG00000021270 | 3252.941309 | 0.043427435 | 0.13893957 | 0.312563473 | 0.754612333 | 0.953048681 | Hsp90aa1 | heat shock protein 90, alpha (cytosolic), c | 3190.7 | 2872 | 3550 | 3923.4 | 2712.4 | 3269.1 |
| 3016 | ENSMUSG00000021271 | 161.075425 | 0.315984282 | 0.183334016 | 1.723544209 | 0.084790158 | 0.429140871 | Zfp839 | zinc finger protein 839 [Source:MGI Symb | 139.74 | 145.96 | 144.9 | 174.48 | 186.5 | 174.87 |
| 3017 | ENSMUSG00000021273 | 1441.086743 | 0.427292229 | 0.128079103 | 3.336158817 | 0.000849446 | 0.01858758 | Fdft1 | farnesyl diphosphate farnesyl transferase | 1267.3 | 1401.1 | 1019.3 | 1699.5 | 1662.1 | 1597.3 |
| 3018 | ENSMUSG00000021275 | 211.184732 | -0.29640231 | 0.189832902 | -1.561385333 | 0.11843286 | 0.504123865 | Tecpr2 | tectonin beta-propeller repeat containing | 202.38 | 235.39 | 261.62 | 220.75 | 183.22 | 163.74 |
| 3019 | ENSMUSG00000021276 | 251.0431171 | -0.020249811 | 0.159496543 | -0.126960813 | 0.898971414 | 0.982798755 | Cinp | cyclin-dependent kinase 2 interacting prot | 256.99 | 240.53 | 260.62 | 242.92 | 232.3 | 272.89 |
| 3020 | ENSMUSG00000021277 | 203.9250644 | 0.014658301 | 0.171908189 | 0.085268195 | 0.932048176 | 0.988420305 | Traf3 | TNF receptor-associated factor 3 [Source: | 194.35 | 191.19 | 223.39 | 211.11 | 195.22 | 208.29 |

| | | | | | | | | | | | | | | | |
|---|---|---|---|---|---|---|---|---|---|---|---|---|---|---|---|
| 3021 | ENSMUSG00000021278 | 1.979171996 | -1.570371184 | 1.781824634 | -0.881327575 | 0.378140545 | NA | Amn | amnionless [Source:MGI Symbol;Acc:MGI | 0.8031 | 6.1675 | 2.0125 | 2.8919 | 0 | 0 |
| 3022 | ENSMUSG00000021279 | 1187.023608 | -0.066497116 | 0.126497134 | -0.525680811 | 0.599109998 | 0.915074771 | Cdc42bpb | CDC42 binding protein kinase beta [Source | 1082.6 | 1262.3 | 1300.1 | 1312.9 | 1095 | 1069.3 |
| 3023 | ENSMUSG00000021280 | 10.18366104 | 0.347430332 | 0.770460902 | 0.450938303 | 0.652034017 | 0.925643431 | Exoc3l4 | exocyst complex component 3-like 4 [Sou | 9.6372 | 2.0558 | 15.094 | 6.7478 | 13.087 | 14.48 |
| 3024 | ENSMUSG00000021281 | 239.3885487 | -0.081180459 | 0.226449009 | -0.358493328 | 0.71997416 | 0.9447719 | Tnfaip2 | tumor necrosis factor, alpha-induced prot | 218.44 | 182.97 | 337.09 | 226.53 | 202.86 | 268.44 |
| 3025 | ENSMUSG00000021282 | 4735.705328 | 0.19270694 | 0.082940226 | 2.32344364 | 0.020155332 | 0.173941958 | Eif5 | eukaryotic translation initiation factor 5 [S | 4340.8 | 4536.2 | 4383.2 | 5063.8 | 5074.6 | 5015.6 |
| 3026 | ENSMUSG00000021285 | 266.9168381 | 0.498126477 | 0.166488188 | 2.991962868 | 0.0027719 | 0.04430139 | Ppp1r13b | protein phosphatase 1, regulatory subunit | 229.69 | 214.83 | 219.36 | 345.1 | 263.93 | 328.58 |
| 3027 | ENSMUSG00000021286 | 874.9494803 | 0.311053082 | 0.111017892 | 2.801828404 | 0.005081389 | 0.067366834 | Zfyve21 | zinc finger, FYVE domain containing 21 [Sc | 772.58 | 770.94 | 799.96 | 1028.6 | 903.03 | 974.61 |
| 3028 | ENSMUSG00000021287 | 121.1285045 | -0.154426007 | 0.258503747 | -0.597384019 | 0.550251021 | 0.895703112 | Xrcc3 | X-ray repair complementing defective rep | 161.42 | 131.57 | 89.556 | 134.96 | 111.24 | 98.018 |
| 3029 | ENSMUSG00000021288 | 255.8313565 | -0.145126259 | 0.19004097 | -0.763657748 | 0.445071226 | 0.852147532 | Klc1 | kinesin light chain 1 [Source:MGI Symbol;. | 280.28 | 236.42 | 289.8 | 301.73 | 201.76 | 225 |
| 3030 | ENSMUSG00000021290 | 1315.658107 | 0.035352015 | 0.118887238 | 0.297357526 | 0.766193567 | 0.955091733 | Atp5mpl | ATP synthase membrane subunit 6.8PL [S | 1352.4 | 1293.1 | 1251.8 | 1171.2 | 1327.3 | 1498.1 |
| 3031 | ENSMUSG00000021294 | 11.55592881 | -0.333899243 | 0.642139543 | -0.519979257 | 0.603078033 | 0.915859383 | Kif26a | kinesin family member 26A [Source:MGI S | 11.243 | 12.335 | 15.094 | 9.6398 | 6.5437 | 14.48 |
| 3032 | ENSMUSG00000021298 | 3.445692414 | -2.48439757 | 1.383350968 | -1.795927157 | 0.072506103 | NA | Gpr132 | G protein-coupled receptor 132 [Source:N | 10.44 | 1.0279 | 6.0375 | 0.964 | 1.0906 | 1.1138 |
| 3033 | ENSMUSG00000021301 | 0 | NA | NA | NA | NA | NA | Hecw1 | HECT, C2 and WW domain containing E3 u | 0 | 0 | 0 | 0 | 0 | 0 |
| 3034 | ENSMUSG00000021302 | 159.8606526 | -0.122369863 | 0.205713908 | -0.594854595 | 0.55194067 | 0.895935901 | Ggps1 | geranylgeranyl diphosphate synthase 1 [S | 176.68 | 189.14 | 133.83 | 148.45 | 142.87 | 168.19 |
| 3035 | ENSMUSG00000021303 | 0 | NA | NA | NA | NA | NA | Gng4 | guanine nucleotide binding protein (G prc | 0 | 0 | 0 | 0 | 0 | 0 |
| 3036 | ENSMUSG00000021306 | 16.78074623 | -1.912843543 | 0.580823408 | -3.293330668 | 0.00099008 | 0.020784486 | Gpr137b | G protein-coupled receptor 137B [Source: | 30.518 | 29.81 | 19.119 | 5.7839 | 6.5437 | 8.9108 |
| 3037 | ENSMUSG00000021311 | 177.561424 | -0.253914491 | 0.184755159 | -1.374329645 | 0.169339367 | 0.5966686 | Mtr | 5-methyltetrahydrofolate-homocysteine r | 204.79 | 202.5 | 172.07 | 167.73 | 170.14 | 148.14 |
| 3038 | ENSMUSG00000021313 | 0 | NA | NA | NA | NA | NA | Ryr2 | ryanodine receptor 2, cardiac [Source:MG | 0 | 0 | 0 | 0 | 0 | 0 |
| 3039 | ENSMUSG00000021314 | 0.17131964 | -0.796629176 | 4.080472857 | -0.195229623 | 0.84521319 | NA | Amph | amphiphysin [Source:MGI Symbol;Acc:MC | 0 | 1.0279 | 0 | 0 | 0 | 0 |
| 3040 | ENSMUSG00000021318 | 5.055082738 | -0.56350983 | 1.006415952 | -0.559917427 | 0.575535762 | NA | Gli3 | GLI-Kruppel family member GLI3 [Source:I | 4.0155 | 6.1675 | 8.05 | 7.7118 | 3.2719 | 1.1138 |
| 3041 | ENSMUSG00000021319 | 0.514011053 | 1.044022102 | 3.082659575 | 0.338675769 | 0.734853997 | NA | Sfrp4 | secreted frizzled-related protein 4 [Source | 0 | 0 | 1.0062 | 0.964 | 0 | 1.1138 |
| 3042 | ENSMUSG00000021322 | 16.17521098 | -2.134160366 | 0.715229165 | -2.983883308 | 0.002846153 | 0.045040215 | Aoah | acyloxyacyl hydrolase [Source:MGI Symbc | 17.668 | 48.312 | 13.081 | 5.7839 | 2.1812 | 10.025 |
| 3043 | ENSMUSG00000021326 | 389.0712334 | 0.174849996 | 0.139628573 | 1.252250824 | 0.210478482 | 0.654419768 | Trim27 | tripartite motif-containing 27 [Source:MG | 327.67 | 380.33 | 389.42 | 411.62 | 416.62 | 408.78 |
| 3044 | ENSMUSG00000021327 | 1192.335767 | -0.076186917 | 0.102780111 | -0.741261286 | 0.45853503 | 0.859322431 | Zkscan3 | zinc finger with KRAB and SCAN domains : | 1176.5 | 1192.4 | 1303.1 | 1173.2 | 1158.2 | 1150.6 |
| 3045 | ENSMUSG00000021335 | 273.2768078 | -0.36322019 | 0.208529498 | -1.741816836 | 0.0815405 | 0.420121054 | Slc17a1 | solute carrier family 17 (sodium phosphat | 244.95 | 310.43 | 368.29 | 297.87 | 188.68 | 229.45 |
| 3046 | ENSMUSG00000021336 | 862.794617 | -0.303695573 | 0.157318817 | -1.930446585 | 0.053551528 | 0.325586084 | Slc17a4 | solute carrier family 17 (sodium phosphat | 1080.2 | 841.86 | 936.81 | 759.61 | 626.01 | 932.29 |
| 3047 | ENSMUSG00000021337 | 0 | NA | NA | NA | NA | NA | Scgn | secretagogin, EF-hand calcium binding prc | 0 | 0 | 0 | 0 | 0 | 0 |
| 3048 | ENSMUSG00000021338 | 472.3708234 | -0.170593436 | 0.154971149 | -1.100807713 | 0.270980354 | 0.726633268 | Carmil1 | capping protein regulator and myosin 1 lir | 467.41 | 508.82 | 525.26 | 463.67 | 510.41 | 358.66 |
| 3049 | ENSMUSG00000021339 | 502.6503761 | 0.066572329 | 0.126986157 | 0.524248708 | 0.600105567 | 0.915217938 | Mrs2 | MRS2 magnesium transporter [Source:MC | 527.64 | 483.12 | 461.87 | 529.22 | 501.68 | 512.37 |
| 3050 | ENSMUSG00000021340 | 3601.974506 | 0.04304975 | 0.127609006 | 0.337356674 | 0.735848042 | 0.948109711 | Gpld1 | glycosylphosphatidylinositol specific phos | 2912 | 3600.8 | 4133.7 | 3756.6 | 3716.8 | 3491.9 |
| 3051 | ENSMUSG00000021342 | 0 | NA | NA | NA | NA | NA | Prl | prolactin [Source:MGI Symbol;Acc:MGI:97 | 0 | 0 | 0 | 0 | 0 | 0 |
| 3052 | ENSMUSG00000021345 | 0 | NA | NA | NA | NA | NA | Prl8a6 | prolactin family 8, subfamily a, member 6 | 0 | 0 | 0 | 0 | 0 | 0 |
| 3053 | ENSMUSG00000021346 | 0 | NA | NA | NA | NA | NA | Prl8a8 | prolactin family 8, subfamily a, member 8 | 0 | 0 | 0 | 0 | 0 | 0 |
| 3054 | ENSMUSG00000021347 | 0 | NA | NA | NA | NA | NA | Prl7b1 | prolactin family 7, subfamily b, member 1 | 0 | 0 | 0 | 0 | 0 | 0 |
| 3055 | ENSMUSG00000021348 | 0 | NA | NA | NA | NA | NA | Prl7d1 | prolactin family 7, subfamily d, member 1 | 0 | 0 | 0 | 0 | 0 | 0 |
| 3056 | ENSMUSG00000021356 | 3.561150536 | -0.968565636 | 1.420401535 | -0.681895656 | 0.495304933 | NA | Irf4 | interferon regulatory factor 4 [Source:MG | 0 | 7.1954 | 7.0437 | 3.8559 | 3.2719 | 0 |
| 3057 | ENSMUSG00000021357 | 223.7660664 | -0.312877628 | 0.164909021 | -1.897274179 | 0.057791761 | 0.341829329 | Exoc2 | exocyst complex component 2 [Source:M | 246.55 | 236.42 | 260.62 | 189.9 | 194.13 | 214.97 |
| 3058 | ENSMUSG00000021359 | 0.346303622 | 2.004339653 | 3.884003423 | 0.516049919 | 0.605819544 | NA | Tfap2a | transcription factor AP-2, alpha [Source:M | 0 | 0 | 0 | 0.964 | 0 | 1.1138 |
| 3059 | ENSMUSG00000021360 | 414.2017198 | 0.120762842 | 0.161979739 | 0.745542888 | 0.455943594 | 0.857604173 | Gcnt2 | glucosaminyl (N-acetyl) transferase 2, I-br | 454.56 | 399.86 | 335.08 | 401.98 | 398.08 | 495.66 |
| 3060 | ENSMUSG00000021361 | 2612.118469 | -0.100315784 | 0.091832717 | -1.092375216 | 0.274668212 | 0.729487777 | Tmem14c | transmembrane protein 14C [Source:MGI | 2625.3 | 2780.5 | 2702.8 | 2396.4 | 2591.3 | 2576.3 |
| 3061 | ENSMUSG00000021362 | 0 | NA | NA | NA | NA | NA | Gcm2 | glial cells missing homolog 2 [Source:MGI | 0 | 0 | 0 | 0 | 0 | 0 |
| 3062 | ENSMUSG00000021363 | 1.19317057 | 1.537463034 | 2.004580332 | 0.766975017 | 0.44309637 | NA | Mak | male germ cell-associated kinase [Source: | 0.8031 | 0 | 1.0062 | 0.964 | 3.2719 | 1.1138 |
| 3063 | ENSMUSG00000021364 | 11737.15022 | 0.469287374 | 0.135151267 | 3.472312064 | 0.000515996 | 0.012751348 | Elovl2 | elongation of very long chain fatty acids (F | 11990 | 9318.1 | 8224 | 13501 | 12573 | 14816 |
| 3064 | ENSMUSG00000021365 | 33.11779356 | -0.336199984 | 0.48055007 | -0.699614889 | 0.484167842 | 0.872431004 | Nedd9 | neural precursor cell expressed, developn | 38.549 | 51.396 | 21.131 | 43.379 | 14.178 | 30.074 |
| 3065 | ENSMUSG00000021366 | 269.5808 | 0.592408254 | 0.171876033 | 3.446718224 | 0.00056744 | 0.013611034 | Hivep1 | human immunodeficiency virus type I enh | 227.28 | 202.5 | 215.34 | 383.66 | 285.74 | 302.97 |
| 3066 | ENSMUSG00000021367 | 6.174401277 | -1.00288778 | 1.038404234 | -0.965797082 | 0.334145729 | 0.775900437 | Edn1 | endothelin 1 [Source:MGI Symbol;Acc:MC | 9.6372 | 1.0279 | 14.087 | 6.7478 | 1.0906 | 4.4554 |
| 3067 | ENSMUSG00000021368 | 101.357824 | -0.223416235 | 0.236481471 | -0.944751542 | 0.344785737 | 0.786033695 | Tbc1d7 | TBC1 domain family, member 7 [Source:M | 93.16 | 106.9 | 127.79 | 90.614 | 89.431 | 100.25 |
| 3068 | ENSMUSG00000021371 | 1126.947824 | -0.074538577 | 0.101084301 | -0.737390243 | 0.460885067 | 0.859821403 | Mcur1 | mitochondrial calcium uniporter regulator | 1162.1 | 1117.3 | 1188.4 | 1070 | 1121.2 | 1102.7 |
| 3069 | ENSMUSG00000021373 | 0.864800165 | 2.10816772 | 2.580439368 | 0.816980142 | 0.413939777 | NA | Cap2 | CAP, adenylate cyclase-associated protein | 0.8031 | 0 | 0 | 0 | 3.2719 | 1.1138 |
| 3070 | ENSMUSG00000021374 | 417.1697411 | 0.048428738 | 0.136857233 | 0.353863196 | 0.72344141 | 0.9447719 | Nup153 | nucleoporin 153 [Source:MGI Symbol;Acc | 431.27 | 371.08 | 427.65 | 425.11 | 405.71 | 442.2 |
| 3071 | ENSMUSG00000021375 | 393.783317 | 0.07969003 | 0.150230065 | 0.53045328 | 0.595797694 | 0.91381309 | Kif13a | kinesin family member 13A [Source:MGI S | 367.82 | 367.99 | 412.56 | 346.07 | 443.88 | 424.37 |
| 3072 | ENSMUSG00000021376 | 1108.98548 | 0.086824045 | 0.162911704 | 0.532951547 | 0.594067121 | 0.913144152 | Tpmt | thiopurine methyltransferase [Source:MG | 1265.7 | 928.21 | 1031.4 | 944.7 | 1410.2 | 1073.7 |
| 3073 | ENSMUSG00000021377 | 680.1726415 | -0.029918833 | 0.126843291 | -0.235872413 | 0.813531676 | 0.966842541 | Dek | DEK proto-oncogene (DNA binding) [Sourc | 619.99 | 733.93 | 708.4 | 635.26 | 729.62 | 653.83 |
| 3074 | ENSMUSG00000021379 | 15.62588931 | -0.388151161 | 0.58503704 | -0.66346425 | 0.507033268 | 0.88185929 | Id4 | inhibitor of DNA binding 4 [Source:MGI Sy | 18.471 | 14.391 | 20.125 | 8.6758 | 9.8156 | 22.277 |
| 3075 | ENSMUSG00000021381 | 0.133850373 | -0.796629176 | 4.080472857 | -0.195229623 | 0.84521319 | NA | Barx1 | BarH-like homeobox 1 [Source:MGI Symb | 0.8031 | 0 | 0 | 0 | 0 | 0 |
| 3076 | ENSMUSG00000021384 | 1.725425169 | -1.125104273 | 1.827651198 | -0.615601201 | 0.538157758 | NA | Susd3 | sushi domain containing 3 [Source:MGI Sy | 4.0155 | 0 | 3.0187 | 0 | 1.0906 | 2.2277 |

| | | | | | | | | | | | | | | | |
|---|---|---|---|---|---|---|---|---|---|---|---|---|---|---|---|
| 3077 | ENSMUSG00000021385 | 165.9908467 | -0.219252804 | 0.186017876 | -1.178665235 | 0.238531504 | 0.687411771 | Ippk | inositol 1,3,4,5,6-pentakisphosphate 2-kin | 179.89 | 175.77 | 180.12 | 155.2 | 167.96 | 137 |
| 3078 | ENSMUSG00000021388 | 0.692375149 | -1.404276945 | 2.772265464 | -0.506544905 | 0.612474175 | NA | Aspn | asporin [Source:MGI Symbol;Acc:MGI:191 | 0 | 1.0279 | 2.0125 | 0 | 0 | 1.1138 |
| 3079 | ENSMUSG00000021390 | 58.09442551 | -0.543825482 | 0.297288228 | -1.82928697 | 0.06735663 | 0.373457298 | Ogn | osteoglycin [Source:MGI Symbol;Acc:MGI | 80.31 | 66.815 | 59.368 | 47.235 | 44.715 | 50.123 |
| 3080 | ENSMUSG00000021391 | 5.970022297 | 0.270909589 | 1.055569574 | 0.256647781 | 0.797450686 | 0.962236316 | Cenpp | centromere protein P [Source:MGI Symbc | 4.0155 | 11.307 | 1.0062 | 10.604 | 1.0906 | 7.7969 |
| 3081 | ENSMUSG00000021392 | 279.321063 | -0.111171254 | 0.156926188 | -0.708430219 | 0.478678132 | 0.870017655 | Nol8 | nucleolar protein 8 [Source:MGI Symbol;A | 272.25 | 301.18 | 296.84 | 243.89 | 295.56 | 266.21 |
| 3082 | ENSMUSG00000021395 | 1207.114785 | 0.119076538 | 0.10610443 | 1.122257932 | 0.261752787 | 0.715688528 | Spin1 | spindlin 1 [Source:MGI Symbol;Acc:MGI:1 | 1072.9 | 1208.8 | 1191.4 | 1287.9 | 1203 | 1278.7 |
| 3083 | ENSMUSG00000021396 | 0.822613314 | -1.749415261 | 2.618121802 | -0.668194757 | 0.504009282 | NA | Nxnl2 | nucleoredoxin-like 2 [Source:MGI Symbol, | 0.8031 | 0 | 3.0187 | 0 | 0 | 1.1138 |
| 3084 | ENSMUSG00000021400 | 1034.341143 | -0.181424007 | 0.13425017 | -1.351387543 | 0.176571322 | 0.607421044 | Wrnip1 | Werner helicase interacting protein 1 [Sou | 1032 | 991.94 | 1274.9 | 1074.8 | 884.49 | 947.88 |
| 3085 | ENSMUSG00000021403 | 0.670829724 | -2.782124174 | 3.401449285 | -0.817923168 | 0.41340106 | NA | Serpinb9b | serine (or cysteine) peptidase inhibitor, cl | 0 | 0 | 4.025 | 0 | 0 | 0 |
| 3086 | ENSMUSG00000021404 | 0 | NA | NA | NA | NA | NA | Serpinb9c | serine (or cysteine) peptidase inhibitor, cl | 0 | 0 | 0 | 0 | 0 | 0 |
| 3087 | ENSMUSG00000021408 | 528.7090546 | 0.186395846 | 0.129230307 | 1.442353966 | 0.149202543 | 0.561319344 | Ripk1 | receptor (TNFRSF)-interacting serine-thre | 470.62 | 510.88 | 503.12 | 602.49 | 510.41 | 574.74 |
| 3088 | ENSMUSG00000021411 | 1302.474549 | 0.210109911 | 0.122040083 | 1.721646746 | 0.08513353 | 0.429948105 | Pxdc1 | PX domain containing 1 [Source:MGI Syml | 1220.7 | 1131.7 | 1270.9 | 1453.7 | 1203 | 1534.9 |
| 3089 | ENSMUSG00000021413 | 1007.386336 | -0.006610138 | 0.128415345 | -0.05147467 | 0.958947285 | 0.992027194 | Prpf4b | pre-mRNA processing factor 4B [Source:N | 952.48 | 1008.4 | 1069.6 | 1158.7 | 967.38 | 887.73 |
| 3090 | ENSMUSG00000021414 | 1.199366721 | -1.417144488 | 2.265583267 | -0.625509779 | 0.531636532 | NA | Fam217a | family with sequence similarity 217, meml | 3.2124 | 2.0558 | 0 | 1.928 | 0 | 0 |
| 3091 | ENSMUSG00000021415 | 0 | NA | NA | NA | NA | NA | 4933417A18 | RIKEN cDNA 4933417A18 gene [Source:M | 0 | 0 | 0 | 0 | 0 | 0 |
| 3092 | ENSMUSG00000021416 | 0.736965982 | -2.935629304 | 2.883220453 | -1.018177192 | 0.308593757 | NA | Eci3 | enoyl-Coenzyme A delta isomerase 3 [Sou | 2.4093 | 0 | 2.0125 | 0 | 0 | 0 |
| 3093 | ENSMUSG00000021417 | 4834.612722 | 0.081503098 | 0.100239105 | 0.813086849 | 0.416168266 | 0.836616789 | Eci2 | enoyl-Coenzyme A delta isomerase 2 [Sou | 4624.3 | 5114.9 | 4356 | 5269.1 | 4930.7 | 4712.7 |
| 3094 | ENSMUSG00000021418 | 130.0810618 | 0.164681535 | 0.208223663 | 0.790887705 | 0.429009524 | 0.842753542 | Rpp40 | ribonuclease P 40 subunit [Source:MGI Sy | 114.84 | 121.29 | 131.82 | 120.5 | 147.23 | 144.8 |
| 3095 | ENSMUSG00000021420 | 545.2835545 | 0.021612703 | 0.126643477 | 0.170657846 | 0.86449281 | 0.977227145 | Fars2 | phenylalanine-tRNA synthetase 2 (mitoch | 544.5 | 570.49 | 508.15 | 510.91 | 556.22 | 581.43 |
| 3096 | ENSMUSG00000021423 | 44.09524664 | -1.774577075 | 0.547151123 | -3.243303359 | 0.001181523 | 0.023604356 | Ly86 | lymphocyte antigen 86 [Source:MGI Symb | 52.202 | 107.93 | 44.275 | 4.8199 | 16.359 | 38.985 |
| 3097 | ENSMUSG00000021427 | 1450.626618 | -0.012436904 | 0.096795151 | -0.128486852 | 0.897763703 | 0.982798755 | Ssr1 | signal sequence receptor, alpha [Source:N | 1486.5 | 1466.8 | 1416.8 | 1428.6 | 1409.1 | 1495.9 |
| 3098 | ENSMUSG00000021428 | 131.2846136 | 0.002815945 | 0.21213911 | 0.013274051 | 0.989409151 | 0.99859176 | Riok1 | RIO kinase 1 [Source:MGI Symbol;Acc:MG | 125.28 | 144.94 | 123.77 | 151.34 | 125.42 | 116.95 |
| 3099 | ENSMUSG00000021431 | 360.5807095 | 0.351411699 | 0.168904908 | 2.080529827 | 0.037476963 | 0.262502077 | Snrnp48 | small nuclear ribonucleoprotein 48 (U11/l | 363.81 | 279.59 | 306.9 | 477.17 | 377.35 | 358.66 |
| 3100 | ENSMUSG00000021432 | 215.5506 | 0.164278735 | 0.191011419 | 0.860046669 | 0.389763317 | 0.819052248 | Slc35b3 | solute carrier family 35, member B3 [Sour | 191.14 | 173.72 | 245.52 | 256.42 | 213.76 | 212.74 |
| 3101 | ENSMUSG00000021439 | 0 | NA | NA | NA | NA | NA | Ctsq | cathepsin Q [Source:MGI Symbol;Acc:MG | 0 | 0 | 0 | 0 | 0 | 0 |
| 3102 | ENSMUSG00000021440 | 0 | NA | NA | NA | NA | NA | Cts7 | cathepsin 7 [Source:MGI Symbol;Acc:MGI | 0 | 0 | 0 | 0 | 0 | 0 |
| 3103 | ENSMUSG00000021441 | 0 | NA | NA | NA | NA | NA | Cts6 | cathepsin 6 [Source:MGI Symbol;Acc:MGI | 0 | 0 | 0 | 0 | 0 | 0 |
| 3104 | ENSMUSG00000021448 | 0 | NA | NA | NA | NA | NA | Shc3 | src homology 2 domain-containing transfc | 0 | 0 | 0 | 0 | 0 | 0 |
| 3105 | ENSMUSG00000021451 | 15.24387704 | -1.386477828 | 0.578447493 | -2.39689487 | 0.016534667 | 0.152621202 | Sema4d | sema domain, immunoglobulin domain (Ig | 23.29 | 26.726 | 16.1 | 7.7118 | 8.7249 | 8.9108 |
| 3106 | ENSMUSG00000021453 | 692.5826839 | -0.428645049 | 1.077566113 | -0.397790023 | 0.690784972 | 0.937606823 | Gadd45g | growth arrest and DNA-damage-inducible | 378.26 | 1597.4 | 408.54 | 420.29 | 107.97 | 1243 |
| 3107 | ENSMUSG00000021456 | 2.978902935 | 1.368373246 | 1.344479072 | 1.017772068 | 0.308786289 | NA | Fbp2 | fructose bisphosphatase 2 [Source:MGI Sy | 0 | 3.0838 | 2.0125 | 2.8919 | 6.5437 | 3.3415 |
| 3108 | ENSMUSG00000021457 | 59.41497716 | -1.135295672 | 0.412447087 | -2.752585016 | 0.005912679 | 0.07450361 | Syk | spleen tyrosine kinase [Source:MGI Symb | 68.264 | 122.32 | 54.337 | 31.811 | 20.722 | 59.034 |
| 3109 | ENSMUSG00000021458 | 439.4579362 | 0.156908038 | 0.139455298 | 1.125149345 | 0.260525754 | 0.71422375 | Aopep | aminopeptidase O [Source:MGI Symbol;A | 423.23 | 376.22 | 446.77 | 429.93 | 501.68 | 458.9 |
| 3110 | ENSMUSG00000021460 | 931.1112505 | 0.073220011 | 0.109721774 | 0.667324342 | 0.504564981 | 0.881510796 | Auh | AU RNA binding protein/enoyl-coenzyme | 887.43 | 924.1 | 910.65 | 883 | 995.73 | 985.75 |
| 3111 | ENSMUSG00000021461 | 104.5162088 | -0.199807496 | 0.24206308 | -0.825435652 | 0.409124301 | 0.833065722 | Fancc | Fanconi anemia, complementation group | 123.68 | 86.345 | 124.77 | 91.578 | 93.793 | 106.93 |
| 3112 | ENSMUSG00000021464 | 0.167707431 | -0.796629176 | 4.080472857 | -0.195229623 | 0.84521319 | NA | Ror2 | receptor tyrosine kinase-like orphan recep | 0 | 0 | 1.0062 | 0 | 0 | 0 |
| 3113 | ENSMUSG00000021466 | 508.2753619 | 0.621886081 | 0.179469032 | 3.465144239 | 0.000529947 | 0.012909222 | Ptch1 | patched 1 [Source:MGI Symbol;Acc:MGI:1 | 340.52 | 401.92 | 459.85 | 713.34 | 464.6 | 669.42 |
| 3114 | ENSMUSG00000021468 | 479.3060653 | -0.077319773 | 0.126296615 | -0.612207802 | 0.54040028 | 0.893324212 | Sptlc1 | serine palmitoyltransferase, long chain ba | 492.3 | 498.54 | 486.02 | 496.45 | 455.88 | 446.65 |
| 3115 | ENSMUSG00000021469 | 0 | NA | NA | NA | NA | NA | Msx2 | msh homeobox 2 [Source:MGI Symbol;Ac | 0 | 0 | 0 | 0 | 0 | 0 |
| 3116 | ENSMUSG00000021470 | 250.4553082 | -0.028959045 | 0.177677553 | -0.162986512 | 0.870529055 | 0.978802475 | Ercc6l2 | excision repair cross-complementing rode | 238.52 | 296.04 | 224.39 | 221.71 | 277.02 | 245.05 |
| 3117 | ENSMUSG00000021474 | 2456.383601 | -0.595554244 | 0.110894756 | -5.370445491 | 7.85E-08 | 8.88E-06 | Sfxn1 | sideroflexin 1 [Source:MGI Symbol;Acc:M | 2653.4 | 3273.9 | 2943.3 | 2090.9 | 1894.4 | 1882.4 |
| 3118 | ENSMUSG00000021476 | 1780.497535 | 0.166288701 | 0.11094283 | 1.498868386 | 0.133907779 | 0.536818769 | Habp4 | hyaluronic acid binding protein 4 [Source: | 1635.9 | 1596.4 | 1801.2 | 1689.9 | 1936.9 | 2022.7 |
| 3119 | ENSMUSG00000021477 | 8694.459378 | 0.157855894 | 0.148769866 | 1.061074381 | 0.2886561 | 0.73951397 | Ctsl | cathepsin L [Source:MGI Symbol;Acc:MGI | 8671.9 | 8269.6 | 7714.9 | 6944.5 | 9263.7 | 11302 |
| 3120 | ENSMUSG00000021478 | 0 | NA | NA | NA | NA | NA | Drd1 | dopamine receptor D1 [Source:MGI Symb | 0 | 0 | 0 | 0 | 0 | 0 |
| 3121 | ENSMUSG00000021481 | 104.6152335 | 0.033993098 | 0.239565052 | 0.141895061 | 0.887162895 | 0.982382091 | Zfp346 | zinc finger protein 346 [Source:MGI Symb | 106.81 | 93.541 | 109.68 | 108.93 | 83.978 | 124.75 |
| 3122 | ENSMUSG00000021482 | 1251.08807 | -0.109747487 | 0.104361909 | -1.051604818 | 0.292980896 | 0.743850082 | Prxl2c | peroxiredoxin like 2C [Source:MGI Symbo | 1218.3 | 1332.2 | 1346.4 | 1220.4 | 1146.2 | 1243 |
| 3123 | ENSMUSG00000021483 | 114.7802649 | 0.001853099 | 0.230866422 | 0.008026714 | 0.993595678 | 0.99859176 | Cdk20 | cyclin-dependent kinase 20 [Source:MGI S | 96.372 | 119.24 | 128.8 | 107 | 103.61 | 133.66 |
| 3124 | ENSMUSG00000021484 | 4753.335557 | 0.072830505 | 0.084454206 | 0.862366817 | 0.388485695 | 0.81838874 | Lman2 | lectin, mannose-binding 2 [Source:MGI Sy | 4628.3 | 4620.5 | 4650.9 | 4682 | 4939.4 | 4998.9 |
| 3125 | ENSMUSG00000021485 | 2.201010822 | -1.044154217 | 1.828885039 | -0.570923921 | 0.568051215 | NA | Mxd3 | Max dimerization protein 3 [Source:MGI S | 4.8186 | 0 | 4.025 | 0 | 4.3625 | 0 |
| 3126 | ENSMUSG00000021486 | 2810.534104 | 0.083922069 | 0.095746532 | 0.876502437 | 0.380756952 | 0.812628154 | Prelid1 | PRELI domain containing 1 [Source:MGI S | 2762.7 | 2629.4 | 2793.3 | 2681.8 | 2973 | 3023 |
| 3127 | ENSMUSG00000021488 | 1078.641 | 0.090033471 | 0.105740576 | 0.851456218 | 0.394515974 | 0.822827574 | Nsd1 | nuclear receptor-binding SET-domain prot | 1003.9 | 1089.6 | 1042.5 | 1166.4 | 1059 | 1110.5 |
| 3128 | ENSMUSG00000021490 | 0 | NA | NA | NA | NA | NA | Slc34a1 | solute carrier family 34 (sodium phosphat | 0 | 0 | 0 | 0 | 0 | 0 |
| 3129 | ENSMUSG00000021492 | 9141.962209 | 0.106913703 | 0.085227044 | 1.254457474 | 0.209675773 | 0.653550636 | F12 | coagulation factor XII (Hageman factor) [S | 8377.2 | 9082.7 | 8951.6 | 9680.3 | 9169.9 | 9590.2 |
| 3130 | ENSMUSG00000021493 | 95.13782506 | -0.162939521 | 0.242402546 | -0.672185684 | 0.501465493 | 0.880974509 | Pdlim7 | PDZ and LIM domain 7 [Source:MGI Symb | 84.326 | 100.74 | 116.72 | 83.866 | 91.612 | 93.563 |
| 3131 | ENSMUSG00000021494 | 458.8576311 | -0.065188283 | 0.130304503 | -0.500276517 | 0.616880387 | 0.917509899 | Ddx41 | DEAD box helicase 41 [Source:MGI Symbc | 479.45 | 444.06 | 484 | 450.18 | 474.42 | 421.03 |
| 3132 | ENSMUSG00000021495 | 1038.551298 | 0.197323185 | 0.161962818 | 1.218323979 | 0.223100881 | 0.668407528 | Fam193b | family with sequence similarity 193, meml | 969.34 | 873.73 | 1060.6 | 1389.1 | 867.04 | 1071.5 |

| | | | | | | | | | | | | | | | |
|---|---|---|---|---|---|---|---|---|---|---|---|---|---|---|---|
| 3133 | ENSMUSG00000021496 | 314.995873 | 0.008442826 | 0.157938251 | 0.053456498 | 0.957368191 | 0.991861398 | Pcbd2 | pterin 4 alpha carbinolamine dehydratase | 288.31 | 291.93 | 362.25 | 304.62 | 305.37 | 337.49 |
| 3134 | ENSMUSG00000021497 | 1508.960034 | 0.09747539 | 0.114080266 | 0.854445682 | 0.392858093 | 0.821390458 | Txndc15 | thioredoxin domain containing 15 [Source | 1325.9 | 1401.1 | 1648.2 | 1608.9 | 1520.3 | 1549.4 |
| 3135 | ENSMUSG00000021499 | 1.322483319 | -0.650659329 | 1.879732149 | -0.346144704 | 0.729233968 | NA | Catsper3 | cation channel, sperm associated 3 [Sourc | 0.8031 | 3.0838 | 1.0062 | 1.928 | 0 | 1.1138 |
| 3136 | ENSMUSG00000021500 | 804.6633138 | 0.030993283 | 0.140297792 | 0.220910696 | 0.825161969 | 0.96943211 | Ddx46 | DEAD box helicase 46 [Source:MGI Symbc | 704.32 | 867.56 | 817.07 | 790.46 | 937.93 | 710.63 |
| 3137 | ENSMUSG00000021501 | 324.18545 | -0.01540464 | 0.14362073 | -0.107259167 | 0.914583377 | 0.98625062 | Caml | calcium modulating ligand [Source:MGI Sy | 327.67 | 317.63 | 332.06 | 298.83 | 337 | 331.93 |
| 3138 | ENSMUSG00000021504 | 184.539432 | 0.030055974 | 0.200764309 | 0.149707757 | 0.880995188 | 0.981490993 | B4galt7 | xylosylprotein beta1,4-galactosyltransfera | 166.24 | 194.28 | 188.17 | 226.53 | 152.69 | 179.33 |
| 3139 | ENSMUSG00000021506 | 0 | NA | NA | NA | NA | NA | Pitx1 | paired-like homeodomain transcription fa | 0 | 0 | 0 | 0 | 0 | 0 |
| 3140 | ENSMUSG00000021508 | 19.88599693 | -0.937365197 | 0.506824695 | -1.849486039 | 0.064387662 | 0.363506301 | Cxcl14 | chemokine (C-X-C motif) ligand 14 [Source | 30.518 | 23.642 | 24.15 | 14.46 | 8.7249 | 17.822 |
| 3141 | ENSMUSG00000021509 | 270.8315466 | 0.201827584 | 0.182302387 | 1.107103355 | 0.268249245 | 0.724039098 | Slc25a48 | solute carrier family 25, member 48 [Sour | 216.84 | 273.43 | 265.65 | 237.14 | 295.56 | 336.38 |
| 3142 | ENSMUSG00000021510 | 148.2790647 | -0.093985885 | 0.220484203 | -0.426270381 | 0.669910838 | 0.930975245 | Zfp729a | zinc finger protein 729a [Source:MGI Sym | 122.87 | 194.28 | 142.89 | 147.49 | 140.69 | 141.46 |
| 3143 | ENSMUSG00000021514 | 246.367779 | 0.000479081 | 0.17489035 | 0.002739323 | 0.997814339 | 0.999680446 | Zfp369 | zinc finger protein 369 [Source:MGI Symb | 228.08 | 268.29 | 243.51 | 282.45 | 246.48 | 209.4 |
| 3144 | ENSMUSG00000021518 | 989.9843558 | -0.057459548 | 0.111652503 | -0.514628393 | 0.606812721 | 0.916376001 | Ptdss1 | phosphatidylserine synthase 1 [Source:M | 938.02 | 1007.4 | 1084.7 | 1002.5 | 924.84 | 982.41 |
| 3145 | ENSMUSG00000021519 | 373.9610264 | 0.063805026 | 0.161475386 | 0.395137785 | 0.692741205 | 0.938623774 | Mterf3 | mitochondrial transcription termination fa | 417.61 | 355.66 | 323 | 375.95 | 435.16 | 336.38 |
| 3146 | ENSMUSG00000021520 | 2134.686161 | -0.05298219 | 0.104566062 | -0.506686289 | 0.612374952 | 0.916441371 | Uqcrb | ubiquinol-cytochrome c reductase binding | 2211.7 | 2248.1 | 2060.8 | 1898.1 | 2154 | 2235.5 |
| 3147 | ENSMUSG00000021532 | 110.3929524 | -0.050498603 | 0.219983373 | -0.229556452 | 0.818436449 | 0.968897682 | Fastkd3 | FAST kinase domains 3 [Source:MGI Symb | 117.25 | 109.99 | 109.68 | 114.71 | 94.884 | 115.84 |
| 3148 | ENSMUSG00000021534 | 0.496077836 | -0.812642266 | 3.444120572 | -0.235950586 | 0.813471014 | NA | 1700001L19I | RIKEN cDNA 1700001L19 gene [Source:M | 0 | 0 | 2.0125 | 0.964 | 0 | 0 |
| 3149 | ENSMUSG00000021536 | 3.677422101 | -0.688481796 | 1.160432188 | -0.593297741 | 0.55298191 | NA | Adcy2 | adenylate cyclase 2 [Source:MGI Symbol;/ | 2.4093 | 9.2513 | 2.0125 | 2.8919 | 3.2719 | 2.2277 |
| 3150 | ENSMUSG00000021537 | 361.9576782 | 0.170927366 | 0.149787713 | 1.141130755 | 0.253815514 | 0.707055046 | Cetn3 | centrin 3 [Source:MGI Symbol;Acc:MGI:1( | 313.21 | 382.39 | 327.03 | 417.4 | 369.72 | 362 |
| 3151 | ENSMUSG00000021538 | 0 | NA | NA | NA | NA | NA | Il9 | interleukin 9 [Source:MGI Symbol;Acc:MG | 0 | 0 | 0 | 0 | 0 | 0 |
| 3152 | ENSMUSG00000021539 | 2484.454881 | 0.022376344 | 0.126678507 | 0.176638837 | 0.859792086 | 0.975604671 | Lect2 | leukocyte cell-derived chemotaxin 2 [Sour | 2776.3 | 2128.8 | 2488.4 | 2211.4 | 2568.4 | 2733.4 |
| 3153 | ENSMUSG00000021540 | 544.8053134 | 0.046961434 | 0.133653321 | 0.351367506 | 0.725312657 | 0.944957869 | Smad5 | SMAD family member 5 [Source:MGI Sym | 503.55 | 593.11 | 512.18 | 600.56 | 514.77 | 544.67 |
| 3154 | ENSMUSG00000021541 | 0 | NA | NA | NA | NA | NA | Trpc7 | transient receptor potential cation channe | 0 | 0 | 0 | 0 | 0 | 0 |
| 3155 | ENSMUSG00000021545 | 0 | NA | NA | NA | NA | NA | 1700067P10 | RIKEN cDNA 1700067P10 gene [Source:M | 0 | 0 | 0 | 0 | 0 | 0 |
| 3156 | ENSMUSG00000021546 | 3914.731215 | 0.094048512 | 0.096421075 | 0.975393728 | 0.329364994 | 0.774371793 | Hnrnpk | heterogeneous nuclear ribonucleoprotein | 3803.5 | 3983.2 | 3574.2 | 3757.6 | 4212 | 4158 |
| 3157 | ENSMUSG00000021548 | 388.8769649 | 0.029999688 | 0.139297002 | 0.215364923 | 0.829482815 | 0.969955221 | Ccnh | cyclin H [Source:MGI Symbol;Acc:MGI:19: | 366.21 | 397.8 | 390.42 | 356.67 | 412.25 | 409.89 |
| 3158 | ENSMUSG00000021549 | 471.8665644 | 0.063567241 | 0.140674044 | 0.451876118 | 0.65135823 | 0.925643431 | Rasa1 | RAS p21 protein activator 1 [Source:MGI S | 424.04 | 524.24 | 436.71 | 463.67 | 516.95 | 465.59 |
| 3159 | ENSMUSG00000021550 | 629.8172721 | -0.030327432 | 0.122044489 | -0.248494887 | 0.803751525 | 0.963596811 | 2210016F16 | RIKEN cDNA 2210016F16 gene [Source:M | 673 | 601.33 | 633.93 | 585.13 | 621.65 | 663.85 |
| 3160 | ENSMUSG00000021552 | 295.5457229 | 0.143739206 | 0.155412867 | 0.924886137 | 0.355025138 | 0.795027651 | Gkap1 | G kinase anchoring protein 1 [Source:MGI | 249.76 | 283.71 | 309.92 | 324.86 | 304.28 | 300.74 |
| 3161 | ENSMUSG00000021553 | 0.301557804 | -1.64335916 | 4.042814367 | -0.406488899 | 0.684383412 | NA | Slc28a3 | solute carrier family 28 (sodium-coupled r | 0.8031 | 0 | 1.0062 | 0 | 0 | 0 |
| 3162 | ENSMUSG00000021555 | 491.1052867 | 0.164337292 | 0.125840859 | 1.30591362 | 0.191581934 | 0.629099068 | Naa35 | N(alpha)-acetyltransferase 35, NatC auxili | 479.45 | 439.95 | 469.92 | 539.83 | 497.32 | 520.17 |
| 3163 | ENSMUSG00000021556 | 22.07462315 | -0.34894176 | 0.489105479 | -0.713428443 | 0.475580678 | 0.869436931 | Golm1 | golgi membrane protein 1 [Source:MGI Sy | 22.487 | 16.447 | 35.219 | 16.388 | 19.631 | 22.277 |
| 3164 | ENSMUSG00000021557 | 177.1046262 | 0.171600499 | 0.199316442 | 0.860945028 | 0.38926832 | 0.818844821 | Agtpbp1 | ATP/GTP binding protein 1 [Source:MGI S | 137.33 | 194.28 | 169.05 | 202.44 | 167.96 | 191.58 |
| 3165 | ENSMUSG00000021559 | 1137.168493 | -0.023230153 | 0.112468429 | -0.206548212 | 0.836362709 | 0.971073158 | Dapk1 | death associated protein kinase 1 [Source | 1135.6 | 1073.1 | 1230.6 | 1191.5 | 1040.4 | 1151.7 |
| 3166 | ENSMUSG00000021565 | 0.583320834 | -1.123770786 | 3.233988906 | -0.34748752 | 0.728225092 | NA | Slc6a19 | solute carrier family 6 (neurotransmitter t | 2.4093 | 0 | 0 | 0 | 1.0906 | 0 |
| 3167 | ENSMUSG00000021566 | 0 | NA | NA | NA | NA | NA | Slc6a19os | solute carrier family 6 (neurotransmitter t | 0 | 0 | 0 | 0 | 0 | 0 |
| 3168 | ENSMUSG00000021567 | 2.888512514 | -2.104609537 | 1.496605989 | -1.406254921 | 0.159648441 | NA | Nkd2 | naked cuticle 2 [Source:MGI Symbol;Acc:I | 0.8031 | 7.1954 | 6.0375 | 0 | 2.1812 | 1.1138 |
| 3169 | ENSMUSG00000021569 | 9.438415212 | -0.189829526 | 0.73344203 | -0.258820082 | 0.795774063 | 0.962236316 | Trip13 | thyroid hormone receptor interactor 13 [S | 12.85 | 4.1117 | 13.081 | 6.7478 | 9.8156 | 10.025 |
| 3170 | ENSMUSG00000021572 | 10.73997727 | -0.191536063 | 0.650640808 | -0.294380648 | 0.768467053 | 0.955856567 | Cep72 | centrosomal protein 72 [Source:MGI Syml | 12.047 | 11.307 | 11.069 | 13.496 | 8.7249 | 7.7969 |
| 3171 | ENSMUSG00000021573 | 370.9489575 | 0.197163641 | 0.158547482 | 1.243562107 | 0.213660769 | 0.658358849 | Tppp | tubulin polymerization promoting protein | 327.67 | 335.1 | 374.32 | 372.1 | 467.88 | 348.63 |
| 3172 | ENSMUSG00000021575 | 1.45266935 | -2.766023049 | 2.035789473 | -1.358697982 | 0.174242312 | NA | Ahrr | aryl-hydrocarbon receptor repressor [Sou | 1.6062 | 5.1396 | 1.0062 | 0.964 | 0 | 0 |
| 3173 | ENSMUSG00000021576 | 1460.620192 | -0.036096415 | 0.10388217 | -0.347474593 | 0.728234803 | 0.946208065 | Pdcd6 | programmed cell death 6 [Source:MGI Syr | 1506.6 | 1496.6 | 1432.9 | 1381.4 | 1566.1 | 1380.1 |
| 3174 | ENSMUSG00000021577 | 10850.69041 | 0.031070424 | 0.093332031 | 0.332902039 | 0.73920823 | 0.948310621 | Sdha | succinate dehydrogenase complex, subun | 10118 | 11490 | 10594 | 10389 | 11547 | 10966 |
| 3175 | ENSMUSG00000021578 | 332.6677629 | 0.122924049 | 0.144426 | 0.851121329 | 0.394701959 | 0.822855164 | Ccdc127 | coiled-coil domain containing 127 [Source | 325.26 | 326.88 | 302.88 | 320.04 | 352.27 | 368.68 |
| 3176 | ENSMUSG00000021579 | 0 | NA | NA | NA | NA | NA | Lrrc14b | leucine rich repeat containing 14B [Source | 0 | 0 | 0 | 0 | 0 | 0 |
| 3177 | ENSMUSG00000021583 | 1724.394965 | -0.054220866 | 0.117520317 | -0.461374402 | 0.644530014 | 0.925643431 | Erap1 | endoplasmic reticulum aminopeptidase 1 | 1537.9 | 1898.6 | 1835.4 | 1710.1 | 1803.9 | 1560.5 |
| 3178 | ENSMUSG00000021585 | 873.451639 | -0.035110357 | 0.107525383 | -0.326530875 | 0.744022734 | 0.949911274 | Cast | calpastatin [Source:MGI Symbol;Acc:MGI: | 872.97 | 861.4 | 917.7 | 830.95 | 886.67 | 871.03 |
| 3179 | ENSMUSG00000021587 | 0.294513347 | 0.165143496 | 4.080472857 | 0.040471657 | 0.967717103 | NA | Pcsk1 | proprotein convertase subtilisin/kexin typ | 0.8031 | 0 | 0 | 0.964 | 0 | 0 |
| 3180 | ENSMUSG00000021589 | 133.802342 | 0.254945987 | 0.221772818 | 1.14958176 | 0.250316174 | 0.702302228 | Rhobtb3 | Rho-related BTB domain containing 3 [Sou | 101.99 | 131.57 | 132.82 | 133.99 | 129.78 | 172.65 |
| 3181 | ENSMUSG00000021590 | 12.00542455 | -0.049549426 | 0.638986263 | -0.077543805 | 0.938190945 | 0.988420305 | Spata9 | spermatogenesis associated 9 [Source:MG | 15.259 | 9.2513 | 12.075 | 13.496 | 15.269 | 6.6831 |
| 3182 | ENSMUSG00000021591 | 767.0666646 | -0.47413654 | 0.127579135 | -3.716411325 | 0.000202073 | 0.006326357 | Glrx | glutaredoxin [Source:MGI Symbol;Acc:MG | 803.1 | 938.49 | 935.81 | 705.63 | 588.93 | 630.44 |
| 3183 | ENSMUSG00000021592 | 151.721234 | -0.077791225 | 0.215811522 | -0.360459091 | 0.718503844 | 0.9447719 | Arsk | arylsulfatase K [Source:MGI Symbol;Acc:N | 134.92 | 176.8 | 155.97 | 130.14 | 179.95 | 132.55 |
| 3184 | ENSMUSG00000021594 | 1510.078372 | 0.823033383 | 0.108246801 | 7.603304459 | 2.89E-14 | 1.61E-11 | Srd5a1 | steroid 5 alpha-reductase 1 [Source:MGI S | 1134 | 1030 | 1106.9 | 1807.5 | 1873.7 | 2108.5 |
| 3185 | ENSMUSG00000021595 | 1421.072565 | 0.172042832 | 0.10436519 | 1.648469491 | 0.099256366 | 0.464258353 | Nsun2 | NOL1/NOP2/Sun domain family member 2 | 1280.9 | 1368.2 | 1360.4 | 1423.8 | 1623.9 | 1469.2 |
| 3186 | ENSMUSG00000021596 | 0.620790101 | -1.233953725 | 2.860353486 | -0.431399032 | 0.666178254 | NA | Mctp1 | multiple C2 domains, transmembrane 1 [S | 1.6062 | 1.0279 | 0 | 0 | 1.0906 | 0 |
| 3187 | ENSMUSG00000021597 | 64.8264745 | 0.383780208 | 0.295514763 | 1.29868371 | 0.194052497 | 0.632584024 | Slf1 | SMC5-SMC6 complex localization factor 1 | 45.777 | 71.954 | 51.318 | 65.55 | 74.162 | 80.197 |
| 3188 | ENSMUSG00000021598 | 0 | NA | NA | NA | NA | NA | Med10 | mediator complex subunit 10 [Source:MG | 0 | 0 | 0 | 0 | 0 | 0 |

| | | | | | | | | | | | | | | | |
|---|---|---|---|---|---|---|---|---|---|---|---|---|---|---|---|
| 3189 | ENSMUSG00000021600 | 0 | NA | NA | NA | NA | NA | 8030423J24I | RIKEN cDNA 8030423J24 gene [Source:MC | 0 | 0 | 0 | 0 | 0 | 0 |
| 3190 | ENSMUSG00000021604 | 0 | NA | NA | NA | NA | NA | Irx4 | Iroquois homeobox 4 [Source:MGI Symbo | 0 | 0 | 0 | 0 | 0 | 0 |
| 3191 | ENSMUSG00000021606 | 1366.218764 | 0.067879652 | 0.110300041 | 0.615409126 | 0.538284566 | 0.891577636 | Ndufs6 | NADH:ubiquinone oxidoreductase core su | 1297.8 | 1400 | 1304.1 | 1283.1 | 1377.5 | 1534.9 |
| 3192 | ENSMUSG00000021607 | 546.7060214 | -0.059069858 | 0.128762424 | -0.458750744 | 0.64641317 | 0.925643431 | Mrpl36 | mitochondrial ribosomal protein L36 [Sou | 599.11 | 522.18 | 551.42 | 515.73 | 512.59 | 579.2 |
| 3193 | ENSMUSG00000021608 | 30.79223679 | -0.749788646 | 0.502420607 | -1.492352494 | 0.135606762 | 0.5381974 | Lpcat1 | lysophosphatidylcholine acyltransferase 1 | 34.533 | 37.005 | 44.275 | 21.208 | 7.6343 | 40.098 |
| 3194 | ENSMUSG00000021609 | 0 | NA | NA | NA | NA | NA | Slc6a3 | solute carrier family 6 (neurotransmitter t | 0 | 0 | 0 | 0 | 0 | 0 |
| 3195 | ENSMUSG00000021610 | 1599.4634 | 0.133134576 | 0.106046276 | 1.25543848 | 0.209319628 | 0.65350014 | Clptm1l | CLPTM1-like [Source:MGI Symbol;Acc:MG | 1660.8 | 1520.3 | 1394.7 | 1635.9 | 1665.4 | 1719.8 |
| 3196 | ENSMUSG00000021611 | 147.7985197 | 0.647557679 | 0.239494785 | 2.703848769 | 0.006854148 | 0.082822382 | Tert | telomerase reverse transcriptase [Source: | 137.33 | 93.541 | 113.71 | 135.92 | 221.4 | 184.9 |
| 3197 | ENSMUSG00000021612 | 0 | NA | NA | NA | NA | NA | Slc6a18 | solute carrier family 6 (neurotransmitter t | 0 | 0 | 0 | 0 | 0 | 0 |
| 3198 | ENSMUSG00000021613 | 5.809758446 | 2.089130168 | 1.108118359 | 1.88529515 | 0.059390001 | 0.346665337 | Hapln1 | hyaluronan and proteoglycan link protein | 1.6062 | 0 | 5.0312 | 3.8559 | 6.5437 | 17.822 |
| 3199 | ENSMUSG00000021614 | 1.805345524 | 0.410335475 | 1.620610909 | 0.253198021 | 0.800115196 | NA | Vcan | versican [Source:MGI Symbol;Acc:MGI:10 | 1.6062 | 2.0558 | 1.0062 | 2.8919 | 3.2719 | 0 |
| 3200 | ENSMUSG00000021615 | 114.938737 | -0.030919411 | 0.261441795 | -0.118264988 | 0.905857698 | 0.983991807 | Xrcc4 | X-ray repair complementing defective rep | 146.16 | 106.9 | 94.587 | 87.722 | 142.87 | 111.38 |
| 3201 | ENSMUSG00000021619 | 88.61675473 | -0.136746231 | 0.288455648 | -0.474063281 | 0.635454782 | 0.923478764 | Atg10 | autophagy related 10 [Source:MGI Symbo | 94.766 | 97.652 | 85.531 | 52.055 | 100.34 | 101.36 |
| 3202 | ENSMUSG00000021620 | 2999.447084 | -0.116658854 | 0.111972122 | -1.041856245 | 0.297478333 | 0.748091545 | Acot12 | acyl-CoA thioesterase 12 [Source:MGI Syn | 3003.6 | 2944 | 3415.2 | 2998.9 | 3083.2 | 2551.8 |
| 3203 | ENSMUSG00000021621 | 145.0396192 | 0.349898677 | 0.207653662 | 1.685010868 | 0.091986475 | 0.446623824 | Zcchc9 | zinc finger, CCHC domain containing 9 [So | 119.66 | 135.69 | 127.79 | 187.01 | 135.24 | 164.85 |
| 3204 | ENSMUSG00000021622 | 0.505131668 | 1.126696199 | 3.421159819 | 0.329331647 | 0.741905026 | NA | Ckmt2 | creatine kinase, mitochondrial 2 [Source:M | 0.8031 | 0 | 0 | 0 | 0 | 2.2277 |
| 3205 | ENSMUSG00000021624 | 14.18706255 | -1.272144215 | 0.641341852 | -1.983566503 | 0.047304186 | 0.301811978 | Cd180 | CD180 antigen [Source:MGI Symbol;Acc:M | 22.487 | 26.726 | 11.069 | 11.568 | 4.3625 | 8.9108 |
| 3206 | ENSMUSG00000021629 | 803.9694908 | 0.311637778 | 0.118874585 | 2.621567756 | 0.008752636 | 0.098272117 | Slc30a5 | solute carrier family 30 (zinc transporter), | 778.21 | 672.26 | 701.35 | 946.63 | 887.76 | 837.61 |
| 3207 | ENSMUSG00000021631 | 1.117074527 | -3.52147603 | 2.24739728 | -1.566913007 | 0.117135027 | NA | Mrps36-ps1 | mitichondrial ribosomal protein S36, pseu | 1.6062 | 3.0838 | 2.0125 | 0 | 0 | 0 |
| 3208 | ENSMUSG00000021635 | 245.0573709 | 0.015989534 | 0.164489815 | 0.097206833 | 0.922562142 | 0.986685804 | Rad17 | RAD17 checkpoint clamp loader compone | 216.84 | 272.4 | 242.5 | 244.85 | 246.48 | 247.27 |
| 3209 | ENSMUSG00000021636 | 139.3976325 | 0.463006064 | 0.220680727 | 2.098081103 | 0.035897982 | 0.254648556 | Marveld2 | MARVEL (membrane-associating) domain | 93.963 | 123.35 | 134.84 | 145.56 | 153.78 | 184.9 |
| 3210 | ENSMUSG00000021638 | 306.4821295 | 0.260607583 | 0.213065034 | 1.223136328 | 0.22127821 | 0.667727967 | Ocln | occludin [Source:MGI Symbol;Acc:MGI:10 | 225.67 | 252.87 | 358.22 | 256.42 | 340.27 | 405.44 |
| 3211 | ENSMUSG00000021639 | 165.6770393 | 0.133081002 | 0.195541186 | 0.680577861 | 0.49613864 | 0.878165817 | Gtf2h2 | general transcription factor II H, polypepti | 142.15 | 152.13 | 180.12 | 162.91 | 194.13 | 162.62 |
| 3212 | ENSMUSG00000021640 | 5.085055653 | -1.926693411 | 1.085207915 | -1.775414079 | 0.075829541 | NA | Naip1 | NLR family, apoptosis inhibitory protein 1 | 12.85 | 7.1954 | 4.025 | 0.964 | 4.3625 | 1.1138 |
| 3213 | ENSMUSG00000021643 | 50.28943663 | 0.206436855 | 0.340185386 | 0.606836343 | 0.543959527 | 0.894702561 | Serf1 | small EDRK-rich factor 1 [Source:MGI Sym | 63.445 | 37.005 | 39.244 | 60.731 | 54.531 | 46.781 |
| 3214 | ENSMUSG00000021645 | 112.5704856 | -0.070959871 | 0.217875419 | -0.325690119 | 0.744658821 | 0.949997329 | Smn1 | survival motor neuron 1 [Source:MGI Sym | 118.06 | 106.9 | 120.75 | 99.29 | 111.24 | 119.18 |
| 3215 | ENSMUSG00000021646 | 1481.526148 | 0.180700349 | 0.123847316 | 1.459057449 | 0.144549296 | 0.55464443 | Mccc2 | methylcrotonoyl-Coenzyme A carboxylase | 1355.6 | 1480.2 | 1330.3 | 1345.7 | 1770.1 | 1607.3 |
| 3216 | ENSMUSG00000021647 | 0 | NA | NA | NA | NA | NA | Cartpt | CART prepropeptide [Source:MGI Symbol, | 0 | 0 | 0 | 0 | 0 | 0 |
| 3217 | ENSMUSG00000021650 | 419.3682283 | 0.078690232 | 0.139125543 | 0.565605928 | 0.571661697 | 0.904651508 | Ptcd2 | pentatricopeptide repeat domain 2 [Sourc | 438.49 | 418.36 | 366.27 | 424.15 | 412.25 | 456.68 |
| 3218 | ENSMUSG00000021660 | 1965.798007 | -0.060613215 | 0.101495997 | -0.597198086 | 0.550375137 | 0.895703112 | Btf3 | basic transcription factor 3 [Source:MGI S | 2129.8 | 1840 | 2050.7 | 1958.8 | 1962 | 1853.4 |
| 3219 | ENSMUSG00000021661 | 235.2044235 | 0.226502401 | 0.166811382 | 1.357835412 | 0.174515915 | 0.604201379 | Ankra2 | ankyrin repeat, family A (RFXANK-like), 2 [ | 201.58 | 206.61 | 242.5 | 255.45 | 237.75 | 267.32 |
| 3220 | ENSMUSG00000021662 | 1.136636081 | 0.517427405 | 1.988767259 | 0.260174942 | 0.794728833 | NA | Arhgef28 | Rho guanine nucleotide exchange factor ( | 0.8031 | 1.0279 | 1.0062 | 2.8919 | 1.0906 | 0 |
| 3221 | ENSMUSG00000021665 | 174.0741335 | -0.158801195 | 0.221889088 | -0.715678254 | 0.474190041 | 0.86899591 | Hexb | hexosaminidase B [Source:MGI Symbol;Ac | 217.64 | 151.1 | 181.12 | 125.32 | 182.13 | 187.13 |
| 3222 | ENSMUSG00000021666 | 993.5819602 | -0.009355547 | 0.108671556 | -0.086090113 | 0.931394783 | 0.988420305 | Gfm2 | G elongation factor, mitochondrial 2 [Sour | 1025.6 | 987.83 | 976.06 | 916.74 | 1033.9 | 1021.4 |
| 3223 | ENSMUSG00000021668 | 92.65408808 | 0.050361829 | 0.233617117 | 0.215574224 | 0.829319649 | 0.969955221 | Polk | polymerase (DNA directed), kappa [Sourc | 85.129 | 98.68 | 89.556 | 97.362 | 90.521 | 94.677 |
| 3224 | ENSMUSG00000021669 | 804.450868 | 0.092976692 | 0.113449529 | 0.819542338 | 0.412477057 | 0.834648594 | Cert1 | ceramide transporter 1 [Source:MGI Symb | 806.31 | 758.6 | 769.78 | 773.11 | 877.95 | 840.95 |
| 3225 | ENSMUSG00000021670 | 845.154281 | 0.448536547 | 0.185217309 | 2.421677273 | 0.015449064 | 0.145384033 | Hmgcr | 3-hydroxy-3-methylglutaryl-Coenzyme A r | 666.57 | 894.29 | 583.62 | 776 | 1221.5 | 928.95 |
| 3226 | ENSMUSG00000021671 | 89.46181379 | 0.39285615 | 0.24913851 | 1.576858395 | 0.114828112 | 0.49756555 | Poc5 | POC5 centriolar protein [Source:MGI Syml | 92.357 | 66.815 | 72.45 | 108.93 | 95.974 | 100.25 |
| 3227 | ENSMUSG00000021675 | 3.185712416 | -1.175100738 | 1.443525849 | -0.814048975 | 0.415616895 | NA | F2rl2 | coagulation factor II (thrombin) receptor-l | 7.2279 | 3.0838 | 3.0187 | 5.7839 | 0 | 0 |
| 3228 | ENSMUSG00000021676 | 7494.430646 | -0.064219776 | 0.103588957 | -0.61994809 | 0.535291963 | 0.889860001 | Iqgap2 | IQ motif containing GTPase activating pro | 7064.1 | 8079.4 | 7841.7 | 7620.2 | 7776.1 | 6585 |
| 3229 | ENSMUSG00000021678 | 1.407401099 | 1.828245829 | 2.025430599 | 0.902645507 | 0.366714066 | NA | F2rl1 | coagulation factor II (thrombin) receptor-l | 0.8031 | 1.0279 | 0 | 0 | 3.2719 | 3.3415 |
| 3230 | ENSMUSG00000021679 | 0 | NA | NA | NA | NA | NA | S100z | S100 calcium binding protein, zeta [Sourc | 0 | 0 | 0 | 0 | 0 | 0 |
| 3231 | ENSMUSG00000021680 | 0 | NA | NA | NA | NA | NA | Crhbp | corticotropin releasing hormone binding p | 0 | 0 | 0 | 0 | 0 | 0 |
| 3232 | ENSMUSG00000021681 | 714.6974934 | -0.043030507 | 0.123351848 | -0.348843634 | 0.727206703 | 0.945907265 | Aggf1 | angiogenic factor with G patch and FHA d | 719.58 | 672.26 | 783.86 | 658.4 | 760.16 | 693.92 |
| 3233 | ENSMUSG00000021684 | 13.04263661 | -0.644911178 | 0.606974188 | -1.062501818 | 0.288007933 | 0.73951397 | Pde8b | phosphodiesterase 8B [Source:MGI Symb | 11.243 | 18.503 | 18.112 | 10.604 | 11.997 | 7.7969 |
| 3234 | ENSMUSG00000021685 | 0 | NA | NA | NA | NA | NA | Otp | orthopedia homeobox [Source:MGI Symb | 0 | 0 | 0 | 0 | 0 | 0 |
| 3235 | ENSMUSG00000021686 | 743.5816932 | 0.155449079 | 0.120351234 | 1.291628459 | 0.196485843 | 0.637356187 | Ap3b1 | adaptor-related protein complex 3, beta 1 | 664.97 | 759.63 | 686.26 | 723.95 | 814.69 | 811.99 |
| 3236 | ENSMUSG00000021687 | 1038.967515 | 0.3250489 | 0.10428797 | 3.116839834 | 0.001828009 | 0.032561965 | Scamp1 | secretory carrier membrane protein 1 [So | 888.23 | 942.6 | 936.81 | 1136.5 | 1123.3 | 1206.3 |
| 3237 | ENSMUSG00000021690 | 476.3815674 | 0.01609536 | 0.135298642 | 0.118961729 | 0.905305676 | 0.983972587 | Jmy | junction-mediating and regulatory protein | 424.04 | 501.62 | 496.08 | 452.11 | 479.87 | 504.57 |
| 3238 | ENSMUSG00000021692 | 66.60254997 | 0.156797554 | 0.273991736 | 0.572271108 | 0.56713832 | 0.902953752 | Dimt1 | DIM1 dimethyladenosine transferase 1-lik | 69.87 | 56.535 | 62.387 | 75.19 | 64.346 | 71.286 |
| 3239 | ENSMUSG00000021693 | 139.1959842 | -0.133458637 | 0.213231596 | -0.625885842 | 0.531389821 | 0.888947136 | Kif2a | kinesin family member 2A [Source:MGI Sy | 150.98 | 161.38 | 124.77 | 151.34 | 131.96 | 114.73 |
| 3240 | ENSMUSG00000021694 | 72.28354122 | 0.392484776 | 0.290936561 | 1.349039033 | 0.177324429 | 0.608561375 | Ercc8 | excision repaiross-complementing rodent | 75.492 | 57.563 | 54.337 | 104.11 | 69.8 | 72.4 |
| 3241 | ENSMUSG00000021696 | 4.461042078 | -0.576725255 | 1.035585746 | -0.556907294 | 0.577590771 | NA | Elovl7 | ELOVL family member 7, elongation of lon | 4.8186 | 4.1117 | 7.0437 | 1.928 | 2.1812 | 6.6831 |
| 3242 | ENSMUSG00000021697 | 1.038626637 | -2.156830076 | 2.453413824 | -0.879113851 | 0.379339548 | NA | Depdc1b | DEP domain containing 1B [Source:MGI Sy | 0 | 4.1117 | 1.0062 | 0 | 0 | 1.1138 |
| 3243 | ENSMUSG00000021699 | 63.38671097 | -0.394024429 | 0.365075151 | -1.079296762 | 0.280455456 | 0.733413074 | Pde4d | phosphodiesterase 4D, cAMP specific [Sou | 93.16 | 88.401 | 34.212 | 58.803 | 55.622 | 50.123 |
| 3244 | ENSMUSG00000021700 | 0.167707431 | -0.796629176 | 4.080472857 | -0.195229623 | 0.84521319 | NA | Rab3c | RAB3C, member RAS oncogene family [So | 0 | 0 | 1.0062 | 0 | 0 | 0 |

| | | | | | | | | | | | | | | | |
|---|---|---|---|---|---|---|---|---|---|---|---|---|---|---|---|
| 3245 | ENSMUSG00000021701 | 132.113455 | 0.228442185 | 0.310711725 | 0.735222284 | 0.46220413 | 0.860650866 | Plk2 | polo like kinase 2 [Source:MGI Symbol;Acc | 150.18 | 60.647 | 153.96 | 172.55 | 143.96 | 111.38 |
| 3246 | ENSMUSG00000021702 | 0 | NA | NA | NA | NA | NA | Thbs4 | thrombospondin 4 [Source:MGI Symbol;A | 0 | 0 | 0 | 0 | 0 | 0 |
| 3247 | ENSMUSG00000021703 | 689.0177152 | 0.077537941 | 0.132874439 | 0.58354294 | 0.559527859 | 0.899423311 | Serinc5 | serine incorporator 5 [Source:MGI Symbo | 689.06 | 596.19 | 725.5 | 642.97 | 780.88 | 699.49 |
| 3248 | ENSMUSG00000021704 | 389.3856672 | 0.296396243 | 0.139960168 | 2.117718539 | 0.034198912 | 0.247603883 | Mtx3 | metaxin 3 [Source:MGI Symbol;Acc:MGI:2 | 335.7 | 324.82 | 388.41 | 437.65 | 424.25 | 425.49 |
| 3249 | ENSMUSG00000021706 | 118.8216235 | -0.094855574 | 0.220581125 | -0.4300258 | 0.667176874 | 0.929709374 | Zfyve16 | zinc finger, FYVE domain containing 16 [Sc | 111.63 | 116.15 | 140.87 | 128.21 | 105.79 | 110.27 |
| 3250 | ENSMUSG00000021707 | 1021.696787 | 0.246387156 | 0.135097918 | 1.823767241 | 0.068187268 | 0.376148227 | Dhfr | dihydrofolate reductase [Source:MGI Sym | 943.65 | 851.12 | 1008.3 | 930.24 | 1153.9 | 1243 |
| 3251 | ENSMUSG00000021708 | 5.166263228 | -0.267065239 | 1.022589068 | -0.261165748 | 0.793964691 | 0.961915077 | Rasgrf2 | RAS protein-specific guanine nucleotide-r | 5.6217 | 8.2233 | 3.0187 | 1.928 | 2.1812 | 10.025 |
| 3252 | ENSMUSG00000021709 | 1656.69399 | 0.006282322 | 0.102330016 | 0.061392755 | 0.951046422 | 0.990883058 | Erbin | Erbb2 interacting protein [Source:MGI Syr | 1635.1 | 1762.9 | 1561.7 | 1680.2 | 1728.6 | 1571.6 |
| 3253 | ENSMUSG00000021710 | 738.1529632 | -0.04520431 | 0.132438831 | -0.341322177 | 0.732861054 | 0.947867718 | Nln | neurolysin (metallopeptidase M3 family) [ | 750.9 | 739.07 | 758.71 | 654.54 | 849.59 | 676.1 |
| 3254 | ENSMUSG00000021711 | 554.0580676 | 0.042580203 | 0.123274985 | 0.345408301 | 0.729787438 | 0.946534338 | Trappc13 | trafficking protein particle complex 13 [So | 563.78 | 556.1 | 517.21 | 535.97 | 595.48 | 555.81 |
| 3255 | ENSMUSG00000021712 | 306.3480668 | -0.003531492 | 0.171799656 | -0.020555873 | 0.983599941 | 0.998580735 | Trim23 | tripartite motif-containing 23 [Source:MG | 249.76 | 326.88 | 344.14 | 273.77 | 326.09 | 317.45 |
| 3256 | ENSMUSG00000021713 | 127.1897964 | -0.36089834 | 0.216130575 | -1.669816218 | 0.094955731 | 0.454630813 | Ppwd1 | peptidylprolyl isomerase domain and WD | 130.1 | 158.3 | 140.87 | 106.04 | 128.69 | 99.132 |
| 3257 | ENSMUSG00000021714 | 15.90163374 | -0.588544341 | 0.608133236 | -0.967788481 | 0.333150018 | 0.77523112 | Cenpk | centromere protein K [Source:MGI Symbc | 26.502 | 19.53 | 11.069 | 8.6758 | 20.722 | 8.9108 |
| 3258 | ENSMUSG00000021715 | 99.24000214 | 0.081521612 | 0.24697859 | 0.330075622 | 0.741342823 | 0.948792385 | Cwc27 | CWC27 spliceosome-associated protein [S | 92.357 | 86.345 | 110.69 | 105.07 | 81.796 | 119.18 |
| 3259 | ENSMUSG00000021716 | 286.3315373 | -0.20296821 | 0.179578509 | -1.130247775 | 0.258371834 | 0.711962318 | Srek1ip1 | splicing regulatory glutamine/lysine-rich p | 330.88 | 351.55 | 236.47 | 253.53 | 271.56 | 274.01 |
| 3260 | ENSMUSG00000021718 | 0 | NA | NA | NA | NA | NA | Nt5el | 5' nucleotidase, ecto-like [Source:MGI Syr | 0 | 0 | 0 | 0 | 0 | 0 |
| 3261 | ENSMUSG00000021719 | 10.69853279 | -0.099994036 | 0.716295959 | -0.139598772 | 0.888977011 | 0.982382091 | Rgs7bp | regulator of G-protein signalling 7 binding | 16.865 | 7.1954 | 9.0562 | 14.46 | 4.3625 | 12.252 |
| 3262 | ENSMUSG00000021720 | 3.998794141 | -1.775669933 | 1.212591491 | -1.464359553 | 0.143095733 | NA | Rnf180 | ring finger protein 180 [Source:MGI Symb | 7.2279 | 8.2233 | 3.0187 | 0 | 2.1812 | 3.3415 |
| 3263 | ENSMUSG00000021721 | 0 | NA | NA | NA | NA | NA | Htr1a | 5-hydroxytryptamine (serotonin) receptor | 0 | 0 | 0 | 0 | 0 | 0 |
| 3264 | ENSMUSG00000021725 | 13.91133267 | -0.279251307 | 0.593931586 | -0.470174198 | 0.638230567 | 0.923699483 | Parp8 | poly (ADP-ribose) polymerase family, men | 19.274 | 10.279 | 16.1 | 13.496 | 8.7249 | 15.594 |
| 3265 | ENSMUSG00000021728 | 3.23710921 | -1.39370912 | 1.260365533 | -1.105797551 | 0.268814155 | NA | Emb | embigin [Source:MGI Symbol;Acc:MGI:95 | 4.8186 | 7.1954 | 2.0125 | 0.964 | 1.0906 | 3.3415 |
| 3266 | ENSMUSG00000021730 | 0 | NA | NA | NA | NA | NA | Hcn1 | hyperpolarization activated cyclic nucleoti | 0 | 0 | 0 | 0 | 0 | 0 |
| 3267 | ENSMUSG00000021731 | 291.1533625 | 0.202778906 | 0.146631537 | 1.382914687 | 0.166691027 | 0.592715132 | Mrps30 | mitochondrial ribosomal protein S30 [Sou | 273.05 | 268.29 | 270.68 | 307.51 | 299.92 | 327.47 |
| 3268 | ENSMUSG00000021732 | 0 | NA | NA | NA | NA | NA | Fgf10 | fibroblast growth factor 10 [Source:MGI S | 0 | 0 | 0 | 0 | 0 | 0 |
| 3269 | ENSMUSG00000021733 | 215.128817 | -0.040477655 | 0.179640111 | -0.225326378 | 0.821725381 | 0.969160361 | Slc4a7 | solute carrier family 4, sodium bicarbonat | 201.58 | 224.09 | 229.42 | 228.46 | 230.12 | 177.1 |
| 3270 | ENSMUSG00000021737 | 1464.510553 | 0.029635982 | 0.104160228 | 0.284523015 | 0.776009599 | 0.95893551 | Psmd6 | proteasome (prosome, macropain) 26S su | 1543.6 | 1346.6 | 1457 | 1428.6 | 1543.2 | 1468 |
| 3271 | ENSMUSG00000021738 | 445.9049777 | 0.144306777 | 0.156139089 | 0.924219411 | 0.355372091 | 0.795424787 | Atxn7 | ataxin 7 [Source:MGI Symbol;Acc:MGI:21 | 420.83 | 453.31 | 397.47 | 540.79 | 479.87 | 383.16 |
| 3272 | ENSMUSG00000021741 | 0 | NA | NA | NA | NA | NA | Gm5457 | predicted pseudogene 5457 [Source:MGI | 0 | 0 | 0 | 0 | 0 | 0 |
| 3273 | ENSMUSG00000021743 | 0 | NA | NA | NA | NA | NA | Fezf2 | Fez family zinc finger 2 [Source:MGI Symb | 0 | 0 | 0 | 0 | 0 | 0 |
| 3274 | ENSMUSG00000021745 | 540.0718661 | 0.186068239 | 0.125225559 | 1.485864715 | 0.137314908 | 0.541266927 | NA | NA | 513.99 | 498.54 | 503.12 | 562.96 | 623.83 | 537.99 |
| 3275 | ENSMUSG00000021747 | 16.02988516 | 0.161499788 | 0.679370578 | 0.237719726 | 0.812098483 | 0.96653525 | Cfap20dc | CFAP20 domain containing [Source:MGI S | 16.865 | 8.2233 | 20.125 | 7.7118 | 8.7249 | 34.529 |
| 3276 | ENSMUSG00000021748 | 3236.793296 | -0.036614492 | 0.099282712 | -0.368790207 | 0.712284105 | 0.944239215 | Pdhb | pyruvate dehydrogenase (lipoamide) beta | 3471 | 3220.5 | 3140.5 | 2967.1 | 3196.6 | 3425.1 |
| 3277 | ENSMUSG00000021749 | 0.556921942 | 2.679043159 | 3.638243063 | 0.736356289 | 0.461513899 | NA | Oit1 | oncoprotein induced transcript 1 [Source: | 0 | 0 | 0 | 0 | 0 | 3.3415 |
| 3278 | ENSMUSG00000021750 | 10.93891417 | -4.324707322 | 1.217612148 | -3.551793836 | 0.000382615 | 0.010379287 | Fam107a | family with sequence similarity 107, meml | 3.2124 | 49.34 | 10.062 | 1.928 | 1.0906 | 0 |
| 3279 | ENSMUSG00000021751 | 4643.771217 | -0.261385323 | 0.120389375 | -2.171166055 | 0.029918622 | 0.228601666 | Acox2 | acyl-Coenzyme A oxidase 2, branched cha | 4380.9 | 5173.5 | 5636 | 3853 | 4142.2 | 4677 |
| 3280 | ENSMUSG00000021752 | 146.0769608 | -0.109893394 | 0.200391512 | -0.548393456 | 0.583421769 | 0.908645447 | Kctd6 | potassium channel tetramerisation domai | 151.79 | 149.05 | 153.96 | 131.1 | 163.59 | 126.98 |
| 3281 | ENSMUSG00000021754 | 174.6086172 | 0.055135958 | 0.195394929 | 0.282177016 | 0.777807781 | 0.959559604 | Map3k1 | mitogen-activated protein kinase kinase k | 153.39 | 190.16 | 171.06 | 200.51 | 181.04 | 151.48 |
| 3282 | ENSMUSG00000021756 | 2449.436639 | 0.045366493 | 0.088321818 | 0.513649895 | 0.607496789 | 0.916376001 | Il6st | interleukin 6 signal transducer [Source:M( | 2414.1 | 2364.2 | 2454.2 | 2463 | 2461.5 | 2539.6 |
| 3283 | ENSMUSG00000021758 | 0.167707431 | -0.796629176 | 4.080472857 | -0.195229623 | 0.84521319 | NA | Ddx4 | DEAD box helicase 4 [Source:MGI Symbol; | 0 | 0 | 1.0062 | 0 | 0 | 0 |
| 3284 | ENSMUSG00000021759 | 61.41419574 | -0.663934586 | 0.310312014 | -2.139570998 | 0.032389453 | 0.24029826 | Plpp1 | phospholipid phosphatase 1 [Source:MGI | 63.445 | 86.345 | 76.475 | 54.947 | 33.809 | 53.465 |
| 3285 | ENSMUSG00000021760 | 13.28735622 | -0.642696534 | 0.649660606 | -0.989280446 | 0.322525948 | 0.769926329 | Gpx8 | glutathione peroxidase 8 (putative) [Sour | 11.243 | 15.419 | 22.137 | 15.424 | 4.3625 | 11.138 |
| 3286 | ENSMUSG00000021763 | 1.957387837 | 0.69362432 | 1.470808191 | 0.471594001 | 0.637216609 | NA | Cspg4b | chondroitin sulfate proteoglycan 4B [Sour | 2.4093 | 1.0279 | 1.0062 | 2.8919 | 2.1812 | 2.2277 |
| 3287 | ENSMUSG00000021764 | 1317.862622 | 0.094527212 | 0.10910126 | 0.866417232 | 0.386261397 | 0.817115886 | Ndufs4 | NADH:ubiquinone oxidoreductase core su | 1284.2 | 1253 | 1286 | 1219.4 | 1423.3 | 1441.3 |
| 3288 | ENSMUSG00000021765 | 28.66902165 | 0.111427298 | 0.408952798 | 0.272469826 | 0.785260789 | 0.960670212 | Fst | follistatin [Source:MGI Symbol;Acc:MGI:9 | 24.896 | 25.698 | 32.2 | 31.811 | 23.994 | 33.415 |
| 3289 | ENSMUSG00000021767 | 205.0086363 | 0.212433342 | 0.177028844 | 1.199992825 | 0.230142127 | 0.676880752 | Kat6b | K(lysine) acetyltransferase 6B [Source:MG | 187.12 | 178.86 | 204.27 | 248.71 | 205.04 | 206.06 |
| 3290 | ENSMUSG00000021768 | 0.486939728 | -0.698472448 | 3.137608162 | -0.222613026 | 0.823836697 | NA | Dusp13 | dual specificity phosphatase 13 [Source:M | 0.8031 | 1.0279 | 0 | 0 | 1.0906 | 0 |
| 3291 | ENSMUSG00000021770 | 568.8840311 | 0.170472221 | 0.138781571 | 1.228349121 | 0.219315939 | 0.664947263 | Samd8 | sterile alpha motif domain containing 8 [S | 494.71 | 589 | 523.25 | 653.58 | 525.68 | 627.09 |
| 3292 | ENSMUSG00000021771 | 1618.122504 | 0.013338087 | 0.117808174 | 0.113218687 | 0.909857181 | 0.985485567 | Vdac2 | voltage-dependent anion channel 2 [Sour | 1623.1 | 1721.8 | 1487.2 | 1628.2 | 1805 | 1443.5 |
| 3293 | ENSMUSG00000021772 | 320.2886648 | 0.116433216 | 0.147187645 | 0.791052922 | 0.42891311 | 0.842753542 | Nkiras1 | NFKB inhibitor interacting Ras-like protein | 286.71 | 316.6 | 318.98 | 310.4 | 333.73 | 355.32 |
| 3294 | ENSMUSG00000021773 | 22.9456591 | 0.312319249 | 0.493475791 | 0.6328968 | 0.526801038 | 0.888334265 | Comtd1 | catechol-O-methyltransferase domain cor | 30.518 | 19.53 | 11.069 | 26.991 | 26.175 | 23.391 |
| 3295 | ENSMUSG00000021774 | 279.4940702 | 0.164099218 | 0.190736531 | 0.860344986 | 0.389598902 | 0.819052248 | Ube2e1 | ubiquitin-conjugating enzyme E2E 1 [Sour | 208 | 246.7 | 337.09 | 305.58 | 295.56 | 284.03 |
| 3296 | ENSMUSG00000021775 | 1650.286793 | 0.729683183 | 0.164310552 | 4.440878404 | 8.96E-06 | 0.000481734 | Nr1d2 | nuclear receptor subfamily 1, group D, me | 1585.3 | 1202.7 | 934.8 | 1876.9 | 2073.3 | 2228.8 |
| 3297 | ENSMUSG00000021779 | 853.2308194 | -0.108615789 | 0.109270551 | -0.99400788 | 0.320219031 | 0.767337321 | Thrb | thyroid hormone receptor beta [Source:M | 841.65 | 898.4 | 916.69 | 830.95 | 840.87 | 790.83 |
| 3298 | ENSMUSG00000021782 | 86.42066001 | 0.082006804 | 0.260736677 | 0.314519632 | 0.753126408 | 0.952989051 | Dlg5 | discs large MAGUK scaffold protein 5 [Sou | 84.326 | 82.233 | 85.531 | 99.29 | 101.43 | 65.717 |
| 3299 | ENSMUSG00000021785 | 808.8617301 | 0.053224709 | 0.108077182 | 0.492469436 | 0.622387523 | 0.918780684 | Ngly1 | N-glycanase 1 [Source:MGI Symbol;Acc:M | 819.97 | 796.64 | 764.75 | 817.45 | 822.33 | 832.04 |
| 3300 | ENSMUSG00000021786 | 983.3668332 | 0.145109931 | 0.121561014 | 1.193720961 | 0.232587138 | 0.679847566 | Oxsm | 3-oxoacyl-ACP synthase, mitochondrial [S | 965.33 | 972.41 | 863.36 | 942.77 | 1153.9 | 1002.5 |

| | | | | | | | | | | | | | | |
|---|---|---|---|---|---|---|---|---|---|---|---|---|---|---|
| 3301 | ENSMUSG00000021789 | 23.32208343 | -0.363646864 | 0.472245071 | -0.770038454 | 0.441277083 | 0.849828766 | Sftpa1 | surfactant associated protein A1 [Source:I | 28.912 | 18.503 | 31.194 | 17.352 | 27.265 | 16.708 |
| 3302 | ENSMUSG00000021790 | 0 | NA | NA | NA | NA | NA | Dydc1 | DPY30 domain containing 1 [Source:MGI S | 0 | 0 | 0 | 0 | 0 | 0 |
| 3303 | ENSMUSG00000021791 | 0 | NA | NA | NA | NA | NA | Dydc2 | DPY30 domain containing 2 [Source:MGI S | 0 | 0 | 0 | 0 | 0 | 0 |
| 3304 | ENSMUSG00000021792 | 1327.995398 | 0.073383744 | 0.149999008 | 0.489228194 | 0.624680151 | 0.919503423 | Prxl2a | peroxiredoxin like 2A [Source:MGI Symbo | 1537.1 | 1302.4 | 1041.5 | 1189.5 | 1500.7 | 1396.8 |
| 3305 | ENSMUSG00000021794 | 21182.24172 | 0.070325124 | 0.08645177 | 0.81346078 | 0.415953924 | 0.836590861 | Glud1 | glutamate dehydrogenase 1 [Source:MGI | 20001 | 21687 | 20310 | 20685 | 21757 | 22653 |
| 3306 | ENSMUSG00000021795 | 21.37438484 | -0.299371606 | 0.469336019 | -0.637861987 | 0.523563524 | 0.888334265 | Sftpd | surfactant associated protein D [Source:M | 20.078 | 26.726 | 24.15 | 23.135 | 17.45 | 16.708 |
| 3307 | ENSMUSG00000021796 | 571.8795871 | 0.125253491 | 0.12305601 | 1.017857565 | 0.30874565 | 0.75728287 | Bmpr1a | bone morphogenetic protein receptor, typ | 511.58 | 544.8 | 585.63 | 609.23 | 605.29 | 574.74 |
| 3308 | ENSMUSG00000021797 | 0 | NA | NA | NA | NA | NA | 9230112D13 | RIKEN cDNA 9230112D13 gene [Source:M | 0 | 0 | 0 | 0 | 0 | 0 |
| 3309 | ENSMUSG00000021798 | 4.076647235 | -0.967898839 | 1.268102133 | -0.763265681 | 0.445304965 | NA | Ldb3 | LIM domain binding 3 [Source:MGI Symbo | 12.047 | 2.0558 | 2.0125 | 2.8919 | 5.4531 | 0 |
| 3310 | ENSMUSG00000021799 | 0 | NA | NA | NA | NA | NA | Opn4 | opsin 4 (melanopsin) [Source:MGI Symbol | 0 | 0 | 0 | 0 | 0 | 0 |
| 3311 | ENSMUSG00000021803 | 0.167707431 | -0.796629176 | 4.080472857 | -0.195229623 | 0.84521319 | NA | Cdhr1 | cadherin-related family member 1 [Source | 0 | 0 | 1.0062 | 0 | 0 | 0 |
| 3312 | ENSMUSG00000021804 | 21.41818584 | 0.204598885 | 0.530234383 | 0.38586499 | 0.69959666 | 0.941362191 | Rgr | retinal G protein coupled receptor [Source | 19.274 | 18.503 | 22.137 | 35.667 | 22.903 | 10.025 |
| 3313 | ENSMUSG00000021806 | 54.93677303 | 0.142020152 | 0.312795573 | 0.454035044 | 0.649803604 | 0.925643431 | Nid2 | nidogen 2 [Source:MGI Symbol;Acc:MGI:1 | 44.974 | 59.619 | 52.325 | 57.839 | 45.806 | 69.058 |
| 3314 | ENSMUSG00000021807 | 1093.688718 | 0.014317127 | 0.104022802 | 0.137634506 | 0.890529283 | 0.982489666 | Rtraf | RNA transcription, translation and transpo | 1127.6 | 1051.6 | 1084.7 | 1046.9 | 1135.3 | 1116.1 |
| 3315 | ENSMUSG00000021809 | 208.6767877 | 0.372969068 | 0.168305141 | 2.216028967 | 0.026689518 | 0.211364326 | Nudt13 | nudix (nucleoside diphosphate linked moi | 184.71 | 166.52 | 194.21 | 237.14 | 235.57 | 233.91 |
| 3316 | ENSMUSG00000021810 | 519.9937335 | 0.013009484 | 0.124818815 | 0.104226947 | 0.91698925 | 0.986374671 | Ecd | ecdysoneless cell cycle regulator [Source:I | 508.36 | 516.01 | 528.28 | 483.92 | 540.95 | 542.44 |
| 3317 | ENSMUSG00000021811 | 130.4780804 | -0.061071433 | 0.222585035 | -0.274373491 | 0.783797613 | 0.960670212 | Dnajc9 | DnaJ heat shock protein family (Hsp40) m | 135.72 | 136.71 | 126.79 | 101.22 | 127.6 | 154.82 |
| 3318 | ENSMUSG00000021814 | 2090.818604 | -0.016279021 | 0.124221617 | -0.131048213 | 0.895737168 | 0.982798755 | Anxa7 | annexin A7 [Source:MGI Symbol;Acc:MGI | 2427.8 | 2041.4 | 1837.4 | 2129.4 | 2202 | 1906.9 |
| 3319 | ENSMUSG00000021815 | 7.017927214 | 1.112538747 | 0.827928046 | 1.343762604 | 0.179025166 | 0.611235187 | Mss51 | MSS51 mitochondrial translational activat | 3.2124 | 4.1117 | 6.0375 | 6.7478 | 13.087 | 8.9108 |
| 3320 | ENSMUSG00000021816 | 578.356004 | -0.085974396 | 0.119931537 | -0.716862289 | 0.473459073 | 0.868952133 | Ppp3cb | protein phosphatase 3, catalytic subunit, I | 559.76 | 621.89 | 605.76 | 562 | 566.03 | 554.69 |
| 3321 | ENSMUSG00000021819 | 643.4529011 | 0.058176423 | 0.164734113 | 0.353153469 | 0.72397339 | 0.944786345 | Zswim8 | zinc finger SWIM-type containing 8 [Sourc | 628.03 | 645.53 | 618.84 | 836.73 | 526.77 | 604.82 |
| 3322 | ENSMUSG00000021820 | 204.2440705 | 0.436504643 | 0.169725085 | 2.571833401 | 0.010116156 | 0.109521017 | Camk2g | calcium/calmodulin-dependent protein ki | 167.85 | 189.14 | 164.02 | 237.14 | 232.3 | 235.02 |
| 3323 | ENSMUSG00000021822 | 20.30514982 | -0.525763404 | 0.497403514 | -1.057015863 | 0.290504348 | 0.740463951 | Plau | plasminogen activator, urokinase [Source: | 28.109 | 25.698 | 18.112 | 21.208 | 11.997 | 16.708 |
| 3324 | ENSMUSG00000021823 | 312.4606449 | -0.081689836 | 0.162608834 | -0.502370225 | 0.615407119 | 0.917509899 | Vcl | vinculin [Source:MGI Symbol;Acc:MGI:989 | 270.65 | 329.96 | 364.26 | 308.47 | 316.28 | 285.14 |
| 3325 | ENSMUSG00000021824 | 2225.369415 | 0.07564161 | 0.115212655 | 0.656539075 | 0.511477334 | 0.883766356 | Ap3m1 | adaptor-related protein complex 3, mu 1 s | 2437.4 | 2040.4 | 2021.5 | 2210.4 | 2147.4 | 2495 |
| 3326 | ENSMUSG00000021830 | 84.76780388 | 0.152752107 | 0.257180132 | 0.593949872 | 0.552545641 | 0.896052884 | Txndc16 | thioredoxin domain containing 16 [Source | 82.72 | 90.457 | 67.418 | 76.154 | 91.612 | 100.25 |
| 3327 | ENSMUSG00000021831 | 276.0920211 | 0.097525308 | 0.162988837 | 0.598355753 | 0.549602583 | 0.895703112 | Ero1a | endoplasmic reticulum oxidoreductase 1 a | 245.75 | 261.09 | 293.82 | 275.7 | 319.55 | 260.64 |
| 3328 | ENSMUSG00000021832 | 1551.533961 | 0.095768792 | 0.100733534 | 0.950714105 | 0.341749526 | 0.783459222 | Psmc6 | proteasome (prosome, macropain) 26S su | 1600.6 | 1455.5 | 1443 | 1580.9 | 1665.4 | 1563.8 |
| 3329 | ENSMUSG00000021835 | 41.23104197 | -0.273453909 | 0.358498337 | -0.762775947 | 0.445597028 | 0.852288235 | Bmp4 | bone morphogenetic protein 4 [Source:M | 44.974 | 45.228 | 45.281 | 43.379 | 25.084 | 43.44 |
| 3330 | ENSMUSG00000021838 | 43.04200251 | -0.521647439 | 0.351814818 | -1.482732995 | 0.138145365 | 0.542971347 | Samd4 | sterile alpha motif domain containing 4 [S | 55.414 | 54.48 | 42.262 | 37.595 | 26.175 | 42.326 |
| 3331 | ENSMUSG00000021840 | 865.4664563 | 0.044536498 | 0.133255554 | 0.334218701 | 0.738214533 | 0.948137969 | Mapk1ip1l | mitogen-activated protein kinase 1 intera | 884.22 | 837.75 | 833.17 | 731.66 | 983.74 | 922.26 |
| 3332 | ENSMUSG00000021843 | 418.650918 | -0.113707895 | 0.150611418 | -0.754975266 | 0.450263823 | 0.855572246 | Ktn1 | kinectin 1 [Source:MGI Symbol;Acc:MGI:1 | 480.26 | 461.54 | 363.25 | 423.19 | 388.26 | 395.41 |
| 3333 | ENSMUSG00000021846 | 553.8387906 | -0.294565638 | 0.136236872 | -2.162157956 | 0.030606003 | 0.231893781 | Peli2 | pellino 2 [Source:MGI Symbol;Acc:MGI:18 | 550.13 | 582.83 | 698.33 | 501.27 | 510.41 | 480.07 |
| 3334 | ENSMUSG00000021848 | 0 | NA | NA | NA | NA | NA | Otx2 | orthodenticle homeobox 2 [Source:MGI S | 0 | 0 | 0 | 0 | 0 | 0 |
| 3335 | ENSMUSG00000021850 | 5.932878734 | -1.236499346 | 0.937540164 | -1.318876134 | 0.187210526 | 0.623632669 | ccdc198 | coiled-coil domain containing 198 [Source | 13.653 | 6.1675 | 5.0312 | 1.928 | 4.3625 | 4.4554 |
| 3336 | ENSMUSG00000021852 | 0 | NA | NA | NA | NA | NA | Slc35f4 | solute carrier family 35, member F4 [Sour | 0 | 0 | 0 | 0 | 0 | 0 |
| 3337 | ENSMUSG00000021866 | 328.7000859 | -0.206094002 | 0.179892187 | -1.145652877 | 0.251938809 | 0.704395904 | Anxa11 | annexin A11 [Source:MGI Symbol;Acc:MG | 301.97 | 321.74 | 433.69 | 343.18 | 303.19 | 268.44 |
| 3338 | ENSMUSG00000021867 | 0 | NA | NA | NA | NA | NA | NA | NA | 0 | 0 | 0 | 0 | 0 | 0 |
| 3339 | ENSMUSG00000021868 | 841.9104615 | -0.238161606 | 0.145752325 | -1.634015827 | 0.102255527 | 0.470577139 | Ppif | peptidylprolyl isomerase F (cyclophilin F) [ | 1050.5 | 870.65 | 811.03 | 665.14 | 883.4 | 770.78 |
| 3340 | ENSMUSG00000021870 | 1229.088223 | -0.196656748 | 0.11333958 | -1.735110968 | 0.082721152 | 0.42416743 | Slmap | sarcolemma associated protein [Source:M | 1293.8 | 1452.4 | 1192.4 | 1167.4 | 1169.1 | 1099.4 |
| 3341 | ENSMUSG00000021871 | 0.181769715 | 1.126924175 | 4.080472857 | 0.276174898 | 0.782413736 | NA | Gm49342 | predicted gene, 49342 [Source:MGI Symb | 0 | 0 | 0 | 0 | 1.0906 | 0 |
| 3342 | ENSMUSG00000021872 | 1.499296716 | 1.249733105 | 1.923011909 | 0.649883185 | 0.51576768 | NA | Rnase10 | ribonuclease, RNase A family, 10 (non-act | 1.6062 | 0 | 1.0062 | 1.928 | 0 | 4.4554 |
| 3343 | ENSMUSG00000021874 | 1.283468866 | 2.785082463 | 2.306011967 | 1.207748487 | 0.227144021 | NA | 4933413J09I | RIKEN cDNA 4933413J09 gene [Source:MC | 0.8031 | 0 | 0 | 5.7839 | 0 | 1.1138 |
| 3344 | ENSMUSG00000021876 | 16936.42886 | 0.575343043 | 0.092399602 | 6.226683151 | 4.76E-10 | 1.06E-07 | Rnase4 | ribonuclease, RNase A family 4 [Source:M | 13262 | 13096 | 14453 | 19106 | 20074 | 21629 |
| 3345 | ENSMUSG00000021877 | 2230.295385 | -0.0438192 | 0.096246971 | -0.455278744 | 0.648908717 | 0.925643431 | Arf4 | ADP-ribosylation factor 4 [Source:MGI Syr | 2431 | 2167.9 | 2192.6 | 2233.5 | 2172.5 | 2184.2 |
| 3346 | ENSMUSG00000021879 | 0.160662974 | 1.126924175 | 4.080472857 | 0.276174898 | 0.782413736 | NA | Dnah12 | dynein, axonemal, heavy chain 12 [Source | 0 | 0 | 0 | 0.964 | 0 | 0 |
| 3347 | ENSMUSG00000021880 | 6.997717663 | -0.558288988 | 0.932924618 | -0.59842883 | 0.549553834 | 0.895703112 | Rnase6 | ribonuclease, RNase A family, 6 [Source:M | 17.668 | 3.0838 | 4.025 | 2.8919 | 7.6343 | 6.6831 |
| 3348 | ENSMUSG00000021884 | 616.5509172 | -0.226296527 | 0.211894257 | -1.067969136 | 0.285534434 | 0.738065506 | Hacl1 | 2-hydroxyacyl-CoA lyase 1 [Source:MGI Sy | 542.09 | 603.39 | 850.28 | 765.4 | 509.32 | 428.83 |
| 3349 | ENSMUSG00000021886 | 10.59615989 | -2.284979048 | 0.894929817 | -2.553249435 | 0.010672304 | 0.113741744 | Gpr65 | G-protein coupled receptor 65 [Source:MG | 9.6372 | 35.977 | 7.0437 | 0.964 | 3.2719 | 6.6831 |
| 3350 | ENSMUSG00000021890 | 815.2467508 | -0.081740868 | 0.117953701 | -0.692991125 | 0.488315121 | 0.874431125 | Eaf1 | ELL associated factor 1 [Source:MGI Symb | 789.45 | 902.51 | 824.11 | 829.98 | 812.51 | 732.91 |
| 3351 | ENSMUSG00000021891 | 203.7303322 | 0.029237473 | 0.177941559 | 0.164309412 | 0.869487572 | 0.978621761 | Mettl6 | methyltransferase like 6 [Source:MGI Sym | 180.7 | 207.64 | 217.35 | 220.75 | 184.31 | 211.63 |
| 3352 | ENSMUSG00000021892 | 1133.30998 | -0.180405889 | 0.122980142 | -1.466951384 | 0.142389281 | 0.55198239 | Sh3bp5 | SH3-domain binding protein 5 (BTK-associ | 1063.3 | 1183.1 | 1366.5 | 1014.1 | 1059 | 1113.8 |
| 3353 | ENSMUSG00000021893 | 712.3101582 | -0.048564136 | 0.124070907 | -0.391422428 | 0.695485017 | 0.939992732 | Capn7 | calpain 7 [Source:MGI Symbol;Acc:MGI:13 | 659.35 | 744.21 | 769.78 | 650.69 | 750.35 | 699.49 |
| 3354 | ENSMUSG00000021895 | 609.4721552 | -0.41151879 | 0.172315058 | -2.388176604 | 0.016932203 | 0.1545157 | Arhgef3 | Rho guanine nucleotide exchange factor ( | 736.44 | 804.86 | 545.38 | 434.75 | 609.66 | 525.73 |
| 3355 | ENSMUSG00000021898 | 0.810359328 | 1.950931649 | 2.89643804 | 0.673562363 | 0.500589588 | NA | Asb14 | ankyrin repeat and SOCS box-containing 1 | 0 | 0 | 1.0062 | 3.8559 | 0 | 0 |
| 3356 | ENSMUSG00000021900 | 1134.446467 | -0.124620313 | 0.116538102 | -1.069352518 | 0.284910854 | 0.737533238 | Btd | biotinidase [Source:MGI Symbol;Acc:MGI: | 1182.2 | 1159.5 | 1209.5 | 1226.2 | 997.92 | 1031.4 |

| | | | | | | | | | | | | | | | |
|---|---|---|---|---|---|---|---|---|---|---|---|---|---|---|---|
| 3357 | ENSMUSG00000021901 | 735.923807 | 0.165149684 | 0.12494666 | 1.321761496 | 0.186247584 | 0.62230226 | Bap1 | Brca1 associated protein 1 [Source:MGI Sy | 767.77 | 684.59 | 627.9 | 751.9 | 755.8 | 827.59 |
| 3358 | ENSMUSG00000021902 | 200.9096196 | 0.106310687 | 0.187368415 | 0.567388517 | 0.570450249 | 0.904602706 | Phf7 | PHD finger protein 7 [Source:MGI Symbol | 198.37 | 208.67 | 173.07 | 185.08 | 195.22 | 245.05 |
| 3359 | ENSMUSG00000021903 | 41.21682157 | -0.96359044 | 0.436621224 | -2.206925331 | 0.027319276 | 0.214171281 | Galnt15 | polypeptide N-acetylgalactosaminyltransf | 81.113 | 37.005 | 45.281 | 38.559 | 15.269 | 30.074 |
| 3360 | ENSMUSG00000021904 | 0.481988923 | 2.492401104 | 3.837077151 | 0.649557203 | 0.515978285 | NA | Sema3g | sema domain, immunoglobulin domain (Ig | 0 | 0 | 0 | 2.8919 | 0 | 0 |
| 3361 | ENSMUSG00000021905 | 252.6640118 | -0.062768721 | 0.16226095 | -0.386838121 | 0.698876054 | 0.940854732 | Dph3 | diphthamine biosynthesis 3 [Source:MGI S | 245.75 | 262.12 | 266.65 | 236.17 | 227.94 | 277.35 |
| 3362 | ENSMUSG00000021906 | 410.622855 | -0.080923917 | 0.130246075 | -0.621315587 | 0.534392 | 0.889858671 | Oxnad1 | oxidoreductase NAD-binding domain cont | 419.22 | 421.45 | 425.64 | 390.41 | 390.44 | 416.58 |
| 3363 | ENSMUSG00000021907 | 0 | NA | NA | NA | NA | NA | Msmb | beta-microseminoprotein [Source:MGI Sy | 0 | 0 | 0 | 0 | 0 | 0 |
| 3364 | ENSMUSG00000021908 | 7.087687294 | -0.494084363 | 0.796157621 | -0.620586112 | 0.534871979 | 0.889858671 | Ncoa4-ps | nuclear receptor coactivator 4, pseudoger | 9.6372 | 7.1954 | 8.05 | 7.7118 | 4.3625 | 5.5692 |
| 3365 | ENSMUSG00000021910 | 2803.778285 | 0.03534726 | 0.107829421 | 0.327807193 | 0.743057449 | 0.949431323 | Nisch | nischarin [Source:MGI Symbol;Acc:MGI:1 | 2676.7 | 2667.4 | 2965.4 | 3146.4 | 2764.7 | 2601.9 |
| 3366 | ENSMUSG00000021911 | 301.3199685 | -0.259946988 | 0.179619634 | -1.447208098 | 0.147838649 | 0.559188866 | Parg | poly (ADP-ribose) glycohydrolase [Source: | 281.89 | 393.69 | 309.92 | 240.99 | 314.1 | 267.32 |
| 3367 | ENSMUSG00000021913 | 0.973672865 | 1.321976834 | 2.461749649 | 0.537007016 | 0.591262767 | NA | Ogdhl | oxoglutarate dehydrogenase-like [Source: | 1.6062 | 0 | 0 | 0.964 | 3.2719 | 0 |
| 3368 | ENSMUSG00000021916 | 132.8850543 | 0.166986383 | 0.205832631 | 0.811272645 | 0.417209116 | 0.836964306 | Glt8d1 | glycosyltransferase 8 domain containing 1 | 133.31 | 120.27 | 121.76 | 141.7 | 124.33 | 155.94 |
| 3369 | ENSMUSG00000021917 | 899.9444289 | 0.243230996 | 0.115111877 | 2.112996523 | 0.034601065 | 0.249237954 | Spcs1 | signal peptidase complex subunit 1 homo | 782.22 | 829.53 | 861.35 | 945.66 | 918.3 | 1062.6 |
| 3370 | ENSMUSG00000021918 | 239.6007755 | -0.266971432 | 0.17599119 | -1.516959066 | 0.129277019 | 0.526260863 | Nek4 | NIMA (never in mitosis gene a)-related ex | 231.29 | 263.15 | 290.8 | 195.69 | 209.4 | 247.27 |
| 3371 | ENSMUSG00000021919 | 0 | NA | NA | NA | NA | NA | Chat | choline acetyltransferase [Source:MGI Syr | 0 | 0 | 0 | 0 | 0 | 0 |
| 3372 | ENSMUSG00000021922 | 23943.46536 | -0.067206138 | 0.097761953 | -0.687446769 | 0.491801234 | 0.876368691 | Itih4 | inter alpha-trypsin inhibitor, heavy chain 4 | 23533 | 25078 | 24893 | 25991 | 22015 | 22150 |
| 3373 | ENSMUSG00000021927 | 0.524408994 | -0.837196744 | 3.377823722 | -0.247850928 | 0.804249749 | NA | Hnrnpa1l2-p | heterogeneous nuclear ribonucleoprotein | 0 | 2.0558 | 0 | 0 | 1.0906 | 0 |
| 3374 | ENSMUSG00000021928 | 1693.437475 | -0.054424801 | 0.130292512 | -0.417712426 | 0.676157386 | 0.932739835 | Ebpl | emopamil binding protein-like [Source:MG | 1914.6 | 1806.1 | 1454 | 1505.7 | 1646.8 | 1833.4 |
| 3375 | ENSMUSG00000021929 | 802.705453 | -0.052280092 | 0.115942897 | -0.450912415 | 0.652052675 | 0.925643431 | Kpna3 | karyopherin (importin) alpha 3 [Source:M | 775.8 | 857.28 | 819.08 | 761.54 | 850.68 | 751.84 |
| 3376 | ENSMUSG00000021930 | 457.5284353 | 0.08522208 | 0.129090423 | 0.660173526 | 0.509142479 | 0.882793133 | Spryd7 | SPRY domain containing 7 [Source:MGI Sy | 476.24 | 425.56 | 429.67 | 478.13 | 473.33 | 462.25 |
| 3377 | ENSMUSG00000021932 | 150.5172818 | -0.280086089 | 0.195904787 | -1.429705177 | 0.152801654 | 0.568739867 | Rnaseh2b | ribonuclease H2, subunit B [Source:MGI S | 170.26 | 178.86 | 145.91 | 128.21 | 142.87 | 137 |
| 3378 | ENSMUSG00000021933 | 0 | NA | NA | NA | NA | NA | Gucy1b2 | guanylate cyclase 1, soluble, beta 2 [Sour | 0 | 0 | 0 | 0 | 0 | 0 |
| 3379 | ENSMUSG00000021936 | 383.1794714 | 0.082640896 | 0.149007228 | 0.554609979 | 0.579161462 | 0.908337213 | Mapk8 | mitogen-activated protein kinase 8 [Sourc | 355.77 | 381.36 | 380.36 | 440.54 | 402.44 | 338.61 |
| 3380 | ENSMUSG00000021938 | 129.6470744 | -0.151794708 | 0.255493837 | -0.594122779 | 0.552429996 | 0.896052884 | Pspc1 | paraspeckle protein 1 [Source:MGI Symbo | 138.94 | 179.89 | 90.562 | 119.53 | 129.78 | 119.18 |
| 3381 | ENSMUSG00000021939 | 9373.984066 | -0.232667488 | 0.088656387 | -2.624373678 | 0.008680849 | 0.097864221 | Ctsb | cathepsin B [Source:MGI Symbol;Acc:MGI | 10395 | 10419 | 9571.4 | 9040.2 | 8275.6 | 8543.2 |
| 3382 | ENSMUSG00000021940 | 0.17131964 | -0.796629176 | 4.080472857 | -0.195229623 | 0.84521319 | NA | Ptpn20 | protein tyrosine phosphatase, non-recept | 0 | 1.0279 | 0 | 0 | 0 | 0 |
| 3383 | ENSMUSG00000021943 | 85.61535283 | -0.994741316 | 0.292611656 | -3.399527311 | 0.000675024 | 0.015516203 | Gdf10 | growth differentiation factor 10 [Source:N | 146.97 | 102.79 | 91.568 | 44.343 | 55.622 | 72.4 |
| 3384 | ENSMUSG00000021944 | 298.5138992 | 0.284418839 | 0.210860107 | 1.348850875 | 0.17738487 | 0.608561375 | Gata4 | GATA binding protein 4 [Source:MGI Symb | 248.96 | 232.31 | 327.03 | 422.22 | 247.57 | 312.99 |
| 3385 | ENSMUSG00000021945 | 448.1158943 | 0.223436643 | 0.126950051 | 1.760035859 | 0.078401727 | 0.410424385 | Zmym2 | zinc finger, MYM-type 2 [Source:MGI Sym | 405.57 | 427.61 | 407.53 | 483.92 | 495.14 | 468.93 |
| 3386 | ENSMUSG00000021947 | 306.2062045 | -0.157319881 | 0.165267998 | -0.951907708 | 0.34114379 | 0.783129922 | Cryl1 | crystallin, lambda 1 [Source:MGI Symbol; | 293.94 | 308.38 | 367.28 | 326.79 | 283.56 | 257.3 |
| 3387 | ENSMUSG00000021948 | 182.4238942 | 0.344101266 | 0.196586337 | 1.750382406 | 0.08005235 | 0.415341728 | Prkcd | protein kinase C, delta [Source:MGI Symb | 136.53 | 162.41 | 184.14 | 219.79 | 175.59 | 216.09 |
| 3388 | ENSMUSG00000021950 | 20.82870846 | -0.821010692 | 0.578662121 | -1.418808425 | 0.155954876 | 0.574855078 | Anxa8 | annexin A8 [Source:MGI Symbol;Acc:MGI | 30.518 | 34.949 | 14.087 | 5.7839 | 21.812 | 17.822 |
| 3389 | ENSMUSG00000021951 | 97.13661893 | -0.317206776 | 0.248702615 | -1.275446081 | 0.202151397 | 0.646741387 | Eef1akmt1 | EEF1A alpha lysine methyltransferase 1 [S | 116.45 | 98.68 | 107.67 | 68.442 | 104.7 | 86.88 |
| 3390 | ENSMUSG00000021952 | 208.7832832 | 0.111903541 | 0.173386902 | 0.645397892 | 0.518669369 | 0.887592015 | Xpo4 | exportin 4 [Source:MGI Symbol;Acc:MGI:1 | 176.68 | 208.67 | 217.35 | 215.93 | 224.67 | 209.4 |
| 3391 | ENSMUSG00000021953 | 0 | NA | NA | NA | NA | NA | Tdh | L-threonine dehydrogenase [Source:MGI S | 0 | 0 | 0 | 0 | 0 | 0 |
| 3392 | ENSMUSG00000021957 | 2468.427611 | -0.175759155 | 0.113003609 | -1.555341084 | 0.119864858 | 0.506640941 | Tkt | transketolase [Source:MGI Symbol;Acc:M | 2873.5 | 2657.2 | 2324.4 | 2376.2 | 2446.3 | 2133 |
| 3393 | ENSMUSG00000021958 | 42.72256578 | -0.071023041 | 0.36311334 | -0.19559469 | 0.844927417 | 0.972803041 | Pinx1 | PIN2/TERF1 interacting, telomerase inhibi | 49.792 | 32.893 | 48.3 | 31.811 | 53.44 | 40.098 |
| 3394 | ENSMUSG00000021959 | 179.8750968 | -0.083110222 | 0.189783185 | -0.437921945 | 0.661442867 | 0.928379255 | Lats2 | large tumor suppressor 2 [Source:MGI Syr | 175.88 | 212.78 | 167.04 | 190.87 | 173.41 | 159.28 |
| 3395 | ENSMUSG00000021961 | 0.836675597 | -0.373395339 | 2.394418521 | -0.155944057 | 0.876077117 | NA | 4930578I06R | RIKEN cDNA 4930578I06 gene [Source:MG | 0.8031 | 0 | 2.0125 | 0 | 1.0906 | 1.1138 |
| 3396 | ENSMUSG00000021962 | 144.3013909 | -0.173679425 | 0.200538158 | -0.866066724 | 0.386453572 | 0.817115886 | Dcp1a | decapping mRNA 1A [Source:MGI Symbol | 154.2 | 151.1 | 153.96 | 153.27 | 136.33 | 116.95 |
| 3397 | ENSMUSG00000021963 | 1379.704104 | -0.034716927 | 0.10491734 | -0.330897895 | 0.740721612 | 0.948792385 | Sap18 | Sin3-associated polypeptide 18 [Source:M | 1370.9 | 1386.7 | 1430.9 | 1250.3 | 1449.4 | 1390.1 |
| 3398 | ENSMUSG00000021965 | 2.274610759 | -0.268465301 | 1.446270839 | -0.185625883 | 0.852738155 | NA | Ska3 | spindle and kinetochore associated compl | 2.4093 | 2.0558 | 3.0187 | 2.8919 | 3.2719 | 0 |
| 3399 | ENSMUSG00000021966 | 0.513958919 | -2.394274821 | 3.756096013 | -0.637437066 | 0.523840191 | NA | Prss52 | protease, serine 52 [Source:MGI Symbol;A | 0 | 3.0838 | 0 | 0 | 0 | 0 |
| 3400 | ENSMUSG00000021967 | 535.6078326 | -0.019758092 | 0.139627185 | -0.141506058 | 0.887470175 | 0.982382091 | Mrpl57 | mitochondrial ribosomal protein L57 [Sou | 600.72 | 541.71 | 474.95 | 576.46 | 492.96 | 526.85 |
| 3401 | ENSMUSG00000021969 | 247.7517166 | 0.054792802 | 0.164152216 | 0.333792642 | 0.738536036 | 0.948137969 | Zdhhc20 | zinc finger, DHHC domain containing 20 [S | 229.69 | 240.53 | 259.61 | 278.59 | 249.75 | 228.34 |
| 3402 | ENSMUSG00000021972 | 361.5674056 | 0.037090006 | 0.140011322 | 0.264907189 | 0.79108096 | 0.961019028 | Hmbox1 | homeobox containing 1 [Source:MGI Sym | 377.46 | 344.35 | 348.16 | 342.21 | 375.17 | 382.05 |
| 3403 | ENSMUSG00000021973 | 783.5094887 | 0.260053314 | 0.157463849 | 1.651511222 | 0.098634231 | 0.463186297 | Micu2 | mitochondrial calcium uptake 2 [Source:N | 799.09 | 743.18 | 595.7 | 707.56 | 1000.1 | 855.43 |
| 3404 | ENSMUSG00000021974 | 11.10495133 | -1.119555312 | 0.716369176 | -1.562818934 | 0.118095189 | 0.503427297 | Fgf9 | fibroblast growth factor 9 [Source:MGI Sy | 24.093 | 12.335 | 9.0562 | 5.7839 | 10.906 | 4.4554 |
| 3405 | ENSMUSG00000021975 | 132.4951067 | 0.017777848 | 0.21217672 | 0.083787931 | 0.933225044 | 0.988420305 | Ints9 | integrator complex subunit 9 [Source:MG | 146.16 | 131.57 | 116.72 | 118.57 | 150.51 | 131.43 |
| 3406 | ENSMUSG00000021977 | 0 | NA | NA | NA | NA | NA | 1700129C05 | RIKEN cDNA 1700129C05 gene [Source:M | 0 | 0 | 0 | 0 | 0 | 0 |
| 3407 | ENSMUSG00000021978 | 549.0960726 | 0.099328344 | 0.129976315 | 0.764203418 | 0.44474603 | 0.851914525 | Extl3 | exostosin-like glycosyltransferase 3 [Sourc | 558.96 | 537.6 | 493.06 | 517.66 | 578.03 | 609.27 |
| 3408 | ENSMUSG00000021981 | 1306.074624 | -0.034949806 | 0.103356761 | -0.338147268 | 0.735252211 | 0.948109711 | Cab39l | calcium binding protein 39-like [Source:M | 1254.4 | 1368.2 | 1343.3 | 1220.4 | 1346.9 | 1303.2 |
| 3409 | ENSMUSG00000021982 | 1147.347605 | -0.015003337 | 0.106239507 | -0.141221823 | 0.887694706 | 0.982382091 | Cdadc1 | cytidine and dCMP deaminase domain co | 1079.4 | 1158.5 | 1222.6 | 1097 | 1158.2 | 1168.4 |
| 3410 | ENSMUSG00000021983 | 0.838717116 | -1.825569927 | 2.582619186 | -0.706867639 | 0.479648736 | NA | Atp8a2 | ATPase, aminophospholipid transporter-li | 0 | 2.0558 | 2.0125 | 0.964 | 0 | 0 |
| 3411 | ENSMUSG00000021986 | 11.9026303 | 0.510650338 | 0.670441977 | 0.761662239 | 0.446261618 | 0.852777619 | Amer2 | APC membrane recruitment 2 [Source:MG | 11.243 | 7.1954 | 11.069 | 23.135 | 7.6343 | 11.138 |
| 3412 | ENSMUSG00000021987 | 461.1218426 | 0.05257876 | 0.135741052 | 0.387346045 | 0.698500042 | 0.940696964 | Mtmr6 | myotubularin related protein 6 [Source:M | 415.2 | 479.01 | 464.88 | 494.52 | 429.7 | 483.41 |

| | | | | | | | | | | | | | | | |
|---|---|---|---|---|---|---|---|---|---|---|---|---|---|---|---|
| 3413 | ENSMUSG00000021990 | 468.6021431 | 0.261821894 | 0.16335038 | 1.602823902 | 0.108973538 | 0.485738292 | Spata13 | spermatogenesis associated 13 [Source:M | 436.89 | 398.83 | 442.75 | 536.94 | 398.08 | 598.13 |
| 3414 | ENSMUSG00000021991 | 0.688736311 | 2.998073904 | 2.697618692 | 1.11137794 | 0.266405704 | NA | Cacna2d3 | calcium channel, voltage-dependent, alph | 0 | 0 | 0 | 1.928 | 1.0906 | 1.1138 |
| 3415 | ENSMUSG00000021993 | 801.0014476 | -0.021880455 | 0.117141941 | -0.186785837 | 0.851828552 | 0.973633287 | Mipep | mitochondrial intermediate peptidase [So | 851.29 | 779.16 | 789.9 | 756.72 | 865.95 | 762.98 |
| 3416 | ENSMUSG00000021994 | 28.12071084 | -0.800083763 | 0.425629085 | -1.879767598 | 0.060139758 | 0.349482308 | Wnt5a | wingless-type MMTV integration site fami | 32.927 | 32.893 | 41.256 | 15.424 | 26.175 | 20.049 |
| 3417 | ENSMUSG00000021996 | 5323.923273 | -0.056163287 | 0.087096334 | -0.644840997 | 0.519030231 | 0.887592015 | Esd | esterase D/formylglutathione hydrolase [S | 5660.3 | 5405.8 | 5215.4 | 5057 | 5380 | 5225 |
| 3418 | ENSMUSG00000021997 | 0.853038123 | -0.516856556 | 2.386032976 | -0.216617524 | 0.828506428 | NA | Lrrc63 | leucine rich repeat containing 63 [Source: | 0 | 1.0279 | 2.0125 | 0.964 | 0 | 1.1138 |
| 3419 | ENSMUSG00000021998 | 1467.86065 | 0.128293174 | 0.094317397 | 1.360228102 | 0.173757752 | 0.604147282 | Lcp1 | lymphocyte cytosolic protein 1 [Source:M | 1411.1 | 1402.1 | 1394.7 | 1525 | 1561.8 | 1512.6 |
| 3420 | ENSMUSG00000021999 | 5358.4576 | 0.101109492 | 0.082871798 | 1.220071168 | 0.222437897 | 0.668407528 | Cpb2 | carboxypeptidase B2 (plasma) [Source:MG | 5021.8 | 5211.5 | 5279.8 | 5621.9 | 5501.1 | 5514.6 |
| 3421 | ENSMUSG00000022000 | 348.6636086 | -0.122158344 | 0.150497865 | -0.811694861 | 0.416966744 | 0.836878476 | Zc3h13 | zinc finger CCCH type containing 13 [Sourc | 373.44 | 373.13 | 344.14 | 380.77 | 309.74 | 310.76 |
| 3422 | ENSMUSG00000022002 | 0 | NA | NA | NA | NA | NA | Erich6b | glutamate rich 6B [Source:MGI Symbol;Ac | 0 | 0 | 0 | 0 | 0 | 0 |
| 3423 | ENSMUSG00000022003 | 948.628883 | 1.378240229 | 0.583485273 | 2.362082288 | 0.018172608 | 0.161653507 | Slc25a30 | solute carrier family 25, member 30 [Sourc | 588.67 | 402.94 | 589.66 | 2306.8 | 557.31 | 1246.4 |
| 3424 | ENSMUSG00000022008 | 378.9806557 | 0.076928936 | 0.149050309 | 0.516127321 | 0.605765488 | 0.916376001 | Gpalpp1 | GPALPP motifs containing 1 [Source:MGI S | 381.47 | 392.66 | 332.06 | 360.53 | 436.25 | 370.91 |
| 3425 | ENSMUSG00000022009 | 171.0125277 | -0.0561467 | 0.187007417 | -0.300237824 | 0.763995756 | 0.954980327 | Nufip1 | nuclear fragile X mental retardation prote | 186.32 | 168.58 | 168.04 | 178.34 | 177.77 | 147.03 |
| 3426 | ENSMUSG00000022010 | 2010.91191 | 0.226602514 | 0.184026968 | 1.23135493 | 0.218190146 | 0.663697152 | Tsc22d1 | TSC22 domain family, member 1 [Source:I | 2239.9 | 1660.1 | 1658.3 | 1534.7 | 2753.8 | 2218.8 |
| 3427 | ENSMUSG00000022012 | 0 | NA | NA | NA | NA | NA | Enox1 | ecto-NOX disulfide-thiol exchanger 1 [Sou | 0 | 0 | 0 | 0 | 0 | 0 |
| 3428 | ENSMUSG00000022013 | 510.8400746 | -0.126946551 | 0.136480408 | -0.930144865 | 0.352296084 | 0.792967718 | Dnajc15 | DnaJ heat shock protein family (Hsp40) m | 494.71 | 548.91 | 556.45 | 446.32 | 547.49 | 471.16 |
| 3429 | ENSMUSG00000022014 | 23.98558324 | -1.359816053 | 0.527344723 | -2.578609387 | 0.009919889 | 0.108043125 | Epsti1 | epithelial stromal interaction 1 (breast) [S | 28.912 | 55.508 | 19.119 | 10.604 | 14.178 | 15.594 |
| 3430 | ENSMUSG00000022015 | 0.17131964 | -0.796629176 | 4.080472857 | -0.195229623 | 0.84521319 | NA | Tnfsf11 | tumor necrosis factor (ligand) superfamily | 0 | 1.0279 | 0 | 0 | 0 | 0 |
| 3431 | ENSMUSG00000022016 | 735.5631108 | -0.003572865 | 0.113482924 | -0.031483721 | 0.974883775 | 0.996336532 | Akap11 | A kinase (PRKA) anchor protein 11 [Source | 721.19 | 782.25 | 706.38 | 740.33 | 749.25 | 713.97 |
| 3432 | ENSMUSG00000022018 | 0.845761573 | -3.112447952 | 2.7647732 | -1.125751636 | 0.260270661 | NA | Rgcc | regulator of cell cycle [Source:MGI Symbo | 0 | 2.0558 | 3.0187 | 0 | 0 | 0 |
| 3433 | ENSMUSG00000022019 | 231.6128895 | 0.05970213 | 0.19007155 | 0.314103451 | 0.753442468 | 0.952989051 | Tdrd3 | tudor domain containing 3 [Source:MGI S | 222.46 | 199.42 | 258.6 | 228.46 | 282.47 | 198.26 |
| 3434 | ENSMUSG00000022020 | 155.9385629 | -0.081636887 | 0.260571291 | -0.313299622 | 0.754053037 | 0.953048681 | Naa16 | N(alpha)-acetyltransferase 16, NatA auxili | 99.585 | 197.36 | 185.15 | 170.62 | 157.05 | 125.86 |
| 3435 | ENSMUSG00000022021 | 1.508201076 | -0.344836786 | 1.925803318 | -0.179061269 | 0.857889593 | NA | Diaph3 | diaphanous related formin 3 [Source:MGI | 4.0155 | 1.0279 | 0 | 2.8919 | 0 | 1.1138 |
| 3436 | ENSMUSG00000022022 | 151.0339481 | 0.109716278 | 0.189173948 | 0.579975619 | 0.561931059 | 0.900565925 | Mtrf1 | mitochondrial translational release factor | 138.94 | 151.1 | 145.91 | 148.45 | 161.41 | 160.39 |
| 3437 | ENSMUSG00000022023 | 322.343226 | 0.043975183 | 0.152314474 | 0.288713097 | 0.772800939 | 0.957899175 | Wbp4 | WW domain binding protein 4 [Source:MG | 301.97 | 350.52 | 299.86 | 304.62 | 319.55 | 357.54 |
| 3438 | ENSMUSG00000022024 | 497.7324615 | 0.217026121 | 0.129529132 | 1.675500472 | 0.093836079 | 0.450983125 | Sugt1 | SGT1, suppressor of G2 allele of SKP1 (S. c | 494.71 | 424.53 | 460.86 | 510.91 | 552.94 | 542.44 |
| 3439 | ENSMUSG00000022025 | 112.4746 | 0.227744771 | 0.248044801 | 0.918159825 | 0.358535198 | 0.798191951 | Cnmd | chondromodulin [Source:MGI Symbol;Acc | 85.129 | 94.568 | 131.82 | 139.78 | 119.97 | 103.59 |
| 3440 | ENSMUSG00000022026 | 0.847770947 | -0.376623742 | 2.576917887 | -0.146152791 | 0.883800772 | NA | Olfm4 | olfactomedin 4 [Source:MGI Symbol;Acc:I | 0.8031 | 2.0558 | 0 | 0 | 0 | 2.2277 |
| 3441 | ENSMUSG00000022031 | 479.7966952 | -0.103994443 | 0.129983225 | -0.800060494 | 0.423675749 | 0.84018348 | Elp3 | elongator acetyltransferase complex subu | 509.17 | 514.99 | 466.9 | 455.96 | 495.14 | 436.63 |
| 3442 | ENSMUSG00000022032 | 352.1298693 | 0.579496708 | 0.2382951 | 2.431844834 | 0.015022141 | 0.142452386 | Scara5 | scavenger receptor class A, member 5 [So | 284.3 | 248.76 | 313.95 | 401.98 | 600.93 | 262.87 |
| 3443 | ENSMUSG00000022033 | 3.663916717 | -0.008681378 | 1.451770898 | -0.005979854 | 0.995228795 | NA | Pbk | PDZ binding kinase [Source:MGI Symbol;A | 8.8341 | 1.0279 | 1.0062 | 0 | 1.0906 | 10.025 |
| 3444 | ENSMUSG00000022034 | 7.99552119 | 0.711103324 | 0.793118162 | 0.896591905 | 0.369936719 | 0.806613237 | Esco2 | establishment of sister chromatid cohesio | 8.8341 | 6.1675 | 3.0187 | 6.7478 | 8.7249 | 14.48 |
| 3445 | ENSMUSG00000022035 | 758.3042288 | 0.368757931 | 0.115387032 | 3.19583512 | 0.001394268 | 0.026817403 | Ccdc25 | coiled-coil domain containing 25 [Source:I | 675.41 | 641.42 | 668.15 | 794.32 | 859.41 | 911.12 |
| 3446 | ENSMUSG00000022037 | 26968.07492 | 0.110441826 | 0.07887061 | 1.400291263 | 0.161426116 | 0.584476975 | Clu | clusterin [Source:MGI Symbol;Acc:MGI:88 | 25674 | 25343 | 26792 | 27882 | 28162 | 27955 |
| 3447 | ENSMUSG00000022039 | 5.028511733 | 1.596454434 | 1.044038935 | 1.529113887 | 0.126236213 | NA | Adam2 | a disintegrin and metallopeptidase domai | 2.4093 | 1.0279 | 4.025 | 2.8919 | 10.906 | 8.9108 |
| 3448 | ENSMUSG00000022040 | 22335.07393 | -0.013036601 | 0.117883939 | -0.110588444 | 0.911942711 | 0.98589875 | Ephx2 | epoxide hydrolase 2, cytoplasmic [Source: | 20012 | 22410 | 24887 | 20981 | 25333 | 20388 |
| 3449 | ENSMUSG00000022041 | 224.4977722 | 0.125810117 | 0.187168107 | 0.672177109 | 0.501470952 | 0.880974509 | Chrna2 | cholinergic receptor, nicotinic, alpha poly | 195.15 | 231.28 | 218.36 | 246.78 | 268.29 | 187.13 |
| 3450 | ENSMUSG00000022043 | 52.64692209 | -0.432687687 | 0.308228104 | -1.40379051 | 0.160381241 | 0.582999313 | Trim35 | tripartite motif-containing 35 [Source:MG | 55.414 | 69.898 | 56.35 | 48.199 | 40.353 | 45.668 |
| 3451 | ENSMUSG00000022044 | 0 | NA | NA | NA | NA | NA | Stmn4 | stathmin-like 4 [Source:MGI Symbol;Acc:N | 0 | 0 | 0 | 0 | 0 | 0 |
| 3452 | ENSMUSG00000022048 | 41.35271334 | -0.928208614 | 0.353776348 | -2.623715856 | 0.008697631 | 0.097878626 | Dpysl2 | dihydropyrimidinase-like 2 [Source:MGI S | 45.777 | 59.619 | 57.356 | 27.955 | 23.994 | 33.415 |
| 3453 | ENSMUSG00000022051 | 1845.567976 | -0.068562692 | 0.09521225 | -0.720103686 | 0.471461158 | 0.867794597 | Bnip3l | BCL2/adenovirus E1B interacting protein 3 | 1801.4 | 1977.7 | 1889.7 | 1763.1 | 1814.8 | 1826.7 |
| 3454 | ENSMUSG00000022052 | 314.8398871 | -0.137157779 | 0.148673833 | -0.922541484 | 0.356246202 | 0.795787977 | Ppp2r2a | protein phosphatase 2, regulatory subunit | 322.85 | 354.63 | 311.94 | 293.05 | 283.56 | 323.01 |
| 3455 | ENSMUSG00000022053 | 0 | NA | NA | NA | NA | NA | Ebf2 | early B cell factor 2 [Source:MGI Symbol;A | 0 | 0 | 0 | 0 | 0 | 0 |
| 3456 | ENSMUSG00000022054 | 0 | NA | NA | NA | NA | NA | Nefm | neurofilament, medium polypeptide [Sour | 0 | 0 | 0 | 0 | 0 | 0 |
| 3457 | ENSMUSG00000022055 | 0 | NA | NA | NA | NA | NA | Nefl | neurofilament, light polypeptide [Source:I | 0 | 0 | 0 | 0 | 0 | 0 |
| 3458 | ENSMUSG00000022056 | 1.059810487 | -2.230732092 | 2.384896422 | -0.935358061 | 0.349603803 | NA | Adam7 | a disintegrin and metallopeptidase domai | 3.2124 | 2.0558 | 0 | 0 | 1.0906 | 0 |
| 3459 | ENSMUSG00000022057 | 14.32483049 | -1.526212196 | 0.63043754 | -2.420877723 | 0.015483084 | 0.145412036 | Adamdec1 | ADAM-like, decysin 1 [Source:MGI Symbo | 20.881 | 30.838 | 12.075 | 6.7478 | 8.7249 | 6.6831 |
| 3460 | ENSMUSG00000022061 | 0 | NA | NA | NA | NA | NA | Nkx3-1 | NK3 homeobox 1 [Source:MGI Symbol;Acc | 0 | 0 | 0 | 0 | 0 | 0 |
| 3461 | ENSMUSG00000022064 | 55.21475051 | 0.172178244 | 0.309141162 | 0.556956709 | 0.577557007 | 0.907877282 | Pibf1 | progesterone immunomodulatory binding | 61.839 | 48.312 | 45.281 | 55.911 | 68.709 | 51.237 |
| 3462 | ENSMUSG00000022066 | 0 | NA | NA | NA | NA | NA | Entpd4b | ectonucleoside triphosphate diphosphohy | 0 | 0 | 0 | 0 | 0 | 0 |
| 3463 | ENSMUSG00000022070 | 41.40374979 | 0.597144417 | 0.366973455 | 1.627214197 | 0.103691614 | 0.474047094 | Bora | bora, aurora kinase A activator [Source:M | 41.761 | 25.698 | 31.194 | 62.659 | 41.443 | 45.668 |
| 3464 | ENSMUSG00000022074 | 5.475807445 | -0.378051478 | 1.085527327 | -0.34826528 | 0.727640966 | 0.945946336 | Tnfrsf10b | tumor necrosis factor receptor superfamil | 2.4093 | 10.279 | 6.0375 | 8.6758 | 5.4531 | 0 |
| 3465 | ENSMUSG00000022075 | 183.45546 | 0.530451139 | 0.185698131 | 2.85652384 | 0.004283078 | 0.060263079 | Rhobtb2 | Rho-related BTB domain containing 2 [Sou | 167.85 | 139.8 | 141.88 | 203.4 | 207.22 | 240.59 |
| 3466 | ENSMUSG00000022076 | 0 | NA | NA | NA | NA | NA | Klhl1 | kelch-like 1 [Source:MGI Symbol;Acc:MGI | 0 | 0 | 0 | 0 | 0 | 0 |
| 3467 | ENSMUSG00000022085 | 0 | NA | NA | NA | NA | NA | Pebp4 | phosphatidylethanolamine binding protei | 0 | 0 | 0 | 0 | 0 | 0 |
| 3468 | ENSMUSG00000022089 | 194.6161419 | -0.018987321 | 0.174122852 | -0.109045541 | 0.91316637 | 0.98625062 | Bin3 | bridging integrator 3 [Source:MGI Symbol | 199.97 | 205.58 | 182.13 | 199.54 | 178.86 | 201.61 |

| | | | | | | | | | | | | | | | |
|---|---|---|---|---|---|---|---|---|---|---|---|---|---|---|---|
| 3469 | ENSMUSG00000022090 | 61.1026275 | -1.670521849 | 0.456121484 | -3.662449386 | 0.000249815 | 0.007496527 | Pdlim2 | PDZ and LIM domain 2 [Source:MGI Symb | 169.45 | 67.843 | 41.256 | 23.135 | 37.081 | 27.846 |
| 3470 | ENSMUSG00000022091 | 543.3483575 | 0.115578205 | 0.200957643 | 0.575137145 | 0.565198548 | 0.901542552 | Sorbs3 | sorbin and SH3 domain containing 3 [Sour | 424.04 | 521.15 | 619.85 | 483.92 | 452.61 | 758.53 |
| 3471 | ENSMUSG00000022092 | 50.34063031 | 0.487806423 | 0.387602833 | 1.258521304 | 0.208203287 | 0.653455953 | Ppp3cc | protein phosphatase 3, catalytic subunit, | 40.958 | 48.312 | 36.225 | 28.919 | 76.343 | 71.286 |
| 3472 | ENSMUSG00000022094 | 4303.939411 | 0.139932842 | 0.112085188 | 1.248450802 | 0.211866015 | 0.656735971 | Slc39a14 | solute carrier family 39 (zinc transporter), | 4061.3 | 4292.6 | 3933.4 | 5159.2 | 4030.9 | 4346.2 |
| 3473 | ENSMUSG00000022095 | 184.4193758 | -0.079902794 | 0.194377195 | -0.411070825 | 0.681020602 | 0.934384556 | Fam160b2 | family with sequence similarity 160, meml | 195.96 | 176.8 | 196.22 | 216.9 | 163.59 | 157.05 |
| 3474 | ENSMUSG00000022096 | 9.146756296 | -1.529418763 | 0.965732176 | -1.583688316 | 0.113264665 | 0.494196113 | Hr | lysine demethylase and nuclear receptor | 27.305 | 6.1675 | 7.0437 | 0 | 5.4531 | 8.9108 |
| 3475 | ENSMUSG00000022097 | 0.469265235 | -2.274672822 | 3.439794491 | -0.661281605 | 0.508431736 | NA | Sftpc | surfactant associated protein C [Source:M | 0.8031 | 0 | 2.0125 | 0 | 0 | 0 |
| 3476 | ENSMUSG00000022098 | 2588.418808 | -0.1934779 | 0.117044515 | -1.653028335 | 0.098325097 | 0.462495676 | Bmp1 | bone morphogenetic protein 1 [Source:M | 2356.3 | 2850.4 | 3080.1 | 2468.7 | 2463.7 | 2311.2 |
| 3477 | ENSMUSG00000022099 | 3.259048115 | -0.520810972 | 1.336898569 | -0.389566557 | 0.696857086 | NA | Dmtn | dematin actin binding protein [Source:MG | 6.4248 | 4.1117 | 1.0062 | 5.7839 | 0 | 2.2277 |
| 3478 | ENSMUSG00000022100 | 1093.569084 | -0.017612465 | 0.108751055 | -0.161952132 | 0.871343549 | 0.979186784 | Xpo7 | exportin 7 [Source:MGI Symbol;Acc:MGI: | 1015.1 | 1130.7 | 1156.2 | 1129.8 | 1048.1 | 1081.5 |
| 3479 | ENSMUSG00000022101 | 0.167707431 | -0.796629176 | 4.080472857 | -0.195229623 | 0.84521319 | NA | Fgf17 | fibroblast growth factor 17 [Source:MGI S | 0 | 0 | 1.0062 | 0 | 0 | 0 |
| 3480 | ENSMUSG00000022102 | 32.59956982 | -0.969021905 | 0.399846127 | -2.423487037 | 0.015372302 | 0.144944573 | Dok2 | docking protein 2 [Source:MGI Symbol;Ac | 36.943 | 54.48 | 38.237 | 24.099 | 22.903 | 18.935 |
| 3481 | ENSMUSG00000022103 | 3.833489803 | -1.705193334 | 1.378579593 | -1.236920481 | 0.216116606 | NA | Gfra2 | glial cell line derived neurotrophic factor f | 3.2124 | 13.363 | 1.0062 | 0.964 | 0 | 4.4554 |
| 3482 | ENSMUSG00000022105 | 164.4441917 | 0.103465441 | 0.20109463 | 0.514511208 | 0.606894627 | 0.916376001 | Rb1 | RB transcriptional corepressor 1 [Source:I | 182.3 | 143.91 | 148.92 | 172.55 | 191.95 | 147.03 |
| 3483 | ENSMUSG00000022106 | 691.0875773 | -0.028696299 | 0.130066348 | -0.220628159 | 0.825381974 | 0.96943211 | Rcbtb2 | regulator of chromosome condensation (I | 762.14 | 670.2 | 660.1 | 607.31 | 737.26 | 709.52 |
| 3484 | ENSMUSG00000022108 | 17641.42841 | -0.081341797 | 0.077553026 | -1.048853933 | 0.294245351 | 0.745164683 | Itm2b | integral membrane protein 2B [Source:MG | 18271 | 18087 | 18058 | 17308 | 17225 | 16899 |
| 3485 | ENSMUSG00000022109 | 170.0252065 | 0.20032993 | 0.198124715 | 1.011130439 | 0.311954003 | 0.759830222 | Med4 | mediator complex subunit 4 [Source:MGI | 167.85 | 151.1 | 155.97 | 215.93 | 175.59 | 153.71 |
| 3486 | ENSMUSG00000022110 | 3089.043614 | 0.061670038 | 0.100241053 | 0.615217379 | 0.538411172 | 0.891577636 | Sucla2 | succinate-Coenzyme A ligase, ADP-formin | 2899.2 | 2982 | 3187.8 | 2904.5 | 3386.4 | 3174.5 |
| 3487 | ENSMUSG00000022111 | 255.7575496 | 0.080118742 | 0.19012227 | 0.421406401 | 0.673458346 | 0.931915649 | Uchl3 | ubiquitin carboxyl-terminal esterase L3 (u | 301.16 | 228.2 | 215.34 | 229.43 | 254.11 | 306.31 |
| 3488 | ENSMUSG00000022112 | 0 | NA | NA | NA | NA | NA | Gpc5 | glypican 5 [Source:MGI Symbol;Acc:MGI:1 | 0 | 0 | 0 | 0 | 0 | 0 |
| 3489 | ENSMUSG00000022113 | 0 | NA | NA | NA | NA | NA | Trim52 | tripartite motif-containing 52 [Source:MG | 0 | 0 | 0 | 0 | 0 | 0 |
| 3490 | ENSMUSG00000022114 | 11.21304974 | -1.105408844 | 0.684113617 | -1.615826403 | 0.106131851 | 0.480498081 | Spry2 | sprouty RTK signaling antagonist 2 [Source | 19.274 | 14.391 | 12.075 | 3.8559 | 6.5437 | 11.138 |
| 3491 | ENSMUSG00000022116 | 0 | NA | NA | NA | NA | NA | 4930449E01 | RIKEN cDNA 4930449E01 gene [Source:M | 0 | 0 | 0 | 0 | 0 | 0 |
| 3492 | ENSMUSG00000022119 | 382.7068778 | -0.064602051 | 0.16787026 | -0.384833213 | 0.70036099 | 0.941626823 | Rbm26 | RNA binding motif protein 26 [Source:MG | 461.78 | 379.3 | 332.06 | 393.3 | 407.89 | 321.9 |
| 3493 | ENSMUSG00000022120 | 18.32283058 | -0.179246907 | 0.540801173 | -0.331446965 | 0.740306895 | 0.948792385 | Obi1 | ORC ubiquitin ligase 1 [Source:MGI Symbo | 13.653 | 30.838 | 14.087 | 18.316 | 17.45 | 15.594 |
| 3494 | ENSMUSG00000022122 | 106.1102209 | -1.146990321 | 0.237057814 | -4.838441311 | 1.31E-06 | 0.000101218 | Ednrb | endothelin receptor type B [Source:MGI S | 147.77 | 137.74 | 152.95 | 64.587 | 54.531 | 79.083 |
| 3495 | ENSMUSG00000022123 | 0 | NA | NA | NA | NA | NA | Scel | sciellin [Source:MGI Symbol;Acc:MGI:189 | 0 | 0 | 0 | 0 | 0 | 0 |
| 3496 | ENSMUSG00000022124 | 766.2459123 | 0.08129369 | 0.111485123 | 0.729188683 | 0.465886253 | 0.863698893 | Fbxl3 | F-box and leucine-rich repeat protein 3 [S | 730.02 | 776.08 | 728.52 | 823.24 | 771.07 | 768.55 |
| 3497 | ENSMUSG00000022125 | 683.4362458 | 0.061348212 | 0.113145908 | 0.542204427 | 0.587677686 | 0.910765033 | Cln5 | ceroid-lipofuscinosis, neuronal 5 [Source: | 670.59 | 654.78 | 681.23 | 695.99 | 672.91 | 725.11 |
| 3498 | ENSMUSG00000022126 | 1.254537109 | -3.692333964 | 2.19368904 | -1.683161969 | 0.092343741 | NA | Acod1 | aconitate decarboxylase 1 [Source:MGI Sy | 2.4093 | 4.1117 | 1.0062 | 0 | 0 | 0 |
| 3499 | ENSMUSG00000022129 | 54.11339507 | 0.988858652 | 0.3220981 | 3.070054286 | 0.002140199 | 0.036657676 | Dct | dopachrome tautomerase [Source:MGI Sy | 44.171 | 32.893 | 31.194 | 59.767 | 70.89 | 85.766 |
| 3500 | ENSMUSG00000022130 | 411.4637527 | 0.167606992 | 0.145593348 | 1.151199522 | 0.249650162 | 0.70183478 | Tgds | TDP-glucose 4,6-dehydratase [Source:MG | 350.96 | 391.64 | 420.61 | 416.44 | 404.62 | 484.52 |
| 3501 | ENSMUSG00000022131 | 593.8562776 | 0.065298875 | 0.131733588 | 0.495688881 | 0.620113933 | 0.917977899 | Gpr180 | G protein-coupled receptor 180 [Source:N | 611.16 | 519.1 | 610.79 | 638.15 | 631.47 | 552.47 |
| 3502 | ENSMUSG00000022132 | 0.181769715 | 1.126924175 | 4.080472857 | 0.276174898 | 0.782413736 | NA | Cldn10 | claudin 10 [Source:MGI Symbol;Acc:MGI: | 0 | 0 | 0 | 0 | 1.0906 | 0 |
| 3503 | ENSMUSG00000022136 | 4888.707344 | 0.110157507 | 0.117650047 | 0.936315023 | 0.349111015 | 0.79040715 | Dnajc3 | DnaJ heat shock protein family (Hsp40) m | 4465.2 | 4574.2 | 5068.5 | 5842.7 | 4458.4 | 4923.2 |
| 3504 | ENSMUSG00000022139 | 669.2420864 | -0.078655671 | 0.155179496 | -0.506868965 | 0.612246763 | 0.916441371 | Mbnl2 | muscleblind like splicing factor 2 [Source:I | 839.24 | 642.45 | 579.6 | 668.04 | 696.91 | 589.22 |
| 3505 | ENSMUSG00000022141 | 710.0007589 | 0.05351622 | 0.130944709 | 0.408693259 | 0.682764781 | 0.935166147 | Nipbl | NIPBL cohesin loading factor [Source:MGI | 688.26 | 752.44 | 650.03 | 716.24 | 809.24 | 643.8 |
| 3506 | ENSMUSG00000022142 | 364.6677483 | 0.089981162 | 0.144151941 | 0.624210544 | 0.532489323 | 0.889143097 | Nup155 | nucleoporin 155 [Source:MGI Symbol;Acc | 367.82 | 316.6 | 375.33 | 392.34 | 381.72 | 354.2 |
| 3507 | ENSMUSG00000022144 | 0 | NA | NA | NA | NA | NA | Gdnf | glial cell line derived neurotrophic factor [ | 0 | 0 | 0 | 0 | 0 | 0 |
| 3508 | ENSMUSG00000022146 | 34.61289613 | -1.093713953 | 0.411625879 | -2.657058287 | 0.007882581 | 0.090992543 | Osmr | oncostatin M receptor [Source:MGI Symb | 44.171 | 48.312 | 49.306 | 32.775 | 14.178 | 18.935 |
| 3509 | ENSMUSG00000022148 | 44.51017353 | -2.082000048 | 0.466307214 | -4.464867764 | 8.01E-06 | 0.000447425 | Fyb | FYN binding protein [Source:MGI Symbol; | 53.808 | 121.29 | 41.256 | 23.135 | 13.087 | 14.48 |
| 3510 | ENSMUSG00000022149 | 8598.212417 | 0.782510158 | 0.1229343 | 6.365271211 | 1.95E-10 | 5.22E-08 | C9 | complement component 9 [Source:MGI S | 6658.5 | 5462.4 | 6843.5 | 9407.5 | 11274 | 11944 |
| 3511 | ENSMUSG00000022150 | 112.7595103 | -0.370709497 | 0.232822854 | -1.592238437 | 0.11133114 | 0.489450944 | Dab2 | disabled 2, mitogen-responsive phosphop | 132.51 | 132.6 | 116.72 | 120.5 | 82.887 | 91.335 |
| 3512 | ENSMUSG00000022151 | 352.3634553 | -0.037496816 | 0.151295343 | -0.247838532 | 0.804259341 | 0.963701061 | Ttc33 | tetratricopeptide repeat domain 33 [Sour | 353.36 | 356.69 | 360.24 | 296.91 | 384.99 | 362 |
| 3513 | ENSMUSG00000022155 | 0 | NA | NA | NA | NA | NA | Mroh2b | maestro heat-like repeat family member 2 | 0 | 0 | 0 | 0 | 0 | 0 |
| 3514 | ENSMUSG00000022156 | 0 | NA | NA | NA | NA | NA | Gzme | granzyme E [Source:MGI Symbol;Acc:MGI | 0 | 0 | 0 | 0 | 0 | 0 |
| 3515 | ENSMUSG00000022157 | 0.335414862 | -1.779047242 | 4.030804179 | -0.441362855 | 0.658950329 | NA | Mcpt8 | mast cell protease 8 [Source:MGI Symbol; | 0 | 0 | 2.0125 | 0 | 0 | 0 |
| 3516 | ENSMUSG00000022159 | 8.465162696 | -0.987237768 | 0.802574332 | -1.230088889 | 0.21866382 | 0.663735712 | Rab2b | RAB2B, member RAS oncogene family [So | 6.4248 | 9.2513 | 18.112 | 4.8199 | 3.2719 | 8.9108 |
| 3517 | ENSMUSG00000022160 | 139.1619831 | 0.072610313 | 0.233858404 | 0.310488362 | 0.756189608 | 0.953085143 | Mettl3 | methyltransferase like 3 [Source:MGI Sym | 119.66 | 160.36 | 127.79 | 178.34 | 136.33 | 112.5 |
| 3518 | ENSMUSG00000022174 | 1072.744807 | 0.023853632 | 0.107954483 | 0.220960088 | 0.82512351 | 0.96943211 | Dad1 | defender against cell death 1 [Source:MG | 1049.7 | 1147.2 | 995.18 | 1090.3 | 1096.1 | 1058.2 |
| 3519 | ENSMUSG00000022175 | 903.3673541 | 0.045891817 | 0.11883019 | 0.386196614 | 0.699351062 | 0.941231391 | Lrp10 | low-density lipoprotein receptor-related p | 934.01 | 783.27 | 948.89 | 893.61 | 927.03 | 933.4 |
| 3520 | ENSMUSG00000022176 | 0.87097134 | 0.683548663 | 2.367771665 | 0.28868859 | 0.772819695 | NA | Rem2 | rad and gem related GTP binding protein : | 0 | 1.0279 | 1.0062 | 0.964 | 0 | 2.2277 |
| 3521 | ENSMUSG00000022177 | 271.2641665 | 0.057395394 | 0.152330173 | 0.37678283 | 0.706335002 | 0.94295036 | Haus4 | HAUS augmin-like complex, subunit 4 [Sou | 269.04 | 259.04 | 269.67 | 294.01 | 258.48 | 277.35 |
| 3522 | ENSMUSG00000022178 | 108.7807592 | 0.636717261 | 0.271469077 | 2.345450423 | 0.019004106 | 0.167504806 | Ajuba | ajuba LIM protein [Source:MGI Symbol;Ac | 93.963 | 60.647 | 100.62 | 145.56 | 97.065 | 154.82 |
| 3523 | ENSMUSG00000022179 | 98.43018324 | 0.11528276 | 0.267661532 | 0.430703504 | 0.66668397 | 0.929596279 | 4931414P19 | RIKEN cDNA 4931414P19 gene [Source:M | 81.916 | 93.541 | 108.67 | 132.06 | 75.253 | 99.132 |
| 3524 | ENSMUSG00000022180 | 30.23507087 | -1.122836715 | 0.44637413 | -2.515461001 | 0.011887683 | 0.121945705 | Slc7a8 | solute carrier family 7 (cationic amino acid | 38.549 | 58.591 | 27.169 | 17.352 | 16.359 | 23.391 |

| | | | | | | | | | | | | | | | |
|---|---|---|---|---|---|---|---|---|---|---|---|---|---|---|---|
| 3525 | ENSMUSG00000022181 | 3795.716903 | 1.077392222 | 0.139425808 | 7.727351464 | 1.10E-14 | 6.91E-12 | C6 | complement component 6 [Source:MGI S | 2825.3 | 2008.6 | 2486.4 | 4471.9 | 5823.9 | 5158.2 |
| 3526 | ENSMUSG00000022184 | 200.3978536 | 0.329353326 | 0.191130528 | 1.723185352 | 0.084855012 | 0.429140871 | Fbxo4 | F-box protein 4 [Source:MGI Symbol;Acc:I | 210.41 | 167.55 | 153.96 | 215.93 | 205.04 | 249.5 |
| 3527 | ENSMUSG00000022185 | 972.628253 | -0.013792691 | 0.102889813 | -0.134053029 | 0.893360642 | 0.982562372 | Acin1 | apoptotic chromatin condensation induce | 954.09 | 991.94 | 986.12 | 962.05 | 987.01 | 954.56 |
| 3528 | ENSMUSG00000022186 | 51.61224987 | -0.575374796 | 0.372922341 | -1.542881006 | 0.12285965 | 0.513267141 | Oxct1 | 3-oxoacid CoA transferase 1 [Source:MGI | 46.58 | 93.541 | 45.281 | 32.775 | 44.715 | 46.781 |
| 3529 | ENSMUSG00000022187 | 0 | NA | NA | NA | NA | NA | Gm5546 | predicted gene 5546 [Source:MGI Symbol | 0 | 0 | 0 | 0 | 0 | 0 |
| 3530 | ENSMUSG00000022191 | 193.1416905 | 0.088610557 | 0.193657208 | 0.457563951 | 0.647265747 | 0.925643431 | Drosha | drosha, ribonuclease type III [Source:MGI | 196.76 | 185.03 | 179.11 | 159.06 | 207.22 | 231.68 |
| 3531 | ENSMUSG00000022193 | 1566.050377 | -0.04654206 | 0.11566593 | -0.402383481 | 0.687401825 | 0.936885029 | Psmb5 | proteasome (prosome, macropain) subun | 1795.7 | 1429.8 | 1546.6 | 1485.5 | 1524.7 | 1614 |
| 3532 | ENSMUSG00000022194 | 548.7382722 | 0.103006432 | 0.139391918 | 0.738969906 | 0.459925272 | 0.859753653 | Pabpn1 | poly(A) binding protein, nuclear 1 [Source | 514.79 | 525.27 | 548.4 | 657.43 | 545.31 | 501.23 |
| 3533 | ENSMUSG00000022195 | 392.9023927 | -0.013549471 | 0.14531915 | -0.093239401 | 0.925713373 | 0.987546253 | 6030458C11 | RIKEN cDNA 6030458C11 gene [Source:M | 418.42 | 412.2 | 353.19 | 380.77 | 428.61 | 364.23 |
| 3534 | ENSMUSG00000022197 | 16.51772036 | 0.52808354 | 0.641537652 | 0.823152839 | 0.410421084 | 0.833241465 | Pdzd2 | PDZ domain containing 2 [Source:MGI Syr | 24.896 | 7.1954 | 8.05 | 12.532 | 16.359 | 30.074 |
| 3535 | ENSMUSG00000022199 | 16.56453402 | -1.251629989 | 0.545058585 | -2.296321942 | 0.02165748 | 0.183082289 | Slc22a17 | solute carrier family 22 (organic cation tra | 23.29 | 20.558 | 26.162 | 10.604 | 7.6343 | 11.138 |
| 3536 | ENSMUSG00000022200 | 1307.210276 | -0.020867898 | 0.101716674 | -0.205157097 | 0.837449388 | 0.971073158 | Golph3 | golgi phosphoprotein 3 [Source:MGI Symb | 1313.1 | 1336.3 | 1301.1 | 1360.2 | 1213.9 | 1318.8 |
| 3537 | ENSMUSG00000022201 | 965.1419314 | -0.122663144 | 0.11561792 | -1.060935399 | 0.288719261 | 0.73951397 | Zfr | zinc finger RNA binding protein [Source:M | 1040.8 | 907.65 | 1069.6 | 933.13 | 963.02 | 876.6 |
| 3538 | ENSMUSG00000022203 | 2.338008318 | 0.297061563 | 1.824465842 | 0.162821115 | 0.870659283 | NA | Efs | embryonal Fyn-associated substrate [Sour | 3.2124 | 0 | 3.0187 | 0 | 0 | 7.7969 |
| 3539 | ENSMUSG00000022204 | 297.062582 | 0.263849106 | 0.155073228 | 1.701448464 | 0.088858807 | 0.439445475 | Ngdn | neuroguidin, EIF4E binding protein [Sourc | 295.54 | 275.48 | 238.48 | 340.28 | 317.37 | 315.22 |
| 3540 | ENSMUSG00000022205 | 783.0725128 | -0.037516646 | 0.141789699 | -0.26459359 | 0.791322559 | 0.961095767 | Sub1 | SUB1 homolog, transcriptional regulator [ | 808.72 | 725.71 | 844.24 | 636.23 | 812.51 | 871.03 |
| 3541 | ENSMUSG00000022206 | 19.11760593 | -1.406929179 | 0.516753214 | -2.722632663 | 0.006476402 | 0.079587668 | Npr3 | natriuretic peptide receptor 3 [Source:MG | 30.518 | 21.586 | 31.194 | 11.568 | 9.8156 | 10.025 |
| 3542 | ENSMUSG00000022208 | 0.133850373 | -0.796629176 | 4.080472857 | -0.195229623 | 0.84521319 | NA | Jph4 | junctophilin 4 [Source:MGI Symbol;Acc:M | 0.8031 | 0 | 0 | 0 | 0 | 0 |
| 3543 | ENSMUSG00000022209 | 0 | NA | NA | NA | NA | NA | Dhrs2 | dehydrogenase/reductase member 2 [Sou | 0 | 0 | 0 | 0 | 0 | 0 |
| 3544 | ENSMUSG00000022210 | 3322.396074 | -0.09309833 | 0.091707299 | -1.015168157 | 0.310025677 | 0.758699916 | Dhrs4 | dehydrogenase/reductase (SDR family) m | 3275.9 | 3464.1 | 3550 | 3350.8 | 3223.9 | 3069.8 |
| 3545 | ENSMUSG00000022211 | 3.877261443 | -0.053015023 | 1.140965301 | -0.046465061 | 0.962939581 | NA | Carmil3 | capping protein regulator and myosin 1 lir | 0.8031 | 5.1396 | 6.0375 | 5.7839 | 3.2719 | 2.2277 |
| 3546 | ENSMUSG00000022212 | 0 | NA | NA | NA | NA | NA | Cpne6 | copine VI [Source:MGI Symbol;Acc:MGI:1 | 0 | 0 | 0 | 0 | 0 | 0 |
| 3547 | ENSMUSG00000022214 | 5004.489217 | 0.432945117 | 0.106457982 | 4.066816876 | 4.77E-05 | 0.002027498 | Dcaf11 | DDB1 and CUL4 associated factor 11 [Sour | 4009.9 | 4028.4 | 4739.4 | 5363.6 | 5709.4 | 6176.3 |
| 3548 | ENSMUSG00000022215 | 891.7821622 | 0.042784368 | 0.378328192 | 0.113087973 | 0.90996081 | 0.98552401 | Fitm1 | fat storage-inducing transmembrane prot | 1237.6 | 814.11 | 583.62 | 936.99 | 1117.9 | 660.51 |
| 3549 | ENSMUSG00000022216 | 1261.64438 | -0.025126204 | 0.105128923 | -0.239003722 | 0.811102697 | 0.966452061 | Psme1 | proteasome (prosome, macropain) activa | 1233.6 | 1352.7 | 1231.6 | 1185.7 | 1310.9 | 1255.3 |
| 3550 | ENSMUSG00000022217 | 467.549293 | 0.219896151 | 0.139911593 | 1.571679279 | 0.11602495 | 0.500730933 | Emc9 | ER membrane protein complex subunit 9 | 422.43 | 384.44 | 489.03 | 485.84 | 526.77 | 496.77 |
| 3551 | ENSMUSG00000022218 | 177.4972585 | 1.114121139 | 0.553714419 | 2.012086195 | 0.044210852 | 0.290666254 | Tgm1 | transglutaminase 1, K polypeptide [Source | 149.38 | 91.485 | 95.593 | 424.15 | 142.87 | 161.51 |
| 3552 | ENSMUSG00000022219 | 1007.757198 | 0.031088131 | 0.108926723 | 0.285404078 | 0.775334584 | 0.95893551 | Cideb | cell death-inducing DNA fragmentation fa | 1052.1 | 935.41 | 1002.2 | 992.9 | 993.55 | 1070.4 |
| 3553 | ENSMUSG00000022220 | 43.42272659 | -0.795528087 | 0.36725988 | -2.166117591 | 0.030302202 | 0.230436937 | Adcy4 | adenylate cyclase 4 [Source:MGI Symbol;/ | 46.58 | 60.647 | 58.362 | 38.559 | 19.631 | 36.757 |
| 3554 | ENSMUSG00000022221 | 13.40897834 | -1.684373196 | 0.672673346 | -2.503998717 | 0.012279848 | 0.124560339 | Ripk3 | receptor-interacting serine-threonine kina | 26.502 | 23.642 | 11.069 | 3.8559 | 9.8156 | 5.5692 |
| 3555 | ENSMUSG00000022223 | 70.77517537 | -0.258264779 | 0.345722496 | -0.747029141 | 0.455045971 | 0.857337687 | Sdr39u1 | short chain dehydrogenase/reductase fan | 87.538 | 93.541 | 50.312 | 79.046 | 76.343 | 37.871 |
| 3556 | ENSMUSG00000022225 | 2.171574289 | -3.41412191 | 1.966839357 | -1.73584177 | 0.082591816 | NA | Cma1 | chymase 1, mast cell [Source:MGI Symbol | 4.0155 | 0 | 8.05 | 0.964 | 0 | 0 |
| 3557 | ENSMUSG00000022226 | 0 | NA | NA | NA | NA | NA | Mcpt2 | mast cell protease 2 [Source:MGI Symbol; | 0 | 0 | 0 | 0 | 0 | 0 |
| 3558 | ENSMUSG00000022227 | 0 | NA | NA | NA | NA | NA | Mcpt1 | mast cell protease 1 [Source:MGI Symbol; | 0 | 0 | 0 | 0 | 0 | 0 |
| 3559 | ENSMUSG00000022228 | 771.238292 | 0.144487843 | 0.119139168 | 1.212765252 | 0.225219584 | 0.672301295 | Zscan26 | zinc finger and SCAN domain containing 2 | 781.42 | 739.07 | 677.2 | 860.83 | 808.15 | 760.76 |
| 3560 | ENSMUSG00000022229 | 0 | NA | NA | NA | NA | NA | Atp12a | ATPase, H+/K+ transporting, nongastric, a | 0 | 0 | 0 | 0 | 0 | 0 |
| 3561 | ENSMUSG00000022231 | 12.99514327 | -0.535599602 | 0.608578362 | -0.880083215 | 0.378814231 | 0.811969331 | Sema5a | sema domain, seven thrombospondin rep | 18.471 | 16.447 | 11.069 | 7.7118 | 10.906 | 13.366 |
| 3562 | ENSMUSG00000022234 | 1957.681655 | 0.015720619 | 0.108871751 | 0.144395758 | 0.885187967 | 0.981795936 | Cct5 | chaperonin containing Tcp1, subunit 5 (ep | 1742.7 | 1981.8 | 2118.1 | 2062.9 | 1885.7 | 1954.8 |
| 3563 | ENSMUSG00000022235 | 3993.632509 | -0.127456774 | 0.101669584 | -1.253637213 | 0.209973899 | 0.653834763 | Cmbl | carboxymethylenebutenolidase-like (Pseu | 4279.7 | 3944.1 | 4284.6 | 3440.4 | 4083.3 | 3929.6 |
| 3564 | ENSMUSG00000022236 | 215.7507009 | 0.287234298 | 0.187546202 | 1.531538867 | 0.125636268 | 0.519794903 | Ropn1l | ropporin 1-like [Source:MGI Symbol;Acc:N | 191.94 | 213.81 | 177.1 | 205.33 | 283.56 | 222.77 |
| 3565 | ENSMUSG00000022237 | 745.6804538 | -0.043961428 | 0.170949706 | -0.25716001 | 0.797055254 | 0.962236316 | Ankrd33b | ankyrin repeat domain 33B [Source:MGI S | 625.62 | 828.5 | 817.07 | 559.11 | 796.15 | 847.64 |
| 3566 | ENSMUSG00000022240 | 0.842381458 | 0.63517692 | 2.579321552 | 0.246257362 | 0.805483014 | NA | Ctnnd2 | catenin (cadherin associated protein), del | 0 | 0 | 2.0125 | 1.928 | 0 | 1.1138 |
| 3567 | ENSMUSG00000022241 | 1762.305757 | -0.190076686 | 0.131197813 | -1.448779373 | 0.147399206 | 0.558402858 | Tars | threonyl-tRNA synthetase [Source:MGI Sy | 1619.9 | 1961.3 | 2054.8 | 1702.4 | 1418.9 | 1816.7 |
| 3568 | ENSMUSG00000022243 | 0.863488199 | 0.685974303 | 2.37316251 | 0.289054922 | 0.772539349 | NA | Slc45a2 | solute carrier family 45, member 2 [Sourc | 0 | 0 | 2.0125 | 0.964 | 1.0906 | 1.1138 |
| 3569 | ENSMUSG00000022244 | 3280.213256 | 0.313903401 | 0.10560706 | 2.972371348 | 0.00295509 | 0.046110081 | Amacr | alpha-methylacyl-CoA racemase [Source:I | 2897.6 | 2792.9 | 3083.1 | 3256.3 | 3956.8 | 3694.6 |
| 3570 | ENSMUSG00000022245 | 0.17131964 | -0.796629176 | 4.080472857 | -0.195229623 | 0.84521319 | NA | Skor1 | SKI family transcriptional corepressor 1 [S | 0 | 1.0279 | 0 | 0 | 0 | 0 |
| 3571 | ENSMUSG00000022246 | 362.170562 | 0.180752072 | 0.162515193 | 1.112216458 | 0.266045093 | 0.722063933 | Rai14 | retinoic acid induced 14 [Source:MGI Sym | 362.2 | 315.57 | 340.11 | 370.17 | 327.19 | 457.79 |
| 3572 | ENSMUSG00000022247 | 310.7083049 | -0.097575749 | 0.149583113 | -0.652317945 | 0.514196086 | 0.885004045 | Brix1 | BRX1, biogenesis of ribosomes [Source:M | 333.29 | 343.32 | 286.78 | 296.91 | 302.1 | 301.85 |
| 3573 | ENSMUSG00000022248 | 105.0393295 | -0.328351583 | 0.24708484 | -1.328902183 | 0.183880243 | 0.618439913 | Rad1 | RAD1 checkpoint DNA exonuclease [Sourc | 138.13 | 108.96 | 103.64 | 111.82 | 76.343 | 91.335 |
| 3574 | ENSMUSG00000022249 | 0 | NA | NA | NA | NA | NA | Ttc23l | tetratricopeptide repeat domain 23-like [S | 0 | 0 | 0 | 0 | 0 | 0 |
| 3575 | ENSMUSG00000022253 | 3544.928933 | 0.164911962 | 0.105242547 | 1.566970465 | 0.117121595 | 0.502643331 | Nadk2 | NAD kinase 2, mitochondrial [Source:MGI | 2986.7 | 3377.7 | 3664.7 | 3785.5 | 3877.1 | 3577.7 |
| 3576 | ENSMUSG00000022255 | 1187.720016 | -0.152258135 | 0.120555206 | -1.262974374 | 0.206598389 | 0.652026861 | Mtdh | metadherin [Source:MGI Symbol;Acc:MGI | 1081.8 | 1388.7 | 1282 | 1132.7 | 1144.1 | 1097.1 |
| 3577 | ENSMUSG00000022257 | 362.7069043 | -0.199128537 | 0.142321847 | -1.399142445 | 0.161770272 | 0.584961303 | Laptm4b | lysosomal-associated protein transmembr | 418.42 | 351.55 | 392.44 | 330.64 | 346.82 | 336.38 |
| 3578 | ENSMUSG00000022261 | 3157.07241 | 0.001689516 | 0.102476382 | 0.016486883 | 0.986845966 | 0.99859176 | Sdc2 | syndecan 2 [Source:MGI Symbol;Acc:MGI | 2879.9 | 3241 | 3345.8 | 3129.1 | 2982.8 | 3363.8 |
| 3579 | ENSMUSG00000022262 | 1.721191773 | 0.739829771 | 1.726422977 | 0.428533321 | 0.66826288 | NA | Dnah5 | dynein, axonemal, heavy chain 5 [Source: | 0.8031 | 3.0838 | 0 | 0.964 | 4.3625 | 1.1138 |
| 3580 | ENSMUSG00000022263 | 330.3476558 | -0.018435921 | 0.152105227 | -0.121205047 | 0.903528627 | 0.983821605 | Trio | triple functional domain (PTPRF interactin | 332.48 | 343.32 | 322 | 364.38 | 285.74 | 334.15 |

| | | | | | | | | | | | | | | | |
|---|---|---|---|---|---|---|---|---|---|---|---|---|---|---|---|
| 3581 | ENSMUSG00000022265 | 796.5945468 | -0.138817031 | 0.129145285 | -1.074890435 | 0.282423785 | 0.735403596 | Ank | progressive ankylosis [Source:MGI Symbo | 736.44 | 908.68 | 860.34 | 736.48 | 707.81 | 829.81 |
| 3582 | ENSMUSG00000022269 | 0 | NA | NA | NA | NA | NA | Marchf11 | membrane associated ring-CH-type finger | 0 | 0 | 0 | 0 | 0 | 0 |
| 3583 | ENSMUSG00000022270 | 1185.275978 | 0.093467369 | 0.348280585 | 0.268368013 | 0.788416058 | 0.960872359 | Retreg1 | reticulophagy regulator 1 [Source:MGI Syı | 1207.9 | 1276.7 | 955.93 | 1045.9 | 839.78 | 1785.5 |
| 3584 | ENSMUSG00000022272 | 577.7431379 | 0.023996347 | 0.147563472 | 0.162617119 | 0.870819908 | 0.97888679 | Myo10 | myosin X [Source:MGI Symbol;Acc:MGI:1( | 505.95 | 609.56 | 604.75 | 670.93 | 503.87 | 571.4 |
| 3585 | ENSMUSG00000022280 | 600.4523536 | -0.122990731 | 0.132948649 | -0.925099518 | 0.354914144 | 0.794902164 | Rnf19a | ring finger protein 19A [Source:MGI Symb | 552.53 | 654.78 | 672.17 | 615.98 | 571.48 | 535.76 |
| 3586 | ENSMUSG00000022283 | 4462.861427 | 0.119162582 | 0.092017442 | 1.294999945 | 0.195320248 | 0.635641153 | Pabpc1 | poly(A) binding protein, cytoplasmic 1 [So | 4275.7 | 4476.6 | 4083.3 | 4439.1 | 4906.7 | 4595.7 |
| 3587 | ENSMUSG00000022285 | 1581.046249 | -0.144636983 | 0.100449438 | -1.43989839 | 0.149896149 | 0.563095827 | Ywhaz | tyrosine 3-monooxygenase/tryptophan 5- | 1741.9 | 1654.9 | 1582.8 | 1462.4 | 1460.3 | 1583.9 |
| 3588 | ENSMUSG00000022286 | 1.700861205 | 0.744792243 | 1.83599023 | 0.405662422 | 0.684990657 | NA | Grhl2 | grainyhead like transcription factor 2 [Sou | 0.8031 | 0 | 3.0187 | 1.928 | 0 | 4.4554 |
| 3589 | ENSMUSG00000022288 | 0 | NA | NA | NA | NA | NA | 4930447A16 | RIKEN cDNA 4930447A16 gene [Source:M | 0 | 0 | 0 | 0 | 0 | 0 |
| 3590 | ENSMUSG00000022292 | 71.2164887 | 0.193101097 | 0.307366116 | 0.628244581 | 0.52984373 | 0.888334265 | Rrm2b | ribonucleotide reductase M2 B (TP53 indu | 66.657 | 89.429 | 43.269 | 71.334 | 73.071 | 83.538 |
| 3591 | ENSMUSG00000022295 | 637.9451829 | 0.212191772 | 0.124743671 | 1.701022353 | 0.08893879 | 0.439498004 | Atp6v1c1 | ATPase, H+ transporting, lysosomal V1 sul | 644.09 | 583.86 | 544.38 | 665.14 | 726.35 | 663.85 |
| 3592 | ENSMUSG00000022296 | 0.133850373 | -0.796629176 | 4.080472857 | -0.195229623 | 0.84521319 | NA | Baalc | brain and acute leukemia, cytoplasmic [Sc | 0.8031 | 0 | 0 | 0 | 0 | 0 |
| 3593 | ENSMUSG00000022297 | 107.9545483 | 0.702814436 | 0.253031889 | 2.777572581 | 0.005476661 | 0.070806769 | Fzd6 | frizzled class receptor 6 [Source:MGI Syml | 93.963 | 57.563 | 94.587 | 144.6 | 117.79 | 139.23 |
| 3594 | ENSMUSG00000022299 | 857.4546642 | -0.315976303 | 0.147628063 | -2.140353913 | 0.032326176 | 0.240270201 | Slc25a32 | solute carrier family 25, member 32 [Sour | 885.82 | 1105 | 862.35 | 705.63 | 898.67 | 687.24 |
| 3595 | ENSMUSG00000022300 | 410.6119533 | -0.23491486 | 0.140322731 | -1.674104102 | 0.094110142 | 0.451490179 | Dcaf13 | DDB1 and CUL4 associated factor 13 [Sou | 462.59 | 448.17 | 420.61 | 353.78 | 420.98 | 357.54 |
| 3596 | ENSMUSG00000022303 | 4.775095863 | 0.83932099 | 1.000260944 | 0.839102031 | 0.401412056 | NA | Dcstamp | dendrocyte expressed seven transmembr | 3.2124 | 4.1117 | 3.0187 | 10.604 | 4.3625 | 3.3415 |
| 3597 | ENSMUSG00000022304 | 6623.625805 | -0.378474618 | 0.138963276 | -2.723558556 | 0.006458277 | 0.079432408 | Dpys | dihydropyrimidinase [Source:MGI Symbol | 6450.5 | 7758.7 | 8255.2 | 6791.2 | 5737.7 | 4748.3 |
| 3598 | ENSMUSG00000022305 | 108.6739119 | 0.008766027 | 0.241717024 | 0.036265659 | 0.971070532 | 0.995010213 | Lrp12 | low density lipoprotein-related protein 12 | 118.06 | 125.41 | 81.506 | 117.61 | 101.43 | 108.04 |
| 3599 | ENSMUSG00000022306 | 10.17738915 | -0.46486327 | 0.694675238 | -0.669180711 | 0.503380209 | 0.881357141 | Zfpm2 | zinc finger protein, multitype 2 [Source:M | 12.85 | 14.391 | 8.05 | 4.8199 | 9.8156 | 11.138 |
| 3600 | ENSMUSG00000022307 | 1082.256386 | 0.117608211 | 0.10835517 | 1.085395472 | 0.277746537 | 0.730821523 | Oxr1 | oxidation resistance 1 [Source:MGI Symbo | 985.41 | 1106 | 1023.4 | 1063.3 | 1160.4 | 1155.1 |
| 3601 | ENSMUSG00000022309 | 12.51083269 | -0.848817462 | 0.693838766 | -1.223364135 | 0.221192193 | 0.66763854 | Angpt1 | angiopoietin 1 [Source:MGI Symbol;Acc:N | 18.471 | 21.586 | 8.05 | 4.8199 | 6.5437 | 15.594 |
| 3602 | ENSMUSG00000022311 | 0.462220778 | -0.685400455 | 3.191475091 | -0.21475977 | 0.82995462 | NA | Csmd3 | CUB and Sushi multiple domains 3 [Source | 0.8031 | 0 | 1.0062 | 0.964 | 0 | 0 |
| 3603 | ENSMUSG00000022312 | 2035.436097 | -0.09329668 | 0.089656689 | -1.040599206 | 0.2980616 | 0.74872577 | Eif3h | eukaryotic translation initiation factor 3, s | 2076.8 | 2087.7 | 2139.3 | 1959.8 | 1965.3 | 1983.8 |
| 3604 | ENSMUSG00000022313 | 71.88427304 | 0.153029106 | 0.311423328 | 0.491386138 | 0.623153367 | 0.919442038 | Utp23 | UTP23 small subunit processome compon | 53.808 | 74.01 | 76.475 | 56.875 | 65.437 | 104.7 |
| 3605 | ENSMUSG00000022314 | 1016.34642 | 0.03858643 | 0.105332231 | 0.366330695 | 0.714118328 | 0.94455746 | Rad21 | RAD21 cohesin complex component [Sour | 1015.9 | 969.33 | 1022.3 | 975.55 | 1057.9 | 1057 |
| 3606 | ENSMUSG00000022315 | 5.259327381 | 0.490765329 | 0.974987444 | 0.503355537 | 0.614714326 | 0.917509899 | Slc30a8 | solute carrier family 30 (zinc transporter), | 4.0155 | 6.1675 | 3.0187 | 10.604 | 2.1812 | 5.5692 |
| 3607 | ENSMUSG00000022321 | 0 | NA | NA | NA | NA | NA | Cdh10 | cadherin 10 [Source:MGI Symbol;Acc:MGI | 0 | 0 | 0 | 0 | 0 | 0 |
| 3608 | ENSMUSG00000022322 | 3.841129646 | 1.678073237 | 1.138170742 | 1.474359844 | 0.14038473 | NA | Shcbp1 | Shc SH2-domain binding protein 1 [Source | 2.4093 | 1.0279 | 2.0125 | 7.7118 | 6.5437 | 3.3415 |
| 3609 | ENSMUSG00000022323 | 11822.28904 | -0.162195758 | 0.099689222 | -1.627013979 | 0.103734129 | 0.474047094 | Rida | reactive intermediate imine deaminase A | 12601 | 11297 | 13560 | 11507 | 10516 | 11453 |
| 3610 | ENSMUSG00000022324 | 17.51650068 | -2.221942875 | 0.640724919 | -3.467857749 | 0.000524625 | 0.012905059 | Matn2 | matrilin 2 [Source:MGI Symbol;Acc:MGI:1 | 41.761 | 29.81 | 15.094 | 9.6398 | 5.4531 | 3.3415 |
| 3611 | ENSMUSG00000022325 | 94.43202244 | 0.016302118 | 0.249562491 | 0.06532279 | 0.947916997 | 0.990677354 | Pop1 | processing of precursor 1, ribonuclease P/ | 73.885 | 101.76 | 106.66 | 101.22 | 86.159 | 96.904 |
| 3612 | ENSMUSG00000022329 | 161.4102538 | 0.228265497 | 0.194269695 | 1.174992823 | 0.239997587 | 0.689857856 | Stk3 | serine/threonine kinase 3 [Source:MGI Sy | 140.54 | 133.63 | 172.07 | 179.3 | 163.59 | 179.33 |
| 3613 | ENSMUSG00000022330 | 0 | NA | NA | NA | NA | NA | Osr2 | odd-skipped related 2 [Source:MGI Symbo | 0 | 0 | 0 | 0 | 0 | 0 |
| 3614 | ENSMUSG00000022332 | 543.1224325 | -0.099105746 | 0.120194302 | -0.824546127 | 0.409629318 | 0.833241465 | Khdrbs3 | KH domain containing, RNA binding, signa | 556.55 | 578.72 | 550.42 | 543.68 | 514.77 | 514.6 |
| 3615 | ENSMUSG00000022335 | 31.24288596 | -0.016291456 | 0.443912663 | -0.036699688 | 0.970724458 | 0.994966572 | Zfat | zinc finger and AT hook domain containin | 36.943 | 27.754 | 29.181 | 17.352 | 29.447 | 46.781 |
| 3616 | ENSMUSG00000022336 | 1511.018612 | -0.031834359 | 0.102088336 | -0.311831504 | 0.75516858 | 0.953085143 | Eif3e | eukaryotic translation initiation factor 3, s | 1454.4 | 1553.2 | 1575.8 | 1425.7 | 1592.3 | 1464.7 |
| 3617 | ENSMUSG00000022337 | 533.9397067 | -0.008220059 | 0.134546249 | -0.061094672 | 0.951283812 | 0.990883058 | Emc2 | ER membrane protein complex subunit 2 | 537.28 | 489.29 | 579.6 | 512.84 | 586.75 | 497.89 |
| 3618 | ENSMUSG00000022338 | 989.4498215 | 0.010258337 | 0.114361389 | 0.089701057 | 0.928524776 | 0.988240056 | Eny2 | ENY2 transcription and export complex 2 | 997.45 | 981.66 | 978.07 | 916.74 | 1101.5 | 961.25 |
| 3619 | ENSMUSG00000022339 | 373.3710664 | 0.1032773 | 0.134499384 | 0.767864486 | 0.442567698 | 0.850670988 | Ebag9 | estrogen receptor-binding fragment-assoc | 363.81 | 361.83 | 354.2 | 379.81 | 376.26 | 404.33 |
| 3620 | ENSMUSG00000022340 | 5.817927321 | -1.50875832 | 1.244352778 | -1.212484391 | 0.225327014 | 0.672301295 | Sybu | syntabulin (syntaxin-interacting) [Source:I | 6.4248 | 19.53 | 0 | 6.7478 | 1.0906 | 1.1138 |
| 3621 | ENSMUSG00000022342 | 0 | NA | NA | NA | NA | NA | Kcnv1 | potassium channel, subfamily V, member | 0 | 0 | 0 | 0 | 0 | 0 |
| 3622 | ENSMUSG00000022346 | 176.3359987 | 2.226394544 | 1.106287565 | 2.012491701 | 0.044168131 | 0.290517436 | Myc | myelocytomatosis oncogene [Source:MGI | 27.305 | 100.74 | 58.362 | 365.35 | 25.084 | 481.18 |
| 3623 | ENSMUSG00000022347 | 1.959713055 | -1.012014342 | 1.893467186 | -0.534476831 | 0.593011675 | NA | A1bg | alpha-1-B glycoprotein [Source:MGI Symb | 4.8186 | 3.0838 | 0 | 3.8559 | 0 | 0 |
| 3624 | ENSMUSG00000022350 | 490.7468026 | 0.122196748 | 0.13887827 | 0.879883861 | 0.378922229 | 0.812043903 | Washc5 | WASH complex subunit 5 [Source:MGI Syr | 491.5 | 468.73 | 448.79 | 448.25 | 570.39 | 516.82 |
| 3625 | ENSMUSG00000022351 | 295.5734324 | 1.618367364 | 0.262019109 | 6.176524187 | 6.55E-10 | 1.44E-07 | Sqle | squalene epoxidase [Source:MGI Symbol; | 172.67 | 177.83 | 84.525 | 350.89 | 387.17 | 600.36 |
| 3626 | ENSMUSG00000022353 | 4203.290138 | 0.347440871 | 0.096915739 | 3.584978827 | 0.000337106 | 0.009541868 | Mtss1 | MTSS I-BAR domain containing 1 [Source: | 4037.2 | 3636.8 | 3423.2 | 4702.3 | 4790 | 4630.2 |
| 3627 | ENSMUSG00000022354 | 1550.124249 | 0.057070396 | 0.101141963 | 0.56426032 | 0.572576982 | 0.905173631 | Ndufb9 | NADH:ubiquinone oxidoreductase subuni | 1519.5 | 1458.6 | 1579.8 | 1514.4 | 1544.3 | 1684.1 |
| 3628 | ENSMUSG00000022357 | 0.517184577 | -0.812115922 | 3.393570547 | -0.239310163 | 0.810865085 | NA | Klhl38 | kelch-like 38 [Source:MGI Symbol;Acc:MG | 0 | 0 | 2.0125 | 0 | 1.0906 | 0 |
| 3629 | ENSMUSG00000022358 | 77.45207434 | -0.25786566 | 0.259071095 | -0.995347088 | 0.319567483 | 0.76641089 | Fbxo32 | F-box protein 32 [Source:MGI Symbol;Acc | 72.279 | 95.596 | 85.531 | 71.334 | 69.8 | 70.172 |
| 3630 | ENSMUSG00000022359 | 96.17138524 | -0.031578009 | 0.233103014 | -0.135468042 | 0.89224183 | 0.982489666 | Wdyhv1 | WDYHV motif containing 1 [Source:MGI S | 96.372 | 98.68 | 96.599 | 94.47 | 83.978 | 106.93 |
| 3631 | ENSMUSG00000022360 | 67.83009505 | -0.192991439 | 0.302196152 | -0.638629704 | 0.523063853 | 0.888334265 | Atad2 | ATPase family, AAA domain containing 2 [ | 65.051 | 70.926 | 81.506 | 75.19 | 71.981 | 42.326 |
| 3632 | ENSMUSG00000022361 | 734.7412473 | 0.066655525 | 0.123862952 | 0.538139325 | 0.590480865 | 0.911690353 | Zhx1 | zinc fingers and homeoboxes 1 [Source:M | 679.42 | 744.21 | 729.53 | 667.07 | 790.7 | 797.51 |
| 3633 | ENSMUSG00000022362 | 0.371281295 | 2.09253362 | 4.031852165 | 0.519000582 | 0.603760335 | NA | Gm29394 | predicted gene 29394 [Source:MGI Symbo | 0 | 0 | 0 | 0 | 0 | 2.2277 |
| 3634 | ENSMUSG00000022364 | 172.0046427 | -0.2517137 | 0.249329137 | -1.009563915 | 0.312704265 | 0.759921531 | Tbc1d31 | TBC1 domain family, member 31 [Source: | 262.61 | 158.3 | 138.86 | 135.92 | 159.23 | 177.1 |
| 3635 | ENSMUSG00000022365 | 2609.37201 | 0.063731322 | 0.099475622 | 0.640672762 | 0.521735314 | 0.888132458 | Derl1 | Der1-like domain family, member 1 [Sourc | 2717.7 | 2551.3 | 2384.8 | 2543.9 | 2636 | 2822.5 |
| 3636 | ENSMUSG00000022366 | 0.167707431 | -0.796629176 | 4.080472857 | -0.195229623 | 0.84521319 | NA | Slc22a22 | solute carrier family 22 (organic cation tra | 0 | 0 | 1.0062 | 0 | 0 | 0 |

| | | | | | | | | | | | | | | | |
|---|---|---|---|---|---|---|---|---|---|---|---|---|---|---|---|
| 3637 | ENSMUSG00000022367 | 0 | NA | NA | NA | NA | NA | Has2 | hyaluronan synthase 2 [Source:MGI Symb | 0 | 0 | 0 | 0 | 0 | 0 |
| 3638 | ENSMUSG00000022369 | 37.1202281 | 0.321830356 | 0.365331171 | 0.88092772 | 0.378356943 | 0.811764373 | Mtbp | Mdm2, transformed 3T3 cell double minu | 37.746 | 34.949 | 26.162 | 42.415 | 45.806 | 35.643 |
| 3639 | ENSMUSG00000022370 | 748.6349644 | 0.042299908 | 0.123535543 | 0.342410829 | 0.73204174 | 0.947494562 | Mrpl13 | mitochondrial ribosomal protein L13 [Sou | 761.34 | 704.12 | 746.63 | 705.63 | 721.99 | 852.09 |
| 3640 | ENSMUSG00000022371 | 657.6137728 | -1.177924757 | 0.17343007 | -6.791929216 | 1.11E-11 | 3.72E-09 | Col14a1 | collagen, type XIV, alpha 1 [Source:MGI Sy | 1118.7 | 923.07 | 694.31 | 452.11 | 363.18 | 394.3 |
| 3641 | ENSMUSG00000022372 | 22.72807903 | -2.522326398 | 0.582060924 | -4.333440531 | 1.47E-05 | 0.000726202 | Sla | src-like adaptor [Source:MGI Symbol;Acc: | 39.352 | 55.508 | 21.131 | 3.8559 | 8.7249 | 7.7969 |
| 3642 | ENSMUSG00000022375 | 0.17131964 | -0.796629176 | 4.080472857 | -0.195229623 | 0.84521319 | NA | Lrrc6 | leucine rich repeat containing 6 (testis) [S | 0 | 1.0279 | 0 | 0 | 0 | 0 |
| 3643 | ENSMUSG00000022376 | 0.472877444 | -2.283655516 | 3.093270275 | -0.738265755 | 0.460352972 | NA | Adcy8 | adenylate cyclase 8 [Source:MGI Symbol;/ | 0.8031 | 1.0279 | 1.0062 | 0 | 0 | 0 |
| 3644 | ENSMUSG00000022377 | 237.1371147 | -0.213022013 | 0.203578474 | -1.046387713 | 0.295382063 | 0.745982248 | Asap1 | ArfGAP with SH3 domain, ankyrin repeat a | 271.45 | 278.57 | 214.33 | 280.52 | 191.95 | 186.01 |
| 3645 | ENSMUSG00000022378 | 116.8251625 | -0.120153934 | 0.213205527 | -0.56355919 | 0.573054167 | 0.905173631 | Cyrib | CYFIP related Rac1 interactor B [Source:M | 110.03 | 132.6 | 122.76 | 111.82 | 111.24 | 112.5 |
| 3646 | ENSMUSG00000022382 | 2.121580398 | 1.198849402 | 1.622652669 | 0.738820713 | 0.460015873 | NA | Wnt7b | wingless-type MMTV integration site fami | 0.8031 | 2.0558 | 1.0062 | 0 | 2.1812 | 6.6831 |
| 3647 | ENSMUSG00000022383 | 2824.839655 | -0.154611555 | 0.106701238 | -1.449013695 | 0.147333758 | 0.558402858 | Ppara | peroxisome proliferator activated recepto | 2800.4 | 3047.8 | 3080.1 | 2459.1 | 2971.9 | 2589.7 |
| 3648 | ENSMUSG00000022385 | 18.34446326 | 0.262219416 | 0.557627379 | 0.470241286 | 0.638182641 | 0.923699483 | Gtse1 | G two S phase expressed protein 1 [Sourc | 27.305 | 10.279 | 12.075 | 16.388 | 25.084 | 18.935 |
| 3649 | ENSMUSG00000022386 | 141.962709 | 0.031248922 | 0.194023234 | 0.161057626 | 0.872048014 | 0.979369708 | Trmu | tRNA 5-methylaminomethyl-2-thiouridyla | 143.76 | 132.6 | 144.9 | 149.42 | 137.42 | 143.69 |
| 3650 | ENSMUSG00000022387 | 515.2527174 | 0.146240448 | 0.136144518 | 1.074155981 | 0.282752778 | 0.735558641 | Brd1 | bromodomain containing 1 [Source:MGI S | 491.5 | 505.74 | 470.92 | 613.09 | 522.41 | 487.86 |
| 3651 | ENSMUSG00000022388 | 21.020649 | -1.116672541 | 0.592936234 | -1.8832928 | 0.059660699 | 0.347816342 | Ttll8 | tubulin tyrosine ligase-like family, membe | 17.668 | 30.838 | 38.237 | 25.063 | 7.6343 | 6.6831 |
| 3652 | ENSMUSG00000022389 | 4394.501068 | 0.376804979 | 0.104651781 | 3.600559646 | 0.000317533 | 0.009130809 | Tef | thyrotroph embryonic factor [Source:MGI | 4228.3 | 3521.6 | 3720.1 | 5011.7 | 5237.1 | 4648.1 |
| 3653 | ENSMUSG00000022390 | 908.5516248 | 0.00391322 | 0.124321806 | 0.031476537 | 0.974889504 | 0.996336532 | Zc3h7b | zinc finger CCCH type containing 7B [Sour | 922.76 | 906.62 | 891.53 | 779.86 | 984.83 | 965.7 |
| 3654 | ENSMUSG00000022391 | 1065.319916 | 0.337945195 | 0.112787266 | 2.996306293 | 0.002732718 | 0.043917814 | Rangap1 | RAN GTPase activating protein 1 [Source:I | 1032 | 858.31 | 931.78 | 1195.3 | 1212.8 | 1161.7 |
| 3655 | ENSMUSG00000022394 | 355.2948122 | -0.107272101 | 0.139583666 | -0.768514719 | 0.44218145 | 0.850193681 | L3mbtl2 | L3MBTL2 polycomb repressive complex 1 | 368.62 | 387.53 | 349.17 | 329.68 | 337 | 359.77 |
| 3656 | ENSMUSG00000022400 | 754.8439924 | 0.115988052 | 0.113457481 | 1.022304136 | 0.306636976 | 0.755610724 | Rbx1 | ring-box 1 [Source:MGI Symbol;Acc:MGI:1 | 728.41 | 751.41 | 693.3 | 741.3 | 804.88 | 809.76 |
| 3657 | ENSMUSG00000022401 | 449.0699396 | 0.082940365 | 0.1494816 | 0.554853344 | 0.578994977 | 0.908337213 | Xpnpep3 | X-prolyl aminopeptidase 3, mitochondrial | 380.67 | 448.17 | 480.98 | 508.98 | 464.6 | 411.01 |
| 3658 | ENSMUSG00000022403 | 3099.564284 | -0.005284802 | 0.089940128 | -0.058759108 | 0.95314398 | 0.9915542 | St13 | suppression of tumorigenicity 13 [Source: | 2989.1 | 3047.8 | 3279.4 | 3073.2 | 3114.8 | 3093.1 |
| 3659 | ENSMUSG00000022404 | 1082.619339 | 0.235398964 | 0.106140864 | 2.217797691 | 0.02656863 | 0.210915839 | Slc25a17 | solute carrier family 25 (mitochondrial car | 980.59 | 1010.4 | 993.16 | 1237.7 | 1183.3 | 1090.5 |
| 3660 | ENSMUSG00000022407 | 285.6450922 | -0.051723685 | 0.169864989 | -0.304498797 | 0.760747909 | 0.95402891 | Adsl | adenylosuccinate lyase [Source:MGI Symb | 289.12 | 284.73 | 297.85 | 227.5 | 321.73 | 292.94 |
| 3661 | ENSMUSG00000022408 | 27.98116435 | -0.47768975 | 0.449823372 | -1.061949601 | 0.288258566 | 0.73951397 | Fam83f | family with sequence similarity 83, memb | 43.368 | 28.782 | 25.156 | 15.424 | 25.084 | 30.074 |
| 3662 | ENSMUSG00000022412 | 558.6382742 | 0.240898827 | 0.132835476 | 1.813512727 | 0.069752777 | 0.381582516 | Mief1 | mitochondrial elongation factor 1 [Source | 492.3 | 547.88 | 496.08 | 558.14 | 580.21 | 677.22 |
| 3663 | ENSMUSG00000022414 | 112.4093667 | 0.357685422 | 0.222577177 | 1.607017516 | 0.108050526 | 0.484761602 | Tab1 | TGF-beta activated kinase 1/MAP3K7 binc | 114.04 | 88.401 | 92.575 | 120.5 | 131.96 | 126.98 |
| 3664 | ENSMUSG00000022415 | 3.947961954 | -0.312342056 | 1.089057179 | -0.28680042 | 0.774265141 | NA | Syngr1 | synaptogyrin 1 [Source:MGI Symbol;Acc:N | 4.0155 | 2.0558 | 7.0437 | 2.8919 | 5.4531 | 2.2277 |
| 3665 | ENSMUSG00000022416 | 0.99233012 | 0.312367548 | 2.428448675 | 0.128628433 | 0.897651667 | NA | Cacna1i | calcium channel, voltage-dependent, alph | 1.6062 | 0 | 1.0062 | 0 | 0 | 3.3415 |
| 3666 | ENSMUSG00000022419 | 4379.940589 | 0.133645272 | 0.106168568 | 1.258802625 | 0.208101632 | 0.653455953 | Deptor | DEP domain containing MTOR-interacting | 3873.4 | 4124 | 4535.1 | 4477.7 | 5019 | 4250.4 |
| 3667 | ENSMUSG00000022420 | 171.2220315 | 0.204945107 | 0.199022045 | 1.029760831 | 0.303122292 | 0.752484788 | Dnal4 | dynein, axonemal, light chain 4 [Source:M | 155.8 | 146.99 | 174.08 | 152.31 | 185.41 | 212.74 |
| 3668 | ENSMUSG00000022421 | 6.642584098 | -0.089739171 | 0.914842979 | -0.09809243 | 0.921858899 | 0.986685804 | Nptxr | neuronal pentraxin receptor [Source:MGI | 6.4248 | 7.1954 | 7.0437 | 12.532 | 1.0906 | 5.5692 |
| 3669 | ENSMUSG00000022422 | 0.910352643 | 1.221794025 | 2.770215377 | 0.441046583 | 0.659179274 | NA | Dscc1 | DNA replication and sister chromatid cohe | 1.6062 | 0 | 0 | 3.8559 | 0 | 0 |
| 3670 | ENSMUSG00000022425 | 2159.75955 | 0.003286439 | 0.163460692 | 0.020105376 | 0.983959312 | 0.998580735 | Enpp2 | ectonucleotide pyrophosphatase/phosph | 2114.6 | 2418.7 | 1938 | 1636.8 | 2736.4 | 2114.1 |
| 3671 | ENSMUSG00000022426 | 1074.843179 | 0.021631895 | 0.108576821 | 0.199231244 | 0.84208186 | 0.971854408 | Josd1 | Josephin domain containing 1 [Source:MG | 1130.8 | 1025.9 | 1043.5 | 1152 | 1063.4 | 1033.6 |
| 3672 | ENSMUSG00000022427 | 852.4615048 | -0.004793847 | 0.115458075 | -0.041520244 | 0.966881154 | 0.994231936 | Tomm22 | translocase of outer mitochondrial memb | 832.01 | 887.09 | 842.23 | 775.04 | 908.49 | 869.91 |
| 3673 | ENSMUSG00000022428 | 149.3432852 | 0.298562531 | 0.203260648 | 1.468865389 | 0.141869304 | 0.551315856 | Cby1 | chibby family member 1, beta catenin ant | 129.3 | 126.43 | 145.91 | 143.63 | 160.32 | 190.47 |
| 3674 | ENSMUSG00000022429 | 0.160662974 | 1.126924175 | 4.080472857 | 0.276174898 | 0.782413736 | NA | Dmc1 | DNA meiotic recombinase 1 [Source:MGI | 0 | 0 | 0 | 0.964 | 0 | 0 |
| 3675 | ENSMUSG00000022431 | 0 | NA | NA | NA | NA | NA | Ribc2 | RIB43A domain with coiled-coils 2 [Source | 0 | 0 | 0 | 0 | 0 | 0 |
| 3676 | ENSMUSG00000022432 | 1.654253829 | 0.876139048 | 1.739479631 | 0.50367882 | 0.614487094 | NA | Smc1b | structural maintenance of chromosomes : | 2.4093 | 0 | 1.0062 | 0.964 | 1.0906 | 4.4554 |
| 3677 | ENSMUSG00000022433 | 97.15497473 | -0.188566728 | 0.233531743 | -0.807456518 | 0.419403516 | 0.837591673 | Csnk1e | casein kinase 1, epsilon [Source:MGI Syml | 110.03 | 105.88 | 94.587 | 97.362 | 94.884 | 80.197 |
| 3678 | ENSMUSG00000022434 | 41.56145144 | 0.399966568 | 0.39953791 | 1.001072883 | 0.316791574 | 0.763467884 | Fam118a | family with sequence similarity 118, meml | 44.171 | 22.614 | 40.25 | 29.883 | 54.531 | 57.92 |
| 3679 | ENSMUSG00000022435 | 0 | NA | NA | NA | NA | NA | Upk3a | uroplakin 3A [Source:MGI Symbol;Acc:MG | 0 | 0 | 0 | 0 | 0 | 0 |
| 3680 | ENSMUSG00000022436 | 6.7914685 | -0.944202851 | 0.855141564 | -1.104148004 | 0.269528941 | 0.725011151 | Sh3bp1 | SH3-domain binding protein 1 [Source:MG | 6.4248 | 11.307 | 9.0562 | 2.8919 | 3.2719 | 7.7969 |
| 3681 | ENSMUSG00000022437 | 1625.927966 | -0.069279684 | 0.096946667 | -0.714616461 | 0.47484607 | 0.869368495 | Samm50 | SAMM50 sorting and assembly machinery | 1657.6 | 1747.5 | 1589.9 | 1568.4 | 1622.8 | 1569.4 |
| 3682 | ENSMUSG00000022438 | 29.61907468 | -0.946607972 | 0.471283287 | -2.008575306 | 0.044582192 | 0.291866583 | Parvb | parvin, beta [Source:MGI Symbol;Acc:MG | 54.611 | 40.089 | 22.137 | 22.171 | 13.087 | 25.618 |
| 3683 | ENSMUSG00000022439 | 14.84202632 | -1.443511241 | 0.659935928 | -2.187350588 | 0.02871694 | 0.222237467 | Parvg | parvin, gamma [Source:MGI Symbol;Acc:N | 15.259 | 34.949 | 15.094 | 11.568 | 3.2719 | 8.9108 |
| 3684 | ENSMUSG00000022440 | 15.66774003 | -0.553800627 | 0.691259454 | -0.801147275 | 0.423046391 | 0.84005258 | C1qtnf6 | C1q and tumor necrosis factor related pro | 16.062 | 21.586 | 18.112 | 3.8559 | 6.5437 | 27.846 |
| 3685 | ENSMUSG00000022441 | 0 | NA | NA | NA | NA | NA | Efcab6 | EF-hand calcium binding domain 6 [Sourc | 0 | 0 | 0 | 0 | 0 | 0 |
| 3686 | ENSMUSG00000022442 | 37.3267531 | 0.096458037 | 0.361996508 | 0.266461236 | 0.789884009 | 0.961019028 | Ttll1 | tubulin tyrosine ligase-like 1 [Source:MGI | 36.14 | 29.81 | 42.262 | 37.595 | 43.625 | 34.529 |
| 3687 | ENSMUSG00000022443 | 1914.367505 | -0.055269268 | 0.112565961 | -0.490994501 | 0.623430337 | 0.919503423 | Myh9 | myosin, heavy polypeptide 9, non-muscle | 1974 | 2002.4 | 1877.7 | 2130.4 | 1786.4 | 1715.3 |
| 3688 | ENSMUSG00000022445 | 21769.78772 | -0.186740781 | 0.112793577 | -1.655597654 | 0.097803326 | 0.460940795 | Cyp2d26 | cytochrome P450, family 2, subfamily d, p | 21525 | 22150 | 25857 | 22267 | 20801 | 18019 |
| 3689 | ENSMUSG00000022449 | 0 | NA | NA | NA | NA | NA | Adamts20 | a disintegrin-like and metallopeptidase (re | 0 | 0 | 0 | 0 | 0 | 0 |
| 3690 | ENSMUSG00000022450 | 2642.38491 | 0.102833302 | 0.09662692 | 1.064230355 | 0.287224359 | 0.73951397 | Ndufa6 | NADH:ubiquinone oxidoreductase subuni | 2525.8 | 2607.8 | 2510.6 | 2540.1 | 2744 | 2926.1 |
| 3691 | ENSMUSG00000022451 | 876.1608074 | -0.016319645 | 0.10595804 | -0.154019883 | 0.877594058 | 0.98072613 | Twf1 | twinfilin actin binding protein 1 [Source:N | 898.67 | 879.9 | 864.36 | 852.16 | 899.76 | 862.12 |
| 3692 | ENSMUSG00000022452 | 1138.314559 | 0.041289897 | 0.106251021 | 0.388607056 | 0.697566846 | 0.940422865 | Smdt1 | single-pass membrane protein with aspar | 1196.6 | 1091.6 | 1076.7 | 1129.8 | 1125.5 | 1209.6 |

| | | | | | | | | | | | | | | | |
|---|---|---|---|---|---|---|---|---|---|---|---|---|---|---|---|
| 3693 | ENSMUSG00000022453 | 479.9768865 | -0.044375602 | 0.132355647 | -0.335275471 | 0.737417295 | 0.948137969 | Naga | N-acetyl galactosaminidase, alpha [Source | 485.88 | 495.46 | 480.98 | 496.45 | 420.98 | 500.12 |
| 3694 | ENSMUSG00000022454 | 0 | NA | NA | NA | NA | NA | Nell2 | NEL-like 2 [Source:MGI Symbol;Acc:MGI:1 | 0 | 0 | 0 | 0 | 0 | 0 |
| 3695 | ENSMUSG00000022455 | 0 | NA | NA | NA | NA | NA | Wbp2nl | WBP2 N-terminal like [Source:MGI Symbo | 0 | 0 | 0 | 0 | 0 | 0 |
| 3696 | ENSMUSG00000022456 | 0.469265235 | -2.274672822 | 3.439794491 | -0.661281605 | 0.508431736 | NA | Septin3 | septin 3 [Source:MGI Symbol;Acc:MGI:13 | 0.8031 | 0 | 2.0125 | 0 | 0 | 0 |
| 3697 | ENSMUSG00000022462 | 2333.850784 | -0.318601341 | 0.189020486 | -1.685538686 | 0.091884688 | 0.446623824 | Slc38a2 | solute carrier family 38, member 2 [Sourc | 2500.1 | 3442.5 | 1829.4 | 2111.1 | 2407 | 1713.1 |
| 3698 | ENSMUSG00000022463 | 1018.970205 | -0.309080243 | 0.134257432 | -2.302146241 | 0.021326926 | 0.181441752 | Srebf2 | sterol regulatory element binding factor 2 | 970.15 | 1275.6 | 1138.1 | 843.48 | 1011 | 875.48 |
| 3699 | ENSMUSG00000022464 | 22178.64542 | 0.236336477 | 0.09709376 | 2.434105711 | 0.014928634 | 0.142198341 | Slc38a4 | solute carrier family 38, member 4 [Sourc | 18915 | 21066 | 21117 | 21906 | 25023 | 25044 |
| 3700 | ENSMUSG00000022466 | 151.8641295 | 0.055214553 | 0.188869966 | 0.292341628 | 0.770025433 | 0.956513901 | Rpap3 | RNA polymerase II associated protein 3 [S | 146.97 | 144.94 | 154.96 | 150.38 | 164.68 | 149.26 |
| 3701 | ENSMUSG00000022468 | 3.433489674 | 0.565657979 | 1.173407734 | 0.482064301 | 0.629760269 | NA | Endou | endonuclease, polyU-specific [Source:MG | 3.2124 | 4.1117 | 1.0062 | 6.7478 | 2.1812 | 3.3415 |
| 3702 | ENSMUSG00000022469 | 38.32408701 | -0.383946796 | 0.360171191 | -1.066011956 | 0.286418237 | 0.738722073 | Rapgef3 | Rap guanine nucleotide exchange factor ( | 50.595 | 44.2 | 35.219 | 32.775 | 37.081 | 30.074 |
| 3703 | ENSMUSG00000022471 | 346.3157798 | 0.026706093 | 0.146782306 | 0.181943547 | 0.855627029 | 0.974158771 | Xrcc6 | X-ray repair complementing defective rep | 349.35 | 308.38 | 371.3 | 338.36 | 371.9 | 338.61 |
| 3704 | ENSMUSG00000022472 | 628.6750425 | 0.285627205 | 0.124535092 | 2.293547952 | 0.021816478 | 0.183889008 | Desi1 | desumoylating isopeptidase 1 [Source:MG | 526.84 | 543.77 | 629.91 | 680.57 | 690.36 | 700.61 |
| 3705 | ENSMUSG00000022474 | 115.1137501 | -0.171625466 | 0.239431291 | -0.716804663 | 0.473494634 | 0.868952133 | Pmm1 | phosphomannomutase 1 [Source:MGI Syr | 137.33 | 103.82 | 124.77 | 132.06 | 104.7 | 87.994 |
| 3706 | ENSMUSG00000022475 | 85.34135295 | -0.580873532 | 0.275116843 | -2.111370306 | 0.034740495 | 0.249868983 | Hdac7 | histone deacetylase 7 [Source:MGI Symbo | 86.735 | 118.21 | 102.64 | 87.722 | 62.165 | 54.578 |
| 3707 | ENSMUSG00000022476 | 190.7290589 | 0.333211084 | 0.195510076 | 1.704316686 | 0.088321939 | 0.438246494 | Polr3h | polymerase (RNA) III (DNA directed) polyp | 166.24 | 173.72 | 166.03 | 166.77 | 239.94 | 231.68 |
| 3708 | ENSMUSG00000022477 | 4248.235824 | -0.370544278 | 0.12850968 | -2.883395857 | 0.003934127 | 0.056579946 | Aco2 | aconitase 2, mitochondrial [Source:MGI S | 4758.4 | 5335.9 | 4278.6 | 3761.4 | 4217.4 | 3137.7 |
| 3709 | ENSMUSG00000022479 | 0.167707431 | -0.796629176 | 4.080472857 | -0.195229623 | 0.84521319 | NA | Vdr | vitamin D (1,25-dihydroxyvitamin D3) rece | 0 | 0 | 1.0062 | 0 | 0 | 0 |
| 3710 | ENSMUSG00000022483 | 0 | NA | NA | NA | NA | NA | Col2a1 | collagen, type II, alpha 1 [Source:MGI Sym | 0 | 0 | 0 | 0 | 0 | 0 |
| 3711 | ENSMUSG00000022484 | 0 | NA | NA | NA | NA | NA | Hoxc10 | homeobox C10 [Source:MGI Symbol;Acc:I | 0 | 0 | 0 | 0 | 0 | 0 |
| 3712 | ENSMUSG00000022485 | 0.824628202 | 0.545567773 | 2.864022412 | 0.19049005 | 0.848925139 | NA | Hoxc5 | homeobox C5 [Source:MGI Symbol;Acc:M | 0 | 2.0558 | 0 | 2.8919 | 0 | 0 |
| 3713 | ENSMUSG00000022487 | 0 | NA | NA | NA | NA | NA | Gtsf1 | gametocyte specific factor 1 [Source:MGI | 0 | 0 | 0 | 0 | 0 | 0 |
| 3714 | ENSMUSG00000022488 | 57.97356761 | -1.518142633 | 0.398915139 | -3.805678159 | 0.000141416 | 0.004734823 | Nckap1l | NCK associated protein 1 like [Source:MG | 85.932 | 116.15 | 55.343 | 16.388 | 28.356 | 45.668 |
| 3715 | ENSMUSG00000022489 | 19.90505833 | -0.34693703 | 0.557356789 | -0.622468474 | 0.533633868 | 0.889143097 | Pde1b | phosphodiesterase 1B, Ca2+-calmodulin c | 19.274 | 25.698 | 22.137 | 27.955 | 6.5437 | 17.822 |
| 3716 | ENSMUSG00000022490 | 0.167707431 | -0.796629176 | 4.080472857 | -0.195229623 | 0.84521319 | NA | Ppp1r1a | protein phosphatase 1, regulatory inhibitc | 0 | 0 | 1.0062 | 0 | 0 | 0 |
| 3717 | ENSMUSG00000022491 | 0 | NA | NA | NA | NA | NA | Glycam1 | glycosylation dependent cell adhesion mo | 0 | 0 | 0 | 0 | 0 | 0 |
| 3718 | ENSMUSG00000022494 | 6.001347906 | -0.756533398 | 0.899922355 | -0.84066519 | 0.400535527 | 0.827266295 | Shisa9 | shisa family member 9 [Source:MGI Symb | 10.44 | 5.1396 | 7.0437 | 6.7478 | 2.1812 | 4.4554 |
| 3719 | ENSMUSG00000022496 | 0.17131964 | -0.796629176 | 4.080472857 | -0.195229623 | 0.84521319 | NA | Tnfrsf17 | tumor necrosis factor receptor superfamil | 0 | 1.0279 | 0 | 0 | 0 | 0 |
| 3720 | ENSMUSG00000022498 | 518.1084524 | 0.02960383 | 0.139332017 | 0.212469688 | 0.831740623 | 0.970533397 | Txndc11 | thioredoxin domain containing 11 [Source | 513.18 | 458.45 | 566.52 | 501.27 | 578.03 | 491.21 |
| 3721 | ENSMUSG00000022500 | 1369.263805 | -0.233445516 | 0.111552499 | -2.092696423 | 0.036376263 | 0.257267344 | Litaf | LPS-induced TN factor [Source:MGI Symb | 1638.3 | 1450.4 | 1349.4 | 1230 | 1303.3 | 1244.2 |
| 3722 | ENSMUSG00000022501 | 0 | NA | NA | NA | NA | NA | Prm1 | protamine 1 [Source:MGI Symbol;Acc:MG | 0 | 0 | 0 | 0 | 0 | 0 |
| 3723 | ENSMUSG00000022503 | 243.743376 | 0.118615367 | 0.164536277 | 0.720907078 | 0.470966688 | 0.867447112 | Nubp1 | nucleotide binding protein 1 [Source:MGI | 249.76 | 245.67 | 205.27 | 243.89 | 265.02 | 252.84 |
| 3724 | ENSMUSG00000022504 | 23.15786915 | -0.810143336 | 0.455163855 | -1.779893827 | 0.075093338 | 0.400645534 | Ciita | class II transactivator [Source:MGI Symbo | 30.518 | 27.754 | 30.187 | 17.352 | 13.087 | 20.049 |
| 3725 | ENSMUSG00000022505 | 127.3070061 | 0.345039073 | 0.264425567 | 1.30486275 | 0.191939587 | 0.629099068 | Emp2 | epithelial membrane protein 2 [Source:M | 72.279 | 114.1 | 150.94 | 147.49 | 129.78 | 149.26 |
| 3726 | ENSMUSG00000022507 | 617.748204 | 0.465630444 | 0.150911176 | 3.085460314 | 0.002032373 | 0.035417168 | 1810013L24 | RIKEN cDNA 1810013L24 gene [Source:M | 569.4 | 481.07 | 505.13 | 658.4 | 630.38 | 862.12 |
| 3727 | ENSMUSG00000022508 | 1030.271074 | -0.593320242 | 0.462422157 | -1.28307053 | 0.199467367 | 0.641417519 | Bcl6 | B cell leukemia/lymphoma 6 [Source:MGI | 1634.3 | 1190.3 | 892.54 | 458.85 | 1223.7 | 781.92 |
| 3728 | ENSMUSG00000022510 | 0.904673413 | -3.227537609 | 2.678727401 | -1.204877214 | 0.228250706 | NA | Trp63 | transformation related protein 63 [Source | 2.4093 | 0 | 3.0187 | 0 | 0 | 0 |
| 3729 | ENSMUSG00000022512 | 822.7888915 | -0.084413395 | 0.220818852 | -0.382274405 | 0.702257838 | 0.942345172 | Cldn1 | claudin 1 [Source:MGI Symbol;Acc:MGI:1 | 561.37 | 793.55 | 1186.4 | 742.26 | 668.55 | 984.64 |
| 3730 | ENSMUSG00000022514 | 3699.174889 | 0.107819492 | 0.119982069 | 0.898630043 | 0.368849751 | 0.806506847 | Il1rap | interleukin 1 receptor accessory protein [ | 3023.7 | 3632.7 | 4028 | 3655.4 | 4109.4 | 3745.9 |
| 3731 | ENSMUSG00000022515 | 246.876794 | 0.318313213 | 0.182760165 | 1.741699091 | 0.081561112 | 0.420121054 | Anks3 | ankyrin repeat and sterile alpha motif dor | 246.55 | 189.14 | 223.39 | 322.93 | 249.75 | 249.5 |
| 3732 | ENSMUSG00000022516 | 187.3218351 | 0.134086144 | 0.217991165 | 0.615098983 | 0.538489353 | 0.891577636 | Nudt16l1 | nudix (nucleoside diphosphate linked moi | 198.37 | 176.8 | 159.99 | 154.24 | 255.2 | 179.33 |
| 3733 | ENSMUSG00000022517 | 1248.481441 | 0.086859131 | 0.103625275 | 0.838204102 | 0.401916084 | 0.82893099 | Mgrn1 | mahogunin, ring finger 1 [Source:MGI Syn | 1172.5 | 1158.5 | 1302.1 | 1279.2 | 1304.4 | 1274.2 |
| 3734 | ENSMUSG00000022518 | 1.150724995 | -1.212126241 | 2.023379121 | -0.599060368 | 0.549132629 | NA | 4930562C15 | RIKEN cDNA 4930562C15 gene [Source:M | 0.8031 | 1.0279 | 3.0187 | 0.964 | 1.0906 | 0 |
| 3735 | ENSMUSG00000022519 | 8.36843835 | -0.348323766 | 0.788887926 | -0.441537707 | 0.65882377 | 0.927599242 | Srl | sarcalumenin [Source:MGI Symbol;Acc:M | 12.85 | 3.0838 | 12.075 | 6.7478 | 6.5437 | 8.9108 |
| 3736 | ENSMUSG00000022521 | 360.528382 | -0.117615786 | 0.149608824 | -0.786155411 | 0.431776445 | 0.84514901 | Crebbp | CREB binding protein [Source:MGI Symbo | 352.56 | 428.64 | 345.14 | 363.42 | 335.91 | 337.49 |
| 3737 | ENSMUSG00000022523 | 5.234499538 | -1.049363309 | 1.087577613 | -0.964862918 | 0.334613477 | 0.776159726 | Fgf12 | fibroblast growth factor 12 [Source:MGI S | 12.047 | 3.0838 | 6.0375 | 5.7839 | 0 | 4.4554 |
| 3738 | ENSMUSG00000022525 | 0 | NA | NA | NA | NA | NA | Plaat1 | phospholipase A and acyltransferase 1 [So | 0 | 0 | 0 | 0 | 0 | 0 |
| 3739 | ENSMUSG00000022526 | 28.68324071 | -0.212287885 | 0.414130461 | -0.512611133 | 0.608223364 | 0.916376001 | Zfp251 | zinc finger protein 251 [Source:MGI Symb | 30.518 | 23.642 | 38.237 | 27.955 | 28.356 | 23.391 |
| 3740 | ENSMUSG00000022528 | 160.6581609 | -0.483641435 | 0.203038115 | -2.382022881 | 0.017217828 | 0.156332651 | Hes1 | hes family bHLH transcription factor 1 [So | 210.41 | 155.22 | 196.22 | 146.52 | 133.06 | 122.52 |
| 3741 | ENSMUSG00000022529 | 235.4248111 | 0.179379876 | 0.178134285 | 1.006992424 | 0.313938415 | 0.760476509 | Zfp263 | zinc finger protein 263 [Source:MGI Symb | 200.78 | 226.14 | 236.47 | 293.05 | 235.57 | 220.54 |
| 3742 | ENSMUSG00000022533 | 3017.00366 | 0.050749618 | 0.117324267 | 0.432558575 | 0.665335489 | 0.929426323 | Atp13a3 | ATPase type 13A3 [Source:MGI Symbol;Ac | 2529.8 | 3322.2 | 3041.9 | 3232.2 | 3026.5 | 2949.5 |
| 3743 | ENSMUSG00000022534 | 3.783705345 | 0.189087247 | 1.244075889 | 0.151990123 | 0.879194726 | NA | Mefv | Mediterranean fever [Source:MGI Symbol | 2.4093 | 6.1675 | 2.0125 | 0 | 6.5437 | 5.5692 |
| 3744 | ENSMUSG00000022536 | 1574.824766 | 0.061927906 | 0.101363093 | 0.610951225 | 0.541231871 | 0.893614211 | Glyr1 | glyoxylate reductase 1 homolog (Arabidop | 1490.6 | 1580.9 | 1551.6 | 1515.4 | 1721 | 1589.5 |
| 3745 | ENSMUSG00000022537 | 30.14919989 | 0.318767278 | 0.405262834 | 0.786569236 | 0.431534074 | 0.844934876 | Tmem44 | transmembrane protein 44 [Source:MGI S | 25.699 | 27.754 | 27.169 | 39.523 | 23.994 | 36.757 |
| 3746 | ENSMUSG00000022538 | 337.3715694 | 0.012116205 | 0.145485756 | 0.083281039 | 0.933628076 | 0.988420305 | Lsg1 | large 60S subunit nuclear export GTPase 1 | 363 | 304.26 | 340.11 | 348 | 340.27 | 328.58 |
| 3747 | ENSMUSG00000022540 | 281.0005839 | -0.16926668 | 0.216066693 | -0.78340015 | 0.433392168 | 0.845264432 | Rogdi | rogdi homolog [Source:MGI Symbol;Acc:N | 330.88 | 238.48 | 322 | 183.16 | 314.1 | 297.4 |
| 3748 | ENSMUSG00000022542 | 0 | NA | NA | NA | NA | NA | Septin12 | septin 12 [Source:MGI Symbol;Acc:MGI:1 | 0 | 0 | 0 | 0 | 0 | 0 |

| | | | | | | | | | | | | | | | |
|---|---|---|---|---|---|---|---|---|---|---|---|---|---|---|---|
| 3749 | ENSMUSG00000022543 | 5.865481757 | -0.272337238 | 0.896245835 | -0.303864439 | 0.761231171 | 0.95402891 | 4930451G09 | RIKEN cDNA 4930451G09 gene [Source:M | 4.0155 | 9.2513 | 6.0375 | 4.8199 | 3.2719 | 7.7969 |
| 3750 | ENSMUSG00000022544 | 152.8348615 | -0.346036832 | 0.188073437 | -1.839902737 | 0.065782518 | 0.368337165 | Eef2kmt | eukaryotic elongation factor 2 lysine meth | 169.45 | 177.83 | 166.03 | 136.88 | 128.69 | 138.12 |
| 3751 | ENSMUSG00000022545 | 143.8492512 | -0.293271582 | 0.235144182 | -1.247198969 | 0.212324551 | 0.656933207 | Ercc4 | excision repair cross-complementing rode | 149.38 | 163.44 | 163.01 | 161.95 | 89.431 | 135.89 |
| 3752 | ENSMUSG00000022546 | 7726.3571 | -0.129543572 | 0.106897947 | -1.211843403 | 0.22557233 | 0.672301295 | Gpt | glutamic pyruvic transaminase, soluble [Sc | 7605.4 | 7854.3 | 8759.4 | 6769.1 | 8206.9 | 7163.1 |
| 3753 | ENSMUSG00000022548 | 0.133850373 | -0.796629176 | 4.080472857 | -0.195229623 | 0.84521319 | NA | Apod | apolipoprotein D [Source:MGI Symbol;Acc | 0.8031 | 0 | 0 | 0 | 0 | 0 |
| 3754 | ENSMUSG00000022550 | 581.5440998 | 0.18679322 | 0.170659516 | 1.09453738 | 0.273719363 | 0.728923893 | Adck5 | aarF domain containing kinase 5 [Source:I | 469.81 | 459.48 | 703.36 | 660.32 | 574.76 | 621.52 |
| 3755 | ENSMUSG00000022551 | 4654.6984 | -0.032350195 | 0.08519417 | -0.379723109 | 0.704150964 | 0.94295036 | Cyc1 | cytochrome c-1 [Source:MGI Symbol;Acc: | 4572.1 | 4751 | 4797.8 | 4497.9 | 4555.5 | 4753.9 |
| 3756 | ENSMUSG00000022552 | 495.7197419 | 0.149105365 | 0.12357318 | 1.206615909 | 0.227580097 | 0.673700979 | Sharpin | SHANK-associated RH domain interacting | 469.81 | 454.34 | 486.02 | 505.12 | 534.4 | 524.62 |
| 3757 | ENSMUSG00000022553 | 417.2235719 | 0.189951804 | 0.134087524 | 1.416625488 | 0.156592452 | 0.575737985 | Maf1 | MAF1 homolog, negative regulator of RN/ | 402.35 | 388.55 | 378.35 | 461.75 | 408.98 | 463.36 |
| 3758 | ENSMUSG00000022554 | 110.1415465 | 0.084530396 | 0.22967475 | 0.368043924 | 0.712840482 | 0.944239215 | Hgh1 | HGH1 homolog [Source:MGI Symbol;Acc:I | 103.6 | 100.74 | 116.72 | 126.28 | 121.06 | 92.449 |
| 3759 | ENSMUSG00000022555 | 431.1176267 | 0.049278132 | 0.130316955 | 0.378140606 | 0.705326145 | 0.94295036 | Dgat1 | diacylglycerol O-acyltransferase 1 [Source | 401.55 | 428.64 | 441.74 | 454.03 | 430.79 | 429.94 |
| 3760 | ENSMUSG00000022556 | 275.3475122 | 0.129753688 | 0.15792766 | 0.821602046 | 0.411303424 | 0.833705705 | Hsf1 | heat shock factor 1 [Source:MGI Symbol;/ | 254.58 | 292.96 | 241.5 | 274.73 | 304.28 | 284.03 |
| 3761 | ENSMUSG00000022557 | 404.7140132 | 0.171240375 | 0.14972215 | 1.143721055 | 0.252739339 | 0.705781321 | Bop1 | block of proliferation 1 [Source:MGI Symb | 344.53 | 371.08 | 427.65 | 475.24 | 417.71 | 392.07 |
| 3762 | ENSMUSG00000022558 | 672.4066204 | -0.079972519 | 0.129313341 | -0.618439818 | 0.536285457 | 0.890560262 | Mroh1 | maestro heat-like repeat family member 1 | 741.26 | 730.85 | 600.73 | 686.35 | 632.56 | 642.69 |
| 3763 | ENSMUSG00000022559 | 331.4283068 | 0.174613545 | 0.151352514 | 1.153687774 | 0.248628198 | 0.700459341 | Fbxl6 | F-box and leucine-rich repeat protein 6 [S | 275.46 | 316.6 | 343.13 | 371.13 | 339.18 | 343.06 |
| 3764 | ENSMUSG00000022560 | 211.6610277 | -0.425454325 | 0.184387798 | -2.307388722 | 0.021033159 | 0.17948296 | Slc52a2 | solute carrier protein 52, member 2 [Sour | 259.4 | 233.34 | 235.46 | 208.22 | 185.41 | 148.14 |
| 3765 | ENSMUSG00000022561 | 356.7501865 | 0.061140509 | 0.140730686 | 0.434450447 | 0.663961372 | 0.929301301 | Gpaa1 | GPI anchor attachment protein 1 [Source: | 339.71 | 369.02 | 339.1 | 369.2 | 339.18 | 384.28 |
| 3766 | ENSMUSG00000022562 | 3301.23783 | -0.029994315 | 0.128297154 | -0.233787846 | 0.815149685 | 0.967689186 | Oplah | 5-oxoprolinase (ATP-hydrolysing) [Source: | 2952.2 | 3316.1 | 3740.2 | 3694 | 3294.8 | 2810.2 |
| 3767 | ENSMUSG00000022564 | 2473.972239 | -0.008376917 | 0.102100304 | -0.082045952 | 0.934610172 | 0.988420305 | Grina | glutamate receptor, ionotropic, N-methyl | 2573.9 | 2240.9 | 2628.3 | 2537.2 | 2487.7 | 2375.8 |
| 3768 | ENSMUSG00000022565 | 1322.341841 | 0.221442893 | 0.113798017 | 1.945929282 | 0.051663235 | 0.319340611 | Plec | plectin [Source:MGI Symbol;Acc:MGI:127 | 1182.2 | 1177 | 1305.1 | 1578 | 1325.1 | 1366.7 |
| 3769 | ENSMUSG00000022568 | 484.7657862 | 0.116068649 | 0.171173335 | 0.678076694 | 0.497723067 | 0.879037876 | Scrib | scribbled planar cell polarity [Source:MGI | 401.55 | 451.26 | 544.38 | 614.05 | 437.34 | 460.02 |
| 3770 | ENSMUSG00000022570 | 498.2612803 | 0.126908605 | 0.137731918 | 0.921417534 | 0.356832478 | 0.796166767 | Gfus | GDP-L-fucose synthase [Source:MGI Symb | 451.34 | 460.51 | 517.21 | 483.92 | 494.05 | 582.54 |
| 3771 | ENSMUSG00000022571 | 612.7417194 | 0.304120403 | 0.127082626 | 2.393091909 | 0.016707055 | 0.153234527 | Pycrl | pyrroline-5-carboxylate reductase-like [So | 534.06 | 502.65 | 608.78 | 696.96 | 645.65 | 688.36 |
| 3772 | ENSMUSG00000022574 | 2342.570856 | 0.076570561 | 0.107731214 | 0.71075558 | 0.477235711 | 0.869688987 | Naprt | nicotinate phosphoribosyltransferase [Sou | 2244.7 | 2144.2 | 2453.2 | 2638.4 | 2233.6 | 2341.3 |
| 3773 | ENSMUSG00000022575 | 623.4114076 | 0.10092249 | 0.14019501 | 0.719872193 | 0.47160369 | 0.867794597 | Gsdmd | gasdermin D [Source:MGI Symbol;Acc:MG | 518.8 | 583.86 | 703.36 | 668.04 | 629.29 | 637.12 |
| 3774 | ENSMUSG00000022577 | 0.133850373 | -0.796629176 | 4.080472857 | -0.195229623 | 0.84521319 | NA | Ly6h | lymphocyte antigen 6 complex, locus H [S | 0.8031 | 0 | 0 | 0 | 0 | 0 |
| 3775 | ENSMUSG00000022579 | 189.8198298 | -0.506603148 | 0.225297315 | -2.248598246 | 0.024538068 | 0.200179705 | Gpihbp1 | GPI-anchored HDL-binding protein 1 [Sour | 216.03 | 232.31 | 220.37 | 178.34 | 102.52 | 189.35 |
| 3776 | ENSMUSG00000022580 | 1.461077907 | -0.762576142 | 1.788015023 | -0.42649314 | 0.669748545 | NA | Rhpn1 | rhophilin, Rho GTPase binding protein 1 [S | 2.4093 | 2.0558 | 1.0062 | 0 | 2.1812 | 1.1138 |
| 3777 | ENSMUSG00000022582 | 0 | NA | NA | NA | NA | NA | Ly6g | lymphocyte antigen 6 complex, locus G [S | 0 | 0 | 0 | 0 | 0 | 0 |
| 3778 | ENSMUSG00000022583 | 0.658052777 | 1.6393978 | 2.811180186 | 0.583170658 | 0.559778421 | NA | Ly6f | lymphocyte antigen 6 complex, locus F [Sc | 0.8031 | 0 | 0 | 0.964 | 2.1812 | 0 |
| 3779 | ENSMUSG00000022584 | 5.453247701 | -1.562071149 | 0.961825151 | -1.624069767 | 0.10436092 | 0.475725288 | Ly6c2 | lymphocyte antigen 6 complex, locus C2 [ | 11.243 | 6.1675 | 7.0437 | 3.8559 | 2.1812 | 2.2277 |
| 3780 | ENSMUSG00000022586 | 0.342639279 | -1.805976596 | 4.028547714 | -0.448294702 | 0.653940524 | NA | Ly6i | lymphocyte antigen 6 complex, locus I [So | 0 | 2.0558 | 0 | 0 | 0 | 0 |
| 3781 | ENSMUSG00000022587 | 2445.396622 | 0.091406506 | 0.142317865 | 0.642270077 | 0.52069784 | 0.887819587 | Ly6e | lymphocyte antigen 6 complex, locus E [Sc | 2745.8 | 2156.6 | 2199.7 | 2247 | 2289.2 | 3034.1 |
| 3782 | ENSMUSG00000022589 | 0 | NA | NA | NA | NA | NA | Cyp11b2 | cytochrome P450, family 11, subfamily b, | 0 | 0 | 0 | 0 | 0 | 0 |
| 3783 | ENSMUSG00000022591 | 0 | NA | NA | NA | NA | NA | Gm9747 | predicted gene 9747 [Source:MGI Symbol | 0 | 0 | 0 | 0 | 0 | 0 |
| 3784 | ENSMUSG00000022594 | 19.19193521 | -0.615287081 | 0.507723904 | -1.211853678 | 0.225568396 | 0.672301295 | Lynx1 | Ly6/neurotoxin 1 [Source:MGI Symbol;Acc | 25.699 | 27.754 | 16.1 | 12.532 | 16.359 | 16.708 |
| 3785 | ENSMUSG00000022595 | 0 | NA | NA | NA | NA | NA | Lypd2 | Ly6/Plaur domain containing 2 [Source:M | 0 | 0 | 0 | 0 | 0 | 0 |
| 3786 | ENSMUSG00000022596 | 0 | NA | NA | NA | NA | NA | Slurp1 | secreted Ly6/Plaur domain containing 1 [S | 0 | 0 | 0 | 0 | 0 | 0 |
| 3787 | ENSMUSG00000022598 | 0 | NA | NA | NA | NA | NA | Psca | prostate stem cell antigen [Source:MGI Sy | 0 | 0 | 0 | 0 | 0 | 0 |
| 3788 | ENSMUSG00000022601 | 202.9189574 | 0.227360596 | 0.181983841 | 1.249344969 | 0.211538927 | 0.6563061 | Zbtb11 | zinc finger and BTB domain containing 11 | 191.94 | 201.47 | 167.04 | 189.9 | 239.94 | 227.22 |
| 3789 | ENSMUSG00000022602 | 1.477544172 | -3.939977216 | 2.089232266 | -1.885849305 | 0.059315265 | NA | Arc | activity regulated cytoskeletal-associated | 4.8186 | 1.0279 | 3.0187 | 0 | 0 | 0 |
| 3790 | ENSMUSG00000022603 | 0 | NA | NA | NA | NA | NA | Mroh4 | maestro heat-like repeat family member 4 | 0 | 0 | 0 | 0 | 0 | 0 |
| 3791 | ENSMUSG00000022604 | 71.65093495 | 0.073132424 | 0.316000927 | 0.231431042 | 0.816979952 | 0.967910123 | Cep97 | centrosomal protein 97 [Source:MGI Syml | 49.792 | 63.731 | 96.599 | 89.65 | 61.075 | 69.058 |
| 3792 | ENSMUSG00000022607 | 362.8312875 | -0.105161017 | 0.149878223 | -0.701643074 | 0.482901782 | 0.871710696 | Ptk2 | PTK2 protein tyrosine kinase 2 [Source:M | 379.06 | 418.36 | 331.05 | 373.06 | 344.64 | 330.81 |
| 3793 | ENSMUSG00000022610 | 47.97378784 | -0.167603398 | 0.332376301 | -0.504257967 | 0.614080111 | 0.917378096 | Mapk12 | mitogen-activated protein kinase 12 [Sour | 49.792 | 41.117 | 61.381 | 47.235 | 37.081 | 51.237 |
| 3794 | ENSMUSG00000022613 | 0.68303045 | 1.69421863 | 2.781340055 | 0.609137537 | 0.542433279 | NA | Miox | myo-inositol oxygenase [Source:MGI Sym | 0.8031 | 0 | 0 | 0 | 2.1812 | 1.1138 |
| 3795 | ENSMUSG00000022614 | 422.424528 | -0.238371722 | 0.147014059 | -1.621421274 | 0.104927321 | 0.47755468 | Lmf2 | lipase maturation factor 2 [Source:MGI Sy | 427.25 | 427.61 | 517.21 | 391.38 | 351.18 | 419.92 |
| 3796 | ENSMUSG00000022615 | 1036.243588 | -0.476278092 | 0.141718411 | -3.36073547 | 0.000777352 | 0.0173275 | Tymp | thymidine phosphorylase [Source:MGI Sy | 1130.8 | 1052.6 | 1433.9 | 832.88 | 800.51 | 966.82 |
| 3797 | ENSMUSG00000022617 | 12.46336838 | 0.150078841 | 0.701002927 | 0.214091604 | 0.830475623 | 0.970274589 | Chkb | choline kinase beta [Source:MGI Symbol;/ | 11.243 | 6.1675 | 18.112 | 17.352 | 17.45 | 4.4554 |
| 3798 | ENSMUSG00000022619 | 0.321325949 | 1.911575114 | 4.048267893 | 0.472195804 | 0.636787034 | NA | Mapk8ip2 | mitogen-activated protein kinase 8 intera | 0 | 0 | 0 | 1.928 | 0 | 0 |
| 3799 | ENSMUSG00000022620 | 1043.980062 | -0.027175419 | 0.136122862 | -0.199638901 | 0.841763002 | 0.971854408 | Arsa | arylsulfatase A [Source:MGI Symbol;Acc:M | 1248 | 1013.5 | 898.58 | 1066.2 | 976.1 | 1061.5 |
| 3800 | ENSMUSG00000022621 | 33.91533218 | -0.263988617 | 0.401645547 | -0.65726763 | 0.511008847 | 0.883679955 | Rabl2 | RAB, member RAS oncogene family-like 2 | 40.958 | 24.67 | 45.281 | 29.883 | 37.081 | 25.618 |
| 3801 | ENSMUSG00000022622 | 0.531944269 | 2.618916564 | 3.285123409 | 0.797204926 | 0.425332026 | NA | Acr | acrosin prepropeptide [Source:MGI Symb | 0 | 0 | 0 | 0.964 | 0 | 2.2277 |
| 3802 | ENSMUSG00000022623 | 68.11472634 | -0.660025245 | 0.318079339 | -2.07503338 | 0.037983441 | 0.26463993 | Shank3 | SH3 and multiple ankyrin repeat domains | 73.885 | 92.513 | 84.525 | 76.154 | 38.172 | 43.44 |
| 3803 | ENSMUSG00000022629 | 961.6455975 | 0.146342683 | 0.115800342 | 1.263750008 | 0.20631977 | 0.65157405 | Kif21a | kinesin family member 21A [Source:MGI S | 978.98 | 926.15 | 833.17 | 1068.1 | 952.11 | 1011.4 |
| 3804 | ENSMUSG00000022634 | 467.3658007 | -0.028995807 | 0.14081027 | -0.205921112 | 0.836852534 | 0.971073158 | Yaf2 | YY1 associated factor 2 [Source:MGI Symb | 476.24 | 466.67 | 473.94 | 529.22 | 449.33 | 408.78 |

| | | | | | | | | | | | | | | | |
|---|---|---|---|---|---|---|---|---|---|---|---|---|---|---|---|
| 3805 | ENSMUSG00000022635 | 469.5289319 | 0.064446265 | 0.133392453 | 0.483132769 | 0.629001468 | 0.92057725 | Zcrb1 | zinc finger CCHC-type and RNA binding mo | 482.66 | 459.48 | 434.7 | 498.38 | 435.16 | 506.8 |
| 3806 | ENSMUSG00000022636 | 788.3388034 | 0.241952983 | 0.126448091 | 1.91345699 | 0.055689575 | 0.332986361 | Alcam | activated leukocyte cell adhesion molecul | 689.86 | 757.58 | 719.46 | 744.19 | 887.76 | 931.17 |
| 3807 | ENSMUSG00000022637 | 131.270315 | -0.246395152 | 0.228955133 | -1.076172211 | 0.281850245 | 0.734802082 | Cblb | Casitas B-lineage lymphoma b [Source:MG | 146.16 | 128.49 | 152.95 | 143.63 | 90.521 | 125.86 |
| 3808 | ENSMUSG00000022639 | 11.98207604 | -0.982753251 | 0.625218565 | -1.571855518 | 0.115984063 | 0.500730933 | Dubr | Dppa2 upstream binding RNA [Source:MG | 15.259 | 14.391 | 18.112 | 8.6758 | 6.5437 | 8.9108 |
| 3809 | ENSMUSG00000022641 | 247.6502825 | -0.024950834 | 0.165535417 | -0.150728068 | 0.880190231 | 0.981359269 | Bbx | bobby sox HMG box containing [Source:M | 224.87 | 261.09 | 263.64 | 221.71 | 261.75 | 252.84 |
| 3810 | ENSMUSG00000022650 | 0 | NA | NA | NA | NA | NA | Retnlb | resistin like beta [Source:MGI Symbol;Acc | 0 | 0 | 0 | 0 | 0 | 0 |
| 3811 | ENSMUSG00000022651 | 0.569258551 | -2.569337407 | 3.18860282 | -0.805787849 | 0.420365185 | NA | Retnlg | resistin like gamma [Source:MGI Symbol;A | 2.4093 | 0 | 1.0062 | 0 | 0 | 0 |
| 3812 | ENSMUSG00000022652 | 0 | NA | NA | NA | NA | NA | Morc1 | microrchidia 1 [Source:MGI Symbol;Acc:M | 0 | 0 | 0 | 0 | 0 | 0 |
| 3813 | ENSMUSG00000022656 | 998.7768453 | 0.12447985 | 0.12692748 | 0.980716316 | 0.326732656 | 0.772451967 | Nectin3 | nectin cell adhesion molecule 3 [Source:M | 820.77 | 956.99 | 1090.8 | 1055.6 | 1038.3 | 1030.3 |
| 3814 | ENSMUSG00000022657 | 0.472877444 | -2.283655516 | 3.093270275 | -0.738265755 | 0.460352972 | NA | Cd96 | CD96 antigen [Source:MGI Symbol;Acc:M | 0.8031 | 1.0279 | 1.0062 | 0 | 0 | 0 |
| 3815 | ENSMUSG00000022658 | 2.424507815 | 2.050475978 | 1.436815582 | 1.427097537 | 0.153551782 | NA | Tagln3 | transgelin 3 [Source:MGI Symbol;Acc:MGI | 0.8031 | 1.0279 | 1.0062 | 2.8919 | 4.3625 | 4.4554 |
| 3816 | ENSMUSG00000022659 | 2.3318748 | -0.778796839 | 1.498249418 | -0.519804533 | 0.60319982 | NA | Gcsam | germinal center associated, signaling and | 0.8031 | 3.0838 | 5.0312 | 2.8919 | 2.1812 | 0 |
| 3817 | ENSMUSG00000022661 | 12.45929885 | -0.890984982 | 0.699182012 | -1.274324807 | 0.202548329 | 0.647482287 | Cd200 | CD200 antigen [Source:MGI Symbol;Acc:M | 7.2279 | 12.335 | 29.181 | 10.604 | 8.7249 | 6.6831 |
| 3818 | ENSMUSG00000022663 | 1677.206823 | -0.040874769 | 0.099536543 | -0.410650879 | 0.681328551 | 0.934452508 | Atg3 | autophagy related 3 [Source:MGI Symbol; | 1727.5 | 1632.3 | 1742.8 | 1639.7 | 1753.7 | 1567.2 |
| 3819 | ENSMUSG00000022664 | 392.9408079 | 0.120142977 | 0.160670236 | 0.747761251 | 0.454604179 | 0.857337687 | Slc35a5 | solute carrier family 35, member A5 [Sour | 452.95 | 355.66 | 319.99 | 416.44 | 389.35 | 423.26 |
| 3820 | ENSMUSG00000022665 | 95.61419485 | -1.139440224 | 0.285510591 | -3.990886009 | 6.58E-05 | 0.002587284 | Ccdc80 | coiled-coil domain containing 80 [Source:I | 178.29 | 123.35 | 92.575 | 62.659 | 58.893 | 57.92 |
| 3821 | ENSMUSG00000022667 | 7.093304262 | -1.576142194 | 0.954828059 | -1.650707873 | 0.098798239 | 0.463365025 | Cd200r1 | CD200 receptor 1 [Source:MGI Symbol;Ac | 14.456 | 9.2513 | 8.05 | 0.964 | 8.7249 | 1.1138 |
| 3822 | ENSMUSG00000022668 | 325.3478758 | 0.089313002 | 0.161377354 | 0.553441977 | 0.579960801 | 0.90863876 | Gtpbp8 | GTP-binding protein 8 (putative) [Source:I | 303.57 | 346.41 | 295.84 | 301.73 | 390.44 | 314.1 |
| 3823 | ENSMUSG00000022671 | 413.3617662 | 0.165263533 | 0.141889207 | 1.164736464 | 0.244125684 | 0.693992511 | Mzt2 | mitotic spindle organizing protein 2 [Sour | 392.72 | 380.33 | 395.45 | 381.74 | 476.6 | 453.33 |
| 3824 | ENSMUSG00000022672 | 168.610369 | -0.275985498 | 0.189119598 | -1.459317285 | 0.1444778 | 0.55464443 | Prkdc | protein kinase, DNA activated, catalytic po | 179.09 | 179.89 | 195.21 | 150.38 | 172.32 | 134.78 |
| 3825 | ENSMUSG00000022673 | 87.43673507 | -0.45689548 | 0.281641351 | -1.622259938 | 0.104747703 | 0.476887243 | Mcm4 | minichromosome maintenance complex c | 118.86 | 85.317 | 98.612 | 51.091 | 89.431 | 81.311 |
| 3826 | ENSMUSG00000022674 | 236.2955517 | -0.080040062 | 0.162645563 | -0.492113408 | 0.622639175 | 0.918870832 | Ube2v2 | ubiquitin-conjugating enzyme E2 variant 2 | 236.11 | 250.81 | 241.5 | 210.15 | 249.75 | 229.45 |
| 3827 | ENSMUSG00000022676 | 99.15900025 | -0.646398297 | 0.28954926 | -2.23242946 | 0.025586593 | 0.206060398 | Snai2 | snail family zinc finger 2 [Source:MGI Sym | 123.68 | 132.6 | 106.66 | 68.442 | 112.33 | 51.237 |
| 3828 | ENSMUSG00000022677 | 274.3879911 | -0.033462144 | 0.167077317 | -0.200279395 | 0.841262077 | 0.971854408 | Cep20 | centrosomal protein 20 [Source:MGI Syml | 305.98 | 261.09 | 264.64 | 235.21 | 304.28 | 275.12 |
| 3829 | ENSMUSG00000022678 | 358.1129959 | 0.031496094 | 0.169620597 | 0.185685551 | 0.85269136 | 0.973781977 | Nde1 | nudE neurodevelopment protein 1 [Sourc | 345.33 | 296.04 | 421.62 | 401.98 | 322.82 | 360.89 |
| 3830 | ENSMUSG00000022679 | 594.5626602 | 0.459315712 | 0.1919078 | 2.393418673 | 0.016692181 | 0.153234527 | Mpv17l | Mpv17 transgene, kidney disease mutant- | 473.03 | 497.51 | 531.3 | 500.3 | 928.12 | 637.12 |
| 3831 | ENSMUSG00000022680 | 1338.211459 | 0.152026115 | 0.098739521 | 1.539668338 | 0.123641218 | 0.515225175 | Pdxdc1 | pyridoxal-dependent decarboxylase doma | 1234.4 | 1293.1 | 1275.9 | 1364 | 1394.9 | 1466.9 |
| 3832 | ENSMUSG00000022681 | 241.8648784 | 0.017065782 | 0.165804802 | 0.102926947 | 0.91802095 | 0.986374671 | Ntan1 | N-terminal Asn amidase [Source:MGI Sym | 220.05 | 253.9 | 247.54 | 224.61 | 236.66 | 268.44 |
| 3833 | ENSMUSG00000022682 | 324.4470348 | 0.180634435 | 0.16316256 | 1.10708262 | 0.268258209 | 0.724039098 | Rrn3 | RRN3 RNA polymerase I transcription fact | 322.85 | 333.05 | 256.59 | 370.17 | 357.72 | 306.31 |
| 3834 | ENSMUSG00000022683 | 0.160662974 | 1.126924175 | 4.080472857 | 0.276174898 | 0.782413736 | NA | Pla2g10 | phospholipase A2, group X [Source:MGI S | 0 | 0 | 0 | 0.964 | 0 | 0 |
| 3835 | ENSMUSG00000022684 | 1428.714207 | 0.15020724 | 0.113564564 | 1.322659415 | 0.185948666 | 0.621903163 | Bfar | bifunctional apoptosis regulator [Source:M | 1483.3 | 1348.6 | 1229.6 | 1393.9 | 1557.4 | 1559.4 |
| 3836 | ENSMUSG00000022685 | 369.3739968 | 0.145838561 | 0.169820323 | 0.858781555 | 0.390461046 | 0.819763149 | Parn | poly(A)-specific ribonuclease (deadenylati | 358.99 | 392.66 | 299.86 | 319.08 | 406.8 | 438.85 |
| 3837 | ENSMUSG00000022686 | 2.093383468 | -1.702435634 | 1.717923104 | -0.990984771 | 0.321693017 | NA | B3gnt5 | UDP-GlcNAc:betaGal beta-1,3-N-acetylglu | 5.6217 | 1.0279 | 3.0187 | 2.8919 | 0 | 0 |
| 3838 | ENSMUSG00000022687 | 5.477262074 | -1.518850553 | 1.123274235 | -1.352163617 | 0.17632298 | 0.607400048 | Boc | biregional cell adhesion molecule-related/ | 16.062 | 6.1675 | 2.0125 | 0.964 | 6.5437 | 1.1138 |
| 3839 | ENSMUSG00000022696 | 0.706721132 | -2.881789029 | 2.959064625 | -0.973885127 | 0.330113579 | NA | Sidt1 | SID1 transmembrane family, member 1 [S | 3.2124 | 1.0279 | 0 | 0 | 0 | 0 |
| 3840 | ENSMUSG00000022698 | 1853.252908 | 0.0942805 | 0.105982269 | 0.88958749 | 0.373687426 | 0.80865857 | Naa50 | N(alpha)-acetyltransferase 50, NatE cataly | 1811.8 | 1885.2 | 1680.4 | 1753.5 | 2042.7 | 1945.9 |
| 3841 | ENSMUSG00000022701 | 92.78708606 | 0.063759411 | 0.272432813 | 0.234037194 | 0.814956104 | 0.967618214 | Ccdc191 | coiled-coil domain containing 191 [Source | 101.99 | 93.541 | 76.475 | 93.506 | 122.15 | 69.058 |
| 3842 | ENSMUSG00000022702 | 83.61153897 | -0.109506668 | 0.267287723 | -0.409695839 | 0.682029082 | 0.934933546 | Hira | histone cell cycle regulator [Source:MGI S | 89.144 | 105.88 | 65.406 | 85.794 | 75.253 | 80.197 |
| 3843 | ENSMUSG00000022704 | 141.206989 | 0.267492453 | 0.199173089 | 1.343015032 | 0.179267107 | 0.611423712 | Qtrt2 | queuine tRNA-ribosyltransferase accessor | 142.15 | 115.13 | 126.79 | 159.06 | 154.87 | 149.26 |
| 3844 | ENSMUSG00000022705 | 0 | NA | NA | NA | NA | NA | Drd3 | dopamine receptor D3 [Source:MGI Symb | 0 | 0 | 0 | 0 | 0 | 0 |
| 3845 | ENSMUSG00000022706 | 341.5793569 | 0.056470723 | 0.151800237 | 0.372006817 | 0.709887774 | 0.944239215 | Mrpl40 | mitochondrial ribosomal protein L40 [Sou | 370.23 | 305.29 | 328.04 | 315.22 | 365.36 | 365.34 |
| 3846 | ENSMUSG00000022707 | 3301.257214 | -0.252924439 | 0.106622772 | -2.372142777 | 0.017685255 | 0.158586872 | Gbe1 | glucan (1,4-alpha-), branching enzyme 1 [S | 3655.7 | 3776.6 | 3337.7 | 3120.4 | 3223.9 | 2693.3 |
| 3847 | ENSMUSG00000022708 | 2152.676522 | -0.18302275 | 0.12381606 | -1.478182637 | 0.1393589 | 0.545633379 | Zbtb20 | zinc finger and BTB domain containing 20 | 2164.4 | 2456.7 | 2245.9 | 1785.3 | 2355.7 | 1908 |
| 3848 | ENSMUSG00000022710 | 1236.091934 | 0.023157679 | 0.105787204 | 0.218908134 | 0.82672161 | 0.969648181 | Usp7 | ubiquitin specific peptidase 7 [Source:MG | 1136.4 | 1259.2 | 1284 | 1261.8 | 1194.2 | 1280.9 |
| 3849 | ENSMUSG00000022711 | 936.6677719 | 0.083306203 | 0.106612849 | 0.781389894 | 0.434573213 | 0.845264432 | Pmm2 | phosphomannomutase 2 [Source:MGI Syr | 872.97 | 946.71 | 909.65 | 951.45 | 940.11 | 999.12 |
| 3850 | ENSMUSG00000022715 | 0 | NA | NA | NA | NA | NA | Tmem114 | transmembrane protein 114 [Source:MGI | 0 | 0 | 0 | 0 | 0 | 0 |
| 3851 | ENSMUSG00000022718 | 119.0614643 | 0.2325268 | 0.231432196 | 1.004729693 | 0.315027026 | 0.761215823 | Dgcr8 | DGCR8, microprocessor complex subunit | | 89.947 | 133.63 | 105.66 | 142.67 | 121.06 | 121.41 |
| 3852 | ENSMUSG00000022721 | 301.4678252 | 0.123881059 | 0.15081241 | 0.821424833 | 0.411404323 | 0.833738738 | Trmt2a | TRM2 tRNA methyltransferase 2A [Source | 290.72 | 277.54 | 296.84 | 283.41 | 323.91 | 336.38 |
| 3853 | ENSMUSG00000022722 | 49.35865518 | 0.254297853 | 0.319373535 | 0.796239591 | 0.42589279 | 0.841545535 | Arl6 | ADP-ribosylation factor-like 6 [Source:MG | 41.761 | 38.033 | 55.343 | 53.019 | 54.531 | 53.465 |
| 3854 | ENSMUSG00000022723 | 233.7070054 | -0.437311345 | 0.277486662 | -1.575972487 | 0.115032144 | 0.498085671 | Crybg3 | beta-gamma crystallin domain containing | 214.43 | 316.6 | 276.72 | 310.4 | 153.78 | 130.32 |
| 3855 | ENSMUSG00000022724 | 203.6790679 | 0.093644126 | 0.179247744 | 0.522428476 | 0.601372034 | 0.915461118 | Riox2 | ribosomal oxygenase 2 [Source:MGI Symb | 187.12 | 211.75 | 192.19 | 181.23 | 219.21 | 230.57 |
| 3856 | ENSMUSG00000022738 | 0.342639279 | -1.805976596 | 4.028547714 | -0.448294702 | 0.653940524 | NA | Gsc2 | goosecoid homebox 2 [Source:MGI Symbo | 0 | 2.0558 | 0 | 0 | 0 | 0 |
| 3857 | ENSMUSG00000022742 | 4586.044802 | -0.306486762 | 0.125769695 | -2.436888807 | 0.014814234 | 0.141621335 | Cpox | coproporphyrinogen oxidase [Source:MGI | 4313.5 | 5329.8 | 5572.6 | 4340.8 | 4386.5 | 3573.2 |
| 3858 | ENSMUSG00000022744 | 360.3361614 | -0.148190001 | 0.15105231 | -0.981050878 | 0.326567652 | 0.772276061 | Cldnd1 | claudin domain containing 1 [Source:MGI | 352.56 | 402.94 | 381.37 | 343.18 | 299.92 | 382.05 |
| 3859 | ENSMUSG00000022747 | 42.71365852 | -0.63282481 | 0.414002968 | -1.528551383 | 0.126375696 | 0.521167236 | St3gal6 | ST3 beta-galactoside alpha-2,3-sialyltrans | 73.082 | 54.48 | 28.175 | 44.343 | 28.356 | 27.846 |
| 3860 | ENSMUSG00000022748 | 70.07089673 | 0.03695435 | 0.267682914 | 0.138052703 | 0.890198765 | 0.982489666 | Cmss1 | cms small ribosomal subunit 1 [Source:MG | 68.264 | 63.731 | 75.468 | 68.442 | 65.437 | 79.083 |

| | | | | | | | | | | | | | | | |
|---|---|---|---|---|---|---|---|---|---|---|---|---|---|---|---|
| 3861 | ENSMUSG00000022749 | 181.1515484 | -0.12380909 | 0.189872142 | -0.652065588 | 0.514358862 | 0.885150099 | Tbc1d23 | TBC1 domain family, member 23 [Source:I | 195.15 | 210.72 | 161 | 184.12 | 178.86 | 157.05 |
| 3862 | ENSMUSG00000022750 | 342.6023269 | 0.123290931 | 0.15296411 | 0.806012152 | 0.420235842 | 0.837906635 | Klhl22 | kelch-like 22 [Source:MGI Symbol;Acc:MG | 338.11 | 303.24 | 342.12 | 346.07 | 320.64 | 405.44 |
| 3863 | ENSMUSG00000022751 | 2781.248167 | 0.351956058 | 0.106220808 | 3.313437972 | 0.000921565 | 0.019682389 | Nit2 | nitrilase family, member 2 [Source:MGI Sy | 2564.3 | 2316.9 | 2448.2 | 2780.1 | 3314.4 | 3263.6 |
| 3864 | ENSMUSG00000022752 | 2739.76783 | 0.140939379 | 0.096111448 | 1.46641614 | 0.142534952 | 0.552122535 | Tomm70a | translocase of outer mitochondrial memb | 2582 | 2762 | 2474.4 | 2837 | 3024.3 | 2759 |
| 3865 | ENSMUSG00000022753 | 0 | NA | NA | NA | NA | NA | Tmem30c | transmembrane protein 30C [Source:MGI | 0 | 0 | 0 | 0 | 0 | 0 |
| 3866 | ENSMUSG00000022754 | 4.157911929 | -0.497418808 | 1.062075146 | -0.468346152 | 0.639537069 | NA | Tmem45a | transmembrane protein 45a [Source:MGI | 6.4248 | 6.1675 | 2.0125 | 4.8199 | 2.1812 | 3.3415 |
| 3867 | ENSMUSG00000022755 | 0.267700746 | -1.497481649 | 4.057001436 | -0.369110456 | 0.712045396 | NA | Adgrg7 | adhesion G protein-coupled receptor G7 [ | 1.6062 | 0 | 0 | 0 | 0 | 0 |
| 3868 | ENSMUSG00000022756 | 11.33051559 | -0.468570958 | 0.69818963 | -0.671122769 | 0.502142323 | 0.88121652 | Slc7a4 | solute carrier family 7 (cationic amino acid | 12.85 | 21.586 | 5.0312 | 8.6758 | 9.8156 | 10.025 |
| 3869 | ENSMUSG00000022757 | 753.3029023 | 0.18925236 | 0.1148593 | 1.647688613 | 0.099416586 | 0.464607916 | Tfg | Trk-fused gene [Source:MGI Symbol;Acc:N | 746.08 | 703.1 | 662.11 | 807.81 | 831.05 | 769.67 |
| 3870 | ENSMUSG00000022758 | 1.29933506 | 0.184946416 | 1.850499756 | 0.099944037 | 0.920388755 | NA | P2rx6 | purinergic receptor P2X, ligand-gated ion | 1.6062 | 1.0279 | 1.0062 | 1.928 | 0 | 2.2277 |
| 3871 | ENSMUSG00000022759 | 0.51014012 | 1.031855138 | 3.09002894 | 0.333930574 | 0.738431948 | NA | Lrrc74b | leucine rich repeat containing 74B [Source | 0 | 0 | 1.0062 | 0.964 | 1.0906 | 0 |
| 3872 | ENSMUSG00000022760 | 302.9805855 | 0.134985072 | 0.151173163 | 0.892916896 | 0.371901678 | 0.807685567 | Thap7 | THAP domain containing 7 [Source:MGI Sy | 263.42 | 290.9 | 312.94 | 333.54 | 313.01 | 304.08 |
| 3873 | ENSMUSG00000022761 | 750.5257593 | 0.041856156 | 0.123497773 | 0.338922356 | 0.734668222 | 0.948109711 | Lztr1 | leucine-zipper-like transcriptional regulato | 809.53 | 672.26 | 736.57 | 814.56 | 733.99 | 736.25 |
| 3874 | ENSMUSG00000022762 | 66.46411804 | 0.069136719 | 0.278998256 | 0.24780341 | 0.804286516 | 0.963701061 | Ncam2 | neural cell adhesion molecule 2 [Source:N | 63.445 | 57.563 | 73.456 | 58.803 | 70.89 | 74.628 |
| 3875 | ENSMUSG00000022763 | 108.1833585 | 0.197609764 | 0.255011273 | 0.774905992 | 0.438395211 | 0.847566655 | Aifm3 | apoptosis-inducing factor, mitochondrion | 82.72 | 93.541 | 126.79 | 136.88 | 115.61 | 93.563 |
| 3876 | ENSMUSG00000022765 | 538.8971003 | 0.251209792 | 0.132335995 | 1.898272584 | 0.057660184 | 0.341802006 | Snap29 | synaptosomal-associated protein 29 [Sour | 538.88 | 461.54 | 474.95 | 589.95 | 539.86 | 628.21 |
| 3877 | ENSMUSG00000022766 | 8136.263065 | 0.221989164 | 0.087163587 | 2.546810782 | 0.010871236 | 0.115286132 | Serpind1 | serine (or cysteine) peptidase inhibitor, cl | 7105 | 7610.7 | 7819.5 | 8465.7 | 8805.7 | 9011 |
| 3878 | ENSMUSG00000022768 | 0.822793274 | -0.447303882 | 2.407293551 | -0.185811939 | 0.852592242 | NA | Ccdc116 | coiled-coil domain containing 116 [Source | 0.8031 | 2.0558 | 0 | 0.964 | 0 | 1.1138 |
| 3879 | ENSMUSG00000022769 | 330.8034402 | 0.742729175 | 0.252307398 | 2.943747115 | 0.003242648 | 0.049318261 | Sdf2l1 | stromal cell-derived factor 2-like 1 [Source | 319.63 | 167.55 | 254.58 | 508.02 | 266.11 | 468.93 |
| 3880 | ENSMUSG00000022770 | 969.9324662 | 0.158589569 | 0.115166361 | 1.377047663 | 0.168497512 | 0.595692028 | Dlg1 | discs large MAGUK scaffold protein 1 [Sou | 905.1 | 982.69 | 862.35 | 1018 | 951.02 | 1100.5 |
| 3881 | ENSMUSG00000022771 | 698.8636648 | 0.047006055 | 0.114319097 | 0.41118288 | 0.680938441 | 0.934384556 | Ppil2 | peptidylprolyl isomerase (cyclophilin)-like | 664.17 | 707.21 | 691.29 | 687.32 | 748.16 | 695.04 |
| 3882 | ENSMUSG00000022772 | 461.2449279 | 0.169049899 | 0.141874579 | 1.191544675 | 0.233439829 | 0.680877937 | Senp5 | SUMO/sentrin specific peptidase 5 [Sourc | 393.52 | 429.67 | 479.98 | 452.11 | 536.58 | 475.61 |
| 3883 | ENSMUSG00000022773 | 8.912879198 | -1.033699202 | 0.741544575 | -1.393981208 | 0.16332328 | 0.587492643 | Ypel1 | yippee like 1 [Source:MGI Symbol;Acc:MG | 13.653 | 10.279 | 12.075 | 8.6758 | 5.4531 | 3.3415 |
| 3884 | ENSMUSG00000022774 | 424.9898424 | -0.03808901 | 0.14802049 | -0.257322551 | 0.796929785 | 0.962236316 | Ncbp2 | nuclear cap binding protein subunit 2 [Sou | 398.34 | 428.64 | 464.88 | 387.52 | 388.26 | 482.29 |
| 3885 | ENSMUSG00000022779 | 411.4899689 | -0.152087482 | 0.145935124 | -1.042158172 | 0.297338352 | 0.748078261 | Top3b | topoisomerase (DNA) III beta [Source:MG | 403.96 | 427.61 | 468.91 | 433.79 | 388.26 | 346.41 |
| 3886 | ENSMUSG00000022780 | 0.439020386 | -2.189747911 | 3.52148864 | -0.621824499 | 0.534057275 | NA | Meltf | melanotransferrin [Source:MGI Symbol;A | 1.6062 | 1.0279 | 0 | 0 | 0 | 0 |
| 3887 | ENSMUSG00000022781 | 572.3676618 | -0.031164293 | 0.137384437 | -0.226840055 | 0.820548116 | 0.969160361 | Pak2 | p21 (RAC1) activated kinase 2 [Source:MG | 533.26 | 614.69 | 587.65 | 489.7 | 609.66 | 599.25 |
| 3888 | ENSMUSG00000022783 | 0.294513347 | 0.165143496 | 4.080472857 | 0.040471657 | 0.967717103 | NA | Spag6l | sperm associated antigen 6-like [Source:N | 0.8031 | 0 | 0 | 0.964 | 0 | 0 |
| 3889 | ENSMUSG00000022787 | 178.045647 | 0.121656499 | 0.186463838 | 0.652440172 | 0.514117256 | 0.885004045 | Wdr53 | WD repeat domain 53 [Source:MGI Symbo | 179.09 | 150.08 | 182.13 | 184.12 | 170.14 | 202.72 |
| 3890 | ENSMUSG00000022788 | 332.0945649 | -0.05711511 | 0.143002414 | -0.399399619 | 0.689598775 | 0.937261276 | Fgd4 | FYVE, RhoGEF and PH domain containing | 356.58 | 319.68 | 339.1 | 309.44 | 339.18 | 328.58 |
| 3891 | ENSMUSG00000022789 | 732.1495782 | 0.124282124 | 0.110686848 | 1.122826483 | 0.261511196 | 0.715569038 | Dnm1l | dynamin 1-like [Source:MGI Symbol;Acc:N | 722.79 | 680.48 | 698.33 | 775.04 | 769.98 | 746.28 |
| 3892 | ENSMUSG00000022790 | 424.7371066 | -0.361448297 | 0.151454438 | -2.38651506 | 0.017008911 | 0.154824975 | Igsf11 | immunoglobulin superfamily, member 11 | 522.02 | 513.96 | 396.46 | 362.46 | 391.53 | 362 |
| 3893 | ENSMUSG00000022791 | 444.8083028 | -0.074544287 | 0.166839541 | -0.446802277 | 0.655017827 | 0.925845583 | Tnk2 | tyrosine kinase, non-receptor, 2 [Source:N | 435.28 | 403.97 | 530.29 | 499.34 | 355.54 | 444.42 |
| 3894 | ENSMUSG00000022792 | 382.8478907 | 0.159993469 | 0.145900623 | 1.096592088 | 0.272819748 | 0.728119873 | Yars2 | tyrosyl-tRNA synthetase 2 (mitochondrial) | 359.79 | 369.02 | 356.21 | 416.44 | 350.09 | 445.54 |
| 3895 | ENSMUSG00000022793 | 15.5902618 | -0.006130978 | 0.625730055 | -0.00979812 | 0.992182356 | 0.99859176 | B4galt4 | UDP-Gal:betaGlcNAc beta 1,4-galactosyltr | 15.259 | 25.698 | 6.0375 | 21.208 | 13.087 | 12.252 |
| 3896 | ENSMUSG00000022797 | 273.1197505 | -0.383810304 | 0.217997359 | -1.760619051 | 0.078302902 | 0.410131696 | Tfrc | transferrin receptor [Source:MGI Symbol; | 350.96 | 359.77 | 216.34 | 197.62 | 287.92 | 226.11 |
| 3897 | ENSMUSG00000022798 | 0 | NA | NA | NA | NA | NA | Tex55 | testis expressed 55 [Source:MGI Symbol;A | 0 | 0 | 0 | 0 | 0 | 0 |
| 3898 | ENSMUSG00000022799 | 73.11738543 | -0.649791131 | 0.26374539 | -2.463706123 | 0.01375088 | 0.134934013 | Arhgap31 | Rho GTPase activating protein 31 [Source: | 89.144 | 88.401 | 90.562 | 63.623 | 50.168 | 56.806 |
| 3899 | ENSMUSG00000022800 | 610.9959993 | -0.000641507 | 0.127702355 | -0.005023457 | 0.995991878 | 0.999003684 | Fyttd1 | forty-two-three domain containing 1 [Sou | 556.55 | 604.42 | 673.18 | 602.49 | 643.46 | 585.88 |
| 3900 | ENSMUSG00000022801 | 71.62932559 | 0.00109204 | 0.326678797 | 0.003342854 | 0.997332793 | 0.999498002 | Lrch3 | leucine-rich repeats and calponin homolo | 61.839 | 77.094 | 76.475 | 105.07 | 45.806 | 63.489 |
| 3901 | ENSMUSG00000022802 | 65.43129183 | 0.509657415 | 0.295487771 | 1.724800363 | 0.084563456 | 0.428715679 | Lmln | leishmanolysin-like (metallopeptidase M8 | 42.564 | 51.396 | 68.425 | 79.046 | 67.618 | 83.538 |
| 3902 | ENSMUSG00000022803 | 1.703723871 | -1.078743695 | 1.724211334 | -0.625644707 | 0.531548008 | NA | Popdc2 | popeye domain containing 2 [Source:MGI | 0.8031 | 4.1117 | 2.0125 | 0 | 2.1812 | 1.1138 |
| 3903 | ENSMUSG00000022805 | 0.82235459 | -1.754619754 | 2.405444547 | -0.729436792 | 0.465734519 | NA | Cfap91 | cilia and flagella associated protein 91 [So | 0.8031 | 1.0279 | 2.0125 | 0 | 1.0906 | 0 |
| 3904 | ENSMUSG00000022807 | 438.9166046 | -0.435244766 | 0.141255213 | -3.081265148 | 0.00206123 | 0.035619632 | Osbpl11 | oxysterol binding protein-like 11 [Source:I | 477.85 | 547.88 | 489.03 | 409.69 | 335.91 | 373.14 |
| 3905 | ENSMUSG00000022808 | 552.2073009 | -0.002634088 | 0.132422543 | -0.01989154 | 0.984129894 | 0.998580735 | Snx4 | sorting nexin 4 [Source:MGI Symbol;Acc:N | 519.61 | 598.25 | 540.35 | 499.34 | 597.66 | 558.04 |
| 3906 | ENSMUSG00000022809 | 2199.925357 | -0.328748826 | 0.109777029 | -2.99469598 | 0.002747185 | 0.044003638 | Nr1i2 | nuclear receptor subfamily 1, group I, mer | 2530.6 | 2577 | 2240.9 | 1982.9 | 2101.6 | 1766.6 |
| 3907 | ENSMUSG00000022811 | 648.3979693 | 0.029465026 | 0.116071262 | 0.253852893 | 0.799609209 | 0.962424862 | Zfp148 | zinc finger protein 148 [Source:MGI Symb | 632.04 | 636.28 | 657.08 | 642.01 | 692.54 | 630.44 |
| 3908 | ENSMUSG00000022812 | 617.7052215 | -0.211627765 | 0.141604794 | -1.494495759 | 0.13504609 | 0.537616739 | Gsk3b | glycogen synthase kinase 3 beta [Source:N | 719.58 | 720.57 | 548.4 | 603.45 | 557.31 | 556.92 |
| 3909 | ENSMUSG00000022814 | 636.7865932 | 0.064019058 | 0.129495355 | 0.494373396 | 0.621042499 | 0.918069106 | Umps | uridine monophosphate synthetase [Sour | 585.46 | 599.28 | 683.24 | 590.92 | 699.09 | 662.74 |
| 3910 | ENSMUSG00000022816 | 66.6599627 | -0.814218316 | 0.321575988 | -2.531962421 | 0.011342615 | 0.118540158 | Fstl1 | follistatin-like 1 [Source:MGI Symbol;Acc:I | 117.25 | 76.066 | 61.381 | 54.947 | 47.987 | 42.326 |
| 3911 | ENSMUSG00000022817 | 1432.139204 | -0.180903155 | 0.111110839 | -1.628132385 | 0.103496818 | 0.473607836 | Itgb5 | integrin beta 5 [Source:MGI Symbol;Acc:N | 1413.5 | 1535.7 | 1617 | 1357.3 | 1228 | 1441.3 |
| 3912 | ENSMUSG00000022818 | 1.588200194 | 0.715132113 | 1.877489912 | 0.380897979 | 0.703278952 | NA | Cyp2ab1 | cytochrome P450, family 2, subfamily ab, | 1.6062 | 0 | 2.0125 | 4.8199 | 1.0906 | 0 |
| 3913 | ENSMUSG00000022820 | 844.4963961 | -0.050883048 | 0.115929455 | -0.43891389 | 0.660723932 | 0.928184201 | Ndufb4 | NADH:ubiquinone oxidoreductase subuni | 890.64 | 872.7 | 814.05 | 793.35 | 791.79 | 904.44 |
| 3914 | ENSMUSG00000022821 | 11122.83252 | 0.165797799 | 0.105876121 | 1.56596027 | 0.117357919 | 0.502643331 | Hgd | homogentisate 1, 2-dioxygenase [Source:I | 11104 | 9710.7 | 10637 | 10593 | 11683 | 13010 |
| 3915 | ENSMUSG00000022822 | 72.04208321 | -0.563831235 | 0.326881839 | -1.724877824 | 0.084549493 | 0.428715679 | Abcc5 | ATP-binding cassette, sub-family C (CFTR/ | 75.492 | 116.15 | 66.412 | 63.623 | 38.172 | 72.4 |
| 3916 | ENSMUSG00000022824 | 0 | NA | NA | NA | NA | NA | Muc13 | mucin 13, epithelial transmembrane [Sour | 0 | 0 | 0 | 0 | 0 | 0 |

| | | | | | | | | | | | | | | | |
|---|---|---|---|---|---|---|---|---|---|---|---|---|---|---|---|
| 3917 | ENSMUSG00000022827 | 238.8893989 | -0.10115632 | 0.164344454 | -0.615514051 | 0.538215293 | 0.891577636 | Rabl3 | RAB, member RAS oncogene family-like 3 | 256.19 | 261.09 | 224.39 | 236.17 | 212.67 | 242.82 |
| 3918 | ENSMUSG00000022828 | 259.2323985 | -0.188454854 | 0.186398439 | -1.011032361 | 0.312000941 | 0.759830222 | Gtf2e1 | general transcription factor II E, polypepti | 252.98 | 267.26 | 307.91 | 190.87 | 284.65 | 251.73 |
| 3919 | ENSMUSG00000022829 | 2.672937875 | -1.454512583 | 1.365552465 | -1.065145881 | 0.286809919 | NA | Stxbp5l | syntaxin binding protein 5-like [Source:M( | 1.6062 | 4.1117 | 6.0375 | 0.964 | 1.0906 | 2.2277 |
| 3920 | ENSMUSG00000022831 | 71.64169778 | -1.496713223 | 0.317958214 | -4.7072639 | 2.51E-06 | 0.000172104 | Hcls1 | hematopoietic cell specific Lyn substrate 1 | 110.03 | 130.55 | 76.475 | 28.919 | 35.99 | 47.895 |
| 3921 | ENSMUSG00000022832 | 0 | NA | NA | NA | NA | NA | Ropn1 | ropporin, rhophilin associated protein 1 [S | 0 | 0 | 0 | 0 | 0 | 0 |
| 3922 | ENSMUSG00000022833 | 24.22383474 | 0.149144312 | 0.456749624 | 0.326534066 | 0.744020319 | 0.949911274 | Ccdc14 | coiled-coil domain containing 14 [Source:I | 24.093 | 18.503 | 26.162 | 20.244 | 21.812 | 34.529 |
| 3923 | ENSMUSG00000022836 | 1805.714068 | 0.015440116 | 0.123458658 | 0.125063044 | 0.900473641 | 0.982850624 | Mylk | myosin, light polypeptide kinase [Source:I | 1782.9 | 1693 | 1912.9 | 1943.4 | 1976.2 | 1526 |
| 3924 | ENSMUSG00000022837 | 58.87620773 | 0.153224395 | 0.294471798 | 0.520336399 | 0.60282913 | 0.91570264 | Iqcb1 | IQ calmodulin-binding motif containing 1 | 54.611 | 55.508 | 57.356 | 73.262 | 51.259 | 61.261 |
| 3925 | ENSMUSG00000022838 | 9.170477637 | -0.03992752 | 0.85970025 | -0.046443536 | 0.962956737 | 0.992956886 | Eaf2 | ELL associated factor 2 [Source:MGI Symb | 18.471 | 8.2233 | 1.0062 | 9.6398 | 6.5437 | 11.138 |
| 3926 | ENSMUSG00000022840 | 23.70067312 | -0.616418255 | 0.453835354 | -1.358242036 | 0.174386896 | 0.604151393 | Adcy5 | adenylate cyclase 5 [Source:MGI Symbol;/ | 28.109 | 23.642 | 34.212 | 15.424 | 18.541 | 22.277 |
| 3927 | ENSMUSG00000022841 | 2483.195492 | 0.168292434 | 0.090613377 | 1.857258153 | 0.063274435 | 0.359877642 | Ap2m1 | adaptor-related protein complex 2, mu 1 s | 2385.2 | 2387.9 | 2241.9 | 2560.3 | 2681.8 | 2642 |
| 3928 | ENSMUSG00000022842 | 0 | NA | NA | NA | NA | NA | Ece2 | endothelin converting enzyme 2 [Source:I | 0 | 0 | 0 | 0 | 0 | 0 |
| 3929 | ENSMUSG00000022843 | 1173.733131 | 0.291622411 | 0.11549722 | 2.524930121 | 0.01157213 | 0.11998515 | Clcn2 | chloride channel, voltage-sensitive 2 [Sou | 1000.7 | 1000.2 | 1166.2 | 1277.3 | 1385.1 | 1213 |
| 3930 | ENSMUSG00000022844 | 878.3831682 | 0.374616781 | 0.138362853 | 2.707495356 | 0.006779302 | 0.082261596 | Pdia5 | protein disulfide isomerase associated 5 [ | 648.91 | 771.97 | 875.43 | 998.68 | 1076.4 | 898.87 |
| 3931 | ENSMUSG00000022847 | 214.5083231 | 0.34255394 | 0.21592029 | 1.586483328 | 0.112629711 | 0.492766469 | Thpo | thrombopoietin [Source:MGI Symbol;Acc: | 178.29 | 199.42 | 190.18 | 273.77 | 163.59 | 281.8 |
| 3932 | ENSMUSG00000022848 | 909.5009471 | -0.146611072 | 0.123307392 | -1.188988509 | 0.234444189 | 0.682322842 | Slc49a4 | solute carrier family 49 member 4 [Source | 992.63 | 932.32 | 940.84 | 760.58 | 861.59 | 969.04 |
| 3933 | ENSMUSG00000022849 | 53.17912943 | -0.456143974 | 0.309494643 | -1.473834797 | 0.140526077 | 0.548157814 | Hspbap1 | Hspb associated protein 1 [Source:MGI Sy | 64.248 | 60.647 | 59.368 | 36.631 | 44.715 | 53.465 |
| 3934 | ENSMUSG00000022853 | 5730.703056 | -0.466214579 | 0.127474611 | -3.657313221 | 0.000254873 | 0.007616696 | Ehhadh | enoyl-Coenzyme A, hydratase/3-hydroxya | 6021.7 | 6821.3 | 7105.1 | 5647.9 | 4706 | 4082.2 |
| 3935 | ENSMUSG00000022855 | 516.2937348 | -0.011083014 | 0.120868913 | -0.091694493 | 0.926940773 | 0.987712228 | Senp2 | SUMO/sentrin specific peptidase 2 [Sourc | 512.38 | 526.29 | 516.2 | 509.94 | 503.87 | 529.08 |
| 3936 | ENSMUSG00000022856 | 270.6322825 | -0.312219929 | 0.159809578 | -1.953699723 | 0.050736739 | 0.316199857 | Tmem41a | transmembrane protein 41a [Source:MGI | 285.9 | 309.4 | 304.89 | 271.84 | 231.21 | 220.54 |
| 3937 | ENSMUSG00000022857 | 0 | NA | NA | NA | NA | NA | Tmprss15 | transmembrane protease, serine 15 [Sour | 0 | 0 | 0 | 0 | 0 | 0 |
| 3938 | ENSMUSG00000022858 | 543.3778505 | -0.268442733 | 0.14738823 | -1.821330871 | 0.068556576 | 0.37732204 | Tra2b | transformer 2 beta [Source:MGI Symbol;A | 582.25 | 571.52 | 626.89 | 406.8 | 567.12 | 505.69 |
| 3939 | ENSMUSG00000022860 | 0 | NA | NA | NA | NA | NA | Chodl | chondrolectin [Source:MGI Symbol;Acc:M | 0 | 0 | 0 | 0 | 0 | 0 |
| 3940 | ENSMUSG00000022861 | 0.328370405 | 0.16514316 | 4.003149536 | 0.041253308 | 0.967093956 | NA | Dgkg | diacylglycerol kinase, gamma [Source:MG | 0 | 0 | 1.0062 | 0.964 | 0 | 0 |
| 3941 | ENSMUSG00000022863 | 26.71534652 | -0.835652316 | 0.428800737 | -1.948812685 | 0.051317798 | 0.318157152 | Btg3 | BTG anti-proliferation factor 3 [Source:M( | 38.549 | 37.005 | 27.169 | 21.208 | 18.541 | 17.822 |
| 3942 | ENSMUSG00000022864 | 233.4072172 | -0.002422992 | 0.206771844 | -0.011718193 | 0.990650449 | 0.99859176 | D16Ertd472e | DNA segment, Chr 16, ERATO Doi 472, exp | 216.84 | 254.92 | 229.42 | 233.28 | 295.56 | 170.42 |
| 3943 | ENSMUSG00000022865 | 949.5751397 | 0.166862776 | 0.112292806 | 1.485961402 | 0.137289331 | 0.541266927 | Cxadr | coxsackie virus and adenovirus receptor [ | 836.03 | 873.73 | 975.05 | 985.19 | 1033.9 | 993.55 |
| 3944 | ENSMUSG00000022867 | 680.8074991 | -0.076889396 | 0.121970372 | -0.630394042 | 0.528436808 | 0.888334265 | Usp25 | ubiquitin specific peptidase 25 [Source:M | 694.68 | 752.44 | 650.03 | 674.78 | 694.72 | 618.18 |
| 3945 | ENSMUSG00000022868 | 106122.5093 | 0.274643445 | 0.100310851 | 2.737923589 | 0.006182844 | 0.076828741 | Ahsg | alpha-2-HS-glycoprotein [Source:MGI Sym | 90490 | 92984 | 104682 | 112365 | 109454 | 126760 |
| 3946 | ENSMUSG00000022871 | 7821.130857 | -0.011488745 | 0.105364825 | -0.109037762 | 0.913172539 | 0.98625062 | Fetub | fetuin beta [Source:MGI Symbol;Acc:MGI: | 7311.4 | 8507 | 7740 | 8352.9 | 8018.2 | 6997.2 |
| 3947 | ENSMUSG00000022875 | 31563.61937 | 0.161156973 | 0.08808526 | 1.829556654 | 0.06731626 | 0.373457298 | Kng1 | kininogen 1 [Source:MGI Symbol;Acc:MGI | 29628 | 30802 | 28979 | 34950 | 34029 | 30994 |
| 3948 | ENSMUSG00000022876 | 3.009715183 | -0.230890534 | 1.241747289 | -0.185940035 | 0.852491787 | NA | Samsn1 | SAM domain, SH3 domain and nuclear loc | 5.6217 | 2.0558 | 2.0125 | 2.8919 | 4.3625 | 1.1138 |
| 3949 | ENSMUSG00000022877 | 15548.74973 | -0.147988849 | 0.108243408 | -1.367185797 | 0.171567076 | 0.599034187 | Hrg | histidine-rich glycoprotein [Source:MGI Sy | 14471 | 16753 | 17814 | 14096 | 16102 | 14057 |
| 3950 | ENSMUSG00000022878 | 0.181769715 | 1.126924175 | 4.080472857 | 0.276174898 | 0.782413736 | NA | Adipoq | adiponectin, C1Q and collagen domain co | 0 | 0 | 0 | 0 | 1.0906 | 0 |
| 3951 | ENSMUSG00000022881 | 49.77591156 | 0.373046215 | 0.338102519 | 1.103352369 | 0.269874173 | 0.725011151 | Rfc4 | replication factor C (activator 1) 4 [Source | 51.399 | 38.033 | 40.25 | 42.415 | 71.981 | 54.578 |
| 3952 | ENSMUSG00000022883 | 53.78924208 | -1.003745702 | 0.350674601 | -2.862327926 | 0.004205415 | 0.059459559 | Robo1 | roundabout guidance receptor 1 [Source: | 75.492 | 92.513 | 47.293 | 33.739 | 43.625 | 30.074 |
| 3953 | ENSMUSG00000022884 | 5874.0596 | 0.31218976 | 0.083607482 | 3.733993069 | 0.000188468 | 0.006024045 | Eif4a2 | eukaryotic translation initiation factor 4A2 | 5384.8 | 5267.1 | 5070.5 | 6485.6 | 6644 | 6392.4 |
| 3954 | ENSMUSG00000022885 | 1361.913642 | 0.223223167 | 0.133768457 | 1.668727977 | 0.095171302 | 0.455057116 | St6gal1 | beta galactoside alpha 2,6 sialyltransferas | 1321.1 | 1366.1 | 1082.7 | 1498 | 1267.3 | 1636.2 |
| 3955 | ENSMUSG00000022886 | 0 | NA | NA | NA | NA | NA | Prl2a1 | prolactin family 2, subfamily a, member 1 | 0 | 0 | 0 | 0 | 0 | 0 |
| 3956 | ENSMUSG00000022887 | 2462.851045 | 0.088423449 | 0.105074674 | 0.841529609 | 0.400051305 | 0.82709119 | Masp1 | mannan-binding lectin serine peptidase 1 | 2358.7 | 2600.6 | 2202.7 | 2384.9 | 2686.2 | 2544 |
| 3957 | ENSMUSG00000022889 | 693.5557011 | 0.043491575 | 0.111338732 | 0.390623953 | 0.696075218 | 0.939992732 | Mrpl39 | mitochondrial ribosomal protein L39 [Sou | 670.59 | 692.82 | 686.26 | 713.34 | 711.08 | 687.24 |
| 3958 | ENSMUSG00000022890 | 3481.785976 | 0.005651544 | 0.087624754 | 0.064497114 | 0.948574405 | 0.990731019 | Atp5j | ATP synthase, H+ transporting, mitochonc | 3361.8 | 3560.7 | 3502.7 | 3412.5 | 3445.3 | 3607.7 |
| 3959 | ENSMUSG00000022892 | 786.0059041 | -0.816210157 | 0.175683591 | -4.645910028 | 3.39E-06 | 0.000217655 | App | amyloid beta (A4) precursor protein [Sour | 894.66 | 1030 | 1084.7 | 747.08 | 443.88 | 515.71 |
| 3960 | ENSMUSG00000022893 | 113.5331464 | -0.923777965 | 0.234711241 | -3.935806231 | 8.29E-05 | 0.003107057 | Adamts1 | a disintegrin-like and metallopeptidase (re | 175.88 | 138.77 | 130.81 | 69.406 | 87.249 | 79.083 |
| 3961 | ENSMUSG00000022894 | 32.15584089 | -0.696776492 | 0.414060584 | -1.682788746 | 0.092415994 | 0.447657382 | Adamts5 | a disintegrin-like and metallopeptidase (re | 40.958 | 41.117 | 37.231 | 23.135 | 34.9 | 15.594 |
| 3962 | ENSMUSG00000022895 | 405.0172182 | -0.431141315 | 0.131491175 | -3.27886122 | 0.001042269 | 0.021567061 | Ets2 | E26 avian leukemia oncogene 2, 3' domair | 465.8 | 471.81 | 457.84 | 356.67 | 330.46 | 347.52 |
| 3963 | ENSMUSG00000022897 | 386.15341 | 0.064759728 | 0.135444493 | 0.478127436 | 0.632559505 | 0.922076476 | Dyrk1a | dual-specificity tyrosine-(Y)-phosphorylati | 375.05 | 387.53 | 370.3 | 421.26 | 375.17 | 387.62 |
| 3964 | ENSMUSG00000022898 | 383.2685913 | 0.132268702 | 0.142860643 | 0.925858227 | 0.354519663 | 0.794633876 | Vps26c | VPS26 endosomal protein sorting factor C | 343.73 | 355.66 | 398.47 | 429.93 | 369.72 | 402.1 |
| 3965 | ENSMUSG00000022899 | 36.7250457 | 0.382235036 | 1.033193502 | 0.369954936 | 0.711416068 | 0.944239215 | Slc15a2 | solute carrier family 15 (H+/peptide transp | 56.217 | 3.0838 | 36.225 | 39.523 | 21.812 | 63.489 |
| 3966 | ENSMUSG00000022900 | 1.168399487 | -0.282366966 | 1.955322271 | -0.144409426 | 0.885177174 | NA | Ildr1 | immunoglobulin-like domain containing re | 0.8031 | 2.0558 | 1.0062 | 0.964 | 2.1812 | 0 |
| 3967 | ENSMUSG00000022901 | 20.02618973 | -1.603853282 | 0.538236723 | -2.97982879 | 0.002884095 | 0.045441781 | Cd86 | CD86 antigen [Source:MGI Symbol;Acc:M( | 35.336 | 24.67 | 30.187 | 5.7839 | 15.269 | 8.9108 |
| 3968 | ENSMUSG00000022902 | 0 | NA | NA | NA | NA | NA | Stfa2 | stefin A2 [Source:MGI Symbol;Acc:MGI:1( | 0 | 0 | 0 | 0 | 0 | 0 |
| 3969 | ENSMUSG00000022905 | 839.489873 | 0.063863141 | 0.125191405 | 0.510124006 | 0.609964588 | 0.916376001 | Kpna1 | karyopherin (importin) alpha 1 [Source:M | 713.15 | 872.7 | 878.45 | 894.57 | 860.5 | 817.56 |
| 3970 | ENSMUSG00000022906 | 1187.881975 | 0.039324626 | 0.104371822 | 0.376774358 | 0.706341298 | 0.94295036 | Parp9 | poly (ADP-ribose) polymerase family, men | 1188.6 | 1148.2 | 1178.3 | 1223.3 | 1268.4 | 1120.5 |
| 3971 | ENSMUSG00000022911 | 114.9312187 | 0.560300312 | 0.240375911 | 2.330933698 | 0.019756856 | 0.172042845 | Arl13b | ADP-ribosylation factor-like 13B [Source:N | 81.113 | 77.094 | 120.75 | 134.96 | 130.87 | 144.8 |
| 3972 | ENSMUSG00000022912 | 1027.326485 | 0.094967678 | 0.102086148 | 0.930269975 | 0.352231319 | 0.792945339 | Pros1 | protein S (alpha) [Source:MGI Symbol;Acc | 1015.1 | 998.11 | 967 | 1052.7 | 1085.2 | 1045.9 |

| | | | | | | | | | | | | | | | |
|---|---|---|---|---|---|---|---|---|---|---|---|---|---|---|---|
| 3973 | ENSMUSG00000022913 | 339.352033 | -0.003362042 | 0.142729766 | -0.023555298 | 0.981207329 | 0.99820894 | Psmg1 | proteasome (prosome, macropain) assem | 348.55 | 340.24 | 330.05 | 320.04 | 367.54 | 329.7 |
| 3974 | ENSMUSG00000022914 | 551.4248477 | 0.137369628 | 0.12679932 | 1.083362497 | 0.278647555 | 0.731329747 | Brwd1 | bromodomain and WD repeat domain con | 501.94 | 518.07 | 556.45 | 624.66 | 562.76 | 544.67 |
| 3975 | ENSMUSG00000022931 | 0 | NA | NA | NA | NA | NA | Krtap15 | keratin associated protein 15 [Source:MG | 0 | 0 | 0 | 0 | 0 | 0 |
| 3976 | ENSMUSG00000022935 | 0 | NA | NA | NA | NA | NA | Grik1 | glutamate receptor, ionotropic, kainate 1 | 0 | 0 | 0 | 0 | 0 | 0 |
| 3977 | ENSMUSG00000022938 | 0 | NA | NA | NA | NA | NA | Fam3b | family with sequence similarity 3, membe | 0 | 0 | 0 | 0 | 0 | 0 |
| 3978 | ENSMUSG00000022940 | 234.1675977 | -0.412557571 | 0.24686657 | -1.671176343 | 0.094686852 | 0.453643796 | Pigp | phosphatidylinositol glycan anchor biosyn | 336.5 | 288.84 | 176.09 | 159.06 | 257.39 | 187.13 |
| 3979 | ENSMUSG00000022941 | 18.27458163 | -1.27557688 | 0.544725151 | -2.341688973 | 0.019196705 | 0.168586 | Ripply3 | ripply transcriptional repressor 3 [Source: | 32.927 | 22.614 | 22.137 | 15.424 | 7.6343 | 8.9108 |
| 3980 | ENSMUSG00000022945 | 29.57346002 | -0.415383664 | 0.415812485 | -0.998968715 | 0.317809847 | 0.764477236 | Chaf1b | chromatin assembly factor 1, subunit B (p | 43.368 | 26.726 | 31.194 | 29.883 | 23.994 | 22.277 |
| 3981 | ENSMUSG00000022946 | 642.724703 | -0.6575808 | 0.186819569 | -3.519871097 | 0.000431757 | 0.011211718 | Dop1b | DOP1 leucine zipper like protein B [Source | 636.86 | 917.93 | 807.01 | 646.83 | 414.43 | 433.29 |
| 3982 | ENSMUSG00000022947 | 9.387352307 | -1.26979769 | 0.757325609 | -1.676686588 | 0.093603785 | 0.450459278 | Cbr3 | carbonyl reductase 3 [Source:MGI Symbol | 18.471 | 11.307 | 10.062 | 7.7118 | 6.5437 | 2.2277 |
| 3983 | ENSMUSG00000022948 | 159.1613134 | -0.086237144 | 0.256160468 | -0.336652819 | 0.736378634 | 0.948109711 | Setd4 | SET domain containing 4 [Source:MGI Sym | 189.53 | 156.24 | 145.91 | 174.48 | 194.13 | 94.677 |
| 3984 | ENSMUSG00000022949 | 0.944152054 | -0.673133018 | 2.474469057 | -0.272031293 | 0.785597958 | NA | Clic6 | chloride intracellular channel 6 [Source:M | 2.4093 | 1.0279 | 0 | 0 | 0 | 2.2277 |
| 3985 | ENSMUSG00000022951 | 406.6227152 | -0.555997001 | 0.173467779 | -3.205188906 | 0.001349739 | 0.026169732 | Rcan1 | regulator of calcineurin 1 [Source:MGI Syr | 519.61 | 548.91 | 383.38 | 342.21 | 280.29 | 365.34 |
| 3986 | ENSMUSG00000022952 | 7.125305504 | -1.057651619 | 0.837426907 | -1.262977831 | 0.206597147 | 0.652026861 | Runx1 | runt related transcription factor 1 [Source | 10.44 | 11.307 | 7.0437 | 2.8919 | 3.2719 | 7.7969 |
| 3987 | ENSMUSG00000022956 | 1298.83382 | -0.050116755 | 0.108654419 | -0.461249119 | 0.644619885 | 0.925643431 | Atp5o | ATP synthase, H+ transporting, mitochonc | 1344.4 | 1223.2 | 1396.7 | 1329.3 | 1310.9 | 1188.5 |
| 3988 | ENSMUSG00000022957 | 427.1133328 | -0.244468865 | 0.161170815 | -1.516830849 | 0.129309396 | 0.526260863 | Itsn1 | intersectin 1 (SH3 domain protein 1A) [So | 408.78 | 506.76 | 474.95 | 419.33 | 318.46 | 434.4 |
| 3989 | ENSMUSG00000022960 | 65.61873732 | 0.223233574 | 0.318110666 | 0.70174816 | 0.482836233 | 0.871710696 | Donson | downstream neighbor of SON [Source:MG | 57.823 | 44.2 | 79.493 | 55.911 | 88.34 | 67.944 |
| 3990 | ENSMUSG00000022961 | 2902.18541 | -0.057239938 | 0.11471604 | -0.498970661 | 0.617800053 | 0.917509899 | Son | Son DNA binding protein [Source:MGI Syr | 2810.9 | 3165 | 2905 | 3190.8 | 2847.6 | 2493.9 |
| 3991 | ENSMUSG00000022962 | 509.9032279 | 0.007152601 | 0.13284666 | 0.053841029 | 0.957061821 | 0.991861398 | Gart | phosphoribosylglycinamide formyltransfe | 456.16 | 533.49 | 537.33 | 523.44 | 530.04 | 478.95 |
| 3992 | ENSMUSG00000022964 | 106.1002752 | 0.009261406 | 0.250967996 | 0.036902736 | 0.970562559 | 0.994966572 | Tmem50b | transmembrane protein 50B [Source:MGI | 94.766 | 118.21 | 104.65 | 115.68 | 77.434 | 125.86 |
| 3993 | ENSMUSG00000022965 | 426.1487968 | 0.090941829 | 0.135307777 | 0.672110882 | 0.50151311 | 0.880974509 | Ifngr2 | interferon gamma receptor 2 [Source:MG | 444.92 | 402.94 | 389.42 | 423.19 | 428.61 | 467.81 |
| 3994 | ENSMUSG00000022967 | 626.1538952 | 0.345510331 | 0.131579708 | 2.625863334 | 0.008642951 | 0.097589107 | Ifnar1 | interferon (alpha and beta) receptor 1 [So | 553.34 | 567.41 | 534.32 | 776 | 609.66 | 716.2 |
| 3995 | ENSMUSG00000022969 | 67.91625801 | -0.533167594 | 0.277021945 | -1.92464028 | 0.054274377 | 0.327913359 | Il10rb | interleukin 10 receptor, beta [Source:MGI | 85.129 | 80.178 | 75.468 | 53.019 | 47.987 | 65.717 |
| 3996 | ENSMUSG00000022971 | 723.444431 | 0.077913032 | 0.112963884 | 0.689716294 | 0.490372617 | 0.875587032 | Ifnar2 | interferon (alpha and beta) receptor 2 [So | 721.99 | 689.73 | 699.34 | 710.45 | 738.35 | 780.8 |
| 3997 | ENSMUSG00000022972 | 134.6989469 | 0.07055159 | 0.209233912 | 0.337190035 | 0.735973649 | 0.948109711 | Cfap298 | cilia and flagella associate protien 298 [So | 118.06 | 146.99 | 129.81 | 152.31 | 138.51 | 122.52 |
| 3998 | ENSMUSG00000022973 | 176.5791692 | -0.01883918 | 0.204642225 | -0.092059103 | 0.926651082 | 0.987712228 | Synj1 | synaptojanin 1 [Source:MGI Symbol;Acc:N | 156.6 | 222.03 | 154.96 | 169.66 | 166.86 | 189.35 |
| 3999 | ENSMUSG00000022974 | 323.5430712 | 0.23119838 | 0.158696529 | 1.456858456 | 0.145155456 | 0.555361204 | Paxbp1 | PAX3 and PAX7 binding protein 1 [Source: | 255.39 | 328.93 | 309.92 | 377.88 | 327.19 | 341.95 |
| 4000 | ENSMUSG00000022978 | 49.30537264 | -0.217181973 | 0.32229205 | -0.673866988 | 0.500395882 | 0.880716237 | Mis18a | MIS18 kinetochore protein A [Source:MGI | 48.186 | 62.703 | 48.3 | 47.235 | 38.172 | 51.237 |
| 4001 | ENSMUSG00000022982 | 14046.17678 | -0.002124916 | 0.084557893 | -0.025129715 | 0.979951499 | 0.99820894 | Sod1 | superoxide dismutase 1, soluble [Source:I | 14027 | 14540 | 13603 | 14652 | 13952 | 13502 |
| 4002 | ENSMUSG00000022983 | 475.8215413 | 0.042122302 | 0.150835948 | 0.279259037 | 0.78004604 | 0.959870443 | Scaf4 | SR-related CTD-associated factor 4 [Sourc | 505.15 | 433.78 | 467.9 | 566.82 | 460.24 | 421.03 |
| 4003 | ENSMUSG00000022986 | 0 | NA | NA | NA | NA | NA | Krt75 | keratin 75 [Source:MGI Symbol;Acc:MGI:1 | 0 | 0 | 0 | 0 | 0 | 0 |
| 4004 | ENSMUSG00000022987 | 4.1781413 | -0.973794781 | 1.082984721 | -0.899176842 | 0.368558474 | NA | Zfp641 | zinc finger protein 641 [Source:MGI Symb | 6.4248 | 4.1117 | 6.0375 | 1.928 | 5.4531 | 1.1138 |
| 4005 | ENSMUSG00000022991 | 0 | NA | NA | NA | NA | NA | Lalba | lactalbumin, alpha [Source:MGI Symbol;A | 0 | 0 | 0 | 0 | 0 | 0 |
| 4006 | ENSMUSG00000022992 | 491.9243254 | 0.24367584 | 0.125058282 | 1.948498228 | 0.051355377 | 0.318253714 | Kansl2 | KAT8 regulatory NSL complex subunit 2 [S | 468.21 | 433.78 | 448.79 | 510.91 | 550.76 | 539.1 |
| 4007 | ENSMUSG00000022993 | 1.060044235 | 0.14943475 | 2.627116342 | 0.056881664 | 0.95463946 | NA | Tex49 | testis expressed 49 [Source:MGI Symbol;A | 0 | 0 | 3.0187 | 0 | 0 | 3.3415 |
| 4008 | ENSMUSG00000022994 | 675.5312984 | 0.163697868 | 0.14986705 | 1.092287248 | 0.274706864 | 0.729487777 | Adcy6 | adenylate cyclase 6 [Source:MGI Symbol;A | 677.82 | 725.71 | 508.15 | 755.76 | 726.35 | 659.4 |
| 4009 | ENSMUSG00000022995 | 4.726861276 | -1.491397284 | 1.018286965 | -1.464613941 | 0.143026276 | NA | Enah | ENAH actin regulator [Source:MGI Symbo | 5.6217 | 9.2513 | 6.0375 | 1.928 | 2.1812 | 3.3415 |
| 4010 | ENSMUSG00000022996 | 0.36353943 | 2.065989254 | 4.034137301 | 0.512126658 | 0.608562368 | NA | Wnt10b | wingless-type MMTV integration site fami | 0 | 0 | 0 | 0 | 2.1812 | 0 |
| 4011 | ENSMUSG00000022997 | 0 | NA | NA | NA | NA | NA | Wnt1 | wingless-type MMTV integration site fami | 0 | 0 | 0 | 0 | 0 | 0 |
| 4012 | ENSMUSG00000022999 | 269.8705784 | 0.065778441 | 0.194163517 | 0.338778582 | 0.734776538 | 0.948109711 | Lmbr1l | limb region 1 like [Source:MGI Symbol;Acc | 233.7 | 231.28 | 327.03 | 323.9 | 268.29 | 235.02 |
| 4013 | ENSMUSG00000023000 | 1.19918676 | -2.469682767 | 2.290014249 | -1.07845738 | 0.280829692 | NA | Dhh | desert hedgehog [Source:MGI Symbol;Acc | 3.2124 | 0 | 3.0187 | 0.964 | 0 | 0 |
| 4014 | ENSMUSG00000023004 | 314.4205589 | -0.087848928 | 0.154993481 | -0.566791115 | 0.570856108 | 0.904651508 | Tuba1b | tubulin, alpha 1B [Source:MGI Symbol;Acc | 354.17 | 322.77 | 294.83 | 332.57 | 278.11 | 304.08 |
| 4015 | ENSMUSG00000023007 | 233.823171 | 0.271374101 | 0.194391565 | 1.396017883 | 0.16270911 | 0.586288131 | Prpf40b | pre-mRNA processing factor 40B [Source: | 209.61 | 191.19 | 235.46 | 319.08 | 217.03 | 230.57 |
| 4016 | ENSMUSG00000023008 | 43.03442019 | -0.990503155 | 0.337233593 | -2.937142606 | 0.003312517 | 0.050222486 | Fmnl3 | formin-like 3 [Source:MGI Symbol;Acc:MG | 57.823 | 54.48 | 59.368 | 26.991 | 28.356 | 31.188 |
| 4017 | ENSMUSG00000023009 | 8.551419851 | -0.409900686 | 0.781693618 | -0.524375121 | 0.600017657 | 0.915217938 | Nckap5l | NCK-associated protein 5-like [Source:MG | 12.85 | 12.335 | 4.025 | 7.7118 | 4.3625 | 10.025 |
| 4018 | ENSMUSG00000023010 | 28881.62751 | -0.030959876 | 0.08182521 | -0.378365977 | 0.705158739 | 0.94295036 | Tmbim6 | transmembrane BAX inhibitor motif conta | 28345 | 29549 | 29682 | 28899 | 29433 | 27383 |
| 4019 | ENSMUSG00000023011 | 0.349477146 | 0.165142768 | 3.911021834 | 0.042224967 | 0.966319359 | NA | Faim2 | Fas apoptotic inhibitory molecule 2 [Sourc | 0 | 0 | 1.0062 | 0 | 1.0906 | 0 |
| 4020 | ENSMUSG00000023013 | 0 | NA | NA | NA | NA | NA | Aqp2 | aquaporin 2 [Source:MGI Symbol;Acc:MG | 0 | 0 | 0 | 0 | 0 | 0 |
| 4021 | ENSMUSG00000023015 | 8.40107747 | -0.142797386 | 0.805464758 | -0.177285703 | 0.85928398 | 0.975487285 | Racgap1 | Rac GTPase-activating protein 1 [Source:N | 16.062 | 7.1954 | 3.0187 | 8.6758 | 6.5437 | 8.9108 |
| 4022 | ENSMUSG00000023017 | 47.63237484 | -0.2611848 | 0.321596114 | -0.812151605 | 0.416704645 | 0.836878476 | Asic1 | acid-sensing (proton-gated) ion channel 1 | 44.974 | 59.619 | 51.318 | 39.523 | 45.806 | 44.554 |
| 4023 | ENSMUSG00000023018 | 102.844116 | -0.849523109 | 0.239567178 | -3.546074702 | 0.000391015 | 0.01049285 | Smarcd1 | SWI/SNF related, matrix associated, actin | 114.04 | 144.94 | 137.86 | 63.623 | 73.071 | 83.538 |
| 4024 | ENSMUSG00000023019 | 7372.94928 | -0.302122803 | 0.105674317 | -2.858999337 | 0.004249797 | 0.059969814 | Gpd1 | glycerol-3-phosphate dehydrogenase 1 (s | 8377.2 | 8918.2 | 7131.3 | 6882.8 | 6491.4 | 6436.9 |
| 4025 | ENSMUSG00000023020 | 621.1446522 | 0.044419406 | 0.132642193 | 0.334881418 | 0.73771454 | 0.948137969 | Cox14 | cytochrome c oxidase assembly protein 1 | 661.76 | 589 | 582.62 | 555.25 | 655.46 | 682.79 |
| 4026 | ENSMUSG00000023021 | 29.21827944 | -0.354747471 | 0.399828835 | -0.887248344 | 0.37494521 | 0.809332934 | Cers5 | ceramide synthase 5 [Source:MGI Symbol | 31.321 | 30.838 | 36.225 | 25.063 | 22.903 | 28.96 |
| 4027 | ENSMUSG00000023022 | 1298.112758 | 0.305945112 | 0.117233336 | 2.609710879 | 0.009061877 | 0.100823842 | Lima1 | LIM domain and actin binding 1 [Source:N | 1202.2 | 1117.3 | 1163.2 | 1528.9 | 1246.6 | 1530.4 |
| 4028 | ENSMUSG00000023025 | 1313.42682 | 0.151048172 | 0.13604791 | 1.110257202 | 0.26688821 | 0.722778036 | Larp4 | La ribonucleoprotein domain family, mem | 1250.4 | 1413.4 | 1070.6 | 1406.4 | 1535.6 | 1204.1 |

| | | | | | | | | | | | | | | | |
|---|---|---|---|---|---|---|---|---|---|---|---|---|---|---|---|
| 4029 | ENSMUSG00000023026 | 117.8447455 | -0.133645581 | 0.22023734 | -0.606825258 | 0.543966884 | 0.894702561 | Dip2b | disco interacting protein 2 homolog B [Sou | 114.04 | 132.6 | 123.77 | 128.21 | 97.065 | 111.38 |
| 4030 | ENSMUSG00000023027 | 472.3143556 | -0.000122846 | 0.145697582 | -0.000843156 | 0.999327259 | 0.999764475 | Atf1 | activating transcription factor 1 [Source:M | 399.94 | 526.29 | 492.05 | 485.84 | 485.33 | 444.42 |
| 4031 | ENSMUSG00000023030 | 520.0108377 | 0.019868221 | 0.139269144 | 0.142660612 | 0.886558226 | 0.982355187 | Slc11a2 | solute carrier family 11 (proton-coupled d | 489.09 | 506.76 | 554.44 | 592.85 | 466.78 | 510.14 |
| 4032 | ENSMUSG00000023031 | 603.0610609 | 0.785626401 | 0.403818502 | 1.945493822 | 0.051715573 | 0.319420348 | Cela1 | chymotrypsin-like elastase family, membe | 505.15 | 276.51 | 546.39 | 508.02 | 777.61 | 1004.7 |
| 4033 | ENSMUSG00000023032 | 10.77226512 | -2.584719815 | 0.818072144 | -3.159525517 | 0.001580263 | 0.029454792 | Slc4a8 | solute carrier family 4 (anion exchanger), | 20.078 | 13.363 | 22.137 | 5.7839 | 3.2719 | 0 |
| 4034 | ENSMUSG00000023033 | 2.036223933 | -0.873374754 | 1.602449542 | -0.545024808 | 0.585736456 | NA | Scn8a | sodium channel, voltage-gated, type VIII, | 0.8031 | 3.0838 | 4.025 | 0.964 | 0 | 3.3415 |
| 4035 | ENSMUSG00000023034 | 17.68774592 | -1.184729404 | 0.541327005 | -2.188565122 | 0.028628463 | 0.221908945 | Nr4a1 | nuclear receptor subfamily 4, group A, me | 28.912 | 23.642 | 21.131 | 12.532 | 6.5437 | 13.366 |
| 4036 | ENSMUSG00000023036 | 1.085201869 | 1.137592316 | 2.627090041 | 0.433023725 | 0.664997534 | NA | Pcdhgc4 | protocadherin gamma subfamily C, 4 [Sou | 0 | 2.0558 | 0 | 0 | 0 | 4.4554 |
| 4037 | ENSMUSG00000023039 | 17.48111888 | 0.025408232 | 0.515317975 | 0.04930593 | 0.960675494 | 0.992515024 | Krt7 | keratin 7 [Source:MGI Symbol;Acc:MGI:96 | 14.456 | 16.447 | 21.131 | 15.424 | 20.722 | 16.708 |
| 4038 | ENSMUSG00000023041 | 0 | NA | NA | NA | NA | NA | Krt6b | keratin 6B [Source:MGI Symbol;Acc:MGI:1 | 0 | 0 | 0 | 0 | 0 | 0 |
| 4039 | ENSMUSG00000023043 | 1733.278643 | 0.310564058 | 0.117638545 | 2.639985544 | 0.008290956 | 0.094351214 | Krt18 | keratin 18 [Source:MGI Symbol;Acc:MGI:9 | 1759.6 | 1418.5 | 1462.1 | 1780.5 | 1970.7 | 2008.3 |
| 4040 | ENSMUSG00000023044 | 944.2367783 | 0.264141509 | 0.217943839 | 1.211970524 | 0.225523664 | 0.672301295 | Csad | cysteine sulfinic acid decarboxylase [Sour | 762.95 | 1191.4 | 620.85 | 1342.8 | 845.23 | 902.21 |
| 4041 | ENSMUSG00000023045 | 738.5365326 | 0.268381134 | 0.126091014 | 2.128471531 | 0.033298008 | 0.243995238 | Soat2 | sterol O-acyltransferase 2 [Source:MGI Sy | 650.51 | 678.43 | 682.23 | 916.74 | 750.35 | 752.96 |
| 4042 | ENSMUSG00000023046 | 55.65268141 | 0.176591103 | 0.306312612 | 0.576506143 | 0.564273119 | 0.901468983 | Igfbp6 | insulin-like growth factor binding protein | 48.989 | 63.731 | 44.275 | 64.587 | 59.984 | 52.351 |
| 4043 | ENSMUSG00000023047 | 7.55626525 | 0.351753937 | 0.868752024 | 0.404895675 | 0.685554198 | 0.935893609 | Amhr2 | anti-Mullerian hormone type 2 receptor [S | 4.8186 | 8.2233 | 7.0437 | 16.388 | 2.1812 | 6.6831 |
| 4044 | ENSMUSG00000023048 | 712.321022 | 0.014562654 | 0.145032351 | 0.100409696 | 0.920019072 | 0.986685804 | Prr13 | proline rich 13 [Source:MGI Symbol;Acc:M | 790.25 | 626 | 708.4 | 623.69 | 698 | 827.59 |
| 4045 | ENSMUSG00000023050 | 23.5688374 | -0.581949731 | 0.463593707 | -1.255301186 | 0.209369445 | 0.65350014 | Map3k12 | mitogen-activated protein kinase kinase k | 28.912 | 25.698 | 30.187 | 19.28 | 25.084 | 12.252 |
| 4046 | ENSMUSG00000023051 | 266.5843606 | 0.28899836 | 0.170278291 | 1.697212006 | 0.089656586 | 0.441714141 | Tarbp2 | TARBP2, RISC loading complex RNA bindir | 261.01 | 258.01 | 200.24 | 269.91 | 316.28 | 294.05 |
| 4047 | ENSMUSG00000023052 | 10.50425203 | -0.427600329 | 0.70028389 | -0.610609975 | 0.541457817 | 0.893717433 | Npff | neuropeptide FF-amide peptide precursor | 8.8341 | 16.447 | 11.069 | 13.496 | 8.7249 | 4.4554 |
| 4048 | ENSMUSG00000023055 | 910.3761673 | 0.101128244 | 0.130961383 | 0.772198961 | 0.439996597 | 0.848765947 | Calcoco1 | calcium binding and coiled coil domain 1 [ | 1010.3 | 813.08 | 811.03 | 1008.3 | 860.5 | 959.02 |
| 4049 | ENSMUSG00000023057 | 3057.241525 | -0.19204148 | 0.135847832 | -1.413651414 | 0.157464277 | 0.576703416 | Fabp2 | fatty acid binding protein 2, intestinal [Sou | 3977.8 | 2966.6 | 2835.6 | 2987.4 | 2969.8 | 2606.4 |
| 4050 | ENSMUSG00000023064 | 0.349477146 | 0.165142768 | 3.911021834 | 0.042224967 | 0.966319359 | NA | Sncg | synuclein, gamma [Source:MGI Symbol;Ac | 0 | 0 | 1.0062 | 0 | 1.0906 | 0 |
| 4051 | ENSMUSG00000023066 | 25.22612471 | -0.314724745 | 0.522444406 | -0.602408105 | 0.546902517 | 0.895518681 | Rttn | rotatin [Source:MGI Symbol;Acc:MGI:217 | 32.927 | 17.475 | 33.206 | 13.496 | 15.269 | 38.985 |
| 4052 | ENSMUSG00000023067 | 361.1177467 | -4.461983043 | 0.634816238 | -7.028779001 | 2.08E-12 | 8.14E-10 | Cdkn1a | cyclin-dependent kinase inhibitor 1A (P21 | 391.91 | 1057.7 | 622.87 | 22.171 | 17.45 | 54.578 |
| 4053 | ENSMUSG00000023068 | 1178.987763 | -0.073435334 | 0.1025253 | -0.71626549 | 0.473827431 | 0.868952133 | Nus1 | NUS1 dehydrodolichyl diphosphate syntha | 1181.4 | 1283.9 | 1162.2 | 1159.7 | 1146.2 | 1140.6 |
| 4054 | ENSMUSG00000023070 | 25250.69744 | 0.230931868 | 0.096957158 | 2.381792881 | 0.017228584 | 0.156332651 | Rgn | regucalcin [Source:MGI Symbol;Acc:MGI:1 | 25280 | 21223 | 23197 | 26538 | 28721 | 26544 |
| 4055 | ENSMUSG00000023072 | 104.9285277 | 0.234112256 | 0.246896159 | 0.94822154 | 0.343016684 | 0.784828638 | Cep89 | centrosomal protein 89 [Source:MGI Syml | 106.81 | 89.429 | 92.575 | 95.434 | 143.96 | 101.36 |
| 4056 | ENSMUSG00000023073 | 546.6662189 | 0.133287555 | 0.235931629 | 0.564941441 | 0.572113595 | 0.905069565 | Slc10a2 | solute carrier family 10, member 2 [Source | 670.59 | 482.09 | 411.55 | 828.06 | 419.89 | 467.81 |
| 4057 | ENSMUSG00000023074 | 223.0329711 | -0.213494786 | 0.169446811 | -1.259951629 | 0.207686812 | 0.652613959 | Mospd1 | motile sperm domain containing 1 [Source | 231.29 | 249.78 | 237.47 | 201.47 | 186.5 | 231.68 |
| 4058 | ENSMUSG00000023075 | 630.2480105 | -0.133387883 | 0.114267526 | -1.167329756 | 0.243077226 | 0.692645587 | Akirin1 | akirin 1 [Source:MGI Symbol;Acc:MGI:191 | 660.95 | 663.01 | 654.06 | 594.77 | 619.47 | 589.22 |
| 4059 | ENSMUSG00000023078 | 14.89321956 | -3.480011172 | 0.88006313 | -3.954274478 | 7.68E-05 | 0.002930291 | Cxcl13 | chemokine (C-X-C motif) ligand 13 [Source | 14.456 | 57.563 | 10.062 | 2.8919 | 3.2719 | 1.1138 |
| 4060 | ENSMUSG00000023079 | 227.6612071 | 0.133791329 | 0.242512019 | 0.551689477 | 0.581161121 | 0.908645447 | Gtf2ird1 | general transcription factor II I repeat don | 197.56 | 213.81 | 240.49 | 294.98 | 141.78 | 277.35 |
| 4061 | ENSMUSG00000023083 | 0 | NA | NA | NA | NA | NA | H2-M10.2 | histocompatibility 2, M region locus 10.2 [ | 0 | 0 | 0 | 0 | 0 | 0 |
| 4062 | ENSMUSG00000023084 | 2.029147333 | 1.292453086 | 1.657037027 | 0.779978398 | 0.43540359 | NA | Lrrc71 | leucine rich repeat containing 71 [Source: | 2.4093 | 1.0279 | 0 | 0.964 | 1.0906 | 6.6831 |
| 4063 | ENSMUSG00000023087 | 531.7670086 | -1.11894462 | 0.495969328 | -2.256076244 | 0.024065857 | 0.197338277 | Noct | nocturnin [Source:MGI Symbol;Acc:MGI:1 | 609.55 | 966.24 | 608.78 | 188.94 | 545.31 | 271.78 |
| 4064 | ENSMUSG00000023088 | 17.67283333 | -0.944906721 | 0.607363464 | -1.555751664 | 0.119767156 | 0.50648835 | Abcc1 | ATP-binding cassette, sub-family C (CFTR/ | 20.881 | 38.033 | 11.069 | 17.352 | 10.906 | 7.7969 |
| 4065 | ENSMUSG00000023089 | 1291.540119 | -0.019075025 | 0.098942279 | -0.192789426 | 0.847123885 | 0.972803041 | Ndufa5 | NADH:ubiquinone oxidoreductase subuni | 1304.2 | 1307.5 | 1288 | 1223.3 | 1318.6 | 1307.7 |
| 4066 | ENSMUSG00000023092 | 72.80776003 | -0.740954662 | 0.269885659 | -2.745439182 | 0.006042997 | 0.075741685 | Fhl1 | four and a half LIM domains 1 [Source:MG | 101.19 | 94.568 | 77.481 | 57.839 | 54.531 | 51.237 |
| 4067 | ENSMUSG00000023093 | 0 | NA | NA | NA | NA | NA | Pate12 | prostate and testis expressed 12 [Source:I | 0 | 0 | 0 | 0 | 0 | 0 |
| 4068 | ENSMUSG00000023094 | 196.2903189 | -0.150749475 | 0.176608676 | -0.853579101 | 0.393338242 | 0.822026346 | Msrb2 | methionine sulfoxide reductase B2 [Sourc | 215.23 | 210.72 | 193.2 | 172.55 | 178.86 | 207.17 |
| 4069 | ENSMUSG00000023104 | 373.4563289 | 0.044216352 | 0.13642236 | 0.324113671 | 0.745851977 | 0.949997329 | Rfc2 | replication factor C (activator 1) 2 [Source | 378.26 | 343.32 | 381.37 | 382.7 | 370.81 | 384.28 |
| 4070 | ENSMUSG00000023106 | 639.2680502 | -0.133558975 | 0.130816534 | -1.020964027 | 0.307271482 | 0.756237058 | Denr | density-regulated protein [Source:MGI Sy | 642.48 | 741.13 | 622.87 | 569.71 | 591.12 | 668.31 |
| 4071 | ENSMUSG00000023110 | 279.0777822 | -0.111029545 | 0.154823435 | -0.717136556 | 0.473289842 | 0.868952133 | Prmt5 | protein arginine N-methyltransferase 5 [S | 285.9 | 290.9 | 292.82 | 270.88 | 293.38 | 240.59 |
| 4072 | ENSMUSG00000023118 | 1234.569911 | 0.00951551 | 0.102178688 | 0.09312617 | 0.925803327 | 0.987546253 | Sympk | symplekin [Source:MGI Symbol;Acc:MGI:1 | 1293 | 1221.2 | 1176.3 | 1195.3 | 1260.8 | 1260.9 |
| 4073 | ENSMUSG00000023120 | 0 | NA | NA | NA | NA | NA | Ldc1 | leucine decarboxylase 1 [Source:MGI Sym | 0 | 0 | 0 | 0 | 0 | 0 |
| 4074 | ENSMUSG00000023122 | 129.209873 | -0.659706591 | 0.261690745 | -2.52093971 | 0.01170419 | 0.120662445 | Sult1c2 | sulfotransferase family, cytosolic, 1C, mer | 145.36 | 191.19 | 137.86 | 67.478 | 129.78 | 103.59 |
| 4075 | ENSMUSG00000023132 | 2.096738607 | -2.322247464 | 1.672606739 | -1.38840016 | 0.165015228 | NA | Gzma | granzyme A [Source:MGI Symbol;Acc:MGI | 2.4093 | 2.0558 | 6.0375 | 0.964 | 0 | 1.1138 |
| 4076 | ENSMUSG00000023140 | 0 | NA | NA | NA | NA | NA | Reg2 | regenerating islet-derived 2 [Source:MGI S | 0 | 0 | 0 | 0 | 0 | 0 |
| 4077 | ENSMUSG00000023143 | 345.6089146 | -0.063395079 | 0.141793116 | -0.447095604 | 0.654806033 | 0.925797231 | Nagpa | N-acetylglucosamine-1-phosphodiester al | 357.38 | 325.85 | 376.34 | 344.14 | 341.36 | 328.58 |
| 4078 | ENSMUSG00000023147 | 58.43352363 | 0.069028215 | 0.30991783 | 0.222730699 | 0.823745107 | 0.96943211 | Get1 | guided entry of tail-anchored proteins fac | 48.989 | 75.038 | 47.293 | 56.875 | 57.803 | 64.603 |
| 4079 | ENSMUSG00000023150 | 3426.996766 | -0.336843443 | 0.125157837 | -2.691349186 | 0.007116367 | 0.084997751 | Ivns1abp | influenza virus NS1A binding protein [Sou | 3574.6 | 4465.3 | 3437.3 | 3362.4 | 2901 | 2821.4 |
| 4080 | ENSMUSG00000023151 | 1.57624605 | 1.108223452 | 1.942426727 | 0.570535525 | 0.568314535 | NA | Lrrc69 | leucine rich repeat containing 69 [Source: | 0 | 1.0279 | 2.0125 | 0.964 | 5.4531 | 0 |
| 4081 | ENSMUSG00000023153 | 1.145199095 | -1.119908671 | 2.303879602 | -0.486096874 | 0.626898484 | NA | Tmem52 | transmembrane protein 52 [Source:MGI S | 1.6062 | 3.0838 | 0 | 0 | 2.1812 | 0 |
| 4082 | ENSMUSG00000023156 | 334.4005127 | 0.031285853 | 0.155285486 | 0.201473133 | 0.840328636 | 0.971838962 | Rpp14 | ribonuclease P 14 subunit [Source:MGI Sy | 345.33 | 364.91 | 281.75 | 329.68 | 327.19 | 357.54 |
| 4083 | ENSMUSG00000023159 | 0 | NA | NA | NA | NA | NA | Psg29 | pregnancy-specific glycoprotein 29 [Sourc | 0 | 0 | 0 | 0 | 0 | 0 |
| 4084 | ENSMUSG00000023165 | 0 | NA | NA | NA | NA | NA | Ssxb2 | synovial sarcoma, X member B2 [Source:M | 0 | 0 | 0 | 0 | 0 | 0 |

| | | | | | | | | | | | | | | |
|---|---|---|---|---|---|---|---|---|---|---|---|---|---|---|
| 4085 | ENSMUSG00000023169 | 11.99025553 | -2.547052095 | 0.740100351 | -3.441495593 | 0.000578508 | 0.013773229 | Slc38a1 | solute carrier family 38, member 1 [Source | 21.684 | 23.642 | 16.1 | 3.8559 | 1.0906 | 5.5692 |
| 4086 | ENSMUSG00000023170 | 252.0437064 | -0.002446681 | 0.169059903 | -0.014472272 | 0.988453201 | 0.99859176 | Gps2 | G protein pathway suppressor 2 [Source:M | 228.08 | 250.81 | 278.73 | 280.52 | 226.85 | 247.27 |
| 4087 | ENSMUSG00000023175 | 6042.381739 | -0.107582836 | 0.095982931 | -1.120853835 | 0.262350084 | 0.716373725 | Bsg | basigin [Source:MGI Symbol;Acc:MGI:882 | 6659.3 | 5985.6 | 6156.2 | 5536.1 | 5646.1 | 6270.9 |
| 4088 | ENSMUSG00000023176 | 6655.489524 | 0.221973027 | 0.101731923 | 2.181940748 | 0.029113906 | 0.223872162 | Cpn2 | carboxypeptidase N, polypeptide 2 [Source | 5533.4 | 6162.4 | 6738.8 | 6881.8 | 7133.7 | 7482.8 |
| 4089 | ENSMUSG00000023185 | 0 | NA | NA | NA | NA | NA | Ceacam14 | carcinoembryonic antigen-related cell adh | 0 | 0 | 0 | 0 | 0 | 0 |
| 4090 | ENSMUSG00000023186 | 561.2484327 | 0.068235326 | 0.132204719 | 0.516133814 | 0.605760953 | 0.916376001 | Vwa5a | von Willebrand factor A domain containin | 596.7 | 557.13 | 489.03 | 537.9 | 605.29 | 581.43 |
| 4091 | ENSMUSG00000023191 | 48.3714459 | -0.690814981 | 0.372116409 | -1.856448585 | 0.063389645 | 0.360000344 | P3h3 | prolyl 3-hydroxylase 3 [Source:MGI Symbo | 71.476 | 44.2 | 63.393 | 41.451 | 21.812 | 47.895 |
| 4092 | ENSMUSG00000023192 | 0 | NA | NA | NA | NA | NA | Grm2 | glutamate receptor, metabotropic 2 [Sour | 0 | 0 | 0 | 0 | 0 | 0 |
| 4093 | ENSMUSG00000023206 | 852.01174 | 0.25597216 | 0.158577971 | 1.614172249 | 0.10649007 | 0.481264827 | Il15ra | interleukin 15 receptor, alpha chain [Sour | 859.32 | 747.3 | 723.49 | 1161.6 | 849.59 | 770.78 |
| 4094 | ENSMUSG00000023210 | 0 | NA | NA | NA | NA | NA | Lcn9 | lipocalin 9 [Source:MGI Symbol;Acc:MGI:1 | 0 | 0 | 0 | 0 | 0 | 0 |
| 4095 | ENSMUSG00000023216 | 0.185640647 | 1.126924175 | 4.080472857 | 0.276174898 | 0.782413736 | NA | Epb42 | erythrocyte membrane protein band 4.2 [ | 0 | 0 | 0 | 0 | 0 | 1.1138 |
| 4096 | ENSMUSG00000023224 | 12868.26995 | 0.143057743 | 0.084949758 | 1.684027676 | 0.09217632 | 0.447243079 | Serping1 | serine (or cysteine) peptidase inhibitor, cl | 12319 | 12215 | 12159 | 14345 | 13143 | 13029 |
| 4097 | ENSMUSG00000023232 | 172.509331 | -0.175031264 | 0.205385545 | -0.852208288 | 0.394098497 | 0.822827574 | Serinc2 | serine incorporator 2 [Source:MGI Symbo | 202.38 | 188.11 | 157.98 | 161.95 | 133.06 | 191.58 |
| 4098 | ENSMUSG00000023235 | 113.6394331 | 0.173708814 | 0.233176178 | 0.744968098 | 0.456291006 | 0.857895893 | Ccl25 | chemokine (C-C motif) ligand 25 [Source:M | 110.83 | 119.24 | 90.562 | 138.81 | 122.15 | 100.25 |
| 4099 | ENSMUSG00000023236 | 0.167707431 | -0.796629176 | 4.080472857 | -0.195229623 | 0.84521319 | NA | Scg5 | secretogranin V [Source:MGI Symbol;Acc: | 0 | 0 | 1.0062 | 0 | 0 | 0 |
| 4100 | ENSMUSG00000023243 | 1365.04043 | -0.502909993 | 0.507618355 | -0.990724601 | 0.321820075 | 0.769262199 | Kcnk5 | potassium channel, subfamily K, member | 1189.4 | 1806.1 | 1806.2 | 537.9 | 1761.3 | 1089.3 |
| 4101 | ENSMUSG00000023247 | 0 | NA | NA | NA | NA | NA | Guca2a | guanylate cyclase activator 2a (guanylin) [ | 0 | 0 | 0 | 0 | 0 | 0 |
| 4102 | ENSMUSG00000023249 | 488.4600667 | 0.119132515 | 0.127667405 | 0.933147463 | 0.350743836 | 0.791943999 | Parp3 | poly (ADP-ribose) polymerase family, men | 479.45 | 447.14 | 477.97 | 506.09 | 478.78 | 541.33 |
| 4103 | ENSMUSG00000023257 | 0 | NA | NA | NA | NA | NA | Cypt1 | cysteine-rich perinuclear theca 1 [Source: | 0 | 0 | 0 | 0 | 0 | 0 |
| 4104 | ENSMUSG00000023259 | 48.76995759 | 0.007589585 | 0.368566722 | 0.020592161 | 0.983570994 | 0.998580735 | Slc26a6 | solute carrier family 26, member 6 [Source | 65.051 | 35.977 | 44.275 | 34.703 | 46.897 | 65.717 |
| 4105 | ENSMUSG00000023262 | 933.6395638 | 0.042287214 | 0.121369018 | 0.348418519 | 0.727525897 | 0.945907265 | Acy1 | aminoacylase 1 [Source:MGI Symbol;Acc:I | 936.42 | 865.51 | 956.94 | 832.88 | 1012.1 | 998 |
| 4106 | ENSMUSG00000023263 | 0 | NA | NA | NA | NA | NA | 9530002B09 | RIKEN cDNA 9530002B09 gene [Source:M | 0 | 0 | 0 | 0 | 0 | 0 |
| 4107 | ENSMUSG00000023266 | 57.75914727 | -0.027659368 | 0.300837513 | -0.09194122 | 0.926744741 | 0.987712228 | Frs3 | fibroblast growth factor receptor substrat | 56.217 | 47.284 | 71.443 | 55.911 | 58.893 | 56.806 |
| 4108 | ENSMUSG00000023267 | 2.425878555 | -0.317361786 | 1.523403258 | -0.208324214 | 0.834975826 | NA | Gabrr2 | gamma-aminobutyric acid (GABA) C recep | 0 | 3.0838 | 5.0312 | 0.964 | 4.3625 | 1.1138 |
| 4109 | ENSMUSG00000023272 | 452.6272787 | 0.659503699 | 0.218573333 | 3.017310895 | 0.002550281 | 0.041917347 | Creld2 | cysteine-rich with EGF-like domains 2 [Sou | 342.92 | 381.36 | 329.04 | 716.24 | 340.27 | 605.93 |
| 4110 | ENSMUSG00000023274 | 41.13578296 | 0.76050591 | 0.476075151 | 1.597449288 | 0.110165593 | 0.488035265 | Cd4 | CD4 antigen [Source:MGI Symbol;Acc:MG | 21.684 | 25.698 | 44.275 | 22.171 | 83.978 | 49.009 |
| 4111 | ENSMUSG00000023277 | 51.33532216 | -0.230779182 | 0.338196541 | -0.682381852 | 0.494997528 | 0.877919732 | Twf2 | twinfilin actin binding protein 2 [Source:M | 69.067 | 55.508 | 41.256 | 39.523 | 43.625 | 59.034 |
| 4112 | ENSMUSG00000023279 | 0 | NA | NA | NA | NA | NA | Bmp15 | bone morphogenetic protein 15 [Source:M | 0 | 0 | 0 | 0 | 0 | 0 |
| 4113 | ENSMUSG00000023284 | 43.61776806 | 0.032740798 | 0.336101309 | 0.097413481 | 0.92239804 | 0.986685804 | Zfp605 | zinc finger protein 605 [Source:MGI Symb | 40.155 | 37.005 | 52.325 | 46.271 | 43.625 | 42.326 |
| 4114 | ENSMUSG00000023286 | 659.5648789 | 0.111627184 | 0.112687933 | 0.990586849 | 0.321887362 | 0.769295903 | Ube2j2 | ubiquitin-conjugating enzyme E2J 2 [Sourc | 640.07 | 637.31 | 624.88 | 701.78 | 678.36 | 674.99 |
| 4115 | ENSMUSG00000023289 | 0 | NA | NA | NA | NA | NA | Sva | seminal vesicle antigen [Source:MGI Symb | 0 | 0 | 0 | 0 | 0 | 0 |
| 4116 | ENSMUSG00000023307 | 1985.476869 | 0.14832838 | 0.103033466 | 1.439613613 | 0.149976746 | 0.563152558 | Marchf5 | membrane associated ring-CH-type finger | 1985.3 | 1954.1 | 1710.6 | 2115 | 2145.2 | 2002.7 |
| 4117 | ENSMUSG00000023328 | 8.137440299 | -0.101113762 | 0.801838427 | -0.126102415 | 0.899650856 | 0.982798755 | Ache | acetylcholinesterase [Source:MGI Symbol | 8.031 | 12.335 | 5.0312 | 13.496 | 4.3625 | 5.5692 |
| 4118 | ENSMUSG00000023330 | 133.5737704 | 0.013953324 | 0.210722722 | 0.066216514 | 0.947205449 | 0.990677354 | Dtwd1 | DTW domain containing 1 [Source:MGI Sy | 138.13 | 132.6 | 127.79 | 123.39 | 159.23 | 120.3 |
| 4119 | ENSMUSG00000023333 | 0 | NA | NA | NA | NA | NA | Gcm1 | glial cells missing homolog 1 [Source:MGI | 0 | 0 | 0 | 0 | 0 | 0 |
| 4120 | ENSMUSG00000023336 | 53.36100002 | -0.82248808 | 0.302947758 | -2.714950214 | 0.006628573 | 0.081044536 | Wfdc1 | WAP four-disulfide core domain 1 [Source | 68.264 | 67.843 | 68.425 | 39.523 | 34.9 | 41.212 |
| 4121 | ENSMUSG00000023341 | 98.73167166 | 0.642905392 | 0.26256023 | 2.448601577 | 0.014341199 | 0.138765904 | Mx2 | MX dynamin-like GTPase 2 [Source:MGI S | 86.735 | 53.452 | 90.562 | 107.97 | 117.79 | 135.89 |
| 4122 | ENSMUSG00000023345 | 49.12822402 | 0.041865422 | 0.356669594 | 0.117378725 | 0.906559942 | 0.984272857 | Poc1a | POC1 centriolar protein A [Source:MGI Sy | 62.642 | 30.838 | 51.318 | 44.343 | 61.075 | 44.554 |
| 4123 | ENSMUSG00000023348 | 274.8124074 | 0.270831245 | 0.162345101 | 1.66824403 | 0.095267294 | 0.455057116 | Trip6 | thyroid hormone receptor interactor 6 [Sc | 281.09 | 228.2 | 237.47 | 325.82 | 293.38 | 282.92 |
| 4124 | ENSMUSG00000023349 | 22.32417682 | -1.476397437 | 0.588957063 | -2.506799782 | 0.01218297 | 0.124007373 | Clec4n | C-type lectin domain family 4, member n [ | 28.912 | 49.34 | 20.125 | 6.7478 | 6.5437 | 22.277 |
| 4125 | ENSMUSG00000023350 | 0 | NA | NA | NA | NA | NA | Phf8-ps | PHD finger protein 8, pseudogene [Source | 0 | 0 | 0 | 0 | 0 | 0 |
| 4126 | ENSMUSG00000023353 | 722.7696813 | 0.175685124 | 0.126229662 | 1.391789545 | 0.163986137 | 0.588267729 | Agap3 | ArfGAP with GTPase domain, ankyrin repe | 658.54 | 704.12 | 674.18 | 791.43 | 670.73 | 837.61 |
| 4127 | ENSMUSG00000023367 | 2368.019657 | -0.365055137 | 0.146200119 | -2.49695513 | 0.012526481 | 0.126529431 | Tmem176a | transmembrane protein 176A [Source:MG | 3021.3 | 2860.7 | 2116.1 | 2355 | 2039.5 | 1815.6 |
| 4128 | ENSMUSG00000023387 | 0 | NA | NA | NA | NA | NA | Kcnk16 | potassium channel, subfamily K, member | 0 | 0 | 0 | 0 | 0 | 0 |
| 4129 | ENSMUSG00000023391 | 0 | NA | NA | NA | NA | NA | Dlx2 | distal-less homeobox 2 [Source:MGI Symb | 0 | 0 | 0 | 0 | 0 | 0 |
| 4130 | ENSMUSG00000023393 | 93.02356433 | 0.523532359 | 0.241453318 | 2.168254982 | 0.030139292 | 0.229802169 | Slc17a9 | solute carrier family 17, member 9 [Source | 83.523 | 69.898 | 75.468 | 124.35 | 106.88 | 98.018 |
| 4131 | ENSMUSG00000023403 | 0 | NA | NA | NA | NA | NA | Stk31 | serine threonine kinase 31 [Source:MGI S | 0 | 0 | 0 | 0 | 0 | 0 |
| 4132 | ENSMUSG00000023411 | 5.284105103 | -0.680590751 | 1.052282371 | -0.64677578 | 0.51777707 | 0.887022094 | Nfatc4 | nuclear factor of activated T cells, cytopla | 2.4093 | 9.2513 | 8.05 | 8.6758 | 1.0906 | 2.2277 |
| 4133 | ENSMUSG00000023433 | 0.267700746 | -1.497481649 | 4.057001436 | -0.369110456 | 0.712045396 | NA | Cela3b | chymotrypsin-like elastase family, membe | 1.6062 | 0 | 0 | 0 | 0 | 0 |
| 4134 | ENSMUSG00000023439 | 0 | NA | NA | NA | NA | NA | Gnb3 | guanine nucleotide binding protein (G pro | 0 | 0 | 0 | 0 | 0 | 0 |
| 4135 | ENSMUSG00000023443 | 0 | NA | NA | NA | NA | NA | Esx1 | extraembryonic, spermatogenesis, homeo | 0 | 0 | 0 | 0 | 0 | 0 |
| 4136 | ENSMUSG00000023452 | 216.4958335 | 0.092061309 | 0.17979432 | 0.5120368 | 0.608625254 | 0.916376001 | Pisd | phosphatidylserine decarboxylase [Source | 207.2 | 217.92 | 204.27 | 264.13 | 208.31 | 197.15 |
| 4137 | ENSMUSG00000023456 | 7423.102317 | 0.043756181 | 0.094689168 | 0.462103348 | 0.644007209 | 0.925643431 | Tpi1 | triosephosphate isomerase 1 [Source:MG | 7192.6 | 7341.4 | 7396.9 | 6822.1 | 7832.8 | 7952.8 |
| 4138 | ENSMUSG00000023460 | 563.6473794 | -0.192288748 | 0.130179345 | -1.477106437 | 0.139647108 | 0.545906967 | Rab12 | RAB12, member RAS oncogene family [So | 660.15 | 571.52 | 570.54 | 492.59 | 524.59 | 562.49 |
| 4139 | ENSMUSG00000023467 | 9.193214908 | 0.428300201 | 0.711727144 | 0.601775841 | 0.547323359 | 0.895518681 | Tulp2 | tubby-like protein 2 [Source:MGI Symbol;. | 7.2279 | 10.279 | 6.0375 | 9.6398 | 14.178 | 7.7969 |
| 4140 | ENSMUSG00000023473 | 0.888749571 | -1.92056428 | 2.547325431 | -0.753953247 | 0.450877296 | NA | Celsr3 | cadherin, EGF LAG seven-pass G-type rece | 3.2124 | 0 | 1.0062 | 0 | 0 | 1.1138 |

| | | | | | | | | | | | | | | | |
|---|---|---|---|---|---|---|---|---|---|---|---|---|---|---|---|
| 4141 | ENSMUSG00000023484 | 16.19838621 | -0.825771012 | 0.592137095 | -1.394560515 | 0.163148409 | 0.587009599 | Prph | peripherin [Source:MGI Symbol;Acc:MGI:9 | 10.44 | 23.642 | 28.175 | 9.6398 | 15.269 | 10.025 |
| 4142 | ENSMUSG00000023495 | 384.0210935 | 0.699209973 | 0.156873582 | 4.457155657 | 8.31E-06 | 0.000455037 | Pcbp4 | poly(rC) binding protein 4 [Source:MGI Sy | 327.67 | 283.71 | 265.65 | 430.9 | 450.43 | 545.78 |
| 4143 | ENSMUSG00000023505 | 35.92054755 | 0.192130524 | 0.370784321 | 0.51817327 | 0.604337383 | 0.916376001 | Cdca3 | cell division cycle associated 3 [Source:MG | 39.352 | 27.754 | 33.206 | 33.739 | 44.715 | 36.757 |
| 4144 | ENSMUSG00000023571 | 14.85146539 | 0.08767994 | 0.565796311 | 0.154967324 | 0.876847078 | 0.980632845 | C1qtnf12 | C1q and tumor necrosis factor related 12 | 12.85 | 13.363 | 17.106 | 18.316 | 17.45 | 10.025 |
| 4145 | ENSMUSG00000023572 | 320.4700945 | 0.065424331 | 0.144450433 | 0.452918897 | 0.650607143 | 0.925643431 | Ccndbp1 | cyclin D-type binding-protein 1 [Source:M | 305.18 | 309.4 | 325.02 | 317.15 | 314.1 | 351.97 |
| 4146 | ENSMUSG00000023577 | 0 | NA | NA | NA | NA | NA | NA | NA | 0 | 0 | 0 | 0 | 0 | 0 |
| 4147 | ENSMUSG00000023707 | 563.1705464 | 0.136648536 | 0.132987889 | 1.027526175 | 0.304172767 | 0.752961647 | Ogfod2 | 2-oxoglutarate and iron-dependent oxyge | 561.37 | 488.26 | 559.47 | 600.56 | 531.13 | 638.23 |
| 4148 | ENSMUSG00000023723 | 382.3325614 | -0.077033485 | 0.135472637 | -0.568627637 | 0.569608866 | 0.90387522 | Mrps23 | mitochondrial ribosomal protein S23 [Sou | 369.43 | 408.08 | 400.49 | 363.42 | 383.9 | 368.68 |
| 4149 | ENSMUSG00000023737 | 2.790734926 | -0.394764107 | 1.376787907 | -0.286728337 | 0.774320338 | NA | 1700082M2 | RIKEN cDNA 1700082M22 gene [Source:M | 6.4248 | 0 | 3.0187 | 2.8919 | 2.1812 | 2.2277 |
| 4150 | ENSMUSG00000023755 | 22.3947203 | 0.642467244 | 0.481033812 | 1.335596849 | 0.18168108 | 0.615095921 | Rhebl1 | Ras homolog enriched in brain like 1 [Sour | 16.865 | 14.391 | 21.131 | 21.208 | 22.903 | 37.871 |
| 4151 | ENSMUSG00000023764 | 273.4759931 | 0.137968807 | 0.191180768 | 0.721666768 | 0.470499378 | 0.867110762 | Sfi1 | Sfi1 homolog, spindle assembly associated | 215.23 | 274.45 | 292.82 | 348.96 | 244.3 | 265.09 |
| 4152 | ENSMUSG00000023781 | 0.513958919 | -2.394274821 | 3.756096013 | -0.637437066 | 0.523840191 | NA | Hes7 | hes family bHLH transcription factor 7 [So | 0 | 3.0838 | 0 | 0 | 0 | 0 |
| 4153 | ENSMUSG00000023791 | 265.1138974 | 0.355924337 | 0.168862434 | 2.107776903 | 0.035050292 | 0.251098275 | Pigx | phosphatidylinositol glycan anchor biosyn | 203.99 | 239.5 | 254.58 | 264.13 | 301.01 | 327.47 |
| 4154 | ENSMUSG00000023795 | 10.92607914 | -0.482782952 | 0.659213924 | -0.732361582 | 0.463947901 | 0.861872906 | Pisd-ps2 | phosphatidylserine decarboxylase, pseudo | 12.85 | 9.2513 | 16.1 | 8.6758 | 11.997 | 6.6831 |
| 4155 | ENSMUSG00000023800 | 245.3115636 | 0.75267796 | 0.197329814 | 3.814314449 | 0.000136561 | 0.004615013 | Tiam2 | T cell lymphoma invasion and metastasis 2 | 184.71 | 143.91 | 219.36 | 304.62 | 262.84 | 356.43 |
| 4156 | ENSMUSG00000023802 | 0 | NA | NA | NA | NA | NA | Nox3 | NADPH oxidase 3 [Source:MGI Symbol;Ac | 0 | 0 | 0 | 0 | 0 | 0 |
| 4157 | ENSMUSG00000023805 | 145.4107797 | 0.334256592 | 0.349661376 | 0.955943709 | 0.339100675 | 0.780887145 | Synj2 | synaptojanin 2 [Source:MGI Symbol;Acc:M | 210.41 | 118.21 | 56.35 | 140.74 | 140.69 | 206.06 |
| 4158 | ENSMUSG00000023806 | 197.3505743 | 0.221347579 | 0.180553995 | 1.22593565 | 0.220222892 | 0.66652101 | Rsph3b | radial spoke 3B homolog (Chlamydomona | 179.89 | 184 | 183.14 | 226.53 | 231.21 | 179.33 |
| 4159 | ENSMUSG00000023809 | 14.71057306 | -1.203365677 | 0.65459961 | -1.838323241 | 0.066014788 | 0.368948184 | Rps6ka2 | ribosomal protein S6 kinase, polypeptide 2 | 20.881 | 28.782 | 12.075 | 14.46 | 8.7249 | 3.3415 |
| 4160 | ENSMUSG00000023826 | 46.56532703 | -0.123141141 | 0.366677142 | -0.335829881 | 0.736999155 | 0.948137969 | Prkn | parkin RBR E3 ubiquitin protein ligase [Sou | 69.87 | 43.173 | 32.2 | 48.199 | 43.625 | 42.326 |
| 4161 | ENSMUSG00000023827 | 17.10173794 | -0.339388804 | 0.523838793 | -0.647887879 | 0.517057471 | 0.886839709 | Agpat4 | 1-acylglycerol-3-phosphate O-acyltransfer | 17.668 | 20.558 | 19.119 | 15.424 | 10.906 | 18.935 |
| 4162 | ENSMUSG00000023828 | 95.88857795 | -0.031209128 | 0.298753509 | -0.104464475 | 0.916800759 | 0.986374671 | Slc22a3 | solute carrier family 22 (organic cation tra | 104.4 | 82.233 | 103.64 | 60.731 | 136.33 | 87.994 |
| 4163 | ENSMUSG00000023829 | 7994.66881 | 0.026054351 | 0.11024923 | 0.23632229 | 0.813182592 | 0.966802988 | Slc22a1 | solute carrier family 22 (organic cation tra | 9157.8 | 7363 | 7245 | 8272.9 | 7880.8 | 8048.6 |
| 4164 | ENSMUSG00000023830 | 713.5472995 | 0.117991647 | 0.133085011 | 0.886588553 | 0.375300459 | 0.809332934 | Igf2r | insulin-like growth factor 2 receptor [Sour | 644.89 | 744.21 | 664.12 | 663.22 | 841.96 | 722.88 |
| 4165 | ENSMUSG00000023832 | 921.6614885 | 0.199776243 | 0.14792553 | 1.350519031 | 0.176849553 | 0.607589534 | Acat2 | acetyl-Coenzyme A acetyltransferase 2 [Sc | 962.92 | 931.29 | 679.22 | 1064.2 | 999.01 | 893.3 |
| 4166 | ENSMUSG00000023845 | 637.0788005 | 0.047494481 | 0.152913396 | 0.310597254 | 0.756106815 | 0.953085143 | Lnpep | leucyl/cystinyl aminopeptidase [Source:M | 548.52 | 739.07 | 593.68 | 739.37 | 628.2 | 573.63 |
| 4167 | ENSMUSG00000023852 | 515.1635141 | 0.081913473 | 0.132460864 | 0.618397549 | 0.536313313 | 0.890560262 | Chd1 | chromodomain helicase DNA binding prot | 452.95 | 537.6 | 512.18 | 556.22 | 494.05 | 537.99 |
| 4168 | ENSMUSG00000023861 | 1854.386775 | -0.334350005 | 0.116549071 | -2.868748769 | 0.00412099 | 0.058613981 | Mpc1 | mitochondrial pyruvate carrier 1 [Source:I | 2208.5 | 1841 | 2155.4 | 1775.6 | 1661 | 1484.8 |
| 4169 | ENSMUSG00000023868 | 6.095222418 | -0.89227348 | 0.874125253 | -1.020761586 | 0.307367408 | 0.756237058 | Pde10a | phosphodiesterase 10A [Source:MGI Sym | 6.4248 | 10.279 | 7.0437 | 2.8919 | 4.3625 | 5.5692 |
| 4170 | ENSMUSG00000023873 | 6.78798388 | -0.539676972 | 0.841935143 | -0.640995897 | 0.521525349 | 0.888064228 | 1700010I14F | RIKEN cDNA 1700010I14 gene [Source:MG | 12.85 | 6.1675 | 5.0312 | 6.7478 | 4.3625 | 5.5692 |
| 4171 | ENSMUSG00000023882 | 38.67777026 | -0.238149811 | 0.362540552 | -0.65689151 | 0.511250677 | 0.883679955 | Zfp54 | zinc finger protein 54 [Source:MGI Symbo | 44.974 | 47.284 | 33.206 | 33.739 | 30.537 | 42.326 |
| 4172 | ENSMUSG00000023883 | 271.4253301 | 0.1817441 | 0.155251096 | 1.170646169 | 0.241741031 | 0.691791181 | Phf10 | PHD finger protein 10 [Source:MGI Symbc | 246.55 | 271.37 | 245.52 | 302.69 | 265.02 | 297.4 |
| 4173 | ENSMUSG00000023885 | 23.11876301 | -0.370692171 | 0.519433443 | -0.7136471 | 0.475445425 | 0.86938592 | Thbs2 | thrombospondin 2 [Source:MGI Symbol;A | 43.368 | 15.419 | 19.119 | 22.171 | 16.359 | 22.277 |
| 4174 | ENSMUSG00000023886 | 100.273102 | -0.514933412 | 0.307139999 | -1.676542992 | 0.093631883 | 0.450459278 | Smoc2 | SPARC related modular calcium binding 2 | 151.79 | 128.49 | 73.456 | 89.65 | 100.34 | 57.92 |
| 4175 | ENSMUSG00000023892 | 66.24335472 | 0.237591603 | 0.291452596 | 0.8151981 | 0.414958925 | 0.836001897 | Zfp51 | zinc finger protein 51 [Source:MGI Symbo | 68.264 | 59.619 | 54.337 | 72.298 | 87.249 | 55.692 |
| 4176 | ENSMUSG00000023902 | 0 | NA | NA | NA | NA | NA | Zscan10 | zinc finger and SCAN domain containing 1 | 0 | 0 | 0 | 0 | 0 | 0 |
| 4177 | ENSMUSG00000023903 | 0.133850373 | -0.796629176 | 4.080472857 | -0.195229623 | 0.84521319 | NA | Mmp25 | matrix metallopeptidase 25 [Source:MGI S | 0.8031 | 0 | 0 | 0 | 0 | 0 |
| 4178 | ENSMUSG00000023904 | 583.2870512 | -0.017227953 | 0.129903708 | -0.132620947 | 0.894493165 | 0.982798755 | Hcfc1r1 | host cell factor C1 regulator 1 (XPO1-depe | 629.63 | 607.5 | 522.24 | 546.58 | 587.84 | 605.93 |
| 4179 | ENSMUSG00000023905 | 125.2305294 | -0.168740441 | 0.233513108 | -0.722616572 | 0.469915484 | 0.866946117 | Tnfrsf12a | tumor necrosis factor receptor superfamil | 123.68 | 141.85 | 132.82 | 150.38 | 107.97 | 94.677 |
| 4180 | ENSMUSG00000023906 | 4.411906453 | -0.543494632 | 1.20408754 | -0.451374683 | 0.651719527 | NA | Cldn6 | claudin 6 [Source:MGI Symbol;Acc:MGI:1 | 6.4248 | 7.1954 | 2.0125 | 1.928 | 0 | 8.9108 |
| 4181 | ENSMUSG00000023908 | 7.487067653 | -0.726697839 | 0.77452261 | -0.938252582 | 0.348114622 | 0.789711685 | Pkmyt1 | protein kinase, membrane associated tyrc | 9.6372 | 10.279 | 8.05 | 4.8199 | 5.4531 | 6.6831 |
| 4182 | ENSMUSG00000023909 | 13.21677394 | -0.763533791 | 0.615932049 | -1.239639651 | 0.215108708 | 0.659728602 | Paqr4 | progestin and adipoQ receptor family mer | 10.44 | 18.503 | 21.131 | 11.568 | 7.6343 | 10.025 |
| 4183 | ENSMUSG00000023911 | 9.128248303 | 0.627129874 | 0.75382934 | 0.831925532 | 0.405450979 | 0.829973745 | Flywch2 | FLYWCH family member 2 [Source:MGI Sy | 6.4248 | 3.0838 | 12.075 | 14.46 | 9.8156 | 8.9108 |
| 4184 | ENSMUSG00000023912 | 23.8557346 | -0.743168627 | 0.524202087 | -1.41771398 | 0.156274287 | 0.574894692 | Slc25a27 | solute carrier family 25, member 27 [Sour | 28.912 | 26.726 | 34.212 | 28.919 | 6.5437 | 17.822 |
| 4185 | ENSMUSG00000023913 | 38.11464329 | -2.031290318 | 0.456982403 | -4.44500774 | 8.79E-06 | 0.000477078 | Pla2g7 | phospholipase A2, group VII (platelet-acti | 60.233 | 90.457 | 33.206 | 18.316 | 11.997 | 14.48 |
| 4186 | ENSMUSG00000023914 | 0.133850373 | -0.796629176 | 4.080472857 | -0.195229623 | 0.84521319 | NA | Mep1a | meprin 1 alpha [Source:MGI Symbol;Acc:I | 0.8031 | 0 | 0 | 0 | 0 | 0 |
| 4187 | ENSMUSG00000023915 | 73.75927299 | -1.578111659 | 0.289204395 | -5.456734713 | 4.85E-08 | 5.89E-06 | Tnfrsf21 | tumor necrosis factor receptor superfamil | 109.22 | 129.52 | 92.575 | 31.811 | 35.99 | 43.44 |
| 4188 | ENSMUSG00000023918 | 0.356960287 | 0.16514264 | 3.880370363 | 0.042558474 | 0.966053499 | NA | Adgrf4 | adhesion G protein-coupled receptor F4 [S | 0 | 1.0279 | 0 | 0 | 0 | 1.1138 |
| 4189 | ENSMUSG00000023919 | 37.89249607 | 0.705920353 | 0.386801908 | 1.825017763 | 0.067998349 | 0.375535746 | Cenpq | centromere protein Q [Source:MGI Symbc | 17.668 | 34.949 | 34.212 | 42.415 | 47.987 | 50.123 |
| 4190 | ENSMUSG00000023921 | 3995.03869 | 0.019944556 | 0.104339285 | 0.191150973 | 0.848407313 | 0.972803041 | Mmut | methylmalonyl-Coenzyme A mutase [Sour | 3646.1 | 4222.7 | 4034 | 3709.4 | 4352.7 | 4005.4 |
| 4191 | ENSMUSG00000023923 | 271.2286344 | 0.041608463 | 0.164908936 | 0.252311757 | 0.800800101 | 0.962424862 | Tbc1d5 | TBC1 domain family, member 5 [Source:M | 299.56 | 249.78 | 251.56 | 245.81 | 297.74 | 282.92 |
| 4192 | ENSMUSG00000023926 | 0.133850373 | -0.796629176 | 4.080472857 | -0.195229623 | 0.84521319 | NA | Rhag | Rhesus blood group-associated A glycopro | 0.8031 | 0 | 0 | 0 | 0 | 0 |
| 4193 | ENSMUSG00000023927 | 7.292269435 | -0.536777256 | 0.802072682 | -0.669237674 | 0.503343878 | 0.881357141 | Satb1 | special AT-rich sequence binding protein 1 | 5.6217 | 12.335 | 8.05 | 6.7478 | 6.5437 | 4.4554 |
| 4194 | ENSMUSG00000023930 | 0.181769715 | 1.126924175 | 4.080472857 | 0.276174898 | 0.782413736 | NA | Crisp2 | cysteine-rich secretory protein 2 [Source:I | 0 | 0 | 0 | 0 | 1.0906 | 0 |
| 4195 | ENSMUSG00000023931 | 0.724629373 | 1.713042495 | 3.014019035 | 0.56835822 | 0.569791754 | NA | Efhb | EF hand domain family, member B [Source | 0 | 0 | 1.0062 | 0 | 0 | 3.3415 |
| 4196 | ENSMUSG00000023932 | 376.3223644 | 0.178183946 | 0.14378025 | 1.239279711 | 0.21524193 | 0.659728602 | Cdc5l | cell division cycle 5-like (S. pombe) [Sourc | 383.08 | 335.1 | 340.11 | 364.38 | 424.25 | 411.01 |

| | | | | | | | | | | | | | | | |
|---|---|---|---|---|---|---|---|---|---|---|---|---|---|---|---|
| 4197 | ENSMUSG00000023935 | 1.343848784 | -0.601579104 | 1.8484757 | -0.325446044 | 0.744843512 | NA | Spats1 | spermatogenesis associated, serine-rich 1 | 0.8031 | 2.0558 | 2.0125 | 0.964 | 0 | 2.2277 |
| 4198 | ENSMUSG00000023938 | 431.814575 | 0.046667547 | 0.134579364 | 0.346765996 | 0.728767125 | 0.946248466 | Aars2 | alanyl-tRNA synthetase 2, mitochondrial [ | 451.34 | 421.45 | 401.49 | 466.57 | 410.07 | 439.97 |
| 4199 | ENSMUSG00000023939 | 982.2291528 | 0.204398054 | 0.113411564 | 1.802268187 | 0.071503221 | 0.38710107 | Mrpl14 | mitochondrial ribosomal protein L14 [Sou | 937.22 | 855.23 | 944.86 | 966.87 | 1075.3 | 1113.8 |
| 4200 | ENSMUSG00000023940 | 2.604835554 | -0.632386176 | 1.443305342 | -0.438151344 | 0.661276577 | NA | Sgo1 | shugoshin 1 [Source:MGI Symbol;Acc:MG | 6.4248 | 1.0279 | 2.0125 | 2.8919 | 3.2719 | 0 |
| 4201 | ENSMUSG00000023942 | 3417.62165 | -0.195185125 | 0.126382196 | -1.544403647 | 0.122490577 | 0.51297445 | Slc29a1 | solute carrier family 29 (nucleoside transp | 3292.7 | 3787.9 | 3865 | 2947.8 | 2852 | 3760.3 |
| 4202 | ENSMUSG00000023943 | 0 | NA | NA | NA | NA | NA | Sult1c1 | sulfotransferase family, cytosolic, 1C, mer | 0 | 0 | 0 | 0 | 0 | 0 |
| 4203 | ENSMUSG00000023944 | 14376.9034 | 0.128699987 | 0.087648941 | 1.468357584 | 0.142007117 | 0.551555032 | Hsp90ab1 | heat shock protein 90 alpha (cytosolic), cla | 14085 | 13406 | 13717 | 15633 | 13989 | 15431 |
| 4204 | ENSMUSG00000023945 | 0 | NA | NA | NA | NA | NA | Slc5a7 | solute carrier family 5 (choline transporte | 0 | 0 | 0 | 0 | 0 | 0 |
| 4205 | ENSMUSG00000023947 | 43.38237462 | -0.347722062 | 0.377512232 | -0.921088199 | 0.357004381 | 0.796166767 | Nfkbie | nuclear factor of kappa light polypeptide g | 55.414 | 28.782 | 61.381 | 38.559 | 32.719 | 43.44 |
| 4206 | ENSMUSG00000023949 | 0 | NA | NA | NA | NA | NA | Tcte1 | t-complex-associated testis expressed 1 [S | 0 | 0 | 0 | 0 | 0 | 0 |
| 4207 | ENSMUSG00000023951 | 2598.985203 | 0.451914939 | 0.134072489 | 3.370676135 | 0.00074984 | 0.016872984 | Vegfa | vascular endothelial growth factor A [Sour | 2129 | 2110.3 | 2347.6 | 3657.3 | 2725.5 | 2624.2 |
| 4208 | ENSMUSG00000023952 | 627.8717816 | 0.375333013 | 0.154110966 | 2.435472459 | 0.014872357 | 0.141989285 | Gtpbp2 | GTP binding protein 2 [Source:MGI Symbo | 646.5 | 445.09 | 547.4 | 781.79 | 688.18 | 658.28 |
| 4209 | ENSMUSG00000023953 | 93.62387994 | 0.075068147 | 0.246846769 | 0.304108283 | 0.761045396 | 0.95402891 | Polh | polymerase (DNA directed), eta (RAD 30 r | 82.72 | 84.289 | 106.66 | 87.722 | 111.24 | 89.108 |
| 4210 | ENSMUSG00000023959 | 86.48736878 | 0.59263511 | 0.30634872 | 1.934511459 | 0.053050275 | 0.323490376 | Clic5 | chloride intracellular channel 5 [Source:M | 76.295 | 60.647 | 69.431 | 62.659 | 138.51 | 111.38 |
| 4211 | ENSMUSG00000023960 | 576.3488149 | 0.184761733 | 0.153341621 | 1.204902697 | 0.228240867 | 0.674708584 | Enpp5 | ectonucleotide pyrophosphatase/phosph | 526.03 | 595.16 | 497.08 | 531.15 | 737.26 | 571.4 |
| 4212 | ENSMUSG00000023961 | 246.7473966 | 0.046260257 | 0.159019066 | 0.290910128 | 0.771120055 | 0.956965056 | Enpp4 | ectonucleotide pyrophosphatase/phosph | 245.75 | 253.9 | 228.42 | 233.28 | 257.39 | 261.75 |
| 4213 | ENSMUSG00000023963 | 688.227254 | -0.756734301 | 0.434849125 | -1.74022266 | 0.081819928 | 0.421154249 | Cyp39a1 | cytochrome P450, family 39, subfamily a, | 555.75 | 1011.5 | 1027.4 | 755.76 | 345.73 | 433.29 |
| 4214 | ENSMUSG00000023964 | 1.357472383 | -0.566420676 | 1.86742493 | -0.303316437 | 0.761648721 | NA | Calcr | calcitonin receptor [Source:MGI Symbol;A | 0.8031 | 1.0279 | 3.0187 | 0 | 2.1812 | 1.1138 |
| 4215 | ENSMUSG00000023965 | 544.1097168 | 0.027813822 | 0.140746777 | 0.197616047 | 0.843345479 | 0.972273952 | Fbxl17 | F-box and leucine-rich repeat protein 17 [ | 554.94 | 486.21 | 575.57 | 604.41 | 477.69 | 565.83 |
| 4216 | ENSMUSG00000023966 | 7.65322901 | 0.29168617 | 0.860874622 | 0.338825379 | 0.734741281 | 0.948109711 | Rsph9 | radial spoke head 9 homolog (Chlamydom | 2.4093 | 14.391 | 4.025 | 9.6398 | 6.5437 | 8.9108 |
| 4217 | ENSMUSG00000023967 | 476.0717646 | 0.074239978 | 0.153474385 | 0.483728784 | 0.628578363 | 0.920328819 | Mrps18a | mitochondrial ribosomal protein S18A [So | 536.47 | 393.69 | 459.85 | 439.57 | 528.95 | 497.89 |
| 4218 | ENSMUSG00000023968 | 22.91863303 | 0.684945653 | 0.484201936 | 1.414586771 | 0.15718969 | 0.576436451 | Crip3 | cysteine-rich protein 3 [Source:MGI Symb | 20.078 | 12.335 | 20.125 | 21.208 | 39.262 | 24.505 |
| 4219 | ENSMUSG00000023971 | 368.8884291 | 0.111200017 | 0.135349975 | 0.821573973 | 0.411319408 | 0.833705705 | Rrp36 | ribosomal RNA processing 36 [Source:MG | 370.23 | 339.21 | 354.2 | 382.7 | 387.17 | 379.82 |
| 4220 | ENSMUSG00000023972 | 6.060310464 | -0.942370357 | 0.908294447 | -1.03751637 | 0.29949527 | 0.750762494 | Ptk7 | PTK7 protein tyrosine kinase 7 [Source:MG | 9.6372 | 10.279 | 4.025 | 5.7839 | 2.1812 | 4.4554 |
| 4221 | ENSMUSG00000023973 | 734.6771926 | -0.043062779 | 0.143661408 | -0.299751891 | 0.764366415 | 0.954980327 | Cnpy3 | canopy FGF signaling regulator 3 [Source:I | 830.41 | 789.44 | 616.83 | 794.32 | 715.45 | 661.62 |
| 4222 | ENSMUSG00000023977 | 1743.221846 | -0.009563765 | 0.136523404 | -0.070052205 | 0.944152108 | 0.990077979 | Ubr2 | ubiquitin protein ligase E3 component n-r | 1623.9 | 1968.5 | 1656.3 | 2048.5 | 1670.8 | 1491.4 |
| 4223 | ENSMUSG00000023978 | 0 | NA | NA | NA | NA | NA | Prph2 | peripherin 2 [Source:MGI Symbol;Acc:MG | 0 | 0 | 0 | 0 | 0 | 0 |
| 4224 | ENSMUSG00000023979 | 4.22675805 | -0.487152862 | 1.012178019 | -0.481291683 | 0.630309207 | NA | Guca1b | guanylate cyclase activator 1B [Source:MG | 5.6217 | 4.1117 | 5.0312 | 2.8919 | 4.3625 | 3.3415 |
| 4225 | ENSMUSG00000023980 | 182.4970391 | 0.045887353 | 0.18885434 | 0.24297749 | 0.808022835 | 0.965249611 | Taf8 | TATA-box binding protein associated facto | 205.59 | 177.83 | 154.96 | 199.54 | 179.95 | 177.1 |
| 4226 | ENSMUSG00000023982 | 18.43699684 | 0.088310758 | 0.54210947 | 0.162902076 | 0.870595536 | 0.978802475 | Guca1a | guanylate cyclase activator 1a (retina) [So | 24.093 | 11.307 | 18.112 | 25.063 | 11.997 | 20.049 |
| 4227 | ENSMUSG00000023984 | 1.004562989 | 0.155751777 | 2.077846764 | 0.07495826 | 0.940247922 | NA | Gm20517 | predicted gene 20517 [Source:MGI Symbo | 0.8031 | 2.0558 | 0 | 0.964 | 1.0906 | 1.1138 |
| 4228 | ENSMUSG00000023987 | 0 | NA | NA | NA | NA | NA | Pgc | progastricsin (pepsinogen C) [Source:MGI | 0 | 0 | 0 | 0 | 0 | 0 |
| 4229 | ENSMUSG00000023988 | 241.7963299 | -0.261479466 | 0.158401252 | -1.650741155 | 0.09879144 | 0.463365025 | Bysl | bystin-like [Source:MGI Symbol;Acc:MGI:1 | 269.04 | 252.87 | 268.67 | 208.22 | 220.3 | 231.68 |
| 4230 | ENSMUSG00000023990 | 206.7018805 | 0.235093425 | 0.173476104 | 1.355191981 | 0.175356399 | 0.605810563 | Tfeb | transcription factor EB [Source:MGI Symb | 201.58 | 165.49 | 202.26 | 224.61 | 226.85 | 219.43 |
| 4231 | ENSMUSG00000023991 | 461.9053434 | 0.074425781 | 0.142533306 | 0.52216414 | 0.601556052 | 0.915461118 | Foxp4 | forkhead box P4 [Source:MGI Symbol;Acc | 435.28 | 400.89 | 514.19 | 498.38 | 450.43 | 472.27 |
| 4232 | ENSMUSG00000023992 | 9.896814709 | -1.32045833 | 0.758051717 | -1.741910613 | 0.081524087 | 0.420121054 | Trem2 | triggering receptor expressed on myeloid | 20.881 | 10.279 | 11.069 | 3.8559 | 3.2719 | 10.025 |
| 4233 | ENSMUSG00000023993 | 1.145948637 | 1.630339718 | 2.307260143 | 0.706612873 | 0.479807087 | NA | Treml1 | triggering receptor expressed on myeloid | 1.6062 | 0 | 0 | 1.928 | 0 | 3.3415 |
| 4234 | ENSMUSG00000023994 | 190.788344 | -0.085507988 | 0.189221599 | -0.45189338 | 0.651345794 | 0.925643431 | Nfya | nuclear transcription factor-Y alpha [Sour | 174.27 | 181.94 | 233.45 | 178.34 | 197.4 | 179.33 |
| 4235 | ENSMUSG00000023995 | 0.371281295 | 2.09253362 | 4.031852165 | 0.519000582 | 0.603760335 | NA | Tspo2 | translocator protein 2 [Source:MGI Symbo | 0 | 0 | 0 | 0 | 0 | 2.2277 |
| 4236 | ENSMUSG00000023999 | 0 | NA | NA | NA | NA | NA | Kif6 | kinesin family member 6 [Source:MGI Sym | 0 | 0 | 0 | 0 | 0 | 0 |
| 4237 | ENSMUSG00000024002 | 818.8310685 | 0.071628057 | 0.111413608 | 0.642902229 | 0.520287543 | 0.887802551 | Brd4 | bromodomain containing 4 [Source:MGI S | 846.47 | 794.58 | 753.68 | 833.84 | 823.42 | 861 |
| 4238 | ENSMUSG00000024006 | 977.7927954 | -0.006340583 | 0.113559897 | -0.055834704 | 0.955473488 | 0.991678478 | Stk38 | serine/threonine kinase 38 [Source:MGI S | 987.01 | 1000.2 | 952.91 | 1029.5 | 876.86 | 1020.3 |
| 4239 | ENSMUSG00000024007 | 133.0515469 | -0.022682026 | 0.26373004 | -0.086004713 | 0.931462671 | 0.988420305 | Ppil1 | peptidylprolyl isomerase (cyclophilin)-like | 128.5 | 103.82 | 170.06 | 141.7 | 91.612 | 162.62 |
| 4240 | ENSMUSG00000024008 | 0.133850373 | -0.796629176 | 4.080472857 | -0.195229623 | 0.84521319 | NA | Cpne5 | copine V [Source:MGI Symbol;Acc:MGI:23 | 0.8031 | 0 | 0 | 0 | 0 | 0 |
| 4241 | ENSMUSG00000024011 | 0.829837731 | -1.7608533 | 2.399900965 | -0.733719152 | 0.463119924 | NA | Pi16 | peptidase inhibitor 16 [Source:MGI Symbo | 0.8031 | 2.0558 | 1.0062 | 0 | 0 | 1.1138 |
| 4242 | ENSMUSG00000024012 | 1473.375895 | 0.034332206 | 0.097047827 | 0.353765841 | 0.723514375 | 0.9447719 | Mtch1 | mitochondrial carrier 1 [Source:MGI Symb | 1417.5 | 1439.1 | 1511.4 | 1486.5 | 1533.4 | 1452.5 |
| 4243 | ENSMUSG00000024013 | 46.31840873 | -1.713478129 | 0.385688969 | -4.442642303 | 8.89E-06 | 0.000479583 | Fgd2 | FYVE, RhoGEF and PH domain containing 2 | 67.461 | 84.289 | 61.381 | 26.027 | 10.906 | 27.846 |
| 4244 | ENSMUSG00000024014 | 474.1090987 | 0.718003936 | 0.183188131 | 3.919489394 | 8.87E-05 | 0.003295748 | Pim1 | proviral integration site 1 [Source:MGI Sym | 309.19 | 308.38 | 458.85 | 667.07 | 490.78 | 610.39 |
| 4245 | ENSMUSG00000024018 | 30.64216515 | -0.33638692 | 0.392045687 | -0.858029895 | 0.390875956 | 0.820195366 | Ccdc167 | coiled-coil domain containing 167 [Source | 35.336 | 35.977 | 31.194 | 24.099 | 31.628 | 25.618 |
| 4246 | ENSMUSG00000024019 | 58.99131273 | 0.094439739 | 0.287297728 | 0.328717318 | 0.742369363 | 0.949056962 | Cmtr1 | cap methyltransferase 1 [Source:MGI Sym | 51.399 | 62.703 | 57.356 | 64.587 | 59.984 | 57.92 |
| 4247 | ENSMUSG00000024026 | 6154.386963 | 0.248086127 | 0.095676718 | 2.592962342 | 0.009515319 | 0.104661272 | Glo1 | glyoxalase 1 [Source:MGI Symbol;Acc:MG | 5384 | 5708 | 5787.9 | 6486.6 | 6284.1 | 7275.6 |
| 4248 | ENSMUSG00000024027 | 0 | NA | NA | NA | NA | NA | Glp1r | glucagon-like peptide 1 receptor [Source:I | 0 | 0 | 0 | 0 | 0 | 0 |
| 4249 | ENSMUSG00000024028 | 0 | NA | NA | NA | NA | NA | Tff2 | trefoil factor 2 (spasmolytic protein 1) [So | 0 | 0 | 0 | 0 | 0 | 0 |
| 4250 | ENSMUSG00000024029 | 51.75867373 | 2.365937593 | 0.415482888 | 5.694428485 | 1.24E-08 | 1.87E-06 | Tff3 | trefoil factor 3, intestinal [Source:MGI Syn | 17.668 | 25.698 | 7.0437 | 72.298 | 70.89 | 116.95 |
| 4251 | ENSMUSG00000024030 | 60.02047986 | -0.819087433 | 0.298950443 | -2.739876969 | 0.006146219 | 0.076570979 | Abcg1 | ATP binding cassette subfamily G member | 81.916 | 79.15 | 68.425 | 34.703 | 45.806 | 50.123 |
| 4252 | ENSMUSG00000024032 | 1.666745423 | 0.787596987 | 1.596783862 | 0.493239571 | 0.621843321 | NA | Tff1 | trefoil factor 1 [Source:MGI Symbol;Acc:M | 1.6062 | 1.0279 | 1.0062 | 1.928 | 1.0906 | 3.3415 |

| | | | | | | | | | | | | | | | |
|---|---|---|---|---|---|---|---|---|---|---|---|---|---|---|---|
| 4253 | ENSMUSG00000024033 | 1.741065311 | -0.452748461 | 2.02834901 | -0.223210334 | 0.823371809 | NA | Rsph1 | radial spoke head 1 homolog (Chlamydom | 0 | 0 | 6.0375 | 0 | 2.1812 | 2.2277 |
| 4254 | ENSMUSG00000024034 | 2.018821526 | 4.538284853 | 2.095621724 | 2.165603077 | 0.030341531 | NA | Tmprss3 | transmembrane protease, serine 3 [Sourc | 0 | 0 | 0 | 0 | 6.5437 | 5.5692 |
| 4255 | ENSMUSG00000024036 | 60.22096339 | 0.309029344 | 0.283816952 | 1.088833281 | 0.276227412 | 0.730038567 | Slc37a1 | solute carrier family 37 (glycerol-3-phosph | 50.595 | 59.619 | 51.318 | 67.478 | 63.256 | 69.058 |
| 4256 | ENSMUSG00000024037 | 279.6937924 | 0.102001179 | 0.154900692 | 0.658494017 | 0.510220741 | 0.883032087 | Wdr4 | WD repeat domain 4 [Source:MGI Symbol | 270.65 | 251.84 | 286.78 | 278.59 | 315.19 | 275.12 |
| 4257 | ENSMUSG00000024038 | 1039.453078 | 0.031481291 | 0.101345768 | 0.310632517 | 0.756080004 | 0.953085143 | Ndufv3 | NADH:ubiquinone oxidoreductase core su | 1003.9 | 1050.5 | 1030.4 | 1063.3 | 1037.2 | 1051.5 |
| 4258 | ENSMUSG00000024039 | 5466.916952 | 0.654873562 | 0.14564899 | 4.496245133 | 6.92E-06 | 0.000403382 | Cbs | cystathionine beta-synthase [Source:MGI | 4905.3 | 3726.2 | 4107.5 | 5666.3 | 6235.1 | 8161.1 |
| 4259 | ENSMUSG00000024041 | 0 | NA | NA | NA | NA | NA | Cryaa | crystallin, alpha A [Source:MGI Symbol;Ac | 0 | 0 | 0 | 0 | 0 | 0 |
| 4260 | ENSMUSG00000024042 | 237.4407896 | -0.507987374 | 0.250152249 | -2.030712804 | 0.042284137 | 0.283193583 | Sik1 | salt inducible kinase 1 [Source:MGI Symbo | 243.34 | 278.57 | 314.95 | 201.47 | 270.47 | 115.84 |
| 4261 | ENSMUSG00000024043 | 10.12915374 | -0.151041924 | 0.759004669 | -0.198999994 | 0.84226275 | 0.971875826 | Arhgap28 | Rho GTPase activating protein 28 [Source: | 14.456 | 12.335 | 5.0312 | 4.8199 | 17.45 | 6.6831 |
| 4262 | ENSMUSG00000024044 | 19.13614944 | -1.581885307 | 0.574372486 | -2.754110519 | 0.005885189 | 0.074343559 | Epb41l3 | erythrocyte membrane protein band 4.1 l | 25.699 | 43.173 | 17.106 | 6.7478 | 8.7249 | 13.366 |
| 4263 | ENSMUSG00000024045 | 667.6241389 | 0.139558682 | 0.122656995 | 1.137796356 | 0.255205527 | 0.708637501 | Akap8 | A kinase (PRKA) anchor protein 8 [Source: | 593.49 | 644.5 | 669.15 | 739.37 | 716.54 | 642.69 |
| 4264 | ENSMUSG00000024048 | 466.45515 | -0.079840938 | 0.141777385 | -0.563142971 | 0.573337533 | 0.905421342 | Myl12a | myosin, light chain 12A, regulatory, non-s | 532.46 | 487.23 | 417.59 | 455 | 428.61 | 477.84 |
| 4265 | ENSMUSG00000024049 | 21.91857568 | -0.138607938 | 0.481331638 | -0.287967645 | 0.773371506 | 0.958177785 | Myom1 | myomesin 1 [Source:MGI Symbol;Acc:MG | 32.124 | 16.447 | 20.125 | 23.135 | 19.631 | 20.049 |
| 4266 | ENSMUSG00000024050 | 313.6250034 | -0.217903312 | 0.161486075 | -1.349362865 | 0.177220443 | 0.608561375 | Wiz | widely-interspaced zinc finger motifs [Sou | 309.19 | 348.46 | 355.2 | 336.43 | 259.57 | 272.89 |
| 4267 | ENSMUSG00000024052 | 6893.997268 | -0.350939524 | 0.132634687 | -2.645910592 | 0.008147135 | 0.093114627 | Lpin2 | lipin 2 [Source:MGI Symbol;Acc:MGI:1891 | 8240.6 | 8465.9 | 6478.2 | 5945.8 | 6953.8 | 5279.6 |
| 4268 | ENSMUSG00000024053 | 21.1613543 | -0.520980866 | 0.49671432 | -1.04885413 | 0.29424526 | 0.745164683 | Emilin2 | elastin microfibril interfacer 2 [Source:MG | 32.927 | 25.698 | 16.1 | 20.244 | 14.178 | 17.822 |
| 4269 | ENSMUSG00000024054 | 230.0694354 | 0.134168433 | 0.178703218 | 0.750789126 | 0.452779575 | 0.856769845 | Smchd1 | SMC hinge domain containing 1 [Source:N | 224.07 | 230.25 | 203.26 | 199.54 | 271.56 | 251.73 |
| 4270 | ENSMUSG00000024055 | 1297.557535 | 0.176339916 | 0.118586477 | 1.487015383 | 0.137010748 | 0.540888479 | Cyp4f13 | cytochrome P450, family 4, subfamily f, po | 1097 | 1333.2 | 1226.6 | 1505.7 | 1324 | 1298.7 |
| 4271 | ENSMUSG00000024056 | 3.814465988 | 0.358560393 | 1.124781522 | 0.318782258 | 0.749891633 | NA | Ndc80 | NDC80 kinetochore complex component [ | 4.8186 | 4.1117 | 1.0062 | 1.928 | 5.4531 | 5.5692 |
| 4272 | ENSMUSG00000024059 | 8.417906976 | -1.113752681 | 0.782963085 | -1.422484281 | 0.154885712 | 0.573104869 | Clip4 | CAP-GLY domain containing linker protein | 8.031 | 17.475 | 9.0562 | 3.8559 | 7.6343 | 4.4554 |
| 4273 | ENSMUSG00000024063 | 90.5827118 | -1.005332806 | 0.313982239 | -3.201877944 | 0.001365348 | 0.026366354 | Lbh | limb-bud and heart [Source:MGI Symbol;A | 121.27 | 145.96 | 95.593 | 65.55 | 33.809 | 81.311 |
| 4274 | ENSMUSG00000024064 | 1.971767618 | -2.285234206 | 1.851155329 | -1.234490791 | 0.217020076 | NA | Galnt14 | polypeptide N-acetylgalactosaminyltransf | 0.8031 | 2.0558 | 7.0437 | 1.928 | 0 | 0 |
| 4275 | ENSMUSG00000024065 | 580.5493194 | -1.124768969 | 0.201396455 | -5.584849902 | 2.34E-08 | 3.19E-06 | Ehd3 | EH-domain containing 3 [Source:MGI Sym | 823.98 | 821.31 | 742.61 | 365.35 | 238.85 | 491.21 |
| 4276 | ENSMUSG00000024066 | 3508.435163 | 0.054446534 | 0.132812663 | 0.409949874 | 0.681842718 | 0.934891751 | Xdh | xanthine dehydrogenase [Source:MGI Syr | 3211.6 | 3468.2 | 3648.6 | 4335 | 3329.7 | 3057.5 |
| 4277 | ENSMUSG00000024067 | 227.9157434 | -0.024833464 | 0.186219006 | -0.133356229 | 0.893911661 | 0.982562372 | Dpy30 | dpy-30, histone methyltransferase comple | 216.03 | 211.75 | 261.62 | 187.98 | 260.66 | 229.45 |
| 4278 | ENSMUSG00000024068 | 651.9935225 | 0.154788049 | 0.124444902 | 1.243827967 | 0.213562885 | 0.658349013 | Spast | spastin [Source:MGI Symbol;Acc:MGI:185 | 579.04 | 594.14 | 678.21 | 651.65 | 683.82 | 725.11 |
| 4279 | ENSMUSG00000024069 | 205.3922621 | 0.035821277 | 0.178754392 | 0.200393828 | 0.841172586 | 0.971854408 | Slc30a6 | solute carrier family 30 (zinc transporter), | 187.12 | 218.95 | 203.26 | 236.17 | 193.04 | 193.81 |
| 4280 | ENSMUSG00000024070 | 2938.511231 | 0.589012627 | 0.145869112 | 4.037953056 | 5.39E-05 | 0.002221919 | Prkd3 | protein kinase D3 [Source:MGI Symbol;Ac | 2084.9 | 2386.8 | 2568.9 | 2738.7 | 3804.1 | 4047.7 |
| 4281 | ENSMUSG00000024072 | 685.8499747 | 0.041847525 | 0.127174505 | 0.329055934 | 0.742113411 | 0.949043964 | Yipf4 | Yip1 domain family, member 4 [Source:M | 608.75 | 697.96 | 721.48 | 639.12 | 741.62 | 706.18 |
| 4282 | ENSMUSG00000024073 | 1395.490356 | -0.048440998 | 0.120435467 | -0.402215386 | 0.68752552 | 0.936885029 | Birc6 | baculoviral IAP repeat-containing 6 [Sourc | 1330.7 | 1497.7 | 1429.9 | 1537.5 | 1375.3 | 1201.8 |
| 4283 | ENSMUSG00000024074 | 310.1901389 | -0.043005402 | 0.160002398 | -0.268779731 | 0.788099192 | 0.960870081 | Crim1 | cysteine rich transmembrane BMP regulat | 300.36 | 333.05 | 311.94 | 351.85 | 297.74 | 266.21 |
| 4284 | ENSMUSG00000024076 | 0.455176322 | 0.955156269 | 3.546674842 | 0.269310357 | 0.787690862 | NA | Vit | vitrin [Source:MGI Symbol;Acc:MGI:19214 | 0.8031 | 0 | 0 | 1.928 | 0 | 0 |
| 4285 | ENSMUSG00000024077 | 282.4860992 | 0.074281645 | 0.151582106 | 0.490042307 | 0.624103962 | 0.919503423 | Strn | striatin, calmodulin binding protein [Sourc | 268.24 | 299.12 | 258.6 | 296.91 | 283.56 | 288.49 |
| 4286 | ENSMUSG00000024078 | 216.5799841 | -0.103074128 | 0.193849228 | -0.531723182 | 0.594917734 | 0.913409341 | Ttc27 | tetratricopeptide repeat domain 27 [Sourc | 200.78 | 254.92 | 217.35 | 177.37 | 253.02 | 196.04 |
| 4287 | ENSMUSG00000024079 | 306.771377 | 0.18935649 | 0.16018864 | 1.182084384 | 0.23717222 | 0.686091798 | Eif2ak2 | eukaryotic translation initiation factor 2-a | 315.62 | 284.73 | 258.6 | 287.27 | 343.54 | 350.86 |
| 4288 | ENSMUSG00000024081 | 595.0627183 | -0.035414901 | 0.121044836 | -0.292576723 | 0.769845708 | 0.956454932 | Cebpz | CCAAT/enhancer binding protein zeta [So | 608.75 | 624.97 | 573.56 | 605.38 | 607.47 | 550.24 |
| 4289 | ENSMUSG00000024082 | 275.7315142 | 0.107123311 | 0.156411031 | 0.684883348 | 0.493417535 | 0.87710702 | Ndufaf7 | NADH:ubiquinone oxidoreductase comple | 273.05 | 267.26 | 255.59 | 254.49 | 299.92 | 304.08 |
| 4290 | ENSMUSG00000024083 | 628.3499793 | 0.042522089 | 0.127348378 | 0.333903655 | 0.738452261 | 0.948137969 | Pja2 | praja ring finger ubiquitin ligase 2 [Source | 586.26 | 679.45 | 591.67 | 587.06 | 671.82 | 653.83 |
| 4291 | ENSMUSG00000024084 | 166.4181909 | -0.054867398 | 0.183999444 | -0.298193283 | 0.765555648 | 0.954980327 | Qpct | glutaminyl-peptide cyclotransferase (gluta | 179.89 | 170.63 | 157.98 | 165.8 | 153.78 | 170.42 |
| 4292 | ENSMUSG00000024085 | 4780.96186 | -0.252695028 | 0.113088748 | -2.234484263 | 0.025451227 | 0.205313187 | Man2a1 | mannosidase 2, alpha 1 [Source:MGI Symb | 4808.2 | 5699.8 | 5089.6 | 4799.6 | 4424.6 | 3863.9 |
| 4293 | ENSMUSG00000024087 | 9.065427045 | -0.372498808 | 0.799760076 | -0.465763196 | 0.641385012 | 0.924019354 | Cyp1b1 | cytochrome P450, family 1, subfamily b, p | 11.243 | 9.2513 | 10.062 | 3.8559 | 3.2719 | 16.708 |
| 4294 | ENSMUSG00000024088 | 0.428363721 | -0.543730382 | 3.624677344 | -0.15000794 | 0.880758351 | NA | 4930583I09F | RIKEN cDNA 4930583I09 gene [Source:MG | 1.6062 | 0 | 0 | 0.964 | 0 | 0 |
| 4295 | ENSMUSG00000024091 | 1552.422447 | -0.082465231 | 0.105951616 | -0.778329154 | 0.436374993 | 0.846530029 | Vapa | vesicle-associated membrane protein, ass | 1745.9 | 1527.5 | 1515.4 | 1453.7 | 1567.2 | 1504.8 |
| 4296 | ENSMUSG00000024095 | 593.2488038 | 0.153510104 | 0.121632405 | 1.262082282 | 0.20691918 | 0.652194904 | Hnrnpll | heterogeneous nuclear ribonucleoprotein | 554.14 | 537.6 | 593.68 | 638.15 | 648.92 | 587 |
| 4297 | ENSMUSG00000024096 | 528.5031676 | 0.077079348 | 0.124161823 | 0.620797489 | 0.534732875 | 0.889858671 | Ralbp1 | ralA binding protein 1 [Source:MGI Symbo | 529.24 | 535.55 | 477.97 | 528.26 | 545.31 | 554.69 |
| 4298 | ENSMUSG00000024097 | 532.1590059 | -0.133235021 | 0.135278453 | -0.984894622 | 0.324675835 | 0.771309263 | Srsf7 | serine and arginine-rich splicing factor 7 [S | 510.77 | 586.94 | 572.55 | 454.03 | 552.94 | 515.71 |
| 4299 | ENSMUSG00000024098 | 1046.885095 | 0.157523197 | 0.105369863 | 1.494954936 | 0.134926203 | 0.537616739 | Twsg1 | twisted gastrulation BMP signaling modul | 1010.3 | 990.91 | 968.01 | 1112.4 | 1159.3 | 1040.3 |
| 4300 | ENSMUSG00000024099 | 2101.708185 | -0.015498933 | 0.095033463 | -0.163089221 | 0.870448187 | 0.978802475 | Ndufv2 | NADH:ubiquinone oxidoreductase core su | 2090.5 | 2184.3 | 2063.8 | 1975.2 | 2127.8 | 2168.7 |
| 4301 | ENSMUSG00000024101 | 321.8479932 | 0.080395065 | 0.169553418 | 0.474157737 | 0.635387429 | 0.923478764 | Washc1 | WASH complex subunit 1 [Source:MGI Syr | 351.76 | 306.32 | 279.74 | 321 | 284.65 | 387.62 |
| 4302 | ENSMUSG00000024104 | 896.5544181 | -0.038171876 | 0.122178788 | -0.312426373 | 0.75471651 | 0.953048681 | Washc2 | WASH complex subunit 2` [Source:MGI Sy | 821.57 | 964.19 | 939.83 | 803.96 | 904.12 | 945.65 |
| 4303 | ENSMUSG00000024105 | 0 | NA | NA | NA | NA | NA | Themis3 | thymocyte selection associated family me | 0 | 0 | 0 | 0 | 0 | 0 |
| 4304 | ENSMUSG00000024107 | 0 | NA | NA | NA | NA | NA | Lhcgr | luteinizing hormone/choriogonadotropin | 0 | 0 | 0 | 0 | 0 | 0 |
| 4305 | ENSMUSG00000024109 | 45.80992265 | -1.193500702 | 0.338720497 | -3.523556188 | 0.000425797 | 0.011136935 | Nrxn1 | neurexin I [Source:MGI Symbol;Acc:MGI:1 | 65.854 | 65.787 | 59.368 | 22.171 | 32.719 | 28.96 |
| 4306 | ENSMUSG00000024112 | 1.058182678 | 2.444751467 | 2.435756566 | 1.003692857 | 0.315526681 | NA | Cacna1h | calcium channel, voltage-dependent, T typ | 0.8031 | 0 | 0 | 0 | 1.0906 | 4.4554 |
| 4307 | ENSMUSG00000024114 | 0.17131964 | -0.796629176 | 4.080472857 | -0.195229623 | 0.84521319 | NA | Prss41 | protease, serine 41 [Source:MGI Symbol;A | 0 | 1.0279 | 0 | 0 | 0 | 0 |
| 4308 | ENSMUSG00000024116 | 0 | NA | NA | NA | NA | NA | Prss21 | protease, serine 21 [Source:MGI Symbol;A | 0 | 0 | 0 | 0 | 0 | 0 |

| | | | | | | | | | | | | | | | |
|---|---|---|---|---|---|---|---|---|---|---|---|---|---|---|---|
| 4309 | ENSMUSG00000024118 | 3151.527249 | -0.029627349 | 0.131059024 | -0.226061113 | 0.821153889 | 0.969160361 | Tedc2 | tubulin epsilon and delta complex 2 [Sour | 2843 | 2839.1 | 3870 | 2941.1 | 3180.2 | 3235.7 |
| 4310 | ENSMUSG00000024120 | 2734.88245 | -0.02616798 | 0.09501335 | -0.275413714 | 0.782998409 | 0.960670212 | Lrpprc | leucine-rich PPR-motif containing [Source | 2719.3 | 2658.2 | 2902 | 2760.8 | 2813.8 | 2555.2 |
| 4311 | ENSMUSG00000024121 | 589.658397 | 0.026016328 | 0.123455147 | 0.210735062 | 0.833094013 | 0.970611245 | Atp6v0c | ATPase, H+ transporting, lysosomal V0 sul | 561.37 | 624.97 | 567.52 | 633.33 | 568.21 | 582.54 |
| 4312 | ENSMUSG00000024122 | 680.6395049 | 0.122453718 | 0.122990236 | 0.995637725 | 0.319426198 | 0.766337562 | Pdpk1 | 3-phosphoinositide dependent protein kir | 614.37 | 719.54 | 621.86 | 689.24 | 692.54 | 746.28 |
| 4313 | ENSMUSG00000024124 | 1.152818647 | -0.434256954 | 2.550829563 | -0.170241462 | 0.864820246 | NA | Prss30 | protease, serine 30 [Source:MGI Symbol;/ | 0 | 0 | 4.025 | 2.8919 | 0 | 0 |
| 4314 | ENSMUSG00000024125 | 0 | NA | NA | NA | NA | NA | Sbpl | spermine binding protein-like [Source:MG | 0 | 0 | 0 | 0 | 0 | 0 |
| 4315 | ENSMUSG00000024127 | 488.9380941 | -0.041259452 | 0.146739972 | -0.281173911 | 0.778577014 | 0.959559604 | Prepl | prolyl endopeptidase-like [Source:MGI Sy | 472.22 | 503.68 | 512.18 | 490.66 | 403.53 | 551.35 |
| 4316 | ENSMUSG00000024128 | 0 | NA | NA | NA | NA | NA | Sbp | spermine binding protein [Source:MGI Sy | 0 | 0 | 0 | 0 | 0 | 0 |
| 4317 | ENSMUSG00000024130 | 811.4071667 | -0.002817811 | 0.156268518 | -0.018031854 | 0.985613441 | 0.99859176 | Abca3 | ATP-binding cassette, sub-family A (ABC1) | 813.54 | 775.05 | 848.26 | 897.46 | 610.75 | 923.38 |
| 4318 | ENSMUSG00000024131 | 524.4533383 | 0.074948625 | 0.177946336 | 0.421186674 | 0.673618774 | 0.931915649 | Slc3a1 | solute carrier family 3, member 1 [Source: | 448.93 | 490.32 | 593.68 | 538.86 | 415.53 | 659.4 |
| 4319 | ENSMUSG00000024132 | 3894.001318 | -0.247586418 | 0.095982934 | -2.579483753 | 0.009894811 | 0.107851206 | Eci1 | enoyl-Coenzyme A delta isomerase 1 [Sou | 4267.7 | 4418 | 3996.8 | 3721 | 3653.6 | 3307 |
| 4320 | ENSMUSG00000024134 | 0 | NA | NA | NA | NA | NA | Six2 | sine oculis-related homeobox 2 [Source:N | 0 | 0 | 0 | 0 | 0 | 0 |
| 4321 | ENSMUSG00000024135 | 410.2626953 | 0.169936439 | 0.148458486 | 1.144673118 | 0.252344591 | 0.705160774 | Srbd1 | S1 RNA binding domain 1 [Source:MGI Sy | 407.17 | 364.91 | 385.39 | 377.88 | 494.05 | 432.17 |
| 4322 | ENSMUSG00000024136 | 12.10859879 | -0.128018527 | 0.630600549 | -0.203010491 | 0.839126832 | 0.971670042 | Dnase1l2 | deoxyribonuclease 1-like 2 [Source:MGI S | 9.6372 | 10.279 | 18.112 | 12.532 | 8.7249 | 13.366 |
| 4323 | ENSMUSG00000024137 | 182.6209585 | 0.266269539 | 0.180408279 | 1.475927491 | 0.139963358 | 0.54655238 | E4f1 | E4F transcription factor 1 [Source:MGI Sy | 161.42 | 158.3 | 178.11 | 218.82 | 191.95 | 187.13 |
| 4324 | ENSMUSG00000024140 | 3666.265317 | -0.100168218 | 0.106323595 | -0.942107141 | 0.346137789 | 0.786987972 | Epas1 | endothelial PAS domain protein 1 [Source | 3660.5 | 4208.3 | 3512.8 | 3790.4 | 3349.3 | 3476.3 |
| 4325 | ENSMUSG00000024142 | 85.58135762 | 0.165318126 | 0.251420584 | 0.657536161 | 0.510836228 | 0.883502953 | Mlst8 | MTOR associated protein, LST8 homolog ( | 87.538 | 75.038 | 79.493 | 106.04 | 79.615 | 85.766 |
| 4326 | ENSMUSG00000024143 | 244.4366318 | -0.13608456 | 0.159767508 | -0.851766184 | 0.394343879 | 0.822827574 | Rhoq | ras homolog family member Q [Source:M | 235.31 | 267.26 | 265.65 | 224.61 | 242.12 | 231.68 |
| 4327 | ENSMUSG00000024145 | 44.10425069 | -0.232784876 | 0.377583268 | -0.61651269 | 0.537556199 | 0.890937649 | Pigf | phosphatidylinositol glycan anchor biosyn | 34.533 | 45.228 | 63.393 | 38.559 | 29.447 | 53.465 |
| 4328 | ENSMUSG00000024146 | 542.0265501 | 0.10123566 | 0.131223244 | 0.77147658 | 0.440424499 | 0.849146888 | Cript | cysteine-rich PDZ-binding protein [Source | 558.96 | 513.96 | 495.07 | 510.91 | 609.66 | 563.61 |
| 4329 | ENSMUSG00000024150 | 3809.75313 | -0.000581517 | 0.093152039 | -0.006242664 | 0.995019107 | 0.998676967 | Mcfd2 | multiple coagulation factor deficiency 2 [S | 3946.4 | 3752.9 | 3731.2 | 3699.7 | 3642.7 | 4085.6 |
| 4330 | ENSMUSG00000024151 | 216.0017663 | 0.237765612 | 0.198451187 | 1.198106269 | 0.230875649 | 0.677910147 | Msh2 | mutS homolog 2 [Source:MGI Symbol;Acc | 188.73 | 165.49 | 240.49 | 225.57 | 203.95 | 271.78 |
| 4331 | ENSMUSG00000024154 | 0 | NA | NA | NA | NA | NA | Gtf2a1l | general transcription factor IIA, 1-like [Sou | 0 | 0 | 0 | 0 | 0 | 0 |
| 4332 | ENSMUSG00000024155 | 154.2803917 | 0.443390473 | 0.204303945 | 2.170249199 | 0.029987973 | 0.22888973 | Meiob | meiosis specific with OB domains [Source: | 132.51 | 128.49 | 130.81 | 143.63 | 190.86 | 199.38 |
| 4333 | ENSMUSG00000024158 | 4989.923406 | 0.340858369 | 0.088141675 | 3.867164649 | 0.000110108 | 0.003913031 | Hagh | hydroxyacyl glutathione hydrolase [Sourc | 4439.5 | 4156.9 | 4612.6 | 5482.1 | 5666.9 | 5581.5 |
| 4334 | ENSMUSG00000024160 | 58.41996381 | -0.180221461 | 0.320987634 | -0.561459203 | 0.574484534 | 0.906163061 | Spsb3 | splA/ryanodine receptor domain and SOC | 46.58 | 59.619 | 80.5 | 63.623 | 54.531 | 45.668 |
| 4335 | ENSMUSG00000024163 | 417.1643504 | 0.02780916 | 0.169807014 | 0.163769208 | 0.869912833 | 0.978765244 | Mapk8ip3 | mitogen-activated protein kinase 8 intera | 392.72 | 407.06 | 440.74 | 522.48 | 399.17 | 340.84 |
| 4336 | ENSMUSG00000024164 | 74540.99834 | 0.038038099 | 0.157093548 | 0.24213661 | 0.808674312 | 0.965682062 | C3 | complement component 3 [Source:MGI S | 57276 | 80015 | 83386 | 91287 | 72859 | 62423 |
| 4337 | ENSMUSG00000024165 | 333.8758531 | 0.415136795 | 0.174157784 | 2.383682114 | 0.017140401 | 0.155792353 | Jpt2 | Jupiter microtubule associated homolog 2 | 296.34 | 244.64 | 316.97 | 353.78 | 333.73 | 457.79 |
| 4338 | ENSMUSG00000024168 | 122.0920209 | -0.82202707 | 0.225347038 | -3.647827277 | 0.000264467 | 0.007854734 | Tmem204 | transmembrane protein 204 [Source:MGI | 140.54 | 164.47 | 163.01 | 86.758 | 73.071 | 104.7 |
| 4339 | ENSMUSG00000024169 | 196.9866329 | -0.110789429 | 0.172429526 | -0.642520055 | 0.520535572 | 0.887802551 | Ift140 | intraflagellar transport 140 [Source:MGI S | 203.18 | 192.22 | 218.36 | 194.72 | 195.22 | 178.22 |
| 4340 | ENSMUSG00000024170 | 104.8861287 | 0.167891116 | 0.22696453 | 0.739724027 | 0.459467466 | 0.859621968 | Telo2 | telomere maintenance 2 [Source:MGI Syn | 110.83 | 89.429 | 95.593 | 104.11 | 109.06 | 120.3 |
| 4341 | ENSMUSG00000024171 | 0 | NA | NA | NA | NA | NA | Prss28 | protease, serine 28 [Source:MGI Symbol;/ | 0 | 0 | 0 | 0 | 0 | 0 |
| 4342 | ENSMUSG00000024172 | 0 | NA | NA | NA | NA | NA | St6gal2 | beta galactoside alpha 2,6 sialyltransferas | 0 | 0 | 0 | 0 | 0 | 0 |
| 4343 | ENSMUSG00000024173 | 0 | NA | NA | NA | NA | NA | Tpsab1 | tryptase alpha/beta 1 [Source:MGI Symbc | 0 | 0 | 0 | 0 | 0 | 0 |
| 4344 | ENSMUSG00000024174 | 52.42536187 | -0.521151805 | 0.318482909 | -1.636357211 | 0.101764849 | 0.469495293 | Pot1b | protection of telomeres 1B [Source:MGI S | 71.476 | 58.591 | 55.343 | 52.055 | 41.443 | 35.643 |
| 4345 | ENSMUSG00000024175 | 0 | NA | NA | NA | NA | NA | Tekt4 | tektin 4 [Source:MGI Symbol;Acc:MGI:191 | 0 | 0 | 0 | 0 | 0 | 0 |
| 4346 | ENSMUSG00000024176 | 0 | NA | NA | NA | NA | NA | Sox8 | SRY (sex determining region Y)-box 8 [Sou | 0 | 0 | 0 | 0 | 0 | 0 |
| 4347 | ENSMUSG00000024177 | 25.18950537 | 0.0259163 | 0.452521358 | 0.057270888 | 0.954329411 | 0.991678478 | Nme4 | NME/NM23 nucleoside diphosphate kinas | 24.896 | 29.81 | 20.125 | 20.244 | 34.9 | 21.163 |
| 4348 | ENSMUSG00000024180 | 651.1495218 | 0.092858242 | 0.122639699 | 0.757162999 | 0.448952215 | 0.854431786 | Pgap6 | post-glycosylphosphatidylinositol attachm | 673 | 613.67 | 602.74 | 619.84 | 690.36 | 707.29 |
| 4349 | ENSMUSG00000024181 | 631.9315774 | 0.007975992 | 0.131885984 | 0.060476422 | 0.951776194 | 0.990883058 | Mrpl28 | mitochondrial ribosomal protein L28 [Sou | 616.78 | 559.19 | 714.43 | 607.31 | 646.74 | 647.14 |
| 4350 | ENSMUSG00000024182 | 401.5816897 | -0.117873151 | 0.137461696 | -0.857498155 | 0.391169634 | 0.820195366 | Axin1 | axin 1 [Source:MGI Symbol;Acc:MGI:1096 | 396.73 | 455.37 | 402.5 | 399.09 | 389.35 | 366.45 |
| 4351 | ENSMUSG00000024184 | 0.545309144 | 2.651974629 | 3.666219369 | 0.72335405 | 0.469462394 | NA | Pdia2 | protein disulfide isomerase associated 2 [! | 0 | 0 | 0 | 0 | 3.2719 | 0 |
| 4352 | ENSMUSG00000024186 | 2.830897027 | 1.260984058 | 1.613897462 | 0.781328484 | 0.434609321 | NA | Rgs11 | regulator of G-protein signaling 11 [Sourc | 0 | 4.1117 | 1.0062 | 9.6398 | 0 | 2.2277 |
| 4353 | ENSMUSG00000024187 | 888.5993657 | 0.00165943 | 0.108383201 | 0.015310764 | 0.987784255 | 0.99859176 | Fam234a | family with sequence similarity 234, meml | 844.86 | 903.54 | 916.69 | 914.81 | 855.04 | 896.64 |
| 4354 | ENSMUSG00000024188 | 529.6362094 | -0.080526529 | 0.131911054 | -0.610460812 | 0.541556595 | 0.89377848 | Luc7l | Luc7-like [Source:MGI Symbol;Acc:MGI:1! | 487.48 | 593.11 | 553.43 | 499.34 | 538.77 | 505.69 |
| 4355 | ENSMUSG00000024190 | 691.7656101 | -1.343530444 | 0.204662649 | -6.564609859 | 5.22E-11 | 1.53E-08 | Dusp1 | dual specificity phosphatase 1 [Source:M | 923.57 | 1158.5 | 896.56 | 508.02 | 412.25 | 251.73 |
| 4356 | ENSMUSG00000024191 | 350.2175784 | 0.107173851 | 0.150648742 | 0.711415507 | 0.476826792 | 0.869671045 | Bnip1 | BCL2/adenovirus E1B interacting protein 1 | 358.99 | 316.6 | 335.08 | 321 | 367.54 | 402.1 |
| 4357 | ENSMUSG00000024193 | 382.5464985 | 0.283162393 | 0.145687075 | 1.943634278 | 0.051939569 | 0.320245495 | Phf1 | PHD finger protein 1 [Source:MGI Symbol | 350.15 | 347.44 | 338.1 | 466.57 | 367.54 | 425.49 |
| 4358 | ENSMUSG00000024194 | 641.1037684 | -0.070402268 | 0.121233818 | -0.580714761 | 0.561432711 | 0.900553224 | Cuta | cutA divalent cation tolerance homolog [S | 656.13 | 647.59 | 667.14 | 682.5 | 611.84 | 581.43 |
| 4359 | ENSMUSG00000024197 | 880.7253283 | -0.069815851 | 0.144240028 | -0.484025498 | 0.628367773 | 0.920328819 | Plin3 | perilipin 3 [Source:MGI Symbol;Acc:MGI:1 | 1040.8 | 871.67 | 791.91 | 757.69 | 836.5 | 985.75 |
| 4360 | ENSMUSG00000024201 | 298.9133051 | 0.076686649 | 0.195400752 | 0.392458309 | 0.69471961 | 0.939895655 | Kdm4b | lysine (K)-specific demethylase 4B [Source | 248.16 | 291.93 | 334.07 | 390.41 | 269.38 | 259.53 |
| 4361 | ENSMUSG00000024205 | 0.818742381 | -1.749431793 | 2.621906314 | -0.667236577 | 0.504621031 | NA | Rpl36-ps2 | ribosomal protein L36, pseudogene 2 [Sou | 0.8031 | 0 | 3.0187 | 0 | 1.0906 | 0 |
| 4362 | ENSMUSG00000024206 | 15.19512201 | -0.156278382 | 0.607630988 | -0.25719291 | 0.797029857 | 0.962236316 | Rfx2 | regulatory factor X, 2 (influences HLA clas | 13.653 | 20.558 | 14.087 | 24.099 | 7.6343 | 11.138 |
| 4363 | ENSMUSG00000024207 | 0 | NA | NA | NA | NA | NA | Acsbg2 | acyl-CoA synthetase bubblegum family me | 0 | 0 | 0 | 0 | 0 | 0 |
| 4364 | ENSMUSG00000024208 | 406.0195579 | -0.254622847 | 0.137719415 | -1.848852222 | 0.064479154 | 0.36373888 | Uqcc2 | ubiquinol-cytochrome c reductase comple | 462.59 | 431.73 | 430.67 | 379.81 | 335.91 | 395.41 |

| | | | | | | | | | | | | | | | |
|---|---|---|---|---|---|---|---|---|---|---|---|---|---|---|---|
| 4365 | ENSMUSG00000024209 | 0.531944269 | 2.618916564 | 3.285123409 | 0.797204926 | 0.425332026 | NA | Acsbg3 | acyl-CoA synthetase bubblegum family me | 0 | 0 | 0 | 0.964 | 0 | 2.2277 |
| 4366 | ENSMUSG00000024210 | 0 | NA | NA | NA | NA | NA | Ip6k3 | inositol hexaphosphate kinase 3 [Source:M | 0 | 0 | 0 | 0 | 0 | 0 |
| 4367 | ENSMUSG00000024211 | 12.83603068 | 1.193397135 | 0.909930127 | 1.311526127 | 0.189680073 | 0.626663447 | Grm8 | glutamate receptor, metabotropic 8 [Sour | 2.4093 | 0 | 21.131 | 11.568 | 19.631 | 22.277 |
| 4368 | ENSMUSG00000024212 | 315.0304342 | 0.164610324 | 0.152755099 | 1.077609362 | 0.281208123 | 0.733710968 | Mllt1 | myeloid/lymphoid or mixed-lineage leuke | 269.04 | 286.79 | 336.09 | 339.32 | 334.82 | 324.13 |
| 4369 | ENSMUSG00000024213 | 440.5431397 | 0.15625076 | 0.143823353 | 1.086407436 | 0.277298773 | 0.730041765 | Nudt3 | nudix (nucleotide diphosphate linked moi | 437.69 | 416.31 | 395.45 | 441.5 | 419.89 | 532.42 |
| 4370 | ENSMUSG00000024215 | 0.160662974 | 1.126924175 | 4.080472857 | 0.276174898 | 0.782413736 | NA | Spdef | SAM pointed domain containing ets transc | 0 | 0 | 0 | 0.964 | 0 | 0 |
| 4371 | ENSMUSG00000024217 | 371.3911797 | 0.0061817 | 0.150426767 | 0.041094416 | 0.967220626 | 0.994231936 | Snrpc | U1 small nuclear ribonucleoprotein C [Sou | 383.88 | 362.85 | 364.26 | 321 | 420.98 | 375.37 |
| 4372 | ENSMUSG00000024218 | 214.9536896 | 0.109108175 | 0.164179952 | 0.664564543 | 0.506329054 | 0.881592135 | Taf11 | TATA-box binding protein associated facto | 210.41 | 202.5 | 207.29 | 211.11 | 232.3 | 226.11 |
| 4373 | ENSMUSG00000024219 | 596.3348593 | -0.282942779 | 0.170429803 | -1.660171955 | 0.096879865 | 0.458530879 | Anks1 | ankyrin repeat and SAM domain containin | 811.13 | 619.83 | 531.3 | 459.82 | 586.75 | 569.17 |
| 4374 | ENSMUSG00000024220 | 403.4208609 | -0.03029328 | 0.137677386 | -0.220030903 | 0.825847088 | 0.96943211 | Zfp523 | zinc finger protein 523 [Source:MGI Symb | 424.04 | 385.47 | 413.57 | 434.75 | 379.54 | 383.16 |
| 4375 | ENSMUSG00000024222 | 2374.438185 | -0.889690843 | 0.137754407 | -6.458529064 | 1.06E-10 | 2.91E-08 | Fkbp5 | FK506 binding protein 5 [Source:MGI Sym | 2898.4 | 3349 | 3006.7 | 1943.4 | 1329.5 | 1719.8 |
| 4376 | ENSMUSG00000024223 | 0 | NA | NA | NA | NA | NA | Armc12 | armadillo repeat containing 12 [Source:M | 0 | 0 | 0 | 0 | 0 | 0 |
| 4377 | ENSMUSG00000024224 | 0 | NA | NA | NA | NA | NA | Clpsl2 | colipase-like 2 [Source:MGI Symbol;Acc:M | 0 | 0 | 0 | 0 | 0 | 0 |
| 4378 | ENSMUSG00000024225 | 0 | NA | NA | NA | NA | NA | Clps | colipase, pancreatic [Source:MGI Symbol; | 0 | 0 | 0 | 0 | 0 | 0 |
| 4379 | ENSMUSG00000024227 | 0 | NA | NA | NA | NA | NA | Pdzph1 | PDZ and pleckstrin homology domains 1 [S | 0 | 0 | 0 | 0 | 0 | 0 |
| 4380 | ENSMUSG00000024228 | 1546.907978 | -0.008096628 | 0.122655965 | -0.06601088 | 0.947369163 | 0.990677354 | Nudt12 | nudix (nucleoside diphosphate linked moi | 1631.9 | 1619 | 1402.7 | 1580 | 1705.7 | 1342.2 |
| 4381 | ENSMUSG00000024231 | 622.1349036 | 0.112523023 | 0.123470925 | 0.911332149 | 0.362120392 | 0.800872989 | Cul2 | cullin 2 [Source:MGI Symbol;Acc:MGI:191 | 577.43 | 606.47 | 609.78 | 605.38 | 709.99 | 623.75 |
| 4382 | ENSMUSG00000024232 | 53.50790624 | -0.519538008 | 0.307568197 | -1.689179874 | 0.091184965 | 0.445423616 | Bambi | BMP and activin membrane-bound inhibit | 53.005 | 68.87 | 67.418 | 42.415 | 41.443 | 47.895 |
| 4383 | ENSMUSG00000024233 | 0 | NA | NA | NA | NA | NA | Lyzl1 | lysozyme-like 1 [Source:MGI Symbol;Acc:M | 0 | 0 | 0 | 0 | 0 | 0 |
| 4384 | ENSMUSG00000024234 | 277.8058517 | 0.022060079 | 0.151008912 | 0.146084615 | 0.883854592 | 0.981769403 | Mtpap | mitochondrial poly(A) polymerase [Source | 285.1 | 263.15 | 278.73 | 290.16 | 286.83 | 262.87 |
| 4385 | ENSMUSG00000024235 | 35.47802461 | -0.786999096 | 0.394192199 | -1.996485721 | 0.045881078 | 0.296229283 | Map3k8 | mitogen-activated protein kinase kinase k | 39.352 | 55.508 | 40.25 | 33.739 | 25.084 | 18.935 |
| 4386 | ENSMUSG00000024236 | 446.2288398 | 0.413499972 | 0.145973227 | 2.832711038 | 0.004615509 | 0.063387888 | Svil | supervillin [Source:MGI Symbol;Acc:MGI:2 | 345.33 | 391.64 | 411.55 | 455 | 570.39 | 503.46 |
| 4387 | ENSMUSG00000024238 | 238.6478377 | 0.222662597 | 0.183710561 | 1.21202938 | 0.225501135 | 0.672301295 | Zeb1 | zinc finger E-box binding homeobox 1 [Sou | 181.5 | 250.81 | 229.42 | 267.02 | 224.67 | 278.46 |
| 4388 | ENSMUSG00000024240 | 236.3145511 | -0.201238177 | 0.187519752 | -1.073157227 | 0.283200578 | 0.735802616 | Epc1 | enhancer of polycomb homolog 1 [Source | 216.03 | 276.51 | 266.65 | 236.17 | 244.3 | 178.22 |
| 4389 | ENSMUSG00000024241 | 765.5054601 | -0.104088874 | 0.115975322 | -0.897508812 | 0.369447473 | 0.80653433 | Sos1 | SOS Ras/Rac guanine nucleotide exchange | 750.9 | 835.7 | 792.92 | 700.81 | 727.44 | 785.26 |
| 4390 | ENSMUSG00000024242 | 518.1373186 | -0.070912979 | 0.129545944 | -0.547396364 | 0.584106453 | 0.908955096 | Map4k3 | mitogen-activated protein kinase kinase k | 500.33 | 580.77 | 512.18 | 510.91 | 525.68 | 478.95 |
| 4391 | ENSMUSG00000024245 | 2.801701033 | -1.089542357 | 1.359652614 | -0.801338773 | 0.42293555 | NA | Tmem178 | transmembrane protein 178 [Source:MGI | 3.2124 | 7.1954 | 1.0062 | 0.964 | 1.0906 | 3.3415 |
| 4392 | ENSMUSG00000024246 | 52.44412395 | -0.124725566 | 0.311647457 | -0.400213648 | 0.688999163 | 0.936908509 | Thumpd2 | THUMP domain containing 2 [Source:MGI | 61.036 | 59.619 | 43.269 | 48.199 | 49.078 | 53.465 |
| 4393 | ENSMUSG00000024247 | 442.6200488 | 0.213768466 | 0.13702194 | 1.560103918 | 0.118735326 | 0.504308497 | Pkdcc | protein kinase domain containing, cytopla | 363 | 444.06 | 423.63 | 476.21 | 478.78 | 470.04 |
| 4394 | ENSMUSG00000024248 | 1869.652187 | 0.110333498 | 0.105547698 | 1.045342537 | 0.295864685 | 0.746382741 | Cox7a2l | cytochrome c oxidase subunit 7A2 like [So | 1632.7 | 1931.5 | 1831.4 | 1860.5 | 1936.9 | 2025 |
| 4395 | ENSMUSG00000024251 | 179.8490766 | -0.000622926 | 0.203959616 | -0.003054165 | 0.997563133 | 0.999498002 | Thada | thyroid adenoma associated [Source:MGI | 160.62 | 193.25 | 186.16 | 170.62 | 220.3 | 148.14 |
| 4396 | ENSMUSG00000024253 | 10.28762903 | -0.473022055 | 0.754927257 | -0.626579648 | 0.530934812 | 0.888794797 | Dync2li1 | dynein cytoplasmic 2 light intermediate ch | 10.44 | 9.2513 | 16.1 | 2.8919 | 16.359 | 6.6831 |
| 4397 | ENSMUSG00000024254 | 2558.868744 | 0.285721163 | 0.114383143 | 2.497930698 | 0.012492061 | 0.12644169 | Abcg8 | ATP binding cassette subfamily G member | 2582 | 1978.7 | 2356.6 | 2818.7 | 2820.3 | 2796.9 |
| 4398 | ENSMUSG00000024256 | 0 | NA | NA | NA | NA | NA | Adcyap1 | adenylate cyclase activating polypeptide 1 | 0 | 0 | 0 | 0 | 0 | 0 |
| 4399 | ENSMUSG00000024258 | 96.45873533 | -0.115815996 | 0.255135676 | -0.453938854 | 0.649872837 | 0.925643431 | Polr2d | polymerase (RNA) II (DNA directed) polyp | 121.27 | 91.485 | 87.543 | 78.082 | 110.15 | 90.221 |
| 4400 | ENSMUSG00000024259 | 906.7557343 | 0.063584425 | 0.104692299 | 0.607345769 | 0.54362147 | 0.894662801 | Slc25a46 | solute carrier family 25, member 46 [Sour | 900.28 | 903.54 | 856.31 | 924.45 | 928.12 | 927.83 |
| 4401 | ENSMUSG00000024260 | 280.4738337 | 0.367618639 | 0.155181957 | 2.368952205 | 0.017838559 | 0.159316366 | Sap130 | Sin3A associated protein [Source:MGI Syn | 248.96 | 240.53 | 245.52 | 343.18 | 321.73 | 282.92 |
| 4402 | ENSMUSG00000024261 | 0.401551119 | -2.072982726 | 4.008251979 | -0.517178744 | 0.605031388 | NA | Syt4 | synaptotagmin IV [Source:MGI Symbol;Ac | 2.4093 | 0 | 0 | 0 | 0 | 0 |
| 4403 | ENSMUSG00000024266 | 0.449470461 | -0.545136533 | 3.562105917 | -0.15303771 | 0.878368537 | NA | Adad2 | adenosine deaminase domain containing | 1.6062 | 0 | 0 | 0 | 1.0906 | 0 |
| 4404 | ENSMUSG00000024268 | 5.733776641 | -0.986824374 | 0.935751326 | -1.054579723 | 0.291617582 | 0.74194489 | Celf4 | CUGBP, Elav-like family member 4 [Source | 5.6217 | 9.2513 | 8.05 | 4.8199 | 1.0906 | 5.5692 |
| 4405 | ENSMUSG00000024269 | 97.91895976 | -0.041032029 | 0.23207713 | -0.176803415 | 0.859662806 | 0.975534508 | Tpgs2 | tubulin polyglutamylase complex subunit | 103.6 | 100.74 | 93.581 | 103.15 | 83.978 | 102.47 |
| 4406 | ENSMUSG00000024270 | 33.24344208 | -0.703697891 | 0.423410975 | -1.66197367 | 0.096518054 | 0.458530879 | Slc39a6 | solute carrier family 39 (metal ion transpo | 39.352 | 45.228 | 39.244 | 34.703 | 13.087 | 27.846 |
| 4407 | ENSMUSG00000024271 | 532.8637912 | -0.123086794 | 0.127218698 | -0.967521256 | 0.333283521 | 0.77523112 | Elp2 | elongator acetyltransferase complex subu | 511.58 | 561.24 | 594.69 | 512.84 | 527.86 | 488.98 |
| 4408 | ENSMUSG00000024273 | 117.0702447 | -0.119474627 | 0.231957102 | -0.515072077 | 0.606502654 | 0.916376001 | 2700062C07 | RIKEN cDNA 2700062C07 gene [Source:M | 124.48 | 121.29 | 119.74 | 106.04 | 90.521 | 140.34 |
| 4409 | ENSMUSG00000024274 | 1.177744186 | -2.412679626 | 2.321486536 | -1.039282197 | 0.298673512 | NA | Zscan30 | zinc finger and SCAN domain containing 3 | 0 | 3.0838 | 3.0187 | 0.964 | 0 | 0 |
| 4410 | ENSMUSG00000024276 | 378.316358 | 0.136379494 | 0.147775903 | 0.922880467 | 0.356069501 | 0.795708016 | Zfp397 | zinc finger protein 397 [Source:MGI Symb | 349.35 | 338.18 | 394.45 | 436.68 | 394.8 | 356.43 |
| 4411 | ENSMUSG00000024277 | 239.4113325 | -0.358861019 | 0.211325266 | -1.69814536 | 0.08948033 | 0.441269518 | Mapre2 | microtubule-associated protein, RP/EB far | 206.4 | 295.01 | 306.9 | 250.63 | 212.67 | 164.85 |
| 4412 | ENSMUSG00000024283 | 847.4068699 | 0.125775388 | 0.110240126 | 1.140922028 | 0.253902372 | 0.707055046 | Wac | WW domain containing adaptor with coile | 790.25 | 848.03 | 793.93 | 923.49 | 886.67 | 842.07 |
| 4413 | ENSMUSG00000024286 | 1270.18485 | 0.018274189 | 0.100252213 | 0.182282149 | 0.855361307 | 0.974158771 | Ccny | cyclin Y [Source:MGI Symbol;Acc:MGI:191 | 1204.7 | 1310.6 | 1271.9 | 1274.4 | 1308.7 | 1250.8 |
| 4414 | ENSMUSG00000024287 | 435.2089901 | -0.00890559 | 0.145299965 | -0.061291068 | 0.951127404 | 0.990883058 | Thoc1 | THO complex 1 [Source:MGI Symbol;Acc:M | 419.22 | 385.47 | 505.13 | 432.83 | 427.52 | 441.08 |
| 4415 | ENSMUSG00000024290 | 706.8561344 | 0.014090791 | 0.121147108 | 0.11631141 | 0.907405743 | 0.984526042 | Rock1 | Rho-associated coiled-coil containing prot | 640.88 | 772.99 | 697.33 | 710.45 | 711.08 | 708.4 |
| 4416 | ENSMUSG00000024292 | 4094.019532 | 0.645173538 | 0.114499093 | 5.634748028 | 1.75E-08 | 2.46E-06 | Cyp4f14 | cytochrome P450, family 4, subfamily f, po | 3025.3 | 3450.7 | 3106.3 | 5714.5 | 4593.7 | 4673.7 |
| 4417 | ENSMUSG00000024293 | 230.1750144 | -0.123859418 | 0.163085311 | -0.759476238 | 0.447567723 | 0.853796936 | Esco1 | establishment of sister chromatid cohesio | 237.72 | 246.7 | 235.46 | 200.51 | 231.21 | 229.45 |
| 4418 | ENSMUSG00000024294 | 1376.786349 | 0.008390164 | 0.11091059 | 0.075647997 | 0.93969915 | 0.988812319 | Mib1 | mindbomb E3 ubiquitin protein ligase 1 [S | 1241.6 | 1405.2 | 1473.1 | 1446 | 1390.5 | 1304.3 |
| 4419 | ENSMUSG00000024295 | 5.381793191 | 0.897113941 | 0.94369718 | 0.950637513 | 0.341788419 | 0.783459222 | Cyp4f41-ps | cytochrome P450, family 4, subfamily f, po | 3.2124 | 2.0558 | 6.0375 | 7.7118 | 4.3625 | 8.9108 |
| 4420 | ENSMUSG00000024298 | 458.1529117 | 0.144622706 | 0.184169422 | 0.785269912 | 0.432295333 | 0.845264432 | Zfp871 | zinc finger protein 871 [Source:MGI Symb | 432.87 | 442 | 431.68 | 636.23 | 431.88 | 374.25 |

| | | | | | | | | | | | | | | | | |
|---|---|---|---|---|---|---|---|---|---|---|---|---|---|---|---|---|
| 4421 | ENSMUSG00000024299 | 123.1866983 | -0.374200734 | 0.266049188 | -1.406509588 | 0.159572859 | 0.581402021 | Adamts10 | a disintegrin-like and metallopeptidase (re | 160.62 | 100.74 | 155.97 | 139.78 | 82.887 | 99.132 |
| 4422 | ENSMUSG00000024300 | 31.94748147 | -0.9311534 | 0.460989797 | -2.019900236 | 0.043393737 | 0.28712123 | Myo1f | myosin IF [Source:MGI Symbol;Acc:MGI:1 | 48.989 | 48.312 | 28.175 | 16.388 | 14.178 | 35.643 |
| 4423 | ENSMUSG00000024301 | 9.556238005 | 0.207253588 | 0.694450283 | 0.298442658 | 0.765365335 | 0.954980327 | Kifc5b | kinesin family member C5B [Source:MGI S | 11.243 | 6.1675 | 9.0562 | 7.7118 | 10.906 | 12.252 |
| 4424 | ENSMUSG00000024302 | 2.035526524 | -1.446736186 | 1.634095103 | -0.885343934 | 0.375971161 | NA | Dtna | dystrobrevin alpha [Source:MGI Symbol;A | 0.8031 | 3.0838 | 5.0312 | 0 | 2.1812 | 1.1138 |
| 4425 | ENSMUSG00000024304 | 1718.975384 | 0.033224867 | 0.139726542 | 0.237784939 | 0.8120479 | 0.96653525 | Cdh2 | cadherin 2 [Source:MGI Symbol;Acc:MGI: | 1473.7 | 1901.6 | 1722.7 | 1499 | 2042.7 | 1674.1 |
| 4426 | ENSMUSG00000024306 | 0 | NA | NA | NA | NA | NA | Ccdc178 | coiled coil domain containing 178 [Source | 0 | 0 | 0 | 0 | 0 | 0 |
| 4427 | ENSMUSG00000024308 | 1141.429779 | 0.028505693 | 0.104540169 | 0.272676939 | 0.785101563 | 0.960670212 | Tapbp | TAP binding protein [Source:MGI Symbol; | 1144.4 | 1188.3 | 1057.6 | 1142.3 | 1135.3 | 1180.7 |
| 4428 | ENSMUSG00000024309 | 590.4824755 | 0.096088404 | 0.116432656 | 0.825270222 | 0.409218194 | 0.833065722 | Pfdn6 | prefoldin subunit 6 [Source:MGI Symbol;A | 566.19 | 576.66 | 569.53 | 591.88 | 626.01 | 612.61 |
| 4429 | ENSMUSG00000024312 | 352.4415971 | 0.150245374 | 0.151774146 | 0.989927319 | 0.322209645 | 0.769684609 | Wdr46 | WD repeat domain 46 [Source:MGI Symbo | 320.44 | 314.54 | 367.28 | 339.32 | 415.53 | 357.54 |
| 4430 | ENSMUSG00000024313 | 1.327718352 | 1.882221139 | 2.032456141 | 0.926082045 | 0.354403344 | NA | Mep1b | meprin 1 beta [Source:MGI Symbol;Acc:M | 1.6062 | 0 | 0 | 1.928 | 1.0906 | 3.3415 |
| 4431 | ENSMUSG00000024317 | 269.202918 | 0.183172676 | 0.165987628 | 1.1035321 | 0.26979616 | 0.725011151 | Rnf138 | ring finger protein 138 [Source:MGI Symb | 269.84 | 230.25 | 255.59 | 267.02 | 263.93 | 328.58 |
| 4432 | ENSMUSG00000024319 | 449.4560336 | 0.008104503 | 0.137594253 | 0.058901461 | 0.953030594 | 0.9915542 | Vps52 | VPS52 GARP complex subunit [Source:MG | 463.39 | 395.75 | 485.01 | 446.32 | 437.34 | 468.93 |
| 4433 | ENSMUSG00000024325 | 332.7050607 | 0.156590359 | 0.165507297 | 0.946123599 | 0.344085549 | 0.785727752 | Ring1 | ring finger protein 1 [Source:MGI Symbol; | 296.34 | 284.73 | 363.25 | 364.38 | 301.01 | 386.5 |
| 4434 | ENSMUSG00000024327 | 36.6810297 | 0.610320028 | 0.390255464 | 1.563898738 | 0.117841351 | 0.502838264 | Slc39a7 | solute carrier family 39 (zinc transporter), | 36.14 | 17.475 | 33.206 | 46.271 | 46.897 | 40.098 |
| 4435 | ENSMUSG00000024330 | 16.60248823 | -0.306415427 | 0.560939025 | -0.546254429 | 0.584891057 | 0.90908382 | Col11a2 | collagen, type XI, alpha 2 [Source:MGI Syr | 14.456 | 19.53 | 21.131 | 13.496 | 8.7249 | 22.277 |
| 4436 | ENSMUSG00000024331 | 817.7463879 | 0.284945799 | 0.135026635 | 2.110293268 | 0.034833103 | 0.250037719 | Dsc2 | desmocollin 2 [Source:MGI Symbol;Acc:M | 811.13 | 645.53 | 754.68 | 1008.3 | 815.78 | 871.03 |
| 4437 | ENSMUSG00000024334 | 1.552482705 | -4.001878574 | 2.003034481 | -1.99790798 | 0.045726637 | NA | H2-Oa | histocompatibility 2, O region alpha locus | 3.2124 | 3.0838 | 3.0187 | 0 | 0 | 0 |
| 4438 | ENSMUSG00000024335 | 1567.148839 | -0.11885895 | 0.107866807 | -1.10190478 | 0.27050307 | 0.725929337 | Brd2 | bromodomain containing 2 [Source:MGI S | 1586.1 | 1700.2 | 1610 | 1656.1 | 1457.1 | 1393.4 |
| 4439 | ENSMUSG00000024338 | 530.7602294 | 0.070509925 | 0.128483404 | 0.548786248 | 0.583152149 | 0.908645447 | Psmb8 | proteasome (prosome, macropain) subun | 556.55 | 484.15 | 512.18 | 561.04 | 563.85 | 506.8 |
| 4440 | ENSMUSG00000024339 | 344.6768652 | 0.024294445 | 0.149312363 | 0.162708862 | 0.870747669 | 0.97888679 | Tap2 | transporter 2, ATP-binding cassette, sub-f | 378.26 | 346.41 | 299.86 | 340.28 | 346.82 | 356.43 |
| 4441 | ENSMUSG00000024340 | 0.342639279 | -1.805976596 | 4.028547714 | -0.448294702 | 0.653940524 | NA | Btnl2 | butyrophilin-like 2 [Source:MGI Symbol;A | 0 | 2.0558 | 0 | 0 | 0 | 0 |
| 4442 | ENSMUSG00000024346 | 419.2025967 | -0.088785299 | 0.139151089 | -0.638049616 | 0.523441383 | 0.888334265 | Pfdn1 | prefoldin 1 [Source:MGI Symbol;Acc:MGI: | 423.23 | 455.37 | 417.59 | 389.45 | 377.35 | 452.22 |
| 4443 | ENSMUSG00000024347 | 0.692116425 | -1.412606127 | 2.773087916 | -0.509398248 | 0.610473104 | NA | Psd2 | pleckstrin and Sec7 domain containing 2 [ | 0 | 2.0558 | 1.0062 | 0 | 1.0906 | 0 |
| 4444 | ENSMUSG00000024349 | 16.96488719 | -0.827830696 | 0.56642291 | -1.461506378 | 0.143876527 | 0.554054867 | Sting1 | stimulator of interferon response cGAMP | 31.321 | 19.53 | 14.087 | 12.532 | 8.7249 | 15.594 |
| 4445 | ENSMUSG00000024350 | 149.2784907 | -0.431169748 | 0.210523062 | -2.048087953 | 0.040551382 | 0.275368634 | Dnajc18 | DnaJ heat shock protein family (Hsp40) m | 150.98 | 184 | 180.12 | 150.38 | 122.15 | 108.04 |
| 4446 | ENSMUSG00000024352 | 44.58867059 | -0.221471336 | 0.377014328 | -0.587434798 | 0.556911729 | 0.898167794 | Spata24 | spermatogenesis associated 24 [Source:M | 36.14 | 69.898 | 38.237 | 38.559 | 50.168 | 34.529 |
| 4447 | ENSMUSG00000024353 | 6.270438854 | -2.318161603 | 0.97842505 | -2.369278673 | 0.017822819 | 0.159316366 | Mzb1 | marginal zone B and B1 cell-specific prote | 16.062 | 6.1675 | 9.0562 | 1.928 | 2.1812 | 2.2277 |
| 4448 | ENSMUSG00000024354 | 1198.024054 | 0.223905434 | 0.183532891 | 1.219974428 | 0.222474569 | 0.668407528 | Slc23a1 | solute carrier family 23 (nucleobase trans| | 1350 | 1142 | 822.1 | 990.97 | 1445.1 | 1438 |
| 4449 | ENSMUSG00000024357 | 912.9227892 | 0.036169481 | 0.110298569 | 0.327923395 | 0.742969584 | 0.949404724 | Sil1 | endoplasmic reticulum chaperone SIL1 ho | 844.06 | 952.88 | 908.64 | 960.12 | 907.39 | 904.44 |
| 4450 | ENSMUSG00000024359 | 10431.24756 | 0.017883644 | 0.103582345 | 0.172651465 | 0.862925396 | 0.976631684 | Hspa9 | heat shock protein 9 [Source:MGI Symbol | 9131.3 | 11236 | 10734 | 9989.7 | 10744 | 10753 |
| 4451 | ENSMUSG00000024360 | 1543.696306 | 0.045443482 | 0.099930357 | 0.454751526 | 0.649288007 | 0.925643431 | Etf1 | eukaryotic translation termination factor : | 1501 | 1564.5 | 1492.3 | 1468.1 | 1594.5 | 1641.8 |
| 4452 | ENSMUSG00000024365 | 20.84430839 | 1.257385719 | 0.540850252 | 2.324831529 | 0.02008097 | 0.173507261 | Cyp21a1 | cytochrome P450, family 21, subfamily a, | 10.44 | 13.363 | 13.081 | 14.46 | 42.534 | 31.188 |
| 4453 | ENSMUSG00000024366 | 0 | NA | NA | NA | NA | NA | Gfra3 | glial cell line derived neurotrophic factor f | 0 | 0 | 0 | 0 | 0 | 0 |
| 4454 | ENSMUSG00000024369 | 403.0017018 | 0.316705595 | 0.152403258 | 2.078076275 | 0.037702334 | 0.263449005 | Nelfe | negative elongation factor complex memk | 412.79 | 318.65 | 344.14 | 435.72 | 416.62 | 490.09 |
| 4455 | ENSMUSG00000024370 | 207.1974797 | 0.070590249 | 0.175550075 | 0.40210891 | 0.687603875 | 0.936885029 | Cdc23 | CDC23 cell division cycle 23 [Source:MGI S | 223.26 | 190.16 | 192.19 | 194.72 | 231.21 | 211.63 |
| 4456 | ENSMUSG00000024371 | 585.3415286 | 0.336829946 | 0.137063686 | 2.45747036 | 0.013991936 | 0.136282395 | C2 | complement component 2 (within H-2S) [ | 496.32 | 513.96 | 542.37 | 689.24 | 714.35 | 555.81 |
| 4457 | ENSMUSG00000024376 | 2.285267424 | -0.700719553 | 1.44809604 | -0.483890249 | 0.628463761 | NA | Epb41l4a | erythrocyte membrane protein band 4.1 l | 2.4093 | 3.0838 | 3.0187 | 1.928 | 3.2719 | 0 |
| 4458 | ENSMUSG00000024378 | 1778.026798 | 1.137376279 | 0.155519967 | 7.313377832 | 2.61E-13 | 1.24E-10 | Stard4 | StAR-related lipid transfer (START) domair | 1389.4 | 1073.1 | 869.4 | 2413.8 | 2231.4 | 2691 |
| 4459 | ENSMUSG00000024379 | 1.665174732 | 0.099506947 | 1.677465077 | 0.059319832 | 0.952697365 | NA | Tslp | thymic stromal lymphopoietin [Source:MC | 0.8031 | 2.0558 | 2.0125 | 2.8919 | 0 | 2.2277 |
| 4460 | ENSMUSG00000024381 | 237.3680833 | 0.034201422 | 0.176123933 | 0.194189517 | 0.846027491 | 0.972803041 | Bin1 | bridging integrator 1 [Source:MGI Symbol | 222.46 | 240.53 | 241.5 | 282.45 | 231.21 | 206.06 |
| 4461 | ENSMUSG00000024382 | 371.3025633 | -0.027310509 | 0.163370733 | -0.167168922 | 0.867237137 | 0.97799142 | Ercc3 | excision repair cross-complementing rode | 393.52 | 380.33 | 350.17 | 340.28 | 446.06 | 317.45 |
| 4462 | ENSMUSG00000024383 | 519.5663656 | 0.020642896 | 0.137528799 | 0.150098716 | 0.880686733 | 0.981448736 | Map3k2 | mitogen-activated protein kinase kinase k | 477.04 | 583.86 | 487.02 | 495.48 | 511.5 | 562.49 |
| 4463 | ENSMUSG00000024384 | 545.8504531 | 0.101635012 | 0.131980944 | 0.77007338 | 0.441256366 | 0.849828766 | Iws1 | IWS1, SUPT6 interacting protein [Source:N | 539.68 | 566.38 | 473.94 | 587.06 | 586.75 | 521.28 |
| 4464 | ENSMUSG00000024386 | 10703.67767 | 0.141912926 | 0.090855369 | 1.56196521 | 0.118296185 | 0.503838639 | Proc | protein C [Source:MGI Symbol;Acc:MGI:9 | 9377 | 10505 | 10652 | 11173 | 11565 | 10950 |
| 4465 | ENSMUSG00000024387 | 1014.999004 | -0.021482245 | 0.108990973 | -0.197101144 | 0.843748388 | 0.97250284 | Csnk2b | casein kinase 2, beta polypeptide [Source | 1034.4 | 1083.4 | 949.89 | 1016 | 1036.1 | 970.16 |
| 4466 | ENSMUSG00000024388 | 0.736965982 | -2.935629304 | 2.883220453 | -1.018177192 | 0.308593757 | NA | Myo7b | myosin VIIB [Source:MGI Symbol;Acc:MGI | 2.4093 | 0 | 2.0125 | 0 | 0 | 0 |
| 4467 | ENSMUSG00000024391 | 2193.641404 | 0.542704938 | 0.147822202 | 3.671335781 | 0.000241286 | 0.007285933 | Apom | apolipoprotein M [Source:MGI Symbol;Ac | 1569.3 | 1764.9 | 2023.6 | 2156.4 | 2487.7 | 3160 |
| 4468 | ENSMUSG00000024392 | 1557.429441 | 0.0083943 | 0.098942638 | 0.08484007 | 0.932388537 | 0.988420305 | Bag6 | BCL2-associated athanogene 6 [Source:M | 1572.5 | 1605.6 | 1480.2 | 1507.7 | 1633.7 | 1544.9 |
| 4469 | ENSMUSG00000024393 | 1775.983758 | 0.03471372 | 0.099639718 | 0.348392393 | 0.727545515 | 0.945907265 | Prrc2a | proline-rich coiled-coil 2A [Source:MGI Sy | 1689.7 | 1684.8 | 1889.7 | 1781.4 | 1764.6 | 1845.6 |
| 4470 | ENSMUSG00000024395 | 1520.922572 | -0.0002014 | 0.116848284 | -0.001723602 | 0.998624765 | 0.999764475 | Lims2 | LIM and senescent cell antigen like domai | 1603.8 | 1348.6 | 1610 | 1545.3 | 1390.5 | 1627.3 |
| 4471 | ENSMUSG00000024397 | 46.84812699 | -1.444770306 | 0.414722461 | -3.48370402 | 0.000494526 | 0.012307964 | Aif1 | allograft inflammatory factor 1 [Source:M | 50.595 | 108.96 | 46.287 | 27.955 | 28.356 | 18.935 |
| 4472 | ENSMUSG00000024399 | 8.986524698 | -2.688278993 | 0.854839995 | -3.144774469 | 0.001662149 | 0.030548062 | Ltb | lymphotoxin B [Source:MGI Symbol;Acc:N | 19.274 | 15.419 | 12.075 | 3.8559 | 2.1812 | 1.1138 |
| 4473 | ENSMUSG00000024400 | 561.9958081 | 0.039458031 | 0.12868851 | 0.306616583 | 0.75913523 | 0.95402891 | Wdr33 | WD repeat domain 33 [Source:MGI Symbo | 505.95 | 581.8 | 575.57 | 530.19 | 574.76 | 603.7 |
| 4474 | ENSMUSG00000024401 | 1.998779642 | -2.254462364 | 1.679887861 | -1.342031463 | 0.179585794 | NA | Tnf | tumor necrosis factor [Source:MGI Symbo | 4.8186 | 3.0838 | 2.0125 | 0.964 | 0 | 1.1138 |
| 4475 | ENSMUSG00000024402 | 0 | NA | NA | NA | NA | NA | Lta | lymphotoxin A [Source:MGI Symbol;Acc:N | 0 | 0 | 0 | 0 | 0 | 0 |
| 4476 | ENSMUSG00000024403 | 29.41831536 | 0.5666383 | 0.406612189 | 1.393559552 | 0.16345065 | 0.587512475 | Atp6v1g2 | ATPase, H+ transporting, lysosomal V1 sul | 27.305 | 23.642 | 20.125 | 40.487 | 35.99 | 28.96 |

| | | | | | | | | | | | | | | | |
|---|---|---|---|---|---|---|---|---|---|---|---|---|---|---|---|
| 4477 | ENSMUSG00000024404 | 1473.47719 | 0.023142102 | 0.10135413 | 0.22832915 | 0.819390365 | 0.969160361 | Riok3 | RIO kinase 3 [Source:MGI Symbol;Acc:MG | 1385.4 | 1571.7 | 1428.9 | 1497.1 | 1486.5 | 1471.4 |
| 4478 | ENSMUSG00000024406 | 0 | NA | NA | NA | NA | NA | Pou5f1 | POU domain, class 5, transcription factor : | 0 | 0 | 0 | 0 | 0 | 0 |
| 4479 | ENSMUSG00000024409 | 0 | NA | NA | NA | NA | NA | Psors1c2 | psoriasis susceptibility 1 candidate 2 (hum | 0 | 0 | 0 | 0 | 0 | 0 |
| 4480 | ENSMUSG00000024410 | 419.3929233 | -0.065836879 | 0.140586297 | -0.468302246 | 0.639568462 | 0.923932929 | Rmc1 | regulator of MON1-CCZ1 [Source:MGI Syn | 411.99 | 434.81 | 440.74 | 432.83 | 437.34 | 358.66 |
| 4481 | ENSMUSG00000024411 | 214.7047321 | -0.511643719 | 0.219939079 | -2.326297452 | 0.020002688 | 0.17340647 | Aqp4 | aquaporin 4 [Source:MGI Symbol;Acc:MG | 291.53 | 287.82 | 177.1 | 149.42 | 194.13 | 188.24 |
| 4482 | ENSMUSG00000024413 | 2724.031662 | 0.183327866 | 0.10138928 | 1.808158272 | 0.070581871 | 0.384228898 | Npc1 | NPC intracellular cholesterol transporter 1 | 2488.8 | 2643.8 | 2520.6 | 2689.5 | 2822.5 | 3178.9 |
| 4483 | ENSMUSG00000024414 | 888.9544265 | 0.015445339 | 0.131567238 | 0.117395026 | 0.906547025 | 0.984272857 | Mrpl27 | mitochondrial ribosomal protein L27 [Sou | 944.45 | 921.01 | 785.88 | 776.97 | 958.65 | 946.77 |
| 4484 | ENSMUSG00000024420 | 5.588804288 | -0.380842851 | 0.89670764 | -0.424712397 | 0.671046345 | 0.93116767 | Zfp521 | zinc finger protein 521 [Source:MGI Symb | 4.8186 | 7.1954 | 7.0437 | 6.7478 | 3.2719 | 4.4554 |
| 4485 | ENSMUSG00000024421 | 156.809359 | 0.061281947 | 0.274287232 | 0.223422529 | 0.823206672 | 0.96943211 | Lama3 | laminin, alpha 3 [Source:MGI Symbol;Acc: | 90.751 | 152.13 | 218.36 | 158.09 | 174.5 | 147.03 |
| 4486 | ENSMUSG00000024422 | 703.4260835 | 0.164972168 | 0.150356387 | 1.09720758 | 0.27255066 | 0.727618887 | Dhx16 | DEAH (Asp-Glu-Ala-His) box polypeptide 1 | 562.97 | 618.81 | 809.02 | 766.36 | 689.27 | 774.12 |
| 4487 | ENSMUSG00000024423 | 204.4352536 | -0.14653913 | 0.181363011 | -0.807987961 | 0.419097513 | 0.837591673 | Impact | impact, RWD domain protein [Source:MG | 214.43 | 224.09 | 206.28 | 218.82 | 162.5 | 200.49 |
| 4488 | ENSMUSG00000024424 | 9641.548895 | 0.265989035 | 0.102354924 | 2.598693098 | 0.00935794 | 0.10348107 | Ttc39c | tetratricopeptide repeat domain 39C [Sou | 8402.1 | 8606.8 | 9257.5 | 10270 | 11691 | 9621.4 |
| 4489 | ENSMUSG00000024425 | 3820.831371 | 0.205202555 | 0.102300723 | 2.005875908 | 0.044869489 | 0.292602477 | Ndfip1 | Nedd4 family interacting protein 1 [Sourc | 3244.5 | 3567.9 | 3837.8 | 4210.7 | 4243.6 | 3820.5 |
| 4490 | ENSMUSG00000024426 | 229.9611012 | 0.237979305 | 0.221468079 | 1.074553526 | 0.282574668 | 0.735497572 | Atat1 | alpha tubulin acetyltransferase 1 [Source: | 218.44 | 164.47 | 250.55 | 319.08 | 232.3 | 194.92 |
| 4491 | ENSMUSG00000024427 | 239.9186908 | 0.374926445 | 0.271516152 | 1.3808624 | 0.167321273 | 0.593441872 | Spry4 | sprouty RTK signaling antagonist 4 [Source | 230.49 | 112.04 | 283.76 | 233.28 | 331.55 | 248.39 |
| 4492 | ENSMUSG00000024429 | 583.3294851 | 0.122916954 | 0.124968231 | 0.983585608 | 0.325319304 | 0.771501263 | Gnl1 | guanine nucleotide binding protein-like 1 | 570.2 | 522.18 | 582.62 | 580.31 | 656.55 | 588.11 |
| 4493 | ENSMUSG00000024430 | 86.13019591 | -0.26152975 | 0.259360322 | -1.00836453 | 0.313279494 | 0.760152431 | Cabyr | calcium-binding tyrosine-(Y)-phosphorylat | 87.538 | 109.99 | 84.525 | 82.902 | 88.34 | 63.489 |
| 4494 | ENSMUSG00000024431 | 1725.837718 | 0.41767534 | 0.106998711 | 3.90355489 | 9.48E-05 | 0.0034713 | Nr3c1 | nuclear receptor subfamily 3, group C, me | 1385.4 | 1492.5 | 1556.7 | 2065.8 | 2055.8 | 1798.9 |
| 4495 | ENSMUSG00000024436 | 257.5598625 | 0.084849821 | 0.166312834 | 0.51018204 | 0.609923934 | 0.916376001 | Mrps18b | mitochondrial ribosomal protein S18B [So | 262.61 | 261.09 | 226.41 | 293.05 | 268.29 | 233.91 |
| 4496 | ENSMUSG00000024437 | 1.029799386 | 0.219173677 | 2.229624068 | 0.098300731 | 0.921693497 | NA | Gm8615 | predicted pseudogene 8615 [Source:MGI | 0.8031 | 1.0279 | 1.0062 | 0 | 0 | 3.3415 |
| 4497 | ENSMUSG00000024440 | 48.86205338 | 0.154075994 | 0.409956594 | 0.375834897 | 0.707039643 | 0.943108897 | Pcdh12 | protocadherin 12 [Source:MGI Symbol;Ac | 46.58 | 27.754 | 64.4 | 55.911 | 28.356 | 70.172 |
| 4498 | ENSMUSG00000024442 | 655.6839067 | -0.013179877 | 0.123018525 | -0.107137333 | 0.91468003 | 0.98625062 | Dele1 | DAP3 binding cell death enhancer 1 [Sour | 693.88 | 623.95 | 657.08 | 596.7 | 667.46 | 695.04 |
| 4499 | ENSMUSG00000024446 | 163.5434058 | 0.048824917 | 0.210529451 | 0.2319149 | 0.816604111 | 0.967767553 | Rpp21 | ribonuclease P 21 subunit [Source:MGI Sy | 139.74 | 159.33 | 184.14 | 199.54 | 157.05 | 141.46 |
| 4500 | ENSMUSG00000024448 | 0 | NA | NA | NA | NA | NA | H2-M10.1 | histocompatibility 2, M region locus 10.1 [ | 0 | 0 | 0 | 0 | 0 | 0 |
| 4501 | ENSMUSG00000024451 | 144.8096927 | -0.770858581 | 0.248103339 | -3.10700607 | 0.001889925 | 0.033417944 | Arap3 | ArfGAP with RhoGAP domain, ankyrin rep | 169.45 | 154.19 | 224.39 | 118.57 | 74.162 | 128.09 |
| 4502 | ENSMUSG00000024454 | 704.2752407 | 0.037955106 | 0.132452889 | 0.286555516 | 0.77445268 | 0.958637874 | Hdac3 | histone deacetylase 3 [Source:MGI Symbo | 778.21 | 631.14 | 674.18 | 651.65 | 777.61 | 712.86 |
| 4503 | ENSMUSG00000024456 | 1026.100109 | -0.160112478 | 0.113250408 | -1.413791629 | 0.157423092 | 0.576703416 | Diaph1 | diaphanous related formin 1 [Source:MGI | 1078.6 | 1084.5 | 1085.7 | 956.27 | 883.4 | 1068.2 |
| 4504 | ENSMUSG00000024457 | 739.5582674 | 0.292337145 | 0.118446445 | 2.468095568 | 0.013583405 | 0.133653312 | Trim26 | tripartite motif-containing 26 [Source:MG | 674.61 | 605.44 | 714.43 | 823.24 | 832.14 | 787.49 |
| 4505 | ENSMUSG00000024459 | 0.513958919 | -2.394274821 | 3.756096013 | -0.637437066 | 0.523840191 | NA | H2-M5 | histocompatibility 2, M region locus 5 [Sou | 0 | 3.0838 | 0 | 0 | 0 | 0 |
| 4506 | ENSMUSG00000024462 | 41.77327517 | -0.76923731 | 0.40290074 | -1.909247698 | 0.056230146 | 0.335386734 | Gabbr1 | gamma-aminobutyric acid (GABA) B recep | 51.399 | 63.731 | 43.269 | 48.199 | 23.994 | 20.049 |
| 4507 | ENSMUSG00000024471 | 0 | NA | NA | NA | NA | NA | Myot | myotilin [Source:MGI Symbol;Acc:MGI:18 | 0 | 0 | 0 | 0 | 0 | 0 |
| 4508 | ENSMUSG00000024472 | 292.7195555 | 0.128794873 | 0.181561559 | 0.709373027 | 0.478093022 | 0.869794311 | Dcp2 | decapping mRNA 2 [Source:MGI Symbol;A | 251.37 | 339.21 | 248.54 | 259.31 | 332.64 | 325.24 |
| 4509 | ENSMUSG00000024474 | 1041.82084 | -0.015408351 | 0.104657636 | -0.147226244 | 0.882953447 | 0.981769403 | Ik | IK cytokine [Source:MGI Symbol;Acc:MGI: | 1032 | 1103 | 1007.3 | 1008.3 | 1060.1 | 1040.3 |
| 4510 | ENSMUSG00000024477 | 173.5861556 | 0.040352332 | 0.190649647 | 0.211656997 | 0.832374639 | 0.970533397 | Pggt1b | protein geranylgeranyltransferase type I, I | 154.2 | 173.72 | 186.16 | 185.08 | 189.77 | 152.6 |
| 4511 | ENSMUSG00000024479 | 911.4246251 | -0.181332615 | 0.116326326 | -1.558826977 | 0.119037337 | 0.504735151 | Mal2 | mal, T cell differentiation protein 2 [Sourc | 995.04 | 1012.5 | 898.58 | 912.89 | 787.43 | 862.12 |
| 4512 | ENSMUSG00000024480 | 597.8841731 | -0.098952434 | 0.125085643 | -0.791077464 | 0.428898789 | 0.842753542 | Ap3s1 | adaptor-related protein complex 3, sigma | 670.59 | 608.53 | 575.57 | 608.27 | 552.94 | 571.4 |
| 4513 | ENSMUSG00000024481 | 2.133426716 | 1.726724247 | 1.714816259 | 1.006944177 | 0.313961601 | NA | Lvrn | laeverin [Source:MGI Symbol;Acc:MGI:19 | 0 | 0 | 3.0187 | 0.964 | 4.3625 | 4.4554 |
| 4514 | ENSMUSG00000024483 | 659.9815607 | 0.142618937 | 0.138845236 | 1.027179186 | 0.304336097 | 0.753237048 | Ankhd1 | ankyrin repeat and KH domain containing | 603.93 | 581.8 | 697.33 | 783.71 | 682.73 | 610.39 |
| 4515 | ENSMUSG00000024485 | 2.938182507 | -0.74406037 | 1.44283098 | -0.515694756 | 0.606067618 | NA | Slc4a9 | solute carrier family 4, sodium bicarbonat | 0.8031 | 8.2233 | 2.0125 | 0 | 4.3625 | 2.2277 |
| 4516 | ENSMUSG00000024486 | 5.386977216 | -0.373686182 | 0.974306378 | -0.383540733 | 0.701318875 | 0.941991169 | Hbegf | heparin-binding EGF-like growth factor [S | 4.0155 | 5.1396 | 9.0562 | 1.928 | 3.2719 | 8.9108 |
| 4517 | ENSMUSG00000024487 | 698.861137 | 0.069431552 | 0.120335106 | 0.57698501 | 0.563949582 | 0.901211606 | Yipf5 | Yip1 domain family, member 5 [Source:M | 695.49 | 644.5 | 706.38 | 773.11 | 665.28 | 708.4 |
| 4518 | ENSMUSG00000024491 | 282.3705547 | -0.089075041 | 0.161573404 | -0.55129767 | 0.581429635 | 0.908645447 | Rbm27 | RNA binding motif protein 27 [Source:MG | 281.89 | 317.63 | 273.7 | 250.63 | 309.74 | 260.64 |
| 4519 | ENSMUSG00000024493 | 540.3273198 | 0.07986935 | 0.133453868 | 0.598479093 | 0.549520305 | 0.895703112 | Lars | leucyl-tRNA synthetase [Source:MGI Symb | 496.32 | 589 | 491.05 | 519.58 | 581.3 | 564.72 |
| 4520 | ENSMUSG00000024497 | 0 | NA | NA | NA | NA | NA | Pou4f3 | POU domain, class 4, transcription factor : | 0 | 0 | 0 | 0 | 0 | 0 |
| 4521 | ENSMUSG00000024498 | 214.2897499 | -0.176861454 | 0.174593056 | -1.012992488 | 0.311063749 | 0.759594595 | Tcerg1 | transcription elongation regulator 1 (CA15 | 212.82 | 247.73 | 222.38 | 221.71 | 201.76 | 179.33 |
| 4522 | ENSMUSG00000024500 | 0.717171207 | -1.535127875 | 3.012718241 | -0.509549102 | 0.61036739 | NA | Ppp2r2b | protein phosphatase 2, regulatory subunit | 3.2124 | 0 | 0 | 0 | 1.0906 | 0 |
| 4523 | ENSMUSG00000024501 | 17.06328015 | 0.003593064 | 0.561104937 | 0.006403552 | 0.99489074 | 0.998676967 | Dpysl3 | dihydropyrimidinase-like 3 [Source:MGI S | 26.502 | 9.2513 | 15.094 | 17.352 | 16.359 | 17.822 |
| 4524 | ENSMUSG00000024502 | 0 | NA | NA | NA | NA | NA | Jakmip2 | janus kinase and microtubule interacting p | 0 | 0 | 0 | 0 | 0 | 0 |
| 4525 | ENSMUSG00000024503 | 0.974396903 | -0.734194653 | 2.427679541 | -0.302426511 | 0.762326948 | NA | Spink1 | serine peptidase inhibitor, Kazal type 1 [S | 1.6062 | 0 | 2.0125 | 0 | 0 | 2.2277 |
| 4526 | ENSMUSG00000024505 | 41.32202418 | -0.085749183 | 0.373391762 | -0.229649369 | 0.818364241 | 0.968897682 | Dtwd2 | DTW domain containing 2 [Source:MGI Sy | 27.305 | 53.452 | 47.293 | 39.523 | 44.715 | 35.643 |
| 4527 | ENSMUSG00000024507 | 11649.68282 | -0.139961527 | 0.093148979 | -1.50255568 | 0.132953659 | 0.533881657 | Hsd17b4 | hydroxysteroid (17-beta) dehydrogenase 4 | 12487 | 12450 | 11707 | 11693 | 11410 | 10152 |
| 4528 | ENSMUSG00000024510 | 0 | NA | NA | NA | NA | NA | Ftmt | ferritin mitochondrial [Source:MGI Symbo | 0 | 0 | 0 | 0 | 0 | 0 |
| 4529 | ENSMUSG00000024511 | 11.31522509 | -1.508377146 | 0.701470747 | -2.150306558 | 0.031530974 | 0.236302595 | Rab27b | RAB27B, member RAS oncogene family [S | 13.653 | 24.67 | 12.075 | 8.6758 | 4.3625 | 4.4554 |
| 4530 | ENSMUSG00000024512 | 0 | NA | NA | NA | NA | NA | Dynap | dynactin associated protein [Source:MGI S | 0 | 0 | 0 | 0 | 0 | 0 |
| 4531 | ENSMUSG00000024513 | 946.4502797 | -0.069501209 | 0.114585481 | -0.606544638 | 0.54415315 | 0.89481775 | Mbd2 | methyl-CpG binding domain protein 2 [So | 936.42 | 945.68 | 1026.4 | 963.01 | 967.38 | 839.84 |
| 4532 | ENSMUSG00000024515 | 525.9776466 | 0.067855026 | 0.129855309 | 0.522543329 | 0.601292087 | 0.915461118 | Smad4 | SMAD family member 4 [Source:MGI Sym | 485.88 | 542.74 | 513.18 | 576.46 | 546.4 | 491.21 |

| | | | | | | | | | | | | | | | |
|---|---|---|---|---|---|---|---|---|---|---|---|---|---|---|---|
| 4533 | ENSMUSG00000024516 | 386.9794706 | 0.073962764 | 0.147732505 | 0.500653283 | 0.616615156 | 0.917509899 | Sec11c | SEC11 homolog C, signal peptidase compl | 416.01 | 366.97 | 347.15 | 371.13 | 379.54 | 441.08 |
| 4534 | ENSMUSG00000024517 | 0 | NA | NA | NA | NA | NA | Grp | gastrin releasing peptide [Source:MGI Syn | 0 | 0 | 0 | 0 | 0 | 0 |
| 4535 | ENSMUSG00000024518 | 0.17131964 | -0.796629176 | 4.080472857 | -0.195229623 | 0.84521319 | NA | Rax | retina and anterior neural fold homeobox | 0 | 1.0279 | 0 | 0 | 0 | 0 |
| 4536 | ENSMUSG00000024519 | 0.17131964 | -0.796629176 | 4.080472857 | -0.195229623 | 0.84521319 | NA | Cplx4 | complexin 4 [Source:MGI Symbol;Acc:MG | 0 | 1.0279 | 0 | 0 | 0 | 0 |
| 4537 | ENSMUSG00000024521 | 8.313596336 | -1.372841802 | 0.830895501 | -1.652243633 | 0.098484895 | 0.462795815 | Pmaip1 | phorbol-12-myristate-13-acetate-induced | 18.471 | 10.279 | 7.0437 | 1.928 | 4.3625 | 7.7969 |
| 4538 | ENSMUSG00000024524 | 80.39576242 | 0.16892479 | 0.272112609 | 0.620790013 | 0.534737795 | 0.889858671 | Gnal | guanine nucleotide binding protein, alpha | 91.554 | 63.731 | 71.443 | 90.614 | 95.974 | 69.058 |
| 4539 | ENSMUSG00000024525 | 99.83703664 | -0.514392468 | 0.236430411 | -2.175661184 | 0.029580605 | 0.226497547 | Impa2 | inositol monophosphatase 2 [Source:MGI | 131.71 | 106.9 | 113.71 | 93.506 | 76.343 | 76.855 |
| 4540 | ENSMUSG00000024526 | 20.03796059 | -2.372466298 | 2.34877659 | -1.010085978 | 0.3124541 | 0.759830222 | Cidea | cell death-inducing DNA fragmentation fa | 84.326 | 16.447 | 0 | 9.6398 | 9.8156 | 0 |
| 4541 | ENSMUSG00000024527 | 1987.093885 | 0.107359862 | 0.093733861 | 1.145369038 | 0.252056318 | 0.704491322 | Afg3l2 | AFG3-like AAA ATPase 2 [Source:MGI Sym | 1955.6 | 1968.5 | 1815.3 | 2061 | 2094 | 2028.3 |
| 4542 | ENSMUSG00000024528 | 92.53352733 | -0.454790789 | 0.28463696 | -1.597792463 | 0.110089173 | 0.487846137 | Srfbp1 | serum response factor binding protein 1 [ | 134.92 | 77.094 | 108.67 | 83.866 | 91.612 | 59.034 |
| 4543 | ENSMUSG00000024529 | 1.816571243 | -4.233540562 | 1.925938062 | -2.198170671 | 0.027936946 | NA | Lox | lysyl oxidase [Source:MGI Symbol;Acc:MG | 4.8186 | 2.0558 | 4.025 | 0 | 0 | 0 |
| 4544 | ENSMUSG00000024530 | 0 | NA | NA | NA | NA | NA | Prelid3a | PRELI domain containing 3A [Source:MGI | 0 | 0 | 0 | 0 | 0 | 0 |
| 4545 | ENSMUSG00000024532 | 0 | NA | NA | NA | NA | NA | 1700034E13 | RIKEN cDNA 1700034E13 gene [Source:M | 0 | 0 | 0 | 0 | 0 | 0 |
| 4546 | ENSMUSG00000024533 | 6.132923779 | -0.395728219 | 0.864493005 | -0.457757572 | 0.647126621 | 0.925643431 | Spire1 | spire type actin nucleation factor 1 [Sourc | 5.6217 | 10.279 | 5.0312 | 4.8199 | 4.3625 | 6.6831 |
| 4547 | ENSMUSG00000024534 | 6.49386128 | -1.80833576 | 1.140079284 | -1.586149126 | 0.112705485 | 0.492822297 | Sncaip | synuclein, alpha interacting protein (synpl | 12.047 | 2.0558 | 16.1 | 0.964 | 0 | 7.7969 |
| 4548 | ENSMUSG00000024535 | 159.1155858 | -0.188033849 | 0.194267693 | -0.967911058 | 0.333088791 | 0.77523112 | Snx24 | sorting nexing 24 [Source:MGI Symbol;Ac | 165.44 | 188.11 | 154.96 | 149.42 | 134.15 | 162.62 |
| 4549 | ENSMUSG00000024537 | 244.3841752 | 0.171412724 | 0.200298012 | 0.855788444 | 0.392114809 | 0.820892834 | Psmg2 | proteasome (prosome, macropain) assem | 263.42 | 215.86 | 209.3 | 198.58 | 316.28 | 262.87 |
| 4550 | ENSMUSG00000024538 | 24.28480984 | -0.863036392 | 0.478719372 | -1.802802315 | 0.071419266 | 0.387039442 | Ppic | peptidylprolyl isomerase C [Source:MGI S | 41.761 | 32.893 | 19.119 | 14.46 | 18.541 | 18.935 |
| 4551 | ENSMUSG00000024539 | 157.9901269 | 0.177906493 | 0.189854984 | 0.937065169 | 0.348725037 | 0.789995175 | Ptpn2 | protein tyrosine phosphatase, non-recept | 159.01 | 134.66 | 150.94 | 179.3 | 161.41 | 162.62 |
| 4552 | ENSMUSG00000024542 | 52.22349373 | 0.535784633 | 0.323568193 | 1.655863108 | 0.097749544 | 0.460837486 | Cep192 | centrosomal protein 192 [Source:MGI Syn | 37.746 | 48.312 | 42.262 | 77.118 | 58.893 | 49.009 |
| 4553 | ENSMUSG00000024544 | 114.420533 | 0.514349068 | 0.243688563 | 2.110682018 | 0.034799653 | 0.250037719 | Ldlrad4 | low density lipoprotein receptor class A d | 115.65 | 68.87 | 97.606 | 146.52 | 129.78 | 128.09 |
| 4554 | ENSMUSG00000024546 | 0 | NA | NA | NA | NA | NA | 4930546C10 | RIKEN cDNA 4930546C10 gene [Source:M | 0 | 0 | 0 | 0 | 0 | 0 |
| 4555 | ENSMUSG00000024548 | 7.245402797 | -0.826422477 | 0.856570039 | -0.964804323 | 0.33464283 | 0.776159726 | Setbp1 | SET binding protein 1 [Source:MGI Symbo | 14.456 | 8.2233 | 5.0312 | 5.7839 | 2.1812 | 7.7969 |
| 4556 | ENSMUSG00000024552 | 0 | NA | NA | NA | NA | NA | Slc14a2 | solute carrier family 14 (urea transporter) | 0 | 0 | 0 | 0 | 0 | 0 |
| 4557 | ENSMUSG00000024553 | 0 | NA | NA | NA | NA | NA | Galr1 | galanin receptor 1 [Source:MGI Symbol;A | 0 | 0 | 0 | 0 | 0 | 0 |
| 4558 | ENSMUSG00000024556 | 23.23863805 | -0.746944218 | 0.477260119 | -1.565067327 | 0.117567124 | 0.502806291 | Me2 | malic enzyme 2, NAD(+)-dependent, mito | 19.274 | 37.005 | 31.194 | 14.46 | 17.45 | 20.049 |
| 4559 | ENSMUSG00000024558 | 28.71254621 | 0.304338826 | 0.41504857 | 0.73326075 | 0.463399409 | 0.861543316 | Mapk4 | mitogen-activated protein kinase 4 [Sourc | 19.274 | 31.865 | 26.162 | 29.883 | 29.447 | 35.643 |
| 4560 | ENSMUSG00000024560 | 650.2394338 | 0.058825719 | 0.126729518 | 0.464183244 | 0.642516465 | 0.924876278 | Cxxc1 | CXXC finger 1 (PHD domain) [Source:MGI | 610.36 | 603.39 | 697.33 | 632.37 | 721.99 | 636 |
| 4561 | ENSMUSG00000024561 | 989.9806682 | 0.314041834 | 0.143352065 | 2.190703252 | 0.028473274 | 0.220942827 | Mbd1 | methyl-CpG binding domain protein 1 [So | 985.41 | 837.75 | 823.11 | 893.61 | 1218.2 | 1181.8 |
| 4562 | ENSMUSG00000024563 | 211.5899678 | 0.394531035 | 0.171975048 | 2.294117897 | 0.021783728 | 0.183720021 | Smad2 | SMAD family member 2 [Source:MGI Sym | 199.97 | 173.72 | 174.08 | 222.68 | 257.39 | 241.7 |
| 4563 | ENSMUSG00000024565 | 0 | NA | NA | NA | NA | NA | Sall3 | spalt like transcription factor 3 [Source:M | 0 | 0 | 0 | 0 | 0 | 0 |
| 4564 | ENSMUSG00000024566 | 637.7075961 | 0.024529826 | 0.119162101 | 0.205852583 | 0.836906065 | 0.971073158 | Atp9b | ATPase, class II, type 9B [Source:MGI Sym | 616.78 | 661.98 | 618.84 | 682.5 | 639.1 | 607.04 |
| 4565 | ENSMUSG00000024570 | 720.6630467 | 0.028967027 | 0.116652677 | 0.24831858 | 0.803887925 | 0.963625663 | Rbfa | ribosome binding factor A [Source:MGI Sy | 698.7 | 737.02 | 704.37 | 676.71 | 777.61 | 729.57 |
| 4566 | ENSMUSG00000024571 | 1114.250681 | -0.047991558 | 0.123357067 | -0.389045874 | 0.697242212 | 0.940268454 | Gm16286 | predicted gene 16286 [Source:MGI Symbo | 1271.3 | 1077.3 | 1048.5 | 1065.2 | 1203 | 1020.3 |
| 4567 | ENSMUSG00000024575 | 0.503122293 | -2.366675966 | 3.785596405 | -0.625179156 | 0.531853481 | NA | Pde6a | phosphodiesterase 6A, cGMP-specific, roc | 0 | 0 | 3.0187 | 0 | 0 | 0 |
| 4568 | ENSMUSG00000024576 | 2173.450585 | 0.038670448 | 0.100627792 | 0.38429192 | 0.700762096 | 0.941778228 | Csnk1a1 | casein kinase 1, alpha 1 [Source:MGI Syml | 2030.2 | 2219.3 | 2183.6 | 2033 | 2299 | 2275.6 |
| 4569 | ENSMUSG00000024578 | 0 | NA | NA | NA | NA | NA | Il17b | interleukin 17B [Source:MGI Symbol;Acc:I | 0 | 0 | 0 | 0 | 0 | 0 |
| 4570 | ENSMUSG00000024579 | 17.52211003 | -0.015359221 | 0.565663967 | -0.027152553 | 0.978338059 | 0.997797092 | Pcyox1l | prenylcysteine oxidase 1 like [Source:MGI | 24.093 | 12.335 | 16.1 | 10.604 | 15.269 | 26.732 |
| 4571 | ENSMUSG00000024580 | 1174.524443 | 0.207045447 | 0.144354298 | 1.434286682 | 0.151490469 | 0.567216709 | Grpel2 | GrpE-like 2, mitochondrial [Source:MGI Sy | 1240.8 | 1117.3 | 912.66 | 1449.8 | 1112.4 | 1214.1 |
| 4572 | ENSMUSG00000024581 | 410.7871748 | 0.158634053 | 0.129222756 | 1.227601533 | 0.219596586 | 0.665321537 | Napg | N-ethylmaleimide sensitive fusion protein | 379.87 | 389.58 | 395.45 | 435.72 | 429.7 | 434.4 |
| 4573 | ENSMUSG00000024583 | 1538.020626 | 0.071807446 | 0.09739274 | 0.737297728 | 0.460941313 | 0.859821403 | Txnl1 | thioredoxin-like 1 [Source:MGI Symbol;Ac | 1560.4 | 1420.6 | 1517.4 | 1558.8 | 1577 | 1593.9 |
| 4574 | ENSMUSG00000024587 | 1625.377364 | -0.003497831 | 0.098133023 | -0.035643768 | 0.971566409 | 0.99516582 | Nars | asparaginyl-tRNA synthetase [Source:MGI | 1687.3 | 1629.2 | 1564.7 | 1578 | 1705.7 | 1587.2 |
| 4575 | ENSMUSG00000024588 | 3300.895956 | 0.208615598 | 0.091966118 | 2.268396258 | 0.023305066 | 0.192840085 | Fech | ferrochelatase [Source:MGI Symbol;Acc:N | 2863.9 | 3148.5 | 3176.7 | 3520.4 | 3473.6 | 3622.2 |
| 4576 | ENSMUSG00000024589 | 746.0642187 | -0.760007806 | 0.213104742 | -3.566358028 | 0.000361977 | 0.010029946 | Nedd4l | neural precursor cell expressed, developn | 709.94 | 967.27 | 1138.1 | 628.51 | 363.18 | 669.42 |
| 4577 | ENSMUSG00000024590 | 106.1344898 | 0.482226131 | 0.259610592 | 1.857497904 | 0.063240349 | 0.359877642 | Lmnb1 | lamin B1 [Source:MGI Symbol;Acc:MGI:96 | 107.62 | 65.787 | 91.568 | 115.68 | 105.79 | 150.37 |
| 4578 | ENSMUSG00000024592 | 240.4817756 | -0.061195115 | 0.168666804 | -0.36281659 | 0.716741898 | 0.9447719 | C330018D20 | RIKEN cDNA C330018D20 gene [Source:M | 261.81 | 254.92 | 219.36 | 214.97 | 232.3 | 259.53 |
| 4579 | ENSMUSG00000024593 | 0 | NA | NA | NA | NA | NA | Megf10 | multiple EGF-like-domains 10 [Source:MG | 0 | 0 | 0 | 0 | 0 | 0 |
| 4580 | ENSMUSG00000024594 | 813.9741027 | 0.018717726 | 0.127177468 | 0.147178003 | 0.882991522 | 0.981769403 | Prrc1 | proline-rich coiled-coil 1 [Source:MGI Sym | 930.8 | 750.38 | 743.61 | 836.73 | 810.33 | 811.99 |
| 4581 | ENSMUSG00000024597 | 243.1366916 | 0.112141066 | 0.167057707 | 0.671271432 | 0.502047631 | 0.881157254 | Slc12a2 | solute carrier family 12, member 2 [Sourc | 220.05 | 222.03 | 259.61 | 275.7 | 249.75 | 231.68 |
| 4582 | ENSMUSG00000024598 | 0 | NA | NA | NA | NA | NA | Fbn2 | fibrillin 2 [Source:MGI Symbol;Acc:MGI:95 | 0 | 0 | 0 | 0 | 0 | 0 |
| 4583 | ENSMUSG00000024600 | 3.472757338 | -0.45258565 | 1.308945446 | -0.345763569 | 0.729520406 | NA | Slc27a6 | solute carrier family 27 (fatty acid transpo | 4.8186 | 4.1117 | 3.0187 | 0 | 1.0906 | 7.7969 |
| 4584 | ENSMUSG00000024601 | 1729.615049 | 0.050464423 | 0.106186969 | 0.475241207 | 0.634615059 | 0.923478764 | Isoc1 | isochorismatase domain containing 1 [Sou | 1539.5 | 1735.1 | 1824.3 | 1691.8 | 1762.4 | 1824.5 |
| 4585 | ENSMUSG00000024603 | 467.8025253 | 0.016804141 | 0.134392175 | 0.12503809 | 0.900493395 | 0.982850624 | Dctn4 | dynactin 4 [Source:MGI Symbol;Acc:MGI: | 481.86 | 479.01 | 433.69 | 439.57 | 456.97 | 515.71 |
| 4586 | ENSMUSG00000024604 | 333.4945122 | -0.139617823 | 0.148723576 | -0.938773982 | 0.3478468 | 0.789340463 | Rbm22 | RNA binding motif protein 22 [Source:MG | 360.59 | 345.38 | 343.13 | 345.1 | 327.19 | 279.57 |
| 4587 | ENSMUSG00000024608 | 4116.072172 | 0.106229238 | 0.098111383 | 1.082741216 | 0.278923304 | 0.731519729 | Rps14 | ribosomal protein S14 [Source:MGI Symbo | 3842 | 4028.4 | 4023 | 3904.1 | 4269.8 | 4629.1 |
| 4588 | ENSMUSG00000024610 | 643.5818148 | -1.22703829 | 0.211997551 | -5.787983327 | 7.12E-09 | 1.14E-06 | Cd74 | CD74 antigen (invariant polypeptide of ma | 900.28 | 1156.4 | 649.03 | 401.01 | 281.38 | 473.38 |

| | | | | | | | | | | | | | | | |
|---|---|---|---|---|---|---|---|---|---|---|---|---|---|---|---|
| 4589 | ENSMUSG00000024613 | 123.3940554 | -0.049376333 | 0.210064291 | -0.235053433 | 0.814167261 | 0.967157956 | Tcof1 | treacle ribosome biogenesis factor 1 [Sour | 119.66 | 124.38 | 132.82 | 133.03 | 109.06 | 121.41 |
| 4590 | ENSMUSG00000024614 | 325.153795 | -0.073549724 | 0.144777813 | -0.508017926 | 0.611440772 | 0.916376001 | Tmx3 | thioredoxin-related transmembrane prote | 338.11 | 349.49 | 312.94 | 336.43 | 303.19 | 310.76 |
| 4591 | ENSMUSG00000024617 | 0.328370405 | 0.16514316 | 4.003149536 | 0.041253308 | 0.967093956 | NA | Camk2a | calcium/calmodulin-dependent protein ki | 0 | 0 | 1.0062 | 0.964 | 0 | 0 |
| 4592 | ENSMUSG00000024619 | 0 | NA | NA | NA | NA | NA | Cdx1 | caudal type homeobox 1 [Source:MGI Syn | 0 | 0 | 0 | 0 | 0 | 0 |
| 4593 | ENSMUSG00000024620 | 82.28267741 | -0.837560905 | 0.301449963 | -2.778440899 | 0.005462045 | 0.070727858 | Pdgfrb | platelet derived growth factor receptor, b | 146.16 | 89.429 | 80.5 | 61.695 | 49.078 | 66.831 |
| 4594 | ENSMUSG00000024621 | 379.8183486 | -1.360576234 | 0.430369761 | -3.16141225 | 0.001570061 | 0.029454792 | Csf1r | colony stimulating factor 1 receptor [Sour | 447.33 | 834.67 | 358.22 | 204.36 | 160.32 | 274.01 |
| 4595 | ENSMUSG00000024622 | 191.2293989 | 0.120236119 | 0.179834883 | 0.66859175 | 0.503755937 | 0.881510796 | Hmgxb3 | HMG box domain containing 3 [Source:M( | 175.08 | 191.19 | 184.14 | 225.57 | 186.5 | 184.9 |
| 4596 | ENSMUSG00000024639 | 383.8912413 | 0.054181678 | 0.151845912 | 0.356820128 | 0.721226467 | 0.9447719 | Gnaq | guanine nucleotide binding protein, alpha | 356.58 | 421.45 | 352.19 | 360.53 | 441.7 | 370.91 |
| 4597 | ENSMUSG00000024640 | 9.568360855 | 0.182907689 | 0.799292346 | 0.228837034 | 0.818995582 | 0.969160361 | Psat1 | phosphoserine aminotransferase 1 [Sourc | 18.471 | 3.0838 | 5.0312 | 7.7118 | 13.087 | 10.025 |
| 4598 | ENSMUSG00000024642 | 176.3402293 | 0.220525395 | 0.183471851 | 1.201957651 | 0.229379936 | 0.675988467 | Tle4 | transducin-like enhancer of split 4 [Source | 152.59 | 162.41 | 174.08 | 195.69 | 202.86 | 170.42 |
| 4599 | ENSMUSG00000024644 | 737.6750457 | -0.177700288 | 0.126528943 | -1.404424032 | 0.160192619 | 0.582606497 | Cndp2 | CNDP dipeptidase 2 (metallopeptidase M2 | 743.67 | 832.61 | 773.8 | 731.66 | 737.26 | 607.04 |
| 4600 | ENSMUSG00000024645 | 368.2056924 | 0.120541137 | 0.139655278 | 0.863133417 | 0.388064116 | 0.817937623 | Timm21 | translocase of inner mitochondrial membr | 346.14 | 374.16 | 338.1 | 358.6 | 404.62 | 387.62 |
| 4601 | ENSMUSG00000024646 | 18024.18146 | -0.129540214 | 0.093414599 | -1.386723432 | 0.165526113 | 0.59115301 | Cyb5a | cytochrome b5 type A (microsomal) [Sour | 18381 | 18569 | 19547 | 15875 | 18618 | 17154 |
| 4602 | ENSMUSG00000024647 | 0 | NA | NA | NA | NA | NA | Cbln2 | cerebellin 2 precursor protein [Source:MG | 0 | 0 | 0 | 0 | 0 | 0 |
| 4603 | ENSMUSG00000024650 | 0.513958919 | -2.394274821 | 3.756096013 | -0.637437066 | 0.523840191 | NA | Slc22a6 | solute carrier family 22 (organic anion tra | 0 | 3.0838 | 0 | 0 | 0 | 0 |
| 4604 | ENSMUSG00000024653 | 0 | NA | NA | NA | NA | NA | Scgb1a1 | secretoglobin, family 1A, member 1 (uterc | 0 | 0 | 0 | 0 | 0 | 0 |
| 4605 | ENSMUSG00000024654 | 306.3678217 | -0.237298336 | 0.169080535 | -1.403463363 | 0.16047871 | 0.583090119 | Asrgl1 | asparaginase like 1 [Source:MGI Symbol;A | 329.27 | 365.94 | 298.85 | 236.17 | 322.82 | 285.14 |
| 4606 | ENSMUSG00000024658 | 0.17131964 | -0.796629176 | 4.080472857 | -0.195229623 | 0.84521319 | NA | Gm9750 | predicted gene 9750 [Source:MGI Symbol | 0 | 1.0279 | 0 | 0 | 0 | 0 |
| 4607 | ENSMUSG00000024659 | 47.79940493 | -0.548974727 | 0.342370072 | -1.603454192 | 0.108834415 | 0.485684837 | Anxa1 | annexin A1 [Source:MGI Symbol;Acc:MGI | 72.279 | 56.535 | 41.256 | 39.523 | 35.99 | 41.212 |
| 4608 | ENSMUSG00000024660 | 22.29807738 | -0.13733305 | 0.466593888 | -0.29433101 | 0.768504979 | 0.955856567 | Incenp | inner centromere protein [Source:MGI Sy | 18.471 | 25.698 | 26.162 | 26.027 | 20.722 | 16.708 |
| 4609 | ENSMUSG00000024661 | 23240.80133 | -0.393612371 | 0.084099942 | -4.680293008 | 2.86E-06 | 0.000190942 | Fth1 | ferritin heavy polypeptide 1 [Source:MGI S | 27182 | 25474 | 26520 | 21084 | 19489 | 19696 |
| 4610 | ENSMUSG00000024663 | 38.30785881 | -0.159893786 | 0.375553385 | -0.425755146 | 0.670286274 | 0.931139134 | Rab3il1 | RAB3A interacting protein (rabin3)-like 1 [ | 41.761 | 46.256 | 33.206 | 35.667 | 26.175 | 46.781 |
| 4611 | ENSMUSG00000024664 | 203.618188 | 0.843450792 | 0.249817587 | 3.376266668 | 0.000734767 | 0.016657782 | Fads3 | fatty acid desaturase 3 [Source:MGI Symb | 203.99 | 108.96 | 122.76 | 199.54 | 286.83 | 299.62 |
| 4612 | ENSMUSG00000024665 | 9254.955281 | 0.34804622 | 0.178849495 | 1.946028531 | 0.051651313 | 0.319340611 | Fads2 | fatty acid desaturase 2 [Source:MGI Symb | 10363 | 8516.3 | 5551.5 | 11780 | 10336 | 8983.2 |
| 4613 | ENSMUSG00000024666 | 27.66538611 | 0.311246527 | 0.461060559 | 0.675066477 | 0.499633531 | 0.880660418 | Tmem138 | transmembrane protein 138 [Source:MGI | 28.109 | 18.503 | 27.169 | 19.28 | 27.265 | 45.668 |
| 4614 | ENSMUSG00000024667 | 66.78233719 | -0.035221078 | 0.274212715 | -0.128444363 | 0.897797325 | 0.982798755 | Tmem216 | transmembrane protein 216 [Source:MGI | 58.626 | 75.038 | 69.431 | 66.514 | 68.709 | 62.375 |
| 4615 | ENSMUSG00000024668 | 464.3180152 | 0.077478519 | 0.132549871 | 0.584523536 | 0.558868135 | 0.899349136 | Sdhaf2 | succinate dehydrogenase complex asseml | 436.89 | 465.65 | 452.81 | 430.9 | 497.32 | 502.34 |
| 4616 | ENSMUSG00000024669 | 1.154595928 | -1.203571421 | 2.021261449 | -0.595455586 | 0.551538979 | NA | Cd5 | CD5 antigen [Source:MGI Symbol;Acc:MG | 0.8031 | 1.0279 | 3.0187 | 0.964 | 0 | 1.1138 |
| 4617 | ENSMUSG00000024670 | 1.852159489 | -0.793632672 | 1.783087598 | -0.445088998 | 0.656255439 | NA | Cd6 | CD6 antigen [Source:MGI Symbol;Acc:MG | 0 | 4.1117 | 3.0187 | 2.8919 | 1.0906 | 0 |
| 4618 | ENSMUSG00000024672 | 5.993197186 | -0.403131679 | 1.071293799 | -0.376303568 | 0.706691227 | 0.94295036 | Ms4a7 | membrane-spanning 4-domains, subfamil | 3.2124 | 13.363 | 4.025 | 8.6758 | 0 | 6.6831 |
| 4619 | ENSMUSG00000024673 | 4.704980905 | -3.614904125 | 1.396000318 | -2.58947228 | 0.009612316 | NA | Ms4a1 | membrane-spanning 4-domains, subfamil | 8.8341 | 5.1396 | 12.075 | 0 | 2.1812 | 0 |
| 4620 | ENSMUSG00000024675 | 3.006954839 | -1.29103205 | 1.300164774 | -0.992975718 | 0.320721788 | NA | Ms4a4c | membrane-spanning 4-domains, subfamil | 5.6217 | 6.1675 | 1.0062 | 1.928 | 1.0906 | 2.2277 |
| 4621 | ENSMUSG00000024677 | 14.34565406 | -0.768520605 | 0.597526168 | -1.286170625 | 0.198383521 | 0.640554426 | Ms4a6b | membrane-spanning 4-domains, subfamil | 14.456 | 26.726 | 13.081 | 8.6758 | 11.997 | 11.138 |
| 4622 | ENSMUSG00000024678 | 56.44710105 | -1.606037854 | 0.341142527 | -4.707820707 | 2.50E-06 | 0.000172104 | Ms4a4d | membrane-spanning 4-domains, subfamil | 82.72 | 91.485 | 80.5 | 23.135 | 19.631 | 41.212 |
| 4623 | ENSMUSG00000024679 | 17.90704988 | -1.247206547 | 0.633193132 | -1.969709532 | 0.04887167 | 0.307874492 | Ms4a6d | membrane-spanning 4-domains, subfamil | 26.502 | 33.921 | 15.094 | 6.7478 | 20.722 | 4.4554 |
| 4624 | ENSMUSG00000024680 | 0.371281295 | 2.09253362 | 4.031852165 | 0.519000582 | 0.603760335 | NA | Ms4a2 | membrane-spanning 4-domains, subfamil | 0 | 0 | 0 | 0 | 0 | 2.2277 |
| 4625 | ENSMUSG00000024681 | 0 | NA | NA | NA | NA | NA | Ms4a3 | membrane-spanning 4-domains, subfamil | 0 | 0 | 0 | 0 | 0 | 0 |
| 4626 | ENSMUSG00000024682 | 0 | NA | NA | NA | NA | NA | Cblif | cobalamin binding intrinsic factor [Source | 0 | 0 | 0 | 0 | 0 | 0 |
| 4627 | ENSMUSG00000024683 | 639.2869839 | -0.109661778 | 0.120180939 | -0.912472308 | 0.361520141 | 0.800421803 | Mrpl16 | mitochondrial ribosomal protein L16 [Sou | 691.47 | 665.06 | 633.93 | 622.73 | 650.01 | 572.52 |
| 4628 | ENSMUSG00000024687 | 998.9556565 | 0.138081047 | 0.105440632 | 1.30956202 | 0.190344044 | 0.627779796 | Osbp | oxysterol binding protein [Source:MGI Syr | 990.23 | 963.16 | 899.58 | 1043 | 1028.5 | 1069.3 |
| 4629 | ENSMUSG00000024691 | 20.38529327 | -1.096502124 | 0.494319512 | -2.218205222 | 0.026540843 | 0.210915839 | Fam111a | family with sequence similarity 111, meml | 31.321 | 28.782 | 23.144 | 11.568 | 16.359 | 11.138 |
| 4630 | ENSMUSG00000024694 | 4003.277694 | 0.525963162 | 0.139059865 | 3.782278683 | 0.000155399 | 0.005155263 | Keg1 | kidney expressed gene 1 [Source:MGI Syn | 3280.7 | 3194.8 | 3369.9 | 5573.7 | 4940.5 | 3660.1 |
| 4631 | ENSMUSG00000024695 | 2395.643967 | 0.046615326 | 0.089876877 | 0.518657602 | 0.603999533 | 0.916322307 | Zfp91 | zinc finger protein 91 [Source:MGI Symbo | 2296.1 | 2431 | 2344.5 | 2494.8 | 2401.5 | 2405.9 |
| 4632 | ENSMUSG00000024696 | 16.33768884 | -0.770019035 | 0.573048442 | -1.343724157 | 0.179037603 | 0.611235187 | Lpxn | leupaxin [Source:MGI Symbol;Acc:MGI:21 | 24.896 | 21.586 | 15.094 | 6.7478 | 17.45 | 12.252 |
| 4633 | ENSMUSG00000024697 | 195.6541491 | 0.467287675 | 0.312694208 | 1.494391844 | 0.135073232 | 0.537616739 | Gna14 | guanine nucleotide binding protein, alpha | 145.36 | 139.8 | 207.29 | 103.15 | 302.1 | 276.23 |
| 4634 | ENSMUSG00000024712 | 937.7234923 | -0.136432088 | 0.107379802 | -1.270556329 | 0.203886531 | 0.649133786 | Rfk | riboflavin kinase [Source:MGI Symbol;Acc | 1014.3 | 990.91 | 940.84 | 933.13 | 859.41 | 887.73 |
| 4635 | ENSMUSG00000024713 | 216.4283605 | -0.28688197 | 0.185777221 | -1.544225758 | 0.122533651 | 0.51297445 | Pcsk5 | proprotein convertase subtilisin/kexin typ | 209.61 | 251.84 | 252.57 | 193.76 | 164.68 | 226.11 |
| 4636 | ENSMUSG00000024725 | 823.5479872 | -0.122997311 | 0.107798306 | -1.140994843 | 0.253872069 | 0.707055046 | Ostf1 | osteoclast stimulating factor 1 [Source:M( | 860.12 | 874.76 | 841.22 | 809.74 | 760.16 | 795.28 |
| 4637 | ENSMUSG00000024726 | 267.8510668 | 0.16206707 | 0.161572786 | 1.003059203 | 0.315832297 | 0.762637454 | Carnmt1 | carnosine N-methyltransferase 1 [Source: | 244.14 | 239.5 | 274.7 | 264.13 | 268.29 | 316.33 |
| 4638 | ENSMUSG00000024727 | 3.288259723 | -0.79482011 | 1.2606124 | -0.630503166 | 0.528365432 | NA | Trpm6 | transient receptor potential cation chann | 2.4093 | 6.1675 | 4.025 | 3.8559 | 3.2719 | 0 |
| 4639 | ENSMUSG00000024728 | 0 | NA | NA | NA | NA | NA | Ms4a19 | membrane-spanning 4-domains, subfamil | 0 | 0 | 0 | 0 | 0 | 0 |
| 4640 | ENSMUSG00000024729 | 0 | NA | NA | NA | NA | NA | Ms4a20 | membrane-spanning 4-domains, subfamil | 0 | 0 | 0 | 0 | 0 | 0 |
| 4641 | ENSMUSG00000024730 | 7.283932473 | -0.830732912 | 0.792894155 | -1.047722331 | 0.294766556 | 0.745164683 | Ms4a8a | membrane-spanning 4-domains, subfamil | 9.6372 | 11.307 | 7.0437 | 5.7839 | 4.3625 | 5.5692 |
| 4642 | ENSMUSG00000024731 | 0 | NA | NA | NA | NA | NA | Ms4a10 | membrane-spanning 4-domains, subfamil | 0 | 0 | 0 | 0 | 0 | 0 |
| 4643 | ENSMUSG00000024732 | 166.1555851 | -0.183766845 | 0.187076068 | -0.982310818 | 0.325946747 | 0.772018857 | Ccdc86 | coiled-coil domain containing 86 [Source:I | 187.12 | 185.03 | 157.98 | 164.84 | 151.6 | 150.37 |
| 4644 | ENSMUSG00000024734 | 0 | NA | NA | NA | NA | NA | Zp1 | zona pellucida glycoprotein 1 [Source:MG | 0 | 0 | 0 | 0 | 0 | 0 |

| | | | | | | | | | | | | | | | |
|---|---|---|---|---|---|---|---|---|---|---|---|---|---|---|---|
| 4645 | ENSMUSG00000024735 | 1873.943178 | 0.032877126 | 0.105173549 | 0.312598805 | 0.754585486 | 0.953048681 | Prpf19 | pre-mRNA processing factor 19 [Source:M | 1997.3 | 1755.7 | 1803.2 | 1801.7 | 1849.7 | 2036.1 |
| 4646 | ENSMUSG00000024736 | 18.62162108 | -1.135522058 | 0.587539855 | -1.932672393 | 0.053276568 | 0.324459908 | Tmem132a | transmembrane protein 132A [Source:MG | 20.078 | 31.865 | 25.156 | 20.244 | 4.3625 | 10.025 |
| 4647 | ENSMUSG00000024737 | 43.72822718 | -1.027721533 | 0.381203965 | -2.695988572 | 0.007018007 | 0.08425311 | Slc15a3 | solute carrier family 15, member 3 [Sourc | 57.02 | 75.038 | 44.275 | 38.559 | 19.631 | 27.846 |
| 4648 | ENSMUSG00000024738 | 0 | NA | NA | NA | NA | NA | Pga5 | pepsinogen 5, group I [Source:MGI Symbo | 0 | 0 | 0 | 0 | 0 | 0 |
| 4649 | ENSMUSG00000024740 | 3741.724281 | 0.080591233 | 0.091307417 | 0.882636219 | 0.377432853 | 0.81030298 | Ddb1 | damage specific DNA binding protein 1 [S | 3516 | 3853.7 | 3543 | 3898.3 | 3944.8 | 3694.6 |
| 4650 | ENSMUSG00000024742 | 5.972590896 | -0.691244253 | 1.00033526 | -0.691012585 | 0.489557633 | 0.875057044 | Fen1 | flap structure specific endonuclease 1 [So | 12.85 | 5.1396 | 4.025 | 2.8919 | 9.8156 | 1.1138 |
| 4651 | ENSMUSG00000024743 | 12.14361283 | -0.300952003 | 0.620617946 | -0.48492314 | 0.627730867 | 0.920328819 | Syt7 | synaptotagmin VII [Source:MGI Symbol;A | 12.85 | 9.2513 | 18.112 | 10.604 | 10.906 | 11.138 |
| 4652 | ENSMUSG00000024747 | 3392.675651 | -0.100356502 | 0.161033743 | -0.623201701 | 0.533151984 | 0.889143097 | Aldh1a7 | aldehyde dehydrogenase family 1, subfam | 3917.5 | 3987.3 | 2626.3 | 3001.8 | 3884.8 | 2938.3 |
| 4653 | ENSMUSG00000024749 | 0.503122293 | -2.366675966 | 3.785596405 | -0.625179156 | 0.531853481 | NA | Tmc1 | transmembrane channel-like gene family | 0 | 0 | 3.0187 | 0 | 0 | 0 |
| 4654 | ENSMUSG00000024750 | 1103.765184 | -0.060950418 | 0.114793712 | -0.530956067 | 0.595449224 | 0.913801334 | Zfand5 | zinc finger, AN1-type domain 5 [Source:M | 1232 | 1117.3 | 1031.4 | 1149.1 | 1050.3 | 1042.6 |
| 4655 | ENSMUSG00000024754 | 166.8180656 | -0.726028313 | 0.20412012 | -3.556867957 | 0.000375303 | 0.010261584 | Cemip2 | cell migration inducing hyaluronidase 2 [S | 204.79 | 216.89 | 202.26 | 133.99 | 97.065 | 145.91 |
| 4656 | ENSMUSG00000024757 | 0 | NA | NA | NA | NA | NA | Slc22a19 | solute carrier family 22 (organic anion tra | 0 | 0 | 0 | 0 | 0 | 0 |
| 4657 | ENSMUSG00000024758 | 1672.300431 | 0.040563916 | 0.098109463 | 0.413455698 | 0.679272774 | 0.934062537 | Rtn3 | reticulon 3 [Source:MGI Symbol;Acc:MGI: | 1593.4 | 1736.2 | 1617 | 1626.2 | 1711.2 | 1749.8 |
| 4658 | ENSMUSG00000024759 | 2162.549595 | 0.118640339 | 0.11319369 | 1.04811796 | 0.294584263 | 0.745164683 | Atl3 | atlastin GTPase 3 [Source:MGI Symbol;Ac | 1958 | 2273.8 | 1989.3 | 2012.8 | 2414.6 | 2326.8 |
| 4659 | ENSMUSG00000024761 | 1.783277137 | 0.023998525 | 1.84120776 | 0.013034121 | 0.989600571 | NA | Gm16437 | predicted pseudogene 16437 [Source:MG | 3.2124 | 1.0279 | 1.0062 | 0 | 5.4531 | 0 |
| 4660 | ENSMUSG00000024764 | 214.1548374 | 0.167157038 | 0.177912003 | 0.939548965 | 0.347448965 | 0.788561404 | Naa40 | N(alpha)-acetyltransferase 40, NatD catal | 232.1 | 187.08 | 185.15 | 210.15 | 242.12 | 228.34 |
| 4661 | ENSMUSG00000024766 | 34.73181014 | -0.256771663 | 0.389255029 | -0.65964893 | 0.509479147 | 0.882793133 | Lipo3 | lipase, member O3 [Source:MGI Symbol;A | 40.155 | 44.2 | 29.181 | 38.559 | 23.994 | 32.301 |
| 4662 | ENSMUSG00000024767 | 656.4478422 | 0.108561678 | 0.119634378 | 0.907445504 | 0.364171255 | 0.802830824 | Otub1 | OTU domain, ubiquitin aldehyde binding 1 | 665.77 | 643.48 | 585.63 | 699.85 | 648.92 | 695.04 |
| 4663 | ENSMUSG00000024768 | 0 | NA | NA | NA | NA | NA | Lipf | lipase, gastric [Source:MGI Symbol;Acc:M | 0 | 0 | 0 | 0 | 0 | 0 |
| 4664 | ENSMUSG00000024769 | 433.4383394 | 0.008989715 | 0.155646239 | 0.057757354 | 0.953941908 | 0.9915542 | Cdc42bpg | CDC42 binding protein kinase gamma (DM | 428.05 | 493.4 | 375.33 | 481.02 | 380.63 | 442.2 |
| 4665 | ENSMUSG00000024770 | 0 | NA | NA | NA | NA | NA | Lipn | lipase, family member N [Source:MGI Sym | 0 | 0 | 0 | 0 | 0 | 0 |
| 4666 | ENSMUSG00000024771 | 0 | NA | NA | NA | NA | NA | Lipk | lipase, family member K [Source:MGI Sym | 0 | 0 | 0 | 0 | 0 | 0 |
| 4667 | ENSMUSG00000024772 | 829.6581892 | -0.147901142 | 0.118016801 | -1.253221077 | 0.210125261 | 0.654024485 | Ehd1 | EH-domain containing 1 [Source:MGI Sym | 864.14 | 904.57 | 847.26 | 729.73 | 762.34 | 869.91 |
| 4668 | ENSMUSG00000024773 | 1070.607851 | 0.133892589 | 0.105448619 | 1.269742458 | 0.204176384 | 0.649416772 | Atg2a | autophagy related 2A [Source:MGI Symbo | 1028.8 | 960.08 | 1073.7 | 1092.2 | 1148.4 | 1120.5 |
| 4669 | ENSMUSG00000024774 | 0.772832414 | -0.17263844 | 2.923794788 | -0.059046018 | 0.952915455 | NA | Ankrd22 | ankyrin repeat domain 22 [Source:MGI Sy | 2.4093 | 0 | 0 | 0 | 0 | 2.2277 |
| 4670 | ENSMUSG00000024776 | 5.978788701 | -1.468683393 | 0.934044026 | -1.572392042 | 0.115859659 | 0.500386415 | Stambpl1 | STAM binding protein like 1 [Source:MGI | 12.047 | 6.1675 | 8.05 | 1.928 | 5.4531 | 2.2277 |
| 4671 | ENSMUSG00000024777 | 223.8838735 | -0.056564312 | 0.164358017 | -0.344153044 | 0.730731193 | 0.946984677 | Ppp2r5b | protein phosphatase 2, regulatory subuni | 210.41 | 236.42 | 238.48 | 221.71 | 225.76 | 210.52 |
| 4672 | ENSMUSG00000024778 | 296.6830061 | -0.313406498 | 0.145073522 | -2.16032873 | 0.03074723 | 0.232720011 | Fas | Fas (TNF receptor superfamily member 6) | 337.3 | 328.93 | 319.99 | 266.06 | 268.29 | 259.53 |
| 4673 | ENSMUSG00000024780 | 1102.333856 | -0.008824041 | 0.113887342 | -0.077480441 | 0.93824135 | 0.988420305 | Cdc37l1 | cell division cycle 37-like 1 [Source:MGI S | 1052.1 | 1107.1 | 1159.2 | 1188.6 | 1112.4 | 994.66 |
| 4674 | ENSMUSG00000024781 | 3524.426833 | -0.241630792 | 0.115720609 | -2.08805323 | 0.036793033 | 0.259024417 | Lipa | lysosomal acid lipase A [Source:MGI Symb | 3945.6 | 4006.8 | 3504.7 | 3528.2 | 3375.5 | 2785.7 |
| 4675 | ENSMUSG00000024782 | 7439.929623 | -0.042244915 | 0.090237224 | -0.468153975 | 0.639674483 | 0.923932929 | Ak3 | adenylate kinase 3 [Source:MGI Symbol;A | 7112.3 | 7634.3 | 7901 | 7333 | 7652.9 | 7006.1 |
| 4676 | ENSMUSG00000024784 | 0.181769715 | 1.126924175 | 4.080472857 | 0.276174898 | 0.782413736 | NA | Gpha2 | glycoprotein hormone alpha 2 [Source:M | 0 | 0 | 0 | 0 | 1.0906 | 0 |
| 4677 | ENSMUSG00000024785 | 2868.861721 | 0.461466927 | 0.121896299 | 3.785733713 | 0.000153256 | 0.005095841 | Rcl1 | RNA terminal phosphate cyclase-like 1 [Sc | 2702.4 | 2114.4 | 2424 | 3718.1 | 3087.5 | 3166.7 |
| 4678 | ENSMUSG00000024786 | 0.615839296 | 1.537131441 | 3.17677869 | 0.483864817 | 0.628481811 | NA | Majin | membrane anchored junction protein [So | 0.8031 | 0 | 0 | 2.8919 | 0 | 0 |
| 4679 | ENSMUSG00000024787 | 222.8835705 | 0.245949569 | 0.169906605 | 1.447557435 | 0.147740863 | 0.559179858 | Snx15 | sorting nexin 15 [Source:MGI Symbol;Acc: | 203.18 | 213.81 | 195.21 | 267.99 | 241.03 | 216.09 |
| 4680 | ENSMUSG00000024789 | 318.8312339 | 0.02052377 | 0.143057363 | 0.143465316 | 0.885922702 | 0.982215695 | Jak2 | Janus kinase 2 [Source:MGI Symbol;Acc:M | 320.44 | 297.07 | 332.06 | 321.97 | 319.55 | 321.9 |
| 4681 | ENSMUSG00000024790 | 160.4581085 | -0.108887203 | 0.188125433 | -0.578801077 | 0.562723405 | 0.900565925 | Sac3d1 | SAC3 domain containing 1 [Source:MGI Sy | 171.06 | 170.63 | 157.98 | 168.7 | 141.78 | 152.6 |
| 4682 | ENSMUSG00000024791 | 2.382449679 | -1.881595796 | 1.677758318 | -1.121493945 | 0.262077667 | NA | Cdca5 | cell division cycle associated 5 [Source:M | 7.2279 | 0 | 4.025 | 1.928 | 0 | 1.1138 |
| 4683 | ENSMUSG00000024792 | 177.1528852 | 0.375586441 | 0.187932498 | 1.998517794 | 0.045660553 | 0.295628575 | Zfpl1 | zinc finger like protein 1 [Source:MGI Sym | 179.09 | 143.91 | 138.86 | 197.62 | 198.49 | 204.95 |
| 4684 | ENSMUSG00000024793 | 3.901275815 | -0.028244375 | 1.167494165 | -0.024192305 | 0.980699216 | NA | Tnfrsf25 | tumor necrosis factor receptor superfamil | 5.6217 | 5.1396 | 1.0062 | 2.8919 | 7.6343 | 1.1138 |
| 4685 | ENSMUSG00000024795 | 14.28520411 | -0.103932264 | 0.680163127 | -0.152804908 | 0.878552127 | 0.980976563 | Kif20b | kinesin family member 20B [Source:MGI S | 21.684 | 18.503 | 4.025 | 9.6398 | 20.722 | 11.138 |
| 4686 | ENSMUSG00000024797 | 20.6267262 | -0.26023016 | 0.50738108 | -0.512888971 | 0.608028988 | 0.916376001 | Vps51 | VPS51 GARP complex subunit [Source:MG | 13.653 | 29.81 | 24.15 | 15.424 | 22.903 | 17.822 |
| 4687 | ENSMUSG00000024798 | 0 | NA | NA | NA | NA | NA | Htr7 | 5-hydroxytryptamine (serotonin) receptor | 0 | 0 | 0 | 0 | 0 | 0 |
| 4688 | ENSMUSG00000024799 | 1725.401342 | 0.890785164 | 0.113745877 | 7.831362197 | 4.83E-15 | 3.17E-12 | Tm7sf2 | transmembrane 7 superfamily member 2 | 1203.9 | 1339.4 | 1083.7 | 2097.6 | 2257.6 | 2370.3 |
| 4689 | ENSMUSG00000024800 | 76.40592441 | -0.28443999 | 0.264190189 | -1.076648573 | 0.281637295 | 0.734379276 | Rpp30 | ribonuclease P/MRP 30 subunit [Source:M | 86.735 | 86.345 | 78.487 | 65.55 | 58.893 | 82.424 |
| 4690 | ENSMUSG00000024803 | 1.099166815 | 0.59718918 | 2.138494465 | 0.279256827 | 0.780047736 | NA | Ankrd1 | ankyrin repeat domain 1 (cardiac muscle) | 1.6062 | 0 | 1.0062 | 2.8919 | 1.0906 | 0 |
| 4691 | ENSMUSG00000024805 | 418.9424968 | 0.265335282 | 0.136029921 | 1.950565578 | 0.051108745 | 0.317405278 | Pcgf5 | polycomb group ring finger 5 [Source:MG | 387.1 | 395.75 | 358.22 | 421.26 | 465.69 | 485.64 |
| 4692 | ENSMUSG00000024806 | 0 | NA | NA | NA | NA | NA | Mlana | melan-A [Source:MGI Symbol;Acc:MGI:10 | 0 | 0 | 0 | 0 | 0 | 0 |
| 4693 | ENSMUSG00000024807 | 991.1987498 | 0.228309866 | 0.132918633 | 1.717666377 | 0.085857486 | 0.431795087 | Syvn1 | synovial apoptosis inhibitor 1, synoviolin [ | 892.25 | 907.65 | 939.83 | 1240.6 | 899.76 | 1067.1 |
| 4694 | ENSMUSG00000024810 | 29.8248199 | -0.449555765 | 0.406856648 | -1.1049488 | 0.269181775 | 0.724979259 | Il33 | interleukin 33 [Source:MGI Symbol;Acc:M | 40.155 | 28.782 | 34.212 | 24.099 | 30.537 | 21.163 |
| 4695 | ENSMUSG00000024811 | 961.9625856 | 0.020159837 | 0.107720123 | 0.18715015 | 0.851542908 | 0.973576526 | Tnks2 | tankyrase, TRF1-interacting ankyrin-relate | 958.9 | 1010.4 | 896.56 | 986.15 | 967.38 | 952.34 |
| 4696 | ENSMUSG00000024812 | 306.2796938 | -0.186288817 | 0.153269663 | -1.215431766 | 0.224201458 | 0.670425862 | Tjp2 | tight junction protein 2 [Source:MGI Symb | 328.47 | 322.77 | 327.03 | 296.91 | 313.01 | 249.5 |
| 4697 | ENSMUSG00000024815 | 0 | NA | NA | NA | NA | NA | Trpd52l3 | tumor protein D52-like 3 [Source:MGI Syn | 0 | 0 | 0 | 0 | 0 | 0 |
| 4698 | ENSMUSG00000024816 | 496.9349903 | 0.064550175 | 0.155901475 | 0.414044671 | 0.678841392 | 0.934052691 | Frmd8 | FERM domain containing 8 [Source:MGI S | 561.37 | 485.18 | 409.54 | 457.89 | 496.23 | 571.4 |
| 4699 | ENSMUSG00000024817 | 575.7934332 | 0.154287391 | 0.122146038 | 1.26313873 | 0.206539327 | 0.652026861 | Uhrf2 | ubiquitin-like, containing PHD and RING fi | 511.58 | 574.61 | 549.41 | 585.13 | 632.56 | 601.48 |
| 4700 | ENSMUSG00000024818 | 587.134788 | 0.087142487 | 0.145735216 | 0.597950788 | 0.54987277 | 0.895703112 | Slc25a45 | solute carrier family 25, member 45 [Sour | 668.18 | 481.07 | 557.46 | 581.28 | 595.48 | 639.35 |

| | | | | | | | | | | | | | | | |
|---|---|---|---|---|---|---|---|---|---|---|---|---|---|---|---|
| 4701 | ENSMUSG00000024824 | 49.40251845 | -0.836425548 | 0.349683311 | -2.391951577 | 0.016759053 | 0.153419587 | Rad9a | RAD9 checkpoint clamp component A [So | 60.233 | 45.228 | 84.525 | 34.703 | 31.628 | 40.098 |
| 4702 | ENSMUSG00000024826 | 434.0038591 | -0.037627688 | 0.148206838 | -0.253886316 | 0.799583387 | 0.962424862 | Dpf2 | D4, zinc and double PHD fingers family 2 [ | 428.86 | 430.7 | 458.85 | 357.64 | 463.51 | 464.47 |
| 4703 | ENSMUSG00000024827 | 2815.045774 | -0.866258707 | 0.17853188 | -4.852123355 | 1.22E-06 | 9.65E-05 | Gldc | glycine decarboxylase [Source:MGI Symbo | 2674.3 | 3526.8 | 4706.2 | 1767 | 1826.8 | 2389.2 |
| 4704 | ENSMUSG00000024829 | 349.8124448 | 0.095672372 | 0.160905279 | 0.594588153 | 0.552118801 | 0.895935901 | Mrpl21 | mitochondrial ribosomal protein L21 [Sou | 385.49 | 338.18 | 289.8 | 326.79 | 362.09 | 396.53 |
| 4705 | ENSMUSG00000024830 | 329.7687644 | -0.015092439 | 0.15984863 | -0.094417068 | 0.924777858 | 0.987546253 | Rps6kb2 | ribosomal protein S6 kinase, polypeptide : | 325.26 | 299.12 | 370.3 | 334.5 | 363.18 | 286.26 |
| 4706 | ENSMUSG00000024831 | 86.65335422 | 0.020972181 | 0.258881309 | 0.081010795 | 0.935433367 | 0.988420305 | Ighmbp2 | immunoglobulin mu binding protein 2 [So | 93.963 | 97.652 | 66.412 | 94.47 | 88.34 | 79.083 |
| 4707 | ENSMUSG00000024833 | 141.0774854 | 0.346710132 | 0.203023159 | 1.707736861 | 0.087685179 | 0.436734997 | Pola2 | polymerase (DNA directed), alpha 2 [Sour | 142.15 | 117.18 | 112.7 | 159.06 | 149.41 | 165.96 |
| 4708 | ENSMUSG00000024835 | 1695.33031 | -0.023947234 | 0.104355191 | -0.22947813 | 0.818497317 | 0.968897682 | Coro1b | coronin, actin binding protein 1B [Source: | 1823 | 1743.3 | 1560.7 | 1622.4 | 1740.6 | 1681.9 |
| 4709 | ENSMUSG00000024837 | 0 | NA | NA | NA | NA | NA | Dmrt1 | doublesex and mab-3 related transcriptio | 0 | 0 | 0 | 0 | 0 | 0 |
| 4710 | ENSMUSG00000024841 | 468.8083778 | 0.016186919 | 0.126922871 | 0.127533508 | 0.898518154 | 0.982798755 | Eif1ad | eukaryotic translation initiation factor 1A | 458.57 | 473.87 | 465.89 | 442.47 | 486.42 | 485.64 |
| 4711 | ENSMUSG00000024842 | 9.111452078 | -0.282593317 | 0.818525576 | -0.345246777 | 0.729908855 | 0.946534338 | Cabp4 | calcium binding protein 4 [Source:MGI Sy | 8.8341 | 7.1954 | 14.087 | 12.532 | 10.906 | 1.1138 |
| 4712 | ENSMUSG00000024843 | 404.0544179 | -0.609798665 | 0.249073276 | -2.448270143 | 0.014354399 | 0.138784774 | Chka | choline kinase alpha [Source:MGI Symbol; | 307.59 | 457.42 | 700.35 | 347.03 | 241.03 | 370.91 |
| 4713 | ENSMUSG00000024844 | 539.9344488 | -0.006346454 | 0.129124345 | -0.049149942 | 0.960799803 | 0.992557454 | Banf1 | BAF nuclear assembly factor 1 [Source:M( | 571.01 | 513.96 | 537.33 | 490.66 | 549.67 | 576.97 |
| 4714 | ENSMUSG00000024845 | 1542.676146 | -0.098484218 | 0.097760085 | -1.007407242 | 0.313739113 | 0.760476509 | Tmem134 | transmembrane protein 134 [Source:MGI | 1638.3 | 1622.1 | 1525.5 | 1524 | 1514.9 | 1431.3 |
| 4715 | ENSMUSG00000024846 | 0.160662974 | 1.126924175 | 4.080472857 | 0.276174898 | 0.782413736 | NA | Cst6 | cystatin E/M [Source:MGI Symbol;Acc:MG | 0 | 0 | 0 | 0.964 | 0 | 0 |
| 4716 | ENSMUSG00000024847 | 441.794784 | -0.052589159 | 0.135040218 | -0.38943331 | 0.696955635 | 0.94023175 | Aip | aryl-hydrocarbon receptor-interacting prc | 454.56 | 438.92 | 455.83 | 429.93 | 394.8 | 476.73 |
| 4717 | ENSMUSG00000024851 | 91.03579131 | -0.404678947 | 0.30560069 | -1.324208223 | 0.185433901 | 0.621059253 | Pitpnm1 | phosphatidylinositol transfer protein, mer | 81.916 | 118.21 | 111.69 | 105.07 | 46.897 | 82.424 |
| 4718 | ENSMUSG00000024853 | 2232.794956 | 0.012961924 | 0.109752263 | 0.118101657 | 0.905987111 | 0.983991807 | Sf3b2 | splicing factor 3b, subunit 2 [Source:MGI ! | 2109.7 | 2299.5 | 2259 | 2016.6 | 2494.2 | 2217.7 |
| 4719 | ENSMUSG00000024854 | 318.5218494 | 0.304337189 | 0.160242639 | 1.899227269 | 0.057534601 | 0.341197403 | Pold4 | polymerase (DNA-directed), delta 4 [Sour | 306.79 | 285.76 | 261.62 | 300.76 | 370.81 | 385.39 |
| 4720 | ENSMUSG00000024855 | 42.41811523 | -0.058848265 | 0.354369649 | -0.166064632 | 0.868106087 | 0.978326885 | Pacs1 | phosphofurin acidic cluster sorting protei | 42.564 | 40.089 | 47.293 | 44.343 | 51.259 | 28.96 |
| 4721 | ENSMUSG00000024856 | 732.1028471 | 0.216602228 | 0.125762142 | 1.722316629 | 0.085012177 | 0.4294852 | Cdk2ap2 | CDK2-associated protein 2 [Source:MGI Sy | 713.15 | 646.56 | 671.17 | 758.65 | 719.81 | 883.28 |
| 4722 | ENSMUSG00000024857 | 1.333010503 | -0.038102961 | 2.187392492 | -0.017419353 | 0.98610207 | NA | Cabp2 | calcium binding protein 2 [Source:MGI Sy | 4.0155 | 0 | 0 | 2.8919 | 1.0906 | 0 |
| 4723 | ENSMUSG00000024858 | 551.3624757 | -0.051474967 | 0.143162934 | -0.35955513 | 0.719179844 | 0.9447719 | Grk2 | G protein-coupled receptor kinase 2 [Sour | 525.23 | 651.7 | 507.15 | 539.83 | 499.5 | 584.77 |
| 4724 | ENSMUSG00000024862 | 192.6145517 | 0.001438948 | 0.181310154 | 0.00793639 | 0.993667744 | 0.99859176 | Klc2 | kinesin light chain 2 [Source:MGI Symbol; | 215.23 | 179.89 | 182.13 | 207.26 | 196.31 | 174.87 |
| 4725 | ENSMUSG00000024863 | 6992.354713 | 0.041032748 | 0.139137491 | 0.29490792 | 0.768064221 | 0.955856567 | Mbl2 | mannose-binding lectin (protein C) 2 [Sou | 7624.7 | 6218.9 | 6835.4 | 8687.4 | 6003.9 | 6583.9 |
| 4726 | ENSMUSG00000024866 | 1638.569079 | 0.26756364 | 0.154467387 | 1.732169129 | 0.083243454 | 0.425303185 | Acy3 | aspartoacylase (aminoacylase) 3 [Source:I | 1789.3 | 1204.7 | 1465.1 | 1495.1 | 1941.3 | 1935.9 |
| 4727 | ENSMUSG00000024867 | 0.52105551 | -0.812028029 | 3.385364865 | -0.239864257 | 0.810435491 | NA | Pip5k1b | phosphatidylinositol-4-phosphate 5-kinas | 0 | 0 | 2.0125 | 0 | 0 | 1.1138 |
| 4728 | ENSMUSG00000024868 | 0.167707431 | -0.796629176 | 4.080472857 | -0.195229623 | 0.84521319 | NA | Dkk1 | dickkopf WNT signaling pathway inhibitor | 0 | 0 | 1.0062 | 0 | 0 | 0 |
| 4729 | ENSMUSG00000024869 | 19.56257394 | 0.198829173 | 0.500011274 | 0.397649379 | 0.690888656 | 0.937606823 | Gm49405 | predicted gene, 49405 [Source:MGI Symb | 16.062 | 15.419 | 23.144 | 16.388 | 19.631 | 26.732 |
| 4730 | ENSMUSG00000024870 | 2705.611685 | 0.025047893 | 0.095078629 | 0.263443985 | 0.792208392 | 0.961434599 | Rab1b | RAB1B, member RAS oncogene family [So | 2855.8 | 2691.1 | 2498.5 | 2752.2 | 2694.9 | 2741.2 |
| 4731 | ENSMUSG00000024871 | 16.97265668 | 0.876604303 | 0.532386763 | 1.646555405 | 0.099649463 | 0.465067707 | Doc2g | double C2, gamma [Source:MGI Symbol;A | 14.456 | 9.2513 | 12.075 | 24.099 | 17.45 | 24.505 |
| 4732 | ENSMUSG00000024873 | 4.985683166 | -0.011214287 | 0.946002901 | -0.01185439 | 0.990541787 | NA | Cnih2 | cornichon family AMPA receptor auxiliary | 4.8186 | 5.1396 | 5.0312 | 3.8559 | 3.2719 | 7.7969 |
| 4733 | ENSMUSG00000024875 | 960.8496164 | 0.03002956 | 0.12120918 | 0.247749885 | 0.804327932 | 0.963701061 | Yif1a | Yip1 interacting factor homolog A (S. cere | 991.83 | 950.82 | 908.64 | 851.19 | 1004.5 | 1058.2 |
| 4734 | ENSMUSG00000024878 | 100.35071 | 0.222556205 | 0.243479896 | 0.914063989 | 0.360683224 | 0.800074801 | Cbwd1 | COBW domain containing 1 [Source:MGI ! | 103.6 | 92.513 | 81.506 | 114.71 | 87.249 | 122.52 |
| 4735 | ENSMUSG00000024883 | 2.89697155 | -0.088147132 | 1.271723963 | -0.069313101 | 0.944740397 | NA | Rin1 | Ras and Rab interactor 1 [Source:MGI Sym | 4.8186 | 2.0558 | 2.0125 | 1.928 | 5.4531 | 1.1138 |
| 4736 | ENSMUSG00000024885 | 26.45219352 | -0.382653561 | 0.463662288 | -0.825285065 | 0.409209769 | 0.833065722 | Aldh3b1 | aldehyde dehydrogenase 3 family, membe | 16.865 | 34.949 | 38.237 | 21.208 | 20.722 | 26.732 |
| 4737 | ENSMUSG00000024887 | 409.9953941 | -0.204420968 | 0.138937527 | -1.471315727 | 0.141205752 | 0.549623251 | Asah2 | N-acylsphingosine amidohydrolase 2 [Sou | 426.45 | 483.12 | 407.53 | 367.28 | 401.35 | 374.25 |
| 4738 | ENSMUSG00000024889 | 187.8785753 | 0.14225961 | 0.177730274 | 0.800424187 | 0.423465072 | 0.84018348 | Rce1 | Ras converting CAAX endopeptidase 1 [So | 175.88 | 181.94 | 178.11 | 198.58 | 176.68 | 216.09 |
| 4739 | ENSMUSG00000024891 | 16.8643672 | -0.633763638 | 0.568652772 | -1.114500219 | 0.265064651 | 0.720853222 | Slc29a2 | solute carrier family 29 (nucleoside transp | 20.078 | 12.335 | 29.181 | 16.388 | 8.7249 | 14.48 |
| 4740 | ENSMUSG00000024892 | 14580.25261 | -0.240116958 | 0.114718839 | -2.0930909 | 0.036341041 | 0.257160869 | Pcx | pyruvate carboxylase [Source:MGI Symbo | 15922 | 17739 | 13712 | 14555 | 13050 | 12503 |
| 4741 | ENSMUSG00000024896 | 847.5668687 | -0.073287772 | 0.116699672 | -0.628003244 | 0.530001815 | 0.888496321 | Minpp1 | multiple inositol polyphosphate histidine | 889.03 | 893.26 | 824.11 | 749.97 | 873.59 | 855.43 |
| 4742 | ENSMUSG00000024897 | 7.656165814 | -0.751376464 | 0.793801595 | -0.946554491 | 0.343865843 | 0.785606627 | Apba1 | amyloid beta (A4) precursor protein bindi | 10.44 | 7.1954 | 11.069 | 2.8919 | 6.5437 | 7.7969 |
| 4743 | ENSMUSG00000024899 | 3478.462534 | 0.032609254 | 0.096837159 | 0.336743189 | 0.736310503 | 0.948109711 | Papss2 | 3'-phosphoadenosine 5'-phosphosulfate s | 3373 | 3245.1 | 3699 | 3335.4 | 3581.6 | 3636.7 |
| 4744 | ENSMUSG00000024900 | 11379.25897 | -0.140597513 | 0.118147872 | -1.190013081 | 0.234041251 | 0.68180718 | Cpt1a | carnitine palmitoyltransferase 1a, liver [Sc | 13407 | 11742 | 10650 | 11061 | 11872 | 9543.4 |
| 4745 | ENSMUSG00000024901 | 35.2682897 | 0.960572541 | 0.381158232 | 2.520141143 | 0.011730779 | 0.120765517 | Peli3 | pellino 3 [Source:MGI Symbol;Acc:MGI:19 | 19.274 | 23.642 | 29.181 | 50.127 | 39.262 | 50.123 |
| 4746 | ENSMUSG00000024902 | 496.2119357 | 0.062747337 | 0.132492804 | 0.473590528 | 0.635791931 | 0.923478764 | Mrpl11 | mitochondrial ribosomal protein L11 [Sou | 492.3 | 498.54 | 464.88 | 460.78 | 557.31 | 503.46 |
| 4747 | ENSMUSG00000024903 | 0 | NA | NA | NA | NA | NA | Lao1 | L-amino acid oxidase 1 [Source:MGI Symb | 0 | 0 | 0 | 0 | 0 | 0 |
| 4748 | ENSMUSG00000024905 | 24.06617827 | -0.372115738 | 0.466232931 | -0.798132678 | 0.4247935 | 0.840504368 | Tesmin | testis expressed metallothionein like [Sou | 22.487 | 38.033 | 21.131 | 23.135 | 22.903 | 16.708 |
| 4749 | ENSMUSG00000024906 | 171.9271188 | -0.159444625 | 0.191823235 | -0.831206007 | 0.405857262 | 0.829973745 | Mus81 | MUS81 structure-specific endonuclease s | 203.18 | 159.33 | 181.12 | 158.09 | 150.51 | 179.33 |
| 4750 | ENSMUSG00000024907 | 0.569258551 | -2.569337407 | 3.18860282 | -0.805787849 | 0.420365185 | NA | Gal | galanin and GMAP prepropeptide [Source | 2.4093 | 0 | 1.0062 | 0 | 0 | 0 |
| 4751 | ENSMUSG00000024908 | 1284.869714 | 0.043610168 | 0.110855413 | 0.39339683 | 0.694026411 | 0.939133502 | Ppp6r3 | protein phosphatase 6, regulatory subuni | 1239.2 | 1337.3 | 1219.6 | 1194.4 | 1417.8 | 1301 |
| 4752 | ENSMUSG00000024909 | 32.60593542 | -0.628433531 | 0.417168766 | -1.506425174 | 0.131958062 | 0.532693668 | Efemp2 | epidermal growth factor-containing fibuli | 43.368 | 41.117 | 34.212 | 26.027 | 15.269 | 35.643 |
| 4753 | ENSMUSG00000024910 | 7.079150382 | -1.849284772 | 0.893387037 | -2.069970456 | 0.038455111 | 0.267025794 | Ctsw | cathepsin W [Source:MGI Symbol;Acc:MG | 12.047 | 9.2513 | 12.075 | 5.7839 | 1.0906 | 2.2277 |
| 4754 | ENSMUSG00000024911 | 321.3171079 | 0.010713582 | 0.163001145 | 0.065727033 | 0.94759515 | 0.990677354 | Fibp | fibroblast growth factor (acidic) intracellu | 351.76 | 307.35 | 300.87 | 329.68 | 365.36 | 272.89 |
| 4755 | ENSMUSG00000024912 | 4.418425317 | 2.244171263 | 1.241558443 | 1.807543798 | 0.070677532 | NA | Fosl1 | fos-like antigen 1 [Source:MGI Symbol;Ac | 1.6062 | 1.0279 | 2.0125 | 16.388 | 4.3625 | 1.1138 |
| 4756 | ENSMUSG00000024913 | 1025.673469 | -0.13663209 | 0.172476047 | -0.792180091 | 0.428255675 | 0.842753542 | Lrp5 | low density lipoprotein receptor-related p | 1033.6 | 1105 | 1083.7 | 885.9 | 747.07 | 1298.7 |

| | | | | | | | | | | | | | | | |
|---|---|---|---|---|---|---|---|---|---|---|---|---|---|---|---|
| 4757 | ENSMUSG00000024914 | 1062.069565 | -0.00296012 | 0.105378866 | -0.02809026 | 0.977590163 | 0.997564159 | Drap1 | Dr1 associated protein 1 (negative cofacto | 1100.3 | 1017.6 | 1070.6 | 1011.2 | 1066.6 | 1106 |
| 4758 | ENSMUSG00000024921 | 2569.020816 | 0.155712875 | 0.136952024 | 1.136988492 | 0.255543096 | 0.708894389 | Smarca2 | SWI/SNF related, matrix associated, actin | 2202.1 | 2394 | 2696.7 | 2965.2 | 2977.4 | 2178.7 |
| 4759 | ENSMUSG00000024922 | 0.342639279 | -1.805976596 | 4.028547714 | -0.448294702 | 0.653940524 | NA | Ovol1 | ovo like zinc finger 1 [Source:MGI Symbol; | 0 | 2.0558 | 0 | 0 | 0 | 0 |
| 4760 | ENSMUSG00000024924 | 56.57557269 | -0.926598219 | 0.411917081 | -2.249477531 | 0.024482131 | 0.199836083 | Vldlr | very low density lipoprotein receptor [Sou | 67.461 | 114.1 | 41.256 | 55.911 | 25.084 | 35.643 |
| 4761 | ENSMUSG00000024925 | 1051.566298 | -0.023937474 | 0.115443197 | -0.207352834 | 0.835734315 | 0.971073158 | Rnaseh2c | ribonuclease H2, subunit C [Source:MGI S | 1039.2 | 994 | 1148.1 | 1098.9 | 955.38 | 1073.7 |
| 4762 | ENSMUSG00000024926 | 184.5626961 | 0.062433807 | 0.213815329 | 0.291998741 | 0.770287587 | 0.956515747 | Kat5 | K(lysine) acetyltransferase 5 [Source:MGI | 216.03 | 148.02 | 177.1 | 206.29 | 149.41 | 210.52 |
| 4763 | ENSMUSG00000024927 | 789.3349369 | 0.105098909 | 0.121260952 | 0.866716839 | 0.386097177 | 0.817115886 | Rela | v-rel reticuloendotheliosis viral oncogene | 739.66 | 739.07 | 802.98 | 788.53 | 757.98 | 907.78 |
| 4764 | ENSMUSG00000024935 | 0.331982614 | 0.16514309 | 3.986751476 | 0.041422971 | 0.966958701 | NA | Slc1a1 | solute carrier family 1 (neuronal/epithelia | 0 | 1.0279 | 0 | 0.964 | 0 | 0 |
| 4765 | ENSMUSG00000024936 | 0 | NA | NA | NA | NA | NA | Kcnk7 | potassium channel, subfamily K, member | 0 | 0 | 0 | 0 | 0 | 0 |
| 4766 | ENSMUSG00000024937 | 122.405464 | -0.404024027 | 0.229579521 | -1.759843497 | 0.078434346 | 0.410446591 | Ehbp1l1 | EH domain binding protein 1-like 1 [Sourc | 134.92 | 158.3 | 124.77 | 84.83 | 107.97 | 123.64 |
| 4767 | ENSMUSG00000024939 | 0 | NA | NA | NA | NA | NA | Fam89b | family with sequence similarity 89, memb | 0 | 0 | 0 | 0 | 0 | 0 |
| 4768 | ENSMUSG00000024940 | 40.6070526 | -0.789109664 | 0.373725785 | -2.111467008 | 0.034732191 | 0.249868983 | Ltbp3 | latent transforming growth factor beta bir | 67.461 | 49.34 | 37.231 | 27.955 | 33.809 | 27.846 |
| 4769 | ENSMUSG00000024941 | 900.8517288 | -0.022833727 | 0.110290445 | -0.207032688 | 0.83598433 | 0.971073158 | Scyl1 | SCY1-like 1 (S. cerevisiae) [Source:MGI Syr | 934.01 | 945.68 | 844.24 | 924.45 | 880.13 | 876.6 |
| 4770 | ENSMUSG00000024942 | 390.8840012 | 0.112910856 | 0.142948029 | 0.789873471 | 0.429601666 | 0.84307112 | Capn1 | calpain 1 [Source:MGI Symbol;Acc:MGI:88 | 424.04 | 351.55 | 350.17 | 402.94 | 411.16 | 405.44 |
| 4771 | ENSMUSG00000024943 | 293.2706733 | 0.16862981 | 0.158076513 | 1.066760695 | 0.286079912 | 0.738507914 | Smc5 | structural maintenance of chromosomes 5 | 249.76 | 266.23 | 312.94 | 306.54 | 295.56 | 328.58 |
| 4772 | ENSMUSG00000024944 | 108.9467515 | 0.231834005 | 0.247816825 | 0.935505507 | 0.349527847 | 0.790905688 | Arl2 | ADP-ribosylation factor-like 2 [Source:MG | 102.8 | 94.568 | 103.64 | 153.27 | 103.61 | 95.791 |
| 4773 | ENSMUSG00000024947 | 452.8944249 | 0.192520769 | 0.136687897 | 1.408469751 | 0.158992018 | 0.580282752 | Men1 | multiple endocrine neoplasia 1 [Source:M | 404.76 | 391.64 | 471.93 | 468.49 | 506.05 | 474.5 |
| 4774 | ENSMUSG00000024948 | 305.4922478 | -0.058091591 | 0.185588582 | -0.31301274 | 0.754270983 | 0.953048681 | Map4k2 | mitogen-activated protein kinase kinase k | 305.18 | 321.74 | 308.92 | 385.59 | 248.66 | 262.87 |
| 4775 | ENSMUSG00000024949 | 1480.469153 | 0.11554097 | 0.110779645 | 1.042980142 | 0.29695749 | 0.747770393 | Sf1 | splicing factor 1 [Source:MGI Symbol;Acc: | 1474.5 | 1443.2 | 1346.4 | 1709.1 | 1451.6 | 1458 |
| 4776 | ENSMUSG00000024952 | 237.4552504 | -0.101472798 | 0.187331082 | -0.541676251 | 0.588041553 | 0.910798643 | NA | NA | 240.13 | 224.09 | 272.69 | 182.19 | 263.93 | 241.7 |
| 4777 | ENSMUSG00000024953 | 4533.834097 | 0.173486361 | 0.110932938 | 1.563885035 | 0.11784457 | 0.502838264 | Prdx5 | peroxiredoxin 5 [Source:MGI Symbol;Acc: | 4442 | 4169.2 | 4171.9 | 4125.8 | 5075.7 | 5218.4 |
| 4778 | ENSMUSG00000024955 | 1055.860162 | 0.139760134 | 0.104443476 | 1.338141343 | 0.180850366 | 0.614186356 | Esrra | estrogen related receptor, alpha [Source:I | 1047.2 | 961.1 | 1005.2 | 1096 | 1147.3 | 1078.2 |
| 4779 | ENSMUSG00000024957 | 5.879757032 | -1.281493204 | 0.917364551 | -1.396929064 | 0.162434904 | 0.585865647 | Kcnk4 | potassium channel, subfamily K, member | 8.8341 | 7.1954 | 9.0562 | 5.7839 | 2.1812 | 2.2277 |
| 4780 | ENSMUSG00000024958 | 383.2836902 | 0.304534549 | 0.150607823 | 2.022036716 | 0.043172559 | 0.286918392 | Gpr137 | G protein-coupled receptor 137 [Source:N | 383.88 | 300.15 | 344.14 | 444.39 | 387.17 | 439.97 |
| 4781 | ENSMUSG00000024959 | 277.5330345 | -0.00536215 | 0.170085769 | -0.03152615 | 0.974849937 | 0.996336532 | Bad | BCL2-associated agonist of cell death [Sou | 285.1 | 265.2 | 283.76 | 290.16 | 225.76 | 315.22 |
| 4782 | ENSMUSG00000024960 | 356.9637677 | 0.133491817 | 0.158371823 | 0.842901314 | 0.399283639 | 0.8263661 | Plcb3 | phospholipase C, beta 3 [Source:MGI Sym | 296.34 | 343.32 | 382.37 | 367.28 | 337 | 415.46 |
| 4783 | ENSMUSG00000024962 | 199.0373685 | -0.212401527 | 0.210057336 | -1.011159768 | 0.311939968 | 0.759830222 | Vegfb | vascular endothelial growth factor B [Sour | 221.66 | 247.73 | 172.07 | 226.53 | 162.5 | 163.74 |
| 4784 | ENSMUSG00000024963 | 227.8132235 | -0.005533425 | 0.180528844 | -0.030651195 | 0.975547714 | 0.996491676 | Dnajc4 | DnaJ heat shock protein family (Hsp40) m | 263.42 | 229.23 | 191.19 | 209.18 | 238.85 | 235.02 |
| 4785 | ENSMUSG00000024965 | 51.51670106 | -1.250507359 | 0.402745584 | -3.104956105 | 0.001903073 | 0.033568345 | Fermt3 | fermitin family member 3 [Source:MGI Sy | 82.72 | 95.596 | 39.244 | 31.811 | 19.631 | 40.098 |
| 4786 | ENSMUSG00000024966 | 1332.534871 | 0.148386727 | 0.135684142 | 1.093618789 | 0.274122205 | 0.729244726 | Stip1 | stress-induced phosphoprotein 1 [Source: | 1271.3 | 1181.1 | 1340.3 | 1599.2 | 1140.8 | 1462.5 |
| 4787 | ENSMUSG00000024968 | 1.640191546 | 0.263594276 | 1.848808724 | 0.142575201 | 0.886625684 | NA | Rcor2 | REST corepressor 2 [Source:MGI Symbol;A | 2.4093 | 0 | 2.0125 | 0.964 | 0 | 4.4554 |
| 4788 | ENSMUSG00000024969 | 332.3365144 | -0.172098937 | 0.149586973 | -1.150494153 | 0.249940401 | 0.702104867 | Mark2 | MAP/microtubule affinity regulating kinas | 350.15 | 330.99 | 375.33 | 321.97 | 279.2 | 336.38 |
| 4789 | ENSMUSG00000024970 | 43.80618732 | 0.036596576 | 0.333213415 | 0.109829239 | 0.912544802 | 0.986248982 | Spindoc | spindlin interactor and repressor of chrom | 39.352 | 42.145 | 48.3 | 41.451 | 40.353 | 51.237 |
| 4790 | ENSMUSG00000024972 | 0 | NA | NA | NA | NA | NA | Lgals12 | lectin, galactose binding, soluble 12 [Sour | 0 | 0 | 0 | 0 | 0 | 0 |
| 4791 | ENSMUSG00000024973 | 0 | NA | NA | NA | NA | NA | Plaat5 | phospholipase A and acyltransferase 5 [Sc | 0 | 0 | 0 | 0 | 0 | 0 |
| 4792 | ENSMUSG00000024974 | 336.8253689 | 0.09185026 | 0.140064387 | 0.65577169 | 0.511971034 | 0.8841969 | Smc3 | structural maintenance of chromosomes 3 | 326.06 | 339.21 | 312.94 | 336.43 | 359.9 | 346.41 |
| 4793 | ENSMUSG00000024975 | 1634.237713 | 0.536779265 | 0.137130383 | 3.914371526 | 9.06E-05 | 0.00334443 | Pdcd4 | programmed cell death 4 [Source:MGI Syr | 1225.5 | 1255.1 | 1520.4 | 1678.3 | 1863.9 | 2262.2 |
| 4794 | ENSMUSG00000024976 | 595.071865 | 0.064592705 | 0.12680529 | 0.509384939 | 0.610482431 | 0.916376001 | Shoc2 | Shoc2, leucine rich repeat scaffold protein | 525.23 | 628.06 | 592.68 | 580.31 | 628.2 | 615.96 |
| 4795 | ENSMUSG00000024978 | 2102.869988 | 0.15237331 | 0.136176426 | 1.118940437 | 0.26316555 | 0.717823918 | Gpam | glycerol-3-phosphate acyltransferase, mit | 2117 | 2243.9 | 1614 | 2001.2 | 2421.2 | 2219.9 |
| 4796 | ENSMUSG00000024979 | 7.679829866 | -0.793364207 | 0.775930361 | -1.022468313 | 0.306559303 | 0.755610724 | Tectb | tectorin beta [Source:MGI Symbol;Acc:MG | 12.85 | 8.2233 | 8.05 | 4.8199 | 5.4531 | 6.6831 |
| 4797 | ENSMUSG00000024981 | 5188.843632 | 0.15282071 | 0.136609946 | 1.118664593 | 0.263283256 | 0.717837703 | Acsl5 | acyl-CoA synthetase long-chain family me | 5798.4 | 5070.7 | 3872 | 5166 | 5461.8 | 5764.1 |
| 4798 | ENSMUSG00000024982 | 1333.418272 | 0.035367343 | 0.101706041 | 0.347740832 | 0.728034828 | 0.946158959 | Zdhhc6 | zinc finger, DHHC domain containing 6 [Sc | 1357.2 | 1257.1 | 1336.3 | 1332.2 | 1420 | 1297.6 |
| 4799 | ENSMUSG00000024983 | 286.0779612 | 0.053830377 | 0.152943007 | 0.35196364 | 0.724865531 | 0.9448775 | Vti1a | vesicle transport through interaction with | 262.61 | 305.29 | 274.7 | 283.41 | 308.64 | 281.8 |
| 4800 | ENSMUSG00000024985 | 265.3499454 | -0.162677587 | 0.188691809 | -0.8621338 | 0.388613893 | 0.818441625 | Tcf7l2 | transcription factor 7 like 2, T cell specific, | 281.09 | 306.32 | 253.57 | 265.09 | 190.86 | 295.17 |
| 4801 | ENSMUSG00000024986 | 1631.118333 | 1.279185138 | 0.682787563 | 1.873474574 | 0.061002881 | 0.352269825 | Hhex | hematopoietically expressed homeobox [S | 695.49 | 1002.2 | 1158.2 | 3104 | 644.56 | 3182.3 |
| 4802 | ENSMUSG00000024987 | 51.14776978 | 0.337880279 | 0.93538339 | 0.361221166 | 0.717934121 | 0.9447719 | Cyp26a1 | cytochrome P450, family 26, subfamily a, | 30.518 | 24.67 | 80.5 | 107.97 | 11.997 | 51.237 |
| 4803 | ENSMUSG00000024989 | 1.737864628 | -3.042880708 | 1.883690737 | -1.615382317 | 0.106227927 | NA | Cep55 | centrosomal protein 55 [Source:MGI Syml | 3.2124 | 4.1117 | 2.0125 | 0 | 1.0906 | 0 |
| 4804 | ENSMUSG00000024990 | 68576.68212 | -0.054086851 | 0.088750885 | -0.609423228 | 0.542243945 | 0.894095573 | Rbp4 | retinol binding protein 4, plasma [Source: | 64630 | 74195 | 70762 | 68493 | 67820 | 65560 |
| 4805 | ENSMUSG00000024991 | 2529.380695 | 0.11249159 | 0.104381885 | 1.077692647 | 0.281170941 | 0.733710968 | Eif3a | eukaryotic translation initiation factor 3, s | 2378 | 2667.4 | 2248 | 2774.3 | 2592.4 | 2516.2 |
| 4806 | ENSMUSG00000024992 | 23.35228129 | 0.23090649 | 0.456903099 | 0.505372999 | 0.613296879 | 0.916813074 | Pde6c | phosphodiesterase 6C, cGMP specific, cor | 20.881 | 15.419 | 28.175 | 26.027 | 23.994 | 25.618 |
| 4807 | ENSMUSG00000024993 | 313.4976012 | 0.002731937 | 0.149917969 | 0.018222881 | 0.985461049 | 0.99859176 | Dennd10 | DENN domain containing 10 [Source:MGI | 322.85 | 302.21 | 313.95 | 290.16 | 303.19 | 348.63 |
| 4808 | ENSMUSG00000024997 | 2182.946367 | -0.072012332 | 0.108262249 | -0.665165674 | 0.505944533 | 0.881592135 | Prdx3 | peroxiredoxin 3 [Source:MGI Symbol;Acc: | 2472.8 | 2233.7 | 2004.4 | 2083.2 | 2207.4 | 2096.3 |
| 4809 | ENSMUSG00000024998 | 310.8200727 | -0.41381816 | 0.195201231 | -2.119956713 | 0.034009696 | 0.246945905 | Plce1 | phospholipase C, epsilon 1 [Source:MGI S | 292.33 | 454.34 | 318.98 | 233.28 | 306.46 | 259.53 |
| 4810 | ENSMUSG00000024999 | 129.2375847 | -0.147925754 | 0.228007379 | -0.648776171 | 0.516483062 | 0.886716042 | Noc3l | NOC3 like DNA replication regulator [Sour | 134.12 | 142.88 | 130.81 | 130.14 | 146.14 | 91.335 |
| 4811 | ENSMUSG00000025001 | 16.9281167 | 0.131813226 | 0.547360057 | 0.240816304 | 0.809697492 | 0.966133025 | Hells | helicase, lymphoid specific [Source:MGI S | 20.881 | 15.419 | 12.075 | 20.244 | 21.812 | 11.138 |
| 4812 | ENSMUSG00000025002 | 16.41892281 | -0.269336767 | 0.561126934 | -0.479992584 | 0.631232667 | 0.921377326 | Cyp2c55 | cytochrome P450, family 2, subfamily c, p | 24.093 | 13.363 | 16.1 | 9.6398 | 15.269 | 20.049 |

| | | | | | | | | | | | | | | | |
|---|---|---|---|---|---|---|---|---|---|---|---|---|---|---|---|
| 4813 | ENSMUSG00000025003 | 46.23350949 | -1.292392964 | 0.343203132 | -3.765679396 | 0.000166097 | 0.005410866 | Cyp2c39 | cytochrome P450, family 2, subfamily c, p | 71.476 | 68.87 | 56.35 | 21.208 | 30.537 | 28.96 |
| 4814 | ENSMUSG00000025004 | 12.29734445 | 1.362359141 | 0.798193725 | 1.706802619 | 0.087858745 | 0.43690328 | Cyp2c40 | cytochrome P450, family 2, subfamily c, p | 1.6062 | 5.1396 | 14.087 | 8.6758 | 13.087 | 31.188 |
| 4815 | ENSMUSG00000025006 | 563.3427036 | 0.079739561 | 0.200096861 | 0.398504806 | 0.690258117 | 0.937454779 | Sorbs1 | sorbin and SH3 domain containing 1 [Sour | 663.36 | 624.97 | 354.2 | 550.43 | 640.19 | 546.9 |
| 4816 | ENSMUSG00000025007 | 10.60982285 | -0.971224651 | 0.723202943 | -1.342948974 | 0.179288497 | 0.611423712 | Aldh18a1 | aldehyde dehydrogenase 18 family, meml | 12.85 | 15.419 | 14.087 | 12.532 | 6.5437 | 2.2277 |
| 4817 | ENSMUSG00000025008 | 11.13839265 | 0.193224882 | 0.71787069 | 0.26916391 | 0.787803551 | 0.960870081 | Tctn3 | tectonic family member 3 [Source:MGI Sy | 8.8341 | 6.1675 | 16.1 | 4.8199 | 13.087 | 17.822 |
| 4818 | ENSMUSG00000025010 | 67.16250281 | -0.040859144 | 0.324524455 | -0.12590467 | 0.899807386 | 0.982798755 | Ccnj | cyclin J [Source:MGI Symbol;Acc:MGI:244 | 50.595 | 96.624 | 57.356 | 54.947 | 63.256 | 80.197 |
| 4819 | ENSMUSG00000025013 | 0 | NA | NA | NA | NA | NA | Tll2 | tolloid-like 2 [Source:MGI Symbol;Acc:MG | 0 | 0 | 0 | 0 | 0 | 0 |
| 4820 | ENSMUSG00000025014 | 44.94084739 | 0.296104793 | 0.356833783 | 0.829811546 | 0.406645342 | 0.830399297 | Dntt | deoxynucleotidyltransferase, terminal [So | 55.414 | 37.005 | 28.175 | 52.055 | 47.987 | 49.009 |
| 4821 | ENSMUSG00000025016 | 3394.564352 | 0.175305856 | 0.089179085 | 1.9657732 | 0.049324815 | 0.309919254 | Tm9sf3 | transmembrane 9 superfamily member 3 | 3085.5 | 3207.1 | 3273.3 | 3485.7 | 3557.6 | 3758.1 |
| 4822 | ENSMUSG00000025017 | 598.195404 | 0.262625907 | 0.142635376 | 1.841239626 | 0.065586451 | 0.367976115 | Pik3ap1 | phosphoinositide-3-kinase adaptor protei | 629.63 | 504.71 | 496.08 | 660.32 | 590.02 | 708.4 |
| 4823 | ENSMUSG00000025019 | 127.5038333 | -0.171725027 | 0.223758345 | -0.767457529 | 0.442809535 | 0.850769625 | Lcor | ligand dependent nuclear receptor corepr | 109.22 | 140.82 | 155.97 | 132.06 | 117.79 | 109.16 |
| 4824 | ENSMUSG00000025020 | 2.153923884 | -2.430521241 | 1.782939597 | -1.363210086 | 0.172816309 | NA | Slit1 | slit guidance ligand 1 [Source:MGI Symbol | 0.8031 | 6.1675 | 4.025 | 1.928 | 0 | 0 |
| 4825 | ENSMUSG00000025024 | 314.725773 | -0.052224876 | 0.149341021 | -0.349702149 | 0.726562241 | 0.945777495 | Smndc1 | survival motor neuron domain containing | 300.36 | 346.41 | 314.95 | 312.33 | 286.83 | 327.47 |
| 4826 | ENSMUSG00000025025 | 540.5388967 | 0.053086431 | 0.128703428 | 0.412470994 | 0.679994236 | 0.934384556 | Mxi1 | MAX interactor 1, dimerization protein [S | 502.74 | 526.29 | 563.5 | 583.21 | 502.78 | 564.72 |
| 4827 | ENSMUSG00000025026 | 179.0872815 | -0.245698948 | 0.175483494 | -1.400125686 | 0.161475685 | 0.584480556 | Add3 | adducin 3 (gamma) [Source:MGI Symbol;A | 190.34 | 199.42 | 193.2 | 161.95 | 160.32 | 169.3 |
| 4828 | ENSMUSG00000025027 | 594.5647149 | -0.065246502 | 0.129783161 | -0.502734723 | 0.615150793 | 0.917509899 | Xpnpep1 | X-prolyl aminopeptidase (aminopeptidase | 648.1 | 604.42 | 570.54 | 535.01 | 642.37 | 566.95 |
| 4829 | ENSMUSG00000025034 | 704.9429522 | 0.026615205 | 0.114435192 | 0.232578847 | 0.816088454 | 0.967767553 | Trim8 | tripartite motif-containing 8 [Source:MGI | 729.22 | 687.68 | 678.21 | 742.26 | 680.55 | 711.75 |
| 4830 | ENSMUSG00000025035 | 200.7390978 | 0.214765386 | 0.173900209 | 1.234992107 | 0.216833442 | 0.662741261 | Arl3 | ADP-ribosylation factor-like 3 [Source:MG | 165.44 | 197.36 | 195.21 | 212.08 | 211.58 | 222.77 |
| 4831 | ENSMUSG00000025036 | 765.1992552 | 0.002486492 | 0.147718232 | 0.01683267 | 0.986570107 | 0.99859176 | Sfxn2 | sideroflexin 2 [Source:MGI Symbol;Acc:M | 759.73 | 733.93 | 800.97 | 935.06 | 714.35 | 647.14 |
| 4832 | ENSMUSG00000025037 | 172.0184173 | -0.364207169 | 0.236760956 | -1.538290665 | 0.123977561 | 0.515924041 | Maoa | monoamine oxidase A [Source:MGI Symb | 161.42 | 230.25 | 190.18 | 192.8 | 149.41 | 108.04 |
| 4833 | ENSMUSG00000025038 | 0 | NA | NA | NA | NA | NA | Efhc2 | EF-hand domain (C-terminal) containing 2 | 0 | 0 | 0 | 0 | 0 | 0 |
| 4834 | ENSMUSG00000025040 | 145.6212652 | 0.002728413 | 0.207315449 | 0.013160685 | 0.989499595 | 0.99859176 | Fundc1 | FUN14 domain containing 1 [Source:MGI | 152.59 | 139.8 | 143.89 | 150.38 | 118.88 | 168.19 |
| 4835 | ENSMUSG00000025041 | 347.230177 | -0.410228756 | 0.155025712 | -2.646198171 | 0.008140211 | 0.093114627 | Nt5c2 | 5'-nucleotidase, cytosolic II [Source:MGI S | 369.43 | 393.69 | 425.64 | 267.02 | 342.45 | 285.14 |
| 4836 | ENSMUSG00000025043 | 0 | NA | NA | NA | NA | NA | Dusp21 | dual specificity phosphatase 21 [Source:M | 0 | 0 | 0 | 0 | 0 | 0 |
| 4837 | ENSMUSG00000025044 | 82.092407 | -0.820529512 | 0.302836809 | -2.709477476 | 0.006738928 | 0.081840349 | Msr1 | macrophage scavenger receptor 1 [Source | 92.357 | 128.49 | 93.581 | 52.055 | 42.534 | 83.538 |
| 4838 | ENSMUSG00000025047 | 452.6279942 | -0.008734111 | 0.142940763 | -0.061103009 | 0.951277173 | 0.990883058 | Pdcd11 | programmed cell death 11 [Source:MGI Sy | 421.63 | 507.79 | 432.69 | 407.76 | 459.15 | 486.75 |
| 4839 | ENSMUSG00000025049 | 89.42438028 | 0.35178488 | 0.242939324 | 1.44803597 | 0.147606991 | 0.558897256 | Taf5 | TATA-box binding protein associated facto | 80.31 | 70.926 | 84.525 | 107 | 105.79 | 87.994 |
| 4840 | ENSMUSG00000025050 | 202.8854163 | 0.110528334 | 0.173150371 | 0.638337268 | 0.523254157 | 0.888334265 | Pcgf6 | polycomb group ring finger 6 [Source:MG | 175.08 | 201.47 | 209.3 | 209.18 | 202.86 | 219.43 |
| 4841 | ENSMUSG00000025051 | 0 | NA | NA | NA | NA | NA | Samt4 | spermatogenesis associated multipass tra | 0 | 0 | 0 | 0 | 0 | 0 |
| 4842 | ENSMUSG00000025056 | 0 | NA | NA | NA | NA | NA | Nr0b1 | nuclear receptor subfamily 0, group B, me | 0 | 0 | 0 | 0 | 0 | 0 |
| 4843 | ENSMUSG00000025058 | 2.972917821 | -1.684922237 | 1.297828273 | -1.298262854 | 0.194197026 | NA | Tasl | TLR adaptor interacting with endolysosom | 6.4248 | 4.1117 | 3.0187 | 0.964 | 1.0906 | 2.2277 |
| 4844 | ENSMUSG00000025059 | 2863.659793 | -0.229364539 | 0.129134453 | -1.776168434 | 0.075705159 | 0.402878781 | Gk | glycerol kinase [Source:MGI Symbol;Acc:N | 3116.8 | 3343.8 | 2812.5 | 2853.4 | 2889 | 2166.4 |
| 4845 | ENSMUSG00000025060 | 940.394985 | -0.178665024 | 0.119437317 | -1.49588946 | 0.134682463 | 0.537616739 | Slk | STE20-like kinase [Source:MGI Symbol;Acc | 955.69 | 999.14 | 1040.5 | 781.79 | 959.74 | 905.56 |
| 4846 | ENSMUSG00000025064 | 3.179324163 | 0.739213293 | 1.451826053 | 0.509161061 | 0.610639335 | NA | Col17a1 | collagen, type XVII, alpha 1 [Source:MGI S | 4.0155 | 1.0279 | 2.0125 | 0 | 10.906 | 1.1138 |
| 4847 | ENSMUSG00000025066 | 625.3365421 | -0.014379611 | 0.135108571 | -0.106430046 | 0.915241155 | 0.98625062 | Sfr1 | SWI5 dependent recombination repair 1 [ | 657.74 | 613.67 | 612.8 | 530.19 | 684.91 | 652.71 |
| 4848 | ENSMUSG00000025068 | 1527.239913 | -0.227026208 | 0.12667914 | -1.792135687 | 0.073111234 | 0.39267764 | Gsto1 | glutathione S-transferase omega 1 [Sourc | 1459.2 | 1639.5 | 1844.4 | 1576.1 | 1366.5 | 1277.6 |
| 4849 | ENSMUSG00000025069 | 4.025059333 | 1.095071351 | 1.15778935 | 0.945829525 | 0.344235544 | NA | Gsto2 | glutathione S-transferase omega 2 [Sourc | 1.6062 | 3.0838 | 3.0187 | 0.964 | 5.4531 | 10.025 |
| 4850 | ENSMUSG00000025075 | 2535.407256 | 0.32955064 | 0.105560172 | 3.121922166 | 0.001796744 | 0.032283367 | Habp2 | hyaluronic acid binding protein 2 [Source: | 2094.5 | 2256.3 | 2390.8 | 2585.4 | 3038.5 | 2847 |
| 4851 | ENSMUSG00000025076 | 655.684445 | 0.081366649 | 0.140131883 | 0.580643372 | 0.561480834 | 0.900553224 | Casp7 | caspase 7 [Source:MGI Symbol;Acc:MGI:1 | 668.18 | 610.58 | 631.92 | 588.99 | 793.97 | 640.46 |
| 4852 | ENSMUSG00000025077 | 302.4215253 | 0.021874485 | 0.155885021 | 0.140324486 | 0.888403618 | 0.982382091 | Dclre1a | DNA cross-link repair 1A [Source:MGI Sym | 331.68 | 304.26 | 263.64 | 290.16 | 317.37 | 307.42 |
| 4853 | ENSMUSG00000025078 | 392.6512918 | 0.084282235 | 0.155430921 | 0.542248827 | 0.587647103 | 0.910765033 | Nhlrc2 | NHL repeat containing 2 [Source:MGI Sym | 382.28 | 392.66 | 369.29 | 468.49 | 406.8 | 336.38 |
| 4854 | ENSMUSG00000025081 | 0 | NA | NA | NA | NA | NA | Tdrd1 | tudor domain containing 1 [Source:MGI S | 0 | 0 | 0 | 0 | 0 | 0 |
| 4855 | ENSMUSG00000025082 | 0 | NA | NA | NA | NA | NA | Vwa2 | von Willebrand factor A domain containin | 0 | 0 | 0 | 0 | 0 | 0 |
| 4856 | ENSMUSG00000025083 | 6.62931029 | -1.139488151 | 0.882775352 | -1.290801956 | 0.196772359 | 0.637822894 | Afap1l2 | actin filament associated protein 1-like 2 [ | 12.047 | 11.307 | 4.025 | 5.7839 | 3.2719 | 3.3415 |
| 4857 | ENSMUSG00000025085 | 320.8329872 | 0.204368095 | 0.180653601 | 1.131270532 | 0.257941243 | 0.711453498 | Ablim1 | actin-binding LIM protein 1 [Source:MGI S | 341.32 | 267.26 | 284.77 | 308.47 | 299.92 | 423.26 |
| 4858 | ENSMUSG00000025086 | 342.4559171 | 0.062988099 | 0.151295118 | 0.416326049 | 0.677171434 | 0.93325821 | Trub1 | TruB pseudouridine (psi) synthase family | 324.45 | 322.77 | 358.22 | 371.13 | 374.08 | 304.08 |
| 4859 | ENSMUSG00000025089 | 4075.773042 | 0.067927278 | 0.129719061 | 0.523649162 | 0.600522581 | 0.91534406 | Gfra1 | glial cell line derived neurotrophic factor f | 3553.7 | 3976 | 4411.4 | 4726.4 | 3491.1 | 4296.1 |
| 4860 | ENSMUSG00000025090 | 0 | NA | NA | NA | NA | NA | Ccdc172 | coiled-coil domain containing 172 [Source | 0 | 0 | 0 | 0 | 0 | 0 |
| 4861 | ENSMUSG00000025091 | 0.181769715 | 1.126924175 | 4.080472857 | 0.276174898 | 0.782413736 | NA | Pnliprp2 | pancreatic lipase-related protein 2 [Sourc | 0 | 0 | 0 | 0 | 1.0906 | 0 |
| 4862 | ENSMUSG00000025092 | 18.77874743 | -0.604509037 | 0.520864849 | -1.160587124 | 0.245809844 | 0.695907924 | Hspa12a | heat shock protein 12A [Source:MGI Symb | 24.093 | 20.558 | 23.144 | 10.604 | 11.997 | 22.277 |
| 4863 | ENSMUSG00000025094 | 29.06903092 | -0.03277292 | 0.413145525 | -0.079325366 | 0.936773831 | 0.988420305 | Slc18a2 | solute carrier family 18 (vesicular monoan | 31.321 | 30.838 | 26.162 | 37.595 | 23.994 | 24.505 |
| 4864 | ENSMUSG00000025102 | 225.9701805 | 0.140978238 | 0.162804586 | 0.865935302 | 0.386525643 | 0.817115886 | 3110040N11 | RIKEN cDNA 3110040N11 gene [Source:M | 209.61 | 216.89 | 218.36 | 232.32 | 223.58 | 255.07 |
| 4865 | ENSMUSG00000025103 | 905.0277388 | 0.107453878 | 0.108635793 | 0.989120395 | 0.32260424 | 0.769926329 | Btbd1 | BTB (POZ) domain containing 1 [Source:M | 826.39 | 861.4 | 926.75 | 924.45 | 947.75 | 943.43 |
| 4866 | ENSMUSG00000025104 | 13.7293566 | -0.438744597 | 0.596931963 | -0.734999337 | 0.462339898 | 0.860650866 | Hdgfl3 | HDGF like 3 [Source:MGI Symbol;Acc:MGI | 12.85 | 17.475 | 17.106 | 10.604 | 7.6343 | 16.708 |
| 4867 | ENSMUSG00000025105 | 10.96052821 | -0.046997913 | 0.644015663 | -0.072976351 | 0.941824936 | 0.989579826 | Bnc1 | basonuclin 1 [Source:MGI Symbol;Acc:MG | 12.047 | 12.335 | 9.0562 | 12.532 | 11.997 | 7.7969 |
| 4868 | ENSMUSG00000025127 | 6246.857981 | 0.1999074 | 0.085832954 | 2.329028551 | 0.019857553 | 0.172383511 | Gcgr | glucagon receptor [Source:MGI Symbol;A | 5844.2 | 5544.6 | 6055.6 | 6838.5 | 6623.3 | 6575 |

| | | | | | | | | | | | | | | | |
|---|---|---|---|---|---|---|---|---|---|---|---|---|---|---|---|
| 4869 | ENSMUSG00000025128 | 1.184762013 | -0.39060714 | 2.10457176 | -0.185599345 | 0.852758967 | NA | Bhlhe22 | basic helix-loop-helix family, member e22 | 0 | 3.0838 | 1.0062 | 1.928 | 1.0906 | 0 |
| 4870 | ENSMUSG00000025129 | 1.031169 | 3.580676128 | 2.33548092 | 1.533164368 | 0.125235361 | NA | Ppp1r27 | protein phosphatase 1, regulatory subunit | 0 | 0 | 0 | 2.8919 | 2.1812 | 1.1138 |
| 4871 | ENSMUSG00000025130 | 24108.12072 | -0.034379456 | 0.101900997 | -0.337380963 | 0.735829734 | 0.948109711 | P4hb | prolyl 4-hydroxylase, beta polypeptide [Sc | 26153 | 24820 | 22213 | 25924 | 23218 | 22320 |
| 4872 | ENSMUSG00000025132 | 743.4201599 | -0.201218957 | 0.131163782 | -1.53410456 | 0.125003932 | 0.518510965 | Arhgdia | Rho GDP dissociation inhibitor (GDI) alpha | 761.34 | 855.23 | 768.77 | 634.3 | 647.83 | 793.06 |
| 4873 | ENSMUSG00000025133 | 372.7642795 | -0.039784214 | 0.152979234 | -0.260062839 | 0.794815302 | 0.962189806 | Ints4 | integrator complex subunit 4 [Source:MGI | 408.78 | 384.44 | 340.11 | 370.17 | 410.07 | 323.01 |
| 4874 | ENSMUSG00000025134 | 282.6276791 | -0.000160602 | 0.175269046 | -0.000916315 | 0.999268886 | 0.999764475 | Alyref | Aly/REF export factor [Source:MGI Symbo | 256.99 | 290.9 | 299.86 | 230.39 | 287.92 | 329.7 |
| 4875 | ENSMUSG00000025135 | 328.0568284 | 0.079183371 | 0.150547758 | 0.525968451 | 0.598910128 | 0.915055389 | Anapc11 | anaphase promoting complex subunit 11 | 326.86 | 328.93 | 300.87 | 312.33 | 320.64 | 378.71 |
| 4876 | ENSMUSG00000025137 | 8429.92623 | 0.23179708 | 0.104671324 | 2.214523245 | 0.026792806 | 0.211649994 | Pcyt2 | phosphate cytidylyltransferase 2, ethanola | 8413.3 | 7714.5 | 7134.3 | 9973.3 | 8868.9 | 8475.2 |
| 4877 | ENSMUSG00000025138 | 946.540591 | 0.281780367 | 0.11608144 | 2.42743687 | 0.015205933 | 0.143938884 | Sirt7 | sirtuin 7 [Source:MGI Symbol;Acc:MGI:23 | 853.7 | 783.27 | 925.75 | 983.26 | 1033.9 | 1099.4 |
| 4878 | ENSMUSG00000025139 | 1321.601677 | 0.042796064 | 0.107863857 | 0.396760006 | 0.691544446 | 0.938056725 | Tollip | toll interacting protein [Source:MGI Symb | 1372.5 | 1339.4 | 1193.4 | 1317.8 | 1415.6 | 1290.9 |
| 4879 | ENSMUSG00000025140 | 4.296817374 | -0.070872974 | 1.076451886 | -0.065839425 | 0.947505667 | NA | Pycr1 | pyrroline-5-carboxylate reductase 1 [Sour | 8.031 | 1.0279 | 4.025 | 3.8559 | 3.2719 | 5.5692 |
| 4880 | ENSMUSG00000025141 | 0 | NA | NA | NA | NA | NA | Myadml2 | myeloid-associated differentiation marker | 0 | 0 | 0 | 0 | 0 | 0 |
| 4881 | ENSMUSG00000025142 | 1638.261475 | 0.071495617 | 0.112837116 | 0.633617906 | 0.526330213 | 0.888334265 | Aspscr1 | alveolar soft part sarcoma chromosome re | 1672.9 | 1460.7 | 1658.3 | 1571.3 | 1613 | 1853.4 |
| 4882 | ENSMUSG00000025144 | 556.2743275 | -0.074110012 | 0.125841182 | -0.588917008 | 0.555916948 | 0.897628185 | Cenpx | centromere protein X [Source:MGI Symbo | 555.75 | 554.05 | 601.73 | 516.69 | 521.32 | 588.11 |
| 4883 | ENSMUSG00000025145 | 193.7021423 | 0.2675097 | 0.18145846 | 1.47422005 | 0.140422353 | 0.548048817 | Lrrc45 | leucine rich repeat containing 45 [Source: | 167.05 | 178.86 | 182.13 | 242.92 | 195.22 | 196.04 |
| 4884 | ENSMUSG00000025147 | 468.4322967 | -0.020161969 | 0.125121049 | -0.161139707 | 0.871983367 | 0.979369708 | Mob2 | MOB kinase activator 2 [Source:MGI Symb | 485.07 | 472.84 | 456.84 | 458.85 | 459.15 | 477.84 |
| 4885 | ENSMUSG00000025150 | 0.840029082 | -0.398118545 | 2.584217466 | -0.154057679 | 0.877564257 | NA | Cbr2 | carbonyl reductase 2 [Source:MGI Symbol | 0.8031 | 2.0558 | 0 | 0 | 2.1812 | 0 |
| 4886 | ENSMUSG00000025151 | 1423.125998 | 0.231777692 | 0.110393895 | 2.099551724 | 0.035768293 | 0.253977709 | Maged1 | MAGE family member D1 [Source:MGI Sy | 1365.3 | 1300.3 | 1261.8 | 1703.3 | 1476.7 | 1431.3 |
| 4887 | ENSMUSG00000025153 | 5185.880996 | 0.016147389 | 0.319822102 | 0.05048866 | 0.959732986 | 0.992148164 | Fasn | fatty acid synthase [Source:MGI Symbol;A | 7016.7 | 5004.9 | 3448.4 | 4882.5 | 6189.3 | 4573.4 |
| 4888 | ENSMUSG00000025154 | 58.74636927 | -1.218669086 | 0.373371562 | -3.263957975 | 0.001098675 | 0.022476987 | Arhgap19 | Rho GTPase activating protein 19 [Source: | 71.476 | 116.15 | 59.368 | 47.235 | 37.081 | 21.163 |
| 4889 | ENSMUSG00000025155 | 1453.845751 | -0.091440885 | 0.122166857 | -0.748491752 | 0.454163598 | 0.857337687 | Dus1l | dihydrouridine synthase 1-like (S. cerevisia | 1379.7 | 1433.9 | 1687.5 | 1558.8 | 1361.1 | 1302.1 |
| 4890 | ENSMUSG00000025156 | 1072.941107 | 0.12054821 | 0.103084207 | 1.169414922 | 0.242236501 | 0.692000002 | Gps1 | G protein pathway suppressor 1 [Source:M | 1027.2 | 984.75 | 1072.7 | 1143.3 | 1101.5 | 1108.3 |
| 4891 | ENSMUSG00000025157 | 157.1853723 | 0.052528546 | 0.215147978 | 0.244150779 | 0.807114046 | 0.964718338 | Zdhhc16 | zinc finger, DHHC domain containing 16 [S | 141.35 | 152.13 | 170.06 | 187.01 | 122.15 | 170.42 |
| 4892 | ENSMUSG00000025158 | 219.7646134 | -0.009386901 | 0.177200788 | -0.052973247 | 0.957753224 | 0.991861398 | Rfng | RFNG O-fucosylpeptide 3-beta-N-acetylglu | 227.28 | 192.22 | 241.5 | 205.33 | 207.22 | 245.05 |
| 4893 | ENSMUSG00000025159 | 513.3121156 | 0.379024901 | 0.129906712 | 2.917669875 | 0.003526575 | 0.052640227 | Mms19 | MMS19 cytosolic iron-sulfur assembly cor | 424.84 | 433.78 | 480.98 | 623.69 | 551.85 | 564.72 |
| 4894 | ENSMUSG00000025161 | 6.362716574 | -0.653407499 | 0.835140511 | -0.782392292 | 0.433984062 | 0.845264432 | Slc16a3 | solute carrier family 16 (monocarboxylic a | 8.031 | 8.2233 | 7.0437 | 3.8559 | 5.4531 | 5.5692 |
| 4895 | ENSMUSG00000025162 | 1217.108546 | -0.119469324 | 0.112807307 | -1.059056616 | 0.289573997 | 0.739999393 | Csnk1d | casein kinase 1, delta [Source:MGI Symbo | 1249.6 | 1315.7 | 1236.7 | 1097 | 1101.5 | 1302.1 |
| 4896 | ENSMUSG00000025163 | 2.581901054 | -1.490476567 | 1.462913831 | -1.018840984 | 0.308278465 | NA | Cd7 | CD7 antigen [Source:MGI Symbol;Acc:MG | 2.4093 | 2.0558 | 7.0437 | 2.8919 | 1.0906 | 0 |
| 4897 | ENSMUSG00000025165 | 8.485135776 | -0.900110059 | 0.752756762 | -1.195751541 | 0.231793531 | 0.67889946 | Sectm1a | secreted and transmembrane 1A [Source: | 8.8341 | 12.335 | 12.075 | 7.7118 | 3.2719 | 6.6831 |
| 4898 | ENSMUSG00000025169 | 118.9868286 | -0.11535511 | 0.215190396 | -0.536060681 | 0.591916613 | 0.912740793 | Ogfod3 | 2-oxoglutarate and iron-dependent oxyge | 126.09 | 114.1 | 130.81 | 105.07 | 128.69 | 109.16 |
| 4899 | ENSMUSG00000025170 | 10.61010496 | 1.377740992 | 0.70574413 | 1.952182006 | 0.0509166 | 0.316755999 | Rab40b | Rab40B, member RAS oncogene family [S | 6.4248 | 5.1396 | 6.0375 | 8.6758 | 22.903 | 14.48 |
| 4900 | ENSMUSG00000025171 | 335.7721928 | -0.104696048 | 0.161740047 | -0.647310609 | 0.517430937 | 0.887022094 | Ubtd1 | ubiquitin domain containing 1 [Source:MC | 358.18 | 376.22 | 309.92 | 374.99 | 289.01 | 306.31 |
| 4901 | ENSMUSG00000025172 | 16.17196135 | -0.851712528 | 0.662046032 | -1.286485359 | 0.198273725 | 0.640554426 | Ankrd2 | ankyrin repeat domain 2 (stretch responsi | 16.865 | 17.475 | 28.175 | 11.568 | 20.722 | 2.2277 |
| 4902 | ENSMUSG00000025173 | 804.3781335 | 0.648328879 | 0.14468115 | 4.481087391 | 7.43E-06 | 0.000424565 | Wdr45b | WD repeat domain 45B [Source:MGI Syml | 691.47 | 613.67 | 574.57 | 1169.3 | 909.58 | 867.68 |
| 4903 | ENSMUSG00000025175 | 418.0564683 | 0.096803447 | 0.149292708 | 0.648413765 | 0.51671737 | 0.886716042 | Fn3k | fructosamine 3 kinase [Source:MGI Symbo | 435.28 | 362.85 | 413.57 | 442.47 | 477.69 | 376.48 |
| 4904 | ENSMUSG00000025176 | 2832.908174 | 0.077788853 | 0.097865709 | 0.794853001 | 0.426699016 | 0.842062628 | Hoga1 | 4-hydroxy-2-oxoglutarate aldolase 1 [Sour | 2823.7 | 2850.4 | 2595.1 | 2846.6 | 3114.8 | 2766.8 |
| 4905 | ENSMUSG00000025178 | 1238.731792 | 0.341006896 | 0.116408601 | 2.929396046 | 0.003396214 | 0.051116377 | Pi4k2a | phosphatidylinositol 4-kinase type 2 alpha | 1063.3 | 1062.9 | 1153.2 | 1411.3 | 1225.9 | 1515.9 |
| 4906 | ENSMUSG00000025184 | 207.6368081 | -0.029088055 | 0.188078394 | -0.154659204 | 0.877089994 | 0.980632845 | R3hcc1l | R3H domain and coiled-coil containing 1 li | 215.23 | 195.3 | 218.36 | 199.54 | 171.23 | 246.16 |
| 4907 | ENSMUSG00000025185 | 9.785495904 | -1.049956136 | 0.802775965 | -1.30790679 | 0.190904926 | 0.628409928 | Loxl4 | lysyl oxidase-like 4 [Source:MGI Symbol;A | 15.259 | 4.1117 | 20.125 | 3.8559 | 10.906 | 4.4554 |
| 4908 | ENSMUSG00000025188 | 178.24296 | 0.045595736 | 0.218577315 | 0.208602327 | 0.834758694 | 0.971073158 | Hps1 | HPS1, biogenesis of lysosomal organelles | 191.14 | 125.41 | 209.3 | 177.37 | 166.86 | 199.38 |
| 4909 | ENSMUSG00000025189 | 6.652562457 | 1.1007501 | 0.9310098 | 1.182318489 | 0.237079352 | 0.685960342 | Cnnm1 | cyclin M1 [Source:MGI Symbol;Acc:MGI:1 | 1.6062 | 9.2513 | 2.0125 | 11.568 | 5.4531 | 10.025 |
| 4910 | ENSMUSG00000025190 | 3534.409947 | 0.003534683 | 0.17462858 | 0.020241148 | 0.983851003 | 0.998580735 | Got1 | glutamic-oxaloacetic transaminase 1, solu | 3910.3 | 3487.7 | 3190.8 | 2654.8 | 3192.2 | 4770.6 |
| 4911 | ENSMUSG00000025192 | 30.96226327 | 0.03316362 | 0.418431208 | 0.079257042 | 0.936828174 | 0.988420305 | Entpd7 | ectonucleoside triphosphate diphosphohy | 20.881 | 37.005 | 34.212 | 31.811 | 23.994 | 37.871 |
| 4912 | ENSMUSG00000025193 | 303.8713326 | 0.084323997 | 0.157596398 | 0.535062975 | 0.592606311 | 0.912925376 | Cutc | cutC copper transporter [Source:MGI Sym | 282.69 | 313.51 | 288.79 | 285.34 | 357.72 | 295.17 |
| 4913 | ENSMUSG00000025194 | 9836.155915 | 0.111761297 | 0.090684101 | 1.232424376 | 0.2177906 | 0.663697152 | Abcc2 | ATP-binding cassette, sub-family C (CFTR/ | 9289.5 | 9926.6 | 9150.8 | 10483 | 10639 | 9527.8 |
| 4914 | ENSMUSG00000025195 | 981.903025 | -0.496708475 | 0.152618813 | -3.254569108 | 0.001135645 | 0.022941303 | Dnmbp | dynamin binding protein [Source:MGI Syn | 1128.4 | 1315.7 | 1004.2 | 842.52 | 942.29 | 658.28 |
| 4915 | ENSMUSG00000025196 | 4871.496046 | 0.208193256 | 0.100823454 | 2.064928818 | 0.038929736 | 0.269932742 | Cpn1 | carboxypeptidase N, polypeptide 1 [Sourc | 4870.8 | 4505.4 | 4185 | 5549.6 | 4862 | 5256.2 |
| 4916 | ENSMUSG00000025197 | 4927.909342 | 0.51937297 | 0.129791254 | 4.001602221 | 6.29E-05 | 0.002506894 | Cyp2c23 | cytochrome P450, family 2, subfamily c, p | 3468.6 | 3883.5 | 4800.8 | 6355.5 | 5766.1 | 5293 |
| 4917 | ENSMUSG00000025198 | 839.6741939 | 0.497154946 | 0.123858124 | 4.013906636 | 5.97E-05 | 0.00240618 | Erlin1 | ER lipid raft associated 1 [Source:MGI Sym | 782.22 | 685.62 | 619.85 | 947.59 | 990.28 | 1012.5 |
| 4918 | ENSMUSG00000025199 | 3116.957163 | 0.161871635 | 0.104773268 | 1.544970747 | 0.122353339 | 0.512759732 | Chuk | conserved helix-loop-helix ubiquitous kina | 2904 | 3042.6 | 2880.9 | 3427.9 | 3533.6 | 2912.7 |
| 4919 | ENSMUSG00000025200 | 244.6556871 | 0.151463005 | 0.171632572 | 0.882484036 | 0.377515109 | 0.81030298 | Cwf19l1 | CWF19-like 1, cell cycle control (S. pombe | 209.61 | 221 | 265.65 | 282.45 | 249.75 | 239.48 |
| 4920 | ENSMUSG00000025201 | 0.319491021 | 0.16514334 | 4.044653141 | 0.040830038 | 0.967431393 | NA | Bloc1s2-ps | biogenesis of lysosomal organelles comple | 0.8031 | 0 | 0 | 0 | 0 | 1.1138 |
| 4921 | ENSMUSG00000025202 | 0.331982614 | 0.16514309 | 3.986751476 | 0.041422971 | 0.966958701 | NA | Scd3 | stearoyl-coenzyme A desaturase 3 [Source | 0 | 1.0279 | 0 | 0.964 | 0 | 0 |
| 4922 | ENSMUSG00000025203 | 69.27408451 | -0.661158207 | 0.289453094 | -2.28416355 | 0.022361921 | 0.187285943 | Scd2 | stearoyl-Coenzyme A desaturase 2 [Sourc | 84.326 | 96.624 | 73.456 | 45.307 | 47.987 | 67.944 |
| 4923 | ENSMUSG00000025204 | 1507.565186 | -0.036029908 | 0.101039479 | -0.356592374 | 0.721396987 | 0.9447719 | Ndufb8 | NADH:ubiquinone oxidoreductase subuni | 1476.9 | 1485.3 | 1617 | 1451.8 | 1450.5 | 1563.8 |
| 4924 | ENSMUSG00000025207 | 5829.963187 | 0.33315092 | 0.090211882 | 3.69298269 | 0.000221639 | 0.006835369 | Sema4g | sema domain, immunoglobulin domain (Ig | 5002.5 | 4963.8 | 5513.2 | 6275.5 | 6587.3 | 6637.4 |

| | | | | | | | | | | | | | | | |
|---|---|---|---|---|---|---|---|---|---|---|---|---|---|---|---|
| 4925 | ENSMUSG00000025208 | 687.048841 | 0.195092688 | 0.12432545 | 1.569209588 | 0.116599111 | 0.501814922 | Mrpl43 | mitochondrial ribosomal protein L43 [Sou | 704.32 | 630.11 | 586.64 | 749.01 | 691.45 | 760.76 |
| 4926 | ENSMUSG00000025209 | 530.2823322 | -0.061127517 | 0.128659291 | -0.475111562 | 0.634707458 | 0.923478764 | Twnk | twinkle mtDNA helicase [Source:MGI Sym | 517.2 | 577.69 | 530.29 | 540.79 | 475.51 | 540.21 |
| 4927 | ENSMUSG00000025212 | 18.10450596 | -0.638005415 | 0.574856269 | -1.109852063 | 0.26706278 | 0.722778036 | Sfxn3 | sideroflexin 3 [Source:MGI Symbol;Acc:M | 25.699 | 10.279 | 30.187 | 19.28 | 9.8156 | 13.366 |
| 4928 | ENSMUSG00000025213 | 31.61437425 | -0.07567792 | 0.400465655 | -0.188974808 | 0.850112561 | 0.973301217 | Kazald1 | Kazal-type serine peptidase inhibitor dom | 36.14 | 34.949 | 26.162 | 32.775 | 22.903 | 36.757 |
| 4929 | ENSMUSG00000025215 | 0 | NA | NA | NA | NA | NA | Tlx1 | T cell leukemia, homeobox 1 [Source:MGI | 0 | 0 | 0 | 0 | 0 | 0 |
| 4930 | ENSMUSG00000025216 | 0 | NA | NA | NA | NA | NA | Lbx1 | ladybird homeobox 1 [Source:MGI Symbo | 0 | 0 | 0 | 0 | 0 | 0 |
| 4931 | ENSMUSG00000025217 | 267.0777862 | 0.115965963 | 0.168073614 | 0.689971262 | 0.49021226 | 0.875501998 | Btrc | beta-transducin repeat containing protein | 257.8 | 232.31 | 278.73 | 261.24 | 319.55 | 252.84 |
| 4932 | ENSMUSG00000025218 | 152.405003 | 0.047681413 | 0.206257869 | 0.231173786 | 0.817179794 | 0.968033459 | Poll | polymerase (DNA directed), lambda [Sour | 142.15 | 166.52 | 140.87 | 134.96 | 146.14 | 183.78 |
| 4933 | ENSMUSG00000025219 | 0 | NA | NA | NA | NA | NA | Fgf8 | fibroblast growth factor 8 [Source:MGI Sy | 0 | 0 | 0 | 0 | 0 | 0 |
| 4934 | ENSMUSG00000025220 | 948.4153675 | 0.180436299 | 0.115360662 | 1.564105955 | 0.117792688 | 0.502838264 | Oga | O-GlcNAcase [Source:MGI Symbol;Acc:M( | 898.67 | 911.76 | 856.31 | 904.21 | 1051.4 | 1068.2 |
| 4935 | ENSMUSG00000025221 | 0.36353943 | 2.065989254 | 4.034137301 | 0.512126658 | 0.608562368 | NA | Kcnip2 | Kv channel-interacting protein 2 [Source:I | 0 | 0 | 0 | 0 | 2.1812 | 0 |
| 4936 | ENSMUSG00000025223 | 726.6089328 | 0.060661376 | 0.115692126 | 0.524334524 | 0.600045888 | 0.915217938 | Ldb1 | LIM domain binding 1 [Source:MGI Symbo | 665.77 | 710.29 | 758.71 | 745.15 | 759.07 | 720.66 |
| 4937 | ENSMUSG00000025224 | 952.9325314 | 0.083879066 | 0.106221563 | 0.78966138 | 0.429725551 | 0.84307112 | Gbf1 | golgi-specific brefeldin A-resistance factor | 946.86 | 897.37 | 930.78 | 942.77 | 971.74 | 1028.1 |
| 4938 | ENSMUSG00000025225 | 156.6415468 | -0.015370194 | 0.190836953 | -0.080540973 | 0.93580701 | 0.988420305 | Nfkb2 | nuclear factor of kappa light polypeptide g | 149.38 | 161.38 | 162.01 | 164.84 | 138.51 | 163.74 |
| 4939 | ENSMUSG00000025226 | 183.8850381 | 0.108279106 | 0.199870184 | 0.541747168 | 0.587992691 | 0.910798643 | Fbxl15 | F-box and leucine-rich repeat protein 15 [ | 192.74 | 146.99 | 191.19 | 221.71 | 165.77 | 184.9 |
| 4940 | ENSMUSG00000025227 | 72.94590683 | 0.062049865 | 0.261859972 | 0.236958191 | 0.812689225 | 0.966802988 | Mfsd13a | major facilitator superfamily domain cont | 70.673 | 71.954 | 71.443 | 71.334 | 67.618 | 84.652 |
| 4941 | ENSMUSG00000025228 | 1015.966865 | 0.155102798 | 0.106728797 | 1.453242255 | 0.146156504 | 0.556855766 | Actr1a | ARP1 actin-related protein 1A, centractin | 1001.5 | 984.75 | 897.57 | 1077.7 | 1090.6 | 1043.7 |
| 4942 | ENSMUSG00000025229 | 32.61463246 | -1.701288361 | 0.599333658 | -2.838633103 | 0.004530722 | 0.062530879 | Pitx3 | paired-like homeodomain transcription fa | 11.243 | 76.066 | 62.387 | 10.604 | 11.997 | 23.391 |
| 4943 | ENSMUSG00000025231 | 97.78696398 | 0.078358938 | 0.232123722 | 0.33757402 | 0.735684223 | 0.948109711 | Sufu | SUFU negative regulator of hedgehog sign | 106.01 | 86.345 | 92.575 | 93.506 | 104.7 | 103.59 |
| 4944 | ENSMUSG00000025232 | 933.22848 | -0.506845544 | 0.125179835 | -4.04893923 | 5.15E-05 | 0.002150801 | Hexa | hexosaminidase A [Source:MGI Symbol;Ac | 1166.1 | 1174.9 | 944.86 | 748.05 | 813.6 | 751.84 |
| 4945 | ENSMUSG00000025234 | 1015.623869 | 0.125070655 | 0.110750021 | 1.129305921 | 0.258768804 | 0.712165697 | Arih1 | ariadne RBR E3 ubiquitin protein ligase 1 [ | 962.12 | 1019.7 | 932.79 | 1003.5 | 1031.7 | 1143.9 |
| 4946 | ENSMUSG00000025235 | 68.65735199 | -0.154303497 | 0.270316596 | -0.570825096 | 0.56811821 | 0.903352921 | Bbs4 | Bardet-Biedl syndrome 4 (human) [Source | 70.673 | 64.759 | 81.506 | 61.695 | 68.709 | 64.603 |
| 4947 | ENSMUSG00000025236 | 350.6281805 | -0.134270589 | 0.169548923 | -0.791928292 | 0.428402489 | 0.842753542 | Adpgk | ADP-dependent glucokinase [Source:MGI | 396.73 | 327.91 | 376.34 | 387.52 | 345.73 | 269.55 |
| 4948 | ENSMUSG00000025237 | 216.9549482 | 0.087235443 | 0.176539928 | 0.494140019 | 0.621207297 | 0.918069106 | Parp6 | poly (ADP-ribose) polymerase family, men | 213.63 | 210.72 | 207.29 | 260.27 | 212.67 | 197.15 |
| 4949 | ENSMUSG00000025239 | 895.3364208 | 0.156587626 | 0.116558466 | 1.343425592 | 0.179134205 | 0.611235187 | Limd1 | LIM domains containing 1 [Source:MGI Sy | 896.26 | 794.58 | 848.26 | 863.72 | 997.92 | 971.27 |
| 4950 | ENSMUSG00000025240 | 664.7274202 | 0.081226791 | 0.138663304 | 0.585784337 | 0.558020451 | 0.899217185 | Sacm1l | SAC1 suppressor of actin mutations 1-like | 557.35 | 660.95 | 720.47 | 627.55 | 748.16 | 673.88 |
| 4951 | ENSMUSG00000025241 | 926.2192685 | 0.09989126 | 0.107835988 | 0.926325815 | 0.354276684 | 0.79449673 | Fyco1 | FYVE and coiled-coil domain containing 1 | 876.18 | 896.34 | 910.65 | 1012.2 | 907.39 | 954.56 |
| 4952 | ENSMUSG00000025243 | 0.267700746 | -1.497481649 | 4.057001436 | -0.369110456 | 0.712045396 | NA | Slc6a20b | solute carrier family 6 (neurotransmitter t | 1.6062 | 0 | 0 | 0 | 0 | 0 |
| 4953 | ENSMUSG00000025245 | 220.8460162 | -0.013902881 | 0.194947088 | -0.071316177 | 0.94314612 | 0.990077979 | Lztfl1 | leucine zipper transcription factor-like 1 [S | 258.6 | 179.89 | 226.41 | 203.4 | 256.3 | 200.49 |
| 4954 | ENSMUSG00000025246 | 773.7366901 | 0.225097985 | 0.113136205 | 1.989619372 | 0.046632881 | 0.299112188 | Tbl1x | transducin (beta)-like 1 X-linked [Source:N | 666.57 | 742.16 | 732.55 | 830.95 | 811.42 | 858.77 |
| 4955 | ENSMUSG00000025255 | 246.8472168 | -0.318782633 | 0.169665475 | -1.878889228 | 0.06025962 | 0.349789984 | Zfhx4 | zinc finger homeodomain 4 [Source:MGI S | 248.96 | 305.29 | 268.67 | 243.89 | 212.67 | 201.61 |
| 4956 | ENSMUSG00000025257 | 45.11311518 | -0.16186667 | 0.361933643 | -0.447227477 | 0.654710824 | 0.925797231 | Ribc1 | RIB43A domain with coiled-coils 1 [Source | 40.958 | 62.703 | 39.244 | 30.847 | 51.259 | 45.668 |
| 4957 | ENSMUSG00000025260 | 4761.239035 | -0.232661927 | 0.135631371 | -1.715399065 | 0.086272087 | 0.43282673 | Hsd17b10 | hydroxysteroid (17-beta) dehydrogenase : | 5869.1 | 5029.6 | 4533.1 | 4132.6 | 5184.8 | 3818.3 |
| 4958 | ENSMUSG00000025261 | 2635.504219 | 0.034180468 | 0.096462438 | 0.354339668 | 0.723084343 | 0.9447719 | Huwe1 | HECT, UBA and WWE domain containing 1 | 2571.5 | 2696.2 | 2545.8 | 2758.9 | 2775.6 | 2464.9 |
| 4959 | ENSMUSG00000025262 | 66.00902994 | 0.349195828 | 0.282432412 | 1.236387232 | 0.21631466 | 0.661699385 | Fam120c | family with sequence similarity 120, meml | 61.036 | 57.563 | 55.343 | 63.623 | 89.431 | 69.058 |
| 4960 | ENSMUSG00000025264 | 199.4243257 | -0.156569774 | 0.182556229 | -0.857652323 | 0.391084474 | 0.820195366 | Tsr2 | TSR2 20S rRNA accumulation [Source:MG | 228.08 | 212.78 | 189.17 | 171.59 | 179.95 | 214.97 |
| 4961 | ENSMUSG00000025265 | 8.698910173 | -2.606183435 | 0.888240528 | -2.93409651 | 0.003345202 | 0.050564445 | Fgd1 | FYVE, RhoGEF and PH domain containing : | 10.44 | 10.279 | 24.15 | 2.8919 | 1.0906 | 3.3415 |
| 4962 | ENSMUSG00000025266 | 308.965327 | -0.171644876 | 0.14866875 | -1.154545767 | 0.248276486 | 0.699604734 | Gnl3l | guanine nucleotide binding protein-like 3 | 304.38 | 322.77 | 355.2 | 285.34 | 303.19 | 282.92 |
| 4963 | ENSMUSG00000025268 | 74.14617889 | 0.08208967 | 0.27419228 | 0.299387238 | 0.764644598 | 0.954980327 | Maged2 | MAGE family member D2 [Source:MGI Syr | 88.341 | 65.787 | 61.381 | 68.442 | 79.615 | 81.311 |
| 4964 | ENSMUSG00000025269 | 141.8982524 | 0.169178407 | 0.224441873 | 0.753773814 | 0.45098505 | 0.855818389 | Apex2 | apurinic/apyrimidinic endonuclease 2 [So | 127.69 | 154.19 | 118.74 | 120.5 | 182.13 | 148.14 |
| 4965 | ENSMUSG00000025270 | 1927.326748 | 0.519896222 | 0.137749126 | 3.774225202 | 0.000160506 | 0.00528828 | Alas2 | aminolevulinic acid synthase 2, erythroid | | 1648 | 1292.1 | 1810.2 | 2027.2 | 2290.3 | 2496.1 |
| 4966 | ENSMUSG00000025271 | 1068.589675 | -0.176981218 | 0.124037054 | -1.426841522 | 0.15362558 | 0.570925073 | Pfkfb1 | 6-phosphofructo-2-kinase/fructose-2,6-bi | 1255.2 | 1138.9 | 1007.3 | 1046.9 | 911.76 | 1051.5 |
| 4967 | ENSMUSG00000025272 | 24.6780039 | -0.919956102 | 0.476634122 | -1.930109618 | 0.053593257 | 0.325702886 | Tro | trophinin [Source:MGI Symbol;Acc:MGI:1! | 28.109 | 23.642 | 45.281 | 20.244 | 18.541 | 12.252 |
| 4968 | ENSMUSG00000025277 | 1003.913624 | -0.335488542 | 0.174226113 | -1.925592769 | 0.054155242 | 0.327604109 | Abhd6 | abhydrolase domain containing 6 [Source | 974.16 | 1190.3 | 1197.4 | 1121.1 | 888.85 | 651.6 |
| 4969 | ENSMUSG00000025278 | 1630.083646 | -0.143289696 | 0.155090298 | -0.923911409 | 0.355532442 | 0.795424787 | Flnb | filamin, beta [Source:MGI Symbol;Acc:MG | 1632.7 | 1985.9 | 1515.4 | 1807.5 | 1630.5 | 1208.5 |
| 4970 | ENSMUSG00000025279 | 334.5759931 | -0.70217444 | 0.183574008 | -3.825021028 | 0.000130761 | 0.004450179 | Dnase1l3 | deoxyribonuclease 1-like 3 [Source:MGI S | 394.32 | 414.25 | 434.7 | 252.56 | 191.95 | 319.67 |
| 4971 | ENSMUSG00000025280 | 340.9523605 | 0.326506953 | 0.163144622 | 2.001334449 | 0.04535636 | 0.294582122 | Polr3a | polymerase (RNA) III (DNA directed) polyp | 314.82 | 310.43 | 282.75 | 448.25 | 366.45 | 323.01 |
| 4972 | ENSMUSG00000025283 | 1078.164479 | -0.121012538 | 0.126161985 | -0.95918385 | 0.337466135 | 0.779481025 | Sat1 | spermidine/spermine N1-acetyl transfera: | 1192.6 | 1225.3 | 951.91 | 1054.6 | 1065.5 | 979.07 |
| 4973 | ENSMUSG00000025287 | 49.25686451 | -0.241105737 | 0.347998491 | -0.692835583 | 0.488412739 | 0.874431125 | Acot9 | acyl-CoA thioesterase 9 [Source:MGI Syml | 73.082 | 48.312 | 38.237 | 44.343 | 41.443 | 50.123 |
| 4974 | ENSMUSG00000025288 | 0 | NA | NA | NA | NA | NA | 4933436I01F | RIKEN cDNA 4933436I01 gene [Source:MC | 0 | 0 | 0 | 0 | 0 | 0 |
| 4975 | ENSMUSG00000025289 | 1259.713812 | 0.261611033 | 0.117150924 | 2.233111129 | 0.025541617 | 0.205927507 | Prdx4 | peroxiredoxin 4 [Source:MGI Symbol;Acc: | 1089.8 | 1085.5 | 1262.8 | 1434.4 | 1242.2 | 1443.5 |
| 4976 | ENSMUSG00000025290 | 3003.827165 | 0.056370173 | 0.098847334 | 0.570275097 | 0.568491129 | 0.903527882 | Rps24 | ribosomal protein S24 [Source:MGI Symbo | 2757 | 3104.3 | 2974.5 | 2864 | 3091.9 | 3231.3 |
| 4977 | ENSMUSG00000025314 | 891.7671541 | -0.076386097 | 0.118195869 | -0.646267064 | 0.518106414 | 0.887376101 | Ptprj | protein tyrosine phosphatase, receptor ty | 929.99 | 994 | 822.1 | 880.11 | 828.87 | 895.53 |
| 4978 | ENSMUSG00000025316 | 193.0433132 | -0.03715135 | 0.226710717 | -0.163871168 | 0.869832565 | 0.978765244 | Banp | BTG3 associated nuclear protein [Source:I | 192.74 | 211.75 | 182.13 | 178.34 | 256.3 | 137 |
| 4979 | ENSMUSG00000025317 | 1673.01347 | 0.07532027 | 0.116116528 | 0.648661064 | 0.516557477 | 0.886716042 | Car5a | carbonic anhydrase 5a, mitochondrial [So | 1688.9 | 1479.2 | 1718.7 | 1524 | 1802.8 | 1824.5 |
| 4980 | ENSMUSG00000025318 | 0 | NA | NA | NA | NA | NA | Jph3 | junctophilin 3 [Source:MGI Symbol;Acc:M | 0 | 0 | 0 | 0 | 0 | 0 |

| | | | | | | | | | | | | | | | |
|---|---|---|---|---|---|---|---|---|---|---|---|---|---|---|---|
| 4981 | ENSMUSG00000025321 | 6.705045884 | -1.915238031 | 0.929083015 | -2.06142831 | 0.039262196 | 0.271078874 | Itgb8 | integrin beta 8 [Source:MGI Symbol;Acc:N | 14.456 | 8.2233 | 9.0562 | 1.928 | 5.4531 | 1.1138 |
| 4982 | ENSMUSG00000025323 | 89.72102933 | 0.644966344 | 0.246448854 | 2.617039335 | 0.008869612 | 0.099142247 | Sp4 | trans-acting transcription factor 4 [Source | 61.036 | 65.787 | 83.518 | 110.86 | 107.97 | 109.16 |
| 4983 | ENSMUSG00000025324 | 2.990360748 | 0.205061354 | 1.197182972 | 0.171286561 | 0.863998447 | NA | Atp10a | ATPase, class V, type 10A [Source:MGI Syr | 3.2124 | 2.0558 | 3.0187 | 1.928 | 3.2719 | 4.4554 |
| 4984 | ENSMUSG00000025326 | 782.7863097 | 0.016630989 | 0.128741136 | 0.129181621 | 0.897213939 | 0.982798755 | Ube3a | ubiquitin protein ligase E3A [Source:MGI ! | 798.28 | 824.39 | 711.41 | 702.74 | 876.86 | 783.03 |
| 4985 | ENSMUSG00000025328 | 0 | NA | NA | NA | NA | NA | Padi3 | peptidyl arginine deiminase, type III [Sour | 0 | 0 | 0 | 0 | 0 | 0 |
| 4986 | ENSMUSG00000025329 | 0.335414862 | -1.779047242 | 4.030804179 | -0.441362855 | 0.658950329 | NA | Padi1 | peptidyl arginine deiminase, type I [Sourc | 0 | 0 | 2.0125 | 0 | 0 | 0 |
| 4987 | ENSMUSG00000025330 | 1.514471015 | -1.671285945 | 1.947671797 | -0.858094237 | 0.390840429 | NA | Padi4 | peptidyl arginine deiminase, type IV [Sour | 0.8031 | 3.0838 | 3.0187 | 0 | 2.1812 | 0 |
| 4988 | ENSMUSG00000025332 | 1034.072912 | 0.015450715 | 0.10808535 | 0.142949207 | 0.886330296 | 0.98229112 | Kdm5c | lysine (K)-specific demethylase 5C [Source | 1019.9 | 1048.5 | 1017.3 | 1050.7 | 961.93 | 1106 |
| 4989 | ENSMUSG00000025333 | 0.503095663 | 2.549494177 | 3.348715477 | 0.761334964 | 0.446457022 | NA | Gpr143 | G protein-coupled receptor 143 [Source:N | 0 | 0 | 0 | 1.928 | 1.0906 | 0 |
| 4990 | ENSMUSG00000025337 | 1073.132118 | -0.017542172 | 0.101024289 | -0.173643113 | 0.862145947 | 0.976334255 | Sbds | SBDS ribosome maturation factor [Source | 1050.5 | 1112.2 | 1076.7 | 1060.4 | 1075.3 | 1063.7 |
| 4991 | ENSMUSG00000025340 | 337.7159576 | -0.542958944 | 0.184058935 | -2.949918971 | 0.003178573 | 0.048599236 | Rabgef1 | RAB guanine nucleotide exchange factor ( | 360.59 | 503.68 | 338.1 | 299.8 | 284.65 | 239.48 |
| 4992 | ENSMUSG00000025347 | 9465.236444 | 0.007446896 | 0.099196668 | 0.075072038 | 0.940157395 | 0.988812319 | Mettl7b | methyltransferase like 7B [Source:MGI Syr | 10236 | 9073.4 | 9011.9 | 9932.8 | 9818.8 | 8718.1 |
| 4993 | ENSMUSG00000025348 | 217.7257787 | -0.044187476 | 0.197242887 | -0.224025699 | 0.822737306 | 0.969219124 | Itga7 | integrin alpha 7 [Source:MGI Symbol;Acc: | 183.91 | 215.86 | 263.64 | 180.26 | 241.03 | 221.65 |
| 4994 | ENSMUSG00000025350 | 234.9916931 | 0.115289789 | 0.176707305 | 0.652433636 | 0.514121471 | 0.885004045 | Rdh5 | retinol dehydrogenase 5 [Source:MGI Sym | 229.69 | 227.17 | 219.36 | 200.51 | 275.93 | 257.3 |
| 4995 | ENSMUSG00000025351 | 70.94070843 | -1.404883699 | 0.298400758 | -4.708043339 | 2.50E-06 | 0.000172104 | Cd63 | CD63 antigen [Source:MGI Symbol;Acc:M | 121.27 | 111.02 | 76.475 | 37.595 | 42.534 | 36.757 |
| 4996 | ENSMUSG00000025352 | 3.030538224 | 0.50964535 | 1.222096226 | 0.41702555 | 0.67665972 | NA | Gdf11 | growth differentiation factor 11 [Source:N | 2.4093 | 2.0558 | 3.0187 | 1.928 | 6.5437 | 2.2277 |
| 4997 | ENSMUSG00000025353 | 312.8909035 | 0.449896306 | 0.167738156 | 2.682134571 | 0.007315403 | 0.086502219 | Ormdl2 | ORM1-like 2 (S. cerevisiae) [Source:MGI S | 263.42 | 223.06 | 306.9 | 362.46 | 326.09 | 395.41 |
| 4998 | ENSMUSG00000025354 | 816.6706571 | 0.120810023 | 0.108458754 | 1.11387988 | 0.265330722 | 0.721244797 | Dnajc14 | DnaJ heat shock protein family (Hsp40) m | 779.01 | 785.33 | 782.86 | 815.53 | 850.68 | 886.62 |
| 4999 | ENSMUSG00000025355 | 904.0138003 | -0.125058441 | 0.121011362 | -1.033443794 | 0.301396261 | 0.752246851 | Mmp19 | matrix metallopeptidase 19 [Source:MGI ! | 934.81 | 878.87 | 1015.3 | 788.53 | 945.57 | 861 |
| 5000 | ENSMUSG00000025357 | 227.8133364 | -0.004851118 | 0.173497938 | -0.027960668 | 0.977693521 | 0.997564159 | Dgka | diacylglycerol kinase, alpha [Source:MGI ! | 240.13 | 198.39 | 245.52 | 209.18 | 248.66 | 225 |
| 5001 | ENSMUSG00000025358 | 252.1996721 | 0.196492337 | 0.189596851 | 1.036369202 | 0.300029933 | 0.751581736 | Cdk2 | cyclin-dependent kinase 2 [Source:MGI Sy | 240.93 | 230.25 | 234.45 | 327.75 | 273.75 | 206.06 |
| 5002 | ENSMUSG00000025359 | 7.698311548 | -0.04807491 | 0.792267744 | -0.060680131 | 0.951613956 | 0.990883058 | Pmel | premelanosome protein [Source:MGI Sym | 11.243 | 5.1396 | 7.0437 | 10.604 | 4.3625 | 7.7969 |
| 5003 | ENSMUSG00000025362 | 2262.324847 | 0.031554017 | 0.093136811 | 0.338792115 | 0.734766341 | 0.948109711 | Rps26 | ribosomal protein S26 [Source:MGI Symbo | 2132.2 | 2284 | 2297.3 | 2288.5 | 2215 | 2356.9 |
| 5004 | ENSMUSG00000025364 | 1307.917353 | -0.016147629 | 0.100713707 | -0.160331988 | 0.872619562 | 0.979419285 | Pa2g4 | proliferation-associated 2G4 [Source:MGI | 1265.7 | 1295.2 | 1385.6 | 1330.3 | 1307.7 | 1263.1 |
| 5005 | ENSMUSG00000025366 | 306.4351411 | -0.147689984 | 0.150151375 | -0.983607271 | 0.325308649 | 0.771501263 | Esyt1 | extended synaptotagmin-like protein 1 [S | 349.35 | 316.6 | 299.86 | 294.98 | 273.75 | 304.08 |
| 5006 | ENSMUSG00000025369 | 2236.631715 | 0.041569913 | 0.105664755 | 0.39341323 | 0.6940143 | 0.939133502 | Smarcc2 | SWI/SNF related, matrix associated, actin | 2121 | 2230.6 | 2263 | 2486.1 | 2268.5 | 2050.6 |
| 5007 | ENSMUSG00000025370 | 0.706721132 | -2.881789029 | 2.959064625 | -0.973885127 | 0.330113579 | NA | Cdh9 | cadherin 9 [Source:MGI Symbol;Acc:MGI: | 3.2124 | 1.0279 | 0 | 0 | 0 | 0 |
| 5008 | ENSMUSG00000025371 | 300.7645704 | 0.239882567 | 0.143671948 | 1.669654866 | 0.094987668 | 0.454633235 | Chmp6 | charged multivesicular body protein 6 [So | 280.28 | 268.29 | 278.73 | 324.86 | 327.19 | 325.24 |
| 5009 | ENSMUSG00000025372 | 908.0808722 | -0.871678049 | 0.137794256 | -6.32593895 | 2.52E-10 | 6.39E-08 | Baiap2 | brain-specific angiogenesis inhibitor 1-ass | 1019.9 | 1215 | 1288 | 552.36 | 688.18 | 685.01 |
| 5010 | ENSMUSG00000025373 | 148.1348449 | -0.002615983 | 0.194779903 | -0.013430454 | 0.98928437 | 0.99859176 | Rnf41 | ring finger protein 41 [Source:MGI Symbo | 150.18 | 143.91 | 150.94 | 162.91 | 148.32 | 132.55 |
| 5011 | ENSMUSG00000025374 | 499.7417989 | 0.064605825 | 0.145732004 | 0.443319404 | 0.657534723 | 0.927303282 | Nabp2 | nucleic acid binding protein 2 [Source:MG | 455.36 | 436.87 | 573.56 | 479.1 | 528.95 | 524.62 |
| 5012 | ENSMUSG00000025375 | 683.6286623 | -0.306036854 | 0.208737529 | -1.466132396 | 0.142612221 | 0.552273941 | Aatk | apoptosis-associated tyrosine kinase [Sou | 955.69 | 675.34 | 635.95 | 628.51 | 785.25 | 421.03 |
| 5013 | ENSMUSG00000025377 | 196.6343157 | -0.022233931 | 0.210028335 | -0.105861578 | 0.915692179 | 0.98625062 | Tepsin | TEPSIN, adaptor related protein complex 4 | 184.71 | 158.3 | 251.56 | 188.94 | 220.3 | 175.99 |
| 5014 | ENSMUSG00000025380 | 0 | NA | NA | NA | NA | NA | Fscn2 | fascin actin-bundling protein 2 [Source:M | 0 | 0 | 0 | 0 | 0 | 0 |
| 5015 | ENSMUSG00000025381 | 1424.482183 | 0.224988976 | 0.11398295 | 1.973882735 | 0.048395078 | 0.30607189 | Cnpy2 | canopy FGF signaling regulator 2 [Source:I | 1328.3 | 1361 | 1250.8 | 1383.3 | 1513.8 | 1709.8 |
| 5016 | ENSMUSG00000025383 | 0.670803094 | 1.59186814 | 2.791527899 | 0.570249769 | 0.568508305 | NA | Il23a | interleukin 23, alpha subunit p19 [Source: | 0 | 0 | 1.0062 | 1.928 | 1.0906 | 0 |
| 5017 | ENSMUSG00000025384 | 116.8622506 | 0.261164094 | 0.215657299 | 1.2110144 | 0.225889886 | 0.6725584 | Faap100 | Fanconi anemia core complex associated | 94.766 | 108.96 | 115.72 | 135.92 | 121.06 | 124.75 |
| 5018 | ENSMUSG00000025386 | 12.32837741 | 0.41955605 | 0.635816626 | 0.659869581 | 0.509337526 | 0.882793133 | Pde6g | phosphodiesterase 6G, cGMP-specific, ro | 15.259 | 8.2233 | 8.05 | 19.28 | 10.906 | 12.252 |
| 5019 | ENSMUSG00000025389 | 0 | NA | NA | NA | NA | NA | Mip | major intrinsic protein of lens fiber [Sourc | 0 | 0 | 0 | 0 | 0 | 0 |
| 5020 | ENSMUSG00000025393 | 23126.89613 | 0.024027994 | 0.089590749 | 0.268197272 | 0.788547475 | 0.960872359 | Atp5b | ATP synthase, H+ transporting mitochond | 21244 | 23729 | 23830 | 22360 | 23939 | 23658 |
| 5021 | ENSMUSG00000025395 | 37.36669711 | -0.226090568 | 0.408045382 | -0.554081917 | 0.579522785 | 0.908350165 | Prim1 | DNA primase, p49 subunit [Source:MGI Sy | 50.595 | 24.67 | 45.281 | 31.811 | 26.175 | 45.668 |
| 5022 | ENSMUSG00000025396 | 3683.80562 | -0.285984901 | 0.30116753 | -0.949587433 | 0.342321927 | 0.784060862 | Hsd17b6 | hydroxysteroid (17-beta) dehydrogenase ( | 2793.2 | 3742.6 | 5607.8 | 3319.9 | 3574 | 3065.3 |
| 5023 | ENSMUSG00000025400 | 0 | NA | NA | NA | NA | NA | Tac2 | tachykinin 2 [Source:MGI Symbol;Acc:MG | 0 | 0 | 0 | 0 | 0 | 0 |
| 5024 | ENSMUSG00000025401 | 0 | NA | NA | NA | NA | NA | Myo1a | myosin IA [Source:MGI Symbol;Acc:MGI:1 | 0 | 0 | 0 | 0 | 0 | 0 |
| 5025 | ENSMUSG00000025402 | 264.5085563 | 0.585582677 | 0.220623186 | 2.654220927 | 0.007949176 | 0.091542106 | Nab2 | Ngfi-A binding protein 2 [Source:MGI Sym | 212.82 | 148.02 | 273.7 | 328.72 | 259.57 | 364.23 |
| 5026 | ENSMUSG00000025403 | 5307.659588 | -0.043493612 | 0.088009801 | -0.494190547 | 0.621171615 | 0.918069106 | Shmt2 | serine hydroxymethyltransferase 2 (mitoc | 5437 | 5151.9 | 5573.6 | 5109.1 | 5461.8 | 5112.5 |
| 5027 | ENSMUSG00000025404 | 1523.411911 | -0.047061874 | 0.104748208 | -0.449285717 | 0.65322556 | 0.925756399 | R3hdm2 | R3H domain containing 2 [Source:MGI Syr | 1489 | 1637.5 | 1518.4 | 1405.5 | 1605.4 | 1484.8 |
| 5028 | ENSMUSG00000025405 | 820.4424709 | 0.108830398 | 0.133680764 | 0.814106646 | 0.415583859 | 0.83614452 | Inhbc | inhibin beta-C [Source:MGI Symbol;Acc:M | 762.95 | 847 | 759.71 | 986.15 | 748.16 | 818.68 |
| 5029 | ENSMUSG00000025407 | 1.335900328 | 0.890498008 | 1.988601539 | 0.447801126 | 0.654296731 | NA | Gli1 | GLI-Kruppel family member GLI1 [Source:I | 0.8031 | 0 | 2.0125 | 1.928 | 3.2719 | 0 |
| 5030 | ENSMUSG00000025408 | 90.81715371 | 0.034707085 | 0.251603781 | 0.137943414 | 0.890285139 | 0.982489666 | Ddit3 | DNA-damage inducible transcript 3 [Sourc | 105.21 | 75.038 | 88.55 | 97.362 | 79.615 | 99.132 |
| 5031 | ENSMUSG00000025409 | 540.9785386 | 0.093032933 | 0.120894276 | 0.769539607 | 0.441573041 | 0.849888552 | Mbd6 | methyl-CpG binding domain protein 6 [So | 508.36 | 521.15 | 541.36 | 558.14 | 538.77 | 578.08 |
| 5032 | ENSMUSG00000025410 | 1952.462475 | 0.058934966 | 0.099689811 | 0.591183442 | 0.554397516 | 0.897460063 | Dctn2 | dynactin 2 [Source:MGI Symbol;Acc:MGI: | 2021.4 | 1866.7 | 1848.5 | 1943.4 | 1912.9 | 2121.9 |
| 5033 | ENSMUSG00000025413 | 365.9812117 | -0.018412545 | 0.138391253 | -0.133047027 | 0.894156189 | 0.982681796 | Ttc4 | tetratricopeptide repeat domain 4 [Sourc | 383.88 | 354.63 | 366.27 | 374.02 | 377.35 | 339.72 |
| 5034 | ENSMUSG00000025417 | 560.9933062 | 0.101682013 | 0.121725676 | 0.835337422 | 0.403527753 | 0.828953151 | Pip4k2c | phosphatidylinositol-5-phosphate 4-kinas | 551.73 | 517.04 | 554.44 | 562.96 | 564.94 | 614.84 |
| 5035 | ENSMUSG00000025418 | 0 | NA | NA | NA | NA | NA | Bsnd | barttin CLCNK type accessory beta subuni | 0 | 0 | 0 | 0 | 0 | 0 |
| 5036 | ENSMUSG00000025420 | 0 | NA | NA | NA | NA | NA | Katnal2 | katanin p60 subunit A-like 2 [Source:MGI | 0 | 0 | 0 | 0 | 0 | 0 |

| | | | | | | | | | | | | | | | |
|---|---|---|---|---|---|---|---|---|---|---|---|---|---|---|---|
| 5037 | ENSMUSG00000025421 | 512.5921014 | -0.172537317 | 0.173310323 | -0.995539753 | 0.31947382 | 0.766337562 | Hdhd2 | haloacid dehalogenase-like hydrolase don | 593.49 | 463.59 | 571.55 | 397.16 | 603.11 | 446.65 |
| 5038 | ENSMUSG00000025422 | 80.44684972 | -0.218601993 | 0.308249782 | -0.709171606 | 0.478217991 | 0.869794311 | Agap2 | ArfGAP with GTPase domain, ankyrin repe | 111.63 | 60.647 | 86.537 | 53.983 | 77.434 | 92.449 |
| 5039 | ENSMUSG00000025423 | 534.6178406 | 0.077615611 | 0.137013402 | 0.566481891 | 0.57106624 | 0.904651508 | Pias2 | protein inhibitor of activated STAT 2 [Sour | 475.44 | 596.19 | 490.04 | 562 | 562.76 | 521.28 |
| 5040 | ENSMUSG00000025425 | 0 | NA | NA | NA | NA | NA | St8sia5 | ST8 alpha-N-acetyl-neuraminide alpha-2,8 | 0 | 0 | 0 | 0 | 0 | 0 |
| 5041 | ENSMUSG00000025427 | 1.748521822 | 1.334906653 | 1.674037031 | 0.797417637 | 0.42520852 | NA | Rnf165 | ring finger protein 165 [Source:MGI Symb | 0 | 1.0279 | 2.0125 | 1.928 | 2.1812 | 3.3415 |
| 5042 | ENSMUSG00000025428 | 20926.18255 | -0.048051921 | 0.08879634 | -0.541147539 | 0.588405893 | 0.910917568 | Atp5a1 | ATP synthase, H+ transporting, mitochonc | 20649 | 22508 | 20667 | 19742 | 21763 | 20227 |
| 5043 | ENSMUSG00000025429 | 112.0139302 | 0.671868298 | 0.269126277 | 2.496479742 | 0.012543283 | 0.126529431 | Pstpip2 | proline-serine-threonine phosphatase-int | 115.65 | 80.178 | 62.387 | 116.64 | 164.68 | 132.55 |
| 5044 | ENSMUSG00000025431 | 0 | NA | NA | NA | NA | NA | Crisp1 | cysteine-rich secretory protein 1 [Source:I | 0 | 0 | 0 | 0 | 0 | 0 |
| 5045 | ENSMUSG00000025432 | 1.922781766 | 4.475799665 | 1.901692444 | 2.353587553 | 0.018593229 | NA | Avil | advillin [Source:MGI Symbol;Acc:MGI:133 | 0 | 0 | 0 | 3.8559 | 5.4531 | 2.2277 |
| 5046 | ENSMUSG00000025433 | 0 | NA | NA | NA | NA | NA | Crisp3 | cysteine-rich secretory protein 3 [Source:I | 0 | 0 | 0 | 0 | 0 | 0 |
| 5047 | ENSMUSG00000025436 | 125.4288262 | 0.334752829 | 0.223291541 | 1.499173803 | 0.133828549 | 0.53673534 | Atp23 | ATP23 metallopeptidase and ATP synthase | 99.585 | 113.07 | 120.75 | 156.16 | 150.51 | 112.5 |
| 5048 | ENSMUSG00000025437 | 405.672417 | 0.176386225 | 0.168598803 | 1.04618907 | 0.295473748 | 0.745982248 | Usp33 | ubiquitin specific peptidase 33 [Source:M | 365.41 | 350.52 | 427.65 | 519.58 | 414.43 | 356.43 |
| 5049 | ENSMUSG00000025439 | 476.5513824 | 0.08745484 | 0.130004616 | 0.672705654 | 0.50113457 | 0.880974509 | Clns1a | chloride channel, nucleotide-sensitive, 1A | 481.86 | 437.89 | 465.89 | 472.35 | 472.24 | 529.08 |
| 5050 | ENSMUSG00000025450 | 21.00011413 | 0.254625122 | 0.538018706 | 0.473264441 | 0.636024528 | 0.923478764 | Gm9752 | predicted gene 9752 [Source:MGI Symbol | 24.896 | 18.503 | 14.087 | 36.631 | 19.631 | 12.252 |
| 5051 | ENSMUSG00000025451 | 1163.715415 | -0.051978463 | 0.103114778 | -0.504083544 | 0.61420267 | 0.917466428 | Paip1 | polyadenylate binding protein-interacting | 1165.3 | 1257.1 | 1132 | 1157.7 | 1147.3 | 1122.8 |
| 5052 | ENSMUSG00000025453 | 18.78661848 | 0.40980625 | 0.49442204 | 0.828859187 | 0.407184092 | 0.83094725 | Nnt | nicotinamide nucleotide transhydrogenas | 16.865 | 13.363 | 18.112 | 20.244 | 19.631 | 24.505 |
| 5053 | ENSMUSG00000025461 | 0.462220778 | -0.685400455 | 3.191475091 | -0.21475977 | 0.82995462 | NA | Scart1 | scavenger receptor family member expres | 0.8031 | 0 | 1.0062 | 0.964 | 0 | 0 |
| 5054 | ENSMUSG00000025464 | 647.464596 | 0.628326045 | 0.178856482 | 3.513018029 | 0.000443047 | 0.011342014 | Paox | polyamine oxidase (exo-N4-amino) [Sourc | 607.95 | 514.99 | 401.49 | 615.98 | 884.49 | 859.89 |
| 5055 | ENSMUSG00000025465 | 5456.855094 | 0.032782556 | 0.094585104 | 0.346593221 | 0.728896939 | 0.946303324 | Echs1 | enoyl Coenzyme A hydratase, short chain, | 5457.9 | 5599.1 | 5126.8 | 5134.1 | 5875.2 | 5548.1 |
| 5056 | ENSMUSG00000025466 | 414.2037898 | 0.170260332 | 0.145176701 | 1.172780003 | 0.240884039 | 0.691298957 | Fuom | fucose mutarotase [Source:MGI Symbol;A | 371.03 | 390.61 | 407.53 | 388.48 | 430.79 | 496.77 |
| 5057 | ENSMUSG00000025467 | 0.688762941 | -1.399623551 | 3.067774249 | -0.456234207 | 0.64822157 | NA | Prap1 | proline-rich acidic protein 1 [Source:MGI S | 0 | 0 | 3.0187 | 0 | 0 | 1.1138 |
| 5058 | ENSMUSG00000025468 | 0.642651897 | 2.906894826 | 3.440535777 | 0.844895974 | 0.398168923 | NA | Caly | calcyon neuron-specific vesicular protein | 0 | 0 | 0 | 3.8559 | 0 | 0 |
| 5059 | ENSMUSG00000025469 | 0.160662974 | 1.126924175 | 4.080472857 | 0.276174898 | 0.782413736 | NA | Msx3 | msh homeobox 3 [Source:MGI Symbol;Ac | 0 | 0 | 0 | 0.964 | 0 | 0 |
| 5060 | ENSMUSG00000025470 | 101.8219014 | 0.08076985 | 0.233829356 | 0.345422197 | 0.729776992 | 0.946534338 | Zfp511 | zinc finger protein 511 [Source:MGI Symb | 101.99 | 85.317 | 109.68 | 115.68 | 104.7 | 93.563 |
| 5061 | ENSMUSG00000025473 | 5.280739476 | 0.01273148 | 0.986185485 | 0.012909823 | 0.989699738 | 0.99859176 | Adam8 | a disintegrin and metallopeptidase domai | 1.6062 | 9.2513 | 5.0312 | 4.8199 | 7.6343 | 3.3415 |
| 5062 | ENSMUSG00000025474 | 164.6524424 | -0.018912229 | 0.199301155 | -0.094892722 | 0.924400038 | 0.987546253 | Tubgcp2 | tubulin, gamma complex associated prote | 191.14 | 165.49 | 139.87 | 153.27 | 178.86 | 159.28 |
| 5063 | ENSMUSG00000025475 | 0.545309144 | 2.651974629 | 3.666219369 | 0.72335405 | 0.469462394 | NA | Adgra1 | adhesion G protein-coupled receptor A1 [ | 0 | 0 | 0 | 0 | 3.2719 | 0 |
| 5064 | ENSMUSG00000025477 | 262.6724065 | 0.030073823 | 0.184025697 | 0.163421867 | 0.870186287 | 0.978802475 | Inpp5a | inositol polyphosphate-5-phosphatase A [ | 253.78 | 272.4 | 253.57 | 258.35 | 211.58 | 326.36 |
| 5065 | ENSMUSG00000025478 | 1.777220956 | -0.71010784 | 1.634042197 | -0.434571299 | 0.663873632 | NA | Dpysl4 | dihydropyrimidinase-like 4 [Source:MGI S | 1.6062 | 2.0558 | 3.0187 | 2.8919 | 1.0906 | 0 |
| 5066 | ENSMUSG00000025479 | 175259.32 | -0.055105057 | 0.102969965 | -0.535156608 | 0.592541569 | 0.912923013 | Cyp2e1 | cytochrome P450, family 2, subfamily e, p | 170141 | 197225 | 168453 | 177011 | 182022 | 156703 |
| 5067 | ENSMUSG00000025480 | 1.477544172 | -3.939977216 | 2.089232266 | -1.885849305 | 0.059315265 | NA | Syce1 | synaptonemal complex central element pr | 4.8186 | 1.0279 | 3.0187 | 0 | 0 | 0 |
| 5068 | ENSMUSG00000025481 | 3336.564592 | 0.179359376 | 0.094569202 | 1.896593952 | 0.057881549 | 0.342132706 | Urah | urate (5-hydroxyiso-) hydrolase [Source:M | 2916.9 | 3121.8 | 3350.8 | 3542.6 | 3612.1 | 3475.2 |
| 5069 | ENSMUSG00000025482 | 0.181769715 | 1.126924175 | 4.080472857 | 0.276174898 | 0.782413736 | NA | Odf3 | outer dense fiber of sperm tails 3 [Source: | 0 | 0 | 0 | 0 | 1.0906 | 0 |
| 5070 | ENSMUSG00000025484 | 373.5029407 | 0.286703186 | 0.149474108 | 1.918079257 | 0.055100962 | 0.330834503 | Bet1l | Bet1 golgi vesicular membrane trafficking | 377.46 | 289.87 | 341.12 | 398.12 | 410.07 | 424.37 |
| 5071 | ENSMUSG00000025485 | 446.6180595 | 0.114920779 | 0.142604248 | 0.805872053 | 0.420316627 | 0.837906635 | Ric8a | RIC8 guanine nucleotide exchange factor A | 387.9 | 408.08 | 491.05 | 451.14 | 454.79 | 486.75 |
| 5072 | ENSMUSG00000025486 | 1849.306162 | 0.147654644 | 0.10907536 | 1.35369385 | 0.175834071 | 0.606476483 | Sirt3 | sirtuin 3 [Source:MGI Symbol;Acc:MGI:19 | 1674.5 | 1709.4 | 1881.7 | 2128.5 | 1911.9 | 1789.9 |
| 5073 | ENSMUSG00000025487 | 1315.993522 | 0.03521878 | 0.101316352 | 0.347611999 | 0.728131594 | 0.946158959 | Psmd13 | proteasome (prosome, macropain) 26S su | 1362.9 | 1297.2 | 1238.7 | 1284 | 1367.6 | 1345.5 |
| 5074 | ENSMUSG00000025488 | 0 | NA | NA | NA | NA | NA | Cox8b | cytochrome c oxidase subunit 8B [Source: | 0 | 0 | 0 | 0 | 0 | 0 |
| 5075 | ENSMUSG00000025489 | 1.277530911 | -0.600203982 | 1.867920928 | -0.321321943 | 0.747966429 | NA | Ifitm5 | interferon induced transmembrane prote | 1.6062 | 1.0279 | 2.0125 | 1.928 | 1.0906 | 0 |
| 5076 | ENSMUSG00000025491 | 74.98769119 | -1.055184467 | 0.292727862 | -3.604660177 | 0.000312562 | 0.009023735 | Ifitm1 | interferon induced transmembrane prote | 118.86 | 105.88 | 78.487 | 37.595 | 54.531 | 54.578 |
| 5077 | ENSMUSG00000025492 | 5753.62655 | 0.048599675 | 0.103234904 | 0.47076786 | 0.637806518 | 0.923478764 | Ifitm3 | interferon induced transmembrane prote | 5951 | 5074.8 | 5942.9 | 5460.9 | 5899.2 | 6193 |
| 5078 | ENSMUSG00000025494 | 308.9609156 | -0.05667824 | 0.169939445 | -0.333520211 | 0.738741636 | 0.948190525 | Sigirr | single immunoglobulin and toll-interleukin | 324.45 | 283.71 | 337.09 | 341.25 | 246.48 | 320.79 |
| 5079 | ENSMUSG00000025495 | 249.2428011 | 0.230526664 | 0.167118952 | 1.379416646 | 0.16776633 | 0.593804353 | Ptdss2 | phosphatidylserine synthase 2 [Source:M | 237.72 | 241.56 | 208.29 | 237.14 | 292.29 | 278.46 |
| 5080 | ENSMUSG00000025496 | 0 | NA | NA | NA | NA | NA | Drd4 | dopamine receptor D4 [Source:MGI Symb | 0 | 0 | 0 | 0 | 0 | 0 |
| 5081 | ENSMUSG00000025497 | 932.8640536 | 0.08283638 | 0.121741065 | 0.680430884 | 0.496231671 | 0.878165817 | Cdhr5 | cadherin-related family member 5 [Source | 825.59 | 929.24 | 963.98 | 892.64 | 1057.9 | 927.83 |
| 5082 | ENSMUSG00000025498 | 630.4204565 | -0.059702038 | 0.130872205 | -0.456185772 | 0.648256396 | 0.925643431 | Irf7 | interferon regulatory factor 7 [Source:MG | 689.86 | 605.44 | 633.93 | 572.6 | 586.75 | 693.92 |
| 5083 | ENSMUSG00000025499 | 751.7521433 | 0.27782953 | 0.113248265 | 2.453278459 | 0.014156071 | 0.137510684 | Hras | Harvey rat sarcoma virus oncogene [Sourc | 645.69 | 671.23 | 722.48 | 831.91 | 802.7 | 836.5 |
| 5084 | ENSMUSG00000025500 | 18.71550733 | 0.410062935 | 0.585056807 | 0.700894221 | 0.483369031 | 0.871960882 | Lmntd2 | lamin tail domain containing 2 [Source:M | 16.865 | 5.1396 | 26.162 | 22.171 | 17.45 | 24.505 |
| 5085 | ENSMUSG00000025503 | 2324.705343 | -0.016549178 | 0.126832431 | -0.130480647 | 0.896186165 | 0.982798755 | Taldo1 | transaldolase 1 [Source:MGI Symbol;Acc:I | 2766.7 | 2265.5 | 1980.3 | 2315.5 | 2352.5 | 2267.8 |
| 5086 | ENSMUSG00000025504 | 492.3599004 | 0.086153936 | 0.188941626 | 0.455981766 | 0.64840309 | 0.925643431 | Eps8l2 | EPS8-like 2 [Source:MGI Symbol;Acc:MGI: | 403.16 | 508.82 | 521.23 | 441.5 | 412.25 | 667.19 |
| 5087 | ENSMUSG00000025505 | 183.2234309 | 0.077559006 | 0.183712135 | 0.422176825 | 0.672895956 | 0.931454408 | Tmem80 | transmembrane protein 80 [Source:MGI S | 196.76 | 162.41 | 175.09 | 181.23 | 176.68 | 207.17 |
| 5088 | ENSMUSG00000025507 | 46.93025027 | 0.192567489 | 0.34114326 | 0.564476898 | 0.572429618 | 0.905173631 | Pidd1 | p53 induced death domain protein 1 [Sou | 32.124 | 46.256 | 53.331 | 47.235 | 44.715 | 57.92 |
| 5089 | ENSMUSG00000025508 | 1797.122812 | -0.014264254 | 0.098229703 | -0.145213245 | 0.884542509 | 0.981795936 | Rplp2 | ribosomal protein, large P2 [Source:MGI S | 1699.4 | 1878 | 1841.4 | 1770.8 | 1733 | 1860.1 |
| 5090 | ENSMUSG00000025509 | 1781.258887 | -0.024382228 | 0.133147753 | -0.183121586 | 0.854702619 | 0.97395562 | Pnpla2 | patatin-like phospholipase domain contai | 2134.6 | 1743.3 | 1509.4 | 1767 | 1668.6 | 1864.6 |
| 5091 | ENSMUSG00000025510 | 818.4991393 | 0.016520932 | 0.120572014 | 0.137021284 | 0.891013972 | 0.982489666 | Cd151 | CD151 antigen [Source:MGI Symbol;Acc:M | 897.87 | 790.47 | 751.66 | 778.89 | 828.87 | 863.23 |
| 5092 | ENSMUSG00000025511 | 1213.111157 | -0.087241197 | 0.199838328 | -0.43655888 | 0.66243129 | 0.9288455 | Tspan4 | tetraspanin 4 [Source:MGI Symbol;Acc:M | 1082.6 | 1310.6 | 1357.4 | 1504.8 | 1281.5 | 741.82 |

| | | | | | | | | | | | | | | | |
|---|---|---|---|---|---|---|---|---|---|---|---|---|---|---|---|
| 5093 | ENSMUSG00000025512 | 480.8761189 | 0.362196516 | 0.138966324 | 2.606361791 | 0.009150975 | 0.101502835 | Chid1 | chitinase domain containing 1 [Source:MG | 419.22 | 406.03 | 436.71 | 483.92 | 526.77 | 612.61 |
| 5094 | ENSMUSG00000025515 | 1.381059326 | -0.664873319 | 1.897391789 | -0.350414355 | 0.726027755 | NA | Muc2 | mucin 2 [Source:MGI Symbol;Acc:MGI:13 | 0 | 4.1117 | 1.0062 | 0.964 | 1.0906 | 1.1138 |
| 5095 | ENSMUSG00000025519 | 0 | NA | NA | NA | NA | NA | Tktl2 | transketolase-like 2 [Source:MGI Symbol;/ | 0 | 0 | 0 | 0 | 0 | 0 |
| 5096 | ENSMUSG00000025521 | 166.9119068 | 0.140555483 | 0.202932717 | 0.692621103 | 0.488547363 | 0.874431125 | Tmem192 | transmembrane protein 192 [Source:MGI | 178.29 | 167.55 | 129.81 | 153.27 | 184.31 | 188.24 |
| 5097 | ENSMUSG00000025525 | 485.223442 | 0.123867999 | 0.135650251 | 0.913142424 | 0.361167641 | 0.800074801 | Apool | apolipoprotein O-like [Source:MGI Symbo | 505.15 | 449.2 | 437.72 | 462.71 | 545.31 | 511.25 |
| 5098 | ENSMUSG00000025527 | 0 | NA | NA | NA | NA | NA | Satl1 | spermidine/spermine N1-acetyl transfera: | 0 | 0 | 0 | 0 | 0 | 0 |
| 5099 | ENSMUSG00000025528 | 0 | NA | NA | NA | NA | NA | 2010106E10 | RIKEN cDNA 2010106E10 gene [Source:M | 0 | 0 | 0 | 0 | 0 | 0 |
| 5100 | ENSMUSG00000025529 | 0 | NA | NA | NA | NA | NA | Zfp711 | zinc finger protein 711 [Source:MGI Symb | 0 | 0 | 0 | 0 | 0 | 0 |
| 5101 | ENSMUSG00000025531 | 284.9957879 | -0.087649813 | 0.153395234 | -0.571398539 | 0.567729519 | 0.903352921 | Chm | choroidermia (RAB escort protein 1) [Sour | 318.03 | 270.34 | 291.81 | 262.2 | 283.56 | 284.03 |
| 5102 | ENSMUSG00000025532 | 657.0887035 | -0.290766204 | 0.136089582 | -2.136579455 | 0.032632212 | 0.241304865 | Crcp | calcitonin gene-related peptide-receptor | 727.61 | 758.6 | 682.23 | 517.66 | 574.76 | 681.67 |
| 5103 | ENSMUSG00000025533 | 16606.0157 | -0.513350354 | 0.099331962 | -5.168027909 | 2.37E-07 | 2.33E-05 | Asl | argininosuccinate lyase [Source:MGI Symb | 17726 | 19802 | 21063 | 14139 | 14136 | 12771 |
| 5104 | ENSMUSG00000025534 | 511.4839165 | -0.307439319 | 0.125903887 | -2.441857253 | 0.014611923 | 0.140429803 | Gusb | glucuronidase, beta [Source:MGI Symbol; | 565.38 | 570.49 | 561.48 | 453.07 | 491.87 | 426.6 |
| 5105 | ENSMUSG00000025537 | 4.080260211 | -0.09003708 | 1.103197794 | -0.08161463 | 0.934953167 | NA | Phkg1 | phosphorylase kinase gamma 1 [Source:M | 2.4093 | 8.2233 | 2.0125 | 1.928 | 5.4531 | 4.4554 |
| 5106 | ENSMUSG00000025538 | 139.4678292 | 0.074853462 | 0.221128398 | 0.338506781 | 0.734981318 | 0.948109711 | Sumf2 | sulfatase modifying factor 2 [Source:MGI | 119.66 | 145.96 | 141.88 | 112.79 | 148.32 | 168.19 |
| 5107 | ENSMUSG00000025544 | 3194.167882 | -0.045386363 | 0.107880806 | -0.420708418 | 0.673968013 | 0.932009833 | Tm9sf2 | transmembrane 9 superfamily member 2 | 3125.7 | 3335.6 | 3273.3 | 3435.6 | 3238 | 2756.8 |
| 5108 | ENSMUSG00000025545 | 775.1689334 | -0.192250196 | 0.111435875 | -1.72520919 | 0.08448978 | 0.428642644 | Clybl | citrate lyase beta like [Source:MGI Symbo | 863.33 | 828.5 | 787.89 | 719.13 | 745.98 | 706.18 |
| 5109 | ENSMUSG00000025551 | 0.301557804 | -1.64335916 | 4.042814367 | -0.406488899 | 0.684383412 | NA | Fgf14 | fibroblast growth factor 14 [Source:MGI S | 0.8031 | 0 | 1.0062 | 0 | 0 | 0 |
| 5110 | ENSMUSG00000025555 | 338.4664952 | 0.211510384 | 0.16144 | 1.310148562 | 0.190145583 | 0.627627958 | Farp1 | FERM, RhoGEF (Arhgef) and pleckstrin do | 289.92 | 340.24 | 310.93 | 306.54 | 412.25 | 370.91 |
| 5111 | ENSMUSG00000025557 | 0.524408994 | -0.837196744 | 3.377823722 | -0.247850928 | 0.804249749 | NA | Slc15a1 | solute carrier family 15 (oligopeptide tran | 0 | 2.0558 | 0 | 0 | 1.0906 | 0 |
| 5112 | ENSMUSG00000025558 | 347.5639357 | -0.148756054 | 0.193359008 | -0.769325698 | 0.441699985 | 0.849906689 | Dock9 | dedicator of cytokinesis 9 [Source:MGI Sy | 304.38 | 417.33 | 375.33 | 313.29 | 415.53 | 259.53 |
| 5113 | ENSMUSG00000025571 | 429.9186089 | 0.022501842 | 0.138061799 | 0.16298384 | 0.870531158 | 0.978802475 | Tnrc6c | trinucleotide repeat containing 6C [Source | 380.67 | 449.2 | 450.8 | 444.39 | 412.25 | 442.2 |
| 5114 | ENSMUSG00000025572 | 202.9506331 | -0.256607167 | 0.186862799 | -1.373238379 | 0.169678252 | 0.596876778 | Tmc6 | transmembrane channel-like gene family | 231.29 | 196.33 | 235.46 | 214.97 | 160.32 | 179.33 |
| 5115 | ENSMUSG00000025573 | 0.321325949 | 1.911575114 | 4.048267893 | 0.472195804 | 0.636787034 | NA | 6030468B19 | RIKEN cDNA 6030468B19 gene [Source:M | 0 | 0 | 0 | 1.928 | 0 | 0 |
| 5116 | ENSMUSG00000025574 | 689.4116469 | 0.219568432 | 0.225181535 | 0.975072992 | 0.329524055 | 0.774441615 | Tk1 | thymidine kinase 1 [Source:MGI Symbol;A | 428.05 | 586.94 | 897.57 | 926.38 | 684.91 | 612.61 |
| 5117 | ENSMUSG00000025575 | 614.7893831 | -0.009349268 | 0.124537543 | -0.075071883 | 0.940157519 | 0.988812319 | Cant1 | calcium activated nucleotidase 1 [Source: | 644.09 | 621.89 | 583.62 | 573.57 | 668.55 | 597.02 |
| 5118 | ENSMUSG00000025576 | 7.61242608 | -1.553579408 | 0.842636071 | -1.843713391 | 0.06522492 | 0.366785337 | Rbfox3 | RNA binding protein, fox-1 homolog (C. el | 17.668 | 8.2233 | 8.05 | 2.8919 | 3.2719 | 5.5692 |
| 5119 | ENSMUSG00000025577 | 192.1704612 | 0.369199955 | 0.19113816 | 1.931586837 | 0.053410522 | 0.325002014 | Cbx2 | chromobox 2 [Source:MGI Symbol;Acc:M | 183.91 | 133.63 | 185.15 | 223.64 | 205.04 | 221.65 |
| 5120 | ENSMUSG00000025578 | 147.6479966 | -0.112882213 | 0.19658487 | -0.574216183 | 0.56582152 | 0.901973059 | Cbx8 | chromobox 8 [Source:MGI Symbol;Acc:M | 151.79 | 169.61 | 138.86 | 135.92 | 151.6 | 138.12 |
| 5121 | ENSMUSG00000025579 | 1391.133037 | -0.042916906 | 0.112289445 | -0.382198933 | 0.702313813 | 0.942345172 | Gaa | glucosidase, alpha, acid [Source:MGI Sym | 1523.5 | 1458.6 | 1252.8 | 1419 | 1322.9 | 1370 |
| 5122 | ENSMUSG00000025580 | 653.3352265 | -0.056847191 | 0.117495813 | -0.48382312 | 0.628511405 | 0.920328819 | Eif4a3 | eukaryotic translation initiation factor 4A3 | 619.19 | 698.98 | 681.23 | 641.05 | 639.1 | 640.46 |
| 5123 | ENSMUSG00000025582 | 2.662249066 | 0.237181228 | 1.329275619 | 0.178428931 | 0.85838613 | NA | Nptx1 | neuronal pentraxin 1 [Source:MGI Symbol | 3.2124 | 1.0279 | 3.0187 | 0.964 | 2.1812 | 5.5692 |
| 5124 | ENSMUSG00000025583 | 777.0146859 | 0.025926369 | 0.114151035 | 0.227123381 | 0.820327804 | 0.969160361 | Rptor | regulatory associated protein of MTOR, co | 748.49 | 784.3 | 777.83 | 795.28 | 829.96 | 726.23 |
| 5125 | ENSMUSG00000025584 | 585.7988012 | 0.130786381 | 0.121000061 | 1.08087863 | 0.279751106 | 0.732834426 | Pde8a | phosphodiesterase 8A [Source:MGI Symb | 569.4 | 589 | 519.22 | 606.34 | 626.01 | 604.82 |
| 5126 | ENSMUSG00000025586 | 98.05775188 | 0.754235322 | 0.300094633 | 2.513324932 | 0.011959912 | 0.122256893 | Cpeb1 | cytoplasmic polyadenylation element binc | 91.554 | 70.926 | 56.35 | 174.48 | 99.246 | 95.791 |
| 5127 | ENSMUSG00000025588 | 42.88196271 | 0.374224789 | 0.350635846 | 1.067274762 | 0.285847781 | 0.738421977 | Nat1 | N-acetyl transferase 1 [Source:MGI Symbo | 42.564 | 41.117 | 28.175 | 45.307 | 57.803 | 42.326 |
| 5128 | ENSMUSG00000025591 | 97.44877032 | -0.134466574 | 0.23749744 | -0.566181153 | 0.571270641 | 0.904651508 | Tma16 | translation machinery associated 16 [Sour | 99.585 | 113.07 | 93.581 | 101.22 | 98.156 | 79.083 |
| 5129 | ENSMUSG00000025592 | 0.535401493 | -2.48596965 | 3.684117979 | -0.674780141 | 0.499815459 | NA | Dach2 | dachshund family transcription factor 2 [S | 3.2124 | 0 | 0 | 0 | 0 | 0 |
| 5130 | ENSMUSG00000025597 | 8.978027946 | -0.379401215 | 0.724867568 | -0.523407629 | 0.600690616 | 0.915429015 | Klhl4 | kelch-like 4 [Source:MGI Symbol;Acc:MGI | 8.031 | 14.391 | 8.05 | 5.7839 | 9.8156 | 7.7969 |
| 5131 | ENSMUSG00000025602 | 11.36504406 | -0.160127873 | 0.705460174 | -0.226983576 | 0.820436513 | 0.969160361 | Zfp202 | zinc finger protein 202 [Source:MGI Symb | 9.6372 | 6.1675 | 20.125 | 5.7839 | 11.997 | 14.48 |
| 5132 | ENSMUSG00000025607 | 520.1729226 | 0.049608115 | 0.123810744 | 0.400676981 | 0.688657961 | 0.936908509 | Copg2 | coatomer protein complex, subunit gamm | 485.88 | 506.76 | 541.36 | 514.76 | 539.86 | 532.42 |
| 5133 | ENSMUSG00000025608 | 54.20153141 | -0.81604342 | 0.327913407 | -2.488594255 | 0.012824924 | 0.128107525 | Podxl | podocalyxin-like [Source:MGI Symbol;Acc | 57.02 | 81.206 | 69.431 | 42.415 | 28.356 | 46.781 |
| 5134 | ENSMUSG00000025609 | 1153.931974 | -0.012909907 | 0.112895372 | -0.114352847 | 0.908958092 | 0.985098284 | Mkln1 | muskelin 1, intracellular mediator contain | 1154.9 | 1226.3 | 1096.8 | 1225.2 | 1179 | 1041.4 |
| 5135 | ENSMUSG00000025610 | 0.181769715 | 1.126924175 | 4.080472857 | 0.276174898 | 0.782413736 | NA | Map3k7cl | Map3k7 C-terminal like [Source:MGI Symb | 0 | 0 | 0 | 0 | 1.0906 | 0 |
| 5136 | ENSMUSG00000025612 | 866.1937753 | 0.540430574 | 0.13280662 | 4.069304482 | 4.72E-05 | 0.002011891 | Bach1 | BTB and CNC homology 1, basic leucine zi | 737.25 | 715.43 | 665.13 | 1182.8 | 1008.8 | 887.73 |
| 5137 | ENSMUSG00000025613 | 2569.671736 | -0.036584003 | 0.090680504 | -0.403438468 | 0.686625693 | 0.936566769 | Cct8 | chaperonin containing Tcp1, subunit 8 (th | 2596.4 | 2685.9 | 2524.7 | 2566.1 | 2602.2 | 2442.7 |
| 5138 | ENSMUSG00000025616 | 557.8650972 | -0.065549286 | 0.125850221 | -0.520851579 | 0.602470169 | 0.915541976 | Usp16 | ubiquitin specific peptidase 16 [Source:M | 566.99 | 583.86 | 560.48 | 508.02 | 597.66 | 530.19 |
| 5139 | ENSMUSG00000025626 | 162.9981979 | -0.08817486 | 0.188659969 | -0.46737451 | 0.640231957 | 0.923932929 | Phf6 | PHD finger protein 6 [Source:MGI Symbol | 165.44 | 159.33 | 179.11 | 150.38 | 175.59 | 148.14 |
| 5140 | ENSMUSG00000025630 | 1104.461401 | -0.180814828 | 0.125674696 | -1.438752857 | 0.150220558 | 0.563917266 | Hprt | hypoxanthine guanine phosphoribosyl tra | 1333.1 | 1069 | 1116.9 | 963.01 | 1127.7 | 1016.9 |
| 5141 | ENSMUSG00000025644 | 12.91472715 | 0.419172148 | 0.668068462 | 0.627438911 | 0.530371568 | 0.888700236 | Gm7628 | predicted gene 7628 [Source:MGI Symbol | 8.8341 | 17.475 | 7.0437 | 23.135 | 7.6343 | 13.366 |
| 5142 | ENSMUSG00000025645 | 176.9748852 | 0.3158516 | 0.182340621 | 1.732206452 | 0.083236811 | 0.425303185 | Ccdc51 | coiled-coil domain containing 51 [Source: | 163.83 | 153.16 | 155.97 | 202.44 | 211.58 | 174.87 |
| 5143 | ENSMUSG00000025646 | 160.7288798 | 0.01185563 | 0.186466506 | 0.063580483 | 0.949304273 | 0.990731019 | Atrip | ATR interacting protein [Source:MGI Symb | 154.2 | 155.22 | 171.06 | 169.66 | 150.51 | 163.74 |
| 5144 | ENSMUSG00000025647 | 504.9317479 | -0.220895313 | 0.128146461 | -1.723772253 | 0.084748965 | 0.429140871 | Shisa5 | shisa family member 5 [Source:MGI Symb | 543.7 | 549.94 | 536.33 | 425.11 | 475.51 | 499 |
| 5145 | ENSMUSG00000025648 | 10.10564988 | 0.07719159 | 0.744656294 | 0.103660697 | 0.917438618 | 0.986374671 | Pfkfb4 | 6-phosphofructo-2-kinase/fructose-2,6-bi | 12.047 | 9.2513 | 8.05 | 5.7839 | 5.4531 | 20.049 |
| 5146 | ENSMUSG00000025650 | 3.7940888 | -0.243959616 | 1.111542902 | -0.219478363 | 0.826277433 | NA | Col7a1 | collagen, type VII, alpha 1 [Source:MGI Sy | 3.2124 | 6.1675 | 3.0187 | 4.8199 | 1.0906 | 4.4554 |
| 5147 | ENSMUSG00000025651 | 5787.462738 | -0.068436099 | 0.086817256 | -0.788277613 | 0.430534339 | 0.843686314 | Uqcrc1 | ubiquinol-cytochrome c reductase core pr | 5763.1 | 5899.2 | 6111.9 | 5450.3 | 5919.9 | 5580.4 |
| 5148 | ENSMUSG00000025652 | 0 | NA | NA | NA | NA | NA | Tmem89 | transmembrane protein 89 [Source:MGI S | 0 | 0 | 0 | 0 | 0 | 0 |

| | | | | | | | | | | | | | | | |
|---|---|---|---|---|---|---|---|---|---|---|---|---|---|---|---|
| 5149 | ENSMUSG00000025656 | 92.06773745 | -0.351593801 | 0.34445028 | -1.020738904 | 0.307378157 | 0.756237058 | Arhgef9 | CDC42 guanine nucleotide exchange facto | 135.72 | 118.21 | 55.343 | 68.442 | 113.42 | 61.261 |
| 5150 | ENSMUSG00000025658 | 0 | NA | NA | NA | NA | NA | Cnksr2 | connector enhancer of kinase suppressor | 0 | 0 | 0 | 0 | 0 | 0 |
| 5151 | ENSMUSG00000025665 | 0.845554982 | -0.525406704 | 2.608531143 | -0.201418605 | 0.840371269 | NA | Rps6ka6 | ribosomal protein S6 kinase polypeptide 6 | 0 | 0 | 3.0187 | 0.964 | 1.0906 | 0 |
| 5152 | ENSMUSG00000025666 | 54.583883 | -1.52456025 | 0.375366054 | -4.061529364 | 4.88E-05 | 0.002067897 | Tmem47 | transmembrane protein 47 [Source:MGI S | 93.16 | 65.787 | 83.518 | 15.424 | 26.175 | 43.44 |
| 5153 | ENSMUSG00000025701 | 0.77082304 | -2.993021951 | 2.843236568 | -1.052681295 | 0.292487084 | NA | Alox5 | arachidonate 5-lipoxygenase [Source:MGI | 1.6062 | 0 | 3.0187 | 0 | 0 | 0 |
| 5154 | ENSMUSG00000025702 | 3141.901361 | 0.119240168 | 0.086634628 | 1.376356897 | 0.168711166 | 0.595783862 | Marchf8 | membrane associated ring-CH-type finger | 2976.3 | 3092 | 2968.4 | 3283.3 | 3198.8 | 3332.6 |
| 5155 | ENSMUSG00000025716 | 0 | NA | NA | NA | NA | NA | Myo3a | myosin IIIA [Source:MGI Symbol;Acc:MGI: | 0 | 0 | 0 | 0 | 0 | 0 |
| 5156 | ENSMUSG00000025722 | 457.9691703 | -0.237427878 | 0.16080065 | -1.476535563 | 0.139800176 | 0.546062582 | Wdr73 | WD repeat domain 73 [Source:MGI Symbo | 583.86 | 463.59 | 438.72 | 456.93 | 446.06 | 358.66 |
| 5157 | ENSMUSG00000025723 | 6.372674405 | -0.629761138 | 0.924450042 | -0.681227876 | 0.495727312 | 0.877965876 | Nmb | neuromedin B [Source:MGI Symbol;Acc:M | 8.8341 | 10.279 | 4.025 | 2.8919 | 2.1812 | 10.025 |
| 5158 | ENSMUSG00000025724 | 903.103537 | -0.017843146 | 0.115875471 | -0.153985529 | 0.877621146 | 0.98072613 | Sec11a | SEC11 homolog A, signal peptidase compl | 890.64 | 862.42 | 973.04 | 893.61 | 834.32 | 964.59 |
| 5159 | ENSMUSG00000025726 | 16.77754992 | 0.457188735 | 0.571342795 | 0.800200404 | 0.423594696 | 0.84018348 | Slc28a1 | solute carrier family 28 (sodium-coupled r | 16.062 | 8.2233 | 18.112 | 23.135 | 23.994 | 11.138 |
| 5160 | ENSMUSG00000025727 | 0 | NA | NA | NA | NA | NA | A930017K11 | RIKEN cDNA A930017K11 gene [Source:M | 0 | 0 | 0 | 0 | 0 | 0 |
| 5161 | ENSMUSG00000025728 | 746.3199366 | -0.008789585 | 0.117260297 | -0.074957895 | 0.940248212 | 0.988812319 | Pigq | phosphatidylinositol glycan anchor biosyn | 795.87 | 707.21 | 741.6 | 709.49 | 731.8 | 791.94 |
| 5162 | ENSMUSG00000025730 | 204.1853502 | 0.279776276 | 0.174878412 | 1.599833125 | 0.109635608 | 0.486451272 | Rab40c | Rab40C, member RAS oncogene family [So | 179.09 | 180.91 | 193.2 | 205.33 | 214.85 | 251.73 |
| 5163 | ENSMUSG00000025731 | 2263.249569 | 0.011362242 | 0.096562111 | 0.117667702 | 0.906330959 | 0.984171996 | Mettl26 | methyltransferase like 26 [Source:MGI Syr | 2235.8 | 2282 | 2245.9 | 2417.7 | 2114.7 | 2283.4 |
| 5164 | ENSMUSG00000025732 | 1184.628642 | -0.390523815 | 0.159463096 | -2.448991805 | 0.014325671 | 0.138765904 | Mcrip2 | MAPK regulated corepressor interacting p | 1186.2 | 1453.5 | 1393.6 | 1167.4 | 1141.9 | 765.21 |
| 5165 | ENSMUSG00000025733 | 442.8139908 | 0.039330428 | 0.137049317 | 0.286980108 | 0.774127551 | 0.958317434 | Rhot2 | ras homolog family member T2 [Source:M | 398.34 | 442 | 470.92 | 472.35 | 417.71 | 455.56 |
| 5166 | ENSMUSG00000025735 | 10.85202923 | -0.226744508 | 0.709701821 | -0.319492639 | 0.749352972 | 0.951342174 | Rhbdl1 | rhomboid like 1 [Source:MGI Symbol;Acc: | 8.8341 | 13.363 | 13.081 | 15.424 | 3.2719 | 11.138 |
| 5167 | ENSMUSG00000025736 | 339.8286418 | 0.182578685 | 0.154585737 | 1.18108364 | 0.237569498 | 0.686377492 | Jmjd8 | jumonji domain containing 8 [Source:MGI | 313.21 | 283.71 | 358.22 | 370.17 | 328.28 | 385.39 |
| 5168 | ENSMUSG00000025737 | 364.7394657 | 0.287812712 | 0.141369679 | 2.035887145 | 0.041761689 | 0.280818721 | Wdr24 | WD repeat domain 24 [Source:MGI Symbo | 335.7 | 300.15 | 349.17 | 383.66 | 419.89 | 399.87 |
| 5169 | ENSMUSG00000025738 | 4.958487873 | 0.094993084 | 0.981067269 | 0.09682627 | 0.922864361 | NA | Fbxl16 | F-box and leucine-rich repeat protein 16 [ | 7.2279 | 4.1117 | 3.0187 | 7.7118 | 5.4531 | 2.2277 |
| 5170 | ENSMUSG00000025739 | 0 | NA | NA | NA | NA | NA | Gng13 | guanine nucleotide binding protein (G pro | 0 | 0 | 0 | 0 | 0 | 0 |
| 5171 | ENSMUSG00000025742 | 320.7213503 | 0.00962458 | 0.168946043 | 0.056968366 | 0.954570395 | 0.991678478 | Prps2 | phosphoribosyl pyrophosphate synthetas | 318.03 | 299.12 | 341.12 | 257.38 | 353.36 | 355.32 |
| 5172 | ENSMUSG00000025743 | 425.3621779 | -1.208069481 | 0.192882673 | -6.263234866 | 3.77E-10 | 8.66E-08 | Sdc3 | syndecan 3 [Source:MGI Symbol;Acc:MGI | 537.28 | 766.83 | 476.96 | 230.39 | 237.75 | 302.97 |
| 5173 | ENSMUSG00000025745 | 8135.624031 | -0.084702536 | 0.109178802 | -0.775814848 | 0.437858316 | 0.847422601 | Hadha | hydroxyacyl-CoA dehydrogenase trifunctio | 7816.6 | 8968.6 | 8339.8 | 8696 | 8034.6 | 6958.2 |
| 5174 | ENSMUSG00000025746 | 0.133850373 | -0.796629176 | 4.080472857 | -0.195229623 | 0.84521319 | NA | Il6 | interleukin 6 [Source:MGI Symbol;Acc:MG | 0.8031 | 0 | 0 | 0 | 0 | 0 |
| 5175 | ENSMUSG00000025747 | 130.4887617 | -0.28424545 | 0.217106693 | -1.309243146 | 0.190452001 | 0.627779796 | Tyms | thymidylate synthase [Source:MGI Symbo | 163.03 | 125.41 | 140.87 | 107 | 135.24 | 111.38 |
| 5176 | ENSMUSG00000025754 | 0 | NA | NA | NA | NA | NA | Agbl1 | ATP/GTP binding protein-like 1 [Source:M | 0 | 0 | 0 | 0 | 0 | 0 |
| 5177 | ENSMUSG00000025757 | 512.4743516 | 0.001346741 | 0.158518552 | 0.008495796 | 0.993221417 | 0.99859176 | Hspa4l | heat shock protein 4 like [Source:MGI Syn | 499.53 | 435.84 | 601.73 | 565.85 | 441.7 | 530.19 |
| 5178 | ENSMUSG00000025758 | 13.84442771 | 0.071508206 | 0.640508231 | 0.111642915 | 0.911106544 | 0.98582838 | Plk4 | polo like kinase 4 [Source:MGI Symbol;Acc | 11.243 | 10.279 | 19.119 | 18.316 | 18.541 | 5.5692 |
| 5179 | ENSMUSG00000025759 | 222.7751258 | 0.136036978 | 0.163899584 | 0.830001973 | 0.406537668 | 0.830399297 | Mfsd8 | major facilitator superfamily domain cont | 221.66 | 210.72 | 204.27 | 235.21 | 246.48 | 218.31 |
| 5180 | ENSMUSG00000025762 | 1616.05779 | 0.148353822 | 0.105894262 | 1.400961855 | 0.16122548 | 0.58429 | Larp1b | La ribonucleoprotein domain family, mem | 1444.8 | 1460.7 | 1694.5 | 1729.4 | 1680.6 | 1686.4 |
| 5181 | ENSMUSG00000025764 | 712.4607416 | 0.636487222 | 0.138706996 | 4.588717508 | 4.46E-06 | 0.000279248 | Jade1 | jade family PHD finger 1 [Source:MGI Sym | 525.23 | 578.72 | 569.53 | 783.71 | 795.06 | 1022.5 |
| 5182 | ENSMUSG00000025766 | 187.208277 | 0.104892465 | 0.199112349 | 0.526800401 | 0.598332204 | 0.915055389 | D3Ertd751e | DNA segment, Chr 3, ERATO Doi 751, expr | 151.79 | 221 | 169.05 | 186.05 | 211.58 | 183.78 |
| 5183 | ENSMUSG00000025774 | 0 | NA | NA | NA | NA | NA | Crisp4 | cysteine-rich secretory protein 4 [Source:I | 0 | 0 | 0 | 0 | 0 | 0 |
| 5184 | ENSMUSG00000025776 | 0 | NA | NA | NA | NA | NA | Crispld1 | cysteine-rich secretory protein LCCL doma | 0 | 0 | 0 | 0 | 0 | 0 |
| 5185 | ENSMUSG00000025777 | 0 | NA | NA | NA | NA | NA | Gdap1 | ganglioside-induced differentiation-associ | 0 | 0 | 0 | 0 | 0 | 0 |
| 5186 | ENSMUSG00000025779 | 32.19367542 | 0.211056747 | 0.477443472 | 0.442055991 | 0.658448689 | 0.927522579 | Ly96 | lymphocyte antigen 96 [Source:MGI Symb | 38.549 | 22.614 | 28.175 | 39.523 | 14.178 | 50.123 |
| 5187 | ENSMUSG00000025780 | 242.2584149 | 0.58925562 | 0.225032738 | 2.61853286 | 0.008830879 | 0.098938285 | Itih5 | inter-alpha (globulin) inhibitor H5 [Source | 156.6 | 164.47 | 259.61 | 244.85 | 270.47 | 357.54 |
| 5188 | ENSMUSG00000025781 | 5071.531706 | -0.014026829 | 0.093675702 | -0.149738173 | 0.88097119 | 0.981490993 | Atp5c1 | ATP synthase, H+ transporting, mitochonc | 5006.5 | 5332.8 | 4948.7 | 4673.4 | 5185.9 | 5281.8 |
| 5189 | ENSMUSG00000025782 | 87.4933832 | -0.204751503 | 0.245629197 | -0.833579662 | 0.40451789 | 0.829271989 | Taf3 | TATA-box binding protein associated facto | 81.113 | 101.76 | 98.612 | 85.794 | 75.253 | 82.424 |
| 5190 | ENSMUSG00000025783 | 0 | NA | NA | NA | NA | NA | 4930412O13 | RIKEN cDNA 4930412O13 gene [Source:M | 0 | 0 | 0 | 0 | 0 | 0 |
| 5191 | ENSMUSG00000025784 | 2.710820084 | -1.601270521 | 1.583923584 | -1.010951878 | 0.312039463 | NA | Clec3b | C-type lectin domain family 3, member b [ | 7.2279 | 0 | 5.0312 | 2.8919 | 0 | 1.1138 |
| 5192 | ENSMUSG00000025785 | 460.0316679 | -0.059506297 | 0.138127949 | -0.430805621 | 0.666609712 | 0.929596279 | Exosc7 | exosome component 7 [Source:MGI Symb | 511.58 | 447.14 | 448.79 | 416.44 | 494.05 | 442.2 |
| 5193 | ENSMUSG00000025786 | 484.1354669 | -0.027794809 | 0.136010824 | -0.204357334 | 0.838074269 | 0.971073158 | Zdhhc3 | zinc finger, DHHC domain containing 3 [Sc | 522.02 | 457.42 | 486.02 | 441.5 | 530.04 | 467.81 |
| 5194 | ENSMUSG00000025787 | 4.599770534 | 0.662697298 | 1.007096137 | 0.658027842 | 0.510520241 | NA | Tgm4 | transglutaminase 4 (prostate) [Source:MG | 1.6062 | 6.1675 | 3.0187 | 5.7839 | 5.4531 | 5.5692 |
| 5195 | ENSMUSG00000025789 | 0.803314871 | 3.228356836 | 3.175155518 | 1.0167555 | 0.309269756 | NA | St8sia2 | ST8 alpha-N-acetyl-neuraminide alpha-2,8 | 0 | 0 | 0 | 4.8199 | 0 | 0 |
| 5196 | ENSMUSG00000025790 | 48.91977756 | -0.092656542 | 0.317395739 | -0.291927492 | 0.770342063 | 0.956515747 | Slco3a1 | solute carrier organic anion transporter fa | 48.989 | 51.396 | 51.318 | 55.911 | 45.806 | 40.098 |
| 5197 | ENSMUSG00000025791 | 4613.043572 | 0.042083534 | 0.096917405 | 0.434220599 | 0.664128256 | 0.929301301 | Pgm1 | phosphoglucomutase 1 [Source:MGI Syml | 4335.9 | 4853.8 | 4448.6 | 4756.3 | 4929.6 | 4354 |
| 5198 | ENSMUSG00000025792 | 5533.606756 | -0.213481692 | 0.12861699 | -1.659824973 | 0.096949668 | 0.458561151 | Slc25a10 | solute carrier family 25 (mitochondrial car | 6530.8 | 5954.7 | 5341.1 | 5547.7 | 5605.8 | 4221.5 |
| 5199 | ENSMUSG00000025793 | 0.824448242 | -0.598621533 | 2.872093747 | -0.208426878 | 0.834895672 | NA | Hgs | HGF-regulated tyrosine kinase substrate [! | 0 | 0 | 3.0187 | 1.928 | 0 | 0 |
| 5200 | ENSMUSG00000025794 | 2842.250652 | -0.127193152 | 0.102489426 | -1.241036829 | 0.214592144 | 0.65927374 | Rpl14 | ribosomal protein L14 [Source:MGI Symbo | 2655.1 | 3156.7 | 3092.2 | 2749.3 | 2756 | 2644.3 |
| 5201 | ENSMUSG00000025795 | 2677.159518 | 0.207634091 | 0.130469886 | 1.59143307 | 0.111512147 | 0.490097749 | Rassf3 | Ras association (RalGDS/AF-6) domain far | 2574.7 | 2537.9 | 2340.5 | 2344.4 | 2929.4 | 3336 |
| 5202 | ENSMUSG00000025804 | 2.367250414 | -1.221000048 | 1.533820264 | -0.796051582 | 0.426002055 | NA | Ccr1 | chemokine (C-C motif) receptor 1 [Source | 0.8031 | 5.1396 | 4.025 | 0.964 | 3.2719 | 0 |
| 5203 | ENSMUSG00000025808 | 0.181769715 | 1.126924175 | 4.080472857 | 0.276174898 | 0.782413736 | NA | Ccdc7a | coiled-coil domain containing 7A [Source: | 0 | 0 | 0 | 0 | 1.0906 | 0 |
| 5204 | ENSMUSG00000025809 | 3171.831529 | 0.066442204 | 0.095717252 | 0.694150763 | 0.487587667 | 0.873910634 | Itgb1 | integrin beta 1 (fibronectin receptor beta) | 3030.9 | 3280.1 | 2985.5 | 3112.7 | 3452.9 | 3168.9 |

| | | | | | | | | | | | | | | | |
|---|---|---|---|---|---|---|---|---|---|---|---|---|---|---|---|
| 5205 | ENSMUSG00000025810 | 1275.106889 | 0.311284241 | 0.135121831 | 2.303730181 | 0.021237795 | 0.180908987 | Nrp1 | neuropilin 1 [Source:MGI Symbol;Acc:MG | 1252 | 1104 | 1057.6 | 1448.9 | 1609.8 | 1178.4 |
| 5206 | ENSMUSG00000025812 | 1390.300144 | 0.263045314 | 0.106958572 | 2.459319604 | 0.013920063 | 0.136099418 | Pard3 | par-3 family cell polarity regulator [Source | 1306.6 | 1333.2 | 1151.1 | 1456.6 | 1518.1 | 1576.1 |
| 5207 | ENSMUSG00000025813 | 798.6641325 | 0.07818905 | 0.11638606 | 0.671807686 | 0.501706135 | 0.880974509 | Homer2 | homer scaffolding protein 2 [Source:MGI : | 785.43 | 782.25 | 763.74 | 863.72 | 856.14 | 740.71 |
| 5208 | ENSMUSG00000025815 | 1262.448204 | 0.081075977 | 0.111743623 | 0.725553502 | 0.468112535 | 0.865229581 | Dhtkd1 | dehydrogenase E1 and transketolase dom | 1171.7 | 1271.5 | 1238.7 | 1352.5 | 1378.5 | 1161.7 |
| 5209 | ENSMUSG00000025816 | 55.20030037 | -0.011488342 | 0.324556091 | -0.035397093 | 0.971763103 | 0.995296808 | Sec61a2 | Sec61, alpha subunit 2 (S. cerevisiae) [Sou | 59.43 | 43.173 | 63.393 | 45.307 | 70.89 | 49.009 |
| 5210 | ENSMUSG00000025817 | 268.9924048 | -0.107317994 | 0.158171257 | -0.678492393 | 0.497459545 | 0.878970787 | Nudt5 | nudix (nucleoside diphosphate linked moi | 305.18 | 270.34 | 260.62 | 240.99 | 263.93 | 272.89 |
| 5211 | ENSMUSG00000025821 | 97.76874153 | 0.106123986 | 0.231348417 | 0.458719308 | 0.646435747 | 0.925643431 | Zfp282 | zinc finger protein 282 [Source:MGI Symb | 88.341 | 104.85 | 89.556 | 103.15 | 93.793 | 106.93 |
| 5212 | ENSMUSG00000025823 | 2431.625652 | 0.276042186 | 0.136894036 | 2.016466128 | 0.043751257 | 0.288610458 | Pdia4 | protein disulfide isomerase associated 4 [! | 2328.2 | 2183.3 | 2088 | 3182.1 | 2156.2 | 2652.1 |
| 5213 | ENSMUSG00000025825 | 2030.680235 | -0.129074845 | 0.106593197 | -1.210910721 | 0.225929623 | 0.6725584 | Iscu | iron-sulfur cluster assembly enzyme [Sour | 2351.5 | 1968.5 | 2042.7 | 1864.3 | 1952.2 | 2004.9 |
| 5214 | ENSMUSG00000025838 | 0 | NA | NA | NA | NA | NA | Pramel6 | PRAME like 6 [Source:MGI Symbol;Acc:M( | 0 | 0 | 0 | 0 | 0 | 0 |
| 5215 | ENSMUSG00000025839 | 0 | NA | NA | NA | NA | NA | Pramel7 | PRAME like 7 [Source:MGI Symbol;Acc:M( | 0 | 0 | 0 | 0 | 0 | 0 |
| 5216 | ENSMUSG00000025854 | 1067.584259 | 0.468848809 | 0.104839118 | 4.472078919 | 7.75E-06 | 0.000435962 | Fam20c | FAM20C, golgi associated secretory pathw | 891.44 | 907.65 | 887.51 | 1189.5 | 1211.7 | 1317.7 |
| 5217 | ENSMUSG00000025855 | 2.47459739 | -0.506488759 | 1.459834581 | -0.346949418 | 0.728629319 | NA | Prkar1b | protein kinase, cAMP dependent regulato | 5.6217 | 2.0558 | 1.0062 | 2.8919 | 3.2719 | 0 |
| 5218 | ENSMUSG00000025856 | 149.2137383 | 0.441593076 | 0.213504636 | 2.068306732 | 0.038611189 | 0.267980921 | Pdgfa | platelet derived growth factor, alpha [Sou | 146.16 | 108.96 | 123.77 | 160.98 | 152.69 | 202.72 |
| 5219 | ENSMUSG00000025857 | 174.9585119 | -0.166923031 | 0.179634881 | -0.929235068 | 0.352767278 | 0.793259753 | Dnaaf5 | dynein, axonemal assembly factor 5 [Sour | 196.76 | 185.03 | 173.07 | 161.95 | 161.41 | 171.53 |
| 5220 | ENSMUSG00000025858 | 517.0684809 | -0.027675732 | 0.143262601 | -0.193181837 | 0.846816563 | 0.972803041 | Get4 | golgi to ER traffic protein 4 [Source:MGI S | 569.4 | 549.94 | 445.77 | 468.49 | 545.31 | 523.51 |
| 5221 | ENSMUSG00000025860 | 1958.426871 | 0.203098152 | 0.09541205 | 2.128642571 | 0.033283844 | 0.243995238 | Xiap | X-linked inhibitor of apoptosis [Source:M( | 1773.2 | 1870.8 | 1818.3 | 1982.9 | 2130 | 2175.3 |
| 5222 | ENSMUSG00000025862 | 1110.236222 | 0.137980367 | 0.118688377 | 1.162543208 | 0.245014884 | 0.695017707 | Stag2 | stromal antigen 2 [Source:MGI Symbol;Ac | 921.16 | 1094.7 | 1157.2 | 1184.7 | 1141.9 | 1161.7 |
| 5223 | ENSMUSG00000025867 | 18.81334877 | -0.633604496 | 0.629595931 | -1.006366885 | 0.314239117 | 0.760822663 | Cplx2 | complexin 2 [Source:MGI Symbol;Acc:MG | 33.73 | 24.67 | 10.062 | 14.46 | 5.4531 | 24.505 |
| 5224 | ENSMUSG00000025868 | 1055.271528 | -0.079080798 | 0.106422555 | -0.743083059 | 0.457431395 | 0.858923498 | Higd2a | HIG1 domain family, member 2A [Source: | 1049.7 | 1058.8 | 1144.1 | 981.33 | 1026.3 | 1071.5 |
| 5225 | ENSMUSG00000025869 | 596.330483 | -0.190638623 | 0.13325008 | -1.430682989 | 0.152521091 | 0.56828054 | Nop16 | NOP16 nucleolar protein [Source:MGI Syn | 560.57 | 653.76 | 693.3 | 530.19 | 542.04 | 598.13 |
| 5226 | ENSMUSG00000025870 | 64.32914759 | -0.523097595 | 0.321542315 | -1.626839052 | 0.103771285 | 0.474047094 | Arl10 | ADP-ribosylation factor-like 10 [Source:M | 55.414 | 84.289 | 88.55 | 67.478 | 51.259 | 38.985 |
| 5227 | ENSMUSG00000025871 | 1623.50044 | -0.335494778 | 0.1257476 | -2.668001441 | 0.007630393 | 0.088790029 | 4833439L19l | RIKEN cDNA 4833439L19 gene [Source:M( | 1537.1 | 1855.4 | 2042.7 | 1360.2 | 1540 | 1405.7 |
| 5228 | ENSMUSG00000025872 | 408.2743749 | 0.086148324 | 0.135041078 | 0.637941616 | 0.523511686 | 0.888334265 | Thoc3 | THO complex 3 [Source:MGI Symbol;Acc:I | 362.2 | 411.17 | 415.58 | 416.44 | 419.89 | 424.37 |
| 5229 | ENSMUSG00000025873 | 886.7323669 | 0.170842478 | 0.111823527 | 1.527786535 | 0.126565547 | 0.521187766 | Faf2 | Fas associated factor family member 2 [Sc | 776.6 | 865.51 | 861.35 | 904.21 | 929.21 | 983.52 |
| 5230 | ENSMUSG00000025875 | 25.24619645 | -0.216424704 | 0.57210658 | -0.37829438 | 0.70521192 | 0.94295036 | Tspan17 | tetraspanin 17 [Source:MGI Symbol;Acc:N | 9.6372 | 38.033 | 34.212 | 37.595 | 14.178 | 17.822 |
| 5231 | ENSMUSG00000025876 | 29.74422114 | -0.945883774 | 0.43022386 | -2.198585114 | 0.027907436 | 0.217720151 | Unc5a | unc-5 netrin receptor A [Source:MGI Syml | 39.352 | 29.81 | 48.3 | 21.208 | 14.178 | 25.618 |
| 5232 | ENSMUSG00000025877 | 35.0340901 | -1.329782998 | 0.4247524 | -3.130725097 | 0.001743753 | 0.031685483 | Hk3 | hexokinase 3 [Source:MGI Symbol;Acc:M( | 47.383 | 69.898 | 33.206 | 22.171 | 15.269 | 22.277 |
| 5233 | ENSMUSG00000025878 | 359.9135664 | 0.2524929 | 0.156991851 | 1.608318509 | 0.107765439 | 0.484241711 | Uimc1 | ubiquitin interaction motif containing 1 [S | 287.51 | 336.13 | 362.25 | 350.89 | 386.08 | 436.63 |
| 5234 | ENSMUSG00000025880 | 212.3474808 | -1.097427843 | 0.223909935 | -4.901202097 | 9.53E-07 | 7.87E-05 | Smad7 | SMAD family member 7 [Source:MGI Sym | 238.52 | 309.4 | 319.99 | 99.29 | 129.78 | 177.1 |
| 5235 | ENSMUSG00000025885 | 553.7794055 | 0.10808772 | 0.121999191 | 0.885970789 | 0.375633268 | 0.809469546 | Myo5b | myosin VB [Source:MGI Symbol;Acc:MGI: | 505.95 | 557.13 | 536.33 | 549.47 | 584.57 | 589.22 |
| 5236 | ENSMUSG00000025887 | 21.77546128 | -0.493788394 | 0.487670488 | -1.012545165 | 0.311277463 | 0.759756535 | Casp12 | caspase 12 [Source:MGI Symbol;Acc:MGI: | 18.471 | 29.81 | 28.175 | 13.496 | 23.994 | 16.708 |
| 5237 | ENSMUSG00000025888 | 25.73564167 | -1.221544258 | 0.438940672 | -2.78293705 | 0.005386926 | 0.069942992 | Casp1 | caspase 1 [Source:MGI Symbol;Acc:MGI:9 | 32.927 | 33.921 | 41.256 | 15.424 | 14.178 | 16.708 |
| 5238 | ENSMUSG00000025889 | 20.66695158 | 0.664275792 | 0.583621163 | 1.138196888 | 0.255038278 | 0.708637501 | Snca | synuclein, alpha [Source:MGI Symbol;Acc | 5.6217 | 16.447 | 26.162 | 18.316 | 21.812 | 35.643 |
| 5239 | ENSMUSG00000025892 | 1.249115476 | 0.348807902 | 2.051884581 | 0.16999392 | 0.865014918 | NA | Gria4 | glutamate receptor, ionotropic, AMPA4 (a | 3.2124 | 0 | 0 | 0.964 | 1.0906 | 2.2277 |
| 5240 | ENSMUSG00000025893 | 51.69079027 | -0.192933795 | 0.339126731 | -0.568913559 | 0.569414803 | 0.903765578 | Kbtbd3 | kelch repeat and BTB (POZ) domain conta | 59.43 | 45.228 | 60.375 | 34.703 | 45.806 | 64.603 |
| 5241 | ENSMUSG00000025894 | 490.3241257 | 0.070230006 | 0.126685384 | 0.554365499 | 0.579328732 | 0.908337213 | Aasdhppt | aminoadipate-semialdehyde dehydrogena | 497.12 | 482.09 | 455.83 | 517.66 | 515.86 | 473.38 |
| 5242 | ENSMUSG00000025898 | 326.5866483 | 0.149607474 | 0.152994627 | 0.977860965 | 0.328143094 | 0.773161429 | Cwf19l2 | CWF19-like 2, cell cycle control (S. pombe | 281.09 | 347.44 | 300.87 | 318.11 | 353.36 | 358.66 |
| 5243 | ENSMUSG00000025899 | 199.7255207 | -0.068855241 | 0.179126761 | -0.384393939 | 0.700686492 | 0.941778228 | Alkbh8 | alkB homolog 8, tRNA methyltransferase [ | 217.64 | 184 | 211.31 | 186.05 | 181.04 | 218.31 |
| 5244 | ENSMUSG00000025900 | 0 | NA | NA | NA | NA | NA | Rp1 | retinitis pigmentosa 1 (human) [Source:M | 0 | 0 | 0 | 0 | 0 | 0 |
| 5245 | ENSMUSG00000025902 | 27.38915294 | 0.071715997 | 0.468068429 | 0.153216907 | 0.878227225 | 0.980901822 | Sox17 | SRY (sex determining region Y)-box 17 [So | 27.305 | 28.782 | 24.15 | 37.595 | 13.087 | 33.415 |
| 5246 | ENSMUSG00000025903 | 2651.556305 | -0.153611858 | 0.096347095 | -1.594358997 | 0.110855651 | 0.489424955 | Lypla1 | lysophospholipase 1 [Source:MGI Symbol; | 2932.1 | 2689 | 2756.1 | 2554.5 | 2626.2 | 2351.3 |
| 5247 | ENSMUSG00000025905 | 0 | NA | NA | NA | NA | NA | Oprk1 | opioid receptor, kappa 1 [Source:MGI Syn | 0 | 0 | 0 | 0 | 0 | 0 |
| 5248 | ENSMUSG00000025907 | 918.1578394 | 0.104763004 | 0.114992319 | 0.911043493 | 0.362272458 | 0.80090868 | Rb1cc1 | RB1-inducible coiled-coil 1 [Source:MGI S | 856.91 | 885.04 | 912.66 | 905.18 | 1049.2 | 899.99 |
| 5249 | ENSMUSG00000025909 | 5.554638914 | -1.829430977 | 0.993852758 | -1.840746492 | 0.065658717 | 0.368238731 | Sntg1 | syntrophin, gamma 1 [Source:MGI Symbo | 8.8341 | 4.1117 | 13.081 | 2.8919 | 2.1812 | 2.2277 |
| 5250 | ENSMUSG00000025911 | 2360.921379 | -0.019520168 | 0.112557321 | -0.173424243 | 0.862317971 | 0.976334255 | Adhfe1 | alcohol dehydrogenase, iron containing, 1 | 2451.9 | 2497.8 | 2180.5 | 2261.5 | 2621.8 | 2151.9 |
| 5251 | ENSMUSG00000025912 | 8.074730009 | -0.584700209 | 0.835969531 | -0.699427655 | 0.48428481 | 0.872521682 | Mybl1 | myeloblastosis oncogene-like 1 [Source:M | 16.865 | 4.1117 | 8.05 | 10.604 | 4.3625 | 4.4554 |
| 5252 | ENSMUSG00000025915 | 198.9132303 | -0.152491146 | 0.209421323 | -0.728154821 | 0.466518819 | 0.863779479 | Sgk3 | serum/glucocorticoid regulated kinase 3 [ | 224.87 | 213.81 | 189.17 | 182.19 | 143.96 | 239.48 |
| 5253 | ENSMUSG00000025916 | 0.133850373 | -0.796629176 | 4.080472857 | -0.195229623 | 0.84521319 | NA | Ppp1r42 | protein phosphatase 1, regulatory subunit | 0.8031 | 0 | 0 | 0 | 0 | 0 |
| 5254 | ENSMUSG00000025917 | 1091.759434 | 0.105307123 | 0.11178849 | 0.942021162 | 0.346181806 | 0.786987972 | Cops5 | COP9 signalosome subunit 5 [Source:MGI | 1094.6 | 1060.8 | 999.2 | 1031.5 | 1161.5 | 1203 |
| 5255 | ENSMUSG00000025920 | 101.1457963 | -0.028159414 | 0.264903975 | -0.106300457 | 0.915343969 | 0.98625062 | Stau2 | staufen double-stranded RNA binding pro | 132.51 | 97.652 | 75.468 | 88.686 | 113.42 | 99.132 |
| 5256 | ENSMUSG00000025921 | 709.2917068 | -0.198138089 | 0.123757365 | -1.60102059 | 0.109372359 | 0.486233674 | Rdh10 | retinol dehydrogenase 10 (all-trans) [Sour | 695.49 | 733.93 | 845.25 | 680.57 | 648.92 | 651.6 |
| 5257 | ENSMUSG00000025922 | 0.267700746 | -1.497481649 | 4.057001436 | -0.369110456 | 0.712045396 | NA | Rpl7-ps8 | ribosomal protein L7, pseudogene 8 [Sour | 1.6062 | 0 | 0 | 0 | 0 | 0 |
| 5258 | ENSMUSG00000025925 | 148.9937504 | 0.008611414 | 0.19637911 | 0.043850965 | 0.965023202 | 0.993480943 | Terf1 | telomeric repeat binding factor 1 [Source: | 159.82 | 141.85 | 143.89 | 160.98 | 154.87 | 132.55 |
| 5259 | ENSMUSG00000025927 | 0 | NA | NA | NA | NA | NA | Tfap2b | transcription factor AP-2 beta [Source:MG | 0 | 0 | 0 | 0 | 0 | 0 |
| 5260 | ENSMUSG00000025929 | 0 | NA | NA | NA | NA | NA | Il17a | interleukin 17A [Source:MGI Symbol;Acc:I | 0 | 0 | 0 | 0 | 0 | 0 |

| | | | | | | | | | | | | | | | |
|---|---|---|---|---|---|---|---|---|---|---|---|---|---|---|---|
| 5261 | ENSMUSG00000025930 | 2.315427997 | -0.173215785 | 1.488547325 | -0.116365656 | 0.907362753 | NA | Msc | musculin [Source:MGI Symbol;Acc:MGI:1 | 3.2124 | 3.0838 | 1.0062 | 0 | 4.3625 | 2.2277 |
| 5262 | ENSMUSG00000025931 | 5.818615657 | -0.243805662 | 0.907431299 | -0.268676717 | 0.788178471 | 0.960870081 | Paqr8 | progestin and adipoQ receptor family mer | 5.6217 | 10.279 | 3.0187 | 3.8559 | 5.4531 | 6.6831 |
| 5263 | ENSMUSG00000025932 | 0 | NA | NA | NA | NA | NA | Eya1 | EYA transcriptional coactivator and phosp | 0 | 0 | 0 | 0 | 0 | 0 |
| 5264 | ENSMUSG00000025933 | 115.1345392 | -0.140940508 | 0.217393643 | -0.648319363 | 0.516778413 | 0.886716042 | Tmem14a | transmembrane protein 14A [Source:MGI | 133.31 | 106.9 | 121.76 | 111.82 | 113.42 | 103.59 |
| 5265 | ENSMUSG00000025934 | 31346.48702 | -0.171248574 | 0.111647004 | -1.533839404 | 0.125069166 | 0.518510965 | Gsta3 | glutathione S-transferase, alpha 3 [Source | 29928 | 33664 | 36024 | 33036 | 28759 | 26668 |
| 5266 | ENSMUSG00000025935 | 1992.626177 | 0.101206042 | 0.105145037 | 0.962537503 | 0.335779674 | 0.777327945 | Tram1 | translocating chain-associating membrane | 1945.9 | 2017.8 | 1805.2 | 2246.1 | 1935.8 | 2004.9 |
| 5267 | ENSMUSG00000025936 | 212.7162995 | -0.38803125 | 0.278573937 | -1.39292015 | 0.163643938 | 0.58776904 | Gm4956 | predicted gene 4956 [Source:MGI Symbol | 146.16 | 240.53 | 337.09 | 138.81 | 239.94 | 173.76 |
| 5268 | ENSMUSG00000025937 | 6522.750927 | 0.114228165 | 0.107688706 | 1.060725575 | 0.288814634 | 0.739603195 | Lactb2 | lactamase, beta 2 [Source:MGI Symbol;Ac | 7048.8 | 6074 | 5669.2 | 6423 | 7183.9 | 6737.6 |
| 5269 | ENSMUSG00000025938 | 0.305170013 | -1.657836477 | 4.041480335 | -0.410205257 | 0.681655385 | NA | Slco5a1 | solute carrier organic anion transporter fa | 0.8031 | 1.0279 | 0 | 0 | 0 | 0 |
| 5270 | ENSMUSG00000025939 | 445.3569053 | 0.011461206 | 0.129180018 | 0.088722742 | 0.929302258 | 0.988405116 | Ube2w | ubiquitin-conjugating enzyme E2W (putat | 466.6 | 436.87 | 426.65 | 431.86 | 463.51 | 446.65 |
| 5271 | ENSMUSG00000025940 | 931.816735 | 0.025480975 | 0.106994704 | 0.238151738 | 0.811763408 | 0.96653525 | Tmem70 | transmembrane protein 70 [Source:MGI S | 937.22 | 932.32 | 900.59 | 891.68 | 945.57 | 983.52 |
| 5272 | ENSMUSG00000025945 | 0 | NA | NA | NA | NA | NA | Crygf | crystallin, gamma F [Source:MGI Symbol;A | 0 | 0 | 0 | 0 | 0 | 0 |
| 5273 | ENSMUSG00000025946 | 0 | NA | NA | NA | NA | NA | Pth2r | parathyroid hormone 2 receptor [Source:I | 0 | 0 | 0 | 0 | 0 | 0 |
| 5274 | ENSMUSG00000025949 | 398.140717 | 0.065195161 | 0.149605001 | 0.435781962 | 0.662994933 | 0.92906822 | Pikfyve | phosphoinositide kinase, FYVE type zinc fi | 381.47 | 430.7 | 355.2 | 388.48 | 374.08 | 458.9 |
| 5275 | ENSMUSG00000025950 | 7778.735115 | 0.017009543 | 0.093679073 | 0.181572497 | 0.855918234 | 0.974266258 | Idh1 | isocitrate dehydrogenase 1 (NADP+), solu | 8153.1 | 7886.2 | 7158.4 | 7702.2 | 8240.7 | 7531.8 |
| 5276 | ENSMUSG00000025952 | 0 | NA | NA | NA | NA | NA | Crygc | crystallin, gamma C [Source:MGI Symbol;A | 0 | 0 | 0 | 0 | 0 | 0 |
| 5277 | ENSMUSG00000025955 | 0 | NA | NA | NA | NA | NA | Akr1cl | aldo-keto reductase family 1, member C-li | 0 | 0 | 0 | 0 | 0 | 0 |
| 5278 | ENSMUSG00000025956 | 193.3423994 | 0.039429419 | 0.179171626 | 0.220065085 | 0.825820466 | 0.96943211 | Mettl21a | methyltransferase like 21A [Source:MGI S | 171.86 | 191.19 | 209.3 | 184.12 | 191.95 | 211.63 |
| 5279 | ENSMUSG00000025958 | 299.4472249 | 0.070224879 | 0.154940042 | 0.453239061 | 0.650376608 | 0.925643431 | Creb1 | cAMP responsive element binding protein | 261.81 | 316.6 | 298.85 | 316.18 | 283.56 | 319.67 |
| 5280 | ENSMUSG00000025959 | 45.50745287 | -0.324805969 | 0.330884601 | -0.981629148 | 0.326282582 | 0.772018857 | Klf7 | Kruppel-like factor 7 (ubiquitous) [Source: | 51.399 | 44.2 | 56.35 | 47.235 | 35.99 | 37.871 |
| 5281 | ENSMUSG00000025961 | 1.011142131 | 1.198131946 | 2.259952339 | 0.530158059 | 0.596002347 | NA | 4933402D24 | RIKEN cDNA 4933402D24 gene [Source:M | 0.8031 | 1.0279 | 0 | 0.964 | 3.2719 | 0 |
| 5282 | ENSMUSG00000025962 | 274.4238784 | 0.188269239 | 0.158300366 | 1.189316511 | 0.23431514 | 0.682322842 | Fastkd2 | FAST kinase domains 2 [Source:MGI Symb | 283.5 | 254.92 | 230.43 | 280.52 | 306.46 | 290.71 |
| 5283 | ENSMUSG00000025963 | 0.133850373 | -0.796629176 | 4.080472857 | -0.195229623 | 0.84521319 | NA | Mdh1b | malate dehydrogenase 1B, NAD (soluble) | 0.8031 | 0 | 0 | 0 | 0 | 0 |
| 5284 | ENSMUSG00000025964 | 148.3665214 | -0.458389281 | 0.243491118 | -1.882570854 | 0.059758549 | 0.348006896 | Adam23 | a disintegrin and metallopeptidase domai | 163.03 | 143.91 | 208.29 | 126.28 | 90.521 | 158.17 |
| 5285 | ENSMUSG00000025967 | 2377.930264 | 0.067438728 | 0.097173744 | 0.694001542 | 0.487681242 | 0.873910634 | Eef1b2 | eukaryotic translation elongation factor 1 | 2265.6 | 2443.4 | 2258 | 2291.4 | 2443 | 2566.3 |
| 5286 | ENSMUSG00000025968 | 2382.099454 | -0.030291219 | 0.105959867 | -0.285874451 | 0.774974282 | 0.95893551 | Ndufs1 | NADH:ubiquinone oxidoreductase core su | 2204.5 | 2549.2 | 2468.3 | 2230.6 | 2552 | 2287.8 |
| 5287 | ENSMUSG00000025969 | 327.4746309 | -0.322543279 | 0.473584991 | -0.681067359 | 0.495828869 | 0.878020172 | Nrp2 | neuropilin 2 [Source:MGI Symbol;Acc:MG | 354.97 | 456.4 | 280.74 | 473.31 | 177.77 | 221.65 |
| 5288 | ENSMUSG00000025971 | 267.2286029 | -0.044233225 | 0.171354169 | -0.258139182 | 0.796299493 | 0.962236316 | Maip1 | matrix AAA peptidase interacting protein | 317.23 | 242.59 | 253.57 | 284.37 | 265.02 | 240.59 |
| 5289 | ENSMUSG00000025977 | 0.867384107 | -1.926017826 | 2.566777421 | -0.750364177 | 0.4530354 | NA | Boll | boule homolog, RNA binding protein [Sou | 3.2124 | 1.0279 | 0 | 0.964 | 0 | 0 |
| 5290 | ENSMUSG00000025978 | 6.778620845 | -1.161956362 | 0.827606186 | -1.403996708 | 0.16031983 | 0.582922579 | Rftn2 | raftlin family member 2 [Source:MGI Syml | 8.8341 | 9.2513 | 10.062 | 4.8199 | 4.3625 | 3.3415 |
| 5291 | ENSMUSG00000025979 | 479.0751515 | 0.167513501 | 0.127528892 | 1.31353373 | 0.189003163 | 0.626143323 | Mob4 | MOB family member 4, phocein [Source:N | 447.33 | 458.45 | 447.78 | 469.46 | 524.59 | 526.85 |
| 5292 | ENSMUSG00000025980 | 9093.115769 | 0.031241067 | 0.100910613 | 0.309591484 | 0.756871633 | 0.953190361 | Hspd1 | heat shock protein 1 (chaperonin) [Source | 8163.5 | 9152.6 | 9669 | 8905.2 | 9870.1 | 8798.3 |
| 5293 | ENSMUSG00000025981 | 748.0438507 | 0.353573496 | 0.189768366 | 1.863184596 | 0.062436299 | 0.356948077 | Coq10b | coenzyme Q10B [Source:MGI Symbol;Acc | 868.96 | 634.23 | 465.89 | 827.09 | 933.57 | 758.53 |
| 5294 | ENSMUSG00000025982 | 2494.767515 | -0.049172734 | 0.114499535 | -0.429457933 | 0.667590001 | 0.930033258 | Sf3b1 | splicing factor 3b, subunit 1 [Source:MGI ! | 2535.4 | 2468 | 2609.2 | 2675 | 2588 | 2092.9 |
| 5295 | ENSMUSG00000025983 | 0 | NA | NA | NA | NA | NA | Ccdc150 | coiled-coil domain containing 150 [Source | 0 | 0 | 0 | 0 | 0 | 0 |
| 5296 | ENSMUSG00000025986 | 49.05993178 | 0.061198105 | 0.351598641 | 0.174056715 | 0.86182089 | 0.976334255 | Slc39a10 | solute carrier family 39 (zinc transporter), | 42.564 | 56.535 | 45.281 | 62.659 | 31.628 | 55.692 |
| 5297 | ENSMUSG00000025991 | 54783.2188 | 0.058632371 | 0.144678965 | 0.405258435 | 0.685287556 | 0.93583845 | Cps1 | carbamoyl-phosphate synthetase 1 [Sourc | 41037 | 56612 | 63363 | 50283 | 60886 | 56519 |
| 5298 | ENSMUSG00000025993 | 2513.328662 | -0.084231705 | 0.133266723 | -0.63205355 | 0.527351886 | 0.888334265 | Slc40a1 | solute carrier family 40 (iron-regulated tra | 2379.6 | 2946 | 2436.1 | 2867.8 | 2168.1 | 2282.3 |
| 5299 | ENSMUSG00000025995 | 215.6198637 | -0.104456662 | 0.171631188 | -0.608611189 | 0.542782186 | 0.894270594 | Wdr75 | WD repeat domain 75 [Source:MGI Symbo | 217.64 | 233.34 | 219.36 | 199.54 | 234.48 | 189.35 |
| 5300 | ENSMUSG00000025997 | 4.595877418 | -0.819090018 | 1.338979788 | -0.611726947 | 0.54071843 | NA | Ikzf2 | IKAROS family zinc finger 2 [Source:MGI S | 10.44 | 7.1954 | 0 | 7.7118 | 0 | 2.2277 |
| 5301 | ENSMUSG00000026000 | 361.419528 | 0.051653273 | 0.164582536 | 0.313844193 | 0.753639377 | 0.952989051 | Lancl1 | LanC (bacterial lantibiotic synthetase com | 346.94 | 385.47 | 332.06 | 304.62 | 432.98 | 366.45 |
| 5302 | ENSMUSG00000026003 | 5231.682337 | -0.224445465 | 0.095295707 | -2.355252642 | 0.018510116 | 0.163849644 | Acadl | acyl-Coenzyme A dehydrogenase, long-ch | 5433 | 6041.1 | 5440.8 | 5139.9 | 4644.9 | 4690.4 |
| 5303 | ENSMUSG00000026004 | 175.1125948 | 0.336930623 | 0.195960438 | 1.719380847 | 0.085545047 | 0.430973029 | Kansl1l | KAT8 regulatory NSL complex subunit 1-lik | 139.74 | 152.13 | 173.07 | 221.71 | 166.86 | 197.15 |
| 5304 | ENSMUSG00000026005 | 581.7070329 | 0.126038383 | 0.122913156 | 1.025426302 | 0.305162083 | 0.75399118 | Rpe | ribulose-5-phosphate-3-epimerase [Sourc | 542.09 | 539.66 | 587.65 | 624.66 | 631.47 | 564.72 |
| 5305 | ENSMUSG00000026009 | 7.012521293 | -0.33201731 | 0.888081132 | -0.373859209 | 0.708509066 | 0.943888782 | Icos | inducible T cell co-stimulator [Source:MGI | 3.2124 | 14.391 | 6.0375 | 9.6398 | 5.4531 | 3.3415 |
| 5306 | ENSMUSG00000026011 | 0.167707431 | -0.796629176 | 4.080472857 | -0.195229623 | 0.84521319 | NA | Ctla4 | cytotoxic T-lymphocyte-associated protei | 0 | 0 | 1.0062 | 0 | 0 | 0 |
| 5307 | ENSMUSG00000026012 | 2.573176126 | -4.72400265 | 1.742078493 | -2.711704822 | 0.006693817 | NA | Cd28 | CD28 antigen [Source:MGI Symbol;Acc:M | 3.2124 | 7.1954 | 5.0312 | 0 | 0 | 0 |
| 5308 | ENSMUSG00000026014 | 829.6473627 | 0.373315814 | 0.141051137 | 2.646670008 | 0.008128863 | 0.093092541 | Raph1 | Ras association (RalGDS/AF-6) and plecks | 819.97 | 638.34 | 709.4 | 993.86 | 801.6 | 1014.7 |
| 5309 | ENSMUSG00000026017 | 82.41351902 | -0.327983456 | 0.249590981 | -1.314083766 | 0.188818017 | 0.625984911 | Carf | calcium response factor [Source:MGI Sym | 85.129 | 96.624 | 93.581 | 69.406 | 81.796 | 67.944 |
| 5310 | ENSMUSG00000026018 | 0.167707431 | -0.796629176 | 4.080472857 | -0.195229623 | 0.84521319 | NA | Ica1l | islet cell autoantigen 1-like [Source:MGI S | 0 | 0 | 1.0062 | 0 | 0 | 0 |
| 5311 | ENSMUSG00000026019 | 288.8862783 | -0.18846388 | 0.155120849 | -1.21494874 | 0.224385643 | 0.670547386 | Wdr12 | WD repeat domain 12 [Source:MGI Symbo | 316.42 | 304.26 | 301.87 | 243.89 | 265.02 | 301.85 |
| 5312 | ENSMUSG00000026020 | 269.5154258 | -0.046637228 | 0.182033779 | -0.256200952 | 0.797795673 | 0.962377049 | Nop58 | NOP58 ribonucleoprotein [Source:MGI Sy | 265.83 | 297.07 | 258.6 | 232.32 | 329.37 | 233.91 |
| 5313 | ENSMUSG00000026021 | 650.7572281 | 0.051082331 | 0.135024958 | 0.378317697 | 0.705194601 | 0.94295036 | Sumo1 | small ubiquitin-like modifier 1 [Source:MG | 660.95 | 600.3 | 656.07 | 640.08 | 761.25 | 585.88 |
| 5314 | ENSMUSG00000026023 | 1.449955927 | 1.315873981 | 2.10518359 | 0.625063765 | 0.531929209 | NA | Cdk15 | cyclin-dependent kinase 15 [Source:MGI S | 2.4093 | 0 | 0 | 1.928 | 4.3625 | 0 |
| 5315 | ENSMUSG00000026024 | 1367.792876 | 0.030613728 | 0.131959713 | 0.231992991 | 0.816543458 | 0.967767553 | Als2 | alsin Rho guanine nucleotide exchange fa | 1289 | 1526.5 | 1245.7 | 1577.1 | 1360 | 1208.5 |
| 5316 | ENSMUSG00000026027 | 795.1072984 | 0.086004641 | 0.123977285 | 0.693712896 | 0.487862277 | 0.873910634 | Stradb | STE20-related kinase adaptor beta [Sourc | 870.56 | 723.65 | 718.46 | 783.71 | 829.96 | 844.29 |

| | | | | | | | | | | | | | | | |
|---|---|---|---|---|---|---|---|---|---|---|---|---|---|---|---|
| 5317 | ENSMUSG00000026028 | 494.3413267 | 0.141202675 | 0.130685597 | 1.080476186 | 0.279930186 | 0.732834426 | Trak2 | trafficking protein, kinesin binding 2 [Sour | 493.91 | 492.37 | 423.63 | 501.27 | 520.22 | 534.65 |
| 5318 | ENSMUSG00000026029 | 694.7103315 | 0.158106132 | 0.125797022 | 1.256835246 | 0.2088133 | 0.653498052 | Casp8 | caspase 8 [Source:MGI Symbol;Acc:MGI:1 | 670.59 | 698.98 | 600.73 | 775.04 | 759.07 | 663.85 |
| 5319 | ENSMUSG00000026031 | 1576.3317 | 0.125230009 | 0.124359859 | 1.006997035 | 0.313936199 | 0.760476509 | Cflar | CASP8 and FADD-like apoptosis regulator | 1294.6 | 1678.6 | 1551.6 | 1502.8 | 1688.3 | 1742.1 |
| 5320 | ENSMUSG00000026032 | 1085.02111 | -0.02563568 | 0.12644649 | -0.202739353 | 0.839338762 | 0.971685662 | Ndufb3 | NADH:ubiquinone oxidoreductase subuni | 1138.8 | 1080.3 | 1063.6 | 912.89 | 1206.2 | 1108.3 |
| 5321 | ENSMUSG00000026034 | 1062.913046 | -0.136621516 | 0.137520894 | -0.993460066 | 0.320485802 | 0.767391055 | Clk1 | CDC-like kinase 1 [Source:MGI Symbol;Acc | 1170.1 | 1154.4 | 1015.3 | 1083.5 | 1126.6 | 827.59 |
| 5322 | ENSMUSG00000026035 | 132.8350195 | 0.01721831 | 0.198680345 | 0.086663376 | 0.930939088 | 0.988420305 | Ppil3 | peptidylprolyl isomerase (cyclophilin)-like | 135.72 | 125.41 | 134.84 | 131.1 | 138.51 | 131.43 |
| 5323 | ENSMUSG00000026036 | 236.9284919 | 0.214604407 | 0.168673968 | 1.272303067 | 0.203265459 | 0.647739134 | Nif3l1 | Ngg1 interacting factor 3-like 1 (S. pombe) | 193.55 | 240.53 | 224.39 | 235.21 | 265.02 | 262.87 |
| 5324 | ENSMUSG00000026037 | 76.51923485 | 0.133428984 | 0.256062527 | 0.521079697 | 0.602311254 | 0.915492852 | Orc2 | origin recognition complex, subunit 2 [Sou | 66.657 | 72.982 | 79.493 | 78.082 | 86.159 | 75.741 |
| 5325 | ENSMUSG00000026039 | 1.148863437 | 0.657647435 | 1.961871891 | 0.335214261 | 0.737463465 | NA | Sgo2a | shugoshin 2A [Source:MGI Symbol;Acc:M | 1.6062 | 1.0279 | 0 | 0.964 | 2.1812 | 1.1138 |
| 5326 | ENSMUSG00000026042 | 35.91042805 | -0.827975726 | 0.431419391 | -1.919189872 | 0.054960309 | 0.330417929 | Col5a2 | collagen, type V, alpha 2 [Source:MGI Sym | 53.808 | 57.563 | 26.162 | 18.316 | 25.084 | 34.529 |
| 5327 | ENSMUSG00000026043 | 326.1997649 | -1.442724096 | 0.499116032 | -2.890558515 | 0.003845579 | 0.055696621 | Col3a1 | collagen, type III, alpha 1 [Source:MGI Syr | 736.44 | 474.9 | 219.36 | 183.16 | 196.31 | 147.03 |
| 5328 | ENSMUSG00000026047 | 93.2619233 | 0.389405227 | 0.242872596 | 1.603331269 | 0.108861536 | 0.485684837 | Poglut2 | protein O-glucosyltransferase 2 [Source:M | 70.673 | 81.206 | 90.562 | 93.506 | 116.7 | 106.93 |
| 5329 | ENSMUSG00000026048 | 323.9993603 | 0.03406893 | 0.144302468 | 0.236093881 | 0.813359822 | 0.966802988 | Ercc5 | excision repair cross-complementing rode | 314.82 | 320.71 | 325.02 | 317.15 | 355.54 | 310.76 |
| 5330 | ENSMUSG00000026049 | 59.92714035 | 0.415517552 | 0.295070713 | 1.40819652 | 0.159072886 | 0.580285052 | Tex30 | testis expressed 30 [Source:MGI Symbol;A | 56.217 | 54.48 | 43.269 | 71.334 | 76.343 | 57.92 |
| 5331 | ENSMUSG00000026051 | 1.524015665 | 1.260818235 | 1.808087234 | 0.697321573 | 0.48560157 | NA | Ecrg4 | ECRG4 augurin precursor [Source:MGI Syr | 1.6062 | 1.0279 | 0 | 0.964 | 1.0906 | 4.4554 |
| 5332 | ENSMUSG00000026058 | 0 | NA | NA | NA | NA | NA | Khdrbs2 | KH domain containing, RNA binding, signa | 0 | 0 | 0 | 0 | 0 | 0 |
| 5333 | ENSMUSG00000026062 | 1.179108286 | 0.594076217 | 1.949629551 | 0.304712357 | 0.760585237 | NA | Slc9a2 | solute carrier family 9 (sodium/hydrogen | 0.8031 | 0 | 2.0125 | 0.964 | 2.1812 | 1.1138 |
| 5334 | ENSMUSG00000026063 | 0 | NA | NA | NA | NA | NA | Dnaaf6 | dynein axonemal assembly factor 6 [Sourc | 0 | 0 | 0 | 0 | 0 | 0 |
| 5335 | ENSMUSG00000026064 | 30.80142298 | -0.203652232 | 0.480569096 | -0.423773051 | 0.671731334 | 0.931191575 | Ptp4a1 | protein tyrosine phosphatase 4a1 [Source | 36.943 | 32.893 | 29.181 | 37.595 | 38.172 | 10.025 |
| 5336 | ENSMUSG00000026065 | 0 | NA | NA | NA | NA | NA | Slc9a4 | solute carrier family 9 (sodium/hydrogen | 0 | 0 | 0 | 0 | 0 | 0 |
| 5337 | ENSMUSG00000026068 | 2.340696537 | -1.722618626 | 1.528581485 | -1.12693935 | 0.259768125 | NA | Il18rap | interleukin 18 receptor accessory protein | 1.6062 | 4.1117 | 5.0312 | 0 | 2.1812 | 1.1138 |
| 5338 | ENSMUSG00000026069 | 0.267700746 | -1.497481649 | 4.057001436 | -0.369110456 | 0.712045396 | NA | Il1rl1 | interleukin 1 receptor-like 1 [Source:MGI | 1.6062 | 0 | 0 | 0 | 0 | 0 |
| 5339 | ENSMUSG00000026070 | 2.538537911 | 0.019245333 | 1.439316376 | 0.013371162 | 0.989331674 | NA | Il18r1 | interleukin 18 receptor 1 [Source:MGI Syn | 2.4093 | 4.1117 | 1.0062 | 0 | 4.3625 | 3.3415 |
| 5340 | ENSMUSG00000026072 | 1289.54355 | -0.644753379 | 0.179704563 | -3.587852016 | 0.000333413 | 0.009479365 | Il1r1 | interleukin 1 receptor, type I [Source:MGI | 1142.8 | 2087.7 | 1489.2 | 918.67 | 1081.9 | 1016.9 |
| 5341 | ENSMUSG00000026073 | 0.449470461 | -0.545136533 | 3.562105917 | -0.15303771 | 0.878368537 | NA | Il1r2 | interleukin 1 receptor, type II [Source:MG | 1.6062 | 0 | 0 | 0 | 1.0906 | 0 |
| 5342 | ENSMUSG00000026074 | 116.4902407 | -0.672918788 | 0.247272163 | -2.721368958 | 0.006501216 | 0.079757066 | Map4k4 | mitogen-activated protein kinase kinase k | 112.43 | 163.44 | 153.96 | 85.794 | 74.162 | 109.16 |
| 5343 | ENSMUSG00000026077 | 0.874144864 | -0.458978284 | 2.368814316 | -0.193758659 | 0.846364859 | NA | Npas2 | neuronal PAS domain protein 2 [Source:M | 0 | 1.0279 | 2.0125 | 0 | 1.0906 | 1.1138 |
| 5344 | ENSMUSG00000026078 | 248.6727474 | -0.257258779 | 0.154716703 | -1.662773147 | 0.096357854 | 0.458386897 | Pdcl3 | phosducin-like 3 [Source:MGI Symbol;Acc | 275.46 | 266.23 | 270.68 | 232.32 | 229.03 | 218.31 |
| 5345 | ENSMUSG00000026080 | 0.887741833 | 3.358069843 | 2.74787414 | 1.222061009 | 0.221684557 | NA | Chst10 | carbohydrate sulfotransferase 10 [Source | 0 | 0 | 0 | 0.964 | 4.3625 | 0 |
| 5346 | ENSMUSG00000026082 | 204.0682782 | 0.043174987 | 0.193582056 | 0.223031969 | 0.823510626 | 0.96943211 | Rev1 | REV1, DNA directed polymerase [Source:N | 191.14 | 184 | 228.42 | 239.07 | 169.05 | 212.74 |
| 5347 | ENSMUSG00000026083 | 1313.355214 | -0.085615383 | 0.11286341 | -0.758575193 | 0.448106717 | 0.854156674 | Eif5b | eukaryotic translation initiation factor 5B | 1365.3 | 1382.5 | 1308.1 | 1127.9 | 1320.7 | 1375.6 |
| 5348 | ENSMUSG00000026085 | 0 | NA | NA | NA | NA | NA | Lyg1 | lysozyme G-like 1 [Source:MGI Symbol;Ac | 0 | 0 | 0 | 0 | 0 | 0 |
| 5349 | ENSMUSG00000026087 | 966.0486266 | 0.034359002 | 0.113254161 | 0.303379597 | 0.761600593 | 0.954216198 | Mrpl30 | mitochondrial ribosomal protein L30 [Sou | 946.05 | 956.99 | 959.96 | 879.15 | 1036.1 | 1018.1 |
| 5350 | ENSMUSG00000026088 | 114.4218546 | 0.300985512 | 0.258148376 | 1.165939981 | 0.243638713 | 0.693425884 | Mitd1 | MIT, microtubule interacting and transpo | 115.65 | 85.317 | 106.66 | 164.84 | 94.884 | 119.18 |
| 5351 | ENSMUSG00000026090 | 2.674972754 | 1.353816897 | 1.325848387 | 1.021094803 | 0.307209525 | NA | Cracdl | capping protein inhibiting regulator of act | 2.4093 | 1.0279 | 1.0062 | 3.8559 | 2.1812 | 5.5692 |
| 5352 | ENSMUSG00000026094 | 69.02061288 | -0.335120283 | 0.27368324 | -1.224482296 | 0.220770341 | 0.667237604 | Stk17b | serine/threonine kinase 17b (apoptosis-in | 76.295 | 84.289 | 70.437 | 61.695 | 52.35 | 69.058 |
| 5353 | ENSMUSG00000026095 | 107.6972581 | 0.166518564 | 0.23929447 | 0.695873014 | 0.486508359 | 0.873586472 | Asnsd1 | asparagine synthetase domain containing | 111.63 | 107.93 | 84.525 | 102.18 | 136.33 | 103.59 |
| 5354 | ENSMUSG00000026096 | 263.4443086 | 0.031759191 | 0.164768259 | 0.192750664 | 0.847154243 | 0.972803041 | Osgepl1 | O-sialoglycoprotein endopeptidase-like 1 | 247.36 | 240.53 | 293.82 | 251.6 | 292.29 | 255.07 |
| 5355 | ENSMUSG00000026097 | 416.8529528 | 0.38209142 | 0.147458789 | 2.591174278 | 0.009564904 | 0.104967201 | Ormdl1 | ORM1-like 1 (S. cerevisiae) [Source:MGI S | 373.44 | 343.32 | 368.29 | 401.01 | 507.14 | 507.91 |
| 5356 | ENSMUSG00000026098 | 96.47967452 | 0.747083603 | 0.290240065 | 2.574019558 | 0.010052459 | 0.109076345 | Pms1 | PMS1 homolog 1, mismatch repair system | 78.704 | 61.675 | 75.468 | 106.04 | 171.23 | 85.766 |
| 5357 | ENSMUSG00000026100 | 0 | NA | NA | NA | NA | NA | Mstn | myostatin [Source:MGI Symbol;Acc:MGI:9 | 0 | 0 | 0 | 0 | 0 | 0 |
| 5358 | ENSMUSG00000026102 | 295.7749608 | 0.067699529 | 0.147192937 | 0.459937348 | 0.645561191 | 0.925643431 | Inpp1 | inositol polyphosphate-1-phosphatase [Sc | 281.89 | 300.15 | 284.77 | 307.51 | 315.19 | 285.14 |
| 5359 | ENSMUSG00000026103 | 85.35471167 | -0.406228167 | 0.272012603 | -1.493416711 | 0.135328141 | 0.538039097 | Gls | glutaminase [Source:MGI Symbol;Acc:MG | 118.86 | 83.261 | 89.556 | 85.794 | 57.803 | 76.855 |
| 5360 | ENSMUSG00000026104 | 343.9492124 | -0.192267941 | 0.155831228 | -1.233821642 | 0.217269373 | 0.663257309 | Stat1 | signal transducer and activator of transcri | 379.06 | 407.06 | 313.95 | 296.91 | 335.91 | 330.81 |
| 5361 | ENSMUSG00000026107 | 219.7879384 | -0.216917342 | 0.183816068 | -1.180078242 | 0.237969097 | 0.686509684 | Nabp1 | nucleic acid binding protein 1 [Source:MG | 267.43 | 250.81 | 190.18 | 209.18 | 202.86 | 198.26 |
| 5362 | ENSMUSG00000026109 | 5.018579106 | 1.353097268 | 1.11675742 | 1.211630426 | 0.225653882 | NA | Tmeff2 | transmembrane protein with EGF-like and | 2.4093 | 0 | 6.0375 | 2.8919 | 7.6343 | 11.138 |
| 5363 | ENSMUSG00000026110 | 14.20739732 | -1.80281623 | 0.606555618 | -2.972219162 | 0.002956556 | 0.046110081 | Mgat4a | mannoside acetylglucosaminyltransferase | 22.487 | 21.586 | 22.137 | 5.7839 | 5.4531 | 7.7969 |
| 5364 | ENSMUSG00000026111 | 571.5336573 | 0.08810782 | 0.118341661 | 0.74452073 | 0.456561505 | 0.858190757 | Unc50 | unc-50 homolog [Source:MGI Symbol;Acc | 544.5 | 576.66 | 541.36 | 593.81 | 575.85 | 597.02 |
| 5365 | ENSMUSG00000026112 | 1947.173663 | 0.028199651 | 0.094414542 | 0.298679108 | 0.7651849 | 0.954980327 | Coa5 | cytochrome C oxidase assembly factor 5 [ | 1832.7 | 1996.2 | 1956.1 | 1917.4 | 1998 | 1982.6 |
| 5366 | ENSMUSG00000026113 | 417.9242949 | 0.584522764 | 0.181258955 | 3.224793851 | 0.001260634 | 0.024875595 | Inpp4a | inositol polyphosphate-4-phosphatase, ty | 308.39 | 342.3 | 353.19 | 573.57 | 364.27 | 565.83 |
| 5367 | ENSMUSG00000026114 | 0 | NA | NA | NA | NA | NA | Cnga3 | cyclic nucleotide gated channel alpha 3 [S | 0 | 0 | 0 | 0 | 0 | 0 |
| 5368 | ENSMUSG00000026116 | 615.513129 | -0.09787708 | 0.123832853 | -0.790396718 | 0.429296118 | 0.842978372 | Tmem131 | transmembrane protein 131 [Source:MGI | 605.54 | 642.45 | 662.11 | 642.97 | 549.67 | 590.34 |
| 5369 | ENSMUSG00000026117 | 280.4980042 | 0.439345712 | 0.206359778 | 2.129027838 | 0.033251957 | 0.243995238 | Zap70 | zeta-chain (TCR) associated protein kinase | 309.19 | 195.3 | 208.29 | 265.09 | 364.27 | 340.84 |
| 5370 | ENSMUSG00000026121 | 37.98051801 | -0.799785101 | 0.378551227 | -2.11275263 | 0.034621946 | 0.249264225 | Sema4c | sema domain, immunoglobulin domain (Ig | 36.14 | 56.535 | 52.325 | 29.883 | 21.812 | 31.188 |
| 5371 | ENSMUSG00000026123 | 253.3362974 | -0.066154045 | 0.162695695 | -0.406612144 | 0.684292876 | 0.935528319 | Plekhb2 | pleckstrin homology domain containing, f | 247.36 | 260.06 | 270.68 | 276.66 | 224.67 | 240.59 |
| 5372 | ENSMUSG00000026124 | 0 | NA | NA | NA | NA | NA | Cfc1 | cripto, FRL-1, cryptic family 1 [Source:MGI | 0 | 0 | 0 | 0 | 0 | 0 |

| | | | | | | | | | | | | | | | |
|---|---|---|---|---|---|---|---|---|---|---|---|---|---|---|---|
| 5373 | ENSMUSG00000026125 | 0 | NA | NA | NA | NA | NA | Prss39 | protease, serine 39 [Source:MGI Symbol;/ | 0 | 0 | 0 | 0 | 0 | 0 |
| 5374 | ENSMUSG00000026126 | 12.54526052 | -0.553147122 | 0.665091581 | -0.831685648 | 0.405586404 | 0.829973745 | Ptpn18 | protein tyrosine phosphatase, non-recept | 23.29 | 6.1675 | 15.094 | 8.6758 | 10.906 | 11.138 |
| 5375 | ENSMUSG00000026127 | 411.3439406 | -0.000161852 | 0.129719093 | -0.001247711 | 0.999004471 | 0.999764475 | Imp4 | IMP4, U3 small nucleolar ribonucleoprote | 403.16 | 423.5 | 407.53 | 403.91 | 411.16 | 418.81 |
| 5376 | ENSMUSG00000026131 | 2236.58921 | -0.147325927 | 0.104223792 | -1.413553705 | 0.157492982 | 0.576703416 | Dst | dystonin [Source:MGI Symbol;Acc:MGI:1C | 2273.6 | 2488.6 | 2291.2 | 2300.1 | 2118 | 1948.1 |
| 5377 | ENSMUSG00000026134 | 36.34311434 | 0.34134466 | 0.369200096 | 0.924551927 | 0.355199029 | 0.795293925 | Prim2 | DNA primase, p58 subunit [Source:MGI Sy | 36.14 | 23.642 | 36.225 | 40.487 | 40.353 | 41.212 |
| 5378 | ENSMUSG00000026135 | 71.48763748 | 0.392812437 | 0.296549912 | 1.324608171 | 0.185301145 | 0.620990675 | Zfp142 | zinc finger protein 142 [Source:MGI Symb | 55.414 | 78.122 | 52.325 | 97.362 | 62.165 | 83.538 |
| 5379 | ENSMUSG00000026141 | 0 | NA | NA | NA | NA | NA | Col19a1 | collagen, type XIX, alpha 1 [Source:MGI Sy | 0 | 0 | 0 | 0 | 0 | 0 |
| 5380 | ENSMUSG00000026142 | 363.9493545 | 0.163970713 | 0.140744028 | 1.16502785 | 0.24400772 | 0.693793525 | Rhbdd1 | rhomboid domain containing 1 [Source:M | 362.2 | 325.85 | 341.12 | 362.46 | 412.25 | 379.82 |
| 5381 | ENSMUSG00000026147 | 0 | NA | NA | NA | NA | NA | Col9a1 | collagen, type IX, alpha 1 [Source:MGI Syr | 0 | 0 | 0 | 0 | 0 | 0 |
| 5382 | ENSMUSG00000026149 | 0.185640647 | 1.126924175 | 4.080472857 | 0.276174898 | 0.782413736 | NA | Tm4sf20 | transmembrane 4 L six family member 20 | 0 | 0 | 0 | 0 | 0 | 1.1138 |
| 5383 | ENSMUSG00000026150 | 557.6430892 | 0.000807122 | 0.128262048 | 0.006292761 | 0.994979137 | 0.998676967 | Mff | mitochondrial fission factor [Source:MGI S | 501.94 | 584.89 | 586.64 | 568.75 | 531.13 | 572.52 |
| 5384 | ENSMUSG00000026153 | 524.8011447 | 0.146652065 | 0.152023419 | 0.964667589 | 0.334711334 | 0.776159726 | Fam135a | family with sequence similarity 135, meml | 455.36 | 548.91 | 491.05 | 569.71 | 627.11 | 456.68 |
| 5385 | ENSMUSG00000026154 | 305.6382298 | -0.028055633 | 0.151400194 | -0.185307771 | 0.852987643 | 0.973813394 | Sdhaf4 | succinate dehydrogenase complex asseml | 317.23 | 297.07 | 310.93 | 271.84 | 330.46 | 306.31 |
| 5386 | ENSMUSG00000026155 | 208.025749 | -0.28803428 | 0.168256042 | -1.711880755 | 0.086918646 | 0.434457217 | Smap1 | small ArfGAP 1 [Source:MGI Symbol;Acc:N | 233.7 | 231.28 | 221.37 | 201.47 | 183.22 | 177.1 |
| 5387 | ENSMUSG00000026156 | 7.114422786 | -0.337785587 | 0.799620382 | -0.422432437 | 0.672709406 | 0.931454408 | B3gat2 | beta-1,3-glucuronyltransferase 2 (glucuro | 5.6217 | 9.2513 | 9.0562 | 7.7118 | 4.3625 | 6.6831 |
| 5388 | ENSMUSG00000026158 | 424.9989705 | 0.342577246 | 0.153131742 | 2.237140661 | 0.025277148 | 0.204593549 | Ogfrl1 | opioid growth factor receptor-like 1 [Sour | 314.82 | 394.72 | 415.58 | 430.9 | 502.78 | 491.21 |
| 5389 | ENSMUSG00000026159 | 415.6814612 | 0.095993547 | 0.142409029 | 0.674069249 | 0.500267289 | 0.880716237 | Agfg1 | ArfGAP with FG repeats 1 [Source:MGI Sy | 372.64 | 421.45 | 412.56 | 472.35 | 429.7 | 385.39 |
| 5390 | ENSMUSG00000026162 | 39.70697228 | 0.249299005 | 0.394411961 | 0.632077699 | 0.527336107 | 0.888334265 | Nhej1 | non-homologous end joining factor 1 [Sou | 38.549 | 31.865 | 38.237 | 36.631 | 28.356 | 64.603 |
| 5391 | ENSMUSG00000026163 | 0 | NA | NA | NA | NA | NA | Sphkap | SPHK1 interactor, AKAP domain containin | 0 | 0 | 0 | 0 | 0 | 0 |
| 5392 | ENSMUSG00000026166 | 0 | NA | NA | NA | NA | NA | Ccl20 | chemokine (C-C motif) ligand 20 [Source:N | 0 | 0 | 0 | 0 | 0 | 0 |
| 5393 | ENSMUSG00000026167 | 0 | NA | NA | NA | NA | NA | Wnt10a | wingless-type MMTV integration site fami | 0 | 0 | 0 | 0 | 0 | 0 |
| 5394 | ENSMUSG00000026170 | 8872.29717 | 0.092312688 | 0.116616438 | 0.791592418 | 0.428598369 | 0.842753542 | Cyp27a1 | cytochrome P450, family 27, subfamily a, | 8580.3 | 9476.4 | 7709.8 | 10146 | 9259.3 | 8062 |
| 5395 | ENSMUSG00000026171 | 248.550981 | -0.114348784 | 0.175522617 | -0.651476068 | 0.514739221 | 0.885488593 | Rnf25 | ring finger protein 25 [Source:MGI Symbo | 252.98 | 266.23 | 256.59 | 270.88 | 251.93 | 192.69 |
| 5396 | ENSMUSG00000026172 | 233.5831401 | 0.097580845 | 0.167153785 | 0.583778853 | 0.559369108 | 0.899423311 | Bcs1l | BCS1-like (yeast) [Source:MGI Symbol;Acc | 219.25 | 224.09 | 233.45 | 212.08 | 248.66 | 263.98 |
| 5397 | ENSMUSG00000026173 | 3.201830933 | -0.564335337 | 1.530807432 | -0.368652075 | 0.712387075 | NA | Plcd4 | phospholipase C, delta 4 [Source:MGI Sym | 3.2124 | 7.1954 | 1.0062 | 0 | 0 | 7.7969 |
| 5398 | ENSMUSG00000026174 | 297.1881053 | 0.057020364 | 0.16511457 | 0.345338175 | 0.729840151 | 0.946534338 | Cnot9 | CCR4-NOT transcription complex, subunit | 248.16 | 324.82 | 301.87 | 310.4 | 273.75 | 324.13 |
| 5399 | ENSMUSG00000026175 | 0.706696157 | 0.149852971 | 3.037999338 | 0.049326203 | 0.960659338 | NA | Vil1 | villin 1 [Source:MGI Symbol;Acc:MGI:9893 | 0 | 0 | 2.0125 | 0 | 0 | 2.2277 |
| 5400 | ENSMUSG00000026176 | 3105.467661 | 0.242032875 | 0.088081869 | 2.747817222 | 0.005999344 | 0.0753251 | Ctdsp1 | CTD (carboxy-terminal domain, RNA polyr | 2774.7 | 2881.3 | 2880.9 | 3239 | 3444.2 | 3412.8 |
| 5401 | ENSMUSG00000026177 | 71.07722468 | -1.448416001 | 0.329838286 | -4.391291316 | 1.13E-05 | 0.000588502 | Slc11a1 | solute carrier family 11 (proton-coupled d | 81.916 | 148.02 | 82.512 | 43.379 | 30.537 | 40.098 |
| 5402 | ENSMUSG00000026179 | 1728.332936 | -0.462302763 | 0.171039313 | -2.702903529 | 0.006873669 | 0.082919727 | Pnkd | paroxysmal nonkinesiogenic dyskinesia [S | 1492.2 | 2154.5 | 2362.7 | 1322.6 | 1278.2 | 1759.9 |
| 5403 | ENSMUSG00000026180 | 0.17131964 | -0.796629176 | 4.080472857 | -0.195229623 | 0.84521319 | NA | Cxcr2 | chemokine (C-X-C motif) receptor 2 [Sour | 0 | 1.0279 | 0 | 0 | 0 | 0 |
| 5404 | ENSMUSG00000026181 | 518.1110703 | 0.000984542 | 0.130951848 | 0.007518353 | 0.994001278 | 0.998618081 | Ppm1f | protein phosphatase 1F (PP2C domain cor | 489.89 | 542.74 | 521.23 | 483.92 | 500.59 | 570.29 |
| 5405 | ENSMUSG00000026182 | 0 | NA | NA | NA | NA | NA | Tnp1 | transition protein 1 [Source:MGI Symbol;/ | 0 | 0 | 0 | 0 | 0 | 0 |
| 5406 | ENSMUSG00000026185 | 373.4918949 | 0.242269837 | 0.168886612 | 1.434511793 | 0.151426266 | 0.567123127 | Igfbp5 | insulin-like growth factor binding protein | 368.62 | 314.54 | 343.13 | 410.65 | 479.87 | 324.13 |
| 5407 | ENSMUSG00000026187 | 458.8963882 | -0.18716222 | 0.147060362 | -1.272689784 | 0.203128144 | 0.647739134 | Xrcc5 | X-ray repair complementing defective rep | 487.48 | 511.9 | 465.89 | 366.31 | 492.96 | 428.83 |
| 5408 | ENSMUSG00000026188 | 1.017313306 | 0.03731332 | 2.211598506 | 0.016871652 | 0.986539008 | NA | Tmem169 | transmembrane protein 169 [Source:MGI | 0 | 2.0558 | 1.0062 | 1.928 | 0 | 1.1138 |
| 5409 | ENSMUSG00000026189 | 4363.447085 | 0.004079263 | 0.106094283 | 0.038449414 | 0.969329364 | 0.994602269 | Pecr | peroxisomal trans-2-enoyl-CoA reductase | 4083 | 4202.1 | 4786.7 | 3984.1 | 4491.2 | 4633.6 |
| 5410 | ENSMUSG00000026192 | 528.5317821 | 0.047696634 | 0.133748278 | 0.356614936 | 0.721380095 | 0.9447719 | Atic | 5-aminoimidazole-4-carboxamide ribonuc | 512.38 | 540.68 | 506.14 | 495.48 | 607.47 | 509.03 |
| 5411 | ENSMUSG00000026193 | 22003.70847 | 0.069145629 | 0.09024493 | 0.766199595 | 0.443557549 | 0.851079912 | Fn1 | fibronectin 1 [Source:MGI Symbol;Acc:MC | 21010 | 22347 | 21074 | 23664 | 23115 | 20813 |
| 5412 | ENSMUSG00000026196 | 3.026104339 | -1.307205043 | 1.238292749 | -1.055651052 | 0.291127668 | NA | Bard1 | BRCA1 associated RING domain 1 [Source | 4.8186 | 3.0838 | 5.0312 | 1.928 | 2.1812 | 1.1138 |
| 5413 | ENSMUSG00000026197 | 538.9366312 | 0.338186495 | 0.133912661 | 2.525425835 | 0.011555817 | 0.119930185 | Zfand2b | zinc finger, AN1 type domain 2B [Source:N | 427.25 | 468.73 | 533.31 | 630.44 | 580.21 | 593.68 |
| 5414 | ENSMUSG00000026198 | 46.2017852 | 0.032371106 | 0.325716158 | 0.099384402 | 0.920833067 | 0.986685804 | Abcb6 | ATP-binding cassette, sub-family B (MDR/ | 41.761 | 46.256 | 49.306 | 53.983 | 45.806 | 40.098 |
| 5415 | ENSMUSG00000026199 | 363.9697958 | 0.231247629 | 0.140742794 | 1.643051292 | 0.100372321 | 0.466511971 | Ankzf1 | ankyrin repeat and zinc finger domain cor | 322.85 | 344.35 | 338.1 | 428.01 | 376.26 | 374.25 |
| 5416 | ENSMUSG00000026200 | 53.4539398 | 0.210484582 | 0.31042679 | 0.678049022 | 0.497740612 | 0.879037876 | Glb1l | galactosidase, beta 1-like [Source:MGI Syr | 48.989 | 56.535 | 43.269 | 62.659 | 46.897 | 62.375 |
| 5417 | ENSMUSG00000026201 | 1363.740792 | 0.102246524 | 0.101788351 | 1.004501236 | 0.315137076 | 0.761215823 | Stk16 | serine/threonine kinase 16 [Source:MGI S | 1354.8 | 1261.3 | 1330.3 | 1496.1 | 1362.2 | 1377.8 |
| 5418 | ENSMUSG00000026202 | 1430.054818 | 0.123480951 | 0.117706991 | 1.049053674 | 0.294153416 | 0.745164683 | Tuba4a | tubulin, alpha 4A [Source:MGI Symbol;Ac | 1260.1 | 1327 | 1520.4 | 1522.1 | 1355.6 | 1595 |
| 5419 | ENSMUSG00000026203 | 1014.035979 | 0.225814686 | 0.120414429 | 1.875312525 | 0.060749741 | 0.352178059 | Dnajb2 | DnaJ heat shock protein family (Hsp40) m | 952.48 | 918.96 | 932.79 | 1102.8 | 958.65 | 1218.5 |
| 5420 | ENSMUSG00000026204 | 1.256424171 | -0.643638749 | 2.022116939 | -0.318299469 | 0.750257787 | NA | Ptprn | protein tyrosine phosphatase, receptor ty | 1.6062 | 1.0279 | 2.0125 | 2.8919 | 0 | 0 |
| 5421 | ENSMUSG00000026205 | 2.204984606 | -0.500668465 | 1.500751 | -0.333611948 | 0.738672401 | NA | Slc23a3 | solute carrier family 23 (nucleobase trans | 1.6062 | 4.1117 | 2.0125 | 0 | 3.2719 | 2.2277 |
| 5422 | ENSMUSG00000026207 | 13.16178208 | -0.371826923 | 0.697708971 | -0.532925529 | 0.594085132 | 0.913144152 | Speg | SPEG complex locus [Source:MGI Symbol; | 12.047 | 19.53 | 13.081 | 15.424 | 2.1812 | 16.708 |
| 5423 | ENSMUSG00000026208 | 54.4328159 | -0.732460741 | 0.306895868 | -2.386675146 | 0.017001507 | 0.154824975 | Des | desmin [Source:MGI Symbol;Acc:MGI:948 | 66.657 | 71.954 | 65.406 | 45.307 | 32.719 | 44.554 |
| 5424 | ENSMUSG00000026209 | 838.4340163 | 0.32788405 | 0.127373449 | 2.574194651 | 0.010047373 | 0.109076345 | Dnpep | aspartyl aminopeptidase [Source:MGI Syn | 839.24 | 735.99 | 654.06 | 915.78 | 898.67 | 986.87 |
| 5425 | ENSMUSG00000026211 | 12.53114175 | -0.5581573 | 0.69835144 | -0.799249873 | 0.424145538 | 0.840450466 | Obsl1 | obscurin-like 1 [Source:MGI Symbol;Acc:N | 14.456 | 17.475 | 13.081 | 20.244 | 4.3625 | 5.5692 |
| 5426 | ENSMUSG00000026213 | 168.059668 | 0.043972835 | 0.198182956 | 0.221880005 | 0.824407296 | 0.96943211 | Stk11ip | serine/threonine kinase 11 interacting prc | 197.56 | 157.27 | 140.87 | 167.73 | 174.5 | 170.42 |
| 5427 | ENSMUSG00000026219 | 2080.633524 | 0.019735384 | 0.101739273 | 0.193980002 | 0.84619154 | 0.972803041 | Trip12 | thyroid hormone receptor interactor 12 [S | 2011.8 | 2152.5 | 2034.6 | 1913.5 | 2215 | 2156.4 |
| 5428 | ENSMUSG00000026220 | 0 | NA | NA | NA | NA | NA | Slc16a14 | solute carrier family 16 (monocarboxylic a | 0 | 0 | 0 | 0 | 0 | 0 |

| | | | | | | | | | | | | | | | |
|---|---|---|---|---|---|---|---|---|---|---|---|---|---|---|---|
| 5429 | ENSMUSG00000026222 | 111.0863518 | -0.689229512 | 0.239512744 | -2.877631894 | 0.004006724 | 0.05726606 | Sp100 | nuclear antigen Sp100 [Source:MGI Symb | 130.1 | 141.85 | 139.87 | 101.22 | 63.256 | 90.221 |
| 5430 | ENSMUSG00000026223 | 605.3771939 | -0.154493662 | 0.139334092 | -1.10880015 | 0.267516406 | 0.72293778 | Itm2c | integral membrane protein 2C [Source:M( | 721.99 | 591.05 | 598.72 | 519.58 | 566.03 | 634.89 |
| 5431 | ENSMUSG00000026224 | 0 | NA | NA | NA | NA | NA | 4933407L21 | RIKEN cDNA 4933407L21 gene [Source:M( | 0 | 0 | 0 | 0 | 0 | 0 |
| 5432 | ENSMUSG00000026226 | 4.282329087 | -0.873791073 | 1.2596791 | -0.693661642 | 0.487894427 | NA | Spata3 | spermatogenesis associated 3 [Source:M( | 1.6062 | 3.0838 | 12.075 | 6.7478 | 2.1812 | 0 |
| 5433 | ENSMUSG00000026227 | 1582.443364 | 0.385028704 | 0.136616116 | 2.81832566 | 0.004827482 | 0.065256728 | 2810459M1 | RIKEN cDNA 2810459M11 gene [Source:M | 1209.5 | 1239.7 | 1669.4 | 1807.5 | 1676.3 | 1892.4 |
| 5434 | ENSMUSG00000026228 | 1.055914579 | 0.129051249 | 2.190846851 | 0.058904733 | 0.953027988 | NA | Htr2b | 5-hydroxytryptamine (serotonin) receptor | 0 | 1.0279 | 2.0125 | 0 | 2.1812 | 1.1138 |
| 5435 | ENSMUSG00000026229 | 1521.907432 | 0.05931361 | 0.106977977 | 0.554446918 | 0.579273024 | 0.908337213 | Psmd1 | proteasome (prosome, macropain) 26S su | 1546.8 | 1484.3 | 1440.9 | 1657.1 | 1596.7 | 1405.7 |
| 5436 | ENSMUSG00000026234 | 3315.894733 | -0.056193154 | 0.11012899 | -0.510248517 | 0.609877366 | 0.916376001 | Ncl | nucleolin [Source:MGI Symbol;Acc:MGI:9 | 3247.7 | 3633.7 | 3260.2 | 3089.5 | 3660.1 | 3004 |
| 5437 | ENSMUSG00000026235 | 3.981376718 | -1.839851654 | 1.22748014 | -1.498885069 | 0.133903451 | NA | Epha4 | Eph receptor A4 [Source:MGI Symbol;Acc | 10.44 | 7.1954 | 1.0062 | 1.928 | 1.0906 | 2.2277 |
| 5438 | ENSMUSG00000026237 | 0 | NA | NA | NA | NA | NA | Nmur1 | neuromedin U receptor 1 [Source:MGI Sy | 0 | 0 | 0 | 0 | 0 | 0 |
| 5439 | ENSMUSG00000026238 | 4643.624976 | 0.059255644 | 0.093603666 | 0.63304833 | 0.526702082 | 0.888334265 | Ptma | prothymosin alpha [Source:MGI Symbol;A | 4353.6 | 4481.7 | 4810.9 | 5032.9 | 4650.4 | 4532.2 |
| 5440 | ENSMUSG00000026239 | 109.8396283 | -0.046249388 | 0.226292594 | -0.204378708 | 0.838057567 | 0.971073158 | Pde6d | phosphodiesterase 6D, cGMP-specific, ro | 126.89 | 108.96 | 98.612 | 112.79 | 97.065 | 114.73 |
| 5441 | ENSMUSG00000026240 | 196.4288225 | 0.092873089 | 0.190573202 | 0.487335511 | 0.626020586 | 0.919638611 | Cops7b | COP9 signalosome subunit 7B [Source:MG | 206.4 | 168.58 | 195.21 | 218.82 | 222.49 | 167.08 |
| 5442 | ENSMUSG00000026241 | 0 | NA | NA | NA | NA | NA | Nppc | natriuretic peptide type C [Source:MGI Sy | 0 | 0 | 0 | 0 | 0 | 0 |
| 5443 | ENSMUSG00000026245 | 542.8603698 | -0.103353599 | 0.125697318 | -0.822241879 | 0.410939248 | 0.833663203 | Farsb | phenylalanyl-tRNA synthetase, beta subu | 519.61 | 599.28 | 568.53 | 504.16 | 539.86 | 525.73 |
| 5444 | ENSMUSG00000026246 | 0.505131668 | 1.126696199 | 3.421159819 | 0.329331647 | 0.741905026 | NA | Alppl2 | alkaline phosphatase, placental-like 2 [So | 0.8031 | 0 | 0 | 0 | 0 | 2.2277 |
| 5445 | ENSMUSG00000026247 | 0 | NA | NA | NA | NA | NA | Ecel1 | endothelin converting enzyme-like 1 [Sou | 0 | 0 | 0 | 0 | 0 | 0 |
| 5446 | ENSMUSG00000026248 | 405.4921884 | -0.071669532 | 0.150886982 | -0.474988172 | 0.634795404 | 0.923478764 | Mrpl44 | mitochondrial ribosomal protein L44 [Sou | 438.49 | 388.55 | 418.6 | 340.28 | 394.8 | 452.22 |
| 5447 | ENSMUSG00000026249 | 466.3916337 | 1.044558942 | 0.418434256 | 2.496351406 | 0.012547823 | 0.126529431 | Serpine2 | serine (or cysteine) peptidase inhibitor, cl | 238.52 | 195.3 | 479.98 | 523.44 | 628.2 | 732.91 |
| 5448 | ENSMUSG00000026251 | 0 | NA | NA | NA | NA | NA | Chrnd | cholinergic receptor, nicotinic, delta polyp | 0 | 0 | 0 | 0 | 0 | 0 |
| 5449 | ENSMUSG00000026253 | 0.556921942 | 2.679043159 | 3.638243063 | 0.736356289 | 0.461513899 | NA | Chrng | cholinergic receptor, nicotinic, gamma po | 0 | 0 | 0 | 0 | 0 | 3.3415 |
| 5450 | ENSMUSG00000026254 | 839.2162484 | 0.097966598 | 0.122273036 | 0.801211788 | 0.423009048 | 0.84005258 | Eif4e2 | eukaryotic translation initiation factor 4E | 820.77 | 839.81 | 770.78 | 768.29 | 887.76 | 947.88 |
| 5451 | ENSMUSG00000026255 | 2.818814527 | -0.77689693 | 1.351610504 | -0.574793498 | 0.565430964 | NA | Efhd1 | EF hand domain containing 1 [Source:MG | 1.6062 | 4.1117 | 5.0312 | 2.8919 | 3.2719 | 0 |
| 5452 | ENSMUSG00000026258 | 0.656740811 | -0.013525372 | 3.112905782 | -0.004344934 | 0.996533255 | NA | Snorc | secondary ossification center associated r | 0 | 0 | 2.0125 | 1.928 | 0 | 0 |
| 5453 | ENSMUSG00000026259 | 878.8294719 | 0.431155073 | 0.115984899 | 3.717338003 | 0.000201333 | 0.006326357 | Ngef | neuronal guanine nucleotide exchange fa | 759.73 | 701.04 | 784.87 | 1079.7 | 1014.3 | 933.4 |
| 5454 | ENSMUSG00000026260 | 2566.888623 | -0.174604897 | 0.098453625 | -1.773473521 | 0.076150276 | 0.403877794 | Ndufa10 | NADH:ubiquinone oxidoreductase subuni | 2912 | 2665.4 | 2587.1 | 2275 | 2473.5 | 2488.3 |
| 5455 | ENSMUSG00000026269 | 1413.948718 | 0.055994512 | 0.09716399 | 0.576288732 | 0.564420038 | 0.901476527 | Rnpepl1 | arginyl aminopeptidase (aminopeptidase | 1370.9 | 1387.7 | 1400.7 | 1378.5 | 1473.4 | 1472.5 |
| 5456 | ENSMUSG00000026270 | 405.1119326 | 0.002658111 | 0.136436833 | 0.019482358 | 0.984456311 | 0.998580735 | Capn10 | calpain 10 [Source:MGI Symbol;Acc:MGI: | 389.5 | 383.41 | 441.74 | 420.29 | 398.08 | 397.64 |
| 5457 | ENSMUSG00000026271 | 5.578664542 | -1.613103411 | 1.011636091 | -1.594549094 | 0.110813104 | 0.489424955 | Gpr35 | G protein-coupled receptor 35 [Source:M | 8.031 | 8.2233 | 9.0562 | 4.8199 | 0 | 3.3415 |
| 5458 | ENSMUSG00000026272 | 4389.334504 | 0.503141112 | 0.143499391 | 3.506224715 | 0.000454511 | 0.011574035 | Agxt | alanine-glyoxylate aminotransferase [Sou | 3854.9 | 3376.7 | 3661.7 | 3944.6 | 5445.5 | 6052.6 |
| 5459 | ENSMUSG00000026273 | 300.9145865 | 0.233114369 | 0.173142087 | 1.346376107 | 0.178181253 | 0.610424831 | Mterf4 | mitochondrial transcription termination fa | 324.45 | 239.5 | 264.64 | 283.41 | 334.82 | 358.66 |
| 5460 | ENSMUSG00000026274 | 10.81500851 | -0.437300722 | 0.685466757 | -0.637960511 | 0.523499386 | 0.888334265 | Pask | PAS domain containing serine/threonine k | 16.865 | 7.1954 | 13.081 | 6.7478 | 7.6343 | 13.366 |
| 5461 | ENSMUSG00000026275 | 406.3191349 | -0.066648767 | 0.14049613 | -0.474381514 | 0.635227873 | 0.923478764 | Ppp1r7 | protein phosphatase 1, regulatory subunit | 402.35 | 443.03 | 401.49 | 355.71 | 420.98 | 414.35 |
| 5462 | ENSMUSG00000026276 | 22.25187684 | 0.31484198 | 0.474496828 | 0.663528103 | 0.506992387 | 0.88185929 | Septin2 | septin 2 [Source:MGI Symbol;Acc:MGI:97 | 20.881 | 20.558 | 18.112 | 28.919 | 29.447 | 15.594 |
| 5463 | ENSMUSG00000026277 | 803.0294583 | 0.188221935 | 0.119497062 | 1.57511768 | 0.115229284 | 0.498557093 | Stk25 | serine/threonine kinase 25 (yeast) [Sourc | 718.78 | 745.24 | 788.9 | 884.93 | 910.67 | 769.67 |
| 5464 | ENSMUSG00000026278 | 185.5817424 | -0.207169864 | 0.226230291 | -0.91574768 | 0.359799251 | 0.799452417 | Bok | BCL2-related ovarian killer [Source:MGI Sy | 186.32 | 166.52 | 243.51 | 141.7 | 153.78 | 221.65 |
| 5465 | ENSMUSG00000026279 | 151.317319 | 0.285586373 | 0.210534301 | 1.356483819 | 0.17494528 | 0.604926735 | Thap4 | THAP domain containing 4 [Source:MGI S | 138.94 | 153.16 | 116.72 | 141.7 | 164.68 | 192.69 |
| 5466 | ENSMUSG00000026280 | 519.3863103 | 0.114152479 | 0.123322431 | 0.925642465 | 0.354631817 | 0.794762101 | Atg4b | autophagy related 4B, cysteine peptidase | 495.51 | 512.93 | 488.03 | 521.51 | 571.48 | 526.85 |
| 5467 | ENSMUSG00000026281 | 298.606826 | -0.1048438 | 0.168854518 | -0.620912019 | 0.534657513 | 0.889858671 | Dtymk | deoxythymidylate kinase [Source:MGI Syn | 362.2 | 261.09 | 303.89 | 266.06 | 301.01 | 297.4 |
| 5468 | ENSMUSG00000026283 | 183.980932 | 0.014635911 | 0.18824052 | 0.077751117 | 0.938026032 | 0.988420305 | Ing5 | inhibitor of growth family, member 5 [Sou | 158.21 | 203.53 | 188.17 | 192.8 | 195.22 | 165.96 |
| 5469 | ENSMUSG00000026285 | 0.572870759 | -2.577107685 | 3.182795569 | -0.809699407 | 0.418112959 | NA | Pdcd1 | programmed cell death 1 [Source:MGI Syr | 2.4093 | 1.0279 | 0 | 0 | 0 | 0 |
| 5470 | ENSMUSG00000026288 | 46.45291792 | -1.153039728 | 0.376809215 | -3.060009369 | 0.002213301 | 0.037530107 | Inpp5d | inositol polyphosphate-5-phosphatase D [ | 50.595 | 93.541 | 48.3 | 28.919 | 26.175 | 31.188 |
| 5471 | ENSMUSG00000026289 | 711.2613531 | 0.128632778 | 0.117532252 | 1.094446637 | 0.273759139 | 0.728923893 | Atg16l1 | autophagy related 16-like 1 (S. cerevisiae) | 703.52 | 709.26 | 625.88 | 774.07 | 725.26 | 729.57 |
| 5472 | ENSMUSG00000026295 | 2273.210435 | 0.151837215 | 0.124437207 | 1.220191442 | 0.22239231 | 0.668407528 | Spp2 | secreted phosphoprotein 2 [Source:MGI S | 2382 | 2144.2 | 1933 | 2064.8 | 2491 | 2624.2 |
| 5473 | ENSMUSG00000026301 | 0.17131964 | -0.796629176 | 4.080472857 | -0.195229623 | 0.84521319 | NA | Iqca | IQ motif containing with AAA domain [Sou | 0 | 1.0279 | 0 | 0 | 0 | 0 |
| 5474 | ENSMUSG00000026303 | 16.87793559 | -0.579616179 | 0.561936754 | -1.031461592 | 0.30232441 | 0.752385931 | Mlph | melanophilin [Source:MGI Symbol;Acc:M( | 24.093 | 16.447 | 20.125 | 17.352 | 6.5437 | 16.708 |
| 5475 | ENSMUSG00000026304 | 423.4257485 | -0.00261855 | 0.150650836 | -0.01738158 | 0.986132204 | 0.99859176 | Rab17 | RAB17, member RAS oncogene family [So | 392.72 | 401.92 | 476.96 | 403.91 | 384.99 | 480.07 |
| 5476 | ENSMUSG00000026305 | 769.6644975 | -0.070856602 | 0.12740918 | -0.556134195 | 0.57811912 | 0.908137944 | Lrrfip1 | leucine rich repeat (in FLII) interacting pro | 753.31 | 811.03 | 800.97 | 661.29 | 745.98 | 845.41 |
| 5477 | ENSMUSG00000026307 | 1691.109709 | -0.013551803 | 0.096888964 | -0.139869415 | 0.888763167 | 0.982382091 | Scly | selenocysteine lyase [Source:MGI Symbol | 1660 | 1756.7 | 1680.4 | 1626.2 | 1653.4 | 1769.9 |
| 5478 | ENSMUSG00000026308 | 0.181769715 | 1.126924175 | 4.080472857 | 0.276174898 | 0.782413736 | NA | Klhl30 | kelch-like 30 [Source:MGI Symbol;Acc:MG | 0 | 0 | 0 | 0 | 1.0906 | 0 |
| 5479 | ENSMUSG00000026309 | 499.4365944 | 0.129392637 | 0.123957724 | 1.043844889 | 0.29655716 | 0.74713095 | Ilkap | integrin-linked kinase-associated serine/th | 492.3 | 454.34 | 484 | 509.94 | 518.04 | 537.99 |
| 5480 | ENSMUSG00000026311 | 148.6328507 | 0.366925496 | 0.209227995 | 1.753711292 | 0.07947999 | 0.413673469 | Asb1 | ankyrin repeat and SOCS box-containing 1 | 127.69 | 122.32 | 139.87 | 198.58 | 139.6 | 163.74 |
| 5481 | ENSMUSG00000026312 | 0 | NA | NA | NA | NA | NA | Cdh7 | cadherin 7, type 2 [Source:MGI Symbol;A | 0 | 0 | 0 | 0 | 0 | 0 |
| 5482 | ENSMUSG00000026313 | 191.6485547 | -0.040293599 | 0.179467942 | -0.224516971 | 0.822355063 | 0.969197142 | Hdac4 | histone deacetylase 4 [Source:MGI Symbo | 191.94 | 195.3 | 196.22 | 214.97 | 181.04 | 170.42 |
| 5483 | ENSMUSG00000026315 | 66.16696511 | -0.082232659 | 0.314037964 | -0.26185579 | 0.793432625 | 0.961915077 | Serpinb8 | serine (or cysteine) peptidase inhibitor, cl | 82.72 | 75.038 | 46.287 | 77.118 | 52.35 | 63.489 |
| 5484 | ENSMUSG00000026317 | 785.058123 | -0.451875727 | 0.127713587 | -3.538196186 | 0.000402871 | 0.010687243 | Cln8 | CLN8 transmembrane ER and ERGIC prote | 936.42 | 903.54 | 880.46 | 615.02 | 764.52 | 610.39 |

| | | | | | | | | | | | | | | | |
|---|---|---|---|---|---|---|---|---|---|---|---|---|---|---|---|
| 5485 | ENSMUSG00000026319 | 365.9569578 | -0.139176069 | 0.148526776 | -0.937043628 | 0.348736117 | 0.789995175 | Relch | RAB11 binding and LisH domain, coiled-cc | 348.55 | 367.99 | 434.7 | 338.36 | 367.54 | 338.61 |
| 5486 | ENSMUSG00000026321 | 9.336848724 | -1.759280753 | 0.814577636 | -2.159745954 | 0.030792342 | 0.232721383 | Tnfrsf11a | tumor necrosis factor receptor superfamil | 17.668 | 14.391 | 11.069 | 2.8919 | 1.0906 | 8.9108 |
| 5487 | ENSMUSG00000026322 | 0 | NA | NA | NA | NA | NA | Htr4 | 5 hydroxytryptamine (serotonin) receptor | 0 | 0 | 0 | 0 | 0 | 0 |
| 5488 | ENSMUSG00000026327 | 0 | NA | NA | NA | NA | NA | Serpinb11 | serine (or cysteine) peptidase inhibitor, cl | 0 | 0 | 0 | 0 | 0 | 0 |
| 5489 | ENSMUSG00000026331 | 0 | NA | NA | NA | NA | NA | Slco6c1 | solute carrier organic anion transporter fa | 0 | 0 | 0 | 0 | 0 | 0 |
| 5490 | ENSMUSG00000026333 | 201.1026767 | -0.683281108 | 0.205487953 | -3.325163829 | 0.000883666 | 0.019076633 | Gin1 | gypsy retrotransposon integrase 1 [Source | 237.72 | 302.21 | 204.27 | 179.3 | 147.23 | 135.89 |
| 5491 | ENSMUSG00000026335 | 306.587524 | -1.114413225 | 0.175901644 | -6.33543384 | 2.37E-10 | 6.11E-08 | Pam | peptidylglycine alpha-amidating monooxy | 504.35 | 402.94 | 350.17 | 184.12 | 195.22 | 202.72 |
| 5492 | ENSMUSG00000026336 | 0 | NA | NA | NA | NA | NA | Slco6d1 | solute carrier organic anion transporter fa | 0 | 0 | 0 | 0 | 0 | 0 |
| 5493 | ENSMUSG00000026339 | 271.5210333 | 0.106253495 | 0.158646917 | 0.669748248 | 0.503018289 | 0.881220881 | Ccdc93 | coiled-coil domain containing 93 [Source:I | 248.96 | 250.81 | 284.77 | 261.24 | 275.93 | 307.42 |
| 5494 | ENSMUSG00000026341 | 1426.154395 | -0.110072149 | 0.104991856 | -1.048387497 | 0.294460112 | 0.745164683 | Actr3 | ARP3 actin-related protein 3 [Source:MGI | 1379.7 | 1605.6 | 1457 | 1340.9 | 1403.6 | 1370 |
| 5495 | ENSMUSG00000026342 | 907.871645 | 0.034804717 | 0.108654963 | 0.32032331 | 0.748723254 | 0.951074124 | Slc35f5 | solute carrier family 35, member F5 [Sour | 921.96 | 910.74 | 857.32 | 870.47 | 946.66 | 940.08 |
| 5496 | ENSMUSG00000026343 | 30.47329253 | -0.023513215 | 0.486231468 | -0.04835807 | 0.961430875 | 0.992661062 | Gpr39 | G protein-coupled receptor 39 [Source:M | 40.155 | 32.893 | 19.119 | 45.307 | 14.178 | 31.188 |
| 5497 | ENSMUSG00000026344 | 0.620790101 | -1.233953725 | 2.860353486 | -0.431399032 | 0.666178254 | NA | Lypd1 | Ly6/Plaur domain containing 1 [Source:M | 1.6062 | 1.0279 | 0 | 0 | 1.0906 | 0 |
| 5498 | ENSMUSG00000026347 | 4.363827452 | 0.421267526 | 1.246051151 | 0.338082049 | 0.735301358 | NA | Tmem163 | transmembrane protein 163 [Source:MGI | 4.0155 | 5.1396 | 2.0125 | 1.928 | 13.087 | 0 |
| 5499 | ENSMUSG00000026348 | 306.9895529 | 0.830901432 | 0.221342699 | 3.753913882 | 0.000174095 | 0.00559579 | Acmsd | amino carboxymuconate semialdehyde de | 218.44 | 198.39 | 245.52 | 270.88 | 375.17 | 533.53 |
| 5500 | ENSMUSG00000026349 | 587.2084341 | 0.188268359 | 0.132720528 | 1.418532328 | 0.156035407 | 0.574894692 | Ccnt2 | cyclin T2 [Source:MGI Symbol;Acc:MGI:19 | 534.06 | 540.68 | 572.55 | 642.01 | 689.27 | 544.67 |
| 5501 | ENSMUSG00000026353 | 1630.238907 | 0.118393787 | 0.103299376 | 1.146122961 | 0.251744278 | 0.704249849 | Ubxn4 | UBX domain protein 4 [Source:MGI Symbo | 1609.4 | 1600.5 | 1480.2 | 1769.9 | 1568.3 | 1753.2 |
| 5502 | ENSMUSG00000026354 | 0 | NA | NA | NA | NA | NA | Lct | lactase [Source:MGI Symbol;Acc:MGI:104 | 0 | 0 | 0 | 0 | 0 | 0 |
| 5503 | ENSMUSG00000026355 | 71.17769793 | -0.312975586 | 0.29625766 | -1.056430357 | 0.290771643 | 0.740574229 | Mcm6 | minichromosome maintenance complex c | 98.782 | 62.703 | 74.462 | 50.127 | 74.162 | 66.831 |
| 5504 | ENSMUSG00000026356 | 1654.320609 | 0.06378171 | 0.110921295 | 0.575017715 | 0.565279316 | 0.901542552 | Dars | aspartyl-tRNA synthetase [Source:MGI Sy | 1630.3 | 1602.5 | 1620.1 | 1597.3 | 1909.7 | 1566.1 |
| 5505 | ENSMUSG00000026357 | 3.149679084 | -1.30803621 | 1.360281816 | -0.961592072 | 0.336254556 | NA | Rgs18 | regulator of G-protein signaling 18 [Sourc | 7.2279 | 4.1117 | 2.0125 | 0 | 1.0906 | 4.4554 |
| 5506 | ENSMUSG00000026358 | 12.47084271 | -2.941808904 | 0.817794 | -3.597249312 | 0.0003216 | 0.00921766 | Rgs1 | regulator of G-protein signaling 1 [Source: | 20.078 | 31.865 | 14.087 | 0 | 5.4531 | 3.3415 |
| 5507 | ENSMUSG00000026360 | 41.80395682 | -0.913875016 | 0.434191362 | -2.10477475 | 0.03531092 | 0.25220602 | Rgs2 | regulator of G-protein signaling 2 [Source: | 46.58 | 72.982 | 44.275 | 26.027 | 14.178 | 46.781 |
| 5508 | ENSMUSG00000026361 | 382.9145348 | -0.175355783 | 0.150333379 | -1.166446101 | 0.243434129 | 0.693217879 | Cdc73 | cell division cycle 73, Paf1/RNA polymeras | 371.03 | 471.81 | 376.34 | 364.38 | 368.63 | 345.29 |
| 5509 | ENSMUSG00000026365 | 15313.43172 | 0.101436335 | 0.138019102 | 0.734944177 | 0.462373492 | 0.860650866 | Cfh | complement component factor h [Source: | 13731 | 15799 | 14798 | 18979 | 16072 | 12502 |
| 5510 | ENSMUSG00000026368 | 3974.553654 | 0.197392767 | 0.106210764 | 1.858500582 | 0.063097961 | 0.359593737 | F13b | coagulation factor XIII, beta subunit [Sour | 3365.8 | 3767.3 | 3976.7 | 3877.1 | 4508.6 | 4351.8 |
| 5511 | ENSMUSG00000026374 | 844.6667451 | 0.041393645 | 0.113418752 | 0.364962976 | 0.715139043 | 0.944595913 | Tsn | translin [Source:MGI Symbol;Acc:MGI:109 | 807.12 | 806.92 | 883.48 | 802.99 | 897.58 | 869.91 |
| 5512 | ENSMUSG00000026377 | 415.3809401 | 0.16971571 | 0.173881393 | 0.976042964 | 0.329043174 | 0.773866742 | Nifk | nucleolar protein interacting with the FHA | 401.55 | 426.59 | 344.14 | 348.96 | 533.31 | 437.74 |
| 5513 | ENSMUSG00000026380 | 109.5731308 | -0.359555058 | 0.221629006 | -1.622328527 | 0.104733024 | 0.476887243 | Tfcp2l1 | transcription factor CP2-like 1 [Source:MG | 125.28 | 127.46 | 116.72 | 99.29 | 83.978 | 104.7 |
| 5514 | ENSMUSG00000026383 | 605.3407531 | 0.496033001 | 0.122786973 | 4.039785238 | 5.35E-05 | 0.002217268 | Epb41l5 | erythrocyte membrane protein band 4.1 l | 458.57 | 522.18 | 527.27 | 729.73 | 691.45 | 702.84 |
| 5515 | ENSMUSG00000026384 | 247.4972247 | 0.01231777 | 0.168175135 | 0.073243706 | 0.941612188 | 0.989428164 | Ptpn4 | protein tyrosine phosphatase, non-recept | 229.69 | 272.4 | 237.47 | 232.32 | 279.2 | 233.91 |
| 5516 | ENSMUSG00000026385 | 10296.24742 | 0.321918897 | 0.126963248 | 2.535528215 | 0.011227788 | 0.11768024 | Dbi | diazepam binding inhibitor [Source:MGI S | 10716 | 8874 | 7864.8 | 10628 | 10921 | 12774 |
| 5517 | ENSMUSG00000026387 | 2.086235272 | 0.364441982 | 1.570103978 | 0.232113279 | 0.816450031 | NA | Sctr | secretin receptor [Source:MGI Symbol;Ac | 2.4093 | 3.0838 | 0 | 4.8199 | 1.0906 | 1.1138 |
| 5518 | ENSMUSG00000026388 | 6.925577898 | -0.11642964 | 0.823481219 | -0.141387122 | 0.887564127 | 0.982382091 | 3110009E18 | RIKEN cDNA 3110009E18 gene [Source:M | 6.4248 | 6.1675 | 9.0562 | 6.7478 | 9.8156 | 3.3415 |
| 5519 | ENSMUSG00000026389 | 469.2649904 | -0.121685548 | 0.161290622 | -0.754448997 | 0.45057966 | 0.855818389 | Steap3 | STEAP family member 3 [Source:MGI Sym | 427.25 | 455.37 | 584.63 | 391.38 | 462.42 | 494.55 |
| 5520 | ENSMUSG00000026390 | 129.9214715 | -3.703773349 | 1.106970989 | -3.345863066 | 0.000820269 | 0.018030957 | Marco | macrophage receptor with collagenous st | 102.8 | 585.91 | 35.219 | 12.532 | 17.45 | 25.618 |
| 5521 | ENSMUSG00000026393 | 1256.340235 | -0.092247125 | 0.104748057 | -0.880657147 | 0.378503418 | 0.811904706 | Nek7 | NIMA (never in mitosis gene a)-related ex | 1212.7 | 1356.9 | 1321.2 | 1261.8 | 1221.5 | 1164 |
| 5522 | ENSMUSG00000026394 | 0 | NA | NA | NA | NA | NA | Atp6v1g3 | ATPase, H+ transporting, lysosomal V1 sul | 0 | 0 | 0 | 0 | 0 | 0 |
| 5523 | ENSMUSG00000026395 | 78.7721263 | -1.818382721 | 0.315846268 | -5.757176532 | 8.55E-09 | 1.36E-06 | Ptprc | protein tyrosine phosphatase, receptor ty | 101.19 | 166.52 | 100.62 | 34.703 | 27.265 | 42.326 |
| 5524 | ENSMUSG00000026398 | 1059.526864 | 0.168621051 | 0.146870687 | 1.148091932 | 0.250930614 | 0.702929347 | Nr5a2 | nuclear receptor subfamily 5, group A, me | 855.3 | 1100.9 | 1037.4 | 1025.7 | 999.01 | 1338.8 |
| 5525 | ENSMUSG00000026399 | 124.235291 | -0.821600706 | 0.230261397 | -3.568121777 | 0.000359549 | 0.009981809 | Cd55 | CD55 molecule, decay accelerating factor | 146.16 | 182.97 | 146.91 | 83.866 | 76.343 | 109.16 |
| 5526 | ENSMUSG00000026401 | 0.133850373 | -0.796629176 | 4.080472857 | -0.195229623 | 0.84521319 | NA | Cd55b | CD55 molecule, decay accelerating factor | 0.8031 | 0 | 0 | 0 | 0 | 0 |
| 5527 | ENSMUSG00000026404 | 104.6567919 | 0.076252448 | 0.250875603 | 0.30394525 | 0.761169603 | 0.95402891 | Ddx59 | DEAD box helicase 59 [Source:MGI Symbo | 85.129 | 105.88 | 114.71 | 81.938 | 118.88 | 121.41 |
| 5528 | ENSMUSG00000026405 | 10845.41695 | 0.083675522 | 0.082333208 | 1.016303432 | 0.309484915 | 0.758143666 | C4bp | complement component 4 binding protei | 10751 | 10587 | 10255 | 10914 | 11579 | 10987 |
| 5529 | ENSMUSG00000026407 | 0 | NA | NA | NA | NA | NA | Cacna1s | calcium channel, voltage-dependent, L typ | 0 | 0 | 0 | 0 | 0 | 0 |
| 5530 | ENSMUSG00000026409 | 230.3568861 | -0.192081743 | 0.165819989 | -1.158375083 | 0.246711003 | 0.696725814 | Pfkfb2 | 6-phosphofructo-2-kinase/fructose-2,6-bi | 240.13 | 239.5 | 257.6 | 218.82 | 233.39 | 192.69 |
| 5531 | ENSMUSG00000026411 | 332.0134039 | 0.109870953 | 0.171428353 | 0.64091471 | 0.521578098 | 0.888064228 | Tmem9 | transmembrane protein 9 [Source:MGI Sy | 319.63 | 332.02 | 305.9 | 274.73 | 411.16 | 348.63 |
| 5532 | ENSMUSG00000026413 | 2.778114618 | 1.756994089 | 1.412372061 | 1.244002298 | 0.213498717 | NA | Pkp1 | plakophilin 1 [Source:MGI Symbol;Acc:MG | 0.8031 | 0 | 3.0187 | 2.8919 | 3.2719 | 6.6831 |
| 5533 | ENSMUSG00000026414 | 4.235274433 | -0.29240038 | 1.196275158 | -0.244425689 | 0.806901148 | NA | Tnnt2 | troponin T2, cardiac [Source:MGI Symbol; | 8.8341 | 2.0558 | 3.0187 | 4.8199 | 0 | 6.6831 |
| 5534 | ENSMUSG00000026415 | 6.074938242 | -0.717174536 | 0.937830619 | -0.764716487 | 0.444440387 | 0.851724159 | Fcamr | Fc receptor, IgA, IgM, high affinity [Source | 10.44 | 2.0558 | 10.062 | 2.8919 | 6.5437 | 4.4554 |
| 5535 | ENSMUSG00000026416 | 0 | NA | NA | NA | NA | NA | Il20 | interleukin 20 [Source:MGI Symbol;Acc:M | 0 | 0 | 0 | 0 | 0 | 0 |
| 5536 | ENSMUSG00000026417 | 31898.3331 | -0.09892232 | 0.13811711 | -0.71622061 | 0.473855139 | 0.868952133 | Pigr | polymeric immunoglobulin receptor [Sour | 25134 | 35998 | 37845 | 31570 | 31350 | 29493 |
| 5537 | ENSMUSG00000026418 | 0 | NA | NA | NA | NA | NA | Tnni1 | troponin I, skeletal, slow 1 [Source:MGI Sy | 0 | 0 | 0 | 0 | 0 | 0 |
| 5538 | ENSMUSG00000026420 | 0.133850373 | -0.796629176 | 4.080472857 | -0.195229623 | 0.84521319 | NA | Il24 | interleukin 24 [Source:MGI Symbol;Acc:M | 0.8031 | 0 | 0 | 0 | 0 | 0 |
| 5539 | ENSMUSG00000026421 | 96.1352738 | -0.577648455 | 0.277598508 | -2.080877377 | 0.037445131 | 0.262483703 | Csrp1 | cysteine and glycine-rich protein 1 [Sourc | 142.95 | 123.35 | 78.487 | 65.55 | 80.706 | 85.766 |
| 5540 | ENSMUSG00000026424 | 0 | NA | NA | NA | NA | NA | Gpr37l1 | G protein-coupled receptor 37-like 1 [Sou | 0 | 0 | 0 | 0 | 0 | 0 |

| | | | | | | | | | | | | | | | |
|---|---|---|---|---|---|---|---|---|---|---|---|---|---|---|---|
| 5541 | ENSMUSG00000026425 | 225.7628977 | -0.005984524 | 0.17375994 | -0.034441335 | 0.972525222 | 0.99573532 | Srgap2 | SLIT-ROBO Rho GTPase activating protein | 214.43 | 244.64 | 220.37 | 255.45 | 220.3 | 199.38 |
| 5542 | ENSMUSG00000026426 | 312.0781193 | 0.156710674 | 0.149526066 | 1.048049201 | 0.294615939 | 0.745164683 | Arl8a | ADP-ribosylation factor-like 8A [Source:M | 321.24 | 284.73 | 278.73 | 323.9 | 313.01 | 350.86 |
| 5543 | ENSMUSG00000026427 | 641.4052694 | -0.147159433 | 0.121557175 | -1.210619054 | 0.226041438 | 0.672590694 | Eif2d | eukaryotic translation initiation factor 2D | 696.29 | 622.92 | 702.36 | 577.42 | 641.28 | 608.16 |
| 5544 | ENSMUSG00000026429 | 13.06654337 | -0.042183673 | 0.651228311 | -0.064775551 | 0.948352708 | 0.990731019 | Ube2t | ubiquitin-conjugating enzyme E2T [Source | 15.259 | 8.2233 | 16.1 | 6.7478 | 10.906 | 21.163 |
| 5545 | ENSMUSG00000026430 | 195.5142271 | 0.203776821 | 0.215219685 | 0.946831703 | 0.343724544 | 0.785513865 | Rassf5 | Ras association (RalGDS/AF-6) domain far | 187.12 | 173.72 | 184.14 | 209.18 | 151.6 | 267.32 |
| 5546 | ENSMUSG00000026432 | 0 | NA | NA | NA | NA | NA | Avpr1b | arginine vasopressin receptor 1B [Source: | 0 | 0 | 0 | 0 | 0 | 0 |
| 5547 | ENSMUSG00000026433 | 247.1886597 | 0.130360653 | 0.15447353 | 0.843902851 | 0.398723695 | 0.82588279 | Rab29 | RAB29, member RAS oncogene family [So | 232.1 | 241.56 | 234.45 | 252.56 | 258.48 | 263.98 |
| 5548 | ENSMUSG00000026434 | 1949.730512 | -0.30852798 | 0.114251089 | -2.70043798 | 0.006924825 | 0.08346722 | Nucks1 | nuclear casein kinase and cyclin-depende | 1917 | 2378.6 | 2177.5 | 1655.1 | 1871.5 | 1698.6 |
| 5549 | ENSMUSG00000026435 | 1201.480897 | -1.657470223 | 0.338771258 | -4.89259399 | 9.95E-07 | 8.09E-05 | Slc45a3 | solute carrier family 45, member 3 [Sourc | 1328.3 | 2341.6 | 1804.2 | 644.9 | 707.81 | 382.05 |
| 5550 | ENSMUSG00000026436 | 756.6046263 | 0.032764414 | 0.127587062 | 0.256800445 | 0.797332827 | 0.962236316 | Elk4 | ELK4, member of ETS oncogene family [So | 713.15 | 833.64 | 698.33 | 837.7 | 737.26 | 719.54 |
| 5551 | ENSMUSG00000026437 | 1143.276117 | 0.035312588 | 0.126357808 | 0.279465027 | 0.779887974 | 0.959870443 | Cdk18 | cyclin-dependent kinase 18 [Source:MGI S | 1172.5 | 1081.4 | 1134 | 1223.3 | 1274.9 | 973.5 |
| 5552 | ENSMUSG00000026439 | 418.5593801 | 0.063960605 | 0.129958344 | 0.492162362 | 0.62260457 | 0.918870832 | Rbbp5 | retinoblastoma binding protein 5, histone | 402.35 | 399.86 | 425.64 | 417.4 | 440.61 | 425.49 |
| 5553 | ENSMUSG00000026442 | 12.84661989 | -0.155438115 | 0.73995624 | -0.210063929 | 0.833617778 | 0.970792812 | Nfasc | neurofascin [Source:MGI Symbol;Acc:MG | 7.2279 | 10.279 | 23.144 | 8.6758 | 4.3625 | 23.391 |
| 5554 | ENSMUSG00000026443 | 0 | NA | NA | NA | NA | NA | Lrrn2 | leucine rich repeat protein 2, neuronal [So | 0 | 0 | 0 | 0 | 0 | 0 |
| 5555 | ENSMUSG00000026447 | 157.5491985 | 0.362113618 | 0.218363817 | 1.658304119 | 0.097256098 | 0.459192119 | Pik3c2b | phosphatidylinositol-4-phosphate 3-kinas | 130.91 | 119.24 | 164.02 | 214.97 | 164.68 | 151.48 |
| 5556 | ENSMUSG00000026450 | 0.167707431 | -0.796629176 | 4.080472857 | -0.195229623 | 0.84521319 | NA | Chit1 | chitinase 1 (chitotriosidase) [Source:MGI S | 0 | 0 | 1.0062 | 0 | 0 | 0 |
| 5557 | ENSMUSG00000026452 | 0.465832987 | -0.698295221 | 3.18348759 | -0.219349126 | 0.826378096 | NA | Syt2 | synaptotagmin II [Source:MGI Symbol;Acc | 0.8031 | 1.0279 | 0 | 0.964 | 0 | 0 |
| 5558 | ENSMUSG00000026455 | 280.8761299 | 0.238804499 | 0.159646156 | 1.495836198 | 0.134696346 | 0.537616739 | Klhl12 | kelch-like 12 [Source:MGI Symbol;Acc:MG | 261.81 | 259.04 | 252.57 | 340.28 | 306.46 | 265.09 |
| 5559 | ENSMUSG00000026456 | 108.2614089 | 0.318138387 | 0.247130794 | 1.287327986 | 0.197979993 | 0.640049759 | Cyb5r1 | cytochrome b5 reductase 1 [Source:MGI S | 80.31 | 90.457 | 118.74 | 120.5 | 100.34 | 139.23 |
| 5560 | ENSMUSG00000026457 | 2013.167549 | 0.099692937 | 0.098121291 | 1.016017375 | 0.309621113 | 0.758272905 | Adipor1 | adiponectin receptor 1 [Source:MGI Symb | 1881.7 | 1991.1 | 1958.2 | 1981.9 | 2032.9 | 2233.3 |
| 5561 | ENSMUSG00000026458 | 13.00429099 | -0.828657259 | 0.686711632 | -1.206703398 | 0.22754639 | 0.673700979 | Ppfia4 | protein tyrosine phosphatase, receptor ty | 11.243 | 29.81 | 9.0562 | 12.532 | 9.8156 | 5.5692 |
| 5562 | ENSMUSG00000026459 | 0 | NA | NA | NA | NA | NA | Myog | myogenin [Source:MGI Symbol;Acc:MGI:9 | 0 | 0 | 0 | 0 | 0 | 0 |
| 5563 | ENSMUSG00000026463 | 42.46016724 | -1.173058048 | 0.380577975 | -3.082306713 | 0.002054031 | 0.035537681 | Atp2b4 | ATPase, Ca++ transporting, plasma memb | 62.642 | 60.647 | 53.331 | 32.775 | 14.178 | 31.188 |
| 5564 | ENSMUSG00000026466 | 718.2334779 | 0.035842301 | 0.116271763 | 0.308263163 | 0.757882094 | 0.953466696 | Tor1aip1 | torsin A interacting protein 1 [Source:MGI | 704.32 | 741.13 | 682.23 | 684.42 | 771.07 | 726.23 |
| 5565 | ENSMUSG00000026468 | 0 | NA | NA | NA | NA | NA | Lhx4 | LIM homeobox protein 4 [Source:MGI Syn | 0 | 0 | 0 | 0 | 0 | 0 |
| 5566 | ENSMUSG00000026469 | 711.2006884 | -0.036842897 | 0.114939211 | -0.320542453 | 0.748557154 | 0.951074124 | Xpr1 | xenotropic and polytropic retrovirus recep | 685.85 | 743.18 | 732.55 | 714.31 | 726.35 | 664.96 |
| 5567 | ENSMUSG00000026470 | 202.2539704 | -0.03815443 | 0.180699401 | -0.211148625 | 0.832771298 | 0.970611245 | Stx6 | syntaxin 6 [Source:MGI Symbol;Acc:MGI:1 | 197.56 | 184 | 233.45 | 209.18 | 181.04 | 208.29 |
| 5568 | ENSMUSG00000026471 | 76.78111172 | 0.173090635 | 0.29709112 | 0.582618004 | 0.560150482 | 0.899475387 | Mr1 | major histocompatibility complex, class I-r | 66.657 | 97.652 | 52.325 | 75.19 | 73.071 | 95.791 |
| 5569 | ENSMUSG00000026473 | 19388.91291 | -0.154365815 | 0.104904088 | -1.471494747 | 0.141157367 | 0.549582815 | Glul | glutamate-ammonia ligase (glutamine syn | 22875 | 19297 | 19102 | 16961 | 18888 | 19210 |
| 5570 | ENSMUSG00000026475 | 4223.035077 | -0.722519933 | 0.63454478 | -1.138642938 | 0.254852112 | 0.708380729 | Rgs16 | regulator of G-protein signaling 16 [Sourc | 7859.2 | 4887.7 | 3029.8 | 2941.1 | 5243.7 | 1376.7 |
| 5571 | ENSMUSG00000026478 | 196.2044305 | -0.519485436 | 0.203477964 | -2.553030446 | 0.010679016 | 0.113741744 | Lamc1 | laminin, gamma 1 [Source:MGI Symbol;Ac | 233.7 | 262.12 | 198.23 | 197.62 | 137.42 | 148.14 |
| 5572 | ENSMUSG00000026479 | 7.411125469 | 0.093451574 | 0.828465159 | 0.11280085 | 0.910188445 | 0.985622944 | Lamc2 | laminin, gamma 2 [Source:MGI Symbol;Ac | 7.2279 | 11.307 | 3.0187 | 9.6398 | 4.3625 | 8.9108 |
| 5573 | ENSMUSG00000026480 | 32.72152786 | -0.90407855 | 0.480324692 | -1.882223763 | 0.05980564 | 0.348100114 | Ncf2 | neutrophil cytosolic factor 2 [Source:MGI | 40.958 | 61.675 | 25.156 | 15.424 | 16.359 | 36.757 |
| 5574 | ENSMUSG00000026482 | 110.0280805 | -0.73101421 | 0.253478266 | -2.883932497 | 0.003927429 | 0.056579946 | Rgl1 | ral guanine nucleotide dissociation stimula | 155 | 138.77 | 117.73 | 75.19 | 65.437 | 108.04 |
| 5575 | ENSMUSG00000026483 | 50.6255237 | -0.944621134 | 0.344560394 | -2.741525579 | 0.00611546 | 0.076332607 | Niban1 | niban apoptosis regulator 1 [Source:MGI S | 68.264 | 76.066 | 55.343 | 28.919 | 27.265 | 47.895 |
| 5576 | ENSMUSG00000026484 | 268.1711404 | -0.023919872 | 0.158042485 | -0.1513509 | 0.879698919 | 0.98128671 | Rnf2 | ring finger protein 2 [Source:MGI Symbol; | 266.63 | 299.12 | 245.52 | 264.13 | 257.39 | 276.23 |
| 5577 | ENSMUSG00000026489 | 3971.168849 | -0.292682347 | 0.165403133 | -1.769509085 | 0.076808956 | 0.405609617 | Coq8a | coenzyme Q8A [Source:MGI Symbol;Acc:M | 3158.6 | 4281.3 | 5679.2 | 3590.8 | 3470.3 | 3646.7 |
| 5578 | ENSMUSG00000026490 | 543.6940696 | -0.066535718 | 0.132083703 | -0.503739039 | 0.614444771 | 0.917509899 | Cdc42bpa | CDC42 binding protein kinase alpha [Sour | 587.07 | 516.01 | 565.51 | 563.93 | 554.03 | 475.61 |
| 5579 | ENSMUSG00000026491 | 1003.075158 | -0.400982164 | 0.178644977 | -2.244575642 | 0.024795388 | 0.201483422 | Ahctf1 | AT hook containing transcription factor 1 | 925.17 | 1220.1 | 1280.9 | 1031.5 | 930.3 | 630.44 |
| 5580 | ENSMUSG00000026492 | 561.2740087 | -0.026714461 | 0.130263275 | -0.205080526 | 0.837509211 | 0.971073158 | Tfb2m | transcription factor B2, mitochondrial [So | 530.85 | 580.77 | 588.65 | 578.39 | 592.21 | 496.77 |
| 5581 | ENSMUSG00000026494 | 9.126330224 | -0.188056023 | 0.717738394 | -0.262011931 | 0.793312244 | 0.961896915 | Kif26b | kinesin family member 26B [Source:MGI S | 8.8341 | 7.1954 | 13.081 | 5.7839 | 8.7249 | 11.138 |
| 5582 | ENSMUSG00000026495 | 107.3324842 | -0.015590306 | 0.232538225 | -0.067044056 | 0.946546631 | 0.990583204 | Efcab2 | EF-hand calcium binding domain 2 [Sourc | 93.16 | 113.07 | 117.73 | 95.434 | 123.24 | 101.36 |
| 5583 | ENSMUSG00000026496 | 722.3669056 | -0.059315525 | 0.116610489 | -0.50866372 | 0.610987957 | 0.916376001 | Parp1 | poly (ADP-ribose) polymerase family, mer | 692.27 | 733.93 | 785.88 | 685.39 | 738.35 | 698.38 |
| 5584 | ENSMUSG00000026497 | 0 | NA | NA | NA | NA | NA | Mixl1 | Mix1 homeobox-like 1 (Xenopus laevis) [S | 0 | 0 | 0 | 0 | 0 | 0 |
| 5585 | ENSMUSG00000026499 | 400.2079042 | -0.103778368 | 0.14935142 | -0.69486027 | 0.487142873 | 0.873910634 | Acbd3 | acyl-Coenzyme A binding domain containi | 375.85 | 465.65 | 402.5 | 345.1 | 411.16 | 400.98 |
| 5586 | ENSMUSG00000026500 | 9.595625589 | 0.614734302 | 0.776893471 | 0.791272323 | 0.428785095 | 0.842753542 | Cox20 | cytochrome c oxidase assembly protein 2( | 1.6062 | 12.335 | 9.0562 | 12.532 | 10.906 | 11.138 |
| 5587 | ENSMUSG00000026502 | 353.7900942 | -0.561238043 | 0.166053163 | -3.379869641 | 0.000725202 | 0.016492648 | Desi2 | desumoylating isopeptidase 2 [Source:MG | 356.58 | 468.73 | 440.74 | 284.37 | 322.82 | 249.5 |
| 5588 | ENSMUSG00000026504 | 176.8472012 | 0.154845824 | 0.182270733 | 0.849537503 | 0.395582271 | 0.823148032 | Sdccag8 | serologically defined colon cancer antigen | 170.26 | 181.94 | 149.93 | 189.9 | 191.95 | 177.1 |
| 5589 | ENSMUSG00000026509 | 677.3385891 | -0.053171399 | 0.137111006 | -0.387798181 | 0.698165392 | 0.940598382 | Capn2 | calpain 2 [Source:MGI Symbol;Acc:MGI:88 | 701.91 | 651.7 | 715.44 | 651.65 | 573.67 | 769.67 |
| 5590 | ENSMUSG00000026510 | 415.2671274 | -0.559781279 | 0.150825007 | -3.711461991 | 0.000206066 | 0.006423563 | Trp53bp2 | transformation related protein 53 binding | 519.61 | 536.57 | 427.65 | 302.69 | 365.36 | 339.72 |
| 5591 | ENSMUSG00000026511 | 1409.811781 | 0.038886206 | 0.099174457 | 0.392099007 | 0.694985061 | 0.939992732 | Srp9 | signal recognition particle 9 [Source:MGI S | 1438.4 | 1343.5 | 1389.6 | 1366 | 1472.3 | 1449.1 |
| 5592 | ENSMUSG00000026514 | 0 | NA | NA | NA | NA | NA | Cnih3 | cornichon family AMPA receptor auxiliary | 0 | 0 | 0 | 0 | 0 | 0 |
| 5593 | ENSMUSG00000026516 | 324.3081214 | -0.043645909 | 0.145334089 | -0.300314327 | 0.763937406 | 0.954980327 | Nvl | nuclear VCP-like [Source:MGI Symbol;Acc | 330.88 | 336.13 | 320.99 | 343.18 | 321.73 | 292.94 |
| 5594 | ENSMUSG00000026519 | 283.0917196 | 0.199985735 | 0.195730638 | 1.021739554 | 0.306904185 | 0.755971071 | Tmem63a | transmembrane protein 63a [Source:MGI | 214.43 | 344.35 | 232.44 | 280.52 | 327.19 | 299.62 |
| 5595 | ENSMUSG00000026520 | 87.09043945 | -0.157114968 | 0.301252829 | -0.521538566 | 0.601991647 | 0.915461118 | Pycr2 | pyrroline-5-carboxylate reductase family, | 97.175 | 95.596 | 82.512 | 79.046 | 51.259 | 116.95 |
| 5596 | ENSMUSG00000026523 | 0 | NA | NA | NA | NA | NA | Wdr64 | WD repeat domain 64 [Source:MGI Symbo | 0 | 0 | 0 | 0 | 0 | 0 |

| | | | | | | | | | | | | | | | |
|---|---|---|---|---|---|---|---|---|---|---|---|---|---|---|---|
| 5597 | ENSMUSG00000026525 | 28.40033654 | 0.579788167 | 0.471366582 | 1.230015425 | 0.218691329 | 0.663735712 | Opn3 | opsin 3 [Source:MGI Symbol;Acc:MGI:133 | 16.865 | 14.391 | 37.231 | 33.739 | 41.443 | 26.732 |
| 5598 | ENSMUSG00000026526 | 4458.241278 | -0.034621009 | 0.093600841 | -0.369879245 | 0.711472467 | 0.944239215 | Fh1 | fumarate hydratase 1 [Source:MGI Symbc | 4662 | 4375.8 | 4495.9 | 4078.6 | 4486.8 | 4650.3 |
| 5599 | ENSMUSG00000026527 | 2.691046067 | 2.859930451 | 1.680378803 | 1.701955801 | 0.088763654 | NA | Rgs7 | regulator of G protein signaling 7 [Source: | 0 | 0 | 2.0125 | 1.928 | 2.1812 | 10.025 |
| 5600 | ENSMUSG00000026531 | 0 | NA | NA | NA | NA | NA | Mptx1 | mucosal pentraxin 1 [Source:MGI Symbol; | 0 | 0 | 0 | 0 | 0 | 0 |
| 5601 | ENSMUSG00000026532 | 1.025928453 | 0.209426832 | 2.06760582 | 0.101289535 | 0.919320623 | NA | Spta1 | spectrin alpha, erythrocytic 1 [Source:MG | 0.8031 | 1.0279 | 1.0062 | 0 | 1.0906 | 2.2277 |
| 5602 | ENSMUSG00000026535 | 0 | NA | NA | NA | NA | NA | Ifi202b | interferon activated gene 202B [Source:M | 0 | 0 | 0 | 0 | 0 | 0 |
| 5603 | ENSMUSG00000026536 | 12.68528616 | -0.940330055 | 0.617642092 | -1.522451379 | 0.127896019 | 0.523750856 | Ifi211 | interferon activated gene 211 [Source:MG | 18.471 | 19.53 | 12.075 | 10.604 | 7.6343 | 7.7969 |
| 5604 | ENSMUSG00000026542 | 846.5232675 | 0.187400269 | 0.195880505 | 0.956707098 | 0.338715114 | 0.780858357 | Apcs | serum amyloid P-component [Source:MG | 966.13 | 877.84 | 530.29 | 1006.4 | 738.35 | 960.13 |
| 5605 | ENSMUSG00000026544 | 259.6385332 | -0.051673137 | 0.160716261 | -0.321517792 | 0.747818032 | 0.951032135 | Dusp23 | dual specificity phosphatase 23 [Source:N | 291.53 | 265.2 | 235.46 | 248.71 | 256.3 | 260.64 |
| 5606 | ENSMUSG00000026546 | 1.216525419 | -1.227051403 | 2.308900077 | -0.531444135 | 0.595111044 | NA | Cfap45 | cilia and flagella associated protein 45 [So | 0 | 4.1117 | 1.0062 | 0 | 2.1812 | 0 |
| 5607 | ENSMUSG00000026547 | 157.7632477 | -0.739082492 | 0.221590457 | -3.335353443 | 0.00085191 | 0.018598477 | Tagln2 | transgelin 2 [Source:MGI Symbol;Acc:MGI | 224.87 | 195.3 | 172.07 | 149.42 | 105.79 | 99.132 |
| 5608 | ENSMUSG00000026548 | 7.846216747 | -1.261890908 | 0.904767883 | -1.394712315 | 0.16310261 | 0.586990831 | Slamf9 | SLAM family member 9 [Source:MGI Symb | 13.653 | 17.475 | 2.0125 | 2.8919 | 4.3625 | 6.6831 |
| 5609 | ENSMUSG00000026553 | 3464.365941 | 0.058044435 | 0.107520447 | 0.539845558 | 0.589303545 | 0.911039598 | Copa | coatomer protein complex subunit alpha [ | 3412.4 | 3729.3 | 3042.9 | 3696.9 | 3605.6 | 3299.2 |
| 5610 | ENSMUSG00000026554 | 2967.946597 | 0.120894427 | 0.088238971 | 1.370079739 | 0.170662012 | 0.597556794 | Dcaf8 | DDB1 and CUL4 associated factor 8 [Sourc | 2798.8 | 2894.6 | 2837.6 | 3002.8 | 3210.8 | 3063.1 |
| 5611 | ENSMUSG00000026556 | 8.415461639 | 0.130945703 | 0.813680833 | 0.160930057 | 0.872148489 | 0.979406502 | Vangl2 | VANGL planar cell polarity 2 [Source:MGI | 12.85 | 7.1954 | 4.025 | 15.424 | 6.5437 | 4.4554 |
| 5612 | ENSMUSG00000026558 | 270.6143368 | 0.150569175 | 0.177436253 | 0.848581801 | 0.396114037 | 0.823764863 | Uck2 | uridine-cytidine kinase 2 [Source:MGI Sym | 257.8 | 210.72 | 300.87 | 287.27 | 257.39 | 309.65 |
| 5613 | ENSMUSG00000026560 | 0 | NA | NA | NA | NA | NA | Fmo9 | flavin containing monooxygenase 9 [Sourc | 0 | 0 | 0 | 0 | 0 | 0 |
| 5614 | ENSMUSG00000026563 | 117.2946491 | 0.129060833 | 0.240650743 | 0.536299335 | 0.59175169 | 0.912678231 | Tada1 | transcriptional adaptor 1 [Source:MGI Syr | 118.86 | 107.93 | 109.68 | 159.06 | 106.88 | 101.36 |
| 5615 | ENSMUSG00000026564 | 0 | NA | NA | NA | NA | NA | Dusp27 | dual specificity phosphatase 27 (putative) | 0 | 0 | 0 | 0 | 0 | 0 |
| 5616 | ENSMUSG00000026565 | 137.3799238 | 0.067816223 | 0.201132085 | 0.337172578 | 0.735986808 | 0.948109711 | Pou2f1 | POU domain, class 2, transcription factor : | 142.15 | 129.52 | 130.81 | 155.2 | 138.51 | 128.09 |
| 5617 | ENSMUSG00000026566 | 21.46610019 | -0.641786469 | 0.528241999 | -1.214947827 | 0.224385991 | 0.670547386 | Mpzl1 | myelin protein zero-like 1 [Source:MGI Sy | 21.684 | 27.754 | 29.181 | 20.244 | 6.5437 | 23.391 |
| 5618 | ENSMUSG00000026567 | 3.412145326 | -0.207469998 | 1.160731942 | -0.178740664 | 0.858141338 | NA | Adcy10 | adenylate cyclase 10 [Source:MGI Symbol | 4.8186 | 3.0838 | 3.0187 | 2.8919 | 1.0906 | 5.5692 |
| 5619 | ENSMUSG00000026568 | 4147.486637 | -0.135834863 | 0.099916393 | -1.359485248 | 0.173992874 | 0.604151393 | Mpc2 | mitochondrial pyruvate carrier 2 [Source:I | 4185 | 4144.6 | 4699.2 | 4172.1 | 3674.3 | 4009.8 |
| 5620 | ENSMUSG00000026571 | 816.9488932 | -0.113712407 | 0.133551093 | -0.851452461 | 0.394518061 | 0.822827574 | Dcaf6 | DDB1 and CUL4 associated factor 6 [Sourc | 799.89 | 890.18 | 858.33 | 871.44 | 658.73 | 823.13 |
| 5621 | ENSMUSG00000026572 | 0 | NA | NA | NA | NA | NA | Tbx19 | T-box 19 [Source:MGI Symbol;Acc:MGI:18 | 0 | 0 | 0 | 0 | 0 | 0 |
| 5622 | ENSMUSG00000026573 | 1.223905709 | -2.475963941 | 2.110557993 | -1.173132389 | 0.24074272 | NA | Xcl1 | chemokine (C motif) ligand 1 [Source:MGI | 3.2124 | 1.0279 | 2.0125 | 0 | 1.0906 | 0 |
| 5623 | ENSMUSG00000026574 | 63.46550312 | -1.14418111 | 0.401836906 | -2.847376862 | 0.004408114 | 0.061484057 | Dpt | dermatopontin [Source:MGI Symbol;Acc:I | 129.3 | 92.513 | 40.25 | 49.163 | 28.356 | 41.212 |
| 5624 | ENSMUSG00000026575 | 98.26284312 | -0.070288344 | 0.261339976 | -0.268953662 | 0.787965341 | 0.960870081 | Nme7 | NME/NM23 family member 7 [Source:MG | 96.372 | 93.541 | 111.69 | 67.478 | 106.88 | 113.61 |
| 5625 | ENSMUSG00000026576 | 925.5054482 | -0.058744964 | 0.173251196 | -0.33907393 | 0.734554037 | 0.948109711 | Atp1b1 | ATPase, Na+/K+ transporting, beta 1 polyp | 787.04 | 1180 | 866.38 | 802.03 | 808.15 | 1109.4 |
| 5626 | ENSMUSG00000026577 | 176.5628735 | -0.090677789 | 0.184023799 | -0.492750335 | 0.622189007 | 0.918675153 | Blzf1 | basic leucine zipper nuclear factor 1 [Sour | 188.73 | 197.36 | 159.99 | 163.88 | 172.32 | 177.1 |
| 5627 | ENSMUSG00000026578 | 94.49446358 | -0.31499161 | 0.2495457 | -1.262260221 | 0.206855165 | 0.652194904 | Ccdc181 | coiled-coil domain containing 181 [Source | 104.4 | 125.41 | 84.525 | 81.938 | 91.612 | 79.083 |
| 5628 | ENSMUSG00000026579 | 9925.971315 | 0.018475394 | 0.098701662 | 0.187184226 | 0.851516192 | 0.973576526 | F5 | coagulation factor V [Source:MGI Symbol; | 9267.8 | 10385 | 9935.7 | 10166 | 10703 | 9097.9 |
| 5629 | ENSMUSG00000026580 | 31.06409068 | 0.648727245 | 0.40404303 | 1.605589498 | 0.108364135 | 0.48522218 | Selp | selectin, platelet [Source:MGI Symbol;Acc | 24.896 | 20.558 | 27.169 | 45.307 | 28.356 | 40.098 |
| 5630 | ENSMUSG00000026581 | 10.11023678 | -1.802239308 | 0.759911285 | -2.371644352 | 0.017709128 | 0.158701872 | Sell | selectin, lymphocyte [Source:MGI Symbol | 9.6372 | 21.586 | 16.1 | 6.7478 | 4.3625 | 2.2277 |
| 5631 | ENSMUSG00000026582 | 1.209480962 | -0.360464025 | 2.290296351 | -0.157387504 | 0.874939461 | NA | Sele | selectin, endothelial cell [Source:MGI Sym | 0 | 4.1117 | 0 | 0.964 | 2.1812 | 0 |
| 5632 | ENSMUSG00000026584 | 520.6447406 | 0.264731975 | 0.159558627 | 1.659151746 | 0.097085218 | 0.458902156 | Scyl3 | SCY1-like 3 (S. cerevisiae) [Source:MGI Syr | 530.85 | 390.61 | 497.08 | 618.87 | 606.38 | 480.07 |
| 5633 | ENSMUSG00000026585 | 102.6942073 | -0.311211422 | 0.25164173 | -1.236724218 | 0.216189485 | 0.66165144 | Kifap3 | kinesin-associated protein 3 [Source:MGI | 116.45 | 96.624 | 127.79 | 76.154 | 115.61 | 83.538 |
| 5634 | ENSMUSG00000026586 | 23.98532552 | 0.194258328 | 0.50254403 | 0.386549868 | 0.699089477 | 0.941054463 | Prrx1 | paired related homeobox 1 [Source:MGI S | 32.124 | 16.447 | 18.112 | 14.46 | 33.809 | 28.96 |
| 5635 | ENSMUSG00000026587 | 1.267080836 | -1.497288836 | 2.029508183 | -0.737759447 | 0.46066064 | NA | Astn1 | astrotactin 1 [Source:MGI Symbol;Acc:MG | 1.6062 | 2.0558 | 2.0125 | 1.928 | 0 | 0 |
| 5636 | ENSMUSG00000026589 | 1948.211878 | 0.114798626 | 0.119641381 | 0.959522738 | 0.337295471 | 0.779335733 | Sec16b | SEC16 homolog B (S. cerevisiae) [Source:N | 1852.8 | 1718.7 | 2041.7 | 2288.5 | 1958.8 | 1828.9 |
| 5637 | ENSMUSG00000026592 | 0.767390791 | -1.69152783 | 2.44773922 | -0.691057207 | 0.489529591 | NA | Tex35 | testis expressed 35 [Source:MGI Symbol;A | 1.6062 | 1.0279 | 1.0062 | 0.964 | 0 | 0 |
| 5638 | ENSMUSG00000026594 | 379.9742514 | 0.041625517 | 0.159073371 | 0.261674951 | 0.793572054 | 0.961915077 | Ralgps2 | Ral GEF with PH domain and SH3 binding r | 393.52 | 367.99 | 362.25 | 451.14 | 319.55 | 385.39 |
| 5639 | ENSMUSG00000026596 | 191.5187786 | -0.076666873 | 0.194243695 | -0.394694266 | 0.693068535 | 0.93867917 | Abl2 | v-abl Abelson murine leukemia viral oncog | 179.89 | 211.75 | 198.23 | 170.62 | 162.5 | 226.11 |
| 5640 | ENSMUSG00000026600 | 39.34724344 | -1.803411209 | 0.416278317 | -4.332224707 | 1.48E-05 | 0.000726202 | Soat1 | sterol O-acyltransferase 1 [Source:MGI Sy | 61.036 | 83.261 | 39.244 | 18.316 | 14.178 | 20.049 |
| 5641 | ENSMUSG00000026601 | 0.167707431 | -0.796629176 | 4.080472857 | -0.195229623 | 0.84521319 | NA | Axdnd1 | axonemal dynein light chain domain conta | 0 | 0 | 1.0062 | 0 | 0 | 0 |
| 5642 | ENSMUSG00000026602 | 0 | NA | NA | NA | NA | NA | Nphs2 | nephrosis 2, podocin [Source:MGI Symbol | 0 | 0 | 0 | 0 | 0 | 0 |
| 5643 | ENSMUSG00000026603 | 191.4368493 | -0.102151628 | 0.20406562 | -0.500582254 | 0.616665155 | 0.917509899 | Smyd2 | SET and MYND domain containing 2 [Sour | 180.7 | 170.63 | 243.51 | 177.37 | 212.67 | 163.74 |
| 5644 | ENSMUSG00000026604 | 33.74040767 | -0.53110841 | 0.398244772 | -1.333623057 | 0.182327423 | 0.61573286 | Ptpn14 | protein tyrosine phosphatase, non-recept | 43.368 | 37.005 | 39.244 | 30.847 | 17.45 | 34.529 |
| 5645 | ENSMUSG00000026605 | 3.678916569 | -1.623680472 | 1.223104756 | -1.327507283 | 0.184340932 | NA | Cenpf | centromere protein F [Source:MGI Symbo | 6.4248 | 4.1117 | 6.0375 | 0 | 3.2719 | 2.2277 |
| 5646 | ENSMUSG00000026608 | 226.7076251 | -0.001813396 | 0.17487225 | -0.010369834 | 0.991726218 | 0.99859176 | Kctd3 | potassium channel tetramerisation domai | 232.9 | 205.58 | 241.5 | 196.65 | 251.93 | 231.68 |
| 5647 | ENSMUSG00000026609 | 2.754118053 | 1.15163846 | 1.558529464 | 0.738926332 | 0.459951732 | NA | Ush2a | usherin [Source:MGI Symbol;Acc:MGI:134 | 4.0155 | 0 | 1.0062 | 4.8199 | 0 | 6.6831 |
| 5648 | ENSMUSG00000026610 | 88.27970434 | -0.924692016 | 0.411877644 | -2.245064835 | 0.024763971 | 0.201358738 | Esrrg | estrogen-related receptor gamma [Source | 112.43 | 133.63 | 100.62 | 31.811 | 118.88 | 32.301 |
| 5649 | ENSMUSG00000026611 | 6.555896356 | -0.179224642 | 0.905622014 | -0.197902258 | 0.843121537 | 0.972273952 | Spata17 | spermatogenesis associated 17 [Source:M | 9.6372 | 9.2513 | 2.0125 | 9.6398 | 5.4531 | 3.3415 |
| 5650 | ENSMUSG00000026614 | 579.1567597 | -0.523945824 | 0.195079341 | -2.685808873 | 0.007235446 | 0.086205517 | Slc30a10 | solute carrier family 30, member 10 [Sour | 500.33 | 643.48 | 906.63 | 496.45 | 407.89 | 520.17 |
| 5651 | ENSMUSG00000026615 | 1993.723408 | 0.111037899 | 0.113105398 | 0.981720596 | 0.326237516 | 0.772018857 | Eprs | glutamyl-prolyl-tRNA synthetase [Source:I | 1891.3 | 1945.8 | 1913.9 | 1957.8 | 2367.7 | 1885.7 |
| 5652 | ENSMUSG00000026616 | 3.434209625 | -0.512378793 | 1.194806874 | -0.42883817 | 0.668040999 | NA | Cr2 | complement receptor 2 [Source:MGI Sym | 4.0155 | 1.0279 | 7.0437 | 1.928 | 4.3625 | 2.2277 |

| | | | | | | | | | | | | | | | |
|---|---|---|---|---|---|---|---|---|---|---|---|---|---|---|---|
| 5653 | ENSMUSG00000026617 | 1302.570259 | -0.047779016 | 0.107018515 | -0.446455605 | 0.655268175 | 0.926018454 | Bpnt1 | 3'(2'), 5'-bisphosphate nucleotidase 1 [Sou | 1258.5 | 1277.7 | 1436.9 | 1304.3 | 1220.4 | 1317.7 |
| 5654 | ENSMUSG00000026618 | 1253.224407 | 0.011344355 | 0.111456084 | 0.101783185 | 0.918928773 | 0.986374671 | Iars2 | isoleucine-tRNA synthetase 2, mitochondı | 1228.7 | 1324 | 1192.4 | 1233.9 | 1375.3 | 1165.1 |
| 5655 | ENSMUSG00000026620 | 16.49383991 | -1.517779438 | 0.611639962 | -2.481491614 | 0.013083379 | 0.129971148 | Mark1 | MAP/microtubule affinity regulating kinas | 24.896 | 19.53 | 29.181 | 15.424 | 4.3625 | 5.5692 |
| 5656 | ENSMUSG00000026621 | 9118.08499 | 0.122878552 | 0.083766333 | 1.466920508 | 0.142397681 | 0.55198239 | Mtarc1 | mitochondrial amidoxime reducing compo | 8752.2 | 8325.1 | 9112.6 | 9495.2 | 9513.5 | 9510 |
| 5657 | ENSMUSG00000026622 | 60.23324465 | 0.201133398 | 0.293371234 | 0.685593455 | 0.49296951 | 0.876713511 | Nek2 | NIMA (never in mitosis gene a)-related ex | 64.248 | 44.2 | 59.368 | 64.587 | 62.165 | 66.831 |
| 5658 | ENSMUSG00000026623 | 5412.294079 | 0.072757972 | 0.101169469 | 0.719169253 | 0.472036641 | 0.867870596 | Lpgat1 | lysophosphatidylglycerol acyltransferase 1 | 5455.5 | 5024.5 | 5347.2 | 5542.9 | 6073.7 | 5030.1 |
| 5659 | ENSMUSG00000026626 | 3827.157788 | -0.044100356 | 0.116606078 | -0.378199461 | 0.705282426 | 0.94295036 | Ppp2r5a | protein phosphatase 2, regulatory subunit | 3563.4 | 4273.1 | 3820.7 | 3328.6 | 4191.2 | 3786 |
| 5660 | ENSMUSG00000026627 | 28.43714085 | -0.316177276 | 0.422601835 | -0.748168251 | 0.454358679 | 0.857337687 | Pacc1 | proton activated chloride channel 1 [Sour | 33.73 | 28.782 | 32.2 | 30.847 | 28.356 | 16.708 |
| 5661 | ENSMUSG00000026628 | 17.9183314 | -2.157564899 | 0.652726185 | -3.305467056 | 0.000948183 | 0.020109255 | Atf3 | activating transcription factor 3 [Source:N | 52.202 | 17.475 | 18.112 | 8.6758 | 4.3625 | 6.6831 |
| 5662 | ENSMUSG00000026630 | 9.086903966 | -0.619941519 | 0.819732385 | -0.75627306 | 0.449485497 | 0.854879451 | Batf3 | basic leucine zipper transcription factor, A | 17.668 | 12.335 | 3.0187 | 11.568 | 4.3625 | 5.5692 |
| 5663 | ENSMUSG00000026632 | 99.2667371 | -0.144905551 | 0.269246099 | -0.538189973 | 0.590445902 | 0.911690353 | Tatdn3 | TatD DNase domain containing 3 [Source: | 110.83 | 69.898 | 131.82 | 94.47 | 88.34 | 100.25 |
| 5664 | ENSMUSG00000026634 | 495.9393504 | 0.000573785 | 0.127527972 | 0.004499285 | 0.996410102 | 0.999154389 | Angel2 | angel homolog 2 [Source:MGI Symbol;Acc | 495.51 | 517.04 | 474.95 | 485.84 | 531.13 | 471.16 |
| 5665 | ENSMUSG00000026637 | 74.86799034 | 0.160755546 | 0.278886867 | 0.576418487 | 0.564332352 | 0.901468983 | Traf5 | TNF receptor-associated factor 5 [Source: | 77.098 | 53.452 | 81.506 | 90.614 | 75.253 | 71.286 |
| 5666 | ENSMUSG00000026638 | 622.3423847 | 0.511213498 | 0.18305309 | 2.792706189 | 0.005226914 | 0.068666748 | Irf6 | interferon regulatory factor 6 [Source:MG | 422.43 | 427.61 | 690.28 | 704.67 | 687.09 | 801.97 |
| 5667 | ENSMUSG00000026639 | 215.7489196 | -0.969533537 | 0.210550355 | -4.604758508 | 4.13E-06 | 0.000260822 | Lamb3 | laminin, beta 3 [Source:MGI Symbol;Acc:N | 300.36 | 342.3 | 214.33 | 151.34 | 161.41 | 124.75 |
| 5668 | ENSMUSG00000026640 | 279.5486086 | 0.04957201 | 0.253151541 | 0.195819506 | 0.844751442 | 0.972803041 | Plxna2 | plexin A2 [Source:MGI Symbol;Acc:MGI:1( | 150.98 | 349.49 | 325.02 | 316.18 | 268.29 | 267.32 |
| 5669 | ENSMUSG00000026641 | 459.4505903 | 0.070217598 | 0.132038287 | 0.531797251 | 0.594866427 | 0.913409341 | Usf1 | upstream transcription factor 1 [Source:N | 438.49 | 420.42 | 486.02 | 472.35 | 449.33 | 490.09 |
| 5670 | ENSMUSG00000026643 | 407.0310163 | -0.053333142 | 0.149644391 | -0.356399204 | 0.721541625 | 0.9447719 | Nmt2 | N-myristoyltransferase 2 [Source:MGI Syn | 415.2 | 430.7 | 398.47 | 458.85 | 395.89 | 343.06 |
| 5671 | ENSMUSG00000026644 | 0.167707431 | -0.796629176 | 4.080472857 | -0.195229623 | 0.84521319 | NA | Acbd7 | acyl-Coenzyme A binding domain containi | 0 | 0 | 1.0062 | 0 | 0 | 0 |
| 5672 | ENSMUSG00000026645 | 0 | NA | NA | NA | NA | NA | Olah | oleoyl-ACP hydrolase [Source:MGI Symbo | 0 | 0 | 0 | 0 | 0 | 0 |
| 5673 | ENSMUSG00000026646 | 22.40438636 | 0.284341325 | 0.470346488 | 0.604535874 | 0.545487426 | 0.89549457 | Suv39h2 | suppressor of variegation 3-9 2 [Source:M | 16.062 | 19.53 | 25.156 | 25.063 | 18.541 | 30.074 |
| 5674 | ENSMUSG00000026648 | 144.4135385 | 0.425937783 | 0.2411955 | 1.765944155 | 0.077405218 | 0.40712138 | Dclre1c | DNA cross-link repair 1C [Source:MGI Sym | 144.56 | 125.41 | 99.618 | 196.65 | 179.95 | 120.3 |
| 5675 | ENSMUSG00000026649 | 42.05828645 | -0.002364059 | 0.365078041 | -0.006475491 | 0.994833342 | 0.998676967 | Cfap126 | cilia and flagella associated protein 126 [S | 48.989 | 27.754 | 49.306 | 42.415 | 35.99 | 47.895 |
| 5676 | ENSMUSG00000026650 | 3.046176711 | 1.151848193 | 1.391290707 | 0.827899006 | 0.407727698 | NA | Meig1 | meiosis expressed gene 1 [Source:MGI Sy | 1.6062 | 3.0838 | 1.0062 | 3.8559 | 8.7249 | 0 |
| 5677 | ENSMUSG00000026655 | 1601.87912 | -0.142040763 | 0.093973089 | -1.511504671 | 0.130659919 | 0.529460957 | Fam107b | family with sequence similarity 107, meml | 1667.2 | 1706.3 | 1668.4 | 1488.4 | 1566.1 | 1514.8 |
| 5678 | ENSMUSG00000026656 | 519.3832909 | -0.514830421 | 0.142959079 | -3.601243274 | 0.000316699 | 0.00912497 | Fcgr2b | Fc receptor, IgG, low affinity IIb [Source:N | 587.07 | 630.11 | 615.82 | 398.12 | 381.72 | 503.46 |
| 5679 | ENSMUSG00000026657 | 73.98119136 | -0.338254315 | 0.272405365 | -1.241731469 | 0.214335655 | 0.658905615 | Frmd4a | FERM domain containing 4A [Source:MGI | 98.782 | 78.122 | 70.437 | 58.803 | 69.8 | 67.944 |
| 5680 | ENSMUSG00000026659 | 168.8878607 | 0.200332994 | 0.189014567 | 1.05988124 | 0.289198631 | 0.7396939 | Dusp12 | dual specificity phosphatase 12 [Source:N | 163.83 | 160.36 | 146.91 | 160.02 | 201.76 | 180.44 |
| 5681 | ENSMUSG00000026662 | 573.4523531 | 0.186862511 | 0.133348389 | 1.401310604 | 0.161121211 | 0.584240557 | Sephs1 | selenophosphate synthetase 1 [Source:M | 562.97 | 570.49 | 474.95 | 568.75 | 658.73 | 604.82 |
| 5682 | ENSMUSG00000026663 | 1452.435054 | 0.160380352 | 0.108970706 | 1.471774911 | 0.141081671 | 0.549435995 | Atf6 | activating transcription factor 6 [Source:N | 1396.6 | 1438.1 | 1280.9 | 1628.2 | 1401.4 | 1569.4 |
| 5683 | ENSMUSG00000026664 | 37189.07926 | 0.165481114 | 0.097034856 | 1.705378058 | 0.088123938 | 0.437714505 | Phyh | phytanoyl-CoA hydroxylase [Source:MGI S | 33496 | 35498 | 36182 | 35254 | 41360 | 41346 |
| 5684 | ENSMUSG00000026667 | 1210.069485 | -0.050474643 | 0.111711528 | -0.451830211 | 0.651391304 | 0.925643431 | Uhmk1 | U2AF homology motif (UHM) kinase 1 [So | 1145.2 | 1337.3 | 1211.5 | 1109.5 | 1218.2 | 1238.6 |
| 5685 | ENSMUSG00000026668 | 0 | NA | NA | NA | NA | NA | Ucma | upper zone of growth plate and cartilage ı | 0 | 0 | 0 | 0 | 0 | 0 |
| 5686 | ENSMUSG00000026669 | 1887.104012 | 0.909951639 | 0.127257647 | 7.150467288 | 8.65E-13 | 3.64E-10 | Mcm10 | minichromosome maintenance 10 replicat | 1441.6 | 1239.7 | 1249.8 | 2084.1 | 2560.8 | 2746.7 |
| 5687 | ENSMUSG00000026670 | 581.4440273 | 0.058797138 | 0.138924407 | 0.423231158 | 0.672126616 | 0.931191575 | Uap1 | UDP-N-acetylglucosamine pyrophosphory | 520.41 | 663.01 | 526.27 | 610.2 | 558.4 | 610.39 |
| 5688 | ENSMUSG00000026672 | 2083.828368 | 0.079560339 | 0.093974017 | 0.846620604 | 0.397206626 | 0.824629917 | Optn | optineurin [Source:MGI Symbol;Acc:MGI: | 2083.2 | 1951 | 2044.7 | 2190.2 | 2188.9 | 2045 |
| 5689 | ENSMUSG00000026674 | 65.12755199 | -0.893651228 | 0.300720738 | -2.971698038 | 0.002961578 | 0.046110081 | Ddr2 | discoidin domain receptor family, membe | 90.751 | 90.457 | 72.45 | 36.631 | 41.443 | 59.034 |
| 5690 | ENSMUSG00000026675 | 865.8273765 | 0.402922452 | 0.119689402 | 3.366400402 | 0.000761561 | 0.017077853 | Hsd17b7 | hydroxysteroid (17-beta) dehydrogenase | 783.02 | 766.83 | 686.26 | 917.71 | 966.29 | 1074.9 |
| 5691 | ENSMUSG00000026676 | 6.290515314 | -0.534334864 | 0.970620482 | -0.550508539 | 0.581970621 | 0.908645447 | Ccdc3 | coiled-coil domain containing 3 [Source:N | 12.047 | 9.2513 | 1.0062 | 7.7118 | 3.2719 | 4.4554 |
| 5692 | ENSMUSG00000026678 | 377.5798548 | -1.443718128 | 0.154015247 | -9.37386497 | 6.99E-21 | 1.26E-17 | Rgs5 | regulator of G-protein signaling 5 [Source: | 579.84 | 586.94 | 489.03 | 185.08 | 199.58 | 225 |
| 5693 | ENSMUSG00000026679 | 0.481988923 | 2.492401104 | 3.837077151 | 0.649557203 | 0.515978285 | NA | Enkur | enkurin, TRPC channel interacting protein | 0 | 0 | 0 | 2.8919 | 0 | 0 |
| 5694 | ENSMUSG00000026683 | 3.567753228 | -0.682208484 | 1.291767758 | -0.528120074 | 0.597415999 | NA | Nuf2 | NUF2, NDC80 kinetochore complex comp | 4.0155 | 8.2233 | 1.0062 | 4.8199 | 0 | 3.3415 |
| 5695 | ENSMUSG00000026686 | 0 | NA | NA | NA | NA | NA | Lmx1a | LIM homeobox transcription factor 1 alph | 0 | 0 | 0 | 0 | 0 | 0 |
| 5696 | ENSMUSG00000026687 | 6294.713497 | 0.094769866 | 0.083831074 | 1.130486123 | 0.258271443 | 0.711956956 | Aldh9a1 | aldehyde dehydrogenase 9, subfamily A1 | 6032.1 | 6109.9 | 6122 | 6329.5 | 6789.1 | 6385.7 |
| 5697 | ENSMUSG00000026688 | 298.1732871 | -0.861138024 | 0.221709257 | -3.884086914 | 0.000102715 | 0.003704918 | Mgst3 | microsomal glutathione S-transferase 3 [S | 347.74 | 439.95 | 367.28 | 293.05 | 148.32 | 192.69 |
| 5698 | ENSMUSG00000026691 | 23.70273962 | -3.656195048 | 0.812256654 | -4.501280513 | 6.75E-06 | 0.000398765 | Fmo3 | flavin containing monooxygenase 3 [Sourc | 7.2279 | 90.457 | 34.212 | 4.8199 | 3.2719 | 2.2277 |
| 5699 | ENSMUSG00000026692 | 164.7396331 | -0.363193132 | 0.212844881 | -1.706374757 | 0.087938327 | 0.437092772 | Fmo4 | flavin containing monooxygenase 4 [Sourc | 217.64 | 195.3 | 142.89 | 158.09 | 133.06 | 141.46 |
| 5700 | ENSMUSG00000026694 | 184.5927137 | 0.328334195 | 0.203571818 | 1.612866643 | 0.106773484 | 0.482164117 | Eef1aknmt | EEF1A lysine and N-terminal methyltransf | 179.09 | 173.72 | 137.86 | 196.65 | 246.48 | 173.76 |
| 5701 | ENSMUSG00000026696 | 215.8582898 | 0.014697188 | 0.177272464 | 0.082907339 | 0.933925219 | 0.988420305 | Vamp4 | vesicle-associated membrane protein 4 [S | 228.08 | 204.56 | 211.31 | 220.75 | 185.41 | 245.05 |
| 5702 | ENSMUSG00000026697 | 0.803102239 | -3.069617558 | 3.178309912 | -0.965801839 | 0.334143348 | NA | Myoc | myocilin [Source:MGI Symbol;Acc:MGI:12 | 4.8186 | 0 | 0 | 0 | 0 | 0 |
| 5703 | ENSMUSG00000026698 | 237.2233601 | 0.431198247 | 0.168709999 | 2.555854712 | 0.010592734 | 0.113239689 | Pigc | phosphatidylinositol glycan anchor biosyn | 202.38 | 190.16 | 213.32 | 260.27 | 248.66 | 308.53 |
| 5704 | ENSMUSG00000026700 | 0 | NA | NA | NA | NA | NA | Tnfsf4 | tumor necrosis factor (ligand) superfamily | 0 | 0 | 0 | 0 | 0 | 0 |
| 5705 | ENSMUSG00000026701 | 10544.82418 | -0.026323264 | 0.087184539 | -0.301925828 | 0.762708606 | 0.95469451 | Prdx6 | peroxiredoxin 6 [Source:MGI Symbol;Acc: | 10446 | 10226 | 11252 | 10536 | 10759 | 10050 |
| 5706 | ENSMUSG00000026705 | 160.0510048 | -0.009560375 | 0.199324462 | -0.047963881 | 0.961745028 | 0.992839918 | Klhl20 | kelch-like 20 [Source:MGI Symbol;Acc:MG | 140.54 | 159.33 | 182.13 | 144.6 | 177.77 | 155.94 |
| 5707 | ENSMUSG00000026707 | 63.31641411 | -0.064573077 | 0.283483177 | -0.227784513 | 0.819813767 | 0.969160361 | Nsun6 | NOL1/NOP2/Sun domain family member 6 | 61.036 | 75.038 | 58.362 | 66.514 | 63.256 | 55.692 |
| 5708 | ENSMUSG00000026708 | 107.3209503 | 0.395140405 | 0.218163928 | 1.811208697 | 0.070108552 | 0.382949435 | Cenpl | centromere protein L [Source:MGI Symbo | 93.16 | 87.373 | 97.606 | 123.39 | 124.33 | 118.07 |

| | | | | | | | | | | | | | | | |
|---|---|---|---|---|---|---|---|---|---|---|---|---|---|---|---|
| 5709 | ENSMUSG00000026709 | 381.8202116 | 0.04631072 | 0.172729921 | 0.268110584 | 0.788614199 | 0.960872359 | Dars2 | aspartyl-tRNA synthetase 2 (mitochondria | 410.39 | 304.26 | 411.55 | 341.25 | 454.79 | 368.68 |
| 5710 | ENSMUSG00000026712 | 400.0204819 | -0.911761053 | 0.191956436 | -4.749833205 | 2.04E-06 | 0.000145775 | Mrc1 | mannose receptor, C type 1 [Source:MGI : | 489.89 | 635.25 | 441.74 | 246.78 | 234.48 | 351.97 |
| 5711 | ENSMUSG00000026715 | 34731.71947 | 0.094824075 | 0.076991287 | 1.231620853 | 0.218090748 | 0.663697152 | Serpinc1 | serine (or cysteine) peptidase inhibitor, cl | 32985 | 33876 | 33911 | 35603 | 36107 | 35908 |
| 5712 | ENSMUSG00000026718 | 264.4521286 | -0.107231975 | 0.172718206 | -0.620849285 | 0.534698792 | 0.889858671 | Stam | signal transducing adaptor molecule (SH3 | 261.81 | 297.07 | 264.64 | 292.09 | 211.58 | 259.53 |
| 5713 | ENSMUSG00000026721 | 666.1856672 | -0.01316459 | 0.121962909 | -0.107939292 | 0.914043849 | 0.98625062 | Rabgap1l | RAB GTPase activating protein 1-like [Sour | 663.36 | 705.15 | 638.97 | 622.73 | 721.99 | 644.92 |
| 5714 | ENSMUSG00000026723 | 52.39624003 | -0.020455014 | 0.338283595 | -0.06046706 | 0.95178365 | 0.990883058 | Trdmt1 | tRNA aspartic acid methyltransferase 1 [S | 55.414 | 46.256 | 56.35 | 37.595 | 71.981 | 46.781 |
| 5715 | ENSMUSG00000026725 | 0.36353943 | 2.065989254 | 4.034137301 | 0.512126658 | 0.608562368 | NA | Tnn | tenascin N [Source:MGI Symbol;Acc:MGI: | 0 | 0 | 0 | 0 | 2.1812 | 0 |
| 5716 | ENSMUSG00000026726 | 2.903090591 | -1.217352591 | 1.276854976 | -0.953399261 | 0.340387816 | NA | Cubn | cubilin (intrinsic factor-cobalamin recepto | 4.0155 | 6.1675 | 2.0125 | 1.928 | 2.1812 | 1.1138 |
| 5717 | ENSMUSG00000026727 | 374.2742752 | -0.020009826 | 0.168537179 | -0.11872648 | 0.905492057 | 0.983972587 | Rsu1 | Ras suppressor protein 1 [Source:MGI Syr | 457.77 | 328.93 | 343.13 | 409.69 | 368.63 | 337.49 |
| 5718 | ENSMUSG00000026728 | 255.5513496 | -0.782945287 | 0.180340449 | -4.341484616 | 1.42E-05 | 0.000708302 | Vim | vimentin [Source:MGI Symbol;Acc:MGI:98 | 344.53 | 356.69 | 268.67 | 203.4 | 197.4 | 162.62 |
| 5719 | ENSMUSG00000026729 | 0.96892865 | 1.129457544 | 2.28221261 | 0.494895848 | 0.620673642 | NA | 4930562F07 | RIKEN cDNA 4930562F07 gene [Source:M | 0.8031 | 1.0279 | 0 | 2.8919 | 1.0906 | 0 |
| 5720 | ENSMUSG00000026730 | 952.8818806 | 0.310441296 | 0.135450125 | 2.291923292 | 0.02191007 | 0.184462894 | Pter | phosphotriesterase related [Source:MGI S | 954.09 | 832.61 | 764.75 | 1127.9 | 905.21 | 1132.8 |
| 5721 | ENSMUSG00000026734 | 0.133850373 | -0.796629176 | 4.080472857 | -0.195229623 | 0.84521319 | NA | 4921504E06 | RIKEN cDNA 4921504E06 gene [Source:M | 0.8031 | 0 | 0 | 0 | 0 | 0 |
| 5722 | ENSMUSG00000026735 | 0 | NA | NA | NA | NA | NA | Ptf1a | pancreas specific transcription factor, 1a [ | 0 | 0 | 0 | 0 | 0 | 0 |
| 5723 | ENSMUSG00000026736 | 1.256159933 | -0.400276649 | 2.054019058 | -0.194874847 | 0.845490926 | NA | 4930426L09I | RIKEN cDNA 4930426L09 gene [Source:M | 3.2124 | 0 | 1.0062 | 0 | 1.0906 | 2.2277 |
| 5724 | ENSMUSG00000026737 | 50.88365649 | -1.018557173 | 0.349282748 | -2.916139373 | 0.003543922 | 0.052790197 | Pip4k2a | phosphatidylinositol-5-phosphate 4-kinas | 77.901 | 76.066 | 50.312 | 34.703 | 23.994 | 42.326 |
| 5725 | ENSMUSG00000026739 | 408.101866 | -0.009567987 | 0.161356346 | -0.059297247 | 0.952715354 | 0.991522443 | Bmi1 | Bmi1 polycomb ring finger oncogene [Sou | 347.74 | 406.03 | 474.95 | 356.67 | 418.8 | 444.42 |
| 5726 | ENSMUSG00000026740 | 587.2626627 | 0.121697868 | 0.119265139 | 1.020397651 | 0.307539907 | 0.756343859 | Dnajc1 | DnaJ heat shock protein family (Hsp40) m | 559.76 | 547.88 | 579.6 | 581.28 | 639.1 | 615.96 |
| 5727 | ENSMUSG00000026743 | 380.1892191 | 0.114757775 | 0.144173644 | 0.795969166 | 0.426049957 | 0.841626138 | Mllt10 | myeloid/lymphoid or mixed-lineage leuke | 355.77 | 361.83 | 378.35 | 444.39 | 362.09 | 378.71 |
| 5728 | ENSMUSG00000026748 | 13.30276277 | -0.515213195 | 0.616580932 | -0.835597029 | 0.403381641 | 0.828953151 | Plxdc2 | plexin domain containing 2 [Source:MGI S | 19.274 | 17.475 | 10.062 | 7.7118 | 15.269 | 10.025 |
| 5729 | ENSMUSG00000026749 | 972.8892187 | 0.133878005 | 0.128522772 | 1.041667576 | 0.297565827 | 0.748091545 | Nek6 | NIMA (never in mitosis gene a)-related ex | 1035.2 | 836.73 | 909.65 | 904.21 | 1064.4 | 1087.1 |
| 5730 | ENSMUSG00000026750 | 1810.601235 | 0.140300553 | 0.103040213 | 1.361609693 | 0.173321094 | 0.603059011 | Psmb7 | proteasome (prosome, macropain) subun | 1856.8 | 1623.1 | 1686.5 | 1805.5 | 1988.2 | 1903.6 |
| 5731 | ENSMUSG00000026751 | 0 | NA | NA | NA | NA | NA | Nr5a1 | nuclear receptor subfamily 5, group A, me | 0 | 0 | 0 | 0 | 0 | 0 |
| 5732 | ENSMUSG00000026753 | 791.3467501 | 0.006188688 | 0.115518157 | 0.053573295 | 0.957275134 | 0.991861398 | Ppp6c | protein phosphatase 6, catalytic subunit [ | 780.62 | 770.94 | 817.07 | 740.33 | 858.32 | 780.8 |
| 5733 | ENSMUSG00000026754 | 305.7629831 | 0.00467685 | 0.150404488 | 0.03109515 | 0.975193658 | 0.996378391 | Golga1 | golgi autoantigen, golgin subfamily a, 1 [S | 299.56 | 323.79 | 292.82 | 317.15 | 323.91 | 277.35 |
| 5734 | ENSMUSG00000026755 | 690.8226795 | 0.117001713 | 0.113456813 | 1.031244492 | 0.302426181 | 0.752385931 | Arpc5l | actin related protein 2/3 complex, subunit | 636.06 | 691.79 | 661.1 | 711.42 | 736.17 | 708.4 |
| 5735 | ENSMUSG00000026761 | 291.0379047 | -0.024624326 | 0.171097829 | -0.14391957 | 0.885563982 | 0.982048364 | Orc4 | origin recognition complex, subunit 4 [Sou | 314.82 | 300.15 | 264.64 | 237.14 | 306.46 | 323.01 |
| 5736 | ENSMUSG00000026764 | 1.279591891 | -3.736153947 | 2.409849227 | -1.55036834 | 0.121053133 | NA | Kif5c | kinesin family member 5C [Source:MGI Sy | 5.6217 | 2.0558 | 0 | 0 | 0 | 0 |
| 5737 | ENSMUSG00000026765 | 0 | NA | NA | NA | NA | NA | Lypd6b | LY6/PLAUR domain containing 6B [Source | 0 | 0 | 0 | 0 | 0 | 0 |
| 5738 | ENSMUSG00000026766 | 1201.44965 | -0.051550102 | 0.130105201 | -0.396218611 | 0.691943763 | 0.938171202 | Mmadhc | methylmalonic aciduria (cobalamin deficie | 1267.3 | 1298.3 | 1103.9 | 1358.2 | 1144.1 | 1037 |
| 5739 | ENSMUSG00000026767 | 593.1478308 | -0.05013677 | 0.118547558 | -0.422925374 | 0.672349709 | 0.931191575 | Mindy3 | MINDY lysine 48 deubiquitinase 3 [Source | 610.36 | 601.33 | 598.72 | 591.88 | 607.47 | 549.13 |
| 5740 | ENSMUSG00000026768 | 60.23783648 | -0.279535523 | 0.37755208 | -0.740389307 | 0.459063805 | 0.859322431 | Itga8 | integrin alpha 8 [Source:MGI Symbol;Acc: | 95.569 | 70.926 | 31.194 | 50.127 | 52.35 | 61.261 |
| 5741 | ENSMUSG00000026770 | 9.978959554 | 0.95467736 | 0.7149173 | 1.335367544 | 0.181756082 | 0.615095921 | Il2ra | interleukin 2 receptor, alpha chain [Sourc | 7.2279 | 3.0838 | 10.062 | 16.388 | 13.087 | 10.025 |
| 5742 | ENSMUSG00000026771 | 190.2515931 | -0.047406108 | 0.197023801 | -0.240611074 | 0.809856567 | 0.966157155 | Spopl | speckle-type BTB/POZ protein-like [Source | 175.88 | 227.17 | 177.1 | 160.98 | 185.41 | 214.97 |
| 5743 | ENSMUSG00000026773 | 364.5304633 | -0.459992306 | 0.460615664 | -0.998646685 | 0.317965876 | 0.764585353 | Pfkfb3 | 6-phosphofructo-2-kinase/fructose-2,6-bi | 564.58 | 499.57 | 202.26 | 375.95 | 254.11 | 290.71 |
| 5744 | ENSMUSG00000026774 | 0 | NA | NA | NA | NA | NA | Potegl | POTE ankyrin domain family, member G lil | 0 | 0 | 0 | 0 | 0 | 0 |
| 5745 | ENSMUSG00000026775 | 1839.202068 | 0.01676815 | 0.103651697 | 0.161773999 | 0.871483829 | 0.979227205 | Yme1l1 | YME1-like 1 (S. cerevisiae) [Source:MGI Sy | 1699.4 | 1797.8 | 1989.3 | 1854.7 | 1912.9 | 1781 |
| 5746 | ENSMUSG00000026778 | 13.33934991 | -8.57E-05 | 0.666372918 | -0.000128667 | 0.999897338 | 0.999897338 | Prkcq | protein kinase C, theta [Source:MGI Symb | 14.456 | 10.279 | 15.094 | 4.8199 | 11.997 | 23.391 |
| 5747 | ENSMUSG00000026779 | 16.81568439 | 0.152318702 | 0.544019453 | 0.279987602 | 0.779487017 | 0.959870443 | Mastl | microtubule associated serine/threonine l | 11.243 | 20.558 | 16.1 | 15.424 | 14.178 | 23.391 |
| 5748 | ENSMUSG00000026781 | 5056.565158 | 0.245829026 | 0.099779285 | 2.463728081 | 0.013750038 | 0.134934013 | Acbd5 | acyl-Coenzyme A binding domain containi | 4433.9 | 4675 | 4771.6 | 5091.7 | 6049.7 | 5317.5 |
| 5749 | ENSMUSG00000026782 | 49.05120784 | 0.293963159 | 0.390310409 | 0.753152241 | 0.451358435 | 0.85613987 | Abi2 | abl interactor 2 [Source:MGI Symbol;Acc:I | 44.974 | 49.34 | 38.237 | 86.758 | 34.9 | 40.098 |
| 5750 | ENSMUSG00000026784 | 45.63529567 | -0.559656903 | 0.36404607 | -1.537324392 | 0.124213891 | 0.516569788 | Pdss1 | prenyl (solanesyl) diphosphate synthase, s | 61.036 | 61.675 | 40.25 | 35.667 | 26.175 | 49.009 |
| 5751 | ENSMUSG00000026785 | 22.26493552 | 0.093412755 | 0.503341487 | 0.185585249 | 0.852770023 | 0.973781977 | Pkn3 | protein kinase N3 [Source:MGI Symbol;Ac | 16.062 | 16.447 | 32.2 | 20.244 | 17.45 | 31.188 |
| 5752 | ENSMUSG00000026786 | 20.47998184 | -1.266789138 | 0.524621223 | -2.414673828 | 0.015749306 | 0.1470999 | Apbb1ip | amyloid beta (A4) precursor protein-bindi | 21.684 | 34.949 | 30.187 | 10.604 | 7.6343 | 17.822 |
| 5753 | ENSMUSG00000026787 | 1.056198278 | -2.228584164 | 2.228987013 | -0.999819268 | 0.317397979 | NA | Gad2 | glutamic acid decarboxylase 2 [Source:MG | 3.2124 | 1.0279 | 1.0062 | 0 | 1.0906 | 0 |
| 5754 | ENSMUSG00000026788 | 481.1984209 | 0.111159421 | 0.128099444 | 0.867758808 | 0.385526384 | 0.816437825 | Zbtb43 | zinc finger and BTB domain containing 43 | 444.12 | 467.7 | 476.96 | 532.12 | 495.14 | 471.16 |
| 5755 | ENSMUSG00000026790 | 213.6415151 | 0.257313648 | 0.197556064 | 1.302484181 | 0.192750922 | 0.630757769 | Odf2 | outer dense fiber of sperm tails 2 [Source: | 212.02 | 171.66 | 200.24 | 278.59 | 184.31 | 235.02 |
| 5756 | ENSMUSG00000026791 | 292.5732146 | 0.115118162 | 0.153257335 | 0.751142916 | 0.45256665 | 0.856769845 | Slc2a8 | solute carrier family 2, (facilitated glucose | 268.24 | 277.54 | 296.84 | 276.66 | 306.46 | 329.7 |
| 5757 | ENSMUSG00000026792 | 204.6525371 | 0.134698977 | 0.183287677 | 0.734904709 | 0.46239753 | 0.860650866 | Lrsam1 | leucine rich repeat and sterile alpha motif | 167.85 | 209.7 | 208.29 | 206.29 | 196.31 | 239.48 |
| 5758 | ENSMUSG00000026796 | 125.7947058 | -0.380136139 | 0.251838186 | -1.509445982 | 0.13118485 | 0.530905895 | Niban2 | niban apoptosis regulator 2 [Source:MGI S | 152.59 | 135.69 | 137.86 | 79.046 | 100.34 | 149.26 |
| 5759 | ENSMUSG00000026797 | 26.26293794 | -0.741850461 | 0.434146086 | -1.708757687 | 0.087495844 | 0.436242634 | Stxbp1 | syntaxin binding protein 1 [Source:MGI Sy | 30.518 | 33.921 | 34.212 | 20.244 | 14.178 | 24.505 |
| 5760 | ENSMUSG00000026798 | 113.5915452 | -0.369346035 | 0.223425155 | -1.653108556 | 0.098308772 | 0.462495676 | Coq4 | coenzyme Q4 [Source:MGI Symbol;Acc:M | 127.69 | 131.57 | 124.77 | 90.614 | 117.79 | 89.108 |
| 5761 | ENSMUSG00000026799 | 82.33675706 | -0.375420031 | 0.255648757 | -1.468499341 | 0.141968635 | 0.551553677 | Med27 | mediator complex subunit 27 [Source:MG | 90.751 | 107.93 | 80.5 | 77.118 | 70.89 | 66.831 |
| 5762 | ENSMUSG00000026803 | 235.2883052 | 0.101088223 | 0.174369843 | 0.579734552 | 0.562093639 | 0.900565925 | Ttf1 | transcription termination factor, RNA poly | 202.38 | 214.83 | 264.64 | 260.27 | 230.12 | 239.48 |
| 5763 | ENSMUSG00000026805 | 14.62489274 | 0.295835873 | 0.562074642 | 0.526328446 | 0.598660022 | 0.915055389 | Barhl1 | BarH like homeobox 1 [Source:MGI Symbo | 12.047 | 10.279 | 17.106 | 16.388 | 17.45 | 14.48 |
| 5764 | ENSMUSG00000026806 | 106.4898696 | 0.249653729 | 0.238558864 | 1.046507868 | 0.295326614 | 0.745982248 | Ddx31 | DEAD/H box helicase 31 [Source:MGI Sym | 86.735 | 94.568 | 110.69 | 104.11 | 103.61 | 139.23 |

| | | | | | | | | | | | | | | | |
|---|---|---|---|---|---|---|---|---|---|---|---|---|---|---|---|
| 5765 | ENSMUSG00000026807 | 0.439020386 | -2.189747911 | 3.52148864 | -0.621824499 | 0.534057275 | NA | Ak8 | adenylate kinase 8 [Source:MGI Symbol;A | 1.6062 | 1.0279 | 0 | 0 | 0 | 0 |
| 5766 | ENSMUSG00000026809 | 1.106237902 | -3.510411771 | 2.527023862 | -1.389148644 | 0.164787555 | NA | Spaca9 | sperm acrosome associated 9 [Source:MG | 1.6062 | 0 | 5.0312 | 0 | 0 | 0 |
| 5767 | ENSMUSG00000026810 | 266.6958636 | 0.160604861 | 0.158511746 | 1.01320479 | 0.310962353 | 0.759594595 | Dpm2 | dolichol-phosphate (beta-D) mannosyltra | 253.78 | 248.76 | 252.57 | 248.71 | 292.29 | 304.08 |
| 5768 | ENSMUSG00000026811 | 1219.697582 | -0.023783789 | 0.116025155 | -0.204988211 | 0.837581335 | 0.971073158 | St6galnac6 | ST6 (alpha-N-acetyl-neuraminyl-2,3-beta-( | 1188.6 | 1112.2 | 1388.6 | 1196.3 | 1210.6 | 1221.9 |
| 5769 | ENSMUSG00000026812 | 491.5803617 | 0.471855247 | 0.151841216 | 3.107557077 | 0.001886406 | 0.033396536 | Tsc1 | TSC complex subunit 1 [Source:MGI Symb | 419.22 | 408.08 | 408.54 | 628.51 | 459.15 | 625.98 |
| 5770 | ENSMUSG00000026814 | 823.3584255 | -0.769448016 | 0.153086146 | -5.026242009 | 5.00E-07 | 4.47E-05 | Eng | endoglin [Source:MGI Symbol;Acc:MGI:95 | 1119.5 | 954.94 | 1038.4 | 595.74 | 489.69 | 741.82 |
| 5771 | ENSMUSG00000026815 | 0.160662974 | 1.126924175 | 4.080472857 | 0.276174898 | 0.782413736 | NA | Gfi1b | growth factor independent 1B [Source:M( | 0 | 0 | 0 | 0.964 | 0 | 0 |
| 5772 | ENSMUSG00000026816 | 126.0151384 | 0.036964472 | 0.223115704 | 0.165674006 | 0.868413504 | 0.978366823 | Gtf3c5 | general transcription factor IIIC, polypepti | 120.47 | 105.88 | 146.91 | 130.14 | 111.24 | 141.46 |
| 5773 | ENSMUSG00000026817 | 4.240185547 | -0.672877264 | 1.412826913 | -0.476263057 | 0.633886981 | NA | Ak1 | adenylate kinase 1 [Source:MGI Symbol;A | 9.6372 | 0 | 6.0375 | 8.6758 | 1.0906 | 0 |
| 5774 | ENSMUSG00000026818 | 0 | NA | NA | NA | NA | NA | Cel | carboxyl ester lipase [Source:MGI Symbol, | 0 | 0 | 0 | 0 | 0 | 0 |
| 5775 | ENSMUSG00000026819 | 2603.633204 | -0.774112351 | 0.142079186 | -5.448457092 | 5.08E-08 | 6.12E-06 | Slc25a25 | solute carrier family 25 (mitochondrial car | 2971.5 | 3926.6 | 2959.4 | 1665.8 | 2215 | 1883.5 |
| 5776 | ENSMUSG00000026820 | 446.004459 | 0.04696175 | 0.129478821 | 0.36269831 | 0.716830262 | 0.9447719 | Ptges2 | prostaglandin E synthase 2 [Source:MGI S | 442.51 | 453.31 | 420.61 | 469.46 | 462.42 | 427.72 |
| 5777 | ENSMUSG00000026821 | 25.06104992 | -0.823241887 | 0.514892785 | -1.598860794 | 0.109851537 | 0.486942272 | Ralgds | ral guanine nucleotide dissociation stimul | 40.155 | 35.977 | 20.125 | 29.883 | 13.087 | 11.138 |
| 5778 | ENSMUSG00000026822 | 140.1523827 | -2.693329362 | 0.366628991 | -7.346198554 | 2.04E-13 | 1.02E-10 | Lcn2 | lipocalin 2 [Source:MGI Symbol;Acc:MGI:( | 295.54 | 319.68 | 112.7 | 28.919 | 27.265 | 56.806 |
| 5779 | ENSMUSG00000026824 | 0 | NA | NA | NA | NA | NA | Kcnj3 | potassium inwardly-rectifying channel, su | 0 | 0 | 0 | 0 | 0 | 0 |
| 5780 | ENSMUSG00000026825 | 3.954365223 | 0.145707812 | 1.078920756 | 0.135049596 | 0.892572662 | NA | Dnm1 | dynamin 1 [Source:MGI Symbol;Acc:MGI: | 3.2124 | 2.0558 | 6.0375 | 5.7839 | 2.1812 | 4.4554 |
| 5781 | ENSMUSG00000026826 | 21.6311738 | 0.115756971 | 0.562504042 | 0.205788691 | 0.836955975 | 0.971073158 | Nr4a2 | nuclear receptor subfamily 4, group A, me | 33.73 | 11.307 | 17.106 | 35.667 | 15.269 | 16.708 |
| 5782 | ENSMUSG00000026827 | 980.5252422 | -0.300412236 | 0.16964568 | -1.770821607 | 0.076590372 | 0.405195002 | Gpd2 | glycerol phosphate dehydrogenase 2, mit | 1252.8 | 1198.6 | 794.93 | 918.67 | 965.2 | 752.96 |
| 5783 | ENSMUSG00000026828 | 0 | NA | NA | NA | NA | NA | Galnt5 | polypeptide N-acetylgalactosaminyltransf | 0 | 0 | 0 | 0 | 0 | 0 |
| 5784 | ENSMUSG00000026829 | 4.487705207 | -1.189069171 | 1.048840482 | -1.133698776 | 0.256920923 | NA | Gbgt1 | globoside alpha-1,3-N-acetylgalactosamin | 5.6217 | 5.1396 | 8.05 | 4.8199 | 2.1812 | 1.1138 |
| 5785 | ENSMUSG00000026830 | 0 | NA | NA | NA | NA | NA | Ermn | ermin, ERM-like protein [Source:MGI Sym | 0 | 0 | 0 | 0 | 0 | 0 |
| 5786 | ENSMUSG00000026831 | 0.824622689 | 0.984749497 | 2.871267556 | 0.342966818 | 0.731623423 | NA | 1700007K13 | RIKEN cDNA 1700007K13 gene [Source:M | 1.6062 | 0 | 0 | 0 | 0 | 3.3415 |
| 5787 | ENSMUSG00000026832 | 8.724318247 | -1.655198979 | 0.869327495 | -1.903999341 | 0.05691027 | 0.338327228 | Cytip | cytohesin 1 interacting protein [Source:M | 11.243 | 16.447 | 12.075 | 3.8559 | 8.7249 | 0 |
| 5788 | ENSMUSG00000026833 | 35.17095474 | -0.322736535 | 0.435521399 | -0.741034852 | 0.458672309 | 0.859322431 | Olfm1 | olfactomedin 1 [Source:MGI Symbol;Acc:I | 53.808 | 26.726 | 36.225 | 20.244 | 28.356 | 45.668 |
| 5789 | ENSMUSG00000026834 | 1.417106607 | -1.652820081 | 2.248611406 | -0.735040335 | 0.46231493 | NA | Acvr1c | activin A receptor, type IC [Source:MGI Sy | 6.4248 | 0 | 0 | 0.964 | 0 | 1.1138 |
| 5790 | ENSMUSG00000026835 | 0.401551119 | -2.072982726 | 4.008251979 | -0.517178744 | 0.605031388 | NA | Fcnb | ficolin B [Source:MGI Symbol;Acc:MGI:13 | 2.4093 | 0 | 0 | 0 | 0 | 0 |
| 5791 | ENSMUSG00000026836 | 231.2455613 | 0.39289899 | 0.183882019 | 2.136690649 | 0.032623161 | 0.241304865 | Acvr1 | activin A receptor, type 1 [Source:MGI Syr | 237.72 | 186.05 | 175.09 | 261.24 | 236.66 | 290.71 |
| 5792 | ENSMUSG00000026837 | 91.67700825 | -0.934175121 | 0.323283952 | -2.889642729 | 0.003856799 | 0.055784738 | Col5a1 | collagen, type V, alpha 1 [Source:MGI Sym | 154.2 | 111.02 | 95.593 | 67.478 | 34.9 | 86.88 |
| 5793 | ENSMUSG00000026839 | 10810.64116 | 0.409759738 | 0.117390889 | 3.49055826 | 0.000482013 | 0.012082892 | Upp2 | uridine phosphorylase 2 [Source:MGI Sym | 10508 | 8803.1 | 8544 | 11305 | 13903 | 11801 |
| 5794 | ENSMUSG00000026840 | 22.02973284 | -0.722485837 | 0.592849135 | -1.21866727 | 0.222970505 | 0.668407528 | Lamc3 | laminin gamma 3 [Source:MGI Symbol;Ac | 45.777 | 14.391 | 22.137 | 28.919 | 9.8156 | 11.138 |
| 5795 | ENSMUSG00000026841 | 0 | NA | NA | NA | NA | NA | Fibcd1 | fibrinogen C domain containing 1 [Source: | 0 | 0 | 0 | 0 | 0 | 0 |
| 5796 | ENSMUSG00000026842 | 598.4803368 | 0.00496473 | 0.13879211 | 0.035770981 | 0.971464972 | 0.99516582 | Abl1 | c-abl oncogene 1, non-receptor tyrosine k | 603.93 | 556.1 | 632.93 | 692.14 | 535.49 | 570.29 |
| 5797 | ENSMUSG00000026843 | 351.2426582 | -0.010668047 | 0.163201737 | -0.065367239 | 0.947881608 | 0.990677354 | Fubp3 | far upstream element (FUSE) binding prot | 335.7 | 396.78 | 326.02 | 397.16 | 356.63 | 295.17 |
| 5798 | ENSMUSG00000026848 | 936.8020366 | -0.112034634 | 0.149209014 | -0.75085701 | 0.452738715 | 0.856769845 | Tor1b | torsin family 1, member B [Source:MGI Sy | 952.48 | 981.66 | 986.12 | 1004.5 | 1007.7 | 688.36 |
| 5799 | ENSMUSG00000026849 | 1160.049503 | -0.071007796 | 0.114089029 | -0.622389347 | 0.533685884 | 0.889143097 | Tor1a | torsin family 1, member A (torsin A) [Sour | 1150 | 1228.4 | 1188.4 | 1208.8 | 1182.2 | 1002.5 |
| 5800 | ENSMUSG00000026851 | 471.0178369 | 0.151295034 | 0.139371825 | 1.085549642 | 0.27767829 | 0.730774888 | BC005624 | cDNA sequence BC005624 [Source:MGI Sy | 388.7 | 495.46 | 455.83 | 481.02 | 503.87 | 501.23 |
| 5801 | ENSMUSG00000026853 | 1392.029795 | -0.676174954 | 0.190900675 | -3.542024949 | 0.000397068 | 0.010631083 | Crat | carnitine acetyltransferase [Source:MGI S | 1788.5 | 1815.3 | 1534.5 | 1463.3 | 1014.3 | 736.25 |
| 5802 | ENSMUSG00000026854 | 150.9598414 | -0.042611862 | 0.212054572 | -0.200947621 | 0.840739531 | 0.971854408 | Usp20 | ubiquitin specific peptidase 20 [Source:M | 150.98 | 132.6 | 176.09 | 152.31 | 170.14 | 123.64 |
| 5803 | ENSMUSG00000026856 | 342.9868328 | 0.312262153 | 0.144639473 | 2.158899956 | 0.030857929 | 0.233016344 | Dolpp1 | dolichyl pyrophosphate phosphatase 1 [S | 289.92 | 305.29 | 323 | 364.38 | 362.09 | 413.24 |
| 5804 | ENSMUSG00000026857 | 157.8856113 | -0.122245405 | 0.194080684 | -0.629868995 | 0.5287803 | 0.888334265 | Ntmt1 | N-terminal Xaa-Pro-Lys N-methyltransfera | 171.06 | 156.24 | 166.03 | 136.88 | 173.41 | 143.69 |
| 5805 | ENSMUSG00000026858 | 2185.890407 | -0.030445529 | 0.120982697 | -0.251651925 | 0.801310119 | 0.96255706 | Miga2 | mitoguardin 2 [Source:MGI Symbol;Acc:M | 2428.6 | 2156.6 | 2040.7 | 2097.6 | 2457.2 | 1934.7 |
| 5806 | ENSMUSG00000026860 | 505.3824522 | 0.013181913 | 0.124418808 | 0.105947915 | 0.915623678 | 0.98625062 | Sh3glb2 | SH3-domain GRB2-like endophilin B2 [Sou | 509.17 | 518.07 | 481.99 | 518.62 | 482.05 | 522.39 |
| 5807 | ENSMUSG00000026864 | 8277.712955 | 0.36048495 | 0.134916242 | 2.671916622 | 0.007541939 | 0.08832923 | Hspa5 | heat shock protein 5 [Source:MGI Symbol | 7007.9 | 7851.2 | 6888.8 | 11122 | 7513.3 | 9282.8 |
| 5808 | ENSMUSG00000026866 | 2919.778075 | 0.277040317 | 0.133336305 | 2.077756073 | 0.037731832 | 0.263521589 | Kynu | kynureninase [Source:MGI Symbol;Acc:M | 2892 | 2338.5 | 2688.7 | 2750.2 | 3074.5 | 3774.8 |
| 5809 | ENSMUSG00000026867 | 826.7187883 | 0.058376791 | 0.117554684 | 0.496592638 | 0.619476346 | 0.917855548 | Gapvd1 | GTPase activating protein and VPS9 doma | 768.57 | 868.59 | 792.92 | 776.97 | 885.58 | 867.68 |
| 5810 | ENSMUSG00000026869 | 612.0519319 | 0.096808638 | 0.120939805 | 0.800469605 | 0.423438767 | 0.84018348 | Psmd5 | proteasome (prosome, macropain) 26S su | 595.1 | 618.81 | 560.48 | 610.2 | 674 | 613.73 |
| 5811 | ENSMUSG00000026870 | 672.2681058 | 0.115883755 | 0.130628236 | 0.887126387 | 0.375010859 | 0.809332934 | Cutal | cutA divalent cation tolerance homolog-lil | 696.29 | 563.3 | 675.19 | 671.89 | 675.09 | 751.84 |
| 5812 | ENSMUSG00000026872 | 103.4820397 | -0.809084416 | 0.258784283 | -3.126482054 | 0.001769113 | 0.031945631 | Zeb2 | zinc finger E-box binding homeobox 2 [Sou | 153.39 | 141.85 | 99.618 | 70.37 | 65.437 | 90.221 |
| 5813 | ENSMUSG00000026873 | 45.31141891 | 0.34987813 | 0.373156391 | 0.93761795 | 0.348440783 | 0.789711685 | Phf19 | PHD finger protein 19 [Source:MGI Symbo | 38.549 | 34.949 | 46.287 | 67.478 | 54.531 | 30.074 |
| 5814 | ENSMUSG00000026874 | 13208.60133 | 0.294974628 | 0.092998866 | 3.17180886 | 0.001514927 | 0.028643008 | Hc | hemolytic complement [Source:MGI Symb | 11223 | 12207 | 12160 | 13864 | 15716 | 14082 |
| 5815 | ENSMUSG00000026875 | 7.054469998 | -0.58691238 | 0.865842685 | -0.677851058 | 0.497866134 | 0.879044893 | Traf1 | TNF receptor-associated factor 1 [Source: | 12.047 | 8.2233 | 5.0312 | 4.8199 | 2.1812 | 10.025 |
| 5816 | ENSMUSG00000026878 | 2750.310376 | -0.013814961 | 0.101574642 | -0.136007968 | 0.891814982 | 0.982489666 | Rab14 | RAB14, member RAS oncogene family [So | 2700.8 | 2835 | 2754.1 | 2471.6 | 2950.1 | 2790.2 |
| 5817 | ENSMUSG00000026879 | 358.7010335 | -0.41588232 | 0.1744817 | -2.383529738 | 0.017147499 | 0.155792353 | Gsn | gelsolin [Source:MGI Symbol;Acc:MGI:95 | 499.53 | 401.92 | 328.04 | 327.75 | 305.37 | 289.6 |
| 5818 | ENSMUSG00000026880 | 945.6898669 | 0.143689381 | 0.141838455 | 1.013049538 | 0.311036499 | 0.759594595 | Stom | stomatin [Source:MGI Symbol;Acc:MGI:95 | 1068.9 | 877.84 | 747.64 | 988.08 | 936.84 | 1054.8 |
| 5819 | ENSMUSG00000026882 | 0 | NA | NA | NA | NA | NA | 4930568D16 | RIKEN cDNA 4930568D16 gene [Source:M | 0 | 0 | 0 | 0 | 0 | 0 |
| 5820 | ENSMUSG00000026883 | 302.9748644 | -0.008792028 | 0.160195175 | -0.054883227 | 0.956231495 | 0.991678478 | Dab2ip | disabled 2 interacting protein [Source:MG | 351.76 | 271.37 | 287.79 | 316.18 | 294.47 | 296.28 |

| | | | | | | | | | | | | | | | |
|---|---|---|---|---|---|---|---|---|---|---|---|---|---|---|---|
| 5821 | ENSMUSG00000026885 | 28.69486758 | 0.802413233 | 0.447696135 | 1.792316644 | 0.073082259 | 0.392667827 | Ttll11 | tubulin tyrosine ligase-like family, membe | 24.896 | 25.698 | 12.075 | 42.415 | 40.353 | 26.732 |
| 5822 | ENSMUSG00000026887 | 475.54696 | 0.004852375 | 0.135898309 | 0.035705924 | 0.971516847 | 0.99516582 | Mrrf | mitochondrial ribosome recycling factor [! | 478.65 | 437.89 | 507.15 | 434.75 | 514.77 | 480.07 |
| 5823 | ENSMUSG00000026888 | 1173.928509 | 0.151691533 | 0.125476451 | 1.208924315 | 0.226691924 | 0.673420584 | Grb14 | growth factor receptor bound protein 14 | | 1260.9 | 1008.4 | 1065.6 | 1117.3 | 1277.1 | 1314.3 |
| 5824 | ENSMUSG00000026889 | 242.0355345 | -0.050552978 | 0.168840574 | -0.299412496 | 0.764625329 | 0.954980327 | Rbm18 | RNA binding motif protein 18 [Source:MG | 244.14 | 225.11 | 269.67 | 240.99 | 260.66 | 211.63 |
| 5825 | ENSMUSG00000026890 | 154.3733672 | -0.747570335 | 0.198181755 | -3.772145097 | 0.00016185 | 0.005308389 | Lhx6 | LIM homeobox protein 6 [Source:MGI Syn | 201.58 | 175.77 | 203.26 | 127.25 | 101.43 | 116.95 |
| 5826 | ENSMUSG00000026893 | 68.5221977 | 0.083397502 | 0.289139943 | 0.288433003 | 0.773015308 | 0.958006461 | Gca | grancalcin [Source:MGI Symbol;Acc:MGI:1 | 64.248 | 78.122 | 57.356 | 68.442 | 86.159 | 56.806 |
| 5827 | ENSMUSG00000026894 | 1.170512601 | -1.184332849 | 2.358814357 | -0.502088198 | 0.615605482 | NA | Morn5 | MORN repeat containing 5 [Source:MGI S | 4.8186 | 0 | 0 | 0 | 1.0906 | 1.1138 |
| 5828 | ENSMUSG00000026895 | 1853.653245 | -0.079308399 | 0.094944452 | -0.835313672 | 0.403541121 | 0.828953151 | Ndufa8 | NADH:ubiquinone oxidoreductase subuni | 1882.5 | 1928.4 | 1902.8 | 1764.1 | 1740.6 | 1903.6 |
| 5829 | ENSMUSG00000026896 | 711.0168258 | 0.166328318 | 0.142722846 | 1.165393782 | 0.243859633 | 0.693748143 | Ifih1 | interferon induced with helicase C domair | 726.81 | 628.06 | 654.06 | 636.23 | 874.68 | 746.28 |
| 5830 | ENSMUSG00000026904 | 2.882889805 | 0.211030814 | 1.500501712 | 0.140640169 | 0.888154213 | NA | Slc4a10 | solute carrier family 4, sodium bicarbonat | 0 | 3.0838 | 5.0312 | 4.8199 | 4.3625 | 0 |
| 5831 | ENSMUSG00000026914 | 808.2317859 | 0.083253268 | 0.12610539 | 0.660188021 | 0.509133179 | 0.882793133 | Psmd14 | proteasome (prosome, macropain) 26S su | 891.44 | 747.3 | 714.43 | 791.43 | 860.5 | 844.29 |
| 5832 | ENSMUSG00000026915 | 846.6230976 | -0.008010262 | 0.12777749 | -0.062689149 | 0.950014039 | 0.990815588 | Strbp | spermatid perinuclear RNA binding protei | 829.6 | 873.73 | 844.24 | 887.82 | 927.03 | 717.32 |
| 5833 | ENSMUSG00000026917 | 652.249766 | 0.134512412 | 0.125612849 | 1.070849142 | 0.284237267 | 0.736651636 | Wdr5 | WD repeat domain 5 [Source:MGI Symbol | 614.37 | 576.66 | 674.18 | 639.12 | 672.91 | 736.25 |
| 5834 | ENSMUSG00000026918 | 889.5203868 | -0.047422807 | 0.120401212 | -0.393873167 | 0.693674682 | 0.939096836 | Brd3 | bromodomain containing 3 [Source:MGI S | 836.83 | 913.82 | 961.97 | 799.14 | 952.11 | 873.25 |
| 5835 | ENSMUSG00000026919 | 0 | NA | NA | NA | NA | NA | Lcn4 | lipocalin 4 [Source:MGI Symbol;Acc:MGI:1 | 0 | 0 | 0 | 0 | 0 | 0 |
| 5836 | ENSMUSG00000026921 | 250.6182103 | -0.32500907 | 0.161351389 | -2.014293596 | 0.043978718 | 0.2895349 | Egfl7 | EGF-like domain 7 [Source:MGI Symbol;Ac | 293.13 | 274.45 | 268.67 | 242.92 | 201.76 | 222.77 |
| 5837 | ENSMUSG00000026922 | 4036.136754 | -0.38394996 | 0.108246385 | -3.547000306 | 0.000389644 | 0.010484566 | Agpat2 | 1-acylglycerol-3-phosphate O-acyltransfer | 4190.6 | 4679.1 | 4842 | 3689.1 | 3697.2 | 3118.8 |
| 5838 | ENSMUSG00000026923 | 261.013808 | -0.46780986 | 0.16084108 | -2.908522249 | 0.003631413 | 0.053773172 | Notch1 | notch 1 [Source:MGI Symbol;Acc:MGI:973 | 270.65 | 315.57 | 323 | 208.22 | 220.3 | 228.34 |
| 5839 | ENSMUSG00000026924 | 1974.410284 | 0.065401061 | 0.094767075 | 0.690124294 | 0.490116027 | 0.875501998 | Sec16a | SEC16 homolog A, endoplasmic reticulum | 1895.3 | 2044.5 | 1849.5 | 2008.9 | 2016.6 | 2031.7 |
| 5840 | ENSMUSG00000026925 | 132.2961693 | -0.357914246 | 0.211972048 | -1.688497377 | 0.091315792 | 0.445789516 | Inpp5e | inositol polyphosphate-5-phosphatase E [ | 126.09 | 152.13 | 168.04 | 113.75 | 111.24 | 122.52 |
| 5841 | ENSMUSG00000026926 | 1088.974085 | -0.041002241 | 0.104623467 | -0.391902908 | 0.695129955 | 0.939992732 | Pmpca | peptidase (mitochondrial processing) alph | 1044 | 1137.9 | 1132 | 1052.7 | 1113.5 | 1053.7 |
| 5842 | ENSMUSG00000026927 | 396.877692 | 0.016587936 | 0.14170359 | 0.117060801 | 0.906811872 | 0.984324722 | Entr1 | endosome associated trafficking regulator | 395.93 | 407.06 | 381.37 | 443.43 | 393.71 | 359.77 |
| 5843 | ENSMUSG00000026928 | 11.29523178 | -1.772129224 | 0.738756785 | -2.398799251 | 0.01644893 | 0.152121049 | Card9 | caspase recruitment domain family, meml | 16.062 | 17.475 | 19.119 | 9.6398 | 4.3625 | 1.1138 |
| 5844 | ENSMUSG00000026930 | 23.16656437 | -0.380059702 | 0.49362337 | -0.769938632 | 0.441336296 | 0.849828766 | Gpsm1 | G-protein signalling modulator 1 (AGS3-lik | 33.73 | 24.67 | 20.125 | 25.063 | 10.906 | 24.505 |
| 5845 | ENSMUSG00000026931 | 0 | NA | NA | NA | NA | NA | 1700019N19 | RIKEN cDNA 1700019N19 gene [Source:M | 0 | 0 | 0 | 0 | 0 | 0 |
| 5846 | ENSMUSG00000026932 | 495.4090366 | 0.026438363 | 0.132440227 | 0.199624873 | 0.841773973 | 0.971854408 | Nacc2 | nucleus accumbens associated 2, BEN anc | 473.83 | 494.43 | 504.13 | 457.89 | 488.6 | 553.58 |
| 5847 | ENSMUSG00000026933 | 162.910651 | -0.041800977 | 0.222955617 | -0.187485642 | 0.85127988 | 0.973568625 | Camsap1 | calmodulin regulated spectrin-associated | 122.07 | 196.33 | 178.11 | 148.45 | 182.13 | 150.37 |
| 5848 | ENSMUSG00000026934 | 0 | NA | NA | NA | NA | NA | Lhx3 | LIM homeobox protein 3 [Source:MGI Syn | 0 | 0 | 0 | 0 | 0 | 0 |
| 5849 | ENSMUSG00000026936 | 0 | NA | NA | NA | NA | NA | Lcn3 | lipocalin 3 [Source:MGI Symbol;Acc:MGI:1 | 0 | 0 | 0 | 0 | 0 | 0 |
| 5850 | ENSMUSG00000026937 | 0 | NA | NA | NA | NA | NA | Lcn5 | lipocalin 5 [Source:MGI Symbol;Acc:MGI:1 | 0 | 0 | 0 | 0 | 0 | 0 |
| 5851 | ENSMUSG00000026938 | 374.6359903 | -0.992824554 | 0.16623568 | -5.972391447 | 2.34E-09 | 4.34E-07 | Fcna | ficolin A [Source:MGI Symbol;Acc:MGI:134 | 461.78 | 470.79 | 563.5 | 243.89 | 211.58 | 296.28 |
| 5852 | ENSMUSG00000026939 | 186.5861547 | -0.205029847 | 0.202150364 | -1.014244263 | 0.310466213 | 0.759154604 | Tmem141 | transmembrane protein 141 [Source:MGI | 204.79 | 167.55 | 227.41 | 204.36 | 148.32 | 167.08 |
| 5853 | ENSMUSG00000026940 | 6.78797046 | 0.743424547 | 0.880748478 | 0.844082693 | 0.398623198 | 0.82588279 | Ccdc183 | coiled-coil domain containing 183 [Source | 3.2124 | 5.1396 | 7.0437 | 15.424 | 5.4531 | 4.4554 |
| 5854 | ENSMUSG00000026941 | 25.81265155 | 0.12671153 | 0.448832085 | 0.282313887 | 0.777702838 | 0.959559604 | Mamdc4 | MAM domain containing 4 [Source:MGI S | 22.487 | 22.614 | 29.181 | 37.595 | 20.722 | 22.277 |
| 5855 | ENSMUSG00000026942 | 200.1944714 | -0.06747253 | 0.181176783 | -0.372412673 | 0.70958562 | 0.944239215 | Traf2 | TNF receptor-associated factor 2 [Source: | 191.14 | 201.47 | 222.38 | 204.36 | 167.96 | 213.86 |
| 5856 | ENSMUSG00000026943 | 3.786136485 | 0.170653403 | 1.238866137 | 0.137749671 | 0.890438262 | NA | Lcn12 | lipocalin 12 [Source:MGI Symbol;Acc:MGI | 1.6062 | 1.0279 | 8.05 | 0.964 | 3.2719 | 7.7969 |
| 5857 | ENSMUSG00000026944 | 688.7872892 | 0.476605689 | 0.123831889 | 3.848812229 | 0.000118692 | 0.00416484 | Abca2 | ATP-binding cassette, sub-family A (ABC1) | 603.93 | 531.43 | 591.67 | 735.52 | 812.51 | 857.66 |
| 5858 | ENSMUSG00000026946 | 399.5662483 | 0.072895891 | 0.143400365 | 0.50833825 | 0.61121615 | 0.916376001 | Nmi | N-myc (and STAT) interactor [Source:MGI | 426.45 | 349.49 | 391.43 | 386.56 | 404.62 | 438.85 |
| 5859 | ENSMUSG00000026950 | 431.2578535 | 0.428931714 | 0.187265791 | 2.2904969 | 0.021992528 | 0.184941818 | Neb | nebulin [Source:MGI Symbol;Acc:MGI:972 | 404.76 | 276.51 | 420.61 | 408.73 | 582.39 | 494.55 |
| 5860 | ENSMUSG00000026955 | 0.342432689 | 1.990653086 | 3.900281263 | 0.51038706 | 0.609780321 | NA | Sapcd2 | suppressor APC domain containing 2 [Sou | 0 | 0 | 0 | 0.964 | 1.0906 | 0 |
| 5861 | ENSMUSG00000026956 | 69.62853578 | -1.191252443 | 0.469258964 | -2.538582181 | 0.011130266 | 0.117069607 | Uap1l1 | UDP-N-acteylglucosamine pyrophosphory | 110.03 | 149.05 | 31.194 | 27.955 | 32.719 | 66.831 |
| 5862 | ENSMUSG00000026958 | 361.6749213 | 0.106503196 | 0.149151533 | 0.714060353 | 0.475189861 | 0.869368495 | Dpp7 | dipeptidylpeptidase 7 [Source:MGI Symbc | 365.41 | 378.27 | 300.87 | 358.6 | 391.53 | 375.37 |
| 5863 | ENSMUSG00000026959 | 0 | NA | NA | NA | NA | NA | Grin1 | glutamate receptor, ionotropic, NMDA1 (z | 0 | 0 | 0 | 0 | 0 | 0 |
| 5864 | ENSMUSG00000026960 | 225.4760391 | 0.276032601 | 0.161991558 | 1.703993739 | 0.088382257 | 0.438393926 | Arl6ip6 | ADP-ribosylation factor-like 6 interacting p | 199.17 | 211.75 | 201.25 | 251.6 | 256.3 | 232.79 |
| 5865 | ENSMUSG00000026961 | 0 | NA | NA | NA | NA | NA | Lrrc26 | leucine rich repeat containing 26 [Source: | 0 | 0 | 0 | 0 | 0 | 0 |
| 5866 | ENSMUSG00000026963 | 0 | NA | NA | NA | NA | NA | Tmem210 | transmembrane protein 210 [Source:MGI | 0 | 0 | 0 | 0 | 0 | 0 |
| 5867 | ENSMUSG00000026965 | 734.6980533 | 0.113029582 | 0.118768199 | 0.951682209 | 0.341258175 | 0.783129922 | Anapc2 | anaphase promoting complex subunit 2 [S | 750.9 | 640.39 | 725.5 | 738.41 | 771.07 | 781.92 |
| 5868 | ENSMUSG00000026966 | 282.5375261 | 0.041851932 | 0.152976311 | 0.273584396 | 0.784404027 | 0.960670212 | Ssna1 | SS nuclear autoantigen 1 [Source:MGI Syn | 268.24 | 280.62 | 286.78 | 295.94 | 259.57 | 304.08 |
| 5869 | ENSMUSG00000026969 | 0.863436065 | -1.832005063 | 2.586421136 | -0.708316615 | 0.478748662 | NA | Fam166a | family with sequence similarity 166, meml | 0 | 3.0838 | 1.0062 | 0 | 1.0906 | 0 |
| 5870 | ENSMUSG00000026970 | 617.3082274 | 0.172781447 | 0.127564567 | 1.354462693 | 0.175588808 | 0.605992966 | Rbms1 | RNA binding motif, single stranded intera | 570.2 | 609.56 | 561.48 | 649.72 | 591.12 | 721.77 |
| 5871 | ENSMUSG00000026971 | 0.824622689 | 0.984749497 | 2.871267556 | 0.342966818 | 0.731623423 | NA | Itgb6 | integrin beta 6 [Source:MGI Symbol;Acc:N | 1.6062 | 0 | 0 | 0 | 0 | 3.3415 |
| 5872 | ENSMUSG00000026972 | 52.10435899 | -0.089199907 | 0.315083583 | -0.283099191 | 0.777100802 | 0.959452497 | Arrdc1 | arrestin domain containing 1 [Source:MGI | 59.43 | 41.117 | 60.375 | 49.163 | 49.078 | 53.465 |
| 5873 | ENSMUSG00000026974 | 194.8119086 | -0.014770387 | 0.188741682 | -0.078257157 | 0.937623496 | 0.988420305 | Zmynd19 | zinc finger, MYND domain containing 19 [! | 187.93 | 222.03 | 178.11 | 221.71 | 188.68 | 170.42 |
| 5874 | ENSMUSG00000026975 | 79.56919429 | -0.133711249 | 0.249638967 | -0.535618499 | 0.592222241 | 0.912923013 | Dph7 | diphthamine biosynthesis 7 [Source:MGI | 81.113 | 85.317 | 83.518 | 80.974 | 77.434 | 69.058 |
| 5875 | ENSMUSG00000026976 | 4.761565514 | 0.783456599 | 0.968598369 | 0.808855996 | 0.418597983 | NA | Pax8 | paired box 8 [Source:MGI Symbol;Acc:MG | 2.4093 | 4.1117 | 4.025 | 4.8199 | 7.6343 | 5.5692 |
| 5876 | ENSMUSG00000026977 | 548.4750494 | 0.224821171 | 0.14128578 | 1.591251233 | 0.111553047 | 0.490128576 | Marchf7 | membrane associated ring-CH-type finger | 502.74 | 529.38 | 486.02 | 656.47 | 619.47 | 496.77 |

| | | | | | | | | | | | | | | | |
|---|---|---|---|---|---|---|---|---|---|---|---|---|---|---|---|
| 5877 | ENSMUSG00000026979 | 65.48801881 | -0.494490676 | 0.338925538 | -1.45899503 | 0.144566476 | 0.55464443 | Psd4 | pleckstrin and Sec7 domain containing 4 [ | 101.99 | 84.289 | 43.269 | 59.767 | 51.259 | 52.351 |
| 5878 | ENSMUSG00000026980 | 49.89179848 | 0.221629166 | 0.342472353 | 0.647144693 | 0.517538302 | 0.887022094 | Ly75 | lymphocyte antigen 75 [Source:MGI Symb | 36.943 | 49.34 | 52.325 | 60.731 | 63.256 | 36.757 |
| 5879 | ENSMUSG00000026981 | 44.97294089 | -0.333873626 | 0.370947326 | -0.900056701 | 0.368090077 | 0.80627882 | Il1rn | interleukin 1 receptor antagonist [Source: | 67.461 | 48.312 | 34.212 | 33.739 | 35.99 | 50.123 |
| 5880 | ENSMUSG00000026983 | 0 | NA | NA | NA | NA | NA | Il1f5 | interleukin 1 family, member 5 (delta) [So | 0 | 0 | 0 | 0 | 0 | 0 |
| 5881 | ENSMUSG00000026984 | 0 | NA | NA | NA | NA | NA | Il1f6 | interleukin 1 family, member 6 [Source:M | 0 | 0 | 0 | 0 | 0 | 0 |
| 5882 | ENSMUSG00000026985 | 0 | NA | NA | NA | NA | NA | Il1f8 | interleukin 1 family, member 8 [Source:M | 0 | 0 | 0 | 0 | 0 | 0 |
| 5883 | ENSMUSG00000026986 | 774.0600512 | -0.286140457 | 0.150782733 | -1.897700427 | 0.057735556 | 0.341829329 | Hnmt | histamine N-methyltransferase [Source:M | 882.61 | 915.87 | 752.67 | 619.84 | 848.5 | 624.87 |
| 5884 | ENSMUSG00000026987 | 241.0385142 | 0.136941843 | 0.174914565 | 0.782907034 | 0.433681707 | 0.845264432 | Baz2b | bromodomain adjacent to zinc finger dom | 212.02 | 259.04 | 218.36 | 259.31 | 279.2 | 218.31 |
| 5885 | ENSMUSG00000026988 | 75.49748111 | 0.436849404 | 0.265771099 | 1.643705451 | 0.100237059 | 0.466182902 | Wdsub1 | WD repeat, SAM and U-box domain conta | 69.067 | 64.759 | 58.362 | 78.082 | 102.52 | 80.197 |
| 5886 | ENSMUSG00000026989 | 0.401551119 | -2.072982726 | 4.008251979 | -0.517178744 | 0.605031388 | NA | Dapl1 | death associated protein-like 1 [Source:M | 2.4093 | 0 | 0 | 0 | 0 | 0 |
| 5887 | ENSMUSG00000026991 | 1603.889273 | 0.082347849 | 0.107885014 | 0.76329275 | 0.445288825 | 0.852147532 | Pkp4 | plakophilin 4 [Source:MGI Symbol;Acc:MC | 1423.1 | 1658 | 1593.9 | 1541.4 | 1689.4 | 1717.5 |
| 5888 | ENSMUSG00000026994 | 3.069015228 | -0.900770308 | 1.218022617 | -0.739534961 | 0.459582219 | NA | Galnt3 | polypeptide N-acetylgalactosaminyltransf | 4.8186 | 3.0838 | 4.025 | 0.964 | 2.1812 | 3.3415 |
| 5889 | ENSMUSG00000026999 | 86.18472643 | -0.142214736 | 0.249795716 | -0.56932416 | 0.569136174 | 0.903735951 | Nup35 | nucleoporin 35 [Source:MGI Symbol;Acc:I | 89.144 | 92.513 | 89.556 | 74.226 | 98.156 | 73.514 |
| 5890 | ENSMUSG00000027001 | 78.70262118 | 0.14242232 | 0.293179187 | 0.485785917 | 0.62711896 | 0.920273791 | Dusp19 | dual specificity phosphatase 19 [Source:M | 77.901 | 59.619 | 86.537 | 62.659 | 76.343 | 109.16 |
| 5891 | ENSMUSG00000027002 | 2033.510321 | -0.009878263 | 0.10385515 | -0.095115777 | 0.924222867 | 0.987482515 | Nckap1 | NCK-associated protein 1 [Source:MGI Syr | 2030.2 | 2169.9 | 1920.9 | 1903.9 | 2182.3 | 1993.8 |
| 5892 | ENSMUSG00000027004 | 19.55847354 | -1.787693042 | 0.588239748 | -3.039055159 | 0.002373214 | 0.039642097 | Frzb | frizzled-related protein [Source:MGI Symb | 36.943 | 29.81 | 24.15 | 6.7478 | 16.359 | 3.3415 |
| 5893 | ENSMUSG00000027006 | 469.418162 | -0.047880269 | 0.128837031 | -0.371634374 | 0.710165093 | 0.944239215 | Dnajc10 | DnaJ heat shock protein family (Hsp40) m | 497.12 | 484.15 | 449.79 | 448.25 | 449.33 | 487.86 |
| 5894 | ENSMUSG00000027007 | 1297.734103 | -0.04295808 | 0.119742843 | -0.3587528 | 0.719780025 | 0.9447719 | Itprid2 | ITPR interacting domain containing 2 [Sou | 1299.4 | 1351.7 | 1301.1 | 1455.6 | 1125.5 | 1253.1 |
| 5895 | ENSMUSG00000027009 | 87.08900912 | -0.776415333 | 0.278293909 | -2.789911344 | 0.005272248 | 0.069074085 | Itga4 | integrin alpha 4 [Source:MGI Symbol;Acc: | 109.22 | 132.6 | 88.55 | 81.938 | 54.531 | 55.692 |
| 5896 | ENSMUSG00000027010 | 339.6349557 | 0.265949141 | 0.152793808 | 1.740575383 | 0.081758036 | 0.420985486 | Slc25a12 | solute carrier family 25 (mitochondrial car | 274.66 | 299.12 | 352.19 | 368.24 | 387.17 | 356.43 |
| 5897 | ENSMUSG00000027011 | 624.2999399 | 0.011736308 | 0.125578583 | 0.093457875 | 0.925539814 | 0.987546253 | Ube2e3 | ubiquitin-conjugating enzyme E2E 3 [Sour | 580.64 | 598.25 | 687.27 | 656.47 | 619.47 | 603.7 |
| 5898 | ENSMUSG00000027012 | 838.0659304 | 0.045388683 | 0.111773506 | 0.406077296 | 0.684685807 | 0.935737593 | Dync1i2 | dynein cytoplasmic 1 intermediate chain 2 | 870.56 | 837.75 | 765.75 | 854.08 | 864.86 | 835.38 |
| 5899 | ENSMUSG00000027014 | 204.9081947 | 0.174471687 | 0.178198672 | 0.979085228 | 0.32753787 | 0.772937193 | Cwc22 | CWC22 spliceosome-associated protein [S | 190.34 | 206.61 | 181.12 | 246.78 | 196.31 | 208.29 |
| 5900 | ENSMUSG00000027015 | 8.446807118 | 0.988307661 | 0.736324661 | 1.34221725 | 0.179525565 | 0.611676727 | Cybrd1 | cytochrome b reductase 1 [Source:MGI Sy | 4.8186 | 7.1954 | 5.0312 | 11.568 | 9.8156 | 12.252 |
| 5901 | ENSMUSG00000027016 | 344.4900727 | -0.149582945 | 0.160662521 | -0.9310382 | 0.351833804 | 0.792889192 | Zfp385b | zinc finger protein 385B [Source:MGI Sym | 371.03 | 410.14 | 305.9 | 341.25 | 295.56 | 343.06 |
| 5902 | ENSMUSG00000027018 | 172.0146932 | -0.010400565 | 0.186006411 | -0.055915087 | 0.955409452 | 0.991678478 | Hat1 | histone aminotransferase 1 [Source:MGI S | 187.12 | 158.3 | 172.07 | 167.73 | 187.59 | 159.28 |
| 5903 | ENSMUSG00000027022 | 0 | NA | NA | NA | NA | NA | Xirp2 | xin actin-binding repeat containing 2 [Sou | 0 | 0 | 0 | 0 | 0 | 0 |
| 5904 | ENSMUSG00000027030 | 33.51679094 | -0.543521726 | 0.393888704 | -1.379886553 | 0.167621577 | 0.593804353 | Stk39 | serine/threonine kinase 39 [Source:MGI S | 41.761 | 37.005 | 40.25 | 20.244 | 25.084 | 36.757 |
| 5905 | ENSMUSG00000027035 | 333.5198431 | -0.286466592 | 0.435129729 | -0.658347553 | 0.510314828 | 0.883032087 | Cers6 | ceramide synthase 6 [Source:MGI Symbol | 427.25 | 439.95 | 232.44 | 402.94 | 178.86 | 319.67 |
| 5906 | ENSMUSG00000027048 | 7042.084005 | 0.590451069 | 0.124970436 | 4.724726011 | 2.30E-06 | 0.000162579 | Abcb11 | ATP-binding cassette, sub-family B (MDR/ | 6097.2 | 4912.4 | 5851.3 | 7796.7 | 7760.8 | 9834.1 |
| 5907 | ENSMUSG00000027067 | 903.1144709 | -0.005226883 | 0.110977382 | -0.047098637 | 0.962434613 | 0.992956886 | Ssrp1 | structure specific recognition protein 1 [S | 838.44 | 910.74 | 965.99 | 906.14 | 906.3 | 891.08 |
| 5908 | ENSMUSG00000027068 | 12.03387401 | -0.369497093 | 0.69041128 | -0.535184032 | 0.592522607 | 0.912923013 | Dhrs9 | dehydrogenase/reductase (SDR family) m | 6.4248 | 11.307 | 23.144 | 12.532 | 6.5437 | 12.252 |
| 5909 | ENSMUSG00000027070 | 29.22350442 | 1.109458182 | 0.545822391 | 2.032635891 | 0.042089323 | 0.282496502 | Lrp2 | low density lipoprotein receptor-related p | 16.062 | 9.2513 | 30.187 | 22.171 | 68.709 | 28.96 |
| 5910 | ENSMUSG00000027071 | 18.83525061 | 0.057300742 | 0.529505022 | 0.108215672 | 0.913824613 | 0.98625062 | P2rx3 | purinergic receptor P2X, ligand-gated ion | 17.668 | 18.503 | 19.119 | 11.568 | 29.447 | 16.708 |
| 5911 | ENSMUSG00000027072 | 0 | NA | NA | NA | NA | NA | Prg3 | proteoglycan 3 [Source:MGI Symbol;Acc:I | 0 | 0 | 0 | 0 | 0 | 0 |
| 5912 | ENSMUSG00000027073 | 0 | NA | NA | NA | NA | NA | Prg2 | proteoglycan 2, bone marrow [Source:MG | 0 | 0 | 0 | 0 | 0 | 0 |
| 5913 | ENSMUSG00000027074 | 400.1778199 | -0.813449491 | 0.177449709 | -4.584112857 | 4.56E-06 | 0.000283021 | Slc43a3 | solute carrier family 43, member 3 [Sourc | 560.57 | 514.99 | 454.82 | 330.64 | 217.03 | 323.01 |
| 5914 | ENSMUSG00000027075 | 227.1454278 | 0.626862497 | 0.234025671 | 2.678605705 | 0.007392939 | 0.087077748 | Slc43a1 | solute carrier family 43, member 1 [Sourc | 197.56 | 113.07 | 224.39 | 240.99 | 268.29 | 318.56 |
| 5915 | ENSMUSG00000027076 | 330.5133814 | 0.192262061 | 0.142989855 | 1.344585328 | 0.178759184 | 0.610813333 | Timm10 | translocase of inner mitochondrial membr | 330.08 | 299.12 | 295.84 | 351.85 | 365.36 | 340.84 |
| 5916 | ENSMUSG00000027077 | 0.167707431 | -0.796629176 | 4.080472857 | -0.195229623 | 0.84521319 | NA | Smtnl1 | smoothelin-like 1 [Source:MGI Symbol;Ac | 0 | 0 | 1.0062 | 0 | 0 | 0 |
| 5917 | ENSMUSG00000027078 | 1293.949907 | 0.152550384 | 0.122824669 | 1.242017459 | 0.21423012 | 0.658874792 | Ube2l6 | ubiquitin-conjugating enzyme E2L 6 [Sour | 1412.7 | 1154.4 | 1107.9 | 1285.9 | 1391.6 | 1411.2 |
| 5918 | ENSMUSG00000027079 | 264.747018 | 0.015612017 | 0.15531663 | 0.100517356 | 0.919933604 | 0.986685804 | Clp1 | CLP1, cleavage and polyadenylation factor | 258.6 | 256.98 | 274.7 | 284.37 | 245.39 | 268.44 |
| 5919 | ENSMUSG00000027080 | 195.7590851 | 0.046204856 | 0.181330571 | 0.254810071 | 0.798869796 | 0.962424862 | Med19 | mediator complex subunit 19 [Source:MG | 203.18 | 179.89 | 194.21 | 180.26 | 188.68 | 228.34 |
| 5920 | ENSMUSG00000027082 | 68.87689055 | -0.430283669 | 0.311902722 | -1.379544449 | 0.167726952 | 0.593804353 | Tfpi | tissue factor pathway inhibitor [Source:M | 83.523 | 84.289 | 69.431 | 66.514 | 35.99 | 73.514 |
| 5921 | ENSMUSG00000027086 | 549.9191304 | 0.103483322 | 0.125550825 | 0.824234505 | 0.409806325 | 0.833241465 | Fastkd1 | FAST kinase domains 1 [Source:MGI Symb | 536.47 | 542.74 | 511.17 | 549.47 | 620.56 | 539.1 |
| 5922 | ENSMUSG00000027087 | 426.9509163 | -0.224854212 | 0.14574947 | -1.542744632 | 0.122892748 | 0.513267141 | Itgav | integrin alpha V [Source:MGI Symbol;Acc: | 434.48 | 485.18 | 461.87 | 443.43 | 394.8 | 341.95 |
| 5923 | ENSMUSG00000027088 | 417.0454084 | 0.171544207 | 0.135507546 | 1.265938411 | 0.205535133 | 0.650907725 | Phospho2 | phosphatase, orphan 2 [Source:MGI Symb | 375.05 | 376.22 | 425.64 | 419.33 | 448.24 | 457.79 |
| 5924 | ENSMUSG00000027091 | 856.5601416 | 0.035814703 | 0.112836438 | 0.3174037 | 0.7509373 | 0.952074518 | Zc3h15 | zinc finger CCCH-type containing 15 [Sour | 800.69 | 911.76 | 826.13 | 882.04 | 833.23 | 885.51 |
| 5925 | ENSMUSG00000027099 | 1031.243564 | 0.024215451 | 0.114237395 | 0.211974821 | 0.832126678 | 0.970533397 | Mtx2 | metaxin 2 [Source:MGI Symbol;Acc:MGI:1 | 1053.7 | 979.61 | 1033.4 | 936.99 | 1120.1 | 1063.7 |
| 5926 | ENSMUSG00000027102 | 0.435408177 | -2.179616019 | 3.531869654 | -0.617128103 | 0.537150234 | NA | Hoxd8 | homeobox D8 [Source:MGI Symbol;Acc:M | 1.6062 | 0 | 1.0062 | 0 | 0 | 0 |
| 5927 | ENSMUSG00000027104 | 468.1504963 | 0.048429084 | 0.127663175 | 0.379350461 | 0.704427632 | 0.94295036 | Atf2 | activating transcription factor 2 [Source:M | 430.46 | 479.01 | 471.93 | 466.57 | 485.33 | 475.61 |
| 5928 | ENSMUSG00000027107 | 0 | NA | NA | NA | NA | NA | Chrna1 | cholinergic receptor, nicotinic, alpha poly | 0 | 0 | 0 | 0 | 0 | 0 |
| 5929 | ENSMUSG00000027108 | 685.7831212 | 0.14756434 | 0.134678852 | 1.095675664 | 0.273220737 | 0.728320076 | Ola1 | Obg-like ATPase 1 [Source:MGI Symbol;Ac | 717.17 | 612.64 | 620.85 | 627.55 | 761.25 | 775.24 |
| 5930 | ENSMUSG00000027109 | 798.7456812 | -0.072003086 | 0.117186812 | -0.614429938 | 0.538931257 | 0.89174748 | Sp3 | trans-acting transcription factor 3 [Source | 752.51 | 856.26 | 848.26 | 785.64 | 815.78 | 734.02 |
| 5931 | ENSMUSG00000027111 | 27.31698379 | -0.587014155 | 0.424819515 | -1.381796584 | 0.167034168 | 0.593319485 | Itga6 | integrin alpha 6 [Source:MGI Symbol;Acc: | 36.14 | 33.921 | 28.175 | 19.28 | 18.541 | 27.846 |
| 5932 | ENSMUSG00000027115 | 8.343356938 | 0.016101413 | 0.736698379 | 0.02185618 | 0.982562679 | 0.998580735 | Kif18a | kinesin family member 18A [Source:MGI S | 9.6372 | 5.1396 | 10.062 | 8.6758 | 7.6343 | 8.9108 |

| | | | | | | | | | | | | | | | |
|---|---|---|---|---|---|---|---|---|---|---|---|---|---|---|---|
| 5933 | ENSMUSG00000027120 | 0 | NA | NA | NA | NA | NA | Fshb | follicle stimulating hormone beta [Source: | 0 | 0 | 0 | 0 | 0 | 0 |
| 5934 | ENSMUSG00000027122 | 229.5063615 | 0.047490166 | 0.166977711 | 0.284410213 | 0.776096034 | 0.95893551 | Arl14ep | ADP-ribosylation factor-like 14 effector pr | 208 | 216.89 | 252.57 | 225.57 | 232.3 | 241.7 |
| 5935 | ENSMUSG00000027130 | 153.9491619 | 0.121179038 | 0.188596473 | 0.642530774 | 0.520528614 | 0.887802551 | Slc12a6 | solute carrier family 12, member 6 [Source | 149.38 | 150.08 | 142.89 | 154.24 | 173.41 | 153.71 |
| 5936 | ENSMUSG00000027131 | 742.6130157 | 0.140082056 | 0.112358315 | 1.24674401 | 0.212491376 | 0.657146732 | Emc4 | ER membrane protein complex subunit 4 [ | 741.26 | 686.65 | 691.29 | 780.82 | 802.7 | 752.96 |
| 5937 | ENSMUSG00000027132 | 96.31991598 | -0.18835368 | 0.241502089 | -0.779925676 | 0.435434624 | 0.845726656 | Katnbl1 | katanin p80 subunit B like 1 [Source:MGI S | 107.62 | 94.568 | 105.66 | 101.22 | 73.071 | 95.791 |
| 5938 | ENSMUSG00000027133 | 746.170651 | 0.023643038 | 0.128284955 | 0.18430094 | 0.853777377 | 0.973825032 | Nop10 | NOP10 ribonucleoprotein [Source:MGI Sy | 737.25 | 718.51 | 763.74 | 676.71 | 719.81 | 861 |
| 5939 | ENSMUSG00000027134 | 7.276887488 | -1.868021835 | 0.875213202 | -2.134362039 | 0.032813156 | 0.242147702 | Lpcat4 | lysophosphatidylcholine acyltransferase 4 | 12.85 | 15.419 | 6.0375 | 3.8559 | 3.2719 | 2.2277 |
| 5940 | ENSMUSG00000027157 | 0 | NA | NA | NA | NA | NA | A26c3 | ANKRD26-like family C, member 3 [Source | 0 | 0 | 0 | 0 | 0 | 0 |
| 5941 | ENSMUSG00000027160 | 38.97284705 | -0.181709743 | 0.426389512 | -0.426159036 | 0.669991965 | 0.930998537 | Ccdc34 | coiled-coil domain containing 34 [Source:I | 50.595 | 34.949 | 38.237 | 17.352 | 40.353 | 52.351 |
| 5942 | ENSMUSG00000027162 | 1033.257268 | -0.071651504 | 0.154362595 | -0.464176596 | 0.642521227 | 0.924876278 | Lin7c | lin-7 homolog C (C. elegans) [Source:MGI | 968.54 | 1226.3 | 983.1 | 1134.6 | 1091.7 | 795.28 |
| 5943 | ENSMUSG00000027163 | 406.8798577 | 0.08832491 | 0.147226699 | 0.599924538 | 0.548556528 | 0.895622715 | Commd9 | COMM domain containing 9 [Source:MGI | 435.28 | 346.41 | 400.49 | 386.56 | 451.52 | 421.03 |
| 5944 | ENSMUSG00000027164 | 361.7890085 | 0.037488674 | 0.16137851 | 0.232302767 | 0.816302864 | 0.967767553 | Traf6 | TNF receptor-associated factor 6 [Source: | 349.35 | 394.72 | 328.04 | 429.93 | 347.91 | 320.79 |
| 5945 | ENSMUSG00000027165 | 230.1232775 | 0.182465528 | 0.166207458 | 1.097817934 | 0.272283999 | 0.727618887 | Iftap | intraflagellar transport associated protein | 207.2 | 229.23 | 210.31 | 221.71 | 265.02 | 247.27 |
| 5946 | ENSMUSG00000027166 | 94.33634092 | 0.147140355 | 0.246659717 | 0.596531762 | 0.550820042 | 0.895703112 | Dnajc24 | DnaJ heat shock protein family (Hsp40) m | 88.341 | 92.513 | 87.543 | 86.758 | 88.34 | 122.52 |
| 5947 | ENSMUSG00000027167 | 138.0049387 | 0.017990787 | 0.198658418 | 0.090561413 | 0.927841094 | 0.987920546 | Elp4 | elongator acetyltransferase complex subu | 140.54 | 140.82 | 129.81 | 130.14 | 135.24 | 151.48 |
| 5948 | ENSMUSG00000027168 | 3.009638074 | -0.160577558 | 1.199222957 | -0.133901337 | 0.893480593 | NA | Pax6 | paired box 6 [Source:MGI Symbol;Acc:MG | 2.4093 | 4.1117 | 3.0187 | 1.928 | 4.3625 | 2.2277 |
| 5949 | ENSMUSG00000027170 | 1117.081802 | -0.001306131 | 0.104459548 | -0.012503705 | 0.990023747 | 0.99859176 | Eif3m | eukaryotic translation initiation factor 3, s | 1061.7 | 1169.8 | 1122 | 1103.8 | 1160.4 | 1084.9 |
| 5950 | ENSMUSG00000027171 | 6.102886408 | -1.521648247 | 0.914311001 | -1.664256741 | 0.096061133 | 0.457754528 | Prrg4 | proline rich Gla (G-carboxyglutamic acid) 4 | 12.85 | 7.1954 | 7.0437 | 2.8919 | 2.1812 | 4.4554 |
| 5951 | ENSMUSG00000027173 | 539.5295178 | 0.752144077 | 0.166038956 | 4.529925346 | 5.90E-06 | 0.000357089 | Depdc7 | DEP domain containing 7 [Source:MGI Syr | 403.96 | 334.07 | 467.9 | 722.02 | 554.03 | 755.19 |
| 5952 | ENSMUSG00000027175 | 77.73531659 | -0.020519387 | 0.287326318 | -0.071414923 | 0.943067533 | 0.990077979 | Tcp11l1 | t-complex 11 like 1 [Source:MGI Symbol;A | 93.16 | 66.815 | 74.462 | 76.154 | 57.803 | 98.018 |
| 5953 | ENSMUSG00000027176 | 176.3810043 | 0.122784339 | 0.181020158 | 0.678290976 | 0.497587219 | 0.879037876 | Cstf3 | cleavage stimulation factor, 3' pre-RNA, su | 175.08 | 153.16 | 178.11 | 177.37 | 194.13 | 180.44 |
| 5954 | ENSMUSG00000027177 | 1562.22514 | -0.177711219 | 0.124577658 | -1.426509549 | 0.153721312 | 0.570987433 | Hipk3 | homeodomain interacting protein kinase 3 | 1444.8 | 1940.7 | 1590.9 | 1493.2 | 1464.7 | 1439.1 |
| 5955 | ENSMUSG00000027180 | 2585.326842 | -0.071311608 | 0.114154412 | -0.624694281 | 0.532171727 | 0.889143097 | Fbxo3 | F-box protein 3 [Source:MGI Symbol;Acc:I | 2390.8 | 2846.3 | 2711.8 | 2553.6 | 2750.5 | 2258.9 |
| 5956 | ENSMUSG00000027184 | 2936.352013 | -0.357302609 | 0.130294763 | -2.742263778 | 0.006101732 | 0.076279561 | Caprin1 | cell cycle associated protein 1 [Source:MG | 2731.4 | 3538.1 | 3625.5 | 2289.4 | 2649.1 | 2784.6 |
| 5957 | ENSMUSG00000027185 | 333.8849885 | -0.101468557 | 0.156546773 | -0.648167672 | 0.516876509 | 0.88673963 | Nat10 | N-acetyltransferase 10 [Source:MGI Symb | 293.94 | 383.41 | 360.24 | 306.54 | 323.91 | 335.27 |
| 5958 | ENSMUSG00000027186 | 2.438885942 | -0.058786538 | 1.410103521 | -0.041689519 | 0.966746209 | NA | Elf5 | E74-like factor 5 [Source:MGI Symbol;Acc | 2.4093 | 3.0838 | 2.0125 | 3.8559 | 3.2719 | 0 |
| 5959 | ENSMUSG00000027187 | 53171.04601 | 0.025422845 | 0.085466158 | 0.297460957 | 0.766114611 | 0.955091733 | Cat | catalase [Source:MGI Symbol;Acc:MGI:88 | 51860 | 53496 | 52751 | 51394 | 57336 | 52189 |
| 5960 | ENSMUSG00000027188 | 58.51841127 | -0.763288321 | 0.313894095 | -2.431674677 | 0.015029199 | 0.142452386 | Pamr1 | peptidase domain containing associated w | 80.31 | 81.206 | 59.368 | 46.271 | 32.719 | 51.237 |
| 5961 | ENSMUSG00000027189 | 1011.788299 | 0.012991698 | 0.107791032 | 0.120526709 | 0.904065923 | 0.983821605 | Trim44 | tripartite motif-containing 44 [Source:MG | 940.43 | 1048.5 | 1033.4 | 982.29 | 1049.2 | 1016.9 |
| 5962 | ENSMUSG00000027193 | 1290.743044 | -0.116328726 | 0.099502649 | -1.169101798 | 0.242362619 | 0.692000002 | Api5 | apoptosis inhibitor 5 [Source:MGI Symbol | 1336.4 | 1376.4 | 1315.2 | 1183.8 | 1259.7 | 1273.1 |
| 5963 | ENSMUSG00000027194 | 521.6661572 | -0.01241819 | 0.163964186 | -0.075737208 | 0.939628173 | 0.988812319 | Ttc17 | tetratricopeptide repeat domain 17 [Sourc | 524.43 | 598.25 | 449.79 | 606.34 | 524.59 | 426.6 |
| 5964 | ENSMUSG00000027195 | 3227.033883 | 0.005912465 | 0.104082287 | 0.05680568 | 0.954699989 | 0.991678478 | Hsd17b12 | hydroxysteroid (17-beta) dehydrogenase 1 | 3514.4 | 3257.5 | 2887.9 | 3111.7 | 3342.7 | 3248 |
| 5965 | ENSMUSG00000027196 | 4.40612823 | -1.7085758 | 1.141795111 | -1.496394392 | 0.13455091 | NA | Alkbh3os1 | alkB homolog 3, opposite strand 1 [Source | 12.047 | 6.1675 | 2.0125 | 2.8919 | 1.0906 | 2.2277 |
| 5966 | ENSMUSG00000027198 | 514.8721633 | -0.074007388 | 0.133097342 | -0.55603956 | 0.578183811 | 0.908137944 | Ext2 | exostosin glycosyltransferase 2 [Source:M | 499.53 | 529.38 | 555.45 | 467.53 | 558.4 | 478.95 |
| 5967 | ENSMUSG00000027199 | 59.98145042 | -0.697069445 | 0.304825454 | -2.286782269 | 0.022208535 | 0.186216956 | Gatm | glycine amidinotransferase (L-arginine:gly | 60.233 | 85.317 | 77.481 | 53.019 | 38.172 | 45.668 |
| 5968 | ENSMUSG00000027200 | 123.1744541 | -1.279505124 | 0.263244456 | -4.860520681 | 1.17E-06 | 9.30E-05 | Sema6d | sema domain, transmembrane domain (TI | 205.59 | 181.94 | 135.84 | 81.938 | 49.078 | 84.652 |
| 5969 | ENSMUSG00000027201 | 135.5241357 | 0.141079039 | 0.240288097 | 0.58712454 | 0.557120067 | 0.898167794 | Myef2 | myelin basic protein expression factor 2, r | 114.04 | 125.41 | 147.92 | 177.37 | 103.61 | 144.8 |
| 5970 | ENSMUSG00000027202 | 0 | NA | NA | NA | NA | NA | Slc12a1 | solute carrier family 12, member 1 [Sourc | 0 | 0 | 0 | 0 | 0 | 0 |
| 5971 | ENSMUSG00000027203 | 182.5112278 | -0.119311104 | 0.195495965 | -0.610299573 | 0.54166338 | 0.893852723 | Dut | deoxyuridine triphosphatase [Source:MGI | 208 | 158.3 | 203.26 | 165.8 | 160.32 | 199.38 |
| 5972 | ENSMUSG00000027204 | 24.43496946 | -0.617648659 | 0.472363335 | -1.307571129 | 0.191018815 | 0.628502306 | Fbn1 | fibrillin 1 [Source:MGI Symbol;Acc:MGI:95 | 26.502 | 33.921 | 28.175 | 10.604 | 22.903 | 24.505 |
| 5973 | ENSMUSG00000027206 | 1144.806691 | 0.067248933 | 0.099614938 | 0.675088836 | 0.499619326 | 0.880660418 | Cops2 | COP9 signalosome subunit 2 [Source:MGI | 1153.3 | 1094.7 | 1105.9 | 1167.4 | 1168.1 | 1179.6 |
| 5974 | ENSMUSG00000027207 | 334.882264 | -0.127428842 | 0.15482905 | -0.823029282 | 0.410491342 | 0.833241465 | Galk2 | galactokinase 2 [Source:MGI Symbol;Acc:I | 338.91 | 355.66 | 354.2 | 283.41 | 370.81 | 306.31 |
| 5975 | ENSMUSG00000027208 | 0.342432689 | 1.990653086 | 3.900281263 | 0.51038706 | 0.609780321 | NA | Fgf7 | fibroblast growth factor 7 [Source:MGI Sy | 0 | 0 | 0 | 0.964 | 1.0906 | 0 |
| 5976 | ENSMUSG00000027209 | 0.503095663 | 2.549494177 | 3.348715477 | 0.761334964 | 0.446457022 | NA | Fam227b | family with sequence similarity 227, meml | 0 | 0 | 0 | 1.928 | 1.0906 | 0 |
| 5977 | ENSMUSG00000027210 | 45.25757524 | -0.386218861 | 0.37217913 | -1.037723049 | 0.299399011 | 0.750651291 | Meis2 | Meis homeobox 2 [Source:MGI Symbol;Ac | 56.217 | 43.173 | 54.337 | 41.451 | 22.903 | 53.465 |
| 5978 | ENSMUSG00000027215 | 1418.485026 | 0.12980772 | 0.102353386 | 1.268230832 | 0.20471553 | 0.650460832 | Cd82 | CD82 antigen [Source:MGI Symbol;Acc:M | 1340.4 | 1351.7 | 1371.5 | 1367.9 | 1533.4 | 1546 |
| 5979 | ENSMUSG00000027217 | 9.711644889 | -1.722115283 | 0.774386625 | -2.223844302 | 0.026158922 | 0.209326659 | Tspan18 | tetraspanin 18 [Source:MGI Symbol;Acc:M | 19.274 | 13.363 | 12.075 | 5.7839 | 1.0906 | 6.6831 |
| 5980 | ENSMUSG00000027219 | 8.558645394 | -1.430577291 | 0.828682017 | -1.726328387 | 0.084288351 | 0.428071177 | Slc28a2 | solute carrier family 28 (sodium-coupled r | 12.85 | 17.475 | 7.0437 | 2.8919 | 2.1812 | 8.9108 |
| 5981 | ENSMUSG00000027220 | 0.167707431 | -0.796629176 | 4.080472857 | -0.195229623 | 0.84521319 | NA | Syt13 | synaptotagmin XIII [Source:MGI Symbol;A | 0 | 0 | 1.0062 | 0 | 0 | 0 |
| 5982 | ENSMUSG00000027221 | 1.786359065 | -0.782159145 | 1.847675767 | -0.423320562 | 0.672061394 | NA | Chst1 | carbohydrate sulfotransferase 1 [Source:M | 0.8031 | 1.0279 | 5.0312 | 3.8559 | 0 | 0 |
| 5983 | ENSMUSG00000027222 | 2002.011938 | 0.037041104 | 0.107741844 | 0.343794965 | 0.731000487 | 0.94705581 | Pex16 | peroxisomal biogenesis factor 16 [Source: | 2167.6 | 1939.7 | 1820.3 | 1970.4 | 2162.7 | 1951.5 |
| 5984 | ENSMUSG00000027223 | 30.59274325 | -0.44218172 | 0.407446802 | -1.085250191 | 0.27781086 | 0.730834142 | Mapk8ip1 | mitogen-activated protein kinase 8 intera | 37.746 | 34.949 | 33.206 | 34.703 | 22.903 | 20.049 |
| 5985 | ENSMUSG00000027224 | 0.133850373 | -0.796629176 | 4.080472857 | -0.195229623 | 0.84521319 | NA | Duoxa1 | dual oxidase maturation factor 1 [Source: | 0.8031 | 0 | 0 | 0 | 0 | 0 |
| 5986 | ENSMUSG00000027225 | 0 | NA | NA | NA | NA | NA | Duoxa2 | dual oxidase maturation factor 2 [Source: | 0 | 0 | 0 | 0 | 0 | 0 |
| 5987 | ENSMUSG00000027227 | 35672.13422 | -0.05574387 | 0.098808068 | -0.564163141 | 0.57264311 | 0.905173631 | Sord | sorbitol dehydrogenase [Source:MGI Sym | 34723 | 37131 | 37230 | 34925 | 38418 | 31605 |
| 5988 | ENSMUSG00000027229 | 0 | NA | NA | NA | NA | NA | Terb2 | telomere repeat binding bouquet formati | 0 | 0 | 0 | 0 | 0 | 0 |

| | | | | | | | | | | | | | | | |
|---|---|---|---|---|---|---|---|---|---|---|---|---|---|---|---|
| 5989 | ENSMUSG00000027230 | 7.940899336 | -1.304898654 | 0.802360195 | -1.626325262 | 0.10388048 | 0.474113351 | Creb3l1 | cAMP responsive element binding protein | 14.456 | 12.335 | 7.0437 | 3.8559 | 3.2719 | 6.6831 |
| 5990 | ENSMUSG00000027233 | 0.160662974 | 1.126924175 | 4.080472857 | 0.276174898 | 0.782413736 | NA | Patl2 | protein associated with topoisomerase II l | 0 | 0 | 0 | 0.964 | 0 | 0 |
| 5991 | ENSMUSG00000027236 | 1039.734023 | 0.029237523 | 0.105349877 | 0.277527828 | 0.781374841 | 0.960079945 | Eif3j1 | eukaryotic translation initiation factor 3, s | 1013.5 | 1010.4 | 1063.6 | 1017 | 1111.3 | 1022.5 |
| 5992 | ENSMUSG00000027238 | 0.335414862 | -1.779047242 | 4.030804179 | -0.441362855 | 0.658950329 | NA | Frmd5 | FERM domain containing 5 [Source:MGI S | 0 | 0 | 2.0125 | 0 | 0 | 0 |
| 5993 | ENSMUSG00000027239 | 3.802342371 | -0.50637119 | 1.096081063 | -0.461983339 | 0.644093268 | NA | Mdk | midkine [Source:MGI Symbol;Acc:MGI:96 | 7.2279 | 2.0558 | 4.025 | 2.8919 | 3.2719 | 3.3415 |
| 5994 | ENSMUSG00000027242 | 35.63705997 | -0.149111602 | 0.372680426 | -0.400105804 | 0.68907859 | 0.936908509 | Wdr76 | WD repeat domain 76 [Source:MGI Symbo | 40.155 | 30.838 | 41.256 | 28.919 | 40.353 | 32.301 |
| 5995 | ENSMUSG00000027243 | 322.0309602 | 0.295462773 | 0.148036238 | 1.995881396 | 0.045946834 | 0.296420611 | Harbi1 | harbinger transposase derived 1 [Source:I | 269.04 | 285.76 | 312.94 | 333.54 | 355.54 | 375.37 |
| 5996 | ENSMUSG00000027244 | 995.6446849 | 0.318151555 | 0.11849881 | 2.68485021 | 0.007256232 | 0.086363086 | Atg13 | autophagy related 13 [Source:MGI Symbo | 881.81 | 927.18 | 850.28 | 1164.5 | 975.01 | 1175.1 |
| 5997 | ENSMUSG00000027245 | 11.81386914 | 0.541661118 | 0.659584387 | 0.821215797 | 0.41152336 | 0.833768578 | Hypk | huntingtin interacting protein K [Source:N | 10.44 | 14.391 | 4.025 | 14.46 | 13.087 | 14.48 |
| 5998 | ENSMUSG00000027246 | 7.577705391 | 0.007568365 | 0.917254615 | 0.008251106 | 0.993416645 | 0.99859176 | Ell3 | elongation factor RNA polymerase II-like 3 | 5.6217 | 1.0279 | 16.1 | 10.604 | 6.5437 | 5.5692 |
| 5999 | ENSMUSG00000027247 | 190.5464241 | -0.291860978 | 0.177038246 | -1.648575855 | 0.099234559 | 0.464258353 | Arhgap1 | Rho GTPase activating protein 1 [Source:N | 201.58 | 212.78 | 215.34 | 186.05 | 153.78 | 173.76 |
| 6000 | ENSMUSG00000027248 | 6705.999865 | 0.214463967 | 0.10171507 | 2.108477791 | 0.034989682 | 0.250788285 | Pdia3 | protein disulfide isomerase associated 3 [ | 6579 | 6119.2 | 5926.8 | 7676.2 | 6454.3 | 7480.6 |
| 6001 | ENSMUSG00000027249 | 34482.58196 | 0.056874908 | 0.088135707 | 0.64531062 | 0.518725912 | 0.887592015 | F2 | coagulation factor II [Source:MGI Symbol; | 31401 | 35124 | 34885 | 36348 | 35062 | 34075 |
| 6002 | ENSMUSG00000027253 | 662.6738287 | 0.071024186 | 0.122834335 | 0.578211181 | 0.563121551 | 0.900821918 | Lrp4 | low density lipoprotein receptor-related p | 629.63 | 675.34 | 634.94 | 728.77 | 693.63 | 613.73 |
| 6003 | ENSMUSG00000027254 | 3.470520256 | 0.262525684 | 1.125772074 | 0.233196124 | 0.815609116 | NA | Map1a | microtubule-associated protein 1 A [Sourc | 2.4093 | 4.1117 | 3.0187 | 5.7839 | 3.2719 | 2.2277 |
| 6004 | ENSMUSG00000027255 | 1183.869463 | 0.174623478 | 0.121382676 | 1.43861944 | 0.150258375 | 0.563917266 | Arfgap2 | ADP-ribosylation factor GTPase activating | 959.71 | 1226.3 | 1152.2 | 1198.2 | 1282.6 | 1284.3 |
| 6005 | ENSMUSG00000027257 | 955.6180549 | 0.303319763 | 0.118736137 | 2.554569907 | 0.010631908 | 0.113490712 | Pacsin3 | protein kinase C and casein kinase substra | 783.02 | 866.53 | 917.7 | 1018 | 999.01 | 1149.5 |
| 6006 | ENSMUSG00000027259 | 205.5939598 | 0.105882206 | 0.185180407 | 0.571778664 | 0.567471934 | 0.903258947 | Adal | adenosine deaminase-like [Source:MGI Sy | 195.96 | 172.69 | 225.4 | 209.18 | 190.86 | 239.48 |
| 6007 | ENSMUSG00000027261 | 6666.154355 | 0.320408907 | 0.093734313 | 3.418266993 | 0.000630213 | 0.014678573 | Hao1 | hydroxyacid oxidase 1, liver [Source:MGI S | 5976.7 | 6077.1 | 5733.6 | 7789.9 | 7642 | 6777.7 |
| 6008 | ENSMUSG00000027263 | 176.7419097 | 0.304263563 | 0.204020439 | 1.491338635 | 0.13587261 | 0.538690549 | Tubgcp4 | tubulin, gamma complex associated prote | 142.95 | 150.08 | 182.13 | 225.57 | 159.23 | 200.49 |
| 6009 | ENSMUSG00000027270 | 0 | NA | NA | NA | NA | NA | Lamp5 | lysosomal-associated membrane protein f | 0 | 0 | 0 | 0 | 0 | 0 |
| 6010 | ENSMUSG00000027272 | 399.9527958 | 0.077413535 | 0.144412609 | 0.536058006 | 0.591918462 | 0.912740793 | Ubr1 | ubiquitin protein ligase E3 component n-r | 366.21 | 429.67 | 372.31 | 406.8 | 448.24 | 376.48 |
| 6011 | ENSMUSG00000027273 | 0 | NA | NA | NA | NA | NA | Snap25 | synaptosomal-associated protein 25 [Sour | 0 | 0 | 0 | 0 | 0 | 0 |
| 6012 | ENSMUSG00000027274 | 198.8711935 | 0.086841176 | 0.178051061 | 0.487731868 | 0.625739776 | 0.919638611 | Mkks | McKusick-Kaufman syndrome [Source:MG | 207.2 | 175.77 | 195.21 | 202.44 | 186.5 | 226.11 |
| 6013 | ENSMUSG00000027276 | 94.07699835 | -0.092736997 | 0.30802853 | -0.301066259 | 0.763363969 | 0.954964232 | Jag1 | jagged 1 [Source:MGI Symbol;Acc:MGI:10 | 99.585 | 94.568 | 97.606 | 136.88 | 55.622 | 80.197 |
| 6014 | ENSMUSG00000027281 | 57.96344269 | -0.163099917 | 0.338544651 | -0.481767816 | 0.629970895 | 0.920740523 | Slx4ip | SLX4 interacting protein [Source:MGI Sym | 82.72 | 47.284 | 53.331 | 61.695 | 39.262 | 63.489 |
| 6015 | ENSMUSG00000027282 | 2470.547437 | 0.057367748 | 0.090055707 | 0.637025124 | 0.524108479 | 0.888334265 | Mtch2 | mitochondrial carrier 2 [Source:MGI Symb | 2421.4 | 2431 | 2412 | 2511.2 | 2622.9 | 2424.8 |
| 6016 | ENSMUSG00000027284 | 149.9385571 | 0.105387301 | 0.204184158 | 0.516138481 | 0.605757693 | 0.916376001 | Cdan1 | congenital dyserythropoietic anemia, type | 142.95 | 143.91 | 146.91 | 181.23 | 128.69 | 155.94 |
| 6017 | ENSMUSG00000027285 | 252.0367244 | 0.01249133 | 0.165071021 | 0.075672457 | 0.93967969 | 0.988812319 | Haus2 | HAUS augmin-like complex, subunit 2 [Sou | 239.32 | 276.51 | 237.47 | 261.24 | 271.56 | 226.11 |
| 6018 | ENSMUSG00000027286 | 194.6836495 | -0.062180778 | 0.198033377 | -0.313991406 | 0.753527566 | 0.952989051 | Lrrc57 | leucine rich repeat containing 57 [Source: | 194.35 | 205.58 | 196.22 | 145.56 | 219.21 | 207.17 |
| 6019 | ENSMUSG00000027287 | 512.8799862 | 0.080466366 | 0.131455935 | 0.612116647 | 0.540460584 | 0.893324212 | Snap23 | synaptosomal-associated protein 23 [Sour | 456.97 | 491.34 | 548.4 | 552.36 | 516.95 | 511.25 |
| 6020 | ENSMUSG00000027288 | 1215.620984 | -0.003386548 | 0.100204806 | -0.033796263 | 0.973039616 | 0.995820834 | Zfp106 | zinc finger protein 106 [Source:MGI Symb | 1191 | 1256.1 | 1204.5 | 1219.4 | 1250.9 | 1171.8 |
| 6021 | ENSMUSG00000027291 | 371.7604214 | 0.190645388 | 0.148949373 | 1.279934145 | 0.200568298 | 0.643866311 | Vps39 | VPS39 HOPS complex subunit [Source:MG | 399.94 | 321.74 | 318.98 | 401.98 | 398.08 | 389.85 |
| 6022 | ENSMUSG00000027293 | 98.21773769 | -0.274196746 | 0.248134331 | -1.10503349 | 0.269145078 | 0.724979259 | Ehd4 | EH-domain containing 4 [Source:MGI Sym | 117.25 | 87.373 | 117.73 | 92.542 | 74.162 | 100.25 |
| 6023 | ENSMUSG00000027296 | 54.46518069 | 0.217372891 | 0.300309409 | 0.723829773 | 0.469170248 | 0.866013077 | Itpka | inositol 1,4,5-trisphosphate 3-kinase A [So | 47.383 | 58.591 | 45.281 | 59.767 | 55.622 | 60.148 |
| 6024 | ENSMUSG00000027297 | 3.561952212 | 0.467032013 | 1.377588564 | 0.339021407 | 0.734593604 | NA | Ltk | leukocyte tyrosine kinase [Source:MGI Syr | 0 | 1.0279 | 8.05 | 6.7478 | 1.0906 | 4.4554 |
| 6025 | ENSMUSG00000027298 | 4.280693582 | -1.331960641 | 1.051777481 | -1.266390148 | 0.205373436 | NA | Tyro3 | TYRO3 protein tyrosine kinase 3 [Source:N | 7.2279 | 4.1117 | 7.0437 | 2.8919 | 2.1812 | 2.2277 |
| 6026 | ENSMUSG00000027300 | 137.2704165 | 0.150929862 | 0.209476108 | 0.720511105 | 0.471210365 | 0.867762102 | Ubox5 | U box domain containing 5 [Source:MGI S | 129.3 | 112.04 | 148.92 | 142.67 | 158.14 | 132.55 |
| 6027 | ENSMUSG00000027301 | 0.167707431 | -0.796629176 | 4.080472857 | -0.195229623 | 0.84521319 | NA | Oxt | oxytocin [Source:MGI Symbol;Acc:MGI:97 | 0 | 0 | 1.0062 | 0 | 0 | 0 |
| 6028 | ENSMUSG00000027303 | 525.5019004 | 0.000617231 | 0.129217609 | 0.004776677 | 0.996188777 | 0.999089896 | Ptpra | protein tyrosine phosphatase, receptor ty | 574.22 | 522.18 | 478.97 | 525.37 | 537.67 | 514.6 |
| 6029 | ENSMUSG00000027304 | 421.5917659 | 0.054677948 | 0.148868198 | 0.367290991 | 0.713401974 | 0.944239215 | Rtf1 | RTF1, Paf1/RNA polymerase II complex co | 403.16 | 398.83 | 438.72 | 396.19 | 501.68 | 390.96 |
| 6030 | ENSMUSG00000027305 | 584.0677836 | 0.044701887 | 0.117440482 | 0.380634397 | 0.703474554 | 0.942540408 | Ndufaf1 | NADH:ubiquinone oxidoreductase comple | 576.63 | 573.58 | 574.57 | 569.71 | 614.02 | 595.91 |
| 6031 | ENSMUSG00000027306 | 15.12622395 | 0.531528161 | 0.577124016 | 0.920994702 | 0.357053193 | 0.796166767 | Nusap1 | nucleolar and spindle associated protein 1 | 13.653 | 12.335 | 11.069 | 16.388 | 26.175 | 11.138 |
| 6032 | ENSMUSG00000027309 | 661.3559337 | 0.012029055 | 0.132855187 | 0.09054261 | 0.927856035 | 0.987920546 | 4930402H24 | RIKEN cDNA 4930402H24 gene [Source:M | 675.41 | 711.32 | 588.65 | 621.77 | 739.44 | 631.55 |
| 6033 | ENSMUSG00000027312 | 1419.332326 | 0.104872219 | 0.126207049 | 0.830953737 | 0.405999765 | 0.830012328 | Atrn | attractin [Source:MGI Symbol;Acc:MGI:13 | 1448.8 | 1442.2 | 1212.5 | 1624.3 | 1505.1 | 1283.1 |
| 6034 | ENSMUSG00000027313 | 133.4513 | 0.110852387 | 0.321507663 | 0.344789254 | 0.730252813 | 0.946534338 | Chac1 | ChaC, cation transport regulator 1 [Source | 107.62 | 153.16 | 124.77 | 193.76 | 64.346 | 157.05 |
| 6035 | ENSMUSG00000027314 | 42.98312845 | -0.790573202 | 0.363035075 | -2.177677189 | 0.029430078 | 0.225823158 | Dll4 | delta like canonical Notch ligand 4 [Source | 65.854 | 49.34 | 48.3 | 41.451 | 23.994 | 28.96 |
| 6036 | ENSMUSG00000027315 | 7.34707108 | -0.835283201 | 0.838084278 | -0.996657763 | 0.318930658 | 0.765645317 | Spint1 | serine protease inhibitor, Kunitz type 1 [S | 12.85 | 10.279 | 5.0312 | 3.8559 | 8.7249 | 3.3415 |
| 6037 | ENSMUSG00000027316 | 1.634491198 | -0.442467688 | 1.583430466 | -0.279436134 | 0.779910144 | NA | Gfra4 | glial cell line derived neurotrophic factor f | 1.6062 | 2.0558 | 2.0125 | 1.928 | 1.0906 | 1.1138 |
| 6038 | ENSMUSG00000027317 | 0 | NA | NA | NA | NA | NA | Ppp1r14d | protein phosphatase 1, regulatory inhibito | 0 | 0 | 0 | 0 | 0 | 0 |
| 6039 | ENSMUSG00000027318 | 12.16737128 | 0.26296948 | 0.71253231 | 0.369063235 | 0.712080592 | 0.944239215 | Adam33 | a disintegrin and metallopeptidase domai | 8.031 | 10.279 | 15.094 | 24.099 | 4.3625 | 11.138 |
| 6040 | ENSMUSG00000027322 | 56.74317199 | -1.997535351 | 0.358217341 | -5.576322304 | 2.46E-08 | 3.32E-06 | Siglec1 | sialic acid binding Ig-like lectin 1, sialoadh | 89.947 | 109.99 | 72.45 | 26.027 | 13.087 | 28.96 |
| 6041 | ENSMUSG00000027323 | 11.9052743 | -0.714121532 | 0.642595146 | -1.111308631 | 0.266435526 | 0.722551261 | Rad51 | RAD51 recombinase [Source:MGI Symbol; | 20.881 | 13.363 | 10.062 | 10.604 | 8.7249 | 7.7969 |
| 6042 | ENSMUSG00000027324 | 61.82884525 | 0.352600193 | 0.286231437 | 1.231870952 | 0.217997294 | 0.663697152 | Rpusd2 | RNA pseudouridylate synthase domain co | 48.186 | 56.535 | 58.362 | 65.55 | 63.256 | 79.083 |
| 6043 | ENSMUSG00000027326 | 4.016294186 | -1.8506079 | 1.197233687 | -1.545736576 | 0.1221682 | NA | Knl1 | kinetochore scaffold 1 [Source:MGI Symb | 5.6217 | 9.2513 | 4.025 | 1.928 | 3.2719 | 0 |
| 6044 | ENSMUSG00000027327 | 114.895372 | -0.021426823 | 0.254670249 | -0.084135555 | 0.932948656 | 0.988420305 | 1700037H04 | RIKEN cDNA 1700037H04 gene [Source:M | 140.54 | 80.178 | 125.78 | 99.29 | 121.06 | 122.52 |

| | | | | | | | | | | | | | | | |
|---|---|---|---|---|---|---|---|---|---|---|---|---|---|---|---|
| 6045 | ENSMUSG00000027329 | 19.07648941 | -0.285217379 | 0.499377104 | -0.571146288 | 0.567900484 | 0.903352921 | Spef1 | sperm flagellar 1 [Source:MGI Symbol;Acc | 20.078 | 26.726 | 16.1 | 17.352 | 15.269 | 18.935 |
| 6046 | ENSMUSG00000027330 | 16.51681539 | -0.667723435 | 0.568306101 | -1.174936242 | 0.240020225 | 0.689857856 | Cdc25b | cell division cycle 25B [Source:MGI Symbc | 16.062 | 22.614 | 22.137 | 10.604 | 7.6343 | 20.049 |
| 6047 | ENSMUSG00000027331 | 11.26531922 | 0.238322798 | 0.708778006 | 0.33624463 | 0.7366864 | 0.948137969 | Knstrn | kinetochore-localized astrin/SPAG5 bindir | 14.456 | 10.279 | 6.0375 | 4.8199 | 14.178 | 17.822 |
| 6048 | ENSMUSG00000027332 | 4300.047855 | 0.146807888 | 0.104073849 | 1.410612652 | 0.158358859 | 0.578994575 | Ivd | isovaleryl coenzyme A dehydrogenase [So | 3757.7 | 4006.8 | 4480.8 | 4434.3 | 4841.3 | 4279.4 |
| 6049 | ENSMUSG00000027333 | 54.7934504 | -0.044309411 | 0.342770599 | -0.129268411 | 0.897145266 | 0.982798755 | Smox | spermine oxidase [Source:MGI Symbol;Ac | 45.777 | 81.206 | 40.25 | 56.875 | 54.531 | 50.123 |
| 6050 | ENSMUSG00000027335 | 0.589026695 | 0.241472922 | 3.228838524 | 0.074786311 | 0.940384733 | NA | Adra1d | adrenergic receptor, alpha 1d [Source:MG | 1.6062 | 0 | 0 | 1.928 | 0 | 0 |
| 6051 | ENSMUSG00000027338 | 0 | NA | NA | NA | NA | NA | Prnd | prion like protein doppel [Source:MGI Syr | 0 | 0 | 0 | 0 | 0 | 0 |
| 6052 | ENSMUSG00000027339 | 27.25471405 | -1.750601079 | 0.537032947 | -3.259764765 | 0.001115047 | 0.022619962 | Rassf2 | Ras association (RalGDS/AF-6) domain far | 36.943 | 66.815 | 22.137 | 6.7478 | 14.178 | 16.708 |
| 6053 | ENSMUSG00000027340 | 544.7977189 | 0.137929981 | 0.163392894 | 0.844161445 | 0.398579196 | 0.82588279 | Slc23a2 | solute carrier family 23 (nucleobase trans | 643.28 | 445.09 | 466.9 | 629.48 | 562.76 | 521.28 |
| 6054 | ENSMUSG00000027341 | 374.6014236 | -0.090394145 | 0.145757193 | -0.620169357 | 0.535146294 | 0.889858671 | Tmem230 | transmembrane protein 230 [Source:MGI | 379.87 | 345.38 | 433.69 | 356.67 | 356.63 | 375.37 |
| 6055 | ENSMUSG00000027342 | 551.1571119 | -0.017073122 | 0.125946812 | -0.135558189 | 0.892170561 | 0.982489666 | Pcna | proliferating cell nuclear antigen [Source:I | 569.4 | 542.74 | 551.42 | 596.7 | 538.77 | 507.91 |
| 6056 | ENSMUSG00000027344 | 2.513639001 | -0.374379339 | 1.44827989 | -0.258499301 | 0.796021589 | NA | Fsip1 | fibrous sheath-interacting protein 1 [Sour | 2.4093 | 1.0279 | 5.0312 | 0 | 3.2719 | 3.3415 |
| 6057 | ENSMUSG00000027345 | 0 | NA | NA | NA | NA | NA | 4921508D12 | RIKEN cDNA 4921508D12 gene [Source:M | 0 | 0 | 0 | 0 | 0 | 0 |
| 6058 | ENSMUSG00000027346 | 4079.184711 | 0.390217736 | 0.325884923 | 1.197409605 | 0.231146942 | 0.678293643 | Gpcpd1 | glycerophosphocholine phosphodiesteras | 5004.1 | 2624.3 | 2963.4 | 3911.8 | 5474.9 | 4496.6 |
| 6059 | ENSMUSG00000027347 | 14.02008165 | -0.832695745 | 0.663617636 | -1.254782422 | 0.209557755 | 0.653550636 | Rasgrp1 | RAS guanyl releasing protein 1 [Source:MG | 11.243 | 26.726 | 16.1 | 12.532 | 14.178 | 3.3415 |
| 6060 | ENSMUSG00000027349 | 264.32129 | -0.07940531 | 0.16283763 | -0.487634888 | 0.625808479 | 0.919638611 | Fam98b | family with sequence similarity 98, membe | 277.07 | 274.45 | 262.63 | 220.75 | 277.02 | 274.01 |
| 6061 | ENSMUSG00000027350 | 0.133850373 | -0.796629176 | 4.080472857 | -0.195229623 | 0.84521319 | NA | Chgb | chromogranin B [Source:MGI Symbol;Acc: | 0.8031 | 0 | 0 | 0 | 0 | 0 |
| 6062 | ENSMUSG00000027351 | 328.6667924 | -0.05466452 | 0.242283242 | -0.225622373 | 0.821495138 | 0.969160361 | Spred1 | sprouty protein with EVH-1 domain 1, rela | 368.62 | 308.38 | 328.04 | 453.07 | 189.77 | 324.13 |
| 6063 | ENSMUSG00000027353 | 51.95099682 | -0.061821972 | 0.332745638 | -0.185793487 | 0.852606713 | 0.973781977 | Mcm8 | minichromosome maintenance 8 homolog | 61.036 | 56.535 | 41.256 | 39.523 | 64.346 | 49.009 |
| 6064 | ENSMUSG00000027355 | 0 | NA | NA | NA | NA | NA | Tmco5 | transmembrane and coiled-coil domains 5 | 0 | 0 | 0 | 0 | 0 | 0 |
| 6065 | ENSMUSG00000027356 | 9.788654205 | -0.457068625 | 0.813063071 | -0.56215642 | 0.574009451 | 0.9059841 | Fermt1 | fermitin family member 1 [Source:MGI Syr | 18.471 | 8.2233 | 7.0437 | 1.928 | 15.269 | 7.7969 |
| 6066 | ENSMUSG00000027357 | 1181.109752 | -0.299823884 | 0.11712721 | -2.55981411 | 0.010472817 | 0.112206532 | Crls1 | cardiolipin synthase 1 [Source:MGI Symbc | 1391 | 1281.8 | 1237.7 | 1182.8 | 1012.1 | 981.3 |
| 6067 | ENSMUSG00000027358 | 158.8887862 | -0.817076186 | 0.280405834 | -2.913905795 | 0.003569376 | 0.053060077 | Bmp2 | bone morphogenetic protein 2 [Source:M | 198.37 | 202.5 | 207.29 | 121.46 | 59.984 | 163.74 |
| 6068 | ENSMUSG00000027359 | 33480.05269 | -0.025254176 | 0.098281313 | -0.256958067 | 0.797211144 | 0.962236316 | Slc27a2 | solute carrier family 27 (fatty acid transpo | 33074 | 33007 | 35239 | 35501 | 34452 | 29607 |
| 6069 | ENSMUSG00000027360 | 191.0822554 | -0.495235755 | 0.538819546 | -0.919112453 | 0.358036756 | 0.797696186 | Hdc | histidine decarboxylase [Source:MGI Syml | 183.91 | 290.9 | 196.22 | 261.24 | 139.6 | 74.628 |
| 6070 | ENSMUSG00000027361 | 146.5725174 | 0.254892291 | 0.21224042 | 1.20096017 | 0.229766652 | 0.676539688 | Gabpb1 | GA repeat binding protein, beta 1 [Source | 130.91 | 127.46 | 142.89 | 181.23 | 125.42 | 171.53 |
| 6071 | ENSMUSG00000027363 | 1055.050376 | 0.037307855 | 0.117956873 | 0.316283856 | 0.751787064 | 0.952398368 | Usp8 | ubiquitin specific peptidase 8 [Source:MG | 1095.4 | 1046.4 | 981.09 | 959.16 | 1076.4 | 1171.8 |
| 6072 | ENSMUSG00000027364 | 2.530517321 | -0.125869405 | 1.345167411 | -0.093571554 | 0.925449507 | NA | Usp50 | ubiquitin specific peptidase 50 [Source:M | 4.8186 | 1.0279 | 2.0125 | 2.8919 | 1.0906 | 3.3415 |
| 6073 | ENSMUSG00000027365 | 985.8919083 | 0.380758613 | 0.108204025 | 3.518895088 | 0.000433348 | 0.011212784 | Trpm7 | transient receptor potential cation channe | 845.67 | 858.31 | 866.38 | 1196.3 | 1093.9 | 1054.8 |
| 6074 | ENSMUSG00000027366 | 3730.793778 | -0.012736815 | 0.112354422 | -0.113362832 | 0.909742906 | 0.985435587 | Sppl2a | signal peptide peptidase like 2A [Source:N | 3516.8 | 3933.8 | 3792.5 | 3955.2 | 3980.8 | 3205.6 |
| 6075 | ENSMUSG00000027367 | 3101.018438 | 0.146997692 | 0.097789291 | 1.503208489 | 0.132785289 | 0.533798339 | Stard7 | START domain containing 7 [Source:MGI S | 2857.4 | 3098.1 | 2873.8 | 3025 | 3470.3 | 3281.4 |
| 6076 | ENSMUSG00000027368 | 4.630634626 | -2.915250514 | 1.290449613 | -2.25909674 | 0.023877368 | NA | Dusp2 | dual specificity phosphatase 2 [Source:MG | 4.0155 | 15.419 | 5.0312 | 0 | 1.0906 | 2.2277 |
| 6077 | ENSMUSG00000027371 | 382.1871784 | -0.169990692 | 0.136018985 | -1.249757096 | 0.211388293 | 0.65612023 | Fahd2a | fumarylacetoacetate hydrolase domain cc | 415.2 | 408.08 | 390.42 | 348 | 383.9 | 347.52 |
| 6078 | ENSMUSG00000027374 | 931.8727362 | 0.069732318 | 0.120301275 | 0.579647371 | 0.562152441 | 0.900565925 | Mrps5 | mitochondrial ribosomal protein S5 [Sour | 948.46 | 799.72 | 979.08 | 914.81 | 996.83 | 952.34 |
| 6079 | ENSMUSG00000027375 | 6.070226171 | -1.347481645 | 0.946879066 | -1.423076815 | 0.154713889 | 0.572785168 | Mal | myelin and lymphocyte protein, T cell diffe | 8.031 | 5.1396 | 13.081 | 5.7839 | 3.2719 | 1.1138 |
| 6080 | ENSMUSG00000027376 | 0 | NA | NA | NA | NA | NA | Prom2 | prominin 2 [Source:MGI Symbol;Acc:MGI: | 0 | 0 | 0 | 0 | 0 | 0 |
| 6081 | ENSMUSG00000027377 | 0 | NA | NA | NA | NA | NA | Mall | mal, T cell differentiation protein-like [Sou | 0 | 0 | 0 | 0 | 0 | 0 |
| 6082 | ENSMUSG00000027378 | 56.69969128 | -0.067954888 | 0.31449588 | -0.216075608 | 0.828928813 | 0.969955221 | Nphp1 | nephronophthisis 1 (juvenile) homolog (h | 57.823 | 60.647 | 55.343 | 39.523 | 57.803 | 69.058 |
| 6083 | ENSMUSG00000027379 | 1.827021318 | -3.132473531 | 1.89602765 | -1.652124394 | 0.098509195 | NA | Bub1 | BUB1, mitotic checkpoint serine/threonin | 4.8186 | 1.0279 | 4.025 | 0 | 1.0906 | 0 |
| 6084 | ENSMUSG00000027380 | 2.462802676 | -4.676689554 | 1.992177992 | -2.347525961 | 0.018898556 | NA | Acoxl | acyl-Coenzyme A oxidase-like [Source:MG | 9.6372 | 5.1396 | 0 | 0 | 0 | 0 |
| 6085 | ENSMUSG00000027381 | 118.2478567 | -0.279544115 | 0.21959596 | -1.27299298 | 0.203020533 | 0.647739134 | Bcl2l11 | BCL2-like 11 (apoptosis facilitator) [Source | 133.31 | 138.77 | 116.72 | 102.18 | 95.974 | 122.52 |
| 6086 | ENSMUSG00000027384 | 85.05873043 | -0.022249328 | 0.249269346 | -0.089258181 | 0.928876728 | 0.988405116 | Ndufaf5 | NADH:ubiquinone oxidoreductase comple | 85.129 | 88.401 | 83.518 | 80.01 | 74.162 | 99.132 |
| 6087 | ENSMUSG00000027386 | 11.20993334 | -1.588555054 | 0.682122563 | -2.328841091 | 0.019867486 | 0.172383511 | Fbln7 | fibulin 7 [Source:MGI Symbol;Acc:MGI:19 | 20.881 | 17.475 | 12.075 | 5.7839 | 4.3625 | 6.6831 |
| 6088 | ENSMUSG00000027387 | 99.89800839 | 0.277832277 | 0.25941323 | 1.07100273 | 0.284168202 | 0.736651636 | Zc3h8 | zinc finger CCCH type containing 8 [Source | 80.31 | 90.457 | 100.62 | 122.43 | 127.6 | 77.969 |
| 6089 | ENSMUSG00000027394 | 159.2522814 | -0.016447083 | 0.18533862 | -0.08874072 | 0.92928797 | 0.988405116 | Ttl | tubulin tyrosine ligase [Source:MGI Symbc | 157.41 | 152.13 | 171.06 | 161.95 | 158.14 | 154.82 |
| 6090 | ENSMUSG00000027395 | 327.928998 | 0.160450863 | 0.152307456 | 1.053466899 | 0.292127057 | 0.742979735 | Polr1b | polymerase (RNA) I polypeptide B [Source | 277.07 | 334.07 | 318.98 | 363.42 | 358.81 | 315.22 |
| 6091 | ENSMUSG00000027397 | 566.2258836 | 0.052898554 | 0.172985042 | 0.305798429 | 0.759758125 | 0.95402891 | Slc20a1 | solute carrier family 20, member 1 [Sourc | 616.78 | 569.47 | 479.98 | 433.79 | 641.28 | 656.05 |
| 6092 | ENSMUSG00000027398 | 11.59088302 | -1.276977433 | 0.708577542 | -1.802170345 | 0.071518609 | 0.38710107 | Il1b | interleukin 1 beta [Source:MGI Symbol;Ac | 22.487 | 17.475 | 9.0562 | 2.8919 | 8.7249 | 8.9108 |
| 6093 | ENSMUSG00000027399 | 25.90683125 | 0.071311276 | 0.44931865 | 0.158709806 | 0.873897512 | 0.979925081 | Il1a | interleukin 1 alpha [Source:MGI Symbol;A | 29.715 | 25.698 | 20.125 | 20.244 | 22.903 | 36.757 |
| 6094 | ENSMUSG00000027400 | 0 | NA | NA | NA | NA | NA | Pdyn | prodynorphin [Source:MGI Symbol;Acc:M | 0 | 0 | 0 | 0 | 0 | 0 |
| 6095 | ENSMUSG00000027401 | 0.331982614 | 0.16514309 | 3.986751476 | 0.041422971 | 0.966958701 | NA | Tgm3 | transglutaminase 3, E polypeptide [Source | 0 | 1.0279 | 0 | 0.964 | 0 | 0 |
| 6096 | ENSMUSG00000027403 | 0 | NA | NA | NA | NA | NA | Tgm6 | transglutaminase 6 [Source:MGI Symbol;A | 0 | 0 | 0 | 0 | 0 | 0 |
| 6097 | ENSMUSG00000027404 | 877.5403416 | 0.064503646 | 0.117575664 | 0.548613918 | 0.583270433 | 0.908645447 | Snrpb | small nuclear ribonucleoprotein B [Source | 772.58 | 869.62 | 932.79 | 911.92 | 911.76 | 866.57 |
| 6098 | ENSMUSG00000027405 | 300.6697458 | -0.099202773 | 0.183541053 | -0.540493645 | 0.588856642 | 0.911039598 | Nop56 | NOP56 ribonucleoprotein [Source:MGI Sy | 286.71 | 351.55 | 295.84 | 355.71 | 279.2 | 235.02 |
| 6099 | ENSMUSG00000027406 | 3581.689918 | -0.029491519 | 0.089334429 | -0.330124891 | 0.741305596 | 0.948792385 | Idh3b | isocitrate dehydrogenase 3 (NAD+) beta [S | 3610.7 | 3514.5 | 3729.1 | 3381.6 | 3675.4 | 3578.8 |
| 6100 | ENSMUSG00000027408 | 18.13935999 | -1.395969309 | 0.566594624 | -2.463788482 | 0.013747721 | 0.134934013 | Cpxm1 | carboxypeptidase X 1 (M14 family) [Sourc | 36.14 | 24.67 | 18.112 | 14.46 | 6.5437 | 8.9108 |

| | | | | | | | | | | | | | | | |
|---|---|---|---|---|---|---|---|---|---|---|---|---|---|---|---|
| 6101 | ENSMUSG00000027409 | 0 | NA | NA | NA | NA | NA | 1700020A23 | RIKEN cDNA 1700020A23 gene [Source:M | 0 | 0 | 0 | 0 | 0 | 0 |
| 6102 | ENSMUSG00000027411 | 438.4470351 | 0.107437064 | 0.152238653 | 0.70571476 | 0.48036554 | 0.870278888 | Vps16 | VSP16 CORVET/HOPS core subunit [Sourc | 452.95 | 402.94 | 410.55 | 508.98 | 478.78 | 376.48 |
| 6103 | ENSMUSG00000027412 | 104.6937389 | 0.053700796 | 0.259850563 | 0.206660302 | 0.836275162 | 0.971073158 | Lpin3 | lipin 3 [Source:MGI Symbol;Acc:MGI:1891 | 121.27 | 95.596 | 90.562 | 78.082 | 112.33 | 130.32 |
| 6104 | ENSMUSG00000027416 | 0 | NA | NA | NA | NA | NA | Otor | otoraplin [Source:MGI Symbol;Acc:MGI:1 | 0 | 0 | 0 | 0 | 0 | 0 |
| 6105 | ENSMUSG00000027419 | 0 | NA | NA | NA | NA | NA | Pcsk2 | proprotein convertase subtilisin/kexin typ | 0 | 0 | 0 | 0 | 0 | 0 |
| 6106 | ENSMUSG00000027420 | 0.496077836 | -0.812642266 | 3.444120572 | -0.235950586 | 0.813471014 | NA | Bfsp1 | beaded filament structural protein 1, in le | 0 | 0 | 2.0125 | 0.964 | 0 | 0 |
| 6107 | ENSMUSG00000027422 | 5107.067914 | 0.142713606 | 0.105333489 | 1.354873998 | 0.175457705 | 0.605810563 | Rrbp1 | ribosome binding protein 1 [Source:MGI S | 4559.2 | 5318.4 | 4688.1 | 5868.7 | 5093.2 | 5114.8 |
| 6108 | ENSMUSG00000027423 | 760.6440086 | -0.188229795 | 0.11379671 | -1.6540882 | 0.098109593 | 0.461932668 | Snx5 | sorting nexin 5 [Source:MGI Symbol;Acc:M | 788.65 | 869.62 | 772.8 | 720.09 | 715.45 | 697.27 |
| 6109 | ENSMUSG00000027424 | 163.0060209 | 0.412559199 | 0.21289314 | 1.937869856 | 0.052639102 | 0.322665669 | Mgme1 | mitochondrial genome maintenance exon | 128.5 | 157.27 | 133.83 | 160.98 | 165.77 | 231.68 |
| 6110 | ENSMUSG00000027425 | 359.0321031 | 0.135877026 | 0.167053932 | 0.813372208 | 0.416004689 | 0.836590861 | Kat14 | lysine acetyltransferase 14 [Source:MGI S | 355.77 | 397.8 | 272.69 | 370.17 | 402.44 | 355.32 |
| 6111 | ENSMUSG00000027427 | 246.7237075 | 0.124994931 | 0.164994381 | 0.757570834 | 0.448707946 | 0.854299293 | Polr3f | polymerase (RNA) III (DNA directed) polyp | 250.57 | 221 | 236.47 | 275.7 | 269.38 | 227.22 |
| 6112 | ENSMUSG00000027428 | 496.546379 | -0.047600772 | 0.129340874 | -0.368025751 | 0.712854033 | 0.944239215 | Rbbp9 | retinoblastoma binding protein 9, serine h | 463.39 | 531.43 | 520.23 | 492.59 | 506.05 | 465.59 |
| 6113 | ENSMUSG00000027429 | 585.6204511 | 0.199476156 | 0.135688101 | 1.470107953 | 0.141532519 | 0.550598803 | Sec23b | SEC23 homolog B, COPII coat complex cor | 522.02 | 560.22 | 554.44 | 708.52 | 622.74 | 545.78 |
| 6114 | ENSMUSG00000027430 | 158.3254182 | -0.049772563 | 0.233229057 | -0.213406357 | 0.831010023 | 0.970505212 | Dtd1 | D-tyrosyl-tRNA deacylase 1 [Source:MGI S | 169.45 | 160.36 | 152.95 | 125.32 | 212.67 | 129.21 |
| 6115 | ENSMUSG00000027431 | 0 | NA | NA | NA | NA | NA | Scp2d1 | SCP2 sterol-binding domain containing 1 [ | 0 | 0 | 0 | 0 | 0 | 0 |
| 6116 | ENSMUSG00000027433 | 736.4195908 | 0.019682202 | 0.131373491 | 0.149818671 | 0.880907678 | 0.981490993 | Xrn2 | 5'-3' exoribonuclease 2 [Source:MGI Symb | 635.25 | 835.7 | 724.5 | 743.23 | 753.62 | 726.23 |
| 6117 | ENSMUSG00000027434 | 0 | NA | NA | NA | NA | NA | Nkx2-2 | NK2 homeobox 2 [Source:MGI Symbol;Ac | 0 | 0 | 0 | 0 | 0 | 0 |
| 6118 | ENSMUSG00000027435 | 41.59114128 | -0.824439633 | 0.3516155 | -2.344719253 | 0.019041411 | 0.167629321 | Cd93 | CD93 antigen [Source:MGI Symbol;Acc:M | 50.595 | 51.396 | 57.356 | 25.063 | 27.265 | 37.871 |
| 6119 | ENSMUSG00000027438 | 39.38829886 | -0.312853846 | 0.36717184 | -0.852063837 | 0.394178662 | 0.822827574 | Napb | N-ethylmaleimide sensitive fusion protein | 46.58 | 43.173 | 41.256 | 40.487 | 41.443 | 23.391 |
| 6120 | ENSMUSG00000027439 | 1048.973041 | 0.251291676 | 0.107351249 | 2.340836085 | 0.019240612 | 0.168766657 | Gzf1 | GDNF-inducible zinc finger protein 1 [Sour | 994.24 | 895.32 | 983.1 | 1095.1 | 1184.4 | 1141.7 |
| 6121 | ENSMUSG00000027442 | 0 | NA | NA | NA | NA | NA | Cst8 | cystatin 8 (cystatin-related epididymal spe | 0 | 0 | 0 | 0 | 0 | 0 |
| 6122 | ENSMUSG00000027443 | 0 | NA | NA | NA | NA | NA | Cst12 | cystatin 12 [Source:MGI Symbol;Acc:MGI: | 0 | 0 | 0 | 0 | 0 | 0 |
| 6123 | ENSMUSG00000027444 | 0 | NA | NA | NA | NA | NA | Cstdc1 | cystatin domain containing 1 [Source:MGI | 0 | 0 | 0 | 0 | 0 | 0 |
| 6124 | ENSMUSG00000027445 | 0 | NA | NA | NA | NA | NA | Cst9 | cystatin 9 [Source:MGI Symbol;Acc:MGI:1 | 0 | 0 | 0 | 0 | 0 | 0 |
| 6125 | ENSMUSG00000027446 | 0 | NA | NA | NA | NA | NA | Cstdc2 | cystatin domain containing 2 [Source:MGI | 0 | 0 | 0 | 0 | 0 | 0 |
| 6126 | ENSMUSG00000027447 | 2686.538004 | -0.122843429 | 0.113754465 | -1.079899839 | 0.280186785 | 0.733108114 | Cst3 | cystatin C [Source:MGI Symbol;Acc:MGI:1 | 2869.5 | 2956.3 | 2577 | 2753.1 | 2246.7 | 2716.7 |
| 6127 | ENSMUSG00000027452 | 2.773085409 | -1.659481454 | 1.605022689 | -1.033930215 | 0.301168788 | NA | Acss1 | acyl-CoA synthetase short-chain family me | 9.6372 | 0 | 3.0187 | 2.8919 | 1.0906 | 0 |
| 6128 | ENSMUSG00000027454 | 74.45794627 | 0.262085782 | 0.305989873 | 0.856517832 | 0.391711414 | 0.820404559 | Gins1 | GINS complex subunit 1 (Psf1 homolog) [S | 60.233 | 48.312 | 94.587 | 68.442 | 90.521 | 84.652 |
| 6129 | ENSMUSG00000027455 | 1028.776343 | 0.031873396 | 0.104410929 | 0.305268774 | 0.760161457 | 0.95402891 | Nsfl1c | NSFL1 (p97) cofactor (p47) [Source:MGI S | 1033.6 | 1023.8 | 994.17 | 986.15 | 1060.1 | 1074.9 |
| 6130 | ENSMUSG00000027456 | 0.794203392 | -0.509190641 | 2.432728345 | -0.209308467 | 0.834207442 | NA | Sdcbp2 | syndecan binding protein (syntenin) 2 [So | 0.8031 | 1.0279 | 1.0062 | 1.928 | 0 | 0 |
| 6131 | ENSMUSG00000027457 | 1.113435689 | -0.265574016 | 2.107410414 | -0.126019125 | 0.899716786 | NA | Snph | syntaphilin [Source:MGI Symbol;Acc:MGI: | 1.6062 | 2.0558 | 0 | 1.928 | 1.0906 | 0 |
| 6132 | ENSMUSG00000027459 | 50.54183823 | -0.148161243 | 0.388109441 | -0.381751195 | 0.702645922 | 0.942421237 | Fam110a | family with sequence similarity 110, meml | 48.186 | 28.782 | 82.512 | 51.091 | 41.443 | 51.237 |
| 6133 | ENSMUSG00000027460 | 1.08158966 | 1.151574547 | 2.421847399 | 0.475494264 | 0.634434721 | NA | Angpt4 | angiopoietin 4 [Source:MGI Symbol;Acc:M | 0 | 1.0279 | 1.0062 | 0 | 0 | 4.4554 |
| 6134 | ENSMUSG00000027463 | 3.742144069 | 0.556125817 | 1.128164266 | 0.492947555 | 0.622049644 | NA | Slc52a3 | solute carrier protein family 52, member 3 | 4.0155 | 1.0279 | 4.025 | 6.7478 | 2.1812 | 4.4554 |
| 6135 | ENSMUSG00000027465 | 981.1122064 | -0.017735932 | 0.131569917 | -0.134802337 | 0.892768158 | 0.982562372 | Tbc1d20 | TBC1 domain family, member 20 [Source:I | 1102.7 | 1016.6 | 841.22 | 987.11 | 895.4 | 1043.7 |
| 6136 | ENSMUSG00000027466 | 1171.696129 | -0.015707534 | 0.104566094 | -0.150216321 | 0.88059395 | 0.981448736 | Rbck1 | RanBP-type and C3HC4-type zinc finger cc | 1208.7 | 1152.3 | 1172.3 | 1087.4 | 1187.7 | 1221.9 |
| 6137 | ENSMUSG00000027468 | 0 | NA | NA | NA | NA | NA | Defb22 | defensin beta 22 [Source:MGI Symbol;Acc | 0 | 0 | 0 | 0 | 0 | 0 |
| 6138 | ENSMUSG00000027469 | 6.154753791 | -1.348009971 | 0.932535824 | -1.445531566 | 0.148308633 | 0.56003169 | Tpx2 | TPX2, microtubule-associated [Source:MG | 15.259 | 5.1396 | 6.0375 | 3.8559 | 2.1812 | 4.4554 |
| 6139 | ENSMUSG00000027470 | 0 | NA | NA | NA | NA | NA | Mylk2 | myosin, light polypeptide kinase 2, skeleta | 0 | 0 | 0 | 0 | 0 | 0 |
| 6140 | ENSMUSG00000027472 | 378.9669626 | -0.019838875 | 0.165465308 | -0.119897489 | 0.904564354 | 0.983865503 | Pdrg1 | p53 and DNA damage regulated 1 [Source | 439.3 | 374.16 | 330.05 | 340.28 | 352.27 | 437.74 |
| 6141 | ENSMUSG00000027474 | 47.25262818 | -0.668237764 | 0.396167906 | -1.686753911 | 0.091650681 | 0.446623824 | Ccm2l | cerebral cavernous malformation 2-like [S | 43.368 | 66.815 | 64.4 | 55.911 | 20.722 | 32.301 |
| 6142 | ENSMUSG00000027475 | 286.3247925 | 0.254291268 | 0.15921082 | 1.597198403 | 0.110221489 | 0.488133378 | Kif3b | kinesin family member 3B [Source:MGI Sy | 247.36 | 284.73 | 251.56 | 298.83 | 286.83 | 348.63 |
| 6143 | ENSMUSG00000027478 | 35.97140623 | -0.240398788 | 0.376371091 | -0.638728089 | 0.522999836 | 0.888334265 | Dnmt3b | DNA methyltransferase 3B [Source:MGI S | 29.715 | 43.173 | 44.275 | 33.739 | 37.081 | 27.846 |
| 6144 | ENSMUSG00000027479 | 491.8858226 | -0.011603114 | 0.124410565 | -0.093264699 | 0.925693276 | 0.987546253 | Mapre1 | microtubule-associated protein, RP/EB far | 485.07 | 501.62 | 495.07 | 485.84 | 514.77 | 468.93 |
| 6145 | ENSMUSG00000027480 | 0 | NA | NA | NA | NA | NA | Sun5 | Sad1 and UNC84 domain containing 5 [So | 0 | 0 | 0 | 0 | 0 | 0 |
| 6146 | ENSMUSG00000027481 | 0 | NA | NA | NA | NA | NA | Bpifb2 | BPI fold containing family B, member 2 [S | 0 | 0 | 0 | 0 | 0 | 0 |
| 6147 | ENSMUSG00000027482 | 0 | NA | NA | NA | NA | NA | Bpifa3 | BPI fold containing family A, member 3 [S | 0 | 0 | 0 | 0 | 0 | 0 |
| 6148 | ENSMUSG00000027483 | 0 | NA | NA | NA | NA | NA | Bpifa1 | BPI fold containing family A, member 1 [S | 0 | 0 | 0 | 0 | 0 | 0 |
| 6149 | ENSMUSG00000027484 | 0 | NA | NA | NA | NA | NA | Bpifa5 | BPI fold containing family A, member 5 [S | 0 | 0 | 0 | 0 | 0 | 0 |
| 6150 | ENSMUSG00000027485 | 0 | NA | NA | NA | NA | NA | Bpifb1 | BPI fold containing family B, member 1 [S | 0 | 0 | 0 | 0 | 0 | 0 |
| 6151 | ENSMUSG00000027487 | 205.2284973 | 0.62775656 | 0.314965544 | 1.993095981 | 0.046250938 | 0.297453787 | Cdk5rap1 | CDK5 regulatory subunit associated protei | 231.29 | 133.63 | 117.73 | 127.25 | 316.28 | 305.19 |
| 6152 | ENSMUSG00000027488 | 413.4881463 | -0.049195441 | 0.13225763 | -0.371966754 | 0.709917603 | 0.944239215 | Snta1 | syntrophin, acidic 1 [Source:MGI Symbol;A | 392.72 | 434.81 | 434.7 | 409.69 | 401.35 | 407.67 |
| 6153 | ENSMUSG00000027489 | 2.083781699 | -0.964914505 | 1.59018311 | -0.606794588 | 0.54398724 | NA | Necab3 | N-terminal EF-hand calcium binding prote | 3.2124 | 0 | 5.0312 | 0.964 | 2.1812 | 1.1138 |
| 6154 | ENSMUSG00000027490 | 79.00073839 | 0.467000469 | 0.286069615 | 1.632471414 | 0.102580216 | 0.470872814 | E2f1 | E2F transcription factor 1 [Source:MGI Sy | 88.341 | 54.48 | 55.343 | 83.866 | 86.159 | 105.82 |
| 6155 | ENSMUSG00000027495 | 2358.465423 | 0.093034664 | 0.100394783 | 0.926688233 | 0.354088429 | 0.79449673 | Fam210b | family with sequence similarity 210, meml | 2227 | 2260.4 | 2359.6 | 2290.4 | 2356.8 | 2656.5 |
| 6156 | ENSMUSG00000027496 | 51.67424667 | -0.155199223 | 0.332713619 | -0.466464893 | 0.640882771 | 0.923932929 | Aurka | aurora kinase A [Source:MGI Symbol;Acc: | 53.005 | 44.2 | 66.412 | 62.659 | 41.443 | 42.326 |

| | | | | | | | | | | | | | | | |
|---|---|---|---|---|---|---|---|---|---|---|---|---|---|---|---|
| 6157 | ENSMUSG00000027498 | 458.6907286 | -0.126677816 | 0.133696672 | -0.947501644 | 0.343383219 | 0.785243795 | Cstf1 | cleavage stimulation factor, 3' pre-RNA, su | 522.82 | 434.81 | 477.97 | 433.79 | 442.79 | 439.97 |
| 6158 | ENSMUSG00000027499 | 13.59450437 | -0.999950403 | 0.591664661 | -1.690062747 | 0.091015951 | 0.44519943 | Pkia | protein kinase inhibitor, alpha [Source:MG | 16.865 | 17.475 | 20.125 | 10.604 | 9.8156 | 6.6831 |
| 6159 | ENSMUSG00000027500 | 0.167707431 | -0.796629176 | 4.080472857 | -0.195229623 | 0.84521319 | NA | Stmn2 | stathmin-like 2 [Source:MGI Symbol;Acc:M | 0 | 0 | 1.0062 | 0 | 0 | 0 |
| 6160 | ENSMUSG00000027502 | 1887.162605 | -0.170015125 | 0.096705973 | -1.75806229 | 0.078736911 | 0.411583189 | Rtf2 | replication termination factor 2 [Source:M | 2029.4 | 2010.6 | 1955.1 | 1885.5 | 1755.9 | 1686.4 |
| 6161 | ENSMUSG00000027505 | 5.708145996 | -2.154732449 | 1.015600022 | -2.121634899 | 0.033868409 | 0.246290937 | Fam209 | family with sequence similarity 209 [Sourc | 13.653 | 9.2513 | 5.0312 | 1.928 | 3.2719 | 1.1138 |
| 6162 | ENSMUSG00000027506 | 307.6979796 | -0.166984052 | 0.149707056 | -1.115405353 | 0.264676757 | 0.720414868 | Tpd52 | tumor protein D52 [Source:MGI Symbol;A | 331.68 | 330.99 | 313.95 | 310.4 | 260.66 | 298.51 |
| 6163 | ENSMUSG00000027508 | 117.7223925 | -0.457113929 | 0.232250277 | -1.96819541 | 0.049045558 | 0.308835417 | Pag1 | phosphoprotein associated with glycosph | 124.48 | 168.58 | 115.72 | 92.542 | 101.43 | 103.59 |
| 6164 | ENSMUSG00000027509 | 330.3724549 | -0.003705107 | 0.146150755 | -0.025351267 | 0.979774782 | 0.99820894 | Rae1 | ribonucleic acid export 1 [Source:MGI Syn | 301.16 | 360.8 | 331.05 | 326.79 | 328.28 | 334.15 |
| 6165 | ENSMUSG00000027510 | 51.58689443 | -0.17976742 | 0.327930968 | -0.548186775 | 0.583563662 | 0.908673141 | Rbm38 | RNA binding motif protein 38 [Source:MG | 44.974 | 53.452 | 66.412 | 58.803 | 46.897 | 38.985 |
| 6166 | ENSMUSG00000027513 | 45503.22639 | -0.651199687 | 0.121761844 | -5.348142468 | 8.89E-08 | 9.81E-06 | Pck1 | phosphoenolpyruvate carboxykinase 1, cy | 53986 | 62712 | 50107 | 31710 | 40689 | 33815 |
| 6167 | ENSMUSG00000027514 | 140.134983 | 0.037745727 | 0.226898525 | 0.166355099 | 0.867877508 | 0.978326885 | Zbp1 | Z-DNA binding protein 1 [Source:MGI Sym | 165.44 | 130.55 | 118.74 | 172.55 | 124.33 | 129.21 |
| 6168 | ENSMUSG00000027517 | 0 | NA | NA | NA | NA | NA | Ankrd60 | ankyrin repeat domain 60 [Source:MGI Sy | 0 | 0 | 0 | 0 | 0 | 0 |
| 6169 | ENSMUSG00000027518 | 0 | NA | NA | NA | NA | NA | 1700021F07 | RIKEN cDNA 1700021F07 gene [Source:MG | 0 | 0 | 0 | 0 | 0 | 0 |
| 6170 | ENSMUSG00000027519 | 681.563921 | 0.134395454 | 0.118688212 | 1.132340376 | 0.257491361 | 0.710872182 | Rab22a | RAB22A, member RAS oncogene family [S | 640.88 | 700.01 | 608.78 | 707.56 | 691.45 | 740.71 |
| 6171 | ENSMUSG00000027520 | 0.342639279 | -1.805976596 | 4.028547714 | -0.448294702 | 0.653940524 | NA | Zdbf2 | zinc finger, DBF-type containing 2 [Source | 0 | 2.0558 | 0 | 0 | 0 | 0 |
| 6172 | ENSMUSG00000027522 | 510.3465906 | 0.040603676 | 0.121600498 | 0.333910441 | 0.738447141 | 0.948137969 | Stx16 | syntaxin 16 [Source:MGI Symbol;Acc:MGI | 514.79 | 489.29 | 505.13 | 513.8 | 530.04 | 509.03 |
| 6173 | ENSMUSG00000027523 | 742.4112514 | -0.022560559 | 0.137275351 | -0.164345304 | 0.869459319 | 0.978621761 | Gnas | GNAS (guanine nucleotide binding proteir | 803.1 | 648.62 | 792.92 | 830.95 | 682.73 | 696.15 |
| 6174 | ENSMUSG00000027524 | 0.401551119 | -2.072982726 | 4.008251979 | -0.517178744 | 0.605031388 | NA | Edn3 | endothelin 3 [Source:MGI Symbol;Acc:MG | 2.4093 | 0 | 0 | 0 | 0 | 0 |
| 6175 | ENSMUSG00000027525 | 0 | NA | NA | NA | NA | NA | Phactr3 | phosphatase and actin regulator 3 [Source | 0 | 0 | 0 | 0 | 0 | 0 |
| 6176 | ENSMUSG00000027528 | 0 | NA | NA | NA | NA | NA | Fabp9 | fatty acid binding protein 9, testis [Source | 0 | 0 | 0 | 0 | 0 | 0 |
| 6177 | ENSMUSG00000027530 | 1.092039735 | 2.444485515 | 2.4136564 | 1.012772785 | 0.311168703 | NA | Fabp12 | fatty acid binding protein 12 [Source:MGI | 0 | 0 | 1.0062 | 0 | 1.0906 | 4.4554 |
| 6178 | ENSMUSG00000027531 | 1325.276689 | 0.196610875 | 0.120079748 | 1.637335846 | 0.101560315 | 0.469189975 | Impa1 | inositol (myo)-1(or 4)-monophosphatase 1 | 1193.4 | 1188.3 | 1323.2 | 1235.8 | 1558.5 | 1452.5 |
| 6179 | ENSMUSG00000027533 | 236.8543548 | 2.430685841 | 0.297881009 | 8.159922139 | 3.35E-16 | 2.55E-13 | Fabp5 | fatty acid binding protein 5, epidermal [So | 76.295 | 93.541 | 52.325 | 221.71 | 399.17 | 578.08 |
| 6180 | ENSMUSG00000027534 | 42.49280431 | -0.418665522 | 0.363565127 | -1.151555774 | 0.249503664 | 0.701784229 | Snx16 | sorting nexin 16 [Source:MGI Symbol;Acc: | 65.051 | 45.228 | 35.219 | 35.667 | 39.262 | 34.529 |
| 6181 | ENSMUSG00000027536 | 46.44335006 | 0.371482117 | 0.466280113 | 0.796693031 | 0.425629332 | 0.841323666 | Chmp4c | charged multivesicular body protein 4C [S | 35.336 | 51.396 | 35.219 | 98.326 | 30.537 | 27.846 |
| 6182 | ENSMUSG00000027540 | 508.0268479 | -0.133766267 | 0.131776329 | -1.015100877 | 0.310057743 | 0.758699916 | Ptpn1 | protein tyrosine phosphatase, non-recept | 523.62 | 574.61 | 497.08 | 518.62 | 484.23 | 449.99 |
| 6183 | ENSMUSG00000027544 | 5.511053278 | -1.343516262 | 1.038934591 | -1.293167322 | 0.195953195 | 0.636484845 | Nfatc2 | nuclear factor of activated T cells, cytopla | 6.4248 | 12.335 | 5.0312 | 4.8199 | 0 | 4.4554 |
| 6184 | ENSMUSG00000027546 | 1265.325629 | -0.077212243 | 0.123453274 | -0.625436981 | 0.531684297 | 0.888947136 | Atp9a | ATPase, class II, type 9A [Source:MGI Sym | 1468.9 | 1309.6 | 1117.9 | 1230 | 1279.3 | 1186.2 |
| 6185 | ENSMUSG00000027547 | 0.167707431 | -0.796629176 | 4.080472857 | -0.195229623 | 0.84521319 | NA | Sall4 | spalt like transcription factor 4 [Source:M | 0 | 0 | 1.0062 | 0 | 0 | 0 |
| 6186 | ENSMUSG00000027550 | 58.09492183 | -0.034960053 | 0.298506763 | -0.117116451 | 0.906767774 | 0.984324722 | Lrrcc1 | leucine rich repeat and coiled-coil domain | 58.626 | 51.396 | 66.412 | 58.803 | 65.437 | 47.895 |
| 6187 | ENSMUSG00000027551 | 155.3291534 | 0.068945021 | 0.200674128 | 0.343567066 | 0.731171896 | 0.94705581 | Zfp64 | zinc finger protein 64 [Source:MGI Symbo | 145.36 | 130.55 | 179.11 | 163.88 | 152.69 | 160.39 |
| 6188 | ENSMUSG00000027552 | 138.4476626 | 0.061555485 | 0.231268606 | 0.266164467 | 0.790112547 | 0.961019028 | E2f5 | E2F transcription factor 5 [Source:MGI Sy | 97.175 | 150.08 | 159.99 | 136.88 | 142.87 | 143.69 |
| 6189 | ENSMUSG00000027555 | 2.906827068 | 0.326987835 | 1.479467432 | 0.221017258 | 0.825078995 | NA | Car13 | carbonic anhydrase 13 [Source:MGI Symb | 1.6062 | 5.1396 | 1.0062 | 0.964 | 8.7249 | 0 |
| 6190 | ENSMUSG00000027556 | 130.7830667 | 0.044498514 | 0.243802597 | 0.182518622 | 0.855175742 | 0.974158771 | Car1 | carbonic anhydrase 1 [Source:MGI Symbo | 101.19 | 136.71 | 148.92 | 126.28 | 165.77 | 105.82 |
| 6191 | ENSMUSG00000027559 | 40780.87344 | -0.195046086 | 0.170786842 | -1.142043984 | 0.253435738 | 0.706571802 | Car3 | carbonic anhydrase 3 [Source:MGI Symbo | 48158 | 48028 | 34413 | 29495 | 48879 | 35712 |
| 6192 | ENSMUSG00000027560 | 0 | NA | NA | NA | NA | NA | Dok5 | docking protein 5 [Source:MGI Symbol;Ac | 0 | 0 | 0 | 0 | 0 | 0 |
| 6193 | ENSMUSG00000027562 | 64.76756839 | -0.172019794 | 0.319782882 | -0.537926837 | 0.59062756 | 0.911722201 | Car2 | carbonic anhydrase 2 [Source:MGI Symbo | 76.295 | 64.759 | 64.4 | 42.415 | 86.159 | 54.578 |
| 6194 | ENSMUSG00000027564 | 0 | NA | NA | NA | NA | NA | Cypt12 | cysteine-rich perinuclear theca 12 [Source | 0 | 0 | 0 | 0 | 0 | 0 |
| 6195 | ENSMUSG00000027566 | 1972.537967 | 0.080347408 | 0.105945204 | 0.758386463 | 0.44821966 | 0.854156674 | Psma7 | proteasome subunit alpha 7 [Source:MGI | 2068 | 1876 | 1807.2 | 1870.1 | 2078.7 | 2135.2 |
| 6196 | ENSMUSG00000027568 | 0 | NA | NA | NA | NA | NA | Ntsr1 | neurotensin receptor 1 [Source:MGI Symb | 0 | 0 | 0 | 0 | 0 | 0 |
| 6197 | ENSMUSG00000027569 | 150.8608712 | 0.117915801 | 0.192857419 | 0.611414387 | 0.540925281 | 0.893466372 | Mrgbp | MRG/MORF4L binding protein [Source:MG | 150.98 | 153.16 | 129.81 | 156.16 | 164.68 | 150.37 |
| 6198 | ENSMUSG00000027570 | 2.758371449 | 0.104917086 | 1.312514018 | 0.079935974 | 0.936288178 | NA | Col9a3 | collagen, type IX, alpha 3 [Source:MGI Syr | 4.8186 | 1.0279 | 2.0125 | 0.964 | 3.2719 | 4.4554 |
| 6199 | ENSMUSG00000027573 | 704.5730221 | 0.076319307 | 0.113148745 | 0.674504231 | 0.499990797 | 0.880716237 | Gid8 | GID complex subunit 8 [Source:MGI Symb | 670.59 | 706.18 | 681.23 | 716.24 | 696.91 | 756.3 |
| 6200 | ENSMUSG00000027574 | 11.73828623 | -0.137361235 | 0.703289968 | -0.195312376 | 0.845148409 | 0.972803041 | Nkain4 | Na+/K+ transporting ATPase interacting 4 | 14.456 | 12.335 | 10.062 | 10.604 | 19.631 | 3.3415 |
| 6201 | ENSMUSG00000027575 | 466.9687596 | 0.150414886 | 0.131505517 | 1.14379145 | 0.252710137 | 0.705781321 | Arfgap1 | ADP-ribosylation factor GTPase activating | 456.16 | 455.37 | 416.59 | 530.19 | 466.78 | 476.73 |
| 6202 | ENSMUSG00000027577 | 20.55362742 | 2.129737616 | 0.605797557 | 3.515592941 | 0.000438773 | 0.011272495 | Chrna4 | cholinergic receptor, nicotinic, alpha poly | 10.44 | 6.1675 | 6.0375 | 13.496 | 38.172 | 49.009 |
| 6203 | ENSMUSG00000027579 | 13.68273835 | -0.157218774 | 0.666064423 | -0.236041393 | 0.81340055 | 0.966802988 | Srms | src-related kinase lacking C-terminal regul | 8.031 | 9.2513 | 26.162 | 15.424 | 7.6343 | 15.594 |
| 6204 | ENSMUSG00000027580 | 1667.263186 | 0.193030493 | 0.166318091 | 1.160610322 | 0.245800406 | 0.695907924 | Helz2 | helicase with zinc finger 2, transcriptional | 1375.7 | 1377.4 | 1915.9 | 2210.4 | 1524.7 | 1599.5 |
| 6205 | ENSMUSG00000027581 | 0 | NA | NA | NA | NA | NA | Stmn3 | stathmin-like 3 [Source:MGI Symbol;Acc:M | 0 | 0 | 0 | 0 | 0 | 0 |
| 6206 | ENSMUSG00000027582 | 354.5315125 | 0.094379529 | 0.137214329 | 0.68782561 | 0.491562607 | 0.876368691 | Zgpat | zinc finger, CCCH-type with G patch doma | 343.73 | 334.07 | 351.18 | 381.74 | 353.36 | 363.11 |
| 6207 | ENSMUSG00000027583 | 91.5716763 | -0.293157848 | 0.257361711 | -1.139088819 | 0.254666111 | 0.708091239 | Zbtb46 | zinc finger and BTB domain containing 46 | 98.782 | 81.206 | 122.76 | 93.506 | 77.434 | 75.741 |
| 6208 | ENSMUSG00000027584 | 0 | NA | NA | NA | NA | NA | Oprl1 | opioid receptor-like 1 [Source:MGI Symbo | 0 | 0 | 0 | 0 | 0 | 0 |
| 6209 | ENSMUSG00000027589 | 1015.027139 | 0.205281264 | 0.107114387 | 1.916467714 | 0.055305589 | 0.331650099 | Pcmtd2 | protein-L-isoaspartate (D-aspartate) O-me | 983.8 | 893.26 | 950.9 | 1068.1 | 1053.5 | 1140.6 |
| 6210 | ENSMUSG00000027593 | 1143.722689 | 0.018056639 | 0.106130629 | 0.170135983 | 0.864903196 | 0.977227145 | Raly | hnRNP-associated with lethal yellow [Sou | 1163.7 | 1107.1 | 1138.1 | 1088.3 | 1127.7 | 1237.5 |
| 6211 | ENSMUSG00000027596 | 0 | NA | NA | NA | NA | NA | a | nonagouti [Source:MGI Symbol;Acc:MGI:8 | 0 | 0 | 0 | 0 | 0 | 0 |
| 6212 | ENSMUSG00000027597 | 50985.3213 | -0.169084475 | 0.131451835 | -1.28628463 | 0.198343744 | 0.640554426 | Ahcy | S-adenosylhomocysteine hydrolase [Sourc | 44031 | 55856 | 62024 | 50662 | 50984 | 42355 |

| | | | | | | | | | | | | | | | |
|---|---|---|---|---|---|---|---|---|---|---|---|---|---|---|---|
| 6213 | ENSMUSG00000027598 | 1433.85682 | 0.016447909 | 0.110518618 | 0.148824777 | 0.881691899 | 0.981676444 | Itch | itchy, E3 ubiquitin protein ligase [Source:I | 1297 | 1404.1 | 1576.8 | 1407.4 | 1486.5 | 1431.3 |
| 6214 | ENSMUSG00000027599 | 854.3324803 | 0.046875832 | 0.112890446 | 0.415232942 | 0.677971386 | 0.93325821 | Armc1 | armadillo repeat containing 1 [Source:MG | 796.68 | 873.73 | 851.28 | 825.17 | 925.93 | 853.2 |
| 6215 | ENSMUSG00000027601 | 1542.125612 | 0.048186151 | 0.11640927 | 0.413937407 | 0.678919948 | 0.934062537 | Mtfr1 | mitochondrial fission regulator 1 [Source:I | 1414.3 | 1665.2 | 1470.1 | 1471 | 1738.4 | 1493.7 |
| 6216 | ENSMUSG00000027602 | 747.5988026 | -0.109899997 | 0.121374525 | -0.905461803 | 0.365220787 | 0.803870323 | Map1lc3a | microtubule-associated protein 1 light cha | 732.43 | 806.92 | 789.9 | 777.93 | 652.19 | 726.23 |
| 6217 | ENSMUSG00000027603 | 3.494630161 | 0.112285609 | 1.523430005 | 0.073705788 | 0.941244493 | NA | Ggt7 | gamma-glutamyltransferase 7 [Source:MG | 0 | 7.1954 | 3.0187 | 9.6398 | 0 | 1.1138 |
| 6218 | ENSMUSG00000027605 | 1935.767222 | 0.605613744 | 0.15350979 | 3.945114806 | 7.98E-05 | 0.003028016 | Acss2 | acyl-CoA synthetase short-chain family me | 1668 | 1593.3 | 1343.3 | 1845.1 | 2301.2 | 2863.7 |
| 6219 | ENSMUSG00000027606 | 0 | NA | NA | NA | NA | NA | Dnajc5b | DnaJ heat shock protein family (Hsp40) m | 0 | 0 | 0 | 0 | 0 | 0 |
| 6220 | ENSMUSG00000027610 | 1262.980411 | 0.105741396 | 0.131949135 | 0.801379987 | 0.422911697 | 0.84005258 | Gss | glutathione synthetase [Source:MGI Symb | 1362.1 | 1198.6 | 1088.8 | 1393.9 | 1122.2 | 1412.4 |
| 6221 | ENSMUSG00000027611 | 8.688805095 | -0.58567723 | 0.846896665 | -0.691556898 | 0.489215638 | 0.874770056 | Procr | protein C receptor, endothelial [Source:M | 12.85 | 15.419 | 3.0187 | 7.7118 | 10.906 | 2.2277 |
| 6222 | ENSMUSG00000027612 | 0 | NA | NA | NA | NA | NA | Mmp24 | matrix metallopeptidase 24 [Source:MGI S | 0 | 0 | 0 | 0 | 0 | 0 |
| 6223 | ENSMUSG00000027613 | 957.9312476 | 0.035894717 | 0.107173266 | 0.334922306 | 0.737683695 | 0.948137969 | Eif6 | eukaryotic translation initiation factor 6 [S | 1003.1 | 902.51 | 931.78 | 989.04 | 951.02 | 970.16 |
| 6224 | ENSMUSG00000027615 | 168.7205671 | -0.134462863 | 0.202378106 | -0.66441408 | 0.506425324 | 0.881592135 | Hps3 | HPS3, biogenesis of lysosomal organelles c | 149.38 | 171.66 | 209.3 | 168.7 | 142.87 | 170.42 |
| 6225 | ENSMUSG00000027618 | 1076.947882 | 0.005749624 | 0.104410467 | 0.055067507 | 0.956084682 | 0.991678478 | Nfs1 | nitrogen fixation gene 1 (S. cerevisiae) [So | 1051.3 | 1109.1 | 1064.6 | 1101.8 | 1115.7 | 1019.2 |
| 6226 | ENSMUSG00000027620 | 2082.512453 | -0.026057544 | 0.100278562 | -0.259851588 | 0.794978256 | 0.962189806 | Rbm39 | RNA binding motif protein 39 [Source:MG | 1956.4 | 2088.7 | 2260 | 2094.7 | 2109.3 | 1986 |
| 6227 | ENSMUSG00000027624 | 29.64745037 | -0.380377395 | 0.456778894 | -0.832738552 | 0.404992197 | 0.829600217 | Epb41l1 | erythrocyte membrane protein band 4.1 l | 42.564 | 35.977 | 22.137 | 37.595 | 22.903 | 16.708 |
| 6228 | ENSMUSG00000027628 | 383.4319518 | -0.173081652 | 0.144511233 | -1.197703794 | 0.231032352 | 0.678094954 | Aar2 | AAR2 splicing factor homolog [Source:MG | 392.72 | 433.78 | 392.44 | 319.08 | 384.99 | 377.59 |
| 6229 | ENSMUSG00000027630 | 743.8030762 | -0.036239963 | 0.114267071 | -0.317151411 | 0.751128717 | 0.95217579 | Tbl1xr1 | transducin (beta)-like 1X-linked receptor 1 | 723.6 | 779.16 | 756.7 | 693.1 | 737.26 | 773.01 |
| 6230 | ENSMUSG00000027634 | 306.5865361 | -0.074705572 | 0.167053637 | -0.447195125 | 0.654734181 | 0.925797231 | Ndrg3 | N-myc downstream regulated gene 3 [Sou | 306.79 | 292.96 | 344.14 | 323.9 | 245.39 | 326.36 |
| 6231 | ENSMUSG00000027635 | 10.25096743 | -0.258423649 | 0.771607744 | -0.334915831 | 0.73768858 | 0.948137969 | Dsn1 | DSN1 homolog, MIS12 kinetochore compl | 16.062 | 13.363 | 4.025 | 12.532 | 3.2719 | 12.252 |
| 6232 | ENSMUSG00000027636 | 2.141405661 | -0.575962194 | 1.475192623 | -0.390431857 | 0.696217235 | NA | Sla2 | Src-like-adaptor 2 [Source:MGI Symbol;Ac | 1.6062 | 5.1396 | 1.0062 | 2.8919 | 1.0906 | 1.1138 |
| 6233 | ENSMUSG00000027637 | 996.6706312 | 0.016129302 | 0.126090123 | 0.127918837 | 0.898213204 | 0.982798755 | Rab5if | RAB5 interacting factor [Source:MGI Symb | 869.76 | 980.63 | 1124 | 1006.4 | 945.57 | 1053.7 |
| 6234 | ENSMUSG00000027639 | 369.3827116 | -0.183288928 | 0.162487767 | -1.128016777 | 0.259312834 | 0.712728614 | Samhd1 | SAM domain and HD domain, 1 [Source:M | 377.46 | 450.23 | 351.18 | 365.35 | 293.38 | 378.71 |
| 6235 | ENSMUSG00000027641 | 26.83431376 | -0.587926591 | 0.440262231 | -1.335400928 | 0.181745161 | 0.615095921 | Rbl1 | RB transcriptional corepressor like 1 [Sour | 37.746 | 30.838 | 28.175 | 27.955 | 21.812 | 14.48 |
| 6236 | ENSMUSG00000027642 | 2461.389995 | 0.111114059 | 0.087713181 | 1.266788616 | 0.205230884 | 0.650728757 | Rpn2 | ribophorin II [Source:MGI Symbol;Acc:MG | 2375.6 | 2393 | 2331.5 | 2589.2 | 2513.9 | 2565.2 |
| 6237 | ENSMUSG00000027643 | 0.17131964 | -0.796629176 | 4.080472857 | -0.195229623 | 0.84521319 | NA | Ghrh | growth hormone releasing hormone [Sour | 0 | 1.0279 | 0 | 0 | 0 | 0 |
| 6238 | ENSMUSG00000027646 | 31.26418451 | -1.14950853 | 0.409369792 | -2.807995494 | 0.004985092 | 0.066565898 | Src | Rous sarcoma oncogene [Source:MGI Sym | 33.73 | 48.312 | 47.293 | 16.388 | 21.812 | 20.049 |
| 6239 | ENSMUSG00000027649 | 294.9283567 | 0.015829451 | 0.152770786 | 0.103615692 | 0.917474334 | 0.986374671 | Ctnnbl1 | catenin, beta like 1 [Source:MGI Symbol;A | 278.68 | 308.38 | 292.82 | 267.02 | 311.92 | 310.76 |
| 6240 | ENSMUSG00000027650 | 229.1501203 | -0.039479631 | 0.175570451 | -0.224864895 | 0.82208438 | 0.969160361 | Tti1 | TELO2 interacting protein 1 [Source:MGI S | 249.76 | 250.81 | 196.22 | 242.92 | 224.67 | 210.52 |
| 6241 | ENSMUSG00000027651 | 500.813495 | 0.038712386 | 0.126322473 | 0.306456843 | 0.759256834 | 0.95402891 | Rprd1b | regulation of nuclear pre-mRNA domain c | 468.21 | 499.57 | 515.2 | 520.55 | 522.41 | 478.95 |
| 6242 | ENSMUSG00000027652 | 351.5530625 | -0.120933665 | 0.142698727 | -0.84747543 | 0.396730176 | 0.824225434 | Ralgapb | Ral GTPase activating protein, beta subun | 354.17 | 401.92 | 343.13 | 341.25 | 340.27 | 328.58 |
| 6243 | ENSMUSG00000027654 | 5.030816422 | -0.693266844 | 1.028149392 | -0.674286101 | 0.500129439 | NA | Fam83d | family with sequence similarity 83, memb | 10.44 | 3.0838 | 5.0312 | 3.8559 | 1.0906 | 6.6831 |
| 6244 | ENSMUSG00000027655 | 121.215966 | 0.479599966 | 0.234100404 | 2.04869346 | 0.040492099 | 0.275224493 | Dhx35 | DEAH (Asp-Glu-Ala-His) box polypeptide 3 | 120.47 | 75.038 | 107.67 | 146.52 | 145.05 | 132.55 |
| 6245 | ENSMUSG00000027656 | 2.118257402 | -0.060295463 | 1.424226128 | -0.042335596 | 0.966231169 | NA | Ccn5 | cellular communication network factor 5 [ | 2.4093 | 3.0838 | 1.0062 | 2.8919 | 1.0906 | 2.2277 |
| 6246 | ENSMUSG00000027660 | 240.8506976 | -0.639859896 | 0.20547661 | -3.114027905 | 0.00184552 | 0.032832144 | Skil | SKI-like [Source:MGI Symbol;Acc:MGI:106 | 209.61 | 345.38 | 326.02 | 187.01 | 181.04 | 196.04 |
| 6247 | ENSMUSG00000027661 | 2.935137697 | -2.856626751 | 1.635210866 | -1.746947021 | 0.080646528 | NA | Slc2a10 | solute carrier family 2 (facilitated glucose | 7.2279 | 7.1954 | 1.0062 | 0 | 2.1812 | 0 |
| 6248 | ENSMUSG00000027663 | 130.5809533 | -0.191718579 | 0.227179671 | -0.843907283 | 0.398721218 | 0.82588279 | Zmat3 | zinc finger matrin type 3 [Source:MGI Sym | 151.79 | 116.15 | 149.93 | 147.49 | 113.42 | 104.7 |
| 6249 | ENSMUSG00000027665 | 692.0814455 | 0.080560456 | 0.120844312 | 0.666646653 | 0.504997862 | 0.881592135 | Pik3ca | phosphatidylinositol-4,5-bisphosphate 3-k | 681.83 | 689.73 | 646.01 | 648.76 | 718.72 | 767.44 |
| 6250 | ENSMUSG00000027667 | 279.2849407 | 0.122428957 | 0.154547952 | 0.792174569 | 0.428258895 | 0.842753542 | Zfp639 | zinc finger protein 639 [Source:MGI Symb | 265.02 | 252.87 | 284.77 | 312.33 | 292.29 | 268.44 |
| 6251 | ENSMUSG00000027668 | 1114.59967 | 0.146234315 | 0.114255678 | 1.279886629 | 0.200585011 | 0.643866311 | Mfn1 | mitofusin 1 [Source:MGI Symbol;Acc:MGI | 1013.5 | 1117.3 | 1043.5 | 1052.7 | 1249.8 | 1210.7 |
| 6252 | ENSMUSG00000027669 | 43.4025492 | -0.150433833 | 0.339727269 | -0.442807648 | 0.657904871 | 0.927303282 | Gnb4 | guanine nucleotide binding protein (G pro | 51.399 | 48.312 | 37.231 | 46.271 | 35.99 | 41.212 |
| 6253 | ENSMUSG00000027670 | 24.65938545 | -0.506567371 | 0.479843094 | -1.055693781 | 0.291108139 | 0.740906421 | Ocstamp | osteoclast stimulatory transmembrane pr | 40.958 | 23.642 | 22.137 | 26.991 | 14.178 | 20.049 |
| 6254 | ENSMUSG00000027671 | 262.9986322 | 0.075249863 | 0.154918584 | 0.48573813 | 0.627152845 | 0.920273791 | Actl6a | actin-like 6A [Source:MGI Symbol;Acc:MG | 260.21 | 249.78 | 258.6 | 287.27 | 273.75 | 248.39 |
| 6255 | ENSMUSG00000027673 | 1390.867272 | -0.024833781 | 0.098000524 | -0.253404577 | 0.799955591 | 0.962424862 | Ndufb5 | NADH:ubiquinone oxidoreductase subuni | 1360.5 | 1379.5 | 1469.1 | 1390.1 | 1389.4 | 1356.7 |
| 6256 | ENSMUSG00000027674 | 0.428363721 | -0.543730382 | 3.624677344 | -0.15000794 | 0.880758351 | NA | Pex5l | peroxisomal biogenesis factor 5-like [Sour | 1.6062 | 0 | 0 | 0.964 | 0 | 0 |
| 6257 | ENSMUSG00000027676 | 26.32698849 | 0.311254712 | 0.491322707 | 0.633503618 | 0.526404819 | 0.888334265 | Ccdc39 | coiled-coil domain containing 39 [Source:I | 29.715 | 25.698 | 15.094 | 42.415 | 29.447 | 15.594 |
| 6258 | ENSMUSG00000027677 | 423.3907007 | 0.04881372 | 0.167095757 | 0.292130219 | 0.770187063 | 0.956515747 | Ttc14 | tetratricopeptide repeat domain 14 [Sourc | 435.28 | 373.13 | 440.74 | 521.51 | 417.71 | 351.97 |
| 6259 | ENSMUSG00000027678 | 341.1126811 | -0.008956776 | 0.155737757 | -0.05751191 | 0.954137419 | 0.991639263 | Ncoa3 | nuclear receptor coactivator 3 [Source:M( | 340.52 | 347.44 | 339.1 | 387.52 | 287.92 | 344.18 |
| 6260 | ENSMUSG00000027679 | 1035.536801 | -0.061286922 | 0.107676573 | -0.569176009 | 0.5692367 | 0.903735951 | Dnajc19 | DnaJ heat shock protein family (Hsp40) m | 1056.9 | 1059.8 | 1055.6 | 959.16 | 1095 | 986.87 |
| 6261 | ENSMUSG00000027680 | 910.4284396 | 0.223564275 | 0.110396907 | 2.025095471 | 0.042857562 | 0.285664413 | Fxr1 | fragile X mental retardation gene 1, autos | 864.94 | 871.67 | 782.86 | 940.84 | 1014.3 | 987.98 |
| 6262 | ENSMUSG00000027684 | 9.2306653 | -0.469667522 | 0.706914285 | -0.664391047 | 0.506440061 | 0.881592135 | Mecom | MDS1 and EVI1 complex locus [Source:M( | 8.8341 | 10.279 | 13.081 | 7.7118 | 5.4531 | 10.025 |
| 6263 | ENSMUSG00000027690 | 9195.831576 | -0.036788675 | 0.124841149 | -0.294683885 | 0.768235375 | 0.955856567 | Slc2a2 | solute carrier family 2 (facilitated glucose | 9096.7 | 10370 | 8472.6 | 8064.6 | 10622 | 8549.9 |
| 6264 | ENSMUSG00000027692 | 20.27299542 | 0.045514067 | 0.471182014 | 0.096595509 | 0.923047623 | 0.987005368 | Tnik | TRAF2 and NCK interacting kinase [Source | 19.274 | 19.53 | 21.131 | 23.135 | 19.631 | 18.935 |
| 6265 | ENSMUSG00000027694 | 2.296182814 | -1.161484606 | 1.402900381 | -0.827916666 | 0.407717696 | NA | Gm8325 | predicted pseudogene 8325 [Source:MGI | 2.4093 | 3.0838 | 4.025 | 0.964 | 2.1812 | 1.1138 |
| 6266 | ENSMUSG00000027695 | 99.24919192 | -0.23903842 | 0.261690196 | -0.913440488 | 0.36101092 | 0.800074801 | Pld1 | phospholipase D1 [Source:MGI Symbol;Ac | 81.916 | 106.9 | 133.83 | 81.938 | 83.978 | 106.93 |
| 6267 | ENSMUSG00000027698 | 1008.688704 | 0.189800757 | 0.138284133 | 1.372541828 | 0.169894827 | 0.596876778 | Nceh1 | neutral cholesterol ester hydrolase 1 [Sou | 1098.6 | 946.71 | 780.85 | 1114.4 | 1111.3 | 1000.2 |
| 6268 | ENSMUSG00000027699 | 24.08798911 | -0.07785156 | 0.467740964 | -0.166441612 | 0.867809429 | 0.978326885 | Ect2 | ect2 oncogene [Source:MGI Symbol;Acc:I | 30.518 | 20.558 | 23.144 | 31.811 | 22.903 | 15.594 |

| | | | | | | | | | | | | | | | |
|---|---|---|---|---|---|---|---|---|---|---|---|---|---|---|---|
| 6269 | ENSMUSG00000027702 | 0.342639279 | -1.805976596 | 4.028547714 | -0.448294702 | 0.653940524 | NA | Lrrc34 | leucine rich repeat containing 34 [Source: | 0 | 2.0558 | 0 | 0 | 0 | 0 |
| 6270 | ENSMUSG00000027703 | 0.133850373 | -0.796629176 | 4.080472857 | -0.195229623 | 0.84521319 | NA | Lrriq4 | leucine-rich repeats and IQ motif containi | 0.8031 | 0 | 0 | 0 | 0 | 0 |
| 6271 | ENSMUSG00000027706 | 3129.192105 | 0.050065079 | 0.090069458 | 0.555849679 | 0.578313621 | 0.908137944 | Sec62 | SEC62 homolog (S. cerevisiae) [Source:MG | 3014 | 3060.1 | 3150.6 | 3071.2 | 3348.2 | 3131 |
| 6272 | ENSMUSG00000027708 | 953.7210595 | 0.079475395 | 0.1249708 | 0.635951714 | 0.524807895 | 0.888334265 | Dcun1d1 | DCN1, defective in cullin neddylation 1, do | 958.1 | 871.67 | 952.91 | 1077.7 | 1014.3 | 847.64 |
| 6273 | ENSMUSG00000027709 | 1118.227559 | -0.033929086 | 0.112159546 | -0.30250734 | 0.762265339 | 0.954414324 | Mccc1 | methylcrotonoyl-Coenzyme A carboxylase | 1167.7 | 1085.5 | 1140.1 | 1028.6 | 1217.1 | 1070.4 |
| 6274 | ENSMUSG00000027710 | 607.6497386 | -0.00228263 | 0.139881144 | -0.016318351 | 0.986980418 | 0.99859176 | Acad9 | acyl-Coenzyme A dehydrogenase family, r | 689.86 | 591.05 | 542.37 | 586.1 | 672.91 | 563.61 |
| 6275 | ENSMUSG00000027712 | 1109.299329 | -0.288923995 | 0.176579996 | -1.636221553 | 0.101793227 | 0.469495293 | Anxa5 | annexin A5 [Source:MGI Symbol;Acc:MGI | 1590.9 | 1184.2 | 883.48 | 958.19 | 961.93 | 1077.1 |
| 6276 | ENSMUSG00000027713 | 13.55603032 | 0.044738911 | 0.605327644 | 0.073908587 | 0.941083123 | 0.989159676 | 1810062G17 | RIKEN cDNA 1810062G17 gene [Source:M | 13.653 | 16.447 | 10.062 | 20.244 | 10.906 | 10.025 |
| 6277 | ENSMUSG00000027714 | 165.1221841 | 0.128070558 | 0.187975437 | 0.681315386 | 0.49567195 | 0.877965876 | Exosc9 | exosome component 9 [Source:MGI Symb | 165.44 | 145.96 | 162.01 | 189.9 | 159.23 | 168.19 |
| 6278 | ENSMUSG00000027715 | 11.01656637 | -0.185120068 | 0.666245064 | -0.277855819 | 0.78112304 | 0.959993512 | Ccna2 | cyclin A2 [Source:MGI Symbol;Acc:MGI:10 | 13.653 | 11.307 | 10.062 | 5.7839 | 15.269 | 10.025 |
| 6279 | ENSMUSG00000027716 | 5.361674498 | 0.186875373 | 1.096141066 | 0.170484784 | 0.8646289 | 0.977227145 | Trpc3 | transient receptor potential cation chann | 4.0155 | 3.0838 | 8.05 | 11.568 | 5.4531 | 0 |
| 6280 | ENSMUSG00000027718 | 0 | NA | NA | NA | NA | NA | Il21 | interleukin 21 [Source:MGI Symbol;Acc:M | 0 | 0 | 0 | 0 | 0 | 0 |
| 6281 | ENSMUSG00000027719 | 0 | NA | NA | NA | NA | NA | Adad1 | adenosine deaminase domain containing | 0 | 0 | 0 | 0 | 0 | 0 |
| 6282 | ENSMUSG00000027720 | 0.401551119 | -2.072982726 | 4.008251979 | -0.517178744 | 0.605031388 | NA | Il2 | interleukin 2 [Source:MGI Symbol;Acc:MG | 2.4093 | 0 | 0 | 0 | 0 | 0 |
| 6283 | ENSMUSG00000027722 | 82.77724721 | -0.134469734 | 0.271506642 | -0.495272356 | 0.620407882 | 0.917977899 | Spata5 | spermatogenesis associated 5 [Source:MG | 97.978 | 76.066 | 85.531 | 73.262 | 100.34 | 63.489 |
| 6284 | ENSMUSG00000027737 | 0 | NA | NA | NA | NA | NA | Slc7a11 | solute carrier family 7 (cationic amino acid | 0 | 0 | 0 | 0 | 0 | 0 |
| 6285 | ENSMUSG00000027739 | 151.9295731 | 0.098484221 | 0.201836782 | 0.487939908 | 0.625592406 | 0.919638611 | Rab33b | RAB33B, member RAS oncogene family [S | 134.12 | 173.72 | 132.82 | 158.09 | 164.68 | 148.14 |
| 6286 | ENSMUSG00000027742 | 446.6118642 | -0.014392482 | 0.132689541 | -0.108467346 | 0.913624982 | 0.98625062 | Cog6 | component of oligomeric golgi complex 6 | 469.81 | 469.76 | 406.52 | 438.61 | 444.97 | 449.99 |
| 6287 | ENSMUSG00000027744 | 0.167707431 | -0.796629176 | 4.080472857 | -0.195229623 | 0.84521319 | NA | Stoml3 | stomatin (Epb7.2)-like 3 [Source:MGI Sym | 0 | 0 | 1.0062 | 0 | 0 | 0 |
| 6288 | ENSMUSG00000027746 | 1059.944227 | 0.005203113 | 0.10478467 | 0.049655293 | 0.960397084 | 0.992456421 | Ufm1 | ubiquitin-fold modifier 1 [Source:MGI Sym | 1101.9 | 1020.7 | 1050.5 | 1014.1 | 1076.4 | 1096 |
| 6289 | ENSMUSG00000027748 | 0.342639279 | -1.805976596 | 4.028547714 | -0.448294702 | 0.653940524 | NA | Trpc4 | transient receptor potential cation chann | 0 | 2.0558 | 0 | 0 | 0 | 0 |
| 6290 | ENSMUSG00000027750 | 106.2321177 | -1.978816683 | 0.272430335 | -7.263569527 | 3.77E-13 | 1.70E-10 | Postn | periostin, osteoblast specific factor [Sourc | 199.17 | 181.94 | 126.79 | 34.703 | 46.897 | 47.895 |
| 6291 | ENSMUSG00000027751 | 265.3890837 | 0.183962959 | 0.17286731 | 1.064185931 | 0.287244479 | 0.73951397 | Supt20 | SPT20 SAGA complex component [Source | 272.25 | 243.62 | 229.42 | 305.58 | 307.55 | 233.91 |
| 6292 | ENSMUSG00000027752 | 167.1973956 | 0.012811274 | 0.195278354 | 0.065605191 | 0.947692156 | 0.990677354 | Exosc8 | exosome component 8 [Source:MGI Symb | 192.74 | 149.05 | 156.97 | 173.52 | 152.69 | 178.22 |
| 6293 | ENSMUSG00000027761 | 9834.387973 | 0.312988013 | 0.091492688 | 3.420907387 | 0.000624126 | 0.014583774 | Aadac | arylacetamide deacetylase [Source:MGI S | 8781.1 | 8380.6 | 9152.8 | 10198 | 10941 | 11553 |
| 6294 | ENSMUSG00000027762 | 496.0955315 | 0.825315164 | 0.171819787 | 4.803376716 | 1.56E-06 | 0.000117529 | Sucnr1 | succinate receptor 1 [Source:MGI Symbol | 354.17 | 364.91 | 354.2 | 468.49 | 724.17 | 710.63 |
| 6295 | ENSMUSG00000027763 | 1386.086446 | -0.221922509 | 0.108884983 | -2.038136964 | 0.041536237 | 0.279562651 | Mbnl1 | muscleblind like splicing factor 1 [Source:I | 1555.6 | 1564.5 | 1357.4 | 1346.7 | 1277.1 | 1215.2 |
| 6296 | ENSMUSG00000027765 | 236.6892028 | 0.314165446 | 0.223390675 | 1.406349867 | 0.159620259 | 0.581402021 | P2ry1 | purinergic receptor P2Y, G-protein couple | 256.19 | 209.7 | 167.04 | 344.14 | 220.3 | 222.77 |
| 6297 | ENSMUSG00000027770 | 828.6005496 | 0.062874209 | 0.121927763 | 0.515667705 | 0.606086514 | 0.916376001 | Dhx36 | DEAH (Asp-Glu-Ala-His) box polypeptide 3 | 818.36 | 852.14 | 761.73 | 922.53 | 858.32 | 758.53 |
| 6298 | ENSMUSG00000027774 | 2031.41609 | -0.200003161 | 0.090687198 | -2.205417809 | 0.027424789 | 0.21476565 | Gfm1 | G elongation factor, mitochondrial 1 [Sour | 2155.5 | 2182.3 | 2178.5 | 1910.6 | 1929.3 | 1832.3 |
| 6299 | ENSMUSG00000027775 | 2146.265575 | 0.141230427 | 0.128390128 | 1.100010019 | 0.271327757 | 0.72702569 | Mfsd1 | major facilitator superfamily domain cont | 1935.5 | 1906.8 | 2281.2 | 1894.2 | 2446.3 | 2413.7 |
| 6300 | ENSMUSG00000027776 | 0.167707431 | -0.796629176 | 4.080472857 | -0.195229623 | 0.84521319 | NA | Il12a | interleukin 12a [Source:MGI Symbol;Acc:M | 0 | 0 | 1.0062 | 0 | 0 | 0 |
| 6301 | ENSMUSG00000027777 | 0.462220778 | -0.685400455 | 3.191475091 | -0.21475977 | 0.82995462 | NA | Schip1 | schwannomin interacting protein 1 [Sourc | 0.8031 | 0 | 1.0062 | 0.964 | 0 | 0 |
| 6302 | ENSMUSG00000027778 | 80.10736866 | -0.186908607 | 0.360077421 | -0.519078942 | 0.603705693 | 0.916106578 | Ift80 | intraflagellar transport 80 [Source:MGI Sy | 75.492 | 79.15 | 101.63 | 107 | 32.719 | 84.652 |
| 6303 | ENSMUSG00000027782 | 903.4025294 | 0.182382411 | 0.116915384 | 1.559952207 | 0.118771176 | 0.504308497 | Kpna4 | karyopherin (importin) alpha 4 [Source:M | 893.85 | 857.28 | 786.88 | 896.5 | 945.57 | 1040.3 |
| 6304 | ENSMUSG00000027784 | 389.4875872 | 0.053811685 | 0.13785336 | 0.390354541 | 0.696274398 | 0.939992732 | Ppm1l | protein phosphatase 1 (formerly 2C)-like [ | 366.21 | 399.86 | 381.37 | 427.04 | 391.53 | 370.91 |
| 6305 | ENSMUSG00000027787 | 576.0804938 | 0.203549113 | 0.120587792 | 1.687974455 | 0.091416133 | 0.445950402 | Nmd3 | NMD3 ribosome export adaptor [Source:M | 513.18 | 522.18 | 571.55 | 612.13 | 629.29 | 608.16 |
| 6306 | ENSMUSG00000027788 | 0 | NA | NA | NA | NA | NA | Otol1 | otolin 1 [Source:MGI Symbol;Acc:MGI:268 | 0 | 0 | 0 | 0 | 0 | 0 |
| 6307 | ENSMUSG00000027790 | 0 | NA | NA | NA | NA | NA | Sis | sucrase isomaltase (alpha-glucosidase) [So | 0 | 0 | 0 | 0 | 0 | 0 |
| 6308 | ENSMUSG00000027792 | 2320.669275 | -0.388660181 | 0.115487361 | -3.365391473 | 0.000764351 | 0.017113896 | Bche | butyrylcholinesterase [Source:MGI Symbo | 2599.6 | 2878.2 | 2417 | 2089.9 | 2161.6 | 1777.7 |
| 6309 | ENSMUSG00000027793 | 0 | NA | NA | NA | NA | NA | Ccna1 | cyclin A1 [Source:MGI Symbol;Acc:MGI:10 | 0 | 0 | 0 | 0 | 0 | 0 |
| 6310 | ENSMUSG00000027794 | 0 | NA | NA | NA | NA | NA | Sohlh2 | spermatogenesis and oogenesis specific b | 0 | 0 | 0 | 0 | 0 | 0 |
| 6311 | ENSMUSG00000027796 | 155.7663023 | 1.225737016 | 0.302339719 | 4.05417131 | 5.03E-05 | 0.002110891 | Smad9 | SMAD family member 9 [Source:MGI Sym | 94.766 | 106.9 | 78.487 | 282.45 | 262.84 | 109.16 |
| 6312 | ENSMUSG00000027797 | 0.133850373 | -0.796629176 | 4.080472857 | -0.195229623 | 0.84521319 | NA | Dclk1 | doublecortin-like kinase 1 [Source:MGI Sy | 0.8031 | 0 | 0 | 0 | 0 | 0 |
| 6313 | ENSMUSG00000027799 | 13.49801288 | -0.116044929 | 0.572004552 | -0.202874136 | 0.83923341 | 0.971685662 | Nbea | neurobeachin [Source:MGI Symbol;Acc:M | 13.653 | 15.419 | 13.081 | 13.496 | 13.087 | 12.252 |
| 6314 | ENSMUSG00000027800 | 200.0034339 | -0.043461015 | 0.224885495 | -0.193258418 | 0.84675659 | 0.972803041 | Tm4sf1 | transmembrane 4 superfamily member 1 | 250.57 | 223.06 | 134.84 | 202.44 | 190.86 | 198.26 |
| 6315 | ENSMUSG00000027801 | 1465.21098 | -0.065453499 | 0.10025629 | -0.65286177 | 0.513845397 | 0.885004045 | Tm4sf4 | transmembrane 4 superfamily member 4 | 1574.9 | 1456.6 | 1463.1 | 1434.4 | 1370.9 | 1491.4 |
| 6316 | ENSMUSG00000027803 | 463.5148921 | -0.209260882 | 0.168791387 | -1.239760429 | 0.215064019 | 0.659728602 | Wwtr1 | WW domain containing transcription regu | 617.59 | 468.73 | 403.5 | 401.01 | 456.97 | 433.29 |
| 6317 | ENSMUSG00000027804 | 703.1640583 | 0.125423195 | 0.124145193 | 1.010294421 | 0.312354253 | 0.759830222 | Ppid | peptidylprolyl isomerase D (cyclophilin D) | 639.27 | 676.37 | 703.36 | 810.71 | 717.63 | 671.65 |
| 6318 | ENSMUSG00000027805 | 48.13801128 | -0.215810476 | 0.324503118 | -0.665049005 | 0.50601915 | 0.881592135 | Pfn2 | profilin 2 [Source:MGI Symbol;Acc:MGI:97 | 52.202 | 54.48 | 48.3 | 35.667 | 44.715 | 53.465 |
| 6319 | ENSMUSG00000027806 | 464.0603222 | 0.125427122 | 0.130635755 | 0.960128582 | 0.336990505 | 0.779072142 | Tsc22d2 | TSC22 domain family, member 2 [Source:I | 443.31 | 469.76 | 418.6 | 480.06 | 459.15 | 513.48 |
| 6320 | ENSMUSG00000027808 | 5532.040162 | 0.10233966 | 0.099021411 | 1.033510419 | 0.301365097 | 0.752246851 | Serp1 | stress-associated endoplasmic reticulum p | 5466.7 | 5310.2 | 5231.5 | 6356.5 | 5454.2 | 5373.2 |
| 6321 | ENSMUSG00000027809 | 5679.01999 | -0.112963608 | 0.089975327 | -1.255495385 | 0.209298982 | 0.65350014 | Etfdh | electron transferring flavoprotein, dehydr | 5620.1 | 6208.6 | 5875.5 | 5246.9 | 5694.1 | 5428.9 |
| 6322 | ENSMUSG00000027810 | 771.9783967 | 0.126580954 | 0.110222223 | 1.148415905 | 0.250796911 | 0.702929347 | Eif2a | eukaryotic translation initiation factor 2A | 723.6 | 755.52 | 735.56 | 801.07 | 839.78 | 776.35 |
| 6323 | ENSMUSG00000027811 | 12.85511441 | -0.965294715 | 0.671398167 | -1.437738086 | 0.150508382 | 0.564307976 | 4930579G24 | RIKEN cDNA 4930579G24 gene [Source:M | 10.44 | 23.642 | 17.106 | 10.604 | 11.997 | 3.3415 |
| 6324 | ENSMUSG00000027820 | 1159.398573 | -0.580846815 | 0.175242434 | -3.31453291 | 0.000917964 | 0.019641165 | Mme | membrane metallo endopeptidase [Sourc | 1190.2 | 1406.2 | 1573.8 | 1028.6 | 1097.2 | 660.51 |

| | | | | | | | | | | | | | | | |
|---|---|---|---|---|---|---|---|---|---|---|---|---|---|---|---|
| 6325 | ENSMUSG00000027822 | 1075.255071 | 0.092697163 | 0.109692344 | 0.84506502 | 0.398074537 | 0.825720652 | Slc33a1 | solute carrier family 33 (acetyl-CoA transp | 1125.9 | 1033.1 | 961.97 | 1083.5 | 1131 | 1116.1 |
| 6326 | ENSMUSG00000027823 | 778.0185965 | 0.025587466 | 0.12218391 | 0.209417637 | 0.834122226 | 0.971005543 | Gmps | guanine monophosphate synthetase [Sou | 736.44 | 774.02 | 803.99 | 857.94 | 792.88 | 702.84 |
| 6327 | ENSMUSG00000027824 | 0.756734126 | -0.422689986 | 2.665959267 | -0.158550804 | 0.874022791 | NA | Vmn2r1 | vomeronasal 2, receptor 1 [Source:MGI Sy | 1.6062 | 0 | 1.0062 | 1.928 | 0 | 0 |
| 6328 | ENSMUSG00000027827 | 18.82203105 | -1.251149454 | 0.537089514 | -2.32949894 | 0.019832649 | 0.17228795 | Kcnab1 | potassium voltage-gated channel, shaker- | 29.715 | 25.698 | 24.15 | 12.532 | 15.269 | 5.5692 |
| 6329 | ENSMUSG00000027828 | 4155.267795 | 0.094308159 | 0.089223193 | 1.056991525 | 0.290515456 | 0.740463951 | Ssr3 | signal sequence receptor, gamma [Source | 4144 | 4020.2 | 3894.2 | 4467.1 | 4080 | 4326.2 |
| 6330 | ENSMUSG00000027829 | 642.6938165 | -0.079184955 | 0.135695939 | -0.583546974 | 0.559525145 | 0.899423311 | Ccnl1 | cyclin L1 [Source:MGI Symbol;Acc:MGI:19 | 712.35 | 664.03 | 604.75 | 711.42 | 592.21 | 571.4 |
| 6331 | ENSMUSG00000027831 | 3.867239025 | 0.100853626 | 1.197475925 | 0.08422184 | 0.932880054 | NA | Veph1 | ventricular zone expressed PH domain-co | 4.0155 | 3.0838 | 4.025 | 0.964 | 1.0906 | 10.025 |
| 6332 | ENSMUSG00000027832 | 0.321325949 | 1.911575114 | 4.048267893 | 0.472195804 | 0.636787034 | NA | Ptx3 | pentraxin related gene [Source:MGI Symb | 0 | 0 | 0 | 1.928 | 0 | 0 |
| 6333 | ENSMUSG00000027833 | 0.36353943 | 2.065989254 | 4.034137301 | 0.512126658 | 0.608562368 | NA | Shox2 | short stature homeobox 2 [Source:MGI Sy | 0 | 0 | 0 | 0 | 2.1812 | 0 |
| 6334 | ENSMUSG00000027834 | 1.878991552 | -1.369551486 | 1.577740219 | -0.868046254 | 0.385369012 | NA | Serpini1 | serine (or cysteine) peptidase inhibitor, cl | 4.0155 | 3.0838 | 1.0062 | 0.964 | 1.0906 | 1.1138 |
| 6335 | ENSMUSG00000027835 | 433.3497566 | 0.043017629 | 0.137652301 | 0.312509336 | 0.754653469 | 0.953048681 | Pdcd10 | programmed cell death 10 [Source:MGI Sy | 445.72 | 423.5 | 410.55 | 394.27 | 449.33 | 476.73 |
| 6336 | ENSMUSG00000027840 | 23.14049527 | 0.362021803 | 0.484672906 | 0.746940461 | 0.455099501 | 0.857337687 | Wnt2b | wingless-type MMTV integration site fami | 15.259 | 17.475 | 28.175 | 24.099 | 34.9 | 18.935 |
| 6337 | ENSMUSG00000027843 | 8.969985711 | -0.944586021 | 0.832835207 | -1.134181183 | 0.256718555 | 0.710192112 | Ptpn22 | protein tyrosine phosphatase, non-recept | 12.047 | 17.475 | 6.0375 | 10.604 | 6.5437 | 1.1138 |
| 6338 | ENSMUSG00000027845 | 123.1137936 | -0.327784187 | 0.237333216 | -1.381113828 | 0.167243965 | 0.593441872 | Dclre1b | DNA cross-link repair 1B [Source:MGI Sym | 158.21 | 144.94 | 107.67 | 108.93 | 127.6 | 91.335 |
| 6339 | ENSMUSG00000027848 | 44.65538194 | -1.087485257 | 0.3587056 | -3.031693001 | 0.002431864 | 0.040430438 | Olfml3 | olfactomedin-like 3 [Source:MGI Symbol;A | 80.31 | 50.368 | 51.318 | 30.847 | 28.356 | 26.732 |
| 6340 | ENSMUSG00000027849 | 0 | NA | NA | NA | NA | NA | Syt6 | synaptotagmin VI [Source:MGI Symbol;Ac | 0 | 0 | 0 | 0 | 0 | 0 |
| 6341 | ENSMUSG00000027852 | 362.9163223 | 0.024022409 | 0.146807772 | 0.163631724 | 0.870021069 | 0.978765244 | Nras | neuroblastoma ras oncogene [Source:MG | 326.06 | 403.97 | 350.17 | 348.96 | 374.08 | 374.25 |
| 6342 | ENSMUSG00000027854 | 277.9841804 | 0.083693376 | 0.180861479 | 0.462748492 | 0.643544655 | 0.925505685 | Sike1 | suppressor of IKBKE 1 [Source:MGI Symbo | 243.34 | 274.45 | 291.81 | 227.5 | 296.65 | 334.15 |
| 6343 | ENSMUSG00000027855 | 0.17131964 | -0.796629176 | 4.080472857 | -0.195229623 | 0.84521319 | NA | Sycp1 | synaptonemal complex protein 1 [Source: | 0 | 1.0279 | 0 | 0 | 0 | 0 |
| 6344 | ENSMUSG00000027857 | 0.160662974 | 1.126924175 | 4.080472857 | 0.276174898 | 0.782413736 | NA | Tshb | thyroid stimulating hormone, beta subuni | 0 | 0 | 0 | 0.964 | 0 | 0 |
| 6345 | ENSMUSG00000027858 | 14.07517327 | -0.537874276 | 0.587886301 | -0.914929085 | 0.36022886 | 0.799598236 | Tspan2 | tetraspanin 2 [Source:MGI Symbol;Acc:M | 14.456 | 22.614 | 13.081 | 14.46 | 9.8156 | 10.025 |
| 6346 | ENSMUSG00000027859 | 35.51226615 | -0.481171407 | 0.414182732 | -1.161737006 | 0.245342309 | 0.695673624 | Ngf | nerve growth factor [Source:MGI Symbol; | 50.595 | 37.005 | 36.225 | 25.063 | 19.631 | 44.554 |
| 6347 | ENSMUSG00000027860 | 135.4527322 | 0.50216631 | 0.237274812 | 2.116391141 | 0.034311555 | 0.247893272 | Vangl1 | VANGL planar cell polarity 1 [Source:MGI | 105.21 | 83.261 | 147.92 | 153.27 | 154.87 | 168.19 |
| 6348 | ENSMUSG00000027861 | 0 | NA | NA | NA | NA | NA | Casq2 | calsequestrin 2 [Source:MGI Symbol;Acc:I | 0 | 0 | 0 | 0 | 0 | 0 |
| 6349 | ENSMUSG00000027863 | 2.715615016 | -2.012165141 | 1.457688029 | -1.380381194 | 0.167469307 | NA | Cd2 | CD2 antigen [Source:MGI Symbol;Acc:MG | 4.8186 | 6.1675 | 2.0125 | 0 | 2.1812 | 1.1138 |
| 6350 | ENSMUSG00000027864 | 68.35382556 | 0.097218849 | 0.291323876 | 0.333713975 | 0.738595403 | 0.948137969 | Ptgfrn | prostaglandin F2 receptor negative regula | 81.916 | 50.368 | 65.406 | 72.298 | 62.165 | 77.969 |
| 6351 | ENSMUSG00000027865 | 450.6588582 | -0.024621956 | 0.159245744 | -0.154616099 | 0.877123978 | 0.980632845 | Gdap2 | ganglioside-induced differentiation-associ | 534.87 | 449.2 | 378.35 | 442.47 | 408.98 | 490.09 |
| 6352 | ENSMUSG00000027867 | 0.185640647 | 1.126924175 | 4.080472857 | 0.276174898 | 0.782413736 | NA | Spag17 | sperm associated antigen 17 [Source:MGI | 0 | 0 | 0 | 0 | 0 | 1.1138 |
| 6353 | ENSMUSG00000027868 | 0 | NA | NA | NA | NA | NA | Tbx15 | T-box 15 [Source:MGI Symbol;Acc:MGI:12 | 0 | 0 | 0 | 0 | 0 | 0 |
| 6354 | ENSMUSG00000027869 | 0 | NA | NA | NA | NA | NA | Hsd3b6 | hydroxy-delta-5-steroid dehydrogenase, 3 | 0 | 0 | 0 | 0 | 0 | 0 |
| 6355 | ENSMUSG00000027870 | 21.41693814 | -2.650694855 | 0.564840604 | -4.692819246 | 2.69E-06 | 0.000182984 | Hao2 | hydroxyacid oxidase 2 [Source:MGI Symbo | 41.761 | 40.089 | 29.181 | 8.6758 | 5.4531 | 3.3415 |
| 6356 | ENSMUSG00000027871 | 0.321325949 | 1.911575114 | 4.048267893 | 0.472195804 | 0.636787034 | NA | Hsd3b1 | hydroxy-delta-5-steroid dehydrogenase, 3 | 0 | 0 | 0 | 1.928 | 0 | 0 |
| 6357 | ENSMUSG00000027875 | 73058.52795 | -0.307564379 | 0.09929554 | -3.097464184 | 0.00195184 | 0.034219892 | Hmgcs2 | 3-hydroxy-3-methylglutaryl-Coenzyme A s | 76701 | 85633 | 80118 | 71312 | 65267 | 59320 |
| 6358 | ENSMUSG00000027876 | 4.494815387 | 1.376320478 | 1.099975606 | 1.251228182 | 0.210851239 | NA | Reg4 | regenerating islet-derived family, member | 2.4093 | 1.0279 | 4.025 | 2.8919 | 4.3625 | 12.252 |
| 6359 | ENSMUSG00000027878 | 594.9241656 | 0.031889934 | 0.119657687 | 0.266509696 | 0.789846693 | 0.961019028 | Notch2 | notch 2 [Source:MGI Symbol;Acc:MGI:97 | 583.86 | 584.89 | 596.7 | 634.3 | 562.76 | 607.04 |
| 6360 | ENSMUSG00000027879 | 968.1469424 | -0.021972545 | 0.117188003 | -0.187498248 | 0.851269998 | 0.973568625 | Sec22b | SEC22 homolog B, vesicle trafficking prote | 1043.2 | 931.29 | 950.9 | 924.45 | 900.85 | 1058.2 |
| 6361 | ENSMUSG00000027880 | 0 | NA | NA | NA | NA | NA | Slc25a54 | solute carrier family 25, member 54 [Sour | 0 | 0 | 0 | 0 | 0 | 0 |
| 6362 | ENSMUSG00000027881 | 787.2037768 | -0.029141017 | 0.132384464 | -0.220124148 | 0.825774469 | 0.96943211 | Prpf38b | PRP38 pre-mRNA processing factor 38 (ye | 754.92 | 886.07 | 745.63 | 860.83 | 787.43 | 688.36 |
| 6363 | ENSMUSG00000027882 | 478.3853245 | -0.057270161 | 0.129668116 | -0.441667256 | 0.658730008 | 0.927599242 | Stxbp3 | syntaxin binding protein 3 [Source:MGI Sy | 456.16 | 501.62 | 506.14 | 443.43 | 495.14 | 467.81 |
| 6364 | ENSMUSG00000027883 | 287.7780074 | 0.113206898 | 0.168806641 | 0.670630592 | 0.502455889 | 0.881220881 | Gpsm2 | G-protein signalling modulator 2 (AGS3-lik | 237.72 | 267.26 | 325.02 | 278.59 | 318.46 | 299.62 |
| 6365 | ENSMUSG00000027884 | 831.7097682 | 0.202832664 | 0.128238863 | 1.581678591 | 0.113722962 | 0.495873456 | Clcc1 | chloride channel CLIC-like 1 [Source:MGI S | 660.15 | 803.83 | 857.32 | 873.36 | 885.58 | 910.01 |
| 6366 | ENSMUSG00000027886 | 0 | NA | NA | NA | NA | NA | 1700013F07 | RIKEN cDNA 1700013F07 gene [Source:M | 0 | 0 | 0 | 0 | 0 | 0 |
| 6367 | ENSMUSG00000027887 | 0.874428563 | -3.186291795 | 2.489702555 | -1.279788137 | 0.200619657 | NA | Sypl2 | synaptophysin-like 2 [Source:MGI Symbol | 3.2124 | 1.0279 | 1.0062 | 0 | 0 | 0 |
| 6368 | ENSMUSG00000027889 | 457.9744921 | 0.313890942 | 0.156123686 | 2.01052736 | 0.044375403 | 0.291350811 | Ampd2 | adenosine monophosphate deaminase 2 [ | 418.42 | 375.19 | 431.68 | 581.28 | 414.43 | 526.85 |
| 6369 | ENSMUSG00000027890 | 569.546703 | 0.108861065 | 0.20611348 | 0.528160823 | 0.597387719 | 0.91454445 | Gstm4 | glutathione S-transferase, mu 4 [Source:M | 768.57 | 465.65 | 408.54 | 601.52 | 516.95 | 656.05 |
| 6370 | ENSMUSG00000027893 | 2485.57968 | -0.131575356 | 0.091148506 | -1.443527288 | 0.148871991 | 0.560897233 | Ahcyl1 | S-adenosylhomocysteine hydrolase-like 1 | 2525 | 2670.5 | 2601.1 | 2282.7 | 2455 | 2379.2 |
| 6371 | ENSMUSG00000027894 | 0 | NA | NA | NA | NA | NA | Slc6a17 | solute carrier family 6 (neurotransmitter t | 0 | 0 | 0 | 0 | 0 | 0 |
| 6372 | ENSMUSG00000027895 | 1.801759945 | -1.275967725 | 1.696123632 | -0.752284622 | 0.451879914 | NA | Kcnc4 | potassium voltage gated channel, Shaw-re | 1.6062 | 1.0279 | 5.0312 | 0.964 | 2.1812 | 0 |
| 6373 | ENSMUSG00000027896 | 32.13579768 | 0.33767554 | 0.386498166 | 0.873679541 | 0.382292796 | 0.813997203 | Slc16a4 | solute carrier family 16 (monocarboxylic a | 27.305 | 31.865 | 26.162 | 42.415 | 30.537 | 34.529 |
| 6374 | ENSMUSG00000027900 | 1193.591609 | -0.209899031 | 0.099889542 | -2.10131138 | 0.035613641 | 0.253377132 | Dram2 | DNA-damage regulated autophagy modul | 1260.1 | 1305.5 | 1275.9 | 1149.1 | 1090.6 | 1080.4 |
| 6375 | ENSMUSG00000027901 | 183.9930355 | -0.18869286 | 0.19776736 | -0.954115279 | 0.340025293 | 0.782393546 | Dennd2d | DENN/MADD domain containing 2D [Sour | 191.94 | 205.58 | 191.19 | 212.08 | 147.23 | 155.94 |
| 6376 | ENSMUSG00000027902 | 0.401551119 | -2.072982726 | 4.008251979 | -0.517178744 | 0.605031388 | NA | Chil6 | chitinase-like 6 [Source:MGI Symbol;Acc:I | 2.4093 | 0 | 0 | 0 | 0 | 0 |
| 6377 | ENSMUSG00000027905 | 200.6674892 | 0.167900613 | 0.173505523 | 0.967696068 | 0.333196183 | 0.77523112 | Ddx20 | DEAD box helicase 20 [Source:MGI Symbo | 172.67 | 208.67 | 186.16 | 212.08 | 206.13 | 218.31 |
| 6378 | ENSMUSG00000027907 | 63.14958963 | -1.019173592 | 0.38061794 | -2.677681437 | 0.007413369 | 0.087176397 | S100a11 | S100 calcium binding protein A11 [Source | 127.69 | 78.122 | 47.293 | 29.883 | 46.897 | 49.009 |
| 6379 | ENSMUSG00000027908 | 0 | NA | NA | NA | NA | NA | Tchhl1 | trichohyalin-like 1 [Source:MGI Symbol;Ac | 0 | 0 | 0 | 0 | 0 | 0 |
| 6380 | ENSMUSG00000027912 | 0 | NA | NA | NA | NA | NA | Lce1m | late cornified envelope 1M [Source:MGI S | 0 | 0 | 0 | 0 | 0 | 0 |

| | | | | | | | | | | | | | | |
|---|---|---|---|---|---|---|---|---|---|---|---|---|---|---|
| 6381 | ENSMUSG00000027913 | 0 | NA | NA | NA | NA | NA | Crct1 | cysteine-rich C-terminal 1 [Source:MGI Sy | 0 | 0 | 0 | 0 | 0 | 0 |
| 6382 | ENSMUSG00000027919 | 0 | NA | NA | NA | NA | NA | Lce1g | late cornified envelope 1G [Source:MGI Sy | 0 | 0 | 0 | 0 | 0 | 0 |
| 6383 | ENSMUSG00000027923 | 0 | NA | NA | NA | NA | NA | Lce1b | late cornified envelope 1B [Source:MGI Sy | 0 | 0 | 0 | 0 | 0 | 0 |
| 6384 | ENSMUSG00000027925 | 0 | NA | NA | NA | NA | NA | Sprr2j-ps | small proline-rich protein 2J, pseudogene | 0 | 0 | 0 | 0 | 0 | 0 |
| 6385 | ENSMUSG00000027927 | 0 | NA | NA | NA | NA | NA | Lelp1 | late cornified envelope-like proline-rich 1 | 0 | 0 | 0 | 0 | 0 | 0 |
| 6386 | ENSMUSG00000027931 | 119.9498248 | -0.412553513 | 0.259416514 | -1.590313225 | 0.111764219 | 0.490460457 | Npr1 | natriuretic peptide receptor 1 [Source:MG | 114.84 | 133.63 | 163.01 | 121.46 | 69.8 | 116.95 |
| 6387 | ENSMUSG00000027932 | 9.599879873 | -0.131483344 | 0.708952472 | -0.185461437 | 0.852867125 | 0.973781977 | Slc27a3 | solute carrier family 27 (fatty acid transpo | 8.8341 | 7.1954 | 14.087 | 7.7118 | 13.087 | 6.6831 |
| 6388 | ENSMUSG00000027933 | 451.0201006 | 0.10168254 | 0.127528229 | 0.79733358 | 0.425257323 | 0.840969906 | Ints3 | integrator complex subunit 3 [Source:MG | 416.81 | 457.42 | 431.68 | 473.31 | 462.42 | 464.47 |
| 6389 | ENSMUSG00000027935 | 98.94354877 | -0.307688254 | 0.249612651 | -1.232662899 | 0.217701559 | 0.663697152 | Rab13 | RAB13, member RAS oncogene family [So | 129.3 | 112.04 | 86.537 | 84.83 | 80.706 | 100.25 |
| 6390 | ENSMUSG00000027936 | 505.705511 | 0.258432473 | 0.142919184 | 1.808242007 | 0.070568843 | 0.384228898 | Crtc2 | CREB regulated transcription coactivator 2 | 418.42 | 446.12 | 518.22 | 613.09 | 509.32 | 529.08 |
| 6391 | ENSMUSG00000027937 | 1003.921773 | 0.069737875 | 0.10751238 | 0.648649711 | 0.516564816 | 0.886716042 | Jtb | jumping translocation breakpoint [Source | 948.46 | 940.54 | 1050.5 | 1049.8 | 1029.5 | 1004.7 |
| 6392 | ENSMUSG00000027938 | 4.216999642 | -0.739121455 | 1.219467096 | -0.606102008 | 0.544447018 | NA | Creb3l4 | cAMP responsive element binding protein | 9.6372 | 3.0838 | 3.0187 | 1.928 | 7.6343 | 0 |
| 6393 | ENSMUSG00000027939 | 0.674674027 | 1.600443604 | 2.78690385 | 0.574272989 | 0.565783085 | NA | Nup210l | nucleoporin 210-like [Source:MGI Symbol | 0 | 0 | 1.0062 | 1.928 | 0 | 1.1138 |
| 6394 | ENSMUSG00000027940 | 893.8591616 | -0.139108934 | 0.126249292 | -1.101859125 | 0.270522921 | 0.725929337 | Tpm3 | tropomyosin 3, gamma [Source:MGI Symb | 829.6 | 998.11 | 984.11 | 864.69 | 771.07 | 915.58 |
| 6395 | ENSMUSG00000027942 | 611.0159904 | 0.136189545 | 0.119973214 | 1.135166258 | 0.256305664 | 0.709512943 | 4933434E20 | RIKEN cDNA 4933434E20 gene [Source:M | 609.55 | 578.72 | 557.46 | 606.34 | 642.37 | 671.65 |
| 6396 | ENSMUSG00000027944 | 913.5655294 | 0.183722151 | 0.118923478 | 1.544877035 | 0.122376009 | 0.512759732 | Hax1 | HCLS1 associated X-1 [Source:MGI Symbo | 841.65 | 830.56 | 894.55 | 1016 | 860.5 | 1038.1 |
| 6397 | ENSMUSG00000027947 | 2107.140534 | -0.781175793 | 0.165822192 | -4.710924303 | 2.47E-06 | 0.000171479 | Il6ra | interleukin 6 receptor, alpha [Source:MGI | 2155.5 | 3179.3 | 2657.5 | 1276.3 | 1919.5 | 1454.7 |
| 6398 | ENSMUSG00000027950 | 62.02247861 | 0.060123956 | 0.316838921 | 0.1897619 | 0.849495712 | 0.9731095 | Chrnb2 | cholinergic receptor, nicotinic, beta polyp | 63.445 | 52.424 | 66.412 | 72.298 | 76.343 | 41.212 |
| 6399 | ENSMUSG00000027951 | 315.9834079 | 0.247267184 | 0.156055137 | 1.584486029 | 0.113083159 | 0.493951959 | Adar | adenosine deaminase, RNA-specific [Sour | 277.07 | 284.73 | 305.9 | 392.34 | 320.64 | 315.22 |
| 6400 | ENSMUSG00000027952 | 288.0192112 | 0.727181774 | 0.217509768 | 3.343214339 | 0.000828139 | 0.018176333 | Pmvk | phosphomevalonate kinase [Source:MGI S | 254.58 | 230.25 | 165.02 | 301.73 | 305.37 | 471.16 |
| 6401 | ENSMUSG00000027953 | 172.3754263 | -0.211766356 | 0.182591382 | -1.159782862 | 0.246137224 | 0.696492381 | Slc50a1 | solute carrier family 50 (sugar transporter | 167.85 | 197.36 | 190.18 | 157.13 | 164.68 | 157.05 |
| 6402 | ENSMUSG00000027954 | 464.5438549 | -0.065458117 | 0.171048634 | -0.382687168 | 0.70195173 | 0.942189108 | Efna1 | ephrin A1 [Source:MGI Symbol;Acc:MGI:1 | 431.27 | 429.67 | 564.5 | 414.51 | 549.67 | 397.64 |
| 6403 | ENSMUSG00000027955 | 2.476872657 | -0.076336198 | 1.336001386 | -0.057137813 | 0.954435415 | NA | Gask1b | golgi associated kinase 1B [Source:MGI Sy | 1.6062 | 2.0558 | 4.025 | 3.8559 | 1.0906 | 2.2277 |
| 6404 | ENSMUSG00000027956 | 79.37527808 | 0.26031978 | 0.273594652 | 0.951479782 | 0.341360877 | 0.783129922 | Tmem144 | transmembrane protein 144 [Source:MGI | 89.947 | 60.647 | 65.406 | 77.118 | 82.887 | 100.25 |
| 6405 | ENSMUSG00000027957 | 1560.52345 | -0.0149405 | 0.096306052 | -0.155135632 | 0.876714392 | 0.980632845 | Slc35a3 | solute carrier family 35 (UDP-N-acetylgluc | 1539.5 | 1616.9 | 1549.6 | 1533.7 | 1614.1 | 1509.3 |
| 6406 | ENSMUSG00000027959 | 13.39502689 | -0.269203621 | 0.588803641 | -0.457204409 | 0.64752413 | 0.925643431 | Sass6 | SAS-6 centriolar assembly protein [Source | 18.471 | 11.307 | 14.087 | 14.46 | 10.906 | 11.138 |
| 6407 | ENSMUSG00000027961 | 26.40714784 | -0.791863129 | 0.550938536 | -1.437298496 | 0.150633196 | 0.564591487 | Lrrc39 | leucine rich repeat containing 39 [Source: | 59.43 | 26.726 | 14.087 | 25.063 | 13.087 | 20.049 |
| 6408 | ENSMUSG00000027962 | 225.6536732 | -2.279930568 | 0.249712729 | -9.130213649 | 6.84E-20 | 8.99E-17 | Vcam1 | vascular cell adhesion molecule 1 [Source | 315.62 | 530.41 | 276.72 | 70.37 | 85.068 | 75.741 |
| 6409 | ENSMUSG00000027963 | 180.3621674 | 0.088632835 | 0.193492826 | 0.458067812 | 0.646903723 | 0.925643431 | Extl2 | exostosin-like glycosyltransferase 2 [Sourc | 167.05 | 177.83 | 180.12 | 214 | 151.6 | 191.58 |
| 6410 | ENSMUSG00000027965 | 110.8360876 | -0.154254795 | 0.245445337 | -0.628469039 | 0.529696722 | 0.888334265 | Olfm3 | olfactomedin 3 [Source:MGI Symbol;Acc:M | 106.81 | 122.32 | 121.76 | 128.21 | 109.06 | 76.855 |
| 6411 | ENSMUSG00000027966 | 0 | NA | NA | NA | NA | NA | Col11a1 | collagen, type XI, alpha 1 [Source:MGI Syr | 0 | 0 | 0 | 0 | 0 | 0 |
| 6412 | ENSMUSG00000027967 | 0 | NA | NA | NA | NA | NA | Neurog2 | neurogenin 2 [Source:MGI Symbol;Acc:MG | 0 | 0 | 0 | 0 | 0 | 0 |
| 6413 | ENSMUSG00000027968 | 147.8641802 | -0.082049439 | 0.218004586 | -0.376365654 | 0.706645076 | 0.94295036 | Larp7 | La ribonucleoprotein domain family, mem | 126.89 | 161.38 | 168.04 | 115.68 | 158.14 | 157.05 |
| 6414 | ENSMUSG00000027971 | 0 | NA | NA | NA | NA | NA | Ndst4 | N-deacetylase/N-sulfotransferase (hepari | 0 | 0 | 0 | 0 | 0 | 0 |
| 6415 | ENSMUSG00000027973 | 0 | NA | NA | NA | NA | NA | 1700006A11 | RIKEN cDNA 1700006A11 gene [Source:M | 0 | 0 | 0 | 0 | 0 | 0 |
| 6416 | ENSMUSG00000027977 | 0.17131964 | -0.796629176 | 4.080472857 | -0.195229623 | 0.84521319 | NA | Ndst3 | N-deacetylase/N-sulfotransferase (hepara | 0 | 1.0279 | 0 | 0 | 0 | 0 |
| 6417 | ENSMUSG00000027978 | 6.629151387 | 0.990599343 | 0.839423363 | 1.180095035 | 0.237962419 | 0.686509684 | Prss12 | protease, serine 12 neurotrypsin (motops | 3.2124 | 5.1396 | 5.0312 | 7.7118 | 11.997 | 6.6831 |
| 6418 | ENSMUSG00000027981 | 293.6383071 | 0.163540831 | 0.156029675 | 1.048139279 | 0.294574441 | 0.745164683 | Rnpc3 | RNA-binding region (RNP1, RRM) containi | 279.48 | 254.92 | 296.84 | 332.57 | 319.55 | 278.46 |
| 6419 | ENSMUSG00000027983 | 483.6450345 | 0.878016348 | 0.214073201 | 4.101477168 | 4.11E-05 | 0.001784488 | Cyp2u1 | cytochrome P450, family 2, subfamily u, p | 362.2 | 259.04 | 400.49 | 433.79 | 652.19 | 794.17 |
| 6420 | ENSMUSG00000027984 | 9613.792507 | -0.301796495 | 0.084421109 | -3.574893759 | 0.00035037 | 0.009859443 | Hadh | hydroxyacyl-Coenzyme A dehydrogenase | 10465 | 10860 | 10522 | 8606.4 | 8998.7 | 8230.2 |
| 6421 | ENSMUSG00000027985 | 1.589951971 | -4.031511759 | 1.994754845 | -2.021056257 | 0.043273942 | NA | Lef1 | lymphoid enhancer binding factor 1 [Sour | 2.4093 | 4.1117 | 3.0187 | 0 | 0 | 0 |
| 6422 | ENSMUSG00000027993 | 510.9213123 | -0.674601556 | 0.146167562 | -4.615261735 | 3.93E-06 | 0.000250157 | Trim2 | tripartite motif-containing 2 [Source:MGI | 567.79 | 688.7 | 628.9 | 408.73 | 337 | 434.4 |
| 6423 | ENSMUSG00000027994 | 1.804053019 | -1.304528934 | 1.646954704 | -0.792085496 | 0.428310827 | NA | Mcub | mitochondrial calcium uniporter dominan | 5.6217 | 1.0279 | 1.0062 | 0.964 | 1.0906 | 1.1138 |
| 6424 | ENSMUSG00000027995 | 27.33255127 | -0.910229764 | 0.443934666 | -2.050368746 | 0.040328461 | 0.274714427 | Tlr2 | toll-like receptor 2 [Source:MGI Symbol;A | 37.746 | 37.005 | 32.2 | 18.316 | 11.997 | 26.732 |
| 6425 | ENSMUSG00000027996 | 0.636972666 | -2.712703855 | 3.077964703 | -0.881330397 | 0.378139018 | NA | Sfrp2 | secreted frizzled-related protein 2 [Source | 0.8031 | 0 | 3.0187 | 0 | 0 | 0 |
| 6426 | ENSMUSG00000027997 | 450.65351 | 0.136009172 | 0.191046919 | 0.711915026 | 0.476517397 | 0.86953366 | Casp6 | caspase 6 [Source:MGI Symbol;Acc:MGI:1 | 468.21 | 477.98 | 341.12 | 357.64 | 484.23 | 574.74 |
| 6427 | ENSMUSG00000027998 | 358.6503135 | -0.147603967 | 0.141800131 | -1.040929693 | 0.29790818 | 0.748600402 | Plrg1 | pleiotropic regulator 1 [Source:MGI Symb | 343.73 | 385.47 | 402.5 | 348.96 | 332.64 | 338.61 |
| 6428 | ENSMUSG00000027999 | 306.9254453 | 0.058958846 | 0.162586894 | 0.362629758 | 0.716881478 | 0.9447719 | Pla2g12a | phospholipase A2, group XIIA [Source:MG | 262.61 | 323.79 | 315.96 | 287.27 | 298.83 | 353.09 |
| 6429 | ENSMUSG00000028001 | 53694.74424 | 0.265686196 | 0.128924045 | 2.060796317 | 0.039322476 | 0.271355101 | Fga | fibrinogen alpha chain [Source:MGI Symb | 41207 | 52549 | 52538 | 54570 | 53196 | 68108 |
| 6430 | ENSMUSG00000028003 | 207.1951778 | -1.271636926 | 0.306441192 | -4.149693188 | 3.33E-05 | 0.001514268 | Lrat | lecithin-retinol acyltransferase (phosphati | 376.65 | 241.56 | 260.62 | 134.96 | 55.622 | 173.76 |
| 6431 | ENSMUSG00000028004 | 0 | NA | NA | NA | NA | NA | Npy2r | neuropeptide Y receptor Y2 [Source:MGI S | 0 | 0 | 0 | 0 | 0 | 0 |
| 6432 | ENSMUSG00000028005 | 55.00651981 | -1.022825792 | 0.319788885 | -3.198440721 | 0.001381729 | 0.026611629 | Gucy1b1 | guanylate cyclase 1, soluble, beta 1 [Sourc | 76.295 | 86.345 | 58.362 | 31.811 | 34.9 | 42.326 |
| 6433 | ENSMUSG00000028007 | 309.5071757 | -0.064366959 | 0.160864523 | -0.400131479 | 0.68905968 | 0.936908509 | Snx7 | sorting nexin 7 [Source:MGI Symbol;Acc:M | 346.14 | 307.35 | 294.83 | 277.63 | 282.47 | 348.63 |
| 6434 | ENSMUSG00000028008 | 182.4579443 | -0.072965886 | 0.182993092 | -0.398735741 | 0.690087929 | 0.937399682 | Asic5 | acid-sensing (proton-gated) ion channel fa | 196.76 | 178.86 | 185.15 | 162.91 | 201.76 | 169.3 |
| 6435 | ENSMUSG00000028009 | 13.31345518 | -0.134762059 | 0.65153734 | -0.206837046 | 0.836137123 | 0.971073158 | 1700061I17F | RIKEN cDNA 1700061I17 gene [Source:MG | 14.456 | 5.1396 | 22.137 | 10.604 | 14.178 | 13.366 |
| 6436 | ENSMUSG00000028010 | 81.17302641 | 0.254320285 | 0.258627277 | 0.983346723 | 0.325436822 | 0.771532239 | Gar1 | GAR1 ribonucleoprotein [Source:MGI Sym | 66.657 | 79.15 | 76.475 | 80.974 | 104.7 | 79.083 |

| | | | | | | | | | | | | | | | |
|---|---|---|---|---|---|---|---|---|---|---|---|---|---|---|---|
| 6437 | ENSMUSG00000028011 | 30835.22185 | -0.103385241 | 0.125873424 | -0.82134288 | 0.411450989 | 0.833738738 | Tdo2 | tryptophan 2,3-dioxygenase [Source:MGI | 27386 | 36118 | 32316 | 26044 | 29813 | 33335 |
| 6438 | ENSMUSG00000028012 | 0 | NA | NA | NA | NA | NA | Rrh | retinal pigment epithelium derived rhodo | 0 | 0 | 0 | 0 | 0 | 0 |
| 6439 | ENSMUSG00000028013 | 868.3548955 | -0.154825579 | 0.109900832 | -1.408775308 | 0.158901619 | 0.580099196 | Ppa2 | pyrophosphatase (inorganic) 2 [Source:M | 908.31 | 886.07 | 949.89 | 775.04 | 836.5 | 854.32 |
| 6440 | ENSMUSG00000028015 | 1206.778195 | -0.140661253 | 0.110929068 | -1.268028806 | 0.204787664 | 0.650460832 | Ctso | cathepsin O [Source:MGI Symbol;Acc:MG | 1264.9 | 1379.5 | 1152.2 | 1108.6 | 1157.1 | 1178.4 |
| 6441 | ENSMUSG00000028016 | 112.1393807 | 0.028461347 | 0.216596728 | 0.131402481 | 0.895456926 | 0.982798755 | Ints12 | integrator complex subunit 12 [Source:M( | 113.24 | 107.93 | 111.69 | 103.15 | 124.33 | 112.5 |
| 6442 | ENSMUSG00000028017 | 16.92449278 | -0.407306038 | 0.542423754 | -0.750900075 | 0.452712795 | 0.856769845 | Egf | epidermal growth factor [Source:MGI Sym | 18.471 | 14.391 | 25.156 | 19.28 | 11.997 | 12.252 |
| 6443 | ENSMUSG00000028018 | 88.8401474 | -0.404236812 | 0.247607068 | -1.632573802 | 0.102558665 | 0.470872814 | Gstcd | glutathione S-transferase, C-terminal dom | 101.19 | 91.485 | 110.69 | 65.55 | 86.159 | 77.969 |
| 6444 | ENSMUSG00000028019 | 46.88700551 | -0.290447448 | 0.343517274 | -0.84551046 | 0.397825895 | 0.82544165 | Pdgfc | platelet-derived growth factor, C polypep | 44.974 | 59.619 | 50.312 | 42.415 | 30.537 | 53.465 |
| 6445 | ENSMUSG00000028020 | 0 | NA | NA | NA | NA | NA | Glrb | glycine receptor, beta subunit [Source:M( | 0 | 0 | 0 | 0 | 0 | 0 |
| 6446 | ENSMUSG00000028023 | 0 | NA | NA | NA | NA | NA | Pitx2 | paired-like homeodomain transcription fa | 0 | 0 | 0 | 0 | 0 | 0 |
| 6447 | ENSMUSG00000028024 | 1540.809983 | -0.100636714 | 0.163829116 | -0.614278564 | 0.539031265 | 0.89174748 | Enpep | glutamyl aminopeptidase [Source:MGI Sy | 1536.3 | 1596.4 | 1652.3 | 1937.6 | 1385.1 | 1137.2 |
| 6448 | ENSMUSG00000028028 | 17.85347411 | -0.782949836 | 0.54450917 | -1.437900184 | 0.150462377 | 0.564307976 | Alpk1 | alpha-kinase 1 [Source:MGI Symbol;Acc:N | 20.078 | 28.782 | 19.119 | 19.28 | 8.7249 | 11.138 |
| 6449 | ENSMUSG00000028029 | 511.1767084 | 0.426368191 | 0.128213066 | 3.325465984 | 0.000882709 | 0.019076633 | Aimp1 | aminoacyl tRNA synthetase complex-inter | 432.07 | 440.98 | 435.7 | 603.45 | 532.22 | 622.64 |
| 6450 | ENSMUSG00000028030 | 243.4941769 | -0.239299153 | 0.159832067 | -1.497191128 | 0.134343533 | 0.537569131 | Tbck | TBC1 domain containing kinase [Source:M | 270.65 | 266.23 | 253.57 | 205.33 | 227.94 | 237.25 |
| 6451 | ENSMUSG00000028031 | 0 | NA | NA | NA | NA | NA | Dkk2 | dickkopf WNT signaling pathway inhibitor | 0 | 0 | 0 | 0 | 0 | 0 |
| 6452 | ENSMUSG00000028032 | 241.1629459 | 0.072180199 | 0.196226465 | 0.367841305 | 0.712991568 | 0.944239215 | Papss1 | 3'-phosphoadenosine 5'-phosphosulfate s | 291.53 | 219.97 | 193.2 | 275.7 | 214.85 | 251.73 |
| 6453 | ENSMUSG00000028033 | 1.327570535 | -1.595323601 | 2.48578983 | -0.641777347 | 0.52101776 | NA | Kcnq5 | potassium voltage-gated channel, subfam | 0 | 0 | 6.0375 | 1.928 | 0 | 0 |
| 6454 | ENSMUSG00000028034 | 622.4612494 | 0.30293009 | 0.151258238 | 2.002734493 | 0.045205794 | 0.294174243 | Fubp1 | far upstream element (FUSE) binding prot | 539.68 | 534.52 | 598.72 | 797.21 | 712.17 | 552.47 |
| 6455 | ENSMUSG00000028035 | 608.8596677 | -0.029077718 | 0.132078247 | -0.220155236 | 0.825750258 | 0.96943211 | Dnajb4 | DnaJ heat shock protein family (Hsp40) m | 603.13 | 574.61 | 667.14 | 570.67 | 674 | 563.61 |
| 6456 | ENSMUSG00000028036 | 2.221914531 | -3.457772302 | 1.951140562 | -1.772180011 | 0.076364681 | NA | Ptgfr | prostaglandin F receptor [Source:MGI Syn | 7.2279 | 5.1396 | 0 | 0.964 | 0 | 0 |
| 6457 | ENSMUSG00000028037 | 69.41000356 | 0.132198652 | 0.304640615 | 0.43394953 | 0.664325091 | 0.929301301 | Ifi44 | interferon-induced protein 44 [Source:M( | 81.113 | 62.703 | 54.337 | 63.623 | 58.893 | 95.791 |
| 6458 | ENSMUSG00000028039 | 1.690152405 | 0.128848849 | 1.67316492 | 0.077009055 | 0.938616341 | NA | Efna3 | ephrin A3 [Source:MGI Symbol;Acc:MGI:1 | 0.8031 | 2.0558 | 2.0125 | 1.928 | 0 | 3.3415 |
| 6459 | ENSMUSG00000028040 | 1.468613182 | 0.608117434 | 1.798340526 | 0.338154774 | 0.735246555 | NA | Efna4 | ephrin A4 [Source:MGI Symbol;Acc:MGI:1 | 2.4093 | 0 | 1.0062 | 0.964 | 1.0906 | 3.3415 |
| 6460 | ENSMUSG00000028041 | 31.28630148 | -0.863796552 | 0.407111707 | -2.121767901 | 0.033857233 | 0.246290937 | Adam15 | a disintegrin and metallopeptidase domai | 44.974 | 39.061 | 37.231 | 27.955 | 22.903 | 15.594 |
| 6461 | ENSMUSG00000028042 | 1666.87009 | 0.449839399 | 0.146534307 | 3.069857211 | 0.002141611 | 0.036657676 | Zbtb7b | zinc finger and BTB domain containing 7B | 1568.5 | 1142 | 1515.4 | 1781.4 | 1741.7 | 2252.2 |
| 6462 | ENSMUSG00000028044 | 47.94398557 | -0.263378336 | 0.355565429 | -0.740730999 | 0.45885656 | 0.859322431 | Cks1b | CDC28 protein kinase 1b [Source:MGI Syn | 64.248 | 34.949 | 57.356 | 37.595 | 54.531 | 38.985 |
| 6463 | ENSMUSG00000028047 | 7.009053042 | 0.215107506 | 0.856926069 | 0.251022245 | 0.801796911 | 0.962796744 | Thbs3 | thrombospondin 3 [Source:MGI Symbol;A | 3.2124 | 10.279 | 6.0375 | 4.8199 | 5.4531 | 12.252 |
| 6464 | ENSMUSG00000028048 | 313.0083824 | -0.050548915 | 0.149445279 | -0.338243642 | 0.73517959 | 0.948109711 | Gba | glucosidase, beta, acid [Source:MGI Symb | 325.26 | 330.99 | 298.85 | 286.3 | 335.91 | 300.74 |
| 6465 | ENSMUSG00000028049 | 575.7342485 | -0.026449016 | 0.118876413 | -0.222491706 | 0.823931129 | 0.96943211 | Scamp3 | secretory carrier membrane protein 3 [So | 582.25 | 559.19 | 601.73 | 588.03 | 554.03 | 569.17 |
| 6466 | ENSMUSG00000028051 | 968.9704163 | 0.046447957 | 0.190879578 | 0.24333644 | 0.807744777 | 0.965081255 | Hcn3 | hyperpolarization-activated, cyclic nucleo | 1108.3 | 889.15 | 862.35 | 1063.3 | 1228 | 662.74 |
| 6467 | ENSMUSG00000028053 | 629.515793 | 0.103833102 | 0.133801594 | 0.77602291 | 0.437735459 | 0.847422601 | Ash1l | ASH1 like histone lysine methyltransferase | 530.85 | 675.34 | 615.82 | 688.28 | 610.75 | 656.05 |
| 6468 | ENSMUSG00000028057 | 184.1095058 | -0.018567978 | 0.173558488 | -0.106983984 | 0.914801686 | 0.98625062 | Rit1 | Ras-like without CAAX 1 [Source:MGI Sym | 189.53 | 182.97 | 183.14 | 175.44 | 188.68 | 184.9 |
| 6469 | ENSMUSG00000028059 | 44.05034573 | -0.732563608 | 0.347108417 | -2.110474917 | 0.034817469 | 0.250037719 | Arhgef2 | rho/rac guanine nucleotide exchange fact | 56.217 | 59.619 | 49.306 | 39.523 | 23.994 | 35.643 |
| 6470 | ENSMUSG00000028060 | 818.1365126 | 0.099074215 | 0.126404417 | 0.783787604 | 0.433164749 | 0.845264432 | Khdc4 | KH domain containing 4, pre-mRNA splicir | 827.2 | 794.58 | 748.65 | 935.06 | 864.86 | 738.48 |
| 6471 | ENSMUSG00000028062 | 1067.874088 | 0.077224834 | 0.117651901 | 0.656384071 | 0.511577037 | 0.883833046 | Lamtor2 | late endosomal/lysosomal adaptor, MAPK | 1011.9 | 1027.9 | 1077.7 | 1006.4 | 1048.1 | 1235.3 |
| 6472 | ENSMUSG00000028063 | 804.5782477 | 0.071098085 | 0.121736785 | 0.58403124 | 0.559199294 | 0.899423311 | Lmna | lamin A [Source:MGI Symbol;Acc:MGI:967 | 840.04 | 791.5 | 721.48 | 755.76 | 835.41 | 883.28 |
| 6473 | ENSMUSG00000028064 | 1426.671313 | 0.211947044 | 0.101017495 | 2.098122152 | 0.035894356 | 0.254648556 | Sema4a | sema domain, immunoglobulin domain (Ig | 1374.1 | 1293.1 | 1298.1 | 1495.1 | 1474.5 | 1625.1 |
| 6474 | ENSMUSG00000028066 | 283.3829933 | -0.047081336 | 0.151526839 | -0.31071285 | 0.756018928 | 0.953085143 | Pmf1 | polyamine-modulated factor 1 [Source:M( | 293.13 | 283.71 | 286.78 | 258.35 | 302.1 | 276.23 |
| 6475 | ENSMUSG00000028068 | 1.717042116 | 0.500813244 | 1.809606202 | 0.276752612 | 0.78197007 | NA | Iqgap3 | IQ motif containing GTPase activating pro | 3.2124 | 0 | 1.0062 | 3.8559 | 0 | 2.2277 |
| 6476 | ENSMUSG00000028069 | 183.9157745 | -0.127560384 | 0.204927929 | -0.622464615 | 0.533636404 | 0.889143097 | Gpatch4 | G patch domain containing 4 [Source:MGI | 182.3 | 194.28 | 199.24 | 136.88 | 217.03 | 173.76 |
| 6477 | ENSMUSG00000028070 | 914.8591029 | -0.07801227 | 0.111470933 | -0.699844057 | 0.484024697 | 0.872431004 | Naxe | NAD(P)HX epimerase [Source:MGI Symbo | 885.02 | 1011.5 | 922.73 | 857.94 | 899.76 | 912.24 |
| 6478 | ENSMUSG00000028071 | 3.495782755 | -4.129065057 | 1.574847919 | -2.621881774 | 0.008744576 | NA | Sh2d2a | SH2 domain containing 2A [Source:MGI S | 5.6217 | 7.1954 | 7.0437 | 0 | 0 | 1.1138 |
| 6479 | ENSMUSG00000028072 | 15.66017731 | 0.064099929 | 0.61909645 | 0.103537872 | 0.917536094 | 0.986374671 | Ntrk1 | neurotrophic tyrosine kinase, receptor, ty | 11.243 | 25.698 | 9.0562 | 10.604 | 23.994 | 13.366 |
| 6480 | ENSMUSG00000028073 | 124.0600795 | -0.833734735 | 0.26426015 | -3.154977155 | 0.001605107 | 0.029802649 | Pear1 | platelet endothelial aggregation receptor | 168.65 | 163.44 | 144.9 | 107 | 53.44 | 106.93 |
| 6481 | ENSMUSG00000028076 | 4571.483451 | -0.046989469 | 0.099080264 | -0.474256603 | 0.635316934 | 0.923478764 | Cd1d1 | CD1d1 antigen [Source:MGI Symbol;Acc:N | 4331.1 | 4847.7 | 4760.5 | 4691.7 | 4672.2 | 4125.7 |
| 6482 | ENSMUSG00000028078 | 12.26790958 | -0.756375681 | 0.732515326 | -1.032573183 | 0.301803683 | 0.752246851 | Dclk2 | doublecortin-like kinase 2 [Source:MGI Sy | 12.047 | 11.307 | 23.144 | 18.316 | 5.4531 | 3.3415 |
| 6483 | ENSMUSG00000028080 | 463.1941324 | 0.162161879 | 0.126268247 | 1.284264915 | 0.199049279 | 0.641213533 | Lrba | LPS-responsive beige-like anchor [Source: | 428.05 | 440.98 | 442.75 | 485.84 | 511.5 | 470.04 |
| 6484 | ENSMUSG00000028081 | 4771.537057 | 0.027962554 | 0.093914872 | 0.297743618 | 0.76589885 | 0.954996635 | Rps3a1 | ribosomal protein S3A1 [Source:MGI Syml | 4341.6 | 4884.7 | 4950.7 | 4692.6 | 4806.4 | 4953.3 |
| 6485 | ENSMUSG00000028082 | 1497.470592 | 0.445848896 | 0.110400084 | 4.038483305 | 5.38E-05 | 0.002221919 | Sh3d19 | SH3 domain protein D19 [Source:MGI Syn | 1215.1 | 1224.3 | 1364.5 | 1640.7 | 1649 | 1891.3 |
| 6486 | ENSMUSG00000028085 | 831.3290758 | -0.160539672 | 0.124953584 | -1.284794456 | 0.19886412 | 0.640976045 | Gatb | glutamyl-tRNA(Gln) amidotransferase, sul | 949.27 | 901.48 | 780.85 | 739.37 | 850.68 | 766.32 |
| 6487 | ENSMUSG00000028086 | 194.8574086 | 0.021082724 | 0.174602264 | 0.120747137 | 0.903891322 | 0.983821605 | Fbxw7 | F-box and WD-40 domain protein 7 [Sourc | 195.96 | 186.05 | 198.23 | 194.72 | 214.85 | 179.33 |
| 6488 | ENSMUSG00000028088 | 21857.36834 | -0.850458615 | 0.134211622 | -6.336698723 | 2.35E-10 | 6.11E-08 | Fmo5 | flavin containing monooxygenase 5 [Sourc | 25548 | 32242 | 26570 | 17290 | 16796 | 12699 |
| 6489 | ENSMUSG00000028089 | 280.7783905 | -0.546120152 | 0.17327576 | -3.151740052 | 0.001623007 | 0.030019402 | Chd1l | chromodomain helicase DNA binding prot | 383.08 | 342.3 | 273.7 | 214 | 241.03 | 230.57 |
| 6490 | ENSMUSG00000028093 | 736.772508 | 0.211729375 | 0.112787055 | 1.877248895 | 0.060483989 | 0.350918739 | Acp6 | acid phosphatase 6, lysophosphatidic [Sou | 673 | 673.29 | 702.36 | 792.39 | 828.87 | 750.73 |
| 6491 | ENSMUSG00000028096 | 400.256124 | 0.162743523 | 0.156220798 | 1.041753245 | 0.297526096 | 0.748091545 | Gpr89 | G protein-coupled receptor 89 [Source:M( | 418.42 | 359.77 | 354.2 | 409.69 | 490.78 | 368.68 |
| 6492 | ENSMUSG00000028098 | 690.675893 | 0.010244473 | 0.126205246 | 0.081173116 | 0.935304279 | 0.988420305 | Rnf115 | ring finger protein 115 [Source:MGI Symb | 640.07 | 704.12 | 720.47 | 618.87 | 719.81 | 740.71 |

| | | | | | | | | | | | | | | | |
|---|---|---|---|---|---|---|---|---|---|---|---|---|---|---|---|
| 6493 | ENSMUSG00000028099 | 394.4035235 | 0.062880161 | 0.142317676 | 0.441829592 | 0.658612523 | 0.927599242 | Polr3c | polymerase (RNA) III (DNA directed) polyp | 349.35 | 401.92 | 406.52 | 374.02 | 402.44 | 432.17 |
| 6494 | ENSMUSG00000028100 | 2.380200682 | 1.52104811 | 1.44764007 | 1.050708765 | 0.293392369 | NA | Nudt17 | nudix (nucleoside diphosphate linked moi | 1.6062 | 2.0558 | 0 | 2.8919 | 3.2719 | 4.4554 |
| 6495 | ENSMUSG00000028101 | 167.6065504 | 0.426173488 | 0.240809632 | 1.769752665 | 0.076768352 | 0.405543261 | Pias3 | protein inhibitor of activated STAT 3 [Sour | 145.36 | 107.93 | 176.09 | 247.74 | 160.32 | 168.19 |
| 6496 | ENSMUSG00000028102 | 398.983095 | 0.228707872 | 0.143899126 | 1.58936248 | 0.11197858 | 0.490954285 | Pex11b | peroxisomal biogenesis factor 11 beta [So | 383.08 | 333.05 | 385.39 | 399.09 | 418.8 | 474.5 |
| 6497 | ENSMUSG00000028104 | 205.1354345 | -0.044062344 | 0.18794639 | -0.234441023 | 0.814642614 | 0.967404825 | Polr3gl | polymerase (RNA) III (DNA directed) polyp | 222.46 | 192.22 | 209.3 | 163.88 | 225.76 | 217.2 |
| 6498 | ENSMUSG00000028106 | 365.5129475 | -0.210051956 | 0.139519317 | -1.505540313 | 0.132185221 | 0.533232592 | Rprd2 | regulation of nuclear pre-mRNA domain c | 387.9 | 417.33 | 371.3 | 348.96 | 347.91 | 319.67 |
| 6499 | ENSMUSG00000028107 | 441.547096 | 0.140353577 | 0.132023076 | 1.063098827 | 0.287737136 | 0.73951397 | Tars2 | threonyl-tRNA synthetase 2, mitochondria | 448.13 | 393.69 | 417.59 | 447.29 | 458.06 | 484.52 |
| 6500 | ENSMUSG00000028108 | 1739.818423 | -1.562212902 | 0.109003102 | -14.33182058 | 1.38E-46 | 2.00E-42 | Ecm1 | extracellular matrix protein 1 [Source:MG | 2764.3 | 2573.9 | 2459.3 | 866.62 | 813.6 | 961.25 |
| 6501 | ENSMUSG00000028109 | 0.167707431 | -0.796629176 | 4.080472857 | -0.195229623 | 0.84521319 | NA | Hormad1 | HORMA domain containing 1 [Source:MG | 0 | 0 | 1.0062 | 0 | 0 | 0 |
| 6502 | ENSMUSG00000028111 | 10.16392143 | -0.58295958 | 0.694684724 | -0.839171439 | 0.401373112 | 0.82816843 | Ctsk | cathepsin K [Source:MGI Symbol;Acc:MGI | 16.062 | 10.279 | 10.062 | 5.7839 | 6.5437 | 12.252 |
| 6503 | ENSMUSG00000028114 | 133.8697894 | 0.080702565 | 0.210141884 | 0.384038457 | 0.700949944 | 0.941805852 | Mettl14 | methyltransferase like 14 [Source:MGI Syr | 132.51 | 137.74 | 119.74 | 116.64 | 142.87 | 153.71 |
| 6504 | ENSMUSG00000028115 | 0.160662974 | 1.126924175 | 4.080472857 | 0.276174898 | 0.782413736 | NA | Bnipl | BCL2/adenovirus E1B 19kD interacting prc | 0 | 0 | 0 | 0.964 | 0 | 0 |
| 6505 | ENSMUSG00000028116 | 0 | NA | NA | NA | NA | NA | Myoz2 | myozenin 2 [Source:MGI Symbol;Acc:MGI | 0 | 0 | 0 | 0 | 0 | 0 |
| 6506 | ENSMUSG00000028121 | 903.9242081 | 0.166312186 | 0.122831148 | 1.353990321 | 0.175739466 | 0.606368233 | Bcar3 | breast cancer anti-estrogen resistance 3 [! | 897.87 | 742.16 | 914.68 | 919.63 | 942.29 | 1006.9 |
| 6507 | ENSMUSG00000028124 | 4497.516401 | -0.007126192 | 0.095031924 | -0.074987348 | 0.940224779 | 0.988812319 | Gclm | glutamate-cysteine ligase, modifier subun | 4280.5 | 4322.4 | 4923.6 | 4447.8 | 4454.1 | 4556.7 |
| 6508 | ENSMUSG00000028125 | 3.431370518 | -1.166167594 | 1.19155305 | -0.978695488 | 0.327730462 | NA | Abca4 | ATP-binding cassette, sub-family A (ABC1) | 4.0155 | 8.2233 | 2.0125 | 1.928 | 2.1812 | 2.2277 |
| 6509 | ENSMUSG00000028126 | 204.6429292 | 0.060791955 | 0.17562311 | 0.346150087 | 0.729229924 | 0.946534338 | Pip5k1a | phosphatidylinositol-4-phosphate 5-kinas | 200.78 | 195.3 | 205.27 | 236.17 | 186.5 | 203.83 |
| 6510 | ENSMUSG00000028127 | 11953.43204 | 0.075528852 | 0.096769213 | 0.780504976 | 0.4350937 | 0.845325456 | Abcd3 | ATP-binding cassette, sub-family D (ALD), | 11186 | 12143 | 11594 | 11918 | 13464 | 11417 |
| 6511 | ENSMUSG00000028128 | 107.1876472 | 0.86671035 | 0.294343183 | 2.944557235 | 0.003234171 | 0.049241108 | F3 | coagulation factor III [Source:MGI Symbol | 109.22 | 47.284 | 70.437 | 163.88 | 129.78 | 122.52 |
| 6512 | ENSMUSG00000028132 | 5328.606163 | -0.291861692 | 0.088821558 | -3.28593304 | 0.001016452 | 0.021093196 | Tlcd4 | TLC domain containing 4 [Source:MGI Sym | 5989.5 | 5875.6 | 5732.6 | 5018.5 | 4850 | 4505.5 |
| 6513 | ENSMUSG00000028133 | 106.2224378 | 0.136817933 | 0.233531244 | 0.585865646 | 0.557965805 | 0.899217185 | Rwdd3 | RWD domain containing 3 [Source:MGI Sy | 91.554 | 106.9 | 105.66 | 126.28 | 115.61 | 91.335 |
| 6514 | ENSMUSG00000028134 | 148.0726613 | 0.161190659 | 0.191567751 | 0.84142899 | 0.40010765 | 0.82709119 | Ptbp2 | polypyrimidine tract binding protein 2 [So | 143.76 | 142.88 | 132.82 | 165.8 | 147.23 | 155.94 |
| 6515 | ENSMUSG00000028136 | 585.136514 | 0.062076392 | 0.122470597 | 0.506867716 | 0.612247639 | 0.916441371 | Snx27 | sorting nexin family member 27 [Source:N | 557.35 | 575.63 | 584.63 | 563.93 | 646.74 | 582.54 |
| 6516 | ENSMUSG00000028137 | 0.663785267 | -1.400021131 | 3.106754789 | -0.450637796 | 0.652250621 | NA | Celf3 | CUGBP, Elav-like family member 3 [Source | 0 | 0 | 3.0187 | 0.964 | 0 | 0 |
| 6517 | ENSMUSG00000028138 | 8846.429104 | -0.003642039 | 0.09775566 | -0.037256552 | 0.970280448 | 0.994966572 | Adh5 | alcohol dehydrogenase 5 (class III), chi po | 8433.4 | 8975.8 | 9163.9 | 8300.8 | 9725 | 8479.7 |
| 6518 | ENSMUSG00000028139 | 0 | NA | NA | NA | NA | NA | Riiad1 | regulatory subunit of type II PKA R-subuni | 0 | 0 | 0 | 0 | 0 | 0 |
| 6519 | ENSMUSG00000028140 | 826.1260648 | 0.142754399 | 0.11345354 | 1.258263067 | 0.208296633 | 0.653455953 | Mrpl9 | mitochondrial ribosomal protein L9 [Sourc | 768.57 | 830.56 | 756.7 | 819.38 | 917.21 | 864.34 |
| 6520 | ENSMUSG00000028141 | 2.406781189 | 0.552529339 | 1.350467613 | 0.409139274 | 0.682437456 | NA | Oaz3 | ornithine decarboxylase antizyme 3 [Sour | 0.8031 | 3.0838 | 2.0125 | 1.928 | 3.2719 | 3.3415 |
| 6521 | ENSMUSG00000028145 | 371.4658209 | -0.154154172 | 0.143550257 | -1.073869006 | 0.282881396 | 0.735634037 | Them4 | thioesterase superfamily member 4 [Sour | 357.38 | 413.22 | 403.5 | 323.9 | 360.99 | 369.8 |
| 6522 | ENSMUSG00000028148 | 1.993843852 | 4.520878464 | 1.938270668 | 2.332428871 | 0.019678141 | NA | Them5 | thioesterase superfamily member 5 [Sour | 0 | 0 | 0 | 0.964 | 6.5437 | 4.4554 |
| 6523 | ENSMUSG00000028149 | 485.2442882 | -0.071802603 | 0.138682432 | -0.517748368 | 0.604633847 | 0.916376001 | Rap1gds1 | RAP1, GTP-GDP dissociation stimulator 1 [ | 445.72 | 563.3 | 484 | 490.66 | 473.33 | 454.45 |
| 6524 | ENSMUSG00000028150 | 2432.618174 | -0.708182213 | 0.316917376 | -2.234595721 | 0.025443902 | 0.205313187 | Rorc | RAR-related orphan receptor gamma [Sou | 2155.5 | 3302.7 | 3596.3 | 2303.9 | 1967.5 | 1269.8 |
| 6525 | ENSMUSG00000028152 | 9.057658321 | -0.201730822 | 0.744009824 | -0.271139997 | 0.78628336 | 0.960870081 | Tspan5 | tetraspanin 5 [Source:MGI Symbol;Acc:M | 13.653 | 9.2513 | 6.0375 | 6.7478 | 13.087 | 5.5692 |
| 6526 | ENSMUSG00000028156 | 804.5628574 | -0.074227288 | 0.111689274 | -0.664587434 | 0.506314409 | 0.881592135 | Eif4e | eukaryotic translation initiation factor 4E | | 836.03 | 824.39 | 815.06 | 762.51 | 839.78 | 749.62 |
| 6527 | ENSMUSG00000028158 | 6424.670231 | -0.111323776 | 0.110193771 | -1.010254711 | 0.312373273 | 0.759830222 | Mttp | microsomal triglyceride transfer protein [ | 6996.6 | 6882.9 | 6138.1 | 6774.8 | 6327.8 | 5427.8 |
| 6528 | ENSMUSG00000028159 | 8.935322231 | -1.095523212 | 0.882059854 | -1.242005525 | 0.214234524 | 0.658874792 | Dapp1 | dual adaptor for phosphotyrosine and 3-p | 4.8186 | 26.726 | 5.0312 | 3.8559 | 8.7249 | 4.4554 |
| 6529 | ENSMUSG00000028161 | 894.7498507 | 0.031331295 | 0.14347912 | 0.218368321 | 0.827142146 | 0.969748238 | Ppp3ca | protein phosphatase 3, catalytic subunit, a | 813.54 | 968.3 | 873.42 | 790.46 | 1085.2 | 837.61 |
| 6530 | ENSMUSG00000028163 | 478.7930615 | -0.101675668 | 0.142669462 | -0.712665956 | 0.476052487 | 0.869483933 | Nfkb1 | nuclear factor of kappa light polypeptide g | 425.64 | 523.21 | 539.35 | 484.88 | 432.98 | 466.7 |
| 6531 | ENSMUSG00000028164 | 385.9088309 | 0.067017171 | 0.155322174 | 0.431472014 | 0.666125197 | 0.929596279 | Manba | mannosidase, beta A, lysosomal [Source:N | 410.39 | 319.68 | 400.49 | 426.08 | 404.62 | 354.2 |
| 6532 | ENSMUSG00000028165 | 770.5442575 | 0.04984968 | 0.11685803 | 0.426583265 | 0.669682889 | 0.930837335 | Cisd2 | CDGSH iron sulfur domain 2 [Source:MGI | 746.08 | 763.74 | 761.73 | 739.37 | 860.5 | 751.84 |
| 6533 | ENSMUSG00000028167 | 308.6239796 | 0.262490116 | 0.200134205 | 1.311570481 | 0.189665099 | 0.626663447 | Bdh2 | 3-hydroxybutyrate dehydrogenase, type 2 | 235.31 | 242.59 | 364.26 | 287.27 | 392.62 | 329.7 |
| 6534 | ENSMUSG00000028172 | 0 | NA | NA | NA | NA | NA | Tacr3 | tachykinin receptor 3 [Source:MGI Symbo | 0 | 0 | 0 | 0 | 0 | 0 |
| 6535 | ENSMUSG00000028173 | 64.07577626 | -0.433345034 | 0.319547526 | -1.356120761 | 0.175060748 | 0.605036718 | Wls | wntless WNT ligand secretion mediator [S | 69.067 | 72.982 | 78.487 | 33.739 | 58.893 | 71.286 |
| 6536 | ENSMUSG00000028174 | 0 | NA | NA | NA | NA | NA | Rpe65 | retinal pigment epithelium 65 [Source:MG | 0 | 0 | 0 | 0 | 0 | 0 |
| 6537 | ENSMUSG00000028175 | 0.185640647 | 1.126924175 | 4.080472857 | 0.276174898 | 0.782413736 | NA | Depdc1a | DEP domain containing 1a [Source:MGI Sy | 0 | 0 | 0 | 0 | 0 | 1.1138 |
| 6538 | ENSMUSG00000028176 | 0 | NA | NA | NA | NA | NA | Lrrc7 | leucine rich repeat containing 7 [Source:N | 0 | 0 | 0 | 0 | 0 | 0 |
| 6539 | ENSMUSG00000028177 | 5.747521427 | 0.290485368 | 1.021301472 | 0.284426662 | 0.776083429 | 0.95893551 | 1810013D15 | RIKEN cDNA 1810013D15 gene [Source:M | 7.2279 | 8.2233 | 0 | 5.7839 | 5.4531 | 7.7969 |
| 6540 | ENSMUSG00000028179 | 5772.702447 | 0.222701124 | 0.152369397 | 1.461586966 | 0.143854428 | 0.554054867 | Cth | cystathionase (cystathionine gamma-lyase | 4732.7 | 4935 | 6316.2 | 4844 | 6643 | 7165.4 |
| 6541 | ENSMUSG00000028180 | 824.1948032 | 0.122638031 | 0.125594232 | 0.976462286 | 0.328835429 | 0.77362974 | Zranb2 | zinc finger, RAN-binding domain containir | 808.72 | 800.75 | 758.71 | 981.33 | 807.06 | 788.6 |
| 6542 | ENSMUSG00000028182 | 29.9603218 | 0.008005324 | 0.397591963 | 0.02013452 | 0.983936062 | 0.998580735 | Lrriq3 | leucine-rich repeats and IQ motif containi | 34.533 | 30.838 | 24.15 | 31.811 | 28.356 | 30.074 |
| 6543 | ENSMUSG00000028184 | 783.2866125 | 0.175862382 | 0.138991663 | 1.265272884 | 0.205773523 | 0.650986055 | Adgrl2 | adhesion G protein-coupled receptor L2 [! | 845.67 | 748.32 | 611.8 | 859.87 | 781.97 | 852.09 |
| 6544 | ENSMUSG00000028185 | 571.7892565 | -0.194885176 | 0.125029778 | -1.558710092 | 0.119065012 | 0.504735151 | Dnase2b | deoxyribonuclease II beta [Source:MGI Sy | 591.89 | 661.98 | 577.58 | 534.04 | 556.22 | 509.03 |
| 6545 | ENSMUSG00000028186 | 45615.65643 | 0.244461979 | 0.09737511 | 2.510518135 | 0.012055413 | 0.123055394 | Uox | urate oxidase [Source:MGI Symbol;Acc:M | 40386 | 40840 | 44053 | 44560 | 52809 | 51045 |
| 6546 | ENSMUSG00000028187 | 342.0753853 | 0.064571204 | 0.141375862 | 0.456734287 | 0.647862043 | 0.925643431 | Rpf1 | ribosome production factor 1 homolog [S | 317.23 | 354.63 | 332.06 | 366.31 | 340.27 | 341.95 |
| 6547 | ENSMUSG00000028188 | 39.47743867 | 0.385507063 | 0.386400743 | 0.997687169 | 0.318431077 | 0.765080913 | Spata1 | spermatogenesis associated 1 [Source:MG | 40.155 | 38.033 | 24.15 | 30.847 | 47.987 | 55.692 |
| 6548 | ENSMUSG00000028189 | 250.2492244 | -0.014542473 | 0.158509644 | -0.091745035 | 0.926900616 | 0.987712228 | Ctbs | chitobiase [Source:MGI Symbol;Acc:MGI: | 258.6 | 265.2 | 230.43 | 238.1 | 255.2 | 253.96 |

| | | | | | | | | | | | | | | | |
|---|---|---|---|---|---|---|---|---|---|---|---|---|---|---|---|
| 6549 | ENSMUSG00000028191 | 230.7383694 | -0.025844561 | 0.176790152 | -0.146187787 | 0.883773147 | 0.981769403 | Bcl10 | B cell leukemia/lymphoma 10 [Source:MG | 246.55 | 249.78 | 202.26 | 259.31 | 212.67 | 213.86 |
| 6550 | ENSMUSG00000028194 | 2737.04156 | -0.057910347 | 0.094639757 | -0.611902957 | 0.540601965 | 0.893324212 | Ddah1 | dimethylarginine dimethylaminohydrolase | 2765.9 | 2985.1 | 2625.3 | 2697.2 | 2714.5 | 2634.2 |
| 6551 | ENSMUSG00000028195 | 112.6201146 | -0.091684775 | 0.244684482 | -0.37470613 | 0.707879033 | 0.943550198 | Ccn1 | cellular communication network factor 1 [ | 116.45 | 90.457 | 141.88 | 126.28 | 97.065 | 103.59 |
| 6552 | ENSMUSG00000028197 | 5.910720945 | -1.201774246 | 0.98229284 | -1.223437856 | 0.221164362 | 0.66763854 | Col24a1 | collagen, type XXIV, alpha 1 [Source:MGI S | 2.4093 | 9.2513 | 13.081 | 1.928 | 5.4531 | 3.3415 |
| 6553 | ENSMUSG00000028199 | 2235.40276 | -0.211745048 | 0.113013819 | -1.873620849 | 0.060982702 | 0.352269825 | Cryz | crystallin, zeta [Source:MGI Symbol;Acc:M | 2534.6 | 2547.2 | 2115.1 | 2035 | 2234.7 | 1945.9 |
| 6554 | ENSMUSG00000028207 | 535.4984117 | 0.075420162 | 0.122713703 | 0.614602606 | 0.538817193 | 0.8916993 | Asph | aspartate-beta-hydroxylase [Source:MGI S | 495.51 | 519.1 | 550.42 | 549.47 | 563.85 | 534.65 |
| 6555 | ENSMUSG00000028211 | 1385.441796 | -1.188943022 | 0.196701464 | -6.044403508 | 1.50E-09 | 3.01E-07 | Trp53inp1 | transformation related protein 53 inducib | 1766 | 2435.1 | 1576.8 | 684.42 | 1141.9 | 708.4 |
| 6556 | ENSMUSG00000028212 | 46.10974333 | 0.071596115 | 0.336069205 | 0.213039798 | 0.831295921 | 0.97052742 | Ccne2 | cyclin E2 [Source:MGI Symbol;Acc:MGI:13 | 51.399 | 37.005 | 46.287 | 46.271 | 56.712 | 38.985 |
| 6557 | ENSMUSG00000028214 | 32.03279116 | -1.580695961 | 0.437011772 | -3.617055793 | 0.000297973 | 0.00865413 | Gem | GTP binding protein (gene overexpressed | 57.823 | 29.81 | 56.35 | 17.352 | 15.269 | 15.594 |
| 6558 | ENSMUSG00000028217 | 0.513958919 | -2.394274821 | 3.756096013 | -0.637437066 | 0.523840191 | NA | Cdh17 | cadherin 17 [Source:MGI Symbol;Acc:MGI | 0 | 3.0838 | 0 | 0 | 0 | 0 |
| 6559 | ENSMUSG00000028218 | 234.7418854 | -0.047585036 | 0.176111436 | -0.270198446 | 0.787007589 | 0.960870081 | Cibar1 | CBY1 interacting BAR domain containing 1 | 275.46 | 228.2 | 211.31 | 218.82 | 254.11 | 220.54 |
| 6560 | ENSMUSG00000028221 | 269.4638073 | -0.163690519 | 0.16338919 | -1.001844245 | 0.316418824 | 0.763155505 | Pip4p2 | phosphatidylinositol-4,5-bisphosphate 4-p | 252.17 | 323.79 | 278.73 | 242.92 | 256.3 | 262.87 |
| 6561 | ENSMUSG00000028222 | 0.945986982 | 0.197573854 | 2.252218024 | 0.087724124 | 0.930095945 | NA | Calb1 | calbindin 1 [Source:MGI Symbol;Acc:MGI: | 1.6062 | 1.0279 | 0 | 1.928 | 0 | 1.1138 |
| 6562 | ENSMUSG00000028223 | 4953.293565 | -0.136559073 | 0.094455791 | -1.445745897 | 0.148248486 | 0.560006817 | Decr1 | 2,4-dienoyl CoA reductase 1, mitochondri | 5325.4 | 5228 | 5009.1 | 4743.7 | 5050.7 | 4362.9 |
| 6563 | ENSMUSG00000028224 | 362.1411659 | 0.185868156 | 0.15374702 | 1.208922005 | 0.226692811 | 0.673420584 | Nbn | nibrin [Source:MGI Symbol;Acc:MGI:1351 | 343.73 | 342.3 | 331.05 | 450.18 | 341.36 | 364.23 |
| 6564 | ENSMUSG00000028226 | 0 | NA | NA | NA | NA | NA | Mmp16 | matrix metallopeptidase 16 [Source:MGI S | 0 | 0 | 0 | 0 | 0 | 0 |
| 6565 | ENSMUSG00000028228 | 998.5621781 | 0.169057637 | 0.10548298 | 1.602700609 | 0.109000769 | 0.485738292 | Cpne3 | copine III [Source:MGI Symbol;Acc:MGI:1 | 907.51 | 924.1 | 989.14 | 1043 | 1090.6 | 1037 |
| 6566 | ENSMUSG00000028229 | 585.1764445 | -0.00169126 | 0.133833832 | -0.012637016 | 0.989917389 | 0.99859176 | Rmdn1 | regulator of microtubule dynamics 1 [Sou | 584.66 | 549.94 | 621.86 | 566.82 | 659.82 | 527.96 |
| 6567 | ENSMUSG00000028232 | 181.5766711 | 0.087489769 | 0.192253087 | 0.455076019 | 0.64905455 | 0.925643431 | Tmem68 | transmembrane protein 68 [Source:MGI S | 179.89 | 193.25 | 154.96 | 178.34 | 215.94 | 167.08 |
| 6568 | ENSMUSG00000028233 | 295.2679208 | 0.041364815 | 0.149977064 | 0.275807609 | 0.782695839 | 0.960670212 | Tgs1 | trimethylguanosine synthase 1 [Source:M | 273.05 | 313.51 | 286.78 | 284.37 | 306.46 | 307.42 |
| 6569 | ENSMUSG00000028234 | 2320.244961 | -0.029830904 | 0.09206889 | -0.324006345 | 0.745933231 | 0.949997329 | Rps20 | ribosomal protein S20 [Source:MGI Symbo | 2282.4 | 2374.5 | 2375.7 | 2202.7 | 2292.5 | 2393.7 |
| 6570 | ENSMUSG00000028236 | 0.730949792 | 3.072456834 | 2.934263219 | 1.04709653 | 0.29505506 | NA | Sdr16c5 | short chain dehydrogenase/reductase fan | 0 | 0 | 0 | 0 | 3.2719 | 1.1138 |
| 6571 | ENSMUSG00000028238 | 26.45654575 | -1.231247076 | 0.434821917 | -2.831612272 | 0.004631397 | 0.063460159 | Atp6v0d2 | ATPase, H+ transporting, lysosomal V0 sul | 40.958 | 39.061 | 31.194 | 14.46 | 16.359 | 16.708 |
| 6572 | ENSMUSG00000028240 | 2514.840553 | 0.070527776 | 0.574879574 | 0.122682696 | 0.902358367 | 0.983821605 | Cyp7a1 | cytochrome P450, family 7, subfamily a, p | 1592.6 | 4151.8 | 1616 | 2516 | 3664.5 | 1548.2 |
| 6573 | ENSMUSG00000028243 | 82.10407173 | -0.229234401 | 0.283718013 | -0.807965623 | 0.419110372 | 0.837591673 | Ubxn2b | UBX domain protein 2B [Source:MGI Syml | 73.082 | 78.122 | 114.71 | 65.55 | 68.709 | 92.449 |
| 6574 | ENSMUSG00000028244 | 0 | NA | NA | NA | NA | NA | 1700025O08 | RIKEN cDNA 1700025O08 gene [Source:M | 0 | 0 | 0 | 0 | 0 | 0 |
| 6575 | ENSMUSG00000028245 | 385.9523978 | 0.136075161 | 0.147330836 | 0.923602726 | 0.355693194 | 0.795525652 | Nsmaf | neutral sphingomyelinase (N-SMase) activ | 367.02 | 409.11 | 327.03 | 386.56 | 388.26 | 437.74 |
| 6576 | ENSMUSG00000028246 | 0 | NA | NA | NA | NA | NA | Faxc | failed axon connections homolog [Source: | 0 | 0 | 0 | 0 | 0 | 0 |
| 6577 | ENSMUSG00000028247 | 482.0574845 | 0.048874086 | 0.140952489 | 0.346741558 | 0.728785486 | 0.946248466 | Coq3 | coenzyme Q3 methyltransferase [Source:I | 464.19 | 475.93 | 480.98 | 418.37 | 509.32 | 543.56 |
| 6578 | ENSMUSG00000028248 | 958.923042 | 0.203328256 | 0.146955672 | 1.383602641 | 0.166480159 | 0.5925379 | Pnisr | PNN interacting serine/arginine-rich [Sour | 921.16 | 863.45 | 890.53 | 1271.5 | 929.21 | 877.71 |
| 6579 | ENSMUSG00000028249 | 1121.014201 | -0.155981371 | 0.104342304 | -1.494900583 | 0.13494039 | 0.537616739 | Sdcbp | syndecan binding protein [Source:MGI Sy | 1202.2 | 1159.5 | 1182.3 | 1029.5 | 1020.8 | 1131.7 |
| 6580 | ENSMUSG00000028251 | 332.9789628 | 0.221179456 | 0.184320701 | 1.199970785 | 0.230150687 | 0.676880752 | Tstd3 | thiosulfate sulfurtransferase (rhodanese)- | 324.45 | 260.06 | 337.09 | 286.3 | 354.45 | 435.51 |
| 6581 | ENSMUSG00000028252 | 407.1988527 | 0.034392954 | 0.149882291 | 0.229466431 | 0.818506409 | 0.968897682 | Ccnc | cyclin C [Source:MGI Symbol;Acc:MGI:185 | 411.19 | 387.53 | 407.53 | 348 | 465.69 | 423.26 |
| 6582 | ENSMUSG00000028255 | 0.181769715 | 1.126924175 | 4.080472857 | 0.276174898 | 0.782413736 | NA | Clca1 | chloride channel accessory 1 [Source:MGI | 0 | 0 | 0 | 0 | 1.0906 | 0 |
| 6583 | ENSMUSG00000028256 | 202.5224405 | -0.098546808 | 0.18587304 | -0.530183444 | 0.595984748 | 0.913847493 | Odf2l | outer dense fiber of sperm tails 2-like [Sou | 216.84 | 216.89 | 194.21 | 174.48 | 233.39 | 179.33 |
| 6584 | ENSMUSG00000028259 | 0 | NA | NA | NA | NA | NA | Fhl5 | four and a half LIM domains 5 [Source:MG | 0 | 0 | 0 | 0 | 0 | 0 |
| 6585 | ENSMUSG00000028261 | 775.7205713 | 0.056544911 | 0.12281406 | 0.460410729 | 0.645221435 | 0.925643431 | Ndufaf4 | NADH:ubiquinone oxidoreductase comple | 813.54 | 758.6 | 708.4 | 739.37 | 870.31 | 764.1 |
| 6586 | ENSMUSG00000028262 | 0.439020386 | -2.189747911 | 3.52148864 | -0.621824499 | 0.534057275 | NA | Clca3a2 | chloride channel accessory 3A2 [Source:M | 1.6062 | 1.0279 | 0 | 0 | 0 | 0 |
| 6587 | ENSMUSG00000028264 | 0.856598198 | -3.132279849 | 3.123122476 | -1.002932121 | 0.315893613 | NA | Spaca1 | sperm acrosome associated 1 [Source:MG | 0 | 5.1396 | 0 | 0 | 0 | 0 |
| 6588 | ENSMUSG00000028266 | 237.8880822 | -0.013018894 | 0.17723864 | -0.073454039 | 0.941444817 | 0.989358531 | Lmo4 | LIM domain only 4 [Source:MGI Symbol;A | 244.95 | 259.04 | 213.32 | 275.7 | 212.67 | 221.65 |
| 6589 | ENSMUSG00000028268 | 189.7363928 | -0.3230311 | 0.182160897 | -1.773328443 | 0.076174299 | 0.403877794 | Gbp3 | guanylate binding protein 3 [Source:MGI S | 226.47 | 207.64 | 198.23 | 170.62 | 148.32 | 187.13 |
| 6590 | ENSMUSG00000028270 | 80.7960645 | -0.630075146 | 0.299147469 | -2.106235925 | 0.035183864 | 0.25151549 | Gbp2 | guanylate binding protein 2 [Source:MGI S | 77.901 | 134.66 | 82.512 | 75.19 | 62.165 | 52.351 |
| 6591 | ENSMUSG00000028271 | 341.8006183 | 0.059280693 | 0.138218539 | 0.428891038 | 0.668002523 | 0.930136128 | Gtf2b | general transcription factor IIB [Source:M | 346.94 | 335.1 | 322 | 348 | 347.91 | 350.86 |
| 6592 | ENSMUSG00000028273 | 906.3523985 | 0.052356447 | 0.110902759 | 0.472093275 | 0.636860213 | 0.923478764 | Pdlim5 | PDZ and LIM domain 5 [Source:MGI Symb | 958.1 | 869.62 | 841.22 | 938.91 | 931.39 | 898.87 |
| 6593 | ENSMUSG00000028274 | 178.8378121 | 0.070156277 | 0.190675875 | 0.367934733 | 0.712921901 | 0.944239215 | Rngtt | RNA guanylyltransferase and 5'-phosphat | 191.94 | 156.24 | 175.09 | 203.4 | 159.23 | 187.13 |
| 6594 | ENSMUSG00000028277 | 654.6711754 | 0.093348291 | 0.118763104 | 0.786004131 | 0.431865067 | 0.84514901 | Ube2j1 | ubiquitin-conjugating enzyme E2J 1 [Sour | 650.51 | 656.84 | 592.68 | 649.72 | 711.08 | 667.19 |
| 6595 | ENSMUSG00000028278 | 26.46515412 | -1.201280773 | 0.443050782 | -2.711383938 | 0.006700299 | 0.08160729 | Rragd | Ras-related GTP binding D [Source:MGI Sy | 30.518 | 42.145 | 38.237 | 19.28 | 16.359 | 12.252 |
| 6596 | ENSMUSG00000028280 | 0 | NA | NA | NA | NA | NA | Gabrr1 | gamma-aminobutyric acid (GABA) C recep | 0 | 0 | 0 | 0 | 0 | 0 |
| 6597 | ENSMUSG00000028282 | 103.7758222 | -0.157077963 | 0.225754976 | -0.695789594 | 0.486560607 | 0.873586472 | Casp8ap2 | caspase 8 associated protein 2 [Source:M | 110.83 | 96.624 | 120.75 | 98.326 | 100.34 | 95.791 |
| 6598 | ENSMUSG00000028284 | 864.3080022 | -0.070337699 | 0.109218012 | -0.6440119 | 0.519567718 | 0.887802551 | Map3k7 | mitogen-activated protein kinase kinase k | 851.29 | 918.96 | 886.5 | 850.23 | 874.68 | 804.2 |
| 6599 | ENSMUSG00000028287 | 0 | NA | NA | NA | NA | NA | 1700009N14 | RIKEN cDNA 1700009N14 gene [Source:M | 0 | 0 | 0 | 0 | 0 | 0 |
| 6600 | ENSMUSG00000028289 | 21.88384761 | -1.132560799 | 0.476400027 | -2.377331518 | 0.017438407 | 0.157348175 | Epha7 | Eph receptor A7 [Source:MGI Symbol;Acc | 28.109 | 32.893 | 29.181 | 12.532 | 17.45 | 11.138 |
| 6601 | ENSMUSG00000028291 | 398.0572056 | 0.012933723 | 0.154887481 | 0.083503989 | 0.933450805 | 0.988420305 | Akirin2 | akirin 2 [Source:MGI Symbol;Acc:MGI:188 | 394.32 | 396.78 | 398.47 | 466.57 | 399.17 | 333.04 |
| 6602 | ENSMUSG00000028292 | 232.7354656 | 0.101337909 | 0.179233658 | 0.565395528 | 0.571804765 | 0.904778922 | Rars2 | arginyl-tRNA synthetase 2, mitochondrial | 209.61 | 215.86 | 248.54 | 237.14 | 279.2 | 206.06 |
| 6603 | ENSMUSG00000028293 | 417.3587395 | 0.076008878 | 0.133065747 | 0.571212948 | 0.567855302 | 0.903352921 | Slc35a1 | solute carrier family 35 (CMP-sialic acid tr | 375.85 | 418.36 | 425.64 | 435.72 | 426.43 | 422.15 |
| 6604 | ENSMUSG00000028294 | 0 | NA | NA | NA | NA | NA | Cfap206 | cilia and flagella associated protein 206 [S | 0 | 0 | 0 | 0 | 0 | 0 |

| | | | | | | | | | | | | | | | |
|---|---|---|---|---|---|---|---|---|---|---|---|---|---|---|---|
| 6605 | ENSMUSG00000028295 | 178.1002921 | -0.04610185 | 0.220613945 | -0.208970696 | 0.834471115 | 0.971073158 | Smim8 | small integral membrane protein 8 [Sourc | 207.2 | 177.83 | 156.97 | 135.92 | 222.49 | 168.19 |
| 6606 | ENSMUSG00000028298 | 0.17131964 | -0.796629176 | 4.080472857 | -0.195229623 | 0.84521319 | NA | Cga | glycoprotein hormones, alpha subunit [So | 0 | 1.0279 | 0 | 0 | 0 | 0 |
| 6607 | ENSMUSG00000028300 | 205.1076699 | -0.275175903 | 0.188641268 | -1.458725902 | 0.144640564 | 0.554723921 | C9orf72 | C9orf72, member of C9orf72-SMCR8 comp | 214.43 | 269.31 | 190.18 | 175.44 | 193.04 | 188.24 |
| 6608 | ENSMUSG00000028307 | 109269.4159 | -0.421031719 | 0.09658473 | -4.359195506 | 1.31E-05 | 0.000669557 | Aldob | aldolase B, fructose-bisphosphate [Source | 135001 | 127716 | 112588 | 94409 | 96213 | 89690 |
| 6609 | ENSMUSG00000028309 | 454.017051 | 0.061845736 | 0.126936621 | 0.487217442 | 0.626104246 | 0.919668103 | Rnf20 | ring finger protein 20 [Source:MGI Symbo | 461.78 | 438.92 | 431.68 | 454.03 | 478.78 | 458.9 |
| 6610 | ENSMUSG00000028310 | 0 | NA | NA | NA | NA | NA | Ppp3r2 | protein phosphatase 3, regulatory subunit | 0 | 0 | 0 | 0 | 0 | 0 |
| 6611 | ENSMUSG00000028312 | 61.11931475 | -0.099492607 | 0.345718669 | -0.287784884 | 0.773511409 | 0.958177785 | Smc2 | structural maintenance of chromosomes 2 | 86.735 | 47.284 | 55.343 | 67.478 | 70.89 | 38.985 |
| 6612 | ENSMUSG00000028314 | 0.401551119 | -2.072982726 | 4.008251979 | -0.517178744 | 0.605031388 | NA | Toporsl | topoisomerase I binding, arginine/serine-r | 2.4093 | 0 | 0 | 0 | 0 | 0 |
| 6613 | ENSMUSG00000028318 | 75.52394175 | 0.088816804 | 0.273345396 | 0.324925188 | 0.745237693 | 0.949997329 | Polr1e | polymerase (RNA) I polypeptide E [Source | 71.476 | 57.563 | 90.562 | 77.118 | 81.796 | 74.628 |
| 6614 | ENSMUSG00000028322 | 237.221916 | -0.264667992 | 0.157451455 | -1.680949796 | 0.092772667 | 0.44871446 | Exosc3 | exosome component 3 [Source:MGI Symb | 265.02 | 261.09 | 250.55 | 217.86 | 210.49 | 218.31 |
| 6615 | ENSMUSG00000028327 | 2163.558517 | 0.257964233 | 0.119308134 | 2.162168041 | 0.030605226 | 0.231893781 | Stra6l | STRA6-like [Source:MGI Symbol;Acc:MGI: | 1949.1 | 1855.4 | 2107.1 | 2261.5 | 2101.6 | 2706.6 |
| 6616 | ENSMUSG00000028328 | 1.238226717 | -1.290915169 | 2.090057231 | -0.617645847 | 0.536808816 | NA | Tmod1 | tropomodulin 1 [Source:MGI Symbol;Acc: | 3.2124 | 0 | 2.0125 | 0 | 1.0906 | 1.1138 |
| 6617 | ENSMUSG00000028329 | 269.6620066 | 0.134967032 | 0.155862183 | 0.865938287 | 0.386524005 | 0.817115886 | Xpa | xeroderma pigmentosum, complementati | 253.78 | 256.98 | 260.62 | 287.27 | 305.37 | 253.96 |
| 6618 | ENSMUSG00000028330 | 443.9819478 | -0.121736411 | 0.159497866 | -0.763247901 | 0.445315566 | 0.852147532 | Ncbp1 | nuclear cap binding protein subunit 1 [Sou | 444.92 | 546.85 | 396.46 | 398.12 | 478.78 | 398.76 |
| 6619 | ENSMUSG00000028331 | 40.86480883 | 0.189733762 | 0.42519241 | 0.446230361 | 0.655430854 | 0.926075335 | Trmo | tRNA methyltransferase O [Source:MGI Sy | 44.974 | 41.117 | 28.175 | 35.667 | 25.084 | 70.172 |
| 6620 | ENSMUSG00000028332 | 0.267700746 | -1.497481649 | 4.057001436 | -0.369110456 | 0.712045396 | NA | Hemgn | hemogen [Source:MGI Symbol;Acc:MGI:2 | 1.6062 | 0 | 0 | 0 | 0 | 0 |
| 6621 | ENSMUSG00000028333 | 876.6008874 | 0.095200578 | 0.113773463 | 0.836755567 | 0.402729978 | 0.828953151 | Anp32b | acidic (leucine-rich) nuclear phosphoprote | 867.35 | 788.41 | 886.5 | 852.16 | 912.85 | 952.34 |
| 6622 | ENSMUSG00000028334 | 257.7249779 | -0.011135303 | 0.174375847 | -0.063858061 | 0.949083246 | 0.990731019 | Nans | N-acetylneuraminic acid synthase (sialic a | 309.19 | 243.62 | 222.38 | 255.45 | 261.75 | 253.96 |
| 6623 | ENSMUSG00000028337 | 10.58155689 | -0.507514169 | 0.654213993 | -0.775761715 | 0.437889693 | 0.847422601 | Coro2a | coronin, actin binding protein 2A [Source: | 12.85 | 13.363 | 11.069 | 9.6398 | 6.5437 | 10.025 |
| 6624 | ENSMUSG00000028339 | 131.9330734 | 0.148756958 | 0.251712229 | 0.590980259 | 0.554533648 | 0.897558119 | Col15a1 | collagen, type XV, alpha 1 [Source:MGI Sy | 166.24 | 90.457 | 117.73 | 129.17 | 127.6 | 160.39 |
| 6625 | ENSMUSG00000028341 | 1.547242687 | -0.125513419 | 2.109179542 | -0.059508172 | 0.952547357 | NA | Nr4a3 | nuclear receptor subfamily 4, group A, me | 0.8031 | 0 | 4.025 | 0 | 0 | 4.4554 |
| 6626 | ENSMUSG00000028343 | 1328.978343 | -0.002362164 | 0.096715497 | -0.024423841 | 0.980514532 | 0.99820894 | Erp44 | endoplasmic reticulum protein 44 [Source | 1314.7 | 1341.4 | 1334.3 | 1322.6 | 1363.3 | 1297.6 |
| 6627 | ENSMUSG00000028344 | 277.6745149 | -0.059070238 | 0.163864034 | -0.36048324 | 0.718485788 | 0.9447719 | Invs | inversin [Source:MGI Symbol;Acc:MGI:13 | 266.63 | 273.43 | 309.92 | 240.03 | 305.37 | 270.66 |
| 6628 | ENSMUSG00000028345 | 126.1277898 | -0.049527414 | 0.222111435 | -0.222984532 | 0.823547545 | 0.96943211 | Tex10 | testis expressed gene 10 [Source:MGI Sym | 131.71 | 114.1 | 138.86 | 121.46 | 103.61 | 147.03 |
| 6629 | ENSMUSG00000028347 | 2.196679788 | -4.500472768 | 1.804468154 | -2.494071596 | 0.012628707 | NA | Tmeff1 | transmembrane protein with EGF-like and | 4.0155 | 5.1396 | 4.025 | 0 | 0 | 0 |
| 6630 | ENSMUSG00000028348 | 0 | NA | NA | NA | NA | NA | Cavin4 | caveolae associated 4 [Source:MGI Symbo | 0 | 0 | 0 | 0 | 0 | 0 |
| 6631 | ENSMUSG00000028351 | 0.167707431 | -0.796629176 | 4.080472857 | -0.195229623 | 0.84521319 | NA | Brinp1 | bone morphogenic protein/retinoic acid in | 0 | 0 | 1.0062 | 0 | 0 | 0 |
| 6632 | ENSMUSG00000028354 | 10.57776118 | 1.502162058 | 0.819668952 | 1.832644818 | 0.066855409 | 0.37192178 | Fmn2 | formin 2 [Source:MGI Symbol;Acc:MGI:18 | 7.2279 | 5.1396 | 4.025 | 2.8919 | 17.45 | 26.732 |
| 6633 | ENSMUSG00000028356 | 39390.45212 | 0.047216199 | 0.086450198 | 0.546166468 | 0.584951513 | 0.90908382 | Ambp | alpha 1 microglobulin/bikunin precursor [ | 37184 | 38718 | 40337 | 40236 | 37939 | 41928 |
| 6634 | ENSMUSG00000028357 | 5.758285441 | 0.332478499 | 1.003956515 | 0.331168227 | 0.740517419 | 0.948792385 | Kif12 | kinesin family member 12 [Source:MGI Sy | 3.2124 | 1.0279 | 11.069 | 3.8559 | 9.8156 | 5.5692 |
| 6635 | ENSMUSG00000028358 | 9.309841616 | -0.009345187 | 0.729812232 | -0.01280492 | 0.989783431 | 0.99859176 | Zfp618 | zinc finger protein 618 [Source:MGI Symb | 8.8341 | 8.2233 | 11.069 | 14.46 | 4.3625 | 8.9108 |
| 6636 | ENSMUSG00000028359 | 75.84791944 | -1.82799118 | 0.338256718 | -5.404153369 | 6.51E-08 | 7.42E-06 | Orm3 | orosomucoid 3 [Source:MGI Symbol;Acc:N | 163.83 | 80.178 | 110.69 | 28.919 | 43.625 | 27.846 |
| 6637 | ENSMUSG00000028360 | 0 | NA | NA | NA | NA | NA | Slc44a5 | solute carrier family 44, member 5 [Source | 0 | 0 | 0 | 0 | 0 | 0 |
| 6638 | ENSMUSG00000028362 | 0.356960287 | 0.16514264 | 3.880370363 | 0.042558474 | 0.966053499 | NA | Tnfsf8 | tumor necrosis factor (ligand) superfamily | 0 | 1.0279 | 0 | 0 | 0 | 1.1138 |
| 6639 | ENSMUSG00000028364 | 4.787936709 | -1.065722585 | 1.143200366 | -0.932227295 | 0.351219072 | NA | Tnc | tenascin C [Source:MGI Symbol;Acc:MGI:1 | 15.259 | 3.0838 | 1.0062 | 3.8559 | 2.1812 | 3.3415 |
| 6640 | ENSMUSG00000028367 | 3554.47123 | -0.054188805 | 0.093811126 | -0.577637294 | 0.563509023 | 0.900821918 | Txn1 | thioredoxin 1 [Source:MGI Symbol;Acc:MG | 3911.1 | 3433.2 | 3517.8 | 3453 | 3503.1 | 3508.6 |
| 6641 | ENSMUSG00000028369 | 6.480003873 | 0.683259064 | 0.846098677 | 0.807540636 | 0.419355072 | 0.837591673 | Svep1 | sushi, von Willebrand factor type A, EGF a | 4.8186 | 3.0838 | 7.0437 | 9.6398 | 8.7249 | 5.5692 |
| 6642 | ENSMUSG00000028370 | 0.453341394 | -0.545391191 | 3.551113277 | -0.153583158 | 0.877938417 | NA | Pappa | pregnancy-associated plasma protein A [S | 1.6062 | 0 | 0 | 0 | 0 | 1.1138 |
| 6643 | ENSMUSG00000028373 | 17.28782261 | -1.430383791 | 0.550395286 | -2.598830016 | 0.009354208 | 0.10348107 | Astn2 | astrotactin 2 [Source:MGI Symbol;Acc:MG | 28.912 | 25.698 | 21.131 | 12.532 | 6.5437 | 8.9108 |
| 6644 | ENSMUSG00000028378 | 51.68287818 | -0.033609051 | 0.369466669 | -0.090966396 | 0.927519292 | 0.987825698 | Ptgr1 | prostaglandin reductase 1 [Source:MGI Sy | 69.87 | 39.061 | 47.293 | 35.667 | 45.806 | 72.4 |
| 6645 | ENSMUSG00000028381 | 349.2423776 | 0.411510217 | 0.171603893 | 2.398023782 | 0.016483795 | 0.152346077 | Ugcg | UDP-glucose ceramide glucosyltransferase | 304.38 | 325.85 | 269.67 | 486.81 | 350.09 | 358.66 |
| 6646 | ENSMUSG00000028382 | 1440.772302 | 0.141584505 | 0.115571309 | 1.225083508 | 0.220543758 | 0.667212909 | Ptbp3 | polypyrimidine tract binding protein 3 [So | 1341.2 | 1393.9 | 1376.5 | 1704.3 | 1484.3 | 1344.4 |
| 6647 | ENSMUSG00000028383 | 1821.752987 | -0.199030637 | 0.129793535 | -1.533440294 | 0.125167407 | 0.518596785 | Hsdl2 | hydroxysteroid dehydrogenase like 2 [Sou | 1941.9 | 1847.2 | 2053.7 | 2030.1 | 1513.8 | 1543.8 |
| 6648 | ENSMUSG00000028385 | 172.6100474 | 0.117310098 | 0.187873366 | 0.624410476 | 0.532358047 | 0.889143097 | Snx30 | sorting nexin family member 30 [Source:M | 171.06 | 167.55 | 157.98 | 179.3 | 157.05 | 202.72 |
| 6649 | ENSMUSG00000028386 | 0 | NA | NA | NA | NA | NA | Slc46a2 | solute carrier family 46, member 2 [Source | 0 | 0 | 0 | 0 | 0 | 0 |
| 6650 | ENSMUSG00000028389 | 5.219417602 | 0.516631099 | 1.019710961 | 0.506644646 | 0.612404176 | 0.916441371 | Zfp37 | zinc finger protein 37 [Source:MGI Symbo | 5.6217 | 5.1396 | 2.0125 | 1.928 | 4.3625 | 12.252 |
| 6651 | ENSMUSG00000028391 | 1.50247024 | 0.551482256 | 1.766361641 | 0.312213673 | 0.754878142 | NA | Wdr31 | WD repeat domain 31 [Source:MGI Symbo | 1.6062 | 0 | 2.0125 | 0.964 | 1.0906 | 3.3415 |
| 6652 | ENSMUSG00000028392 | 0.859823856 | -1.825223727 | 2.563005556 | -0.712141931 | 0.476376891 | NA | Bspry | B-box and SPRY domain containing [Sourc | 0 | 2.0558 | 2.0125 | 0 | 1.0906 | 0 |
| 6653 | ENSMUSG00000028393 | 2483.347745 | 0.055515139 | 0.095843645 | 0.57922608 | 0.562436635 | 0.900565925 | Alad | aminolevulinate, delta-, dehydratase [Sou | 2462.3 | 2420.7 | 2423 | 2400.3 | 2727.6 | 2466.1 |
| 6654 | ENSMUSG00000028394 | 414.7536702 | 0.022509678 | 0.135767607 | 0.165795648 | 0.868317771 | 0.978366823 | Pole3 | polymerase (DNA directed), epsilon 3 (p17 | 406.37 | 380.33 | 447.78 | 407.76 | 429.7 | 416.58 |
| 6655 | ENSMUSG00000028396 | 0 | NA | NA | NA | NA | NA | 2310002L09 | RIKEN cDNA 2310002L09 gene [Source:MG | 0 | 0 | 0 | 0 | 0 | 0 |
| 6656 | ENSMUSG00000028397 | 165.8333017 | 0.118823454 | 0.188829776 | 0.629262271 | 0.529177367 | 0.888334265 | Kdm4c | lysine (K)-specific demethylase 4C [Source | 162.23 | 164.47 | 149.93 | 151.34 | 182.13 | 184.9 |
| 6657 | ENSMUSG00000028398 | 534.8230651 | 0.016202642 | 0.129234848 | 0.125373631 | 0.900227763 | 0.982805511 | Dmac1 | distal membrane arm assembly complex 1 | 571.01 | 484.15 | 539.35 | 519.58 | 525.68 | 569.17 |
| 6658 | ENSMUSG00000028399 | 2978.474033 | 0.046937339 | 0.129845 | 0.361487454 | 0.717735082 | 0.9447719 | Ptprd | protein tyrosine phosphatase, receptor ty | 2503.3 | 3088.9 | 3199.9 | 3330.5 | 3150.8 | 2597.5 |
| 6659 | ENSMUSG00000028402 | 413.2766538 | 0.44879819 | 0.159044811 | 2.821834854 | 0.004774976 | 0.064850002 | Mpdz | multiple PDZ domain crumbs cell polarity | 348.55 | 391.64 | 307.91 | 401.01 | 511.5 | 519.05 |
| 6660 | ENSMUSG00000028403 | 246.5102998 | 0.182200767 | 0.174821849 | 1.042208213 | 0.297315156 | 0.748078261 | Zdhhc21 | zinc finger, DHHC domain containing 21 [S | 235.31 | 251.84 | 205.27 | 223.64 | 290.1 | 272.89 |

| | | | | | | | | | | | | | | | |
|---|---|---|---|---|---|---|---|---|---|---|---|---|---|---|---|
| 6661 | ENSMUSG00000028405 | 4913.621817 | -0.069387929 | 0.10953832 | -0.633458031 | 0.52643458 | 0.888334265 | Aco1 | aconitase 1 [Source:MGI Symbol;Acc:MGI | 4617 | 5381.1 | 5098.6 | 5017.5 | 5127 | 4240.4 |
| 6662 | ENSMUSG00000028407 | 75.64888642 | -0.153331332 | 0.262952952 | -0.58311318 | 0.559817112 | 0.899475387 | Smim27 | small integral membrane protein 27 [Sour | 92.357 | 68.87 | 77.481 | 75.19 | 68.709 | 71.286 |
| 6663 | ENSMUSG00000028409 | 868.3538022 | 0.064529111 | 0.108861951 | 0.59276093 | 0.553341158 | 0.896251569 | Smu1 | smu-1 suppressor of mec-8 and unc-52 hc | 870.56 | 850.09 | 826.13 | 912.89 | 912.85 | 837.61 |
| 6664 | ENSMUSG00000028410 | 5023.212609 | -0.003360565 | 0.087506502 | -0.038403606 | 0.969365886 | 0.994602269 | Dnaja1 | DnaJ heat shock protein family (Hsp40) m | 4940.7 | 5078.9 | 5068.5 | 5204.5 | 5111.7 | 4735 |
| 6665 | ENSMUSG00000028411 | 236.1069213 | 0.222019942 | 0.186566878 | 1.19002871 | 0.234035108 | 0.68180718 | Aptx | aprataxin [Source:MGI Symbol;Acc:MGI:1 | 236.11 | 227.17 | 190.18 | 236.17 | 306.46 | 220.54 |
| 6666 | ENSMUSG00000028412 | 659.4119275 | -0.154063302 | 0.129097024 | -1.193391585 | 0.232716049 | 0.67995822 | Slc44a1 | solute carrier family 44, member 1 [Sourc | 628.03 | 734.96 | 721.48 | 618.87 | 572.57 | 680.56 |
| 6667 | ENSMUSG00000028413 | 944.8965425 | 0.339703117 | 0.197032919 | 1.724093206 | 0.084691018 | 0.429140871 | B4galt1 | UDP-Gal:betaGlcNAc beta 1,4- galactosylt | 933.2 | 757.58 | 812.04 | 1477.8 | 775.43 | 913.35 |
| 6668 | ENSMUSG00000028414 | 266.6207801 | 0.010615551 | 0.17425649 | 0.060919116 | 0.951423625 | 0.990883058 | Fktn | fukutin [Source:MGI Symbol;Acc:MGI:217 | 289.92 | 249.78 | 256.59 | 247.74 | 319.55 | 236.13 |
| 6669 | ENSMUSG00000028415 | 0 | NA | NA | NA | NA | NA | Spink4 | serine peptidase inhibitor, Kazal type 4 [S | 0 | 0 | 0 | 0 | 0 | 0 |
| 6670 | ENSMUSG00000028416 | 1793.911863 | -0.067496099 | 0.099153501 | -0.680723308 | 0.496046585 | 0.878165817 | Bag1 | BCL2-associated athanogene 1 [Source:M | 1729.1 | 1823.5 | 1956.1 | 1807.5 | 1729.7 | 1717.5 |
| 6671 | ENSMUSG00000028417 | 0.17131964 | -0.796629176 | 4.080472857 | -0.195229623 | 0.84521319 | NA | Tal2 | T cell acute lymphocytic leukemia 2 [Sour | 0 | 1.0279 | 0 | 0 | 0 | 0 |
| 6672 | ENSMUSG00000028419 | 1005.65638 | 0.21546358 | 0.106590644 | 2.021411743 | 0.04323716 | 0.286918392 | Chmp5 | charged multivesicular body protein 5 [So | 923.57 | 931.29 | 936.81 | 1007.4 | 1125.5 | 1109.4 |
| 6673 | ENSMUSG00000028420 | 1343.507703 | -0.050268954 | 0.10764673 | -0.466980779 | 0.64051363 | 0.923932929 | Tmem38b | transmembrane protein 38B [Source:MGI | 1353.2 | 1479.2 | 1268.9 | 1374.6 | 1259.7 | 1325.5 |
| 6674 | ENSMUSG00000028423 | 824.3291619 | 0.401833867 | 0.121005644 | 3.320786157 | 0.000897643 | 0.019349488 | Nfx1 | nuclear transcription factor, X-box binding | 766.16 | 737.02 | 626.89 | 949.52 | 960.83 | 905.56 |
| 6675 | ENSMUSG00000028426 | 3280.161698 | 0.032156095 | 0.086914878 | 0.369972276 | 0.711403148 | 0.944239215 | Rad23b | RAD23 homolog B, nucleotide excision rep | 3258.2 | 3309.9 | 3162.6 | 3284.3 | 3416.9 | 3249.1 |
| 6676 | ENSMUSG00000028427 | 0.94232264 | -2.048339122 | 2.477551956 | -0.8267593 | 0.408373503 | NA | Aqp7 | aquaporin 7 [Source:MGI Symbol;Acc:MG | 1.6062 | 3.0838 | 0 | 0.964 | 0 | 0 |
| 6677 | ENSMUSG00000028430 | 446.1858399 | -0.103162549 | 0.127421359 | -0.809617395 | 0.418160107 | 0.837575303 | Nol6 | nucleolar protein family 6 (RNA-associate | 459.37 | 449.2 | 477.97 | 436.68 | 438.43 | 415.46 |
| 6678 | ENSMUSG00000028431 | 205.0236708 | -0.048168663 | 0.183133394 | -0.263025009 | 0.792531303 | 0.961523605 | Elp1 | elongator complex protein 1 [Source:MGI | 205.59 | 219.97 | 200.24 | 224.61 | 213.76 | 165.96 |
| 6679 | ENSMUSG00000028433 | 335.8935156 | -0.084583068 | 0.139775246 | -0.605136248 | 0.545088471 | 0.895484962 | Ubap2 | ubiquitin-associated protein 2 [Source:MC | 346.94 | 345.38 | 345.14 | 343.18 | 321.73 | 312.99 |
| 6680 | ENSMUSG00000028434 | 2029.33423 | 0.381217714 | 0.11598256 | 3.286853774 | 0.001013134 | 0.021080742 | Epb41l4b | erythrocyte membrane protein band 4.1 l | 1601.4 | 1740.3 | 1947.1 | 2061 | 2405.9 | 2420.4 |
| 6681 | ENSMUSG00000028435 | 3.220121938 | 1.507950346 | 1.551847413 | 0.971713026 | 0.331193335 | NA | Aqp3 | aquaporin 3 [Source:MGI Symbol;Acc:MG | 0 | 3.0838 | 2.0125 | 0 | 11.997 | 2.2277 |
| 6682 | ENSMUSG00000028436 | 554.9169943 | 0.092300729 | 0.135298074 | 0.682202833 | 0.495110704 | 0.877919732 | Dcaf12 | DDB1 and CUL4 associated factor 12 [Sou | 566.99 | 525.27 | 518.22 | 516.69 | 551.85 | 650.48 |
| 6683 | ENSMUSG00000028437 | 856.7066098 | 0.083045871 | 0.116197724 | 0.714694475 | 0.474797852 | 0.869368495 | Ubap1 | ubiquitin-associated protein 1 [Source:MC | 880.2 | 807.94 | 807.01 | 847.34 | 836.5 | 961.25 |
| 6684 | ENSMUSG00000028438 | 10.95901404 | 0.716763145 | 0.710545972 | 1.008749852 | 0.313094617 | 0.760114938 | Kif24 | kinesin family member 24 [Source:MGI Sy | 9.6372 | 9.2513 | 6.0375 | 22.171 | 13.087 | 5.5692 |
| 6685 | ENSMUSG00000028439 | 77.3684852 | 0.192125214 | 0.30231231 | 0.635518991 | 0.525089985 | 0.888334265 | Fam219a | family with sequence similarity 219, meml | 89.947 | 68.87 | 57.356 | 80.01 | 59.984 | 108.04 |
| 6686 | ENSMUSG00000028441 | 0.48903338 | 0.95637552 | 3.456690196 | 0.276673773 | 0.782030612 | NA | 1110017D15 | RIKEN cDNA 1110017D15 gene [Source:M | 0 | 0 | 1.0062 | 1.928 | 0 | 0 |
| 6687 | ENSMUSG00000028443 | 173.4021609 | 0.387169599 | 0.206536849 | 1.874578803 | 0.060850692 | 0.352269825 | Nudt2 | nudix (nucleoside diphosphate linked moi | 182.3 | 129.52 | 137.86 | 169.66 | 207.22 | 213.86 |
| 6688 | ENSMUSG00000028444 | 42.10987143 | -0.815087494 | 0.3775898 | -2.158658669 | 0.030876658 | 0.233016344 | Cntfr | ciliary neurotrophic factor receptor [Sour | 54.611 | 49.34 | 57.356 | 40.487 | 17.45 | 33.415 |
| 6689 | ENSMUSG00000028445 | 307.4347229 | 1.261947776 | 0.599106685 | 2.106382399 | 0.035171149 | 0.25151549 | Enho | energy homeostasis associated [Source:M | 141.35 | 95.596 | 305.9 | 248.71 | 448.24 | 604.82 |
| 6690 | ENSMUSG00000028447 | 521.9112162 | 0.092327507 | 0.128866183 | 0.71646032 | 0.47370716 | 0.868952133 | Dctn3 | dynactin 3 [Source:MGI Symbol;Acc:MGI: | 546.91 | 465.65 | 502.12 | 520.55 | 564.94 | 531.3 |
| 6691 | ENSMUSG00000028451 | 0.160662974 | 1.126924175 | 4.080472857 | 0.276174898 | 0.782413736 | NA | 1700022I11F | RIKEN cDNA 1700022I11 gene [Source:MC | 0 | 0 | 0 | 0.964 | 0 | 0 |
| 6692 | ENSMUSG00000028452 | 5206.578019 | 0.118165496 | 0.092874931 | 1.272307767 | 0.20326379 | 0.647739134 | Vcp | valosin containing protein [Source:MGI Sy | 5386.4 | 4922.7 | 4670 | 5425.3 | 5508.7 | 5326.4 |
| 6693 | ENSMUSG00000028453 | 54.72539685 | -0.042554752 | 0.304886326 | -0.139575796 | 0.888995165 | 0.982382091 | Fancg | Fanconi anemia, complementation group | 53.005 | 56.535 | 57.356 | 65.55 | 46.897 | 49.009 |
| 6694 | ENSMUSG00000028454 | 280.416161 | 0.155414642 | 0.170206393 | 0.913095207 | 0.361192471 | 0.800074801 | Pigo | phosphatidylinositol glycan anchor biosyn | 240.13 | 242.59 | 313.95 | 319.08 | 270.47 | 296.28 |
| 6695 | ENSMUSG00000028455 | 879.8258912 | 0.083668552 | 0.124918972 | 0.669782584 | 0.502996397 | 0.881220881 | Stoml2 | stomatin (Epb7.2)-like 2 [Source:MGI Sym | 854.5 | 745.24 | 962.98 | 886.86 | 920.48 | 908.9 |
| 6696 | ENSMUSG00000028456 | 122.7237175 | 0.632870325 | 0.240108104 | 2.635772448 | 0.0083946 | 0.095354881 | Unc13b | unc-13 homolog B [Source:MGI Symbol;A | 74.689 | 114.1 | 100.62 | 153.27 | 123.24 | 170.42 |
| 6697 | ENSMUSG00000028457 | 12.23733145 | 0.132922185 | 0.643137608 | 0.206677674 | 0.836261595 | 0.971073158 | Atp8b5 | ATPase, class I, type 8B, member 5 [Sourc | 9.6372 | 10.279 | 15.094 | 9.6398 | 8.7249 | 20.049 |
| 6698 | ENSMUSG00000028458 | 185.5182202 | -0.089204067 | 0.201955524 | -0.441701547 | 0.65870519 | 0.927599242 | Tesk1 | testis specific protein kinase 1 [Source:MC | 191.94 | 192.22 | 190.18 | 225.57 | 147.23 | 165.96 |
| 6699 | ENSMUSG00000028459 | 23.11603506 | -1.049451247 | 0.475292128 | -2.208013107 | 0.02724336 | 0.213715335 | Cd72 | CD72 antigen [Source:MGI Symbol;Acc:M | 31.321 | 40.089 | 22.137 | 15.424 | 16.359 | 13.366 |
| 6700 | ENSMUSG00000028460 | 0.167707431 | -0.796629176 | 4.080472857 | -0.195229623 | 0.84521319 | NA | Sit1 | suppression inducing transmembrane ada | 0 | 0 | 1.0062 | 0 | 0 | 0 |
| 6701 | ENSMUSG00000028461 | 1097.902504 | -0.175838041 | 0.112141915 | -1.567995706 | 0.116882135 | 0.50225288 | Ccdc107 | coiled-coil domain containing 107 [Source | 1102.7 | 1133.8 | 1257.8 | 979.4 | 1008.8 | 1104.9 |
| 6702 | ENSMUSG00000028463 | 3.692932761 | 0.105058947 | 1.142726336 | 0.091937101 | 0.926748014 | NA | Car9 | carbonic anhydrase 9 [Source:MGI Symbo | 1.6062 | 5.1396 | 4.025 | 4.8199 | 5.4531 | 1.1138 |
| 6703 | ENSMUSG00000028464 | 243.1477459 | -0.991945091 | 0.287026377 | -3.455937056 | 0.000548383 | 0.013219695 | Tpm2 | tropomyosin 2, beta [Source:MGI Symbol; | 421.63 | 273.43 | 275.71 | 190.87 | 215.94 | 81.311 |
| 6704 | ENSMUSG00000028465 | 1395.817054 | -0.185190353 | 0.106869501 | -1.732864389 | 0.083119775 | 0.425083636 | Tln1 | talin 1 [Source:MGI Symbol;Acc:MGI:1099 | 1475.3 | 1558.3 | 1422.8 | 1361.1 | 1363.3 | 1194 |
| 6705 | ENSMUSG00000028466 | 690.9051099 | 0.173975265 | 0.111316194 | 1.562892676 | 0.118077841 | 0.503427297 | Creb3 | cAMP responsive element binding protein | 651.32 | 646.56 | 650.03 | 741.3 | 712.17 | 744.05 |
| 6706 | ENSMUSG00000028467 | 228.6967727 | 0.408934214 | 0.168802077 | 2.422566247 | 0.015411316 | 0.145123221 | Gba2 | glucosidase beta 2 [Source:MGI Symbol;A | 198.37 | 209.7 | 181.12 | 234.25 | 279.2 | 269.55 |
| 6707 | ENSMUSG00000028468 | 720.7349847 | 0.193970946 | 0.12422368 | 1.561465134 | 0.118414044 | 0.504123865 | Rgp1 | RAB6A GEF compex partner 1 [Source:MG | 652.12 | 732.91 | 631.92 | 705.63 | 778.7 | 823.13 |
| 6708 | ENSMUSG00000028469 | 1263.430747 | 0.364016618 | 0.145957717 | 2.493986778 | 0.012631725 | 0.127143542 | Npr2 | natriuretic peptide receptor 2 [Source:MG | 987.01 | 1016.6 | 1311.1 | 1197.3 | 1572.7 | 1495.9 |
| 6709 | ENSMUSG00000028470 | 1685.706532 | -0.014493332 | 0.097899896 | -0.148042365 | 0.882309334 | 0.981769403 | Hint2 | histidine triad nucleotide binding protein | 1675.3 | 1700.2 | 1706.6 | 1575.1 | 1688.3 | 1768.8 |
| 6710 | ENSMUSG00000028475 | 5.021426757 | -0.902462276 | 0.950211845 | -0.949748502 | 0.342240059 | NA | Spaar | small regulatory polypeptide of amino aci | 6.4248 | 6.1675 | 7.0437 | 3.8559 | 2.1812 | 4.4554 |
| 6711 | ENSMUSG00000028476 | 81.69672529 | 0.42942486 | 0.292758061 | 1.466825057 | 0.142423651 | 0.55198239 | Reck | reversion-inducing-cysteine-rich protein v | 81.916 | 46.256 | 80.5 | 112.79 | 79.615 | 89.108 |
| 6712 | ENSMUSG00000028478 | 1791.693465 | 0.030594088 | 0.092148596 | 0.332008186 | 0.739883078 | 0.948393197 | Clta | clathrin, light polypeptide (Lca) [Source:M | 1745.1 | 1782.4 | 1791.1 | 1858.5 | 1790.8 | 1782.2 |
| 6713 | ENSMUSG00000028479 | 3790.515786 | 0.107345165 | 0.102015681 | 1.052241817 | 0.292688618 | 0.743754072 | Gne | glucosamine (UDP-N-acetyl)-2-epimerase/ | 4025.1 | 3596.7 | 3325.6 | 4055.5 | 3815 | 3925.2 |
| 6714 | ENSMUSG00000028480 | 6.856614676 | -0.93313171 | 1.000083435 | -0.933053861 | 0.35079216 | 0.791943999 | Glipr2 | GLI pathogenesis-related 2 [Source:MGI S | 4.8186 | 11.307 | 11.069 | 10.604 | 0 | 3.3415 |
| 6715 | ENSMUSG00000028483 | 116.8394266 | 0.255185023 | 0.218324915 | 1.168831429 | 0.242471555 | 0.692000002 | Snapc3 | small nuclear RNA activating complex, pol | 109.22 | 94.568 | 115.72 | 131.1 | 113.42 | 137 |
| 6716 | ENSMUSG00000028484 | 360.0209796 | -0.013618991 | 0.144116304 | -0.094500003 | 0.92471198 | 0.987546253 | Psip1 | PC4 and SFRS1 interacting protein 1 [Sour | 375.85 | 327.91 | 381.37 | 379.81 | 358.81 | 336.38 |

| | | | | | | | | | | | | | | | |
|---|---|---|---|---|---|---|---|---|---|---|---|---|---|---|---|
| 6717 | ENSMUSG00000028487 | 1.590514924 | 0.432082731 | 2.100429761 | 0.205711583 | 0.83701621 | NA | Bnc2 | basonuclin 2 [Source:MGI Symbol;Acc:MG | 0 | 3.0838 | 1.0062 | 0 | 5.4531 | 0 |
| 6718 | ENSMUSG00000028488 | 0.335414862 | -1.779047242 | 4.030804179 | -0.441362855 | 0.658950329 | NA | Sh3gl2 | SH3-domain GRB2-like 2 [Source:MGI Sym | 0 | 0 | 2.0125 | 0 | 0 | 0 |
| 6719 | ENSMUSG00000028492 | 0 | NA | NA | NA | NA | NA | Saxo1 | stabilizer of axonemal microtubules 1 [Sou | 0 | 0 | 0 | 0 | 0 | 0 |
| 6720 | ENSMUSG00000028494 | 9531.962064 | -0.149928542 | 0.14925708 | -1.004498698 | 0.315138298 | 0.761215823 | Plin2 | perilipin 2 [Source:MGI Symbol;Acc:MGI:8 | 12705 | 9115.6 | 8258.2 | 9156.8 | 9896.3 | 8059.8 |
| 6721 | ENSMUSG00000028495 | 3760.521578 | 0.01236295 | 0.108737447 | 0.113695427 | 0.909479238 | 0.985399467 | Rps6 | ribosomal protein S6 [Source:MGI Symbol | 3320 | 4005.8 | 3908.3 | 3515.6 | 3774.6 | 4038.8 |
| 6722 | ENSMUSG00000028496 | 53.49279304 | 0.199496584 | 0.310912972 | 0.641647669 | 0.521101974 | 0.887887809 | Mllt3 | myeloid/lymphoid or mixed-lineage leuke | 53.808 | 38.033 | 57.356 | 54.947 | 58.893 | 57.92 |
| 6723 | ENSMUSG00000028497 | 21.46775094 | -1.14896653 | 0.575535887 | -1.996342115 | 0.045896697 | 0.296229283 | Hacd4 | 3-hydroxyacyl-CoA dehydratase 4 [Source | 42.564 | 35.977 | 10.062 | 11.568 | 15.269 | 13.366 |
| 6724 | ENSMUSG00000028514 | 1603.353125 | 0.007849117 | 0.106562538 | 0.07365738 | 0.941283013 | 0.989297885 | Usp24 | ubiquitin specific peptidase 24 [Source:M | 1554 | 1644.7 | 1598.9 | 1627.2 | 1739.5 | 1455.8 |
| 6725 | ENSMUSG00000028517 | 3906.595541 | -0.076020295 | 0.126792049 | -0.599566732 | 0.548795024 | 0.895703112 | Plpp3 | phospholipid phosphatase 3 [Source:MGI | 4163.3 | 4360.4 | 3503.7 | 3273.7 | 4307.9 | 3830.5 |
| 6726 | ENSMUSG00000028518 | 1438.58414 | -0.143961895 | 0.140078143 | -1.027725611 | 0.304078917 | 0.752961647 | Prkaa2 | protein kinase, AMP-activated, alpha 2 ca | 1342.8 | 1751.6 | 1437.9 | 1472 | 1480 | 1147.3 |
| 6727 | ENSMUSG00000028519 | 3.147172251 | -0.236426567 | 1.417246935 | -0.166821011 | 0.867510886 | NA | Dab1 | disabled 1 [Source:MGI Symbol;Acc:MGI:1 | 8.031 | 1.0279 | 1.0062 | 0 | 4.3625 | 4.4554 |
| 6728 | ENSMUSG00000028520 | 0 | NA | NA | NA | NA | NA | 4921539E11 | RIKEN cDNA 4921539E11 gene [Source:M | 0 | 0 | 0 | 0 | 0 | 0 |
| 6729 | ENSMUSG00000028521 | 2384.171062 | 0.147657878 | 0.111266715 | 1.327062433 | 0.184488031 | 0.619137675 | Slc35d1 | solute carrier family 35 (UDP-glucuronic a | 2225.4 | 2437.2 | 2125.2 | 2705.9 | 2583.7 | 2227.7 |
| 6730 | ENSMUSG00000028522 | 605.1270869 | 0.081176699 | 0.139377337 | 0.582423947 | 0.560281155 | 0.899475387 | Mier1 | MEIR1 treanscription regulator [Source:M | 511.58 | 596.19 | 657.08 | 562.96 | 639.1 | 663.85 |
| 6731 | ENSMUSG00000028523 | 0 | NA | NA | NA | NA | NA | Dynlt5 | dynein light chain Tctex-type 5 [Source:M | 0 | 0 | 0 | 0 | 0 | 0 |
| 6732 | ENSMUSG00000028524 | 3.654926046 | 0.026128482 | 1.16917038 | 0.022347882 | 0.982170454 | NA | Sgip1 | SH3-domain GRB2-like (endophilin) intera | 4.8186 | 2.0558 | 4.025 | 7.7118 | 1.0906 | 2.2277 |
| 6733 | ENSMUSG00000028525 | 136.2427256 | 1.162813034 | 0.305340969 | 3.808244394 | 0.000139957 | 0.00470776 | Pde4b | phosphodiesterase 4B, cAMP specific [Sou | 87.538 | 76.066 | 88.55 | 167.73 | 107.97 | 289.6 |
| 6734 | ENSMUSG00000028527 | 2829.891189 | -0.097427343 | 0.146428848 | -0.665356207 | 0.505822688 | 0.881592135 | Ak4 | adenylate kinase 4 [Source:MGI Symbol;A | 2415.7 | 3027.2 | 3334.7 | 2829.3 | 3121.3 | 2251.1 |
| 6735 | ENSMUSG00000028528 | 4.562275764 | -0.408813532 | 1.01251325 | -0.403761167 | 0.686388355 | NA | Dnajc6 | DnaJ heat shock protein family (Hsp40) m | 2.4093 | 8.2233 | 5.0312 | 2.8919 | 4.3625 | 4.4554 |
| 6736 | ENSMUSG00000028530 | 1762.379931 | 0.101635977 | 0.099376968 | 1.022731712 | 0.306434713 | 0.755586718 | Jak1 | Janus kinase 1 [Source:MGI Symbol;Acc:M | 1599 | 1715.6 | 1787.1 | 1771.8 | 1905.3 | 1795.5 |
| 6737 | ENSMUSG00000028532 | 14.74963205 | -2.174857185 | 0.705584799 | -3.082346995 | 0.002053753 | 0.035537681 | Cachd1 | cache domain containing 1 [Source:MGI S | 33.73 | 11.307 | 27.169 | 1.928 | 5.4531 | 8.9108 |
| 6738 | ENSMUSG00000028533 | 0 | NA | NA | NA | NA | NA | Izumo3 | IZUMO family member 3 [Source:MGI Sym | 0 | 0 | 0 | 0 | 0 | 0 |
| 6739 | ENSMUSG00000028536 | 235.9776982 | 0.353251844 | 0.199150373 | 1.773794541 | 0.076097141 | 0.403877794 | 2610528J11I | RIKEN cDNA 2610528J11 gene [Source:M | 202.38 | 164.47 | 254.58 | 236.17 | 250.84 | 307.42 |
| 6740 | ENSMUSG00000028538 | 1658.690178 | 0.262396419 | 0.123277183 | 2.128507589 | 0.033295021 | 0.243995238 | St3gal3 | ST3 beta-galactoside alpha-2,3-sialyltrans | 1756.4 | 1396.9 | 1369.5 | 1757.3 | 1742.8 | 1929.2 |
| 6741 | ENSMUSG00000028539 | 1.208395577 | 1.546058676 | 2.154712835 | 0.717524234 | 0.473050689 | NA | Artn | artemin [Source:MGI Symbol;Acc:MGI:13 | 0.8031 | 1.0279 | 0 | 0.964 | 0 | 4.4554 |
| 6742 | ENSMUSG00000028540 | 74.5721853 | -0.335864543 | 0.271511994 | -1.237015495 | 0.21608133 | 0.66165144 | Dph2 | DPH2 homolog [Source:MGI Symbol;Acc:M | 98.782 | 70.926 | 79.493 | 61.695 | 74.162 | 62.375 |
| 6743 | ENSMUSG00000028541 | 4.420275319 | -0.011751813 | 1.087896419 | -0.010802327 | 0.991381158 | NA | B4galt2 | UDP-Gal:betaGlcNAc beta 1,4- galactosylt | 7.2279 | 1.0279 | 5.0312 | 7.7118 | 2.1812 | 3.3415 |
| 6744 | ENSMUSG00000028542 | 664.1015993 | -0.320090187 | 0.172742993 | -1.852985069 | 0.063884499 | 0.361795376 | Slc6a9 | solute carrier family 6 (neurotransmitter t | 803.91 | 727.77 | 681.23 | 720.09 | 620.56 | 431.06 |
| 6745 | ENSMUSG00000028544 | 0 | NA | NA | NA | NA | NA | Slc5a9 | solute carrier family 5 (sodium/glucose co | 0 | 0 | 0 | 0 | 0 | 0 |
| 6746 | ENSMUSG00000028545 | 0.979791906 | -2.106318065 | 2.490808728 | -0.845636215 | 0.397755716 | NA | Bend5 | BEN domain containing 5 [Source:MGI Syr | 0.8031 | 4.1117 | 0 | 0.964 | 0 | 0 |
| 6747 | ENSMUSG00000028546 | 0 | NA | NA | NA | NA | NA | Elavl4 | ELAV like RNA binding protein 4 [Source:M | 0 | 0 | 0 | 0 | 0 | 0 |
| 6748 | ENSMUSG00000028549 | 40.90908854 | 0.359660746 | 0.433647526 | 0.829384983 | 0.406886596 | 0.830690236 | Itgb3bp | integrin beta 3 binding protein (beta3-enc | 36.14 | 30.838 | 40.25 | 19.28 | 63.256 | 55.692 |
| 6749 | ENSMUSG00000028550 | 298.9393929 | -0.158454583 | 0.147913836 | -1.071262751 | 0.284051303 | 0.736651636 | Atg4c | autophagy related 4C, cysteine peptidase | 314.82 | 307.35 | 324.01 | 294.98 | 260.66 | 291.83 |
| 6750 | ENSMUSG00000028551 | 93.98576808 | 0.217609556 | 0.268014991 | 0.81193054 | 0.416831489 | 0.836878476 | Cdkn2c | cyclin dependent kinase inhibitor 2C [Sou | 88.341 | 61.675 | 110.69 | 107.97 | 89.431 | 105.82 |
| 6751 | ENSMUSG00000028552 | 844.9158277 | 0.105886972 | 0.115897016 | 0.913629839 | 0.360911382 | 0.800074801 | Eps15 | epidermal growth factor receptor pathwa | 818.36 | 869.62 | 753.68 | 852.16 | 931.39 | 844.29 |
| 6752 | ENSMUSG00000028553 | 10507.63426 | -0.324903065 | 0.122556953 | -2.651037387 | 0.008024496 | 0.092335965 | Angptl3 | angiopoietin-like 3 [Source:MGI Symbol;A | 10595 | 13558 | 10905 | 8873.4 | 10485 | 8628.9 |
| 6753 | ENSMUSG00000028555 | 33.84072829 | -0.943974425 | 0.525238307 | -1.797230728 | 0.072298998 | 0.389617254 | Ttc39a | tetratricopeptide repeat domain 39A [Sou | 73.082 | 41.117 | 19.119 | 19.28 | 37.081 | 13.366 |
| 6754 | ENSMUSG00000028556 | 164.3024974 | -0.217616833 | 0.22120006 | -0.983800965 | 0.325213385 | 0.771501263 | Dock7 | dedicator of cytokinesis 7 [Source:MGI Sy | 173.47 | 210.72 | 145.91 | 145.56 | 185.41 | 124.75 |
| 6755 | ENSMUSG00000028557 | 1839.233221 | 0.00517457 | 0.099525828 | 0.051992233 | 0.958534882 | 0.992004045 | Rnf11 | ring finger protein 11 [Source:MGI Symbo | 1948.3 | 1761.9 | 1797.2 | 1924.1 | 1850.8 | 1753.2 |
| 6756 | ENSMUSG00000028558 | 0.133850373 | -0.796629176 | 4.080472857 | -0.195229623 | 0.84521319 | NA | Calr4 | calreticulin 4 [Source:MGI Symbol;Acc:M | 0.8031 | 0 | 0 | 0 | 0 | 0 |
| 6757 | ENSMUSG00000028559 | 2595.284274 | 0.119043134 | 0.096713012 | 1.230890573 | 0.218363794 | 0.663735712 | Osbpl9 | oxysterol binding protein-like 9 [Source:M | 2496.8 | 2598.6 | 2369.7 | 2758.9 | 2819.2 | 2528.4 |
| 6758 | ENSMUSG00000028560 | 380.2935909 | 0.183570521 | 0.145167955 | 1.264538862 | 0.206036681 | 0.651217828 | Usp1 | ubiquitin specific peptidase 1 [Source:MG | 363 | 351.55 | 353.19 | 354.74 | 447.15 | 412.12 |
| 6759 | ENSMUSG00000028563 | 321.8465676 | 0.043270996 | 0.160188382 | 0.270125686 | 0.787063562 | 0.960870081 | Tm2d1 | TM2 domain containing 1 [Source:MGI Sy | 273.86 | 344.35 | 334.07 | 358.6 | 323.91 | 296.28 |
| 6760 | ENSMUSG00000028565 | 795.0015542 | -0.007893127 | 0.117147754 | -0.06737754 | 0.946281149 | 0.990448697 | Nfia | nuclear factor I/A [Source:MGI Symbol;Ac | 783.02 | 848.03 | 760.72 | 801.07 | 733.99 | 843.18 |
| 6761 | ENSMUSG00000028567 | 536.3187335 | 0.060390186 | 0.131415403 | 0.459536587 | 0.645848885 | 0.925643431 | Txndc12 | thioredoxin domain containing 12 (endop | 518.8 | 471.81 | 584.63 | 543.68 | 540.95 | 558.04 |
| 6762 | ENSMUSG00000028568 | 546.2987635 | 0.144323851 | 0.122018546 | 1.182802573 | 0.236887401 | 0.685662114 | Btf3l4 | basic transcription factor 3-like 4 [Source: | 540.49 | 517.04 | 499.1 | 575.49 | 597.66 | 548.01 |
| 6763 | ENSMUSG00000028571 | 0 | NA | NA | NA | NA | NA | Cyp2j13 | cytochrome P450, family 2, subfamily j, po | 0 | 0 | 0 | 0 | 0 | 0 |
| 6764 | ENSMUSG00000028572 | 1361.525557 | 0.137270679 | 0.104482997 | 1.313808787 | 0.18891056 | 0.626123359 | Hook1 | hook microtubule tethering protein 1 [Sou | 1200.6 | 1395.9 | 1295 | 1437.3 | 1413.4 | 1426.8 |
| 6765 | ENSMUSG00000028573 | 1914.983373 | 0.112254836 | 0.094914205 | 1.182697955 | 0.236928875 | 0.685662114 | Fggy | FGGY carbohydrate kinase domain contai | 1771.6 | 1909.9 | 1840.4 | 1924.1 | 2063.4 | 1980.4 |
| 6766 | ENSMUSG00000028575 | 0.267700746 | -1.497481649 | 4.057001436 | -0.369110456 | 0.712045396 | NA | Eqtn | equatorin, sperm acrosome associated [So | 1.6062 | 0 | 0 | 0 | 0 | 0 |
| 6767 | ENSMUSG00000028576 | 64.11686248 | -0.140259938 | 0.309707594 | -0.452878589 | 0.650636169 | 0.925643431 | Ift74 | intraflagellar transport 74 [Source:MGI Sy | 51.399 | 86.345 | 64.4 | 62.659 | 70.89 | 49.009 |
| 6768 | ENSMUSG00000028577 | 829.0983199 | 0.060046704 | 0.109835715 | 0.546695611 | 0.58458787 | 0.908955096 | Plaa | phospholipase A2, activating protein [Sou | 829.6 | 835.7 | 769.78 | 819.38 | 872.49 | 847.64 |
| 6769 | ENSMUSG00000028578 | 88.96260643 | 0.16049972 | 0.240394161 | 0.667652323 | 0.50435555 | 0.881510796 | Caap1 | caspase activity and apoptosis inhibitor 1 | 84.326 | 89.429 | 78.487 | 104.11 | 89.431 | 87.994 |
| 6770 | ENSMUSG00000028580 | 772.8895693 | 0.11511673 | 0.124680085 | 0.923296852 | 0.355852528 | 0.795525652 | Pum1 | pumilio RNA-binding family member 1 [So | 816.75 | 722.63 | 686.26 | 853.12 | 822.33 | 736.25 |
| 6771 | ENSMUSG00000028581 | 137.249628 | -1.14667347 | 0.236584228 | -4.846787456 | 1.25E-06 | 9.86E-05 | Laptm5 | lysosomal-associated protein transmembr | 175.08 | 230.25 | 162.01 | 83.866 | 68.709 | 103.59 |
| 6772 | ENSMUSG00000028582 | 221.77805 | -0.354877978 | 0.198103686 | -1.791374936 | 0.073233148 | 0.392921239 | Cc2d1b | coiled-coil and C2 domain containing 1B [S | 259.4 | 231.28 | 256.59 | 240.03 | 195.22 | 148.14 |

| | | | | | | | | | | | | | | | |
|---|---|---|---|---|---|---|---|---|---|---|---|---|---|---|---|
| 6773 | ENSMUSG00000028583 | 10.49705169 | 0.235789517 | 0.714765365 | 0.329883803 | 0.741487763 | 0.948792385 | Pdpn | podoplanin [Source:MGI Symbol;Acc:MGI | 9.6372 | 12.335 | 7.0437 | 17.352 | 4.3625 | 12.252 |
| 6774 | ENSMUSG00000028584 | 0 | NA | NA | NA | NA | NA | Lrrc38 | leucine rich repeat containing 38 [Source: | 0 | 0 | 0 | 0 | 0 | 0 |
| 6775 | ENSMUSG00000028587 | 1.596583247 | -1.035272675 | 1.872224846 | -0.552963858 | 0.580288158 | NA | Orc1 | origin recognition complex, subunit 1 [Sou | 2.4093 | 0 | 4.025 | 0.964 | 2.1812 | 0 |
| 6776 | ENSMUSG00000028589 | 0 | NA | NA | NA | NA | NA | 1700012P22 | RIKEN cDNA 1700012P22 gene [Source:M | 0 | 0 | 0 | 0 | 0 | 0 |
| 6777 | ENSMUSG00000028590 | 0 | NA | NA | NA | NA | NA | Pramel13os | PRAME like 13, opposite strand [Source:M | 0 | 0 | 0 | 0 | 0 | 0 |
| 6778 | ENSMUSG00000028591 | 0 | NA | NA | NA | NA | NA | Pramel13 | PRAME like 13 [Source:MGI Symbol;Acc:M | 0 | 0 | 0 | 0 | 0 | 0 |
| 6779 | ENSMUSG00000028593 | 0 | NA | NA | NA | NA | NA | Aadacl4fm1 | AADACL4 family member 1 [Source:MGI S | 0 | 0 | 0 | 0 | 0 | 0 |
| 6780 | ENSMUSG00000028597 | 28.18722099 | -0.23033574 | 0.436016361 | -0.52827316 | 0.597309758 | 0.914521894 | Gpx7 | glutathione peroxidase 7 [Source:MGI Syr | 35.336 | 31.865 | 24.15 | 34.703 | 17.45 | 25.618 |
| 6781 | ENSMUSG00000028599 | 691.6431598 | -0.487556486 | 0.173362853 | -2.812346919 | 0.004918143 | 0.066064903 | Tnfrsf1b | tumor necrosis factor receptor superfamil | 789.45 | 974.47 | 658.08 | 496.45 | 703.45 | 527.96 |
| 6782 | ENSMUSG00000028600 | 81.03108338 | -0.301240102 | 0.315504025 | -0.954790046 | 0.339683882 | 0.781856727 | Podn | podocan [Source:MGI Symbol;Acc:MGI:26 | 110.03 | 97.652 | 60.375 | 66.514 | 98.156 | 53.465 |
| 6783 | ENSMUSG00000028601 | 3961.810175 | 0.262019414 | 0.086885035 | 3.015702448 | 0.002563847 | 0.04206803 | Echdc2 | enoyl Coenzyme A hydratase domain cont | 3709.5 | 3589.5 | 3509.8 | 4455.5 | 4178.2 | 4328.4 |
| 6784 | ENSMUSG00000028602 | 1.006604507 | -1.038622879 | 2.695419007 | -0.385328914 | 0.699993742 | NA | Tnfrsf8 | tumor necrosis factor receptor superfamil | 0 | 4.1117 | 0 | 1.928 | 0 | 0 |
| 6785 | ENSMUSG00000028603 | 100678.6686 | 0.149988612 | 0.089898951 | 1.668413375 | 0.095233695 | 0.455057116 | Scp2 | sterol carrier protein 2, liver [Source:MGI | 100234 | 94930 | 91185 | 104813 | 112931 | 99979 |
| 6786 | ENSMUSG00000028607 | 3699.304756 | -0.2307222 | 0.10871255 | -2.122314309 | 0.033811353 | 0.246247434 | Cpt2 | carnitine palmitoyltransferase 2 [Source:N | 3781 | 3872.2 | 4330.9 | 3368.1 | 3757.2 | 3086.5 |
| 6787 | ENSMUSG00000028608 | 427.9345131 | 0.010108611 | 0.149763913 | 0.067496974 | 0.946186071 | 0.990420851 | Czib | CXXC motif containing zinc binding proteir | 470.62 | 402.94 | 404.51 | 373.06 | 480.96 | 435.51 |
| 6788 | ENSMUSG00000028609 | 228.7439544 | -0.08793037 | 0.16897411 | -0.520377765 | 0.602800304 | 0.91570264 | Magoh | mago homolog, exon junction complex co | 253.78 | 248.76 | 204.27 | 222.68 | 224.67 | 218.31 |
| 6789 | ENSMUSG00000028610 | 0 | NA | NA | NA | NA | NA | Dmrtb1 | DMRT-like family B with proline-rich C-ter | 0 | 0 | 0 | 0 | 0 | 0 |
| 6790 | ENSMUSG00000028612 | 0 | NA | NA | NA | NA | NA | Lrp8os1 | low density lipoprotein receptor-related p | 0 | 0 | 0 | 0 | 0 | 0 |
| 6791 | ENSMUSG00000028613 | 1.313397344 | 0.961420072 | 1.854794958 | 0.518343048 | 0.604218943 | NA | Lrp8 | low density lipoprotein receptor-related p | 1.6062 | 1.0279 | 0 | 1.928 | 1.0906 | 2.2277 |
| 6792 | ENSMUSG00000028614 | 211.8586272 | 0.184090318 | 0.173069088 | 1.063681101 | 0.287473189 | 0.73951397 | Ndc1 | NDC1 transmembrane nucleoporin [Sourc | 190.34 | 204.56 | 200.24 | 209.18 | 255.2 | 211.63 |
| 6793 | ENSMUSG00000028617 | 304.9518688 | 0.385446077 | 0.154914428 | 2.488121726 | 0.012841977 | 0.128189342 | Lrrc42 | leucine rich repeat containing 42 [Source: | 281.89 | 283.71 | 227.41 | 348.96 | 342.45 | 345.29 |
| 6794 | ENSMUSG00000028618 | 4136.26458 | -0.043574666 | 0.090301286 | -0.482547564 | 0.629417019 | 0.92057725 | Tmem59 | transmembrane protein 59 [Source:MGI S | 3984.2 | 4338.8 | 4274.5 | 4207.8 | 4113.8 | 3898.5 |
| 6795 | ENSMUSG00000028619 | 296.8657132 | 0.189079016 | 0.159507958 | 1.18538923 | 0.235863588 | 0.684222009 | Tceanc2 | transcription elongation factor A (SII) N-te | 273.86 | 271.37 | 286.78 | 278.59 | 363.18 | 307.42 |
| 6796 | ENSMUSG00000028621 | 183.0566808 | 0.066801272 | 0.181169273 | 0.368722967 | 0.712334228 | 0.944239215 | Cyb5rl | cytochrome b5 reductase-like [Source:MG | 198.37 | 160.36 | 177.1 | 182.19 | 185.41 | 194.92 |
| 6797 | ENSMUSG00000028622 | 769.5490459 | 0.005216449 | 0.116538355 | 0.044761649 | 0.964297294 | 0.993347771 | Mrpl37 | mitochondrial ribosomal protein L37 [Sou | 758.93 | 792.52 | 752.67 | 722.98 | 750.35 | 839.84 |
| 6798 | ENSMUSG00000028626 | 0.181769715 | 1.126924175 | 4.080472857 | 0.276174898 | 0.782413736 | NA | Col9a2 | collagen, type IX, alpha 2 [Source:MGI Syr | 0 | 0 | 0 | 0 | 1.0906 | 0 |
| 6799 | ENSMUSG00000028629 | 59.65229061 | 0.167808369 | 0.309877622 | 0.541531099 | 0.588141568 | 0.910800818 | Exo5 | exonuclease 5 [Source:MGI Symbol;Acc:M | 58.626 | 48.312 | 61.381 | 55.911 | 51.259 | 82.424 |
| 6800 | ENSMUSG00000028630 | 1157.594683 | 0.125950196 | 0.120384676 | 1.046231136 | 0.295454331 | 0.745982248 | Dyrk2 | dual-specificity tyrosine-(Y)-phosphorylati | 991.83 | 1100.9 | 1229.6 | 1223.3 | 1117.9 | 1282 |
| 6801 | ENSMUSG00000028631 | 5.569607152 | -0.738250432 | 1.085435966 | -0.680141856 | 0.496414644 | 0.878165817 | Kcnq4 | potassium voltage-gated channel, subfam | 4.8186 | 1.0279 | 15.094 | 4.8199 | 6.5437 | 1.1138 |
| 6802 | ENSMUSG00000028633 | 132.2512627 | -0.474636742 | 0.293869795 | -1.615125986 | 0.106283415 | 0.480851832 | Ctps | cytidine 5'-triphosphate synthase [Source | 220.05 | 145.96 | 94.587 | 89.65 | 136.33 | 106.93 |
| 6803 | ENSMUSG00000028634 | 11.1473336 | -1.232912485 | 0.679984189 | -1.813148755 | 0.069808881 | 0.38174505 | Hivep3 | human immunodeficiency virus type I enh | 18.471 | 10.279 | 18.112 | 6.7478 | 4.3625 | 8.9108 |
| 6804 | ENSMUSG00000028635 | 0.167707431 | -0.796629176 | 4.080472857 | -0.195229623 | 0.84521319 | NA | Edn2 | endothelin 2 [Source:MGI Symbol;Acc:MG | 0 | 0 | 1.0062 | 0 | 0 | 0 |
| 6805 | ENSMUSG00000028636 | 581.0627609 | 0.126550106 | 0.13599615 | 0.930541823 | 0.35209062 | 0.792945339 | Ppcs | phosphopantothenoylcysteine synthetase | 606.34 | 554.05 | 505.13 | 550.43 | 597.66 | 672.76 |
| 6806 | ENSMUSG00000028637 | 120.2370162 | 0.051009911 | 0.228425072 | 0.223311349 | 0.823293195 | 0.96943211 | Ccdc30 | coiled-coil domain containing 30 [Source:I | 120.47 | 97.652 | 135.84 | 104.11 | 134.15 | 129.21 |
| 6807 | ENSMUSG00000028639 | 4278.026379 | 0.00695065 | 0.08736564 | 0.079558166 | 0.936588669 | 0.988420305 | Ybx1 | Y box protein 1 [Source:MGI Symbol;Acc:I | 4046.8 | 4336.8 | 4420.4 | 4256 | 4272 | 4336.2 |
| 6808 | ENSMUSG00000028640 | 0 | NA | NA | NA | NA | NA | Tfap2c | transcription factor AP-2, gamma [Source: | 0 | 0 | 0 | 0 | 0 | 0 |
| 6809 | ENSMUSG00000028641 | 269.3434602 | -0.093340786 | 0.185953688 | -0.501957165 | 0.615697653 | 0.917509899 | P3h1 | prolyl 3-hydroxylase 1 [Source:MGI Symbo | 246.55 | 256.98 | 331.05 | 267.02 | 215.94 | 298.51 |
| 6810 | ENSMUSG00000028642 | 1.643803754 | 0.255028028 | 1.738937021 | 0.146657426 | 0.883402424 | NA | Tmem269 | transmembrane protein 269 [Source:MGI | 2.4093 | 1.0279 | 1.0062 | 0.964 | 0 | 4.4554 |
| 6811 | ENSMUSG00000028643 | 94.29962991 | -0.312637918 | 0.266382076 | -1.173644721 | 0.24053736 | 0.690715184 | Svbp | small vasohibin binding protein [Source:M | 95.569 | 131.57 | 86.537 | 88.686 | 66.528 | 96.904 |
| 6812 | ENSMUSG00000028644 | 1.449082645 | 1.182143685 | 1.823582063 | 0.648253626 | 0.516820922 | NA | Ermap | erythroblast membrane-associated protei | 1.6062 | 1.0279 | 0 | 3.8559 | 1.0906 | 1.1138 |
| 6813 | ENSMUSG00000028645 | 227.202555 | -0.156321438 | 0.24054932 | -0.649851921 | 0.515787877 | 0.886238519 | Slc2a1 | solute carrier family 2 (facilitated glucose | 162.23 | 223.06 | 334.07 | 232.32 | 184.31 | 227.22 |
| 6814 | ENSMUSG00000028646 | 747.4016844 | 0.041802114 | 0.116863959 | 0.357698939 | 0.720568627 | 0.9447719 | Rragc | Ras-related GTP binding C [Source:MGI Sy | 760.54 | 769.91 | 679.22 | 787.57 | 723.08 | 764.1 |
| 6815 | ENSMUSG00000028647 | 250.0351578 | -0.060931218 | 0.170229515 | -0.357935683 | 0.720391447 | 0.9447719 | Mycbp | MYC binding protein [Source:MGI Symbol | 262.61 | 267.26 | 235.46 | 208.22 | 270.47 | 256.18 |
| 6816 | ENSMUSG00000028648 | 1151.289427 | -0.05818762 | 0.128266351 | -0.453646802 | 0.650083062 | 0.925643431 | Ndufs5 | NADH:ubiquinone oxidoreductase core su | 1338.8 | 1180 | 1003.2 | 1083.5 | 1104.8 | 1197.4 |
| 6817 | ENSMUSG00000028649 | 1233.845708 | -0.172274239 | 0.122951386 | -1.401157356 | 0.161167023 | 0.584240557 | Macf1 | microtubule-actin crosslinking factor 1 [Sc | 1322.7 | 1399 | 1201.5 | 1313.9 | 1117.9 | 1048.1 |
| 6818 | ENSMUSG00000028651 | 196.6444288 | -0.078413715 | 0.172312217 | -0.455067647 | 0.649060573 | 0.925643431 | Ppie | peptidylprolyl isomerase E (cyclophilin E) | | 207.2 | 208.67 | 190.18 | 203.4 | 179.95 | 190.47 |
| 6819 | ENSMUSG00000028653 | 304.2411833 | 0.229609159 | 0.155000114 | 1.481348329 | 0.138513776 | 0.543828246 | Trit1 | tRNA isopentenyltransferase 1 [Source:M | 281.09 | 286.79 | 272.69 | 351.85 | 349 | 284.03 |
| 6820 | ENSMUSG00000028654 | 395.9415355 | -0.376400283 | 0.136021166 | -2.767218471 | 0.005653685 | 0.072431271 | Mycl | v-myc avian myelocytomatosis viral oncog | 429.66 | 431.73 | 480.98 | 356.67 | 342.45 | 334.15 |
| 6821 | ENSMUSG00000028655 | 3141.010127 | -1.471825865 | 0.393368256 | -3.741597958 | 0.000182854 | 0.005864298 | Mfsd2a | major facilitator superfamily domain cont | 4529.5 | 5371.9 | 3950.5 | 1283.1 | 2521.5 | 1189.6 |
| 6822 | ENSMUSG00000028656 | 1365.642982 | -0.028231196 | 0.099158608 | -0.284707468 | 0.775868269 | 0.95893551 | Cap1 | CAP, adenylate cyclase-associated proteir | 1443.2 | 1346.6 | 1346.4 | 1320.6 | 1393.8 | 1343.3 |
| 6823 | ENSMUSG00000028657 | 418.9903101 | -0.189382767 | 0.139410647 | -1.358452677 | 0.174320089 | 0.604151393 | Ppt1 | palmitoyl-protein thioesterase 1 [Source:I | 464.19 | 422.47 | 451.8 | 348.96 | 416.62 | 409.89 |
| 6824 | ENSMUSG00000028661 | 0 | NA | NA | NA | NA | NA | Epha8 | Eph receptor A8 [Source:MGI Symbol;Acc | 0 | 0 | 0 | 0 | 0 | 0 |
| 6825 | ENSMUSG00000028664 | 8.299977484 | -1.972849396 | 0.952073695 | -2.072160386 | 0.038250486 | 0.266030047 | Ephb2 | Eph receptor B2 [Source:MGI Symbol;Acc: | 26.502 | 8.2233 | 5.0312 | 6.7478 | 2.1812 | 1.1138 |
| 6826 | ENSMUSG00000028668 | 623.5826849 | -0.057583707 | 0.115272872 | -0.499542568 | 0.617397206 | 0.917509899 | Eloa | elongin A [Source:MGI Symbol;Acc:MGI:1 | 637.66 | 634.23 | 635.95 | 596.7 | 599.84 | 637.12 |
| 6827 | ENSMUSG00000028669 | 511.7224434 | 0.131860062 | 0.127769307 | 1.032016725 | 0.302064282 | 0.752248241 | Pithd1 | PITH (C-terminal proteasome-interacting c | 462.59 | 493.4 | 509.16 | 495.48 | 560.58 | 549.13 |
| 6828 | ENSMUSG00000028670 | 951.8556686 | 0.04863303 | 0.106330808 | 0.457374785 | 0.647401685 | 0.925643431 | Lypla2 | lysophospholipase 2 [Source:MGI Symbol, | 924.37 | 941.57 | 941.84 | 995.79 | 910.67 | 996.89 |

| | | | | | | | | | | | | | | | |
|---|---|---|---|---|---|---|---|---|---|---|---|---|---|---|---|
| 6829 | ENSMUSG00000028671 | 190.744432 | 0.290362389 | 0.316527008 | 0.917338432 | 0.358965324 | 0.798534982 | Gale | galactose-4-epimerase, UDP [Source:MGI | 289.92 | 112.04 | 111.69 | 203.4 | 171.23 | 256.18 |
| 6830 | ENSMUSG00000028672 | 5449.433367 | -0.244492745 | 0.094234287 | -2.594520031 | 0.009472309 | 0.104426435 | Hmgcl | 3-hydroxy-3-methylglutaryl-Coenzyme A l | 6361.4 | 5932.1 | 5435.7 | 5008.8 | 5036.5 | 4922.1 |
| 6831 | ENSMUSG00000028673 | 1589.677419 | -0.035323539 | 0.092730529 | -0.380926745 | 0.703257607 | 0.942506832 | Fuca1 | fucosidase, alpha-L- 1, tissue [Source:MGI | 1603.8 | 1608.7 | 1615 | 1573.2 | 1585.8 | 1551.6 |
| 6832 | ENSMUSG00000028675 | 886.7246361 | 0.064734252 | 0.113887801 | 0.568403742 | 0.56976085 | 0.904017215 | Pnrc2 | proline-rich nuclear receptor coactivator 2 | 879.4 | 858.31 | 862.35 | 885.9 | 841.96 | 992.43 |
| 6833 | ENSMUSG00000028676 | 656.7308511 | -0.118089819 | 0.124100536 | -0.951565744 | 0.341317262 | 0.783129922 | Srsf10 | serine and arginine-rich splicing factor 10 | 616.78 | 731.88 | 703.36 | 659.36 | 607.47 | 621.52 |
| 6834 | ENSMUSG00000028677 | 451.3542283 | -0.056175594 | 0.157707192 | -0.356201854 | 0.721689404 | 0.9447719 | Rnf220 | ring finger protein 220 [Source:MGI Symb | 469.81 | 471.81 | 438.72 | 459.82 | 353.36 | 514.6 |
| 6835 | ENSMUSG00000028678 | 4.150320961 | -0.167268353 | 1.202762849 | -0.139070103 | 0.889394753 | NA | Kif2c | kinesin family member 2C [Source:MGI Sy | 4.0155 | 4.1117 | 5.0312 | 1.928 | 9.8156 | 0 |
| 6836 | ENSMUSG00000028680 | 842.0371326 | 0.831470505 | 0.799735715 | 1.039681597 | 0.298487853 | 0.749276 | Plk3 | polo like kinase 3 [Source:MGI Symbol;Acc | 621.6 | 893.26 | 302.88 | 1781.4 | 285.74 | 1167.3 |
| 6837 | ENSMUSG00000028681 | 14.37762298 | 0.578479746 | 0.697054792 | 0.829891355 | 0.406600213 | 0.830399297 | Ptch2 | patched 2 [Source:MGI Symbol;Acc:MGI:1 | 24.093 | 4.1117 | 6.0375 | 21.208 | 17.45 | 13.366 |
| 6838 | ENSMUSG00000028683 | 195.4493233 | 0.00169126 | 0.206815108 | 0.008177641 | 0.99347526 | 0.99859176 | Eif2b3 | eukaryotic translation initiation factor 2B, | 189.53 | 245.67 | 150.94 | 188.94 | 210.49 | 187.13 |
| 6839 | ENSMUSG00000028684 | 1632.120996 | -0.008623633 | 0.100090818 | -0.086158083 | 0.931340752 | 0.988420305 | Urod | uroporphyrinogen decarboxylase [Source: | 1587.7 | 1578.9 | 1744.8 | 1629.1 | 1686.1 | 1566.1 |
| 6840 | ENSMUSG00000028687 | 8.455119463 | 0.107580996 | 0.741240199 | 0.145136484 | 0.884603113 | 0.981795936 | Mutyh | mutY DNA glycosylase [Source:MGI Symbo | 7.2279 | 5.1396 | 12.075 | 8.6758 | 9.8156 | 7.7969 |
| 6841 | ENSMUSG00000028688 | 85.16293782 | -0.247053884 | 0.240148353 | -1.028755273 | 0.303594689 | 0.752690022 | Toe1 | target of EGR1, member 1 (nuclear) [Sour | 93.16 | 93.541 | 90.562 | 76.154 | 80.706 | 76.855 |
| 6842 | ENSMUSG00000028689 | 88.7181367 | 0.242170215 | 0.235116255 | 1.03000201 | 0.303009062 | 0.752484788 | Ccdc163 | coiled-coil domain containing 163 [Source | 81.113 | 82.233 | 80.5 | 94.47 | 95.974 | 98.018 |
| 6843 | ENSMUSG00000028690 | 318.7572661 | -0.026396368 | 0.166109453 | -0.158909485 | 0.873740188 | 0.979925081 | Mmachc | methylmalonic aciduria cblC type, with ho | 335.7 | 311.46 | 316.97 | 255.45 | 358.81 | 334.15 |
| 6844 | ENSMUSG00000028691 | 14437.5733 | 0.105717627 | 0.084080732 | 1.25733477 | 0.208632439 | 0.653455953 | Prdx1 | peroxiredoxin 1 [Source:MGI Symbol;Acc: | 14602 | 13491 | 13632 | 14516 | 15311 | 15074 |
| 6845 | ENSMUSG00000028692 | 7565.874145 | -0.079256285 | 0.09694972 | -0.817498851 | 0.413643406 | 0.835385466 | Akr1a1 | aldo-keto reductase family 1, member A1 | 8323.4 | 7721.7 | 7274.1 | 6860.6 | 7779.4 | 7436 |
| 6846 | ENSMUSG00000028693 | 319.7509412 | 0.077486902 | 0.157775584 | 0.491120996 | 0.623340873 | 0.919503423 | Nasp | nuclear autoantigenic sperm protein (hist | 307.59 | 319.68 | 305.9 | 287.27 | 380.63 | 317.45 |
| 6847 | ENSMUSG00000028696 | 241.4990055 | 0.123819558 | 0.1755701 | 0.70524285 | 0.480659119 | 0.870278888 | Ipp | IAP promoted placental gene [Source:MG | 259.4 | 204.56 | 228.42 | 218.82 | 270.47 | 267.32 |
| 6848 | ENSMUSG00000028698 | 31.73571158 | 0.163638028 | 0.414648535 | 0.394642725 | 0.693106577 | 0.93867917 | Pik3r3 | phosphoinositide-3-kinase regulatory sub | 38.549 | 26.726 | 24.15 | 25.063 | 43.625 | 32.301 |
| 6849 | ENSMUSG00000028699 | 0 | NA | NA | NA | NA | NA | Tspan1 | tetraspanin 1 [Source:MGI Symbol;Acc:M | 0 | 0 | 0 | 0 | 0 | 0 |
| 6850 | ENSMUSG00000028700 | 392.0195695 | 0.166990177 | 0.13318711 | 1.253801343 | 0.209914221 | 0.653789683 | Pomgnt1 | protein O-linked mannose beta 1,2-N-acet | 360.59 | 388.55 | 359.23 | 420.29 | 403.53 | 419.92 |
| 6851 | ENSMUSG00000028701 | 5.966634236 | 0.415569013 | 0.928088637 | 0.447768668 | 0.654320158 | 0.925797231 | Lurap1 | leucine rich adaptor protein 1 [Source:MG | 3.2124 | 3.0838 | 9.0562 | 3.8559 | 5.4531 | 11.138 |
| 6852 | ENSMUSG00000028702 | 3.927764726 | -0.668479301 | 1.163863949 | -0.574362065 | 0.565722818 | NA | Rad54l | RAD54 like (S. cerevisiae) [Source:MGI Syr | 2.4093 | 6.1675 | 6.0375 | 6.7478 | 1.0906 | 1.1138 |
| 6853 | ENSMUSG00000028703 | 307.7832236 | 0.00994804 | 0.146326293 | 0.06798532 | 0.945797321 | 0.990309554 | Lrrc41 | leucine rich repeat containing 41 [Source: | 312.41 | 289.87 | 317.97 | 324.86 | 308.64 | 292.94 |
| 6854 | ENSMUSG00000028706 | 387.6789249 | 0.295273633 | 0.136277689 | 2.166705602 | 0.030257309 | 0.230381759 | Nsun4 | NOL1/NOP2/Sun domain family, member | 346.94 | 360.8 | 337.09 | 455.96 | 422.07 | 403.21 |
| 6855 | ENSMUSG00000028707 | 0 | NA | NA | NA | NA | NA | Dmbx1 | diencephalon/mesencephalon homeobox | 0 | 0 | 0 | 0 | 0 | 0 |
| 6856 | ENSMUSG00000028708 | 332.0700666 | -0.051026571 | 0.151310755 | -0.337230299 | 0.735943298 | 0.948109711 | Mknk1 | MAP kinase-interacting serine/threonine k | 345.33 | 359.77 | 308.92 | 344.14 | 343.54 | 290.71 |
| 6857 | ENSMUSG00000028709 | 259.1066536 | 0.048257394 | 0.15786038 | 0.305696682 | 0.7598356 | 0.95402891 | Mob3c | MOB kinase activator 3C [Source:MGI Sym | 264.22 | 240.53 | 259.61 | 285.34 | 244.3 | 260.64 |
| 6858 | ENSMUSG00000028710 | 271.5368727 | 0.062947909 | 0.151130674 | 0.416513123 | 0.677034567 | 0.933164472 | Atpaf1 | ATP synthase mitochondrial F1 complex a | 274.66 | 270.34 | 251.56 | 271.84 | 287.92 | 272.89 |
| 6859 | ENSMUSG00000028712 | 97.98531022 | -0.749242465 | 0.274163668 | -2.732829154 | 0.006279289 | 0.077627045 | Cyp4a31 | cytochrome P450, family 4, subfamily a, p | 114.84 | 133.63 | 120.75 | 98.326 | 49.078 | 71.286 |
| 6860 | ENSMUSG00000028713 | 165.1579274 | -0.406536779 | 0.269257921 | -1.509841485 | 0.131083877 | 0.53064573 | Cyp4b1 | cytochrome P450, family 4, subfamily b, p | 150.98 | 217.92 | 196.22 | 143.63 | 86.159 | 196.04 |
| 6861 | ENSMUSG00000028715 | 8034.53038 | -2.743982052 | 1.091132218 | -2.51480252 | 0.011909907 | 0.122087102 | Cyp4a14 | cytochrome P450, family 4, subfamily a, p | 9797.8 | 21508 | 10640 | 5035.8 | 675.09 | 550.24 |
| 6862 | ENSMUSG00000028716 | 23.40118364 | 0.942136907 | 0.536391596 | 1.756434876 | 0.079014182 | 0.412857069 | Pdzk1ip1 | PDZK1 interacting protein 1 [Source:MGI S | 14.456 | 16.447 | 17.106 | 19.28 | 18.541 | 54.578 |
| 6863 | ENSMUSG00000028717 | 14.18939024 | -0.238232895 | 0.6569219 | -0.362650256 | 0.716866163 | 0.9447719 | Tal1 | T cell acute lymphocytic leukemia 1 [Sourc | 21.684 | 10.279 | 14.087 | 20.244 | 4.3625 | 14.48 |
| 6864 | ENSMUSG00000028718 | 1.479611194 | -0.378768569 | 2.116041761 | -0.178998627 | 0.85793878 | NA | Stil | Scl/Tal1 interrupting locus [Source:MGI Sy | 4.0155 | 0 | 1.0062 | 3.8559 | 0 | 0 |
| 6865 | ENSMUSG00000028719 | 1984.913151 | -0.095887394 | 0.100617213 | -0.952991955 | 0.340594147 | 0.783094143 | Cmpk1 | cytidine monophosphate (UMP-CMP) kina | 2183.6 | 2017.8 | 1950.1 | 1881.7 | 2036.2 | 1840.1 |
| 6866 | ENSMUSG00000028729 | 447.2734413 | -0.177126763 | 0.132394617 | -1.337869831 | 0.180938873 | 0.61434269 | Ebna1bp2 | EBNA1 binding protein 2 [Source:MGI Sym | 456.97 | 492.37 | 474.95 | 405.83 | 455.88 | 397.64 |
| 6867 | ENSMUSG00000028730 | 0 | NA | NA | NA | NA | NA | Cfap57 | cilia and flagella associated protein 57 [So | 0 | 0 | 0 | 0 | 0 | 0 |
| 6868 | ENSMUSG00000028736 | 0 | NA | NA | NA | NA | NA | Pax7 | paired box 7 [Source:MGI Symbol;Acc:MG | 0 | 0 | 0 | 0 | 0 | 0 |
| 6869 | ENSMUSG00000028737 | 4943.687318 | 0.096647287 | 0.091549133 | 1.055687629 | 0.291110951 | 0.740906421 | Aldh4a1 | aldehyde dehydrogenase 4 family, membe | 4691.7 | 4792.2 | 4850.1 | 4724.5 | 5339.7 | 5264 |
| 6870 | ENSMUSG00000028738 | 0.48903338 | 0.95637552 | 3.456690196 | 0.276673773 | 0.782030612 | NA | Tas1r2 | taste receptor, type 1, member 2 [Source: | 0 | 0 | 1.0062 | 1.928 | 0 | 0 |
| 6871 | ENSMUSG00000028741 | 363.2525057 | 0.073124828 | 0.147118639 | 0.497046655 | 0.619156152 | 0.917569117 | Mrto4 | mRNA turnover 4, ribosome maturation fa | 342.92 | 373.13 | 346.15 | 349.92 | 420.98 | 346.41 |
| 6872 | ENSMUSG00000028743 | 3343.998935 | 0.01714643 | 0.091902153 | 0.186572671 | 0.851995696 | 0.973709367 | Akr7a5 | aldo-keto reductase family 7, member A5 | 3410 | 3116.6 | 3445.4 | 3426 | 3258.8 | 3407.2 |
| 6873 | ENSMUSG00000028744 | 261.4469473 | 0.269504091 | 0.168459651 | 1.599813899 | 0.109639875 | 0.486451272 | Slc66a1 | solute carrier family 66 member 1 [Source | 244.95 | 227.17 | 238.48 | 244.85 | 284.65 | 328.58 |
| 6874 | ENSMUSG00000028745 | 1489.083454 | -0.060665298 | 0.097390603 | -0.622907095 | 0.533345576 | 0.889143097 | Capzb | capping protein (actin filament) muscle Z- | 1562.8 | 1542.9 | 1455 | 1451.8 | 1499.6 | 1422.4 |
| 6875 | ENSMUSG00000028747 | 0 | NA | NA | NA | NA | NA | Htr6 | 5-hydroxytryptamine (serotonin) receptor | 0 | 0 | 0 | 0 | 0 | 0 |
| 6876 | ENSMUSG00000028749 | 0 | NA | NA | NA | NA | NA | Pla2g2f | phospholipase A2, group IIF [Source:MGI | 0 | 0 | 0 | 0 | 0 | 0 |
| 6877 | ENSMUSG00000028750 | 0.181769715 | 1.126924175 | 4.080472857 | 0.276174898 | 0.782413736 | NA | Pla2g2c | phospholipase A2, group IIC [Source:MGI | 0 | 0 | 0 | 0 | 1.0906 | 0 |
| 6878 | ENSMUSG00000028751 | 0 | NA | NA | NA | NA | NA | Pla2g2e | phospholipase A2, group IIE [Source:MGI | 0 | 0 | 0 | 0 | 0 | 0 |
| 6879 | ENSMUSG00000028753 | 0 | NA | NA | NA | NA | NA | Vwa5b1 | von Willebrand factor A domain containin | 0 | 0 | 0 | 0 | 0 | 0 |
| 6880 | ENSMUSG00000028755 | 165.304185 | -0.335976057 | 0.250998669 | -1.338557126 | 0.180714891 | 0.614014609 | Cda | cytidine deaminase [Source:MGI Symbol;A | 187.12 | 173.72 | 193.2 | 208.22 | 99.246 | 130.32 |
| 6881 | ENSMUSG00000028756 | 4208.286173 | 0.043841174 | 0.093808126 | 0.467349434 | 0.640249894 | 0.923932929 | Pink1 | PTEN induced putative kinase 1 [Source:M | 4099.8 | 4271 | 4063.2 | 4596.2 | 4010.2 | 4209.2 |
| 6882 | ENSMUSG00000028757 | 1352.329653 | 0.186298968 | 0.113907427 | 1.635529592 | 0.101938076 | 0.469863715 | Ddost | dolichyl-di-phosphooligosaccharide-prote | 1296.2 | 1327 | 1172.3 | 1527.9 | 1290.2 | 1500.3 |
| 6883 | ENSMUSG00000028758 | 0.335414862 | -1.779047242 | 4.030804179 | -0.441362855 | 0.658950329 | NA | Kif17 | kinesin family member 17 [Source:MGI Sy | 0 | 0 | 2.0125 | 0 | 0 | 0 |
| 6884 | ENSMUSG00000028759 | 1906.814915 | 0.013216542 | 0.098829843 | 0.133730275 | 0.893615864 | 0.982562372 | Hp1bp3 | heterochromatin protein 1, binding protei | 1906.6 | 1904.7 | 1883.7 | 2070.6 | 1849.7 | 1825.6 |

| | | | | | | | | | | | | | | | |
|---|---|---|---|---|---|---|---|---|---|---|---|---|---|---|---|
| 6885 | ENSMUSG00000028760 | 913.8246197 | -0.003089249 | 0.110523365 | -0.027951096 | 0.977701156 | 0.997564159 | Eif4g3 | eukaryotic translation initiation factor 4 g | 972.56 | 893.26 | 877.45 | 873.36 | 911.76 | 954.56 |
| 6886 | ENSMUSG00000028763 | 220.0607156 | -0.657063692 | 0.225348354 | -2.915768762 | 0.003548134 | 0.052798565 | Hspg2 | perlecan (heparan sulfate proteoglycan 2) | 280.28 | 279.59 | 248.54 | 216.9 | 110.15 | 184.9 |
| 6887 | ENSMUSG00000028766 | 435.1562885 | -0.422591004 | 0.157549116 | -2.682281021 | 0.007312201 | 0.086502219 | Alpl | alkaline phosphatase, liver/bone/kidney [ | 540.49 | 489.29 | 465.89 | 433.79 | 375.17 | 306.31 |
| 6888 | ENSMUSG00000028771 | 387.7621268 | 0.16190921 | 0.166506449 | 0.972390022 | 0.330856554 | 0.774747658 | Ptpn12 | protein tyrosine phosphatase, non-recept | 347.74 | 384.44 | 366.27 | 431.86 | 320.64 | 475.61 |
| 6889 | ENSMUSG00000028772 | 255.1990986 | -0.063546674 | 0.187772333 | -0.338424054 | 0.73504365 | 0.948109711 | Zcchc17 | zinc finger, CCHC domain containing 17 [S | 217.64 | 262.12 | 302.88 | 207.26 | 262.84 | 278.46 |
| 6890 | ENSMUSG00000028773 | 0.342432689 | 1.990653086 | 3.900281263 | 0.51038706 | 0.609780321 | NA | Fabp3 | fatty acid binding protein 3, muscle and h | 0 | 0 | 0 | 0.964 | 1.0906 | 0 |
| 6891 | ENSMUSG00000028776 | 244.7508504 | -0.421184882 | 0.212823006 | -1.979038312 | 0.047811693 | 0.303663969 | Tinagl1 | tubulointerstitial nephritis antigen-like 1 [ | 277.87 | 293.98 | 268.67 | 203.4 | 147.23 | 277.35 |
| 6892 | ENSMUSG00000028777 | 0 | NA | NA | NA | NA | NA | Gnat3 | guanine nucleotide binding protein, alpha | 0 | 0 | 0 | 0 | 0 | 0 |
| 6893 | ENSMUSG00000028778 | 0 | NA | NA | NA | NA | NA | Hcrtr1 | hypocretin (orexin) receptor 1 [Source:M | 0 | 0 | 0 | 0 | 0 | 0 |
| 6894 | ENSMUSG00000028779 | 601.5314267 | 0.060776915 | 0.134230475 | 0.452780299 | 0.650706951 | 0.925643431 | Pef1 | penta-EF hand domain containing 1 [Sour | 639.27 | 546.85 | 579.6 | 612.13 | 549.67 | 681.67 |
| 6895 | ENSMUSG00000028780 | 28.01996946 | 0.148530993 | 0.419702698 | 0.353895731 | 0.723417027 | 0.9447719 | Sema3c | sema domain, immunoglobulin domain (Ig | 20.881 | 34.949 | 24.15 | 30.847 | 29.447 | 27.846 |
| 6896 | ENSMUSG00000028782 | 0.305170013 | -1.657836477 | 4.041480335 | -0.410205257 | 0.681655385 | NA | Adgrb2 | adhesion G protein-coupled receptor B2 [ | 0.8031 | 1.0279 | 0 | 0 | 0 | 0 |
| 6897 | ENSMUSG00000028784 | 0 | NA | NA | NA | NA | NA | Spocd1 | SPOC domain containing 1 [Source:MGI Sy | 0 | 0 | 0 | 0 | 0 | 0 |
| 6898 | ENSMUSG00000028785 | 0.806689472 | -0.274753491 | 2.612595279 | -0.105164965 | 0.91624491 | NA | Hpca | hippocalcin [Source:MGI Symbol;Acc:MGI | 1.6062 | 0 | 1.0062 | 0 | 0 | 2.2277 |
| 6899 | ENSMUSG00000028786 | 0.686901383 | 1.702364833 | 2.776876575 | 0.613050233 | 0.539843125 | NA | Tmem54 | transmembrane protein 54 [Source:MGI S | 0.8031 | 0 | 0 | 0 | 1.0906 | 2.2277 |
| 6900 | ENSMUSG00000028788 | 2776.793444 | 0.060376473 | 0.086894407 | 0.694825769 | 0.487164497 | 0.873910634 | Ptp4a2 | protein tyrosine phosphatase 4a2 [Source | 2740.2 | 2721.9 | 2693.7 | 2799.4 | 2908.7 | 2796.9 |
| 6901 | ENSMUSG00000028789 | 41.27144979 | 1.202916125 | 0.411143165 | 2.925784076 | 0.003435893 | 0.051606188 | Azin2 | antizyme inhibitor 2 [Source:MGI Symbol; | 28.109 | 13.363 | 33.206 | 39.523 | 63.256 | 70.172 |
| 6902 | ENSMUSG00000028790 | 573.6754657 | -0.060986607 | 0.136405498 | -0.447097865 | 0.654804401 | 0.925797231 | Khdrbs1 | KH domain containing, RNA binding, signa | 565.38 | 631.14 | 560.48 | 485.84 | 594.39 | 604.82 |
| 6903 | ENSMUSG00000028792 | 6955.923518 | 0.225590545 | 0.121474697 | 1.857099052 | 0.063297063 | 0.359877642 | Ak2 | adenylate kinase 2 [Source:MGI Symbol;A | 7096.2 | 6305.2 | 5837.2 | 7625.1 | 8432.7 | 6439.1 |
| 6904 | ENSMUSG00000028793 | 711.5907312 | 0.569572325 | 0.122972214 | 4.631715605 | 3.63E-06 | 0.000232095 | Rnf19b | ring finger protein 19B [Source:MGI Symb | 555.75 | 579.75 | 583.62 | 864.69 | 761.25 | 924.49 |
| 6905 | ENSMUSG00000028794 | 0.331982614 | 0.16514309 | 3.986751476 | 0.041422971 | 0.966958701 | NA | A3galt2 | alpha 1,3-galactosyltransferase 2 (isoglob | 0 | 1.0279 | 0 | 0.964 | 0 | 0 |
| 6906 | ENSMUSG00000028795 | 56.59764317 | -0.117034869 | 0.327775268 | -0.357058267 | 0.721048186 | 0.9447719 | Ccdc28b | coiled coil domain containing 28B [Source | 56.217 | 51.396 | 69.431 | 72.298 | 51.259 | 38.985 |
| 6907 | ENSMUSG00000028796 | 568.4494036 | -0.155698964 | 0.13352607 | -1.166056666 | 0.243591535 | 0.693425884 | Phc2 | polyhomeotic 2 [Source:MGI Symbol;Acc: | 616.78 | 571.52 | 608.78 | 554.29 | 467.88 | 591.45 |
| 6908 | ENSMUSG00000028797 | 1136.251081 | 0.049847544 | 0.112631432 | 0.44257223 | 0.658075175 | 0.927303282 | Tmem234 | transmembrane protein 234 [Source:MGI | 1080.2 | 1054.6 | 1215.5 | 1194.4 | 1073.2 | 1199.6 |
| 6909 | ENSMUSG00000028798 | 1261.075957 | 0.08044065 | 0.098598976 | 0.815836559 | 0.414593621 | 0.836001897 | Eif3i | eukaryotic translation initiation factor 3, s | 1179 | 1235.6 | 1263.8 | 1293.7 | 1293.5 | 1301 |
| 6910 | ENSMUSG00000028799 | 133.0140568 | 0.021172002 | 0.205093724 | 0.103230861 | 0.917779748 | 0.986374671 | Zfp362 | zinc finger protein 362 [Source:MGI Symb | 136.53 | 134.66 | 124.77 | 133.99 | 118.88 | 149.26 |
| 6911 | ENSMUSG00000028800 | 464.5872638 | -0.033004376 | 0.152611495 | -0.216264027 | 0.828781948 | 0.969955221 | Hdac1 | histone deacetylase 1 [Source:MGI Symbo | 420.83 | 485.18 | 504.13 | 423.19 | 539.86 | 414.35 |
| 6912 | ENSMUSG00000028801 | 2.116273531 | 1.322494987 | 1.566253494 | 0.844368419 | 0.398463564 | NA | Stpg1 | sperm tail PG rich repeat containing 1 [So | 1.6062 | 0 | 2.0125 | 5.7839 | 2.1812 | 1.1138 |
| 6913 | ENSMUSG00000028803 | 200.0915693 | 0.068288245 | 0.227122063 | 0.300667597 | 0.76366798 | 0.954980327 | Nipal3 | NIPA-like domain containing 3 [Source:M | 187.12 | 150.08 | 248.54 | 166.77 | 196.31 | 251.73 |
| 6914 | ENSMUSG00000028804 | 0.824448242 | -0.598621533 | 2.872093747 | -0.208426878 | 0.834895672 | NA | Csmd2 | CUB and Sushi multiple domains 2 [Source | 0 | 0 | 3.0187 | 1.928 | 0 | 0 |
| 6915 | ENSMUSG00000028807 | 25.8721642 | -0.358698818 | 0.447588287 | -0.801403495 | 0.422898092 | 0.84005258 | Zbtb8a | zinc finger and BTB domain containing 8a | 23.29 | 25.698 | 38.237 | 17.352 | 27.265 | 23.391 |
| 6916 | ENSMUSG00000028809 | 782.046835 | 0.04287081 | 0.116872301 | 0.366817541 | 0.713755123 | 0.944425938 | Srrm1 | serine/arginine repetitive matrix 1 [Source | 708.34 | 819.25 | 784.87 | 816.49 | 808.15 | 755.19 |
| 6917 | ENSMUSG00000028811 | 521.8277337 | -0.00302185 | 0.128244671 | -0.023563162 | 0.981201056 | 0.99820894 | Yars | tyrosyl-tRNA synthetase [Source:MGI Sym | 505.95 | 544.8 | 517.21 | 568.75 | 489.69 | 504.57 |
| 6918 | ENSMUSG00000028813 | 0 | NA | NA | NA | NA | NA | CK137956 | cDNA sequence CK137956 [Source:MGI Sy | 0 | 0 | 0 | 0 | 0 | 0 |
| 6919 | ENSMUSG00000028820 | 1062.981924 | -0.388917328 | 0.135259813 | -2.875335383 | 0.004035985 | 0.057627338 | Sfpq | splicing factor proline/glutamine rich (pol | 1045.6 | 1410.3 | 1161.2 | 897.46 | 1002.3 | 861 |
| 6920 | ENSMUSG00000028821 | 686.4605425 | 0.055192907 | 0.118959028 | 0.463965689 | 0.642672328 | 0.924936573 | Syf2 | SYF2 homolog, RNA splicing factor (S. cere | 681.03 | 653.76 | 685.25 | 727.8 | 638.01 | 732.91 |
| 6921 | ENSMUSG00000028822 | 2093.779765 | -0.004529784 | 0.109904164 | -0.041215759 | 0.96712389 | 0.994231936 | Tmem50a | transmembrane protein 50A [Source:MGI | 2002.1 | 2030.1 | 2259 | 1957.8 | 2318.7 | 1994.9 |
| 6922 | ENSMUSG00000028825 | 0.795774083 | 0.899657507 | 2.433208733 | 0.369741196 | 0.711575334 | NA | Rhd | Rh blood group, D antigen [Source:MGI Sy | 1.6062 | 0 | 0 | 0.964 | 1.0906 | 1.1138 |
| 6923 | ENSMUSG00000028826 | 973.755684 | 0.080177217 | 0.133175006 | 0.602044027 | 0.547144832 | 0.895518681 | Maco1 | macoilin 1 [Source:MGI Symbol;Acc:MGI: | 905.1 | 999.14 | 936.81 | 1076.8 | 1098.3 | 826.47 |
| 6924 | ENSMUSG00000028830 | 550.8667084 | 0.059356332 | 0.129032702 | 0.46000999 | 0.64550905 | 0.925643431 | AU040320 | expressed sequence AU040320 [Source:M | 558.16 | 497.51 | 562.49 | 562 | 520.22 | 604.82 |
| 6925 | ENSMUSG00000028832 | 91.81459748 | -0.191409086 | 0.255269788 | -0.749830555 | 0.453356764 | 0.857073339 | Stmn1 | stathmin 1 [Source:MGI Symbol;Acc:MGI: | 106.01 | 98.68 | 88.55 | 73.262 | 76.343 | 108.04 |
| 6926 | ENSMUSG00000028833 | 231.8361468 | -0.167377315 | 0.16947778 | -0.987606251 | 0.323345524 | 0.770475661 | Ncdn | neurochondrin [Source:MGI Symbol;Acc:M | 232.1 | 246.7 | 257.6 | 240.99 | 190.86 | 222.77 |
| 6927 | ENSMUSG00000028834 | 0 | NA | NA | NA | NA | NA | Trim63 | tripartite motif-containing 63 [Source:MG | 0 | 0 | 0 | 0 | 0 | 0 |
| 6928 | ENSMUSG00000028836 | 3.717957055 | 0.75348777 | 1.223002684 | 0.616096579 | 0.53783078 | NA | Slc30a2 | solute carrier family 30 (zinc transporter), | 3.2124 | 0 | 5.0312 | 1.928 | 5.4531 | 6.6831 |
| 6929 | ENSMUSG00000028837 | 948.8908612 | 0.023562691 | 0.108340232 | 0.217487913 | 0.827828126 | 0.969794868 | Psmb2 | proteasome (prosome, macropain) subun | 998.26 | 905.6 | 918.7 | 942.77 | 995.73 | 932.29 |
| 6930 | ENSMUSG00000028838 | 461.8037883 | -0.298571598 | 0.163248824 | -1.82893567 | 0.067409246 | 0.373457298 | Extl1 | exostosin-like glycosyltransferase 1 [Sourc | 576.63 | 445.09 | 505.13 | 338.36 | 467.88 | 437.74 |
| 6931 | ENSMUSG00000028840 | 4.303506536 | 0.190293241 | 1.159192331 | 0.164160196 | 0.869605035 | NA | Zfp593 | zinc finger protein 593 [Source:MGI Symb | 0 | 5.1396 | 7.0437 | 4.8199 | 4.3625 | 4.4554 |
| 6932 | ENSMUSG00000028841 | 2.1341491 | 0.960342201 | 1.53492285 | 0.625661544 | 0.531536962 | NA | Cnksr1 | connector enhancer of kinase suppressor | 3.2124 | 1.0279 | 0 | 1.928 | 2.1812 | 4.4554 |
| 6933 | ENSMUSG00000028842 | 221.3203103 | -0.05319338 | 0.196785923 | -0.2703109 | 0.786921081 | 0.960870081 | Ago3 | argonaute RISC catalytic subunit 3 [Source | 221.66 | 194.28 | 260.62 | 240.99 | 238.85 | 171.53 |
| 6934 | ENSMUSG00000028843 | 57.80354451 | -0.976247937 | 0.302824741 | -3.223805077 | 0.001264995 | 0.024927635 | Sh3bgrl3 | SH3 domain binding glutamic acid-rich pro | 72.279 | 83.261 | 74.462 | 39.523 | 31.628 | 45.668 |
| 6935 | ENSMUSG00000028845 | 2.737886434 | -0.231456743 | 1.386362172 | -0.166952581 | 0.86740736 | NA | Tekt2 | tektin 2 [Source:MGI Symbol;Acc:MGI:134 | 0.8031 | 2.0558 | 6.0375 | 0.964 | 5.4531 | 1.1138 |
| 6936 | ENSMUSG00000028847 | 626.0906661 | 0.158409692 | 0.120900415 | 1.310249363 | 0.190111491 | 0.627627958 | Trappc3 | trafficking protein particle complex 3 [Sou | 629.63 | 582.83 | 562.49 | 699.85 | 641.28 | 640.46 |
| 6937 | ENSMUSG00000028848 | 195.7519676 | 0.147888887 | 0.176695677 | 0.836969472 | 0.402609728 | 0.828953151 | Gpn2 | GPN-loop GTPase 2 [Source:MGI Symbol;A | 188.73 | 182.97 | 185.15 | 191.83 | 193.04 | 232.79 |
| 6938 | ENSMUSG00000028849 | 824.2026745 | 0.099500482 | 0.121608669 | 0.818202208 | 0.413241735 | 0.8353299 | Map7d1 | MAP7 domain containing 1 [Source:MGI S | 777.4 | 779.16 | 831.16 | 884.93 | 752.53 | 920.04 |
| 6939 | ENSMUSG00000028850 | 105.0599099 | 0.152112181 | 0.24157051 | 0.62968026 | 0.5289038 | 0.888334265 | Gpatch3 | G patch domain containing 3 [Source:MGI | 122.87 | 88.401 | 86.537 | 101.22 | 121.06 | 110.27 |
| 6940 | ENSMUSG00000028851 | 1115.788088 | 0.105550845 | 0.103469837 | 1.020112213 | 0.307675245 | 0.756343859 | Nudc | nudC nuclear distribution protein [Source | 1078.6 | 1032 | 1113.9 | 1113.4 | 1153.9 | 1203 |

| | | | | | | | | | | | | | | | |
|---|---|---|---|---|---|---|---|---|---|---|---|---|---|---|---|
| 6941 | ENSMUSG00000028854 | 169.8399755 | 0.046612801 | 0.190791995 | 0.244312141 | 0.806989081 | 0.964718338 | Slc9a1 | solute carrier family 9 (sodium/hydrogen | 146.97 | 167.55 | 187.16 | 167.73 | 162.5 | 187.13 |
| 6942 | ENSMUSG00000028857 | 393.3001411 | -0.118577079 | 0.13952532 | -0.849860653 | 0.395402564 | 0.823010891 | Tmem222 | transmembrane protein 222 [Source:MGI | 421.63 | 419.39 | 387.4 | 391.38 | 399.17 | 340.84 |
| 6943 | ENSMUSG00000028859 | 25.76335953 | -1.420420428 | 0.548520703 | -2.589547524 | 0.009610216 | 0.105304666 | Csf3r | colony stimulating factor 3 receptor (gran | 24.896 | 59.619 | 28.175 | 16.388 | 5.4531 | 20.049 |
| 6944 | ENSMUSG00000028860 | 217.1204528 | 0.589891595 | 0.195168532 | 3.022472887 | 0.002507185 | 0.04139718 | Sytl1 | synaptotagmin-like 1 [Source:MGI Symbol | 161.42 | 162.41 | 196.22 | 223.64 | 319.55 | 239.48 |
| 6945 | ENSMUSG00000028861 | 485.452849 | 0.060829952 | 0.135087542 | 0.450300235 | 0.652493969 | 0.925756399 | Mrps15 | mitochondrial ribosomal protein S15 [Sou | 477.85 | 458.45 | 489.03 | 480.06 | 451.52 | 555.81 |
| 6946 | ENSMUSG00000028862 | 57.00932266 | -0.668000537 | 0.332141141 | -2.011194802 | 0.044304885 | 0.291019914 | Map3k6 | mitogen-activated protein kinase kinase k | 69.067 | 50.368 | 90.562 | 46.271 | 51.259 | 34.529 |
| 6947 | ENSMUSG00000028863 | 192.8861171 | 0.074430869 | 0.184744142 | 0.402886219 | 0.68703193 | 0.936679219 | Meaf6 | MYST/Esa1-associated factor 6 [Source:M | 201.58 | 191.19 | 171.06 | 225.57 | 193.04 | 174.87 |
| 6948 | ENSMUSG00000028864 | 94.89439165 | -0.858289975 | 0.264328364 | -3.247059688 | 0.001166039 | 0.023424435 | Hgf | hepatocyte growth factor [Source:MGI Sy | 138.94 | 117.18 | 110.69 | 73.262 | 47.987 | 81.311 |
| 6949 | ENSMUSG00000028865 | 0.335414862 | -1.779047242 | 4.030804179 | -0.441362855 | 0.658950329 | NA | Cd164l2 | CD164 sialomucin-like 2 [Source:MGI Sym | 0 | 0 | 2.0125 | 0 | 0 | 0 |
| 6950 | ENSMUSG00000028868 | 518.6313352 | -0.09287329 | 0.135242405 | -0.686717231 | 0.492260937 | 0.876368691 | Wasf2 | WASP family, member 2 [Source:MGI Sym | 558.96 | 570.49 | 475.95 | 482.95 | 479.87 | 543.56 |
| 6951 | ENSMUSG00000028869 | 472.6757455 | -0.069063718 | 0.145676781 | -0.474088718 | 0.635436643 | 0.923478764 | Gnl2 | guanine nucleotide binding protein-like 2 | 529.24 | 490.32 | 431.68 | 442.47 | 521.32 | 421.03 |
| 6952 | ENSMUSG00000028871 | 15.35895841 | 0.24694557 | 0.561435554 | 0.439846689 | 0.660048149 | 0.927787797 | Rspo1 | R-spondin 1 [Source:MGI Symbol;Acc:MG | 13.653 | 11.307 | 17.106 | 11.568 | 21.812 | 16.708 |
| 6953 | ENSMUSG00000028873 | 17.37115914 | 0.2132651 | 0.509058499 | 0.41894026 | 0.675259793 | 0.932446243 | Cdca8 | cell division cycle associated 8 [Source:M( | 17.668 | 14.391 | 16.1 | 16.388 | 19.631 | 20.049 |
| 6954 | ENSMUSG00000028874 | 34.12602137 | -0.986833453 | 0.389659163 | -2.532555488 | 0.011323445 | 0.118425382 | Fgr | FGR proto-oncogene, Src family tyrosine k | 45.777 | 47.284 | 43.269 | 29.883 | 20.722 | 17.822 |
| 6955 | ENSMUSG00000028876 | 0 | NA | NA | NA | NA | NA | Epha10 | Eph receptor A10 [Source:MGI Symbol;Ac | 0 | 0 | 0 | 0 | 0 | 0 |
| 6956 | ENSMUSG00000028878 | 428.7732892 | 0.318027534 | 0.143525328 | 2.215828651 | 0.026703239 | 0.211364326 | Fam76a | family with sequence similarity 76, membe | 377.46 | 425.56 | 342.12 | 464.64 | 447.15 | 515.71 |
| 6957 | ENSMUSG00000028879 | 416.9051805 | 0.010246763 | 0.131965459 | 0.077647314 | 0.938108605 | 0.988420305 | Stx12 | syntaxin 12 [Source:MGI Symbol;Acc:MGI | 420.02 | 436.87 | 389.42 | 423.19 | 423.16 | 408.78 |
| 6958 | ENSMUSG00000028882 | 174.4970679 | 0.077824029 | 0.203956064 | 0.381572516 | 0.702778473 | 0.942506832 | Ppp1r8 | protein phosphatase 1, regulatory subunit | 145.36 | 198.39 | 166.03 | 169.66 | 158.14 | 209.4 |
| 6959 | ENSMUSG00000028883 | 2.029204452 | -1.568501427 | 1.613733801 | -0.971970362 | 0.331065293 | NA | Sema3a | sema domain, immunoglobulin domain (Ig | 4.0155 | 4.1117 | 1.0062 | 1.928 | 0 | 1.1138 |
| 6960 | ENSMUSG00000028884 | 113.1479239 | -0.2800127 | 0.222068368 | -1.26093015 | 0.207334014 | 0.652364421 | Rpa2 | replication protein A2 [Source:MGI Symbo | 131.71 | 134.66 | 105.66 | 96.398 | 106.88 | 103.59 |
| 6961 | ENSMUSG00000028885 | 9.455241398 | -0.334164753 | 0.704782811 | -0.474138625 | 0.635401057 | 0.923478764 | Smpdl3b | sphingomyelin phosphodiesterase, acid-lil | 15.259 | 7.1954 | 9.0562 | 8.6758 | 7.6343 | 8.9108 |
| 6962 | ENSMUSG00000028886 | 357.0726991 | 0.058303379 | 0.149171995 | 0.390846679 | 0.695910569 | 0.939992732 | Eya3 | EYA transcriptional coactivator and phosp | 374.25 | 340.24 | 334.07 | 320.04 | 379.54 | 394.3 |
| 6963 | ENSMUSG00000028889 | 197.0839891 | -0.155170554 | 0.1717445 | -0.903496492 | 0.366262449 | 0.804498111 | Yrdc | yrdC domain containing (E.coli) [Source:M | 203.99 | 217.92 | 201.25 | 189.9 | 173.41 | 196.04 |
| 6964 | ENSMUSG00000028890 | 309.100949 | 0.171111179 | 0.163309661 | 1.047771321 | 0.294743979 | 0.745164683 | Mtf1 | metal response element binding transcrip | 275.46 | 282.68 | 313.95 | 273.77 | 350.09 | 358.66 |
| 6965 | ENSMUSG00000028893 | 214.6772214 | -0.806929215 | 0.187233031 | -4.309758871 | 1.63E-05 | 0.000795922 | Sesn2 | sestrin 2 [Source:MGI Symbol;Acc:MGI:26 | 236.92 | 261.09 | 322 | 154.24 | 170.14 | 143.69 |
| 6966 | ENSMUSG00000028894 | 751.8802721 | -0.048773781 | 0.133518757 | -0.365295349 | 0.71489095 | 0.944595913 | Inpp5b | inositol polyphosphate-5-phosphatase B [ | 845.67 | 783.27 | 664.12 | 764.43 | 669.64 | 784.15 |
| 6967 | ENSMUSG00000028896 | 134.0855775 | -0.503125241 | 0.20547801 | -2.448560023 | 0.014342853 | 0.138765904 | Rcc1 | regulator of chromosome condensation 1 | 146.16 | 177.83 | 147.92 | 108.93 | 114.51 | 109.16 |
| 6968 | ENSMUSG00000028898 | 367.5169637 | 0.065644446 | 0.162525061 | 0.40390353 | 0.68628366 | 0.936489198 | Trnau1ap | tRNA selenocysteine 1 associated protein | 311.6 | 362.85 | 403.5 | 342.21 | 434.07 | 350.86 |
| 6969 | ENSMUSG00000028899 | 249.5844465 | 0.089158521 | 0.156060604 | 0.571307035 | 0.567791534 | 0.903352921 | Taf12 | TATA-box binding protein associated facto | 234.51 | 240.53 | 250.55 | 243.89 | 258.48 | 269.55 |
| 6970 | ENSMUSG00000028901 | 102.3828273 | -0.008478069 | 0.247492697 | -0.034255835 | 0.972673143 | 0.99573532 | Gmeb1 | glucocorticoid modulatory element bindir | 84.326 | 130.55 | 93.581 | 96.398 | 102.52 | 106.93 |
| 6971 | ENSMUSG00000028902 | 268.2249978 | 0.00240161 | 0.154665858 | 0.015527734 | 0.987611159 | 0.99859176 | Sf3a3 | splicing factor 3a, subunit 3 [Source:MGI S | 273.86 | 268.29 | 261.62 | 265.09 | 248.66 | 291.83 |
| 6972 | ENSMUSG00000028906 | 2692.211107 | 0.00220481 | 0.093398356 | 0.023606517 | 0.981166474 | 0.99820894 | Epb41 | erythrocyte membrane protein band 4.1 [ | 2712.9 | 2606.8 | 2751.1 | 2792.6 | 2761.4 | 2528.4 |
| 6973 | ENSMUSG00000028907 | 305.1670672 | 0.022470856 | 0.155397951 | 0.144602011 | 0.885025109 | 0.981795936 | Utp11 | UTP11 small subunit processome compon | 316.42 | 300.15 | 291.81 | 324.86 | 331.55 | 266.21 |
| 6974 | ENSMUSG00000028909 | 12.33598043 | 0.163524889 | 0.612798562 | 0.266849336 | 0.789585167 | 0.961009963 | Ptpru | protein tyrosine phosphatase, receptor ty | 14.456 | 8.2233 | 12.075 | 10.604 | 14.178 | 14.48 |
| 6975 | ENSMUSG00000028910 | 468.1672471 | 0.033541942 | 0.137755105 | 0.243489651 | 0.807626101 | 0.965081255 | Mecr | mitochondrial trans-2-enoyl-CoA reductas | 403.96 | 488.26 | 497.08 | 481.99 | 477.69 | 460.02 |
| 6976 | ENSMUSG00000028911 | 282.0756066 | -0.245361647 | 0.163968581 | -1.496394278 | 0.13455094 | 0.537616739 | Srsf4 | serine and arginine-rich splicing factor 4 [S | 261.01 | 328.93 | 329.04 | 268.95 | 263.93 | 240.59 |
| 6977 | ENSMUSG00000028914 | 425.2690489 | 0.491735079 | 0.142955671 | 3.439773149 | 0.000582202 | 0.013773229 | Casp9 | caspase 9 [Source:MGI Symbol;Acc:MGI:1 | 346.14 | 318.65 | 395.45 | 461.75 | 502.78 | 526.85 |
| 6978 | ENSMUSG00000028917 | 271.1756897 | -0.126079415 | 0.164084407 | -0.768381453 | 0.442260596 | 0.850193681 | Plekhm2 | pleckstrin homology domain containing, fa | 297.95 | 302.21 | 248.54 | 255.45 | 237.75 | 285.14 |
| 6979 | ENSMUSG00000028919 | 1267.325439 | -0.042893517 | 0.175917371 | -0.243827638 | 0.807364314 | 0.964861393 | Arhgef19 | Rho guanine nucleotide exchange factor ( | 1056.9 | 1288 | 1514.4 | 1284 | 932.48 | 1528.2 |
| 6980 | ENSMUSG00000028920 | 235.4506443 | -0.202574047 | 0.172060829 | -1.177339715 | 0.239059941 | 0.688386022 | Fbxo42 | F-box protein 42 [Source:MGI Symbol;Acc | 223.26 | 273.43 | 259.61 | 214 | 199.58 | 242.82 |
| 6981 | ENSMUSG00000028923 | 136.9209592 | -0.152184763 | 0.215261186 | -0.706977256 | 0.479580612 | 0.870236354 | Necap2 | NECAP endocytosis associated 2 [Source:I | 167.85 | 124.38 | 139.87 | 144.6 | 114.51 | 130.32 |
| 6982 | ENSMUSG00000028926 | 15.44518115 | -1.348242315 | 0.610594074 | -2.208082869 | 0.027238497 | 0.213715335 | Cdk14 | cyclin-dependent kinase 14 [Source:MGI S | 25.699 | 28.782 | 12.075 | 9.6398 | 10.906 | 5.5692 |
| 6983 | ENSMUSG00000028927 | 3.00716143 | -2.227883015 | 1.468301122 | -1.517320243 | 0.12918585 | NA | Padi2 | peptidyl arginine deiminase, type II [Sourc | 5.6217 | 8.2233 | 1.0062 | 0.964 | 0 | 2.2277 |
| 6984 | ENSMUSG00000028931 | 6.795592415 | -1.107269378 | 0.888804279 | -1.245796633 | 0.212839066 | 0.657720288 | Kcnab2 | potassium voltage-gated channel, shaker- | 10.44 | 8.2233 | 9.0562 | 0.964 | 7.6343 | 4.4554 |
| 6985 | ENSMUSG00000028932 | 1283.617655 | 0.119840053 | 0.110618401 | 1.08336454 | 0.278646648 | 0.731329747 | Psmc2 | proteasome (prosome, macropain) 26S su | 1318.7 | 1262.3 | 1108.9 | 1287.9 | 1330.6 | 1393.4 |
| 6986 | ENSMUSG00000028933 | 35.52802671 | -0.481194358 | 0.40975921 | -1.174334453 | 0.240261088 | 0.690333012 | Xrcc2 | X-ray repair complementing defective rep | 38.549 | 41.117 | 44.275 | 16.388 | 31.628 | 41.212 |
| 6987 | ENSMUSG00000028936 | 1886.51327 | 0.034421243 | 0.118303857 | 0.290956218 | 0.771084805 | 0.956965056 | Rpl22 | ribosomal protein L22 [Source:MGI Symbo | 1694.5 | 2083.6 | 1814.3 | 1721.7 | 1952.2 | 2052.8 |
| 6988 | ENSMUSG00000028937 | 415.9471651 | -0.067114853 | 0.142981252 | -0.469396174 | 0.638786483 | 0.923932929 | Acot7 | acyl-CoA thioesterase 7 [Source:MGI Syml | 441.71 | 393.69 | 441.74 | 455.96 | 384.99 | 377.59 |
| 6989 | ENSMUSG00000028938 | 0 | NA | NA | NA | NA | NA | Galntl5 | UDP-N-acetyl-alpha-D-galactosamine:poly | 0 | 0 | 0 | 0 | 0 | 0 |
| 6990 | ENSMUSG00000028940 | 0 | NA | NA | NA | NA | NA | Hes2 | hes family bHLH transcription factor 2 [So | 0 | 0 | 0 | 0 | 0 | 0 |
| 6991 | ENSMUSG00000028943 | 20.36529342 | -0.315446784 | 0.519408141 | -0.607319676 | 0.543638783 | 0.894662801 | Espn | espin [Source:MGI Symbol;Acc:MGI:1861( | 16.865 | 27.754 | 23.144 | 13.496 | 13.087 | 27.846 |
| 6992 | ENSMUSG00000028944 | 716.0031051 | -0.016387863 | 0.136413416 | -0.120133804 | 0.904377155 | 0.98383695 | Prkag2 | protein kinase, AMP-activated, gamma 2 r | 701.91 | 674.31 | 784.87 | 818.42 | 630.38 | 686.13 |
| 6993 | ENSMUSG00000028945 | 1002.173091 | 0.001120342 | 0.109818467 | 0.010201759 | 0.991860315 | 0.99859176 | Rheb | Ras homolog enriched in brain [Source:M | 1006.3 | 958.02 | 1040.5 | 936.99 | 1067.7 | 1003.6 |
| 6994 | ENSMUSG00000028946 | 0 | NA | NA | NA | NA | NA | Hes3 | hes family bHLH transcription factor 3 [So | 0 | 0 | 0 | 0 | 0 | 0 |
| 6995 | ENSMUSG00000028948 | 334.1617652 | -0.01939643 | 0.156468456 | -0.123963833 | 0.901343911 | 0.98333371 | Nol9 | nucleolar protein 9 [Source:MGI Symbol;A | 352.56 | 335.1 | 320.99 | 301.73 | 387.17 | 307.42 |
| 6996 | ENSMUSG00000028949 | 34.53220241 | -0.165082051 | 0.366169359 | -0.450835241 | 0.6521083 | 0.925643431 | Smarcd3 | SWI/SNF related, matrix associated, actin | 35.336 | 38.033 | 36.225 | 33.739 | 34.9 | 28.96 |

| | | | | | | | | | | | | | | | |
|---|---|---|---|---|---|---|---|---|---|---|---|---|---|---|---|
| 6997 | ENSMUSG00000028950 | 2.805758855 | 0.186815713 | 1.420833876 | 0.13148315 | 0.895393116 | NA | Tas1r1 | taste receptor, type 1, member 1 [Source: | 4.8186 | 1.0279 | 2.0125 | 6.7478 | 0 | 2.2277 |
| 6998 | ENSMUSG00000028952 | 383.6484318 | 0.429191975 | 0.143748591 | 2.985712574 | 0.002829184 | 0.044869869 | Zbtb48 | zinc finger and BTB domain containing 48 | 345.33 | 286.79 | 348.16 | 421.26 | 453.7 | 446.65 |
| 6999 | ENSMUSG00000028953 | 736.3989027 | 0.086136158 | 0.110351292 | 0.780563204 | 0.435059441 | 0.845325456 | Abcf2 | ATP-binding cassette, sub-family F (GCN2( | 697.9 | 705.15 | 740.6 | 768.29 | 759.07 | 747.39 |
| 7000 | ENSMUSG00000028954 | 449.7008242 | 0.004169977 | 0.13254926 | 0.031459827 | 0.97490283 | 0.996336532 | Nub1 | negative regulator of ubiquitin-like protei | 432.07 | 443.03 | 471.93 | 414.51 | 474.42 | 462.25 |
| 7001 | ENSMUSG00000028955 | 179.0758433 | -0.08283855 | 0.186910243 | -0.443199626 | 0.657621349 | 0.927303282 | Vamp3 | vesicle-associated membrane protein 3 [S | 202.38 | 178.86 | 171.06 | 181.23 | 152.69 | 188.24 |
| 7002 | ENSMUSG00000028957 | 773.3140664 | 0.654260936 | 0.165985332 | 3.941679227 | 8.09E-05 | 0.003047728 | Per3 | period circadian clock 3 [Source:MGI Syml | 740.46 | 504.71 | 556.45 | 1085.4 | 958.65 | 794.17 |
| 7003 | ENSMUSG00000028958 | 174.6043853 | 0.479499252 | 0.179628998 | 2.669386657 | 0.007598992 | 0.088781756 | Tmub1 | transmembrane and ubiquitin-like domair | 134.92 | 153.16 | 149.93 | 206.29 | 203.95 | 199.38 |
| 7004 | ENSMUSG00000028959 | 840.7373197 | 0.254933768 | 0.110959662 | 2.297535542 | 0.021588238 | 0.182976706 | Fastk | Fas-activated serine/threonine kinase [So | 791.86 | 752.44 | 755.69 | 974.58 | 891.04 | 878.82 |
| 7005 | ENSMUSG00000028960 | 955.364317 | 0.015641441 | 0.11935348 | 0.131051406 | 0.895734642 | 0.982798755 | Ube4b | ubiquitination factor E4B [Source:MGI Syr | 962.92 | 892.23 | 995.18 | 940.84 | 1065.5 | 875.48 |
| 7006 | ENSMUSG00000028961 | 551.8583637 | -0.10682324 | 0.168697151 | -0.633224917 | 0.526586776 | 0.888334265 | Pgd | phosphogluconate dehydrogenase [Sourc | 685.85 | 581.8 | 447.78 | 461.75 | 571.48 | 562.49 |
| 7007 | ENSMUSG00000028962 | 474.997234 | 0.289455794 | 0.133676209 | 2.165350116 | 0.030360883 | 0.230640658 | Slc4a2 | solute carrier family 4 (anion exchanger), | 419.22 | 394.72 | 468.91 | 543.68 | 531.13 | 492.32 |
| 7008 | ENSMUSG00000028963 | 0.85435009 | 0.823372753 | 2.383559862 | 0.345438252 | 0.729764924 | NA | Uts2 | urotensin 2 [Source:MGI Symbol;Acc:MGI | 0.8031 | 1.0279 | 0 | 0 | 2.1812 | 1.1138 |
| 7009 | ENSMUSG00000028964 | 3141.285537 | -0.107841374 | 0.093142115 | -1.15781538 | 0.246939386 | 0.696923175 | Park7 | Parkinson disease (autosomal recessive, e | 3258.2 | 3172.2 | 3344.8 | 2831.2 | 3060.3 | 3181.1 |
| 7010 | ENSMUSG00000028965 | 0.315620088 | 0.165143421 | 4.063300985 | 0.040642675 | 0.967580763 | NA | Tnfrsf9 | tumor necrosis factor receptor superfamil | 0.8031 | 0 | 0 | 0 | 1.0906 | 0 |
| 7011 | ENSMUSG00000028967 | 21293.9535 | -0.427802162 | 0.139719795 | -3.061857932 | 0.002199678 | 0.037342895 | Errfi1 | ERBB receptor feedback inhibitor 1 [Sourc | 21383 | 27374 | 24527 | 14547 | 21148 | 18784 |
| 7012 | ENSMUSG00000028969 | 364.6294693 | 0.119509871 | 0.148571657 | 0.804392126 | 0.421170547 | 0.838715738 | Cdk5 | cyclin-dependent kinase 5 [Source:MGI Sy | 345.33 | 317.63 | 385.39 | 351.85 | 414.43 | 373.14 |
| 7013 | ENSMUSG00000028970 | 28.17942123 | -0.151763398 | 0.416681574 | -0.364219125 | 0.715694387 | 0.9447719 | Abcb1b | ATP-binding cassette, sub-family B (MDR/ | 28.109 | 23.642 | 37.231 | 26.027 | 23.994 | 30.074 |
| 7014 | ENSMUSG00000028971 | 0.160662974 | 1.126924175 | 4.080472857 | 0.276174898 | 0.782413736 | NA | Cort | cortistatin [Source:MGI Symbol;Acc:MGI:: | 0 | 0 | 0 | 0.964 | 0 | 0 |
| 7015 | ENSMUSG00000028972 | 0.335414862 | -1.779047242 | 4.030804179 | -0.441362855 | 0.658950329 | NA | Car6 | carbonic anhydrase 6 [Source:MGI Symbo | 0 | 0 | 2.0125 | 0 | 0 | 0 |
| 7016 | ENSMUSG00000028973 | 694.5654902 | 0.120416713 | 0.142656943 | 0.844099916 | 0.398613575 | 0.82588279 | Abcb8 | ATP-binding cassette, sub-family B (MDR/ | 762.14 | 635.25 | 597.71 | 628.51 | 785.25 | 758.53 |
| 7017 | ENSMUSG00000028974 | 418.9480519 | 0.211408955 | 0.130977136 | 1.614090523 | 0.106507793 | 0.481264827 | Dffa | DNA fragmentation factor, alpha subunit | | 376.65 | 405 | 383.38 | 434.75 | 444.97 | 468.93 |
| 7018 | ENSMUSG00000028975 | 791.3659285 | -0.019849578 | 0.115855913 | -0.171329865 | 0.863964399 | 0.976966701 | Pex14 | peroxisomal biogenesis factor 14 [Source: | 861.73 | 789.44 | 738.58 | 806.85 | 784.15 | 767.44 |
| 7019 | ENSMUSG00000028976 | 205.32991 | 0.758011427 | 0.194853387 | 3.890162956 | 0.000100177 | 0.003649771 | Slc2a5 | solute carrier family 2 (facilitated glucose | 172.67 | 118.21 | 166.03 | 235.21 | 278.11 | 261.75 |
| 7020 | ENSMUSG00000028977 | 14.37859149 | 0.151216428 | 0.64019803 | 0.236202583 | 0.813275475 | 0.966802988 | Casz1 | castor zinc finger 1 [Source:MGI Symbol;A | 18.471 | 10.279 | 12.075 | 22.171 | 5.4531 | 17.822 |
| 7021 | ENSMUSG00000028978 | 32.48490448 | -0.169181483 | 0.497579145 | -0.340009191 | 0.733849606 | 0.948020741 | Nos3 | nitric oxide synthase 3, endothelial cell [S( | 30.518 | 26.726 | 46.287 | 54.947 | 15.269 | 21.163 |
| 7022 | ENSMUSG00000028979 | 3138.247326 | 0.101815883 | 0.106302514 | 0.957793738 | 0.338166778 | 0.780725343 | Masp2 | mannan-binding lectin serine peptidase 2 | 2819.7 | 3026.2 | 3238.1 | 3372 | 3447.4 | 2926.1 |
| 7023 | ENSMUSG00000028980 | 8539.831329 | 0.28974926 | 0.099101515 | 2.923762157 | 0.003458289 | 0.051888682 | H6pd | hexose-6-phosphate dehydrogenase (gluc | 7593.3 | 7384.6 | 8078.1 | 10022 | 8455.6 | 9704.9 |
| 7024 | ENSMUSG00000028982 | 389.7246193 | -0.147494491 | 0.161537098 | -0.913068841 | 0.361206336 | 0.800074801 | Slc25a33 | solute carrier family 25, member 33 [Sour | 377.46 | 390.61 | 461.87 | 434.75 | 323.91 | 349.75 |
| 7025 | ENSMUSG00000028986 | 328.5963254 | 0.16652257 | 0.161617392 | 1.030350557 | 0.302845474 | 0.752484788 | Klhl7 | kelch-like 7 [Source:MGI Symbol;Acc:MGI | 364.61 | 282.68 | 280.74 | 359.56 | 362.09 | 321.9 |
| 7026 | ENSMUSG00000028988 | 175.2324407 | 0.209031287 | 0.209633964 | 0.997125096 | 0.318703793 | 0.7652277 | Ctnnbip1 | catenin beta interacting protein 1 [Source | 128.5 | 176.8 | 183.14 | 182.19 | 217.03 | 163.74 |
| 7027 | ENSMUSG00000028989 | 15.01690277 | -0.419806126 | 0.556850151 | -0.753894249 | 0.450912724 | 0.855818389 | Angptl7 | angiopoietin-like 7 [Source:MGI Symbol;A | 16.062 | 19.53 | 16.1 | 16.388 | 11.997 | 10.025 |
| 7028 | ENSMUSG00000028990 | 94.02213114 | -0.173722496 | 0.250850052 | -0.692535218 | 0.488601277 | 0.874431125 | Lzic | leucine zipper and CTNNBIP1 domain con | 123.68 | 84.289 | 90.562 | 92.542 | 85.068 | 87.994 |
| 7029 | ENSMUSG00000028991 | 945.9706368 | -0.062826836 | 0.13346485 | -0.470736944 | 0.637828598 | 0.923478764 | Mtor | mechanistic target of rapamycin kinase [S | 1032.8 | 1055.7 | 811.03 | 959.16 | 970.65 | 846.52 |
| 7030 | ENSMUSG00000028992 | 296.6383451 | 0.020819644 | 0.172246164 | 0.120871453 | 0.903792854 | 0.983821605 | Nmnat1 | nicotinamide nucleotide adenylyltransfer | 333.29 | 304.26 | 245.52 | 313.29 | 322.82 | 260.64 |
| 7031 | ENSMUSG00000028995 | 123.7316161 | -0.635995225 | 0.328494946 | -1.936088314 | 0.052856886 | 0.32299197 | Fam126a | family with sequence similarity 126, meml | 191.94 | 192.22 | 67.418 | 108.93 | 89.431 | 92.449 |
| 7032 | ENSMUSG00000028996 | 0.315620088 | 0.165143421 | 4.063300985 | 0.040642675 | 0.967580763 | NA | Rbp7 | retinol binding protein 7, cellular [Source: | 0.8031 | 0 | 0 | 0 | 1.0906 | 0 |
| 7033 | ENSMUSG00000028998 | 598.6542186 | 0.100726031 | 0.123009906 | 0.818844878 | 0.412874922 | 0.835103185 | Tomm7 | translocase of outer mitochondrial memb | 609.55 | 560.22 | 562.49 | 576.46 | 624.92 | 658.28 |
| 7034 | ENSMUSG00000028999 | 241.7485505 | 0.06411614 | 0.162078289 | 0.395587467 | 0.692409384 | 0.93834998 | Rint1 | RAD50 interactor 1 [Source:MGI Symbol;A | 238.52 | 219.97 | 250.55 | 250.63 | 227.94 | 262.87 |
| 7035 | ENSMUSG00000029001 | 74.1063933 | -0.188370504 | 0.279346926 | -0.674324601 | 0.500104967 | 0.880716237 | Fbxo44 | F-box protein 44 [Source:MGI Symbol;Acc | 66.657 | 69.898 | 100.62 | 75.19 | 65.437 | 66.831 |
| 7036 | ENSMUSG00000029003 | 230.039496 | -0.133575558 | 0.215324156 | -0.620346365 | 0.535029776 | 0.889858671 | Mad2l2 | MAD2 mitotic arrest deficient-like 2 [Sour | 201.58 | 244.64 | 275.71 | 163.88 | 214.85 | 279.57 |
| 7037 | ENSMUSG00000029004 | 565.870373 | 0.02950207 | 0.143117639 | 0.206138603 | 0.836682645 | 0.971073158 | Kmt2e | lysine (K)-specific methyltransferase 2E [S | 515.59 | 648.62 | 517.21 | 628.51 | 556.22 | 529.08 |
| 7038 | ENSMUSG00000029005 | 0.476283062 | 1.031501528 | 3.159856949 | 0.326439312 | 0.744091998 | NA | Draxin | dorsal inhibitory axon guidance protein [S | 0.8031 | 0 | 0 | 0.964 | 1.0906 | 0 |
| 7039 | ENSMUSG00000029007 | 59.5546434 | -0.751294896 | 0.302156638 | -2.486441807 | 0.012902767 | 0.128577932 | Agtrap | angiotensin II, type I receptor-associated | 82.72 | 70.926 | 70.437 | 48.199 | 33.809 | 51.237 |
| 7040 | ENSMUSG00000029009 | 163.5636636 | -1.410565001 | 0.235366505 | -5.993057504 | 2.06E-09 | 3.92E-07 | Mthfr | methylenetetrahydrofolate reductase [So | 178.29 | 250.81 | 284.77 | 100.25 | 95.974 | 71.286 |
| 7041 | ENSMUSG00000029012 | 325.7977238 | 0.072839049 | 0.156122947 | 0.466549281 | 0.640822382 | 0.923932929 | Orc5 | origin recognition complex, subunit 5 [Sou | 360.59 | 299.12 | 291.81 | 305.58 | 346.82 | 350.86 |
| 7042 | ENSMUSG00000029014 | 343.8060876 | 0.013289908 | 0.149085642 | 0.089142779 | 0.92896844 | 0.988405116 | Dnajc2 | DnaJ heat shock protein family (Hsp40) m | 336.5 | 369.02 | 320.99 | 324.86 | 327.19 | 384.28 |
| 7043 | ENSMUSG00000029015 | 0.524202404 | 2.600864256 | 3.301637836 | 0.787749713 | 0.430843121 | NA | Slc26a5 | solute carrier family 26, member 5 [Sourc | 0 | 0 | 0 | 0.964 | 2.1812 | 0 |
| 7044 | ENSMUSG00000029016 | 40.51832293 | 0.448001937 | 0.345998239 | 1.294809875 | 0.195385825 | 0.635641153 | Clcn6 | chloride channel, voltage-sensitive 6 [Sou | 33.73 | 38.033 | 31.194 | 53.019 | 40.353 | 46.781 |
| 7045 | ENSMUSG00000029017 | 2419.433887 | -0.033361287 | 0.091745648 | -0.363628011 | 0.716135807 | 0.9447719 | Pmpcb | peptidase (mitochondrial processing) beta | 2566.7 | 2401.2 | 2373.7 | 2450.4 | 2373.2 | 2351.3 |
| 7046 | ENSMUSG00000029019 | 0 | NA | NA | NA | NA | NA | Nppb | natriuretic peptide type B [Source:MGI Sy | 0 | 0 | 0 | 0 | 0 | 0 |
| 7047 | ENSMUSG00000029020 | 2659.805465 | -0.1351237 | 0.107374013 | -1.25843951 | 0.20823285 | 0.653455953 | Mfn2 | mitofusin 2 [Source:MGI Symbol;Acc:MGI | 2726.5 | 2954.2 | 2671.6 | 2283.7 | 2783.3 | 2539.6 |
| 7048 | ENSMUSG00000029022 | 322.7848025 | 0.210356318 | 0.144812601 | 1.452610588 | 0.146331905 | 0.556855766 | Miip | migration and invasion inhibitory protein | 295.54 | 302.21 | 299.86 | 317.15 | 357.72 | 364.23 |
| 7049 | ENSMUSG00000029026 | 0 | NA | NA | NA | NA | NA | Trp73 | transformation related protein 73 [Source | 0 | 0 | 0 | 0 | 0 | 0 |
| 7050 | ENSMUSG00000029027 | 62.92045873 | 0.145166684 | 0.309882496 | 0.468457193 | 0.639457677 | 0.923932929 | Dffb | DNA fragmentation factor, beta subunit [S | 66.657 | 47.284 | 65.406 | 85.794 | 57.803 | 54.578 |
| 7051 | ENSMUSG00000029028 | 318.8154627 | 0.172099232 | 0.14630237 | 1.176325658 | 0.239464766 | 0.688924491 | Lrrc47 | leucine rich repeat containing 47 [Source: | 287.51 | 296.04 | 315.96 | 321.97 | 327.19 | 364.23 |
| 7052 | ENSMUSG00000029029 | 136.7016077 | 0.07636175 | 0.212047444 | 0.360116342 | 0.718760132 | 0.9447719 | Wrap73 | WD repeat containing, antisense to Trp73 | 114.04 | 134.66 | 150.94 | 135.92 | 127.6 | 157.05 |

| | | | | | | | | | | | | | | | |
|---|---|---|---|---|---|---|---|---|---|---|---|---|---|---|---|
| 7053 | ENSMUSG00000029030 | 1006.272788 | 0.033151762 | 0.105385685 | 0.31457557 | 0.75308393 | 0.952989051 | Tprgl | transformation related protein 63 regulat | 955.69 | 1002.2 | 1026.4 | 975.55 | 1023 | 1054.8 |
| 7054 | ENSMUSG00000029032 | 131.5293587 | 0.398446641 | 0.204426076 | 1.949098906 | 0.051283614 | 0.31808156 | Arhgef16 | Rho guanine nucleotide exchange factor ( | 106.01 | 116.15 | 118.74 | 163.88 | 139.6 | 144.8 |
| 7055 | ENSMUSG00000029033 | 80.85311872 | -0.114483771 | 0.274625499 | -0.416872328 | 0.676771795 | 0.932891189 | Acap3 | ArfGAP with coiled-coil, ankyrin repeat an | 94.766 | 71.954 | 85.531 | 98.326 | 63.256 | 71.286 |
| 7056 | ENSMUSG00000029034 | 244.2706777 | 0.097022488 | 0.189893865 | 0.510930083 | 0.609400015 | 0.916376001 | Ints11 | integrator complex subunit 11 [Source:M( | 208.81 | 231.28 | 268.67 | 269.91 | 199.58 | 287.37 |
| 7057 | ENSMUSG00000029036 | 1339.821305 | 0.023937677 | 0.097719348 | 0.244963537 | 0.806484664 | 0.964686916 | Atad3a | ATPase family, AAA domain containing 3A | 1348.4 | 1342.5 | 1295 | 1343.8 | 1394.9 | 1314.3 |
| 7058 | ENSMUSG00000029038 | 585.7425021 | 0.134129987 | 0.124475308 | 1.077563011 | 0.281228817 | 0.733710968 | Ssu72 | Ssu72 RNA polymerase II CTD phosphatas | 541.29 | 592.08 | 542.37 | 584.17 | 663.1 | 591.45 |
| 7059 | ENSMUSG00000029044 | 0 | NA | NA | NA | NA | NA | 4930584F24 | RIKEN cDNA 4930584F24 gene [Source:M( | 0 | 0 | 0 | 0 | 0 | 0 |
| 7060 | ENSMUSG00000029047 | 136.6289646 | 0.063611746 | 0.209063297 | 0.304270273 | 0.760921991 | 0.95402891 | Pex10 | peroxisomal biogenesis factor 10 [Source: | 144.56 | 116.15 | 139.87 | 145.56 | 122.15 | 151.48 |
| 7061 | ENSMUSG00000029048 | 1921.453428 | 0.032410943 | 0.097137248 | 0.333661323 | 0.738635138 | 0.948137969 | Rer1 | retention in endoplasmic reticulum sortin | 1959.6 | 1974.6 | 1765 | 1932.8 | 1960.9 | 1935.9 |
| 7062 | ENSMUSG00000029049 | 42.05097811 | 0.294417769 | 0.368922577 | 0.798047577 | 0.424842881 | 0.840504368 | Morn1 | MORN repeat containing 1 [Source:MGI S | 35.336 | 39.061 | 39.244 | 61.695 | 46.897 | 30.074 |
| 7063 | ENSMUSG00000029050 | 501.4677797 | -0.130702356 | 0.152483271 | -0.857158661 | 0.391357205 | 0.820365651 | Ski | ski sarcoma viral oncogene homolog (avia | 501.14 | 544.8 | 527.27 | 535.97 | 382.81 | 516.82 |
| 7064 | ENSMUSG00000029053 | 655.7874387 | 0.282729508 | 0.138562136 | 2.040452872 | 0.04130524 | 0.278526339 | Prkcz | protein kinase C, zeta [Source:MGI Symbc | 664.97 | 560.22 | 548.4 | 629.48 | 778.7 | 752.96 |
| 7065 | ENSMUSG00000029054 | 8.214741369 | 0.278854074 | 0.887236095 | 0.314295232 | 0.753296819 | 0.952989051 | Gabrd | gamma-aminobutyric acid (GABA) A recep | 12.047 | 2.0558 | 8.05 | 9.6398 | 15.269 | 2.2277 |
| 7066 | ENSMUSG00000029055 | 2.120088471 | 0.148644442 | 2.137000357 | 0.069557518 | 0.94454585 | NA | Plch2 | phospholipase C, eta 2 [Source:MGI Symb | 0 | 0 | 6.0375 | 0 | 0 | 6.6831 |
| 7067 | ENSMUSG00000029056 | 251.2996652 | 0.334464998 | 0.173390544 | 1.928969077 | 0.053734701 | 0.326014563 | Pank4 | pantothenate kinase 4 [Source:MGI Symb | 199.17 | 226.14 | 242.5 | 310.4 | 290.1 | 239.48 |
| 7068 | ENSMUSG00000029059 | 222.5253666 | -0.49894751 | 0.224372968 | -2.223741633 | 0.026165832 | 0.209326659 | Prxl2b | peroxiredoxin like 2B [Source:MGI Symbo | 218.44 | 235.39 | 329.04 | 238.1 | 153.78 | 160.39 |
| 7069 | ENSMUSG00000029060 | 535.81084 | 0.17999883 | 0.122813464 | 1.465627819 | 0.142749708 | 0.552480247 | Mib2 | mindbomb E3 ubiquitin protein ligase 2 [S | 497.12 | 477.98 | 532.3 | 572.6 | 582.39 | 552.47 |
| 7070 | ENSMUSG00000029061 | 16.62909568 | -0.277319663 | 0.601213187 | -0.461266767 | 0.644607225 | 0.925643431 | Mmp23 | matrix metallopeptidase 23 [Source:MGI S | 28.109 | 16.447 | 10.062 | 23.135 | 11.997 | 10.025 |
| 7071 | ENSMUSG00000029062 | 564.4443197 | 0.143246362 | 0.14175768 | 1.010501593 | 0.312255037 | 0.759830222 | Cdk11b | cyclin-dependent kinase 11B [Source:MGI | 592.69 | 525.27 | 490.04 | 531.15 | 574.76 | 672.76 |
| 7072 | ENSMUSG00000029063 | 4666.428302 | -0.036819624 | 0.098790538 | -0.372703947 | 0.709368798 | 0.944239215 | Nadk | NAD kinase [Source:MGI Symbol;Acc:MGI | 5012.2 | 4850.7 | 4314.8 | 4872.9 | 4528.2 | 4419.7 |
| 7073 | ENSMUSG00000029064 | 1176.921968 | -0.017152674 | 0.11463911 | -0.149623227 | 0.881061882 | 0.981490993 | Gnb1 | guanine nucleotide binding protein (G prc | 1275.3 | 1190.3 | 1084.7 | 1093.2 | 1158.2 | 1259.8 |
| 7074 | ENSMUSG00000029066 | 730.4186631 | 0.061717338 | 0.130311494 | 0.473613924 | 0.635775245 | 0.923478764 | Mrpl20 | mitochondrial ribosomal protein L20 [Sou | 703.52 | 731.88 | 708.4 | 639.12 | 831.05 | 768.55 |
| 7075 | ENSMUSG00000029068 | 2208.320542 | 0.196293097 | 0.091690853 | 2.140814397 | 0.032289009 | 0.240179886 | Ccnl2 | cyclin L2 [Source:MGI Symbol;Acc:MGI:19 | 2094.5 | 2066.1 | 2014.5 | 2454.3 | 2339.4 | 2281.2 |
| 7076 | ENSMUSG00000029070 | 86.62839066 | -0.587787788 | 0.262687074 | -2.237596922 | 0.025247352 | 0.204466801 | Mxra8 | matrix-remodelling associated 8 [Source:N | 97.978 | 96.624 | 117.73 | 77.118 | 52.35 | 77.969 |
| 7077 | ENSMUSG00000029071 | 1468.021602 | -0.010908371 | 0.110565471 | -0.09865983 | 0.921408365 | 0.986685804 | Dvl1 | dishevelled segment polarity protein 1 [Sc | 1355.6 | 1505.9 | 1560.7 | 1572.2 | 1355.6 | 1458 |
| 7078 | ENSMUSG00000029072 | 4.090056038 | 0.155306788 | 1.25660445 | 0.123592422 | 0.901637993 | NA | Tas1r3 | taste receptor, type 1, member 3 [Source: | 1.6062 | 4.1117 | 6.0375 | 10.604 | 2.1812 | 0 |
| 7079 | ENSMUSG00000029073 | 341.4337723 | 0.112808282 | 0.140090185 | 0.805254715 | 0.420672709 | 0.837913304 | Cptp | ceramide-1-phosphate transfer protein [S | 324.45 | 325.85 | 334.07 | 359.56 | 332.64 | 372.02 |
| 7080 | ENSMUSG00000029074 | 1.038420047 | 0.084391034 | 2.062151212 | 0.040923786 | 0.967356655 | NA | Ttll10 | tubulin tyrosine ligase-like family, membe | 0 | 2.0558 | 1.0062 | 0.964 | 1.0906 | 1.1138 |
| 7081 | ENSMUSG00000029075 | 4.400817145 | 1.042984454 | 1.061062448 | 0.982962365 | 0.325625962 | NA | Tnfrsf4 | tumor necrosis factor receptor superfamil | 1.6062 | 3.0838 | 4.025 | 7.7118 | 2.1812 | 7.7969 |
| 7082 | ENSMUSG00000029076 | 2271.782456 | 0.099653609 | 0.095614182 | 1.042247146 | 0.29729711 | 0.748078261 | Sdf4 | stromal cell derived factor 4 [Source:MGI | 2227 | 2255.3 | 2097 | 2214.3 | 2417.9 | 2419.3 |
| 7083 | ENSMUSG00000029082 | 15.8878645 | -0.453586616 | 0.545992646 | -0.830755909 | 0.406111535 | 0.830012328 | Bst1 | bone marrow stromal cell antigen 1 [Sour | 18.471 | 15.419 | 21.131 | 10.604 | 17.45 | 12.252 |
| 7084 | ENSMUSG00000029084 | 124.7705323 | -1.099656002 | 0.267159403 | -4.116104431 | 3.85E-05 | 0.001694046 | Cd38 | CD38 antigen [Source:MGI Symbol;Acc:M( | 142.95 | 200.44 | 167.04 | 72.298 | 55.622 | 110.27 |
| 7085 | ENSMUSG00000029086 | 7.984444623 | -0.776388093 | 0.913325864 | -0.850066908 | 0.395287888 | 0.822933206 | Prom1 | prominin 1 [Source:MGI Symbol;Acc:MGI: | 8.031 | 7.1954 | 15.094 | 8.6758 | 0 | 8.9108 |
| 7086 | ENSMUSG00000029088 | 6.593831765 | 1.569205736 | 0.955861814 | 1.64166589 | 0.100659264 | 0.46725495 | Kcnip4 | Kv channel interacting protein 4 [Source:N | 4.8186 | 3.0838 | 2.0125 | 15.424 | 11.997 | 2.2277 |
| 7087 | ENSMUSG00000029089 | 98.58899474 | 0.181088262 | 0.238914227 | 0.757963494 | 0.448472838 | 0.854238022 | 5730480H06 | RIKEN cDNA 5730480H06 gene [Source:M | 100.39 | 83.261 | 93.581 | 120.5 | 104.7 | 89.108 |
| 7088 | ENSMUSG00000029090 | 707.4723112 | 0.069655568 | 0.125076012 | 0.556905894 | 0.577591728 | 0.907877282 | Adgra3 | adhesion G protein-coupled receptor A3 [ | 682.64 | 746.27 | 642.99 | 790.46 | 670.73 | 711.75 |
| 7089 | ENSMUSG00000029092 | 0.185640647 | 1.126924175 | 4.080472857 | 0.276174898 | 0.782413736 | NA | D5Ertd615e | DNA segment, Chr 5, ERATO Doi 615, expr | 0 | 0 | 0 | 0 | 0 | 1.1138 |
| 7090 | ENSMUSG00000029093 | 1.447092732 | -1.704069753 | 2.053523113 | -0.829827404 | 0.406636375 | NA | Sorcs2 | sortilin-related VPS10 domain containing | 5.6217 | 0 | 1.0062 | 0.964 | 1.0906 | 0 |
| 7091 | ENSMUSG00000029094 | 14.2341267 | -0.458882992 | 0.578826284 | -0.792781884 | 0.427904913 | 0.842753542 | Afap1 | actin filament associated protein 1 [Sourc | 16.865 | 14.391 | 18.112 | 10.604 | 8.7249 | 16.708 |
| 7092 | ENSMUSG00000029095 | 44.65101636 | -0.906446621 | 0.336800546 | -2.691345458 | 0.007116446 | 0.084997751 | Ablim2 | actin-binding LIM protein 2 [Source:MGI S | 53.005 | 56.535 | 65.406 | 35.667 | 29.447 | 27.846 |
| 7093 | ENSMUSG00000029096 | 44.77317912 | -0.925026486 | 0.401592211 | -2.303397478 | 0.02125649 | 0.180961662 | Htra3 | HtrA serine peptidase 3 [Source:MGI Syml | 77.901 | 66.815 | 31.194 | 37.595 | 26.175 | 28.96 |
| 7094 | ENSMUSG00000029097 | 71.43034036 | 0.083511623 | 0.269190424 | 0.310232517 | 0.756384145 | 0.953085143 | Trmt44 | tRNA methyltransferase 44 [Source:MGI S | 61.839 | 81.206 | 65.406 | 79.046 | 69.8 | 71.286 |
| 7095 | ENSMUSG00000029098 | 649.4035202 | 0.254358087 | 0.122527903 | 2.075919691 | 0.037901379 | 0.26432283 | Acox3 | acyl-Coenzyme A oxidase 3, pristanoyl [So | 595.1 | 604.42 | 577.58 | 722.02 | 759.07 | 638.23 |
| 7096 | ENSMUSG00000029101 | 143.676283 | -0.001107338 | 0.215969283 | -0.005127294 | 0.99590903 | 0.999003684 | Rgs12 | regulator of G-protein signaling 12 [Sourc | 170.26 | 121.29 | 138.86 | 135.92 | 130.87 | 164.85 |
| 7097 | ENSMUSG00000029102 | 5091.300583 | -0.063604149 | 0.099508366 | -0.639183938 | 0.52270328 | 0.888334265 | Hgfac | hepatocyte growth factor activator [Sourc | 5065.2 | 5120.1 | 5426.7 | 5498.5 | 4665.7 | 4771.7 |
| 7098 | ENSMUSG00000029103 | 727.7767066 | 0.035174426 | 0.114741047 | 0.306554863 | 0.759182214 | 0.95402891 | Lrpap1 | low density lipoprotein receptor-related p | 742.87 | 738.05 | 675.19 | 712.38 | 729.62 | 768.55 |
| 7099 | ENSMUSG00000029104 | 389.0461117 | -0.094052844 | 0.133030486 | -0.707002182 | 0.479565122 | 0.870236354 | Htt | huntingtin [Source:MGI Symbol;Acc:MGI:! | 411.99 | 391.64 | 401.49 | 386.56 | 381.72 | 360.89 |
| 7100 | ENSMUSG00000029106 | 619.8782599 | 0.057108261 | 0.134909066 | 0.423309291 | 0.672069617 | 0.931191575 | Add1 | adducin 1 (alpha) [Source:MGI Symbol;Ac | 604.74 | 678.43 | 539.35 | 579.35 | 640.19 | 677.22 |
| 7101 | ENSMUSG00000029108 | 16.06917969 | -2.025369726 | 0.603636754 | -3.355279005 | 0.00079285 | 0.017588618 | Pcdh7 | protocadherin 7 [Source:MGI Symbol;Acc | 26.502 | 24.67 | 26.162 | 5.7839 | 3.2719 | 10.025 |
| 7102 | ENSMUSG00000029110 | 1538.495029 | 0.140535479 | 0.11845969 | 1.186356977 | 0.235481354 | 0.683799474 | Rnf4 | ring finger protein 4 [Source:MGI Symbol; | 1426.3 | 1557.3 | 1406.7 | 1394.9 | 1748.3 | 1697.5 |
| 7103 | ENSMUSG00000029111 | 180.5567251 | -0.1920846 | 0.197239833 | -0.973863122 | 0.330124506 | 0.774747658 | Nelfa | negative elongation factor complex memb | 192.74 | 168.58 | 216.34 | 166.77 | 195.22 | 143.69 |
| 7104 | ENSMUSG00000029112 | 0 | NA | NA | NA | NA | NA | Nkx1-1 | NK1 homeobox 1 [Source:MGI Symbol;Ac | 0 | 0 | 0 | 0 | 0 | 0 |
| 7105 | ENSMUSG00000029119 | 756.9539752 | -0.047283121 | 0.116267932 | -0.406673796 | 0.684247589 | 0.935528319 | Man2b2 | mannosidase 2, alpha B2 [Source:MGI Syr | 735.64 | 761.69 | 811.03 | 721.06 | 798.33 | 713.97 |
| 7106 | ENSMUSG00000029120 | 0.167707431 | -0.796629176 | 4.080472857 | -0.195229623 | 0.84521319 | NA | Ppp2r2c | protein phosphatase 2, regulatory subunit | 0 | 0 | 1.0062 | 0 | 0 | 0 |
| 7107 | ENSMUSG00000029121 | 0.160662974 | 1.126924175 | 4.080472857 | 0.276174898 | 0.782413736 | NA | Crmp1 | collapsin response mediator protein 1 [So | 0 | 0 | 0 | 0.964 | 0 | 0 |
| 7108 | ENSMUSG00000029122 | 19.06078264 | -0.48249504 | 0.554911468 | -0.869499131 | 0.384574183 | 0.816437825 | Evc | EvC ciliary complex subunit 1 [Source:MGI | 24.896 | 27.754 | 14.087 | 22.171 | 7.6343 | 17.822 |

| | | | | | | | | | | | | | | | |
|---|---|---|---|---|---|---|---|---|---|---|---|---|---|---|---|
| 7109 | ENSMUSG00000029123 | 0 | NA | NA | NA | NA | NA | Stk32b | serine/threonine kinase 32B [Source:MGI | 0 | 0 | 0 | 0 | 0 | 0 |
| 7110 | ENSMUSG00000029125 | 252.8426559 | -0.086433761 | 0.164254529 | -0.526218434 | 0.598736448 | 0.915055389 | Stx18 | syntaxin 18 [Source:MGI Symbol;Acc:MGI | 233.7 | 265.2 | 282.75 | 239.07 | 230.12 | 266.21 |
| 7111 | ENSMUSG00000029126 | 6.605155948 | -0.421130332 | 0.815176209 | -0.516612638 | 0.605426592 | 0.916376001 | Nsg1 | neuron specific gene family member 1 [Sc | 6.4248 | 8.2233 | 8.05 | 4.8199 | 6.5437 | 5.5692 |
| 7112 | ENSMUSG00000029127 | 45.64161731 | 0.311241691 | 0.362979459 | 0.857463646 | 0.391188698 | 0.820195366 | Zbtb49 | zinc finger and BTB domain containing 49 | 36.14 | 43.173 | 43.269 | 65.55 | 54.531 | 31.188 |
| 7113 | ENSMUSG00000029128 | 266.939076 | 0.001681897 | 0.157444902 | 0.010682451 | 0.9914768 | 0.99859176 | Rab28 | RAB28, member RAS oncogene family [So | 284.3 | 245.67 | 269.67 | 247.74 | 282.47 | 271.78 |
| 7114 | ENSMUSG00000029130 | 12.72853478 | 0.919208531 | 0.660733201 | 1.391194705 | 0.164166392 | 0.588768336 | Rnf32 | ring finger protein 32 [Source:MGI Symbo | 7.2279 | 9.2513 | 10.062 | 20.244 | 22.903 | 6.6831 |
| 7115 | ENSMUSG00000029131 | 1543.766462 | 0.001892521 | 0.130988593 | 0.014447985 | 0.988472577 | 0.99859176 | Dnajb6 | DnaJ heat shock protein family (Hsp40) m | 1442.4 | 1583 | 1604 | 1642.6 | 1726.4 | 1264.2 |
| 7116 | ENSMUSG00000029134 | 0.349477146 | 0.165142768 | 3.911021834 | 0.042224967 | 0.966319359 | NA | Plb1 | phospholipase B1 [Source:MGI Symbol;Ac | 0 | 0 | 1.0062 | 0 | 1.0906 | 0 |
| 7117 | ENSMUSG00000029135 | 173.6724129 | -0.288799868 | 0.212601352 | -1.358410309 | 0.174333525 | 0.604151393 | Fosl2 | fos-like antigen 2 [Source:MGI Symbol;Acc | 180.7 | 167.55 | 225.4 | 191.83 | 141.78 | 134.78 |
| 7118 | ENSMUSG00000029136 | 226.6477465 | -0.000599862 | 0.186768927 | -0.003211785 | 0.997437371 | 0.999498002 | Rbks | ribokinase [Source:MGI Symbol;Acc:MGI:1 | 183.11 | 226.14 | 271.69 | 229.43 | 231.21 | 218.31 |
| 7119 | ENSMUSG00000029138 | 0 | NA | NA | NA | NA | NA | 4930548H24 | RIKEN cDNA 4930548H24 gene [Source:M | 0 | 0 | 0 | 0 | 0 | 0 |
| 7120 | ENSMUSG00000029141 | 251.6369281 | -0.057443657 | 0.163863795 | -0.350557348 | 0.725920459 | 0.945409105 | Slc4a1ap | solute carrier family 4 (anion exchanger), | 244.14 | 271.37 | 254.58 | 233.28 | 278.11 | 228.34 |
| 7121 | ENSMUSG00000029145 | 594.2250299 | 0.202766302 | 0.131089032 | 1.546783116 | 0.121915553 | 0.511776306 | Eif2b4 | eukaryotic translation initiation factor 2B, | 610.36 | 563.3 | 483 | 633.33 | 627.11 | 648.26 |
| 7122 | ENSMUSG00000029146 | 335.0770075 | -0.121799485 | 0.150885106 | -0.807233323 | 0.419532071 | 0.837591673 | Snx17 | sorting nexin 17 [Source:MGI Symbol;Acc: | 314.01 | 370.05 | 364.26 | 326.79 | 292.29 | 343.06 |
| 7123 | ENSMUSG00000029147 | 783.1448505 | 0.047657538 | 0.112463183 | 0.423761244 | 0.671739945 | 0.931191575 | Ppm1g | protein phosphatase 1G (formerly 2C), ma | 753.31 | 821.31 | 736.57 | 819.38 | 784.15 | 784.15 |
| 7124 | ENSMUSG00000029148 | 863.5973323 | 0.106098387 | 0.109604956 | 0.968007206 | 0.333040771 | 0.77523112 | Nrbp1 | nuclear receptor binding protein 1 [Sourc | 825.59 | 822.33 | 847.26 | 846.37 | 891.04 | 948.99 |
| 7125 | ENSMUSG00000029149 | 16.35339343 | 0.34868018 | 0.559562855 | 0.623129605 | 0.533199357 | 0.889143097 | Krtcap3 | keratinocyte associated protein 3 [Source | 8.8341 | 14.391 | 20.125 | 18.316 | 14.178 | 22.277 |
| 7126 | ENSMUSG00000029151 | 7.283293769 | 1.311205551 | 0.896357598 | 1.462815236 | 0.143517943 | 0.553373526 | Slc30a3 | solute carrier family 30 (zinc transporter), | 1.6062 | 2.0558 | 9.0562 | 14.46 | 8.7249 | 7.7969 |
| 7127 | ENSMUSG00000029152 | 1802.273897 | 0.028634427 | 0.103382555 | 0.276975419 | 0.781798981 | 0.960090038 | Ociad1 | OCIA domain containing 1 [Source:MGI Sy | 1848.7 | 1786.5 | 1716.7 | 1660.9 | 1918.4 | 1882.4 |
| 7128 | ENSMUSG00000029153 | 877.3110978 | 0.067385587 | 0.109110492 | 0.617590353 | 0.536845405 | 0.890579628 | Ociad2 | OCIA domain containing 2 [Source:MGI Sy | 895.46 | 843.92 | 830.15 | 855.05 | 925.93 | 913.35 |
| 7129 | ENSMUSG00000029154 | 0 | NA | NA | NA | NA | NA | Cwh43 | cell wall biogenesis 43 C-terminal homolo | 0 | 0 | 0 | 0 | 0 | 0 |
| 7130 | ENSMUSG00000029155 | 0 | NA | NA | NA | NA | NA | Spata18 | spermatogenesis associated 18 [Source:M | 0 | 0 | 0 | 0 | 0 | 0 |
| 7131 | ENSMUSG00000029156 | 27.38498418 | -0.46225191 | 0.509706116 | -0.906898888 | 0.36446027 | 0.802897466 | Sgcb | sarcoglycan, beta (dystrophin-associated | 33.73 | 38.033 | 23.144 | 8.6758 | 25.084 | 35.643 |
| 7132 | ENSMUSG00000029158 | 0 | NA | NA | NA | NA | NA | Yipf7 | Yip1 domain family, member 7 [Source:M | 0 | 0 | 0 | 0 | 0 | 0 |
| 7133 | ENSMUSG00000029161 | 64.11872885 | -0.61486463 | 0.487993119 | -1.259986271 | 0.207674315 | 0.652613959 | Cgref1 | cell growth regulator with EF hand domai | 142.15 | 44.2 | 46.287 | 82.902 | 46.897 | 22.277 |
| 7134 | ENSMUSG00000029162 | 13181.04645 | -0.282928782 | 0.082154462 | -3.443863844 | 0.000573464 | 0.013709799 | Khk | ketohexokinase [Source:MGI Symbol;Acc: | 14731 | 14413 | 14264 | 12285 | 11397 | 11996 |
| 7135 | ENSMUSG00000029163 | 224.7227758 | -0.799245345 | 0.220404165 | -3.626271522 | 0.000287543 | 0.008385125 | Emilin1 | elastin microfibril interfacer 1 [Source:MG | 357.38 | 239.5 | 258.6 | 158.09 | 128.69 | 206.06 |
| 7136 | ENSMUSG00000029165 | 101.8637832 | 0.360036095 | 0.243713717 | 1.477291062 | 0.139597633 | 0.545861088 | Agbl5 | ATP/GTP binding protein-like 5 [Source:M | 69.067 | 100.74 | 98.612 | 123.39 | 106.88 | 112.5 |
| 7137 | ENSMUSG00000029166 | 690.5210327 | -0.087592343 | 0.143252335 | -0.611454906 | 0.540898463 | 0.893466372 | Mapre3 | microtubule-associated protein, RP/EB fa | 811.13 | 658.9 | 664.12 | 771.18 | 612.93 | 624.87 |
| 7138 | ENSMUSG00000029167 | 194.3422418 | -0.612412497 | 0.203477562 | -3.009729876 | 0.002614801 | 0.042638653 | Ppargc1a | peroxisome proliferative activated recept | 199.97 | 296.04 | 209.3 | 146.52 | 151.6 | 162.62 |
| 7139 | ENSMUSG00000029168 | 0.520796785 | -0.820103173 | 3.072141206 | -0.266948398 | 0.789508892 | NA | Dpysl5 | dihydropyrimidinase-like 5 [Source:MGI S | 0 | 1.0279 | 1.0062 | 0 | 1.0906 | 0 |
| 7140 | ENSMUSG00000029169 | 1052.607089 | 0.000798596 | 0.116370134 | 0.006862551 | 0.994524519 | 0.998676967 | Dhx15 | DEAH (Asp-Glu-Ala-His) box polypeptide 1 | 944.45 | 1104 | 1109.9 | 1077.7 | 1096.1 | 983.52 |
| 7141 | ENSMUSG00000029171 | 72.42259784 | -0.126840992 | 0.265468357 | -0.477800794 | 0.632791996 | 0.922107984 | Pgm2 | phosphoglucomutase 2 [Source:MGI Syml | 81.113 | 72.982 | 72.45 | 62.659 | 79.615 | 65.717 |
| 7142 | ENSMUSG00000029173 | 1595.804858 | 0.108894222 | 0.107403101 | 1.013883413 | 0.310638387 | 0.759349946 | Sepsecs | Sep (O-phosphoserine) tRNA:Sec (selenoc | 1547.6 | 1514.1 | 1544.6 | 1548.1 | 1841 | 1579.4 |
| 7143 | ENSMUSG00000029174 | 89.56962234 | -0.637018437 | 0.237571904 | -2.681371096 | 0.007332116 | 0.086502219 | Tbc1d1 | TBC1 domain family, member 1 [Source:M | 112.43 | 106.9 | 107.67 | 70.37 | 66.528 | 73.514 |
| 7144 | ENSMUSG00000029175 | 384.3951228 | -0.113981273 | 0.149674418 | -0.761528086 | 0.44634171 | 0.852777619 | Slc35f6 | solute carrier family 35, member F6 [Sour | 415.2 | 384.44 | 399.48 | 427.04 | 330.46 | 349.75 |
| 7145 | ENSMUSG00000029176 | 299.8504141 | -0.190691885 | 0.156419757 | -1.219103576 | 0.222804883 | 0.668407528 | Anapc4 | anaphase promoting complex subunit 4 [S | 344.53 | 335.1 | 278.73 | 268.95 | 295.56 | 276.23 |
| 7146 | ENSMUSG00000029177 | 97.24858093 | 0.207150738 | 0.304044231 | 0.681317772 | 0.49567044 | 0.877965876 | Cenpa | centromere protein A [Source:MGI Symbo | 77.098 | 67.843 | 125.78 | 72.298 | 110.15 | 130.32 |
| 7147 | ENSMUSG00000029178 | 942.4725758 | -0.477850719 | 0.142706278 | -3.348491213 | 0.000812529 | 0.017915264 | Klf3 | Kruppel-like factor 3 (basic) [Source:MGI S | 1044.8 | 1202.7 | 1043.5 | 658.4 | 933.57 | 771.89 |
| 7148 | ENSMUSG00000029179 | 107.7241406 | -0.151401359 | 0.25529733 | -0.593039334 | 0.553154829 | 0.896245536 | Zcchc4 | zinc finger, CCHC domain containing 4 [So | 111.63 | 125.41 | 103.64 | 135.92 | 83.978 | 85.766 |
| 7149 | ENSMUSG00000029182 | 0.781659665 | -3.010437285 | 2.830777252 | -1.063466679 | 0.287570368 | NA | Fam166c | family with sequence similarity 166, meml | 1.6062 | 3.0838 | 0 | 0 | 0 | 0 |
| 7150 | ENSMUSG00000029184 | 0 | NA | NA | NA | NA | NA | Olfr109 | olfactory receptor 109 [Source:MGI Symb | 0 | 0 | 0 | 0 | 0 | 0 |
| 7151 | ENSMUSG00000029185 | 554.0324528 | 0.416121895 | 0.171075895 | 2.43238181 | 0.014999886 | 0.142361123 | Fam114a1 | family with sequence similarity 114, meml | 566.19 | 422.47 | 433.69 | 520.55 | 621.65 | 759.64 |
| 7152 | ENSMUSG00000029186 | 728.97611 | 0.016151477 | 0.130554828 | 0.123714128 | 0.901541625 | 0.983401061 | Pi4k2b | phosphatidylinositol 4-kinase type 2 beta | 770.98 | 677.4 | 726.51 | 834.8 | 693.63 | 670.53 |
| 7153 | ENSMUSG00000029188 | 4.239870302 | -5.456176079 | 1.550062914 | -3.519970725 | 0.000431594 | NA | Slc34a2 | solute carrier family 34 (sodium phosphat | 11.243 | 5.1396 | 9.0562 | 0 | 0 | 0 |
| 7154 | ENSMUSG00000029189 | 227.5284161 | 0.803793635 | 0.226954731 | 3.541647412 | 0.000397637 | 0.010631083 | Sel1l3 | sel-1 suppressor of lin-12-like 3 (C. elegan | 218.44 | 132.6 | 144.9 | 280.52 | 233.39 | 355.32 |
| 7155 | ENSMUSG00000029190 | 1107.95168 | 0.171403327 | 0.110670294 | 1.548774474 | 0.121435944 | 0.510744254 | D5Ertd579e | DNA segment, Chr 5, ERATO Doi 579, expr | 1139.6 | 978.58 | 1007.3 | 1171.2 | 1216 | 1135 |
| 7156 | ENSMUSG00000029191 | 489.7042046 | 0.064437119 | 0.155217975 | 0.41513954 | 0.678039755 | 0.93325821 | Rfc1 | replication factor C (activator 1) 1 [Source | 459.37 | 493.4 | 483 | 397.16 | 558.4 | 546.9 |
| 7157 | ENSMUSG00000029192 | 389.195036 | 0.0610093 | 0.151215602 | 0.403459032 | 0.686610567 | 0.936566769 | Tbc1d14 | TBC1 domain family, member 14 [Source:I | 342.92 | 393.69 | 406.52 | 362.46 | 377.35 | 452.22 |
| 7158 | ENSMUSG00000029193 | 0 | NA | NA | NA | NA | NA | Cckar | cholecystokinin A receptor [Source:MGI S | 0 | 0 | 0 | 0 | 0 | 0 |
| 7159 | ENSMUSG00000029195 | 543.4343997 | 0.271723816 | 0.149536681 | 1.817104773 | 0.069201072 | 0.379983891 | Klb | klotho beta [Source:MGI Symbol;Acc:MGI | 514.79 | 481.07 | 480.98 | 603.45 | 487.51 | 692.81 |
| 7160 | ENSMUSG00000029196 | 545.8365267 | -0.062549259 | 0.142572695 | -0.438718364 | 0.660865619 | 0.928184201 | Tada2b | transcriptional adaptor 2B [Source:MGI Sy | 471.42 | 638.34 | 564.5 | 543.68 | 521.32 | 535.76 |
| 7161 | ENSMUSG00000029198 | 2374.889065 | -0.052406573 | 0.10203005 | -0.513638606 | 0.607504683 | 0.916376001 | Grpel1 | GrpE-like 1, mitochondrial [Source:MGI Sy | 2420.6 | 2463.9 | 2368.7 | 2101.5 | 2488.8 | 2405.9 |
| 7162 | ENSMUSG00000029199 | 611.7743899 | 0.022884774 | 0.116155381 | 0.197018629 | 0.84381296 | 0.97250284 | Lias | lipoic acid synthetase [Source:MGI Symbo | 586.26 | 626 | 608.78 | 607.31 | 610.75 | 631.55 |
| 7163 | ENSMUSG00000029201 | 1943.695807 | -0.125081434 | 0.139299904 | -0.897929074 | 0.369223364 | 0.80653433 | Ugdh | UDP-glucose dehydrogenase [Source:MGI | 2360.3 | 1995.2 | 1726.7 | 1707.2 | 2148.5 | 1724.2 |
| 7164 | ENSMUSG00000029202 | 881.9443635 | 0.062566867 | 0.106162016 | 0.589352662 | 0.555624725 | 0.897628185 | Pds5a | PDS5 cohesin associated factor A [Source: | 838.44 | 888.12 | 862.35 | 908.07 | 875.77 | 918.92 |

| | | | | | | | | | | | | | | | |
|---|---|---|---|---|---|---|---|---|---|---|---|---|---|---|---|
| 7165 | ENSMUSG00000029203 | 758.36577 | -0.076574673 | 0.11481466 | -0.666941594 | 0.504809441 | 0.881592135 | Ube2k | ubiquitin-conjugating enzyme E2K [Source | 734.84 | 768.88 | 832.16 | 719.13 | 765.61 | 729.57 |
| 7166 | ENSMUSG00000029204 | 4.497197195 | -2.946490795 | 1.264948028 | -2.329337435 | 0.019841197 | NA | Rhoh | ras homolog family member H [Source:M( | 9.6372 | 10.279 | 4.025 | 1.928 | 0 | 1.1138 |
| 7167 | ENSMUSG00000029205 | 0 | NA | NA | NA | NA | NA | Chrna9 | cholinergic receptor, nicotinic, alpha poly| | 0 | 0 | 0 | 0 | 0 | 0 |
| 7168 | ENSMUSG00000029206 | 6.019644653 | 0.451580496 | 0.902360935 | 0.500443313 | 0.616762962 | 0.917509899 | Nsun7 | NOL1/NOP2/Sun domain family, member | 3.2124 | 3.0838 | 9.0562 | 8.6758 | 7.6343 | 4.4554 |
| 7169 | ENSMUSG00000029207 | 151.0111045 | -0.134299253 | 0.209352543 | -0.641498074 | 0.521199132 | 0.887887809 | Apbb2 | amyloid beta (A4) precursor protein-bindi | 178.29 | 171.66 | 123.77 | 143.63 | 146.14 | 142.57 |
| 7170 | ENSMUSG00000029208 | 337.2618245 | 0.076220563 | 0.154446957 | 0.493506407 | 0.621654814 | 0.918448951 | Guf1 | GUF1 homolog, GTPase [Source:MGI Symb | 333.29 | 339.21 | 312.94 | 379.81 | 364.27 | 294.05 |
| 7171 | ENSMUSG00000029209 | 35.11420416 | 0.08989173 | 0.406430444 | 0.221173713 | 0.824957177 | 0.96943211 | Gnpda2 | glucosamine-6-phosphate deaminase 2 [S | 32.124 | 23.642 | 46.287 | 32.775 | 46.897 | 28.96 |
| 7172 | ENSMUSG00000029211 | 0 | NA | NA | NA | NA | NA | Gabra4 | gamma-aminobutyric acid (GABA) A recep | 0 | 0 | 0 | 0 | 0 | 0 |
| 7173 | ENSMUSG00000029212 | 0.472877444 | -2.283655516 | 3.093270275 | -0.738265755 | 0.460352972 | NA | Gabrb1 | gamma-aminobutyric acid (GABA) A recep | 0.8031 | 1.0279 | 1.0062 | 0 | 0 | 0 |
| 7174 | ENSMUSG00000029213 | 411.1644325 | -0.096459799 | 0.136390067 | -0.707234783 | 0.479420586 | 0.870236354 | Commd8 | COMM domain containing 8 [Source:MGI | 411.99 | 454.34 | 408.54 | 383.66 | 427.52 | 380.93 |
| 7175 | ENSMUSG00000029217 | 348.8752061 | -0.318687965 | 0.160265821 | -1.988496132 | 0.046756846 | 0.299774391 | Tec | tec protein tyrosine kinase [Source:MGI S | 330.88 | 428.64 | 403.5 | 346.07 | 290.1 | 294.05 |
| 7176 | ENSMUSG00000029219 | 0 | NA | NA | NA | NA | NA | Slc10a4 | solute carrier family 10 (sodium/bile acid | 0 | 0 | 0 | 0 | 0 | 0 |
| 7177 | ENSMUSG00000029221 | 1409.420914 | 0.018526947 | 0.096977146 | 0.191044464 | 0.848490758 | 0.972803041 | Slc30a9 | solute carrier family 30 (zinc transporter), | 1432.7 | 1361 | 1406.7 | 1376.6 | 1452.7 | 1426.8 |
| 7178 | ENSMUSG00000029223 | 0.513958919 | -2.394274821 | 3.756096013 | -0.637437066 | 0.523840191 | NA | Uchl1 | ubiquitin carboxy-terminal hydrolase L1 [S | 0 | 3.0838 | 0 | 0 | 0 | 0 |
| 7179 | ENSMUSG00000029227 | 556.1039796 | -0.012936889 | 0.126833001 | -0.101999389 | 0.91875716 | 0.986374671 | Fip1l1 | FIP1 like 1 (S. cerevisiae) [Source:MGI Sym | 538.08 | 603.39 | 534.32 | 515.73 | 571.48 | 573.63 |
| 7180 | ENSMUSG00000029228 | 11.22107079 | 1.738542797 | 0.703467549 | 2.471390186 | 0.013458888 | 0.132518282 | Lnx1 | ligand of numb-protein X 1 [Source:MGI S | 7.2279 | 2.0558 | 6.0375 | 14.46 | 15.269 | 22.277 |
| 7181 | ENSMUSG00000029229 | 357.2463046 | -0.113048428 | 0.142464414 | -0.793520469 | 0.427474648 | 0.842415355 | Chic2 | cysteine-rich hydrophobic domain 2 [Sour | 367.82 | 373.13 | 373.32 | 378.84 | 319.55 | 330.81 |
| 7182 | ENSMUSG00000029231 | 82.52934615 | -1.181393765 | 0.286619579 | -4.121818089 | 3.76E-05 | 0.001672902 | Pdgfra | platelet derived growth factor receptor, a | 139.74 | 94.568 | 108.67 | 38.559 | 51.259 | 62.375 |
| 7183 | ENSMUSG00000029233 | 186.3565014 | -0.120079365 | 0.186105372 | -0.645222452 | 0.518783038 | 0.887592015 | Srd5a3 | steroid 5 alpha-reductase 3 [Source:MGI S | 214.43 | 194.28 | 173.07 | 174.48 | 162.5 | 199.38 |
| 7184 | ENSMUSG00000029234 | 171.0437889 | -0.016000538 | 0.22317149 | -0.071696158 | 0.942843714 | 0.990074885 | Tmem165 | transmembrane protein 165 [Source:MGI | 174.27 | 140.82 | 201.25 | 207.26 | 171.23 | 131.43 |
| 7185 | ENSMUSG00000029235 | 0 | NA | NA | NA | NA | NA | Pdcl2 | phosducin-like 2 [Source:MGI Symbol;Acc | 0 | 0 | 0 | 0 | 0 | 0 |
| 7186 | ENSMUSG00000029236 | 0 | NA | NA | NA | NA | NA | Nmu | neuromedin U [Source:MGI Symbol;Acc:N | 0 | 0 | 0 | 0 | 0 | 0 |
| 7187 | ENSMUSG00000029238 | 689.8137034 | -0.347031643 | 0.16872689 | -2.056765485 | 0.03970879 | 0.272849375 | Clock | circadian locomotor output cycles kaput [ | 621.6 | 821.31 | 874.43 | 504.16 | 588.93 | 728.45 |
| 7188 | ENSMUSG00000029245 | 0 | NA | NA | NA | NA | NA | Epha5 | Eph receptor A5 [Source:MGI Symbol;Acc | 0 | 0 | 0 | 0 | 0 | 0 |
| 7189 | ENSMUSG00000029246 | 440.4273671 | 0.584480885 | 0.143436343 | 4.074845142 | 4.60E-05 | 0.001976245 | Ppat | phosphoribosyl pyrophosphate amidotrar | 366.21 | 326.88 | 363.25 | 460.78 | 554.03 | 571.4 |
| 7190 | ENSMUSG00000029247 | 3822.528322 | 0.211016632 | 0.116665547 | 1.808731355 | 0.070492749 | 0.384032816 | Paics | phosphoribosylaminoimidazole carboxyla | 3047 | 3649.1 | 3935.4 | 3779.8 | 4252.3 | 4271.6 |
| 7191 | ENSMUSG00000029248 | 0 | NA | NA | NA | NA | NA | Thegl | theg spermatid protein like [Source:MGI S | 0 | 0 | 0 | 0 | 0 | 0 |
| 7192 | ENSMUSG00000029249 | 390.7281559 | 0.000892477 | 0.163183722 | 0.005469152 | 0.99563627 | 0.999003684 | Rest | RE1-silencing transcription factor [Source: | 359.79 | 365.94 | 446.77 | 412.58 | 435.16 | 324.13 |
| 7193 | ENSMUSG00000029250 | 1019.1083 | -0.129576741 | 0.144543807 | -0.896453082 | 0.370010828 | 0.806613237 | Polr2b | polymerase (RNA) II (DNA directed) polyp | 948.46 | 1106 | 1141.1 | 1009.3 | 1120.1 | 789.72 |
| 7194 | ENSMUSG00000029253 | 84.6822472 | -0.050809862 | 0.248111435 | -0.20478646 | 0.837738966 | 0.971073158 | Cenpc1 | centromere protein C1 [Source:MGI Symb | 94.766 | 74.01 | 89.556 | 87.722 | 79.615 | 82.424 |
| 7195 | ENSMUSG00000029254 | 144.1359916 | -0.4678352 | 0.27228678 | -1.718170821 | 0.085765462 | 0.431795087 | Stap1 | signal transducing adaptor family membe | 231.29 | 138.77 | 130.81 | 88.686 | 150.51 | 124.75 |
| 7196 | ENSMUSG00000029255 | 0 | NA | NA | NA | NA | NA | Gnrhr | gonadotropin releasing hormone recepto | 0 | 0 | 0 | 0 | 0 | 0 |
| 7197 | ENSMUSG00000029260 | 9262.873105 | 0.165532595 | 0.111421898 | 1.485637906 | 0.137374923 | 0.541266927 | Ugt2b34 | UDP glucuronosyltransferase 2 family, pol | 7815.8 | 8726 | 9655.9 | 10047 | 10515 | 8818.3 |
| 7198 | ENSMUSG00000029263 | 325.1093864 | 0.111682441 | 0.153455981 | 0.727781613 | 0.466747282 | 0.863779479 | Pigg | phosphatidylinositol glycan anchor biosyn | 293.13 | 346.41 | 298.85 | 362.46 | 345.73 | 304.08 |
| 7199 | ENSMUSG00000029265 | 474.2679836 | -0.139245713 | 0.128120195 | -1.086836569 | 0.277109043 | 0.730041765 | Dr1 | down-regulator of transcription 1 [Source | 489.89 | 484.15 | 517.21 | 428.97 | 480.96 | 444.42 |
| 7200 | ENSMUSG00000029267 | 124.8583019 | 0.023118353 | 0.23600845 | 0.097955617 | 0.921967536 | 0.986685804 | Mtf2 | metal response element binding transcrip | 122.87 | 95.596 | 152.95 | 115.68 | 142.87 | 119.18 |
| 7201 | ENSMUSG00000029268 | 0 | NA | NA | NA | NA | NA | Ugt2a2 | UDP glucuronosyltransferase 2 family, pol | 0 | 0 | 0 | 0 | 0 | 0 |
| 7202 | ENSMUSG00000029269 | 851.8601502 | -0.097138873 | 0.166172714 | -0.584565724 | 0.55883976 | 0.899349136 | Sult1b1 | sulfotransferase family 1B, member 1 [Sou | 858.52 | 870.65 | 911.66 | 606.34 | 1020.8 | 843.18 |
| 7203 | ENSMUSG00000029270 | 118.7795821 | -0.027540301 | 0.246169468 | -0.111875373 | 0.910922224 | 0.985827089 | Dipk1a | divergent protein kinase domain 1A [Sour | 95.569 | 127.46 | 136.85 | 93.506 | 115.61 | 143.69 |
| 7204 | ENSMUSG00000029272 | 10.43327306 | -1.374262615 | 1.986687602 | -0.691735638 | 0.489103362 | 0.874770056 | Sult1e1 | sulfotransferase family 1E, member 1 [Sou | 13.653 | 14.391 | 17.106 | 0 | 17.45 | 0 |
| 7205 | ENSMUSG00000029273 | 4171.43891 | -0.701070763 | 0.134595155 | -5.208737003 | 1.90E-07 | 1.94E-05 | Sult1d1 | sulfotransferase family 1D, member 1 [So | 4298.2 | 5639.2 | 5559.5 | 2792.6 | 3643.8 | 3095.4 |
| 7206 | ENSMUSG00000029275 | 1.013648964 | -2.157154189 | 2.470907524 | -0.873021013 | 0.382651625 | NA | Gfi1 | growth factor independent 1 transcriptio | 0 | 4.1117 | 1.0062 | 0.964 | 0 | 0 |
| 7207 | ENSMUSG00000029276 | 44.47195392 | 0.063765786 | 0.337687462 | 0.188830778 | 0.850225448 | 0.973301217 | Glmn | glomulin, FKBP associated protein [Source | 36.943 | 52.424 | 41.256 | 42.415 | 41.443 | 52.351 |
| 7208 | ENSMUSG00000029279 | 34.49809628 | 0.156549927 | 0.402038191 | 0.389390685 | 0.696987162 | 0.94023175 | Brdt | bromodomain, testis-specific [Source:MG | 24.093 | 30.838 | 43.269 | 39.523 | 42.534 | 26.732 |
| 7209 | ENSMUSG00000029280 | 0 | NA | NA | NA | NA | NA | Smr3a | submaxillary gland androgen regulated pr | 0 | 0 | 0 | 0 | 0 | 0 |
| 7210 | ENSMUSG00000029281 | 0 | NA | NA | NA | NA | NA | Smr2 | submaxillary gland androgen regulated pr | 0 | 0 | 0 | 0 | 0 | 0 |
| 7211 | ENSMUSG00000029282 | 0 | NA | NA | NA | NA | NA | Amtn | amelotin [Source:MGI Symbol;Acc:MGI:19 | 0 | 0 | 0 | 0 | 0 | 0 |
| 7212 | ENSMUSG00000029283 | 3.2128309 | 0.55737552 | 1.254601979 | 0.444264818 | 0.656851134 | NA | Cdc7 | cell division cycle 7 (S. cerevisiae) [Source | 0.8031 | 2.0558 | 5.0312 | 4.8199 | 5.4531 | 1.1138 |
| 7213 | ENSMUSG00000029286 | 0 | NA | NA | NA | NA | NA | Enam | enamelin [Source:MGI Symbol;Acc:MGI:1 | 0 | 0 | 0 | 0 | 0 | 0 |
| 7214 | ENSMUSG00000029287 | 159.2956751 | -0.268317493 | 0.195712126 | -1.37098042 | 0.17038106 | 0.597556794 | Tgfbr3 | transforming growth factor, beta recepto | 183.11 | 161.38 | 177.1 | 124.35 | 148.32 | 161.51 |
| 7215 | ENSMUSG00000029288 | 0 | NA | NA | NA | NA | NA | Ambn | ameloblastin [Source:MGI Symbol;Acc:M( | 0 | 0 | 0 | 0 | 0 | 0 |
| 7216 | ENSMUSG00000029290 | 248.1808946 | -0.074623029 | 0.163260052 | -0.457080763 | 0.647612997 | 0.925643431 | Zfp326 | zinc finger protein 326 [Source:MGI Symb | 227.28 | 270.34 | 266.65 | 235.21 | 232.3 | 257.3 |
| 7217 | ENSMUSG00000029291 | 1081.976933 | 0.381139009 | 0.122120692 | 3.121002684 | 0.001802364 | 0.03231892 | Rufy3 | RUN and FYVE domain containing 3 [Sour | 850.49 | 924.1 | 1046.5 | 1323.5 | 1133.2 | 1214.1 |
| 7218 | ENSMUSG00000029298 | 42.70970287 | -0.551421776 | 0.388918263 | -1.417834616 | 0.156239055 | 0.574894692 | Gbp9 | guanylate-binding protein 9 [Source:MGI | 44.171 | 70.926 | 37.231 | 29.883 | 27.265 | 46.781 |
| 7219 | ENSMUSG00000029299 | 24.46694605 | -1.598194136 | 0.564242126 | -2.832461565 | 0.004619112 | 0.063387888 | Abcg3 | ATP binding cassette subfamily G membe | 34.533 | 55.508 | 20.125 | 6.7478 | 8.7249 | 21.163 |
| 7220 | ENSMUSG00000029304 | 401.2073201 | -0.224549472 | 0.200439397 | -1.120286109 | 0.262591859 | 0.716898576 | Spp1 | secreted phosphoprotein 1 [Source:MGI S | 555.75 | 408.08 | 332.06 | 345.1 | 317.37 | 448.88 |

| | | | | | | | | | | | | | | | |
|---|---|---|---|---|---|---|---|---|---|---|---|---|---|---|---|
| 7221 | ENSMUSG00000029306 | 0 | NA | NA | NA | NA | NA | Ibsp | integrin binding sialoprotein [Source:MGI | 0 | 0 | 0 | 0 | 0 | 0 |
| 7222 | ENSMUSG00000029307 | 0 | NA | NA | NA | NA | NA | Dmp1 | dentin matrix protein 1 [Source:MGI Symb | 0 | 0 | 0 | 0 | 0 | 0 |
| 7223 | ENSMUSG00000029309 | 71.07837888 | -1.530800173 | 0.292001427 | -5.242440734 | 1.58E-07 | 1.65E-05 | Sparcl1 | SPARC-like 1 [Source:MGI Symbol;Acc:MG | 112.43 | 113.07 | 91.568 | 43.379 | 38.172 | 27.846 |
| 7224 | ENSMUSG00000029310 | 642.6541068 | -0.241943733 | 0.118664211 | -2.038893877 | 0.04146062 | 0.279183617 | Nudt9 | nudix (nucleoside diphosphate linked moi | 687.46 | 698.98 | 702.36 | 544.65 | 599.84 | 622.64 |
| 7225 | ENSMUSG00000029311 | 4392.581946 | -0.23646473 | 0.111023189 | -2.129867921 | 0.033182519 | 0.243878028 | Hsd17b11 | hydroxysteroid (17-beta) dehydrogenase | 4281.3 | 5119 | 4856.1 | 4012.1 | 4426.8 | 3660.1 |
| 7226 | ENSMUSG00000029312 | 64.15207674 | 0.165162493 | 0.275253191 | 0.600038434 | 0.548480622 | 0.895622715 | Klhl8 | kelch-like 8 [Source:MGI Symbol;Acc:MGI | 63.445 | 59.619 | 58.362 | 71.334 | 70.89 | 61.261 |
| 7227 | ENSMUSG00000029313 | 605.6368089 | -0.007192822 | 0.142601133 | -0.050440143 | 0.959771648 | 0.992148164 | Aff1 | AF4/FMR2 family, member 1 [Source:MGI | 589.48 | 639.36 | 593.68 | 714.31 | 530.04 | 566.95 |
| 7228 | ENSMUSG00000029314 | 161.2813327 | -0.877737526 | 0.244806383 | -3.585435622 | 0.000336516 | 0.009541868 | Gpat3 | glycerol-3-phosphate acyltransferase 3 [S | 251.37 | 203.53 | 172.07 | 153.27 | 89.431 | 98.018 |
| 7229 | ENSMUSG00000029319 | 893.4353855 | -0.034772557 | 0.104728821 | -0.332024719 | 0.739870594 | 0.948393197 | Coq2 | coenzyme Q2 4-hydroxybenzoate polypre | 900.28 | 916.9 | 895.56 | 884.93 | 901.94 | 861 |
| 7230 | ENSMUSG00000029320 | 0 | NA | NA | NA | NA | NA | 1700016H13 | RIKEN cDNA 1700016H13 gene [Source:M | 0 | 0 | 0 | 0 | 0 | 0 |
| 7231 | ENSMUSG00000029321 | 5.205083243 | -3.103399781 | 1.279975734 | -2.424577044 | 0.015326231 | 0.14470442 | Slc10a6 | solute carrier family 10 (sodium/bile acid | 2.4093 | 16.447 | 9.0562 | 0 | 1.0906 | 2.2277 |
| 7232 | ENSMUSG00000029322 | 82.53154884 | -0.464842761 | 0.267726176 | -1.736261906 | 0.082517536 | 0.423554602 | Plac8 | placenta-specific 8 [Source:MGI Symbol;A | 100.39 | 102.79 | 83.518 | 53.019 | 74.162 | 81.311 |
| 7233 | ENSMUSG00000029326 | 92.55357723 | -0.160562579 | 0.245486905 | -0.654057613 | 0.513074688 | 0.884860213 | Enoph1 | enolase-phosphatase 1 [Source:MGI Symb | 85.129 | 102.79 | 105.66 | 97.362 | 74.162 | 90.221 |
| 7234 | ENSMUSG00000029328 | 1043.622517 | 0.367167917 | 0.126899245 | 2.893381401 | 0.003811182 | 0.055346321 | Hnrnpdl | heterogeneous nuclear ribonucleoprotein | 913.13 | 944.66 | 877.45 | 1290.8 | 1243.3 | 992.43 |
| 7235 | ENSMUSG00000029330 | 2.724700991 | -3.746809028 | 1.690662092 | -2.21617853 | 0.026679278 | NA | Cds1 | CDP-diacylglycerol synthase 1 [Source:MG | 4.0155 | 8.2233 | 3.0187 | 0 | 1.0906 | 0 |
| 7236 | ENSMUSG00000029331 | 0 | NA | NA | NA | NA | NA | 4930522N08 | RIKEN cDNA 4930522N08 gene [Source:M | 0 | 0 | 0 | 0 | 0 | 0 |
| 7237 | ENSMUSG00000029334 | 6.980456531 | 0.688744801 | 1.004181362 | 0.685876902 | 0.492790737 | 0.876660964 | Prkg2 | protein kinase, cGMP-dependent, type II | 0.8031 | 13.363 | 2.0125 | 4.8199 | 13.087 | 7.7969 |
| 7238 | ENSMUSG00000029335 | 0.465832987 | -0.698295221 | 3.18348759 | -0.219349126 | 0.826378096 | NA | Bmp3 | bone morphogenetic protein 3 [Source:M | 0.8031 | 1.0279 | 0 | 0.964 | 0 | 0 |
| 7239 | ENSMUSG00000029337 | 0 | NA | NA | NA | NA | NA | Fgf5 | fibroblast growth factor 5 [Source:MGI Sy | 0 | 0 | 0 | 0 | 0 | 0 |
| 7240 | ENSMUSG00000029338 | 542.8628306 | 0.07916764 | 0.150533475 | 0.525913855 | 0.598948062 | 0.915055389 | Antxr2 | anthrax toxin receptor 2 [Source:MGI Sym | 575.82 | 467.7 | 539.35 | 496.45 | 655.46 | 522.39 |
| 7241 | ENSMUSG00000029343 | 0 | NA | NA | NA | NA | NA | Crybb1 | crystallin, beta B1 [Source:MGI Symbol;Ac | 0 | 0 | 0 | 0 | 0 | 0 |
| 7242 | ENSMUSG00000029344 | 611.7769909 | -0.368691298 | 0.155375373 | -2.372906915 | 0.017648711 | 0.15855339 | Tpst2 | protein-tyrosine sulfotransferase 2 [Sourc | 549.32 | 750.38 | 769.78 | 515.73 | 496.23 | 589.22 |
| 7243 | ENSMUSG00000029345 | 403.5535247 | 0.105260844 | 0.142657527 | 0.737856925 | 0.460601397 | 0.859817816 | Tfip11 | tuftelin interacting protein 11 [Source:MG | 354.17 | 396.78 | 416.59 | 445.36 | 428.61 | 379.82 |
| 7244 | ENSMUSG00000029346 | 91.08482521 | 0.015137604 | 0.25036201 | 0.060462863 | 0.951786992 | 0.990883058 | Srrd | SRR1 domain containing [Source:MGI Sym | 82.72 | 95.596 | 93.581 | 85.794 | 77.434 | 111.38 |
| 7245 | ENSMUSG00000029348 | 0.371281295 | 2.09253362 | 4.031852165 | 0.519000582 | 0.603760335 | NA | Asphd2 | aspartate beta-hydroxylase domain conta | 0 | 0 | 0 | 0 | 0 | 2.2277 |
| 7246 | ENSMUSG00000029352 | 8.146034231 | 1.102408149 | 0.840410842 | 1.311749081 | 0.189604811 | 0.626663447 | Crybb3 | crystallin, beta B3 [Source:MGI Symbol;Ac | 7.2279 | 4.1117 | 4.025 | 3.8559 | 19.631 | 10.025 |
| 7247 | ENSMUSG00000029359 | 4.228826198 | -1.584020008 | 1.07448484 | -1.474213455 | 0.140424128 | NA | Tesc | tescalcin [Source:MGI Symbol;Acc:MGI:19 | 4.8186 | 6.1675 | 8.05 | 1.928 | 2.1812 | 2.2277 |
| 7248 | ENSMUSG00000029360 | 1.632920508 | -1.194424845 | 1.858045464 | -0.642839407 | 0.52032831 | NA | Gm9754 | predicted gene 9754 [Source:MGI Symbol | 0.8031 | 3.0838 | 3.0187 | 2.8919 | 0 | 0 |
| 7249 | ENSMUSG00000029361 | 0.401551119 | -2.072982726 | 4.008251979 | -0.517178744 | 0.605031388 | NA | Nos1 | nitric oxide synthase 1, neuronal [Source: | 2.4093 | 0 | 0 | 0 | 0 | 0 |
| 7250 | ENSMUSG00000029363 | 168.4353541 | 0.049463331 | 0.190452546 | 0.259714725 | 0.795083833 | 0.962189806 | Rfc5 | replication factor C (activator 1) 5 [Source | 179.89 | 156.24 | 159.99 | 165.8 | 154.87 | 193.81 |
| 7251 | ENSMUSG00000029364 | 875.4779316 | -0.033029802 | 0.108058794 | -0.305665097 | 0.759859651 | 0.95402891 | Wsb2 | WD repeat and SOCS box-containing 2 [Sc | 881.81 | 872.7 | 901.6 | 825.17 | 861.59 | 910.01 |
| 7252 | ENSMUSG00000029366 | 116.339472 | -0.085208171 | 0.214600207 | -0.397055397 | 0.69132661 | 0.937937162 | Dck | deoxycytidine kinase [Source:MGI Symbol | 116.45 | 126.43 | 116.72 | 123.39 | 101.43 | 113.61 |
| 7253 | ENSMUSG00000029368 | 1250468.502 | -0.262167726 | 0.102471729 | -2.558439573 | 0.010514309 | 0.112567704 | Alb | albumin [Source:MGI Symbol;Acc:MGI:87 | 1E+06 | 1E+06 | 1E+06 | 1E+06 | 1E+06 | 1E+06 |
| 7254 | ENSMUSG00000029369 | 3363.050028 | 0.064389281 | 0.103166341 | 0.624130703 | 0.532541752 | 0.889143097 | Afm | afamin [Source:MGI Symbol;Acc:MGI:242 | 2934.5 | 3336.6 | 3594.3 | 3408.6 | 3475.8 | 3428.4 |
| 7255 | ENSMUSG00000029370 | 166.6076362 | -0.01125324 | 0.194416117 | -0.05788224 | 0.95384243 | 0.9915542 | Rassf6 | Ras association (RalGDS/AF-6) domain far | 182.3 | 142.88 | 176.09 | 158.09 | 183.22 | 157.05 |
| 7256 | ENSMUSG00000029371 | 0.185640647 | 1.126924175 | 4.080472857 | 0.276174898 | 0.782413736 | NA | Cxcl5 | chemokine (C-X-C motif) ligand 5 [Source: | 0 | 0 | 0 | 0 | 0 | 1.1138 |
| 7257 | ENSMUSG00000029372 | 3.010412591 | 0.085379122 | 1.240805719 | 0.06880942 | 0.94514132 | NA | Ppbp | pro-platelet basic protein [Source:MGI Sy | 5.6217 | 2.0558 | 1.0062 | 3.8559 | 2.1812 | 3.3415 |
| 7258 | ENSMUSG00000029373 | 8.120349347 | -0.784747832 | 0.748810691 | -1.047992291 | 0.294642159 | 0.745164683 | Pf4 | platelet factor 4 [Source:MGI Symbol;Acc: | 10.44 | 12.335 | 8.05 | 5.7839 | 6.5437 | 5.5692 |
| 7259 | ENSMUSG00000029375 | 0 | NA | NA | NA | NA | NA | Cxcl15 | chemokine (C-X-C motif) ligand 15 [Source | 0 | 0 | 0 | 0 | 0 | 0 |
| 7260 | ENSMUSG00000029376 | 63.1329973 | -0.370193097 | 0.283895262 | -1.303977722 | 0.192241178 | 0.629944813 | Mthfd2l | methylenetetrahydrofolate dehydrogenas | 80.31 | 64.759 | 68.425 | 61.695 | 51.259 | 52.351 |
| 7261 | ENSMUSG00000029377 | 2.403607137 | -2.506259432 | 1.645587804 | -1.523017748 | 0.127754265 | NA | Ereg | epiregulin [Source:MGI Symbol;Acc:MGI:1 | 4.0155 | 7.1954 | 1.0062 | 0 | 1.0906 | 1.1138 |
| 7262 | ENSMUSG00000029378 | 0.185640647 | 1.126924175 | 4.080472857 | 0.276174898 | 0.782413736 | NA | Areg | amphiregulin [Source:MGI Symbol;Acc:MG | 0 | 0 | 0 | 0 | 0 | 1.1138 |
| 7263 | ENSMUSG00000029379 | 0 | NA | NA | NA | NA | NA | Cxcl3 | chemokine (C-X-C motif) ligand 3 [Source: | 0 | 0 | 0 | 0 | 0 | 0 |
| 7264 | ENSMUSG00000029380 | 83.91389464 | 0.363784919 | 0.248903491 | 1.461550084 | 0.143864541 | 0.554054867 | Cxcl1 | chemokine (C-X-C motif) ligand 1 [Source: | 73.885 | 67.843 | 78.487 | 98.326 | 102.52 | 82.424 |
| 7265 | ENSMUSG00000029381 | 139.618897 | 0.206958068 | 0.225530522 | 0.917649932 | 0.358802168 | 0.798417624 | Shroom3 | shroom family member 3 [Source:MGI Syr | 114.04 | 125.41 | 149.93 | 166.77 | 115.61 | 165.96 |
| 7266 | ENSMUSG00000029384 | 1.104376344 | -1.045764527 | 2.161210491 | -0.483879072 | 0.628471694 | NA | 2010109A12 | RIKEN cDNA 2010109A12 gene [Source:M | 2.4093 | 0 | 2.0125 | 0 | 1.0906 | 1.1138 |
| 7267 | ENSMUSG00000029385 | 381.0195578 | -0.806763528 | 0.175600723 | -4.5943064 | 4.34E-06 | 0.00027305 | Ccng2 | cyclin G2 [Source:MGI Symbol;Acc:MGI:1( | 372.64 | 579.75 | 503.12 | 278.59 | 282.47 | 269.55 |
| 7268 | ENSMUSG00000029386 | 36.1861373 | 0.119415968 | 0.394100196 | 0.303009156 | 0.761882884 | 0.954306538 | NA | NA | 24.896 | 38.033 | 41.256 | 27.955 | 37.081 | 47.895 |
| 7269 | ENSMUSG00000029387 | 260.2894865 | 0.145410727 | 0.15376371 | 0.945676499 | 0.344313613 | 0.785808801 | Gtf2h3 | general transcription factor IIH, polypepti | 256.19 | 243.62 | 241.5 | 275.7 | 258.48 | 286.26 |
| 7270 | ENSMUSG00000029388 | 450.013502 | -0.002286404 | 0.134601772 | -0.016986433 | 0.986447439 | 0.99859176 | Eif2b1 | eukaryotic translation initiation factor 2B, | 449.74 | 446.12 | 454.82 | 408.73 | 495.14 | 445.54 |
| 7271 | ENSMUSG00000029389 | 114.16008 | 0.004453743 | 0.234089578 | 0.019025808 | 0.984820517 | 0.99859176 | Ddx55 | DEAD box helicase 55 [Source:MGI Symbc | 107.62 | 108.96 | 125.78 | 133.99 | 89.431 | 119.18 |
| 7272 | ENSMUSG00000029390 | 2356.881283 | 0.058956522 | 0.08748591 | 0.673897342 | 0.500376582 | 0.880716237 | Tmed2 | transmembrane p24 trafficking protein 2 | 2311.3 | 2299.5 | 2315.4 | 2399.3 | 2427.7 | 2388.1 |
| 7273 | ENSMUSG00000029392 | 38.40902645 | 0.225716706 | 0.369575753 | 0.610745441 | 0.541368118 | 0.89367136 | Rilpl1 | Rab interacting lysosomal protein-like 1 [S | 27.305 | 33.921 | 45.281 | 43.379 | 34.9 | 45.668 |
| 7274 | ENSMUSG00000029394 | 299.1198802 | 0.003290886 | 0.171982088 | 0.019135052 | 0.984733369 | 0.99859176 | Cdk2ap1 | CDK2 (cyclin-dependent kinase 2)-associa | 296.34 | 272.4 | 327.03 | 253.53 | 290.1 | 355.32 |
| 7275 | ENSMUSG00000029397 | 560.3568691 | -0.095611307 | 0.123084432 | -0.776794472 | 0.437280039 | 0.84703609 | Rchy1 | ring finger and CHY zinc finger domain cor | 587.87 | 602.36 | 546.39 | 535.01 | 572.57 | 517.94 |
| 7276 | ENSMUSG00000029401 | 183.3272884 | -0.098079592 | 0.192724806 | -0.508910055 | 0.610815272 | 0.916376001 | Rilpl2 | Rab interacting lysosomal protein-like 2 [S | 216.03 | 196.33 | 155.97 | 188.94 | 174.5 | 168.19 |

| | | | | | | | | | | | | | | | |
|---|---|---|---|---|---|---|---|---|---|---|---|---|---|---|---|
| 7277 | ENSMUSG00000029402 | 84.00391036 | -0.004137353 | 0.274790743 | -0.015056376 | 0.987987204 | 0.99859176 | Snrnp35 | small nuclear ribonucleoprotein 35 (U11/U | 93.963 | 90.457 | 67.418 | 63.623 | 89.431 | 99.132 |
| 7278 | ENSMUSG00000029403 | 117.2760495 | 0.18461523 | 0.215985753 | 0.854756516 | 0.392685956 | 0.821375223 | Cdkl2 | cyclin-dependent kinase-like 2 (CDC2-rela | 116.45 | 105.88 | 106.66 | 117.61 | 115.61 | 141.46 |
| 7279 | ENSMUSG00000029404 | 606.6593465 | 0.095966657 | 0.116621825 | 0.822887631 | 0.410571897 | 0.833241465 | Arl6ip4 | ADP-ribosylation factor-like 6 interacting p | 611.96 | 565.35 | 581.61 | 625.62 | 622.74 | 632.66 |
| 7280 | ENSMUSG00000029405 | 593.2829881 | -0.081069185 | 0.123821815 | -0.654724579 | 0.512645097 | 0.884860213 | G3bp2 | GTPase activating protein (SH3 domain) b | 619.19 | 594.14 | 615.82 | 526.33 | 620.56 | 583.65 |
| 7281 | ENSMUSG00000029406 | 966.6895539 | -0.041055926 | 0.129663203 | -0.316635136 | 0.751520472 | 0.952255112 | Pitpnm2 | phosphatidylinositol transfer protein, mer | 1004.7 | 905.6 | 1031.4 | 1068.1 | 816.87 | 973.5 |
| 7282 | ENSMUSG00000029407 | 1877.721445 | 0.097715381 | 0.109242459 | 0.894481705 | 0.371064214 | 0.807214774 | Uso1 | USO1 vesicle docking factor [Source:MGI S | 1947.5 | 1858.5 | 1636.2 | 2074.5 | 1911.9 | 1837.8 |
| 7283 | ENSMUSG00000029408 | 79.60025664 | -0.02667876 | 0.288020572 | -0.092627968 | 0.926199124 | 0.98758158 | Abcb9 | ATP-binding cassette, sub-family B (MDR/ | 100.39 | 64.759 | 75.468 | 86.758 | 58.893 | 91.335 |
| 7284 | ENSMUSG00000029409 | 0 | NA | NA | NA | NA | NA | U90926 | cDNA sequence U90926 [Source:MGI Sym | 0 | 0 | 0 | 0 | 0 | 0 |
| 7285 | ENSMUSG00000029410 | 1.193894609 | -0.207782015 | 2.285126421 | -0.090928018 | 0.927549788 | NA | Ppef2 | protein phosphatase, EF hand calcium-bir | 0.8031 | 0 | 3.0187 | 0 | 0 | 3.3415 |
| 7286 | ENSMUSG00000029413 | 81.29916215 | -0.297204452 | 0.266801049 | -1.113955336 | 0.265298348 | 0.721244797 | Naaa | N-acylethanolamine acid amidase [Source | 72.279 | 97.652 | 99.618 | 83.866 | 70.89 | 63.489 |
| 7287 | ENSMUSG00000029414 | 5.96702002 | -0.355901001 | 0.943225557 | -0.377323322 | 0.705933342 | 0.94295036 | Kntc1 | kinetochore associated 1 [Source:MGI Syr | 12.85 | 2.0558 | 5.0312 | 4.8199 | 4.3625 | 6.6831 |
| 7288 | ENSMUSG00000029415 | 213.8098227 | 0.076975316 | 0.167949851 | 0.458323218 | 0.646720246 | 0.925643431 | Sdad1 | SDA1 domain containing 1 [Source:MGI Sy | 203.18 | 216.89 | 204.27 | 214 | 205.04 | 239.48 |
| 7289 | ENSMUSG00000029416 | 481.3361745 | -0.522728136 | 0.164691054 | -3.17399229 | 0.001503577 | 0.028465622 | Slc15a4 | solute carrier family 15, member 4 [Source | 465 | 605.44 | 632.93 | 362.46 | 462.42 | 359.77 |
| 7290 | ENSMUSG00000029417 | 171.4896143 | -0.923639911 | 0.27225991 | -3.392493262 | 0.000692596 | 0.015825772 | Cxcl9 | chemokine (C-X-C motif) ligand 9 [Source: | 227.28 | 291.93 | 154.96 | 150.38 | 78.525 | 125.86 |
| 7291 | ENSMUSG00000029419 | 0 | NA | NA | NA | NA | NA | Ajm1 | apical junction component 1 [Source:MGI | 0 | 0 | 0 | 0 | 0 | 0 |
| 7292 | ENSMUSG00000029420 | 0 | NA | NA | NA | NA | NA | Rimbp2 | RIMS binding protein 2 [Source:MGI Symb | 0 | 0 | 0 | 0 | 0 | 0 |
| 7293 | ENSMUSG00000029422 | 598.9796916 | 0.084906341 | 0.131089344 | 0.64769827 | 0.517180123 | 0.886944902 | Rsrc2 | arginine/serine-rich coiled-coil 2 [Source:I | 538.88 | 645.53 | 560.48 | 618.87 | 659.82 | 570.29 |
| 7294 | ENSMUSG00000029423 | 0 | NA | NA | NA | NA | NA | Piwil1 | piwi-like RNA-mediated gene silencing 1 [S | 0 | 0 | 0 | 0 | 0 | 0 |
| 7295 | ENSMUSG00000029426 | 3708.168252 | 0.027069737 | 0.105021127 | 0.257755153 | 0.796595878 | 0.962236316 | Scarb2 | scavenger receptor class B, member 2 [So | 3338.5 | 4036.6 | 3646.6 | 3899.3 | 3505.2 | 3822.7 |
| 7296 | ENSMUSG00000029427 | 337.8818715 | 0.138942764 | 0.143045352 | 0.971319668 | 0.331389117 | 0.774747658 | Zcchc8 | zinc finger, CCHC domain containing 8 [So | 306.79 | 313.51 | 345.14 | 373.06 | 345.73 | 343.06 |
| 7297 | ENSMUSG00000029428 | 352.324647 | -0.151740236 | 0.162567602 | -0.933397762 | 0.350614635 | 0.791943999 | Stx2 | syntaxin 2 [Source:MGI Symbol;Acc:MGI:1 | 342.92 | 366.97 | 403.5 | 382.7 | 277.02 | 340.84 |
| 7298 | ENSMUSG00000029430 | 1538.618924 | 0.063457647 | 0.100463579 | 0.63164828 | 0.527616731 | 0.888334265 | Ran | RAN, member RAS oncogene family [Sour | 1606.2 | 1423.7 | 1483.2 | 1534.7 | 1592.3 | 1591.7 |
| 7299 | ENSMUSG00000029431 | 0 | NA | NA | NA | NA | NA | B3gnt4 | UDP-GlcNAc:betaGal beta-1,3-N-acetylglu | 0 | 0 | 0 | 0 | 0 | 0 |
| 7300 | ENSMUSG00000029432 | 266.3509612 | -0.128486978 | 0.16280194 | -0.789222648 | 0.429981887 | 0.843289222 | Nipsnap2 | nipsnap homolog 2 [Source:MGI Symbol;A | 276.27 | 304.26 | 254.58 | 282.45 | 238.85 | 241.7 |
| 7301 | ENSMUSG00000029433 | 55.4506822 | 0.04303763 | 0.303328061 | 0.141884763 | 0.88717103 | 0.982382091 | Diablo | diablo, IAP-binding mitochondrial protein | 52.202 | 54.48 | 57.356 | 59.767 | 64.346 | 44.554 |
| 7302 | ENSMUSG00000029434 | 486.6166571 | 0.062593765 | 0.141177527 | 0.44336918 | 0.657498725 | 0.927303282 | Vps33a | VPS33A CORVET/HOPS core subunit [Sour | 436.08 | 548.91 | 443.75 | 476.21 | 521.32 | 493.43 |
| 7303 | ENSMUSG00000029436 | 9.009409247 | -1.557881511 | 0.748709159 | -2.080756582 | 0.037456192 | 0.262483703 | Mmp17 | matrix metallopeptidase 17 [Source:MGI S | 16.865 | 12.335 | 11.069 | 3.8559 | 4.3625 | 5.5692 |
| 7304 | ENSMUSG00000029437 | 0 | NA | NA | NA | NA | NA | Il31 | interleukin 31 [Source:MGI Symbol;Acc:M | 0 | 0 | 0 | 0 | 0 | 0 |
| 7305 | ENSMUSG00000029438 | 290.9161892 | 0.079498215 | 0.155292816 | 0.511924613 | 0.608703772 | 0.916376001 | Bcl7a | B cell CLL/lymphoma 7A [Source:MGI Sym | 261.01 | 278.57 | 309.92 | 322.93 | 287.92 | 285.14 |
| 7306 | ENSMUSG00000029439 | 426.5609696 | 0.056511306 | 0.157131772 | 0.359642771 | 0.719114294 | 0.9447719 | Sfswap | splicing factor SWAP [Source:MGI Symbol | 457.77 | 379.3 | 417.59 | 504.16 | 432.98 | 367.57 |
| 7307 | ENSMUSG00000029440 | 880.1325078 | 0.065522528 | 0.116752939 | 0.561206668 | 0.574656658 | 0.906163061 | Psmd9 | proteasome (prosome, macropain) 26S su | 819.97 | 948.77 | 812.04 | 865.65 | 895.4 | 938.97 |
| 7308 | ENSMUSG00000029442 | 10.08335234 | -0.884941052 | 0.875370378 | -1.010933285 | 0.312048362 | 0.759830222 | Wdr66 | WD repeat domain 66 [Source:MGI Symbo | 16.062 | 7.1954 | 16.1 | 14.46 | 0 | 6.6831 |
| 7309 | ENSMUSG00000029445 | 41735.9908 | -0.069833401 | 0.094701345 | -0.737406644 | 0.460875096 | 0.859821403 | Hpd | 4-hydroxyphenylpyruvic acid dioxygenase | 39215 | 42927 | 46096 | 38719 | 41201 | 42257 |
| 7310 | ENSMUSG00000029446 | 84.37690359 | 0.278547423 | 0.246612102 | 1.12949616 | 0.258688588 | 0.712165697 | Psph | phosphoserine phosphatase [Source:MGI | 85.129 | 66.815 | 76.475 | 91.578 | 92.703 | 93.563 |
| 7311 | ENSMUSG00000029447 | 2163.241104 | 0.027571027 | 0.096566634 | 0.285512978 | 0.775251163 | 0.95893551 | Cct6a | chaperonin containing Tcp1, subunit 6a (z | 2092.1 | 2041.4 | 2294.2 | 2130.4 | 2228.1 | 2193.2 |
| 7312 | ENSMUSG00000029449 | 5.238210511 | -1.200162433 | 0.974085729 | -1.232091178 | 0.217915026 | 0.663697152 | Rhof | ras homolog family member F (in filopodia | 9.6372 | 5.1396 | 7.0437 | 1.928 | 5.4531 | 2.2277 |
| 7313 | ENSMUSG00000029451 | 0 | NA | NA | NA | NA | NA | Pramel27 | PRAME like 27 [Source:MGI Symbol;Acc:M | 0 | 0 | 0 | 0 | 0 | 0 |
| 7314 | ENSMUSG00000029452 | 8.245260644 | -0.816242673 | 0.979258042 | -0.833531754 | 0.404544897 | 0.829271989 | Tmem116 | transmembrane protein 116 [Source:MGI | 14.456 | 5.1396 | 12.075 | 14.46 | 0 | 3.3415 |
| 7315 | ENSMUSG00000029454 | 74.58898409 | 0.152365446 | 0.315239853 | 0.483331801 | 0.628860163 | 0.92053774 | Mapkapk5 | MAP kinase-activated protein kinase 5 [Sc | 48.989 | 81.206 | 82.512 | 97.362 | 82.887 | 54.578 |
| 7316 | ENSMUSG00000029455 | 48304.19499 | -0.158542465 | 0.09735004 | -1.628581395 | 0.103401665 | 0.473591415 | Aldh2 | aldehyde dehydrogenase 2, mitochondria | 51933 | 49450 | 51485 | 50833 | 43662 | 42462 |
| 7317 | ENSMUSG00000029456 | 795.2383157 | -0.182955556 | 0.141711406 | -1.291043259 | 0.196688678 | 0.637822894 | Acad10 | acyl-Coenzyme A dehydrogenase family, r | 772.58 | 956.99 | 808.01 | 747.08 | 847.41 | 639.35 |
| 7318 | ENSMUSG00000029458 | 3288.507764 | 0.148656128 | 0.104858577 | 1.417682109 | 0.156283596 | 0.574894692 | Brap | BRCA1 associated protein [Source:MGI Sy | 2999.6 | 2920.3 | 3438.3 | 3538.8 | 3214.1 | 3620 |
| 7319 | ENSMUSG00000029461 | 608.6352013 | 0.132341191 | 0.12082163 | 1.095343531 | 0.273366163 | 0.728364449 | Fam168a | family with sequence similarity 168, meml | 554.14 | 590.02 | 598.72 | 647.79 | 667.46 | 593.68 |
| 7320 | ENSMUSG00000029462 | 675.0718367 | 0.005024658 | 0.119066027 | 0.042200605 | 0.96633878 | 0.993964167 | Vps29 | VPS29 retromer complex component [Sou | 706.73 | 656.84 | 657.08 | 626.59 | 691.45 | 711.75 |
| 7321 | ENSMUSG00000029463 | 32.90778766 | -0.101339002 | 0.404579537 | -0.250479802 | 0.802216323 | 0.963085732 | Fam216a | family with sequence similarity 216, meml | 40.155 | 33.921 | 28.175 | 43.379 | 25.084 | 26.732 |
| 7322 | ENSMUSG00000029464 | 298.630406 | 0.048568544 | 0.160998671 | 0.301670463 | 0.762903287 | 0.95469451 | Gpn3 | GPN-loop GTPase 3 [Source:MGI Symbol;A | 293.94 | 270.34 | 315.96 | 262.2 | 315.19 | 334.15 |
| 7323 | ENSMUSG00000029465 | 1005.914603 | -0.066785274 | 0.108282259 | -0.616770234 | 0.537386288 | 0.890758111 | Arpc3 | actin related protein 2/3 complex, subunit | 1068.1 | 1038.2 | 981.09 | 1021.8 | 921.57 | 1004.7 |
| 7324 | ENSMUSG00000029466 | 425.2088546 | -0.066801171 | 0.135176418 | -0.494177697 | 0.621180689 | 0.918069106 | Anapc7 | anaphase promoting complex subunit 7 [S | 403.96 | 448.17 | 453.82 | 428.01 | 430.79 | 386.5 |
| 7325 | ENSMUSG00000029467 | 2605.181375 | -0.183361701 | 0.153602386 | -1.193742536 | 0.232578696 | 0.679847566 | Atp2a2 | ATPase, Ca++ transporting, cardiac muscle | 2288 | 3013.9 | 3011.7 | 3015.3 | 2120.2 | 2182 |
| 7326 | ENSMUSG00000029468 | 30.60722264 | -0.57522602 | 0.44026649 | -1.306540548 | 0.191368799 | 0.629099068 | P2rx7 | purinergic receptor P2X, ligand-gated ion | 31.321 | 46.256 | 32.2 | 16.388 | 20.722 | 36.757 |
| 7327 | ENSMUSG00000029469 | 41.31206595 | 0.041834985 | 0.343944676 | 0.121632891 | 0.903189764 | 0.983821605 | Ift81 | intraflagellar transport 81 [Source:MGI Sy | 46.58 | 37.005 | 38.237 | 35.667 | 44.715 | 45.668 |
| 7328 | ENSMUSG00000029470 | 1039.394951 | 0.286854316 | 0.149036859 | 1.924720621 | 0.054264319 | 0.327913359 | P2rx4 | purinergic receptor P2X, ligand-gated ion | 1039.2 | 928.21 | 840.21 | 930.24 | 1144.1 | 1354.4 |
| 7329 | ENSMUSG00000029471 | 230.3070514 | 0.187176575 | 0.160309753 | 1.167593184 | 0.242970901 | 0.692615513 | Camkk2 | calcium/calmodulin-dependent protein ki | 224.07 | 202.5 | 219.36 | 248.71 | 239.94 | 247.27 |
| 7330 | ENSMUSG00000029472 | 1157.050575 | -0.150059517 | 0.109340655 | -1.372403668 | 0.169937809 | 0.596876778 | Anapc5 | anaphase-promoting complex subunit 5 [S | 1228.7 | 1205.7 | 1217.6 | 1171.2 | 1127.7 | 991.32 |
| 7331 | ENSMUSG00000029474 | 156.6509475 | 0.046325365 | 0.186446175 | 0.248465086 | 0.80377458 | 0.963596811 | Rnf34 | ring finger protein 34 [Source:MGI Symbo | 159.01 | 153.16 | 149.93 | 150.38 | 159.23 | 168.19 |
| 7332 | ENSMUSG00000029475 | 132.6459891 | -0.207626798 | 0.217286682 | -0.955543141 | 0.339303099 | 0.781219101 | Kdm2b | lysine (K)-specific demethylase 2B [Source | 138.13 | 133.63 | 154.96 | 133.99 | 98.156 | 137 |

| | | | | | | | | | | | | | | | |
|---|---|---|---|---|---|---|---|---|---|---|---|---|---|---|---|
| 7333 | ENSMUSG00000029477 | 0.513958919 | -2.394274821 | 3.756096013 | -0.637437066 | 0.523840191 | NA | Morn3 | MORN repeat containing 3 [Source:MGI S | 0 | 3.0838 | 0 | 0 | 0 | 0 |
| 7334 | ENSMUSG00000029478 | 875.1765894 | 0.186863607 | 0.118368346 | 1.578661976 | 0.114413611 | 0.497223548 | Ncor2 | nuclear receptor co-repressor 2 [Source:N | 823.98 | 820.28 | 812.04 | 1030.5 | 839.78 | 924.49 |
| 7335 | ENSMUSG00000029480 | 206.7557904 | -0.067808134 | 0.18276187 | -0.371019041 | 0.71062335 | 0.944239215 | Dhx37 | DEAH (Asp-Glu-Ala-His) box polypeptide 3 | 206.4 | 224.09 | 204.27 | 191.83 | 175.59 | 238.36 |
| 7336 | ENSMUSG00000029482 | 1866.254486 | 0.35076681 | 0.51224615 | 0.684762219 | 0.49349398 | 0.87710702 | Aacs | acetoacetyl-CoA synthetase [Source:MGI : | 2349.1 | 1684.8 | 887.51 | 2831.2 | 2203 | 1241.9 |
| 7337 | ENSMUSG00000029484 | 47.80427137 | -1.122890544 | 0.334872859 | -3.353184685 | 0.000798874 | 0.017668059 | Anxa3 | annexin A3 [Source:MGI Symbol;Acc:MGI | 72.279 | 69.898 | 54.337 | 31.811 | 25.084 | 33.415 |
| 7338 | ENSMUSG00000029486 | 580.7343557 | 0.142085043 | 0.133995381 | 1.060372689 | 0.288975082 | 0.739603195 | Mrpl1 | mitochondrial ribosomal protein L1 [Sourc | 551.73 | 590.02 | 514.19 | 535.97 | 659.82 | 632.66 |
| 7339 | ENSMUSG00000029490 | 46.24863546 | 0.596145313 | 0.338286421 | 1.762250203 | 0.078027033 | 0.409203412 | Mfsd7a | major facilitator superfamily domain cont | 44.171 | 32.893 | 33.206 | 64.587 | 44.715 | 57.92 |
| 7340 | ENSMUSG00000029491 | 0 | NA | NA | NA | NA | NA | Pde6b | phosphodiesterase 6B, cGMP, rod recepto | 0 | 0 | 0 | 0 | 0 | 0 |
| 7341 | ENSMUSG00000029499 | 1792.487379 | 0.142592653 | 0.114623421 | 1.244009747 | 0.213495976 | 0.658283051 | Pxmp2 | peroxisomal membrane protein 2 [Source | 1644.8 | 1638.5 | 1828.3 | 1680.2 | 2089.6 | 1873.5 |
| 7342 | ENSMUSG00000029500 | 126.0917047 | -0.113296267 | 0.209309198 | -0.541286613 | 0.588310046 | 0.910917568 | Pgam5 | phosphoglycerate mutase family member | 126.89 | 132.6 | 133.83 | 128.21 | 104.7 | 130.32 |
| 7343 | ENSMUSG00000029501 | 522.5885949 | -0.020711869 | 0.128583089 | -0.16107771 | 0.872032196 | 0.979369708 | Ankle2 | ankyrin repeat and LEM domain containin | 497.92 | 577.69 | 504.13 | 535.01 | 499.5 | 521.28 |
| 7344 | ENSMUSG00000029502 | 498.2102808 | -0.036103125 | 0.138276806 | -0.261093135 | 0.794020686 | 0.961915077 | Golga3 | golgi autoantigen, golgin subfamily a, 3 [S | 466.6 | 551.99 | 495.07 | 456.93 | 546.4 | 472.27 |
| 7345 | ENSMUSG00000029503 | 0.49081066 | -0.698504554 | 3.1295234 | -0.22319838 | 0.823381113 | NA | P2rx2 | purinergic receptor P2X, ligand-gated ion | 0.8031 | 1.0279 | 0 | 0 | 0 | 1.1138 |
| 7346 | ENSMUSG00000029504 | 212.8512525 | -0.01418976 | 0.188493597 | -0.075279796 | 0.939992096 | 0.988812319 | Ddx51 | DEAD box helicase 51 [Source:MGI Symbo | 216.84 | 202.5 | 222.38 | 213.04 | 251.93 | 170.42 |
| 7347 | ENSMUSG00000029505 | 639.4267769 | -0.066373643 | 0.124255731 | -0.534169675 | 0.593224148 | 0.913011006 | Ep400 | E1A binding protein p400 [Source:MGI Sy | 630.44 | 698.98 | 632.93 | 576.46 | 674 | 623.75 |
| 7348 | ENSMUSG00000029507 | 296.3690864 | -0.095745022 | 0.167863901 | -0.570372912 | 0.568424798 | 0.903527882 | Pus1 | pseudouridine synthase 1 [Source:MGI Sy | 342.12 | 291.93 | 283.76 | 266.06 | 334.82 | 259.53 |
| 7349 | ENSMUSG00000029510 | 1.329089092 | -2.591198613 | 2.080256964 | -1.24561468 | 0.21290589 | NA | Gpc2 | glypican 2 (cerebroglycan) [Source:MGI Sy | 0.8031 | 2.0558 | 4.025 | 0 | 1.0906 | 0 |
| 7350 | ENSMUSG00000029512 | 2089.542752 | -0.090195256 | 0.123513379 | -0.730246846 | 0.465239312 | 0.863163342 | Ulk1 | unc-51 like kinase 1 [Source:MGI Symbol;/ | 2237.4 | 2331.3 | 1894.8 | 1822.9 | 1967.5 | 2283.4 |
| 7351 | ENSMUSG00000029513 | 448.0916642 | 0.155952408 | 0.130281858 | 1.197038561 | 0.231291525 | 0.678442632 | Prkab1 | protein kinase, AMP-activated, beta 1 nor | 445.72 | 394.72 | 430.67 | 470.42 | 459.15 | 487.86 |
| 7352 | ENSMUSG00000029516 | 13.90407622 | -0.901140719 | 0.754531673 | -1.194304693 | 0.232358802 | 0.679781092 | Cit | citron [Source:MGI Symbol;Acc:MGI:1053 | 16.865 | 24.67 | 13.081 | 22.171 | 2.1812 | 4.4554 |
| 7353 | ENSMUSG00000029517 | 0 | NA | NA | NA | NA | NA | Ankrd7 | ankyrin repeat domain 7 [Source:MGI Sym | 0 | 0 | 0 | 0 | 0 | 0 |
| 7354 | ENSMUSG00000029518 | 318.5960786 | 0.116099914 | 0.150716299 | 0.770320888 | 0.441109569 | 0.849828766 | Rab35 | RAB35, member RAS oncogene family [So | 297.15 | 301.18 | 318.98 | 319.08 | 304.28 | 370.91 |
| 7355 | ENSMUSG00000029521 | 47.4506193 | 0.381137098 | 0.328552337 | 1.160049875 | 0.2460285 | 0.696390652 | Chek2 | checkpoint kinase 2 [Source:MGI Symbol;. | 48.186 | 31.865 | 43.269 | 51.091 | 51.259 | 59.034 |
| 7356 | ENSMUSG00000029522 | 0 | NA | NA | NA | NA | NA | Pla2g1b | phospholipase A2, group IB, pancreas [So | 0 | 0 | 0 | 0 | 0 | 0 |
| 7357 | ENSMUSG00000029524 | 240.0371309 | 0.098007505 | 0.166319216 | 0.589273491 | 0.555677824 | 0.897628185 | Sirt4 | sirtuin 4 [Source:MGI Symbol;Acc:MGI:19 | 244.14 | 201.47 | 249.55 | 238.1 | 254.11 | 252.84 |
| 7358 | ENSMUSG00000029526 | 0 | NA | NA | NA | NA | NA | 1700123K08 | RIKEN cDNA 1700123K08 gene [Source:M | 0 | 0 | 0 | 0 | 0 | 0 |
| 7359 | ENSMUSG00000029528 | 840.0099001 | 0.020672605 | 0.128637402 | 0.16070447 | 0.872326169 | 0.979419285 | Pxn | paxillin [Source:MGI Symbol;Acc:MGI:108 | 774.99 | 838.78 | 888.51 | 817.45 | 759.07 | 961.25 |
| 7360 | ENSMUSG00000029530 | 4.401484963 | -0.293265409 | 1.063924129 | -0.27564504 | 0.782820713 | NA | Ccr9 | chemokine (C-C motif) receptor 9 [Source | 2.4093 | 3.0838 | 9.0562 | 1.928 | 4.3625 | 5.5692 |
| 7361 | ENSMUSG00000029534 | 121.455 | -0.17193737 | 0.22153969 | -0.776101881 | 0.437688833 | 0.847422601 | St7 | suppression of tumorigenicity 7 [Source:N | 128.5 | 127.46 | 129.81 | 101.22 | 102.52 | 139.23 |
| 7362 | ENSMUSG00000029535 | 333.5008018 | 0.038614795 | 0.149860304 | 0.257671938 | 0.796660105 | 0.962236316 | Triap1 | TP53 regulated inhibitor of apoptosis 1 [S | 371.03 | 300.15 | 314.95 | 332.57 | 337 | 345.29 |
| 7363 | ENSMUSG00000029536 | 423.4300353 | -0.044891385 | 0.150191935 | -0.298893448 | 0.765021347 | 0.954980327 | Gatc | glutamyl-tRNA(Gln) amidotransferase, sul | 442.51 | 482.09 | 365.27 | 404.87 | 450.43 | 395.41 |
| 7364 | ENSMUSG00000029538 | 441.3853313 | 0.105961633 | 0.132757989 | 0.798156355 | 0.424779761 | 0.840504368 | Srsf9 | serine and arginine-rich splicing factor 9 [ | 420.02 | 421.45 | 433.69 | 416.44 | 474.42 | 482.29 |
| 7365 | ENSMUSG00000029541 | 0 | NA | NA | NA | NA | NA | Cyp2w1 | cytochrome P450, family 2, subfamily w, p | 0 | 0 | 0 | 0 | 0 | 0 |
| 7366 | ENSMUSG00000029544 | 1.277763006 | 1.830198481 | 2.0837712 | 0.878310671 | 0.379775147 | NA | Cabp1 | calcium binding protein 1 [Source:MGI Sy | 1.6062 | 0 | 0 | 3.8559 | 1.0906 | 1.1138 |
| 7367 | ENSMUSG00000029545 | 4217.608928 | -0.017511069 | 0.085412423 | -0.205017819 | 0.837558203 | 0.971073158 | Acads | acyl-Coenzyme A dehydrogenase, short ch | 4240.4 | 4298.8 | 4190 | 4027.5 | 4276.3 | 4272.7 |
| 7368 | ENSMUSG00000029546 | 0 | NA | NA | NA | NA | NA | Uncx | UNC homeobox [Source:MGI Symbol;Acc: | 0 | 0 | 0 | 0 | 0 | 0 |
| 7369 | ENSMUSG00000029547 | 434.7300994 | -0.138926792 | 0.130278749 | -1.066381074 | 0.286251413 | 0.73856672 | Ints1 | integrator complex subunit 1 [Source:MG | 447.33 | 448.17 | 471.93 | 437.65 | 407.89 | 395.41 |
| 7370 | ENSMUSG00000029550 | 300.3604427 | -0.035253653 | 0.151383197 | -0.232876922 | 0.815856979 | 0.967767553 | Sppl3 | signal peptide peptidase 3 [Source:MGI Sy | 282.69 | 328.93 | 300.87 | 293.05 | 280.29 | 316.33 |
| 7371 | ENSMUSG00000029551 | 156.134814 | -0.018016858 | 0.192911812 | -0.093394273 | 0.92559034 | 0.987546253 | Psmg3 | proteasome (prosome, macropain) assem | 162.23 | 138.77 | 170.06 | 150.38 | 149.41 | 165.96 |
| 7372 | ENSMUSG00000029552 | 15.14721585 | 0.50096372 | 0.662923523 | 0.755688556 | 0.449835947 | 0.855321038 | Tes | testin LIM domain protein [Source:MGI Sy | 12.047 | 14.391 | 11.069 | 5.7839 | 14.178 | 33.415 |
| 7373 | ENSMUSG00000029553 | 16.32096264 | -1.382772279 | 0.572676526 | -2.414578239 | 0.015753439 | 0.1470999 | Tfec | transcription factor EC [Source:MGI Symb | 24.093 | 22.614 | 24.15 | 11.568 | 4.3625 | 11.138 |
| 7374 | ENSMUSG00000029554 | 168.629523 | -0.331974853 | 0.195811825 | -1.695376946 | 0.090003936 | 0.442644317 | Mad1l1 | MAD1 mitotic arrest deficient 1-like 1 [Sou | 171.86 | 177.83 | 214.33 | 141.7 | 169.05 | 137 |
| 7375 | ENSMUSG00000029556 | 596.0889124 | -0.097515103 | 0.140007114 | -0.696501059 | 0.486115094 | 0.873111725 | Hnf1a | HNF1 homeobox A [Source:MGI Symbol;A | 566.19 | 607.5 | 676.2 | 652.61 | 509.32 | 564.72 |
| 7376 | ENSMUSG00000029557 | 186.6863391 | -0.183289294 | 0.176447454 | -1.038775505 | 0.298909156 | 0.750073219 | Mrm2 | mitochondrial rRNA methyltransferase 2 [ | 199.17 | 198.39 | 198.23 | 187.98 | 158.14 | 178.22 |
| 7377 | ENSMUSG00000029559 | 268.1955151 | 0.285111051 | 0.164919869 | 1.728785339 | 0.083847521 | 0.426710495 | 2210016L21 | RIKEN cDNA 2210016L21 gene [Source:M | 232.1 | 235.39 | 257.6 | 254.49 | 301.01 | 328.58 |
| 7378 | ENSMUSG00000029560 | 337.642625 | -0.08407871 | 0.141704234 | -0.593339437 | 0.552954011 | 0.896226672 | Snx8 | sorting nexin 8 [Source:MGI Symbol;Acc:N | 350.96 | 345.38 | 346.15 | 339.32 | 302.1 | 341.95 |
| 7379 | ENSMUSG00000029561 | 48.85102072 | -0.855580312 | 0.32345195 | -2.645154284 | 0.008165368 | 0.093215374 | Oasl2 | 2'-5' oligoadenylate synthetase-like 2 [Sou | 70.673 | 53.452 | 64.4 | 31.811 | 34.9 | 37.871 |
| 7380 | ENSMUSG00000029563 | 96.58048023 | -0.332512419 | 0.250512554 | -1.327328368 | 0.184400084 | 0.619137675 | Foxp2 | forkhead box P2 [Source:MGI Symbol;Acc | 113.24 | 112.04 | 97.606 | 88.686 | 65.437 | 102.47 |
| 7381 | ENSMUSG00000029564 | 0 | NA | NA | NA | NA | NA | 4930519G04 | RIKEN cDNA 4930519G04 gene [Source:M | 0 | 0 | 0 | 0 | 0 | 0 |
| 7382 | ENSMUSG00000029569 | 388.2947797 | -0.055534064 | 0.146611424 | -0.378784019 | 0.704848256 | 0.94295036 | Tmem168 | transmembrane protein 168 [Source:MGI | 445.72 | 393.69 | 347.15 | 390.41 | 374.08 | 378.71 |
| 7383 | ENSMUSG00000029570 | 50.26978059 | -1.18579944 | 0.323621107 | -3.664159764 | 0.000248152 | 0.007477643 | Lfng | LFNG O-fucosylpeptide 3-beta-N-acetylglu | 69.87 | 75.038 | 64.4 | 25.063 | 32.719 | 34.529 |
| 7384 | ENSMUSG00000029571 | 5126.594486 | -0.209303481 | 0.106171327 | -1.971374818 | 0.048681019 | 0.307208666 | Tmem106b | transmembrane protein 106B [Source:MG | 5390.4 | 6114.1 | 4990 | 5058 | 4702.7 | 4504.4 |
| 7385 | ENSMUSG00000029575 | 379.2143166 | 0.456179723 | 0.160968627 | 2.833966671 | 0.004597413 | 0.063210054 | Mmab | methylmalonic aciduria (cobalamin deficie | 352.56 | 323.79 | 281.75 | 370.17 | 459.15 | 487.86 |
| 7386 | ENSMUSG00000029576 | 0.17131964 | -0.796629176 | 4.080472857 | -0.195229623 | 0.84521319 | NA | Radil | Ras association and DIL domains [Source:I | 0 | 1.0279 | 0 | 0 | 0 | 0 |
| 7387 | ENSMUSG00000029577 | 1276.933564 | 0.135884829 | 0.10895366 | 1.247180026 | 0.212331495 | 0.656933207 | Ube3b | ubiquitin protein ligase E3B [Source:MGI S | 1207.1 | 1222.2 | 1220.6 | 1200.2 | 1399.3 | 1412.4 |
| 7388 | ENSMUSG00000029578 | 1751.164867 | 0.182415968 | 0.096325604 | 1.893743315 | 0.058259089 | 0.342880809 | Wipi2 | WD repeat domain, phosphoinositide inte | 1611 | 1622.1 | 1688.5 | 1764.1 | 1894.4 | 1926.9 |

| | | | | | | | | | | | | | | | |
|---|---|---|---|---|---|---|---|---|---|---|---|---|---|---|---|
| 7389 | ENSMUSG00000029580 | 5825.189311 | -0.527630229 | 0.088541961 | -5.959098048 | 2.54E-09 | 4.60E-07 | Actb | actin, beta [Source:MGI Symbol;Acc:MGI: | 7256.8 | 6891.2 | 6487.3 | 4791.9 | 4679.8 | 4844.1 |
| 7390 | ENSMUSG00000029581 | 37.27313484 | -0.603388155 | 0.390120143 | -1.54667265 | 0.121942201 | 0.511776306 | Fscn1 | fascin actin-bundling protein 1 [Source:M | 59.43 | 37.005 | 38.237 | 33.739 | 21.812 | 33.415 |
| 7391 | ENSMUSG00000029586 | 0 | NA | NA | NA | NA | NA | Spdye4b | speedy/RINGO cell cycle regulator family, | 0 | 0 | 0 | 0 | 0 | 0 |
| 7392 | ENSMUSG00000029587 | 197.4786971 | 0.198790504 | 0.188075358 | 1.056972621 | 0.290524083 | 0.740463951 | Zfp12 | zinc finger protein 12 [Source:MGI Symbo | 170.26 | 214.83 | 167.04 | 211.11 | 233.39 | 188.24 |
| 7393 | ENSMUSG00000029591 | 170.1324634 | 0.569127776 | 0.29734794 | 1.914012846 | 0.055618514 | 0.332908719 | Ung | uracil DNA glycosylase [Source:MGI Symb | 208 | 91.485 | 110.69 | 258.35 | 142.87 | 209.4 |
| 7394 | ENSMUSG00000029592 | 299.6208719 | 0.20042675 | 0.14987346 | 1.337306481 | 0.181122616 | 0.614561135 | Usp30 | ubiquitin specific peptidase 30 [Source:M | 264.22 | 284.73 | 287.79 | 311.36 | 302.1 | 347.52 |
| 7395 | ENSMUSG00000029594 | 166.0290661 | -0.265334449 | 0.186369929 | -1.423697758 | 0.154533983 | 0.572594786 | Rbm19 | RNA binding motif protein 19 [Source:MG | 177.49 | 196.33 | 170.06 | 152.31 | 139.6 | 160.39 |
| 7396 | ENSMUSG00000029595 | 0 | NA | NA | NA | NA | NA | Lhx5 | LIM homeobox protein 5 [Source:MGI Syn | 0 | 0 | 0 | 0 | 0 | 0 |
| 7397 | ENSMUSG00000029596 | 234.9007229 | -0.009978016 | 0.189712231 | -0.052595534 | 0.958054176 | 0.991861398 | Sdsl | serine dehydratase-like [Source:MGI Syml | 221.66 | 243.62 | 241.5 | 184.12 | 234.48 | 284.03 |
| 7398 | ENSMUSG00000029597 | 2818.939825 | -0.543380419 | 0.164460662 | -3.30401455 | 0.000953109 | 0.020184138 | Sds | serine dehydratase [Source:MGI Symbol;A | 2954.6 | 4005.8 | 3070.1 | 1816.1 | 2188.9 | 2878.2 |
| 7399 | ENSMUSG00000029598 | 1270.598634 | -0.231626992 | 0.109976327 | -2.106153186 | 0.035191048 | 0.25151549 | Plbd2 | phospholipase B domain containing 2 [Sou | 1233.6 | 1427.8 | 1457 | 1177 | 1191 | 1137.2 |
| 7400 | ENSMUSG00000029599 | 532.0632676 | 0.163235889 | 0.130162619 | 1.25409192 | 0.209808597 | 0.653601453 | Ddx54 | DEAD box helicase 54 [Source:MGI Symbc | 457.77 | 494.43 | 554.44 | 551.4 | 554.03 | 580.31 |
| 7401 | ENSMUSG00000029600 | 81.57731592 | 0.208450078 | 0.246172115 | 0.846765599 | 0.397125787 | 0.824612828 | Rita1 | RBPJ interacting and tubulin associated 1 | 75.492 | 76.066 | 75.468 | 82.902 | 94.884 | 84.652 |
| 7402 | ENSMUSG00000029601 | 5.799969419 | 0.644848358 | 0.963437073 | 0.669320681 | 0.503290937 | 0.881357141 | Iqcd | IQ motif containing D [Source:MGI Symbo | 6.4248 | 5.1396 | 2.0125 | 13.496 | 3.2719 | 4.4554 |
| 7403 | ENSMUSG00000029602 | 1.514986808 | -1.005766966 | 1.909033141 | -0.526846257 | 0.598300357 | NA | Rasal1 | RAS protein activator like 1 (GAP1 like) [Sc | 4.0155 | 2.0558 | 0 | 1.928 | 1.0906 | 0 |
| 7404 | ENSMUSG00000029603 | 4.702731909 | -0.923654042 | 1.044705205 | -0.884128879 | 0.376626645 | NA | Dtx1 | deltex 1, E3 ubiquitin ligase [Source:MGI S | 3.2124 | 7.1954 | 8.05 | 0.964 | 5.4531 | 3.3415 |
| 7405 | ENSMUSG00000029605 | 25.28340109 | 0.883953187 | 0.470326169 | 1.87944717 | 0.060183461 | 0.349595814 | Oas1b | 2'-5' oligoadenylate synthetase 1B [Source | 21.684 | 17.475 | 14.087 | 42.415 | 35.99 | 20.049 |
| 7406 | ENSMUSG00000029607 | 7.802667153 | 0.57706204 | 0.769366061 | 0.750048735 | 0.453225354 | 0.857073339 | Ankrd61 | ankyrin repeat domain 61 [Source:MGI Sy | 5.6217 | 8.2233 | 5.0312 | 12.532 | 8.7249 | 6.6831 |
| 7407 | ENSMUSG00000029608 | 0 | NA | NA | NA | NA | NA | Rph3a | rabphilin 3A [Source:MGI Symbol;Acc:MG | 0 | 0 | 0 | 0 | 0 | 0 |
| 7408 | ENSMUSG00000029610 | 91.21461131 | 0.038961775 | 0.284102802 | 0.137139708 | 0.890920366 | 0.982489666 | Aimp2 | aminoacyl tRNA synthetase complex-inter | 94.766 | 86.345 | 88.55 | 73.262 | 131.96 | 72.4 |
| 7409 | ENSMUSG00000029613 | 865.6939028 | 0.028327533 | 0.108308293 | 0.26154537 | 0.793671966 | 0.961915077 | Eif2ak1 | eukaryotic translation initiation factor 2 al | 835.23 | 873.73 | 863.36 | 917.71 | 838.69 | 865.46 |
| 7410 | ENSMUSG00000029614 | 4713.803627 | 0.014903065 | 0.086088548 | 0.173113206 | 0.862562444 | 0.976334255 | Rpl6 | ribosomal protein L6 [Source:MGI Symbol | 4699 | 4887.7 | 4481.8 | 4773.6 | 4719.1 | 4721.6 |
| 7411 | ENSMUSG00000029616 | 673.7009097 | 0.123395928 | 0.126348489 | 0.976631609 | 0.328751565 | 0.77362974 | Erp29 | endoplasmic reticulum protein 29 [Source | 648.1 | 659.92 | 626.89 | 740.33 | 615.11 | 751.84 |
| 7412 | ENSMUSG00000029617 | 618.1257071 | -0.014774493 | 0.129080523 | -0.114459506 | 0.908873545 | 0.985098284 | Ccz1 | CCZ1 vacuolar protein trafficking and biog | 652.92 | 642.45 | 567.52 | 557.18 | 629.29 | 659.4 |
| 7413 | ENSMUSG00000029618 | 0 | NA | NA | NA | NA | NA | Ocm | oncomodulin [Source:MGI Symbol;Acc:M | 0 | 0 | 0 | 0 | 0 | 0 |
| 7414 | ENSMUSG00000029620 | 0 | NA | NA | NA | NA | NA | 1700018F24 | RIKEN cDNA 1700018F24 gene [Source:M | 0 | 0 | 0 | 0 | 0 | 0 |
| 7415 | ENSMUSG00000029621 | 982.7246045 | 0.018704341 | 0.109057877 | 0.171508391 | 0.863824033 | 0.976954351 | Arpc1a | actin related protein 2/3 complex, subunit | 981.39 | 1006.3 | 941.84 | 1057.5 | 933.57 | 975.73 |
| 7416 | ENSMUSG00000029622 | 664.5894404 | -0.048020325 | 0.130642633 | -0.367570093 | 0.713193818 | 0.944239215 | Arpc1b | actin related protein 2/3 complex, subunit | 704.32 | 698.98 | 623.87 | 722.98 | 580.21 | 657.17 |
| 7417 | ENSMUSG00000029623 | 1008.846193 | 0.044186344 | 0.11402287 | 0.387521768 | 0.698369973 | 0.940696964 | Pdap1 | PDGFA associated protein 1 [Source:MGI S | 950.07 | 971.38 | 1058.6 | 939.88 | 1038.3 | 1094.9 |
| 7418 | ENSMUSG00000029624 | 234.780566 | 0.085719625 | 0.166695941 | 0.514227428 | 0.607092994 | 0.916376001 | Ptcd1 | pentatricopeptide repeat domain 1 [Sourc | 240.93 | 204.56 | 237.47 | 232.32 | 262.84 | 230.57 |
| 7419 | ENSMUSG00000029625 | 376.6540108 | 0.398655588 | 0.155599813 | 2.562056981 | 0.010405425 | 0.111804391 | Cpsf4 | cleavage and polyadenylation specific fact | 281.89 | 320.71 | 373.32 | 449.21 | 446.06 | 388.73 |
| 7420 | ENSMUSG00000029627 | 131.7185842 | 0.391425404 | 0.249896564 | 1.566349686 | 0.117266775 | 0.502643331 | Zkscan14 | zinc finger with KRAB and SCAN domains 1 | 97.978 | 87.373 | 156.97 | 164.84 | 146.14 | 137 |
| 7421 | ENSMUSG00000029629 | 464.6478701 | -0.288889481 | 0.140832455 | -2.051299041 | 0.040237834 | 0.274604506 | Phf14 | PHD finger protein 14 [Source:MGI Symbc | 509.17 | 560.22 | 463.88 | 422.22 | 453.7 | 378.71 |
| 7422 | ENSMUSG00000029630 | 9714.224233 | -0.447307273 | 0.128744078 | -3.474391056 | 0.000512014 | 0.01268112 | Cyp3a25 | cytochrome P450, family 3, subfamily a, p | 11413 | 10302 | 11910 | 8125.4 | 9710.9 | 6824.5 |
| 7423 | ENSMUSG00000029632 | 5085.500673 | 0.048293608 | 0.087968654 | 0.54898655 | 0.58301468 | 0.908645447 | Ndufa4 | Ndufa4, mitochondrial complex associate | 5048.3 | 5076.9 | 4875.3 | 4953.9 | 5099.7 | 5458.9 |
| 7424 | ENSMUSG00000029633 | 0.349477146 | 0.165142768 | 3.911021834 | 0.042224967 | 0.966319359 | NA | Gm5578 | predicted pseudogene 5578 [Source:MGI | 0 | 0 | 1.0062 | 0 | 1.0906 | 0 |
| 7425 | ENSMUSG00000029634 | 674.8009121 | 0.242749085 | 0.118170536 | 2.054226827 | 0.039953743 | 0.273655693 | Rnf6 | ring finger protein (C3H2C3 type) 6 [Sourc | 613.57 | 620.86 | 619.85 | 705.63 | 694.72 | 794.17 |
| 7426 | ENSMUSG00000029635 | 597.0911237 | 0.368477378 | 0.116383742 | 3.166055415 | 0.001545214 | 0.029101522 | Cdk8 | cyclin-dependent kinase 8 [Source:MGI Sy | 536.47 | 503.68 | 523.25 | 675.75 | 675.09 | 668.31 |
| 7427 | ENSMUSG00000029636 | 1.182255181 | 2.639215885 | 2.317584413 | 1.138778752 | 0.254795446 | NA | Wasf3 | WASP family, member 3 [Source:MGI Sym | 0.8031 | 0 | 0 | 1.928 | 4.3625 | 0 |
| 7428 | ENSMUSG00000029638 | 21.26118229 | -0.404842772 | 0.498901885 | -0.811467714 | 0.417097127 | 0.836878476 | Glcci1 | glucocorticoid induced transcript 1 [Sourc | 24.093 | 21.586 | 27.169 | 25.063 | 19.631 | 10.025 |
| 7429 | ENSMUSG00000029640 | 397.593124 | -0.008091203 | 0.147430321 | -0.05488154 | 0.956232838 | 0.991678478 | Usp12 | ubiquitin specific peptidase 12 [Source:M | 369.43 | 453.31 | 374.32 | 426.08 | 392.62 | 369.8 |
| 7430 | ENSMUSG00000029641 | 2.053898425 | -0.351790157 | 1.497411951 | -0.234932115 | 0.814261423 | NA | Rasl11a | RAS-like, family 11, member A [Source:MC | 0.8031 | 4.1117 | 2.0125 | 0.964 | 1.0906 | 3.3415 |
| 7431 | ENSMUSG00000029642 | 905.7286897 | 0.101465429 | 0.105017175 | 0.966179384 | 0.333954427 | 0.775900437 | Polr1d | polymerase (RNA) I polypeptide D [Source | 885.02 | 841.86 | 894.55 | 936.99 | 930.3 | 945.65 |
| 7432 | ENSMUSG00000029644 | 0 | NA | NA | NA | NA | NA | Pdx1 | pancreatic and duodenal homeobox 1 [So | 0 | 0 | 0 | 0 | 0 | 0 |
| 7433 | ENSMUSG00000029646 | 0 | NA | NA | NA | NA | NA | Cdx2 | caudal type homeobox 2 [Source:MGI Syn | 0 | 0 | 0 | 0 | 0 | 0 |
| 7434 | ENSMUSG00000029647 | 322.4857144 | -0.065521071 | 0.159065776 | -0.411911804 | 0.680404068 | 0.934384556 | Pan3 | PAN3 poly(A) specific ribonuclease subun | 288.31 | 352.58 | 349.17 | 300.76 | 352.27 | 291.83 |
| 7435 | ENSMUSG00000029648 | 96.03306139 | -0.479535076 | 0.273912371 | -1.750687907 | 0.079999684 | 0.415341728 | Flt1 | FMS-like tyrosine kinase 1 [Source:MGI Sy | 98.782 | 130.55 | 106.66 | 95.434 | 53.44 | 91.335 |
| 7436 | ENSMUSG00000029649 | 2076.953726 | 0.115700458 | 0.10433879 | 1.108892084 | 0.267476739 | 0.72293778 | Pomp | proteasome maturation protein [Source:N | 2098.5 | 1957.2 | 1923.9 | 1963.6 | 2214 | 2304.5 |
| 7437 | ENSMUSG00000029650 | 1099.79285 | 0.076741356 | 0.138173151 | 0.555399917 | 0.578621149 | 0.908137944 | Slc46a3 | solute carrier family 46, member 3 [Sourc | 1085.8 | 969.33 | 1156.2 | 1061.3 | 1340.4 | 985.75 |
| 7438 | ENSMUSG00000029651 | 13.67791694 | 0.30502629 | 0.669754541 | 0.455429969 | 0.648799939 | 0.925643431 | Mtus2 | microtubule associated tumor suppressor | 19.274 | 3.0838 | 14.087 | 12.532 | 15.269 | 17.822 |
| 7439 | ENSMUSG00000029655 | 790.6473862 | -0.063683652 | 0.114949357 | -0.554014862 | 0.579568674 | 0.908350165 | N4bp2l2 | NEDD4 binding protein 2-like 2 [Source:M | 841.65 | 781.22 | 800.97 | 768.29 | 828.87 | 722.88 |
| 7440 | ENSMUSG00000029656 | 4744.468125 | 0.998484758 | 0.175423948 | 5.691838386 | 1.26E-08 | 1.87E-06 | C8b | complement component 8, beta polypept | 2223.8 | 3541.2 | 3732.2 | 5206.4 | 7347.5 | 6415.7 |
| 7441 | ENSMUSG00000029657 | 835.1411963 | 0.326355584 | 0.146536005 | 2.227135812 | 0.025938198 | 0.208275484 | Hsph1 | heat shock 105kDa/110kDa protein 1 [Sou | 671.39 | 730.85 | 822.1 | 1045 | 755.8 | 985.75 |
| 7442 | ENSMUSG00000029658 | 0.17131964 | -0.796629176 | 4.080472857 | -0.195229623 | 0.84521319 | NA | Wdr95 | WD40 repeat domain 95 [Source:MGI Sym | 0 | 1.0279 | 0 | 0 | 0 | 0 |
| 7443 | ENSMUSG00000029659 | 1.995115828 | 0.655771166 | 1.467070164 | 0.446993731 | 0.654879586 | NA | Medag | mesenteric estrogen dependent adipogen | 1.6062 | 1.0279 | 2.0125 | 2.8919 | 1.0906 | 3.3415 |
| 7444 | ENSMUSG00000029660 | 0 | NA | NA | NA | NA | NA | Tex26 | testis expressed 26 [Source:MGI Symbol;A | 0 | 0 | 0 | 0 | 0 | 0 |

| | | | | | | | | | | | | | | | |
|---|---|---|---|---|---|---|---|---|---|---|---|---|---|---|---|
| 7445 | ENSMUSG00000029661 | 186.0421339 | -1.316499336 | 0.307714277 | -4.278317372 | 1.88E-05 | 0.000910949 | Col1a2 | collagen, type I, alpha 2 [Source:MGI Sym | 405.57 | 251.84 | 138.86 | 127.25 | 102.52 | 90.221 |
| 7446 | ENSMUSG00000029663 | 8.783527137 | -0.233445588 | 0.839520634 | -0.278070101 | 0.780958547 | 0.959993512 | Gngt1 | guanine nucleotide binding protein (G prc | 20.078 | 4.1117 | 4.025 | 5.7839 | 10.906 | 7.7969 |
| 7447 | ENSMUSG00000029664 | 804.8096287 | 0.114876938 | 0.156168469 | 0.735596236 | 0.461976454 | 0.860650866 | Tfpi2 | tissue factor pathway inhibitor 2 [Source:I | 949.27 | 652.73 | 714.43 | 748.05 | 834.32 | 930.06 |
| 7448 | ENSMUSG00000029669 | 3447.865194 | 0.165246144 | 0.087327155 | 1.892265282 | 0.058455644 | 0.343085466 | Tspan12 | tetraspanin 12 [Source:MGI Symbol;Acc:N | 3281.5 | 3345.9 | 3124.4 | 3633.2 | 3720.1 | 3582.1 |
| 7449 | ENSMUSG00000029670 | 106.9448459 | -0.129355482 | 0.234052049 | -0.552678273 | 0.580483733 | 0.908645447 | Ing3 | inhibitor of growth family, member 3 [Sou | 114.84 | 102.79 | 117.73 | 113.75 | 111.24 | 81.311 |
| 7450 | ENSMUSG00000029671 | 0 | NA | NA | NA | NA | NA | Wnt16 | wingless-type MMTV integration site fami | 0 | 0 | 0 | 0 | 0 | 0 |
| 7451 | ENSMUSG00000029672 | 520.4626177 | 0.174334686 | 0.130428285 | 1.336632509 | 0.181342622 | 0.614703465 | Fam3c | family with sequence similarity 3, membe | 485.88 | 514.99 | 466.9 | 604.41 | 511.5 | 539.1 |
| 7452 | ENSMUSG00000029673 | 3.859631387 | -1.456127778 | 1.138027078 | -1.279519448 | 0.200714196 | NA | Auts2 | autism susceptibility candidate 2 [Source: | 8.8341 | 4.1117 | 4.025 | 2.8919 | 2.1812 | 1.1138 |
| 7453 | ENSMUSG00000029674 | 39.36144249 | -1.092647595 | 0.412333796 | -2.64991035 | 0.008051313 | 0.092428861 | Limk1 | LIM-domain containing, protein kinase [Sc | 68.264 | 52.424 | 40.25 | 35.667 | 25.084 | 14.48 |
| 7454 | ENSMUSG00000029675 | 26.46969304 | -0.486143356 | 0.471797369 | -1.030407095 | 0.302818944 | 0.752484788 | Eln | elastin [Source:MGI Symbol;Acc:MGI:953: | 37.746 | 30.838 | 24.15 | 31.811 | 11.997 | 22.277 |
| 7455 | ENSMUSG00000029676 | 89.7264095 | -0.089473218 | 0.240911449 | -0.371394629 | 0.710343625 | 0.944239215 | Pot1a | protection of telomeres 1A [Source:MGI S | 91.554 | 87.373 | 98.612 | 88.686 | 76.343 | 95.791 |
| 7456 | ENSMUSG00000029678 | 0.799645015 | 0.918084596 | 2.625685367 | 0.34965522 | 0.726597464 | NA | Hyal5 | hyaluronoglucosaminidase 5 [Source:MGI | 1.6062 | 0 | 0 | 0.964 | 0 | 2.2277 |
| 7457 | ENSMUSG00000029679 | 0.349477146 | 0.165142768 | 3.911021834 | 0.042224967 | 0.966319359 | NA | Hyal6 | hyaluronoglucosaminidase 6 [Source:MGI | 0 | 0 | 1.0062 | 0 | 1.0906 | 0 |
| 7458 | ENSMUSG00000029680 | 0.167707431 | -0.796629176 | 4.080472857 | -0.195229623 | 0.84521319 | NA | Hyal4 | hyaluronoglucosaminidase 4 [Source:MGI | 0 | 0 | 1.0062 | 0 | 0 | 0 |
| 7459 | ENSMUSG00000029681 | 226.9850086 | -0.007676053 | 0.16270869 | -0.047176665 | 0.962372426 | 0.992956886 | Bcl7b | B cell CLL/lymphoma 7B [Source:MGI Sym | 240.93 | 228.2 | 213.32 | 225.57 | 235.57 | 218.31 |
| 7460 | ENSMUSG00000029682 | 0 | NA | NA | NA | NA | NA | Spam1 | sperm adhesion molecule 1 [Source:MGI ! | 0 | 0 | 0 | 0 | 0 | 0 |
| 7461 | ENSMUSG00000029683 | 40.61564384 | -1.047104511 | 0.56975192 | -1.837825329 | 0.066088148 | 0.369154172 | Lmod2 | leiomodin 2 (cardiac) [Source:MGI Symbo | 54.611 | 34.949 | 74.462 | 9.6398 | 58.893 | 11.138 |
| 7462 | ENSMUSG00000029684 | 1376.551625 | -0.022407732 | 0.101646448 | -0.220447758 | 0.825522455 | 0.96943211 | Wasl | WASP like actin nucleation promoting fact | 1427.9 | 1331.2 | 1401.7 | 1289.8 | 1429.8 | 1378.9 |
| 7463 | ENSMUSG00000029685 | 0.845761573 | -3.112447952 | 2.7647732 | -1.125751636 | 0.260270661 | NA | Asb15 | ankyrin repeat and SOCS box-containing 1 | 0 | 2.0558 | 3.0187 | 0 | 0 | 0 |
| 7464 | ENSMUSG00000029686 | 1114.700613 | 0.016666483 | 0.106737966 | 0.156143908 | 0.875919589 | 0.980632845 | Cul1 | cullin 1 [Source:MGI Symbol;Acc:MGI:134 | 1028 | 1175.9 | 1122 | 1128.8 | 1138.6 | 1094.9 |
| 7465 | ENSMUSG00000029687 | 63.61317957 | 0.336787643 | 0.296104008 | 1.137396437 | 0.255372596 | 0.708829251 | Ezh2 | enhancer of zeste 2 polycomb repressive ( | 57.823 | 61.675 | 49.306 | 84.83 | 54.531 | 73.514 |
| 7466 | ENSMUSG00000029695 | 6286.703762 | -0.054180144 | 0.114828276 | -0.47183626 | 0.637043667 | 0.923478764 | Aass | aminoadipate-semialdehyde synthase [So | 5637.8 | 6246.7 | 7330.5 | 5759.8 | 6232.9 | 6512.6 |
| 7467 | ENSMUSG00000029697 | 0 | NA | NA | NA | NA | NA | Fezf1 | Fez family zinc finger 1 [Source:MGI Symb | 0 | 0 | 0 | 0 | 0 | 0 |
| 7468 | ENSMUSG00000029699 | 68.22513991 | 0.303366753 | 0.278217227 | 1.090395286 | 0.275539057 | 0.730038567 | Ssc4d | scavenger receptor cysteine rich family, 4 | 64.248 | 52.424 | 66.412 | 76.154 | 64.346 | 85.766 |
| 7469 | ENSMUSG00000029700 | 2.878661384 | 0.405302053 | 1.422632953 | 0.28489573 | 0.775724028 | NA | Slc13a1 | solute carrier family 13 (sodium/sulfate sy | 2.4093 | 3.0838 | 2.0125 | 8.6758 | 1.0906 | 0 |
| 7470 | ENSMUSG00000029701 | 444.00064 | -0.111052185 | 0.146276177 | -0.759195294 | 0.447735741 | 0.853796936 | Rbm28 | RNA binding motif protein 28 [Source:MG | 420.02 | 501.62 | 462.87 | 484.88 | 398.08 | 396.53 |
| 7471 | ENSMUSG00000029703 | 119.4754473 | 0.050180634 | 0.22811743 | 0.219977202 | 0.82588891 | 0.96943211 | Lrwd1 | leucine-rich repeats and WD repeat doma | 96.372 | 138.77 | 117.73 | 127.25 | 128.69 | 108.04 |
| 7472 | ENSMUSG00000029705 | 493.8344272 | 0.01077439 | 0.130114904 | 0.08280673 | 0.934005218 | 0.988420305 | Cux1 | cut-like homeobox 1 [Source:MGI Symbol | 474.63 | 475.93 | 525.26 | 457.89 | 518.04 | 511.25 |
| 7473 | ENSMUSG00000029706 | 0 | NA | NA | NA | NA | NA | Pax4 | paired box 4 [Source:MGI Symbol;Acc:MG | 0 | 0 | 0 | 0 | 0 | 0 |
| 7474 | ENSMUSG00000029707 | 0 | NA | NA | NA | NA | NA | Fscn3 | fascin actin-bundling protein 3 [Source:M | 0 | 0 | 0 | 0 | 0 | 0 |
| 7475 | ENSMUSG00000029708 | 292.4299179 | 0.146668118 | 0.180159375 | 0.814102057 | 0.415586488 | 0.83614452 | Gcc1 | golgi coiled coil 1 [Source:MGI Symbol;Acc | 235.31 | 282.68 | 314.95 | 251.6 | 337 | 333.04 |
| 7476 | ENSMUSG00000029710 | 356.8352591 | 0.025515883 | 0.176210728 | 0.144803233 | 0.88486623 | 0.981795936 | Ephb4 | Eph receptor B4 [Source:MGI Symbol;Acc | 291.53 | 360.8 | 409.54 | 368.24 | 301.01 | 409.89 |
| 7477 | ENSMUSG00000029711 | 0 | NA | NA | NA | NA | NA | Epo | erythropoietin [Source:MGI Symbol;Acc:N | 0 | 0 | 0 | 0 | 0 | 0 |
| 7478 | ENSMUSG00000029712 | 1.66287449 | 0.783544442 | 1.575943233 | 0.497190778 | 0.619054524 | NA | Actl6b | actin-like 6B [Source:MGI Symbol;Acc:MG | 1.6062 | 1.0279 | 1.0062 | 1.928 | 2.1812 | 2.2277 |
| 7479 | ENSMUSG00000029713 | 1754.647514 | 0.06352195 | 0.09637314 | 0.659125042 | 0.509815476 | 0.882793133 | Gnb2 | guanine nucleotide binding protein (G prc | 1705 | 1715.6 | 1727.7 | 1799.7 | 1885.7 | 1694.2 |
| 7480 | ENSMUSG00000029714 | 547.8060423 | 0.064745613 | 0.13281545 | 0.487485553 | 0.625914279 | 0.919638611 | Gigyf1 | GRB10 interacting GYF protein 1 [Source:I | 527.64 | 521.15 | 558.47 | 617.91 | 567.12 | 494.55 |
| 7481 | ENSMUSG00000029715 | 88.18081288 | 0.233319776 | 0.263710031 | 0.884758822 | 0.376286723 | 0.809731241 | Pop7 | processing of precursor 7, ribonuclease P | 95.569 | 64.759 | 82.512 | 104.11 | 77.434 | 104.7 |
| 7482 | ENSMUSG00000029716 | 7927.740636 | 0.173933964 | 0.097030908 | 1.792562468 | 0.073042912 | 0.392602261 | Tfr2 | transferrin receptor 2 [Source:MGI Symbc | 6934 | 7562.4 | 7856.8 | 8996.8 | 8366.1 | 7850.4 |
| 7483 | ENSMUSG00000029718 | 99.78249715 | -0.537889633 | 0.293715132 | -1.831331021 | 0.06705115 | 0.372627026 | Pcolce | procollagen C-endopeptidase enhancer pı | 171.86 | 89.429 | 92.575 | 84.83 | 70.89 | 89.108 |
| 7484 | ENSMUSG00000029720 | 7.124070947 | -0.390836062 | 0.887016841 | -0.440618536 | 0.659489181 | 0.927787797 | Gm20605 | predicted gene 20605 [Source:MGI Symbc | 12.047 | 5.1396 | 7.0437 | 8.6758 | 8.7249 | 1.1138 |
| 7485 | ENSMUSG00000029722 | 636.3042334 | 0.279732656 | 0.144428453 | 1.936825122 | 0.052766724 | 0.32296796 | Agfg2 | ArfGAP with FG repeats 2 [Source:MGI Sy | 497.92 | 587.97 | 639.97 | 775.04 | 610.75 | 706.18 |
| 7486 | ENSMUSG00000029723 | 198.1661696 | 0.123375896 | 0.178480303 | 0.69125777 | 0.489403565 | 0.874889775 | Tsc22d4 | TSC22 domain family, member 4 [Source:I | 195.96 | 175.77 | 197.22 | 214 | 181.04 | 225 |
| 7487 | ENSMUSG00000029725 | 252.7779561 | 0.126429979 | 0.160086125 | 0.789762255 | 0.429666626 | 0.84307112 | Ppp1r35 | protein phosphatase 1, regulatory subunit | 247.36 | 215.86 | 261.62 | 258.35 | 263.93 | 269.55 |
| 7488 | ENSMUSG00000029726 | 499.4694854 | 0.106412315 | 0.124000945 | 0.858157291 | 0.390805616 | 0.820195366 | Mepce | methylphosphate capping enzyme [Source | 480.26 | 485.18 | 477.97 | 531.15 | 534.4 | 487.86 |
| 7489 | ENSMUSG00000029727 | 3370.434913 | -0.226605383 | 0.140617095 | -1.611506642 | 0.107069341 | 0.483009289 | Cyp3a13 | cytochrome P450, family 3, subfamily a, p | 2993.2 | 3698.4 | 4214.2 | 3567.7 | 3004.7 | 2744.5 |
| 7490 | ENSMUSG00000029729 | 779.5226051 | -0.181177431 | 0.145010158 | -1.249411995 | 0.211514423 | 0.6563061 | Zkscan1 | zinc finger with KRAB and SCAN domains : | 728.41 | 859.34 | 897.57 | 616.95 | 841.96 | 732.91 |
| 7491 | ENSMUSG00000029730 | 221.5156046 | -0.086863033 | 0.181836928 | -0.477697433 | 0.632865572 | 0.922107984 | Mcm7 | minichromosome maintenance complex c | 207.2 | 250.81 | 227.41 | 242.92 | 220.3 | 180.44 |
| 7492 | ENSMUSG00000029735 | 428.4784257 | -0.156831523 | 0.144787501 | -1.083184125 | 0.278726704 | 0.731404763 | Tpk1 | thiamine pyrophosphokinase [Source:MG | 490.7 | 437.89 | 425.64 | 365.35 | 455.88 | 395.41 |
| 7493 | ENSMUSG00000029736 | 0.160662974 | 1.126924175 | 4.080472857 | 0.276174898 | 0.782413736 | NA | Nobox | NOBOX oogenesis homeobox [Source:MG | 0 | 0 | 0 | 0.964 | 0 | 0 |
| 7494 | ENSMUSG00000029752 | 7.725551447 | 2.010608348 | 0.903148745 | 2.226220606 | 0.025999408 | 0.20857207 | Asns | asparagine synthetase [Source:MGI Symb | 4.0155 | 4.1117 | 1.0062 | 10.604 | 5.4531 | 21.163 |
| 7495 | ENSMUSG00000029754 | 0 | NA | NA | NA | NA | NA | Dlx6 | distal-less homeobox 6 [Source:MGI Symb | 0 | 0 | 0 | 0 | 0 | 0 |
| 7496 | ENSMUSG00000029755 | 0 | NA | NA | NA | NA | NA | Dlx5 | distal-less homeobox 5 [Source:MGI Symb | 0 | 0 | 0 | 0 | 0 | 0 |
| 7497 | ENSMUSG00000029757 | 0.583320834 | -1.123770786 | 3.233988906 | -0.34748752 | 0.728225092 | NA | Dync1i1 | dynein cytoplasmic 1 intermediate chain 1 | 2.4093 | 0 | 0 | 0 | 1.0906 | 0 |
| 7498 | ENSMUSG00000029759 | 1183.041026 | -0.062427586 | 0.109528197 | -0.569968171 | 0.568699287 | 0.903527882 | Pon3 | paraoxonase 3 [Source:MGI Symbol;Acc:N | 1109.9 | 1200.6 | 1316.2 | 1130.7 | 1172.4 | 1168.4 |
| 7499 | ENSMUSG00000029761 | 3215.38768 | 0.375465296 | 0.124413155 | 3.017890637 | 0.002545407 | 0.041884839 | Cald1 | caldesmon 1 [Source:MGI Symbol;Acc:MG | 3208.4 | 2631.5 | 2555.9 | 3198.5 | 3697.2 | 4000.9 |
| 7500 | ENSMUSG00000029762 | 26.08853358 | -0.321321175 | 0.461120261 | -0.696827275 | 0.485910895 | 0.872998689 | Akr1b8 | aldo-keto reductase family 1, member B8 | 20.881 | 30.838 | 35.219 | 14.46 | 26.175 | 28.96 |

| | | | | | | | | | | | | | | | |
|---|---|---|---|---|---|---|---|---|---|---|---|---|---|---|---|
| 7501 | ENSMUSG00000029763 | 365.2103175 | 0.070659384 | 0.195109995 | 0.362151534 | 0.717238795 | 0.9447719 | Exoc4 | exocyst complex component 4 [Source:M( | 450.54 | 335.1 | 281.75 | 312.33 | 439.52 | 372.02 |
| 7502 | ENSMUSG00000029765 | 43.00200093 | -0.687384405 | 0.403015999 | -1.705600788 | 0.088082432 | 0.437658641 | Plxna4 | plexin A4 [Source:MGI Symbol;Acc:MGI:2 | 68.264 | 62.703 | 28.175 | 40.487 | 30.537 | 27.846 |
| 7503 | ENSMUSG00000029766 | 0.160662974 | 1.126924175 | 4.080472857 | 0.276174898 | 0.782413736 | NA | 1700012A03 | RIKEN cDNA 1700012A03 gene [Source:M | 0 | 0 | 0 | 0.964 | 0 | 0 |
| 7504 | ENSMUSG00000029767 | 1552.861936 | 0.067758104 | 0.109846372 | 0.616844262 | 0.537337454 | 0.890758111 | Calu | calumenin [Source:MGI Symbol;Acc:MGI: | 1502.6 | 1534.7 | 1511.4 | 1475.9 | 1510.5 | 1782.2 |
| 7505 | ENSMUSG00000029769 | 1.816546267 | -1.992111541 | 1.871326098 | -1.064545374 | 0.287081711 | NA | Ccdc136 | coiled-coil domain containing 136 [Source | 1.6062 | 1.0279 | 6.0375 | 0 | 0 | 2.2277 |
| 7506 | ENSMUSG00000029771 | 244.1216453 | 1.014798781 | 0.222403139 | 4.562879765 | 5.05E-06 | 0.000307935 | Irf5 | interferon regulatory factor 5 [Source:MG | 128.5 | 130.55 | 226.41 | 289.19 | 337 | 353.09 |
| 7507 | ENSMUSG00000029772 | 349.5435206 | -0.103687905 | 0.15413221 | -0.67272055 | 0.501125091 | 0.880974509 | Ahcyl2 | S-adenosylhomocysteine hydrolase-like 2 | 386.29 | 384.44 | 314.95 | 313.29 | 370.81 | 327.47 |
| 7508 | ENSMUSG00000029775 | 1128.600309 | 0.010925844 | 0.101446015 | 0.107701065 | 0.914232825 | 0.98625062 | Klhdc10 | kelch domain containing 10 [Source:MGI S | 1086.6 | 1171.8 | 1114.9 | 1111.5 | 1147.3 | 1139.5 |
| 7509 | ENSMUSG00000029776 | 7170.684211 | -0.024118897 | 0.096396324 | -0.250205571 | 0.802428377 | 0.963260357 | Hibadh | 3-hydroxyisobutyrate dehydrogenase [Sou | 7668.8 | 6815.1 | 7206.7 | 6886.7 | 7676.9 | 6769.9 |
| 7510 | ENSMUSG00000029777 | 693.7918692 | 0.065808173 | 0.123687038 | 0.532053916 | 0.594688654 | 0.913409341 | Gars | glycyl-tRNA synthetase [Source:MGI Symb | 630.44 | 711.32 | 692.3 | 643.94 | 731.8 | 752.96 |
| 7511 | ENSMUSG00000029778 | 15.65376179 | -0.800322188 | 0.57265627 | -1.397561208 | 0.162244874 | 0.585529252 | Adcyap1r1 | adenylate cyclase activating polypeptide 1 | 16.865 | 20.558 | 22.137 | 6.7478 | 10.906 | 16.708 |
| 7512 | ENSMUSG00000029780 | 382.0039283 | 0.328442803 | 0.150451697 | 2.183044852 | 0.029032507 | 0.223715404 | Nt5c3 | 5'-nucleotidase, cytosolic III [Source:MGI S | 361.4 | 329.96 | 324.01 | 404.87 | 382.81 | 488.98 |
| 7513 | ENSMUSG00000029781 | 528.7982185 | -0.0694879 | 0.132141904 | -0.525858169 | 0.598986755 | 0.915055389 | Fkbp9 | FK506 binding protein 9 [Source:MGI Sym | 510.77 | 606.47 | 508.15 | 536.94 | 513.68 | 496.77 |
| 7514 | ENSMUSG00000029782 | 161.7298647 | 0.187466547 | 0.189333507 | 0.990139301 | 0.322106036 | 0.769593967 | Tmem209 | transmembrane protein 209 [Source:MGI | 147.77 | 141.85 | 164.02 | 160.98 | 188.68 | 167.08 |
| 7515 | ENSMUSG00000029784 | 1.648373522 | -0.40726801 | 1.719057758 | -0.236913511 | 0.812723888 | NA | Ssmem1 | serine-rich single-pass membrane protein | 1.6062 | 0 | 4.025 | 0.964 | 2.1812 | 1.1138 |
| 7516 | ENSMUSG00000029787 | 362.5089613 | 0.172195584 | 0.145424622 | 1.184088236 | 0.236378136 | 0.685294058 | Avl9 | AVL9 cell migration associated [Source:M( | 319.63 | 360.8 | 343.13 | 419.33 | 349 | 383.16 |
| 7517 | ENSMUSG00000029788 | 0 | NA | NA | NA | NA | NA | Cpa5 | carboxypeptidase A5 [Source:MGI Symbol | 0 | 0 | 0 | 0 | 0 | 0 |
| 7518 | ENSMUSG00000029790 | 52.43192576 | 0.118400352 | 0.364030319 | 0.3252486 | 0.744992929 | 0.949997329 | Cep41 | centrosomal protein 41 [Source:MGI Syml | 48.989 | 45.228 | 56.35 | 34.703 | 46.897 | 82.424 |
| 7519 | ENSMUSG00000029797 | 33.18990543 | 0.445189236 | 0.470962745 | 0.945274845 | 0.344518577 | 0.785874854 | Sspo | SCO-spondin [Source:MGI Symbol;Acc:M( | 36.14 | 22.614 | 25.156 | 17.352 | 58.893 | 38.985 |
| 7520 | ENSMUSG00000029798 | 196.2052307 | -0.023363862 | 0.19063642 | -0.122557179 | 0.902457765 | 0.983821605 | Herc6 | hect domain and RLD 6 [Source:MGI Symb | 222.46 | 174.75 | 195.21 | 165.8 | 199.58 | 219.43 |
| 7521 | ENSMUSG00000029802 | 1771.4425 | -0.089050569 | 0.134843409 | -0.660399862 | 0.50899726 | 0.882793133 | Abcg2 | ATP binding cassette subfamily G member | 1860.8 | 1690.9 | 1926 | 1538.5 | 2078.7 | 1533.8 |
| 7522 | ENSMUSG00000029804 | 105.7470256 | -0.044508069 | 0.250355407 | -0.177779539 | 0.858896117 | 0.975439379 | Herc3 | hect domain and RLD 3 [Source:MGI Symb | 98.782 | 124.38 | 99.618 | 131.1 | 97.065 | 83.538 |
| 7523 | ENSMUSG00000029810 | 2483.561931 | -0.264544327 | 0.118869525 | -2.225501666 | 0.026047579 | 0.208842677 | Tmem176b | transmembrane protein 176B [Source:MG | 2918.5 | 2941.9 | 2271.1 | 2315.5 | 2241.2 | 2213.2 |
| 7524 | ENSMUSG00000029811 | 0.167707431 | -0.796629176 | 4.080472857 | -0.195229623 | 0.84521319 | NA | Aoc1 | amine oxidase, copper-containing 1 [Sour | 0 | 0 | 1.0062 | 0 | 0 | 0 |
| 7525 | ENSMUSG00000029813 | 0 | NA | NA | NA | NA | NA | Doxl1 | diamine oxidase-like protein 1 [Source:M( | 0 | 0 | 0 | 0 | 0 | 0 |
| 7526 | ENSMUSG00000029814 | 3.70413185 | -0.751673017 | 1.10685422 | -0.679107513 | 0.497069743 | NA | Igf2bp3 | insulin-like growth factor 2 mRNA binding | 4.8186 | 5.1396 | 4.025 | 3.8559 | 3.2719 | 1.1138 |
| 7527 | ENSMUSG00000029815 | 351.187394 | 0.056285484 | 0.147347325 | 0.381991895 | 0.702467377 | 0.942405009 | Malsu1 | mitochondrial assembly of ribosomal large | 384.69 | 307.35 | 340.11 | 358.6 | 357.72 | 358.66 |
| 7528 | ENSMUSG00000029816 | 7.078834011 | -2.246081355 | 0.893390351 | -2.514109708 | 0.011933331 | 0.122154064 | Gpnmb | glycoprotein (transmembrane) nmb [Sour | 13.653 | 11.307 | 10.062 | 1.928 | 2.1812 | 3.3415 |
| 7529 | ENSMUSG00000029817 | 326.2029294 | -0.158321122 | 0.14454687 | -1.095292634 | 0.273388454 | 0.728364449 | Tra2a | transformer 2 alpha [Source:MGI Symbol; | 330.88 | 346.41 | 355.2 | 310.4 | 285.74 | 328.58 |
| 7530 | ENSMUSG00000029819 | 0 | NA | NA | NA | NA | NA | Npy | neuropeptide Y [Source:MGI Symbol;Acc: | 0 | 0 | 0 | 0 | 0 | 0 |
| 7531 | ENSMUSG00000029821 | 189.9049308 | -0.142441479 | 0.201542782 | -0.706755545 | 0.479718405 | 0.870278888 | Gsdme | gasdermin E [Source:MGI Symbol;Acc:MG | 206.4 | 235.39 | 155.97 | 183.16 | 163.59 | 194.92 |
| 7532 | ENSMUSG00000029822 | 332.1094492 | -0.627016229 | 0.725143211 | -0.864679169 | 0.387214906 | 0.817618864 | Osbpl3 | oxysterol binding protein-like 3 [Source:M | 517.2 | 476.95 | 215.34 | 294.01 | 410.07 | 79.083 |
| 7533 | ENSMUSG00000029823 | 2068.464742 | 0.082324837 | 0.120702479 | 0.682047607 | 0.495208848 | 0.877919732 | Luc7l2 | LUC7-like 2 (S. cerevisiae) [Source:MGI Sy | 1998.9 | 2128.8 | 1901.8 | 2456.2 | 2047.1 | 1877.9 |
| 7534 | ENSMUSG00000029826 | 870.4071048 | 0.11014462 | 0.117868471 | 0.934470598 | 0.350061196 | 0.791509322 | Zc3hav1 | zinc finger CCCH type, antiviral 1 [Source:I | 847.27 | 822.33 | 841.22 | 813.6 | 992.46 | 905.56 |
| 7535 | ENSMUSG00000029828 | 11.58295435 | 0.663696279 | 0.68162291 | 0.973700076 | 0.33020548 | 0.774747658 | 4921507P07 | RIKEN cDNA 4921507P07 gene [Source:M | 4.8186 | 8.2233 | 14.087 | 19.28 | 14.178 | 8.9108 |
| 7536 | ENSMUSG00000029829 | 0 | NA | NA | NA | NA | NA | Tmem213 | transmembrane protein 213 [Source:MGI | 0 | 0 | 0 | 0 | 0 | 0 |
| 7537 | ENSMUSG00000029830 | 0 | NA | NA | NA | NA | NA | Svopl | SV2 related protein homolog (rat)-like [So | 0 | 0 | 0 | 0 | 0 | 0 |
| 7538 | ENSMUSG00000029831 | 0 | NA | NA | NA | NA | NA | Npvf | neuropeptide VF precursor [Source:MGI S | 0 | 0 | 0 | 0 | 0 | 0 |
| 7539 | ENSMUSG00000029832 | 4.757131629 | -0.344474205 | 0.952393456 | -0.361693167 | 0.717581334 | NA | Nfe2l3 | nuclear factor, erythroid derived 2, like 3 [ | 4.8186 | 5.1396 | 6.0375 | 4.8199 | 3.2719 | 4.4554 |
| 7540 | ENSMUSG00000029833 | 534.9746957 | -0.218734666 | 0.238221088 | -0.918200262 | 0.358514032 | 0.798191951 | Trim24 | tripartite motif-containing 24 [Source:MG | 804.71 | 504.71 | 416.59 | 664.18 | 371.9 | 447.77 |
| 7541 | ENSMUSG00000029836 | 420.5963114 | 0.099110577 | 0.137301608 | 0.721845709 | 0.470389343 | 0.867110762 | Cbx3 | chromobox 3 [Source:MGI Symbol;Acc:M( | 417.61 | 386.5 | 414.57 | 474.28 | 432.98 | 397.64 |
| 7542 | ENSMUSG00000029837 | 0 | NA | NA | NA | NA | NA | 1700111E14 | RIKEN cDNA 1700111E14 gene [Source:M( | 0 | 0 | 0 | 0 | 0 | 0 |
| 7543 | ENSMUSG00000029838 | 0.535401493 | -2.48596965 | 3.684117979 | -0.674780141 | 0.499815459 | NA | Ptn | pleiotrophin [Source:MGI Symbol;Acc:MG | 3.2124 | 0 | 0 | 0 | 0 | 0 |
| 7544 | ENSMUSG00000029840 | 2182.370957 | 0.00229479 | 0.100243551 | 0.022892147 | 0.981736304 | 0.998368917 | Mtpn | myotrophin [Source:MGI Symbol;Acc:MGI | 1998.1 | 2279.9 | 2265.1 | 2157.4 | 2266.3 | 2127.4 |
| 7545 | ENSMUSG00000029843 | 7.897047858 | 0.385816777 | 0.801135261 | 0.481587561 | 0.630098964 | 0.920766965 | Slc13a4 | solute carrier family 13 (sodium/sulfate sy | 7.2279 | 7.1954 | 6.0375 | 3.8559 | 15.269 | 7.7969 |
| 7546 | ENSMUSG00000029844 | 1.363694037 | 0.592025543 | 2.168456282 | 0.273017053 | 0.784840107 | NA | Hoxa1 | homeobox A1 [Source:MGI Symbol;Acc:M | 3.2124 | 0 | 0 | 3.8559 | 0 | 1.1138 |
| 7547 | ENSMUSG00000029847 | 0.167707431 | -0.796629176 | 4.080472857 | -0.195229623 | 0.84521319 | NA | Slc23a4 | solute carrier family 23 member 4 [Source | 0 | 0 | 1.0062 | 0 | 0 | 0 |
| 7548 | ENSMUSG00000029848 | 0.17131964 | -0.796629176 | 4.080472857 | -0.195229623 | 0.84521319 | NA | Stra8 | stimulated by retinoic acid gene 8 [Source | 0 | 1.0279 | 0 | 0 | 0 | 0 |
| 7549 | ENSMUSG00000029851 | 8.861470489 | -0.393324971 | 0.719448631 | -0.546703342 | 0.584582558 | 0.908955096 | Tcaf2 | TRPM8 channel-associated factor 2 [Sourc | 12.85 | 8.2233 | 9.0562 | 8.6758 | 5.4531 | 8.9108 |
| 7550 | ENSMUSG00000029859 | 794.939844 | 0.502973325 | 0.113504296 | 4.431315311 | 9.37E-06 | 0.00050174 | Epha1 | Eph receptor A1 [Source:MGI Symbol;Acc | 705.12 | 624.97 | 641.98 | 907.1 | 929.21 | 961.25 |
| 7551 | ENSMUSG00000029860 | 674.7855721 | 0.17849558 | 0.128649306 | 1.387458556 | 0.16530198 | 0.590790176 | Zyx | zyxin [Source:MGI Symbol;Acc:MGI:10307 | 656.13 | 576.66 | 666.13 | 735.52 | 642.37 | 771.89 |
| 7552 | ENSMUSG00000029861 | 0 | NA | NA | NA | NA | NA | Fam131b | family with sequence similarity 131, meml | 0 | 0 | 0 | 0 | 0 | 0 |
| 7553 | ENSMUSG00000029862 | 0.36353943 | 2.065989254 | 4.034137301 | 0.512126658 | 0.608562368 | NA | Clcn1 | chloride channel, voltage-sensitive 1 [Sou | 0 | 0 | 0 | 0 | 2.1812 | 0 |
| 7554 | ENSMUSG00000029863 | 78.74566768 | -0.124740025 | 0.269673679 | -0.462559141 | 0.643680402 | 0.925505685 | Casp2 | caspase 2 [Source:MGI Symbol;Acc:MGI:9 | 69.067 | 78.122 | 99.618 | 80.01 | 64.346 | 81.311 |
| 7555 | ENSMUSG00000029864 | 2813.462916 | 0.039378957 | 0.093677892 | 0.420365535 | 0.674218441 | 0.932114677 | Gstk1 | glutathione S-transferase kappa 1 [Source | 2952.2 | 2662.3 | 2709.8 | 2925.7 | 2758.2 | 2872.6 |
| 7556 | ENSMUSG00000029865 | 0 | NA | NA | NA | NA | NA | Sval1 | seminal vesicle antigen-like 1 [Source:MG | 0 | 0 | 0 | 0 | 0 | 0 |

| | | | | | | | | | | | | | | | |
|---|---|---|---|---|---|---|---|---|---|---|---|---|---|---|---|
| 7557 | ENSMUSG00000029866 | 0 | NA | NA | NA | NA | NA | Kel | Kell blood group [Source:MGI Symbol;Acc | 0 | 0 | 0 | 0 | 0 | 0 |
| 7558 | ENSMUSG00000029867 | 0 | NA | NA | NA | NA | NA | Llcfc1 | LLLL and CFNLAS motif containing 1 [Sourc | 0 | 0 | 0 | 0 | 0 | 0 |
| 7559 | ENSMUSG00000029868 | 0.181769715 | 1.126924175 | 4.080472857 | 0.276174898 | 0.782413736 | NA | Trpv6 | transient receptor potential cation channe | 0 | 0 | 0 | 0 | 1.0906 | 0 |
| 7560 | ENSMUSG00000029869 | 17.723492 | -0.698570317 | 0.535141188 | -1.305394413 | 0.19175858 | 0.629099068 | Ephb6 | Eph receptor B6 [Source:MGI Symbol;Acc | 23.29 | 15.419 | 27.169 | 17.352 | 13.087 | 10.025 |
| 7561 | ENSMUSG00000029875 | 0 | NA | NA | NA | NA | NA | Ccdc184 | coiled-coil domain containing 184 [Source | 0 | 0 | 0 | 0 | 0 | 0 |
| 7562 | ENSMUSG00000029878 | 0 | NA | NA | NA | NA | NA | Dbpht2 | DNA binding protein with his-thr domain [ | 0 | 0 | 0 | 0 | 0 | 0 |
| 7563 | ENSMUSG00000029882 | 0 | NA | NA | NA | NA | NA | 2210010C04 | RIKEN cDNA 2210010C04 gene [Source:M | 0 | 0 | 0 | 0 | 0 | 0 |
| 7564 | ENSMUSG00000029883 | 0 | NA | NA | NA | NA | NA | 1700074P13 | RIKEN cDNA 1700074P13 gene [Source:M | 0 | 0 | 0 | 0 | 0 | 0 |
| 7565 | ENSMUSG00000029885 | 0 | NA | NA | NA | NA | NA | Moxd2 | monooxygenase, DBH-like 2 [Source:MGI | 0 | 0 | 0 | 0 | 0 | 0 |
| 7566 | ENSMUSG00000029909 | 0 | NA | NA | NA | NA | NA | Prss37 | protease, serine 37 [Source:MGI Symbol;A | 0 | 0 | 0 | 0 | 0 | 0 |
| 7567 | ENSMUSG00000029910 | 69.20721596 | -0.213896288 | 0.310140964 | -0.689674415 | 0.490398959 | 0.875587032 | Mad2l1 | MAD2 mitotic arrest deficient-like 1 [Sourc | 100.39 | 60.647 | 61.381 | 63.623 | 52.35 | 76.855 |
| 7568 | ENSMUSG00000029911 | 415.9876308 | 0.222816354 | 0.132123678 | 1.686422579 | 0.091714435 | 0.446623824 | Ssbp1 | single-stranded DNA binding protein 1 [Sc | 381.47 | 403.97 | 366.27 | 431.86 | 465.69 | 446.65 |
| 7569 | ENSMUSG00000029913 | 1.431356019 | -0.928509842 | 1.937491009 | -0.47923311 | 0.631772802 | NA | Prdm5 | PR domain containing 5 [Source:MGI Sym | 1.6062 | 3.0838 | 1.0062 | 2.8919 | 0 | 0 |
| 7570 | ENSMUSG00000029915 | 1.676063492 | 1.348842178 | 1.846556211 | 0.730463644 | 0.465106828 | NA | Clec5a | C-type lectin domain family 5, member a [ | 0.8031 | 2.0558 | 0 | 3.8559 | 0 | 3.3415 |
| 7571 | ENSMUSG00000029916 | 114.7982471 | -0.304882972 | 0.216502833 | -1.408217012 | 0.15906682 | 0.580285052 | Agk | acylglycerol kinase [Source:MGI Symbol;A | 117.25 | 136.71 | 126.79 | 95.434 | 111.24 | 101.36 |
| 7572 | ENSMUSG00000029917 | 0 | NA | NA | NA | NA | NA | Qrfprl | pyroglutamylated RFamide peptide recept | 0 | 0 | 0 | 0 | 0 | 0 |
| 7573 | ENSMUSG00000029918 | 365.4521624 | 0.187710181 | 0.169057518 | 1.110333234 | 0.266855457 | 0.722778036 | Mrps33 | mitochondrial ribosomal protein S33 [Sou | 340.52 | 371.08 | 312.94 | 307.51 | 434.07 | 426.6 |
| 7574 | ENSMUSG00000029919 | 13.56666797 | -1.283609447 | 0.736039533 | -1.7439409 | 0.081169396 | 0.418998622 | Hpgds | hematopoietic prostaglandin D synthase [ | 21.684 | 29.81 | 6.0375 | 2.8919 | 8.7249 | 12.252 |
| 7575 | ENSMUSG00000029920 | 274.5866798 | 0.089892825 | 0.164928934 | 0.54503975 | 0.585726179 | 0.90955446 | Smarcad1 | SWI/SNF-related, matrix-associated actin- | 241.73 | 300.15 | 256.59 | 263.17 | 314.1 | 271.78 |
| 7576 | ENSMUSG00000029922 | 482.2179402 | 0.009245598 | 0.145857411 | 0.063387919 | 0.949457607 | 0.990731019 | Mkrn1 | makorin, ring finger protein, 1 [Source:MG | 515.59 | 450.23 | 474.95 | 409.69 | 509.32 | 533.53 |
| 7577 | ENSMUSG00000029923 | 4.457351105 | -0.360755075 | 0.994547369 | -0.362732924 | 0.716804403 | NA | Rab19 | RAB19, member RAS oncogene family [So | 4.8186 | 6.1675 | 4.025 | 2.8919 | 3.2719 | 5.5692 |
| 7578 | ENSMUSG00000029924 | 423.4756392 | 0.137492099 | 0.144290967 | 0.952880844 | 0.340650447 | 0.783094143 | Slc37a3 | solute carrier family 37 (glycerol-3-phosph | 404.76 | 402.94 | 402.5 | 481.02 | 377.35 | 472.27 |
| 7579 | ENSMUSG00000029925 | 20.96947247 | -1.02289441 | 0.55236685 | -1.851838882 | 0.064048963 | 0.362159575 | Tbxas1 | thromboxane A synthase 1, platelet [Sour | 20.078 | 41.117 | 23.144 | 11.568 | 7.6343 | 22.277 |
| 7580 | ENSMUSG00000029992 | 354.725388 | 0.093391877 | 0.148632751 | 0.628339826 | 0.529781347 | 0.888334265 | Gfpt1 | glutamine fructose-6-phosphate transami | 338.11 | 364.91 | 327.03 | 385.59 | 320.64 | 392.07 |
| 7581 | ENSMUSG00000029993 | 663.9740521 | 0.087304071 | 0.114418887 | 0.763021496 | 0.445450576 | 0.852147532 | Nfu1 | NFU1 iron-sulfur cluster scaffold [Source:I | 619.19 | 646.56 | 666.13 | 663.22 | 690.36 | 698.38 |
| 7582 | ENSMUSG00000029994 | 913.0972648 | -0.107244359 | 0.115352854 | -0.929707021 | 0.352522798 | 0.793231138 | Anxa4 | annexin A4 [Source:MGI Symbol;Acc:MGI | 987.01 | 1004.3 | 849.27 | 862.76 | 898.67 | 876.6 |
| 7583 | ENSMUSG00000029998 | 3751.541808 | 0.006580112 | 0.09695248 | 0.067869459 | 0.945889551 | 0.990319806 | Pcyox1 | prenylcysteine oxidase 1 [Source:MGI Sym | 3752.1 | 3742.6 | 3734.2 | 3705.5 | 4100.7 | 3474.1 |
| 7584 | ENSMUSG00000029999 | 347.8746139 | 0.290741697 | 0.154126991 | 1.886377555 | 0.059244095 | 0.346085052 | Tgfa | transforming growth factor alpha [Source | 289.12 | 318.65 | 331.05 | 333.54 | 387.17 | 427.72 |
| 7585 | ENSMUSG00000030000 | 0 | NA | NA | NA | NA | NA | Add2 | adducin 2 (beta) [Source:MGI Symbol;Acc | 0 | 0 | 0 | 0 | 0 | 0 |
| 7586 | ENSMUSG00000030001 | 0 | NA | NA | NA | NA | NA | Figla | folliculogenesis specific basic helix-loop-h | 0 | 0 | 0 | 0 | 0 | 0 |
| 7587 | ENSMUSG00000030002 | 921.5451601 | 0.150667211 | 0.118789338 | 1.268356353 | 0.204670721 | 0.650460832 | Dusp11 | dual specificity phosphatase 11 (RNA/RNF | 813.54 | 910.74 | 897.57 | 1070 | 909.58 | 927.83 |
| 7588 | ENSMUSG00000030004 | 295.9836187 | 0.268558066 | 0.519250685 | 0.517203104 | 0.605014385 | 0.916376001 | Nat8 | N-acetyltransferase 8 (GCN5-related) [Sou | 165.44 | 203.53 | 436.71 | 237.14 | 462.42 | 270.66 |
| 7589 | ENSMUSG00000030007 | 1684.498759 | 0.047073548 | 0.09801389 | 0.48027426 | 0.631032389 | 0.921293275 | Cct7 | chaperonin containing Tcp1, subunit 7 (et | 1727.5 | 1582 | 1661.3 | 1775.6 | 1719.9 | 1640.7 |
| 7590 | ENSMUSG00000030008 | 102.7276017 | -0.153907794 | 0.225791784 | -0.681635937 | 0.495469185 | 0.877965876 | Pradc1 | protease-associated domain containing 1 | 105.21 | 106.9 | 112.7 | 102.18 | 103.61 | 85.766 |
| 7591 | ENSMUSG00000030016 | 532.6500172 | -0.173705046 | 0.13208951 | -1.315055566 | 0.188491229 | 0.625737234 | Zfp638 | zinc finger protein 638 [Source:MGI Symb | 567.79 | 565.35 | 560.48 | 461.75 | 566.03 | 474.5 |
| 7592 | ENSMUSG00000030017 | 0 | NA | NA | NA | NA | NA | Reg3g | regenerating islet-derived 3 gamma [Sourc | 0 | 0 | 0 | 0 | 0 | 0 |
| 7593 | ENSMUSG00000030019 | 248.8098958 | -0.048732275 | 0.170160333 | -0.286390335 | 0.774579176 | 0.958712409 | Fbxl14 | F-box and leucine-rich repeat protein 14 [ | 227.28 | 286.79 | 245.52 | 246.78 | 221.4 | 265.09 |
| 7594 | ENSMUSG00000030020 | 15.97277469 | -1.083553658 | 0.617862044 | -1.753714552 | 0.079479431 | 0.413673469 | Prickle2 | prickle planar cell polarity protein 2 [Sourc | 26.502 | 25.698 | 13.081 | 17.352 | 7.6343 | 5.5692 |
| 7595 | ENSMUSG00000030022 | 11.50975989 | -0.35175007 | 0.640408826 | -0.549258623 | 0.582827978 | 0.908645447 | Adamts9 | a disintegrin-like and metallopeptidase (re | 15.259 | 13.363 | 10.062 | 10.604 | 13.087 | 6.6831 |
| 7596 | ENSMUSG00000030029 | 547.6821258 | 0.328160137 | 0.143960658 | 2.279512615 | 0.022636611 | 0.188541979 | Lrig1 | leucine-rich repeats and immunoglobulin- | 419.22 | 503.68 | 535.32 | 642.01 | 540.95 | 644.92 |
| 7597 | ENSMUSG00000030030 | 50.52419328 | 0.203025067 | 0.374844671 | 0.541624525 | 0.588077193 | 0.910798643 | 1700003E16 | RIKEN cDNA 1700003E16 gene [Source:M | 32.927 | 34.949 | 73.456 | 62.659 | 51.259 | 47.895 |
| 7598 | ENSMUSG00000030031 | 25.90247834 | -0.123407656 | 0.464098321 | -0.265908429 | 0.790309733 | 0.961019028 | Kbtbd8 | kelch repeat and BTB (POZ) domain conta | 36.943 | 27.754 | 16.1 | 26.027 | 19.631 | 28.96 |
| 7599 | ENSMUSG00000030032 | 14.30842487 | 0.631000035 | 0.580015157 | 1.087902664 | 0.27663808 | 0.730038567 | Wdr54 | WD repeat domain 54 [Source:MGI Symbo | 10.44 | 8.2233 | 15.094 | 21.208 | 14.178 | 16.708 |
| 7600 | ENSMUSG00000030034 | 335.6196862 | 0.424230544 | 0.146060504 | 2.904485008 | 0.003678578 | 0.054127116 | Ino80b | INO80 complex subunit B [Source:MGI Sy | 279.48 | 272.4 | 307.91 | 364.38 | 374.08 | 415.46 |
| 7601 | ENSMUSG00000030035 | 663.3869902 | 0.178181706 | 0.152108676 | 1.171410534 | 0.2414338 | 0.691388521 | Wbp1 | WW domain binding protein 1 [Source:M | 650.51 | 580.77 | 634.94 | 554.29 | 764.52 | 795.28 |
| 7602 | ENSMUSG00000030036 | 702.4130465 | 0.291554755 | 0.143908502 | 2.025973113 | 0.04276754 | 0.285195804 | Mogs | mannosyl-oligosaccharide glucosidase [So | 523.62 | 673.29 | 699.34 | 760.58 | 708.9 | 848.75 |
| 7603 | ENSMUSG00000030037 | 0 | NA | NA | NA | NA | NA | Mrpl53 | mitochondrial ribosomal protein L53 [Sou | 0 | 0 | 0 | 0 | 0 | 0 |
| 7604 | ENSMUSG00000030041 | 0.815516723 | -3.06336425 | 2.557025255 | -1.19801877 | 0.23090971 | NA | M1ap | meiosis 1 associated protein [Source:MGI | 0.8031 | 3.0838 | 1.0062 | 0 | 0 | 0 |
| 7605 | ENSMUSG00000030042 | 614.4626325 | -0.359534348 | 0.139152787 | -2.583738029 | 0.0097736 | 0.106771413 | Pole4 | polymerase (DNA-directed), epsilon 4 (p1 | 708.34 | 775.05 | 588.65 | 563.93 | 501.68 | 549.13 |
| 7606 | ENSMUSG00000030043 | 0 | NA | NA | NA | NA | NA | Tacr1 | tachykinin receptor 1 [Source:MGI Symbo | 0 | 0 | 0 | 0 | 0 | 0 |
| 7607 | ENSMUSG00000030045 | 219.4324954 | 0.082780917 | 0.189550798 | 0.436721543 | 0.662313303 | 0.9288455 | Mrpl19 | mitochondrial ribosomal protein L19 [Sou | 222.46 | 177.83 | 238.48 | 194.72 | 223.58 | 259.53 |
| 7608 | ENSMUSG00000030046 | 54.83621019 | -1.464989921 | 0.328670925 | -4.45731524 | 8.30E-06 | 0.000455037 | Bmp10 | bone morphogenetic protein 10 [Source:M | 92.357 | 66.815 | 82.512 | 36.631 | 25.084 | 25.618 |
| 7609 | ENSMUSG00000030047 | 16.39621536 | -1.756195322 | 0.631413671 | -2.781370444 | 0.005412993 | 0.070155495 | Arhgap25 | Rho GTPase activating protein 25 [Source: | 27.305 | 27.754 | 21.131 | 14.46 | 3.2719 | 4.4554 |
| 7610 | ENSMUSG00000030048 | 0.321325949 | 1.911575114 | 4.048267893 | 0.472195804 | 0.636787034 | NA | Gkn3 | gastrokine 3 [Source:MGI Symbol;Acc:MG | 0 | 0 | 0 | 1.928 | 0 | 0 |
| 7611 | ENSMUSG00000030049 | 0 | NA | NA | NA | NA | NA | Gkn2 | gastrokine 2 [Source:MGI Symbol;Acc:MG | 0 | 0 | 0 | 0 | 0 | 0 |
| 7612 | ENSMUSG00000030050 | 0 | NA | NA | NA | NA | NA | Gkn1 | gastrokine 1 [Source:MGI Symbol;Acc:MG | 0 | 0 | 0 | 0 | 0 | 0 |

| | | | | | | | | | | | | | | | |
|---|---|---|---|---|---|---|---|---|---|---|---|---|---|---|---|
| 7613 | ENSMUSG00000030051 | 71.85861896 | -0.014566758 | 0.279057837 | -0.05219978 | 0.958369508 | 0.991903869 | Aplf | aprataxin and PNKP like factor [Source:MG | 73.082 | 81.206 | 62.387 | 74.226 | 56.712 | 83.538 |
| 7614 | ENSMUSG00000030054 | 1.497203064 | 0.640740167 | 1.941800609 | 0.329972173 | 0.741420988 | NA | Gp9 | glycoprotein 9 (platelet) [Source:MGI Sym | 2.4093 | 1.0279 | 0 | 0 | 1.0906 | 4.4554 |
| 7615 | ENSMUSG00000030055 | 271.6641752 | -0.058801572 | 0.158261506 | -0.371546901 | 0.71023023 | 0.944239215 | Rab43 | RAB43, member RAS oncogene family [So | 277.87 | 275.48 | 278.73 | 298.83 | 258.48 | 240.59 |
| 7616 | ENSMUSG00000030056 | 204.2880893 | 0.019244107 | 0.173870624 | 0.110680611 | 0.911869622 | 0.98589875 | Isy1 | ISY1 splicing factor homolog [Source:MGI | 188.73 | 215.86 | 204.27 | 186.05 | 220.3 | 210.52 |
| 7617 | ENSMUSG00000030057 | 5665.955327 | -0.111695808 | 0.096211587 | -1.160939252 | 0.245666605 | 0.695774774 | Cnbp | cellular nucleic acid binding protein [Sourc | 5386.4 | 6079.1 | 6191.4 | 5456.1 | 5681 | 5201.7 |
| 7618 | ENSMUSG00000030058 | 1009.377358 | 0.003433236 | 0.118987411 | 0.028853777 | 0.976981211 | 0.997251675 | Copg1 | coatomer protein complex, subunit gamm | 995.04 | 996.05 | 1034.4 | 1131.7 | 996.83 | 902.21 |
| 7619 | ENSMUSG00000030059 | 464.0874156 | -0.191250386 | 0.134232087 | -1.424773988 | 0.154222544 | 0.57167475 | Tmf1 | TATA element modulatory factor 1 [Sourc | 456.97 | 529.38 | 499.1 | 464.64 | 410.07 | 424.37 |
| 7620 | ENSMUSG00000030060 | 122.6065409 | 0.294170351 | 0.216803856 | 1.356850178 | 0.17482882 | 0.604813215 | Hmces | 5-hydroxymethylcytosine (hmC) binding, E | 108.42 | 118.21 | 103.64 | 115.68 | 152.69 | 137 |
| 7621 | ENSMUSG00000030061 | 687.8475362 | -0.036615445 | 0.134311959 | -0.27261493 | 0.785149234 | 0.960670212 | Uba3 | ubiquitin-like modifier activating enzyme : | 769.37 | 709.26 | 609.78 | 620.8 | 682.73 | 735.14 |
| 7622 | ENSMUSG00000030062 | 2014.398962 | 0.088665053 | 0.106638073 | 0.831457756 | 0.405715083 | 0.829973745 | Rpn1 | ribophorin I [Source:MGI Symbol;Acc:MG | 1892.9 | 1923.2 | 2042.7 | 2303.9 | 1954.4 | 1969.3 |
| 7623 | ENSMUSG00000030064 | 738.5215399 | -0.377022788 | 0.437901561 | -0.860976121 | 0.389251195 | 0.818844821 | Frmd4b | FERM domain containing 4B [Source:MGI | 950.87 | 842.89 | 709.4 | 402.94 | 1038.3 | 486.75 |
| 7624 | ENSMUSG00000030067 | 288.7456906 | 0.180537339 | 0.149581234 | 1.206951793 | 0.22745071 | 0.673700979 | Foxp1 | forkhead box P1 [Source:MGI Symbol;Acc | 259.4 | 260.06 | 292.82 | 300.76 | 307.55 | 311.88 |
| 7625 | ENSMUSG00000030068 | 0.469265235 | -2.274672822 | 3.439794491 | -0.661281605 | 0.508431736 | NA | Gm20696 | predicted gene 20696 [Source:MGI Symbo | 0.8031 | 0 | 2.0125 | 0 | 0 | 0 |
| 7626 | ENSMUSG00000030069 | 0 | NA | NA | NA | NA | NA | Prok2 | prokineticin 2 [Source:MGI Symbol;Acc:M | 0 | 0 | 0 | 0 | 0 | 0 |
| 7627 | ENSMUSG00000030074 | 0 | NA | NA | NA | NA | NA | Gxylt2 | glucoside xylosyltransferase 2 [Source:MG | 0 | 0 | 0 | 0 | 0 | 0 |
| 7628 | ENSMUSG00000030075 | 0 | NA | NA | NA | NA | NA | Cntn3 | contactin 3 [Source:MGI Symbol;Acc:MGI | 0 | 0 | 0 | 0 | 0 | 0 |
| 7629 | ENSMUSG00000030077 | 0.160662974 | 1.126924175 | 4.080472857 | 0.276174898 | 0.782413736 | NA | Chl1 | cell adhesion molecule L1-like [Source:MG | 0 | 0 | 0 | 0.964 | 0 | 0 |
| 7630 | ENSMUSG00000030079 | 301.6336832 | -0.036249231 | 0.147570679 | -0.245639791 | 0.805961084 | 0.964428432 | Ruvbl1 | RuvB-like protein 1 [Source:MGI Symbol;A | 310.8 | 318.65 | 286.78 | 303.65 | 281.38 | 308.53 |
| 7631 | ENSMUSG00000030082 | 1735.774187 | 0.220563342 | 0.110689631 | 1.99262876 | 0.046302113 | 0.297518334 | Sec61a1 | Sec61 alpha 1 subunit (S. cerevisiae) [Sour | 1517.9 | 1626.2 | 1667.3 | 2051.3 | 1679.6 | 1872.4 |
| 7632 | ENSMUSG00000030083 | 628.1796647 | 0.217368775 | 0.135274632 | 1.606870196 | 0.108082846 | 0.484761602 | Abtb1 | ankyrin repeat and BTB (POZ) domain cor | 656.94 | 507.79 | 576.58 | 632.37 | 691.45 | 703.95 |
| 7633 | ENSMUSG00000030084 | 158.3101279 | -0.014317956 | 0.232190379 | -0.061664725 | 0.950829832 | 0.990883058 | Plxna1 | plexin A1 [Source:MGI Symbol;Acc:MGI:1( | 172.67 | 148.02 | 156.97 | 210.15 | 142.87 | 119.18 |
| 7634 | ENSMUSG00000030086 | 62.28211253 | -0.432970012 | 0.289892246 | -1.493554996 | 0.13529197 | 0.538039097 | Chchd6 | coiled-coil-helix-coiled-coil-helix domain c | 73.082 | 79.15 | 62.387 | 50.127 | 62.165 | 46.781 |
| 7635 | ENSMUSG00000030087 | 3870.894064 | -0.292727656 | 0.152534768 | -1.919088078 | 0.054973188 | 0.330417929 | Klf15 | Kruppel-like factor 15 [Source:MGI Symbo | 3958.5 | 4765.4 | 4062.2 | 2599.8 | 4072.4 | 3767 |
| 7636 | ENSMUSG00000030088 | 19177.54597 | 0.030856046 | 0.112112249 | 0.275224577 | 0.783143706 | 0.960670212 | Aldh1l1 | aldehyde dehydrogenase 1 family, membe | 17552 | 19344 | 20021 | 16979 | 21891 | 19278 |
| 7637 | ENSMUSG00000030089 | 92.70627268 | -0.412868575 | 0.260011092 | -1.587888319 | 0.112311595 | 0.491966964 | Slc41a3 | solute carrier family 41, member 3 [Sourc | 102.8 | 126.43 | 88.55 | 83.866 | 63.256 | 91.335 |
| 7638 | ENSMUSG00000030091 | 316.9358364 | 0.396398223 | 0.180060988 | 2.201466442 | 0.02770302 | 0.216475678 | Nup210 | nucleoporin 210 [Source:MGI Symbol;Acc | 232.1 | 264.17 | 325.02 | 298.83 | 386.08 | 395.41 |
| 7639 | ENSMUSG00000030092 | 0 | NA | NA | NA | NA | NA | Cntn6 | contactin 6 [Source:MGI Symbol;Acc:MGI | 0 | 0 | 0 | 0 | 0 | 0 |
| 7640 | ENSMUSG00000030093 | 0 | NA | NA | NA | NA | NA | Wnt7a | wingless-type MMTV integration site fami | 0 | 0 | 0 | 0 | 0 | 0 |
| 7641 | ENSMUSG00000030094 | 468.5132289 | 0.126820835 | 0.132871479 | 0.954462432 | 0.339849617 | 0.782113741 | Xpc | xeroderma pigmentosum, complementati | 444.92 | 423.5 | 474.95 | 450.18 | 495.14 | 522.39 |
| 7642 | ENSMUSG00000030095 | 189.1385535 | -0.155782656 | 0.18106141 | -0.860385747 | 0.38957644 | 0.819052248 | Tmem43 | transmembrane protein 43 [Source:MGI S | 186.32 | 211.75 | 200.24 | 181.23 | 158.14 | 197.15 |
| 7643 | ENSMUSG00000030096 | 1557.523763 | 0.330641175 | 0.119861153 | 2.758534921 | 0.00580611 | 0.073795762 | Slc6a6 | solute carrier family 6 (neurotransmitter t | 1256.1 | 1393.9 | 1491.3 | 1946.3 | 1582.5 | 1675.2 |
| 7644 | ENSMUSG00000030098 | 1.136894805 | 0.527420275 | 2.120428083 | 0.248732923 | 0.80356738 | NA | Grip2 | glutamate receptor interacting protein 2 [ | 0.8031 | 0 | 2.0125 | 2.8919 | 0 | 1.1138 |
| 7645 | ENSMUSG00000030101 | 656.6170058 | -0.097265348 | 0.140459807 | -0.692478158 | 0.488637098 | 0.874431125 | Sumf1 | sulfatase modifying factor 1 [Source:MGI : | 619.19 | 758.6 | 659.09 | 615.02 | 723.08 | 564.72 |
| 7646 | ENSMUSG00000030102 | 1189.245991 | -0.144342408 | 0.162440983 | -0.888583687 | 0.374226862 | 0.809332734 | Itpr1 | inositol 1,4,5-trisphosphate receptor 1 [Sc | 1082.6 | 1273.6 | 1391.6 | 1434.4 | 1013.2 | 940.08 |
| 7647 | ENSMUSG00000030103 | 1956.054082 | -0.055698793 | 0.165427708 | -0.336695674 | 0.736346325 | 0.948109711 | Bhlhe40 | basic helix-loop-helix family, member e40 | 2310.5 | 1864.6 | 1805.2 | 1726.5 | 2477.9 | 1551.6 |
| 7648 | ENSMUSG00000030104 | 2982.14831 | 0.044563537 | 0.096139466 | 0.463530105 | 0.642984441 | 0.925293697 | Edem1 | ER degradation enhancer, mannosidase a | 2781.1 | 2990.2 | 3037.9 | 3078 | 2831.2 | 3174.5 |
| 7649 | ENSMUSG00000030105 | 1006.833473 | 0.070974689 | 0.109044729 | 0.650876847 | 0.515125988 | 0.885627279 | Arl8b | ADP-ribosylation factor-like 8B [Source:M | 937.22 | 986.8 | 1022.3 | 984.22 | 1008.8 | 1101.6 |
| 7650 | ENSMUSG00000030107 | 204.0652543 | 0.292366732 | 0.222383539 | 1.314695923 | 0.188612117 | 0.625851266 | Usp18 | ubiquitin specific peptidase 18 [Source:M | 238.52 | 177.83 | 132.82 | 195.69 | 234.48 | 245.05 |
| 7651 | ENSMUSG00000030108 | 3664.601978 | 0.089410494 | 0.130346427 | 0.685945105 | 0.492747726 | 0.876660964 | Slc6a13 | solute carrier family 6 (neurotransmitter t | 3618.8 | 3378.8 | 3656.7 | 4599.1 | 3402.7 | 3331.5 |
| 7652 | ENSMUSG00000030109 | 1297.628289 | 0.25514258 | 0.110837568 | 2.301950364 | 0.021337972 | 0.181441752 | Slc6a12 | solute carrier family 6 (neurotransmitter t | 1169.3 | 1237.6 | 1142.1 | 1268.6 | 1475.6 | 1492.6 |
| 7653 | ENSMUSG00000030110 | 0 | NA | NA | NA | NA | NA | Ret | ret proto-oncogene [Source:MGI Symbol;, | 0 | 0 | 0 | 0 | 0 | 0 |
| 7654 | ENSMUSG00000030111 | 1.183185809 | -1.148989598 | 2.104741745 | -0.545905264 | 0.58513106 | NA | A2m | alpha-2-macroglobulin [Source:MGI Symb | 0.8031 | 2.0558 | 2.0125 | 0 | 0 | 2.2277 |
| 7655 | ENSMUSG00000030114 | 0.342639279 | -1.805976596 | 4.028547714 | -0.448294702 | 0.653940524 | NA | Klrg1 | killer cell lectin-like receptor subfamily G, | 0 | 2.0558 | 0 | 0 | 0 | 0 |
| 7656 | ENSMUSG00000030116 | 0.487198452 | -0.685759572 | 3.136894395 | -0.218610985 | 0.826953095 | NA | Mfap5 | microfibrillar associated protein 5 [Source | 0.8031 | 0 | 1.0062 | 0 | 0 | 1.1138 |
| 7657 | ENSMUSG00000030117 | 0.401551119 | -2.072982726 | 4.008251979 | -0.517178744 | 0.605031388 | NA | Gdf3 | growth differentiation factor 3 [Source:M | 2.4093 | 0 | 0 | 0 | 0 | 0 |
| 7658 | ENSMUSG00000030120 | 2488.122505 | 0.058392198 | 0.099456351 | 0.587113821 | 0.557127266 | 0.898167794 | Mlf2 | myeloid leukemia factor 2 [Source:MGI Sy | 2312.1 | 2329.3 | 2672.6 | 2531.4 | 2521.5 | 2561.8 |
| 7659 | ENSMUSG00000030122 | 13835.29901 | 0.066344006 | 0.092844145 | 0.714573937 | 0.474872354 | 0.869368495 | Ptms | parathymosin [Source:MGI Symbol;Acc:M | 13717 | 12394 | 14441 | 14079 | 13764 | 14618 |
| 7660 | ENSMUSG00000030123 | 285.2167791 | -0.754306983 | 0.194565944 | -3.876870573 | 0.000105809 | 0.003807004 | Plxnd1 | plexin D1 [Source:MGI Symbol;Acc:MGI:2 | 324.45 | 351.55 | 398.47 | 209.18 | 160.32 | 267.32 |
| 7661 | ENSMUSG00000030124 | 38.66399165 | -0.725064653 | 0.376076739 | -1.927969955 | 0.053858863 | 0.326357185 | Lag3 | lymphocyte-activation gene 3 [Source:MG | 36.14 | 45.228 | 63.393 | 29.883 | 27.265 | 30.074 |
| 7662 | ENSMUSG00000030125 | 0 | NA | NA | NA | NA | NA | Lrrc23 | leucine rich repeat containing 23 [Source: | 0 | 0 | 0 | 0 | 0 | 0 |
| 7663 | ENSMUSG00000030126 | 633.8389197 | -0.305200447 | 0.177494416 | -1.719493233 | 0.085524599 | 0.430973029 | Tmcc1 | transmembrane and coiled coil domains 1 | 667.38 | 780.19 | 655.07 | 627.55 | 669.64 | 403.21 |
| 7664 | ENSMUSG00000030127 | 1164.163574 | 0.145532847 | 0.099149569 | 1.46781119 | 0.142155517 | 0.551920354 | Cops7a | COP9 signalosome subunit 7A [Source:MG | 1097.8 | 1094.7 | 1124 | 1214.6 | 1254.2 | 1199.6 |
| 7665 | ENSMUSG00000030131 | 7901.189428 | 0.671753852 | 0.382666399 | 1.755455544 | 0.079181418 | 0.413309288 | Mug2 | murinoglobulin 2 [Source:MGI Symbol;Acc | 8050.3 | 4656.5 | 5575.6 | 11198 | 12192 | 5735.2 |
| 7666 | ENSMUSG00000030134 | 0.556921942 | 2.679043159 | 3.638243063 | 0.736356289 | 0.461513899 | NA | Rasgef1a | RasGEF domain family, member 1A [Sourc | 0 | 0 | 0 | 0 | 0 | 3.3415 |
| 7667 | ENSMUSG00000030137 | 3.546104064 | -0.08266918 | 1.296363056 | -0.063770083 | 0.9491533 | NA | Tuba8 | tubulin, alpha 8 [Source:MGI Symbol;Acc: | 0.8031 | 9.2513 | 1.0062 | 5.7839 | 1.0906 | 3.3415 |
| 7668 | ENSMUSG00000030138 | 442.2180509 | -0.032251579 | 0.152661962 | -0.211261391 | 0.832683309 | 0.970611245 | Bms1 | BMS1, ribosome biogenesis factor [Source | 388.7 | 488.26 | 464.88 | 381.74 | 486.42 | 443.31 |

| | | | | | | | | | | | | | | | |
|---|---|---|---|---|---|---|---|---|---|---|---|---|---|---|---|
| 7669 | ENSMUSG00000030142 | 0.535401493 | -2.48596965 | 3.684117979 | -0.674780141 | 0.499815459 | NA | Clec4e | C-type lectin domain family 4, member e [ | 3.2124 | 0 | 0 | 0 | 0 | 0 |
| 7670 | ENSMUSG00000030143 | 0 | NA | NA | NA | NA | NA | Gm8882 | predicted gene 8882 [Source:MGI Symbol | 0 | 0 | 0 | 0 | 0 | 0 |
| 7671 | ENSMUSG00000030144 | 1.566520013 | -0.876242964 | 1.911025802 | -0.458519693 | 0.646579118 | NA | Clec4d | C-type lectin domain family 4, member d [ | 0 | 2.0558 | 4.025 | 0 | 1.0906 | 2.2277 |
| 7672 | ENSMUSG00000030145 | 4.911964775 | -0.114386571 | 1.097327389 | -0.104241061 | 0.916978049 | NA | Zfp248 | zinc finger protein 248 [Source:MGI Symb | 7.2279 | 3.0838 | 5.0312 | 8.6758 | 5.4531 | 0 |
| 7673 | ENSMUSG00000030147 | 1.256521509 | 0.543933952 | 2.487612764 | 0.218657003 | 0.826917245 | NA | Clec4b1 | C-type lectin domain family 4, member b1 | 0 | 3.0838 | 0 | 0 | 0 | 4.4554 |
| 7674 | ENSMUSG00000030148 | 13.14138565 | -2.226787937 | 0.755428525 | -2.947714925 | 0.003201321 | 0.048843788 | Clec4a2 | C-type lectin domain family 4, member a2 | 20.881 | 33.921 | 10.062 | 2.8919 | 2.1812 | 8.9108 |
| 7675 | ENSMUSG00000030149 | 9.754061402 | -1.269626488 | 0.764513893 | -1.660697732 | 0.096774169 | 0.458530879 | Klrk1 | killer cell lectin-like receptor subfamily K, | 16.865 | 8.2233 | 16.1 | 1.928 | 8.7249 | 6.6831 |
| 7676 | ENSMUSG00000030154 | 0.17131964 | -0.796629176 | 4.080472857 | -0.195229623 | 0.84521319 | NA | Klrb1f | killer cell lectin-like receptor subfamily B r | 0 | 1.0279 | 0 | 0 | 0 | 0 |
| 7677 | ENSMUSG00000030155 | 0 | NA | NA | NA | NA | NA | Clec2e | C-type lectin domain family 2, member e [ | 0 | 0 | 0 | 0 | 0 | 0 |
| 7678 | ENSMUSG00000030156 | 2.237813397 | -2.43564826 | 1.612029887 | -1.510920039 | 0.130808825 | NA | Cd69 | CD69 antigen [Source:MGI Symbol;Acc:M | 3.2124 | 4.1117 | 4.025 | 0.964 | 0 | 1.1138 |
| 7679 | ENSMUSG00000030157 | 1762.711571 | 0.347595045 | 0.168403665 | 2.064058667 | 0.039012154 | 0.270232919 | Clec2d | C-type lectin domain family 2, member d [ | 1146.8 | 1764.9 | 1743.8 | 1961.7 | 1655.6 | 2303.4 |
| 7680 | ENSMUSG00000030158 | 1.476096323 | 0.614272012 | 1.947304907 | 0.315447268 | 0.752422083 | NA | Clec12b | C-type lectin domain family 12, member B | 2.4093 | 1.0279 | 0 | 0.964 | 0 | 4.4554 |
| 7681 | ENSMUSG00000030159 | 101.8555636 | -1.258090977 | 0.305099027 | -4.123549615 | 3.73E-05 | 0.00167065 | Clec1b | C-type lectin domain family 1, member b [ | 127.69 | 204.56 | 98.612 | 51.091 | 53.44 | 75.741 |
| 7682 | ENSMUSG00000030160 | 0 | NA | NA | NA | NA | NA | Tmem52b | transmembrane protein 52B [Source:MGI | 0 | 0 | 0 | 0 | 0 | 0 |
| 7683 | ENSMUSG00000030161 | 6498.11064 | 0.133088208 | 0.12383687 | 1.074705837 | 0.282506449 | 0.735452355 | Gabarapl1 | gamma-aminobutyric acid (GABA) A recep | 6821.6 | 6373.1 | 5399.5 | 6100.1 | 6514.3 | 7780.2 |
| 7684 | ENSMUSG00000030162 | 0.503122293 | -2.366675966 | 3.785596405 | -0.625179156 | 0.531853481 | NA | Olr1 | oxidized low density lipoprotein (lectin-lik | 0 | 0 | 3.0187 | 0 | 0 | 0 |
| 7685 | ENSMUSG00000030165 | 1.891794003 | -0.288063386 | 1.710028191 | -0.168455343 | 0.866225073 | NA | Klrd1 | killer cell lectin-like receptor, subfamily D, | 3.2124 | 0 | 3.0187 | 2.8919 | 0 | 2.2277 |
| 7686 | ENSMUSG00000030166 | 97.65175054 | 0.480951256 | 0.233245359 | 2.06199711 | 0.039208011 | 0.270953021 | Rad52 | RAD52 homolog, DNA repair protein [Sou | 77.098 | 86.345 | 81.506 | 127.25 | 111.24 | 102.47 |
| 7687 | ENSMUSG00000030167 | 1.627214648 | -1.880408741 | 1.778601828 | -1.057239856 | 0.290402134 | NA | Klrc1 | killer cell lectin-like receptor subfamily C, | 1.6062 | 3.0838 | 3.0187 | 0.964 | 1.0906 | 0 |
| 7688 | ENSMUSG00000030168 | 12243.72143 | 0.039107608 | 0.129784904 | 0.30132632 | 0.763165673 | 0.954864872 | Adipor2 | adiponectin receptor 2 [Source:MGI Symb | 13833 | 12504 | 9895.4 | 12863 | 13279 | 11087 |
| 7689 | ENSMUSG00000030170 | 260.4370997 | -0.224100961 | 0.16504691 | -1.357801614 | 0.174526643 | 0.604201379 | Wnt5b | wingless-type MMTV integration site fami | 270.65 | 286.79 | 284.77 | 243.89 | 270.47 | 206.06 |
| 7690 | ENSMUSG00000030172 | 429.9757441 | -0.099904983 | 0.161845668 | -0.617285493 | 0.537046434 | 0.890705151 | Erc1 | ELKS/RAB6-interacting/CAST family memb | 442.51 | 407.06 | 485.01 | 404.87 | 496.23 | 344.18 |
| 7691 | ENSMUSG00000030173 | 0 | NA | NA | NA | NA | NA | Klra5 | killer cell lectin-like receptor, subfamily A, | 0 | 0 | 0 | 0 | 0 | 0 |
| 7692 | ENSMUSG00000030177 | 61.01091516 | 0.201725198 | 0.290592536 | 0.694185753 | 0.487565726 | 0.873910634 | Ccdc77 | coiled-coil domain containing 77 [Source:I | 63.445 | 47.284 | 59.368 | 71.334 | 57.803 | 66.831 |
| 7693 | ENSMUSG00000030178 | 0.620790101 | -1.233953725 | 2.860353486 | -0.431399032 | 0.666178254 | NA | Klra13-ps | killer cell lectin-like receptor subfamily A, | 1.6062 | 1.0279 | 0 | 0 | 1.0906 | 0 |
| 7694 | ENSMUSG00000030180 | 523.5454264 | 0.115941251 | 0.1255697 | 0.923321877 | 0.35583949 | 0.795525652 | Kdm5a | lysine (K)-specific demethylase 5A [Source | 484.27 | 538.63 | 485.01 | 535.01 | 570.39 | 527.96 |
| 7695 | ENSMUSG00000030187 | 20.39217989 | -1.3817675 | 0.537114468 | -2.572575459 | 0.010094495 | 0.109368369 | Klra2 | killer cell lectin-like receptor, subfamily A, | 32.124 | 39.061 | 17.106 | 8.6758 | 10.906 | 14.48 |
| 7696 | ENSMUSG00000030188 | 32.39000172 | 0.085770725 | 0.408099595 | 0.210171062 | 0.833534164 | 0.97078977 | Magohb | mago homolog B, exon junction complex | 32.927 | 37.005 | 24.15 | 24.099 | 32.719 | 43.44 |
| 7697 | ENSMUSG00000030189 | 469.7796502 | -0.242166156 | 0.137157147 | -1.765610915 | 0.077461148 | 0.40712138 | Ybx3 | Y box protein 3 [Source:MGI Symbol;Acc:I | 497.12 | 504.71 | 526.27 | 480.06 | 382.81 | 427.72 |
| 7698 | ENSMUSG00000030194 | 0 | NA | NA | NA | NA | NA | Tas2r116 | taste receptor, type 2, member 116 [Sour | 0 | 0 | 0 | 0 | 0 | 0 |
| 7699 | ENSMUSG00000030196 | 0 | NA | NA | NA | NA | NA | Tas2r103 | taste receptor, type 2, member 103 [Sour | 0 | 0 | 0 | 0 | 0 | 0 |
| 7700 | ENSMUSG00000030199 | 283.010732 | -0.038928131 | 0.151655209 | -0.25668839 | 0.797419335 | 0.962236316 | Etv6 | ets variant 6 [Source:MGI Symbol;Acc:MG | 294.74 | 293.98 | 271.69 | 281.48 | 296.65 | 259.53 |
| 7701 | ENSMUSG00000030200 | 14.15983501 | -0.110092342 | 0.762546664 | -0.144374564 | 0.885204701 | 0.981795936 | Bcl2l14 | BCL2-like 14 (apoptosis facilitator) [Source | 30.518 | 5.1396 | 8.05 | 4.8199 | 15.269 | 21.163 |
| 7702 | ENSMUSG00000030201 | 1412.843584 | 0.065622387 | 0.109338953 | 0.60017391 | 0.54839034 | 0.895622715 | Lrp6 | low density lipoprotein receptor-related p | 1334 | 1435 | 1374.5 | 1597.3 | 1379.6 | 1356.7 |
| 7703 | ENSMUSG00000030203 | 663.5545016 | -0.126880586 | 0.13423206 | -0.945233102 | 0.344539883 | 0.785874854 | Dusp16 | dual specificity phosphatase 16 [Source:M | 642.48 | 653.76 | 781.85 | 570.67 | 660.91 | 671.65 |
| 7704 | ENSMUSG00000030204 | 559.9807552 | 0.067417616 | 0.124918055 | 0.53969473 | 0.589407575 | 0.911039845 | Ddx47 | DEAD box helicase 47 [Source:MGI Symbo | 571.81 | 571.52 | 497.08 | 584.17 | 561.67 | 573.63 |
| 7705 | ENSMUSG00000030205 | 0.185640647 | 1.126924175 | 4.080472857 | 0.276174898 | 0.782413736 | NA | Gprc5d | G protein-coupled receptor, family C, grou | 0 | 0 | 0 | 0 | 0 | 1.1138 |
| 7706 | ENSMUSG00000030206 | 0.929831046 | -1.988873913 | 2.481927043 | -0.801342617 | 0.422933325 | NA | Gsg1 | germ cell associated 1 [Source:MGI Symbo | 2.4093 | 2.0558 | 0 | 0 | 0 | 1.1138 |
| 7707 | ENSMUSG00000030207 | 931.083932 | -0.235945976 | 0.159524618 | -1.479056832 | 0.139125126 | 0.545066664 | Fam234b | family with sequence similarity 234, meml | 761.34 | 1048.5 | 1212.5 | 827.09 | 913.94 | 823.13 |
| 7708 | ENSMUSG00000030208 | 20.03166338 | -0.973941387 | 0.569076734 | -1.711441233 | 0.086999691 | 0.434517794 | Emp1 | epithelial membrane protein 1 [Source:M | 37.746 | 27.754 | 14.087 | 17.352 | 6.5437 | 16.708 |
| 7709 | ENSMUSG00000030209 | 1.199289611 | -1.270320185 | 2.16443215 | -0.58690691 | 0.557266229 | NA | Grin2b | glutamate receptor, ionotropic, NMDA2B | 0 | 4.1117 | 1.0062 | 0.964 | 0 | 1.1138 |
| 7710 | ENSMUSG00000030213 | 472.8700909 | -0.117389477 | 0.135825304 | -0.864268099 | 0.38744063 | 0.817618864 | Atf7ip | activating transcription factor 7 interactin | 495.51 | 452.28 | 528.28 | 451.14 | 418.8 | 491.21 |
| 7711 | ENSMUSG00000030214 | 862.2124429 | 0.017615606 | 0.125940632 | 0.1398723 | 0.888760887 | 0.982382091 | Plbd1 | phospholipase B domain containing 1 [So | 949.27 | 866.53 | 753.68 | 814.56 | 870.31 | 918.92 |
| 7712 | ENSMUSG00000030216 | 592.0029087 | -0.07350614 | 0.123200602 | -0.596637833 | 0.550749206 | 0.895703112 | Wbp11 | WW domain binding protein 11 [Source:M | 636.86 | 619.83 | 564.5 | 611.16 | 563.85 | 555.81 |
| 7713 | ENSMUSG00000030217 | 45.11182964 | -0.920455817 | 0.332513462 | -2.768176097 | 0.005637099 | 0.072346936 | Art4 | ADP-ribosyltransferase 4 [Source:MGI Syn | 60.233 | 53.452 | 63.393 | 30.847 | 34.9 | 27.846 |
| 7714 | ENSMUSG00000030218 | 17.94060285 | -1.070686644 | 0.593185343 | -1.804978252 | 0.071078085 | 0.385913448 | Mgp | matrix Gla protein [Source:MGI Symbol;A | 22.487 | 20.558 | 30.187 | 21.208 | 7.6343 | 5.5692 |
| 7715 | ENSMUSG00000030219 | 1.168941911 | -2.435929877 | 2.172449565 | -1.121282591 | 0.262167593 | NA | Erp27 | endoplasmic reticulum protein 27 [Source | 4.0155 | 1.0279 | 1.0062 | 0.964 | 0 | 0 |
| 7716 | ENSMUSG00000030220 | 44.98144481 | -0.718702828 | 0.363992882 | -1.974496932 | 0.048325266 | 0.305821368 | Arhgdib | Rho, GDP dissociation inhibitor (GDI) beta | 73.885 | 50.368 | 43.269 | 32.775 | 27.265 | 42.326 |
| 7717 | ENSMUSG00000030222 | 41.49504584 | -0.876820549 | 0.364597855 | -2.404897717 | 0.016176993 | 0.149989759 | Rerg | RAS-like, estrogen-regulated, growth-inhil | 59.43 | 56.535 | 45.281 | 32.775 | 34.9 | 20.049 |
| 7718 | ENSMUSG00000030223 | 3.608266537 | -0.463059347 | 1.202734594 | -0.385005428 | 0.700233394 | NA | Ptpro | protein tyrosine phosphatase, receptor ty | 6.4248 | 5.1396 | 1.0062 | 5.7839 | 2.1812 | 1.1138 |
| 7719 | ENSMUSG00000030224 | 1569.640021 | -0.052166995 | 0.104022204 | -0.501498654 | 0.616020225 | 0.917509899 | Strap | serine/threonine kinase receptor associat | 1533.1 | 1712.5 | 1548.6 | 1460.4 | 1628.3 | 1534.9 |
| 7720 | ENSMUSG00000030225 | 1507.93596 | -0.116677843 | 0.103308172 | -1.129415424 | 0.258722629 | 0.712165697 | Dera | deoxyribose-phosphate aldolase (putative | 1643.1 | 1489.5 | 1572.8 | 1351.5 | 1512.7 | 1478.1 |
| 7721 | ENSMUSG00000030226 | 0 | NA | NA | NA | NA | NA | Lmo3 | LIM domain only 3 [Source:MGI Symbol;A | 0 | 0 | 0 | 0 | 0 | 0 |
| 7722 | ENSMUSG00000030228 | 61.88157623 | -0.396370852 | 0.308392611 | -1.285279991 | 0.19869446 | 0.640976045 | Pik3c2g | phosphatidylinositol-4-phosphate 3-kinas | 84.326 | 70.926 | 55.343 | 46.271 | 66.528 | 47.895 |
| 7723 | ENSMUSG00000030230 | 0 | NA | NA | NA | NA | NA | Plcz1 | phospholipase C, zeta 1 [Source:MGI Sym | 0 | 0 | 0 | 0 | 0 | 0 |
| 7724 | ENSMUSG00000030231 | 569.292956 | 0.147167802 | 0.139786491 | 1.052804182 | 0.292430748 | 0.743376015 | Plekha5 | pleckstrin homology domain containing, f | 543.7 | 534.52 | 543.37 | 701.78 | 536.58 | 555.81 |

| | | | | | | | | | | | | | | |
|---|---|---|---|---|---|---|---|---|---|---|---|---|---|---|
| 7725 | ENSMUSG00000030232 | 559.4530368 | -0.129934008 | 0.134662784 | -0.964884315 | 0.334602758 | 0.776159726 | Aebp2 | AE binding protein 2 [Source:MGI Symbol; | 572.61 | 651.7 | 530.29 | 578.39 | 518.04 | 505.69 |
| 7726 | ENSMUSG00000030235 | 1.80219863 | -0.674544505 | 1.531097253 | -0.440562808 | 0.659529533 | NA | Slco1c1 | solute carrier organic anion transporter fa | 1.6062 | 2.0558 | 3.0187 | 1.928 | 1.0906 | 1.1138 |
| 7727 | ENSMUSG00000030236 | 25797.8004 | 0.242317177 | 0.089322249 | 2.712842308 | 0.006670884 | 0.081424194 | Slco1b2 | solute carrier organic anion transporter fa | 22995 | 23897 | 24017 | 26862 | 30238 | 26777 |
| 7728 | ENSMUSG00000030237 | 942.6522244 | -1.598268885 | 0.520751987 | -3.069155618 | 0.002146647 | 0.036657743 | Slco1a4 | solute carrier organic anion transporter fa | 1217.5 | 2225.4 | 809.02 | 660.32 | 386.08 | 357.54 |
| 7729 | ENSMUSG00000030243 | 82.10856318 | 0.25333186 | 0.283263687 | 0.894332283 | 0.371144132 | 0.807214774 | Recql | RecQ protein-like [Source:MGI Symbol;Ac | 79.507 | 51.396 | 93.581 | 80.974 | 101.43 | 85.766 |
| 7730 | ENSMUSG00000030244 | 5798.104503 | 0.14460619 | 0.116133596 | 1.245171035 | 0.213068887 | 0.657724029 | Gys2 | glycogen synthase 2 [Source:MGI Symbol; | 5601.6 | 5736.8 | 5185.2 | 6375.7 | 6743.3 | 5146 |
| 7731 | ENSMUSG00000030245 | 51.56807393 | -0.359364727 | 0.328555824 | -1.093770678 | 0.274055567 | 0.729244726 | Golt1b | golgi transport 1B [Source:MGI Symbol;Ac | 50.595 | 75.038 | 48.3 | 39.523 | 44.715 | 51.237 |
| 7732 | ENSMUSG00000030246 | 28.10542451 | -0.211895486 | 0.411286923 | -0.515201126 | 0.606412483 | 0.916376001 | Ldhb | lactate dehydrogenase B [Source:MGI Syn | 35.336 | 27.754 | 27.169 | 22.171 | 28.356 | 27.846 |
| 7733 | ENSMUSG00000030247 | 47.36545177 | -0.664343933 | 0.32860231 | -2.021726299 | 0.043204635 | 0.286918392 | Kcnj8 | potassium inwardly-rectifying channel, su | 68.264 | 57.563 | 48.3 | 39.523 | 34.9 | 35.643 |
| 7734 | ENSMUSG00000030249 | 322.9411825 | -1.047010318 | 0.189449894 | -5.526581692 | 3.27E-08 | 4.14E-06 | Abcc9 | ATP-binding cassette, sub-family C (CFTR/ | 512.38 | 396.78 | 395.45 | 169.66 | 260.66 | 202.72 |
| 7735 | ENSMUSG00000030254 | 29.58686162 | -0.183127432 | 0.494503182 | -0.370326094 | 0.711139534 | 0.944239215 | Rad18 | RAD18 E3 ubiquitin protein ligase [Source | 24.896 | 52.424 | 17.106 | 21.208 | 22.903 | 38.985 |
| 7736 | ENSMUSG00000030255 | 25.51730369 | -0.323697781 | 0.503731459 | -0.642599891 | 0.520483754 | 0.887802551 | Sspn | sarcospan [Source:MGI Symbol;Acc:MGI:1 | 20.881 | 34.949 | 29.181 | 9.6398 | 27.265 | 31.188 |
| 7737 | ENSMUSG00000030256 | 77.81241048 | 2.186778486 | 0.429169559 | 5.095371843 | 3.48E-07 | 3.19E-05 | Bhlhe41 | basic helix-loop-helix family, member e41 | 42.564 | 25.698 | 15.094 | 79.046 | 86.159 | 218.31 |
| 7738 | ENSMUSG00000030257 | 16.30681964 | 1.378008241 | 0.587612306 | 2.345097654 | 0.019022097 | 0.167561271 | Srgap3 | SLIT-ROBO Rho GTPase activating protein | 8.8341 | 8.2233 | 10.062 | 15.424 | 18.541 | 36.757 |
| 7739 | ENSMUSG00000030259 | 652.1501632 | 0.3910447 | 0.145349082 | 2.690383005 | 0.007137006 | 0.085172977 | Rassf8 | Ras association (RalGDS/AF-6) domain far | 477.85 | 558.16 | 658.08 | 752.87 | 674 | 791.94 |
| 7740 | ENSMUSG00000030263 | 31.3409266 | -0.886753771 | 0.414732219 | -2.138135716 | 0.03250573 | 0.240985585 | Lrmp | lymphoid-restricted membrane protein [S | 38.549 | 53.452 | 30.187 | 25.063 | 19.631 | 21.163 |
| 7741 | ENSMUSG00000030264 | 308.3343402 | 0.114946583 | 0.175848497 | 0.65366827 | 0.51332555 | 0.885004045 | Thumpd3 | THUMP domain containing 3 [Source:MGI | 308.39 | 322.77 | 256.59 | 284.37 | 387.17 | 290.71 |
| 7742 | ENSMUSG00000030265 | 580.9428992 | -0.17781826 | 0.134851489 | -1.318622887 | 0.187295218 | 0.623771135 | Kras | Kirsten rat sarcoma viral oncogene homol | 538.08 | 658.9 | 654.06 | 532.12 | 584.57 | 517.94 |
| 7743 | ENSMUSG00000030268 | 0 | NA | NA | NA | NA | NA | Bcat1 | branched chain aminotransferase 1, cytos | 0 | 0 | 0 | 0 | 0 | 0 |
| 7744 | ENSMUSG00000030269 | 442.9598854 | 0.295227458 | 0.138849018 | 2.126248082 | 0.033482608 | 0.24471574 | Mtmr14 | myotubularin related protein 14 [Source:I | 372.64 | 389.58 | 431.68 | 496.45 | 442.79 | 524.62 |
| 7745 | ENSMUSG00000030270 | 0 | NA | NA | NA | NA | NA | Cpne9 | copine family member IX [Source:MGI Syr | 0 | 0 | 0 | 0 | 0 | 0 |
| 7746 | ENSMUSG00000030271 | 84.28286495 | -0.032320884 | 0.253749721 | -0.127373083 | 0.89864512 | 0.982798755 | Ogg1 | 8-oxoguanine DNA-glycosylase 1 [Source:I | 85.129 | 71.954 | 98.612 | 84.83 | 89.431 | 75.741 |
| 7747 | ENSMUSG00000030272 | 202.1560443 | 0.02263907 | 0.17316001 | 0.130740754 | 0.895980392 | 0.982798755 | Camk1 | calcium/calmodulin-dependent protein ki | 189.53 | 199.42 | 213.32 | 222.68 | 191.95 | 196.04 |
| 7748 | ENSMUSG00000030275 | 1424.504227 | -0.18049514 | 0.10289389 | -1.754187147 | 0.079398445 | 0.413580628 | Etnk1 | ethanolamine kinase 1 [Source:MGI Symb | 1429.5 | 1603.6 | 1508.4 | 1365 | 1270.6 | 1370 |
| 7749 | ENSMUSG00000030276 | 16.21207954 | 0.411753035 | 0.654332726 | 0.629271651 | 0.529171227 | 0.888334265 | Ttll3 | tubulin tyrosine ligase-like family, membe | 18.471 | 8.2233 | 15.094 | 35.667 | 10.906 | 8.9108 |
| 7750 | ENSMUSG00000030278 | 473.0312943 | -1.840272984 | 0.814052066 | -2.260633024 | 0.023781991 | 0.19577844 | Cidec | cell death-inducing DFFA-like effector c [S | 1358 | 679.45 | 181.12 | 293.05 | 201.76 | 124.75 |
| 7751 | ENSMUSG00000030279 | 98.79052207 | 0.030526599 | 0.249529365 | 0.1223367 | 0.902632368 | 0.983821605 | C2cd5 | C2 calcium-dependent domain containing | 91.554 | 115.13 | 86.537 | 88.686 | 89.431 | 121.41 |
| 7752 | ENSMUSG00000030281 | 635.2673375 | -0.073312319 | 0.121672434 | -0.602538448 | 0.546815778 | 0.895509898 | Il17rc | interleukin 17 receptor C [Source:MGI Syr | 627.22 | 691.79 | 635.95 | 643.94 | 640.19 | 572.52 |
| 7753 | ENSMUSG00000030282 | 1050.925252 | 0.105758361 | 0.109493557 | 0.965886612 | 0.334100922 | 0.775900437 | Cmas | cytidine monophospho-N-acetylneuramin | 1085.8 | 1022.8 | 927.76 | 1081.6 | 1096.1 | 1091.6 |
| 7754 | ENSMUSG00000030283 | 2.028999516 | -0.867672368 | 1.663116493 | -0.521714727 | 0.601868969 | NA | St8sia1 | ST8 alpha-N-acetyl-neuraminide alpha-2,8 | 0.8031 | 1.0279 | 6.0375 | 0.964 | 0 | 3.3415 |
| 7755 | ENSMUSG00000030284 | 847.6303369 | 0.272228156 | 0.140312154 | 1.94016091 | 0.052360138 | 0.321586849 | Creld1 | cysteine-rich with EGF-like domains 1 [Sou | 874.58 | 677.4 | 750.66 | 974.58 | 799.42 | 1009.1 |
| 7756 | ENSMUSG00000030286 | 1001.478364 | 0.083327365 | 0.103002348 | 0.808985103 | 0.418523715 | 0.837575303 | Emc3 | ER membrane protein complex subunit 3 [ | 1010.3 | 952.88 | 953.92 | 1029.5 | 1021.9 | 1040.3 |
| 7757 | ENSMUSG00000030287 | 680.0000037 | -0.070882568 | 0.118828963 | -0.596509184 | 0.55083512 | 0.895703112 | Itpr2 | inositol 1,4,5-triphosphate receptor 2 [Sou | 652.12 | 723.65 | 715.44 | 701.78 | 655.46 | 631.55 |
| 7758 | ENSMUSG00000030291 | 226.5820588 | 0.01385313 | 0.161099748 | 0.085991011 | 0.931473563 | 0.988420305 | Med21 | mediator complex subunit 21 [Source:MG | 224.87 | 236.42 | 215.34 | 234.25 | 221.4 | 227.22 |
| 7759 | ENSMUSG00000030292 | 0 | NA | NA | NA | NA | NA | Smco2 | single-pass membrane protein with coiled | 0 | 0 | 0 | 0 | 0 | 0 |
| 7760 | ENSMUSG00000030298 | 1015.283601 | -0.075935652 | 0.108694519 | -0.698615285 | 0.484792489 | 0.87256358 | Sec13 | SEC13 homolog, nuclear pore and COPII c | 1127.6 | 998.11 | 999.2 | 976.51 | 999.01 | 991.32 |
| 7761 | ENSMUSG00000030301 | 491.4773258 | -0.06761864 | 0.134743027 | -0.501834054 | 0.615784256 | 0.917509899 | Ccdc91 | coiled-coil domain containing 91 [Source:I | 547.72 | 447.14 | 513.18 | 464.64 | 501.68 | 474.5 |
| 7762 | ENSMUSG00000030302 | 26.20766482 | -0.36214027 | 0.570151003 | -0.635165542 | 0.525320454 | 0.888334265 | Atp2b2 | ATPase, Ca++ transporting, plasma memb | 31.321 | 39.061 | 18.112 | 29.883 | 5.4531 | 33.415 |
| 7763 | ENSMUSG00000030303 | 22.49995215 | 0.349356495 | 0.573765569 | 0.608883686 | 0.542601538 | 0.894095573 | Far2 | fatty acyl CoA reductase 2 [Source:MGI Sy | 25.699 | 23.642 | 10.062 | 40.487 | 25.084 | 10.025 |
| 7764 | ENSMUSG00000030304 | 686.2957147 | 0.169445592 | 0.128795863 | 1.315613626 | 0.188303758 | 0.625258391 | Ergic2 | ERGIC and golgi 2 [Source:MGI Symbol;Ac | 601.52 | 683.57 | 654.06 | 760.58 | 778.7 | 639.35 |
| 7765 | ENSMUSG00000030306 | 39.75686485 | -0.317330691 | 0.371507255 | -0.854170914 | 0.393010297 | 0.821459672 | Tmtc1 | transmembrane and tetratricopeptide rep | 40.155 | 43.173 | 49.306 | 46.271 | 23.994 | 35.643 |
| 7766 | ENSMUSG00000030307 | 0 | NA | NA | NA | NA | NA | Slc6a11 | solute carrier family 6 (neurotransmitter t | 0 | 0 | 0 | 0 | 0 | 0 |
| 7767 | ENSMUSG00000030309 | 37.3792761 | 0.522068645 | 0.395644194 | 1.319540772 | 0.186988389 | 0.623323359 | Caprin2 | caprin family member 2 [Source:MGI Sym | 28.109 | 27.754 | 36.225 | 36.631 | 63.256 | 32.301 |
| 7768 | ENSMUSG00000030310 | 0.133850373 | -0.796629176 | 4.080472857 | -0.195229623 | 0.84521319 | NA | Slc6a1 | solute carrier family 6 (neurotransmitter t | 0.8031 | 0 | 0 | 0 | 0 | 0 |
| 7769 | ENSMUSG00000030313 | 1401.764098 | 0.105892583 | 0.109322364 | 0.9686269 | 0.332731377 | 0.77523112 | Dennd5b | DENN/MADD domain containing 5B [Sour | 1308.3 | 1498.7 | 1244.7 | 1474.9 | 1451.6 | 1432.4 |
| 7770 | ENSMUSG00000030314 | 813.5379692 | -0.0330567 | 0.108592346 | -0.304410957 | 0.760814821 | 0.95402891 | Atg7 | autophagy related 7 [Source:MGI Symbol; | 837.64 | 828.5 | 801.98 | 783.71 | 791.79 | 837.61 |
| 7771 | ENSMUSG00000030315 | 497.9284956 | 0.091916605 | 0.141216847 | 0.650889794 | 0.51511763 | 0.885627279 | Vgll4 | vestigial like family member 4 [Source:MG | 513.18 | 496.48 | 436.71 | 581.28 | 480.96 | 478.95 |
| 7772 | ENSMUSG00000030316 | 57.33656512 | -0.067108668 | 0.30223212 | -0.222043467 | 0.824280046 | 0.96943211 | Tamm41 | TAM41 mitochondrial translocator asseml | 69.067 | 55.508 | 51.318 | 64.587 | 54.531 | 49.009 |
| 7773 | ENSMUSG00000030317 | 0.503122293 | -2.366675966 | 3.785596405 | -0.625179156 | 0.531853481 | NA | Timp4 | tissue inhibitor of metalloproteinase 4 [So | 0 | 0 | 3.0187 | 0 | 0 | 0 |
| 7774 | ENSMUSG00000030319 | 2.312804064 | -1.207519777 | 1.482067309 | -0.814753668 | 0.415213326 | NA | Cand2 | cullin-associated and neddylation-dissocia | 1.6062 | 3.0838 | 5.0312 | 1.928 | 0 | 2.2277 |
| 7775 | ENSMUSG00000030321 | 2.942577288 | -2.952867063 | 1.597682631 | -1.848218792 | 0.064570698 | NA | Efcab12 | EF-hand calcium binding domain 12 [Sour | 5.6217 | 2.0558 | 8.05 | 1.928 | 0 | 0 |
| 7776 | ENSMUSG00000030322 | 16.02117245 | -0.338120424 | 0.549794766 | -0.614993894 | 0.538558753 | 0.891577636 | Mbd4 | methyl-CpG binding domain protein 4 [So | 16.062 | 16.447 | 21.131 | 10.604 | 19.631 | 12.252 |
| 7777 | ENSMUSG00000030323 | 71.37023833 | 0.16611237 | 0.321157937 | 0.517229534 | 0.604995937 | 0.916376001 | Ift122 | intraflagellar transport 122 [Source:MGI S | 81.916 | 43.173 | 76.475 | 84.83 | 87.249 | 54.578 |
| 7778 | ENSMUSG00000030324 | 0.160662974 | 1.126924175 | 4.080472857 | 0.276174898 | 0.782413736 | NA | Rho | rhodopsin [Source:MGI Symbol;Acc:MGI:9 | 0 | 0 | 0 | 0.964 | 0 | 0 |
| 7779 | ENSMUSG00000030325 | 1.933955351 | -1.364704768 | 1.603715128 | -0.850964579 | 0.39478903 | NA | Klrb1c | killer cell lectin-like receptor subfamily B r | 3.2124 | 3.0838 | 2.0125 | 0 | 2.1812 | 1.1138 |
| 7780 | ENSMUSG00000030327 | 895.2761281 | 0.113545212 | 0.117171654 | 0.969050183 | 0.332520151 | 0.77523112 | Necap1 | NECAP endocytosis associated 1 [Source:I | 889.84 | 775.05 | 914.68 | 918.67 | 892.13 | 981.3 |

| | | | | | | | | | | | | | | | |
|---|---|---|---|---|---|---|---|---|---|---|---|---|---|---|---|
| 7781 | ENSMUSG00000030329 | 2.569923838 | -2.739120955 | 1.854547871 | -1.47697506 | 0.139682323 | NA | Pianp | PILR alpha associated neural protein [Sou | 3.2124 | 10.279 | 0 | 1.928 | 0 | 0 |
| 7782 | ENSMUSG00000030330 | 12.97152291 | 0.119157403 | 0.642203581 | 0.185544594 | 0.852801906 | 0.973781977 | Ing4 | inhibitor of growth family, member 4 [Sou | 12.047 | 8.2233 | 17.106 | 18.316 | 6.5437 | 15.594 |
| 7783 | ENSMUSG00000030335 | 937.0037187 | 0.003709919 | 0.11389536 | 0.032573049 | 0.974015062 | 0.996324688 | Mrpl51 | mitochondrial ribosomal protein L51 [Sou | 950.07 | 951.85 | 904.61 | 853.12 | 952.11 | 1010.3 |
| 7784 | ENSMUSG00000030336 | 5.378962139 | -0.500624402 | 1.059569277 | -0.47247916 | 0.636584813 | 0.923478764 | Cd27 | CD27 antigen [Source:MGI Symbol;Acc:M | 4.8186 | 4.1117 | 10.062 | 7.7118 | 0 | 5.5692 |
| 7785 | ENSMUSG00000030337 | 105.9150658 | 0.309463513 | 0.253701197 | 1.219795239 | 0.222542507 | 0.668407528 | Vamp1 | vesicle-associated membrane protein 1 [S | 110.03 | 68.87 | 104.65 | 134.96 | 114.51 | 102.47 |
| 7786 | ENSMUSG00000030339 | 1897.017515 | 0.197507568 | 0.10190476 | 1.938158402 | 0.0526039 | 0.322665669 | Ltbr | lymphotoxin B receptor [Source:MGI Sym | 1892.1 | 1740.3 | 1668.4 | 1967.5 | 1965.3 | 2148.6 |
| 7787 | ENSMUSG00000030340 | 574.7697528 | 0.216562898 | 0.18529894 | 1.168721731 | 0.242515764 | 0.692000002 | Scnn1a | sodium channel, nonvoltage-gated 1 alph | 383.88 | 571.52 | 640.98 | 597.67 | 714.35 | 540.21 |
| 7788 | ENSMUSG00000030341 | 1328.894401 | 0.09131797 | 0.130151991 | 0.701625604 | 0.48291268 | 0.871710696 | Tnfrsf1a | tumor necrosis factor receptor superfami | 1373.3 | 1258.2 | 1227.6 | 1180.9 | 1329.5 | 1603.9 |
| 7789 | ENSMUSG00000030342 | 125.8851431 | 0.303822382 | 0.244671835 | 1.241754621 | 0.21432711 | 0.658905615 | Cd9 | CD9 antigen [Source:MGI Symbol;Acc:MG | 138.13 | 104.85 | 94.587 | 157.13 | 105.79 | 154.82 |
| 7790 | ENSMUSG00000030344 | 0 | NA | NA | NA | NA | NA | Akap3 | A kinase (PRKA) anchor protein 3 [Source: | 0 | 0 | 0 | 0 | 0 | 0 |
| 7791 | ENSMUSG00000030345 | 0 | NA | NA | NA | NA | NA | Dyrk4 | dual-specificity tyrosine-(Y)-phosphorylati | 0 | 0 | 0 | 0 | 0 | 0 |
| 7792 | ENSMUSG00000030346 | 1.885777285 | -2.095794515 | 1.817350062 | -1.15321454 | 0.248822337 | NA | Rad51ap1 | RAD51 associated protein 1 [Source:MGI S | 4.0155 | 4.1117 | 1.0062 | 0 | 2.1812 | 0 |
| 7793 | ENSMUSG00000030347 | 162.1485911 | -0.185220458 | 0.256541883 | -0.721989158 | 0.470301143 | 0.867104619 | D6Wsu163e | DNA segment, Chr 6, Wayne State Univers | 188.73 | 154.19 | 174.08 | 104.11 | 218.12 | 133.66 |
| 7794 | ENSMUSG00000030350 | 0 | NA | NA | NA | NA | NA | Prmt8 | protein arginine N-methyltransferase 8 [S | 0 | 0 | 0 | 0 | 0 | 0 |
| 7795 | ENSMUSG00000030351 | 0 | NA | NA | NA | NA | NA | Tspan11 | tetraspanin 11 [Source:MGI Symbol;Acc:M | 0 | 0 | 0 | 0 | 0 | 0 |
| 7796 | ENSMUSG00000030352 | 448.4213102 | 0.319529062 | 0.132131154 | 2.418271944 | 0.015594417 | 0.145989013 | Tspan9 | tetraspanin 9 [Source:MGI Symbol;Acc:M | 410.39 | 380.33 | 405.52 | 465.6 | 494.05 | 534.65 |
| 7797 | ENSMUSG00000030353 | 4.488526884 | 0.037263384 | 1.019639141 | 0.036545659 | 0.970847272 | NA | Tead4 | TEA domain family member 4 [Source:MG | 3.2124 | 3.0838 | 7.0437 | 4.8199 | 6.5437 | 2.2277 |
| 7798 | ENSMUSG00000030357 | 3572.147831 | 0.116653466 | 0.101661776 | 1.147466331 | 0.251188941 | 0.7034263 | Fkbp4 | FK506 binding protein 4 [Source:MGI Sym | 3317.6 | 3218.4 | 3748.3 | 3937.8 | 3517.2 | 3693.5 |
| 7799 | ENSMUSG00000030359 | 64285.85609 | 0.236720596 | 0.106667349 | 2.219241386 | 0.026470307 | 0.210708529 | Pzp | PZP, alpha-2-macroglobulin like [Source:M | 52360 | 64616 | 60097 | 72257 | 72713 | 63673 |
| 7800 | ENSMUSG00000030361 | 0 | NA | NA | NA | NA | NA | Klrb1a | killer cell lectin-like receptor subfamily B r | 0 | 0 | 0 | 0 | 0 | 0 |
| 7801 | ENSMUSG00000030363 | 0 | NA | NA | NA | NA | NA | Clec2j | C-type lectin domain family 2, member J [ | 0 | 0 | 0 | 0 | 0 | 0 |
| 7802 | ENSMUSG00000030364 | 150.1811117 | 2.198011693 | 0.216898746 | 10.13381467 | 3.91E-24 | 1.13E-20 | Clec2h | C-type lectin domain family 2, member h [ | 61.839 | 47.284 | 51.318 | 221.71 | 267.2 | 251.73 |
| 7803 | ENSMUSG00000030365 | 5.448318013 | -1.767398549 | 1.104337267 | -1.600415563 | 0.109506425 | 0.486396098 | Clec2i | C-type lectin domain family 2, member i [ | 12.047 | 3.0838 | 10.062 | 1.928 | 0 | 5.5692 |
| 7804 | ENSMUSG00000030366 | 0 | NA | NA | NA | NA | NA | Ceacam12 | carcinoembryonic antigen-related cell adh | 0 | 0 | 0 | 0 | 0 | 0 |
| 7805 | ENSMUSG00000030368 | 0 | NA | NA | NA | NA | NA | Ceacam11 | carcinoembryonic antigen-related cell adh | 0 | 0 | 0 | 0 | 0 | 0 |
| 7806 | ENSMUSG00000030373 | 0 | NA | NA | NA | NA | NA | Psg28 | pregnancy-specific glycoprotein 28 [Sourc | 0 | 0 | 0 | 0 | 0 | 0 |
| 7807 | ENSMUSG00000030374 | 528.7126817 | 0.076929227 | 0.136504924 | 0.56356375 | 0.573051063 | 0.905173631 | Strn4 | striatin, calmodulin binding protein 4 [Sou | 480.26 | 530.41 | 534.32 | 607.31 | 484.23 | 535.76 |
| 7808 | ENSMUSG00000030376 | 0.267700746 | -1.497481649 | 4.057001436 | -0.369110456 | 0.712045396 | NA | Slc8a2 | solute carrier family 8 (sodium/calcium ex | 1.6062 | 0 | 0 | 0 | 0 | 0 |
| 7809 | ENSMUSG00000030378 | 18854.32328 | 0.288208567 | 0.103867117 | 2.774781615 | 0.005523879 | 0.071146377 | Sult2a8 | sulfotransferase family 2A, dehydroepian | 15965 | 16732 | 18235 | 19843 | 23094 | 19256 |
| 7810 | ENSMUSG00000030380 | 12.49393725 | -0.012570341 | 0.726520368 | -0.017302118 | 0.986195596 | 0.99859176 | Mzf1 | myeloid zinc finger 1 [Source:MGI Symbol | 10.44 | 13.363 | 14.087 | 26.027 | 4.3625 | 6.6831 |
| 7811 | ENSMUSG00000030382 | 17049.16772 | 0.03354866 | 0.119853774 | 0.279913257 | 0.779544056 | 0.959870443 | Slc27a5 | solute carrier family 27 (fatty acid transpo | 14635 | 17607 | 18313 | 18531 | 18234 | 14976 |
| 7812 | ENSMUSG00000030385 | 0 | NA | NA | NA | NA | NA | 2900092C05 | RIKEN cDNA 2900092C05 gene [Source:M | 0 | 0 | 0 | 0 | 0 | 0 |
| 7813 | ENSMUSG00000030386 | 139.7361647 | 0.051358643 | 0.209247876 | 0.245444037 | 0.806112635 | 0.964428432 | Zfp606 | zinc finger protein 606 [Source:MGI Symb | 133.31 | 151.1 | 127.79 | 161.95 | 143.96 | 120.3 |
| 7814 | ENSMUSG00000030393 | 6.535038739 | -0.402414655 | 0.887766239 | -0.453288982 | 0.650340665 | 0.925643431 | Zik1 | zinc finger protein interacting with K prot | 12.047 | 3.0838 | 7.0437 | 3.8559 | 8.7249 | 4.4554 |
| 7815 | ENSMUSG00000030397 | 257.8288317 | 0.101193272 | 0.151634208 | 0.667351209 | 0.504547823 | 0.881510796 | Mark4 | MAP/microtubule affinity regulating kinas | 244.14 | 249.78 | 252.57 | 267.02 | 265.02 | 268.44 |
| 7816 | ENSMUSG00000030399 | 0 | NA | NA | NA | NA | NA | Ckm | creatine kinase, muscle [Source:MGI Syml | 0 | 0 | 0 | 0 | 0 | 0 |
| 7817 | ENSMUSG00000030400 | 125.7780838 | 0.008869693 | 0.20772346 | 0.042699522 | 0.96594106 | 0.993767088 | Ercc2 | excision repair cross-complementing rode | 126.09 | 120.27 | 129.81 | 129.17 | 112.33 | 137 |
| 7818 | ENSMUSG00000030401 | 5.55356343 | 0.037470076 | 1.074971812 | 0.034856798 | 0.97219393 | 0.995667563 | Rtn2 | reticulon 2 (Z-band associated protein) [S | 6.4248 | 2.0558 | 8.05 | 13.496 | 2.1812 | 1.1138 |
| 7819 | ENSMUSG00000030402 | 0.51034671 | -2.381743948 | 3.331081356 | -0.715006238 | 0.474605189 | NA | Ppm1n | protein phosphatase, Mg2+/Mn2+ depen | 0 | 2.0558 | 1.0062 | 0 | 0 | 0 |
| 7820 | ENSMUSG00000030403 | 255.2234412 | -0.217906982 | 0.193523938 | -1.125994976 | 0.260167646 | 0.713728596 | Vasp | vasodilator-stimulated phosphoprotein [S | 244.14 | 332.02 | 247.54 | 228.46 | 198.49 | 280.69 |
| 7821 | ENSMUSG00000030406 | 0 | NA | NA | NA | NA | NA | Gipr | gastric inhibitory polypeptide receptor [S | 0 | 0 | 0 | 0 | 0 | 0 |
| 7822 | ENSMUSG00000030407 | 196.4691501 | -0.014619489 | 0.175413046 | -0.083343226 | 0.93357863 | 0.988420305 | Qpctl | glutaminyl-peptide cyclotransferase-like [ | 190.34 | 188.11 | 214.33 | 208.22 | 198.49 | 179.33 |
| 7823 | ENSMUSG00000030409 | 243.0652727 | -1.292378194 | 0.212678335 | -6.076680036 | 1.23E-09 | 2.50E-07 | Dmpk | dystrophia myotonica-protein kinase [Sou | 294.74 | 438.92 | 302.88 | 171.59 | 121.06 | 129.21 |
| 7824 | ENSMUSG00000030410 | 400.1857373 | 0.281836311 | 0.137418971 | 2.050927235 | 0.040274033 | 0.274645742 | Dmwd | dystrophia myotonica-containing WD rep | 359.79 | 359.77 | 364.26 | 456.93 | 395.89 | 464.47 |
| 7825 | ENSMUSG00000030411 | 12.2344691 | 0.449838255 | 0.665531516 | 0.675908269 | 0.499098888 | 0.880361745 | Nova2 | NOVA alternative splicing regulator 2 [Sou | 4.8186 | 11.307 | 15.094 | 12.532 | 19.631 | 10.025 |
| 7826 | ENSMUSG00000030413 | 2.73401906 | -2.82732167 | 1.649430404 | -1.714120015 | 0.086506687 | NA | Pglyrp1 | peptidoglycan recognition protein 1 [Sour | 3.2124 | 9.2513 | 2.0125 | 1.928 | 0 | 0 |
| 7827 | ENSMUSG00000030417 | 382.5489984 | 0.035219703 | 0.139283569 | 0.2528633 | 0.800373851 | 0.962424862 | Pdcd5 | programmed cell death 5 [Source:MGI Syr | 379.06 | 378.27 | 376.34 | 397.16 | 349 | 415.46 |
| 7828 | ENSMUSG00000030421 | 685.8174219 | 0.149814288 | 0.116729132 | 1.283435297 | 0.199339614 | 0.641417519 | Uri1 | URI1, prefoldin-like chaperone [Source:M | 648.1 | 666.09 | 636.95 | 753.83 | 741.62 | 668.31 |
| 7829 | ENSMUSG00000030423 | 283.5151923 | 0.33152335 | 0.157894698 | 2.099648398 | 0.035759782 | 0.253977709 | Pop4 | processing of precursor 4, ribonuclease P/ | 263.42 | 240.53 | 248.54 | 281.48 | 317.37 | 349.75 |
| 7830 | ENSMUSG00000030424 | 44.22143501 | -0.248818517 | 0.389060157 | -0.63953739 | 0.522473398 | 0.888334265 | Zfp939 | zinc finger protein 939 [Source:MGI Symb | 36.943 | 45.228 | 62.387 | 57.839 | 26.175 | 36.757 |
| 7831 | ENSMUSG00000030427 | 0.692116425 | -1.412606127 | 2.773087916 | -0.509398248 | 0.610473104 | NA | Lilra6 | leukocyte immunoglobulin-like receptor, s | 0 | 2.0558 | 1.0062 | 0 | 1.0906 | 0 |
| 7832 | ENSMUSG00000030428 | 19.01264325 | -0.526913923 | 0.525706532 | -1.002296702 | 0.316200315 | 0.762889301 | Ttyh1 | tweety family member 1 [Source:MGI Sym | 26.502 | 17.475 | 23.144 | 11.568 | 11.997 | 23.391 |
| 7833 | ENSMUSG00000030431 | 519.2941543 | 0.155142927 | 0.142640315 | 1.087651318 | 0.276749066 | 0.730041765 | Tmem238 | transmembrane protein 238 [Source:MGI | 450.54 | 454.34 | 569.53 | 505.12 | 570.39 | 565.83 |
| 7834 | ENSMUSG00000030432 | 2783.754945 | -0.033758786 | 0.110728158 | -0.304879869 | 0.760457648 | 0.95402891 | Rpl28 | ribosomal protein L28 [Source:MGI Symbo | 2546.6 | 3196.8 | 2706.8 | 2777.2 | 2643.7 | 2831.4 |
| 7835 | ENSMUSG00000030433 | 0 | NA | NA | NA | NA | NA | Sbk2 | SH3-binding domain kinase family, memb | 0 | 0 | 0 | 0 | 0 | 0 |
| 7836 | ENSMUSG00000030435 | 1017.093871 | 0.017709152 | 0.106472622 | 0.16632587 | 0.867900508 | 0.978326885 | U2af2 | U2 small nuclear ribonucleoprotein auxili | 970.95 | 1078.3 | 984.11 | 1047.8 | 1005.6 | 1015.8 |

| | | | | | | | | | | | | | | | |
|---|---|---|---|---|---|---|---|---|---|---|---|---|---|---|---|
| 7837 | ENSMUSG00000030443 | 1.384982393 | 0.827058939 | 1.972217336 | 0.419354867 | 0.674956803 | NA | Zfp583 | zinc finger protein 583 [Source:MGI Symb | 0 | 1.0279 | 2.0125 | 1.928 | 0 | 3.3415 |
| 7838 | ENSMUSG00000030446 | 46.16463705 | 0.437873394 | 0.340051861 | 1.28766651 | 0.197862076 | 0.639811551 | Zfp273 | zinc finger protein 273 [Source:MGI Symb | 34.533 | 32.893 | 50.312 | 51.091 | 46.897 | 61.261 |
| 7839 | ENSMUSG00000030447 | 749.4831915 | -0.121864233 | 0.124079704 | -0.98214477 | 0.326028532 | 0.772018857 | Cyfip1 | cytoplasmic FMR1 interacting protein 1 [S | 767.77 | 733.93 | 841.22 | 653.58 | 781.97 | 718.43 |
| 7840 | ENSMUSG00000030450 | 0 | NA | NA | NA | NA | NA | Oca2 | oculocutaneous albinism II [Source:MGI S | 0 | 0 | 0 | 0 | 0 | 0 |
| 7841 | ENSMUSG00000030451 | 717.1763383 | -0.011093569 | 0.121954922 | -0.090964502 | 0.927520797 | 0.987825698 | Herc2 | HECT and RLD domain containing E3 ubiqu | 676.21 | 724.68 | 759.71 | 746.12 | 647.83 | 748.5 |
| 7842 | ENSMUSG00000030452 | 371.8686547 | -0.002341031 | 0.164022234 | -0.014272643 | 0.988612465 | 0.99859176 | Nipa2 | non imprinted in Prader-Willi/Angelman s | 358.99 | 352.58 | 404.51 | 304.62 | 435.16 | 375.37 |
| 7843 | ENSMUSG00000030463 | 0 | NA | NA | NA | NA | NA | 4933421I07R | RIKEN cDNA 4933421I07 gene [Source:MC | 0 | 0 | 0 | 0 | 0 | 0 |
| 7844 | ENSMUSG00000030465 | 885.5642976 | 0.037909452 | 0.142067077 | 0.266841924 | 0.789590874 | 0.961009963 | Psd3 | pleckstrin and Sec7 domain containing 3 [ | 807.92 | 944.66 | 870.4 | 988.08 | 976.1 | 726.23 |
| 7845 | ENSMUSG00000030468 | 2.952787742 | 1.41259024 | 1.281348136 | 1.102425016 | 0.270276941 | NA | Siglecg | sialic acid binding Ig-like lectin G [Source:I | 0.8031 | 3.0838 | 1.0062 | 2.8919 | 4.3625 | 5.5692 |
| 7846 | ENSMUSG00000030469 | 58.14059333 | 0.236578833 | 0.308406123 | 0.767101607 | 0.443021107 | 0.850769625 | Zfp719 | zinc finger protein 719 [Source:MGI Symb | 45.777 | 54.48 | 60.375 | 78.082 | 58.893 | 51.237 |
| 7847 | ENSMUSG00000030470 | 712.2556384 | -0.253155284 | 0.187841457 | -1.347707198 | 0.177752576 | 0.609100512 | Csrp3 | cysteine and glycine-rich protein 3 [Sourc | 1037.6 | 721.6 | 563.5 | 696.96 | 642.37 | 611.5 |
| 7848 | ENSMUSG00000030471 | 118.0330988 | -0.032348274 | 0.257844049 | -0.125456742 | 0.900161969 | 0.982805511 | Zdhhc13 | zinc finger, DHHC domain containing 13 [S | 120.47 | 121.29 | 116.72 | 154.24 | 78.525 | 116.95 |
| 7849 | ENSMUSG00000030472 | 0 | NA | NA | NA | NA | NA | Ceacam18 | carcinoembryonic antigen-related cell adh | 0 | 0 | 0 | 0 | 0 | 0 |
| 7850 | ENSMUSG00000030474 | 35.37684921 | -1.832608992 | 0.518532655 | -3.53422099 | 0.000408979 | 0.010774994 | Siglece | sialic acid binding Ig-like lectin E [Source:N | 56.217 | 84.289 | 25.156 | 14.46 | 7.6343 | 24.505 |
| 7851 | ENSMUSG00000030483 | 126.2873501 | -3.829390628 | 0.353402113 | -10.83578871 | 2.33E-27 | 1.12E-23 | Cyp2b10 | cytochrome P450, family 2, subfamily b, p | 169.45 | 186.05 | 352.19 | 12.532 | 17.45 | 20.049 |
| 7852 | ENSMUSG00000030484 | 0 | NA | NA | NA | NA | NA | Lypd5 | Ly6/Plaur domain containing 5 [Source:M | 0 | 0 | 0 | 0 | 0 | 0 |
| 7853 | ENSMUSG00000030486 | 35.46840321 | 0.265079477 | 0.376007236 | 0.70498504 | 0.480819546 | 0.870278888 | Zfp108 | zinc finger protein 108 [Source:MGI Symb | 29.715 | 25.698 | 41.256 | 35.667 | 39.262 | 41.212 |
| 7854 | ENSMUSG00000030491 | 0.133850373 | -0.796629176 | 4.080472857 | -0.195229623 | 0.84521319 | NA | Tdrd12 | tudor domain containing 12 [Source:MGI | 0.8031 | 0 | 0 | 0 | 0 | 0 |
| 7855 | ENSMUSG00000030492 | 0.133850373 | -0.796629176 | 4.080472857 | -0.195229623 | 0.84521319 | NA | Slc7a9 | solute carrier family 7 (cationic amino acid | 0.8031 | 0 | 0 | 0 | 0 | 0 |
| 7856 | ENSMUSG00000030493 | 44.70646305 | -0.523093778 | 0.364103886 | -1.436660793 | 0.1508144 | 0.565124219 | Faap24 | Fanconi anemia core complex associated | 48.186 | 50.368 | 59.368 | 23.135 | 38.172 | 49.009 |
| 7857 | ENSMUSG00000030494 | 298.8995507 | -0.208585042 | 0.195513631 | -1.066856775 | 0.286036516 | 0.738507914 | Rhpn2 | rhophilin, Rho GTPase binding protein 2 [S | 349.35 | 237.45 | 374.32 | 295.94 | 234.48 | 301.85 |
| 7858 | ENSMUSG00000030495 | 0 | NA | NA | NA | NA | NA | Slc7a10 | solute carrier family 7 (cationic amino acid | 0 | 0 | 0 | 0 | 0 | 0 |
| 7859 | ENSMUSG00000030498 | 733.9178043 | 0.10880459 | 0.121817433 | 0.893177503 | 0.371762124 | 0.807685567 | Gas2 | growth arrest specific 2 [Source:MGI Syml | 734.04 | 725.71 | 658.08 | 696.96 | 766.7 | 822.02 |
| 7860 | ENSMUSG00000030499 | 189.7578033 | 0.298338764 | 0.203572453 | 1.465516383 | 0.142780086 | 0.552480247 | Kctd15 | potassium channel tetramerisation domai | 147.77 | 196.33 | 167.04 | 211.11 | 170.14 | 246.16 |
| 7861 | ENSMUSG00000030500 | 0 | NA | NA | NA | NA | NA | Slc17a6 | solute carrier family 17 (sodium-depender | 0 | 0 | 0 | 0 | 0 | 0 |
| 7862 | ENSMUSG00000030505 | 115.5605756 | -0.258202426 | 0.23907367 | -1.080011975 | 0.280136847 | 0.733108114 | Prmt3 | protein arginine N-methyltransferase 3 [S | 103.6 | 149.05 | 125.78 | 124.35 | 99.246 | 91.335 |
| 7863 | ENSMUSG00000030507 | 0 | NA | NA | NA | NA | NA | Dbx1 | developing brain homeobox 1 [Source:MC | 0 | 0 | 0 | 0 | 0 | 0 |
| 7864 | ENSMUSG00000030509 | 240.5198819 | 0.269922453 | 0.162552269 | 1.660527135 | 0.096808454 | 0.458530879 | Asb7 | ankyrin repeat and SOCS box-containing 7 | 230.49 | 216.89 | 206.28 | 245.81 | 257.39 | 286.26 |
| 7865 | ENSMUSG00000030510 | 0.133850373 | -0.796629176 | 4.080472857 | -0.195229623 | 0.84521319 | NA | Cers3 | ceramide synthase 3 [Source:MGI Symbol | 0.8031 | 0 | 0 | 0 | 0 | 0 |
| 7866 | ENSMUSG00000030512 | 193.2989032 | 0.122778332 | 0.181567869 | 0.676211781 | 0.498906194 | 0.880236544 | Snrpa1 | small nuclear ribonucleoprotein polypepti | 200.78 | 177.83 | 176.09 | 188.94 | 229.03 | 187.13 |
| 7867 | ENSMUSG00000030513 | 2654.76478 | -0.244246544 | 0.115017746 | -2.123555308 | 0.033707346 | 0.245744069 | Pcsk6 | proprotein convertase subtilisin/kexin typ | 2814.1 | 2907 | 2917.1 | 2714.6 | 2499.7 | 2076.2 |
| 7868 | ENSMUSG00000030515 | 297.9928354 | -0.050669828 | 0.162166906 | -0.312454801 | 0.754694908 | 0.953048681 | Tarsl2 | threonyl-tRNA synthetase-like 2 [Source:N | 293.94 | 332.02 | 283.76 | 272.81 | 337 | 268.44 |
| 7869 | ENSMUSG00000030516 | 637.7706229 | 0.04678346 | 0.12906607 | 0.36247683 | 0.716995735 | 0.9447719 | Tjp1 | tight junction protein 1 [Source:MGI Symb | 569.4 | 675.34 | 638.97 | 707.56 | 622.74 | 612.61 |
| 7870 | ENSMUSG00000030518 | 1.555836718 | 1.905225419 | 1.802547931 | 1.056962418 | 0.29052874 | NA | Fam189a1 | family with sequence similarity 189, meml | 0 | 1.0279 | 1.0062 | 2.8919 | 2.1812 | 2.2277 |
| 7871 | ENSMUSG00000030519 | 3.842649329 | -0.741869465 | 1.12598732 | -0.658861296 | 0.509984841 | NA | Apba2 | amyloid beta (A4) precursor protein-bindi | 3.2124 | 6.1675 | 5.0312 | 0.964 | 5.4531 | 2.2277 |
| 7872 | ENSMUSG00000030521 | 209.996807 | -0.238906636 | 0.174876926 | -1.366141556 | 0.171894537 | 0.599537638 | Mphosph10 | M-phase phosphoprotein 10 (U3 small nu | 225.67 | 237.45 | 219.36 | 216.9 | 170.14 | 190.47 |
| 7873 | ENSMUSG00000030522 | 754.3847452 | -0.19400345 | 0.125627705 | -1.544272818 | 0.122522255 | 0.51297445 | Mtmr10 | myotubularin related protein 10 [Source:I | 779.01 | 885.04 | 751.66 | 722.02 | 750.35 | 638.23 |
| 7874 | ENSMUSG00000030523 | 0.922347905 | -1.985378883 | 2.314342092 | -0.857858866 | 0.3909704 | NA | Trpm1 | transient receptor potential cation channe | 2.4093 | 1.0279 | 1.0062 | 0 | 1.0906 | 0 |
| 7875 | ENSMUSG00000030525 | 1.883896265 | 0.294826654 | 1.940862371 | 0.151904977 | 0.879261883 | NA | Chrna7 | cholinergic receptor, nicotinic, alpha poly | 0 | 5.1396 | 0 | 2.8919 | 3.2719 | 0 |
| 7876 | ENSMUSG00000030527 | 195.608944 | 0.021254467 | 0.199349572 | 0.106619078 | 0.915091183 | 0.98625062 | Crtc3 | CREB regulated transcription coactivator 3 | 188.73 | 230.25 | 164.02 | 225.57 | 169.05 | 196.04 |
| 7877 | ENSMUSG00000030528 | 251.5516207 | 0.374610711 | 0.280398469 | 1.335994139 | 0.181551189 | 0.615095921 | Blm | Bloom syndrome, RecQ like helicase [Sour | 272.25 | 181.94 | 202.26 | 198.58 | 449.33 | 204.95 |
| 7878 | ENSMUSG00000030530 | 5532.29575 | -0.080173313 | 0.099406502 | -0.806519803 | 0.419943194 | 0.837906635 | Furin | furin (paired basic amino acid cleaving en | 5588.8 | 5995.8 | 5473 | 5094.6 | 5124.8 | 5916.7 |
| 7879 | ENSMUSG00000030532 | 12.20773367 | -1.091775275 | 0.648148573 | -1.684452177 | 0.092094314 | 0.44699737 | Hddc3 | HD domain containing 3 [Source:MGI Sym | 19.274 | 11.307 | 19.119 | 4.8199 | 9.8156 | 8.9108 |
| 7880 | ENSMUSG00000030533 | 292.0778306 | -0.237052542 | 0.148451217 | -1.596837982 | 0.110301827 | 0.488191174 | Unc45a | unc-45 myosin chaperone A [Source:MGI | 326.06 | 317.63 | 303.89 | 251.6 | 274.84 | 278.46 |
| 7881 | ENSMUSG00000030534 | 116.8102098 | 0.284735324 | 0.222500228 | 1.279708012 | 0.200647846 | 0.643925103 | Vps33b | vacuolar protein sorting 33B [Source:MGI | 97.175 | 120.27 | 98.612 | 115.68 | 124.33 | 144.8 |
| 7882 | ENSMUSG00000030536 | 109.0022466 | -1.272688808 | 0.249728598 | -5.096287808 | 3.46E-07 | 3.19E-05 | Iqgap1 | IQ motif containing GTPase activating prot | 129.3 | 184 | 149.93 | 76.154 | 56.712 | 57.92 |
| 7883 | ENSMUSG00000030538 | 986.4134021 | 0.010713789 | 0.112699145 | 0.095065401 | 0.924262879 | 0.987482515 | Cib1 | calcium and integrin binding 1 (calmyrin) [ | 1051.3 | 960.08 | 935.81 | 962.05 | 945.57 | 1063.7 |
| 7884 | ENSMUSG00000030539 | 94.55370795 | -0.260693699 | 0.247849576 | -1.051822252 | 0.292881108 | 0.743850082 | Sema4b | sema domain, immunoglobulin domain (Ig | 118.06 | 88.401 | 102.64 | 96.398 | 89.431 | 72.4 |
| 7885 | ENSMUSG00000030541 | 1889.721406 | -0.477770363 | 0.097570426 | -4.896671889 | 9.75E-07 | 8.01E-05 | Idh2 | isocitrate dehydrogenase 2 (NADP+), mitc | 2196.5 | 2299.5 | 2104.1 | 1655.1 | 1513.8 | 1569.4 |
| 7886 | ENSMUSG00000030543 | 5.77337327 | -0.401400871 | 1.053520676 | -0.381009011 | 0.703196562 | 0.942506832 | Mesp2 | mesoderm posterior 2 [Source:MGI Symb | 14.456 | 1.0279 | 4.025 | 1.928 | 7.6343 | 5.5692 |
| 7887 | ENSMUSG00000030544 | 1.318405796 | 0.858286845 | 1.874838376 | 0.457792446 | 0.647101564 | NA | Mesp1 | mesoderm posterior 1 [Source:MGI Symb | 0.8031 | 1.0279 | 1.0062 | 2.8919 | 2.1812 | 0 |
| 7888 | ENSMUSG00000030545 | 3881.173633 | -0.729904895 | 0.182938719 | -3.98988742 | 6.61E-05 | 0.002591159 | Pex11a | peroxisomal biogenesis factor 11 alpha [S | 5023.4 | 5537.4 | 3967.6 | 3477.1 | 3308.9 | 1972.6 |
| 7889 | ENSMUSG00000030546 | 0 | NA | NA | NA | NA | NA | Plin1 | perilipin 1 [Source:MGI Symbol;Acc:MGI:1 | 0 | 0 | 0 | 0 | 0 | 0 |
| 7890 | ENSMUSG00000030549 | 0 | NA | NA | NA | NA | NA | Rhcg | Rhesus blood group-associated C glycopro | 0 | 0 | 0 | 0 | 0 | 0 |
| 7891 | ENSMUSG00000030551 | 577.9179002 | 0.290095682 | 0.142089225 | 2.041644477 | 0.04118681 | 0.278116723 | Nr2f2 | nuclear receptor subfamily 2, group F, me | 529.24 | 507.79 | 522.24 | 562 | 594.39 | 751.84 |
| 7892 | ENSMUSG00000030553 | 4.067258099 | 0.545969019 | 1.133706869 | 0.481578646 | 0.630105299 | NA | Pgpep1l | pyroglutamyl-peptidase I-like [Source:MG | 4.8186 | 1.0279 | 4.025 | 5.7839 | 7.6343 | 1.1138 |

| | | | | | | | | | | | | | | | |
|---|---|---|---|---|---|---|---|---|---|---|---|---|---|---|---|
| 7893 | ENSMUSG00000030554 | 33.12924876 | -0.251906389 | 0.40189625 | -0.626794574 | 0.5307939 | 0.888794797 | Synm | synemin, intermediate filament protein [S | 32.124 | 32.893 | 43.269 | 36.631 | 33.809 | 20.049 |
| 7894 | ENSMUSG00000030555 | 1502.458942 | -0.455194736 | 0.141587338 | -3.214939587 | 0.00130472 | 0.025467567 | Ttc23 | tetratricopeptide repeat domain 23 [Sour | 1930.7 | 1829.7 | 1452 | 1328.4 | 1403.6 | 1070.4 |
| 7895 | ENSMUSG00000030556 | 690.6091087 | -0.426925715 | 0.14798924 | -2.884842938 | 0.00391609 | 0.056472904 | Lrrc28 | leucine rich repeat containing 28 [Source: | 917.95 | 789.44 | 668.15 | 612.13 | 635.83 | 520.17 |
| 7896 | ENSMUSG00000030557 | 494.6278183 | -0.030094841 | 0.141918352 | -0.212057428 | 0.832062232 | 0.970533397 | Mef2a | myocyte enhancer factor 2A [Source:MGI | 509.97 | 553.02 | 436.71 | 527.3 | 490.78 | 449.99 |
| 7897 | ENSMUSG00000030559 | 7.132996292 | 0.717797231 | 0.858738924 | 0.835873641 | 0.403225992 | 0.828953151 | Rab38 | RAB38, member RAS oncogene family [So | 4.0155 | 6.1675 | 6.0375 | 7.7118 | 3.2719 | 15.594 |
| 7898 | ENSMUSG00000030560 | 1183.221094 | -0.17315528 | 0.1638605 | -1.056723732 | 0.290637691 | 0.740463951 | Ctsc | cathepsin C [Source:MGI Symbol;Acc:MGI | 1017.5 | 1366.1 | 1380.6 | 1357.3 | 905.21 | 1072.6 |
| 7899 | ENSMUSG00000030562 | 1307.375075 | 0.423666251 | 0.123546253 | 3.429211664 | 0.000605337 | 0.014236745 | Nox4 | NADPH oxidase 4 [Source:MGI Symbol;Ac | 960.51 | 1116.3 | 1274.9 | 1469.1 | 1548.7 | 1474.7 |
| 7900 | ENSMUSG00000030577 | 13.02612289 | -1.741224095 | 0.685382345 | -2.540514953 | 0.011068936 | 0.116691757 | Cd22 | CD22 antigen [Source:MGI Symbol;Acc:M | 11.243 | 32.893 | 16.1 | 5.7839 | 5.4531 | 6.6831 |
| 7901 | ENSMUSG00000030579 | 89.66511602 | -1.160866186 | 0.311267461 | -3.729481336 | 0.000191874 | 0.006099495 | Tyrobp | TYRO protein tyrosine kinase binding prot | 111.63 | 152.13 | 107.67 | 40.487 | 42.534 | 83.538 |
| 7902 | ENSMUSG00000030583 | 148.1337739 | 0.458310103 | 0.200479041 | 2.286074891 | 0.022249878 | 0.186455522 | Sipa1l3 | signal-induced proliferation-associated 1 l | 122.87 | 142.88 | 108.67 | 162.91 | 181.04 | 170.42 |
| 7903 | ENSMUSG00000030584 | 2.607698977 | 1.628011946 | 1.529200433 | 1.064616457 | 0.28704953 | NA | Dpf1 | D4, zinc and double PHD fingers family 1 [ | 0.8031 | 1.0279 | 2.0125 | 2.8919 | 0 | 8.9108 |
| 7904 | ENSMUSG00000030587 | 182.7605327 | 0.449553388 | 0.243242783 | 1.848167425 | 0.064578126 | 0.364013257 | 2200002D01 | RIKEN cDNA 2200002D01 gene [Source:M | 160.62 | 116.15 | 186.16 | 149.42 | 223.58 | 260.64 |
| 7905 | ENSMUSG00000030588 | 418.8327537 | 0.231788674 | 0.140727619 | 1.647073093 | 0.099543023 | 0.464748317 | Yif1b | Yip1 interacting factor homolog B (S. cere | 347.74 | 424.53 | 384.39 | 459.82 | 424.25 | 472.27 |
| 7906 | ENSMUSG00000030589 | 3.121154017 | -1.114861913 | 1.549400915 | -0.719543859 | 0.471805889 | NA | Rasgrp4 | RAS guanyl releasing protein 4 [Source:M | 0.8031 | 3.0838 | 9.0562 | 5.7839 | 0 | 0 |
| 7907 | ENSMUSG00000030590 | 79.46715845 | 0.425193261 | 0.309579722 | 1.373453204 | 0.1696115 | 0.596876778 | Fam98c | family with sequence similarity 98, memb | 53.808 | 71.954 | 78.487 | 127.25 | 80.706 | 64.603 |
| 7908 | ENSMUSG00000030591 | 1960.824869 | 0.076903219 | 0.092148855 | 0.834554257 | 0.403968726 | 0.829030031 | Psmd8 | proteasome (prosome, macropain) 26S su | 1940.3 | 1927.3 | 1857.5 | 1955.9 | 2067.8 | 2016.1 |
| 7909 | ENSMUSG00000030592 | 1.220137628 | -1.233564006 | 2.542691384 | -0.485141065 | 0.627576284 | NA | Ryr1 | ryanodine receptor 1, skeletal muscle [Sou | 0 | 5.1396 | 0 | 0 | 2.1812 | 0 |
| 7910 | ENSMUSG00000030595 | 367.3237694 | 0.062863276 | 0.170289831 | 0.369154611 | 0.712012485 | 0.944239215 | Nfkbib | nuclear factor of kappa light polypeptide g | 408.78 | 359.77 | 308.92 | 406.8 | 307.55 | 412.12 |
| 7911 | ENSMUSG00000030598 | 18.93536549 | 0.657006819 | 0.522166385 | 1.258232696 | 0.208307614 | 0.653455953 | Fbxo17 | F-box protein 17 [Source:MGI Symbol;Acc | 8.8341 | 14.391 | 21.131 | 24.099 | 23.994 | 21.163 |
| 7912 | ENSMUSG00000030600 | 5.400335451 | 1.202036511 | 1.072106758 | 1.121191058 | 0.262206545 | 0.716133499 | Lrfn1 | leucine rich repeat and fibronectin type III | 5.6217 | 0 | 4.025 | 2.8919 | 8.7249 | 11.138 |
| 7913 | ENSMUSG00000030602 | 214.5297348 | 0.21410638 | 0.181907556 | 1.177006526 | 0.239192902 | 0.688631793 | Pak4 | p21 (RAC1) activated kinase 4 [Source:MG | 191.14 | 197.36 | 207.29 | 210.15 | 206.13 | 275.12 |
| 7914 | ENSMUSG00000030603 | 1331.898054 | -0.035484671 | 0.120730314 | -0.293916827 | 0.76882146 | 0.956003576 | Psmc4 | proteasome (prosome, macropain) 26S su | 1543.6 | 1291.1 | 1208.5 | 1271.5 | 1297.8 | 1378.9 |
| 7915 | ENSMUSG00000030604 | 114.726051 | 0.384665426 | 0.229775531 | 1.67409221 | 0.094112479 | 0.451490179 | Zfp626 | zinc finger protein 626 [Source:MGI Symb | 112.43 | 83.261 | 102.64 | 145.56 | 131.96 | 112.5 |
| 7916 | ENSMUSG00000030605 | 158.2715271 | -0.926365714 | 0.201057486 | -4.60746691 | 4.08E-06 | 0.000258578 | Mfge8 | milk fat globule EGF and factor V/VIII dom | 221.66 | 204.56 | 196.22 | 125.32 | 91.612 | 110.27 |
| 7917 | ENSMUSG00000030606 | 0 | NA | NA | NA | NA | NA | Hapln3 | hyaluronan and proteoglycan link protein | 0 | 0 | 0 | 0 | 0 | 0 |
| 7918 | ENSMUSG00000030607 | 0 | NA | NA | NA | NA | NA | Acan | aggrecan [Source:MGI Symbol;Acc:MGI:9 | 0 | 0 | 0 | 0 | 0 | 0 |
| 7919 | ENSMUSG00000030609 | 263.9816415 | -0.058977378 | 0.163686677 | -0.36030653 | 0.718617917 | 0.9447719 | Aen | apoptosis enhancing nuclease [Source:M | 284.3 | 238.48 | 284.77 | 245.81 | 245.39 | 285.14 |
| 7920 | ENSMUSG00000030610 | 183.4976122 | -0.306198585 | 0.204851811 | -1.494732137 | 0.134984364 | 0.537616739 | Det1 | de-etiolated homolog 1 (Arabidopsis) [Sou | 246.55 | 203.53 | 157.98 | 167.73 | 159.23 | 165.96 |
| 7921 | ENSMUSG00000030611 | 264.3056049 | 0.0218331 | 0.152403422 | 0.143258595 | 0.886085955 | 0.982222482 | Mrps11 | mitochondrial ribosomal protein S11 [Sou | 272.25 | 249.78 | 264.64 | 267.02 | 274.84 | 257.3 |
| 7922 | ENSMUSG00000030612 | 368.9155351 | 0.105903196 | 0.14624503 | 0.724149029 | 0.468974247 | 0.865872289 | Mrpl46 | mitochondrial ribosomal protein L46 [Sou | 353.36 | 343.32 | 369.29 | 374.02 | 343.54 | 429.94 |
| 7923 | ENSMUSG00000030613 | 119.8531153 | -0.095261764 | 0.209461431 | -0.454793819 | 0.649257577 | 0.925643431 | Ccdc90b | coiled-coil domain containing 90B [Source | 133.31 | 114.1 | 123.77 | 117.61 | 115.61 | 114.73 |
| 7924 | ENSMUSG00000030614 | 557.9122033 | 0.07249378 | 0.141594961 | 0.511979941 | 0.608665049 | 0.916376001 | Tmem126b | transmembrane protein 126B [Source:MG | 485.07 | 520.13 | 626.89 | 525.37 | 607.47 | 582.54 |
| 7925 | ENSMUSG00000030615 | 571.3021919 | -0.038857536 | 0.117978926 | -0.329359976 | 0.741883616 | 0.948939213 | Tmem126a | transmembrane protein 126A [Source:MG | 573.41 | 582.83 | 580.6 | 542.72 | 581.3 | 566.95 |
| 7926 | ENSMUSG00000030616 | 8.079596537 | -0.655239627 | 0.835535343 | -0.784215333 | 0.43291377 | 0.845264432 | Sytl2 | synaptotagmin-like 2 [Source:MGI Symbol | 19.274 | 6.1675 | 4.025 | 5.7839 | 6.5437 | 6.6831 |
| 7927 | ENSMUSG00000030617 | 0 | NA | NA | NA | NA | NA | Ccdc83 | coiled-coil domain containing 83 [Source:I | 0 | 0 | 0 | 0 | 0 | 0 |
| 7928 | ENSMUSG00000030619 | 295.3432425 | 0.221897006 | 0.160434999 | 1.383096005 | 0.16663543 | 0.592715132 | Eed | embryonic ectoderm development [Sourc | 277.07 | 266.23 | 274.7 | 319.08 | 362.09 | 272.89 |
| 7929 | ENSMUSG00000030621 | 1.919686477 | -0.786714795 | 1.516205739 | -0.518870741 | 0.603850883 | NA | Me3 | malic enzyme 3, NADP(+)-dependent, mit | 3.2124 | 1.0279 | 3.0187 | 0.964 | 2.1812 | 1.1138 |
| 7930 | ENSMUSG00000030623 | 0 | NA | NA | NA | NA | NA | Prss23os | protease, serine 23, opposite strand [Sour | 0 | 0 | 0 | 0 | 0 | 0 |
| 7931 | ENSMUSG00000030629 | 2310.007837 | 0.003724523 | 0.092685203 | 0.040184653 | 0.967945913 | 0.994231936 | Zfand6 | zinc finger, AN1-type domain 6 [Source:M | 2243.1 | 2323.1 | 2355.6 | 2343.4 | 2395 | 2199.8 |
| 7932 | ENSMUSG00000030630 | 11953.11011 | 0.004952783 | 0.083954733 | 0.058993498 | 0.952957287 | 0.9915542 | Fah | fumarylacetoacetate hydrolase [Source:M | 12285 | 11634 | 11878 | 11573 | 12518 | 11831 |
| 7933 | ENSMUSG00000030636 | 0 | NA | NA | NA | NA | NA | 1700010L04I | RIKEN cDNA 1700010L04 gene [Source:M | 0 | 0 | 0 | 0 | 0 | 0 |
| 7934 | ENSMUSG00000030638 | 0 | NA | NA | NA | NA | NA | Sh3gl3 | SH3-domain GRB2-like 3 [Source:MGI Sym | 0 | 0 | 0 | 0 | 0 | 0 |
| 7935 | ENSMUSG00000030641 | 14.34922105 | -0.372818331 | 0.595480886 | -0.626079426 | 0.531262846 | 0.888877582 | Ddias | DNA damage-induced apoptosis suppress | 20.078 | 13.363 | 15.094 | 15.424 | 6.5437 | 15.594 |
| 7936 | ENSMUSG00000030643 | 429.9909996 | -0.669884206 | 0.174739043 | -3.833626384 | 0.000126268 | 0.00434447 | Rab30 | RAB30, member RAS oncogene family [So | 611.96 | 443.03 | 529.28 | 392.34 | 331.55 | 271.78 |
| 7937 | ENSMUSG00000030647 | 959.1372858 | 0.042855007 | 0.120118991 | 0.356771285 | 0.721263034 | 0.9447719 | Ndufc2 | NADH:ubiquinone oxidoreductase subuni | 958.9 | 965.21 | 909.65 | 866.62 | 969.56 | 1084.9 |
| 7938 | ENSMUSG00000030649 | 71.23065134 | -0.171598144 | 0.287978575 | -0.595871217 | 0.551261263 | 0.89589637 | Anapc15 | anaphase promoting complex C subunit 1 | 56.217 | 76.066 | 94.587 | 69.406 | 67.618 | 63.489 |
| 7939 | ENSMUSG00000030650 | 0 | NA | NA | NA | NA | NA | Tmc5 | transmembrane channel-like gene family ! | 0 | 0 | 0 | 0 | 0 | 0 |
| 7940 | ENSMUSG00000030651 | 3.981074683 | -4.318237734 | 1.587475229 | -2.720192199 | 0.006524398 | NA | Art2b | ADP-ribosyltransferase 2b [Source:MGI Sy | 2.4093 | 11.307 | 9.0562 | 0 | 0 | 1.1138 |
| 7941 | ENSMUSG00000030652 | 489.6500092 | -0.001197464 | 0.161963007 | -0.007393444 | 0.994100939 | 0.998618081 | Coq7 | demethyl-Q 7 [Source:MGI Symbol;Acc:M | 554.14 | 498.54 | 415.58 | 418.37 | 479.87 | 571.4 |
| 7942 | ENSMUSG00000030653 | 1.187987671 | 0.476455097 | 2.259281564 | 0.210887879 | 0.832974763 | NA | Gm45837 | predicted gene 45837 [Source:MGI Symbo | 0 | 0 | 3.0187 | 1.928 | 2.1812 | 0 |
| 7943 | ENSMUSG00000030654 | 4696.97984 | 0.157359797 | 0.099545214 | 1.580787169 | 0.113926709 | 0.496336122 | Arl6ip1 | ADP-ribosylation factor-like 6 interacting p | 4752.8 | 4525.9 | 4043.1 | 4703.2 | 5191.3 | 4965.5 |
| 7944 | ENSMUSG00000030655 | 1022.919609 | -0.064075681 | 0.113270765 | -0.565685952 | 0.571607286 | 0.904651508 | Smg1 | SMG1 homolog, phosphatidylinositol 3-kin | 983 | 1134.8 | 1020.3 | 1065.2 | 968.47 | 965.7 |
| 7945 | ENSMUSG00000030657 | 8.206435125 | -0.108356284 | 0.800382902 | -0.135380558 | 0.892310995 | 0.982489666 | Xylt1 | xylosyltransferase 1 [Source:MGI Symbol;. | 15.259 | 4.1117 | 6.0375 | 10.604 | 6.5437 | 6.6831 |
| 7946 | ENSMUSG00000030659 | 23.29491345 | 0.212062663 | 0.580447946 | 0.365343119 | 0.714855296 | 0.944595913 | Nucb2 | nucleobindin 2 [Source:MGI Symbol;Acc:N | 32.927 | 20.558 | 11.069 | 30.847 | 8.7249 | 35.643 |
| 7947 | ENSMUSG00000030660 | 585.1213382 | -0.137698594 | 0.140164929 | -0.982404054 | 0.32590083 | 0.772018857 | Pik3c2a | phosphatidylinositol-4-phosphate 3-kinas | 531.65 | 701.04 | 607.77 | 595.74 | 538.77 | 535.76 |
| 7948 | ENSMUSG00000030662 | 1113.456103 | 0.261145636 | 0.118191446 | 2.209513832 | 0.027138921 | 0.21354341 | Ipo5 | importin 5 [Source:MGI Symbol;Acc:MGI:: | 1017.5 | 1093.7 | 927.76 | 1212.7 | 1318.6 | 1110.5 |

| | | | | | | | | | | | | | | | |
|---|---|---|---|---|---|---|---|---|---|---|---|---|---|---|---|
| 7949 | ENSMUSG00000030663 | 724.1056538 | -0.003195615 | 0.118545469 | -0.026956873 | 0.978494132 | 0.997797092 | 1110004F10 | RIKEN cDNA 1110004F10 gene [Source:M( | 735.64 | 686.65 | 751.66 | 667.07 | 736.17 | 767.44 |
| 7950 | ENSMUSG00000030664 | 29.20100102 | -0.326651836 | 0.444558076 | -0.734778769 | 0.462474239 | 0.860682887 | Sox6os | SRY (sex determining region Y)-box 6, opp | 40.155 | 39.061 | 18.112 | 25.063 | 30.537 | 22.277 |
| 7951 | ENSMUSG00000030666 | 0 | NA | NA | NA | NA | NA | Calcb | calcitonin-related polypeptide, beta [Sour | 0 | 0 | 0 | 0 | 0 | 0 |
| 7952 | ENSMUSG00000030669 | 0 | NA | NA | NA | NA | NA | Calca | calcitonin/calcitonin-related polypeptide, | 0 | 0 | 0 | 0 | 0 | 0 |
| 7953 | ENSMUSG00000030670 | 196.1840093 | 0.525872011 | 0.17097838 | 3.075663776 | 0.002100345 | 0.036165948 | Cyp2r1 | cytochrome P450, family 2, subfamily r, po | 159.01 | 160.36 | 163.01 | 221.71 | 226.85 | 246.16 |
| 7954 | ENSMUSG00000030671 | 1811.563512 | -0.102202708 | 0.109459252 | -0.933705516 | 0.350455819 | 0.791877747 | Pde3b | phosphodiesterase 3B, cGMP-inhibited [S | 1715.4 | 2008.6 | 1904.8 | 1865.3 | 1762.4 | 1612.8 |
| 7955 | ENSMUSG00000030672 | 13.16528125 | 0.19606676 | 0.583909947 | 0.335782532 | 0.737034864 | 0.948137969 | Mylpf | myosin light chain, phosphorylatable, fast | 10.44 | 13.363 | 13.081 | 13.496 | 16.359 | 12.252 |
| 7956 | ENSMUSG00000030674 | 3983.547675 | 0.120019104 | 0.109482586 | 1.096239209 | 0.272974105 | 0.728199457 | Qprt | quinolinate phosphoribosyltransferase [So | 3989 | 3814.6 | 3648.6 | 3602.4 | 4370.1 | 4476.5 |
| 7957 | ENSMUSG00000030677 | 12.73478134 | 0.502162244 | 0.653465091 | 0.768460705 | 0.442213527 | 0.850193681 | Kif22 | kinesin family member 22 [Source:MGI Sy | 11.243 | 13.363 | 7.0437 | 19.28 | 6.5437 | 18.935 |
| 7958 | ENSMUSG00000030678 | 1225.644945 | 0.063006137 | 0.116057519 | 0.542887162 | 0.587207496 | 0.910765033 | Maz | MYC-associated zinc finger protein (purine | 1200.6 | 1134.8 | 1261.8 | 1388.1 | 1131 | 1237.5 |
| 7959 | ENSMUSG00000030680 | 0 | NA | NA | NA | NA | NA | Pagr1a | PAXIP1 associated glutamate rich protein | 0 | 0 | 0 | 0 | 0 | 0 |
| 7960 | ENSMUSG00000030681 | 613.2499727 | -0.205727182 | 0.150121474 | -1.370404756 | 0.170560588 | 0.597556794 | Mvp | major vault protein [Source:MGI Symbol;A | 712.35 | 733.93 | 524.25 | 600.56 | 569.3 | 539.1 |
| 7961 | ENSMUSG00000030682 | 787.3612754 | 0.149450326 | 0.109662098 | 1.362825703 | 0.172937449 | 0.601945568 | Cdipt | CDP-diacylglycerol--inositol 3-phosphatidy | 762.95 | 724.68 | 751.66 | 805.89 | 816.87 | 862.12 |
| 7962 | ENSMUSG00000030683 | 8.01764835 | 0.024379151 | 0.769506484 | 0.031681541 | 0.974726016 | 0.996336532 | Sez6l2 | seizure related 6 homolog like 2 [Source:N | 5.6217 | 9.2513 | 9.0562 | 8.6758 | 4.3625 | 11.138 |
| 7963 | ENSMUSG00000030685 | 61.53359613 | -0.021669972 | 0.290013275 | -0.074720621 | 0.940437 | 0.988839835 | Kctd13 | potassium channel tetramerisation domai | 66.657 | 58.591 | 60.375 | 50.127 | 62.165 | 71.286 |
| 7964 | ENSMUSG00000030688 | 20511.17312 | 0.034984603 | 0.084441009 | 0.414308208 | 0.678648404 | 0.933875988 | Stard10 | START domain containing 10 [Source:MGI | 19651 | 19859 | 21277 | 19968 | 21074 | 21238 |
| 7965 | ENSMUSG00000030689 | 115.3803117 | -0.06696851 | 0.227408561 | -0.294485437 | 0.76838699 | 0.955856567 | Ino80e | INO80 complex subunit E [Source:MGI Syr | 104.4 | 136.71 | 113.71 | 129.17 | 104.7 | 103.59 |
| 7966 | ENSMUSG00000030691 | 463.3624431 | 0.037201301 | 0.129438159 | 0.287405979 | 0.773801484 | 0.958261993 | Fchsd2 | FCH and double SH3 domains 2 [Source:M | 424.04 | 467.7 | 480.98 | 462.71 | 461.33 | 483.41 |
| 7967 | ENSMUSG00000030693 | 0 | NA | NA | NA | NA | NA | Klk10 | kallikrein related-peptidase 10 [Source:M( | 0 | 0 | 0 | 0 | 0 | 0 |
| 7968 | ENSMUSG00000030695 | 1521.867006 | -0.089200301 | 0.121603901 | -0.733531577 | 0.463234276 | 0.861432317 | Aldoa | aldolase A, fructose-bisphosphate [Source | 1753.2 | 1620 | 1332.3 | 1436.3 | 1470.2 | 1519.3 |
| 7969 | ENSMUSG00000030697 | 616.8999302 | 0.165590278 | 0.128068967 | 1.292977384 | 0.196018881 | 0.636555253 | Ppp4c | protein phosphatase 4, catalytic subunit [ | 595.1 | 570.49 | 578.59 | 642.01 | 586.75 | 728.45 |
| 7970 | ENSMUSG00000030699 | 8.636895817 | -0.796908351 | 0.787692325 | -1.011700034 | 0.3116815 | 0.759830222 | Tbx6 | T-box 6 [Source:MGI Symbol;Acc:MGI:102 | 5.6217 | 10.279 | 17.106 | 6.7478 | 8.7249 | 3.3415 |
| 7971 | ENSMUSG00000030701 | 485.7377935 | 0.117886308 | 0.231403833 | 0.509439735 | 0.61044403 | 0.916376001 | Plekhb1 | pleckstrin homology domain containing, fa | 350.15 | 448.17 | 600.73 | 692.14 | 465.69 | 357.54 |
| 7972 | ENSMUSG00000030703 | 3.189889232 | 0.282262628 | 1.186789104 | 0.237837226 | 0.812007345 | NA | Gdpd3 | glycerophosphodiester phosphodiesterase | 1.6062 | 2.0558 | 5.0312 | 3.8559 | 4.3625 | 2.2277 |
| 7973 | ENSMUSG00000030704 | 932.3754221 | 0.118926462 | 0.105314094 | 1.129254948 | 0.2587903 | 0.712165697 | Rab6a | RAB6A, member RAS oncogene family [So | 917.95 | 874.76 | 888.51 | 939.88 | 968.47 | 1004.7 |
| 7974 | ENSMUSG00000030706 | 509.0666443 | 0.021578442 | 0.133489854 | 0.161648552 | 0.871582621 | 0.979227205 | Mrpl48 | mitochondrial ribosomal protein L48 [Sou | 528.44 | 486.21 | 500.1 | 454.03 | 540.95 | 544.67 |
| 7975 | ENSMUSG00000030707 | 68.81771931 | -1.353756446 | 0.333807862 | -4.055495995 | 5.00E-05 | 0.002109637 | Coro1a | coronin, actin binding protein 1A [Source: | 114.04 | 97.652 | 84.525 | 26.027 | 31.628 | 59.034 |
| 7976 | ENSMUSG00000030708 | 2.079165311 | -1.670228279 | 1.732849949 | -0.963862035 | 0.3351151 | NA | Dnajb13 | DnaJ heat shock protein family (Hsp40) m | 2.4093 | 6.1675 | 1.0062 | 2.8919 | 0 | 0 |
| 7977 | ENSMUSG00000030711 | 6974.084205 | -0.238587888 | 0.087655098 | -2.721894026 | 0.006490895 | 0.079698057 | Sult1a1 | sulfotransferase family 1A, phenol-prefer | 7719.4 | 7330.1 | 7599.2 | 6723.7 | 6415 | 6057.1 |
| 7978 | ENSMUSG00000030713 | 0 | NA | NA | NA | NA | NA | Klk7 | kallikrein related-peptidase 7 (chymotrypt | 0 | 0 | 0 | 0 | 0 | 0 |
| 7979 | ENSMUSG00000030714 | 210.9769064 | 0.202361059 | 0.184339139 | 1.097765021 | 0.272307109 | 0.727618887 | Sgf29 | SAGA complex associated factor 29 [Sourc | 182.3 | 219.97 | 187.16 | 263.17 | 208.31 | 204.95 |
| 7980 | ENSMUSG00000030717 | 11.46418907 | -0.395162984 | 0.653457253 | -0.604726601 | 0.54536067 | 0.89549457 | Nupr1 | nuclear protein transcription regulator 1 [ | 13.653 | 8.2233 | 17.106 | 7.7118 | 8.7249 | 13.366 |
| 7981 | ENSMUSG00000030718 | 447.3995737 | 0.131447847 | 0.135098875 | 0.972975137 | 0.330565659 | 0.774747658 | Ppme1 | protein phosphatase methylesterase 1 [So | 418.42 | 446.12 | 416.59 | 457.89 | 430.79 | 514.6 |
| 7982 | ENSMUSG00000030720 | 215.9541629 | 0.161500348 | 0.181524256 | 0.889690179 | 0.373632269 | 0.80865857 | Cln3 | ceroid lipofuscinosis, neuronal 3, juvenile | 237.72 | 191.19 | 182.13 | 241.96 | 236.66 | 206.06 |
| 7983 | ENSMUSG00000030722 | 79.34104722 | -0.334808054 | 0.284607582 | -1.176384871 | 0.239441114 | 0.688924491 | Nfatc2ip | nuclear factor of activated T cells, cytopla | 70.673 | 87.373 | 107.67 | 60.731 | 88.34 | 61.261 |
| 7984 | ENSMUSG00000030724 | 10.16746904 | -3.766091359 | 0.951468722 | -3.958187246 | 7.55E-05 | 0.002897433 | Cd19 | CD19 antigen [Source:MGI Symbol;Acc:M( | 19.274 | 18.503 | 19.119 | 1.928 | 2.1812 | 0 |
| 7985 | ENSMUSG00000030725 | 80.4651417 | -0.328481453 | 0.248516998 | -1.321766542 | 0.186245903 | 0.62230226 | Lipt2 | lipoyl(octanoyl) transferase 2 (putative) [S | 94.766 | 89.429 | 84.525 | 75.19 | 68.709 | 70.172 |
| 7986 | ENSMUSG00000030726 | 173.2933931 | 0.264157193 | 0.198786852 | 1.328846403 | 0.183898649 | 0.618439913 | Pold3 | polymerase (DNA-directed), delta 3, acces | 167.85 | 143.91 | 159.99 | 161.95 | 179.95 | 226.11 |
| 7987 | ENSMUSG00000030727 | 212.7642404 | 0.229033135 | 0.221099818 | 1.035881154 | 0.300257591 | 0.751875664 | Rabep2 | rabaptin, RAB GTPase binding effector pro | 211.22 | 161.38 | 215.34 | 296.91 | 172.32 | 219.43 |
| 7988 | ENSMUSG00000030729 | 113.2511317 | 0.182198431 | 0.234792758 | 0.775996811 | 0.437750869 | 0.847422601 | Pgm2l1 | phosphoglucomutase 2-like 1 [Source:MG | 101.99 | 117.18 | 99.618 | 141.7 | 124.33 | 94.677 |
| 7989 | ENSMUSG00000030730 | 1.339512537 | 0.884662434 | 1.869203337 | 0.473283145 | 0.636011186 | NA | Atp2a1 | ATPase, Ca++ transporting, cardiac muscle | 0.8031 | 1.0279 | 1.0062 | 1.928 | 3.2719 | 0 |
| 7990 | ENSMUSG00000030731 | 124.5261372 | -0.254967801 | 0.221709264 | -1.150009685 | 0.250139882 | 0.702302228 | Syt3 | synaptotagmin III [Source:MGI Symbol;Ac | 137.33 | 133.63 | 135.84 | 133.99 | 90.521 | 115.84 |
| 7991 | ENSMUSG00000030732 | 4.30384184 | -1.651702192 | 1.073979807 | -1.537926674 | 0.124066544 | NA | Chrdl2 | chordin-like 2 [Source:MGI Symbol;Acc:M | 6.4248 | 6.1675 | 7.0437 | 2.8919 | 2.1812 | 1.1138 |
| 7992 | ENSMUSG00000030733 | 937.5791941 | 0.136845404 | 0.108179355 | 1.264986321 | 0.205876231 | 0.651085207 | Sh2b1 | SH2B adaptor protein 1 [Source:MGI Syml | 881.81 | 913.82 | 884.49 | 1045 | 932.48 | 967.93 |
| 7993 | ENSMUSG00000030735 | 9.292037643 | 0.070480478 | 0.799958292 | 0.088105191 | 0.929793071 | 0.988405116 | Gm9755 | predicted pseudogene 9755 [Source:MGI | 12.047 | 3.0838 | 12.075 | 16.388 | 4.3625 | 7.7969 |
| 7994 | ENSMUSG00000030737 | 3191.150211 | -0.027472592 | 0.095479417 | -0.287733136 | 0.773551023 | 0.958177785 | Slco2b1 | solute carrier organic anion transporter fa | 3447.7 | 3188.6 | 3026.8 | 3052.9 | 3159.5 | 3271.4 |
| 7995 | ENSMUSG00000030738 | 2022.648172 | 0.005961165 | 0.092784112 | 0.064247693 | 0.948773003 | 0.990731019 | Eif3c | eukaryotic translation initiation factor 3, s | 1994.9 | 2092.8 | 1968.2 | 2073.5 | 2043.8 | 1962.6 |
| 7996 | ENSMUSG00000030739 | 457.6232164 | 0.21827457 | 0.157999623 | 1.381487913 | 0.167128992 | 0.593361251 | Myh14 | myosin, heavy polypeptide 14 [Source:MG | 461.78 | 388.55 | 417.59 | 399.09 | 550.76 | 527.96 |
| 7997 | ENSMUSG00000030741 | 587.8864819 | 0.210590972 | 0.125357107 | 1.679928471 | 0.092971234 | 0.449293659 | Spns1 | spinster homolog 1 [Source:MGI Symbol;A | 595.1 | 517.04 | 522.24 | 652.61 | 598.75 | 641.57 |
| 7998 | ENSMUSG00000030742 | 5.138766787 | -1.975745113 | 1.046337205 | -1.888248936 | 0.058992539 | 0.345173174 | Lat | linker for activation of T cells [Source:MGI | 6.4248 | 5.1396 | 13.081 | 2.8919 | 2.1812 | 1.1138 |
| 7999 | ENSMUSG00000030744 | 4493.630513 | 0.003512303 | 0.094512192 | 0.037162437 | 0.970355489 | 0.994966572 | Rps3 | ribosomal protein S3 [Source:MGI Symbol | 4193.8 | 4521.8 | 4749.5 | 4336 | 4423.5 | 4737.2 |
| 8000 | ENSMUSG00000030745 | 4.254197352 | -1.110679667 | 1.149597121 | -0.966146876 | 0.333970691 | NA | Il21r | interleukin 21 receptor [Source:MGI Symb | 6.4248 | 2.0558 | 9.0562 | 5.7839 | 1.0906 | 1.1138 |
| 8001 | ENSMUSG00000030747 | 15088.88824 | 0.024208647 | 0.083476346 | 0.290006068 | 0.771811595 | 0.957088727 | Dgat2 | diacylglycerol O-acyltransferase 2 [Source | 15181 | 14871 | 14834 | 14504 | 15940 | 15203 |
| 8002 | ENSMUSG00000030748 | 567.3468595 | -0.392761098 | 0.139777179 | -2.809908617 | 0.004955557 | 0.066429262 | Il4ra | interleukin 4 receptor, alpha [Source:MGI | 550.93 | 673.29 | 709.4 | 522.48 | 465.69 | 482.29 |
| 8003 | ENSMUSG00000030750 | 196.4382211 | 0.141366332 | 0.183527206 | 0.770274529 | 0.441137062 | 0.849828766 | Nsmce1 | NSE1 homolog, SMC5-SMC6 complex com | 189.53 | 205.58 | 165.02 | 182.19 | 224.67 | 211.63 |
| 8004 | ENSMUSG00000030751 | 1528.383897 | 0.050437419 | 0.115476858 | 0.436775123 | 0.662274442 | 0.9288455 | Psma1 | proteasome subunit alpha 1 [Source:MGI | 1686.5 | 1406.2 | 1410.8 | 1541.4 | 1464.7 | 1660.7 |

| | | | | | | | | | | | | | | | |
|---|---|---|---|---|---|---|---|---|---|---|---|---|---|---|---|
| 8005 | ENSMUSG00000030752 | 56.48390102 | 0.279389219 | 0.291846243 | 0.957316481 | 0.338407539 | 0.780780576 | Kdm8 | lysine (K)-specific demethylase 8 [Source:I | 47.383 | 53.452 | 52.325 | 57.839 | 61.075 | 66.831 |
| 8006 | ENSMUSG00000030753 | 704.7367564 | -0.157444014 | 0.142873766 | -1.101979875 | 0.270470421 | 0.725929337 | Thap12 | THAP domain containing 12 [Source:MGI ! | 665.77 | 840.84 | 723.49 | 632.37 | 766.7 | 599.25 |
| 8007 | ENSMUSG00000030754 | 2075.723085 | -0.018582958 | 0.102549488 | -0.181209663 | 0.856203011 | 0.974319798 | Copb1 | coatomer protein complex, subunit beta 1 | 2215.8 | 2134 | 1916.9 | 2074.5 | 2152.9 | 1960.4 |
| 8008 | ENSMUSG00000030757 | 2.191445284 | -1.148462182 | 1.696862069 | -0.676815283 | 0.49852316 | NA | Zkscan2 | zinc finger with KRAB and SCAN domains : | 0 | 4.1117 | 5.0312 | 2.8919 | 0 | 1.1138 |
| 8009 | ENSMUSG00000030759 | 33.5700284 | -1.165445185 | 0.413266248 | -2.820083158 | 0.004801121 | 0.065021918 | Far1 | fatty acyl CoA reductase 1 [Source:MGI Sy | 48.989 | 50.368 | 40.25 | 29.883 | 17.45 | 14.48 |
| 8010 | ENSMUSG00000030760 | 214.2397471 | 0.012915009 | 0.18794762 | 0.068716 | 0.945215682 | 0.990309554 | Acer3 | alkaline ceramidase 3 [Source:MGI Symbc | 210.41 | 244.64 | 185.15 | 238.1 | 182.13 | 225 |
| 8011 | ENSMUSG00000030761 | 188.8208668 | -0.189553357 | 0.258561766 | -0.733106675 | 0.46349337 | 0.861581814 | Myo7a | myosin VIIA [Source:MGI Symbol;Acc:MG | 158.21 | 208.67 | 237.47 | 217.86 | 106.88 | 203.83 |
| 8012 | ENSMUSG00000030762 | 1928.997538 | -0.055566264 | 0.165303877 | -0.336146164 | 0.736760648 | 0.948137969 | Aqp8 | aquaporin 8 [Source:MGI Symbol;Acc:MG | 2511.3 | 1724.8 | 1660.3 | 1669.6 | 1763.5 | 2244.4 |
| 8013 | ENSMUSG00000030763 | 169.3290771 | -0.091996548 | 0.186248504 | -0.493945163 | 0.621344908 | 0.918178662 | Lcmt1 | leucine carboxyl methyltransferase 1 [Sou | 185.52 | 157.27 | 181.12 | 163.88 | 175.59 | 152.6 |
| 8014 | ENSMUSG00000030766 | 186.623718 | -0.300989362 | 0.178811757 | -1.683275013 | 0.092321865 | 0.447351243 | Arhgap17 | Rho GTPase activating protein 17 [Source: | 202.38 | 208.67 | 207.29 | 179.3 | 147.23 | 174.87 |
| 8015 | ENSMUSG00000030768 | 28.1173979 | 0.214949539 | 0.42417108 | 0.50675199 | 0.612328846 | 0.916441371 | Disp1 | dispatched RND transporter family memb | 22.487 | 23.642 | 32.2 | 38.559 | 25.084 | 26.732 |
| 8016 | ENSMUSG00000030769 | 0.781711799 | -0.350673833 | 2.438876442 | -0.143784994 | 0.885670253 | NA | Slc5a11 | solute carrier family 5 (sodium/glucose co | 1.6062 | 0 | 1.0062 | 0.964 | 0 | 1.1138 |
| 8017 | ENSMUSG00000030770 | 1706.844806 | -0.183071212 | 0.096877929 | -1.889710225 | 0.058796726 | 0.344306009 | Parva | parvin, alpha [Source:MGI Symbol;Acc:M( | 1884.9 | 1844.1 | 1715.6 | 1626.2 | 1558.5 | 1611.7 |
| 8018 | ENSMUSG00000030772 | 24.16302738 | -0.351731052 | 0.493806025 | -0.712285866 | 0.476287775 | 0.86953366 | Dkk3 | dickkopf WNT signaling pathway inhibitor | 40.155 | 21.586 | 19.119 | 14.46 | 21.812 | 27.846 |
| 8019 | ENSMUSG00000030774 | 19.84928296 | -0.989334169 | 0.539791141 | -1.832809197 | 0.066830951 | 0.37192178 | Pak1 | p21 (RAC1) activated kinase 1 [Source:MG | 23.29 | 30.838 | 25.156 | 15.424 | 5.4531 | 18.935 |
| 8020 | ENSMUSG00000030775 | 0.17131964 | -0.796629176 | 4.080472857 | -0.195229623 | 0.84521319 | NA | Trat1 | T cell receptor associated transmembrane | 0 | 1.0279 | 0 | 0 | 0 | 0 |
| 8021 | ENSMUSG00000030779 | 811.8102197 | 0.041543426 | 0.134975512 | 0.307784912 | 0.758246003 | 0.953675669 | Rbbp6 | retinoblastoma binding protein 6, ubiquiti | 701.11 | 907.65 | 792.92 | 867.58 | 737.26 | 864.34 |
| 8022 | ENSMUSG00000030780 | 288.1912737 | -0.009949093 | 0.150359321 | -0.066168779 | 0.947243453 | 0.990677354 | Rusf1 | RUS family member 1 [Source:MGI Symbc | 306.79 | 270.34 | 289.8 | 273.77 | 297.74 | 290.71 |
| 8023 | ENSMUSG00000030781 | 0 | NA | NA | NA | NA | NA | Slc5a2 | solute carrier family 5 (sodium/glucose co | 0 | 0 | 0 | 0 | 0 | 0 |
| 8024 | ENSMUSG00000030782 | 19.0630628 | -0.4700718 | 0.507363887 | -0.926498343 | 0.354187058 | 0.79449673 | Tgfb1i1 | transforming growth factor beta 1 induce | 22.487 | 24.67 | 19.119 | 10.604 | 17.45 | 20.049 |
| 8025 | ENSMUSG00000030785 | 0.133850373 | -0.796629176 | 4.080472857 | -0.195229623 | 0.84521319 | NA | Cox6a2 | cytochrome c oxidase subunit 6A2 [Sourc | 0.8031 | 0 | 0 | 0 | 0 | 0 |
| 8026 | ENSMUSG00000030786 | 6.865948359 | -2.177532834 | 0.980987838 | -2.219734791 | 0.026436776 | 0.210562513 | Itgam | integrin alpha M [Source:MGI Symbol;Acc | 3.2124 | 17.475 | 13.081 | 1.928 | 3.2719 | 2.2277 |
| 8027 | ENSMUSG00000030787 | 112.5740221 | -0.939223214 | 0.273066182 | -3.439544247 | 0.000582695 | 0.013773229 | Lyve1 | lymphatic vessel endothelial hyaluronan r | 157.41 | 173.72 | 112.7 | 79.046 | 53.44 | 99.132 |
| 8028 | ENSMUSG00000030788 | 959.5329842 | 0.079576257 | 0.141766878 | 0.561317694 | 0.574580981 | 0.906163061 | Rnf141 | ring finger protein 141 [Source:MGI Symb | 876.18 | 989.88 | 932.79 | 821.31 | 961.93 | 1175.1 |
| 8029 | ENSMUSG00000030789 | 8.279306108 | -1.56010744 | 0.798054284 | -1.954888873 | 0.050596188 | 0.315985866 | Itgax | integrin alpha X [Source:MGI Symbol;Acc: | 17.668 | 11.307 | 8.05 | 3.8559 | 5.4531 | 3.3415 |
| 8030 | ENSMUSG00000030790 | 19.96738981 | -1.562488538 | 0.561230928 | -2.78403855 | 0.005368665 | 0.069842592 | Adm | adrenomedullin [Source:MGI Symbol;Acc: | 34.533 | 30.838 | 24.15 | 12.532 | 3.2719 | 14.48 |
| 8031 | ENSMUSG00000030792 | 0 | NA | NA | NA | NA | NA | Dkkl1 | dickkopf-like 1 [Source:MGI Symbol;Acc:N | 0 | 0 | 0 | 0 | 0 | 0 |
| 8032 | ENSMUSG00000030793 | 18.18458631 | -0.142891269 | 0.551765037 | -0.258971227 | 0.795657441 | 0.962236316 | Pycard | PYD and CARD domain containing [Source | 14.456 | 30.838 | 12.075 | 15.424 | 20.722 | 15.594 |
| 8033 | ENSMUSG00000030795 | 1237.801553 | -0.118632827 | 0.106959701 | -1.10913573 | 0.267371633 | 0.72293778 | Fus | fused in sarcoma [Source:MGI Symbol;Acc | 1261.7 | 1388.7 | 1215.5 | 1130.7 | 1216 | 1214.1 |
| 8034 | ENSMUSG00000030796 | 7.023819136 | -0.162310537 | 0.812737986 | -0.19970832 | 0.841708707 | 0.971854408 | Tead2 | TEA domain family member 2 [Source:MG | 8.031 | 6.1675 | 8.05 | 7.7118 | 3.2719 | 8.9108 |
| 8035 | ENSMUSG00000030798 | 31.38574959 | -1.457228302 | 0.442040635 | -3.296593538 | 0.00097865 | 0.020634397 | Cd37 | CD37 antigen [Source:MGI Symbol;Acc:M | 51.399 | 53.452 | 33.206 | 19.28 | 9.8156 | 21.163 |
| 8036 | ENSMUSG00000030800 | 66.24029478 | 0.81560314 | 0.317867652 | 2.565857631 | 0.010292106 | 0.111093302 | Prss8 | protease, serine 8 (prostasin) [Source:MG | 48.989 | 57.563 | 37.231 | 62.659 | 79.615 | 111.38 |
| 8037 | ENSMUSG00000030801 | 318.8333347 | 0.22169939 | 0.143801109 | 1.54170848 | 0.12314445 | 0.513662281 | Kat8 | K(lysine) acetyltransferase 8 [Source:MGI | 299.56 | 292.96 | 290.8 | 367.28 | 329.37 | 333.04 |
| 8038 | ENSMUSG00000030802 | 287.4810337 | -0.181615897 | 0.161799196 | -1.122477135 | 0.261659624 | 0.715612553 | Bckdk | branched chain ketoacid dehydrogenase k | 302.77 | 328.93 | 284.77 | 245.81 | 308.64 | 253.96 |
| 8039 | ENSMUSG00000030804 | 0.133850373 | -0.796629176 | 4.080472857 | -0.195229623 | 0.84521319 | NA | Gm21974 | predicted gene 21974 [Source:MGI Symbc | 0.8031 | 0 | 0 | 0 | 0 | 0 |
| 8040 | ENSMUSG00000030805 | 431.1013043 | 0.25901779 | 0.172251373 | 1.503719737 | 0.132653545 | 0.533565316 | Stx4a | syntaxin 4A (placental) [Source:MGI Symb | 426.45 | 400.89 | 349.17 | 392.34 | 431.88 | 585.88 |
| 8041 | ENSMUSG00000030806 | 90.35040528 | -0.197835528 | 0.236932713 | -0.834986128 | 0.403725519 | 0.828953151 | Stx1b | syntaxin 1B [Source:MGI Symbol;Acc:MGI | 87.538 | 99.708 | 102.64 | 85.794 | 82.887 | 83.538 |
| 8042 | ENSMUSG00000030811 | 100.9153619 | 0.069867332 | 0.256389347 | 0.272504819 | 0.785233887 | 0.960670212 | Fbxl19 | F-box and leucine-rich repeat protein 19 [ | 94.766 | 82.233 | 118.74 | 120.5 | 109.06 | 80.197 |
| 8043 | ENSMUSG00000030814 | 247.668945 | -0.038181122 | 0.172626015 | -0.221178263 | 0.824953634 | 0.96943211 | Bcl7c | B cell CLL/lymphoma 7C [Source:MGI Sym | 249.76 | 282.68 | 220.37 | 240.03 | 221.4 | 271.78 |
| 8044 | ENSMUSG00000030815 | 214.148607 | 0.234233434 | 0.165003401 | 1.419567307 | 0.155733689 | 0.574732247 | Phkg2 | phosphorylase kinase, gamma 2 (testis) [S | 204.79 | 186.05 | 199.24 | 228.46 | 225.76 | 240.59 |
| 8045 | ENSMUSG00000030816 | 619.0274453 | 0.137763377 | 0.115580537 | 1.191925394 | 0.2332905 | 0.680673483 | Rnf40 | ring finger protein 40 [Source:MGI Symbo | 572.61 | 600.3 | 595.7 | 628.51 | 657.64 | 659.4 |
| 8046 | ENSMUSG00000030822 | 578.5752566 | 0.204255092 | 0.156297491 | 1.306835385 | 0.191268625 | 0.629099068 | Prr14 | proline rich 14 [Source:MGI Symbol;Acc:N | 476.24 | 487.23 | 650.03 | 595.74 | 564.94 | 697.27 |
| 8047 | ENSMUSG00000030823 | 40.92643129 | -0.222390443 | 0.372742613 | -0.596632732 | 0.550752613 | 0.895703112 | 9130019O22 | RIKEN cDNA 9130019O22 gene [Source:M | 33.73 | 49.34 | 49.306 | 34.703 | 28.356 | 50.123 |
| 8048 | ENSMUSG00000030824 | 2762.359483 | -0.031904045 | 0.100363966 | -0.317883463 | 0.750573337 | 0.952074518 | Nucb1 | nucleobindin 1 [Source:MGI Symbol;Acc:N | 2926.5 | 2829.9 | 2622.3 | 2913.1 | 2537.9 | 2744.5 |
| 8049 | ENSMUSG00000030825 | 0.589026695 | 0.241472922 | 3.228838524 | 0.074786311 | 0.940384733 | NA | Hsd17b14 | hydroxysteroid (17-beta) dehydrogenase : | 1.6062 | 0 | 0 | 1.928 | 0 | 0 |
| 8050 | ENSMUSG00000030826 | 591.7252748 | 0.168669994 | 0.142602401 | 1.182799117 | 0.236888771 | 0.685662114 | Bcat2 | branched chain aminotransferase 2, mito | 546.91 | 534.52 | 589.66 | 564.89 | 573.67 | 740.71 |
| 8051 | ENSMUSG00000030827 | 76.29384073 | -1.846407857 | 0.413223836 | -4.468299494 | 7.88E-06 | 0.000442015 | Fgf21 | fibroblast growth factor 21 [Source:MGI S | 202.38 | 84.289 | 71.443 | 41.451 | 39.262 | 18.935 |
| 8052 | ENSMUSG00000030830 | 91.1178418 | -1.155786222 | 0.336029068 | -3.439542384 | 0.000582699 | 0.013773229 | Itgal | integrin alpha L [Source:MGI Symbol;Acc:I | 83.523 | 192.22 | 101.63 | 41.451 | 62.165 | 65.717 |
| 8053 | ENSMUSG00000030834 | 3824.712142 | 0.191715303 | 0.099380189 | 1.929109871 | 0.053717224 | 0.326014563 | Abcc6 | ATP-binding cassette, sub-family C (CFTR/ | 3594.7 | 3475.4 | 3643.6 | 4499.8 | 3913.1 | 3821.6 |
| 8054 | ENSMUSG00000030835 | 640.8382987 | 0.009409897 | 0.125590298 | 0.074925354 | 0.940274104 | 0.988812319 | Nomo1 | nodal modulator 1 [Source:MGI Symbol;A | 600.72 | 712.35 | 603.75 | 637.19 | 623.83 | 667.19 |
| 8055 | ENSMUSG00000030838 | 0.181769715 | 1.126924175 | 4.080472857 | 0.276174898 | 0.782413736 | NA | Ush1c | USH1 protein network component harmo | 0 | 0 | 0 | 0 | 1.0906 | 0 |
| 8056 | ENSMUSG00000030839 | 61.02477518 | -0.271334 | 0.346463864 | -0.783152383 | 0.433537634 | 0.845264432 | Sergef | secretion regulating guanine nucleotide e: | 50.595 | 87.373 | 62.387 | 40.487 | 78.525 | 46.781 |
| 8057 | ENSMUSG00000030842 | 700.3096728 | -0.09175044 | 0.123760027 | -0.74135763 | 0.458476628 | 0.859322431 | Lamtor1 | late endosomal/lysosomal adaptor, MAPK | 717.17 | 726.74 | 723.49 | 648.76 | 623.83 | 761.87 |
| 8058 | ENSMUSG00000030844 | 17.82298228 | -0.949580729 | 0.57924679 | -1.639337057 | 0.101143082 | 0.468438535 | Rgs10 | regulator of G-protein signalling 10 [Sourc | 22.487 | 23.642 | 24.15 | 6.7478 | 7.6343 | 22.277 |
| 8059 | ENSMUSG00000030846 | 977.7508489 | 0.209129719 | 0.119479203 | 1.750344098 | 0.080058956 | 0.415341728 | Tial1 | Tia1 cytotoxic granule-associated RNA bin | 856.11 | 986.8 | 879.46 | 1153.9 | 1002.3 | 987.98 |
| 8060 | ENSMUSG00000030847 | 380.3868931 | -0.318343009 | 0.147536785 | -2.157719581 | 0.030949641 | 0.233275457 | Bag3 | BCL2-associated athanogene 3 [Source:M | 425.64 | 388.55 | 451.8 | 303.65 | 375.17 | 337.49 |

| | | | | | | | | | | | | | | | |
|---|---|---|---|---|---|---|---|---|---|---|---|---|---|---|---|
| 8061 | ENSMUSG00000030849 | 396.5448968 | 0.198440814 | 0.182536382 | 1.087130206 | 0.27697927 | 0.730041765 | Fgfr2 | fibroblast growth factor receptor 2 [Sourc | 402.35 | 323.79 | 381.37 | 445.36 | 317.37 | 509.03 |
| 8062 | ENSMUSG00000030850 | 555.503239 | 0.086549603 | 0.136664495 | 0.63329984 | 0.526537858 | 0.888334265 | Ate1 | arginyltransferase 1 [Source:MGI Symbol; | 522.02 | 612.64 | 481.99 | 540.79 | 605.29 | 570.29 |
| 8063 | ENSMUSG00000030851 | 0 | NA | NA | NA | NA | NA | Ldhc | lactate dehydrogenase C [Source:MGI Syn | 0 | 0 | 0 | 0 | 0 | 0 |
| 8064 | ENSMUSG00000030852 | 439.3423857 | -0.775674663 | 0.160811891 | -4.823490721 | 1.41E-06 | 0.000107389 | Tacc2 | transforming, acidic coiled-coil containing | 542.09 | 659.92 | 461.87 | 300.76 | 325 | 346.41 |
| 8065 | ENSMUSG00000030854 | 0 | NA | NA | NA | NA | NA | Ptpn5 | protein tyrosine phosphatase, non-recept | 0 | 0 | 0 | 0 | 0 | 0 |
| 8066 | ENSMUSG00000030858 | 0 | NA | NA | NA | NA | NA | Fam24b | family with sequence similarity 24 membe | 0 | 0 | 0 | 0 | 0 | 0 |
| 8067 | ENSMUSG00000030859 | 0.371281295 | 2.09253362 | 4.031852165 | 0.519000582 | 0.603760335 | NA | Fam24a | family with sequence similarity 24, memb | 0 | 0 | 0 | 0 | 0 | 2.2277 |
| 8068 | ENSMUSG00000030861 | 2920.305182 | 0.29906501 | 0.106277923 | 2.813989976 | 0.004893075 | 0.065864706 | Acadsb | acyl-Coenzyme A dehydrogenase, short/b | 2478.4 | 2464.9 | 2913.1 | 3053.9 | 3410.4 | 3201.2 |
| 8069 | ENSMUSG00000030862 | 3.404744807 | -0.357705509 | 1.291054896 | -0.277064523 | 0.781730562 | NA | Cpxm2 | carboxypeptidase X 2 (M14 family) [Sourc | 6.4248 | 2.0558 | 3.0187 | 6.7478 | 2.1812 | 0 |
| 8070 | ENSMUSG00000030865 | 22.21942215 | -0.649423227 | 0.628771389 | -1.032844749 | 0.301676558 | 0.752246851 | Chp2 | calcineurin-like EF hand protein 2 [Source | 28.109 | 37.005 | 16.1 | 7.7118 | 7.6343 | 36.757 |
| 8071 | ENSMUSG00000030866 | 0.663785267 | -1.400021131 | 3.106754789 | -0.450637796 | 0.652250621 | NA | Ern2 | endoplasmic reticulum (ER) to nucleus sig | 0 | 0 | 3.0187 | 0.964 | 0 | 0 |
| 8072 | ENSMUSG00000030867 | 5.140537428 | -0.302228286 | 1.017266796 | -0.297098349 | 0.766391425 | 0.955199101 | Plk1 | polo like kinase 1 [Source:MGI Symbol;Ac | 8.8341 | 1.0279 | 7.0437 | 2.8919 | 4.3625 | 6.6831 |
| 8073 | ENSMUSG00000030868 | 338.2744978 | 0.036954425 | 0.143646545 | 0.257259406 | 0.796978527 | 0.962236316 | Dctn5 | dynactin 5 [Source:MGI Symbol;Acc:MGI: | 339.71 | 332.02 | 330.05 | 352.82 | 310.83 | 364.23 |
| 8074 | ENSMUSG00000030869 | 1468.77939 | -0.040596343 | 0.094914656 | -0.427714169 | 0.668859233 | 0.930586759 | Ndufab1 | NADH:ubiquinone oxidoreductase subuni | 1511.4 | 1499.7 | 1457 | 1458.5 | 1464.7 | 1421.3 |
| 8075 | ENSMUSG00000030870 | 556.6437554 | 0.099179761 | 0.122995141 | 0.806371377 | 0.420028746 | 0.837906635 | Ubfd1 | ubiquitin family domain containing 1 [Sou | 571.81 | 521.15 | 519.22 | 601.52 | 572.57 | 553.58 |
| 8076 | ENSMUSG00000030871 | 194.684394 | 0.155961171 | 0.175320026 | 0.889579902 | 0.373691502 | 0.80865857 | Ears2 | glutamyl-tRNA synthetase 2, mitochondri | 190.34 | 182.97 | 179.11 | 213.04 | 183.22 | 219.43 |
| 8077 | ENSMUSG00000030872 | 348.752194 | -0.577142882 | 0.17783284 | -3.245423515 | 0.001172761 | 0.023494196 | Gga2 | golgi associated, gamma adaptin ear cont | 387.1 | 447.14 | 419.6 | 332.57 | 294.47 | 211.63 |
| 8078 | ENSMUSG00000030873 | 0 | NA | NA | NA | NA | NA | Scnn1b | sodium channel, nonvoltage-gated 1 beta | 0 | 0 | 0 | 0 | 0 | 0 |
| 8079 | ENSMUSG00000030876 | 734.9318476 | -0.04215113 | 0.135075181 | -0.312056808 | 0.754997351 | 0.953085143 | Mettl9 | methyltransferase like 9 [Source:MGI Sym | 874.58 | 687.68 | 673.18 | 700.81 | 744.89 | 728.45 |
| 8080 | ENSMUSG00000030877 | 0 | NA | NA | NA | NA | NA | Mfsd13b | major facilitator superfamily domain cont | 0 | 0 | 0 | 0 | 0 | 0 |
| 8081 | ENSMUSG00000030878 | 12.53808326 | -0.189469748 | 0.70399204 | -0.269136208 | 0.787824868 | 0.960870081 | Cdr2 | cerebellar degeneration-related 2 [Source | 17.668 | 4.1117 | 18.112 | 6.7478 | 17.45 | 11.138 |
| 8082 | ENSMUSG00000030879 | 1086.162527 | -0.117476216 | 0.106288665 | -1.105256295 | 0.26904855 | 0.724979259 | Mrpl17 | mitochondrial ribosomal protein L17 [Sou | 1167.7 | 1126.6 | 1095.8 | 973.62 | 1089.5 | 1063.7 |
| 8083 | ENSMUSG00000030880 | 309.9817117 | 0.104526012 | 0.146334504 | 0.714295055 | 0.475044749 | 0.869368495 | Polr3e | polymerase (RNA) III (DNA directed) polyp | 293.13 | 319.68 | 283.76 | 331.61 | 306.46 | 325.24 |
| 8084 | ENSMUSG00000030881 | 273.5023069 | 0.142371821 | 0.156539764 | 0.909493008 | 0.363089947 | 0.802036193 | Arfip2 | ADP-ribosylation factor interacting protei | 278.68 | 244.64 | 256.59 | 312.33 | 278.11 | 270.66 |
| 8085 | ENSMUSG00000030882 | 21.91700493 | 0.591049937 | 0.474987909 | 1.244347332 | 0.213371758 | 0.658040322 | Dnhd1 | dynein heavy chain domain 1 [Source:MG | 20.881 | 15.419 | 16.1 | 31.811 | 28.356 | 18.935 |
| 8086 | ENSMUSG00000030884 | 4610.103066 | -0.040428014 | 0.089150113 | -0.453482475 | 0.65020136 | 0.925643431 | Uqcrc2 | ubiquinol cytochrome c reductase core pr | 4740.7 | 4723.3 | 4560.3 | 4676.3 | 4692.9 | 4267.1 |
| 8087 | ENSMUSG00000030887 | 3.686662822 | 0.560578957 | 1.154314656 | 0.48563791 | 0.627223913 | NA | Pdzd9 | PDZ domain containing 9 [Source:MGI Syr | 4.8186 | 3.0838 | 1.0062 | 7.7118 | 3.2719 | 2.2277 |
| 8088 | ENSMUSG00000030888 | 203.6777748 | 0.097370537 | 0.17052421 | 0.571007111 | 0.567994822 | 0.903352921 | Rrp8 | ribosomal RNA processing 8 [Source:MGI | 192.74 | 197.36 | 200.24 | 196.65 | 230.12 | 204.95 |
| 8089 | ENSMUSG00000030889 | 0.331982614 | 0.16514309 | 3.986751476 | 0.041422971 | 0.966958701 | NA | Vwa3a | von Willebrand factor A domain containin | 0 | 1.0279 | 0 | 0.964 | 0 | 0 |
| 8090 | ENSMUSG00000030890 | 14.50855343 | 0.600932254 | 0.633090945 | 0.949203678 | 0.342517031 | 0.784247367 | Ilk | integrin linked kinase [Source:MGI Symbo | 10.44 | 6.1675 | 18.112 | 26.991 | 13.087 | 12.252 |
| 8091 | ENSMUSG00000030894 | 1598.815219 | -0.090802972 | 0.105742979 | -0.858713959 | 0.390498347 | 0.819763149 | Tpp1 | tripeptidyl peptidase I [Source:MGI Symbo | 1709.8 | 1646.7 | 1590.9 | 1621.4 | 1621.7 | 1402.3 |
| 8092 | ENSMUSG00000030895 | 59978.10462 | -0.016641995 | 0.103652551 | -0.160555575 | 0.872443447 | 0.979419285 | Hpx | hemopexin [Source:MGI Symbol;Acc:MGI | 65273 | 63210 | 52489 | 61799 | 58680 | 58419 |
| 8093 | ENSMUSG00000030897 | 0.622883753 | 0.113321311 | 2.884614835 | 0.039284729 | 0.968663382 | NA | Cnga4 | cyclic nucleotide gated channel alpha 4 [S | 0.8031 | 0 | 1.0062 | 1.928 | 0 | 0 |
| 8094 | ENSMUSG00000030898 | 0 | NA | NA | NA | NA | NA | Cckbr | cholecystokinin B receptor [Source:MGI S | 0 | 0 | 0 | 0 | 0 | 0 |
| 8095 | ENSMUSG00000030905 | 39.31262797 | 0.097757843 | 0.361780872 | 0.270212857 | 0.786996502 | 0.960870081 | Crym | crystallin, mu [Source:MGI Symbol;Acc:M | 41.761 | 27.754 | 44.275 | 39.523 | 45.806 | 36.757 |
| 8096 | ENSMUSG00000030909 | 1243.697286 | 0.492318543 | 0.15367737 | 3.20358517 | 0.001357279 | 0.026260602 | Anks4b | ankyrin repeat and sterile alpha motif dor | 1146 | 1045.4 | 907.63 | 1135.6 | 1533.4 | 1694.2 |
| 8097 | ENSMUSG00000030911 | 0.17131964 | -0.796629176 | 4.080472857 | -0.195229623 | 0.84521319 | NA | Zp2 | zona pellucida glycoprotein 2 [Source:MG | 0 | 1.0279 | 0 | 0 | 0 | 0 |
| 8098 | ENSMUSG00000030917 | 30.17029205 | -0.524372027 | 0.478420498 | -1.096048412 | 0.273057589 | 0.728287842 | Tmem159 | transmembrane protein 159 [Source:MGI | 27.305 | 41.117 | 38.237 | 13.496 | 18.541 | 42.326 |
| 8099 | ENSMUSG00000030921 | 78.88257975 | -0.509552585 | 0.292424719 | -1.742508593 | 0.081419489 | 0.419989833 | Trim30a | tripartite motif-containing 30A [Source:M | 95.569 | 93.541 | 88.55 | 53.019 | 50.168 | 92.449 |
| 8100 | ENSMUSG00000030922 | 51.75445759 | -0.588966926 | 0.370226832 | -1.59082723 | 0.111648463 | 0.490292648 | Lyrm1 | LYR motif containing 1 [Source:MGI Symb | 86.735 | 53.452 | 46.287 | 56.875 | 35.99 | 31.188 |
| 8101 | ENSMUSG00000030924 | 85.42290252 | -0.192285076 | 0.249706305 | -0.770044938 | 0.441273236 | 0.849828766 | Rexo5 | RNA exonuclease 5 [Source:MGI Symbol;A | 92.357 | 91.485 | 89.556 | 82.902 | 90.521 | 65.717 |
| 8102 | ENSMUSG00000030929 | 112.6847758 | -0.384292351 | 0.23059388 | -1.666533177 | 0.095607266 | 0.456089543 | Eri2 | exoribonuclease 2 [Source:MGI Symbol;A | 111.63 | 149.05 | 122.76 | 110.86 | 92.703 | 89.108 |
| 8103 | ENSMUSG00000030930 | 143.2723105 | -1.058077234 | 0.215181142 | -4.917146655 | 8.78E-07 | 7.38E-05 | Chst15 | carbohydrate sulfotransferase 15 [Source | 212.82 | 192.22 | 175.09 | 81.938 | 85.068 | 112.5 |
| 8104 | ENSMUSG00000030934 | 15406.18363 | 0.221265664 | 0.1355068 | 1.632874977 | 0.102495294 | 0.470781813 | Oat | ornithine aminotransferase [Source:MGI S | 16807 | 12159 | 13713 | 14089 | 17785 | 17885 |
| 8105 | ENSMUSG00000030935 | 2220.228519 | -0.178110578 | 0.128635637 | -1.384613021 | 0.166170828 | 0.5925379 | Acsm3 | acyl-CoA synthetase medium-chain family | 2084.1 | 2707.5 | 2281.2 | 2169.9 | 2257.6 | 1821.1 |
| 8106 | ENSMUSG00000030942 | 420.4844757 | 0.010198233 | 0.144243051 | 0.070701726 | 0.943635147 | 0.990077979 | Thumpd1 | THUMP domain containing 1 [Source:MGI | 389.5 | 432.75 | 434.7 | 375.95 | 416.62 | 473.38 |
| 8107 | ENSMUSG00000030945 | 10.53609616 | 2.108568347 | 0.762734938 | 2.764483759 | 0.005701295 | 0.07278334 | Acsm2 | acyl-CoA synthetase medium-chain family | 5.6217 | 2.0558 | 4.025 | 10.604 | 14.178 | 26.732 |
| 8108 | ENSMUSG00000030946 | 976.1250643 | 0.187019238 | 0.158484446 | 1.180047901 | 0.237981164 | 0.686509684 | Lhpp | phospholysine phosphohistidine inorgani | 1135.6 | 886.07 | 715.44 | 960.12 | 1108.1 | 1051.5 |
| 8109 | ENSMUSG00000030954 | 0 | NA | NA | NA | NA | NA | Gp2 | glycoprotein 2 (zymogen granule membra | 0 | 0 | 0 | 0 | 0 | 0 |
| 8110 | ENSMUSG00000030956 | 257.283711 | -0.202164326 | 0.198463234 | -1.018648753 | 0.30836975 | 0.757131228 | Fam53b | family with sequence similarity 53, memb | 354.97 | 223.06 | 246.53 | 218.82 | 251.93 | 248.39 |
| 8111 | ENSMUSG00000030960 | 53.91672567 | -0.16009448 | 0.306062537 | -0.523077674 | 0.600920201 | 0.915461118 | Eef1akmt2 | EEF1A lysine methyltransferase 2 [Source: | 65.854 | 50.368 | 54.337 | 54.947 | 53.44 | 44.554 |
| 8112 | ENSMUSG00000030963 | 0 | NA | NA | NA | NA | NA | Umod | uromodulin [Source:MGI Symbol;Acc:MGI | 0 | 0 | 0 | 0 | 0 | 0 |
| 8113 | ENSMUSG00000030965 | 424.3446209 | 0.097724155 | 0.139946656 | 0.698295749 | 0.484992257 | 0.87256358 | Abraxas2 | BRISC complex subunit [Source:MGI Symb | 394.32 | 430.7 | 405.52 | 469.46 | 387.17 | 458.9 |
| 8114 | ENSMUSG00000030966 | 130.6758241 | 0.241605828 | 0.200787562 | 1.20329081 | 0.228863804 | 0.67545951 | Trim21 | tripartite motif-containing 21 [Source:MG | 120.47 | 117.18 | 121.76 | 151.34 | 137.42 | 135.89 |
| 8115 | ENSMUSG00000030967 | 1110.480484 | 0.045617194 | 0.116768083 | 0.390664926 | 0.696044927 | 0.939992732 | Zranb1 | zinc finger, RAN-binding domain containir | 1109.9 | 1183.1 | 986.12 | 1190.5 | 1147.3 | 1045.9 |
| 8116 | ENSMUSG00000030968 | 136.7432314 | 1.144821547 | 0.259852093 | 4.405666056 | 1.05E-05 | 0.000558741 | Pdilt | protein disulfide isomerase-like, testis exp | 104.4 | 85.317 | 65.406 | 234.25 | 196.31 | 134.78 |

| | | | | | | | | | | | | | | | |
|---|---|---|---|---|---|---|---|---|---|---|---|---|---|---|---|
| 8117 | ENSMUSG00000030970 | 15.70816098 | -1.118386488 | 0.566315395 | -1.974847404 | 0.048285468 | 0.305779774 | Ctbp2 | C-terminal binding protein 2 [Source:MGI | 21.684 | 22.614 | 20.125 | 7.7118 | 7.6343 | 14.48 |
| 8118 | ENSMUSG00000030972 | 3295.318895 | -0.015525996 | 0.1378986 | -0.112589945 | 0.910355657 | 0.985650811 | Acsm5 | acyl-CoA synthetase medium-chain family | 3183.5 | 3672.8 | 3084.1 | 3763.4 | 3455.1 | 2613.1 |
| 8119 | ENSMUSG00000030976 | 0 | NA | NA | NA | NA | NA | Tex36 | testis expressed 36 [Source:MGI Symbol;A | 0 | 0 | 0 | 0 | 0 | 0 |
| 8120 | ENSMUSG00000030978 | 218.5520025 | -0.044350221 | 0.178937897 | -0.247852587 | 0.804248466 | 0.963701061 | Rrm1 | ribonucleotide reductase M1 [Source:MG | 244.95 | 217.92 | 202.26 | 201.47 | 247.57 | 197.15 |
| 8121 | ENSMUSG00000030979 | 425.6202484 | 0.074115922 | 0.171235742 | 0.43282974 | 0.665138466 | 0.929426323 | Uros | uroporphyrinogen III synthase [Source:M( | 435.28 | 369.02 | 438.72 | 335.46 | 494.05 | 481.18 |
| 8122 | ENSMUSG00000030980 | 540.9812471 | -0.023354015 | 0.133303138 | -0.175194787 | 0.860926581 | 0.9759732 | Knop1 | lysine rich nucleolar protein 1 [Source:MG | 487.48 | 584.89 | 564.5 | 518.62 | 578.03 | 512.37 |
| 8123 | ENSMUSG00000030981 | 0.51014012 | 1.031855138 | 3.09002894 | 0.333930574 | 0.738431948 | NA | Mmp21 | matrix metallopeptidase 21 [Source:MGI S | 0 | 0 | 1.0062 | 0.964 | 1.0906 | 0 |
| 8124 | ENSMUSG00000030982 | 364.9315179 | 0.097568715 | 0.139605004 | 0.698891241 | 0.484620002 | 0.87256358 | Vps35l | VPS35 endosomal protein sorting factor li | 371.03 | 328.93 | 357.22 | 363.42 | 398.08 | 370.91 |
| 8125 | ENSMUSG00000030983 | 1067.889592 | 0.159209748 | 0.11429619 | 1.39295761 | 0.16363261 | 0.58776904 | Bccip | BRCA2 and CDKN1A interacting protein [S | 1056.9 | 989.88 | 979.08 | 1030.5 | 1113.5 | 1237.5 |
| 8126 | ENSMUSG00000030986 | 587.5836033 | 0.032856072 | 0.133726899 | 0.245695312 | 0.805918101 | 0.964428432 | Dhx32 | DEAH (Asp-Glu-Ala-His) box polypeptide 3 | 530.85 | 604.42 | 607.77 | 544.65 | 664.19 | 573.63 |
| 8127 | ENSMUSG00000030987 | 700.0785893 | 0.08301072 | 0.116624374 | 0.711778478 | 0.476601962 | 0.86953366 | Stim1 | stromal interaction molecule 1 [Source:M | 669.79 | 680.48 | 690.28 | 777.93 | 693.63 | 688.36 |
| 8128 | ENSMUSG00000030990 | 303.0364597 | 0.186025091 | 0.173492742 | 1.072235579 | 0.283614234 | 0.736605908 | Pgap2 | post-GPI attachment to proteins 2 [Source | 299.56 | 274.45 | 275.71 | 259.31 | 381.72 | 327.47 |
| 8129 | ENSMUSG00000030994 | 0.185640647 | 1.126924175 | 4.080472857 | 0.276174898 | 0.782413736 | NA | D7Ertd443e | DNA segment, Chr 7, ERATO Doi 443, expr | 0 | 0 | 0 | 0 | 0 | 1.1138 |
| 8130 | ENSMUSG00000030996 | 0 | NA | NA | NA | NA | NA | Art1 | ADP-ribosyltransferase 1 [Source:MGI Syn | 0 | 0 | 0 | 0 | 0 | 0 |
| 8131 | ENSMUSG00000031004 | 26.56555576 | -0.542999253 | 0.522151247 | -1.039927139 | 0.298373753 | 0.74911959 | Mki67 | antigen identified by monoclonal antibody | 47.383 | 21.586 | 25.156 | 15.424 | 13.087 | 36.757 |
| 8132 | ENSMUSG00000031007 | 632.9839247 | -0.17947097 | 0.125860848 | -1.425947571 | 0.153883477 | 0.571097853 | Atp6ap2 | ATPase, H+ transporting, lysosomal access | 685.05 | 716.46 | 615.82 | 640.08 | 579.12 | 561.38 |
| 8133 | ENSMUSG00000031010 | 1215.765903 | 0.079805145 | 0.116540069 | 0.684787179 | 0.493478227 | 0.87710702 | Usp9x | ubiquitin specific peptidase 9, X chromoso | 1045.6 | 1319.8 | 1182.3 | 1248.4 | 1252 | 1246.4 |
| 8134 | ENSMUSG00000031012 | 415.0906499 | -0.207471802 | 0.156810489 | -1.3230735 | 0.185810936 | 0.621690813 | Cask | calcium/calmodulin-dependent serine prc | 399.94 | 473.87 | 461.87 | 416.44 | 420.98 | 317.45 |
| 8135 | ENSMUSG00000031015 | 124.3691147 | -0.274415421 | 0.206017498 | -1.332000549 | 0.182860008 | 0.61680331 | Swap70 | SWA-70 protein [Source:MGI Symbol;Acc: | 130.1 | 133.63 | 144.9 | 113.75 | 106.88 | 116.95 |
| 8136 | ENSMUSG00000031016 | 474.072406 | 0.376156734 | 0.197295247 | 1.906567637 | 0.056576599 | 0.337175086 | Wee1 | WEE 1 homolog 1 (S. pombe) [Source:MG | 455.36 | 414.25 | 368.29 | 629.48 | 617.29 | 359.77 |
| 8137 | ENSMUSG00000031021 | 389.8657433 | -0.020072222 | 0.142455254 | -0.140901944 | 0.887947407 | 0.982382091 | Tmem9b | TMEM9 domain family, member B [Source | 383.88 | 415.28 | 378.35 | 357.64 | 372.99 | 431.06 |
| 8138 | ENSMUSG00000031022 | 1.216318829 | 0.500682538 | 2.087023804 | 0.239902648 | 0.810405728 | NA | BC051019 | cDNA sequence BC051019 [Source:MGI Sy | 0 | 2.0558 | 1.0062 | 0.964 | 3.2719 | 0 |
| 8139 | ENSMUSG00000031023 | 354.0844675 | -0.089496096 | 0.143554766 | -0.623428245 | 0.533003142 | 0.889143097 | Akip1 | A kinase (PRKA) interacting protein 1 [Sou | 379.06 | 335.1 | 380.36 | 319.08 | 353.36 | 357.54 |
| 8140 | ENSMUSG00000031024 | 394.855336 | -0.973657391 | 0.175684259 | -5.542086686 | 2.99E-08 | 3.87E-06 | Denn2b | DENN domain containing 2B [Source:MGI | 490.7 | 462.56 | 616.83 | 283.41 | 212.67 | 302.97 |
| 8141 | ENSMUSG00000031026 | 2.208501132 | -1.666613533 | 1.764125691 | -0.944724938 | 0.344799322 | NA | Trim66 | tripartite motif-containing 66 [Source:MG | 0 | 2.0558 | 8.05 | 0.964 | 2.1812 | 0 |
| 8142 | ENSMUSG00000031027 | 0 | NA | NA | NA | NA | NA | Stk33 | serine/threonine kinase 33 [Source:MGI S | 0 | 0 | 0 | 0 | 0 | 0 |
| 8143 | ENSMUSG00000031028 | 0 | NA | NA | NA | NA | NA | Tub | tubby bipartite transcription factor [Sourc | 0 | 0 | 0 | 0 | 0 | 0 |
| 8144 | ENSMUSG00000031029 | 1516.214472 | -0.027933628 | 0.107545218 | -0.259738449 | 0.795065532 | 0.962189806 | Eif3f | eukaryotic translation initiation factor 3, s | 1470.5 | 1601.5 | 1520.4 | 1362.1 | 1540 | 1602.8 |
| 8145 | ENSMUSG00000031059 | 2460.285424 | 0.142919446 | 0.098380998 | 1.45271393 | 0.146303198 | 0.556855766 | Ndufb11 | NADH:ubiquinone oxidoreductase subuni | 2258.3 | 2371.4 | 2385.8 | 2448.5 | 2499.7 | 2798 |
| 8146 | ENSMUSG00000031060 | 434.8604876 | 0.191971607 | 0.144608464 | 1.32752677 | 0.18433449 | 0.619137675 | Rbm10 | RNA binding motif protein 10 [Source:MG | 379.87 | 382.39 | 455.83 | 442.47 | 436.25 | 512.37 |
| 8147 | ENSMUSG00000031065 | 558.6436802 | 0.10609637 | 0.13122569 | 0.80850304 | 0.418801057 | 0.837591673 | Cdk16 | cyclin-dependent kinase 16 [Source:MGI S | 501.94 | 558.16 | 555.45 | 642.97 | 544.22 | 549.13 |
| 8148 | ENSMUSG00000031066 | 11.70139176 | -0.478815223 | 0.617184033 | -0.775806239 | 0.4378634 | 0.847422601 | Usp11 | ubiquitin specific peptidase 11 [Source:M( | 14.456 | 12.335 | 14.087 | 10.604 | 9.8156 | 8.9108 |
| 8149 | ENSMUSG00000031068 | 1170.272089 | 0.006664121 | 0.104231395 | 0.063935828 | 0.949021324 | 0.990731019 | Glrx3 | glutaredoxin 3 [Source:MGI Symbol;Acc:N | 1133.2 | 1253 | 1116.9 | 1165.4 | 1174.6 | 1178.4 |
| 8150 | ENSMUSG00000031070 | 2.545050432 | -0.624984375 | 1.432633703 | -0.43624855 | 0.662656407 | NA | Mrgprf | MAS-related GPR, member F [Source:MGI | 3.2124 | 4.1117 | 2.0125 | 4.8199 | 0 | 1.1138 |
| 8151 | ENSMUSG00000031072 | 267.5259794 | 0.291048266 | 0.172125235 | 1.690909913 | 0.09085401 | 0.444639662 | LTO1 | ABCE maturation factor [Source:MGI Sym | 244.95 | 246.7 | 230.43 | 326.79 | 238.85 | 317.45 |
| 8152 | ENSMUSG00000031073 | 0 | NA | NA | NA | NA | NA | Fgf15 | fibroblast growth factor 15 [Source:MGI S | 0 | 0 | 0 | 0 | 0 | 0 |
| 8153 | ENSMUSG00000031074 | 0 | NA | NA | NA | NA | NA | Fgf3 | fibroblast growth factor 3 [Source:MGI Sy | 0 | 0 | 0 | 0 | 0 | 0 |
| 8154 | ENSMUSG00000031075 | 28.89040217 | -0.654063078 | 0.437095465 | -1.496384957 | 0.134553367 | 0.537616739 | Ano1 | anoctamin 1, calcium activated chloride cl | 32.927 | 30.838 | 42.262 | 21.208 | 31.628 | 14.48 |
| 8155 | ENSMUSG00000031077 | 210.6855922 | 0.004761257 | 0.169975745 | 0.028011391 | 0.977653066 | 0.997564159 | Fadd | Fas (TNFRSF6)-associated via death doma | 211.22 | 198.39 | 221.37 | 209.18 | 229.03 | 194.92 |
| 8156 | ENSMUSG00000031078 | 887.7607373 | 0.098829464 | 0.118376756 | 0.83487221 | 0.403789663 | 0.828953151 | Cttn | cortactin [Source:MGI Symbol;Acc:MGI:9! | 770.98 | 890.18 | 911.66 | 866.62 | 928.12 | 959.02 |
| 8157 | ENSMUSG00000031079 | 2.732313375 | -2.119027265 | 1.46393342 | -1.447488824 | 0.147760064 | NA | Zfp300 | zinc finger protein 300 [Source:MGI Symb | 7.2279 | 4.1117 | 2.0125 | 1.928 | 0 | 1.1138 |
| 8158 | ENSMUSG00000031085 | 0 | NA | NA | NA | NA | NA | Acte1 | actin, epsilon 1 [Source:MGI Symbol;Acc:I | 0 | 0 | 0 | 0 | 0 | 0 |
| 8159 | ENSMUSG00000031089 | 0.501260735 | 1.111687294 | 3.109072805 | 0.357562323 | 0.720670879 | NA | Slc6a14 | solute carrier family 6 (neurotransmitter t | 0.8031 | 0 | 0 | 0 | 1.0906 | 1.1138 |
| 8160 | ENSMUSG00000031090 | 475.9840566 | 0.172215405 | 0.142443509 | 1.209008442 | 0.226659602 | 0.673420584 | Nadsyn1 | NAD synthetase 1 [Source:MGI Symbol;Ac | 489.09 | 455.37 | 397.47 | 473.31 | 559.49 | 481.18 |
| 8161 | ENSMUSG00000031093 | 13.14801799 | -0.568572432 | 0.629205475 | -0.903635545 | 0.366188687 | 0.804458258 | Dock11 | dedicator of cytokinesis 11 [Source:MGI S | 14.456 | 21.586 | 11.069 | 9.6398 | 6.5437 | 15.594 |
| 8162 | ENSMUSG00000031095 | 451.9115923 | 0.193645923 | 0.128670219 | 1.504978573 | 0.132329586 | 0.533283479 | Cul4b | cullin 4B [Source:MGI Symbol;Acc:MGI:19 | 408.78 | 439.95 | 416.59 | 487.77 | 500.59 | 457.79 |
| 8163 | ENSMUSG00000031097 | 0.133850373 | -0.796629176 | 4.080472857 | -0.195229623 | 0.84521319 | NA | Tnni2 | troponin I, skeletal, fast 2 [Source:MGI Syı | 0.8031 | 0 | 0 | 0 | 0 | 0 |
| 8164 | ENSMUSG00000031098 | 0.17131964 | -0.796629176 | 4.080472857 | -0.195229623 | 0.84521319 | NA | Syt8 | synaptotagmin VIII [Source:MGI Symbol;A | 0 | 1.0279 | 0 | 0 | 0 | 0 |
| 8165 | ENSMUSG00000031099 | 1.609946696 | 0.310181083 | 1.891928032 | 0.163949726 | 0.869770721 | NA | Smarca1 | SWI/SNF related, matrix associated, actin | 3.2124 | 1.0279 | 0 | 0.964 | 0 | 4.4554 |
| 8166 | ENSMUSG00000031101 | 8.905502995 | -0.688503958 | 0.836547894 | -0.823029933 | 0.410490971 | 0.833241465 | Sash3 | SAM and SH3 domain containing 3 [Sourc | 9.6372 | 16.447 | 7.0437 | 12.532 | 1.0906 | 6.6831 |
| 8167 | ENSMUSG00000031103 | 20.09994874 | -1.191634847 | 0.500446309 | -2.38114424 | 0.017258953 | 0.156510025 | Elf4 | E74-like factor 4 (ets domain transcription | 28.912 | 30.838 | 24.15 | 13.496 | 8.7249 | 14.48 |
| 8168 | ENSMUSG00000031104 | 1.219931038 | 0.487877328 | 2.270901307 | 0.214838631 | 0.829893133 | NA | Rab33a | RAB33A, member RAS oncogene family [S | 0 | 3.0838 | 0 | 0.964 | 3.2719 | 0 |
| 8169 | ENSMUSG00000031105 | 39.76039831 | 0.787609378 | 0.385438145 | 2.043413161 | 0.041011556 | 0.277451423 | Slc25a14 | solute carrier family 25 (mitochondrial car | 16.865 | 33.921 | 37.231 | 49.163 | 51.259 | 50.123 |
| 8170 | ENSMUSG00000031107 | 17.66563559 | 0.92411623 | 0.604324064 | 1.52917331 | 0.126221485 | 0.521094669 | Rbmx2 | RNA binding motif protein, X-linked 2 [Sou | 16.062 | 6.1675 | 14.087 | 15.424 | 15.269 | 38.985 |
| 8171 | ENSMUSG00000031109 | 224.9223664 | 0.153136155 | 0.165472707 | 0.925446607 | 0.354733645 | 0.794867148 | Enox2 | ecto-NOX disulfide-thiol exchanger 2 [Sou | 208.81 | 205.58 | 224.39 | 215.93 | 248.66 | 246.16 |
| 8172 | ENSMUSG00000031111 | 1.068948596 | -2.273572368 | 2.191117856 | -1.037631254 | 0.299441762 | NA | Igsf1 | immunoglobulin superfamily, member 1 [! | 2.4093 | 1.0279 | 2.0125 | 0.964 | 0 | 0 |

| | | | | | | | | | | | | | | | |
|---|---|---|---|---|---|---|---|---|---|---|---|---|---|---|---|
| 8173 | ENSMUSG00000031112 | 9.56919273 | -0.681527878 | 0.736544615 | -0.925304271 | 0.354807658 | 0.794902164 | Stk26 | serine/threonine kinase 26 [Source:MGI S | 16.865 | 13.363 | 5.0312 | 6.7478 | 8.7249 | 6.6831 |
| 8174 | ENSMUSG00000031118 | 0 | NA | NA | NA | NA | NA | Magea14 | MAGE family member A14 [Source:MGI Sy | 0 | 0 | 0 | 0 | 0 | 0 |
| 8175 | ENSMUSG00000031119 | 986.7312778 | -0.294190548 | 0.118056071 | -2.491956126 | 0.012704173 | 0.127461989 | Gpc4 | glypican 4 [Source:MGI Symbol;Acc:MGI:1 | 1058.5 | 1142 | 1061.6 | 988.08 | 810.33 | 859.89 |
| 8176 | ENSMUSG00000031125 | 2.069926006 | -3.317193035 | 1.811059714 | -1.831630956 | 0.067006422 | NA | 3830403N18 | RIKEN cDNA 3830403N18 gene [Source:M | 3.2124 | 2.0558 | 6.0375 | 0 | 0 | 1.1138 |
| 8177 | ENSMUSG00000031129 | 31.13497568 | -0.536535437 | 0.397280205 | -1.350521446 | 0.176848779 | 0.607589534 | Slc9a9 | solute carrier family 9 (sodium/hydrogen | 39.352 | 41.117 | 30.187 | 29.883 | 23.994 | 22.277 |
| 8178 | ENSMUSG00000031130 | 0.181769715 | 1.126924175 | 4.080472857 | 0.276174898 | 0.782413736 | NA | Brs3 | bombesin-like receptor 3 [Source:MGI Syr | 0 | 0 | 0 | 0 | 1.0906 | 0 |
| 8179 | ENSMUSG00000031131 | 0 | NA | NA | NA | NA | NA | Vgll1 | vestigial like family member 1 [Source:MG | 0 | 0 | 0 | 0 | 0 | 0 |
| 8180 | ENSMUSG00000031132 | 0 | NA | NA | NA | NA | NA | Cd40lg | CD40 ligand [Source:MGI Symbol;Acc:MG | 0 | 0 | 0 | 0 | 0 | 0 |
| 8181 | ENSMUSG00000031133 | 42.61499479 | -1.088625234 | 0.381636799 | -2.852516419 | 0.004337457 | 0.060746309 | Arhgef6 | Rac/Cdc42 guanine nucleotide exchange f | 77.901 | 56.535 | 39.244 | 26.991 | 31.628 | 23.391 |
| 8182 | ENSMUSG00000031134 | 278.4346939 | -0.143772962 | 0.155698276 | -0.923407544 | 0.355794862 | 0.795525652 | Rbmx | RNA binding motif protein, X chromosome | 266.63 | 298.1 | 312.94 | 281.48 | 251.93 | 259.53 |
| 8183 | ENSMUSG00000031137 | 0.17131964 | -0.796629176 | 4.080472857 | -0.195229623 | 0.84521319 | NA | Fgf13 | fibroblast growth factor 13 [Source:MGI S | 0 | 1.0279 | 0 | 0 | 0 | 0 |
| 8184 | ENSMUSG00000031138 | 4271.627516 | -0.038828246 | 0.086472277 | -0.449025367 | 0.653413358 | 0.925756399 | F9 | coagulation factor IX [Source:MGI Symbol | 4421.1 | 4307 | 4259.4 | 4391.9 | 4177.1 | 4073.3 |
| 8185 | ENSMUSG00000031139 | 0.569258551 | -2.569337407 | 3.18860282 | -0.805787849 | 0.420365185 | NA | Mcf2 | mcf.2 transforming sequence [Source:MG | 2.4093 | 0 | 1.0062 | 0 | 0 | 0 |
| 8186 | ENSMUSG00000031142 | 0.877550482 | 2.072039088 | 2.361583201 | 0.877394066 | 0.380272635 | NA | Cacna1f | calcium channel, voltage-dependent, alph | 0 | 0 | 1.0062 | 0.964 | 2.1812 | 1.1138 |
| 8187 | ENSMUSG00000031143 | 0 | NA | NA | NA | NA | NA | Ccdc22 | coiled-coil domain containing 22 [Source:I | 0 | 0 | 0 | 0 | 0 | 0 |
| 8188 | ENSMUSG00000031144 | 7.108198051 | -0.169512753 | 0.892125406 | -0.190010005 | 0.84930129 | 0.973071229 | Syp | synaptophysin [Source:MGI Symbol;Acc:N | 7.2279 | 12.335 | 3.0187 | 6.7478 | 2.1812 | 11.138 |
| 8189 | ENSMUSG00000031145 | 67.35943751 | 0.114829788 | 0.287243778 | 0.399764229 | 0.689330179 | 0.937074409 | Prickle3 | prickle planar cell polarity protein 3 [Sour | 56.217 | 67.843 | 70.437 | 83.866 | 55.622 | 70.172 |
| 8190 | ENSMUSG00000031146 | 18.53795912 | 0.196422618 | 0.551481032 | 0.356172936 | 0.721711059 | 0.9447719 | Plp2 | proteolipid protein 2 [Source:MGI Symbol | 28.109 | 13.363 | 10.062 | 22.171 | 16.359 | 21.163 |
| 8191 | ENSMUSG00000031147 | 334.5116962 | 0.245540361 | 0.192300113 | 1.276860206 | 0.201651606 | 0.645999739 | Magix | MAGI family member, X-linked [Source:M | 245.75 | 287.82 | 385.39 | 344.14 | 317.37 | 426.6 |
| 8192 | ENSMUSG00000031148 | 345.4483452 | -0.214409888 | 0.142125353 | -1.508597048 | 0.131401789 | 0.531189345 | Gpkow | G patch domain and KOW motifs [Source: | 342.92 | 386.5 | 384.39 | 321 | 330.46 | 307.42 |
| 8193 | ENSMUSG00000031149 | 26.83961754 | -0.483854865 | 0.43916654 | -1.101757126 | 0.270567274 | 0.725929337 | Praf2 | PRA1 domain family 2 [Source:MGI Symbo | 25.699 | 34.949 | 33.206 | 16.388 | 20.722 | 30.074 |
| 8194 | ENSMUSG00000031150 | 11.99809503 | -1.075198188 | 0.753525793 | -1.426889694 | 0.153611692 | 0.570925073 | Ccdc120 | coiled-coil domain containing 120 [Source | 23.29 | 12.335 | 13.081 | 7.7118 | 1.0906 | 14.48 |
| 8195 | ENSMUSG00000031153 | 412.3981151 | -0.016390237 | 0.141789875 | -0.115595253 | 0.907973325 | 0.984639154 | Gripap1 | GRIP1 associated protein 1 [Source:MGI S | 398.34 | 398.83 | 447.78 | 446.32 | 414.43 | 368.68 |
| 8196 | ENSMUSG00000031154 | 903.4221531 | 0.0970584 | 0.115638968 | 0.83932261 | 0.401288299 | 0.82816843 | Otud5 | OTU domain containing 5 [Source:MGI Sy | 908.31 | 823.36 | 887.51 | 1013.1 | 865.95 | 922.26 |
| 8197 | ENSMUSG00000031155 | 64.50674179 | 0.311933938 | 0.298688899 | 1.044343929 | 0.296326295 | 0.746961229 | Pim2 | proviral integration site 2 [Source:MGI Sy | 44.171 | 63.731 | 65.406 | 84.83 | 66.528 | 62.375 |
| 8198 | ENSMUSG00000031156 | 216.3965365 | 0.229188354 | 0.167012777 | 1.372280355 | 0.169976179 | 0.596876778 | Slc35a2 | solute carrier family 35 (UDP-galactose tra | 191.94 | 216.89 | 189.17 | 233.28 | 243.21 | 223.88 |
| 8199 | ENSMUSG00000031157 | 411.7229085 | -0.144341893 | 0.140189146 | -1.029622457 | 0.303187269 | 0.752484788 | Pqbp1 | polyglutamine binding protein 1 [Source:N | 441.71 | 458.45 | 396.46 | 373.06 | 427.52 | 373.14 |
| 8200 | ENSMUSG00000031158 | 743.4149102 | 0.123676183 | 0.119224476 | 1.037338872 | 0.299577956 | 0.750839638 | Timm17b | translocase of inner mitochondrial membr | 677.82 | 707.21 | 749.65 | 710.45 | 823.42 | 791.94 |
| 8201 | ENSMUSG00000031160 | 0 | NA | NA | NA | NA | NA | Eras | ES cell-expressed Ras [Source:MGI Symbo | 0 | 0 | 0 | 0 | 0 | 0 |
| 8202 | ENSMUSG00000031161 | 288.8985067 | 0.080973425 | 0.154590508 | 0.523792996 | 0.600422526 | 0.91534406 | Hdac6 | histone deacetylase 6 [Source:MGI Symbo | 274.66 | 263.15 | 304.89 | 316.18 | 272.65 | 301.85 |
| 8203 | ENSMUSG00000031162 | 0.371281295 | 2.09253362 | 4.031852165 | 0.519000582 | 0.603760335 | NA | Gata1 | GATA binding protein 1 [Source:MGI Sym | 0 | 0 | 0 | 0 | 0 | 2.2277 |
| 8204 | ENSMUSG00000031163 | 1.008278937 | -3.39371293 | 2.386458974 | -1.42207051 | 0.155005783 | NA | Glod5 | glyoxalase domain containing 5 [Source:M | 4.0155 | 1.0279 | 1.0062 | 0 | 0 | 0 |
| 8205 | ENSMUSG00000031165 | 16.35660031 | -2.010226423 | 0.628014173 | -3.20092525 | 0.001369871 | 0.026418412 | Was | Wiskott-Aldrich syndrome [Source:MGI Sy | 28.912 | 30.838 | 19.119 | 11.568 | 4.3625 | 3.3415 |
| 8206 | ENSMUSG00000031166 | 530.310969 | 0.202005913 | 0.126713802 | 1.594190293 | 0.11089342 | 0.489424955 | Wdr13 | WD repeat domain 13 [Source:MGI Symbo | 487.48 | 511.9 | 480.98 | 616.95 | 538.77 | 545.78 |
| 8207 | ENSMUSG00000031167 | 564.1284891 | -0.289128235 | 0.198875744 | -1.453813469 | 0.145998028 | 0.556855766 | Rbm3 | RNA binding motif (RNP1, RRM) protein 3 | 511.58 | 863.45 | 487.02 | 488.74 | 559.49 | 474.5 |
| 8208 | ENSMUSG00000031168 | 3209.380261 | 0.144204421 | 0.087976902 | 1.639116842 | 0.101188928 | 0.46850085 | Ebp | phenylalkylamine Ca2+ antagonist (emopa | 3040.5 | 3026.2 | 3080.1 | 3220.6 | 3415.8 | 3473 |
| 8209 | ENSMUSG00000031169 | 39.37721705 | -0.306467115 | 0.431042289 | -0.71099083 | 0.477089918 | 0.869671045 | Porcn | porcupine O-acyltransferase [Source:MGI | 34.533 | 49.34 | 47.293 | 58.803 | 22.903 | 23.391 |
| 8210 | ENSMUSG00000031170 | 0.17131964 | -0.796629176 | 4.080472857 | -0.195229623 | 0.84521319 | NA | Slc38a5 | solute carrier family 38, member 5 [Sourc | 0 | 1.0279 | 0 | 0 | 0 | 0 |
| 8211 | ENSMUSG00000031171 | 339.7918631 | -0.002653641 | 0.146731989 | -0.018084952 | 0.985571083 | 0.99859176 | Ftsj1 | FtsJ RNA methyltransferase homolog 1 (E. | 329.27 | 325.85 | 365.27 | 326.79 | 371.9 | 319.67 |
| 8212 | ENSMUSG00000031173 | 10702.13983 | 0.109805704 | 0.089683545 | 1.224368457 | 0.220813263 | 0.667237604 | Otc | ornithine transcarbamylase [Source:MGI S | 10611 | 9863.9 | 10409 | 10430 | 11754 | 11145 |
| 8213 | ENSMUSG00000031174 | 38.05764692 | -0.050713164 | 0.379511554 | -0.133627457 | 0.893697171 | 0.982562372 | Rpgr | retinitis pigmentosa GTPase regulator [So | 45.777 | 39.061 | 31.194 | 37.595 | 47.987 | 26.732 |
| 8214 | ENSMUSG00000031176 | 1071.430049 | 0.071797384 | 0.114007936 | 0.629757775 | 0.528853076 | 0.888334265 | Dynlt3 | dynein light chain Tctex-type 3 [Source:M | 1077.8 | 1037.2 | 1018.3 | 1001.6 | 1081.9 | 1211.9 |
| 8215 | ENSMUSG00000031179 | 0 | NA | NA | NA | NA | NA | Magea9 | MAGE family member A9 [Source:MGI Syr | 0 | 0 | 0 | 0 | 0 | 0 |
| 8216 | ENSMUSG00000031181 | 0 | NA | NA | NA | NA | NA | Ctag2 | cancer/testis antigen 2 [Source:MGI Symb | 0 | 0 | 0 | 0 | 0 | 0 |
| 8217 | ENSMUSG00000031182 | 0 | NA | NA | NA | NA | NA | 4930447F04 | RIKEN cDNA 4930447F04 gene [Source:M | 0 | 0 | 0 | 0 | 0 | 0 |
| 8218 | ENSMUSG00000031189 | 0.160662974 | 1.126924175 | 4.080472857 | 0.276174898 | 0.782413736 | NA | Aff2 | AF4/FMR2 family, member 2 [Source:MGI | 0 | 0 | 0 | 0.964 | 0 | 0 |
| 8219 | ENSMUSG00000031194 | 0 | NA | NA | NA | NA | NA | 4931400O07 | RIKEN cDNA 4931400O07 gene [Source:M | 0 | 0 | 0 | 0 | 0 | 0 |
| 8220 | ENSMUSG00000031196 | 134.6606161 | -0.425250781 | 0.20777319 | -2.046706707 | 0.04068689 | 0.276029635 | F8 | coagulation factor VIII [Source:MGI Symbo | 134.12 | 164.47 | 165.02 | 120.5 | 105.79 | 118.07 |
| 8221 | ENSMUSG00000031197 | 210.5449081 | -0.195137898 | 0.174348436 | -1.119240886 | 0.263037387 | 0.717708502 | Vbp1 | von Hippel-Lindau binding protein 1 [Sour | 206.4 | 216.89 | 251.56 | 209.18 | 184.31 | 194.92 |
| 8222 | ENSMUSG00000031198 | 135.4423755 | 0.17164315 | 0.211869363 | 0.810136716 | 0.417861604 | 0.837344173 | Fundc2 | FUN14 domain containing 2 [Source:MGI | 119.66 | 130.55 | 131.82 | 118.57 | 148.32 | 163.74 |
| 8223 | ENSMUSG00000031200 | 85.9134273 | -0.004876464 | 0.249889472 | -0.019514483 | 0.984430683 | 0.998580735 | Mtcp1 | mature T cell proliferation 1 [Source:MGI | 78.704 | 86.345 | 93.581 | 99.29 | 80.706 | 76.855 |
| 8224 | ENSMUSG00000031201 | 429.0703983 | 0.094880581 | 0.131371581 | 0.722230641 | 0.470152687 | 0.867051954 | Brcc3 | BRCA1/BRCA2-containing complex, subun | 413.6 | 440.98 | 390.42 | 442.47 | 454.79 | 432.17 |
| 8225 | ENSMUSG00000031202 | 0.185640647 | 1.126924175 | 4.080472857 | 0.276174898 | 0.782413736 | NA | Rab39b | RAB39B, member RAS oncogene family [S | 0 | 0 | 0 | 0 | 0 | 1.1138 |
| 8226 | ENSMUSG00000031204 | 0 | NA | NA | NA | NA | NA | Asb12 | ankyrin repeat and SOCS box-containing 1 | 0 | 0 | 0 | 0 | 0 | 0 |
| 8227 | ENSMUSG00000031207 | 234.839817 | -1.122942549 | 0.209263171 | -5.366173807 | 8.04E-08 | 9.02E-06 | Msn | moesin [Source:MGI Symbol;Acc:MGI:971 | 317.23 | 392.66 | 255.59 | 138.81 | 125.42 | 179.33 |
| 8228 | ENSMUSG00000031209 | 14.90753267 | -1.257375914 | 0.603501817 | -2.08346666 | 0.037208708 | 0.261128941 | Heph | hephaestin [Source:MGI Symbol;Acc:MGI | 27.305 | 18.503 | 17.106 | 7.7118 | 5.4531 | 13.366 |

| | | | | | | | | | | | | | | | |
|---|---|---|---|---|---|---|---|---|---|---|---|---|---|---|---|
| 8229 | ENSMUSG00000031210 | 0 | NA | NA | NA | NA | NA | Gpr165 | G protein-coupled receptor 165 [Source:N | 0 | 0 | 0 | 0 | 0 | 0 |
| 8230 | ENSMUSG00000031212 | 0 | NA | NA | NA | NA | NA | Pgr15l | G protein-coupled receptor 15-like [Sourc | 0 | 0 | 0 | 0 | 0 | 0 |
| 8231 | ENSMUSG00000031214 | 33.49919664 | -0.519249599 | 0.450933923 | -1.151498197 | 0.249527337 | 0.701784229 | Ophn1 | oligophrenin 1 [Source:MGI Symbol;Acc:N | 60.233 | 22.614 | 35.219 | 29.883 | 19.631 | 33.415 |
| 8232 | ENSMUSG00000031216 | 128.0605401 | -0.466215605 | 0.270106993 | -1.726040481 | 0.08434013 | 0.428183799 | Stard8 | START domain containing 8 [Source:MGI S | 145.36 | 126.43 | 174.08 | 123.39 | 65.437 | 133.66 |
| 8233 | ENSMUSG00000031217 | 307.6889492 | -0.265793881 | 0.201793601 | -1.317157133 | 0.187785959 | 0.624686319 | Efnb1 | ephrin B1 [Source:MGI Symbol;Acc:MGI:1 | 254.58 | 320.71 | 433.69 | 319.08 | 255.2 | 262.87 |
| 8234 | ENSMUSG00000031220 | 46.66055399 | -0.064113872 | 0.346298957 | -0.18514024 | 0.85311904 | 0.973825032 | Awat2 | acyl-CoA wax alcohol acyltransferase 2 [Sc | 54.611 | 31.865 | 56.35 | 42.415 | 50.168 | 44.554 |
| 8235 | ENSMUSG00000031221 | 932.4765847 | 0.067573484 | 0.106579563 | 0.634019157 | 0.52606832 | 0.888334265 | Igbp1 | immunoglobulin (CD79A) binding protein | 874.58 | 937.46 | 920.71 | 981.33 | 965.2 | 915.58 |
| 8236 | ENSMUSG00000031224 | 0 | NA | NA | NA | NA | NA | Magee2 | MAGE family member E2 [Source:MGI Syr | 0 | 0 | 0 | 0 | 0 | 0 |
| 8237 | ENSMUSG00000031226 | 114.0918356 | -0.013969138 | 0.239333287 | -0.058366882 | 0.953456394 | 0.9915542 | Pbdc1 | polysaccharide biosynthesis domain conta | 126.09 | 91.485 | 125.78 | 97.362 | 109.06 | 134.78 |
| 8238 | ENSMUSG00000031227 | 41.39504264 | 0.109883995 | 0.376724729 | 0.291682457 | 0.770529423 | 0.956515747 | Magee1 | MAGE family member E1 [Source:MGI Syr | 38.549 | 28.782 | 52.325 | 54.947 | 40.353 | 33.415 |
| 8239 | ENSMUSG00000031229 | 879.7505123 | 0.103077687 | 0.132043512 | 0.780634244 | 0.435017645 | 0.845325456 | Atrx | ATRX, chromatin remodeler [Source:MGI | 805.51 | 905.6 | 835.18 | 1035.3 | 917.21 | 779.69 |
| 8240 | ENSMUSG00000031230 | 0 | NA | NA | NA | NA | NA | Fgf16 | fibroblast growth factor 16 [Source:MGI S | 0 | 0 | 0 | 0 | 0 | 0 |
| 8241 | ENSMUSG00000031231 | 3051.068986 | 0.042183343 | 0.102510253 | 0.411503648 | 0.680703267 | 0.934384556 | Cox7b | cytochrome c oxidase subunit 7B [Source: | 3121.7 | 3042.6 | 2853.7 | 2807.1 | 3145.3 | 3336 |
| 8242 | ENSMUSG00000031232 | 1406.876081 | 0.066443825 | 0.108383224 | 0.613045286 | 0.539846396 | 0.892687582 | Magt1 | magnesium transporter 1 [Source:MGI Syr | 1295.4 | 1418.5 | 1410.8 | 1536.6 | 1315.3 | 1464.7 |
| 8243 | ENSMUSG00000031233 | 0 | NA | NA | NA | NA | NA | Pgk2 | phosphoglycerate kinase 2 [Source:MGI S | 0 | 0 | 0 | 0 | 0 | 0 |
| 8244 | ENSMUSG00000031239 | 13.48110135 | -0.617051859 | 0.63415502 | -0.97303 | 0.330538392 | 0.774747658 | Itm2a | integral membrane protein 2A [Source:M( | 23.29 | 12.335 | 13.081 | 6.7478 | 8.7249 | 16.708 |
| 8245 | ENSMUSG00000031241 | 0 | NA | NA | NA | NA | NA | Tbx22 | T-box 22 [Source:MGI Symbol;Acc:MGI:23 | 0 | 0 | 0 | 0 | 0 | 0 |
| 8246 | ENSMUSG00000031242 | 330.4721717 | -0.019742315 | 0.154342556 | -0.127912321 | 0.89821836 | 0.982798755 | 2610002M0( | RIKEN cDNA 2610002M06 gene [Source:M | 364.61 | 329.96 | 302.88 | 310.4 | 366.45 | 308.53 |
| 8247 | ENSMUSG00000031245 | 811.4416855 | 0.146299697 | 0.134769962 | 1.085551227 | 0.277677588 | 0.730774888 | Hmgn5 | high-mobility group nucleosome binding c | 718.78 | 829.53 | 762.73 | 771.18 | 792.88 | 993.55 |
| 8248 | ENSMUSG00000031246 | 2276.569737 | -0.057906633 | 0.098985855 | -0.58499907 | 0.558548342 | 0.899348015 | Sh3bgrl | SH3-binding domain glutamic acid-rich pro | 2422.2 | 2414.6 | 2129.2 | 2169.9 | 2290.3 | 2233.3 |
| 8249 | ENSMUSG00000031250 | 0 | NA | NA | NA | NA | NA | Tnmd | tenomodulin [Source:MGI Symbol;Acc:M( | 0 | 0 | 0 | 0 | 0 | 0 |
| 8250 | ENSMUSG00000031253 | 28.87097033 | -0.638222476 | 0.44751595 | -1.426144645 | 0.153826594 | 0.571085179 | Srpx2 | sushi-repeat-containing protein, X-linked | 47.383 | 27.754 | 30.187 | 26.991 | 14.178 | 26.732 |
| 8251 | ENSMUSG00000031255 | 65.54853732 | -0.041186207 | 0.317838779 | -0.129582069 | 0.896897091 | 0.982798755 | Sytl4 | synaptotagmin-like 4 [Source:MGI Symbol | 89.144 | 61.675 | 48.3 | 74.226 | 68.709 | 51.237 |
| 8252 | ENSMUSG00000031256 | 255.914882 | -0.154328448 | 0.15628542 | -0.987478216 | 0.323408257 | 0.770475661 | Cstf2 | cleavage stimulation factor, 3' pre-RNA su | 256.19 | 261.09 | 291.81 | 244.85 | 244.3 | 237.25 |
| 8253 | ENSMUSG00000031257 | 0.531944269 | 2.618916564 | 3.285123409 | 0.797204926 | 0.425332026 | NA | Nox1 | NADPH oxidase 1 [Source:MGI Symbol;Ac | 0 | 0 | 0 | 0.964 | 0 | 2.2277 |
| 8254 | ENSMUSG00000031258 | 2.1777838 | -0.088202879 | 1.74430227 | -0.050566281 | 0.959671132 | NA | Xkrx | X-linked Kx blood group related, X-linked | 5.6217 | 1.0279 | 0 | 0.964 | 5.4531 | 0 |
| 8255 | ENSMUSG00000031262 | 1.82348533 | -0.119783331 | 1.631350139 | -0.073425887 | 0.941467219 | NA | Cenpi | centromere protein I [Source:MGI Symbol | 1.6062 | 4.1117 | 0 | 1.928 | 2.1812 | 1.1138 |
| 8256 | ENSMUSG00000031264 | 17.17262309 | -1.165516085 | 0.593984027 | -1.962201057 | 0.049739081 | 0.31211543 | Btk | Bruton agammaglobulinemia tyrosine kina | 23.29 | 29.81 | 18.112 | 9.6398 | 4.3625 | 17.822 |
| 8257 | ENSMUSG00000031266 | 92.39364518 | 0.014763994 | 0.268100995 | 0.055068777 | 0.956083671 | 0.991678478 | Gla | galactosidase, alpha [Source:MGI Symbol; | 95.569 | 76.066 | 103.64 | 67.478 | 105.79 | 105.82 |
| 8258 | ENSMUSG00000031270 | 2.475244319 | -1.29905859 | 1.449842486 | -0.895999809 | 0.370252865 | NA | 4930513O0( | RIKEN cDNA 4930513O06 gene [Source:M | 2.4093 | 4.1117 | 4.025 | 0.964 | 0 | 3.3415 |
| 8259 | ENSMUSG00000031271 | 1155.845209 | -0.189919735 | 0.671184216 | -0.282962159 | 0.777205845 | 0.959467933 | Serpina7 | serine (or cysteine) peptidase inhibitor, cl | 1937.1 | 1307.5 | 450.8 | 1650.3 | 684.91 | 904.44 |
| 8260 | ENSMUSG00000031273 | 5.071405424 | -1.179261056 | 1.158531886 | -1.017892619 | 0.308728989 | NA | Col4a6 | collagen, type IV, alpha 6 [Source:MGI Syr | 12.85 | 8.2233 | 0 | 3.8559 | 3.2719 | 2.2277 |
| 8261 | ENSMUSG00000031274 | 22.24313825 | -0.273259996 | 0.53974912 | -0.506272241 | 0.612665548 | 0.916495447 | Col4a5 | collagen, type IV, alpha 5 [Source:MGI Syr | 27.305 | 21.586 | 24.15 | 26.027 | 6.5437 | 27.846 |
| 8262 | ENSMUSG00000031278 | 855.3459806 | 0.13599952 | 0.135020923 | 1.007247745 | 0.313815734 | 0.760476509 | Acsl4 | acyl-CoA synthetase long-chain family me | 805.51 | 919.99 | 720.47 | 990.97 | 895.4 | 799.74 |
| 8263 | ENSMUSG00000031283 | 8.53885755 | -0.92861086 | 0.760721759 | -1.22069712 | 0.222200719 | 0.668407528 | Chrdl1 | chordin-like 1 [Source:MGI Symbol;Acc:M | 15.259 | 9.2513 | 9.0562 | 7.7118 | 3.2719 | 6.6831 |
| 8264 | ENSMUSG00000031284 | 1.134335839 | 1.615183059 | 2.312890309 | 0.698339672 | 0.484964795 | NA | Pak3 | p21 (RAC1) activated kinase 3 [Source:MG | 1.6062 | 0 | 0 | 1.928 | 3.2719 | 0 |
| 8265 | ENSMUSG00000031285 | 0 | NA | NA | NA | NA | NA | Dcx | doublecortin [Source:MGI Symbol;Acc:M( | 0 | 0 | 0 | 0 | 0 | 0 |
| 8266 | ENSMUSG00000031286 | 87.26128975 | -0.400414179 | 0.274377008 | -1.459357626 | 0.144466702 | 0.55464443 | Glt28d2 | glycosyltransferase 28 domain containing | 97.978 | 108.96 | 90.562 | 52.055 | 92.703 | 81.311 |
| 8267 | ENSMUSG00000031289 | 4.631286233 | 1.447751176 | 1.075835029 | 1.345699979 | 0.178399293 | NA | Il13ra2 | interleukin 13 receptor, alpha 2 [Source:N | 2.4093 | 3.0838 | 2.0125 | 12.532 | 2.1812 | 5.5692 |
| 8268 | ENSMUSG00000031290 | 0.988723425 | -0.017012424 | 2.431983019 | -0.006995289 | 0.994418612 | NA | Lrch2 | leucine-rich repeats and calponin homolo | 0 | 1.0279 | 2.0125 | 2.8919 | 0 | 0 |
| 8269 | ENSMUSG00000031292 | 111.2830635 | 0.010583799 | 0.241756979 | 0.043778668 | 0.965080831 | 0.993480943 | Cdkl5 | cyclin-dependent kinase-like 5 [Source:M( | 121.27 | 118.21 | 92.575 | 89.65 | 112.33 | 133.66 |
| 8270 | ENSMUSG00000031293 | 0 | NA | NA | NA | NA | NA | Rs1 | retinoschisis (X-linked, juvenile) 1 (human | 0 | 0 | 0 | 0 | 0 | 0 |
| 8271 | ENSMUSG00000031294 | 0 | NA | NA | NA | NA | NA | D630029K05 | RIKEN cDNA D630029K05 gene [Source:M | 0 | 0 | 0 | 0 | 0 | 0 |
| 8272 | ENSMUSG00000031295 | 459.1412272 | 0.239722573 | 0.128498972 | 1.865560233 | 0.062102918 | 0.355990107 | Phka2 | phosphorylase kinase alpha 2 [Source:MG | 440.1 | 417.33 | 405.52 | 512.84 | 471.15 | 507.91 |
| 8273 | ENSMUSG00000031297 | 0.506966596 | 2.558866737 | 3.339834942 | 0.766165628 | 0.443577757 | NA | Slc7a3 | solute carrier family 7 (cationic amino acic | 0 | 0 | 0 | 1.928 | 0 | 1.1138 |
| 8274 | ENSMUSG00000031298 | 32.7753253 | -0.494196679 | 0.43328619 | -1.140577962 | 0.254045593 | 0.707252596 | Adgrg2 | adhesion G protein-coupled receptor G2 [ | 49.792 | 40.089 | 25.156 | 37.595 | 25.084 | 18.935 |
| 8275 | ENSMUSG00000031299 | 2262.811924 | -0.197527356 | 0.104319133 | -1.893491164 | 0.058292582 | 0.342880809 | Pdha1 | pyruvate dehydrogenase E1 alpha 1 [Sour | 2443 | 2456.7 | 2353.6 | 2332.8 | 2050.4 | 1940.3 |
| 8276 | ENSMUSG00000031302 | 0.133850373 | -0.796629176 | 4.080472857 | -0.195229623 | 0.84521319 | NA | Nlgn3 | neuroligin 3 [Source:MGI Symbol;Acc:MG | 0.8031 | 0 | 0 | 0 | 0 | 0 |
| 8277 | ENSMUSG00000031303 | 28.92504529 | -0.227905086 | 0.446497147 | -0.510428987 | 0.609750954 | 0.916376001 | Map3k15 | mitogen-activated protein kinase kinase k | 32.927 | 19.53 | 41.256 | 34.703 | 25.084 | 20.049 |
| 8278 | ENSMUSG00000031304 | 10.42463507 | -1.5436584 | 0.725838087 | -2.126725544 | 0.033442894 | 0.244672745 | Il2rg | interleukin 2 receptor, gamma chain [Sour | 12.85 | 22.614 | 11.069 | 3.8559 | 4.3625 | 7.7969 |
| 8279 | ENSMUSG00000031309 | 210.7302771 | -0.10529036 | 0.172886984 | -0.609012646 | 0.542516057 | 0.894095573 | Rps6ka3 | ribosomal protein S6 kinase polypeptide 3 | 191.94 | 237.45 | 226.41 | 198.58 | 203.95 | 206.06 |
| 8280 | ENSMUSG00000031310 | 243.5979152 | 0.136733874 | 0.182486124 | 0.749283676 | 0.453686245 | 0.857120931 | Zmym3 | zinc finger, MYM-type 3 [Source:MGI Sym | 207.2 | 237.45 | 252.57 | 300.76 | 249.75 | 213.86 |
| 8281 | ENSMUSG00000031311 | 1791.831007 | 0.171501258 | 0.102262383 | 1.67707082 | 0.093528634 | 0.450459278 | Nono | non-POU-domain-containing, octamer bir | 1673.7 | 1797.8 | 1584.8 | 1833.5 | 2012.2 | 1849 |
| 8282 | ENSMUSG00000031312 | 0.831466069 | 1.993315099 | 2.60242648 | 0.765944827 | 0.443709132 | NA | Itgb1bp2 | integrin beta 1 binding protein 2 [Source:I | 0 | 0 | 1.0062 | 2.8919 | 1.0906 | 0 |
| 8283 | ENSMUSG00000031314 | 261.4359201 | 0.071177543 | 0.151481551 | 0.469875983 | 0.638443624 | 0.923906811 | Taf1 | TATA-box binding protein associated facto | 255.39 | 250.81 | 258.6 | 259.31 | 269.38 | 275.12 |
| 8284 | ENSMUSG00000031320 | 3476.592766 | 0.017380761 | 0.095137882 | 0.182690221 | 0.855041089 | 0.974158771 | Rps4x | ribosomal protein S4, X-linked [Source:M( | 3192.3 | 3685.1 | 3490.7 | 3428.9 | 3569.6 | 3493 |

| | | | | | | | | | | | | | | | |
|---|---|---|---|---|---|---|---|---|---|---|---|---|---|---|---|
| 8285 | ENSMUSG00000031323 | 0.167707431 | -0.796629176 | 4.080472857 | -0.195229623 | 0.84521319 | NA | Dmrtc1a | DMRT-like family C1a [Source:MGI Symbo | 0 | 0 | 1.0062 | 0 | 0 | 0 |
| 8286 | ENSMUSG00000031325 | 0.36353943 | 2.065989254 | 4.034137301 | 0.512126658 | 0.608562368 | NA | 4930519F16 | RIKEN cDNA 4930519F16 gene [Source:M( | 0 | 0 | 0 | 0 | 2.1812 | 0 |
| 8287 | ENSMUSG00000031326 | 0 | NA | NA | NA | NA | NA | Cdx4 | caudal type homeobox 4 [Source:MGI Syn | 0 | 0 | 0 | 0 | 0 | 0 |
| 8288 | ENSMUSG00000031327 | 60.07894559 | 0.40275282 | 0.459256789 | 0.876966502 | 0.380504833 | 0.812628154 | Chic1 | cysteine-rich hydrophobic domain 1 [Sour | 46.58 | 35.977 | 72.45 | 22.171 | 75.253 | 108.04 |
| 8289 | ENSMUSG00000031328 | 227.9409527 | -0.72644185 | 0.206603601 | -3.516114181 | 0.000437913 | 0.011270406 | Flna | filamin, alpha [Source:MGI Symbol;Acc:M | 299.56 | 315.57 | 237.47 | 194.72 | 126.51 | 193.81 |
| 8290 | ENSMUSG00000031329 | 0.371281295 | 2.09253362 | 4.031852165 | 0.519000582 | 0.603760335 | NA | Tsx | testis specific X-linked gene [Source:MGI S | 0 | 0 | 0 | 0 | 0 | 2.2277 |
| 8291 | ENSMUSG00000031330 | 0 | NA | NA | NA | NA | NA | Zcchc13 | zinc finger, CCHC domain containing 13 [S | 0 | 0 | 0 | 0 | 0 | 0 |
| 8292 | ENSMUSG00000031333 | 903.0986512 | 0.015568502 | 0.114118899 | 0.136423523 | 0.891486479 | 0.982489666 | Abcb7 | ATP-binding cassette, sub-family B (MDR/ | 873.78 | 929.24 | 891.53 | 841.55 | 989.19 | 893.3 |
| 8293 | ENSMUSG00000031337 | 323.8317206 | 0.028904199 | 0.149571741 | 0.19324639 | 0.846766009 | 0.972803041 | Mtm1 | X-linked myotubular myopathy gene 1 [So | 354.97 | 308.38 | 297.85 | 341.25 | 309.74 | 330.81 |
| 8294 | ENSMUSG00000031340 | 2.037825641 | 2.398671016 | 1.687764115 | 1.421212239 | 0.155255068 | NA | Gabre | gamma-aminobutyric acid (GABA) A recep | 0 | 1.0279 | 1.0062 | 5.7839 | 2.1812 | 2.2277 |
| 8295 | ENSMUSG00000031342 | 8.302875244 | -1.411675375 | 0.776267207 | -1.818543102 | 0.068981165 | 0.379081905 | Gpm6b | glycoprotein m6b [Source:MGI Symbol;Ac | 8.8341 | 14.391 | 13.081 | 5.7839 | 3.2719 | 4.4554 |
| 8296 | ENSMUSG00000031343 | 33.37238864 | -1.110721762 | 0.401737948 | -2.764791742 | 0.005695915 | 0.072778897 | Gabra3 | gamma-aminobutyric acid (GABA) A recep | 46.58 | 46.256 | 44.275 | 28.919 | 15.269 | 18.935 |
| 8297 | ENSMUSG00000031344 | 2.95906817 | -0.248988864 | 1.344243524 | -0.185226009 | 0.85305177 | NA | Gabrq | gamma-aminobutyric acid (GABA) A recep | 1.6062 | 4.1117 | 4.025 | 5.7839 | 0 | 2.2277 |
| 8298 | ENSMUSG00000031347 | 179.5038439 | -0.044989132 | 0.208035916 | -0.216256564 | 0.828787765 | 0.969955221 | Cetn2 | centrin 2 [Source:MGI Symbol;Acc:MGI:1: | 208.81 | 146.99 | 190.18 | 149.42 | 176.68 | 204.95 |
| 8299 | ENSMUSG00000031349 | 493.8615567 | 1.253799311 | 0.408651515 | 3.068138168 | 0.00215397 | 0.036696139 | Nsdhl | NAD(P) dependent steroid dehydrogenase | 360.59 | 349.49 | 165.02 | 508.02 | 703.45 | 876.6 |
| 8300 | ENSMUSG00000031351 | 8.946557062 | -0.736920415 | 0.800368759 | -0.920726111 | 0.35719344 | 0.79630794 | Zfp185 | zinc finger protein 185 [Source:MGI Symb | 3.2124 | 13.363 | 17.106 | 6.7478 | 5.4531 | 7.7969 |
| 8301 | ENSMUSG00000031352 | 250.7344663 | -0.037996664 | 0.155680448 | -0.244068313 | 0.807177913 | 0.964718338 | Hccs | holocytochrome c synthetase [Source:MG | 248.96 | 251.84 | 261.62 | 258.35 | 250.84 | 232.79 |
| 8302 | ENSMUSG00000031353 | 962.5079021 | 0.153249057 | 0.112322811 | 1.364362726 | 0.172453433 | 0.600762634 | Rbbp7 | retinoblastoma binding protein 7, chroma | 889.03 | 929.24 | 915.68 | 919.63 | 1065.5 | 1055.9 |
| 8303 | ENSMUSG00000031354 | 0.181769715 | 1.126924175 | 4.080472857 | 0.276174898 | 0.782413736 | NA | Amelx | amelogenin, X-linked [Source:MGI Symbo | 0 | 0 | 0 | 0 | 1.0906 | 0 |
| 8304 | ENSMUSG00000031355 | 104.5744763 | 0.229044244 | 0.229191066 | 0.999359389 | 0.317620625 | 0.764257918 | Arhgap6 | Rho GTPase activating protein 6 [Source:N | 93.963 | 98.68 | 96.599 | 131.1 | 107.97 | 99.132 |
| 8305 | ENSMUSG00000031357 | 1274.122576 | -0.128397045 | 0.106649331 | -1.203917963 | 0.228621288 | 0.675041782 | Syap1 | synapse associated protein 1 [Source:MGI | 1363.7 | 1374.3 | 1253.8 | 1163.5 | 1307.7 | 1181.8 |
| 8306 | ENSMUSG00000031358 | 553.0340335 | 0.153434527 | 0.121963255 | 1.258038962 | 0.208377666 | 0.653455953 | Msl3 | MSL complex subunit 3 [Source:MGI Symb | 530.05 | 527.32 | 513.18 | 560.07 | 564.94 | 622.64 |
| 8307 | ENSMUSG00000031360 | 140.0214514 | -0.096489629 | 0.217823613 | -0.442971391 | 0.657786428 | 0.927303282 | Ctps2 | cytidine 5'-triphosphate synthase 2 [Sourc | 175.88 | 130.55 | 126.79 | 122.43 | 135.24 | 149.26 |
| 8308 | ENSMUSG00000031362 | 1.176089219 | -2.379463953 | 2.371229009 | -1.003472859 | 0.315632766 | NA | Xlr4c | X-linked lymphocyte-regulated 4C [Source | 0.8031 | 5.1396 | 0 | 0 | 0 | 1.1138 |
| 8309 | ENSMUSG00000031364 | 0 | NA | NA | NA | NA | NA | Grpr | gastrin releasing peptide receptor [Source | 0 | 0 | 0 | 0 | 0 | 0 |
| 8310 | ENSMUSG00000031365 | 230.3265502 | 0.421844347 | 0.180369059 | 2.338784435 | 0.019346593 | 0.1693571 | Zfp275 | zinc finger protein 275 [Source:MGI Symb | 191.94 | 168.58 | 230.43 | 288.23 | 241.03 | 261.75 |
| 8311 | ENSMUSG00000031367 | 29.22692153 | -1.434806218 | 0.436117281 | -3.289954979 | 0.001002034 | 0.02092365 | Ap1s2 | adaptor-related protein complex 1, sigma | 47.383 | 50.368 | 30.187 | 15.424 | 14.178 | 17.822 |
| 8312 | ENSMUSG00000031370 | 168.428797 | 0.063741067 | 0.179895718 | 0.354322313 | 0.723097347 | 0.9447719 | Zrsr2 | zinc finger (CCCH type), RNA binding moti | 159.82 | 161.38 | 173.07 | 173.52 | 170.14 | 172.65 |
| 8313 | ENSMUSG00000031371 | 181.955781 | -0.084029584 | 0.237549042 | -0.35373573 | 0.723536943 | 0.9447719 | Haus7 | HAUS augmin-like complex, subunit 7 [Sou | 237.72 | 199.42 | 123.77 | 158.09 | 175.59 | 197.15 |
| 8314 | ENSMUSG00000031372 | 0.367410362 | 2.078255025 | 3.800070157 | 0.54689912 | 0.58444804 | NA | Trex2 | three prime repair exonuclease 2 [Source: | 0 | 0 | 0 | 0 | 1.0906 | 1.1138 |
| 8315 | ENSMUSG00000031373 | 43.23101763 | 0.339285985 | 0.33746081 | 1.005408554 | 0.314700163 | 0.761014653 | Car5b | carbonic anhydrase 5b, mitochondrial [So | 40.155 | 41.117 | 33.206 | 51.091 | 40.353 | 53.465 |
| 8316 | ENSMUSG00000031374 | 0 | NA | NA | NA | NA | NA | Zfp92 | zinc finger protein 92 [Source:MGI Symbo | 0 | 0 | 0 | 0 | 0 | 0 |
| 8317 | ENSMUSG00000031375 | 1107.843206 | -1.000425625 | 0.147577895 | -6.778966612 | 1.21E-11 | 3.98E-09 | Bgn | biglycan [Source:MGI Symbol;Acc:MGI:88 | 1768.4 | 1432.9 | 1229.6 | 758.65 | 655.46 | 801.97 |
| 8318 | ENSMUSG00000031376 | 0 | NA | NA | NA | NA | NA | Atp2b3 | ATPase, Ca++ transporting, plasma memb | 0 | 0 | 0 | 0 | 0 | 0 |
| 8319 | ENSMUSG00000031377 | 0.80281854 | -0.292476728 | 2.419692615 | -0.120873505 | 0.903791228 | NA | Bmx | BMX non-receptor tyrosine kinase [Source | 1.6062 | 0 | 1.0062 | 0 | 1.0906 | 1.1138 |
| 8320 | ENSMUSG00000031378 | 578.9040216 | -0.158489734 | 0.129722128 | -1.221763292 | 0.221797154 | 0.667949912 | Abcd1 | ATP-binding cassette, sub-family D (ALD), | 660.15 | 551.99 | 618.84 | 517.66 | 584.57 | 540.21 |
| 8321 | ENSMUSG00000031379 | 258.1824471 | 0.32537217 | 0.200729121 | 1.6209515 | 0.105028041 | 0.477712447 | Pir | pirin [Source:MGI Symbol;Acc:MGI:19169 | 296.34 | 211.75 | 178.11 | 261.24 | 305.37 | 296.28 |
| 8322 | ENSMUSG00000031380 | 10.48478977 | -0.172948672 | 0.740158978 | -0.233664223 | 0.815245665 | 0.967723701 | Vegfd | vascular endothelial growth factor D [Sou | 4.0155 | 15.419 | 14.087 | 9.6398 | 14.178 | 5.5692 |
| 8323 | ENSMUSG00000031381 | 97.26618424 | 0.200789695 | 0.238306577 | 0.842568837 | 0.399469626 | 0.826481 | Piga | phosphatidylinositol glycan anchor biosyn | 86.735 | 78.122 | 106.66 | 99.29 | 102.52 | 110.27 |
| 8324 | ENSMUSG00000031382 | 0.133850373 | -0.796629176 | 4.080472857 | -0.195229623 | 0.84521319 | NA | Asb11 | ankyrin repeat and SOCS box-containing 1 | 0.8031 | 0 | 0 | 0 | 0 | 0 |
| 8325 | ENSMUSG00000031383 | 1.168916935 | -0.258095295 | 2.108876362 | -0.122385219 | 0.902593944 | NA | Dusp9 | dual specificity phosphatase 9 [Source:M( | 0.8031 | 0 | 3.0187 | 0.964 | 0 | 2.2277 |
| 8326 | ENSMUSG00000031384 | 0 | NA | NA | NA | NA | NA | Asb9 | ankyrin repeat and SOCS box-containing 9 | 0 | 0 | 0 | 0 | 0 | 0 |
| 8327 | ENSMUSG00000031385 | 2.628629088 | -1.542449981 | 1.5943627 | -0.967439831 | 0.333324206 | NA | Plxnb3 | plexin B3 [Source:MGI Symbol;Acc:MGI:2: | 5.6217 | 6.1675 | 0 | 2.8919 | 1.0906 | 0 |
| 8328 | ENSMUSG00000031386 | 667.7678992 | 0.03537317 | 0.130650337 | 0.270746875 | 0.786585722 | 0.960870081 | Hcfc1 | host cell factor C1 [Source:MGI Symbol;Ac | 595.9 | 741.13 | 642.99 | 712.38 | 686 | 628.21 |
| 8329 | ENSMUSG00000031387 | 35.23992934 | -0.627668935 | 0.381296564 | -1.646143697 | 0.099734178 | 0.465190309 | Renbp | renin binding protein [Source:MGI Symbo | 41.761 | 53.452 | 33.206 | 27.955 | 29.447 | 25.618 |
| 8330 | ENSMUSG00000031388 | 226.3622948 | -0.006651061 | 0.162573675 | -0.040911057 | 0.967366803 | 0.994231936 | Naa10 | N(alpha)-acetyltransferase 10, NatA cataly | 240.13 | 215.86 | 224.39 | 229.43 | 233.39 | 214.97 |
| 8331 | ENSMUSG00000031389 | 72.49245975 | -0.308824058 | 0.279029265 | -1.106780172 | 0.268388983 | 0.724039098 | Arhgap4 | Rho GTPase activating protein 4 [Source:N | 65.051 | 95.596 | 80.5 | 69.406 | 68.709 | 55.692 |
| 8332 | ENSMUSG00000031390 | 1.617689988 | -0.994256779 | 2.025800961 | -0.490796874 | 0.623570121 | NA | Avpr2 | arginine vasopressin receptor 2 [Source:M | 2.4093 | 0 | 4.025 | 0 | 3.2719 | 0 |
| 8333 | ENSMUSG00000031391 | 1.213687728 | -0.49359056 | 2.07197653 | -0.238222081 | 0.811708852 | NA | L1cam | L1 cell adhesion molecule [Source:MGI Sy | 3.2124 | 1.0279 | 0 | 1.928 | 0 | 1.1138 |
| 8334 | ENSMUSG00000031392 | 1213.275119 | 0.081052996 | 0.098852242 | 0.819940897 | 0.412249801 | 0.834424863 | Irak1 | interleukin-1 receptor-associated kinase 1 | 1176.5 | 1154.4 | 1206.5 | 1221.4 | 1242.2 | 1278.7 |
| 8335 | ENSMUSG00000031393 | 383.4314911 | 0.031622393 | 0.142598121 | 0.221758838 | 0.824501625 | 0.96943211 | Mecp2 | methyl CpG binding protein 2 [Source:MG | 368.62 | 385.47 | 384.39 | 434.75 | 365.36 | 362 |
| 8336 | ENSMUSG00000031394 | 0 | NA | NA | NA | NA | NA | Opn1mw | opsin 1 (cone pigments), medium-wave-se | 0 | 0 | 0 | 0 | 0 | 0 |
| 8337 | ENSMUSG00000031397 | 0 | NA | NA | NA | NA | NA | Tktl1 | transketolase-like 1 [Source:MGI Symbol;/ | 0 | 0 | 0 | 0 | 0 | 0 |
| 8338 | ENSMUSG00000031398 | 6.779163797 | -0.380878004 | 0.843859894 | -0.451352182 | 0.651735742 | 0.925643431 | Plxna3 | plexin A3 [Source:MGI Symbol;Acc:MGI:1( | 8.8341 | 4.1117 | 10.062 | 7.7118 | 3.2719 | 6.6831 |
| 8339 | ENSMUSG00000031399 | 309.3613785 | -0.047073483 | 0.146567752 | -0.321172172 | 0.74807992 | 0.951074124 | Fam3a | family with sequence similarity 3, membe | 315.62 | 306.32 | 320.99 | 285.34 | 328.28 | 299.62 |
| 8340 | ENSMUSG00000031400 | 57.61901985 | -0.340850067 | 0.334706937 | -1.018353758 | 0.308509869 | 0.757218183 | G6pdx | glucose-6-phosphate dehydrogenase X-lir | 69.067 | 76.066 | 48.3 | 68.442 | 38.172 | 45.668 |

| | | | | | | | | | | | | | | | |
|---|---|---|---|---|---|---|---|---|---|---|---|---|---|---|---|
| 8341 | ENSMUSG00000031402 | 629.3983167 | -0.227276565 | 0.13298228 | -1.709074066 | 0.087437231 | 0.43616036 | Mpp1 | membrane protein, palmitoylated [Source | 689.06 | 734.96 | 612.8 | 596.7 | 624.92 | 517.94 |
| 8342 | ENSMUSG00000031403 | 289.4710083 | 0.248770395 | 0.152191345 | 1.634589632 | 0.102135103 | 0.47032223 | Dkc1 | dyskeratosis congenita 1, dyskerin [Source | 262.61 | 262.12 | 268.67 | 283.41 | 337 | 323.01 |
| 8343 | ENSMUSG00000031409 | 0 | NA | NA | NA | NA | NA | Tceal6 | transcription elongation factor A (SII)-like | 0 | 0 | 0 | 0 | 0 | 0 |
| 8344 | ENSMUSG00000031410 | 0.679159517 | 1.687875387 | 3.079592583 | 0.548083989 | 0.583634234 | NA | Nxf7 | nuclear RNA export factor 7 [Source:MGI | 0.8031 | 0 | 0 | 0 | 3.2719 | 0 |
| 8345 | ENSMUSG00000031411 | 0 | NA | NA | NA | NA | NA | Pramex1 | PRAME like, X-linked 1 [Source:MGI Symb | 0 | 0 | 0 | 0 | 0 | 0 |
| 8346 | ENSMUSG00000031422 | 1354.605302 | 0.13675682 | 0.121016421 | 1.130068291 | 0.25844745 | 0.712035031 | Morf4l2 | mortality factor 4 like 2 [Source:MGI Symb | 1430.3 | 1135.8 | 1304.1 | 1468.1 | 1300 | 1489.2 |
| 8347 | ENSMUSG00000031424 | 0 | NA | NA | NA | NA | NA | Kir3dl1 | killer cell immunoglobulin-like receptor, th | 0 | 0 | 0 | 0 | 0 | 0 |
| 8348 | ENSMUSG00000031425 | 32.55115248 | 0.578824003 | 0.396207669 | 1.460910649 | 0.144039964 | 0.554282225 | Plp1 | proteolipid protein (myelin) 1 [Source:MG | 27.305 | 23.642 | 27.169 | 28.919 | 39.262 | 49.009 |
| 8349 | ENSMUSG00000031428 | 3.126897914 | -0.629558392 | 1.348725992 | -0.466780055 | 0.640657247 | NA | Zcchc18 | zinc finger, CCHC domain containing 18 [S | 3.2124 | 7.1954 | 1.0062 | 2.8919 | 0 | 4.4554 |
| 8350 | ENSMUSG00000031429 | 196.4559438 | 0.211885559 | 0.181551259 | 1.167083944 | 0.243176471 | 0.6927919 | Psmd10 | proteasome (prosome, macropain) 26S su | 171.06 | 182.97 | 192.19 | 189.9 | 242.12 | 200.49 |
| 8351 | ENSMUSG00000031430 | 0 | NA | NA | NA | NA | NA | Vsig1 | V-set and immunoglobulin domain contai | 0 | 0 | 0 | 0 | 0 | 0 |
| 8352 | ENSMUSG00000031431 | 2826.105761 | -1.696714922 | 0.518490278 | -3.272414147 | 0.001066333 | 0.022018741 | Tsc22d3 | TSC22 domain family, member 3 [Source:I | 3189.1 | 5254.7 | 4515 | 813.6 | 2210.7 | 973.5 |
| 8353 | ENSMUSG00000031432 | 585.8415105 | -0.157740367 | 0.124814138 | -1.263802074 | 0.206301077 | 0.65157405 | Prps1 | phosphoribosyl pyrophosphate synthetas | 590.28 | 638.34 | 625.88 | 603.45 | 520.22 | 536.87 |
| 8354 | ENSMUSG00000031433 | 50.33698696 | -0.29638485 | 0.327428704 | -0.905188966 | 0.365365286 | 0.803870323 | Rbm41 | RNA binding motif protein 41 [Source:MG | 46.58 | 68.87 | 51.318 | 49.163 | 37.081 | 49.009 |
| 8355 | ENSMUSG00000031434 | 42.54680058 | -1.20498259 | 0.392900439 | -3.066890414 | 0.002162981 | 0.036806309 | Morc4 | microrchidia 4 [Source:MGI Symbol;Acc:M | 81.113 | 57.563 | 39.244 | 29.883 | 19.631 | 27.846 |
| 8356 | ENSMUSG00000031438 | 2109.749609 | -0.070731318 | 0.101686414 | -0.695582779 | 0.486690154 | 0.873681495 | Rnf128 | ring finger protein 128 [Source:MGI Symb | 1986.1 | 2271.7 | 2227.8 | 2055.2 | 2152.9 | 1964.8 |
| 8357 | ENSMUSG00000031441 | 199.0302855 | 0.722708105 | 0.268736221 | 2.689284319 | 0.00716054 | 0.085383392 | Atp11a | ATPase, class VI, type 11A [Source:MGI Sy | 159.82 | 120.27 | 170.06 | 206.29 | 167.96 | 369.8 |
| 8358 | ENSMUSG00000031442 | 2.462086334 | 0.299758663 | 1.425245847 | 0.210320671 | 0.833417403 | NA | Mcf2l | mcf.2 transforming sequence-like [Source | 1.6062 | 2.0558 | 3.0187 | 4.8199 | 3.2719 | 0 |
| 8359 | ENSMUSG00000031443 | 1852.925264 | 0.221494317 | 0.135764973 | 1.631454059 | 0.102794546 | 0.471407836 | F7 | coagulation factor VII [Source:MGI Symbo | 1435.9 | 1625.1 | 2072.9 | 1899 | 2035.1 | 2049.5 |
| 8360 | ENSMUSG00000031444 | 6947.421194 | 0.191085119 | 0.094193095 | 2.028653151 | 0.042493631 | 0.283892783 | F10 | coagulation factor X [Source:MGI Symbol; | 5979.9 | 6473.8 | 7011.5 | 7371.5 | 7260.2 | 7587.5 |
| 8361 | ENSMUSG00000031445 | 7307.561443 | 0.128794611 | 0.100909079 | 1.276343139 | 0.201834247 | 0.646227482 | Proz | protein Z, vitamin K-dependent plasma gly | 6284.3 | 7170.8 | 7491.5 | 7787 | 7935.3 | 7176.5 |
| 8362 | ENSMUSG00000031446 | 2709.102149 | -0.09239076 | 0.112420598 | -0.821831247 | 0.411172947 | 0.833705705 | Cul4a | cullin 4A [Source:MGI Symbol;Acc:MGI:19 | 2628.6 | 3021.1 | 2739 | 2712.6 | 2842.2 | 2311.2 |
| 8363 | ENSMUSG00000031447 | 11665.31952 | -0.033303668 | 0.080904019 | -0.411644179 | 0.680600245 | 0.934384556 | Lamp1 | lysosomal-associated membrane protein 1 | 11684 | 11702 | 12015 | 11360 | 11928 | 11304 |
| 8364 | ENSMUSG00000031448 | 0 | NA | NA | NA | NA | NA | Adprhl1 | ADP-ribosylhydrolase like 1 [Source:MGI S | 0 | 0 | 0 | 0 | 0 | 0 |
| 8365 | ENSMUSG00000031449 | 0 | NA | NA | NA | NA | NA | Atp4b | ATPase, H+/K+ exchanging, beta polypept | 0 | 0 | 0 | 0 | 0 | 0 |
| 8366 | ENSMUSG00000031450 | 0 | NA | NA | NA | NA | NA | Grk1 | G protein-coupled receptor kinase 1 [Sour | 0 | 0 | 0 | 0 | 0 | 0 |
| 8367 | ENSMUSG00000031451 | 678.9272029 | -0.611816178 | 0.171647477 | -3.564376173 | 0.000364723 | 0.010067457 | Gas6 | growth arrest specific 6 [Source:MGI Syml | 917.95 | 862.42 | 682.23 | 649.72 | 418.8 | 542.44 |
| 8368 | ENSMUSG00000031452 | 0 | NA | NA | NA | NA | NA | 1700029H14 | RIKEN cDNA 1700029H14 gene [Source:M | 0 | 0 | 0 | 0 | 0 | 0 |
| 8369 | ENSMUSG00000031453 | 257.2517352 | 0.253992479 | 0.189908524 | 1.337446437 | 0.181076955 | 0.614561135 | Rasa3 | RAS p21 protein activator 3 [Source:MGI S | 246.55 | 197.36 | 259.61 | 267.99 | 233.39 | 338.61 |
| 8370 | ENSMUSG00000031458 | 26.13946136 | -0.177816572 | 0.427373281 | -0.416068528 | 0.677359859 | 0.93325821 | Coprs | coordinator of PRMT5, differentiation stin | 22.487 | 26.726 | 34.212 | 26.027 | 23.994 | 23.391 |
| 8371 | ENSMUSG00000031461 | 0 | NA | NA | NA | NA | NA | Myom2 | myomesin 2 [Source:MGI Symbol;Acc:MG | 0 | 0 | 0 | 0 | 0 | 0 |
| 8372 | ENSMUSG00000031465 | 14.31242564 | -0.291350929 | 0.583777121 | -0.499079047 | 0.617723698 | 0.917509899 | Angpt2 | angiopoietin 2 [Source:MGI Symbol;Acc:M | 16.865 | 10.279 | 20.125 | 15.424 | 9.8156 | 13.366 |
| 8373 | ENSMUSG00000031467 | 184.5419639 | -0.180277829 | 0.208863712 | -0.863136193 | 0.388062591 | 0.817937623 | Agpat5 | 1-acylglycerol-3-phosphate O-acyltransfer | 205.59 | 165.49 | 216.34 | 133.99 | 188.68 | 197.15 |
| 8374 | ENSMUSG00000031471 | 0 | NA | NA | NA | NA | NA | Defb8 | defensin beta 8 [Source:MGI Symbol;Acc: | 0 | 0 | 0 | 0 | 0 | 0 |
| 8375 | ENSMUSG00000031478 | 37.70225726 | -0.138588725 | 0.361865429 | -0.382984154 | 0.701731514 | 0.942189108 | Nek3 | NIMA (never in mitosis gene a)-related ex | 44.974 | 41.117 | 32.2 | 31.811 | 34.9 | 41.212 |
| 8376 | ENSMUSG00000031479 | 622.6697606 | -0.129448524 | 0.137295476 | -0.942846242 | 0.345759555 | 0.786899236 | Vps36 | vacuolar protein sorting 36 [Source:MGI S | 629.63 | 686.65 | 634.94 | 509.94 | 598.75 | 676.1 |
| 8377 | ENSMUSG00000031480 | 52.16554324 | -0.384106534 | 0.32334682 | -1.187908803 | 0.234869341 | 0.682844252 | Thsd1 | thrombospondin, type I, domain 1 [Source | 58.626 | 48.312 | 70.437 | 53.983 | 37.081 | 44.554 |
| 8378 | ENSMUSG00000031481 | 10.27380459 | -0.181397754 | 0.701378753 | -0.258630238 | 0.79592055 | 0.962236316 | Tpte | transmembrane phosphatase with tensin | 6.4248 | 10.279 | 16.1 | 6.7478 | 8.7249 | 13.366 |
| 8379 | ENSMUSG00000031482 | 5194.652267 | 0.028188962 | 0.145603516 | 0.193600836 | 0.846488443 | 0.972803041 | Slc25a15 | solute carrier family 25 (mitochondrial car | 4580.9 | 5211.5 | 5639 | 4136.4 | 5260.1 | 6340 |
| 8380 | ENSMUSG00000031483 | 788.4591802 | 0.054098161 | 0.114806934 | 0.471209873 | 0.637490869 | 0.923478764 | Erlin2 | ER lipid raft associated 2 [Source:MGI Sym | 750.9 | 775.05 | 794.93 | 745.15 | 859.41 | 805.31 |
| 8381 | ENSMUSG00000031485 | 1794.785283 | 0.025284394 | 0.095250679 | 0.265451069 | 0.790662 | 0.961019028 | Plpbp | pyridoxal phosphate binding protein [Sou | 1836.7 | 1683.7 | 1816.3 | 1834.4 | 1785.3 | 1812.2 |
| 8382 | ENSMUSG00000031486 | 68.71594346 | -0.626996453 | 0.310932208 | -2.016505328 | 0.043747162 | 0.288610458 | Adgra2 | adhesion G protein-coupled receptor A2 [ | 99.585 | 77.094 | 73.456 | 62.659 | 34.9 | 64.603 |
| 8383 | ENSMUSG00000031487 | 106.5523383 | 0.125921818 | 0.233200831 | 0.539971566 | 0.589216642 | 0.911039598 | Brf2 | BRF2, RNA polymerase III transcription ini | 92.357 | 96.624 | 116.72 | 95.434 | 113.42 | 124.75 |
| 8384 | ENSMUSG00000031488 | 190.2477001 | 0.12719769 | 0.182141814 | 0.69834426 | 0.484961926 | 0.87256358 | Rab11fip1 | RAB11 family interacting protein 1 (class I | 171.86 | 165.49 | 208.29 | 187.01 | 207.22 | 201.61 |
| 8385 | ENSMUSG00000031489 | 132.5596933 | 0.947580577 | 0.279007899 | 3.396250005 | 0.000683159 | 0.015634836 | Adrb3 | adrenergic receptor, beta 3 [Source:MGI S | 86.735 | 69.898 | 114.71 | 115.68 | 230.12 | 178.22 |
| 8386 | ENSMUSG00000031490 | 898.4842514 | 0.038326549 | 0.114752483 | 0.333993195 | 0.738384693 | 0.948137969 | Eif4ebp1 | eukaryotic translation initiation factor 4E l | 962.92 | 855.23 | 840.21 | 879.15 | 891.04 | 962.36 |
| 8387 | ENSMUSG00000031491 | 0 | NA | NA | NA | NA | NA | Chrna6 | cholinergic receptor, nicotinic, alpha poly | 0 | 0 | 0 | 0 | 0 | 0 |
| 8388 | ENSMUSG00000031492 | 0 | NA | NA | NA | NA | NA | Chrnb3 | cholinergic receptor, nicotinic, beta polyp | 0 | 0 | 0 | 0 | 0 | 0 |
| 8389 | ENSMUSG00000031493 | 0.815310133 | -0.44746426 | 2.258708535 | -0.198106242 | 0.842961941 | NA | Ggn | gametogenetin [Source:MGI Symbol;Acc:I | 0.8031 | 1.0279 | 1.0062 | 0.964 | 1.0906 | 0 |
| 8390 | ENSMUSG00000031494 | 1.928772452 | -2.187117985 | 1.685911891 | -1.297290799 | 0.194531149 | NA | Cd209a | CD209a antigen [Source:MGI Symbol;Acc: | 2.4093 | 3.0838 | 4.025 | 0.964 | 1.0906 | 0 |
| 8391 | ENSMUSG00000031495 | 0.133850373 | -0.796629176 | 4.080472857 | -0.195229623 | 0.84521319 | NA | Cd209d | CD209d antigen [Source:MGI Symbol;Acc: | 0.8031 | 0 | 0 | 0 | 0 | 0 |
| 8392 | ENSMUSG00000031497 | 4.160440178 | -0.485247292 | 1.068147194 | -0.454288786 | 0.649620986 | NA | Tnfsf13b | tumor necrosis factor (ligand) superfamily | 6.4248 | 3.0838 | 5.0312 | 3.8559 | 5.4531 | 1.1138 |
| 8393 | ENSMUSG00000031502 | 332.4325561 | -1.240474067 | 0.50830795 | -2.440398711 | 0.01467106 | 0.14062439 | Col4a1 | collagen, type IV, alpha 1 [Source:MGI Syr | 780.62 | 371.08 | 249.55 | 210.15 | 154.87 | 228.34 |
| 8394 | ENSMUSG00000031503 | 177.051016 | -0.614179515 | 0.313843905 | -1.956958556 | 0.050352341 | 0.315279767 | Col4a2 | collagen, type IV, alpha 2 [Source:MGI Syr | 320.44 | 200.44 | 120.75 | 107.97 | 117.79 | 194.92 |
| 8395 | ENSMUSG00000031504 | 138.1673025 | 0.205947711 | 0.214245134 | 0.961271361 | 0.336415744 | 0.778173248 | Rab20 | RAB20, member RAS oncogene family [So | 148.57 | 107.93 | 127.79 | 144.6 | 134.15 | 165.96 |
| 8396 | ENSMUSG00000031505 | 1056.998432 | 0.058715728 | 0.115070519 | 0.510258654 | 0.609870265 | 0.916376001 | Naxd | NAD(P)HX dehydratase [Source:MGI Symb | 1034.4 | 979.61 | 1091.8 | 990.01 | 1065.5 | 1180.7 |

| | | | | | | | | | | | | | | | |
|---|---|---|---|---|---|---|---|---|---|---|---|---|---|---|---|
| 8397 | ENSMUSG00000031506 | 4.661602387 | -0.963766203 | 1.138723641 | -0.846356542 | 0.397353874 | NA | Ptpn7 | protein tyrosine phosphatase, non-recept | 3.2124 | 11.307 | 4.025 | 3.8559 | 0 | 5.5692 |
| 8398 | ENSMUSG00000031508 | 205.0500474 | 0.135567389 | 0.185297085 | 0.731621813 | 0.464399431 | 0.86249016 | Ankrd10 | ankyrin repeat domain 10 [Source:MGI Sy | 191.14 | 188.11 | 207.29 | 234.25 | 174.5 | 235.02 |
| 8399 | ENSMUSG00000031509 | 0 | NA | NA | NA | NA | NA | 1700016D06 | RIKEN cDNA 1700016D06 gene [Source:M | 0 | 0 | 0 | 0 | 0 | 0 |
| 8400 | ENSMUSG00000031510 | 0 | NA | NA | NA | NA | NA | 1700128E19 | RIKEN cDNA 1700128E19 gene [Source:M | 0 | 0 | 0 | 0 | 0 | 0 |
| 8401 | ENSMUSG00000031511 | 434.571903 | -0.050687854 | 0.150969829 | -0.335748239 | 0.737060726 | 0.948137969 | Arhgef7 | Rho guanine nucleotide exchange factor ( | 466.6 | 393.69 | 465.89 | 434.75 | 367.54 | 478.95 |
| 8402 | ENSMUSG00000031512 | 0.160662974 | 1.126924175 | 4.080472857 | 0.276174898 | 0.782413736 | NA | Tex29 | testis expressed 29 [Source:MGI Symbol;A | 0 | 0 | 0 | 0.964 | 0 | 0 |
| 8403 | ENSMUSG00000031513 | 263.2616726 | -0.169226689 | 0.17759634 | -0.952872619 | 0.340654615 | 0.783094143 | Leprotl1 | leptin receptor overlapping transcript-like | 256.19 | 292.96 | 286.78 | 214.97 | 227.94 | 300.74 |
| 8404 | ENSMUSG00000031516 | 192.7074059 | 0.075814775 | 0.181592547 | 0.417499374 | 0.676313182 | 0.932771138 | Dctn6 | dynactin 6 [Source:MGI Symbol;Acc:MGI: | 204.79 | 168.58 | 189.17 | 199.54 | 215.94 | 178.22 |
| 8405 | ENSMUSG00000031517 | 55.83285681 | 0.103486287 | 0.387117013 | 0.267325599 | 0.789218479 | 0.960994657 | Gpm6a | glycoprotein m6a [Source:MGI Symbol;Ac | 51.399 | 71.954 | 38.237 | 54.947 | 86.159 | 32.301 |
| 8406 | ENSMUSG00000031518 | 0 | NA | NA | NA | NA | NA | Spata4 | spermatogenesis associated 4 [Source:MC | 0 | 0 | 0 | 0 | 0 | 0 |
| 8407 | ENSMUSG00000031519 | 2.605480829 | -1.048300662 | 1.476815896 | -0.70983842 | 0.477804341 | NA | Asb5 | ankyrin repeat and SOCs box-containing 5 | 6.4248 | 4.1117 | 0 | 2.8919 | 1.0906 | 1.1138 |
| 8408 | ENSMUSG00000031520 | 60.62862376 | 0.069435476 | 0.321511713 | 0.215965617 | 0.829014549 | 0.969955221 | Vegfc | vascular endothelial growth factor C [Sour | 75.492 | 42.145 | 59.368 | 59.767 | 51.259 | 75.741 |
| 8409 | ENSMUSG00000031521 | 363.9904305 | -0.125834811 | 0.137551077 | -0.914822433 | 0.360284856 | 0.799598236 | Aga | aspartylglucosaminidase [Source:MGI Sym | 400.75 | 363.88 | 374.32 | 336.43 | 358.81 | 349.75 |
| 8410 | ENSMUSG00000031523 | 601.889762 | 0.217186911 | 0.139766596 | 1.553925737 | 0.120202132 | 0.507621499 | Dlc1 | deleted in liver cancer 1 [Source:MGI Sym | 546.91 | 480.04 | 642.99 | 639.12 | 618.38 | 683.9 |
| 8411 | ENSMUSG00000031527 | 213.3201482 | 0.338684147 | 0.171891627 | 1.970335333 | 0.048799952 | 0.307687718 | Eri1 | exoribonuclease 1 [Source:MGI Symbol;A | 180.7 | 190.16 | 194.21 | 210.15 | 255.2 | 249.5 |
| 8412 | ENSMUSG00000031529 | 604.9839305 | 0.071947288 | 0.137198721 | 0.524402033 | 0.599998943 | 0.915217938 | Tnks | tankyrase, TRF1-interacting ankyrin-relate | 611.16 | 644.5 | 514.19 | 657.43 | 643.46 | 559.15 |
| 8413 | ENSMUSG00000031530 | 33.07222008 | 0.301431738 | 0.502232221 | 0.600183989 | 0.548383623 | 0.895622715 | Dusp4 | dual specificity phosphatase 4 [Source:MC | 41.761 | 25.698 | 21.131 | 36.631 | 14.178 | 59.034 |
| 8414 | ENSMUSG00000031532 | 2158.357843 | 0.036811587 | 0.103712677 | 0.35493816 | 0.72263592 | 0.9447719 | Saraf | store-operated calcium entry-associated r | 2038.3 | 2154.5 | 2200.7 | 2304.9 | 1974 | 2277.8 |
| 8415 | ENSMUSG00000031533 | 533.7033059 | -0.111817889 | 0.128544383 | -0.869877679 | 0.384367255 | 0.816437825 | Mrps31 | mitochondrial ribosomal protein S31 [Sou | 562.97 | 551.99 | 547.4 | 462.71 | 520.22 | 556.92 |
| 8416 | ENSMUSG00000031534 | 356.7458545 | -0.020199055 | 0.167453728 | -0.120624696 | 0.903988307 | 0.983821605 | Smim19 | small integral membrane protein 19 [Sour | 348.55 | 344.35 | 384.39 | 289.19 | 424.25 | 349.75 |
| 8417 | ENSMUSG00000031535 | 13.01100643 | 0.217061574 | 0.670816945 | 0.323577953 | 0.746257581 | 0.949997329 | Dkk4 | dickkopf WNT signaling pathway inhibitor | 13.653 | 6.1675 | 16.1 | 5.7839 | 18.541 | 17.822 |
| 8418 | ENSMUSG00000031536 | 310.3817527 | 0.232904483 | 0.174903297 | 1.331618597 | 0.182985551 | 0.61680331 | Polb | polymerase (DNA directed), beta [Source: | 317.23 | 256.98 | 280.74 | 271.84 | 349 | 386.5 |
| 8419 | ENSMUSG00000031537 | 1036.501917 | 0.05444283 | 0.106626088 | 0.510595773 | 0.609634136 | 0.916376001 | Ikbkb | inhibitor of kappaB kinase beta [Source:M | 1014.3 | 955.96 | 1080.7 | 1079.7 | 1050.3 | 1038.1 |
| 8420 | ENSMUSG00000031538 | 7.780422364 | -0.324165389 | 0.803275612 | -0.403554377 | 0.686540441 | 0.936566769 | Plat | plasminogen activator, tissue [Source:MG | 9.6372 | 12.335 | 4.025 | 9.6398 | 4.3625 | 6.6831 |
| 8421 | ENSMUSG00000031539 | 27.66473526 | -0.73926679 | 0.417094976 | -1.772418352 | 0.076325138 | 0.404235368 | Ap3m2 | adaptor-related protein complex 3, mu 2 s | 38.549 | 33.921 | 31.194 | 19.28 | 18.541 | 24.505 |
| 8422 | ENSMUSG00000031540 | 407.5141582 | 0.196975554 | 0.139274544 | 1.414296882 | 0.157274752 | 0.576488093 | Kat6a | K(lysine) acetyltransferase 6A [Source:MG | 354.17 | 400.89 | 384.39 | 400.05 | 468.97 | 436.63 |
| 8423 | ENSMUSG00000031543 | 1.497235207 | -1.708541429 | 1.853711467 | -0.921686821 | 0.356691957 | NA | Ank1 | ankyrin 1, erythroid [Source:MGI Symbol;. | 0.8031 | 3.0838 | 3.0187 | 0.964 | 0 | 1.1138 |
| 8424 | ENSMUSG00000031545 | 3963.013507 | -0.230527909 | 0.107024938 | -2.153964431 | 0.031242961 | 0.234752309 | Gpat4 | glycerol-3-phosphate acyltransferase 4 [So | 3968.1 | 4643.1 | 4227.2 | 3861.7 | 3792.1 | 3285.8 |
| 8425 | ENSMUSG00000031546 | 334.4577521 | 0.061014635 | 0.143460204 | 0.425307038 | 0.670612863 | 0.93116767 | Gins4 | GINS complex subunit 4 (Sld5 homolog) [S | 342.12 | 306.32 | 333.07 | 321.97 | 345.73 | 357.54 |
| 8426 | ENSMUSG00000031548 | 3.871275203 | -0.172847907 | 1.261957373 | -0.136968103 | 0.891056008 | NA | Sfrp1 | secreted frizzled-related protein 1 [Source | 7.2279 | 3.0838 | 2.0125 | 8.6758 | 0 | 2.2277 |
| 8427 | ENSMUSG00000031549 | 2633.363765 | -0.097271609 | 0.124470309 | -0.781484434 | 0.434517628 | 0.845264432 | Ido2 | indoleamine 2,3-dioxygenase 2 [Source:M | 2450.3 | 2534.8 | 3181.7 | 2404.2 | 2787.6 | 2441.5 |
| 8428 | ENSMUSG00000031551 | 0.17131964 | -0.796629176 | 4.080472857 | -0.195229623 | 0.84521319 | NA | Ido1 | indoleamine 2,3-dioxygenase 1 [Source:M | 0 | 1.0279 | 0 | 0 | 0 | 0 |
| 8429 | ENSMUSG00000031552 | 0 | NA | NA | NA | NA | NA | Adam18 | a disintegrin and metallopeptidase domai | 0 | 0 | 0 | 0 | 0 | 0 |
| 8430 | ENSMUSG00000031553 | 0 | NA | NA | NA | NA | NA | Adam3 | a disintegrin and metallopeptidase domai | 0 | 0 | 0 | 0 | 0 | 0 |
| 8431 | ENSMUSG00000031554 | 0 | NA | NA | NA | NA | NA | Adam5 | a disintegrin and metallopeptidase domai | 0 | 0 | 0 | 0 | 0 | 0 |
| 8432 | ENSMUSG00000031555 | 1445.391144 | 0.058414208 | 0.102096074 | 0.572149405 | 0.567220761 | 0.902958517 | Adam9 | a disintegrin and metallopeptidase domai | 1342 | 1488.4 | 1418.8 | 1464.3 | 1546.5 | 1412.4 |
| 8433 | ENSMUSG00000031556 | 904.9652297 | 0.194187956 | 0.106841586 | 1.817531563 | 0.06913576 | 0.379787175 | Tm2d2 | TM2 domain containing 2 [Source:MGI Sy | 839.24 | 830.56 | 862.35 | 919.63 | 1002.3 | 975.73 |
| 8434 | ENSMUSG00000031557 | 25.09521549 | -1.144656483 | 0.489185544 | -2.339922954 | 0.019287718 | 0.169077308 | Plekha2 | pleckstrin homology domain-containing, f | 40.155 | 41.117 | 22.137 | 9.6398 | 16.359 | 21.163 |
| 8435 | ENSMUSG00000031558 | 1.338639255 | 0.752904846 | 2.022714537 | 0.372224964 | 0.709725361 | NA | Slit2 | slit guidance ligand 2 [Source:MGI Symbol | 0 | 2.0558 | 1.0062 | 3.8559 | 0 | 1.1138 |
| 8436 | ENSMUSG00000031559 | 0 | NA | NA | NA | NA | NA | 4930555F03 | RIKEN cDNA 4930555F03 gene [Source:M | 0 | 0 | 0 | 0 | 0 | 0 |
| 8437 | ENSMUSG00000031561 | 414.2375096 | -0.20630191 | 0.221397334 | -0.9318175 | 0.351430849 | 0.792511048 | Tenm3 | teneurin transmembrane protein 3 [Sourc | 516.39 | 543.77 | 270.68 | 345.1 | 431.88 | 377.59 |
| 8438 | ENSMUSG00000031562 | 1.506341172 | 0.559890583 | 1.913681709 | 0.292572469 | 0.76984896 | NA | Dctd | dCMP deaminase [Source:MGI Symbol;Ac | 1.6062 | 0 | 2.0125 | 0.964 | 0 | 4.4554 |
| 8439 | ENSMUSG00000031563 | 292.2610632 | -0.132713067 | 0.188707765 | -0.70327295 | 0.481885665 | 0.871358202 | Wwc2 | WW, C2 and coiled-coil domain containin | 248.16 | 338.18 | 332.06 | 328.72 | 278.11 | 228.34 |
| 8440 | ENSMUSG00000031565 | 104.9246746 | 0.955384218 | 0.252000649 | 3.791197451 | 0.000149923 | 0.004996502 | Fgfr1 | fibroblast growth factor receptor 1 [Sourc | 66.657 | 76.066 | 71.443 | 106.04 | 171.23 | 138.12 |
| 8441 | ENSMUSG00000031568 | 466.2370747 | 0.012781108 | 0.13563769 | 0.094229765 | 0.924926641 | 0.987546253 | Rwdd4a | RWD domain containing 4A [Source:MGI S | 447.33 | 494.43 | 450.8 | 435.72 | 518.04 | 451.11 |
| 8442 | ENSMUSG00000031570 | 263.6591334 | -0.340211356 | 0.234701035 | -1.449552001 | 0.147183489 | 0.558351692 | Plpp5 | phospholipid phosphatase 5 [Source:MGI | 392.72 | 222.03 | 268.67 | 290.16 | 175.59 | 232.79 |
| 8443 | ENSMUSG00000031574 | 53.48810696 | 0.003898289 | 0.337603792 | 0.011546935 | 0.990787083 | 0.99859176 | Star | steroidogenic acute regulatory protein [Sc | 34.533 | 65.787 | 60.375 | 50.127 | 59.984 | 50.123 |
| 8444 | ENSMUSG00000031575 | 378.7687678 | -0.074694576 | 0.146896415 | -0.508484676 | 0.611113483 | 0.916376001 | Ash2l | ASH2 like histone lysine methyltransferase | 399.94 | 394.72 | 370.3 | 321 | 405.71 | 380.93 |
| 8445 | ENSMUSG00000031576 | 0.321325949 | 1.911575114 | 4.048267893 | 0.472195804 | 0.636787034 | NA | Kcnu1 | potassium channel, subfamily U, member | 0 | 0 | 0 | 1.928 | 0 | 0 |
| 8446 | ENSMUSG00000031577 | 173.2518699 | 0.007902163 | 0.191437127 | 0.04127811 | 0.967074184 | 0.994231936 | Tti2 | TELO2 interacting protein 2 [Source:MGI S | 179.09 | 162.41 | 177.1 | 201.47 | 167.96 | 151.48 |
| 8447 | ENSMUSG00000031578 | 84.84494358 | -0.16265668 | 0.248561893 | -0.654391057 | 0.512859895 | 0.884860213 | Mak16 | MAK16 homolog [Source:MGI Symbol;Acc | 82.72 | 103.82 | 82.512 | 79.046 | 77.434 | 83.538 |
| 8448 | ENSMUSG00000031583 | 239.538867 | -0.040340946 | 0.163538811 | -0.246675063 | 0.805159707 | 0.96404377 | Wrn | Werner syndrome RecQ like helicase [Sou | 230.49 | 247.73 | 250.55 | 221.71 | 261.75 | 225 |
| 8449 | ENSMUSG00000031584 | 1105.089083 | -0.015271989 | 0.1098472 | -0.139029386 | 0.889426928 | 0.982382091 | Gsr | glutathione reductase [Source:MGI Symbo | 1061.7 | 1193.4 | 1078.7 | 1156.8 | 1033.9 | 1106 |
| 8450 | ENSMUSG00000031585 | 186.3357924 | 0.087744696 | 0.180545121 | 0.48599871 | 0.626968081 | 0.920273791 | Gtf2e2 | general transcription factor II E, polypepti | 159.01 | 194.28 | 189.17 | 183.16 | 194.13 | 198.26 |
| 8451 | ENSMUSG00000031586 | 541.4229993 | -0.43723981 | 0.140425549 | -3.113677065 | 0.001847716 | 0.032832144 | Rbpms | RNA binding protein gene with multiple sp | 607.15 | 566.38 | 695.32 | 483.92 | 406.8 | 488.98 |
| 8452 | ENSMUSG00000031590 | 215.7521021 | 0.026660819 | 0.172157564 | 0.154862895 | 0.876929406 | 0.980632845 | Frg1 | FSHD region gene 1 [Source:MGI Symbol;/ | 221.66 | 210.72 | 208.29 | 191.83 | 220.3 | 241.7 |

| | | | | | | | | | | | | | | |
|---|---|---|---|---|---|---|---|---|---|---|---|---|---|---|
| 8453 | ENSMUSG00000031591 | 455.1169777 | -0.146938659 | 0.142205368 | -1.033284891 | 0.301470596 | 0.752246851 | Asah1 | N-acylsphingosine amidohydrolase 1 [Sou | 518 | 508.82 | 407.53 | 436.68 | 428.61 | 431.06 |
| 8454 | ENSMUSG00000031592 | 331.2411613 | 0.02421277 | 0.177083051 | 0.136731152 | 0.891243305 | 0.982489666 | Pcm1 | pericentriolar material 1 [Source:MGI Sym | 338.91 | 335.1 | 311.94 | 403.91 | 339.18 | 258.41 |
| 8455 | ENSMUSG00000031594 | 3537.237111 | -0.073452617 | 0.166715001 | -0.44058793 | 0.659511343 | 0.927787797 | Fgl1 | fibrinogen-like protein 1 [Source:MGI Sym | 4024.3 | 3781.7 | 3076.1 | 4365.9 | 2579.3 | 3396.1 |
| 8456 | ENSMUSG00000031595 | 5.128878238 | -0.500201736 | 0.986857492 | -0.506863189 | 0.612250816 | 0.916441371 | Pdgfrl | platelet-derived growth factor receptor-lil | 4.8186 | 6.1675 | 7.0437 | 3.8559 | 1.0906 | 7.7969 |
| 8457 | ENSMUSG00000031596 | 9142.045059 | 0.176071898 | 0.150882236 | 1.166949163 | 0.2432309 | 0.692810504 | Slc7a2 | solute carrier family 7 (cationic amino acic | 7533.1 | 9321.2 | 8900.2 | 7447.7 | 9818.8 | 11831 |
| 8458 | ENSMUSG00000031600 | 1046.118017 | 0.132618481 | 0.115355043 | 1.149654818 | 0.250286071 | 0.702302228 | Vps37a | vacuolar protein sorting 37A [Source:MGI | 910.72 | 1096.8 | 988.13 | 1141.3 | 1097.2 | 1042.6 |
| 8459 | ENSMUSG00000031601 | 441.0808077 | 0.104185749 | 0.135957308 | 0.766312232 | 0.443490542 | 0.851079912 | Cnot7 | CCR4-NOT transcription complex, subunit | 430.46 | 463.59 | 381.37 | 452.11 | 468.97 | 449.99 |
| 8460 | ENSMUSG00000031603 | 0 | NA | NA | NA | NA | NA | Fgf20 | fibroblast growth factor 20 [Source:MGI S | 0 | 0 | 0 | 0 | 0 | 0 |
| 8461 | ENSMUSG00000031604 | 1074.810443 | 1.039880092 | 0.208793987 | 4.980412072 | 6.34E-07 | 5.46E-05 | Msmo1 | methylsterol monoxygenase 1 [Source:M( | 791.06 | 812.06 | 506.14 | 974.58 | 1721 | 1644 |
| 8462 | ENSMUSG00000031605 | 785.5651396 | 0.37243072 | 0.14483903 | 2.571342271 | 0.010130515 | 0.109594442 | Klhl2 | kelch-like 2, Mayven [Source:MGI Symbol, | 664.17 | 586.94 | 802.98 | 827.09 | 996.83 | 835.38 |
| 8463 | ENSMUSG00000031608 | 41.25366614 | -0.10368606 | 0.351955853 | -0.294599619 | 0.768299754 | 0.955856567 | Galnt7 | polypeptide N-acetylgalactosaminyltransf | 45.777 | 44.2 | 38.237 | 45.307 | 29.447 | 44.554 |
| 8464 | ENSMUSG00000031609 | 65.37456974 | -0.190941484 | 0.292395994 | -0.653023597 | 0.513741066 | 0.885004045 | Sap30 | sin3 associated polypeptide [Source:MGI : | 60.233 | 62.703 | 86.537 | 70.37 | 56.712 | 55.692 |
| 8465 | ENSMUSG00000031610 | 0 | NA | NA | NA | NA | NA | Scrg1 | scrapie responsive gene 1 [Source:MGI Sy | 0 | 0 | 0 | 0 | 0 | 0 |
| 8466 | ENSMUSG00000031613 | 2315.303159 | -0.202739467 | 0.104433333 | -1.94132908 | 0.052218374 | 0.321125242 | Hpgd | hydroxyprostaglandin dehydrogenase 15 ( | 2272.8 | 2595.5 | 2565.9 | 2122.7 | 2307.7 | 2027.2 |
| 8467 | ENSMUSG00000031616 | 49.3947536 | -1.040156846 | 0.351348622 | -2.960469405 | 0.003071706 | 0.047432745 | Ednra | endothelin receptor type A [Source:MGI S | 65.854 | 65.787 | 67.418 | 22.171 | 28.356 | 46.781 |
| 8468 | ENSMUSG00000031617 | 298.3977763 | 0.03731927 | 0.185051609 | 0.201669522 | 0.840175091 | 0.971838962 | Tmem184c | transmembrane protein 184C [Source:MG | 334.89 | 302.21 | 245.52 | 251.6 | 360.99 | 295.17 |
| 8469 | ENSMUSG00000031618 | 287.632013 | 0.562253786 | 0.18893568 | 2.97590051 | 0.002921296 | 0.045927856 | Nr3c2 | nuclear receptor subfamily 3, group C, me | 240.13 | 266.23 | 190.18 | 300.76 | 416.62 | 311.88 |
| 8470 | ENSMUSG00000031620 | 0 | NA | NA | NA | NA | NA | Iqcm | IQ motif containing M [Source:MGI Symbo | 0 | 0 | 0 | 0 | 0 | 0 |
| 8471 | ENSMUSG00000031621 | 0 | NA | NA | NA | NA | NA | Isx | intestine specific homeobox [Source:MGI | 0 | 0 | 0 | 0 | 0 | 0 |
| 8472 | ENSMUSG00000031622 | 672.7902818 | 0.075199413 | 0.132733753 | 0.566543257 | 0.571024536 | 0.904651508 | Sin3b | transcriptional regulator, SIN3B (yeast) [S | 597.51 | 739.07 | 629.91 | 681.53 | 639.1 | 749.62 |
| 8473 | ENSMUSG00000031626 | 689.1031726 | 0.001205008 | 0.151705135 | 0.007943096 | 0.993662393 | 0.99859176 | Sorbs2 | sorbin and SH3 domain containing 2 [Sour | 542.09 | 734.96 | 790.91 | 717.2 | 652.19 | 697.27 |
| 8474 | ENSMUSG00000031627 | 1271.589636 | -0.127077296 | 0.104043961 | -1.221380796 | 0.221941875 | 0.667949912 | Irf2 | interferon regulatory factor 2 [Source:MG | 1368.5 | 1282.8 | 1330.3 | 1133.6 | 1237.9 | 1276.5 |
| 8475 | ENSMUSG00000031628 | 326.7735706 | 0.319034318 | 0.15876559 | 2.009467657 | 0.044487561 | 0.291685757 | Casp3 | caspase 3 [Source:MGI Symbol;Acc:MGI:1 | 287.51 | 253.9 | 331.05 | 377.88 | 329.37 | 380.93 |
| 8476 | ENSMUSG00000031629 | 6.453396208 | 0.341628843 | 0.899222216 | 0.379915929 | 0.704007823 | 0.94295036 | Cenpu | centromere protein U [Source:MGI Symbo | 8.8341 | 6.1675 | 2.0125 | 9.6398 | 8.7249 | 3.3415 |
| 8477 | ENSMUSG00000031631 | 263.7825369 | 0.122673094 | 0.156312144 | 0.784795672 | 0.432573378 | 0.845264432 | Cfap97 | cilia and flagella associated protein 97 [So | 247.36 | 240.53 | 269.67 | 255.45 | 290.1 | 279.57 |
| 8478 | ENSMUSG00000031633 | 61.82570248 | -0.973050977 | 0.327520051 | -2.970966126 | 0.002968645 | 0.046170414 | Slc25a4 | solute carrier family 25 (mitochondrial car | 98.782 | 86.345 | 60.375 | 41.451 | 30.537 | 53.465 |
| 8479 | ENSMUSG00000031634 | 703.2066075 | 0.08558532 | 0.118275072 | 0.723612494 | 0.469303669 | 0.866148815 | Ufsp2 | UFM1-specific peptidase 2 [Source:MGI S | 667.38 | 714.4 | 665.13 | 672.86 | 773.25 | 726.23 |
| 8480 | ENSMUSG00000031635 | 0.167707431 | -0.796629176 | 4.080472857 | -0.195229623 | 0.84521319 | NA | Anxa10 | annexin A10 [Source:MGI Symbol;Acc:MG | 0 | 0 | 1.0062 | 0 | 0 | 0 |
| 8481 | ENSMUSG00000031636 | 2.006056193 | -0.936667107 | 1.585330403 | -0.590834002 | 0.55463165 | NA | Pdlim3 | PDZ and LIM domain 3 [Source:MGI Symb | 4.8186 | 2.0558 | 1.0062 | 1.928 | 0 | 2.2277 |
| 8482 | ENSMUSG00000031637 | 22.87579136 | -0.18358825 | 0.49336105 | -0.372117438 | 0.709805413 | 0.944239215 | Lrp2bp | Lrp2 binding protein [Source:MGI Symbol, | 32.124 | 26.726 | 14.087 | 27.955 | 18.541 | 17.822 |
| 8483 | ENSMUSG00000031639 | 108.0610789 | 0.099366965 | 0.233826838 | 0.424959622 | 0.67086611 | 0.93116767 | Tlr3 | toll-like receptor 3 [Source:MGI Symbol;A | 120.47 | 103.82 | 88.55 | 127.25 | 104.7 | 103.59 |
| 8484 | ENSMUSG00000031640 | 0.874403588 | -0.441200908 | 2.817192576 | -0.156610134 | 0.875552115 | NA | Gm45753 | predicted gene 45753 [Source:MGI Symbo | 0 | 0 | 3.0187 | 0 | 0 | 2.2277 |
| 8485 | ENSMUSG00000031641 | 610.7229923 | 0.038388851 | 0.123325032 | 0.311281907 | 0.75558632 | 0.953085143 | Cbr4 | carbonyl reductase 4 [Source:MGI Symbol | 583.05 | 634.23 | 590.67 | 590.92 | 672.91 | 592.56 |
| 8486 | ENSMUSG00000031642 | 54.9384085 | 0.067000163 | 0.345760515 | 0.193776212 | 0.846351114 | 0.972803041 | Sh3rf1 | SH3 domain containing ring finger 1 [Sour | 69.067 | 42.145 | 49.306 | 53.019 | 40.353 | 75.741 |
| 8487 | ENSMUSG00000031644 | 191.8718841 | 0.250588636 | 0.199384754 | 1.256809413 | 0.208822657 | 0.653498052 | Nek1 | NIMA (never in mitosis gene a)-related ex | 183.91 | 149.05 | 192.19 | 173.52 | 245.39 | 207.17 |
| 8488 | ENSMUSG00000031645 | 1465.224125 | 0.897822257 | 0.178819927 | 5.020817723 | 5.15E-07 | 4.54E-05 | F11 | coagulation factor XI [Source:MGI Symbol | 1117.1 | 999.14 | 952.91 | 1292.7 | 2104.9 | 2324.6 |
| 8489 | ENSMUSG00000031647 | 9.318540512 | -0.904990744 | 0.753686978 | -1.200751466 | 0.229847623 | 0.676539688 | Mfap3l | microfibrillar-associated protein 3-like [So | 11.243 | 10.279 | 15.094 | 11.568 | 3.2719 | 4.4554 |
| 8490 | ENSMUSG00000031651 | 0 | NA | NA | NA | NA | NA | Triml1 | tripartite motif family-like 1 [Source:MGI S | 0 | 0 | 0 | 0 | 0 | 0 |
| 8491 | ENSMUSG00000031652 | 297.715048 | 0.123517471 | 0.149062912 | 0.828626446 | 0.407315819 | 0.83094725 | N4bp1 | NEDD4 binding protein 1 [Source:MGI Syn | 289.92 | 298.1 | 266.65 | 299.8 | 301.01 | 330.81 |
| 8492 | ENSMUSG00000031654 | 0 | NA | NA | NA | NA | NA | Cbln1 | cerebellin 1 precursor protein [Source:MG | 0 | 0 | 0 | 0 | 0 | 0 |
| 8493 | ENSMUSG00000031657 | 429.4259308 | -0.002914373 | 0.146276597 | -0.01992371 | 0.984104231 | 0.998580735 | Heatr3 | HEAT repeat containing 3 [Source:MGI Syı | 414.4 | 457.42 | 417.59 | 379.81 | 491.87 | 415.46 |
| 8494 | ENSMUSG00000031659 | 39.1123649 | -1.306313617 | 0.366031954 | -3.568851306 | 0.00035855 | 0.009973582 | Adcy7 | adenylate cyclase 7 [Source:MGI Symbol;/ | 53.005 | 64.759 | 49.306 | 20.244 | 25.084 | 22.277 |
| 8495 | ENSMUSG00000031660 | 638.0256394 | 0.066984665 | 0.127756153 | 0.524316548 | 0.600058389 | 0.915217938 | Brd7 | bromodomain containing 7 [Source:MGI S | 672.2 | 611.61 | 584.63 | 604.41 | 639.1 | 716.2 |
| 8496 | ENSMUSG00000031661 | 90.33589057 | -0.614295686 | 0.296633673 | -2.070889932 | 0.038369083 | 0.266556398 | Nkd1 | naked cuticle 1 [Source:MGI Symbol;Acc:I | 91.554 | 85.317 | 150.94 | 60.731 | 63.256 | 90.221 |
| 8497 | ENSMUSG00000031662 | 6.923747655 | -2.762079505 | 0.988073762 | -2.795418328 | 0.00518326 | 0.068217168 | Snx20 | sorting nexin 20 [Source:MGI Symbol;Acc: | 9.6372 | 16.447 | 10.062 | 0.964 | 1.0906 | 3.3415 |
| 8498 | ENSMUSG00000031665 | 421.9608085 | -0.449357646 | 0.142549273 | -3.152297014 | 0.001619914 | 0.03000056 | Sall1 | spalt like transcription factor 1 [Source:M | 447.33 | 480.04 | 534.32 | 335.46 | 390.44 | 344.18 |
| 8499 | ENSMUSG00000031666 | 693.4983966 | 0.172903379 | 0.129947923 | 1.33055901 | 0.183334157 | 0.617546635 | Rbl2 | RB transcriptional corepressor like 2 [Sour | 611.16 | 607.5 | 737.58 | 710.45 | 702.36 | 791.94 |
| 8500 | ENSMUSG00000031667 | 196.1941209 | 0.081807441 | 0.177776852 | 0.46016925 | 0.645394741 | 0.925643431 | Aktip | AKT interacting protein [Source:MGI Symk | 187.12 | 214.83 | 170.06 | 198.58 | 208.31 | 198.26 |
| 8501 | ENSMUSG00000031668 | 246.4187941 | 0.079907387 | 0.174651068 | 0.457525898 | 0.647293092 | 0.925643431 | Eif2ak3 | eukaryotic translation initiation factor 2 al | 240.93 | 261.09 | 216.34 | 220.75 | 247.57 | 291.83 |
| 8502 | ENSMUSG00000031669 | 11.63655478 | -0.243852249 | 0.675980827 | -0.360738411 | 0.718295008 | 0.9447719 | Gins3 | GINS complex subunit 3 (Psf3 homolog) [S | 19.274 | 9.2513 | 9.0562 | 5.7839 | 13.087 | 13.366 |
| 8503 | ENSMUSG00000031671 | 307.5625349 | 0.042109035 | 0.152980382 | 0.275257745 | 0.783118226 | 0.960670212 | Setd6 | SET domain containing 6 [Source:MGI Syn | 280.28 | 325.85 | 303.89 | 335.46 | 283.56 | 316.33 |
| 8504 | ENSMUSG00000031672 | 10266.83004 | -0.151242409 | 0.086701288 | -1.74440786 | 0.081087995 | 0.41872787 | Got2 | glutamatic-oxaloacetic transaminase 2, m | 10775 | 10990 | 10648 | 9818.1 | 10231 | 9138 |
| 8505 | ENSMUSG00000031673 | 2.776595533 | -0.69915904 | 1.274993354 | -0.54836289 | 0.583442753 | NA | Cdh11 | cadherin 11 [Source:MGI Symbol;Acc:MG | 3.2124 | 2.0558 | 5.0312 | 1.928 | 1.0906 | 3.3415 |
| 8506 | ENSMUSG00000031681 | 255.0942139 | 0.202467213 | 0.161322576 | 1.255045742 | 0.209462155 | 0.653550636 | Smad1 | SMAD family member 1 [Source:MGI Sym | 262.61 | 230.25 | 218.36 | 288.23 | 270.47 | 260.64 |
| 8507 | ENSMUSG00000031682 | 0.133850373 | -0.796629176 | 4.080472857 | -0.195229623 | 0.84521319 | NA | 1700011L22I | RIKEN cDNA 1700011L22 gene [Source:M( | 0.8031 | 0 | 0 | 0 | 0 | 0 |
| 8508 | ENSMUSG00000031683 | 547.0944949 | 0.083117129 | 0.13027909 | 0.637992858 | 0.523478329 | 0.888334265 | Lsm6 | LSM6 homolog, U6 small nuclear RNA and | 510.77 | 504.71 | 578.59 | 533.08 | 610.75 | 544.67 |

| | | | | | | | | | | | | | | |
|---|---|---|---|---|---|---|---|---|---|---|---|---|---|---|
| 8509 | ENSMUSG00000031684 | 134.6882152 | 0.122033349 | 0.204311706 | 0.597290051 | 0.550313746 | 0.895703112 | Slc10a7 | solute carrier family 10 (sodium/bile acid | 126.89 | 117.18 | 142.89 | 138.81 | 130.87 | 151.48 |
| 8510 | ENSMUSG00000031688 | 0 | NA | NA | NA | NA | NA | Pou4f2 | POU domain, class 4, transcription factor | 0 | 0 | 0 | 0 | 0 | 0 |
| 8511 | ENSMUSG00000031691 | 570.4093402 | 0.202753932 | 0.136351792 | 1.486991319 | 0.137017104 | 0.540888479 | Tnpo2 | transportin 2 (importin 3, karyopherin bet | 480.26 | 600.3 | 511.17 | 572.6 | 599.84 | 658.28 |
| 8512 | ENSMUSG00000031696 | 1751.865467 | 0.070563217 | 0.104893962 | 0.672710004 | 0.501131802 | 0.880974509 | Vps35 | VPS35 retromer complex component [Sou | 1751.6 | 1707.4 | 1667.3 | 1674.4 | 1975.1 | 1735.4 |
| 8513 | ENSMUSG00000031697 | 69.99394753 | 0.107205707 | 0.327582736 | 0.327262993 | 0.74346898 | 0.949707262 | Orc6 | origin recognition complex, subunit 6 [Sou | 71.476 | 88.401 | 42.262 | 69.406 | 91.612 | 56.806 |
| 8514 | ENSMUSG00000031698 | 7.212724224 | -1.686979613 | 0.943713181 | -1.787597807 | 0.073840919 | 0.395275744 | Mylk3 | myosin light chain kinase 3 [Source:MGI S | 4.8186 | 10.279 | 18.112 | 6.7478 | 1.0906 | 2.2277 |
| 8515 | ENSMUSG00000031700 | 13510.45116 | -0.358170033 | 0.101386748 | -3.532710538 | 0.000411323 | 0.010817036 | Gpt2 | glutamic pyruvate transaminase (alanine | 15475 | 16196 | 13865 | 10825 | 12518 | 12183 |
| 8516 | ENSMUSG00000031701 | 2487.603078 | -0.074754635 | 0.102833791 | -0.726946218 | 0.467258903 | 0.864254244 | Dnaja2 | DnaJ heat shock protein family (Hsp40) m | 2417.3 | 2590.4 | 2649.4 | 2470.7 | 2592.4 | 2205.4 |
| 8517 | ENSMUSG00000031703 | 1026.924404 | 0.042734123 | 0.102949565 | 0.415097652 | 0.678070418 | 0.93325821 | Itfg1 | integrin alpha FG-GAP repeat containing 1 | 1032.8 | 1033.1 | 969.01 | 1036.3 | 1059 | 1031.4 |
| 8518 | ENSMUSG00000031706 | 178.2678835 | 0.239398148 | 0.186467013 | 1.283863268 | 0.199189801 | 0.641237767 | Rfx1 | regulatory factor X, 1 (influences HLA clas | 170.26 | 166.52 | 153.96 | 220.75 | 182.13 | 175.99 |
| 8519 | ENSMUSG00000031708 | 2353.566668 | 0.095465588 | 0.097713159 | 0.976998269 | 0.328570009 | 0.77362974 | Tecr | trans-2,3-enoyl-CoA reductase [Source:M | 2390 | 2341.6 | 2095 | 2482.2 | 2372.1 | 2440.4 |
| 8520 | ENSMUSG00000031709 | 18.23558546 | -1.593689231 | 0.570629642 | -2.792860928 | 0.005224415 | 0.068666748 | Tbc1d9 | TBC1 domain family, member 9 [Source:M | 28.109 | 38.033 | 16.1 | 10.604 | 6.5437 | 10.025 |
| 8521 | ENSMUSG00000031710 | 0 | NA | NA | NA | NA | NA | Ucp1 | uncoupling protein 1 (mitochondrial, prot | 0 | 0 | 0 | 0 | 0 | 0 |
| 8522 | ENSMUSG00000031711 | 256.1469025 | 0.024475422 | 0.164994686 | 0.14834067 | 0.882073921 | 0.981738667 | Zfp330 | zinc finger protein 330 [Source:MGI Symb | 236.92 | 281.65 | 243.51 | 236.17 | 283.56 | 255.07 |
| 8523 | ENSMUSG00000031712 | 24.26329437 | 0.666706917 | 0.495327872 | 1.345991119 | 0.178305381 | 0.610642232 | Il15 | interleukin 15 [Source:MGI Symbol;Acc:M | 27.305 | 16.447 | 12.075 | 19.28 | 38.172 | 32.301 |
| 8524 | ENSMUSG00000031714 | 286.2057227 | 0.066226446 | 0.193251851 | 0.342695014 | 0.731827914 | 0.947387358 | Gab1 | growth factor receptor bound protein 2-a | 273.86 | 252.87 | 312.94 | 372.1 | 271.56 | 233.91 |
| 8525 | ENSMUSG00000031715 | 742.1371612 | -0.084282854 | 0.126279301 | -0.66743206 | 0.504496193 | 0.881510796 | Smarca5 | SWI/SNF related, matrix associated, actin | 691.47 | 828.5 | 772.8 | 763.47 | 740.53 | 656.05 |
| 8526 | ENSMUSG00000031722 | 62093.27425 | -0.314473966 | 0.10875003 | -2.891713822 | 0.003831468 | 0.055585105 | Hp | haptoglobin [Source:MGI Symbol;Acc:MG | 71314 | 73734 | 61455 | 61101 | 54412 | 50544 |
| 8527 | ENSMUSG00000031723 | 227.1149601 | 0.348269987 | 0.180797135 | 1.926302578 | 0.054066604 | 0.327341714 | Txnl4b | thioredoxin-like 4B [Source:MGI Symbol;A | 206.4 | 190.16 | 202.26 | 209.18 | 261.75 | 292.94 |
| 8528 | ENSMUSG00000031725 | 8510.039457 | 0.164496388 | 0.112844366 | 1.457727965 | 0.14491554 | 0.555361204 | Ces1f | carboxylesterase 1F [Source:MGI Symbol; | 7324.3 | 8396 | 8357.9 | 10024 | 9078.3 | 7879.3 |
| 8529 | ENSMUSG00000031727 | 0 | NA | NA | NA | NA | NA | Pmfbp1 | polyamine modulated factor 1 binding pro | 0 | 0 | 0 | 0 | 0 | 0 |
| 8530 | ENSMUSG00000031728 | 36.7912874 | 0.280566323 | 0.408508832 | 0.686806014 | 0.492204979 | 0.876368691 | Zfp821 | zinc finger protein 821 [Source:MGI Symb | 24.896 | 25.698 | 49.306 | 42.415 | 30.537 | 47.895 |
| 8531 | ENSMUSG00000031729 | 1040.329103 | 0.089490688 | 0.106559264 | 0.839820817 | 0.40100886 | 0.827897196 | Ist1 | increased sodium tolerance 1 homolog (y | 1009.5 | 967.27 | 1047.5 | 1079.7 | 1123.3 | 1014.7 |
| 8532 | ENSMUSG00000031730 | 190.5868048 | -0.035323607 | 0.17147752 | -0.205995557 | 0.836794382 | 0.971073158 | Dhodh | dihydroorotate dehydrogenase [Source:M | 199.17 | 194.28 | 185.15 | 187.01 | 194.13 | 183.78 |
| 8533 | ENSMUSG00000031731 | 943.2097188 | -0.023343806 | 0.114004002 | -0.204763041 | 0.837757264 | 0.971073158 | Ap1g1 | adaptor protein complex AP-1, gamma 1 s | 908.31 | 1027.9 | 916.69 | 920.6 | 888.85 | 996.89 |
| 8534 | ENSMUSG00000031732 | 71.40757621 | 0.113920756 | 0.310620925 | 0.366751714 | 0.713804228 | 0.944425938 | Phlpp2 | PH domain and leucine rich repeat protei | 61.036 | 65.787 | 79.493 | 104.11 | 54.531 | 63.489 |
| 8535 | ENSMUSG00000031734 | 14.34312994 | 0.385587472 | 0.620911813 | 0.621001991 | 0.534598313 | 0.889858671 | Irx3 | Iroquois related homeobox 3 [Source:MG | 8.031 | 10.279 | 19.119 | 15.424 | 9.8156 | 23.391 |
| 8536 | ENSMUSG00000031736 | 1.370661385 | 0.064653576 | 1.948658061 | 0.033178513 | 0.973532233 | NA | Crnde | colorectal neoplasia differentially express | 0 | 2.0558 | 2.0125 | 1.928 | 0 | 2.2277 |
| 8537 | ENSMUSG00000031737 | 0.133850373 | -0.796629176 | 4.080472857 | -0.195229623 | 0.84521319 | NA | Irx5 | Iroquois homeobox 5 [Source:MGI Symbo | 0.8031 | 0 | 0 | 0 | 0 | 0 |
| 8538 | ENSMUSG00000031738 | 0 | NA | NA | NA | NA | NA | Irx6 | Iroquois homeobox 6 [Source:MGI Symbo | 0 | 0 | 0 | 0 | 0 | 0 |
| 8539 | ENSMUSG00000031740 | 33.7717349 | -0.379181618 | 0.407755366 | -0.929924286 | 0.352410287 | 0.793101353 | Mmp2 | matrix metallopeptidase 2 [Source:MGI Sy | 47.383 | 37.005 | 30.187 | 38.559 | 29.447 | 20.049 |
| 8540 | ENSMUSG00000031748 | 6.105776232 | -1.285101178 | 0.891460517 | -1.44156825 | 0.149424211 | 0.561953141 | Gnao1 | guanine nucleotide binding protein, alpha | 9.6372 | 7.1954 | 9.0562 | 1.928 | 4.3625 | 4.4554 |
| 8541 | ENSMUSG00000031749 | 43.28048757 | -0.677437294 | 0.339826307 | -1.993481023 | 0.0462088 | 0.297429405 | St3gal2 | ST3 beta-galactoside alpha-2,3-sialyltrans | 57.823 | 58.591 | 43.269 | 32.775 | 33.809 | 33.415 |
| 8542 | ENSMUSG00000031750 | 36.06815045 | -1.301906517 | 0.378515312 | -3.439508194 | 0.000582772 | 0.013773229 | Il34 | interleukin 34 [Source:MGI Symbol;Acc:M | 48.186 | 48.312 | 57.356 | 17.352 | 21.812 | 23.391 |
| 8543 | ENSMUSG00000031751 | 6132.07136 | 0.070173835 | 0.087767447 | 0.799542854 | 0.423975708 | 0.840450466 | Amfr | autocrine motility factor receptor [Source | 5661.1 | 6112 | 6177.3 | 6510.7 | 6174 | 6157.3 |
| 8544 | ENSMUSG00000031753 | 1386.652648 | -0.001853164 | 0.10846267 | -0.017085726 | 0.986368226 | 0.99859176 | Cog4 | component of oligomeric golgi complex 4 | 1452.8 | 1404.1 | 1305.1 | 1364 | 1501.8 | 1292.1 |
| 8545 | ENSMUSG00000031754 | 476.3752832 | 0.153112143 | 0.144734399 | 1.05788357 | 0.290108527 | 0.740058153 | Nudt21 | nudix (nucleoside diphosphate linked moi | 394.32 | 457.42 | 502.12 | 453.07 | 530.04 | 521.28 |
| 8546 | ENSMUSG00000031755 | 60.51680606 | -0.251881808 | 0.33201278 | -0.758650941 | 0.448061391 | 0.854156674 | Bbs2 | Bardet-Biedl syndrome 2 (human) [Source | 47.383 | 80.178 | 70.437 | 69.406 | 56.712 | 38.985 |
| 8547 | ENSMUSG00000031756 | 7.736501304 | -0.252110772 | 0.867463806 | -0.29062973 | 0.77133452 | 0.956965056 | Cenpn | centromere protein N [Source:MGI Symbo | 4.0155 | 8.2233 | 13.081 | 5.7839 | 13.087 | 2.2277 |
| 8548 | ENSMUSG00000031757 | 0 | NA | NA | NA | NA | NA | Mt4 | metallothionein 4 [Source:MGI Symbol;Ac | 0 | 0 | 0 | 0 | 0 | 0 |
| 8549 | ENSMUSG00000031758 | 20.46642456 | -0.053754815 | 0.476361393 | -0.112844609 | 0.910153751 | 0.985622944 | Cdyl2 | chromodomain protein, Y chromosome-lil | 16.865 | 23.642 | 22.137 | 18.316 | 22.903 | 18.935 |
| 8550 | ENSMUSG00000031760 | 0 | NA | NA | NA | NA | NA | Mt3 | metallothionein 3 [Source:MGI Symbol;Ac | 0 | 0 | 0 | 0 | 0 | 0 |
| 8551 | ENSMUSG00000031762 | 7.487950007 | -1.575952967 | 0.912662014 | -1.726765159 | 0.084209848 | 0.427953408 | Mt2 | metallothionein 2 [Source:MGI Symbol;Ac | 11.243 | 18.503 | 4.025 | 6.7478 | 2.1812 | 2.2277 |
| 8552 | ENSMUSG00000031765 | 3127.708545 | -1.418127832 | 0.520489195 | -2.724605707 | 0.006437832 | 0.079266903 | Mt1 | metallothionein 1 [Source:MGI Symbol;Ac | 4947.1 | 5550.8 | 3158.6 | 2780.1 | 1382.9 | 946.77 |
| 8553 | ENSMUSG00000031766 | 11.18611981 | -1.985548059 | 0.706029098 | -2.812275109 | 0.004919241 | 0.066064903 | Slc12a3 | solute carrier family 12, member 3 [Sourc | 20.078 | 17.475 | 16.1 | 5.7839 | 5.4531 | 2.2277 |
| 8554 | ENSMUSG00000031767 | 9402.525768 | 1.369423714 | 0.166782643 | 8.210828732 | 2.20E-16 | 1.87E-13 | Nudt7 | nudix (nucleoside diphosphate linked moi | 6892.2 | 4506.4 | 4340.9 | 11265 | 15087 | 14324 |
| 8555 | ENSMUSG00000031770 | 9587.5244 | -0.664634009 | 0.166911235 | -3.981960888 | 6.83E-05 | 0.002664691 | Herpud1 | homocysteine-inducible, endoplasmic reti | 10644 | 13167 | 11462 | 6204.2 | 9953 | 6095 |
| 8556 | ENSMUSG00000031772 | 0 | NA | NA | NA | NA | NA | Cntnap4 | contactin associated protein-like 4 [Sourc | 0 | 0 | 0 | 0 | 0 | 0 |
| 8557 | ENSMUSG00000031774 | 411.9339828 | 0.177278389 | 0.151154908 | 1.172825888 | 0.240865635 | 0.691298957 | Psme3ip1 | proteasome activator subunit 3 interactin | 411.19 | 389.58 | 358.22 | 382.7 | 425.34 | 504.57 |
| 8558 | ENSMUSG00000031775 | 60.51598471 | 0.074359375 | 0.291074899 | 0.255464747 | 0.798364168 | 0.962424862 | Pllp | plasma membrane proteolipid [Source:M | 65.854 | 50.368 | 60.375 | 58.803 | 70.89 | 56.806 |
| 8559 | ENSMUSG00000031776 | 117.7685182 | 0.347467166 | 0.227257384 | 1.528958754 | 0.126274669 | 0.521094669 | Arl2bp | ADP-ribosylation factor-like 2 binding prot | 106.01 | 121.29 | 83.518 | 121.46 | 141.78 | 132.55 |
| 8560 | ENSMUSG00000031778 | 11.12206287 | -1.583291621 | 0.720798713 | -2.196579422 | 0.028050497 | 0.21848271 | Cx3cl1 | chemokine (C-X3-C motif) ligand 1 [Source | 26.502 | 14.391 | 9.0562 | 5.7839 | 6.5437 | 4.4554 |
| 8561 | ENSMUSG00000031779 | 1.495425783 | 1.239937803 | 1.780032402 | 0.696581591 | 0.48606468 | NA | Ccl22 | chemokine (C-C motif) ligand 22 [Source:N | 1.6062 | 0 | 1.0062 | 1.928 | 1.0906 | 3.3415 |
| 8562 | ENSMUSG00000031780 | 0.319491021 | 0.16514334 | 4.044653141 | 0.040830038 | 0.967431393 | NA | Ccl17 | chemokine (C-C motif) ligand 17 [Source:N | 0.8031 | 0 | 0 | 0 | 0 | 1.1138 |
| 8563 | ENSMUSG00000031781 | 576.7094037 | -0.08708095 | 0.125851304 | -0.691935223 | 0.48897801 | 0.874706988 | Ciapin1 | cytokine induced apoptosis inhibitor 1 [So | 651.32 | 568.44 | 561.48 | 544.65 | 552.94 | 581.43 |
| 8564 | ENSMUSG00000031782 | 2272.297876 | 0.034662023 | 0.093634797 | 0.370183134 | 0.711246043 | 0.944239215 | Coq9 | coenzyme Q9 [Source:MGI Symbol;Acc:M | 2337.8 | 2234.7 | 2161.4 | 2212.3 | 2384.1 | 2303.4 |

| | | | | | | | | | | | | | | | |
|---|---|---|---|---|---|---|---|---|---|---|---|---|---|---|---|
| 8565 | ENSMUSG00000031783 | 428.4030725 | 0.188908462 | 0.135440681 | 1.394768993 | 0.163085512 | 0.586990831 | Polr2c | polymerase (RNA) II (DNA directed) polyp | 387.1 | 413.22 | 401.49 | 489.7 | 414.43 | 464.47 |
| 8566 | ENSMUSG00000031785 | 26.16994532 | -0.224414817 | 0.434566942 | -0.516410235 | 0.605567919 | 0.916376001 | Adgrg1 | adhesion G protein-coupled receptor G1 [ | 26.502 | 29.81 | 28.175 | 17.352 | 23.994 | 31.188 |
| 8567 | ENSMUSG00000031786 | 0.160662974 | 1.126924175 | 4.080472857 | 0.276174898 | 0.782413736 | NA | Drc7 | dynein regulatory complex subunit 7 [Sou | 0 | 0 | 0 | 0.964 | 0 | 0 |
| 8568 | ENSMUSG00000031787 | 47.32503308 | 0.068986124 | 0.346917546 | 0.198854526 | 0.842376542 | 0.971875826 | Katnb1 | katanin p80 (WD40-containing) subunit B | 54.611 | 49.34 | 34.212 | 36.631 | 52.35 | 56.806 |
| 8569 | ENSMUSG00000031788 | 199.4689797 | 0.215869697 | 0.236763806 | 0.911751253 | 0.361899677 | 0.800872989 | Kifc3 | kinesin family member C3 [Source:MGI Sy | 155.8 | 158.3 | 240.49 | 277.63 | 191.95 | 172.65 |
| 8570 | ENSMUSG00000031789 | 0.931665974 | -0.888641078 | 2.464021547 | -0.360646634 | 0.718363623 | NA | Cngb1 | cyclic nucleotide gated channel beta 1 [So | 1.6062 | 2.0558 | 0 | 1.928 | 0 | 0 |
| 8571 | ENSMUSG00000031790 | 1169.809886 | 0.360099881 | 0.109442381 | 3.290314747 | 0.001000754 | 0.02092365 | Mmp15 | matrix metallopeptidase 15 [Source:MGI S | 982.19 | 1089.6 | 1002.2 | 1241.6 | 1414.5 | 1288.7 |
| 8572 | ENSMUSG00000031791 | 54.06749829 | 0.312315242 | 0.321343544 | 0.97190452 | 0.331098051 | 0.774747658 | Tmem38a | transmembrane protein 38A [Source:MGI | 44.171 | 45.228 | 55.343 | 55.911 | 46.897 | 76.855 |
| 8573 | ENSMUSG00000031792 | 190.8147292 | -0.003480955 | 0.191123067 | -0.01821316 | 0.985468804 | 0.99859176 | Usb1 | U6 snRNA biogenesis 1 [Source:MGI Symb | 194.35 | 177.83 | 201.25 | 221.71 | 157.05 | 192.69 |
| 8574 | ENSMUSG00000031796 | 526.3666517 | 0.489424087 | 0.129590926 | 3.776684851 | 0.00015893 | 0.005248306 | Cfap20 | cilia and flagella associated protein 20 [So | 430.46 | 452.28 | 431.68 | 672.86 | 562.76 | 608.16 |
| 8575 | ENSMUSG00000031799 | 222.8802904 | -0.159518589 | 0.212773453 | -0.749710958 | 0.453428807 | 0.857073339 | Tpm4 | tropomyosin 4 [Source:MGI Symbol;Acc:N | 289.12 | 195.3 | 220.37 | 214.97 | 164.68 | 252.84 |
| 8576 | ENSMUSG00000031802 | 0 | NA | NA | NA | NA | NA | Phxr4 | per-hexamer repeat gene 4 [Source:MGI S | 0 | 0 | 0 | 0 | 0 | 0 |
| 8577 | ENSMUSG00000031803 | 24.42750196 | -0.272495674 | 0.455833376 | -0.59779667 | 0.549975613 | 0.895703112 | B3gnt3 | UDP-GlcNAc:betaGal beta-1,3-N-acetylglu | 19.274 | 35.977 | 25.156 | 23.135 | 19.631 | 23.391 |
| 8578 | ENSMUSG00000031805 | 275.8387112 | -0.160901839 | 0.181588193 | -0.886080952 | 0.375573907 | 0.809462224 | Jak3 | Janus kinase 3 [Source:MGI Symbol;Acc:M | 302.77 | 281.65 | 289.8 | 326.79 | 229.03 | 225 |
| 8579 | ENSMUSG00000031807 | 1102.43285 | -0.034896671 | 0.106111085 | -0.328869227 | 0.742254535 | 0.949043964 | Pgls | 6-phosphogluconolactonase [Source:MGI | 1178.2 | 1042.3 | 1126 | 1077.7 | 1122.2 | 1068.2 |
| 8580 | ENSMUSG00000031808 | 258.3846717 | -0.123376366 | 0.209902942 | -0.587778165 | 0.556681202 | 0.898167794 | Slc27a1 | solute carrier family 27 (fatty acid transpo | 254.58 | 237.45 | 316.97 | 304.62 | 258.48 | 178.22 |
| 8581 | ENSMUSG00000031809 | 0.321325949 | 1.911575114 | 4.048267893 | 0.472195804 | 0.636787034 | NA | 1700018B08 | RIKEN cDNA 1700018B08 gene [Source:M | 0 | 0 | 0 | 1.928 | 0 | 0 |
| 8582 | ENSMUSG00000031812 | 2625.58125 | -0.073479548 | 0.114251622 | -0.643137899 | 0.520134625 | 0.887802551 | Map1lc3b | microtubule-associated protein 1 light cha | 2716.9 | 2728.1 | 2631.3 | 2301 | 2423.4 | 2952.8 |
| 8583 | ENSMUSG00000031813 | 996.7879389 | 0.026349862 | 0.109188687 | 0.241324106 | 0.809303929 | 0.965982178 | Mvb12a | multivesicular body subunit 12A [Source:N | 1016.7 | 926.15 | 1019.3 | 959.16 | 1000.1 | 1059.3 |
| 8584 | ENSMUSG00000031816 | 205.388772 | -0.239210302 | 0.17452736 | -1.370617779 | 0.170494138 | 0.597556794 | Mthfsd | methenyltetrahydrofolate synthetase don | 217.64 | 210.72 | 238.48 | 169.66 | 189.77 | 206.06 |
| 8585 | ENSMUSG00000031818 | 7440.709648 | -0.072549842 | 0.081761449 | -0.887335566 | 0.374898263 | 0.809332934 | Cox4i1 | cytochrome c oxidase subunit 4I1 [Source | 7579.7 | 7700.1 | 7603.2 | 7065 | 7207.9 | 7488.4 |
| 8586 | ENSMUSG00000031819 | 447.8354258 | -0.228377857 | 0.169717916 | -1.345631989 | 0.17842123 | 0.610812939 | Emc8 | ER membrane protein complex subunit 8 [ | 499.53 | 545.82 | 403.5 | 333.54 | 462.42 | 442.2 |
| 8587 | ENSMUSG00000031820 | 868.0568425 | 0.068586007 | 0.110191973 | 0.622422893 | 0.533663831 | 0.889143097 | Babam1 | BRISC and BRCA1 A complex member 1 [S | 836.83 | 836.73 | 868.39 | 840.59 | 879.04 | 946.77 |
| 8588 | ENSMUSG00000031821 | 9.338201509 | 0.514084119 | 0.802840202 | 0.640331809 | 0.521956905 | 0.888290937 | Gins2 | GINS complex subunit 2 (Psf2 homolog) [S | 13.653 | 5.1396 | 4.025 | 5.7839 | 19.631 | 7.7969 |
| 8589 | ENSMUSG00000031822 | 193.07039 | 0.05363969 | 0.222656966 | 0.240907306 | 0.809626959 | 0.966128565 | Gse1 | genetic suppressor element 1, coiled-coil | 165.44 | 169.61 | 233.45 | 173.52 | 164.68 | 251.73 |
| 8590 | ENSMUSG00000031823 | 451.9063497 | 0.111936263 | 0.140221067 | 0.798284207 | 0.424705581 | 0.840504368 | Zdhhc7 | zinc finger, DHHC domain containing 7 [Sc | 398.34 | 445.09 | 459.85 | 417.4 | 497.32 | 493.43 |
| 8591 | ENSMUSG00000031824 | 66.3749932 | -0.466173157 | 0.31190486 | -1.494600494 | 0.135018737 | 0.537616739 | 6430548M08 | RIKEN cDNA 6430548M08 gene [Source:M | 81.113 | 82.233 | 67.418 | 49.163 | 40.353 | 77.969 |
| 8592 | ENSMUSG00000031825 | 21.0655876 | -0.635802276 | 0.512187772 | -1.241346067 | 0.214477934 | 0.659202897 | Crispld2 | cysteine-rich secretory protein LCCL doma | 23.29 | 26.726 | 27.169 | 26.027 | 9.8156 | 13.366 |
| 8593 | ENSMUSG00000031826 | 1508.963487 | 0.004252748 | 0.097810938 | 0.043479263 | 0.965319494 | 0.993480943 | Usp10 | ubiquitin specific peptidase 10 [Source:M | 1552.4 | 1498.7 | 1469.1 | 1576.1 | 1507.2 | 1450.2 |
| 8594 | ENSMUSG00000031827 | 160.1081826 | -0.61528704 | 0.219083022 | -2.808465181 | 0.004977827 | 0.066542775 | Cotl1 | coactosin-like 1 (Dictyostelium) [Source:M | 212.82 | 210.72 | 156.97 | 102.18 | 128.69 | 149.26 |
| 8595 | ENSMUSG00000031828 | 123.5043776 | -0.191751155 | 0.243933447 | -0.786079799 | 0.431820738 | 0.84514901 | Klhl36 | kelch-like 36 [Source:MGI Symbol;Acc:MG | 118.06 | 136.71 | 140.87 | 138.81 | 80.706 | 125.86 |
| 8596 | ENSMUSG00000031831 | 1.020693421 | -3.383091396 | 2.599252854 | -1.301563021 | 0.193065808 | NA | Dnaaf1 | dynein, axonemal assembly factor 1 [Sour | 0 | 4.1117 | 2.0125 | 0 | 0 | 0 |
| 8597 | ENSMUSG00000031832 | 37.005792 | -0.329466972 | 0.375159623 | -0.878204773 | 0.379832602 | 0.812628154 | Taf1c | TATA-box binding protein associated facto | 32.124 | 53.452 | 38.237 | 29.883 | 33.809 | 34.529 |
| 8598 | ENSMUSG00000031833 | 749.7199308 | 0.03081914 | 0.129791337 | 0.237451438 | 0.812306589 | 0.96653525 | Mast3 | microtubule associated serine/threonine l | 747.69 | 687.68 | 788.9 | 658.4 | 809.24 | 806.42 |
| 8599 | ENSMUSG00000031834 | 487.7571869 | -0.057810743 | 0.131678828 | -0.439028382 | 0.660640972 | 0.928170084 | Pik3r2 | phosphoinositide-3-kinase regulatory sub | 461.78 | 523.21 | 508.15 | 477.17 | 444.97 | 511.25 |
| 8600 | ENSMUSG00000031835 | 1156.922357 | 0.047418017 | 0.10386162 | 0.456549945 | 0.647994564 | 0.925643431 | Mbtps1 | membrane-bound transcription factor pe | 1112.3 | 1104 | 1197.4 | 1144.2 | 1152.8 | 1230.8 |
| 8601 | ENSMUSG00000031837 | 0.813971537 | 1.950992709 | 2.892144212 | 0.674583481 | 0.499940431 | NA | Necab2 | N-terminal EF-hand calcium binding prote | 0 | 1.0279 | 0 | 3.8559 | 0 | 0 |
| 8602 | ENSMUSG00000031838 | 221.7031279 | -0.223579602 | 0.216866648 | -1.030954295 | 0.302562254 | 0.752385931 | Ifi30 | interferon gamma inducible protein 30 [Sc | 195.15 | 297.07 | 224.39 | 158.09 | 211.58 | 243.93 |
| 8603 | ENSMUSG00000031839 | 1277.180098 | -0.081590727 | 0.103856939 | -0.785606892 | 0.432097826 | 0.845261421 | Hsbp1 | heat shock factor binding protein 1 [Sourc | 1399.8 | 1263.3 | 1275.9 | 1259.9 | 1188.8 | 1275.4 |
| 8604 | ENSMUSG00000031840 | 74.19475645 | 0.589134867 | 0.271969727 | 2.166178103 | 0.03029758 | 0.230436937 | Rab3a | RAB3A, member RAS oncogene family [So | 48.186 | 63.731 | 66.412 | 98.326 | 89.431 | 79.083 |
| 8605 | ENSMUSG00000031841 | 29.66593663 | -0.41165926 | 0.41642568 | -0.988553973 | 0.322881415 | 0.769995293 | Cdh13 | cadherin 13 [Source:MGI Symbol;Acc:MG | 43.368 | 33.921 | 24.15 | 26.991 | 26.175 | 23.391 |
| 8606 | ENSMUSG00000031842 | 290.1553603 | 0.188181743 | 0.188433293 | 0.998665043 | 0.31795698 | 0.764585353 | Pde4c | phosphodiesterase 4C, cAMP specific [Sou | 214.43 | 281.65 | 318.98 | 355.71 | 267.2 | 302.97 |
| 8607 | ENSMUSG00000031843 | 48.25040181 | -0.057348313 | 0.322962049 | -0.17756982 | 0.859060828 | 0.975487285 | Mphosph6 | M phase phosphoprotein 6 [Source:MGI S | 53.808 | 53.452 | 40.25 | 48.199 | 41.443 | 52.351 |
| 8608 | ENSMUSG00000031844 | 2370.188774 | 0.617920006 | 0.123474982 | 5.004414615 | 5.60E-07 | 4.88E-05 | Hsd17b2 | hydroxysteroid (17-beta) dehydrogenase 2 | 1979.6 | 1717.7 | 1911.9 | 2478.4 | 2841.1 | 3292.5 |
| 8609 | ENSMUSG00000031845 | 116.0830416 | 0.025016698 | 0.268755936 | 0.093083332 | 0.925837359 | 0.987546253 | Bco1 | beta-carotene oxygenase 1 [Source:MGI S | 127.69 | 123.35 | 93.581 | 86.758 | 103.61 | 161.51 |
| 8610 | ENSMUSG00000031847 | 9.968254842 | 1.036864948 | 0.717784812 | 1.444534532 | 0.148588673 | 0.560454158 | 1700030J22 | RIKEN cDNA 1700030J22 gene [Source:MG | 6.4248 | 8.2233 | 5.0312 | 20.244 | 7.6343 | 12.252 |
| 8611 | ENSMUSG00000031848 | 514.7480006 | 0.089629854 | 0.129211069 | 0.693670087 | 0.487889129 | 0.873910634 | Lsm4 | LSM4 homolog, U6 small nuclear RNA and | 517.2 | 501.62 | 476.96 | 520.55 | 492.96 | 579.2 |
| 8612 | ENSMUSG00000031849 | 0.688504217 | -1.404357195 | 2.777143853 | -0.505683994 | 0.613078506 | NA | Comp | cartilage oligomeric matrix protein [Sourc | 0 | 1.0279 | 2.0125 | 0 | 1.0906 | 0 |
| 8613 | ENSMUSG00000031851 | 154.3912231 | -0.114710002 | 0.201564884 | -0.569097155 | 0.569290209 | 0.903735951 | Ntpcr | nucleoside-triphosphatase, cancer-relatec | 170.26 | 158.3 | 152.95 | 153.27 | 169.05 | 122.52 |
| 8614 | ENSMUSG00000031853 | 126.0855512 | 0.452212312 | 0.262725664 | 1.721233871 | 0.085208394 | 0.430025893 | Map3k21 | mitogen-activated protein kinase kinase k | 81.113 | 113.07 | 125.78 | 143.63 | 105.79 | 187.13 |
| 8615 | ENSMUSG00000031858 | 601.9711969 | 0.192577468 | 0.127414897 | 1.511420344 | 0.130681389 | 0.529460957 | Mau2 | MAU2 sister chromatid cohesion factor [S | 559.76 | 545.82 | 580.6 | 715.27 | 593.3 | 617.07 |
| 8616 | ENSMUSG00000031860 | 9.087319858 | 0.426881636 | 0.761298851 | 0.56072807 | 0.574982928 | 0.906337519 | Pbx4 | pre B cell leukemia homeobox 4 [Source:N | 8.031 | 7.1954 | 8.05 | 13.496 | 3.2719 | 14.48 |
| 8617 | ENSMUSG00000031861 | 23.74168618 | 1.214373154 | 0.536258517 | 2.264529358 | 0.023541576 | 0.194241505 | Lpar2 | lysophosphatidic acid receptor 2 [Source:I | 18.471 | 5.1396 | 19.119 | 43.379 | 21.812 | 34.529 |
| 8618 | ENSMUSG00000031862 | 817.6261283 | -0.011338507 | 0.116106017 | -0.097656495 | 0.922205062 | 0.986685804 | Atp13a1 | ATPase type 13A1 [Source:MGI Symbol;Ac | 815.15 | 852.14 | 795.94 | 885.9 | 756.89 | 799.74 |
| 8619 | ENSMUSG00000031864 | 108.6808381 | 0.078919946 | 0.230732543 | 0.34204081 | 0.73232018 | 0.947515796 | Ints10 | integrator complex subunit 10 [Source:MG | 97.978 | 91.485 | 127.79 | 108.93 | 114.51 | 111.38 |
| 8620 | ENSMUSG00000031865 | 653.373867 | -0.008651979 | 0.119557714 | -0.072366549 | 0.942310205 | 0.989802092 | Dctn1 | dynactin 1 [Source:MGI Symbol;Acc:MGI:1 | 648.91 | 632.17 | 685.25 | 682.5 | 603.11 | 668.31 |

| | | | | | | | | | | | | | | | |
|---|---|---|---|---|---|---|---|---|---|---|---|---|---|---|---|
| 8621 | ENSMUSG00000031870 | 0.133850373 | -0.796629176 | 4.080472857 | -0.195229623 | 0.84521319 | NA | Pgr | progesterone receptor [Source:MGI Symb | 0.8031 | 0 | 0 | 0 | 0 | 0 |
| 8622 | ENSMUSG00000031871 | 183.1699674 | -0.868203723 | 0.238659114 | -3.63784021 | 0.000274934 | 0.008113038 | Cdh5 | cadherin 5 [Source:MGI Symbol;Acc:MGI: | 220.85 | 297.07 | 192.19 | 129.17 | 94.884 | 164.85 |
| 8623 | ENSMUSG00000031872 | 0 | NA | NA | NA | NA | NA | Bean1 | brain expressed, associated with Nedd4, 1 | 0 | 0 | 0 | 0 | 0 | 0 |
| 8624 | ENSMUSG00000031875 | 84.3876386 | -0.918616169 | 0.278519588 | -3.298210284 | 0.000973032 | 0.020545898 | Cmtm3 | CKLF-like MARVEL transmembrane domai | 104.4 | 134.66 | 92.575 | 73.262 | 49.078 | 52.351 |
| 8625 | ENSMUSG00000031876 | 0 | NA | NA | NA | NA | NA | Gm45711 | predicted gene 45711 [Source:MGI Symbo | 0 | 0 | 0 | 0 | 0 | 0 |
| 8626 | ENSMUSG00000031877 | 894.4812326 | -0.177553432 | 0.108991028 | -1.629064651 | 0.103299333 | 0.473422547 | Ces2g | carboxylesterase 2G [Source:MGI Symbol; | 984.6 | 923.07 | 939.83 | 797.21 | 881.22 | 840.95 |
| 8627 | ENSMUSG00000031878 | 398.3273599 | 0.041185926 | 0.144848689 | 0.284337582 | 0.776151688 | 0.95893551 | Nae1 | NEDD8 activating enzyme E1 subunit 1 [Sc | 440.9 | 369.02 | 367.28 | 416.44 | 422.07 | 374.25 |
| 8628 | ENSMUSG00000031879 | 203.6563571 | 0.277139345 | 0.172150865 | 1.609863217 | 0.107427721 | 0.48315751 | Ciao2b | cytosolic iron-sulfur assembly component | 191.14 | 188.11 | 173.07 | 223.64 | 241.03 | 204.95 |
| 8629 | ENSMUSG00000031880 | 2.337909564 | -1.758177114 | 1.610422893 | -1.091748708 | 0.274943569 | NA | Rrad | Ras-related associated with diabetes [Sou | 1.6062 | 8.2233 | 1.0062 | 0.964 | 0 | 2.2277 |
| 8630 | ENSMUSG00000031881 | 0 | NA | NA | NA | NA | NA | Cdh16 | cadherin 16 [Source:MGI Symbol;Acc:MGI | 0 | 0 | 0 | 0 | 0 | 0 |
| 8631 | ENSMUSG00000031883 | 34.90541187 | 0.021720008 | 0.372388484 | 0.058326207 | 0.953488793 | 0.9915542 | Car7 | carbonic anhydrase 7 [Source:MGI Symbo | 37.746 | 27.754 | 38.237 | 31.811 | 34.9 | 38.985 |
| 8632 | ENSMUSG00000031884 | 3.645188395 | 2.755254916 | 1.281626292 | 2.14981148 | 0.03157013 | NA | Ces2d-ps | carboxylesterase 2D, pseudogene [Source | 0.8031 | 1.0279 | 1.0062 | 5.7839 | 5.4531 | 7.7969 |
| 8633 | ENSMUSG00000031885 | 276.5387114 | 0.246949307 | 0.162795902 | 1.516925821 | 0.129285414 | 0.526260863 | Cbfb | core binding factor beta [Source:MGI Sym | 265.02 | 268.29 | 225.4 | 310.4 | 323.91 | 266.21 |
| 8634 | ENSMUSG00000031886 | 4320.930567 | -0.068009447 | 0.105392351 | -0.645297748 | 0.518734252 | 0.887592015 | Ces2e | carboxylesterase 2E [Source:MGI Symbol; | 4890.9 | 4173.3 | 4203.1 | 4390 | 4382.1 | 3886.2 |
| 8635 | ENSMUSG00000031887 | 268.8748828 | 0.281971477 | 0.168425919 | 1.674157267 | 0.094099695 | 0.451490179 | Tradd | TNFRSF1A-associated via death domain [S | 258.6 | 203.53 | 265.65 | 309.44 | 305.37 | 270.66 |
| 8636 | ENSMUSG00000031889 | 946.6007934 | 0.139925669 | 0.135701377 | 1.031129325 | 0.302480178 | 0.752385931 | D230025D16 | RIKEN cDNA D230025D16 gene [Source:M | 872.17 | 948.77 | 882.48 | 1163.5 | 972.83 | 839.84 |
| 8637 | ENSMUSG00000031891 | 0.321325949 | 1.911575114 | 4.048267893 | 0.472195804 | 0.636787034 | NA | Hsd11b2 | hydroxysteroid 11-beta dehydrogenase 2 | 0 | 0 | 0 | 1.928 | 0 | 0 |
| 8638 | ENSMUSG00000031893 | 0 | NA | NA | NA | NA | NA | Tsnaxip1 | translin-associated factor X (Tsnax) intera | 0 | 0 | 0 | 0 | 0 | 0 |
| 8639 | ENSMUSG00000031896 | 3.19193075 | -0.077533648 | 1.243537991 | -0.06234924 | 0.950284717 | NA | Ctrl | chymotrypsin-like [Source:MGI Symbol;Ac | 0.8031 | 4.1117 | 5.0312 | 4.8199 | 3.2719 | 1.1138 |
| 8640 | ENSMUSG00000031897 | 723.0642111 | 0.064209122 | 0.116992275 | 0.54883215 | 0.583120645 | 0.908645447 | Psmb10 | proteasome (prosome, macropain) subun | 700.31 | 712.35 | 708.4 | 730.7 | 800.51 | 686.13 |
| 8641 | ENSMUSG00000031898 | 0.160662974 | 1.126924175 | 4.080472857 | 0.276174898 | 0.782413736 | NA | Dpep3 | dipeptidase 3 [Source:MGI Symbol;Acc:M | 0 | 0 | 0 | 0.964 | 0 | 0 |
| 8642 | ENSMUSG00000031901 | 125.2660015 | 0.141305945 | 0.223976315 | 0.63089682 | 0.528107991 | 0.888334265 | Dus2 | dihydrouridine synthase 2 [Source:MGI Sy | 130.1 | 105.88 | 120.75 | 106.04 | 140.69 | 148.14 |
| 8643 | ENSMUSG00000031902 | 638.230123 | -0.111790331 | 0.118013623 | -0.947266327 | 0.343503085 | 0.785243795 | Nfatc3 | nuclear factor of activated T cells, cytopla | 691.47 | 619.83 | 677.2 | 626.59 | 618.38 | 595.91 |
| 8644 | ENSMUSG00000031903 | 366.8663384 | -0.034691927 | 0.173079956 | -0.200438736 | 0.841137467 | 0.971854408 | Pla2g15 | phospholipase A2, group XV [Source:MGI | 428.86 | 350.52 | 333.07 | 292.09 | 405.71 | 390.96 |
| 8645 | ENSMUSG00000031904 | 5.238133402 | -1.165856858 | 0.980092679 | -1.18953736 | 0.234228278 | 0.682214623 | Slc7a6 | solute carrier family 7 (cationic amino acid | 6.4248 | 7.1954 | 8.05 | 0.964 | 5.4531 | 3.3415 |
| 8646 | ENSMUSG00000031906 | 150.2527769 | -0.694406293 | 0.275082167 | -2.524359543 | 0.011590931 | 0.119989074 | Smpd3 | sphingomyelin phosphodiesterase 3, neut | 201.58 | 141.85 | 213.32 | 87.722 | 169.05 | 87.994 |
| 8647 | ENSMUSG00000031907 | 4.744149217 | -0.974781187 | 0.983667402 | -0.99096624 | 0.321702066 | NA | Zfp90 | zinc finger protein 90 [Source:MGI Symbo | 5.6217 | 6.1675 | 7.0437 | 1.928 | 4.3625 | 3.3415 |
| 8648 | ENSMUSG00000031910 | 9.679277495 | 0.313177793 | 0.707970812 | 0.442359752 | 0.658228899 | 0.927303282 | Has3 | hyaluronan synthase 3 [Source:MGI Symb | 5.6217 | 9.2513 | 11.069 | 6.7478 | 10.906 | 14.48 |
| 8649 | ENSMUSG00000031913 | 360.2604574 | 0.010342506 | 0.136375499 | 0.075838445 | 0.93954763 | 0.988812319 | Vps4a | vacuolar protein sorting 4A [Source:MGI S | 358.99 | 367.99 | 350.17 | 376.92 | 356.63 | 350.86 |
| 8650 | ENSMUSG00000031916 | 483.2796216 | -0.117456514 | 0.148680711 | -0.789991607 | 0.42953267 | 0.84307112 | Cog8 | component of oligomeric golgi complex 8 | 499.53 | 523.21 | 487.02 | 552.36 | 420.98 | 416.58 |
| 8651 | ENSMUSG00000031917 | 229.9680581 | 0.20146042 | 0.176847167 | 1.139178099 | 0.254628878 | 0.708091239 | Nip7 | NIP7, nucleolar pre-rRNA processing prote | 228.88 | 222.03 | 190.18 | 215.93 | 243.21 | 279.57 |
| 8652 | ENSMUSG00000031918 | 473.300533 | 0.06977599 | 0.132818246 | 0.525349429 | 0.599340302 | 0.915117178 | Mtmr2 | myotubularin related protein 2 [Source:M | 469.01 | 443.03 | 472.93 | 442.47 | 528.95 | 483.41 |
| 8653 | ENSMUSG00000031919 | 0.167707431 | -0.796629176 | 4.080472857 | -0.195229623 | 0.84521319 | NA | Tmed6 | transmembrane p24 trafficking protein 6 [ | 0 | 0 | 1.0062 | 0 | 0 | 0 |
| 8654 | ENSMUSG00000031921 | 179.8337954 | 0.222627938 | 0.177775595 | 1.252297528 | 0.210461469 | 0.654419768 | Terf2 | telomeric repeat binding factor 2 [Source: | 171.06 | 168.58 | 157.98 | 180.26 | 202.86 | 198.26 |
| 8655 | ENSMUSG00000031922 | 244.3807309 | 0.067170138 | 0.164459991 | 0.40842844 | 0.682959157 | 0.935181411 | Cep57 | centrosomal protein 57 [Source:MGI Syml | 256.19 | 221 | 238.48 | 256.42 | 225.76 | 268.44 |
| 8656 | ENSMUSG00000031924 | 6376.213896 | 0.287746133 | 0.110664375 | 2.600169501 | 0.009317772 | 0.103194686 | Cyb5b | cytochrome b5 type B [Source:MGI Symbo | 6512.4 | 5603.2 | 5109.7 | 6706.4 | 6880.7 | 7444.9 |
| 8657 | ENSMUSG00000031925 | 20.18716232 | -0.561441953 | 0.491162644 | -1.143087651 | 0.253002201 | 0.706179821 | Maml2 | mastermind like transcriptional coactivato | 19.274 | 28.782 | 24.15 | 12.532 | 17.45 | 18.935 |
| 8658 | ENSMUSG00000031927 | 0 | NA | NA | NA | NA | NA | 1700012B09 | RIKEN cDNA 1700012B09 gene [Source:M | 0 | 0 | 0 | 0 | 0 | 0 |
| 8659 | ENSMUSG00000031928 | 117.923405 | 0.109388937 | 0.21769575 | 0.502485406 | 0.615326116 | 0.917509899 | Mre11a | MRE11A homolog A, double strand break | 117.25 | 109.99 | 112.7 | 104.11 | 127.6 | 135.89 |
| 8660 | ENSMUSG00000031930 | 354.5812705 | -0.162353467 | 0.155105834 | -1.046727015 | 0.2952255 | 0.745876268 | Wwp2 | WW domain containing E3 ubiquitin prote | 409.58 | 359.77 | 353.19 | 306.54 | 313.01 | 385.39 |
| 8661 | ENSMUSG00000031931 | 146.7856189 | 0.222582015 | 0.222469912 | 1.000503905 | 0.317066709 | 0.763688233 | Ankrd49 | ankyrin repeat domain 49 [Source:MGI Sy | 145.36 | 146.99 | 113.71 | 136.88 | 196.31 | 141.46 |
| 8662 | ENSMUSG00000031932 | 0 | NA | NA | NA | NA | NA | Gpr83 | G protein-coupled receptor 83 [Source:M | 0 | 0 | 0 | 0 | 0 | 0 |
| 8663 | ENSMUSG00000031933 | 0.476489653 | -2.293616034 | 3.421506946 | -0.670352587 | 0.502633051 | NA | Izumo1r | IZUMO1 receptor, JUNO [Source:MGI Sym | 0.8031 | 2.0558 | 0 | 0 | 0 | 0 |
| 8664 | ENSMUSG00000031934 | 6.867720416 | -0.348965571 | 0.859568432 | -0.405977648 | 0.684759024 | 0.935737593 | Panx1 | pannexin 1 [Source:MGI Symbol;Acc:MGI: | 8.8341 | 3.0838 | 11.069 | 3.8559 | 5.4531 | 8.9108 |
| 8665 | ENSMUSG00000031935 | 238.6878476 | 0.14376227 | 0.164188684 | 0.875591831 | 0.381251968 | 0.8130053 | Med17 | mediator complex subunit 17 [Source:MG | 216.84 | 228.2 | 235.46 | 241.96 | 231.21 | 278.46 |
| 8666 | ENSMUSG00000031936 | 0 | NA | NA | NA | NA | NA | Hephl1 | hephaestin-like 1 [Source:MGI Symbol;Ac | 0 | 0 | 0 | 0 | 0 | 0 |
| 8667 | ENSMUSG00000031937 | 0.965523032 | -0.944628791 | 2.267086424 | -0.416670834 | 0.67691919 | NA | Vstm5 | V-set and transmembrane domain contair | 0.8031 | 2.0558 | 1.0062 | 1.928 | 0 | 0 |
| 8668 | ENSMUSG00000031938 | 4864.912706 | -0.263817104 | 0.114940859 | -2.295242144 | 0.021719251 | 0.183390105 | 4931406C07 | RIKEN cDNA 4931406C07 gene [Source:M | 5751 | 5181.7 | 4992 | 4389 | 4990.7 | 3885.1 |
| 8669 | ENSMUSG00000031939 | 198.5089388 | 0.236015342 | 0.178167863 | 1.324679645 | 0.185277427 | 0.620990675 | Taf1d | TATA-box binding protein associated facto | 205.59 | 175.77 | 165.02 | 224.61 | 202.86 | 217.2 |
| 8670 | ENSMUSG00000031948 | 894.2958646 | 0.151328975 | 0.111253013 | 1.360223608 | 0.173759174 | 0.604147282 | Kars | lysyl-tRNA synthetase [Source:MGI Symbo | 799.09 | 886.07 | 858.33 | 990.97 | 933.57 | 897.76 |
| 8671 | ENSMUSG00000031949 | 102.8618189 | 0.101404747 | 0.23904569 | 0.424206549 | 0.671415185 | 0.931191575 | Adat1 | adenosine deaminase, tRNA-specific 1 [Sc | 85.932 | 91.485 | 120.75 | 110.86 | 111.24 | 96.904 |
| 8672 | ENSMUSG00000031950 | 900.906772 | 0.087125745 | 0.110777925 | 0.786490134 | 0.431580397 | 0.844934876 | Gabarapl2 | gamma-aminobutyric acid (GABA) A recep | 862.53 | 854.2 | 904.61 | 928.31 | 988.1 | 867.68 |
| 8673 | ENSMUSG00000031951 | 4.02679805 | -1.133795045 | 1.142742876 | -0.992169865 | 0.32111467 | NA | Tmem231 | transmembrane protein 231 [Source:MGI | 2.4093 | 4.1117 | 10.062 | 0.964 | 3.2719 | 3.3415 |
| 8674 | ENSMUSG00000031952 | 0 | NA | NA | NA | NA | NA | Chst5 | carbohydrate (N-acetylglucosamine 6-O) s | 0 | 0 | 0 | 0 | 0 | 0 |
| 8675 | ENSMUSG00000031953 | 205.5558421 | -0.111006118 | 0.199214568 | -0.55721888 | 0.577377892 | 0.907838603 | Tmem170 | transmembrane protein 170 [Source:MGI | 162.23 | 248.76 | 230.43 | 207.26 | 190.86 | 193.81 |
| 8676 | ENSMUSG00000031954 | 632.9700161 | 0.064487034 | 0.123879329 | 0.520563313 | 0.602671012 | 0.915657359 | Cfdp1 | craniofacial development protein 1 [Sourc | 652.92 | 624.97 | 577.58 | 595.74 | 681.64 | 664.96 |

| | | | | | | | | | | | | | | | |
|---|---|---|---|---|---|---|---|---|---|---|---|---|---|---|---|
| 8677 | ENSMUSG00000031955 | 781.3735916 | 0.294943981 | 0.115129992 | 2.561834468 | 0.010412093 | 0.111804391 | Bcar1 | breast cancer anti-estrogen resistance 1 [! | 751.7 | 668.15 | 684.25 | 818.42 | 875.77 | 889.96 |
| 8678 | ENSMUSG00000031957 | 0 | NA | NA | NA | NA | NA | Ctrb1 | chymotrypsinogen B1 [Source:MGI Symbo | 0 | 0 | 0 | 0 | 0 | 0 |
| 8679 | ENSMUSG00000031958 | 3905.405022 | 0.080683049 | 0.097361302 | 0.828697313 | 0.407275707 | 0.83094725 | Ldhd | lactate dehydrogenase D [Source:MGI Syr | 4147.2 | 3581.3 | 3658.7 | 4002.4 | 4144.3 | 3898.5 |
| 8680 | ENSMUSG00000031959 | 188.3333833 | 0.168387083 | 0.183202852 | 0.919129159 | 0.358028019 | 0.797696186 | Wdr59 | WD repeat domain 59 [Source:MGI Symbo | 175.88 | 177.83 | 178.11 | 182.19 | 184.31 | 231.68 |
| 8681 | ENSMUSG00000031960 | 3490.639427 | 0.180078641 | 0.087740772 | 2.052394063 | 0.040131381 | 0.274113751 | Aars | alanyl-tRNA synthetase [Source:MGI Syml | 3320.8 | 3288.3 | 3209.9 | 3723.8 | 3844.4 | 3556.5 |
| 8682 | ENSMUSG00000031962 | 7.392734106 | 0.082046953 | 0.853257195 | 0.096157352 | 0.923395602 | 0.98705827 | Cdh15 | cadherin 15 [Source:MGI Symbol;Acc:MG | 2.4093 | 11.307 | 8.05 | 11.568 | 5.4531 | 5.5692 |
| 8683 | ENSMUSG00000031963 | 5.874878061 | -1.660578607 | 1.009426448 | -1.645071427 | 0.099955082 | 0.465620067 | Bmper | BMP-binding endothelial regulator [Sourc | 6.4248 | 11.307 | 9.0562 | 2.8919 | 0 | 5.5692 |
| 8684 | ENSMUSG00000031965 | 51.2583381 | -1.369642111 | 0.336555949 | -4.069582238 | 4.71E-05 | 0.002011891 | Tbx20 | T-box 20 [Source:MGI Symbol;Acc:MGI:18 | 76.295 | 80.178 | 65.406 | 33.739 | 19.631 | 32.301 |
| 8685 | ENSMUSG00000031966 | 0.167707431 | -0.796629176 | 4.080472857 | -0.195229623 | 0.84521319 | NA | Glb1l3 | galactosidase, beta 1 like 3 [Source:MGI S | 0 | 0 | 1.0062 | 0 | 0 | 0 |
| 8686 | ENSMUSG00000031967 | 1510.528239 | 0.027667759 | 0.099281893 | 0.278678799 | 0.780491334 | 0.959905354 | Afg3l1 | AFG3-like AAA ATPase 1 [Source:MGI Sym | 1510.6 | 1490.5 | 1486.2 | 1425.7 | 1561.8 | 1588.3 |
| 8687 | ENSMUSG00000031969 | 1287.931223 | 0.177339088 | 0.105562435 | 1.679945028 | 0.092968012 | 0.449293659 | Acad8 | acyl-Coenzyme A dehydrogenase family, r | 1189.4 | 1291.1 | 1146.1 | 1313.9 | 1346.9 | 1440.2 |
| 8688 | ENSMUSG00000031970 | 7.214673171 | -0.2759853 | 0.823486631 | -0.335142417 | 0.737517657 | 0.948137969 | Dbndd1 | dysbindin (dystrobrevin binding protein 1) | 10.44 | 9.2513 | 4.025 | 9.6398 | 4.3625 | 5.5692 |
| 8689 | ENSMUSG00000031971 | 6.286125257 | 0.414146153 | 0.907819635 | 0.456198717 | 0.648247088 | 0.925643431 | Ccsap | centriole, cilia and spindle associated prot | 4.0155 | 3.0838 | 9.0562 | 3.8559 | 5.4531 | 12.252 |
| 8690 | ENSMUSG00000031972 | 0 | NA | NA | NA | NA | NA | Acta1 | actin, alpha 1, skeletal muscle [Source:MG | 0 | 0 | 0 | 0 | 0 | 0 |
| 8691 | ENSMUSG00000031974 | 622.332738 | 0.375133084 | 0.135838954 | 2.761601679 | 0.00575186 | 0.073364117 | Abcb10 | ATP-binding cassette, sub-family B (MDR/ | 478.65 | 518.07 | 629.91 | 705.63 | 707.81 | 693.92 |
| 8692 | ENSMUSG00000031976 | 135.9496552 | 0.161574073 | 0.212572777 | 0.760088264 | 0.447201827 | 0.853566538 | Urb2 | URB2 ribosome biogenesis 2 homolog (S. | 118.06 | 115.13 | 151.94 | 132.06 | 157.05 | 141.46 |
| 8693 | ENSMUSG00000031979 | 393.7391705 | 0.178360977 | 0.144726329 | 1.23240172 | 0.217799059 | 0.663697152 | Cog2 | component of oligomeric golgi complex 2 | 375.85 | 324.82 | 407.53 | 432.83 | 394.8 | 426.6 |
| 8694 | ENSMUSG00000031980 | 11373.32136 | -0.371527588 | 0.10866119 | -3.419137855 | 0.000628199 | 0.014655273 | Agt | angiotensinogen (serpin peptidase inhibit | 11983 | 13055 | 13453 | 11281 | 9108.8 | 9358.5 |
| 8695 | ENSMUSG00000031981 | 0.331982614 | 0.16514309 | 3.986751476 | 0.041422971 | 0.966958701 | NA | Capn9 | calpain 9 [Source:MGI Symbol;Acc:MGI:19 | 0 | 1.0279 | 0 | 0.964 | 0 | 0 |
| 8696 | ENSMUSG00000031982 | 34.43972414 | -0.147268017 | 0.377539921 | -0.390072703 | 0.696482787 | 0.939992732 | Arv1 | ARV1 homolog, fatty acid homeostasis mc | 38.549 | 31.865 | 38.237 | 38.559 | 33.809 | 25.618 |
| 8697 | ENSMUSG00000031983 | 383.7052852 | 0.230118986 | 0.146307743 | 1.572842157 | 0.115755371 | 0.500151317 | 2310022B05 | RIKEN cDNA 2310022B05 gene [Source:M | 389.5 | 318.65 | 350.17 | 380.77 | 423.16 | 439.97 |
| 8698 | ENSMUSG00000031984 | 601.6296811 | 0.168878016 | 0.128110675 | 1.31821971 | 0.187430109 | 0.623933049 | 2810004N23 | RIKEN cDNA 2810004N23 gene [Source:M | 554.14 | 619.83 | 525.26 | 589.95 | 647.83 | 672.76 |
| 8699 | ENSMUSG00000031985 | 751.2590506 | -0.028851042 | 0.115903083 | -0.24892385 | 0.803419686 | 0.963596811 | Gnpat | glyceronephosphate O-acyltransferase [So | 715.56 | 775.05 | 785.88 | 695.99 | 775.43 | 759.64 |
| 8700 | ENSMUSG00000031986 | 199.1169207 | -0.093390841 | 0.173054102 | -0.539662685 | 0.589429678 | 0.911039845 | Sprtn | SprT-like N-terminal domain [Source:MGI | 222.46 | 202.5 | 191.19 | 184.12 | 202.86 | 191.58 |
| 8701 | ENSMUSG00000031987 | 812.6321003 | -0.162002918 | 0.111166402 | -1.457301079 | 0.145033289 | 0.555361204 | Egln1 | egl-9 family hypoxia-inducible factor 1 [So | 833.62 | 855.23 | 886.5 | 807.81 | 728.53 | 764.1 |
| 8702 | ENSMUSG00000031988 | 616.0855519 | 0.134215647 | 0.123739636 | 1.084661728 | 0.278071505 | 0.730997543 | Vps26b | VPS26 retromer complex component B [So | 534.06 | 600.3 | 628.9 | 642.01 | 666.37 | 624.87 |
| 8703 | ENSMUSG00000031990 | 11.70612149 | -0.96775029 | 0.729248043 | -1.327052296 | 0.184491384 | 0.619137675 | Jam3 | junction adhesion molecule 3 [Source:MG | 12.047 | 18.503 | 16.1 | 11.568 | 10.906 | 1.1138 |
| 8704 | ENSMUSG00000031991 | 0 | NA | NA | NA | NA | NA | Spata19 | spermatogenesis associated 19 [Source:M | 0 | 0 | 0 | 0 | 0 | 0 |
| 8705 | ENSMUSG00000031993 | 704.8893688 | -0.096496722 | 0.13641848 | -0.707358139 | 0.479343944 | 0.870236354 | Snx19 | sorting nexin 19 [Source:MGI Symbol;Acc: | 803.1 | 726.74 | 654.06 | 593.81 | 718.72 | 732.91 |
| 8706 | ENSMUSG00000031994 | 1.170512601 | -1.184332849 | 2.358814357 | -0.502088198 | 0.615605482 | NA | Adamts8 | a disintegrin-like and metallopeptidase (re | 4.8186 | 0 | 0 | 0 | 1.0906 | 1.1138 |
| 8707 | ENSMUSG00000031995 | 7.118190339 | -1.637252636 | 0.844890534 | -1.937828122 | 0.052644196 | 0.322665669 | St14 | suppression of tumorigenicity 14 (colon ca | 12.047 | 9.2513 | 11.069 | 4.8199 | 2.1812 | 3.3415 |
| 8708 | ENSMUSG00000031996 | 5842.351348 | -0.239002894 | 0.112331828 | -2.127650718 | 0.033366054 | 0.244357773 | Aplp2 | amyloid beta (A4) precursor-like protein 2 | 6371 | 6549.9 | 6055.6 | 5908.2 | 5596 | 4573.4 |
| 8709 | ENSMUSG00000031997 | 0 | NA | NA | NA | NA | NA | Trpc6 | transient receptor potential cation channe | 0 | 0 | 0 | 0 | 0 | 0 |
| 8710 | ENSMUSG00000031998 | 0 | NA | NA | NA | NA | NA | 1700128F08 | RIKEN cDNA 1700128F08 gene [Source:MG | 0 | 0 | 0 | 0 | 0 | 0 |
| 8711 | ENSMUSG00000032000 | 258.2347097 | 0.350094294 | 0.160016082 | 2.187869429 | 0.028679115 | 0.222176208 | Birc3 | baculoviral IAP repeat-containing 3 [Sourc | 224.87 | 211.75 | 244.52 | 289.19 | 267.2 | 311.88 |
| 8712 | ENSMUSG00000032002 | 532.051757 | 0.019341905 | 0.125028701 | 0.154699719 | 0.877058052 | 0.980632845 | Dcun1d5 | DCN1, defective in cullin neddylation 1, do | 516.39 | 566.38 | 503.12 | 546.58 | 548.58 | 511.25 |
| 8713 | ENSMUSG00000032006 | 18.72187794 | 0.038593687 | 0.492564445 | 0.078352564 | 0.937547606 | 0.988420305 | Pdgfd | platelet-derived growth factor, D polypep | 16.865 | 16.447 | 22.137 | 18.316 | 19.631 | 18.935 |
| 8714 | ENSMUSG00000032009 | 604.4968362 | 0.30544123 | 0.163700456 | 1.865854478 | 0.062061728 | 0.355990107 | Sesn3 | sestrin 3 [Source:MGI Symbol;Acc:MGI:19 | 540.49 | 457.42 | 623.87 | 613.09 | 584.57 | 807.54 |
| 8715 | ENSMUSG00000032010 | 1141.595877 | -0.310322449 | 0.153086934 | -2.027099515 | 0.042652237 | 0.284558094 | Usp2 | ubiquitin specific peptidase 2 [Source:MG | 1289 | 1431.9 | 1070.6 | 952.41 | 1232.4 | 873.25 |
| 8716 | ENSMUSG00000032011 | 7.805744165 | -1.627380547 | 0.896166289 | -1.815935912 | 0.069380204 | 0.380407607 | Thy1 | thymus cell antigen 1, theta [Source:MGI | 8.031 | 13.363 | 14.087 | 5.7839 | 0 | 5.5692 |
| 8717 | ENSMUSG00000032012 | 1048.608912 | -0.320009286 | 0.124640195 | -2.567464576 | 0.010244526 | 0.110745006 | Nectin1 | nectin cell adhesion molecule 1 [Source:M | 1029.6 | 1290 | 1174.3 | 881.08 | 900.85 | 1015.8 |
| 8718 | ENSMUSG00000032013 | 0 | NA | NA | NA | NA | NA | Trim29 | tripartite motif-containing 29 [Source:MG | 0 | 0 | 0 | 0 | 0 | 0 |
| 8719 | ENSMUSG00000032014 | 5809.304531 | 0.226866018 | 0.159877541 | 1.41899867 | 0.155899404 | 0.574797088 | Oaf | out at first homolog [Source:MGI Symbol; | 5020.2 | 5205.4 | 5834.2 | 4471.9 | 6793.5 | 7530.7 |
| 8720 | ENSMUSG00000032015 | 0.513958919 | -2.394274821 | 3.756096013 | -0.637437066 | 0.523840191 | NA | Pou2f3 | POU domain, class 2, transcription factor 3 | 0 | 3.0838 | 0 | 0 | 0 | 0 |
| 8721 | ENSMUSG00000032017 | 1.816571243 | -4.233540562 | 1.925938062 | -2.198170671 | 0.027936946 | NA | Grik4 | glutamate receptor, ionotropic, kainate 4 | 4.8186 | 2.0558 | 4.025 | 0 | 0 | 0 |
| 8722 | ENSMUSG00000032018 | 5695.538125 | 0.570125684 | 0.151220203 | 3.770168758 | 0.000163137 | 0.005338499 | Sc5d | sterol-C5-desaturase [Source:MGI Symbol | 5267.5 | 4566 | 3918.3 | 5425.3 | 6780.4 | 8215.7 |
| 8723 | ENSMUSG00000032020 | 11.30378175 | -0.635902643 | 0.657300927 | -0.967445225 | 0.333321511 | 0.77523112 | Ubash3b | ubiquitin associated and SH3 domain cont | 13.653 | 18.503 | 9.0562 | 6.7478 | 8.7249 | 11.138 |
| 8724 | ENSMUSG00000032021 | 0.133850373 | -0.796629176 | 4.080472857 | -0.195229623 | 0.84521319 | NA | Crtam | cytotoxic and regulatory T cell molecule [S | 0.8031 | 0 | 0 | 0 | 0 | 0 |
| 8725 | ENSMUSG00000032023 | 0.133850373 | -0.796629176 | 4.080472857 | -0.195229623 | 0.84521319 | NA | Jhy | junctional cadherin complex regulator [So | 0.8031 | 0 | 0 | 0 | 0 | 0 |
| 8726 | ENSMUSG00000032024 | 13.60063676 | -0.213278876 | 0.591741511 | -0.360425747 | 0.718528776 | 0.9447719 | Clmp | CXADR-like membrane protein [Source:MG | 19.274 | 11.307 | 13.081 | 12.532 | 9.8156 | 15.594 |
| 8727 | ENSMUSG00000032026 | 1270.846841 | -0.072737254 | 0.110412617 | -0.658776653 | 0.510039202 | 0.882863453 | Rexo2 | RNA exonuclease 2 [Source:MGI Symbol;A | 1423.9 | 1246.9 | 1236.7 | 1238.7 | 1173.5 | 1305.4 |
| 8728 | ENSMUSG00000032028 | 608.4572424 | 0.503905323 | 0.130066858 | 3.874202311 | 0.000106975 | 0.0038299 | Nxpe2 | neurexophilin and PC-esterase domain far | 501.94 | 492.37 | 515.2 | 660.32 | 808.15 | 672.76 |
| 8729 | ENSMUSG00000032030 | 420.9612548 | -0.067790551 | 0.136318606 | -0.497294928 | 0.618981088 | 0.917509899 | Cul5 | cullin 5 [Source:MGI Symbol;Acc:MGI:192 | 416.81 | 463.59 | 412.56 | 414.51 | 436.25 | 382.05 |
| 8730 | ENSMUSG00000032033 | 1.643397213 | -2.961425256 | 1.984843834 | -1.492019274 | 0.135694093 | NA | Barx2 | BarH-like homeobox 2 [Source:MGI Symbo | 0.8031 | 2.0558 | 6.0375 | 0.964 | 0 | 0 |
| 8731 | ENSMUSG00000032034 | 1.793267111 | -1.985540124 | 1.765104342 | -1.124885411 | 0.260637595 | NA | Kcnj5 | potassium inwardly-rectifying channel, su | 2.4093 | 5.1396 | 1.0062 | 0 | 1.0906 | 1.1138 |
| 8732 | ENSMUSG00000032035 | 219.3140044 | -0.652111198 | 0.176625272 | -3.692060551 | 0.000222444 | 0.006845611 | Ets1 | E26 avian leukemia oncogene 1, 5' domair | 256.99 | 270.34 | 276.72 | 161.95 | 150.51 | 199.38 |

| | | | | | | | | | | | | | | | |
|---|---|---|---|---|---|---|---|---|---|---|---|---|---|---|---|
| 8733 | ENSMUSG00000032036 | 2.662306185 | -2.069641035 | 1.437172571 | -1.440078302 | 0.149845247 | NA | Kirrel3 | kirre like nephrin family adhesion molecul | 4.8186 | 4.1117 | 4.025 | 1.928 | 1.0906 | 0 |
| 8734 | ENSMUSG00000032038 | 1620.695486 | -0.016038554 | 0.100948265 | -0.158878943 | 0.873764251 | 0.979925081 | St3gal4 | ST3 beta-galactoside alpha-2,3-sialyltransf | 1671.3 | 1668.3 | 1548.6 | 1510.6 | 1644.7 | 1680.8 |
| 8735 | ENSMUSG00000032040 | 286.9578621 | 0.266150686 | 0.153132812 | 1.738038263 | 0.082204072 | 0.422379999 | Dcps | decapping enzyme, scavenger [Source:MC | 248.16 | 279.59 | 254.58 | 333.54 | 316.28 | 289.6 |
| 8736 | ENSMUSG00000032041 | 348.0643399 | 0.156013649 | 0.179810745 | 0.867654762 | 0.385583358 | 0.816437825 | Tirap | toll-interleukin 1 receptor (TIR) domain-cc | 281.09 | 286.79 | 420.61 | 379.81 | 340.27 | 379.82 |
| 8737 | ENSMUSG00000032042 | 3348.521843 | 0.255264259 | 0.087881055 | 2.904656268 | 0.003676566 | 0.054127116 | Srpr | signal recognition particle receptor ('dock | 2991.6 | 3174.2 | 2993.6 | 3551.3 | 3653.6 | 3726.9 |
| 8738 | ENSMUSG00000032044 | 130.7153274 | -0.002048695 | 0.218116133 | -0.009392679 | 0.992505837 | 0.99859176 | Rpusd4 | RNA pseudouridylate synthase domain co | 114.04 | 138.77 | 139.87 | 123.39 | 114.51 | 153.71 |
| 8739 | ENSMUSG00000032046 | 528.3922104 | 0.071147761 | 0.150899779 | 0.471490159 | 0.637290745 | 0.923478764 | Abhd12 | abhydrolase domain containing 12 [Sourc | 530.85 | 473.87 | 540.35 | 445.36 | 558.4 | 621.52 |
| 8740 | ENSMUSG00000032047 | 11698.05197 | -0.233455438 | 0.093624444 | -2.493530826 | 0.01264796 | 0.127218425 | Acat1 | acetyl-Coenzyme A acetyltransferase 1 [Sc | 12252 | 13737 | 11939 | 10957 | 11034 | 10270 |
| 8741 | ENSMUSG00000032048 | 0 | NA | NA | NA | NA | NA | 4930510E17 | RIKEN cDNA 4930510E17 gene [Source:M | 0 | 0 | 0 | 0 | 0 | 0 |
| 8742 | ENSMUSG00000032050 | 3914.278108 | 0.227916269 | 0.107663211 | 2.116937324 | 0.034265167 | 0.247805691 | Rdx | radixin [Source:MGI Symbol;Acc:MGI:978 | 3307.2 | 3661.4 | 3848.9 | 3784.6 | 4413.7 | 4469.9 |
| 8743 | ENSMUSG00000032051 | 2907.900643 | 0.176348269 | 0.11130555 | 1.584361868 | 0.113111394 | 0.493951959 | Fdx1 | ferredoxin 1 [Source:MGI Symbol;Acc:MG | 2802 | 2547.2 | 2841.6 | 3049.1 | 3449.6 | 2757.9 |
| 8744 | ENSMUSG00000032053 | 2.61366446 | -1.984198213 | 1.478448286 | -1.342081581 | 0.179569545 | NA | Pou2af1 | POU domain, class 2, associating factor 1 | | 2.4093 | 3.0838 | 7.0437 | 0.964 | 2.1812 | 0 |
| 8745 | ENSMUSG00000032056 | 0 | NA | NA | NA | NA | NA | Btg4 | BTG anti-proliferation factor 4 [Source:M( | 0 | 0 | 0 | 0 | 0 | 0 |
| 8746 | ENSMUSG00000032057 | 0 | NA | NA | NA | NA | NA | Hoatz | HOATZ cilia and flagella associated protei | 0 | 0 | 0 | 0 | 0 | 0 |
| 8747 | ENSMUSG00000032058 | 518.5082885 | -0.351861699 | 0.150287914 | -2.341250795 | 0.019219252 | 0.168681589 | Ppp2r1b | protein phosphatase 2, regulatory subunit | 633.65 | 635.25 | 474.95 | 453.07 | 486.42 | 427.72 |
| 8748 | ENSMUSG00000032059 | 294.166602 | 0.119576591 | 0.158543186 | 0.754220942 | 0.450716565 | 0.855818389 | Alg9 | asparagine-linked glycosylation 9 (alpha 1 | 296.34 | 270.34 | 278.73 | 284.37 | 351.18 | 284.03 |
| 8749 | ENSMUSG00000032060 | 42.63989409 | 0.170772832 | 0.395758947 | 0.431507191 | 0.666099625 | 0.929596279 | Cryab | crystallin, alpha B [Source:MGI Symbol;Ac | 61.839 | 32.893 | 25.156 | 51.091 | 42.534 | 42.326 |
| 8750 | ENSMUSG00000032062 | 270.1971425 | 0.34072525 | 0.171476903 | 1.987003756 | 0.046921981 | 0.300126868 | 2310030G06 | RIKEN cDNA 2310030G06 gene [Source:M | 285.9 | 219.97 | 208.29 | 300.76 | 298.83 | 307.42 |
| 8751 | ENSMUSG00000032064 | 359.1768967 | 0.445655723 | 0.197128668 | 2.260735218 | 0.023775658 | 0.19577844 | Dixdc1 | DIX domain containing 1 [Source:MGI Sym | 306.79 | 232.31 | 373.32 | 452.11 | 327.19 | 463.36 |
| 8752 | ENSMUSG00000032065 | 29.94479858 | 0.235343013 | 0.458946899 | 0.5127892 | 0.608098785 | 0.916376001 | Tex12 | testis expressed 12 [Source:MGI Symbol;A | 20.881 | 18.503 | 43.269 | 31.811 | 23.994 | 41.212 |
| 8753 | ENSMUSG00000032066 | 382.0488391 | 0.156163427 | 0.150853655 | 1.035198166 | 0.300576375 | 0.751875664 | Bco2 | beta-carotene oxygenase 2 [Source:MGI S | 335.7 | 369.02 | 379.35 | 344.14 | 430.79 | 433.29 |
| 8754 | ENSMUSG00000032067 | 751.926311 | 0.002652158 | 0.119383908 | 0.022215373 | 0.982276155 | 0.998580735 | Pts | 6-pyruvoyl-tetrahydropterin synthase [So | 796.68 | 751.41 | 705.38 | 786.61 | 779.79 | 691.7 |
| 8755 | ENSMUSG00000032068 | 40.09975524 | 0.125582467 | 0.368635481 | 0.340668419 | 0.733353216 | 0.947915087 | Plet1 | placenta expressed transcript 1 [Source:M | 50.595 | 35.977 | 28.175 | 45.307 | 35.99 | 44.554 |
| 8756 | ENSMUSG00000032076 | 442.3383757 | -0.142578657 | 0.159344352 | -0.894783247 | 0.370902968 | 0.807214774 | Cadm1 | cell adhesion molecule 1 [Source:MGI Syn | 437.69 | 482.09 | 473.94 | 507.05 | 404.62 | 348.63 |
| 8757 | ENSMUSG00000032077 | 91.82498822 | 0.112450614 | 0.24954083 | 0.450630119 | 0.652256155 | 0.925650807 | Bud13 | BUD13 homolog [Source:MGI Symbol;Acc | 74.689 | 104.85 | 85.531 | 90.614 | 88.34 | 106.93 |
| 8758 | ENSMUSG00000032078 | 726.4150733 | 0.004985345 | 0.138453242 | 0.036007429 | 0.971276435 | 0.995150695 | Zpr1 | ZPR1 zinc finger [Source:MGI Symbol;Acc: | 619.99 | 830.56 | 726.51 | 777.93 | 729.62 | 673.88 |
| 8759 | ENSMUSG00000032079 | 22378.03267 | -0.381213274 | 0.134063601 | -2.843525541 | 0.004461743 | 0.06187407 | Apoa5 | apolipoprotein A-V [Source:MGI Symbol;A | 22591 | 26014 | 27350 | 23247 | 18969 | 16097 |
| 8760 | ENSMUSG00000032080 | 8680.701973 | -1.0071535 | 0.400727278 | -2.513314057 | 0.011960281 | 0.122256893 | Apoa4 | apolipoprotein A-IV [Source:MGI Symbol; | 17838 | 9783.7 | 7158.4 | 5969 | 5761.7 | 5573.7 |
| 8761 | ENSMUSG00000032081 | 35983.74466 | -0.055058745 | 0.100920188 | -0.545567207 | 0.585363471 | 0.909202452 | Apoc3 | apolipoprotein C-III [Source:MGI Symbol; | 37178 | 32750 | 40082 | 34398 | 34297 | 37198 |
| 8762 | ENSMUSG00000032083 | 157602.5844 | 0.163454262 | 0.094844893 | 1.723384957 | 0.084818933 | 0.429140871 | Apoa1 | apolipoprotein A-I [Source:MGI Symbol;A | 144849 | 162891 | 138312 | 161232 | 164088 | 174243 |
| 8763 | ENSMUSG00000032085 | 6.548025776 | 0.637963858 | 0.911707064 | 0.699746534 | 0.48408561 | 0.872431004 | Tagln | transgelin [Source:MGI Symbol;Acc:MGI:1 | 3.2124 | 10.279 | 2.0125 | 10.604 | 8.7249 | 4.4554 |
| 8764 | ENSMUSG00000032086 | 505.9290842 | -0.117366888 | 0.140188105 | -0.83721003 | 0.402474521 | 0.828953151 | Bace1 | beta-site APP cleaving enzyme 1 [Source:I | 477.04 | 497.51 | 605.76 | 506.09 | 462.42 | 486.75 |
| 8765 | ENSMUSG00000032087 | 0.335414862 | -1.779047242 | 4.030804179 | -0.441362855 | 0.658950329 | NA | Dscaml1 | DS cell adhesion molecule like 1 [Source:N | 0 | 0 | 2.0125 | 0 | 0 | 0 |
| 8766 | ENSMUSG00000032089 | 35.96286206 | -1.036596147 | 0.432630262 | -2.39603245 | 0.016573623 | 0.152883216 | Il10ra | interleukin 10 receptor, alpha [Source:MC | 57.02 | 50.368 | 37.231 | 11.568 | 26.175 | 33.415 |
| 8767 | ENSMUSG00000032091 | 4.2796537 | 0.458511146 | 1.04490757 | 0.438805459 | 0.660802504 | NA | Tmprss4 | transmembrane protease, serine 4 [Sourc | 5.6217 | 2.0558 | 3.0187 | 2.8919 | 7.6343 | 4.4554 |
| 8768 | ENSMUSG00000032092 | 355.7348101 | -0.10095877 | 0.142862929 | -0.706682769 | 0.47976364 | 0.870278888 | Mpzl2 | myelin protein zero-like 2 [Source:MGI Sy | 371.84 | 360.8 | 371.3 | 308.47 | 355.54 | 366.45 |
| 8769 | ENSMUSG00000032093 | 5.131877315 | -0.961965542 | 0.990253628 | -0.971433494 | 0.331332456 | 0.774747658 | Cd3e | CD3 antigen, epsilon polypeptide [Source: | 3.2124 | 6.1675 | 11.069 | 4.8199 | 2.1812 | 3.3415 |
| 8770 | ENSMUSG00000032094 | 5.150036584 | -1.633600027 | 1.036442424 | -1.576160902 | 0.114988726 | 0.498085671 | Cd3d | CD3 antigen, delta polypeptide [Source:M | 8.031 | 7.1954 | 8.05 | 0.964 | 1.0906 | 5.5692 |
| 8771 | ENSMUSG00000032096 | 3095.589251 | 0.135120712 | 0.088526699 | 1.526327235 | 0.126928389 | 0.522005181 | Arcn1 | archain 1 [Source:MGI Symbol;Acc:MGI:2 | 2896.8 | 3088.9 | 2866.8 | 3196.6 | 3260.9 | 3263.6 |
| 8772 | ENSMUSG00000032097 | 2279.760775 | -0.019698804 | 0.110811395 | -0.177768755 | 0.858904586 | 0.975439379 | Ddx6 | DEAD (Asp-Glu-Ala-Asp) box polypeptide | 2034.3 | 2568.8 | 2284.2 | 2197.9 | 2301.2 | 2292.3 |
| 8773 | ENSMUSG00000032098 | 44.8323235 | -0.649690068 | 0.397878094 | -1.632887255 | 0.102492711 | 0.470781813 | Treh | trehalase (brush-border membrane glycop | 73.082 | 59.619 | 31.194 | 26.991 | 45.806 | 32.301 |
| 8774 | ENSMUSG00000032099 | 0 | NA | NA | NA | NA | NA | Pate4 | prostate and testis expressed 4 [Source:M | 0 | 0 | 0 | 0 | 0 | 0 |
| 8775 | ENSMUSG00000032101 | 1.935816909 | -3.210126137 | 1.8298281 | -1.75433208 | 0.079373622 | NA | Ddx25 | DEAD box helicase 25 [Source:MGI Symbc | 2.4093 | 3.0838 | 5.0312 | 0 | 1.0906 | 0 |
| 8776 | ENSMUSG00000032103 | 0 | NA | NA | NA | NA | NA | Pus3 | pseudouridine synthase 3 [Source:MGI Sy | 0 | 0 | 0 | 0 | 0 | 0 |
| 8777 | ENSMUSG00000032105 | 0.913882229 | 2.129306995 | 2.79764157 | 0.761107862 | 0.446592645 | NA | Pdzd3 | PDZ domain containing 3 [Source:MGI Syr | 0 | 1.0279 | 0 | 0 | 0 | 4.4554 |
| 8778 | ENSMUSG00000032108 | 0 | NA | NA | NA | NA | NA | Pate6 | prostate and testis expressed 6 [Source:M | 0 | 0 | 0 | 0 | 0 | 0 |
| 8779 | ENSMUSG00000032109 | 186.5749927 | 0.239127455 | 0.188704898 | 1.267203226 | 0.205082632 | 0.650728757 | Nlrx1 | NLR family member X1 [Source:MGI Symb | 188.73 | 143.91 | 180.12 | 187.01 | 220.3 | 199.38 |
| 8780 | ENSMUSG00000032110 | 0.583320834 | -1.123770786 | 3.233988906 | -0.34748752 | 0.728225092 | NA | Acrv1 | acrosomal vesicle protein 1 [Source:MGI S | 2.4093 | 0 | 0 | 0 | 1.0906 | 0 |
| 8781 | ENSMUSG00000032112 | 283.8833976 | 0.240594497 | 0.150282812 | 1.600944871 | 0.10938913 | 0.486233674 | Trappc4 | trafficking protein particle complex 4 [Sou | 265.02 | 259.04 | 256.59 | 309.44 | 285.74 | 327.47 |
| 8782 | ENSMUSG00000032113 | 3.94871701 | 0.353617651 | 1.147818762 | 0.308077949 | 0.75802302 | NA | Chek1 | checkpoint kinase 1 [Source:MGI Symbol; | 2.4093 | 1.0279 | 7.0437 | 7.7118 | 3.2719 | 2.2277 |
| 8783 | ENSMUSG00000032114 | 5960.291631 | -0.22302674 | 0.092614869 | -2.408109449 | 0.016035373 | 0.148963156 | Slc37a4 | solute carrier family 37 (glucose-6-phosph | 6190.3 | 6373.1 | 6697.6 | 5532.3 | 5834.8 | 5133.7 |
| 8784 | ENSMUSG00000032115 | 1521.597718 | 0.695216151 | 0.180540999 | 3.850738356 | 0.000117762 | 0.004144315 | Hyou1 | hypoxia up-regulated 1 [Source:MGI Symb | 1023.2 | 1315.7 | 1148.1 | 2399.3 | 1320.7 | 1922.5 |
| 8785 | ENSMUSG00000032116 | 1438.982757 | 0.180100052 | 0.112980897 | 1.594075262 | 0.110919179 | 0.489424955 | Stt3a | STT3, subunit of the oligosaccharyltransfe | 1289 | 1513.1 | 1246.7 | 1583.8 | 1542.1 | 1459.1 |
| 8786 | ENSMUSG00000032118 | 1.843048011 | -3.123341269 | 1.909007933 | -1.636107014 | 0.101817193 | NA | Fez1 | fasciculation and elongation protein zeta | 0.8031 | 5.1396 | 4.025 | 0 | 1.0906 | 0 |
| 8787 | ENSMUSG00000032119 | 157.5975991 | 0.078730097 | 0.19248475 | 0.409019922 | 0.682525041 | 0.93502957 | Hinfp | histone H4 transcription factor [Source:M | 149.38 | 167.55 | 142.89 | 149.42 | 158.14 | 178.22 |
| 8788 | ENSMUSG00000032120 | 998.5812293 | 0.115354405 | 0.135502426 | 0.851308776 | 0.394597851 | 0.822855164 | C2cd2l | C2 calcium-dependent domain containing | 1017.5 | 900.46 | 956.94 | 946.63 | 933.57 | 1236.4 |

| | | | | | | | | | | | | | | | |
|---|---|---|---|---|---|---|---|---|---|---|---|---|---|---|---|
| 8789 | ENSMUSG00000032121 | 33.06619332 | 0.567206698 | 0.391478154 | 1.448884676 | 0.147369791 | 0.558402858 | Tmem218 | transmembrane protein 218 [Source:MGI | 28.109 | 22.614 | 29.181 | 43.379 | 29.447 | 45.668 |
| 8790 | ENSMUSG00000032122 | 25.93576934 | -1.030857782 | 0.453047756 | -2.27538437 | 0.022882882 | 0.190326629 | Slc37a2 | solute carrier family 37 (glycerol-3-phospl | 26.502 | 38.033 | 40.25 | 22.171 | 14.178 | 14.48 |
| 8791 | ENSMUSG00000032123 | 416.5370035 | 0.263048697 | 0.138145897 | 1.904136885 | 0.056892359 | 0.338327228 | Dpagt1 | dolichyl-phosphate (UDP-N-acetylglucosa | 389.5 | 406.03 | 340.11 | 438.61 | 448.24 | 476.73 |
| 8792 | ENSMUSG00000032125 | 124.1846314 | -1.016110216 | 0.257745781 | -3.942296218 | 8.07E-05 | 0.003047728 | Robo4 | roundabout guidance receptor 4 [Source: | 198.37 | 168.58 | 131.82 | 105.07 | 61.075 | 80.197 |
| 8793 | ENSMUSG00000032126 | 902.7705374 | 0.218902509 | 0.123791603 | 1.768314677 | 0.077008312 | 0.406191172 | Hmbs | hydroxymethylbilane synthase [Source:M | 908.31 | 847 | 746.63 | 910.96 | 945.57 | 1058.2 |
| 8794 | ENSMUSG00000032127 | 516.0231988 | 0.082674057 | 0.137481346 | 0.601347455 | 0.547608588 | 0.895600915 | Vps11 | VPS11, CORVET/HOPS core subunit [Sourc | 499.53 | 550.96 | 453.82 | 583.21 | 495.14 | 513.48 |
| 8795 | ENSMUSG00000032128 | 6.534015985 | -0.976294197 | 0.978016566 | -0.998238916 | 0.31816352 | 0.764585353 | Robo3 | roundabout guidance receptor 3 [Source: | 12.85 | 5.1396 | 8.05 | 7.7118 | 5.4531 | 0 |
| 8796 | ENSMUSG00000032131 | 0.267700746 | -1.497481649 | 4.057001436 | -0.369110456 | 0.712045396 | NA | Abcg4 | ATP binding cassette subfamily G member | 1.6062 | 0 | 0 | 0 | 0 | 0 |
| 8797 | ENSMUSG00000032135 | 43.41911282 | -1.072730279 | 0.387955833 | -2.765083516 | 0.005690822 | 0.072778121 | Mcam | melanoma cell adhesion molecule [Source | 48.989 | 83.261 | 44.275 | 22.171 | 26.175 | 35.643 |
| 8798 | ENSMUSG00000032171 | 316.5496374 | 0.169615072 | 0.163208747 | 1.039252338 | 0.298687395 | 0.749646795 | Pin1 | protein (peptidyl-prolyl cis/trans isomeras | 251.37 | 316.6 | 327.03 | 348 | 354.45 | 301.85 |
| 8799 | ENSMUSG00000032172 | 7.24093979 | 0.908936451 | 0.832128459 | 1.092303047 | 0.274699922 | 0.729487777 | Olfm2 | olfactomedin 2 [Source:MGI Symbol;Acc:I | 4.0155 | 3.0838 | 8.05 | 9.6398 | 13.087 | 5.5692 |
| 8800 | ENSMUSG00000032174 | 1.28666902 | -0.696779623 | 2.228977613 | -0.312600548 | 0.754584162 | NA | Icam5 | intercellular adhesion molecule 5, telence | 0.8031 | 0 | 4.025 | 2.8919 | 0 | 0 |
| 8801 | ENSMUSG00000032175 | 435.088484 | 0.234362247 | 0.136052598 | 1.722585602 | 0.08496349 | 0.429396285 | Tyk2 | tyrosine kinase 2 [Source:MGI Symbol;Acc | 413.6 | 391.64 | 393.44 | 425.11 | 476.6 | 510.14 |
| 8802 | ENSMUSG00000032177 | 50.04528991 | -0.305483524 | 0.374162511 | -0.816446102 | 0.414245038 | 0.835831451 | Pde4a | phosphodiesterase 4A, cAMP specific [Sou | 58.626 | 44.2 | 63.393 | 65.55 | 26.175 | 42.326 |
| 8803 | ENSMUSG00000032178 | 269.5102112 | 0.124268883 | 0.160788403 | 0.772872173 | 0.439598034 | 0.84844279 | Ilf3 | interleukin enhancer binding factor 3 [Sou | 240.93 | 292.96 | 240.49 | 294.98 | 275.93 | 271.78 |
| 8804 | ENSMUSG00000032179 | 54.33716288 | -0.940609835 | 0.34910743 | -2.694327749 | 0.007053077 | 0.08445009 | Bmp5 | bone morphogenetic protein 5 [Source:M | 74.689 | 68.87 | 70.437 | 27.955 | 27.265 | 56.806 |
| 8805 | ENSMUSG00000032180 | 253.6980387 | 0.17950572 | 0.154993602 | 1.158149228 | 0.246803144 | 0.696810594 | Tmed1 | transmembrane p24 trafficking protein 1 [ | 232.1 | 229.23 | 252.57 | 267.02 | 263.93 | 277.35 |
| 8806 | ENSMUSG00000032181 | 0 | NA | NA | NA | NA | NA | Scg3 | secretogranin III [Source:MGI Symbol;Acc | 0 | 0 | 0 | 0 | 0 | 0 |
| 8807 | ENSMUSG00000032182 | 208.8490384 | 0.091394112 | 0.16663775 | 0.548459828 | 0.583376206 | 0.908645447 | Yipf2 | Yip1 domain family, member 2 [Source:M | 191.14 | 210.72 | 205.27 | 222.68 | 208.31 | 214.97 |
| 8808 | ENSMUSG00000032184 | 8.484322476 | 0.373051804 | 0.737242252 | 0.5060098 | 0.61284977 | 0.916676222 | Lysmd2 | LysM, putative peptidoglycan-binding, do | 8.8341 | 7.1954 | 6.0375 | 6.7478 | 8.7249 | 13.366 |
| 8809 | ENSMUSG00000032185 | 341.4122887 | -0.188044603 | 0.14257522 | -1.318915049 | 0.187197515 | 0.623632669 | Carm1 | coactivator-associated arginine methyltra | 350.96 | 375.19 | 365.27 | 332.57 | 331.55 | 292.94 |
| 8810 | ENSMUSG00000032186 | 0.866661723 | -0.464185162 | 2.825297661 | -0.16429602 | 0.869498114 | NA | Tmod2 | tropomodulin 2 [Source:MGI Symbol;Acc: | 0 | 0 | 3.0187 | 0 | 2.1812 | 0 |
| 8811 | ENSMUSG00000032187 | 1200.933381 | -0.60797498 | 0.170281804 | -3.570404853 | 0.00035643 | 0.009958034 | Smarca4 | SWI/SNF related, matrix associated, actin | 1892.9 | 1366.1 | 1090.8 | 968.8 | 985.92 | 901.1 |
| 8812 | ENSMUSG00000032191 | 0 | NA | NA | NA | NA | NA | Bcl2l10 | Bcl2-like 10 [Source:MGI Symbol;Acc:MGI | 0 | 0 | 0 | 0 | 0 | 0 |
| 8813 | ENSMUSG00000032192 | 9.213858576 | -0.527947201 | 0.773171169 | -0.682833533 | 0.494712038 | 0.877869576 | Gnb5 | guanine nucleotide binding protein (G prc | 11.243 | 6.1675 | 15.094 | 2.8919 | 7.6343 | 12.252 |
| 8814 | ENSMUSG00000032193 | 2301.966294 | 0.024592623 | 0.189534014 | 0.129753083 | 0.896761784 | 0.982798755 | Ldlr | low density lipoprotein receptor [Source:I | 2905.6 | 2206.9 | 1732.8 | 1733.2 | 2857.4 | 2375.8 |
| 8815 | ENSMUSG00000032194 | 828.8502421 | 0.278960726 | 0.147161569 | 1.895608533 | 0.058011828 | 0.342343159 | Kank2 | KN motif and ankyrin repeat domains 2 [S | 695.49 | 789.44 | 761.73 | 746.12 | 893.22 | 1087.1 |
| 8816 | ENSMUSG00000032198 | 255.631103 | 0.041267427 | 0.18890436 | 0.218456718 | 0.827073278 | 0.969748238 | Dock6 | dedicator of cytokinesis 6 [Source:MGI Sy | 279.48 | 252.87 | 223.39 | 291.12 | 200.67 | 286.26 |
| 8817 | ENSMUSG00000032199 | 1368.84425 | 0.053203854 | 0.12154347 | 0.437735191 | 0.661578256 | 0.928404763 | Polr2m | polymerase (RNA) II (DNA directed) polyp | 1160.5 | 1384.6 | 1487.2 | 1408.4 | 1463.6 | 1308.8 |
| 8818 | ENSMUSG00000032202 | 33.58761884 | -1.041736743 | 0.421444416 | -2.471824761 | 0.013442539 | 0.132518282 | Rab27a | RAB27A, member RAS oncogene family [S | 32.927 | 52.424 | 50.312 | 18.316 | 16.359 | 31.188 |
| 8819 | ENSMUSG00000032204 | 9019.52838 | 0.267898049 | 0.115849155 | 2.312473045 | 0.020751629 | 0.177549827 | Aqp9 | aquaporin 9 [Source:MGI Symbol;Acc:MG | 8806.8 | 8513.2 | 7231.9 | 8667.1 | 10765 | 10133 |
| 8820 | ENSMUSG00000032207 | 3156.100156 | 0.241293414 | 0.105443953 | 2.288357055 | 0.022116736 | 0.185770311 | Lipc | lipase, hepatic [Source:MGI Symbol;Acc:N | 2615.7 | 2892.6 | 3171.7 | 3560.9 | 3223.9 | 3471.9 |
| 8821 | ENSMUSG00000032212 | 427.3943172 | -0.07781814 | 0.176829702 | -0.440073917 | 0.659883572 | 0.927787797 | Sltm | SAFB-like, transcription modulator [Sourc | 343.73 | 533.49 | 440.74 | 424.15 | 459.15 | 363.11 |
| 8822 | ENSMUSG00000032215 | 298.2493814 | -0.055174297 | 0.165346892 | -0.333688143 | 0.738614898 | 0.948137969 | Rsl24d1 | ribosomal L24 domain containing 1 [Sourc | 278.68 | 351.55 | 281.75 | 260.27 | 305.37 | 311.88 |
| 8823 | ENSMUSG00000032216 | 4689.975807 | 0.04524444 | 0.09105387 | 0.496897495 | 0.619261338 | 0.91763098 | Nedd4 | neural precursor cell expressed, developn | 4414.7 | 4848.7 | 4586.5 | 4583.7 | 5007 | 4699.3 |
| 8824 | ENSMUSG00000032217 | 362.5153259 | 0.115617897 | 0.140572726 | 0.822477447 | 0.410805217 | 0.833598017 | Rnf111 | ring finger 111 [Source:MGI Symbol;Acc:N | 334.09 | 338.18 | 372.31 | 401.01 | 369.72 | 359.77 |
| 8825 | ENSMUSG00000032218 | 10.69620563 | -0.728763448 | 0.708747093 | -1.028241886 | 0.30383606 | 0.75290128 | Ccnb2 | cyclin B2 [Source:MGI Symbol;Acc:MGI:88 | 14.456 | 19.53 | 6.0375 | 8.6758 | 5.4531 | 10.025 |
| 8826 | ENSMUSG00000032220 | 871.24739 | -0.006100461 | 0.140978233 | -0.043272362 | 0.965484422 | 0.993576285 | Myo1e | myosin IE [Source:MGI Symbol;Acc:MGI:1 | 732.43 | 895.32 | 993.16 | 958.19 | 789.61 | 858.77 |
| 8827 | ENSMUSG00000032221 | 3.689340015 | -0.405292498 | 1.120590168 | -0.361677721 | 0.717592877 | NA | Mns1 | meiosis-specific nuclear structural protein | 6.4248 | 3.0838 | 3.0187 | 1.928 | 5.4531 | 2.2277 |
| 8828 | ENSMUSG00000032224 | 43.33678384 | 0.758436948 | 0.428840419 | 1.768576178 | 0.076964629 | 0.406191172 | Fam81a | family with sequence similarity 81, membe | 21.684 | 26.726 | 48.3 | 29.883 | 63.256 | 70.172 |
| 8829 | ENSMUSG00000032226 | 0 | NA | NA | NA | NA | NA | Gcnt3 | glucosaminyl (N-acetyl) transferase 3, mu | 0 | 0 | 0 | 0 | 0 | 0 |
| 8830 | ENSMUSG00000032228 | 368.2258851 | 0.158951484 | 0.157559342 | 1.00883567 | 0.313053451 | 0.760114938 | Tcf12 | transcription factor 12 [Source:MGI Symb | 303.57 | 389.58 | 351.18 | 343.18 | 426.43 | 395.41 |
| 8831 | ENSMUSG00000032231 | 180.7361799 | -1.226759332 | 0.319144115 | -3.843903978 | 0.000121092 | 0.00421029 | Anxa2 | annexin A2 [Source:MGI Symbol;Acc:MGI | 413.6 | 198.39 | 146.91 | 81.938 | 121.06 | 122.52 |
| 8832 | ENSMUSG00000032232 | 2007.123303 | -0.000665087 | 0.108002241 | -0.006158084 | 0.995086591 | 0.998676967 | Cgnl1 | cingulin-like 1 [Source:MGI Symbol;Acc:M | 1953.9 | 2074.3 | 1995.4 | 2108.2 | 2137.6 | 1773.2 |
| 8833 | ENSMUSG00000032235 | 226.4695539 | -0.028872785 | 0.184467385 | -0.156519727 | 0.875623371 | 0.980566196 | Ice2 | interactor of little elongation complex ELL | 203.99 | 259.04 | 223.39 | 195.69 | 261.75 | 214.97 |
| 8834 | ENSMUSG00000032238 | 1249.204534 | 0.118739959 | 0.107743774 | 1.102058657 | 0.270436172 | 0.725929337 | Rora | RAR-related orphan receptor alpha [Sourc | 1135.6 | 1278.7 | 1180.3 | 1373.7 | 1221.5 | 1305.4 |
| 8835 | ENSMUSG00000032239 | 876.4478412 | 0.057509105 | 0.11368799 | 0.505850305 | 0.612961741 | 0.916748901 | Rp9 | retinitis pigmentosa 9 (human) [Source:M | 789.45 | 890.18 | 898.58 | 926.38 | 898.67 | 855.43 |
| 8836 | ENSMUSG00000032243 | 1.302792284 | -2.617024899 | 2.117274403 | -1.236034827 | 0.216445618 | NA | Itga11 | integrin alpha 11 [Source:MGI Symbol;Acc | 4.8186 | 1.0279 | 1.0062 | 0.964 | 0 | 0 |
| 8837 | ENSMUSG00000032244 | 351.3612728 | -0.158976575 | 0.144670259 | -1.098889131 | 0.271816428 | 0.727526428 | Fem1b | fem 1 homolog b [Source:MGI Symbol;Acc | 367.02 | 385.47 | 359.23 | 297.87 | 356.63 | 341.95 |
| 8838 | ENSMUSG00000032245 | 71.15743073 | 0.099281982 | 0.399908491 | 0.248261751 | 0.803931891 | 0.963625663 | Cln6 | ceroid-lipofuscinosis, neuronal 6 [Source:I | 100.39 | 68.87 | 36.225 | 51.091 | 54.531 | 115.84 |
| 8839 | ENSMUSG00000032246 | 8.591463099 | 0.356950935 | 0.771096125 | 0.462913667 | 0.643426251 | 0.925477092 | Calml4 | calmodulin-like 4 [Source:MGI Symbol;Acc | 7.2279 | 11.307 | 4.025 | 5.7839 | 8.7249 | 14.48 |
| 8840 | ENSMUSG00000032249 | 2066.861427 | -0.056502864 | 0.146804557 | -0.384884944 | 0.700322661 | 0.941626823 | Anp32a | acidic (leucine-rich) nuclear phosphoprote | 1786.9 | 2103.1 | 2432.1 | 1664.8 | 2087.4 | 2326.8 |
| 8841 | ENSMUSG00000032251 | 36.33271255 | 0.224393569 | 0.372881502 | 0.601782517 | 0.547318914 | 0.895518681 | Irak1bp1 | interleukin-1 receptor-associated kinase 1 | 30.518 | 27.754 | 42.262 | 33.739 | 43.625 | 40.098 |
| 8842 | ENSMUSG00000032252 | 437.1704822 | -0.218650347 | 0.153663269 | -1.422918752 | 0.15475971 | 0.572785168 | Glce | glucuronyl C5-epimerase [Source:MGI Syr | 421.63 | 532.46 | 456.84 | 369.2 | 379.54 | 463.36 |
| 8843 | ENSMUSG00000032253 | 262.7139059 | -0.007915617 | 0.171807898 | -0.046072485 | 0.963252477 | 0.993120238 | Phip | pleckstrin homology domain interacting p | 240.13 | 311.46 | 239.49 | 279.55 | 262.84 | 242.82 |
| 8844 | ENSMUSG00000032254 | 4.235966338 | -1.219524958 | 1.193013564 | -1.022222207 | 0.306675743 | NA | Kif23 | kinesin family member 23 [Source:MGI Sy | 6.4248 | 9.2513 | 2.0125 | 0 | 3.2719 | 4.4554 |

| | | | | | | | | | | | | | | | |
|---|---|---|---|---|---|---|---|---|---|---|---|---|---|---|---|
| 8845 | ENSMUSG00000032257 | 0 | NA | NA | NA | NA | NA | Ankk1 | ankyrin repeat and kinase domain contain | 0 | 0 | 0 | 0 | 0 | 0 |
| 8846 | ENSMUSG00000032258 | 2.820565178 | -0.022253972 | 1.29160034 | -0.017229766 | 0.986253316 | NA | Lca5 | Leber congenital amaurosis 5 (human) [Sc | 2.4093 | 5.1396 | 1.0062 | 2.8919 | 4.3625 | 1.1138 |
| 8847 | ENSMUSG00000032259 | 0 | NA | NA | NA | NA | NA | Drd2 | dopamine receptor D2 [Source:MGI Symb | 0 | 0 | 0 | 0 | 0 | 0 |
| 8848 | ENSMUSG00000032261 | 885.2593095 | 0.100216923 | 0.134355945 | 0.745906134 | 0.45572412 | 0.857604173 | Sh3bgrl2 | SH3 domain binding glutamic acid-rich pro | 960.51 | 884.01 | 718.46 | 982.29 | 901.94 | 864.34 |
| 8849 | ENSMUSG00000032262 | 0.342639279 | -1.805976596 | 4.028547714 | -0.448294702 | 0.653940524 | NA | Elovl4 | elongation of very long chain fatty acids (F | 0 | 2.0558 | 0 | 0 | 0 | 0 |
| 8850 | ENSMUSG00000032263 | 1668.343115 | -0.095371853 | 0.101449714 | -0.940089919 | 0.347171439 | 0.788352979 | Bckdhb | branched chain ketoacid dehydrogenase E | 1653.6 | 1759.8 | 1756.9 | 1494.2 | 1691.5 | 1654.1 |
| 8851 | ENSMUSG00000032264 | 232.1241839 | -0.142079264 | 0.170137191 | -0.835086453 | 0.403669034 | 0.828953151 | Zw10 | zw10 kinetochore protein [Source:MGI Sy | 218.44 | 272.4 | 240.49 | 231.35 | 203.95 | 226.11 |
| 8852 | ENSMUSG00000032265 | 193.9257214 | -0.319950615 | 0.22072881 | -1.449519046 | 0.147192685 | 0.558351692 | Tent5a | terminal nucleotidyltransferase 5A [Sourc | 215.23 | 184 | 246.53 | 161.95 | 131.96 | 223.88 |
| 8853 | ENSMUSG00000032267 | 146.726304 | 0.272504912 | 0.198481428 | 1.372949171 | 0.169768149 | 0.596876778 | Usp28 | ubiquitin specific peptidase 28 [Source:M | 133.31 | 125.41 | 139.87 | 150.38 | 182.13 | 149.26 |
| 8854 | ENSMUSG00000032268 | 0.181769715 | 1.126924175 | 4.080472857 | 0.276174898 | 0.782413736 | NA | Tmprss5 | transmembrane protease, serine 5 (spines | 0 | 0 | 0 | 0 | 1.0906 | 0 |
| 8855 | ENSMUSG00000032269 | 0 | NA | NA | NA | NA | NA | Htr3a | 5-hydroxytryptamine (serotonin) receptor | 0 | 0 | 0 | 0 | 0 | 0 |
| 8856 | ENSMUSG00000032271 | 2680.954756 | -0.140820912 | 0.312369695 | -0.450814896 | 0.652122964 | 0.925643431 | Nnmt | nicotinamide N-methyltransferase [Source | 2765.9 | 2227.5 | 3441.4 | 1790.1 | 2518.2 | 3342.6 |
| 8857 | ENSMUSG00000032274 | 0 | NA | NA | NA | NA | NA | Cyp19a1 | cytochrome P450, family 19, subfamily a, | 0 | 0 | 0 | 0 | 0 | 0 |
| 8858 | ENSMUSG00000032278 | 0 | NA | NA | NA | NA | NA | Paqr5 | progestin and adipoQ receptor family mer | 0 | 0 | 0 | 0 | 0 | 0 |
| 8859 | ENSMUSG00000032279 | 741.9361276 | -0.178619753 | 0.145023834 | -1.231657919 | 0.218076895 | 0.663697152 | Idh3a | isocitrate dehydrogenase 3 (NAD+) alpha | 844.86 | 871.67 | 646.01 | 624.66 | 745.98 | 718.43 |
| 8860 | ENSMUSG00000032280 | 414.7035305 | 0.051913665 | 0.138161484 | 0.375746291 | 0.70710552 | 0.943108897 | Tle3 | transducin-like enhancer of split 3 [Source | 429.66 | 373.13 | 418.6 | 430.9 | 444.97 | 390.96 |
| 8861 | ENSMUSG00000032281 | 0.678054141 | -2.792799308 | 2.990673428 | -0.933836266 | 0.350388359 | NA | Acsbg1 | acyl-CoA synthetase bubblegum family me | 0 | 2.0558 | 2.0125 | 0 | 0 | 0 |
| 8862 | ENSMUSG00000032285 | 47.67354218 | 0.157913479 | 0.333816042 | 0.473055394 | 0.63617366 | 0.923478764 | Dnaja4 | DnaJ heat shock protein family (Hsp40) m | 32.927 | 48.312 | 54.337 | 50.127 | 47.987 | 52.351 |
| 8863 | ENSMUSG00000032288 | 785.6055622 | 0.301432678 | 0.109676651 | 2.748376021 | 0.005989128 | 0.075262161 | Imp3 | IMP3, U3 small nucleolar ribonucleoprote | 723.6 | 697.96 | 689.28 | 832.88 | 887.76 | 882.16 |
| 8864 | ENSMUSG00000032289 | 17.5184767 | 0.12948366 | 0.5301262 | 0.244250634 | 0.807036714 | 0.964718338 | Thsd4 | thrombospondin, type I, domain containir | 22.487 | 13.363 | 14.087 | 15.424 | 16.359 | 23.391 |
| 8865 | ENSMUSG00000032290 | 273.8954399 | -0.297128444 | 0.158462949 | -1.875065725 | 0.060783682 | 0.352233645 | Ptpn9 | protein tyrosine phosphatase, non-recept | 273.05 | 323.79 | 309.92 | 260.27 | 227.94 | 248.39 |
| 8866 | ENSMUSG00000032291 | 0 | NA | NA | NA | NA | NA | Crabp1 | cellular retinoic acid binding protein I [Sou | 0 | 0 | 0 | 0 | 0 | 0 |
| 8867 | ENSMUSG00000032292 | 0 | NA | NA | NA | NA | NA | Nr2e3 | nuclear receptor subfamily 2, group E, me | 0 | 0 | 0 | 0 | 0 | 0 |
| 8868 | ENSMUSG00000032293 | 1928.338195 | 0.280110321 | 0.0966525 | 2.898117689 | 0.003754097 | 0.054737155 | Ireb2 | iron responsive element binding protein 2 | 1762.8 | 1788.6 | 1673.4 | 2045.6 | 2236.9 | 2062.8 |
| 8869 | ENSMUSG00000032294 | 190.7604837 | -1.050186654 | 0.205001331 | -5.122828477 | 3.01E-07 | 2.81E-05 | Pkm | pyruvate kinase, muscle [Source:MGI Sym | 274.66 | 278.57 | 218.36 | 123.39 | 100.34 | 149.26 |
| 8870 | ENSMUSG00000032295 | 573.069327 | 0.107541481 | 0.133555932 | 0.805216805 | 0.42069458 | 0.837913304 | Man2c1 | mannosidase, alpha, class 2C, member 1 [ | 549.32 | 593.11 | 513.18 | 625.62 | 632.56 | 524.62 |
| 8871 | ENSMUSG00000032297 | 0 | NA | NA | NA | NA | NA | Celf6 | CUGBP, Elav-like family member 6 [Source | 0 | 0 | 0 | 0 | 0 | 0 |
| 8872 | ENSMUSG00000032298 | 127.9325055 | 0.092822298 | 0.221000658 | 0.420009146 | 0.674478773 | 0.932114677 | Neil1 | nei endonuclease VIII-like 1 (E. coli) [Sour | 118.86 | 102.79 | 149.93 | 136.88 | 123.24 | 135.89 |
| 8873 | ENSMUSG00000032299 | 394.4579593 | 0.133872853 | 0.149752099 | 0.893963117 | 0.371341626 | 0.807214774 | Commd4 | COMM domain containing 4 [Source:MGI | 373.44 | 334.07 | 420.61 | 381.74 | 402.44 | 454.45 |
| 8874 | ENSMUSG00000032300 | 192.7581397 | -0.905182728 | 0.176420658 | -5.130820487 | 2.88E-07 | 2.73E-05 | 1700017B05 | RIKEN cDNA 1700017B05 gene [Source:M | 259.4 | 259.04 | 235.46 | 136.88 | 125.42 | 140.34 |
| 8875 | ENSMUSG00000032301 | 2102.480195 | 0.169562702 | 0.1027503 | 1.650240454 | 0.098893766 | 0.463570426 | Psma4 | proteasome subunit alpha 4 [Source:MGI | 2029.4 | 1923.2 | 1983.3 | 2011.8 | 2350.3 | 2316.8 |
| 8876 | ENSMUSG00000032303 | 0 | NA | NA | NA | NA | NA | Chrna3 | cholinergic receptor, nicotinic, alpha poly | 0 | 0 | 0 | 0 | 0 | 0 |
| 8877 | ENSMUSG00000032305 | 274.328295 | 0.171291024 | 0.171159517 | 1.00076833 | 0.316938824 | 0.763524842 | Fam219b | family with sequence similarity 219, meml | 293.13 | 234.37 | 245.52 | 256.42 | 286.83 | 329.7 |
| 8878 | ENSMUSG00000032306 | 181.3767223 | -0.300738015 | 0.19859572 | -1.514322742 | 0.129944003 | 0.527655829 | Mpi | mannose phosphate isomerase [Source:M | 180.7 | 192.22 | 228.42 | 187.98 | 136.33 | 162.62 |
| 8879 | ENSMUSG00000032307 | 562.7303037 | 0.067537514 | 0.122879931 | 0.549622009 | 0.58257866 | 0.908645447 | Ube2q2 | ubiquitin-conjugating enzyme E2Q family | 540.49 | 531.43 | 576.58 | 551.4 | 562.76 | 613.73 |
| 8880 | ENSMUSG00000032308 | 209.7239464 | 0.497552082 | 0.178810804 | 2.782561636 | 0.005393162 | 0.069961162 | Ulk3 | unc-51-like kinase 3 [Source:MGI Symbol; | 170.26 | 164.47 | 187.16 | 256.42 | 210.49 | 269.55 |
| 8881 | ENSMUSG00000032309 | 839.7099076 | 0.081717942 | 0.12882853 | 0.63431557 | 0.525874897 | 0.888334265 | Fbxo22 | F-box protein 22 [Source:MGI Symbol;Acc | 736.44 | 857.28 | 854.3 | 765.4 | 872.49 | 952.34 |
| 8882 | ENSMUSG00000032310 | 7064.328557 | 0.503673025 | 0.322365344 | 1.562429193 | 0.118186914 | 0.503669867 | Cyp1a2 | cytochrome P450, family 1, subfamily a, p | 4273.3 | 6168.5 | 7089 | 5423.3 | 9795.9 | 9635.9 |
| 8883 | ENSMUSG00000032311 | 308.8912871 | -0.534119991 | 0.212874634 | -2.50908237 | 0.012104525 | 0.123460168 | Nrg4 | neuregulin 4 [Source:MGI Symbol;Acc:MG | 376.65 | 419.39 | 300.87 | 313.29 | 269.38 | 173.76 |
| 8884 | ENSMUSG00000032312 | 1193.374085 | 0.132223476 | 0.114896472 | 1.150805357 | 0.24981232 | 0.702104867 | Csk | c-src tyrosine kinase [Source:MGI Symbol; | 1035.2 | 1178 | 1204.5 | 1332.2 | 1150.6 | 1259.8 |
| 8885 | ENSMUSG00000032313 | 13.37507895 | 0.934091012 | 0.614781022 | 1.519388169 | 0.128664818 | 0.524858587 | Tmem266 | transmembrane protein 266 [Source:MGI | 6.4248 | 7.1954 | 14.087 | 18.316 | 14.178 | 20.049 |
| 8886 | ENSMUSG00000032314 | 7567.557916 | -0.011716258 | 0.086547028 | -0.13537447 | 0.892315808 | 0.982489666 | Etfa | electron transferring flavoprotein, alpha p | 7434.3 | 7927.3 | 7433.1 | 7196.1 | 7635.4 | 7779.1 |
| 8887 | ENSMUSG00000032315 | 11.58704605 | 0.471416519 | 0.704058655 | 0.669569951 | 0.503131975 | 0.881242539 | Cyp1a1 | cytochrome P450, family 1, subfamily a, p | 8.8341 | 7.1954 | 13.081 | 11.568 | 5.4531 | 23.391 |
| 8888 | ENSMUSG00000032316 | 629.1082289 | -0.109292808 | 0.14305002 | -0.764018124 | 0.444856442 | 0.852013186 | Clk3 | CDC-like kinase 3 [Source:MGI Symbol;Acc | 655.33 | 633.2 | 671.17 | 721.06 | 570.39 | 523.51 |
| 8889 | ENSMUSG00000032318 | 0.267700746 | -1.497481649 | 4.057001436 | -0.369110456 | 0.712045396 | NA | Isl2 | insulin related protein 2 (islet 2) [Source:M | 1.6062 | 0 | 0 | 0 | 0 | 0 |
| 8890 | ENSMUSG00000032320 | 94.21677812 | -0.01589055 | 0.264279443 | -0.060127832 | 0.952053824 | 0.991040411 | Rcn2 | reticulocalbin 2 [Source:MGI Symbol;Acc:I | 78.704 | 106.9 | 98.612 | 69.406 | 102.52 | 109.16 |
| 8891 | ENSMUSG00000032322 | 7.060738811 | 0.27658738 | 0.819652329 | 0.337444755 | 0.735781651 | 0.948109711 | Pstpip1 | proline-serine-threonine phosphatase-int | 8.8341 | 7.1954 | 3.0187 | 6.7478 | 6.5437 | 10.025 |
| 8892 | ENSMUSG00000032323 | 0 | NA | NA | NA | NA | NA | Cyp11a1 | cytochrome P450, family 11, subfamily a, | 0 | 0 | 0 | 0 | 0 | 0 |
| 8893 | ENSMUSG00000032324 | 54.50394232 | -0.580934567 | 0.349992263 | -1.659849739 | 0.096944685 | 0.458561151 | Tspan3 | tetraspanin 3 [Source:MGI Symbol;Acc:M | 91.554 | 57.563 | 46.287 | 35.667 | 44.715 | 51.237 |
| 8894 | ENSMUSG00000032327 | 0 | NA | NA | NA | NA | NA | Stra6 | stimulated by retinoic acid gene 6 [Source | 0 | 0 | 0 | 0 | 0 | 0 |
| 8895 | ENSMUSG00000032328 | 3746.932318 | 0.231017875 | 0.097222018 | 2.376188846 | 0.017492508 | 0.157512348 | Tmem30a | transmembrane protein 30A [Source:MGI | 3262.2 | 3636.8 | 3445.4 | 4314.8 | 4029.8 | 3792.6 |
| 8896 | ENSMUSG00000032329 | 251.8599868 | -0.127978813 | 0.16536505 | -0.773916935 | 0.438979914 | 0.848057363 | Hmg20a | high mobility group 20A [Source:MGI Sym | 239.32 | 277.54 | 272.69 | 243.89 | 214.85 | 262.87 |
| 8897 | ENSMUSG00000032330 | 2286.694376 | 0.159231976 | 0.09151267 | 1.739999231 | 0.081859153 | 0.421206257 | Cox7a2 | cytochrome c oxidase subunit 7A2 [Sourc | 2251.9 | 2085.6 | 2143.3 | 2367.5 | 2413.5 | 2458.3 |
| 8898 | ENSMUSG00000032332 | 30.69771175 | -0.095767776 | 0.420655307 | -0.227663302 | 0.819908004 | 0.969160361 | Col12a1 | collagen, type XII, alpha 1 [Source:MGI Sy | 40.155 | 29.81 | 25.156 | 39.523 | 27.265 | 22.277 |
| 8899 | ENSMUSG00000032333 | 99.08375146 | -0.029577045 | 0.234399715 | -0.126182088 | 0.89958779 | 0.982798755 | Stoml1 | stomatin-like 1 [Source:MGI Symbol;Acc:M | 114.84 | 89.429 | 95.593 | 96.398 | 105.79 | 92.449 |
| 8900 | ENSMUSG00000032334 | 30.89859715 | 0.204749072 | 0.478830139 | 0.427602725 | 0.668940382 | 0.930610146 | Loxl1 | lysyl oxidase-like 1 [Source:MGI Symbol;A | 48.186 | 16.447 | 21.131 | 43.379 | 26.175 | 30.074 |

| | | | | | | | | | | | | | | | |
|---|---|---|---|---|---|---|---|---|---|---|---|---|---|---|---|
| 8901 | ENSMUSG00000032336 | 801.830053 | -0.082003605 | 0.119360873 | -0.687022499 | 0.492068551 | 0.876368691 | Nptn | neuroplastin [Source:MGI Symbol;Acc:MG | 860.93 | 852.14 | 760.72 | 810.71 | 812.51 | 713.97 |
| 8902 | ENSMUSG00000032338 | 0.185640647 | 1.126924175 | 4.080472857 | 0.276174898 | 0.782413736 | NA | Hcn4 | hyperpolarization-activated, cyclic nucleo | 0 | 0 | 0 | 0 | 0 | 1.1138 |
| 8903 | ENSMUSG00000032340 | 305.8119079 | 0.047234896 | 0.150918391 | 0.312983036 | 0.754293551 | 0.953048681 | Neo1 | neogenin [Source:MGI Symbol;Acc:MGI:1 | 285.9 | 308.38 | 308.92 | 343.18 | 295.56 | 292.94 |
| 8904 | ENSMUSG00000032342 | 297.5283961 | 0.118437513 | 0.151769813 | 0.780375954 | 0.435169618 | 0.845325456 | Mto1 | mitochondrial tRNA translation optimizati | 288.31 | 305.29 | 262.63 | 330.64 | 306.46 | 291.83 |
| 8905 | ENSMUSG00000032343 | 0 | NA | NA | NA | NA | NA | Impg1 | interphotoreceptor matrix proteoglycan 1 | 0 | 0 | 0 | 0 | 0 | 0 |
| 8906 | ENSMUSG00000032344 | 5.670554057 | 0.289126749 | 0.899695286 | 0.321360747 | 0.747937026 | 0.951047134 | Cgas | cyclic GMP-AMP synthase [Source:MGI Sy | 3.2124 | 7.1954 | 5.0312 | 8.6758 | 5.4531 | 4.4554 |
| 8907 | ENSMUSG00000032346 | 0 | NA | NA | NA | NA | NA | Ooep | oocyte expressed protein [Source:MGI Sy | 0 | 0 | 0 | 0 | 0 | 0 |
| 8908 | ENSMUSG00000032348 | 861.2292187 | -0.169771539 | 0.151729526 | -1.118909046 | 0.263178944 | 0.717823918 | Gsta4 | glutathione S-transferase, alpha 4 [Source | 783.02 | 885.04 | 1068.6 | 856.98 | 896.49 | 677.22 |
| 8909 | ENSMUSG00000032349 | 8696.092749 | -0.128983954 | 0.409833434 | -0.314722869 | 0.752972079 | 0.952989051 | Elovl5 | ELOVL family member 5, elongation of lon | 14065 | 8144.2 | 5044.3 | 8133.1 | 7210.1 | 9580.2 |
| 8910 | ENSMUSG00000032350 | 4091.162813 | -0.171997669 | 0.112020541 | -1.535411875 | 0.124682689 | 0.51807251 | Gclc | glutamate-cysteine ligase, catalytic subun | 3814.7 | 4609.2 | 4582.4 | 4211.6 | 3665.6 | 3663.4 |
| 8911 | ENSMUSG00000032352 | 7.825469439 | -0.00954734 | 0.807275477 | -0.01182662 | 0.990563943 | 0.99859176 | Lrrc1 | leucine rich repeat containing 1 [Source:M | 11.243 | 3.0838 | 9.0562 | 4.8199 | 8.7249 | 10.025 |
| 8912 | ENSMUSG00000032353 | 194.483094 | 0.297544807 | 0.19142583 | 1.554360801 | 0.120098378 | 0.507331466 | Tmed3 | transmembrane p24 trafficking protein 3 | | 173.47 | 193.25 | 156.97 | 230.39 | 176.68 | 236.13 |
| 8913 | ENSMUSG00000032355 | 0 | NA | NA | NA | NA | NA | Mlip | muscular LMNA-interacting protein [Sourc | 0 | 0 | 0 | 0 | 0 | 0 |
| 8914 | ENSMUSG00000032356 | 0 | NA | NA | NA | NA | NA | Rasgrf1 | RAS protein-specific guanine nucleotide-r | 0 | 0 | 0 | 0 | 0 | 0 |
| 8915 | ENSMUSG00000032357 | 0.185640647 | 1.126924175 | 4.080472857 | 0.276174898 | 0.782413736 | NA | Tinag | tubulointerstitial nephritis antigen [Source | 0 | 0 | 0 | 0 | 0 | 1.1138 |
| 8916 | ENSMUSG00000032358 | 0.185640647 | 1.126924175 | 4.080472857 | 0.276174898 | 0.782413736 | NA | Fam83b | family with sequence similarity 83, memb | 0 | 0 | 0 | 0 | 0 | 1.1138 |
| 8917 | ENSMUSG00000032359 | 4471.97204 | 0.105039193 | 0.10820322 | 0.970758484 | 0.331668561 | 0.774747658 | Ctsh | cathepsin H [Source:MGI Symbol;Acc:MGI | 3736.8 | 4474.5 | 4718.3 | 4749.5 | 4540.2 | 4612.4 |
| 8918 | ENSMUSG00000032360 | 10.64516563 | 0.873094538 | 0.849302192 | 1.028013994 | 0.303943245 | 0.752961647 | Hcrtr2 | hypocretin (orexin) receptor 2 [Source:MG | 7.2279 | 8.2233 | 7.0437 | 1.928 | 30.537 | 8.9108 |
| 8919 | ENSMUSG00000032363 | 103.8512603 | -0.087017127 | 0.318142357 | -0.273516321 | 0.784456348 | 0.960670212 | Adamts7 | a disintegrin-like and metallopeptidase (re | 58.626 | 107.93 | 154.96 | 97.362 | 86.159 | 118.07 |
| 8920 | ENSMUSG00000032366 | 453.4168586 | -0.061962249 | 0.135685165 | -0.456661925 | 0.647914062 | 0.925643431 | Tpm1 | tropomyosin 1, alpha [Source:MGI Symbo | 485.07 | 474.9 | 428.66 | 403.91 | 464.6 | 463.36 |
| 8921 | ENSMUSG00000032368 | 1.668734807 | -1.163316616 | 1.922224918 | -0.605192767 | 0.545050922 | NA | Zic1 | zinc finger protein of the cerebellum 1 [Sc | 0.8031 | 6.1675 | 0 | 1.928 | 0 | 1.1138 |
| 8922 | ENSMUSG00000032369 | 461.9162716 | -0.131024001 | 0.179366103 | -0.730483626 | 0.465094618 | 0.863116315 | Plscr1 | phospholipid scramblase 1 [Source:MGI S | 611.16 | 408.08 | 428.66 | 504.16 | 435.16 | 384.28 |
| 8923 | ENSMUSG00000032370 | 1135.784591 | 0.006465725 | 0.128978001 | 0.050130451 | 0.960018434 | 0.992261442 | Lactb | lactamase, beta [Source:MGI Symbol;Acc: | 959.71 | 1216 | 1225.6 | 1172.2 | 1197.5 | 1043.7 |
| 8924 | ENSMUSG00000032372 | 967.0518655 | -0.467892632 | 0.14095356 | -3.319480764 | 0.00090185 | 0.019361951 | Plscr2 | phospholipid scramblase 2 [Source:MGI S | 1316.3 | 989.88 | 1060.6 | 850.23 | 873.59 | 711.75 |
| 8925 | ENSMUSG00000032373 | 0 | NA | NA | NA | NA | NA | Car12 | carbonic anhydrase 12 [Source:MGI Symb | 0 | 0 | 0 | 0 | 0 | 0 |
| 8926 | ENSMUSG00000032374 | 11.04658214 | -1.190228784 | 0.758457285 | -1.56927596 | 0.116583651 | 0.501814922 | Plod2 | procollagen lysine, 2-oxoglutarate 5-dioxy | 22.487 | 13.363 | 10.062 | 4.8199 | 2.1812 | 13.366 |
| 8927 | ENSMUSG00000032375 | 27.77280776 | 0.114974775 | 0.483990632 | 0.237555786 | 0.812225647 | 0.96653525 | Aph1b | aph1 homolog B, gamma secretase subun | 28.912 | 39.061 | 12.075 | 34.703 | 21.812 | 30.074 |
| 8928 | ENSMUSG00000032376 | 536.5047537 | 0.111807186 | 0.129546704 | 0.863064688 | 0.388101902 | 0.817937623 | Usp3 | ubiquitin specific peptidase 3 [Source:MG | 540.49 | 479.01 | 527.27 | 572.6 | 509.32 | 590.34 |
| 8929 | ENSMUSG00000032377 | 66.94347465 | -1.884442537 | 0.306254922 | -6.153182856 | 7.59E-10 | 1.64E-07 | Plscr4 | phospholipid scramblase 4 [Source:MGI S | 101.99 | 123.35 | 90.562 | 24.099 | 33.809 | 27.846 |
| 8930 | ENSMUSG00000032380 | 226.3015911 | -0.072526156 | 0.220897431 | -0.328325033 | 0.742665919 | 0.949206972 | Dapk2 | death-associated protein kinase 2 [Source | 246.55 | 255.95 | 193.2 | 218.82 | 158.14 | 285.14 |
| 8931 | ENSMUSG00000032381 | 709.0488495 | 0.009619255 | 0.120469115 | 0.079848308 | 0.936357902 | 0.988420305 | Ciao2a | cytosolic iron-sulfur assembly component | 673.8 | 688.7 | 757.7 | 688.28 | 678.36 | 767.44 |
| 8932 | ENSMUSG00000032382 | 323.4344662 | -0.032554658 | 0.153465978 | -0.212129477 | 0.832006024 | 0.970533397 | Snx1 | sorting nexin 1 [Source:MGI Symbol;Acc:M | 328.47 | 312.49 | 340.11 | 316.18 | 282.47 | 360.89 |
| 8933 | ENSMUSG00000032383 | 1465.577482 | 0.28992347 | 0.102629923 | 2.824940937 | 0.004728933 | 0.064285047 | Ppib | peptidylprolyl isomerase B [Source:MGI S | 1351.6 | 1235.6 | 1368.5 | 1566.5 | 1570.5 | 1700.8 |
| 8934 | ENSMUSG00000032384 | 81.658754 | 0.130216648 | 0.266214427 | 0.489141966 | 0.624741193 | 0.919503423 | Csnk1g1 | casein kinase 1, gamma 1 [Source:MGI Sy | 78.704 | 75.038 | 80.5 | 107.97 | 70.89 | 76.855 |
| 8935 | ENSMUSG00000032386 | 1107.639536 | 0.265428911 | 0.100165919 | 2.649892448 | 0.00805174 | 0.092428861 | Trip4 | thyroid hormone receptor interactor 4 [Sc | 1025.6 | 985.77 | 1006.2 | 1183.8 | 1221.5 | 1223 |
| 8936 | ENSMUSG00000032387 | 1000.137421 | -0.107665792 | 0.112686575 | -0.955444718 | 0.339352849 | 0.781219101 | Rbpms2 | RNA binding protein with multiple splicing | 967.74 | 1079.3 | 1066.6 | 1035.3 | 911.76 | 940.08 |
| 8937 | ENSMUSG00000032388 | 460.2844987 | 0.036198472 | 0.133151907 | 0.271858456 | 0.785730856 | 0.960775321 | Spg21 | SPG21, maspardin [Source:MGI Symbol;Ac | 466.6 | 459.48 | 436.71 | 422.22 | 477.69 | 499 |
| 8938 | ENSMUSG00000032392 | 706.4173895 | -0.342299118 | 0.167497542 | -2.043606815 | 0.040992405 | 0.277451423 | Parp16 | poly (ADP-ribose) polymerase family, men | 652.92 | 792.52 | 924.74 | 588.99 | 754.71 | 524.62 |
| 8939 | ENSMUSG00000032393 | 903.4354135 | -0.078070706 | 0.109228565 | -0.71474624 | 0.47476586 | 0.869368495 | Dpp8 | dipeptidylpeptidase 8 [Source:MGI Symbo | 901.08 | 983.72 | 899.58 | 913.85 | 871.4 | 850.98 |
| 8940 | ENSMUSG00000032394 | 0 | NA | NA | NA | NA | NA | Igdcc3 | immunoglobulin superfamily, DCC subclas | 0 | 0 | 0 | 0 | 0 | 0 |
| 8941 | ENSMUSG00000032396 | 416.0785448 | 0.155762353 | 0.139428553 | 1.117148168 | 0.263930978 | 0.71906153 | Dis3l | DIS3 like exosome 3'-5' exoribonuclease [S | 382.28 | 363.88 | 434.7 | 415.47 | 463.51 | 436.63 |
| 8942 | ENSMUSG00000032397 | 79.47153323 | 0.030914347 | 0.271983204 | 0.113662705 | 0.909505178 | 0.985399467 | Tipin | timeless interacting protein [Source:MGI S | 84.326 | 81.206 | 70.437 | 97.362 | 61.075 | 82.424 |
| 8943 | ENSMUSG00000032398 | 276.3576514 | -0.079252971 | 0.184593295 | -0.429338299 | 0.667677049 | 0.930033258 | Snapc5 | small nuclear RNA activating complex, pol | 276.27 | 309.4 | 265.65 | 211.11 | 270.47 | 325.24 |
| 8944 | ENSMUSG00000032399 | 8430.721827 | 0.088793795 | 0.094467472 | 0.939940422 | 0.347248122 | 0.788352979 | Rpl4 | ribosomal protein L4 [Source:MGI Symbol | 7579.7 | 8635.5 | 8299.5 | 8160.1 | 8942 | 8967.6 |
| 8945 | ENSMUSG00000032400 | 14.28547994 | -0.512443476 | 0.62902758 | -0.814659789 | 0.415267076 | 0.836001897 | Zwilch | zwilch kinetochore protein [Source:MGI S | 12.85 | 12.335 | 25.156 | 7.7118 | 8.7249 | 18.935 |
| 8946 | ENSMUSG00000032401 | 0 | NA | NA | NA | NA | NA | Lctl | lactase-like [Source:MGI Symbol;Acc:MGI | 0 | 0 | 0 | 0 | 0 | 0 |
| 8947 | ENSMUSG00000032402 | 215.6230326 | 0.135084973 | 0.230771595 | 0.585362218 | 0.558304188 | 0.899286756 | Smad3 | SMAD family member 3 [Source:MGI Sym | 169.45 | 207.64 | 239.49 | 164.84 | 211.58 | 300.74 |
| 8948 | ENSMUSG00000032403 | 79.51849488 | 0.27855559 | 0.267674155 | 1.040651793 | 0.298037184 | 0.74872577 | 2300009A05 | RIKEN cDNA 2300009A05 gene [Source:M | 85.932 | 60.647 | 68.425 | 77.118 | 100.34 | 84.652 |
| 8949 | ENSMUSG00000032405 | 291.9207401 | 0.065770279 | 0.152733012 | 0.43062255 | 0.666742842 | 0.929596279 | Pias1 | protein inhibitor of activated STAT 1 [Sour | 313.21 | 270.34 | 271.69 | 307.51 | 282.47 | 306.31 |
| 8950 | ENSMUSG00000032407 | 560.6378184 | -0.096457567 | 0.124460058 | -0.775008216 | 0.438334805 | 0.847566655 | U2surp | U2 snRNP-associated SURP domain contai | 615.98 | 571.52 | 550.42 | 575.49 | 520.22 | 530.19 |
| 8951 | ENSMUSG00000032409 | 257.5570196 | 0.031733347 | 0.176307778 | 0.179988353 | 0.857161712 | 0.974809103 | Atr | ataxia telangiectasia and Rad3 related [So | 227.28 | 292.96 | 244.52 | 254.49 | 296.65 | 229.45 |
| 8952 | ENSMUSG00000032410 | 329.0301812 | 0.074454703 | 0.147997676 | 0.50308022 | 0.614907873 | 0.917509899 | Xrn1 | 5'-3' exoribonuclease 1 [Source:MGI Symb | 289.92 | 334.07 | 338.1 | 319.08 | 356.63 | 336.38 |
| 8953 | ENSMUSG00000032411 | 399.7963184 | 0.309426763 | 0.16070923 | 1.925382648 | 0.054181505 | 0.327625956 | Tfdp2 | transcription factor Dp 2 [Source:MGI Sym | 324.45 | 318.65 | 428.66 | 451.14 | 400.26 | 475.61 |
| 8954 | ENSMUSG00000032412 | 485.2942477 | -0.183727447 | 0.142542807 | -1.288928229 | 0.197423039 | 0.639249348 | Atp1b3 | ATPase, Na+/K+ transporting, beta 3 poly | 471.42 | 542.74 | 535.32 | 497.41 | 393.71 | 471.16 |
| 8955 | ENSMUSG00000032413 | 49.77242895 | 0.097510427 | 0.394917141 | 0.246913635 | 0.804975064 | 0.963917487 | Rasa2 | RAS p21 protein activator 2 [Source:MGI S | 40.958 | 55.508 | 48.3 | 78.082 | 50.168 | 25.618 |
| 8956 | ENSMUSG00000032415 | 35.84324806 | 0.131136362 | 0.406969443 | 0.322226557 | 0.747281067 | 0.950714518 | Ube2cbp | ubiquitin-conjugating enzyme E2C binding | 24.896 | 32.893 | 45.281 | 48.199 | 38.172 | 25.618 |

| | | | | | | | | | | | | | | | |
|---|---|---|---|---|---|---|---|---|---|---|---|---|---|---|---|
| 8957 | ENSMUSG00000032417 | 8.179339063 | -0.385165649 | 0.75790774 | -0.508195956 | 0.611315927 | 0.916376001 | Rwdd2a | RWD domain containing 2A [Source:MGI S | 6.4248 | 13.363 | 8.05 | 5.7839 | 6.5437 | 8.9108 |
| 8958 | ENSMUSG00000032418 | 3135.783679 | -0.146928726 | 0.185114914 | -0.793716306 | 0.427360605 | 0.842374461 | Me1 | malic enzyme 1, NADP(+)-dependent, cyto | 4316.7 | 3152.6 | 2416 | 3575.4 | 2416.8 | 2937.2 |
| 8959 | ENSMUSG00000032419 | 0 | NA | NA | NA | NA | NA | Tbx18 | T-box18 [Source:MGI Symbol;Acc:MGI:19 | 0 | 0 | 0 | 0 | 0 | 0 |
| 8960 | ENSMUSG00000032420 | 97.79479538 | -1.128959977 | 0.289214871 | -3.903533637 | 9.48E-05 | 0.0034713 | Nt5e | 5' nucleotidase, ecto [Source:MGI Symbol | 156.6 | 129.52 | 116.72 | 80.01 | 37.081 | 66.831 |
| 8961 | ENSMUSG00000032422 | 347.054483 | 0.071070051 | 0.147369801 | 0.482256543 | 0.629623714 | 0.92057725 | Snx14 | sorting nexin 14 [Source:MGI Symbol;Acc: | 318.83 | 353.6 | 343.13 | 321 | 390.44 | 355.32 |
| 8962 | ENSMUSG00000032423 | 713.3012378 | -0.056813514 | 0.126520805 | -0.449044832 | 0.653399316 | 0.925756399 | Syncrip | synaptotagmin binding, cytoplasmic RNA i | 680.23 | 810 | 692.3 | 670.93 | 754.71 | 671.65 |
| 8963 | ENSMUSG00000032425 | 108.7083012 | 0.074825466 | 0.233505794 | 0.320443722 | 0.748631986 | 0.951074124 | Zfp949 | zinc finger protein 949 [Source:MGI Symb | 109.22 | 96.624 | 111.69 | 107 | 134.15 | 93.563 |
| 8964 | ENSMUSG00000032431 | 63.33572554 | -0.149232323 | 0.309801766 | -0.481702623 | 0.630017212 | 0.920740523 | Crtap | cartilage associated protein [Source:MGI S | 60.233 | 82.233 | 57.356 | 52.055 | 50.168 | 77.969 |
| 8965 | ENSMUSG00000032434 | 2168.169807 | 0.422489929 | 0.0962119 | 4.391244015 | 1.13E-05 | 0.000588502 | Cmtm6 | CKLF-like MARVEL transmembrane domai | 1778.9 | 1809.1 | 1971.2 | 2404.2 | 2567.3 | 2478.3 |
| 8966 | ENSMUSG00000032435 | 340.9581094 | 0.07266262 | 0.144929296 | 0.501365995 | 0.616113567 | 0.917509899 | Dync1li1 | dynein cytoplasmic 1 light intermediate ch | 331.68 | 321.74 | 344.14 | 384.63 | 327.19 | 336.38 |
| 8967 | ENSMUSG00000032436 | 25.74546582 | -0.876215522 | 0.471698994 | -1.857573439 | 0.063229613 | 0.359877642 | Cmtm7 | CKLF-like MARVEL transmembrane domai | 21.684 | 40.089 | 38.237 | 12.532 | 18.541 | 23.391 |
| 8968 | ENSMUSG00000032437 | 1439.091014 | -0.088712536 | 0.098865435 | -0.897305877 | 0.369555722 | 0.80653433 | Stt3b | STT3, subunit of the oligosaccharyltransfe | 1415.9 | 1540.8 | 1494.3 | 1440.2 | 1361.1 | 1382.3 |
| 8969 | ENSMUSG00000032440 | 646.0995514 | -0.443491968 | 0.162236763 | -2.733609565 | 0.006264428 | 0.077509566 | Tgfbr2 | transforming growth factor, beta receptor | 885.02 | 739.07 | 608.78 | 516.69 | 635.83 | 491.21 |
| 8970 | ENSMUSG00000032443 | 0.836675597 | -0.373395339 | 2.394418521 | -0.155944057 | 0.876077117 | NA | Zcwpw2 | zinc finger, CW type with PWWP domain 2 | 0.8031 | 0 | 2.0125 | 0 | 1.0906 | 1.1138 |
| 8971 | ENSMUSG00000032446 | 1.048817988 | -0.908682365 | 2.659333108 | -0.341695579 | 0.73258 | NA | Eomes | eomesodermin [Source:MGI Symbol;Acc:I | 0 | 4.1117 | 0 | 0 | 2.1812 | 0 |
| 8972 | ENSMUSG00000032449 | 47.6245819 | -1.031757423 | 0.345093883 | -2.989787626 | 0.002791715 | 0.044426743 | Slc25a36 | solute carrier family 25, member 36 [Sour | 57.823 | 77.094 | 57.356 | 39.523 | 29.447 | 24.505 |
| 8973 | ENSMUSG00000032451 | 0 | NA | NA | NA | NA | NA | Trim42 | tripartite motif-containing 42 [Source:MG | 0 | 0 | 0 | 0 | 0 | 0 |
| 8974 | ENSMUSG00000032452 | 1.82194127 | -3.123453192 | 1.914091973 | -1.631819806 | 0.102717452 | NA | Clstn2 | calsyntenin 2 [Source:MGI Symbol;Acc:MG | 0.8031 | 5.1396 | 4.025 | 0.964 | 0 | 0 |
| 8975 | ENSMUSG00000032454 | 0.17131964 | -0.796629176 | 4.080472857 | -0.195229623 | 0.84521319 | NA | Rbp2 | retinol binding protein 2, cellular [Source: | 0 | 1.0279 | 0 | 0 | 0 | 0 |
| 8976 | ENSMUSG00000032456 | 151.863253 | 0.155756623 | 0.236634714 | 0.658215444 | 0.510399703 | 0.883064748 | Nmnat3 | nicotinamide nucleotide adenylyltransfera | 159.82 | 128.49 | 141.88 | 115.68 | 158.14 | 207.17 |
| 8977 | ENSMUSG00000032458 | 2402.891624 | 0.035123537 | 0.093333149 | 0.376324359 | 0.706675772 | 0.94295036 | Copb2 | coatomer protein complex, subunit beta 2 | 2488.8 | 2336.5 | 2295.2 | 2517.9 | 2433.2 | 2345.8 |
| 8978 | ENSMUSG00000032459 | 346.581704 | 0.194034397 | 0.144938687 | 1.33873434 | 0.180657173 | 0.613962723 | Mrps22 | mitochondrial ribosomal protein S22 [Sou | 335.7 | 330.99 | 302.88 | 356.67 | 404.62 | 348.63 |
| 8979 | ENSMUSG00000032462 | 442.9811922 | -0.243252301 | 0.15967188 | -1.523451102 | 0.127645885 | 0.523319186 | Pik3cb | phosphatidylinositol-4,5-bisphosphate 3-k | 497.12 | 517.04 | 426.65 | 418.37 | 466.78 | 331.93 |
| 8980 | ENSMUSG00000032463 | 136.8755794 | 0.240639579 | 0.209849711 | 1.146723424 | 0.251495946 | 0.703994757 | Faim | Fas apoptotic inhibitory molecule [Source | 143.76 | 123.35 | 108.67 | 135.92 | 161.41 | 148.14 |
| 8981 | ENSMUSG00000032468 | 318.4961154 | -0.047505117 | 0.143490649 | -0.331067685 | 0.74059336 | 0.948792385 | Armc8 | armadillo repeat containing 8 [Source:MG | 303.57 | 332.02 | 336.09 | 312.33 | 319.55 | 307.42 |
| 8982 | ENSMUSG00000032469 | 221.3355297 | 0.147302728 | 0.207840403 | 0.708729994 | 0.478492048 | 0.869898062 | Dbr1 | debranching RNA lariats 1 [Source:MGI Sy | 265.83 | 182.97 | 180.12 | 201.47 | 273.75 | 223.88 |
| 8983 | ENSMUSG00000032470 | 109.5624755 | -0.035752553 | 0.230210212 | -0.155303939 | 0.87658171 | 0.980632845 | Mras | muscle and microspikes RAS [Source:MGI | 112.43 | 91.485 | 128.8 | 111.82 | 100.34 | 112.5 |
| 8984 | ENSMUSG00000032473 | 0.734820724 | 3.078876299 | 2.902525826 | 1.06075759 | 0.28880008 | NA | Cldn18 | claudin 18 [Source:MGI Symbol;Acc:MGI:1 | 0 | 0 | 0 | 0 | 2.1812 | 2.2277 |
| 8985 | ENSMUSG00000032475 | 314.6206458 | 0.067319484 | 0.156634959 | 0.429785817 | 0.667351451 | 0.929830593 | Nck1 | non-catalytic region of tyrosine kinase ada | 266.63 | 320.71 | 335.08 | 302.69 | 319.55 | 343.06 |
| 8986 | ENSMUSG00000032477 | 69.48298598 | 0.462738096 | 0.340906721 | 1.35737452 | 0.17466224 | 0.604526117 | Cdc25a | cell division cycle 25A [Source:MGI Symbo | 74.689 | 35.977 | 64.4 | 106.04 | 56.712 | 79.083 |
| 8987 | ENSMUSG00000032478 | 143.0996017 | 0.424104036 | 0.201369162 | 2.106102207 | 0.035195475 | 0.25151549 | Nme6 | NME/NM23 nucleoside diphosphate kinas | 122.87 | 118.21 | 125.78 | 181.23 | 169.05 | 141.46 |
| 8988 | ENSMUSG00000032479 | 595.2353686 | -0.097580698 | 0.125567059 | -0.777120201 | 0.437087857 | 0.846852749 | Map4 | microtubule-associated protein 4 [Source | 610.36 | 668.15 | 567.52 | 561.04 | 557.31 | 607.04 |
| 8989 | ENSMUSG00000032480 | 515.4911485 | 0.048348841 | 0.132524554 | 0.364829302 | 0.715238831 | 0.944595913 | Dhx30 | DEAH (Asp-Glu-Ala-His) box polypeptide 3 | 496.32 | 540.68 | 484 | 536.94 | 563.85 | 471.16 |
| 8990 | ENSMUSG00000032481 | 447.6582455 | 0.019038405 | 0.13347118 | 0.142640569 | 0.886574056 | 0.982355187 | Smarcc1 | SWI/SNF related, matrix associated, actin | 416.81 | 440.98 | 476.96 | 464.64 | 420.98 | 465.59 |
| 8991 | ENSMUSG00000032482 | 11.47118957 | 2.306364029 | 0.768239299 | 3.002142734 | 0.002680864 | 0.043485382 | Cspg5 | chondroitin sulfate proteoglycan 5 [Sourc | 7.2279 | 2.0558 | 2.0125 | 28.919 | 16.359 | 12.252 |
| 8992 | ENSMUSG00000032484 | 0 | NA | NA | NA | NA | NA | Ngp | neutrophilic granule protein [Source:MGI | 0 | 0 | 0 | 0 | 0 | 0 |
| 8993 | ENSMUSG00000032485 | 1900.045 | 0.389173325 | 0.124856937 | 3.116953978 | 0.001827301 | 0.032561965 | Scap | SREBF chaperone [Source:MGI Symbol;Ac | 1582.1 | 1648.8 | 1704.6 | 1836.4 | 2125.6 | 2502.8 |
| 8994 | ENSMUSG00000032487 | 0 | NA | NA | NA | NA | NA | Ptgs2 | prostaglandin-endoperoxide synthase 2 [S | 0 | 0 | 0 | 0 | 0 | 0 |
| 8995 | ENSMUSG00000032489 | 8.98298568 | -0.298838333 | 0.719832894 | -0.415149593 | 0.678032396 | 0.93325821 | Kif9 | kinesin family member 9 [Source:MGI Sym | 10.44 | 7.1954 | 12.075 | 7.7118 | 10.906 | 5.5692 |
| 8996 | ENSMUSG00000032491 | 12.49922095 | 0.305080145 | 0.618936938 | 0.492909902 | 0.622076251 | 0.918602439 | Nradd | neurotrophin receptor associated death d | 12.047 | 13.363 | 8.05 | 10.604 | 11.997 | 18.935 |
| 8997 | ENSMUSG00000032492 | 141.2555587 | -0.592585497 | 0.205014665 | -2.890454189 | 0.003846856 | 0.055696621 | Pth1r | parathyroid hormone 1 receptor [Source:I | 184.71 | 155.22 | 169.05 | 99.29 | 114.51 | 124.75 |
| 8998 | ENSMUSG00000032493 | 0 | NA | NA | NA | NA | NA | Prss44 | protease, serine 44 [Source:MGI Symbol;A | 0 | 0 | 0 | 0 | 0 | 0 |
| 8999 | ENSMUSG00000032494 | 0 | NA | NA | NA | NA | NA | Tdgf1 | teratocarcinoma-derived growth factor 1 | 0 | 0 | 0 | 0 | 0 | 0 |
| 9000 | ENSMUSG00000032495 | 0.167707431 | -0.796629176 | 4.080472857 | -0.195229623 | 0.84521319 | NA | Lrrc2 | leucine rich repeat containing 2 [Source:M | 0 | 0 | 1.0062 | 0 | 0 | 0 |
| 9001 | ENSMUSG00000032496 | 0.133850373 | -0.796629176 | 4.080472857 | -0.195229623 | 0.84521319 | NA | Ltf | lactotransferrin [Source:MGI Symbol;Acc: | 0.8031 | 0 | 0 | 0 | 0 | 0 |
| 9002 | ENSMUSG00000032497 | 633.0017545 | -0.03324847 | 0.129162778 | -0.257415263 | 0.796858222 | 0.962236316 | Lrrfip2 | leucine rich repeat (in FLII) interacting pro | 674.61 | 686.65 | 559.47 | 651.65 | 608.57 | 617.07 |
| 9003 | ENSMUSG00000032498 | 253.0547654 | 0.161742147 | 0.16660231 | 0.970827756 | 0.331634058 | 0.774747658 | Mlh1 | mutL homolog 1 [Source:MGI Symbol;Acc | 236.11 | 217.92 | 262.63 | 262.2 | 296.65 | 242.82 |
| 9004 | ENSMUSG00000032500 | 255.0889051 | 0.682000682 | 0.213877803 | 3.188739884 | 0.001428944 | 0.027375159 | Dclk3 | doublecortin-like kinase 3 [Source:MGI Sy | 158.21 | 174.75 | 255.59 | 354.74 | 249.75 | 337.49 |
| 9005 | ENSMUSG00000032501 | 980.9775998 | -0.205161586 | 0.149352911 | -1.373669818 | 0.169544211 | 0.596876778 | Trib1 | tribbles pseudokinase 1 [Source:MGI Sym | 1071.3 | 1092.7 | 987.13 | 767.33 | 1134.2 | 833.16 |
| 9006 | ENSMUSG00000032502 | 0 | NA | NA | NA | NA | NA | Stac | src homology three (SH3) and cysteine ricl | 0 | 0 | 0 | 0 | 0 | 0 |
| 9007 | ENSMUSG00000032503 | 0 | NA | NA | NA | NA | NA | Arpp21 | cyclic AMP-regulated phosphoprotein, 21 | 0 | 0 | 0 | 0 | 0 | 0 |
| 9008 | ENSMUSG00000032504 | 1998.544051 | -0.091750844 | 0.092783179 | -0.988873689 | 0.322724945 | 0.769926329 | Pdcd6ip | programmed cell death 6 interacting prote | 2053.5 | 2145.3 | 1987.3 | 1899 | 1942.4 | 1963.7 |
| 9009 | ENSMUSG00000032507 | 2.607864222 | 1.05621716 | 1.832501946 | 0.57637983 | 0.564358475 | NA | Fbxl2 | F-box and leucine-rich repeat protein 2 [S | 4.0155 | 1.0279 | 0 | 10.604 | 0 | 0 |
| 9010 | ENSMUSG00000032508 | 243.3661901 | -0.005825151 | 0.185241399 | -0.031446268 | 0.974913643 | 0.996336532 | Myd88 | myeloid differentiation primary response | 264.22 | 210.72 | 256.59 | 279.55 | 198.49 | 250.61 |
| 9011 | ENSMUSG00000032511 | 0 | NA | NA | NA | NA | NA | Scn5a | sodium channel, voltage-gated, type V, alp | 0 | 0 | 0 | 0 | 0 | 0 |
| 9012 | ENSMUSG00000032512 | 408.9606487 | 0.158095324 | 0.13794822 | 1.146048309 | 0.251775164 | 0.704249849 | Wdr48 | WD repeat domain 48 [Source:MGI Symbo | 373.44 | 407.06 | 379.35 | 420.29 | 401.35 | 472.27 |

| | | | | | | | | | | | | | | | |
|---|---|---|---|---|---|---|---|---|---|---|---|---|---|---|---|
| 9013 | ENSMUSG00000032513 | 751.9005196 | 0.135405569 | 0.114818045 | 1.179305648 | 0.23827649 | 0.687224557 | Gorasp1 | golgi reassembly stacking protein 1 [Sourc | 742.87 | 693.84 | 712.42 | 756.72 | 761.25 | 844.29 |
| 9014 | ENSMUSG00000032514 | 1.054859682 | -0.240938279 | 2.360206831 | -0.102083544 | 0.918690362 | NA | Ttc21a | tetratricopeptide repeat domain 21A [Sou | 2.4093 | 1.0279 | 0 | 2.8919 | 0 | 0 |
| 9015 | ENSMUSG00000032515 | 222.038474 | 0.450016838 | 0.240210199 | 1.873429352 | 0.06100912 | 0.352269825 | Csrnp1 | cysteine-serine-rich nuclear protein 1 [Sou | 170.26 | 213.81 | 179.11 | 238.1 | 173.41 | 357.54 |
| 9016 | ENSMUSG00000032517 | 0.133850373 | -0.796629176 | 4.080472857 | -0.195229623 | 0.84521319 | NA | Mobp | myelin-associated oligodendrocytic basic | 0.8031 | 0 | 0 | 0 | 0 | 0 |
| 9017 | ENSMUSG00000032518 | 6241.635647 | 0.152202607 | 0.10538448 | 1.444260167 | 0.148665806 | 0.560558451 | Rpsa | ribosomal protein SA [Source:MGI Symbo | 5586.4 | 6161.3 | 5990.2 | 5789.7 | 6836 | 7086.3 |
| 9018 | ENSMUSG00000032519 | 443.287615 | 0.318752062 | 0.150534532 | 2.117468051 | 0.034220144 | 0.247603883 | Slc25a38 | solute carrier family 25, member 38 [Sour | 452.95 | 335.1 | 394.45 | 508.98 | 455.88 | 512.37 |
| 9019 | ENSMUSG00000032523 | 0 | NA | NA | NA | NA | NA | Hhatl | hedgehog acyltransferase-like [Source:MG | 0 | 0 | 0 | 0 | 0 | 0 |
| 9020 | ENSMUSG00000032525 | 509.3139309 | 0.281400808 | 0.144830033 | 1.942972755 | 0.05201945 | 0.320310485 | Nktr | natural killer tumor recognition sequence | 453.75 | 465.65 | 460.86 | 654.54 | 537.67 | 483.41 |
| 9021 | ENSMUSG00000032526 | 540.9634948 | 0.254193859 | 0.138594936 | 1.834077532 | 0.066642486 | 0.371533109 | Ss18l2 | SS18, nBAF chromatin remodeling comple | 522.82 | 497.51 | 458.85 | 508.02 | 630.38 | 628.21 |
| 9022 | ENSMUSG00000032527 | 4317.361494 | 0.131389495 | 0.089340071 | 1.470667005 | 0.141381193 | 0.55015808 | Pccb | propionyl Coenzyme A carboxylase, beta p | 3877.4 | 4278.2 | 4208.1 | 4428.5 | 4620.9 | 4491 |
| 9023 | ENSMUSG00000032528 | 576.0284785 | -1.430730875 | 0.138509923 | -10.32944677 | 5.19E-25 | 1.88E-21 | Vipr1 | vasoactive intestinal peptide receptor 1 [S | 813.54 | 840.84 | 866.38 | 300.76 | 272.65 | 362 |
| 9024 | ENSMUSG00000032530 | 0.68166635 | -2.79976525 | 3.012029182 | -0.92952793 | 0.352615558 | NA | Lyzl4 | lysozyme-like 4 [Source:MGI Symbol;Acc:I | 0 | 3.0838 | 1.0062 | 0 | 0 | 0 |
| 9025 | ENSMUSG00000032531 | 830.4017001 | -0.256647047 | 0.165995082 | -1.546112352 | 0.122077436 | 0.511805228 | Amotl2 | angiomotin-like 2 [Source:MGI Symbol;Ac | 718.78 | 942.6 | 1052.5 | 888.79 | 748.16 | 631.55 |
| 9026 | ENSMUSG00000032532 | 0 | NA | NA | NA | NA | NA | Cck | cholecystokinin [Source:MGI Symbol;Acc: | 0 | 0 | 0 | 0 | 0 | 0 |
| 9027 | ENSMUSG00000032534 | 211.6067524 | -0.037639021 | 0.170620788 | -0.220600439 | 0.825403559 | 0.96943211 | Cep63 | centrosomal protein 63 [Source:MGI Symb | 211.22 | 205.58 | 226.41 | 214.97 | 187.59 | 223.88 |
| 9028 | ENSMUSG00000032536 | 1571.553901 | 0.019957222 | 0.110816445 | 0.180092599 | 0.857079873 | 0.974809103 | Trak1 | trafficking protein, kinesin binding 1 [Sour | 1721.9 | 1556.3 | 1402.7 | 1587.7 | 1529 | 1631.8 |
| 9029 | ENSMUSG00000032537 | 2.824572522 | 0.239899004 | 1.367375703 | 0.175444834 | 0.860730114 | NA | Ephb1 | Eph receptor B1 [Source:MGI Symbol;Acc | 0.8031 | 1.0279 | 6.0375 | 5.7839 | 2.1812 | 1.1138 |
| 9030 | ENSMUSG00000032540 | 461.5588158 | 0.091193772 | 0.124819665 | 0.730604202 | 0.465020945 | 0.863116315 | Abhd5 | abhydrolase domain containing 5 [Source | 436.08 | 453.31 | 451.8 | 479.1 | 467.88 | 481.18 |
| 9031 | ENSMUSG00000032547 | 810.3813482 | -0.141182847 | 0.143313868 | -0.985130388 | 0.324560028 | 0.771309263 | Ryk | receptor-like tyrosine kinase [Source:MGI | 705.93 | 847 | 998.19 | 761.54 | 824.51 | 725.11 |
| 9032 | ENSMUSG00000032548 | 626.1057389 | 0.267213526 | 0.193381636 | 1.381793698 | 0.167035055 | 0.593319485 | Slco2a1 | solute carrier organic anion transporter fa | 441.71 | 533.49 | 730.53 | 697.92 | 538.77 | 814.22 |
| 9033 | ENSMUSG00000032549 | 17.92303734 | -1.058670824 | 0.559072253 | -1.893620759 | 0.058275366 | 0.342880809 | Rab6b | RAB6B, member RAS oncogene family [So | 17.668 | 34.949 | 20.125 | 11.568 | 7.6343 | 15.594 |
| 9034 | ENSMUSG00000032551 | 222.9220913 | 0.466392457 | 0.18309463 | 2.547275453 | 0.01085677 | 0.115286132 | 1110059G10 | RIKEN cDNA 1110059G10 gene [Source:M | 201.58 | 185.03 | 175.09 | 308.47 | 230.12 | 237.25 |
| 9035 | ENSMUSG00000032553 | 206.0850568 | 0.121394128 | 0.167307217 | 0.725576163 | 0.468098638 | 0.865229581 | Srprb | signal recognition particle receptor, B sub | 192.74 | 203.53 | 196.22 | 221.71 | 201.76 | 220.54 |
| 9036 | ENSMUSG00000032554 | 11263.98681 | 0.041282512 | 0.10280138 | 0.401575468 | 0.687996487 | 0.936885029 | Trf | transferrin [Source:MGI Symbol;Acc:MGI: | 11805 | 10021 | 11482 | 12257 | 11443 | 10576 |
| 9037 | ENSMUSG00000032555 | 217.3563438 | 0.08086404 | 0.169239051 | 0.477809581 | 0.632785741 | 0.922107984 | Topbp1 | topoisomerase (DNA) II binding protein 1 | 233.7 | 199.42 | 200.24 | 234.25 | 213.76 | 222.77 |
| 9038 | ENSMUSG00000032556 | 0 | NA | NA | NA | NA | NA | Bfsp2 | beaded filament structural protein 2, phal | 0 | 0 | 0 | 0 | 0 | 0 |
| 9039 | ENSMUSG00000032557 | 551.5467 | 0.225559692 | 0.135566454 | 1.663831167 | 0.096146174 | 0.457754528 | Uba5 | ubiquitin-like modifier activating enzyme ! | 544.5 | 537.6 | 442.75 | 581.28 | 566.03 | 637.12 |
| 9040 | ENSMUSG00000032558 | 122.8019588 | 0.209312741 | 0.21559653 | 0.970853942 | 0.331621017 | 0.774747658 | Nphp3 | nephronophthisis 3 (adolescent) [Source:I | 100.39 | 118.21 | 123.77 | 146.52 | 126.51 | 121.41 |
| 9041 | ENSMUSG00000032560 | 561.106492 | 0.144163965 | 0.13269549 | 1.08642702 | 0.277290113 | 0.730041765 | Dnajc13 | DnaJ heat shock protein family (Hsp40) m | 509.97 | 538.63 | 550.42 | 520.55 | 595.48 | 651.6 |
| 9042 | ENSMUSG00000032561 | 111.1180956 | 0.40654488 | 0.240809079 | 1.688245649 | 0.091364084 | 0.445846864 | Acpp | acid phosphatase, prostate [Source:MGI S | 87.538 | 87.373 | 111.69 | 100.25 | 143.96 | 135.89 |
| 9043 | ENSMUSG00000032562 | 1526.537286 | -0.200676404 | 0.1020473 | -1.966503802 | 0.049240444 | 0.309658163 | Gnai2 | guanine nucleotide binding protein (G pro | 1745.1 | 1605.6 | 1545.6 | 1373.7 | 1416.7 | 1472.5 |
| 9044 | ENSMUSG00000032563 | 452.52247 | -0.027787799 | 0.137873829 | -0.201545134 | 0.840272342 | 0.971838962 | Mrpl3 | mitochondrial ribosomal protein L3 [Sourc | 426.45 | 457.42 | 487.02 | 420.29 | 496.23 | 427.72 |
| 9045 | ENSMUSG00000032564 | 0 | NA | NA | NA | NA | NA | Cpne4 | copine IV [Source:MGI Symbol;Acc:MGI:1 | 0 | 0 | 0 | 0 | 0 | 0 |
| 9046 | ENSMUSG00000032565 | 179.9135763 | 0.390297775 | 0.211713564 | 1.843518039 | 0.06525341 | 0.366785337 | Nudt16 | nudix (nucleoside diphosphate linked moi | 164.64 | 139.8 | 162.01 | 154.24 | 213.76 | 245.05 |
| 9047 | ENSMUSG00000032566 | 0 | NA | NA | NA | NA | NA | Nudt16l2 | nudix (nucleoside diphosphate linked moi | 0 | 0 | 0 | 0 | 0 | 0 |
| 9048 | ENSMUSG00000032567 | 60.62158951 | 0.214742031 | 0.296023087 | 0.725423253 | 0.468192412 | 0.865229581 | Aste1 | asteroid homolog 1 [Source:MGI Symbol;/ | 67.461 | 48.312 | 52.325 | 72.298 | 66.528 | 56.806 |
| 9049 | ENSMUSG00000032570 | 688.7654683 | -0.030677156 | 0.12457025 | -0.246263905 | 0.805477949 | 0.964280173 | Atp2c1 | ATPase, Ca++-sequestering [Source:MGI S | 636.86 | 698.98 | 753.68 | 732.62 | 653.28 | 657.17 |
| 9050 | ENSMUSG00000032571 | 381.4368091 | 0.005961551 | 0.134769 | 0.044235326 | 0.964716824 | 0.993480943 | Pik3r4 | phosphoinositide-3-kinase regulatory sub | 379.87 | 394.72 | 367.28 | 371.13 | 400.26 | 375.37 |
| 9051 | ENSMUSG00000032572 | 0.167707431 | -0.796629176 | 4.080472857 | -0.195229623 | 0.84521319 | NA | Col6a4 | collagen, type VI, alpha 4 [Source:MGI Syr | 0 | 0 | 1.0062 | 0 | 0 | 0 |
| 9052 | ENSMUSG00000032575 | 777.8088406 | 0.771718002 | 0.19035974 | 4.053997987 | 5.03E-05 | 0.002110891 | Manf | mesencephalic astrocyte-derived neurotro | 591.08 | 616.75 | 516.2 | 1185.7 | 652.19 | 1104.9 |
| 9053 | ENSMUSG00000032577 | 83.22401834 | 0.831451043 | 0.301154041 | 2.760882903 | 0.005764534 | 0.073396321 | Mapkapk3 | mitogen-activated protein kinase-activate | 78.704 | 41.117 | 59.368 | 134.96 | 90.521 | 94.677 |
| 9054 | ENSMUSG00000032578 | 304.370222 | 3.01665836 | 1.050317767 | 2.872138751 | 0.004077039 | 0.058098812 | Cish | cytokine inducible SH2-containing protein | 32.927 | 95.596 | 72.45 | 844.44 | 52.35 | 728.45 |
| 9055 | ENSMUSG00000032579 | 351.6655249 | 0.229866748 | 0.151872615 | 1.51354968 | 0.130140092 | 0.528303758 | Hemk1 | HemK methyltransferase family member 1 | 347.74 | 296.04 | 327.03 | 397.16 | 408.98 | 333.04 |
| 9056 | ENSMUSG00000032580 | 630.041617 | 0.105716019 | 0.131567422 | 0.80351213 | 0.421678789 | 0.838959049 | Rbm5 | RNA binding motif protein 5 [Source:MGI | 554.14 | 627.03 | 640.98 | 715.27 | 585.66 | 657.17 |
| 9057 | ENSMUSG00000032582 | 436.7140112 | 0.093655138 | 0.133365276 | 0.702245299 | 0.4825262 | 0.871710696 | Rbm6 | RNA binding motif protein 6 [Source:MGI | 395.13 | 456.4 | 416.59 | 441.5 | 439.52 | 471.16 |
| 9058 | ENSMUSG00000032583 | 393.0127999 | 0.04016076 | 0.132423734 | 0.303274637 | 0.761680573 | 0.954216198 | Mon1a | MON1 homolog A, secretory traffciking as | 393.52 | 394.72 | 374.32 | 400.05 | 410.07 | 385.39 |
| 9059 | ENSMUSG00000032584 | 7.551560498 | 0.126818769 | 0.844511607 | 0.150168177 | 0.880631932 | 0.981448736 | Mst1r | macrophage stimulating 1 receptor (c-met | 6.4248 | 12.335 | 3.0187 | 12.532 | 6.5437 | 4.4554 |
| 9060 | ENSMUSG00000032586 | 3.425510201 | -0.200044455 | 1.12671525 | -0.177546594 | 0.85907907 | NA | Traip | TRAF-interacting protein [Source:MGI Syn | 4.8186 | 3.0838 | 3.0187 | 1.928 | 4.3625 | 3.3415 |
| 9061 | ENSMUSG00000032589 | 1.429055777 | -0.12521544 | 1.898409786 | -0.065958067 | 0.94741121 | NA | Bsn | bassoon [Source:MGI Symbol;Acc:MGI:12 | 2.4093 | 2.0558 | 0 | 1.928 | 2.1812 | 0 |
| 9062 | ENSMUSG00000032590 | 1394.447725 | -0.015276547 | 0.128822444 | -0.118586069 | 0.905603303 | 0.983972587 | Apeh | acylpeptide hydrolase [Source:MGI Symbo | 1220.7 | 1428.8 | 1557.7 | 1506.7 | 1429.8 | 1223 |
| 9063 | ENSMUSG00000032591 | 2142.647749 | 0.347558579 | 0.091192595 | 3.811258783 | 0.000138261 | 0.004661553 | Mst1 | macrophage stimulating 1 (hepatocyte gro | 1874.4 | 1831.7 | 1951.1 | 2359.8 | 2447.3 | 2391.4 |
| 9064 | ENSMUSG00000032593 | 49.00848653 | 0.303233421 | 0.352483142 | 0.860277798 | 0.389635928 | 0.819052248 | Amigo3 | adhesion molecule with Ig like domain 3 [! | 61.839 | 40.089 | 29.181 | 54.947 | 54.531 | 53.465 |
| 9065 | ENSMUSG00000032594 | 1198.667563 | 0.161410388 | 0.103472617 | 1.559933371 | 0.118775627 | 0.504308497 | Ip6k1 | inositol hexaphosphate kinase 1 [Source:I | 1136.4 | 1153.3 | 1105.9 | 1349.6 | 1214.9 | 1231.9 |
| 9066 | ENSMUSG00000032595 | 0.503122293 | -2.366675966 | 3.785596405 | -0.625179156 | 0.531853481 | NA | Cdhr4 | cadherin-related family member 4 [Source | 0 | 0 | 3.0187 | 0 | 0 | 0 |
| 9067 | ENSMUSG00000032596 | 206.7361859 | -0.215911266 | 0.191783052 | -1.125809939 | 0.260245976 | 0.713728596 | Uba7 | ubiquitin-like modifier activating enzyme | 262.61 | 225.11 | 178.11 | 190.87 | 183.22 | 200.49 |
| 9068 | ENSMUSG00000032598 | 82.14355983 | -0.155929272 | 0.277957561 | -0.560982303 | 0.574809602 | 0.906163061 | Nckipsd | NCK interacting protein with SH3 domain | 108.42 | 84.289 | 66.412 | 67.478 | 90.521 | 75.741 |

| | | | | | | | | | | | | | | | |
|---|---|---|---|---|---|---|---|---|---|---|---|---|---|---|---|
| 9069 | ENSMUSG00000032599 | 863.0832638 | -0.697254529 | 0.184407389 | -3.781055271 | 0.000156165 | 0.005168811 | Ip6k2 | inositol hexaphosphate kinase 2 [Source:M | 996.65 | 1234.5 | 972.03 | 608.27 | 873.59 | 493.43 |
| 9070 | ENSMUSG00000032601 | 1570.832818 | -0.08516668 | 0.116485537 | -0.731135227 | 0.464696559 | 0.862709669 | Prkar2a | protein kinase, cAMP dependent regulato | 1431.9 | 1746.4 | 1674.4 | 1494.2 | 1649 | 1429.1 |
| 9071 | ENSMUSG00000032602 | 3121.475129 | -0.272624821 | 0.101549543 | -2.684648438 | 0.007260613 | 0.086363086 | Slc25a20 | solute carrier family 25 (mitochondrial car | 3355.4 | 3509.3 | 3383 | 3021.1 | 2928.3 | 2531.8 |
| 9072 | ENSMUSG00000032604 | 1004.208352 | 0.187185432 | 0.109805726 | 1.704696448 | 0.088251053 | 0.438115648 | Qars | glutaminyl-tRNA synthetase [Source:MGI | 988.62 | 919.99 | 907.63 | 996.75 | 1090.6 | 1121.6 |
| 9073 | ENSMUSG00000032606 | 272.8796491 | -0.134236533 | 0.163832652 | -0.819351523 | 0.412585884 | 0.834752026 | Nicn1 | nicolin 1 [Source:MGI Symbol;Acc:MGI:19 | 281.89 | 282.68 | 291.81 | 239.07 | 237.75 | 304.08 |
| 9074 | ENSMUSG00000032607 | 775.0357978 | 0.064884405 | 0.136862997 | 0.474082889 | 0.6354408 | 0.923478764 | Amt | aminomethyltransferase [Source:MGI Syn | 823.98 | 737.02 | 710.41 | 692.14 | 769.98 | 916.69 |
| 9075 | ENSMUSG00000032609 | 195.566601 | 0.107411217 | 0.17754444 | 0.604982148 | 0.545190858 | 0.895484962 | Klhdc8b | kelch domain containing 8B [Source:MGI S | 180.7 | 188.11 | 196.22 | 196.65 | 227.94 | 183.78 |
| 9076 | ENSMUSG00000032611 | 5.362478218 | 0.865821248 | 1.025069857 | 0.844646092 | 0.398308466 | 0.825882163 | 1700102P08 | RIKEN cDNA 1700102P08 gene [Source:M | 3.2124 | 5.1396 | 3.0187 | 0.964 | 9.8156 | 10.025 |
| 9077 | ENSMUSG00000032612 | 1458.499102 | 0.085292984 | 0.104653551 | 0.815003248 | 0.415070451 | 0.836001897 | Usp4 | ubiquitin specific peptidase 4 (proto-onco | 1386.2 | 1380.5 | 1479.2 | 1388.1 | 1597.8 | 1519.3 |
| 9078 | ENSMUSG00000032615 | 173.1881307 | -0.130345753 | 0.198422125 | -0.656911384 | 0.511237898 | 0.883679955 | Nt5m | 5',3'-nucleotidase, mitochondrial [Source: | 191.94 | 155.22 | 195.21 | 139.78 | 183.22 | 173.76 |
| 9079 | ENSMUSG00000032621 | 438.7390776 | 0.030239994 | 0.142066796 | 0.212857578 | 0.831438052 | 0.970533397 | Srek1 | splicing regulatory glutamine/lysine-rich p | 426.45 | 467.7 | 408.54 | 442.47 | 491.87 | 395.41 |
| 9080 | ENSMUSG00000032623 | 0 | NA | NA | NA | NA | NA | Oas1d | 2'-5' oligoadenylate synthetase 1D [Sourc | 0 | 0 | 0 | 0 | 0 | 0 |
| 9081 | ENSMUSG00000032624 | 294.8950477 | -0.297959948 | 0.158277249 | -1.882519131 | 0.059765565 | 0.348006896 | Eml4 | echinoderm microtubule associated prote | 318.03 | 346.41 | 311.94 | 293.05 | 269.38 | 230.57 |
| 9082 | ENSMUSG00000032625 | 5.41611338 | -0.692082547 | 0.952797623 | -0.726368885 | 0.467612661 | 0.864682885 | Thsd7a | thrombospondin, type I, domain containir | 8.8341 | 8.2233 | 3.0187 | 5.7839 | 2.1812 | 4.4554 |
| 9083 | ENSMUSG00000032633 | 627.8792109 | 0.322710736 | 0.149846542 | 2.153608163 | 0.031270914 | 0.234840339 | Flcn | folliculin [Source:MGI Symbol;Acc:MGI:24 | 600.72 | 628.06 | 444.76 | 729.73 | 648.92 | 715.09 |
| 9084 | ENSMUSG00000032637 | 1010.923483 | 0.233858974 | 0.14319532 | 1.633146772 | 0.102438132 | 0.470781813 | Atxn2l | ataxin 2-like [Source:MGI Symbol;Acc:MG | 938.02 | 901.48 | 948.89 | 1343.8 | 954.29 | 979.07 |
| 9085 | ENSMUSG00000032640 | 55.9896455 | -0.931516566 | 0.311569717 | -2.989753226 | 0.002792029 | 0.044426743 | Chsy1 | chondroitin sulfate synthase 1 [Source:M( | 78.704 | 84.289 | 57.356 | 39.523 | 37.081 | 38.985 |
| 9086 | ENSMUSG00000032641 | 79.85669916 | 0.500639442 | 0.274388892 | 1.82456162 | 0.06806721 | 0.375772565 | Gpr19 | G protein-coupled receptor 19 [Source:M | 69.87 | 62.703 | 65.406 | 68.442 | 105.79 | 106.93 |
| 9087 | ENSMUSG00000032643 | 4.375781296 | -1.053097062 | 1.08713906 | -0.96868662 | 0.33270157 | NA | Fhl3 | four and a half LIM domains 3 [Source:MC | 6.4248 | 8.2233 | 3.0187 | 1.928 | 1.0906 | 5.5692 |
| 9088 | ENSMUSG00000032648 | 8.796031112 | -1.107632858 | 0.760988244 | -1.455519014 | 0.14552563 | 0.556332213 | Pygm | muscle glycogen phosphorylase [Source:M | 13.653 | 12.335 | 10.062 | 6.7478 | 2.1812 | 7.7969 |
| 9089 | ENSMUSG00000032649 | 4.998402766 | 0.197609579 | 0.988691609 | 0.199869784 | 0.841582422 | NA | Colgalt2 | collagen beta(1-O)galactosyltransferase 2 | 4.8186 | 2.0558 | 7.0437 | 2.8919 | 8.7249 | 4.4554 |
| 9090 | ENSMUSG00000032652 | 1257.607496 | 0.091865212 | 0.113420064 | 0.809955565 | 0.417965715 | 0.837436778 | Crebl2 | cAMP responsive element binding protein | 1123.5 | 1268.5 | 1261.8 | 1305.2 | 1400.4 | 1186.2 |
| 9091 | ENSMUSG00000032656 | 28.01375277 | 0.17476959 | 0.42698642 | 0.409309481 | 0.682312559 | 0.93502957 | Marchf3 | membrane associated ring-CH-type finger | 28.109 | 23.642 | 27.169 | 32.775 | 19.631 | 36.757 |
| 9092 | ENSMUSG00000032657 | 13.98692365 | -0.439284903 | 0.610720066 | -0.719290108 | 0.471962189 | 0.867870596 | Fam189b | family with sequence similarity 189, meml | 12.85 | 16.447 | 19.119 | 14.46 | 5.4531 | 15.594 |
| 9093 | ENSMUSG00000032661 | 9.806455844 | -1.608125651 | 0.764176513 | -2.104390312 | 0.035344414 | 0.25220602 | Oas3 | 2'-5' oligoadenylate synthetase 3 [Source: | 12.85 | 18.503 | 13.081 | 6.7478 | 6.5437 | 1.1138 |
| 9094 | ENSMUSG00000032666 | 38.7714731 | -0.94242155 | 0.387318794 | -2.433193444 | 0.014966303 | 0.142229042 | 1700025G04 | RIKEN cDNA 1700025G04 gene [Source:M | 54.611 | 52.424 | 46.287 | 38.559 | 21.812 | 18.935 |
| 9095 | ENSMUSG00000032667 | 876.4461991 | 0.155707812 | 0.111530988 | 1.396094615 | 0.162686005 | 0.586288131 | Pon2 | paraoxonase 2 [Source:MGI Symbol;Acc:M | 841.65 | 815.14 | 830.15 | 877.22 | 895.4 | 999.12 |
| 9096 | ENSMUSG00000032671 | 5.346250148 | 0.059557002 | 1.032170842 | 0.057700721 | 0.953987019 | 0.9915542 | A930018P22 | RIKEN cDNA A930018P22 gene [Source:M | 6.4248 | 5.1396 | 4.025 | 0.964 | 3.2719 | 12.252 |
| 9097 | ENSMUSG00000032673 | 297.8778349 | 0.176075961 | 0.168584619 | 1.044436686 | 0.296283398 | 0.746961229 | Prorsd1 | prolyl-tRNA synthetase domain containing | 316.42 | 267.26 | 254.58 | 305.58 | 279.2 | 364.23 |
| 9098 | ENSMUSG00000032679 | 1627.018506 | 0.002847128 | 0.105113803 | 0.027086145 | 0.978391025 | 0.997797092 | Cd59a | CD59a antigen [Source:MGI Symbol;Acc:M | 1737.9 | 1589.2 | 1547.6 | 1513.4 | 1666.5 | 1707.5 |
| 9099 | ENSMUSG00000032680 | 0.181769715 | 1.126924175 | 4.080472857 | 0.276174898 | 0.782413736 | NA | 6820408C15 | RIKEN cDNA 6820408C15 gene [Source:M | 0 | 0 | 0 | 0 | 1.0906 | 0 |
| 9100 | ENSMUSG00000032688 | 121.3315309 | 0.125984874 | 0.212142134 | 0.59387012 | 0.552598986 | 0.896052884 | Malt1 | MALT1 paracaspase [Source:MGI Symbol; | 104.4 | 115.13 | 128.8 | 119.53 | 128.69 | 131.43 |
| 9101 | ENSMUSG00000032690 | 30.44424214 | -0.812222345 | 0.4225038 | -1.922402462 | 0.054555136 | 0.32878562 | Oas2 | 2'-5' oligoadenylate synthetase 2 [Source: | 39.352 | 39.061 | 38.237 | 31.811 | 15.269 | 18.935 |
| 9102 | ENSMUSG00000032691 | 6.034663666 | -1.950293609 | 0.950290122 | -2.052313881 | 0.040139168 | 0.274113751 | Nlrp3 | NLR family, pyrin domain containing 3 [So | 6.4248 | 9.2513 | 13.081 | 1.928 | 2.1812 | 3.3415 |
| 9103 | ENSMUSG00000032698 | 60.88700443 | -0.272340689 | 0.321773291 | -0.846374441 | 0.397343892 | 0.824796506 | Lmo2 | LIM domain only 2 [Source:MGI Symbol;A | 72.279 | 68.87 | 58.362 | 45.307 | 42.534 | 77.969 |
| 9104 | ENSMUSG00000032702 | 1081.063369 | 0.327573227 | 0.140251564 | 2.335611927 | 0.019511476 | 0.170316231 | Kank1 | KN motif and ankyrin repeat domains 1 [S | 950.87 | 817.19 | 1107.9 | 1046.9 | 1279.3 | 1284.3 |
| 9105 | ENSMUSG00000032705 | 249.5910946 | -0.046406454 | 0.171908186 | -0.269949066 | 0.787199439 | 0.960870081 | Exd2 | exonuclease 3'-5' domain containing 2 [Sc | 236.11 | 242.59 | 282.75 | 272.81 | 212.67 | 250.61 |
| 9106 | ENSMUSG00000032712 | 456.0951687 | 0.492174616 | 0.243511768 | 2.021153307 | 0.043263898 | 0.286918392 | Resf1 | retroelement silencing factor 1 [Source:M | 334.09 | 460.51 | 343.13 | 643.94 | 297.74 | 657.17 |
| 9107 | ENSMUSG00000032714 | 38.80968419 | -0.885801498 | 0.378872785 | -2.337991888 | 0.019387669 | 0.169440027 | Syde1 | synapse defective 1, Rho GTPase, homolo | 50.595 | 44.2 | 56.35 | 30.847 | 17.45 | 33.415 |
| 9108 | ENSMUSG00000032715 | 159.1842429 | 0.258083656 | 0.324720001 | 0.794788295 | 0.426736661 | 0.842062628 | Trib3 | tribbles pseudokinase 3 [Source:MGI Sym | 165.44 | 152.13 | 116.72 | 90.614 | 145.05 | 285.14 |
| 9109 | ENSMUSG00000032717 | 7.59034836 | -0.162112132 | 0.867773605 | -0.186813854 | 0.851806584 | 0.973633287 | Mdfi | MyoD family inhibitor [Source:MGI Symbo | 8.8341 | 9.2513 | 6.0375 | 12.532 | 1.0906 | 7.7969 |
| 9110 | ENSMUSG00000032718 | 14.60667363 | 0.270836732 | 0.602486775 | 0.449531414 | 0.653048352 | 0.925756399 | Mansc1 | MANSC domain containing 1 [Source:MGI | 19.274 | 6.1675 | 14.087 | 18.316 | 13.087 | 16.708 |
| 9111 | ENSMUSG00000032719 | 1.284755329 | -0.613014061 | 2.01101397 | -0.304828346 | 0.760496892 | NA | Sbspon | somatomedin B and thrombospondin, typ | 1.6062 | 3.0838 | 0 | 1.928 | 1.0906 | 0 |
| 9112 | ENSMUSG00000032724 | 213.8158166 | -0.579451616 | 0.234106873 | -2.475158502 | 0.013317706 | 0.131756016 | Abtb2 | ankyrin repeat and BTB (POZ) domain con | 166.24 | 292.96 | 309.92 | 154.24 | 167.96 | 191.58 |
| 9113 | ENSMUSG00000032725 | 17.79442454 | -1.161949318 | 0.558594823 | -2.080129049 | 0.037513698 | 0.2626322 | Folr2 | folate receptor 2 (fetal) [Source:MGI Syml | 20.881 | 24.67 | 28.175 | 8.6758 | 6.5437 | 17.822 |
| 9114 | ENSMUSG00000032726 | 0.267700746 | -1.497481649 | 4.057001436 | -0.369110456 | 0.712045396 | NA | Bmp8a | bone morphogenetic protein 8a [Source:M | 1.6062 | 0 | 0 | 0 | 0 | 0 |
| 9115 | ENSMUSG00000032727 | 257.5746627 | -0.048972145 | 0.175328914 | -0.279315856 | 0.780002439 | 0.959870443 | Mier3 | MIER family member 3 [Source:MGI Symb | 242.54 | 273.43 | 270.68 | 292.09 | 208.31 | 258.41 |
| 9116 | ENSMUSG00000032733 | 311.8315898 | -0.044486865 | 0.176147505 | -0.252554616 | 0.800612405 | 0.962424862 | Snx33 | sorting nexin 33 [Source:MGI Symbol;Acc: | 280.28 | 296.04 | 374.32 | 335.46 | 256.3 | 328.58 |
| 9117 | ENSMUSG00000032735 | 458.3369418 | -0.08849225 | 0.154439142 | -0.572991078 | 0.566650735 | 0.902528221 | Ablim3 | actin binding LIM protein family, member | 392.72 | 472.84 | 552.43 | 428.97 | 430.79 | 472.27 |
| 9118 | ENSMUSG00000032737 | 686.0142474 | 0.034348429 | 0.119141319 | 0.28829989 | 0.773117192 | 0.958050639 | Inppl1 | inositol polyphosphate phosphatase-like 1 | 703.52 | 650.67 | 679.22 | 730.7 | 638.01 | 713.97 |
| 9119 | ENSMUSG00000032739 | 0.342432689 | 1.990653086 | 3.900281263 | 0.51038706 | 0.609780321 | NA | Pram1 | PML-RAR alpha-regulated adaptor molecu | 0 | 0 | 0 | 0.964 | 1.0906 | 0 |
| 9120 | ENSMUSG00000032740 | 18.29795806 | -0.197387198 | 0.498633225 | -0.39585649 | 0.6922109 | 0.938256814 | Ccdc88a | coiled coil domain containing 88A [Source | 20.078 | 18.503 | 20.125 | 20.244 | 16.359 | 14.48 |
| 9121 | ENSMUSG00000032741 | 1328.932615 | -0.057980145 | 0.102498487 | -0.565668299 | 0.571619289 | 0.904651508 | Tpcn1 | two pore channel 1 [Source:MGI Symbol;A | 1368.5 | 1281.8 | 1416.8 | 1352.5 | 1307.7 | 1246.4 |
| 9122 | ENSMUSG00000032743 | 321.7331604 | 0.003403107 | 0.146792465 | 0.023183117 | 0.981504206 | 0.998323671 | Katnip | katanin interacting protein [Source:MGI S | 318.03 | 308.38 | 338.1 | 349.92 | 313.01 | 302.97 |
| 9123 | ENSMUSG00000032744 | 16.21871293 | -0.86520055 | 0.567673245 | -1.524117188 | 0.12747944 | 0.523285632 | Heyl | hairy/enhancer-of-split related with YRPW | 24.093 | 23.642 | 15.094 | 12.532 | 15.269 | 6.6831 |
| 9124 | ENSMUSG00000032745 | 612.687782 | 0.163264914 | 0.117995024 | 1.383659317 | 0.166462796 | 0.5925379 | Gpbp1 | GC-rich promoter binding protein 1 [Sourc | 578.23 | 602.36 | 553.43 | 632.37 | 634.74 | 674.99 |

| | | | | | | | | | | | | | | | |
|---|---|---|---|---|---|---|---|---|---|---|---|---|---|---|---|
| 9125 | ENSMUSG00000032750 | 1.51140123 | -3.968382626 | 2.069908584 | -1.917177723 | 0.055215357 | NA | Gab3 | growth factor receptor bound protein 2-a | 4.0155 | 1.0279 | 4.025 | 0 | 0 | 0 |
| 9126 | ENSMUSG00000032754 | 1394.230201 | 0.215638262 | 0.134126133 | 1.60772742 | 0.10789489 | 0.48458257 | Slc8b1 | solute carrier family 8 (sodium/lithium/ca | 1361.3 | 1199.6 | 1309.1 | 1483.6 | 1254.2 | 1757.6 |
| 9127 | ENSMUSG00000032757 | 987.8721168 | -0.03066012 | 0.120491371 | -0.25445905 | 0.799140936 | 0.962424862 | Bet1 | Bet1 golgi vesicular membrane trafficking | 1129.2 | 921.01 | 943.86 | 1019.9 | 954.29 | 959.02 |
| 9128 | ENSMUSG00000032758 | 0 | NA | NA | NA | NA | NA | Kap | kidney androgen regulated protein [Sourc | 0 | 0 | 0 | 0 | 0 | 0 |
| 9129 | ENSMUSG00000032763 | 474.705808 | 0.024646124 | 0.141909672 | 0.173674732 | 0.862121096 | 0.976334255 | Ilvbl | ilvB (bacterial acetolactate synthase)-like | 431.27 | 462.56 | 519.22 | 525.37 | 479.87 | 429.94 |
| 9130 | ENSMUSG00000032766 | 59.75556411 | -0.878726377 | 0.332766718 | -2.640667857 | 0.008274279 | 0.094235571 | Gng11 | guanine nucleotide binding protein (G prc | 68.264 | 100.74 | 63.393 | 44.343 | 29.447 | 52.351 |
| 9131 | ENSMUSG00000032769 | 0 | NA | NA | NA | NA | NA | Trpa1 | transient receptor potential cation chann | 0 | 0 | 0 | 0 | 0 | 0 |
| 9132 | ENSMUSG00000032773 | 6.449515903 | 0.339523937 | 0.976635884 | 0.347646388 | 0.728105764 | 0.946158959 | Chrm1 | cholinergic receptor, muscarinic 1, CNS [S | 8.8341 | 1.0279 | 7.0437 | 1.928 | 8.7249 | 11.138 |
| 9133 | ENSMUSG00000032776 | 244.109195 | 0.237278639 | 0.215000874 | 1.10361709 | 0.269759275 | 0.725011151 | Mctp2 | multiple C2 domains, transmembrane 2 [S | 223.26 | 291.93 | 156.97 | 238.1 | 275.93 | 278.46 |
| 9134 | ENSMUSG00000032777 | 1019.113093 | -0.197880545 | 0.11104749 | -1.78194522 | 0.074758165 | 0.399358478 | Gtf3c1 | general transcription factor III C 1 [Source | 1061.7 | 1060.8 | 1145.1 | 1028.6 | 908.49 | 910.01 |
| 9135 | ENSMUSG00000032782 | 157.4298282 | 0.136894134 | 0.1858983 | 0.736392608 | 0.461491802 | 0.860071824 | Cntrob | centrobin, centrosomal BRCA2 interacting | 155.8 | 143.91 | 149.93 | 161.95 | 159.23 | 173.76 |
| 9136 | ENSMUSG00000032783 | 0.342639279 | -1.805976596 | 4.028547714 | -0.448294702 | 0.653940524 | NA | Troap | trophinin associated protein [Source:MGI | 0 | 2.0558 | 0 | 0 | 0 | 0 |
| 9137 | ENSMUSG00000032786 | 10383.05473 | 0.052124788 | 0.451937893 | 0.115336175 | 0.908178666 | 0.984699867 | Alas1 | aminolevulinic acid synthase 1 [Source:M | 9200.3 | 13923 | 7463.3 | 14934 | 10786 | 5991.4 |
| 9138 | ENSMUSG00000032788 | 845.7466948 | -0.029740183 | 0.166989066 | -0.178096589 | 0.858647121 | 0.975439379 | Pdxk | pyridoxal (pyridoxine, vitamin B6) kinase [ | 793.47 | 819.25 | 950.9 | 776 | 1069.9 | 664.96 |
| 9139 | ENSMUSG00000032796 | 53.14651197 | -1.496539799 | 0.332983159 | -4.494340808 | 6.98E-06 | 0.000404554 | Lama1 | laminin, alpha 1 [Source:MGI Symbol;Acc: | 89.947 | 82.233 | 63.393 | 33.739 | 26.175 | 23.391 |
| 9140 | ENSMUSG00000032802 | 709.1902688 | -0.206861137 | 0.147610593 | -1.401397645 | 0.161095196 | 0.584240557 | Srxn1 | sulfiredoxin 1 homolog (S. cerevisiae) [Sou | 900.28 | 684.59 | 693.3 | 601.52 | 634.74 | 740.71 |
| 9141 | ENSMUSG00000032803 | 2585.083174 | 0.11903001 | 0.091819592 | 1.29634653 | 0.194856125 | 0.634631612 | Cdv3 | carnitine deficiency-associated gene expr | 2523.3 | 2549.2 | 2362.7 | 2678.9 | 2782.2 | 2614.2 |
| 9142 | ENSMUSG00000032806 | 73.13436065 | -0.374679903 | 0.313296818 | -1.195926296 | 0.231725323 | 0.67889946 | Slc10a3 | solute carrier family 10 (sodium/bile acid | 110.83 | 83.261 | 53.331 | 72.298 | 56.712 | 62.375 |
| 9143 | ENSMUSG00000032807 | 0 | NA | NA | NA | NA | NA | Alox12b | arachidonate 12-lipoxygenase, 12R type [S | 0 | 0 | 0 | 0 | 0 | 0 |
| 9144 | ENSMUSG00000032808 | 949.8943213 | -1.133556355 | 0.542901197 | -2.087960686 | 0.036801381 | 0.259024417 | Cyp2c38 | cytochrome P450, family 2, subfamily c, p | 1500.2 | 1340.4 | 1074.7 | 1057.5 | 400.26 | 326.36 |
| 9145 | ENSMUSG00000032812 | 1140.352583 | 0.371773976 | 0.130863196 | 2.840936088 | 0.004498133 | 0.062259321 | Arap1 | ArfGAP with RhoGAP domain, ankyrin rep | 958.9 | 946.71 | 1076.7 | 1133.6 | 1226.9 | 1499.2 |
| 9146 | ENSMUSG00000032815 | 28.6844688 | -0.337427096 | 0.544948558 | -0.619190732 | 0.535790717 | 0.890255822 | Fanca | Fanconi anemia, complementation group | 28.109 | 39.061 | 29.181 | 38.559 | 31.628 | 5.5692 |
| 9147 | ENSMUSG00000032816 | 1.823537464 | 0.992305125 | 1.523144579 | 0.651484526 | 0.514733762 | NA | Igdcc4 | immunoglobulin superfamily, DCC subclas | 1.6062 | 1.0279 | 1.0062 | 2.8919 | 2.1812 | 2.2277 |
| 9148 | ENSMUSG00000032818 | 0 | NA | NA | NA | NA | NA | Loxhd1 | lipoxygenase homology domains 1 [Sourc | 0 | 0 | 0 | 0 | 0 | 0 |
| 9149 | ENSMUSG00000032826 | 27.60217811 | 0.34971983 | 0.420835667 | 0.83101281 | 0.405966393 | 0.830012328 | Ank2 | ankyrin 2, brain [Source:MGI Symbol;Acc: | 24.896 | 27.754 | 20.125 | 29.883 | 25.084 | 37.871 |
| 9150 | ENSMUSG00000032827 | 247.7278867 | 0.24096472 | 0.274423699 | 0.878075476 | 0.379902761 | 0.812628154 | Ppp1r9a | protein phosphatase 1, regulatory subunit | 293.13 | 128.49 | 258.6 | 187.01 | 321.73 | 297.4 |
| 9151 | ENSMUSG00000032834 | 245.7669767 | 0.094788112 | 0.15587526 | 0.60810235 | 0.543119594 | 0.894344786 | Pwp2 | PWP2 periodic tryptophan protein homol | 227.28 | 244.64 | 241.5 | 255.45 | 259.57 | 246.16 |
| 9152 | ENSMUSG00000032839 | 2.718199486 | -0.338614449 | 1.353029183 | -0.250263965 | 0.802383221 | NA | Trpc1 | transient receptor potential cation chann | 4.0155 | 3.0838 | 2.0125 | 3.8559 | 0 | 3.3415 |
| 9153 | ENSMUSG00000032840 | 197.3540798 | 0.020904591 | 0.196807651 | 0.106218386 | 0.915409084 | 0.98625062 | Spring1 | SREBF pathway regulator in golgi 1 [Sourc | 179.09 | 205.58 | 203.26 | 190.87 | 162.5 | 242.82 |
| 9154 | ENSMUSG00000032841 | 12.25847254 | -0.037805951 | 0.632702682 | -0.059753107 | 0.952352274 | 0.991279741 | Prr5l | proline rich 5 like [Source:MGI Symbol;Ac | 12.85 | 16.447 | 8.05 | 16.388 | 10.906 | 8.9108 |
| 9155 | ENSMUSG00000032842 | 257.6351685 | -0.027591219 | 0.210008659 | -0.131381341 | 0.895473648 | 0.982798755 | Abcc10 | ATP-binding cassette, sub-family C (CFTR/ | 236.92 | 235.39 | 308.92 | 329.68 | 237.75 | 197.15 |
| 9156 | ENSMUSG00000032845 | 0 | NA | NA | NA | NA | NA | Alpk2 | alpha-kinase 2 [Source:MGI Symbol;Acc:M | 0 | 0 | 0 | 0 | 0 | 0 |
| 9157 | ENSMUSG00000032846 | 41.72462053 | -0.252264743 | 0.346478085 | -0.728082826 | 0.466562886 | 0.863779479 | Zswim6 | zinc finger SWIM-type containing 6 [Sourc | 43.368 | 52.424 | 40.25 | 33.739 | 34.9 | 45.668 |
| 9158 | ENSMUSG00000032849 | 19.95697966 | -0.353424785 | 0.497979995 | -0.709716833 | 0.477879751 | 0.869794311 | Abcc4 | ATP-binding cassette, sub-family C (CFTR/ | 23.29 | 28.782 | 15.094 | 17.352 | 19.631 | 15.594 |
| 9159 | ENSMUSG00000032850 | 0.469265235 | -2.274672822 | 3.439794491 | -0.661281605 | 0.508431736 | NA | Rnft2 | ring finger protein, transmembrane 2 [Sou | 0.8031 | 0 | 2.0125 | 0 | 0 | 0 |
| 9160 | ENSMUSG00000032852 | 0 | NA | NA | NA | NA | NA | Rspo4 | R-spondin 4 [Source:MGI Symbol;Acc:MG | 0 | 0 | 0 | 0 | 0 | 0 |
| 9161 | ENSMUSG00000032854 | 1.124144488 | 0.633232731 | 1.971100216 | 0.321258517 | 0.74801449 | NA | Ugt8a | UDP galactosyltransferase 8A [Source:MG | 1.6062 | 0 | 1.0062 | 1.928 | 1.0906 | 1.1138 |
| 9162 | ENSMUSG00000032855 | 440.773873 | -0.031088046 | 0.157444646 | -0.197453817 | 0.843472419 | 0.972342797 | Pkd1 | polycystin 1, transient receptor poteintial | 372.64 | 497.51 | 467.9 | 484.88 | 432.98 | 388.73 |
| 9163 | ENSMUSG00000032860 | 421.6730336 | 0.280943064 | 0.151243413 | 1.857555696 | 0.063232135 | 0.359877642 | P2ry2 | purinergic receptor P2Y, G-protein couple | 438.49 | 335.1 | 367.28 | 439.57 | 442.79 | 506.8 |
| 9164 | ENSMUSG00000032864 | 0.36353943 | 2.065989254 | 4.034137301 | 0.512126658 | 0.608562368 | NA | Rag2 | recombination activating gene 2 [Source:I | 0 | 0 | 0 | 0 | 2.1812 | 0 |
| 9165 | ENSMUSG00000032867 | 731.6669642 | 0.106426398 | 0.119881455 | 0.887763649 | 0.3746679 | 0.809332934 | Fbxw8 | F-box and WD-40 domain protein 8 [Sourc | 715.56 | 693.84 | 704.37 | 733.59 | 838.69 | 703.95 |
| 9166 | ENSMUSG00000032869 | 651.2885545 | -0.007253786 | 0.131548217 | -0.055141653 | 0.956025612 | 0.991678478 | Psmf1 | proteasome (prosome, macropain) inhibit | 735.64 | 578.72 | 642.99 | 619.84 | 651.1 | 679.44 |
| 9167 | ENSMUSG00000032870 | 945.710427 | -0.030134108 | 0.119265279 | -0.252664547 | 0.800527446 | 0.962424862 | Smap2 | small ArfGAP 2 [Source:MGI Symbol;Acc:N | 990.23 | 970.35 | 905.62 | 922.53 | 846.32 | 1039.2 |
| 9168 | ENSMUSG00000032872 | 543.0263024 | 0.141924239 | 0.13792787 | 1.028974342 | 0.303491732 | 0.752690022 | Cyb5r4 | cytochrome b5 reductase 4 [Source:MGI S | 465 | 551.99 | 532.3 | 506.09 | 583.48 | 619.3 |
| 9169 | ENSMUSG00000032875 | 45.48234611 | -1.035140411 | 0.36213 | -2.858477371 | 0.004256795 | 0.060010018 | Arhgef17 | Rho guanine nucleotide exchange factor ( | 57.823 | 74.01 | 51.318 | 20.244 | 31.628 | 37.871 |
| 9170 | ENSMUSG00000032878 | 0 | NA | NA | NA | NA | NA | Ccdc85a | coiled-coil domain containing 85A [Source | 0 | 0 | 0 | 0 | 0 | 0 |
| 9171 | ENSMUSG00000032883 | 277.8173022 | 0.276782562 | 0.16662278 | 1.661132777 | 0.096686783 | 0.458530879 | Acsl3 | acyl-CoA synthetase long-chain family me | 213.63 | 273.43 | 267.66 | 306.54 | 327.19 | 278.46 |
| 9172 | ENSMUSG00000032889 | 0 | NA | NA | NA | NA | NA | Gm6685 | predicted pseudogene 6685 [Source:MGI | 0 | 0 | 0 | 0 | 0 | 0 |
| 9173 | ENSMUSG00000032890 | 21.75890182 | -1.171339765 | 0.47889716 | -2.445910862 | 0.014448672 | 0.13941667 | Rims3 | regulating synaptic membrane exocytosis | 27.305 | 33.921 | 29.181 | 12.532 | 10.906 | 16.708 |
| 9174 | ENSMUSG00000032892 | 0 | NA | NA | NA | NA | NA | Rangrf | RAN guanine nucleotide release factor [Sc | 0 | 0 | 0 | 0 | 0 | 0 |
| 9175 | ENSMUSG00000032894 | 0 | NA | NA | NA | NA | NA | 1700031F05 | RIKEN cDNA 1700031F05 gene [Source:M | 0 | 0 | 0 | 0 | 0 | 0 |
| 9176 | ENSMUSG00000032897 | 372.0461942 | -0.11692374 | 0.147915913 | -0.790474385 | 0.429250776 | 0.842978372 | Nfyc | nuclear transcription factor-Y gamma [Sou | 399.14 | 358.74 | 403.5 | 374.02 | 384.99 | 311.88 |
| 9177 | ENSMUSG00000032898 | 1611.592246 | 0.548711082 | 0.155999849 | 3.517382125 | 0.000435826 | 0.01125676 | Fbxo21 | F-box protein 21 [Source:MGI Symbol;Acc | 1060.9 | 1402.1 | 1464.1 | 1616.6 | 1820.2 | 2305.7 |
| 9178 | ENSMUSG00000032899 | 0.321325949 | 1.911575114 | 4.048267893 | 0.472195804 | 0.636787034 | NA | Styk1 | serine/threonine/tyrosine kinase 1 [Sourc | 0 | 0 | 0 | 1.928 | 0 | 0 |
| 9179 | ENSMUSG00000032900 | 0.160662974 | 1.126924175 | 4.080472857 | 0.276174898 | 0.782413736 | NA | Tex43 | testis expressed 43 [Source:MGI Symbol;A | 0 | 0 | 0 | 0.964 | 0 | 0 |
| 9180 | ENSMUSG00000032902 | 869.2517805 | -0.27513571 | 0.117710944 | -2.337384279 | 0.019419212 | 0.169613212 | Slc16a1 | solute carrier family 16 (monocarboxylic a | 860.93 | 1011.5 | 984.11 | 778.89 | 752.53 | 827.59 |

| | | | | | | | | | | | | | | | |
|---|---|---|---|---|---|---|---|---|---|---|---|---|---|---|---|
| 9181 | ENSMUSG00000032905 | 616.1715424 | -0.020362156 | 0.118752409 | -0.171467309 | 0.863856333 | 0.976954351 | Atg12 | autophagy related 12 [Source:MGI Symbo | 640.88 | 616.75 | 603.75 | 630.44 | 573.67 | 631.55 |
| 9182 | ENSMUSG00000032908 | 2.177758824 | 0.932080269 | 1.537333436 | 0.606296752 | 0.544317715 | NA | Sgpp2 | sphingosine-1-phosphate phosphatase 2 [ | 2.4093 | 0 | 2.0125 | 0.964 | 5.4531 | 2.2277 |
| 9183 | ENSMUSG00000032911 | 0.514011053 | 1.044022102 | 3.082659575 | 0.338675769 | 0.734853997 | NA | Cspg4 | chondroitin sulfate proteoglycan 4 [Sourc | 0 | 0 | 1.0062 | 0.964 | 0 | 1.1138 |
| 9184 | ENSMUSG00000032913 | 134.701928 | -0.046823333 | 0.207349566 | -0.22581833 | 0.82134272 | 0.969160361 | Lrig2 | leucine-rich repeats and immunoglobulin- | 117.25 | 143.91 | 149.93 | 129.17 | 127.6 | 140.34 |
| 9185 | ENSMUSG00000032915 | 42.75560435 | -1.481606204 | 0.507079181 | -2.921843882 | 0.003479659 | 0.052101231 | Adgre4 | adhesion G protein-coupled receptor E4 [ | 42.564 | 109.99 | 36.225 | 10.604 | 35.99 | 21.163 |
| 9186 | ENSMUSG00000032921 | 0 | NA | NA | NA | NA | NA | Odf4 | outer dense fiber of sperm tails 4 [Source: | 0 | 0 | 0 | 0 | 0 | 0 |
| 9187 | ENSMUSG00000032925 | 5.874927414 | 0.288347423 | 1.058362906 | 0.272446645 | 0.785278611 | 0.960670212 | Itgbl1 | integrin, beta-like 1 [Source:MGI Symbol;/ | 9.6372 | 6.1675 | 0 | 10.604 | 3.2719 | 5.5692 |
| 9188 | ENSMUSG00000032932 | 480.3864192 | 0.158074343 | 0.136307902 | 1.159685833 | 0.24617674 | 0.696492381 | Hspa13 | heat shock protein 70 family, member 13 | 457.77 | 447.14 | 457.84 | 566.82 | 451.52 | 501.23 |
| 9189 | ENSMUSG00000032936 | 0.133850373 | -0.796629176 | 4.080472857 | -0.195229623 | 0.84521319 | NA | Camkv | CaM kinase-like vesicle-associated [Sourc | 0.8031 | 0 | 0 | 0 | 0 | 0 |
| 9190 | ENSMUSG00000032937 | 0 | NA | NA | NA | NA | NA | Fshr | follicle stimulating hormone receptor [Sou | 0 | 0 | 0 | 0 | 0 | 0 |
| 9191 | ENSMUSG00000032939 | 94.56399521 | -0.254708922 | 0.274445774 | -0.928084694 | 0.353363638 | 0.793888112 | Nup93 | nucleoporin 93 [Source:MGI Symbol;Acc:I | 115.65 | 125.41 | 67.418 | 84.83 | 89.431 | 84.652 |
| 9192 | ENSMUSG00000032940 | 0 | NA | NA | NA | NA | NA | Rbm11 | RNA binding motif protein 11 [Source:MG | 0 | 0 | 0 | 0 | 0 | 0 |
| 9193 | ENSMUSG00000032942 | 0.898940922 | -0.606261109 | 2.77650979 | -0.218353672 | 0.827153559 | NA | Ucp3 | uncoupling protein 3 (mitochondrial, prot | 3.2124 | 0 | 0 | 0 | 2.1812 | 0 |
| 9194 | ENSMUSG00000032946 | 312.7224193 | -0.376866535 | 0.202133915 | -1.864439898 | 0.062259954 | 0.356503555 | Rasgrp2 | RAS, guanyl releasing protein 2 [Source:M | 452.95 | 259.04 | 347.15 | 276.66 | 298.83 | 241.7 |
| 9195 | ENSMUSG00000032948 | 0 | NA | NA | NA | NA | NA | Lipi | lipase, member I [Source:MGI Symbol;Acc | 0 | 0 | 0 | 0 | 0 | 0 |
| 9196 | ENSMUSG00000032952 | 36.16467707 | -0.206974232 | 0.360810846 | -0.573636393 | 0.56621388 | 0.902076435 | Ap4b1 | adaptor-related protein complex AP-4, be | 40.958 | 39.061 | 36.225 | 34.703 | 37.081 | 28.96 |
| 9197 | ENSMUSG00000032959 | 3763.055007 | -0.136763774 | 0.088648821 | -1.542759078 | 0.122889242 | 0.513267141 | Pebp1 | phosphatidylethanolamine binding protei | 4139.2 | 3770.4 | 3913.3 | 3585 | 3528.2 | 3642.3 |
| 9198 | ENSMUSG00000032965 | 11.61517507 | -0.619321795 | 0.653241285 | -0.948075097 | 0.343091225 | 0.784828638 | Ift57 | intraflagellar transport 57 [Source:MGI Sy | 13.653 | 17.475 | 11.069 | 8.6758 | 5.4531 | 13.366 |
| 9199 | ENSMUSG00000032966 | 717.732896 | -0.165953626 | 0.122993126 | -1.349291875 | 0.177243234 | 0.608561375 | Fkbp1a | FK506 binding protein 1a [Source:MGI Syr | 782.22 | 801.78 | 692.3 | 643.94 | 654.37 | 731.8 |
| 9200 | ENSMUSG00000032968 | 1.934478312 | -1.529593741 | 1.722984031 | -0.887758513 | 0.374670664 | NA | Inha | inhibin alpha [Source:MGI Symbol;Acc:MO | 1.6062 | 3.0838 | 4.025 | 2.8919 | 0 | 0 |
| 9201 | ENSMUSG00000032977 | 481.5450286 | 0.110626168 | 0.149488198 | 0.740032784 | 0.459280102 | 0.859442977 | Fam207a | family with sequence similarity 207, meml | 432.07 | 416.31 | 541.36 | 516.69 | 450.43 | 532.42 |
| 9202 | ENSMUSG00000032978 | 0 | NA | NA | NA | NA | NA | Guca2b | guanylate cyclase activator 2b (retina) [So | 0 | 0 | 0 | 0 | 0 | 0 |
| 9203 | ENSMUSG00000032985 | 12.51727751 | 0.225080805 | 0.650917871 | 0.345789869 | 0.729500639 | 0.946534338 | 5730522E02 | RIKEN cDNA 5730522E02 gene [Source:M | 10.44 | 6.1675 | 18.112 | 18.316 | 9.8156 | 12.252 |
| 9204 | ENSMUSG00000032987 | 0 | NA | NA | NA | NA | NA | Olfr281 | olfactory receptor 281 [Source:MGI Symb | 0 | 0 | 0 | 0 | 0 | 0 |
| 9205 | ENSMUSG00000032988 | 0.455176322 | 0.955156269 | 3.546674842 | 0.269310357 | 0.787690862 | NA | Slc16a8 | solute carrier family 16 (monocarboxylic a | 0.8031 | 0 | 0 | 1.928 | 0 | 0 |
| 9206 | ENSMUSG00000032997 | 61.0650163 | -0.570940125 | 0.291518727 | -1.95850239 | 0.050171091 | 0.314553384 | Chpf | chondroitin polymerizing factor [Source:N | 78.704 | 69.898 | 70.437 | 56.875 | 40.353 | 50.123 |
| 9207 | ENSMUSG00000032998 | 256.2501092 | -0.016982672 | 0.174017541 | -0.097591723 | 0.922256497 | 0.986685804 | Foxj3 | forkhead box J3 [Source:MGI Symbol;Acc: | 254.58 | 289.87 | 229.42 | 293.05 | 236.66 | 233.91 |
| 9208 | ENSMUSG00000032999 | 0.167707431 | -0.796629176 | 4.080472857 | -0.195229623 | 0.84521319 | NA | Nlrp4f | NLR family, pyrin domain containing 4F [S | 0 | 0 | 1.0062 | 0 | 0 | 0 |
| 9209 | ENSMUSG00000033004 | 543.4263554 | -0.242087908 | 0.143607044 | -1.685766253 | 0.09184083 | 0.446623824 | Mycbp2 | MYC binding protein 2, E3 ubiquitin prote | 547.72 | 682.54 | 537.33 | 520.55 | 521.32 | 451.11 |
| 9210 | ENSMUSG00000033006 | 0 | NA | NA | NA | NA | NA | Sox10 | SRY (sex determining region Y)-box 10 [So | 0 | 0 | 0 | 0 | 0 | 0 |
| 9211 | ENSMUSG00000033007 | 0 | NA | NA | NA | NA | NA | Asic4 | acid-sensing (proton-gated) ion channel fa | 0 | 0 | 0 | 0 | 0 | 0 |
| 9212 | ENSMUSG00000033009 | 287.1097995 | -0.025703551 | 0.170862208 | -0.150434386 | 0.880421913 | 0.981448736 | Ogfod1 | 2-oxoglutarate and iron-dependent oxyge | 252.17 | 277.54 | 340.11 | 297.87 | 301.01 | 253.96 |
| 9213 | ENSMUSG00000033014 | 336.7981286 | -0.159114734 | 0.158215514 | -1.005683518 | 0.314567834 | 0.761014653 | Trim33 | tripartite motif-containing 33 [Source:MG | 329.27 | 390.61 | 347.15 | 354.74 | 325 | 274.01 |
| 9214 | ENSMUSG00000033015 | 0 | NA | NA | NA | NA | NA | 4933427E13 | RIKEN cDNA 4933427E13 gene [Source:M | 0 | 0 | 0 | 0 | 0 | 0 |
| 9215 | ENSMUSG00000033016 | 221.7026627 | 0.211672071 | 0.168348174 | 1.257347001 | 0.208628011 | 0.653455953 | Nfatc1 | nuclear factor of activated T cells, cytopla | 205.59 | 192.22 | 218.36 | 226.53 | 225.76 | 261.75 |
| 9216 | ENSMUSG00000033020 | 442.467192 | -0.002541081 | 0.150243077 | -0.016913133 | 0.986505916 | 0.99859176 | Polr2f | polymerase (RNA) II (DNA directed) polyp | 498.73 | 395.75 | 432.69 | 389.45 | 454.79 | 483.41 |
| 9217 | ENSMUSG00000033021 | 447.4410373 | 0.091024686 | 0.141536424 | 0.643118449 | 0.520147244 | 0.887802551 | Gmppa | GDP-mannose pyrophosphorylase A [Sour | 419.22 | 439.95 | 441.74 | 523.44 | 451.52 | 408.78 |
| 9218 | ENSMUSG00000033022 | 27575.01481 | -0.37868998 | 0.131384977 | -2.882292859 | 0.003947926 | 0.056705863 | Cdo1 | cysteine dioxygenase 1, cytosolic [Source: | 28048 | 35382 | 30090 | 20266 | 23650 | 28014 |
| 9219 | ENSMUSG00000033024 | 0 | NA | NA | NA | NA | NA | Klra9 | killer cell lectin-like receptor subfamily A, | 0 | 0 | 0 | 0 | 0 | 0 |
| 9220 | ENSMUSG00000033027 | 0 | NA | NA | NA | NA | NA | Klrc3 | killer cell lectin-like receptor subfamily C, | 0 | 0 | 0 | 0 | 0 | 0 |
| 9221 | ENSMUSG00000033029 | 3.404408974 | -0.304786079 | 1.241178549 | -0.245561833 | 0.806021438 | NA | 1700088E04 | RIKEN cDNA 1700088E04 gene [Source:M | 3.2124 | 5.1396 | 3.0187 | 5.7839 | 3.2719 | 0 |
| 9222 | ENSMUSG00000033031 | 21.282417 | -0.090365703 | 0.539550408 | -0.167483337 | 0.866989757 | 0.977930657 | Cip2a | cell proliferation regulating inhibitor of pr | 36.943 | 17.475 | 11.069 | 19.28 | 23.994 | 18.935 |
| 9223 | ENSMUSG00000033032 | 46.1280452 | -0.333299507 | 0.343369134 | -0.970674048 | 0.331710619 | 0.774747658 | Afap1l1 | actin filament associated protein 1-like 1 [ | 44.974 | 43.173 | 66.412 | 47.235 | 35.99 | 38.985 |
| 9224 | ENSMUSG00000033033 | 18.43861499 | -1.033320831 | 0.533490352 | -1.936906313 | 0.052756797 | 0.32296796 | Calhm2 | calcium homeostasis modulator family me | 19.274 | 30.838 | 24.15 | 7.7118 | 14.178 | 14.48 |
| 9225 | ENSMUSG00000033036 | 0.315620088 | 0.165143421 | 4.063300985 | 0.040642675 | 0.967580763 | NA | Gm7879 | predicted pseudogene 7879 [Source:MGI | 0.8031 | 0 | 0 | 0 | 1.0906 | 0 |
| 9226 | ENSMUSG00000033039 | 176.3111045 | 0.223846656 | 0.203933624 | 1.09764467 | 0.272359679 | 0.727618887 | Micall1 | microtubule associated monooxygenase, | 170.26 | 138.77 | 179.11 | 220.75 | 193.04 | 155.94 |
| 9227 | ENSMUSG00000033044 | 0 | NA | NA | NA | NA | NA | Dhrs7c | dehydrogenase/reductase (SDR family) m | 0 | 0 | 0 | 0 | 0 | 0 |
| 9228 | ENSMUSG00000033047 | 1369.861776 | 0.017773023 | 0.098101756 | 0.181169266 | 0.856234718 | 0.974319798 | Eif3l | eukaryotic translation initiation factor 3, s | 1318.7 | 1369.2 | 1396.7 | 1344.7 | 1426.5 | 1363.3 |
| 9229 | ENSMUSG00000033053 | 0 | NA | NA | NA | NA | NA | 1700028P14 | RIKEN cDNA 1700028P14 gene [Source:M | 0 | 0 | 0 | 0 | 0 | 0 |
| 9230 | ENSMUSG00000033054 | 126.0682233 | 0.090801888 | 0.210130672 | 0.432121055 | 0.665653433 | 0.929596279 | Npat | nuclear protein in the AT region [Source:N | 116.45 | 111.02 | 138.86 | 123.39 | 134.15 | 132.55 |
| 9231 | ENSMUSG00000033055 | 162.8030916 | 0.041976888 | 0.204283404 | 0.205483594 | 0.837194315 | 0.971073158 | Ankrd54 | ankyrin repeat domain 54 [Source:MGI Sy | 148.57 | 137.74 | 195.21 | 167.73 | 152.69 | 174.87 |
| 9232 | ENSMUSG00000033059 | 51.41009705 | -0.936935926 | 0.338789312 | -2.765541572 | 0.005682836 | 0.072740302 | Pygb | brain glycogen phosphorylase [Source:MG | 65.854 | 86.345 | 50.312 | 29.883 | 37.081 | 38.985 |
| 9233 | ENSMUSG00000033060 | 501.4718334 | 0.037151149 | 0.144698168 | 0.256749268 | 0.797372336 | 0.962236316 | Lmo7 | LIM domain only 7 [Source:MGI Symbol;A | 536.47 | 503.68 | 444.76 | 548.5 | 537.67 | 437.74 |
| 9234 | ENSMUSG00000033061 | 0 | NA | NA | NA | NA | NA | Resp18 | regulated endocrine-specific protein 18 [S | 0 | 0 | 0 | 0 | 0 | 0 |
| 9235 | ENSMUSG00000033063 | 0.534859069 | 1.101323472 | 3.351640271 | 0.328592386 | 0.742463804 | NA | Cntnap3 | contactin associated protein-like 3 [Sourc | 0 | 1.0279 | 0 | 0 | 2.1812 | 0 |
| 9236 | ENSMUSG00000033065 | 338.5346257 | 0.101972486 | 0.14795541 | 0.689210932 | 0.490690538 | 0.875999499 | Pfkm | phosphofructokinase, muscle [Source:MG | 299.56 | 324.82 | 356.21 | 373.06 | 350.09 | 327.47 |

| | | | | | | | | | | | | | | | |
|---|---|---|---|---|---|---|---|---|---|---|---|---|---|---|---|
| 9237 | ENSMUSG00000033066 | 55.04396153 | 0.186925421 | 0.366496852 | 0.510032815 | 0.610028472 | 0.916376001 | Gas7 | growth arrest specific 7 [Source:MGI Syml | 42.564 | 52.424 | 59.368 | 36.631 | 50.168 | 89.108 |
| 9238 | ENSMUSG00000033068 | 116.5671276 | 0.014057857 | 0.222393862 | 0.063211532 | 0.949598062 | 0.990731019 | Entpd6 | ectonucleoside triphosphate diphosphohy | 130.91 | 100.74 | 115.72 | 105.07 | 118.88 | 128.09 |
| 9239 | ENSMUSG00000033075 | 154.428649 | -0.100818008 | 0.197508087 | -0.510450025 | 0.609736218 | 0.916376001 | Senp1 | SUMO1/sentrin specific peptidase 1 [Sour | 149.38 | 173.72 | 156.97 | 166.77 | 149.41 | 130.32 |
| 9240 | ENSMUSG00000033080 | 0 | NA | NA | NA | NA | NA | Vsx1 | visual system homeobox 1 [Source:MGI Sy | 0 | 0 | 0 | 0 | 0 | 0 |
| 9241 | ENSMUSG00000033082 | 20.87512601 | -0.465543355 | 0.516113924 | -0.902016656 | 0.36704802 | 0.805239112 | Clec1a | C-type lectin domain family 1, member a [ | 25.699 | 29.81 | 17.106 | 18.316 | 9.8156 | 24.505 |
| 9242 | ENSMUSG00000033083 | 89.60076144 | 0.085706954 | 0.25379466 | 0.337701961 | 0.735587797 | 0.948109711 | Tbc1d4 | TBC1 domain family, member 4 [Source:M | 76.295 | 94.568 | 90.562 | 109.89 | 89.431 | 76.855 |
| 9243 | ENSMUSG00000033088 | 337.0285259 | 0.215662287 | 0.149264879 | 1.44482941 | 0.148505808 | 0.560454158 | Triobp | TRIO and F-actin binding protein [Source:I | 316.42 | 292.96 | 326.02 | 358.6 | 327.19 | 400.98 |
| 9244 | ENSMUSG00000033096 | 732.4470427 | -0.118810703 | 0.111975011 | -1.061046588 | 0.28866873 | 0.73951397 | Apmap | adipocyte plasma membrane associated p | 742.87 | 758.6 | 786.88 | 730.7 | 678.36 | 697.27 |
| 9245 | ENSMUSG00000033099 | 199.6091765 | -0.031836968 | 0.169950646 | -0.187330668 | 0.851401379 | 0.973568625 | Nol12 | nucleolar protein 12 [Source:MGI Symbol; | 191.94 | 208.67 | 205.27 | 208.22 | 190.86 | 192.69 |
| 9246 | ENSMUSG00000033102 | 623.2508921 | 0.274248967 | 0.127569977 | 2.149792395 | 0.03157164 | 0.236342058 | Cdc14b | CDC14 cell division cycle 14B [Source:MGI | 527.64 | 539.66 | 625.88 | 692.14 | 639.1 | 715.09 |
| 9247 | ENSMUSG00000033105 | 464.5914994 | 0.686654059 | 0.165069753 | 4.15978121 | 3.19E-05 | 0.001456931 | Lss | lanosterol synthase [Source:MGI Symbol;/ | 434.48 | 353.6 | 278.73 | 576.46 | 558.4 | 585.88 |
| 9248 | ENSMUSG00000033106 | 396.4235966 | 0.017792558 | 0.140151979 | 0.126951882 | 0.898978483 | 0.982798755 | Slc7a6os | solute carrier family 7, member 6 opposite | 358.99 | 401.92 | 421.62 | 387.52 | 425.34 | 383.16 |
| 9249 | ENSMUSG00000033107 | 838.4344173 | -0.069831484 | 0.165776697 | -0.421238242 | 0.673581122 | 0.931915649 | Rnf125 | ring finger protein 125 [Source:MGI Symb | 713.96 | 745.24 | 1117.9 | 856.98 | 820.14 | 776.35 |
| 9250 | ENSMUSG00000033111 | 205.7011525 | -0.047522335 | 0.186476834 | -0.254843106 | 0.79884428 | 0.962424862 | 3830406C13 | RIKEN cDNA 3830406C13 gene [Source:M | 195.96 | 212.78 | 218.36 | 167.73 | 236.66 | 202.72 |
| 9251 | ENSMUSG00000033114 | 564.4116225 | 0.244578134 | 0.143545828 | 1.703833109 | 0.088412271 | 0.438393926 | Slc35d2 | solute carrier family 35, member D2 [Sour | 520.41 | 442 | 587.65 | 629.48 | 650.01 | 556.92 |
| 9252 | ENSMUSG00000033122 | 0.181769715 | 1.126924175 | 4.080472857 | 0.276174898 | 0.782413736 | NA | Hsd17b3 | hydroxysteroid (17-beta) dehydrogenase : | 0 | 0 | 0 | 0 | 1.0906 | 0 |
| 9253 | ENSMUSG00000033124 | 508.3735269 | 0.303023803 | 0.134975313 | 2.245031312 | 0.024766122 | 0.201358738 | Atg9a | autophagy related 9A [Source:MGI Symbo | 422.43 | 442 | 502.12 | 609.23 | 542.04 | 532.42 |
| 9254 | ENSMUSG00000033126 | 84.53703814 | 0.03679711 | 0.261243727 | 0.140853565 | 0.887985626 | 0.982382091 | Ybey | ybeY metallopeptidase [Source:MGI Symb | 79.507 | 76.066 | 94.587 | 67.478 | 93.793 | 95.791 |
| 9255 | ENSMUSG00000033128 | 642.7930657 | 0.111681075 | 0.121731759 | 0.91743581 | 0.358914315 | 0.798534982 | Gga1 | golgi associated, gamma adaptin ear conta | 646.5 | 610.58 | 595.7 | 615.98 | 672.91 | 715.09 |
| 9256 | ENSMUSG00000033147 | 248.5744752 | -0.197485726 | 0.190280444 | -1.037866643 | 0.299332145 | 0.750613756 | Slc22a15 | solute carrier family 22 (organic anion/cat | 316.42 | 259.04 | 220.37 | 226.53 | 201.76 | 267.32 |
| 9257 | ENSMUSG00000033149 | 907.4607275 | 0.161746639 | 0.111019135 | 1.456925772 | 0.145136871 | 0.555361204 | Phldb2 | pleckstrin homology like domain, family B | 887.43 | 896.34 | 785.88 | 971.69 | 947.75 | 955.68 |
| 9258 | ENSMUSG00000033152 | 3.142898865 | -1.057230039 | 1.301940219 | -0.812041923 | 0.416767576 | NA | Podxl2 | podocalyxin-like 2 [Source:MGI Symbol;A | 5.6217 | 5.1396 | 2.0125 | 3.8559 | 0 | 2.2277 |
| 9259 | ENSMUSG00000033156 | 0 | NA | NA | NA | NA | NA | Cst10 | cystatin 10 (chondrocytes) [Source:MGI Sy | 0 | 0 | 0 | 0 | 0 | 0 |
| 9260 | ENSMUSG00000033157 | 94.56774262 | 0.117358115 | 0.314015032 | 0.373734066 | 0.708602178 | 0.943888782 | Abhd10 | abhydrolase domain containing 10 [Sourc | 115.65 | 82.233 | 73.456 | 56.875 | 117.79 | 121.41 |
| 9261 | ENSMUSG00000033159 | 1338.295746 | 0.244184563 | 0.101178739 | 2.413397965 | 0.015804552 | 0.147481965 | Cnppd1 | cyclin Pas1/PHO80 domain containing 1 [S | 1252.8 | 1169.8 | 1252.8 | 1440.2 | 1393.8 | 1520.4 |
| 9262 | ENSMUSG00000033161 | 9035.23664 | -0.03426852 | 0.105063595 | -0.32616931 | 0.74429626 | 0.949997329 | Atp1a1 | ATPase, Na+/K+ transporting, alpha 1 poly | 8340.2 | 10064 | 9024 | 8527.3 | 9694.5 | 8561 |
| 9263 | ENSMUSG00000033166 | 135.7127998 | 0.001553376 | 0.20360311 | 0.007629432 | 0.993912653 | 0.998618081 | Dis3 | DIS3 homolog, exosome endoribonucleas | 125.28 | 153.16 | 128.8 | 134.96 | 142.87 | 129.21 |
| 9264 | ENSMUSG00000033170 | 315.5493573 | 0.232139122 | 0.234779413 | 0.988754163 | 0.322783435 | 0.769926329 | Card10 | caspase recruitment domain family, meml | 289.12 | 237.45 | 344.14 | 378.84 | 211.58 | 432.17 |
| 9265 | ENSMUSG00000033174 | 6327.133104 | -0.315842821 | 0.139211778 | -2.268793815 | 0.023280867 | 0.192750123 | Mgll | monoglyceride lipase [Source:MGI Symbo | 7241.6 | 7381.5 | 6428.9 | 6799.9 | 5621 | 4489.9 |
| 9266 | ENSMUSG00000033177 | 2.702658108 | -1.001936689 | 1.455879218 | -0.68820042 | 0.49132658 | NA | Tmprss7 | transmembrane serine protease 7 [Source | 5.6217 | 4.1117 | 1.0062 | 0 | 4.3625 | 1.1138 |
| 9267 | ENSMUSG00000033182 | 6.676780449 | 0.069022074 | 0.949716636 | 0.072676493 | 0.942063555 | 0.989758645 | Kbtbd12 | kelch repeat and BTB (POZ) domain conta | 11.243 | 2.0558 | 6.0375 | 1.928 | 6.5437 | 12.252 |
| 9268 | ENSMUSG00000033184 | 2148.214868 | 0.0256744 | 0.091241052 | 0.281390882 | 0.778410611 | 0.959559604 | Tmed7 | transmembrane p24 trafficking protein 7 [ | 2050.3 | 2134 | 2203.7 | 2171.8 | 2145.2 | 2184.2 |
| 9269 | ENSMUSG00000033186 | 392.615201 | 0.140365396 | 0.140121513 | 1.001740506 | 0.316468938 | 0.763155505 | Mzt1 | mitotic spindle organizing protein 1 [Sour | 348.55 | 366.97 | 405.52 | 401.98 | 440.61 | 392.07 |
| 9270 | ENSMUSG00000033187 | 1.20269523 | 0.466011887 | 2.092262081 | 0.222731125 | 0.823744776 | NA | BC016579 | cDNA sequence, BC016579 [Source:MGI S | 0 | 3.0838 | 0 | 1.928 | 1.0906 | 1.1138 |
| 9271 | ENSMUSG00000033191 | 151.6904026 | -0.36259223 | 0.219069681 | -1.655145653 | 0.097894956 | 0.46122236 | Tie1 | tyrosine kinase with immunoglobulin-like | 137.33 | 175.77 | 199.24 | 126.28 | 116.7 | 154.82 |
| 9272 | ENSMUSG00000033192 | 17.37633415 | -1.11053226 | 0.665508465 | -1.668697423 | 0.09517736 | 0.455057116 | Lpcat2 | lysophosphatidylcholine acyltransferase 2 | 23.29 | 40.089 | 8.05 | 17.352 | 5.4531 | 10.025 |
| 9273 | ENSMUSG00000033196 | 0 | NA | NA | NA | NA | NA | Myh2 | myosin, heavy polypeptide 2, skeletal mus | 0 | 0 | 0 | 0 | 0 | 0 |
| 9274 | ENSMUSG00000033200 | 0.133850373 | -0.796629176 | 4.080472857 | -0.195229623 | 0.84521319 | NA | Tpsg1 | tryptase gamma 1 [Source:MGI Symbol;A | 0.8031 | 0 | 0 | 0 | 0 | 0 |
| 9275 | ENSMUSG00000033207 | 44.76585745 | -1.020684302 | 0.363772994 | -2.805827587 | 0.005018754 | 0.066818229 | Mamdc2 | MAM domain containing 2 [Source:MGI S | 69.067 | 42.145 | 68.425 | 26.027 | 26.175 | 36.757 |
| 9276 | ENSMUSG00000033208 | 0 | NA | NA | NA | NA | NA | S100b | S100 protein, beta polypeptide, neural [S | 0 | 0 | 0 | 0 | 0 | 0 |
| 9277 | ENSMUSG00000033209 | 23.76571706 | -0.642454116 | 0.486225202 | -1.321309781 | 0.186398094 | 0.622503355 | Ttc28 | tetratricopeptide repeat domain 28 [Sour | 29.715 | 28.782 | 28.175 | 10.604 | 16.359 | 28.96 |
| 9278 | ENSMUSG00000033210 | 0 | NA | NA | NA | NA | NA | Slc9c1 | solute carrier family 9, subfamily C (Na+-t | 0 | 0 | 0 | 0 | 0 | 0 |
| 9279 | ENSMUSG00000033213 | 0.342639279 | -1.805976596 | 4.028547714 | -0.448294702 | 0.653940524 | NA | AA467197 | expressed sequence AA467197 [Source:M | 0 | 2.0558 | 0 | 0 | 0 | 0 |
| 9280 | ENSMUSG00000033214 | 0 | NA | NA | NA | NA | NA | Slitrk5 | SLIT and NTRK-like family, member 5 [Sou | 0 | 0 | 0 | 0 | 0 | 0 |
| 9281 | ENSMUSG00000033216 | 481.7600917 | -0.071122096 | 0.134988661 | -0.526874596 | 0.598280677 | 0.915055389 | Eefsec | eukaryotic elongation factor, selenocystei | 488.29 | 508.82 | 484 | 470.42 | 520.22 | 418.81 |
| 9282 | ENSMUSG00000033219 | 0 | NA | NA | NA | NA | NA | Gm9758 | predicted gene 9758 [Source:MGI Symbol | 0 | 0 | 0 | 0 | 0 | 0 |
| 9283 | ENSMUSG00000033220 | 57.96420765 | -1.776195938 | 0.405638839 | -4.378762015 | 1.19E-05 | 0.000616556 | Rac2 | Rac family small GTPase 2 [Source:MGI Sy | 57.823 | 146.99 | 64.4 | 21.208 | 26.175 | 31.188 |
| 9284 | ENSMUSG00000033222 | 88.18461337 | -0.321839144 | 0.251844899 | -1.277925996 | 0.201275521 | 0.645509786 | Ttf2 | transcription termination factor, RNA poly | 83.523 | 115.13 | 95.593 | 77.118 | 71.981 | 85.766 |
| 9285 | ENSMUSG00000033227 | 0.751028265 | -1.614680879 | 2.682726358 | -0.601880573 | 0.547253636 | NA | Wnt6 | wingless-type MMTV integration site fami | 2.4093 | 0 | 1.0062 | 0 | 1.0906 | 0 |
| 9286 | ENSMUSG00000033228 | 1112.67344 | 0.067959909 | 0.117195046 | 0.579887213 | 0.56199068 | 0.900565925 | Scaf11 | SR-related CTD-associated factor 11 [Sour | 1077 | 1163.6 | 1019.3 | 1172.2 | 1223.7 | 1020.3 |
| 9287 | ENSMUSG00000033233 | 20.1990266 | 0.593894661 | 0.491381284 | 1.208622877 | 0.226807763 | 0.67343689 | Trim45 | tripartite motif-containing 45 [Source:MG | 17.668 | 15.419 | 15.094 | 22.171 | 18.541 | 32.301 |
| 9288 | ENSMUSG00000033237 | 437.8425409 | 0.107091118 | 0.133379999 | 0.802902372 | 0.422031165 | 0.839188723 | Arid2 | AT rich interactive domain 2 (ARID, RFX-lik | 402.35 | 457.42 | 405.52 | 454.03 | 473.33 | 434.4 |
| 9289 | ENSMUSG00000033249 | 189.3961018 | 0.407953729 | 0.175058676 | 2.330382807 | 0.019785928 | 0.172127966 | Hsf4 | heat shock transcription factor 4 [Source:I | 154.2 | 163.44 | 171.06 | 215.93 | 229.03 | 202.72 |
| 9290 | ENSMUSG00000033253 | 262.2009245 | 0.109580082 | 0.184301821 | 0.594568634 | 0.552131851 | 0.895935901 | Szt2 | SZT2 subunit of KICSTOR complex [Source | 208.81 | 276.51 | 272.69 | 319.08 | 239.94 | 256.18 |
| 9291 | ENSMUSG00000033255 | 0.319491021 | 0.16514334 | 4.044653141 | 0.040830038 | 0.967431393 | NA | Gm5134 | predicted gene 5134 [Source:MGI Symbol | 0.8031 | 0 | 0 | 0 | 0 | 1.1138 |
| 9292 | ENSMUSG00000033256 | 192.3415168 | 0.606932426 | 0.1821028 | 3.332911016 | 0.000859424 | 0.018720952 | Shf | Src homology 2 domain containing F [Sour | 143.76 | 140.82 | 173.07 | 234.25 | 213.76 | 248.39 |

| | | | | | | | | | | | | | | | |
|---|---|---|---|---|---|---|---|---|---|---|---|---|---|---|---|
| 9293 | ENSMUSG00000033257 | 392.2697869 | 0.079285461 | 0.134537238 | 0.589319821 | 0.555646751 | 0.897628185 | Ttll4 | tubulin tyrosine ligase-like family, membe | 373.44 | 402.94 | 368.29 | 398.12 | 420.98 | 389.85 |
| 9294 | ENSMUSG00000033268 | 0 | NA | NA | NA | NA | NA | Duox1 | dual oxidase 1 [Source:MGI Symbol;Acc:M | 0 | 0 | 0 | 0 | 0 | 0 |
| 9295 | ENSMUSG00000033272 | 295.2463962 | -0.05294949 | 0.163553929 | -0.323743308 | 0.74613238 | 0.949997329 | Slc35a4 | solute carrier family 35, member A4 [Sour | 307.59 | 311.46 | 282.75 | 290.16 | 246.48 | 333.04 |
| 9296 | ENSMUSG00000033276 | 13.54723941 | -0.461111014 | 0.674804001 | -0.683325845 | 0.494400966 | 0.877473722 | Stk36 | serine/threonine kinase 36 [Source:MGI S | 12.85 | 7.1954 | 27.169 | 16.388 | 6.5437 | 11.138 |
| 9297 | ENSMUSG00000033278 | 25.45913356 | -0.560701995 | 0.465025924 | -1.205743521 | 0.227916399 | 0.674239914 | Ptprm | protein tyrosine phosphatase, receptor ty | 32.927 | 37.005 | 21.131 | 24.099 | 13.087 | 24.505 |
| 9298 | ENSMUSG00000033282 | 53.09010948 | -0.124201897 | 0.344103564 | -0.360943362 | 0.718141788 | 0.9447719 | Rpgrip1l | Rpgrip1-like [Source:MGI Symbol;Acc:MG | 34.533 | 62.703 | 69.431 | 54.947 | 51.259 | 45.668 |
| 9299 | ENSMUSG00000033285 | 287.588474 | -0.034616421 | 0.160844839 | -0.215216239 | 0.829598731 | 0.969955221 | Wdr3 | WD repeat domain 3 [Source:MGI Symbol | 268.24 | 327.91 | 277.72 | 298.83 | 296.65 | 256.18 |
| 9300 | ENSMUSG00000033287 | 76.93371357 | -0.410231944 | 0.294924761 | -1.390971523 | 0.164234062 | 0.588865016 | Kctd17 | potassium channel tetramerisation domai | 110.83 | 75.038 | 77.481 | 84.83 | 61.075 | 52.351 |
| 9301 | ENSMUSG00000033294 | 188.9550468 | -0.032233502 | 0.178997522 | -0.180077922 | 0.857091395 | 0.974809103 | Noc4l | NOC4 like [Source:MGI Symbol;Acc:MGI:2 | 184.71 | 212.78 | 176.09 | 197.62 | 184.31 | 178.22 |
| 9302 | ENSMUSG00000033295 | 2614.822116 | 0.329073059 | 0.122748662 | 2.68086881 | 0.00734313 | 0.086561556 | Ptprf | protein tyrosine phosphatase, receptor ty | 2170 | 2206.9 | 2577 | 2778.2 | 2615.3 | 3341.5 |
| 9303 | ENSMUSG00000033306 | 920.8908344 | -0.198444195 | 0.134274863 | -1.477895346 | 0.139435792 | 0.545670806 | Lpp | LIM domain containing preferred transloc | 910.72 | 1062.9 | 979.08 | 788.53 | 1004.5 | 779.69 |
| 9304 | ENSMUSG00000033307 | 1858.214979 | -0.22193382 | 0.106952236 | -2.075074137 | 0.037979664 | 0.26463993 | Mif | macrophage migration inhibitory factor (g | 1856.8 | 2026 | 2121.2 | 1829.6 | 1579.2 | 1736.5 |
| 9305 | ENSMUSG00000033308 | 10473.6969 | -0.067861431 | 0.103279941 | -0.657063032 | 0.511140388 | 0.883679955 | Dpyd | dihydropyrimidine dehydrogenase [Sourc | 9548.9 | 11249 | 11365 | 10691 | 10480 | 9510 |
| 9306 | ENSMUSG00000033313 | 77.88714926 | 0.527304756 | 0.270144937 | 1.951932771 | 0.050946187 | 0.316803804 | Fbxl8 | F-box and leucine-rich repeat protein 8 [S | 67.461 | 57.563 | 66.412 | 107 | 71.981 | 96.904 |
| 9307 | ENSMUSG00000033316 | 1.186152743 | -0.224486212 | 2.101231109 | -0.106835565 | 0.914919432 | NA | Galnt9 | polypeptide N-acetylgalactosaminyltransf | 0.8031 | 0 | 3.0187 | 0 | 2.1812 | 1.1138 |
| 9308 | ENSMUSG00000033318 | 1338.543597 | 0.216194666 | 0.116555824 | 1.854859402 | 0.063616307 | 0.360558879 | Gstt2 | glutathione S-transferase, theta 2 [Source | 1086.6 | 1265.4 | 1364.5 | 1376.6 | 1466.9 | 1471.4 |
| 9309 | ENSMUSG00000033319 | 294.2607365 | -0.099230618 | 0.149049212 | -0.66575741 | 0.505566172 | 0.881592135 | Fem1c | fem 1 homolog c [Source:MGI Symbol;Acc | 294.74 | 308.38 | 309.92 | 267.02 | 279.2 | 306.31 |
| 9310 | ENSMUSG00000033323 | 198.2134752 | 0.070267656 | 0.177436017 | 0.396016868 | 0.692092584 | 0.938256814 | Ctdp1 | CTD (carboxy-terminal domain, RNA polyr | 191.14 | 212.78 | 176.09 | 187.01 | 203.95 | 218.31 |
| 9311 | ENSMUSG00000033326 | 480.2339056 | 0.08229618 | 0.139433139 | 0.590219661 | 0.555043392 | 0.897628185 | Kdm4a | lysine (K)-specific demethylase 4A [Source | 524.43 | 466.67 | 407.53 | 487.77 | 507.14 | 487.86 |
| 9312 | ENSMUSG00000033327 | 233.0990063 | -0.579593596 | 0.232211273 | -2.495975276 | 0.012561136 | 0.126529431 | Tnxb | tenascin XB [Source:MGI Symbol;Acc:MGI | 354.17 | 274.45 | 209.3 | 238.1 | 177.77 | 144.8 |
| 9313 | ENSMUSG00000033335 | 939.7887882 | 0.080642601 | 0.111786434 | 0.721398819 | 0.470664173 | 0.867110762 | Dnm2 | dynamin 2 [Source:MGI Symbol;Acc:MGI: | 958.9 | 870.65 | 910.65 | 1007.4 | 895.4 | 995.78 |
| 9314 | ENSMUSG00000033342 | 4.69821926 | -2.124403721 | 1.090455922 | -1.948179362 | 0.051393505 | NA | Plppr5 | phospholipid phosphatase related 5 [Sour | 5.6217 | 11.307 | 6.0375 | 1.928 | 2.1812 | 1.1138 |
| 9315 | ENSMUSG00000033343 | 0 | NA | NA | NA | NA | NA | Magea4 | MAGE family member A4 [Source:MGI Syr | 0 | 0 | 0 | 0 | 0 | 0 |
| 9316 | ENSMUSG00000033350 | 22.87542728 | -0.538077782 | 0.5004221 | -1.075247839 | 0.282263782 | 0.735230549 | Chst2 | carbohydrate sulfotransferase 2 [Source:N | 31.321 | 23.642 | 26.162 | 17.352 | 9.8156 | 28.96 |
| 9317 | ENSMUSG00000033352 | 712.7213357 | -0.055237055 | 0.111062536 | -0.497350927 | 0.618941604 | 0.917509899 | Map2k4 | mitogen-activated protein kinase kinase 4 | 743.67 | 731.88 | 703.36 | 705.63 | 704.54 | 687.24 |
| 9318 | ENSMUSG00000033355 | 280.3580798 | 0.392924352 | 0.211037306 | 1.861871532 | 0.062621201 | 0.357440034 | Rtp4 | receptor transporter protein 4 [Source:M( | 285.1 | 170.63 | 270.68 | 314.26 | 266.11 | 375.37 |
| 9319 | ENSMUSG00000033356 | 41.54151344 | 0.067288338 | 0.379223114 | 0.177437332 | 0.859164886 | 0.975487285 | Pus7l | pseudouridylate synthase 7-like [Source:N | 40.958 | 53.452 | 27.169 | 31.811 | 49.078 | 46.781 |
| 9320 | ENSMUSG00000033361 | 10.9118028 | -0.841709452 | 0.699719307 | -1.202924435 | 0.229005563 | 0.67545951 | Prrg3 | proline rich Gla (G-carboxyglutamic acid) 3 | 18.471 | 10.279 | 13.081 | 4.8199 | 5.4531 | 13.366 |
| 9321 | ENSMUSG00000033364 | 111.8868047 | 0.178970817 | 0.23225919 | 0.770565062 | 0.440964777 | 0.849828766 | Usp37 | ubiquitin specific peptidase 37 [Source:M | 108.42 | 112.04 | 94.587 | 133.99 | 127.6 | 94.677 |
| 9322 | ENSMUSG00000033365 | 253.7892485 | 0.013636018 | 0.155789541 | 0.087528456 | 0.930251467 | 0.988420305 | Ipo13 | importin 13 [Source:MGI Symbol;Acc:MGI | 245.75 | 250.81 | 261.62 | 271.84 | 243.21 | 249.5 |
| 9323 | ENSMUSG00000033368 | 1.277737502 | -2.558404199 | 2.061944144 | -1.240772795 | 0.214689694 | NA | Trim69 | tripartite motif-containing 69 [Source:MG | 1.6062 | 3.0838 | 2.0125 | 0.964 | 0 | 0 |
| 9324 | ENSMUSG00000033373 | 62.70216984 | -0.027871406 | 0.291568213 | -0.095591375 | 0.923845116 | 0.987327897 | Fntb | farnesyltransferase, CAAX box, beta [Sour | 71.476 | 60.647 | 57.356 | 50.127 | 70.89 | 65.717 |
| 9325 | ENSMUSG00000033377 | 811.1263478 | 0.099143876 | 0.152932846 | 0.648283732 | 0.516801454 | 0.886716042 | Palmd | palmdelphin [Source:MGI Symbol;Acc:MG | 954.09 | 756.55 | 637.96 | 869.51 | 881.22 | 767.44 |
| 9326 | ENSMUSG00000033379 | 1282.853375 | 0.068515285 | 0.098329266 | 0.696794439 | 0.485931447 | 0.872998689 | Atp6v0b | ATPase, H+ transporting, lysosomal V0 sul | 1297 | 1207.8 | 1251.8 | 1311 | 1319.6 | 1309.9 |
| 9327 | ENSMUSG00000033382 | 917.0727755 | -0.014974681 | 0.11911996 | -0.125710928 | 0.899960751 | 0.982798755 | Trappc8 | trafficking protein particle complex 8 [Sou | 827.2 | 992.97 | 946.88 | 969.76 | 880.13 | 885.51 |
| 9328 | ENSMUSG00000033383 | 0 | NA | NA | NA | NA | NA | Rtp1 | receptor transporter protein 1 [Source:M( | 0 | 0 | 0 | 0 | 0 | 0 |
| 9329 | ENSMUSG00000033386 | 451.2823654 | 0.245073009 | 0.142178505 | 1.723699439 | 0.084762116 | 0.429140871 | Frrs1 | ferric-chelate reductase 1 [Source:MGI Sy | 410.39 | 465.65 | 363.25 | 485.84 | 515.86 | 466.7 |
| 9330 | ENSMUSG00000033389 | 14.99672852 | 0.071257616 | 0.662856394 | 0.107500835 | 0.914391663 | 0.98625062 | Arhgap44 | Rho GTPase activating protein 44 [Source: | 27.305 | 7.1954 | 9.0562 | 13.496 | 22.903 | 10.025 |
| 9331 | ENSMUSG00000033392 | 189.021799 | -0.296433062 | 0.190160012 | -1.558861186 | 0.119029239 | 0.504735151 | Clasp2 | CLIP associating protein 2 [Source:MGI Sy | 188.73 | 209.7 | 227.41 | 187.98 | 178.86 | 141.46 |
| 9332 | ENSMUSG00000033396 | 293.6528955 | 0.110812585 | 0.153070216 | 0.723933033 | 0.469106849 | 0.866006568 | Spg11 | SPG11, spatacsin vesicle trafficking associ | 298.75 | 253.9 | 293.82 | 287.27 | 315.19 | 312.99 |
| 9333 | ENSMUSG00000033400 | 1157.903927 | 0.118902382 | 0.151464939 | 0.785015875 | 0.432444261 | 0.845264432 | Agl | amylo-1,6-glucosidase, 4-alpha-glucanotra | 1056.9 | 1237.6 | 1037.4 | 1455.6 | 1196.4 | 963.47 |
| 9334 | ENSMUSG00000033405 | 21.8322699 | 0.102009468 | 0.468403565 | 0.217781152 | 0.827599631 | 0.969794868 | Nudt15 | nudix (nucleoside diphosphate linked moi | 17.668 | 21.586 | 24.15 | 27.955 | 21.812 | 17.822 |
| 9335 | ENSMUSG00000033409 | 0 | NA | NA | NA | NA | NA | Syce1l | synaptonemal complex central element p | 0 | 0 | 0 | 0 | 0 | 0 |
| 9336 | ENSMUSG00000033411 | 141.6984574 | 0.050596335 | 0.212648796 | 0.237933797 | 0.811932442 | 0.96653525 | Ctdspl2 | CTD (carboxy-terminal domain, RNA polyr | 140.54 | 133.63 | 143.89 | 176.41 | 126.51 | 129.21 |
| 9337 | ENSMUSG00000033416 | 3615.83079 | 0.026389683 | 0.092581914 | 0.285041453 | 0.775612384 | 0.95893551 | Gucd1 | guanylyl cyclase domain containing 1 [Sou | 3524.8 | 3473.3 | 3750.3 | 3655.4 | 3455.1 | 3836.1 |
| 9338 | ENSMUSG00000033417 | 1051.227109 | 0.010243615 | 0.101626838 | 0.100796361 | 0.919712116 | 0.986685804 | Cacul1 | CDK2 associated, cullin domain 1 [Source: | 1015.1 | 1075.2 | 1052.5 | 1045 | 1050.3 | 1069.3 |
| 9339 | ENSMUSG00000033419 | 0 | NA | NA | NA | NA | NA | Snap91 | synaptosomal-associated protein 91 [Sour | 0 | 0 | 0 | 0 | 0 | 0 |
| 9340 | ENSMUSG00000033420 | 19.27428505 | -0.436875608 | 0.503619854 | -0.867470979 | 0.385684006 | 0.816530074 | Antxr1 | anthrax toxin receptor 1 [Source:MGI Sym | 25.699 | 25.698 | 15.094 | 18.316 | 16.359 | 14.48 |
| 9341 | ENSMUSG00000033423 | 462.3283273 | -0.061636534 | 0.134708573 | -0.457554654 | 0.647272428 | 0.925643431 | Eri3 | exoribonuclease 3 [Source:MGI Symbol;A | 477.04 | 440.98 | 498.09 | 427.04 | 435.16 | 495.66 |
| 9342 | ENSMUSG00000033427 | 6536.228047 | 0.125821136 | 0.107355254 | 1.172007252 | 0.241194145 | 0.691388521 | Upb1 | ureidopropionase, beta [Source:MGI Sym | 5620.1 | 6651.7 | 6483.2 | 6293.8 | 7457.6 | 6710.9 |
| 9343 | ENSMUSG00000033429 | 854.1661937 | 0.002055334 | 0.110825976 | 0.018545599 | 0.985203601 | 0.99859176 | Mcee | methylmalonyl CoA epimerase [Source:M | 820.77 | 894.29 | 846.25 | 861.8 | 892.13 | 809.76 |
| 9344 | ENSMUSG00000033430 | 341.9363914 | -0.07883259 | 0.143233682 | -0.550377461 | 0.582060505 | 0.908645447 | Terf2ip | telomeric repeat binding factor 2, interact | 325.26 | 354.63 | 374.32 | 324.86 | 323.91 | 348.63 |
| 9345 | ENSMUSG00000033434 | 100.3764765 | 0.113122405 | 0.247484902 | 0.457088106 | 0.647607719 | 0.925643431 | Gtpbp6 | GTP binding protein 6 (putative) [Source:N | 113.24 | 86.345 | 89.556 | 120.5 | 107.97 | 84.652 |
| 9346 | ENSMUSG00000033436 | 23.93912529 | -0.234402776 | 0.479852098 | -0.488489636 | 0.625203065 | 0.919558383 | Armcx2 | armadillo repeat containing, X-linked 2 [S | 20.078 | 18.503 | 39.244 | 25.063 | 21.812 | 18.935 |
| 9347 | ENSMUSG00000033439 | 55.34906235 | 0.119417905 | 0.335243369 | 0.35621258 | 0.721681372 | 0.9447719 | Trmt13 | tRNA methyltransferase 13 [Source:MGI S | 57.823 | 42.145 | 59.368 | 78.082 | 52.35 | 42.326 |
| 9348 | ENSMUSG00000033444 | 331.5855794 | 0.041522355 | 0.147301602 | 0.281886647 | 0.778030429 | 0.959559604 | Specc1l | sperm antigen with calponin homology ar | 305.98 | 355.66 | 318.98 | 312.33 | 346.82 | 349.75 |

| | | | | | | | | | | | | | | | |
|---|---|---|---|---|---|---|---|---|---|---|---|---|---|---|---|
| 9349 | ENSMUSG00000033446 | 313.6190633 | 0.199568537 | 0.154167547 | 1.294491226 | 0.195495798 | 0.635855908 | Lpar6 | lysophosphatidic acid receptor 6 [Source:I | 297.95 | 262.12 | 315.96 | 363.42 | 333.73 | 308.53 |
| 9350 | ENSMUSG00000033450 | 4.686786951 | -0.104823514 | 1.068663876 | -0.098088386 | 0.92186211 | NA | Tagap | T cell activation Rho GTPase activating prc | 2.4093 | 9.2513 | 3.0187 | 5.7839 | 6.5437 | 1.1138 |
| 9351 | ENSMUSG00000033453 | 14.07899793 | -1.206676823 | 0.709292193 | -1.701240807 | 0.088897778 | 0.439445475 | Adamts15 | a disintegrin-like and metallopeptidase (re | 26.502 | 16.447 | 16.1 | 15.424 | 1.0906 | 8.9108 |
| 9352 | ENSMUSG00000033454 | 202.2504072 | 0.0983798 | 0.177802273 | 0.553310137 | 0.580051061 | 0.908645447 | Zbtb1 | zinc finger and BTB domain containing 1 [! | 197.56 | 195.3 | 193.2 | 216.9 | 178.86 | 231.68 |
| 9353 | ENSMUSG00000033458 | 113.4497684 | 0.014836703 | 0.233052406 | 0.06366252 | 0.949238949 | 0.990731019 | Fan1 | FANCD2/FANCI-associated nuclease 1 [So | 96.372 | 115.13 | 127.79 | 131.1 | 114.51 | 95.791 |
| 9354 | ENSMUSG00000033460 | 19.97901113 | -0.491200576 | 0.520935691 | -0.942919799 | 0.345721927 | 0.786899236 | Armcx1 | armadillo repeat containing, X-linked 1 [Sc | 22.487 | 19.53 | 28.175 | 23.135 | 8.7249 | 17.822 |
| 9355 | ENSMUSG00000033467 | 119.1879385 | 0.104950015 | 0.225405805 | 0.465604756 | 0.641498438 | 0.924019354 | Crlf2 | cytokine receptor-like factor 2 [Source:MC | 115.65 | 116.15 | 112.7 | 124.35 | 99.246 | 147.03 |
| 9356 | ENSMUSG00000033470 | 2.580246086 | -1.441009965 | 1.342823356 | -1.073119528 | 0.28321749 | NA | Cysltr2 | cysteinyl leukotriene receptor 2 [Source:N | 3.2124 | 4.1117 | 4.025 | 1.928 | 1.0906 | 1.1138 |
| 9357 | ENSMUSG00000033475 | 0.428363721 | -0.543730382 | 3.624677344 | -0.15000794 | 0.880758351 | NA | Tomm6 | translocase of outer mitochondrial memb | 1.6062 | 0 | 0 | 0.964 | 0 | 0 |
| 9358 | ENSMUSG00000033478 | 977.2023122 | 0.213372832 | 0.120326821 | 1.773277404 | 0.076182751 | 0.403877794 | Fam160b1 | family with sequence similarity 160, meml | 944.45 | 944.66 | 825.12 | 947.59 | 1129.9 | 1071.5 |
| 9359 | ENSMUSG00000033486 | 56.03746067 | -0.162382564 | 0.329614154 | -0.49264439 | 0.622263877 | 0.918691918 | Catsper2 | cation channel, sperm associated 2 [Sourc | 63.445 | 47.284 | 66.412 | 36.631 | 55.622 | 66.831 |
| 9360 | ENSMUSG00000033487 | 1012.489087 | 0.019286643 | 0.106049548 | 0.181864451 | 0.855689103 | 0.974158771 | Fndc3a | fibronectin type III domain containing 3A | 954.89 | 1000.2 | 1062.6 | 997.72 | 1042.6 | 1016.9 |
| 9361 | ENSMUSG00000033488 | 476.3612862 | -0.012182384 | 0.145138368 | -0.083936346 | 0.933107041 | 0.988420305 | Cryzl2 | crystallin zeta like 2 [Source:MGI Symbol;/ | 523.62 | 413.22 | 497.08 | 428.01 | 501.68 | 494.55 |
| 9362 | ENSMUSG00000033491 | 0.342639279 | -1.805976596 | 4.028547714 | -0.448294702 | 0.653940524 | NA | Prss35 | protease, serine 35 [Source:MGI Symbol;/ | 0 | 2.0558 | 0 | 0 | 0 | 0 |
| 9363 | ENSMUSG00000033498 | 0.160662974 | 1.126924175 | 4.080472857 | 0.276174898 | 0.782413736 | NA | Strc | stereocilin [Source:MGI Symbol;Acc:MGI:1 | 0 | 0 | 0 | 0.964 | 0 | 0 |
| 9364 | ENSMUSG00000033499 | 1716.654203 | 0.18300401 | 0.114678442 | 1.595801333 | 0.110533156 | 0.488765381 | Larp4b | La ribonucleoprotein domain family, mem | 1614.2 | 1729 | 1481.2 | 1954.9 | 1895.5 | 1625.1 |
| 9365 | ENSMUSG00000033501 | 0.51034671 | -2.381743948 | 3.331081356 | -0.715006238 | 0.474605189 | NA | Crygs | crystallin, gamma S [Source:MGI Symbol;/ | 0 | 2.0558 | 1.0062 | 0 | 0 | 0 |
| 9366 | ENSMUSG00000033502 | 38.79914865 | -0.214406737 | 0.351980813 | -0.609143253 | 0.542429491 | 0.894095573 | Cdc14a | CDC14 cell division cycle 14A [Source:MGI | 41.761 | 41.117 | 42.262 | 41.451 | 29.447 | 36.757 |
| 9367 | ENSMUSG00000033508 | 0.6996517 | 1.656076801 | 2.758183935 | 0.600422901 | 0.548224429 | NA | Asprv1 | aspartic peptidase, retroviral-like 1 [Sourc | 0 | 0 | 1.0062 | 0.964 | 0 | 2.2277 |
| 9368 | ENSMUSG00000033510 | 0 | NA | NA | NA | NA | NA | Otud7a | OTU domain containing 7A [Source:MGI S | 0 | 0 | 0 | 0 | 0 | 0 |
| 9369 | ENSMUSG00000033520 | 0.685278559 | -2.809870862 | 3.376578027 | -0.832165239 | 0.405315682 | NA | Idi2 | isopentenyl-diphosphate delta isomerase | 0 | 4.1117 | 0 | 0 | 0 | 0 |
| 9370 | ENSMUSG00000033526 | 77.92812832 | 0.017859424 | 0.256496512 | 0.069628329 | 0.944489488 | 0.990077979 | Ppip5k1 | diphosphoinositol pentakisphosphate kina | 81.916 | 74.01 | 76.475 | 89.65 | 70.89 | 74.628 |
| 9371 | ENSMUSG00000033530 | 296.5416611 | -0.101575021 | 0.157440453 | -0.645164685 | 0.518820469 | 0.887592015 | Ttc7b | tetratricopeptide repeat domain 7B [Sour | 286.71 | 340.24 | 294.83 | 315.22 | 270.47 | 271.78 |
| 9372 | ENSMUSG00000033533 | 6785.753811 | 0.187193078 | 0.100343837 | 1.86551645 | 0.062109049 | 0.355990107 | Acsm1 | acyl-CoA synthetase medium-chain family | 6322 | 6403.9 | 6313.2 | 7643.4 | 7645.2 | 6386.8 |
| 9373 | ENSMUSG00000033538 | 14.28499683 | -0.31277807 | 0.60894722 | -0.513637405 | 0.607505523 | 0.916376001 | Casp4 | caspase 4, apoptosis-related cysteine pep | 20.078 | 11.307 | 16.1 | 18.316 | 6.5437 | 13.366 |
| 9374 | ENSMUSG00000033540 | 94.25966818 | -0.032725768 | 0.230422006 | -0.142025359 | 0.887059975 | 0.982382091 | Idua | iduronidase, alpha-L [Source:MGI Symbol, | 98.782 | 93.541 | 93.581 | 93.506 | 98.156 | 87.994 |
| 9375 | ENSMUSG00000033542 | 460.8839779 | 0.026681761 | 0.172138416 | 0.155001783 | 0.876819912 | 0.980632845 | Arhgef5 | Rho guanine nucleotide exchange factor ( | 448.13 | 498.54 | 423.63 | 522.48 | 347.91 | 524.62 |
| 9376 | ENSMUSG00000033543 | 600.9875687 | -0.029832118 | 0.126212721 | -0.236363801 | 0.813150383 | 0.966802988 | Gtf2a2 | general transcription factor II A, 2 [Source | 590.28 | 613.67 | 617.83 | 577.42 | 659.82 | 546.9 |
| 9377 | ENSMUSG00000033544 | 2.664651393 | -1.100779192 | 1.512700818 | -0.727691279 | 0.46680259 | NA | Angptl1 | angiopoietin-like 1 [Source:MGI Symbol;A | 8.8341 | 1.0279 | 1.0062 | 2.8919 | 0 | 2.2277 |
| 9378 | ENSMUSG00000033545 | 763.6910676 | 0.218523077 | 0.14123757 | 1.547202182 | 0.1218145 | 0.511642546 | Znrf1 | zinc and ring finger 1 [Source:MGI Symbol | 633.65 | 691.79 | 793.93 | 944.7 | 788.52 | 729.57 |
| 9379 | ENSMUSG00000033554 | 153.3398959 | -0.142380667 | 0.212252715 | -0.670807284 | 0.502343307 | 0.881220881 | Dph5 | diphthamide biosynthesis 5 [Source:MGI S | 152.59 | 176.8 | 153.96 | 172.55 | 149.41 | 114.73 |
| 9380 | ENSMUSG00000033557 | 460.339586 | 0.06605336 | 0.144867777 | 0.455956193 | 0.64842148 | 0.925643431 | Fam20b | FAM20B, glycosaminoglycan xylosylkinase | 412.79 | 440.98 | 496.08 | 466.57 | 419.89 | 525.73 |
| 9381 | ENSMUSG00000033565 | 471.1590851 | 0.169424401 | 0.140816438 | 1.203157839 | 0.228915247 | 0.67545951 | Rbfox2 | RNA binding protein, fox-1 homolog (C. el | 427.25 | 495.46 | 408.54 | 524.4 | 521.32 | 449.99 |
| 9382 | ENSMUSG00000033569 | 0 | NA | NA | NA | NA | NA | Adgrb3 | adhesion G protein-coupled receptor B3 [ | 0 | 0 | 0 | 0 | 0 | 0 |
| 9383 | ENSMUSG00000033576 | 0 | NA | NA | NA | NA | NA | Apol6 | apolipoprotein L 6 [Source:MGI Symbol;A | 0 | 0 | 0 | 0 | 0 | 0 |
| 9384 | ENSMUSG00000033577 | 1095.236876 | -0.007600506 | 0.122891303 | -0.061847383 | 0.950684369 | 0.990883058 | Myo6 | myosin VI [Source:MGI Symbol;Acc:MGI:1 | 1071.3 | 1144.1 | 1079.7 | 1152 | 1191 | 933.4 |
| 9385 | ENSMUSG00000033578 | 0.335414862 | -1.779047242 | 4.030804179 | -0.441362855 | 0.658950329 | NA | Tmem35a | transmembrane protein 35A [Source:MGI | 0 | 0 | 2.0125 | 0 | 0 | 0 |
| 9386 | ENSMUSG00000033579 | 0.353348078 | 0.165142701 | 3.895042568 | 0.042398176 | 0.966181282 | NA | Fa2h | fatty acid 2-hydroxylase [Source:MGI Sym | 0 | 0 | 1.0062 | 0 | 0 | 1.1138 |
| 9387 | ENSMUSG00000033581 | 34.52702768 | -0.097236944 | 0.398117243 | -0.24424198 | 0.807043416 | 0.964718338 | Igf2bp2 | insulin-like growth factor 2 mRNA binding | 32.927 | 44.2 | 30.187 | 42.415 | 22.903 | 34.529 |
| 9388 | ENSMUSG00000033585 | 10.3779268 | -1.066668061 | 0.715157355 | -1.491515195 | 0.135826285 | 0.538686971 | Ndn | necdin, MAGE family member [Source:MC | 13.653 | 13.363 | 15.094 | 4.8199 | 11.997 | 3.3415 |
| 9389 | ENSMUSG00000033589 | 149.779944 | -0.294174409 | 0.242375734 | -1.2137123 | 0.224857606 | 0.671416269 | Reep4 | receptor accessory protein 4 [Source:MGI | 152.59 | 132.6 | 210.31 | 160.02 | 140.69 | 102.47 |
| 9390 | ENSMUSG00000033590 | 4.080699783 | -0.926093106 | 1.081038278 | -0.856670041 | 0.391627265 | NA | Myo5c | myosin VC [Source:MGI Symbol;Acc:MGI:1 | 8.8341 | 3.0838 | 4.025 | 1.928 | 3.2719 | 3.3415 |
| 9391 | ENSMUSG00000033594 | 955.5542208 | -1.098822013 | 0.188242019 | -5.837283404 | 5.31E-09 | 9.03E-07 | Spata2l | spermatogenesis associated 2-like [Source | 1421.5 | 1416.5 | 1069.6 | 474.28 | 820.14 | 531.3 |
| 9392 | ENSMUSG00000033595 | 2.630355651 | -1.512329909 | 1.547547236 | -0.977243133 | 0.328448799 | NA | Lgi3 | leucine-rich repeat LGI family, member 3 [ | 9.6372 | 0 | 2.0125 | 1.928 | 1.0906 | 1.1138 |
| 9393 | ENSMUSG00000033596 | 383.399886 | 0.265785447 | 0.159576949 | 1.665562909 | 0.095800504 | 0.456860696 | Rfwd3 | ring finger and WD repeat domain 3 [Sour | 327.67 | 305.29 | 411.55 | 383.66 | 414.43 | 457.79 |
| 9394 | ENSMUSG00000033597 | 2.737270222 | -0.021978048 | 1.273093666 | -0.017263497 | 0.986226407 | NA | Caskin1 | CASK interacting protein 1 [Source:MGI Sy | 3.2124 | 3.0838 | 2.0125 | 4.8199 | 2.1812 | 1.1138 |
| 9395 | ENSMUSG00000033610 | 5779.076898 | -0.023452642 | 0.087230306 | -0.268858877 | 0.788038283 | 0.960870081 | Pank1 | pantothenate kinase 1 [Source:MGI Symb | 6089.9 | 5670 | 5717.5 | 5762.7 | 5925.3 | 5509.1 |
| 9396 | ENSMUSG00000033615 | 0 | NA | NA | NA | NA | NA | Cplx1 | complexin 1 [Source:MGI Symbol;Acc:MG | 0 | 0 | 0 | 0 | 0 | 0 |
| 9397 | ENSMUSG00000033618 | 317.9292538 | -0.303970745 | 0.166866532 | -1.821639972 | 0.068509632 | 0.37732204 | Map3k13 | mitogen-activated protein kinase kinase k | 381.47 | 337.16 | 335.08 | 294.98 | 326.09 | 232.79 |
| 9398 | ENSMUSG00000033623 | 456.870463 | 0.215169507 | 0.127641731 | 1.685730088 | 0.091847799 | 0.446623824 | Pcgf3 | polycomb group ring finger 3 [Source:MG | 428.86 | 401.92 | 437.72 | 495.48 | 503.87 | 473.38 |
| 9399 | ENSMUSG00000033624 | 423.0766451 | 0.035318005 | 0.148516719 | 0.23780491 | 0.81203241 | 0.96653525 | Pdpr | pyruvate dehydrogenase phosphatase reg | 351.76 | 457.42 | 445.77 | 439.57 | 435.16 | 408.78 |
| 9400 | ENSMUSG00000033628 | 395.3349803 | 0.109270986 | 0.136023107 | 0.803326641 | 0.421785963 | 0.838966521 | Pik3c3 | phosphatidylinositol 3-kinase catalytic sut | 357.38 | 391.64 | 392.44 | 393.3 | 435.16 | 402.1 |
| 9401 | ENSMUSG00000033629 | 3660.818596 | 0.046174376 | 0.104219215 | 0.443050502 | 0.657729207 | 0.927303282 | Hacd3 | 3-hydroxyacyl-CoA dehydratase 3 [Source | 3446.1 | 3567.9 | 3792.5 | 3373.9 | 4089.8 | 3694.6 |
| 9402 | ENSMUSG00000033632 | 45.51933938 | -0.003571713 | 0.389486376 | -0.009170317 | 0.992683248 | 0.99859176 | AW554918 | expressed sequence AW554918 [Source:N | 44.171 | 57.563 | 35.219 | 54.947 | 56.712 | 24.505 |
| 9403 | ENSMUSG00000033633 | 0 | NA | NA | NA | NA | NA | Clec18a | C-type lectin domain family 18, member A | 0 | 0 | 0 | 0 | 0 | 0 |
| 9404 | ENSMUSG00000033634 | 23840.92045 | 0.392190297 | 0.132750995 | 2.954330372 | 0.003133484 | 0.048164407 | Nat8f2 | N-acetyltransferase 8 (GCN5-related) fam | 21392 | 19946 | 20521 | 21342 | 31893 | 27951 |

| | | | | | | | | | | | | | | | |
|---|---|---|---|---|---|---|---|---|---|---|---|---|---|---|---|
| 9405 | ENSMUSG00000033644 | 5.440607403 | -1.53687662 | 0.98224077 | -1.564663845 | 0.11766175 | 0.502838264 | Piwil2 | piwi-like RNA-mediated gene silencing 2 [! | 4.0155 | 11.307 | 9.0562 | 3.8559 | 2.1812 | 2.2277 |
| 9406 | ENSMUSG00000033653 | 141.1074129 | -0.057800801 | 0.207232148 | -0.27891812 | 0.780307662 | 0.959892424 | Vps8 | VPS8 CORVET complex subunit [Source:M | 138.94 | 165.49 | 127.79 | 153.27 | 133.06 | 128.09 |
| 9407 | ENSMUSG00000033658 | 411.4743115 | 0.491747632 | 0.147675135 | 3.32992845 | 0.000868683 | 0.018865813 | Ddx19b | DEAD box helicase 19b [Source:MGI Symb | 311.6 | 340.24 | 375.33 | 538.86 | 442.79 | 460.02 |
| 9408 | ENSMUSG00000033669 | 30.79816977 | 0.327118316 | 0.400180419 | 0.817427091 | 0.413684399 | 0.835385466 | Zfp7 | zinc finger protein 7 [Source:MGI Symbol; | 23.29 | 23.642 | 35.219 | 37.595 | 31.628 | 33.415 |
| 9409 | ENSMUSG00000033671 | 539.8697161 | 0.025504288 | 0.144082726 | 0.177011424 | 0.859499416 | 0.975502162 | Cep350 | centrosomal protein 350 [Source:MGI Syn | 471.42 | 624.97 | 510.17 | 582.24 | 520.22 | 530.19 |
| 9410 | ENSMUSG00000033676 | 25.54690652 | 0.508888648 | 0.43521732 | 1.169274806 | 0.242292931 | 0.692000002 | Gabrb3 | gamma-aminobutyric acid (GABA) A recep | 22.487 | 17.475 | 23.144 | 25.063 | 28.356 | 36.757 |
| 9411 | ENSMUSG00000033684 | 6435.637796 | 0.230375314 | 0.103098679 | 2.234512763 | 0.025449354 | 0.205313187 | Qsox1 | quiescin Q6 sulfhydryl oxidase 1 [Source:N | 5441.8 | 5976.3 | 6352.4 | 7560.5 | 6476.1 | 6806.7 |
| 9412 | ENSMUSG00000033685 | 386.2093125 | -1.218376246 | 0.176419517 | -6.906130716 | 4.98E-12 | 1.90E-09 | Ucp2 | uncoupling protein 2 (mitochondrial, prot | 612.77 | 592.08 | 415.58 | 239.07 | 210.49 | 247.27 |
| 9413 | ENSMUSG00000033688 | 12387.41941 | 0.192103305 | 0.091577445 | 2.097714183 | 0.035930401 | 0.25475359 | 1300017J02I | RIKEN cDNA 1300017J02 gene [Source:MC | 10872 | 12189 | 11632 | 13014 | 13996 | 12622 |
| 9414 | ENSMUSG00000033697 | 86.29063208 | -0.016372975 | 0.25074825 | -0.065296468 | 0.947937954 | 0.990677354 | Arhgap39 | Rho GTPase activating protein 39 [Source: | 93.16 | 96.624 | 70.437 | 86.758 | 88.34 | 82.424 |
| 9415 | ENSMUSG00000033701 | 628.1558999 | 0.018008151 | 0.123089101 | 0.146301749 | 0.883683185 | 0.981769403 | Acbd6 | acyl-Coenzyme A binding domain containi | 599.92 | 649.64 | 623.87 | 655.5 | 666.37 | 573.63 |
| 9416 | ENSMUSG00000033703 | 256.0814419 | -0.011344022 | 0.159089489 | -0.071305917 | 0.943154286 | 0.990077979 | Fcsk | fucose kinase [Source:MGI Symbol;Acc:M | 244.14 | 263.15 | 264.64 | 277.63 | 253.02 | 233.91 |
| 9417 | ENSMUSG00000033705 | 60.89627769 | -1.171187572 | 0.335359528 | -3.492334274 | 0.000478819 | 0.012044578 | Stard9 | START domain containing 9 [Source:MGI S | 72.279 | 101.76 | 79.493 | 51.091 | 23.994 | 36.757 |
| 9418 | ENSMUSG00000033706 | 192.3956689 | -0.105843582 | 0.178027882 | -0.594533734 | 0.552155187 | 0.895935901 | Smyd5 | SET and MYND domain containing 5 [Sour | 181.5 | 215.86 | 201.25 | 176.41 | 178.86 | 200.49 |
| 9419 | ENSMUSG00000033707 | 1.156610816 | 0.369776097 | 2.544046901 | 0.14534956 | 0.884434887 | NA | Lrrc24 | leucine rich repeat containing 24 [Source: | 0 | 3.0838 | 0 | 3.8559 | 0 | 0 |
| 9420 | ENSMUSG00000033712 | 377.8096219 | -0.081005181 | 0.14101951 | -0.57442535 | 0.565680002 | 0.901895454 | Ccar2 | cell cycle activator and apoptosis regulato | 400.75 | 381.36 | 382.37 | 329.68 | 380.63 | 392.07 |
| 9421 | ENSMUSG00000033713 | 1013.523411 | 0.02743429 | 0.106487857 | 0.257628343 | 0.796693753 | 0.962236316 | Foxn3 | forkhead box N3 [Source:MGI Symbol;Acc | 979.78 | 1052.6 | 980.08 | 1071 | 1018.6 | 979.07 |
| 9422 | ENSMUSG00000033715 | 9807.877027 | 0.27326862 | 0.16619431 | 1.644271812 | 0.100120069 | 0.465788575 | Akr1c14 | aldo-keto reductase family 1, member C1 | 6478.6 | 9064.2 | 11104 | 10163 | 12452 | 9585.7 |
| 9423 | ENSMUSG00000033717 | 0.845761573 | -3.112447952 | 2.7647732 | -1.125751636 | 0.260270661 | NA | Adra2a | adrenergic receptor, alpha 2a [Source:MG | 0 | 2.0558 | 3.0187 | 0 | 0 | 0 |
| 9424 | ENSMUSG00000033720 | 741.6858288 | 0.026903038 | 0.124671772 | 0.21579093 | 0.829150718 | 0.969955221 | Sfxn5 | sideroflexin 5 [Source:MGI Symbol;Acc:M | 770.18 | 654.78 | 778.83 | 745.15 | 798.33 | 702.84 |
| 9425 | ENSMUSG00000033721 | 12.0288762 | -0.363908344 | 0.649778809 | -0.560049572 | 0.575445625 | 0.906770403 | Vav3 | vav 3 oncogene [Source:MGI Symbol;Acc: | 20.078 | 10.279 | 10.062 | 9.6398 | 7.6343 | 14.48 |
| 9426 | ENSMUSG00000033722 | 5.626436595 | -0.804027251 | 1.059501175 | -0.758873393 | 0.447928297 | 0.854051527 | BC034090 | cDNA sequence BC034090 [Source:MGI Sy | 2.4093 | 4.1117 | 15.094 | 7.7118 | 1.0906 | 3.3415 |
| 9427 | ENSMUSG00000033726 | 0 | NA | NA | NA | NA | NA | Emx1 | empty spiracles homeobox 1 [Source:MGI | 0 | 0 | 0 | 0 | 0 | 0 |
| 9428 | ENSMUSG00000033728 | 243.5417483 | -0.057958082 | 0.160504487 | -0.361099448 | 0.718025106 | 0.9447719 | Lrrc14 | leucine rich repeat containing 14 [Source: | 244.14 | 233.34 | 267.66 | 224.61 | 247.57 | 243.93 |
| 9429 | ENSMUSG00000033730 | 0.669251866 | -2.807208097 | 3.391018549 | -0.827836255 | 0.407763239 | NA | Egr3 | early growth response 3 [Source:MGI Sym | 4.0155 | 0 | 0 | 0 | 0 | 0 |
| 9430 | ENSMUSG00000033731 | 0 | NA | NA | NA | NA | NA | 3300002A11 | RIKEN cDNA 3300002A11 gene [Source:M | 0 | 0 | 0 | 0 | 0 | 0 |
| 9431 | ENSMUSG00000033732 | 501.3046368 | -0.131911049 | 0.136835812 | -0.964009691 | 0.335041067 | 0.776159726 | Sf3b3 | splicing factor 3b, subunit 3 [Source:MGI S | 467.41 | 591.05 | 515.2 | 488.74 | 480.96 | 464.47 |
| 9432 | ENSMUSG00000033735 | 2637.045942 | -0.032599183 | 0.091647222 | -0.3557029 | 0.722063073 | 0.9447719 | Spr | sepiapterin reductase [Source:MGI Symbo | 2617.3 | 2604.7 | 2778.2 | 2501.5 | 2632.8 | 2687.7 |
| 9433 | ENSMUSG00000033737 | 0.36353943 | 2.065989254 | 4.034137301 | 0.512126658 | 0.608562368 | NA | Fndc3c1 | fibronectin type III domain containing 3C1 | 0 | 0 | 0 | 0 | 2.1812 | 0 |
| 9434 | ENSMUSG00000033739 | 82.47354222 | 0.302477853 | 0.247492439 | 1.222170075 | 0.221643318 | 0.667949912 | Fkbpl | FK506 binding protein-like [Source:MGI Sy | 77.901 | 71.954 | 71.443 | 83.866 | 89.431 | 100.25 |
| 9435 | ENSMUSG00000033740 | 0 | NA | NA | NA | NA | NA | St18 | suppression of tumorigenicity 18 [Source: | 0 | 0 | 0 | 0 | 0 | 0 |
| 9436 | ENSMUSG00000033751 | 468.973867 | -0.100263346 | 0.12740964 | -0.786936893 | 0.431318809 | 0.844880181 | Gadd45gip1 | growth arrest and DNA-damage-inducible | 480.26 | 465.65 | 510.17 | 470.42 | 436.25 | 451.11 |
| 9437 | ENSMUSG00000033752 | 113.4722127 | 0.233719059 | 0.218203576 | 1.07110554 | 0.284121978 | 0.736651636 | Mnd1 | meiotic nuclear divisions 1 [Source:MGI Sy | 105.21 | 108.96 | 98.612 | 117.61 | 112.33 | 138.12 |
| 9438 | ENSMUSG00000033760 | 226.5906228 | 0.143537864 | 0.161313181 | 0.889808653 | 0.37356864 | 0.80865857 | Rbm4b | RNA binding motif protein 4B [Source:MG | 208.81 | 228.2 | 209.3 | 244.85 | 234.48 | 233.91 |
| 9439 | ENSMUSG00000033762 | 51.3998899 | 0.387043629 | 0.321775762 | 1.202836495 | 0.229039599 | 0.67545951 | Recql4 | RecQ protein-like 4 [Source:MGI Symbol;A | 44.171 | 35.977 | 53.331 | 48.199 | 64.346 | 62.375 |
| 9440 | ENSMUSG00000033763 | 41.06586801 | -0.533465808 | 0.357918683 | -1.490466504 | 0.136101618 | 0.538744881 | Mtss2 | MTSS I-BAR domain containing 2 [Source: | 40.155 | 57.563 | 48.3 | 38.559 | 26.175 | 35.643 |
| 9441 | ENSMUSG00000033765 | 0 | NA | NA | NA | NA | NA | Calm4 | calmodulin 4 [Source:MGI Symbol;Acc:MC | 0 | 0 | 0 | 0 | 0 | 0 |
| 9442 | ENSMUSG00000033767 | 500.9437301 | 0.283815264 | 0.140476577 | 2.020374289 | 0.043344578 | 0.287075127 | Tmem131l | transmembrane 131 like [Source:MGI Sym | 416.01 | 471.81 | 468.91 | 623.69 | 499.5 | 525.73 |
| 9443 | ENSMUSG00000033768 | 30.54035896 | -0.398280976 | 0.448785999 | -0.887463016 | 0.374829669 | 0.809332934 | Nrxn2 | neurexin II [Source:MGI Symbol;Acc:MGI: | 28.109 | 44.2 | 32.2 | 33.739 | 31.628 | 13.366 |
| 9444 | ENSMUSG00000033769 | 98.24777546 | 0.209605101 | 0.24205332 | 0.865945985 | 0.386519784 | 0.817115886 | Exoc6b | exocyst complex component 6B [Source:N | 89.144 | 86.345 | 97.606 | 84.83 | 110.15 | 121.41 |
| 9445 | ENSMUSG00000033770 | 0 | NA | NA | NA | NA | NA | Clcnka | chloride channel, voltage-sensitive Ka [So | 0 | 0 | 0 | 0 | 0 | 0 |
| 9446 | ENSMUSG00000033773 | 185.9115435 | 0.225450747 | 0.205794771 | 1.095512511 | 0.273292168 | 0.728364449 | Rpap2 | RNA polymerase II associated protein 2 [S | 203.99 | 154.19 | 154.96 | 163.88 | 227.94 | 210.52 |
| 9447 | ENSMUSG00000033774 | 0.371281295 | 2.09253362 | 4.031852165 | 0.519000582 | 0.603760335 | NA | Npbwr1 | neuropeptides B/W receptor 1 [Source:M | 0 | 0 | 0 | 0 | 0 | 2.2277 |
| 9448 | ENSMUSG00000033777 | 15.94512553 | -1.990655203 | 0.616529145 | -3.22880957 | 0.001243066 | 0.024562448 | Tlr13 | toll-like receptor 13 [Source:MGI Symbol;/ | 27.305 | 29.81 | 19.119 | 2.8919 | 7.6343 | 8.9108 |
| 9449 | ENSMUSG00000033781 | 673.9610039 | 0.190853102 | 0.118369001 | 1.612357122 | 0.10688425 | 0.482381339 | Asb13 | ankyrin repeat and SOCS box-containing 1 | 626.42 | 649.64 | 612.8 | 770.22 | 671.82 | 712.86 |
| 9450 | ENSMUSG00000033788 | 43.25812235 | -0.81776478 | 0.358480265 | -2.281198883 | 0.022536681 | 0.187879281 | Dysf | dysferlin [Source:MGI Symbol;Acc:MGI:13 | 54.611 | 58.591 | 52.325 | 29.883 | 21.812 | 42.326 |
| 9451 | ENSMUSG00000033790 | 92.87336607 | -0.000275429 | 0.245695332 | -0.001121018 | 0.999105557 | 0.999764475 | Tubgcp5 | tubulin, gamma complex associated prote | 82.72 | 97.652 | 98.612 | 98.326 | 76.343 | 103.59 |
| 9452 | ENSMUSG00000033792 | 41.62978996 | -0.152333779 | 0.36361558 | -0.418941837 | 0.67525864 | 0.932446243 | Atp7a | ATPase, Cu++ transporting, alpha polypep | 48.989 | 44.2 | 38.237 | 41.451 | 50.168 | 26.732 |
| 9453 | ENSMUSG00000033793 | 1186.761816 | 0.020143129 | 0.111043872 | 0.181397933 | 0.856055241 | 0.974319798 | Atp6v1h | ATPase, H+ transporting, lysosomal V1 sul | 1242.4 | 1133.8 | 1159.2 | 1283.1 | 1211.7 | 1090.5 |
| 9454 | ENSMUSG00000033794 | 0 | NA | NA | NA | NA | NA | Lpcat2b | lysophosphatidylcholine acyltransferase 2 | 0 | 0 | 0 | 0 | 0 | 0 |
| 9455 | ENSMUSG00000033799 | 202.4473067 | 0.139205105 | 0.173337331 | 0.803087852 | 0.421923959 | 0.839090903 | Tasor2 | transcription activation suppressor family | 179.09 | 190.16 | 209.3 | 219.79 | 220.3 | 196.04 |
| 9456 | ENSMUSG00000033805 | 0.795980673 | -1.697257209 | 2.614316861 | -0.649216334 | 0.516198555 | NA | Ephx4 | epoxide hydrolase 4 [Source:MGI Symbol; | 1.6062 | 2.0558 | 0 | 0 | 0 | 1.1138 |
| 9457 | ENSMUSG00000033808 | 165.5899063 | 0.076799323 | 0.18926737 | 0.405771597 | 0.68491043 | 0.935737593 | Tmem87a | transmembrane protein 87A [Source:MGI | 141.35 | 170.63 | 172.07 | 165.8 | 179.95 | 163.74 |
| 9458 | ENSMUSG00000033809 | 178.0317608 | 0.012945844 | 0.182096082 | 0.071093477 | 0.94332336 | 0.990077979 | Alg3 | asparagine-linked glycosylation 3 (alpha-1 | 187.93 | 156.24 | 187.16 | 175.44 | 184.31 | 177.1 |
| 9459 | ENSMUSG00000033813 | 853.8109291 | 0.030320808 | 0.115575479 | 0.262346373 | 0.793054413 | 0.961851842 | Tcea1 | transcription elongation factor A (SII) 1 [Sc | 775.8 | 837.75 | 921.72 | 846.37 | 875.77 | 865.46 |
| 9460 | ENSMUSG00000033819 | 684.2943162 | -0.07070802 | 0.126218457 | -0.560203489 | 0.575340647 | 0.906703762 | Ppp1r16a | protein phosphatase 1, regulatory subuni | 715.56 | 624.97 | 762.73 | 706.6 | 656.55 | 639.35 |

| | | | | | | | | | | | | | | | |
|---|---|---|---|---|---|---|---|---|---|---|---|---|---|---|---|
| 9461 | ENSMUSG00000033825 | 2.107368642 | -1.026713842 | 1.438822086 | -0.713579429 | 0.475487282 | NA | Tpsb2 | tryptase beta 2 [Source:MGI Symbol;Acc:I | 2.4093 | 3.0838 | 3.0187 | 1.928 | 1.0906 | 1.1138 |
| 9462 | ENSMUSG00000033826 | 5.241104423 | -0.580773589 | 0.936558991 | -0.620114264 | 0.535182562 | 0.889858671 | Dnah8 | dynein, axonemal, heavy chain 8 [Source: | 5.6217 | 8.2233 | 5.0312 | 4.8199 | 2.1812 | 5.5692 |
| 9463 | ENSMUSG00000033831 | 61454.78165 | 0.387767594 | 0.114076099 | 3.39920104 | 0.00067583 | 0.015516203 | Fgb | fibrinogen beta chain [Source:MGI Symbo | 47989 | 53700 | 58047 | 63784 | 65981 | 79227 |
| 9464 | ENSMUSG00000033834 | 0 | NA | NA | NA | NA | NA | Tpbpa | trophoblast specific protein alpha [Source | 0 | 0 | 0 | 0 | 0 | 0 |
| 9465 | ENSMUSG00000033837 | 0.321325949 | 1.911575114 | 4.048267893 | 0.472195804 | 0.636787034 | NA | Foxh1 | forkhead box H1 [Source:MGI Symbol;Acc | 0 | 0 | 0 | 1.928 | 0 | 0 |
| 9466 | ENSMUSG00000033845 | 963.1569554 | -0.0868286 | 0.112429482 | -0.772293877 | 0.439940391 | 0.848765947 | Mrpl15 | mitochondrial ribosomal protein L15 [Sou | 974.97 | 999.14 | 1002.2 | 889.75 | 1027.4 | 885.51 |
| 9467 | ENSMUSG00000033847 | 0 | NA | NA | NA | NA | NA | Pla2g4c | phospholipase A2, group IVC (cytosolic, ca | 0 | 0 | 0 | 0 | 0 | 0 |
| 9468 | ENSMUSG00000033849 | 0.688942901 | 0.052211695 | 2.79596103 | 0.018673971 | 0.985101193 | NA | B3galt2 | UDP-Gal:betaGlcNAc beta 1,3-galactosyltr | 0 | 2.0558 | 0 | 0.964 | 0 | 1.1138 |
| 9469 | ENSMUSG00000033850 | 0 | NA | NA | NA | NA | NA | Olfr1015 | olfactory receptor 1015 [Source:MGI Symb | 0 | 0 | 0 | 0 | 0 | 0 |
| 9470 | ENSMUSG00000033852 | 0 | NA | NA | NA | NA | NA | Gm28042 | predicted gene, 28042 [Source:MGI Symb | 0 | 0 | 0 | 0 | 0 | 0 |
| 9471 | ENSMUSG00000033854 | 79.15961677 | 0.74865773 | 0.266953268 | 2.804452384 | 0.005040213 | 0.066943658 | Kcnk10 | potassium channel, subfamily K, member | 59.43 | 46.256 | 71.443 | 97.362 | 105.79 | 94.677 |
| 9472 | ENSMUSG00000033855 | 85.13092939 | -0.77719704 | 0.243472364 | -3.192136581 | 0.001412245 | 0.027091137 | Ston1 | stonin 1 [Source:MGI Symbol;Acc:MGI:19 | 110.03 | 103.82 | 108.67 | 62.659 | 64.346 | 61.261 |
| 9473 | ENSMUSG00000033857 | 171.8954563 | -0.135939541 | 0.188183374 | -0.722378059 | 0.470062073 | 0.867051954 | Engase | endo-beta-N-acetylglucosaminidase [Sour | 167.85 | 189.14 | 183.14 | 153.27 | 186.5 | 151.48 |
| 9474 | ENSMUSG00000033860 | 44965.89056 | 0.334520125 | 0.116201926 | 2.878782964 | 0.00399213 | 0.05711391 | Fgg | fibrinogen gamma chain [Source:MGI Sym | 36577 | 39534 | 43217 | 46325 | 46078 | 58065 |
| 9475 | ENSMUSG00000033862 | 276.1983056 | 0.053165815 | 0.163681495 | 0.324812621 | 0.745322892 | 0.949997329 | Cdk10 | cyclin-dependent kinase 10 [Source:MGI S | 236.92 | 265.2 | 311.94 | 285.34 | 272.65 | 285.14 |
| 9476 | ENSMUSG00000033863 | 1838.949775 | 0.42373823 | 0.11410394 | 3.713616098 | 0.000204319 | 0.006382862 | Klf9 | Kruppel-like factor 9 [Source:MGI Symbol; | 1521.9 | 1531.6 | 1659.3 | 2140 | 1853 | 2327.9 |
| 9477 | ENSMUSG00000033871 | 104.4266782 | 0.62238028 | 0.231985658 | 2.682839473 | 0.007300002 | 0.086502219 | Ppargc1b | peroxisome proliferative activated recepto | 85.932 | 77.094 | 83.518 | 127.25 | 107.97 | 144.8 |
| 9478 | ENSMUSG00000033872 | 0.185640647 | 1.126924175 | 4.080472857 | 0.276174898 | 0.782413736 | NA | Best4-ps | bestrophin 4, pseudogene [Source:MGI Sy | 0 | 0 | 0 | 0 | 0 | 1.1138 |
| 9479 | ENSMUSG00000033880 | 1050.107918 | -0.258868729 | 0.128991709 | -2.006863312 | 0.044764219 | 0.292311362 | Lgals3bp | lectin, galactoside-binding, soluble, 3 bind | 1334.8 | 1068 | 1028.4 | 998.68 | 961.93 | 908.9 |
| 9480 | ENSMUSG00000033882 | 1.580607272 | -1.771860989 | 1.909966014 | -0.927692418 | 0.353567143 | NA | Rbm46 | RNA binding motif protein 46 [Source:MG | 3.2124 | 3.0838 | 1.0062 | 0 | 2.1812 | 0 |
| 9481 | ENSMUSG00000033883 | 84.83598117 | -0.020259803 | 0.252336617 | -0.080288795 | 0.936007569 | 0.988420305 | Zfp267 | zinc finger protein 267 [Source:MGI Symb | 96.372 | 81.206 | 78.487 | 86.758 | 93.793 | 72.4 |
| 9482 | ENSMUSG00000033885 | 470.4504903 | -0.095146195 | 0.134122502 | -0.709397702 | 0.478077713 | 0.869794311 | Pxk | PX domain containing serine/threonine ki | 501.94 | 448.17 | 507.15 | 435.72 | 434.07 | 495.66 |
| 9483 | ENSMUSG00000033898 | 10526.08998 | 0.319601646 | 0.098459309 | 3.246027718 | 0.001170274 | 0.023476907 | Cfhr2 | complement factor H-related 2 [Source:M | 8862.2 | 9221.5 | 10011 | 10664 | 12362 | 12035 |
| 9484 | ENSMUSG00000033900 | 13.39652675 | 0.071906003 | 0.647903313 | 0.110982614 | 0.911630133 | 0.98589875 | Map9 | microtubule-associated protein 9 [Source | 4.8186 | 19.53 | 15.094 | 14.46 | 11.997 | 14.48 |
| 9485 | ENSMUSG00000033902 | 92.32349958 | -0.151534234 | 0.239057932 | -0.633880802 | 0.526158615 | 0.888334265 | Mapkbp1 | mitogen-activated protein kinase binding | 105.21 | 94.568 | 91.568 | 91.578 | 76.343 | 94.677 |
| 9486 | ENSMUSG00000033904 | 144.8708133 | 0.243912698 | 0.253030051 | 0.963967311 | 0.335062315 | 0.776159726 | Ccp110 | centriolar coiled coil protein 110 [Source:I | 122.87 | 148.02 | 127.79 | 217.86 | 112.33 | 140.34 |
| 9487 | ENSMUSG00000033906 | 0.160662974 | 1.126924175 | 4.080472857 | 0.276174898 | 0.782413736 | NA | Zdhhc15 | zinc finger, DHHC domain containing 15 [S | 0 | 0 | 0 | 0.964 | 0 | 0 |
| 9488 | ENSMUSG00000033909 | 341.355307 | 0.115056307 | 0.157083914 | 0.732451231 | 0.463893199 | 0.861872906 | Usp36 | ubiquitin specific peptidase 36 [Source:M | 308.39 | 330.99 | 344.14 | 353.78 | 304.28 | 406.55 |
| 9489 | ENSMUSG00000033910 | 59.67021687 | -1.330106316 | 0.306125889 | -4.344965143 | 1.39E-05 | 0.000704477 | Gucy1a1 | guanylate cyclase 1, soluble, alpha 1 [Sou | 97.175 | 88.401 | 70.437 | 33.739 | 37.081 | 31.188 |
| 9490 | ENSMUSG00000033916 | 1315.104407 | -0.057413138 | 0.101475926 | -0.565780873 | 0.571542751 | 0.904651508 | Chmp2a | charged multivesicular body protein 2A [S | 1381.3 | 1296.2 | 1345.3 | 1233.9 | 1273.8 | 1360 |
| 9491 | ENSMUSG00000033917 | 1476.135717 | -0.407967492 | 0.164624053 | -2.478176698 | 0.013205573 | 0.130993706 | Gde1 | glycerophosphodiester phosphodiesterase | 1549.2 | 2003.4 | 1499.3 | 1583.8 | 1198.6 | 1022.5 |
| 9492 | ENSMUSG00000033918 | 483.1589333 | -0.137156415 | 0.137850631 | -0.994963995 | 0.319753776 | 0.76673054 | Parl | presenilin associated, rhomboid-like [Sour | 460.98 | 539.66 | 518.22 | 438.61 | 509.32 | 432.17 |
| 9493 | ENSMUSG00000033931 | 222.4851152 | -0.062104333 | 0.173677949 | -0.357583292 | 0.720655184 | 0.9447719 | Rbm34 | RNA binding motif protein 34 [Source:MG | 229.69 | 248.76 | 203.26 | 207.26 | 242.12 | 203.83 |
| 9494 | ENSMUSG00000033933 | 227.1040372 | 0.158429269 | 0.172618269 | 0.917801286 | 0.358722909 | 0.79836408 | Vhl | von Hippel-Lindau tumor suppressor [Sou | 193.55 | 248.76 | 202.26 | 242.92 | 231.21 | 243.93 |
| 9495 | ENSMUSG00000033938 | 1234.34834 | -0.095417732 | 0.110519763 | -0.863354476 | 0.387942601 | 0.817937623 | Ndufb7 | NADH:ubiquinone oxidoreductase subuni | 1287.4 | 1235.6 | 1302.1 | 1167.4 | 1098.3 | 1315.4 |
| 9496 | ENSMUSG00000033940 | 889.2887876 | -0.041692911 | 0.116141248 | -0.358984525 | 0.719606666 | 0.9447719 | Brk1 | BRICK1, SCAR/WAVE actin-nucleating com | 858.52 | 990.91 | 857.32 | 847.34 | 860.5 | 921.15 |
| 9497 | ENSMUSG00000033943 | 464.7638814 | 0.228246047 | 0.156938173 | 1.454369208 | 0.145843972 | 0.556690797 | Mga | MAX gene associated [Source:MGI Symbo | 428.86 | 388.55 | 466.9 | 535.01 | 407.89 | 561.38 |
| 9498 | ENSMUSG00000033948 | 67.06922457 | 0.096118342 | 0.273204836 | 0.351817866 | 0.724974859 | 0.9448775 | Zswim5 | zinc finger SWIM-type containing 5 [Sourc | 60.233 | 72.982 | 61.381 | 65.55 | 66.528 | 75.741 |
| 9499 | ENSMUSG00000033949 | 3.060115853 | -3.924235894 | 1.633373358 | -2.402534531 | 0.016281897 | NA | Trim36 | tripartite motif-containing 36 [Source:MG | 4.0155 | 8.2233 | 5.0312 | 0 | 1.0906 | 0 |
| 9500 | ENSMUSG00000033952 | 7.309855563 | -0.46721941 | 0.936558347 | -0.498868448 | 0.617872063 | 0.917509899 | Aspm | abnormal spindle microtubule assembly [S | 8.031 | 11.307 | 6.0375 | 0.964 | 14.178 | 3.3415 |
| 9501 | ENSMUSG00000033953 | 707.4075753 | -0.133178075 | 0.122850423 | -1.084066883 | 0.278335145 | 0.73110838 | Ppp3r1 | protein phosphatase 3, regulatory subunit | 682.64 | 798.69 | 739.59 | 669.96 | 720.9 | 632.66 |
| 9502 | ENSMUSG00000033955 | 1387.762494 | -0.076168615 | 0.119737708 | -0.636128888 | 0.524692418 | 0.888334265 | Tnks1bp1 | tankyrase 1 binding protein 1 [Source:MG | 1442.4 | 1377.4 | 1454 | 1486.5 | 1411.3 | 1155.1 |
| 9503 | ENSMUSG00000033960 | 80.7876753 | -0.531235164 | 0.265391922 | -2.00170058 | 0.045316944 | 0.294458346 | Jcad | junctional cadherin 5 associated [Source:I | 85.932 | 118.21 | 82.512 | 63.623 | 67.618 | 66.831 |
| 9504 | ENSMUSG00000033961 | 62.91431717 | 0.112748145 | 0.317468894 | 0.355147061 | 0.722479422 | 0.9447719 | Zfp446 | zinc finger protein 446 [Source:MGI Symb | 58.626 | 64.759 | 58.362 | 85.794 | 67.618 | 42.326 |
| 9505 | ENSMUSG00000033963 | 0 | NA | NA | NA | NA | NA | Fancd2os | Fancd2 opposite strand [Source:MGI Symb | 0 | 0 | 0 | 0 | 0 | 0 |
| 9506 | ENSMUSG00000033964 | 369.9231398 | 0.00684662 | 0.141595108 | 0.048353504 | 0.961434514 | 0.992661062 | Zbtb41 | zinc finger and BTB domain containing 41 | 338.91 | 397.8 | 371.3 | 386.56 | 372.99 | 351.97 |
| 9507 | ENSMUSG00000033965 | 2930.933919 | 0.108977457 | 0.100086545 | 1.088832237 | 0.276227873 | 0.730038567 | Slc16a2 | solute carrier family 16 (monocarboxylic a | 2816.5 | 2838.1 | 2805.4 | 2734.8 | 3209.7 | 3181.1 |
| 9508 | ENSMUSG00000033966 | 11.68716756 | -1.095007236 | 0.653166965 | -1.676458385 | 0.093648441 | 0.450459278 | Cdkl4 | cyclin-dependent kinase-like 4 [Source:M | 16.062 | 19.53 | 12.075 | 4.8199 | 8.7249 | 8.9108 |
| 9509 | ENSMUSG00000033967 | 3.121942124 | 3.198127935 | 1.674589881 | 1.909797719 | 0.056159263 | NA | Rnf225 | ring finger protein 225 [Source:MGI Symb | 0.8031 | 0 | 1.0062 | 5.7839 | 0 | 11.138 |
| 9510 | ENSMUSG00000033970 | 86.20334343 | 0.39898664 | 0.303227824 | 1.315798248 | 0.188241767 | 0.625244593 | Rfc3 | replication factor C (activator 1) 3 [Source | 54.611 | 87.373 | 81.506 | 83.866 | 135.24 | 74.628 |
| 9511 | ENSMUSG00000033972 | 175.3465739 | -0.011921822 | 0.211672691 | -0.056321966 | 0.95508532 | 0.991678478 | Zfp944 | zinc finger protein 944 [Source:MGI Symb | 173.47 | 196.33 | 157.98 | 132.06 | 201.76 | 190.47 |
| 9512 | ENSMUSG00000033981 | 0 | NA | NA | NA | NA | NA | Gria2 | glutamate receptor, ionotropic, AMPA2 (a | 0 | 0 | 0 | 0 | 0 | 0 |
| 9513 | ENSMUSG00000033983 | 100.0642716 | 0.244189389 | 0.240234983 | 1.016460575 | 0.309410112 | 0.758141261 | Coil | coilin [Source:MGI Symbol;Acc:MGI:10484 | 103.6 | 81.206 | 89.556 | 99.29 | 127.6 | 99.132 |
| 9514 | ENSMUSG00000033985 | 241.3792676 | 0.126327085 | 0.173290866 | 0.728988713 | 0.466008567 | 0.863779479 | Tesk2 | testis-specific kinase 2 [Source:MGI Symb | 197.56 | 252.87 | 242.5 | 227.5 | 267.2 | 260.64 |
| 9515 | ENSMUSG00000033987 | 6.744007833 | -0.134072356 | 0.90200518 | -0.148638122 | 0.88183919 | 0.981676444 | Dnah17 | dynein, axonemal, heavy chain 17 [Source | 8.8341 | 7.1954 | 5.0312 | 3.8559 | 2.1812 | 13.366 |
| 9516 | ENSMUSG00000033991 | 355.2638542 | 0.037368365 | 0.139106731 | 0.268630891 | 0.788213738 | 0.960870081 | Ttc37 | tetratricopeptide repeat domain 37 [Sour | 332.48 | 361.83 | 358.22 | 366.31 | 371.9 | 340.84 |

| | | | | | | | | | | | | | | | |
|---|---|---|---|---|---|---|---|---|---|---|---|---|---|---|---|
| 9517 | ENSMUSG00000033998 | 3.160336278 | 0.718126908 | 1.251897724 | 0.573630652 | 0.566217765 | NA | Kcnk1 | potassium channel, subfamily K, member | 4.0155 | 1.0279 | 2.0125 | 1.928 | 2.1812 | 7.7969 |
| 9518 | ENSMUSG00000034000 | 0 | NA | NA | NA | NA | NA | Neu4 | sialidase 4 [Source:MGI Symbol;Acc:MGI:2 | 0 | 0 | 0 | 0 | 0 | 0 |
| 9519 | ENSMUSG00000034006 | 1864.398704 | 0.132369776 | 0.118899396 | 1.113292241 | 0.265582938 | 0.721565724 | Slc66a2 | solute carrier family 66 member 2 [Source | 1994.9 | 1646.7 | 1693.5 | 1894.2 | 1799.5 | 2157.5 |
| 9520 | ENSMUSG00000034007 | 103.1044283 | 0.360289235 | 0.300682833 | 1.198236799 | 0.230824843 | 0.677910147 | Scaper | S phase cyclin A-associated protein in the | 84.326 | 100.74 | 85.531 | 64.587 | 131.96 | 151.48 |
| 9521 | ENSMUSG00000034009 | 0 | NA | NA | NA | NA | NA | Rxfp1 | relaxin/insulin-like family peptide recepto | 0 | 0 | 0 | 0 | 0 | 0 |
| 9522 | ENSMUSG00000034021 | 364.4640568 | 0.118504603 | 0.162724991 | 0.728250786 | 0.466460082 | 0.863779479 | Pds5b | PDS5 cohesin associated factor B [Source: | 304.38 | 376.22 | 368.29 | 329.68 | 438.43 | 369.8 |
| 9523 | ENSMUSG00000034022 | 844.1490628 | 0.090290472 | 0.13073893 | 0.690616578 | 0.489806527 | 0.875278016 | Cpsf1 | cleavage and polyadenylation specific fact | 783.02 | 865.51 | 806 | 1008.3 | 821.24 | 780.8 |
| 9524 | ENSMUSG00000034023 | 4.539204614 | -0.212834086 | 0.975270605 | -0.218230802 | 0.827249287 | NA | Fancd2 | Fanconi anemia, complementation group | 6.4248 | 4.1117 | 4.025 | 3.8559 | 4.3625 | 4.4554 |
| 9525 | ENSMUSG00000034024 | 1933.283678 | 0.017321882 | 0.097434023 | 0.177780628 | 0.858895262 | 0.975439379 | Cct2 | chaperonin containing Tcp1, subunit 2 (be | 2002.1 | 1905.8 | 1856.5 | 1933.7 | 2051.5 | 1850.1 |
| 9526 | ENSMUSG00000034028 | 1.274125293 | -2.554690053 | 2.062976523 | -1.238351491 | 0.215585759 | NA | Cd226 | CD226 antigen [Source:MGI Symbol;Acc:N | 1.6062 | 2.0558 | 3.0187 | 0.964 | 0 | 0 |
| 9527 | ENSMUSG00000034031 | 0 | NA | NA | NA | NA | NA | Ccdc182 | coiled-coil domain containing 182 [Source | 0 | 0 | 0 | 0 | 0 | 0 |
| 9528 | ENSMUSG00000034032 | 189.3005602 | -0.041396843 | 0.196622744 | -0.210539444 | 0.833246669 | 0.970611245 | Rpap1 | RNA polymerase II associated protein 1 [S | 182.3 | 194.28 | 200.24 | 227.5 | 177.77 | 153.71 |
| 9529 | ENSMUSG00000034035 | 73.34872564 | 0.150832949 | 0.270194146 | 0.558239145 | 0.576681094 | 0.907752116 | Ccdc17 | coiled-coil domain containing 17 [Source:I | 71.476 | 67.843 | 69.431 | 93.506 | 65.437 | 72.4 |
| 9530 | ENSMUSG00000034037 | 59.346772 | -0.559274965 | 0.300084583 | -1.863724416 | 0.062360415 | 0.356831368 | Fgd5 | FYVE, RhoGEF and PH domain containing ! | 73.082 | 65.787 | 73.456 | 58.803 | 38.172 | 46.781 |
| 9531 | ENSMUSG00000034039 | 0 | NA | NA | NA | NA | NA | Prss29 | protease, serine 29 [Source:MGI Symbol;A | 0 | 0 | 0 | 0 | 0 | 0 |
| 9532 | ENSMUSG00000034040 | 0 | NA | NA | NA | NA | NA | Galnt17 | polypeptide N-acetylgalactosaminyltransf | 0 | 0 | 0 | 0 | 0 | 0 |
| 9533 | ENSMUSG00000034041 | 17.62984469 | -0.726845574 | 0.589361876 | -1.233275519 | 0.217472988 | 0.663473804 | Lyl1 | lymphoblastomic leukemia 1 [Source:MGI | 24.896 | 25.698 | 15.094 | 4.8199 | 17.45 | 17.822 |
| 9534 | ENSMUSG00000034042 | 946.0495957 | -0.013446037 | 0.106695501 | -0.126022531 | 0.899714089 | 0.982798755 | Gpbp1l1 | GC-rich promoter binding protein 1-like 1 | 951.68 | 899.43 | 1000.2 | 942.77 | 949.93 | 932.29 |
| 9535 | ENSMUSG00000034055 | 120.2902861 | 0.078836861 | 0.21448682 | 0.367560397 | 0.713201049 | 0.944239215 | Phka1 | phosphorylase kinase alpha 1 [Source:MG | 126.09 | 101.76 | 122.76 | 116.64 | 131.96 | 122.52 |
| 9536 | ENSMUSG00000034057 | 0 | NA | NA | NA | NA | NA | Myrfl | myelin regulatory factor-like [Source:MGI | 0 | 0 | 0 | 0 | 0 | 0 |
| 9537 | ENSMUSG00000034059 | 2.829807026 | -1.215375179 | 1.351479774 | -0.89929217 | 0.368497057 | NA | Ypel4 | yippee like 4 [Source:MGI Symbol;Acc:MG | 4.8186 | 2.0558 | 5.0312 | 2.8919 | 2.1812 | 0 |
| 9538 | ENSMUSG00000034063 | 3.766909648 | -0.120959941 | 1.226830295 | -0.098595496 | 0.921459447 | NA | 4930590J08I | RIKEN cDNA 4930590J08 gene [Source:MC | 1.6062 | 6.1675 | 4.025 | 0.964 | 8.7249 | 1.1138 |
| 9539 | ENSMUSG00000034064 | 189.8646634 | -0.34996347 | 0.196407719 | -1.781821364 | 0.074778367 | 0.399358478 | Poglut1 | protein O-glucosyltransferase 1 [Source:N | 190.34 | 198.39 | 249.55 | 141.7 | 183.22 | 175.99 |
| 9540 | ENSMUSG00000034066 | 257.5147477 | 0.319588928 | 0.182931946 | 1.747037272 | 0.080630873 | 0.417261162 | Farp2 | FERM, RhoGEF and pleckstrin domain pro | 181.5 | 261.09 | 245.52 | 257.38 | 297.74 | 301.85 |
| 9541 | ENSMUSG00000034071 | 22.53633495 | 0.049681044 | 0.459652264 | 0.108083974 | 0.91392908 | 0.98625062 | Zfp551 | zinc fingr protein 551 [Source:MGI Symbo | 20.881 | 20.558 | 25.156 | 28.919 | 18.541 | 21.163 |
| 9542 | ENSMUSG00000034075 | 1445.020051 | -0.033384954 | 0.107496499 | -0.310567828 | 0.756129188 | 0.953085143 | Zdhhc5 | zinc finger, DHHC domain containing 5 [Sc | 1370.9 | 1527.5 | 1488.2 | 1515.4 | 1453.8 | 1314.3 |
| 9543 | ENSMUSG00000034083 | 151.5992769 | 0.188901774 | 0.199416692 | 0.947271623 | 0.343500387 | 0.785243795 | Ccdc174 | coiled-coil domain containing 174 [Source | 123.68 | 156.24 | 145.91 | 177.37 | 152.69 | 153.71 |
| 9544 | ENSMUSG00000034087 | 0 | NA | NA | NA | NA | NA | Nlrp4b | NLR family, pyrin domain containing 4B [S | 0 | 0 | 0 | 0 | 0 | 0 |
| 9545 | ENSMUSG00000034088 | 6756.54743 | 0.137134422 | 0.091090506 | 1.505474367 | 0.132202162 | 0.533232592 | Hdlbp | high density lipoprotein (HDL) binding pro | 6548.5 | 6713.3 | 6045.5 | 7417.8 | 7049.8 | 6764.4 |
| 9546 | ENSMUSG00000034098 | 0 | NA | NA | NA | NA | NA | Fstl5 | follistatin-like 5 [Source:MGI Symbol;Acc:I | 0 | 0 | 0 | 0 | 0 | 0 |
| 9547 | ENSMUSG00000034101 | 369.0414458 | -0.042349385 | 0.137279854 | -0.308489439 | 0.757709935 | 0.953466696 | Ctnnd1 | catenin (cadherin associated protein), del | 377.46 | 387.53 | 358.22 | 351.85 | 384.99 | 354.2 |
| 9548 | ENSMUSG00000034105 | 228.8414113 | 0.162018184 | 0.179643495 | 0.901887285 | 0.367116746 | 0.805239112 | Meak7 | MTOR associated protein , eak-7 homolog | 208 | 195.3 | 244.52 | 218.82 | 227.94 | 278.46 |
| 9549 | ENSMUSG00000034107 | 0 | NA | NA | NA | NA | NA | Ano7 | anoctamin 7 [Source:MGI Symbol;Acc:MG | 0 | 0 | 0 | 0 | 0 | 0 |
| 9550 | ENSMUSG00000034108 | 3894.81005 | 0.194084984 | 0.084505274 | 2.296720373 | 0.021634727 | 0.183082289 | Ccs | copper chaperone for superoxide dismuta | 3696.7 | 3524.7 | 3677.8 | 4154.7 | 4182.5 | 4132.4 |
| 9551 | ENSMUSG00000034109 | 1603.924358 | -0.076605693 | 0.108349526 | -0.707023793 | 0.479551692 | 0.870236354 | Golim4 | golgi integral membrane protein 4 [Source | 1517.1 | 1819.4 | 1604 | 1529.8 | 1621.7 | 1531.5 |
| 9552 | ENSMUSG00000034110 | 55.22798163 | 0.23707276 | 0.305396826 | 0.776277746 | 0.43758501 | 0.847422601 | Kctd7 | potassium channel tetramerisation domai | 48.186 | 56.535 | 47.293 | 48.199 | 65.437 | 65.717 |
| 9553 | ENSMUSG00000034111 | 79.67178892 | 0.167722481 | 0.267666421 | 0.626610091 | 0.530914852 | 0.888794797 | Tmed8 | transmembrane p24 trafficking protein 8 [ | 57.823 | 86.345 | 81.506 | 83.866 | 90.521 | 77.969 |
| 9554 | ENSMUSG00000034112 | 0 | NA | NA | NA | NA | NA | Atp2c2 | ATPase, Ca++ transporting, type 2C, meml | 0 | 0 | 0 | 0 | 0 | 0 |
| 9555 | ENSMUSG00000034115 | 0 | NA | NA | NA | NA | NA | Scn11a | sodium channel, voltage-gated, type XI, al | 0 | 0 | 0 | 0 | 0 | 0 |
| 9556 | ENSMUSG00000034116 | 36.03373774 | -1.385138646 | 0.425854654 | -3.252608924 | 0.001143508 | 0.023067913 | Vav1 | vav 1 oncogene [Source:MGI Symbol;Acc: | 39.352 | 70.926 | 46.287 | 23.135 | 11.997 | 24.505 |
| 9557 | ENSMUSG00000034117 | 0 | NA | NA | NA | NA | NA | Ptgdr2 | prostaglandin D2 receptor 2 [Source:MGI | 0 | 0 | 0 | 0 | 0 | 0 |
| 9558 | ENSMUSG00000034118 | 459.7296756 | 0.046912328 | 0.169108002 | 0.277410458 | 0.781464952 | 0.960079945 | Tpst1 | protein-tyrosine sulfotransferase 1 [Sourc | 346.94 | 464.62 | 546.39 | 461.75 | 484.23 | 454.45 |
| 9559 | ENSMUSG00000034120 | 1284.483952 | -0.079003395 | 0.131868254 | -0.599108521 | 0.54910052 | 0.895703112 | Srsf2 | serine and arginine-rich splicing factor 2 [! | 1227.1 | 1306.5 | 1426.9 | 1456.6 | 1212.8 | 1077.1 |
| 9560 | ENSMUSG00000034121 | 39.87699804 | -0.065048753 | 0.379255091 | -0.171517151 | 0.863817145 | 0.976954351 | Mks1 | MKS transition zone complex subunit 1 [S | 48.186 | 26.726 | 47.293 | 45.307 | 30.537 | 41.212 |
| 9561 | ENSMUSG00000034122 | 0 | NA | NA | NA | NA | NA | Nlrp4d | NLR family, pyrin domain containing 4D [S | 0 | 0 | 0 | 0 | 0 | 0 |
| 9562 | ENSMUSG00000034126 | 382.2085713 | 0.335638906 | 0.161597053 | 2.0770113 | 0.037800516 | 0.263873875 | Pomt2 | protein-O-mannosyltransferase 2 [Source | 312.41 | 336.13 | 366.27 | 497.41 | 355.54 | 425.49 |
| 9563 | ENSMUSG00000034127 | 8.643620224 | 0.992989453 | 0.771773898 | 1.286632595 | 0.198222377 | 0.64054702 | Tspan8 | tetraspanin 8 [Source:MGI Symbol;Acc:M | 8.031 | 7.1954 | 2.0125 | 12.532 | 8.7249 | 13.366 |
| 9564 | ENSMUSG00000034135 | 924.9405809 | -0.057961869 | 0.112647344 | -0.514542709 | 0.606872609 | 0.916376001 | Sik3 | SIK family kinase 3 [Source:MGI Symbol;A | 885.02 | 1015.6 | 930.78 | 917.71 | 861.59 | 938.97 |
| 9565 | ENSMUSG00000034139 | 0 | NA | NA | NA | NA | NA | Serpini2 | serine (or cysteine) peptidase inhibitor, cl | 0 | 0 | 0 | 0 | 0 | 0 |
| 9566 | ENSMUSG00000034145 | 0.74256259 | 3.09446134 | 3.273353887 | 0.945348852 | 0.344480805 | NA | Tmem63c | transmembrane protein 63c [Source:MGI | 0 | 0 | 0 | 0 | 0 | 4.4554 |
| 9567 | ENSMUSG00000034151 | 0 | NA | NA | NA | NA | NA | Zbbx | zinc finger, B-box domain containing [Sou | 0 | 0 | 0 | 0 | 0 | 0 |
| 9568 | ENSMUSG00000034152 | 493.3499582 | 0.037244682 | 0.127645235 | 0.291782783 | 0.77045271 | 0.956515747 | Exoc3 | exocyst complex component 3 [Source:M | 500.33 | 480.04 | 479.98 | 467.53 | 537.67 | 494.55 |
| 9569 | ENSMUSG00000034154 | 312.7850644 | 0.211710929 | 0.144243753 | 1.467730319 | 0.142177492 | 0.551920354 | Ino80 | INO80 complex subunit [Source:MGI Syml | 305.98 | 272.4 | 290.8 | 337.39 | 332.64 | 337.49 |
| 9570 | ENSMUSG00000034156 | 5.581811965 | 0.144055238 | 0.907307211 | 0.158772284 | 0.873848286 | 0.979925081 | Tspoap1 | TSPO associated protein 1 [Source:MGI Sy | 4.8186 | 4.1117 | 7.0437 | 8.6758 | 3.2719 | 5.5692 |
| 9571 | ENSMUSG00000034157 | 1169.82625 | -0.165446877 | 0.114016388 | -1.451079791 | 0.146757643 | 0.55769976 | Cipc | CLOCK interacting protein, circadian [Sour | 1355.6 | 1194.4 | 1159.2 | 1067.1 | 1184.4 | 1058.2 |
| 9572 | ENSMUSG00000034158 | 1195.112403 | -0.203443712 | 0.133351446 | -1.525620592 | 0.127104381 | 0.522431874 | Lrrc58 | leucine rich repeat containing 58 [Source: | 1154.1 | 1446.3 | 1238.7 | 1181.8 | 1198.6 | 951.22 |

| | | | | | | | | | | | | | | | |
|---|---|---|---|---|---|---|---|---|---|---|---|---|---|---|---|
| 9573 | ENSMUSG00000034159 | 1.279591891 | -3.736153947 | 2.409849227 | -1.55036834 | 0.121053133 | NA | Mab21l4 | mab-21-like 4 [Source:MGI Symbol;Acc:M | 5.6217 | 2.0558 | 0 | 0 | 0 | 0 |
| 9574 | ENSMUSG00000034160 | 1653.604853 | 0.071744708 | 0.14354436 | 0.499808613 | 0.617209845 | 0.917509899 | Ogt | O-linked N-acetylglucosamine (GlcNAc) tr | 1627.9 | 1595.3 | 1615 | 1957.8 | 1825.7 | 1299.9 |
| 9575 | ENSMUSG00000034161 | 3.620082367 | 0.471669182 | 1.196820155 | 0.394101971 | 0.693505756 | NA | Scx | scleraxis [Source:MGI Symbol;Acc:MGI:10 | 4.0155 | 0 | 5.0312 | 3.8559 | 4.3625 | 4.4554 |
| 9576 | ENSMUSG00000034163 | 695.9259435 | 0.306372145 | 0.144675675 | 2.117647937 | 0.034204895 | 0.247603883 | Zfc3h1 | zinc finger, C3H1-type containing [Source: | 588.67 | 646.56 | 632.93 | 922.53 | 663.1 | 721.77 |
| 9577 | ENSMUSG00000034164 | 2.810833928 | 1.120404662 | 1.409845222 | 0.794700471 | 0.426787758 | NA | Emid1 | EMI domain containing 1 [Source:MGI Syr | 3.2124 | 0 | 2.0125 | 2.8919 | 7.6343 | 1.1138 |
| 9578 | ENSMUSG00000034165 | 664.8584367 | -0.05963249 | 0.12234538 | -0.487411047 | 0.625967066 | 0.919638611 | Ccnd3 | cyclin D3 [Source:MGI Symbol;Acc:MGI:8 | 696.29 | 615.72 | 723.49 | 660.32 | 620.56 | 672.76 |
| 9579 | ENSMUSG00000034168 | 776.2154536 | 0.335937687 | 0.178370273 | 1.883372614 | 0.059649889 | 0.347816342 | Irf2bpl | interferon regulatory factor 2 binding pro | 610.36 | 655.81 | 792.92 | 844.44 | 653.28 | 1100.5 |
| 9580 | ENSMUSG00000034171 | 3097.675225 | 0.032174368 | 0.08545395 | 0.376511181 | 0.706536905 | 0.94295036 | Faah | fatty acid amide hydrolase [Source:MGI S | 3086.3 | 3064.2 | 3038.9 | 3174.4 | 3070.1 | 3152.2 |
| 9581 | ENSMUSG00000034173 | 163.5949719 | 0.263953359 | 0.254902757 | 1.035506101 | 0.300432619 | 0.751875664 | Zbed5 | zinc finger, BED type containing 5 [Source | 210.41 | 111.02 | 123.77 | 211.11 | 155.96 | 169.3 |
| 9582 | ENSMUSG00000034175 | 380.2608053 | 0.267260948 | 0.151351611 | 1.765828234 | 0.07742467 | 0.40712138 | Rhbdd3 | rhomboid domain containing 3 [Source:M | 400.75 | 318.65 | 314.95 | 428.01 | 393.71 | 425.49 |
| 9583 | ENSMUSG00000034177 | 397.7942753 | 0.828675788 | 0.201845604 | 4.105493359 | 4.03E-05 | 0.001768346 | Rnf43 | ring finger protein 43 [Source:MGI Symbo | 317.23 | 300.15 | 241.5 | 399.09 | 448.24 | 680.56 |
| 9584 | ENSMUSG00000034185 | 0 | NA | NA | NA | NA | NA | 6430628N08 | RIKEN cDNA 6430628N08 gene [Source:M | 0 | 0 | 0 | 0 | 0 | 0 |
| 9585 | ENSMUSG00000034187 | 273.3384652 | -0.073687883 | 0.16485465 | -0.446986985 | 0.654884457 | 0.925797231 | Nsf | N-ethylmaleimide sensitive fusion protein | 248.96 | 321.74 | 270.68 | 249.67 | 268.29 | 280.69 |
| 9586 | ENSMUSG00000034189 | 190.7224429 | 0.077504942 | 0.178774778 | 0.433533986 | 0.664626881 | 0.929301301 | Hsdl1 | hydroxysteroid dehydrogenase like 1 [Sou | 194.35 | 177.83 | 184.14 | 183.16 | 184.31 | 220.54 |
| 9587 | ENSMUSG00000034190 | 841.5349513 | 0.190623226 | 0.115715513 | 1.647343738 | 0.099487413 | 0.464748317 | Chmp7 | charged multivesicular body protein 7 [So | 800.69 | 778.13 | 778.83 | 886.86 | 824.51 | 980.18 |
| 9588 | ENSMUSG00000034192 | 59.48711329 | 0.025725496 | 0.327228148 | 0.07861639 | 0.93733775 | 0.988420305 | Lsm3 | LSM3 homolog, U6 small nuclear RNA and | 69.067 | 38.033 | 69.431 | 54.947 | 71.981 | 53.465 |
| 9589 | ENSMUSG00000034194 | 229.4371838 | 0.003644038 | 0.172774078 | 0.021091346 | 0.983172788 | 0.998580735 | R3hcc1 | R3H domain and coiled-coil containing 1 [ | 201.58 | 252.87 | 233.45 | 214 | 250.84 | 223.88 |
| 9590 | ENSMUSG00000034201 | 355.5993564 | -0.072004828 | 0.153913115 | -0.467827757 | 0.639907769 | 0.923932929 | Gas2l1 | growth arrest-specific 2 like 1 [Source:MG | 386.29 | 328.93 | 378.35 | 393.3 | 333.73 | 312.99 |
| 9591 | ENSMUSG00000034203 | 252.6967967 | 0.121771206 | 0.164220954 | 0.741508334 | 0.458385281 | 0.859322431 | Chchd4 | coiled-coil-helix-coiled-coil-helix domain c | 240.93 | 254.92 | 230.43 | 267.02 | 291.2 | 231.68 |
| 9592 | ENSMUSG00000034205 | 24.15576078 | -0.861686663 | 0.496774528 | -1.734562892 | 0.082818257 | 0.424179628 | Loxl2 | lysyl oxidase-like 2 [Source:MGI Symbol;A | 47.383 | 25.698 | 20.125 | 16.388 | 14.178 | 21.163 |
| 9593 | ENSMUSG00000034206 | 2.13260504 | -0.98964569 | 1.534730285 | -0.644833623 | 0.51903501 | NA | Polq | polymerase (DNA directed), theta [Source | 2.4093 | 2.0558 | 4.025 | 0.964 | 0 | 3.3415 |
| 9594 | ENSMUSG00000034209 | 1.209843426 | -3.647354608 | 2.210792771 | -1.649794886 | 0.098984895 | NA | Rasl10a | RAS-like, family 10, member A [Source:M | 3.2124 | 1.0279 | 3.0187 | 0 | 0 | 0 |
| 9595 | ENSMUSG00000034210 | 380.3345637 | 0.065262839 | 0.143741271 | 0.454029931 | 0.649807284 | 0.925643431 | Efcab14 | EF-hand calcium binding domain 14 [Sour | 368.62 | 355.66 | 390.42 | 347.03 | 395.89 | 424.37 |
| 9596 | ENSMUSG00000034211 | 741.5220824 | -0.061620984 | 0.119477144 | -0.515755411 | 0.606025248 | 0.916376001 | Mrps17 | mitochondrial ribosomal protein S17 [Sou | 709.94 | 793.55 | 768.77 | 671.89 | 776.52 | 728.45 |
| 9597 | ENSMUSG00000034212 | 2.044638003 | 0.061889169 | 1.570441361 | 0.039408774 | 0.968564485 | NA | Ankmy1 | ankyrin repeat and MYND domain contair | 0 | 2.0558 | 4.025 | 2.8919 | 2.1812 | 1.1138 |
| 9598 | ENSMUSG00000034216 | 321.4904241 | 0.003831232 | 0.156490205 | 0.024482251 | 0.980467941 | 0.99820894 | Vps18 | VPS18 CORVET/HOPS core subunit [Sourc | 323.65 | 319.68 | 319.99 | 343.18 | 270.47 | 351.97 |
| 9599 | ENSMUSG00000034218 | 175.2452096 | 0.067940631 | 0.190856046 | 0.355978405 | 0.721856738 | 0.9447719 | Atm | ataxia telangiectasia mutated [Source:MG | 161.42 | 197.36 | 154.96 | 183.16 | 191.95 | 162.62 |
| 9600 | ENSMUSG00000034219 | 0.305170013 | -1.657836477 | 4.041480335 | -0.410205257 | 0.681655385 | NA | Septin14 | septin 14 [Source:MGI Symbol;Acc:MGI:1 | 0.8031 | 1.0279 | 0 | 0 | 0 | 0 |
| 9601 | ENSMUSG00000034220 | 1015.462862 | -0.652273036 | 0.186610688 | -3.495368052 | 0.000473408 | 0.011929233 | Gpc1 | glypican 1 [Source:MGI Symbol;Acc:MGI:1 | 1541.2 | 1123.5 | 1058.6 | 873.36 | 928.12 | 568.06 |
| 9602 | ENSMUSG00000034224 | 0 | NA | NA | NA | NA | NA | Slc38a8 | solute carrier family 38, member 8 [Sourc | 0 | 0 | 0 | 0 | 0 | 0 |
| 9603 | ENSMUSG00000034226 | 0.339027071 | -1.79174149 | 3.919585585 | -0.457125237 | 0.647581032 | NA | Rhov | ras homolog family member V [Source:M | 0 | 1.0279 | 1.0062 | 0 | 0 | 0 |
| 9604 | ENSMUSG00000034227 | 4.880796165 | -0.695494088 | 1.035388389 | -0.671722897 | 0.501760122 | NA | Foxj1 | forkhead box J1 [Source:MGI Symbol;Acc: | 4.0155 | 7.1954 | 7.0437 | 7.7118 | 1.0906 | 2.2277 |
| 9605 | ENSMUSG00000034235 | 63.70290348 | -0.019690595 | 0.282535846 | -0.069692378 | 0.944438508 | 0.990077979 | Usp54 | ubiquitin specific peptidase 54 [Source:M | 63.445 | 60.647 | 68.425 | 69.406 | 52.35 | 67.944 |
| 9606 | ENSMUSG00000034239 | 0 | NA | NA | NA | NA | NA | Gm884 | predicted gene 884 [Source:MGI Symbol;/ | 0 | 0 | 0 | 0 | 0 | 0 |
| 9607 | ENSMUSG00000034243 | 1344.089754 | 0.150973038 | 0.102141869 | 1.47807201 | 0.139388505 | 0.545633379 | Golgb1 | golgi autoantigen, golgin subfamily b, mac | 1301 | 1292.1 | 1228.6 | 1497.1 | 1406.9 | 1338.8 |
| 9608 | ENSMUSG00000034245 | 957.4761453 | 0.488115288 | 0.108148286 | 4.513389023 | 6.38E-06 | 0.000381323 | Hdac11 | histone deacetylase 11 [Source:MGI Symb | 844.86 | 790.47 | 754.68 | 1086.4 | 1120.1 | 1148.4 |
| 9609 | ENSMUSG00000034247 | 387.2302406 | -0.020722796 | 0.146222332 | -0.141721145 | 0.887300272 | 0.982382091 | Plekhm1 | pleckstrin homology domain containing, f | 415.2 | 360.8 | 393.44 | 382.7 | 344.64 | 426.6 |
| 9610 | ENSMUSG00000034248 | 821.6414005 | 0.119403142 | 0.125805729 | 0.94910735 | 0.342566017 | 0.784247367 | Slc25a37 | solute carrier family 25, member 37 [Sour | 774.99 | 753.46 | 835.18 | 923.49 | 898.67 | 744.05 |
| 9611 | ENSMUSG00000034252 | 775.8454754 | 0.051496077 | 0.113989774 | 0.451760501 | 0.651441528 | 0.925643431 | Senp6 | SUMO/sentrin specific peptidase 6 [Sourc | 756.52 | 794.58 | 734.56 | 742.26 | 792.88 | 834.27 |
| 9612 | ENSMUSG00000034254 | 223.974712 | 0.095145796 | 0.190407108 | 0.499696658 | 0.617288686 | 0.917509899 | Agpat1 | 1-acylglycerol-3-phosphate O-acyltransfer | 241.73 | 217.92 | 190.18 | 279.55 | 203.95 | 210.52 |
| 9613 | ENSMUSG00000034255 | 31.78596098 | -1.111861188 | 0.417265403 | -2.664637857 | 0.007707127 | 0.089323626 | Arhgap27 | Rho GTPase activating protein 27 [Source: | 37.746 | 50.368 | 42.262 | 18.316 | 14.178 | 27.846 |
| 9614 | ENSMUSG00000034258 | 82.06835463 | -0.193082447 | 0.303684307 | -0.635799884 | 0.524906863 | 0.888334265 | Flvcr2 | feline leukemia virus subgroup C cellular r | 97.175 | 67.843 | 97.606 | 91.578 | 47.987 | 90.221 |
| 9615 | ENSMUSG00000034259 | 345.4199995 | -0.244587949 | 0.152408845 | -1.604814657 | 0.108534599 | 0.485347914 | Exosc4 | exosome component 4 [Source:MGI Symb | 359.79 | 376.22 | 388.41 | 330.64 | 347.91 | 269.55 |
| 9616 | ENSMUSG00000034263 | 185.6284237 | -0.174687013 | 0.188015888 | -0.929107717 | 0.352833266 | 0.793259753 | Ints14 | integrator complex subunit 14 [Source:M | 174.27 | 213.81 | 203.26 | 189.9 | 181.04 | 151.48 |
| 9617 | ENSMUSG00000034265 | 79.0685126 | -0.207402068 | 0.288529475 | -0.718824544 | 0.472249033 | 0.868150739 | Zdhhc14 | zinc finger, DHHC domain containing 14 [S | 65.854 | 74.01 | 114.71 | 72.298 | 80.706 | 66.831 |
| 9618 | ENSMUSG00000034266 | 4.629056767 | -2.919468993 | 1.359227938 | -2.147887718 | 0.031722676 | NA | Batf | basic leucine zipper transcription factor, A | 8.031 | 15.419 | 1.0062 | 0 | 1.0906 | 2.2277 |
| 9619 | ENSMUSG00000034269 | 599.6189726 | 0.080731349 | 0.125118053 | 0.645241412 | 0.518770754 | 0.887592015 | Setd5 | SET domain containing 5 [Source:MGI Sym | 589.48 | 587.97 | 571.55 | 676.71 | 563.85 | 608.16 |
| 9620 | ENSMUSG00000034271 | 19.76375114 | -2.112304938 | 0.573845584 | -3.680964004 | 0.000232354 | 0.007054905 | Jdp2 | Jun dimerization protein 2 [Source:MGI Sy | 36.943 | 40.089 | 19.119 | 4.8199 | 9.8156 | 7.7969 |
| 9621 | ENSMUSG00000034274 | 281.1418586 | -0.173844815 | 0.158551643 | -1.096455461 | 0.272879504 | 0.728119873 | Thoc5 | THO complex 5 [Source:MGI Symbol;Acc:I | 315.62 | 313.51 | 264.64 | 254.49 | 285.74 | 252.84 |
| 9622 | ENSMUSG00000034275 | 1.156404226 | 2.563524626 | 2.366885343 | 1.083079345 | 0.278773206 | NA | Igsf9b | immunoglobulin superfamily, member 9B | 0 | 1.0279 | 0 | 4.8199 | 1.0906 | 0 |
| 9623 | ENSMUSG00000034278 | 52.79910292 | 0.221923158 | 0.352653405 | 0.629295377 | 0.529155697 | 0.888334265 | Dnajc17 | DnaJ heat shock protein family (Hsp40) m | 64.248 | 43.173 | 38.237 | 46.271 | 46.897 | 77.969 |
| 9624 | ENSMUSG00000034282 | 5.200869838 | -0.676202645 | 1.140649617 | -0.592822402 | 0.553300014 | 0.896251569 | Evpl | envoplakin [Source:MGI Symbol;Acc:MGI: | 0.8031 | 14.391 | 4.025 | 0.964 | 5.4531 | 5.5692 |
| 9625 | ENSMUSG00000034285 | 4974.172771 | 0.030119688 | 0.089268966 | 0.337403799 | 0.735812521 | 0.948109711 | Nipsnap1 | nipsnap homolog 1 [Source:MGI Symbol;/ | 5079.6 | 4627.7 | 5058.4 | 4885.4 | 5021.2 | 5172.7 |
| 9626 | ENSMUSG00000034290 | 1248.977064 | 0.068181872 | 0.106899675 | 0.637811779 | 0.523596211 | 0.888334265 | Nek9 | NIMA (never in mitosis gene a)-related ex | 1142 | 1277.7 | 1239.7 | 1308.1 | 1196.4 | 1329.9 |
| 9627 | ENSMUSG00000034292 | 187.1281512 | 0.058434805 | 0.198099453 | 0.294977112 | 0.768011364 | 0.955856567 | Traf3ip1 | TRAF3 interacting protein 1 [Source:MGI S | 150.98 | 215.86 | 184.14 | 207.26 | 195.22 | 169.3 |
| 9628 | ENSMUSG00000034295 | 41.62096519 | -0.447360399 | 0.338354548 | -1.32216458 | 0.186113353 | 0.622270813 | Fhod3 | formin homology 2 domain containing 3 [ | 47.383 | 47.284 | 49.306 | 30.847 | 39.262 | 35.643 |

| | | | | | | | | | | | | | | | |
|---|---|---|---|---|---|---|---|---|---|---|---|---|---|---|---|
| 9629 | ENSMUSG00000034297 | 517.3450876 | -0.172134459 | 0.128393443 | -1.340679517 | 0.180024528 | 0.612244244 | Med13 | mediator complex subunit 13 [Source:MG | 537.28 | 600.3 | 507.15 | 482.95 | 491.87 | 484.52 |
| 9630 | ENSMUSG00000034300 | 287.8790206 | -0.183438784 | 0.161399393 | -1.136551882 | 0.255725665 | 0.708918985 | Fam53c | family with sequence similarity 53, memb | 265.83 | 337.16 | 315.96 | 251.6 | 270.47 | 286.26 |
| 9631 | ENSMUSG00000034303 | 18.91972012 | -1.398531807 | 0.552806467 | -2.529875988 | 0.011410285 | 0.11916127 | Ccdc15 | coiled-coil domain containing 15 [Source:I | 21.684 | 21.586 | 39.244 | 14.46 | 7.6343 | 8.9108 |
| 9632 | ENSMUSG00000034308 | 2977.009375 | 0.162062288 | 0.11079586 | 1.462710682 | 0.143546562 | 0.553373526 | Sdr42e1 | short chain dehydrogenase/reductase fan | 2960.2 | 2458.8 | 3009.7 | 2984.5 | 3098.4 | 3350.4 |
| 9633 | ENSMUSG00000034310 | 0.185640647 | 1.126924175 | 4.080472857 | 0.276174898 | 0.782413736 | NA | Tmem132d | transmembrane protein 132D [Source:MG | 0 | 0 | 0 | 0 | 0 | 1.1138 |
| 9634 | ENSMUSG00000034311 | 2.623306219 | -0.106310505 | 1.611451643 | -0.065971887 | 0.947400208 | NA | Kif4 | kinesin family member 4 [Source:MGI Syn | 4.0155 | 3.0838 | 1.0062 | 0 | 7.6343 | 0 |
| 9635 | ENSMUSG00000034312 | 527.4664418 | -0.051039253 | 0.151982075 | -0.335824166 | 0.737003466 | 0.948137969 | Iqsec1 | IQ motif and Sec7 domain 1 [Source:MGI S | 503.55 | 533.49 | 574.57 | 622.73 | 451.52 | 478.95 |
| 9636 | ENSMUSG00000034317 | 3.281958767 | -1.861148399 | 1.328362847 | -1.401084352 | 0.16118885 | NA | Trim59 | tripartite motif-containing 59 [Source:MG | 7.2279 | 5.1396 | 3.0187 | 0.964 | 0 | 3.3415 |
| 9637 | ENSMUSG00000034320 | 110.5993673 | 0.356979335 | 0.234662535 | 1.521245541 | 0.128198231 | 0.524098138 | Slc26a2 | solute carrier family 26 (sulfate transporte | 116.45 | 93.541 | 80.5 | 133.03 | 128.69 | 111.38 |
| 9638 | ENSMUSG00000034321 | 168.5455768 | -0.06398083 | 0.182048018 | -0.351450296 | 0.725250555 | 0.944957869 | Exosc1 | exosome component 1 [Source:MGI Symb | 183.91 | 167.55 | 165.02 | 162.91 | 159.23 | 172.65 |
| 9639 | ENSMUSG00000034324 | 0 | NA | NA | NA | NA | NA | Tmem132c | transmembrane protein 132C [Source:MG | 0 | 0 | 0 | 0 | 0 | 0 |
| 9640 | ENSMUSG00000034327 | 173.2340633 | 0.324136385 | 0.184610538 | 1.755784851 | 0.079125152 | 0.413164692 | Kctd9 | potassium channel tetramerisation domai | 149.38 | 161.38 | 150.94 | 193.76 | 172.32 | 211.63 |
| 9641 | ENSMUSG00000034329 | 41.15812398 | 0.483829711 | 0.345981446 | 1.398426756 | 0.161984953 | 0.585444868 | Brip1 | BRCA1 interacting protein C-terminal helic | 32.927 | 31.865 | 38.237 | 48.199 | 55.622 | 40.098 |
| 9642 | ENSMUSG00000034330 | 17.11021953 | -2.809510867 | 0.716330219 | -3.922088991 | 8.78E-05 | 0.003272463 | Plcg2 | phospholipase C, gamma 2 [Source:MGI S | 38.549 | 37.005 | 14.087 | 1.928 | 2.1812 | 8.9108 |
| 9643 | ENSMUSG00000034333 | 63.21963452 | -0.20984762 | 0.288671663 | -0.726942221 | 0.467261351 | 0.864254244 | Zbed4 | zinc finger, BED type containing 4 [Source | 59.43 | 71.954 | 72.45 | 66.514 | 61.075 | 47.895 |
| 9644 | ENSMUSG00000034334 | 95.06535837 | -0.119886125 | 0.241658983 | -0.496096291 | 0.619826476 | 0.917977899 | Fam151b | family with sequence similarity 151, meml | 103.6 | 101.76 | 91.568 | 93.506 | 75.253 | 104.7 |
| 9645 | ENSMUSG00000034336 | 0 | NA | NA | NA | NA | NA | Ina | internexin neuronal intermediate filament | 0 | 0 | 0 | 0 | 0 | 0 |
| 9646 | ENSMUSG00000034339 | 0 | NA | NA | NA | NA | NA | Cstdc7 | cystatin domain containing 7 [Source:MGI | 0 | 0 | 0 | 0 | 0 | 0 |
| 9647 | ENSMUSG00000034341 | 1895.645964 | -0.01038685 | 0.101666094 | -0.102166316 | 0.918624664 | 0.986374671 | Wbp2 | WW domain binding protein 2 [Source:MG | 2030.2 | 1860.5 | 1816.3 | 2010.9 | 1814.8 | 1841.2 |
| 9648 | ENSMUSG00000034342 | 184.6047161 | -0.299416311 | 0.193527298 | -1.547152853 | 0.121826392 | 0.511642546 | Cbl | Casitas B-lineage lymphoma [Source:MGI | 196.76 | 213.81 | 201.25 | 200.51 | 150.51 | 144.8 |
| 9649 | ENSMUSG00000034343 | 676.7920955 | -0.075680264 | 0.113249948 | -0.668258705 | 0.503968468 | 0.881510796 | Ube2f | ubiquitin-conjugating enzyme E2F (putativ | 713.15 | 690.76 | 679.22 | 641.05 | 681.64 | 654.94 |
| 9650 | ENSMUSG00000034345 | 585.8656975 | 0.036715719 | 0.140481629 | 0.261356015 | 0.793817973 | 0.961915077 | Gtf2h5 | general transcription factor IIH, polypepti | 669.79 | 555.08 | 509.16 | 586.1 | 631.47 | 563.61 |
| 9651 | ENSMUSG00000034349 | 190.182998 | 0.127845017 | 0.195510026 | 0.653905168 | 0.513172904 | 0.884899009 | Smc4 | structural maintenance of chromosomes 4 | 220.05 | 163.44 | 161 | 207.26 | 179.95 | 209.4 |
| 9652 | ENSMUSG00000034353 | 65.6794889 | -1.371535928 | 0.291912827 | -4.698443509 | 2.62E-06 | 0.000178857 | Ramp1 | receptor (calcitonin) activity modifying pro | 97.978 | 88.401 | 97.606 | 30.847 | 44.715 | 34.529 |
| 9653 | ENSMUSG00000034354 | 632.9515546 | 0.01762313 | 0.125183785 | 0.140778053 | 0.888045282 | 0.982382091 | Mtmr3 | myotubularin related protein 3 [Source:M | 564.58 | 651.7 | 672.17 | 660.32 | 612.93 | 636 |
| 9654 | ENSMUSG00000034359 | 0 | NA | NA | NA | NA | NA | Skint2 | selection and upkeep of intraepithelial T c | 0 | 0 | 0 | 0 | 0 | 0 |
| 9655 | ENSMUSG00000034361 | 195.4198037 | -0.042081007 | 0.218893551 | -0.192244163 | 0.847550954 | 0.972803041 | Cpne2 | copine II [Source:MGI Symbol;Acc:MGI:23 | 168.65 | 206.61 | 220.37 | 232.32 | 138.51 | 206.06 |
| 9656 | ENSMUSG00000034362 | 0.342639279 | -1.805976596 | 4.028547714 | -0.448294702 | 0.653940524 | NA | Csta1 | cystatin A1 [Source:MGI Symbol;Acc:MGI: | 0 | 2.0558 | 0 | 0 | 0 | 0 |
| 9657 | ENSMUSG00000034371 | 11883.39237 | -0.509510884 | 0.108711361 | -4.68682278 | 2.77E-06 | 0.000186672 | Tkfc | triokinase, FMN cyclase [Source:MGI Sym | 14917 | 13128 | 13837 | 10954 | 9774.1 | 8691.3 |
| 9658 | ENSMUSG00000034377 | 614.7462038 | 0.083344886 | 0.119572916 | 0.697021439 | 0.485789377 | 0.872998689 | Tulp4 | tubby like protein 4 [Source:MGI Symbol;A | 583.05 | 624.97 | 583.62 | 669.96 | 603.11 | 623.75 |
| 9659 | ENSMUSG00000034379 | 48.95594203 | 0.499062448 | 0.364452964 | 1.369346655 | 0.170890941 | 0.597622477 | Wdr5b | WD repeat domain 5B [Source:MGI Symbo | 41.761 | 28.782 | 51.318 | 76.154 | 55.622 | 40.098 |
| 9660 | ENSMUSG00000034382 | 394.7413512 | 0.077049088 | 0.160705311 | 0.479443321 | 0.631623281 | 0.921549084 | AI661453 | expressed sequence AI661453 [Source:MG | 378.26 | 347.44 | 426.65 | 382.7 | 354.45 | 478.95 |
| 9661 | ENSMUSG00000034384 | 0 | NA | NA | NA | NA | NA | Barhl2 | BarH like homeobox 2 [Source:MGI Symbo | 0 | 0 | 0 | 0 | 0 | 0 |
| 9662 | ENSMUSG00000034387 | 0 | NA | NA | NA | NA | NA | Ssu2 | ssu-2 homolog (C. elegans) [Source:MGI S | 0 | 0 | 0 | 0 | 0 | 0 |
| 9663 | ENSMUSG00000034390 | 489.2857318 | -0.092759309 | 0.16078838 | -0.576903065 | 0.56400494 | 0.901211606 | Cmip | c-Maf inducing protein [Source:MGI Symb | 534.06 | 500.6 | 480.98 | 587.06 | 425.34 | 407.67 |
| 9664 | ENSMUSG00000034391 | 1.127783327 | -2.316944858 | 2.16487464 | -1.070244353 | 0.284509335 | NA | Fbxo15 | F-box protein 15 [Source:MGI Symbol;Acc | 1.6062 | 1.0279 | 3.0187 | 0 | 0 | 1.1138 |
| 9665 | ENSMUSG00000034394 | 0.353348078 | 0.165142701 | 3.895042568 | 0.042398176 | 0.966181282 | NA | Lif | leukemia inhibitory factor [Source:MGI Sy | 0 | 0 | 1.0062 | 0 | 0 | 1.1138 |
| 9666 | ENSMUSG00000034401 | 14.0173665 | -0.724499198 | 0.638077904 | -1.135440035 | 0.256190993 | 0.709331265 | Spata6 | spermatogenesis associated 6 [Source:MG | 13.653 | 26.726 | 12.075 | 10.604 | 5.4531 | 15.594 |
| 9667 | ENSMUSG00000034402 | 0 | NA | NA | NA | NA | NA | Kcnh5 | potassium voltage-gated channel, subfam | 0 | 0 | 0 | 0 | 0 | 0 |
| 9668 | ENSMUSG00000034403 | 1279.300192 | 0.188630786 | 0.107105728 | 1.761164319 | 0.078210595 | 0.409868858 | Pja1 | praja ring finger ubiquitin ligase 1 [Source | 1262.5 | 1154.4 | 1169.3 | 1271.5 | 1454.9 | 1363.3 |
| 9669 | ENSMUSG00000034412 | 265.8281536 | 0.422571408 | 0.182871705 | 2.310753365 | 0.020846482 | 0.177994989 | Tbc1d10a | TBC1 domain family, member 10a [Source | 235.31 | 230.25 | 215.34 | 258.35 | 277.02 | 378.71 |
| 9670 | ENSMUSG00000034413 | 8.6190597 | -0.312293163 | 0.790774274 | -0.394920742 | 0.692901382 | 0.938679052 | Neurl1b | neuralized E3 ubiquitin protein ligase 1B [ | 10.44 | 3.0838 | 15.094 | 7.7118 | 9.8156 | 5.5692 |
| 9671 | ENSMUSG00000034416 | 1.521824675 | -0.314308103 | 1.896441304 | -0.16573574 | 0.868364918 | NA | Pkd1l2 | polycystic kidney disease 1 like 2 [Source:I | 4.0155 | 0 | 1.0062 | 1.928 | 2.1812 | 0 |
| 9672 | ENSMUSG00000034422 | 516.971657 | 0.121700401 | 0.133526315 | 0.911433833 | 0.362066834 | 0.800872989 | Parp14 | poly (ADP-ribose) polymerase family, men | 501.94 | 520.13 | 462.87 | 481.99 | 580.21 | 554.69 |
| 9673 | ENSMUSG00000034424 | 1717.272032 | -0.021436181 | 0.093420549 | -0.229458951 | 0.818512222 | 0.968897682 | Gcsh | glycine cleavage system protein H (aminor | 1741.9 | 1719.7 | 1728.7 | 1761.2 | 1701.4 | 1650.7 |
| 9674 | ENSMUSG00000034427 | 14.3991389 | -0.24917431 | 0.610847016 | -0.407916063 | 0.6833353 | 0.935342271 | Myo15b | myosin XVB [Source:MGI Symbol;Acc:MGI | 9.6372 | 14.391 | 23.144 | 18.316 | 11.997 | 8.9108 |
| 9675 | ENSMUSG00000034429 | 456.2783601 | 0.438320908 | 0.158796082 | 2.760275341 | 0.005775266 | 0.073468297 | Zfp707 | zinc finger protein 707 [Source:MGI Symb | 353.36 | 376.22 | 433.69 | 575.49 | 428.61 | 570.29 |
| 9676 | ENSMUSG00000034430 | 281.7419925 | -0.018985206 | 0.166139064 | -0.114272982 | 0.909021399 | 0.985098284 | Zxdc | ZXD family zinc finger C [Source:MGI Symb | 288.31 | 315.57 | 246.53 | 250.63 | 305.37 | 284.03 |
| 9677 | ENSMUSG00000034432 | 454.0400705 | 0.070557457 | 0.134491482 | 0.524623981 | 0.599844612 | 0.915217938 | Cops8 | COP9 signalosome subunit 8 [Source:MGI | 433.68 | 440.98 | 453.82 | 423.19 | 461.33 | 511.25 |
| 9678 | ENSMUSG00000034435 | 1000.289896 | 0.095211003 | 0.126752981 | 0.751153959 | 0.452560005 | 0.856769845 | Tmem30b | transmembrane protein 30B [Source:MGI | 848.88 | 1002.2 | 1052.5 | 1138.5 | 975.01 | 984.64 |
| 9679 | ENSMUSG00000034437 | 2.837213058 | -1.149592003 | 1.294892422 | -0.887789582 | 0.374653948 | NA | Gm9761 | predicted gene 9761 [Source:MGI Symbol | 1.6062 | 5.1396 | 5.0312 | 1.928 | 1.0906 | 2.2277 |
| 9680 | ENSMUSG00000034438 | 22.82469985 | -0.691115581 | 0.487442455 | -1.417840351 | 0.156237381 | 0.574894692 | Gbp8 | guanylate-binding protein 8 [Source:MGI S | 26.502 | 39.061 | 19.119 | 20.244 | 13.087 | 18.935 |
| 9681 | ENSMUSG00000034442 | 127.1759046 | 0.065040134 | 0.221565608 | 0.293547971 | 0.769103338 | 0.956045284 | Trmt5 | TRM5 tRNA methyltransferase 5 [Source:I | 110.03 | 114.1 | 148.92 | 117.61 | 127.6 | 144.8 |
| 9682 | ENSMUSG00000034445 | 184.5010615 | 0.238740442 | 0.177981046 | 1.341381271 | 0.179796695 | 0.612025777 | Cyb561a3 | cytochrome b561 family, member A3 [Sou | 179.09 | 154.19 | 174.08 | 192.8 | 195.22 | 211.63 |
| 9683 | ENSMUSG00000034449 | 508.949344 | 0.164152786 | 0.135879269 | 1.208078224 | 0.227017175 | 0.67343689 | Dhrs11 | dehydrogenase/reductase (SDR family) m | 444.92 | 449.2 | 546.39 | 523.44 | 556.22 | 533.53 |
| 9684 | ENSMUSG00000034450 | 11094.48348 | 0.391943377 | 0.101544006 | 3.859837661 | 0.000113462 | 0.004011279 | Gulo | gulonolactone (L-) oxidase [Source:MGI Sy | 10093 | 9047.7 | 9647.9 | 11254 | 13480 | 13044 |

| | | | | | | | | | | | | | | | |
|---|---|---|---|---|---|---|---|---|---|---|---|---|---|---|---|
| 9685 | ENSMUSG00000034452 | 0.884336215 | 0.707502939 | 2.570766407 | 0.275210901 | 0.783154212 | NA | Slc24a1 | solute carrier family 24 (sodium/potassiur | 0 | 1.0279 | 1.0062 | 0 | 3.2719 | 0 |
| 9686 | ENSMUSG00000034453 | 293.9916048 | -0.171438014 | 0.175887728 | -0.9747014 | 0.329708398 | 0.774655633 | Polr3b | polymerase (RNA) III (DNA directed) polyp | 311.6 | 299.12 | 323 | 223.64 | 335.91 | 270.66 |
| 9687 | ENSMUSG00000034456 | 8670.824592 | -0.198004077 | 0.103292659 | -1.91692303 | 0.055247711 | 0.331563872 | Uroc1 | urocanase domain containing 1 [Source:M | 8743.4 | 9008.7 | 10044 | 8893.7 | 7764.1 | 7570.8 |
| 9688 | ENSMUSG00000034457 | 22.63205242 | -0.662198831 | 0.524690994 | -1.262073943 | 0.20692218 | 0.652194904 | Eda2r | ectodysplasin A2 receptor [Source:MGI Sy | 43.368 | 13.363 | 26.162 | 15.424 | 18.541 | 18.935 |
| 9689 | ENSMUSG00000034459 | 251.8425859 | 0.135326933 | 0.237376054 | 0.570095133 | 0.568613176 | 0.903527882 | Ifit1 | interferon-induced protein with tetratrico | 326.86 | 171.66 | 220.37 | 225.57 | 330.46 | 236.13 |
| 9690 | ENSMUSG00000034460 | 0 | NA | NA | NA | NA | NA | Six4 | sine oculis-related homeobox 4 [Source:M | 0 | 0 | 0 | 0 | 0 | 0 |
| 9691 | ENSMUSG00000034462 | 377.6143435 | -0.045771002 | 0.150942142 | -0.303235405 | 0.761710469 | 0.954216198 | Pkd2 | polycystin 2, transient receptor potential | 420.83 | 349.49 | 380.36 | 415.47 | 364.27 | 335.27 |
| 9692 | ENSMUSG00000034463 | 30.91820324 | -0.163620069 | 0.432289828 | -0.378496227 | 0.705061996 | 0.94295036 | Scara3 | scavenger receptor class A, member 3 [So | 36.14 | 32.893 | 29.181 | 44.343 | 22.903 | 20.049 |
| 9693 | ENSMUSG00000034467 | 0.319491021 | 0.16514334 | 4.044653141 | 0.040830038 | 0.967431393 | NA | Dynlrb2 | dynein light chain roadblock-type 2 [Sourc | 0.8031 | 0 | 0 | 0 | 0 | 1.1138 |
| 9694 | ENSMUSG00000034471 | 259.1520908 | -0.220431181 | 0.158171206 | -1.393623952 | 0.163431192 | 0.587512475 | Caskin2 | CASK-interacting protein 2 [Source:MGI Sy | 258.6 | 297.07 | 281.75 | 254.49 | 225.76 | 237.25 |
| 9695 | ENSMUSG00000034472 | 0.449470461 | -0.545136533 | 3.562105917 | -0.15303771 | 0.878368537 | NA | Rasd2 | RASD family, member 2 [Source:MGI Syml | 1.6062 | 0 | 0 | 0 | 1.0906 | 0 |
| 9696 | ENSMUSG00000034473 | 264.7135179 | 0.06828607 | 0.161390929 | 0.423109715 | 0.672215215 | 0.931191575 | Sec22a | SEC22 homolog A, vesicle trafficking prote | 273.05 | 263.15 | 238.48 | 242.92 | 295.56 | 275.12 |
| 9697 | ENSMUSG00000034480 | 253.5389964 | 0.169782438 | 0.171882205 | 0.987783685 | 0.323258599 | 0.770475661 | Diaph2 | diaphanous related formin 2 [Source:MGI | 251.37 | 239.5 | 224.39 | 232.32 | 313.01 | 260.64 |
| 9698 | ENSMUSG00000034482 | 0 | NA | NA | NA | NA | NA | Ly6g5c | lymphocyte antigen 6 complex, locus G5C | 0 | 0 | 0 | 0 | 0 | 0 |
| 9699 | ENSMUSG00000034484 | 673.7633169 | -0.172004662 | 0.122906695 | -1.399473488 | 0.161671043 | 0.584894158 | Snx2 | sorting nexin 2 [Source:MGI Symbol;Acc:N | 729.22 | 760.66 | 652.05 | 669.96 | 632.56 | 598.13 |
| 9700 | ENSMUSG00000034485 | 249.00118 | 0.133054884 | 0.169701666 | 0.784051724 | 0.433009761 | 0.845264432 | Uaca | uveal autoantigen with coiled-coil domain | 236.92 | 262.12 | 213.32 | 234.25 | 290.1 | 257.3 |
| 9701 | ENSMUSG00000034486 | 0 | NA | NA | NA | NA | NA | Gbx2 | gastrulation brain homeobox 2 [Source:M | 0 | 0 | 0 | 0 | 0 | 0 |
| 9702 | ENSMUSG00000034487 | 149.6328329 | -0.255323872 | 0.218197918 | -1.170148067 | 0.241941388 | 0.691791181 | Poglut3 | protein O-glucosyltransferase 3 [Source:M | 193.55 | 154.19 | 139.87 | 115.68 | 135.24 | 159.28 |
| 9703 | ENSMUSG00000034488 | 0.181769715 | 1.126924175 | 4.080472857 | 0.276174898 | 0.782413736 | NA | Edil3 | EGF-like repeats and discoidin I-like doma | 0 | 0 | 0 | 0 | 1.0906 | 0 |
| 9704 | ENSMUSG00000034493 | 1.93086059 | -1.333059531 | 1.915656195 | -0.695876189 | 0.48650637 | NA | 4930556J24I | RIKEN cDNA 4930556J24 gene [Source:MG | 3.2124 | 0 | 5.0312 | 0 | 0 | 3.3415 |
| 9705 | ENSMUSG00000034501 | 151.6770267 | 0.238540317 | 0.203482637 | 1.172288313 | 0.241081322 | 0.691388521 | Pcnx4 | pecanex homolog 4 [Source:MGI Symbol;A | 121.27 | 131.57 | 165.02 | 158.09 | 159.23 | 174.87 |
| 9706 | ENSMUSG00000034509 | 108.8601801 | 0.381272239 | 0.245460252 | 1.553295232 | 0.120352619 | 0.507960396 | Mad2l1bp | MAD2L1 binding protein [Source:MGI Syn | 109.22 | 89.429 | 84.525 | 121.46 | 98.156 | 150.37 |
| 9707 | ENSMUSG00000034518 | 128.3587896 | 0.398539919 | 0.218588557 | 1.82324237 | 0.068266691 | 0.376317774 | Hmgxb4 | HMG box domain containing 4 [Source:MG | 119.66 | 87.373 | 124.77 | 140.74 | 147.23 | 150.37 |
| 9708 | ENSMUSG00000034520 | 3.085506469 | -3.96636637 | 1.660162988 | -2.389142751 | 0.016887739 | NA | Gjc1 | gap junction protein, gamma 1 [Source:M | 10.44 | 3.0838 | 4.025 | 0.964 | 0 | 0 |
| 9709 | ENSMUSG00000034522 | 570.1321866 | -0.013257815 | 0.187729256 | -0.070621996 | 0.943698604 | 0.990077979 | Zfp395 | zinc finger protein 395 [Source:MGI Symb | 484.27 | 459.48 | 774.81 | 522.48 | 566.03 | 613.73 |
| 9710 | ENSMUSG00000034525 | 378.1135621 | -0.178542259 | 0.157002736 | -1.137192022 | 0.255458021 | 0.708894389 | Ice1 | interactor of little elongation complex ELL | 358.18 | 439.95 | 407.53 | 405.83 | 313.01 | 344.18 |
| 9711 | ENSMUSG00000034528 | 9139.938146 | -0.310868226 | 0.108966482 | -2.852879347 | 0.004332507 | 0.060746309 | Hsd17b13 | hydroxysteroid (17-beta) dehydrogenase | 8987.5 | 11363 | 10014 | 8382.8 | 8482.8 | 7609.8 |
| 9712 | ENSMUSG00000034532 | 0.856598198 | -3.132279849 | 3.123122476 | -1.002932121 | 0.315893613 | NA | Fbxo16 | F-box protein 16 [Source:MGI Symbol;Acc | 0 | 5.1396 | 0 | 0 | 0 | 0 |
| 9713 | ENSMUSG00000034533 | 0.133850373 | -0.796629176 | 4.080472857 | -0.195229623 | 0.84521319 | NA | Scn10a | sodium channel, voltage-gated, type X, alp | 0.8031 | 0 | 0 | 0 | 0 | 0 |
| 9714 | ENSMUSG00000034538 | 15.72045233 | -0.152780272 | 0.567236244 | -0.26934152 | 0.787666884 | 0.960870081 | Zfp418 | zinc finger protein 418 [Source:MGI Symb | 15.259 | 15.419 | 19.119 | 19.28 | 17.45 | 7.7969 |
| 9715 | ENSMUSG00000034543 | 318.6321554 | 0.132535009 | 0.162382281 | 0.816191325 | 0.414390717 | 0.835831451 | Morc2a | microrchidia 2A [Source:MGI Symbol;Acc: | 301.97 | 308.38 | 301.87 | 338.36 | 384.99 | 276.23 |
| 9716 | ENSMUSG00000034544 | 393.8823789 | 0.041370754 | 0.14012982 | 0.295231624 | 0.767816946 | 0.95578799 | Rsrc1 | arginine/serine-rich coiled-coil 1 [Source:I | 393.52 | 405 | 366.27 | 406.8 | 429.7 | 362 |
| 9717 | ENSMUSG00000034551 | 0.856470372 | -1.816219066 | 2.859255434 | -0.635207 | 0.525293418 | NA | Hdx | highly divergent homeobox [Source:MGI S | 0 | 0 | 4.025 | 0 | 0 | 1.1138 |
| 9718 | ENSMUSG00000034552 | 0 | NA | NA | NA | NA | NA | Zswim2 | zinc finger SWIM-type containing 2 [Sourc | 0 | 0 | 0 | 0 | 0 | 0 |
| 9719 | ENSMUSG00000034555 | 0 | NA | NA | NA | NA | NA | Tex16 | testis expressed gene 16 [Source:MGI Sym | 0 | 0 | 0 | 0 | 0 | 0 |
| 9720 | ENSMUSG00000034557 | 596.4484272 | -0.013078949 | 0.139109166 | -0.094019318 | 0.925093811 | 0.987546253 | Zfyve9 | zinc finger, FYVE domain containing 9 [Sou | 528.44 | 650.67 | 618.84 | 535.01 | 658.73 | 587 |
| 9721 | ENSMUSG00000034560 | 479.8580487 | 0.115717009 | 0.132034566 | 0.876414508 | 0.380804734 | 0.812628154 | Washc4 | WASH complex subunit 4 [Source:MGI Syr | 424.04 | 475.93 | 483 | 534.04 | 480.96 | 481.18 |
| 9722 | ENSMUSG00000034563 | 1.728933638 | 0.747891267 | 1.690952491 | 0.442289935 | 0.658279414 | NA | Ccpg1 | cell cycle progression 1 [Source:MGI Symb | 0.8031 | 3.0838 | 0 | 0.964 | 2.1812 | 3.3415 |
| 9723 | ENSMUSG00000034566 | 3518.296671 | -0.116566844 | 0.085031104 | -1.370872987 | 0.170414554 | 0.597556794 | Atp5h | ATP synthase, H+ transporting, mitochonc | 3675.8 | 3580.2 | 3725.1 | 3430.8 | 3334 | 3363.8 |
| 9724 | ENSMUSG00000034570 | 0.167707431 | -0.796629176 | 4.080472857 | -0.195229623 | 0.84521319 | NA | Inpp5j | inositol polyphosphate 5-phosphatase J [S | 0 | 0 | 1.0062 | 0 | 0 | 0 |
| 9725 | ENSMUSG00000034573 | 28.68909244 | -0.384560378 | 0.406668808 | -0.945635293 | 0.344334637 | 0.785808801 | Ptpn13 | protein tyrosine phosphatase, non-recept | 31.321 | 38.033 | 28.175 | 25.063 | 27.265 | 22.277 |
| 9726 | ENSMUSG00000034574 | 1181.765922 | 0.406549438 | 0.15666719 | 2.594987748 | 0.009459429 | 0.104363983 | Daam1 | dishevelled associated activator of morph | 1085.8 | 907.65 | 1055.6 | 1491.3 | 1534.5 | 1015.8 |
| 9727 | ENSMUSG00000034575 | 407.5013063 | 0.17488693 | 0.152173442 | 1.149260525 | 0.25044857 | 0.702302228 | Tent4a | terminal nucleotidyltransferase 4A [Sourc | 347.74 | 374.16 | 426.65 | 376.92 | 441.7 | 477.84 |
| 9728 | ENSMUSG00000034579 | 0.68166635 | -2.79976525 | 3.012029182 | -0.92952793 | 0.352615558 | NA | Pla2g3 | phospholipase A2, group III [Source:MGI S | 0 | 3.0838 | 1.0062 | 0 | 0 | 0 |
| 9729 | ENSMUSG00000034583 | 0 | NA | NA | NA | NA | NA | Olfr1347 | olfactory receptor 1347 [Source:MGI Sym | 0 | 0 | 0 | 0 | 0 | 0 |
| 9730 | ENSMUSG00000034584 | 237.6754514 | 0.431320291 | 0.170597925 | 2.528285685 | 0.011462104 | 0.11952983 | Exph5 | exophilin 5 [Source:MGI Symbol;Acc:MGI: | 196.76 | 181.94 | 228.42 | 251.6 | 295.56 | 271.78 |
| 9731 | ENSMUSG00000034586 | 17.09755726 | -0.634764361 | 0.581550117 | -1.091504142 | 0.275051111 | 0.729768468 | Hid1 | HID1 domain containing [Source:MGI Sym | 21.684 | 18.503 | 22.137 | 12.532 | 5.4531 | 22.277 |
| 9732 | ENSMUSG00000034587 | 11.23314287 | 0.36102222 | 0.710419304 | 0.508181883 | 0.611325795 | 0.916376001 | 8430429K09 | RIKEN cDNA 8430429K09 gene [Source:M | 8.031 | 14.391 | 7.0437 | 4.8199 | 14.178 | 18.935 |
| 9733 | ENSMUSG00000034591 | 352.7443903 | 0.5125368 | 0.181368931 | 2.825934935 | 0.004714284 | 0.06420853 | Slc41a2 | solute carrier family 41, member 2 [Sourc | 252.98 | 254.92 | 365.27 | 475.24 | 389.35 | 378.71 |
| 9734 | ENSMUSG00000034593 | 9.644629249 | -0.491587483 | 0.689481837 | -0.712981049 | 0.475857484 | 0.86947987 | Myo5a | myosin VA [Source:MGI Symbol;Acc:MGI: | 10.44 | 14.391 | 9.0562 | 9.6398 | 6.5437 | 7.7969 |
| 9735 | ENSMUSG00000034595 | 50.01630432 | -1.494458756 | 0.416548722 | -3.587716579 | 0.000333587 | 0.009479365 | Ppp1r18 | protein phosphatase 1, regulatory subunit | 54.611 | 99.708 | 67.418 | 31.811 | 10.906 | 35.643 |
| 9736 | ENSMUSG00000034601 | 56.13009697 | 0.501028137 | 0.318446964 | 1.573348764 | 0.115638084 | 0.49987724 | 2700049A03 | RIKEN cDNA 2700049A03 gene [Source:M | 40.155 | 39.061 | 60.375 | 63.623 | 56.712 | 76.855 |
| 9737 | ENSMUSG00000034602 | 1295.665371 | 0.018453516 | 0.105513572 | 0.174892341 | 0.861164229 | 0.976013118 | Mon2 | MON2 homolog, regulator of endosome t | 1213.5 | 1314.7 | 1335.3 | 1391 | 1256.4 | 1263.1 |
| 9738 | ENSMUSG00000034607 | 0.160662974 | 1.126924175 | 4.080472857 | 0.276174898 | 0.782413736 | NA | Pof1b | premature ovarian failure 1B [Source:MGI | 0 | 0 | 0 | 0.964 | 0 | 0 |
| 9739 | ENSMUSG00000034610 | 306.4810445 | -0.030259066 | 0.190309653 | -0.158999115 | 0.873669571 | 0.979925081 | Tut4 | terminal uridylyl transferase 4 [Source:MG | 294.74 | 327.91 | 306.9 | 319.08 | 369.72 | 220.54 |
| 9740 | ENSMUSG00000034612 | 3.175180247 | 0.964995413 | 1.450043957 | 0.665493904 | 0.505734641 | NA | Chst11 | carbohydrate sulfotransferase 11 [Source | 2.4093 | 0 | 4.025 | 4.8199 | 0 | 7.7969 |

| | | | | | | | | | | | | | | | |
|---|---|---|---|---|---|---|---|---|---|---|---|---|---|---|---|
| 9741 | ENSMUSG00000034613 | 42.6388866 | 0.069577327 | 0.378918441 | 0.183620852 | 0.854310904 | 0.973825032 | Ppm1h | protein phosphatase 1H (PP2C domain co | 48.186 | 51.396 | 25.156 | 45.307 | 51.259 | 34.529 |
| 9742 | ENSMUSG00000034614 | 16.01301598 | 0.036483086 | 0.53286291 | 0.068466176 | 0.945414544 | 0.990309554 | Pik3ip1 | phosphoinositide-3-kinase interacting pro | 16.865 | 13.363 | 17.106 | 13.496 | 18.541 | 16.708 |
| 9743 | ENSMUSG00000034616 | 246.4629517 | 0.12807055 | 0.171787082 | 0.745519097 | 0.455957971 | 0.857604173 | Ssh3 | slingshot protein phosphatase 3 [Source:I | 250.57 | 229.23 | 226.41 | 259.31 | 219.21 | 294.05 |
| 9744 | ENSMUSG00000034617 | 130.6678218 | -0.164214502 | 0.212289369 | -0.773540865 | 0.439202354 | 0.848147242 | Mtrr | 5-methyltetrahydrofolate-homocysteine r | 149.38 | 129.52 | 134.84 | 109.89 | 116.7 | 143.69 |
| 9745 | ENSMUSG00000034620 | 360.6873811 | 0.038918513 | 0.140389497 | 0.277218127 | 0.781612622 | 0.960079945 | Rxylt1 | ribitol xylosyltransferase 1 [Source:MGI Sy | 347.74 | 370.05 | 350.17 | 384.63 | 376.26 | 335.27 |
| 9746 | ENSMUSG00000034621 | 323.1051466 | 0.172476779 | 0.158251752 | 1.089888593 | 0.275762222 | 0.730038567 | Gpatch8 | G patch domain containing 8 [Source:MGI | 273.05 | 286.79 | 352.19 | 339.32 | 364.27 | 323.01 |
| 9747 | ENSMUSG00000034623 | 0 | NA | NA | NA | NA | NA | Prss55 | protease, serine 55 [Source:MGI Symbol;A | 0 | 0 | 0 | 0 | 0 | 0 |
| 9748 | ENSMUSG00000034634 | 24.73551332 | -1.381247204 | 0.548795364 | -2.516871122 | 0.011840213 | 0.121544955 | Ly6d | lymphocyte antigen 6 complex, locus D [S | 61.036 | 28.782 | 17.106 | 10.604 | 14.178 | 16.708 |
| 9749 | ENSMUSG00000034636 | 1066.968421 | 0.179806081 | 0.122121446 | 1.472354675 | 0.140925126 | 0.548974151 | Zyg11b | zyg-ll family member B, cell cycle regulato | 984.6 | 1050.5 | 965.99 | 982.29 | 1244.4 | 1174 |
| 9750 | ENSMUSG00000034639 | 84.69342954 | -0.011607415 | 0.265601093 | -0.043702437 | 0.965141597 | 0.993480943 | Setmar | SET domain without mariner transposase | 77.901 | 108.96 | 68.425 | 80.974 | 87.249 | 84.652 |
| 9751 | ENSMUSG00000034640 | 127.8039602 | 0.07656299 | 0.258906056 | 0.295717263 | 0.767446013 | 0.95578799 | Tiparp | TCDD-inducible poly(ADP-ribose) polymer | 163.03 | 104.85 | 104.65 | 136.88 | 99.246 | 158.17 |
| 9752 | ENSMUSG00000034641 | 61.08435473 | -1.400252402 | 0.365774579 | -3.828184033 | 0.000129092 | 0.00441416 | Cd300ld | CD300 molecule like family member d [So | 101.19 | 95.596 | 68.425 | 16.388 | 40.353 | 44.554 |
| 9753 | ENSMUSG00000034645 | 118.0980846 | 0.706862303 | 0.269431958 | 2.623528061 | 0.008702428 | 0.097878626 | Zyg11a | zyg-11 family member A, cell cycle regulat | 93.16 | 107.93 | 68.425 | 196.65 | 123.24 | 119.18 |
| 9754 | ENSMUSG00000034647 | 156.9851118 | 0.120094315 | 0.2286802 | 0.525162715 | 0.599470082 | 0.915117178 | Ankrd12 | ankyrin repeat domain 12 [Source:MGI Sy | 173.47 | 153.16 | 124.77 | 208.22 | 134.15 | 148.14 |
| 9755 | ENSMUSG00000034648 | 0.339027071 | -1.79174149 | 3.919585585 | -0.457125237 | 0.647581032 | NA | Lrrn1 | leucine rich repeat protein 1, neuronal [S | 0 | 1.0279 | 1.0062 | 0 | 0 | 0 |
| 9756 | ENSMUSG00000034652 | 50.00494066 | -1.530799215 | 0.373310541 | -4.10060539 | 4.12E-05 | 0.001784488 | Cd300a | CD300A molecule [Source:MGI Symbol;Ac | 72.279 | 93.541 | 57.356 | 33.739 | 15.269 | 27.846 |
| 9757 | ENSMUSG00000034653 | 181.7085358 | 0.124024264 | 0.192512111 | 0.644241355 | 0.519418937 | 0.887731952 | Ythdc2 | YTH domain containing 2 [Source:MGI Syr | 146.97 | 202.5 | 173.07 | 197.62 | 195.22 | 174.87 |
| 9758 | ENSMUSG00000034656 | 14.01858503 | 0.329967882 | 0.632262338 | 0.521884449 | 0.601750786 | 0.915461118 | Cacna1a | calcium channel, voltage-dependent, P/Q | 12.047 | 5.1396 | 20.125 | 19.28 | 15.269 | 12.252 |
| 9759 | ENSMUSG00000034659 | 495.1410248 | -0.178828169 | 0.156379951 | -1.143549211 | 0.252810635 | 0.705781321 | Tmem109 | transmembrane protein 109 [Source:MGI | 599.92 | 529.38 | 447.78 | 511.87 | 483.14 | 398.76 |
| 9760 | ENSMUSG00000034660 | 0 | NA | NA | NA | NA | NA | Galp | galanin-like peptide [Source:MGI Symbol; | 0 | 0 | 0 | 0 | 0 | 0 |
| 9761 | ENSMUSG00000034663 | 192.6181161 | 0.010018273 | 0.176724173 | 0.056688755 | 0.954793132 | 0.991678478 | Bmp2k | BMP2 inducible kinase [Source:MGI Symb | 199.97 | 191.19 | 184.14 | 172.55 | 200.67 | 207.17 |
| 9762 | ENSMUSG00000034664 | 58.31402498 | -0.127110255 | 0.305426098 | -0.416173522 | 0.677283033 | 0.93325821 | Itga2b | integrin alpha 2b [Source:MGI Symbol;Acc | 70.673 | 45.228 | 66.412 | 53.983 | 53.44 | 60.148 |
| 9763 | ENSMUSG00000034667 | 2061.828568 | -0.11039846 | 0.111697844 | -0.988366969 | 0.322972958 | 0.769995293 | Xpot | exportin, tRNA (nuclear export receptor fc | 2319.4 | 2150.4 | 1952.1 | 2160.3 | 1957.7 | 1831.2 |
| 9764 | ENSMUSG00000034673 | 331.9909748 | 0.235188082 | 0.143030161 | 1.644325086 | 0.10010907 | 0.465788575 | Pbx2 | pre B cell leukemia homeobox 2 [Source:I | 288.31 | 302.21 | 325.02 | 370.17 | 363.18 | 343.06 |
| 9765 | ENSMUSG00000034674 | 234.6618062 | -0.018698273 | 0.179077531 | -0.104414399 | 0.916840496 | 0.986374671 | Tdg | thymine DNA glycosylase [Source:MGI Syr | 231.29 | 254.92 | 222.38 | 220.75 | 277.02 | 201.61 |
| 9766 | ENSMUSG00000034675 | 17.78032822 | -0.746043404 | 0.560164238 | -1.331829764 | 0.182916135 | 0.61680331 | Dbn1 | drebrin 1 [Source:MGI Symbol;Acc:MGI:1 | 32.124 | 14.391 | 20.125 | 13.496 | 8.7249 | 17.822 |
| 9767 | ENSMUSG00000034677 | 0 | NA | NA | NA | NA | NA | Gpr142 | G protein-coupled receptor 142 [Source:I | 0 | 0 | 0 | 0 | 0 | 0 |
| 9768 | ENSMUSG00000034681 | 359.2199111 | -0.034528387 | 0.147167586 | -0.23461951 | 0.814504066 | 0.967404825 | Rnps1 | RNA binding protein with serine rich doma | 378.26 | 382.39 | 330.05 | 386.56 | 326.09 | 351.97 |
| 9769 | ENSMUSG00000034683 | 0 | NA | NA | NA | NA | NA | Ppp1r1c | protein phosphatase 1, regulatory inhibitc | 0 | 0 | 0 | 0 | 0 | 0 |
| 9770 | ENSMUSG00000034684 | 74.60302662 | -0.18571705 | 0.270133507 | -0.687500978 | 0.491767085 | 0.876368691 | Sema3f | sema domain, immunoglobulin domain (Ig | 69.87 | 71.954 | 96.599 | 71.334 | 64.346 | 73.514 |
| 9771 | ENSMUSG00000034685 | 27.36893107 | -0.898126711 | 0.442887326 | -2.027889846 | 0.042571492 | 0.284150464 | Fam171a2 | family with sequence similarity 171, meml | 38.549 | 31.865 | 36.225 | 14.46 | 15.269 | 27.846 |
| 9772 | ENSMUSG00000034686 | 3.763250281 | -1.400413478 | 1.132768605 | -1.236274974 | 0.21635637 | NA | Prr7 | proline rich 7 (synaptic) [Source:MGI Symb | 7.2279 | 5.1396 | 4.025 | 2.8919 | 2.1812 | 1.1138 |
| 9773 | ENSMUSG00000034687 | 76.84283558 | -0.016227333 | 0.285687866 | -0.056800916 | 0.954703784 | 0.991678478 | Fras1 | Fraser extracellular matrix complex subun | 93.16 | 65.787 | 72.45 | 71.334 | 97.065 | 61.261 |
| 9774 | ENSMUSG00000034689 | 0 | NA | NA | NA | NA | NA | Spertl | spermatid associated like [Source:MGI Syr | 0 | 0 | 0 | 0 | 0 | 0 |
| 9775 | ENSMUSG00000034690 | 0 | NA | NA | NA | NA | NA | Nlrp4c | NLR family, pyrin domain containing 4C [S | 0 | 0 | 0 | 0 | 0 | 0 |
| 9776 | ENSMUSG00000034701 | 0 | NA | NA | NA | NA | NA | Neurod1 | neurogenic differentiation 1 [Source:MGI | 0 | 0 | 0 | 0 | 0 | 0 |
| 9777 | ENSMUSG00000034706 | 0 | NA | NA | NA | NA | NA | Dnai2 | dynein axonemal intermediate chain 2 [Sc | 0 | 0 | 0 | 0 | 0 | 0 |
| 9778 | ENSMUSG00000034707 | 3114.200063 | -0.005998834 | 0.095440029 | -0.062854486 | 0.949882378 | 0.990776571 | Gns | glucosamine (N-acetyl)-6-sulfatase [Sourc | 3314.4 | 3096.1 | 2950.3 | 3017.3 | 3268.6 | 3038.6 |
| 9779 | ENSMUSG00000034708 | 3578.092959 | -0.479848046 | 0.109063508 | -4.399712207 | 1.08E-05 | 0.000570116 | Grn | granulin [Source:MGI Symbol;Acc:MGI:95 | 4553.6 | 4318.3 | 3630.5 | 3020.1 | 3071.2 | 2874.8 |
| 9780 | ENSMUSG00000034709 | 338.8069935 | -0.004538375 | 0.147293986 | -0.030811679 | 0.975419726 | 0.996491676 | Ppp1r21 | protein phosphatase 1, regulatory subunit | 330.88 | 349.49 | 338.1 | 362.46 | 298.83 | 353.09 |
| 9781 | ENSMUSG00000034714 | 888.3870579 | 0.193397658 | 0.115165715 | 1.679298895 | 0.093093806 | 0.449435518 | Ttyh2 | tweety family member 2 [Source:MGI Sym | 870.56 | 818.22 | 796.95 | 892.64 | 918.3 | 1033.6 |
| 9782 | ENSMUSG00000034723 | 449.8700092 | 0.09659564 | 0.132680015 | 0.728034589 | 0.466592413 | 0.863779479 | Tmx4 | thioredoxin-related transmembrane prote | 414.4 | 459.48 | 430.67 | 432.83 | 495.14 | 466.7 |
| 9783 | ENSMUSG00000034724 | 865.2803102 | 0.084161488 | 0.118539317 | 0.709987962 | 0.477711601 | 0.869794311 | Cnot6l | CCR4-NOT transcription complex, subunit | 796.68 | 802.8 | 920.71 | 829.02 | 933.57 | 908.9 |
| 9784 | ENSMUSG00000034729 | 618.4496199 | 0.072937799 | 0.123732655 | 0.589478976 | 0.555540012 | 0.897628185 | Mrps10 | mitochondrial ribosomal protein S10 [Sou | 609.55 | 555.08 | 644 | 668.04 | 632.56 | 601.48 |
| 9785 | ENSMUSG00000034730 | 0 | NA | NA | NA | NA | NA | Adgrb1 | adhesion G protein-coupled receptor B1 [ | 0 | 0 | 0 | 0 | 0 | 0 |
| 9786 | ENSMUSG00000034731 | 64.44000371 | 0.100817643 | 0.325076143 | 0.310135471 | 0.756457939 | 0.953085143 | Dgkh | diacylglycerol kinase, eta [Source:MGI Syr | 48.989 | 59.619 | 78.487 | 85.794 | 45.806 | 67.944 |
| 9787 | ENSMUSG00000034732 | 0 | NA | NA | NA | NA | NA | Pabpc5 | poly(A) binding protein, cytoplasmic 5 [So | 0 | 0 | 0 | 0 | 0 | 0 |
| 9788 | ENSMUSG00000034738 | 317.5643162 | 0.190333862 | 0.175564095 | 1.084127493 | 0.278308275 | 0.73110838 | Nostrin | nitric oxide synthase trafficker [Source:M( | 344.53 | 233.34 | 310.93 | 315.22 | 330.46 | 370.91 |
| 9789 | ENSMUSG00000034739 | 0 | NA | NA | NA | NA | NA | Mfrp | membrane frizzled-related protein [Sourc | 0 | 0 | 0 | 0 | 0 | 0 |
| 9790 | ENSMUSG00000034744 | 271.6790806 | -0.218198093 | 0.166297431 | -1.312095394 | 0.18948795 | 0.626663447 | Nagk | N-acetylglucosamine kinase [Source:MGI : | 289.92 | 311.46 | 274.7 | 213.04 | 279.2 | 261.75 |
| 9791 | ENSMUSG00000034748 | 59.371453 | -0.146772445 | 0.299525013 | -0.490017322 | 0.624121641 | 0.919503423 | Sirt6 | sirtuin 6 [Source:MGI Symbol;Acc:MGI:13 | 72.279 | 51.396 | 63.393 | 65.55 | 51.259 | 52.351 |
| 9792 | ENSMUSG00000034751 | 67.74342206 | 0.354708833 | 0.276648825 | 1.282162802 | 0.199785542 | 0.642000421 | Mast4 | microtubule associated serine/threonine I | 59.43 | 65.787 | 53.331 | 86.758 | 66.528 | 74.628 |
| 9793 | ENSMUSG00000034755 | 1.533049506 | -1.663064797 | 2.170614423 | -0.76617237 | 0.443573746 | NA | Pcdh11x | protocadherin 11 X-linked [Source:MGI Sy | 0.8031 | 6.1675 | 0 | 0 | 0 | 2.2277 |
| 9794 | ENSMUSG00000034757 | 529.1308304 | 0.134439458 | 0.124856581 | 1.076751079 | 0.281591486 | 0.734379276 | Tmub2 | transmembrane and ubiquitin-like domair | 508.36 | 466.67 | 538.34 | 557.18 | 556.22 | 548.01 |
| 9795 | ENSMUSG00000034758 | 28.88897826 | -0.494011309 | 0.469230201 | -1.05281226 | 0.292427045 | 0.743376015 | Tle6 | transducin-like enhancer of split 6 [Source | 18.471 | 38.033 | 45.281 | 30.847 | 23.994 | 16.708 |
| 9796 | ENSMUSG00000034761 | 303.7835721 | 0.291589269 | 0.146813163 | 1.986124832 | 0.047019465 | 0.300361156 | Map4k5 | mitogen-activated protein kinase kinase k | 253.78 | 281.65 | 284.77 | 340.28 | 338.09 | 324.13 |

| | | | | | | | | | | | | | | | |
|---|---|---|---|---|---|---|---|---|---|---|---|---|---|---|---|
| 9797 | ENSMUSG00000034762 | 1.108247276 | -1.030184547 | 2.320402292 | -0.443968079 | 0.657065663 | NA | Glis1 | GLIS family zinc finger 1 [Source:MGI Sym | 2.4093 | 0 | 2.0125 | 0 | 0 | 2.2277 |
| 9798 | ENSMUSG00000034764 | 0.160662974 | 1.126924175 | 4.080472857 | 0.276174898 | 0.782413736 | NA | 1700006J14I | RIKEN cDNA 1700006J14 gene [Source:MG | 0 | 0 | 0 | 0.964 | 0 | 0 |
| 9799 | ENSMUSG00000034765 | 14.16916306 | -0.004884488 | 0.614362946 | -0.007950493 | 0.993656492 | 0.99859176 | Dusp5 | dual specificity phosphatase 5 [Source:MG | 11.243 | 16.447 | 15.094 | 21.208 | 6.5437 | 14.48 |
| 9800 | ENSMUSG00000034768 | 2.713904885 | 2.866898116 | 1.818136056 | 1.576833652 | 0.114833806 | NA | Asb16 | ankyrin repeat and SOCS box-containing 1 | 0 | 0 | 2.0125 | 0 | 9.8156 | 4.4554 |
| 9801 | ENSMUSG00000034771 | 186.002366 | 0.552012521 | 0.230609857 | 2.393707395 | 0.016679049 | 0.153234527 | Tle2 | transducin-like enhancer of split 2 [Source | 113.24 | 161.38 | 179.11 | 278.59 | 184.31 | 199.38 |
| 9802 | ENSMUSG00000034773 | 3.232540869 | -1.804233941 | 1.430108251 | -1.261606554 | 0.207090397 | NA | Hrob | homologous recombination factor with OI | 4.8186 | 7.1954 | 3.0187 | 0 | 4.3625 | 0 |
| 9803 | ENSMUSG00000034774 | 317.0890726 | -0.832900362 | 0.267891262 | -3.109098653 | 0.00187659 | 0.033263486 | Dsg1c | desmoglein 1 gamma [Source:MGI Symbo | 261.81 | 548.91 | 408.54 | 242.92 | 295.56 | 144.8 |
| 9804 | ENSMUSG00000034777 | 0 | NA | NA | NA | NA | NA | Vax2 | ventral anterior homeobox 2 [Source:MGI | 0 | 0 | 0 | 0 | 0 | 0 |
| 9805 | ENSMUSG00000034780 | 545.8271546 | 0.488230523 | 0.219794136 | 2.221308227 | 0.026330092 | 0.210051502 | B3galt1 | UDP-Gal:betaGlcNAc beta 1,3-galactosyltr | 515.59 | 440.98 | 406.52 | 816.49 | 709.99 | 385.39 |
| 9806 | ENSMUSG00000034781 | 1254.933793 | 0.005336761 | 0.098823151 | 0.054003145 | 0.956932659 | 0.991861398 | Gna11 | guanine nucleotide binding protein, alpha | 1273.7 | 1243.8 | 1239.7 | 1208.8 | 1278.2 | 1285.4 |
| 9807 | ENSMUSG00000034783 | 22.62539403 | 0.184706806 | 0.489128244 | 0.377624494 | 0.705709566 | 0.94295036 | Cd207 | CD207 antigen [Source:MGI Symbol;Acc:N | 17.668 | 30.838 | 15.094 | 20.244 | 20.722 | 31.188 |
| 9808 | ENSMUSG00000034785 | 5028.28825 | 0.101556777 | 0.115073623 | 0.882537412 | 0.377486258 | 0.81030298 | Dio1 | deiodinase, iodothyronine, type I [Source: | 5587.2 | 4254.6 | 4710.2 | 4937.5 | 5352.8 | 5327.5 |
| 9809 | ENSMUSG00000034786 | 26.30890684 | -0.934100912 | 0.501246478 | -1.863556062 | 0.062384073 | 0.356831368 | Gpsm3 | G-protein signalling modulator 3 (AGS3-lik | 51.399 | 35.977 | 16.1 | 20.244 | 18.541 | 15.594 |
| 9810 | ENSMUSG00000034789 | 171.299035 | 0.26844203 | 0.196332814 | 1.367280511 | 0.171537398 | 0.599034187 | Rab24 | RAB24, member RAS oncogene family [So | 160.62 | 155.22 | 149.93 | 165.8 | 171.23 | 225 |
| 9811 | ENSMUSG00000034792 | 3.608448152 | -0.4037061 | 1.136187516 | -0.355316437 | 0.722352543 | NA | Gna15 | guanine nucleotide binding protein, alpha | 3.2124 | 6.1675 | 3.0187 | 4.8199 | 1.0906 | 3.3415 |
| 9812 | ENSMUSG00000034793 | 112.4247169 | 0.035477296 | 0.216700328 | 0.163715933 | 0.869954774 | 0.978765244 | G6pc3 | glucose 6 phosphatase, catalytic, 3 [Sourc | 111.63 | 111.02 | 110.69 | 125.32 | 113.42 | 102.47 |
| 9813 | ENSMUSG00000034795 | 85.02991391 | -0.310609978 | 0.273441238 | -1.13592953 | 0.255986058 | 0.708998869 | Ccdc122 | coiled-coil domain containing 122 [Source | 116.45 | 71.954 | 93.581 | 78.082 | 64.346 | 85.766 |
| 9814 | ENSMUSG00000034796 | 0.353089354 | 0.165142706 | 3.896102244 | 0.042386646 | 0.966190474 | NA | Cpne7 | copine VII [Source:MGI Symbol;Acc:MGI:2 | 0 | 1.0279 | 0 | 0 | 1.0906 | 0 |
| 9815 | ENSMUSG00000034799 | 45.7107852 | 0.018449139 | 0.334206631 | 0.055202791 | 0.955976906 | 0.991678478 | Unc13a | unc-13 homolog A [Source:MGI Symbol;A | 40.958 | 38.033 | 57.356 | 45.307 | 44.715 | 47.895 |
| 9816 | ENSMUSG00000034800 | 98.02456207 | 0.147135896 | 0.240791305 | 0.611051533 | 0.541165465 | 0.893614211 | Zfp661 | zinc finger protein 661 [Source:MGI Symb | 91.554 | 105.88 | 81.506 | 87.722 | 112.33 | 109.16 |
| 9817 | ENSMUSG00000034801 | 265.598905 | 0.20355395 | 0.160668911 | 1.266915601 | 0.20518547 | 0.650728757 | Sos2 | SOS Ras/Rho guanine nucleotide exchang | 237.72 | 248.76 | 254.58 | 291.12 | 311.92 | 249.5 |
| 9818 | ENSMUSG00000034807 | 1149.615367 | -0.094037375 | 0.109927923 | -0.855445757 | 0.392304421 | 0.821170933 | Colgalt1 | collagen beta(1-O)galactosyltransferase 1 | 1146 | 1159.5 | 1255.8 | 1064.2 | 1207.3 | 1064.8 |
| 9819 | ENSMUSG00000034810 | 3.022226238 | -0.895003168 | 1.466268153 | -0.610395286 | 0.54159999 | NA | Scn7a | sodium channel, voltage-gated, type VII, a | 9.6372 | 2.0558 | 0 | 0.964 | 4.3625 | 1.1138 |
| 9820 | ENSMUSG00000034813 | 5.736661252 | 0.320216446 | 0.949963077 | 0.337083044 | 0.736054299 | 0.948109711 | Grip1 | glutamate receptor interacting protein 1 [ | 3.2124 | 3.0838 | 9.0562 | 4.8199 | 10.906 | 3.3415 |
| 9821 | ENSMUSG00000034818 | 0.160662974 | 1.126924175 | 4.080472857 | 0.276174898 | 0.782413736 | NA | Celf5 | CUGBP, Elav-like family member 5 [Source | 0 | 0 | 0 | 0.964 | 0 | 0 |
| 9822 | ENSMUSG00000034820 | 460.7293958 | 0.08445622 | 0.147182821 | 0.573818465 | 0.566090652 | 0.902076435 | Cpsf7 | cleavage and polyadenylation specific fact | 409.58 | 460.51 | 472.93 | 546.58 | 451.52 | 423.26 |
| 9823 | ENSMUSG00000034825 | 1.368851432 | -2.657829223 | 2.207244444 | -1.204139048 | 0.22853584 | NA | Nrip3 | nuclear receptor interacting protein 3 [So | 4.0155 | 3.0838 | 0 | 0 | 0 | 1.1138 |
| 9824 | ENSMUSG00000034826 | 370.4062157 | 0.011356367 | 0.136941375 | 0.082928678 | 0.933908251 | 0.988420305 | Nup54 | nucleoporin 54 [Source:MGI Symbol;Acc:I | 388.7 | 344.35 | 373.32 | 372.1 | 369.72 | 374.25 |
| 9825 | ENSMUSG00000034829 | 1.363461943 | -1.65198086 | 2.004705142 | -0.82405179 | 0.409910132 | NA | Nxnl1 | nucleoredoxin-like 1 [Source:MGI Symbol, | 3.2124 | 1.0279 | 2.0125 | 1.928 | 0 | 0 |
| 9826 | ENSMUSG00000034832 | 448.4455626 | 0.297879657 | 0.140763581 | 2.116169937 | 0.034330357 | 0.247905284 | Tet3 | tet methylcytosine dioxygenase 3 [Source | 368.62 | 420.42 | 418.6 | 502.23 | 441.7 | 539.1 |
| 9827 | ENSMUSG00000034833 | 0.644197084 | -2.725354879 | 2.762261801 | -0.986638876 | 0.323819705 | NA | Tespa1 | thymocyte expressed, positive selection a | 0.8031 | 2.0558 | 1.0062 | 0 | 0 | 0 |
| 9828 | ENSMUSG00000034837 | 606.9249304 | 1.289813293 | 0.419874143 | 3.071904555 | 0.002126977 | 0.036494186 | Gnat1 | guanine nucleotide binding protein, alpha | 554.14 | 241.56 | 260.62 | 720.09 | 810.33 | 1054.8 |
| 9829 | ENSMUSG00000034839 | 0.356960287 | 0.16514264 | 3.880370363 | 0.042558474 | 0.966053499 | NA | Larp6 | La ribonucleoprotein domain family, mem | 0 | 1.0279 | 0 | 0 | 0 | 1.1138 |
| 9830 | ENSMUSG00000034842 | 21.82019909 | -0.729588941 | 0.531115918 | -1.373690595 | 0.169537758 | 0.596876778 | Art3 | ADP-ribosyltransferase 3 [Source:MGI Syn | 30.518 | 27.754 | 23.144 | 9.6398 | 10.906 | 28.96 |
| 9831 | ENSMUSG00000034845 | 645.0202679 | -1.431897259 | 0.167101424 | -8.569030856 | 1.04E-17 | 1.08E-14 | Plvap | plasmalemma vesicle associated protein [ | 1086.6 | 908.68 | 828.14 | 380.77 | 268.29 | 397.64 |
| 9832 | ENSMUSG00000034848 | 71.92672051 | -0.007406485 | 0.290099622 | -0.025530833 | 0.979631555 | 0.99820894 | Ttc21b | tetratricopeptide repeat domain 21B [Sou | 72.279 | 70.926 | 73.456 | 90.614 | 74.162 | 50.123 |
| 9833 | ENSMUSG00000034850 | 972.7481985 | 0.084210635 | 0.113702977 | 0.740619441 | 0.458924217 | 0.859322431 | Tmem127 | transmembrane protein 127 [Source:MGI | 848.88 | 1002.2 | 983.1 | 983.26 | 1011 | 1008 |
| 9834 | ENSMUSG00000034853 | 247.1037237 | -0.480678289 | 0.201329946 | -2.387515121 | 0.016962705 | 0.154696447 | Acot11 | acyl-CoA thioesterase 11 [Source:MGI Syn | 360.59 | 228.2 | 273.7 | 178.34 | 227.94 | 213.86 |
| 9835 | ENSMUSG00000034854 | 148.3310921 | 0.012797412 | 0.204426253 | 0.062601605 | 0.950083751 | 0.990815588 | Mfsd12 | major facilitator superfamily domain cont | 140.54 | 171.66 | 130.81 | 134.96 | 150.51 | 161.51 |
| 9836 | ENSMUSG00000034855 | 32.87195517 | -0.199245725 | 0.396928899 | -0.501968302 | 0.615689819 | 0.917509899 | Cxcl10 | chemokine (C-X-C motif) ligand 10 [Source | 44.171 | 34.949 | 26.162 | 34.703 | 31.628 | 25.618 |
| 9837 | ENSMUSG00000034858 | 2829.767003 | -0.540298463 | 0.159052396 | -3.396984107 | 0.000681329 | 0.015617664 | Fam214a | family with sequence similarity 214, meml | 2961 | 3623.4 | 3475.6 | 1683.1 | 2700.4 | 2535.1 |
| 9838 | ENSMUSG00000034863 | 83.31931241 | 0.111766073 | 0.290994965 | 0.384082499 | 0.700917303 | 0.941805852 | Ano8 | anoctamin 8 [Source:MGI Symbol;Acc:MG | 61.036 | 88.401 | 91.568 | 106.04 | 91.612 | 61.261 |
| 9839 | ENSMUSG00000034867 | 474.3497724 | 0.188847478 | 0.125350777 | 1.506552114 | 0.131925499 | 0.532693668 | Ankrd27 | ankyrin repeat domain 27 (VPS9 domain) | 436.89 | 462.56 | 430.67 | 494.52 | 520.22 | 501.23 |
| 9840 | ENSMUSG00000034868 | 24.96551612 | -0.073733096 | 0.452030029 | -0.163115483 | 0.87042751 | 0.978802475 | Myl12b | myosin, light chain 12B, regulatory [Sourc | 33.73 | 22.614 | 20.125 | 19.28 | 25.084 | 28.96 |
| 9841 | ENSMUSG00000034871 | 0 | NA | NA | NA | NA | NA | Fam151a | family with sequence simliarity 151, meml | 0 | 0 | 0 | 0 | 0 | 0 |
| 9842 | ENSMUSG00000034872 | 5.844092204 | -0.645990072 | 0.93440924 | -0.691335279 | 0.489354867 | 0.874889775 | Gipc3 | GIPC PDZ domain containing family, meml | 7.2279 | 3.0838 | 11.069 | 4.8199 | 2.1812 | 6.6831 |
| 9843 | ENSMUSG00000034875 | 760.9104962 | -0.070298904 | 0.122163717 | -0.575448308 | 0.564988141 | 0.901542552 | Nudt19 | nudix (nucleoside diphosphate linked moi | 706.73 | 797.66 | 835.18 | 774.07 | 762.34 | 689.47 |
| 9844 | ENSMUSG00000034880 | 739.8050476 | 0.062334802 | 0.116953503 | 0.532987902 | 0.594041955 | 0.913144152 | Mrpl34 | mitochondrial ribosomal protein L34 [Sou | 706.73 | 689.73 | 774.81 | 713.34 | 765.61 | 788.6 |
| 9845 | ENSMUSG00000034881 | 11.4589019 | -0.863174861 | 0.638613575 | -1.35163876 | 0.176490905 | 0.607400048 | Tbxa2r | thromboxane A2 receptor [Source:MGI Sy | 12.85 | 14.391 | 17.106 | 6.7478 | 7.6343 | 10.025 |
| 9846 | ENSMUSG00000034883 | 1.059810487 | -2.230732092 | 2.384896422 | -0.935358061 | 0.349603803 | NA | Lrr1 | leucine rich repeat protein 1 [Source:MGI | 3.2124 | 2.0558 | 0 | 0 | 1.0906 | 0 |
| 9847 | ENSMUSG00000034889 | 278.7483188 | -0.209553566 | 0.164390821 | -1.2747279 | 0.202405568 | 0.647300167 | Cactin | cactin, spliceosome C complex subunit [Sc | 274.66 | 272.4 | 350.17 | 258.35 | 257.39 | 259.53 |
| 9848 | ENSMUSG00000034891 | 0 | NA | NA | NA | NA | NA | Sncb | synuclein, beta [Source:MGI Symbol;Acc:I | 0 | 0 | 0 | 0 | 0 | 0 |
| 9849 | ENSMUSG00000034892 | 1412.495927 | 0.022221915 | 0.141364957 | 0.157195357 | 0.875090887 | 0.980503106 | Rps29 | ribosomal protein S29 [Source:MGI Symb | 1221.5 | 1637.5 | 1346.4 | 1284 | 1336 | 1649.6 |
| 9850 | ENSMUSG00000034893 | 310.3427603 | -0.057751665 | 0.153059775 | -0.377314451 | 0.705939934 | 0.94295036 | Cog3 | component of oligomeric golgi complex 3 | 281.09 | 349.49 | 319.99 | 315.22 | 296.65 | 299.62 |
| 9851 | ENSMUSG00000034898 | 8.72258646 | -0.240779683 | 0.738605424 | -0.325992302 | 0.744430179 | 0.949997329 | Filip1 | filamin A interacting protein 1 [Source:MG | 12.047 | 7.1954 | 9.0562 | 8.6758 | 10.906 | 4.4554 |
| 9852 | ENSMUSG00000034902 | 487.5217504 | -0.130909862 | 0.160241871 | -0.816951656 | 0.413956056 | 0.835653928 | Pip5k1c | phosphatidylinositol-4-phosphate 5-kinas | 437.69 | 470.79 | 620.85 | 439.57 | 444.97 | 511.25 |

| | | | | | | | | | | | | | | | |
|---|---|---|---|---|---|---|---|---|---|---|---|---|---|---|---|
| 9853 | ENSMUSG00000034903 | 890.6102957 | 0.048522205 | 0.132671956 | 0.365730686 | 0.714566046 | 0.944595913 | Cobll1 | Cobl-like 1 [Source:MGI Symbol;Acc:MGI: | 912.32 | 928.21 | 786.88 | 1042.1 | 833.23 | 840.95 |
| 9854 | ENSMUSG00000034906 | 7.030294011 | -0.591373725 | 0.803030522 | -0.736427457 | 0.4614706 | 0.860071824 | Ncaph | non-SMC condensin I complex, subunit H | 8.031 | 11.307 | 6.0375 | 5.7839 | 5.4531 | 5.5692 |
| 9855 | ENSMUSG00000034908 | 2440.205243 | -0.000152946 | 0.10689836 | -0.001430763 | 0.998858416 | 0.999764475 | Sidt2 | SID1 transmembrane family, member 2 [S | 2534.6 | 2174 | 2612.2 | 2570.9 | 2404.8 | 2344.6 |
| 9856 | ENSMUSG00000034910 | 3.011932803 | -0.434143513 | 1.31981705 | -0.328942192 | 0.742199383 | NA | Pygo1 | pygopus 1 [Source:MGI Symbol;Acc:MGI: | 3.2124 | 3.0838 | 4.025 | 0 | 2.1812 | 5.5692 |
| 9857 | ENSMUSG00000034911 | 211.564027 | -0.142394838 | 0.234109266 | -0.608240932 | 0.543027691 | 0.894344786 | Ushbp1 | USH1 protein network component harmo | 217.64 | 208.67 | 240.49 | 286.3 | 159.23 | 157.05 |
| 9858 | ENSMUSG00000034912 | 0 | NA | NA | NA | NA | NA | Mdga2 | MAM domain containing glycosylphospha | 0 | 0 | 0 | 0 | 0 | 0 |
| 9859 | ENSMUSG00000034913 | 0 | NA | NA | NA | NA | NA | Cby2 | chibby family member 2 [Source:MGI Sym | 0 | 0 | 0 | 0 | 0 | 0 |
| 9860 | ENSMUSG00000034917 | 500.3690674 | -0.346207234 | 0.171616491 | -2.0173308 | 0.043661003 | 0.288482134 | Tjp3 | tight junction protein 3 [Source:MGI Symb | 533.26 | 484.15 | 663.12 | 455.96 | 353.36 | 512.37 |
| 9861 | ENSMUSG00000034918 | 1.570209331 | -0.999497949 | 1.663052205 | -0.601002149 | 0.547838555 | NA | Cdhr2 | cadherin-related family member 2 [Source | 3.2124 | 1.0279 | 2.0125 | 0.964 | 1.0906 | 1.1138 |
| 9862 | ENSMUSG00000034919 | 0.795774083 | 0.899657507 | 2.433208733 | 0.369741196 | 0.711575334 | NA | Ttc22 | tetratricopeptide repeat domain 22 [Sour | 1.6062 | 0 | 0 | 0.964 | 1.0906 | 1.1138 |
| 9863 | ENSMUSG00000034923 | 0 | NA | NA | NA | NA | NA | Ly6g6f | lymphocyte antigen 6 complex, locus G6F | 0 | 0 | 0 | 0 | 0 | 0 |
| 9864 | ENSMUSG00000034926 | 9623.515868 | 0.040294959 | 0.112510007 | 0.358145553 | 0.720234391 | 0.9447719 | Dhcr24 | 24-dehydrocholesterol reductase [Source | 9711.1 | 10647 | 8109.3 | 10182 | 9637.8 | 9453.2 |
| 9865 | ENSMUSG00000034928 | 1058.14941 | 0.056352583 | 0.105641323 | 0.533433138 | 0.593733783 | 0.913106373 | Rnf44 | ring finger protein 44 [Source:MGI Symbo | 1022.3 | 1064.9 | 1025.4 | 1082.5 | 1015.4 | 1138.3 |
| 9866 | ENSMUSG00000034930 | 102.3147606 | 0.751433435 | 0.240585507 | 3.123352869 | 0.001788032 | 0.032166791 | Rtkn | rhotekin [Source:MGI Symbol;Acc:MGI:10 | 70.673 | 76.066 | 82.512 | 154.24 | 112.33 | 118.07 |
| 9867 | ENSMUSG00000034931 | 301.0666981 | -0.114322432 | 0.158822468 | -0.719812715 | 0.471640315 | 0.867794597 | Dhx8 | DEAH (Asp-Glu-Ala-His) box polypeptide 8 | 281.09 | 360.8 | 297.85 | 297.87 | 279.2 | 289.6 |
| 9868 | ENSMUSG00000034932 | 384.7392075 | 0.012578799 | 0.150237483 | 0.083726106 | 0.9332742 | 0.988420305 | Mrpl54 | mitochondrial ribosomal protein L54 [Sou | 404.76 | 378.27 | 365.27 | 347.03 | 366.45 | 446.65 |
| 9869 | ENSMUSG00000034936 | 539.5142574 | -1.082602389 | 0.444077267 | -2.437869421 | 0.01477411 | 0.141505508 | Arl4d | ADP-ribosylation factor-like 4D [Source:M | 606.34 | 935.41 | 657.08 | 260.27 | 550.76 | 227.22 |
| 9870 | ENSMUSG00000034940 | 339.5058405 | 0.052312539 | 0.149785282 | 0.349250192 | 0.726901488 | 0.945902846 | Synrg | synergin, gamma [Source:MGI Symbol;Acc | 346.14 | 344.35 | 308.92 | 309.44 | 378.44 | 349.75 |
| 9871 | ENSMUSG00000034947 | 131.9032759 | -0.34730738 | 0.215938089 | -1.608365534 | 0.107755146 | 0.484241711 | Tmem106a | transmembrane protein 106A [Source:MG | 160.62 | 146.99 | 134.84 | 95.434 | 124.33 | 129.21 |
| 9872 | ENSMUSG00000034949 | 18.53330958 | 0.062048217 | 0.531529205 | 0.116735293 | 0.907069822 | 0.984383096 | Zfr2 | zinc finger RNA binding protein 2 [Source: | 25.699 | 12.335 | 16.1 | 16.388 | 25.084 | 15.594 |
| 9873 | ENSMUSG00000034951 | 406.831693 | 0.021008936 | 0.138083835 | 0.152146238 | 0.879071596 | 0.981233798 | Cog7 | component of oligomeric golgi complex 7 | 400.75 | 400.89 | 409.54 | 368.24 | 443.88 | 417.69 |
| 9874 | ENSMUSG00000034957 | 12474.27378 | 0.426220413 | 0.095887131 | 4.44502209 | 8.79E-06 | 0.000477078 | Cebpa | CCAAT/enhancer binding protein (C/EBP), | 11145 | 9757 | 11031 | 14726 | 13381 | 14805 |
| 9875 | ENSMUSG00000034958 | 0 | NA | NA | NA | NA | NA | Atcay | ataxia, cerebellar, Cayman type [Source:M | 0 | 0 | 0 | 0 | 0 | 0 |
| 9876 | ENSMUSG00000034959 | 9.047263939 | -0.689454047 | 0.794178258 | -0.868135132 | 0.38532036 | 0.816437825 | Rubcnl | RUN and cysteine rich domain containing | 16.062 | 14.391 | 3.0187 | 8.6758 | 5.4531 | 6.6831 |
| 9877 | ENSMUSG00000034968 | 3.330345377 | -0.432070464 | 1.266425276 | -0.341173279 | 0.732973138 | NA | Lbx2 | ladybird homeobox 2 [Source:MGI Symbo | 2.4093 | 1.0279 | 8.05 | 1.928 | 5.4531 | 1.1138 |
| 9878 | ENSMUSG00000034973 | 327.0165983 | 0.031206322 | 0.155848706 | 0.200234717 | 0.841297017 | 0.971854408 | Dop1a | DOP1 leucine zipper like protein A [Source | 331.68 | 343.32 | 295.84 | 371.13 | 327.19 | 292.94 |
| 9879 | ENSMUSG00000034974 | 495.7868368 | 0.035060189 | 0.139661357 | 0.251037151 | 0.801785386 | 0.962796744 | Dapk3 | death-associated protein kinase 3 [Source | 497.12 | 428.64 | 543.37 | 502.23 | 533.31 | 470.04 |
| 9880 | ENSMUSG00000034981 | 3.531440943 | -1.280837323 | 1.328024954 | -0.964467813 | 0.334811438 | NA | Parm1 | prostate androgen-regulated mucin-like p | 8.8341 | 6.1675 | 0 | 2.8919 | 2.1812 | 1.1138 |
| 9881 | ENSMUSG00000034987 | 0.94941923 | -0.804713261 | 2.270652885 | -0.354397304 | 0.723041155 | NA | Hrh2 | histamine receptor H2 [Source:MGI Symb | 1.6062 | 0 | 2.0125 | 0.964 | 0 | 1.1138 |
| 9882 | ENSMUSG00000034990 | 0.535117793 | 1.111049431 | 3.044213722 | 0.364970903 | 0.715133126 | NA | Otoa | otoancorin [Source:MGI Symbol;Acc:MGI: | 0 | 0 | 1.0062 | 0 | 1.0906 | 1.1138 |
| 9883 | ENSMUSG00000034993 | 540.5661341 | -0.293085874 | 0.123256703 | -2.37784938 | 0.017413937 | 0.157225462 | Vat1 | vesicle amine transport 1 [Source:MGI Syr | 605.54 | 589 | 591.67 | 520.55 | 475.51 | 461.13 |
| 9884 | ENSMUSG00000034994 | 28239.13189 | 0.160171871 | 0.105239039 | 1.521981504 | 0.128013715 | 0.524051388 | Eef2 | eukaryotic translation elongation factor 2 | 23104 | 29099 | 27819 | 30590 | 29915 | 28909 |
| 9885 | ENSMUSG00000034997 | 0.990320745 | -2.084832509 | 2.480930004 | -0.84034314 | 0.40071602 | NA | Htr2a | 5-hydroxytryptamine (serotonin) receptor | 0.8031 | 0 | 4.025 | 0 | 0 | 1.1138 |
| 9886 | ENSMUSG00000034998 | 306.5107965 | -0.070982125 | 0.16618019 | -0.427139506 | 0.669277719 | 0.930837335 | Foxn2 | forkhead box N2 [Source:MGI Symbol;Acc | 315.62 | 301.18 | 325.02 | 283.41 | 256.3 | 357.54 |
| 9887 | ENSMUSG00000035000 | 2457.491959 | -0.042691547 | 0.104341068 | -0.409153828 | 0.682426776 | 0.93502957 | Dpp4 | dipeptidylpeptidase 4 [Source:MGI Symbo | 2306.5 | 2598.6 | 2577 | 2266.3 | 2621.8 | 2374.7 |
| 9888 | ENSMUSG00000035004 | 30.6015385 | -1.61858536 | 0.561950138 | -2.88030067 | 0.003972961 | 0.056970643 | Igsf6 | immunoglobulin superfamily, member 6 [ | 28.109 | 90.457 | 20.125 | 17.352 | 13.087 | 14.48 |
| 9889 | ENSMUSG00000035007 | 262.831958 | 0.06501655 | 0.153624053 | 0.423218556 | 0.67213581 | 0.931191575 | Rundc1 | RUN domain containing 1 [Source:MGI Sy | 246.55 | 250.81 | 273.7 | 275.7 | 259.57 | 270.66 |
| 9890 | ENSMUSG00000035011 | 1098.715393 | 0.13487947 | 0.12459603 | 1.082534247 | 0.279015207 | 0.731519729 | Zbtb7a | zinc finger and BTB domain containing 7a | 954.89 | 1056.7 | 1132 | 1246.4 | 1194.2 | 1008 |
| 9891 | ENSMUSG00000035020 | 0 | NA | NA | NA | NA | NA | Epgn | epithelial mitogen [Source:MGI Symbol;A | 0 | 0 | 0 | 0 | 0 | 0 |
| 9892 | ENSMUSG00000035021 | 186.302305 | -0.21909803 | 0.214862348 | -1.01971347 | 0.307864372 | 0.756343859 | Baz1a | bromodomain adjacent to zinc finger dom | 161.42 | 244.64 | 196.22 | 203.4 | 145.05 | 167.08 |
| 9893 | ENSMUSG00000035024 | 230.4756091 | -0.004372671 | 0.184736077 | -0.023669825 | 0.981115975 | 0.99820894 | Ncapd3 | non-SMC condensin II complex, subunit D | 249.76 | 242.59 | 200.24 | 268.95 | 196.31 | 225 |
| 9894 | ENSMUSG00000035027 | 2619.241367 | -0.014363841 | 0.097887952 | -0.146737582 | 0.883339154 | 0.981769403 | Map2k2 | mitogen-activated protein kinase kinase 2 | 2407.7 | 2743.5 | 2747 | 2637.4 | 2545.5 | 2634.2 |
| 9895 | ENSMUSG00000035031 | 11676.76282 | 0.630181372 | 0.105465654 | 5.975228409 | 2.30E-09 | 4.32E-07 | C8a | complement component 8, alpha polypep | 8108.1 | 9700.5 | 9692.1 | 13947 | 15245 | 13368 |
| 9896 | ENSMUSG00000035032 | 0.328370405 | 0.16514316 | 4.003149536 | 0.041253308 | 0.967093956 | NA | Nek11 | NIMA (never in mitosis gene a)-related ex | 0 | 0 | 1.0062 | 0.964 | 0 | 0 |
| 9897 | ENSMUSG00000035033 | 0 | NA | NA | NA | NA | NA | Tbr1 | T-box brain transcription factor 1 [Source: | 0 | 0 | 0 | 0 | 0 | 0 |
| 9898 | ENSMUSG00000035041 | 7787.25403 | 0.155276897 | 0.092115125 | 1.685682959 | 0.091856881 | 0.446623824 | Creb3l3 | cAMP responsive element binding protein | 7384.5 | 6904.5 | 7816.5 | 8106.1 | 8618.1 | 7893.8 |
| 9899 | ENSMUSG00000035042 | 16.69781091 | -2.922561661 | 0.708578229 | -4.124543406 | 3.71E-05 | 0.00166862 | Ccl5 | chemokine (C-C motif) ligand 5 [Source:M | 28.109 | 45.228 | 15.094 | 2.8919 | 2.1812 | 6.6831 |
| 9900 | ENSMUSG00000035045 | 3.449562459 | -0.827454045 | 1.193725345 | -0.693169537 | 0.488203163 | NA | Zc3h12b | zinc finger CCCH-type containing 12B [Sou | 4.0155 | 7.1954 | 2.0125 | 1.928 | 1.0906 | 4.4554 |
| 9901 | ENSMUSG00000035047 | 66.76716988 | 0.199745136 | 0.288689969 | 0.691901891 | 0.488998943 | 0.874706988 | Kri1 | KRI1 homolog [Source:MGI Symbol;Acc:M | 66.657 | 50.368 | 69.431 | 80.974 | 75.253 | 57.92 |
| 9902 | ENSMUSG00000035048 | 666.6839224 | 0.002448319 | 0.128999753 | 0.018979255 | 0.984857654 | 0.99859176 | Anapc13 | anaphase promoting complex subunit 13 | 651.32 | 673.29 | 673.18 | 583.21 | 676.18 | 742.93 |
| 9903 | ENSMUSG00000035049 | 155.2196761 | -0.345477531 | 0.227538943 | -1.518322647 | 0.128933081 | 0.52561671 | Rrp12 | ribosomal RNA processing 12 homolog [S | 142.95 | 204.56 | 174.08 | 128.21 | 170.14 | 111.38 |
| 9904 | ENSMUSG00000035051 | 178.2692249 | 0.3235535 | 0.198230386 | 1.632209397 | 0.102635382 | 0.470976574 | Dhx57 | DEAH (Asp-Glu-Ala-Asp/His) box polypept | 179.09 | 132.6 | 163.01 | 214 | 210.49 | 170.42 |
| 9905 | ENSMUSG00000035057 | 0 | NA | NA | NA | NA | NA | Mgat4d | MGAT4 family, member C [Source:MGI Sy | 0 | 0 | 0 | 0 | 0 | 0 |
| 9906 | ENSMUSG00000035062 | 16.58791265 | 0.292873099 | 0.551137341 | 0.531397671 | 0.595143235 | 0.913525602 | Zc4h2 | zinc finger, C4H2 domain containing [Sour | 19.274 | 11.307 | 14.087 | 25.063 | 13.087 | 16.708 |
| 9907 | ENSMUSG00000035064 | 204.3784809 | 0.377882978 | 0.18953219 | 1.993766745 | 0.046177551 | 0.297429405 | Eef2k | eukaryotic elongation factor-2 kinase [Sou | 195.15 | 156.24 | 181.12 | 200.51 | 270.47 | 222.77 |
| 9908 | ENSMUSG00000035067 | 35.08324734 | -0.078819645 | 0.397510737 | -0.198283059 | 0.842823605 | 0.972137211 | Xkr6 | X-linked Kx blood group related 6 [Source | 32.124 | 26.726 | 49.306 | 32.775 | 28.356 | 41.212 |

| | | | | | | | | | | | | | | | |
|---|---|---|---|---|---|---|---|---|---|---|---|---|---|---|---|
| 9909 | ENSMUSG00000035069 | 274.9400322 | 0.118843755 | 0.17288283 | 0.687423704 | 0.491815765 | 0.876368691 | Oma1 | OMA1 zinc metallopeptidase [Source:MGI | 267.43 | 278.57 | 244.52 | 251.6 | 343.54 | 263.98 |
| 9910 | ENSMUSG00000035078 | 464.0413582 | -0.113132431 | 0.130293281 | -0.868290593 | 0.385235271 | 0.816437825 | Mtmr9 | myotubularin related protein 9 [Source:M | 501.94 | 488.26 | 455.83 | 419.33 | 472.24 | 446.65 |
| 9911 | ENSMUSG00000035085 | 2.30701944 | -1.168923237 | 1.458542889 | -0.8014322 | 0.42288148 | NA | 1700020L24I | RIKEN cDNA 1700020L24 gene [Source:MG | 2.4093 | 6.1675 | 1.0062 | 0.964 | 2.1812 | 1.1138 |
| 9912 | ENSMUSG00000035086 | 797.742743 | 0.095887222 | 0.124048184 | 0.772983678 | 0.43953204 | 0.84844279 | Becn1 | beclin 1, autophagy related [Source:MGI S | 712.35 | 758.6 | 843.23 | 800.1 | 771.07 | 901.1 |
| 9913 | ENSMUSG00000035093 | 1324.085772 | -0.155940056 | 0.113186454 | -1.377727193 | 0.168287532 | 0.595160122 | Secisbp2l | SECIS binding protein 2-like [Source:MGI S | 1497.8 | 1461.7 | 1226.6 | 1267.6 | 1244.4 | 1246.4 |
| 9914 | ENSMUSG00000035095 | 8.616654292 | 0.419220635 | 0.732588167 | 0.572245982 | 0.56715534 | 0.902953752 | Fam167a | family with sequence similarity 167, meml | 8.8341 | 5.1396 | 8.05 | 8.6758 | 7.6343 | 13.366 |
| 9915 | ENSMUSG00000035104 | 2353.898044 | 0.53766642 | 0.124103039 | 4.332419439 | 1.47E-05 | 0.000726202 | Eva1a | eva-1 homolog A (C. elegans) [Source:MG | 1913.8 | 1975.7 | 1870.6 | 2373.3 | 2730.9 | 3259.1 |
| 9916 | ENSMUSG00000035105 | 333.2097212 | 0.344796893 | 0.158471319 | 2.17576843 | 0.02957258 | 0.226497547 | Egln3 | egl-9 family hypoxia-inducible factor 3 [So | 290.72 | 289.87 | 299.86 | 345.1 | 333.73 | 439.97 |
| 9917 | ENSMUSG00000035107 | 26.8540856 | -0.419202093 | 0.436580101 | -0.960195145 | 0.33695701 | 0.779072142 | Dcbld2 | discoidin, CUB and LCCL domain containin | 36.943 | 28.782 | 26.162 | 17.352 | 21.812 | 30.074 |
| 9918 | ENSMUSG00000035109 | 1.612112483 | 3.090316284 | 1.984167877 | 1.557487307 | 0.119354832 | NA | Shc4 | SHC (Src homology 2 domain containing) f | 0 | 1.0279 | 0 | 0.964 | 5.4531 | 2.2277 |
| 9919 | ENSMUSG00000035112 | 36.19002703 | 1.14175388 | 0.390590918 | 2.923144978 | 0.003465151 | 0.051937771 | Wnk4 | WNK lysine deficient protein kinase 4 [Sou | 20.078 | 18.503 | 29.181 | 43.379 | 43.625 | 62.375 |
| 9920 | ENSMUSG00000035121 | 21.59513514 | 0.760880549 | 0.471075047 | 1.615200282 | 0.106267329 | 0.480851832 | Neil2 | nei like 2 (E. coli) [Source:MGI Symbol;Acc | 18.471 | 17.475 | 12.075 | 30.847 | 25.084 | 25.618 |
| 9921 | ENSMUSG00000035125 | 61.64998807 | 0.25080909 | 0.295700535 | 0.848186121 | 0.396334325 | 0.823764863 | Gcfc2 | GC-rich sequence DNA binding factor 2 [S | 47.383 | 58.591 | 63.393 | 76.154 | 69.8 | 54.578 |
| 9922 | ENSMUSG00000035126 | 12.62052789 | 0.581244249 | 0.623791861 | 0.931791973 | 0.351444044 | 0.792511048 | Dnai4 | dynein axonemal intermediate chain 4 [Sc | 12.85 | 11.307 | 6.0375 | 12.532 | 19.631 | 13.366 |
| 9923 | ENSMUSG00000035129 | 0.665794642 | 1.656189793 | 2.801711575 | 0.591135007 | 0.554429965 | NA | Gm6781 | predicted gene 6781 [Source:MGI Symbol | 0.8031 | 0 | 0 | 0.964 | 0 | 2.2277 |
| 9924 | ENSMUSG00000035131 | 0 | NA | NA | NA | NA | NA | Brinp3 | bone morphogenetic protein/retinoic acic | 0 | 0 | 0 | 0 | 0 | 0 |
| 9925 | ENSMUSG00000035133 | 1980.511853 | -0.15270404 | 0.121502404 | -1.256798516 | 0.208826604 | 0.653498052 | Arhgap5 | Rho GTPase activating protein 5 [Source:N | 1905 | 2392 | 1960.2 | 1983.9 | 1942.4 | 1699.7 |
| 9926 | ENSMUSG00000035139 | 335.2218009 | 0.12919108 | 0.143632816 | 0.899453786 | 0.368411001 | 0.80627882 | Secisbp2 | SECIS binding protein 2 [Source:MGI Symb | 311.6 | 333.05 | 315.96 | 325.82 | 376.26 | 348.63 |
| 9927 | ENSMUSG00000035142 | 225.2544998 | -0.22614011 | 0.194785445 | -1.160970263 | 0.245653993 | 0.695774774 | Nubpl | nucleotide binding protein-like [Source:M | 232.9 | 244.64 | 250.55 | 155.2 | 242.12 | 226.11 |
| 9928 | ENSMUSG00000035148 | 0.17131964 | -0.796629176 | 4.080472857 | -0.195229623 | 0.84521319 | NA | Gpr33 | G protein-coupled receptor 33 [Source:M | 0 | 1.0279 | 0 | 0 | 0 | 0 |
| 9929 | ENSMUSG00000035150 | 776.7276645 | 0.129055965 | 0.113345742 | 1.138604437 | 0.254868177 | 0.708380729 | Eif2s3x | eukaryotic translation initiation factor 2, s | 690.67 | 748.32 | 787.89 | 815.53 | 805.97 | 811.99 |
| 9930 | ENSMUSG00000035151 | 387.2869041 | 0.23356385 | 0.141604848 | 1.649405741 | 0.099064539 | 0.463861929 | Elmod2 | ELMO/CED-12 domain containing 2 [Sourc | 346.94 | 332.02 | 389.42 | 434.75 | 432.98 | 387.62 |
| 9931 | ENSMUSG00000035152 | 843.8831508 | 0.011057361 | 0.122169176 | 0.090508597 | 0.927883063 | 0.987920546 | Ap2b1 | adaptor-related protein complex 2, beta 1 | 839.24 | 897.37 | 784.87 | 768.29 | 928.12 | 845.41 |
| 9932 | ENSMUSG00000035158 | 65.28337144 | -0.223289358 | 0.33153806 | -0.673495399 | 0.500632175 | 0.880917734 | Mitf | melanogenesis associated transcription fa | 60.233 | 92.513 | 58.362 | 57.839 | 41.443 | 81.311 |
| 9933 | ENSMUSG00000035161 | 299.2229246 | -0.051254155 | 0.157711168 | -0.324987478 | 0.745190549 | 0.949997329 | Ints6 | integrator complex subunit 6 [Source:MG | 279.48 | 339.21 | 295.84 | 320.04 | 290.1 | 270.66 |
| 9934 | ENSMUSG00000035164 | 123.421183 | 0.02430839 | 0.205490674 | 0.118294369 | 0.905834419 | 0.983991807 | Zc3h12c | zinc finger CCCH type containing 12C [Sou | 122.87 | 123.35 | 120.75 | 115.68 | 129.78 | 128.09 |
| 9935 | ENSMUSG00000035165 | 5.654784144 | -1.623623921 | 1.056001408 | -1.537520602 | 0.124165874 | 0.516518608 | Kcne3 | potassium voltage-gated channel, Isk-rela | 7.2279 | 16.447 | 2.0125 | 3.8559 | 3.2719 | 1.1138 |
| 9936 | ENSMUSG00000035168 | 139.254101 | -0.35011355 | 0.234848633 | -1.49080514 | 0.136012662 | 0.538690549 | Tanc1 | tetratricopeptide repeat, ankyrin repeat a | 179.89 | 167.55 | 120.75 | 145.56 | 99.246 | 122.52 |
| 9937 | ENSMUSG00000035171 | 472.3786612 | 0.021796994 | 0.155763356 | 0.1399366 | 0.888710083 | 0.982382091 | 1110059E24 | RIKEN cDNA 1110059E24 gene [Source:M | 534.06 | 450.23 | 421.62 | 506.09 | 522.41 | 399.87 |
| 9938 | ENSMUSG00000035172 | 263.3529509 | -0.00412265 | 0.189858672 | -0.02171431 | 0.982675849 | 0.998580735 | Plekhh3 | pleckstrin homology domain containing, f | 277.87 | 226.14 | 286.78 | 260.27 | 210.49 | 318.56 |
| 9939 | ENSMUSG00000035173 | 212.8284066 | 0.128923044 | 0.190958949 | 0.675134863 | 0.499590086 | 0.880660418 | Ccdc186 | coiled-coil domain containing 186 [Source | 213.63 | 203.53 | 192.19 | 191.83 | 200.67 | 275.12 |
| 9940 | ENSMUSG00000035177 | 0 | NA | NA | NA | NA | NA | Nlrp2 | NLR family, pyrin domain containing 2 [So | 0 | 0 | 0 | 0 | 0 | 0 |
| 9941 | ENSMUSG00000035179 | 0.328370405 | 0.16514316 | 4.003149536 | 0.041253308 | 0.967093956 | NA | Ppp1r32 | protein phosphatase 1, regulatory subunit | 0 | 0 | 1.0062 | 0.964 | 0 | 0 |
| 9942 | ENSMUSG00000035181 | 258.6929482 | 0.088148962 | 0.164657314 | 0.535347987 | 0.592409249 | 0.912923013 | Heatr5a | HEAT repeat containing 5A [Source:MGI S | 256.99 | 267.26 | 228.42 | 297.87 | 243.21 | 258.41 |
| 9943 | ENSMUSG00000035183 | 11.86229234 | -0.209096657 | 0.638030099 | -0.327722245 | 0.743121682 | 0.949431323 | Slc24a5 | solute carrier family 24, member 5 [Sourc | 12.85 | 13.363 | 12.075 | 16.388 | 9.8156 | 6.6831 |
| 9944 | ENSMUSG00000035184 | 200.1503513 | -0.595959203 | 0.200319819 | -2.975038653 | 0.002929516 | 0.046007084 | Fam124a | family with sequence similarity 124, meml | 277.87 | 243.62 | 201.25 | 187.01 | 136.33 | 154.82 |
| 9945 | ENSMUSG00000035186 | 19.65010291 | -1.020453529 | 0.537804838 | -1.897442078 | 0.057769617 | 0.341829329 | Ubd | ubiquitin D [Source:MGI Symbol;Acc:MGI: | 32.927 | 30.838 | 15.094 | 11.568 | 17.45 | 10.025 |
| 9946 | ENSMUSG00000035187 | 0 | NA | NA | NA | NA | NA | Nkx6-1 | NK6 homeobox 1 [Source:MGI Symbol;Ac | 0 | 0 | 0 | 0 | 0 | 0 |
| 9947 | ENSMUSG00000035189 | 0.636972666 | -2.712703855 | 3.077964703 | -0.881330397 | 0.378139018 | NA | Ano4 | anoctamin 4 [Source:MGI Symbol;Acc:MG | 0.8031 | 0 | 3.0187 | 0 | 0 | 0 |
| 9948 | ENSMUSG00000035191 | 0 | NA | NA | NA | NA | NA | Rfpl4 | ret finger protein-like 4 [Source:MGI Symb | 0 | 0 | 0 | 0 | 0 | 0 |
| 9949 | ENSMUSG00000035198 | 567.1451672 | -0.151330923 | 0.122684036 | -1.23350135 | 0.217388773 | 0.663356797 | Tubg1 | tubulin, gamma 1 [Source:MGI Symbol;Ac | 585.46 | 563.3 | 641.98 | 541.76 | 524.59 | 545.78 |
| 9950 | ENSMUSG00000035199 | 572.9857921 | -0.138805752 | 0.126682946 | -1.095694062 | 0.273212682 | 0.728320076 | Arl6ip5 | ADP-ribosylation factor-like 6 interacting p | 643.28 | 568.44 | 589.66 | 582.24 | 547.49 | 506.8 |
| 9951 | ENSMUSG00000035200 | 0 | NA | NA | NA | NA | NA | Chrnb4 | cholinergic receptor, nicotinic, beta polyp | 0 | 0 | 0 | 0 | 0 | 0 |
| 9952 | ENSMUSG00000035201 | 0 | NA | NA | NA | NA | NA | Pramel17 | PRAME like 17 [Source:MGI Symbol;Acc:M | 0 | 0 | 0 | 0 | 0 | 0 |
| 9953 | ENSMUSG00000035202 | 8967.379013 | 0.687859589 | 0.990079749 | 0.6947517 | 0.487210922 | 0.873910634 | Lars2 | leucyl-tRNA synthetase, mitochondrial [Sc | 5783.9 | 8286 | 6537.6 | 4388 | 26036 | 2772.4 |
| 9954 | ENSMUSG00000035203 | 3360.573408 | -0.115278237 | 0.088038274 | -1.309410451 | 0.190395353 | 0.627779796 | Epn1 | epsin 1 [Source:MGI Symbol;Acc:MGI:133 | 3348.1 | 3522.7 | 3614.4 | 3266 | 3250 | 3162.2 |
| 9955 | ENSMUSG00000035206 | 373.6858037 | -0.115033632 | 0.15371773 | -0.748343293 | 0.454253117 | 0.857337687 | Sppl2b | signal peptide peptidase like 2B [Source:N | 363 | 351.55 | 451.8 | 384.63 | 340.27 | 350.86 |
| 9956 | ENSMUSG00000035208 | 20.83632763 | -0.738966026 | 0.485163995 | -1.523126269 | 0.127727118 | 0.523389132 | Slfn8 | schlafen 8 [Source:MGI Symbol;Acc:MGI:2 | 26.502 | 22.614 | 29.181 | 20.244 | 11.997 | 14.48 |
| 9957 | ENSMUSG00000035211 | 0.428363721 | -0.543730382 | 3.624677344 | -0.15000794 | 0.880758351 | NA | Xrra1 | X-ray radiation resistance associated 1 [Sc | 1.6062 | 0 | 0 | 0.964 | 0 | 0 |
| 9958 | ENSMUSG00000035212 | 266.3988286 | -0.066497299 | 0.165702453 | -0.40130546 | 0.688195244 | 0.936908509 | Leprot | leptin receptor overlapping transcript [Sou | 296.34 | 279.59 | 241.5 | 282.45 | 232.3 | 266.21 |
| 9959 | ENSMUSG00000035215 | 213.2271064 | -0.112346489 | 0.177177784 | -0.634089031 | 0.526022721 | 0.888334265 | Lsm7 | LSM7 homolog, U6 small nuclear RNA and | 221.66 | 225.11 | 217.35 | 173.52 | 231.21 | 210.52 |
| 9960 | ENSMUSG00000035226 | 1.190203636 | 0.585913931 | 1.945514715 | 0.301161398 | 0.763291424 | NA | Rims4 | regulating synaptic membrane exocytosis | 0.8031 | 2.0558 | 0 | 0.964 | 1.0906 | 2.2277 |
| 9961 | ENSMUSG00000035227 | 972.1540455 | 0.051431634 | 0.115282947 | 0.446133932 | 0.655500503 | 0.926075335 | Spcs2 | signal peptidase complex subunit 2 homol | 867.35 | 997.08 | 1001.2 | 1012.2 | 927.03 | 1028.1 |
| 9962 | ENSMUSG00000035228 | 0.428363721 | -0.543730382 | 3.624677344 | -0.15000794 | 0.880758351 | NA | Ccdc106 | coiled-coil domain containing 106 [Source | 1.6062 | 0 | 0 | 0.964 | 0 | 0 |
| 9963 | ENSMUSG00000035232 | 4.294763034 | -1.350658803 | 1.124953613 | -1.200635109 | 0.229892776 | NA | Pdk3 | pyruvate dehydrogenase kinase, isoenzym | 2.4093 | 5.1396 | 11.069 | 3.8559 | 2.1812 | 1.1138 |
| 9964 | ENSMUSG00000035234 | 116.0075505 | 0.073812524 | 0.229324875 | 0.321868806 | 0.747552086 | 0.950808422 | Abraxas1 | BRCA1 A complex subunit [Source:MGI Sy | 138.13 | 99.708 | 100.62 | 118.57 | 126.51 | 112.5 |

| | | | | | | | | | | | | | | | |
|---|---|---|---|---|---|---|---|---|---|---|---|---|---|---|---|
| 9965 | ENSMUSG00000035235 | 32.97960418 | -0.178664773 | 0.395761169 | -0.45144594 | 0.651668179 | 0.925643431 | Trim13 | tripartite motif-containing 13 [Source:MG | 37.746 | 37.005 | 30.187 | 26.991 | 41.443 | 24.505 |
| 9966 | ENSMUSG00000035236 | 71.67774711 | 0.265629643 | 0.284845057 | 0.932540818 | 0.351057102 | 0.792398553 | Scai | suppressor of cancer cell invasion [Source | 50.595 | 65.787 | 79.493 | 87.722 | 78.525 | 67.944 |
| 9967 | ENSMUSG00000035237 | 7144.260737 | -0.096048358 | 0.10701541 | -0.897518946 | 0.369442068 | 0.80653433 | Lcat | lecithin cholesterol acyltransferase [Sourc | 6617.6 | 7494.5 | 8035.9 | 7478.5 | 6804.4 | 6434.7 |
| 9968 | ENSMUSG00000035238 | 0 | NA | NA | NA | NA | NA | Kcnk15 | potassium channel, subfamily K, member | 0 | 0 | 0 | 0 | 0 | 0 |
| 9969 | ENSMUSG00000035239 | 27.00372102 | -1.29093497 | 0.518110907 | -2.49161898 | 0.012716237 | 0.127461989 | Neu3 | neuraminidase 3 [Source:MGI Symbol;Acc | 28.109 | 26.726 | 60.375 | 20.244 | 7.6343 | 18.935 |
| 9970 | ENSMUSG00000035242 | 5730.720668 | 0.225226998 | 0.098463306 | 2.287420637 | 0.022171283 | 0.186120392 | Oaz1 | ornithine decarboxylase antizyme 1 [Sour | 5155.9 | 5122.1 | 5574.6 | 5840.7 | 5918.8 | 6772.2 |
| 9971 | ENSMUSG00000035245 | 101.4515531 | 0.121605484 | 0.242963828 | 0.500508593 | 0.616717007 | 0.917509899 | Eogt | EGF domain-specific O-linked N-acetylgluc | 80.31 | 106.9 | 104.65 | 98.326 | 94.884 | 123.64 |
| 9972 | ENSMUSG00000035246 | 1.546750214 | -1.732713903 | 2.087872254 | -0.829894597 | 0.40659838 | NA | Pcyt1b | phosphate cytidylyltransferase 1, choline, | 4.0155 | 3.0838 | 0 | 0 | 2.1812 | 0 |
| 9973 | ENSMUSG00000035247 | 2921.070356 | 0.024296818 | 0.126899839 | 0.191464531 | 0.848161667 | 0.972803041 | Hectd1 | HECT domain E3 ubiquitin protein ligase 1 | 2418.9 | 3130 | 3142.5 | 3159.9 | 3033 | 2642 |
| 9974 | ENSMUSG00000035248 | 1137.436603 | 0.22651568 | 0.09990043 | 2.267414474 | 0.023364918 | 0.193224801 | Tut7 | terminal uridylyl transferase 7 [Source:MC | 1056.9 | 1043.3 | 1044.5 | 1212.7 | 1199.7 | 1267.6 |
| 9975 | ENSMUSG00000035258 | 58.95070794 | -0.970235239 | 0.313095196 | -3.098850613 | 0.001942729 | 0.034101504 | Abi3bp | ABI family member 3 binding protein [Sou | 81.113 | 90.457 | 62.387 | 32.775 | 47.987 | 38.985 |
| 9976 | ENSMUSG00000035262 | 1.716680779 | -3.00402501 | 2.103677169 | -1.427987647 | 0.153295414 | NA | Amh | anti-Mullerian hormone [Source:MGI Sym | 0 | 6.1675 | 3.0187 | 0 | 0 | 1.1138 |
| 9977 | ENSMUSG00000035266 | 79.55312032 | 0.237187914 | 0.261337037 | 0.907593948 | 0.364092793 | 0.802784204 | Helq | helicase, POLQ-like [Source:MGI Symbol;A | 69.87 | 77.094 | 72.45 | 98.326 | 90.521 | 69.058 |
| 9978 | ENSMUSG00000035268 | 333.8787471 | -0.075710839 | 0.182684354 | -0.414435269 | 0.678555364 | 0.933836802 | Pkig | protein kinase inhibitor, gamma [Source:N | 326.06 | 335.1 | 366.27 | 245.81 | 396.99 | 333.04 |
| 9979 | ENSMUSG00000035270 | 101.4289452 | -0.335519645 | 0.23446919 | -1.430975406 | 0.152437264 | 0.568249894 | Impg2 | interphotoreceptor matrix proteoglycan 2 | 131.71 | 97.652 | 109.68 | 87.722 | 92.703 | 89.108 |
| 9980 | ENSMUSG00000035273 | 55.4945235 | -1.375684364 | 0.324726389 | -4.236441542 | 2.27E-05 | 0.001076928 | Hpse | heparanase [Source:MGI Symbol;Acc:MGI | 98.782 | 64.759 | 76.475 | 27.955 | 33.809 | 31.188 |
| 9981 | ENSMUSG00000035274 | 1.002521471 | 1.366653481 | 2.416683574 | 0.565507829 | 0.571728401 | NA | Tpbg | trophoblast glycoprotein [Source:MGI Syn | 1.6062 | 0 | 0 | 0 | 2.1812 | 2.2277 |
| 9982 | ENSMUSG00000035275 | 3.755916611 | -0.402885734 | 1.123118879 | -0.358720471 | 0.719804212 | NA | Raver2 | ribonucleoprotein, PTB-binding 2 [Source: | 5.6217 | 3.0838 | 4.025 | 0.964 | 3.2719 | 5.5692 |
| 9983 | ENSMUSG00000035277 | 0 | NA | NA | NA | NA | NA | Arx | aristaless related homeobox [Source:MGI | 0 | 0 | 0 | 0 | 0 | 0 |
| 9984 | ENSMUSG00000035278 | 339.6715759 | 0.246131061 | 0.168581501 | 1.460012272 | 0.144286701 | 0.554600809 | Plekhj1 | pleckstrin homology domain containing, fa | 261.81 | 291.93 | 379.35 | 367.28 | 352.27 | 385.39 |
| 9985 | ENSMUSG00000035279 | 2.571882495 | -1.017859161 | 1.57761233 | -0.645189659 | 0.518804287 | NA | Ssc5d | scavenger receptor cysteine rich family, 5 | 7.2279 | 3.0838 | 0 | 2.8919 | 0 | 2.2277 |
| 9986 | ENSMUSG00000035283 | 13.61312279 | -1.340932706 | 0.847656054 | -1.581930194 | 0.113665507 | 0.495795505 | Adrb1 | adrenergic receptor, beta 1 [Source:MGI S | 10.44 | 32.893 | 15.094 | 0 | 6.5437 | 16.708 |
| 9987 | ENSMUSG00000035284 | 577.76597 | 0.062861656 | 0.129074087 | 0.487019954 | 0.62624419 | 0.919780257 | Vps13c | vacuolar protein sorting 13C [Source:MGI | 520.41 | 605.44 | 570.54 | 591.88 | 633.65 | 544.67 |
| 9988 | ENSMUSG00000035285 | 5.862004304 | -0.022013788 | 0.892287266 | -0.024671189 | 0.980317236 | 0.99820894 | Nat14 | N-acetyltransferase 14 [Source:MGI Symb | 5.6217 | 5.1396 | 7.0437 | 9.6398 | 3.2719 | 4.4554 |
| 9989 | ENSMUSG00000035293 | 86.84242009 | 0.118108413 | 0.253511392 | 0.465889964 | 0.641294265 | 0.924019354 | G2e3 | G2/M-phase specific E3 ubiquitin ligase [S | 93.16 | 82.233 | 74.462 | 107 | 82.887 | 81.311 |
| 9990 | ENSMUSG00000035295 | 4.581856779 | 0.015521232 | 1.004513949 | 0.015451485 | 0.987671989 | NA | Wdr38 | WD repeat domain 38 [Source:MGI Symbo | 6.4248 | 5.1396 | 2.0125 | 2.8919 | 5.4531 | 5.5692 |
| 9991 | ENSMUSG00000035296 | 0 | NA | NA | NA | NA | NA | Sgcg | sarcoglycan, gamma (dystrophin-associate | 0 | 0 | 0 | 0 | 0 | 0 |
| 9992 | ENSMUSG00000035297 | 568.1684001 | 0.016697424 | 0.12635651 | 0.132145338 | 0.894869335 | 0.982798755 | Cops4 | COP9 signalosome subunit 4 [Source:MGI | 521.21 | 603.39 | 570.54 | 542.72 | 604.2 | 566.95 |
| 9993 | ENSMUSG00000035298 | 2.163131934 | -1.776184496 | 1.890800391 | -0.93938234 | 0.347534477 | NA | Klhl35 | kelch-like 35 [Source:MGI Symbol;Acc:MG | 8.031 | 2.0558 | 0 | 2.8919 | 0 | 0 |
| 9994 | ENSMUSG00000035299 | 13.11459401 | 0.036367284 | 0.68475013 | 0.053110299 | 0.957644026 | 0.991861398 | Mid1 | midline 1 [Source:MGI Symbol;Acc:MGI:1 | 13.653 | 7.1954 | 18.112 | 23.135 | 5.4531 | 11.138 |
| 9995 | ENSMUSG00000035305 | 68.6978284 | 0.03077314 | 0.287011237 | 0.107219287 | 0.914615015 | 0.98625062 | Ror1 | receptor tyrosine kinase-like orphan recep | 70.673 | 53.452 | 79.493 | 60.731 | 66.528 | 81.311 |
| 9996 | ENSMUSG00000035310 | 162.7919323 | 0.283062325 | 0.198256394 | 1.42775887 | 0.153361275 | 0.570529188 | NA | NA | 145.36 | 132.6 | 163.01 | 206.29 | 166.86 | 162.62 |
| 9997 | ENSMUSG00000035311 | 242.1435364 | 0.078019133 | 0.162093545 | 0.481321656 | 0.630287908 | 0.920950025 | Gnptab | N-acetylglucosamine-1-phosphate transfe | 247.36 | 238.48 | 220.37 | 227.5 | 256.3 | 262.87 |
| 9998 | ENSMUSG00000035314 | 11.31021769 | -0.12429808 | 0.674504229 | -0.184280654 | 0.85379329 | 0.973825032 | Gdpd5 | glycerophosphodiester phosphodiesteras | 8.031 | 16.447 | 11.069 | 13.496 | 5.4531 | 13.366 |
| 9999 | ENSMUSG00000035325 | 2256.760944 | -0.08557324 | 0.113917406 | -0.751186693 | 0.452540307 | 0.856769845 | Sec31a | Sec31 homolog A (S. cerevisiae) [Source:N | 2443.8 | 2448.5 | 2078.9 | 2378.1 | 2206.3 | 1984.9 |
| 10000 | ENSMUSG00000035329 | 279.0346198 | -0.024817482 | 0.19323258 | -0.12843322 | 0.897806143 | 0.982798755 | Fbxo33 | F-box protein 33 [Source:MGI Symbol;Acc | 216.03 | 330.99 | 297.85 | 234.25 | 299.92 | 295.17 |
| 10001 | ENSMUSG00000035337 | 1.107988552 | -1.056291367 | 2.031653537 | -0.519917076 | 0.603121373 | NA | Uchl4 | ubiquitin carboxyl-terminal esterase L4 [S | 2.4093 | 1.0279 | 1.0062 | 0 | 1.0906 | 1.1138 |
| 10002 | ENSMUSG00000035342 | 216.8852961 | -0.14358353 | 0.17568345 | -0.817285463 | 0.413765312 | 0.835385466 | Lzts2 | leucine zipper, putative tumor suppressor | 204.79 | 228.2 | 250.55 | 214 | 183.22 | 220.54 |
| 10003 | ENSMUSG00000035351 | 92.44256566 | 0.179385875 | 0.23755679 | 0.755128382 | 0.450171955 | 0.855539645 | Nup37 | nucleoporin 37 [Source:MGI Symbol;Acc:I | 89.144 | 78.122 | 92.575 | 88.686 | 101.43 | 104.7 |
| 10004 | ENSMUSG00000035352 | 0.331982614 | 0.16514309 | 3.986751476 | 0.041422971 | 0.966958701 | NA | Ccl12 | chemokine (C-C motif) ligand 12 [Source:N | 0 | 1.0279 | 0 | 0.964 | 0 | 0 |
| 10005 | ENSMUSG00000035354 | 744.5189849 | 0.085437644 | 0.114445285 | 0.746537039 | 0.455343066 | 0.857339509 | Uvrag | UV radiation resistance associated gene [S | 755.72 | 743.18 | 668.15 | 776.97 | 762.34 | 760.76 |
| 10006 | ENSMUSG00000035355 | 0.678054141 | -2.792799308 | 2.990673428 | -0.933836266 | 0.350388359 | NA | Kcnh4 | potassium voltage-gated channel, subfam | 0 | 2.0558 | 2.0125 | 0 | 0 | 0 |
| 10007 | ENSMUSG00000035356 | 139.6283897 | 0.015734388 | 0.199544771 | 0.078851416 | 0.937150807 | 0.988420305 | Nfkbiz | nuclear factor of kappa light polypeptide g | 146.16 | 131.57 | 138.86 | 151.34 | 143.96 | 125.86 |
| 10008 | ENSMUSG00000035357 | 182.0995027 | -0.778984426 | 0.221969974 | -3.509413505 | 0.000449096 | 0.011476548 | Pdzrn3 | PDZ domain containing RING finger 3 [Sou | 234.51 | 289.87 | 166.03 | 138.81 | 133.06 | 130.32 |
| 10009 | ENSMUSG00000035364 | 4.988035542 | 0.161348971 | 1.019907091 | 0.158199676 | 0.874299459 | NA | NA | NA | 4.0155 | 3.0838 | 7.0437 | 5.7839 | 1.0906 | 8.9108 |
| 10010 | ENSMUSG00000035365 | 2.011066299 | -1.469429503 | 1.703075104 | -0.862809573 | 0.388242175 | NA | Parpbp | PARP1 binding protein [Source:MGI Symb | 0.8031 | 1.0279 | 7.0437 | 0.964 | 0 | 2.2277 |
| 10011 | ENSMUSG00000035367 | 236.6094071 | -0.120324886 | 0.164997248 | -0.729253897 | 0.465846368 | 0.863698893 | Rmi1 | RecQ mediated genome instability 1 [Sour | 248.16 | 241.56 | 249.55 | 216.9 | 255.2 | 208.29 |
| 10012 | ENSMUSG00000035370 | 5.946131425 | 0.377705211 | 0.937169444 | 0.403027663 | 0.686927874 | 0.936625639 | Gm49322 | predicted gene, 49322 [Source:MGI Symb | 6.4248 | 4.1117 | 5.0312 | 13.496 | 3.2719 | 3.3415 |
| 10013 | ENSMUSG00000035371 | 0.167707431 | -0.796629176 | 4.080472857 | -0.195229623 | 0.84521319 | NA | Ssx9 | synovial sarcoma, X 9 [Source:MGI Symbo | 0 | 0 | 1.0062 | 0 | 0 | 0 |
| 10014 | ENSMUSG00000035372 | 1468.010228 | -0.667389776 | 0.34305083 | -1.945454487 | 0.051720302 | 0.319420348 | 1810055G02 | RIKEN cDNA 1810055G02 gene [Source:M | 2229.4 | 1932.5 | 1242.7 | 997.72 | 1524.7 | 881.05 |
| 10015 | ENSMUSG00000035373 | 0.535401493 | -2.48596965 | 3.684117979 | -0.674780141 | 0.499815459 | NA | Ccl7 | chemokine (C-C motif) ligand 7 [Source:M | 3.2124 | 0 | 0 | 0 | 0 | 0 |
| 10016 | ENSMUSG00000035376 | 983.6015999 | -0.01591787 | 0.132935772 | -0.119741059 | 0.904688274 | 0.983865503 | Hacd2 | 3-hydroxyacyl-CoA dehydratase 2 [Source | 1130 | 950.82 | 884.49 | 870.47 | 1008.8 | 1057 |
| 10017 | ENSMUSG00000035378 | 93.0484261 | 0.143899379 | 0.235556539 | 0.610891041 | 0.541271716 | 0.893614211 | Shq1 | SHQ1 homolog (S. cerevisiae) [Source:MG | 87.538 | 87.373 | 90.562 | 109.89 | 92.703 | 90.221 |
| 10018 | ENSMUSG00000035382 | 622.157923 | 0.140220335 | 0.120526404 | 1.163399308 | 0.24466753 | 0.694577264 | Pcsk7 | proprotein convertase subtilisin/kexin typ | 553.34 | 618.81 | 604.75 | 684.42 | 645.65 | 625.98 |
| 10019 | ENSMUSG00000035383 | 0.267700746 | -1.497481649 | 4.057001436 | -0.369110456 | 0.712045396 | NA | Pmch | pro-melanin-concentrating hormone [Sou | 1.6062 | 0 | 0 | 0 | 0 | 0 |
| 10020 | ENSMUSG00000035385 | 3.697733556 | -2.205556477 | 1.415439347 | -1.558213343 | 0.119182683 | NA | Ccl2 | chemokine (C-C motif) ligand 2 [Source:M | 11.243 | 2.0558 | 5.0312 | 3.8559 | 0 | 0 |

| | | | | | | | | | | | | | | | |
|---|---|---|---|---|---|---|---|---|---|---|---|---|---|---|---|
| 10021 | ENSMUSG00000035387 | 0 | NA | NA | NA | NA | NA | 4930415L06I | RIKEN cDNA 4930415L06 gene [Source:MG | 0 | 0 | 0 | 0 | 0 | 0 |
| 10022 | ENSMUSG00000035390 | 4.326731904 | -0.111191495 | 1.030450766 | -0.107905685 | 0.914070508 | NA | Brsk1 | BR serine/threonine kinase 1 [Source:MGI | 2.4093 | 5.1396 | 6.0375 | 5.7839 | 4.3625 | 2.2277 |
| 10023 | ENSMUSG00000035392 | 375.0601253 | -0.236838625 | 0.135606135 | -1.746518508 | 0.080720894 | 0.417428319 | Dennd1a | DENN/MADD domain containing 1A [Sour | 422.43 | 403.97 | 390.42 | 333.54 | 342.45 | 357.54 |
| 10024 | ENSMUSG00000035394 | 1.190203636 | 0.585913931 | 1.945514715 | 0.301161398 | 0.763291424 | NA | Cfap53 | cilia and flagella associated protein 53 [So | 0.8031 | 2.0558 | 0 | 0.964 | 1.0906 | 2.2277 |
| 10025 | ENSMUSG00000035395 | 0 | NA | NA | NA | NA | NA | Dcaf8l | DDB1 and CUL4 associated factor 8 like [S | 0 | 0 | 0 | 0 | 0 | 0 |
| 10026 | ENSMUSG00000035397 | 84.8580594 | 0.505832806 | 0.278509696 | 1.816212552 | 0.069337773 | 0.380319133 | Klf16 | Kruppel-like factor 16 [Source:MGI Symbc | 73.082 | 68.87 | 68.425 | 110.86 | 67.618 | 120.3 |
| 10027 | ENSMUSG00000035399 | 232.4352802 | -0.043388739 | 0.178311089 | -0.24333169 | 0.807748457 | 0.965081255 | Oser1 | oxidative stress responsive serine rich 1 [S | 228.88 | 202.5 | 276.72 | 246.78 | 220.3 | 219.43 |
| 10028 | ENSMUSG00000035401 | 224.7364546 | 0.092087846 | 0.167325895 | 0.550350234 | 0.582079176 | 0.908645447 | Emsy | EMSY, BRCA2-interacting transcriptional r | 207.2 | 226.14 | 219.36 | 215.93 | 222.49 | 257.3 |
| 10029 | ENSMUSG00000035403 | 4.963652965 | 0.566955218 | 1.109248605 | 0.511116458 | 0.609269513 | NA | Crb2 | crumbs family member 2 [Source:MGI Syr | 0 | 4.1117 | 8.05 | 7.7118 | 5.4531 | 4.4554 |
| 10030 | ENSMUSG00000035407 | 1.198824297 | 0.453633123 | 2.254176062 | 0.201241212 | 0.840509968 | NA | Kank4 | KN motif and ankyrin repeat domains 4 [S | 0 | 3.0838 | 0 | 1.928 | 2.1812 | 0 |
| 10031 | ENSMUSG00000035413 | 191.1587253 | -0.461750908 | 0.218340938 | -2.114815998 | 0.034445633 | 0.24861359 | Tmem98 | transmembrane protein 98 [Source:MGI S | 245.75 | 250.81 | 168.04 | 184.12 | 170.14 | 128.09 |
| 10032 | ENSMUSG00000035420 | 0 | NA | NA | NA | NA | NA | Fam170a | family with sequence similarity 170, meml | 0 | 0 | 0 | 0 | 0 | 0 |
| 10033 | ENSMUSG00000035427 | 0 | NA | NA | NA | NA | NA | Mageb4 | MAGE family member B4 [Source:MGI Syr | 0 | 0 | 0 | 0 | 0 | 0 |
| 10034 | ENSMUSG00000035429 | 0.476489653 | -2.293616034 | 3.421506946 | -0.670352587 | 0.502633051 | NA | Ptprh | protein tyrosine phosphatase, receptor ty | 0.8031 | 2.0558 | 0 | 0 | 0 | 0 |
| 10035 | ENSMUSG00000035431 | 0 | NA | NA | NA | NA | NA | Sstr1 | somatostatin receptor 1 [Source:MGI Sym | 0 | 0 | 0 | 0 | 0 | 0 |
| 10036 | ENSMUSG00000035435 | 0.133850373 | -0.796629176 | 4.080472857 | -0.195229623 | 0.84521319 | NA | Abca17 | ATP-binding cassette, sub-family A (ABC1) | 0.8031 | 0 | 0 | 0 | 0 | 0 |
| 10037 | ENSMUSG00000035437 | 372.4600832 | -0.116938058 | 0.141394764 | -0.827032446 | 0.408218671 | 0.832151413 | Rabgap1 | RAB GTPase activating protein 1 [Source:N | 393.52 | 363.88 | 405.52 | 386.56 | 353.36 | 331.93 |
| 10038 | ENSMUSG00000035439 | 77.94124047 | -0.229545113 | 0.318801993 | -0.720024085 | 0.471510167 | 0.867794597 | Haus8 | 4HAUS augmin-like complex, subunit 8 [Sc | 124.48 | 67.843 | 59.368 | 68.442 | 81.796 | 65.717 |
| 10039 | ENSMUSG00000035441 | 290.400822 | -0.196446625 | 0.167163902 | -1.175173726 | 0.23992522 | 0.689857856 | Myo1d | myosin ID [Source:MGI Symbol;Acc:MGI:1 | 321.24 | 319.68 | 289.8 | 298.83 | 291.2 | 221.65 |
| 10040 | ENSMUSG00000035443 | 303.0112614 | 0.025204324 | 0.169249129 | 0.148918488 | 0.881617953 | 0.981676444 | Thyn1 | thymocyte nuclear protein 1 [Source:MGI | 258.6 | 338.18 | 304.89 | 294.01 | 273.75 | 348.63 |
| 10041 | ENSMUSG00000035448 | 0.913882229 | 2.129306995 | 2.79764157 | 0.761107862 | 0.446592645 | NA | Ccr3 | chemokine (C-C motif) receptor 3 [Source | 0 | 1.0279 | 0 | 0 | 0 | 4.4554 |
| 10042 | ENSMUSG00000035451 | 506.401409 | 0.554895937 | 0.20569 | 2.697729284 | 0.006981419 | 0.084057311 | Foxa1 | forkhead box A1 [Source:MGI Symbol;Acc | 336.5 | 413.22 | 481.99 | 752.87 | 412.25 | 641.57 |
| 10043 | ENSMUSG00000035454 | 0 | NA | NA | NA | NA | NA | Samt3 | spermatogenesis associated multipass tra | 0 | 0 | 0 | 0 | 0 | 0 |
| 10044 | ENSMUSG00000035455 | 5.979443778 | 0.805337009 | 1.208923139 | 0.666160638 | 0.50530843 | 0.881592135 | Fignl1 | fidgetin-like 1 [Source:MGI Symbol;Acc:M | 8.8341 | 1.0279 | 3.0187 | 0 | 18.541 | 4.4554 |
| 10045 | ENSMUSG00000035456 | 0 | NA | NA | NA | NA | NA | Prdm8 | PR domain containing 8 [Source:MGI Sym | 0 | 0 | 0 | 0 | 0 | 0 |
| 10046 | ENSMUSG00000035458 | 2.112455858 | -1.646283536 | 1.776649521 | -0.926622565 | 0.354122535 | NA | Tnni3 | troponin I, cardiac 3 [Source:MGI Symbol; | 1.6062 | 0 | 8.05 | 1.928 | 1.0906 | 0 |
| 10047 | ENSMUSG00000035459 | 507.3825464 | -0.484367067 | 0.161269801 | -3.00345796 | 0.002669304 | 0.043380693 | Stab2 | stabilin 2 [Source:MGI Symbol;Acc:MGI:2 | 610.36 | 621.89 | 542.37 | 380.77 | 363.18 | 525.73 |
| 10048 | ENSMUSG00000035469 | 1365.289287 | 0.249619459 | 0.101477934 | 2.459839788 | 0.013899905 | 0.136027217 | Rcbtb1 | regulator of chromosome condensation (I | 1183.8 | 1319.8 | 1239.7 | 1483.6 | 1525.8 | 1439.1 |
| 10049 | ENSMUSG00000035472 | 282.5994859 | 0.131480701 | 0.16243864 | 0.809417642 | 0.418274958 | 0.837575303 | Slc25a21 | solute carrier family 25 (mitochondrial oxo | 281.89 | 249.78 | 276.72 | 255.45 | 304.28 | 327.47 |
| 10050 | ENSMUSG00000035473 | 3265.168565 | -0.088307759 | 0.094868094 | -0.930847823 | 0.351932288 | 0.792889192 | Galm | galactose mutarotase [Source:MGI Symbo | 3169.8 | 3359.2 | 3567.1 | 3179.2 | 3290.4 | 3025.2 |
| 10051 | ENSMUSG00000035476 | 149.9785586 | 0.12608229 | 0.198384205 | 0.635546012 | 0.525072368 | 0.888334265 | Tab3 | TGF-beta activated kinase 1/MAP3K7 binc | 152.59 | 144.94 | 132.82 | 177.37 | 141.78 | 150.37 |
| 10052 | ENSMUSG00000035478 | 998.8866602 | -0.049002624 | 0.130625768 | -0.3751375 | 0.707558209 | 0.943361388 | Mbd3 | methyl-CpG binding domain protein 3 [So | 958.9 | 1085.5 | 1004.2 | 1092.2 | 827.78 | 1024.7 |
| 10053 | ENSMUSG00000035486 | 48.94679327 | 0.081391427 | 0.377199768 | 0.215778042 | 0.829160765 | 0.969955221 | Plk5 | polo like kinase 5 [Source:MGI Symbol;Acc | 47.383 | 42.145 | 53.331 | 59.767 | 65.437 | 25.618 |
| 10054 | ENSMUSG00000035491 | 0 | NA | NA | NA | NA | NA | Fthl17a | ferritin, heavy polypeptide-like 17, memb | 0 | 0 | 0 | 0 | 0 | 0 |
| 10055 | ENSMUSG00000035493 | 595.39136 | -1.232051811 | 0.150430383 | -8.190179292 | 2.61E-16 | 2.10E-13 | Tgfbi | transforming growth factor, beta induced | 985.41 | 754.49 | 764.75 | 345.1 | 327.19 | 395.41 |
| 10056 | ENSMUSG00000035495 | 215.053858 | 0.020867004 | 0.164970584 | 0.12648924 | 0.899344665 | 0.982798755 | Tstd2 | thiosulfate sulfurtransferase (rhodanese)- | 209.61 | 214.83 | 216.34 | 226.53 | 221.4 | 201.61 |
| 10057 | ENSMUSG00000035498 | 126.4949762 | -0.658012442 | 0.255764436 | -2.572728456 | 0.010090034 | 0.109368369 | Cdcp1 | CUB domain containing protein 1 [Source: | 146.97 | 148.02 | 169.05 | 67.478 | 93.793 | 133.66 |
| 10058 | ENSMUSG00000035504 | 13161.48896 | -0.169466091 | 0.092914158 | -1.823899553 | 0.068167259 | 0.376148227 | Reep6 | receptor accessory protein 6 [Source:MGI | 14987 | 13776 | 13037 | 12414 | 12924 | 11831 |
| 10059 | ENSMUSG00000035505 | 187.6877581 | 0.0834204 | 0.182006949 | 0.458336349 | 0.646710813 | 0.925643431 | Cox18 | cytochrome c oxidase assembly protein 18 | 199.97 | 162.41 | 184.14 | 207.26 | 194.13 | 178.22 |
| 10060 | ENSMUSG00000035506 | 3.089428647 | -0.606378063 | 1.341373011 | -0.452057749 | 0.651227381 | NA | Slc12a8 | solute carrier family 12 (potassium/chloric | 4.0155 | 6.1675 | 1.0062 | 2.8919 | 0 | 4.4554 |
| 10061 | ENSMUSG00000035509 | 81.35823092 | 0.43954332 | 0.260564057 | 1.686891605 | 0.091624197 | 0.446623824 | Fbxl21 | F-box and leucine-rich repeat protein 21 [ | 70.673 | 63.731 | 72.45 | 75.19 | 102.52 | 103.59 |
| 10062 | ENSMUSG00000035513 | 33.09865057 | -0.704742594 | 0.422583188 | -1.667701447 | 0.095375008 | 0.455281886 | Ntng2 | netrin G2 [Source:MGI Symbol;Acc:MGI:2 | 44.974 | 39.061 | 39.244 | 36.631 | 14.178 | 24.505 |
| 10063 | ENSMUSG00000035517 | 529.6759202 | 0.248781791 | 0.138893084 | 1.79117479 | 0.07326525 | 0.392921239 | Tdrd7 | tudor domain containing 7 [Source:MGI S | 461.78 | 483.12 | 507.15 | 494.52 | 595.48 | 636 |
| 10064 | ENSMUSG00000035521 | 435.3499502 | -0.299980446 | 0.129467084 | -2.317040257 | 0.020501539 | 0.176403485 | Gnptg | N-acetylglucosamine-1-phosphotransfera | 455.36 | 494.43 | 492.05 | 392.34 | 396.99 | 380.93 |
| 10065 | ENSMUSG00000035522 | 0 | NA | NA | NA | NA | NA | Tsga8 | testis specific gene A8 [Source:MGI Symbo | 0 | 0 | 0 | 0 | 0 | 0 |
| 10066 | ENSMUSG00000035523 | 0 | NA | NA | NA | NA | NA | Vmn1r172 | vomeronasal 1 receptor 172 [Source:MGI | 0 | 0 | 0 | 0 | 0 | 0 |
| 10067 | ENSMUSG00000035528 | 0 | NA | NA | NA | NA | NA | Npffr2 | neuropeptide FF receptor 2 [Source:MGI S | 0 | 0 | 0 | 0 | 0 | 0 |
| 10068 | ENSMUSG00000035529 | 140.2871822 | 0.334810627 | 0.197834389 | 1.692378304 | 0.090573866 | 0.444540347 | Prdm4 | PR domain containing 4 [Source:MGI Sym | 130.1 | 125.41 | 116.72 | 169.66 | 147.23 | 152.6 |
| 10069 | ENSMUSG00000035530 | 6492.291293 | -0.321072578 | 0.113826747 | -2.820712944 | 0.004791706 | 0.064976213 | Eif1 | eukaryotic translation initiation factor 1 [S | 6960.5 | 7581.9 | 7092 | 4889.3 | 6298.3 | 6131.7 |
| 10070 | ENSMUSG00000035539 | 0.335414862 | -1.779047242 | 4.030804179 | -0.441362855 | 0.658950329 | NA | Ccdc180 | coiled-coil domain containing 180 [Source | 0 | 0 | 2.0125 | 0 | 0 | 0 |
| 10071 | ENSMUSG00000035540 | 106533.5909 | 0.039226581 | 0.081065481 | 0.483887593 | 0.628465646 | 0.920328819 | Gc | vitamin D binding protein [Source:MGI Sy | 103660 | 104722 | 106875 | 113008 | 106805 | 104131 |
| 10072 | ENSMUSG00000035545 | 1250.426774 | 0.239144272 | 0.174502624 | 1.370433678 | 0.170551565 | 0.597556794 | Leng8 | leukocyte receptor cluster (LRC) member | 1230.4 | 1028.9 | 1182.3 | 1814.2 | 1093.9 | 1152.8 |
| 10073 | ENSMUSG00000035547 | 38.60963263 | 0.01951137 | 0.350588087 | 0.055653261 | 0.955618034 | 0.991678478 | Capn5 | calpain 5 [Source:MGI Symbol;Acc:MGI:1 | 32.927 | 40.089 | 42.262 | 41.451 | 38.172 | 36.757 |
| 10074 | ENSMUSG00000035551 | 0 | NA | NA | NA | NA | NA | Igfbpl1 | insulin-like growth factor binding protein- | 0 | 0 | 0 | 0 | 0 | 0 |
| 10075 | ENSMUSG00000035557 | 0 | NA | NA | NA | NA | NA | Krt17 | keratin 17 [Source:MGI Symbol;Acc:MGI:9 | 0 | 0 | 0 | 0 | 0 | 0 |
| 10076 | ENSMUSG00000035559 | 292.5482674 | 0.104869861 | 0.146485014 | 0.715908463 | 0.474047872 | 0.86899591 | Mpv17l2 | MPV17 mitochondrial membrane protein- | 285.1 | 280.62 | 279.74 | 289.19 | 303.19 | 317.45 |

| | | | | | | | | | | | | | | | |
|---|---|---|---|---|---|---|---|---|---|---|---|---|---|---|---|
| 10077 | ENSMUSG00000035560 | 0 | NA | NA | NA | NA | NA | Wdr20rt | WD repeat domain 20, retrogene [Source: | 0 | 0 | 0 | 0 | 0 | 0 |
| 10078 | ENSMUSG00000035561 | 1054.042372 | -0.166285299 | 0.174338215 | -0.953808656 | 0.340180507 | 0.78262619 | Aldh1b1 | aldehyde dehydrogenase 1 family, membe | 1215.9 | 991.94 | 1135 | 724.91 | 1264 | 992.43 |
| 10079 | ENSMUSG00000035566 | 15.47865077 | -0.958769234 | 0.598211318 | -1.602726672 | 0.108995012 | 0.485738292 | Pcdh17 | protocadherin 17 [Source:MGI Symbol;Ac | 18.471 | 30.838 | 12.075 | 10.604 | 13.087 | 7.7969 |
| 10080 | ENSMUSG00000035569 | 970.5394209 | -0.070585426 | 0.119947176 | -0.588470931 | 0.55621624 | 0.897828342 | Ankrd11 | ankyrin repeat domain 11 [Source:MGI Sy | 985.41 | 1048.5 | 949.89 | 1061.3 | 869.22 | 908.9 |
| 10081 | ENSMUSG00000035572 | 465.2045297 | 0.032874856 | 0.130082989 | 0.252722177 | 0.800482909 | 0.962424862 | Dcaf10 | DDB1 and CUL4 associated factor 10 [Sou | 432.07 | 497.51 | 450.8 | 474.28 | 478.78 | 457.79 |
| 10082 | ENSMUSG00000035575 | 655.9044193 | 0.128446497 | 0.12895538 | 0.996053806 | 0.319224003 | 0.766222366 | Utp6 | UTP6 small subunit processome compone | 587.07 | 632.17 | 661.1 | 625.62 | 765.61 | 663.85 |
| 10083 | ENSMUSG00000035576 | 0.636972666 | -2.712703855 | 3.077964703 | -0.881330397 | 0.378139018 | NA | L3mbtl1 | L3MBTL1 histone methyl-lysine binding pr | 0.8031 | 0 | 3.0187 | 0 | 0 | 0 |
| 10084 | ENSMUSG00000035578 | 36.55036034 | -0.142928184 | 0.381041937 | -0.375098303 | 0.707587359 | 0.943361388 | Iqcg | IQ motif containing G [Source:MGI Symbo | 45.777 | 39.061 | 30.187 | 40.487 | 38.172 | 25.618 |
| 10085 | ENSMUSG00000035580 | 0 | NA | NA | NA | NA | NA | Kcnh8 | potassium voltage-gated channel, subfam | 0 | 0 | 0 | 0 | 0 | 0 |
| 10086 | ENSMUSG00000035582 | 0 | NA | NA | NA | NA | NA | Gdpd4 | glycerophosphodiester phosphodiesteras | 0 | 0 | 0 | 0 | 0 | 0 |
| 10087 | ENSMUSG00000035585 | 725.2097866 | 0.012099598 | 0.112174241 | 0.107864314 | 0.914103325 | 0.98625062 | Tsen34 | tRNA splicing endonuclease subunit 34 [S | 709.94 | 719.54 | 737.58 | 760.58 | 726.35 | 697.27 |
| 10088 | ENSMUSG00000035592 | 0 | NA | NA | NA | NA | NA | Krt33a | keratin 33A [Source:MGI Symbol;Acc:MGI | 0 | 0 | 0 | 0 | 0 | 0 |
| 10089 | ENSMUSG00000035594 | 0 | NA | NA | NA | NA | NA | Chrna5 | cholinergic receptor, nicotinic, alpha poly | 0 | 0 | 0 | 0 | 0 | 0 |
| 10090 | ENSMUSG00000035595 | 74.74217447 | -0.107899129 | 0.268361926 | -0.402065713 | 0.687635666 | 0.936885029 | Fam174c | family with sequence similarity 174, meml | 81.916 | 85.317 | 65.406 | 77.118 | 77.434 | 61.261 |
| 10091 | ENSMUSG00000035596 | 666.6269441 | 0.161643791 | 0.126433028 | 1.2784934 | 0.201075511 | 0.645135365 | Mboat7 | membrane bound O-acyltransferase doma | 626.42 | 583.86 | 678.21 | 772.15 | 666.37 | 672.76 |
| 10092 | ENSMUSG00000035597 | 390.3095663 | -0.086992688 | 0.149372825 | -0.582386308 | 0.560306502 | 0.899475387 | Prpf39 | pre-mRNA processing factor 39 [Source:M | 392.72 | 387.53 | 426.65 | 419.33 | 392.62 | 323.01 |
| 10093 | ENSMUSG00000035601 | 65.79590263 | 0.122915275 | 0.284103426 | 0.432642705 | 0.66527436 | 0.929426323 | Trmt10b | tRNA methyltransferase 10B [Source:MGI | 66.657 | 72.982 | 49.306 | 71.334 | 65.437 | 69.058 |
| 10094 | ENSMUSG00000035606 | 0.160662974 | 1.126924175 | 4.080472857 | 0.276174898 | 0.782413736 | NA | Ky | kyphoscoliosis peptidase [Source:MGI Syr | 0 | 0 | 0 | 0.964 | 0 | 0 |
| 10095 | ENSMUSG00000035614 | 276.2660136 | 0.238861243 | 0.150012522 | 1.592275358 | 0.111322847 | 0.489450944 | Togaram1 | TOG array regulator of axonemal microtul | 252.17 | 251.84 | 256.59 | 315.22 | 297.74 | 284.03 |
| 10096 | ENSMUSG00000035615 | 0.606727817 | -2.651277273 | 2.811524265 | -0.943003518 | 0.345679103 | NA | Frmpd1 | FERM and PDZ domain containing 1 [Sour | 1.6062 | 1.0279 | 1.0062 | 0 | 0 | 0 |
| 10097 | ENSMUSG00000035620 | 174.6047605 | -0.158186985 | 0.199895141 | -0.791349827 | 0.428739879 | 0.842753542 | Ric8b | RIC8 guanine nucleotide exchange factor | 151.79 | 196.33 | 205.27 | 179.3 | 170.14 | 144.8 |
| 10098 | ENSMUSG00000035621 | 863.4279422 | -0.093257758 | 0.115762461 | -0.805595843 | 0.420475923 | 0.837906635 | Midn | midnolin [Source:MGI Symbol;Acc:MGI:1 | 902.69 | 940.54 | 830.15 | 781.79 | 834.32 | 891.08 |
| 10099 | ENSMUSG00000035623 | 258.5387779 | 0.136306345 | 0.158030286 | 0.862533055 | 0.388394252 | 0.818315288 | Rsf1 | remodeling and spacing factor 1 [Source:! | 243.34 | 253.9 | 241.5 | 243.89 | 287.92 | 280.69 |
| 10100 | ENSMUSG00000035626 | 0 | NA | NA | NA | NA | NA | Olfr1509 | olfactory receptor 1509 [Source:MGI Sym | 0 | 0 | 0 | 0 | 0 | 0 |
| 10101 | ENSMUSG00000035629 | 410.3280275 | 0.002690415 | 0.14850328 | 0.018116871 | 0.985545619 | 0.99859176 | Rubcn | RUN domain and cysteine-rich domain co | 395.13 | 413.22 | 421.62 | 409.69 | 352.27 | 470.04 |
| 10102 | ENSMUSG00000035632 | 285.9398809 | -0.193388603 | 0.152576059 | -1.267489831 | 0.204980198 | 0.650619832 | Cnot3 | CCR4-NOT transcription complex, subunit | 295.54 | 295.01 | 325.02 | 274.73 | 280.29 | 245.05 |
| 10103 | ENSMUSG00000035637 | 4624.345104 | 0.014243001 | 0.09322012 | 0.152788911 | 0.878564743 | 0.980976563 | Grhpr | glyoxylate reductase/hydroxypyruvate rec | 4648.4 | 4456 | 4699.2 | 4342.7 | 4622 | 4977.8 |
| 10104 | ENSMUSG00000035638 | 0 | NA | NA | NA | NA | NA | Muc20 | mucin 20 [Source:MGI Symbol;Acc:MGI:2 | 0 | 0 | 0 | 0 | 0 | 0 |
| 10105 | ENSMUSG00000035640 | 27.85870796 | -0.204912435 | 0.440985715 | -0.4646691 | 0.64216844 | 0.924834503 | Cbarp | calcium channel, voltage-dependent, beta | 34.533 | 24.67 | 30.187 | 25.063 | 35.99 | 16.708 |
| 10106 | ENSMUSG00000035642 | 395.4329482 | 0.105971802 | 0.157794593 | 0.671580692 | 0.501850673 | 0.881001421 | Aamdc | adipogenesis associated Mth938 domain | 399.94 | 343.32 | 398.47 | 343.18 | 420.98 | 466.7 |
| 10107 | ENSMUSG00000035649 | 309.111909 | 0.049055257 | 0.15825133 | 0.309983222 | 0.756573716 | 0.953085143 | Zcchc7 | zinc finger, CCHC domain containing 7 [So | 305.98 | 334.07 | 271.69 | 321.97 | 340.27 | 280.69 |
| 10108 | ENSMUSG00000035651 | 0 | NA | NA | NA | NA | NA | 4930480E11 | RIKEN cDNA 4930480E11 gene [Source:M | 0 | 0 | 0 | 0 | 0 | 0 |
| 10109 | ENSMUSG00000035653 | 0 | NA | NA | NA | NA | NA | Lrfn5 | leucine rich repeat and fibronectin type II | 0 | 0 | 0 | 0 | 0 | 0 |
| 10110 | ENSMUSG00000035666 | 564.8163604 | 0.304797055 | 0.142685744 | 2.136142318 | 0.032667815 | 0.241444702 | Gtf3c4 | general transcription factor IIIC, polypepti | 532.46 | 451.26 | 531.3 | 540.79 | 635.83 | 697.27 |
| 10111 | ENSMUSG00000035671 | 140.1016579 | 0.327112421 | 0.233939914 | 1.398275374 | 0.16203039 | 0.585462793 | Zswim4 | zinc finger SWIM-type containing 4 [Sourc | 128.5 | 92.513 | 151.94 | 182.19 | 141.78 | 143.69 |
| 10112 | ENSMUSG00000035673 | 573.5964289 | 0.112920443 | 0.14053506 | 0.803503719 | 0.421683648 | 0.838959049 | Sbno2 | strawberry notch 2 [Source:MGI Symbol;A | 535.67 | 509.85 | 608.78 | 675.75 | 579.12 | 532.42 |
| 10113 | ENSMUSG00000035674 | 1228.85923 | -0.029184634 | 0.098838649 | -0.295275526 | 0.767783411 | 0.95578799 | Ndufa3 | NADH:ubiquinone oxidoreductase subuni | 1236.8 | 1221.2 | 1265.9 | 1210.8 | 1184.4 | 1254.2 |
| 10114 | ENSMUSG00000035678 | 0.160662974 | 1.126924175 | 4.080472857 | 0.276174898 | 0.782413736 | NA | Tnfsf9 | tumor necrosis factor (ligand) superfamily | 0 | 0 | 0 | 0.964 | 0 | 0 |
| 10115 | ENSMUSG00000035681 | 0 | NA | NA | NA | NA | NA | Kcnc2 | potassium voltage gated channel, Shaw-re | 0 | 0 | 0 | 0 | 0 | 0 |
| 10116 | ENSMUSG00000035683 | 0.472877444 | -2.283655516 | 3.093270275 | -0.738265755 | 0.460352972 | NA | Melk | maternal embryonic leucine zipper kinase | 0.8031 | 1.0279 | 1.0062 | 0 | 0 | 0 |
| 10117 | ENSMUSG00000035686 | 16594.51181 | -0.683111442 | 0.440437396 | -1.5509842 | 0.12090547 | 0.509634296 | Thrsp | thyroid hormone responsive [Source:MGI | 28782 | 20253 | 12319 | 12270 | 17321 | 8622.3 |
| 10118 | ENSMUSG00000035692 | 161.5668194 | 0.323100446 | 0.243360726 | 1.327660592 | 0.184290258 | 0.619137675 | Isg15 | ISG15 ubiquitin-like modifier [Source:MGI | 181.5 | 117.18 | 130.81 | 134.96 | 179.95 | 225 |
| 10119 | ENSMUSG00000035694 | 0.167707431 | -0.796629176 | 4.080472857 | -0.195229623 | 0.84521319 | NA | Caps2 | calcyphosphine 2 [Source:MGI Symbol;Ac | 0 | 0 | 1.0062 | 0 | 0 | 0 |
| 10120 | ENSMUSG00000035696 | 210.5565401 | 0.366578316 | 0.175201418 | 2.092325056 | 0.036409448 | 0.257267344 | Rnf38 | ring finger protein 38 [Source:MGI Symbo | 187.12 | 170.63 | 194.21 | 260.27 | 209.4 | 241.7 |
| 10121 | ENSMUSG00000035697 | 130.5332811 | -0.535627508 | 0.261130852 | -2.051184312 | 0.040249001 | 0.274604506 | Arhgap45 | Rho GTPase activating protein 45 [Source: | 186.32 | 119.24 | 157.98 | 139.78 | 78.525 | 101.36 |
| 10122 | ENSMUSG00000035699 | 3.292286528 | -1.160196146 | 1.274044965 | -0.910639873 | 0.362485154 | NA | Slc51a | solute carrier family 51, alpha subunit [So | 5.6217 | 1.0279 | 7.0437 | 3.8559 | 1.0906 | 1.1138 |
| 10123 | ENSMUSG00000035704 | 79.43278842 | 0.227446828 | 0.297137293 | 0.765460387 | 0.443997446 | 0.851432536 | Alg8 | asparagine-linked glycosylation 8 (alpha-1 | 93.963 | 67.843 | 57.356 | 92.542 | 102.52 | 62.375 |
| 10124 | ENSMUSG00000035711 | 26.00453654 | -0.905568013 | 0.475233125 | -1.905523763 | 0.056712021 | 0.337842948 | Dok3 | docking protein 3 [Source:MGI Symbol;Ac | 39.352 | 33.921 | 28.175 | 12.532 | 13.087 | 28.96 |
| 10125 | ENSMUSG00000035713 | 28.75195104 | -0.220915019 | 0.433312759 | -0.509828095 | 0.610171901 | 0.916376001 | Usp35 | ubiquitin specific peptidase 35 [Source:M | 28.109 | 30.838 | 34.212 | 38.559 | 19.631 | 21.163 |
| 10126 | ENSMUSG00000035722 | 139.9923321 | -0.122715666 | 0.240310119 | -0.510655424 | 0.609592359 | 0.916376001 | Abca7 | ATP-binding cassette, sub-family A (ABC1) | 129.3 | 159.33 | 149.93 | 170.62 | 94.884 | 135.89 |
| 10127 | ENSMUSG00000035725 | 58.48055948 | -0.491767412 | 0.314421746 | -1.564037537 | 0.117808753 | 0.502838264 | Prkx | protein kinase, X-linked [Source:MGI Sym | 50.595 | 86.345 | 68.425 | 46.271 | 46.897 | 52.351 |
| 10128 | ENSMUSG00000035726 | 407.034709 | 0.038733169 | 0.136383833 | 0.284001174 | 0.776409481 | 0.959090164 | Supt16 | SPT16, facilitates chromatin remodeling su | 391.91 | 406.03 | 406.52 | 374.99 | 440.61 | 422.15 |
| 10129 | ENSMUSG00000035735 | 10.33880754 | -1.322679328 | 0.701157267 | -1.88642319 | 0.05923795 | 0.346085052 | Dagla | diacylglycerol lipase, alpha [Source:MGI S | 20.881 | 11.307 | 12.075 | 6.7478 | 5.4531 | 5.5692 |
| 10130 | ENSMUSG00000035745 | 20.87858772 | -0.37338536 | 0.599819309 | -0.6224964 | 0.53361551 | 0.889143097 | Grin3b | glutamate receptor, ionotropic, NMDA3B | 32.124 | 8.2233 | 30.187 | 19.28 | 8.7249 | 26.732 |
| 10131 | ENSMUSG00000035754 | 1372.284734 | 0.007109019 | 0.112145849 | 0.063390833 | 0.949455287 | 0.990731019 | Wdr18 | WD repeat domain 18 [Source:MGI Symbo | 1493.8 | 1221.2 | 1390.6 | 1372.7 | 1422.2 | 1333.3 |
| 10132 | ENSMUSG00000035757 | 1114.173993 | 0.353675712 | 0.122578452 | 2.885300846 | 0.003910398 | 0.056472904 | Selenoo | selenoprotein O [Source:MGI Symbol;Acc | 934.01 | 934.38 | 1067.6 | 1407.4 | 1187.7 | 1153.9 |

| | | | | | | | | | | | | | | | |
|---|---|---|---|---|---|---|---|---|---|---|---|---|---|---|---|
| 10133 | ENSMUSG00000035759 | 131.8413618 | 0.115638573 | 0.214108752 | 0.540092697 | 0.589133107 | 0.911039598 | Bbs10 | Bardet-Biedl syndrome 10 (human) [Sourc | 115.65 | 136.71 | 127.79 | 144.6 | 152.69 | 113.61 |
| 10134 | ENSMUSG00000035762 | 154.3700101 | 0.277223245 | 0.220906052 | 1.254937303 | 0.209501521 | 0.653550636 | Tmem161b | transmembrane protein 161B [Source:MG | 123.68 | 180.91 | 114.71 | 178.34 | 157.05 | 171.53 |
| 10135 | ENSMUSG00000035764 | 205.0114126 | 0.056966758 | 0.176515853 | 0.322728849 | 0.746900602 | 0.950481287 | Fbxo45 | F-box protein 45 [Source:MGI Symbol;Acc | 194.35 | 184 | 224.39 | 190.87 | 217.03 | 219.43 |
| 10136 | ENSMUSG00000035765 | 595.891069 | -0.001995639 | 0.155107374 | -0.01286618 | 0.989734557 | 0.99859176 | Dym | dymeclin [Source:MGI Symbol;Acc:MGI:1 | 660.15 | 525.27 | 602.74 | 572.6 | 708.9 | 505.69 |
| 10137 | ENSMUSG00000035769 | 3018.261694 | -0.006092356 | 0.106257154 | -0.057335958 | 0.954277577 | 0.991678478 | Xylb | xylulokinase homolog (H. influenzae) [Sou | 3088.7 | 3124.9 | 2860.8 | 3209.1 | 3156.2 | 2669.9 |
| 10138 | ENSMUSG00000035770 | 410.5676767 | 0.052790656 | 0.148863388 | 0.354624845 | 0.722870661 | 0.9447719 | Dync1li2 | dynein, cytoplasmic 1 light intermediate c | 423.23 | 388.55 | 397.47 | 467.53 | 354.45 | 432.17 |
| 10139 | ENSMUSG00000035772 | 725.9972199 | -0.056965073 | 0.124036409 | -0.459260905 | 0.646046819 | 0.925643431 | Mrps2 | mitochondrial ribosomal protein S2 [Sourc | 733.23 | 709.26 | 777.83 | 631.41 | 757.98 | 746.28 |
| 10140 | ENSMUSG00000035773 | 17.73051625 | -0.244111356 | 0.533706966 | -0.457388364 | 0.647391926 | 0.925643431 | Kiss1r | KISS1 receptor [Source:MGI Symbol;Acc:M | 20.881 | 22.614 | 14.087 | 12.532 | 22.903 | 13.366 |
| 10141 | ENSMUSG00000035775 | 0 | NA | NA | NA | NA | NA | Krt20 | keratin 20 [Source:MGI Symbol;Acc:MGI:1 | 0 | 0 | 0 | 0 | 0 | 0 |
| 10142 | ENSMUSG00000035776 | 495.4642272 | 0.062552659 | 0.138669711 | 0.451091002 | 0.651923963 | 0.925643431 | Cd99l2 | CD99 antigen-like 2 [Source:MGI Symbol;A | 477.85 | 549.94 | 426.65 | 508.02 | 519.13 | 491.21 |
| 10143 | ENSMUSG00000035778 | 23.84414905 | -0.855107948 | 0.490912622 | -1.74187403 | 0.081530489 | 0.420121054 | Ggta1 | glycoprotein galactosyltransferase alpha 1 | 28.912 | 35.977 | 27.169 | 14.46 | 9.8156 | 26.732 |
| 10144 | ENSMUSG00000035780 | 6385.401113 | 0.085843158 | 0.11637473 | 0.737644309 | 0.460730623 | 0.859821403 | Ugt2a3 | UDP glucuronosyltransferase 2 family, pol | 5640.2 | 5849.9 | 7097 | 6257.2 | 7240.6 | 6227.5 |
| 10145 | ENSMUSG00000035781 | 656.496252 | -0.081816599 | 0.112371853 | -0.728088007 | 0.466559715 | 0.863779479 | R3hdm4 | R3H domain containing 4 [Source:MGI Syr | 677.02 | 666.09 | 682.23 | 643.94 | 631.47 | 638.23 |
| 10146 | ENSMUSG00000035783 | 11.32784698 | 0.008619077 | 0.659336057 | 0.013072359 | 0.989570064 | 0.99859176 | Acta2 | actin, alpha 2, smooth muscle, aorta [Sou | 9.6372 | 12.335 | 12.075 | 17.352 | 6.5437 | 10.025 |
| 10147 | ENSMUSG00000035785 | 0.74256259 | 3.09446134 | 3.273353887 | 0.945348852 | 0.344480805 | NA | Cmtm2b | CKLF-like MARVEL transmembrane domai | 0 | 0 | 0 | 0 | 0 | 4.4554 |
| 10148 | ENSMUSG00000035790 | 96.7456049 | 0.160396648 | 0.245000019 | 0.654680145 | 0.512673711 | 0.884860213 | Cep19 | centrosomal protein 19 [Source:MGI Symb | 77.098 | 107.93 | 89.556 | 105.07 | 89.431 | 111.38 |
| 10149 | ENSMUSG00000035798 | 233.7320029 | 0.117108562 | 0.176200275 | 0.664633256 | 0.506285093 | 0.881592135 | Zdhhc17 | zinc finger, DHHC domain containing 17 [S | 208 | 224.09 | 241.5 | 283.41 | 215.94 | 229.45 |
| 10150 | ENSMUSG00000035799 | 15.68057385 | -0.883775434 | 0.636223146 | -1.389096639 | 0.164803366 | 0.590028684 | Twist1 | twist basic helix-loop-helix transcription fa | 23.29 | 15.419 | 22.137 | 7.7118 | 4.3625 | 21.163 |
| 10151 | ENSMUSG00000035804 | 0 | NA | NA | NA | NA | NA | Ins1 | insulin I [Source:MGI Symbol;Acc:MGI:965 | 0 | 0 | 0 | 0 | 0 | 0 |
| 10152 | ENSMUSG00000035805 | 0 | NA | NA | NA | NA | NA | Mlc1 | megalencephalic leukoencephalopathy wi | 0 | 0 | 0 | 0 | 0 | 0 |
| 10153 | ENSMUSG00000035811 | 2670.247355 | 0.470411794 | 0.105477972 | 4.459810771 | 8.20E-06 | 0.000455037 | Ugt2b35 | UDP glucuronosyltransferase 2 family, pol | 2431 | 2238.8 | 2044.7 | 2905.4 | 3280.6 | 3121 |
| 10154 | ENSMUSG00000035818 | 0.863488199 | 0.685974303 | 2.37316251 | 0.289054922 | 0.772539349 | NA | Plekhs1 | pleckstrin homology domain containing, fa | 0 | 0 | 2.0125 | 0.964 | 1.0906 | 1.1138 |
| 10155 | ENSMUSG00000035824 | 451.6076157 | 0.012374556 | 0.131794678 | 0.093892681 | 0.925194408 | 0.987546253 | Tk2 | thymidine kinase 2, mitochondrial [Source | 477.85 | 423.5 | 446.77 | 427.04 | 472.24 | 462.25 |
| 10156 | ENSMUSG00000035828 | 2247.689088 | -0.311801626 | 0.157266689 | -1.982629812 | 0.047408795 | 0.302152946 | Pim3 | proviral integration site 3 [Source:MGI Syr | 2137.9 | 2901.8 | 2430.1 | 1945.3 | 2459.3 | 1611.7 |
| 10157 | ENSMUSG00000035829 | 1.770273837 | 0.688912014 | 1.858592431 | 0.370663306 | 0.710888324 | NA | Ppp1r26 | protein phosphatase 1, regulatory subunit | 0 | 4.1117 | 0 | 0.964 | 1.0906 | 4.4554 |
| 10158 | ENSMUSG00000035831 | 0 | NA | NA | NA | NA | NA | Krt25 | keratin 25 [Source:MGI Symbol;Acc:MGI:1 | 0 | 0 | 0 | 0 | 0 | 0 |
| 10159 | ENSMUSG00000035834 | 150.8529324 | 0.268877529 | 0.191388259 | 1.404879951 | 0.160056979 | 0.582406075 | Polr3g | polymerase (RNA) III (DNA directed) polyp | 146.97 | 124.38 | 138.86 | 168.7 | 163.59 | 162.62 |
| 10160 | ENSMUSG00000035835 | 26.91673198 | 0.131751605 | 0.461457062 | 0.285512167 | 0.775251785 | 0.95893551 | Plppr3 | phospholipid phosphatase related 3 [Sour | 19.274 | 28.782 | 29.181 | 28.919 | 16.359 | 38.985 |
| 10161 | ENSMUSG00000035836 | 9403.969722 | 0.637177201 | 0.117133095 | 5.439770904 | 5.33E-08 | 6.27E-06 | Ugt2b1 | UDP glucuronosyltransferase 2 family, pol | 7469.7 | 6536.5 | 8074.1 | 10077 | 12662 | 11604 |
| 10162 | ENSMUSG00000035840 | 549.553678 | -0.301962986 | 0.144592003 | -2.088379581 | 0.036763608 | 0.259010631 | Lysmd3 | LysM, putative peptidoglycan-binding, do | 559.76 | 601.33 | 660.1 | 494.52 | 560.58 | 421.03 |
| 10163 | ENSMUSG00000035842 | 87.99459116 | 0.183380538 | 0.281047664 | 0.652489103 | 0.5140857 | 0.885004045 | Ddx11 | DEAD/H box helicase 11 [Source:MGI Sym | 85.129 | 61.675 | 100.62 | 118.57 | 82.887 | 79.083 |
| 10164 | ENSMUSG00000035845 | 119.6046759 | 0.059564006 | 0.213008719 | 0.279631776 | 0.779760025 | 0.959870443 | Alg12 | asparagine-linked glycosylation 12 (alpha- | 106.81 | 113.07 | 131.82 | 123.39 | 117.79 | 124.75 |
| 10165 | ENSMUSG00000035847 | 208.366509 | -0.3832038 | 0.183473485 | -2.088605887 | 0.036743215 | 0.259010631 | Ids | iduronate 2-sulfatase [Source:MGI Symbo | 214.43 | 240.53 | 253.57 | 211.11 | 169.05 | 161.51 |
| 10166 | ENSMUSG00000035849 | 1.799025105 | -0.131635114 | 1.599985855 | -0.082272673 | 0.934429884 | NA | Krt222 | keratin 222 [Source:MGI Symbol;Acc:MGI | 1.6062 | 2.0558 | 2.0125 | 2.8919 | 0 | 2.2277 |
| 10167 | ENSMUSG00000035851 | 530.1051969 | -0.164160055 | 0.134384956 | -1.221565719 | 0.221871899 | 0.667949912 | Ythdc1 | YTH domain containing 1 [Source:MGI Syr | 494.71 | 613.67 | 573.56 | 523.44 | 492.96 | 482.29 |
| 10168 | ENSMUSG00000035852 | 2.836560615 | -0.364735835 | 1.325473205 | -0.275174054 | 0.78318252 | NA | Misp | mitotic spindle positioning [Source:MGI S | 6.4248 | 1.0279 | 2.0125 | 0.964 | 4.3625 | 2.2277 |
| 10169 | ENSMUSG00000035860 | 0 | NA | NA | NA | NA | NA | Cdhr3 | cadherin-related family member 3 [Source | 0 | 0 | 0 | 0 | 0 | 0 |
| 10170 | ENSMUSG00000035861 | 0 | NA | NA | NA | NA | NA | Tmprss11b | transmembrane protease, serine 11B [Sou | 0 | 0 | 0 | 0 | 0 | 0 |
| 10171 | ENSMUSG00000035863 | 30.65844567 | -0.148172676 | 0.451579293 | -0.328121059 | 0.742820132 | 0.949297614 | Palm | paralemmin [Source:MGI Symbol;Acc:MG | 33.73 | 32.893 | 30.187 | 39.523 | 13.087 | 34.529 |
| 10172 | ENSMUSG00000035864 | 197.3506674 | 0.351373415 | 0.209043335 | 1.680863992 | 0.092789335 | 0.44871446 | Syt1 | synaptotagmin I [Source:MGI Symbol;Acc | 190.34 | 144.94 | 184.14 | 169.66 | 238.85 | 256.18 |
| 10173 | ENSMUSG00000035868 | 93.19257069 | -0.079202026 | 0.25207615 | -0.314198808 | 0.753370048 | 0.952989051 | Zfp983 | zinc finger protein 983 [Source:MGI Symb | 110.03 | 77.094 | 99.618 | 84.83 | 82.887 | 104.7 |
| 10174 | ENSMUSG00000035873 | 163.9181809 | 0.020827591 | 0.236330936 | 0.088128923 | 0.929774209 | 0.988405116 | Pawr | PRKC, apoptosis, WT1, regulator [Source:I | 157.41 | 164.47 | 167.04 | 226.53 | 122.15 | 145.91 |
| 10175 | ENSMUSG00000035875 | 6591.482913 | 0.378256735 | 0.110831993 | 3.412883986 | 0.000642793 | 0.014923528 | AI182371 | expressed sequence AI182371 [Source:MG | 5191.3 | 5661.8 | 6344.4 | 6709.3 | 7664.9 | 7977.3 |
| 10176 | ENSMUSG00000035877 | 1302.624281 | -0.138372204 | 0.126938887 | -1.090069458 | 0.275682549 | 0.730038567 | Zhx3 | zinc fingers and homeoboxes 3 [Source:M | 1157.3 | 1571.7 | 1367.5 | 1218.5 | 1243.3 | 1257.5 |
| 10177 | ENSMUSG00000035878 | 2461.204872 | 0.057236122 | 0.0943216 | 0.606818821 | 0.543971156 | 0.894702561 | Hykk | hydroxylysine kinase 1 [Source:MGI Symb | 2468.7 | 2321 | 2447.2 | 2545.9 | 2617.5 | 2366.9 |
| 10178 | ENSMUSG00000035885 | 4279.75984 | 0.000729877 | 0.08993661 | 0.008115459 | 0.993524872 | 0.99859176 | Cox8a | cytochrome c oxidase subunit 8A [Source: | 4242 | 4465.3 | 4128.6 | 4171.1 | 4171.6 | 4499.9 |
| 10179 | ENSMUSG00000035890 | 497.1265241 | -0.061230925 | 0.152168776 | -0.402388236 | 0.687398327 | 0.936885029 | Rnf126 | ring finger protein 126 [Source:MGI Symb | 480.26 | 548.91 | 495.07 | 578.39 | 461.33 | 418.81 |
| 10180 | ENSMUSG00000035891 | 45.44383593 | -0.823370605 | 0.366862523 | -2.244357364 | 0.024809417 | 0.20148423 | Cerk | ceramide kinase [Source:MGI Symbol;Acc | 57.02 | 77.094 | 40.25 | 37.595 | 26.175 | 34.529 |
| 10181 | ENSMUSG00000035896 | 0 | NA | NA | NA | NA | NA | Rnase1 | ribonuclease, RNase A family, 1 (pancreat | 0 | 0 | 0 | 0 | 0 | 0 |
| 10182 | ENSMUSG00000035898 | 144.4654957 | -0.026546311 | 0.197759662 | -0.134235217 | 0.893216579 | 0.982562372 | Uba6 | ubiquitin-like modifier activating enzyme 6 | 152.59 | 148.02 | 136.85 | 158.09 | 129.78 | 141.46 |
| 10183 | ENSMUSG00000035900 | 97.00847496 | 0.19042729 | 0.254742197 | 0.747529433 | 0.454744043 | 0.857337687 | Gramd4 | GRAM domain containing 4 [Source:MGI S | 94.766 | 76.066 | 100.62 | 88.686 | 92.703 | 129.21 |
| 10184 | ENSMUSG00000035901 | 544.5677769 | 0.070625226 | 0.122939718 | 0.574470379 | 0.565649539 | 0.901895454 | Dennd5a | DENN/MADD domain containing 5A [Sour | 524.43 | 527.32 | 542.37 | 586.1 | 519.13 | 568.06 |
| 10185 | ENSMUSG00000035910 | 18.49103048 | -0.082829458 | 0.539960272 | -0.15339917 | 0.878083499 | 0.980817031 | Dcdc2a | doublecortin domain containing 2a [Sourc | 10.44 | 27.754 | 19.119 | 18.316 | 15.269 | 20.049 |
| 10186 | ENSMUSG00000035914 | 26.85686474 | 0.828619731 | 0.448548171 | 1.847337217 | 0.064698282 | 0.364548482 | Cd276 | CD276 antigen [Source:MGI Symbol;Acc:M | 28.109 | 14.391 | 15.094 | 30.847 | 38.172 | 34.529 |
| 10187 | ENSMUSG00000035916 | 1.156404226 | 2.563524626 | 2.366885343 | 1.083079345 | 0.278773206 | NA | Ptprq | protein tyrosine phosphatase, receptor ty | 0 | 1.0279 | 0 | 4.8199 | 1.0906 | 0 |
| 10188 | ENSMUSG00000035919 | 85.25793355 | 0.01137616 | 0.273476831 | 0.041598259 | 0.966818961 | 0.994231936 | Bbs9 | Bardet-Biedl syndrome 9 (human) [Source | 73.082 | 81.206 | 100.62 | 80.974 | 67.618 | 108.04 |

| | | | | | | | | | | | | | | | |
|---|---|---|---|---|---|---|---|---|---|---|---|---|---|---|---|
| 10189 | ENSMUSG00000035923 | 0 | NA | NA | NA | NA | NA | Myf6 | myogenic factor 6 [Source:MGI Symbol;Ac | 0 | 0 | 0 | 0 | 0 | 0 |
| 10190 | ENSMUSG00000035929 | 643.8932062 | -0.285325641 | 0.127083877 | -2.24517576 | 0.024756852 | 0.201358738 | H2-Q4 | histocompatibility 2, Q region locus 4 [Sou | 743.67 | 721.6 | 657.08 | 638.15 | 568.21 | 534.65 |
| 10191 | ENSMUSG00000035930 | 9.612725345 | -0.664710584 | 0.7249936 | -0.916850278 | 0.359221102 | 0.798981089 | Chst4 | carbohydrate sulfotransferase 4 [Source:N | 12.85 | 14.391 | 8.05 | 4.8199 | 11.997 | 5.5692 |
| 10192 | ENSMUSG00000035932 | 0 | NA | NA | NA | NA | NA | Olfr750 | olfactory receptor 750 [Source:MGI Symb | 0 | 0 | 0 | 0 | 0 | 0 |
| 10193 | ENSMUSG00000035933 | 314.5647189 | -0.094220564 | 0.177413749 | -0.531078142 | 0.595364631 | 0.913768465 | Cog5 | component of oligomeric golgi complex 5 | 334.89 | 291.93 | 347.15 | 263.17 | 377.35 | 272.89 |
| 10194 | ENSMUSG00000035934 | 15.47191331 | -0.78140909 | 0.567129584 | -1.37783165 | 0.168255272 | 0.595160122 | Pknox2 | Pbx/knotted 1 homeobox 2 [Source:MGI S | 16.865 | 23.642 | 18.112 | 7.7118 | 10.906 | 15.594 |
| 10195 | ENSMUSG00000035936 | 1145.119596 | -0.13008147 | 0.10451203 | -1.24465547 | 0.213258421 | 0.657933491 | Aldh5a1 | aldhehyde dehydrogenase family 5, subfa | 1170.9 | 1197.5 | 1221.6 | 1041.1 | 1167 | 1072.6 |
| 10196 | ENSMUSG00000035941 | 1040.221214 | 0.056131134 | 0.107205611 | 0.523583917 | 0.60056797 | 0.91534406 | Ibtk | inhibitor of Bruton agammaglobulinemia t | 1071.3 | 1005.3 | 983.1 | 1112.4 | 1063.4 | 1005.8 |
| 10197 | ENSMUSG00000035944 | 2074.671986 | 0.126488374 | 0.117912291 | 1.072732726 | 0.283391053 | 0.736165263 | Ttc38 | tetratricopeptide repeat domain 38 [Sour | 1884.9 | 1857.4 | 2208.7 | 1919.3 | 2257.6 | 2320.1 |
| 10198 | ENSMUSG00000035946 | 0 | NA | NA | NA | NA | NA | Gsx2 | GS homeobox 2 [Source:MGI Symbol;Acc: | 0 | 0 | 0 | 0 | 0 | 0 |
| 10199 | ENSMUSG00000035948 | 425.7756172 | -0.728189873 | 0.16695639 | -4.36155737 | 1.29E-05 | 0.000664726 | Acss3 | acyl-CoA synthetase short-chain family me | 535.67 | 555.08 | 503.12 | 379.81 | 334.82 | 246.16 |
| 10200 | ENSMUSG00000035949 | 769.7406713 | 0.190019005 | 0.118379596 | 1.605166867 | 0.108457087 | 0.48522218 | Fbxw2 | F-box and WD-40 domain protein 2 [Sourc | 682.64 | 731.88 | 743.61 | 833.84 | 876.86 | 749.62 |
| 10201 | ENSMUSG00000035951 | 0.17131964 | -0.796629176 | 4.080472857 | -0.195229623 | 0.84521319 | NA | Ascl3 | achaete-scute family bHLH transcription fa | 0 | 1.0279 | 0 | 0 | 0 | 0 |
| 10202 | ENSMUSG00000035953 | 354.9884416 | 0.37754755 | 0.149413361 | 2.526866063 | 0.011508538 | 0.119817569 | Pip4p1 | phosphatidylinositol-4,5-bisphosphate 4-p | 297.15 | 329.96 | 299.86 | 431.86 | 351.18 | 419.92 |
| 10203 | ENSMUSG00000035954 | 512.6003543 | -0.099819487 | 0.197068488 | -0.506521807 | 0.612490385 | 0.916441371 | Dock4 | dedicator of cytokinesis 4 [Source:MGI Sy | 487.48 | 618.81 | 485.01 | 464.64 | 656.55 | 363.11 |
| 10204 | ENSMUSG00000035958 | 408.0496796 | 0.188852263 | 0.134375677 | 1.405405111 | 0.159900848 | 0.581984365 | Tdp2 | tyrosyl-DNA phosphodiesterase 2 [Source | 410.39 | 360.8 | 372.31 | 440.54 | 422.07 | 442.2 |
| 10205 | ENSMUSG00000035960 | 105.3891841 | 0.078494217 | 0.237475167 | 0.330536527 | 0.740994598 | 0.948792385 | Apex1 | apurinic/apyrimidinic endonuclease 1 [So | 97.175 | 94.568 | 115.72 | 96.398 | 98.156 | 130.32 |
| 10206 | ENSMUSG00000035963 | 0 | NA | NA | NA | NA | NA | Odf3l2 | outer dense fiber of sperm tails 3-like 2 [S | 0 | 0 | 0 | 0 | 0 | 0 |
| 10207 | ENSMUSG00000035964 | 0 | NA | NA | NA | NA | NA | Tmem59l | transmembrane protein 59-like [Source:M | 0 | 0 | 0 | 0 | 0 | 0 |
| 10208 | ENSMUSG00000035967 | 56.10871629 | -0.789378981 | 0.404070452 | -1.953567697 | 0.050752364 | 0.316199857 | Ints6l | integrator complex subunit 6 like [Source: | 53.808 | 92.513 | 67.418 | 54.947 | 51.259 | 16.708 |
| 10209 | ENSMUSG00000035969 | 734.6982106 | -0.702050513 | 0.147237392 | -4.768153682 | 1.86E-06 | 0.00013379 | Rusc2 | RUN and SH3 domain containing 2 [Sourc | 785.43 | 995.02 | 949.89 | 502.23 | 656.55 | 519.05 |
| 10210 | ENSMUSG00000035983 | 4.048859268 | -0.639954925 | 1.044387674 | -0.612756106 | 0.540037618 | NA | Gm7008 | predicted gene 7008 [Source:MGI Symbol | 5.6217 | 4.1117 | 5.0312 | 2.8919 | 2.1812 | 4.4554 |
| 10211 | ENSMUSG00000035984 | 1.977485773 | -2.162101321 | 2.226406456 | -0.971117073 | 0.331489983 | NA | Nme5 | NME/NM23 family member 5 [Source:MG | 9.6372 | 0 | 0 | 0 | 0 | 2.2277 |
| 10212 | ENSMUSG00000035992 | 1027.814515 | 0.182087995 | 0.115901306 | 1.571060771 | 0.116168534 | 0.500730933 | Fnip1 | folliculin interacting protein 1 [Source:MG | 1044.8 | 927.18 | 916.69 | 1186.7 | 1059 | 1032.5 |
| 10213 | ENSMUSG00000036002 | 345.5123993 | 0.236620093 | 0.143196604 | 1.652414147 | 0.098450154 | 0.462782912 | Fam214b | family with sequence similarity 214, meml | 338.11 | 296.04 | 316.97 | 372.1 | 354.45 | 395.41 |
| 10214 | ENSMUSG00000036006 | 95.74357694 | 0.541327808 | 0.304951514 | 1.775127465 | 0.075876843 | 0.403486271 | Ripor2 | RHO family interacting cell polarization re | 100.39 | 49.34 | 83.518 | 91.578 | 150.51 | 99.132 |
| 10215 | ENSMUSG00000036009 | 34.66784839 | 0.110105113 | 0.39436385 | 0.279196769 | 0.780093823 | 0.959870443 | Mettl25 | methyltransferase like 25 [Source:MGI Syr | 45.777 | 25.698 | 28.175 | 36.631 | 31.628 | 40.098 |
| 10216 | ENSMUSG00000036013 | 0 | NA | NA | NA | NA | NA | Fam122c | family with sequence similarity 122, meml | 0 | 0 | 0 | 0 | 0 | 0 |
| 10217 | ENSMUSG00000036019 | 289.5038838 | -0.181684702 | 0.16418083 | -1.106613373 | 0.268461123 | 0.724039098 | Tmtc2 | transmembrane and tetratricopeptide rep | 296.34 | 272.4 | 354.2 | 249.67 | 275.93 | 288.49 |
| 10218 | ENSMUSG00000036022 | 47.67398941 | -0.003462373 | 0.379180092 | -0.009131211 | 0.992714449 | 0.99859176 | Fam122b | family with sequence similarity 122, meml | 41.761 | 32.893 | 68.425 | 33.739 | 49.078 | 60.148 |
| 10219 | ENSMUSG00000036023 | 134.8227291 | -0.104383476 | 0.216312843 | -0.482557923 | 0.629409661 | 0.92057725 | Parp2 | poly (ADP-ribose) polymerase family, men | 140.54 | 120.27 | 157.98 | 117.61 | 121.06 | 151.48 |
| 10220 | ENSMUSG00000036026 | 1047.152175 | 0.288782862 | 0.123937299 | 2.330072258 | 0.019802332 | 0.172127966 | Tmem63b | transmembrane protein 63b [Source:MGI | 836.83 | 985.77 | 1007.3 | 1270.5 | 1074.3 | 1108.3 |
| 10221 | ENSMUSG00000036027 | 38.45552583 | 0.471166184 | 0.440805732 | 1.068874901 | 0.285126043 | 0.737769696 | 1810046K07 | RIKEN cDNA 1810046K07 gene [Source:M | 34.533 | 23.642 | 38.237 | 22.171 | 68.709 | 43.44 |
| 10222 | ENSMUSG00000036030 | 1.755797845 | -1.954451883 | 1.763849358 | -1.108060546 | 0.267835668 | NA | Prtg | protogenin [Source:MGI Symbol;Acc:MGI: | 3.2124 | 4.1117 | 1.0062 | 0 | 1.0906 | 1.1138 |
| 10223 | ENSMUSG00000036036 | 6.2620494 | -0.791121765 | 0.92601079 | -0.854333205 | 0.392920394 | 0.821390458 | Zfp57 | zinc finger protein 57 [Source:MGI Symbo | 10.44 | 10.279 | 3.0187 | 3.8559 | 2.1812 | 7.7969 |
| 10224 | ENSMUSG00000036040 | 156.8783041 | -1.094204729 | 0.264116321 | -4.142889489 | 3.43E-05 | 0.001555026 | Adamtsl2 | ADAMTS-like 2 [Source:MGI Symbol;Acc:N | 306.79 | 175.77 | 157.98 | 109.89 | 87.249 | 103.59 |
| 10225 | ENSMUSG00000036045 | 0 | NA | NA | NA | NA | NA | Gm12221 | predicted gene 12221 [Source:MGI Symbo | 0 | 0 | 0 | 0 | 0 | 0 |
| 10226 | ENSMUSG00000036046 | 1145.635133 | -0.052483366 | 0.111093213 | -0.47242639 | 0.636622471 | 0.923478764 | 5031439G07 | RIKEN cDNA 5031439G07 gene [Source:M | 1130 | 1163.6 | 1205.5 | 1018 | 1207.3 | 1149.5 |
| 10227 | ENSMUSG00000036052 | 66.53795763 | 0.107811936 | 0.283010068 | 0.380947353 | 0.703242314 | 0.942506832 | Dnajb5 | DnaJ heat shock protein family (Hsp40) m | 55.414 | 64.759 | 72.45 | 80.974 | 63.256 | 62.375 |
| 10228 | ENSMUSG00000036053 | 103.3579781 | -0.653453053 | 0.225789029 | -2.894086815 | 0.00380263 | 0.05527763 | Fmnl2 | formin-like 2 [Source:MGI Symbol;Acc:MC | 122.07 | 119.24 | 137.86 | 80.01 | 77.434 | 83.538 |
| 10229 | ENSMUSG00000036054 | 180.944644 | 0.027011143 | 0.221114286 | 0.122159196 | 0.902772941 | 0.983821605 | Sugp2 | SURP and G patch domain containing 2 [S | 168.65 | 172.69 | 197.22 | 241.96 | 159.23 | 145.91 |
| 10230 | ENSMUSG00000036057 | 282.3053609 | -0.118307856 | 0.149304303 | -0.792394147 | 0.428130891 | 0.842753542 | Ptpn23 | protein tyrosine phosphatase, non-recept | 281.89 | 308.38 | 291.81 | 279.55 | 271.56 | 260.64 |
| 10231 | ENSMUSG00000036061 | 195.6655413 | -0.136823336 | 0.181418043 | -0.754188138 | 0.45073626 | 0.855818389 | Smug1 | single-strand selective monofunctional ur | 213.63 | 179.89 | 221.37 | 205.33 | 176.68 | 177.1 |
| 10232 | ENSMUSG00000036062 | 4.066565904 | -1.244090438 | 1.092573389 | -1.138679059 | 0.25483704 | NA | Phf24 | PHD finger protein 24 [Source:MGI Symbo | 4.0155 | 7.1954 | 6.0375 | 3.8559 | 2.1812 | 1.1138 |
| 10233 | ENSMUSG00000036067 | 3.490153116 | -1.927447306 | 1.350785392 | -1.42690861 | 0.153606239 | NA | Slc2a6 | solute carrier family 2 (facilitated glucose | 3.2124 | 11.307 | 2.0125 | 0 | 2.1812 | 2.2277 |
| 10234 | ENSMUSG00000036073 | 550.4233715 | 0.128599156 | 0.14522845 | 0.885495615 | 0.375889384 | 0.809690069 | Galt | galactose-1-phosphate uridyl transferase | 533.26 | 493.4 | 550.42 | 522.48 | 521.32 | 681.67 |
| 10235 | ENSMUSG00000036078 | 1453.623389 | 0.078652186 | 0.122165825 | 0.643814963 | 0.51969543 | 0.887802551 | Sigmar1 | sigma non-opioid intracellular receptor 1 | | 1319.5 | 1415.4 | 1507.4 | 1431.5 | 1338.2 | 1709.8 |
| 10236 | ENSMUSG00000036083 | 1396.408751 | 0.103268413 | 0.136708729 | 0.755390044 | 0.450014985 | 0.855479371 | Slc17a3 | solute carrier family 17 (sodium phosphat | 1215.1 | 1208.8 | 1616 | 1401.6 | 1584.7 | 1352.2 |
| 10237 | ENSMUSG00000036086 | 91.70033022 | 0.260364959 | 0.255869871 | 1.017567869 | 0.308883363 | 0.757462943 | Zranb3 | zinc finger, RAN-binding domain containir | 94.766 | 61.675 | 93.581 | 105.07 | 95.974 | 99.132 |
| 10238 | ENSMUSG00000036087 | 684.3110941 | -0.428902725 | 0.133368573 | -3.215920484 | 0.001300269 | 0.025414979 | Slain2 | SLAIN motif family, member 2 [Source:MC | 693.88 | 878.87 | 783.86 | 562 | 632.56 | 554.69 |
| 10239 | ENSMUSG00000036091 | 4.092647527 | 0.104540167 | 1.346318205 | 0.077648929 | 0.93810732 | NA | Hyal3 | hyaluronoglucosaminidase 3 [Source:MGI | 5.6217 | 6.1675 | 0 | 3.8559 | 0 | 8.9108 |
| 10240 | ENSMUSG00000036093 | 2007.750652 | 0.192403588 | 0.111303221 | 1.728643487 | 0.083872922 | 0.426710495 | Arl5a | ADP-ribosylation factor-like 5A [Source:M | 1835.1 | 1897.5 | 1890.7 | 2379.1 | 2159.4 | 1884.6 |
| 10241 | ENSMUSG00000036095 | 0 | NA | NA | NA | NA | NA | Dgkb | diacylglycerol kinase, beta [Source:MGI Sy | 0 | 0 | 0 | 0 | 0 | 0 |
| 10242 | ENSMUSG00000036097 | 283.00599 | 0.151284478 | 0.152190073 | 0.994049577 | 0.320198732 | 0.767337321 | Slf2 | SMC5-SMC6 complex localization factor 2 | 267.43 | 287.82 | 249.55 | 310.4 | 299.92 | 282.92 |
| 10243 | ENSMUSG00000036098 | 456.8835377 | 0.157863319 | 0.148674917 | 1.061801963 | 0.288325599 | 0.73951397 | Myrf | myelin regulatory factor [Source:MGI Sym | 371.84 | 496.48 | 428.66 | 507.05 | 446.06 | 491.21 |
| 10244 | ENSMUSG00000036099 | 279.0326934 | 0.338856767 | 0.154336515 | 2.195570936 | 0.028122668 | 0.218809186 | Vezt | vezatin, adherens junctions transmembra | 224.07 | 268.29 | 247.54 | 308.47 | 320.64 | 305.19 |

| | | | | | | | | | | | | | | |
|---|---|---|---|---|---|---|---|---|---|---|---|---|---|---|
| 10245 | ENSMUSG00000036103 | 932.4143313 | -0.219316421 | 0.136214365 | -1.610082911 | 0.107379757 | 0.48315751 | Colec12 | collectin sub-family member 12 [Source:M | 882.61 | 1011.5 | 1116.9 | 959.16 | 873.59 | 750.73 |
| 10246 | ENSMUSG00000036104 | 823.788753 | 0.258909103 | 0.117029177 | 2.212346603 | 0.026942725 | 0.212718111 | Rab3gap1 | RAB3 GTPase activating protein subunit 1 | 741.26 | 720.57 | 787.89 | 817.45 | 948.84 | 926.72 |
| 10247 | ENSMUSG00000036106 | 125.6218986 | 0.044645718 | 0.218967327 | 0.203892145 | 0.838437783 | 0.97126094 | Prr5 | proline rich 5 (renal) [Source:MGI Symbol, | 133.31 | 105.88 | 131.82 | 142.67 | 129.78 | 110.27 |
| 10248 | ENSMUSG00000036109 | 2.446859373 | -1.982979018 | 1.78344671 | -1.111880163 | 0.266189679 | NA | Mbnl3 | muscleblind like splicing factor 3 [Source:I | 5.6217 | 6.1675 | 0 | 2.8919 | 0 | 0 |
| 10249 | ENSMUSG00000036110 | 4574.285303 | 0.27044465 | 0.16668658 | 1.622474052 | 0.104701885 | 0.476887243 | Slc17a2 | solute carrier family 17 (sodium phosphat | 3465.4 | 4207.3 | 4769.6 | 6495.3 | 4415.9 | 4092.3 |
| 10250 | ENSMUSG00000036111 | 0.956437057 | 1.282470111 | 2.271719443 | 0.56453719 | 0.572388597 | NA | Lmo1 | LIM domain only 1 [Source:MGI Symbol;A | 1.6062 | 0 | 0 | 1.928 | 1.0906 | 1.1138 |
| 10251 | ENSMUSG00000036112 | 1109.792443 | -0.016809026 | 0.120771325 | -0.1391806 | 0.889307438 | 0.982382091 | Metap2 | methionine aminopeptidase 2 [Source:MC | 975.77 | 1218.1 | 1156.2 | 1082.5 | 1169.1 | 1057 |
| 10252 | ENSMUSG00000036114 | 288.7045164 | -0.15451195 | 0.149743493 | -1.031844167 | 0.302145125 | 0.752320036 | Rpp25l | ribonuclease P/MRP 25 subunit-like [Sour | 310 | 297.07 | 304.89 | 252.56 | 278.11 | 289.6 |
| 10253 | ENSMUSG00000036117 | 0 | NA | NA | NA | NA | NA | Il5 | interleukin 5 [Source:MGI Symbol;Acc:MG | 0 | 0 | 0 | 0 | 0 | 0 |
| 10254 | ENSMUSG00000036120 | 159.4585844 | -0.296923456 | 0.202144747 | -1.468865551 | 0.14186926 | 0.551315856 | Rfxank | regulatory factor X-associated ankyrin-cor | 158.21 | 162.41 | 207.29 | 154.24 | 128.69 | 145.91 |
| 10255 | ENSMUSG00000036123 | 0 | NA | NA | NA | NA | NA | Slc9a3 | solute carrier family 9 (sodium/hydrogen | 0 | 0 | 0 | 0 | 0 | 0 |
| 10256 | ENSMUSG00000036131 | 0 | NA | NA | NA | NA | NA | Frmd7 | FERM domain containing 7 [Source:MGI S | 0 | 0 | 0 | 0 | 0 | 0 |
| 10257 | ENSMUSG00000036136 | 28.274459 | 0.038088216 | 0.439020187 | 0.086757322 | 0.930864412 | 0.988420305 | Fam110c | family with sequence similarity 110, meml | 20.881 | 25.698 | 37.231 | 25.063 | 22.903 | 37.871 |
| 10258 | ENSMUSG00000036138 | 6388.115326 | -0.013634723 | 0.08863359 | -0.153832463 | 0.877741837 | 0.98072613 | Acaa1a | acetyl-Coenzyme A acyltransferase 1A [So | 6372.6 | 6670.2 | 6212.6 | 6421.1 | 6635.3 | 6017 |
| 10259 | ENSMUSG00000036139 | 0 | NA | NA | NA | NA | NA | Hoxc9 | homeobox C9 [Source:MGI Symbol;Acc:M | 0 | 0 | 0 | 0 | 0 | 0 |
| 10260 | ENSMUSG00000036144 | 0.267700746 | -1.497481649 | 4.057001436 | -0.369110456 | 0.712045396 | NA | Meox2 | mesenchyme homeobox 2 [Source:MGI Sy | 1.6062 | 0 | 0 | 0 | 0 | 0 |
| 10261 | ENSMUSG00000036151 | 521.8786299 | -0.279198555 | 0.192331954 | -1.451649342 | 0.146599132 | 0.55726934 | Tm6sf2 | transmembrane 6 superfamily member 2 | 644.09 | 485.18 | 587.65 | 606.34 | 343.54 | 464.47 |
| 10262 | ENSMUSG00000036155 | 182.4271871 | -0.124912188 | 0.196279577 | -0.636399312 | 0.524516189 | 0.888334265 | Mgat5 | mannoside acetylglucosaminyltransferase | 182.3 | 187.08 | 201.25 | 146.52 | 165.77 | 211.63 |
| 10263 | ENSMUSG00000036158 | 38.42216919 | -0.108424535 | 0.361834089 | -0.299652625 | 0.764442139 | 0.954980327 | Prickle1 | prickle planar cell polarity protein 1 [Sour | 34.533 | 38.033 | 47.293 | 43.379 | 30.537 | 36.757 |
| 10264 | ENSMUSG00000036160 | 267.4091354 | 0.088689594 | 0.167344122 | 0.529983325 | 0.596123492 | 0.913894721 | Surf6 | surfeit gene 6 [Source:MGI Symbol;Acc:M | 258.6 | 282.68 | 236.47 | 278.59 | 308.64 | 239.48 |
| 10265 | ENSMUSG00000036162 | 0 | NA | NA | NA | NA | NA | Fam219aos | family with sequence similarity 219, meml | 0 | 0 | 0 | 0 | 0 | 0 |
| 10266 | ENSMUSG00000036167 | 321.8664741 | -0.017800866 | 0.160470372 | -0.110929299 | 0.911672412 | 0.98589875 | Pphln1 | periphilin 1 [Source:MGI Symbol;Acc:MGI | 343.73 | 342.3 | 284.77 | 279.55 | 352.27 | 328.58 |
| 10267 | ENSMUSG00000036168 | 6.586441574 | 0.553389156 | 0.833253006 | 0.664131005 | 0.506606467 | 0.881669588 | Ccdc38 | coiled-coil domain containing 38 [Source:I | 4.8186 | 5.1396 | 6.0375 | 4.8199 | 10.906 | 7.7969 |
| 10268 | ENSMUSG00000036169 | 0 | NA | NA | NA | NA | NA | Sostdc1 | sclerostin domain containing 1 [Source:M | 0 | 0 | 0 | 0 | 0 | 0 |
| 10269 | ENSMUSG00000036172 | 1.688270857 | -2.021375987 | 2.131042748 | -0.948538451 | 0.342855408 | NA | Cd200r3 | CD200 receptor 3 [Source:MGI Symbol;Ac | 0 | 7.1954 | 1.0062 | 1.928 | 0 | 0 |
| 10270 | ENSMUSG00000036180 | 761.2361958 | -0.03641445 | 0.12030743 | -0.302678316 | 0.762135025 | 0.954414324 | Gatad2a | GATA zinc finger domain containing 2A [S | 764.55 | 817.19 | 731.54 | 822.27 | 705.63 | 726.23 |
| 10271 | ENSMUSG00000036181 | 1654.241681 | -0.277819027 | 0.393885691 | -0.705329066 | 0.480605476 | 0.870278888 | H1f2 | H1.2 linker histone, cluster member [Sour | 1425.5 | 1182.1 | 2831.6 | 1376.6 | 1628.3 | 1481.4 |
| 10272 | ENSMUSG00000036185 | 1.809216456 | 0.415211641 | 1.519791261 | 0.273203072 | 0.784697119 | NA | Sapcd1 | suppressor APC domain containing 1 [Sou | 1.6062 | 2.0558 | 1.0062 | 2.8919 | 2.1812 | 1.1138 |
| 10273 | ENSMUSG00000036186 | 2.050052996 | 2.500818252 | 1.660010363 | 1.506507615 | 0.131936914 | NA | Dipk1b | divergent protein kinase domain 1B [Sour | 0.8031 | 1.0279 | 0 | 3.8559 | 3.2719 | 3.3415 |
| 10274 | ENSMUSG00000036188 | 168.3071498 | -0.025227866 | 0.188142005 | -0.134089491 | 0.89333181 | 0.982562372 | Ankmy2 | ankyrin repeat and MYND domain contair | 179.89 | 146.99 | 182.13 | 166.77 | 161.41 | 172.65 |
| 10275 | ENSMUSG00000036192 | 0 | NA | NA | NA | NA | NA | Rorb | RAR-related orphan receptor beta [Source | 0 | 0 | 0 | 0 | 0 | 0 |
| 10276 | ENSMUSG00000036196 | 3.512837476 | -0.609779514 | 1.156848124 | -0.527104208 | 0.598121225 | NA | Slc26a8 | solute carrier family 26, member 8 [Sourc | 5.6217 | 2.0558 | 5.0312 | 2.8919 | 4.3625 | 1.1138 |
| 10277 | ENSMUSG00000036197 | 290.4030842 | 0.119446428 | 0.156052642 | 0.76542394 | 0.444019142 | 0.851432536 | Gxylt1 | glucoside xylosyltransferase 1 [Source:MG | 254.58 | 266.23 | 314.95 | 310.4 | 297.74 | 298.51 |
| 10278 | ENSMUSG00000036198 | 0.305170013 | -1.657836477 | 4.041480335 | -0.410205257 | 0.681655385 | NA | Arhgap36 | Rho GTPase activating protein 36 [Source: | 0.8031 | 1.0279 | 0 | 0 | 0 | 0 |
| 10279 | ENSMUSG00000036199 | 2142.99725 | 0.070410677 | 0.099765781 | 0.705759794 | 0.480337529 | 0.870278888 | Ndufa13 | NADH:ubiquinone oxidoreductase subuni | 2170.8 | 2031.2 | 2068.8 | 2033 | 2212.9 | 2341.3 |
| 10280 | ENSMUSG00000036202 | 224.3528241 | -0.083651834 | 0.171118811 | -0.488852355 | 0.624946229 | 0.919503423 | Rif1 | replication timing regulatory factor 1 [Sou | 211.22 | 242.59 | 239.49 | 233.28 | 226.85 | 192.69 |
| 10281 | ENSMUSG00000036206 | 31.99900201 | 0.463535471 | 0.390861369 | 1.18593319 | 0.235648685 | 0.683802256 | Sh3bp4 | SH3-domain binding protein 4 [Source:MC | 24.896 | 27.754 | 28.175 | 40.487 | 28.356 | 42.326 |
| 10282 | ENSMUSG00000036208 | 97.94788721 | 0.212664619 | 0.252811269 | 0.841199129 | 0.400236388 | 0.827239085 | Nepro | nucleolus and neural progenitor protein [! | 96.372 | 87.373 | 88.55 | 118.57 | 119.97 | 76.855 |
| 10283 | ENSMUSG00000036211 | 0.62992821 | -1.312600165 | 2.851194982 | -0.460368433 | 0.645251789 | NA | H1f6 | H1.6 linker histone, cluster member [Sour | 0.8031 | 0 | 2.0125 | 0.964 | 0 | 0 |
| 10284 | ENSMUSG00000036214 | 182.0472247 | 0.02275506 | 0.179775044 | 0.126575189 | 0.899276636 | 0.982798755 | Polr1has | RNA polymerase I subunit H, antisense [Sc | 194.35 | 178.86 | 168.04 | 169.66 | 188.68 | 192.69 |
| 10285 | ENSMUSG00000036216 | 1207.237604 | -0.671067622 | 0.148772184 | -4.510706272 | 6.46E-06 | 0.000384588 | Leap2 | liver-expressed antimicrobial peptide 2 [S | 1489.8 | 1515.2 | 1443 | 704.67 | 1036.1 | 1054.8 |
| 10286 | ENSMUSG00000036218 | 0 | NA | NA | NA | NA | NA | Pdzrn4 | PDZ domain containing RING finger 4 [Sou | 0 | 0 | 0 | 0 | 0 | 0 |
| 10287 | ENSMUSG00000036223 | 1.457904383 | -0.085543281 | 1.777067292 | -0.048137333 | 0.961606793 | NA | Ska1 | spindle and kinetochore associated compl | 2.4093 | 2.0558 | 0 | 0.964 | 1.0906 | 2.2277 |
| 10288 | ENSMUSG00000036225 | 18.40650829 | 0.247803612 | 0.517278223 | 0.479052859 | 0.631901025 | 0.921724125 | Kctd1 | potassium channel tetramerisation domai | 16.865 | 20.558 | 13.081 | 19.28 | 26.175 | 14.48 |
| 10289 | ENSMUSG00000036231 | 0 | NA | NA | NA | NA | NA | Agr3 | anterior gradient 3 [Source:MGI Symbol;A | 0 | 0 | 0 | 0 | 0 | 0 |
| 10290 | ENSMUSG00000036241 | 2168.637214 | 0.049986196 | 0.102298207 | 0.48863218 | 0.625102126 | 0.919503423 | Ube2r2 | ubiquitin-conjugating enzyme E2R 2 [Sour | 1966.8 | 2093.9 | 2333.5 | 2185.3 | 2180.1 | 2252.2 |
| 10291 | ENSMUSG00000036242 | 3.464503538 | -0.5659049 | 1.314542899 | -0.430495575 | 0.666835185 | NA | Armh4 | armadillo-like helical domain containing 4 | 3.2124 | 8.2233 | 1.0062 | 2.8919 | 5.4531 | 0 |
| 10292 | ENSMUSG00000036244 | 0.321325949 | 1.911575114 | 4.048267893 | 0.472195804 | 0.636787034 | NA | Tbc1d21 | TBC1 domain family, member 21 [Source:I | 0 | 0 | 0 | 1.928 | 0 | 0 |
| 10293 | ENSMUSG00000036246 | 83.28433282 | 0.069258944 | 0.273760187 | 0.252991293 | 0.800274941 | 0.962424862 | Gmip | Gem-interacting protein [Source:MGI Sym | 75.492 | 74.01 | 94.587 | 95.434 | 62.165 | 98.018 |
| 10294 | ENSMUSG00000036249 | 110.8606584 | 0.052094237 | 0.23715597 | 0.219662348 | 0.82613413 | 0.96948417 | Rbm43 | RNA binding motif protein 43 [Source:MG | 95.569 | 131.57 | 99.618 | 98.326 | 128.69 | 111.38 |
| 10295 | ENSMUSG00000036251 | 0.959817171 | -2.042882424 | 2.2694873 | -0.900151512 | 0.368039626 | NA | Trpm8 | transient receptor potential cation channe | 1.6062 | 2.0558 | 1.0062 | 0 | 1.0906 | 0 |
| 10296 | ENSMUSG00000036256 | 1050.870332 | -0.963426906 | 0.172345797 | -5.590080656 | 2.27E-08 | 3.13E-06 | Igfbp7 | insulin-like growth factor binding protein | 1656 | 1263.3 | 1247.7 | 723.95 | 539.86 | 874.37 |
| 10297 | ENSMUSG00000036257 | 2957.838287 | 0.143366778 | 0.097993151 | 1.463028558 | 0.143459565 | 0.553333105 | Pnpla8 | patatin-like phospholipase domain contai | 2842.2 | 2936.8 | 2653.5 | 2929.5 | 3322 | 3063.1 |
| 10298 | ENSMUSG00000036264 | 0.160662974 | 1.126924175 | 4.080472857 | 0.276174898 | 0.782413736 | NA | Fstl4 | follistatin-like 4 [Source:MGI Symbol;Acc:I | 0 | 0 | 0 | 0.964 | 0 | 0 |
| 10299 | ENSMUSG00000036270 | 371.4713701 | -0.004478161 | 0.147709255 | -0.0303174 | 0.97581392 | 0.996622832 | Edc4 | enhancer of mRNA decapping 4 [Source:N | 368.62 | 359.77 | 388.41 | 423.19 | 343.54 | 345.29 |
| 10300 | ENSMUSG00000036273 | 25.11984698 | -0.664320327 | 0.439801488 | -1.510500408 | 0.130915785 | 0.530113639 | Lrrk2 | leucine-rich repeat kinase 2 [Source:MGI ! | 27.305 | 31.865 | 33.206 | 16.388 | 17.45 | 24.505 |

| | | | | | | | | | | | | | | | |
|---|---|---|---|---|---|---|---|---|---|---|---|---|---|---|---|
| 10301 | ENSMUSG00000036275 | 2249.93963 | 0.22232571 | 0.098772731 | 2.250881464 | 0.024393046 | 0.19944659 | 9530068E07 | RIKEN cDNA 9530068E07 gene [Source:M | 2056.7 | 1975.7 | 2198.6 | 2499.6 | 2271.8 | 2497.2 |
| 10302 | ENSMUSG00000036278 | 1324.926559 | -0.073794688 | 0.111746528 | -0.660375668 | 0.509012782 | 0.882793133 | Macrod1 | mono-ADP ribosylhydrolase 1 [Source:MG | 1288.2 | 1295.2 | 1493.3 | 1229.1 | 1378.5 | 1265.3 |
| 10303 | ENSMUSG00000036281 | 130.5662923 | 0.204445473 | 0.2121677 | 0.963603194 | 0.335244905 | 0.776458335 | Snapc4 | small nuclear RNA activating complex, pol | 129.3 | 105.88 | 128.8 | 156.16 | 138.51 | 124.75 |
| 10304 | ENSMUSG00000036282 | 559.7369821 | -0.012436732 | 0.127808873 | -0.097307265 | 0.922482387 | 0.986685804 | Naa30 | N(alpha)-acetyltransferase 30, NatC cataly | 526.03 | 596.19 | 564.5 | 520.55 | 602.02 | 549.13 |
| 10305 | ENSMUSG00000036285 | 283.9940919 | -0.047468946 | 0.176927408 | -0.268296171 | 0.788471354 | 0.960872359 | Noa1 | nitric oxide associated 1 [Source:MGI Sym | 318.03 | 259.04 | 288.79 | 305.58 | 308.64 | 223.88 |
| 10306 | ENSMUSG00000036291 | 390.29389 | 0.008063718 | 0.143279493 | 0.056279641 | 0.955119037 | 0.991678478 | Ap5m1 | adaptor-related protein complex 5, mu 1 s | 363 | 402.94 | 402.5 | 413.55 | 412.25 | 347.52 |
| 10307 | ENSMUSG00000036292 | 771.0336336 | 0.171830352 | 0.122662231 | 1.40084157 | 0.161261455 | 0.58429 | Gramd1c | GRAM domain containing 1C [Source:MGI | 783.83 | 655.81 | 734.56 | 760.58 | 860.5 | 830.93 |
| 10308 | ENSMUSG00000036295 | 13.82381798 | -0.972994897 | 0.655521634 | -1.48430631 | 0.137727677 | 0.541918695 | Lrrn3 | leucine rich repeat protein 3, neuronal [So | 20.078 | 21.586 | 13.081 | 3.8559 | 7.6343 | 16.708 |
| 10309 | ENSMUSG00000036298 | 3.886496122 | -0.617154328 | 1.147234169 | -0.537949745 | 0.590611744 | NA | Slc2a13 | solute carrier family 2 (facilitated glucose | 8.031 | 1.0279 | 5.0312 | 4.8199 | 2.1812 | 2.2277 |
| 10310 | ENSMUSG00000036299 | 1028.821499 | 0.087821239 | 0.112310763 | 0.78194856 | 0.434244805 | 0.845264432 | BC031181 | cDNA sequence BC031181 [Source:MGI Sy | 1046.4 | 948.77 | 996.18 | 994.83 | 1035 | 1151.7 |
| 10311 | ENSMUSG00000036304 | 9.296608566 | 0.255325606 | 0.819047827 | 0.311734672 | 0.755242175 | 0.953085143 | Zdhhc23 | zinc finger, DHHC domain containing 23 [S | 4.0155 | 16.447 | 5.0312 | 3.8559 | 14.178 | 12.252 |
| 10312 | ENSMUSG00000036305 | 3.009690207 | 0.501921708 | 1.21366154 | 0.413559869 | 0.679196468 | NA | Rpl39-ps | ribosomal protein L39, pseudogene [Sour | 2.4093 | 1.0279 | 4.025 | 2.8919 | 4.3625 | 3.3415 |
| 10313 | ENSMUSG00000036306 | 7.588445297 | -0.113341464 | 0.900057458 | -0.12592692 | 0.899789773 | 0.982798755 | Lzts1 | leucine zipper, putative tumor suppressor | 7.2279 | 11.307 | 5.0312 | 0.964 | 7.6343 | 13.366 |
| 10314 | ENSMUSG00000036309 | 3964.738356 | -0.156401775 | 0.087752428 | -1.782307087 | 0.074699168 | 0.399278921 | Skp1 | S-phase kinase-associated protein 1 [Sour | 4366.5 | 4077.8 | 4093.4 | 3774 | 3836.8 | 3640 |
| 10315 | ENSMUSG00000036315 | 144.5013926 | 0.042648575 | 0.209041269 | 0.204019881 | 0.838337963 | 0.971223091 | Polr1h | RNA polymerase I subunit H [Source:MGI | 131.71 | 132.6 | 163.01 | 149.42 | 125.42 | 164.85 |
| 10316 | ENSMUSG00000036322 | 0 | NA | NA | NA | NA | NA | H2-Ea | histocompatibility 2, class II antigen E alph | 0 | 0 | 0 | 0 | 0 | 0 |
| 10317 | ENSMUSG00000036323 | 1683.199916 | 0.217375479 | 0.094962868 | 2.289057645 | 0.022076003 | 0.185535916 | Srp72 | signal recognition particle 72 [Source:MGI | 1490.6 | 1599.4 | 1580.8 | 1830.6 | 1826.8 | 1771 |
| 10318 | ENSMUSG00000036327 | 145.4316424 | -0.317931504 | 0.21015433 | -1.51284774 | 0.130318339 | 0.528582293 | Qsox2 | quiescin Q6 sulfhydryl oxidase 2 [Source:M | 136.53 | 193.25 | 154.96 | 132.06 | 123.24 | 132.55 |
| 10319 | ENSMUSG00000036330 | 78.73058047 | -0.330428829 | 0.263451945 | -1.254228086 | 0.209759114 | 0.653588071 | Slc18a1 | solute carrier family 18 (vesicular monoan | 90.751 | 98.68 | 73.456 | 60.731 | 75.253 | 73.514 |
| 10320 | ENSMUSG00000036333 | 1030.284971 | -0.054865745 | 0.119331306 | -0.459776622 | 0.645676565 | 0.925643431 | Kidins220 | kinase D-interacting substrate 220 [Source | 955.69 | 1166.7 | 1028.4 | 1020.9 | 1064.4 | 945.65 |
| 10321 | ENSMUSG00000036334 | 14.26833278 | -1.230945563 | 0.663073919 | -1.856422832 | 0.063393313 | 0.360000344 | Igsf10 | immunoglobulin superfamily, member 10 | 31.321 | 12.335 | 16.1 | 3.8559 | 13.087 | 8.9108 |
| 10322 | ENSMUSG00000036339 | 523.897356 | 0.151288615 | 0.13080252 | 1.156618504 | 0.247428259 | 0.697758303 | Tmem260 | transmembrane protein 260 [Source:MGI | 447.33 | 495.46 | 547.4 | 545.61 | 555.12 | 552.47 |
| 10323 | ENSMUSG00000036352 | 382.4414793 | 0.220343752 | 0.140662378 | 1.566472536 | 0.117238033 | 0.502643331 | Ubac1 | ubiquitin associated domain containing 1 | 327.67 | 386.5 | 346.15 | 391.38 | 428.61 | 414.35 |
| 10324 | ENSMUSG00000036353 | 13.28501639 | -1.53794555 | 0.724768122 | -2.121982883 | 0.033839175 | 0.246290937 | P2ry12 | purinergic receptor P2Y, G-protein couple | 11.243 | 33.921 | 14.087 | 2.8919 | 11.997 | 5.5692 |
| 10325 | ENSMUSG00000036356 | 9.414056423 | -0.373697128 | 0.732006192 | -0.510510884 | 0.609693591 | 0.916376001 | Csgalnact1 | chondroitin sulfate N-acetylgalactosaminy | 6.4248 | 16.447 | 9.0562 | 5.7839 | 7.6343 | 11.138 |
| 10326 | ENSMUSG00000036357 | 0 | NA | NA | NA | NA | NA | Gpr101 | G protein-coupled receptor 101 [Source:N | 0 | 0 | 0 | 0 | 0 | 0 |
| 10327 | ENSMUSG00000036362 | 15.98264938 | -2.029244388 | 0.674051517 | -3.010518244 | 0.002608023 | 0.042624228 | P2ry13 | purinergic receptor P2Y, G-protein couple | 24.896 | 41.117 | 11.069 | 6.7478 | 8.7249 | 3.3415 |
| 10328 | ENSMUSG00000036368 | 893.6002396 | 0.161067718 | 0.11080348 | 1.453634105 | 0.146047776 | 0.556855766 | Rmdn2 | regulator of microtubule dynamics 2 [Sou | 814.35 | 873.73 | 843.23 | 888.79 | 937.93 | 1003.6 |
| 10329 | ENSMUSG00000036371 | 3365.21025 | 0.029053569 | 0.105910576 | 0.274321702 | 0.783837408 | 0.960670212 | Serbp1 | serpine1 mRNA binding protein 1 [Source | 3069.5 | 3288.3 | 3636.6 | 3132.9 | 3608.9 | 3455.1 |
| 10330 | ENSMUSG00000036372 | 390.8456157 | 0.24259559 | 0.154420752 | 1.571003816 | 0.116181762 | 0.500730933 | Tmem258 | transmembrane protein 258 [Source:MGI | 386.29 | 338.18 | 349.17 | 434.75 | 357.72 | 478.95 |
| 10331 | ENSMUSG00000036376 | 233.9440804 | -0.049897907 | 0.167253169 | -0.298337591 | 0.765445517 | 0.954980327 | Abt1 | activator of basal transcription 1 [Source:I | 260.21 | 215.86 | 237.47 | 239.07 | 211.58 | 239.48 |
| 10332 | ENSMUSG00000036377 | 0.784885323 | -1.683590003 | 2.624742731 | -0.641430485 | 0.521243032 | NA | Cracd | capping protein inhibiting regulator of act | 1.6062 | 0 | 2.0125 | 0 | 1.0906 | 0 |
| 10333 | ENSMUSG00000036381 | 10.76456039 | -1.251215294 | 0.751971214 | -1.663913817 | 0.096129653 | 0.457754528 | P2ry14 | purinergic receptor P2Y, G-protein couple | 20.881 | 14.391 | 10.062 | 4.8199 | 2.1812 | 12.252 |
| 10334 | ENSMUSG00000036390 | 106.2360198 | 1.399058658 | 0.322136117 | 4.343066745 | 1.41E-05 | 0.000708118 | Gadd45a | growth arrest and DNA-damage-inducible | 85.129 | 27.754 | 61.381 | 155.2 | 131.96 | 175.99 |
| 10335 | ENSMUSG00000036391 | 2688.090453 | -0.047653306 | 0.10859553 | -0.438814614 | 0.66079587 | 0.928184201 | Sec24a | Sec24 related gene family, member A (S. c | 2819.7 | 2745.6 | 2632.3 | 2766.6 | 2850.9 | 2313.5 |
| 10336 | ENSMUSG00000036395 | 0.160662974 | 1.126924175 | 4.080472857 | 0.276174898 | 0.782413736 | NA | Glb1l2 | galactosidase, beta 1-like 2 [Source:MGI S | 0 | 0 | 0 | 0.964 | 0 | 0 |
| 10337 | ENSMUSG00000036398 | 438.1549537 | 0.157848986 | 0.149370608 | 1.056760678 | 0.290620824 | 0.740463951 | Ppp1r11 | protein phosphatase 1, regulatory inhibito | 451.34 | 386.5 | 403.5 | 393.3 | 488.6 | 505.69 |
| 10338 | ENSMUSG00000036401 | 0 | NA | NA | NA | NA | NA | Glt6d1 | glycosyltransferase 6 domain containing 1 | 0 | 0 | 0 | 0 | 0 | 0 |
| 10339 | ENSMUSG00000036402 | 1015.577273 | 0.004824265 | 0.103709309 | 0.046517182 | 0.96289804 | 0.992956886 | Gng12 | guanine nucleotide binding protein (G pro | 986.21 | 1061.8 | 994.17 | 1034.3 | 1006.6 | 1010.3 |
| 10340 | ENSMUSG00000036403 | 72.78567688 | -0.248473926 | 0.282228397 | -0.880400163 | 0.378642568 | 0.811969331 | Cep135 | centrosomal protein 135 [Source:MGI Syn | 77.901 | 78.122 | 81.506 | 86.758 | 55.622 | 56.806 |
| 10341 | ENSMUSG00000036411 | 5.089649498 | -0.53816246 | 1.073600733 | -0.501268715 | 0.61618202 | 0.917509899 | 9530077C05 | RIKEN cDNA 9530077C05 gene [Source:M | 12.85 | 4.1117 | 1.0062 | 4.8199 | 2.1812 | 5.5692 |
| 10342 | ENSMUSG00000036412 | 0 | NA | NA | NA | NA | NA | Arsi | arylsulfatase i [Source:MGI Symbol;Acc:M | 0 | 0 | 0 | 0 | 0 | 0 |
| 10343 | ENSMUSG00000036422 | 0 | NA | NA | NA | NA | NA | Pcdh8 | protocadherin 8 [Source:MGI Symbol;Acc | 0 | 0 | 0 | 0 | 0 | 0 |
| 10344 | ENSMUSG00000036427 | 2697.775197 | -0.225006048 | 0.105527413 | -2.132204726 | 0.03299002 | 0.242957052 | Gpi1 | glucose-6-phosphate isomerase 1 [Source | 3198 | 2819.6 | 2704.8 | 2601.8 | 2324.1 | 2538.5 |
| 10345 | ENSMUSG00000036430 | 194.3815366 | -0.132182954 | 0.185394255 | -0.712983006 | 0.475856273 | 0.86947987 | Tbcc | tubulin-specific chaperone C [Source:MGI | 219.25 | 171.66 | 218.36 | 175.44 | 178.86 | 202.72 |
| 10346 | ENSMUSG00000036432 | 293.8882186 | -0.200783058 | 0.200919818 | -0.999319331 | 0.317640024 | 0.764257918 | Siah2 | siah E3 ubiquitin protein ligase 2 [Source:I | 263.42 | 313.51 | 366.27 | 236.17 | 352.27 | 231.68 |
| 10347 | ENSMUSG00000036435 | 198.812113 | -0.010040296 | 0.183228694 | -0.054796527 | 0.956300568 | 0.991678478 | Exoc1 | exocyst complex component 1 [Source:M | 167.85 | 216.89 | 214.33 | 186.05 | 205.04 | 202.72 |
| 10348 | ENSMUSG00000036437 | 2.703878479 | -0.722810538 | 1.364929442 | -0.529558903 | 0.596417795 | NA | Npy1r | neuropeptide Y receptor Y1 [Source:MGI S | 4.0155 | 4.1117 | 2.0125 | 3.8559 | 0 | 2.2277 |
| 10349 | ENSMUSG00000036438 | 1341.358049 | 0.053809546 | 0.137999515 | 0.389925617 | 0.696591551 | 0.939992732 | Calm2 | calmodulin 2 [Source:MGI Symbol;Acc:MG | 1289 | 1378.4 | 1280.9 | 1158.7 | 1282.6 | 1658.5 |
| 10350 | ENSMUSG00000036442 | 355.8619709 | 0.085205808 | 0.141386271 | 0.602645557 | 0.546744507 | 0.89549457 | Thap11 | THAP domain containing 11 [Source:MGI S | 325.26 | 369.02 | 342.12 | 351.85 | 388.26 | 358.66 |
| 10351 | ENSMUSG00000036446 | 97.73137813 | -1.090644593 | 0.258162735 | -4.224639904 | 2.39E-05 | 0.001123886 | Lum | lumican [Source:MGI Symbol;Acc:MGI:10 | 158.21 | 102.79 | 137.86 | 68.442 | 56.712 | 62.375 |
| 10352 | ENSMUSG00000036449 | 6.218286645 | -0.472640635 | 0.855848884 | -0.552247768 | 0.580778611 | 0.908645447 | Lcn8 | lipocalin 8 [Source:MGI Symbol;Acc:MGI:2 | 10.44 | 5.1396 | 6.0375 | 5.7839 | 5.4531 | 4.4554 |
| 10353 | ENSMUSG00000036450 | 540.3260391 | 0.115645896 | 0.13629743 | 0.848481854 | 0.396169674 | 0.823764863 | Hif1an | hypoxia-inducible factor 1, alpha subunit i | 553.34 | 499.57 | 502.12 | 504.16 | 633.65 | 549.13 |
| 10354 | ENSMUSG00000036452 | 552.1324705 | 0.846154432 | 0.148937529 | 5.681270765 | 1.34E-08 | 1.97E-06 | Arhgap26 | Rho GTPase activating protein 26 [Source: | 431.27 | 329.96 | 421.62 | 629.48 | 726.35 | 774.12 |
| 10355 | ENSMUSG00000036459 | 29.94264134 | -0.297691756 | 0.471207077 | -0.631764187 | 0.527540978 | 0.888334265 | Wtip | WT1-interacting protein [Source:MGI Sym | 28.109 | 31.865 | 39.244 | 32.775 | 10.906 | 36.757 |
| 10356 | ENSMUSG00000036461 | 279.5580251 | -0.09605517 | 0.160423018 | -0.598761766 | 0.549331762 | 0.895703112 | Elf1 | E74-like factor 1 [Source:MGI Symbol;Acc | 279.48 | 322.77 | 264.64 | 270.88 | 291.2 | 248.39 |

| | | | | | | | | | | | | | | | |
|---|---|---|---|---|---|---|---|---|---|---|---|---|---|---|---|
| 10357 | ENSMUSG00000036463 | 0 | NA | NA | NA | NA | NA | 4930544G11 | RIKEN cDNA 4930544G11 gene [Source:M | 0 | 0 | 0 | 0 | 0 | 0 |
| 10358 | ENSMUSG00000036466 | 7.145667977 | -0.986793859 | 0.953589099 | -1.034820826 | 0.300752595 | 0.751875664 | Megf11 | multiple EGF-like-domains 11 [Source:MG | 15.259 | 7.1954 | 6.0375 | 6.7478 | 7.6343 | 0 |
| 10359 | ENSMUSG00000036469 | 21.44317244 | -1.87910897 | 0.624336262 | -3.009770669 | 0.00261445 | 0.042638653 | Marchf1 | membrane associated ring-CH-type finger | 30.518 | 56.535 | 14.087 | 8.6758 | 4.3625 | 14.48 |
| 10360 | ENSMUSG00000036473 | 473.3236154 | -0.061338694 | 0.141854236 | -0.432406503 | 0.665445993 | 0.929426323 | Tbc1d24 | TBC1 domain family, member 24 [Source: | 538.88 | 471.81 | 438.72 | 465.6 | 502.78 | 422.15 |
| 10361 | ENSMUSG00000036478 | 899.4292694 | -1.008761734 | 0.17326995 | -5.821908145 | 5.82E-09 | 9.63E-07 | Btg1 | BTG anti-proliferation factor 1 [Source:M( | 859.32 | 1274.6 | 1472.1 | 621.77 | 560.58 | 608.16 |
| 10362 | ENSMUSG00000036480 | 0 | NA | NA | NA | NA | NA | Prss56 | protease, serine 56 [Source:MGI Symbol;A | 0 | 0 | 0 | 0 | 0 | 0 |
| 10363 | ENSMUSG00000036492 | 128.6396865 | 0.025683026 | 0.213602243 | 0.120237625 | 0.904294914 | 0.98383695 | Rnf39 | ring finger protein 39 [Source:MGI Symbo | 119.66 | 112.04 | 150.94 | 130.14 | 126.51 | 132.55 |
| 10364 | ENSMUSG00000036499 | 689.7040182 | 0.219849208 | 0.137443687 | 1.59955843 | 0.109696577 | 0.486553599 | Eea1 | early endosome antigen 1 [Source:MGI Sy | 553.34 | 697.96 | 662.11 | 811.67 | 747.07 | 666.08 |
| 10365 | ENSMUSG00000036500 | 0 | NA | NA | NA | NA | NA | Akp3 | alkaline phosphatase 3, intestine, not Mn | 0 | 0 | 0 | 0 | 0 | 0 |
| 10366 | ENSMUSG00000036501 | 237.2476934 | -0.35821337 | 0.222423972 | -1.610498031 | 0.107289175 | 0.48315751 | Fam13b | family with sequence similarity 13, memb | 237.72 | 253.9 | 308.92 | 271.84 | 145.05 | 206.06 |
| 10367 | ENSMUSG00000036502 | 2.405237129 | -1.233999931 | 1.552071828 | -0.795066252 | 0.426574965 | NA | Tmem255a | transmembrane protein 255A [Source:MG | 0 | 4.1117 | 6.0375 | 0.964 | 1.0906 | 2.2277 |
| 10368 | ENSMUSG00000036503 | 3093.941213 | 0.07486314 | 0.096722433 | 0.77399976 | 0.438930933 | 0.848057363 | Rnf13 | ring finger protein 13 [Source:MGI Symbo | 3026.1 | 3100.2 | 2914.1 | 2945 | 3401.6 | 3176.7 |
| 10369 | ENSMUSG00000036504 | 5.682968013 | -0.991927021 | 1.02129323 | -0.971246056 | 0.331425764 | 0.774747658 | Phpt1 | phosphohistidine phosphatase 1 [Source: | 2.4093 | 14.391 | 6.0375 | 5.7839 | 4.3625 | 1.1138 |
| 10370 | ENSMUSG00000036510 | 0 | NA | NA | NA | NA | NA | Cdh8 | cadherin 8 [Source:MGI Symbol;Acc:MGI: | 0 | 0 | 0 | 0 | 0 | 0 |
| 10371 | ENSMUSG00000036513 | 260.1155071 | 0.066493793 | 0.190586747 | 0.34888991 | 0.72717196 | 0.945907265 | Commd2 | COMM domain containing 2 [Source:MGI | 271.45 | 242.59 | 247.54 | 209.18 | 332.64 | 257.3 |
| 10372 | ENSMUSG00000036521 | 0 | NA | NA | NA | NA | NA | Scgb2b2 | secretoglobin, family 2B, member 2 [Sour | 0 | 0 | 0 | 0 | 0 | 0 |
| 10373 | ENSMUSG00000036523 | 3.569319831 | -0.867536977 | 1.275880097 | -0.679951807 | 0.496534977 | NA | Greb1 | gene regulated by estrogen in breast canc | 5.6217 | 4.1117 | 4.025 | 0 | 6.5437 | 1.1138 |
| 10374 | ENSMUSG00000036526 | 5.381584128 | -2.337725822 | 1.162203941 | -2.011459211 | 0.044276975 | 0.290968723 | Card11 | caspase recruitment domain family, meml | 6.4248 | 18.503 | 2.0125 | 0.964 | 3.2719 | 1.1138 |
| 10375 | ENSMUSG00000036528 | 453.4722575 | 0.098081559 | 0.132939697 | 0.737789848 | 0.460642164 | 0.859817816 | Ppfibp2 | PTPRF interacting protein, binding protein | 456.97 | 400.89 | 455.83 | 466.57 | 447.15 | 493.43 |
| 10376 | ENSMUSG00000036529 | 822.1163886 | 0.0805863 | 0.115334706 | 0.69871683 | 0.484729014 | 0.87256358 | Sbf1 | SET binding factor 1 [Source:MGI Symbol; | 766.96 | 778.13 | 853.3 | 904.21 | 811.42 | 818.68 |
| 10377 | ENSMUSG00000036533 | 15.85958353 | -0.653261578 | 0.554096778 | -1.17896657 | 0.238411488 | 0.687339796 | Cdc42ep3 | CDC42 effector protein (Rho GTPase bindi | 22.487 | 15.419 | 20.125 | 10.604 | 9.8156 | 16.708 |
| 10378 | ENSMUSG00000036534 | 467.7622051 | 0.206901653 | 0.150745141 | 1.372526182 | 0.169899694 | 0.596876778 | Slc38a7 | solute carrier family 38, member 7 [Sourc | 504.35 | 418.36 | 379.35 | 547.54 | 462.42 | 494.55 |
| 10379 | ENSMUSG00000036537 | 30.61852034 | 0.24357925 | 0.416771404 | 0.58444329 | 0.558922109 | 0.899349136 | Rnf113a1 | ring finger protein 113A1 [Source:MGI Syr | 22.487 | 22.614 | 39.244 | 37.595 | 28.356 | 33.415 |
| 10380 | ENSMUSG00000036545 | 85.45054881 | -1.41765555 | 0.356283895 | -3.979005419 | 6.92E-05 | 0.002683563 | Adamts2 | a disintegrin-like and metallopeptidase (re | 160.62 | 111.02 | 101.63 | 67.478 | 20.722 | 51.237 |
| 10381 | ENSMUSG00000036550 | 1571.343988 | 0.090005028 | 0.100139031 | 0.898800666 | 0.368758845 | 0.806506847 | Cnot1 | CCR4-NOT transcription complex, subunit | 1428.7 | 1547 | 1591.9 | 1563.6 | 1626.1 | 1670.8 |
| 10382 | ENSMUSG00000036551 | 0.267700746 | -1.497481649 | 4.057001436 | -0.369110456 | 0.712045396 | NA | Akap14 | A kinase (PRKA) anchor protein 14 [Sourc | 1.6062 | 0 | 0 | 0 | 0 | 0 |
| 10383 | ENSMUSG00000036552 | 143.6748542 | -0.097794648 | 0.200112222 | -0.488699026 | 0.625054794 | 0.919503423 | Ermard | ER membrane associated RNA degradatio | 146.97 | 143.91 | 154.96 | 146.52 | 150.51 | 119.18 |
| 10384 | ENSMUSG00000036553 | 68.74824423 | -0.536978246 | 0.306118932 | -1.754149088 | 0.079404964 | 0.413580628 | Sh3tc1 | SH3 domain and tetratricopeptide repeats | 93.963 | 66.815 | 83.518 | 73.262 | 40.353 | 54.578 |
| 10385 | ENSMUSG00000036555 | 159.1160941 | 0.051653603 | 0.218744992 | 0.236136161 | 0.813327015 | 0.966802988 | Iqce | IQ motif containing E [Source:MGI Symbo | 142.15 | 127.46 | 199.24 | 146.52 | 175.59 | 163.74 |
| 10386 | ENSMUSG00000036557 | 0 | NA | NA | NA | NA | NA | Stpg4 | sperm tail PG rich repeat containing 4 [So | 0 | 0 | 0 | 0 | 0 | 0 |
| 10387 | ENSMUSG00000036560 | 1.816054323 | 0.995151601 | 1.621281033 | 0.613805738 | 0.539343705 | NA | Lgi4 | leucine-rich repeat LGI family, member 4 | 1.6062 | 0 | 2.0125 | 2.8919 | 3.2719 | 1.1138 |
| 10388 | ENSMUSG00000036561 | 649.6625837 | 0.067148154 | 0.127847309 | 0.525221486 | 0.59942923 | 0.915117178 | Ppp6r2 | protein phosphatase 6, regulatory subunit | 582.25 | 617.78 | 704.37 | 675.75 | 620.56 | 697.27 |
| 10389 | ENSMUSG00000036564 | 1.569873498 | -0.891123148 | 1.910581038 | -0.466414735 | 0.640918667 | NA | Ndrg4 | N-myc downstream regulated gene 4 [Sou | 0 | 4.1117 | 2.0125 | 0 | 2.1812 | 1.1138 |
| 10390 | ENSMUSG00000036565 | 120.5042951 | -0.095934726 | 0.210332945 | -0.456108889 | 0.648311679 | 0.925643431 | Ttyh3 | tweety family member 3 [Source:MGI Sym | 124.48 | 125.41 | 123.77 | 124.35 | 103.61 | 121.41 |
| 10391 | ENSMUSG00000036568 | 270.9493292 | 0.178412421 | 0.160401886 | 1.112283813 | 0.266016142 | 0.722063933 | Bicral | BRD4 interacting chromatin remodeling co | 277.87 | 231.28 | 252.57 | 262.2 | 299.92 | 301.85 |
| 10392 | ENSMUSG00000036570 | 3449.928201 | 0.01550466 | 0.08788609 | 0.176417684 | 0.859965813 | 0.97570916 | Fxyd1 | FXYD domain-containing ion transport reg | 3420.4 | 3335.6 | 3538 | 3426 | 3386.4 | 3593.3 |
| 10393 | ENSMUSG00000036572 | 254.586788 | 0.007532543 | 0.16173626 | 0.046572999 | 0.962853552 | 0.992956886 | Upf3b | UPF3 regulator of nonsense transcripts ho | 229.69 | 263.15 | 269.67 | 269.91 | 235.57 | 259.53 |
| 10394 | ENSMUSG00000036574 | 0 | NA | NA | NA | NA | NA | Tex44 | testis expressed 44 [Source:MGI Symbol;A | 0 | 0 | 0 | 0 | 0 | 0 |
| 10395 | ENSMUSG00000036578 | 2.737212575 | -0.640471344 | 1.315141345 | -0.486998106 | 0.626259673 | NA | Fxyd7 | FXYD domain-containing ion transport reg | 4.8186 | 4.1117 | 1.0062 | 0.964 | 2.1812 | 3.3415 |
| 10396 | ENSMUSG00000036580 | 705.7437613 | -0.453328737 | 0.130148514 | -3.483164896 | 0.000495523 | 0.012307964 | Spg20 | spastic paraplegia 20, spartin (Troyer sync | 770.18 | 878.87 | 798.96 | 639.12 | 519.13 | 628.21 |
| 10397 | ENSMUSG00000036585 | 3538.132658 | -0.612905568 | 0.148458795 | -4.128455775 | 3.65E-05 | 0.001645596 | Fgf1 | fibroblast growth factor 1 [Source:MGI Sy | 3360.2 | 4659.6 | 4817.9 | 3141.6 | 2876 | 2373.6 |
| 10398 | ENSMUSG00000036586 | 0 | NA | NA | NA | NA | NA | Grifin | galectin-related inter-fiber protein [Sourc | 0 | 0 | 0 | 0 | 0 | 0 |
| 10399 | ENSMUSG00000036587 | 1.704421279 | -0.452606536 | 1.733726794 | -0.261059896 | 0.794046318 | NA | Fut7 | fucosyltransferase 7 [Source:MGI Symbol; | 0.8031 | 4.1117 | 1.0062 | 0.964 | 0 | 3.3415 |
| 10400 | ENSMUSG00000036591 | 724.6000027 | 0.175825555 | 0.147984651 | 1.188133727 | 0.234780728 | 0.682729836 | Arhgap21 | Rho GTPase activating protein 21 [Source: | 716.37 | 686.65 | 638.97 | 912.89 | 764.52 | 628.21 |
| 10401 | ENSMUSG00000036594 | 239.9978385 | -1.273305069 | 0.249244892 | -5.108650606 | 3.24E-07 | 3.01E-05 | H2-Aa | histocompatibility 2, class II antigen A, alp | 322.04 | 448.17 | 248.54 | 150.38 | 95.974 | 174.87 |
| 10402 | ENSMUSG00000036596 | 12.2068878 | -1.562631456 | 0.76138908 | -2.052342879 | 0.040136352 | 0.274113751 | Cpz | carboxypeptidase Z [Source:MGI Symbol;A | 20.078 | 26.726 | 8.05 | 9.6398 | 7.6343 | 1.1138 |
| 10403 | ENSMUSG00000036598 | 0 | NA | NA | NA | NA | NA | Ccdc113 | coiled-coil domain containing 113 [Source | 0 | 0 | 0 | 0 | 0 | 0 |
| 10404 | ENSMUSG00000036599 | 11.46485179 | 0.409625124 | 0.687675499 | 0.595666305 | 0.551398172 | 0.89589637 | Chst12 | carbohydrate sulfotransferase 12 [Source | 12.047 | 13.363 | 4.025 | 10.604 | 9.8156 | 18.935 |
| 10405 | ENSMUSG00000036602 | 0 | NA | NA | NA | NA | NA | Alx1 | ALX homeobox 1 [Source:MGI Symbol;Acc | 0 | 0 | 0 | 0 | 0 | 0 |
| 10406 | ENSMUSG00000036606 | 3906.450379 | 0.274740437 | 0.100466543 | 2.734646073 | 0.006244738 | 0.077332099 | Plxnb2 | plexin B2 [Source:MGI Symbol;Acc:MGI:2 | 3343.3 | 3417.8 | 3845.9 | 3994.7 | 4353.7 | 4483.2 |
| 10407 | ENSMUSG00000036611 | 381.4227853 | 0.724949733 | 0.236428497 | 3.066253612 | 0.002167594 | 0.036841452 | Eepd1 | endonuclease/exonuclease/phosphatase | 414.4 | 240.53 | 206.28 | 482.95 | 374.08 | 570.29 |
| 10408 | ENSMUSG00000036613 | 174.5019617 | 0.057232042 | 0.216012194 | 0.264948202 | 0.791049365 | 0.961019028 | Eipr1 | EARP complex and GARP complex interact | 194.35 | 144.94 | 173.07 | 148.45 | 223.58 | 162.62 |
| 10409 | ENSMUSG00000036615 | 197.2421052 | 0.293597937 | 0.169320346 | 1.733979079 | 0.082921795 | 0.424559592 | Rfxap | regulatory factor X-associated protein [So | 171.06 | 175.77 | 185.15 | 221.71 | 218.12 | 211.63 |
| 10410 | ENSMUSG00000036617 | 126.3591017 | 0.357926169 | 0.247019356 | 1.448980253 | 0.147343097 | 0.558402858 | Etl4 | enhancer trap locus 4 [Source:MGI Symbo | 109.22 | 83.261 | 139.87 | 154.24 | 114.51 | 157.05 |
| 10411 | ENSMUSG00000036620 | 616.0938776 | -0.035595769 | 0.118430643 | -0.300562152 | 0.763748396 | 0.954980327 | Mgat4b | mannoside acetylglucosaminyltransferase | 600.72 | 658.9 | 611.8 | 602.49 | 591.12 | 631.55 |
| 10412 | ENSMUSG00000036622 | 364.8840055 | 0.022798757 | 0.150425888 | 0.151561391 | 0.879532887 | 0.98128671 | Atp13a2 | ATPase type 13A2 [Source:MGI Symbol;Ac | 338.11 | 394.72 | 354.2 | 414.51 | 341.36 | 346.41 |

| | | | | | | | | | | | | | | | |
|---|---|---|---|---|---|---|---|---|---|---|---|---|---|---|---|
| 10413 | ENSMUSG00000036632 | 672.6401275 | 0.028189909 | 0.118200068 | 0.238493168 | 0.811498614 | 0.966527993 | Alg5 | asparagine-linked glycosylation 5 (dolichy | 685.05 | 677.4 | 635.95 | 726.84 | 645.65 | 664.96 |
| 10414 | ENSMUSG00000036634 | 3.16553133 | -1.818041301 | 1.314589439 | -1.382972696 | 0.166673238 | NA | Mag | myelin-associated glycoprotein [Source:M | 1.6062 | 8.2233 | 5.0312 | 1.928 | 1.0906 | 1.1138 |
| 10415 | ENSMUSG00000036636 | 954.4525277 | 0.017344143 | 0.137703559 | 0.125952757 | 0.899769321 | 0.982798755 | Clcn7 | chloride channel, voltage-sensitive 7 [Sou | 1101.1 | 935.41 | 808.01 | 872.4 | 1024.1 | 985.75 |
| 10416 | ENSMUSG00000036639 | 335.6175942 | -0.006580425 | 0.148277542 | -0.044379108 | 0.964602214 | 0.993480943 | Nudt1 | nudix (nucleoside diphosphate linked moi | 369.43 | 323.79 | 314.95 | 314.26 | 333.73 | 357.54 |
| 10417 | ENSMUSG00000036641 | 24.8588751 | -0.612121594 | 0.474724495 | -1.28942492 | 0.197250402 | 0.638920846 | Ccdc148 | coiled-coil domain containing 148 [Source | 32.927 | 39.061 | 18.112 | 19.28 | 15.269 | 24.505 |
| 10418 | ENSMUSG00000036644 | 1037.155487 | 0.027976073 | 0.105716581 | 0.26463278 | 0.791292366 | 0.961095767 | Tbc1d9b | TBC1 domain family, member 9B [Source: | 966.94 | 1054.6 | 1060.6 | 1059.4 | 1013.2 | 1068.2 |
| 10419 | ENSMUSG00000036646 | 769.2633053 | 0.072623946 | 0.113309802 | 0.640932598 | 0.521566475 | 0.888064228 | Man1b1 | mannosidase, alpha, class 1B, member 1 [ | 702.71 | 777.11 | 770.78 | 797.21 | 809.24 | 758.53 |
| 10420 | ENSMUSG00000036647 | 0 | NA | NA | NA | NA | NA | Olfr6 | olfactory receptor 6 [Source:MGI Symbol; | 0 | 0 | 0 | 0 | 0 | 0 |
| 10421 | ENSMUSG00000036655 | 527.8987465 | -1.109754371 | 0.145320689 | -7.636588958 | 2.23E-14 | 1.29E-11 | Colec11 | collectin sub-family member 11 [Source:N | 819.16 | 688.7 | 656.07 | 347.03 | 296.65 | 359.77 |
| 10422 | ENSMUSG00000036658 | 0 | NA | NA | NA | NA | NA | Olfr11 | olfactory receptor 11 [Source:MGI Symbo | 0 | 0 | 0 | 0 | 0 | 0 |
| 10423 | ENSMUSG00000036661 | 63.48590618 | 0.508482599 | 0.326769934 | 1.556087468 | 0.119687295 | 0.506334317 | Dennd3 | DENN/MADD domain containing 3 [Source | 41.761 | 63.731 | 52.325 | 98.326 | 51.259 | 73.514 |
| 10424 | ENSMUSG00000036667 | 38.05672859 | -0.120790538 | 0.378285803 | -0.319310261 | 0.749491253 | 0.951356173 | Tcaf1 | TRPM8 channel-associated factor 1 [Sourc | 42.564 | 39.061 | 37.231 | 37.595 | 23.994 | 47.895 |
| 10425 | ENSMUSG00000036672 | 25.73518618 | 0.238990022 | 0.493797731 | 0.483983637 | 0.628397482 | 0.920328819 | Cenpt | centromere protein T [Source:MGI Symbo | 27.305 | 13.363 | 30.187 | 40.487 | 17.45 | 25.618 |
| 10426 | ENSMUSG00000036676 | 168.1466084 | -0.136011005 | 0.206960824 | -0.65718237 | 0.51106366 | 0.883679955 | Tmtc3 | transmembrane and tetratricopeptide rep | 159.82 | 189.14 | 180.12 | 195.69 | 152.69 | 131.43 |
| 10427 | ENSMUSG00000036678 | 183.4035284 | -0.049455033 | 0.196142227 | -0.252138633 | 0.80093391 | 0.962424862 | Aaas | achalasia, adrenocortical insufficiency, ala | 203.18 | 164.47 | 191.19 | 153.27 | 176.68 | 211.63 |
| 10428 | ENSMUSG00000036686 | 216.3678387 | -0.010012933 | 0.188890599 | -0.053009165 | 0.957724605 | 0.991861398 | Cc2d1a | coiled-coil and C2 domain containing 1A [ | 216.84 | 212.78 | 222.38 | 265.09 | 200.67 | 180.44 |
| 10429 | ENSMUSG00000036687 | 311.0069461 | 0.240397717 | 0.219812125 | 1.093650851 | 0.274108138 | 0.729244726 | Tmem184a | transmembrane protein 184a [Source:MG | 393.52 | 240.53 | 220.37 | 382.7 | 280.29 | 348.63 |
| 10430 | ENSMUSG00000036693 | 272.4315382 | -0.19121422 | 0.159624248 | -1.197902089 | 0.230955137 | 0.678005905 | Nop14 | NOP14 nucleolar protein [Source:MGI Syn | 289.12 | 324.82 | 257.6 | 258.35 | 255.2 | 249.5 |
| 10431 | ENSMUSG00000036698 | 431.762253 | -0.026808425 | 0.14291495 | -0.187583072 | 0.851203497 | 0.973568625 | Ago2 | argonaute RISC catalytic subunit 2 [Source | 466.6 | 446.12 | 394.45 | 461.75 | 382.81 | 438.85 |
| 10432 | ENSMUSG00000036699 | 0 | NA | NA | NA | NA | NA | Zcchc12 | zinc finger, CCHC domain containing 12 [S | 0 | 0 | 0 | 0 | 0 | 0 |
| 10433 | ENSMUSG00000036707 | 1389.541486 | -0.00322961 | 0.103962254 | -0.031065221 | 0.975217526 | 0.996378391 | Cab39 | calcium binding protein 39 [Source:MGI S | 1309.9 | 1373.3 | 1491.3 | 1444 | 1364.4 | 1354.4 |
| 10434 | ENSMUSG00000036712 | 497.6462888 | 0.134783082 | 0.144153617 | 0.934996182 | 0.349790267 | 0.791144084 | Cyld | CYLD lysine 63 deubiquitinase [Source:MG | 465 | 417.33 | 541.36 | 562.96 | 518.04 | 481.18 |
| 10435 | ENSMUSG00000036718 | 19.98433165 | 0.449489091 | 0.537622114 | 0.836068829 | 0.403116183 | 0.828953151 | Micall2 | MICAL-like 2 [Source:MGI Symbol;Acc:MG | 24.093 | 7.1954 | 19.119 | 23.135 | 19.631 | 26.732 |
| 10436 | ENSMUSG00000036721 | 109.5844726 | 0.021512362 | 0.230799304 | 0.093208089 | 0.925738248 | 0.987546253 | Zscan12 | zinc finger and SCAN domain containing 1 | 105.21 | 92.513 | 128.8 | 120.5 | 105.79 | 104.7 |
| 10437 | ENSMUSG00000036731 | 0 | NA | NA | NA | NA | NA | Cysrt1 | cysteine rich tail 1 [Source:MGI Symbol;A | 0 | 0 | 0 | 0 | 0 | 0 |
| 10438 | ENSMUSG00000036733 | 0.663965228 | -0.037580459 | 3.10181903 | -0.012115619 | 0.990333371 | NA | Rbm42 | RNA binding motif protein 42 [Source:MG | 0 | 2.0558 | 0 | 1.928 | 0 | 0 |
| 10439 | ENSMUSG00000036737 | 563.2568266 | 0.155195467 | 0.129231781 | 1.200907904 | 0.229786928 | 0.676539688 | Oxsr1 | oxidative-stress responsive 1 [Source:MGI | 478.65 | 535.55 | 585.63 | 588.03 | 581.3 | 610.39 |
| 10440 | ENSMUSG00000036743 | 15.13060509 | 0.312277382 | 0.625610131 | 0.49915653 | 0.617669116 | 0.917509899 | Psma8 | proteasome subunit alpha 8 [Source:MGI | 16.865 | 13.363 | 10.062 | 10.604 | 9.8156 | 30.074 |
| 10441 | ENSMUSG00000036744 | 1.006392403 | 1.374439583 | 2.441002378 | 0.563063599 | 0.573391578 | NA | Olfr701 | olfactory receptor 701 [Source:MGI Symb | 1.6062 | 0 | 0 | 0 | 1.0906 | 3.3415 |
| 10442 | ENSMUSG00000036745 | 11.62926653 | -1.279814816 | 0.730370353 | -1.752281991 | 0.079725331 | 0.414529629 | Ttll7 | tubulin tyrosine ligase-like family, membe | 16.865 | 10.279 | 22.137 | 3.8559 | 3.2719 | 13.366 |
| 10443 | ENSMUSG00000036748 | 255.6911848 | -0.018975365 | 0.159365947 | -0.11906788 | 0.905221577 | 0.983972587 | Cuedc2 | CUE domain containing 2 [Source:MGI Syr | 243.34 | 285.76 | 243.51 | 261.24 | 253.02 | 247.27 |
| 10444 | ENSMUSG00000036749 | 0 | NA | NA | NA | NA | NA | Pramel5 | PRAME like 5 [Source:MGI Symbol;Acc:M | 0 | 0 | 0 | 0 | 0 | 0 |
| 10445 | ENSMUSG00000036751 | 2934.761378 | -0.029454862 | 0.088626172 | -0.332349474 | 0.739625386 | 0.948393197 | Cox6b1 | cytochrome c oxidase, subunit 6B1 [Sourc | 3025.3 | 3011.8 | 2856.7 | 2915.1 | 2829.1 | 2970.6 |
| 10446 | ENSMUSG00000036752 | 1062.46287 | -0.075114557 | 0.116539454 | -0.644541863 | 0.51922412 | 0.887713673 | Tubb4b | tubulin, beta 4B class IVB [Source:MGI Syr | 1214.3 | 1055.7 | 999.2 | 1045 | 991.37 | 1069.3 |
| 10447 | ENSMUSG00000036760 | 0 | NA | NA | NA | NA | NA | Kcnk9 | potassium channel, subfamily K, member | 0 | 0 | 0 | 0 | 0 | 0 |
| 10448 | ENSMUSG00000036764 | 127.0330186 | -0.19629912 | 0.244366972 | -0.803296446 | 0.421803411 | 0.838966521 | Dnajc12 | DnaJ heat shock protein family (Hsp40) m | 106.81 | 144.94 | 155.97 | 130.14 | 135.24 | 89.108 |
| 10449 | ENSMUSG00000036766 | 0 | NA | NA | NA | NA | NA | Dner | delta/notch-like EGF repeat containing [S | 0 | 0 | 0 | 0 | 0 | 0 |
| 10450 | ENSMUSG00000036768 | 5.748225237 | -0.333981766 | 0.975003582 | -0.342544142 | 0.73194143 | 0.947449512 | Kif15 | kinesin family member 15 [Source:MGI Sy | 12.047 | 4.1117 | 3.0187 | 8.6758 | 2.1812 | 4.4554 |
| 10451 | ENSMUSG00000036769 | 60.73054968 | 0.294373278 | 0.292457693 | 1.006549956 | 0.314151094 | 0.760800449 | Wdr44 | WD repeat domain 44 [Source:MGI Symbo | 44.171 | 56.535 | 63.393 | 71.334 | 64.346 | 64.603 |
| 10452 | ENSMUSG00000036770 | 0 | NA | NA | NA | NA | NA | Stpg3 | sperm tail PG rich repeat containing 3 [So | 0 | 0 | 0 | 0 | 0 | 0 |
| 10453 | ENSMUSG00000036775 | 6119.940327 | -0.080870102 | 0.100380918 | -0.805632223 | 0.42045494 | 0.837906635 | Decr2 | 2-4-dienoyl-Coenzyme A reductase 2, per | 6771.8 | 6415.2 | 5686.3 | 5917.9 | 6260.1 | 5668.4 |
| 10454 | ENSMUSG00000036777 | 9.356352931 | -0.739971722 | 0.733731198 | -1.00850519 | 0.313211997 | 0.760152431 | Anln | anillin, actin binding protein [Source:MGI | 17.668 | 9.2513 | 8.05 | 5.7839 | 9.8156 | 5.5692 |
| 10455 | ENSMUSG00000036779 | 839.7300985 | 0.006444469 | 0.113851199 | 0.056604312 | 0.954860399 | 0.991678478 | Tent4b | terminal nucleotidyltransferase 4B [Sourc | 833.62 | 860.37 | 820.09 | 881.08 | 874.68 | 768.55 |
| 10456 | ENSMUSG00000036781 | 2959.023965 | -0.073391386 | 0.087933958 | -0.834619388 | 0.403932043 | 0.829030031 | Rps27l | ribosomal protein S27-like [Source:MGI Sy | 3118.4 | 2978.9 | 3004.6 | 2811.9 | 2965.4 | 2874.8 |
| 10457 | ENSMUSG00000036782 | 89.35186012 | -0.202776576 | 0.24775315 | -0.818462151 | 0.413093346 | 0.835261854 | Klhl13 | kelch-like 13 [Source:MGI Symbol;Acc:MG | 89.947 | 87.373 | 109.68 | 85.794 | 71.981 | 91.335 |
| 10458 | ENSMUSG00000036790 | 0 | NA | NA | NA | NA | NA | Slitrk2 | SLIT and NTRK-like family, member 2 [Sou | 0 | 0 | 0 | 0 | 0 | 0 |
| 10459 | ENSMUSG00000036792 | 112.5090622 | -0.042609833 | 0.230704744 | -0.184694223 | 0.853468878 | 0.973825032 | Mbd5 | methyl-CpG binding domain protein 5 [So | 127.69 | 114.1 | 100.62 | 122.43 | 118.88 | 91.335 |
| 10460 | ENSMUSG00000036800 | 0 | NA | NA | NA | NA | NA | Fam135b | family with sequence similarity 135, meml | 0 | 0 | 0 | 0 | 0 | 0 |
| 10461 | ENSMUSG00000036805 | 16.0547302 | 0.240545804 | 0.548902659 | 0.438230349 | 0.66121931 | 0.928341045 | Noxa1 | NADPH oxidase activator 1 [Source:MGI S | 12.85 | 11.307 | 20.125 | 21.208 | 16.359 | 14.48 |
| 10462 | ENSMUSG00000036810 | 288.4771087 | 0.139064235 | 0.154390296 | 0.900731705 | 0.367730987 | 0.805888029 | Cnep1r1 | CTD nuclear envelope phosphatase 1 regu | 272.25 | 292.96 | 258.6 | 303.65 | 275.93 | 327.47 |
| 10463 | ENSMUSG00000036813 | 1920.142084 | 0.172005275 | 0.099387307 | 1.730656368 | 0.083513072 | 0.426078684 | Entpd8 | ectonucleoside triphosphate diphosphohy | 1776.5 | 1707.4 | 1934 | 2094.7 | 1955.5 | 2052.8 |
| 10464 | ENSMUSG00000036814 | 0 | NA | NA | NA | NA | NA | Slc6a20a | solute carrier family 6 (neurotransmitter t | 0 | 0 | 0 | 0 | 0 | 0 |
| 10465 | ENSMUSG00000036815 | 0 | NA | NA | NA | NA | NA | Dpp10 | dipeptidylpeptidase 10 [Source:MGI Symb | 0 | 0 | 0 | 0 | 0 | 0 |
| 10466 | ENSMUSG00000036816 | 0 | NA | NA | NA | NA | NA | Atoh7 | atonal bHLH transcription factor 7 [Source | 0 | 0 | 0 | 0 | 0 | 0 |
| 10467 | ENSMUSG00000036817 | 304.1056899 | 0.431323439 | 0.161846285 | 2.665019083 | 0.007698395 | 0.089301824 | Sun1 | Sad1 and UNC84 domain containing 1 [So | 267.43 | 230.25 | 278.73 | 327.75 | 322.82 | 397.64 |
| 10468 | ENSMUSG00000036819 | 210.8238783 | 0.71657755 | 0.19967681 | 3.58868689 | 0.000332348 | 0.009479365 | Jmjd4 | jumonji domain containing 4 [Source:MGI | 155.8 | 156.24 | 167.04 | 325.82 | 208.31 | 251.73 |

| | | | | | | | | | | | | | | |
|---|---|---|---|---|---|---|---|---|---|---|---|---|---|---|
| 10469 | ENSMUSG00000036820 | 455.0868881 | 0.067413139 | 0.13519943 | 0.498619992 | 0.618047118 | 0.917509899 | Amdhd2 | amidohydrolase domain containing 2 [Sou | 457.77 | 450.23 | 424.64 | 418.37 | 501.68 | 477.84 |
| 10470 | ENSMUSG00000036822 | 509.4769326 | 0.196859724 | 0.131862023 | 1.492922065 | 0.135457588 | 0.538110014 | Topors | topoisomerase I binding, arginine/serine-r | 452.95 | 482.09 | 490.04 | 573.57 | 572.57 | 485.64 |
| 10471 | ENSMUSG00000036825 | 176.5692215 | 0.020777048 | 0.209785522 | 0.099039474 | 0.921106929 | 0.986685804 | Ssx2ip | synovial sarcoma, X 2 interacting protein [ | 144.56 | 192.22 | 190.18 | 214.97 | 154.87 | 162.62 |
| 10472 | ENSMUSG00000036826 | 0 | NA | NA | NA | NA | NA | Igflr1 | IGF-like family receptor 1 [Source:MGI Syr | 0 | 0 | 0 | 0 | 0 | 0 |
| 10473 | ENSMUSG00000036832 | 4.011746592 | 1.705832118 | 1.282856378 | 1.329714025 | 0.183612512 | NA | Lpar3 | lysophosphatidic acid receptor 3 [Source:I | 1.6062 | 0 | 4.025 | 2.8919 | 2.1812 | 13.366 |
| 10474 | ENSMUSG00000036833 | 2237.487835 | 0.333624317 | 0.152837262 | 2.182872897 | 0.029045171 | 0.223715404 | Pnpla7 | patatin-like phospholipase domain contai | 2452.7 | 1840 | 1645.2 | 2375.2 | 2230.3 | 2881.5 |
| 10475 | ENSMUSG00000036834 | 0.428363721 | -0.543730382 | 3.624677344 | -0.15000794 | 0.880758351 | NA | Plch1 | phospholipase C, eta 1 [Source:MGI Symb | 1.6062 | 0 | 0 | 0.964 | 0 | 0 |
| 10476 | ENSMUSG00000036835 | 735.5693096 | 0.135986375 | 0.113715446 | 1.195847884 | 0.231755926 | 0.67889946 | Psenen | presenilin enhancer gamma secretase sub | 715.56 | 728.79 | 658.08 | 757.69 | 756.89 | 796.4 |
| 10477 | ENSMUSG00000036840 | 137.7369744 | -0.049810309 | 0.219464214 | -0.226963242 | 0.820452325 | 0.969160361 | Siah1a | siah E3 ubiquitin protein ligase 1A [Source | 159.82 | 133.63 | 126.79 | 159.06 | 111.24 | 135.89 |
| 10478 | ENSMUSG00000036845 | 91.5211551 | 0.110927003 | 0.246855243 | 0.449360532 | 0.653171598 | 0.925756399 | Lin37 | lin-37 homolog (C. elegans) [Source:MGI S | 101.99 | 80.178 | 81.506 | 98.326 | 104.7 | 82.424 |
| 10479 | ENSMUSG00000036850 | 444.2221875 | 0.122439713 | 0.130531191 | 0.93801115 | 0.34823868 | 0.789711685 | Mrpl41 | mitochondrial ribosomal protein L41 [Sou | 428.86 | 413.22 | 433.69 | 443.43 | 448.24 | 497.89 |
| 10480 | ENSMUSG00000036853 | 0 | NA | NA | NA | NA | NA | Mcoln3 | mucolipin 3 [Source:MGI Symbol;Acc:MGI | 0 | 0 | 0 | 0 | 0 | 0 |
| 10481 | ENSMUSG00000036854 | 261.9282327 | 0.09512764 | 0.173151342 | 0.549390144 | 0.582737737 | 0.908645447 | Hspb6 | heat shock protein, alpha-crystallin-relate | 269.84 | 250.81 | 239.49 | 321.97 | 238.85 | 250.61 |
| 10482 | ENSMUSG00000036855 | 0 | NA | NA | NA | NA | NA | Gjd4 | gap junction protein, delta 4 [Source:MGI | 0 | 0 | 0 | 0 | 0 | 0 |
| 10483 | ENSMUSG00000036856 | 36.93531765 | -0.485754076 | 0.378926861 | -1.281920408 | 0.199870568 | 0.642000421 | Wnt4 | wingless-type MMTV integration site fami | 53.808 | 37.005 | 38.237 | 31.811 | 23.994 | 36.757 |
| 10484 | ENSMUSG00000036858 | 0.267700746 | -1.497481649 | 4.057001436 | -0.369110456 | 0.712045396 | NA | Ptcra | pre T cell antigen receptor alpha [Source:I | 1.6062 | 0 | 0 | 0 | 0 | 0 |
| 10485 | ENSMUSG00000036860 | 327.9646598 | 0.158679152 | 0.15403882 | 1.03012443 | 0.302951598 | 0.752484788 | Mrpl55 | mitochondrial ribosomal protein L55 [Sou | 338.91 | 269.31 | 320.99 | 348 | 366.45 | 324.13 |
| 10486 | ENSMUSG00000036862 | 28.57123674 | -0.531181256 | 0.426254768 | -1.246159095 | 0.212705993 | 0.657529571 | Dchs1 | dachsous cadherin related 1 [Source:MGI | 44.171 | 28.782 | 28.175 | 24.099 | 27.265 | 18.935 |
| 10487 | ENSMUSG00000036863 | 227.0909607 | -0.371738203 | 0.17798643 | -2.088576098 | 0.036745899 | 0.259010631 | Syde2 | synapse defective 1, Rho GTPase, homolo | 230.49 | 302.21 | 236.47 | 203.4 | 202.86 | 187.13 |
| 10488 | ENSMUSG00000036864 | 50.57982823 | 0.363369773 | 0.33118004 | 1.09719708 | 0.27255525 | 0.727618887 | Proser3 | proline and serine rich 3 [Source:MGI Sym | 38.549 | 34.949 | 59.368 | 55.911 | 53.44 | 61.261 |
| 10489 | ENSMUSG00000036867 | 138.2957159 | -0.059181367 | 0.277359691 | -0.213374074 | 0.831035201 | 0.970505212 | Smad6 | SMAD family member 6 [Source:MGI Sym | 136.53 | 129.52 | 156.97 | 99.29 | 102.52 | 204.95 |
| 10490 | ENSMUSG00000036872 | 3.984815008 | -0.185532652 | 1.079452149 | -0.17187668 | 0.86353448 | NA | Abcc12 | ATP-binding cassette, sub-family C (CFTR/ | 5.6217 | 4.1117 | 3.0187 | 6.7478 | 2.1812 | 2.2277 |
| 10491 | ENSMUSG00000036873 | 118.001272 | -0.258382211 | 0.217425642 | -1.188370465 | 0.234687487 | 0.682729239 | 2410004B18 | RIKEN cDNA 2410004B18 gene [Source:M | 124.48 | 139.8 | 121.76 | 121.46 | 103.61 | 96.904 |
| 10492 | ENSMUSG00000036875 | 225.5683465 | 0.18949381 | 0.160347937 | 1.181766435 | 0.23729839 | 0.686182309 | Dna2 | DNA replication helicase/nuclease 2 [Sour | 211.22 | 217.92 | 203.26 | 244.85 | 235.57 | 240.59 |
| 10493 | ENSMUSG00000036879 | 412.0777337 | 0.272701293 | 0.13600444 | 2.005091106 | 0.044953308 | 0.293016967 | Phkb | phosphorylase kinase beta [Source:MGI S | 395.93 | 355.66 | 367.28 | 424.15 | 447.15 | 482.29 |
| 10494 | ENSMUSG00000036880 | 21912.85708 | 0.047664319 | 0.086298046 | 0.55232211 | 0.580727685 | 0.908645447 | Acaa2 | acetyl-Coenzyme A acyltransferase 2 (mit | 20268 | 22193 | 22194 | 22971 | 22381 | 21472 |
| 10495 | ENSMUSG00000036882 | 6.08864879 | -1.133786138 | 0.923054642 | -1.228297964 | 0.219335135 | 0.664947263 | Arhgap33 | Rho GTPase activating protein 33 [Source: | 4.8186 | 13.363 | 7.0437 | 5.7839 | 2.1812 | 3.3415 |
| 10496 | ENSMUSG00000036885 | 728.4008669 | -0.267395296 | 0.216668411 | -1.234122201 | 0.217157372 | 0.663070346 | Arhgef26 | Rho guanine nucleotide exchange factor ( | 829.6 | 953.91 | 603.75 | 680.57 | 865.95 | 436.63 |
| 10497 | ENSMUSG00000036887 | 498.0967887 | -1.298360052 | 0.451893724 | -2.873153536 | 0.004063966 | 0.057969624 | C1qa | complement component 1, q subcompon | 593.49 | 1031 | 500.1 | 223.64 | 214.85 | 425.49 |
| 10498 | ENSMUSG00000036890 | 108.2095615 | 0.175935531 | 0.236961418 | 0.742464881 | 0.457805722 | 0.859309158 | Gtdc1 | glycosyltransferase-like domain containin | 120.47 | 94.568 | 89.556 | 132.06 | 111.24 | 101.36 |
| 10499 | ENSMUSG00000036892 | 5285.645299 | 0.011268463 | 0.0895932 | 0.125773643 | 0.899911106 | 0.982798755 | Prodh2 | proline dehydrogenase (oxidase) 2 [Sourc | 5026.6 | 5191 | 5578.6 | 5496.6 | 5233.9 | 5187.2 |
| 10500 | ENSMUSG00000036893 | 519.3223066 | 0.160888302 | 0.137216094 | 1.172517717 | 0.240989263 | 0.691388521 | Ehmt1 | euchromatic histone methyltransferase 1 | 423.23 | 505.74 | 543.37 | 552.36 | 538.77 | 552.47 |
| 10501 | ENSMUSG00000036894 | 23.50826656 | -0.454208094 | 0.578087263 | -0.785708531 | 0.432038264 | 0.845259226 | Rap2b | RAP2B, member of RAS oncogene family [ | 23.29 | 42.145 | 16.1 | 17.352 | 7.6343 | 34.529 |
| 10502 | ENSMUSG00000036896 | 402.7824742 | -1.612699937 | 0.464967286 | -3.468415921 | 0.000523536 | 0.01290022 | C1qc | complement component 1, q subcompon | 511.58 | 878.87 | 430.67 | 170.62 | 129.78 | 295.17 |
| 10503 | ENSMUSG00000036898 | 91.21204393 | -0.003816108 | 0.239866772 | -0.01590928 | 0.987306766 | 0.99859176 | Zfp157 | zinc finger protein 157 [Source:MGI Symb | 89.947 | 95.596 | 88.55 | 91.578 | 102.52 | 79.083 |
| 10504 | ENSMUSG00000036899 | 0 | NA | NA | NA | NA | NA | Trpv5 | transient receptor potential cation chann | 0 | 0 | 0 | 0 | 0 | 0 |
| 10505 | ENSMUSG00000036902 | 12.29010237 | 0.287065232 | 0.77014786 | 0.372740413 | 0.709341655 | 0.944239215 | Neto2 | neuropilin (NRP) and tolloid (TLL)-like 2 [S | 20.881 | 1.0279 | 11.069 | 15.424 | 13.087 | 12.252 |
| 10506 | ENSMUSG00000036904 | 476.2342151 | 0.970830792 | 0.182211798 | 5.328034737 | 9.93E-08 | 1.09E-05 | Fzd8 | frizzled class receptor 8 [Source:MGI Sym | 360.59 | 302.21 | 301.87 | 665.14 | 464.6 | 762.98 |
| 10507 | ENSMUSG00000036905 | 420.8283743 | -1.366407202 | 0.237632882 | -5.750076295 | 8.92E-09 | 1.40E-06 | C1qb | complement component 1, q subcompon | 532.46 | 850.09 | 436.71 | 214 | 185.41 | 306.31 |
| 10508 | ENSMUSG00000036907 | 0 | NA | NA | NA | NA | NA | C1ql2 | complement component 1, q subcompon | 0 | 0 | 0 | 0 | 0 | 0 |
| 10509 | ENSMUSG00000036908 | 676.4797623 | -0.133605781 | 0.125195924 | -1.067173571 | 0.285893465 | 0.738421977 | Unc93b1 | unc-93 homolog B1, TLR signaling regulat | 707.53 | 730.85 | 684.25 | 573.57 | 658.73 | 703.95 |
| 10510 | ENSMUSG00000036912 | 0.342639279 | -1.805976596 | 4.028547714 | -0.448294702 | 0.653940524 | NA | Piwil4 | piwi-like RNA-mediated gene silencing 4 [ | 0 | 2.0558 | 0 | 0 | 0 | 0 |
| 10511 | ENSMUSG00000036913 | 0.647809292 | -2.733819264 | 3.061311498 | -0.893022244 | 0.371845261 | NA | Trim67 | tripartite motif-containing 67 [Source:MG | 0.8031 | 3.0838 | 0 | 0 | 0 | 0 |
| 10512 | ENSMUSG00000036915 | 0 | NA | NA | NA | NA | NA | Kirrel2 | kirre like nephrin family adhesion molecul | 0 | 0 | 0 | 0 | 0 | 0 |
| 10513 | ENSMUSG00000036916 | 104.4393681 | 0.125861155 | 0.23199467 | 0.542517443 | 0.587462094 | 0.910765033 | Zfp280c | zinc finger protein 280C [Source:MGI Sym | 98.782 | 92.513 | 108.67 | 127.25 | 102.52 | 96.904 |
| 10514 | ENSMUSG00000036918 | 1357.407831 | 0.488361423 | 0.112654735 | 4.335027913 | 1.46E-05 | 0.000724401 | Ttc7 | tetratricopeptide repeat domain 7 [Sourc | 1234.4 | 1055.7 | 1097.8 | 1514.4 | 1526.9 | 1715.3 |
| 10515 | ENSMUSG00000036921 | 0 | NA | NA | NA | NA | NA | Tex46 | testis expressed 46 [Source:MGI Symbol;A | 0 | 0 | 0 | 0 | 0 | 0 |
| 10516 | ENSMUSG00000036923 | 4.388766789 | -1.055233039 | 1.113910541 | -0.947322967 | 0.343474231 | NA | Stox1 | storkhead box 1 [Source:MGI Symbol;Acc | 1.6062 | 5.1396 | 11.069 | 1.928 | 4.3625 | 2.2277 |
| 10517 | ENSMUSG00000036924 | 0 | NA | NA | NA | NA | NA | Cst13 | cystatin 13 [Source:MGI Symbol;Acc:MGI: | 0 | 0 | 0 | 0 | 0 | 0 |
| 10518 | ENSMUSG00000036925 | 0 | NA | NA | NA | NA | NA | Mucl2 | mucin-like 2 [Source:MGI Symbol;Acc:MG | 0 | 0 | 0 | 0 | 0 | 0 |
| 10519 | ENSMUSG00000036928 | 1.407975666 | -2.72347108 | 1.994677936 | -1.36536883 | 0.172137156 | NA | Stag3 | stromal antigen 3 [Source:MGI Symbol;Ac | 2.4093 | 2.0558 | 3.0187 | 0.964 | 0 | 0 |
| 10520 | ENSMUSG00000036931 | 4.417411237 | 0.019627578 | 1.162300797 | 0.016886831 | 0.986526898 | NA | Nfkbid | nuclear factor of kappa light polypeptide g | 4.0155 | 7.1954 | 2.0125 | 7.7118 | 0 | 5.5692 |
| 10521 | ENSMUSG00000036932 | 2324.60647 | 0.012163322 | 0.099046521 | 0.122804128 | 0.902262206 | 0.983821605 | Aifm1 | apoptosis-inducing factor, mitochondrion | 2243.1 | 2381.7 | 2319.4 | 2143.9 | 2462.6 | 2397 |
| 10522 | ENSMUSG00000036934 | 142.1207123 | 0.161637004 | 0.19918414 | 0.811495353 | 0.417081261 | 0.836878476 | 4921524J17I | RIKEN cDNA 4921524J17 gene [Source:M | 142.15 | 130.55 | 129.81 | 163.88 | 152.69 | 133.66 |
| 10523 | ENSMUSG00000036938 | 0 | NA | NA | NA | NA | NA | Try5 | trypsin 5 [Source:MGI Symbol;Acc:MGI:1 | 0 | 0 | 0 | 0 | 0 | 0 |
| 10524 | ENSMUSG00000036940 | 253.3418049 | -0.124524703 | 0.166630841 | -0.747308854 | 0.454877149 | 0.857337687 | Kdm1a | lysine (K)-specific demethylase 1A [Source | 260.21 | 280.62 | 252.57 | 271.84 | 244.3 | 210.52 |

| | | | | | | | | | | | | | | | |
|---|---|---|---|---|---|---|---|---|---|---|---|---|---|---|---|
| 10525 | ENSMUSG00000036941 | 223.1139584 | 0.125420886 | 0.167829391 | 0.747311812 | 0.454875364 | 0.857337687 | Elac1 | elaC ribonuclease Z 1 [Source:MGI Symbo | 225.67 | 211.75 | 202.26 | 212.08 | 253.02 | 233.91 |
| 10526 | ENSMUSG00000036943 | 42.95307851 | -0.808800061 | 0.361685516 | -2.236196984 | 0.025338871 | 0.204853713 | Rab8b | RAB8B, member RAS oncogene family [So | 57.02 | 63.731 | 43.269 | 31.811 | 22.903 | 38.985 |
| 10527 | ENSMUSG00000036944 | 25.94592101 | -0.161838745 | 0.443534106 | -0.364884556 | 0.715197583 | 0.944595913 | Tmem71 | transmembrane protein 71 [Source:MGI S | 27.305 | 27.754 | 27.169 | 26.991 | 15.269 | 31.188 |
| 10528 | ENSMUSG00000036948 | 263.0530726 | 0.06594944 | 0.162532602 | 0.405761301 | 0.684917996 | 0.935737593 | Map11 | microtubule associated protein 11 [Source | 265.83 | 249.78 | 255.59 | 285.34 | 290.1 | 231.68 |
| 10529 | ENSMUSG00000036949 | 1.106417862 | -2.321245515 | 2.151108314 | -1.07909281 | 0.280546356 | NA | Slc39a12 | solute carrier family 39 (zinc transporter), | 1.6062 | 2.0558 | 2.0125 | 0.964 | 0 | 0 |
| 10530 | ENSMUSG00000036951 | 0 | NA | NA | NA | NA | NA | Aadacl2fm1 | AADACL2 family member 1 [Source:MGI S | 0 | 0 | 0 | 0 | 0 | 0 |
| 10531 | ENSMUSG00000036955 | 287.6450514 | -0.088345124 | 0.164709592 | -0.536369027 | 0.591703533 | 0.912678231 | Kifbp | kinesin family binding protein [Source:MG | 304.38 | 328.93 | 255.59 | 250.63 | 292.29 | 294.05 |
| 10532 | ENSMUSG00000036957 | 169.3964474 | -1.126461142 | 0.233162211 | -4.831233749 | 1.36E-06 | 0.000104394 | Lrfn3 | leucine rich repeat and fibronectin type II | 187.12 | 203.53 | 306.9 | 114.71 | 91.612 | 112.5 |
| 10533 | ENSMUSG00000036958 | 0 | NA | NA | NA | NA | NA | Cst11 | cystatin 11 [Source:MGI Symbol;Acc:MGI: | 0 | 0 | 0 | 0 | 0 | 0 |
| 10534 | ENSMUSG00000036959 | 149.9934907 | 0.467943079 | 0.239293611 | 1.955518481 | 0.050521904 | 0.315931181 | Bcorl1 | BCL6 co-repressor-like 1 [Source:MGI Sym | 126.89 | 108.96 | 141.88 | 199.54 | 119.97 | 202.72 |
| 10535 | ENSMUSG00000036960 | 0 | NA | NA | NA | NA | NA | Clca2 | chloride channel accessory 2 [Source:MGI | 0 | 0 | 0 | 0 | 0 | 0 |
| 10536 | ENSMUSG00000036961 | 0.160662974 | 1.126924175 | 4.080472857 | 0.276174898 | 0.782413736 | NA | Wnt8b | wingless-type MMTV integration site fami | 0 | 0 | 0 | 0.964 | 0 | 0 |
| 10537 | ENSMUSG00000036962 | 0 | NA | NA | NA | NA | NA | Cfap221 | cilia and flagella associated protein 221 [S | 0 | 0 | 0 | 0 | 0 | 0 |
| 10538 | ENSMUSG00000036964 | 3.612080351 | 1.066231675 | 1.251933468 | 0.851668002 | 0.394398385 | NA | Trim17 | tripartite motif-containing 17 [Source:MG | 4.8186 | 2.0558 | 0 | 4.8199 | 2.1812 | 7.7969 |
| 10539 | ENSMUSG00000036966 | 363.604023 | -0.060499238 | 0.147413481 | -0.410405059 | 0.681508835 | 0.934522544 | Spryd3 | SPRY domain containing 3 [Source:MGI Sy | 363.81 | 372.11 | 377.34 | 316.18 | 350.09 | 402.1 |
| 10540 | ENSMUSG00000036968 | 79.04694358 | -0.281819186 | 0.263053675 | -1.071337194 | 0.284017842 | 0.736651636 | Cnpy4 | canopy FGF signaling regulator 4 [Source:I | 90.751 | 75.038 | 94.587 | 83.866 | 65.437 | 64.603 |
| 10541 | ENSMUSG00000036972 | 0 | NA | NA | NA | NA | NA | Zic4 | zinc finger protein of the cerebellum 4 [Sc | 0 | 0 | 0 | 0 | 0 | 0 |
| 10542 | ENSMUSG00000036975 | 197.232783 | -0.092505985 | 0.196448164 | -0.470892593 | 0.637717438 | 0.923478764 | Tmem177 | transmembrane protein 177 [Source:MGI | 218.44 | 188.11 | 203.26 | 153.27 | 190.86 | 229.45 |
| 10543 | ENSMUSG00000036977 | 52.83208387 | 0.185267841 | 0.363529362 | 0.509636526 | 0.61030613 | 0.916376001 | Anapc10 | anaphase promoting complex subunit 10 [ | 57.823 | 30.838 | 59.368 | 53.983 | 40.353 | 74.628 |
| 10544 | ENSMUSG00000036980 | 171.5159337 | 0.134835083 | 0.181783479 | 0.741734531 | 0.458248194 | 0.859322431 | Taf6 | TATA-box binding protein associated facto | 153.39 | 159.33 | 178.11 | 182.19 | 173.41 | 182.67 |
| 10545 | ENSMUSG00000036983 | 137.1705651 | 0.0386777 | 0.195612925 | 0.197725689 | 0.84325969 | 0.972273952 | Tfb1m | transcription factor B1, mitochondrial [So | 137.33 | 136.71 | 131.82 | 133.99 | 145.05 | 138.12 |
| 10546 | ENSMUSG00000036985 | 1687.242273 | 0.03363698 | 0.09916002 | 0.339219172 | 0.734444627 | 0.948109711 | Zdhhc9 | zinc finger, DHHC domain containing 9 [Sc | 1576.5 | 1658 | 1769 | 1695.6 | 1761.3 | 1663 |
| 10547 | ENSMUSG00000036986 | 313.7287597 | 0.107512085 | 0.149469774 | 0.719289807 | 0.471962375 | 0.867870596 | Pml | promyelocytic leukemia [Source:MGI Sym | 312.41 | 272.4 | 320.99 | 320.04 | 343.54 | 312.99 |
| 10548 | ENSMUSG00000036989 | 241.9043815 | 0.107520045 | 0.200880189 | 0.535244641 | 0.592480701 | 0.912923013 | Trim3 | tripartite motif-containing 3 [Source:MGI | 225.67 | 222.03 | 251.56 | 309.44 | 184.31 | 258.41 |
| 10549 | ENSMUSG00000036990 | 735.5060614 | -0.004766674 | 0.129318514 | -0.036859954 | 0.97059667 | 0.994966572 | Otud4 | OTU domain containing 4 [Source:MGI Sy | 721.99 | 746.27 | 742.61 | 786.61 | 790.7 | 624.87 |
| 10550 | ENSMUSG00000036992 | 140.3668855 | 0.044302867 | 0.21331431 | 0.20768821 | 0.835472424 | 0.971073158 | Nxt1 | NTF2-related export protein 1 [Source:MC | 116.45 | 149.05 | 149.93 | 160.98 | 124.33 | 141.46 |
| 10551 | ENSMUSG00000036995 | 187.4250645 | 0.397870067 | 0.279860122 | 1.421674745 | 0.155120695 | 0.573534185 | Asap3 | ArfGAP with SH3 domain, ankyrin repeat a | 129.3 | 115.13 | 241.5 | 279.55 | 189.77 | 169.3 |
| 10552 | ENSMUSG00000037001 | 90.41786082 | 0.404402059 | 0.250514775 | 1.614284264 | 0.106465782 | 0.481264827 | Zfp39 | zinc finger protein 39 [Source:MGI Symbo | 76.295 | 64.759 | 92.575 | 113.75 | 94.884 | 100.25 |
| 10553 | ENSMUSG00000037003 | 1064.516857 | 0.603756184 | 0.134929567 | 4.474602534 | 7.66E-06 | 0.000432528 | Tns2 | tensin 2 [Source:MGI Symbol;Acc:MGI:23 | 893.05 | 818.22 | 823.11 | 1331.3 | 1059 | 1462.5 |
| 10554 | ENSMUSG00000037005 | 1.14027492 | -2.363583184 | 2.163739702 | -1.092360223 | 0.274674799 | NA | Xpnpep2 | X-prolyl aminopeptidase (aminopeptidase | 0.8031 | 2.0558 | 3.0187 | 0.964 | 0 | 0 |
| 10555 | ENSMUSG00000037007 | 153.6932403 | 0.204771622 | 0.239657195 | 0.854435529 | 0.392863717 | 0.821390458 | Zfp113 | zinc finger protein 113 [Source:MGI Symb | 171.86 | 137.74 | 118.74 | 213.04 | 152.69 | 128.09 |
| 10556 | ENSMUSG00000037010 | 0.952979305 | -3.293912175 | 2.649451307 | -1.243243145 | 0.213778248 | NA | Apln | apelin [Source:MGI Symbol;Acc:MGI:1353 | 1.6062 | 4.1117 | 0 | 0 | 0 | 0 |
| 10557 | ENSMUSG00000037012 | 30.23155039 | -0.811550825 | 0.410137179 | -1.978730208 | 0.04784639 | 0.303663969 | Hk1 | hexokinase 1 [Source:MGI Symbol;Acc:MC | 35.336 | 44.2 | 36.225 | 26.991 | 16.359 | 22.277 |
| 10558 | ENSMUSG00000037013 | 595.7634327 | 0.104956665 | 0.13417402 | 0.782242825 | 0.434071881 | 0.845264432 | Ss18 | SS18, nBAF chromatin remodeling comple | 507.56 | 607.5 | 607.77 | 562 | 631.47 | 658.28 |
| 10559 | ENSMUSG00000037014 | 0 | NA | NA | NA | NA | NA | Sstr4 | somatostatin receptor 4 [Source:MGI Sym | 0 | 0 | 0 | 0 | 0 | 0 |
| 10560 | ENSMUSG00000037016 | 1.959223891 | -4.328304717 | 1.90474694 | -2.272377829 | 0.0230637 | NA | Frem2 | Fras1 related extracellular matrix protein 2 | 1.6062 | 4.1117 | 6.0375 | 0 | 0 | 0 |
| 10561 | ENSMUSG00000037017 | 239.8231092 | 0.023888917 | 0.174430343 | 0.136953906 | 0.891067229 | 0.982489666 | Zscan21 | zinc finger and SCAN domain containing 2 | 277.07 | 233.34 | 202.26 | 239.07 | 239.94 | 247.27 |
| 10562 | ENSMUSG00000037020 | 63.53401538 | 0.38180351 | 0.317812578 | 1.201348016 | 0.229616232 | 0.676409202 | Wdr62 | WD repeat domain 62 [Source:MGI Symbo | 75.492 | 48.312 | 41.256 | 82.902 | 71.981 | 61.261 |
| 10563 | ENSMUSG00000037022 | 492.6586225 | -0.17719145 | 0.134525018 | -1.317163552 | 0.187783808 | 0.624686319 | Mmaa | methylmalonic aciduria (cobalamin deficie | 513.18 | 497.51 | 558.47 | 492.59 | 480.96 | 413.24 |
| 10564 | ENSMUSG00000037025 | 705.2385981 | 0.594986082 | 0.399031644 | 1.491074933 | 0.135941823 | 0.538690549 | Foxa2 | forkhead box A2 [Source:MGI Symbol;Acc | 453.75 | 543.77 | 688.27 | 1021.8 | 466.78 | 1057 |
| 10565 | ENSMUSG00000037028 | 0 | NA | NA | NA | NA | NA | Pramel18 | PRAME like 18 [Source:MGI Symbol;Acc:M | 0 | 0 | 0 | 0 | 0 | 0 |
| 10566 | ENSMUSG00000037029 | 305.8834221 | 0.226963772 | 0.152092985 | 1.492269825 | 0.135628424 | 0.5381974 | Zfp146 | zinc finger protein 146 [Source:MGI Symb | 297.95 | 274.45 | 272.69 | 315.22 | 366.45 | 308.53 |
| 10567 | ENSMUSG00000037031 | 36.20328901 | -0.795379853 | 0.420355544 | -1.89215978 | 0.058469695 | 0.343085466 | Tspan15 | tetraspanin 15 [Source:MGI Symbol;Acc:M | 28.912 | 68.87 | 40.25 | 24.099 | 28.356 | 26.732 |
| 10568 | ENSMUSG00000037032 | 10.96420138 | -0.027250844 | 0.716316734 | -0.038043009 | 0.96965339 | 0.994674947 | Apbb1 | amyloid beta (A4) precursor protein-bindi | 4.0155 | 10.279 | 19.119 | 12.532 | 9.8156 | 10.025 |
| 10569 | ENSMUSG00000037033 | 0 | NA | NA | NA | NA | NA | Clca3b | chloride channel accessory 3B [Source:MC | 0 | 0 | 0 | 0 | 0 | 0 |
| 10570 | ENSMUSG00000037034 | 0 | NA | NA | NA | NA | NA | Pax1 | paired box 1 [Source:MGI Symbol;Acc:MG | 0 | 0 | 0 | 0 | 0 | 0 |
| 10571 | ENSMUSG00000037035 | 24.53610497 | -0.997292783 | 0.496342511 | -2.00928343 | 0.044507084 | 0.291685757 | Inhbb | inhibin beta-B [Source:MGI Symbol;Acc:M | 33.73 | 34.949 | 29.181 | 10.604 | 10.906 | 27.846 |
| 10572 | ENSMUSG00000037049 | 792.313965 | 0.089882843 | 0.116897484 | 0.768903141 | 0.441950813 | 0.850050074 | Smpd1 | sphingomyelin phosphodiesterase 1, acid | 786.24 | 707.21 | 809.02 | 811.67 | 776.52 | 863.23 |
| 10573 | ENSMUSG00000037053 | 26478.5561 | 0.033723848 | 0.07828852 | 0.430763635 | 0.666640244 | 0.929596279 | Azgp1 | alpha-2-glycoprotein 1, zinc [Source:MGI S | 25588 | 26716 | 26204 | 26402 | 27144 | 26817 |
| 10574 | ENSMUSG00000037058 | 1343.368185 | 0.06540189 | 0.101112256 | 0.646824558 | 0.517745496 | 0.887022094 | Paip2 | polyadenylate-binding protein-interacting | 1360.5 | 1256.1 | 1321.2 | 1309.1 | 1422.2 | 1391.2 |
| 10575 | ENSMUSG00000037060 | 9.301763389 | -0.76569598 | 0.740331067 | -1.034261581 | 0.301013892 | 0.7521219 | Cavin3 | caveolae associated 3 [Source:MGI Symbo | 8.8341 | 15.419 | 11.069 | 10.604 | 6.5437 | 3.3415 |
| 10576 | ENSMUSG00000037062 | 1160.038543 | -0.148898326 | 0.100264216 | -1.485059502 | 0.137528063 | 0.541722739 | Sh3glb1 | SH3-domain GRB2-like B1 (endophilin) [Sc | 1258.5 | 1228.4 | 1172.3 | 1098 | 1099.3 | 1103.8 |
| 10577 | ENSMUSG00000037064 | 0 | NA | NA | NA | NA | NA | Gm5229 | predicted gene 5229 [Source:MGI Symbol | 0 | 0 | 0 | 0 | 0 | 0 |
| 10578 | ENSMUSG00000037070 | 208.2656103 | -0.094511264 | 0.172959103 | -0.546437059 | 0.584765542 | 0.909076612 | Rbmxl1 | RNA binding motif protein, X-linked like-1 | 221.66 | 211.75 | 211.31 | 177.37 | 219.21 | 208.29 |
| 10579 | ENSMUSG00000037071 | 107207.0154 | 0.273845608 | 0.379836943 | 0.720955698 | 0.470936772 | 0.867447112 | Scd1 | stearoyl-Coenzyme A desaturase 1 [Sourc | 152291 | 71537 | 67359 | 139818 | 108992 | 103244 |
| 10580 | ENSMUSG00000037072 | 3178.54652 | 0.063362926 | 0.092460077 | 0.685300385 | 0.493154389 | 0.876934483 | Selenof | selenoprotein F [Source:MGI Symbol;Acc: | 3304 | 2979.9 | 3040.9 | 3144.5 | 3282.8 | 3319.3 |

| | | | | | | | | | | | | | | | |
|---|---|---|---|---|---|---|---|---|---|---|---|---|---|---|---|
| 10581 | ENSMUSG00000037075 | 559.4152983 | -0.019582934 | 0.124891495 | -0.156799582 | 0.875402802 | 0.980525066 | Rnf139 | ring finger protein 139 [Source:MGI Symb | 577.43 | 516.01 | 595.7 | 537.9 | 574.76 | 554.69 |
| 10582 | ENSMUSG00000037085 | 101.9903316 | 0.150871575 | 0.237465445 | 0.635341177 | 0.525205924 | 0.888334265 | Trmt12 | tRNA methyltransferase 12 [Source:MGI S | 97.978 | 82.233 | 109.68 | 104.11 | 122.15 | 95.791 |
| 10583 | ENSMUSG00000037086 | 0 | NA | NA | NA | NA | NA | Prr32 | proline rich 32 [Source:MGI Symbol;Acc:M | 0 | 0 | 0 | 0 | 0 | 0 |
| 10584 | ENSMUSG00000037089 | 935.0040408 | 0.025773503 | 0.129625562 | 0.198830405 | 0.842395411 | 0.971875826 | Slc35b2 | solute carrier family 35, member B2 [Sour | 938.83 | 950.82 | 889.52 | 814.56 | 1087.3 | 928.95 |
| 10585 | ENSMUSG00000037095 | 7941.045269 | -0.055185016 | 0.103606766 | -0.532639109 | 0.594283424 | 0.913144152 | Lrg1 | leucine-rich alpha-2-glycoprotein 1 [Sourc | 7735.5 | 9063.2 | 7480.4 | 7501.7 | 7638.7 | 8226.9 |
| 10586 | ENSMUSG00000037096 | 18.9347652 | 0.608914811 | 0.542437315 | 1.122553323 | 0.261627249 | 0.715612553 | Gm9762 | predicted pseudogene 9762 [Source:MGI | 9.6372 | 16.447 | 19.119 | 21.208 | 32.719 | 14.48 |
| 10587 | ENSMUSG00000037098 | 318.0400541 | 0.117383148 | 0.149966244 | 0.782730465 | 0.433785408 | 0.845264432 | Rab11fip3 | RAB11 family interacting protein 3 (class I | 291.53 | 303.24 | 320.99 | 346.07 | 295.56 | 350.86 |
| 10588 | ENSMUSG00000037101 | 0 | NA | NA | NA | NA | NA | Ttc29 | tetratricopeptide repeat domain 29 [Sour | 0 | 0 | 0 | 0 | 0 | 0 |
| 10589 | ENSMUSG00000037103 | 79.57600184 | 0.136259204 | 0.255425731 | 0.533459192 | 0.593715753 | 0.913106373 | Dcaf15 | DDB1 and CUL4 associated factor 15 [Sou | 81.916 | 78.122 | 67.418 | 93.506 | 78.525 | 77.969 |
| 10590 | ENSMUSG00000037104 | 104.100918 | 0.254406863 | 0.256341867 | 0.99245147 | 0.320977342 | 0.768136378 | Socs5 | suppressor of cytokine signaling 5 [Source | 102.8 | 72.982 | 108.67 | 106.04 | 94.884 | 139.23 |
| 10591 | ENSMUSG00000037106 | 0 | NA | NA | NA | NA | NA | Fer1l6 | fer-1-like 6 (C. elegans) [Source:MGI Symb | 0 | 0 | 0 | 0 | 0 | 0 |
| 10592 | ENSMUSG00000037108 | 67.71920674 | -0.199047605 | 0.284256329 | -0.700239837 | 0.483777537 | 0.872271041 | Zcwpw1 | zinc finger, CW type with PWWP domain 1 | 69.87 | 79.15 | 68.425 | 75.19 | 49.078 | 64.603 |
| 10593 | ENSMUSG00000037110 | 884.2397176 | 0.070256377 | 0.124046639 | 0.566370662 | 0.571141834 | 0.904651508 | Ralgapa2 | Ral GTPase activating protein, alpha subu | 901.08 | 780.19 | 906.63 | 958.19 | 809.24 | 950.11 |
| 10594 | ENSMUSG00000037111 | 123.9739017 | -0.217023787 | 0.222586226 | -0.97500996 | 0.32955532 | 0.774441615 | Setd7 | SET domain containing (lysine methyltrans | 126.89 | 149.05 | 123.77 | 100.25 | 106.88 | 137 |
| 10595 | ENSMUSG00000037112 | 137.532821 | -0.219310292 | 0.230225813 | -0.952587762 | 0.34079898 | 0.783129922 | Sik2 | salt inducible kinase 2 [Source:MGI Symbo | 113.24 | 161.38 | 170.06 | 133.99 | 139.6 | 106.93 |
| 10596 | ENSMUSG00000037119 | 1904.902677 | 0.134938394 | 0.094653411 | 1.425605182 | 0.15398234 | 0.57119049 | Fam91a1 | family with sequence similarity 91, memb | 1744.3 | 1824.6 | 1879.7 | 2032.1 | 2029.6 | 1919.2 |
| 10597 | ENSMUSG00000037124 | 0 | NA | NA | NA | NA | NA | Trim58 | tripartite motif-containing 58 [Source:MG | 0 | 0 | 0 | 0 | 0 | 0 |
| 10598 | ENSMUSG00000037126 | 23.67887056 | 0.040558533 | 0.486429506 | 0.083380085 | 0.933549323 | 0.988420305 | Psd | pleckstrin and Sec7 domain containing [Sc | 14.456 | 20.558 | 35.219 | 21.208 | 28.356 | 22.277 |
| 10599 | ENSMUSG00000037129 | 0 | NA | NA | NA | NA | NA | Tmprss13 | transmembrane protease, serine 13 [Sour | 0 | 0 | 0 | 0 | 0 | 0 |
| 10600 | ENSMUSG00000037130 | 0.545309144 | 2.651974629 | 3.666219369 | 0.72335405 | 0.469462394 | NA | H2-M10.6 | histocompatibility 2, M region locus 10.6 [ | 0 | 0 | 0 | 0 | 3.2719 | 0 |
| 10601 | ENSMUSG00000037134 | 274.3323183 | 0.073598523 | 0.178627609 | 0.412022102 | 0.680323223 | 0.934384556 | Prmt9 | protein arginine methyltransferase 9 [Sou | 261.81 | 251.84 | 287.79 | 221.71 | 304.28 | 318.56 |
| 10602 | ENSMUSG00000037138 | 4.102252376 | -0.938216208 | 1.050303903 | -0.893280702 | 0.37170687 | NA | Aff3 | AF4/FMR2 family, member 3 [Source:MGI | 4.0155 | 5.1396 | 7.0437 | 2.8919 | 2.1812 | 3.3415 |
| 10603 | ENSMUSG00000037139 | 24.9187871 | -1.680548068 | 0.544769264 | -3.084880476 | 0.00203634 | 0.03544235 | Myom3 | myomesin family, member 3 [Source:MGI | 43.368 | 38.033 | 32.2 | 2.8919 | 18.541 | 14.48 |
| 10604 | ENSMUSG00000037140 | 0.167707431 | -0.796629176 | 4.080472857 | -0.195229623 | 0.84521319 | NA | Tas2r108 | taste receptor, type 2, member 108 [Sour | 0 | 0 | 1.0062 | 0 | 0 | 0 |
| 10605 | ENSMUSG00000037143 | 0.528279927 | -0.837071592 | 3.369085264 | -0.248456636 | 0.803781118 | NA | Cfap61 | cilia and flagella associated protein 61 [So | 0 | 2.0558 | 0 | 0 | 0 | 1.1138 |
| 10606 | ENSMUSG00000037145 | 0 | NA | NA | NA | NA | NA | Lypd8l | LY6/PLAUR domain containing 8 like [Sour | 0 | 0 | 0 | 0 | 0 | 0 |
| 10607 | ENSMUSG00000037148 | 35.03279386 | -0.220306728 | 0.413251257 | -0.533106008 | 0.5939602 | 0.913144152 | Arhgap10 | Rho GTPase activating protein 10 [Source: | 46.58 | 31.865 | 34.212 | 20.244 | 31.628 | 45.668 |
| 10608 | ENSMUSG00000037149 | 1166.326444 | 0.052389137 | 0.102972909 | 0.508766219 | 0.6109161 | 0.916376001 | Ddx1 | DEAD box helicase 1 [Source:MGI Symbol; | 1196.6 | 1145.1 | 1092.8 | 1145.2 | 1198.6 | 1219.7 |
| 10609 | ENSMUSG00000037151 | 141.3728281 | -0.165690217 | 0.205275721 | -0.807159347 | 0.419574684 | 0.837591673 | Lrrc20 | leucine rich repeat containing 20 [Source: | 130.91 | 143.91 | 174.08 | 135.92 | 130.87 | 132.55 |
| 10610 | ENSMUSG00000037152 | 679.443164 | 0.079943071 | 0.131489454 | 0.607980861 | 0.543200168 | 0.894348005 | Ndufc1 | NADH:ubiquinone oxidoreductase subuni | 615.98 | 686.65 | 679.22 | 625.62 | 678.36 | 790.83 |
| 10611 | ENSMUSG00000037157 | 76.99079515 | -1.034101823 | 0.328672479 | -3.146298794 | 0.00165351 | 0.030466709 | Il22ra1 | interleukin 22 receptor, alpha 1 [Source:M | 68.264 | 103.82 | 138.86 | 62.659 | 35.99 | 52.351 |
| 10612 | ENSMUSG00000037159 | 0 | NA | NA | NA | NA | NA | Wee2 | WEE1 homolog 2 (S. pombe) [Source:MGI | 0 | 0 | 0 | 0 | 0 | 0 |
| 10613 | ENSMUSG00000037161 | 0 | NA | NA | NA | NA | NA | Mgarp | mitochondria localized glutamic acid rich | 0 | 0 | 0 | 0 | 0 | 0 |
| 10614 | ENSMUSG00000037166 | 32.17403766 | -0.184493616 | 0.421157308 | -0.438063432 | 0.661340302 | 0.928341045 | Ppp1r14a | protein phosphatase 1, regulatory inhibito | 38.549 | 29.81 | 34.212 | 31.811 | 17.45 | 41.212 |
| 10615 | ENSMUSG00000037167 | 0.535401493 | -2.48596965 | 3.684117979 | -0.674780141 | 0.499815459 | NA | Spaca5 | sperm acrosome associated 5 [Source:MG | 3.2124 | 0 | 0 | 0 | 0 | 0 |
| 10616 | ENSMUSG00000037169 | 40.54759963 | -0.458622632 | 0.3510891 | -1.306285591 | 0.191455455 | 0.629099068 | Mycn | v-myc avian myelocytomatosis viral relate | 48.989 | 53.452 | 38.237 | 29.883 | 37.081 | 35.643 |
| 10617 | ENSMUSG00000037171 | 14.83257906 | -0.555696016 | 0.592979458 | -0.937125239 | 0.34869414 | 0.789995175 | Nodal | nodal [Source:MGI Symbol;Acc:MGI:9735 | 22.487 | 16.447 | 14.087 | 18.316 | 7.6343 | 10.025 |
| 10618 | ENSMUSG00000037172 | 273.2672696 | -0.136980786 | 0.180027432 | -0.760888408 | 0.446723723 | 0.853104294 | Dennd11 | DENN domain containing 11 [Source:MGI | 226.47 | 308.38 | 325.02 | 281.48 | 244.3 | 253.96 |
| 10619 | ENSMUSG00000037174 | 443.0982512 | -0.053814691 | 0.140535896 | -0.382924878 | 0.701775465 | 0.942189108 | Elf2 | E74-like factor 2 [Source:MGI Symbol;Acc | 423.23 | 502.65 | 428.66 | 433.79 | 402.44 | 467.81 |
| 10620 | ENSMUSG00000037185 | 3.52997452 | -0.416310654 | 1.232441913 | -0.337793326 | 0.735518939 | NA | Krt80 | keratin 80 [Source:MGI Symbol;Acc:MGI:1 | 8.031 | 1.0279 | 3.0187 | 5.7839 | 1.0906 | 2.2277 |
| 10621 | ENSMUSG00000037188 | 3.860141667 | -1.104046848 | 1.314871713 | -0.839661267 | 0.401098337 | NA | Grhl3 | grainyhead like transcription factor 3 [Sou | 13.653 | 1.0279 | 1.0062 | 1.928 | 1.0906 | 4.4554 |
| 10622 | ENSMUSG00000037190 | 138.3340517 | 0.022254093 | 0.210271012 | 0.105835289 | 0.915713037 | 0.98625062 | Cyb561d2 | cytochrome b-561 domain containing 2 [S | 163.03 | 121.29 | 126.79 | 136.88 | 147.23 | 134.78 |
| 10623 | ENSMUSG00000037196 | 2.874261976 | 0.621783239 | 1.375946128 | 0.451895046 | 0.651344594 | NA | Pacrg | PARK2 co-regulated [Source:MGI Symbol; | 5.6217 | 1.0279 | 0 | 2.8919 | 4.3625 | 3.3415 |
| 10624 | ENSMUSG00000037197 | 702.4763008 | 0.06053002 | 0.116115516 | 0.5212914 | 0.602163791 | 0.915461118 | Rbm17 | RNA binding motif protein 17 [Source:MG | 646.5 | 692.82 | 724.5 | 713.34 | 692.54 | 745.16 |
| 10625 | ENSMUSG00000037202 | 2.326917953 | -4.585908496 | 1.774559963 | -2.584251077 | 0.009759072 | NA | Prf1 | perforin 1 (pore forming protein) [Source: | 4.8186 | 4.1117 | 5.0312 | 0 | 0 | 0 |
| 10626 | ENSMUSG00000037204 | 729.4214847 | 0.043018138 | 0.138810272 | 0.309906013 | 0.75663243 | 0.953085143 | Atg101 | autophagy related 101 [Source:MGI Symb | 685.05 | 634.23 | 836.19 | 678.64 | 743.8 | 798.63 |
| 10627 | ENSMUSG00000037206 | 52.84190251 | -1.075901954 | 0.370982008 | -2.900145913 | 0.00372989 | 0.054645736 | Islr | immunoglobulin superfamily containing le | 96.372 | 78.122 | 40.25 | 31.811 | 37.081 | 33.415 |
| 10628 | ENSMUSG00000037210 | 256.596589 | 0.166567451 | 0.16754241 | 0.994180824 | 0.320134842 | 0.767337321 | Fam193a | family with sequence homology 193, mem | 228.88 | 246.7 | 250.55 | 303.65 | 278.11 | 231.68 |
| 10629 | ENSMUSG00000037211 | 63.05774716 | -1.483662814 | 0.366774506 | -4.045163412 | 5.23E-05 | 0.002179464 | Spry1 | sprouty RTK signaling antagonist 1 [Source | 112.43 | 101.76 | 64.4 | 36.631 | 17.45 | 45.668 |
| 10630 | ENSMUSG00000037214 | 128.189128 | 0.141344315 | 0.204636471 | 0.690709305 | 0.489748241 | 0.875278016 | Thap1 | THAP domain containing, apoptosis assoc | 114.84 | 127.46 | 123.77 | 141.7 | 123.24 | 138.12 |
| 10631 | ENSMUSG00000037216 | 121.8039222 | 0.109347792 | 0.213796737 | 0.511456788 | 0.609031239 | 0.916376001 | Lipt1 | lipoyltransferase 1 [Source:MGI Symbol;A | 128.5 | 105.88 | 116.72 | 117.61 | 139.6 | 122.52 |
| 10632 | ENSMUSG00000037217 | 2.935712792 | -1.784506585 | 1.454279616 | -1.227072541 | 0.219795327 | NA | Syn1 | synapsin I [Source:MGI Symbol;Acc:MGI:9 | 5.6217 | 4.1117 | 4.025 | 3.8559 | 0 | 0 |
| 10633 | ENSMUSG00000037221 | 467.8173948 | -0.02887167 | 0.126539072 | -0.228164072 | 0.819518691 | 0.969160361 | Mospd3 | motile sperm domain containing 3 [Sourc | 485.07 | 462.56 | 469.92 | 487.77 | 448.24 | 453.33 |
| 10634 | ENSMUSG00000037224 | 0.428363721 | -0.543730382 | 3.624677344 | -0.15000794 | 0.880758351 | NA | Zfyve28 | zinc finger, FYVE domain containing 28 [Sc | 1.6062 | 0 | 0 | 0.964 | 0 | 0 |
| 10635 | ENSMUSG00000037225 | 35.61293076 | -0.011421482 | 0.431429177 | -0.026473597 | 0.978879593 | 0.997992136 | Fgf2 | fibroblast growth factor 2 [Source:MGI Sy | 43.368 | 24.67 | 39.244 | 48.199 | 39.262 | 18.935 |
| 10636 | ENSMUSG00000037234 | 1204.83759 | 0.110911459 | 0.137356489 | 0.807471565 | 0.41939485 | 0.837591673 | Hook3 | hook microtubule tethering protein 3 [Sou | 1146 | 1205.7 | 1125 | 1488.4 | 1229.1 | 1034.8 |

| | | | | | | | | | | | | | | | |
|---|---|---|---|---|---|---|---|---|---|---|---|---|---|---|---|
| 10637 | ENSMUSG00000037235 | 1616.241873 | 0.157800139 | 0.111560475 | 1.4144807 | 0.15722081 | 0.576436451 | Mxd4 | Max dimerization protein 4 [Source:MGI S | 1657.6 | 1388.7 | 1536.5 | 1763.1 | 1572.7 | 1778.8 |
| 10638 | ENSMUSG00000037236 | 1021.163961 | 0.027561306 | 0.121784657 | 0.226311811 | 0.820958913 | 0.969160361 | Matr3 | matrin 3 [Source:MGI Symbol;Acc:MGI:12 | 932.4 | 1060.8 | 1041.5 | 957.23 | 1158.2 | 976.84 |
| 10639 | ENSMUSG00000037239 | 8.570756479 | -0.428259531 | 0.743005702 | -0.576387947 | 0.564352989 | 0.901468983 | Spred3 | sprouty-related EVH1 domain containing 3 | 7.2279 | 12.335 | 10.062 | 9.6398 | 4.3625 | 7.7969 |
| 10640 | ENSMUSG00000037242 | 1009.332254 | -0.322328509 | 0.133751957 | -2.409897522 | 0.015957002 | 0.148616919 | Clic4 | chloride intracellular channel 4 (mitochon | 945.25 | 1208.8 | 1211.5 | 823.24 | 977.19 | 889.96 |
| 10641 | ENSMUSG00000037243 | 111.3350953 | 0.260355965 | 0.230214557 | 1.130927462 | 0.258085623 | 0.711580339 | Zfp692 | zinc finger protein 692 [Source:MGI Symb | 101.99 | 82.233 | 119.74 | 128.21 | 118.88 | 116.95 |
| 10642 | ENSMUSG00000037246 | 0 | NA | NA | NA | NA | NA | H2-M10.5 | histocompatibility 2, M region locus 10.5 [ | 0 | 0 | 0 | 0 | 0 | 0 |
| 10643 | ENSMUSG00000037247 | 0 | NA | NA | NA | NA | NA | Pldi | polymorphic derived intron containing [Sc | 0 | 0 | 0 | 0 | 0 | 0 |
| 10644 | ENSMUSG00000037251 | 147.9337546 | 0.177471221 | 0.257118429 | 0.69023143 | 0.490048661 | 0.875501956 | Pomk | protein-O-mannose kinase [Source:MGI S | 131.71 | 127.46 | 156.97 | 107 | 146.14 | 218.31 |
| 10645 | ENSMUSG00000037253 | 332.579454 | 0.205909731 | 0.141669158 | 1.453454897 | 0.146097494 | 0.556855766 | Mex3c | mex3 RNA binding family member C [Sour | 304.38 | 323.79 | 298.85 | 374.02 | 342.45 | 351.97 |
| 10646 | ENSMUSG00000037254 | 15477.25411 | 0.155403358 | 0.100491422 | 1.546434067 | 0.121999772 | 0.511776306 | Itih2 | inter-alpha trypsin inhibitor, heavy chain 2 | 13124 | 15565 | 15247 | 17381 | 15850 | 15697 |
| 10647 | ENSMUSG00000037257 | 624.2362301 | 0.124526762 | 0.129139424 | 0.964281537 | 0.334904795 | 0.776159726 | Aagab | alpha- and gamma-adaptin binding protei | 580.64 | 587.97 | 622.87 | 570.67 | 686 | 697.27 |
| 10648 | ENSMUSG00000037259 | 11.50748592 | 0.487974442 | 0.650951803 | 0.749632215 | 0.453476243 | 0.857073339 | Dzank1 | double zinc ribbon and ankyrin repeat dor | 6.4248 | 14.391 | 8.05 | 11.568 | 16.359 | 12.252 |
| 10649 | ENSMUSG00000037260 | 1105.223373 | -0.053536964 | 0.120340552 | -0.444878831 | 0.656407321 | 0.92672284 | Hgsnat | heparan-alpha-glucosaminide N-acetyltra | 1014.3 | 1185.2 | 1178.3 | 1011.2 | 1196.4 | 1045.9 |
| 10650 | ENSMUSG00000037262 | 200.2478425 | 0.166230673 | 0.185187406 | 0.897634872 | 0.369380241 | 0.80653433 | Kin | Kin17 DNA and RNA binding protein [Sour | 213.63 | 181.94 | 170.06 | 212.08 | 236.66 | 187.13 |
| 10651 | ENSMUSG00000037263 | 2.289783632 | -1.191369508 | 1.52702254 | -0.780191174 | 0.435278357 | NA | Aldh3b3 | aldehyde dehydrogenase 3 family, membe | 2.4093 | 6.1675 | 1.0062 | 1.928 | 0 | 2.2277 |
| 10652 | ENSMUSG00000037266 | 7005.187266 | -0.051850623 | 0.108431467 | -0.478187973 | 0.632516422 | 0.922076476 | Rsrp1 | arginine/serine rich protein 1 [Source:MG | 7379.7 | 6696.9 | 7316.4 | 7006.2 | 7602.7 | 6029.2 |
| 10653 | ENSMUSG00000037270 | 712.724082 | -0.398401522 | 0.164079359 | -2.428102626 | 0.015178048 | 0.143769012 | 4932438A13 | RIKEN cDNA 4932438A13 gene [Source:M | 636.06 | 905.6 | 891.53 | 696.96 | 624.92 | 521.28 |
| 10654 | ENSMUSG00000037275 | 264.0064067 | 0.054891363 | 0.155152295 | 0.353790209 | 0.723496111 | 0.9447719 | Gemin5 | gem nuclear organelle associated protein | 263.42 | 239.5 | 273.7 | 256.42 | 278.11 | 272.89 |
| 10655 | ENSMUSG00000037278 | 1201.088156 | 0.348159527 | 0.159948574 | 2.17669666 | 0.029503208 | 0.226164801 | Tmem97 | transmembrane protein 97 [Source:MGI S | 1219.1 | 1148.2 | 801.98 | 1165.4 | 1501.8 | 1370 |
| 10656 | ENSMUSG00000037279 | 0.160662974 | 1.126924175 | 4.080472857 | 0.276174898 | 0.782413736 | NA | Ovol2 | ovo like zinc finger 2 [Source:MGI Symbol; | 0 | 0 | 0 | 0.964 | 0 | 0 |
| 10657 | ENSMUSG00000037280 | 13.44510027 | -1.98376672 | 0.765225572 | -2.592394704 | 0.009531035 | 0.104754479 | Galnt6 | polypeptide N-acetylgalactosaminyltransf | 17.668 | 33.921 | 13.081 | 11.568 | 1.0906 | 3.3415 |
| 10658 | ENSMUSG00000037286 | 408.6055907 | 0.057703875 | 0.135234612 | 0.426694576 | 0.669601802 | 0.930837335 | Stag1 | stromal antigen 1 [Source:MGI Symbol;Ac | 379.06 | 408.08 | 414.57 | 418.37 | 389.35 | 442.2 |
| 10659 | ENSMUSG00000037287 | 1397.070567 | 0.565460758 | 0.172807533 | 3.27219971 | 0.001067142 | 0.022018741 | Tbcel | tubulin folding cofactor E-like [Source:MG | 1205.5 | 1056.7 | 1116.9 | 1295.6 | 1475.6 | 2232.1 |
| 10660 | ENSMUSG00000037295 | 722.0893218 | 0.163712686 | 0.118688461 | 1.379347949 | 0.167787499 | 0.593804353 | Ldlrap1 | low density lipoprotein receptor adaptor p | 641.68 | 657.87 | 744.62 | 776.97 | 736.17 | 775.24 |
| 10661 | ENSMUSG00000037296 | 245.5406474 | 0.111881436 | 0.169456462 | 0.660237057 | 0.509101715 | 0.882793133 | Lsm1 | LSM1 homolog, mRNA degradation associ | 261.81 | 223.06 | 222.38 | 241.96 | 236.66 | 287.37 |
| 10662 | ENSMUSG00000037300 | 475.2924561 | 0.148032099 | 0.130120446 | 1.137654405 | 0.255264819 | 0.708666093 | Ttc13 | tetratricopeptide repeat domain 13 [Sour | 452.15 | 486.21 | 414.57 | 508.98 | 487.51 | 502.34 |
| 10663 | ENSMUSG00000037306 | 34.85167042 | -0.731983386 | 0.388818903 | -1.882581789 | 0.059757066 | 0.348006896 | Man1c1 | mannosidase, alpha, class 1C, member 1 [ | 48.989 | 47.284 | 34.212 | 28.919 | 19.631 | 30.074 |
| 10664 | ENSMUSG00000037307 | 0 | NA | NA | NA | NA | NA | Banf2 | BANF family member 2 [Source:MGI Symb | 0 | 0 | 0 | 0 | 0 | 0 |
| 10665 | ENSMUSG00000037313 | 25.76182917 | -0.176330016 | 0.456311933 | -0.386424293 | 0.699182461 | 0.941092046 | Tacc3 | transforming, acidic coiled-coil containing | 36.14 | 23.642 | 22.137 | 31.811 | 17.45 | 23.391 |
| 10666 | ENSMUSG00000037315 | 56.99554634 | 0.122304651 | 0.293342468 | 0.416934691 | 0.676726178 | 0.932891189 | Jade3 | jade family PHD finger 3 [Source:MGI Sym | 48.989 | 56.535 | 58.362 | 57.839 | 54.531 | 65.717 |
| 10667 | ENSMUSG00000037316 | 636.4547807 | 0.072911028 | 0.119978409 | 0.607701238 | 0.543385642 | 0.894449746 | Bag4 | BCL2-associated athanogene 4 [Source:M | 609.55 | 672.26 | 579.6 | 653.58 | 654.37 | 649.37 |
| 10668 | ENSMUSG00000037318 | 5.375405924 | -0.058290488 | 1.099534626 | -0.053013781 | 0.957720927 | 0.991861398 | Traf3ip3 | TRAF3 interacting protein 3 [Source:MGI S | 6.4248 | 3.0838 | 7.0437 | 13.496 | 1.0906 | 1.1138 |
| 10669 | ENSMUSG00000037321 | 700.2060027 | 0.329948123 | 0.131636788 | 2.506503903 | 0.012193171 | 0.124023929 | Tap1 | transporter 1, ATP-binding cassette, sub-f | 681.83 | 562.27 | 615.82 | 695.99 | 828.87 | 816.45 |
| 10670 | ENSMUSG00000037325 | 62.93885815 | 0.06301725 | 0.32622581 | 0.193170644 | 0.846825329 | 0.972803041 | Bbs7 | Bardet-Biedl syndrome 7 (human) [Source | 73.082 | 42.145 | 69.431 | 83.866 | 54.531 | 54.578 |
| 10671 | ENSMUSG00000037326 | 272.1949305 | -0.239545316 | 0.168493747 | -1.421686683 | 0.155117228 | 0.573534185 | Capn15 | calpain 15 [Source:MGI Symbol;Acc:MGI:1 | 284.3 | 285.76 | 314.95 | 294.01 | 223.58 | 230.57 |
| 10672 | ENSMUSG00000037331 | 2280.759034 | 0.084805286 | 0.099081644 | 0.855913196 | 0.392045795 | 0.820867166 | Larp1 | La ribonucleoprotein domain family, mem | 2070.4 | 2261.4 | 2310.3 | 2349.2 | 2222.7 | 2470.5 |
| 10673 | ENSMUSG00000037334 | 0 | NA | NA | NA | NA | NA | H2-M1 | histocompatibility 2, M region locus 1 [Sou | 0 | 0 | 0 | 0 | 0 | 0 |
| 10674 | ENSMUSG00000037335 | 0.906399088 | 2.120308852 | 2.545194173 | 0.833063691 | 0.40480881 | NA | Hand1 | heart and neural crest derivatives express | 0 | 0 | 1.0062 | 0 | 1.0906 | 3.3415 |
| 10675 | ENSMUSG00000037336 | 12.57755921 | 0.14725181 | 0.653508605 | 0.225324975 | 0.821726472 | 0.969160361 | Mfsd2b | major facilitator superfamily domain cont | 5.6217 | 12.335 | 18.112 | 17.352 | 10.906 | 11.138 |
| 10676 | ENSMUSG00000037337 | 14.11557845 | -0.812306416 | 0.579990939 | -1.400550182 | 0.161348627 | 0.58445944 | Map4k1 | mitogen-activated protein kinase kinase k | 15.259 | 22.614 | 16.1 | 8.6758 | 10.906 | 11.138 |
| 10677 | ENSMUSG00000037339 | 495.5483508 | 0.144176043 | 0.12914894 | 1.116354826 | 0.264270282 | 0.719492304 | Fam53a | family with sequence similarity 53, membe | 503.55 | 440.98 | 466.9 | 490.66 | 537.67 | 533.53 |
| 10678 | ENSMUSG00000037341 | 4.203293949 | 1.108503806 | 1.186531167 | 0.934239097 | 0.350180572 | NA | Slc9a7 | solute carrier family 9 (sodium/hydrogen e | 4.8186 | 0 | 3.0187 | 1.928 | 6.5437 | 8.9108 |
| 10679 | ENSMUSG00000037343 | 302.7322716 | -0.038048107 | 0.150739467 | -0.252409725 | 0.800724384 | 0.962424862 | Taf2 | TATA-box binding protein associated facto | 336.5 | 303.24 | 279.74 | 294.01 | 298.83 | 304.08 |
| 10680 | ENSMUSG00000037344 | 286.0766818 | 0.012668225 | 0.164038009 | 0.077227376 | 0.938442664 | 0.988467426 | Slc12a9 | solute carrier family 12 (potassium/chlorid | 301.16 | 291.93 | 261.62 | 329.68 | 278.11 | 253.96 |
| 10681 | ENSMUSG00000037346 | 0 | NA | NA | NA | NA | NA | Hrh4 | histamine receptor H4 [Source:MGI Symb | 0 | 0 | 0 | 0 | 0 | 0 |
| 10682 | ENSMUSG00000037347 | 11.62575439 | -0.371290453 | 0.642612206 | -0.57778307 | 0.563410587 | 0.900821918 | Chst7 | carbohydrate (N-acetylglucosamino) sulfo | 16.865 | 11.307 | 11.069 | 10.604 | 6.5437 | 13.366 |
| 10683 | ENSMUSG00000037348 | 2053.203846 | -0.0061369 | 0.151399605 | -0.040534451 | 0.967667042 | 0.994231936 | Paqr7 | progestin and adipoQ receptor family mer | 2371.6 | 2038.4 | 1761.9 | 2028.2 | 2450.6 | 1668.5 |
| 10684 | ENSMUSG00000037349 | 138.1256152 | 0.165575032 | 0.209202366 | 0.791458699 | 0.428676368 | 0.842753542 | Nudt22 | nudix (nucleoside diphosphate linked moi | 120.47 | 135.69 | 134.84 | 155.2 | 121.06 | 161.51 |
| 10685 | ENSMUSG00000037351 | 1613.689302 | -0.004849134 | 0.097078333 | -0.049950737 | 0.960161645 | 0.992338552 | Actr1b | ARP1 actin-related protein 1B, centractin | 1650.4 | 1584 | 1615 | 1698.5 | 1577 | 1557.2 |
| 10686 | ENSMUSG00000037353 | 592.876522 | 0.13506459 | 0.116975298 | 1.154641979 | 0.248237069 | 0.699604734 | Letmd1 | LETM1 domain containing 1 [Source:MGI S | 566.99 | 556.1 | 572.55 | 643.94 | 616.2 | 601.48 |
| 10687 | ENSMUSG00000037355 | 268.1931292 | -0.122784054 | 0.189822047 | -0.646837686 | 0.517736999 | 0.887022094 | Uvssa | UV stimulated scaffold protein A [Source:I | 284.3 | 261.09 | 293.82 | 295.94 | 284.65 | 189.35 |
| 10688 | ENSMUSG00000037358 | 0.353348078 | 0.165142701 | 3.895042568 | 0.042398176 | 0.966181282 | NA | Dipk2b | divergent protein kinase domain 2B [Sour | 0 | 0 | 1.0062 | 0 | 0 | 1.1138 |
| 10689 | ENSMUSG00000037361 | 444.6616178 | -0.006907822 | 0.152592339 | -0.04526978 | 0.963892275 | 0.993245949 | Sf3b6 | splicing factor 3B, subunit 6 [Source:MGI S | 429.66 | 406.03 | 501.11 | 391.38 | 431.88 | 507.91 |
| 10690 | ENSMUSG00000037362 | 0.185640647 | 1.126924175 | 4.080472857 | 0.276174898 | 0.782413736 | NA | Ccn3 | cellular communication network factor 3 [ | 0 | 0 | 0 | 0 | 0 | 1.1138 |
| 10691 | ENSMUSG00000037363 | 49.8373266 | -0.09098873 | 0.333079314 | -0.273174364 | 0.784719185 | 0.960670212 | Letm2 | leucine zipper-EF-hand containing transm | 63.445 | 43.173 | 47.293 | 47.235 | 58.893 | 38.985 |
| 10692 | ENSMUSG00000037364 | 737.9756864 | -0.058581427 | 0.126964354 | -0.461400583 | 0.644511233 | 0.925643431 | Srrt | serrate RNA effector molecule homolog (A | 691.47 | 762.72 | 806 | 797.21 | 659.82 | 710.63 |

| | | | | | | | | | | | | | | | |
|---|---|---|---|---|---|---|---|---|---|---|---|---|---|---|---|
| 10693 | ENSMUSG00000037366 | 2861.311049 | -0.133605219 | 0.114195153 | -1.169972767 | 0.242011928 | 0.691791181 | Pafah2 | platelet-activating factor acetylhydrolase | 3156.2 | 2918.3 | 2905 | 2345.4 | 2990.5 | 2852.6 |
| 10694 | ENSMUSG00000037369 | 255.2911536 | 0.014920785 | 0.175297315 | 0.085117017 | 0.932168361 | 0.988420305 | Kdm6a | lysine (K)-specific demethylase 6A [Source | 224.87 | 248.76 | 288.79 | 248.71 | 292.29 | 228.34 |
| 10695 | ENSMUSG00000037370 | 556.1440148 | -0.087195356 | 0.130606655 | -0.667618013 | 0.504377456 | 0.881510796 | Enpp1 | ectonucleotide pyrophosphatase/phosph | 506.76 | 606.47 | 606.77 | 551.4 | 545.31 | 520.17 |
| 10696 | ENSMUSG00000037373 | 995.9344467 | -0.023353823 | 0.105178033 | -0.222040875 | 0.824282063 | 0.96943211 | Ctbp1 | C-terminal binding protein 1 [Source:MGI | 985.41 | 1060.8 | 965.99 | 979.4 | 982.65 | 1001.3 |
| 10697 | ENSMUSG00000037375 | 26.43902377 | 0.229681624 | 0.477742417 | 0.480764563 | 0.630683839 | 0.921020101 | Hhat | hedgehog acyltransferase [Source:MGI Sy | 22.487 | 21.586 | 29.181 | 43.379 | 15.269 | 26.732 |
| 10698 | ENSMUSG00000037376 | 206.9281727 | 0.086426499 | 0.178929253 | 0.483020507 | 0.629081176 | 0.92057725 | Trmt6 | tRNA methyltransferase 6 [Source:MGI Sy | 191.94 | 215.86 | 194.21 | 182.19 | 229.03 | 228.34 |
| 10699 | ENSMUSG00000037379 | 68.29530342 | 0.024732183 | 0.30448972 | 0.081225017 | 0.935263004 | 0.988420305 | Spon2 | spondin 2, extracellular matrix protein [Sc | 64.248 | 54.48 | 84.525 | 76.154 | 50.168 | 80.197 |
| 10700 | ENSMUSG00000037386 | 1.439764576 | 0.572545823 | 1.780903968 | 0.321491688 | 0.74783781 | NA | Rims2 | regulating synaptic membrane exocytosis | 2.4093 | 0 | 1.0062 | 1.928 | 2.1812 | 1.1138 |
| 10701 | ENSMUSG00000037390 | 0 | NA | NA | NA | NA | NA | Muc3 | mucin 3, intestinal [Source:MGI Symbol;A | 0 | 0 | 0 | 0 | 0 | 0 |
| 10702 | ENSMUSG00000037393 | 0 | NA | NA | NA | NA | NA | Nmur2 | neuromedin U receptor 2 [Source:MGI Syı | 0 | 0 | 0 | 0 | 0 | 0 |
| 10703 | ENSMUSG00000037395 | 132.0082838 | 0.04279627 | 0.241969599 | 0.17686631 | 0.859613402 | 0.975534508 | Rcor3 | REST corepressor 3 [Source:MGI Symbol;A | 119.66 | 114.1 | 156.97 | 169.66 | 105.79 | 125.86 |
| 10704 | ENSMUSG00000037400 | 916.3123007 | -0.030090571 | 0.110723249 | -0.271763803 | 0.785803639 | 0.960775321 | Atp11b | ATPase, class VI, type 11B [Source:MGI Sy | 875.38 | 968.3 | 934.8 | 920.6 | 944.48 | 854.32 |
| 10705 | ENSMUSG00000037405 | 82.81008788 | -0.482918743 | 0.251652862 | -1.918987681 | 0.054985893 | 0.330417929 | Icam1 | intercellular adhesion molecule 1 [Source | 95.569 | 100.74 | 93.581 | 79.046 | 59.984 | 67.944 |
| 10706 | ENSMUSG00000037406 | 8.002550174 | 2.250638132 | 1.106725699 | 2.033600678 | 0.041991873 | 0.281973285 | Htra4 | HtrA serine peptidase 4 [Source:MGI Syml | 3.2124 | 1.0279 | 4.025 | 0 | 16.359 | 23.391 |
| 10707 | ENSMUSG00000037408 | 67.88778785 | -0.224271254 | 0.308142053 | -0.727817743 | 0.466725162 | 0.863779479 | Cnnm4 | cyclin M4 [Source:MGI Symbol;Acc:MGI:2 | 55.414 | 62.703 | 101.63 | 59.767 | 65.437 | 62.375 |
| 10708 | ENSMUSG00000037410 | 487.8975967 | -0.084298929 | 0.182108126 | -0.462905917 | 0.643431806 | 0.925477092 | Tbc1d2b | TBC1 domain family, member 2B [Source: | 460.18 | 602.36 | 443.75 | 393.3 | 431.88 | 595.91 |
| 10709 | ENSMUSG00000037411 | 23.82891777 | -1.242806226 | 0.473710387 | -2.62355705 | 0.008701687 | 0.097878626 | Serpine1 | serine (or cysteine) peptidase inhibitor, cl | 36.14 | 37.005 | 27.169 | 9.6398 | 18.541 | 14.48 |
| 10710 | ENSMUSG00000037415 | 1674.528472 | 0.056035647 | 0.123742469 | 0.452840867 | 0.650663334 | 0.925643431 | Ranbp10 | RAN binding protein 10 [Source:MGI Syml | 1460 | 1558.3 | 1908.8 | 1768.9 | 1592.3 | 1758.8 |
| 10711 | ENSMUSG00000037416 | 427.9642051 | -0.119968073 | 0.180535657 | -0.66451179 | 0.506362805 | 0.881592135 | Dmxl1 | Dmx-like 1 [Source:MGI Symbol;Acc:MGI: | 393.52 | 529.38 | 415.58 | 489.7 | 417.71 | 321.9 |
| 10712 | ENSMUSG00000037418 | 0.636946036 | 1.591416711 | 2.83781177 | 0.560790087 | 0.574940645 | NA | Best1 | bestrophin 1 [Source:MGI Symbol;Acc:MG | 0.8031 | 0 | 0 | 1.928 | 1.0906 | 0 |
| 10713 | ENSMUSG00000037419 | 71.51259868 | -0.490092175 | 0.306327229 | -1.599897524 | 0.109621319 | 0.486451272 | Endod1 | endonuclease domain containing 1 [Sourc | 99.585 | 81.206 | 69.431 | 49.163 | 81.796 | 47.895 |
| 10714 | ENSMUSG00000037424 | 0 | NA | NA | NA | NA | NA | Taar9 | trace amine-associated receptor 9 [Source | 0 | 0 | 0 | 0 | 0 | 0 |
| 10715 | ENSMUSG00000037426 | 215.4032971 | 0.240155466 | 0.170712181 | 1.406785764 | 0.159490925 | 0.581370147 | Depdc5 | DEP domain containing 5 [Source:MGI Syr | 186.32 | 208.67 | 198.23 | 257.38 | 226.85 | 214.97 |
| 10716 | ENSMUSG00000037428 | 0.17131964 | -0.796629176 | 4.080472857 | -0.195229623 | 0.84521319 | NA | Vgf | VGF nerve growth factor inducible [Sourc | 0 | 1.0279 | 0 | 0 | 0 | 0 |
| 10717 | ENSMUSG00000037432 | 66.23074602 | 0.54801286 | 0.303701182 | 1.80444757 | 0.071161171 | 0.386047033 | Fer1l5 | fer-1-like 5 (C. elegans) [Source:MGI Symb | 53.005 | 41.117 | 67.418 | 93.506 | 63.256 | 79.083 |
| 10718 | ENSMUSG00000037434 | 948.1415847 | 0.095153691 | 0.11265636 | 0.84463665 | 0.39831374 | 0.825882163 | Slc30a1 | solute carrier family 30 (zinc transporter), | 910.72 | 849.06 | 991.15 | 1018.9 | 948.84 | 970.16 |
| 10719 | ENSMUSG00000037437 | 1.03119563 | 0.094051261 | 2.229235409 | 0.042189919 | 0.966347299 | NA | Adam32 | a disintegrin and metallopeptidase domai | 0 | 0 | 3.0187 | 0.964 | 1.0906 | 1.1138 |
| 10720 | ENSMUSG00000037438 | 1.374681561 | 1.793959471 | 2.085650988 | 0.860143658 | 0.389709858 | NA | Uqcrh-ps1 | ubiquinol-cytochrome c reductase hinge p | 0.8031 | 1.0279 | 0 | 0.964 | 5.4531 | 0 |
| 10721 | ENSMUSG00000037440 | 2054.593451 | -0.546100836 | 0.183734498 | -2.972228094 | 0.00295647 | 0.046110081 | Vnn1 | vanin 1 [Source:MGI Symbol;Acc:MGI:108 | 2761.9 | 2584.2 | 1971.2 | 2176.7 | 1621.7 | 1211.9 |
| 10722 | ENSMUSG00000037443 | 3333.216197 | -0.179751784 | 0.171845583 | -1.04600759 | 0.295557528 | 0.74606354 | Cep85 | centrosomal protein 85 [Source:MGI Syml | 4396.2 | 3528.8 | 2695.7 | 3264 | 2481.2 | 3633.4 |
| 10723 | ENSMUSG00000037446 | 0 | NA | NA | NA | NA | NA | Tulp1 | tubby like protein 1 [Source:MGI Symbol;A | 0 | 0 | 0 | 0 | 0 | 0 |
| 10724 | ENSMUSG00000037447 | 35.77194348 | 0.652665165 | 0.413259948 | 1.579309023 | 0.114265193 | 0.496868018 | Arid5a | AT rich interactive domain 5A (MRF1-like) | 25.699 | 27.754 | 30.187 | 56.875 | 23.994 | 50.123 |
| 10725 | ENSMUSG00000037451 | 0.503302254 | -0.837877584 | 3.426698643 | -0.244514523 | 0.806832356 | NA | Slc22a20 | solute carrier family 22 (organic anion trai | 0 | 2.0558 | 0 | 0.964 | 0 | 0 |
| 10726 | ENSMUSG00000037455 | 83.53606013 | -0.315810394 | 0.242753942 | -1.300948569 | 0.19327606 | 0.631476153 | Slc18b1 | solute carrier family 18, subfamily B, mem | 94.766 | 90.457 | 92.575 | 71.334 | 76.343 | 75.741 |
| 10727 | ENSMUSG00000037458 | 2124.598094 | 0.295716603 | 0.121478322 | 2.434315835 | 0.01491997 | 0.142198341 | Azin1 | antizyme inhibitor 1 [Source:MGI Symbol; | 2146.7 | 1669.3 | 1904.8 | 2123.6 | 2512.8 | 2390.3 |
| 10728 | ENSMUSG00000037461 | 338.2039839 | 0.206648277 | 0.1656354 | 1.247609372 | 0.212174145 | 0.656735971 | Ints7 | integrator complex subunit 7 [Source:MG | 333.29 | 278.57 | 330.05 | 379.81 | 407.89 | 299.62 |
| 10729 | ENSMUSG00000037463 | 0.462220778 | -0.685400455 | 3.191475091 | -0.21475977 | 0.82995462 | NA | Fbxo27 | F-box protein 27 [Source:MGI Symbol;Acc | 0.8031 | 0 | 1.0062 | 0.964 | 0 | 0 |
| 10730 | ENSMUSG00000037465 | 631.4656791 | 0.113316732 | 0.199744725 | 0.567307757 | 0.570505108 | 0.904602706 | Klf10 | Kruppel-like factor 10 [Source:MGI Symbc | 815.15 | 485.18 | 518.22 | 636.23 | 539.86 | 794.17 |
| 10731 | ENSMUSG00000037466 | 4.11765213 | 0.805458991 | 1.051088709 | 0.766309241 | 0.443492321 | NA | Tedc1 | tubulin epsilon and delta complex 1 [Sour | 4.8186 | 2.0558 | 2.0125 | 4.8199 | 6.5437 | 4.4554 |
| 10732 | ENSMUSG00000037469 | 0 | NA | NA | NA | NA | NA | Acp7 | acid phosphatase 7, tartrate resistant [Sou | 0 | 0 | 0 | 0 | 0 | 0 |
| 10733 | ENSMUSG00000037470 | 1084.942865 | 0.132618955 | 0.111597481 | 1.188368716 | 0.234688176 | 0.682729239 | Uggt1 | UDP-glucose glycoprotein glucosyltransfe | 1035.2 | 997.08 | 1073.7 | 1239.7 | 1053.5 | 1110.5 |
| 10734 | ENSMUSG00000037474 | 8.436183483 | 0.174624554 | 0.807656389 | 0.216211444 | 0.828822933 | 0.969955221 | Dtl | denticleless E3 ubiquitin protein ligase [Sc | 9.6372 | 2.0558 | 12.075 | 12.532 | 7.6343 | 6.6831 |
| 10735 | ENSMUSG00000037475 | 508.2388478 | 0.056513211 | 0.133030302 | 0.424814573 | 0.670971854 | 0.93116767 | Thoc2 | THO complex 2 [Source:MGI Symbol;Acc:I | 470.62 | 518.07 | 507.15 | 559.11 | 531.13 | 463.36 |
| 10736 | ENSMUSG00000037477 | 1.32696881 | -0.53124189 | 2.17162591 | -0.244628639 | 0.806743987 | NA | Tbx10 | T-box 10 [Source:MGI Symbol;Acc:MGI:12 | 1.6062 | 3.0838 | 0 | 0 | 3.2719 | 0 |
| 10737 | ENSMUSG00000037482 | 0 | NA | NA | NA | NA | NA | Erv3 | endogenous retroviral sequence 3 [Source | 0 | 0 | 0 | 0 | 0 | 0 |
| 10738 | ENSMUSG00000037486 | 476.205341 | 0.124523658 | 0.138786225 | 0.897233553 | 0.369594305 | 0.80653433 | Asxl2 | ASXL transcriptional regulator 2 [Source:N | 454.56 | 451.26 | 461.87 | 561.04 | 490.78 | 437.74 |
| 10739 | ENSMUSG00000037487 | 2126.99282 | -0.062355522 | 0.149108396 | -0.418189206 | 0.675808787 | 0.932630311 | Ubr5 | ubiquitin protein ligase E3 component n-r | 1934.7 | 2435.1 | 2150.3 | 2271.1 | 2362.3 | 1608.4 |
| 10740 | ENSMUSG00000037490 | 1.391768125 | 0.099984466 | 1.825726943 | 0.054764195 | 0.956326326 | NA | Slc2a12 | solute carrier family 2 (facilitated glucose | 0 | 2.0558 | 2.0125 | 0.964 | 1.0906 | 2.2277 |
| 10741 | ENSMUSG00000037492 | 0.305170013 | -1.657836477 | 4.041480335 | -0.410205257 | 0.681655385 | NA | Zmat4 | zinc finger, matrin type 4 [Source:MGI Syr | 0.8031 | 1.0279 | 0 | 0 | 0 | 0 |
| 10742 | ENSMUSG00000037493 | 79.73165382 | 0.258182524 | 0.268566234 | 0.961336502 | 0.336383 | 0.778173248 | Cib2 | calcium and integrin binding family memb | 84.326 | 55.508 | 77.481 | 78.082 | 89.431 | 93.563 |
| 10743 | ENSMUSG00000037499 | 119.7409864 | -0.01575357 | 0.275858204 | -0.057107493 | 0.954459568 | 0.991678478 | Nenf | neuron derived neurotrophic factor [Sour | 109.22 | 154.19 | 97.606 | 85.794 | 111.24 | 160.39 |
| 10744 | ENSMUSG00000037503 | 1095.437299 | 0.062216045 | 0.116799224 | 0.532675155 | 0.594258467 | 0.913144152 | Fam168b | family with sequence similarity 168, meml | 1028 | 1126.6 | 1060.6 | 992.9 | 1156.1 | 1208.5 |
| 10745 | ENSMUSG00000037509 | 2.610877486 | -2.01617492 | 1.480706014 | -1.361630803 | 0.173314429 | NA | Arhgef4 | Rho guanine nucleotide exchange factor ( | 2.4093 | 7.1954 | 3.0187 | 1.928 | 0 | 1.1138 |
| 10746 | ENSMUSG00000037514 | 383.1688733 | 0.009370568 | 0.144770747 | 0.064726947 | 0.948391407 | 0.990731019 | Pank2 | pantothenate kinase 2 [Source:MGI Symb | 384.69 | 401.92 | 359.23 | 383.66 | 425.34 | 344.18 |
| 10747 | ENSMUSG00000037519 | 561.472256 | 0.097818559 | 0.137054739 | 0.713718912 | 0.475401009 | 0.86938592 | Ppfia1 | protein tyrosine phosphatase, receptor ty | 492.3 | 575.63 | 560.48 | 605.38 | 626.01 | 509.03 |
| 10748 | ENSMUSG00000037523 | 1649.621885 | 0.26353904 | 0.113789321 | 2.316026126 | 0.020556842 | 0.176557791 | Mavs | mitochondrial antiviral signaling protein [S | 1665.6 | 1394.9 | 1435.9 | 1749.6 | 1702.5 | 1949.2 |

| | | | | | | | | | | | | | | | |
|---|---|---|---|---|---|---|---|---|---|---|---|---|---|---|---|
| 10749 | ENSMUSG00000037525 | 71.22161414 | 0.434941666 | 0.264456144 | 1.644664628 | 0.100038991 | 0.4657776 | Bcdin3d | BCDIN3 domain containing [Source:MGI S | 53.808 | 63.731 | 64.4 | 80.01 | 79.615 | 85.766 |
| 10750 | ENSMUSG00000037526 | 412.1909676 | 0.311627917 | 0.15100857 | 2.063643911 | 0.039051491 | 0.270232919 | Atg14 | autophagy related 14 [Source:MGI Symbo | 417.61 | 338.18 | 347.15 | 508.02 | 415.53 | 446.65 |
| 10751 | ENSMUSG00000037529 | 0 | NA | NA | NA | NA | NA | Prss40 | protease, serine 40 [Source:MGI Symbol;A | 0 | 0 | 0 | 0 | 0 | 0 |
| 10752 | ENSMUSG00000037531 | 258.8675761 | 0.050671298 | 0.156146649 | 0.324510952 | 0.745551232 | 0.949997329 | Mrpl47 | mitochondrial ribosomal protein L47 [Sou | 256.99 | 262.12 | 243.51 | 244.85 | 263.93 | 281.8 |
| 10753 | ENSMUSG00000037533 | 321.3819836 | 0.225101304 | 0.145914278 | 1.542695531 | 0.122904667 | 0.513267141 | Rapgef6 | Rap guanine nucleotide exchange factor ( | 278.68 | 301.18 | 309.92 | 368.24 | 326.09 | 344.18 |
| 10754 | ENSMUSG00000037535 | 0.562214094 | -1.122836648 | 3.276281221 | -0.34271681 | 0.731811515 | NA | 1700021A07 | RIKEN cDNA 1700021A07 gene [Source:M | 2.4093 | 0 | 0 | 0.964 | 0 | 0 |
| 10755 | ENSMUSG00000037536 | 418.2794771 | -0.043403292 | 0.163782159 | -0.26500623 | 0.791004663 | 0.961019028 | Fbxo34 | F-box protein 34 [Source:MGI Symbol;Acc | 471.42 | 435.84 | 365.27 | 340.28 | 460.24 | 436.63 |
| 10756 | ENSMUSG00000037537 | 0 | NA | NA | NA | NA | NA | H2-M11 | histocompatibility 2, M region locus 11 [S | 0 | 0 | 0 | 0 | 0 | 0 |
| 10757 | ENSMUSG00000037541 | 139.5853915 | 0.119520682 | 0.22777316 | 0.524735582 | 0.599767018 | 0.915217938 | Shank2 | SH3 and multiple ankyrin repeat domains | 119.66 | 117.18 | 165.02 | 156.16 | 161.41 | 118.07 |
| 10758 | ENSMUSG00000037542 | 6460.518521 | 0.106913666 | 0.088052794 | 1.214199575 | 0.224671523 | 0.670999154 | Aldh8a1 | aldehyde dehydrogenase 8 family, membe | 5947.8 | 6235.3 | 6481.2 | 6536.7 | 7014.9 | 6547.2 |
| 10759 | ENSMUSG00000037544 | 2.486267835 | 0.160132045 | 1.508514452 | 0.106152145 | 0.915461639 | NA | Dlgap5 | DLG associated protein 5 [Source:MGI Syr | 4.0155 | 1.0279 | 2.0125 | 6.7478 | 0 | 1.1138 |
| 10760 | ENSMUSG00000037548 | 6.090479092 | -1.765868572 | 0.952722585 | -1.853497125 | 0.063811138 | 0.361521464 | H2-DMb2 | histocompatibility 2, class II, locus Mb2 [S | 12.047 | 5.1396 | 11.069 | 3.8559 | 1.0906 | 3.3415 |
| 10761 | ENSMUSG00000037552 | 98.23835246 | -0.213069089 | 0.251161808 | -0.848333952 | 0.396252014 | 0.823764863 | Plekhg2 | pleckstrin homology domain containing, f | 101.19 | 106.9 | 108.67 | 107 | 66.528 | 99.132 |
| 10762 | ENSMUSG00000037553 | 500.4278194 | 0.04559506 | 0.126764375 | 0.359683542 | 0.719083801 | 0.9447719 | Zdhhc18 | zinc finger, DHHC domain containing 18 [S | 464.19 | 516.01 | 498.09 | 525.37 | 480.96 | 517.94 |
| 10763 | ENSMUSG00000037563 | 3190.025607 | 0.013389123 | 0.10184717 | 0.131462886 | 0.895409144 | 0.982798755 | Rps16 | ribosomal protein S16 [Source:MGI Symbo | 2917.7 | 3475.4 | 3133.4 | 3070.3 | 3211.9 | 3331.5 |
| 10764 | ENSMUSG00000037568 | 0.874376958 | 3.338720136 | 2.469103386 | 1.352199407 | 0.176311533 | NA | Vash2 | vasohibin 2 [Source:MGI Symbol;Acc:MGI | 0 | 0 | 0 | 1.928 | 1.0906 | 2.2277 |
| 10765 | ENSMUSG00000037570 | 158.6351445 | 0.24787768 | 0.190223311 | 1.303087826 | 0.192544779 | 0.63052317 | Mcrs1 | microspherule protein 1 [Source:MGI Sym | 157.41 | 131.57 | 145.91 | 184.12 | 167.96 | 164.85 |
| 10766 | ENSMUSG00000037572 | 23.22635861 | -0.211065324 | 0.463091389 | -0.455774667 | 0.648552023 | 0.925643431 | Wdhd1 | WD repeat and HMG-box DNA binding pro | 28.109 | 17.475 | 29.181 | 26.027 | 19.631 | 18.935 |
| 10767 | ENSMUSG00000037573 | 2366.704281 | -0.036253996 | 0.098002962 | -0.369927554 | 0.711436471 | 0.944239215 | Tob1 | transducer of ErbB-2.1 [Source:MGI Symb | 2325.8 | 2415.6 | 2448.2 | 2323.2 | 2178 | 2509.5 |
| 10768 | ENSMUSG00000037577 | 0.913882229 | 2.129306995 | 2.79764157 | 0.761107862 | 0.446592645 | NA | Ephx3 | epoxide hydrolase 3 [Source:MGI Symbol; | 0 | 1.0279 | 0 | 0 | 0 | 4.4554 |
| 10769 | ENSMUSG00000037578 | 0 | NA | NA | NA | NA | NA | Pkd2l1 | polycystic kidney disease 2-like 1 [Source: | 0 | 0 | 0 | 0 | 0 | 0 |
| 10770 | ENSMUSG00000037579 | 0.17131964 | -0.796629176 | 4.080472857 | -0.195229623 | 0.84521319 | NA | Kcnh3 | potassium voltage-gated channel, subfam | 0 | 1.0279 | 0 | 0 | 0 | 0 |
| 10771 | ENSMUSG00000037580 | 3657.194648 | 0.352033561 | 0.101553095 | 3.466497623 | 0.000527286 | 0.012909222 | Gch1 | GTP cyclohydrolase 1 [Source:MGI Symbo | 3292.7 | 2892.6 | 3453.4 | 3948.5 | 4082.2 | 4273.8 |
| 10772 | ENSMUSG00000037583 | 450.6573618 | -0.3354752 | 0.634332618 | -0.528863234 | 0.596900328 | 0.91428215 | Nr0b2 | nuclear receptor subfamily 0, group B, me | 487.48 | 312.49 | 708.4 | 227.5 | 725.26 | 242.82 |
| 10773 | ENSMUSG00000037593 | 3.686967638 | -0.724654797 | 1.217964184 | -0.594972173 | 0.551862072 | NA | Rskr | ribosomal protein S6 kinase related [Sour | 9.6372 | 1.0279 | 3.0187 | 2.8919 | 1.0906 | 4.4554 |
| 10774 | ENSMUSG00000037594 | 20.90869771 | -0.327870598 | 0.515343445 | -0.636217654 | 0.524634568 | 0.888334265 | Clba1 | clathrin binding box of aftiphilin containin | 24.896 | 30.838 | 14.087 | 20.244 | 11.997 | 23.391 |
| 10775 | ENSMUSG00000037600 | 102.5327216 | 0.301344966 | 0.242073072 | 1.24485125 | 0.213186433 | 0.657889604 | Kdf1 | keratinocyte differentiation factor 1 [Sour | 100.39 | 75.038 | 99.618 | 95.434 | 119.97 | 124.75 |
| 10776 | ENSMUSG00000037601 | 309.0077772 | 0.115421841 | 0.157025333 | 0.735052361 | 0.462307606 | 0.860650866 | Nme1 | NME/NM23 nucleoside diphosphate kinas | 323.65 | 276.51 | 288.79 | 290.16 | 316.28 | 358.66 |
| 10777 | ENSMUSG00000037603 | 0 | NA | NA | NA | NA | NA | Gm5519 | predicted pseudogene 5519 [Source:MGI | 0 | 0 | 0 | 0 | 0 | 0 |
| 10778 | ENSMUSG00000037605 | 1.839667896 | -0.711099816 | 1.548660795 | -0.4591708 | 0.646111518 | NA | Adgrl3 | adhesion G protein-coupled receptor L3 [S | 0.8031 | 3.0838 | 3.0187 | 1.928 | 1.0906 | 1.1138 |
| 10779 | ENSMUSG00000037606 | 194.5509482 | -0.565067478 | 0.560447618 | -1.008243161 | 0.313337742 | 0.760152431 | Osbpl5 | oxysterol binding protein-like 5 [Source:M | 227.28 | 262.12 | 207.29 | 212.08 | 202.86 | 55.692 |
| 10780 | ENSMUSG00000037608 | 873.3940686 | 0.101017739 | 0.117864806 | 0.857064485 | 0.391409247 | 0.820365651 | Bclaf1 | BCL2-associated transcription factor 1 [So | 783.83 | 921.01 | 824.11 | 849.26 | 951.02 | 911.12 |
| 10781 | ENSMUSG00000037610 | 1.211103787 | 2.67104802 | 2.127358732 | 1.255570102 | 0.209271877 | NA | Kcnmb2 | potassium large conductance calcium-acti | 0.8031 | 0 | 0 | 0.964 | 3.2719 | 2.2277 |
| 10782 | ENSMUSG00000037613 | 6.403356643 | -2.667321835 | 1.122597241 | -2.376027427 | 0.017500162 | 0.157512348 | Tnfrsf23 | tumor necrosis factor receptor superfamil | 20.881 | 9.2513 | 3.0187 | 1.928 | 0 | 3.3415 |
| 10783 | ENSMUSG00000037617 | 18.08891067 | -0.366672139 | 0.518167783 | -0.707632067 | 0.479173774 | 0.870169438 | Spag1 | sperm associated antigen 1 [Source:MGI S | 14.456 | 26.726 | 20.125 | 16.388 | 16.359 | 14.48 |
| 10784 | ENSMUSG00000037621 | 271.9296421 | -0.923257301 | 0.240695494 | -3.835789712 | 0.000125161 | 0.004328686 | Atoh8 | atonal bHLH transcription factor 8 [Source | 295.54 | 275.48 | 497.08 | 157.13 | 166.86 | 239.48 |
| 10785 | ENSMUSG00000037622 | 2569.321192 | 0.141135322 | 0.107240402 | 1.316064846 | 0.188152279 | 0.625185978 | Wdtc1 | WD and tetratricopeptide repeats 1 [Sour | 2372.4 | 2423.8 | 2534.7 | 2504.4 | 2552 | 3028.5 |
| 10786 | ENSMUSG00000037624 | 0.803102239 | -3.069617558 | 3.178309912 | -0.965801839 | 0.334143348 | NA | Kcnk2 | potassium channel, subfamily K, member | 4.8186 | 0 | 0 | 0 | 0 | 0 |
| 10787 | ENSMUSG00000037625 | 0 | NA | NA | NA | NA | NA | Cldn11 | claudin 11 [Source:MGI Symbol;Acc:MGI:1 | 0 | 0 | 0 | 0 | 0 | 0 |
| 10788 | ENSMUSG00000037627 | 0 | NA | NA | NA | NA | NA | Rgs22 | regulator of G-protein signalling 22 [Sourc | 0 | 0 | 0 | 0 | 0 | 0 |
| 10789 | ENSMUSG00000037628 | 3.513540398 | -0.419240679 | 1.237975094 | -0.338650334 | 0.73487316 | NA | Cdkn3 | cyclin-dependent kinase inhibitor 3 [Sourc | 4.0155 | 4.1117 | 4.025 | 6.7478 | 2.1812 | 0 |
| 10790 | ENSMUSG00000037636 | 0.462220778 | -0.685400455 | 3.191475091 | -0.21475977 | 0.82995462 | NA | Slc25a43 | solute carrier family 25, member 43 [Sour | 0.8031 | 0 | 1.0062 | 0.964 | 0 | 0 |
| 10791 | ENSMUSG00000037638 | 95.74181438 | 0.465560804 | 0.263270535 | 1.768374131 | 0.076998378 | 0.406191172 | Zbtb42 | zinc finger and BTB domain containing 42 | 60.233 | 80.178 | 101.63 | 126.28 | 101.43 | 104.7 |
| 10792 | ENSMUSG00000037640 | 161.8501997 | -0.024289766 | 0.201873111 | -0.120321949 | 0.904228118 | 0.98383695 | Zfp60 | zinc finger protein 60 [Source:MGI Symbo | 147.77 | 169.61 | 173.07 | 190.87 | 148.32 | 141.46 |
| 10793 | ENSMUSG00000037643 | 335.8959234 | -0.167153666 | 0.148674324 | -1.124294107 | 0.260888277 | 0.714835715 | Prkci | protein kinase C, iota [Source:MGI Symbo | 336.5 | 392.66 | 337.09 | 303.65 | 340.27 | 305.19 |
| 10794 | ENSMUSG00000037646 | 702.3396853 | -0.018934705 | 0.128237332 | -0.147653607 | 0.882616146 | 0.981769403 | Vps13b | vacuolar protein sorting 13B [Source:MGI | 666.57 | 756.55 | 698.33 | 691.17 | 775.43 | 625.98 |
| 10795 | ENSMUSG00000037649 | 30.5428002 | -1.651249551 | 0.443299559 | -3.724906821 | 0.000195387 | 0.006197551 | H2-DMa | histocompatibility 2, class II, locus DMa [S | 48.186 | 57.563 | 33.206 | 14.46 | 10.906 | 18.935 |
| 10796 | ENSMUSG00000037652 | 421.1865457 | 0.088406821 | 0.13380209 | 0.660728248 | 0.508786605 | 0.882793133 | Phc3 | polyhomeotic 3 [Source:MGI Symbol;Acc: | 399.14 | 426.59 | 399.48 | 439.57 | 460.24 | 402.1 |
| 10797 | ENSMUSG00000037653 | 0 | NA | NA | NA | NA | NA | Kctd8 | potassium channel tetramerisation domai | 0 | 0 | 0 | 0 | 0 | 0 |
| 10798 | ENSMUSG00000037656 | 1567.110383 | 0.064049929 | 0.116406412 | 0.550226812 | 0.582163816 | 0.908645447 | Slc20a2 | solute carrier family 20, member 2 [Sourc | 1553.2 | 1621 | 1421.8 | 1412.2 | 1650.1 | 1744.3 |
| 10799 | ENSMUSG00000037660 | 0 | NA | NA | NA | NA | NA | Gdf7 | growth differentiation factor 7 [Source:M | 0 | 0 | 0 | 0 | 0 | 0 |
| 10800 | ENSMUSG00000037661 | 6.915825231 | -0.121031911 | 0.793170282 | -0.152592594 | 0.878719565 | 0.980997205 | Gpr160 | G protein-coupled receptor 160 [Source:N | 6.4248 | 7.1954 | 8.05 | 7.7118 | 6.5437 | 5.5692 |
| 10801 | ENSMUSG00000037664 | 29.04209221 | -0.826792236 | 0.406387641 | -2.034491584 | 0.041902056 | 0.281500855 | Cdkn1c | cyclin-dependent kinase inhibitor 1C (P57 | 35.336 | 35.977 | 40.25 | 24.099 | 18.541 | 20.049 |
| 10802 | ENSMUSG00000037669 | 1048.82883 | 0.057937538 | 0.11174827 | 0.518464741 | 0.604134054 | 0.916322307 | Ldah | lipid droplet associated hydrolase [Source | 1023.2 | 1053.6 | 1007.3 | 1147.1 | 1093.9 | 967.93 |
| 10803 | ENSMUSG00000037674 | 191.9372385 | 0.215218452 | 0.174889444 | 1.23059716 | 0.218473568 | 0.663735712 | Rfx7 | regulatory factor X, 7 [Source:MGI Symbo | 173.47 | 170.63 | 189.17 | 222.68 | 197.4 | 198.26 |
| 10804 | ENSMUSG00000037679 | 955.2271602 | 0.242856899 | 0.116553444 | 2.0836527 | 0.03719177 | 0.261128941 | Inf2 | inverted formin, FH2 and WH2 domain co | 912.32 | 843.92 | 868.39 | 1068.1 | 927.03 | 1111.6 |

| | | | | | | | | | | | | | | | |
|---|---|---|---|---|---|---|---|---|---|---|---|---|---|---|---|
| 10805 | ENSMUSG00000037681 | 2.642872986 | -1.004766823 | 1.370925625 | -0.732911257 | 0.463612557 | NA | Esyt3 | extended synaptotagmin-like protein 3 [S | 2.4093 | 7.1954 | 1.0062 | 1.928 | 1.0906 | 2.2277 |
| 10806 | ENSMUSG00000037683 | 0 | NA | NA | NA | NA | NA | Armc3 | armadillo repeat containing 3 [Source:MG | 0 | 0 | 0 | 0 | 0 | 0 |
| 10807 | ENSMUSG00000037685 | 64.34544881 | -0.860433082 | 0.343319667 | -2.506215533 | 0.012203121 | 0.124037905 | Atp8a1 | ATPase, aminophospholipid transporter ( | 72.279 | 119.24 | 57.356 | 34.703 | 51.259 | 51.237 |
| 10808 | ENSMUSG00000037686 | 4923.064031 | 0.349767798 | 0.105342619 | 3.320287646 | 0.000899247 | 0.01935523 | Aspg | asparaginase [Source:MGI Symbol;Acc:M | 4414.7 | 3938 | 4634.8 | 5729.9 | 5014.7 | 5806.5 |
| 10809 | ENSMUSG00000037689 | 0.181769715 | 1.126924175 | 4.080472857 | 0.276174898 | 0.782413736 | NA | Tmem247 | transmembrane protein 247 [Source:MGI | 0 | 0 | 0 | 0 | 1.0906 | 0 |
| 10810 | ENSMUSG00000037692 | 304.7527542 | -0.257828701 | 0.152936429 | -1.685855374 | 0.091823659 | 0.446623824 | Ahdc1 | AT hook, DNA binding motif, containing 1 | 305.98 | 342.3 | 348.16 | 294.98 | 250.84 | 286.26 |
| 10811 | ENSMUSG00000037697 | 251.923347 | -0.182611546 | 0.165829093 | -1.10120331 | 0.270808182 | 0.726306238 | Ddhd1 | DDHD domain containing 1 [Source:MGI S | 261.01 | 279.59 | 263.64 | 269.91 | 225.76 | 211.63 |
| 10812 | ENSMUSG00000037703 | 601.0059747 | 0.279628029 | 0.15578253 | 1.794989649 | 0.072655349 | 0.390954974 | Lzts3 | leucine zipper, putative tumor suppressor | 505.95 | 491.34 | 631.92 | 678.64 | 549.67 | 748.5 |
| 10813 | ENSMUSG00000037705 | 0.346303622 | 2.004339653 | 3.884003423 | 0.516049919 | 0.605819544 | NA | Tecta | tectorin alpha [Source:MGI Symbol;Acc:M | 0 | 0 | 0 | 0.964 | 0 | 1.1138 |
| 10814 | ENSMUSG00000037706 | 4062.499082 | -0.322637951 | 0.130542366 | -2.471519109 | 0.013454036 | 0.132518282 | Cd81 | CD81 antigen [Source:MGI Symbol;Acc:M | 5134.2 | 4789.1 | 3620.5 | 3642.9 | 3784.4 | 3403.9 |
| 10815 | ENSMUSG00000037708 | 0 | NA | NA | NA | NA | NA | Spag6 | sperm associated antigen 6 [Source:MGI S | 0 | 0 | 0 | 0 | 0 | 0 |
| 10816 | ENSMUSG00000037709 | 456.1983471 | -0.085929481 | 0.226390739 | -0.379562705 | 0.70427005 | 0.94295036 | Fam13a | family with sequence similarity 13, memb | 432.87 | 474.9 | 501.11 | 312.33 | 358.81 | 657.17 |
| 10817 | ENSMUSG00000037710 | 3287.75646 | 0.074733322 | 0.095463269 | 0.782848973 | 0.433715805 | 0.845264432 | Cisd1 | CDGSH iron sulfur domain 1 [Source:MGI | 3238.1 | 3208.1 | 3160.6 | 3092.4 | 3556.5 | 3470.7 |
| 10818 | ENSMUSG00000037712 | 1679.257454 | 0.318018548 | 0.134491663 | 2.36459674 | 0.018049713 | 0.160758032 | Fermt2 | fermitin family member 2 [Source:MGI Sy | 1293.8 | 1471 | 1720.7 | 1701.4 | 1765.7 | 2123 |
| 10819 | ENSMUSG00000037716 | 0 | NA | NA | NA | NA | NA | Ccdc33 | coiled-coil domain containing 33 [Source: | 0 | 0 | 0 | 0 | 0 | 0 |
| 10820 | ENSMUSG00000037720 | 1447.018517 | 0.030924135 | 0.099570545 | 0.310575135 | 0.756123633 | 0.953085143 | Tmem33 | transmembrane protein 33 [Source:MGI S | 1402.2 | 1380.5 | 1512.4 | 1509.6 | 1447.3 | 1430.2 |
| 10821 | ENSMUSG00000037722 | 25.51777156 | 0.423885719 | 0.497736554 | 0.851626658 | 0.394421339 | 0.822827574 | Gnpnat1 | glucosamine-phosphate N-acetyltransfera | 34.533 | 13.363 | 17.106 | 29.883 | 38.172 | 20.049 |
| 10822 | ENSMUSG00000037725 | 6.311462084 | 0.149404063 | 0.970656477 | 0.153920637 | 0.877672312 | 0.98072613 | Ckap2 | cytoskeleton associated protein 2 [Source | 13.653 | 2.0558 | 2.0125 | 5.7839 | 5.4531 | 8.9108 |
| 10823 | ENSMUSG00000037727 | 0 | NA | NA | NA | NA | NA | Avp | arginine vasopressin [Source:MGI Symbol | 0 | 0 | 0 | 0 | 0 | 0 |
| 10824 | ENSMUSG00000037730 | 181.714242 | -0.140123877 | 0.18823371 | -0.74441436 | 0.456625834 | 0.858190757 | Mynn | myoneurin [Source:MGI Symbol;Acc:MGI: | 183.91 | 213.81 | 174.08 | 170.62 | 193.04 | 154.82 |
| 10825 | ENSMUSG00000037731 | 41.93663714 | -1.525725382 | 0.39869277 | -3.826819791 | 0.000129809 | 0.004428218 | Themis2 | thymocyte selection associated family me | 61.036 | 82.233 | 43.269 | 15.424 | 21.812 | 27.846 |
| 10826 | ENSMUSG00000037735 | 0.335414862 | -1.779047242 | 4.030804179 | -0.441362855 | 0.658950329 | NA | 2810032G03 | RIKEN cDNA 2810032G03 gene [Source:M | 0 | 0 | 2.0125 | 0 | 0 | 0 |
| 10827 | ENSMUSG00000037736 | 9.380740662 | 0.236512315 | 0.744875911 | 0.317519081 | 0.750849763 | 0.952074518 | Limch1 | LIM and calponin homology domains 1 [S | 10.44 | 12.335 | 3.0187 | 10.604 | 7.6343 | 12.252 |
| 10828 | ENSMUSG00000037737 | 0 | NA | NA | NA | NA | NA | Actrt3 | actin related protein T3 [Source:MGI Sym | 0 | 0 | 0 | 0 | 0 | 0 |
| 10829 | ENSMUSG00000037738 | 3.486154885 | 0.689660205 | 1.273622761 | 0.541494881 | 0.588166525 | NA | Nek5 | NIMA (never in mitosis gene a)-related ex | 0 | 2.0558 | 6.0375 | 2.8919 | 4.3625 | 5.5692 |
| 10830 | ENSMUSG00000037740 | 488.9141475 | 0.16865106 | 0.136038543 | 1.239729982 | 0.215075284 | 0.659728602 | Mrps26 | mitochondrial ribosomal protein S26 [Sou | 432.07 | 487.23 | 461.87 | 462.71 | 562.76 | 526.85 |
| 10831 | ENSMUSG00000037742 | 70187.1818 | 0.088847683 | 0.088621366 | 1.002553744 | 0.316076223 | 0.762717132 | Eef1a1 | eukaryotic translation elongation factor 1 | 62827 | 72173 | 69081 | 71718 | 73935 | 71388 |
| 10832 | ENSMUSG00000037747 | 0 | NA | NA | NA | NA | NA | Phyhipl | phytanoyl-CoA hydroxylase interacting pr | 0 | 0 | 0 | 0 | 0 | 0 |
| 10833 | ENSMUSG00000037750 | 282.7584746 | -0.309898111 | 0.153149395 | -2.023502032 | 0.043021414 | 0.28636067 | Fam222b | family with sequence similarity 222, mem | 309.19 | 299.12 | 331.05 | 267.02 | 258.48 | 231.68 |
| 10834 | ENSMUSG00000037752 | 182.0012042 | 0.224964271 | 0.188521406 | 1.193308896 | 0.23274842 | 0.67995822 | Xkr8 | X-linked Kx blood group related 8 [Source | 150.18 | 160.36 | 193.2 | 178.34 | 207.22 | 202.72 |
| 10835 | ENSMUSG00000037754 | 29.71682635 | -0.263657492 | 0.456474048 | -0.577595797 | 0.563537046 | 0.900821918 | Ppp1r16b | protein phosphatase 1, regulatory subuni | 28.912 | 50.368 | 18.112 | 27.955 | 23.994 | 28.96 |
| 10836 | ENSMUSG00000037759 | 18.10859768 | -1.39618651 | 0.544754414 | -2.562965025 | 0.01037825 | 0.111734379 | Ptger2 | prostaglandin E receptor 2 (subtype EP2) | 21.684 | 27.754 | 29.181 | 5.7839 | 11.997 | 12.252 |
| 10837 | ENSMUSG00000037761 | 160.2370717 | 0.006899798 | 0.233424619 | 0.029559 | 0.976418764 | 0.9970485 | Actr5 | ARP5 actin-related protein 5 [Source:MGI | 150.98 | 154.19 | 175.09 | 207.26 | 161.41 | 112.5 |
| 10838 | ENSMUSG00000037762 | 22.41605505 | -1.534768828 | 0.481908952 | -3.184769285 | 0.001448695 | 0.027643691 | Slc16a9 | solute carrier family 16 (monocarboxylic a | 36.943 | 32.893 | 30.187 | 13.496 | 8.7249 | 12.252 |
| 10839 | ENSMUSG00000037771 | 0 | NA | NA | NA | NA | NA | Slc32a1 | solute carrier family 32 (GABA vesicular tr | 0 | 0 | 0 | 0 | 0 | 0 |
| 10840 | ENSMUSG00000037772 | 448.4026888 | -0.147183731 | 0.157552384 | -0.934189171 | 0.350206321 | 0.791589971 | Mrpl23 | mitochondrial ribosomal protein L23 [Sou | 517.2 | 464.62 | 430.67 | 354.74 | 427.52 | 495.66 |
| 10841 | ENSMUSG00000037773 | 370.7541571 | 0.11742125 | 0.147768895 | 0.794627653 | 0.426830128 | 0.842132174 | Pced1a | PC-esterase domain containing 1A [Sourc | 388.7 | 318.65 | 359.23 | 406.8 | 399.17 | 351.97 |
| 10842 | ENSMUSG00000037780 | 3343.812016 | 0.182922607 | 0.09300974 | 1.966703776 | 0.049217371 | 0.309647699 | Mbl1 | mannose-binding lectin (protein A) 1 [Sou | 3141.7 | 3142.3 | 3112.3 | 3630.3 | 3295.8 | 3740.3 |
| 10843 | ENSMUSG00000037784 | 10.64277009 | -1.477397163 | 0.712469949 | -2.073627339 | 0.038113934 | 0.265293524 | Dzip1l | DAZ interacting protein 1-like [Source:MG | 14.456 | 15.419 | 17.106 | 5.7839 | 2.1812 | 8.9108 |
| 10844 | ENSMUSG00000037787 | 343.7247425 | -0.089652667 | 0.155677884 | -0.575885698 | 0.564692443 | 0.901542552 | Coa8 | cytochrome c oxidase assembly factor 8 [ | 365.41 | 398.83 | 298.85 | 334.5 | 323.91 | 340.84 |
| 10845 | ENSMUSG00000037788 | 53.11794895 | -0.534843698 | 0.315062178 | -1.697581416 | 0.089586793 | 0.441643958 | Vopp1 | vesicular, overexpressed in cancer, prosu | 68.264 | 58.591 | 61.381 | 36.631 | 39.262 | 54.578 |
| 10846 | ENSMUSG00000037790 | 0 | NA | NA | NA | NA | NA | Defb7 | defensin beta 7 [Source:MGI Symbol;Acc: | 0 | 0 | 0 | 0 | 0 | 0 |
| 10847 | ENSMUSG00000037791 | 482.9642188 | -0.015811942 | 0.132931787 | -0.118947788 | 0.905316721 | 0.983972587 | Phf12 | PHD finger protein 12 [Source:MGI Symbo | 479.45 | 476.95 | 501.11 | 529.22 | 474.42 | 436.63 |
| 10848 | ENSMUSG00000037795 | 710.05316 | 0.008916706 | 0.128845171 | 0.069204813 | 0.944826592 | 0.990287814 | N4bp2 | NEDD4 binding protein 2 [Source:MGI Syr | 662.56 | 668.15 | 792.92 | 653.58 | 756.89 | 726.23 |
| 10849 | ENSMUSG00000037797 | 2220.732812 | 0.640178078 | 0.108638706 | 5.89272554 | 3.80E-09 | 6.62E-07 | Adh4 | alcohol dehydrogenase 4 (class II), pi poly | 1741.1 | 1597.4 | 1869.6 | 2857.2 | 2796.3 | 2462.7 |
| 10850 | ENSMUSG00000037798 | 75113.52688 | -0.448376315 | 0.090932015 | -4.930896067 | 8.19E-07 | 6.96E-05 | Mat1a | methionine adenosyltransferase I, alpha [ | 80590 | 92163 | 87327 | 64708 | 65583 | 60310 |
| 10851 | ENSMUSG00000037801 | 3.105171402 | 1.320303835 | 1.258896979 | 1.048778301 | 0.294280167 | NA | Iqch | IQ motif containing H [Source:MGI Symbo | 3.2124 | 1.0279 | 1.0062 | 6.7478 | 2.1812 | 4.4554 |
| 10852 | ENSMUSG00000037805 | 2706.025193 | 0.13369445 | 0.101082722 | 1.322624156 | 0.185960397 | 0.621903163 | Rpl10a | ribosomal protein L10A [Source:MGI Sym | 2371.6 | 2720.9 | 2650.4 | 2655.8 | 2937 | 2900.4 |
| 10853 | ENSMUSG00000037808 | 178.5555943 | 0.128374323 | 0.216500489 | 0.592951657 | 0.553213506 | 0.896245536 | Fam76b | family with sequence similarity 76, memb | 159.01 | 162.41 | 191.19 | 236.17 | 178.86 | 143.69 |
| 10854 | ENSMUSG00000037813 | 17.51151676 | -0.086344522 | 0.523807222 | -0.164840267 | 0.869069709 | 0.978605211 | D630003M2 | RIKEN cDNA D630003M21 gene [Source:N | 14.456 | 21.586 | 18.112 | 14.46 | 14.178 | 22.277 |
| 10855 | ENSMUSG00000037815 | 2107.374702 | -0.156610198 | 0.107972535 | -1.450463291 | 0.146929369 | 0.558084661 | Ctnna1 | catenin (cadherin associated protein), alp | 2199.7 | 2310.8 | 2154.4 | 1899 | 2231.4 | 1849 |
| 10856 | ENSMUSG00000037816 | 14.49929882 | -0.808914469 | 0.686755797 | -1.177877891 | 0.23884529 | 0.68804188 | Fbxw17 | F-box and WD-40 domain protein 17 [Sou | 12.047 | 17.475 | 26.162 | 20.244 | 3.2719 | 7.7969 |
| 10857 | ENSMUSG00000037818 | 463.5726186 | 0.328464268 | 0.144732299 | 2.269460727 | 0.023240323 | 0.192524645 | Abhd18 | abhydrolase domain containing 18 [Sourc | 424.04 | 388.55 | 419.6 | 438.61 | 569.3 | 541.33 |
| 10858 | ENSMUSG00000037820 | 1130.763577 | -0.251573006 | 0.122536747 | -2.053041325 | 0.04006857 | 0.274075154 | Tgm2 | transglutaminase 2, C polypeptide [Sourc | 1304.2 | 1285.9 | 1096.8 | 1070 | 922.66 | 1104.9 |
| 10859 | ENSMUSG00000037822 | 1086.739412 | 0.048822501 | 0.118624407 | 0.41157214 | 0.680653055 | 0.934384556 | Smim14 | small integral membrane protein 14 [Sour | 1206.3 | 1016.6 | 981.09 | 1150 | 1097.2 | 1069.3 |
| 10860 | ENSMUSG00000037824 | 498.7727854 | -0.106046856 | 0.138982328 | -0.763024029 | 0.445449066 | 0.852147532 | Tspan14 | tetraspanin 14 [Source:MGI Symbol;Acc:N | 505.15 | 495.46 | 551.42 | 540.79 | 426.43 | 473.38 |

| | | | | | | | | | | | | | | | |
|---|---|---|---|---|---|---|---|---|---|---|---|---|---|---|---|
| 10861 | ENSMUSG00000037826 | 1656.673869 | 0.247800111 | 0.141478735 | 1.75150075 | 0.079859692 | 0.415052314 | Ppm1k | protein phosphatase 1K (PP2C domain co | 1362.9 | 1298.3 | 1883.7 | 1759.3 | 1865 | 1771 |
| 10862 | ENSMUSG00000037827 | 33.15979013 | 0.500211457 | 0.438243632 | 1.141400401 | 0.253703338 | 0.706909089 | Gm5884 | predicted pseudogene 5884 [Source:MGI | 26.502 | 18.503 | 37.231 | 25.063 | 37.081 | 54.578 |
| 10863 | ENSMUSG00000037833 | 1.035014429 | -2.151020407 | 2.418909279 | -0.889252204 | 0.373867552 | NA | Sh2d4b | SH2 domain containing 4B [Source:MGI Sy | 0 | 3.0838 | 2.0125 | 0 | 0 | 1.1138 |
| 10864 | ENSMUSG00000037843 | 0.160662974 | 1.126924175 | 4.080472857 | 0.276174898 | 0.782413736 | NA | Vstm2l | V-set and transmembrane domain contai | 0 | 0 | 0 | 0.964 | 0 | 0 |
| 10865 | ENSMUSG00000037845 | 58.14180326 | 0.462317155 | 0.301618938 | 1.532785569 | 0.125328697 | 0.51896773 | Fdxacb1 | ferredoxin-fold anticodon binding domain | 48.989 | 46.256 | 51.318 | 55.911 | 82.887 | 63.489 |
| 10866 | ENSMUSG00000037846 | 1.020977649 | 2.347692013 | 2.415115044 | 0.972082891 | 0.331009313 | NA | Rtkn2 | rhotekin 2 [Source:MGI Symbol;Acc:MGI:2 | 0 | 0 | 1.0062 | 2.8919 | 0 | 2.2277 |
| 10867 | ENSMUSG00000037847 | 1112.678397 | 0.11434517 | 0.195456728 | 0.585015267 | 0.558537451 | 0.899348015 | Nmrk1 | nicotinamide riboside kinase 1 [Source:M | 1163.7 | 1038.2 | 1004.2 | 1421.9 | 1320.7 | 727.34 |
| 10868 | ENSMUSG00000037849 | 7.083997448 | -1.607515253 | 0.835430232 | -1.924176539 | 0.054332459 | 0.328127217 | Ifi206 | interferon activated gene 206 [Source:MG | 9.6372 | 12.335 | 10.062 | 3.8559 | 3.2719 | 3.3415 |
| 10869 | ENSMUSG00000037851 | 522.7697205 | 0.001309583 | 0.141627386 | 0.00924668 | 0.992622322 | 0.99859176 | Iars | isoleucine-tRNA synthetase [Source:MGI S | 463.39 | 585.91 | 519.22 | 517.66 | 571.48 | 478.95 |
| 10870 | ENSMUSG00000037852 | 21.82222939 | -0.322300912 | 0.507491658 | -0.635086127 | 0.525372245 | 0.888334265 | Cpe | carboxypeptidase E [Source:MGI Symbol;A | 24.896 | 32.893 | 15.094 | 26.027 | 13.087 | 18.935 |
| 10871 | ENSMUSG00000037855 | 0 | NA | NA | NA | NA | NA | Zfp365 | zinc finger protein 365 [Source:MGI Symb | 0 | 0 | 0 | 0 | 0 | 0 |
| 10872 | ENSMUSG00000037857 | 576.0423406 | -0.000308759 | 0.138226668 | -0.002233712 | 0.998217757 | 0.999764475 | Nufip2 | nuclear fragile X mental retardation prote | 515.59 | 641.42 | 572.55 | 611.16 | 600.93 | 514.6 |
| 10873 | ENSMUSG00000037860 | 14.86429954 | -0.959057319 | 0.669441951 | -1.432622079 | 0.151965869 | 0.567340036 | Aim2 | absent in melanoma 2 [Source:MGI Symb | 16.865 | 29.81 | 12.075 | 2.8919 | 14.178 | 13.366 |
| 10874 | ENSMUSG00000037868 | 1.559707122 | -4.007492068 | 2.057955009 | -1.947317629 | 0.051496666 | NA | Egr2 | early growth response 2 [Source:MGI Sym | 3.2124 | 5.1396 | 1.0062 | 0 | 0 | 0 |
| 10875 | ENSMUSG00000037872 | 7.155859556 | -0.643341895 | 0.825283812 | -0.779540185 | 0.435661575 | 0.845940264 | Ackr1 | atypical chemokine receptor 1 (Duffy bloc | 12.85 | 7.1954 | 6.0375 | 5.7839 | 3.2719 | 7.7969 |
| 10876 | ENSMUSG00000037876 | 682.7958026 | -0.061992458 | 0.131332018 | -0.472028516 | 0.636906435 | 0.923478764 | Jmjd1c | jumonji domain containing 1C [Source:MG | 624.01 | 789.44 | 680.22 | 699.85 | 677.27 | 625.98 |
| 10877 | ENSMUSG00000037884 | 0 | NA | NA | NA | NA | NA | 1700017G19 | RIKEN cDNA 1700017G19 gene [Source:M | 0 | 0 | 0 | 0 | 0 | 0 |
| 10878 | ENSMUSG00000037885 | 476.1807673 | 0.095082304 | 0.135485512 | 0.701789459 | 0.482810474 | 0.871710696 | Stk35 | serine/threonine kinase 35 [Source:MGI S | 512.38 | 426.59 | 441.74 | 508.02 | 502.78 | 465.59 |
| 10879 | ENSMUSG00000037887 | 9.228233922 | -0.076737532 | 0.743200965 | -0.103252734 | 0.917762388 | 0.986374671 | Dusp8 | dual specificity phosphatase 8 [Source:M | 16.062 | 6.1675 | 6.0375 | 10.604 | 9.8156 | 6.6831 |
| 10880 | ENSMUSG00000037890 | 57.70197539 | 0.388390365 | 0.306826002 | 1.265832628 | 0.205573011 | 0.650907725 | Wdr19 | WD repeat domain 19 [Source:MGI Symb | 50.595 | 40.089 | 59.368 | 77.118 | 58.893 | 60.148 |
| 10881 | ENSMUSG00000037892 | 6.226467195 | -0.313916421 | 0.897844239 | -0.349633497 | 0.726613768 | 0.945777495 | Pcdh18 | protocadherin 18 [Source:MGI Symbol;Ac | 10.44 | 6.1675 | 4.025 | 6.7478 | 2.1812 | 7.7969 |
| 10882 | ENSMUSG00000037894 | 1065.053921 | 0.057564338 | 0.101880675 | 0.56501724 | 0.572062038 | 0.905069565 | H2az1 | H2A.Z variant histone 1 [Source:MGI Sym | 1034.4 | 1016.6 | 1080.7 | 1109.5 | 1076.4 | 1072.6 |
| 10883 | ENSMUSG00000037896 | 378.4056529 | 0.050256472 | 0.153080159 | 0.328301673 | 0.74268358 | 0.949206972 | Rcor1 | REST corepressor 1 [Source:MGI Symbol;A | 318.03 | 387.53 | 410.55 | 361.49 | 376.26 | 416.58 |
| 10884 | ENSMUSG00000037902 | 167.3490703 | -1.088150009 | 0.272389856 | -3.994825745 | 6.47E-05 | 0.00255157 | Sirpa | signal-regulatory protein alpha [Source:M | 203.99 | 318.65 | 159.99 | 81.938 | 102.52 | 137 |
| 10885 | ENSMUSG00000037904 | 68.38846425 | 0.2700311 | 0.268150312 | 1.00701393 | 0.31392808 | 0.760476509 | Ankrd9 | ankyrin repeat domain 9 [Source:MGI Sym | 65.051 | 56.535 | 64.4 | 80.01 | 74.162 | 70.172 |
| 10886 | ENSMUSG00000037905 | 636.745448 | 0.073062441 | 0.132481011 | 0.551493685 | 0.581295295 | 0.908645447 | Bri3bp | Bri3 binding protein [Source:MGI Symbol; | 628.83 | 562.27 | 670.16 | 610.2 | 620.56 | 728.45 |
| 10887 | ENSMUSG00000037907 | 48.01864499 | 0.10446382 | 0.392377476 | 0.266232968 | 0.790059794 | 0.961019028 | Ankrd13b | ankyrin repeat domain 13b [Source:MGI S | 52.202 | 52.424 | 34.212 | 54.947 | 68.709 | 25.618 |
| 10888 | ENSMUSG00000037910 | 0 | NA | NA | NA | NA | NA | 1700018B24 | RIKEN cDNA 1700018B24 gene [Source:M | 0 | 0 | 0 | 0 | 0 | 0 |
| 10889 | ENSMUSG00000037913 | 4.504137913 | -2.048155047 | 1.136343815 | -1.80240788 | 0.071481256 | NA | Tmem156 | transmembrane protein 156 [Source:MGI | 6.4248 | 12.335 | 3.0187 | 1.928 | 1.0906 | 2.2277 |
| 10890 | ENSMUSG00000037916 | 3136.149384 | -0.024984727 | 0.091202664 | -0.273947332 | 0.784125097 | 0.960670212 | Ndufv1 | NADH:ubiquinone oxidoreductase core su | 3093.5 | 3282.1 | 3114.3 | 3000.9 | 3271.9 | 3054.2 |
| 10891 | ENSMUSG00000037921 | 114.2314747 | 0.276456049 | 0.221097075 | 1.250383115 | 0.21115963 | 0.655691903 | Ddx60 | DExD/H box helicase 60 [Source:MGI Sym | 109.22 | 96.624 | 103.64 | 107 | 135.24 | 133.66 |
| 10892 | ENSMUSG00000037922 | 16.48976993 | -1.155788149 | 0.558006348 | -2.071281361 | 0.03833251 | 0.266430286 | Bank1 | B cell scaffold protein with ankyrin repeat | 26.502 | 18.503 | 23.144 | 8.6758 | 7.6343 | 14.48 |
| 10893 | ENSMUSG00000037924 | 1.479611194 | -0.378768569 | 2.116041761 | -0.178998627 | 0.85793878 | NA | Olfr16 | olfactory receptor 16 [Source:MGI Symbo | 4.0155 | 0 | 1.0062 | 3.8559 | 0 | 0 |
| 10894 | ENSMUSG00000037926 | 499.9766776 | 0.111552286 | 0.174549514 | 0.639086775 | 0.522766483 | 0.888334265 | Ssh2 | slingshot protein phosphatase 2 [Source:M | 468.21 | 484.15 | 490.04 | 549.47 | 629.29 | 378.71 |
| 10895 | ENSMUSG00000037933 | 305.6988412 | 0.120544357 | 0.169551207 | 0.710961361 | 0.47710818 | 0.869671045 | Bicd2 | BICD cargo adaptor 2 [Source:MGI Symbo | 323.65 | 273.43 | 281.75 | 378.84 | 282.47 | 294.05 |
| 10896 | ENSMUSG00000037935 | 533.7322822 | 0.06599296 | 0.134501819 | 0.49064734 | 0.623675898 | 0.919503423 | Smarce1 | SWI/SNF related, matrix associated, actin | 525.23 | 547.88 | 491.05 | 511.87 | 510.41 | 615.96 |
| 10897 | ENSMUSG00000037936 | 3712.841807 | -0.106393346 | 0.116461202 | -0.913551841 | 0.360952382 | 0.800074801 | Scarb1 | scavenger receptor class B, member 1 [So | 3439.7 | 3749.8 | 4360.1 | 3369.1 | 3437.6 | 3920.7 |
| 10898 | ENSMUSG00000037938 | 190.9526762 | 0.166953872 | 0.187149723 | 0.892087194 | 0.372346198 | 0.807923104 | Chchd5 | coiled-coil-helix-coiled-coil-helix domain c | 204.79 | 149.05 | 185.15 | 196.65 | 200.67 | 209.4 |
| 10899 | ENSMUSG00000037940 | 6.264014696 | -1.793102717 | 0.986872769 | -1.816954296 | 0.069224112 | 0.379983891 | Inpp4b | inositol polyphosphate-4-phosphatase, ty | 12.85 | 8.2233 | 8.05 | 2.8919 | 0 | 5.5692 |
| 10900 | ENSMUSG00000037942 | 6471.623075 | 0.207579155 | 0.106323507 | 1.952335479 | 0.050898388 | 0.316755999 | Crp | C-reactive protein, pentraxin-related [Sou | 6174.3 | 5908.5 | 5936.8 | 6557.9 | 6352.9 | 7899.4 |
| 10901 | ENSMUSG00000037944 | 2.304882238 | -1.752931151 | 1.613517193 | -1.086403764 | 0.277300397 | NA | Ccr7 | chemokine (C-C motif) receptor 7 [Source | 1.6062 | 1.0279 | 8.05 | 0.964 | 2.1812 | 0 |
| 10902 | ENSMUSG00000037946 | 14.31871773 | -1.003706526 | 0.654417537 | -1.533740265 | 0.125093564 | 0.518510965 | Fgd3 | FYVE, RhoGEF and PH domain containing | 10.44 | 33.921 | 13.081 | 8.6758 | 11.997 | 7.7969 |
| 10903 | ENSMUSG00000037949 | 166.5981774 | 0.214282566 | 0.19363585 | 1.106626516 | 0.268455438 | 0.724039098 | Ano10 | anoctamin 10 [Source:MGI Symbol;Acc:M | 158.21 | 149.05 | 154.96 | 150.38 | 186.5 | 200.49 |
| 10904 | ENSMUSG00000037953 | 17.49083984 | -1.531768895 | 0.667480799 | -2.294850872 | 0.021741672 | 0.183472313 | A4gnt | alpha-1,4-N-acetylglucosaminyltransfera | 51.399 | 13.363 | 13.081 | 10.604 | 9.8156 | 6.6831 |
| 10905 | ENSMUSG00000037957 | 341.9036228 | 0.124409076 | 0.146313803 | 0.850289399 | 0.395164209 | 0.822933206 | Wdr20 | WD repeat domain 20 [Source:MGI Symb | 342.12 | 321.74 | 316.97 | 324.86 | 391.53 | 354.2 |
| 10906 | ENSMUSG00000037958 | 205.9181676 | -0.237565711 | 0.185929763 | -1.277717493 | 0.201349055 | 0.645602468 | Nsrp1 | nuclear speckle regulatory protein 1 [Sou | 214.43 | 231.28 | 223.39 | 212.08 | 202.86 | 151.48 |
| 10907 | ENSMUSG00000037960 | 160.7676463 | -0.051740618 | 0.188598354 | -0.274342893 | 0.783821124 | 0.960670212 | Card19 | caspase recruitment domain family, mem | 151.79 | 167.55 | 172.07 | 167.73 | 143.96 | 161.51 |
| 10908 | ENSMUSG00000037962 | 1.007736513 | -0.832948274 | 2.237007798 | -0.372349294 | 0.709632802 | NA | Rflna | refilin A [Source:MGI Symbol;Acc:MGI:19 | 0.8031 | 2.0558 | 1.0062 | 0 | 2.1812 | 0 |
| 10909 | ENSMUSG00000037965 | 625.0498252 | 0.002784614 | 0.164917838 | 0.016884858 | 0.986528473 | 0.99859176 | Zc3h7a | zinc finger CCCH type containing 7 A [Sou | 586.26 | 629.09 | 659.09 | 747.08 | 658.73 | 470.04 |
| 10910 | ENSMUSG00000037966 | 2573.189434 | 0.116266788 | 0.096956269 | 1.199167305 | 0.230462898 | 0.677523445 | Ninj1 | ninjurin 1 [Source:MGI Symbol;Acc:MGI:1 | 2418.1 | 2420.7 | 2569.9 | 2669.3 | 2505.2 | 2855.9 |
| 10911 | ENSMUSG00000037971 | 558.2938565 | 0.199736747 | 0.125408669 | 1.592686919 | 0.111230443 | 0.489450944 | 1110032A03 | RIKEN cDNA 1110032A03 gene [Source:M | 568.6 | 492.37 | 497.08 | 588.03 | 593.3 | 610.39 |
| 10912 | ENSMUSG00000037972 | 18.82220603 | -0.755668253 | 0.502513996 | -1.503775533 | 0.132639173 | 0.533565316 | Snn | stannin [Source:MGI Symbol;Acc:MGI:127 | 28.109 | 21.586 | 21.131 | 13.496 | 16.359 | 12.252 |
| 10913 | ENSMUSG00000037973 | 0 | NA | NA | NA | NA | NA | Itprid1 | ITPR interacting domain containing 1 [Sou | 0 | 0 | 0 | 0 | 0 | 0 |
| 10914 | ENSMUSG00000037974 | 0 | NA | NA | NA | NA | NA | Muc5ac | mucin 5, subtypes A and C, tracheobronc | 0 | 0 | 0 | 0 | 0 | 0 |
| 10915 | ENSMUSG00000037977 | 24.18716145 | 0.757547722 | 0.460251376 | 1.645943417 | 0.099775409 | 0.465232597 | 6430571L13 | RIKEN cDNA 6430571L13 gene [Source:M | 18.471 | 16.447 | 19.119 | 41.451 | 22.903 | 26.732 |
| 10916 | ENSMUSG00000037979 | 1.91264202 | -0.274392484 | 1.488100162 | -0.184391139 | 0.85370662 | NA | Ccdc92 | coiled-coil domain containing 92 [Source: | 3.2124 | 1.0279 | 2.0125 | 1.928 | 2.1812 | 1.1138 |

| | | | | | | | | | | | | | | | |
|---|---|---|---|---|---|---|---|---|---|---|---|---|---|---|---|
| 10917 | ENSMUSG00000037982 | 5.191997383 | -0.143988892 | 0.925455681 | -0.15558702 | 0.876358557 | 0.980632845 | Gm9725 | predicted gene 9725 [Source:MGI Symbol | 3.2124 | 6.1675 | 7.0437 | 4.8199 | 5.4531 | 4.4554 |
| 10918 | ENSMUSG00000037984 | 0 | NA | NA | NA | NA | NA | Neurod6 | neurogenic differentiation 6 [Source:MGI | 0 | 0 | 0 | 0 | 0 | 0 |
| 10919 | ENSMUSG00000037989 | 4.207739448 | -0.218377496 | 1.102301624 | -0.198110473 | 0.842958631 | NA | Wnk2 | WNK lysine deficient protein kinase 2 [Sou | 6.4248 | 1.0279 | 6.0375 | 2.8919 | 2.1812 | 6.6831 |
| 10920 | ENSMUSG00000037990 | 0.744190399 | -2.947713985 | 2.874421214 | -1.025498271 | 0.305128141 | NA | Sh3rf3 | SH3 domain containing ring finger 3 [Sour | 2.4093 | 2.0558 | 0 | 0 | 0 | 0 |
| 10921 | ENSMUSG00000037991 | 1.63273338 | -1.101533746 | 1.808437309 | -0.609108063 | 0.542452814 | NA | Rmi2 | RecQ mediated genome instability 2 [Sour | 5.6217 | 0 | 1.0062 | 0.964 | 1.0906 | 1.1138 |
| 10922 | ENSMUSG00000037992 | 461.2276063 | 0.013841235 | 0.154108086 | 0.089815109 | 0.928434141 | 0.988240056 | Rara | retinoic acid receptor, alpha [Source:MGI | 454.56 | 393.69 | 528.28 | 409.69 | 477.69 | 503.46 |
| 10923 | ENSMUSG00000037993 | 387.2063016 | -0.008471686 | 0.138464695 | -0.061183004 | 0.951213465 | 0.990883058 | Dhx38 | DEAH (Asp-Glu-Ala-His) box polypeptide 3 | 390.31 | 415.28 | 359.23 | 367.28 | 403.53 | 387.62 |
| 10924 | ENSMUSG00000037994 | 1.589538791 | -0.401591289 | 1.826131279 | -0.219913701 | 0.825938365 | NA | Slc9b2 | solute carrier family 9, subfamily B (NHA2 | 2.4093 | 0 | 3.0187 | 1.928 | 2.1812 | 0 |
| 10925 | ENSMUSG00000037995 | 8.959417311 | -0.016803713 | 0.728253721 | -0.023073982 | 0.98159126 | 0.998323671 | Igsf9 | immunoglobulin superfamily, member 9 [ | 9.6372 | 9.2513 | 8.05 | 4.8199 | 13.087 | 8.9108 |
| 10926 | ENSMUSG00000037996 | 0 | NA | NA | NA | NA | NA | Slc24a2 | solute carrier family 24 (sodium/potassiur | 0 | 0 | 0 | 0 | 0 | 0 |
| 10927 | ENSMUSG00000037997 | 101.4255915 | 0.050437285 | 0.27429323 | 0.183880896 | 0.854106892 | 0.973825032 | Parp11 | poly (ADP-ribose) polymerase family, men | 108.42 | 101.76 | 88.55 | 104.11 | 68.709 | 137 |
| 10928 | ENSMUSG00000037999 | 256.5442483 | -0.038348852 | 0.178142926 | -0.215270137 | 0.829556711 | 0.969955221 | Arap2 | ArfGAP with RhoGAP domain, ankyrin rep | 270.65 | 291.93 | 217.35 | 272.81 | 219.21 | 267.32 |
| 10929 | ENSMUSG00000038000 | 218.9068206 | 0.060007043 | 0.16995035 | 0.353085727 | 0.724024173 | 0.944786345 | Acd | adrenocortical dysplasia [Source:MGI Sym | 208 | 200.44 | 234.45 | 205.33 | 226.85 | 238.36 |
| 10930 | ENSMUSG00000038002 | 273.0965638 | 0.088596314 | 0.17568899 | 0.504279259 | 0.614065151 | 0.917378096 | Cramp1l | cramped chromatin regulator homolog 1 | 235.31 | 247.73 | 311.94 | 314.26 | 248.66 | 280.69 |
| 10931 | ENSMUSG00000038005 | 115.1061092 | 0.200139556 | 0.23424289 | 0.854410374 | 0.39287765 | 0.821390458 | Hpf1 | histone PARylation factor 1 [Source:MGI S | 113.24 | 119.24 | 88.55 | 107 | 116.7 | 145.91 |
| 10932 | ENSMUSG00000038007 | 185.2842309 | -0.889381895 | 0.255312474 | -3.483503487 | 0.000494897 | 0.012307964 | Acer2 | alkaline ceramidase 2 [Source:MGI Symbo | 285.1 | 275.48 | 161 | 128.21 | 97.065 | 164.85 |
| 10933 | ENSMUSG00000038009 | 2144.58437 | 0.248989521 | 0.093025555 | 2.676571203 | 0.007437976 | 0.087394707 | Dnajc22 | DnaJ heat shock protein family (Hsp40) m | 2021.4 | 1969.5 | 1888.7 | 2370.4 | 2378.6 | 2238.8 |
| 10934 | ENSMUSG00000038010 | 88.76289673 | 0.14076216 | 0.259192389 | 0.543079835 | 0.587074836 | 0.910708969 | Ccdc138 | coiled-coil domain containing 138 [Source | 73.885 | 72.982 | 106.66 | 89.65 | 102.52 | 86.88 |
| 10935 | ENSMUSG00000038011 | 0.935330317 | 1.257565026 | 2.487133718 | 0.505628232 | 0.613117658 | NA | Dnah10 | dynein, axonemal, heavy chain 10 [Source | 1.6062 | 0 | 0 | 2.8919 | 0 | 1.1138 |
| 10936 | ENSMUSG00000038013 | 226.1958861 | 0.076416416 | 0.172605642 | 0.442722585 | 0.657966405 | 0.927303282 | Wipf2 | WAS/WASL interacting protein family, me | 203.99 | 250.81 | 206.28 | 236.17 | 213.76 | 246.16 |
| 10937 | ENSMUSG00000038014 | 2911.517952 | 0.006365086 | 0.096277127 | 0.066112126 | 0.947288556 | 0.990677354 | Fam120a | family with sequence similarity 120, meml | 2750.6 | 3051.9 | 2913.1 | 2743.5 | 3046.1 | 2963.9 |
| 10938 | ENSMUSG00000038015 | 0 | NA | NA | NA | NA | NA | Prm2 | protamine 2 [Source:MGI Symbol;Acc:MG | 0 | 0 | 0 | 0 | 0 | 0 |
| 10939 | ENSMUSG00000038020 | 6.776513234 | -0.284321312 | 0.995254683 | -0.28567694 | 0.775125568 | 0.95893551 | Rapgefl1 | Rap guanine nucleotide exchange factor ( | 7.2279 | 4.1117 | 11.069 | 11.568 | 0 | 6.6831 |
| 10940 | ENSMUSG00000038022 | 57.51383597 | 0.25589376 | 0.323755493 | 0.790392024 | 0.429298859 | 0.842978372 | Mindy4 | MINDY lysine 48 deubiquitinase 4 [Source | 48.989 | 41.117 | 67.418 | 74.226 | 65.437 | 47.895 |
| 10941 | ENSMUSG00000038023 | 742.1203824 | 0.114862968 | 0.118298438 | 0.970959301 | 0.331568546 | 0.774747658 | Atp6v0a2 | ATPase, H+ transporting, lysosomal V0 sul | 668.18 | 748.32 | 722.48 | 825.17 | 762.34 | 726.23 |
| 10942 | ENSMUSG00000038024 | 493.6761604 | 0.049284643 | 0.125898289 | 0.391463964 | 0.69545432 | 0.939992732 | Dennd4c | DENN/MADD domain containing 4C [Sour | 459.37 | 503.68 | 493.06 | 486.81 | 524.59 | 494.55 |
| 10943 | ENSMUSG00000038025 | 434.6263982 | -0.079907494 | 0.136537851 | -0.585240601 | 0.558385948 | 0.899286756 | Phf2 | PHD finger protein 2 [Source:MGI Symbol, | 405.57 | 482.09 | 452.81 | 401.98 | 425.34 | 439.97 |
| 10944 | ENSMUSG00000038026 | 1.54179941 | -0.399207299 | 1.874986936 | -0.212912043 | 0.831395569 | NA | Kcnj9 | potassium inwardly-rectifying channel, su | 3.2124 | 2.0558 | 0 | 2.8919 | 1.0906 | 0 |
| 10945 | ENSMUSG00000038028 | 189.3602405 | 0.112983692 | 0.185225342 | 0.609979666 | 0.541875277 | 0.894000766 | Tigar | Trp53 induced glycolysis regulatory phosp | 194.35 | 170.63 | 180.12 | 167.73 | 206.13 | 217.2 |
| 10946 | ENSMUSG00000038034 | 161.0993136 | 0.319700079 | 0.223126836 | 1.432817694 | 0.151909943 | 0.567340036 | Igsf8 | immunoglobulin superfamily, member 8 [ | 123.68 | 156.24 | 150.94 | 224.61 | 139.6 | 171.53 |
| 10947 | ENSMUSG00000038037 | 12.43091302 | 1.534253644 | 0.809605403 | 1.895063495 | 0.05808399 | 0.342489537 | Socs1 | suppressor of cytokine signaling 1 [Source | 8.8341 | 4.1117 | 6.0375 | 14.46 | 3.2719 | 37.871 |
| 10948 | ENSMUSG00000038039 | 501.3483985 | 0.106032523 | 0.127948713 | 0.828711133 | 0.407267885 | 0.83094725 | Gcc2 | GRIP and coiled-coil domain containing 2 | 442.51 | 516.01 | 491.05 | 514.76 | 519.13 | 524.62 |
| 10949 | ENSMUSG00000038042 | 18.35926309 | -0.608087267 | 0.521880091 | -1.165185792 | 0.243943795 | 0.693748143 | Ptpdc1 | protein tyrosine phosphatase domain con | 21.684 | 20.558 | 24.15 | 10.604 | 11.997 | 21.163 |
| 10950 | ENSMUSG00000038044 | 0 | NA | NA | NA | NA | NA | Cct8l1 | chaperonin containing TCP1, subunit 8 (th | 0 | 0 | 0 | 0 | 0 | 0 |
| 10951 | ENSMUSG00000038045 | 1.035246523 | 1.060038137 | 2.384517192 | 0.444550427 | 0.656644679 | NA | Sult6b1 | sulfotransferase family, cytosolic, 6B, mer | 0 | 2.0558 | 0 | 1.928 | 0 | 2.2277 |
| 10952 | ENSMUSG00000038046 | 146.2974077 | 0.235078585 | 0.199963085 | 1.175609911 | 0.239750793 | 0.689551695 | Mrm3 | mitochondrial rRNA methyltransferase 3 [ | 149.38 | 124.38 | 128.8 | 143.63 | 172.32 | 159.28 |
| 10953 | ENSMUSG00000038047 | 84.47501317 | 0.423054583 | 0.264984882 | 1.59652347 | 0.11037197 | 0.488351231 | Haus6 | HAUS augmin-like complex, subunit 6 [Sou | 68.264 | 70.926 | 77.481 | 94.47 | 119.97 | 75.741 |
| 10954 | ENSMUSG00000038048 | 0 | NA | NA | NA | NA | NA | Cntnap5c | contactin associated protein-like 5C [Sour | 0 | 0 | 0 | 0 | 0 | 0 |
| 10955 | ENSMUSG00000038055 | 619.0578049 | 0.037734308 | 0.162473418 | 0.23224912 | 0.816344528 | 0.967767553 | Dexi | dexamethasone-induced transcript [Sourc | 594.3 | 505.74 | 732.55 | 577.42 | 573.67 | 730.68 |
| 10956 | ENSMUSG00000038056 | 531.066863 | 0.168482507 | 0.140616194 | 1.198172854 | 0.230849731 | 0.677910147 | Kmt2c | lysine (K)-specific methyltransferase 2C [S | 452.15 | 562.27 | 487.02 | 594.77 | 587.84 | 502.34 |
| 10957 | ENSMUSG00000038057 | 6.080197891 | 1.399854846 | 0.893769862 | 1.56623635 | 0.117293296 | 0.502643331 | Dbil5 | diazepam binding inhibitor-like 5 [Source: | 4.8186 | 2.0558 | 3.0187 | 6.7478 | 9.8156 | 10.025 |
| 10958 | ENSMUSG00000038058 | 268.8702335 | 0.098364886 | 0.171316937 | 0.57416907 | 0.565853397 | 0.901973059 | Nod1 | nucleotide-binding oligomerization domai | 240.13 | 244.64 | 294.83 | 294.01 | 237.75 | 301.85 |
| 10959 | ENSMUSG00000038059 | 21.73583838 | -1.066576106 | 0.498520009 | -2.13948505 | 0.032396405 | 0.24029826 | Smim3 | small integral membrane protein 3 [Sourc | 19.274 | 35.977 | 33.206 | 14.46 | 16.359 | 11.138 |
| 10960 | ENSMUSG00000038060 | 28.49902616 | -0.340173294 | 0.48689332 | -0.698660836 | 0.484764015 | 0.87256358 | Dlec1 | deleted in lung and esophageal cancer 1 [ | 34.533 | 20.558 | 40.25 | 16.388 | 41.443 | 17.822 |
| 10961 | ENSMUSG00000038064 | 0 | NA | NA | NA | NA | NA | Cldn22 | claudin 22 [Source:MGI Symbol;Acc:MGI: | 0 | 0 | 0 | 0 | 0 | 0 |
| 10962 | ENSMUSG00000038065 | 43.82679767 | 0.370643924 | 0.339589237 | 1.091447796 | 0.275075891 | 0.729768468 | Mturn | maturin, neural progenitor differentiation | 30.518 | 40.089 | 44.275 | 43.379 | 52.35 | 52.351 |
| 10963 | ENSMUSG00000038067 | 0 | NA | NA | NA | NA | NA | Csf3 | colony stimulating factor 3 (granulocyte) [ | 0 | 0 | 0 | 0 | 0 | 0 |
| 10964 | ENSMUSG00000038068 | 945.3533943 | -0.458332219 | 0.121603125 | -3.76908257 | 0.000163849 | 0.005349678 | Rnf144b | ring finger protein 144B [Source:MGI Sym | 1168.5 | 1140 | 974.04 | 813.6 | 840.87 | 735.14 |
| 10965 | ENSMUSG00000038069 | 270.390786 | 0.191163436 | 0.158703144 | 1.20453465 | 0.228382998 | 0.674837933 | Cdkn2aip | CDKN2A interacting protein [Source:MGI S | 248.16 | 246.7 | 262.63 | 274.73 | 323.91 | 266.21 |
| 10966 | ENSMUSG00000038070 | 10.12708349 | -0.520485723 | 0.697171332 | -0.74656788 | 0.455324443 | 0.857339509 | Cntln | centlein, centrosomal protein [Source:MG | 10.44 | 15.419 | 10.062 | 11.568 | 4.3625 | 8.9108 |
| 10967 | ENSMUSG00000038071 | 0 | NA | NA | NA | NA | NA | Npy6r | neuropeptide Y receptor Y6 [Source:MGI S | 0 | 0 | 0 | 0 | 0 | 0 |
| 10968 | ENSMUSG00000038072 | 273.0284666 | 0.119477159 | 0.152820455 | 0.78181392 | 0.434323939 | 0.845264432 | Galnt11 | polypeptide N-acetylgalactosaminyltransf | 270.65 | 241.56 | 272.69 | 281.48 | 295.56 | 276.23 |
| 10969 | ENSMUSG00000038074 | 108.2409603 | -0.154904651 | 0.229344665 | -0.675422954 | 0.499407086 | 0.880660418 | Fkbp14 | FK506 binding protein 14 [Source:MGI Syr | 106.81 | 103.82 | 131.82 | 109.89 | 106.88 | 90.221 |
| 10970 | ENSMUSG00000038077 | 0 | NA | NA | NA | NA | NA | Kcna6 | potassium voltage-gated channel, shaker- | 0 | 0 | 0 | 0 | 0 | 0 |
| 10971 | ENSMUSG00000038079 | 70.26695439 | 0.394026231 | 0.296188284 | 1.330323488 | 0.183411712 | 0.617664027 | Tmem237 | transmembrane protein 237 [Source:MGI | 69.067 | 44.2 | 68.425 | 63.623 | 90.521 | 85.766 |
| 10972 | ENSMUSG00000038080 | 677.5138625 | -0.052413464 | 0.115869503 | -0.452349089 | 0.651017518 | 0.925643431 | Kdm1b | lysine (K)-specific demethylase 1B [Source | 685.85 | 703.1 | 680.22 | 627.55 | 705.63 | 662.74 |

| | | | | | | | | | | | | | | | |
|---|---|---|---|---|---|---|---|---|---|---|---|---|---|---|---|
| 10973 | ENSMUSG00000038084 | 1925.076081 | -0.072258041 | 0.105742411 | -0.683340214 | 0.494391889 | 0.877473722 | Opa1 | OPA1, mitochondrial dynamin like GTPase | 1842.3 | 2027.1 | 2051.7 | 1943.4 | 1975.1 | 1710.9 |
| 10974 | ENSMUSG00000038085 | 10.66860402 | -0.963695311 | 0.765898117 | -1.25825523 | 0.208299467 | 0.653455953 | Cnbd2 | cyclic nucleotide binding domain containi | 12.85 | 19.53 | 10.062 | 11.568 | 1.0906 | 8.9108 |
| 10975 | ENSMUSG00000038086 | 1.238640426 | 1.454958615 | 2.28972858 | 0.635428421 | 0.525149037 | NA | Hspb2 | heat shock protein 2 [Source:MGI Symbol | 0 | 0 | 2.0125 | 0.964 | 0 | 4.4554 |
| 10976 | ENSMUSG00000038092 | 3222.67487 | 2.678911241 | 0.472747879 | 5.666680616 | 1.46E-08 | 2.11E-06 | Hsd3b5 | hydroxy-delta-5-steroid dehydrogenase, 3 | 1097 | 422.47 | 1091.8 | 3990.9 | 6726.9 | 6007 |
| 10977 | ENSMUSG00000038094 | 0.133850373 | -0.796629176 | 4.080472857 | -0.195229623 | 0.84521319 | NA | Atp13a4 | ATPase type 13A4 [Source:MGI Symbol;Ac | 0.8031 | 0 | 0 | 0 | 0 | 0 |
| 10978 | ENSMUSG00000038095 | 895.9631719 | 0.174924304 | 0.107517408 | 1.626939359 | 0.103749978 | 0.474047094 | Sbno1 | strawberry notch 1 [Source:MGI Symbol;A | 882.61 | 835.7 | 806 | 927.35 | 969.56 | 954.56 |
| 10979 | ENSMUSG00000038102 | 878.7385429 | -0.039867196 | 0.116988409 | -0.340779023 | 0.733269943 | 0.947915087 | Trappc11 | trafficking protein particle complex 11 [So | 906.7 | 895.32 | 870.4 | 848.3 | 958.65 | 793.06 |
| 10980 | ENSMUSG00000038112 | 1.301169988 | 0.83530979 | 1.913806454 | 0.436465134 | 0.662499291 | NA | AW551984 | expressed sequence AW551984 [Source:N | 0.8031 | 1.0279 | 1.0062 | 3.8559 | 0 | 1.1138 |
| 10981 | ENSMUSG00000038115 | 1.379560231 | -1.495251236 | 2.030912621 | -0.736245972 | 0.46158102 | NA | Ano2 | anoctamin 2 [Source:MGI Symbol;Acc:MG | 4.0155 | 1.0279 | 1.0062 | 0 | 0 | 2.2277 |
| 10982 | ENSMUSG00000038116 | 155.170715 | 0.06012659 | 0.245204802 | 0.245209675 | 0.806294085 | 0.964538719 | Phf20 | PHD finger protein 20 [Source:MGI Symbo | 108.42 | 203.53 | 144.9 | 182.19 | 149.41 | 142.57 |
| 10983 | ENSMUSG00000038119 | 30.33071509 | -0.028430767 | 0.417311198 | -0.068128456 | 0.945683379 | 0.990309554 | Cdon | cell adhesion molecule-related/down-reg | 36.943 | 29.81 | 25.156 | 40.487 | 25.084 | 24.505 |
| 10984 | ENSMUSG00000038121 | 1720.618921 | 0.081333197 | 0.095593971 | 0.850819314 | 0.394869731 | 0.822933206 | Fam210a | family with sequence similarity 210, meml | 1672.1 | 1595.3 | 1748.9 | 1752.5 | 1798.4 | 1756.5 |
| 10985 | ENSMUSG00000038122 | 165.8838668 | -0.157938142 | 0.202803459 | -0.778774398 | 0.436112622 | 0.84624805 | Tbc1d32 | TBC1 domain family, member 32 [Source:I | 147.77 | 195.3 | 182.13 | 139.78 | 179.95 | 150.37 |
| 10986 | ENSMUSG00000038126 | 72.7622076 | 0.730065919 | 0.262163242 | 2.78477606 | 0.00535647 | 0.069798186 | Mphosph9 | M-phase phosphoprotein 9 [Source:MGI S | 51.399 | 56.535 | 56.35 | 83.866 | 95.974 | 92.449 |
| 10987 | ENSMUSG00000038127 | 951.1010157 | 0.04445759 | 0.109178898 | 0.407199476 | 0.683861486 | 0.935528319 | Ccdc50 | coiled-coil domain containing 50 [Source:I | 876.99 | 977.55 | 955.93 | 1003.5 | 931.39 | 961.25 |
| 10988 | ENSMUSG00000038128 | 18.60799939 | 1.272292136 | 0.573353416 | 2.219036461 | 0.026484244 | 0.210708529 | Camk4 | calcium/calmodulin-dependent protein ki | 6.4248 | 18.503 | 8.05 | 28.919 | 17.45 | 32.301 |
| 10989 | ENSMUSG00000038132 | 1.611574505 | -2.954982165 | 2.117944162 | -1.395212498 | 0.162951769 | NA | Rbm24 | RNA binding motif protein 24 [Source:MG | 5.6217 | 3.0838 | 0 | 0.964 | 0 | 0 |
| 10990 | ENSMUSG00000038135 | 44.58435397 | 1.029935802 | 0.442902915 | 2.32542114 | 0.020049452 | 0.173442151 | Crygn | crystallin, gamma N [Source:MGI Symbol;. | 32.124 | 11.307 | 44.275 | 43.379 | 79.615 | 56.806 |
| 10991 | ENSMUSG00000038141 | 79.25439704 | 0.281229905 | 0.264630383 | 1.062727196 | 0.287905684 | 0.73951397 | Tmem181a | transmembrane protein 181A [Source:MG | 72.279 | 77.094 | 65.406 | 97.362 | 68.709 | 94.677 |
| 10992 | ENSMUSG00000038143 | 9.657353465 | 0.539292322 | 0.700628272 | 0.769726749 | 0.441461999 | 0.849828766 | Stox2 | storkhead box 2 [Source:MGI Symbol;Acc | 6.4248 | 10.279 | 7.0437 | 15.424 | 7.6343 | 11.138 |
| 10993 | ENSMUSG00000038145 | 996.034393 | 0.021956792 | 0.118207975 | 0.185747128 | 0.852643068 | 0.973781977 | Snrk | SNF related kinase [Source:MGI Symbol;A | 1082.6 | 969.33 | 912.66 | 1023.7 | 1061.2 | 926.72 |
| 10994 | ENSMUSG00000038146 | 13.04610372 | 0.045377316 | 0.64845706 | 0.069977364 | 0.944211676 | 0.990077979 | Notch3 | notch 3 [Source:MGI Symbol;Acc:MGI:994 | 15.259 | 10.279 | 13.081 | 23.135 | 8.7249 | 7.7969 |
| 10995 | ENSMUSG00000038147 | 41.71996589 | -1.132988466 | 0.382454486 | -2.962413853 | 0.003052372 | 0.047319947 | Cd84 | CD84 antigen [Source:MGI Symbol;Acc:M | 42.564 | 79.15 | 50.312 | 23.135 | 25.084 | 30.074 |
| 10996 | ENSMUSG00000038148 | 0 | NA | NA | NA | NA | NA | Cldn16 | claudin 16 [Source:MGI Symbol;Acc:MGI:: | 0 | 0 | 0 | 0 | 0 | 0 |
| 10997 | ENSMUSG00000038150 | 2599.785689 | -0.25025645 | 0.093309076 | -2.68201617 | 0.007317992 | 0.086502219 | Ormdl3 | ORM1-like 3 (S. cerevisiae) [Source:MGI S | 2910.4 | 2736.3 | 2827.5 | 2513.1 | 2301.2 | 2310.1 |
| 10998 | ENSMUSG00000038151 | 3.520552183 | -2.043009025 | 1.310623115 | -1.558807411 | 0.11904197 | NA | Prdm1 | PR domain containing 1, with ZNF domain | 8.8341 | 6.1675 | 2.0125 | 1.928 | 2.1812 | 0 |
| 10999 | ENSMUSG00000038152 | 16.48845813 | -0.051075754 | 0.568452568 | -0.089850512 | 0.928406008 | 0.988240056 | 5033430I15R | RIKEN cDNA 5033430I15 gene [Source:MG | 12.85 | 16.447 | 21.131 | 16.388 | 8.7249 | 23.391 |
| 11000 | ENSMUSG00000038155 | 58.93603617 | -1.280604199 | 0.865283353 | -1.479982476 | 0.138877923 | 0.544666563 | Gstp2 | glutathione S-transferase, pi 2 [Source:MG | 165.44 | 63.731 | 21.131 | 18.316 | 35.99 | 49.009 |
| 11001 | ENSMUSG00000038156 | 9.519913126 | -0.514293836 | 0.749341916 | -0.686327329 | 0.492506719 | 0.876429279 | Spon1 | spondin 1, (f-spondin) extracellular matrix | 7.2279 | 10.279 | 16.1 | 5.7839 | 4.3625 | 13.366 |
| 11002 | ENSMUSG00000038160 | 1211.298004 | 0.018974195 | 0.121310596 | 0.156410035 | 0.875709827 | 0.980566196 | Atg5 | autophagy related 5 [Source:MGI Symbol; | 1260.9 | 1066 | 1282 | 1098.9 | 1285.8 | 1274.2 |
| 11003 | ENSMUSG00000038165 | 0 | NA | NA | NA | NA | NA | 1700022A21 | RIKEN cDNA 1700022A21 gene [Source:M | 0 | 0 | 0 | 0 | 0 | 0 |
| 11004 | ENSMUSG00000038167 | 418.6944034 | 0.561402742 | 0.162810996 | 3.448186886 | 0.000564363 | 0.013559721 | Plekhg6 | pleckstrin homology domain containing, fa | 346.14 | 299.12 | 369.29 | 520.55 | 407.89 | 569.17 |
| 11005 | ENSMUSG00000038168 | 11.56413102 | -0.285473496 | 0.708062169 | -0.40317575 | 0.686818938 | 0.936582996 | P3h2 | prolyl 3-hydroxylase 2 [Source:MGI Symbo | 5.6217 | 16.447 | 16.1 | 5.7839 | 8.7249 | 16.708 |
| 11006 | ENSMUSG00000038170 | 1690.41564 | 0.059306166 | 0.095249748 | 0.622638557 | 0.533522068 | 0.889143097 | Pde4dip | phosphodiesterase 4D interacting protein | 1623.1 | 1630.3 | 1713.6 | 1665.8 | 1776.6 | 1733.1 |
| 11007 | ENSMUSG00000038172 | 607.3569659 | 0.296634171 | 0.127591093 | 2.324881484 | 0.020078298 | 0.173507261 | Ttc39b | tetratricopeptide repeat domain 39B [Sou | 572.61 | 546.85 | 515.2 | 628.51 | 741.62 | 639.35 |
| 11008 | ENSMUSG00000038173 | 38.80897461 | -0.083667502 | 0.397339942 | -0.210569069 | 0.833223551 | 0.970611245 | Enpp6 | ectonucleotide pyrophosphatase/phosph | 32.124 | 55.508 | 32.2 | 26.027 | 46.897 | 40.098 |
| 11009 | ENSMUSG00000038174 | 824.9830672 | 0.101674627 | 0.141129649 | 0.720434206 | 0.471257696 | 0.867762102 | Fam126b | family with sequence similarity 126, meml | 897.87 | 775.05 | 714.43 | 939.88 | 898.67 | 724 |
| 11010 | ENSMUSG00000038175 | 141.7668861 | -0.864425188 | 0.260419151 | -3.319361052 | 0.000902237 | 0.019361951 | Mylip | myosin regulatory light chain interacting p | 187.12 | 201.47 | 159.99 | 81.938 | 74.162 | 145.91 |
| 11011 | ENSMUSG00000038178 | 109.5680764 | -0.48690362 | 0.259779414 | -1.874296399 | 0.060889584 | 0.352269825 | Slc43a2 | solute carrier family 43, member 2 [Sourc | 138.94 | 134.66 | 109.68 | 82.902 | 68.709 | 122.52 |
| 11012 | ENSMUSG00000038179 | 3.552684332 | -2.50493085 | 1.397361801 | -1.792614373 | 0.073034607 | NA | Slamf7 | SLAM family member 7 [Source:MGI Symb | 0.8031 | 11.307 | 6.0375 | 0.964 | 1.0906 | 1.1138 |
| 11013 | ENSMUSG00000038180 | 36.18787881 | -0.362122966 | 0.362696437 | -0.998418869 | 0.318076287 | 0.764585353 | Spag4 | sperm associated antigen 4 [Source:MGI S | 37.746 | 41.117 | 43.269 | 29.883 | 28.356 | 36.757 |
| 11014 | ENSMUSG00000038181 | 260.4492782 | 0.104883566 | 0.184047962 | 0.569870839 | 0.568765305 | 0.903527882 | Chpf2 | chondroitin polymerizing factor 2 [Source | 248.16 | 259.04 | 246.53 | 336.43 | 248.66 | 223.88 |
| 11015 | ENSMUSG00000038187 | 36.29034394 | 0.053514213 | 0.398510169 | 0.13428569 | 0.893176668 | 0.982562372 | Btbd10 | BTB (POZ) domain containing 10 [Source:I | 43.368 | 30.838 | 32.2 | 23.135 | 42.534 | 45.668 |
| 11016 | ENSMUSG00000038188 | 195.3772235 | 0.136712699 | 0.180388304 | 0.757880061 | 0.448522788 | 0.854238022 | Scarf1 | scavenger receptor class F, member 1 [So | 188.73 | 161.38 | 208.29 | 214.97 | 202.86 | 196.04 |
| 11017 | ENSMUSG00000038192 | 0 | NA | NA | NA | NA | NA | Cer1 | cerberus 1, DAN family BMP antagonist [S | 0 | 0 | 0 | 0 | 0 | 0 |
| 11018 | ENSMUSG00000038193 | 105.4989308 | -1.318030011 | 0.231480287 | -5.69391902 | 1.24E-08 | 1.87E-06 | Hand2 | heart and neural crest derivatives express | 147.77 | 145.96 | 157.98 | 58.803 | 54.531 | 67.944 |
| 11019 | ENSMUSG00000038195 | 474.4083073 | 0.124896335 | 0.14043884 | 0.889329011 | 0.373826284 | 0.808708253 | Rilp | Rab interacting lysosomal protein [Source | 437.69 | 428.64 | 496.08 | 545.61 | 496.23 | 442.2 |
| 11020 | ENSMUSG00000038199 | 0 | NA | NA | NA | NA | NA | Iqca1l | IQ motif containing with AAA domain 1 lik | 0 | 0 | 0 | 0 | 0 | 0 |
| 11021 | ENSMUSG00000038201 | 0.185640647 | 1.126924175 | 4.080472857 | 0.276174898 | 0.782413736 | NA | Kcna7 | potassium voltage-gated channel, shaker- | 0 | 0 | 0 | 0 | 0 | 1.1138 |
| 11022 | ENSMUSG00000038203 | 0 | NA | NA | NA | NA | NA | Hoxa13 | homeobox A13 [Source:MGI Symbol;Acc:I | 0 | 0 | 0 | 0 | 0 | 0 |
| 11023 | ENSMUSG00000038204 | 0.315620088 | 0.165143421 | 4.063300985 | 0.040642675 | 0.967580763 | NA | Asb10 | ankyrin repeat and SOCS box-containing 1 | 0.8031 | 0 | 0 | 0 | 1.0906 | 0 |
| 11024 | ENSMUSG00000038205 | 34.86272545 | -0.741107212 | 0.386350149 | -1.918226805 | 0.055082259 | 0.330834503 | Prkab2 | protein kinase, AMP-activated, beta 2 nor | 52.202 | 41.117 | 37.231 | 20.244 | 30.537 | 27.846 |
| 11025 | ENSMUSG00000038206 | 1074.356347 | -0.019898529 | 0.102994215 | -0.193200449 | 0.846801987 | 0.972803041 | Fbxo8 | F-box protein 8 [Source:MGI Symbol;Acc:I | 1041.6 | 1111.2 | 1092.8 | 1042.1 | 1052.4 | 1106 |
| 11026 | ENSMUSG00000038208 | 29.71023721 | -0.152156539 | 0.443741914 | -0.342894224 | 0.731678038 | 0.947387358 | Pgap3 | post-GPI attachment to proteins 3 [Source | 32.927 | 27.754 | 33.206 | 35.667 | 14.178 | 34.529 |
| 11027 | ENSMUSG00000038209 | 0 | NA | NA | NA | NA | NA | Itln1 | intelectin 1 (galactofuranose binding) [Sou | 0 | 0 | 0 | 0 | 0 | 0 |
| 11028 | ENSMUSG00000038210 | 0 | NA | NA | NA | NA | NA | Hoxa11 | homeobox A11 [Source:MGI Symbol;Acc:I | 0 | 0 | 0 | 0 | 0 | 0 |

| | | | | | | | | | | | | | | | |
|---|---|---|---|---|---|---|---|---|---|---|---|---|---|---|---|
| 11029 | ENSMUSG00000038212 | 789.0600581 | 0.051710667 | 0.109608987 | 0.47177397 | 0.637088132 | 0.923478764 | Mfsd14b | major facilitator superfamily domain cont | 747.69 | 785.33 | 791.91 | 776.97 | 804.88 | 827.59 |
| 11030 | ENSMUSG00000038213 | 410.998893 | 0.156646946 | 0.149615542 | 1.046996483 | 0.2951012 | 0.745692481 | Tapbpl | TAP binding protein-like [Source:MGI Sym | 432.07 | 390.61 | 343.13 | 476.21 | 430.79 | 393.19 |
| 11031 | ENSMUSG00000038214 | 118.0305859 | -0.068260655 | 0.230970962 | -0.29553782 | 0.767583065 | 0.95578799 | Bend3 | BEN domain containing 3 [Source:MGI Syr | 126.89 | 99.708 | 135.84 | 127.25 | 95.974 | 122.52 |
| 11032 | ENSMUSG00000038215 | 169.1520346 | -0.017023989 | 0.183366051 | -0.092841551 | 0.926029441 | 0.987546253 | Cep44 | centrosomal protein 44 [Source:MGI Syml | 180.7 | 172.69 | 156.97 | 178.34 | 163.59 | 162.62 |
| 11033 | ENSMUSG00000038216 | 0 | NA | NA | NA | NA | NA | Pnmt | phenylethanolamine-N-methyltransferase | 0 | 0 | 0 | 0 | 0 | 0 |
| 11034 | ENSMUSG00000038217 | 2096.752267 | 0.096580764 | 0.152370658 | 0.633854082 | 0.526176055 | 0.888334265 | Tlcd2 | TLC domain containing 2 [Source:MGI Sym | 2336.2 | 1740.3 | 2001.4 | 1707.2 | 2496.4 | 2299 |
| 11035 | ENSMUSG00000038218 | 0 | NA | NA | NA | NA | NA | Gm9765 | predicted gene 9765 [Source:MGI Symbol | 0 | 0 | 0 | 0 | 0 | 0 |
| 11036 | ENSMUSG00000038224 | 10506.25954 | 0.355531718 | 0.117743456 | 3.019545465 | 0.002531543 | 0.041751696 | Serpinf2 | serine (or cysteine) peptidase inhibitor, cl | 8883.1 | 8760.9 | 10010 | 10418 | 11299 | 13667 |
| 11037 | ENSMUSG00000038225 | 69.64316016 | 0.4817961 | 0.328843927 | 1.465120866 | 0.142887945 | 0.552601934 | Primpol | primase and polymerase (DNA-directed) [ | 81.113 | 41.117 | 51.318 | 66.514 | 71.981 | 105.82 |
| 11038 | ENSMUSG00000038227 | 0 | NA | NA | NA | NA | NA | Hoxa9 | homeobox A9 [Source:MGI Symbol;Acc:M | 0 | 0 | 0 | 0 | 0 | 0 |
| 11039 | ENSMUSG00000038233 | 98.72580898 | -0.163597072 | 0.296152879 | -0.5524075 | 0.580669193 | 0.908645447 | Gask1a | golgi associated kinase 1A [Source:MGI Sy | 78.704 | 91.485 | 142.89 | 77.118 | 78.525 | 123.64 |
| 11040 | ENSMUSG00000038235 | 1215.304581 | 0.118344907 | 0.099391165 | 1.19069846 | 0.233771983 | 0.681342761 | F11r | F11 receptor [Source:MGI Symbol;Acc:MC | 1143.6 | 1155.4 | 1197.4 | 1226.2 | 1274.9 | 1294.3 |
| 11041 | ENSMUSG00000038236 | 0 | NA | NA | NA | NA | NA | Hoxa7 | homeobox A7 [Source:MGI Symbol;Acc:M | 0 | 0 | 0 | 0 | 0 | 0 |
| 11042 | ENSMUSG00000038239 | 0 | NA | NA | NA | NA | NA | Hrc | histidine rich calcium binding protein [Sou | 0 | 0 | 0 | 0 | 0 | 0 |
| 11043 | ENSMUSG00000038240 | 201.7486474 | -0.028110548 | 0.170620367 | -0.164754938 | 0.869136874 | 0.978605211 | Pdss2 | prenyl (solanesyl) diphosphate synthase, s | 193.55 | 198.39 | 219.36 | 193.76 | 209.4 | 196.04 |
| 11044 | ENSMUSG00000038241 | 259.994714 | 0.094141714 | 0.156771252 | 0.600503681 | 0.548170609 | 0.895600915 | Cep250 | centrosomal protein 250 [Source:MGI Syn | 267.43 | 233.34 | 253.57 | 284.37 | 262.84 | 258.41 |
| 11045 | ENSMUSG00000038242 | 0 | NA | NA | NA | NA | NA | Aox4 | aldehyde oxidase 4 [Source:MGI Symbol;A | 0 | 0 | 0 | 0 | 0 | 0 |
| 11046 | ENSMUSG00000038244 | 316.5845451 | -0.154407974 | 0.216935758 | -0.711768199 | 0.476608328 | 0.86953366 | Mical2 | microtubule associated monooxygenase, | 252.98 | 322.77 | 425.64 | 332.57 | 222.49 | 343.06 |
| 11047 | ENSMUSG00000038246 | 0.492645588 | 0.9565034 | 3.447648281 | 0.277436479 | 0.781444974 | NA | Fam50b | family with sequence similarity 50, memb | 0 | 1.0279 | 0 | 1.928 | 0 | 0 |
| 11048 | ENSMUSG00000038248 | 1.95039664 | -2.125345478 | 1.817712029 | -1.169242127 | 0.242306093 | NA | Sobp | sine oculis binding protein [Source:MGI Sy | 2.4093 | 1.0279 | 6.0375 | 0 | 0 | 2.2277 |
| 11049 | ENSMUSG00000038250 | 494.4945812 | -0.21480588 | 0.13877141 | -1.547911632 | 0.121643574 | 0.511468794 | Usp38 | ubiquitin specific peptidase 38 [Source:M | 540.49 | 516.01 | 536.33 | 392.34 | 499.5 | 482.29 |
| 11050 | ENSMUSG00000038252 | 69.83966767 | 0.067508052 | 0.334571953 | 0.201774391 | 0.840093103 | 0.971838962 | Ncapd2 | non-SMC condensin I complex, subunit D2 | 73.885 | 83.261 | 47.293 | 67.478 | 47.987 | 99.132 |
| 11051 | ENSMUSG00000038253 | 5.000526607 | -1.374485166 | 1.050738001 | -1.308114073 | 0.190834621 | NA | Hoxa5 | homeobox A5 [Source:MGI Symbol;Acc:M | 6.4248 | 8.2233 | 7.0437 | 3.8559 | 0 | 4.4554 |
| 11052 | ENSMUSG00000038255 | 0 | NA | NA | NA | NA | NA | Neurod2 | neurogenic differentiation 2 [Source:MGI | 0 | 0 | 0 | 0 | 0 | 0 |
| 11053 | ENSMUSG00000038256 | 113.6750924 | -0.046112289 | 0.229098864 | -0.201276812 | 0.840482133 | 0.971838962 | Bcl9 | B cell CLL/lymphoma 9 [Source:MGI Symb | 105.21 | 102.79 | 138.86 | 122.43 | 103.61 | 109.16 |
| 11054 | ENSMUSG00000038257 | 1.299392708 | 1.568460528 | 2.48138834 | 0.632089908 | 0.527328129 | NA | Glra3 | glycine receptor, alpha 3 subunit [Source: | 0 | 0 | 2.0125 | 5.7839 | 0 | 0 |
| 11055 | ENSMUSG00000038259 | 0 | NA | NA | NA | NA | NA | Gdf5 | growth differentiation factor 5 [Source:M | 0 | 0 | 0 | 0 | 0 | 0 |
| 11056 | ENSMUSG00000038260 | 31.18957833 | -0.129724311 | 0.426551968 | -0.30412311 | 0.761034101 | 0.95402891 | Trpm4 | transient receptor potential cation chann | 38.549 | 24.67 | 34.212 | 19.28 | 40.353 | 30.074 |
| 11057 | ENSMUSG00000038264 | 1.238665401 | -0.445718538 | 2.061328571 | -0.216228768 | 0.82880943 | NA | Sema7a | sema domain, immunoglobulin domain (Ig | 3.2124 | 1.0279 | 0 | 0.964 | 0 | 2.2277 |
| 11058 | ENSMUSG00000038267 | 1967.027648 | 0.23277067 | 0.137497531 | 1.692907996 | 0.090472981 | 0.444346757 | Slc22a23 | solute carrier family 22, member 23 [Sour | 1925.8 | 1998.3 | 1501.3 | 1993.5 | 2446.3 | 1937 |
| 11059 | ENSMUSG00000038268 | 75.04297296 | -0.146740708 | 0.280911119 | -0.522374152 | 0.601409849 | 0.915461118 | Ovca2 | candidate tumor suppressor in ovarian ca | 83.523 | 78.122 | 74.462 | 51.091 | 83.978 | 79.083 |
| 11060 | ENSMUSG00000038271 | 159.367196 | 0.229126838 | 0.227050607 | 1.009144353 | 0.312905409 | 0.760010721 | Iffo1 | intermediate filament family orphan 1 [So | 150.98 | 136.71 | 152.95 | 221.71 | 166.86 | 126.98 |
| 11061 | ENSMUSG00000038274 | 3538.776314 | 0.006198772 | 0.10455319 | 0.059288217 | 0.952722546 | 0.991522443 | Fau | Finkel-Biskis-Reilly murine sarcoma virus ( | 3252.6 | 3773.5 | 3568.1 | 3407.7 | 3360.2 | 3870.6 |
| 11062 | ENSMUSG00000038276 | 1.813087389 | 0.418784123 | 1.519329652 | 0.275637432 | 0.782826556 | NA | Asic3 | acid-sensing (proton-gated) ion channel 3 | 1.6062 | 2.0558 | 1.0062 | 2.8919 | 1.0906 | 2.2277 |
| 11063 | ENSMUSG00000038279 | 339.2772075 | -0.187845635 | 0.150488662 | -1.248237792 | 0.211943988 | 0.656735971 | Nop2 | NOP2 nucleolar protein [Source:MGI Syml | 355.77 | 382.39 | 346.15 | 323.9 | 349 | 278.46 |
| 11064 | ENSMUSG00000038280 | 268.096294 | 0.122512409 | 0.150896107 | 0.811899074 | 0.416849546 | 0.836878476 | Ostm1 | osteopetrosis associated transmembrane | 251.37 | 268.29 | 250.55 | 271.84 | 281.38 | 285.14 |
| 11065 | ENSMUSG00000038286 | 3387.983698 | -0.148037869 | 0.087635463 | -1.689246157 | 0.091172267 | 0.445423616 | Bphl | biphenyl hydrolase-like (serine hydrolase, | 3564.2 | 3600.8 | 3519.8 | 3154.1 | 3361.3 | 3127.7 |
| 11066 | ENSMUSG00000038290 | 586.3907935 | -0.191457014 | 0.120044498 | -1.594883711 | 0.110738243 | 0.489424955 | Smg6 | Smg-6 homolog, nonsense mediated mRN | 594.3 | 633.2 | 649.03 | 571.64 | 533.31 | 536.87 |
| 11067 | ENSMUSG00000038291 | 286.9957651 | 0.305112445 | 0.153547566 | 1.987087469 | 0.046912705 | 0.300126868 | Snx25 | sorting nexin 25 [Source:MGI Symbol;Acc: | 239.32 | 245.67 | 285.77 | 322.93 | 310.83 | 317.45 |
| 11068 | ENSMUSG00000038292 | 0.401551119 | -2.072982726 | 4.008251979 | -0.517178744 | 0.605031388 | NA | Kash5 | KASH domain containing 5 [Source:MGI Sy | 2.4093 | 0 | 0 | 0 | 0 | 0 |
| 11069 | ENSMUSG00000038295 | 2.77225787 | -1.075590256 | 1.84121023 | -0.58417569 | 0.559102115 | NA | Atg9b | autophagy related 9B [Source:MGI Symbo | 0 | 11.307 | 0 | 0.964 | 4.3625 | 0 |
| 11070 | ENSMUSG00000038296 | 10.72106151 | 0.019096154 | 0.65963837 | 0.028949429 | 0.976904924 | 0.997244182 | Galnt18 | polypeptide N-acetylgalactosaminyltransf | 9.6372 | 12.335 | 10.062 | 13.496 | 6.5437 | 12.252 |
| 11071 | ENSMUSG00000038298 | 632.9585899 | 0.285394561 | 0.122039213 | 2.338548028 | 0.019358837 | 0.1693571 | Pdzk1 | PDZ domain containing 1 [Source:MGI Syr | 552.53 | 609.56 | 550.42 | 744.19 | 680.55 | 660.51 |
| 11072 | ENSMUSG00000038299 | 543.5197331 | -0.01248806 | 0.156425718 | -0.079833801 | 0.936369441 | 0.988420305 | Wdr36 | WD repeat domain 36 [Source:MGI Symbo | 451.34 | 613.67 | 573.56 | 511.87 | 619.47 | 491.21 |
| 11073 | ENSMUSG00000038300 | 0 | NA | NA | NA | NA | NA | Pth2 | parathyroid hormone 2 [Source:MGI Syml | 0 | 0 | 0 | 0 | 0 | 0 |
| 11074 | ENSMUSG00000038301 | 818.6039833 | -0.030330377 | 0.154576248 | -0.196216286 | 0.844440882 | 0.972803041 | Snx10 | sorting nexin 10 [Source:MGI Symbol;Acc: | 1022.3 | 694.87 | 762.73 | 765.4 | 891.04 | 775.24 |
| 11075 | ENSMUSG00000038302 | 283.2767859 | -0.063529293 | 0.152965597 | -0.415317524 | 0.677909474 | 0.93325821 | Afg1l | AFG1 like ATPase [Source:MGI Symbol;Ac | 308.39 | 289.87 | 269.67 | 266.06 | 269.38 | 296.28 |
| 11076 | ENSMUSG00000038304 | 5.370242486 | -0.609560106 | 1.200511989 | -0.50775012 | 0.611628595 | 0.916376001 | Cd160 | CD160 antigen [Source:MGI Symbol;Acc:N | 10.44 | 2.0558 | 7.0437 | 11.568 | 0 | 1.1138 |
| 11077 | ENSMUSG00000038305 | 15.48869663 | -0.22573088 | 0.610503019 | -0.369745722 | 0.711571961 | 0.944239215 | Spats2l | spermatogenesis associated, serine-rich 2 | 23.29 | 16.447 | 10.062 | 6.7478 | 17.45 | 18.935 |
| 11078 | ENSMUSG00000038311 | 0 | NA | NA | NA | NA | NA | 2410017I17F | RIKEN cDNA 2410017I17 gene [Source:MC | 0 | 0 | 0 | 0 | 0 | 0 |
| 11079 | ENSMUSG00000038312 | 799.8987619 | 0.346819313 | 0.111688224 | 3.105245128 | 0.001901214 | 0.033568345 | Edem2 | ER degradation enhancer, mannosidase a | 707.53 | 686.65 | 718.46 | 915.78 | 838.69 | 932.29 |
| 11080 | ENSMUSG00000038319 | 0.428363721 | -0.543730382 | 3.624677344 | -0.15000794 | 0.880758351 | NA | Kcnh2 | potassium voltage-gated channel, subfam | 1.6062 | 0 | 0 | 0.964 | 0 | 0 |
| 11081 | ENSMUSG00000038323 | 548.4960926 | 0.020166373 | 0.132319955 | 0.152406134 | 0.878866619 | 0.981085651 | 1700066M2: | RIKEN cDNA 1700066M21 gene [Source:M | 518 | 522.18 | 593.68 | 507.05 | 548.58 | 601.48 |
| 11082 | ENSMUSG00000038324 | 1886.453497 | 0.092671546 | 0.095442721 | 0.970965039 | 0.331565688 | 0.774747658 | Trpc4ap | transient receptor potential cation chann | 1796.5 | 1817.4 | 1864.6 | 2045.6 | 1943.5 | 1851.2 |
| 11083 | ENSMUSG00000038327 | 0 | NA | NA | NA | NA | NA | Serpinb9f | serine (or cysteine) peptidase inhibitor, cl | 0 | 0 | 0 | 0 | 0 | 0 |
| 11084 | ENSMUSG00000038330 | 0 | NA | NA | NA | NA | NA | Pramel32 | PRAME like 32 [Source:MGI Symbol;Acc:M | 0 | 0 | 0 | 0 | 0 | 0 |

| | | | | | | | | | | | | | | | |
|---|---|---|---|---|---|---|---|---|---|---|---|---|---|---|---|
| 11085 | ENSMUSG00000038331 | 21.11262686 | -0.334050485 | 0.545259872 | -0.612644543 | 0.540111399 | 0.893023694 | Satb2 | special AT-rich sequence binding protein 2 | 17.668 | 26.726 | 26.162 | 9.6398 | 14.178 | 32.301 |
| 11086 | ENSMUSG00000038332 | 220.5016519 | 0.087767596 | 0.161786444 | 0.542490423 | 0.587480703 | 0.910765033 | Sesn1 | sestrin 1 [Source:MGI Symbol;Acc:MGI:21 | 220.05 | 216.89 | 204.27 | 225.57 | 230.12 | 226.11 |
| 11087 | ENSMUSG00000038335 | 190.7405822 | -0.039690043 | 0.188082359 | -0.211024802 | 0.832867918 | 0.970611245 | Tsr1 | TSR1 20S rRNA accumulation [Source:MG | 170.26 | 211.75 | 198.23 | 161.95 | 201.76 | 200.49 |
| 11088 | ENSMUSG00000038342 | 148.9220061 | -0.354969277 | 0.212568693 | -1.669903844 | 0.09493839 | 0.454630813 | Mlxip | MLX interacting protein [Source:MGI Sym | 157.41 | 184 | 159.99 | 120.5 | 111.24 | 160.39 |
| 11089 | ENSMUSG00000038344 | 569.2774683 | 0.171568916 | 0.118395353 | 1.44911867 | 0.147304445 | 0.558402858 | Txlng | taxilin gamma [Source:MGI Symbol;Acc:M | 549.32 | 530.41 | 526.27 | 589.95 | 624.92 | 594.79 |
| 11090 | ENSMUSG00000038346 | 433.4853375 | 0.135158317 | 0.138879721 | 0.973204119 | 0.330451864 | 0.774747658 | Zfp384 | zinc finger protein 384 [Source:MGI Symb | 389.5 | 420.42 | 430.67 | 505.12 | 429.7 | 425.49 |
| 11091 | ENSMUSG00000038347 | 17.55257741 | 0.666706125 | 0.516443541 | 1.290956459 | 0.196718776 | 0.637822894 | Tcte2 | t-complex-associated testis expressed 2 [S | 15.259 | 12.335 | 13.081 | 26.027 | 17.45 | 21.163 |
| 11092 | ENSMUSG00000038349 | 12.37652724 | -0.566960131 | 0.666717743 | -0.850375045 | 0.395116606 | 0.822933206 | Plcl1 | phospholipase C-like 1 [Source:MGI Symb | 12.85 | 10.279 | 21.131 | 6.7478 | 6.5437 | 16.708 |
| 11093 | ENSMUSG00000038351 | 145.23953 | -0.4465829 | 0.211263496 | -2.113866843 | 0.034526642 | 0.249073989 | Sgsm2 | small G protein signaling modulator 2 [Sou | 140.54 | 163.44 | 199.24 | 130.14 | 117.79 | 120.3 |
| 11094 | ENSMUSG00000038352 | 2.227027489 | -3.420490785 | 1.976943475 | -1.730191494 | 0.083596068 | NA | Arl5c | ADP-ribosylation factor-like 5C [Source:M | 0 | 8.2233 | 4.025 | 0 | 0 | 1.1138 |
| 11095 | ENSMUSG00000038354 | 0.636946036 | 1.591416711 | 2.83781177 | 0.560790087 | 0.574940645 | NA | Ankrd35 | ankyrin repeat domain 35 [Source:MGI Sy | 0.8031 | 0 | 0 | 1.928 | 1.0906 | 0 |
| 11096 | ENSMUSG00000038357 | 1.345760821 | -0.129104346 | 2.177075009 | -0.059301745 | 0.952711771 | NA | Camp | cathelicidin antimicrobial peptide [Source | 3.2124 | 0 | 1.0062 | 3.8559 | 0 | 0 |
| 11097 | ENSMUSG00000038365 | 112.1650653 | 0.022274584 | 0.219120723 | 0.101654392 | 0.919031004 | 0.986374671 | Fbxo25 | F-box protein 25 [Source:MGI Symbol;Acc | 117.25 | 113.07 | 103.64 | 126.28 | 104.7 | 108.04 |
| 11098 | ENSMUSG00000038366 | 1620.418617 | 0.111536136 | 0.15330817 | 0.727528981 | 0.466901968 | 0.86392095 | Lasp1 | LIM and SH3 protein 1 [Source:MGI Symb | 1278.5 | 1415.4 | 1980.3 | 1602.1 | 1627.2 | 1818.9 |
| 11099 | ENSMUSG00000038368 | 162.5229665 | -0.181031146 | 0.190090889 | -0.952339939 | 0.340924609 | 0.783129922 | Focad | focadhesin [Source:MGI Symbol;Acc:MGI: | 178.29 | 184 | 155.97 | 163.88 | 153.78 | 139.23 |
| 11100 | ENSMUSG00000038369 | 396.9970189 | 0.23042162 | 0.15964317 | 1.443354079 | 0.148920753 | 0.560934836 | Ncoa6 | nuclear receptor coactivator 6 [Source:MG | 386.29 | 407.06 | 302.88 | 478.13 | 414.43 | 393.19 |
| 11101 | ENSMUSG00000038370 | 173.8104646 | -1.14077808 | 0.277918771 | -4.10471764 | 4.05E-05 | 0.00176893 | Pcp4l1 | Purkinje cell protein 4-like 1 [Source:MGI | 139.74 | 314.54 | 263.64 | 98.326 | 133.06 | 93.563 |
| 11102 | ENSMUSG00000038371 | 523.9752596 | -0.12095812 | 0.153333038 | -0.788858824 | 0.430194524 | 0.84347751 | Sbf2 | SET binding factor 2 [Source:MGI Symbol; | 510.77 | 615.72 | 511.17 | 427.04 | 583.48 | 495.66 |
| 11103 | ENSMUSG00000038372 | 23.35820187 | -0.112262904 | 0.529385378 | -0.212062722 | 0.832058102 | 0.970533397 | Gmds | GDP-mannose 4, 6-dehydratase [Source:M | 28.912 | 16.447 | 27.169 | 14.46 | 14.178 | 38.985 |
| 11104 | ENSMUSG00000038374 | 576.2914387 | 0.005518375 | 0.124548309 | 0.044307101 | 0.964659611 | 0.993480943 | Rbm8a | RNA binding motif protein 8a [Source:MG | 579.84 | 539.66 | 605.76 | 566.82 | 547.49 | 618.18 |
| 11105 | ENSMUSG00000038375 | 6778.237962 | -0.517578882 | 0.129504763 | -3.996601128 | 6.43E-05 | 0.002539437 | Trp53inp2 | transformation related protein 53 inducib | 6797.5 | 7947.9 | 9199.1 | 5202.6 | 6341.9 | 5180.5 |
| 11106 | ENSMUSG00000038379 | 2.101379082 | -0.364329998 | 1.684075909 | -0.216338228 | 0.828724113 | NA | Ttk | Ttk protein kinase [Source:MGI Symbol;Ac | 0 | 3.0838 | 4.025 | 0 | 3.2719 | 2.2277 |
| 11107 | ENSMUSG00000038383 | 249.7178939 | -0.15496383 | 0.169669007 | -0.913330208 | 0.361068899 | 0.800074801 | Pigu | phosphatidylinositol glycan anchor biosyn | 253.78 | 305.29 | 230.43 | 230.39 | 234.48 | 243.93 |
| 11108 | ENSMUSG00000038384 | 420.5917112 | 0.307041368 | 0.150468992 | 2.0405624 | 0.041294343 | 0.278526339 | Setd1b | SET domain containing 1B [Source:MGI Sy | 322.04 | 377.25 | 429.67 | 462.71 | 437.34 | 494.55 |
| 11109 | ENSMUSG00000038387 | 378.8679545 | 0.132847254 | 0.14132425 | 0.940017398 | 0.347208637 | 0.788352979 | Rras | related RAS viral (r-ras) oncogene [Source | 340.52 | 358.74 | 385.39 | 393.3 | 367.54 | 427.72 |
| 11110 | ENSMUSG00000038388 | 1285.224425 | -0.107334426 | 0.110348551 | -0.972685419 | 0.330709674 | 0.774747658 | Mpp6 | membrane protein, palmitoylated 6 (MAG | 1219.1 | 1432.9 | 1348.4 | 1289.8 | 1167 | 1254.2 |
| 11111 | ENSMUSG00000038390 | 26.250148 | 0.241260102 | 0.458967967 | 0.525657823 | 0.599125974 | 0.915074771 | Gpr162 | G protein-coupled receptor 162 [Source:M | 18.471 | 19.53 | 34.212 | 20.244 | 31.628 | 33.415 |
| 11112 | ENSMUSG00000038393 | 1982.502327 | -0.183699486 | 0.499978206 | -0.367414986 | 0.713309495 | 0.944239215 | Txnip | thioredoxin interacting protein [Source:M | 2745 | 1980.8 | 1599.9 | 2929.5 | 1676.3 | 963.47 |
| 11113 | ENSMUSG00000038398 | 240.734168 | 0.001598837 | 0.176215637 | 0.009073183 | 0.992760747 | 0.99859176 | Upf3a | UPF3 regulator of nonsense transcripts hc | 281.89 | 212.78 | 226.41 | 253.53 | 220.3 | 249.5 |
| 11114 | ENSMUSG00000038400 | 39.82508998 | -1.019330898 | 0.372142416 | -2.739088196 | 0.006160984 | 0.076688879 | Pmepa1 | prostate transmembrane protein, androge | 58.626 | 45.228 | 56.35 | 34.703 | 23.994 | 20.049 |
| 11115 | ENSMUSG00000038402 | 0 | NA | NA | NA | NA | NA | Foxf2 | forkhead box F2 [Source:MGI Symbol;Acc | 0 | 0 | 0 | 0 | 0 | 0 |
| 11116 | ENSMUSG00000038403 | 1968.347406 | 0.075140708 | 0.123827583 | 0.606817204 | 0.543972229 | 0.894702561 | Hjv | hemojuvelin BMP co-receptor [Source:MG | 2117.8 | 1614.9 | 2017.5 | 2046.5 | 1872.6 | 2140.8 |
| 11117 | ENSMUSG00000038406 | 825.0848651 | -0.006072273 | 0.134538572 | -0.045134067 | 0.964000448 | 0.993245949 | Scaf1 | SR-related CTD-associated factor 1 [Sourc | 739.66 | 839.81 | 902.6 | 940.84 | 760.16 | 767.44 |
| 11118 | ENSMUSG00000038408 | 0 | NA | NA | NA | NA | NA | 1700018A04 | RIKEN cDNA 1700018A04 gene [Source:M | 0 | 0 | 0 | 0 | 0 | 0 |
| 11119 | ENSMUSG00000038412 | 1156.050815 | 0.1096833 | 0.109031706 | 1.005976185 | 0.314427026 | 0.760895517 | Higd1a | HIG1 domain family, member 1A [Source: | 1166.1 | 1078.3 | 1090.8 | 1106.6 | 1279.3 | 1215.2 |
| 11120 | ENSMUSG00000038415 | 499.0932768 | 2.041926736 | 0.860788608 | 2.372158177 | 0.017684518 | 0.158586872 | Foxq1 | forkhead box Q1 [Source:MGI Symbol;Acc | 114.04 | 251.84 | 219.36 | 874.33 | 151.6 | 1383.4 |
| 11121 | ENSMUSG00000038416 | 433.5560672 | 0.125275238 | 0.140736505 | 0.890140321 | 0.373390545 | 0.80865857 | Cdc16 | CDC16 cell division cycle 16 [Source:MGI S | 418.42 | 375.19 | 450.8 | 485.84 | 415.53 | 455.56 |
| 11122 | ENSMUSG00000038417 | 332.0803977 | 0.057793071 | 0.151737739 | 0.380874735 | 0.7032962 | 0.942506832 | Fig4 | FIG4 phosphoinositide 5-phosphatase [So | 305.18 | 327.91 | 343.13 | 297.87 | 367.54 | 350.86 |
| 11123 | ENSMUSG00000038418 | 141.1590753 | 0.255247316 | 0.313247578 | 0.814842105 | 0.415162696 | 0.836001897 | Egr1 | early growth response 1 [Source:MGI Sym | 149.38 | 86.345 | 149.93 | 116.64 | 238.85 | 105.82 |
| 11124 | ENSMUSG00000038421 | 2.515964219 | -1.921103131 | 1.413459368 | -1.359149881 | 0.174099099 | NA | Fcrla | Fc receptor-like A [Source:MGI Symbol;Ac | 4.8186 | 3.0838 | 4.025 | 0.964 | 1.0906 | 1.1138 |
| 11125 | ENSMUSG00000038422 | 382.1801365 | -0.114698302 | 0.150015644 | -0.764575605 | 0.4445243 | 0.851724159 | Hdhd3 | haloacid dehalogenase-like hydrolase don | 408.78 | 363.88 | 418.6 | 317.15 | 393.71 | 390.96 |
| 11126 | ENSMUSG00000038425 | 118.2908397 | 0.570024606 | 0.237300643 | 2.402119945 | 0.016300363 | 0.150940106 | Poli | polymerase (DNA directed), iota [Source:I | 90.751 | 91.485 | 103.64 | 144.6 | 171.23 | 108.04 |
| 11127 | ENSMUSG00000038429 | 980.7262626 | 0.076425328 | 0.11182606 | 0.683430394 | 0.49433492 | 0.877473722 | Usp5 | ubiquitin specific peptidase 5 (isopeptidas | 995.04 | 911.76 | 956.94 | 998.68 | 1084.1 | 937.86 |
| 11128 | ENSMUSG00000038437 | 586.9212821 | -0.014117758 | 0.127297641 | -0.110903533 | 0.911692844 | 0.98589875 | Mllt6 | myeloid/lymphoid or mixed-lineage leuke | 534.87 | 623.95 | 611.8 | 615.02 | 585.66 | 550.24 |
| 11129 | ENSMUSG00000038446 | 536.7296528 | 0.080024028 | 0.121440399 | 0.658957223 | 0.509923238 | 0.882852507 | Cdc40 | cell division cycle 40 [Source:MGI Symbol; | 521.21 | 511.9 | 532.3 | 537.9 | 581.3 | 535.76 |
| 11130 | ENSMUSG00000038451 | 309.2300781 | 0.368725047 | 0.146259298 | 2.521036619 | 0.011700967 | 0.120662445 | Spsb2 | splA/ryanodine receptor domain and SOC | 283.5 | 272.4 | 253.57 | 355.71 | 331.55 | 358.66 |
| 11131 | ENSMUSG00000038453 | 2.566333274 | -0.394259243 | 1.585441669 | -0.248674708 | 0.803612415 | NA | Srcin1 | SRC kinase signaling inhibitor 1 [Source:M | 1.6062 | 3.0838 | 4.025 | 0 | 0 | 6.6831 |
| 11132 | ENSMUSG00000038456 | 53.54099342 | -0.910983976 | 0.347522502 | -2.621366876 | 0.008757796 | 0.098272117 | Dennd2a | DENN/MADD domain containing 2A [Sour | 81.113 | 64.759 | 63.393 | 27.955 | 29.447 | 54.578 |
| 11133 | ENSMUSG00000038457 | 1.755668363 | -0.670584633 | 2.006833331 | -0.334150636 | 0.738265892 | NA | Tmem255b | transmembrane protein 255B [Source:MG | 6.4248 | 0 | 0 | 1.928 | 2.1812 | 0 |
| 11134 | ENSMUSG00000038459 | 1383.649714 | -0.059758105 | 0.110423381 | -0.541172573 | 0.58838864 | 0.910917568 | Abhd17c | abhydrolase domain containing 17C [Sour | 1548.4 | 1315.7 | 1371.5 | 1352.5 | 1284.7 | 1429.1 |
| 11135 | ENSMUSG00000038462 | 5104.32367 | -0.088774098 | 0.083950194 | -1.057461508 | 0.290301013 | 0.740288056 | Uqcrfs1 | ubiquinol-cytochrome c reductase, Rieske | 5303.7 | 5143.7 | 5336.1 | 4843 | 5105.2 | 4894.2 |
| 11136 | ENSMUSG00000038463 | 1.975347684 | -0.276641231 | 1.49287429 | -0.185307787 | 0.85298763 | NA | Olfml2b | olfactomedin-like 2B [Source:MGI Symbol | 2.4093 | 1.0279 | 3.0187 | 0.964 | 1.0906 | 3.3415 |
| 11137 | ENSMUSG00000038467 | 1016.693575 | -0.019295366 | 0.104155896 | -0.185254667 | 0.853029293 | 0.973813394 | Chmp4b | charged multivesicular body protein 4B [S | 1019.1 | 1037.2 | 1014.3 | 1022.8 | 958.65 | 1048.1 |
| 11138 | ENSMUSG00000038473 | 44.50956253 | 0.05367503 | 0.376355149 | 0.142618031 | 0.886591856 | 0.982355187 | Nos1ap | nitric oxide synthase 1 (neuronal) adaptor | 40.958 | 38.033 | 52.325 | 61.695 | 27.265 | 46.781 |
| 11139 | ENSMUSG00000038481 | 163.4162524 | 0.335878359 | 0.19231251 | 1.746523718 | 0.080719989 | 0.417428319 | Cdk19 | cyclin-dependent kinase 19 [Source:MGI S | 138.13 | 130.55 | 165.02 | 189.9 | 187.59 | 169.3 |
| 11140 | ENSMUSG00000038482 | 484.006514 | 0.046216117 | 0.126982691 | 0.363956041 | 0.715890835 | 0.9447719 | Tfdp1 | transcription factor Dp 1 [Source:MGI Sym | 481.06 | 449.2 | 498.09 | 469.46 | 502.78 | 503.46 |

| | | | | | | | | | | | | | | | |
|---|---|---|---|---|---|---|---|---|---|---|---|---|---|---|---|
| 11141 | ENSMUSG00000038485 | 318.4997562 | 0.101466203 | 0.145353282 | 0.698066132 | 0.485135837 | 0.872653246 | Socs7 | suppressor of cytokine signaling 7 [Source | 301.97 | 300.15 | 319.99 | 339.32 | 304.28 | 345.29 |
| 11142 | ENSMUSG00000038486 | 1.395225349 | -2.690668074 | 2.005635346 | -1.341553977 | 0.179740657 | NA | Sv2a | synaptic vesicle glycoprotein 2 a [Source:N | 3.2124 | 2.0558 | 2.0125 | 0 | 1.0906 | 0 |
| 11143 | ENSMUSG00000038489 | 644.1522584 | -0.00116796 | 0.119828001 | -0.009746971 | 0.992223166 | 0.99859176 | Polr2l | polymerase (RNA) II (DNA directed) polyp | 693.88 | 603.39 | 634.94 | 625.62 | 654.37 | 652.71 |
| 11144 | ENSMUSG00000038491 | 0.133850373 | -0.796629176 | 4.080472857 | -0.195229623 | 0.84521319 | NA | Cdk19os | cyclin-dependent kinase 19, opposite stra | 0.8031 | 0 | 0 | 0 | 0 | 0 |
| 11145 | ENSMUSG00000038495 | 430.5999931 | -0.098082073 | 0.129675287 | -0.75636673 | 0.44942935 | 0.854879451 | Otud7b | OTU domain containing 7B [Source:MGI S | 449.74 | 457.42 | 428.66 | 430.9 | 398.08 | 418.81 |
| 11146 | ENSMUSG00000038496 | 0.795980673 | -1.697257209 | 2.614316861 | -0.649216334 | 0.516198555 | NA | Slc19a3 | solute carrier family 19, member 3 [Source | 1.6062 | 2.0558 | 0 | 0 | 0 | 1.1138 |
| 11147 | ENSMUSG00000038497 | 121.0347446 | -0.110380197 | 0.213565856 | -0.51684384 | 0.605265174 | 0.916376001 | Tmco3 | transmembrane and coiled-coil domains 3 | 118.86 | 139.8 | 118.74 | 126.28 | 115.61 | 106.93 |
| 11148 | ENSMUSG00000038498 | 0 | NA | NA | NA | NA | NA | Catsper1 | cation channel, sperm associated 1 [Sourc | 0 | 0 | 0 | 0 | 0 | 0 |
| 11149 | ENSMUSG00000038500 | 156.4309186 | 0.271527637 | 0.193771046 | 1.401280755 | 0.161130134 | 0.584240557 | Prr3 | proline-rich polypeptide 3 [Source:MGI Sy | 142.95 | 129.52 | 152.95 | 186.05 | 173.41 | 153.71 |
| 11150 | ENSMUSG00000038502 | 759.5949678 | 0.000841237 | 0.14200018 | 0.005924198 | 0.995273201 | 0.998794948 | Ptov1 | prostate tumor over expressed gene 1 [So | 795.87 | 793.55 | 689.28 | 902.28 | 686 | 690.58 |
| 11151 | ENSMUSG00000038503 | 359.3245968 | -0.131269173 | 0.146809405 | -0.894146891 | 0.371243304 | 0.807214774 | Mesd | mesoderm development LRP chaperone [ | 350.96 | 363.88 | 412.56 | 348 | 314.1 | 366.45 |
| 11152 | ENSMUSG00000038506 | 287.9880166 | 0.09931725 | 0.150400258 | 0.660352923 | 0.509027375 | 0.882793133 | Dcun1d2 | DCN1, defective in cullin neddylation 1, do | 266.63 | 298.1 | 269.67 | 288.23 | 291.2 | 314.1 |
| 11153 | ENSMUSG00000038507 | 682.8501398 | 0.184269071 | 0.145093569 | 1.270001649 | 0.204084043 | 0.649416772 | Parp12 | poly (ADP-ribose) polymerase family, men | 630.44 | 613.67 | 673.18 | 599.59 | 851.77 | 728.45 |
| 11154 | ENSMUSG00000038508 | 131.1737062 | -0.907155554 | 0.369110165 | -2.457682397 | 0.013983678 | 0.136282395 | Gdf15 | growth differentiation factor 15 [Source:N | 305.18 | 107.93 | 99.618 | 89.65 | 115.61 | 69.058 |
| 11155 | ENSMUSG00000038510 | 139.092778 | 0.128198651 | 0.206816062 | 0.619867964 | 0.535344719 | 0.889860001 | Rpf2 | ribosome production factor 2 homolog [So | 134.12 | 132.6 | 131.82 | 136.88 | 127.6 | 171.53 |
| 11156 | ENSMUSG00000038515 | 445.5110242 | 0.426071923 | 0.135961633 | 3.133765858 | 0.001725785 | 0.031437978 | Grtp1 | GH regulated TBC protein 1 [Source:MGI S | 371.84 | 350.52 | 418.6 | 537.9 | 491.87 | 502.34 |
| 11157 | ENSMUSG00000038517 | 233.0876868 | -0.494651623 | 0.194162689 | -2.547614197 | 0.010846235 | 0.11526814 | Tbkbp1 | TBK1 binding protein 1 [Source:MGI Symb | 289.12 | 243.62 | 285.77 | 242.92 | 175.59 | 161.51 |
| 11158 | ENSMUSG00000038518 | 203.0567033 | 0.184121279 | 0.178655759 | 1.030592464 | 0.302731971 | 0.752484788 | Jarid2 | jumonji, AT rich interactive domain 2 [Sou | 196.76 | 176.8 | 196.22 | 190.87 | 213.76 | 243.93 |
| 11159 | ENSMUSG00000038520 | 641.6027725 | 0.156796762 | 0.131260625 | 1.194545291 | 0.232264735 | 0.67964336 | Tbc1d17 | TBC1 domain family, member 17 [Source:I | 654.53 | 596.19 | 569.53 | 759.61 | 629.29 | 640.46 |
| 11160 | ENSMUSG00000038521 | 8214.247005 | 0.032987935 | 0.093760799 | 0.351830776 | 0.724965177 | 0.9448775 | C1s1 | complement component 1, s subcompone | 7656.8 | 8146.2 | 8559.1 | 8369.3 | 8780.6 | 7773.5 |
| 11161 | ENSMUSG00000038522 | 1203.887958 | 0.053285064 | 0.122665132 | 0.43439454 | 0.664001962 | 0.929301301 | Mfsd4b1 | major facilitator superfamily domain cont | 1095.4 | 1093.7 | 1356.4 | 1205 | 1304.4 | 1168.4 |
| 11162 | ENSMUSG00000038523 | 1.886268103 | -0.236130036 | 1.606040561 | -0.147026197 | 0.883111343 | NA | 1700003F12 | RIKEN cDNA 1700003F12 gene [Source:M | 4.0155 | 2.0558 | 0 | 1.928 | 1.0906 | 2.2277 |
| 11163 | ENSMUSG00000038524 | 9.784988585 | -0.143688719 | 0.709245737 | -0.202593702 | 0.839452613 | 0.971736102 | Fchsd1 | FCH and double SH3 domains 1 [Source:M | 9.6372 | 7.1954 | 14.087 | 13.496 | 8.7249 | 5.5692 |
| 11164 | ENSMUSG00000038525 | 141.200121 | -0.161155267 | 0.211158049 | -0.763197363 | 0.445345701 | 0.852147532 | Armc10 | armadillo repeat containing 10 [Source:M | 149.38 | 164.47 | 133.83 | 157.13 | 124.33 | 118.07 |
| 11165 | ENSMUSG00000038526 | 598.1847741 | -0.318626286 | 0.150096441 | -2.1228104 | 0.033769743 | 0.246068294 | Car14 | carbonic anhydrase 14 [Source:MGI Symb | 532.46 | 701.04 | 759.71 | 544.65 | 533.31 | 517.94 |
| 11166 | ENSMUSG00000038527 | 3277.597126 | 0.166803834 | 0.100321537 | 1.66269217 | 0.09637407 | 0.458386897 | C1rl | complement component 1, r subcompone | 3010.8 | 3000.5 | 3253.2 | 3174.4 | 3500.9 | 3725.8 |
| 11167 | ENSMUSG00000038528 | 3.431326969 | -1.229591605 | 1.267669621 | -0.969962192 | 0.332065338 | NA | Mfsd4b5 | major facilitator superfamily domain cont | 2.4093 | 2.0558 | 10.062 | 3.8559 | 1.0906 | 1.1138 |
| 11168 | ENSMUSG00000038530 | 19.9597928 | -1.956373239 | 0.550793487 | -3.551917887 | 0.000382434 | 0.010379287 | Rgs4 | regulator of G-protein signaling 4 [Source: | 36.943 | 30.838 | 27.169 | 3.8559 | 9.8156 | 11.138 |
| 11169 | ENSMUSG00000038533 | 224.4691533 | 0.184746768 | 0.163158922 | 1.132311773 | 0.257503382 | 0.710872182 | Cbfa2t2 | CBFA2/RUNX1 translocation partner 2 [So | 203.18 | 227.17 | 200.24 | 240.03 | 234.48 | 241.7 |
| 11170 | ENSMUSG00000038534 | 34.47746098 | -0.353755306 | 0.430826029 | -0.821109408 | 0.411583952 | 0.833774549 | Osbpl7 | oxysterol binding protein-like 7 [Source:M | 40.958 | 41.117 | 34.212 | 43.379 | 32.719 | 14.48 |
| 11171 | ENSMUSG00000038535 | 158.96259 | 0.145421686 | 0.202052188 | 0.719723392 | 0.471695321 | 0.867794597 | Zfp280d | zinc finger protein 280D [Source:MGI Sym | 165.44 | 141.85 | 144.9 | 148.45 | 154.87 | 198.26 |
| 11172 | ENSMUSG00000038537 | 0 | NA | NA | NA | NA | NA | Mc3r | melanocortin 3 receptor [Source:MGI Sym | 0 | 0 | 0 | 0 | 0 | 0 |
| 11173 | ENSMUSG00000038538 | 371.6488951 | 0.095115666 | 0.145572292 | 0.653391279 | 0.513504059 | 0.885004045 | Ubn2 | ubinuclein 2 [Source:MGI Symbol;Acc:MG | 335.7 | 363.88 | 379.35 | 400.05 | 410.07 | 340.84 |
| 11174 | ENSMUSG00000038539 | 3876.935491 | 0.021563536 | 0.140991536 | 0.152942063 | 0.878443964 | 0.980976563 | Atf5 | activating transcription factor 5 [Source:N | 4355.2 | 3633.7 | 3554.1 | 4080.5 | 3114.8 | 4523.3 |
| 11175 | ENSMUSG00000038540 | 0.501260735 | 1.111687294 | 3.109072805 | 0.357562323 | 0.720670879 | NA | Tmc3 | transmembrane channel-like gene family | 0.8031 | 0 | 0 | 0 | 1.0906 | 1.1138 |
| 11176 | ENSMUSG00000038541 | 44.52719045 | 0.234821356 | 0.333123487 | 0.704907835 | 0.480867594 | 0.870278888 | Srd5a2 | steroid 5 alpha-reductase 2 [Source:MGI S | 48.186 | 34.949 | 39.244 | 43.379 | 50.168 | 51.237 |
| 11177 | ENSMUSG00000038542 | 248.7061703 | 0.016974363 | 0.161356401 | 0.105197955 | 0.916218733 | 0.986374671 | Pcid2 | PCI domain containing 2 [Source:MGI Sym | 238.52 | 264.17 | 239.49 | 267.02 | 226.85 | 256.18 |
| 11178 | ENSMUSG00000038543 | 3.877383756 | -0.241469629 | 1.110553833 | -0.217431719 | 0.827871914 | NA | BC028528 | cDNA sequence BC028528 [Source:MGI Sy | 2.4093 | 8.2233 | 2.0125 | 2.8919 | 3.2719 | 4.4554 |
| 11179 | ENSMUSG00000038544 | 89.36606148 | 0.100662158 | 0.243817405 | 0.412858786 | 0.679710078 | 0.934260934 | Inip | INTS3 and NABP interacting protein [Sour | 82.72 | 85.317 | 90.562 | 78.082 | 98.156 | 101.36 |
| 11180 | ENSMUSG00000038545 | 150.8066853 | -0.142634026 | 0.212101678 | -0.672479482 | 0.501278498 | 0.880974509 | Cul7 | cullin 7 [Source:MGI Symbol;Acc:MGI:191 | 189.53 | 137.74 | 146.91 | 145.56 | 159.23 | 125.86 |
| 11181 | ENSMUSG00000038546 | 705.3566332 | 0.049765504 | 0.129468893 | 0.384381939 | 0.700695385 | 0.941778228 | Ranbp9 | RAN binding protein 9 [Source:MGI Symb | 732.43 | 644.5 | 702.36 | 739.37 | 784.15 | 629.32 |
| 11182 | ENSMUSG00000038550 | 345.2707812 | 0.068175154 | 0.238362888 | 0.28601413 | 0.7748673 | 0.958904913 | Ciart | circadian associated repressor of transcrip | 432.07 | 365.94 | 212.32 | 277.63 | 440.61 | 343.06 |
| 11183 | ENSMUSG00000038552 | 728.529092 | 0.122943305 | 0.140703102 | 0.873778212 | 0.382239048 | 0.813997203 | Fndc4 | fibronectin type III domain containing 4 [S | 672.2 | 644.5 | 775.81 | 742.26 | 660.91 | 875.48 |
| 11184 | ENSMUSG00000038555 | 1.72823623 | 0.164878706 | 1.724917247 | 0.095586444 | 0.923849032 | NA | Reep2 | receptor accessory protein 2 [Source:MGI | 0.8031 | 3.0838 | 1.0062 | 0 | 4.3625 | 1.1138 |
| 11185 | ENSMUSG00000038560 | 0 | NA | NA | NA | NA | NA | Sp6 | trans-acting transcription factor 6 [Source | 0 | 0 | 0 | 0 | 0 | 0 |
| 11186 | ENSMUSG00000038563 | 229.3248381 | 0.063811292 | 0.174599245 | 0.365472896 | 0.714758436 | 0.944595913 | Efl1 | elongation factor like GTPase 1 [Source:M | 238.52 | 242.59 | 191.19 | 217.86 | 254.11 | 231.68 |
| 11187 | ENSMUSG00000038564 | 344.5640962 | -0.022879545 | 0.143610819 | -0.159316307 | 0.873419673 | 0.979925081 | Ift172 | intraflagellar transport 172 [Source:MGI S | 337.3 | 369.02 | 336.09 | 358.6 | 352.27 | 314.1 |
| 11188 | ENSMUSG00000038567 | 0 | NA | NA | NA | NA | NA | Cyp24a1 | cytochrome P450, family 24, subfamily a, | 0 | 0 | 0 | 0 | 0 | 0 |
| 11189 | ENSMUSG00000038569 | 40.54259892 | 0.045156633 | 0.364526597 | 0.123877472 | 0.90141229 | 0.983334141 | Rad9b | RAD9 checkpoint clamp component B [So | 51.399 | 27.754 | 40.25 | 43.379 | 39.262 | 41.212 |
| 11190 | ENSMUSG00000038570 | 7.280697743 | -0.174959145 | 0.848076464 | -0.206301145 | 0.836555684 | 0.971073158 | Saxo2 | stabilizer of axonemal microtubules 2 [Sou | 8.8341 | 5.1396 | 9.0562 | 1.928 | 9.8156 | 8.9108 |
| 11191 | ENSMUSG00000038572 | 0 | NA | NA | NA | NA | NA | Bpifb5 | BPI fold containing family B, member 5 [So | 0 | 0 | 0 | 0 | 0 | 0 |
| 11192 | ENSMUSG00000038576 | 552.4398821 | 1.18504804 | 0.395075473 | 2.999548494 | 0.002703801 | 0.043695838 | Susd4 | sushi domain containing 4 [Source:MGI Sy | 337.3 | 230.25 | 444.76 | 485.84 | 846.32 | 970.16 |
| 11193 | ENSMUSG00000038578 | 166.825975 | -0.168592075 | 0.250702623 | -0.672478307 | 0.501279246 | 0.880974509 | Susd1 | sushi domain containing 1 [Source:MGI Sy | 184.71 | 191.19 | 153.96 | 173.52 | 200.67 | 96.904 |
| 11194 | ENSMUSG00000038580 | 0.185640647 | 1.126924175 | 4.080472857 | 0.276174898 | 0.782413736 | NA | Sct | secretin [Source:MGI Symbol;Acc:MGI:99 | 0 | 0 | 0 | 0 | 0 | 1.1138 |
| 11195 | ENSMUSG00000038582 | 475.8108659 | 0.065576299 | 0.158869178 | 0.412769171 | 0.679775741 | 0.934260934 | Pptc7 | PTC7 protein phosphatase homolog [Sour | 462.59 | 534.52 | 397.47 | 413.55 | 535.49 | 511.25 |
| 11196 | ENSMUSG00000038583 | 1.259856419 | -1.497590767 | 2.241077249 | -0.668245937 | 0.503976617 | NA | Pln | phospholamban [Source:MGI Symbol;Acc | 1.6062 | 0 | 4.025 | 1.928 | 0 | 0 |

| | | | | | | | | | | | | | | | |
|---|---|---|---|---|---|---|---|---|---|---|---|---|---|---|---|
| 11197 | ENSMUSG00000038587 | 38.39041717 | -0.398700431 | 0.387252431 | -1.029562113 | 0.303215608 | 0.752484788 | Akap12 | A kinase (PRKA) anchor protein (gravin) 1: | 40.958 | 39.061 | 51.318 | 47.235 | 27.265 | 24.505 |
| 11198 | ENSMUSG00000038591 | 298.6303126 | -1.249738149 | 0.204081324 | -6.12372618 | 9.14E-10 | 1.92E-07 | Colec10 | collectin sub-family member 10 [Source:N | 409.58 | 416.31 | 434.7 | 137.85 | 149.41 | 243.93 |
| 11199 | ENSMUSG00000038593 | 51.59483636 | 0.203418481 | 0.354592526 | 0.573668271 | 0.566192303 | 0.902076435 | Tctn1 | tectonic family member 1 [Source:MGI Sy | 47.383 | 52.424 | 44.275 | 70.37 | 31.628 | 63.489 |
| 11200 | ENSMUSG00000038594 | 702.9154556 | -0.626477856 | 0.191880939 | -3.264930119 | 0.001094911 | 0.022431715 | Cep85l | centrosomal protein 85-like [Source:MGI ! | 808.72 | 1079.3 | 671.17 | 437.65 | 686 | 534.65 |
| 11201 | ENSMUSG00000038598 | 0 | NA | NA | NA | NA | NA | Shoc1 | shortage in chiasmata 1 [Source:MGI Sym | 0 | 0 | 0 | 0 | 0 | 0 |
| 11202 | ENSMUSG00000038599 | 88.49782628 | 1.196758116 | 0.808438611 | 1.480332705 | 0.138784479 | 0.544447711 | Capn8 | calpain 8 [Source:MGI Symbol;Acc:MGI:2: | 88.341 | 11.307 | 61.381 | 71.334 | 151.6 | 147.03 |
| 11203 | ENSMUSG00000038600 | 8.437586158 | 0.894390646 | 0.785937224 | 1.137992474 | 0.255123625 | 0.708637501 | Atp6v0a4 | ATPase, H+ transporting, lysosomal V0 sul | 6.4248 | 8.2233 | 3.0187 | 7.7118 | 17.45 | 7.7969 |
| 11204 | ENSMUSG00000038602 | 1.125921769 | -0.10056922 | 2.125718962 | -0.047310685 | 0.962265612 | NA | Slc35f1 | solute carrier family 35, member F1 [Sour | 2.4093 | 1.0279 | 0 | 0 | 1.0906 | 2.2277 |
| 11205 | ENSMUSG00000038604 | 229.6481679 | 0.032974894 | 0.160463917 | 0.205497253 | 0.837183644 | 0.971073158 | Ripor1 | RHO family interacting cell polarization re | 229.69 | 225.11 | 226.41 | 243.89 | 234.48 | 218.31 |
| 11206 | ENSMUSG00000038605 | 20.07162113 | 0.172678224 | 0.528802332 | 0.326545883 | 0.74401138 | 0.949911274 | Samd10 | sterile alpha motif domain containing 10 [ | 15.259 | 11.307 | 30.187 | 25.063 | 17.45 | 21.163 |
| 11207 | ENSMUSG00000038607 | 1.177744186 | -2.412679626 | 2.321486536 | -1.039282197 | 0.298673512 | NA | Gng10 | guanine nucleotide binding protein (G prc | 0 | 3.0838 | 3.0187 | 0.964 | 0 | 0 |
| 11208 | ENSMUSG00000038608 | 40.58611081 | -1.964537139 | 0.416854364 | -4.712766159 | 2.44E-06 | 0.000170757 | Dock10 | dedicator of cytokinesis 10 [Source:MGI S | 72.279 | 78.122 | 43.269 | 14.46 | 11.997 | 23.391 |
| 11209 | ENSMUSG00000038611 | 374.9571462 | -0.022171284 | 0.135349621 | -0.163807505 | 0.869882683 | 0.978765244 | Phrf1 | PHD and ring finger domains 1 [Source:M( | 377.46 | 394.72 | 361.24 | 361.49 | 382.81 | 372.02 |
| 11210 | ENSMUSG00000038612 | 1889.849568 | 0.279630462 | 0.100442482 | 2.783985978 | 0.005369536 | 0.069842592 | Mcl1 | myeloid cell leukemia sequence 1 [Source | 1807.8 | 1611.8 | 1701.6 | 2157.4 | 1957.7 | 2102.9 |
| 11211 | ENSMUSG00000038615 | 2837.72939 | -0.168075702 | 0.090729301 | -1.852496386 | 0.063954576 | 0.362050484 | Nfe2l1 | nuclear factor, erythroid derived 2,-like 1 | 3071.9 | 3052.9 | 2882.9 | 2563.2 | 2725.5 | 2730 |
| 11212 | ENSMUSG00000038618 | 369.9207067 | 0.571236848 | 0.16208556 | 3.524292042 | 0.000424616 | 0.011126169 | Rassf7 | Ras association (RalGDS/AF-6) domain far | 304.38 | 282.68 | 305.9 | 488.74 | 355.54 | 482.29 |
| 11213 | ENSMUSG00000038619 | 757.0950256 | 0.064459642 | 0.13943183 | 0.462302203 | 0.64386462 | 0.925547393 | Ensa | endosulfine alpha [Source:MGI Symbol;A( | 836.83 | 746.27 | 635.95 | 695.03 | 781.97 | 846.52 |
| 11214 | ENSMUSG00000038622 | 123.9634542 | -0.175781137 | 0.224920689 | -0.781524984 | 0.434493788 | 0.845264432 | Med30 | mediator complex subunit 30 [Source:MG | 131.71 | 129.52 | 132.82 | 89.65 | 130.87 | 129.21 |
| 11215 | ENSMUSG00000038623 | 16.79195137 | -1.597607293 | 0.568721455 | -2.809120841 | 0.0049677 | 0.066530377 | Tm6sf1 | transmembrane 6 superfamily member 1 | 28.912 | 27.754 | 19.119 | 10.604 | 5.4531 | 8.9108 |
| 11216 | ENSMUSG00000038624 | 0.503095663 | 2.549494177 | 3.348715477 | 0.761334964 | 0.446457022 | NA | Nepn | nephrocan [Source:MGI Symbol;Acc:MGI: | 0 | 0 | 0 | 1.928 | 1.0906 | 0 |
| 11217 | ENSMUSG00000038628 | 284.4803298 | 0.195741714 | 0.168345529 | 1.162737826 | 0.24493589 | 0.695017707 | Polr3k | polymerase (RNA) III (DNA directed) polyp | 291.53 | 236.42 | 266.65 | 267.02 | 297.74 | 347.52 |
| 11218 | ENSMUSG00000038630 | 0.185640647 | 1.126924175 | 4.080472857 | 0.276174898 | 0.782413736 | NA | Zkscan16 | zinc finger with KRAB and SCAN domains : | 0 | 0 | 0 | 0 | 0 | 1.1138 |
| 11219 | ENSMUSG00000038633 | 917.8066327 | -0.044295257 | 0.110493222 | -0.400886643 | 0.688503584 | 0.936908509 | Degs1 | delta(4)-desaturase, sphingolipid 1 [Sourc | 868.96 | 970.35 | 956.94 | 877.22 | 889.94 | 943.43 |
| 11220 | ENSMUSG00000038637 | 120.8062693 | 0.211496768 | 0.2133198 | 0.991453993 | 0.321463947 | 0.76866499 | Lrrc56 | leucine rich repeat containing 56 [Source: | 103.6 | 109.99 | 122.76 | 141.7 | 127.6 | 119.18 |
| 11221 | ENSMUSG00000038641 | 4659.567816 | -0.322374312 | 0.1255985 | -2.566705104 | 0.010266989 | 0.110904947 | Akr1d1 | aldo-keto reductase family 1, member D1 | 4418.7 | 5281.4 | 5835.2 | 4132.6 | 4589.3 | 3700.2 |
| 11222 | ENSMUSG00000038642 | 279.9622558 | -1.513655739 | 0.25453059 | -5.946851971 | 2.73E-09 | 4.88E-07 | Ctss | cathepsin S [Source:MGI Symbol;Acc:MGI | 385.49 | 566.38 | 291.81 | 115.68 | 122.15 | 198.26 |
| 11223 | ENSMUSG00000038644 | 71.4907197 | 0.221533463 | 0.271696679 | 0.815370522 | 0.414860253 | 0.836001897 | Pold1 | polymerase (DNA directed), delta 1, catal | 74.689 | 55.508 | 67.418 | 69.406 | 85.068 | 76.855 |
| 11224 | ENSMUSG00000038646 | 633.9511433 | -0.073741799 | 0.123779609 | -0.595750785 | 0.551341726 | 0.89589637 | Ramac | RNA guanine-7 methyltransferase activati | 651.32 | 637.31 | 661.1 | 553.32 | 642.37 | 658.28 |
| 11225 | ENSMUSG00000038648 | 265.8509798 | -0.063870932 | 0.180724069 | -0.353416854 | 0.723775952 | 0.944786345 | Creb3l2 | cAMP responsive element binding protein | 218.44 | 274.45 | 323 | 247.74 | 254.11 | 277.35 |
| 11226 | ENSMUSG00000038650 | 2132.507292 | -0.018264599 | 0.09528206 | -0.191689805 | 0.847985192 | 0.972803041 | Rnh1 | ribonuclease/angiogenin inhibitor 1 [Sour | 2209.3 | 2059.9 | 2168.5 | 2152.6 | 2197.6 | 2007.1 |
| 11227 | ENSMUSG00000038651 | 0.167707431 | -0.796629176 | 4.080472857 | -0.195229623 | 0.84521319 | NA | Sycp2l | synaptonemal complex protein 2-like [Sou | 0 | 0 | 1.0062 | 0 | 0 | 0 |
| 11228 | ENSMUSG00000038656 | 0 | NA | NA | NA | NA | NA | Cyp3a16 | cytochrome P450, family 3, subfamily a, p | 0 | 0 | 0 | 0 | 0 | 0 |
| 11229 | ENSMUSG00000038658 | 586.1835508 | -0.015918473 | 0.137256276 | -0.115976286 | 0.907671336 | 0.984592636 | Ric1 | RAB6A GEF complex partner 1 [Source:M( | 518 | 590.02 | 661.1 | 570.67 | 629.29 | 548.01 |
| 11230 | ENSMUSG00000038663 | 6.984927894 | -1.064336573 | 0.873415702 | -1.218591067 | 0.222999441 | 0.668407528 | Fsd2 | fibronectin type III and SPRY domain cont | 12.047 | 9.2513 | 7.0437 | 4.8199 | 7.6343 | 1.1138 |
| 11231 | ENSMUSG00000038664 | 603.6816495 | -0.215383105 | 0.151713844 | -1.419666784 | 0.155704713 | 0.574732247 | Herc1 | HECT and RLD domain containing E3 ubiqu | 567.79 | 757.58 | 621.86 | 601.52 | 594.39 | 478.95 |
| 11232 | ENSMUSG00000038665 | 0 | NA | NA | NA | NA | NA | Dgki | diacylglycerol kinase, iota [Source:MGI Sy | 0 | 0 | 0 | 0 | 0 | 0 |
| 11233 | ENSMUSG00000038668 | 21.17428766 | -0.617721801 | 0.471690608 | -1.309591056 | 0.190334216 | 0.627779796 | Lpar1 | lysophosphatidic acid receptor 1 [Source:I | 28.109 | 25.698 | 23.144 | 19.28 | 17.45 | 13.366 |
| 11234 | ENSMUSG00000038670 | 1.265323017 | -2.571964304 | 2.315565711 | -1.110728273 | 0.266685328 | NA | Mybpc2 | myosin binding protein C, fast-type [Sourc | 5.6217 | 0 | 1.0062 | 0.964 | 0 | 0 |
| 11235 | ENSMUSG00000038671 | 279.9737964 | 0.144478597 | 0.153565991 | 0.940824173 | 0.346794971 | 0.78794258 | Arfrp1 | ADP-ribosylation factor related protein 1 [ | 273.86 | 256.98 | 266.65 | 283.41 | 275.93 | 323.01 |
| 11236 | ENSMUSG00000038676 | 0 | NA | NA | NA | NA | NA | Ucn | urocortin [Source:MGI Symbol;Acc:MGI:1 | 0 | 0 | 0 | 0 | 0 | 0 |
| 11237 | ENSMUSG00000038677 | 1.507865243 | -0.22394421 | 1.802042691 | -0.124272422 | 0.901099582 | NA | Scube3 | signal peptide, CUB domain, EGF-like 3 [S( | 0.8031 | 4.1117 | 0 | 1.928 | 1.0906 | 1.1138 |
| 11238 | ENSMUSG00000038679 | 6.222209421 | -2.23914636 | 1.035727974 | -2.161905844 | 0.030625434 | 0.231919521 | Trps1 | transcriptional repressor GATA binding 1 [ | 7.2279 | 15.419 | 8.05 | 0 | 2.1812 | 4.4554 |
| 11239 | ENSMUSG00000038683 | 724.7193033 | 0.067606819 | 0.118680296 | 0.56965496 | 0.568911744 | 0.903661263 | Pak1ip1 | PAK1 interacting protein 1 [Source:MGI Sy | 721.19 | 741.13 | 661.1 | 773.11 | 763.43 | 688.36 |
| 11240 | ENSMUSG00000038685 | 235.5675798 | 0.024425231 | 0.159180043 | 0.153444055 | 0.878048106 | 0.980817031 | Rtel1 | regulator of telomere elongation helicase | 245.75 | 232.31 | 222.38 | 238.1 | 244.3 | 230.57 |
| 11241 | ENSMUSG00000038690 | 3111.989691 | 0.059680206 | 0.089355663 | 0.667895062 | 0.50420058 | 0.881510796 | Atp5j2 | ATP synthase, H+ transporting, mitochonc | 3088.7 | 2909 | 3144.5 | 3098.2 | 3184.6 | 3246.9 |
| 11242 | ENSMUSG00000038691 | 0 | NA | NA | NA | NA | NA | Mbd3l1 | methyl-CpG binding domain protein 3-like | 0 | 0 | 0 | 0 | 0 | 0 |
| 11243 | ENSMUSG00000038692 | 28.02573543 | -0.850946003 | 0.507114307 | -1.678016162 | 0.093343944 | 0.450090286 | Hoxb4 | homeobox B4 [Source:MGI Symbol;Acc:M | 19.274 | 38.033 | 51.318 | 30.847 | 13.087 | 15.594 |
| 11244 | ENSMUSG00000038695 | 771.2025397 | 0.352502506 | 0.15499104 | 2.274341191 | 0.02294548 | 0.190737602 | Josd2 | Josephin domain containing 2 [Source:M( | 769.37 | 531.43 | 730.53 | 886.86 | 766.7 | 942.31 |
| 11245 | ENSMUSG00000038696 | 768.9062845 | 0.102526884 | 0.121612323 | 0.843063282 | 0.399193053 | 0.8263661 | Mapkap1 | mitogen-activated protein kinase associat | 752.51 | 799.72 | 672.17 | 761.54 | 776.52 | 850.98 |
| 11246 | ENSMUSG00000038697 | 245.2753834 | 0.007719256 | 0.160283495 | 0.048160018 | 0.961588714 | 0.992749404 | Taf5l | TATA-box binding protein associated facto | 226.47 | 266.23 | 241.5 | 238.1 | 245.39 | 253.96 |
| 11247 | ENSMUSG00000038700 | 6.615203569 | -0.644194692 | 0.892891966 | -0.721469916 | 0.470620444 | 0.867110762 | Hoxb5 | homeobox B5 [Source:MGI Symbol;Acc:M | 4.0155 | 11.307 | 9.0562 | 8.6758 | 2.1812 | 4.4554 |
| 11248 | ENSMUSG00000038702 | 14.02664027 | -1.349811476 | 0.655818554 | -2.058208734 | 0.039570101 | 0.27228446 | Dsel | dermatan sulfate epimerase-like [Source:I | 16.865 | 25.698 | 18.112 | 12.532 | 8.7249 | 2.2277 |
| 11249 | ENSMUSG00000038704 | 2287.174772 | 0.067562148 | 0.106949662 | 0.631719132 | 0.527570424 | 0.888334265 | Aspdh | aspartate dehydrogenase domain contain | 2285.6 | 1983.9 | 2431.1 | 2392.6 | 2267.4 | 2362.5 |
| 11250 | ENSMUSG00000038705 | 204.6626603 | 0.112577215 | 0.191708618 | 0.587230852 | 0.557048674 | 0.898167794 | Gmeb2 | glucocorticoid modulatory element bindir | 203.99 | 187.08 | 199.24 | 252.56 | 170.14 | 214.97 |
| 11251 | ENSMUSG00000038708 | 1485.756363 | 0.105672029 | 0.099761765 | 1.059243776 | 0.289488773 | 0.739912637 | Golga4 | golgi autoantigen, golgin subfamily a, 4 [S( | 1372.5 | 1440.1 | 1482.2 | 1565.5 | 1461.4 | 1592.8 |
| 11252 | ENSMUSG00000038709 | 0 | NA | NA | NA | NA | NA | Txndc8 | thioredoxin domain containing 8 [Source:I | 0 | 0 | 0 | 0 | 0 | 0 |

| | | | | | | | | | | | | | | | |
|---|---|---|---|---|---|---|---|---|---|---|---|---|---|---|---|
| 11253 | ENSMUSG00000038712 | 1856.585816 | 0.042061454 | 0.096799411 | 0.434521799 | 0.663909569 | 0.929301301 | Mindy1 | MINDY lysine 48 deubiquitinase 1 [Source | 1882.5 | 1761.9 | 1843.4 | 1842.2 | 1819.2 | 1990.4 |
| 11254 | ENSMUSG00000038717 | 2077.765407 | 0.034132007 | 0.095654781 | 0.356824896 | 0.721222897 | 0.9447719 | Atp5l | ATP synthase, H+ transporting, mitochonc | 2130.6 | 1998.3 | 2029.6 | 2030.1 | 2054.7 | 2223.2 |
| 11255 | ENSMUSG00000038718 | 47.05872344 | -0.249405438 | 0.3397305 | -0.734127311 | 0.462871149 | 0.860978434 | Pbx3 | pre B cell leukemia homeobox 3 [Source:N | 40.958 | 49.34 | 63.393 | 47.235 | 46.897 | 34.529 |
| 11256 | ENSMUSG00000038721 | 1.165329702 | -2.428778977 | 2.321199838 | -1.04634635 | 0.295401153 | NA | Hoxb7 | homeobox B7 [Source:MGI Symbol;Acc:M | 4.0155 | 0 | 2.0125 | 0.964 | 0 | 0 |
| 11257 | ENSMUSG00000038722 | 588.0416881 | 0.058905583 | 0.125066097 | 0.470995614 | 0.637643867 | 0.923478764 | Bud31 | BUD31 homolog [Source:MGI Symbol;Acc | 607.95 | 535.55 | 583.62 | 558.14 | 630.38 | 612.61 |
| 11258 | ENSMUSG00000038725 | 22.90452195 | 0.251990817 | 0.505059504 | 0.498932928 | 0.617826636 | 0.917509899 | Pkhd1l1 | polycystic kidney and hepatic disease 1-lik | 24.896 | 22.614 | 15.094 | 22.171 | 38.172 | 14.48 |
| 11259 | ENSMUSG00000038729 | 0 | NA | NA | NA | NA | NA | Pakap | paralemmin A kinase anchor protein [Sou | 0 | 0 | 0 | 0 | 0 | 0 |
| 11260 | ENSMUSG00000038732 | 1.923764 | -2.106737264 | 1.758021002 | -1.198357278 | 0.230777957 | NA | Mboat1 | membrane bound O-acyltransferase dom | 3.2124 | 3.0838 | 3.0187 | 0 | 0 | 2.2277 |
| 11261 | ENSMUSG00000038733 | 1801.082663 | 0.03302982 | 0.101081634 | 0.326763806 | 0.743846537 | 0.949911274 | Wdr26 | WD repeat domain 26 [Source:MGI Symb | 1717.8 | 1932.5 | 1691.5 | 1778.5 | 1862.8 | 1823.4 |
| 11262 | ENSMUSG00000038736 | 263.0516027 | 0.116039301 | 0.172934025 | 0.671003296 | 0.50221843 | 0.881220881 | Nudcd1 | NudC domain containing 1 [Source:MGI S | 252.17 | 237.45 | 267.66 | 255.45 | 326.09 | 239.48 |
| 11263 | ENSMUSG00000038738 | 1.22362201 | -0.289417703 | 2.087230194 | -0.138661133 | 0.889717933 | NA | Shank1 | SH3 and multiple ankyrin repeat domains | 0 | 1.0279 | 3.0187 | 0 | 2.1812 | 1.1138 |
| 11264 | ENSMUSG00000038740 | 19.86393904 | -1.433440053 | 0.516090531 | -2.777497294 | 0.00547793 | 0.070806769 | Mvb12b | multivesicular body subunit 12B [Source:N | 32.124 | 30.838 | 24.15 | 14.46 | 9.8156 | 7.7969 |
| 11265 | ENSMUSG00000038742 | 215.1439076 | -1.021850736 | 0.18774146 | -5.442861358 | 5.24E-08 | 6.26E-06 | Angptl6 | angiopoietin-like 6 [Source:MGI Symbol;A | 334.09 | 273.43 | 256.59 | 131.1 | 134.15 | 161.51 |
| 11266 | ENSMUSG00000038745 | 3247.916177 | 0.379522724 | 0.109475669 | 3.466731243 | 0.000526828 | 0.012909222 | Nlrp6 | NLR family, pyrin domain containing 6 [So | 2513.7 | 2820.6 | 3136.5 | 3565.8 | 3901.1 | 3549.8 |
| 11267 | ENSMUSG00000038750 | 0 | NA | NA | NA | NA | NA | Omt2b | oocyte maturation, beta [Source:MGI Sym | 0 | 0 | 0 | 0 | 0 | 0 |
| 11268 | ENSMUSG00000038751 | 45.94869902 | 0.407908206 | 0.462244258 | 0.882451645 | 0.377532618 | 0.81030298 | Ptk6 | PTK6 protein tyrosine kinase 6 [Source:M | 17.668 | 27.754 | 73.456 | 53.019 | 42.534 | 61.261 |
| 11269 | ENSMUSG00000038754 | 1552.82778 | -0.919564682 | 0.560194 | -1.641511122 | 0.10069136 | 0.46725495 | Elovl3 | elongation of very long chain fatty acids (I | 963.72 | 2636.6 | 2494.5 | 642.97 | 911.76 | 1667.4 |
| 11270 | ENSMUSG00000038756 | 0.267700746 | -1.497481649 | 4.057001436 | -0.369110456 | 0.712045396 | NA | Ttll6 | tubulin tyrosine ligase-like family, membe | 1.6062 | 0 | 0 | 0 | 0 | 0 |
| 11271 | ENSMUSG00000038759 | 200.6636573 | -0.118764125 | 0.191682897 | -0.619586444 | 0.535530094 | 0.890027263 | Nup205 | nucleoporin 205 [Source:MGI Symbol;Acc | 208 | 218.95 | 200.24 | 215.93 | 210.49 | 150.37 |
| 11272 | ENSMUSG00000038760 | 0 | NA | NA | NA | NA | NA | Trhr | thyrotropin releasing hormone receptor [ | 0 | 0 | 0 | 0 | 0 | 0 |
| 11273 | ENSMUSG00000038762 | 1445.188727 | -0.042830761 | 0.097330714 | -0.440053909 | 0.659898063 | 0.927787797 | Abcf1 | ATP-binding cassette, sub-family F (GCN2 | 1472.9 | 1462.7 | 1464.1 | 1393.9 | 1495.2 | 1382.3 |
| 11274 | ENSMUSG00000038763 | 4.39137458 | -0.47214008 | 1.13999439 | -0.414160003 | 0.678756931 | NA | Alpk3 | alpha-kinase 3 [Source:MGI Symbol;Acc:N | 7.2279 | 4.1117 | 4.025 | 7.7118 | 3.2719 | 0 |
| 11275 | ENSMUSG00000038764 | 632.025735 | 0.543022601 | 0.146553372 | 3.705289022 | 0.00021115 | 0.006551645 | Ptpn3 | protein tyrosine phosphatase, non-recept | 573.41 | 418.36 | 550.42 | 726.84 | 708.9 | 814.22 |
| 11276 | ENSMUSG00000038765 | 0 | NA | NA | NA | NA | NA | Lmx1b | LIM homeobox transcription factor 1 beta | 0 | 0 | 0 | 0 | 0 | 0 |
| 11277 | ENSMUSG00000038766 | 551.9858132 | 0.103486876 | 0.129476417 | 0.799272013 | 0.424132704 | 0.840450466 | Gabpb2 | GA repeat binding protein, beta 2 [Source | 478.65 | 579.75 | 539.35 | 590.92 | 551.85 | 571.4 |
| 11278 | ENSMUSG00000038768 | 237.3714419 | -0.614850446 | 0.204889269 | -3.000891405 | 0.002691905 | 0.04355226 | 9130409I23F | RIKEN cDNA 9130409I23 gene [Source:MC | 358.99 | 267.26 | 234.45 | 170.62 | 222.49 | 170.42 |
| 11279 | ENSMUSG00000038770 | 0 | NA | NA | NA | NA | NA | Kpna7 | karyopherin alpha 7 (importin alpha 8) [Sc | 0 | 0 | 0 | 0 | 0 | 0 |
| 11280 | ENSMUSG00000038773 | 534.5132384 | -0.047228456 | 0.13807377 | -0.342052339 | 0.732311504 | 0.947515796 | Kdm3b | KDM3B lysine (K)-specific demethylase 3B | 505.15 | 569.47 | 556.45 | 599.59 | 490.78 | 485.64 |
| 11281 | ENSMUSG00000038774 | 484.7004985 | 0.219693468 | 0.137859116 | 1.593608565 | 0.111023735 | 0.489438373 | Ascc3 | activating signal cointegrator 1 complex su | 412.79 | 479.01 | 452.81 | 549.47 | 550.76 | 463.36 |
| 11282 | ENSMUSG00000038775 | 8.672857394 | -0.544266204 | 0.778672322 | -0.698966932 | 0.484572697 | 0.87256358 | Vill | villin-like [Source:MGI Symbol;Acc:MGI:12 | 14.456 | 8.2233 | 8.05 | 5.7839 | 3.2719 | 12.252 |
| 11283 | ENSMUSG00000038776 | 5393.940438 | -0.354919772 | 0.122107623 | -2.906614364 | 0.003653633 | 0.054018047 | Ephx1 | epoxide hydrolase 1, microsomal [Source: | 6166.2 | 5667.9 | 6329.3 | 5617.1 | 4297 | 4286.1 |
| 11284 | ENSMUSG00000038777 | 31.0533992 | -0.709338079 | 0.471992684 | -1.502858208 | 0.132875612 | 0.533805738 | Sema6c | sema domain, transmembrane domain (T | 49.792 | 26.726 | 39.244 | 38.559 | 14.178 | 17.822 |
| 11285 | ENSMUSG00000038780 | 423.2578695 | 0.197885005 | 0.151901546 | 1.302718832 | 0.19267077 | 0.630757769 | Smurf1 | SMAD specific E3 ubiquitin protein ligase | 436.08 | 381.36 | 365.27 | 519.58 | 435.16 | 402.1 |
| 11286 | ENSMUSG00000038781 | 889.0168431 | 0.205807544 | 0.110939415 | 1.855134564 | 0.063577013 | 0.360558879 | Stap2 | signal transducing adaptor family member | 854.5 | 760.66 | 861.35 | 937.95 | 970.65 | 948.99 |
| 11287 | ENSMUSG00000038782 | 5.374070787 | 0.517346144 | 0.975777252 | 0.530188773 | 0.595981054 | 0.913847493 | 1700028J19F | RIKEN cDNA 1700028J19 gene [Source:MC | 8.031 | 1.0279 | 4.025 | 4.8199 | 6.5437 | 7.7969 |
| 11288 | ENSMUSG00000038784 | 171.6975168 | 0.000833205 | 0.192677802 | 0.004324345 | 0.996549683 | 0.999154389 | Cnot4 | CCR4-NOT transcription complex, subunit | 179.89 | 181.94 | 152.95 | 168.7 | 195.22 | 151.48 |
| 11289 | ENSMUSG00000038791 | 0 | NA | NA | NA | NA | NA | Scgb3a2 | secretoglobin, family 3A, member 2 [Sour | 0 | 0 | 0 | 0 | 0 | 0 |
| 11290 | ENSMUSG00000038793 | 12.12227584 | 0.483281895 | 0.67742741 | 0.713407647 | 0.475593542 | 0.869436931 | Lefty1 | left right determination factor 1 [Source:N | 8.031 | 16.447 | 6.0375 | 20.244 | 14.178 | 7.7969 |
| 11291 | ENSMUSG00000038797 | 56.31293043 | -0.08814466 | 0.320609801 | -0.274928151 | 0.783371439 | 0.960670212 | Zscan2 | zinc finger and SCAN domain containing 2 | 61.036 | 63.731 | 49.306 | 58.803 | 38.172 | 66.831 |
| 11292 | ENSMUSG00000038801 | 0.853270218 | 3.307793827 | 2.742080224 | 1.206308188 | 0.227698682 | NA | Scgb1c1 | secretoglobin, family 1C, member 1 [Sour | 0 | 0 | 0 | 2.8919 | 0 | 2.2277 |
| 11293 | ENSMUSG00000038803 | 484.8898633 | 0.037412361 | 0.134135802 | 0.278914058 | 0.78031078 | 0.959892424 | Ost4 | oligosaccharyltransferase complex subuni | 525.23 | 443.03 | 466.9 | 513.8 | 459.15 | 501.23 |
| 11294 | ENSMUSG00000038805 | 0 | NA | NA | NA | NA | NA | Six3 | sine oculis-related homeobox 3 [Source:N | 0 | 0 | 0 | 0 | 0 | 0 |
| 11295 | ENSMUSG00000038806 | 222.5964886 | 0.03403995 | 0.166106368 | 0.20492863 | 0.837627886 | 0.971073158 | Sde2 | SDE2 telomere maintenance homolog (S. | 236.11 | 224.09 | 199.24 | 218.82 | 231.21 | 226.11 |
| 11296 | ENSMUSG00000038807 | 9.348570715 | -0.10940489 | 0.705294499 | -0.155119443 | 0.876727154 | 0.980632845 | Rap1gap2 | RAP1 GTPase activating protein 2 [Source: | 8.8341 | 12.335 | 8.05 | 12.532 | 6.5437 | 7.7969 |
| 11297 | ENSMUSG00000038811 | 50.95089336 | -0.869499191 | 0.373140348 | -2.33022024 | 0.019794514 | 0.172127966 | Gngt2 | guanine nucleotide binding protein (G prc | 57.823 | 77.094 | 62.387 | 22.171 | 30.537 | 55.692 |
| 11298 | ENSMUSG00000038812 | 450.4503747 | -0.034746463 | 0.130817581 | -0.265610043 | 0.79053955 | 0.961019028 | Trmt112 | tRNA methyltransferase 11-2 [Source:MG | 445.72 | 457.42 | 464.88 | 456.93 | 468.97 | 408.78 |
| 11299 | ENSMUSG00000038816 | 94.32793402 | 0.128308038 | 0.247730396 | 0.517934174 | 0.604504197 | 0.916376001 | Ctnnal1 | catenin (cadherin associated protein), alp | 74.689 | 95.596 | 100.62 | 100.25 | 110.15 | 84.652 |
| 11300 | ENSMUSG00000038822 | 67.05599329 | 0.077743555 | 0.28445585 | 0.273306226 | 0.78461783 | 0.960670212 | Hace1 | HECT domain and ankyrin repeat contain | 72.279 | 59.619 | 63.393 | 55.911 | 68.709 | 82.424 |
| 11301 | ENSMUSG00000038827 | 68.11279036 | 0.047224703 | 0.302045257 | 0.156349759 | 0.875757336 | 0.980566196 | Abitram | actin binding transcription modulator [Sou | 48.989 | 64.759 | 87.543 | 60.731 | 69.8 | 76.855 |
| 11302 | ENSMUSG00000038828 | 958.1967852 | 0.040796806 | 0.117488109 | 0.347241996 | 0.728409523 | 0.946208446 | Tmem214 | transmembrane protein 214 [Source:MGI | 974.16 | 930.27 | 929.77 | 1043 | 1013.2 | 858.77 |
| 11303 | ENSMUSG00000038831 | 75.07581209 | 0.364837572 | 0.295186653 | 1.235955514 | 0.2164751 | 0.661953589 | Ralgps1 | Ral GEF with PH domain and SH3 binding | 75.492 | 75.038 | 46.287 | 100.25 | 67.618 | 85.766 |
| 11304 | ENSMUSG00000038836 | 127.6822534 | 0.040803485 | 0.213374747 | 0.191229213 | 0.848346017 | 0.972803041 | Agbl3 | ATP/GTP binding protein-like 3 [Source:M | 124.48 | 130.55 | 122.76 | 133.03 | 147.23 | 108.04 |
| 11305 | ENSMUSG00000038838 | 407.0371582 | 0.047326489 | 0.136376157 | 0.347029055 | 0.728569491 | 0.946248466 | Vars2 | valyl-tRNA synthetase 2, mitochondrial [S | 410.39 | 362.85 | 427.65 | 420.29 | 412.25 | 408.78 |
| 11306 | ENSMUSG00000038840 | 0.160662974 | 1.126924175 | 4.080472857 | 0.276174898 | 0.782413736 | NA | Birc7 | baculoviral IAP repeat-containing 7 (livin) | 0 | 0 | 0 | 0.964 | 0 | 0 |
| 11307 | ENSMUSG00000038843 | 10.79077322 | -0.528324704 | 0.682629188 | -0.773955631 | 0.43895703 | 0.848057363 | Gcnt1 | glucosaminyl (N-acetyl) transferase 1, core | 17.668 | 12.335 | 8.05 | 5.7839 | 11.997 | 8.9108 |
| 11308 | ENSMUSG00000038844 | 291.9477075 | -0.02906825 | 0.173566574 | -0.167476081 | 0.866995466 | 0.977930657 | Kif16b | kinesin family member 16B [Source:MGI S | 315.62 | 329.96 | 238.48 | 259.31 | 306.46 | 301.85 |

| | | | | | | | | | | | | | | | |
|---|---|---|---|---|---|---|---|---|---|---|---|---|---|---|---|
| 11309 | ENSMUSG00000038845 | 2173.985431 | -0.057682375 | 0.099644927 | -0.578879189 | 0.562670694 | 0.900565925 | Phb | prohibitin [Source:MGI Symbol;Acc:MGI:9 | 2359.5 | 2236.7 | 2054.8 | 2043.6 | 2158.3 | 2190.9 |
| 11310 | ENSMUSG00000038848 | 571.542124 | 0.079875875 | 0.12301311 | 0.649328151 | 0.516126294 | 0.886609349 | Ythdf1 | YTH N6-methyladenosine RNA binding pro | 525.23 | 562.27 | 580.6 | 623.69 | 564.94 | 572.52 |
| 11311 | ENSMUSG00000038855 | 89.84985785 | -0.40051337 | 0.26009143 | -1.539894525 | 0.123586066 | 0.515143761 | Itpkb | inositol 1,4,5-trisphosphate 3-kinase B [Sc | 113.24 | 97.652 | 95.593 | 79.046 | 58.893 | 94.677 |
| 11312 | ENSMUSG00000038859 | 755.429458 | -0.432537942 | 0.128163763 | -3.374884843 | 0.000738466 | 0.016715446 | Baiap2l1 | BAI1-associated protein 2-like 1 [Source:N | 809.53 | 982.69 | 812.04 | 650.69 | 677.27 | 600.36 |
| 11313 | ENSMUSG00000038860 | 3.854101041 | 1.240897639 | 1.234814711 | 1.004926186 | 0.314932394 | NA | Garnl3 | GTPase activating RANGAP domain-like 3 | 0.8031 | 5.1396 | 1.0062 | 1.928 | 10.906 | 3.3415 |
| 11314 | ENSMUSG00000038861 | 475.5432419 | 0.117256726 | 0.130362506 | 0.899466647 | 0.368404153 | 0.80627882 | Pi4kb | phosphatidylinositol 4-kinase beta [Source | 424.84 | 449.2 | 495.07 | 479.1 | 506.05 | 499 |
| 11315 | ENSMUSG00000038866 | 568.9926181 | 0.502968845 | 0.202899226 | 2.478909623 | 0.013178469 | 0.13082593 | Zcchc2 | zinc finger, CCHC domain containing 2 [So | 429.66 | 504.71 | 478.97 | 877.22 | 440.61 | 682.79 |
| 11316 | ENSMUSG00000038871 | 274.2027862 | 0.335198275 | 0.156999714 | 2.135024747 | 0.032758989 | 0.24187137 | Bpgm | 2,3-bisphosphoglycerate mutase [Source: | 218.44 | 255.95 | 253.57 | 292.09 | 299.92 | 325.24 |
| 11317 | ENSMUSG00000038872 | 134.1121554 | -0.004382302 | 0.199799764 | -0.021933468 | 0.982501027 | 0.998580735 | Zfhx3 | zinc finger homeobox 3 [Source:MGI Symb | 127.69 | 131.57 | 143.89 | 136.88 | 126.51 | 138.12 |
| 11318 | ENSMUSG00000038876 | 113.3849948 | 0.242861648 | 0.217493436 | 1.11663898 | 0.264148718 | 0.719492304 | Rnf146 | ring finger protein 146 [Source:MGI Symb | 96.372 | 105.88 | 109.68 | 130.14 | 110.15 | 128.09 |
| 11319 | ENSMUSG00000038879 | 1.447454308 | -0.79447158 | 1.928813837 | -0.411896454 | 0.68041532 | NA | Nipal2 | NIPA-like domain containing 2 [Source:MG | 2.4093 | 3.0838 | 0 | 0.964 | 0 | 2.2277 |
| 11320 | ENSMUSG00000038880 | 583.1131192 | 0.018720942 | 0.126627835 | 0.147842232 | 0.88246728 | 0.981769403 | Mrps34 | mitochondrial ribosomal protein S34 [Sou | 562.97 | 581.8 | 593.68 | 598.63 | 634.74 | 526.85 |
| 11321 | ENSMUSG00000038883 | 0 | NA | NA | NA | NA | NA | Prl3a1 | prolactin family 3, subfamily a, member 1 | 0 | 0 | 0 | 0 | 0 | 0 |
| 11322 | ENSMUSG00000038884 | 807.4950626 | 0.053225125 | 0.149104196 | 0.356965974 | 0.721117279 | 0.9447719 | Shfl | shiftless antiviral inhibitor of ribosomal fra | 761.34 | 739.07 | 878.45 | 992.9 | 699.09 | 774.12 |
| 11323 | ENSMUSG00000038886 | 532.8693726 | 0.019843944 | 0.133443816 | 0.148706359 | 0.881785343 | 0.981676444 | Man2a2 | mannosidase 2, alpha 2 [Source:MGI Symb | 548.52 | 527.32 | 512.18 | 605.38 | 511.5 | 492.32 |
| 11324 | ENSMUSG00000038888 | 117.5289756 | -0.049728691 | 0.225074452 | -0.220943297 | 0.825136584 | 0.96943211 | Ctu1 | cytosolic thiouridylase subunit 1 [Source:N | 132.51 | 109.99 | 115.72 | 111.82 | 99.246 | 135.89 |
| 11325 | ENSMUSG00000038891 | 0 | NA | NA | NA | NA | NA | Prl3b1 | prolactin family 3, subfamily b, member 1 | 0 | 0 | 0 | 0 | 0 | 0 |
| 11326 | ENSMUSG00000038893 | 24.80900242 | -0.046635126 | 0.451027551 | -0.103397511 | 0.917647488 | 0.986374671 | Fam117a | family with sequence similarity 117, meml | 28.912 | 16.447 | 30.187 | 26.991 | 21.812 | 24.505 |
| 11327 | ENSMUSG00000038894 | 319.1540072 | 0.520225869 | 0.214681895 | 2.423240528 | 0.015382738 | 0.144948483 | Irs2 | insulin receptor substrate 2 [Source:MGI S | 181.5 | 319.68 | 286.78 | 404.87 | 298.83 | 423.26 |
| 11328 | ENSMUSG00000038895 | 87.15289679 | 0.133322369 | 0.2653636 | 0.502413928 | 0.615376384 | 0.917509899 | Zfp653 | zinc finger protein 653 [Source:MGI Symb | 73.082 | 76.066 | 100.62 | 103.15 | 71.981 | 98.018 |
| 11329 | ENSMUSG00000038900 | 2397.259161 | 0.033666399 | 0.107870553 | 0.312099995 | 0.754964531 | 0.953085143 | Rpl12 | ribosomal protein L12 [Source:MGI Symbo | 2276 | 2455.7 | 2375.7 | 2148.7 | 2492.1 | 2635.4 |
| 11330 | ENSMUSG00000038902 | 242.4020837 | 0.019911739 | 0.184409322 | 0.107975772 | 0.914014911 | 0.98625062 | Pogz | pogo transposable element with ZNF dom | 220.85 | 211.75 | 289.8 | 240.99 | 218.12 | 272.89 |
| 11331 | ENSMUSG00000038903 | 17.92678062 | 0.052480589 | 0.67136301 | 0.078170213 | 0.937692656 | 0.988420305 | Ccdc68 | coiled-coil domain containing 68 [Source:I | 20.078 | 27.754 | 5.0312 | 26.991 | 6.5437 | 21.163 |
| 11332 | ENSMUSG00000038909 | 363.1121615 | 0.208312624 | 0.141370773 | 1.473519735 | 0.140610947 | 0.548229787 | Kat7 | K(lysine) acetyltransferase 7 [Source:MGI | 336.5 | 351.55 | 323 | 400.05 | 412.25 | 355.32 |
| 11333 | ENSMUSG00000038910 | 280.5789951 | -0.244462751 | 0.185678228 | -1.316593517 | 0.187974913 | 0.624893591 | Plcl2 | phospholipase C-like 2 [Source:MGI Symb | 324.45 | 358.74 | 229.42 | 251.6 | 251.93 | 267.32 |
| 11334 | ENSMUSG00000038914 | 935.0283101 | 0.018911485 | 0.130626878 | 0.144774838 | 0.88488865 | 0.981795936 | Dido1 | death inducer-obliterator 1 [Source:MGI S | 950.07 | 904.57 | 932.79 | 1089.3 | 927.03 | 806.42 |
| 11335 | ENSMUSG00000038916 | 0 | NA | NA | NA | NA | NA | Soga3 | SOGA family member 3 [Source:MGI Symb | 0 | 0 | 0 | 0 | 0 | 0 |
| 11336 | ENSMUSG00000038917 | 3.19682301 | 1.988638763 | 1.600460861 | 1.242541328 | 0.214036902 | NA | 3930402G23 | RIKEN cDNA 3930402G23 gene [Source:M | 0.8031 | 3.0838 | 0 | 1.928 | 0 | 13.366 |
| 11337 | ENSMUSG00000038925 | 0 | NA | NA | NA | NA | NA | E330034G19 | RIKEN cDNA E330034G19 gene [Source:M | 0 | 0 | 0 | 0 | 0 | 0 |
| 11338 | ENSMUSG00000038930 | 45.5256862 | 0.353842062 | 0.372610675 | 0.949629428 | 0.34230058 | 0.784060862 | Rccd1 | RCC1 domain containing 1 [Source:MGI Sy | 38.549 | 45.228 | 36.225 | 56.875 | 29.447 | 66.831 |
| 11339 | ENSMUSG00000038932 | 0 | NA | NA | NA | NA | NA | Tcfl5 | transcription factor-like 5 (basic helix-loop | 0 | 0 | 0 | 0 | 0 | 0 |
| 11340 | ENSMUSG00000038936 | 27.65540199 | 0.041431753 | 0.449047173 | 0.092265926 | 0.92648676 | 0.987712228 | Sccpdh | saccharopine dehydrogenase (putative) [S | 20.881 | 34.949 | 26.162 | 30.847 | 17.45 | 35.643 |
| 11341 | ENSMUSG00000038943 | 9.915633001 | -0.857383164 | 0.701600063 | -1.222039747 | 0.221692596 | 0.667949912 | Prc1 | protein regulator of cytokinesis 1 [Source: | 8.8341 | 18.503 | 11.069 | 6.7478 | 6.5437 | 7.7969 |
| 11342 | ENSMUSG00000038949 | 312.3921159 | -0.754303334 | 0.168530531 | -4.475766678 | 7.61E-06 | 0.000431864 | Cnst | consortin, connexin sorting protein [Sourc | 377.46 | 438.92 | 360.24 | 219.79 | 203.95 | 274.01 |
| 11343 | ENSMUSG00000038954 | 77.54477291 | 0.13205607 | 0.283138906 | 0.466400299 | 0.640928998 | 0.923932929 | Supt3 | SPT3, SAGA and STAGA complex compone | 96.372 | 66.815 | 58.362 | 90.614 | 80.706 | 72.4 |
| 11344 | ENSMUSG00000038957 | 355.2806786 | 0.23418875 | 0.152574655 | 1.534912533 | 0.124805315 | 0.518284258 | Edc3 | enhancer of mRNA decapping 3 [Source:N | 314.01 | 377.25 | 288.79 | 400.05 | 378.44 | 373.14 |
| 11345 | ENSMUSG00000038963 | 5.167820329 | -2.322527861 | 1.056025105 | -2.19931122 | 0.0278558 | 0.217463141 | Slco4a1 | solute carrier organic anion transporter fa | 9.6372 | 8.2233 | 8.05 | 2.8919 | 1.0906 | 1.1138 |
| 11346 | ENSMUSG00000038965 | 1220.671421 | -0.084050397 | 0.104330575 | -0.805616158 | 0.420464206 | 0.837906635 | Ube2l3 | ubiquitin-conjugating enzyme E2L 3 [Sour | 1300.2 | 1211.9 | 1255.8 | 1166.4 | 1127.7 | 1262 |
| 11347 | ENSMUSG00000038967 | 2531.897743 | -0.026595112 | 0.111448237 | -0.238631964 | 0.811390977 | 0.966527993 | Pdk2 | pyruvate dehydrogenase kinase, isoenzym | 2838.2 | 2418.7 | 2406.9 | 2342.5 | 2449.5 | 2735.6 |
| 11348 | ENSMUSG00000038968 | 0 | NA | NA | NA | NA | NA | Klk1b16 | kallikrein 1-related peptidase b16 [Source | 0 | 0 | 0 | 0 | 0 | 0 |
| 11349 | ENSMUSG00000038970 | 344.7799475 | -0.037592532 | 0.14346105 | -0.262039992 | 0.79329061 | 0.961896915 | Lmtk2 | lemur tyrosine kinase 2 [Source:MGI Symb | 342.92 | 370.05 | 335.08 | 348.96 | 313.01 | 358.66 |
| 11350 | ENSMUSG00000038973 | 0.698954292 | 0.126053087 | 3.048972385 | 0.04134281 | 0.967022605 | NA | Cldnd2 | claudin domain containing 2 [Source:MGI | 0 | 0 | 2.0125 | 0 | 2.1812 | 0 |
| 11351 | ENSMUSG00000038975 | 713.9831649 | 0.147901268 | 0.125515809 | 1.17834772 | 0.238658011 | 0.687639336 | Rabggtb | Rab geranylgeranyl transferase, b subunit | 664.97 | 742.16 | 625.88 | 808.78 | 747.07 | 695.04 |
| 11352 | ENSMUSG00000038976 | 127.3219773 | -0.937788824 | 0.233960492 | -4.008321297 | 6.12E-05 | 0.002443373 | Ppp1r9b | protein phosphatase 1, regulatory subunit | 179.09 | 161.38 | 161 | 79.046 | 69.8 | 113.61 |
| 11353 | ENSMUSG00000038980 | 0.371281295 | 2.09253362 | 4.031852165 | 0.519000582 | 0.603760335 | NA | Rbbp8nl | RBBP8 N-terminal like [Source:MGI Symbo | 0 | 0 | 0 | 0 | 0 | 2.2277 |
| 11354 | ENSMUSG00000038982 | 512.5288894 | 0.014206582 | 0.124004365 | 0.114565174 | 0.908789785 | 0.985098284 | Bloc1s5 | biogenesis of lysosomal organelles comple | 516.39 | 521.15 | 492.05 | 491.63 | 511.5 | 542.44 |
| 11355 | ENSMUSG00000038984 | 11.11920844 | -0.45408291 | 0.652485096 | -0.695928402 | 0.48647367 | 0.873586472 | Tspyl5 | testis-specific protein, Y-encoded-like 5 [S | 11.243 | 12.335 | 15.094 | 11.568 | 10.906 | 5.5692 |
| 11356 | ENSMUSG00000038987 | 4.911881624 | -1.583489554 | 1.023200004 | -1.547585562 | 0.12172211 | NA | Cfap157 | cilia and flagella associated protein 157 [S | 8.8341 | 7.1954 | 6.0375 | 1.928 | 4.3625 | 1.1138 |
| 11357 | ENSMUSG00000038990 | 548.6204662 | 0.119604784 | 0.145328837 | 0.822994159 | 0.410511315 | 0.833241465 | Cables2 | CDK5 and Abl enzyme substrate 2 [Source | 510.77 | 481.07 | 586.64 | 658.4 | 521.32 | 533.53 |
| 11358 | ENSMUSG00000038991 | 2991.75418 | 0.243279962 | 0.110732958 | 2.196996863 | 0.02802067 | 0.21848271 | Txndc5 | thioredoxin domain containing 5 [Source: | 2893.6 | 2619.1 | 2707.8 | 3679.5 | 3028.6 | 3021.9 |
| 11359 | ENSMUSG00000038994 | 0 | NA | NA | NA | NA | NA | H1f9 | H1.9 linker histone [Source:MGI Symbol;A | 0 | 0 | 0 | 0 | 0 | 0 |
| 11360 | ENSMUSG00000038997 | 0 | NA | NA | NA | NA | NA | Asb17 | ankyrin repeat and SOCS box-containing 1 | 0 | 0 | 0 | 0 | 0 | 0 |
| 11361 | ENSMUSG00000039000 | 887.794201 | 0.042034761 | 0.116601649 | 0.360498855 | 0.718474113 | 0.9447719 | Ube3c | ubiquitin protein ligase E3C [Source:MGI S | 893.85 | 864.48 | 865.37 | 812.63 | 982.65 | 907.78 |
| 11362 | ENSMUSG00000039001 | 1625.805446 | 0.094002354 | 0.134627121 | 0.698242323 | 0.485025662 | 0.87256358 | Rps21 | ribosomal protein S21 [Source:MGI Symbo | 1460 | 1726.9 | 1531.5 | 1454.6 | 1586.8 | 1994.9 |
| 11363 | ENSMUSG00000039004 | 57.10854065 | -0.184582651 | 0.376050439 | -0.490845461 | 0.623535754 | 0.919503423 | Bmp6 | bone morphogenetic protein 6 [Source:M | 54.611 | 42.145 | 85.531 | 53.019 | 32.719 | 74.628 |
| 11364 | ENSMUSG00000039005 | 14.4630371 | -1.260842257 | 0.637877019 | -1.976622798 | 0.048084283 | 0.304906213 | Tlr4 | toll-like receptor 4 [Source:MGI Symbol;A | 23.29 | 26.726 | 11.069 | 5.7839 | 6.5437 | 13.366 |

| | | | | | | | | | | | | | | | |
|---|---|---|---|---|---|---|---|---|---|---|---|---|---|---|---|
| 11365 | ENSMUSG00000039007 | 1072.368522 | -0.112925357 | 0.114759068 | -0.984021208 | 0.325105085 | 0.771501263 | Cpq | carboxypeptidase Q [Source:MGI Symbol; | 1045.6 | 1169.8 | 1129 | 1132.7 | 989.19 | 967.93 |
| 11366 | ENSMUSG00000039013 | 0.610340026 | -2.657561537 | 3.094276798 | -0.858863544 | 0.390415805 | NA | Siglecf | sialic acid binding Ig-like lectin F [Source:N | 1.6062 | 2.0558 | 0 | 0 | 0 | 0 |
| 11367 | ENSMUSG00000039016 | 591.1234557 | 0.033275543 | 0.130985175 | 0.25404053 | 0.799464246 | 0.962424862 | Timm8b | translocase of inner mitochondrial membr | 561.37 | 565.35 | 625.88 | 530.19 | 640.19 | 623.75 |
| 11368 | ENSMUSG00000039018 | 411.1959581 | -0.006772853 | 0.136452026 | -0.049635417 | 0.960412923 | 0.992456421 | Mtg1 | mitochondrial ribosome-associated GTPas | 392.72 | 411.17 | 432.69 | 384.63 | 443.88 | 402.1 |
| 11369 | ENSMUSG00000039021 | 3.297644413 | 0.279796601 | 1.338317009 | 0.209066013 | 0.834396706 | NA | Ttc16 | tetratricopeptide repeat domain 16 [Sourc | 0.8031 | 6.1675 | 2.0125 | 0.964 | 8.7249 | 1.1138 |
| 11370 | ENSMUSG00000039031 | 186.6076901 | 0.285222879 | 0.200960859 | 1.419295677 | 0.155812832 | 0.574770926 | Arhgap18 | Rho GTPase activating protein 18 [Source: | 209.61 | 146.99 | 146.91 | 191.83 | 212.67 | 211.63 |
| 11371 | ENSMUSG00000039032 | 0 | NA | NA | NA | NA | NA | Tsga13 | testis specific gene A13 [Source:MGI Symb | 0 | 0 | 0 | 0 | 0 | 0 |
| 11372 | ENSMUSG00000039033 | 127.6691422 | 0.190666094 | 0.227099912 | 0.839569209 | 0.40114997 | 0.828062389 | Tasp1 | taspase, threonine aspartase 1 [Source:M | 108.42 | 100.74 | 148.92 | 143.63 | 141.78 | 122.52 |
| 11373 | ENSMUSG00000039037 | 0.17131964 | -0.796629176 | 4.080472857 | -0.195229623 | 0.84521319 | NA | St6galnac5 | ST6 (alpha-N-acetyl-neuraminyl-2,3-beta-| | 0 | 1.0279 | 0 | 0 | 0 | 0 |
| 11374 | ENSMUSG00000039041 | 1338.056118 | 0.020202227 | 0.108558652 | 0.186095039 | 0.852370232 | 0.973781977 | Adrm1 | adhesion regulating molecule 1 [Source:M | 1381.3 | 1328.1 | 1276.9 | 1475.9 | 1261.8 | 1304.3 |
| 11375 | ENSMUSG00000039043 | 164.0335884 | -0.120899385 | 0.204911783 | -0.590006991 | 0.555185962 | 0.897628185 | Arpin | actin-related protein 2/3 complex inhibito | 175.88 | 151.1 | 185.15 | 127.25 | 178.86 | 165.96 |
| 11376 | ENSMUSG00000039046 | 341.9655032 | -0.056736388 | 0.179165398 | -0.316670452 | 0.751493671 | 0.952255112 | Usp6nl | USP6 N-terminal like [Source:MGI Symbol | 355.77 | 370.05 | 320.99 | 421.26 | 310.83 | 272.89 |
| 11377 | ENSMUSG00000039047 | 706.2865232 | -0.036914311 | 0.128111118 | -0.288142913 | 0.773237345 | 0.958117447 | Pigk | phosphatidylinositol glycan anchor biosyn | 768.57 | 667.12 | 709.4 | 689.24 | 630.38 | 773.01 |
| 11378 | ENSMUSG00000039048 | 678.6540157 | 0.174752384 | 0.113283135 | 1.54261606 | 0.122923959 | 0.513267141 | Foxred1 | FAD-dependent oxidoreductase domain c | 662.56 | 628.06 | 621.86 | 736.48 | 704.54 | 718.43 |
| 11379 | ENSMUSG00000039050 | 616.453607 | 0.21163705 | 0.122001572 | 1.734707562 | 0.082792616 | 0.424179628 | Osbpl2 | oxysterol binding protein-like 2 [Source:M | 606.34 | 520.13 | 586.64 | 648.76 | 668.55 | 668.31 |
| 11380 | ENSMUSG00000039055 | 2.695951139 | -0.980921351 | 1.542107593 | -0.636091382 | 0.524716862 | NA | Eme1 | essential meiotic structure-specific endon | 5.6217 | 0 | 5.0312 | 0 | 2.1812 | 3.3415 |
| 11381 | ENSMUSG00000039057 | 0 | NA | NA | NA | NA | NA | Myo16 | myosin XVI [Source:MGI Symbol;Acc:MGI: | 0 | 0 | 0 | 0 | 0 | 0 |
| 11382 | ENSMUSG00000039058 | 2.258688572 | -0.110194109 | 1.578820231 | -0.069795222 | 0.94435665 | NA | Ak5 | adenylate kinase 5 [Source:MGI Symbol;A | 0 | 1.0279 | 6.0375 | 0.964 | 2.1812 | 3.3415 |
| 11383 | ENSMUSG00000039059 | 0 | NA | NA | NA | NA | NA | Hrh3 | histamine receptor H3 [Source:MGI Symb | 0 | 0 | 0 | 0 | 0 | 0 |
| 11384 | ENSMUSG00000039062 | 2198.197653 | 0.053640766 | 0.117289102 | 0.457338025 | 0.647428102 | 0.925643431 | Anpep | alanyl (membrane) aminopeptidase [Sour | 1999.7 | 2157.6 | 2315.4 | 2167 | 2537.9 | 2011.6 |
| 11385 | ENSMUSG00000039063 | 1619.965993 | 0.035041429 | 0.100091584 | 0.350093664 | 0.726268406 | 0.945691954 | Echdc3 | enoyl Coenzyme A hydratase domain cont | 1575.7 | 1608.7 | 1616 | 1517.3 | 1706.8 | 1695.3 |
| 11386 | ENSMUSG00000039065 | 324.0566457 | -0.027521882 | 0.143256515 | -0.192116094 | 0.847651269 | 0.972803041 | Atpsckmt | ATP synthase C subunit lysine N-methyltra | 334.89 | 317.63 | 329.04 | 340.28 | 320.64 | 301.85 |
| 11387 | ENSMUSG00000039067 | 1115.505136 | 0.069011266 | 0.106388037 | 0.648675059 | 0.516548429 | 0.886716042 | Psmd7 | proteasome (prosome, macropain) 26S su | 1128.4 | 1063.9 | 1073.7 | 1144.2 | 1069.9 | 1213 |
| 11388 | ENSMUSG00000039068 | 455.8272516 | -0.039001286 | 0.144107121 | -0.270640935 | 0.786667209 | 0.960870081 | Zzz3 | zinc finger, ZZ domain containing 3 [Sourc | 430.46 | 532.46 | 423.63 | 446.32 | 476.6 | 425.49 |
| 11389 | ENSMUSG00000039069 | 316.9632844 | 0.230561931 | 0.15280451 | 1.508868628 | 0.131332358 | 0.531175668 | Mtg2 | mitochondrial ribosome associated GTPas | 273.86 | 275.48 | 326.02 | 339.32 | 320.64 | 366.45 |
| 11390 | ENSMUSG00000039070 | 0 | NA | NA | NA | NA | NA | Cpa4 | carboxypeptidase A4 [Source:MGI Symbol | 0 | 0 | 0 | 0 | 0 | 0 |
| 11391 | ENSMUSG00000039079 | 0 | NA | NA | NA | NA | NA | Trhr2 | thyrotropin releasing hormone receptor 2 | 0 | 0 | 0 | 0 | 0 | 0 |
| 11392 | ENSMUSG00000039081 | 33.60244211 | -0.203455052 | 0.476798247 | -0.42671099 | 0.669589845 | 0.930837335 | Zfp503 | zinc finger protein 503 [Source:MGI Symb | 49.792 | 21.586 | 36.225 | 30.847 | 15.269 | 47.895 |
| 11393 | ENSMUSG00000039084 | 46.70429643 | -0.106787627 | 0.405266688 | -0.263499642 | 0.792165499 | 0.961434599 | Chad | chondroadherin [Source:MGI Symbol;Acc | 52.202 | 28.782 | 64.4 | 56.875 | 52.35 | 25.618 |
| 11394 | ENSMUSG00000039086 | 88.42043089 | 0.405102609 | 0.256157443 | 1.581459454 | 0.113773023 | 0.495873456 | Ss18l1 | SS18, nBAF chromatin remodeling comple | 91.554 | 69.898 | 66.412 | 111.82 | 87.249 | 103.59 |
| 11395 | ENSMUSG00000039087 | 951.3183891 | -0.058425229 | 0.105896859 | -0.551718245 | 0.581141407 | 0.908645447 | Rreb1 | ras responsive element binding protein 1 | 1020.7 | 937.46 | 952.91 | 942.77 | 913.94 | 940.08 |
| 11396 | ENSMUSG00000039089 | 118.8166997 | 0.204079054 | 0.211303361 | 0.965810731 | 0.334138898 | 0.775900437 | L3mbtl3 | L3MBTL3 histone methyl-lysine binding pr | 103.6 | 115.13 | 112.7 | 123.39 | 119.97 | 138.12 |
| 11397 | ENSMUSG00000039092 | 0.133850373 | -0.796629176 | 4.080472857 | -0.195229623 | 0.84521319 | NA | Sptlc3 | serine palmitoyltransferase, long chain ba | 0.8031 | 0 | 0 | 0 | 0 | 0 |
| 11398 | ENSMUSG00000039095 | 1.391871864 | -2.68210978 | 2.193998595 | -1.222475614 | 0.22152782 | NA | En2 | engrailed 2 [Source:MGI Symbol;Acc:MGI: | 3.2124 | 0 | 4.025 | 0 | 0 | 1.1138 |
| 11399 | ENSMUSG00000039096 | 178.9163033 | -0.054374459 | 0.19372378 | -0.280680355 | 0.778955578 | 0.959828458 | Rsad1 | radical S-adenosyl methionine domain cor | 203.18 | 176.8 | 166.03 | 148.45 | 194.13 | 184.9 |
| 11400 | ENSMUSG00000039097 | 0 | NA | NA | NA | NA | NA | Rln1 | relaxin 1 [Source:MGI Symbol;Acc:MGI:97 | 0 | 0 | 0 | 0 | 0 | 0 |
| 11401 | ENSMUSG00000039098 | 0 | NA | NA | NA | NA | NA | Gm9767 | predicted gene 9767 [Source:MGI Symbol | 0 | 0 | 0 | 0 | 0 | 0 |
| 11402 | ENSMUSG00000039099 | 42.46634621 | -0.031419497 | 0.392368171 | -0.080076569 | 0.936176358 | 0.988420305 | Wdr93 | WD repeat domain 93 [Source:MGI Symbo | 43.368 | 50.368 | 35.219 | 50.127 | 53.44 | 22.277 |
| 11403 | ENSMUSG00000039100 | 3314.507082 | 0.102258478 | 0.092782084 | 1.10213603 | 0.270402538 | 0.725929337 | Marchf6 | membrane associated ring-CH-type finger | 3112 | 3246.2 | 3233.1 | 3249.6 | 3659 | 3387.2 |
| 11404 | ENSMUSG00000039103 | 12.96767167 | -0.298003668 | 0.624239414 | -0.477386819 | 0.633086699 | 0.922107984 | Nexn | nexilin [Source:MGI Symbol;Acc:MGI:191 | 10.44 | 16.447 | 16.1 | 11.568 | 6.5437 | 16.708 |
| 11405 | ENSMUSG00000039105 | 884.2121734 | 0.050187106 | 0.108317936 | 0.463331445 | 0.64312681 | 0.925364325 | Atp6v1g1 | ATPase, H+ transporting, lysosomal V1 sul | 885.02 | 895.32 | 826.13 | 920.6 | 864.86 | 913.35 |
| 11406 | ENSMUSG00000039106 | 0 | NA | NA | NA | NA | NA | Htr5a | 5-hydroxytryptamine (serotonin) receptor | 0 | 0 | 0 | 0 | 0 | 0 |
| 11407 | ENSMUSG00000039108 | 492.7280054 | 0.086661747 | 0.129839876 | 0.66745094 | 0.504484137 | 0.881510796 | Lsm14b | LSM family member 14B [Source:MGI Sym | 480.26 | 452.28 | 501.11 | 508.02 | 471.15 | 543.56 |
| 11408 | ENSMUSG00000039109 | 1.827460002 | -2.101153165 | 1.731039385 | -1.213810144 | 0.224820232 | NA | F13a1 | coagulation factor XIII, A1 subunit [Source | 4.8186 | 2.0558 | 2.0125 | 0.964 | 0 | 1.1138 |
| 11409 | ENSMUSG00000039110 | 1.28271381 | 0.267922475 | 1.87956843 | 0.142544677 | 0.886649792 | NA | Mycbpap | MYCBP associated protein [Source:MGI Sy | 2.4093 | 1.0279 | 0 | 0.964 | 2.1812 | 1.1138 |
| 11410 | ENSMUSG00000039114 | 2653.513888 | 0.257264387 | 0.119912795 | 2.145429002 | 0.031918562 | 0.238097001 | Nrn1 | neuritin 1 [Source:MGI Symbol;Acc:MGI:1 | 2415.7 | 2255.3 | 2581 | 2676 | 2639.3 | 3353.8 |
| 11411 | ENSMUSG00000039115 | 139.367679 | -0.797758175 | 0.216404411 | -3.68642289 | 0.000227428 | 0.006969329 | Itga9 | integrin alpha 9 [Source:MGI Symbol;Acc: | 174.27 | 197.36 | 158.99 | 92.542 | 90.521 | 122.52 |
| 11412 | ENSMUSG00000039116 | 14.12592366 | 0.054408681 | 0.58877602 | 0.092409812 | 0.926372444 | 0.987693574 | Adgrg6 | adhesion G protein-coupled receptor G6 [ | 12.047 | 20.558 | 9.0562 | 14.46 | 15.269 | 13.366 |
| 11413 | ENSMUSG00000039117 | 298.6081281 | 0.073742904 | 0.164889403 | 0.447226458 | 0.65471156 | 0.925797231 | Taf4 | TATA-box binding protein associated facto | 306.79 | 282.68 | 283.76 | 355.71 | 303.19 | 259.53 |
| 11414 | ENSMUSG00000039126 | 5.480982875 | -1.061173094 | 1.272328962 | -0.83403988 | 0.404258512 | 0.829271989 | Prune2 | prune homolog 2 [Source:MGI Symbol;Ac | 8.8341 | 13.363 | 0 | 2.8919 | 0 | 7.7969 |
| 11415 | ENSMUSG00000039128 | 827.8094174 | -0.031765119 | 0.106417864 | -0.298494234 | 0.765325976 | 0.954980327 | Cdc123 | cell division cycle 123 [Source:MGI Symbo | 816.75 | 852.14 | 842.23 | 820.35 | 824.51 | 810.88 |
| 11416 | ENSMUSG00000039130 | 160.8997606 | -0.27369634 | 0.199802592 | -1.369833783 | 0.170738794 | 0.597556794 | Zc3hc1 | zinc finger, C3HC type 1 [Source:MGI Sym | 156.6 | 172.69 | 199.24 | 137.85 | 133.06 | 165.96 |
| 11417 | ENSMUSG00000039131 | 3.02501677 | -3.935546141 | 1.842622616 | -2.135839486 | 0.032692499 | NA | Gipc2 | GIPC PDZ domain containing family, meml | 12.047 | 5.1396 | 0 | 0.964 | 0 | 0 |
| 11418 | ENSMUSG00000039133 | 0 | NA | NA | NA | NA | NA | 9330171B17 | RIKEN cDNA 9330171B17 gene [Source:M | 0 | 0 | 0 | 0 | 0 | 0 |
| 11419 | ENSMUSG00000039137 | 29.70699181 | 0.236657349 | 0.486076609 | 0.486872532 | 0.626348666 | 0.919840299 | Whrn | whirlin [Source:MGI Symbol;Acc:MGI:268 | 16.062 | 39.061 | 27.169 | 44.343 | 34.9 | 16.708 |
| 11420 | ENSMUSG00000039138 | 0 | NA | NA | NA | NA | NA | 4930591A17 | RIKEN cDNA 4930591A17 gene [Source:M | 0 | 0 | 0 | 0 | 0 | 0 |

| | | | | | | | | | | | | | | | |
|---|---|---|---|---|---|---|---|---|---|---|---|---|---|---|---|
| 11421 | ENSMUSG00000039145 | 713.1523526 | 0.821525454 | 0.213694954 | 3.844383963 | 0.000120856 | 0.00421029 | Camk1d | calcium/calmodulin-dependent protein ki | 533.26 | 479.01 | 533.31 | 658.4 | 739.44 | 1335.5 |
| 11422 | ENSMUSG00000039146 | 0 | NA | NA | NA | NA | NA | Ifi44l | interferon-induced protein 44 like [Source | 0 | 0 | 0 | 0 | 0 | 0 |
| 11423 | ENSMUSG00000039148 | 610.6868807 | 0.003428186 | 0.116870711 | 0.029333149 | 0.97659889 | 0.997072516 | Sart1 | squamous cell carcinoma antigen recogni: | 622.4 | 594.14 | 612.8 | 588.03 | 610.75 | 636 |
| 11424 | ENSMUSG00000039153 | 1.446756899 | -1.598585954 | 1.96261736 | -0.814517382 | 0.415348619 | NA | Runx2 | runt related transcription factor 2 [Source | 2.4093 | 3.0838 | 1.0062 | 0 | 2.1812 | 0 |
| 11425 | ENSMUSG00000039154 | 2.183755553 | -0.192054085 | 1.714467931 | -0.112019643 | 0.910807832 | NA | Shd | src homology 2 domain-containing transfo | 0 | 1.0279 | 6.0375 | 3.8559 | 2.1812 | 0 |
| 11426 | ENSMUSG00000039155 | 0 | NA | NA | NA | NA | NA | Cdh26 | cadherin-like 26 [Source:MGI Symbol;Acc: | 0 | 0 | 0 | 0 | 0 | 0 |
| 11427 | ENSMUSG00000039156 | 397.2609616 | 0.203869093 | 0.133165375 | 1.530946706 | 0.125782565 | 0.520102634 | Stim2 | stromal interaction molecule 2 [Source:M | 378.26 | 373.13 | 356.21 | 429.93 | 441.7 | 404.33 |
| 11428 | ENSMUSG00000039157 | 972.9609474 | -0.114823057 | 0.153746598 | -0.74683316 | 0.455164277 | 0.857337687 | Fam102a | family with sequence similarity 102, meml | 1192.6 | 1002.2 | 839.21 | 934.09 | 1069.9 | 799.74 |
| 11429 | ENSMUSG00000039158 | 32.10354793 | -1.077979165 | 0.435026327 | -2.477963049 | 0.013213483 | 0.130993706 | Akna | AT-hook transcription factor [Source:MGI | 53.005 | 38.033 | 39.244 | 11.568 | 21.812 | 28.96 |
| 11430 | ENSMUSG00000039159 | 2011.274624 | 0.209654355 | 0.102032521 | 2.054779711 | 0.039900287 | 0.273576354 | Ube2h | ubiquitin-conjugating enzyme E2H [Source | 1944.3 | 1717.7 | 1933 | 2035.9 | 2233.6 | 2203.2 |
| 11431 | ENSMUSG00000039163 | 315.8854906 | -0.001200782 | 0.146014068 | -0.008223741 | 0.993438478 | 0.99859176 | Cmc1 | COX assembly mitochondrial protein 1 [Sc | 314.01 | 327.91 | 305.9 | 293.05 | 337 | 317.45 |
| 11432 | ENSMUSG00000039164 | 99.08671776 | 0.190945122 | 0.23331405 | 0.818403873 | 0.413126611 | 0.835261854 | Naif1 | nuclear apoptosis inducing factor 1 [Sourc | 99.585 | 91.485 | 86.537 | 119.53 | 93.793 | 103.59 |
| 11433 | ENSMUSG00000039166 | 132.7607783 | 0.044912554 | 0.205016086 | 0.219068441 | 0.826596734 | 0.969648181 | Akap7 | A kinase (PRKA) anchor protein 7 [Source: | 132.51 | 143.91 | 115.72 | 137.85 | 139.6 | 126.98 |
| 11434 | ENSMUSG00000039167 | 115.7445765 | -0.451306534 | 0.24527338 | -1.840014329 | 0.065766134 | 0.368337165 | Adgrl4 | adhesion G protein-coupled receptor L4 [! | 117.25 | 148.02 | 135.84 | 86.758 | 79.615 | 126.98 |
| 11435 | ENSMUSG00000039168 | 4475.994685 | -0.100477971 | 0.093068427 | -1.079613933 | 0.280314134 | 0.73330867 | Dap | death-associated protein [Source:MGI Syr | 4946.3 | 4344 | 4603.6 | 4175 | 4445.4 | 4341.8 |
| 11436 | ENSMUSG00000039174 | 0 | NA | NA | NA | NA | NA | 1700003H04 | RIKEN cDNA 1700003H04 gene [Source:M | 0 | 0 | 0 | 0 | 0 | 0 |
| 11437 | ENSMUSG00000039176 | 930.0728085 | 0.122168827 | 0.105210941 | 1.161179867 | 0.245568761 | 0.695774774 | Polg | polymerase (DNA directed), gamma [Sour | 926.78 | 858.31 | 886.5 | 972.65 | 978.28 | 957.91 |
| 11438 | ENSMUSG00000039178 | 40.02434667 | 0.032744055 | 0.379593741 | 0.086260786 | 0.93125911 | 0.988420305 | Tbc1d19 | TBC1 domain family, member 19 [Source:I | 36.14 | 54.48 | 28.175 | 36.631 | 49.078 | 35.643 |
| 11439 | ENSMUSG00000039179 | 0.455176322 | 0.955156269 | 3.546674842 | 0.269310357 | 0.787690862 | NA | Tekt5 | tektin 5 [Source:MGI Symbol;Acc:MGI:191 | 0.8031 | 0 | 0 | 1.928 | 0 | 0 |
| 11440 | ENSMUSG00000039182 | 272.5562777 | 0.064693247 | 0.162779217 | 0.397429405 | 0.691050835 | 0.937657013 | AW209491 | expressed sequence AW209491 [Source:N | 282.69 | 276.51 | 239.49 | 250.63 | 307.55 | 278.46 |
| 11441 | ENSMUSG00000039183 | 474.3149503 | 0.180536391 | 0.137907521 | 1.309112006 | 0.190496413 | 0.627783121 | Nubp2 | nucleotide binding protein 2 [Source:MGI | 407.17 | 429.67 | 498.09 | 533.08 | 474.42 | 503.46 |
| 11442 | ENSMUSG00000039187 | 2.525514382 | -0.195898988 | 1.438902948 | -0.136144684 | 0.891706903 | NA | Fanci | Fanconi anemia, complementation group | 4.0155 | 3.0838 | 1.0062 | 4.8199 | 0 | 2.2277 |
| 11443 | ENSMUSG00000039191 | 246.2553427 | 0.017749567 | 0.159287926 | 0.111430714 | 0.911274805 | 0.985839849 | Rbpj | recombination signal binding protein for i | 257.8 | 239.5 | 236.47 | 241.96 | 234.48 | 267.32 |
| 11444 | ENSMUSG00000039193 | 9.988174482 | -0.165773183 | 0.693074054 | -0.239185383 | 0.810961837 | 0.966452061 | Nlrc4 | NLR family, CARD domain containing 4 [Sc | 6.4248 | 13.363 | 12.075 | 11.568 | 9.8156 | 6.6831 |
| 11445 | ENSMUSG00000039194 | 0 | NA | NA | NA | NA | NA | Rlbp1 | retinaldehyde binding protein 1 [Source:N | 0 | 0 | 0 | 0 | 0 | 0 |
| 11446 | ENSMUSG00000039195 | 283.7533228 | -0.047838008 | 0.16874108 | -0.283499479 | 0.776793981 | 0.959295521 | 1110008P14 | RIKEN cDNA 1110008P14 gene [Source:M | 325.26 | 237.45 | 301.87 | 276.66 | 266.11 | 295.17 |
| 11447 | ENSMUSG00000039196 | 8215.764356 | 0.390674489 | 0.10992718 | 3.553938982 | 0.000379507 | 0.010356968 | Orm1 | orosomucoid 1 [Source:MGI Symbol;Acc:I | 7965.2 | 6774 | 6588.9 | 8477.2 | 9448 | 10041 |
| 11448 | ENSMUSG00000039197 | 16143.22645 | -0.060214146 | 0.081600282 | -0.737915906 | 0.460565553 | 0.859817816 | Adk | adenosine kinase [Source:MGI Symbol;Ac | 16794 | 16461 | 16185 | 16150 | 16122 | 15147 |
| 11449 | ENSMUSG00000039198 | 0 | NA | NA | NA | NA | NA | Ptchd3 | patched domain containing 3 [Source:MG | 0 | 0 | 0 | 0 | 0 | 0 |
| 11450 | ENSMUSG00000039199 | 44.97499447 | -0.50915376 | 0.347894225 | -1.463530362 | 0.143322312 | 0.553098698 | Zdhhc1 | zinc finger, DHHC domain containing 1 [Sc | 40.958 | 53.452 | 64.4 | 39.523 | 41.443 | 30.074 |
| 11451 | ENSMUSG00000039200 | 0 | NA | NA | NA | NA | NA | Atf7ip2 | activating transcription factor 7 interactin | 0 | 0 | 0 | 0 | 0 | 0 |
| 11452 | ENSMUSG00000039201 | 68.36229028 | -0.393782232 | 0.278088365 | -1.416032752 | 0.156765915 | 0.575786484 | Tbc1d25 | TBC1 domain family, member 25 [Source:I | 67.461 | 86.345 | 79.493 | 64.587 | 62.165 | 50.123 |
| 11453 | ENSMUSG00000039202 | 1741.900354 | -0.719804276 | 0.181430637 | -3.967379972 | 7.27E-05 | 0.002795364 | Abhd2 | abhydrolase domain containing 2 [Source | 2451.1 | 2358 | 1694.5 | 1687.9 | 1261.8 | 998 |
| 11454 | ENSMUSG00000039205 | 314.3813327 | -0.004883627 | 0.148099152 | -0.032975391 | 0.973694212 | 0.996075612 | Ciz1 | CDKN1A interacting zinc finger protein 1 [: | 331.68 | 293.98 | 318.98 | 333.54 | 316.28 | 291.83 |
| 11455 | ENSMUSG00000039206 | 567.2153472 | 0.082052749 | 0.137587335 | 0.596368473 | 0.550929097 | 0.895729712 | Daglb | diacylglycerol lipase, beta [Source:MGI Sy | 615.98 | 503.68 | 532.3 | 539.83 | 644.56 | 566.95 |
| 11456 | ENSMUSG00000039208 | 39.36510382 | -0.413947208 | 0.379911411 | -1.089588774 | 0.27589433 | 0.730038567 | Metrnl | meteorin, glial cell differentiation regulatc | 40.155 | 50.368 | 44.275 | 22.171 | 45.806 | 33.415 |
| 11457 | ENSMUSG00000039209 | 0.160662974 | 1.126924175 | 4.080472857 | 0.276174898 | 0.782413736 | NA | Rpl39l | ribosomal protein L39-like [Source:MGI Sy | 0 | 0 | 0 | 0.964 | 0 | 0 |
| 11458 | ENSMUSG00000039210 | 273.1742967 | 0.050847725 | 0.184637385 | 0.275392359 | 0.783014814 | 0.960670212 | Gpatch2 | G patch domain containing 2 [Source:MGI | 318.83 | 268.29 | 217.35 | 301.73 | 241.03 | 291.83 |
| 11459 | ENSMUSG00000039215 | 0.267700746 | -1.497481649 | 4.057001436 | -0.369110456 | 0.712045396 | NA | Svs1 | seminal vesicle secretory protein 1 [Sourc | 1.6062 | 0 | 0 | 0 | 0 | 0 |
| 11460 | ENSMUSG00000039217 | 174.682894 | 0.226209584 | 0.199804799 | 1.132152906 | 0.257570156 | 0.710872182 | Il18 | interleukin 18 [Source:MGI Symbol;Acc:M | 173.47 | 140.82 | 168.04 | 161.95 | 181.04 | 222.77 |
| 11461 | ENSMUSG00000039218 | 3282.271897 | 0.158427744 | 0.142813853 | 1.109330365 | 0.267287691 | 0.722896252 | Srrm2 | serine/arginine repetitive matrix 2 [Source | 3013.2 | 3140.3 | 3154.6 | 4361 | 3151.9 | 2872.6 |
| 11462 | ENSMUSG00000039219 | 390.9779899 | 0.01082876 | 0.141742515 | 0.076397401 | 0.939102938 | 0.98880286 | Arid4b | AT rich interactive domain 4B (RBP1-like) | 352.56 | 409.11 | 407.53 | 383.66 | 420.98 | 372.02 |
| 11463 | ENSMUSG00000039220 | 659.884053 | -0.847667175 | 0.156250127 | -5.425065512 | 5.79E-08 | 6.76E-06 | Ppp1r10 | protein phosphatase 1, regulatory subunit | 721.99 | 943.63 | 879.46 | 399.09 | 555.12 | 460.02 |
| 11464 | ENSMUSG00000039221 | 260.5637494 | 0.39146033 | 0.21602523 | 1.812104679 | 0.069970024 | 0.38248164 | Rpl22l1 | ribosomal protein L22 like 1 [Source:MGI ! | 244.95 | 229.23 | 201.25 | 199.54 | 363.18 | 325.24 |
| 11465 | ENSMUSG00000039224 | 0 | NA | NA | NA | NA | NA | D1Pas1 | DNA segment, Chr 1, Pasteur Institute 1 [S | 0 | 0 | 0 | 0 | 0 | 0 |
| 11466 | ENSMUSG00000039230 | 496.4408632 | 0.027082232 | 0.138673028 | 0.195295597 | 0.845161544 | 0.972803041 | Tbcd | tubulin-specific chaperone d [Source:MGI | 542.09 | 456.4 | 475.95 | 486.81 | 554.03 | 463.36 |
| 11467 | ENSMUSG00000039231 | 106.0981165 | 0.078554677 | 0.239912232 | 0.327430895 | 0.743342003 | 0.949628928 | Suv39h1 | suppressor of variegation 3-9 1 [Source:M | 103.6 | 81.206 | 124.77 | 108.93 | 115.61 | 102.47 |
| 11468 | ENSMUSG00000039232 | 7.7583099 | -1.456042846 | 0.806980527 | -1.804309765 | 0.07118276 | 0.386047033 | Stx11 | syntaxin 11 [Source:MGI Symbol;Acc:MGI | 12.85 | 9.2513 | 12.075 | 5.7839 | 4.3625 | 2.2277 |
| 11469 | ENSMUSG00000039233 | 259.1077958 | 0.106357758 | 0.17069998 | 0.623068372 | 0.533239593 | 0.889143097 | Tbce | tubulin-specific chaperone E [Source:MGI | 278.68 | 243.62 | 225.4 | 235.21 | 297.74 | 274.01 |
| 11470 | ENSMUSG00000039234 | 1642.432621 | -0.169118717 | 0.119103293 | -1.419933172 | 0.155627138 | 0.574732247 | Sec24d | Sec24 related gene family, member D (S. c | 1753.2 | 1726.9 | 1736.8 | 1795.9 | 1444 | 1397.9 |
| 11471 | ENSMUSG00000039236 | 38.9684383 | -0.428440295 | 0.359477017 | -1.191843356 | 0.233322672 | 0.680673483 | Isg20 | interferon-stimulated protein [Source:MG | 42.564 | 42.145 | 49.306 | 26.027 | 40.353 | 33.415 |
| 11472 | ENSMUSG00000039238 | 177.7306086 | -0.573976126 | 0.242228457 | -2.369565216 | 0.017809014 | 0.159300914 | Zfp750 | zinc finger protein 750 [Source:MGI Symb | 186.32 | 223.06 | 228.42 | 109.89 | 202.86 | 115.84 |
| 11473 | ENSMUSG00000039239 | 8.504948886 | -0.301278659 | 0.750029491 | -0.401689084 | 0.687912859 | 0.936885029 | Tgfb2 | transforming growth factor, beta 2 [Sourc | 12.85 | 8.2233 | 7.0437 | 9.6398 | 4.3625 | 8.9108 |
| 11474 | ENSMUSG00000039242 | 350.9541008 | -0.006661019 | 0.164342821 | -0.040531244 | 0.967669598 | 0.994231936 | B3galnt2 | UDP-GalNAc:betaGlcNAc beta 1,3-galacto | 301.16 | 405 | 350.17 | 362.46 | 380.63 | 306.31 |
| 11475 | ENSMUSG00000039244 | 334.4560629 | 0.034128957 | 0.15119476 | 0.22572844 | 0.821412637 | 0.969160361 | E130309D02 | RIKEN cDNA E130309D02 gene [Source:M | 338.11 | 317.63 | 336.09 | 374.02 | 345.73 | 295.17 |
| 11476 | ENSMUSG00000039246 | 293.1527706 | -0.154360592 | 0.18058791 | -0.85476703 | 0.392680134 | 0.821375223 | Lyplal1 | lysophospholipase-like 1 [Source:MGI Syn | 326.86 | 280.62 | 317.97 | 215.93 | 292.29 | 325.24 |

| | | | | | | | | | | | | | | | |
|---|---|---|---|---|---|---|---|---|---|---|---|---|---|---|---|
| 11477 | ENSMUSG00000039252 | 2.543830062 | -0.914350285 | 1.476673896 | -0.619195807 | 0.535787374 | NA | Lgi2 | leucine-rich repeat LGI family, member 2 [ | 4.8186 | 4.1117 | 1.0062 | 0.964 | 4.3625 | 0 |
| 11478 | ENSMUSG00000039253 | 217.5341724 | 0.099124715 | 0.169693643 | 0.584139236 | 0.559126639 | 0.899423311 | Fn3krp | fructosamine 3 kinase related protein [Sou | 201.58 | 193.25 | 235.46 | 220.75 | 222.49 | 231.68 |
| 11479 | ENSMUSG00000039254 | 247.961363 | -0.161256548 | 0.162342916 | -0.993308191 | 0.320559787 | 0.767391055 | Pomt1 | protein-O-mannosyltransferase 1 [Source | 288.31 | 260.06 | 236.47 | 231.35 | 242.12 | 229.45 |
| 11480 | ENSMUSG00000039257 | 0 | NA | NA | NA | NA | NA | Vstm2b | V-set and transmembrane domain contair | 0 | 0 | 0 | 0 | 0 | 0 |
| 11481 | ENSMUSG00000039262 | 939.1773313 | 0.080866917 | 0.10725157 | 0.753992852 | 0.450853514 | 0.855818389 | Prrc2b | proline-rich coiled-coil 2B [Source:MGI Sy | 892.25 | 938.49 | 908.64 | 1020.9 | 930.3 | 944.54 |
| 11482 | ENSMUSG00000039263 | 1330.067978 | 0.056296788 | 0.108712754 | 0.517848972 | 0.604563647 | 0.916376001 | Npepl1 | aminopeptidase-like 1 [Source:MGI Symbo | 1260.1 | 1410.3 | 1242.7 | 1376.6 | 1425.4 | 1265.3 |
| 11483 | ENSMUSG00000039264 | 18.57879686 | -1.85113533 | 0.589090772 | -3.142360091 | 0.001675918 | 0.030723037 | Gimap3 | GTPase, IMAP family member 3 [Source:M | 31.321 | 27.754 | 28.175 | 8.6758 | 2.1812 | 13.366 |
| 11484 | ENSMUSG00000039269 | 0 | NA | NA | NA | NA | NA | 2300002M2: | RIKEN cDNA 2300002M23 gene [Source:M | 0 | 0 | 0 | 0 | 0 | 0 |
| 11485 | ENSMUSG00000039270 | 569.6024769 | -0.351158802 | 0.162021748 | -2.167355969 | 0.030207722 | 0.230202576 | Megf9 | multiple EGF-like-domains 9 [Source:MGI | 661.76 | 757.58 | 496.08 | 472.35 | 491.87 | 537.99 |
| 11486 | ENSMUSG00000039275 | 332.2923741 | 0.212202324 | 0.148091207 | 1.432916431 | 0.151881721 | 0.567340036 | Foxk2 | forkhead box K2 [Source:MGI Symbol;Acc | 284.3 | 337.16 | 302.88 | 355.71 | 378.44 | 335.27 |
| 11487 | ENSMUSG00000039278 | 0 | NA | NA | NA | NA | NA | Pcsk1n | proprotein convertase subtilisin/kexin typ | 0 | 0 | 0 | 0 | 0 | 0 |
| 11488 | ENSMUSG00000039285 | 914.8903899 | 0.067910404 | 0.107148183 | 0.633798935 | 0.526212048 | 0.888334265 | Azi2 | 5-azacytidine induced gene 2 [Source:MG | 914.73 | 919.99 | 845.25 | 950.48 | 946.66 | 912.24 |
| 11489 | ENSMUSG00000039286 | 427.7182505 | -0.080217476 | 0.184074755 | -0.435787492 | 0.66299092 | 0.92906822 | Fndc3b | fibronectin type III domain containing 3B [ | 538.08 | 445.09 | 335.08 | 470.42 | 357.72 | 419.92 |
| 11490 | ENSMUSG00000039294 | 157.586616 | -0.147562953 | 0.194353436 | -0.759250548 | 0.447702693 | 0.853796936 | Cybc1 | cytochrome b 245 chaperone 1 [Source:M | 158.21 | 178.86 | 159.99 | 149.42 | 131.96 | 167.08 |
| 11491 | ENSMUSG00000039296 | 0 | NA | NA | NA | NA | NA | Spdye4a | speedy/RINGO cell cycle regulator family, | 0 | 0 | 0 | 0 | 0 | 0 |
| 11492 | ENSMUSG00000039298 | 76.84942006 | 0.227294464 | 0.321033427 | 0.708008714 | 0.478939846 | 0.870169438 | Cdk5rap2 | CDK5 regulatory subunit associated protei | 61.839 | 79.15 | 71.443 | 69.406 | 55.622 | 123.64 |
| 11493 | ENSMUSG00000039304 | 128.1390702 | 0.15806224 | 0.221359789 | 0.714051276 | 0.475195473 | 0.869368495 | Tnfsf10 | tumor necrosis factor (ligand) superfamily | 125.28 | 102.79 | 134.84 | 113.75 | 140.69 | 151.48 |
| 11494 | ENSMUSG00000039307 | 131.8585068 | 0.29452695 | 0.238320742 | 1.235842705 | 0.216517038 | 0.661953589 | Hexdc | hexosaminidase (glycosyl hydrolase family | 85.932 | 130.55 | 139.87 | 164.84 | 137.42 | 132.55 |
| 11495 | ENSMUSG00000039308 | 471.4505358 | 0.122743717 | 0.143128461 | 0.857577283 | 0.391125923 | 0.820195366 | Ndst2 | N-deacetylase/N-sulfotransferase (hepara | 431.27 | 437.89 | 486.02 | 558.14 | 479.87 | 435.51 |
| 11496 | ENSMUSG00000039313 | 0 | NA | NA | NA | NA | NA | Minar1 | membrane integral NOTCH2 associated re | 0 | 0 | 0 | 0 | 0 | 0 |
| 11497 | ENSMUSG00000039315 | 0.305170013 | -1.657836477 | 4.041480335 | -0.410205257 | 0.681655385 | NA | Clnk | cytokine-dependent hematopoietic cell lir | 0.8031 | 1.0279 | 0 | 0 | 0 | 0 |
| 11498 | ENSMUSG00000039316 | 14.37600851 | -1.562078508 | 0.639064128 | -2.444322002 | 0.014512467 | 0.139752549 | Rftn1 | raftlin lipid raft linker 1 [Source:MGI Symb | 28.912 | 14.391 | 21.131 | 9.6398 | 3.2719 | 8.9108 |
| 11499 | ENSMUSG00000039318 | 266.9195605 | 0.056284662 | 0.157831334 | 0.356612728 | 0.721381748 | 0.9447719 | Rab3gap2 | RAB3 GTPase activating protein subunit 2 | 264.22 | 281.65 | 239.49 | 286.3 | 277.02 | 252.84 |
| 11500 | ENSMUSG00000039321 | 0 | NA | NA | NA | NA | NA | Uts2r | urotensin 2 receptor [Source:MGI Symbol | 0 | 0 | 0 | 0 | 0 | 0 |
| 11501 | ENSMUSG00000039323 | 7165.525411 | 0.419782506 | 0.319253299 | 1.314888549 | 0.188547362 | 0.625779955 | Igfbp2 | insulin-like growth factor binding protein : | 3709.5 | 6059.6 | 8622.5 | 8080.1 | 7853.5 | 8667.9 |
| 11502 | ENSMUSG00000039328 | 11.46837532 | -1.628920298 | 0.667473238 | -2.440427877 | 0.014669875 | 0.14062439 | Rnf122 | ring finger protein 122 [Source:MGI Symb | 18.471 | 14.391 | 19.119 | 5.7839 | 4.3625 | 6.6831 |
| 11503 | ENSMUSG00000039329 | 0 | NA | NA | NA | NA | NA | Tex19.1 | testis expressed gene 19.1 [Source:MGI Sy | 0 | 0 | 0 | 0 | 0 | 0 |
| 11504 | ENSMUSG00000039330 | 0.401551119 | -2.072982726 | 4.008251979 | -0.517178744 | 0.605031388 | NA | Tsga10ip | testis specific 10 interacting protein [Sour | 2.4093 | 0 | 0 | 0 | 0 | 0 |
| 11505 | ENSMUSG00000039335 | 0 | NA | NA | NA | NA | NA | Spata16 | spermatogenesis associated 16 [Source:M | 0 | 0 | 0 | 0 | 0 | 0 |
| 11506 | ENSMUSG00000039337 | 0 | NA | NA | NA | NA | NA | Tex19.2 | testis expressed gene 19.2 [Source:MGI Sy | 0 | 0 | 0 | 0 | 0 | 0 |
| 11507 | ENSMUSG00000039339 | 0 | NA | NA | NA | NA | NA | Mfsd4b2 | major facilitator superfamily domain cont | 0 | 0 | 0 | 0 | 0 | 0 |
| 11508 | ENSMUSG00000039342 | 0 | NA | NA | NA | NA | NA | Ankar | ankyrin and armadillo repeat containing [S | 0 | 0 | 0 | 0 | 0 | 0 |
| 11509 | ENSMUSG00000039345 | 138.2199234 | 0.230933995 | 0.2002231 | 1.153383374 | 0.248753063 | 0.70053822 | Mettl22 | methyltransferase like 22 [Source:MGI Syr | 126.09 | 135.69 | 119.74 | 145.56 | 138.51 | 163.74 |
| 11510 | ENSMUSG00000039347 | 16.54394617 | -0.379354467 | 0.59029231 | -0.642655275 | 0.520447808 | 0.887802551 | Atp6v0e2 | ATPase, H+ transporting, lysosomal V0 sul | 19.274 | 19.53 | 17.106 | 5.7839 | 14.178 | 23.391 |
| 11511 | ENSMUSG00000039349 | 3758.889033 | 0.107868684 | 0.122033004 | 0.883930414 | 0.376733778 | 0.810151258 | C130074G19 | RIKEN cDNA C130074G19 gene [Source:M | 3285.5 | 3488.8 | 4081.3 | 3595.6 | 3660.1 | 4442 |
| 11512 | ENSMUSG00000039354 | 228.0871985 | 0.066669981 | 0.165882231 | 0.401911531 | 0.687749136 | 0.936885029 | Smarcal1 | SWI/SNF related matrix associated, actin c | 245.75 | 207.64 | 214.33 | 223.64 | 241.03 | 236.13 |
| 11513 | ENSMUSG00000039356 | 197.5702633 | -0.151083266 | 0.169289119 | -0.892457041 | 0.372148009 | 0.807785387 | Exosc2 | exosome component 2 [Source:MGI Symb | 205.59 | 211.75 | 206.28 | 178.34 | 195.22 | 188.24 |
| 11514 | ENSMUSG00000039357 | 116.932494 | 0.018421293 | 0.260844817 | 0.070621655 | 0.943698875 | 0.990077979 | Fut11 | fucosyltransferase 11 [Source:MGI Symbo | 113.24 | 85.317 | 149.93 | 115.68 | 95.974 | 141.46 |
| 11515 | ENSMUSG00000039358 | 0 | NA | NA | NA | NA | NA | Drd5 | dopamine receptor D5 [Source:MGI Symb | 0 | 0 | 0 | 0 | 0 | 0 |
| 11516 | ENSMUSG00000039361 | 2096.164291 | -0.241177521 | 0.09941755 | -2.425904882 | 0.015270272 | 0.144358963 | Picalm | phosphatidylinositol binding clathrin asse | 2339.4 | 2374.5 | 2099 | 1985.8 | 1931.5 | 1846.8 |
| 11517 | ENSMUSG00000039364 | 0.486939728 | -0.698472448 | 3.137608162 | -0.222613026 | 0.823836697 | NA | Sectm1b | secreted and transmembrane 1B [Source: | 0.8031 | 1.0279 | 0 | 0 | 1.0906 | 0 |
| 11518 | ENSMUSG00000039367 | 1883.783791 | 0.129507164 | 0.108497144 | 1.193645832 | 0.232616538 | 0.679847566 | Sec24c | Sec24 related gene family, member C (S. c | 1928.2 | 1864.6 | 1604 | 1914.5 | 2068.9 | 1922.5 |
| 11519 | ENSMUSG00000039372 | 0 | NA | NA | NA | NA | NA | Marchf4 | membrane associated ring-CH-type finger | 0 | 0 | 0 | 0 | 0 | 0 |
| 11520 | ENSMUSG00000039375 | 0 | NA | NA | NA | NA | NA | Wdr17 | WD repeat domain 17 [Source:MGI Symbo | 0 | 0 | 0 | 0 | 0 | 0 |
| 11521 | ENSMUSG00000039376 | 5.564828252 | 0.741079597 | 0.932379429 | 0.794826198 | 0.426714609 | 0.842062628 | Synpo2l | synaptopodin 2-like [Source:MGI Symbol; | 2.4093 | 7.1954 | 3.0187 | 8.6758 | 7.6343 | 4.4554 |
| 11522 | ENSMUSG00000039377 | 60.4977074 | -1.356631354 | 0.389067922 | -3.486875371 | 0.000488699 | 0.012229313 | Hlx | H2.0-like homeobox [Source:MGI Symbol; | 85.932 | 109.99 | 65.406 | 46.271 | 14.178 | 41.212 |
| 11523 | ENSMUSG00000039382 | 350.0113067 | 0.084574431 | 0.15383403 | 0.549777126 | 0.58247225 | 0.908645447 | Wdr45 | WD repeat domain 45 [Source:MGI Symbo | 371.84 | 343.32 | 303.89 | 389.45 | 371.9 | 319.67 |
| 11524 | ENSMUSG00000039384 | 47.46015723 | 0.117362163 | 0.33552186 | 0.349789917 | 0.726496367 | 0.945777495 | Dusp10 | dual specificity phosphatase 10 [Source:M | 48.186 | 35.977 | 52.325 | 51.091 | 39.262 | 57.92 |
| 11525 | ENSMUSG00000039385 | 3.318499059 | -0.712948128 | 1.181130495 | -0.603615038 | 0.546099613 | NA | Cdh6 | cadherin 6 [Source:MGI Symbol;Acc:MGI: | 3.2124 | 3.0838 | 6.0375 | 0.964 | 3.2719 | 3.3415 |
| 11526 | ENSMUSG00000039391 | 0 | NA | NA | NA | NA | NA | Ccdc81 | coiled-coil domain containing 81 [Source:I | 0 | 0 | 0 | 0 | 0 | 0 |
| 11527 | ENSMUSG00000039395 | 738.3821404 | -0.64527526 | 0.183602149 | -3.514529997 | 0.000440533 | 0.011297638 | Mreg | melanoregulin [Source:MGI Symbol;Acc:M | 644.89 | 986.8 | 1071.7 | 549.47 | 672.91 | 504.57 |
| 11528 | ENSMUSG00000039396 | 1.381163065 | -3.83556022 | 2.086882674 | -1.837937641 | 0.066071595 | NA | Neil3 | nei like 3 (E. coli) [Source:MGI Symbol;Acc | 3.2124 | 2.0558 | 3.0187 | 0 | 0 | 0 |
| 11529 | ENSMUSG00000039405 | 37.11921113 | -0.632240303 | 0.422318528 | -1.497069776 | 0.134375103 | 0.537569131 | Prss23 | protease, serine 23 [Source:MGI Symbol;A | 66.657 | 43.173 | 25.156 | 26.027 | 31.628 | 30.074 |
| 11530 | ENSMUSG00000039410 | 5.143871451 | 0.057432135 | 0.922389588 | 0.062264509 | 0.950352191 | 0.990815588 | Prdm16 | PR domain containing 16 [Source:MGI Syr | 4.0155 | 4.1117 | 7.0437 | 5.7839 | 5.4531 | 4.4554 |
| 11531 | ENSMUSG00000039414 | 243.2266106 | 0.012901244 | 0.183113857 | 0.070454767 | 0.943831702 | 0.990077979 | Heatr5b | HEAT repeat containing 5B [Source:MGI S | 219.25 | 284.73 | 223.39 | 282.45 | 230.12 | 219.43 |
| 11532 | ENSMUSG00000039419 | 0 | NA | NA | NA | NA | NA | Cntnap2 | contactin associated protein-like 2 [Sourc | 0 | 0 | 0 | 0 | 0 | 0 |

| | | | | | | | | | | | | | | | |
|---|---|---|---|---|---|---|---|---|---|---|---|---|---|---|---|
| 11533 | ENSMUSG00000039427 | 290.6242143 | 0.046490067 | 0.155868073 | 0.298265488 | 0.765500543 | 0.954980327 | Alg1 | asparagine-linked glycosylation 1 (beta-1,4 | 281.09 | 258.01 | 318.98 | 310.4 | 289.01 | 286.26 |
| 11534 | ENSMUSG00000039428 | 1699.041646 | -0.255497366 | 0.11749979 | -2.174449553 | 0.02967139 | 0.22707248 | Tmem135 | transmembrane protein 135 [Source:MGI | 1984.5 | 1964.4 | 1597.9 | 1549.1 | 1635.9 | 1462.5 |
| 11535 | ENSMUSG00000039431 | 36.8916743 | 0.001204785 | 0.385539887 | 0.003124928 | 0.997506672 | 0.999498002 | Mtmr7 | myotubularin related protein 7 [Source:M | 27.305 | 44.2 | 39.244 | 28.919 | 34.9 | 46.781 |
| 11536 | ENSMUSG00000039438 | 2496.447575 | 0.033568673 | 0.108650873 | 0.308959077 | 0.757352657 | 0.953381099 | Ttc36 | tetratricopeptide repeat domain 36 [Sour | 2356.3 | 2331.3 | 2714.8 | 2425.4 | 2751.6 | 2399.2 |
| 11537 | ENSMUSG00000039449 | 671.1405274 | 0.154573324 | 0.113064093 | 1.367130101 | 0.17158453 | 0.599034187 | Prpf18 | pre-mRNA processing factor 18 [Source:M | 635.25 | 635.25 | 634.94 | 683.46 | 733.99 | 703.95 |
| 11538 | ENSMUSG00000039450 | 2085.933448 | -0.028230025 | 0.121324679 | -0.232681641 | 0.816008627 | 0.967767553 | Dcxr | dicarbonyl L-xylulose reductase [Source:M | 2002.1 | 1994.2 | 2322.4 | 1813.2 | 2321.9 | 2061.7 |
| 11539 | ENSMUSG00000039452 | 5.432168119 | -0.694820623 | 1.009952763 | -0.687973387 | 0.491469541 | 0.876368691 | Snx22 | sorting nexin 22 [Source:MGI Symbol;Acc: | 8.031 | 2.0558 | 10.062 | 5.7839 | 1.0906 | 5.5692 |
| 11540 | ENSMUSG00000039456 | 307.0917498 | -0.424179431 | 0.190472209 | -2.226988566 | 0.025948038 | 0.208275484 | Morc3 | microrchidia 3 [Source:MGI Symbol;Acc:M | 355.77 | 391.64 | 308.92 | 305.58 | 286.83 | 193.81 |
| 11541 | ENSMUSG00000039457 | 207.2081409 | -0.079877557 | 0.217575439 | -0.367125799 | 0.713525184 | 0.94429556 | Ppl | periplakin [Source:MGI Symbol;Acc:MGI:1 | 163.83 | 235.39 | 240.49 | 223.64 | 153.78 | 226.11 |
| 11542 | ENSMUSG00000039458 | 135.4075206 | -0.223608514 | 0.216167255 | -1.034423617 | 0.300938168 | 0.7521219 | Mtmr12 | myotubularin related protein 12 [Source:M | 159.01 | 126.43 | 151.94 | 131.1 | 103.61 | 140.34 |
| 11543 | ENSMUSG00000039461 | 422.1639077 | -0.047638651 | 0.15094495 | -0.315602814 | 0.752304001 | 0.952830567 | Tcta | T cell leukemia translocation altered gene | 465 | 388.55 | 432.69 | 364.38 | 408.98 | 473.38 |
| 11544 | ENSMUSG00000039462 | 1.077718728 | 1.149008664 | 2.221862456 | 0.517137621 | 0.605060093 | NA | Col10a1 | collagen, type X, alpha 1 [Source:MGI Sym | 0 | 1.0279 | 1.0062 | 0 | 1.0906 | 3.3415 |
| 11545 | ENSMUSG00000039463 | 576.2988064 | 0.107719304 | 0.121823508 | 0.884224284 | 0.376575152 | 0.809930558 | Slc9a8 | solute carrier family 9 (sodium/hydrogen e | 562.17 | 572.55 | 529.28 | 576.46 | 580.21 | 637.12 |
| 11546 | ENSMUSG00000039470 | 10.02409062 | -0.013895516 | 0.781686286 | -0.017776333 | 0.985817285 | 0.99859176 | Zdhhc2 | zinc finger, DHHC domain containing 2 [Sc | 4.0155 | 6.1675 | 20.125 | 6.7478 | 14.178 | 8.9108 |
| 11547 | ENSMUSG00000039473 | 561.4215721 | 0.050920821 | 0.140785955 | 0.36168964 | 0.71758397 | 0.9447719 | Ubn1 | ubinuclein 1 [Source:MGI Symbol;Acc:MG | 492.3 | 616.75 | 546.39 | 582.24 | 509.32 | 621.52 |
| 11548 | ENSMUSG00000039474 | 136.9849126 | 0.209563092 | 0.211802779 | 0.989425599 | 0.322454955 | 0.76988915 | Wfs1 | wolframin ER transmembrane glycoprotei | 123.68 | 131.57 | 125.78 | 133.99 | 129.78 | 177.1 |
| 11549 | ENSMUSG00000039476 | 0 | NA | NA | NA | NA | NA | Prrx2 | paired related homeobox 2 [Source:MGI S | 0 | 0 | 0 | 0 | 0 | 0 |
| 11550 | ENSMUSG00000039477 | 657.7687601 | 0.160720896 | 0.131060191 | 1.226313607 | 0.220080683 | 0.666369478 | Tnrc18 | trinucleotide repeat containing 18 [Source | 557.35 | 644.5 | 662.11 | 639.12 | 680.55 | 762.98 |
| 11551 | ENSMUSG00000039478 | 55.47981355 | 0.00327494 | 0.304345016 | 0.010760616 | 0.991414436 | 0.99859176 | Micu3 | mitochondrial calcium uptake family, men | 47.383 | 64.759 | 54.337 | 54.947 | 49.078 | 62.375 |
| 11552 | ENSMUSG00000039480 | 417.3043415 | -0.025522605 | 0.13728354 | -0.185911621 | 0.852514069 | 0.973781977 | Nt5dc1 | 5'-nucleotidase domain containing 1 [Sour | 448.93 | 386.5 | 426.65 | 386.56 | 429.7 | 425.49 |
| 11553 | ENSMUSG00000039481 | 40.22969484 | -0.061910479 | 0.407389875 | -0.151968625 | 0.879211681 | 0.981243654 | Nrtn | neurturin [Source:MGI Symbol;Acc:MGI:1 | 23.29 | 42.145 | 58.362 | 48.199 | 37.081 | 32.301 |
| 11554 | ENSMUSG00000039483 | 172.0641976 | -0.040052248 | 0.192604165 | -0.207951099 | 0.83526715 | 0.971073158 | Asb6 | ankyrin repeat and SOCS box-containing 6 | 163.83 | 184 | 176.09 | 191.83 | 142.87 | 173.76 |
| 11555 | ENSMUSG00000039485 | 23.8120648 | 0.899284566 | 0.588986644 | 1.526833545 | 0.126802408 | 0.521635389 | Tspyl4 | TSPY-like 4 [Source:MGI Symbol;Acc:MGI: | 5.6217 | 11.307 | 33.206 | 22.171 | 33.809 | 36.757 |
| 11556 | ENSMUSG00000039488 | 0 | NA | NA | NA | NA | NA | Cntn5 | contactin 5 [Source:MGI Symbol;Acc:MGI | 0 | 0 | 0 | 0 | 0 | 0 |
| 11557 | ENSMUSG00000039492 | 0.96342938 | -2.048320316 | 2.460986955 | -0.832316608 | 0.405230258 | NA | Ccdc27 | coiled-coil domain containing 27 [Source:I | 1.6062 | 3.0838 | 0 | 0 | 1.0906 | 0 |
| 11558 | ENSMUSG00000039496 | 11.72873351 | 0.380986648 | 0.683705473 | 0.557237966 | 0.577364853 | 0.907838603 | Cdnf | cerebral dopamine neurotrophic factor [S | 16.062 | 10.279 | 4.025 | 12.532 | 17.45 | 10.025 |
| 11559 | ENSMUSG00000039497 | 85.97398555 | -1.347595147 | 0.259969386 | -5.183668616 | 2.18E-07 | 2.16E-05 | Dse | dermatan sulfate epimerase [Source:MGI | 122.87 | 133.63 | 113.71 | 46.271 | 41.443 | 57.92 |
| 11560 | ENSMUSG00000039501 | 917.1782629 | -0.004904799 | 0.132975181 | -0.036885074 | 0.970576641 | 0.994966572 | Znfx1 | zinc finger, NFX1-type containing 1 [Sourc | 1054.5 | 882.98 | 818.08 | 1009.3 | 858.32 | 879.94 |
| 11561 | ENSMUSG00000039508 | 0 | NA | NA | NA | NA | NA | Calhm4 | calcium homeostasis modulator family me | 0 | 0 | 0 | 0 | 0 | 0 |
| 11562 | ENSMUSG00000039509 | 184.5801807 | 0.219094677 | 0.184403182 | 1.188128506 | 0.234782785 | 0.682729836 | Nup133 | nucleoporin 133 [Source:MGI Symbol;Acc | 176.68 | 176.8 | 157.98 | 183.16 | 226.85 | 186.01 |
| 11563 | ENSMUSG00000039512 | 557.7249892 | 0.060351431 | 0.128838762 | 0.468426037 | 0.639479952 | 0.923932929 | Uhrf1bp1 | UHRF1 (ICBP90) binding protein 1 [Source | 592.69 | 502.65 | 542.37 | 606.34 | 542.04 | 560.26 |
| 11564 | ENSMUSG00000039515 | 848.8255858 | -0.089229668 | 0.12175481 | -0.732863599 | 0.463641627 | 0.861746657 | Ptpa | protein phosphatase 2 protein activator [S | 766.96 | 945.68 | 913.67 | 807.81 | 822.33 | 836.5 |
| 11565 | ENSMUSG00000039518 | 0 | NA | NA | NA | NA | NA | Cdsn | corneodesmosin [Source:MGI Symbol;Acc | 0 | 0 | 0 | 0 | 0 | 0 |
| 11566 | ENSMUSG00000039519 | 4087.746594 | 1.420882474 | 0.322801644 | 4.401720072 | 1.07E-05 | 0.000566926 | Cyp7b1 | cytochrome P450, family 7, subfamily b, p | 2063.2 | 1933.5 | 2672.6 | 3617.8 | 7478.4 | 6761 |
| 11567 | ENSMUSG00000039521 | 1.661407539 | -2.014491774 | 2.044004678 | -0.985561235 | 0.324348469 | NA | Foxp3 | forkhead box P3 [Source:MGI Symbol;Acc | 4.0155 | 0 | 4.025 | 1.928 | 0 | 0 |
| 11568 | ENSMUSG00000039523 | 216.9421208 | -0.091070039 | 0.170550438 | -0.533977164 | 0.593357334 | 0.913011006 | Cep104 | centrosomal protein 104 [Source:MGI Syn | 231.29 | 200.44 | 239.49 | 216.9 | 195.22 | 218.31 |
| 11569 | ENSMUSG00000039529 | 654.4054383 | 0.195449648 | 0.137074994 | 1.425859243 | 0.153908976 | 0.571097853 | Atp8b1 | ATPase, class I, type 8B, member 1 [Sourc | 546.11 | 622.92 | 662.11 | 687.32 | 623.83 | 784.15 |
| 11570 | ENSMUSG00000039530 | 124.3620409 | 0.10545567 | 0.219367281 | 0.480726522 | 0.630710879 | 0.921020101 | NA | NA | 128.5 | 101.76 | 128.8 | 123.39 | 116.7 | 147.03 |
| 11571 | ENSMUSG00000039531 | 93.64425592 | -0.080978668 | 0.252899614 | -0.320200838 | 0.748816088 | 0.951074124 | Zup1 | zinc finger containing ubiquitin peptidase | 81.916 | 90.457 | 116.72 | 86.758 | 104.7 | 81.311 |
| 11572 | ENSMUSG00000039533 | 54.51251932 | -0.444584256 | 0.315557748 | -1.40888398 | 0.158869477 | 0.580099196 | Mmd2 | monocyte to macrophage differentiation- | 73.082 | 63.731 | 51.318 | 38.559 | 45.806 | 54.578 |
| 11573 | ENSMUSG00000039536 | 1360.835751 | -0.051449921 | 0.103480725 | -0.497193281 | 0.619052759 | 0.917509899 | Stau1 | staufen double-stranded RNA binding pro | 1388.6 | 1465.8 | 1301.1 | 1360.2 | 1276 | 1373.4 |
| 11574 | ENSMUSG00000039539 | 0 | NA | NA | NA | NA | NA | Sgcz | sarcoglycan zeta [Source:MGI Symbol;Acc | 0 | 0 | 0 | 0 | 0 | 0 |
| 11575 | ENSMUSG00000039540 | 0 | NA | NA | NA | NA | NA | 4921524L21I | RIKEN cDNA 4921524L21 gene [Source:MG | 0 | 0 | 0 | 0 | 0 | 0 |
| 11576 | ENSMUSG00000039542 | 2.04752066 | 1.223483741 | 1.515685623 | 0.807214717 | 0.419542788 | NA | Ncam1 | neural cell adhesion molecule 1 [Source:M | 1.6062 | 1.0279 | 1.0062 | 0.964 | 5.4531 | 2.2277 |
| 11577 | ENSMUSG00000039543 | 1.076225146 | -0.183775317 | 2.150113873 | -0.085472365 | 0.931885865 | NA | Cfap70 | cilia and flagella associated protein 70 [So | 2.4093 | 0 | 1.0062 | 1.928 | 0 | 1.1138 |
| 11578 | ENSMUSG00000039545 | 0.699599566 | -1.425530384 | 3.053137229 | -0.466906751 | 0.640566595 | NA | Flicr | Foxp3 regulating long intergenic noncodir | 0 | 3.0838 | 0 | 0 | 0 | 1.1138 |
| 11579 | ENSMUSG00000039546 | 0.435408177 | -2.179616019 | 3.531869654 | -0.617128103 | 0.537150234 | NA | Ajap1 | adherens junction associated protein 1 [S | 1.6062 | 0 | 1.0062 | 0 | 0 | 0 |
| 11580 | ENSMUSG00000039552 | 7.229070998 | -1.849171254 | 0.920763775 | -2.008301482 | 0.044611264 | 0.291866583 | Rsph4a | radial spoke head 4 homolog A (Chlamydo | 10.44 | 19.53 | 4.025 | 3.8559 | 2.1812 | 3.3415 |
| 11581 | ENSMUSG00000039555 | 0 | NA | NA | NA | NA | NA | Cylc2 | cylicin, basic protein of sperm head cytosk | 0 | 0 | 0 | 0 | 0 | 0 |
| 11582 | ENSMUSG00000039556 | 7.847134484 | 0.308890486 | 0.973363331 | 0.317343459 | 0.750983005 | 0.952074518 | Ppp1r3f | protein phosphatase 1, regulatory subunit | 0.8031 | 14.391 | 6.0375 | 12.532 | 2.1812 | 11.138 |
| 11583 | ENSMUSG00000039563 | 16.9482218 | -0.228926889 | 0.541354291 | -0.422878127 | 0.672384182 | 0.931191575 | 2210406O10 | RIKEN cDNA 2210406O10 gene [Source:M | 23.29 | 15.419 | 16.1 | 18.316 | 18.541 | 10.025 |
| 11584 | ENSMUSG00000039568 | 389.7213316 | -0.06400579 | 0.134740937 | -0.475028537 | 0.634766633 | 0.923478764 | Ubald1 | UBA-like domain containing 1 [Source:MG | 397.54 | 394.72 | 402.5 | 358.6 | 379.54 | 405.44 |
| 11585 | ENSMUSG00000039577 | 0.811697924 | -0.441734208 | 2.416070595 | -0.182831664 | 0.854930103 | NA | Nphp4 | nephronophthisis 4 (juvenile) homolog (h | 0.8031 | 0 | 2.0125 | 0.964 | 1.0906 | 0 |
| 11586 | ENSMUSG00000039578 | 0.808472266 | -1.769372749 | 2.633653003 | -0.671832146 | 0.501690561 | NA | Ccser1 | coiled-coil serine rich 1 [Source:MGI Symb | 0.8031 | 3.0838 | 0 | 0.964 | 0 | 0 |
| 11587 | ENSMUSG00000039579 | 0.342432689 | 1.990653086 | 3.900281263 | 0.51038706 | 0.609780321 | NA | Grin3a | glutamate receptor ionotropic, NMDA3A [ | 0 | 0 | 0 | 0.964 | 1.0906 | 0 |
| 11588 | ENSMUSG00000039585 | 67.35482467 | -1.019257979 | 0.309291317 | -3.295462641 | 0.000982598 | 0.020687471 | Myo9a | myosin IXa [Source:MGI Symbol;Acc:MGI: | 79.507 | 115.13 | 76.475 | 55.911 | 40.353 | 36.757 |

| | | | | | | | | | | | | | | | |
|---|---|---|---|---|---|---|---|---|---|---|---|---|---|---|---|
| 11589 | ENSMUSG00000039599 | 469.9499513 | 0.196128442 | 0.130122698 | 1.507257726 | 0.131744609 | 0.53212902 | Fam149b | family with sequence similarity 149, meml | 439.3 | 419.39 | 455.83 | 538.86 | 494.05 | 472.27 |
| 11590 | ENSMUSG00000039601 | 169.2903736 | -1.351639485 | 0.265884556 | -5.083557713 | 3.70E-07 | 3.35E-05 | Rcan2 | regulator of calcineurin 2 [Source:MGI Syr | 257.8 | 272.4 | 200.24 | 128.21 | 102.52 | 54.578 |
| 11591 | ENSMUSG00000039607 | 43.11247312 | -0.581673325 | 0.33596829 | -1.731334005 | 0.08339221 | 0.425868616 | Rbms3 | RNA binding motif, single stranded interac | 48.186 | 56.535 | 50.312 | 30.847 | 33.809 | 38.985 |
| 11592 | ENSMUSG00000039608 | 0 | NA | NA | NA | NA | NA | Olfr303 | olfactory receptor 303 [Source:MGI Symb | 0 | 0 | 0 | 0 | 0 | 0 |
| 11593 | ENSMUSG00000039611 | 6.21978574 | -0.306164276 | 0.849320552 | -0.360481417 | 0.718487151 | 0.9447719 | Pgap4 | post-GPI attachment to proteins GalNAc t | 6.4248 | 8.2233 | 6.0375 | 6.7478 | 6.5437 | 3.3415 |
| 11594 | ENSMUSG00000039615 | 470.7088125 | 0.081174963 | 0.1313415 | 0.618045047 | 0.536545645 | 0.890560262 | Stub1 | STIP1 homology and U-Box containing prc | 415.2 | 474.9 | 483 | 472.35 | 483.14 | 495.66 |
| 11595 | ENSMUSG00000039616 | 873.4811398 | 0.050961065 | 0.116031366 | 0.439200766 | 0.66051607 | 0.928166095 | Mocos | molybdenum cofactor sulfurase [Source:M | 785.43 | 878.87 | 910.65 | 876.26 | 849.59 | 940.08 |
| 11596 | ENSMUSG00000039617 | 0.767390791 | -1.69152783 | 2.44773922 | -0.691057207 | 0.489529591 | NA | Gm7488 | predicted gene 7488 [Source:MGI Symbol | 1.6062 | 1.0279 | 1.0062 | 0.964 | 0 | 0 |
| 11597 | ENSMUSG00000039620 | 90.17169237 | -0.299835885 | 0.370030867 | -0.810299659 | 0.417767971 | 0.837344173 | Trmt9b | tRNA methyltransferase 9B [Source:MGI S | 64.248 | 86.345 | 147.92 | 50.127 | 66.528 | 125.86 |
| 11598 | ENSMUSG00000039621 | 225.3187599 | 0.136243383 | 0.213200369 | 0.63903915 | 0.522797464 | 0.888334265 | Prex1 | phosphatidylinositol-3,4,5-trisphosphate- | 228.08 | 225.11 | 190.18 | 172.55 | 303.19 | 232.79 |
| 11599 | ENSMUSG00000039623 | 268.3679676 | 0.350087464 | 0.16665913 | 2.100619773 | 0.035674357 | 0.253684316 | Ap5z1 | adaptor-related protein complex 5, zeta 1 | 268.24 | 208.67 | 230.43 | 328.72 | 289.01 | 285.14 |
| 11600 | ENSMUSG00000039628 | 18.741288 | 0.988405542 | 0.521211701 | 1.896360997 | 0.057912326 | 0.342174787 | Hs3st6 | heparan sulfate (glucosamine) 3-O-sulfotr | 15.259 | 9.2513 | 13.081 | 31.811 | 18.541 | 24.505 |
| 11601 | ENSMUSG00000039629 | 2.323712285 | -1.134178715 | 1.679488462 | -0.675312001 | 0.499477561 | NA | Strip2 | striatin interacting protein 2 [Source:MGI | 6.4248 | 2.0558 | 1.0062 | 0 | 0 | 4.4554 |
| 11602 | ENSMUSG00000039630 | 4009.552144 | 0.020454787 | 0.092422376 | 0.221318564 | 0.824844396 | 0.96943211 | Hnrnpu | heterogeneous nuclear ribonucleoprotein | 3824.4 | 3950.3 | 4169.9 | 4195.2 | 4116 | 3801.5 |
| 11603 | ENSMUSG00000039632 | 118.7895475 | -0.032758928 | 0.226050971 | -0.144918325 | 0.884775358 | 0.981795936 | Odad3 | outer dynein arm docking complex subuni | 98.782 | 131.57 | 130.81 | 131.1 | 107.97 | 112.5 |
| 11604 | ENSMUSG00000039633 | 173.5443594 | 0.730010072 | 0.224494803 | 3.251790513 | 0.001146805 | 0.023102213 | Lonrf1 | LON peptidase N-terminal domain and rin | 136.53 | 136.71 | 118.74 | 280.52 | 205.04 | 163.74 |
| 11605 | ENSMUSG00000039634 | 114.0575346 | 0.0098527 | 0.218615339 | 0.045068656 | 0.964052585 | 0.993245949 | Zfp189 | zinc finger protein 189 [Source:MGI Symb | 118.86 | 114.1 | 107.67 | 100.25 | 126.51 | 116.95 |
| 11606 | ENSMUSG00000039637 | 358.2075546 | -0.418929103 | 0.145205067 | -2.88508598 | 0.003913068 | 0.056472904 | Coro7 | coronin 7 [Source:MGI Symbol;Acc:MGI:1 | 384.69 | 401.92 | 443.75 | 332.57 | 293.38 | 292.94 |
| 11607 | ENSMUSG00000039639 | 0 | NA | NA | NA | NA | NA | Kcne1 | potassium voltage-gated channel, Isk-rela | 0 | 0 | 0 | 0 | 0 | 0 |
| 11608 | ENSMUSG00000039640 | 1399.113115 | 0.053607349 | 0.105580217 | 0.507740467 | 0.611635365 | 0.916376001 | Mrpl12 | mitochondrial ribosomal protein L12 [Sou | 1354.8 | 1425.7 | 1338.3 | 1303.3 | 1480 | 1492.6 |
| 11609 | ENSMUSG00000039646 | 112.8001482 | -0.75290968 | 0.25547646 | -2.947080446 | 0.003207897 | 0.048892548 | Vasn | vasorin [Source:MGI Symbol;Acc:MGI:217 | 117.25 | 134.66 | 173.07 | 78.082 | 105.79 | 67.944 |
| 11610 | ENSMUSG00000039648 | 2030.191592 | 0.606413741 | 0.153358341 | 3.954227302 | 7.68E-05 | 0.002930291 | Kyat1 | kynurenine aminotransferase 1 [Source:M | 1765.2 | 1192.4 | 1870.6 | 2518.9 | 2249.9 | 2584.1 |
| 11611 | ENSMUSG00000039652 | 240.6017956 | 0.008368464 | 0.161833954 | 0.051710187 | 0.95875962 | 0.992027194 | Cpeb3 | cytoplasmic polyadenylation element binc | 233.7 | 252.87 | 233.45 | 246.78 | 257.39 | 219.43 |
| 11612 | ENSMUSG00000039653 | 5489.704221 | -0.123378747 | 0.104392701 | -1.181871394 | 0.237256734 | 0.686182309 | Baat | bile acid-Coenzyme A: amino acid N-acyltr | 5057.1 | 5948.6 | 6168.3 | 5001.1 | 5436.7 | 5326.4 |
| 11613 | ENSMUSG00000039656 | 719.5441779 | 0.114685555 | 0.11883396 | 0.965090747 | 0.334499361 | 0.776159726 | Rxrb | retinoid X receptor beta [Source:MGI Sym | 712.35 | 675.34 | 684.25 | 686.35 | 751.44 | 807.54 |
| 11614 | ENSMUSG00000039660 | 308.3014367 | 0.08627484 | 0.172271433 | 0.500807585 | 0.616506547 | 0.917509899 | Spout1 | SPOUT domain containing methyltransfera | 317.23 | 274.45 | 305.9 | 385.59 | 275.93 | 290.71 |
| 11615 | ENSMUSG00000039661 | 0.185640647 | 1.126924175 | 4.080472857 | 0.276174898 | 0.782413736 | NA | Dusp26 | dual specificity phosphatase 26 (putative) | 0 | 0 | 0 | 0 | 0 | 1.1138 |
| 11616 | ENSMUSG00000039662 | 836.0918264 | -0.036365737 | 0.122510862 | -0.296836841 | 0.766591078 | 0.955283307 | Icmt | isoprenylcysteine carboxyl methyltransfer | 811.13 | 871.67 | 858.33 | 931.2 | 764.52 | 779.69 |
| 11617 | ENSMUSG00000039670 | 323.1018025 | 0.072697628 | 0.161500784 | 0.450137929 | 0.65261099 | 0.925756399 | Oxld1 | oxidoreductase like domain containing 1 [ | 359.79 | 297.07 | 286.78 | 293.05 | 356.63 | 345.29 |
| 11618 | ENSMUSG00000039671 | 475.8734832 | -0.04643427 | 0.139587891 | -0.332652563 | 0.739396561 | 0.948310621 | Zmynd8 | zinc finger, MYND-type containing 8 [Sour | 507.56 | 445.09 | 497.08 | 434.75 | 442.79 | 527.96 |
| 11619 | ENSMUSG00000039672 | 0.562214094 | -1.122836648 | 3.276281221 | -0.34271681 | 0.731811515 | NA | Kcne2 | potassium voltage-gated channel, Isk-rela | 2.4093 | 0 | 0 | 0.964 | 0 | 0 |
| 11620 | ENSMUSG00000039676 | 0.895690289 | 0.713065541 | 2.535541507 | 0.281228108 | 0.778535447 | NA | Capsl | calcyphosine-like [Source:MGI Symbol;Ac | 0 | 2.0558 | 0 | 0 | 1.0906 | 2.2277 |
| 11621 | ENSMUSG00000039678 | 737.6468171 | 0.160581528 | 0.123380509 | 1.301514554 | 0.193082386 | 0.631271165 | Tbc1d13 | TBC1 domain family, member 13 [Source:I | 709.94 | 644.5 | 735.56 | 833.84 | 705.63 | 796.4 |
| 11622 | ENSMUSG00000039680 | 108.0745834 | -0.273666344 | 0.235005372 | -1.164511017 | 0.244216981 | 0.694115626 | Mrps6 | mitochondrial ribosomal protein S6 [Sourc | 101.19 | 111.02 | 142.89 | 90.614 | 103.61 | 99.132 |
| 11623 | ENSMUSG00000039682 | 6406.694963 | 0.163250514 | 0.099495049 | 1.640790335 | 0.100840945 | 0.46735231 | Lap3 | leucine aminopeptidase 3 [Source:MGI Sy | 6212.8 | 5806.7 | 6112.9 | 6183.9 | 6682.2 | 7441.6 |
| 11624 | ENSMUSG00000039683 | 1.600840731 | -1.845954437 | 1.813816175 | -1.017718589 | 0.308811711 | NA | Sdk1 | sidekick cell adhesion molecule 1 [Source: | 2.4093 | 4.1117 | 1.0062 | 0.964 | 0 | 1.1138 |
| 11625 | ENSMUSG00000039684 | 0.643990493 | 0.179747576 | 2.633236711 | 0.068261078 | 0.945577807 | NA | Gm5422 | predicted pseudogene 5422 [Source:MGI | 0.8031 | 0 | 1.0062 | 0.964 | 1.0906 | 0 |
| 11626 | ENSMUSG00000039686 | 479.0345304 | 0.09919448 | 0.125318982 | 0.791535961 | 0.4286313 | 0.842753542 | Zer1 | zyg-11 related, cell cycle regulator [Source | 444.12 | 481.07 | 462.87 | 490.66 | 484.23 | 511.25 |
| 11627 | ENSMUSG00000039691 | 0 | NA | NA | NA | NA | NA | Tspan10 | tetraspanin 10 [Source:MGI Symbol;Acc:M | 0 | 0 | 0 | 0 | 0 | 0 |
| 11628 | ENSMUSG00000039693 | 3.472814985 | 0.028643179 | 1.144200776 | 0.02503335 | 0.980028362 | NA | Msantd3 | Myb/SANT-like DNA-binding domain cont | 3.2124 | 3.0838 | 4.025 | 3.8559 | 1.0906 | 5.5692 |
| 11629 | ENSMUSG00000039697 | 14.37453071 | -0.768124564 | 0.578422263 | -1.327965075 | 0.184189645 | 0.619137675 | Ncoa7 | nuclear receptor coactivator 7 [Source:MC | 17.668 | 16.447 | 20.125 | 7.7118 | 9.8156 | 14.48 |
| 11630 | ENSMUSG00000039699 | 4.958876078 | -0.250944375 | 0.945154129 | -0.265506299 | 0.790619458 | NA | Batf2 | basic leucine zipper transcription factor, A | 4.0155 | 7.1954 | 5.0312 | 5.7839 | 3.2719 | 4.4554 |
| 11631 | ENSMUSG00000039701 | 157.1661232 | -0.183598195 | 0.200418458 | -0.916074284 | 0.359627935 | 0.799452417 | Usp53 | ubiquitin specific peptidase 53 [Source:M | 146.97 | 182.97 | 172.07 | 155.2 | 126.51 | 159.28 |
| 11632 | ENSMUSG00000039703 | 1340.732159 | 0.147745207 | 0.101552788 | 1.454861159 | 0.145707703 | 0.556690797 | Nploc4 | NPL4 homolog, ubiquitin recognition facto | 1292.2 | 1294.1 | 1229.6 | 1382.3 | 1350.2 | 1495.9 |
| 11633 | ENSMUSG00000039704 | 1319.715291 | 0.058193842 | 0.110605141 | 0.526140482 | 0.598790603 | 0.915055389 | Lmbrd2 | LMBR1 domain containing 2 [Source:MGI | 1174.9 | 1422.6 | 1283 | 1343.8 | 1379.6 | 1314.3 |
| 11634 | ENSMUSG00000039706 | 33.58893223 | -1.07419702 | 0.408275304 | -2.631060484 | 0.008511889 | 0.096334864 | Ldb2 | LIM domain binding 2 [Source:MGI Symbo | 32.927 | 51.396 | 52.325 | 16.388 | 23.994 | 24.505 |
| 11635 | ENSMUSG00000039710 | 0 | NA | NA | NA | NA | NA | Slc7a12 | solute carrier family 7 (cationic amino acic | 0 | 0 | 0 | 0 | 0 | 0 |
| 11636 | ENSMUSG00000039713 | 168.0581027 | 0.41097515 | 0.203659548 | 2.01795179 | 0.043596283 | 0.288197729 | Plekhg5 | pleckstrin homology domain containing, fa | 153.39 | 126.43 | 152.95 | 216.9 | 155.96 | 202.72 |
| 11637 | ENSMUSG00000039714 | 0.321325949 | 1.911575114 | 4.048267893 | 0.472195804 | 0.636787034 | NA | Cplx3 | complexin 3 [Source:MGI Symbol;Acc:MG | 0 | 0 | 0 | 1.928 | 0 | 0 |
| 11638 | ENSMUSG00000039715 | 120.50141 | 0.130158088 | 0.215779644 | 0.603199105 | 0.546376243 | 0.89549457 | Dync2i2 | dynein 2 intermediate chain 2 [Source:MC | 104.4 | 127.46 | 113.71 | 125.32 | 138.51 | 113.61 |
| 11639 | ENSMUSG00000039716 | 0.428363721 | -0.543730382 | 3.624677344 | -0.15000794 | 0.880758351 | NA | Dock3 | dedicator of cyto-kinesis 3 [Source:MGI Sy | 1.6062 | 0 | 0 | 0.964 | 0 | 0 |
| 11640 | ENSMUSG00000039717 | 0 | NA | NA | NA | NA | NA | Ralyl | RALY RNA binding protein-like [Source:MC | 0 | 0 | 0 | 0 | 0 | 0 |
| 11641 | ENSMUSG00000039720 | 0.832804665 | -0.381337792 | 2.590357773 | -0.147214333 | 0.882962848 | NA | Got1l1 | glutamic-oxaloacetic transaminase 1-like 1 | 0.8031 | 0 | 2.0125 | 0 | 2.1812 | 0 |
| 11642 | ENSMUSG00000039725 | 157.0146944 | -0.069842745 | 0.203794129 | -0.342712253 | 0.731814944 | 0.947387358 | Trp53rka | transformation related protein 53 regulati | 139.74 | 192.22 | 150.94 | 150.38 | 143.96 | 164.85 |
| 11643 | ENSMUSG00000039728 | 0 | NA | NA | NA | NA | NA | Slc6a5 | solute carrier family 6 (neurotransmitter t | 0 | 0 | 0 | 0 | 0 | 0 |
| 11644 | ENSMUSG00000039735 | 374.2640963 | 0.185496198 | 0.148191336 | 1.251734428 | 0.21066665 | 0.654864052 | Fnbp1l | formin binding protein 1-like [Source:MGI | 320.44 | 334.07 | 396.46 | 370.17 | 408.98 | 415.46 |

| | | | | | | | | | | | | | | | |
|---|---|---|---|---|---|---|---|---|---|---|---|---|---|---|---|
| 11645 | ENSMUSG00000039737 | 132.6447914 | 0.095427907 | 0.227681191 | 0.419129515 | 0.675121481 | 0.932446243 | Prkrip1 | Prkr interacting protein 1 (IL11 inducible) | 143.76 | 141.85 | 98.612 | 120.5 | 136.33 | 154.82 |
| 11646 | ENSMUSG00000039738 | 51.21835087 | -0.139229913 | 0.347551748 | -0.400601964 | 0.6887132 | 0.936908509 | Slx4 | SLX4 structure-specific endonuclease subu | 42.564 | 56.535 | 62.387 | 60.731 | 31.628 | 53.465 |
| 11647 | ENSMUSG00000039740 | 219.0266548 | -0.371659649 | 0.192053424 | -1.935188869 | 0.052967123 | 0.323119557 | Alg2 | asparagine-linked glycosylation 2 (alpha-1 | 240.93 | 295.01 | 205.27 | 175.44 | 215.94 | 181.56 |
| 11648 | ENSMUSG00000039741 | 260.3723456 | 0.242320596 | 0.206006793 | 1.176274781 | 0.23948509 | 0.688924491 | Bahcc1 | BAH domain and coiled-coil containing 1 [ | 228.88 | 182.97 | 303.89 | 290.16 | 236.66 | 319.67 |
| 11649 | ENSMUSG00000039742 | 0 | NA | NA | NA | NA | NA | Fam71f1 | family with sequence similarity 71, memb | 0 | 0 | 0 | 0 | 0 | 0 |
| 11650 | ENSMUSG00000039745 | 448.3568491 | -0.156560283 | 0.158662303 | -0.986751612 | 0.323764421 | 0.770868138 | Htatip2 | HIV-1 Tat interactive protein 2 [Source:M( | 551.73 | 466.67 | 398.47 | 411.62 | 388.26 | 473.38 |
| 11651 | ENSMUSG00000039747 | 5.023074556 | -0.699001945 | 0.97799982 | -0.714726047 | 0.47477834 | NA | Orai2 | ORAI calcium release-activated calcium m | 10.44 | 3.0838 | 5.0312 | 3.8559 | 3.2719 | 4.4554 |
| 11652 | ENSMUSG00000039748 | 2.554538861 | -0.491798685 | 1.356074862 | -0.362663374 | 0.716856363 | NA | Exo1 | exonuclease 1 [Source:MGI Symbol;Acc:M | 4.8186 | 2.0558 | 2.0125 | 0.964 | 4.3625 | 1.1138 |
| 11653 | ENSMUSG00000039753 | 512.7521525 | -0.206885018 | 0.127220592 | -1.62619129 | 0.103908969 | 0.474113351 | Fbxl5 | F-box and leucine-rich repeat protein 5 [S | 559.76 | 576.66 | 512.18 | 503.2 | 460.24 | 464.47 |
| 11654 | ENSMUSG00000039754 | 61.4556302 | -0.155643769 | 0.299864877 | -0.519046346 | 0.603728422 | 0.916106578 | Alkbh4 | alkB homolog 4, lysine demethylase [Sour | 60.233 | 76.066 | 58.362 | 70.37 | 46.897 | 56.806 |
| 11655 | ENSMUSG00000039756 | 393.3549677 | -0.061676405 | 0.134370133 | -0.459003829 | 0.646231416 | 0.925643431 | Dnttip2 | deoxynucleotidyltransferase, terminal, int | 405.57 | 422.47 | 377.34 | 393.3 | 386.08 | 375.37 |
| 11656 | ENSMUSG00000039759 | 140.1196905 | -0.109865216 | 0.20030721 | -0.548483582 | 0.583359899 | 0.908645447 | Thap3 | THAP domain containing, apoptosis assoc | 151.79 | 127.46 | 156.97 | 137.85 | 136.33 | 130.32 |
| 11657 | ENSMUSG00000039760 | 0 | NA | NA | NA | NA | NA | Il22ra2 | interleukin 22 receptor, alpha 2 [Source:M | 0 | 0 | 0 | 0 | 0 | 0 |
| 11658 | ENSMUSG00000039763 | 380.2453415 | 0.134560901 | 0.149903938 | 0.897647534 | 0.369373489 | 0.80653433 | Dnajc28 | DnaJ heat shock protein family (Hsp40) m | 355.77 | 328.93 | 402.5 | 359.56 | 398.08 | 436.63 |
| 11659 | ENSMUSG00000039765 | 76.53527611 | -0.225015573 | 0.275665023 | -0.816264503 | 0.414348872 | 0.835831451 | Cc2d2a | coiled-coil and C2 domain containing 2A [ | 70.673 | 94.568 | 82.512 | 72.298 | 55.622 | 83.538 |
| 11660 | ENSMUSG00000039768 | 1089.423959 | 0.057707146 | 0.113620328 | 0.507894553 | 0.611527295 | 0.916376001 | Dnajc11 | DnaJ heat shock protein family (Hsp40) m | 1095.4 | 1154.4 | 952.91 | 1104.7 | 1135.3 | 1093.8 |
| 11661 | ENSMUSG00000039770 | 1039.321686 | 0.255728505 | 0.120043064 | 2.130306384 | 0.033146327 | 0.243735878 | Ypel5 | yippee like 5 [Source:MGI Symbol;Acc:MG | 1077 | 857.28 | 906.63 | 1149.1 | 1127.7 | 1118.3 |
| 11662 | ENSMUSG00000039771 | 260.1086047 | 0.187105067 | 0.159716028 | 1.171485846 | 0.241403544 | 0.691388521 | Polr2j | polymerase (RNA) II (DNA directed) polyp | 231.29 | 261.09 | 237.47 | 253.53 | 299.92 | 277.35 |
| 11663 | ENSMUSG00000039774 | 0.94782191 | 1.090542322 | 2.508664729 | 0.43471027 | 0.663772743 | NA | Galnt12 | polypeptide N-acetylgalactosaminyltransf | 0.8031 | 1.0279 | 0 | 3.8559 | 0 | 0 |
| 11664 | ENSMUSG00000039775 | 0 | NA | NA | NA | NA | NA | Defb3 | defensin beta 3 [Source:MGI Symbol;Acc: | 0 | 0 | 0 | 0 | 0 | 0 |
| 11665 | ENSMUSG00000039781 | 93.98039912 | 0.26116305 | 0.253906597 | 1.028579222 | 0.303677446 | 0.75273701 | Cep131 | centrosomal protein 131 [Source:MGI Syn | 77.098 | 89.429 | 90.562 | 128.21 | 87.249 | 91.335 |
| 11666 | ENSMUSG00000039782 | 1323.703404 | -1.027706018 | 0.170153028 | -6.039892628 | 1.54E-09 | 3.06E-07 | Cpeb2 | cytoplasmic polyadenylation element binc | 1830.3 | 1996.2 | 1501.3 | 696.96 | 1125.5 | 791.94 |
| 11667 | ENSMUSG00000039783 | 7727.201137 | 0.078212445 | 0.096067383 | 0.814141524 | 0.41556388 | 0.83614452 | Kmo | kynurenine 3-monooxygenase (kynurenin | 7482.5 | 7282.8 | 7787.3 | 7270.3 | 8593 | 7947.3 |
| 11668 | ENSMUSG00000039785 | 0 | NA | NA | NA | NA | NA | Defb5 | defensin beta 5 [Source:MGI Symbol;Acc: | 0 | 0 | 0 | 0 | 0 | 0 |
| 11669 | ENSMUSG00000039787 | 4.021322629 | -0.373161532 | 1.061610325 | -0.351505184 | 0.725209384 | NA | Cercam | cerebral endothelial cell adhesion molecu | 6.4248 | 4.1117 | 3.0187 | 2.8919 | 5.4531 | 2.2277 |
| 11670 | ENSMUSG00000039789 | 96.60130441 | 0.108581482 | 0.240796466 | 0.450926393 | 0.6520426 | 0.925643431 | Zfp597 | zinc finger protein 597 [Source:MGI Symb | 81.113 | 93.541 | 104.65 | 102.18 | 111.24 | 86.88 |
| 11671 | ENSMUSG00000039795 | 457.8725885 | 0.121471774 | 0.140315988 | 0.865701595 | 0.386653825 | 0.817267416 | Zfand1 | zinc finger, AN1-type domain 1 [Source:M | 443.31 | 448.17 | 423.63 | 411.62 | 512.59 | 507.91 |
| 11672 | ENSMUSG00000039798 | 0 | NA | NA | NA | NA | NA | 2600006K01 | RIKEN cDNA 2600006K01 gene [Source:M | 0 | 0 | 0 | 0 | 0 | 0 |
| 11673 | ENSMUSG00000039801 | 95.64978517 | 0.293785939 | 0.252967426 | 1.161358768 | 0.245496029 | 0.695774774 | Cplane1 | ciliogenesis and planar polarity effector 1 | 94.766 | 75.038 | 87.543 | 93.506 | 91.612 | 131.43 |
| 11674 | ENSMUSG00000039804 | 230.8029157 | -0.284090191 | 0.169411591 | -1.676922988 | 0.093557542 | 0.450459278 | Ncoa5 | nuclear receptor coactivator 5 [Source:M( | 220.85 | 276.51 | 263.64 | 208.22 | 202.86 | 212.74 |
| 11675 | ENSMUSG00000039809 | 541.6465713 | 0.189926737 | 0.152741947 | 1.243448447 | 0.213702627 | 0.658358849 | Gabbr2 | gamma-aminobutyric acid (GABA) B recep | 538.88 | 475.93 | 502.12 | 463.67 | 651.1 | 618.18 |
| 11676 | ENSMUSG00000039810 | 105.3972922 | -0.133238892 | 0.24362598 | -0.546899357 | 0.584447878 | 0.908955096 | Zc3h10 | zinc finger CCCH type containing 10 [Sour | 100.39 | 122.32 | 108.67 | 123.39 | 80.706 | 96.904 |
| 11677 | ENSMUSG00000039813 | 23.20091588 | 0.386294745 | 0.483132093 | 0.799563413 | 0.423963792 | 0.840450466 | Tbc1d2 | TBC1 domain family, member 2 [Source:M | 18.471 | 27.754 | 14.087 | 19.28 | 25.084 | 34.529 |
| 11678 | ENSMUSG00000039814 | 0.804680097 | -3.048447205 | 2.843448763 | -1.072095001 | 0.283677364 | NA | Xkr5 | X-linked Kx blood group related 5 [Source | 0.8031 | 0 | 4.025 | 0 | 0 | 0 |
| 11679 | ENSMUSG00000039824 | 28.85212076 | -0.01426303 | 0.424077796 | -0.033633051 | 0.973169766 | 0.995820834 | Myl6b | myosin, light polypeptide 6B [Source:MGI | 27.305 | 25.698 | 34.212 | 38.559 | 26.175 | 21.163 |
| 11680 | ENSMUSG00000039826 | 218.8970286 | -0.273674684 | 0.168978153 | -1.619586197 | 0.105321197 | 0.478186913 | Trub2 | TruB pseudouridine (psi) synthase family | 224.87 | 246.7 | 247.54 | 195.69 | 217.03 | 181.56 |
| 11681 | ENSMUSG00000039828 | 163.1495752 | -0.067078583 | 0.185335603 | -0.361930366 | 0.717404067 | 0.9447719 | Wdr70 | WD repeat domain 70 [Source:MGI Symbo | 165.44 | 171.66 | 164.02 | 173.52 | 148.32 | 155.94 |
| 11682 | ENSMUSG00000039830 | 0 | NA | NA | NA | NA | NA | Olig2 | oligodendrocyte transcription factor 2 [So | 0 | 0 | 0 | 0 | 0 | 0 |
| 11683 | ENSMUSG00000039831 | 1128.88826 | 0.527053605 | 0.156829069 | 3.360688231 | 0.000777485 | 0.0173275 | Arhgap29 | Rho GTPase activating protein 29 [Source: | 869.76 | 823.36 | 1082.7 | 1624.3 | 1117.9 | 1255.3 |
| 11684 | ENSMUSG00000039834 | 154.6943884 | -0.019826284 | 0.209789359 | -0.094505671 | 0.924707478 | 0.987546253 | Zfp335 | zinc finger protein 335 [Source:MGI Symb | 144.56 | 165.49 | 157.98 | 187.01 | 146.14 | 126.98 |
| 11685 | ENSMUSG00000039835 | 135.712323 | 0.033838005 | 0.212779377 | 0.159028595 | 0.873646345 | 0.979925081 | Nhsl1 | NHS-like 1 [Source:MGI Symbol;Acc:MGI: | 137.33 | 108.96 | 155.97 | 139.78 | 135.24 | 137 |
| 11686 | ENSMUSG00000039838 | 0.167707431 | -0.796629176 | 4.080472857 | -0.195229623 | 0.84521319 | NA | Slc45a1 | solute carrier family 45, member 1 [Sourc | 0 | 0 | 1.0062 | 0 | 0 | 0 |
| 11687 | ENSMUSG00000039840 | 428.7520195 | -0.025823458 | 0.138172704 | -0.186892618 | 0.851744827 | 0.973633287 | Epg5 | ectopic P-granules autophagy protein 5 h | 415.2 | 425.56 | 456.84 | 383.66 | 462.42 | 428.83 |
| 11688 | ENSMUSG00000039841 | 182.9028937 | 0.048249422 | 0.18046765 | 0.267357736 | 0.789193737 | 0.960994657 | Zfp800 | zinc finger protein 800 [Source:MGI Symb | 182.3 | 171.66 | 185.15 | 164.84 | 196.31 | 197.15 |
| 11689 | ENSMUSG00000039842 | 48.80585513 | 0.030145171 | 0.368799434 | 0.081738659 | 0.934854536 | 0.988420305 | Mcph1 | microcephaly, primary autosomal recessiv | 57.02 | 49.34 | 38.237 | 39.523 | 73.071 | 35.643 |
| 11690 | ENSMUSG00000039844 | 1017.251289 | 0.055857376 | 0.103742321 | 0.538424196 | 0.590284226 | 0.911690353 | Rapgef1 | Rap guanine nucleotide exchange factor ( | 1028.8 | 1011.5 | 951.91 | 1020.9 | 1054.6 | 1035.9 |
| 11691 | ENSMUSG00000039849 | 471.9958058 | -0.082271779 | 0.128339872 | -0.641046131 | 0.521492712 | 0.888064228 | Pcif1 | phosphorylated CTD interacting factor 1 [ | 466.6 | 475.93 | 514.19 | 465.6 | 435.16 | 474.5 |
| 11692 | ENSMUSG00000039850 | 135.036835 | 0.151235994 | 0.207636394 | 0.728369395 | 0.466387493 | 0.863779479 | Endov | endonuclease V [Source:MGI Symbol;Acc: | 110.03 | 135.69 | 138.86 | 151.34 | 142.87 | 131.43 |
| 11693 | ENSMUSG00000039851 | 0.614004368 | 0.315622926 | 2.898299649 | 0.108899342 | 0.913282328 | NA | 4932438H23 | RIKEN cDNA 4932438H23 gene [Source:M | 1.6062 | 0 | 0 | 0.964 | 0 | 1.1138 |
| 11694 | ENSMUSG00000039852 | 658.5943164 | -0.054943676 | 0.129749175 | -0.423460698 | 0.671959167 | 0.931191575 | Rere | arginine glutamic acid dipeptide (RE) repe | 743.67 | 678.43 | 590.67 | 676.71 | 622.74 | 639.35 |
| 11695 | ENSMUSG00000039853 | 347.9801121 | 0.487944673 | 0.207834026 | 2.347761254 | 0.018886623 | 0.166685225 | Trim14 | tripartite motif-containing 14 [Source:MG | 252.98 | 215.86 | 400.49 | 376.92 | 438.43 | 403.21 |
| 11696 | ENSMUSG00000039860 | 0 | NA | NA | NA | NA | NA | Srrm3 | serine/arginine repetitive matrix 3 [Source | 0 | 0 | 0 | 0 | 0 | 0 |
| 11697 | ENSMUSG00000039865 | 113.3382006 | -0.193128998 | 0.251846786 | -0.766851152 | 0.443170019 | 0.850769625 | Slc44a3 | solute carrier family 44, member 3 [Sourc | 124.48 | 139.8 | 98.612 | 113.75 | 125.42 | 77.969 |
| 11698 | ENSMUSG00000039873 | 31.04413867 | 0.229019847 | 0.460401507 | 0.497435052 | 0.618882293 | 0.917509899 | Neurl2 | neuralized E3 ubiquitin protein ligase 2 [S | 33.73 | 28.782 | 23.144 | 38.559 | 15.269 | 46.781 |
| 11699 | ENSMUSG00000039878 | 31.13058997 | -0.576042107 | 0.461252541 | -1.248864896 | 0.211714493 | 0.656595245 | Slc39a5 | solute carrier family 39 (metal ion transpo | 35.336 | 56.535 | 20.125 | 31.811 | 21.812 | 21.163 |
| 11700 | ENSMUSG00000039879 | 909.0240081 | 0.105071458 | 0.114844287 | 0.914903657 | 0.36024221 | 0.799598236 | Heca | hdc homolog, cell cycle regulator [Source: | 805.51 | 956.99 | 866.38 | 953.37 | 911.76 | 960.13 |

| | | | | | | | | | | | | | | | |
|---|---|---|---|---|---|---|---|---|---|---|---|---|---|---|---|
| 11701 | ENSMUSG00000039883 | 1.019142721 | 1.225935691 | 2.256581771 | 0.543271113 | 0.58694315 | NA | Lrrc17 | leucine rich repeat containing 17 [Source: | 0.8031 | 0 | 1.0062 | 0.964 | 0 | 3.3415 |
| 11702 | ENSMUSG00000039886 | 1619.769949 | -0.106471447 | 0.12256926 | -0.868663539 | 0.38503119 | 0.816437825 | Tmem120a | transmembrane protein 120A [Source:MG | 1920.2 | 1628.2 | 1488.2 | 1436.3 | 1627.2 | 1618.4 |
| 11703 | ENSMUSG00000039887 | 223.3072616 | 0.050747754 | 0.175821256 | 0.288632644 | 0.772862512 | 0.957899175 | Alg14 | asparagine-linked glycosylation 14 [Source | 212.02 | 202.5 | 243.51 | 214 | 209.4 | 258.41 |
| 11704 | ENSMUSG00000039891 | 0.346303622 | 2.004339653 | 3.884003423 | 0.516049919 | 0.605819544 | NA | Txlnb | taxilin beta [Source:MGI Symbol;Acc:MGI: | 0 | 0 | 0 | 0.964 | 0 | 1.1138 |
| 11705 | ENSMUSG00000039899 | 56.08049932 | -0.694544492 | 0.37541081 | -1.850091881 | 0.064300308 | 0.36329674 | Fgl2 | fibrinogen-like protein 2 [Source:MGI Sym | 71.476 | 93.541 | 43.269 | 59.767 | 29.447 | 38.985 |
| 11706 | ENSMUSG00000039901 | 118.6172181 | 0.145190849 | 0.226658988 | 0.640569563 | 0.52180238 | 0.888132458 | Armh3 | armadillo-like helical domain containing 3 | 108.42 | 124.38 | 105.66 | 149.42 | 106.88 | 116.95 |
| 11707 | ENSMUSG00000039903 | 4.877074175 | -1.573183865 | 1.025240776 | -1.534453079 | 0.124918228 | NA | Eva1c | eva-1 homolog C (C. elegans) [Source:MG | 5.6217 | 10.279 | 6.0375 | 2.8919 | 1.0906 | 3.3415 |
| 11708 | ENSMUSG00000039904 | 0 | NA | NA | NA | NA | NA | Gpr37 | G protein-coupled receptor 37 [Source:M | 0 | 0 | 0 | 0 | 0 | 0 |
| 11709 | ENSMUSG00000039908 | 125.3292507 | 0.192311739 | 0.215021881 | 0.894382183 | 0.371117442 | 0.807214774 | Slc26a11 | solute carrier family 26, member 11 [Sour | 134.12 | 101.76 | 114.71 | 144.6 | 128.69 | 128.09 |
| 11710 | ENSMUSG00000039910 | 538.2401224 | 0.099812981 | 0.17703259 | 0.563811337 | 0.572882535 | 0.905173631 | Cited2 | Cbp/p300-interacting transactivator, with | 474.63 | 446.12 | 637.96 | 473.31 | 521.32 | 676.1 |
| 11711 | ENSMUSG00000039911 | 13.71432573 | -2.213013073 | 0.674621582 | -3.28037693 | 0.001036685 | 0.021482248 | Spsb1 | splA/ryanodine receptor domain and SOC | 16.062 | 22.614 | 29.181 | 6.7478 | 5.4531 | 2.2277 |
| 11712 | ENSMUSG00000039913 | 0 | NA | NA | NA | NA | NA | Pak5 | p21 (RAC1) activated kinase 5 [Source:MG | 0 | 0 | 0 | 0 | 0 | 0 |
| 11713 | ENSMUSG00000039914 | 549.720363 | 0.249304144 | 0.14300243 | 1.743355995 | 0.08127145 | 0.419375759 | Coq10a | coenzyme Q10A [Source:MGI Symbol;Acc | 568.6 | 516.01 | 421.62 | 624.66 | 569.3 | 598.13 |
| 11714 | ENSMUSG00000039917 | 480.1148391 | -0.342117274 | 0.155056645 | -2.20640189 | 0.027355873 | 0.214342007 | Rhbdd2 | rhomboid domain containing 2 [Source:M | 533.26 | 606.47 | 470.92 | 434.75 | 475.51 | 359.77 |
| 11715 | ENSMUSG00000039929 | 100.3185735 | -0.213815487 | 0.282681581 | -0.756382804 | 0.449419715 | 0.854879451 | Urb1 | URB1 ribosome biogenesis 1 homolog (S. | 119.66 | 123.35 | 80.5 | 125.32 | 81.796 | 71.286 |
| 11716 | ENSMUSG00000039934 | 472.9960112 | 0.506519053 | 0.14885566 | 3.402753073 | 0.000667105 | 0.01536467 | Gsap | gamma-secretase activating protein [Sour | 399.14 | 364.91 | 407.53 | 464.64 | 579.12 | 622.64 |
| 11717 | ENSMUSG00000039936 | 39.31210692 | -0.828087285 | 0.374354585 | -2.212039919 | 0.026963907 | 0.212769203 | Pik3cd | phosphatidylinositol-4,5-bisphosphate 3-k | 49.792 | 60.647 | 40.25 | 21.208 | 29.447 | 34.529 |
| 11718 | ENSMUSG00000039942 | 5.181707278 | -1.248653779 | 1.161688393 | -1.074861199 | 0.282436876 | 0.735403596 | Ptger4 | prostaglandin E receptor 4 (subtype EP4) | | 1.6062 | 14.391 | 6.0375 | 5.7839 | 3.2719 | 0 |
| 11719 | ENSMUSG00000039943 | 22.95993264 | -0.070204398 | 0.492909092 | -0.142428693 | 0.8867414 | 0.982382091 | Plcb4 | phospholipase C, beta 4 [Source:MGI Sym | 21.684 | 22.614 | 26.162 | 16.388 | 15.269 | 35.643 |
| 11720 | ENSMUSG00000039952 | 1550.9602 | -0.110360077 | 0.126772943 | -0.870533368 | 0.384008992 | 0.816413186 | Dag1 | dystroglycan 1 [Source:MGI Symbol;Acc:M | 1435.1 | 1884.2 | 1512.4 | 1444 | 1604.3 | 1425.7 |
| 11721 | ENSMUSG00000039953 | 54.83955178 | -0.600636633 | 0.327154734 | -1.835940522 | 0.066366454 | 0.370435957 | Clstn1 | calsyntenin 1 [Source:MGI Symbol;Acc:M | 61.839 | 66.815 | 69.431 | 30.847 | 58.893 | 41.212 |
| 11722 | ENSMUSG00000039954 | 0 | NA | NA | NA | NA | NA | Stk32a | serine/threonine kinase 32A [Source:MGI | 0 | 0 | 0 | 0 | 0 | 0 |
| 11723 | ENSMUSG00000039956 | 727.9388792 | -0.329264295 | 0.123661201 | -2.662632195 | 0.007753211 | 0.089713954 | Mrap | melanocortin 2 receptor accessory protei | 825.59 | 835.7 | 769.78 | 580.31 | 646.74 | 709.52 |
| 11724 | ENSMUSG00000039958 | 762.7274943 | 0.025932304 | 0.214751987 | 0.120754665 | 0.903885359 | 0.983821605 | Etfbkmt | electron transfer flavoprotein beta subuni | 683.44 | 889.15 | 696.32 | 1110.5 | 647.83 | 549.13 |
| 11725 | ENSMUSG00000039959 | 66.58719174 | -0.887421452 | 0.331809096 | -2.674494049 | 0.00748421 | 0.087866575 | Hip1 | huntingtin interacting protein 1 [Source:M | 73.082 | 106.9 | 79.493 | 48.199 | 28.356 | 63.489 |
| 11726 | ENSMUSG00000039960 | 1366.051905 | 0.316632351 | 0.139738806 | 2.265887047 | 0.0234583 | 0.1938862 | Rhou | ras homolog family member U [Source:M | 1177.3 | 1242.8 | 1229.6 | 1284 | 1406.9 | 1855.7 |
| 11727 | ENSMUSG00000039962 | 0 | NA | NA | NA | NA | NA | Olfr906 | olfactory receptor 906 [Source:MGI Symb | 0 | 0 | 0 | 0 | 0 | 0 |
| 11728 | ENSMUSG00000039963 | 2.42652383 | -0.759308984 | 1.513160495 | -0.501803336 | 0.615805867 | NA | Ccdc40 | coiled-coil domain containing 40 [Source: | 0 | 6.1675 | 3.0187 | 0.964 | 2.1812 | 2.2277 |
| 11729 | ENSMUSG00000039967 | 307.248839 | 0.17408751 | 0.173535023 | 1.003183722 | 0.315772226 | 0.762619715 | Zfp292 | zinc finger protein 292 [Source:MGI Symb | 273.05 | 305.29 | 288.79 | 388.48 | 325 | 262.87 |
| 11730 | ENSMUSG00000039968 | 209.7736618 | 0.043366354 | 0.16691618 | 0.259809168 | 0.795010979 | 0.962189806 | Rsbn1l | round spermatid basic protein 1-like [Sour | 212.82 | 204.56 | 202.26 | 211.11 | 200.67 | 227.22 |
| 11731 | ENSMUSG00000039976 | 692.4129214 | 0.114230623 | 0.121445314 | 0.940593084 | 0.346913427 | 0.788087924 | Tbc1d16 | TBC1 domain family, member 16 [Source: | 693.08 | 609.56 | 692.3 | 767.33 | 683.82 | 708.4 |
| 11732 | ENSMUSG00000039977 | 0.956643647 | -0.817327529 | 2.26666353 | -0.360586174 | 0.718408827 | NA | Deup1 | deuterosome assembly protein 1 [Source: | 1.6062 | 2.0558 | 0 | 0.964 | 0 | 1.1138 |
| 11733 | ENSMUSG00000039981 | 92.7907329 | -0.141800744 | 0.278467691 | -0.509217941 | 0.610599469 | 0.916376001 | Zc3h12d | zinc finger CCCH type containing 12D [Sou | 89.947 | 77.094 | 124.77 | 67.478 | 90.521 | 106.93 |
| 11734 | ENSMUSG00000039982 | 143.1869239 | -0.557979103 | 0.285367806 | -1.955298005 | 0.050547907 | 0.315957183 | Dtx4 | deltex 4, E3 ubiquitin ligase [Source:MGI S | 111.63 | 181.94 | 218.36 | 105.07 | 85.068 | 157.05 |
| 11735 | ENSMUSG00000039983 | 191.1767463 | -0.082098996 | 0.218066155 | -0.376486651 | 0.706555137 | 0.94295036 | Ccdc32 | coiled-coil domain containing 32 [Source: | 209.61 | 192.22 | 187.16 | 129.17 | 206.13 | 222.77 |
| 11736 | ENSMUSG00000039985 | 18.39537098 | -0.05930161 | 0.568214002 | -0.104364922 | 0.916879759 | 0.986374671 | Sinhcaf | SIN3-HDAC complex associated factor [So | 25.699 | 15.419 | 15.094 | 23.135 | 7.6343 | 23.391 |
| 11737 | ENSMUSG00000039987 | 145.9333781 | 0.349327149 | 0.196103872 | 1.781337332 | 0.074857359 | 0.399533891 | Phtf2 | putative homeodomain transcription fact | 126.09 | 125.41 | 133.83 | 176.41 | 167.96 | 145.91 |
| 11738 | ENSMUSG00000039988 | 604.5555603 | -0.030968651 | 0.140432713 | -0.220523055 | 0.825463819 | 0.96943211 | Ankrd13c | ankyrin repeat domain 13c [Source:MGI S | 610.36 | 549.94 | 672.17 | 533.08 | 584.57 | 677.22 |
| 11739 | ENSMUSG00000039989 | 292.1419423 | -0.03148783 | 0.20303875 | -0.155082862 | 0.876755992 | 0.980632845 | Cbx4 | chromobox 4 [Source:MGI Symbol;Acc:M | 263.42 | 260.06 | 362.25 | 220.75 | 297.74 | 348.63 |
| 11740 | ENSMUSG00000039990 | 394.0388432 | 0.09561659 | 0.139211533 | 0.686843886 | 0.492181111 | 0.876368691 | Edrf1 | erythroid differentiation regulatory factor | 347.74 | 382.39 | 413.57 | 409.69 | 419.89 | 390.96 |
| 11741 | ENSMUSG00000039994 | 130.5521361 | -0.079973215 | 0.201725399 | -0.39644594 | 0.691776081 | 0.938106998 | Timeless | timeless circadian clock 1 [Source:MGI Sy | 130.1 | 135.69 | 136.85 | 126.28 | 136.33 | 118.07 |
| 11742 | ENSMUSG00000039997 | 100.5451102 | -0.487333811 | 0.227650535 | -2.140710148 | 0.03229742 | 0.240179886 | Ifi203 | interferon activated gene 203 [Source:MG | 126.09 | 116.15 | 109.68 | 84.83 | 78.525 | 87.994 |
| 11743 | ENSMUSG00000040003 | 1.869963225 | 0.705654062 | 1.764931387 | 0.399819544 | 0.689289435 | NA | Magi2 | membrane associated guanylate kinase, W | 3.2124 | 0 | 1.0062 | 4.8199 | 2.1812 | 0 |
| 11744 | ENSMUSG00000040006 | 945.8719484 | -0.145661434 | 0.152158122 | -0.95730305 | 0.338414316 | 0.780780576 | Ginm1 | glycoprotein integral membrane 1 [Source | 1230.4 | 847 | 901.6 | 828.06 | 979.38 | 888.85 |
| 11745 | ENSMUSG00000040007 | 303.1595427 | 0.022671006 | 0.158217073 | 0.143290518 | 0.886060744 | 0.982222482 | Bahd1 | bromo adjacent homology domain contai | 306.79 | 267.26 | 328.04 | 304.62 | 278.11 | 334.15 |
| 11746 | ENSMUSG00000040009 | 0.48903338 | 0.95637552 | 3.456690196 | 0.276673773 | 0.782030612 | NA | Gnaz | guanine nucleotide binding protein, alpha | 0 | 0 | 1.0062 | 1.928 | 0 | 0 |
| 11747 | ENSMUSG00000040010 | 26.68666956 | -0.257451497 | 0.426665066 | -0.603404209 | 0.546239823 | 0.89549457 | Slc7a5 | solute carrier family 7 (cationic amino acid | 32.124 | 25.698 | 29.181 | 21.208 | 20.722 | 31.188 |
| 11748 | ENSMUSG00000040013 | 0 | NA | NA | NA | NA | NA | Fkbp6 | FK506 binding protein 6 [Source:MGI Sym | 0 | 0 | 0 | 0 | 0 | 0 |
| 11749 | ENSMUSG00000040016 | 0.589026695 | 0.241472922 | 3.228838524 | 0.074786311 | 0.940384733 | NA | Ptger3 | prostaglandin E receptor 3 (subtype EP3) | | 1.6062 | 0 | 0 | 1.928 | 0 | 0 |
| 11750 | ENSMUSG00000040017 | 9465.062929 | -0.198110306 | 0.150278783 | -1.318285272 | 0.187408169 | 0.623933049 | Saa4 | serum amyloid A 4 [Source:MGI Symbol;A | 8439 | 10921 | 10982 | 6825.9 | 10089 | 9533.4 |
| 11751 | ENSMUSG00000040018 | 566.6657116 | -0.213896978 | 0.119231678 | -1.793960981 | 0.072819397 | 0.391632053 | Cox15 | cytochrome c oxidase assembly protein 1! | 589.48 | 616.75 | 619.85 | 519.58 | 545.31 | 509.03 |
| 11752 | ENSMUSG00000040021 | 575.9246718 | 0.052533673 | 0.123797374 | 0.424352083 | 0.671309061 | 0.931191575 | Lats1 | large tumor suppressor [Source:MGI Syml | 560.57 | 608.53 | 527.27 | 572.6 | 611.84 | 574.74 |
| 11753 | ENSMUSG00000040022 | 221.3612398 | 0.027146457 | 0.182444893 | 0.148792637 | 0.881717261 | 0.981676444 | Rab11fip2 | RAB11 family interacting protein 2 (class I | 236.92 | 235.39 | 185.15 | 214.97 | 253.02 | 202.72 |
| 11754 | ENSMUSG00000040025 | 443.0369241 | -0.363412768 | 0.138065025 | -2.632185585 | 0.008483751 | 0.096227639 | Ythdf2 | YTH N6-methyladenosine RNA binding pro | 460.98 | 554.05 | 480.98 | 371.13 | 406.8 | 384.28 |
| 11755 | ENSMUSG00000040026 | 111.9622029 | -1.119736276 | 0.327718333 | -3.416764225 | 0.000633701 | 0.014736104 | Saa3 | serum amyloid A 3 [Source:MGI Symbol;A | 97.175 | 225.11 | 137.86 | 56.875 | 55.622 | 99.132 |
| 11756 | ENSMUSG00000040028 | 486.9159294 | -0.038816095 | 0.127000425 | -0.305637525 | 0.759880646 | 0.95402891 | Elavl1 | ELAV (embryonic lethal, abnormal vision)- | 520.41 | 501.62 | 457.84 | 476.21 | 484.23 | 481.18 |

| | | | | | | | | | | | | | | | |
|---|---|---|---|---|---|---|---|---|---|---|---|---|---|---|---|
| 11757 | ENSMUSG00000040029 | 1133.858405 | 0.159226057 | 0.106617874 | 1.493427417 | 0.135325341 | 0.538039097 | Ipo8 | importin 8 [Source:MGI Symbol;Acc:MGI:: | 1097 | 1087.5 | 1029.4 | 1214.6 | 1261.8 | 1112.7 |
| 11758 | ENSMUSG00000040033 | 1089.278475 | 0.261241395 | 0.109325697 | 2.389569903 | 0.016868114 | 0.154222752 | Stat2 | signal transducer and activator of transcri | 1006.3 | 913.82 | 1052.5 | 1221.4 | 1133.2 | 1208.5 |
| 11759 | ENSMUSG00000040034 | 58.8931675 | 0.116779622 | 0.305562644 | 0.382178988 | 0.702328606 | 0.942345172 | Nup43 | nucleoporin 43 [Source:MGI Symbol;Acc:I | 53.808 | 45.228 | 70.437 | 55.911 | 57.803 | 70.172 |
| 11760 | ENSMUSG00000040035 | 42.04584616 | -0.880011926 | 0.367868733 | -2.392190062 | 0.016748167 | 0.153417027 | Disp2 | dispatched RND tramsporter family meml | 48.989 | 48.312 | 66.412 | 34.703 | 33.809 | 20.049 |
| 11761 | ENSMUSG00000040037 | 0.702825224 | 0.139451984 | 2.778672966 | 0.050186541 | 0.959973737 | NA | Negr1 | neuronal growth regulator 1 [Source:MGI | 0 | 0 | 2.0125 | 0 | 1.0906 | 1.1138 |
| 11762 | ENSMUSG00000040040 | 99.52577828 | 0.0114849 | 0.245408459 | 0.046799119 | 0.962673331 | 0.992956886 | Ift88 | intraflagellar transport 88 [Source:MGI Sy | 89.144 | 107.93 | 100.62 | 99.29 | 119.97 | 80.197 |
| 11763 | ENSMUSG00000040043 | 206.0789204 | -0.224678751 | 0.170692511 | -1.316277728 | 0.188080843 | 0.625092215 | Rbms2 | RNA binding motif, single stranded intera | 207.2 | 227.17 | 232.44 | 199.54 | 194.13 | 175.99 |
| 11764 | ENSMUSG00000040044 | 382.2500799 | 0.048807336 | 0.134338504 | 0.363316061 | 0.716368797 | 0.9447719 | Orc3 | origin recognition complex, subunit 3 [Sou | 380.67 | 375.19 | 371.3 | 379.81 | 411.16 | 375.37 |
| 11765 | ENSMUSG00000040046 | 0 | NA | NA | NA | NA | NA | Tph1 | tryptophan hydroxylase 1 [Source:MGI Sy | 0 | 0 | 0 | 0 | 0 | 0 |
| 11766 | ENSMUSG00000040048 | 2085.618179 | -0.107528813 | 0.105295684 | -1.021208164 | 0.307155825 | 0.756237058 | Ndufb10 | NADH:ubiquinone oxidoreductase subuni | 1998.9 | 2261.4 | 2229.8 | 1845.1 | 2071.1 | 2107.4 |
| 11767 | ENSMUSG00000040054 | 725.2576124 | 0.183331047 | 0.125307787 | 1.46304592 | 0.143454814 | 0.553333105 | Baz2a | bromodomain adjacent to zinc finger dom | 653.73 | 681.51 | 703.36 | 848.3 | 682.73 | 781.92 |
| 11768 | ENSMUSG00000040055 | 0.506966596 | 2.558866737 | 3.339834942 | 0.766165628 | 0.443577757 | NA | Gjb6 | gap junction protein, beta 6 [Source:MGI : | 0 | 0 | 0 | 1.928 | 0 | 1.1138 |
| 11769 | ENSMUSG00000040061 | 15.89855083 | -1.627460766 | 0.574066211 | -2.834970487 | 0.004582992 | 0.063131805 | Plcb2 | phospholipase C, beta 2 [Source:MGI Sym | 24.093 | 27.754 | 20.125 | 5.7839 | 8.7249 | 8.9108 |
| 11770 | ENSMUSG00000040065 | 0 | NA | NA | NA | NA | NA | Pfpl | pore forming protein-like [Source:MGI Syr | 0 | 0 | 0 | 0 | 0 | 0 |
| 11771 | ENSMUSG00000040078 | 123.7966241 | -0.026338839 | 0.230922793 | -0.114059069 | 0.909190969 | 0.985206635 | Ptges3-ps | prostaglandin E synthase 3, pseudogene [: | 118.86 | 104.85 | 150.94 | 106.04 | 140.69 | 121.41 |
| 11772 | ENSMUSG00000040084 | 13.04246782 | -0.17261912 | 0.621823641 | -0.277601411 | 0.781318349 | 0.960079945 | Bub1b | BUB1B, mitotic checkpoint serine/threoni | 20.881 | 11.307 | 9.0562 | 10.604 | 15.269 | 11.138 |
| 11773 | ENSMUSG00000040086 | 0 | NA | NA | NA | NA | NA | Tnni3k | TNNI3 interacting kinase [Source:MGI Syn | 0 | 0 | 0 | 0 | 0 | 0 |
| 11774 | ENSMUSG00000040093 | 351.1937819 | 1.20344013 | 0.267456256 | 4.499577423 | 6.81E-06 | 0.000400339 | Bmf | BCL2 modifying factor [Source:MGI Symbo | 182.3 | 187.08 | 268.67 | 361.49 | 344.64 | 762.98 |
| 11775 | ENSMUSG00000040097 | 223.7688598 | -0.064109679 | 0.21364091 | -0.300081473 | 0.76411501 | 0.954980327 | Flywch1 | FLYWCH-type zinc finger 1 [Source:MGI Sy | 159.01 | 283.71 | 244.52 | 218.82 | 212.67 | 223.88 |
| 11776 | ENSMUSG00000040102 | 326.6957287 | 0.060866687 | 0.156924407 | 0.387872658 | 0.698110273 | 0.940598382 | Klhl42 | kelch-like 42 [Source:MGI Symbol;Acc:MG | 342.12 | 308.38 | 307.91 | 287.27 | 341.36 | 373.14 |
| 11777 | ENSMUSG00000040105 | 259.8893183 | 0.077122462 | 0.167981724 | 0.45911222 | 0.646153582 | 0.925643431 | Plpp6 | phospholipid phosphatase 6 [Source:MGI | 240.13 | 261.09 | 257.6 | 248.71 | 239.94 | 311.88 |
| 11778 | ENSMUSG00000040111 | 23.78786588 | -0.91745306 | 0.464244503 | -1.976228159 | 0.048128942 | 0.30505566 | Gramd1b | GRAM domain containing 1B [Source:MGI | 36.14 | 34.949 | 22.137 | 16.388 | 14.178 | 18.935 |
| 11779 | ENSMUSG00000040112 | 591.5903422 | 0.073712981 | 0.130799464 | 0.563557213 | 0.573055513 | 0.905173631 | Mrps35 | mitochondrial ribosomal protein S35 [Sou | 557.35 | 562.27 | 609.78 | 564.89 | 682.73 | 572.52 |
| 11780 | ENSMUSG00000040113 | 0 | NA | NA | NA | NA | NA | Mettl11b | methyltransferase like 11B [Source:MGI S | 0 | 0 | 0 | 0 | 0 | 0 |
| 11781 | ENSMUSG00000040118 | 12.419442 | -0.861154217 | 0.65637919 | -1.311976721 | 0.189527989 | 0.626663447 | Cacna2d1 | calcium channel, voltage-dependent, alph | 10.44 | 22.614 | 15.094 | 7.7118 | 13.087 | 5.5692 |
| 11782 | ENSMUSG00000040121 | 3.206194111 | -0.28472836 | 1.191354525 | -0.238995492 | 0.811109079 | NA | Rep15 | RAB15 effector protein [Source:MGI Symb | 2.4093 | 4.1117 | 4.025 | 0.964 | 3.2719 | 4.4554 |
| 11783 | ENSMUSG00000040123 | 710.7801082 | 0.207221149 | 0.116185463 | 1.78353766 | 0.074498823 | 0.398650011 | Zmym5 | zinc finger, MYM-type 5 [Source:MGI Sym | 615.98 | 669.17 | 695.32 | 785.64 | 763.43 | 735.14 |
| 11784 | ENSMUSG00000040124 | 75.140307 | 0.358315298 | 0.277165368 | 1.292785244 | 0.196085345 | 0.636628156 | Gorab | golgin, RAB6-interacting [Source:MGI Sym | 49.792 | 69.898 | 78.487 | 89.65 | 86.159 | 76.855 |
| 11785 | ENSMUSG00000040125 | 0 | NA | NA | NA | NA | NA | Gpr26 | G protein-coupled receptor 26 [Source:M | 0 | 0 | 0 | 0 | 0 | 0 |
| 11786 | ENSMUSG00000040127 | 1849.383417 | 0.232859765 | 0.170614017 | 1.364833728 | 0.172305315 | 0.600396752 | Sdr9c7 | 4short chain dehydrogenase/reductase fa | 1452 | 1560.4 | 2089 | 1550.1 | 2010 | 2434.9 |
| 11787 | ENSMUSG00000040128 | 884.0495817 | -0.676263673 | 0.167618472 | -4.034541445 | 5.47E-05 | 0.002248044 | Pnrc1 | proline-rich nuclear receptor coactivator : | 925.17 | 1144.1 | 1194.4 | 727.8 | 802.7 | 510.14 |
| 11788 | ENSMUSG00000040132 | 0 | NA | NA | NA | NA | NA | Svs2 | seminal vesicle secretory protein 2 [Sourc | 0 | 0 | 0 | 0 | 0 | 0 |
| 11789 | ENSMUSG00000040133 | 0.342639279 | -1.805976596 | 4.028547714 | -0.448294702 | 0.653940524 | NA | Gpr176 | G protein-coupled receptor 176 [Source:N | 0 | 2.0558 | 0 | 0 | 0 | 0 |
| 11790 | ENSMUSG00000040134 | 23590.90218 | 0.155511064 | 0.084266398 | 1.845469462 | 0.064969277 | 0.365648101 | Rdh7 | retinol dehydrogenase 7 [Source:MGI Sym | 21049 | 22987 | 22927 | 24616 | 24467 | 25499 |
| 11791 | ENSMUSG00000040136 | 0 | NA | NA | NA | NA | NA | Abcc8 | ATP-binding cassette, sub-family C (CFTR/ | 0 | 0 | 0 | 0 | 0 | 0 |
| 11792 | ENSMUSG00000040138 | 0 | NA | NA | NA | NA | NA | Ndp | Norrie disease (pseudoglioma) (human) [S | 0 | 0 | 0 | 0 | 0 | 0 |
| 11793 | ENSMUSG00000040139 | 201.5809292 | 0.214556672 | 0.174833117 | 1.227208412 | 0.219744269 | 0.66562955 | 9430038I01F | RIKEN cDNA 9430038I01 gene [Source:MC | 187.12 | 167.55 | 205.27 | 228.46 | 207.22 | 213.86 |
| 11794 | ENSMUSG00000040140 | 0 | NA | NA | NA | NA | NA | Tdrd6 | tudor domain containing 6 [Source:MGI S | 0 | 0 | 0 | 0 | 0 | 0 |
| 11795 | ENSMUSG00000040146 | 577.8261355 | 0.242948564 | 0.157512173 | 1.54241135 | 0.122973665 | 0.513267141 | Rgl3 | ral guanine nucleotide dissociation stimul | 497.92 | 440.98 | 649.03 | 598.63 | 599.84 | 680.56 |
| 11796 | ENSMUSG00000040147 | 4691.123638 | -0.237863612 | 0.094159495 | -2.526177647 | 0.011531116 | 0.119817569 | Maob | monoamine oxidase B [Source:MGI Symbo | 4995.3 | 4865.1 | 5371.3 | 4526.8 | 4351.6 | 4036.6 |
| 11797 | ENSMUSG00000040148 | 0 | NA | NA | NA | NA | NA | Hmx3 | H6 homeobox 3 [Source:MGI Symbol;Acc: | 0 | 0 | 0 | 0 | 0 | 0 |
| 11798 | ENSMUSG00000040151 | 265.510608 | -0.215465939 | 0.170421066 | -1.264315172 | 0.206116926 | 0.651217828 | Hs2st1 | heparan sulfate 2-O-sulfotransferase 1 [Sc | 270.65 | 308.38 | 276.72 | 219.79 | 227.94 | 289.6 |
| 11799 | ENSMUSG00000040152 | 15.05371195 | -2.117903993 | 0.670918229 | -3.156724471 | 0.00159552 | 0.029700908 | Thbs1 | thrombospondin 1 [Source:MGI Symbol;A | 38.549 | 21.586 | 13.081 | 3.8559 | 5.4531 | 7.7969 |
| 11800 | ENSMUSG00000040154 | 0 | NA | NA | NA | NA | NA | Wfdc5 | WAP four-disulfide core domain 5 [Source | 0 | 0 | 0 | 0 | 0 | 0 |
| 11801 | ENSMUSG00000040158 | 148.1279657 | -0.11583483 | 0.196728035 | -0.588806927 | 0.555990798 | 0.897628185 | Tax1bp3 | Tax1 (human T cell leukemia virus type I) k | 148.57 | 145.96 | 168.04 | 155.2 | 141.78 | 129.21 |
| 11802 | ENSMUSG00000040163 | 0.342639279 | -1.805976596 | 4.028547714 | -0.448294702 | 0.653940524 | NA | 1700034J05I | RIKEN cDNA 1700034J05 gene [Source:MC | 0 | 2.0558 | 0 | 0 | 0 | 0 |
| 11803 | ENSMUSG00000040164 | 0 | NA | NA | NA | NA | NA | Kcns1 | K+ voltage-gated channel, subfamily S, 1 [ | 0 | 0 | 0 | 0 | 0 | 0 |
| 11804 | ENSMUSG00000040165 | 0.160662974 | 1.126924175 | 4.080472857 | 0.276174898 | 0.782413736 | NA | Cd209c | CD209c antigen [Source:MGI Symbol;Acc: | 0 | 0 | 0 | 0.964 | 0 | 0 |
| 11805 | ENSMUSG00000040167 | 16.90022723 | -0.235134832 | 0.550909436 | -0.426812133 | 0.669516169 | 0.930837335 | Ikzf5 | IKAROS family zinc finger 5 [Source:MGI S | 15.259 | 16.447 | 23.144 | 14.46 | 9.8156 | 22.277 |
| 11806 | ENSMUSG00000040170 | 860.0982398 | -1.20107034 | 0.44245373 | -2.714567101 | 0.006636245 | 0.08106981 | Fmo2 | flavin containing monooxygenase 2 [Sourc | 962.12 | 1590.2 | 1044.5 | 761.54 | 509.32 | 292.94 |
| 11807 | ENSMUSG00000040174 | 265.0546875 | -0.160623111 | 0.1641225 | -0.978678186 | 0.327739014 | 0.772937193 | Alkbh3 | alkB homolog 3, alpha-ketoglutarate-depe | 308.39 | 286.79 | 243.51 | 237.14 | 266.11 | 248.39 |
| 11808 | ENSMUSG00000040177 | 51.03006148 | 0.20656239 | 0.342708159 | 0.602735548 | 0.546684629 | 0.89549457 | 2310057M21 | RIKEN cDNA 2310057M21 gene [Source:M | 43.368 | 57.563 | 41.256 | 50.127 | 40.353 | 73.514 |
| 11809 | ENSMUSG00000040181 | 5184.996116 | -0.737601415 | 0.173367504 | -4.254554043 | 2.09E-05 | 0.001006551 | Fmo1 | flavin containing monooxygenase 1 [Sourc | 5730.1 | 6831.5 | 6886.7 | 4764 | 4257.8 | 2639.8 |
| 11810 | ENSMUSG00000040183 | 4.262888721 | -0.076459105 | 1.007632606 | -0.075879943 | 0.939514614 | NA | Ankrd6 | ankyrin repeat domain 6 [Source:MGI Sym | 4.0155 | 5.1396 | 4.025 | 5.7839 | 3.2719 | 3.3415 |
| 11811 | ENSMUSG00000040187 | 23.50591891 | 0.707314427 | 0.447270573 | 1.581401661 | 0.113786228 | 0.495873456 | Arntl2 | aryl hydrocarbon receptor nuclear translo | 16.062 | 16.447 | 21.131 | 28.919 | 26.175 | 32.301 |
| 11812 | ENSMUSG00000040188 | 320.8842556 | -0.017619246 | 0.14825094 | -0.11884745 | 0.905396215 | 0.983972587 | Scamp2 | secretory carrier membrane protein 2 [So | 342.92 | 317.63 | 307.91 | 346.07 | 295.56 | 315.22 |

| | | | | | | | | | | | | | | | |
|---|---|---|---|---|---|---|---|---|---|---|---|---|---|---|---|
| 11813 | ENSMUSG00000040189 | 2.358629754 | -1.158259516 | 1.471008772 | -0.787391305 | 0.431052835 | NA | Odad1 | outer dynein arm docking complex subuni | 1.6062 | 4.1117 | 4.025 | 0 | 2.1812 | 2.2277 |
| 11814 | ENSMUSG00000040195 | 106.0108866 | 0.505711471 | 0.245010821 | 2.064037288 | 0.039014181 | 0.270232919 | Nemp1 | nuclear envelope integral membrane prot | 86.735 | 90.457 | 85.531 | 113.75 | 101.43 | 158.17 |
| 11815 | ENSMUSG00000040197 | 0 | NA | NA | NA | NA | NA | Cd209e | CD209e antigen [Source:MGI Symbol;Acc: | 0 | 0 | 0 | 0 | 0 | 0 |
| 11816 | ENSMUSG00000040204 | 5.499408865 | 0.618628781 | 0.916133809 | 0.675260289 | 0.499510409 | 0.880660418 | Pclaf | PCNA clamp associated factor [Source:MC | 4.8186 | 6.1675 | 2.0125 | 6.7478 | 5.4531 | 7.7969 |
| 11817 | ENSMUSG00000040205 | 0 | NA | NA | NA | NA | NA | Cuzd1 | CUB and zona pellucida-like domains 1 [Sc | 0 | 0 | 0 | 0 | 0 | 0 |
| 11818 | ENSMUSG00000040209 | 159.5812735 | -0.42404609 | 0.214110128 | -1.980504587 | 0.047646858 | 0.303195847 | Zfp704 | zinc finger protein 704 [Source:MGI Symb | 219.25 | 176.8 | 151.94 | 136.88 | 117.79 | 154.82 |
| 11819 | ENSMUSG00000040212 | 29.05406099 | -0.719881361 | 0.444251819 | -1.62043537 | 0.105138787 | 0.477915593 | Emp3 | epithelial membrane protein 3 [Source:M | 44.974 | 37.005 | 26.162 | 14.46 | 29.447 | 22.277 |
| 11820 | ENSMUSG00000040213 | 3152.260901 | 0.478453159 | 0.125687563 | 3.806686578 | 0.000140841 | 0.004726508 | Kyat3 | kynurenine aminotransferase 3 [Source:M | 2409.3 | 2585.2 | 2910.1 | 4280.1 | 3286 | 3442.9 |
| 11821 | ENSMUSG00000040217 | 0 | NA | NA | NA | NA | NA | Gm6369 | predicted gene 6369 [Source:MGI Symbol | 0 | 0 | 0 | 0 | 0 | 0 |
| 11822 | ENSMUSG00000040219 | 4.068781869 | -1.214261558 | 1.110708769 | -1.09323127 | 0.274292271 | NA | Ttc12 | tetratricopeptide repeat domain 12 [Sour | 4.8186 | 9.2513 | 3.0187 | 2.8919 | 1.0906 | 3.3415 |
| 11823 | ENSMUSG00000040220 | 54.45685495 | -0.466064654 | 0.319143683 | -1.460359952 | 0.144191173 | 0.554380949 | Gas8 | growth arrest specific 8 [Source:MGI Syml | 51.399 | 70.926 | 67.418 | 42.415 | 56.712 | 37.871 |
| 11824 | ENSMUSG00000040225 | 733.9111969 | -0.179080024 | 0.127714039 | -1.402195293 | 0.160856937 | 0.583943402 | Prrc2c | proline-rich coiled-coil 2C [Source:MGI Sy | 742.07 | 828.5 | 768.77 | 754.79 | 705.63 | 603.7 |
| 11825 | ENSMUSG00000040229 | 0.942116049 | 0.190003924 | 2.254635356 | 0.084272573 | 0.932839718 | NA | Gpr34 | G protein-coupled receptor 34 [Source:M | 1.6062 | 1.0279 | 0 | 1.928 | 1.0906 | 0 |
| 11826 | ENSMUSG00000040231 | 0.660353019 | -0.01500701 | 2.83315355 | -0.005296928 | 0.995773683 | NA | Syngr4 | synaptogyrin 4 [Source:MGI Symbol;Acc:N | 0 | 1.0279 | 1.0062 | 1.928 | 0 | 0 |
| 11827 | ENSMUSG00000040234 | 1560.826407 | 0.075206224 | 0.09905767 | 0.759216559 | 0.447723022 | 0.853796936 | Tm7sf3 | transmembrane 7 superfamily member 3 | 1563.6 | 1488.4 | 1507.4 | 1509.6 | 1617.4 | 1678.6 |
| 11828 | ENSMUSG00000040236 | 495.5498224 | 0.012600764 | 0.122039009 | 0.103251935 | 0.917763022 | 0.986374671 | Trappc5 | trafficking protein particle complex 5 [Sou | 498.73 | 484.15 | 497.08 | 493.56 | 491.87 | 507.91 |
| 11829 | ENSMUSG00000040242 | 883.3698579 | 0.134572918 | 0.117189704 | 1.148333968 | 0.250830722 | 0.702929347 | Fgfr1op2 | FGFR1 oncogene partner 2 [Source:MGI S | 866.55 | 869.62 | 789.9 | 902.28 | 860.5 | 1011.4 |
| 11830 | ENSMUSG00000040247 | 8.673446977 | -0.272433213 | 0.756688875 | -0.360033327 | 0.718822211 | 0.9447719 | Tbc1d10c | TBC1 domain family, member 10c [Source | 12.047 | 7.1954 | 9.0562 | 3.8559 | 7.6343 | 12.252 |
| 11831 | ENSMUSG00000040249 | 12488.55386 | 0.312364791 | 0.085471443 | 3.65461 | 0.000257573 | 0.007665718 | Lrp1 | low density lipoprotein receptor-related p | 10848 | 10884 | 11695 | 14345 | 13812 | 13348 |
| 11832 | ENSMUSG00000040250 | 310.7661611 | -0.028737276 | 0.142713416 | -0.201363524 | 0.840414335 | 0.971838962 | Ints13 | integrator complex subunit 13 [Source:MC | 301.16 | 320.71 | 319.99 | 305.58 | 309.74 | 307.42 |
| 11833 | ENSMUSG00000040253 | 332.343196 | 0.12519158 | 0.214448906 | 0.583782788 | 0.55936646 | 0.899423311 | Gbp7 | guanylate binding protein 7 [Source:MGI S | 420.83 | 318.65 | 213.32 | 350.89 | 376.26 | 314.1 |
| 11834 | ENSMUSG00000040254 | 44.56156135 | -0.94115744 | 0.378052291 | -2.48949011 | 0.012792648 | 0.127961865 | Sema3d | sema domain, immunoglobulin domain (Ig | 73.885 | 61.675 | 40.25 | 38.559 | 21.812 | 31.188 |
| 11835 | ENSMUSG00000040258 | 1.034369154 | -0.882584563 | 2.669352151 | -0.330636242 | 0.740919267 | NA | Nxph4 | neurexophilin 4 [Source:MGI Symbol;Acc: | 0 | 0 | 4.025 | 0 | 2.1812 | 0 |
| 11836 | ENSMUSG00000040260 | 55.53202715 | -0.781236738 | 0.305777969 | -2.554915061 | 0.010621371 | 0.113461975 | Daam2 | dishevelled associated activator of morph | 76.295 | 71.954 | 62.387 | 45.307 | 33.809 | 43.44 |
| 11837 | ENSMUSG00000040263 | 288.002757 | 0.057168999 | 0.161275172 | 0.354481089 | 0.722978374 | 0.9447719 | Klhdc4 | kelch domain containing 4 [Source:MGI Sy | 312.41 | 250.81 | 282.75 | 277.63 | 280.29 | 324.13 |
| 11838 | ENSMUSG00000040264 | 17.71192844 | -1.576870434 | 0.641222176 | -2.459163912 | 0.013926102 | 0.136099418 | Gbp2b | guanylate binding protein 2b [Source:MGI | 11.243 | 46.256 | 22.137 | 6.7478 | 7.6343 | 12.252 |
| 11839 | ENSMUSG00000040265 | 7.184167694 | 0.374340972 | 0.823211112 | 0.454732652 | 0.649301587 | 0.925643431 | Dnm3 | dynamin 3 [Source:MGI Symbol;Acc:MGI: | 5.6217 | 3.0838 | 10.062 | 6.7478 | 10.906 | 6.6831 |
| 11840 | ENSMUSG00000040268 | 227.0421921 | -0.440092417 | 0.179269723 | -2.454917711 | 0.014091684 | 0.136977231 | Plekha1 | pleckstrin homology domain containing, f | 282.69 | 286.79 | 214.33 | 186.05 | 194.13 | 198.26 |
| 11841 | ENSMUSG00000040269 | 445.1379203 | -0.042139746 | 0.144255853 | -0.292118102 | 0.770196327 | 0.956515747 | Mrps28 | mitochondrial ribosomal protein S28 [Sou | 489.09 | 452.28 | 412.56 | 406.8 | 414.43 | 495.66 |
| 11842 | ENSMUSG00000040270 | 128.1677984 | 0.417419954 | 0.302143422 | 1.381529181 | 0.167116312 | 0.593361251 | Bach2 | BTB and CNC homology, basic leucine zipp | 157.41 | 88.401 | 82.512 | 117.61 | 206.13 | 116.95 |
| 11843 | ENSMUSG00000040272 | 82.77337008 | -0.515934917 | 0.260961005 | -1.977057519 | 0.048035129 | 0.304728116 | Accs | 1-aminocyclopropane-1-carboxylate synth | 118.06 | 84.289 | 89.556 | 71.334 | 64.346 | 69.058 |
| 11844 | ENSMUSG00000040274 | 525.1305354 | -0.105156876 | 0.175125058 | -0.600467328 | 0.548194829 | 0.895600915 | Cdk6 | cyclin-dependent kinase 6 [Source:MGI Sy | 676.21 | 530.41 | 424.64 | 441.5 | 531.13 | 546.9 |
| 11845 | ENSMUSG00000040276 | 2.085946059 | 1.771224257 | 1.530148286 | 1.157550724 | 0.247047428 | NA | Pacsin1 | protein kinase C and casein kinase substra | 0.8031 | 1.0279 | 1.0062 | 1.928 | 2.1812 | 5.5692 |
| 11846 | ENSMUSG00000040280 | 47.92142604 | 0.112894263 | 0.331724906 | 0.340324952 | 0.733611829 | 0.947915087 | Ndufa4l2 | Ndufa4, mitochondrial complex associate | 45.777 | 45.228 | 47.293 | 56.875 | 56.712 | 35.643 |
| 11847 | ENSMUSG00000040282 | 67.67088775 | -0.00420356 | 0.318894495 | -0.013181665 | 0.989482858 | 0.99859176 | Cdin1 | CDAN1 interacting nuclease 1 [Source:MG | 94.766 | 53.452 | 54.337 | 55.911 | 79.615 | 67.944 |
| 11848 | ENSMUSG00000040283 | 82.21161379 | -0.83960783 | 0.269209587 | -3.118788741 | 0.001815961 | 0.032507502 | Btnl9 | butyrophilin-like 9 [Source:MGI Symbol;A | 89.144 | 116.15 | 111.69 | 70.37 | 47.987 | 57.92 |
| 11849 | ENSMUSG00000040284 | 0 | NA | NA | NA | NA | NA | Gzmg | granzyme G [Source:MGI Symbol;Acc:MG | 0 | 0 | 0 | 0 | 0 | 0 |
| 11850 | ENSMUSG00000040287 | 6.368359803 | 0.3964819 | 0.94351571 | 0.420217593 | 0.674326504 | 0.932114677 | Stac3 | SH3 and cysteine rich domain 3 [Source:M | 7.2279 | 3.0838 | 6.0375 | 1.928 | 5.4531 | 14.48 |
| 11851 | ENSMUSG00000040289 | 6.138210417 | -0.732549966 | 0.877489989 | -0.8348243 | 0.403816642 | 0.828953151 | Hey1 | hairy/enhancer-of-split related with YRPW | 9.6372 | 8.2233 | 5.0312 | 2.8919 | 4.3625 | 6.6831 |
| 11852 | ENSMUSG00000040296 | 440.1515492 | 0.239515301 | 0.140648502 | 1.702935316 | 0.088580174 | 0.438625004 | Ddx58 | DEAD/H box helicase 58 [Source:MGI Sym | 417.61 | 432.75 | 360.24 | 452.11 | 459.15 | 519.05 |
| 11853 | ENSMUSG00000040297 | 327.7607128 | -0.11146885 | 0.153134768 | -0.727913406 | 0.466666596 | 0.863779479 | Suco | SUN domain containing ossification factor | 304.38 | 388.55 | 329.04 | 314.26 | 318.46 | 311.88 |
| 11854 | ENSMUSG00000040298 | 0.17131964 | -0.796629176 | 4.080472857 | -0.195229623 | 0.84521319 | NA | Btbd16 | BTB (POZ) domain containing 16 [Source:I | 0 | 1.0279 | 0 | 0 | 0 | 0 |
| 11855 | ENSMUSG00000040302 | 112.3276301 | -0.024227224 | 0.227214736 | -0.106626993 | 0.915084903 | 0.98625062 | Rbm48 | RNA binding motif protein 48 [Source:MG | 106.01 | 109.99 | 123.77 | 96.398 | 130.87 | 106.93 |
| 11856 | ENSMUSG00000040310 | 0.717584917 | 3.048786435 | 2.951298099 | 1.033032358 | 0.301588756 | NA | Alx4 | aristaless-like homeobox 4 [Source:MGI S | 0 | 0 | 0 | 0.964 | 0 | 3.3415 |
| 11857 | ENSMUSG00000040312 | 51.98539148 | 0.483627722 | 0.333019581 | 1.452250109 | 0.146432074 | 0.556855766 | Cchcr1 | coiled-coil alpha-helical rod protein 1 [Sou | 49.792 | 33.921 | 46.287 | 77.118 | 47.987 | 56.806 |
| 11858 | ENSMUSG00000040314 | 0 | NA | NA | NA | NA | NA | Ctsg | cathepsin G [Source:MGI Symbol;Acc:MG | 0 | 0 | 0 | 0 | 0 | 0 |
| 11859 | ENSMUSG00000040321 | 144.374751 | -0.255254856 | 0.235353418 | -1.0845598 | 0.278116668 | 0.730997543 | Zfp770 | zinc finger protein 770 [Source:MGI Symb | 183.11 | 145.96 | 141.88 | 128.21 | 167.96 | 99.132 |
| 11860 | ENSMUSG00000040322 | 19.53958571 | -0.863225901 | 0.538323315 | -1.603545447 | 0.108814284 | 0.485684837 | Slc25a24 | solute carrier family 25 (mitochondrial car | 25.699 | 22.614 | 27.169 | 9.6398 | 8.7249 | 23.391 |
| 11861 | ENSMUSG00000040323 | 5.72649345 | 0.119861384 | 0.898717062 | 0.133369432 | 0.893901219 | 0.982562372 | Gm15429 | predicted pseudogene 15429 [Source:MG | 7.2279 | 5.1396 | 4.025 | 5.7839 | 3.2719 | 8.9108 |
| 11862 | ENSMUSG00000040325 | 511.94531 | 0.125676855 | 0.131828467 | 0.953336245 | 0.340419733 | 0.783052007 | Dcaf1 | DDB1 and CUL4 associated factor 1 [Sourc | 490.7 | 531.43 | 446.77 | 525.37 | 508.23 | 569.17 |
| 11863 | ENSMUSG00000040327 | 215.88266 | -0.051378397 | 0.173534314 | -0.296070532 | 0.767176218 | 0.955600828 | Cul9 | cullin 9 [Source:MGI Symbol;Acc:MGI:192 | 245.75 | 208.67 | 204.27 | 215.93 | 225.76 | 194.92 |
| 11864 | ENSMUSG00000040328 | 1.568870735 | 0.164657267 | 1.753435172 | 0.093905535 | 0.925184197 | NA | Olfr56 | olfactory receptor 56 [Source:MGI Symbo | 2.4093 | 1.0279 | 1.0062 | 3.8559 | 0 | 1.1138 |
| 11865 | ENSMUSG00000040329 | 3.040423692 | -0.649538282 | 1.483682319 | -0.437787977 | 0.661539987 | NA | Il7 | interleukin 7 [Source:MGI Symbol;Acc:MG | 8.031 | 3.0838 | 0 | 3.8559 | 3.2719 | 0 |
| 11866 | ENSMUSG00000040331 | 799.9922329 | -0.037427462 | 0.121405162 | -0.308285591 | 0.757865029 | 0.953466696 | Nsmce4a | NSE4 homolog A, SMC5-SMC6 complex cc | 739.66 | 880.93 | 811.03 | 736.48 | 833.23 | 798.63 |
| 11867 | ENSMUSG00000040339 | 26.15046368 | -0.932819785 | 0.455024138 | -2.050044616 | 0.040360077 | 0.274714427 | Fam102b | family with sequence similarity 102, meml | 38.549 | 41.117 | 23.144 | 14.46 | 21.812 | 17.822 |
| 11868 | ENSMUSG00000040340 | 1.481719333 | 0.62921854 | 2.133055425 | 0.294984618 | 0.76800563 | NA | Tex45 | testis expressed 45 [Source:MGI Symbol;A | 2.4093 | 1.0279 | 0 | 0 | 5.4531 | 0 |

| | | | | | | | | | | | | | | | |
|---|---|---|---|---|---|---|---|---|---|---|---|---|---|---|---|
| 11869 | ENSMUSG00000040345 | 31.16430438 | -1.321045403 | 0.448977758 | -2.942340413 | 0.003257416 | 0.04949083 | Arhgap9 | Rho GTPase activating protein 9 [Source:N | 42.564 | 55.508 | 35.219 | 10.604 | 16.359 | 26.732 |
| 11870 | ENSMUSG00000040350 | 415.5832817 | -0.352473961 | 0.151147168 | -2.33199183 | 0.019701121 | 0.171867922 | Trim7 | tripartite motif-containing 7 [Source:MGI | 481.06 | 476.95 | 440.74 | 406.8 | 384.99 | 302.97 |
| 11871 | ENSMUSG00000040351 | 454.5607713 | -0.088814538 | 0.143986502 | -0.616825442 | 0.537349869 | 0.890758111 | Ankib1 | ankyrin repeat and IBR domain containing | 445.72 | 543.77 | 416.59 | 439.57 | 439.52 | 442.2 |
| 11872 | ENSMUSG00000040354 | 1035.132288 | 0.087386102 | 0.119933834 | 0.728619262 | 0.466234593 | 0.863779479 | Mars1 | methionine-tRNA synthetase 1 [Source:M | 1026.4 | 1019.7 | 965.99 | 1210.8 | 1004.5 | 983.52 |
| 11873 | ENSMUSG00000040356 | 1263.709925 | 0.045281808 | 0.100662364 | 0.449838509 | 0.652826888 | 0.925756399 | Skiv2l | superkiller viralicidic activity 2-like (S. cere | 1227.9 | 1232.5 | 1271.9 | 1348.6 | 1271.7 | 1229.7 |
| 11874 | ENSMUSG00000040359 | 557.9407024 | 0.096384058 | 0.141748694 | 0.679964341 | 0.49652704 | 0.878181353 | Ufl1 | UFM1 specific ligase 1 [Source:MGI Symbo | 542.9 | 575.63 | 500.1 | 657.43 | 572.57 | 499 |
| 11875 | ENSMUSG00000040363 | 233.7841218 | -0.08165874 | 0.182225664 | -0.448118767 | 0.654067485 | 0.925797231 | Bcor | BCL6 interacting corepressor [Source:MGI | 229.69 | 221 | 270.68 | 234.25 | 188.68 | 258.41 |
| 11876 | ENSMUSG00000040364 | 0.133850373 | -0.796629176 | 4.080472857 | -0.195229623 | 0.84521319 | NA | Sec1 | secretory blood group 1 [Source:MGI Sym | 0.8031 | 0 | 0 | 0 | 0 | 0 |
| 11877 | ENSMUSG00000040365 | 775.1191861 | 0.163946588 | 0.117634211 | 1.393698199 | 0.16340876 | 0.587512475 | Trim41 | tripartite motif-containing 41 [Source:MG | 749.29 | 677.4 | 766.76 | 868.54 | 819.05 | 769.67 |
| 11878 | ENSMUSG00000040367 | 5.643839682 | -0.365623229 | 0.912167857 | -0.40082889 | 0.688546107 | 0.936908509 | Lrrd1 | leucine rich repeats and death domain co | 8.8341 | 3.0838 | 7.0437 | 3.8559 | 4.3625 | 6.6831 |
| 11879 | ENSMUSG00000040370 | 1147.351625 | -0.162324666 | 0.115924333 | -1.400263969 | 0.161434286 | 0.584476975 | Etfrf1 | electron transfer flavoprotein regulatory f | 1208.7 | 1197.5 | 1228.6 | 971.69 | 1212.8 | 1064.8 |
| 11880 | ENSMUSG00000040372 | 1.018186588 | 0.19712111 | 2.235934965 | 0.088160484 | 0.929749124 | NA | Gpr63 | G protein-coupled receptor 63 [Source:M | 0.8031 | 1.0279 | 1.0062 | 0 | 3.2719 | 0 |
| 11881 | ENSMUSG00000040373 | 0 | NA | NA | NA | NA | NA | Cacng5 | calcium channel, voltage-dependent, gam | 0 | 0 | 0 | 0 | 0 | 0 |
| 11882 | ENSMUSG00000040374 | 983.4141336 | 0.100723711 | 0.114872196 | 0.876832819 | 0.38057745 | 0.812628154 | Pex2 | peroxisomal biogenesis factor 2 [Source:N | 927.58 | 1038.2 | 881.47 | 959.16 | 1043.7 | 1050.4 |
| 11883 | ENSMUSG00000040380 | 10.78560188 | -0.032017154 | 0.685471021 | -0.046708254 | 0.962745752 | 0.992956886 | Cbln3 | cerebellin 3 precursor protein [Source:MG | 10.44 | 12.335 | 10.062 | 15.424 | 11.997 | 4.4554 |
| 11884 | ENSMUSG00000040383 | 375.822991 | -0.210703569 | 0.153258167 | -1.374827669 | 0.169184878 | 0.596414836 | Aqr | aquarius [Source:MGI Symbol;Acc:MGI:12 | 385.49 | 415.28 | 409.54 | 371.13 | 383.9 | 289.6 |
| 11885 | ENSMUSG00000040385 | 1848.166612 | -0.003165671 | 0.098065149 | -0.032281301 | 0.974247721 | 0.996336532 | Ppp1ca | protein phosphatase 1 catalytic subunit al | 1826.3 | 1855.4 | 1868.6 | 1763.1 | 1791.9 | 1983.8 |
| 11886 | ENSMUSG00000040387 | 1.107988552 | -1.056291367 | 2.031653537 | -0.519917076 | 0.603121373 | NA | Klhl32 | kelch-like 32 [Source:MGI Symbol;Acc:MG | 2.4093 | 1.0279 | 1.0062 | 0 | 1.0906 | 1.1138 |
| 11887 | ENSMUSG00000040389 | 31.25582205 | -0.472677951 | 0.437680333 | -1.079961597 | 0.280159282 | 0.733108114 | Wdr47 | WD repeat domain 47 [Source:MGI Symbo | 32.927 | 34.949 | 41.256 | 27.955 | 37.081 | 13.366 |
| 11888 | ENSMUSG00000040390 | 342.5440439 | 0.060720545 | 0.145148388 | 0.418334268 | 0.675702738 | 0.932630311 | Map3k10 | mitogen-activated protein kinase kinase k | 367.82 | 314.54 | 323 | 358.6 | 332.64 | 358.66 |
| 11889 | ENSMUSG00000040396 | 512.4735402 | 0.33288667 | 0.131875551 | 2.52424856 | 0.011594592 | 0.119989074 | Abhd13 | abhydrolase domain containing 13 [Sourc | 434.48 | 429.67 | 497.08 | 572.6 | 607.47 | 533.53 |
| 11890 | ENSMUSG00000040405 | 0.710308366 | 0.13927508 | 2.769785472 | 0.050283707 | 0.959896308 | NA | Havcr1 | hepatitis A virus cellular receptor 1 [Sourc | 0 | 1.0279 | 1.0062 | 0 | 0 | 2.2277 |
| 11891 | ENSMUSG00000040407 | 744.0983742 | 0.038405955 | 0.13373567 | 0.287178097 | 0.773975956 | 0.958293805 | Akap9 | A kinase (PRKA) anchor protein (yotiao) 9 | 661.76 | 784.3 | 757.7 | 784.68 | 822.33 | 653.83 |
| 11892 | ENSMUSG00000040410 | 395.531344 | 0.124505524 | 0.156418029 | 0.795979368 | 0.426044028 | 0.841626138 | Fbxl4 | F-box and leucine-rich repeat protein 4 [S | 354.17 | 362.85 | 418.6 | 384.63 | 483.14 | 369.8 |
| 11893 | ENSMUSG00000040412 | 12.75717953 | -0.374309148 | 0.655796755 | -0.570770053 | 0.568155526 | 0.903352921 | Elapor1 | endosome-lysosome associated apoptosis | 13.653 | 22.614 | 7.0437 | 14.46 | 7.6343 | 11.138 |
| 11894 | ENSMUSG00000040413 | 1389.647694 | 0.070119261 | 0.110962408 | 0.631919066 | 0.527439764 | 0.888334265 | Timd2 | T cell immunoglobulin and mucin domain | 1417.5 | 1221.2 | 1427.9 | 1335.1 | 1456 | 1480.3 |
| 11895 | ENSMUSG00000040414 | 345.5902219 | 0.017307822 | 0.143263318 | 0.120811263 | 0.903840529 | 0.983821605 | Slc25a28 | solute carrier family 25, member 28 [Sour | 358.99 | 344.35 | 327.03 | 353.78 | 369.72 | 319.67 |
| 11896 | ENSMUSG00000040415 | 44.36741857 | -0.391847738 | 0.373276266 | -1.049752621 | 0.293831863 | 0.745164683 | Dtx3 | deltex 3, E3 ubiquitin ligase [Source:MGI S | 58.626 | 45.228 | 47.293 | 52.055 | 22.903 | 40.098 |
| 11897 | ENSMUSG00000040420 | 19.40209749 | -0.120506152 | 0.595493376 | -0.202363547 | 0.839632524 | 0.971801133 | Cdh18 | cadherin 18 [Source:MGI Symbol;Acc:MG | 25.699 | 22.614 | 12.075 | 7.7118 | 32.719 | 15.594 |
| 11898 | ENSMUSG00000040423 | 527.1283377 | -0.072660211 | 0.145275125 | -0.500155905 | 0.616965304 | 0.917509899 | Rc3h1 | RING CCCH (C3H) domains 1 [Source:MGI | 547.72 | 574.61 | 499.1 | 531.15 | 578.03 | 432.17 |
| 11899 | ENSMUSG00000040424 | 0.133850373 | -0.796629176 | 4.080472857 | -0.195229623 | 0.84521319 | NA | Hipk4 | homeodomain interacting protein kinase | 0.8031 | 0 | 0 | 0 | 0 | 0 |
| 11900 | ENSMUSG00000040428 | 18.90130524 | -0.506861393 | 0.499580587 | -1.014573837 | 0.310309016 | 0.759065371 | Plekha4 | pleckstrin homology domain containing, f | 24.896 | 21.586 | 20.125 | 19.28 | 15.269 | 12.252 |
| 11901 | ENSMUSG00000040429 | 30.99993173 | -0.042974116 | 0.393180841 | -0.109298603 | 0.912965655 | 0.98625062 | Mterf1a | mitochondrial transcription termination fa | 27.305 | 39.061 | 28.175 | 30.847 | 30.537 | 30.074 |
| 11902 | ENSMUSG00000040430 | 453.8961345 | 0.203902767 | 0.162093395 | 1.257933839 | 0.208415684 | 0.653455953 | Pitpnc1 | phosphatidylinositol transfer protein, cyto | 489.09 | 374.16 | 401.49 | 513.8 | 404.62 | 540.21 |
| 11903 | ENSMUSG00000040432 | 50.37569615 | 0.073416878 | 0.379059943 | 0.193681446 | 0.846425321 | 0.972803041 | Ltb4r2 | leukotriene B4 receptor 2 [Source:MGI Sy | 28.912 | 62.703 | 56.35 | 69.406 | 41.443 | 43.44 |
| 11904 | ENSMUSG00000040433 | 366.1302312 | -0.013898781 | 0.135939418 | -0.102242465 | 0.918564222 | 0.986374671 | Zbtb38 | zinc finger and BTB domain containing 38 | 378.26 | 371.08 | 354.2 | 371.13 | 350.09 | 372.02 |
| 11905 | ENSMUSG00000040434 | 12.44382599 | -0.780314313 | 0.61916587 | -1.260267 | 0.20757306 | 0.652593069 | Large2 | LARGE xylosyl- and glucuronyltransferase | 17.668 | 11.307 | 18.112 | 7.7118 | 8.7249 | 11.138 |
| 11906 | ENSMUSG00000040435 | 163.9035251 | -0.286063731 | 0.215618248 | -1.32671392 | 0.184603335 | 0.619369666 | Ppp1r15a | protein phosphatase 1, regulatory subunit | 199.17 | 193.25 | 147.92 | 171.59 | 117.79 | 153.71 |
| 11907 | ENSMUSG00000040441 | 64.052007 | -0.40592993 | 0.332913581 | -1.219325236 | 0.222720774 | 0.668407528 | Slc26a10 | solute carrier family 26, member 10 [Sour | 65.051 | 76.066 | 78.487 | 80.01 | 50.168 | 34.529 |
| 11908 | ENSMUSG00000040446 | 200.2647124 | -0.014714449 | 0.224139321 | -0.065648674 | 0.947657536 | 0.990677354 | Rprd1a | regulation of nuclear pre-mRNA domain c | 157.41 | 190.16 | 256.59 | 173.52 | 178.86 | 245.05 |
| 11909 | ENSMUSG00000040447 | 803.4864957 | 0.272206889 | 0.141699773 | 1.921011469 | 0.054730262 | 0.329566405 | Spns2 | spinster homolog 2 [Source:MGI Symbol;A | 742.07 | 621.89 | 819.08 | 808.78 | 828.87 | 1000.2 |
| 11910 | ENSMUSG00000040451 | 241.3261474 | -0.007859209 | 0.198348413 | -0.039623251 | 0.96839349 | 0.994245784 | Sgms1 | sphingomyelin synthase 1 [Source:MGI Sy | 295.54 | 242.59 | 187.16 | 259.31 | 209.4 | 253.96 |
| 11911 | ENSMUSG00000040452 | 0 | NA | NA | NA | NA | NA | Cdh12 | cadherin 12 [Source:MGI Symbol;Acc:MGI | 0 | 0 | 0 | 0 | 0 | 0 |
| 11912 | ENSMUSG00000040455 | 286.0373383 | 0.110805836 | 0.156230158 | 0.709247416 | 0.478170953 | 0.869794311 | Usp45 | ubiquitin specific petidase 45 [Source:MG | 278.68 | 251.84 | 294.83 | 318.11 | 302.1 | 270.66 |
| 11913 | ENSMUSG00000040456 | 0 | NA | NA | NA | NA | NA | H2ap | H2A.P histone [Source:MGI Symbol;Acc:N | 0 | 0 | 0 | 0 | 0 | 0 |
| 11914 | ENSMUSG00000040459 | 637.5826981 | 0.011029455 | 0.13760383 | 0.080153693 | 0.936115019 | 0.988420305 | Arglu1 | arginine and glutamate rich 1 [Source:MG | 682.64 | 586.94 | 635.95 | 723.95 | 639.1 | 556.92 |
| 11915 | ENSMUSG00000040462 | 4324.667658 | 0.123229546 | 0.088826319 | 1.387308931 | 0.165347581 | 0.590807167 | Os9 | amplified in osteosarcoma [Source:MGI S | 3935.2 | 4257.6 | 4228.2 | 4428.5 | 4682 | 4416.4 |
| 11916 | ENSMUSG00000040463 | 854.9308905 | -0.120295807 | 0.123458009 | -0.974386414 | 0.329864712 | 0.774747658 | Mybbp1a | MYB binding protein (P160) 1a [Source:M | 859.32 | 937.46 | 875.43 | 827.09 | 907.39 | 722.88 |
| 11917 | ENSMUSG00000040464 | 464.9848218 | 0.055990585 | 0.146806465 | 0.381390461 | 0.702913538 | 0.942506832 | Gtpbp10 | GTP-binding protein 10 (putative) [Source | 473.03 | 446.12 | 447.78 | 396.19 | 531.13 | 495.66 |
| 11918 | ENSMUSG00000040466 | 2039.399209 | -0.337614961 | 0.108754918 | -3.104365006 | 0.001906879 | 0.03359452 | Blvrb | biliverdin reductase B (flavin reductase (N | 2384.4 | 2045.6 | 2400.9 | 1928 | 1771.2 | 1706.4 |
| 11919 | ENSMUSG00000040467 | 0.160662974 | 1.126924175 | 4.080472857 | 0.276174898 | 0.782413736 | NA | 4921522P10 | RIKEN cDNA 4921522P10 gene [Source:M | 0 | 0 | 0 | 0.964 | 0 | 0 |
| 11920 | ENSMUSG00000040471 | 438.2130866 | -0.05561199 | 0.188190097 | -0.295509653 | 0.76760458 | 0.95578799 | Ggt6 | gamma-glutamyltransferase 6 [Source:MG | 391.11 | 363.88 | 585.63 | 472.35 | 372.99 | 443.31 |
| 11921 | ENSMUSG00000040472 | 600.6623608 | 0.146469223 | 0.142055075 | 1.031073498 | 0.302506355 | 0.752385931 | Rabggta | Rab geranylgeranyl transferase, a subunit | 575.02 | 535.55 | 600.73 | 743.23 | 576.94 | 572.52 |
| 11922 | ENSMUSG00000040473 | 2.87450071 | -1.240235791 | 1.328301199 | -0.933700724 | 0.350458292 | NA | Cfap69 | cilia and flagella associated protein 69 [So | 4.0155 | 5.1396 | 3.0187 | 2.8919 | 2.1812 | 0 |
| 11923 | ENSMUSG00000040478 | 0 | NA | NA | NA | NA | NA | Prdm13 | PR domain containing 13 [Source:MGI Syr | 0 | 0 | 0 | 0 | 0 | 0 |
| 11924 | ENSMUSG00000040479 | 707.8326841 | 0.03541654 | 0.118103817 | 0.299876334 | 0.764271487 | 0.954980327 | Dgkz | diacylglycerol kinase zeta [Source:MGI Syr | 685.05 | 683.57 | 729.53 | 773.11 | 672.91 | 702.84 |

| | | | | | | | | | | | | | | | |
|---|---|---|---|---|---|---|---|---|---|---|---|---|---|---|---|
| 11925 | ENSMUSG00000040481 | 468.4742462 | -0.034580117 | 0.129981934 | -0.266037869 | 0.790210044 | 0.961019028 | Bptf | bromodomain PHD finger transcription fa | 444.92 | 510.88 | 466.9 | 451.14 | 458.06 | 478.95 |
| 11926 | ENSMUSG00000040482 | 355.0966312 | 0.172126079 | 0.137393652 | 1.252794992 | 0.210280325 | 0.654366311 | Dxo | decapping exoribonuclease [Source:MGI S | 326.86 | 324.82 | 350.17 | 369.2 | 371.9 | 387.62 |
| 11927 | ENSMUSG00000040483 | 106.6758101 | 0.326606721 | 0.24143753 | 1.352758711 | 0.176132727 | 0.607001136 | Xaf1 | XIAP associated factor 1 [Source:MGI Sym | 114.84 | 78.122 | 90.562 | 133.99 | 115.61 | 106.93 |
| 11928 | ENSMUSG00000040485 | 0 | NA | NA | NA | NA | NA | Lrrc52 | leucine rich repeat containing 52 [Source: | 0 | 0 | 0 | 0 | 0 | 0 |
| 11929 | ENSMUSG00000040488 | 581.5523817 | -0.682396326 | 0.139436891 | -4.893943913 | 9.88E-07 | 8.08E-05 | Ltbp4 | latent transforming growth factor beta bi | 744.48 | 766.83 | 638.97 | 502.23 | 403.53 | 433.29 |
| 11930 | ENSMUSG00000040489 | 0.668968166 | 0.251312211 | 2.604595067 | 0.096488016 | 0.923132991 | NA | Sox30 | SRY (sex determining region Y)-box 30 [So | 0.8031 | 0 | 1.0062 | 0 | 1.0906 | 1.1138 |
| 11931 | ENSMUSG00000040490 | 0.852779399 | -0.534702824 | 2.386795691 | -0.224025385 | 0.822737551 | NA | Lrfn2 | leucine rich repeat and fibronectin type II | 0 | 2.0558 | 1.0062 | 0.964 | 1.0906 | 0 |
| 11932 | ENSMUSG00000040495 | 0.185640647 | 1.126924175 | 4.080472857 | 0.276174898 | 0.782413736 | NA | Chrm4 | cholinergic receptor, muscarinic 4 [Source | 0 | 0 | 0 | 0 | 0 | 1.1138 |
| 11933 | ENSMUSG00000040498 | 9.051650954 | 0.075584092 | 0.787375706 | 0.09599495 | 0.923524583 | 0.98705827 | Igsf23 | immunoglobulin superfamily, member 23 | 3.2124 | 9.2513 | 14.087 | 5.7839 | 14.178 | 7.7969 |
| 11934 | ENSMUSG00000040502 | 10.54700407 | -0.671175669 | 0.664340262 | -1.010289014 | 0.312356843 | 0.759830222 | Marchf9 | membrane associated ring-CH-type finger | 10.44 | 12.335 | 16.1 | 6.7478 | 7.6343 | 10.025 |
| 11935 | ENSMUSG00000040505 | 2107.838575 | 0.061705908 | 0.123128131 | 0.501151996 | 0.616264155 | 0.917509899 | Abcg5 | ATP binding cassette subfamily G membe | 2304.9 | 1734.1 | 2148.3 | 2246.1 | 2198.7 | 2014.9 |
| 11936 | ENSMUSG00000040506 | 465.6688539 | -0.149309778 | 0.130050863 | -1.148087558 | 0.25093242 | 0.702929347 | Ambra1 | autophagy/beclin 1 regulator 1 [Source:M | 485.07 | 521.15 | 462.87 | 420.29 | 462.42 | 442.2 |
| 11937 | ENSMUSG00000040511 | 81.67889428 | 0.311760825 | 0.315433866 | 0.98835559 | 0.322978529 | 0.769995293 | Pvr | poliovirus receptor [Source:MGI Symbol;A | 70.673 | 87.373 | 60.375 | 61.695 | 78.525 | 131.43 |
| 11938 | ENSMUSG00000040514 | 0 | NA | NA | NA | NA | NA | Tex47 | testis expressed 47 [Source:MGI Symbol;A | 0 | 0 | 0 | 0 | 0 | 0 |
| 11939 | ENSMUSG00000040520 | 454.0734213 | 0.0308368 | 0.14034823 | 0.219716344 | 0.826092075 | 0.96948417 | Manea | mannosidase, endo-alpha [Source:MGI Sy | 440.9 | 458.45 | 447.78 | 395.23 | 486.42 | 495.66 |
| 11940 | ENSMUSG00000040521 | 278.2347685 | -0.080791026 | 0.155850394 | -0.518388337 | 0.604187351 | 0.916322307 | Tsfm | Ts translation elongation factor, mitochon | 268.24 | 301.18 | 288.79 | 256.42 | 257.39 | 297.4 |
| 11941 | ENSMUSG00000040522 | 8.256499434 | -2.093439095 | 0.828238827 | -2.527579035 | 0.011485197 | 0.119684359 | Tlr8 | toll-like receptor 8 [Source:MGI Symbol;A | 13.653 | 17.475 | 9.0562 | 3.8559 | 3.2719 | 2.2277 |
| 11942 | ENSMUSG00000040524 | 344.1359024 | 0.50156688 | 0.199486065 | 2.514295323 | 0.011927051 | 0.122154064 | Zfp609 | zinc finger protein 609 [Source:MGI Symb | 262.61 | 309.4 | 283.76 | 533.08 | 372.99 | 302.97 |
| 11943 | ENSMUSG00000040525 | 387.0706512 | 0.099698002 | 0.157546953 | 0.632814535 | 0.526854764 | 0.888334265 | Cblc | Casitas B-lineage lymphoma c [Source:MG | 418.42 | 303.24 | 398.47 | 386.56 | 399.17 | 416.58 |
| 11944 | ENSMUSG00000040528 | 8.324363322 | -0.711187227 | 0.789547504 | -0.900752929 | 0.3677197 | 0.805888029 | Milr1 | mast cell immunoglobulin like receptor 1 [ | 9.6372 | 12.335 | 9.0562 | 5.7839 | 10.906 | 2.2277 |
| 11945 | ENSMUSG00000040532 | 428.4227069 | -0.067605912 | 0.137761416 | -0.49074635 | 0.623605859 | 0.919503423 | Abhd11 | abhydrolase domain containing 11 [Sourc | 411.19 | 453.31 | 451.8 | 455.96 | 382.81 | 415.46 |
| 11946 | ENSMUSG00000040533 | 0 | NA | NA | NA | NA | NA | Matn1 | matrilin 1, cartilage matrix protein [Source | 0 | 0 | 0 | 0 | 0 | 0 |
| 11947 | ENSMUSG00000040536 | 762.8093419 | 0.281965333 | 0.125148947 | 2.253037996 | 0.024256753 | 0.198556695 | Necab1 | N-terminal EF-hand calcium binding prote | 732.43 | 647.59 | 684.25 | 749.97 | 917.21 | 845.41 |
| 11948 | ENSMUSG00000040537 | 4.70890143 | -1.003481056 | 1.000520547 | -1.002958968 | 0.315880659 | NA | Adam22 | a disintegrin and metallopeptidase domai | 5.6217 | 9.2513 | 4.025 | 3.8559 | 3.2719 | 2.2277 |
| 11949 | ENSMUSG00000040540 | 1.373834909 | -0.654567451 | 1.865508732 | -0.350878792 | 0.725679283 | NA | Ctdsp2-ps | CTD (carboxy-terminal domain, RNA polyr | 0 | 2.0558 | 3.0187 | 0.964 | 1.0906 | 1.1138 |
| 11950 | ENSMUSG00000040541 | 0 | NA | NA | NA | NA | NA | Tmem225 | transmembrane protein 225 [Source:MGI | 0 | 0 | 0 | 0 | 0 | 0 |
| 11951 | ENSMUSG00000040543 | 1.734691104 | -1.996525895 | 1.771958162 | -1.126734219 | 0.25985487 | NA | Pitpnm3 | PITPNM family member 3 [Source:MGI Sy | 3.2124 | 4.1117 | 1.0062 | 0.964 | 0 | 1.1138 |
| 11952 | ENSMUSG00000040548 | 2400.705349 | -0.01229311 | 0.104069685 | -0.118123828 | 0.905969544 | 0.983991807 | Tex2 | testis expressed gene 2 [Source:MGI Symb | 2295.3 | 2417.7 | 2519.6 | 2170.9 | 2423.4 | 2577.4 |
| 11953 | ENSMUSG00000040549 | 219.1423525 | 0.049470078 | 0.166124553 | 0.29778908 | 0.76586415 | 0.954996635 | Ckap5 | cytoskeleton associated protein 5 [Source | 228.08 | 210.72 | 207.29 | 240.03 | 213.76 | 214.97 |
| 11954 | ENSMUSG00000040550 | 690.3909825 | -0.015759519 | 0.129302685 | -0.12188083 | 0.902993398 | 0.983821605 | Otud6b | OTU domain containing 6B [Source:MGI S | 604.74 | 744.21 | 734.56 | 664.18 | 726.35 | 668.31 |
| 11955 | ENSMUSG00000040552 | 31.20520951 | -1.196899719 | 0.446358943 | -2.681473596 | 0.00732987 | 0.086502219 | C3ar1 | complement component 3a receptor 1 [S | 34.533 | 49.34 | 46.287 | 9.6398 | 21.812 | 25.618 |
| 11956 | ENSMUSG00000040554 | 0 | NA | NA | NA | NA | NA | Aipl1 | aryl hydrocarbon receptor-interacting pro | 0 | 0 | 0 | 0 | 0 | 0 |
| 11957 | ENSMUSG00000040557 | 197.420688 | 0.442365903 | 0.1964757 | 2.251504396 | 0.024353609 | 0.199236763 | Mettl27 | methyltransferase like 27 [Source:MGI Sy | 187.12 | 166.52 | 147.92 | 223.64 | 189.77 | 269.55 |
| 11958 | ENSMUSG00000040560 | 414.1527403 | -0.034799819 | 0.148010183 | -0.235117738 | 0.814117351 | 0.967157956 | Wdr7 | WD repeat domain 7 [Source:MGI Symbol | 419.22 | 479.01 | 359.23 | 402.94 | 405.71 | 418.81 |
| 11959 | ENSMUSG00000040562 | 643.3507523 | -0.029179389 | 0.218755508 | -0.133388134 | 0.893886429 | 0.982562372 | Gstm2 | glutathione S-transferase, mu 2 [Source:M | 901.08 | 455.37 | 591.67 | 589.95 | 527.86 | 794.17 |
| 11960 | ENSMUSG00000040563 | 103.8282096 | 0.3538953 | 0.239649231 | 1.476722034 | 0.139750163 | 0.546014685 | Plppr2 | phospholipid phosphatase related 2 [Sour | 105.21 | 74.01 | 93.581 | 102.18 | 123.24 | 124.75 |
| 11961 | ENSMUSG00000040564 | 67022.38071 | -0.072438119 | 0.082306104 | -0.880106276 | 0.378801739 | 0.811969331 | Apoc1 | apolipoprotein C-I [Source:MGI Symbol;A | 68682 | 65524 | 71908 | 66153 | 64079 | 65788 |
| 11962 | ENSMUSG00000040565 | 510.5959997 | -0.179507251 | 0.169182371 | -1.061028109 | 0.288677127 | 0.73951397 | Btaf1 | B-TFIID TATA-box binding protein associat | 540.49 | 583.86 | 503.12 | 523.44 | 555.12 | 357.54 |
| 11963 | ENSMUSG00000040569 | 0 | NA | NA | NA | NA | NA | Slc26a7 | solute carrier family 26, member 7 [Sourc | 0 | 0 | 0 | 0 | 0 | 0 |
| 11964 | ENSMUSG00000040570 | 5.572291392 | 0.664583789 | 1.065760671 | 0.623576951 | 0.532905452 | 0.889143097 | Rundc3b | RUN domain containing 3B [Source:MGI S | 4.8186 | 4.1117 | 4.025 | 11.568 | 0 | 8.9108 |
| 11965 | ENSMUSG00000040576 | 0 | NA | NA | NA | NA | NA | Tmem270 | transmembrane protein 270 [Source:MGI | 0 | 0 | 0 | 0 | 0 | 0 |
| 11966 | ENSMUSG00000040583 | 26.3677911 | -6.199073006 | 1.125136718 | -5.50961755 | 3.60E-08 | 4.52E-06 | Cyp2b13 | cytochrome P450, family 2, subfamily b, p | 28.109 | 51.396 | 76.475 | 0 | 0 | 2.2277 |
| 11967 | ENSMUSG00000040584 | 102.495108 | -1.310564837 | 0.337351174 | -3.88486817 | 0.000102385 | 0.003702257 | Abcb1a | ATP-binding cassette, sub-family B (MDR/ | 115.65 | 208.67 | 114.71 | 85.794 | 55.622 | 34.529 |
| 11968 | ENSMUSG00000040586 | 67.91679761 | -0.186942393 | 0.325900874 | -0.573617341 | 0.566226774 | 0.902076435 | Ofd1 | OFD1, centriole and centriolar satellite pr | 96.372 | 75.038 | 45.281 | 76.154 | 55.622 | 59.034 |
| 11969 | ENSMUSG00000040591 | 27.95619437 | -0.262928712 | 0.419624835 | -0.626580436 | 0.530934296 | 0.888794797 | 1110051M2( | RIKEN cDNA 1110051M20 gene [Source:M | 35.336 | 25.698 | 30.187 | 20.244 | 25.084 | 31.188 |
| 11970 | ENSMUSG00000040592 | 11.80708898 | -0.872723667 | 0.714463435 | -1.221509211 | 0.22189328 | 0.667949912 | Cd79b | CD79B antigen [Source:MGI Symbol;Acc:N | 19.274 | 17.475 | 9.0562 | 10.604 | 2.1812 | 12.252 |
| 11971 | ENSMUSG00000040594 | 1.38402626 | 0.084723649 | 1.970541585 | 0.042995108 | 0.965705433 | NA | Ranbp17 | RAN binding protein 17 [Source:MGI Sym | 0 | 2.0558 | 2.0125 | 0.964 | 3.2719 | 0 |
| 11972 | ENSMUSG00000040596 | 159.8157675 | 0.382326903 | 0.233521971 | 1.637220263 | 0.101584455 | 0.469189975 | Pogk | pogo transposable element with KRAB do | 122.87 | 160.36 | 133.83 | 229.43 | 131.96 | 180.44 |
| 11973 | ENSMUSG00000040599 | 326.3640646 | 0.300199317 | 0.163173157 | 1.839759203 | 0.065803597 | 0.368337165 | Mis12 | MIS12 kinetochore complex component [ | 293.13 | 296.04 | 287.79 | 293.05 | 386.08 | 402.1 |
| 11974 | ENSMUSG00000040600 | 0 | NA | NA | NA | NA | NA | Eps8l3 | EPS8-like 3 [Source:MGI Symbol;Acc:MGI: | 0 | 0 | 0 | 0 | 0 | 0 |
| 11975 | ENSMUSG00000040601 | 0 | NA | NA | NA | NA | NA | Nlrp4a | NLR family, pyrin domain containing 4A [S | 0 | 0 | 0 | 0 | 0 | 0 |
| 11976 | ENSMUSG00000040605 | 84.81856856 | -0.083576112 | 0.253061614 | -0.330259934 | 0.741203564 | 0.948792385 | Bace2 | beta-site APP-cleaving enzyme 2 [Source:I | 99.585 | 81.206 | 80.5 | 71.334 | 90.521 | 85.766 |
| 11977 | ENSMUSG00000040606 | 12.49864932 | 0.261662779 | 0.651996851 | 0.401325219 | 0.688180699 | 0.936908509 | Kazn | kazrin, periplakin interacting protein [Sou | 12.85 | 10.279 | 11.069 | 23.135 | 7.6343 | 10.025 |
| 11978 | ENSMUSG00000040610 | 0 | NA | NA | NA | NA | NA | Tlx3 | T cell leukemia, homeobox 3 [Source:MGI | 0 | 0 | 0 | 0 | 0 | 0 |
| 11979 | ENSMUSG00000040612 | 794.9658028 | -1.257451124 | 0.199298872 | -6.309374021 | 2.80E-10 | 6.75E-08 | Ildr2 | immunoglobulin-like domain containing re | 942.04 | 1333.2 | 1088.8 | 522.48 | 573.67 | 309.65 |
| 11980 | ENSMUSG00000040613 | 264.7263398 | 0.073143887 | 0.190742354 | 0.383469562 | 0.701371635 | 0.941991169 | Apobec1 | apolipoprotein B mRNA editing enzyme, c | 202.38 | 290.9 | 281.75 | 282.45 | 229.03 | 301.85 |

| | | | | | | | | | | | | | | | |
|---|---|---|---|---|---|---|---|---|---|---|---|---|---|---|---|
| 11981 | ENSMUSG00000040614 | 0 | NA | NA | NA | NA | NA | Nlrp9c | NLR family, pyrin domain containing 9C [S | 0 | 0 | 0 | 0 | 0 | 0 |
| 11982 | ENSMUSG00000040616 | 69.66566515 | 0.554693135 | 0.324040415 | 1.711802324 | 0.086933104 | 0.434457217 | Tmem51 | transmembrane protein 51 [Source:MGI S | 52.202 | 59.619 | 57.356 | 59.767 | 66.528 | 122.52 |
| 11983 | ENSMUSG00000040618 | 63.2347362 | 0.056632139 | 0.33847287 | 0.167316627 | 0.867120922 | 0.977936609 | Pck2 | phosphoenolpyruvate carboxykinase 2 (m | 40.155 | 57.563 | 88.55 | 52.055 | 69.8 | 71.286 |
| 11984 | ENSMUSG00000040620 | 346.7510721 | 0.015696045 | 0.167499482 | 0.09370802 | 0.925341099 | 0.987546253 | Dhx33 | DEAH (Asp-Glu-Ala-His) box polypeptide 3 | 361.4 | 391.64 | 281.75 | 387.52 | 317.37 | 340.84 |
| 11985 | ENSMUSG00000040621 | 17.29348501 | -0.151021987 | 0.559854755 | -0.269752083 | 0.787350988 | 0.960870081 | Gemin8 | gem nuclear organelle associated protein | 24.896 | 17.475 | 12.075 | 16.388 | 22.903 | 10.025 |
| 11986 | ENSMUSG00000040624 | 31.05632939 | -0.35162508 | 0.432897917 | -0.812258656 | 0.416643229 | 0.836878476 | Plekhg1 | pleckstrin homology domain containing, f | 52.202 | 25.698 | 26.162 | 26.027 | 26.175 | 30.074 |
| 11987 | ENSMUSG00000040627 | 0 | NA | NA | NA | NA | NA | Aicda | activation-induced cytidine deaminase [S | 0 | 0 | 0 | 0 | 0 | 0 |
| 11988 | ENSMUSG00000040629 | 0 | NA | NA | NA | NA | NA | Mael | maelstrom spermatogenic transposon sile | 0 | 0 | 0 | 0 | 0 | 0 |
| 11989 | ENSMUSG00000040631 | 11.10800241 | 0.043193136 | 0.645390224 | 0.066925612 | 0.946640923 | 0.990610209 | Dok4 | docking protein 4 [Source:MGI Symbol;Ac | 10.44 | 11.307 | 11.069 | 10.604 | 7.6343 | 15.594 |
| 11990 | ENSMUSG00000040632 | 0 | NA | NA | NA | NA | NA | Nrl | neural retina leucine zipper gene [Source: | 0 | 0 | 0 | 0 | 0 | 0 |
| 11991 | ENSMUSG00000040640 | 119.4735143 | 0.16191246 | 0.216586071 | 0.747566357 | 0.454721763 | 0.857337687 | Erc2 | ELKS/RAB6-interacting/CAST family memb | 125.28 | 103.82 | 108.67 | 114.71 | 139.6 | 124.75 |
| 11992 | ENSMUSG00000040648 | 1080.383822 | -0.420000427 | 0.128564439 | -3.266847587 | 0.001087522 | 0.022311943 | Ppip5k2 | diphosphoinositol pentakisphosphate kin | 1150.8 | 1433.9 | 1126 | 990.97 | 911.76 | 868.8 |
| 11993 | ENSMUSG00000040649 | 0.315620088 | 0.165143421 | 4.063300985 | 0.040642675 | 0.967580763 | NA | Rimklb | ribosomal modification protein rimK-like f | 0.8031 | 0 | 0 | 0 | 1.0906 | 0 |
| 11994 | ENSMUSG00000040650 | 0 | NA | NA | NA | NA | NA | Cyp2b23 | cytochrome P450, family 2, subfamily b, p | 0 | 0 | 0 | 0 | 0 | 0 |
| 11995 | ENSMUSG00000040651 | 479.1390796 | 0.147821006 | 0.136275459 | 1.084722124 | 0.278044746 | 0.730997543 | Tasor | transcription activation suppressor [Sourc | 474.63 | 418.36 | 470.92 | 551.4 | 499.5 | 460.02 |
| 11996 | ENSMUSG00000040652 | 582.2225231 | 0.114449525 | 0.124034519 | 0.922723176 | 0.356151485 | 0.795708016 | Oaz2 | ornithine decarboxylase antizyme 2 [Sour | 547.72 | 549.94 | 579.6 | 576.46 | 579.12 | 660.51 |
| 11997 | ENSMUSG00000040653 | 0 | NA | NA | NA | NA | NA | Ppp1r14c | protein phosphatase 1, regulatory inhibit | 0 | 0 | 0 | 0 | 0 | 0 |
| 11998 | ENSMUSG00000040657 | 0 | NA | NA | NA | NA | NA | 1700063H04 | RIKEN cDNA 1700063H04 gene [Source:M | 0 | 0 | 0 | 0 | 0 | 0 |
| 11999 | ENSMUSG00000040658 | 255.6556497 | -0.322549975 | 0.179426729 | -1.797669592 | 0.072229383 | 0.389401088 | Dnph1 | 2'-deoxynucleoside 5'-phosphate N-hydro | 279.48 | 242.59 | 330.05 | 207.26 | 258.48 | 216.09 |
| 12000 | ENSMUSG00000040659 | 1275.748773 | 0.188252894 | 0.10217431 | 1.842467975 | 0.065406727 | 0.367252677 | Efhd2 | EF hand domain containing 2 [Source:MG | 1231.2 | 1129.7 | 1216.5 | 1373.7 | 1301.1 | 1402.3 |
| 12001 | ENSMUSG00000040660 | 1021.698504 | -4.607191134 | 0.409316224 | -11.2558234 | 2.17E-29 | 1.57E-25 | Cyp2b9 | cytochrome P450, family 2, subfamily b, p | 1847.9 | 2049.7 | 1991.4 | 104.11 | 99.246 | 37.871 |
| 12002 | ENSMUSG00000040661 | 909.9127905 | -0.518960665 | 0.191448457 | -2.710706961 | 0.006713993 | 0.08160729 | Rad54l2 | RAD54 like 2 (S. cerevisiae) [Source:MGI S | 1036.8 | 1325 | 854.3 | 824.2 | 882.31 | 536.87 |
| 12003 | ENSMUSG00000040663 | 9.609093664 | 0.048509646 | 0.711802305 | 0.068150448 | 0.945665872 | 0.990309554 | Clcf1 | cardiotrophin-like cytokine factor 1 [Sourc | 12.047 | 5.1396 | 11.069 | 10.604 | 6.5437 | 12.252 |
| 12004 | ENSMUSG00000040666 | 0.688942901 | 0.052211695 | 2.79596103 | 0.018673971 | 0.985101193 | NA | Sh3bgr | SH3-binding domain glutamic acid-rich pr | 0 | 2.0558 | 0 | 0.964 | 0 | 1.1138 |
| 12005 | ENSMUSG00000040667 | 491.9964955 | 0.191620782 | 0.143127844 | 1.338808553 | 0.180633006 | 0.613962723 | Nup88 | nucleoporin 88 [Source:MGI Symbol;Acc:I | 436.89 | 500.6 | 440.74 | 477.17 | 600.93 | 495.66 |
| 12006 | ENSMUSG00000040669 | 53.10926662 | 0.479972726 | 0.357830057 | 1.341342677 | 0.17980922 | 0.612025777 | Phc1 | polyhomeotic 1 [Source:MGI Symbol;Acc: | 53.808 | 35.977 | 43.269 | 89.65 | 44.715 | 51.237 |
| 12007 | ENSMUSG00000040675 | 27.83806522 | -0.626042396 | 0.500275898 | -1.251394277 | 0.210790664 | 0.654968028 | Mthfd1l | methylenetetrahydrofolate dehydrogenas | 31.321 | 45.228 | 25.156 | 35.667 | 19.631 | 10.025 |
| 12008 | ENSMUSG00000040680 | 0.724629373 | 1.713042495 | 3.014019035 | 0.56835822 | 0.569791754 | NA | Kremen2 | kringle containing transmembrane protei | 0 | 0 | 1.0062 | 0 | 0 | 3.3415 |
| 12009 | ENSMUSG00000040681 | 884.3108475 | 0.140673184 | 0.124739448 | 1.127736147 | 0.259431367 | 0.712728614 | Hmgn1 | high mobility group nucleosomal binding | 749.29 | 856.26 | 918.7 | 848.3 | 956.47 | 976.84 |
| 12010 | ENSMUSG00000040687 | 457.915099 | 0.357123655 | 0.142873804 | 2.499574072 | 0.01243427 | 0.125944877 | Madd | MAP-kinase activating death domain [Sou | 358.99 | 413.22 | 433.69 | 570.67 | 487.51 | 483.41 |
| 12011 | ENSMUSG00000040688 | 366.6154801 | -0.177627855 | 0.151791151 | -1.170212183 | 0.241915591 | 0.691791181 | Tbl3 | transducin (beta)-like 3 [Source:MGI Symb | 381.47 | 420.42 | 366.27 | 380.77 | 353.36 | 297.4 |
| 12012 | ENSMUSG00000040690 | 58.87639717 | -1.154870885 | 0.405123802 | -2.850661645 | 0.004362837 | 0.060970118 | Col16a1 | collagen, type XVI, alpha 1 [Source:MGI Sy | 110.83 | 89.429 | 43.269 | 31.811 | 54.531 | 23.391 |
| 12013 | ENSMUSG00000040693 | 0.531944269 | 2.618916564 | 3.285123409 | 0.797204926 | 0.425332026 | NA | Slco4c1 | solute carrier organic anion transporter fa | 0 | 0 | 0 | 0.964 | 0 | 2.2277 |
| 12014 | ENSMUSG00000040694 | 0.849605875 | 0.613471193 | 2.572643751 | 0.238459442 | 0.811524769 | NA | Apobec2 | apolipoprotein B mRNA editing enzyme, c | 0 | 2.0558 | 0 | 1.928 | 0 | 1.1138 |
| 12015 | ENSMUSG00000040697 | 311.8698287 | 0.166694851 | 0.143048877 | 1.165299966 | 0.243897593 | 0.693748143 | Dnajc16 | DnaJ heat shock protein family (Hsp40) m | 294.74 | 291.93 | 294.83 | 321 | 346.82 | 321.9 |
| 12016 | ENSMUSG00000040699 | 379.1210339 | 0.077538403 | 0.193778436 | 0.400139482 | 0.689053786 | 0.936908509 | Limd2 | LIM domain containing 2 [Source:MGI Syn | 329.27 | 352.58 | 424.64 | 280.52 | 418.8 | 468.93 |
| 12017 | ENSMUSG00000040701 | 127.9259451 | -0.122316048 | 0.217905993 | -0.561324844 | 0.574576108 | 0.906163061 | Ap1g2 | adaptor protein complex AP-1, gamma 2 s | 138.94 | 129.52 | 131.82 | 145.56 | 101.43 | 120.3 |
| 12018 | ENSMUSG00000040703 | 16.97611118 | 0.169679138 | 0.649733194 | 0.261152023 | 0.793975275 | 0.961915077 | Cyp2s1 | cytochrome P450, family 2, subfamily s, p | 31.321 | 9.2513 | 7.0437 | 22.171 | 10.906 | 21.163 |
| 12019 | ENSMUSG00000040705 | 0 | NA | NA | NA | NA | NA | A930016O22 | RIKEN cDNA A930016O22 gene [Source:M | 0 | 0 | 0 | 0 | 0 | 0 |
| 12020 | ENSMUSG00000040706 | 3546.066665 | 0.072109523 | 0.106961629 | 0.674162541 | 0.500207981 | 0.880716237 | Agmat | agmatine ureohydrolase (agmatinase) [So | 3494.3 | 3212.2 | 3664.7 | 3266 | 3693.9 | 3945.2 |
| 12021 | ENSMUSG00000040710 | 8.119088986 | -1.697347155 | 0.789603234 | -2.149620319 | 0.03158526 | 0.236342058 | St8sia4 | ST8 alpha-N-acetyl-neuraminide alpha-2,8 | 12.85 | 13.363 | 11.069 | 4.8199 | 3.2719 | 3.3415 |
| 12022 | ENSMUSG00000040711 | 19.5094958 | -0.052199806 | 0.563247032 | -0.092676574 | 0.926160508 | 0.98758158 | Sh3pxd2b | SH3 and PX domains 2B [Source:MGI Sym | 8.8341 | 31.865 | 19.119 | 15.424 | 23.994 | 17.822 |
| 12023 | ENSMUSG00000040712 | 508.5310748 | -0.16335286 | 0.143230221 | -1.140491571 | 0.254081563 | 0.707252596 | Camta2 | calmodulin binding transcription activator | 505.95 | 521.15 | 585.63 | 535.01 | 413.34 | 490.09 |
| 12024 | ENSMUSG00000040713 | 6494.137012 | -0.075527106 | 0.091869036 | -0.822117102 | 0.411010253 | 0.833663203 | Creg1 | cellular repressor of E1A-stimulated gene | 6652.9 | 6511.9 | 6828.4 | 6823 | 5964.6 | 6184.1 |
| 12025 | ENSMUSG00000040714 | 3.068911489 | 0.149414242 | 1.214255074 | 0.123050128 | 0.902067403 | NA | Klc3 | kinesin light chain 3 [Source:MGI Symbol; | 1.6062 | 5.1396 | 2.0125 | 1.928 | 3.2719 | 4.4554 |
| 12026 | ENSMUSG00000040715 | 0 | NA | NA | NA | NA | NA | Rsc1a1 | regulatory solute carrier protein, family 1, | 0 | 0 | 0 | 0 | 0 | 0 |
| 12027 | ENSMUSG00000040717 | 9.079598353 | -0.80250432 | 0.742688795 | -1.080539152 | 0.279902162 | 0.732834426 | Il17rd | interleukin 17 receptor D [Source:MGI Syr | 15.259 | 9.2513 | 10.062 | 6.7478 | 9.8156 | 3.3415 |
| 12028 | ENSMUSG00000040720 | 363.7562632 | -0.052277268 | 0.140147777 | -0.373015319 | 0.709137042 | 0.944237849 | Virma | vir like m6A methyltransferase associated | 378.26 | 369.02 | 363.25 | 336.43 | 345.73 | 389.85 |
| 12029 | ENSMUSG00000040721 | 153.2542619 | 0.265482229 | 0.241665524 | 1.098552352 | 0.271963371 | 0.727618887 | Zfhx2 | zinc finger homeobox 2 [Source:MGI Syml | 130.91 | 104.85 | 182.13 | 193.76 | 135.24 | 172.65 |
| 12030 | ENSMUSG00000040722 | 293.8779039 | 0.056758792 | 0.211234156 | 0.268700824 | 0.788159917 | 0.960870081 | Scamp5 | secretory carrier membrane protein 5 [So | 375.05 | 225.11 | 262.63 | 239.07 | 323.91 | 337.49 |
| 12031 | ENSMUSG00000040723 | 65.39906246 | -1.380913797 | 0.390328047 | -3.537828775 | 0.000403432 | 0.010687243 | Rcsd1 | RCSD domain containing 1 [Source:MGI Sy | 91.554 | 129.52 | 62.387 | 34.703 | 18.541 | 55.692 |
| 12032 | ENSMUSG00000040724 | 30.23889334 | -1.743031681 | 0.492338782 | -3.540309525 | 0.000399658 | 0.010645771 | Kcna2 | potassium voltage-gated channel, shaker- | 57.02 | 57.563 | 25.156 | 17.352 | 7.6343 | 16.708 |
| 12033 | ENSMUSG00000040725 | 1214.419703 | 0.100632613 | 0.10891755 | 0.923933869 | 0.355520748 | 0.795424787 | Hnrnpul1 | heterogeneous nuclear ribonucleoprotein | 1165.3 | 1217.1 | 1134 | 1277.3 | 1147.3 | 1345.5 |
| 12034 | ENSMUSG00000040726 | 0 | NA | NA | NA | NA | NA | Hesx1 | homeobox gene expressed in ES cells [Sou | 0 | 0 | 0 | 0 | 0 | 0 |
| 12035 | ENSMUSG00000040728 | 0.994165048 | 1.185966287 | 2.243927044 | 0.528522659 | 0.597136625 | NA | Esrp1 | epithelial splicing regulatory protein 1 [So | 0.8031 | 0 | 1.0062 | 1.928 | 0 | 2.2277 |
| 12036 | ENSMUSG00000040729 | 2.805088077 | -1.237186471 | 1.429825179 | -0.865271146 | 0.386889984 | NA | Cep126 | centrosomal protein 126 [Source:MGI Syn | 4.8186 | 1.0279 | 6.0375 | 3.8559 | 1.0906 | 0 |

| | | | | | | | | | | | | | | | |
|---|---|---|---|---|---|---|---|---|---|---|---|---|---|---|---|
| 12037 | ENSMUSG00000040731 | 2990.443554 | 0.107080654 | 0.098918328 | 1.082515814 | 0.279023392 | 0.731519729 | Eif4h | eukaryotic translation initiation factor 4H | 2702.4 | 3089.9 | 2846.7 | 2916 | 3204.2 | 3183.4 |
| 12038 | ENSMUSG00000040732 | 87.11280511 | -0.536428279 | 0.266104391 | -2.015856547 | 0.043814978 | 0.288851344 | Erg | ETS transcription factor [Source:MGI Syml | 112.43 | 114.1 | 82.512 | 64.587 | 62.165 | 86.88 |
| 12039 | ENSMUSG00000040734 | 77.67720879 | 0.239082306 | 0.267936422 | 0.892309841 | 0.372226881 | 0.807785387 | Ppp1r13l | protein phosphatase 1, regulatory subunit | 63.445 | 74.01 | 76.475 | 80.974 | 69.8 | 101.36 |
| 12040 | ENSMUSG00000040738 | 302.4188926 | 0.32006609 | 0.151005677 | 2.119563295 | 0.034042891 | 0.247046246 | Ints8 | integrator complex subunit 8 [Source:MG | 242.54 | 280.62 | 284.77 | 350.89 | 330.46 | 325.24 |
| 12041 | ENSMUSG00000040740 | 186.9234125 | -0.13975384 | 0.197136587 | -0.708918838 | 0.478374844 | 0.869794311 | Slc25a34 | solute carrier family 25, member 34 [Sour | 183.11 | 164.47 | 240.49 | 177.37 | 172.32 | 183.78 |
| 12042 | ENSMUSG00000040746 | 835.6094476 | 0.060494933 | 0.132690924 | 0.455908593 | 0.64845571 | 0.925643431 | Rnf167 | ring finger protein 167 [Source:MGI Symb | 949.27 | 716.46 | 786.88 | 821.31 | 840.87 | 898.87 |
| 12043 | ENSMUSG00000040747 | 63.27368297 | -1.66151146 | 0.382678068 | -4.341799546 | 1.41E-05 | 0.000708302 | Cd53 | CD53 antigen [Source:MGI Symbol;Acc:M | 85.129 | 141.85 | 61.381 | 26.027 | 21.812 | 43.44 |
| 12044 | ENSMUSG00000040749 | 18.12337445 | 0.278337795 | 0.571797393 | 0.486776958 | 0.626416401 | 0.919846378 | Siah1b | siah E3 ubiquitin protein ligase 1B [Source | 14.456 | 22.614 | 12.075 | 16.388 | 10.906 | 32.301 |
| 12045 | ENSMUSG00000040751 | 3.852381995 | -0.810231655 | 1.144359391 | -0.708022026 | 0.478931579 | NA | Lat2 | linker for activation of T cells family, mem | 5.6217 | 1.0279 | 8.05 | 2.8919 | 2.1812 | 3.3415 |
| 12046 | ENSMUSG00000040752 | 0 | NA | NA | NA | NA | NA | Myh6 | myosin, heavy polypeptide 6, cardiac mus | 0 | 0 | 0 | 0 | 0 | 0 |
| 12047 | ENSMUSG00000040759 | 0 | NA | NA | NA | NA | NA | Cmtm5 | CKLF-like MARVEL transmembrane domai | 0 | 0 | 0 | 0 | 0 | 0 |
| 12048 | ENSMUSG00000040760 | 252.6858293 | 0.083194916 | 0.200184314 | 0.415591585 | 0.677708885 | 0.93325821 | Appl1 | adaptor protein, phosphotyrosine interac | 231.29 | 253.9 | 250.55 | 186.05 | 283.56 | 310.76 |
| 12049 | ENSMUSG00000040761 | 468.0494609 | 0.081984197 | 0.149263506 | 0.549258148 | 0.582828305 | 0.908645447 | Spen | spen family transcription repressor [Sourc | 423.23 | 436.87 | 505.13 | 547.54 | 418.8 | 476.73 |
| 12050 | ENSMUSG00000040767 | 32.70397098 | 0.445068331 | 0.401125936 | 1.109547628 | 0.26719401 | 0.722778036 | Snrnp25 | small nuclear ribonucleoprotein 25 (U11/U | 24.093 | 28.782 | 30.187 | 26.027 | 40.353 | 46.781 |
| 12051 | ENSMUSG00000040770 | 0 | NA | NA | NA | NA | NA | Il25 | interleukin 25 [Source:MGI Symbol;Acc:M | 0 | 0 | 0 | 0 | 0 | 0 |
| 12052 | ENSMUSG00000040771 | 504.596846 | 0.152147281 | 0.134391212 | 1.132122252 | 0.257583041 | 0.710872182 | Oard1 | O-acyl-ADP-ribose deacylase 1 [Source:M | 469.81 | 481.07 | 483 | 508.02 | 485.33 | 600.36 |
| 12053 | ENSMUSG00000040774 | 1110.568033 | -0.189542876 | 0.126936558 | -1.493209518 | 0.135382351 | 0.538068113 | Cept1 | choline/ethanolaminephosphotransferase | 1111.5 | 1292.1 | 1148.1 | 1185.7 | 933.57 | 992.43 |
| 12054 | ENSMUSG00000040775 | 0.267700746 | -1.497481649 | 4.057001436 | -0.369110456 | 0.712045396 | NA | Gm9772 | predicted gene 9772 [Source:MGI Symbol | 1.6062 | 0 | 0 | 0 | 0 | 0 |
| 12055 | ENSMUSG00000040782 | 1047.79026 | 0.036289145 | 0.116760275 | 0.310800445 | 0.755952331 | 0.953085143 | Cop1 | COP1, E3 ubiquitin ligase [Source:MGI Sym | 1003.9 | 979.61 | 1121 | 1099.9 | 1121.2 | 961.25 |
| 12056 | ENSMUSG00000040785 | 358.0709879 | -0.220124704 | 0.170204749 | -1.293293549 | 0.195909551 | 0.636484845 | Ttc3 | tetratricopeptide repeat domain 3 [Source | 409.58 | 440.98 | 304.89 | 301.73 | 334.82 | 356.43 |
| 12057 | ENSMUSG00000040794 | 10.96568636 | 0.383410376 | 0.686990448 | 0.558101466 | 0.5767751 | 0.907752116 | C1qtnf4 | C1q and tumor necrosis factor related pro | 7.2279 | 11.307 | 10.062 | 10.604 | 6.5437 | 20.049 |
| 12058 | ENSMUSG00000040795 | 126.4516975 | -0.109763193 | 0.216798506 | -0.506291277 | 0.612652185 | 0.916495447 | Iqcc | IQ motif containing C [Source:MGI Symbo | 130.91 | 121.29 | 141.88 | 137.85 | 125.42 | 101.36 |
| 12059 | ENSMUSG00000040797 | 0.506734502 | -2.372847156 | 3.338060886 | -0.710845978 | 0.477179685 | NA | Iqsec3 | IQ motif and Sec7 domain 3 [Source:MGI S | 0 | 1.0279 | 2.0125 | 0 | 0 | 0 |
| 12060 | ENSMUSG00000040808 | 1.765602645 | -0.538112305 | 1.961752587 | -0.27430182 | 0.783852686 | NA | S100g | S100 calcium binding protein G [Source:M | 3.2124 | 0 | 3.0187 | 0 | 4.3625 | 0 |
| 12061 | ENSMUSG00000040809 | 1.827280042 | -3.128059463 | 2.039372122 | -1.53383457 | 0.125070356 | NA | Chil3 | chitinase-like 3 [Source:MGI Symbol;Acc:M | 4.8186 | 0 | 5.0312 | 0 | 0 | 1.1138 |
| 12062 | ENSMUSG00000040811 | 239.1167418 | 0.017817802 | 0.167036226 | 0.106670284 | 0.915050558 | 0.98625062 | Eml2 | echinoderm microtubule associated prote | 232.1 | 229.23 | 251.56 | 235.21 | 215.94 | 270.66 |
| 12063 | ENSMUSG00000040812 | 0.465832987 | -0.698295221 | 3.18348759 | -0.219349126 | 0.826378096 | NA | Agbl2 | ATP/GTP binding protein-like 2 [Source:M | 0.8031 | 1.0279 | 0 | 0.964 | 0 | 0 |
| 12064 | ENSMUSG00000040813 | 1536.933659 | 0.054350916 | 0.106289611 | 0.511347395 | 0.609107823 | 0.916376001 | Tex264 | testis expressed gene 264 [Source:MGI Sy | 1606.2 | 1444.2 | 1472.1 | 1462.4 | 1680.6 | 1556 |
| 12065 | ENSMUSG00000040818 | 378.764191 | -0.084412248 | 0.14910643 | -0.566120774 | 0.571311682 | 0.904651508 | Dennd6a | DENN/MADD domain containing 6A [Sour | 358.99 | 396.78 | 414.57 | 385.59 | 396.99 | 319.67 |
| 12066 | ENSMUSG00000040820 | 295.0661813 | 0.00243288 | 0.178106684 | 0.013659676 | 0.989101494 | 0.99859176 | Hlcs | holocarboxylase synthetase (biotin- [prop | 316.42 | 330.99 | 236.47 | 263.17 | 332.64 | 290.71 |
| 12067 | ENSMUSG00000040822 | 410.817781 | -0.072485344 | 0.134130689 | -0.540408348 | 0.588915452 | 0.911039598 | 1700123O20 | RIKEN cDNA 1700123O20 gene [Source:M | 411.19 | 435.84 | 416.59 | 403.91 | 371.9 | 425.49 |
| 12068 | ENSMUSG00000040824 | 524.5110191 | 0.158648113 | 0.133754062 | 1.186118093 | 0.235575666 | 0.683799474 | Snrpd2 | small nuclear ribonucleoprotein D2 [Sourc | 495.51 | 469.76 | 521.23 | 499.34 | 547.49 | 613.73 |
| 12069 | ENSMUSG00000040828 | 17.25560325 | 0.319715162 | 0.646773928 | 0.494322898 | 0.621078156 | 0.918069106 | Catsperd | cation channel sperm associated auxiliary | 18.471 | 12.335 | 15.094 | 15.424 | 5.4531 | 36.757 |
| 12070 | ENSMUSG00000040829 | 56.86244927 | -0.441027668 | 0.377804067 | -1.167344946 | 0.243071094 | 0.692645587 | Zmynd15 | zinc finger, MYND-type containing 15 [Sou | 69.87 | 56.535 | 70.437 | 78.082 | 27.265 | 38.985 |
| 12071 | ENSMUSG00000040836 | 3.770346871 | -1.076005261 | 1.127415888 | -0.954399589 | 0.339881414 | NA | Gpr161 | G protein-coupled receptor 161 [Source:M | 7.2279 | 2.0558 | 6.0375 | 2.8919 | 2.1812 | 2.2277 |
| 12072 | ENSMUSG00000040838 | 0.301557804 | -1.64335916 | 4.042814367 | -0.406488899 | 0.684383412 | NA | Gm11639 | predicted gene 11639 [Source:MGI Symbo | 0.8031 | 0 | 1.0062 | 0 | 0 | 0 |
| 12073 | ENSMUSG00000040840 | 24.8105311 | 0.260782061 | 0.45056225 | 0.578792522 | 0.562729178 | 0.900565925 | 4930579G18 | RIKEN cDNA 4930579G18 gene [Source:M | 24.093 | 19.53 | 24.15 | 34.703 | 18.541 | 27.846 |
| 12074 | ENSMUSG00000040841 | 83.01590547 | 0.225779257 | 0.294197405 | 0.767441363 | 0.442819144 | 0.850769625 | Six5 | sine oculis-related homeobox 5 [Source:M | 59.43 | 71.954 | 98.612 | 93.506 | 65.437 | 109.16 |
| 12075 | ENSMUSG00000040842 | 1467.656644 | 0.055532582 | 0.109638916 | 0.506504291 | 0.612502678 | 0.916441371 | Szrd1 | SUZ RNA binding domain containing 1 [So | 1344.4 | 1457.6 | 1516.4 | 1380.4 | 1487.6 | 1619.5 |
| 12076 | ENSMUSG00000040843 | 354.7968344 | -0.038726483 | 0.138896153 | -0.278816098 | 0.78038596 | 0.959903268 | Tiprl | TIP41, TOR signalling pathway regulator-li | 358.18 | 372.11 | 348.16 | 329.68 | 366.45 | 354.2 |
| 12077 | ENSMUSG00000040848 | 1371.3657 | 0.117116043 | 0.114121335 | 1.026241437 | 0.304777794 | 0.753556583 | Sft2d2 | SFT2 domain containing 2 [Source:MGI Sy | 1273.7 | 1270.5 | 1402.7 | 1327.4 | 1362.2 | 1591.7 |
| 12078 | ENSMUSG00000040850 | 2095.772183 | 0.080223546 | 0.101357936 | 0.791487567 | 0.428659528 | 0.842753542 | Psme4 | proteasome (prosome, macropain) activat | 1904.2 | 2083.6 | 2125.2 | 2009.9 | 2259.8 | 2192 |
| 12079 | ENSMUSG00000040852 | 20.19018329 | -0.64285829 | 0.49710569 | -1.293202439 | 0.195941052 | 0.636484845 | Plekhh2 | pleckstrin homology domain containing, f | 24.896 | 31.865 | 17.106 | 16.388 | 14.178 | 16.708 |
| 12080 | ENSMUSG00000040855 | 818.5627102 | 0.120357581 | 0.118824289 | 1.012903864 | 0.311106083 | 0.759594595 | Reps2 | RALBP1 associated Eps domain containing | 763.75 | 778.13 | 811.03 | 763.47 | 913.94 | 881.05 |
| 12081 | ENSMUSG00000040856 | 0.703263909 | 1.656064937 | 2.753745348 | 0.601386377 | 0.54758267 | NA | Dlk1 | delta like non-canonical Notch ligand 1 [S | 0 | 1.0279 | 0 | 0.964 | 0 | 2.2277 |
| 12082 | ENSMUSG00000040857 | 723.8389746 | 0.461490114 | 0.116283155 | 3.968675563 | 7.23E-05 | 0.002787626 | Erf | Ets2 repressor factor [Source:MGI Symbol | 583.05 | 610.58 | 633.93 | 834.8 | 790.7 | 889.96 |
| 12083 | ENSMUSG00000040859 | 1260.620224 | 0.028992571 | 0.107418792 | 0.269902228 | 0.787235473 | 0.960870081 | Bsdc1 | BSD domain containing 1 [Source:MGI Syr | 1311.5 | 1238.6 | 1193.4 | 1331.3 | 1173.5 | 1315.4 |
| 12084 | ENSMUSG00000040860 | 36.36844279 | -0.263708042 | 0.387116103 | -0.68121176 | 0.495737508 | 0.877965876 | Crocc | ciliary rootlet coiled-coil, rootletin [Source | 28.912 | 39.061 | 51.318 | 31.811 | 39.262 | 27.846 |
| 12085 | ENSMUSG00000040865 | 309.0051702 | -0.079307111 | 0.157696164 | -0.502910843 | 0.615026958 | 0.917509899 | Ino80d | INO80 complex subunit D [Source:MGI Sy | 293.13 | 345.38 | 314.95 | 338.36 | 274.84 | 287.37 |
| 12086 | ENSMUSG00000040866 | 0 | NA | NA | NA | NA | NA | Rsph6a | radial spoke head 6 homolog A (Chlamydo | 0 | 0 | 0 | 0 | 0 | 0 |
| 12087 | ENSMUSG00000040867 | 0 | NA | NA | NA | NA | NA | Begain | brain-enriched guanylate kinase-associate | 0 | 0 | 0 | 0 | 0 | 0 |
| 12088 | ENSMUSG00000040875 | 1.268212842 | -1.340829108 | 2.218250673 | -0.604453376 | 0.545542258 | NA | Osbpl10 | oxysterol binding protein-like 10 [Source:I | 2.4093 | 0 | 3.0187 | 0 | 2.1812 | 0 |
| 12089 | ENSMUSG00000040877 | 70.60487199 | -0.358874658 | 0.2770011 | -1.295571239 | 0.195123243 | 0.63535853 | Wdr25 | WD repeat domain 25 [Source:MGI Symbo | 88.341 | 66.815 | 82.512 | 54.947 | 71.981 | 59.034 |
| 12090 | ENSMUSG00000040883 | 7092.821361 | 0.112804086 | 0.083380403 | 1.352884878 | 0.176092411 | 0.607001136 | Tmem205 | transmembrane protein 205 [Source:MGI | 7072.9 | 6631.1 | 6741.8 | 7221.2 | 7372.6 | 7517.3 |
| 12091 | ENSMUSG00000040888 | 619.744443 | -0.016371659 | 0.13421165 | -0.121983892 | 0.902911775 | 0.983821605 | Gfer | growth factor, augmenter of liver regener | 531.65 | 676.37 | 663.12 | 630.44 | 600.93 | 615.96 |
| 12092 | ENSMUSG00000040891 | 980.6901316 | 0.499086019 | 0.146504335 | 3.406629691 | 0.000657703 | 0.01519651 | Foxa3 | forkhead box A3 [Source:MGI Symbol;Acc | 644.09 | 951.85 | 844.24 | 1226.2 | 1092.8 | 1125 |

| | | | | | | | | | | | | | | | |
|---|---|---|---|---|---|---|---|---|---|---|---|---|---|---|---|
| 12093 | ENSMUSG00000040896 | 2.581971762 | 0.323790554 | 1.330431546 | 0.243372577 | 0.807716785 | NA | Kcnd3 | potassium voltage-gated channel, Shal-rel | 0.8031 | 4.1117 | 2.0125 | 1.928 | 2.1812 | 4.4554 |
| 12094 | ENSMUSG00000040899 | 2.867921567 | -1.685143673 | 1.378075961 | -1.222823502 | 0.221396367 | NA | Ccr6 | chemokine (C-C motif) receptor 6 [Source | 4.0155 | 6.1675 | 3.0187 | 2.8919 | 0 | 1.1138 |
| 12095 | ENSMUSG00000040901 | 0 | NA | NA | NA | NA | NA | Kcnk18 | potassium channel, subfamily K, member | 0 | 0 | 0 | 0 | 0 | 0 |
| 12096 | ENSMUSG00000040904 | 1.841581588 | -0.783093515 | 2.023706327 | -0.386960056 | 0.698785779 | NA | Gm21988 | predicted gene 21988 [Source:MGI Symbo | 0 | 0 | 7.0437 | 2.8919 | 0 | 1.1138 |
| 12097 | ENSMUSG00000040907 | 1.552508209 | 0.243045542 | 1.75637778 | 0.138378853 | 0.889941009 | NA | Atp1a3 | ATPase, Na+/K+ transporting, alpha 3 poly | 3.2124 | 0 | 1.0062 | 2.8919 | 1.0906 | 1.1138 |
| 12098 | ENSMUSG00000040913 | 264.8659433 | -0.019765515 | 0.186354075 | -0.106064305 | 0.915531332 | 0.98625062 | Fbxw4 | F-box and WD-40 domain protein 4 [Sourc | 272.25 | 234.37 | 292.82 | 214 | 318.46 | 257.3 |
| 12099 | ENSMUSG00000040918 | 1524.885944 | 0.245784842 | 0.134101855 | 1.832822093 | 0.066829033 | 0.37192178 | Slc19a2 | solute carrier family 19 (thiamine transpo | 1243.2 | 1510 | 1432.9 | 1398.7 | 1665.4 | 1899.1 |
| 12100 | ENSMUSG00000040919 | 0 | NA | NA | NA | NA | NA | 4930505A04 | RIKEN cDNA 4930505A04 gene [Source:M | 0 | 0 | 0 | 0 | 0 | 0 |
| 12101 | ENSMUSG00000040928 | 265.8491386 | 0.45702349 | 0.17128923 | 2.668139092 | 0.007627267 | 0.088790029 | S100pbp | S100P binding protein [Source:MGI Symb | 198.37 | 236.42 | 238.48 | 346.07 | 266.11 | 309.65 |
| 12102 | ENSMUSG00000040929 | 47.13554102 | -0.3391438 | 0.328578566 | -1.032154361 | 0.301999811 | 0.752248241 | Rfx3 | regulatory factor X, 3 (influences HLA clas | 51.399 | 46.256 | 60.375 | 45.307 | 33.809 | 45.668 |
| 12103 | ENSMUSG00000040935 | 0 | NA | NA | NA | NA | NA | Padi6 | peptidyl arginine deiminase, type VI [Sour | 0 | 0 | 0 | 0 | 0 | 0 |
| 12104 | ENSMUSG00000040936 | 1.866758683 | -1.280213463 | 1.790252286 | -0.715102264 | 0.474545855 | NA | Ulk4 | unc-51-like kinase 4 [Source:MGI Symbol; | 4.8186 | 1.0279 | 2.0125 | 0 | 0 | 3.3415 |
| 12105 | ENSMUSG00000040938 | 401.7818788 | -0.154812269 | 0.165923004 | -0.933036801 | 0.350800968 | 0.791943999 | Slc16a11 | solute carrier family 16 (monocarboxylic a | 469.01 | 378.27 | 422.62 | 443.43 | 309.74 | 387.62 |
| 12106 | ENSMUSG00000040940 | 702.5592599 | -0.118490503 | 0.132897406 | -0.891593793 | 0.372610698 | 0.808133324 | Arhgef1 | Rho guanine nucleotide exchange factor ( | 665.77 | 730.85 | 798.96 | 757.69 | 622.74 | 639.35 |
| 12107 | ENSMUSG00000040943 | 191.2991315 | -0.048055752 | 0.290958124 | -0.165163811 | 0.868815049 | 0.97847745 | Tet2 | tet methylcytosine dioxygenase 2 [Source | 185.52 | 203.53 | 195.21 | 306.54 | 118.88 | 138.12 |
| 12108 | ENSMUSG00000040945 | 612.4303306 | 0.180832294 | 0.120316399 | 1.502972959 | 0.132846017 | 0.533805738 | Rcc2 | regulator of chromosome condensation 2 | 534.06 | 598.25 | 590.67 | 637.19 | 676.18 | 638.23 |
| 12109 | ENSMUSG00000040950 | 0.428363721 | -0.543730382 | 3.624677344 | -0.15000794 | 0.880758351 | NA | Mgl2 | macrophage galactose N-acetyl-galactosa | 1.6062 | 0 | 0 | 0.964 | 0 | 0 |
| 12110 | ENSMUSG00000040952 | 2187.435379 | -0.029942045 | 0.099133765 | -0.3020368 | 0.76262401 | 0.95469451 | Rps19 | ribosomal protein S19 [Source:MGI Symb | 2178.8 | 2389.9 | 2061.8 | 2121.7 | 2171.4 | 2201 |
| 12111 | ENSMUSG00000040957 | 22.69214257 | -1.012539635 | 0.515976315 | -1.962376192 | 0.049718703 | 0.31211543 | Cables1 | CDK5 and Abl enzyme substrate 1 [Source | 28.912 | 38.033 | 24.15 | 17.352 | 6.5437 | 21.163 |
| 12112 | ENSMUSG00000040963 | 3511.676845 | 0.042124889 | 0.105571066 | 0.399019263 | 0.68987901 | 0.937261276 | Asgr2 | asialoglycoprotein receptor 2 [Source:MG | 3116 | 3540.1 | 3726.1 | 3427.9 | 3827 | 3432.9 |
| 12113 | ENSMUSG00000040964 | 508.8313129 | 0.250989208 | 0.160903677 | 1.559872418 | 0.118790034 | 0.504308497 | Arhgef10l | Rho guanine nucleotide exchange factor ( | 444.12 | 421.45 | 529.28 | 655.5 | 462.42 | 540.21 |
| 12114 | ENSMUSG00000040966 | 0.342639279 | -1.805976596 | 4.028547714 | -0.448294702 | 0.653940524 | NA | Slc22a2 | solute carrier family 22 (organic cation tra | 0 | 2.0558 | 0 | 0 | 0 | 0 |
| 12115 | ENSMUSG00000040969 | 0.435408177 | -2.179616019 | 3.531869654 | -0.617128103 | 0.537150234 | NA | Arhgef38 | Rho guanine nucleotide exchange factor ( | 1.6062 | 0 | 1.0062 | 0 | 0 | 0 |
| 12116 | ENSMUSG00000040972 | 1.771515096 | -1.247833515 | 1.649051971 | -0.756697507 | 0.449231109 | NA | Igsf21 | immunoglobulin superfamily, member 21 | 2.4093 | 2.0558 | 3.0187 | 0.964 | 2.1812 | 0 |
| 12117 | ENSMUSG00000040978 | 29.12610448 | -0.374652117 | 0.463989832 | -0.807457602 | 0.419402892 | 0.837591673 | Gm11992 | predicted gene 11992 [Source:MGI Symbo | 51.399 | 23.642 | 23.144 | 19.28 | 29.447 | 27.846 |
| 12118 | ENSMUSG00000040985 | 9.542211952 | -0.049850668 | 0.733372282 | -0.067974573 | 0.945805875 | 0.990309554 | Sun3 | Sad1 and UNC84 domain containing 3 [So | 8.8341 | 11.307 | 9.0562 | 12.532 | 3.2719 | 12.252 |
| 12119 | ENSMUSG00000040987 | 21.58874764 | -0.777403634 | 0.507652526 | -1.531369576 | 0.125678079 | 0.519819197 | Mill2 | MHC I like leukocyte 2 [Source:MGI Symb | 28.912 | 33.921 | 19.119 | 22.171 | 9.8156 | 15.594 |
| 12120 | ENSMUSG00000040990 | 21.27772598 | -0.831054232 | 0.533714235 | -1.557114608 | 0.119443277 | 0.505745774 | Sh3kbp1 | SH3-domain kinase binding protein 1 [Sou | 30.518 | 31.865 | 19.119 | 9.6398 | 10.906 | 25.618 |
| 12121 | ENSMUSG00000040997 | 719.5145675 | -0.210587274 | 0.124965135 | -1.685168226 | 0.09195612 | 0.446623824 | Abhd4 | abhydrolase domain containing 4 [Source | 824.79 | 779.16 | 711.41 | 690.21 | 603.11 | 708.4 |
| 12122 | ENSMUSG00000040998 | 7.19338618 | -2.278551647 | 0.946187534 | -2.408139575 | 0.01603405 | 0.148963156 | Npnt | nephronectin [Source:MGI Symbol;Acc:M | 7.2279 | 20.558 | 8.05 | 2.8919 | 1.0906 | 3.3415 |
| 12123 | ENSMUSG00000041000 | 15.0244968 | -0.370729366 | 0.662500953 | -0.559590691 | 0.575758656 | 0.907164837 | Trim62 | tripartite motif-containing 62 [Source:MG | 18.471 | 10.279 | 22.137 | 19.28 | 3.2719 | 16.708 |
| 12124 | ENSMUSG00000041009 | 0 | NA | NA | NA | NA | NA | Iqcf4 | IQ motif containing F4 [Source:MGI Symb | 0 | 0 | 0 | 0 | 0 | 0 |
| 12125 | ENSMUSG00000041012 | 994.6960177 | 0.341757215 | 0.112723667 | 3.031814215 | 0.002430888 | 0.040430438 | Cmtm8 | CKLF-like MARVEL transmembrane domai | 891.44 | 866.53 | 873.42 | 1007.4 | 1133.2 | 1196.3 |
| 12126 | ENSMUSG00000041014 | 0 | NA | NA | NA | NA | NA | Nrg3 | neuregulin 3 [Source:MGI Symbol;Acc:MG | 0 | 0 | 0 | 0 | 0 | 0 |
| 12127 | ENSMUSG00000041020 | 0.160662974 | 1.126924175 | 4.080472857 | 0.276174898 | 0.782413736 | NA | Map7d2 | MAP7 domain containing 2 [Source:MGI S | 0 | 0 | 0 | 0.964 | 0 | 0 |
| 12128 | ENSMUSG00000041025 | 344.2941638 | 0.177273443 | 0.172291422 | 1.028916242 | 0.303519035 | 0.752690022 | Iffo2 | intermediate filament family orphan 2 [So | 345.33 | 298.1 | 325.02 | 292.09 | 369.72 | 435.51 |
| 12129 | ENSMUSG00000041028 | 7145.011704 | -0.089731823 | 0.084990621 | -1.055785003 | 0.291066452 | 0.740906421 | Ghitm | growth hormone inducible transmembrar | 7587.7 | 7426.7 | 7086 | 6674.6 | 7089 | 7006.1 |
| 12130 | ENSMUSG00000041035 | 120.1832696 | 0.043730704 | 0.229359671 | 0.190664313 | 0.8487886 | 0.972888368 | Gm17018 | predicted gene 17018 [Source:MGI Symbo | 122.07 | 103.82 | 128.8 | 107 | 110.15 | 149.26 |
| 12131 | ENSMUSG00000041037 | 314.401273 | -0.041895692 | 0.16285245 | -0.257261659 | 0.796976788 | 0.962236316 | Irgq | immunity-related GTPase family, Q [Sourc | 308.39 | 283.71 | 365.27 | 335.46 | 318.46 | 275.12 |
| 12132 | ENSMUSG00000041040 | 124.2283931 | -0.053019221 | 0.204275534 | -0.259547583 | 0.795212774 | 0.962236316 | Fam117b | family with sequence similarity 117, meml | 120.47 | 128.49 | 130.81 | 125.32 | 118.88 | 121.41 |
| 12133 | ENSMUSG00000041044 | 307.9567409 | 0.13163069 | 0.160515554 | 0.820049436 | 0.412187926 | 0.834424863 | Lrit1 | leucine-rich repeat, immunoglobulin-like a | 253.78 | 334.07 | 294.83 | 322.93 | 340.27 | 301.85 |
| 12134 | ENSMUSG00000041046 | 3.435189317 | -1.935999368 | 1.336782725 | -1.448252833 | 0.147546354 | NA | Ramp3 | receptor (calcitonin) activity modifying pr | 4.0155 | 11.307 | 1.0062 | 0.964 | 1.0906 | 2.2277 |
| 12135 | ENSMUSG00000041052 | 0.545309144 | 2.651974629 | 3.666219369 | 0.72335405 | 0.469462394 | NA | Slc7a13 | solute carrier family 7, (cationic amino aci | 0 | 0 | 0 | 0 | 3.2719 | 0 |
| 12136 | ENSMUSG00000041057 | 407.5344256 | -0.03805817 | 0.150269815 | -0.253265567 | 0.800063002 | 0.962424862 | Wdr43 | WD repeat domain 43 [Source:MGI Symb | 417.61 | 428.64 | 392.44 | 393.3 | 465.69 | 347.52 |
| 12137 | ENSMUSG00000041058 | 2129.187788 | -0.166814256 | 0.120564706 | -1.383607705 | 0.166478608 | 0.5925379 | Wwp1 | WW domain containing E3 ubiquitin prote | 2106.5 | 2468 | 2182.5 | 1959.8 | 2277.2 | 1781 |
| 12138 | ENSMUSG00000041062 | 0 | NA | NA | NA | NA | NA | Mslnl | mesothelin-like [Source:MGI Symbol;Acc:I | 0 | 0 | 0 | 0 | 0 | 0 |
| 12139 | ENSMUSG00000041064 | 1.172011696 | -0.290561157 | 2.108054195 | -0.137833818 | 0.890371757 | NA | Pif1 | PIF1 5'-to-3' DNA helicase [Source:MGI Sy | 0.8031 | 3.0838 | 0 | 0.964 | 2.1812 | 0 |
| 12140 | ENSMUSG00000041068 | 0 | NA | NA | NA | NA | NA | 4930596D02 | RIKEN cDNA 4930596D02 gene [Source:M | 0 | 0 | 0 | 0 | 0 | 0 |
| 12141 | ENSMUSG00000041073 | 1.939687841 | -3.210118107 | 1.828993089 | -1.755128615 | 0.079237311 | NA | Nacad | NAC alpha domain containing [Source:MG | 2.4093 | 3.0838 | 5.0312 | 0 | 0 | 1.1138 |
| 12142 | ENSMUSG00000041075 | 484.3545986 | 0.879819332 | 0.237479062 | 3.704829074 | 0.000211533 | 0.006551645 | Fzd7 | frizzled class receptor 7 [Source:MGI Syml | 344.53 | 348.46 | 330.05 | 588.99 | 375.17 | 918.92 |
| 12143 | ENSMUSG00000041078 | 33.11768538 | -0.599122211 | 0.457565077 | -1.309370495 | 0.190408881 | 0.627779796 | Grid1 | glutamate receptor, ionotropic, delta 1 [S | 26.502 | 39.061 | 54.337 | 25.063 | 39.262 | 14.48 |
| 12144 | ENSMUSG00000041079 | 175.0258615 | 0.3639036 | 0.222330642 | 1.636767639 | 0.10167903 | 0.469365216 | Rwdd2b | RWD domain containing 2B [Source:MGI S | 184.71 | 127.46 | 145.91 | 151.34 | 225.76 | 214.97 |
| 12145 | ENSMUSG00000041084 | 577.3102617 | 0.210377978 | 0.1236557 | 1.701320506 | 0.088882819 | 0.439445475 | Ostc | oligosaccharyltransferase complex subuni | 542.09 | 528.35 | 535.32 | 630.44 | 566.03 | 661.62 |
| 12146 | ENSMUSG00000041096 | 67.99129791 | 0.154863116 | 0.270521235 | 0.572461959 | 0.56700905 | 0.90291962 | Tspyl2 | TSPY-like 2 [Source:MGI Symbol;Acc:MGI: | 65.854 | 69.898 | 57.356 | 77.118 | 70.89 | 66.831 |
| 12147 | ENSMUSG00000041112 | 19.90460348 | -1.224849007 | 0.493564657 | -2.481638404 | 0.013077991 | 0.129971148 | Elmo1 | engulfment and cell motility 1 [Source:MG | 29.715 | 25.698 | 28.175 | 11.568 | 10.906 | 13.366 |
| 12148 | ENSMUSG00000041115 | 280.4348792 | -0.165060031 | 0.155191379 | -1.063590204 | 0.287514382 | 0.73951397 | Iqsec2 | IQ motif and Sec7 domain 2 [Source:MGI S | 311.6 | 283.71 | 293.82 | 272.81 | 237.75 | 282.92 |

| | | | | | | | | | | | | | | | |
|---|---|---|---|---|---|---|---|---|---|---|---|---|---|---|---|
| 12149 | ENSMUSG00000041117 | 8.976974704 | -0.505094465 | 0.718498268 | -0.702986337 | 0.482064264 | 0.87157219 | Ccdc8 | coiled-coil domain containing 8 [Source:M | 7.2279 | 13.363 | 11.069 | 6.7478 | 6.5437 | 8.9108 |
| 12150 | ENSMUSG00000041119 | 269.2143448 | -0.17994966 | 0.474300291 | -0.379400273 | 0.704390648 | 0.94295036 | Pde9a | phosphodiesterase 9A [Source:MGI Symb | 171.06 | 277.54 | 409.54 | 244.85 | 160.32 | 351.97 |
| 12151 | ENSMUSG00000041120 | 28.91156554 | 0.000175997 | 0.470448765 | 0.000374104 | 0.999701508 | 0.999897338 | Nbl1 | NBL1, DAN family BMP antagonist [Source | 12.85 | 38.033 | 36.225 | 27.955 | 29.447 | 28.96 |
| 12152 | ENSMUSG00000041124 | 402.448773 | 0.046367903 | 0.14641887 | 0.31667983 | 0.751486555 | 0.952255112 | Msantd4 | Myb/SANT-like DNA-binding domain cont | 399.94 | 414.25 | 373.32 | 379.81 | 378.44 | 468.93 |
| 12153 | ENSMUSG00000041126 | 568.0248689 | 0.082634173 | 0.125059335 | 0.660759739 | 0.508766406 | 0.882793133 | H2az2 | H2A.Z histone variant 2 [Source:MGI Symb | 526.84 | 545.82 | 582.62 | 540.79 | 617.29 | 594.79 |
| 12154 | ENSMUSG00000041130 | 1185.055973 | 0.360897148 | 0.133387895 | 2.705621435 | 0.006817672 | 0.0825195 | Zfp598 | zinc finger protein 598 [Source:MGI Symb | 1072.1 | 1079.3 | 961.97 | 1581.9 | 1193.1 | 1221.9 |
| 12155 | ENSMUSG00000041132 | 1600.153275 | -0.341306777 | 0.11181628 | -3.052388951 | 0.002270277 | 0.038271898 | N4bp2l1 | NEDD4 binding protein 2-like 1 [Source:M | 1742.7 | 1793.7 | 1829.4 | 1417 | 1253.1 | 1565 |
| 12156 | ENSMUSG00000041133 | 1024.864451 | 0.053981233 | 0.116873481 | 0.461877515 | 0.64416916 | 0.925643431 | Smc1a | structural maintenance of chromosomes 1 | 948.46 | 1050.5 | 1018.3 | 967.83 | 1158.2 | 1005.8 |
| 12157 | ENSMUSG00000041134 | 22.4236712 | -0.101029203 | 0.548014068 | -0.184355127 | 0.85373487 | 0.973825032 | Cyyr1 | cysteine and tyrosine-rich protein 1 [Sour | 24.093 | 11.307 | 34.212 | 25.063 | 10.906 | 28.96 |
| 12158 | ENSMUSG00000041135 | 145.2906719 | 0.787872924 | 0.234323221 | 3.362333956 | 0.000772866 | 0.017277794 | Ripk2 | receptor (TNFRSF)-interacting serine-thre | 127.69 | 93.541 | 97.606 | 161.95 | 157.05 | 233.91 |
| 12159 | ENSMUSG00000041138 | 0 | NA | NA | NA | NA | NA | Nme8 | NME/NM23 family member 8 [Source:MG | 0 | 0 | 0 | 0 | 0 | 0 |
| 12160 | ENSMUSG00000041141 | 0 | NA | NA | NA | NA | NA | Pnmal1 | PNMA-like 1 [Source:MGI Symbol;Acc:MG | 0 | 0 | 0 | 0 | 0 | 0 |
| 12161 | ENSMUSG00000041143 | 396.5688224 | 0.135107057 | 0.151409046 | 0.892331474 | 0.372215289 | 0.807785387 | Tmco4 | transmembrane and coiled-coil domains 4 | 402.35 | 379.3 | 352.19 | 428.97 | 463.51 | 353.09 |
| 12162 | ENSMUSG00000041144 | 0.572870759 | -2.577107685 | 3.182795569 | -0.809699407 | 0.418112959 | NA | Dnah7b | dynein, axonemal, heavy chain 7B [Source | 2.4093 | 1.0279 | 0 | 0 | 0 | 0 |
| 12163 | ENSMUSG00000041147 | 171.3945401 | -0.107316196 | 0.216148597 | -0.496492678 | 0.619546852 | 0.91786599 | Brca2 | breast cancer 2, early onset [Source:MGI S | 167.05 | 168.58 | 198.23 | 189.9 | 184.31 | 120.3 |
| 12164 | ENSMUSG00000041153 | 51.03366499 | 0.51882878 | 0.308908686 | 1.679553873 | 0.093044149 | 0.449345765 | Osgin2 | oxidative stress induced growth inhibitor f | 43.368 | 41.117 | 41.256 | 54.947 | 68.709 | 56.806 |
| 12165 | ENSMUSG00000041161 | 114.928871 | 0.019564866 | 0.220443249 | 0.088752391 | 0.929278695 | 0.988405116 | Otud3 | OTU domain containing 3 [Source:MGI Sy | 118.86 | 112.04 | 111.69 | 133.03 | 100.34 | 113.61 |
| 12166 | ENSMUSG00000041164 | 602.4898409 | -0.005260563 | 0.120580529 | -0.04362697 | 0.965201753 | 0.993480943 | Zmiz2 | zinc finger, MIZ-type containing 2 [Source | 612.77 | 604.42 | 593.68 | 617.91 | 631.47 | 554.69 |
| 12167 | ENSMUSG00000041165 | 0 | NA | NA | NA | NA | NA | Spem1 | sperm maturation 1 [Source:MGI Symbol; | 0 | 0 | 0 | 0 | 0 | 0 |
| 12168 | ENSMUSG00000041168 | 3328.081513 | -0.084490999 | 0.087621731 | -0.964269918 | 0.334910618 | 0.776159726 | Lonp1 | lon peptidase 1, mitochondrial [Source:M | 3318.4 | 3479.5 | 3479.6 | 3321.9 | 3235.9 | 3133.2 |
| 12169 | ENSMUSG00000041180 | 34.81877447 | -0.135178749 | 0.377870323 | -0.357738463 | 0.720539046 | 0.9447719 | Hectd2 | HECT domain E3 ubiquitin protein ligase 2 | 31.321 | 37.005 | 41.256 | 36.631 | 37.081 | 25.618 |
| 12170 | ENSMUSG00000041187 | 109.9996862 | -0.060348713 | 0.233716741 | -0.258213054 | 0.796242484 | 0.962236316 | Prkd2 | protein kinase D2 [Source:MGI Symbol;Ac | 112.43 | 119.24 | 105.66 | 132.06 | 98.156 | 92.449 |
| 12171 | ENSMUSG00000041189 | 5.813925531 | 0.972179287 | 0.986258996 | 0.985724126 | 0.324268507 | 0.771164039 | Chrnb1 | cholinergic receptor, nicotinic, beta polyp | 0.8031 | 3.0838 | 8.05 | 8.6758 | 9.8156 | 4.4554 |
| 12172 | ENSMUSG00000041193 | 0.342639279 | -1.805976596 | 4.028547714 | -0.448294702 | 0.653940524 | NA | Pla2g5 | phospholipase A2, group V [Source:MGI S | 0 | 2.0558 | 0 | 0 | 0 | 0 |
| 12173 | ENSMUSG00000041199 | 29.7251453 | -0.081770629 | 0.521688017 | -0.156742395 | 0.875447874 | 0.980525066 | Rpusd1 | RNA pseudouridylate synthase domain co | 36.14 | 17.475 | 38.237 | 49.163 | 25.084 | 12.252 |
| 12174 | ENSMUSG00000041202 | 0 | NA | NA | NA | NA | NA | Pla2g2d | phospholipase A2, group IID [Source:MGI | 0 | 0 | 0 | 0 | 0 | 0 |
| 12175 | ENSMUSG00000041203 | 1017.060812 | 0.029813526 | 0.114402562 | 0.260601909 | 0.794399518 | 0.962046762 | Trir | telomerase RNA component interacting R | 1021.5 | 945.68 | 1052.5 | 1086.4 | 932.48 | 1063.7 |
| 12176 | ENSMUSG00000041205 | 1.256521509 | 0.543933952 | 2.487612764 | 0.218657003 | 0.826917245 | NA | Map6d1 | MAP6 domain containing 1 [Source:MGI S | 0 | 3.0838 | 0 | 0 | 0 | 4.4554 |
| 12177 | ENSMUSG00000041215 | 200.8724415 | 0.044517803 | 0.191627219 | 0.232314612 | 0.816293664 | 0.967767553 | Yeats2 | YEATS domain containing 2 [Source:MGI S | 163.03 | 224.09 | 207.29 | 228.46 | 194.13 | 188.24 |
| 12178 | ENSMUSG00000041216 | 0 | NA | NA | NA | NA | NA | Clvs1 | clavesin 1 [Source:MGI Symbol;Acc:MGI:1 | 0 | 0 | 0 | 0 | 0 | 0 |
| 12179 | ENSMUSG00000041219 | 26.88771642 | -0.79184282 | 0.483003346 | -1.639414773 | 0.101126907 | 0.468438535 | Arhgap11a | Rho GTPase activating protein 11A [Sourc | 48.186 | 20.558 | 33.206 | 17.352 | 14.178 | 27.846 |
| 12180 | ENSMUSG00000041220 | 1778.309523 | -0.107354728 | 0.364640898 | -0.294412197 | 0.768442948 | 0.955856567 | Elovl6 | ELOVL family member 6, elongation of lon | 2653.4 | 1657 | 1222.6 | 2079.3 | 1673 | 1384.5 |
| 12181 | ENSMUSG00000041225 | 434.4183898 | 0.397605056 | 0.136580323 | 2.911144497 | 0.003601074 | 0.053476324 | Arhgap12 | Rho GTPase activating protein 12 [Source: | 350.15 | 387.53 | 387.4 | 464.64 | 477.69 | 539.1 |
| 12182 | ENSMUSG00000041229 | 185.9041325 | 0.445065745 | 0.231416628 | 1.923222841 | 0.05445207 | 0.328575194 | Phf8 | PHD finger protein 8 [Source:MGI Symbol | 175.88 | 167.55 | 128.8 | 233.28 | 260.66 | 149.26 |
| 12183 | ENSMUSG00000041231 | 541.8420287 | 0.050524275 | 0.124609538 | 0.405460739 | 0.685138872 | 0.935774188 | Ublcp1 | ubiquitin-like domain containing CTD pho | 521.21 | 562.27 | 514.19 | 580.31 | 555.12 | 517.94 |
| 12184 | ENSMUSG00000041235 | 488.6312514 | 0.105379855 | 0.137549858 | 0.766121147 | 0.443604221 | 0.851079912 | Chd7 | chromodomain helicase DNA binding prot | 462.59 | 507.79 | 442.75 | 558.14 | 504.96 | 455.56 |
| 12185 | ENSMUSG00000041236 | 1051.0298 | -0.016239754 | 0.118565327 | -0.13696883 | 0.891055433 | 0.982489666 | Vps41 | VPS41 HOPS complex subunit [Source:MG | 1132.4 | 1082.4 | 954.93 | 970.73 | 1129.9 | 1035.9 |
| 12186 | ENSMUSG00000041237 | 8801.060021 | -0.210851449 | 0.12354031 | -1.706742106 | 0.087869997 | 0.43690328 | Pklr | pyruvate kinase liver and red blood cell [S | 10030 | 9040.5 | 9258.5 | 8337.4 | 9396.8 | 6743.2 |
| 12187 | ENSMUSG00000041238 | 85.79398025 | -0.616197517 | 0.247599917 | -2.488682247 | 0.01282175 | 0.128107525 | Rbbp8 | retinoblastoma binding protein 8, endonu | 102.8 | 116.15 | 92.575 | 66.514 | 65.437 | 71.286 |
| 12188 | ENSMUSG00000041241 | 1033.644767 | 0.052865369 | 0.118279132 | 0.446954317 | 0.654908044 | 0.925797231 | Mul1 | mitochondrial ubiquitin ligase activator of | 1059.3 | 1093.7 | 890.53 | 1020.9 | 1044.8 | 1092.7 |
| 12189 | ENSMUSG00000041245 | 0.822586684 | 2.037574431 | 2.401368532 | 0.84850551 | 0.396156505 | NA | Wnk3 | WNK lysine deficient protein kinase 3 [Sou | 0.8031 | 0 | 0 | 1.928 | 1.0906 | 1.1138 |
| 12190 | ENSMUSG00000041247 | 0 | NA | NA | NA | NA | NA | Lamp3 | lysosomal-associated membrane protein 3 | 0 | 0 | 0 | 0 | 0 | 0 |
| 12191 | ENSMUSG00000041248 | 0 | NA | NA | NA | NA | NA | Kcnj1 | potassium inwardly-rectifying channel, su | 0 | 0 | 0 | 0 | 0 | 0 |
| 12192 | ENSMUSG00000041255 | 0 | NA | NA | NA | NA | NA | Tmco5b | transmembrane and coiled-coil domains 5 | 0 | 0 | 0 | 0 | 0 | 0 |
| 12193 | ENSMUSG00000041258 | 315.575 | -0.032989447 | 0.172430344 | -0.191320429 | 0.848274557 | 0.972803041 | Zfp236 | zinc finger protein 236 [Source:MGI Symb | 258.6 | 372.11 | 328.04 | 326.79 | 325 | 282.92 |
| 12194 | ENSMUSG00000041261 | 1708.093913 | 0.180929212 | 0.128015903 | 1.413333874 | 0.157557579 | 0.57679393 | Car8 | carbonic anhydrase 8 [Source:MGI Symbo | 1480.1 | 1474 | 1849.5 | 1721.7 | 1688.3 | 2035 |
| 12195 | ENSMUSG00000041263 | 40.53464705 | -0.452046072 | 0.375742426 | -1.203074343 | 0.228947553 | 0.67545951 | Rusc1 | RUN and SH3 domain containing 1 [Sourc | 42.564 | 63.731 | 34.212 | 29.883 | 32.719 | 40.098 |
| 12196 | ENSMUSG00000041264 | 322.8599156 | 0.176789701 | 0.15898126 | 1.11201598 | 0.26613128 | 0.722063933 | Uspl1 | ubiquitin specific peptidase like 1 [Source | 277.07 | 302.21 | 330.05 | 294.98 | 368.63 | 364.23 |
| 12197 | ENSMUSG00000041268 | 585.3156395 | 0.086273248 | 0.13070335 | 0.660069142 | 0.50920946 | 0.882793133 | Dmxl2 | Dmx-like 2 [Source:MGI Symbol;Acc:MGI: | 536.47 | 629.09 | 538.34 | 594.77 | 563.85 | 649.37 |
| 12198 | ENSMUSG00000041272 | 112.978861 | -0.374879841 | 0.293728637 | -1.276279507 | 0.201856732 | 0.646227482 | Tox | thymocyte selection-associated high mob | 160.62 | 122.32 | 99.618 | 110.86 | 125.42 | 59.034 |
| 12199 | ENSMUSG00000041278 | 323.4886509 | 0.042850731 | 0.149123169 | 0.287351265 | 0.773843373 | 0.958261993 | Ttc1 | tetratricopeptide repeat domain 1 [Sourc | 329.27 | 321.74 | 304.89 | 327.75 | 360.99 | 296.28 |
| 12200 | ENSMUSG00000041287 | 1.295205404 | 0.168345827 | 1.972095081 | 0.08536395 | 0.931972052 | NA | Sox15 | SRY (sex determining region Y)-box 15 [So | 1.6062 | 2.0558 | 0 | 1.928 | 2.1812 | 0 |
| 12201 | ENSMUSG00000041293 | 54.14768668 | 2.543147596 | 0.381204538 | 6.671346591 | 2.53E-11 | 7.80E-09 | Adgrf1 | adhesion G protein-coupled receptor F1 [ | 10.44 | 15.419 | 22.137 | 66.514 | 111.24 | 99.132 |
| 12202 | ENSMUSG00000041297 | 563.9105012 | 0.135269359 | 0.131809211 | 1.026251186 | 0.3047732 | 0.753556583 | Cdk13 | cyclin-dependent kinase 13 [Source:MGI S | 482.66 | 604.42 | 526.27 | 576.46 | 593.3 | 600.36 |
| 12203 | ENSMUSG00000041298 | 109.5225719 | 0.203444709 | 0.228473149 | 0.890453474 | 0.37322244 | 0.808611351 | Katnal1 | katanin p60 subunit A-like 1 [Source:MGI | 98.782 | 111.02 | 95.593 | 108.93 | 104.7 | 138.12 |
| 12204 | ENSMUSG00000041301 | 2.086261902 | -1.05519918 | 1.529757687 | -0.68978191 | 0.490331346 | NA | Cftr | cystic fibrosis transmembrane conductanc | 2.4093 | 3.0838 | 3.0187 | 2.8919 | 0 | 1.1138 |

| | | | | | | | | | | | | | | | |
|---|---|---|---|---|---|---|---|---|---|---|---|---|---|---|---|
| 12205 | ENSMUSG00000041303 | 296.416108 | 0.043369015 | 0.152944758 | 0.283559996 | 0.776747598 | 0.959295521 | Gtf3c3 | general transcription factor IIIC, polypepti | 275.46 | 283.71 | 316.97 | 291.12 | 326.09 | 285.14 |
| 12206 | ENSMUSG00000041308 | 156.1208305 | 0.001373258 | 0.203815744 | 0.006737741 | 0.994624101 | 0.998676967 | Sntb2 | syntrophin, basic 2 [Source:MGI Symbol;A | 138.94 | 176.8 | 152.95 | 166.77 | 131.96 | 169.3 |
| 12207 | ENSMUSG00000041309 | 0.305170013 | -1.657836477 | 4.041480335 | -0.410205257 | 0.681655385 | NA | Nkx6-2 | NK6 homeobox 2 [Source:MGI Symbol;Ac | 0.8031 | 1.0279 | 0 | 0 | 0 | 0 |
| 12208 | ENSMUSG00000041313 | 9.403684344 | -0.984852123 | 0.713173602 | -1.380943042 | 0.167296474 | 0.593441872 | Slc7a1 | solute carrier family 7 (cationic amino acid | 16.062 | 9.2513 | 12.075 | 5.7839 | 5.4531 | 7.7969 |
| 12209 | ENSMUSG00000041319 | 66.61594507 | 0.272098335 | 0.333984106 | 0.814704445 | 0.415241508 | 0.836001897 | Thoc6 | THO complex 6 [Source:MGI Symbol;Acc:I | 60.233 | 46.256 | 74.462 | 61.695 | 104.7 | 52.351 |
| 12210 | ENSMUSG00000041323 | 3.990051695 | 0.414580617 | 1.200390893 | 0.345371344 | 0.729815217 | NA | Ak7 | adenylate kinase 7 [Source:MGI Symbol;A | 3.2124 | 0 | 7.0437 | 4.8199 | 2.1812 | 6.6831 |
| 12211 | ENSMUSG00000041324 | 153.9411317 | 0.3830939 | 0.258707416 | 1.480799841 | 0.138659919 | 0.544254293 | Inhba | inhibin beta-A [Source:MGI Symbol;Acc:M | 156.6 | 135.69 | 108.67 | 244.85 | 134.15 | 143.69 |
| 12212 | ENSMUSG00000041328 | 489.253845 | 0.107960142 | 0.131378588 | 0.821748381 | 0.411220118 | 0.833705705 | Pcf11 | PCF11 cleavage and polyadenylation factc | 430.46 | 490.32 | 493.06 | 528.26 | 476.6 | 516.82 |
| 12213 | ENSMUSG00000041329 | 4.821774305 | -0.669778115 | 1.020998146 | -0.656003263 | 0.511822025 | NA | Atp1b2 | ATPase, Na+/K+ transporting, beta 2 polyp | 9.6372 | 4.1117 | 4.025 | 6.7478 | 2.1812 | 2.2277 |
| 12214 | ENSMUSG00000041333 | 0 | NA | NA | NA | NA | NA | Mup4 | major urinary protein 4 [Source:MGI Symb | 0 | 0 | 0 | 0 | 0 | 0 |
| 12215 | ENSMUSG00000041341 | 596.9337255 | -0.141474082 | 0.130193179 | -1.086647419 | 0.27719266 | 0.730041765 | Atg2b | autophagy related 2B [Source:MGI Symbc | 589.48 | 673.29 | 615.82 | 527.3 | 546.4 | 629.32 |
| 12216 | ENSMUSG00000041343 | 23.4614424 | 0.008110379 | 0.519092378 | 0.015624153 | 0.987534237 | 0.99859176 | Ankrd42 | ankyrin repeat domain 42 [Source:MGI Sy | 20.078 | 37.005 | 13.081 | 15.424 | 23.994 | 31.188 |
| 12217 | ENSMUSG00000041346 | 157.0398737 | 0.048274833 | 0.190505509 | 0.253403868 | 0.799956139 | 0.962424862 | Wrap53 | WD repeat containing, antisense to Trp53 | 163.83 | 144.94 | 153.96 | 146.52 | 159.23 | 173.76 |
| 12218 | ENSMUSG00000041347 | 0.17131964 | -0.796629176 | 4.080472857 | -0.195229623 | 0.84521319 | NA | Bdkrb1 | bradykinin receptor, beta 1 [Source:MGI S | 0 | 1.0279 | 0 | 0 | 0 | 0 |
| 12219 | ENSMUSG00000041351 | 163.5825432 | -0.281254908 | 0.202422022 | -1.389448171 | 0.164696511 | 0.589893637 | Rap1gap | Rap1 GTPase-activating protein [Source:M | 202.38 | 155.22 | 180.12 | 135.92 | 136.33 | 171.53 |
| 12220 | ENSMUSG00000041353 | 105.8378578 | 0.323796921 | 0.232912247 | 1.390209937 | 0.164465137 | 0.589361417 | Tmem29 | transmembrane protein 29 [Source:MGI S | 81.113 | 98.68 | 102.64 | 113.75 | 134.15 | 104.7 |
| 12221 | ENSMUSG00000041354 | 368.7893631 | -0.123328447 | 0.145152597 | -0.849646851 | 0.395521456 | 0.823139905 | Rgl2 | ral guanine nucleotide dissociation stimul | 425.64 | 353.6 | 373.32 | 331.61 | 360.99 | 367.57 |
| 12222 | ENSMUSG00000041355 | 1143.705768 | 0.218081517 | 0.099765931 | 2.185931763 | 0.028820598 | 0.22274774 | Ssr2 | signal sequence receptor, beta [Source:M | 1050.5 | 1085.5 | 1036.4 | 1226.2 | 1209.5 | 1254.2 |
| 12223 | ENSMUSG00000041358 | 0 | NA | NA | NA | NA | NA | Nutm1 | NUT midline carcinoma, family member 1 | 0 | 0 | 0 | 0 | 0 | 0 |
| 12224 | ENSMUSG00000041359 | 0 | NA | NA | NA | NA | NA | Tcl1 | T cell lymphoma breakpoint 1 [Source:MG | 0 | 0 | 0 | 0 | 0 | 0 |
| 12225 | ENSMUSG00000041360 | 444.4468398 | 0.141478145 | 0.14508909 | 0.975112228 | 0.329504594 | 0.774441615 | Pum3 | pumilio RNA-binding family member 3 [So | 440.1 | 423.5 | 403.5 | 398.12 | 518.04 | 483.41 |
| 12226 | ENSMUSG00000041361 | 11.70122895 | -0.605575928 | 0.712360584 | -0.850097468 | 0.395270899 | 0.822933206 | Myzap | myocardial zonula adherens protein [Sour | 9.6372 | 25.698 | 7.0437 | 5.7839 | 10.906 | 11.138 |
| 12227 | ENSMUSG00000041362 | 748.0184637 | 0.088475255 | 0.142051711 | 0.622838361 | 0.533390747 | 0.889143097 | Shtn1 | shootin 1 [Source:MGI Symbol;Acc:MGI:1 | 700.31 | 723.65 | 750.66 | 621.77 | 846.32 | 845.41 |
| 12228 | ENSMUSG00000041372 | 74.6680577 | 0.574205458 | 0.337852118 | 1.699576316 | 0.089210648 | 0.440239788 | B4galnt3 | beta-1,4-N-acetyl-galactosaminyl transfer | 51.399 | 38.033 | 90.562 | 67.478 | 102.52 | 98.018 |
| 12229 | ENSMUSG00000041375 | 351.4167481 | 0.159113378 | 0.156305701 | 1.017962725 | 0.30869567 | 0.75728287 | Ccdc9 | coiled-coil domain containing 9 [Source:M | 305.98 | 333.05 | 358.22 | 428.01 | 344.64 | 338.61 |
| 12230 | ENSMUSG00000041377 | 0 | NA | NA | NA | NA | NA | Ninj2 | ninjurin 2 [Source:MGI Symbol;Acc:MGI:1 | 0 | 0 | 0 | 0 | 0 | 0 |
| 12231 | ENSMUSG00000041378 | 71.91994795 | -0.141914431 | 0.281427584 | -0.504266244 | 0.614074295 | 0.917378096 | Cldn5 | claudin 5 [Source:MGI Symbol;Acc:MGI:1: | 78.704 | 69.898 | 77.481 | 60.731 | 56.712 | 87.994 |
| 12232 | ENSMUSG00000041380 | 0 | NA | NA | NA | NA | NA | Htr2c | 5-hydroxytryptamine (serotonin) receptor | 0 | 0 | 0 | 0 | 0 | 0 |
| 12233 | ENSMUSG00000041390 | 491.6252112 | -0.141540751 | 0.132216948 | -1.070518967 | 0.284385776 | 0.736709833 | Mdfic | MyoD family inhibitor domain containing | 546.11 | 530.41 | 469.92 | 441.5 | 496.23 | 465.59 |
| 12234 | ENSMUSG00000041396 | 43.28610944 | -0.483313147 | 0.35002954 | -1.380778169 | 0.167347178 | 0.593441872 | Mettl18 | methyltransferase like 18 [Source:MGI Syı | 42.564 | 55.508 | 53.331 | 27.955 | 44.715 | 35.643 |
| 12235 | ENSMUSG00000041399 | 0 | NA | NA | NA | NA | NA | 1700013G24 | RIKEN cDNA 1700013G24 gene [Source:M | 0 | 0 | 0 | 0 | 0 | 0 |
| 12236 | ENSMUSG00000041406 | 4.567903748 | -0.975854489 | 1.027868441 | -0.949396294 | 0.342419095 | NA | BC055324 | cDNA sequence BC055324 [Source:MGI Sy | 8.031 | 5.1396 | 5.0312 | 4.8199 | 3.2719 | 1.1138 |
| 12237 | ENSMUSG00000041408 | 1389.102633 | 0.027737919 | 0.105780997 | 0.262220247 | 0.793151644 | 0.961863451 | Wapl | WAPL cohesin release factor [Source:MGI | 1252.8 | 1471 | 1404.7 | 1404.5 | 1399.3 | 1402.3 |
| 12238 | ENSMUSG00000041415 | 755.6892147 | -0.000501484 | 0.10951341 | -0.004579199 | 0.99634634 | 0.999154389 | Dicer1 | dicer 1, ribonuclease type III [Source:MGI | 746.89 | 780.19 | 740.6 | 755.76 | 768.89 | 741.82 |
| 12239 | ENSMUSG00000041417 | 1801.821433 | -0.215583565 | 0.156017235 | -1.381793267 | 0.167035187 | 0.593319485 | Pik3r1 | phosphoinositide-3-kinase regulatory sub | 2009.4 | 2284 | 1515.4 | 1753.5 | 1850.8 | 1397.9 |
| 12240 | ENSMUSG00000041420 | 8.608942616 | -0.543968147 | 0.785346124 | -0.69264765 | 0.488530699 | 0.874431125 | Meis3 | Meis homeobox 3 [Source:MGI Symbol;Ac | 11.243 | 12.335 | 7.0437 | 7.7118 | 2.1812 | 11.138 |
| 12241 | ENSMUSG00000041423 | 0.439020386 | -2.189747911 | 3.52148864 | -0.621824499 | 0.534057275 | NA | Paqr6 | progestin and adipoQ receptor family mer | 1.6062 | 1.0279 | 0 | 0 | 0 | 0 |
| 12242 | ENSMUSG00000041426 | 1928.345552 | -0.006734822 | 0.111496944 | -0.060403647 | 0.951834154 | 0.990883058 | Hibch | 3-hydroxyisobutyryl-Coenzyme A hydrolas | 1967.6 | 1926.3 | 1903.8 | 1747.7 | 2181.2 | 1843.4 |
| 12243 | ENSMUSG00000041429 | 104.446536 | -0.006037918 | 0.231782508 | -0.026049929 | 0.979217514 | 0.998055255 | Nthl1 | nth (endonuclease III)-like 1 (E.coli) [Sourc | 98.782 | 102.79 | 112.7 | 113.75 | 86.159 | 112.5 |
| 12244 | ENSMUSG00000041431 | 6.498344228 | -0.844107207 | 0.869012245 | -0.971340981 | 0.331378508 | 0.774747658 | Ccnb1 | cyclin B1 [Source:MGI Symbol;Acc:MGI:88 | 9.6372 | 9.2513 | 6.0375 | 1.928 | 5.4531 | 6.6831 |
| 12245 | ENSMUSG00000041438 | 248.9278 | -0.063237803 | 0.155959606 | -0.405475525 | 0.685128006 | 0.935774188 | Utp4 | UTP4 small subunit processome compone | 242.54 | 263.15 | 257.6 | 233.28 | 250.84 | 246.16 |
| 12246 | ENSMUSG00000041439 | 222.2793749 | 0.214769102 | 0.218308676 | 0.983786382 | 0.325220556 | 0.771501263 | Mfsd6 | major facilitator superfamily domain cont | 175.08 | 174.75 | 267.66 | 214.97 | 209.4 | 291.83 |
| 12247 | ENSMUSG00000041440 | 3.918621104 | -0.557643651 | 1.057098536 | -0.527522868 | 0.597830541 | NA | Gk5 | glycerol kinase 5 (putative) [Source:MGI S | 4.8186 | 5.1396 | 4.025 | 2.8919 | 2.1812 | 4.4554 |
| 12248 | ENSMUSG00000041444 | 233.1731492 | -0.0613406 | 0.193201536 | -0.317495407 | 0.750867724 | 0.952074518 | Arhgap32 | Rho GTPase activating protein 32 [Source: | 208 | 279.59 | 227.41 | 227.5 | 268.29 | 188.24 |
| 12249 | ENSMUSG00000041445 | 89.51951572 | -0.463815844 | 0.256142756 | -1.810770885 | 0.070176325 | 0.383030326 | Mmrn2 | multimerin 2 [Source:MGI Symbol;Acc:MC | 93.963 | 101.76 | 115.72 | 73.262 | 61.075 | 91.335 |
| 12250 | ENSMUSG00000041449 | 3.310498469 | -0.722237425 | 1.310680415 | -0.551040068 | 0.581606208 | NA | Serpina3h | serine (or cysteine) peptidase inhibitor, cl | 3.2124 | 4.1117 | 5.0312 | 0.964 | 6.5437 | 0 |
| 12251 | ENSMUSG00000041453 | 2963.595701 | -0.07343352 | 0.090496895 | -0.811447951 | 0.417108472 | 0.836878476 | Rpl21 | ribosomal protein L21 [Source:MGI Symbo | 2896.8 | 3157.8 | 3063 | 2809 | 2952.3 | 2902.7 |
| 12252 | ENSMUSG00000041459 | 1321.085726 | 0.243969028 | 0.120282967 | 2.028292403 | 0.042530414 | 0.284007344 | Tardbp | TAR DNA binding protein [Source:MGI Syr | 1156.5 | 1085.5 | 1387.6 | 1495.1 | 1388.4 | 1413.5 |
| 12253 | ENSMUSG00000041460 | 0.181769715 | 1.126924175 | 4.080472857 | 0.276174898 | 0.782413736 | NA | Cacna2d4 | calcium channel, voltage-dependent, alph | 0 | 0 | 0 | 0 | 1.0906 | 0 |
| 12254 | ENSMUSG00000041468 | 20.63048803 | 0.268689936 | 0.529040176 | 0.507881911 | 0.611536161 | 0.916376001 | Gpr12 | G-protein coupled receptor 12 [Source:M | 21.684 | 16.447 | 18.112 | 36.631 | 13.087 | 17.822 |
| 12255 | ENSMUSG00000041471 | 1062.021019 | -0.83085382 | 0.161203194 | -5.154077903 | 2.55E-07 | 2.44E-05 | Shld2 | shieldin complex subunit 2 [Source:MGI S | 1137.2 | 1711.5 | 1230.6 | 687.32 | 868.13 | 737.36 |
| 12256 | ENSMUSG00000041476 | 0 | NA | NA | NA | NA | NA | Smpx | small muscle protein, X-linked [Source:MC | 0 | 0 | 0 | 0 | 0 | 0 |
| 12257 | ENSMUSG00000041477 | 54.8629122 | -0.092921079 | 0.313379484 | -0.296512961 | 0.76683837 | 0.955295274 | Dcp1b | decapping mRNA 1B [Source:MGI Symbol | 49.792 | 72.982 | 47.293 | 51.091 | 53.44 | 54.578 |
| 12258 | ENSMUSG00000041479 | 14.64022043 | -0.91060315 | 0.563563767 | -1.615794349 | 0.106138784 | 0.480498081 | Syt15 | synaptotagmin XV [Source:MGI Symbol;Ac | 21.684 | 17.475 | 18.112 | 9.6398 | 10.906 | 10.025 |
| 12259 | ENSMUSG00000041481 | 99.3602937 | -0.932426548 | 0.290117245 | -3.21396458 | 0.001309158 | 0.025519763 | Serpina3g | serine (or cysteine) peptidase inhibitor, cl | 126.89 | 170.63 | 93.581 | 63.623 | 53.44 | 87.994 |
| 12260 | ENSMUSG00000041482 | 10.07206625 | -2.030249428 | 0.805289638 | -2.521141875 | 0.011697468 | 0.120662445 | Piezo2 | piezo-type mechanosensitive ion channel | 8.8341 | 17.475 | 22.137 | 0.964 | 5.4531 | 5.5692 |

| | | | | | | | | | | | | | | | |
|---|---|---|---|---|---|---|---|---|---|---|---|---|---|---|---|
| 12261 | ENSMUSG00000041483 | 480.5428073 | 0.032230351 | 0.129147518 | 0.249562292 | 0.802925863 | 0.963537682 | Zfp281 | zinc finger protein 281 [Source:MGI Symb | 467.41 | 469.76 | 488.03 | 445.36 | 512.59 | 500.12 |
| 12262 | ENSMUSG00000041488 | 143.2257946 | -0.300602632 | 0.203451041 | -1.477518282 | 0.139536761 | 0.545861088 | Stx3 | syntaxin 3 [Source:MGI Symbol;Acc:MGI:1 | 155.8 | 179.89 | 138.86 | 136.88 | 126.51 | 121.41 |
| 12263 | ENSMUSG00000041491 | 52.43831902 | 0.271529944 | 0.301184959 | 0.901538859 | 0.367301882 | 0.805313692 | Cep78 | centrosomal protein 78 [Source:MGI Syml | 44.974 | 47.284 | 50.312 | 53.019 | 58.893 | 60.148 |
| 12264 | ENSMUSG00000041498 | 0.981621321 | -0.747528389 | 2.423078149 | -0.308503623 | 0.757699144 | NA | Kif14 | kinesin family member 14 [Source:MGI Sy | 1.6062 | 2.0558 | 0 | 0 | 0 | 2.2277 |
| 12265 | ENSMUSG00000041505 | 0 | NA | NA | NA | NA | NA | Gm3402 | predicted gene 3402 [Source:MGI Symbol | 0 | 0 | 0 | 0 | 0 | 0 |
| 12266 | ENSMUSG00000041506 | 165.3500947 | 0.527376015 | 0.188504104 | 2.797689835 | 0.005146951 | 0.067924728 | Rrp9 | ribosomal RNA processing 9, U3 small nuc | 131.71 | 134.66 | 139.87 | 173.52 | 198.49 | 213.86 |
| 12267 | ENSMUSG00000041512 | 0 | NA | NA | NA | NA | NA | 4930569F06 | RIKEN cDNA 4930569F06 gene [Source:M( | 0 | 0 | 0 | 0 | 0 | 0 |
| 12268 | ENSMUSG00000041515 | 78.47621812 | -0.590015971 | 0.294302247 | -2.004796012 | 0.044984859 | 0.293090541 | Irf8 | interferon regulatory factor 8 [Source:MG | 81.113 | 105.88 | 96.599 | 85.794 | 46.897 | 54.578 |
| 12269 | ENSMUSG00000041523 | 0 | NA | NA | NA | NA | NA | Upk2 | uroplakin 2 [Source:MGI Symbol;Acc:MGI | 0 | 0 | 0 | 0 | 0 | 0 |
| 12270 | ENSMUSG00000041528 | 1136.085141 | 0.078550116 | 0.131032925 | 0.599468538 | 0.548860484 | 0.895703112 | Rnf123 | ring finger protein 123 [Source:MGI Symb | 1228.7 | 1028.9 | 1057.6 | 1320.6 | 1164.8 | 1015.8 |
| 12271 | ENSMUSG00000041530 | 446.8251942 | 0.285334707 | 0.148404499 | 1.922682328 | 0.054519958 | 0.328710574 | Ago1 | argonaute RISC catalytic subunit 1 [Source | 345.33 | 403.97 | 459.85 | 469.46 | 476.6 | 525.73 |
| 12272 | ENSMUSG00000041534 | 0 | NA | NA | NA | NA | NA | Rbp3 | retinol binding protein 3, interstitial [Sour | 0 | 0 | 0 | 0 | 0 | 0 |
| 12273 | ENSMUSG00000041536 | 0.503122293 | -2.366675966 | 3.785596405 | -0.625179156 | 0.531853481 | NA | Serpina3a | serine (or cysteine) peptidase inhibitor, cl | 0 | 0 | 3.0187 | 0 | 0 | 0 |
| 12274 | ENSMUSG00000041538 | 7.109030336 | -1.768827716 | 1.001066279 | -1.766943661 | 0.077237664 | 0.406981993 | H2-Ob | histocompatibility 2, O region beta locus [ | 5.6217 | 7.1954 | 20.125 | 0.964 | 7.6343 | 1.1138 |
| 12275 | ENSMUSG00000041540 | 240.8800754 | 0.229609743 | 0.181442611 | 1.265467583 | 0.205703762 | 0.650907725 | Sox5 | SRY (sex determining region Y)-box 5 [Sou | 199.97 | 250.81 | 215.34 | 295.94 | 218.12 | 265.09 |
| 12276 | ENSMUSG00000041544 | 0 | NA | NA | NA | NA | NA | Disp3 | dispatched RND transporter family memb | 0 | 0 | 0 | 0 | 0 | 0 |
| 12277 | ENSMUSG00000041548 | 3088.964151 | -0.03842354 | 0.112484982 | -0.34158818 | 0.732660834 | 0.947722168 | Hspb8 | heat shock protein 8 [Source:MGI Symbol | 3380.3 | 3018 | 2990.6 | 2870.7 | 2824.7 | 3449.6 |
| 12278 | ENSMUSG00000041550 | 3.051443587 | -0.858592818 | 1.285310797 | -0.66800405 | 0.504131007 | NA | Serpina5 | serine (or cysteine) peptidase inhibitor, cl | 1.6062 | 6.1675 | 4.025 | 0.964 | 1.0906 | 4.4554 |
| 12279 | ENSMUSG00000041552 | 0 | NA | NA | NA | NA | NA | Ptchd1 | patched domain containing 1 [Source:MG | 0 | 0 | 0 | 0 | 0 | 0 |
| 12280 | ENSMUSG00000041556 | 2.706038751 | 0.575011924 | 1.296426854 | 0.443535956 | 0.657378118 | NA | Fbxo2 | F-box protein 2 [Source:MGI Symbol;Acc:I | 2.4093 | 2.0558 | 2.0125 | 0.964 | 5.4531 | 3.3415 |
| 12281 | ENSMUSG00000041559 | 54.8949949 | -0.474137094 | 0.328410245 | -1.443734174 | 0.148813765 | 0.560823943 | Fmod | fibromodulin [Source:MGI Symbol;Acc:M( | 72.279 | 71.954 | 47.293 | 52.055 | 51.259 | 34.529 |
| 12282 | ENSMUSG00000041560 | 1720.73852 | 0.244511698 | 0.098747269 | 2.47613631 | 0.013281286 | 0.131539526 | Nop53 | NOP53 ribosome biogenesis factor [Sourc | 1510.6 | 1591.2 | 1624.1 | 1755.4 | 1919.5 | 1923.6 |
| 12283 | ENSMUSG00000041565 | 0 | NA | NA | NA | NA | NA | L3mbtl4 | L3MBTL4 histone methyl-lysine binding pr | 0 | 0 | 0 | 0 | 0 | 0 |
| 12284 | ENSMUSG00000041566 | 4.778031779 | 0.077009919 | 0.953332769 | 0.080779683 | 0.935617166 | NA | Tssk1 | testis-specific serine kinase 1 [Source:MG | 4.8186 | 3.0838 | 6.0375 | 4.8199 | 5.4531 | 4.4554 |
| 12285 | ENSMUSG00000041567 | 1986.559975 | 0.677370814 | 0.187520643 | 3.612246654 | 0.000303556 | 0.008781257 | Serpina12 | serine (or cysteine) peptidase inhibitor, cl | 1921 | 1347.6 | 1315.2 | 1716.8 | 2800.7 | 2818 |
| 12286 | ENSMUSG00000041570 | 311.1030333 | 0.116536673 | 0.146719632 | 0.794281391 | 0.427031635 | 0.842220356 | Camsap2 | calmodulin regulated spectrin-associated | 285.9 | 299.12 | 310.93 | 321.97 | 345.73 | 302.97 |
| 12287 | ENSMUSG00000041571 | 1098.30408 | -0.105710249 | 0.112965422 | -0.935775276 | 0.349388904 | 0.790730887 | Selenow | selenoprotein W [Source:MGI Symbol;Acc | 1129.2 | 1076.2 | 1209.5 | 964.94 | 1092.8 | 1117.2 |
| 12288 | ENSMUSG00000041577 | 455.0057343 | -0.954419169 | 0.164421438 | -5.804712465 | 6.45E-09 | 1.05E-06 | Prelp | proline arginine-rich end leucine-rich repe | 664.17 | 523.21 | 612.8 | 310.4 | 254.11 | 365.34 |
| 12289 | ENSMUSG00000041578 | 0 | NA | NA | NA | NA | NA | Crx | cone-rod homeobox [Source:MGI Symbol | 0 | 0 | 0 | 0 | 0 | 0 |
| 12290 | ENSMUSG00000041583 | 0 | NA | NA | NA | NA | NA | Obox6 | oocyte specific homeobox 6 [Source:MGI | 0 | 0 | 0 | 0 | 0 | 0 |
| 12291 | ENSMUSG00000041592 | 10.06851388 | -0.997360802 | 0.790020936 | -1.262448572 | 0.206787421 | 0.652194904 | Sdk2 | sidekick cell adhesion molecule 2 [Source: | 16.062 | 10.279 | 14.087 | 14.46 | 2.1812 | 3.3415 |
| 12292 | ENSMUSG00000041594 | 95.45354687 | 0.208640678 | 0.243719729 | 0.85606807 | 0.391960129 | 0.820806617 | Tmtc4 | transmembrane and tetratricopeptide rep | 77.098 | 86.345 | 102.64 | 104.11 | 113.42 | 89.108 |
| 12293 | ENSMUSG00000041596 | 12.66507633 | 1.088667742 | 0.723438648 | 1.50485151 | 0.132362257 | 0.533283479 | Nlrp5-ps | NLR family, pyrin domain containing 5, ps( | 7.2279 | 5.1396 | 12.075 | 32.775 | 7.6343 | 11.138 |
| 12294 | ENSMUSG00000041598 | 1098.443985 | -0.112660482 | 0.132743491 | -0.848708143 | 0.396043714 | 0.823764863 | Cdc42ep4 | CDC42 effector protein (Rho GTPase bindi | 1195 | 1130.7 | 1096.8 | 865.65 | 1143 | 1159.5 |
| 12295 | ENSMUSG00000041605 | 0.346303622 | 2.004339653 | 3.884003423 | 0.516049919 | 0.605819544 | NA | Inava | innate immunity activator [Source:MGI Sy | 0 | 0 | 0 | 0.964 | 0 | 1.1138 |
| 12296 | ENSMUSG00000041607 | 16.96741099 | -0.245017214 | 0.531891176 | -0.460652902 | 0.64504765 | 0.925643431 | Mbp | myelin basic protein [Source:MGI Symbol; | 22.487 | 15.419 | 17.106 | 11.568 | 19.631 | 15.594 |
| 12297 | ENSMUSG00000041608 | 0.501260735 | 1.111687294 | 3.109072805 | 0.357562323 | 0.720670879 | NA | Entpd3 | ectonucleoside triphosphate diphosphohy | 0.8031 | 0 | 0 | 0 | 1.0906 | 1.1138 |
| 12298 | ENSMUSG00000041609 | 52.23013712 | -0.277506554 | 0.310295013 | -0.89433134 | 0.371144637 | 0.807214774 | Bicdl1 | BICD family like cargo adaptor 1 [Source:N | 49.792 | 59.619 | 62.387 | 43.379 | 43.625 | 54.578 |
| 12299 | ENSMUSG00000041616 | 0 | NA | NA | NA | NA | NA | Nppa | natriuretic peptide type A [Source:MGI Sy | 0 | 0 | 0 | 0 | 0 | 0 |
| 12300 | ENSMUSG00000041617 | 6.286931918 | 0.014994287 | 0.91573351 | 0.016374072 | 0.986935964 | 0.99859176 | Ccdc74a | coiled-coil domain containing 74A [Source | 5.6217 | 3.0838 | 10.062 | 6.7478 | 2.1812 | 10.025 |
| 12301 | ENSMUSG00000041620 | 0 | NA | NA | NA | NA | NA | Mmp1b | matrix metallopeptidase 1b (interstitial co | 0 | 0 | 0 | 0 | 0 | 0 |
| 12302 | ENSMUSG00000041623 | 82.49777811 | 0.454376524 | 0.251726247 | 1.8050423 | 0.071068063 | 0.385913448 | D11Wsu47e | DNA segment, Chr 11, Wayne State Unive | 75.492 | 58.591 | 74.462 | 92.542 | 100.34 | 93.563 |
| 12303 | ENSMUSG00000041624 | 0.301557804 | -1.64335916 | 4.042814367 | -0.406488899 | 0.684383412 | NA | Gucy1a2 | guanylate cyclase 1, soluble, alpha 2 [Sour | 0.8031 | 0 | 1.0062 | 0 | 0 | 0 |
| 12304 | ENSMUSG00000041625 | 1105.331936 | 0.085562177 | 0.118288285 | 0.723336019 | 0.469473469 | 0.866351653 | Ggact | gamma-glutamylamine cyclotransferase [S | 1151.6 | 1007.4 | 1058.6 | 1254.1 | 1125.5 | 1034.8 |
| 12305 | ENSMUSG00000041629 | 349.8873763 | -0.014157067 | 0.143550704 | -0.098620674 | 0.921439455 | 0.986685804 | Fam104a | family with sequence similarity 104, meml | 365.41 | 340.24 | 349.17 | 374.02 | 316.28 | 354.2 |
| 12306 | ENSMUSG00000041632 | 388.7878372 | 0.202336455 | 0.151688817 | 1.333891703 | 0.182239351 | 0.615722956 | Mrps27 | mitochondrial ribosomal protein S27 [Sou | 396.73 | 370.05 | 316.97 | 387.52 | 396.99 | 464.47 |
| 12307 | ENSMUSG00000041633 | 28.43826597 | -0.844525219 | 0.441150024 | -1.914371924 | 0.05557265 | 0.332837604 | Kctd12b | potassium channel tetramerisation domai | 47.383 | 27.754 | 34.212 | 19.28 | 15.269 | 26.732 |
| 12308 | ENSMUSG00000041638 | 995.001934 | 0.157095824 | 0.11653182 | 1.348093801 | 0.177628215 | 0.608962907 | Gcn1 | GCN1 activator of EIF2AK4 [Source:MGI S) | 876.18 | 977.55 | 970.02 | 1081.6 | 1116.8 | 947.88 |
| 12309 | ENSMUSG00000041642 | 26.67320103 | -1.219843708 | 0.465952034 | -2.617959832 | 0.008845722 | 0.098951677 | Kif21b | kinesin family member 21B [Source:MGI S | 31.321 | 48.312 | 32.2 | 9.6398 | 19.631 | 18.935 |
| 12310 | ENSMUSG00000041644 | 0 | NA | NA | NA | NA | NA | Slc5a12 | solute carrier family 5 (sodium/glucose co | 0 | 0 | 0 | 0 | 0 | 0 |
| 12311 | ENSMUSG00000041645 | 640.6147377 | -0.073969021 | 0.123101186 | -0.600879839 | 0.547920023 | 0.895600915 | Ddx24 | DEAD box helicase 24 [Source:MGI Symbc | 693.88 | 632.17 | 644 | 573.57 | 669.64 | 630.44 |
| 12312 | ENSMUSG00000041649 | 0.638982041 | 0.404325377 | 3.14176733 | 0.128693609 | 0.897600092 | NA | Klf8 | Kruppel-like factor 8 [Source:MGI Symbol; | 1.6062 | 0 | 0 | 0 | 0 | 2.2277 |
| 12313 | ENSMUSG00000041650 | 1765.970169 | 0.062269752 | 0.097076219 | 0.641452181 | 0.52122894 | 0.887887809 | Pcca | propionyl-Coenzyme A carboxylase, alpha | 1802.2 | 1682.7 | 1697.5 | 1714.9 | 1857.3 | 1841.2 |
| 12314 | ENSMUSG00000041653 | 7.263574149 | 0.658903481 | 0.888437495 | 0.741643036 | 0.458303642 | 0.859322431 | Pnpla3 | patatin-like phospholipase domain contai | 9.6372 | 5.1396 | 2.0125 | 13.496 | 3.2719 | 10.025 |
| 12315 | ENSMUSG00000041654 | 446.3503287 | 0.343494082 | 0.157661475 | 2.178681145 | 0.029355363 | 0.225489101 | Slc39a11 | solute carrier family 39 (metal ion transpo | 348.55 | 450.23 | 382.37 | 500.3 | 429.7 | 566.95 |
| 12316 | ENSMUSG00000041658 | 0.305170013 | -1.657836477 | 4.041480335 | -0.410205257 | 0.681655385 | NA | Rragb | Ras-related GTP binding B [Source:MGI Sy | 0.8031 | 1.0279 | 0 | 0 | 0 | 0 |

| | | | | | | | | | | | | | | | |
|---|---|---|---|---|---|---|---|---|---|---|---|---|---|---|---|
| 12317 | ENSMUSG00000041660 | 3974.213679 | -0.087656281 | 0.09487515 | -0.923911909 | 0.355532182 | 0.795424787 | Bbox1 | butyrobetaine (gamma), 2-oxoglutarate d | 4070.1 | 3984.2 | 4230.3 | 3741.2 | 4169.4 | 3650.1 |
| 12318 | ENSMUSG00000041669 | 0 | NA | NA | NA | NA | NA | Prima1 | proline rich membrane anchor 1 [Source:I | 0 | 0 | 0 | 0 | 0 | 0 |
| 12319 | ENSMUSG00000041670 | 0 | NA | NA | NA | NA | NA | Rims1 | regulating synaptic membrane exocytosis | 0 | 0 | 0 | 0 | 0 | 0 |
| 12320 | ENSMUSG00000041671 | 113.5819325 | 0.048068211 | 0.21733351 | 0.221172572 | 0.824958065 | 0.96943211 | Pyroxd1 | pyridine nucleotide-disulphide oxidoredu | 113.24 | 103.82 | 117.73 | 103.15 | 122.15 | 121.41 |
| 12321 | ENSMUSG00000041673 | 13.44621598 | 0.05432194 | 0.639666269 | 0.084922314 | 0.932323152 | 0.988420305 | Lrrc18 | leucine rich repeat containing 18 [Source: | 12.047 | 19.53 | 8.05 | 12.532 | 20.722 | 7.7969 |
| 12322 | ENSMUSG00000041674 | 0.915303448 | -0.756900117 | 2.316458517 | -0.326748833 | 0.743857862 | NA | BC006965 | cDNA sequence BC006965 [Source:MGI Sy | 2.4093 | 1.0279 | 0 | 0.964 | 1.0906 | 0 |
| 12323 | ENSMUSG00000041679 | 80.641121 | 0.130695454 | 0.265441187 | 0.492370666 | 0.622457333 | 0.918789964 | Lrrc29 | leucine rich repeat containing 29 [Source: | 89.947 | 81.206 | 59.368 | 78.082 | 87.249 | 87.994 |
| 12324 | ENSMUSG00000041681 | 0 | NA | NA | NA | NA | NA | Iapp | islet amyloid polypeptide [Source:MGI Syr | 0 | 0 | 0 | 0 | 0 | 0 |
| 12325 | ENSMUSG00000041684 | 265.9948858 | 0.043685317 | 0.159630632 | 0.273665 | 0.784342077 | 0.960670212 | Bivm | basic, immunoglobulin-like variable motif | 273.86 | 246.7 | 264.64 | 242.92 | 271.56 | 296.28 |
| 12326 | ENSMUSG00000041685 | 730.8166238 | 0.059961409 | 0.11258089 | 0.532607344 | 0.594305417 | 0.913144152 | Fcho2 | FCH domain only 2 [Source:MGI Symbol;A | 699.5 | 735.99 | 711.41 | 722.02 | 730.71 | 785.26 |
| 12327 | ENSMUSG00000041688 | 118.5797479 | -0.313134913 | 0.228108564 | -1.372745101 | 0.169831603 | 0.596876778 | Amot | angiomotin [Source:MGI Symbol;Acc:MGI | 124.48 | 149.05 | 120.75 | 104.11 | 88.34 | 124.75 |
| 12328 | ENSMUSG00000041695 | 9.484528927 | -0.610509573 | 0.762705338 | -0.800452734 | 0.423448538 | 0.84018348 | Kcnj2 | potassium inwardly-rectifying channel, su | 8.8341 | 17.475 | 8.05 | 4.8199 | 4.3625 | 13.366 |
| 12329 | ENSMUSG00000041696 | 0.17131964 | -0.796629176 | 4.080472857 | -0.195229623 | 0.84521319 | NA | Rasl12 | RAS-like, family 12 [Source:MGI Symbol;A | 0 | 1.0279 | 0 | 0 | 0 | 0 |
| 12330 | ENSMUSG00000041697 | 6825.642522 | 0.011710894 | 0.094411087 | 0.124041516 | 0.901282403 | 0.98333371 | Cox6a1 | cytochrome c oxidase subunit 6A1 [Sourc | 7029.6 | 6656.8 | 6706.6 | 6770 | 6363.8 | 7427.1 |
| 12331 | ENSMUSG00000041698 | 8507.661457 | 0.997193915 | 0.108241804 | 9.212650518 | 3.18E-20 | 4.60E-17 | Slco1a1 | solute carrier organic anion transporter fa | 6395.9 | 5076.9 | 5562.5 | 11675 | 10617 | 11719 |
| 12332 | ENSMUSG00000041700 | 0 | NA | NA | NA | NA | NA | Lhfpl1 | lipoma HMGIC fusion partner-like 1 [Sour | 0 | 0 | 0 | 0 | 0 | 0 |
| 12333 | ENSMUSG00000041702 | 363.5627674 | 0.241946298 | 0.156828242 | 1.542746986 | 0.122892177 | 0.513267141 | Btbd7 | BTB (POZ) domain containing 7 [Source:M | 275.46 | 370.05 | 355.2 | 406.8 | 378.44 | 395.41 |
| 12334 | ENSMUSG00000041703 | 0.321325949 | 1.911575114 | 4.048267893 | 0.472195804 | 0.636787034 | NA | Zic5 | zinc finger protein of the cerebellum 5 [Sc | 0 | 0 | 0 | 1.928 | 0 | 0 |
| 12335 | ENSMUSG00000041707 | 8.445759458 | -2.140261132 | 0.841595207 | -2.54310043 | 0.010987364 | 0.116085638 | Tmem273 | transmembrane protein 273 [Source:MGI | 8.8341 | 19.53 | 13.081 | 4.8199 | 2.1812 | 2.2277 |
| 12336 | ENSMUSG00000041708 | 0.840571505 | -3.133009828 | 2.792076153 | -1.122107585 | 0.261816699 | NA | Mpped1 | metallophosphoesterase domain containi | 4.0155 | 1.0279 | 0 | 0 | 0 | 0 |
| 12337 | ENSMUSG00000041710 | 0 | NA | NA | NA | NA | NA | Trpc5 | transient receptor potential cation channe | 0 | 0 | 0 | 0 | 0 | 0 |
| 12338 | ENSMUSG00000041712 | 398.8929159 | 0.037050052 | 0.144292874 | 0.256769795 | 0.797356489 | 0.962236316 | Ubr7 | ubiquitin protein ligase E3 component n-r | 359.79 | 411.17 | 410.55 | 368.24 | 398.08 | 445.54 |
| 12339 | ENSMUSG00000041716 | 25.31361356 | 0.087556428 | 0.456624228 | 0.191747225 | 0.847940212 | 0.972803041 | Gm20604 | predicted gene 20604 [Source:MGI Symbo | 33.73 | 20.558 | 19.119 | 29.883 | 19.631 | 28.96 |
| 12340 | ENSMUSG00000041718 | 122.942909 | 0.038154647 | 0.224226622 | 0.170161094 | 0.864883448 | 0.977227145 | Alg13 | asparagine-linked glycosylation 13 [Source | 133.31 | 123.35 | 106.66 | 104.11 | 125.42 | 144.8 |
| 12341 | ENSMUSG00000041720 | 812.1122025 | -0.018736139 | 0.133982949 | -0.13983973 | 0.888786621 | 0.982382091 | Pi4ka | phosphatidylinositol 4-kinase alpha [Sour | 742.07 | 905.6 | 806 | 901.32 | 807.06 | 710.63 |
| 12342 | ENSMUSG00000041722 | 0 | NA | NA | NA | NA | NA | Khdc1c | KH domain containing 1C [Source:MGI Syr | 0 | 0 | 0 | 0 | 0 | 0 |
| 12343 | ENSMUSG00000041729 | 19.33850694 | 0.103396124 | 0.658522698 | 0.15701224 | 0.875235201 | 0.980512892 | Coro2b | coronin, actin binding protein, 2B [Source | 24.093 | 25.698 | 6.0375 | 20.244 | 6.5437 | 33.415 |
| 12344 | ENSMUSG00000041730 | 0.133850373 | -0.796629176 | 4.080472857 | -0.195229623 | 0.84521319 | NA | Prrxl1 | paired related homeobox protein-like 1 [S | 0.8031 | 0 | 0 | 0 | 0 | 0 |
| 12345 | ENSMUSG00000041731 | 8.297250101 | -0.957667526 | 0.871882043 | -1.098391157 | 0.272033722 | 0.727618887 | Pgm5 | phosphoglucomutase 5 [Source:MGI Syml | 5.6217 | 6.1675 | 21.131 | 4.8199 | 9.8156 | 2.2277 |
| 12346 | ENSMUSG00000041733 | 576.219308 | -0.059353801 | 0.119019728 | -0.49868876 | 0.617998664 | 0.917509899 | Coq5 | coenzyme Q5 methyltransferase [Source:I | 616.78 | 561.24 | 585.63 | 563.93 | 561.67 | 568.06 |
| 12347 | ENSMUSG00000041734 | 22.55563537 | -1.383133049 | 0.495963535 | -2.78877972 | 0.005290704 | 0.069144802 | Kirrel | kirre like nephrin family adhesion molecul | 33.73 | 40.089 | 24.15 | 16.388 | 8.7249 | 12.252 |
| 12348 | ENSMUSG00000041735 | 0 | NA | NA | NA | NA | NA | AAdacl4fm3 | AADACL4 family member 3 [Source:MGI S | 0 | 0 | 0 | 0 | 0 | 0 |
| 12349 | ENSMUSG00000041736 | 1617.322335 | -0.063873096 | 0.09688056 | -0.659297343 | 0.509704848 | 0.882793133 | Tspo | translocator protein [Source:MGI Symbol; | 1690.5 | 1658 | 1611 | 1662.9 | 1540 | 1541.6 |
| 12350 | ENSMUSG00000041737 | 4.61053041 | -2.074181778 | 1.104088684 | -1.878636931 | 0.060294085 | NA | Tmem45b | transmembrane protein 45b [Source:MGI | 8.031 | 9.2513 | 5.0312 | 0.964 | 3.2719 | 1.1138 |
| 12351 | ENSMUSG00000041740 | 3005.597952 | -0.049248888 | 0.096869037 | -0.508406918 | 0.611168002 | 0.916376001 | Rnf10 | ring finger protein 10 [Source:MGI Symbo | 2875.9 | 3189.6 | 3106.3 | 3031.7 | 2762.5 | 3067.5 |
| 12352 | ENSMUSG00000041741 | 21.87854498 | -0.880612453 | 0.530867818 | -1.658816796 | 0.097152715 | 0.45892125 | Pde3a | phosphodiesterase 3A, cGMP inhibited [S | 36.943 | 33.921 | 14.087 | 14.46 | 20.722 | 11.138 |
| 12353 | ENSMUSG00000041747 | 184.1459081 | -0.042282389 | 0.194181706 | -0.217746513 | 0.827626621 | 0.969794868 | Utp15 | UTP15 small subunit processome compon | 224.07 | 169.61 | 166.03 | 181.23 | 170.14 | 193.81 |
| 12354 | ENSMUSG00000041750 | 5.176558618 | 0.427165542 | 1.022192229 | 0.417891596 | 0.676026377 | 0.932739835 | Cd1d2 | CD1d2 antigen [Source:MGI Symbol;Acc:M | 8.031 | 2.0558 | 3.0187 | 4.8199 | 10.906 | 2.2277 |
| 12355 | ENSMUSG00000041754 | 0.167707431 | -0.796629176 | 4.080472857 | -0.195229623 | 0.84521319 | NA | Trem3 | triggering receptor expressed on myeloid | 0 | 0 | 1.0062 | 0 | 0 | 0 |
| 12356 | ENSMUSG00000041757 | 1310.231135 | 0.155039456 | 0.151307574 | 1.024664208 | 0.305521657 | 0.754621797 | Plekha6 | pleckstrin homology domain containing, f | 1179.8 | 1152.3 | 1387.6 | 1326.4 | 1122.2 | 1693 |
| 12357 | ENSMUSG00000041762 | 861.0858495 | 0.030726658 | 0.15651752 | 0.196314498 | 0.844364014 | 0.972803041 | Gpr155 | G protein-coupled receptor 155 [Source:N | 746.89 | 873.73 | 935.81 | 826.13 | 1065.5 | 718.43 |
| 12358 | ENSMUSG00000041763 | 718.1532975 | 0.119649284 | 0.117208027 | 1.020828406 | 0.307335744 | 0.756237058 | Tpp2 | tripeptidyl peptidase II [Source:MGI Symb | 654.53 | 709.26 | 702.36 | 787.57 | 760.16 | 695.04 |
| 12359 | ENSMUSG00000041765 | 521.6626465 | 0.068798418 | 0.121522149 | 0.566138918 | 0.571299349 | 0.904651508 | Ubac2 | ubiquitin associated domain containing 2 | 486.68 | 521.15 | 520.23 | 527.3 | 534.4 | 540.21 |
| 12360 | ENSMUSG00000041769 | 706.4418672 | -0.00852514 | 0.129515157 | -0.06582349 | 0.947518354 | 0.990677354 | Ppp2r2d | protein phosphatase 2, regulatory subunit | 770.18 | 663.01 | 692.3 | 789.5 | 659.82 | 663.85 |
| 12361 | ENSMUSG00000041771 | 0 | NA | NA | NA | NA | NA | Slc24a4 | solute carrier family 24 (sodium/potassiur | 0 | 0 | 0 | 0 | 0 | 0 |
| 12362 | ENSMUSG00000041773 | 180.1729489 | -1.211094283 | 0.203249691 | -5.95865253 | 2.54E-09 | 4.60E-07 | Enc1 | ectodermal-neural cortex 1 [Source:MGI S | 295.54 | 240.53 | 218.36 | 103.15 | 124.33 | 99.132 |
| 12363 | ENSMUSG00000041774 | 3.981327296 | 1.330994957 | 1.194566758 | 1.114207262 | 0.265190281 | NA | Ydjc | YdjC homolog (bacterial) [Source:MGI Syn | 0.8031 | 1.0279 | 5.0312 | 4.8199 | 2.1812 | 10.025 |
| 12364 | ENSMUSG00000041775 | 170.8823952 | 0.11087067 | 0.18426445 | 0.601693219 | 0.547378364 | 0.895518681 | Mapk1ip1 | mitogen-activated protein kinase 1 intera | 173.47 | 166.52 | 152.95 | 192.8 | 164.68 | 174.87 |
| 12365 | ENSMUSG00000041777 | 327.9661618 | 0.243770529 | 0.146136897 | 1.668097062 | 0.09529646 | 0.455057116 | Cir1 | corepressor interacting with RBPJ, 1 [Sour | 320.44 | 293.98 | 285.77 | 335.46 | 350.09 | 382.05 |
| 12366 | ENSMUSG00000041779 | 419.6734065 | -0.15263154 | 0.138117148 | -1.105087545 | 0.269121657 | 0.724979259 | Tram2 | translocating chain-associating membrane | 431.27 | 415.28 | 478.97 | 380.77 | 429.7 | 382.05 |
| 12367 | ENSMUSG00000041781 | 450.0515334 | 0.016017155 | 0.138630455 | 0.115538501 | 0.908018305 | 0.984639154 | Cpsf2 | cleavage and polyadenylation specific fact | 403.96 | 497.51 | 441.74 | 428.97 | 456.97 | 471.16 |
| 12368 | ENSMUSG00000041782 | 202.4736368 | -0.146934934 | 0.243231912 | -0.604093983 | 0.54578116 | 0.89549457 | Lad1 | ladinin [Source:MGI Symbol;Acc:MGI:109 | 242.54 | 137.74 | 257.6 | 193.76 | 154.87 | 228.34 |
| 12369 | ENSMUSG00000041789 | 0.952566124 | 1.285634123 | 2.448533191 | 0.525062975 | 0.599539413 | NA | 2700046A07 | RIKEN cDNA 2700046A07 gene [Source:M | 1.6062 | 0 | 0 | 1.928 | 2.1812 | 0 |
| 12370 | ENSMUSG00000041791 | 0 | NA | NA | NA | NA | NA | Capza3 | capping protein (actin filament) muscle Z- | 0 | 0 | 0 | 0 | 0 | 0 |
| 12371 | ENSMUSG00000041794 | 1.897189006 | -0.921505127 | 1.752291123 | -0.525885862 | 0.598967513 | NA | Myrip | myosin VIIA and Rab interacting protein [S | 2.4093 | 4.1117 | 1.0062 | 3.8559 | 0 | 0 |
| 12372 | ENSMUSG00000041797 | 84.77905916 | -0.900363782 | 0.283050617 | -3.180928527 | 0.001468038 | 0.027902376 | Abca9 | ATP-binding cassette, sub-family A (ABC1) | 106.01 | 131.57 | 93.581 | 51.091 | 78.525 | 47.895 |

| | | | | | | | | | | | | | | | |
|---|---|---|---|---|---|---|---|---|---|---|---|---|---|---|---|
| 12373 | ENSMUSG00000041798 | 4500.240162 | 0.101685904 | 0.121165672 | 0.8392303 | 0.401340087 | 0.82816843 | Gck | glucokinase [Source:MGI Symbol;Acc:MGI | 5018.6 | 3805.4 | 4200.1 | 5086.9 | 4349.4 | 4541.1 |
| 12374 | ENSMUSG00000041801 | 27.32886406 | -0.598931106 | 0.461691896 | -1.297252803 | 0.194544218 | 0.633957463 | Phlda3 | pleckstrin homology like domain, family A | 39.352 | 32.893 | 26.162 | 11.568 | 27.265 | 26.732 |
| 12375 | ENSMUSG00000041805 | 0 | NA | NA | NA | NA | NA | Pramel1 | PRAME like 1 [Source:MGI Symbol;Acc:MG | 0 | 0 | 0 | 0 | 0 | 0 |
| 12376 | ENSMUSG00000041809 | 3.194115998 | 0.324191162 | 1.330780252 | 0.243609838 | 0.807533008 | NA | Efhc1 | EF-hand domain (C-terminal) containing 1 | 2.4093 | 3.0838 | 3.0187 | 1.928 | 8.7249 | 0 |
| 12377 | ENSMUSG00000041815 | 614.3143749 | 0.041387586 | 0.123041802 | 0.33637012 | 0.736591779 | 0.948137969 | Poldip3 | polymerase (DNA-directed), delta interact | 568.6 | 593.11 | 655.07 | 594.77 | 622.74 | 651.6 |
| 12378 | ENSMUSG00000041817 | 7.169075727 | -0.629903611 | 0.822955196 | -0.765416652 | 0.44402348 | 0.851432536 | Fam169a | family with sequence similarity 169, meml | 4.8186 | 13.363 | 8.05 | 5.7839 | 6.5437 | 4.4554 |
| 12379 | ENSMUSG00000041827 | 214.3157942 | 0.12785409 | 0.231851551 | 0.55144807 | 0.581326556 | 0.908645447 | Oasl1 | 2'-5' oligoadenylate synthetase-like 1 [Sou | 255.39 | 173.72 | 184.14 | 157.13 | 270.47 | 245.05 |
| 12380 | ENSMUSG00000041828 | 2283.876439 | 0.141974189 | 0.193020989 | 0.735537569 | 0.462012168 | 0.860650866 | Abca8a | ATP-binding cassette, sub-family A (ABC1) | 1432.7 | 2200.8 | 2882.9 | 2619.1 | 2517.1 | 2050.6 |
| 12381 | ENSMUSG00000041831 | 0.674441933 | -2.786326546 | 3.022203109 | -0.921952114 | 0.356553554 | NA | Sytl3 | synaptotagmin-like 3 [Source:MGI Symbol | 0 | 1.0279 | 3.0187 | 0 | 0 | 0 |
| 12382 | ENSMUSG00000041836 | 26.02165951 | 0.717332943 | 0.437910551 | 1.638080977 | 0.101404803 | 0.46908328 | Ptpre | protein tyrosine phosphatase, receptor ty | 18.471 | 22.614 | 18.112 | 39.523 | 23.994 | 33.415 |
| 12383 | ENSMUSG00000041837 | 158.4259416 | 0.002079217 | 0.211936806 | 0.00981055 | 0.992172439 | 0.99859176 | Pdcd7 | programmed cell death 7 [Source:MGI Syr | 129.3 | 154.19 | 192.19 | 171.59 | 141.78 | 161.51 |
| 12384 | ENSMUSG00000041840 | 111.2116206 | 0.080464415 | 0.216898678 | 0.370976973 | 0.710654683 | 0.944239215 | Haus1 | HAUS augmin-like complex, subunit 1 [Sou | 111.63 | 107.93 | 104.65 | 111.82 | 125.42 | 105.82 |
| 12385 | ENSMUSG00000041841 | 1520.838336 | 0.024672886 | 0.117540459 | 0.209909728 | 0.83373813 | 0.970792812 | Rpl37 | ribosomal protein L37 [Source:MGI Symbo | 1336.4 | 1646.7 | 1541.6 | 1523.1 | 1424.3 | 1652.9 |
| 12386 | ENSMUSG00000041842 | 4.07182644 | 0.791141726 | 1.074271148 | 0.736445103 | 0.461459865 | NA | Fhdc1 | FH2 domain containing 1 [Source:MGI Syr | 4.8186 | 1.0279 | 3.0187 | 6.7478 | 4.3625 | 4.4554 |
| 12387 | ENSMUSG00000041845 | 327.5288311 | -0.044753883 | 0.218881309 | -0.204466443 | 0.837989013 | 0.971073158 | Rhod | ras homolog family member D [Source:MG | 394.32 | 274.45 | 329.04 | 424.15 | 225.76 | 317.45 |
| 12388 | ENSMUSG00000041846 | 1113.381883 | 0.081035501 | 0.110631989 | 0.732478031 | 0.463876847 | 0.861872906 | Ppp4r3a | protein phosphatase 4 regulatory subunit | 1036 | 1080.3 | 1131 | 1205.9 | 1183.3 | 1043.7 |
| 12389 | ENSMUSG00000041849 | 288.00439 | 0.187955588 | 0.156409866 | 1.201686266 | 0.229485104 | 0.676160633 | Card6 | caspase recruitment domain family, meml | 278.68 | 256.98 | 271.69 | 294.01 | 281.38 | 345.29 |
| 12390 | ENSMUSG00000041852 | 499.8578783 | 0.066005378 | 0.142673997 | 0.462630745 | 0.643629067 | 0.925505685 | Tcf20 | transcription factor 20 [Source:MGI Symb | 466.6 | 443.03 | 555.45 | 461.75 | 537.67 | 534.65 |
| 12391 | ENSMUSG00000041857 | 0 | NA | NA | NA | NA | NA | Oosp1 | oocyte secreted protein 1 [Source:MGI Sy | 0 | 0 | 0 | 0 | 0 | 0 |
| 12392 | ENSMUSG00000041859 | 72.18356616 | -0.456970606 | 0.312236675 | -1.463539176 | 0.143319902 | 0.553098698 | Mcm3 | minichromosome maintenance complex c | 118.86 | 61.675 | 69.431 | 60.731 | 57.803 | 64.603 |
| 12393 | ENSMUSG00000041870 | 431.5860651 | -0.322853912 | 0.143231726 | -2.25406703 | 0.024191951 | 0.198138383 | Ankrd13a | ankyrin repeat domain 13a [Source:MGI S | 462.59 | 494.43 | 483 | 439.57 | 346.82 | 363.11 |
| 12394 | ENSMUSG00000041872 | 0 | NA | NA | NA | NA | NA | Il17f | interleukin 17F [Source:MGI Symbol;Acc:M | 0 | 0 | 0 | 0 | 0 | 0 |
| 12395 | ENSMUSG00000041878 | 6.428968126 | -0.157301756 | 0.917587746 | -0.171429661 | 0.863885934 | 0.976954351 | 8430432A02 | RIKEN cDNA 8430432A02 gene [Source:M | 7.2279 | 6.1675 | 7.0437 | 11.568 | 5.4531 | 1.1138 |
| 12396 | ENSMUSG00000041879 | 458.3908897 | -0.02299735 | 0.135592852 | -0.16960592 | 0.865320067 | 0.977353541 | Ipo9 | importin 9 [Source:MGI Symbol;Acc:MGI: | 503.55 | 429.67 | 451.8 | 423.19 | 478.78 | 463.36 |
| 12397 | ENSMUSG00000041881 | 1306.340624 | -0.031432343 | 0.100413285 | -0.313029727 | 0.754258077 | 0.953048681 | Ndufa7 | NADH:ubiquinone oxidoreductase subuni | 1364.5 | 1305.5 | 1291 | 1261.8 | 1257.5 | 1357.8 |
| 12398 | ENSMUSG00000041885 | 0 | NA | NA | NA | NA | NA | Olfr680-ps1 | olfactory receptor 680, pseudogene 1 [Sou | 0 | 0 | 0 | 0 | 0 | 0 |
| 12399 | ENSMUSG00000041886 | 1.461775315 | -0.076374232 | 1.915478993 | -0.039872133 | 0.968195068 | NA | Macc1 | metastasis associated in colon cancer 1 [S | 2.4093 | 2.0558 | 0 | 0.964 | 0 | 3.3415 |
| 12400 | ENSMUSG00000041889 | 1.113642279 | -2.331062147 | 2.36105891 | -0.987295208 | 0.323497939 | NA | Shisa4 | shisa family member 4 [Source:MGI Symb | 1.6062 | 4.1117 | 0 | 0.964 | 0 | 0 |
| 12401 | ENSMUSG00000041890 | 248.432577 | -0.075510808 | 0.161427034 | -0.467770521 | 0.639948704 | 0.923932929 | Git2 | GIT ArfGAP 2 [Source:MGI Symbol;Acc:MG | 257.8 | 232.31 | 274.7 | 247.74 | 251.93 | 226.11 |
| 12402 | ENSMUSG00000041891 | 4655.344999 | -0.166955572 | 0.111706495 | -1.494591442 | 0.135021101 | 0.537616739 | Lman1 | lectin, mannose-binding, 1 [Source:MGI S | 5143.1 | 4983.3 | 4645.8 | 4019.8 | 5028.8 | 4111.2 |
| 12403 | ENSMUSG00000041895 | 465.4511387 | 0.028011217 | 0.130495446 | 0.214652834 | 0.830037998 | 0.9702546 | Wipi1 | WD repeat domain, phosphoinositide inte | 494.71 | 428.64 | 458.85 | 480.06 | 452.61 | 477.84 |
| 12404 | ENSMUSG00000041907 | 0.160662974 | 1.126924175 | 4.080472857 | 0.276174898 | 0.782413736 | NA | Gpr45 | G protein-coupled receptor 45 [Source:M | 0 | 0 | 0 | 0.964 | 0 | 0 |
| 12405 | ENSMUSG00000041911 | 0.160662974 | 1.126924175 | 4.080472857 | 0.276174898 | 0.782413736 | NA | Dlx1 | distal-less homeobox 1 [Source:MGI Symb | 0 | 0 | 0 | 0.964 | 0 | 0 |
| 12406 | ENSMUSG00000041912 | 24.00631291 | 0.625847411 | 0.487191511 | 1.284602454 | 0.198931241 | 0.640976045 | Tdrkh | tudor and KH domain containing protein [ | 29.715 | 13.363 | 13.081 | 25.063 | 31.628 | 31.188 |
| 12407 | ENSMUSG00000041915 | 775.2467522 | 0.104988581 | 0.118908336 | 0.882937097 | 0.377270259 | 0.81030298 | Ammecr1l | AMME chromosomal region gene 1-like [S | 695.49 | 782.25 | 763.74 | 753.83 | 789.61 | 866.57 |
| 12408 | ENSMUSG00000041920 | 121.5112814 | -0.443912949 | 0.274082427 | -1.619633019 | 0.105311132 | 0.478186913 | Slc16a6 | solute carrier family 16 (monocarboxylic a | 162.23 | 154.19 | 103.64 | 116.64 | 67.618 | 124.75 |
| 12409 | ENSMUSG00000041921 | 696.0318758 | 0.062371001 | 0.118336094 | 0.527066591 | 0.598147346 | 0.915029425 | Metap1d | methionyl aminopeptidase type 1D (mitoc | 700.31 | 694.87 | 647.02 | 662.25 | 715.45 | 756.3 |
| 12410 | ENSMUSG00000041923 | 0 | NA | NA | NA | NA | NA | Nol4 | nucleolar protein 4 [Source:MGI Symbol;A | 0 | 0 | 0 | 0 | 0 | 0 |
| 12411 | ENSMUSG00000041926 | 540.3982158 | 0.028935956 | 0.128381751 | 0.225389949 | 0.82167593 | 0.969160361 | Rnpep | arginyl aminopeptidase (aminopeptidase | 527.64 | 517.04 | 560.48 | 550.43 | 590.02 | 496.77 |
| 12412 | ENSMUSG00000041930 | 53.92847451 | -0.018114965 | 0.437442373 | -0.041411088 | 0.966968173 | 0.994231936 | Fam222a | family with sequence similarity 222, meml | 20.078 | 57.563 | 85.531 | 41.451 | 63.256 | 55.692 |
| 12413 | ENSMUSG00000041935 | 677.3965375 | 0.248076393 | 0.116821853 | 2.123544421 | 0.033708257 | 0.245744069 | AW549877 | expressed sequence AW549877 [Source:N | 608.75 | 589 | 660.1 | 717.2 | 726.35 | 762.98 |
| 12414 | ENSMUSG00000041936 | 771.7144518 | 0.251582109 | 0.131309411 | 1.915948797 | 0.055371613 | 0.331889018 | Agrn | agrin [Source:MGI Symbol;Acc:MGI:87961 | 641.68 | 682.54 | 790.91 | 927.35 | 809.24 | 778.58 |
| 12415 | ENSMUSG00000041939 | 188.4869215 | 0.749049802 | 0.218685055 | 3.425244588 | 0.000614246 | 0.014399444 | Mvk | mevalonate kinase [Source:MGI Symbol;A | 140.54 | 152.13 | 128.8 | 176.41 | 232.3 | 300.74 |
| 12416 | ENSMUSG00000041945 | 15.42544318 | 0.128660831 | 0.56110143 | 0.229300486 | 0.818635376 | 0.968964163 | Mfsd9 | major facilitator superfamily domain cont | 12.85 | 11.307 | 20.125 | 16.388 | 19.631 | 12.252 |
| 12417 | ENSMUSG00000041949 | 146.486606 | -0.072747895 | 0.202603486 | -0.359065364 | 0.719546192 | 0.9447719 | Tango6 | transport and golgi organization 6 [Source | 136.53 | 163.44 | 150.94 | 140.74 | 161.41 | 125.86 |
| 12418 | ENSMUSG00000041954 | 18.1974869 | 0.1625067 | 0.582012462 | 0.279215156 | 0.780079713 | 0.959870443 | Tnfrsf18 | tumor necrosis factor receptor superfamil | 15.259 | 10.279 | 26.162 | 28.919 | 18.541 | 10.025 |
| 12419 | ENSMUSG00000041957 | 701.839913 | 0.062184517 | 0.139029941 | 0.447274282 | 0.654677034 | 0.925797231 | Pkp2 | plakophilin 2 [Source:MGI Symbol;Acc:MG | 690.67 | 657.87 | 712.42 | 849.26 | 634.74 | 666.08 |
| 12420 | ENSMUSG00000041958 | 372.6870877 | 0.120238903 | 0.154320455 | 0.779150782 | 0.435890899 | 0.84624805 | Pigs | phosphatidylinositol glycan anchor biosyn | 325.26 | 363.88 | 382.37 | 336.43 | 390.44 | 437.74 |
| 12421 | ENSMUSG00000041959 | 1586.667862 | -0.304815074 | 0.180228627 | -1.691268915 | 0.090785454 | 0.444602826 | S100a10 | S100 calcium binding protein A10 (calpact | 2206.1 | 1489.5 | 1563.7 | 1188.6 | 1247.7 | 1824.5 |
| 12422 | ENSMUSG00000041961 | 229.7896744 | 0.074726477 | 0.178679831 | 0.41821439 | 0.675790375 | 0.932630311 | Znrf3 | zinc and ring finger 3 [Source:MGI Symbol | 205.59 | 268.29 | 198.23 | 244.85 | 231.21 | 230.57 |
| 12423 | ENSMUSG00000041966 | 290.0564772 | 0.291360302 | 0.152324791 | 1.91275694 | 0.055779176 | 0.333350319 | Dcaf17 | DDB1 and CUL4 associated factor 17 [Sour | 274.66 | 248.76 | 258.6 | 300.76 | 347.91 | 309.65 |
| 12424 | ENSMUSG00000041974 | 69.63846718 | 0.208137414 | 0.287278843 | 0.72451355 | 0.468750512 | 0.865790755 | Spidr | scaffolding protein involved in DNA repair | 53.808 | 61.675 | 78.487 | 60.731 | 80.706 | 82.424 |
| 12425 | ENSMUSG00000041975 | 206.6181426 | 0.192863405 | 0.202351438 | 0.953111119 | 0.340533773 | 0.783094143 | Mettl8 | methyltransferase like 8 [Source:MGI Sym | 187.12 | 177.83 | 213.32 | 185.08 | 279.2 | 197.15 |
| 12426 | ENSMUSG00000041977 | 479.6720753 | 0.132636631 | 0.151031677 | 0.878204051 | 0.379832994 | 0.812628154 | Arhgef11 | Rho guanine nucleotide exchange factor ( | 390.31 | 502.65 | 480.98 | 509.94 | 442.79 | 551.35 |
| 12427 | ENSMUSG00000041984 | 0 | NA | NA | NA | NA | NA | Rptn | repetin [Source:MGI Symbol;Acc:MGI:109 | 0 | 0 | 0 | 0 | 0 | 0 |
| 12428 | ENSMUSG00000041986 | 0 | NA | NA | NA | NA | NA | Elmod1 | ELMO/CED-12 domain containing 1 [Sourc | 0 | 0 | 0 | 0 | 0 | 0 |

| | | | | | | | | | | | | | | | |
|---|---|---|---|---|---|---|---|---|---|---|---|---|---|---|---|
| 12429 | ENSMUSG00000041991 | 0 | NA | NA | NA | NA | NA | Hrnr | hornerin [Source:MGI Symbol;Acc:MGI:30 | 0 | 0 | 0 | 0 | 0 | 0 |
| 12430 | ENSMUSG00000041992 | 109.4259717 | -1.183111644 | 0.25095097 | -4.714513128 | 2.42E-06 | 0.00017012 | Rapgef5 | Rap guanine nucleotide exchange factor ( | 163.03 | 172.69 | 119.74 | 56.875 | 79.615 | 64.603 |
| 12431 | ENSMUSG00000041995 | 63.76029518 | -0.343552426 | 0.280466351 | -1.224932777 | 0.220600549 | 0.667237604 | Zbed3 | zinc finger, BED type containing 3 [Source | 74.689 | 68.87 | 70.437 | 62.659 | 47.987 | 57.92 |
| 12432 | ENSMUSG00000041997 | 792.4255257 | 0.07912974 | 0.112252774 | 0.704924584 | 0.48085717 | 0.870278888 | Tlk1 | tousled-like kinase 1 [Source:MGI Symbol, | 721.99 | 800.75 | 789.9 | 787.57 | 823.42 | 830.93 |
| 12433 | ENSMUSG00000042002 | 0 | NA | NA | NA | NA | NA | Foxn4 | forkhead box N4 [Source:MGI Symbol;Acc | 0 | 0 | 0 | 0 | 0 | 0 |
| 12434 | ENSMUSG00000042010 | 1408.675156 | -0.343781274 | 0.444069864 | -0.774160332 | 0.438835983 | 0.848057363 | Acacb | acetyl-Coenzyme A carboxylase beta [Sou | 2378 | 1546 | 802.98 | 1505.7 | 1071 | 1148.4 |
| 12435 | ENSMUSG00000042015 | 296.6581985 | 0.000639709 | 0.170801723 | 0.003745331 | 0.997011665 | 0.999360826 | Wdr41 | WD repeat domain 41 [Source:MGI Symbo | 314.82 | 324.82 | 249.55 | 271.84 | 279.2 | 339.72 |
| 12436 | ENSMUSG00000042029 | 22.41482624 | -0.25853706 | 0.587064117 | -0.440389819 | 0.659654797 | 0.927787797 | Ncapg2 | non-SMC condensin II complex, subunit G | 35.336 | 21.586 | 16.1 | 19.28 | 6.5437 | 35.643 |
| 12437 | ENSMUSG00000042031 | 0 | NA | NA | NA | NA | NA | Lce3b | late cornified envelope 3B [Source:MGI Sy | 0 | 0 | 0 | 0 | 0 | 0 |
| 12438 | ENSMUSG00000042032 | 788.8954642 | 0.13623022 | 0.112019584 | 1.216128602 | 0.223935937 | 0.669770345 | Mat2b | methionine adenosyltransferase II, beta [! | 799.89 | 723.65 | 730.53 | 808.78 | 848.5 | 822.02 |
| 12439 | ENSMUSG00000042035 | 3.296927004 | -0.119189289 | 1.152374946 | -0.103429261 | 0.91762229 | NA | Igsf3 | immunoglobulin superfamily, member 3 [ | 3.2124 | 2.0558 | 5.0312 | 2.8919 | 4.3625 | 2.2277 |
| 12440 | ENSMUSG00000042041 | 546.5436038 | -0.705178618 | 0.457581648 | -1.541098996 | 0.123292694 | 0.514069049 | 2010003K11 | RIKEN cDNA 2010003K11 gene [Source:M | 1121.9 | 507.79 | 402.5 | 422.22 | 443.88 | 380.93 |
| 12441 | ENSMUSG00000042042 | 206.575914 | -0.034107907 | 0.192334368 | -0.177336517 | 0.859244069 | 0.975487285 | Csgalnact2 | chondroitin sulfate N-acetylgalactosaminy | 208.81 | 234.37 | 184.14 | 214.97 | 231.21 | 165.96 |
| 12442 | ENSMUSG00000042043 | 902.9421498 | 0.141769816 | 0.132084765 | 1.073324509 | 0.283125542 | 0.735802616 | Tbca | tubulin cofactor A [Source:MGI Symbol;A | 848.08 | 870.65 | 856.31 | 787.57 | 993.55 | 1061.5 |
| 12443 | ENSMUSG00000042045 | 0 | NA | NA | NA | NA | NA | Sln | sarcolipin [Source:MGI Symbol;Acc:MGI:1 | 0 | 0 | 0 | 0 | 0 | 0 |
| 12444 | ENSMUSG00000042046 | 225.5366257 | -0.037664988 | 0.180159902 | -0.209064211 | 0.834398113 | 0.971073158 | Dstyk | dual serine/threonine and tyrosine protei | 187.12 | 240.53 | 258.6 | 222.68 | 215.94 | 228.34 |
| 12445 | ENSMUSG00000042050 | 85.87118652 | 0.216203024 | 0.254876117 | 0.848267098 | 0.396289237 | 0.823764863 | Dync2i1 | dynein 2 intermediate chain 1 [Source:MG | 86.735 | 79.15 | 72.45 | 104.11 | 98.156 | 74.628 |
| 12446 | ENSMUSG00000042055 | 706.2074388 | -0.088148169 | 0.124984555 | -0.705272492 | 0.480640676 | 0.870278888 | Wdr11 | WD repeat domain 11 [Source:MGI Symbo | 667.38 | 710.29 | 806 | 642.97 | 709.99 | 700.61 |
| 12447 | ENSMUSG00000042063 | 207.3494703 | 0.086182266 | 0.183733727 | 0.469060678 | 0.639026265 | 0.923932929 | Zfp386 | zinc finger protein 386 (Kruppel-like) [Sou | 215.23 | 192.22 | 195.21 | 176.41 | 235.57 | 229.45 |
| 12448 | ENSMUSG00000042064 | 1.413442265 | -3.880278124 | 2.342991152 | -1.656121544 | 0.097697207 | NA | Myo3b | myosin IIIB [Source:MGI Symbol;Acc:MGI: | 6.4248 | 2.0558 | 0 | 0 | 0 | 0 |
| 12449 | ENSMUSG00000042066 | 49.30686583 | -0.703072528 | 0.343670976 | -2.045772197 | 0.040778789 | 0.276523394 | Tmcc2 | transmembrane and coiled-coil domains 2 | 57.02 | 75.038 | 51.318 | 36.631 | 47.987 | 27.846 |
| 12450 | ENSMUSG00000042073 | 2377.749667 | -0.035183927 | 0.096130397 | -0.366002096 | 0.714363511 | 0.944595913 | Abhd14b | abhydrolase domain containing 14b [Sour | 2255.9 | 2401.2 | 2563.9 | 2311.6 | 2405.9 | 2327.9 |
| 12451 | ENSMUSG00000042078 | 0 | NA | NA | NA | NA | NA | Svop | SV2 related protein [Source:MGI Symbol;/ | 0 | 0 | 0 | 0 | 0 | 0 |
| 12452 | ENSMUSG00000042079 | 1746.817465 | 0.035658184 | 0.101329939 | 0.351901759 | 0.72491194 | 0.9448775 | Hnrnpf | heterogeneous nuclear ribonucleoprotein | 1721 | 1625.1 | 1829.4 | 1735.2 | 1867.1 | 1703.1 |
| 12453 | ENSMUSG00000042082 | 292.3082353 | -0.184373571 | 0.152337623 | -1.210295708 | 0.226165444 | 0.672682908 | Arsb | arylsulfatase B [Source:MGI Symbol;Acc:N | 293.13 | 296.04 | 344.14 | 280.52 | 270.47 | 269.55 |
| 12454 | ENSMUSG00000042087 | 36.04417852 | -0.41054858 | 0.44493394 | -0.922718054 | 0.356154155 | 0.795708016 | 4933440N22 | RIKEN cDNA 4933440N22 gene [Source:M | 36.14 | 45.228 | 42.262 | 35.667 | 44.715 | 12.252 |
| 12455 | ENSMUSG00000042092 | 0 | NA | NA | NA | NA | NA | Lce1c | late cornified envelope 1C [Source:MGI Sy | 0 | 0 | 0 | 0 | 0 | 0 |
| 12456 | ENSMUSG00000042096 | 0 | NA | NA | NA | NA | NA | Dao | D-amino acid oxidase [Source:MGI Symbo | 0 | 0 | 0 | 0 | 0 | 0 |
| 12457 | ENSMUSG00000042097 | 1.463610243 | 0.484975761 | 1.887111255 | 0.25699373 | 0.797183613 | NA | Zfp239 | zinc finger protein 239 [Source:MGI Symb | 1.6062 | 2.0558 | 0 | 2.8919 | 0 | 2.2277 |
| 12458 | ENSMUSG00000042099 | 114.2360838 | -0.591940446 | 0.242854962 | -2.437423724 | 0.014792335 | 0.141505508 | Kank3 | KN motif and ankyrin repeat domains 3 [S | 124.48 | 164.47 | 123.77 | 111.82 | 81.796 | 79.083 |
| 12459 | ENSMUSG00000042102 | 10287.36554 | 0.076681627 | 0.116222655 | 0.659782098 | 0.509393673 | 0.882793133 | Dmgdh | dimethylglycine dehydrogenase precursor | 9626 | 10460 | 9955.8 | 9284.1 | 12345 | 10054 |
| 12460 | ENSMUSG00000042104 | 33.21281437 | 0.21118157 | 0.37999867 | 0.55574292 | 0.578386611 | 0.908137944 | Uggt2 | UDP-glucose glycoprotein glucosyltransfe | 35.336 | 24.67 | 32.2 | 37.595 | 32.719 | 36.757 |
| 12461 | ENSMUSG00000042105 | 466.8025166 | 0.096619661 | 0.137848827 | 0.700910284 | 0.483359006 | 0.871960882 | Inpp5f | inositol polyphosphate-5-phosphatase F [ | 404.76 | 501.62 | 447.78 | 457.89 | 486.42 | 502.34 |
| 12462 | ENSMUSG00000042106 | 4.991465607 | -1.386533323 | 1.0183322 | -1.361572699 | 0.173332776 | NA | Inka1 | inka box actin regulator 1 [Source:MGI Sy | 10.44 | 7.1954 | 4.025 | 3.8559 | 1.0906 | 3.3415 |
| 12463 | ENSMUSG00000042109 | 3.345930305 | 0.142252212 | 1.1907846 | 0.11946091 | 0.904910208 | NA | Csdc2 | cold shock domain containing C2, RNA bir | 2.4093 | 6.1675 | 1.0062 | 3.8559 | 2.1812 | 4.4554 |
| 12464 | ENSMUSG00000042111 | 242.2799924 | -0.029378946 | 0.165578414 | -0.17743222 | 0.859168901 | 0.975487285 | Ccdc115 | coiled-coil domain containing 115 [Source | 265.83 | 244.64 | 223.39 | 247.74 | 218.12 | 253.96 |
| 12465 | ENSMUSG00000042115 | 53.12667828 | -1.038905961 | 0.372480865 | -2.789152568 | 0.005284616 | 0.069144802 | Klhdc8a | kelch domain containing 8A [Source:MGI ! | 44.171 | 82.233 | 88.55 | 45.307 | 25.084 | 33.415 |
| 12466 | ENSMUSG00000042116 | 12.69670494 | 0.050221257 | 0.771237514 | 0.065117756 | 0.948080243 | 0.990682895 | Vwa1 | von Willebrand factor A domain containin | 16.062 | 2.0558 | 19.119 | 5.7839 | 11.997 | 21.163 |
| 12467 | ENSMUSG00000042118 | 4448.249586 | -0.192740665 | 0.109408076 | -1.761667621 | 0.078125472 | 0.409571156 | Bhmt2 | betaine-homocysteine methyltransferase | 4451.6 | 5034.7 | 4748.5 | 3721.9 | 4643.9 | 4088.9 |
| 12468 | ENSMUSG00000042121 | 242.8353549 | -0.128670097 | 0.161517396 | -0.796633058 | 0.425664172 | 0.841323666 | Ssh1 | slingshot protein phosphatase 1 [Source:N | 254.58 | 266.23 | 240.49 | 252.56 | 217.03 | 226.11 |
| 12469 | ENSMUSG00000042124 | 0 | NA | NA | NA | NA | NA | Lce1f | late cornified envelope 1F [Source:MGI Sy | 0 | 0 | 0 | 0 | 0 | 0 |
| 12470 | ENSMUSG00000042129 | 166.7729306 | -0.455501836 | 0.296628953 | -1.535594659 | 0.124637825 | 0.518034914 | Rassf4 | Ras association (RalGDS/AF-6) domain far | 216.03 | 234.37 | 127.79 | 113.75 | 97.065 | 211.63 |
| 12471 | ENSMUSG00000042133 | 574.454149 | 0.1647575 | 0.124459111 | 1.323788183 | 0.185573402 | 0.621183449 | Ppig | peptidyl-prolyl isomerase G (cyclophilin G | 529.24 | 569.47 | 526.27 | 565.85 | 652.19 | 603.7 |
| 12472 | ENSMUSG00000042138 | 103.593641 | 0.368659944 | 0.241005131 | 1.529676745 | 0.126096763 | 0.521094669 | Msantd2 | Myb/SANT-like DNA-binding domain cont | 80.31 | 109.99 | 81.506 | 129.17 | 102.52 | 118.07 |
| 12473 | ENSMUSG00000042148 | 268.127737 | 0.031175866 | 0.205542483 | 0.151676023 | 0.879442469 | 0.98128671 | Cox10 | heme A:farnesyltransferase cytochrome c | 280.28 | 328.93 | 186.16 | 246.78 | 277.02 | 289.6 |
| 12474 | ENSMUSG00000042155 | 7.734784691 | -0.922652042 | 0.950669514 | -0.970528694 | 0.331783029 | 0.774747658 | Klhl23 | kelch-like 23 [Source:MGI Symbol;Acc:MG | 11.243 | 8.2233 | 11.069 | 12.532 | 0 | 3.3415 |
| 12475 | ENSMUSG00000042156 | 3.988421703 | -0.110469136 | 1.089157803 | -0.101426199 | 0.91921214 | NA | Dzip1 | DAZ interacting protein 1 [Source:MGI Syr | 7.2279 | 3.0838 | 2.0125 | 3.8559 | 2.1812 | 5.5692 |
| 12476 | ENSMUSG00000042157 | 0 | NA | NA | NA | NA | NA | Sprr2i | small proline-rich protein 2I [Source:MGI ! | 0 | 0 | 0 | 0 | 0 | 0 |
| 12477 | ENSMUSG00000042165 | 9.665943108 | 0.04733102 | 0.686566313 | 0.068938745 | 0.945038378 | 0.990309554 | Gm9774 | predicted pseudogene 9774 [Source:MGI | 7.2279 | 12.335 | 9.0562 | 10.604 | 7.6343 | 11.138 |
| 12478 | ENSMUSG00000042167 | 491.8017094 | 0.192773293 | 0.128980233 | 1.494595629 | 0.135020007 | 0.537616739 | Tent2 | terminal nucleotidyltransferase 2 [Source: | 480.26 | 427.61 | 468.91 | 549.47 | 531.13 | 493.43 |
| 12479 | ENSMUSG00000042178 | 176.2058758 | -0.113887311 | 0.192021174 | -0.593097672 | 0.553115788 | 0.896245536 | Armc5 | armadillo repeat containing 5 [Source:MG | 175.08 | 186.05 | 188.17 | 144.6 | 198.49 | 164.85 |
| 12480 | ENSMUSG00000042179 | 0 | NA | NA | NA | NA | NA | Pnliprp1 | pancreatic lipase related protein 1 [Source | 0 | 0 | 0 | 0 | 0 | 0 |
| 12481 | ENSMUSG00000042182 | 1.604014255 | -2.909372143 | 1.929202543 | -1.508069826 | 0.131536657 | NA | Bend6 | BEN domain containing 6 [Source:MGI Syr | 2.4093 | 4.1117 | 2.0125 | 0 | 1.0906 | 0 |
| 12482 | ENSMUSG00000042184 | 0.181769715 | 1.126924175 | 4.080472857 | 0.276174898 | 0.782413736 | NA | 1700069L16 | RIKEN cDNA 1700069L16 gene [Source:M | 0 | 0 | 0 | 0 | 1.0906 | 0 |
| 12483 | ENSMUSG00000042185 | 224.2123213 | 0.253959113 | 0.188016157 | 1.350730262 | 0.176781855 | 0.607589534 | Nfrkb | nuclear factor related to kappa B binding | 179.89 | 231.28 | 203.26 | 263.17 | 268.29 | 199.38 |
| 12484 | ENSMUSG00000042189 | 0 | NA | NA | NA | NA | NA | Tekt3 | tektin 3 [Source:MGI Symbol;Acc:MGI:191 | 0 | 0 | 0 | 0 | 0 | 0 |

| | | | | | | | | | | | | | | | |
|---|---|---|---|---|---|---|---|---|---|---|---|---|---|---|---|
| 12485 | ENSMUSG00000042190 | 23.73847213 | -1.677578377 | 0.517022081 | -3.244693869 | 0.00117577 | 0.023521895 | Cmklr1 | chemokine-like receptor 1 [Source:MGI Sy | 40.958 | 46.256 | 21.131 | 8.6758 | 9.8156 | 15.594 |
| 12486 | ENSMUSG00000042195 | 26.83704825 | -0.228966824 | 0.663327277 | -0.345179269 | 0.729959603 | 0.946534338 | Slc35f2 | solute carrier family 35, member F2 [Sour | 46.58 | 35.977 | 4.025 | 10.604 | 35.99 | 27.846 |
| 12487 | ENSMUSG00000042197 | 239.4595579 | -0.091513983 | 0.164047845 | -0.557849347 | 0.576947263 | 0.907752116 | Zfp451 | zinc finger protein 451 [Source:MGI Symb | 235.31 | 256.98 | 249.55 | 256.42 | 225.76 | 212.74 |
| 12488 | ENSMUSG00000042198 | 341.4572475 | 0.069880054 | 0.153617377 | 0.454896802 | 0.649183484 | 0.925643431 | Chchd7 | coiled-coil-helix-coiled-coil-helix domain c | 304.38 | 318.65 | 377.34 | 376.92 | 322.82 | 348.63 |
| 12489 | ENSMUSG00000042200 | 0.17131964 | -0.796629176 | 4.080472857 | -0.195229623 | 0.84521319 | NA | Cdrt4 | CMT1A duplicated region transcript 4 [Sou | 0 | 1.0279 | 0 | 0 | 0 | 0 |
| 12490 | ENSMUSG00000042202 | 1045.404667 | -0.130579219 | 0.134809042 | -0.968623592 | 0.332733028 | 0.77523112 | Slc35e2 | solute carrier family 35, member E2 [Sour | 1096.2 | 1060.8 | 1120 | 811.67 | 1121.2 | 1062.6 |
| 12491 | ENSMUSG00000042203 | 82.91712559 | 0.588788728 | 0.265692329 | 2.216054671 | 0.026687758 | 0.211364326 | Tbc1d22b | TBC1 domain family, member 22B [Source | 72.279 | 59.619 | 66.412 | 91.578 | 83.978 | 123.64 |
| 12492 | ENSMUSG00000042207 | 404.0723525 | 0.524690214 | 0.141425556 | 3.710009905 | 0.000207251 | 0.006446625 | Kdm5b | lysine (K)-specific demethylase 5B [Source | 343.73 | 308.38 | 342.12 | 526.33 | 444.97 | 458.9 |
| 12493 | ENSMUSG00000042208 | 342.9698002 | 0.118979743 | 0.154728791 | 0.768956717 | 0.441919006 | 0.850050074 | 0610010F05 | RIKEN cDNA 0610010F05 gene [Source:M | 301.97 | 319.68 | 365.27 | 343.18 | 400.26 | 327.47 |
| 12494 | ENSMUSG00000042210 | 181.6300019 | 0.24663344 | 0.183337038 | 1.345246128 | 0.178545764 | 0.610813333 | Abhd14a | abhydrolase domain containing 14A [Sour | 167.05 | 148.02 | 183.14 | 187.98 | 190.86 | 212.74 |
| 12495 | ENSMUSG00000042211 | 491.2769566 | 0.127856595 | 0.148559154 | 0.86064434 | 0.389433957 | 0.819052248 | Fbxo38 | F-box protein 38 [Source:MGI Symbol;Acc | 400.75 | 491.34 | 517.21 | 484.88 | 480.96 | 572.52 |
| 12496 | ENSMUSG00000042212 | 0 | NA | NA | NA | NA | NA | Sprr2d | small proline-rich protein 2D [Source:MGI | 0 | 0 | 0 | 0 | 0 | 0 |
| 12497 | ENSMUSG00000042213 | 321.8247437 | 0.024443461 | 0.176322346 | 0.138629397 | 0.889743012 | 0.982384956 | Zfand4 | zinc finger, AN1-type domain 4 [Source:M | 292.33 | 381.36 | 283.76 | 295.94 | 297.74 | 379.82 |
| 12498 | ENSMUSG00000042215 | 35.39372377 | -0.188448894 | 0.384299208 | -0.490370238 | 0.623871933 | 0.919503423 | Bag2 | BCL2-associated athanogene 2 [Source:M | 36.943 | 29.81 | 46.287 | 28.919 | 41.443 | 28.96 |
| 12499 | ENSMUSG00000042216 | 31.34311524 | 0.748688906 | 0.460767595 | 1.624873178 | 0.104189584 | 0.475225645 | Sgsm1 | small G protein signaling modulator 1 [Sou | 22.487 | 18.503 | 29.181 | 41.451 | 19.631 | 56.806 |
| 12500 | ENSMUSG00000042219 | 0 | NA | NA | NA | NA | NA | Olfr631 | olfactory receptor 631 [Source:MGI Symb | 0 | 0 | 0 | 0 | 0 | 0 |
| 12501 | ENSMUSG00000042225 | 31.83777781 | 0.398943277 | 0.393841812 | 1.012953082 | 0.311082572 | 0.759594595 | Ammecr1 | Alport syndrome, mental retardation, mid | 25.699 | 23.642 | 33.206 | 42.415 | 35.99 | 30.074 |
| 12502 | ENSMUSG00000042228 | 456.5436809 | -0.092257238 | 0.138215753 | -0.667487141 | 0.50446102 | 0.881510796 | Lyn | LYN proto-oncogene, Src family tyrosine k | 433.68 | 522.18 | 457.84 | 409.69 | 456.97 | 458.9 |
| 12503 | ENSMUSG00000042229 | 274.3465334 | 0.160694128 | 0.150372851 | 1.068637901 | 0.285232864 | 0.737769696 | Rabif | RAB interacting factor [Source:MGI Symbo | 271.45 | 259.04 | 246.53 | 294.98 | 292.29 | 281.8 |
| 12504 | ENSMUSG00000042240 | 0 | NA | NA | NA | NA | NA | Crybb2 | crystallin, beta B2 [Source:MGI Symbol;Ac | 0 | 0 | 0 | 0 | 0 | 0 |
| 12505 | ENSMUSG00000042243 | 0 | NA | NA | NA | NA | NA | BC051665 | cDNA sequence BC051665 [Source:MGI Sy | 0 | 0 | 0 | 0 | 0 | 0 |
| 12506 | ENSMUSG00000042244 | 0 | NA | NA | NA | NA | NA | Pglyrp3 | peptidoglycan recognition protein 3 [Sour | 0 | 0 | 0 | 0 | 0 | 0 |
| 12507 | ENSMUSG00000042246 | 133.6561279 | -1.218801638 | 0.266063657 | -4.580864788 | 4.63E-06 | 0.000286225 | Tmc7 | transmembrane channel-like gene family | 167.05 | 260.06 | 133.83 | 72.298 | 80.706 | 87.994 |
| 12508 | ENSMUSG00000042248 | 6634.133448 | -0.742317142 | 0.453554244 | -1.636666733 | 0.101700124 | 0.469365216 | Cyp2c37 | cytochrome P450, family 2. subfamily c, p | 5267.5 | 8426.9 | 11219 | 7288.6 | 3503.1 | 4100.1 |
| 12509 | ENSMUSG00000042249 | 71.73900489 | -0.168263589 | 0.378591546 | -0.44444624 | 0.656719988 | 0.9268505 | Grk3 | G protein-coupled receptor kinase 3 [Sour | 95.569 | 98.68 | 33.206 | 62.659 | 53.44 | 86.88 |
| 12510 | ENSMUSG00000042250 | 0 | NA | NA | NA | NA | NA | Pglyrp4 | peptidoglycan recognition protein 4 [Sour | 0 | 0 | 0 | 0 | 0 | 0 |
| 12511 | ENSMUSG00000042251 | 1976.360701 | 0.102427932 | 0.097628347 | 1.049161796 | 0.294103659 | 0.745164683 | Pm20d1 | peptidase M20 domain containing 1 [Sour | 1872.8 | 1906.8 | 1939 | 1962.7 | 2199.8 | 1977.1 |
| 12512 | ENSMUSG00000042254 | 3.013903613 | -2.984672678 | 1.579013989 | -1.890212942 | 0.058729486 | NA | Cilp | cartilage intermediate layer protein, nucle | 4.0155 | 3.0838 | 9.0562 | 1.928 | 0 | 0 |
| 12513 | ENSMUSG00000042256 | 0 | NA | NA | NA | NA | NA | Ptchd4 | patched domain containing 4 [Source:MG | 0 | 0 | 0 | 0 | 0 | 0 |
| 12514 | ENSMUSG00000042258 | 0.133850373 | -0.796629176 | 4.080472857 | -0.195229623 | 0.84521319 | NA | Isl1 | ISL1 transcription factor, LIM/homeodom | 0.8031 | 0 | 0 | 0 | 0 | 0 |
| 12515 | ENSMUSG00000042262 | 0 | NA | NA | NA | NA | NA | Ccr8 | chemokine (C-C motif) receptor 8 [Source | 0 | 0 | 0 | 0 | 0 | 0 |
| 12516 | ENSMUSG00000042265 | 0.17131964 | -0.796629176 | 4.080472857 | -0.195229623 | 0.84521319 | NA | Trem1 | triggering receptor expressed on myeloid | 0 | 1.0279 | 0 | 0 | 0 | 0 |
| 12517 | ENSMUSG00000042268 | 0 | NA | NA | NA | NA | NA | Slc26a9 | solute carrier family 26, member 9 [Sourc | 0 | 0 | 0 | 0 | 0 | 0 |
| 12518 | ENSMUSG00000042269 | 0 | NA | NA | NA | NA | NA | Cibar2 | CBY1 interacting BAR domain containing 2 | 0 | 0 | 0 | 0 | 0 | 0 |
| 12519 | ENSMUSG00000042271 | 636.1473016 | -0.102345048 | 0.117804236 | -0.868772226 | 0.384971727 | 0.816437825 | Nxt2 | nuclear transport factor 2-like export fact | 641.68 | 681.51 | 653.05 | 594.77 | 652.19 | 593.68 |
| 12520 | ENSMUSG00000042272 | 34.537472 | 0.054931804 | 0.394546948 | 0.139227548 | 0.88927034 | 0.982382091 | Sestd1 | SEC14 and spectrin domains 1 [Source:MC | 25.699 | 31.865 | 44.275 | 34.703 | 28.356 | 42.326 |
| 12521 | ENSMUSG00000042275 | 68.42080246 | 0.191036803 | 0.292718611 | 0.652629507 | 0.513995158 | 0.885004045 | Pelo | pelota mRNA surveillance and ribosome r | 79.507 | 58.591 | 53.331 | 86.758 | 62.165 | 70.172 |
| 12522 | ENSMUSG00000042279 | 0 | NA | NA | NA | NA | NA | H1f8 | H1.8 linker histone [Source:MGI Symbol;A | 0 | 0 | 0 | 0 | 0 | 0 |
| 12523 | ENSMUSG00000042282 | 0 | NA | NA | NA | NA | NA | Gucy2f | guanylate cyclase 2f [Source:MGI Symbol; | 0 | 0 | 0 | 0 | 0 | 0 |
| 12524 | ENSMUSG00000042284 | 473.0126814 | -0.246462754 | 0.131836563 | -1.869456765 | 0.061559293 | 0.354584001 | Itga1 | integrin alpha 1 [Source:MGI Symbol;Acc: | 485.07 | 542.74 | 512.18 | 403.91 | 429.7 | 464.47 |
| 12525 | ENSMUSG00000042286 | 556.4023565 | -0.468497573 | 0.181179647 | -2.585817895 | 0.009714824 | 0.106289877 | Stab1 | stabilin 1 [Source:MGI Symbol;Acc:MGI:2 | 642.48 | 658.9 | 636.95 | 543.68 | 320.64 | 535.76 |
| 12526 | ENSMUSG00000042289 | 21.8639681 | 0.065987903 | 0.513532509 | 0.128498005 | 0.897754877 | 0.982798755 | Hsd3b7 | hydroxy-delta-5-steroid dehydrogenase, 3 | 21.684 | 13.363 | 29.181 | 32.775 | 16.359 | 17.822 |
| 12527 | ENSMUSG00000042292 | 109.0850648 | -0.343708726 | 0.262799792 | -1.307872902 | 0.190916422 | 0.628409928 | Mrtfa | myocardin related transcription factor A [ | 125.28 | 131.57 | 109.68 | 126.28 | 95.974 | 65.717 |
| 12528 | ENSMUSG00000042293 | 395.1882951 | -0.069092448 | 0.134830938 | -0.51243764 | 0.608344754 | 0.916376001 | Gm5617 | predicted gene 5617 [Source:MGI Symbol | 427.25 | 395.75 | 390.42 | 383.66 | 369.72 | 404.33 |
| 12529 | ENSMUSG00000042298 | 384.3624029 | 0.026854716 | 0.144760389 | 0.185511494 | 0.852827866 | 0.973781977 | Ttc19 | tetratricopeptide repeat domain 19 [Sour | 409.58 | 382.39 | 350.17 | 409.69 | 405.71 | 348.63 |
| 12530 | ENSMUSG00000042302 | 161.7502631 | 0.106400711 | 0.190328214 | 0.559038036 | 0.576135762 | 0.907462448 | Ehbp1 | EH domain binding protein 1 [Source:MGI | 153.39 | 151.1 | 163.01 | 176.41 | 146.14 | 180.44 |
| 12531 | ENSMUSG00000042303 | 157.6541894 | -0.020266858 | 0.204289502 | -0.099206556 | 0.92097427 | 0.986685804 | Sgsm3 | small G protein signaling modulator 3 [Sou | 164.64 | 148.02 | 163.01 | 125.32 | 173.41 | 171.53 |
| 12532 | ENSMUSG00000042305 | 2495.003335 | 0.190806951 | 0.095563069 | 1.996659925 | 0.045862138 | 0.296229283 | Tmem183a | transmembrane protein 183A [Source:MG | 2278.4 | 2209 | 2503.5 | 2626.8 | 2709.1 | 2643.2 |
| 12533 | ENSMUSG00000042306 | 0.428363721 | -0.543730382 | 3.624677344 | -0.15000794 | 0.880758351 | NA | S100a14 | S100 calcium binding protein A14 [Source | 1.6062 | 0 | 0 | 0.964 | 0 | 0 |
| 12534 | ENSMUSG00000042308 | 173.5123136 | 0.157741673 | 0.194289707 | 0.811888984 | 0.416855336 | 0.836878476 | Setd1a | SET domain containing 1A [Source:MGI Sy | 175.88 | 162.41 | 153.96 | 215.93 | 163.59 | 169.3 |
| 12535 | ENSMUSG00000042312 | 377.8811944 | -0.094661924 | 0.172627387 | -0.548359828 | 0.583444855 | 0.908645447 | S100a13 | S100 calcium binding protein A13 [Source | 301.97 | 410.14 | 459.85 | 367.28 | 384.99 | 343.06 |
| 12536 | ENSMUSG00000042320 | 54.78089324 | -0.133071936 | 0.310487198 | -0.428590732 | 0.668221091 | 0.930146267 | Prox2 | prospero homeobox 2 [Source:MGI Symb | 60.233 | 50.368 | 61.381 | 58.803 | 57.803 | 40.098 |
| 12537 | ENSMUSG00000042323 | 717.2811144 | -0.109383394 | 0.123581704 | -0.885109935 | 0.376097341 | 0.809731241 | Pbrm1 | polybromo 1 [Source:MGI Symbol;Acc:MC | 720.38 | 799.72 | 713.43 | 658.4 | 760.16 | 651.6 |
| 12538 | ENSMUSG00000042328 | 270.3704459 | 0.258595264 | 0.160987437 | 1.606307102 | 0.108206451 | 0.4851397 | Hps4 | HPS4, biogenesis of lysosomal organelles | 242.54 | 226.14 | 269.67 | 271.84 | 289.01 | 323.01 |
| 12539 | ENSMUSG00000042331 | 29.9510955 | -0.276878886 | 0.524089595 | -0.528304489 | 0.597288017 | 0.914521894 | Specc1 | sperm antigen with calponin homology ar | 26.502 | 39.061 | 33.206 | 44.343 | 7.6343 | 28.96 |
| 12540 | ENSMUSG00000042333 | 70.12857908 | -0.029350741 | 0.29278493 | -0.100246761 | 0.920148423 | 0.986685804 | Tnfrsf14 | tumor necrosis factor receptor superfamil | 62.642 | 61.675 | 88.55 | 80.974 | 54.531 | 72.4 |

| | | | | | | | | | | | | | | | |
|---|---|---|---|---|---|---|---|---|---|---|---|---|---|---|---|
| 12541 | ENSMUSG00000042340 | 149.3553122 | -0.328124172 | 0.214600259 | -1.529001754 | 0.126264009 | 0.521094669 | Ctf1 | cardiotrophin 1 [Source:MGI Symbol;Acc: | 159.01 | 157.27 | 183.14 | 160.02 | 130.87 | 105.82 |
| 12542 | ENSMUSG00000042345 | 0.774435248 | -2.999089927 | 2.565713408 | -1.168910728 | 0.242439601 | NA | Ubash3a | ubiquitin associated and SH3 domain cont | 1.6062 | 1.0279 | 2.0125 | 0 | 0 | 0 |
| 12543 | ENSMUSG00000042348 | 197.9374097 | -0.084272507 | 0.184869262 | -0.45584921 | 0.648498415 | 0.925643431 | Arl15 | ADP-ribosylation factor-like 15 [Source:M | 175.88 | 213.81 | 222.38 | 214 | 177.77 | 183.78 |
| 12544 | ENSMUSG00000042349 | 286.0525047 | -0.156482502 | 0.193488259 | -0.808744171 | 0.418662315 | 0.837591673 | Ikbke | inhibitor of kappaB kinase epsilon [Source | 357.38 | 272.4 | 274.7 | 329.68 | 215.94 | 266.21 |
| 12545 | ENSMUSG00000042350 | 955.1321766 | 0.021368131 | 0.112394618 | 0.190117032 | 0.849217424 | 0.973071229 | Arel1 | apoptosis resistant E3 ubiquitin protein lig | 921.96 | 906.62 | 1016.3 | 1014.1 | 968.47 | 903.33 |
| 12546 | ENSMUSG00000042351 | 5.854846208 | -0.559349614 | 1.003397396 | -0.557455716 | 0.577216108 | 0.907838603 | Grap2 | GRB2-related adaptor protein 2 [Source:N | 5.6217 | 6.1675 | 9.0562 | 0.964 | 2.1812 | 11.138 |
| 12547 | ENSMUSG00000042353 | 0.867306998 | -1.831940351 | 2.582928097 | -0.709249457 | 0.478169687 | NA | Frem3 | Fras1 related extracellular matrix protein 3 | 0 | 3.0838 | 1.0062 | 0 | 0 | 1.1138 |
| 12548 | ENSMUSG00000042354 | 436.2690763 | 0.07466197 | 0.165394804 | 0.451416659 | 0.651689279 | 0.925643431 | Gnl3 | guanine nucleotide binding protein-like 3 | 515.59 | 412.2 | 346.15 | 471.39 | 411.16 | 461.13 |
| 12549 | ENSMUSG00000042357 | 0.315620088 | 0.165143421 | 4.063300985 | 0.040642675 | 0.967580763 | NA | Gjb5 | gap junction protein, beta 5 [Source:MGI | 0.8031 | 0 | 0 | 0 | 1.0906 | 0 |
| 12550 | ENSMUSG00000042359 | 1.655804529 | -1.834171817 | 1.778544603 | -1.031276817 | 0.302411027 | NA | Osbpl6 | oxysterol binding protein-like 6 [Source:M | 1.6062 | 4.1117 | 2.0125 | 0 | 1.0906 | 1.1138 |
| 12551 | ENSMUSG00000042360 | 0.328370405 | 0.16514316 | 4.003149536 | 0.041253308 | 0.967093956 | NA | 4930433N12 | RIKEN cDNA 4930433N12 gene [Source:M | 0 | 0 | 1.0062 | 0.964 | 0 | 0 |
| 12552 | ENSMUSG00000042363 | 384.1953103 | -0.868669694 | 0.165631282 | -5.244599234 | 1.57E-07 | 1.64E-05 | Lgalsl | lectin, galactoside binding-like [Source:M | 425.64 | 499.57 | 565.51 | 315.22 | 250.84 | 248.39 |
| 12553 | ENSMUSG00000042364 | 531.9144809 | -0.310960902 | 0.152728702 | -2.03603447 | 0.041746894 | 0.280818721 | Snx18 | sorting nexin 18 [Source:MGI Symbol;Acc: | 556.55 | 652.73 | 557.46 | 396.19 | 552.94 | 475.61 |
| 12554 | ENSMUSG00000042367 | 0 | NA | NA | NA | NA | NA | Gjb3 | gap junction protein, beta 3 [Source:MGI | 0 | 0 | 0 | 0 | 0 | 0 |
| 12555 | ENSMUSG00000042369 | 217.3087003 | -0.063843716 | 0.172205768 | -0.370740869 | 0.710830547 | 0.944239215 | Rbm45 | RNA binding motif protein 45 [Source:MG | 205.59 | 209.7 | 251.56 | 222.68 | 210.49 | 203.83 |
| 12556 | ENSMUSG00000042371 | 0.17131964 | -0.796629176 | 4.080472857 | -0.195229623 | 0.84521319 | NA | Slc5a10 | solute carrier family 5 (sodium/glucose co | 0 | 1.0279 | 0 | 0 | 0 | 0 |
| 12557 | ENSMUSG00000042372 | 0 | NA | NA | NA | NA | NA | Dmrt3 | doublesex and mab-3 related transcriptio | 0 | 0 | 0 | 0 | 0 | 0 |
| 12558 | ENSMUSG00000042377 | 40.84316343 | 0.408527102 | 0.384655419 | 1.062059916 | 0.288208486 | 0.73951397 | Fam83g | family with sequence similarity 83, memb | 40.155 | 31.865 | 33.206 | 56.875 | 27.265 | 55.692 |
| 12559 | ENSMUSG00000042379 | 18.44141723 | -0.794977285 | 0.576143906 | -1.379824168 | 0.167640789 | 0.593804353 | Esm1 | endothelial cell-specific molecule 1 [Sourc | 24.093 | 33.921 | 12.075 | 8.6758 | 19.631 | 12.252 |
| 12560 | ENSMUSG00000042380 | 295.0278986 | -0.001811917 | 0.154675608 | -0.011714302 | 0.990653553 | 0.99859176 | Smim12 | small integral membrane protein 12 [Sour | 297.15 | 274.45 | 313.95 | 298.83 | 266.11 | 319.67 |
| 12561 | ENSMUSG00000042385 | 0.587191767 | -1.12394062 | 3.226482889 | -0.348348545 | 0.72757844 | NA | Gzmk | granzyme K [Source:MGI Symbol;Acc:MGI | 2.4093 | 0 | 0 | 0 | 0 | 1.1138 |
| 12562 | ENSMUSG00000042386 | 0 | NA | NA | NA | NA | NA | Tex13b | testis expressed 13B [Source:MGI Symbol | 0 | 0 | 0 | 0 | 0 | 0 |
| 12563 | ENSMUSG00000042388 | 0.36353943 | 2.065989254 | 4.034137301 | 0.512126658 | 0.608562368 | NA | Dlgap3 | DLG associated protein 3 [Source:MGI Syr | 0 | 0 | 0 | 0 | 2.1812 | 0 |
| 12564 | ENSMUSG00000042389 | 129.0213178 | 0.246039938 | 0.215897052 | 1.139616944 | 0.254445923 | 0.707887253 | Tsen2 | tRNA splicing endonuclease subunit 2 [So | 138.13 | 104.85 | 110.69 | 148.45 | 146.14 | 125.86 |
| 12565 | ENSMUSG00000042390 | 369.7152069 | -0.08042936 | 0.148787583 | -0.540565002 | 0.588807446 | 0.911039598 | Gatad2b | GATA zinc finger domain containing 2B [S | 373.44 | 432.75 | 334.07 | 360.53 | 357.72 | 359.77 |
| 12566 | ENSMUSG00000042396 | 791.129531 | 0.036444393 | 0.113577705 | 0.320876292 | 0.748304141 | 0.951074124 | Rbm7 | RNA binding motif protein 7 [Source:MGI | 757.33 | 822.33 | 763.74 | 759.61 | 797.24 | 846.52 |
| 12567 | ENSMUSG00000042401 | 0 | NA | NA | NA | NA | NA | Crtac1 | cartilage acidic protein 1 [Source:MGI Sym | 0 | 0 | 0 | 0 | 0 | 0 |
| 12568 | ENSMUSG00000042404 | 150.9666159 | 0.017718694 | 0.20443807 | 0.086670225 | 0.930933644 | 0.988420305 | Dennd4b | DENN/MADD domain containing 4B [Sour | 124.48 | 156.24 | 170.06 | 156.16 | 140.69 | 158.17 |
| 12569 | ENSMUSG00000042406 | 1201.387687 | 0.096424518 | 0.112218067 | 0.859260192 | 0.390196981 | 0.819725365 | Atf4 | activating transcription factor 4 [Source:N | 1085 | 1256.1 | 1144.1 | 1333.2 | 1220.4 | 1169.5 |
| 12570 | ENSMUSG00000042408 | 144.8014721 | 0.225418827 | 0.211792059 | 1.064340318 | 0.28717456 | 0.73951397 | Zmym6 | zinc finger, MYM-type 6 [Source:MGI Sym | 123.68 | 123.35 | 153.96 | 175.44 | 130.87 | 161.51 |
| 12571 | ENSMUSG00000042410 | 251.5484531 | -0.18512965 | 0.165300585 | -1.119957623 | 0.262731821 | 0.717145321 | Agps | alkylglycerone phosphate synthase [Sourc | 265.83 | 256.98 | 279.74 | 204.36 | 259.57 | 242.82 |
| 12572 | ENSMUSG00000042414 | 0.267700746 | -1.497481649 | 4.057001436 | -0.369110456 | 0.712045396 | NA | Prdm14 | PR domain containing 14 [Source:MGI Syr | 1.6062 | 0 | 0 | 0 | 0 | 0 |
| 12573 | ENSMUSG00000042417 | 14.13858048 | 0.34205883 | 0.679482409 | 0.503410869 | 0.614675432 | 0.917509899 | Ccno | cyclin O [Source:MGI Symbol;Acc:MGI:214 | 3.2124 | 12.335 | 22.137 | 17.352 | 13.087 | 16.708 |
| 12574 | ENSMUSG00000042419 | 98.08990751 | 0.109393897 | 0.231046358 | 0.473471636 | 0.635876732 | 0.923478764 | Nfkbil1 | nuclear factor of kappa light polypeptide g | 85.932 | 106.9 | 90.562 | 101.22 | 100.34 | 103.59 |
| 12575 | ENSMUSG00000042423 | 458.7671121 | 0.060439256 | 0.142923062 | 0.422879665 | 0.67238306 | 0.931191575 | Fbrs | fibrosin [Source:MGI Symbol;Acc:MGI:104 | 426.45 | 490.32 | 431.68 | 525.37 | 419.89 | 458.9 |
| 12576 | ENSMUSG00000042425 | 0 | NA | NA | NA | NA | NA | Frmpd3 | FERM and PDZ domain containing 3 [Sour | 0 | 0 | 0 | 0 | 0 | 0 |
| 12577 | ENSMUSG00000042426 | 202.9806416 | -0.051979792 | 0.175299272 | -0.296520295 | 0.76683277 | 0.955295274 | Dhx29 | DEAH (Asp-Glu-Ala-His) box polypeptide 2 | 217.64 | 204.56 | 197.22 | 182.19 | 223.58 | 192.69 |
| 12578 | ENSMUSG00000042428 | 8.013369222 | -0.410676723 | 0.779778262 | -0.526658337 | 0.598430873 | 0.915055389 | Mgat3 | mannoside acetylglucosaminyltransferase | 7.2279 | 11.307 | 9.0562 | 10.604 | 6.5437 | 3.3415 |
| 12579 | ENSMUSG00000042429 | 577.5873272 | 0.044710223 | 0.180889753 | 0.247168357 | 0.804777933 | 0.963787787 | Adora1 | adenosine A1 receptor [Source:MGI Symb | 676.21 | 562.27 | 465.89 | 451.14 | 621.65 | 688.36 |
| 12580 | ENSMUSG00000042433 | 0 | NA | NA | NA | NA | NA | Dnaaf6b | dynein axonemal assembly factor 6B [Sou | 0 | 0 | 0 | 0 | 0 | 0 |
| 12581 | ENSMUSG00000042436 | 16.51686216 | -1.371291283 | 0.685768981 | -1.999640287 | 0.04553912 | 0.295238834 | Mfap4 | microfibrillar-associated protein 4 [Source | 44.974 | 15.419 | 11.069 | 14.46 | 8.7249 | 4.4554 |
| 12582 | ENSMUSG00000042439 | 44.30650738 | -0.086433093 | 0.405176137 | -0.213322268 | 0.831075606 | 0.970505212 | Zfp532 | zinc finger protein 532 [Source:MGI Symb | 53.005 | 43.173 | 40.25 | 21.208 | 44.715 | 63.489 |
| 12583 | ENSMUSG00000042444 | 258.4721159 | 0.005328059 | 0.17069088 | 0.031214666 | 0.975098344 | 0.996378391 | Mindy2 | MINDY lysine 48 deubiquitinase 2 [Source | 245.75 | 269.31 | 258.6 | 215.93 | 268.29 | 292.94 |
| 12584 | ENSMUSG00000042446 | 160.6760115 | 0.071124307 | 0.212814569 | 0.334207886 | 0.738222694 | 0.948137969 | Zmym4 | zinc finger, MYM-type 4 [Source:MGI Sym | 138.94 | 174.75 | 156.97 | 166.77 | 196.31 | 130.32 |
| 12585 | ENSMUSG00000042447 | 237.9599124 | -0.00024909 | 0.17279964 | -0.001441494 | 0.998849855 | 0.999764475 | Mios | meiosis regulator for oocyte development | 223.26 | 274.45 | 216.34 | 217.86 | 253.02 | 242.82 |
| 12586 | ENSMUSG00000042448 | 0 | NA | NA | NA | NA | NA | Hoxd1 | homeobox D1 [Source:MGI Symbol;Acc:M | 0 | 0 | 0 | 0 | 0 | 0 |
| 12587 | ENSMUSG00000042451 | 0.545309144 | 2.651974629 | 3.666219369 | 0.72335405 | 0.469462394 | NA | Mybph | myosin binding protein H [Source:MGI Syr | 0 | 0 | 0 | 0 | 3.2719 | 0 |
| 12588 | ENSMUSG00000042453 | 614.3765643 | -0.891657384 | 0.135873543 | -6.562406227 | 5.29E-11 | 1.53E-08 | Reln | reelin [Source:MGI Symbol;Acc:MGI:1030 | 893.85 | 712.35 | 787.89 | 407.76 | 417.71 | 466.7 |
| 12589 | ENSMUSG00000042459 | 0 | NA | NA | NA | NA | NA | Bpifa2 | BPI fold containing family A, member 2 [S | 0 | 0 | 0 | 0 | 0 | 0 |
| 12590 | ENSMUSG00000042460 | 1001.577168 | -0.065425637 | 0.11377989 | -0.575019335 | 0.56527822 | 0.901542552 | C1galt1 | core 1 synthase, glycoprotein-N-acetylgal | 940.43 | 1080.3 | 1053.5 | 1045 | 941.2 | 948.99 |
| 12591 | ENSMUSG00000042462 | 487.1261455 | 0.199118662 | 0.124423908 | 1.600324768 | 0.109526555 | 0.486396098 | Dctpp1 | dCTP pyrophosphatase 1 [Source:MGI Syr | 433.68 | 470.79 | 456.84 | 532.12 | 515.86 | 513.48 |
| 12592 | ENSMUSG00000042472 | 344.4905577 | -0.114460466 | 0.141196873 | -0.810644485 | 0.417569862 | 0.837107483 | Zfp410 | zinc finger protein 410 [Source:MGI Symb | 374.25 | 367.99 | 332.06 | 340.28 | 330.46 | 321.9 |
| 12593 | ENSMUSG00000042473 | 416.1272305 | 0.052630262 | 0.153111408 | 0.343738344 | 0.731043072 | 0.94705581 | Tbc1d8b | TBC1 domain family, member 8B [Source: | 344.53 | 459.48 | 422.62 | 424.15 | 391.53 | 454.45 |
| 12594 | ENSMUSG00000042474 | 3.158563982 | -1.463956485 | 1.319605235 | -1.109389722 | 0.267262094 | NA | Fcmr | Fc fragment of IgM receptor [Source:MGI | 4.8186 | 6.1675 | 3.0187 | 3.8559 | 1.0906 | 0 |
| 12595 | ENSMUSG00000042476 | 6118.184496 | 0.020596796 | 0.095486933 | 0.215702768 | 0.829219444 | 0.969955221 | Abcb4 | ATP-binding cassette, sub-family B (MDR/ | 6583.8 | 6068.8 | 5569.6 | 6143.4 | 6284.1 | 6059.3 |
| 12596 | ENSMUSG00000042477 | 0 | NA | NA | NA | NA | NA | Tfap2e | transcription factor AP-2, epsilon [Source: | 0 | 0 | 0 | 0 | 0 | 0 |

| | | | | | | | | | | | | | | | |
|---|---|---|---|---|---|---|---|---|---|---|---|---|---|---|---|
| 12597 | ENSMUSG00000042485 | 41.68268935 | -0.2807833 | 0.404210807 | -0.694645703 | 0.487277363 | 0.873910634 | Mustn1 | musculoskeletal, embryonic nuclear prote | 63.445 | 32.893 | 40.25 | 24.099 | 38.172 | 51.237 |
| 12598 | ENSMUSG00000042487 | 208.6748807 | -0.613888396 | 0.205956223 | -2.980674174 | 0.002876146 | 0.045365954 | Leo1 | Leo1, Paf1/RNA polymerase II complex co | 192.74 | 258.01 | 306.9 | 150.38 | 164.68 | 179.33 |
| 12599 | ENSMUSG00000042489 | 1.379094917 | 0.028286433 | 1.992516577 | 0.014196335 | 0.988673344 | NA | Clspn | claspin [Source:MGI Symbol;Acc:MGI:244 | 4.0155 | 0 | 0 | 0.964 | 2.1812 | 1.1138 |
| 12600 | ENSMUSG00000042492 | 226.9191188 | 0.013856423 | 0.168213319 | 0.082374112 | 0.934349221 | 0.988420305 | Tbc1d10b | TBC1 domain family, member 10b [Source | 206.4 | 219.97 | 251.56 | 230.39 | 215.94 | 237.25 |
| 12601 | ENSMUSG00000042496 | 98.42245232 | -0.144809047 | 0.24842727 | -0.582903186 | 0.559958476 | 0.899475387 | Prdm10 | PR domain containing 10 [Source:MGI Syr | 113.24 | 117.18 | 79.493 | 96.398 | 83.978 | 100.25 |
| 12602 | ENSMUSG00000042498 | 0.531944269 | 2.618916564 | 3.285123409 | 0.797204926 | 0.425332026 | NA | Radx | RPA1 related single stranded DNA binding | 0 | 0 | 0 | 0.964 | 0 | 2.2277 |
| 12603 | ENSMUSG00000042499 | 0 | NA | NA | NA | NA | NA | Hoxd11 | homeobox D11 [Source:MGI Symbol;Acc:I | 0 | 0 | 0 | 0 | 0 | 0 |
| 12604 | ENSMUSG00000042500 | 87.23952402 | 0.001690141 | 0.256842435 | 0.006580459 | 0.994749591 | 0.998676967 | Ago4 | argonaute RISC catalytic subunit 4 [Source | 85.129 | 80.178 | 96.599 | 104.11 | 87.249 | 70.172 |
| 12605 | ENSMUSG00000042501 | 0 | NA | NA | NA | NA | NA | Cpa6 | carboxypeptidase A6 [Source:MGI Symbol | 0 | 0 | 0 | 0 | 0 | 0 |
| 12606 | ENSMUSG00000042502 | 630.4929127 | 0.236235894 | 0.123309653 | 1.915793994 | 0.055391323 | 0.331889018 | Cd2bp2 | CD2 cytoplasmic tail binding protein 2 [So | 538.88 | 589 | 609.78 | 671.89 | 730.71 | 642.69 |
| 12607 | ENSMUSG00000042505 | 133.9639871 | 0.16799837 | 0.211544712 | 0.794150645 | 0.427107737 | 0.842220356 | Sdhaf3 | succinate dehydrogenase complex asseml | 121.27 | 119.24 | 137.86 | 121.46 | 162.5 | 141.46 |
| 12608 | ENSMUSG00000042506 | 590.3472608 | 0.26420263 | 0.140502855 | 1.880407552 | 0.060052555 | 0.349115819 | Usp22 | ubiquitin specific peptidase 22 [Source:M | 609.55 | 539.66 | 458.85 | 621.77 | 620.56 | 691.7 |
| 12609 | ENSMUSG00000042507 | 245.6029909 | -0.056717026 | 0.162285472 | -0.349489241 | 0.726722047 | 0.945777495 | Mideas | mitotic deacetylase associated SANT dom | 232.1 | 254.92 | 264.64 | 235.21 | 261.75 | 225 |
| 12610 | ENSMUSG00000042508 | 386.0682708 | 0.046712646 | 0.138434481 | 0.337435047 | 0.735788968 | 0.948109711 | Dmtf1 | cyclin D binding myb-like transcription fac | 383.08 | 397.8 | 358.22 | 370.17 | 384.99 | 422.15 |
| 12611 | ENSMUSG00000042510 | 58.6343336 | -0.981428185 | 0.312369363 | -3.141883618 | 0.001678647 | 0.030734123 | AA986860 | expressed sequence AA986860 [Source:M | 85.129 | 71.954 | 76.475 | 42.415 | 47.987 | 27.846 |
| 12612 | ENSMUSG00000042514 | 0.685278559 | -2.809870862 | 3.376578027 | -0.832165239 | 0.405315682 | NA | Klhl14 | kelch-like 14 [Source:MGI Symbol;Acc:MG | 0 | 4.1117 | 0 | 0 | 0 | 0 |
| 12613 | ENSMUSG00000042515 | 1.613177868 | 0.743173382 | 1.721904591 | 0.431599629 | 0.666032428 | NA | Pwwp3b | PWWP domain containing 3B [Source:MG | 1.6062 | 0 | 2.0125 | 3.8559 | 1.0906 | 1.1138 |
| 12614 | ENSMUSG00000042520 | 762.3865904 | -0.051979949 | 0.112383937 | -0.462521159 | 0.643707632 | 0.925505685 | Ubap2l | ubiquitin-associated protein 2-like [Source | 795.87 | 770.94 | 761.73 | 792.39 | 740.53 | 712.86 |
| 12615 | ENSMUSG00000042523 | 23.20341822 | -0.468627732 | 0.493366491 | -0.949857236 | 0.342184798 | 0.783995077 | Dnal1 | dynein, axonemal, light chain 1 [Source:M | 37.746 | 20.558 | 22.137 | 12.532 | 25.084 | 21.163 |
| 12616 | ENSMUSG00000042524 | 2188.17259 | 0.062780314 | 0.100870559 | 0.62238491 | 0.5336888 | 0.889143097 | Sun2 | Sad1 and UNC84 domain containing 2 [So | 2156.3 | 2022.9 | 2242.9 | 2385.8 | 2230.3 | 2090.7 |
| 12617 | ENSMUSG00000042525 | 0 | NA | NA | NA | NA | NA | 4933428M09 | RIKEN cDNA 4933428M09 gene [Source:M | 0 | 0 | 0 | 0 | 0 | 0 |
| 12618 | ENSMUSG00000042529 | 0 | NA | NA | NA | NA | NA | Kcnj12 | potassium inwardly-rectifying channel, su | 0 | 0 | 0 | 0 | 0 | 0 |
| 12619 | ENSMUSG00000042532 | 1.192033051 | 1.693401922 | 2.323696699 | 0.728753422 | 0.466152508 | NA | Golga7b | golgi autoantigen, golgin subfamily a, 7B [ | 1.6062 | 0 | 0 | 0 | 1.0906 | 4.4554 |
| 12620 | ENSMUSG00000042535 | 404.1300519 | 0.071289766 | 0.135494674 | 0.526144418 | 0.598787869 | 0.915055389 | Gtpbp1 | GTP binding protein 1 [Source:MGI Symbo | 424.84 | 385.47 | 371.3 | 401.01 | 414.43 | 427.72 |
| 12621 | ENSMUSG00000042540 | 2.223801593 | -1.273618493 | 1.692502795 | -0.752505991 | 0.451746829 | NA | Acot5 | acyl-CoA thioesterase 5 [Source:MGI Syml | 6.4248 | 2.0558 | 1.0062 | 3.8559 | 0 | 0 |
| 12622 | ENSMUSG00000042541 | 759.5229809 | 0.007300456 | 0.142138297 | 0.05136164 | 0.959037352 | 0.992027194 | Sem1 | SEM1, 26S proteasome complex subunit [ | 759.73 | 804.86 | 707.39 | 637.19 | 756.89 | 891.08 |
| 12623 | ENSMUSG00000042548 | 347.0790561 | 0.220819355 | 0.13998438 | 1.577457113 | 0.114690383 | 0.497415802 | Asxl1 | ASXL transcriptional regulator 1 [Source:M | 342.92 | 308.38 | 309.92 | 381.74 | 369.72 | 369.8 |
| 12624 | ENSMUSG00000042549 | 28.96115474 | 0.922566741 | 0.447541068 | 2.061412474 | 0.039263706 | 0.271078874 | Map2k3os | mitogen-activated protein kinase kinase 3 | 14.456 | 26.726 | 19.119 | 40.487 | 25.084 | 47.895 |
| 12625 | ENSMUSG00000042554 | 0 | NA | NA | NA | NA | NA | Zp3r | zona pellucida 3 receptor [Source:MGI Syr | 0 | 0 | 0 | 0 | 0 | 0 |
| 12626 | ENSMUSG00000042557 | 413.1293043 | 0.47246612 | 0.148198279 | 3.188067528 | 0.001432271 | 0.0274026 | Sin3a | transcriptional regulator, SIN3A (yeast) [S | 344.53 | 326.88 | 366.27 | 414.51 | 488.6 | 537.99 |
| 12627 | ENSMUSG00000042558 | 289.3630523 | 0.090187799 | 0.153498152 | 0.587549738 | 0.556834556 | 0.898167794 | Adprs | ADP-ribosylserine hydrolase [Source:MGI | 306.79 | 255.95 | 277.72 | 308.47 | 302.1 | 285.14 |
| 12628 | ENSMUSG00000042564 | 22.97297323 | 0.435553422 | 0.500966628 | 0.869426021 | 0.384614156 | 0.816437825 | Fam227a | family with sequence similarity 227, meml | 16.062 | 16.447 | 26.162 | 26.027 | 15.269 | 37.871 |
| 12629 | ENSMUSG00000042567 | 0 | NA | NA | NA | NA | NA | Nek10 | NIMA (never in mitosis gene a)- related ki | 0 | 0 | 0 | 0 | 0 | 0 |
| 12630 | ENSMUSG00000042569 | 524.0869856 | -0.046513017 | 0.156613768 | -0.296991877 | 0.76647271 | 0.955218101 | Dhrs7b | dehydrogenase/reductase (SDR family) m | 628.83 | 496.48 | 470.92 | 470.42 | 591.12 | 486.75 |
| 12631 | ENSMUSG00000042570 | 137.6517983 | 0.228833491 | 0.205333464 | 1.114448108 | 0.265086994 | 0.720853222 | Mier2 | MIER family member 2 [Source:MGI Symb | 120.47 | 128.49 | 131.82 | 170.62 | 133.06 | 141.46 |
| 12632 | ENSMUSG00000042572 | 755.9851682 | 0.043915008 | 0.112270273 | 0.391154374 | 0.695683132 | 0.939992732 | Ube2q1 | ubiquitin-conjugating enzyme E2Q family | 724.4 | 726.74 | 782.86 | 796.25 | 743.8 | 761.87 |
| 12633 | ENSMUSG00000042579 | 127.2582677 | 0.087985492 | 0.238475971 | 0.368949088 | 0.712165674 | 0.944239215 | 4632404H12 | RIKEN cDNA 4632404H12 gene [Source:M | 123.68 | 116.15 | 130.81 | 172.55 | 113.42 | 106.93 |
| 12634 | ENSMUSG00000042581 | 0 | NA | NA | NA | NA | NA | Thsd7b | thrombospondin, type I, domain containir | 0 | 0 | 0 | 0 | 0 | 0 |
| 12635 | ENSMUSG00000042589 | 9.434365985 | -0.390113463 | 0.922106463 | -0.423067703 | 0.672245866 | 0.931191575 | Cux2 | cut-like homeobox 2 [Source:MGI Symbol | 24.896 | 1.0279 | 6.0375 | 12.532 | 6.5437 | 5.5692 |
| 12636 | ENSMUSG00000042590 | 305.3096481 | 0.126684481 | 0.155732716 | 0.813473778 | 0.415946474 | 0.836590861 | Ipo11 | importin 11 [Source:MGI Symbol;Acc:MGI | 316.42 | 284.73 | 273.7 | 287.27 | 351.18 | 318.56 |
| 12637 | ENSMUSG00000042594 | 477.4537044 | 0.123824288 | 0.13304097 | 0.930722982 | 0.351996878 | 0.792911205 | Sh2b3 | SH2B adaptor protein 3 [Source:MGI Syml | 460.18 | 418.36 | 492.05 | 482.95 | 479.87 | 531.3 |
| 12638 | ENSMUSG00000042595 | 175.6054647 | 0.060833723 | 0.184494321 | 0.329732223 | 0.741602303 | 0.948831112 | Fam199x | family with sequence similarity 199, X-link | 163.83 | 187.08 | 165.02 | 175.44 | 197.4 | 164.85 |
| 12639 | ENSMUSG00000042596 | 0 | NA | NA | NA | NA | NA | Tfap2d | transcription factor AP-2, delta [Source:M | 0 | 0 | 0 | 0 | 0 | 0 |
| 12640 | ENSMUSG00000042599 | 380.2494191 | 0.067009271 | 0.143109397 | 0.468238094 | 0.639614333 | 0.923932929 | Kdm7a | lysine (K)-specific demethylase 7A [Source | 349.35 | 418.36 | 347.15 | 395.23 | 389.35 | 382.05 |
| 12641 | ENSMUSG00000042604 | 0 | NA | NA | NA | NA | NA | Kcna4 | potassium voltage-gated channel, shaker- | 0 | 0 | 0 | 0 | 0 | 0 |
| 12642 | ENSMUSG00000042605 | 1141.456642 | 0.251523724 | 0.12271096 | 2.049725015 | 0.040391273 | 0.274797444 | Atxn2 | ataxin 2 [Source:MGI Symbol;Acc:MGI:12 | 1076.2 | 996.05 | 1054.5 | 1368.8 | 1065.5 | 1287.6 |
| 12643 | ENSMUSG00000042606 | 79.94174406 | 0.214479025 | 0.2626881 | 0.816477886 | 0.414226866 | 0.835831451 | Hirip3 | HIRA interacting protein 3 [Source:MGI Sy | 66.657 | 88.401 | 67.418 | 97.362 | 79.615 | 80.197 |
| 12644 | ENSMUSG00000042607 | 2.060967857 | -2.243284682 | 1.696417045 | -1.322366271 | 0.186046215 | NA | Asb4 | ankyrin repeat and SOCS box-containing 4 | 4.0155 | 5.1396 | 1.0062 | 0 | 1.0906 | 1.1138 |
| 12645 | ENSMUSG00000042608 | 597.3645406 | -0.249897718 | 0.14395675 | -1.735922201 | 0.082577592 | 0.423697158 | Stk40 | serine/threonine kinase 40 [Source:MGI S | 624.01 | 629.09 | 694.31 | 587.06 | 448.24 | 601.48 |
| 12646 | ENSMUSG00000042613 | 193.9173494 | -0.213139232 | 0.178418691 | -1.194601481 | 0.23224277 | 0.67964336 | Pbxip1 | pre B cell leukemia transcription factor int | 194.35 | 221 | 209.3 | 160.98 | 196.31 | 181.56 |
| 12647 | ENSMUSG00000042616 | 3.657298423 | 0.350950241 | 1.23713258 | 0.28368038 | 0.776655331 | NA | Oscp1 | organic solute carrier partner 1 [Source:M | 1.6062 | 4.1117 | 4.025 | 6.7478 | 5.4531 | 0 |
| 12648 | ENSMUSG00000042622 | 158.1579711 | 0.268964881 | 0.277407913 | 0.969564561 | 0.332263585 | 0.775138789 | Maff | v-maf musculoaponeurotic fibrosarcoma | 208 | 113.07 | 108.67 | 219.79 | 166.86 | 132.55 |
| 12649 | ENSMUSG00000042625 | 835.5345719 | 0.128006588 | 0.122238002 | 1.047191433 | 0.295011296 | 0.745595559 | Safb2 | scaffold attachment factor B2 [Source:MG | 828.8 | 780.19 | 786.88 | 984.22 | 826.69 | 806.42 |
| 12650 | ENSMUSG00000042626 | 1045.512709 | 0.233792031 | 0.106868062 | 2.187669792 | 0.028693664 | 0.222176208 | Shc1 | src homology 2 domain-containing transfo | 1032 | 925.13 | 924.74 | 1120.1 | 1152.8 | 1118.3 |
| 12651 | ENSMUSG00000042628 | 1737.532282 | -0.09848791 | 0.129204745 | -0.762262325 | 0.445903455 | 0.85243822 | Zfyve1 | zinc finger, FYVE domain containing 1 [Sou | 1656 | 1708.4 | 2027.6 | 1935.7 | 1571.6 | 1526 |
| 12652 | ENSMUSG00000042631 | 0 | NA | NA | NA | NA | NA | Xkr7 | X-linked Kx blood group related 7 [Source | 0 | 0 | 0 | 0 | 0 | 0 |

| | | | | | | | | | | | | | | | |
|---|---|---|---|---|---|---|---|---|---|---|---|---|---|---|---|
| 12653 | ENSMUSG00000042632 | 1070.281105 | -0.070050181 | 0.166523789 | -0.420661705 | 0.674002129 | 0.932009833 | Pla2g6 | phospholipase A2, group VI [Source:MGI S | 1227.1 | 1045.4 | 1016.3 | 1211.7 | 1162.6 | 758.53 |
| 12654 | ENSMUSG00000042638 | 7.171674145 | -0.309111173 | 0.829511872 | -0.372642253 | 0.709414721 | 0.944239215 | Gucy2c | guanylate cyclase 2c [Source:MGI Symbol, | 10.44 | 7.1954 | 6.0375 | 3.8559 | 4.3625 | 11.138 |
| 12655 | ENSMUSG00000042641 | 0.462220778 | -0.685400455 | 3.191475091 | -0.21475977 | 0.82995462 | NA | Rgsl1 | regulator of G-protein signaling like 1 [Sou | 0.8031 | 0 | 1.0062 | 0.964 | 0 | 0 |
| 12656 | ENSMUSG00000042642 | 691.4252587 | 0.20153422 | 0.11161646 | 1.805595874 | 0.070981488 | 0.385759452 | Flad1 | flavin adenine dinucleotide synthetase 1 [ | 644.89 | 658.9 | 625.88 | 742.26 | 748.16 | 728.45 |
| 12657 | ENSMUSG00000042644 | 23.03507565 | -0.249575499 | 0.470543742 | -0.530398083 | 0.595835955 | 0.91381309 | Itpr3 | inositol 1,4,5-triphosphate receptor 3 [Sou | 28.109 | 27.754 | 19.119 | 19.28 | 28.356 | 15.594 |
| 12658 | ENSMUSG00000042647 | 287.5963328 | 0.231087287 | 0.199891254 | 1.156065021 | 0.247654562 | 0.698260349 | Acad12 | acyl-Coenzyme A dehydrogenase family, r | 310.8 | 293.98 | 188.17 | 273.77 | 339.18 | 319.67 |
| 12659 | ENSMUSG00000042650 | 1791.605441 | 0.098651126 | 0.09856655 | 1.00085806 | 0.316895435 | 0.763524842 | Alkbh5 | alkB homolog 5, RNA demethylase [Source | 1668 | 1838.9 | 1684.5 | 1797.8 | 1829 | 1931.4 |
| 12660 | ENSMUSG00000042655 | 0 | NA | NA | NA | NA | NA | Shisal2b | shisa like 2B [Source:MGI Symbol;Acc:MG | 0 | 0 | 0 | 0 | 0 | 0 |
| 12661 | ENSMUSG00000042659 | 222.2281291 | -0.182329042 | 0.171279053 | -1.064514534 | 0.287095674 | 0.73951397 | Arrdc4 | arrestin domain containing 4 [Source:MGI | 248.96 | 213.81 | 245.52 | 201.47 | 193.04 | 230.57 |
| 12662 | ENSMUSG00000042660 | 271.1802806 | -0.001953246 | 0.187084082 | -0.010440474 | 0.991669858 | 0.99859176 | Wdr55 | WD repeat domain 55 [Source:MGI Symbo | 228.88 | 241.56 | 344.14 | 258.35 | 286.83 | 267.32 |
| 12663 | ENSMUSG00000042662 | 0 | NA | NA | NA | NA | NA | Dusp15 | dual specificity phosphatase-like 15 [Sourc | 0 | 0 | 0 | 0 | 0 | 0 |
| 12664 | ENSMUSG00000042670 | 337.7310912 | 0.139763503 | 0.154593175 | 0.904072923 | 0.365956735 | 0.804235439 | Immp1l | IMP1 inner mitochondrial membrane pep | 311.6 | 349.49 | 302.88 | 320.04 | 340.27 | 402.1 |
| 12665 | ENSMUSG00000042671 | 3.635177364 | -1.006879696 | 1.236422496 | -0.814349221 | 0.41544492 | NA | Rgs8 | regulator of G-protein signaling 8 [Source: | 10.44 | 1.0279 | 3.0187 | 2.8919 | 1.0906 | 3.3415 |
| 12666 | ENSMUSG00000042672 | 0.339027071 | -1.79174149 | 3.919585585 | -0.457125237 | 0.647581032 | NA | Dcst1 | DC-STAMP domain containing 1 [Source:N | 0 | 1.0279 | 1.0062 | 0 | 0 | 0 |
| 12667 | ENSMUSG00000042675 | 1085.673289 | 0.125637748 | 0.123630204 | 1.016238293 | 0.309515926 | 0.758143666 | Ypel3 | yippee like 3 [Source:MGI Symbol;Acc:MG | 1064.1 | 919.99 | 1131 | 1181.8 | 1020.8 | 1196.3 |
| 12668 | ENSMUSG00000042677 | 63.37054831 | -0.358188795 | 0.31509763 | -1.136754965 | 0.255640735 | 0.708918985 | Zc3h12a | zinc finger CCCH type containing 12A [Sou | 60.233 | 84.289 | 69.431 | 64.587 | 37.081 | 64.603 |
| 12669 | ENSMUSG00000042678 | 0.267700746 | -1.497481649 | 4.057001436 | -0.369110456 | 0.712045396 | NA | Myo15 | myosin XV [Source:MGI Symbol;Acc:MGI:1 | 1.6062 | 0 | 0 | 0 | 0 | 0 |
| 12670 | ENSMUSG00000042680 | 568.1839758 | -0.261268023 | 0.136671492 | -1.911649742 | 0.055921135 | 0.333956768 | Garem1 | GRB2 associated regulator of MAPK1 subt | 556.55 | 692.82 | 609.78 | 510.91 | 478.78 | 560.26 |
| 12671 | ENSMUSG00000042682 | 1115.862804 | 0.252272906 | 0.138870512 | 1.816605289 | 0.069277573 | 0.380133087 | Selenok | selenoprotein K [Source:MGI Symbol;Acc: | 1161.3 | 1013.5 | 879.46 | 1171.2 | 1085.2 | 1384.5 |
| 12672 | ENSMUSG00000042684 | 6.97251341 | -1.069259962 | 0.912487807 | -1.171807396 | 0.241274392 | 0.691388521 | Npl | N-acetylneuraminate pyruvate lyase [Sour | 16.062 | 6.1675 | 6.0375 | 4.8199 | 7.6343 | 1.1138 |
| 12673 | ENSMUSG00000042686 | 44.15817823 | 1.148244437 | 0.458535462 | 2.504156237 | 0.012274382 | 0.124560339 | Jph1 | junctophilin 1 [Source:MGI Symbol;Acc:M | 16.865 | 30.838 | 35.219 | 100.25 | 30.537 | 51.237 |
| 12674 | ENSMUSG00000042688 | 601.3336816 | 0.109314832 | 0.124929311 | 0.875013493 | 0.381566564 | 0.8130053 | Mapk6 | mitogen-activated protein kinase 6 [Sourc | 609.55 | 578.72 | 546.39 | 571.64 | 645.65 | 656.05 |
| 12675 | ENSMUSG00000042694 | 63.43398808 | -0.243494126 | 0.295776654 | -0.823236459 | 0.410373539 | 0.833241465 | Stn1 | STN1, CST complex subunit [Source:MGI S | 65.854 | 64.759 | 75.468 | 44.343 | 58.893 | 71.286 |
| 12676 | ENSMUSG00000042699 | 1053.297992 | -0.098460514 | 0.158107588 | -0.622743757 | 0.533452923 | 0.889143097 | Dhx9 | DEAH (Asp-Glu-Ala-His) box polypeptide 9 | 897.07 | 1226.3 | 1146.1 | 1195.3 | 1026.3 | 828.7 |
| 12677 | ENSMUSG00000042700 | 545.4430286 | 0.088595167 | 0.126959554 | 0.697821973 | 0.485288536 | 0.872797142 | Sipa1l1 | signal-induced proliferation-associated 1 l | 533.26 | 486.21 | 566.52 | 569.71 | 533.31 | 583.65 |
| 12678 | ENSMUSG00000042705 | 108.2119064 | -0.000495044 | 0.237027371 | -0.00208855 | 0.998333579 | 0.999764475 | Commd10 | COMM domain containing 10 [Source:MG | 107.62 | 122.32 | 94.587 | 97.362 | 97.065 | 130.32 |
| 12679 | ENSMUSG00000042707 | 1.97374485 | 0.797404686 | 1.650514433 | 0.48312494 | 0.629007027 | NA | Dnali1 | dynein, axonemal, light intermediate poly | 3.2124 | 0 | 1.0062 | 0.964 | 1.0906 | 5.5692 |
| 12680 | ENSMUSG00000042708 | 0 | NA | NA | NA | NA | NA | Shcbp1l | Shc SH2-domain binding protein 1-like [So | 0 | 0 | 0 | 0 | 0 | 0 |
| 12681 | ENSMUSG00000042709 | 373.591683 | 0.059827344 | 0.139740139 | 0.428132854 | 0.668554399 | 0.930350163 | Atpaf2 | ATP synthase mitochondrial F1 complex a | 335.7 | 375.19 | 387.4 | 391.38 | 364.27 | 387.62 |
| 12682 | ENSMUSG00000042712 | 201.5646958 | -0.314889911 | 0.207873652 | -1.514813964 | 0.129819523 | 0.527586234 | Tceal9 | transcription elongation factor A like 9 [Sc | 275.46 | 211.75 | 182.13 | 156.16 | 175.59 | 208.29 |
| 12683 | ENSMUSG00000042717 | 0.185640647 | 1.126924175 | 4.080472857 | 0.276174898 | 0.782413736 | NA | Ppp1r3a | protein phosphatase 1, regulatory subunit | 0 | 0 | 0 | 0 | 0 | 1.1138 |
| 12684 | ENSMUSG00000042719 | 317.9753768 | 0.08235324 | 0.157832897 | 0.521774877 | 0.601827083 | 0.915461118 | Naa25 | N(alpha)-acetyltransferase 25, NatB auxili | 267.43 | 330.99 | 329.04 | 315.22 | 356.63 | 308.53 |
| 12685 | ENSMUSG00000042724 | 4.782605634 | 0.215833028 | 0.993668375 | 0.21720831 | 0.828046009 | NA | Map3k9 | mitogen-activated protein kinase kinase k | 3.2124 | 5.1396 | 5.0312 | 8.6758 | 2.1812 | 4.4554 |
| 12686 | ENSMUSG00000042726 | 905.3813298 | 0.233407692 | 0.121197114 | 1.925851904 | 0.054122868 | 0.327545259 | Trafd1 | TRAF type zinc finger domain containing 1 | 813.54 | 833.64 | 850.28 | 1045 | 851.77 | 1038.1 |
| 12687 | ENSMUSG00000042729 | 1.72470113 | 0.045957583 | 1.852394656 | 0.024809823 | 0.980206656 | NA | Wdr74 | WD repeat domain 74 [Source:MGI Symbo | 4.0155 | 0 | 1.0062 | 0.964 | 4.3625 | 0 |
| 12688 | ENSMUSG00000042734 | 0 | NA | NA | NA | NA | NA | Ttc9 | tetratricopeptide repeat domain 9 [Source | 0 | 0 | 0 | 0 | 0 | 0 |
| 12689 | ENSMUSG00000042737 | 428.0273664 | 0.07017536 | 0.140933834 | 0.497931247 | 0.618532502 | 0.917509899 | Dpm3 | dolichyl-phosphate mannosyltransferase p | 440.9 | 366.97 | 444.76 | 462.71 | 436.25 | 416.58 |
| 12690 | ENSMUSG00000042742 | 192.1793897 | 0.009145524 | 0.185362131 | 0.049338687 | 0.960649389 | 0.992515024 | Bmt2 | base methyltransferase of 25S rRNA 2 [So | 169.45 | 200.44 | 205.27 | 192.8 | 170.14 | 214.97 |
| 12691 | ENSMUSG00000042743 | 70.30431568 | 0.373209346 | 0.302509342 | 1.233711803 | 0.217310314 | 0.663257309 | Sgtb | small glutamine-rich tetratricopeptide rep | 57.02 | 44.2 | 82.512 | 68.442 | 88.34 | 81.311 |
| 12692 | ENSMUSG00000042744 | 543.3420989 | 0.054397551 | 0.144927089 | 0.375344259 | 0.707404454 | 0.943292895 | Hectd4 | HECT domain E3 ubiquitin protein ligase 4 | 526.03 | 580.77 | 493.06 | 613.09 | 466.78 | 580.31 |
| 12693 | ENSMUSG00000042745 | 115.3728727 | -0.58354633 | 0.288677831 | -2.021444902 | 0.043233731 | 0.286918392 | Id1 | inhibitor of DNA binding 1, HLH protein [S | 127.69 | 113.07 | 175.09 | 131.1 | 81.796 | 63.489 |
| 12694 | ENSMUSG00000042747 | 650.4127915 | 0.08709717 | 0.142843132 | 0.609739987 | 0.542034062 | 0.894071846 | Krtcap2 | keratinocyte associated protein 2 [Source | 732.43 | 589 | 569.53 | 654.54 | 612.93 | 744.05 |
| 12695 | ENSMUSG00000042750 | 0.472877444 | -2.283655516 | 3.093270275 | -0.738265755 | 0.460352972 | NA | Bex2 | brain expressed X-linked 2 [Source:MGI Sy | 0.8031 | 1.0279 | 1.0062 | 0 | 0 | 0 |
| 12696 | ENSMUSG00000042751 | 5.378768382 | 0.120381724 | 0.934524227 | 0.128816055 | 0.897503201 | 0.982798755 | Nmnat2 | nicotinamide nucleotide adenylyltransfera | 7.2279 | 5.1396 | 3.0187 | 4.8199 | 8.7249 | 3.3415 |
| 12697 | ENSMUSG00000042757 | 6.1469137 | -0.838394221 | 1.003366729 | -0.835581046 | 0.403390635 | 0.828953151 | Tmem108 | transmembrane protein 108 [Source:MGI | 10.44 | 9.2513 | 4.025 | 7.7118 | 5.4531 | 0 |
| 12698 | ENSMUSG00000042759 | 16.15497591 | -0.885429332 | 0.552554376 | -1.602429318 | 0.109060706 | 0.485818926 | Apobr | apolipoprotein B receptor [Source:MGI Sy | 16.062 | 25.698 | 21.131 | 8.6758 | 11.997 | 13.366 |
| 12699 | ENSMUSG00000042761 | 0.167707431 | -0.796629176 | 4.080472857 | -0.195229623 | 0.84521319 | NA | Mrap2 | melanocortin 2 receptor accessory protei | 0 | 0 | 1.0062 | 0 | 0 | 0 |
| 12700 | ENSMUSG00000042763 | 0.167707431 | -0.796629176 | 4.080472857 | -0.195229623 | 0.84521319 | NA | Maneal | mannosidase, endo-alpha-like [Source:MG | 0 | 0 | 1.0062 | 0 | 0 | 0 |
| 12701 | ENSMUSG00000042766 | 2.347973088 | -0.69676879 | 1.371449581 | -0.508052793 | 0.61141632 | NA | Trim46 | tripartite motif-containing 46 [Source:MG | 1.6062 | 3.0838 | 4.025 | 0.964 | 2.1812 | 2.2277 |
| 12702 | ENSMUSG00000042770 | 4874.970229 | -0.022374358 | 0.09904983 | -0.225889919 | 0.821287039 | 0.969160361 | Hebp1 | heme binding protein 1 [Source:MGI Syml | 4713.4 | 4836.4 | 5188.2 | 4394.8 | 5019 | 5098.1 |
| 12703 | ENSMUSG00000042772 | 538.2732728 | 0.039292802 | 0.131479026 | 0.298852247 | 0.765052784 | 0.954980327 | Smg7 | Smg-7 homolog, nonsense mediated mRN | 526.84 | 569.47 | 497.08 | 582.24 | 560.58 | 493.43 |
| 12704 | ENSMUSG00000042774 | 0 | NA | NA | NA | NA | NA | Olfr1353 | olfactory receptor 1353 [Source:MGI Syml | 0 | 0 | 0 | 0 | 0 | 0 |
| 12705 | ENSMUSG00000042784 | 5.802588965 | -0.256232135 | 0.904142762 | -0.283397872 | 0.776871859 | 0.959295521 | Muc1 | mucin 1, transmembrane [Source:MGI Syr | 9.6372 | 6.1675 | 3.0187 | 3.8559 | 5.4531 | 6.6831 |
| 12706 | ENSMUSG00000042787 | 25.35599145 | -0.644797 | 0.44969044 | -1.433868596 | 0.151609765 | 0.567340036 | Exog | endo/exonuclease (5'-3'), endonuclease G | 21.684 | 38.033 | 33.206 | 17.352 | 21.812 | 20.049 |
| 12707 | ENSMUSG00000042788 | 1.298844242 | -2.558331636 | 2.053310462 | -1.24595461 | 0.212781059 | NA | Fam166b | family with sequence similarity 166, meml | 1.6062 | 3.0838 | 2.0125 | 0 | 1.0906 | 0 |
| 12708 | ENSMUSG00000042790 | 316.4927697 | 0.010133293 | 0.16637586 | 0.060906027 | 0.95143405 | 0.990883058 | Rnf214 | ring finger protein 214 [Source:MGI Symb | 269.84 | 351.55 | 326.02 | 359.56 | 289.01 | 302.97 |

| | | | | | | | | | | | | | | | |
|---|---|---|---|---|---|---|---|---|---|---|---|---|---|---|---|
| 12709 | ENSMUSG00000042793 | 0.647809292 | -2.733819264 | 3.061311498 | -0.893022244 | 0.371845261 | NA | Lgr6 | leucine-rich repeat-containing G protein-c | 0.8031 | 3.0838 | 0 | 0 | 0 | 0 |
| 12710 | ENSMUSG00000042796 | 0 | NA | NA | NA | NA | NA | Olfr1032 | olfactory receptor 1032 [Source:MGI Syml | 0 | 0 | 0 | 0 | 0 | 0 |
| 12711 | ENSMUSG00000042797 | 1181.415582 | 0.170131063 | 0.1164524 | 1.460949389 | 0.144029331 | 0.554282225 | Aqp11 | aquaporin 11 [Source:MGI Symbol;Acc:M( | 1207.9 | 1025.9 | 1100.8 | 1253.2 | 1351.3 | 1149.5 |
| 12712 | ENSMUSG00000042800 | 4.807868661 | -0.380930794 | 1.109381379 | -0.343372263 | 0.731318424 | NA | Spata46 | spermatogenesis associated 46 [Source:M | 3.2124 | 4.1117 | 9.0562 | 5.7839 | 0 | 6.6831 |
| 12713 | ENSMUSG00000042801 | 0 | NA | NA | NA | NA | NA | Olfr769 | olfactory receptor 769 [Source:MGI Symb | 0 | 0 | 0 | 0 | 0 | 0 |
| 12714 | ENSMUSG00000042804 | 11.62181992 | -0.856407102 | 0.673121448 | -1.272292102 | 0.203269354 | 0.647739134 | Gpr153 | G protein-coupled receptor 153 [Source:N | 13.653 | 10.279 | 21.131 | 12.532 | 5.4531 | 6.6831 |
| 12715 | ENSMUSG00000042807 | 31.78252517 | -0.889748171 | 0.400080028 | -2.223925486 | 0.026153458 | 0.209326659 | Hecw2 | HECT, C2 and WW domain containing E3 u | 35.336 | 44.2 | 44.275 | 18.316 | 20.722 | 27.846 |
| 12716 | ENSMUSG00000042808 | 0.319491021 | 0.16514334 | 4.044653141 | 0.040830038 | 0.967431393 | NA | Gpx2 | glutathione peroxidase 2 [Source:MGI Syr | 0.8031 | 0 | 0 | 0 | 0 | 1.1138 |
| 12717 | ENSMUSG00000042810 | 292.3070308 | 0.182755077 | 0.167782487 | 1.0892381 | 0.276048902 | 0.730038567 | Krba1 | KRAB-A domain containing 1 [Source:MGI | 277.07 | 261.09 | 283.76 | 367.28 | 265.02 | 299.62 |
| 12718 | ENSMUSG00000042812 | 39.43609985 | -1.176416386 | 0.365655446 | -3.217281182 | 0.001294117 | 0.025328969 | Foxf1 | forkhead box F1 [Source:MGI Symbol;Acc | 51.399 | 47.284 | 65.406 | 24.099 | 27.265 | 21.163 |
| 12719 | ENSMUSG00000042814 | 79.9341569 | 0.166706041 | 0.247839342 | 0.672637522 | 0.501177924 | 0.880974509 | Mcts2 | malignant T cell amplified sequence 2 [Sou | 80.31 | 72.982 | 72.45 | 81.938 | 86.159 | 85.766 |
| 12720 | ENSMUSG00000042816 | 8.149766349 | 3.999694908 | 1.086563873 | 3.681049046 | 0.000232276 | 0.007054905 | Gpr151 | G protein-coupled receptor 151 [Source:N | 0 | 2.0558 | 1.0062 | 18.316 | 15.269 | 12.252 |
| 12721 | ENSMUSG00000042817 | 2.542699711 | -1.34979786 | 1.419931018 | -0.950608053 | 0.341803379 | NA | Flt3 | FMS-like tyrosine kinase 3 [Source:MGI Sy | 0.8031 | 5.1396 | 5.0312 | 0.964 | 1.0906 | 2.2277 |
| 12722 | ENSMUSG00000042821 | 11.07455718 | 0.273858198 | 0.694647767 | 0.394240378 | 0.693403577 | 0.938905574 | Snai1 | snail family zinc finger 1 [Source:MGI Sym | 13.653 | 12.335 | 4.025 | 14.46 | 14.178 | 7.7969 |
| 12723 | ENSMUSG00000042826 | 12.03273431 | -0.908935414 | 0.642411647 | -1.414880037 | 0.157103672 | 0.576436451 | Fgf11 | fibroblast growth factor 11 [Source:MGI S | 13.653 | 16.447 | 17.106 | 10.604 | 4.3625 | 10.025 |
| 12724 | ENSMUSG00000042828 | 19.01455431 | 0.075159844 | 0.58951899 | 0.127493508 | 0.898549811 | 0.982798755 | Trim72 | tripartite motif-containing 72 [Source:MG | 19.274 | 8.2233 | 28.175 | 30.847 | 13.087 | 14.48 |
| 12725 | ENSMUSG00000042831 | 112.7612527 | 0.041625651 | 0.246721715 | 0.168714988 | 0.866020829 | 0.977767277 | Alkbh6 | alkB homolog 6 [Source:MGI Symbol;Acc:I | 121.27 | 103.82 | 107.67 | 87.722 | 109.06 | 147.03 |
| 12726 | ENSMUSG00000042834 | 899.1014029 | 0.600155116 | 0.42508336 | 1.411852764 | 0.157993319 | 0.578096473 | Nrep | neuronal regeneration related protein [So | 894.66 | 654.78 | 594.69 | 1415.1 | 1267.3 | 568.06 |
| 12727 | ENSMUSG00000042842 | 56.20805569 | -0.723383749 | 0.335440069 | -2.156521582 | 0.03104296 | 0.233856969 | Serpinb6b | serine (or cysteine) peptidase inhibitor, cl | 94.766 | 63.731 | 51.318 | 49.163 | 38.172 | 40.098 |
| 12728 | ENSMUSG00000042845 | 0.717584917 | 3.048786435 | 2.951298099 | 1.033032358 | 0.301588756 | NA | Wfdc12 | WAP four-disulfide core domain 12 [Sourc | 0 | 0 | 0 | 0.964 | 0 | 3.3415 |
| 12729 | ENSMUSG00000042846 | 8.255282304 | 0.62339041 | 0.83371492 | 0.747726106 | 0.454625381 | 0.857337687 | Lrrtm3 | leucine rich repeat transmembrane neuro | 2.4093 | 4.1117 | 13.081 | 6.7478 | 9.8156 | 13.366 |
| 12730 | ENSMUSG00000042848 | 0 | NA | NA | NA | NA | NA | Vmn1r226 | vomeronasal 1 receptor 226 [Source:MGI | 0 | 0 | 0 | 0 | 0 | 0 |
| 12731 | ENSMUSG00000042849 | 0 | NA | NA | NA | NA | NA | Olfr1414 | olfactory receptor 1414 [Source:MGI Syml | 0 | 0 | 0 | 0 | 0 | 0 |
| 12732 | ENSMUSG00000042851 | 56.74121343 | 0.30626811 | 0.296143716 | 1.034187434 | 0.301048548 | 0.7521219 | Zc3h6 | zinc finger CCCH type containing 6 [Source | 46.58 | 56.535 | 49.306 | 65.55 | 54.531 | 67.944 |
| 12733 | ENSMUSG00000042854 | 44.72547637 | 0.084908658 | 0.347595027 | 0.244274664 | 0.807018105 | 0.964718338 | Trp53rkb | transformation related protein 53 regulati | 41.761 | 35.977 | 52.325 | 34.703 | 52.35 | 51.237 |
| 12734 | ENSMUSG00000042857 | 24.31867707 | -0.099369188 | 0.438785979 | -0.226463909 | 0.820840626 | 0.969160361 | Gm9776 | predicted gene 9776 [Source:MGI Symbol | 25.699 | 20.558 | 29.181 | 23.135 | 26.175 | 21.163 |
| 12735 | ENSMUSG00000042861 | 0 | NA | NA | NA | NA | NA | Kcna10 | potassium voltage-gated channel, shaker- | 0 | 0 | 0 | 0 | 0 | 0 |
| 12736 | ENSMUSG00000042863 | 0.133850373 | -0.796629176 | 4.080472857 | -0.195229623 | 0.84521319 | NA | Olfr1026 | olfactory receptor 1026 [Source:MGI Syml | 0.8031 | 0 | 0 | 0 | 0 | 0 |
| 12737 | ENSMUSG00000042869 | 0 | NA | NA | NA | NA | NA | Olfr1370 | olfactory receptor 1370 [Source:MGI Syml | 0 | 0 | 0 | 0 | 0 | 0 |
| 12738 | ENSMUSG00000042870 | 320.915468 | -0.015951977 | 0.148358978 | -0.107522827 | 0.914374217 | 0.98625062 | Tom1 | target of myb1 trafficking protein [Source | 310.8 | 358.74 | 298.85 | 319.08 | 323.91 | 314.1 |
| 12739 | ENSMUSG00000042873 | 0.167707431 | -0.796629176 | 4.080472857 | -0.195229623 | 0.84521319 | NA | Lhfpl4 | lipoma HMGIC fusion partner-like protein | 0 | 0 | 1.0062 | 0 | 0 | 0 |
| 12740 | ENSMUSG00000042874 | 0 | NA | NA | NA | NA | NA | D930007J09 | RIKEN cDNA D930007J09 gene [Source:M( | 0 | 0 | 0 | 0 | 0 | 0 |
| 12741 | ENSMUSG00000042888 | 0 | NA | NA | NA | NA | NA | Prr30 | proline rich 30 [Source:MGI Symbol;Acc:N | 0 | 0 | 0 | 0 | 0 | 0 |
| 12742 | ENSMUSG00000042894 | 0 | NA | NA | NA | NA | NA | Olfr1260 | olfactory receptor 1260 [Source:MGI Syml | 0 | 0 | 0 | 0 | 0 | 0 |
| 12743 | ENSMUSG00000042895 | 0 | NA | NA | NA | NA | NA | Abra | actin-binding Rho activating protein [Sour | 0 | 0 | 0 | 0 | 0 | 0 |
| 12744 | ENSMUSG00000042901 | 174.0858923 | -0.166519142 | 0.191322585 | -0.870357997 | 0.384104793 | 0.816413186 | Aida | axin interactor, dorsalization associated [S | 192.74 | 202.5 | 156.97 | 165.8 | 150.51 | 175.99 |
| 12745 | ENSMUSG00000042903 | 691.7259948 | 0.156209005 | 0.123345172 | 1.26643793 | 0.205356339 | 0.650805014 | Foxo4 | forkhead box O4 [Source:MGI Symbol;Acc | 598.31 | 656.84 | 708.4 | 694.06 | 775.43 | 717.32 |
| 12746 | ENSMUSG00000042909 | 0 | NA | NA | NA | NA | NA | Olfr648 | olfactory receptor 648 [Source:MGI Symb | 0 | 0 | 0 | 0 | 0 | 0 |
| 12747 | ENSMUSG00000042918 | 2.330329613 | 0.863967558 | 1.681310098 | 0.513865681 | 0.607345903 | NA | Mamstr | MEF2 activating motif and SAP domain co | 0 | 1.0279 | 4.025 | 6.7478 | 2.1812 | 0 |
| 12748 | ENSMUSG00000042938 | 1.216784143 | -1.211426352 | 2.119017616 | -0.571692441 | 0.567530357 | NA | Gm14117 | predicted gene 14117 [Source:MGI Symbo | 0 | 3.0838 | 2.0125 | 0 | 1.0906 | 1.1138 |
| 12749 | ENSMUSG00000042940 | 0 | NA | NA | NA | NA | NA | 9230106D20 | RIKEN cDNA 9230106D20 gene [Source:M | 0 | 0 | 0 | 0 | 0 | 0 |
| 12750 | ENSMUSG00000042942 | 4.115175486 | -0.449619639 | 1.19413257 | -0.376524056 | 0.706527335 | NA | Greb1l | growth regulation by estrogen in breast ca | 8.031 | 6.1675 | 0 | 3.8559 | 2.1812 | 4.4554 |
| 12751 | ENSMUSG00000042961 | 5.642603937 | 0.157383321 | 0.89947249 | 0.174972913 | 0.861100918 | 0.976013118 | Egflam | EGF-like, fibronectin type III and laminin G | 4.8186 | 7.1954 | 4.025 | 6.7478 | 3.2719 | 7.7969 |
| 12752 | ENSMUSG00000042962 | 0.68303045 | 1.69421863 | 2.781340055 | 0.609137537 | 0.542433279 | NA | Gm5436 | predicted pseudogene 5436 [Source:MGI | 0.8031 | 0 | 0 | 0 | 2.1812 | 1.1138 |
| 12753 | ENSMUSG00000042976 | 0 | NA | NA | NA | NA | NA | 9930038B18 | RIKEN cDNA 9930038B18 gene [Source:M | 0 | 0 | 0 | 0 | 0 | 0 |
| 12754 | ENSMUSG00000042978 | 1193.761256 | -0.168731911 | 0.167163203 | -1.009384293 | 0.312790368 | 0.759921531 | Sbk1 | SH3-binding kinase 1 [Source:MGI Symbol | 1540.4 | 1164.6 | 1083.7 | 897.46 | 1344.7 | 1131.7 |
| 12755 | ENSMUSG00000042985 | 86.72057697 | 0.268067142 | 0.245854419 | 1.090349086 | 0.2755594 | 0.730038567 | Upk3b | uroplakin 3B [Source:MGI Symbol;Acc:MG | 79.507 | 79.15 | 77.481 | 99.29 | 104.7 | 80.197 |
| 12756 | ENSMUSG00000042988 | 561.7783287 | 0.214923236 | 0.153779394 | 1.397607515 | 0.16223096 | 0.585529252 | Notum | notum palmitoleoyl-protein carboxylester | 526.84 | 442 | 590.67 | 562 | 549.67 | 699.49 |
| 12757 | ENSMUSG00000042992 | 211.2771394 | -0.070241409 | 0.18076002 | -0.388589296 | 0.697579986 | 0.940422865 | Borcs5 | BLOC-1 related complex subunit 5 [Source | 248.96 | 186.05 | 213.32 | 192.8 | 212.67 | 213.86 |
| 12758 | ENSMUSG00000042993 | 1.270693045 | -1.50280405 | 2.054132149 | -0.731600472 | 0.46441246 | NA | Ifnk | interferon kappa [Source:MGI Symbol;Acc | 1.6062 | 3.0838 | 1.0062 | 1.928 | 0 | 0 |
| 12759 | ENSMUSG00000042997 | 370.9239222 | -0.071108272 | 0.136396497 | -0.52133503 | 0.602133402 | 0.915461118 | Nhlrc3 | NHL repeat containing 3 [Source:MGI Sym | 391.11 | 379.3 | 369.29 | 346.07 | 357.72 | 382.05 |
| 12760 | ENSMUSG00000043003 | 0.838717116 | -1.825569927 | 2.582619186 | -0.706867639 | 0.479648736 | NA | Rasef | RAS and EF hand domain containing [Sour | 0 | 2.0558 | 2.0125 | 0.964 | 0 | 0 |
| 12761 | ENSMUSG00000043004 | 10.67494698 | -1.367961573 | 0.691200612 | -1.979109319 | 0.047803699 | 0.303663969 | Gng2 | guanine nucleotide binding protein (G pro | 13.653 | 16.447 | 16.1 | 5.7839 | 8.7249 | 3.3415 |
| 12762 | ENSMUSG00000043008 | 23.6681791 | -1.125034488 | 0.470092593 | -2.393218919 | 0.016701273 | 0.153234527 | Klhl6 | kelch-like 6 [Source:MGI Symbol;Acc:MGI | 36.14 | 28.782 | 32.2 | 9.6398 | 18.541 | 16.708 |
| 12763 | ENSMUSG00000043013 | 372.7866909 | 3.244810157 | 0.805937223 | 4.026132636 | 5.67E-05 | 0.002323325 | Onecut1 | one cut domain, family member 1 [Source | 88.341 | 59.619 | 65.406 | 1401.6 | 199.58 | 422.15 |
| 12764 | ENSMUSG00000043015 | 10.97874897 | 0.365334704 | 0.699895514 | 0.521984635 | 0.601681029 | 0.915461118 | Nemp2 | nuclear envelope integral membrane prot | 5.6217 | 6.1675 | 17.106 | 11.568 | 9.8156 | 15.594 |

| | | | | | | | | | | | | | | | |
|---|---|---|---|---|---|---|---|---|---|---|---|---|---|---|---|
| 12765 | ENSMUSG00000043017 | 1.775179438 | -0.065077508 | 1.62655559 | -0.040009397 | 0.968085635 | NA | Ptgir | prostaglandin I receptor (IP) [Source:MGI | 2.4093 | 0 | 3.0187 | 1.928 | 2.1812 | 1.1138 |
| 12766 | ENSMUSG00000043019 | 841.8868481 | 0.008858343 | 0.11774072 | 0.075236017 | 0.940026928 | 0.988812319 | Edem3 | ER degradation enhancer, mannosidase al | 817.56 | 921.01 | 779.84 | 854.08 | 876.86 | 801.97 |
| 12767 | ENSMUSG00000043020 | 5.274864671 | 0.479129491 | 0.993424386 | 0.482300916 | 0.629592197 | 0.92057725 | Dnai3 | dynein axonemal intermediate chain 3 [Sc | 3.2124 | 2.0558 | 8.05 | 10.604 | 3.2719 | 4.4554 |
| 12768 | ENSMUSG00000043024 | 0.831492699 | -1.816502146 | 2.887260166 | -0.629143909 | 0.529254847 | NA | Gm16433 | predicted gene 16433 [Source:MGI Symbo | 0 | 0 | 4.025 | 0.964 | 0 | 0 |
| 12769 | ENSMUSG00000043029 | 0.767390791 | -1.69152783 | 2.44773922 | -0.691057207 | 0.489529591 | NA | Trpv3 | transient receptor potential cation chann | 1.6062 | 1.0279 | 1.0062 | 0.964 | 0 | 0 |
| 12770 | ENSMUSG00000043036 | 0 | NA | NA | NA | NA | NA | Ccdc63 | coiled-coil domain containing 63 [Source:I | 0 | 0 | 0 | 0 | 0 | 0 |
| 12771 | ENSMUSG00000043050 | 0 | NA | NA | NA | NA | NA | Tnp2 | transition protein 2 [Source:MGI Symbol;A | 0 | 0 | 0 | 0 | 0 | 0 |
| 12772 | ENSMUSG00000043051 | 2.01467134 | -0.839818341 | 1.652159903 | -0.508315411 | 0.611232164 | NA | Disc1 | disrupted in schizophrenia 1 [Source:MGI | 5.6217 | 1.0279 | 1.0062 | 0 | 1.0906 | 3.3415 |
| 12773 | ENSMUSG00000043059 | 153.3221996 | -0.004216298 | 0.196175687 | -0.02149246 | 0.982852818 | 0.998580735 | Zfp513 | zinc finger protein 513 [Source:MGI Symb | 155 | 135.69 | 170.06 | 164.84 | 142.87 | 151.48 |
| 12774 | ENSMUSG00000043060 | 0 | NA | NA | NA | NA | NA | Fscb | fibrous sheath CABYR binding protein [Sou | 0 | 0 | 0 | 0 | 0 | 0 |
| 12775 | ENSMUSG00000043061 | 297.9472418 | 0.007580874 | 0.15640712 | 0.048468857 | 0.961342584 | 0.992661062 | Tmem18 | transmembrane protein 18 [Source:MGI S | 288.31 | 307.35 | 295.84 | 278.59 | 341.36 | 276.23 |
| 12776 | ENSMUSG00000043065 | 51.20130948 | -0.318737737 | 0.326351097 | -0.976671258 | 0.328731929 | 0.77362974 | Spice1 | spindle and centriole associated protein 1 | 67.461 | 53.452 | 49.306 | 45.307 | 35.99 | 55.692 |
| 12777 | ENSMUSG00000043066 | 0 | NA | NA | NA | NA | NA | Vmn1r66 | vomeronasal 1 receptor 66 [Source:MGI S | 0 | 0 | 0 | 0 | 0 | 0 |
| 12778 | ENSMUSG00000043067 | 1500.833647 | -0.217337188 | 0.119390875 | -1.820383585 | 0.06870061 | 0.377683629 | Dpy19l1 | dpy-19-like 1 (C. elegans) [Source:MGI Syr | 1484.9 | 1673.5 | 1682.4 | 1206.9 | 1463.6 | 1493.7 |
| 12779 | ENSMUSG00000043068 | 81.72628398 | 0.301324348 | 0.316419538 | 0.952293748 | 0.340948028 | 0.783129922 | Fam89a | family with sequence similarity 89, membe | 65.051 | 70.926 | 83.518 | 59.767 | 77.434 | 133.66 |
| 12780 | ENSMUSG00000043073 | 0 | NA | NA | NA | NA | NA | Usp17le | ubiquitin specific peptidase 17-like E [Sou | 0 | 0 | 0 | 0 | 0 | 0 |
| 12781 | ENSMUSG00000043079 | 99.1624334 | -0.882595093 | 0.326025089 | -2.707138571 | 0.006786592 | 0.082281032 | Synpo | synaptopodin [Source:MGI Symbol;Acc:M | 188.73 | 122.32 | 74.462 | 82.902 | 71.981 | 54.578 |
| 12782 | ENSMUSG00000043085 | 263.3899728 | 0.015856819 | 0.154530999 | 0.102612545 | 0.918270486 | 0.986374671 | Tmem82 | transmembrane protein 82 [Source:MGI S | 250.57 | 252.87 | 282.75 | 272.81 | 258.48 | 262.87 |
| 12783 | ENSMUSG00000043087 | 0 | NA | NA | NA | NA | NA | Olfr855 | olfactory receptor 855 [Source:MGI Symb | 0 | 0 | 0 | 0 | 0 | 0 |
| 12784 | ENSMUSG00000043088 | 8.413576232 | -0.282230195 | 0.736099535 | -0.383413087 | 0.701413502 | 0.941991169 | Il17re | interleukin 17 receptor E [Source:MGI Syn | 10.44 | 9.2513 | 8.05 | 10.604 | 5.4531 | 6.6831 |
| 12785 | ENSMUSG00000043089 | 0 | NA | NA | NA | NA | NA | Mmp1a | matrix metallopeptidase 1a (interstitial co | 0 | 0 | 0 | 0 | 0 | 0 |
| 12786 | ENSMUSG00000043090 | 191.8145109 | -0.085661937 | 0.181861365 | -0.471028781 | 0.637620182 | 0.923478764 | Zfp866 | zinc finger protein 866 [Source:MGI Symb | 224.07 | 172.69 | 195.21 | 190.87 | 186.5 | 181.56 |
| 12787 | ENSMUSG00000043091 | 603.2960376 | -0.017411304 | 0.126967072 | -0.137132434 | 0.890926116 | 0.982489666 | Tuba1c | tubulin, alpha 1C [Source:MGI Symbol;Acc | 657.74 | 614.69 | 547.4 | 573.57 | 626.01 | 600.36 |
| 12788 | ENSMUSG00000043099 | 42.35212178 | -0.664125873 | 0.391827546 | -1.694944318 | 0.090085984 | 0.442897238 | Hic1 | hypermethylated in cancer 1 [Source:MGI | 67.461 | 38.033 | 50.312 | 45.307 | 21.812 | 31.188 |
| 12789 | ENSMUSG00000043102 | 2.878372171 | 1.383647897 | 1.302980174 | 1.061910169 | 0.288276468 | NA | Qrfp | pyroglutamylated RFamide peptide [Sourc | 0.8031 | 1.0279 | 3.0187 | 5.7839 | 2.1812 | 4.4554 |
| 12790 | ENSMUSG00000043110 | 27.55750533 | 0.033022715 | 0.42382106 | 0.077916645 | 0.937894359 | 0.988420305 | Lrrn4 | leucine rich repeat neuronal 4 [Source:MG | 25.699 | 31.865 | 24.15 | 23.135 | 35.99 | 24.505 |
| 12791 | ENSMUSG00000043119 | 0 | NA | NA | NA | NA | NA | Olfr448 | olfactory receptor 448 [Source:MGI Symb | 0 | 0 | 0 | 0 | 0 | 0 |
| 12792 | ENSMUSG00000043122 | 1.824027754 | -1.372520812 | 1.800673256 | -0.762226466 | 0.445924853 | NA | A530016L24 | RIKEN cDNA A530016L24 gene [Source:MG | 4.8186 | 3.0838 | 0 | 1.928 | 0 | 1.1138 |
| 12793 | ENSMUSG00000043126 | 0.133850373 | -0.796629176 | 4.080472857 | -0.195229623 | 0.84521319 | NA | D830039M1 | RIKEN cDNA D830039M14 gene [Source:M | 0.8031 | 0 | 0 | 0 | 0 | 0 |
| 12794 | ENSMUSG00000043131 | 882.9397475 | 0.038652505 | 0.121665863 | 0.317693919 | 0.750717124 | 0.952074518 | Mob1a | MOB kinase activator 1A [Source:MGI Syn | 878.59 | 953.91 | 780.85 | 903.25 | 836.5 | 944.54 |
| 12795 | ENSMUSG00000043140 | 321.0752806 | 0.076341247 | 0.155992481 | 0.489390553 | 0.624565223 | 0.919503423 | Tmem186 | transmembrane protein 186 [Source:MGI | 350.96 | 285.76 | 299.86 | 298.83 | 344.64 | 346.41 |
| 12796 | ENSMUSG00000043144 | 0 | NA | NA | NA | NA | NA | Aqp6 | aquaporin 6 [Source:MGI Symbol;Acc:MG | 0 | 0 | 0 | 0 | 0 | 0 |
| 12797 | ENSMUSG00000043145 | 0 | NA | NA | NA | NA | NA | Gm11292 | predicted gene 11292 [Source:MGI Symbo | 0 | 0 | 0 | 0 | 0 | 0 |
| 12798 | ENSMUSG00000043153 | 46.83280611 | 0.003554279 | 0.328673201 | 0.01081402 | 0.991371828 | 0.99859176 | Crppa | CDP-L-ribitol pyrophosphorylase A [Sourc | 45.777 | 50.368 | 44.275 | 48.199 | 55.622 | 36.757 |
| 12799 | ENSMUSG00000043154 | 301.9132558 | -0.083926189 | 0.15629909 | -0.536958906 | 0.591296 | 0.912461895 | Ppp2r3a | protein phosphatase 2, regulatory subunit | 280.28 | 350.52 | 301.87 | 287.27 | 309.74 | 281.8 |
| 12800 | ENSMUSG00000043155 | 7.245948301 | -1.644610579 | 0.866726422 | -1.897496763 | 0.057762406 | 0.341829329 | Hpdl | 4-hydroxyphenylpyruvate dioxygenase-lik | 13.653 | 5.1396 | 14.087 | 2.8919 | 4.3625 | 3.3415 |
| 12801 | ENSMUSG00000043157 | 4.094349125 | -0.571167267 | 1.192263398 | -0.479061311 | 0.631895012 | NA | Arl11 | ADP-ribosylation factor-like 11 [Source:M | 2.4093 | 8.2233 | 4.025 | 0 | 5.4531 | 4.4554 |
| 12802 | ENSMUSG00000043162 | 365.0701837 | -0.015277038 | 0.145892168 | -0.104714584 | 0.916602289 | 0.986374671 | Pyurf | Pigy upstream reading frame [Source:MGI | 340.52 | 348.46 | 412.56 | 371.13 | 345.73 | 372.02 |
| 12803 | ENSMUSG00000043164 | 0 | NA | NA | NA | NA | NA | Tmem212 | transmembrane protein 212 [Source:MGI | 0 | 0 | 0 | 0 | 0 | 0 |
| 12804 | ENSMUSG00000043165 | 24.19409468 | 0.00081856 | 0.516921645 | 0.001583529 | 0.998736527 | 0.999764475 | Lor | loricrin [Source:MGI Symbol;Acc:MGI:968 | 24.093 | 19.53 | 29.181 | 41.451 | 13.087 | 17.822 |
| 12805 | ENSMUSG00000043168 | 0.503122293 | -2.366675966 | 3.785596405 | -0.625179156 | 0.531853481 | NA | 4930426D05 | RIKEN cDNA 4930426D05 gene [Source:M | 0 | 0 | 3.0187 | 0 | 0 | 0 |
| 12806 | ENSMUSG00000043183 | 96.97826758 | 0.044834931 | 0.302829309 | 0.148053475 | 0.882300566 | 0.981769403 | Simc1 | SUMO-interacting motifs containing 1 [So | 92.357 | 100.74 | 93.581 | 110.86 | 130.87 | 53.465 |
| 12807 | ENSMUSG00000043190 | 223.9723316 | 0.370797468 | 0.161144591 | 2.301023358 | 0.021390311 | 0.181779939 | Rfesd | Rieske (Fe-S) domain containing [Source:M | 202.38 | 189.14 | 194.21 | 247.74 | 250.84 | 259.53 |
| 12808 | ENSMUSG00000043192 | 3.097404033 | -0.314470817 | 1.19104509 | -0.264029313 | 0.791757331 | NA | Gpi-ps | glucose-6-phosphate isomerase, pseudog | 3.2124 | 4.1117 | 3.0187 | 3.8559 | 3.2719 | 1.1138 |
| 12809 | ENSMUSG00000043207 | 1154.49804 | -0.087294442 | 0.104942603 | -0.831830346 | 0.405504712 | 0.829973745 | Zmpste24 | zinc metallopeptidase, STE24 [Source:MG | 1153.3 | 1150.2 | 1264.8 | 1084.5 | 1164.8 | 1109.4 |
| 12810 | ENSMUSG00000043219 | 0 | NA | NA | NA | NA | NA | Hoxa6 | homeobox A6 [Source:MGI Symbol;Acc:M | 0 | 0 | 0 | 0 | 0 | 0 |
| 12811 | ENSMUSG00000043223 | 0 | NA | NA | NA | NA | NA | Gm4835 | predicted pseudogene 4835 [Source:MGI | 0 | 0 | 0 | 0 | 0 | 0 |
| 12812 | ENSMUSG00000043226 | 0 | NA | NA | NA | NA | NA | Olfr1009 | olfactory receptor 1009 [Source:MGI Symb | 0 | 0 | 0 | 0 | 0 | 0 |
| 12813 | ENSMUSG00000043230 | 0.17131964 | -0.796629176 | 4.080472857 | -0.195229623 | 0.84521319 | NA | Fam124b | family with sequence similarity 124, meml | 0 | 1.0279 | 0 | 0 | 0 | 0 |
| 12814 | ENSMUSG00000043241 | 323.2389381 | 0.262805822 | 0.155114057 | 1.694274691 | 0.090213098 | 0.44322087 | Upf2 | UPF2 regulator of nonsense transcripts hc | 256.19 | 298.1 | 328.04 | 335.46 | 371.9 | 349.75 |
| 12815 | ENSMUSG00000043243 | 6.441844066 | -0.521725742 | 0.969480775 | -0.538149652 | 0.590473736 | 0.911690353 | Niban3 | niban apoptosis regulator 3 [Source:MGI S | 1.6062 | 7.1954 | 14.087 | 5.7839 | 2.1812 | 7.7969 |
| 12816 | ENSMUSG00000043251 | 35.91844254 | -0.379207197 | 0.435648733 | -0.870442557 | 0.384058598 | 0.816413186 | Exoc3l | exocyst complex component 3-like [Sourc | 46.58 | 47.284 | 28.175 | 48.199 | 18.541 | 26.732 |
| 12817 | ENSMUSG00000043252 | 916.5590353 | -0.328719754 | 0.12859196 | -2.556300991 | 0.010579157 | 0.113178195 | Tmem64 | transmembrane protein 64 [Source:MGI S | 1069.7 | 1092.7 | 899.58 | 899.39 | 796.15 | 741.82 |
| 12818 | ENSMUSG00000043257 | 171.7346748 | -0.189870519 | 0.188019031 | -1.009847344 | 0.312568433 | 0.759830222 | Pigv | phosphatidylinositol glycan anchor biosyn | 180.7 | 176.8 | 191.19 | 137.85 | 171.23 | 172.65 |
| 12819 | ENSMUSG00000043259 | 2.009384702 | 0.162838868 | 1.466055496 | 0.111072786 | 0.911558629 | NA | Fam13c | family with sequence similarity 13, membe | 1.6062 | 3.0838 | 1.0062 | 1.928 | 1.0906 | 3.3415 |
| 12820 | ENSMUSG00000043262 | 263.698036 | 0.166612586 | 0.163390852 | 1.01971796 | 0.307862241 | 0.756343859 | Uevld | UEV and lactate/malate dehyrogenase do | 282.69 | 242.59 | 219.36 | 276.66 | 284.65 | 276.23 |

| | | | | | | | | | | | | | | | |
|---|---|---|---|---|---|---|---|---|---|---|---|---|---|---|---|
| 12821 | ENSMUSG00000043263 | 18.99554742 | -1.068987571 | 0.519252366 | -2.058705248 | 0.039522483 | 0.272086243 | Ifi209 | interferon activated gene 209 [Source:MG | 23.29 | 29.81 | 24.15 | 13.496 | 7.6343 | 15.594 |
| 12822 | ENSMUSG00000043267 | 0 | NA | NA | NA | NA | NA | Olfr1031 | olfactory receptor 1031 [Source:MGI Syml | 0 | 0 | 0 | 0 | 0 | 0 |
| 12823 | ENSMUSG00000043274 | 0 | NA | NA | NA | NA | NA | Olfr1123 | olfactory receptor 1123 [Source:MGI Syml | 0 | 0 | 0 | 0 | 0 | 0 |
| 12824 | ENSMUSG00000043279 | 489.3847024 | 0.154760752 | 0.133605532 | 1.158340902 | 0.246724946 | 0.696725814 | Trim56 | tripartite motif-containing 56 [Source:MG | 444.92 | 453.31 | 491.05 | 466.57 | 521.32 | 559.15 |
| 12825 | ENSMUSG00000043282 | 2.819123731 | 0.196420479 | 1.395576181 | 0.140745079 | 0.888071332 | NA | Teddm1b | transmembrane epididymal protein 1B [So | 4.8186 | 1.0279 | 2.0125 | 5.7839 | 3.2719 | 0 |
| 12826 | ENSMUSG00000043284 | 520.463309 | 0.386479878 | 0.14760887 | 2.618270018 | 0.008837684 | 0.098938285 | Tmem11 | transmembrane protein 11 [Source:MGI S | 538.08 | 423.5 | 390.42 | 586.1 | 596.57 | 588.11 |
| 12827 | ENSMUSG00000043286 | 0.36353943 | 2.065989254 | 4.034137301 | 0.512126658 | 0.608562368 | NA | Pnpla1 | patatin-like phospholipase domain contai | 0 | 0 | 0 | 0 | 2.1812 | 0 |
| 12828 | ENSMUSG00000043289 | 0 | NA | NA | NA | NA | NA | Mei4 | meiotic double-stranded break formation | 0 | 0 | 0 | 0 | 0 | 0 |
| 12829 | ENSMUSG00000043290 | 0.371281295 | 2.09253362 | 4.031852165 | 0.519000582 | 0.603760335 | NA | Zfp784 | zinc finger protein 784 [Source:MGI Symb | 0 | 0 | 0 | 0 | 0 | 2.2277 |
| 12830 | ENSMUSG00000043295 | 0 | NA | NA | NA | NA | NA | Gm16373 | predicted pseudogene 16373 [Source:MG | 0 | 0 | 0 | 0 | 0 | 0 |
| 12831 | ENSMUSG00000043298 | 0 | NA | NA | NA | NA | NA | Smco3 | single-pass membrane protein with coiled | 0 | 0 | 0 | 0 | 0 | 0 |
| 12832 | ENSMUSG00000043300 | 12.51396155 | -0.700715634 | 0.641485554 | -1.092332678 | 0.274686902 | 0.729487777 | B3galnt1 | UDP-GalNAc:betaGlcNAc beta 1,3-galacto | 12.85 | 21.586 | 12.075 | 7.7118 | 14.178 | 6.6831 |
| 12833 | ENSMUSG00000043301 | 0 | NA | NA | NA | NA | NA | Kcnj6 | potassium inwardly-rectifying channel, su | 0 | 0 | 0 | 0 | 0 | 0 |
| 12834 | ENSMUSG00000043308 | 0 | NA | NA | NA | NA | NA | Vmn1r75 | vomeronasal 1 receptor 75 [Source:MGI S | 0 | 0 | 0 | 0 | 0 | 0 |
| 12835 | ENSMUSG00000043310 | 0 | NA | NA | NA | NA | NA | Olfr571 | olfactory receptor 571 [Source:MGI Symb | 0 | 0 | 0 | 0 | 0 | 0 |
| 12836 | ENSMUSG00000043311 | 177.6079784 | 0.10507653 | 0.203799641 | 0.515587417 | 0.6061426 | 0.916376001 | D17H6S53E | DNA segment, Chr 17, human D6S53E [So | 195.96 | 141.85 | 175.09 | 184.12 | 211.58 | 157.05 |
| 12837 | ENSMUSG00000043312 | 0 | NA | NA | NA | NA | NA | Olfr131 | olfactory receptor 131 [Source:MGI Symb | 0 | 0 | 0 | 0 | 0 | 0 |
| 12838 | ENSMUSG00000043313 | 1.031479329 | -2.220845979 | 2.403244684 | -0.924103148 | 0.355432614 | NA | Pcdhb19 | protocadherin beta 19 [Source:MGI Symb | 3.2124 | 0 | 2.0125 | 0.964 | 0 | 0 |
| 12839 | ENSMUSG00000043314 | 0 | NA | NA | NA | NA | NA | Olfr30 | olfactory receptor 30 [Source:MGI Symbo | 0 | 0 | 0 | 0 | 0 | 0 |
| 12840 | ENSMUSG00000043319 | 0 | NA | NA | NA | NA | NA | Cox8c | cytochrome c oxidase subunit 8C [Source: | 0 | 0 | 0 | 0 | 0 | 0 |
| 12841 | ENSMUSG00000043323 | 476.2110881 | 0.025868944 | 0.127274882 | 0.203252545 | 0.838937645 | 0.971535186 | Fbrsl1 | fibrosin-like 1 [Source:MGI Symbol;Acc:M | 491.5 | 454.34 | 469.92 | 507.05 | 473.33 | 461.13 |
| 12842 | ENSMUSG00000043329 | 0 | NA | NA | NA | NA | NA | Gm8849 | predicted gene 8849 [Source:MGI Symbol | 0 | 0 | 0 | 0 | 0 | 0 |
| 12843 | ENSMUSG00000043331 | 0 | NA | NA | NA | NA | NA | Olfr975 | olfactory receptor 975 [Source:MGI Symb | 0 | 0 | 0 | 0 | 0 | 0 |
| 12844 | ENSMUSG00000043333 | 0 | NA | NA | NA | NA | NA | Rhbdl2 | rhomboid like 2 [Source:MGI Symbol;Acc: | 0 | 0 | 0 | 0 | 0 | 0 |
| 12845 | ENSMUSG00000043336 | 39.29214038 | -1.005137151 | 0.427605289 | -2.350619082 | 0.01874221 | 0.165701295 | Filip1l | filamin A interacting protein 1-like [Source | 40.958 | 83.261 | 33.206 | 22.171 | 30.537 | 25.618 |
| 12846 | ENSMUSG00000043340 | 5.435315013 | -0.328048115 | 0.952056413 | -0.344567938 | 0.730419213 | 0.946665188 | 6530409C15 | RIKEN cDNA 6530409C15 gene [Source:M | 8.031 | 2.0558 | 8.05 | 6.7478 | 3.2719 | 4.4554 |
| 12847 | ENSMUSG00000043342 | 0 | NA | NA | NA | NA | NA | Hoxd9 | homeobox D9 [Source:MGI Symbol;Acc:M | 0 | 0 | 0 | 0 | 0 | 0 |
| 12848 | ENSMUSG00000043346 | 1.35357482 | 1.770076822 | 2.050987331 | 0.863036448 | 0.388117427 | NA | Gm6741 | predicted gene 6741 [Source:MGI Symbol | 0.8031 | 1.0279 | 0 | 1.928 | 4.3625 | 0 |
| 12849 | ENSMUSG00000043354 | 0 | NA | NA | NA | NA | NA | Olfr577 | olfactory receptor 577 [Source:MGI Symb | 0 | 0 | 0 | 0 | 0 | 0 |
| 12850 | ENSMUSG00000043357 | 0 | NA | NA | NA | NA | NA | Olfr187 | olfactory receptor 187 [Source:MGI Symb | 0 | 0 | 0 | 0 | 0 | 0 |
| 12851 | ENSMUSG00000043366 | 0 | NA | NA | NA | NA | NA | Olfr78 | olfactory receptor 78 [Source:MGI Symbo | 0 | 0 | 0 | 0 | 0 | 0 |
| 12852 | ENSMUSG00000043372 | 37.63446001 | 0.10889267 | 0.386439123 | 0.281784797 | 0.778108529 | 0.959559604 | Hexim2 | hexamethylene bis-acetamide inducible 2 | 26.502 | 43.173 | 39.244 | 39.523 | 28.356 | 49.009 |
| 12853 | ENSMUSG00000043383 | 0 | NA | NA | NA | NA | NA | Olfr1342 | olfactory receptor 1342 [Source:MGI Syml | 0 | 0 | 0 | 0 | 0 | 0 |
| 12854 | ENSMUSG00000043384 | 102.0787728 | 0.140991156 | 0.287314046 | 0.490721418 | 0.623623496 | 0.919503423 | Gprasp1 | G protein-coupled receptor associated so | 126.09 | 94.568 | 70.437 | 141.7 | 88.34 | 91.335 |
| 12855 | ENSMUSG00000043385 | 0 | NA | NA | NA | NA | NA | Olfr267 | olfactory receptor 267 [Source:MGI Symb | 0 | 0 | 0 | 0 | 0 | 0 |
| 12856 | ENSMUSG00000043388 | 0 | NA | NA | NA | NA | NA | Tmem130 | transmembrane protein 130 [Source:MGI | 0 | 0 | 0 | 0 | 0 | 0 |
| 12857 | ENSMUSG00000043391 | 101.7060713 | -0.371299047 | 0.261319626 | -1.420861694 | 0.155356971 | 0.574114264 | 2510009E07 | RIKEN cDNA 2510009E07 gene [Source:M | 118.06 | 119.24 | 106.66 | 83.866 | 64.346 | 118.07 |
| 12858 | ENSMUSG00000043398 | 44.06600306 | -0.744952247 | 0.384729212 | -1.936302791 | 0.052830627 | 0.32296796 | Gpr135 | G protein-coupled receptor 135 [Source:M | 60.233 | 50.368 | 55.343 | 51.091 | 25.084 | 22.277 |
| 12859 | ENSMUSG00000043410 | 0 | NA | NA | NA | NA | NA | Hfm1 | HFM1, ATP-dependent DNA helicase hom | 0 | 0 | 0 | 0 | 0 | 0 |
| 12860 | ENSMUSG00000043411 | 596.9343363 | 0.093344562 | 0.121734414 | 0.766788605 | 0.443207213 | 0.850769625 | Usp48 | ubiquitin specific peptidase 48 [Source:M | 542.09 | 591.05 | 600.73 | 650.69 | 591.12 | 605.93 |
| 12861 | ENSMUSG00000043415 | 179.8308187 | -0.32653026 | 0.185706818 | -1.758310559 | 0.078694682 | 0.411511165 | Otud1 | OTU domain containing 1 [Source:MGI Sy | 172.67 | 210.72 | 217.35 | 154.24 | 162.5 | 161.51 |
| 12862 | ENSMUSG00000043418 | 142.4349258 | 0.284004849 | 0.233482383 | 1.216386632 | 0.223837674 | 0.669614916 | Lrit2 | leucine-rich repeat, immunoglobulin-like a | 92.357 | 139.8 | 153.96 | 143.63 | 174.5 | 150.37 |
| 12863 | ENSMUSG00000043419 | 14.49712896 | -0.060750401 | 0.606432796 | -0.100176641 | 0.92020409 | 0.986685804 | Rnf227 | ring finger protein 227 [Source:MGI Symb | 12.85 | 16.447 | 15.094 | 11.568 | 7.6343 | 23.391 |
| 12864 | ENSMUSG00000043421 | 48.31170885 | -0.905974006 | 0.319401082 | -2.836477569 | 0.004561418 | 0.062894522 | Hilpda | hypoxia inducible lipid droplet associated | 61.036 | 59.619 | 68.425 | 34.703 | 34.9 | 31.188 |
| 12865 | ENSMUSG00000043424 | 15.17754997 | -0.085501583 | 0.609303172 | -0.140326831 | 0.888401766 | 0.982382091 | Eif3j2 | eukaryotic translation initiation factor 3, s | 11.243 | 18.503 | 17.106 | 7.7118 | 11.997 | 24.505 |
| 12866 | ENSMUSG00000043429 | 0 | NA | NA | NA | NA | NA | Ccdc185 | coiled-coil domain containing 185 [Source | 0 | 0 | 0 | 0 | 0 | 0 |
| 12867 | ENSMUSG00000043430 | 0 | NA | NA | NA | NA | NA | Psapl1 | prosaposin-like 1 [Source:MGI Symbol;Acc | 0 | 0 | 0 | 0 | 0 | 0 |
| 12868 | ENSMUSG00000043432 | 102.8834538 | -0.015007397 | 0.247945249 | -0.060527061 | 0.951735864 | 0.990883058 | Leng9 | leukocyte receptor cluster (LRC) member | 110.03 | 106.9 | 93.581 | 126.28 | 101.43 | 79.083 |
| 12869 | ENSMUSG00000043439 | 97.0497419 | 0.538312208 | 0.263945201 | 2.039484735 | 0.041401673 | 0.278916536 | Epop | elongin BC and polycomb repressive comp | 81.113 | 79.15 | 77.481 | 146.52 | 115.61 | 82.424 |
| 12870 | ENSMUSG00000043441 | 0 | NA | NA | NA | NA | NA | Gpr149 | G protein-coupled receptor 149 [Source:M | 0 | 0 | 0 | 0 | 0 | 0 |
| 12871 | ENSMUSG00000043445 | 426.5750733 | 0.092672899 | 0.134975914 | 0.686588413 | 0.492342132 | 0.876368691 | Pgp | phosphoglycolate phosphatase [Source:M | 427.25 | 408.08 | 403.5 | 481.99 | 423.16 | 415.46 |
| 12872 | ENSMUSG00000043448 | 15.78711833 | -0.217524868 | 0.583794668 | -0.372605095 | 0.70944238 | 0.944239215 | Gjc2 | gap junction protein, gamma 2 [Source:M | 27.305 | 11.307 | 12.075 | 15.424 | 16.359 | 12.252 |
| 12873 | ENSMUSG00000043453 | 0 | NA | NA | NA | NA | NA | Magea10 | MAGE family member A10 [Source:MGI Sy | 0 | 0 | 0 | 0 | 0 | 0 |
| 12874 | ENSMUSG00000043456 | 0 | NA | NA | NA | NA | NA | Zfp536 | zinc finger protein 536 [Source:MGI Symb | 0 | 0 | 0 | 0 | 0 | 0 |
| 12875 | ENSMUSG00000043458 | 0 | NA | NA | NA | NA | NA | Pcdhb12 | protocadherin beta 12 [Source:MGI Symb | 0 | 0 | 0 | 0 | 0 | 0 |
| 12876 | ENSMUSG00000043460 | 0 | NA | NA | NA | NA | NA | Elfn2 | leucine rich repeat and fibronectin type III | 0 | 0 | 0 | 0 | 0 | 0 |

| | | | | | | | | | | | | | | | |
|---|---|---|---|---|---|---|---|---|---|---|---|---|---|---|---|
| 12877 | ENSMUSG00000043461 | 0 | NA | NA | NA | NA | NA | Sptssb | serine palmitoyltransferase, small subunit | 0 | 0 | 0 | 0 | 0 | 0 |
| 12878 | ENSMUSG00000043463 | 0 | NA | NA | NA | NA | NA | Rab9b | RAB9B, member RAS oncogene family [So | 0 | 0 | 0 | 0 | 0 | 0 |
| 12879 | ENSMUSG00000043467 | 182.8515333 | -0.019062226 | 0.204656605 | -0.093142492 | 0.92579036 | 0.987546253 | Zbtb37 | zinc finger and BTB domain containing 37 | 187.93 | 175.77 | 188.17 | 158.09 | 231.21 | 155.94 |
| 12880 | ENSMUSG00000043468 | 0.513958919 | -2.394274821 | 3.756096013 | -0.637437066 | 0.523840191 | NA | Adam30 | a disintegrin and metallopeptidase domai | 0 | 3.0838 | 0 | 0 | 0 | 0 |
| 12881 | ENSMUSG00000043472 | 0 | NA | NA | NA | NA | NA | Lce3d | late cornified envelope 3D [Source:MGI Sy | 0 | 0 | 0 | 0 | 0 | 0 |
| 12882 | ENSMUSG00000043483 | 18.9748738 | -0.059463015 | 0.510511734 | -0.116477273 | 0.907274297 | 0.984526042 | Gm6863 | predicted gene 6863 [Source:MGI Symbol | 14.456 | 26.726 | 17.106 | 20.244 | 15.269 | 20.049 |
| 12883 | ENSMUSG00000043484 | 0 | NA | NA | NA | NA | NA | Gm5867 | predicted gene 5867 [Source:MGI Symbol | 0 | 0 | 0 | 0 | 0 | 0 |
| 12884 | ENSMUSG00000043485 | 0 | NA | NA | NA | NA | NA | Krt34 | keratin 34 [Source:MGI Symbol;Acc:MGI:1 | 0 | 0 | 0 | 0 | 0 | 0 |
| 12885 | ENSMUSG00000043487 | 41.96983086 | -0.401246665 | 0.361902294 | -1.10871545 | 0.267552955 | 0.72293778 | Acot6 | acyl-CoA thioesterase 6 [Source:MGI Syml | 61.839 | 35.977 | 45.281 | 41.451 | 31.628 | 35.643 |
| 12886 | ENSMUSG00000043488 | 1.700420866 | 0.570186631 | 1.819321624 | 0.313406175 | 0.753972094 | NA | Frmd8os | FERM domain containing 8, opposite strar | 4.0155 | 0 | 0 | 2.8919 | 2.1812 | 1.1138 |
| 12887 | ENSMUSG00000043496 | 4.803034428 | -0.369659881 | 0.968453835 | -0.381701087 | 0.702683094 | NA | Tril | TLR4 interactor with leucine-rich repeats [ | 8.031 | 4.1117 | 4.025 | 3.8559 | 5.4531 | 3.3415 |
| 12888 | ENSMUSG00000043498 | 0 | NA | NA | NA | NA | NA | NA | NA | 0 | 0 | 0 | 0 | 0 | 0 |
| 12889 | ENSMUSG00000043501 | 3.223228842 | 0.242624395 | 1.333569364 | 0.181936089 | 0.855632882 | NA | Lgals2 | lectin, galactose-binding, soluble 2 [Sourc | 0.8031 | 4.1117 | 4.025 | 3.8559 | 6.5437 | 0 |
| 12890 | ENSMUSG00000043505 | 9.030092946 | -0.626658478 | 0.80799523 | -0.775572002 | 0.438001737 | 0.847526036 | Gimap5 | GTPase, IMAP family member 5 [Source:N | 9.6372 | 14.391 | 9.0562 | 14.46 | 2.1812 | 4.4554 |
| 12891 | ENSMUSG00000043510 | 83.2668781 | 0.23233033 | 0.261050912 | 0.889980915 | 0.373476134 | 0.80865857 | Hscb | HscB iron-sulfur cluster co-chaperone [So | 85.932 | 64.759 | 78.487 | 74.226 | 97.065 | 99.132 |
| 12892 | ENSMUSG00000043518 | 42.48615536 | 0.21159591 | 0.395247599 | 0.535350274 | 0.592407669 | 0.912923013 | Rai2 | retinoic acid induced 2 [Source:MGI Symb | 36.943 | 28.782 | 52.325 | 39.523 | 31.628 | 65.717 |
| 12893 | ENSMUSG00000043522 | 0 | NA | NA | NA | NA | NA | A430060F13 | RIKEN cDNA A430060F13 gene [Source:M | 0 | 0 | 0 | 0 | 0 | 0 |
| 12894 | ENSMUSG00000043529 | 0 | NA | NA | NA | NA | NA | Olfr266 | olfactory receptor 266 [Source:MGI Symb | 0 | 0 | 0 | 0 | 0 | 0 |
| 12895 | ENSMUSG00000043531 | 0 | NA | NA | NA | NA | NA | Sorcs1 | sortilin-related VPS10 domain containing | 0 | 0 | 0 | 0 | 0 | 0 |
| 12896 | ENSMUSG00000043535 | 406.4773007 | 0.209685181 | 0.140435705 | 1.493104479 | 0.13540984 | 0.538068113 | Setx | senataxin [Source:MGI Symbol;Acc:MGI:2 | 392.72 | 392.66 | 345.14 | 427.04 | 406.8 | 474.5 |
| 12897 | ENSMUSG00000043537 | 0 | NA | NA | NA | NA | NA | Vmn1r225 | vomeronasal 1 receptor 225 [Source:MGI | 0 | 0 | 0 | 0 | 0 | 0 |
| 12898 | ENSMUSG00000043541 | 0.469265235 | -2.274672822 | 3.439794491 | -0.661281605 | 0.508431736 | NA | Casc1 | cancer susceptibility candidate 1 [Source: | 0.8031 | 0 | 2.0125 | 0 | 0 | 0 |
| 12899 | ENSMUSG00000043542 | 7.994084728 | -0.14398419 | 0.78332067 | -0.183812576 | 0.85416049 | 0.973825032 | Zc2hc1a | zinc finger, C2HC-type containing 1A [Sou | 12.85 | 8.2233 | 4.025 | 9.6398 | 6.5437 | 6.6831 |
| 12900 | ENSMUSG00000043549 | 0 | NA | NA | NA | NA | NA | Fam90a1b | family with sequence similarity 90, memb | 0 | 0 | 0 | 0 | 0 | 0 |
| 12901 | ENSMUSG00000043556 | 7.355905489 | -0.582844203 | 0.833899033 | -0.698938576 | 0.484590418 | 0.87256358 | Fbxl7 | F-box and leucine-rich repeat protein 7 [S | 11.243 | 5.1396 | 10.062 | 7.7118 | 2.1812 | 7.7969 |
| 12902 | ENSMUSG00000043557 | 1.045748203 | -3.439391613 | 2.321767015 | -1.481368109 | 0.138508508 | NA | Mdga1 | MAM domain containing glycosylphospha | 3.2124 | 2.0558 | 1.0062 | 0 | 0 | 0 |
| 12903 | ENSMUSG00000043569 | 0 | NA | NA | NA | NA | NA | Cldn34c4 | claudin 34C4 [Source:MGI Symbol;Acc:MC | 0 | 0 | 0 | 0 | 0 | 0 |
| 12904 | ENSMUSG00000043572 | 73.64081876 | 0.024012274 | 0.286420134 | 0.083835843 | 0.933186949 | 0.988420305 | Pars2 | prolyl-tRNA synthetase (mitochondrial)(pu | 73.082 | 65.787 | 80.5 | 91.578 | 77.434 | 53.465 |
| 12905 | ENSMUSG00000043583 | 0 | NA | NA | NA | NA | NA | 4930470P17 | RIKEN cDNA 4930470P17 gene [Source:M | 0 | 0 | 0 | 0 | 0 | 0 |
| 12906 | ENSMUSG00000043587 | 102.162579 | 0.472571194 | 0.246969655 | 1.913478774 | 0.055686788 | 0.332986361 | Pxylp1 | 2-phosphoxylose phosphatase 1 [Source:I | 67.461 | 83.261 | 106.66 | 126.28 | 111.24 | 118.07 |
| 12907 | ENSMUSG00000043592 | 0.17131964 | -0.796629176 | 4.080472857 | -0.195229623 | 0.84521319 | NA | Unc5cl | unc-5 family C-terminal like [Source:MGI S | 0 | 1.0279 | 0 | 0 | 0 | 0 |
| 12908 | ENSMUSG00000043602 | 38.75214146 | 0.108190335 | 0.349338646 | 0.309700446 | 0.756788763 | 0.953168989 | Zfp3 | zinc finger protein 3 [Source:MGI Symbol; | 42.564 | 35.977 | 33.206 | 41.451 | 41.443 | 37.871 |
| 12909 | ENSMUSG00000043605 | 0 | NA | NA | NA | NA | NA | Olfr13 | olfactory receptor 13 [Source:MGI Symbo | 0 | 0 | 0 | 0 | 0 | 0 |
| 12910 | ENSMUSG00000043613 | 0 | NA | NA | NA | NA | NA | Mmp3 | matrix metallopeptidase 3 [Source:MGI Sy | 0 | 0 | 0 | 0 | 0 | 0 |
| 12911 | ENSMUSG00000043614 | 7.960585158 | -0.208908436 | 0.769695622 | -0.271416947 | 0.786070369 | 0.960870081 | Vps37d | vacuolar protein sorting 37D [Source:MGI | 11.243 | 8.2233 | 6.0375 | 5.7839 | 10.906 | 5.5692 |
| 12912 | ENSMUSG00000043618 | 0.465832987 | -0.698295221 | 3.18348759 | -0.219349126 | 0.826378096 | NA | Eif5al3-ps | eukaryotic translation initiation factor 5A- | 0.8031 | 1.0279 | 0 | 0.964 | 0 | 0 |
| 12913 | ENSMUSG00000043621 | 1.718303603 | -0.415412219 | 1.696679303 | -0.244838384 | 0.806581572 | NA | Ubxn10 | UBX domain protein 10 [Source:MGI Syml | 0.8031 | 2.0558 | 3.0187 | 0 | 1.0906 | 3.3415 |
| 12914 | ENSMUSG00000043623 | 0.983276288 | -0.866716944 | 2.277998595 | -0.380472993 | 0.70359434 | NA | 4933412A08 | RIKEN cDNA 4933412A08 gene [Source:M | 0.8031 | 0 | 3.0187 | 0.964 | 0 | 1.1138 |
| 12915 | ENSMUSG00000043629 | 12.45883279 | 0.250305346 | 0.632295201 | 0.395867857 | 0.692202515 | 0.938256814 | 1700019D03 | RIKEN cDNA 1700019D03 gene [Source:M | 13.653 | 8.2233 | 12.075 | 7.7118 | 15.269 | 17.822 |
| 12916 | ENSMUSG00000043631 | 11.37279619 | -0.528770013 | 0.707499865 | -0.747378252 | 0.454835269 | 0.857337687 | Ecm2 | extracellular matrix protein 2, female orga | 16.062 | 5.1396 | 19.119 | 12.532 | 9.8156 | 5.5692 |
| 12917 | ENSMUSG00000043633 | 0.925960114 | -1.988843075 | 2.485090455 | -0.800310134 | 0.423531133 | NA | Fam221b | family with sequence similarity 221, meml | 2.4093 | 2.0558 | 0 | 0 | 1.0906 | 0 |
| 12918 | ENSMUSG00000043635 | 4.154718414 | -1.676813404 | 1.333365569 | -1.257579648 | 0.208543817 | NA | Adamts3 | a disintegrin-like and metallopeptidase (re | 4.8186 | 11.307 | 3.0187 | 5.7839 | 0 | 0 |
| 12919 | ENSMUSG00000043639 | 55.72560465 | 0.24283761 | 0.316532954 | 0.767179551 | 0.442974769 | 0.850769625 | Rbm20 | RNA binding motif protein 20 [Source:MG | 49.792 | 45.228 | 58.362 | 76.154 | 46.897 | 57.92 |
| 12920 | ENSMUSG00000043644 | 7.813155851 | -0.506141851 | 0.965399308 | -0.524282385 | 0.600082147 | 0.915217938 | 0610009L18I | RIKEN cDNA 0610009L18 gene [Source:M( | 16.062 | 9.2513 | 2.0125 | 2.8919 | 2.1812 | 14.48 |
| 12921 | ENSMUSG00000043648 | 24.08054055 | 0.766828894 | 0.466995268 | 1.642048532 | 0.100579947 | 0.46725495 | Pld6 | phospholipase D family, member 6 [Sourc | 16.865 | 18.503 | 18.112 | 27.955 | 20.722 | 42.326 |
| 12922 | ENSMUSG00000043659 | 0 | NA | NA | NA | NA | NA | Npsr1 | neuropeptide S receptor 1 [Source:MGI Sy | 0 | 0 | 0 | 0 | 0 | 0 |
| 12923 | ENSMUSG00000043661 | 1.249115476 | 0.348807902 | 2.051884581 | 0.16999392 | 0.865014918 | NA | Gm9785 | predicted pseudogene 9785 [Source:MGI | 3.2124 | 0 | 0 | 0.964 | 1.0906 | 2.2277 |
| 12924 | ENSMUSG00000043664 | 4.795659112 | -0.652165529 | 1.127006435 | -0.578670634 | 0.562811434 | NA | Tmem221 | transmembrane protein 221 [Source:MGI | 10.44 | 4.1117 | 3.0187 | 6.7478 | 0 | 4.4554 |
| 12925 | ENSMUSG00000043668 | 0.465832987 | -0.698295221 | 3.18348759 | -0.219349126 | 0.826378096 | NA | Tox3 | TOX high mobility group box family memb | 0.8031 | 1.0279 | 0 | 0.964 | 0 | 0 |
| 12926 | ENSMUSG00000043670 | 0 | NA | NA | NA | NA | NA | Diras1 | DIRAS family, GTP-binding RAS-like 1 [Sou | 0 | 0 | 0 | 0 | 0 | 0 |
| 12927 | ENSMUSG00000043671 | 277.4283967 | 1.028695602 | 0.163860686 | 6.277867049 | 3.43E-10 | 8.14E-08 | Dpy19l3 | dpy-19-like 3 (C. elegans) [Source:MGI Syr | 155.8 | 206.61 | 186.16 | 365.35 | 369.72 | 380.93 |
| 12928 | ENSMUSG00000043673 | 2.101946481 | -3.358108752 | 1.810171328 | -1.855133103 | 0.063577222 | NA | Kcns3 | potassium voltage-gated channel, delayec | 6.4248 | 3.0838 | 2.0125 | 0 | 1.0906 | 0 |
| 12929 | ENSMUSG00000043681 | 210.8118932 | 0.56254696 | 0.197676446 | 2.84579661 | 0.004430048 | 0.061582136 | Fam25c | family with sequence similarity 25, memb | 183.91 | 140.82 | 185.15 | 211.11 | 246.48 | 297.4 |
| 12930 | ENSMUSG00000043683 | 889.427801 | -0.050853287 | 0.108945851 | -0.466775801 | 0.640660291 | 0.923932929 | Fem1a | fem 1 homolog a [Source:MGI Symbol;Acc | 889.03 | 911.76 | 914.68 | 859.87 | 930.3 | 830.93 |
| 12931 | ENSMUSG00000043687 | 11.76599843 | -0.366823068 | 0.68730168 | -0.533714785 | 0.593538879 | 0.913011006 | 1190005I06F | RIKEN cDNA 1190005I06 gene [Source:M( | 19.274 | 6.1675 | 14.087 | 6.7478 | 8.7249 | 15.594 |
| 12932 | ENSMUSG00000043692 | 0 | NA | NA | NA | NA | NA | Olfr399 | olfactory receptor 399 [Source:MGI Symb | 0 | 0 | 0 | 0 | 0 | 0 |

| | | | | | | | | | | | | | | | |
|---|---|---|---|---|---|---|---|---|---|---|---|---|---|---|---|
| 12933 | ENSMUSG00000043698 | 0 | NA | NA | NA | NA | NA | Olfr62 | olfactory receptor 62 [Source:MGI Symbo | 0 | 0 | 0 | 0 | 0 | 0 |
| 12934 | ENSMUSG00000043702 | 251.8112509 | 0.0431901 | 0.161976569 | 0.266644124 | 0.789743179 | 0.961019028 | Pde12 | phosphodiesterase 12 [Source:MGI Symb | 230.49 | 279.59 | 234.45 | 251.6 | 255.2 | 259.53 |
| 12935 | ENSMUSG00000043705 | 0 | NA | NA | NA | NA | NA | Capn13 | calpain 13 [Source:MGI Symbol;Acc:MGI: | 0 | 0 | 0 | 0 | 0 | 0 |
| 12936 | ENSMUSG00000043715 | 0 | NA | NA | NA | NA | NA | Olfr1326-ps1 | olfactory receptor 1326, pseudogene 1 [S | 0 | 0 | 0 | 0 | 0 | 0 |
| 12937 | ENSMUSG00000043716 | 5105.956505 | -0.031264862 | 0.085433657 | -0.365954856 | 0.714398762 | 0.944595913 | Rpl7 | ribosomal protein L7 [Source:MGI Symbol | 4947.9 | 5365.7 | 5171.1 | 5067.6 | 4980.9 | 5102.5 |
| 12938 | ENSMUSG00000043719 | 8.523807518 | 0.065531899 | 0.762306636 | 0.085965274 | 0.931494022 | 0.988420305 | Col6a6 | collagen, type VI, alpha 6 [Source:MGI Syr | 13.653 | 4.1117 | 7.0437 | 8.6758 | 7.6343 | 10.025 |
| 12939 | ENSMUSG00000043727 | 0 | NA | NA | NA | NA | NA | F830045P16 | RIKEN cDNA F830045P16 gene [Source:M | 0 | 0 | 0 | 0 | 0 | 0 |
| 12940 | ENSMUSG00000043733 | 1350.643715 | 0.022591232 | 0.103674717 | 0.217904936 | 0.827503181 | 0.969794868 | Ptpn11 | protein tyrosine phosphatase, non-recept | 1276.9 | 1437 | 1307.1 | 1373.7 | 1400.4 | 1308.8 |
| 12941 | ENSMUSG00000043740 | 17.48704213 | -1.901258064 | 0.58374612 | -3.256994779 | 0.001125985 | 0.022809871 | B430306N03 | RIKEN cDNA B430306N03 gene [Source:M | 17.668 | 38.033 | 27.169 | 7.7118 | 6.5437 | 7.7969 |
| 12942 | ENSMUSG00000043747 | 0.835284868 | -0.623834031 | 2.861223713 | -0.218030498 | 0.827405349 | NA | 1520401A03 | RIKEN cDNA 1520401A03 gene [Source:M | 0 | 3.0838 | 0 | 1.928 | 0 | 0 |
| 12943 | ENSMUSG00000043753 | 10.69275012 | 0.372119919 | 0.688803131 | 0.540241329 | 0.589030614 | 0.911039598 | Dmrta1 | doublesex and mab-3 related transcriptio | 13.653 | 5.1396 | 9.0562 | 15.424 | 13.087 | 7.7969 |
| 12944 | ENSMUSG00000043760 | 99.92931367 | 0.048641345 | 0.252347791 | 0.192755183 | 0.847150704 | 0.972803041 | Pkhd1 | polycystic kidney and hepatic disease 1 [S | 115.65 | 77.094 | 101.63 | 106.04 | 114.51 | 84.652 |
| 12945 | ENSMUSG00000043770 | 0 | NA | NA | NA | NA | NA | Gm12481 | predicted gene 12481 [Source:MGI Symb | 0 | 0 | 0 | 0 | 0 | 0 |
| 12946 | ENSMUSG00000043773 | 6.703753548 | 0.6228343 | 0.817935664 | 0.761471015 | 0.446375785 | 0.852777619 | 1700048O20 | RIKEN cDNA 1700048O20 gene [Source:M | 5.6217 | 5.1396 | 5.0312 | 6.7478 | 6.5437 | 11.138 |
| 12947 | ENSMUSG00000043782 | 0.863771898 | -1.920093103 | 2.569535405 | -0.747253025 | 0.454910842 | NA | Bicdl2 | BICD family like cargo adaptor 2 [Source:N | 3.2124 | 0 | 1.0062 | 0.964 | 0 | 0 |
| 12948 | ENSMUSG00000043787 | 0 | NA | NA | NA | NA | NA | Defb12 | defensin beta 12 [Source:MGI Symbol;Acc | 0 | 0 | 0 | 0 | 0 | 0 |
| 12949 | ENSMUSG00000043789 | 701.9859117 | 0.269686697 | 0.161758774 | 1.667215261 | 0.095471609 | 0.455592661 | Vwce | von Willebrand factor C and EGF domains | 538.08 | 707.21 | 666.13 | 929.27 | 728.53 | 642.69 |
| 12950 | ENSMUSG00000043794 | 4.750139306 | 0.271104 | 0.983925484 | 0.275533061 | 0.782906729 | NA | D830025C05 | RIKEN cDNA D830025C05 gene [Source:M | 4.8186 | 2.0558 | 6.0375 | 6.7478 | 3.2719 | 5.5692 |
| 12951 | ENSMUSG00000043795 | 1.873628693 | -3.18161861 | 1.892492825 | -1.681178691 | 0.092728211 | NA | Prr33 | proline rich 33 [Source:MGI Symbol;Acc:N | 3.2124 | 1.0279 | 6.0375 | 0.964 | 0 | 0 |
| 12952 | ENSMUSG00000043800 | 0 | NA | NA | NA | NA | NA | Kifc5c-ps | kinesin family member C5C, pseudogene [ | 0 | 0 | 0 | 0 | 0 | 0 |
| 12953 | ENSMUSG00000043801 | 28.81107749 | 0.658115628 | 0.410368988 | 1.60371677 | 0.108776498 | 0.485684837 | Oaz1-ps | ornithine decarboxylase antizyme 1, pseu | 18.471 | 25.698 | 23.144 | 40.487 | 30.537 | 34.529 |
| 12954 | ENSMUSG00000043807 | 1.643397213 | -2.961425256 | 1.984843834 | -1.492019274 | 0.135694093 | NA | Ly6g5b | lymphocyte antigen 6 complex, locus G5B | 0.8031 | 2.0558 | 6.0375 | 0.964 | 0 | 0 |
| 12955 | ENSMUSG00000043811 | 0.185640647 | 1.126924175 | 4.080472857 | 0.276174898 | 0.782413736 | NA | Rtn4r | reticulon 4 receptor [Source:MGI Symbol; | 0 | 0 | 0 | 0 | 0 | 1.1138 |
| 12956 | ENSMUSG00000043822 | 17.06504425 | 0.256545529 | 0.524536356 | 0.489090081 | 0.624777923 | 0.919503423 | Adamtsl5 | ADAMTS-like 5 [Source:MGI Symbol;Acc:N | 11.243 | 18.503 | 17.106 | 20.244 | 16.359 | 18.935 |
| 12957 | ENSMUSG00000043827 | 0 | NA | NA | NA | NA | NA | Olfr94 | olfactory receptor 94 [Source:MGI Symbo | 0 | 0 | 0 | 0 | 0 | 0 |
| 12958 | ENSMUSG00000043831 | 159.1070136 | 0.25702298 | 0.205355306 | 1.251601363 | 0.210715158 | 0.654874096 | Lysmd4 | LysM, putative peptidoglycan-binding, do | 149.38 | 122.32 | 163.01 | 174.48 | 149.41 | 196.04 |
| 12959 | ENSMUSG00000043832 | 19.20569671 | -1.00503339 | 0.679519193 | -1.479036059 | 0.139130678 | 0.545066664 | Clec4a3 | C-type lectin domain family 4, member a3 | 10.44 | 55.508 | 11.069 | 10.604 | 10.906 | 16.708 |
| 12960 | ENSMUSG00000043833 | 54.5140049 | 0.335152621 | 0.354242427 | 0.9461109 | 0.344092025 | 0.785727752 | 2900005J15R | RIKEN cDNA 2900005J15 gene [Source:M | 51.399 | 37.005 | 56.35 | 79.046 | 66.528 | 36.757 |
| 12961 | ENSMUSG00000043843 | 0 | NA | NA | NA | NA | NA | Tmem145 | transmembrane protein 145 [Source:MGI | 0 | 0 | 0 | 0 | 0 | 0 |
| 12962 | ENSMUSG00000043850 | 0 | NA | NA | NA | NA | NA | Clrn1 | clarin 1 [Source:MGI Symbol;Acc:MGI:238 | 0 | 0 | 0 | 0 | 0 | 0 |
| 12963 | ENSMUSG00000043855 | 0 | NA | NA | NA | NA | NA | Olfr479 | olfactory receptor 479 [Source:MGI Symb | 0 | 0 | 0 | 0 | 0 | 0 |
| 12964 | ENSMUSG00000043857 | 0 | NA | NA | NA | NA | NA | Mgat5b | mannoside acetylglucosaminyltransferase | 0 | 0 | 0 | 0 | 0 | 0 |
| 12965 | ENSMUSG00000043859 | 0 | NA | NA | NA | NA | NA | 1700049L16R | RIKEN cDNA 1700049L16 gene [Source:M | 0 | 0 | 0 | 0 | 0 | 0 |
| 12966 | ENSMUSG00000043865 | 0 | NA | NA | NA | NA | NA | Tas2r118 | taste receptor, type 2, member 118 [Sour | 0 | 0 | 0 | 0 | 0 | 0 |
| 12967 | ENSMUSG00000043866 | 1105.346654 | 0.00257418 | 0.105124048 | 0.024487071 | 0.980464097 | 0.99820894 | Taf10 | TATA-box binding protein associated fact | 1143.6 | 1026.9 | 1142.1 | 1117.3 | 1090.6 | 1111.6 |
| 12968 | ENSMUSG00000043870 | 0.48903338 | 0.95637552 | 3.456690196 | 0.276673773 | 0.782030612 | NA | Gm5809 | predicted pseudogene 5809 [Source:MGI | 0 | 0 | 1.0062 | 1.928 | 0 | 0 |
| 12969 | ENSMUSG00000043872 | 25.05284612 | 0.211671926 | 0.437633186 | 0.483674302 | 0.628617034 | 0.920328819 | Zmym1 | zinc finger, MYM domain containing 1 [So | 20.881 | 21.586 | 27.169 | 21.208 | 31.628 | 27.846 |
| 12970 | ENSMUSG00000043873 | 0.301557804 | -1.64335916 | 4.042814367 | -0.406488899 | 0.684383412 | NA | Chil5 | chitinase-like 5 [Source:MGI Symbol;Acc:N | 0.8031 | 0 | 1.0062 | 0 | 0 | 0 |
| 12971 | ENSMUSG00000043880 | 0 | NA | NA | NA | NA | NA | Olfr323 | olfactory receptor 323 [Source:MGI Symb | 0 | 0 | 0 | 0 | 0 | 0 |
| 12972 | ENSMUSG00000043881 | 256.9846392 | -0.105640454 | 0.157623358 | -0.670208116 | 0.502725131 | 0.881220881 | Kbtbd7 | kelch repeat and BTB (POZ) domain conta | 260.21 | 268.29 | 270.68 | 236.17 | 272.65 | 233.91 |
| 12973 | ENSMUSG00000043885 | 157.7125872 | 0.128371297 | 0.186927999 | 0.686741943 | 0.492245361 | 0.876368691 | Slc36a4 | solute carrier family 36 (proton/amino aci | 159.01 | 151.1 | 141.88 | 173.52 | 158.14 | 162.62 |
| 12974 | ENSMUSG00000043889 | 0.822793274 | -0.447303882 | 2.407293551 | -0.185811939 | 0.852592242 | NA | Gm8399 | predicted gene 8399 [Source:MGI Symbol | 0.8031 | 2.0558 | 0 | 0.964 | 0 | 1.1138 |
| 12975 | ENSMUSG00000043892 | 0 | NA | NA | NA | NA | NA | Olfr1018 | olfactory receptor 1018 [Source:MGI Sym | 0 | 0 | 0 | 0 | 0 | 0 |
| 12976 | ENSMUSG00000043895 | 112.3103264 | 0.022258281 | 0.222482181 | 0.100045232 | 0.920308416 | 0.986685804 | S1pr2 | sphingosine-1-phosphate receptor 2 [Sou | 124.48 | 102.79 | 106.66 | 113.75 | 101.43 | 124.75 |
| 12977 | ENSMUSG00000043897 | 0 | NA | NA | NA | NA | NA | Vmn2r2 | vomeronasal 2, receptor 2 [Source:MGI S | 0 | 0 | 0 | 0 | 0 | 0 |
| 12978 | ENSMUSG00000043903 | 0.938530471 | -3.274726771 | 2.66105904 | -1.230610341 | 0.218468636 | NA | Zfp469 | zinc finger protein 469 [Source:MGI Symb | 1.6062 | 0 | 4.025 | 0 | 0 | 0 |
| 12979 | ENSMUSG00000043909 | 105.9888366 | 0.452882663 | 0.233709409 | 1.937802441 | 0.05264733 | 0.322665669 | Trp53bp1 | transformation related protein 53 binding | 82.72 | 79.15 | 106.66 | 113.75 | 118.88 | 134.78 |
| 12980 | ENSMUSG00000043911 | 0 | NA | NA | NA | NA | NA | Olfr922 | olfactory receptor 922 [Source:MGI Symb | 0 | 0 | 0 | 0 | 0 | 0 |
| 12981 | ENSMUSG00000043913 | 0 | NA | NA | NA | NA | NA | Ccdc60 | coiled-coil domain containing 60 [Source: | 0 | 0 | 0 | 0 | 0 | 0 |
| 12982 | ENSMUSG00000043923 | 219.0892791 | 0.032780101 | 0.24557524 | 0.133482925 | 0.893811468 | 0.982562372 | Ccdc84 | coiled-coil domain containing 84 [Source: | 234.51 | 202.5 | 213.32 | 322.93 | 188.68 | 152.6 |
| 12983 | ENSMUSG00000043924 | 12.78817768 | 0.404005011 | 0.64236575 | 0.628932989 | 0.529392928 | 0.888334265 | Ncmap | noncompact myelin associated protein [S | 14.456 | 12.335 | 6.0375 | 8.6758 | 19.631 | 15.594 |
| 12984 | ENSMUSG00000043925 | 0 | NA | NA | NA | NA | NA | Olfr544 | olfactory receptor 544 [Source:MGI Symb | 0 | 0 | 0 | 0 | 0 | 0 |
| 12985 | ENSMUSG00000043929 | 82.51132869 | -0.152175577 | 0.254280574 | -0.598455377 | 0.549536125 | 0.895703112 | Klhl15 | kelch-like 15 [Source:MGI Symbol;Acc:MG | 86.735 | 94.568 | 79.493 | 86.758 | 81.796 | 65.717 |
| 12986 | ENSMUSG00000043931 | 2.732236266 | -2.054537401 | 1.438243813 | -1.428504251 | 0.153146772 | NA | Gimap7 | GTPase, IMAP family member 7 [Source:N | 4.0155 | 6.1675 | 3.0187 | 0.964 | 0 | 2.2277 |
| 12987 | ENSMUSG00000043932 | 1.117074527 | -3.52147603 | 2.24739728 | -1.566913007 | 0.117135027 | NA | Klri2 | killer cell lectin-like receptor family I mem | 1.6062 | 3.0838 | 2.0125 | 0 | 0 | 0 |
| 12988 | ENSMUSG00000043939 | 0.906682787 | -0.583665192 | 2.768799895 | -0.210800785 | 0.833042726 | NA | A530064D06 | RIKEN cDNA A530064D06 gene [Source:M | 3.2124 | 0 | 0 | 0 | 0 | 2.2277 |

| | | | | | | | | | | | | | | | |
|---|---|---|---|---|---|---|---|---|---|---|---|---|---|---|---|
| 12989 | ENSMUSG00000043940 | 892.7116096 | -0.202352544 | 0.118839144 | -1.7027432 | 0.088616137 | 0.438652909 | Wdfy3 | WD repeat and FYVE domain containing 3 | 952.48 | 1043.3 | 870.4 | 879.15 | 823.42 | 787.49 |
| 12990 | ENSMUSG00000043943 | 21.66751424 | -0.555695646 | 0.50864301 | -1.09250621 | 0.274610663 | 0.729487777 | Naalad2 | N-acetylated alpha-linked acidic dipeptida | 27.305 | 34.949 | 15.094 | 16.388 | 22.903 | 13.366 |
| 12991 | ENSMUSG00000043945 | 0 | NA | NA | NA | NA | NA | Adam6a | a disintegrin and metallopeptidase domai | 0 | 0 | 0 | 0 | 0 | 0 |
| 12992 | ENSMUSG00000043948 | 0 | NA | NA | NA | NA | NA | Olfr691 | olfactory receptor 691 [Source:MGI Symb | 0 | 0 | 0 | 0 | 0 | 0 |
| 12993 | ENSMUSG00000043953 | 18.38159254 | -0.060721969 | 0.58598743 | -0.10362333 | 0.917468273 | 0.986374671 | Ccrl2 | chemokine (C-C motif) receptor-like 2 [Sou | 22.487 | 15.419 | 18.112 | 6.7478 | 17.45 | 30.074 |
| 12994 | ENSMUSG00000043962 | 1265.876912 | -0.023417067 | 0.103026256 | -0.227292228 | 0.820196516 | 0.969160361 | Thrap3 | thyroid hormone receptor associated prot | 1214.3 | 1317.8 | 1297 | 1243.5 | 1319.6 | 1203 |
| 12995 | ENSMUSG00000043964 | 334.7369584 | -0.012543726 | 0.154782742 | -0.08104086 | 0.935409458 | 0.988420305 | Orai3 | ORAI calcium release-activated calcium m | 312.41 | 325.85 | 370.3 | 301.73 | 325 | 373.14 |
| 12996 | ENSMUSG00000043969 | 0 | NA | NA | NA | NA | NA | Emx2 | empty spiracles homeobox 2 [Source:MGI | 0 | 0 | 0 | 0 | 0 | 0 |
| 12997 | ENSMUSG00000043972 | 0 | NA | NA | NA | NA | NA | Opn5 | opsin 5 [Source:MGI Symbol;Acc:MGI:266 | 0 | 0 | 0 | 0 | 0 | 0 |
| 12998 | ENSMUSG00000043982 | 0 | NA | NA | NA | NA | NA | Krtap19-4 | keratin associated protein 19-4 [Source:M | 0 | 0 | 0 | 0 | 0 | 0 |
| 12999 | ENSMUSG00000043986 | 0 | NA | NA | NA | NA | NA | Spata31d1d | spermatogenesis associated 31 subfamily | 0 | 0 | 0 | 0 | 0 | 0 |
| 13000 | ENSMUSG00000043987 | 255.5890094 | -0.104002796 | 0.159994054 | -0.650041635 | 0.515665328 | 0.886238519 | Cep164 | centrosomal protein 164 [Source:MGI Syn | 244.14 | 262.12 | 288.79 | 260.27 | 244.3 | 233.91 |
| 13001 | ENSMUSG00000043991 | 248.7814281 | -0.088543432 | 0.173018619 | -0.511756668 | 0.608821321 | 0.916376001 | Pura | purine rich element binding protein A [Sou | 232.1 | 247.73 | 289.8 | 241.96 | 212.67 | 268.44 |
| 13002 | ENSMUSG00000043993 | 112.6485902 | 0.334555121 | 0.273206407 | 1.224550784 | 0.220744521 | 0.667237604 | 2900052L18 | RIKEN cDNA 2900052L18 gene [Source:M | 100.39 | 71.954 | 126.79 | 157.13 | 94.884 | 124.75 |
| 13003 | ENSMUSG00000043998 | 1066.731807 | -0.082425868 | 0.108500213 | -0.759683925 | 0.44744354 | 0.85368993 | Mgat2 | mannoside acetylglucosaminyltransferase | 1036 | 1093.7 | 1162.2 | 1002.5 | 1009.9 | 1096 |
| 13004 | ENSMUSG00000043999 | 0.160662974 | 1.126924175 | 4.080472857 | 0.276174898 | 0.782413736 | NA | Gpr75 | G protein-coupled receptor 75 [Source:M | 0 | 0 | 0 | 0.964 | 0 | 0 |
| 13005 | ENSMUSG00000044005 | 7436.291701 | 0.148982817 | 0.118506089 | 1.257174359 | 0.208690506 | 0.653496315 | Gls2 | glutaminase 2 (liver, mitochondrial) [Sour | 6416.8 | 6584.8 | 8156.6 | 7048.6 | 8181.8 | 8229.1 |
| 13006 | ENSMUSG00000044006 | 0.614004368 | 0.315622926 | 2.898299649 | 0.108899342 | 0.913282328 | NA | Cilp2 | cartilage intermediate layer protein 2 [Sou | 1.6062 | 0 | 0 | 0.964 | 0 | 1.1138 |
| 13007 | ENSMUSG00000044014 | 0 | NA | NA | NA | NA | NA | Npy5r | neuropeptide Y receptor Y5 [Source:MGI S | 0 | 0 | 0 | 0 | 0 | 0 |
| 13008 | ENSMUSG00000044017 | 18.62851183 | -0.157633 | 0.678394452 | -0.232361865 | 0.816256966 | 0.967767553 | Adgrd1 | adhesion G protein-coupled receptor D1 [ | 27.305 | 18.503 | 13.081 | 21.208 | 29.447 | 2.2277 |
| 13009 | ENSMUSG00000044018 | 851.3730091 | 0.172371269 | 0.117686415 | 1.464665806 | 0.143012118 | 0.552624717 | Mrpl50 | mitochondrial ribosomal protein L50 [Sou | 779.01 | 824.39 | 797.95 | 806.85 | 949.93 | 950.11 |
| 13010 | ENSMUSG00000044021 | 0 | NA | NA | NA | NA | NA | NA | NA | 0 | 0 | 0 | 0 | 0 | 0 |
| 13011 | ENSMUSG00000044022 | 0 | NA | NA | NA | NA | NA | Pcdhb21 | protocadherin beta 21 [Source:MGI Symb | 0 | 0 | 0 | 0 | 0 | 0 |
| 13012 | ENSMUSG00000044024 | 3.254113452 | 0.008031834 | 1.201216531 | 0.006686417 | 0.994665051 | NA | Rell2 | RELT-like 2 [Source:MGI Symbol;Acc:MGI: | 1.6062 | 4.1117 | 4.025 | 0.964 | 4.3625 | 4.4554 |
| 13013 | ENSMUSG00000044025 | 0 | NA | NA | NA | NA | NA | Olfr790 | olfactory receptor 790 [Source:MGI Symb | 0 | 0 | 0 | 0 | 0 | 0 |
| 13014 | ENSMUSG00000044026 | 547.892702 | -0.906940091 | 0.171279989 | -5.295073281 | 1.19E-07 | 1.28E-05 | Slc35g1 | solute carrier family 35, member G1 [Sour | 679.42 | 902.51 | 562.49 | 398.12 | 382.81 | 362 |
| 13015 | ENSMUSG00000044029 | 0 | NA | NA | NA | NA | NA | Olfr178 | olfactory receptor 178 [Source:MGI Symb | 0 | 0 | 0 | 0 | 0 | 0 |
| 13016 | ENSMUSG00000044030 | 385.1442533 | 0.022605558 | 0.155263364 | 0.145594929 | 0.884241172 | 0.981769403 | Irf2bp1 | interferon regulatory factor 2 binding prot | 409.58 | 347.44 | 389.42 | 445.36 | 335.91 | 383.16 |
| 13017 | ENSMUSG00000044033 | 262.1703097 | -0.097663611 | 0.17536613 | -0.556912619 | 0.577587133 | 0.907877282 | Ccdc141 | coiled-coil domain containing 141 [Source | 279.48 | 264.17 | 269.67 | 287.27 | 201.76 | 270.66 |
| 13018 | ENSMUSG00000044034 | 2.293365652 | 3.655610874 | 1.784528181 | 2.048502743 | 0.040510764 | NA | Npb | neuropeptide B [Source:MGI Symbol;Acc: | 0 | 0 | 1.0062 | 2.8919 | 7.6343 | 2.2277 |
| 13019 | ENSMUSG00000044037 | 674.2825813 | -0.189096774 | 0.208199574 | -0.908247651 | 0.363747394 | 0.802534854 | Als2cl | ALS2 C-terminal like [Source:MGI Symbol; | 632.84 | 728.79 | 794.93 | 897.46 | 454.79 | 536.87 |
| 13020 | ENSMUSG00000044039 | 0 | NA | NA | NA | NA | NA | Olfr1288 | olfactory receptor 1288 [Source:MGI Sym | 0 | 0 | 0 | 0 | 0 | 0 |
| 13021 | ENSMUSG00000044040 | 0 | NA | NA | NA | NA | NA | Olfr1497 | olfactory receptor 1497 [Source:MGI Sym | 0 | 0 | 0 | 0 | 0 | 0 |
| 13022 | ENSMUSG00000044041 | 0 | NA | NA | NA | NA | NA | Krt13 | keratin 13 [Source:MGI Symbol;Acc:MGI:1 | 0 | 0 | 0 | 0 | 0 | 0 |
| 13023 | ENSMUSG00000044042 | 85.41544511 | 0.96494692 | 0.311866068 | 3.094106795 | 0.001974064 | 0.034525833 | Fmn1 | formin 1 [Source:MGI Symbol;Acc:MGI:10 | 65.051 | 76.066 | 32.2 | 93.506 | 127.6 | 118.07 |
| 13024 | ENSMUSG00000044043 | 2.814735578 | 0.67830071 | 1.384834885 | 0.489806198 | 0.624271045 | NA | Pcdhb14 | protocadherin beta 14 [Source:MGI Symb | 2.4093 | 3.0838 | 1.0062 | 4.8199 | 0 | 5.5692 |
| 13025 | ENSMUSG00000044052 | 6.898304957 | -0.322961594 | 0.903119374 | -0.357606761 | 0.720637618 | 0.9447719 | Ccr10 | chemokine (C-C motif) receptor 10 [Sourc | 12.85 | 2.0558 | 8.05 | 9.6398 | 5.4531 | 3.3415 |
| 13026 | ENSMUSG00000044055 | 0.133850373 | -0.796629176 | 4.080472857 | -0.195229623 | 0.84521319 | NA | Otos | otospiralin [Source:MGI Symbol;Acc:MGI: | 0.8031 | 0 | 0 | 0 | 0 | 0 |
| 13027 | ENSMUSG00000044056 | 1.299180075 | -2.610228666 | 2.254841034 | -1.157610947 | 0.24702284 | NA | Efcab9 | EF-hand calcium binding domain 9 [Sourc | 4.8186 | 0 | 2.0125 | 0.964 | 0 | 0 |
| 13028 | ENSMUSG00000044058 | 0 | NA | NA | NA | NA | NA | Mthfr-ps1 | 5,10-methylenetetrahydrofolate reductas | 0 | 0 | 0 | 0 | 0 | 0 |
| 13029 | ENSMUSG00000044061 | 0 | NA | NA | NA | NA | NA | Olfr225 | olfactory receptor 225 [Source:MGI Symb | 0 | 0 | 0 | 0 | 0 | 0 |
| 13030 | ENSMUSG00000044062 | 0 | NA | NA | NA | NA | NA | Plekhd1os | pleckstrin homology domain containing, f | 0 | 0 | 0 | 0 | 0 | 0 |
| 13031 | ENSMUSG00000044065 | 0 | NA | NA | NA | NA | NA | Gm9788 | predicted pseudogene 9788 [Source:MGI | 0 | 0 | 0 | 0 | 0 | 0 |
| 13032 | ENSMUSG00000044066 | 270.9401298 | 0.185120088 | 0.153237506 | 1.208059915 | 0.227024217 | 0.67343689 | Cep68 | centrosomal protein 68 [Source:MGI Sym | 249.76 | 242.59 | 268.67 | 304.62 | 273.75 | 286.26 |
| 13033 | ENSMUSG00000044067 | 1.059268063 | 0.100102635 | 2.627854962 | 0.038092907 | 0.969613606 | NA | Gpr22 | G protein-coupled receptor 22 [Source:M | 0 | 3.0838 | 0 | 0 | 3.2719 | 0 |
| 13034 | ENSMUSG00000044068 | 302.3463499 | -0.090823218 | 0.150696665 | -0.602688971 | 0.54671562 | 0.89549457 | Zrsr1 | zinc finger (CCCH type), RNA binding moti | 297.15 | 311.46 | 327.03 | 273.77 | 320.64 | 284.03 |
| 13035 | ENSMUSG00000044071 | 0.836675597 | -0.373395339 | 2.394418521 | -0.155944057 | 0.876077117 | NA | Tafa2 | TAFA chemokine like family member 2 [Sc | 0.8031 | 0 | 2.0125 | 0 | 1.0906 | 1.1138 |
| 13036 | ENSMUSG00000044072 | 15.32644841 | 1.090994749 | 0.630249479 | 1.731052203 | 0.083442454 | 0.425868616 | Eml6 | echinoderm microtubule associated prote | 16.865 | 4.1117 | 8.05 | 23.135 | 14.178 | 25.618 |
| 13037 | ENSMUSG00000044080 | 1648.218907 | -0.082297959 | 0.092371258 | -0.890947682 | 0.372957238 | 0.808400044 | S100a1 | S100 calcium binding protein A1 [Source:I | 1697 | 1705.3 | 1683.4 | 1607.9 | 1618.5 | 1577.2 |
| 13038 | ENSMUSG00000044081 | 8.766707232 | 0.285042582 | 0.7303036 | 0.390306966 | 0.696309573 | 0.939992732 | Zfp85os | zinc finger protein 85, opposite strand [So | 6.4248 | 8.2233 | 9.0562 | 5.7839 | 13.087 | 10.025 |
| 13039 | ENSMUSG00000044083 | 0 | NA | NA | NA | NA | NA | Efcab8 | EF-hand calcium binding domain 8 [Sourc | 0 | 0 | 0 | 0 | 0 | 0 |
| 13040 | ENSMUSG00000044084 | 0 | NA | NA | NA | NA | NA | Spem2 | SPEM family member 2 [Source:MGI Symb | 0 | 0 | 0 | 0 | 0 | 0 |
| 13041 | ENSMUSG00000044086 | 4.235068382 | 0.703746856 | 1.069975183 | 0.657722597 | 0.5107164 | NA | Lmod3 | leiomodin 3 (fetal) [Source:MGI Symbol;A | 1.6062 | 5.1396 | 3.0187 | 5.7839 | 7.6343 | 2.2277 |
| 13042 | ENSMUSG00000044092 | 3.73597402 | -1.316841996 | 1.154304592 | -1.140809804 | 0.25394908 | NA | C130050O18 | RIKEN cDNA C130050O18 gene [Source:M | 4.8186 | 3.0838 | 8.05 | 0.964 | 3.2719 | 2.2277 |
| 13043 | ENSMUSG00000044098 | 108.835848 | 0.012225441 | 0.262634865 | 0.046549191 | 0.962872527 | 0.992956886 | Rsbn1 | rosbin, round spermatid basic protein 1 [S | 86.735 | 132.6 | 106.66 | 139.78 | 99.246 | 87.994 |
| 13044 | ENSMUSG00000044103 | 2.077790485 | 0.05279525 | 1.623176971 | 0.032525874 | 0.974052682 | NA | Il1f9 | interleukin 1 family, member 9 [Source:M | 4.0155 | 1.0279 | 1.0062 | 0.964 | 5.4531 | 0 |

| | | | | | | | | | | | | | | | |
|---|---|---|---|---|---|---|---|---|---|---|---|---|---|---|---|
| 13045 | ENSMUSG00000044106 | 0 | NA | NA | NA | NA | NA | Olfr868 | olfactory receptor 868 [Source:MGI Symb | 0 | 0 | 0 | 0 | 0 | 0 |
| 13046 | ENSMUSG00000044117 | 10.08165528 | -0.034906851 | 0.692006128 | -0.050442979 | 0.959769388 | 0.992148164 | Bmerb1 | bMERB domain containing 1 [Source:MGI | 7.2279 | 13.363 | 10.062 | 6.7478 | 14.178 | 8.9108 |
| 13047 | ENSMUSG00000044120 | 0 | NA | NA | NA | NA | NA | Olfr683 | olfactory receptor 683 [Source:MGI Symb | 0 | 0 | 0 | 0 | 0 | 0 |
| 13048 | ENSMUSG00000044121 | 0 | NA | NA | NA | NA | NA | 5430402E10 | RIKEN cDNA 5430402E10 gene [Source:M | 0 | 0 | 0 | 0 | 0 | 0 |
| 13049 | ENSMUSG00000044122 | 59.70147 | -0.04553487 | 0.43715752 | -0.104161242 | 0.917041391 | 0.986374671 | Proca1 | protein interacting with cyclin A1 [Source: | 82.72 | 75.038 | 24.15 | 82.902 | 59.984 | 33.415 |
| 13050 | ENSMUSG00000044125 | 10.45519493 | -0.163522409 | 0.719969434 | -0.227124099 | 0.820327246 | 0.969160361 | 9530080O11 | RIKEN cDNA 9530080O11 gene [Source:M | 16.865 | 7.1954 | 9.0562 | 16.388 | 6.5437 | 6.6831 |
| 13051 | ENSMUSG00000044134 | 190.7299604 | 0.736641268 | 0.218765953 | 3.367257364 | 0.000759198 | 0.017051304 | Pheta1 | PH domain containing endocytic traffickin | 130.1 | 126.43 | 173.07 | 264.13 | 178.86 | 271.78 |
| 13052 | ENSMUSG00000044139 | 18.18081077 | -0.118701453 | 0.652480186 | -0.18192346 | 0.855642793 | 0.974158771 | Prss53 | protease, serine 53 [Source:MGI Symbol;/ | 18.471 | 4.1117 | 34.212 | 20.244 | 11.997 | 20.049 |
| 13053 | ENSMUSG00000044141 | 0 | NA | NA | NA | NA | NA | E130201H02 | RIKEN cDNA E130201H02 gene [Source:M | 0 | 0 | 0 | 0 | 0 | 0 |
| 13054 | ENSMUSG00000044145 | 25.17811615 | 0.21063422 | 0.451633132 | 0.466383455 | 0.640941052 | 0.923932929 | 1810024B03 | RIKEN cDNA 1810024B03 gene [Source:M | 19.274 | 28.782 | 22.137 | 28.919 | 18.541 | 33.415 |
| 13055 | ENSMUSG00000044147 | 554.2505567 | 0.05589947 | 0.124988285 | 0.447237675 | 0.654703462 | 0.925797231 | Arf6 | ADP-ribosylation factor 6 [Source:MGI Syr | 521.21 | 575.63 | 534.32 | 586.1 | 524.59 | 583.65 |
| 13056 | ENSMUSG00000044148 | 121.5135267 | -0.271573363 | 0.226864226 | -1.197074426 | 0.231277547 | 0.678442632 | 1810030O07 | RIKEN cDNA 1810030O07 gene [Source:M | 152.59 | 108.96 | 136.85 | 116.64 | 95.974 | 118.07 |
| 13057 | ENSMUSG00000044149 | 70.44529716 | -0.060123994 | 0.28740359 | -0.209197086 | 0.834294387 | 0.971073158 | Nkrf | NF-kappaB repressing factor [Source:MGI | 85.932 | 71.954 | 57.356 | 58.803 | 81.796 | 66.831 |
| 13058 | ENSMUSG00000044150 | 40.87944309 | 0.092468012 | 0.351732878 | 0.262892718 | 0.792633269 | 0.961557293 | Bclaf3 | Bclaf1 and Thrap3 family member 3 [Sour | 48.186 | 33.921 | 36.225 | 36.631 | 47.987 | 42.326 |
| 13059 | ENSMUSG00000044155 | 123.1687765 | -0.00226084 | 0.208720752 | -0.010831891 | 0.991357571 | 0.99859176 | Lsm8 | LSM8 homolog, U6 small nuclear RNA ass | 127.69 | 116.15 | 125.78 | 123.39 | 112.33 | 133.66 |
| 13060 | ENSMUSG00000044156 | 1.627473372 | -1.870384392 | 1.7879862 | -1.046084356 | 0.295522088 | NA | Hepacam2 | HEPACAM family member 2 [Source:MGI : | 1.6062 | 2.0558 | 4.025 | 0.964 | 0 | 1.1138 |
| 13061 | ENSMUSG00000044162 | 0.267700746 | -1.497481649 | 4.057001436 | -0.369110456 | 0.712045396 | NA | Tnip3 | TNFAIP3 interacting protein 3 [Source:MG | 1.6062 | 0 | 0 | 0 | 0 | 0 |
| 13062 | ENSMUSG00000044164 | 0 | NA | NA | NA | NA | NA | Rnf182 | ring finger protein 182 [Source:MGI Symb | 0 | 0 | 0 | 0 | 0 | 0 |
| 13063 | ENSMUSG00000044165 | 0.998650538 | 1.362383175 | 2.445753587 | 0.557040244 | 0.577499933 | NA | Bcl2l15 | BCLl2-like 15 [Source:MGI Symbol;Acc:MG | 1.6062 | 0 | 0 | 0 | 3.2719 | 1.1138 |
| 13064 | ENSMUSG00000044167 | 901.9099436 | -0.596672109 | 0.13714632 | -4.350624282 | 1.36E-05 | 0.000691371 | Foxo1 | forkhead box O1 [Source:MGI Symbol;Acc | 1088.2 | 1157.4 | 1011.3 | 640.08 | 853.95 | 660.51 |
| 13065 | ENSMUSG00000044170 | 0 | NA | NA | NA | NA | NA | Olfr1384 | olfactory receptor 1384 [Source:MGI Sym | 0 | 0 | 0 | 0 | 0 | 0 |
| 13066 | ENSMUSG00000044172 | 0.17131964 | -0.796629176 | 4.080472857 | -0.195229623 | 0.84521319 | NA | Ptx4 | pentraxin 4 [Source:MGI Symbol;Acc:MGI | 0 | 1.0279 | 0 | 0 | 0 | 0 |
| 13067 | ENSMUSG00000044176 | 0.698954292 | 0.126053087 | 3.048972385 | 0.04134281 | 0.967022605 | NA | Spink10 | serine peptidase inhibitor, Kazal type 10 [! | 0 | 0 | 2.0125 | 0 | 2.1812 | 0 |
| 13068 | ENSMUSG00000044177 | 2.242092297 | -1.082932787 | 1.626413984 | -0.665840799 | 0.505512864 | NA | Wfikkn2 | WAP, follistatin/kazal, immunoglobulin, ku | 4.0155 | 2.0558 | 3.0187 | 0 | 4.3625 | 0 |
| 13069 | ENSMUSG00000044186 | 11.31454321 | 0.266914608 | 0.636293044 | 0.419483775 | 0.67486261 | 0.932315978 | Nkx2-6 | NK2 homeobox 6 [Source:MGI Symbol;Ac | 10.44 | 9.2513 | 11.069 | 10.604 | 9.8156 | 16.708 |
| 13070 | ENSMUSG00000044197 | 1975.502227 | 1.142401364 | 0.150385919 | 7.596464968 | 3.04E-14 | 1.63E-11 | Gpr146 | G protein-coupled receptor 146 [Source:N | 1441.6 | 1057.7 | 1194.4 | 2660.6 | 2264.1 | 3234.6 |
| 13071 | ENSMUSG00000044199 | 3.919965213 | -1.143647206 | 1.099582869 | -1.040073685 | 0.298305669 | NA | S1pr4 | sphingosine-1-phosphate receptor 4 [Sou | 4.0155 | 7.1954 | 5.0312 | 2.8919 | 3.2719 | 1.1138 |
| 13072 | ENSMUSG00000044201 | 0.729921525 | -1.61423457 | 2.706831497 | -0.596355766 | 0.550937584 | NA | Cdc25c | cell division cycle 25C [Source:MGI Symbo | 2.4093 | 0 | 1.0062 | 0.964 | 0 | 0 |
| 13073 | ENSMUSG00000044205 | 0 | NA | NA | NA | NA | NA | Olfr983 | olfactory receptor 983 [Source:MGI Symb | 0 | 0 | 0 | 0 | 0 | 0 |
| 13074 | ENSMUSG00000044206 | 185.0584051 | -1.690324775 | 0.275346433 | -6.13890202 | 8.31E-10 | 1.77E-07 | Vsig4 | V-set and immunoglobulin domain contai | 246.55 | 385.47 | 215.34 | 68.442 | 69.8 | 124.75 |
| 13075 | ENSMUSG00000044211 | 2.662687751 | 0.601091865 | 1.347945625 | 0.445931834 | 0.655646486 | NA | Gm7887 | predicted gene 7887 [Source:MGI Symbol | 3.2124 | 2.0558 | 1.0062 | 1.928 | 1.0906 | 6.6831 |
| 13076 | ENSMUSG00000044213 | 0 | NA | NA | NA | NA | NA | Olfr1094 | olfactory receptor 1094 [Source:MGI Sym | 0 | 0 | 0 | 0 | 0 | 0 |
| 13077 | ENSMUSG00000044216 | 0 | NA | NA | NA | NA | NA | Kcnj4 | potassium inwardly-rectifying channel, su | 0 | 0 | 0 | 0 | 0 | 0 |
| 13078 | ENSMUSG00000044217 | 0.371281295 | 2.09253362 | 4.031852165 | 0.519000582 | 0.603760335 | NA | Aqp5 | aquaporin 5 [Source:MGI Symbol;Acc:MG | 0 | 0 | 0 | 0 | 0 | 2.2277 |
| 13079 | ENSMUSG00000044220 | 0 | NA | NA | NA | NA | NA | Nkx2-3 | NK2 homeobox 3 [Source:MGI Symbol;Ac | 0 | 0 | 0 | 0 | 0 | 0 |
| 13080 | ENSMUSG00000044221 | 2165.709523 | 0.122280826 | 0.102839578 | 1.189044423 | 0.234422187 | 0.682322842 | Grsf1 | G-rich RNA sequence binding factor 1 [Sou | 2024.6 | 2175.1 | 2022.6 | 2199.8 | 2463.7 | 2108.5 |
| 13081 | ENSMUSG00000044222 | 0 | NA | NA | NA | NA | NA | Defb13 | defensin beta 13 [Source:MGI Symbol;Acc | 0 | 0 | 0 | 0 | 0 | 0 |
| 13082 | ENSMUSG00000044224 | 480.4178161 | -0.008280471 | 0.158196686 | -0.052342886 | 0.958255482 | 0.991891687 | Dnajc21 | DnaJ heat shock protein family (Hsp40) m | 382.28 | 559.19 | 505.13 | 472.35 | 466.78 | 496.77 |
| 13083 | ENSMUSG00000044227 | 0 | NA | NA | NA | NA | NA | Gm9789 | predicted gene 9789 [Source:MGI Symbol | 0 | 0 | 0 | 0 | 0 | 0 |
| 13084 | ENSMUSG00000044229 | 6.947696463 | -0.452197408 | 0.875813285 | -0.516317137 | 0.605632929 | 0.916376001 | Nxpe4 | neurexophilin and PC-esterase domain far | 8.8341 | 9.2513 | 6.0375 | 8.6758 | 1.0906 | 7.7969 |
| 13085 | ENSMUSG00000044231 | 46.83430466 | 0.08160822 | 0.324115485 | 0.251787475 | 0.801205339 | 0.96255706 | Nhlrc1 | NHL repeat containing 1 [Source:MGI Sym | 53.005 | 39.061 | 44.275 | 52.055 | 44.715 | 47.895 |
| 13086 | ENSMUSG00000044243 | 0.695728634 | -1.425610641 | 3.05889997 | -0.46605337 | 0.641177299 | NA | Bhlha9 | basic helix-loop-helix family, member a9 [ | 0 | 3.0838 | 0 | 0 | 1.0906 | 0 |
| 13087 | ENSMUSG00000044244 | 10.16485698 | -0.007851552 | 0.6980707 | -0.011247502 | 0.991025981 | 0.99859176 | Il20rb | interleukin 20 receptor beta [Source:MGI | 7.2279 | 16.447 | 7.0437 | 11.568 | 10.906 | 7.7969 |
| 13088 | ENSMUSG00000044248 | 0 | NA | NA | NA | NA | NA | Vmn1r45 | vomeronasal 1 receptor 45 [Source:MGI S | 0 | 0 | 0 | 0 | 0 | 0 |
| 13089 | ENSMUSG00000044249 | 0 | NA | NA | NA | NA | NA | Defb29 | defensin beta 29 [Source:MGI Symbol;Acc | 0 | 0 | 0 | 0 | 0 | 0 |
| 13090 | ENSMUSG00000044250 | 51.25171109 | -0.433244283 | 0.340197418 | -1.27350844 | 0.202837679 | 0.647682949 | Pced1b | PC-esterase domain containing 1B [Source | 57.823 | 69.898 | 49.306 | 58.803 | 33.809 | 37.871 |
| 13091 | ENSMUSG00000044252 | 2843.784558 | 0.191372785 | 0.101123914 | 1.892458242 | 0.058429952 | 0.343085466 | Osbpl1a | oxysterol binding protein-like 1A [Source:I | 2853.4 | 2548.2 | 2562.9 | 2826.4 | 3046.1 | 3225.7 |
| 13092 | ENSMUSG00000044254 | 277.1793839 | 1.335681141 | 0.203660256 | 6.558378981 | 5.44E-11 | 1.54E-08 | Pcsk9 | proprotein convertase subtilisin/kexin typ | 189.53 | 175.77 | 105.66 | 377.88 | 417.71 | 396.53 |
| 13093 | ENSMUSG00000044258 | 19.5691414 | -0.489540347 | 0.494428358 | -0.990113813 | 0.322118493 | 0.769593967 | Ctla2a | cytotoxic T lymphocyte-associated proteir | 24.896 | 23.642 | 20.125 | 21.208 | 14.178 | 13.366 |
| 13094 | ENSMUSG00000044265 | 0 | NA | NA | NA | NA | NA | Olfm5 | olfactomedin 5 [Source:MGI Symbol;Acc:I | 0 | 0 | 0 | 0 | 0 | 0 |
| 13095 | ENSMUSG00000044268 | 1.359513901 | -1.534879831 | 2.048575836 | -0.749242378 | 0.453711131 | NA | Gm4895 | predicted gene 4895 [Source:MGI Symbol | 0 | 3.0838 | 3.0187 | 0.964 | 1.0906 | 0 |
| 13096 | ENSMUSG00000044276 | 0.294513347 | 0.165143496 | 4.080472857 | 0.040471657 | 0.967717103 | NA | 4933427E11 | RIKEN cDNA 4933427E11 gene [Source:M | 0.8031 | 0 | 0 | 0.964 | 0 | 0 |
| 13097 | ENSMUSG00000044279 | 240.7519562 | 0.433440404 | 0.179782206 | 2.410919379 | 0.015912365 | 0.148296684 | Crb3 | crumbs family member 3 [Source:MGI Syr | 220.05 | 178.86 | 215.34 | 294.98 | 231.21 | 304.08 |
| 13098 | ENSMUSG00000044285 | 27.71728338 | 0.210838083 | 0.555737402 | 0.379384369 | 0.704402456 | 0.94295036 | Ubb-ps | ubiquitin B, pseudogene [Source:MGI Sym | 47.383 | 10.279 | 19.119 | 44.343 | 22.903 | 22.277 |
| 13099 | ENSMUSG00000044286 | 0 | NA | NA | NA | NA | NA | Olfr221 | olfactory receptor 221 [Source:MGI Symb | 0 | 0 | 0 | 0 | 0 | 0 |
| 13100 | ENSMUSG00000044287 | 0 | NA | NA | NA | NA | NA | Nrn1l | neuritin 1-like [Source:MGI Symbol;Acc:M | 0 | 0 | 0 | 0 | 0 | 0 |

| | | | | | | | | | | | | | | | |
|---|---|---|---|---|---|---|---|---|---|---|---|---|---|---|---|
| 13101 | ENSMUSG00000044288 | 0 | NA | NA | NA | NA | NA | Cnr1 | cannabinoid receptor 1 (brain) [Source:M | 0 | 0 | 0 | 0 | 0 | 0 |
| 13102 | ENSMUSG00000044292 | 0 | NA | NA | NA | NA | NA | Olfr978 | olfactory receptor 978 [Source:MGI Symb | 0 | 0 | 0 | 0 | 0 | 0 |
| 13103 | ENSMUSG00000044293 | 0 | NA | NA | NA | NA | NA | Olfr794 | olfactory receptor 794 [Source:MGI Symb | 0 | 0 | 0 | 0 | 0 | 0 |
| 13104 | ENSMUSG00000044294 | 0 | NA | NA | NA | NA | NA | Krt84 | keratin 84 [Source:MGI Symbol;Acc:MGI:9 | 0 | 0 | 0 | 0 | 0 | 0 |
| 13105 | ENSMUSG00000044296 | 6.459976476 | -0.717173528 | 0.896247141 | -0.800196168 | 0.42359715 | 0.84018348 | Zfp879 | zinc finger protein 879 [Source:MGI Symb | 8.8341 | 8.2233 | 7.0437 | 4.8199 | 8.7249 | 1.1138 |
| 13106 | ENSMUSG00000044303 | 1.61446433 | -1.805419496 | 1.867585336 | -0.966713253 | 0.3336874 | NA | Cdkn2a | cyclin dependent kinase inhibitor 2A [Sou | 2.4093 | 3.0838 | 2.0125 | 0 | 2.1812 | 0 |
| 13107 | ENSMUSG00000044306 | 0 | NA | NA | NA | NA | NA | 4930500M09 | RIKEN cDNA 4930500M09 gene [Source:M | 0 | 0 | 0 | 0 | 0 | 0 |
| 13108 | ENSMUSG00000044308 | 2390.748746 | 0.023580182 | 0.103671137 | 0.22745176 | 0.820072476 | 0.969160361 | Ubr3 | ubiquitin protein ligase E3 component n-r | 2255.9 | 2496.8 | 2361.7 | 2383.9 | 2621.8 | 2224.3 |
| 13109 | ENSMUSG00000044309 | 0.935098222 | -2.037664976 | 2.285197225 | -0.891680138 | 0.372564403 | NA | Apol7c | apolipoprotein L 7c [Source:MGI Symbol;A | 1.6062 | 1.0279 | 2.0125 | 0.964 | 0 | 0 |
| 13110 | ENSMUSG00000044312 | 0 | NA | NA | NA | NA | NA | Neurog3 | neurogenin 3 [Source:MGI Symbol;Acc:M | 0 | 0 | 0 | 0 | 0 | 0 |
| 13111 | ENSMUSG00000044313 | 1.68083985 | -0.568684001 | 1.835637312 | -0.30980194 | 0.756711576 | NA | Mab21l3 | mab-21-like 3 [Source:MGI Symbol;Acc:M | 0 | 3.0838 | 3.0187 | 2.8919 | 1.0906 | 0 |
| 13112 | ENSMUSG00000044317 | 11.91379974 | -1.148034799 | 0.753365781 | -1.523874362 | 0.127540099 | 0.523285632 | Gpr4 | G protein-coupled receptor 4 [Source:MG | 9.6372 | 20.558 | 19.119 | 7.7118 | 1.0906 | 13.366 |
| 13113 | ENSMUSG00000044320 | 0 | NA | NA | NA | NA | NA | 1700001O22 | RIKEN cDNA 1700001O22 gene [Source:M | 0 | 0 | 0 | 0 | 0 | 0 |
| 13114 | ENSMUSG00000044322 | 0.301557804 | -1.64335916 | 4.042814367 | -0.406488899 | 0.684383412 | NA | Dsc1 | desmocollin 1 [Source:MGI Symbol;Acc:M | 0.8031 | 0 | 1.0062 | 0 | 0 | 0 |
| 13115 | ENSMUSG00000044328 | 82.64890949 | 0.177385215 | 0.277872081 | 0.638370054 | 0.52323282 | 0.888334265 | Trp53i13 | transformation related protein 53 inducib | 98.782 | 69.898 | 63.393 | 83.866 | 75.253 | 104.7 |
| 13116 | ENSMUSG00000044330 | 3.581837156 | 0.896889782 | 1.204266697 | 0.744760097 | 0.456416761 | NA | Gm9790 | predicted gene 9790 [Source:MGI Symbol | 2.4093 | 2.0558 | 3.0187 | 2.8919 | 1.0906 | 10.025 |
| 13117 | ENSMUSG00000044337 | 14.88666129 | -0.299760748 | 0.688525739 | -0.435366075 | 0.663296731 | 0.9291936 | Ackr3 | atypical chemokine receptor 3 [Source:M | 21.684 | 22.614 | 5.0312 | 20.244 | 14.178 | 5.5692 |
| 13118 | ENSMUSG00000044338 | 0.729921525 | -1.61423457 | 2.706831497 | -0.596355766 | 0.550937584 | NA | Aplnr | apelin receptor [Source:MGI Symbol;Acc:I | 2.4093 | 0 | 1.0062 | 0.964 | 0 | 0 |
| 13119 | ENSMUSG00000044339 | 64.3126562 | 0.323511004 | 0.273958109 | 1.180877634 | 0.237651337 | 0.686377492 | Alkbh2 | alkB homolog 2, alpha-ketoglutarate-depe | 59.43 | 56.535 | 55.343 | 72.298 | 66.528 | 75.741 |
| 13120 | ENSMUSG00000044340 | 468.4551494 | 0.026433396 | 0.128093342 | 0.206360425 | 0.836509382 | 0.971073158 | Phlpp1 | PH domain and leucine rich repeat protein | 440.9 | 474.9 | 476.96 | 459.82 | 459.15 | 499 |
| 13121 | ENSMUSG00000044345 | 1468.358427 | -0.136475922 | 0.124714143 | -1.09430991 | 0.27381908 | 0.728923893 | Marveld1 | MARVEL (membrane-associating) domain | 1472.9 | 1462.7 | 1678.4 | 1464.3 | 1536.7 | 1195.2 |
| 13122 | ENSMUSG00000044348 | 32.31894566 | -0.186375419 | 0.391203695 | -0.476415282 | 0.63377855 | 0.922787693 | Slc25a53 | solute carrier family 25, member 53 [Sour | 31.321 | 32.893 | 39.244 | 37.595 | 28.356 | 24.505 |
| 13123 | ENSMUSG00000044349 | 66.81638873 | 0.360187428 | 0.352953111 | 1.020496539 | 0.30749303 | 0.756343859 | Snhg11 | small nucleolar RNA host gene 11 [Source | 83.523 | 43.173 | 48.3 | 73.262 | 102.52 | 50.123 |
| 13124 | ENSMUSG00000044350 | 95.24818836 | -0.146932248 | 0.279204823 | -0.526252541 | 0.598712753 | 0.915055389 | Lacc1 | laccase domain containing 1 [Source:MGI | 124.48 | 107.93 | 67.418 | 90.614 | 76.343 | 104.7 |
| 13125 | ENSMUSG00000044352 | 5.316342558 | -0.353176298 | 1.030459586 | -0.342736681 | 0.731796565 | 0.947387358 | Sowaha | sosondowah ankyrin repeat domain famil | 9.6372 | 2.0558 | 6.0375 | 0.964 | 7.6343 | 5.5692 |
| 13126 | ENSMUSG00000044359 | 166.6520736 | 0.165861183 | 0.215490616 | 0.769690977 | 0.441483224 | 0.849828766 | P2ry4 | pyrimidinergic receptor P2Y, G-protein co | 120.47 | 176.8 | 175.09 | 202.44 | 162.5 | 162.62 |
| 13127 | ENSMUSG00000044361 | 258.8530221 | -0.003439801 | 0.197129821 | -0.017449421 | 0.986078083 | 0.99859176 | BC024139 | cDNA sequence BC024139 [Source:MGI Sy | 189.53 | 293.98 | 294.83 | 237.14 | 278.11 | 259.53 |
| 13128 | ENSMUSG00000044362 | 4.541453251 | 0.678987794 | 0.99691136 | 0.68109144 | 0.495813633 | NA | Ccdc89 | coiled-coil domain containing 89 [Source:I | 2.4093 | 4.1117 | 4.025 | 6.7478 | 3.2719 | 6.6831 |
| 13129 | ENSMUSG00000044364 | 0.335414862 | -1.779047242 | 4.030804179 | -0.441362855 | 0.658950329 | NA | Tmem74b | transmembrane protein 74B [Source:MGI | 0 | 0 | 2.0125 | 0 | 0 | 0 |
| 13130 | ENSMUSG00000044365 | 4.04333941 | 1.121619374 | 1.078024269 | 1.040439818 | 0.298135611 | NA | Cxxc4 | CXXC finger 4 [Source:MGI Symbol;Acc:M | 1.6062 | 4.1117 | 2.0125 | 7.7118 | 4.3625 | 4.4554 |
| 13131 | ENSMUSG00000044367 | 57.10756085 | -0.807442987 | 0.342145525 | -2.359940222 | 0.01827788 | 0.162390207 | Slc16a13 | solute carrier family 16 (monocarboxylic a | 81.916 | 70.926 | 65.406 | 53.019 | 47.987 | 23.391 |
| 13132 | ENSMUSG00000044375 | 0 | NA | NA | NA | NA | NA | Pcare | photoreceptor cilium actin regulator [Sou | 0 | 0 | 0 | 0 | 0 | 0 |
| 13133 | ENSMUSG00000044377 | 0.315620088 | 0.165143421 | 4.063300985 | 0.040642675 | 0.967580763 | NA | Gm13080 | predicted gene 13080 [Source:MGI Symbo | 0.8031 | 0 | 0 | 0 | 1.0906 | 0 |
| 13134 | ENSMUSG00000044378 | 129.0054822 | 0.447447774 | 0.231074617 | 1.936377866 | 0.052821438 | 0.32296796 | Slc15a5 | solute carrier family 15, member 5 [Sourc | 129.3 | 99.708 | 97.606 | 121.46 | 175.59 | 150.37 |
| 13135 | ENSMUSG00000044387 | 0.970525971 | -0.749213202 | 2.255458623 | -0.332177764 | 0.739755033 | NA | 2410080I02R | RIKEN cDNA 2410080I02 gene [Source:MC | 1.6062 | 0 | 2.0125 | 0 | 1.0906 | 1.1138 |
| 13136 | ENSMUSG00000044390 | 1.793583482 | -0.786477115 | 1.793018796 | -0.438632945 | 0.660927521 | NA | Tigd3 | tigger transposable element derived 3 [So | 0.8031 | 3.0838 | 3.0187 | 3.8559 | 0 | 0 |
| 13137 | ENSMUSG00000044393 | 1864.450887 | 0.091440616 | 0.127641748 | 0.716384863 | 0.473753739 | 0.868952133 | Dsg2 | desmoglein 2 [Source:MGI Symbol;Acc:M | 1563.6 | 1913 | 1941 | 1962.7 | 2125.6 | 1680.8 |
| 13138 | ENSMUSG00000044400 | 0.36353943 | 2.065989254 | 4.034137301 | 0.512126658 | 0.608562368 | NA | Sowahd | sosondowah ankyrin repeat domain famil | 0 | 0 | 0 | 0 | 2.1812 | 0 |
| 13139 | ENSMUSG00000044405 | 0.133850373 | -0.796629176 | 4.080472857 | -0.195229623 | 0.84521319 | NA | Adig | adipogenin [Source:MGI Symbol;Acc:MGI | 0.8031 | 0 | 0 | 0 | 0 | 0 |
| 13140 | ENSMUSG00000044408 | 846.7271829 | 0.167734268 | 0.120007204 | 1.397701662 | 0.162202675 | 0.585529252 | Sptssa | serine palmitoyltransferase, small subunit | 742.07 | 837.75 | 813.05 | 816.49 | 900.85 | 970.16 |
| 13141 | ENSMUSG00000044424 | 7.301531273 | -0.25956445 | 0.800389749 | -0.324297569 | 0.74571276 | 0.949997329 | Gm9493 | predicted gene 9493 [Source:MGI Symbol | 9.6372 | 4.1117 | 10.062 | 6.7478 | 5.4531 | 7.7969 |
| 13142 | ENSMUSG00000044429 | 0 | NA | NA | NA | NA | NA | Cryga | crystallin, gamma A [Source:MGI Symbol;A | 0 | 0 | 0 | 0 | 0 | 0 |
| 13143 | ENSMUSG00000044430 | 0 | NA | NA | NA | NA | NA | Klk12 | kallikrein related-peptidase 12 [Source:M | 0 | 0 | 0 | 0 | 0 | 0 |
| 13144 | ENSMUSG00000044433 | 254.2674647 | -0.031709224 | 0.171691398 | -0.184687321 | 0.853474292 | 0.973825032 | Camsap3 | calmodulin regulated spectrin-associated | 239.32 | 261.09 | 271.69 | 293.05 | 241.03 | 219.43 |
| 13145 | ENSMUSG00000044434 | 0.614004368 | 0.315622926 | 2.898299649 | 0.108899342 | 0.913282328 | NA | Gm9791 | predicted pseudogene 9791 [Source:MGI | 1.6062 | 0 | 0 | 0.964 | 0 | 1.1138 |
| 13146 | ENSMUSG00000044441 | 1.15844023 | 1.480569095 | 2.137934749 | 0.692523051 | 0.488608915 | NA | Olfr1442 | olfactory receptor 1442 [Source:MGI Sym | 0.8031 | 1.0279 | 0 | 2.8919 | 0 | 2.2277 |
| 13147 | ENSMUSG00000044442 | 179.4183222 | 0.061118622 | 0.203541963 | 0.300275292 | 0.763967178 | 0.954980327 | N6amt1 | N-6 adenine-specific DNA methyltransfera | 181.5 | 171.66 | 173.07 | 141.7 | 219.21 | 189.35 |
| 13148 | ENSMUSG00000044444 | 0 | NA | NA | NA | NA | NA | Pfn3 | profilin 3 [Source:MGI Symbol;Acc:MGI:2 | 0 | 0 | 0 | 0 | 0 | 0 |
| 13149 | ENSMUSG00000044447 | 450.8600744 | 0.106459517 | 0.138763048 | 0.767203654 | 0.442960441 | 0.850769625 | Dock5 | dedicator of cytokinesis 5 [Source:MGI Sy | 409.58 | 457.42 | 436.71 | 514.76 | 467.88 | 418.81 |
| 13150 | ENSMUSG00000044452 | 163.9166543 | 0.250602312 | 0.202163454 | 1.239602443 | 0.215122477 | 0.659728602 | Zfp507 | zinc finger protein 507 [Source:MGI Symb | 139.74 | 152.13 | 157.98 | 212.08 | 172.32 | 149.26 |
| 13151 | ENSMUSG00000044453 | 0.133850373 | -0.796629176 | 4.080472857 | -0.195229623 | 0.84521319 | NA | Ffar1 | free fatty acid receptor 1 [Source:MGI Syr | 0.8031 | 0 | 0 | 0 | 0 | 0 |
| 13152 | ENSMUSG00000044454 | 0 | NA | NA | NA | NA | NA | Olfr867 | olfactory receptor 867 [Source:MGI Symb | 0 | 0 | 0 | 0 | 0 | 0 |
| 13153 | ENSMUSG00000044456 | 428.1106483 | -0.043396131 | 0.129395612 | -0.335375601 | 0.737341771 | 0.948137969 | Rin3 | Ras and Rab interactor 3 [Source:MGI Syn | 422.43 | 437.89 | 443.75 | 436.68 | 403.53 | 424.37 |
| 13154 | ENSMUSG00000044457 | 0 | NA | NA | NA | NA | NA | Stk-ps2 | serine/threonine kinase 2 [Source:MGI Sy | 0 | 0 | 0 | 0 | 0 | 0 |
| 13155 | ENSMUSG00000044461 | 3.725497315 | -0.734142813 | 1.080634579 | -0.679362689 | 0.496908085 | NA | Shisa2 | shisa family member 2 [Source:MGI Symb | 4.8186 | 4.1117 | 5.0312 | 2.8919 | 3.2719 | 2.2277 |
| 13156 | ENSMUSG00000044465 | 444.4320252 | 0.227867772 | 0.130148653 | 1.750826966 | 0.07997572 | 0.415341728 | Fam160a2 | family with sequence similarity 160, meml | 428.86 | 382.39 | 416.59 | 486.81 | 488.6 | 463.36 |

| | | | | | | | | | | | | | | | |
|---|---|---|---|---|---|---|---|---|---|---|---|---|---|---|---|
| 13157 | ENSMUSG00000044468 | 333.8244915 | -0.583268929 | 0.216373492 | -2.69565797 | 0.007024976 | 0.08425311 | Tent5c | terminal nucleotidyltransferase 5C [Source | 296.34 | 443.03 | 461.87 | 193.76 | 323.91 | 284.03 |
| 13158 | ENSMUSG00000044469 | 769.782729 | -0.183351018 | 0.118222253 | -1.550901069 | 0.120925394 | 0.509634296 | Tnfaip8l1 | tumor necrosis factor, alpha-induced prot | 840.85 | 771.97 | 842.23 | 664.18 | 773.25 | 726.23 |
| 13159 | ENSMUSG00000044471 | 12.08212446 | -0.419842689 | 0.661744998 | -0.634447847 | 0.525788592 | 0.888334265 | Lncpint | long non-protein coding RNA, Trp53 induc | 12.047 | 20.558 | 9.0562 | 15.424 | 8.7249 | 6.6831 |
| 13160 | ENSMUSG00000044475 | 262.2961598 | -0.021021019 | 0.160934981 | -0.130618088 | 0.896077434 | 0.982798755 | Ascc1 | activating signal cointegrator 1 complex su | 262.61 | 251.84 | 277.72 | 230.39 | 268.29 | 282.92 |
| 13161 | ENSMUSG00000044477 | 1563.273439 | 0.081504023 | 0.106507105 | 0.765244937 | 0.444125706 | 0.851512043 | Zfand3 | zinc finger, AN1-type domain 3 [Source:M | 1660 | 1456.6 | 1439.9 | 1686 | 1543.2 | 1593.9 |
| 13162 | ENSMUSG00000044485 | 0 | NA | NA | NA | NA | NA | Klk1b11 | kallikrein 1-related peptidase b11 [Source | 0 | 0 | 0 | 0 | 0 | 0 |
| 13163 | ENSMUSG00000044487 | 0 | NA | NA | NA | NA | NA | Olfr1112 | olfactory receptor 1112 [Source:MGI Symb | 0 | 0 | 0 | 0 | 0 | 0 |
| 13164 | ENSMUSG00000044496 | 768.4960094 | -0.061743043 | 0.112837628 | -0.547184871 | 0.584251729 | 0.908955096 | 2510039O18 | RIKEN cDNA 2510039O18 gene [Source:M | 828.8 | 755.52 | 769.78 | 751.9 | 724.17 | 780.8 |
| 13165 | ENSMUSG00000044499 | 0 | NA | NA | NA | NA | NA | Hs3st5 | heparan sulfate (glucosamine) 3-O-sulfotr | 0 | 0 | 0 | 0 | 0 | 0 |
| 13166 | ENSMUSG00000044501 | 94.42987379 | 0.142771934 | 0.231771807 | 0.616002161 | 0.537893094 | 0.891189657 | Zfp758 | zinc finger protein 758 [Source:MGI Symb | 93.16 | 94.568 | 81.506 | 101.22 | 100.34 | 95.791 |
| 13167 | ENSMUSG00000044502 | 513.667562 | -0.008775749 | 0.128543929 | -0.068270426 | 0.945570365 | 0.990309554 | Bod1 | biorientation of chromosomes in cell divis | 502.74 | 537.6 | 506.14 | 559.11 | 490.78 | 485.64 |
| 13168 | ENSMUSG00000044505 | 62.4695085 | -0.953646199 | 0.339677312 | -2.80750632 | 0.00499267 | 0.066565898 | Lingo4 | leucine rich repeat and Ig domain contain | 60.233 | 97.652 | 89.556 | 39.523 | 58.893 | 28.96 |
| 13169 | ENSMUSG00000044515 | 0 | NA | NA | NA | NA | NA | Gm8385 | predicted gene 8385 [Source:MGI Symbol | 0 | 0 | 0 | 0 | 0 | 0 |
| 13170 | ENSMUSG00000044518 | 0 | NA | NA | NA | NA | NA | Foxe3 | forkhead box E3 [Source:MGI Symbol;Acc | 0 | 0 | 0 | 0 | 0 | 0 |
| 13171 | ENSMUSG00000044519 | 0.17131964 | -0.796629176 | 4.080472857 | -0.195229623 | 0.84521319 | NA | Zfp488 | zinc finger protein 488 [Source:MGI Symb | 0 | 1.0279 | 0 | 0 | 0 | 0 |
| 13172 | ENSMUSG00000044522 | 1.728288363 | 1.41789506 | 1.713683695 | 0.827396015 | 0.408012639 | NA | A730020M07 | RIKEN cDNA A730020M07 gene [Source:M | 0.8031 | 0 | 2.0125 | 0.964 | 4.3625 | 2.2277 |
| 13173 | ENSMUSG00000044526 | 0 | NA | NA | NA | NA | NA | Znrf4 | zinc and ring finger 4 [Source:MGI Symbol | 0 | 0 | 0 | 0 | 0 | 0 |
| 13174 | ENSMUSG00000044528 | 0 | NA | NA | NA | NA | NA | Tram1l1 | translocation associated membrane prote | 0 | 0 | 0 | 0 | 0 | 0 |
| 13175 | ENSMUSG00000044533 | 10831.0263 | 0.070332922 | 0.099912634 | 0.70394423 | 0.481467506 | 0.871037649 | Rps2 | ribosomal protein S2 [Source:MGI Symbol | 10401 | 10699 | 10601 | 9820 | 11486 | 11979 |
| 13176 | ENSMUSG00000044534 | 33.22209772 | 0.025202689 | 0.429285345 | 0.058708477 | 0.953184307 | 0.9915542 | Ackr2 | atypical chemokine receptor 2 [Source:MG | 39.352 | 32.893 | 26.162 | 18.316 | 43.625 | 38.985 |
| 13177 | ENSMUSG00000044537 | 0 | NA | NA | NA | NA | NA | Olfr779 | olfactory receptor 779 [Source:MGI Symb | 0 | 0 | 0 | 0 | 0 | 0 |
| 13178 | ENSMUSG00000044538 | 0 | NA | NA | NA | NA | NA | Obox3-ps8 | oocyte specific homeobox 3, pseudogene | 0 | 0 | 0 | 0 | 0 | 0 |
| 13179 | ENSMUSG00000044542 | 0 | NA | NA | NA | NA | NA | Prop1 | paired like homeodomain factor 1 [Source | 0 | 0 | 0 | 0 | 0 | 0 |
| 13180 | ENSMUSG00000044544 | 0.167707431 | -0.796629176 | 4.080472857 | -0.195229623 | 0.84521319 | NA | 4921513I03R | RIKEN cDNA 4921513I03 gene [Source:MG | 0 | 0 | 1.0062 | 0 | 0 | 0 |
| 13181 | ENSMUSG00000044548 | 51.72989245 | 0.55440363 | 0.453586513 | 1.222266568 | 0.221606838 | 0.667949912 | Dact1 | dishevelled-binding antagonist of beta-cat | 27.305 | 47.284 | 51.318 | 49.163 | 27.265 | 108.04 |
| 13182 | ENSMUSG00000044550 | 0 | NA | NA | NA | NA | NA | Tceal3 | transcription elongation factor A (SII)-like | 0 | 0 | 0 | 0 | 0 | 0 |
| 13183 | ENSMUSG00000044551 | 5.750412378 | -0.892807391 | 1.04222716 | -0.856634163 | 0.391647099 | 0.820404559 | 9930012K11 | RIKEN cDNA 9930012K11 gene [Source:M | 4.0155 | 11.307 | 7.0437 | 0 | 5.4531 | 6.6831 |
| 13184 | ENSMUSG00000044556 | 0.294513347 | 0.165143496 | 4.080472857 | 0.040471657 | 0.967717103 | NA | Tex38 | testis expressed 38 [Source:MGI Symbol;A | 0.8031 | 0 | 0 | 0.964 | 0 | 0 |
| 13185 | ENSMUSG00000044560 | 0 | NA | NA | NA | NA | NA | Olfr1302 | olfactory receptor 1302 [Source:MGI Sym | 0 | 0 | 0 | 0 | 0 | 0 |
| 13186 | ENSMUSG00000044562 | 37.1145837 | -0.638002104 | 0.38602248 | -1.65275894 | 0.098379934 | 0.462603173 | Rasip1 | Ras interacting protein 1 [Source:MGI Sym | 35.336 | 41.117 | 59.368 | 32.775 | 22.903 | 31.188 |
| 13187 | ENSMUSG00000044566 | 13.3016072 | -0.148304314 | 0.603022093 | -0.245935125 | 0.805732453 | 0.964425206 | Cage1 | cancer antigen 1 [Source:MGI Symbol;Acc | 18.471 | 8.2233 | 15.094 | 11.568 | 13.087 | 13.366 |
| 13188 | ENSMUSG00000044573 | 1323.930485 | 0.102707349 | 0.099343462 | 1.033861183 | 0.301201064 | 0.7521219 | Acp1 | acid phosphatase 1, soluble [Source:MGI | 1237.6 | 1255.1 | 1338.3 | 1375.6 | 1398.2 | 1338.8 |
| 13189 | ENSMUSG00000044574 | 0 | NA | NA | NA | NA | NA | 5031434C07 | RIKEN cDNA 5031434C07 gene [Source:M | 0 | 0 | 0 | 0 | 0 | 0 |
| 13190 | ENSMUSG00000044576 | 0.335414862 | -1.779047242 | 4.030804179 | -0.441362855 | 0.658950329 | NA | Garem2 | GRB2 associated regulator of MAPK1 subt | 0 | 0 | 2.0125 | 0 | 0 | 0 |
| 13191 | ENSMUSG00000044581 | 0 | NA | NA | NA | NA | NA | 4932415D10 | RIKEN cDNA 4932415D10 gene [Source:M | 0 | 0 | 0 | 0 | 0 | 0 |
| 13192 | ENSMUSG00000044583 | 18.06976878 | -1.200101487 | 0.749361313 | -1.601499125 | 0.109266414 | 0.486233674 | Tlr7 | toll-like receptor 7 [Source:MGI Symbol;A | 27.305 | 32.893 | 15.094 | 0.964 | 6.5437 | 25.618 |
| 13193 | ENSMUSG00000044594 | 0 | NA | NA | NA | NA | NA | Serpinb3a | serine (or cysteine) peptidase inhibitor, cl | 0 | 0 | 0 | 0 | 0 | 0 |
| 13194 | ENSMUSG00000044595 | 9.240133739 | -1.040036548 | 0.734778392 | -1.415442478 | 0.156938802 | 0.575986511 | Dnd1 | DND microRNA-mediated repression inhib | 8.8341 | 13.363 | 15.094 | 3.8559 | 8.7249 | 5.5692 |
| 13195 | ENSMUSG00000044597 | 0 | NA | NA | NA | NA | NA | Mycs | myc-like oncogene, s-myc protein [Source | 0 | 0 | 0 | 0 | 0 | 0 |
| 13196 | ENSMUSG00000044600 | 1051.54135 | 0.06047722 | 0.13027788 | 0.4642171 | 0.642492211 | 0.924876278 | Smim7 | small integral membrane protein 7 [Sourc | 1081.8 | 999.14 | 1006.2 | 915.78 | 1067.7 | 1238.6 |
| 13197 | ENSMUSG00000044609 | 0.181769715 | 1.126924175 | 4.080472857 | 0.276174898 | 0.782413736 | NA | Gm9294 | predicted pseudogene 9294 [Source:MGI | 0 | 0 | 0 | 0 | 1.0906 | 0 |
| 13198 | ENSMUSG00000044617 | 213.3394411 | 0.070020499 | 0.18801618 | 0.372417412 | 0.709582092 | 0.944239215 | Zbtb39 | zinc finger and BTB domain containing 39 | 240.93 | 186.05 | 197.22 | 253.53 | 199.58 | 202.72 |
| 13199 | ENSMUSG00000044624 | 0 | NA | NA | NA | NA | NA | Gm4922 | predicted gene 4922 [Source:MGI Symbol | 0 | 0 | 0 | 0 | 0 | 0 |
| 13200 | ENSMUSG00000044626 | 0.692607243 | 3.004183657 | 2.957557172 | 1.015765202 | 0.309741211 | NA | Liph | lipase, member H [Source:MGI Symbol;Ac | 0 | 0 | 0 | 1.928 | 0 | 2.2277 |
| 13201 | ENSMUSG00000044627 | 1717.152342 | -0.017839188 | 0.102525634 | -0.173997348 | 0.861867546 | 0.976334255 | Swi5 | SWI5 recombination repair homolog (yeas | 1694.5 | 1745.4 | 1742.8 | 1553.9 | 1793 | 1773.2 |
| 13202 | ENSMUSG00000044628 | 0.514011053 | 1.044022102 | 3.082659575 | 0.338675769 | 0.734853997 | NA | Rnf208 | ring finger protein 208 [Source:MGI Symb | 0 | 0 | 1.0062 | 0.964 | 0 | 1.1138 |
| 13203 | ENSMUSG00000044629 | 8.988988301 | -0.146000125 | 0.748107881 | -0.195159185 | 0.84526833 | 0.972803041 | Cnrip1 | cannabinoid receptor interacting protein | 12.85 | 10.279 | 5.0312 | 4.8199 | 9.8156 | 11.138 |
| 13204 | ENSMUSG00000044633 | 0.315620088 | 0.165143421 | 4.063300985 | 0.040642675 | 0.967580763 | NA | B530045E10 | RIKEN cDNA B530045E10 gene [Source:M | 0.8031 | 0 | 0 | 0 | 1.0906 | 0 |
| 13205 | ENSMUSG00000044636 | 43.59272122 | 0.085262068 | 0.339683523 | 0.251004427 | 0.801810686 | 0.962796744 | Csrnp2 | cysteine-serine-rich nuclear protein 2 [Sou | 37.746 | 45.228 | 44.275 | 54.947 | 39.262 | 40.098 |
| 13206 | ENSMUSG00000044639 | 0 | NA | NA | NA | NA | NA | Gm11507 | predicted gene 11507 [Source:MGI Symbo | 0 | 0 | 0 | 0 | 0 | 0 |
| 13207 | ENSMUSG00000044641 | 107.1100015 | 0.012929394 | 0.245069244 | 0.052758126 | 0.957924626 | 0.991861398 | Pard6b | par-6 family cell polarity regulator beta [So | 110.83 | 109.99 | 98.612 | 80.974 | 130.87 | 111.38 |
| 13208 | ENSMUSG00000044645 | 0.815516723 | -3.06336425 | 2.557025255 | -1.19801877 | 0.23090971 | NA | Gm7334 | predicted gene 7334 [Source:MGI Symbol | 0.8031 | 3.0838 | 1.0062 | 0 | 0 | 0 |
| 13209 | ENSMUSG00000044646 | 43.96768653 | 0.281874303 | 0.353730959 | 0.796860709 | 0.425531932 | 0.841292218 | Zbtb7c | zinc finger and BTB domain containing 7C | 37.746 | 38.033 | 43.269 | 41.451 | 65.437 | 37.871 |
| 13210 | ENSMUSG00000044647 | 0 | NA | NA | NA | NA | NA | Csrnp3 | cysteine-serine-rich nuclear protein 3 [Sou | 0 | 0 | 0 | 0 | 0 | 0 |
| 13211 | ENSMUSG00000044649 | 0 | NA | NA | NA | NA | NA | Krtap4-2 | keratin associated protein 4-2 [Source:MG | 0 | 0 | 0 | 0 | 0 | 0 |
| 13212 | ENSMUSG00000044664 | 0 | NA | NA | NA | NA | NA | Prss42 | protease, serine 42 [Source:MGI Symbol;A | 0 | 0 | 0 | 0 | 0 | 0 |

| | | | | | | | | | | | | | | | |
|---|---|---|---|---|---|---|---|---|---|---|---|---|---|---|---|
| 13213 | ENSMUSG00000044667 | 0.524667718 | -0.819976748 | 3.064824146 | -0.267544469 | 0.78904998 | NA | Plppr4 | phospholipid phosphatase related 4 [Sour | 0 | 1.0279 | 1.0062 | 0 | 0 | 1.1138 |
| 13214 | ENSMUSG00000044674 | 244.320639 | 0.089182625 | 0.159827523 | 0.557992912 | 0.576849224 | 0.907752116 | Fzd1 | frizzled class receptor 1 [Source:MGI Syml | 247.36 | 232.31 | 230.43 | 252.56 | 270.47 | 232.79 |
| 13215 | ENSMUSG00000044676 | 180.0692264 | -0.074584611 | 0.20918818 | -0.356543141 | 0.72143385 | 0.9447719 | Zfp612 | zinc finger protein 612 [Source:MGI Symb | 163.83 | 176.8 | 214.33 | 206.29 | 181.04 | 138.12 |
| 13216 | ENSMUSG00000044678 | 0.133850373 | -0.796629176 | 4.080472857 | -0.195229623 | 0.84521319 | NA | Ly6k | lymphocyte antigen 6 complex, locus K [S | 0.8031 | 0 | 0 | 0 | 0 | 0 |
| 13217 | ENSMUSG00000044681 | 0 | NA | NA | NA | NA | NA | Cnpy1 | canopy FGF signaling regulator 1 [Source:I | 0 | 0 | 0 | 0 | 0 | 0 |
| 13218 | ENSMUSG00000044689 | 0 | NA | NA | NA | NA | NA | Gm13749 | predicted gene 13749 [Source:MGI Symbo | 0 | 0 | 0 | 0 | 0 | 0 |
| 13219 | ENSMUSG00000044694 | 1.878759987 | 0.969656089 | 1.612585343 | 0.601305285 | 0.54763667 | NA | 2010007H06 | RIKEN cDNA 2010007H06 gene [Source:M | 0.8031 | 0 | 3.0187 | 1.928 | 2.1812 | 3.3415 |
| 13220 | ENSMUSG00000044697 | 0 | NA | NA | NA | NA | NA | Gm5479 | predicted gene 5479 [Source:MGI Symbol | 0 | 0 | 0 | 0 | 0 | 0 |
| 13221 | ENSMUSG00000044700 | 438.530669 | 0.192142416 | 0.138675663 | 1.385552534 | 0.165883581 | 0.591991145 | Tmem201 | transmembrane protein 201 [Source:MGI | 386.29 | 400.89 | 441.74 | 508.02 | 426.43 | 467.81 |
| 13222 | ENSMUSG00000044701 | 0.621048825 | -1.224891493 | 2.859518283 | -0.428355888 | 0.668392035 | NA | Il27 | interleukin 27 [Source:MGI Symbol;Acc:M | 1.6062 | 0 | 1.0062 | 0 | 0 | 1.1138 |
| 13223 | ENSMUSG00000044702 | 7.210647253 | -0.716504635 | 0.856457802 | -0.836590703 | 0.402822673 | 0.828953151 | Palb2 | partner and localizer of BRCA2 [Source:M | 13.653 | 8.2233 | 5.0312 | 8.6758 | 5.4531 | 2.2277 |
| 13224 | ENSMUSG00000044703 | 2.215977105 | -1.052150141 | 1.507034885 | -0.698159115 | 0.485077692 | NA | Phf11a | PHD finger protein 11A [Source:MGI Symt | 4.8186 | 2.0558 | 2.0125 | 0 | 2.1812 | 2.2277 |
| 13225 | ENSMUSG00000044705 | 0 | NA | NA | NA | NA | NA | Olfr670 | olfactory receptor 670 [Source:MGI Symb | 0 | 0 | 0 | 0 | 0 | 0 |
| 13226 | ENSMUSG00000044707 | 0.339027071 | -1.79174149 | 3.919585585 | -0.457125237 | 0.647581032 | NA | Ccnjl | cyclin J-like [Source:MGI Symbol;Acc:MGI | 0 | 1.0279 | 1.0062 | 0 | 0 | 0 |
| 13227 | ENSMUSG00000044708 | 9.056221797 | -2.426911511 | 0.857253315 | -2.831031934 | 0.004639809 | 0.063491197 | Kcnj10 | potassium inwardly-rectifying channel, su | 12.047 | 24.67 | 9.0562 | 1.928 | 2.1812 | 4.4554 |
| 13228 | ENSMUSG00000044709 | 395.7650896 | 0.154607952 | 0.145675874 | 1.061314742 | 0.288546889 | 0.73951397 | Gemin7 | gem nuclear organelle associated protein | 390.31 | 357.72 | 375.33 | 420.29 | 365.36 | 465.59 |
| 13229 | ENSMUSG00000044712 | 129.1693998 | -0.021398596 | 0.228423336 | -0.093679554 | 0.925363712 | 0.987546253 | Slc38a6 | solute carrier family 38, member 6 [Sourc | 141.35 | 145.96 | 102.64 | 110.86 | 147.23 | 126.98 |
| 13230 | ENSMUSG00000044715 | 228.5785891 | -0.117178063 | 0.200170462 | -0.585391383 | 0.558284581 | 0.899286756 | Gskip | GSK3B interacting protein [Source:MGI Sy | 265.02 | 224.09 | 223.39 | 163.88 | 235.57 | 259.53 |
| 13231 | ENSMUSG00000044716 | 1.346789088 | 2.851836319 | 2.053945616 | 1.388467298 | 0.164994796 | NA | Dok7 | docking protein 7 [Source:MGI Symbol;Ac | 0.8031 | 0 | 0 | 2.8919 | 3.2719 | 1.1138 |
| 13232 | ENSMUSG00000044719 | 0.916590439 | 3.39841529 | 2.684534563 | 1.265923463 | 0.205540485 | NA | E230025N22 | Riken cDNA E230025N22 gene [Source:M | 0 | 0 | 0 | 0 | 3.2719 | 2.2277 |
| 13233 | ENSMUSG00000044724 | 0.294513347 | 0.165143496 | 4.080472857 | 0.040471657 | 0.967717103 | NA | Gpr152 | G protein-coupled receptor 152 [Source:N | 0.8031 | 0 | 0 | 0.964 | 0 | 0 |
| 13234 | ENSMUSG00000044726 | 0.346303622 | 2.004339653 | 3.884003423 | 0.516049919 | 0.605819544 | NA | Erich5 | glutamate rich 5 [Source:MGI Symbol;Acc | 0 | 0 | 0 | 0.964 | 0 | 1.1138 |
| 13235 | ENSMUSG00000044730 | 100.3000578 | -0.03728462 | 0.24587269 | -0.151641975 | 0.879469325 | 0.98128671 | 9930104L06 | RIKEN cDNA 9930104L06 gene [Source:M | 98.782 | 88.401 | 117.73 | 107 | 79.615 | 110.27 |
| 13236 | ENSMUSG00000044734 | 73.01086917 | -1.255469814 | 0.390015165 | -3.219028198 | 0.001286258 | 0.02520927 | Serpinb1a | serine (or cysteine) peptidase inhibitor, cl | 61.839 | 78.122 | 169.05 | 47.235 | 28.356 | 53.465 |
| 13237 | ENSMUSG00000044737 | 0 | NA | NA | NA | NA | NA | Klk14 | kallikrein related-peptidase 14 [Source:M | 0 | 0 | 0 | 0 | 0 | 0 |
| 13238 | ENSMUSG00000044743 | 0 | NA | NA | NA | NA | NA | Defb10 | defensin beta 10 [Source:MGI Symbol;Acc | 0 | 0 | 0 | 0 | 0 | 0 |
| 13239 | ENSMUSG00000044748 | 31.0907313 | -0.334912451 | 0.410859017 | -0.81515176 | 0.414985447 | 0.836001897 | Defb1 | defensin beta 1 [Source:MGI Symbol;Acc: | 41.761 | 28.782 | 33.206 | 23.135 | 22.903 | 36.757 |
| 13240 | ENSMUSG00000044749 | 3805.755116 | 0.027149664 | 0.102488243 | 0.264905154 | 0.791082528 | 0.961019028 | Abca6 | ATP-binding cassette, sub-family A (ABC1) | 3344.1 | 4003.7 | 3963.6 | 3822.2 | 3932.8 | 3768.1 |
| 13241 | ENSMUSG00000044751 | 6.024361937 | 0.007249882 | 0.883100072 | 0.008209582 | 0.993449775 | 0.99859176 | Atp5pb-ps | ATP synthase peripheral stalk-membrane | 4.8186 | 6.1675 | 7.0437 | 4.8199 | 3.2719 | 10.025 |
| 13242 | ENSMUSG00000044757 | 0.181769715 | 1.126924175 | 4.080472857 | 0.276174898 | 0.782413736 | NA | Gm6430 | predicted gene 6430 [Source:MGI Symbol | 0 | 0 | 0 | 0 | 1.0906 | 0 |
| 13243 | ENSMUSG00000044763 | 335.1334228 | -0.158099775 | 0.156614591 | -1.009483051 | 0.312743026 | 0.759921531 | Trmt10c | tRNA methyltransferase 10C, mitochondri | 317.23 | 371.08 | 372.31 | 280.52 | 354.45 | 315.22 |
| 13244 | ENSMUSG00000044768 | 340.2313635 | -0.177302306 | 0.158365932 | -1.119573528 | 0.262895542 | 0.717456814 | Macir | macrophage immunometabolism regulatc | 320.44 | 381.36 | 382.37 | 339.32 | 274.84 | 343.06 |
| 13245 | ENSMUSG00000044770 | 1.672244164 | -2.018254233 | 1.885226569 | -1.070563224 | 0.284365867 | NA | Scml4 | Scm polycomb group protein like 4 [Sourc | 4.0155 | 3.0838 | 1.0062 | 1.928 | 0 | 0 |
| 13246 | ENSMUSG00000044772 | 0 | NA | NA | NA | NA | NA | Sntn | sentan, cilia apical structure protein [Sour | 0 | 0 | 0 | 0 | 0 | 0 |
| 13247 | ENSMUSG00000044783 | 2297.649246 | 0.370887474 | 0.117711585 | 3.150815394 | 0.001628154 | 0.030076136 | Hjurp | Holliday junction recognition protein [Sou | 2270.4 | 1777.3 | 1962.2 | 2444.6 | 2797.4 | 2534 |
| 13248 | ENSMUSG00000044786 | 1215.493303 | -0.275979225 | 0.166165458 | -1.660869999 | 0.096739559 | 0.458530879 | Zfp36 | zinc finger protein 36 [Source:MGI Symbo | 1256.9 | 1565.5 | 1172.3 | 1088.3 | 1354.5 | 855.43 |
| 13249 | ENSMUSG00000044787 | 0 | NA | NA | NA | NA | NA | Spata32 | spermatogenesis associated 32 [Source:M | 0 | 0 | 0 | 0 | 0 | 0 |
| 13250 | ENSMUSG00000044788 | 3037.048631 | 0.388594692 | 0.102127739 | 3.80498673 | 0.000141812 | 0.004737103 | Fads6 | fatty acid desaturase domain family, mem | 2535.4 | 2700.3 | 2655.5 | 3088.6 | 3595.8 | 3646.7 |
| 13251 | ENSMUSG00000044791 | 682.7196665 | 0.045182222 | 0.122441011 | 0.369012163 | 0.712118659 | 0.944239215 | Setd2 | SET domain containing 2 [Source:MGI Syn | 633.65 | 712.35 | 671.17 | 736.48 | 708.9 | 633.78 |
| 13252 | ENSMUSG00000044792 | 1034.162213 | 0.03370483 | 0.105037519 | 0.320883721 | 0.748298511 | 0.951074124 | Isca1 | iron-sulfur cluster assembly 1 [Source:MG | 1024.8 | 1013.5 | 1027.4 | 981.33 | 1076.4 | 1081.5 |
| 13253 | ENSMUSG00000044795 | 306.6731546 | 0.134027864 | 0.156071291 | 0.858760528 | 0.390472649 | 0.819763149 | Cyb5d1 | cytochrome b5 domain containing 1 [Sour | 291.53 | 272.4 | 312.94 | 283.41 | 351.18 | 328.58 |
| 13254 | ENSMUSG00000044798 | 0 | NA | NA | NA | NA | NA | Olfr923 | olfactory receptor 923 [Source:MGI Symb | 0 | 0 | 0 | 0 | 0 | 0 |
| 13255 | ENSMUSG00000044801 | 0 | NA | NA | NA | NA | NA | Olfr159 | olfactory receptor 159 [Source:MGI Symb | 0 | 0 | 0 | 0 | 0 | 0 |
| 13256 | ENSMUSG00000044807 | 10.54331862 | -0.607988616 | 0.678219905 | -0.896447615 | 0.370013746 | 0.806613237 | Zfp354c | zinc finger protein 354C [Source:MGI Sym | 8.8341 | 16.447 | 13.081 | 10.604 | 8.7249 | 5.5692 |
| 13257 | ENSMUSG00000044811 | 35.80456268 | -1.682380169 | 0.476496974 | -3.530725821 | 0.000414421 | 0.010878742 | Cd300c2 | CD300C molecule 2 [Source:MGI Symbol;/ | 52.202 | 84.289 | 27.169 | 12.532 | 16.359 | 22.277 |
| 13258 | ENSMUSG00000044813 | 962.2651128 | -0.361109141 | 0.131519481 | -2.745670354 | 0.006038741 | 0.075741685 | Shb | src homology 2 domain-containing transfc | 1048 | 1188.3 | 1009.3 | 720.09 | 882.31 | 925.6 |
| 13259 | ENSMUSG00000044814 | 0 | NA | NA | NA | NA | NA | Olfr543 | olfactory receptor 543 [Source:MGI Symb | 0 | 0 | 0 | 0 | 0 | 0 |
| 13260 | ENSMUSG00000044816 | 0.569258551 | -2.569337407 | 3.18860282 | -0.805787849 | 0.420365185 | NA | D630023F18 | RIKEN cDNA D630023F18 gene [Source:M | 2.4093 | 0 | 1.0062 | 0 | 0 | 0 |
| 13261 | ENSMUSG00000044819 | 0 | NA | NA | NA | NA | NA | Oxgr1 | oxoglutarate (alpha-ketoglutarate) recept | 0 | 0 | 0 | 0 | 0 | 0 |
| 13262 | ENSMUSG00000044820 | 9.588814114 | 0.564126043 | 0.697374039 | 0.808928941 | 0.418556021 | 0.837575303 | AY074887 | cDNA sequence AY074887 [Source:MGI Sy | 8.031 | 5.1396 | 10.062 | 14.46 | 9.8156 | 10.025 |
| 13263 | ENSMUSG00000044824 | 0 | NA | NA | NA | NA | NA | Olfr545 | olfactory receptor 545 [Source:MGI Symb | 0 | 0 | 0 | 0 | 0 | 0 |
| 13264 | ENSMUSG00000044827 | 13.47845294 | -1.707998061 | 0.748367653 | -2.282298086 | 0.022471748 | 0.187679993 | Tlr1 | toll-like receptor 1 [Source:MGI Symbol;A | 16.865 | 39.061 | 6.0375 | 6.7478 | 4.3625 | 7.7969 |
| 13265 | ENSMUSG00000044835 | 0 | NA | NA | NA | NA | NA | Ankrd45 | ankyrin repeat domain 45 [Source:MGI Sy | 0 | 0 | 0 | 0 | 0 | 0 |
| 13266 | ENSMUSG00000044847 | 79.58028462 | -0.173281165 | 0.267885516 | -0.646847832 | 0.517730432 | 0.887022094 | Lsm11 | U7 snRNP-specific Sm-like protein LSM11 | 86.735 | 69.898 | 96.599 | 88.686 | 67.618 | 67.944 |
| 13267 | ENSMUSG00000044854 | 19.86240193 | -0.645393914 | 0.511806825 | -1.261010761 | 0.20730497 | 0.652364421 | 1700056E22 | RIKEN cDNA 1700056E22 gene [Source:M | 29.715 | 25.698 | 17.106 | 13.496 | 11.997 | 21.163 |
| 13268 | ENSMUSG00000044857 | 572.6938758 | 0.188338223 | 0.137405514 | 1.370674414 | 0.170476474 | 0.597556794 | Lemd2 | LEM domain containing 2 [Source:MGI Syı | 487.48 | 514.99 | 604.75 | 667.07 | 569.3 | 592.56 |

| | | | | | | | | | | | | | | | |
|---|---|---|---|---|---|---|---|---|---|---|---|---|---|---|---|
| 13269 | ENSMUSG00000044860 | 0.267700746 | -1.497481649 | 4.057001436 | -0.369110456 | 0.712045396 | NA | Gm1123 | predicted gene 1123 [Source:MGI Symbol | 1.6062 | 0 | 0 | 0 | 0 | 0 |
| 13270 | ENSMUSG00000044863 | 0 | NA | NA | NA | NA | NA | Defb36 | defensin beta 36 [Source:MGI Symbol;Acc | 0 | 0 | 0 | 0 | 0 | 0 |
| 13271 | ENSMUSG00000044864 | 187.3903596 | 0.179523396 | 0.190981092 | 0.940006122 | 0.347214421 | 0.788352979 | Ankrd50 | ankyrin repeat domain 50 [Source:MGI Sy | 194.35 | 146.99 | 185.15 | 183.16 | 193.04 | 221.65 |
| 13272 | ENSMUSG00000044867 | 3.412376892 | -1.540282279 | 1.335553142 | -1.153291644 | 0.248790699 | NA | Gimap1os | GTPase, IMAP family member 1, opposite | 8.031 | 6.1675 | 1.0062 | 1.928 | 0 | 3.3415 |
| 13273 | ENSMUSG00000044876 | 521.9541251 | 0.001214725 | 0.13971771 | 0.00869414 | 0.993063167 | 0.99859176 | Zfp444 | zinc finger protein 444 [Source:MGI Symb | 529.24 | 450.23 | 585.63 | 537.9 | 492.96 | 535.76 |
| 13274 | ENSMUSG00000044881 | 118.9120625 | 0.168823496 | 0.221352672 | 0.762690119 | 0.445648225 | 0.852288235 | Coa4 | cytochrome c oxidase assembly factor 4 [S | 123.68 | 107.93 | 103.64 | 107 | 130.87 | 140.34 |
| 13275 | ENSMUSG00000044894 | 7126.752877 | 0.012315346 | 0.083825803 | 0.146915929 | 0.883198379 | 0.981769403 | Uqcrq | ubiquinol-cytochrome c reductase, compl | 7378.1 | 7039.2 | 6870.6 | 7035.1 | 7120.6 | 7316.8 |
| 13276 | ENSMUSG00000044897 | 0 | NA | NA | NA | NA | NA | Olfr821 | olfactory receptor 821 [Source:MGI Symb | 0 | 0 | 0 | 0 | 0 | 0 |
| 13277 | ENSMUSG00000044899 | 0 | NA | NA | NA | NA | NA | Olfr649 | olfactory receptor 649 [Source:MGI Symb | 0 | 0 | 0 | 0 | 0 | 0 |
| 13278 | ENSMUSG00000044903 | 0.160662974 | 1.126924175 | 4.080472857 | 0.276174898 | 0.782413736 | NA | Psg22 | pregnancy-specific glycoprotein 22 [Sourc | 0 | 0 | 0 | 0.964 | 0 | 0 |
| 13279 | ENSMUSG00000044906 | 39.10708068 | -1.525666733 | 0.406093645 | -3.756933293 | 0.000172008 | 0.005553412 | 4930503L19I | RIKEN cDNA 4930503L19 gene [Source:M( | 74.689 | 39.061 | 60.375 | 23.135 | 22.903 | 14.48 |
| 13280 | ENSMUSG00000044912 | 0 | NA | NA | NA | NA | NA | Syt16 | synaptotagmin XVI [Source:MGI Symbol;A | 0 | 0 | 0 | 0 | 0 | 0 |
| 13281 | ENSMUSG00000044916 | 30.67823295 | 0.022303193 | 0.410029792 | 0.054394079 | 0.956621196 | 0.991798415 | 1700029I15F | RIKEN cDNA 1700029I15 gene [Source:M( | 40.155 | 22.614 | 28.175 | 26.991 | 32.719 | 33.415 |
| 13282 | ENSMUSG00000044921 | 1.395716167 | -0.052563455 | 1.945055679 | -0.027024139 | 0.978440481 | NA | Rassf9 | Ras association (RalGDS/AF-6) domain far | 3.2124 | 0 | 1.0062 | 1.928 | 0 | 2.2277 |
| 13283 | ENSMUSG00000044923 | 1.356081653 | -0.715969176 | 2.025878503 | -0.353411705 | 0.723779812 | NA | Olfr1030 | olfactory receptor 1030 [Source:MGI Syml | 0 | 4.1117 | 1.0062 | 1.928 | 1.0906 | 0 |
| 13284 | ENSMUSG00000044927 | 3.008268989 | 1.392856744 | 1.400779515 | 0.994344027 | 0.320055408 | NA | H1f10 | H1.10 linker histone [Source:MGI Symbol; | 0 | 1.0279 | 4.025 | 1.928 | 3.2719 | 7.7969 |
| 13285 | ENSMUSG00000044933 | 0.487198452 | -0.685759572 | 3.136894395 | -0.218610985 | 0.826953095 | NA | Sstr3 | somatostatin receptor 3 [Source:MGI Sym | 0.8031 | 0 | 1.0062 | 0 | 0 | 1.1138 |
| 13286 | ENSMUSG00000044934 | 339.3335793 | 0.183979528 | 0.142845693 | 1.287959929 | 0.197759912 | 0.639767249 | Zfp367 | zinc finger protein 367 [Source:MGI Symb | 326.86 | 289.87 | 336.09 | 355.71 | 359.9 | 367.57 |
| 13287 | ENSMUSG00000044937 | 41.96694506 | 0.515059345 | 0.373124998 | 1.380393559 | 0.167465502 | 0.593681623 | Ttc41 | tetratricopeptide repeat domain 41 [Sour | 39.352 | 29.81 | 34.212 | 43.379 | 35.99 | 69.058 |
| 13288 | ENSMUSG00000044938 | 7.854730099 | 0.488555293 | 0.809104184 | 0.603822477 | 0.545961674 | 0.89549457 | Klhl31 | kelch-like 31 [Source:MGI Symbol;Acc:MG | 11.243 | 4.1117 | 4.025 | 6.7478 | 7.6343 | 13.366 |
| 13289 | ENSMUSG00000044948 | 17.9916738 | -1.063412928 | 0.588795913 | -1.80608069 | 0.070905737 | 0.385556607 | Cfap43 | cilia and flagella associated protein 43 [So | 16.865 | 42.145 | 14.087 | 10.604 | 11.997 | 12.252 |
| 13290 | ENSMUSG00000044949 | 5.819893675 | 0.155407751 | 1.006336676 | 0.154429183 | 0.877271345 | 0.980667187 | Ubtd2 | ubiquitin domain containing 2 [Source:M( | 2.4093 | 9.2513 | 5.0312 | 11.568 | 1.0906 | 5.5692 |
| 13291 | ENSMUSG00000044950 | 126.5855184 | 0.323815116 | 0.216992117 | 1.492289769 | 0.135623198 | 0.5381974 | Pwwp2a | PWWP domain containing 2A [Source:MG | 97.978 | 121.29 | 118.74 | 159.06 | 124.33 | 138.12 |
| 13292 | ENSMUSG00000044951 | 0 | NA | NA | NA | NA | NA | Mylk4 | myosin light chain kinase family, member | 0 | 0 | 0 | 0 | 0 | 0 |
| 13293 | ENSMUSG00000044952 | 184.3779918 | 0.22697419 | 0.182372282 | 1.244565169 | 0.21329163 | 0.657933491 | Kctd21 | potassium channel tetramerisation domai | 173.47 | 153.16 | 183.14 | 214.97 | 181.04 | 200.49 |
| 13294 | ENSMUSG00000044957 | 0.79190315 | 0.90205007 | 2.632846219 | 0.342614036 | 0.731888841 | NA | Pp2d1 | protein phosphatase 2C-like domain conta | 1.6062 | 0 | 0 | 0.964 | 2.1812 | 0 |
| 13295 | ENSMUSG00000044966 | 1.416487074 | 0.12239615 | 1.960786118 | 0.062421979 | 0.950226792 | NA | Fbxo48 | F-box protein 48 [Source:MGI Symbol;Acc | 0 | 3.0838 | 1.0062 | 0 | 2.1812 | 2.2277 |
| 13296 | ENSMUSG00000044968 | 16.7883347 | -0.60472494 | 0.555211381 | -1.089179655 | 0.276074669 | 0.730038567 | Napepld | N-acyl phosphatidylethanolamine phosph | 24.093 | 16.447 | 20.125 | 13.496 | 7.6343 | 18.935 |
| 13297 | ENSMUSG00000044976 | 1.245244544 | 0.343065865 | 2.05333203 | 0.167077638 | 0.867308961 | NA | Wdr72 | WD repeat domain 72 [Source:MGI Symb | 3.2124 | 0 | 0 | 0.964 | 2.1812 | 1.1138 |
| 13298 | ENSMUSG00000044982 | 129.6263233 | 0.019757725 | 0.227263267 | 0.086937607 | 0.930721107 | 0.988420305 | Sft2d3 | SFT2 domain containing 3 [Source:MGI Sy | 123.68 | 111.02 | 151.94 | 153.27 | 128.69 | 109.16 |
| 13299 | ENSMUSG00000044985 | 0 | NA | NA | NA | NA | NA | Olfr124 | olfactory receptor 124 [Source:MGI Symb | 0 | 0 | 0 | 0 | 0 | 0 |
| 13300 | ENSMUSG00000044986 | 5139.934786 | 0.094813322 | 0.097098061 | 0.976469781 | 0.328831716 | 0.77362974 | Tst | thiosulfate sulfurtransferase, mitochondri | 5112.5 | 4608.2 | 5192.2 | 5358.8 | 5606.9 | 4961.1 |
| 13301 | ENSMUSG00000044988 | 0 | NA | NA | NA | NA | NA | Ucn3 | urocortin 3 [Source:MGI Symbol;Acc:MGI: | 0 | 0 | 0 | 0 | 0 | 0 |
| 13302 | ENSMUSG00000044991 | 135.5549137 | -0.144853229 | 0.225583149 | -0.642127881 | 0.520790154 | 0.887872323 | Shld1 | shieldin complex subunit 1 [Source:MGI S | 171.86 | 140.82 | 113.71 | 120.5 | 146.14 | 120.3 |
| 13303 | ENSMUSG00000044994 | 0 | NA | NA | NA | NA | NA | Olfr1426 | olfactory receptor 1426 [Source:MGI Syml | 0 | 0 | 0 | 0 | 0 | 0 |
| 13304 | ENSMUSG00000044997 | 0 | NA | NA | NA | NA | NA | E130304I02F | RIKEN cDNA E130304I02 gene [Source:M( | 0 | 0 | 0 | 0 | 0 | 0 |
| 13305 | ENSMUSG00000045004 | 0.534859069 | 1.101323472 | 3.351640271 | 0.328592386 | 0.742463804 | NA | Spata21 | spermatogenesis associated 21 [Source:M | 0 | 1.0279 | 0 | 0 | 2.1812 | 0 |
| 13306 | ENSMUSG00000045005 | 153.670277 | 0.167166389 | 0.251902677 | 0.663614974 | 0.506936771 | 0.88185929 | Fzd5 | frizzled class receptor 5 [Source:MGI Syml | 106.01 | 127.46 | 201.25 | 136.88 | 177.77 | 172.65 |
| 13307 | ENSMUSG00000045007 | 1.453594766 | -0.760720938 | 1.926474833 | -0.394877174 | 0.692933537 | NA | Tubg2 | tubulin, gamma 2 [Source:MGI Symbol;Ac | 2.4093 | 1.0279 | 2.0125 | 0 | 3.2719 | 0 |
| 13308 | ENSMUSG00000045008 | 0.133850373 | -0.796629176 | 4.080472857 | -0.195229623 | 0.84521319 | NA | 9030612E09 | RIKEN cDNA 9030612E09 gene [Source:M( | 0.8031 | 0 | 0 | 0 | 0 | 0 |
| 13309 | ENSMUSG00000045009 | 4.04098152 | 0.966418352 | 1.162897412 | 0.831043514 | 0.405949048 | NA | Prrt3 | proline-rich transmembrane protein 3 [So | 4.0155 | 4.1117 | 0 | 2.8919 | 6.5437 | 6.6831 |
| 13310 | ENSMUSG00000045010 | 5.489066336 | -4.810967548 | 3.048295474 | -1.578248431 | 0.114508548 | 0.497223548 | Gm4779 | predicted gene 4779 [Source:MGI Symbol | 0 | 30.838 | 1.0062 | 0 | 1.0906 | 0 |
| 13311 | ENSMUSG00000045013 | 0 | NA | NA | NA | NA | NA | Olfr711 | olfactory receptor 711 [Source:MGI Symb | 0 | 0 | 0 | 0 | 0 | 0 |
| 13312 | ENSMUSG00000045019 | 0 | NA | NA | NA | NA | NA | Acer1 | alkaline ceramidase 1 [Source:MGI Symbo | 0 | 0 | 0 | 0 | 0 | 0 |
| 13313 | ENSMUSG00000045022 | 0 | NA | NA | NA | NA | NA | H2bl1 | H2B.L histone variant 1 [Source:MGI Symb | 0 | 0 | 0 | 0 | 0 | 0 |
| 13314 | ENSMUSG00000045027 | 0 | NA | NA | NA | NA | NA | Prss22 | protease, serine 22 [Source:MGI Symbol;A | 0 | 0 | 0 | 0 | 0 | 0 |
| 13315 | ENSMUSG00000045030 | 0.133850373 | -0.796629176 | 4.080472857 | -0.195229623 | 0.84521319 | NA | Olfr1443 | olfactory receptor 1443 [Source:MGI Syml | 0.8031 | 0 | 0 | 0 | 0 | 0 |
| 13316 | ENSMUSG00000045031 | 1.284575368 | -1.441415965 | 1.931383639 | -0.74631261 | 0.455478596 | NA | Cetn4 | centrin 4 [Source:MGI Symbol;Acc:MGI:2( | 1.6062 | 1.0279 | 3.0187 | 0.964 | 1.0906 | 0 |
| 13317 | ENSMUSG00000045034 | 0 | NA | NA | NA | NA | NA | Ankrd34b | ankyrin repeat domain 34B [Source:MGI S | 0 | 0 | 0 | 0 | 0 | 0 |
| 13318 | ENSMUSG00000045036 | 0 | NA | NA | NA | NA | NA | Tmem232 | transmembrane protein 232 [Source:MGI | 0 | 0 | 0 | 0 | 0 | 0 |
| 13319 | ENSMUSG00000045038 | 297.7873487 | -0.111243338 | 0.159263604 | -0.698485623 | 0.484873546 | 0.87256358 | Prkce | protein kinase C, epsilon [Source:MGI Syn | 330.88 | 321.74 | 274.7 | 274.73 | 318.46 | 266.21 |
| 13320 | ENSMUSG00000045039 | 215.251965 | -0.056165319 | 0.200617871 | -0.279961695 | 0.779506893 | 0.959870443 | Megf8 | multiple EGF-like-domains 8 [Source:MGI | 215.23 | 182.97 | 260.62 | 253.53 | 187.59 | 191.58 |
| 13321 | ENSMUSG00000045045 | 81.53035589 | -0.696028791 | 0.30844024 | -2.256608251 | 0.024032564 | 0.197338277 | Lrfn4 | leucine rich repeat and fibronectin type II | 110.83 | 82.233 | 109.68 | 80.974 | 68.709 | 36.757 |
| 13322 | ENSMUSG00000045048 | 0 | NA | NA | NA | NA | NA | Gm9795 | predicted pseudogene 9795 [Source:MGI | 0 | 0 | 0 | 0 | 0 | 0 |
| 13323 | ENSMUSG00000045052 | 0 | NA | NA | NA | NA | NA | Prlhr | prolactin releasing hormone receptor [Sou | 0 | 0 | 0 | 0 | 0 | 0 |
| 13324 | ENSMUSG00000045053 | 0 | NA | NA | NA | NA | NA | Kcng3 | potassium voltage-gated channel, subfam | 0 | 0 | 0 | 0 | 0 | 0 |

| | | | | | | | | | | | | | | | |
|---|---|---|---|---|---|---|---|---|---|---|---|---|---|---|---|
| 13325 | ENSMUSG00000045055 | 2.230639698 | -3.423046393 | 1.991814356 | -1.718556944 | 0.085695077 | NA | Rpsa-ps2 | ribosomal protein SA, pseudogene 2 [Sour | 0 | 9.2513 | 3.0187 | 0 | 0 | 1.1138 |
| 13326 | ENSMUSG00000045062 | 2.020352225 | 0.687206082 | 1.706356834 | 0.402732927 | 0.687144708 | NA | Pcdhb7 | protocadherin beta 7 [Source:MGI Symbo | 1.6062 | 0 | 3.0187 | 1.928 | 0 | 5.5692 |
| 13327 | ENSMUSG00000045064 | 100.9201636 | 0.631954278 | 0.282510381 | 2.236924095 | 0.025291301 | 0.204593616 | Zc2hc1c | zinc finger, C2HC-type containing 1C [Sour | 85.932 | 62.703 | 88.55 | 119.53 | 85.068 | 163.74 |
| 13328 | ENSMUSG00000045065 | 0 | NA | NA | NA | NA | NA | 9930022D16 | RIKEN cDNA 9930022D16 gene [Source:M | 0 | 0 | 0 | 0 | 0 | 0 |
| 13329 | ENSMUSG00000045071 | 98.23523137 | 0.144104793 | 0.226294901 | 0.636800887 | 0.524254548 | 0.888334265 | E130308A19 | RIKEN cDNA E130308A19 gene [Source:M | 98.782 | 88.401 | 92.575 | 100.25 | 104.7 | 104.7 |
| 13330 | ENSMUSG00000045075 | 1.266900876 | -2.548032382 | 2.280137557 | -1.117490642 | 0.263784599 | NA | Gm9796 | predicted gene 9796 [Source:MGI Symbol | 1.6062 | 0 | 5.0312 | 0.964 | 0 | 0 |
| 13331 | ENSMUSG00000045078 | 411.6661392 | 0.013755114 | 0.143172337 | 0.096073828 | 0.923461937 | 0.98705827 | Rnf216 | ring finger protein 216 [Source:MGI Symb | 371.03 | 425.56 | 433.69 | 453.07 | 405.71 | 380.93 |
| 13332 | ENSMUSG00000045083 | 0 | NA | NA | NA | NA | NA | Lingo2 | leucine rich repeat and Ig domain contain | 0 | 0 | 0 | 0 | 0 | 0 |
| 13333 | ENSMUSG00000045087 | 175.7783031 | 0.792635462 | 0.253754586 | 3.123630094 | 0.001786349 | 0.032166791 | S1pr5 | sphingosine-1-phosphate receptor 5 [Sou | 107.62 | 90.457 | 188.17 | 196.65 | 232.3 | 239.48 |
| 13334 | ENSMUSG00000045091 | 0 | NA | NA | NA | NA | NA | Aqp12 | aquaporin 12 [Source:MGI Symbol;Acc:M | 0 | 0 | 0 | 0 | 0 | 0 |
| 13335 | ENSMUSG00000045092 | 700.0742308 | 0.430858497 | 0.159722789 | 2.697539282 | 0.006985404 | 0.084057311 | S1pr1 | sphingosine-1-phosphate receptor 1 [Sou | 487.48 | 649.64 | 653.05 | 833.84 | 664.19 | 912.24 |
| 13336 | ENSMUSG00000045094 | 217.1797727 | -0.169907305 | 0.180256814 | -0.942584646 | 0.345893397 | 0.786899236 | Arhgef37 | Rho guanine nucleotide exchange factor ( | 193.55 | 232.31 | 264.64 | 200.51 | 211.58 | 200.49 |
| 13337 | ENSMUSG00000045095 | 420.0061688 | 0.053245115 | 0.15182859 | 0.350692282 | 0.725819216 | 0.945362372 | Magi1 | membrane associated guanylate kinase, W | 452.95 | 433.78 | 349.17 | 395.23 | 467.88 | 421.03 |
| 13338 | ENSMUSG00000045098 | 489.5730095 | 0.090288386 | 0.133577819 | 0.675923493 | 0.499089221 | 0.880361745 | Kmt5b | lysine methyltransferase 5B [Source:MGI S | 444.12 | 520.13 | 458.85 | 480.06 | 492.96 | 541.33 |
| 13339 | ENSMUSG00000045100 | 181.0532134 | 0.035624039 | 0.184167278 | 0.193433056 | 0.846619829 | 0.972803041 | Slc25a26 | solute carrier family 25 (mitochondrial car | 159.01 | 181.94 | 196.22 | 197.62 | 177.77 | 173.76 |
| 13340 | ENSMUSG00000045102 | 1.613177868 | 0.743173382 | 1.721904591 | 0.431599629 | 0.666032428 | NA | Poln | DNA polymerase N [Source:MGI Symbol;A | 1.6062 | 0 | 2.0125 | 3.8559 | 1.0906 | 1.1138 |
| 13341 | ENSMUSG00000045103 | 569.720066 | -0.127911655 | 0.155438486 | -0.822908524 | 0.410560015 | 0.833241465 | Dmd | dystrophin, muscular dystrophy [Source:N | 628.03 | 677.4 | 478.97 | 505.12 | 605.29 | 523.51 |
| 13342 | ENSMUSG00000045104 | 0 | NA | NA | NA | NA | NA | Ldhb-ps | lactate dehydrogenase B, pseudogene [Sc | 0 | 0 | 0 | 0 | 0 | 0 |
| 13343 | ENSMUSG00000045106 | 28.34861946 | 0.178601348 | 0.403579175 | 0.442543518 | 0.658095947 | 0.927303282 | Ccdc73 | coiled-coil domain containing 73 [Source:I | 28.912 | 22.614 | 28.175 | 30.847 | 28.356 | 31.188 |
| 13344 | ENSMUSG00000045107 | 216.9711623 | -0.059524141 | 0.179171527 | -0.33221875 | 0.739724087 | 0.948393197 | Saysd1 | SAYSVFN motif domain containing 1 [Sour | 216.84 | 197.36 | 250.55 | 237.14 | 206.13 | 193.81 |
| 13345 | ENSMUSG00000045109 | 0 | NA | NA | NA | NA | NA | Krtap4-7 | keratin associated protein 4-7 [Source:MG | 0 | 0 | 0 | 0 | 0 | 0 |
| 13346 | ENSMUSG00000045111 | 0 | NA | NA | NA | NA | NA | Taar6 | trace amine-associated receptor 6 [Source | 0 | 0 | 0 | 0 | 0 | 0 |
| 13347 | ENSMUSG00000045114 | 0.167707431 | -0.796629176 | 4.080472857 | -0.195229623 | 0.84521319 | NA | Prrt2 | proline-rich transmembrane protein 2 [So | 0 | 0 | 1.0062 | 0 | 0 | 0 |
| 13348 | ENSMUSG00000045126 | 0 | NA | NA | NA | NA | NA | Olfr1445 | olfactory receptor 1445 [Source:MGI Syml | 0 | 0 | 0 | 0 | 0 | 0 |
| 13349 | ENSMUSG00000045128 | 3970.867664 | -0.017143967 | 0.084351053 | -0.203245435 | 0.838943202 | 0.971535186 | Rpl18a | ribosomal protein L18A [Source:MGI Syml | 3883 | 4034.6 | 4066.2 | 3907 | 3952.4 | 3982 |
| 13350 | ENSMUSG00000045132 | 0 | NA | NA | NA | NA | NA | Olfr620 | olfactory receptor 620 [Source:MGI Symb | 0 | 0 | 0 | 0 | 0 | 0 |
| 13351 | ENSMUSG00000045136 | 1.045257913 | 1.114000467 | 2.212392902 | 0.50352741 | 0.614593514 | NA | Tubb2b | tubulin, beta 2B class IIB [Source:MGI Sym | 0 | 0 | 2.0125 | 0.964 | 2.1812 | 1.1138 |
| 13352 | ENSMUSG00000045140 | 83.9784141 | 0.14756286 | 0.245674891 | 0.600642825 | 0.548077908 | 0.895600915 | Pigw | phosphatidylinositol glycan anchor biosyn | 79.507 | 76.066 | 83.518 | 91.578 | 78.525 | 94.677 |
| 13353 | ENSMUSG00000045148 | 0 | NA | NA | NA | NA | NA | Olfr1255 | olfactory receptor 1255 [Source:MGI Syml | 0 | 0 | 0 | 0 | 0 | 0 |
| 13354 | ENSMUSG00000045150 | 0 | NA | NA | NA | NA | NA | Olfr1161 | olfactory receptor 1161 [Source:MGI Syml | 0 | 0 | 0 | 0 | 0 | 0 |
| 13355 | ENSMUSG00000045160 | 811.7404979 | -0.022997899 | 0.117030804 | -0.196511502 | 0.844209831 | 0.972803041 | Bola3 | bolA-like 3 (E. coli) [Source:MGI Symbol;A | 862.53 | 814.11 | 776.82 | 740.33 | 822.33 | 854.32 |
| 13356 | ENSMUSG00000045165 | 6.689620028 | -1.323194144 | 0.848822718 | -1.558858071 | 0.119029976 | 0.504735151 | AI467606 | expressed sequence AI467606 [Source:MG | 10.44 | 7.1954 | 11.069 | 4.8199 | 3.2719 | 3.3415 |
| 13357 | ENSMUSG00000045174 | 0 | NA | NA | NA | NA | NA | Amer3 | APC membrane recruitment 3 [Source:MG | 0 | 0 | 0 | 0 | 0 | 0 |
| 13358 | ENSMUSG00000045176 | 123.5038012 | 0.086953354 | 0.225899628 | 0.384920305 | 0.700296461 | 0.941626823 | Borcs6 | BLOC-1 related complex subunit 6 [Source | 128.5 | 116.15 | 114.71 | 146.52 | 99.246 | 135.89 |
| 13359 | ENSMUSG00000045178 | 0 | NA | NA | NA | NA | NA | 4933411O13 | RIKEN cDNA 4933411O13 gene [Source:M | 0 | 0 | 0 | 0 | 0 | 0 |
| 13360 | ENSMUSG00000045179 | 0 | NA | NA | NA | NA | NA | Sox3 | SRY (sex determining region Y)-box 3 [Sou | 0 | 0 | 0 | 0 | 0 | 0 |
| 13361 | ENSMUSG00000045180 | 290.8268971 | 0.197974675 | 0.174220841 | 1.136343242 | 0.25581294 | 0.708918985 | Shroom2 | shroom family member 2 [Source:MGI Syr | 275.46 | 262.12 | 275.71 | 381.74 | 275.93 | 274.01 |
| 13362 | ENSMUSG00000045193 | 372.7843334 | 0.094928297 | 0.173316411 | 0.547716727 | 0.583886425 | 0.908955096 | Cirbp | cold inducible RNA binding protein [Sourc | 320.44 | 364.91 | 397.47 | 464.64 | 376.26 | 312.99 |
| 13363 | ENSMUSG00000045201 | 0 | NA | NA | NA | NA | NA | Lrrc3b | leucine rich repeat containing 3B [Source: | 0 | 0 | 0 | 0 | 0 | 0 |
| 13364 | ENSMUSG00000045202 | 0 | NA | NA | NA | NA | NA | Olfr123 | olfactory receptor 123 [Source:MGI Symb | 0 | 0 | 0 | 0 | 0 | 0 |
| 13365 | ENSMUSG00000045204 | 0 | NA | NA | NA | NA | NA | Olfr835 | olfactory receptor 835 [Source:MGI Symb | 0 | 0 | 0 | 0 | 0 | 0 |
| 13366 | ENSMUSG00000045205 | 220.0346412 | -0.221636418 | 0.188333194 | -1.176831413 | 0.239262802 | 0.688695954 | Dpy19l4 | dpy-19-like 4 (C. elegans) [Source:MGI Syr | 224.07 | 286.79 | 200.24 | 212.08 | 185.41 | 211.63 |
| 13367 | ENSMUSG00000045210 | 683.9877024 | 0.105057971 | 0.113085089 | 0.929017007 | 0.352880273 | 0.793259753 | Vcpip1 | valosin containing protein (p97)/p47 comp | 644.09 | 663.01 | 670.16 | 680.57 | 716.54 | 729.57 |
| 13368 | ENSMUSG00000045211 | 193.0687368 | 0.212250014 | 0.215425781 | 0.985258188 | 0.324497265 | 0.771309263 | Nudt18 | nudix (nucleoside diphosphate linked moi | 235.31 | 155.22 | 144.9 | 190.87 | 211.58 | 220.54 |
| 13369 | ENSMUSG00000045215 | 0.17131964 | -0.796629176 | 4.080472857 | -0.195229623 | 0.84521319 | NA | Asxl3 | ASXL transcriptional regulator 3 [Source:N | 0 | 1.0279 | 0 | 0 | 0 | 0 |
| 13370 | ENSMUSG00000045216 | 2809.248078 | 0.090240915 | 0.128427975 | 0.702657777 | 0.482269047 | 0.871593699 | Hs6st1 | heparan sulfate 6-O-sulfotransferase 1 [Sc | 2871.1 | 2915.2 | 2376.7 | 2517.9 | 2852 | 3322.6 |
| 13371 | ENSMUSG00000045217 | 0 | NA | NA | NA | NA | NA | Ppp1r2-ps9 | protein phosphatase 1, regulatory (inhibit | 0 | 0 | 0 | 0 | 0 | 0 |
| 13372 | ENSMUSG00000045225 | 0 | NA | NA | NA | NA | NA | Olfr1152 | olfactory receptor 1152 [Source:MGI Syml | 0 | 0 | 0 | 0 | 0 | 0 |
| 13373 | ENSMUSG00000045231 | 5.524185461 | 0.947102919 | 0.93092124 | 1.017382436 | 0.308971534 | 0.757462943 | BC106179 | cDNA sequence BC106179 [Source:MGI Sy | 3.2124 | 6.1675 | 2.0125 | 9.6398 | 6.5437 | 5.5692 |
| 13374 | ENSMUSG00000045232 | 0.698954292 | 0.126053087 | 3.048972385 | 0.04134281 | 0.967022605 | NA | Rln3 | relaxin 3 [Source:MGI Symbol;Acc:MGI:21 | 0 | 0 | 2.0125 | 0 | 2.1812 | 0 |
| 13375 | ENSMUSG00000045236 | 0 | NA | NA | NA | NA | NA | Krtap5-4 | keratin associated protein 5-4 [Source:MG | 0 | 0 | 0 | 0 | 0 | 0 |
| 13376 | ENSMUSG00000045237 | 205.9247338 | 0.39669356 | 0.211704722 | 1.873805912 | 0.060957181 | 0.352269825 | Eola1 | endothelium and lymphocyte associated A | 176.68 | 145.96 | 210.31 | 193.76 | 218.12 | 290.71 |
| 13377 | ENSMUSG00000045238 | 0 | NA | NA | NA | NA | NA | A730035I17I | RIKEN cDNA A730035I17 gene [Source:MG | 0 | 0 | 0 | 0 | 0 | 0 |
| 13378 | ENSMUSG00000045246 | 0 | NA | NA | NA | NA | NA | Kcng4 | potassium voltage-gated channel, subfam | 0 | 0 | 0 | 0 | 0 | 0 |
| 13379 | ENSMUSG00000045248 | 94.82754498 | -0.105754176 | 0.279220466 | -0.378747939 | 0.704875051 | 0.94295036 | Med26 | mediator complex subunit 26 [Source:MG | 85.129 | 96.624 | 113.71 | 116.64 | 61.075 | 95.791 |
| 13380 | ENSMUSG00000045251 | 94.46159122 | 0.323679224 | 0.255129723 | 1.268684886 | 0.204553475 | 0.650398212 | Zfp688 | zinc finger protein 688 [Source:MGI Symb | 71.476 | 74.01 | 106.66 | 112.79 | 93.793 | 108.04 |

| | | | | | | | | | | | | | | | |
|---|---|---|---|---|---|---|---|---|---|---|---|---|---|---|---|
| 13381 | ENSMUSG00000045252 | 298.224159 | -0.124838044 | 0.162113382 | -0.770066248 | 0.441260596 | 0.849828766 | Zfp574 | zinc finger protein 574 [Source:MGI Symb | 326.86 | 273.43 | 333.07 | 306.54 | 298.83 | 250.61 |
| 13382 | ENSMUSG00000045257 | 53.60736799 | -0.459209918 | 0.315243392 | -1.456683721 | 0.145203705 | 0.555361204 | Morn2 | MORN repeat containing 2 [Source:MGI S | 61.839 | 68.87 | 55.343 | 39.523 | 39.262 | 56.806 |
| 13383 | ENSMUSG00000045259 | 0.294513347 | 0.165143496 | 4.080472857 | 0.040471657 | 0.967717103 | NA | Klhdc9 | kelch domain containing 9 [Source:MGI Sy | 0.8031 | 0 | 0 | 0.964 | 0 | 0 |
| 13384 | ENSMUSG00000045267 | 0 | NA | NA | NA | NA | NA | Tas2r119 | taste receptor, type 2, member 119 [Sour | 0 | 0 | 0 | 0 | 0 | 0 |
| 13385 | ENSMUSG00000045268 | 105.2316474 | 0.133718628 | 0.21990409 | 0.608077039 | 0.54313638 | 0.894344786 | Zfp691 | zinc finger protein 691 [Source:MGI Symb | 103.6 | 101.76 | 95.593 | 109.89 | 104.7 | 115.84 |
| 13386 | ENSMUSG00000045273 | 2.399736204 | -2.510937729 | 1.717938497 | -1.461599314 | 0.143851043 | NA | Cenph | centromere protein H [Source:MGI Symbo | 4.0155 | 7.1954 | 1.0062 | 0 | 2.1812 | 0 |
| 13387 | ENSMUSG00000045275 | 0.538730002 | 1.11098254 | 3.037664833 | 0.365735722 | 0.714562288 | NA | Lca5l | Leber congenital amaurosis 5-like [Source | 0 | 1.0279 | 0 | 0 | 1.0906 | 1.1138 |
| 13388 | ENSMUSG00000045281 | 0 | NA | NA | NA | NA | NA | Gpr20 | G protein-coupled receptor 20 [Source:M | 0 | 0 | 0 | 0 | 0 | 0 |
| 13389 | ENSMUSG00000045282 | 1172.855733 | 0.173157476 | 0.131494434 | 1.316842625 | 0.187891382 | 0.624893296 | Tmem86b | transmembrane protein 86B [Source:MGI | 1105.1 | 1002.2 | 1199.4 | 1053.6 | 1297.8 | 1378.9 |
| 13390 | ENSMUSG00000045284 | 74.90658154 | 0.042234569 | 0.312754522 | 0.135040636 | 0.892579746 | 0.982512308 | Dcaf12l1 | DDB1 and CUL4 associated factor 12-like 1 | 92.357 | 80.178 | 48.3 | 59.767 | 74.162 | 94.677 |
| 13391 | ENSMUSG00000045287 | 174.0063265 | 0.024835766 | 0.249044447 | 0.099724231 | 0.920563263 | 0.986685804 | Rtn4rl1 | reticulon 4 receptor-like 1 [Source:MGI Sy | 114.04 | 173.72 | 230.43 | 163.88 | 158.14 | 203.83 |
| 13392 | ENSMUSG00000045288 | 3.447423603 | -0.915202129 | 1.164792726 | -0.78572102 | 0.432030946 | NA | Ush1g | USH1 protein network component sans [S | 6.4248 | 3.0838 | 4.025 | 3.8559 | 2.1812 | 1.1138 |
| 13393 | ENSMUSG00000045291 | 0 | NA | NA | NA | NA | NA | E330021D16 | RIKEN cDNA E330021D16 gene [Source:M | 0 | 0 | 0 | 0 | 0 | 0 |
| 13394 | ENSMUSG00000045294 | 2518.62265 | 0.793127773 | 0.153731018 | 5.159191572 | 2.48E-07 | 2.41E-05 | Insig1 | insulin induced gene 1 [Source:MGI Symb | 2075.2 | 1845.1 | 1608 | 2627.8 | 3990.6 | 2965.1 |
| 13395 | ENSMUSG00000045302 | 1573.221861 | 0.097504519 | 0.09362035 | 1.041488508 | 0.297648884 | 0.748091545 | Preb | prolactin regulatory element binding [Sou | 1495.4 | 1527.5 | 1537.5 | 1605 | 1658.8 | 1615.1 |
| 13396 | ENSMUSG00000045306 | 0 | NA | NA | NA | NA | NA | Olfr734 | olfactory receptor 734 [Source:MGI Symb | 0 | 0 | 0 | 0 | 0 | 0 |
| 13397 | ENSMUSG00000045312 | 68.32186497 | -0.03073991 | 0.319780796 | -0.096128069 | 0.923418859 | 0.98705827 | Lhfpl2 | lipoma HMGIC fusion partner-like 2 [Sour | 81.113 | 45.228 | 80.5 | 69.406 | 51.259 | 82.424 |
| 13398 | ENSMUSG00000045314 | 101.8303287 | -0.103219576 | 0.234645877 | -0.439895121 | 0.66001307 | 0.927787797 | Sowahb | sosondowah ankyrin repeat domain famil | 96.372 | 98.68 | 121.76 | 108.93 | 88.34 | 96.904 |
| 13399 | ENSMUSG00000045316 | 1210.348791 | 0.17242788 | 0.119226956 | 1.446215571 | 0.148116748 | 0.559801578 | Fahd1 | fumarylacetoacetate hydrolase domain cc | 1110.7 | 1113.2 | 1191.4 | 1421.9 | 1304.4 | 1120.5 |
| 13400 | ENSMUSG00000045318 | 0 | NA | NA | NA | NA | NA | Adra2c | adrenergic receptor, alpha 2c [Source:MG | 0 | 0 | 0 | 0 | 0 | 0 |
| 13401 | ENSMUSG00000045319 | 353.3008983 | -0.790843471 | 0.205617227 | -3.846192667 | 0.000119967 | 0.004191326 | Proser2 | proline and serine rich 2 [Source:MGI Sym | 373.44 | 463.59 | 506.14 | 194.72 | 343.54 | 238.36 |
| 13402 | ENSMUSG00000045322 | 1.932616755 | -0.336864596 | 1.497862992 | -0.224896802 | 0.822059557 | NA | Tlr9 | toll-like receptor 9 [Source:MGI Symbol;A | 2.4093 | 3.0838 | 1.0062 | 2.8919 | 1.0906 | 1.1138 |
| 13403 | ENSMUSG00000045326 | 0 | NA | NA | NA | NA | NA | Fndc7 | fibronectin type III domain containing 7 [S | 0 | 0 | 0 | 0 | 0 | 0 |
| 13404 | ENSMUSG00000045327 | 16.30982756 | 0.727488186 | 0.54533827 | 1.334012715 | 0.182199689 | 0.615722956 | 6330549D23 | RIKEN cDNA 6330549D23 gene [Source:M | 10.44 | 17.475 | 9.0562 | 21.208 | 19.631 | 20.049 |
| 13405 | ENSMUSG00000045328 | 4.242545481 | -0.943713882 | 1.186938841 | -0.795082147 | 0.426565719 | NA | Cenpe | centromere protein E [Source:MGI Symbo | 6.4248 | 8.2233 | 2.0125 | 0 | 5.4531 | 3.3415 |
| 13406 | ENSMUSG00000045330 | 0 | NA | NA | NA | NA | NA | Magec2 | MAGE family member C2 [Source:MGI Syr | 0 | 0 | 0 | 0 | 0 | 0 |
| 13407 | ENSMUSG00000045331 | 0 | NA | NA | NA | NA | NA | 2310079G19 | RIKEN cDNA 2310079G19 gene [Source:M | 0 | 0 | 0 | 0 | 0 | 0 |
| 13408 | ENSMUSG00000045333 | 13.41817297 | -0.8384067 | 0.659549369 | -1.271181113 | 0.203664224 | 0.64863958 | Zfp423 | zinc finger protein 423 [Source:MGI Symb | 24.093 | 18.503 | 9.0562 | 13.496 | 10.906 | 4.4554 |
| 13409 | ENSMUSG00000045336 | 0 | NA | NA | NA | NA | NA | Hsfy2 | heat shock transcription factor, Y-linked 2 | 0 | 0 | 0 | 0 | 0 | 0 |
| 13410 | ENSMUSG00000045337 | 0 | NA | NA | NA | NA | NA | Defb11 | defensin beta 11 [Source:MGI Symbol;Acc | 0 | 0 | 0 | 0 | 0 | 0 |
| 13411 | ENSMUSG00000045339 | 0 | NA | NA | NA | NA | NA | A930104D05 | RIKEN cDNA A930104D05 gene [Source:M | 0 | 0 | 0 | 0 | 0 | 0 |
| 13412 | ENSMUSG00000045340 | 0 | NA | NA | NA | NA | NA | Vmn1r70 | vomeronasal 1 receptor 70 [Source:MGI S | 0 | 0 | 0 | 0 | 0 | 0 |
| 13413 | ENSMUSG00000045341 | 0 | NA | NA | NA | NA | NA | Olfr167 | olfactory receptor 167 [Source:MGI Symb | 0 | 0 | 0 | 0 | 0 | 0 |
| 13414 | ENSMUSG00000045345 | 5.493429793 | 0.334902814 | 0.990875479 | 0.337986782 | 0.735373149 | 0.948109711 | 9530056K15 | RIKEN cDNA 9530056K15 gene [Source:M | 6.4248 | 5.1396 | 3.0187 | 10.604 | 1.0906 | 6.6831 |
| 13415 | ENSMUSG00000045348 | 2.860587897 | -0.332179554 | 1.291933542 | -0.257118144 | 0.797087571 | NA | Nyap1 | neuronal tyrosine-phosphorylated phospl | 2.4093 | 4.1117 | 3.0187 | 0.964 | 1.0906 | 5.5692 |
| 13416 | ENSMUSG00000045349 | 2.633348855 | 2.83378612 | 1.6064032 | 1.764056571 | 0.077722455 | NA | Sh2d5 | SH2 domain containing 5 [Source:MGI Syr | 0 | 0 | 2.0125 | 3.8559 | 4.3625 | 5.5692 |
| 13417 | ENSMUSG00000045350 | 6.306529474 | 0.159933192 | 0.986098202 | 0.162187895 | 0.871157892 | 0.979054215 | Fam186a | family with sequence similarity 186, meml | 4.8186 | 6.1675 | 7.0437 | 15.424 | 3.2719 | 1.1138 |
| 13418 | ENSMUSG00000045362 | 5.555385914 | -0.197186531 | 0.926994331 | -0.212716005 | 0.831548482 | 0.970533397 | Tnfrsf26 | tumor necrosis factor receptor superfamil | 5.6217 | 8.2233 | 4.025 | 7.7118 | 2.1812 | 5.5692 |
| 13419 | ENSMUSG00000045364 | 0 | NA | NA | NA | NA | NA | Ifne | interferon epsilon [Source:MGI Symbol;Ac | 0 | 0 | 0 | 0 | 0 | 0 |
| 13420 | ENSMUSG00000045374 | 876.8602918 | 0.124094618 | 0.145179421 | 0.854767279 | 0.392679996 | 0.821375223 | Wdr81 | WD repeat domain 81 [Source:MGI Symbo | 822.38 | 703.1 | 992.16 | 947.59 | 816.87 | 979.07 |
| 13421 | ENSMUSG00000045377 | 24.54089159 | -1.335273408 | 0.454204349 | -2.939807625 | 0.003284161 | 0.049844808 | Tmem88 | transmembrane protein 88 [Source:MGI S | 39.352 | 34.949 | 31.194 | 16.388 | 11.997 | 13.366 |
| 13422 | ENSMUSG00000045378 | 0 | NA | NA | NA | NA | NA | Pxt1 | peroxisomal, testis specific 1 [Source:MGI | 0 | 0 | 0 | 0 | 0 | 0 |
| 13423 | ENSMUSG00000045381 | 0.17131964 | -0.796629176 | 4.080472857 | -0.195229623 | 0.84521319 | NA | Olfr433 | olfactory receptor 433 [Source:MGI Symb | 0 | 1.0279 | 0 | 0 | 0 | 0 |
| 13424 | ENSMUSG00000045382 | 11.41118721 | -2.421431235 | 0.792805658 | -3.054255743 | 0.002256197 | 0.038078911 | Cxcr4 | chemokine (C-X-C motif) receptor 4 [Sour | 13.653 | 30.838 | 13.081 | 0.964 | 4.3625 | 5.5692 |
| 13425 | ENSMUSG00000045391 | 0.319491021 | 0.16514334 | 4.044653141 | 0.040830038 | 0.967431393 | NA | 1700120B22 | RIKEN cDNA 1700120B22 gene [Source:M | 0.8031 | 0 | 0 | 0 | 0 | 1.1138 |
| 13426 | ENSMUSG00000045392 | 201.4750414 | -0.077620915 | 0.190549992 | -0.40735197 | 0.683749498 | 0.935528319 | Olfr1033 | olfactory receptor 1033 [Source:MGI Syml | 225.67 | 215.86 | 179.11 | 214 | 211.58 | 162.62 |
| 13427 | ENSMUSG00000045394 | 17.99235907 | -0.545629171 | 0.582771914 | -0.936265386 | 0.349136565 | 0.79040715 | Epcam | epithelial cell adhesion molecule [Source: | 17.668 | 10.279 | 36.225 | 17.352 | 14.178 | 12.252 |
| 13428 | ENSMUSG00000045395 | 0 | NA | NA | NA | NA | NA | Olfr1499 | olfactory receptor 1499 [Source:MGI Syml | 0 | 0 | 0 | 0 | 0 | 0 |
| 13429 | ENSMUSG00000045404 | 8.097955326 | -3.838955649 | 1.118173269 | -3.433238617 | 0.000596417 | 0.014072716 | Kcnk13 | potassium channel, subfamily K, member | 9.6372 | 26.726 | 9.0562 | 0.964 | 1.0906 | 1.1138 |
| 13430 | ENSMUSG00000045409 | 299.3299836 | 0.185015061 | 0.159708865 | 1.158452041 | 0.246679612 | 0.696725814 | Trim39 | tripartite motif-containing 39 [Source:MG | 270.65 | 289.87 | 280.74 | 368.24 | 285.74 | 300.74 |
| 13431 | ENSMUSG00000045410 | 881.2495024 | 0.142328006 | 0.14072531 | 1.011388825 | 0.311830368 | 0.759830222 | Akr1e1 | aldo-keto reductase family 1, member E1 | 878.59 | 788.41 | 845.25 | 758.65 | 1073.2 | 943.43 |
| 13432 | ENSMUSG00000045411 | 192.3022802 | 0.449452291 | 0.195509447 | 2.298877618 | 0.02151189 | 0.182705797 | 2410002F23 | RIKEN cDNA 2410002F23 gene [Source:M | 164.64 | 169.61 | 153.96 | 270.88 | 188.68 | 206.06 |
| 13433 | ENSMUSG00000045414 | 895.4276975 | 0.215936226 | 0.169536155 | 1.273688355 | 0.202773885 | 0.647682949 | Dipk2a | divergent protein kinase domain 2A [Sour | 853.7 | 841.86 | 790.91 | 1263.8 | 862.68 | 759.64 |
| 13434 | ENSMUSG00000045417 | 0 | NA | NA | NA | NA | NA | Vmn1r230 | vomeronasal 1 receptor 230 [Source:MGI | 0 | 0 | 0 | 0 | 0 | 0 |
| 13435 | ENSMUSG00000045421 | 0 | NA | NA | NA | NA | NA | Olfr1390 | olfactory receptor 1390 [Source:MGI Syml | 0 | 0 | 0 | 0 | 0 | 0 |
| 13436 | ENSMUSG00000045427 | 956.4788906 | 0.074427182 | 0.115444137 | 0.644703005 | 0.519119669 | 0.887640015 | Hnrnph2 | heterogeneous nuclear ribonucleoprotein | 899.47 | 949.8 | 945.87 | 891.68 | 977.19 | 1074.9 |

| | | | | | | | | | | | | | | | |
|---|---|---|---|---|---|---|---|---|---|---|---|---|---|---|---|
| 13437 | ENSMUSG00000045435 | 160.7225274 | 0.20794179 | 0.195936803 | 1.06126969 | 0.288567356 | 0.73951397 | Tmem60 | transmembrane protein 60 [Source:MGI S | 163.03 | 141.85 | 141.88 | 148.45 | 188.68 | 180.44 |
| 13438 | ENSMUSG00000045438 | 723.1899757 | 0.064777052 | 0.119974218 | 0.539924772 | 0.589248913 | 0.911039598 | Cox19 | cytochrome c oxidase assembly protein 1! | 770.98 | 659.92 | 689.28 | 775.04 | 714.35 | 729.57 |
| 13439 | ENSMUSG00000045440 | 0 | NA | NA | NA | NA | NA | Insm2 | insulinoma-associated 2 [Source:MGI Sym | 0 | 0 | 0 | 0 | 0 | 0 |
| 13440 | ENSMUSG00000045441 | 653.5101147 | 0.376309165 | 0.149805053 | 2.511992476 | 0.012005165 | 0.122629028 | Gprin3 | GPRIN family member 3 [Source:MGI Sym | 607.15 | 555.08 | 544.38 | 892.64 | 694.72 | 627.09 |
| 13441 | ENSMUSG00000045455 | 2.686766939 | 0.890021341 | 1.269721055 | 0.700958165 | 0.483329124 | NA | Gm9797 | predicted pseudogene 9797 [Source:MGI | 1.6062 | 2.0558 | 2.0125 | 3.8559 | 4.3625 | 2.2277 |
| 13442 | ENSMUSG00000045463 | 0 | NA | NA | NA | NA | NA | Slco6b1 | solute carrier organic anion transporter fa | 0 | 0 | 0 | 0 | 0 | 0 |
| 13443 | ENSMUSG00000045464 | 31.51828889 | -0.036936965 | 0.415014812 | -0.089001557 | 0.929080672 | 0.988405116 | 2810002D19 | RIKEN cDNA 2810002D19 gene [Source:M | 38.549 | 24.67 | 32.2 | 24.099 | 27.265 | 42.326 |
| 13444 | ENSMUSG00000045466 | 93.74671738 | 0.484769034 | 0.242789995 | 1.996659841 | 0.045862147 | 0.296229283 | Zfp956 | zinc finger protein 956 [Source:MGI Symb | 85.932 | 63.731 | 84.525 | 116.64 | 103.61 | 108.04 |
| 13445 | ENSMUSG00000045467 | 18.3966416 | -0.19613297 | 0.547618081 | -0.358156491 | 0.720226205 | 0.9447719 | Ttll13 | tubulin tyrosine ligase-like family, membe | 10.44 | 22.614 | 26.162 | 19.28 | 19.631 | 12.252 |
| 13446 | ENSMUSG00000045471 | 0 | NA | NA | NA | NA | NA | Hcrt | hypocretin [Source:MGI Symbol;Acc:MGI: | 0 | 0 | 0 | 0 | 0 | 0 |
| 13447 | ENSMUSG00000045474 | 0 | NA | NA | NA | NA | NA | Olfr1368 | olfactory receptor 1368 [Source:MGI Sym | 0 | 0 | 0 | 0 | 0 | 0 |
| 13448 | ENSMUSG00000045475 | 0 | NA | NA | NA | NA | NA | Lce3c | late cornified envelope 3C [Source:MGI Sy | 0 | 0 | 0 | 0 | 0 | 0 |
| 13449 | ENSMUSG00000045479 | 0 | NA | NA | NA | NA | NA | Olfr459 | olfactory receptor 459 [Source:MGI Symb | 0 | 0 | 0 | 0 | 0 | 0 |
| 13450 | ENSMUSG00000045482 | 578.5073767 | -0.085721385 | 0.156108136 | -0.549115427 | 0.582926239 | 0.908645447 | Trrap | transformation/transcription domain-asso | 575.82 | 653.76 | 558.47 | 650.69 | 585.66 | 446.65 |
| 13451 | ENSMUSG00000045493 | 0 | NA | NA | NA | NA | NA | Bhlhe23 | basic helix-loop-helix family, member e23 | 0 | 0 | 0 | 0 | 0 | 0 |
| 13452 | ENSMUSG00000045498 | 0.371281295 | 2.09253362 | 4.031852165 | 0.519000582 | 0.603760335 | NA | Pcdhb3 | protocadherin beta 3 [Source:MGI Symbo | 0 | 0 | 0 | 0 | 0 | 2.2277 |
| 13453 | ENSMUSG00000045502 | 3.191024796 | -1.437605492 | 1.312081305 | -1.095667994 | 0.273224094 | NA | Hcar2 | hydroxycarboxylic acid receptor 2 [Source | 4.8186 | 7.1954 | 2.0125 | 2.8919 | 0 | 2.2277 |
| 13454 | ENSMUSG00000045503 | 425.1164895 | -0.008596433 | 0.161292805 | -0.053297065 | 0.957495219 | 0.991861398 | Sys1 | SYS1 Golgi-localized integral membrane pr | 440.9 | 385.47 | 451.8 | 346.07 | 430.79 | 495.66 |
| 13455 | ENSMUSG00000045506 | 1.660838485 | 1.346556011 | 1.746883686 | 0.770833239 | 0.440805782 | NA | A930002H24 | RIKEN cDNA A930002H24 gene [Source:M | 0.8031 | 1.0279 | 1.0062 | 3.8559 | 3.2719 | 0 |
| 13456 | ENSMUSG00000045508 | 0 | NA | NA | NA | NA | NA | Olfr1367 | olfactory receptor 1367 [Source:MGI Sym | 0 | 0 | 0 | 0 | 0 | 0 |
| 13457 | ENSMUSG00000045509 | 0.267700746 | -1.497481649 | 4.057001436 | -0.369110456 | 0.712045396 | NA | Gpr150 | G protein-coupled receptor 150 [Source:N | 1.6062 | 0 | 0 | 0 | 0 | 0 |
| 13458 | ENSMUSG00000045514 | 0 | NA | NA | NA | NA | NA | Olfr460 | olfactory receptor 460 [Source:MGI Symb | 0 | 0 | 0 | 0 | 0 | 0 |
| 13459 | ENSMUSG00000045515 | 0 | NA | NA | NA | NA | NA | Pou3f3 | POU domain, class 3, transcription factor : | 0 | 0 | 0 | 0 | 0 | 0 |
| 13460 | ENSMUSG00000045518 | 0 | NA | NA | NA | NA | NA | Onecut3 | one cut domain, family member 3 [Source | 0 | 0 | 0 | 0 | 0 | 0 |
| 13461 | ENSMUSG00000045519 | 68.03140995 | -0.303611437 | 0.28691216 | -1.058203448 | 0.2899627 | 0.740058153 | Zfp560 | zinc finger protein 560 [Source:MGI Symb | 57.02 | 85.317 | 83.518 | 57.839 | 64.346 | 60.148 |
| 13462 | ENSMUSG00000045521 | 0.17131964 | -0.796629176 | 4.080472857 | -0.195229623 | 0.84521319 | NA | Tssk2 | testis-specific serine kinase 2 [Source:MG | 0 | 1.0279 | 0 | 0 | 0 | 0 |
| 13463 | ENSMUSG00000045528 | 0 | NA | NA | NA | NA | NA | Olfr891 | olfactory receptor 891 [Source:MGI Symb | 0 | 0 | 0 | 0 | 0 | 0 |
| 13464 | ENSMUSG00000045532 | 0 | NA | NA | NA | NA | NA | C1ql1 | complement component 1, q subcompon | 0 | 0 | 0 | 0 | 0 | 0 |
| 13465 | ENSMUSG00000045534 | 0.503122293 | -2.366675966 | 3.785596405 | -0.625179156 | 0.531853481 | NA | Kcna5 | potassium voltage-gated channel, shaker- | 0 | 0 | 3.0187 | 0 | 0 | 0 |
| 13466 | ENSMUSG00000045538 | 336.5114285 | 0.122560465 | 0.148158439 | 0.827225679 | 0.408109159 | 0.832096262 | Ddx28 | DEAD box helicase 28 [Source:MGI Symbo | 342.92 | 289.87 | 333.07 | 325.82 | 364.27 | 363.11 |
| 13467 | ENSMUSG00000045539 | 0 | NA | NA | NA | NA | NA | Sprr3 | small proline-rich protein 3 [Source:MGI S | 0 | 0 | 0 | 0 | 0 | 0 |
| 13468 | ENSMUSG00000045540 | 0 | NA | NA | NA | NA | NA | Olfr600 | olfactory receptor 600 [Source:MGI Symb | 0 | 0 | 0 | 0 | 0 | 0 |
| 13469 | ENSMUSG00000045545 | 0.17131964 | -0.796629176 | 4.080472857 | -0.195229623 | 0.84521319 | NA | Krt14 | keratin 14 [Source:MGI Symbol;Acc:MGI:! | 0 | 1.0279 | 0 | 0 | 0 | 0 |
| 13470 | ENSMUSG00000045551 | 19.2285411 | -2.245423788 | 0.665475658 | -3.37416367 | 0.000740403 | 0.016733115 | Fpr1 | formyl peptide receptor 1 [Source:MGI Sy | 21.684 | 55.508 | 18.112 | 6.7478 | 2.1812 | 11.138 |
| 13471 | ENSMUSG00000045555 | 1.90508177 | -0.16483056 | 1.645020183 | -0.100199719 | 0.920185769 | NA | Mettl24 | methyltransferase like 24 [Source:MGI Syı | 0 | 2.0558 | 4.025 | 0.964 | 3.2719 | 1.1138 |
| 13472 | ENSMUSG00000045559 | 0 | NA | NA | NA | NA | NA | Olfr827 | olfactory receptor 827 [Source:MGI Symb | 0 | 0 | 0 | 0 | 0 | 0 |
| 13473 | ENSMUSG00000045566 | 0 | NA | NA | NA | NA | NA | Sprr4 | small proline-rich protein 4 [Source:MGI S | 0 | 0 | 0 | 0 | 0 | 0 |
| 13474 | ENSMUSG00000045569 | 0 | NA | NA | NA | NA | NA | Mc2r | melanocortin 2 receptor [Source:MGI Sym | 0 | 0 | 0 | 0 | 0 | 0 |
| 13475 | ENSMUSG00000045573 | 1.006397917 | 1.009706429 | 2.42879401 | 0.415723369 | 0.677612439 | NA | Penk | preproenkephalin [Source:MGI Symbol;Ac | 0 | 2.0558 | 0 | 2.8919 | 1.0906 | 0 |
| 13476 | ENSMUSG00000045575 | 0 | NA | NA | NA | NA | NA | Vmn1r233 | vomeronasal 1 receptor 233 [Source:MGI | 0 | 0 | 0 | 0 | 0 | 0 |
| 13477 | ENSMUSG00000045576 | 402.9370869 | -0.166516567 | 0.140725455 | -1.183272537 | 0.236701153 | 0.685551758 | St7l | suppression of tumorigenicity 7-like [Sour | 380.67 | 439.95 | 458.85 | 393.3 | 380.63 | 364.23 |
| 13478 | ENSMUSG00000045581 | 0 | NA | NA | NA | NA | NA | Olfr710 | olfactory receptor 710 [Source:MGI Symb | 0 | 0 | 0 | 0 | 0 | 0 |
| 13479 | ENSMUSG00000045584 | 0 | NA | NA | NA | NA | NA | Olfr610 | olfactory receptor 610 [Source:MGI Symb | 0 | 0 | 0 | 0 | 0 | 0 |
| 13480 | ENSMUSG00000045587 | 0 | NA | NA | NA | NA | NA | Lypd10 | Ly6/PLAUR domain containing 10 [Source: | 0 | 0 | 0 | 0 | 0 | 0 |
| 13481 | ENSMUSG00000045589 | 0 | NA | NA | NA | NA | NA | Frrs1l | ferric-chelate reductase 1 like [Source:MG | 0 | 0 | 0 | 0 | 0 | 0 |
| 13482 | ENSMUSG00000045591 | 0 | NA | NA | NA | NA | NA | Olig3 | oligodendrocyte transcription factor 3 [So | 0 | 0 | 0 | 0 | 0 | 0 |
| 13483 | ENSMUSG00000045594 | 229.7134277 | -0.344739144 | 0.171720627 | -2.007558151 | 0.044690265 | 0.291960252 | Glb1 | galactosidase, beta 1 [Source:MGI Symbol | 232.9 | 243.62 | 294.83 | 194.72 | 205.04 | 207.17 |
| 13484 | ENSMUSG00000045598 | 116.994134 | 0.044722606 | 0.217297609 | 0.205812691 | 0.836937227 | 0.971073158 | Zfp553 | zinc finger protein 553 [Source:MGI Symb | 106.01 | 112.04 | 127.79 | 120.5 | 128.69 | 106.93 |
| 13485 | ENSMUSG00000045608 | 0 | NA | NA | NA | NA | NA | Dbx2 | developing brain homeobox 2 [Source:MG | 0 | 0 | 0 | 0 | 0 | 0 |
| 13486 | ENSMUSG00000045609 | 0 | NA | NA | NA | NA | NA | Gm5455 | predicted pseudogene 5455 [Source:MGI | 0 | 0 | 0 | 0 | 0 | 0 |
| 13487 | ENSMUSG00000045613 | 0 | NA | NA | NA | NA | NA | Chrm2 | cholinergic receptor, muscarinic 2, cardiac | 0 | 0 | 0 | 0 | 0 | 0 |
| 13488 | ENSMUSG00000045620 | 0 | NA | NA | NA | NA | NA | Odf3l1 | outer dense fiber of sperm tails 3-like 1 [S | 0 | 0 | 0 | 0 | 0 | 0 |
| 13489 | ENSMUSG00000045624 | 283.3409427 | -0.214887856 | 0.155774659 | -1.379478908 | 0.167747145 | 0.593804353 | Esf1 | ESF1 nucleolar pre-rRNA processing prote | 306.79 | 302.21 | 303.89 | 240.99 | 295.56 | 250.61 |
| 13490 | ENSMUSG00000045625 | 3.938563695 | 0.238345212 | 1.216086195 | 0.195993683 | 0.84461511 | NA | Pigz | phosphatidylinositol glycan anchor biosyn | 5.6217 | 5.1396 | 0 | 2.8919 | 2.1812 | 7.7969 |
| 13491 | ENSMUSG00000045629 | 16.33347304 | 0.544760857 | 0.53652919 | 1.01534244 | 0.309942621 | 0.758686885 | Sh3tc2 | SH3 domain and tetratricopeptide repeats | 14.456 | 13.363 | 12.075 | 25.063 | 17.45 | 15.594 |
| 13492 | ENSMUSG00000045631 | 0 | NA | NA | NA | NA | NA | Tmprss12 | transmembrane (C-terminal) protease, se | 0 | 0 | 0 | 0 | 0 | 0 |

| | | | | | | | | | | | | | | | |
|---|---|---|---|---|---|---|---|---|---|---|---|---|---|---|---|
| 13493 | ENSMUSG00000045636 | 2941.448925 | 0.050462267 | 0.101368996 | 0.497807697 | 0.618619591 | 0.917509899 | Mtus1 | mitochondrial tumor suppressor 1 [Source | 2830.9 | 3097.1 | 2743 | 3182.1 | 3024.3 | 2771.2 |
| 13494 | ENSMUSG00000045639 | 351.4902087 | 0.04217157 | 0.145220954 | 0.290395901 | 0.771513379 | 0.956965056 | Zfp629 | zinc finger protein 629 [Source:MGI Symb | 341.32 | 317.63 | 380.36 | 374.02 | 339.18 | 356.43 |
| 13495 | ENSMUSG00000045648 | 0 | NA | NA | NA | NA | NA | Vwc2l | von Willebrand factor C domain-containin | 0 | 0 | 0 | 0 | 0 | 0 |
| 13496 | ENSMUSG00000045655 | 0 | NA | NA | NA | NA | NA | Fam216b | family with sequence similarity 216, meml | 0 | 0 | 0 | 0 | 0 | 0 |
| 13497 | ENSMUSG00000045657 | 0.301557804 | -1.64335916 | 4.042814367 | -0.406488899 | 0.684383412 | NA | Pcdhb10 | protocadherin beta 10 [Source:MGI Symb | 0.8031 | 0 | 1.0062 | 0 | 0 | 0 |
| 13498 | ENSMUSG00000045658 | 1421.104151 | -0.009532102 | 0.113011367 | -0.084346404 | 0.932781019 | 0.988420305 | Pid1 | phosphotyrosine interaction domain cont | 1390.2 | 1392.8 | 1494.3 | 1369.8 | 1295.7 | 1583.9 |
| 13499 | ENSMUSG00000045659 | 193.1241301 | 0.590571786 | 0.211144299 | 2.797005592 | 0.005157864 | 0.067944759 | Plekha7 | pleckstrin homology domain containing, fa | 146.97 | 151.1 | 165.02 | 299.8 | 188.68 | 207.17 |
| 13500 | ENSMUSG00000045662 | 0 | NA | NA | NA | NA | NA | Henmt1 | HEN1 methyltransferase homolog 1 (Arab | 0 | 0 | 0 | 0 | 0 | 0 |
| 13501 | ENSMUSG00000045664 | 32.3222336 | -0.691188012 | 0.452308937 | -1.528132556 | 0.12647963 | 0.521187766 | Cdc42ep2 | CDC42 effector protein (Rho GTPase bindi | 46.58 | 22.614 | 50.312 | 20.244 | 18.541 | 35.643 |
| 13502 | ENSMUSG00000045665 | 372.9101992 | -0.017868569 | 0.139159001 | -0.128403975 | 0.897829285 | 0.982798755 | Mfsd5 | major facilitator superfamily domain cont | 374.25 | 382.39 | 369.29 | 391.38 | 337 | 383.16 |
| 13503 | ENSMUSG00000045667 | 13.72564043 | -0.306337914 | 0.586584752 | -0.522239818 | 0.601503365 | 0.915461118 | Smtnl2 | smoothelin-like 2 [Source:MGI Symbol;Ac | 12.047 | 18.503 | 15.094 | 12.532 | 15.269 | 8.9108 |
| 13504 | ENSMUSG00000045671 | 229.5205282 | 0.346898356 | 0.231750559 | 1.496860928 | 0.134429449 | 0.537569131 | Spred2 | sprouty-related EVH1 domain containing 2 | 255.39 | 130.55 | 219.36 | 264.13 | 219.21 | 288.49 |
| 13505 | ENSMUSG00000045672 | 1776.257435 | -0.393958036 | 0.493428538 | -0.798409507 | 0.424632888 | 0.840504368 | Col27a1 | collagen, type XXVII, alpha 1 [Source:MGI | 1198.2 | 2141.2 | 2712.8 | 2011.8 | 1755.9 | 837.61 |
| 13506 | ENSMUSG00000045678 | 0 | NA | NA | NA | NA | NA | Olfr1489 | olfactory receptor 1489 [Source:MGI Syml | 0 | 0 | 0 | 0 | 0 | 0 |
| 13507 | ENSMUSG00000045679 | 11.17828487 | -0.065535537 | 0.710792211 | -0.092200697 | 0.926538585 | 0.987712228 | Pqlc3 | PQ loop repeat containing [Source:MGI Sy | 8.031 | 16.447 | 10.062 | 19.28 | 5.4531 | 7.7969 |
| 13508 | ENSMUSG00000045680 | 71.59308547 | -0.833529052 | 0.30935489 | -2.694410461 | 0.007051327 | 0.08445009 | Tcf21 | transcription factor 21 [Source:MGI Symb | 101.19 | 95.596 | 78.487 | 63.623 | 31.628 | 59.034 |
| 13509 | ENSMUSG00000045684 | 0.160662974 | 1.126924175 | 4.080472857 | 0.276174898 | 0.782413736 | NA | Lcn6 | lipocalin 6 [Source:MGI Symbol;Acc:MGI:3 | 0 | 0 | 0 | 0.964 | 0 | 0 |
| 13510 | ENSMUSG00000045689 | 0.744190399 | -2.947713985 | 2.874421214 | -1.025498271 | 0.305128141 | NA | Pcdhb4 | protocadherin beta 4 [Source:MGI Symbo | 2.4093 | 2.0558 | 0 | 0 | 0 | 0 |
| 13511 | ENSMUSG00000045690 | 50.54749212 | -0.311086044 | 0.341341638 | -0.911362722 | 0.362104288 | 0.800872989 | Wdr89 | WD repeat domain 89 [Source:MGI Symbo | 48.186 | 50.368 | 69.431 | 38.559 | 59.984 | 36.757 |
| 13512 | ENSMUSG00000045691 | 190.2826864 | 0.27706432 | 0.179632283 | 1.542397146 | 0.122977115 | 0.513267141 | Thtpa | thiamine triphosphatase [Source:MGI Syn | 173.47 | 159.33 | 183.14 | 191.83 | 231.21 | 202.72 |
| 13513 | ENSMUSG00000045693 | 0 | NA | NA | NA | NA | NA | Nlrp4e | NLR family, pyrin domain containing 4E [S | 0 | 0 | 0 | 0 | 0 | 0 |
| 13514 | ENSMUSG00000045694 | 0 | NA | NA | NA | NA | NA | Gm4997 | predicted gene 4997 [Source:MGI Symbol | 0 | 0 | 0 | 0 | 0 | 0 |
| 13515 | ENSMUSG00000045699 | 0 | NA | NA | NA | NA | NA | Gm21411 | predicted gene, 21411 [Source:MGI Symb | 0 | 0 | 0 | 0 | 0 | 0 |
| 13516 | ENSMUSG00000045708 | 0 | NA | NA | NA | NA | NA | Olfr447 | olfactory receptor 447 [Source:MGI Symb | 0 | 0 | 0 | 0 | 0 | 0 |
| 13517 | ENSMUSG00000045709 | 0.167707431 | -0.796629176 | 4.080472857 | -0.195229623 | 0.84521319 | NA | Smkr-ps | smal lysine rich protein 1, pseudogene [Sc | 0 | 0 | 1.0062 | 0 | 0 | 0 |
| 13518 | ENSMUSG00000045725 | 4.590374589 | -0.262107829 | 1.041387533 | -0.251690961 | 0.801279944 | NA | Prr15 | proline rich 15 [Source:MGI Symbol;Acc:M | 8.8341 | 2.0558 | 4.025 | 3.8559 | 6.5437 | 2.2277 |
| 13519 | ENSMUSG00000045730 | 15.22016365 | -2.400051646 | 0.673815281 | -3.561883668 | 0.000368204 | 0.010124897 | Adrb2 | adrenergic receptor, beta 2 [Source:MGI S | 32.927 | 30.838 | 13.081 | 6.7478 | 3.2719 | 4.4554 |
| 13520 | ENSMUSG00000045731 | 0 | NA | NA | NA | NA | NA | Pnoc | prepronociceptin [Source:MGI Symbol;Ac | 0 | 0 | 0 | 0 | 0 | 0 |
| 13521 | ENSMUSG00000045733 | 18.2636409 | -0.419559849 | 0.513927564 | -0.816379346 | 0.414283206 | 0.835831451 | Sprn | shadow of prion protein [Source:MGI Sym | 20.078 | 23.642 | 19.119 | 20.244 | 10.906 | 15.594 |
| 13522 | ENSMUSG00000045744 | 0 | NA | NA | NA | NA | NA | Bricd5 | BRICHOS domain containing 5 [Source:MC | 0 | 0 | 0 | 0 | 0 | 0 |
| 13523 | ENSMUSG00000045746 | 7.076533938 | 0.917066289 | 0.859987869 | 1.066371192 | 0.286255879 | 0.73856672 | B230317F23 | RIKEN cDNA B230317F23 gene [Source:M | 1.6062 | 8.2233 | 5.0312 | 7.7118 | 7.6343 | 12.252 |
| 13524 | ENSMUSG00000045751 | 35.21154407 | 0.078766262 | 0.36498176 | 0.215808764 | 0.829136817 | 0.969955221 | Mms22l | MMS22-like, DNA repair protein [Source:M | 29.715 | 34.949 | 38.237 | 35.667 | 38.172 | 34.529 |
| 13525 | ENSMUSG00000045752 | 269.4614403 | -0.10822079 | 0.153243235 | -0.706202723 | 0.480062077 | 0.870278888 | Tssc4 | tumor-suppressing subchromosomal trans | 301.16 | 271.37 | 265.65 | 257.38 | 265.02 | 256.18 |
| 13526 | ENSMUSG00000045757 | 40.23632213 | -0.199589401 | 0.346595697 | -0.575856545 | 0.56471215 | 0.901542552 | Zfp764 | zinc finger protein 764 [Source:MGI Symb | 42.564 | 48.312 | 38.237 | 38.559 | 41.443 | 32.301 |
| 13527 | ENSMUSG00000045761 | 0 | NA | NA | NA | NA | NA | Togaram2 | TOG array regulator of axonemal microtul | 0 | 0 | 0 | 0 | 0 | 0 |
| 13528 | ENSMUSG00000045763 | 18.99105673 | -1.01123881 | 0.542866381 | -1.862776632 | 0.062493699 | 0.356994022 | Basp1 | brain abundant, membrane attached sign | 19.274 | 32.893 | 24.15 | 13.496 | 17.45 | 6.6831 |
| 13529 | ENSMUSG00000045767 | 865.7522936 | 0.043557234 | 0.106715515 | 0.408162146 | 0.683154637 | 0.935211432 | B230219D22 | RIKEN cDNA B230219D22 gene [Source:M | 822.38 | 881.95 | 854.3 | 888.79 | 861.59 | 885.51 |
| 13530 | ENSMUSG00000045775 | 354.0996478 | -0.848150152 | 0.464594945 | -1.825569051 | 0.067915201 | 0.375219813 | Slc16a5 | solute carrier family 16 (monocarboxylic a | 660.95 | 409.11 | 295.84 | 369.2 | 173.41 | 216.09 |
| 13531 | ENSMUSG00000045776 | 237.9799599 | -0.424182161 | 0.48381086 | -0.876752047 | 0.38062133 | 0.812628154 | Lrtm1 | leucine-rich repeats and transmembrane | 233.7 | 318.65 | 265.65 | 96.398 | 314.1 | 199.38 |
| 13532 | ENSMUSG00000045777 | 7.041227964 | 0.324617734 | 1.114155546 | 0.291357643 | 0.770777806 | 0.956709018 | Ifitm10 | interferon induced transmembrane prote | 10.44 | 7.1954 | 1.0062 | 5.7839 | 0 | 17.822 |
| 13533 | ENSMUSG00000045780 | 0 | NA | NA | NA | NA | NA | Olfr624 | olfactory receptor 624 [Source:MGI Symb | 0 | 0 | 0 | 0 | 0 | 0 |
| 13534 | ENSMUSG00000045790 | 22.64780684 | 0.285561853 | 0.472731316 | 0.604067983 | 0.545798445 | 0.89549457 | Ccdc149 | coiled-coil domain containing 149 [Source | 17.668 | 19.53 | 24.15 | 26.991 | 16.359 | 31.188 |
| 13535 | ENSMUSG00000045792 | 0 | NA | NA | NA | NA | NA | Olfr578 | olfactory receptor 578 [Source:MGI Symb | 0 | 0 | 0 | 0 | 0 | 0 |
| 13536 | ENSMUSG00000045794 | 0 | NA | NA | NA | NA | NA | Lkaaear1 | LKAAEAR motif containing 1 (IKAAEAR mu | 0 | 0 | 0 | 0 | 0 | 0 |
| 13537 | ENSMUSG00000045795 | 773.8912877 | 0.230085587 | 0.127960924 | 1.798092564 | 0.07216234 | 0.389315961 | Whamm | WAS protein homolog associated with act | 746.89 | 608.53 | 780.85 | 826.13 | 828.87 | 852.09 |
| 13538 | ENSMUSG00000045797 | 0 | NA | NA | NA | NA | NA | 4930402K13 | RIKEN cDNA 4930402K13 gene [Source:M | 0 | 0 | 0 | 0 | 0 | 0 |
| 13539 | ENSMUSG00000045799 | 15.38138807 | -0.004234053 | 0.601231822 | -0.007042297 | 0.994381107 | 0.998676967 | Gm9800 | predicted gene 9800 [Source:MGI Symbol | 9.6372 | 19.53 | 17.106 | 10.604 | 10.906 | 24.505 |
| 13540 | ENSMUSG00000045802 | 0 | NA | NA | NA | NA | NA | Hsf3 | heat shock transcription factor 3 [Source:I | 0 | 0 | 0 | 0 | 0 | 0 |
| 13541 | ENSMUSG00000045812 | 0 | NA | NA | NA | NA | NA | Olfr984 | olfactory receptor 984 [Source:MGI Symb | 0 | 0 | 0 | 0 | 0 | 0 |
| 13542 | ENSMUSG00000045813 | 0 | NA | NA | NA | NA | NA | Gm9801 | predicted gene 9801 [Source:MGI Symbol | 0 | 0 | 0 | 0 | 0 | 0 |
| 13543 | ENSMUSG00000045815 | 0 | NA | NA | NA | NA | NA | 1700101I19F | RIKEN cDNA 1700101I19 gene [Source:MC | 0 | 0 | 0 | 0 | 0 | 0 |
| 13544 | ENSMUSG00000045817 | 1210.757098 | -0.065911295 | 0.176185715 | -0.374101245 | 0.708328993 | 0.943888782 | Zfp36l2 | zinc finger protein 36, C3H type-like 2 [Sou | 1191 | 1347.6 | 1177.3 | 1401.6 | 807.06 | 1340 |
| 13545 | ENSMUSG00000045822 | 31.4712503 | -0.067416564 | 0.404279456 | -0.166757333 | 0.867560992 | 0.978204099 | Zswim3 | zinc finger SWIM-type containing 3 [Sourc | 30.518 | 29.81 | 36.225 | 24.099 | 41.443 | 26.732 |
| 13546 | ENSMUSG00000045824 | 0 | NA | NA | NA | NA | NA | Olfr574 | olfactory receptor 574 [Source:MGI Symb | 0 | 0 | 0 | 0 | 0 | 0 |
| 13547 | ENSMUSG00000045826 | 8.246155281 | -1.70269394 | 0.829385745 | -2.052957807 | 0.04007667 | 0.274075154 | Ptprcap | protein tyrosine phosphatase, receptor ty | 10.44 | 11.307 | 16.1 | 3.8559 | 1.0906 | 6.6831 |
| 13548 | ENSMUSG00000045827 | 176.0073696 | -0.145894962 | 0.23183665 | -0.629300684 | 0.529152224 | 0.888334265 | Serpinb9 | serine (or cysteine) peptidase inhibitor, cl | 220.05 | 166.52 | 167.04 | 117.61 | 183.22 | 201.61 |

| | | | | | | | | | | | | | | | |
|---|---|---|---|---|---|---|---|---|---|---|---|---|---|---|---|
| 13549 | ENSMUSG00000045835 | 0 | NA | NA | NA | NA | NA | Hdgfl1 | HDGF like 1 [Source:MGI Symbol;Acc:MGI | 0 | 0 | 0 | 0 | 0 | 0 |
| 13550 | ENSMUSG00000045838 | 43.50252553 | -0.821083217 | 0.413292653 | -1.986687184 | 0.046957073 | 0.300126868 | Ccdc9b | coiled-coil domain containing 9B [Source: | 64.248 | 53.452 | 49.306 | 52.055 | 17.45 | 24.505 |
| 13551 | ENSMUSG00000045842 | 0 | NA | NA | NA | NA | NA | Tgif2-ps2 | TGFB-induced factor homeobox 2, pseudo | 0 | 0 | 0 | 0 | 0 | 0 |
| 13552 | ENSMUSG00000045854 | 155.5902988 | 0.006298236 | 0.239630595 | 0.026283104 | 0.979031532 | 0.998055255 | Lyrm2 | LYR motif containing 2 [Source:MGI Symb | 146.97 | 160.36 | 157.98 | 102.18 | 175.59 | 190.47 |
| 13553 | ENSMUSG00000045867 | 323.5039314 | -0.156889315 | 0.157135259 | -0.998434826 | 0.318068553 | 0.764585353 | Cradd | CASP2 and RIPK1 domain containing adap | 383.88 | 321.74 | 316.97 | 308.47 | 333.73 | 276.23 |
| 13554 | ENSMUSG00000045868 | 42.95901764 | -0.04693034 | 0.353542107 | -0.132743282 | 0.894396412 | 0.982798755 | Gvin1 | GTPase, very large interferon inducible 1 [ | 41.761 | 37.005 | 52.325 | 45.307 | 50.168 | 31.188 |
| 13555 | ENSMUSG00000045871 | 0 | NA | NA | NA | NA | NA | Slitrk6 | SLIT and NTRK-like family, member 6 [Sou | 0 | 0 | 0 | 0 | 0 | 0 |
| 13556 | ENSMUSG00000045875 | 192.9264124 | -0.422404396 | 0.172751664 | -2.445153849 | 0.014479036 | 0.139616521 | Adra1a | adrenergic receptor, alpha 1a [Source:MG | 219.25 | 235.39 | 208.29 | 162.91 | 166.86 | 164.85 |
| 13557 | ENSMUSG00000045876 | 0.638982041 | 0.404325377 | 3.14176733 | 0.128693609 | 0.897600092 | NA | Pcdhb8 | protocadherin beta 8 [Source:MGI Symbo | 1.6062 | 0 | 0 | 0 | 0 | 2.2277 |
| 13558 | ENSMUSG00000045877 | 0.294513347 | 0.165143496 | 4.080472857 | 0.040471657 | 0.967717103 | NA | 4933415A04 | RIKEN cDNA 4933415A04 gene [Source:M | 0.8031 | 0 | 0 | 0.964 | 0 | 0 |
| 13559 | ENSMUSG00000045883 | 0 | NA | NA | NA | NA | NA | Olfr1461 | olfactory receptor 1461 [Source:MGI Sym | 0 | 0 | 0 | 0 | 0 | 0 |
| 13560 | ENSMUSG00000045886 | 12.36212083 | 0.088997513 | 0.628394837 | 0.141626741 | 0.887374843 | 0.982382091 | Gm9803 | predicted gene 9803 [Source:MGI Symbol | 14.456 | 9.2513 | 12.075 | 9.6398 | 9.8156 | 18.935 |
| 13561 | ENSMUSG00000045896 | 411.6622234 | 0.319878188 | 0.137650353 | 2.323845748 | 0.020133763 | 0.17385955 | Paip2b | poly(A) binding protein interacting protei | 332.48 | 383.41 | 383.38 | 443.43 | 444.97 | 482.29 |
| 13562 | ENSMUSG00000045903 | 0 | NA | NA | NA | NA | NA | Npas4 | neuronal PAS domain protein 4 [Source:M | 0 | 0 | 0 | 0 | 0 | 0 |
| 13563 | ENSMUSG00000045912 | 2.362551165 | 0.058997824 | 1.453486794 | 0.040590547 | 0.96762232 | NA | C2cd4c | C2 calcium-dependent domain containing | 4.8186 | 2.0558 | 0 | 2.8919 | 2.1812 | 2.2277 |
| 13564 | ENSMUSG00000045915 | 0 | NA | NA | NA | NA | NA | Ccdc42 | coiled-coil domain containing 42 [Source: | 0 | 0 | 0 | 0 | 0 | 0 |
| 13565 | ENSMUSG00000045917 | 136.6014194 | 0.303167065 | 0.238198924 | 1.272747418 | 0.203107685 | 0.647739134 | Tmem268 | transmembrane protein 268 [Source:MGI | 106.01 | 131.57 | 129.81 | 153.27 | 188.68 | 110.27 |
| 13566 | ENSMUSG00000045928 | 0 | NA | NA | NA | NA | NA | 4933440M0: | RIKEN cDNA 4933440M02 gene [Source:M | 0 | 0 | 0 | 0 | 0 | 0 |
| 13567 | ENSMUSG00000045929 | 0 | NA | NA | NA | NA | NA | Gm20715 | predicted gene 20715 [Source:MGI Symbo | 0 | 0 | 0 | 0 | 0 | 0 |
| 13568 | ENSMUSG00000045930 | 150.3172018 | -0.354340612 | 0.293630526 | -1.206756725 | 0.227525846 | 0.673700979 | Clec14a | C-type lectin domain family 14, member a | 179.09 | 166.52 | 159.99 | 105.07 | 80.706 | 210.52 |
| 13569 | ENSMUSG00000045932 | 54.46095913 | -0.195099753 | 0.3793515 | -0.514298094 | 0.607043594 | 0.916376001 | Ifit2 | interferon-induced protein with tetratrico | 68.264 | 45.228 | 60.375 | 25.063 | 64.346 | 63.489 |
| 13570 | ENSMUSG00000045934 | 9.528194644 | -0.351193835 | 0.718588767 | -0.488727143 | 0.625034885 | 0.919503423 | Mtmr11 | myotubularin related protein 11 [Source:M | 9.6372 | 16.447 | 6.0375 | 9.6398 | 8.7249 | 6.6831 |
| 13571 | ENSMUSG00000045942 | 11.86091147 | -0.506578714 | 0.651573324 | -0.777470003 | 0.436881526 | 0.846852749 | BC049762 | cDNA sequence BC049762 [Source:MGI Sy | 19.274 | 10.279 | 12.075 | 8.6758 | 14.178 | 6.6831 |
| 13572 | ENSMUSG00000045948 | 583.089115 | -0.027114296 | 0.133997058 | -0.202349933 | 0.839643167 | 0.971801133 | Mrps12 | mitochondrial ribosomal protein S12 [Sou | 590.28 | 543.77 | 630.92 | 508.98 | 605.29 | 619.3 |
| 13573 | ENSMUSG00000045952 | 0 | NA | NA | NA | NA | NA | Gm5548 | predicted pseudogene 5548 [Source:MGI | 0 | 0 | 0 | 0 | 0 | 0 |
| 13574 | ENSMUSG00000045954 | 133.2887445 | -0.923508994 | 0.278742948 | -3.313120571 | 0.000922612 | 0.019682389 | Cavin2 | caveolae associated 2 [Source:MGI Symbo | 183.91 | 200.44 | 138.86 | 75.19 | 65.437 | 135.89 |
| 13575 | ENSMUSG00000045962 | 1142.393311 | 0.013102958 | 0.103118142 | 0.127067437 | 0.898887024 | 0.982798755 | Wnk1 | WNK lysine deficient protein kinase 1 [Sou | 1092.2 | 1158.5 | 1161.2 | 1117.3 | 1121.2 | 1204.1 |
| 13576 | ENSMUSG00000045967 | 0 | NA | NA | NA | NA | NA | Gpr158 | G protein-coupled receptor 158 [Source:M | 0 | 0 | 0 | 0 | 0 | 0 |
| 13577 | ENSMUSG00000045968 | 58.65094953 | 0.432954564 | 0.297226979 | 1.45664625 | 0.145214054 | 0.555361204 | Teddm2 | transmembrane epididymal family membe | 47.383 | 43.173 | 59.368 | 75.19 | 61.075 | 65.717 |
| 13578 | ENSMUSG00000045969 | 330.7387201 | 0.439641253 | 0.194177806 | 2.264116905 | 0.023566925 | 0.194339801 | Ing1 | inhibitor of growth family, member 1 [Sou | 205.59 | 274.45 | 363.25 | 374.02 | 381.72 | 385.39 |
| 13579 | ENSMUSG00000045973 | 1725.689865 | -0.718194614 | 0.118025069 | -6.085102256 | 1.16E-09 | 2.41E-07 | Slc25a51 | solute carrier family 25, member 51 [Sour | 2041.5 | 2394 | 2004.4 | 1248.4 | 1445.1 | 1220.8 |
| 13580 | ENSMUSG00000045975 | 2654.385686 | -0.078263555 | 0.096985392 | -0.806962297 | 0.419688205 | 0.837639051 | C2cd2 | C2 calcium-dependent domain containing | 2538.6 | 2941.9 | 2699.8 | 2592.1 | 2555.3 | 2598.6 |
| 13581 | ENSMUSG00000045980 | 201.6294864 | 0.06221144 | 0.180411939 | 0.34482995 | 0.730222216 | 0.946534338 | Tmem104 | transmembrane protein 104 [Source:MGI | 203.18 | 199.42 | 189.17 | 205.33 | 235.57 | 177.1 |
| 13582 | ENSMUSG00000045983 | 9398.644248 | 0.052178586 | 0.088673171 | 0.588437129 | 0.556238922 | 0.897828342 | Eif4g1 | eukaryotic translation initiation factor 4, g | 9073.4 | 9392.1 | 9221.2 | 9722.7 | 10081 | 8901.8 |
| 13583 | ENSMUSG00000045989 | 0 | NA | NA | NA | NA | NA | 4930451I11F | RIKEN cDNA 4930451I11 gene [Source:MG | 0 | 0 | 0 | 0 | 0 | 0 |
| 13584 | ENSMUSG00000045991 | 1456.536706 | 0.434616558 | 0.137549618 | 3.159707481 | 0.001579276 | 0.029454792 | Onecut2 | one cut domain, family member 2 [Source | 1230.4 | 1364 | 1123 | 2000.3 | 1529 | 1492.6 |
| 13585 | ENSMUSG00000045994 | 0 | NA | NA | NA | NA | NA | B3gat1 | beta-1,3-glucuronyltransferase 1 (glucuro | 0 | 0 | 0 | 0 | 0 | 0 |
| 13586 | ENSMUSG00000045996 | 169.0947776 | 0.034244318 | 0.259229188 | 0.13210055 | 0.894904761 | 0.982798755 | Polr2k | polymerase (RNA) II (DNA directed) polyp | 171.06 | 166.52 | 163.01 | 100.25 | 186.5 | 227.22 |
| 13587 | ENSMUSG00000046000 | 0 | NA | NA | NA | NA | NA | Naa11 | N(alpha)-acetyltransferase 11, NatA cataly | 0 | 0 | 0 | 0 | 0 | 0 |
| 13588 | ENSMUSG00000046005 | 16.58978265 | -0.198213274 | 0.530611324 | -0.373556434 | 0.708734352 | 0.94393496 | D830044D21 | RIKEN cDNA D830044D21 gene [Source:M | 22.487 | 15.419 | 15.094 | 13.496 | 17.45 | 15.594 |
| 13589 | ENSMUSG00000046006 | 0.167707431 | -0.796629176 | 4.080472857 | -0.195229623 | 0.84521319 | NA | Gapt | Grb2-binding adaptor, transmembrane [S | 0 | 0 | 1.0062 | 0 | 0 | 0 |
| 13590 | ENSMUSG00000046008 | 0 | NA | NA | NA | NA | NA | Pnlip | pancreatic lipase [Source:MGI Symbol;Acc | 0 | 0 | 0 | 0 | 0 | 0 |
| 13591 | ENSMUSG00000046010 | 149.2384565 | 0.086035009 | 0.198159851 | 0.434169728 | 0.664165194 | 0.929301301 | Zfp830 | zinc finger protein 830 [Source:MGI Symb | 133.31 | 162.41 | 138.86 | 143.63 | 167.96 | 149.26 |
| 13592 | ENSMUSG00000046016 | 0 | NA | NA | NA | NA | NA | Olfr1364 | olfactory receptor 1364 [Source:MGI Sym | 0 | 0 | 0 | 0 | 0 | 0 |
| 13593 | ENSMUSG00000046020 | 265.9660474 | -0.077356857 | 0.163404043 | -0.473408466 | 0.635921791 | 0.923478764 | Pofut1 | protein O-fucosyltransferase 1 [Source:M | 256.99 | 307.35 | 255.59 | 278.59 | 237.75 | 259.53 |
| 13594 | ENSMUSG00000046027 | 3168.044767 | -0.129128452 | 0.099050074 | -1.303668404 | 0.192346667 | 0.630147721 | Stard5 | StAR-related lipid transfer (START) domai | 3137.7 | 3296.5 | 3495.7 | 3005.7 | 3244.6 | 2828 |
| 13595 | ENSMUSG00000046031 | 29.32636157 | -0.623698836 | 0.448479077 | -1.390697734 | 0.164317105 | 0.589016756 | Calhm6 | calcium homeostasis modulator family me | 36.14 | 45.228 | 25.156 | 16.388 | 19.631 | 33.415 |
| 13596 | ENSMUSG00000046032 | 171.8480509 | -0.232433501 | 0.190590298 | -1.219545296 | 0.222637295 | 0.668407528 | Snx12 | sorting nexin 12 [Source:MGI Symbol;Acc: | 189.53 | 206.61 | 161 | 169.66 | 147.23 | 157.05 |
| 13597 | ENSMUSG00000046034 | 924.4695813 | 0.202239932 | 0.117964639 | 1.714411489 | 0.08645318 | 0.433134324 | Otulin | OTU deubiquitinase with linear linkage sp | 942.04 | 868.59 | 767.76 | 1007.4 | 960.83 | 1000.2 |
| 13598 | ENSMUSG00000046041 | 0 | NA | NA | NA | NA | NA | Olfr803 | olfactory receptor 803 [Source:MGI Symb | 0 | 0 | 0 | 0 | 0 | 0 |
| 13599 | ENSMUSG00000046049 | 0 | NA | NA | NA | NA | NA | Rp1l1 | retinitis pigmentosa 1 homolog like 1 [Sou | 0 | 0 | 0 | 0 | 0 | 0 |
| 13600 | ENSMUSG00000046056 | 3.247385367 | 0.845993078 | 1.298664207 | 0.651433275 | 0.514766837 | NA | Sbsn | suprabasin [Source:MGI Symbol;Acc:MGI: | 0 | 2.0558 | 5.0312 | 5.7839 | 3.2719 | 3.3415 |
| 13601 | ENSMUSG00000046057 | 0 | NA | NA | NA | NA | NA | Gm15428 | predicted pseudogene 15428 [Source:MG | 0 | 0 | 0 | 0 | 0 | 0 |
| 13602 | ENSMUSG00000046058 | 6.85394237 | -0.497709496 | 0.859398364 | -0.579137123 | 0.562496653 | 0.900565925 | Eid2 | EP300 interacting inhibitor of differentiati | 4.8186 | 8.2233 | 11.069 | 3.8559 | 9.8156 | 3.3415 |
| 13603 | ENSMUSG00000046062 | 2392.56041 | 0.106915118 | 0.101838127 | 1.049853534 | 0.293785458 | 0.745164683 | Ppp1r15b | protein phosphatase 1, regulatory subuni | 2174.8 | 2499.9 | 2237.9 | 2411.9 | 2630.6 | 2400.3 |
| 13604 | ENSMUSG00000046070 | 2456.083508 | 0.358224128 | 0.122093483 | 2.934015138 | 0.003346079 | 0.050564445 | Igfals | insulin-like growth factor binding protein, | 2247.1 | 1914 | 2297.3 | 3105.9 | 2740.7 | 2431.5 |

| | | | | | | | | | | | | | | | |
|---|---|---|---|---|---|---|---|---|---|---|---|---|---|---|---|
| 13605 | ENSMUSG00000046079 | 735.2330348 | 0.028840779 | 0.137076489 | 0.210399164 | 0.833356145 | 0.970660596 | Lrrc8d | leucine rich repeat containing 8D [Source: | 766.16 | 624.97 | 791.91 | 742.26 | 669.64 | 816.45 |
| 13606 | ENSMUSG00000046080 | 34.59322948 | -0.753141416 | 0.416281922 | -1.80921 | 0.070418385 | 0.383772237 | Clec9a | C-type lectin domain family 9, member a [ | 48.186 | 40.089 | 42.262 | 39.523 | 17.45 | 20.049 |
| 13607 | ENSMUSG00000046082 | 0 | NA | NA | NA | NA | NA | Tmem174 | transmembrane protein 174 [Source:MGI | 0 | 0 | 0 | 0 | 0 | 0 |
| 13608 | ENSMUSG00000046085 | 17.81163138 | 0.301581423 | 0.659812062 | 0.457071702 | 0.647619509 | 0.925643431 | 4931422A03 | RIKEN cDNA 4931422A03 gene [Source:M | 17.668 | 5.1396 | 25.156 | 34.703 | 14.178 | 10.025 |
| 13609 | ENSMUSG00000046093 | 0 | NA | NA | NA | NA | NA | Hpcal4 | hippocalcin-like 4 [Source:MGI Symbol;Ac | 0 | 0 | 0 | 0 | 0 | 0 |
| 13610 | ENSMUSG00000046095 | 0 | NA | NA | NA | NA | NA | Krt32 | keratin 32 [Source:MGI Symbol;Acc:MGI:1 | 0 | 0 | 0 | 0 | 0 | 0 |
| 13611 | ENSMUSG00000046096 | 357.1743944 | -0.125982519 | 0.165479862 | -0.761316312 | 0.44646816 | 0.85284145 | Mosmo | modulator of smoothened [Source:MGI Sy | 301.97 | 405 | 412.56 | 366.31 | 311.92 | 345.29 |
| 13612 | ENSMUSG00000046101 | 0.549180077 | 2.659271622 | 3.249668912 | 0.818320787 | 0.41317404 | NA | Mcmdc2 | minichromosome maintenance domain cc | 0 | 0 | 0 | 0 | 2.1812 | 1.1138 |
| 13613 | ENSMUSG00000046108 | 0 | NA | NA | NA | NA | NA | Il17c | interleukin 17C [Source:MGI Symbol;Acc:I | 0 | 0 | 0 | 0 | 0 | 0 |
| 13614 | ENSMUSG00000046109 | 0 | NA | NA | NA | NA | NA | AB041806 | hypothetical protein, MNCb-2457 [Source | 0 | 0 | 0 | 0 | 0 | 0 |
| 13615 | ENSMUSG00000046110 | 0 | NA | NA | NA | NA | NA | Serinc4 | serine incorporator 4 [Source:MGI Symbo | 0 | 0 | 0 | 0 | 0 | 0 |
| 13616 | ENSMUSG00000046111 | 90.3833795 | 0.471308913 | 0.241079828 | 1.954991077 | 0.050584124 | 0.315985866 | Cep295 | centrosomal protein 295 [Source:MGI Syn | 77.901 | 70.926 | 78.487 | 117.61 | 93.793 | 103.59 |
| 13617 | ENSMUSG00000046119 | 0 | NA | NA | NA | NA | NA | 1110008E08 | RIKEN cDNA 1110008E08 gene [Source:M | 0 | 0 | 0 | 0 | 0 | 0 |
| 13618 | ENSMUSG00000046130 | 1.541646079 | -3.992862386 | 2.258766145 | -1.76771836 | 0.077107999 | NA | Vmn1r184 | vomeronasal 1 receptor, 184 [Source:MGI | 3.2124 | 0 | 6.0375 | 0 | 0 | 0 |
| 13619 | ENSMUSG00000046133 | 0 | NA | NA | NA | NA | NA | C130073F10 | RIKEN cDNA C130073F10 gene [Source:M | 0 | 0 | 0 | 0 | 0 | 0 |
| 13620 | ENSMUSG00000046138 | 309.3424405 | 0.12976156 | 0.152644991 | 0.850087242 | 0.395276584 | 0.822933206 | 9930021J03I | RIKEN cDNA 9930021J03 gene [Source:M | 299.56 | 309.4 | 277.72 | 354.74 | 323.91 | 290.71 |
| 13621 | ENSMUSG00000046139 | 292.6468938 | -0.189622094 | 0.170363824 | -1.113042013 | 0.265690386 | 0.721565724 | Patl1 | protein associated with topoisomerase II I | 295.54 | 323.79 | 316.97 | 304.62 | 296.65 | 218.31 |
| 13622 | ENSMUSG00000046145 | 2.56778551 | 0.479881734 | 1.429054934 | 0.33580356 | 0.737019006 | NA | 1700018M1 | RIKEN cDNA 1700018M17 gene [Source:M | 2.4093 | 2.0558 | 2.0125 | 6.7478 | 2.1812 | 0 |
| 13623 | ENSMUSG00000046150 | 0 | NA | NA | NA | NA | NA | Olfr918 | olfactory receptor 918 [Source:MGI Symb | 0 | 0 | 0 | 0 | 0 | 0 |
| 13624 | ENSMUSG00000046152 | 16.95826438 | -0.428272051 | 0.58625991 | -0.730515669 | 0.465075039 | 0.863116315 | Fut10 | fucosyltransferase 10 [Source:MGI Symbo | 13.653 | 25.698 | 19.119 | 14.46 | 6.5437 | 22.277 |
| 13625 | ENSMUSG00000046157 | 20.97043643 | -1.334681164 | 0.49841981 | -2.677825273 | 0.007410186 | 0.087176397 | Tmem229b | transmembrane protein 229B [Source:MG | 24.093 | 34.949 | 31.194 | 13.496 | 8.7249 | 13.366 |
| 13626 | ENSMUSG00000046159 | 20.68529721 | 0.337279782 | 0.503480398 | 0.669896551 | 0.502923739 | 0.881220881 | Chrm3 | cholinergic receptor, muscarinic 3, cardiac | 15.259 | 15.419 | 24.15 | 17.352 | 19.631 | 32.301 |
| 13627 | ENSMUSG00000046160 | 262.8813735 | -0.083469898 | 0.227892745 | -0.366268343 | 0.714164849 | 0.94455746 | Olig1 | oligodendrocyte transcription factor 1 [So | 346.94 | 287.82 | 176.09 | 268.95 | 280.29 | 217.2 |
| 13628 | ENSMUSG00000046167 | 3.564900063 | 3.480044868 | 1.616200111 | 2.153226474 | 0.031300884 | NA | Gldn | gliomedin [Source:MGI Symbol;Acc:MGI:2 | 1.6062 | 0 | 0 | 0.964 | 5.4531 | 13.366 |
| 13629 | ENSMUSG00000046168 | 5.84230634 | -0.819969108 | 0.924671215 | -0.886768286 | 0.375203665 | 0.809332934 | Kcnrg | potassium channel regulator [Source:MGI | 8.031 | 11.307 | 3.0187 | 3.8559 | 3.2719 | 5.5692 |
| 13630 | ENSMUSG00000046169 | 24.41471488 | -0.29745463 | 0.443902981 | -0.670089282 | 0.502800877 | 0.881220881 | Adamts6 | a disintegrin-like and metallopeptidase (re | 29.715 | 25.698 | 25.156 | 17.352 | 20.722 | 27.846 |
| 13631 | ENSMUSG00000046173 | 0 | NA | NA | NA | NA | NA | Pabpc6 | poly(A) binding protein, cytoplasmic 6 [So | 0 | 0 | 0 | 0 | 0 | 0 |
| 13632 | ENSMUSG00000046178 | 3.461719635 | 0.030575743 | 1.164924404 | 0.026246976 | 0.979060347 | NA | Nxph1 | neurexophilin 1 [Source:MGI Symbol;Acc: | 3.2124 | 1.0279 | 6.0375 | 3.8559 | 2.1812 | 4.4554 |
| 13633 | ENSMUSG00000046179 | 246.7077233 | 0.933155408 | 0.735958834 | 1.267945115 | 0.204817552 | 0.650460832 | E2f8 | E2F transcription factor 8 [Source:MGI Syr | 210.41 | 169.61 | 128.8 | 667.07 | 143.96 | 160.39 |
| 13634 | ENSMUSG00000046180 | 0 | NA | NA | NA | NA | NA | Magea13 | MAGE family member A13 [Source:MGI Sy | 0 | 0 | 0 | 0 | 0 | 0 |
| 13635 | ENSMUSG00000046182 | 5.361740819 | -0.387217727 | 0.926985243 | -0.41771725 | 0.676153858 | 0.932739835 | Gsg1l | GSG1-like [Source:MGI Symbol;Acc:MGI:2 | 4.0155 | 6.1675 | 8.05 | 2.8919 | 4.3625 | 6.6831 |
| 13636 | ENSMUSG00000046185 | 158.0803812 | 0.094170581 | 0.206196979 | 0.456702037 | 0.647885226 | 0.925643431 | Zfp84 | zinc finger protein 84 [Source:MGI Symbo | 140.54 | 150.08 | 168.04 | 137.85 | 157.05 | 194.92 |
| 13637 | ENSMUSG00000046186 | 0.301557804 | -1.64335916 | 4.042814367 | -0.406488899 | 0.684383412 | NA | Cd109 | CD109 antigen [Source:MGI Symbol;Acc:N | 0.8031 | 0 | 1.0062 | 0 | 0 | 0 |
| 13638 | ENSMUSG00000046191 | 2.279534934 | 0.216897638 | 1.619752515 | 0.133907888 | 0.893475413 | NA | Pcdhb20 | protocadherin beta 20 [Source:MGI Symb | 3.2124 | 3.0838 | 0 | 1.928 | 5.4531 | 0 |
| 13639 | ENSMUSG00000046192 | 0.839822492 | 2.066357436 | 2.600192809 | 0.794693928 | 0.426791566 | NA | Iqub | IQ motif and ubiquitin domain containing | 0.8031 | 0 | 0 | 0.964 | 3.2719 | 0 |
| 13640 | ENSMUSG00000046196 | 1.060044235 | 0.14943475 | 2.627116342 | 0.056881664 | 0.95463946 | NA | Ttc39d | tetratricopeptide repeat domain 39D [Sou | 0 | 0 | 3.0187 | 0 | 0 | 3.3415 |
| 13641 | ENSMUSG00000046201 | 370.4910396 | -0.075105215 | 0.134548026 | -0.558203766 | 0.57670525 | 0.907752116 | Scaf8 | SR-related CTD-associated factor 8 [Sourc | 373.44 | 392.66 | 374.32 | 352.82 | 359.9 | 369.8 |
| 13642 | ENSMUSG00000046203 | 0 | NA | NA | NA | NA | NA | Sprr2g | small proline-rich protein 2G [Source:MGI | 0 | 0 | 0 | 0 | 0 | 0 |
| 13643 | ENSMUSG00000046204 | 0 | NA | NA | NA | NA | NA | Pnma2 | paraneoplastic antigen MA2 [Source:MGI | 0 | 0 | 0 | 0 | 0 | 0 |
| 13644 | ENSMUSG00000046207 | 11.56607189 | -0.384419644 | 0.661095557 | -0.581488773 | 0.560911083 | 0.900146222 | Pik3r6 | phosphoinositide-3-kinase regulatory sub | 12.85 | 18.503 | 8.05 | 13.496 | 9.8156 | 6.6831 |
| 13645 | ENSMUSG00000046210 | 0 | NA | NA | NA | NA | NA | Olfr735 | olfactory receptor 735 [Source:MGI Symb | 0 | 0 | 0 | 0 | 0 | 0 |
| 13646 | ENSMUSG00000046213 | 1.878062578 | 0.421017775 | 1.679595165 | 0.250666222 | 0.802072178 | NA | Cym | chymosin [Source:MGI Symbol;Acc:MGI:2 | 0.8031 | 0 | 4.025 | 0.964 | 4.3625 | 1.1138 |
| 13647 | ENSMUSG00000046215 | 0 | NA | NA | NA | NA | NA | Rprml | reprimo-like [Source:MGI Symbol;Acc:MG | 0 | 0 | 0 | 0 | 0 | 0 |
| 13648 | ENSMUSG00000046219 | 0 | NA | NA | NA | NA | NA | C230012O17 | RIKEN cDNA C230012O17 gene [Source:M | 0 | 0 | 0 | 0 | 0 | 0 |
| 13649 | ENSMUSG00000046223 | 7.840594566 | 0.43919602 | 0.845975897 | 0.519159023 | 0.603649852 | 0.916106578 | Plaur | plasminogen activator, urokinase recepto | 9.6372 | 9.2513 | 1.0062 | 10.604 | 7.6343 | 8.9108 |
| 13650 | ENSMUSG00000046229 | 594.7039867 | -0.224090477 | 0.129537553 | -1.729926743 | 0.083643364 | 0.426388452 | Scand1 | SCAN domain-containing 1 [Source:MGI S | 705.12 | 575.63 | 640.98 | 571.64 | 523.5 | 551.35 |
| 13651 | ENSMUSG00000046230 | 414.0027027 | -0.006026291 | 0.224862101 | -0.026799942 | 0.978619299 | 0.997797092 | Vps13a | vacuolar protein sorting 13A [Source:MGI | 370.23 | 469.76 | 405.52 | 515.73 | 475.51 | 247.27 |
| 13652 | ENSMUSG00000046240 | 0 | NA | NA | NA | NA | NA | Hepacam | hepatocyte cell adhesion molecule [Sourc | 0 | 0 | 0 | 0 | 0 | 0 |
| 13653 | ENSMUSG00000046242 | 1.610852122 | -1.802134023 | 1.868246277 | -0.964612667 | 0.334738853 | NA | Nme9 | NME/NM23 family member 9 [Source:MG | 2.4093 | 2.0558 | 3.0187 | 0 | 2.1812 | 0 |
| 13654 | ENSMUSG00000046245 | 18.98818518 | -1.296016635 | 0.570712005 | -2.270876773 | 0.023154439 | 0.192033143 | Pilra | paired immunoglobin-like type 2 receptor | 21.684 | 42.145 | 17.106 | 8.6758 | 8.7249 | 15.594 |
| 13655 | ENSMUSG00000046248 | 0 | NA | NA | NA | NA | NA | Krtap5-3 | keratin associated protein 5-3 [Source:MG | 0 | 0 | 0 | 0 | 0 | 0 |
| 13656 | ENSMUSG00000046258 | 0 | NA | NA | NA | NA | NA | Adam29 | a disintegrin and metallopeptidase domai | 0 | 0 | 0 | 0 | 0 | 0 |
| 13657 | ENSMUSG00000046259 | 0 | NA | NA | NA | NA | NA | Sprr2h | small proline-rich protein 2H [Source:MGI | 0 | 0 | 0 | 0 | 0 | 0 |
| 13658 | ENSMUSG00000046262 | 0 | NA | NA | NA | NA | NA | Pramel29 | PRAME like 29 [Source:MGI Symbol;Acc:M | 0 | 0 | 0 | 0 | 0 | 0 |
| 13659 | ENSMUSG00000046269 | 22.99708735 | -0.064168225 | 0.495530781 | -0.129493923 | 0.896966833 | 0.982798755 | Usp27x | ubiquitin specific peptidase 27, X chromos | 24.896 | 20.558 | 25.156 | 30.847 | 10.906 | 25.618 |
| 13660 | ENSMUSG00000046272 | 0 | NA | NA | NA | NA | NA | Olfr1444 | olfactory receptor 1444 [Source:MGI Syml | 0 | 0 | 0 | 0 | 0 | 0 |

| | | | | | | | | | | | | | | | |
|---|---|---|---|---|---|---|---|---|---|---|---|---|---|---|---|
| 13661 | ENSMUSG00000046275 | 10.31239869 | -0.070348153 | 0.711471053 | -0.098877042 | 0.921235897 | 0.986685804 | Trarg1 | trafficking regulator of GLUT4 (SLC2A4) 1 [ | 11.243 | 8.2233 | 12.075 | 3.8559 | 11.997 | 14.48 |
| 13662 | ENSMUSG00000046280 | 31.35518308 | -0.504534077 | 0.4052089 | -1.245120917 | 0.213087306 | 0.657724029 | She | src homology 2 domain-containing transfo | 32.927 | 39.061 | 38.237 | 18.316 | 26.175 | 33.415 |
| 13663 | ENSMUSG00000046282 | 0.17131964 | -0.796629176 | 4.080472857 | -0.195229623 | 0.84521319 | NA | Adam20 | a disintegrin and metallopeptidase domai | 0 | 1.0279 | 0 | 0 | 0 | 0 |
| 13664 | ENSMUSG00000046287 | 0 | NA | NA | NA | NA | NA | Pnma3 | paraneoplastic antigen MA3 [Source:MGI | 0 | 0 | 0 | 0 | 0 | 0 |
| 13665 | ENSMUSG00000046295 | 2.70870697 | -1.156480633 | 1.410993723 | -0.819621387 | 0.412431977 | NA | Ankle1 | ankyrin repeat and LEM domain containin | 3.2124 | 2.0558 | 6.0375 | 3.8559 | 1.0906 | 0 |
| 13666 | ENSMUSG00000046300 | 0 | NA | NA | NA | NA | NA | Olfr1412 | olfactory receptor 1412 [Source:MGI Syml | 0 | 0 | 0 | 0 | 0 | 0 |
| 13667 | ENSMUSG00000046311 | 266.6601596 | 0.139150106 | 0.153702857 | 0.90531893 | 0.365296451 | 0.803870323 | Zfp62 | zinc finger protein 62 [Source:MGI Symbo | 258.6 | 260.06 | 242.5 | 270.88 | 268.29 | 299.62 |
| 13668 | ENSMUSG00000046312 | 951.2679112 | 0.266877354 | 0.130157324 | 2.050421334 | 0.040323333 | 0.274714427 | Myorg | myogenesis regulating glycosidase (putati | 870.56 | 943.63 | 775.81 | 910.96 | 1145.1 | 1061.5 |
| 13669 | ENSMUSG00000046314 | 274.4573937 | 0.306655476 | 0.18099019 | 1.694320978 | 0.090204306 | 0.44322087 | Stxbp6 | syntaxin binding protein 6 (amisyn) [Sourc | 281.89 | 205.58 | 247.54 | 257.38 | 341.36 | 312.99 |
| 13670 | ENSMUSG00000046317 | 0.133850373 | -0.796629176 | 4.080472857 | -0.195229623 | 0.84521319 | NA | BC107364 | cDNA sequence BC107364 [Source:MGI Sy | 0.8031 | 0 | 0 | 0 | 0 | 0 |
| 13671 | ENSMUSG00000046318 | 59.35819368 | -1.746720895 | 0.31919048 | -5.472346465 | 4.44E-08 | 5.44E-06 | Ccbe1 | collagen and calcium binding EGF domain: | 97.175 | 104.85 | 72.45 | 29.883 | 26.175 | 25.618 |
| 13672 | ENSMUSG00000046321 | 0 | NA | NA | NA | NA | NA | Hs3st2 | heparan sulfate (glucosamine) 3-O-sulfotr | 0 | 0 | 0 | 0 | 0 | 0 |
| 13673 | ENSMUSG00000046323 | 0.17131964 | -0.796629176 | 4.080472857 | -0.195229623 | 0.84521319 | NA | Dppa3 | developmental pluripotency-associated 3 | 0 | 1.0279 | 0 | 0 | 0 | 0 |
| 13674 | ENSMUSG00000046324 | 848.0276302 | -0.227673806 | 0.155457632 | -1.464539263 | 0.143046663 | 0.552624717 | Ermp1 | endoplasmic reticulum metallopeptidase : | 1034.4 | 999.14 | 710.41 | 803.96 | 845.23 | 695.04 |
| 13675 | ENSMUSG00000046329 | 4607.945649 | 0.117500864 | 0.090002869 | 1.305523532 | 0.19171464 | 0.629099068 | Slc25a23 | solute carrier family 25 (mitochondrial car | 4464.4 | 4329.6 | 4467.7 | 5112 | 4703.8 | 4570.1 |
| 13676 | ENSMUSG00000046330 | 2429.801901 | 0.039071299 | 0.103066095 | 0.379089738 | 0.704621227 | 0.94295036 | Rpl37a | ribosomal protein L37a [Source:MGI Symb | 2281.6 | 2554.4 | 2355.6 | 2534.3 | 2254.3 | 2598.6 |
| 13677 | ENSMUSG00000046334 | 0 | NA | NA | NA | NA | NA | Gm6195 | predicted pseudogene 6195 [Source:MGI | 0 | 0 | 0 | 0 | 0 | 0 |
| 13678 | ENSMUSG00000046337 | 0 | NA | NA | NA | NA | NA | Fam178b | family with sequence similarity 178, meml | 0 | 0 | 0 | 0 | 0 | 0 |
| 13679 | ENSMUSG00000046338 | 38.2698931 | -0.465452664 | 0.385668218 | -1.206873271 | 0.227480953 | 0.673700979 | Gpat2 | glycerol-3-phosphate acyltransferase 2, m | 62.642 | 33.921 | 36.225 | 31.811 | 32.719 | 32.301 |
| 13680 | ENSMUSG00000046341 | 1.02048683 | -0.947758949 | 2.234944437 | -0.424063763 | 0.671519312 | NA | Gm11223 | predicted gene 11223 [Source:MGI Symbc | 0 | 2.0558 | 2.0125 | 0.964 | 1.0906 | 0 |
| 13681 | ENSMUSG00000046345 | 0.342639279 | -1.805976596 | 4.028547714 | -0.448294702 | 0.653940524 | NA | Smco1 | single-pass membrane protein with coiled | 0 | 2.0558 | 0 | 0 | 0 | 0 |
| 13682 | ENSMUSG00000046351 | 188.407947 | 0.271907165 | 0.178808338 | 1.520662672 | 0.128344511 | 0.524399718 | Zfp322a | zinc finger protein 322A [Source:MGI Sym | 163.83 | 158.3 | 190.18 | 210.15 | 194.13 | 213.86 |
| 13683 | ENSMUSG00000046352 | 4077.817606 | 0.505122221 | 0.11935337 | 4.232157167 | 2.31E-05 | 0.001090503 | Gjb2 | gap junction protein, beta 2 [Source:MGI | 3809.1 | 2939.8 | 3362.9 | 4889.3 | 4325.4 | 5140.4 |
| 13684 | ENSMUSG00000046354 | 0 | NA | NA | NA | NA | NA | Defb14 | defensin beta 14 [Source:MGI Symbol;Acc | 0 | 0 | 0 | 0 | 0 | 0 |
| 13685 | ENSMUSG00000046364 | 3293.891586 | 0.124157271 | 0.097598976 | 1.272116526 | 0.20333172 | 0.647795155 | Rpl27a | ribosomal protein L27A [Source:MGI Syml | 2996.4 | 3267.8 | 3192.8 | 3244.7 | 3352.6 | 3709.1 |
| 13686 | ENSMUSG00000046367 | 0.160662974 | 1.126924175 | 4.080472857 | 0.276174898 | 0.782413736 | NA | Mgat4e | MGAT4 family, member E [Source:MGI Sy | 0 | 0 | 0 | 0.964 | 0 | 0 |
| 13687 | ENSMUSG00000046372 | 0.321325949 | 1.911575114 | 4.048267893 | 0.472195804 | 0.636787034 | NA | Gm5590 | predicted gene 5590 [Source:MGI Symbol | 0 | 0 | 0 | 1.928 | 0 | 0 |
| 13688 | ENSMUSG00000046378 | 1.681971856 | -0.45628717 | 1.737711254 | -0.262579395 | 0.792874783 | NA | Asphd1 | aspartate beta-hydroxylase domain conta | 0.8031 | 1.0279 | 4.025 | 0.964 | 3.2719 | 0 |
| 13689 | ENSMUSG00000046380 | 23.60560172 | 0.109782818 | 0.444061376 | 0.247224424 | 0.804734545 | 0.963787787 | Jrk | jerky [Source:MGI Symbol;Acc:MGI:10621 | 27.305 | 18.503 | 22.137 | 24.099 | 25.084 | 24.505 |
| 13690 | ENSMUSG00000046387 | 2.042880184 | -0.446897916 | 1.554642031 | -0.287460333 | 0.77375987 | NA | Pcdhb17 | protocadherin beta 17 [Source:MGI Symb | 4.0155 | 0 | 3.0187 | 1.928 | 2.1812 | 1.1138 |
| 13691 | ENSMUSG00000046388 | 0 | NA | NA | NA | NA | NA | Gm9806 | predicted gene 9806 [Source:MGI Symbol | 0 | 0 | 0 | 0 | 0 | 0 |
| 13692 | ENSMUSG00000046396 | 0 | NA | NA | NA | NA | NA | Olfr609 | olfactory receptor 609 [Source:MGI Symb | 0 | 0 | 0 | 0 | 0 | 0 |
| 13693 | ENSMUSG00000046402 | 114.6584823 | -0.111797392 | 0.304150887 | -0.367572139 | 0.713192292 | 0.944239215 | Rbp1 | retinol binding protein 1, cellular [Source: | 159.01 | 118.21 | 79.493 | 108.93 | 74.162 | 148.14 |
| 13694 | ENSMUSG00000046404 | 235.8740346 | -0.478975641 | 0.181033315 | -2.645787281 | 0.008150105 | 0.093114627 | Yod1 | YOD1 deubiquitinase [Source:MGI Symbo | 236.92 | 318.65 | 268.67 | 174.48 | 211.58 | 204.95 |
| 13695 | ENSMUSG00000046408 | 22.4267782 | 0.404814221 | 0.453159415 | 0.893315261 | 0.371688368 | 0.807685567 | 1700067K01 | RIKEN cDNA 1700067K01 gene [Source:M | 19.274 | 19.53 | 19.119 | 26.027 | 29.447 | 21.163 |
| 13696 | ENSMUSG00000046410 | 23.35252931 | -1.106373941 | 0.493729989 | -2.240848166 | 0.025035912 | 0.202995271 | Kcnk6 | potassium inwardly-rectifying channel, su | 35.336 | 40.089 | 20.125 | 11.568 | 19.631 | 13.366 |
| 13697 | ENSMUSG00000046411 | 0 | NA | NA | NA | NA | NA | Rps4l-ps | ribosomal protein S4-like, pseudogene [Sc | 0 | 0 | 0 | 0 | 0 | 0 |
| 13698 | ENSMUSG00000046413 | 0.453341394 | -0.545391191 | 3.551113277 | -0.153583158 | 0.877938417 | NA | Irx3os | iroquois homeobox 3, opposite strand [Sc | 1.6062 | 0 | 0 | 0 | 0 | 1.1138 |
| 13699 | ENSMUSG00000046415 | 9.992191408 | -1.659935691 | 0.892538952 | -1.859790755 | 0.062915136 | 0.358834592 | B430212C06 | RIKEN cDNA B430212C06 gene [Source:M | 24.093 | 16.447 | 5.0312 | 6.7478 | 7.6343 | 0 |
| 13700 | ENSMUSG00000046417 | 4.734545794 | -0.507667055 | 1.015832651 | -0.499754615 | 0.617247871 | NA | Lrrc75a | leucine rich repeat containing 75A [Source | 6.4248 | 6.1675 | 4.025 | 1.928 | 7.6343 | 2.2277 |
| 13701 | ENSMUSG00000046431 | 0 | NA | NA | NA | NA | NA | Olfr518 | olfactory receptor 518 [Source:MGI Symb | 0 | 0 | 0 | 0 | 0 | 0 |
| 13702 | ENSMUSG00000046432 | 7.42761285 | -0.701656436 | 0.870276318 | -0.806245582 | 0.420101261 | 0.837906635 | Bex3 | brain expressed X-linked 3 [Source:MGI Sy | 11.243 | 8.2233 | 8.05 | 4.8199 | 1.0906 | 11.138 |
| 13703 | ENSMUSG00000046434 | 887.200067 | 0.138998132 | 0.11021613 | 1.261141466 | 0.207257883 | 0.652364421 | Hnrnpa1 | heterogeneous nuclear ribonucleoprotein | 791.06 | 886.07 | 857.32 | 938.91 | 953.2 | 896.64 |
| 13704 | ENSMUSG00000046435 | 0 | NA | NA | NA | NA | NA | Pramel24 | PRAME like 24 [Source:MGI Symbol;Acc:M | 0 | 0 | 0 | 0 | 0 | 0 |
| 13705 | ENSMUSG00000046438 | 0 | NA | NA | NA | NA | NA | Scgb2b24 | secretoglobin, family 2B, member 24 [Sou | 0 | 0 | 0 | 0 | 0 | 0 |
| 13706 | ENSMUSG00000046440 | 0.346303622 | 2.004339653 | 3.884003423 | 0.516049919 | 0.605819544 | NA | Gm5564 | predicted gene 5564 [Source:MGI Symbol | 0 | 0 | 0 | 0.964 | 0 | 1.1138 |
| 13707 | ENSMUSG00000046441 | 46.07022481 | 0.108287692 | 0.334843055 | 0.32339835 | 0.746393579 | 0.950047651 | Cmtr2 | cap methyltransferase 2 [Source:MGI Sym | 33.73 | 48.312 | 51.318 | 48.199 | 43.625 | 51.237 |
| 13708 | ENSMUSG00000046442 | 1.818381195 | -1.336872336 | 1.738895199 | -0.768805582 | 0.442008734 | NA | Ppm1e | protein phosphatase 1E (PP2C domain cor | 0.8031 | 1.0279 | 6.0375 | 1.928 | 0 | 1.1138 |
| 13709 | ENSMUSG00000046447 | 2890.180309 | 0.204362982 | 0.097931622 | 2.086792587 | 0.036906888 | 0.259514449 | Camk2n1 | calcium/calmodulin-dependent protein ki | 2639.8 | 2601.7 | 2816.5 | 3211 | 2849.8 | 3222.4 |
| 13710 | ENSMUSG00000046449 | 0 | NA | NA | NA | NA | NA | Nexmif | neurite extension and migration factor [Sc | 0 | 0 | 0 | 0 | 0 | 0 |
| 13711 | ENSMUSG00000046450 | 0 | NA | NA | NA | NA | NA | Olfr71 | olfactory receptor 71 [Source:MGI Symbo | 0 | 0 | 0 | 0 | 0 | 0 |
| 13712 | ENSMUSG00000046456 | 1.353988001 | -1.451133495 | 2.240453263 | -0.647696392 | 0.517181338 | NA | Tmem150b | transmembrane protein 150B [Source:MG | 0.8031 | 5.1396 | 0 | 0 | 2.1812 | 0 |
| 13713 | ENSMUSG00000046460 | 0 | NA | NA | NA | NA | NA | Sh2d7 | SH2 domain containing 7 [Source:MGI Syr | 0 | 0 | 0 | 0 | 0 | 0 |
| 13714 | ENSMUSG00000046463 | 0.505131668 | 1.126696199 | 3.421159819 | 0.329331647 | 0.741905026 | NA | 5930403N24 | RIKEN cDNA 5930403N24 gene [Source:M | 0.8031 | 0 | 0 | 0 | 0 | 2.2277 |
| 13715 | ENSMUSG00000046470 | 70.62508162 | -0.529920629 | 0.313847089 | -1.68846756 | 0.091321511 | 0.445789516 | Sox18 | SRY (sex determining region Y)-box 18 [So | 86.735 | 74.01 | 89.556 | 61.695 | 34.9 | 76.855 |
| 13716 | ENSMUSG00000046474 | 0 | NA | NA | NA | NA | NA | Krtap4-16 | keratin associated protein 4-16 [Source:M | 0 | 0 | 0 | 0 | 0 | 0 |

| | | | | | | | | | | | | | | | |
|---|---|---|---|---|---|---|---|---|---|---|---|---|---|---|---|
| 13717 | ENSMUSG00000046480 | 3.880887599 | -0.45548396 | 1.217924378 | -0.373983778 | 0.708416385 | NA | Scn4b | sodium channel, type IV, beta [Source:MG | 7.2279 | 3.0838 | 3.0187 | 0 | 3.2719 | 6.6831 |
| 13718 | ENSMUSG00000046486 | 0 | NA | NA | NA | NA | NA | Olfr231 | olfactory receptor 231 [Source:MGI Symb | 0 | 0 | 0 | 0 | 0 | 0 |
| 13719 | ENSMUSG00000046487 | 0 | NA | NA | NA | NA | NA | Mospd4 | motile sperm domain containing 4 [Source | 0 | 0 | 0 | 0 | 0 | 0 |
| 13720 | ENSMUSG00000046490 | 0.339027071 | -1.79174149 | 3.919585585 | -0.457125237 | 0.647581032 | NA | Rnf222 | ring finger protein 222 [Source:MGI Symb | 0 | 1.0279 | 1.0062 | 0 | 0 | 0 |
| 13721 | ENSMUSG00000046491 | 14.22997409 | -1.221083547 | 0.689822647 | -1.770141286 | 0.076703607 | 0.405497433 | C1qtnf2 | C1q and tumor necrosis factor related pro | 35.336 | 11.307 | 13.081 | 13.496 | 4.3625 | 7.7969 |
| 13722 | ENSMUSG00000046493 | 0 | NA | NA | NA | NA | NA | Olfr1352 | olfactory receptor 1352 [Source:MGI Syml | 0 | 0 | 0 | 0 | 0 | 0 |
| 13723 | ENSMUSG00000046500 | 0 | NA | NA | NA | NA | NA | Tafa4 | TAFA chemokine like family member 4 [So | 0 | 0 | 0 | 0 | 0 | 0 |
| 13724 | ENSMUSG00000046516 | 57.63141054 | -0.190256485 | 0.329259506 | -0.577831409 | 0.563377947 | 0.900821918 | Cox17 | cytochrome c oxidase assembly protein 1 | 71.476 | 58.591 | 54.337 | 74.226 | 39.262 | 47.895 |
| 13725 | ENSMUSG00000046518 | 0 | NA | NA | NA | NA | NA | Ferd3l | Fer3 like bHLH transcription factor [Source | 0 | 0 | 0 | 0 | 0 | 0 |
| 13726 | ENSMUSG00000046519 | 106.3298332 | -0.199950977 | 0.234727134 | -0.851844323 | 0.394300502 | 0.822827574 | Golph3l | golgi phosphoprotein 3-like [Source:MGI S | 114.84 | 115.13 | 110.69 | 78.082 | 113.42 | 105.82 |
| 13727 | ENSMUSG00000046523 | 0.160662974 | 1.126924175 | 4.080472857 | 0.276174898 | 0.782413736 | NA | Kctd4 | potassium channel tetramerisation domai | 0 | 0 | 0 | 0.964 | 0 | 0 |
| 13728 | ENSMUSG00000046532 | 234.3253327 | -0.315930579 | 0.188891155 | -1.672553586 | 0.094415211 | 0.452792309 | Ar | androgen receptor [Source:MGI Symbol;A | 258.6 | 265.2 | 255.59 | 182.19 | 263.93 | 180.44 |
| 13729 | ENSMUSG00000046541 | 73.03515295 | 0.258812843 | 0.273940905 | 0.944776185 | 0.344773152 | 0.786033695 | Zfp526 | zinc finger protein 526 [Source:MGI Symb | 63.445 | 62.703 | 73.456 | 82.902 | 63.256 | 92.449 |
| 13730 | ENSMUSG00000046546 | 75.2715405 | -0.456415811 | 0.294538224 | -1.549597889 | 0.12123806 | 0.510505764 | Fam43a | family with sequence similarity 43, memb | 91.554 | 81.206 | 88.55 | 74.226 | 40.353 | 75.741 |
| 13731 | ENSMUSG00000046550 | 1.488355822 | -1.619254888 | 1.926476882 | -0.840526509 | 0.400613245 | NA | Spin2c | spindlin family, member 2C [Source:MGI S | 1.6062 | 3.0838 | 2.0125 | 0 | 0 | 2.2277 |
| 13732 | ENSMUSG00000046556 | 135.6376977 | -0.074937911 | 0.198050356 | -0.378378068 | 0.705149758 | 0.94295036 | Zfp319 | zinc finger protein 319 [Source:MGI Symb | 142.95 | 138.77 | 135.84 | 141.7 | 128.69 | 125.86 |
| 13733 | ENSMUSG00000046561 | 0 | NA | NA | NA | NA | NA | Arsj | arylsulfatase J [Source:MGI Symbol;Acc:M | 0 | 0 | 0 | 0 | 0 | 0 |
| 13734 | ENSMUSG00000046562 | 639.9254737 | 0.00994413 | 0.123492734 | 0.080524009 | 0.935820502 | 0.988420305 | Unc119b | unc-119 lipid binding chaperone B [Source | 588.67 | 632.17 | 693.3 | 670.93 | 614.02 | 640.46 |
| 13735 | ENSMUSG00000046567 | 69.01705788 | -0.122742954 | 0.292290356 | -0.41993501 | 0.674532932 | 0.932114677 | 4930430F08 | RIKEN cDNA 4930430F08 gene [Source:M | 57.823 | 80.178 | 78.487 | 81.938 | 59.984 | 55.692 |
| 13736 | ENSMUSG00000046572 | 21.80231659 | 0.406620535 | 0.470536926 | 0.86416286 | 0.387498432 | 0.817618864 | Zfp518b | zinc finger protein 518B [Source:MGI Sym | 22.487 | 19.53 | 14.087 | 24.099 | 29.447 | 21.163 |
| 13737 | ENSMUSG00000046573 | 164.9248665 | -0.146563602 | 0.222825749 | -0.657749845 | 0.510698887 | 0.883371049 | Lyrm4 | LYR motif containing 4 [Source:MGI Symb | 138.94 | 182.97 | 198.23 | 128.21 | 191.95 | 149.26 |
| 13738 | ENSMUSG00000046574 | 228.1628205 | 0.096644475 | 0.226944307 | 0.425851064 | 0.670216375 | 0.931131462 | Prr12 | proline rich 12 [Source:MGI Symbol;Acc:M | 174.27 | 239.5 | 248.54 | 261.24 | 162.5 | 282.92 |
| 13739 | ENSMUSG00000046580 | 1.144992505 | 0.656673711 | 2.118104369 | 0.310028968 | 0.756538928 | NA | Gm7862 | predicted gene 7862 [Source:MGI Symbol | 1.6062 | 1.0279 | 0 | 0.964 | 3.2719 | 0 |
| 13740 | ENSMUSG00000046585 | 0 | NA | NA | NA | NA | NA | Cfap58 | cilia and flagella associated protein 58 [So | 0 | 0 | 0 | 0 | 0 | 0 |
| 13741 | ENSMUSG00000046589 | 2.027344548 | -0.808545912 | 1.694727033 | -0.477095069 | 0.633294426 | NA | Lrrc8e | leucine rich repeat containing 8 family, m | 1.6062 | 3.0838 | 3.0187 | 0 | 0 | 4.4554 |
| 13742 | ENSMUSG00000046590 | 0 | NA | NA | NA | NA | NA | Olfr1165-ps | olfactory receptor 1165, pseudogene [Sou | 0 | 0 | 0 | 0 | 0 | 0 |
| 13743 | ENSMUSG00000046591 | 0.858221022 | 0.830865716 | 2.38087913 | 0.348974337 | 0.727108576 | NA | Ticrr | TOPBP1-interacting checkpoint and replic | 0.8031 | 1.0279 | 0 | 0 | 1.0906 | 2.2277 |
| 13744 | ENSMUSG00000046593 | 1.038910865 | 3.589047152 | 2.558249368 | 1.402930925 | 0.160637438 | NA | Tmem215 | transmembrane protein 215 [Source:MGI | 0 | 0 | 0 | 2.8919 | 0 | 3.3415 |
| 13745 | ENSMUSG00000046598 | 6118.834218 | -0.080273036 | 0.108312493 | -0.741124444 | 0.45861799 | 0.859322431 | Bdh1 | 3-hydroxybutyrate dehydrogenase, type 1 | 7102.6 | 6240.5 | 5522.3 | 5851.3 | 6007.1 | 5989.1 |
| 13746 | ENSMUSG00000046603 | 411.8016307 | 0.15489817 | 0.161131431 | 0.961315669 | 0.336393472 | 0.778173248 | Tcaim | T cell activation inhibitor, mitochondrial [ | 349.35 | 431.73 | 389.42 | 514.76 | 404.62 | 380.93 |
| 13747 | ENSMUSG00000046605 | 57.26895164 | 0.481945772 | 0.330303469 | 1.45909994 | 0.144537603 | 0.55464443 | B3gntl1 | UDP-GlcNAc:betaGal beta-1,3-N-acetylglu | 61.839 | 30.838 | 50.312 | 76.154 | 65.437 | 59.034 |
| 13748 | ENSMUSG00000046607 | 3.059909263 | -2.266537259 | 1.381344901 | -1.640819217 | 0.100834948 | NA | Hrk | harakiri, BCL2 interacting protein (contain | 4.0155 | 6.1675 | 5.0312 | 0.964 | 2.1812 | 0 |
| 13749 | ENSMUSG00000046610 | 0 | NA | NA | NA | NA | NA | Oacyl | O-acyltransferase like [Source:MGI Symbo | 0 | 0 | 0 | 0 | 0 | 0 |
| 13750 | ENSMUSG00000046613 | 0 | NA | NA | NA | NA | NA | Vwa5b2 | von Willebrand factor A domain containin | 0 | 0 | 0 | 0 | 0 | 0 |
| 13751 | ENSMUSG00000046615 | 0 | NA | NA | NA | NA | NA | Actrt1 | actin-related protein T1 [Source:MGI Sym | 0 | 0 | 0 | 0 | 0 | 0 |
| 13752 | ENSMUSG00000046618 | 2.070048847 | 0.723003262 | 1.626432696 | 0.444533158 | 0.656657162 | NA | Olfml2a | olfactomedin-like 2A [Source:MGI Symbol | 1.6062 | 1.0279 | 2.0125 | 0 | 1.0906 | 6.6831 |
| 13753 | ENSMUSG00000046620 | 0 | NA | NA | NA | NA | NA | Rps11-ps4 | ribosomal protein S11, pseudogene 4 [Sou | 0 | 0 | 0 | 0 | 0 | 0 |
| 13754 | ENSMUSG00000046623 | 0 | NA | NA | NA | NA | NA | Gjb4 | gap junction protein, beta 4 [Source:MGI | 0 | 0 | 0 | 0 | 0 | 0 |
| 13755 | ENSMUSG00000046634 | 0 | NA | NA | NA | NA | NA | Pkd1l1 | polycystic kidney disease 1 like 1 [Source:I | 0 | 0 | 0 | 0 | 0 | 0 |
| 13756 | ENSMUSG00000046636 | 0 | NA | NA | NA | NA | NA | Gm7729 | predicted gene 7729 [Source:MGI Symbol | 0 | 0 | 0 | 0 | 0 | 0 |
| 13757 | ENSMUSG00000046637 | 0.505131668 | 1.126696199 | 3.421159819 | 0.329331647 | 0.741905026 | NA | Ttc34 | tetratricopeptide repeat domain 34 [Sour | 0.8031 | 0 | 0 | 0 | 0 | 2.2277 |
| 13758 | ENSMUSG00000046643 | 0 | NA | NA | NA | NA | NA | Olfr218 | olfactory receptor 218 [Source:MGI Symb | 0 | 0 | 0 | 0 | 0 | 0 |
| 13759 | ENSMUSG00000046650 | 0 | NA | NA | NA | NA | NA | Olfr1440 | olfactory receptor 1440 [Source:MGI Syml | 0 | 0 | 0 | 0 | 0 | 0 |
| 13760 | ENSMUSG00000046652 | 0 | NA | NA | NA | NA | NA | Tas2r143 | taste receptor, type 2, member 143 [Sour | 0 | 0 | 0 | 0 | 0 | 0 |
| 13761 | ENSMUSG00000046658 | 118.6914563 | 0.094426245 | 0.256682091 | 0.367872354 | 0.712968415 | 0.944239215 | Zfp316 | zinc finger protein 316 [Source:MGI Symb | 87.538 | 141.85 | 115.72 | 133.99 | 92.703 | 140.34 |
| 13762 | ENSMUSG00000046667 | 118.6992258 | -0.513973608 | 0.243644346 | -2.109524054 | 0.034899372 | 0.250389145 | Rbm12b1 | RNA binding motif protein 12 B1 [Source:I | 121.27 | 165.49 | 132.82 | 120.5 | 77.434 | 94.677 |
| 13763 | ENSMUSG00000046668 | 849.3526751 | 0.307445155 | 0.158174677 | 1.94370655 | 0.051930848 | 0.320245495 | Cxxc5 | CXXC finger 5 [Source:MGI Symbol;Acc:M | 702.71 | 679.45 | 895.56 | 899.39 | 791.79 | 1127.2 |
| 13764 | ENSMUSG00000046671 | 1510.588547 | 0.019203042 | 0.095674589 | 0.200712045 | 0.840923741 | 0.971854408 | Mtfr1l | mitochondrial fission regulator 1-like [Sou | 1474.5 | 1503.8 | 1523.5 | 1499.9 | 1577 | 1484.8 |
| 13765 | ENSMUSG00000046675 | 141.380464 | 0.033087184 | 0.239206845 | 0.138320389 | 0.889987212 | 0.982489666 | Tmem251 | transmembrane protein 251 [Source:MGI | 175.88 | 105.88 | 136.85 | 143.63 | 166.86 | 119.18 |
| 13766 | ENSMUSG00000046676 | 0 | NA | NA | NA | NA | NA | Lce1l | late cornified envelope 1L [Source:MGI Sy | 0 | 0 | 0 | 0 | 0 | 0 |
| 13767 | ENSMUSG00000046678 | 0 | NA | NA | NA | NA | NA | Olfr981 | olfactory receptor 981 [Source:MGI Symb | 0 | 0 | 0 | 0 | 0 | 0 |
| 13768 | ENSMUSG00000046679 | 226.7633485 | 0.171275962 | 0.16339437 | 1.048236615 | 0.294529605 | 0.745164683 | C87436 | expressed sequence C87436 [Source:MGI | 208 | 203.53 | 228.42 | 233.28 | 233.39 | 253.96 |
| 13769 | ENSMUSG00000046683 | 1.124092354 | -1.17552103 | 2.158249136 | -0.544664196 | 0.585984494 | NA | 0610025J13I | RIKEN cDNA 0610025J13 gene [Source:M | 1.6062 | 3.0838 | 0 | 0.964 | 1.0906 | 0 |
| 13770 | ENSMUSG00000046687 | 187.9441439 | 0.002134526 | 0.213441583 | 0.010000514 | 0.992020877 | 0.99859176 | Gm5424 | predicted gene 5424 [Source:MGI Symbol | 174.27 | 210.72 | 178.11 | 137.85 | 203.95 | 222.77 |
| 13771 | ENSMUSG00000046688 | 228.0221946 | 0.531521446 | 0.288339856 | 1.84338528 | 0.065272778 | 0.366785337 | Tifa | TRAF-interacting protein with forkhead-as | 216.03 | 130.55 | 212.32 | 239.07 | 162.5 | 407.67 |
| 13772 | ENSMUSG00000046691 | 604.2454449 | 0.059355223 | 0.125425839 | 0.473229626 | 0.636049364 | 0.923478764 | Chtf8 | CTF8, chromosome transmission fidelity fa | 547.72 | 600.3 | 627.9 | 586.1 | 663.1 | 600.36 |

| | | | | | | | | | | | | | | | |
|---|---|---|---|---|---|---|---|---|---|---|---|---|---|---|---|
| 13773 | ENSMUSG00000046694 | 1.913210186 | -0.724126197 | 1.834322115 | -0.394765015 | 0.693016317 | NA | Tent5b | terminal nucleotidyltransferase 5B [Sourc | 0 | 6.1675 | 1.0062 | 0.964 | 0 | 3.3415 |
| 13774 | ENSMUSG00000046697 | 0 | NA | NA | NA | NA | NA | Enpp7 | ectonucleotide pyrophosphatase/phosph | 0 | 0 | 0 | 0 | 0 | 0 |
| 13775 | ENSMUSG00000046699 | 0.305170013 | -1.657836477 | 4.041480335 | -0.410205257 | 0.681655385 | NA | Slitrk4 | SLIT and NTRK-like family, member 4 [Sou | 0.8031 | 1.0279 | 0 | 0 | 0 | 0 |
| 13776 | ENSMUSG00000046707 | 568.25506 | 0.247307387 | 0.118086577 | 2.094288728 | 0.036234268 | 0.256530813 | Csnk2a2 | casein kinase 2, alpha prime polypeptide [ | 522.82 | 502.65 | 533.31 | 606.34 | 616.2 | 628.21 |
| 13777 | ENSMUSG00000046709 | 0 | NA | NA | NA | NA | NA | Mapk10 | mitogen-activated protein kinase 10 [Sour | 0 | 0 | 0 | 0 | 0 | 0 |
| 13778 | ENSMUSG00000046711 | 350.7667498 | 0.210431524 | 0.157475211 | 1.336283485 | 0.181456633 | 0.614945814 | Hmga1 | high mobility group AT-hook 1 [Source:MC | 363 | 282.68 | 329.04 | 365.35 | 345.73 | 418.81 |
| 13779 | ENSMUSG00000046714 | 1.373912018 | -0.790726746 | 1.9618327 | -0.403055136 | 0.686907664 | NA | Foxc2 | forkhead box C2 [Source:MGI Symbol;Acc | 3.2124 | 0 | 2.0125 | 1.928 | 1.0906 | 0 |
| 13780 | ENSMUSG00000046716 | 0 | NA | NA | NA | NA | NA | Vmn1r67 | vomeronasal 1 receptor 67 [Source:MGI S | 0 | 0 | 0 | 0 | 0 | 0 |
| 13781 | ENSMUSG00000046717 | 0 | NA | NA | NA | NA | NA | Igbp1b | immunoglobulin (CD79A) binding protein | 0 | 0 | 0 | 0 | 0 | 0 |
| 13782 | ENSMUSG00000046718 | 2415.353252 | 0.058714042 | 0.115127836 | 0.50998997 | 0.610058489 | 0.916376001 | Bst2 | bone marrow stromal cell antigen 2 [Sour | 2462.3 | 2127.8 | 2507.6 | 2362.7 | 2277.2 | 2754.5 |
| 13783 | ENSMUSG00000046719 | 0.824622689 | 0.984749497 | 2.871267556 | 0.342966818 | 0.731623423 | NA | Nxph3 | neurexophilin 3 [Source:MGI Symbol;Acc: | 1.6062 | 0 | 0 | 0 | 0 | 3.3415 |
| 13784 | ENSMUSG00000046721 | 11.90967447 | -0.4636625 | 0.683055275 | -0.678806704 | 0.497260345 | 0.878825085 | Rpl14-ps1 | ribosomal protein L14, pseudogene 1 [Sou | 8.031 | 20.558 | 13.081 | 15.424 | 5.4531 | 8.9108 |
| 13785 | ENSMUSG00000046722 | 352.5317256 | -0.011639624 | 0.14202794 | -0.081953059 | 0.934684041 | 0.988420305 | Cdc42se1 | CDC42 small effector 1 [Source:MGI Symb | 342.12 | 357.72 | 362.25 | 353.78 | 321.73 | 377.59 |
| 13786 | ENSMUSG00000046723 | 0.505131668 | 1.126696199 | 3.421159819 | 0.329331647 | 0.741905026 | NA | Adam24 | a disintegrin and metallopeptidase domai | 0.8031 | 0 | 0 | 0 | 0 | 2.2277 |
| 13787 | ENSMUSG00000046727 | 98.60315768 | 0.399593063 | 0.233018724 | 1.714853879 | 0.08637202 | 0.433027696 | Cystm1 | cysteine-rich transmembrane module con | 93.16 | 76.066 | 85.531 | 111.82 | 102.52 | 122.52 |
| 13788 | ENSMUSG00000046731 | 33.16149244 | -0.247912597 | 0.391876798 | -0.632628924 | 0.526975994 | 0.888334265 | Kctd11 | potassium channel tetramerisation domai | 41.761 | 37.005 | 29.181 | 34.703 | 22.903 | 33.415 |
| 13789 | ENSMUSG00000046733 | 1.83440072 | -0.65028449 | 1.557672604 | -0.417471867 | 0.676333298 | NA | Gprc5a | G protein-coupled receptor, family C, grou | 1.6062 | 4.1117 | 1.0062 | 0.964 | 1.0906 | 2.2277 |
| 13790 | ENSMUSG00000046743 | 14.06646886 | 0.068123887 | 0.589261083 | 0.115609005 | 0.907962426 | 0.984639154 | Fat4 | FAT atypical cadherin 4 [Source:MGI Symb | 13.653 | 18.503 | 9.0562 | 14.46 | 10.906 | 17.822 |
| 13791 | ENSMUSG00000046747 | 0.647809292 | -2.733819264 | 3.061311498 | -0.893022244 | 0.371845261 | NA | Gm9812 | predicted gene 9812 [Source:MGI Symbol | 0.8031 | 3.0838 | 0 | 0 | 0 | 0 |
| 13792 | ENSMUSG00000046748 | 0.328370405 | 0.16514316 | 4.003149536 | 0.041253308 | 0.967093956 | NA | Tmem45a2 | transmembrane protein 45A2 [Source:MG | 0 | 0 | 1.0062 | 0.964 | 0 | 0 |
| 13793 | ENSMUSG00000046750 | 0 | NA | NA | NA | NA | NA | Selenov | selenoprotein V [Source:MGI Symbol;Acc: | 0 | 0 | 0 | 0 | 0 | 0 |
| 13794 | ENSMUSG00000046753 | 49.0456422 | 0.00304425 | 0.347472491 | 0.008761126 | 0.993009722 | 0.99859176 | Ccdc66 | coiled-coil domain containing 66 [Source:I | 40.155 | 54.48 | 52.325 | 32.775 | 61.075 | 53.465 |
| 13795 | ENSMUSG00000046755 | 0 | NA | NA | NA | NA | NA | Kif2b | kinesin family member 2B [Source:MGI Sy | 0 | 0 | 0 | 0 | 0 | 0 |
| 13796 | ENSMUSG00000046756 | 1339.044333 | -0.001237594 | 0.101790169 | -0.012158288 | 0.990299329 | 0.99859176 | Mrps7 | mitchondrial ribosomal protein S7 [Source | 1393.4 | 1345.5 | 1278.9 | 1272.5 | 1386.2 | 1357.8 |
| 13797 | ENSMUSG00000046761 | 338.9172448 | 0.153591756 | 0.145182828 | 1.057919574 | 0.29009211 | 0.740058153 | Fam83h | family with sequence similarity 83, memb | 348.55 | 305.29 | 307.91 | 335.46 | 364.27 | 372.02 |
| 13798 | ENSMUSG00000046764 | 0 | NA | NA | NA | NA | NA | A530053G22 | RIKEN cDNA A530053G22 gene [Source:M | 0 | 0 | 0 | 0 | 0 | 0 |
| 13799 | ENSMUSG00000046768 | 30.96052528 | -0.429802327 | 0.399141084 | -1.076818059 | 0.281561556 | 0.734379276 | Rhoj | ras homolog family member J [Source:MG | 39.352 | 31.865 | 35.219 | 24.099 | 21.812 | 33.415 |
| 13800 | ENSMUSG00000046774 | 1.199237478 | -3.618525487 | 2.801611289 | -1.291587274 | 0.196500113 | NA | 8030474K03 | RIKEN cDNA 8030474K03 gene [Source:M | 0 | 7.1954 | 0 | 0 | 0 | 0 |
| 13801 | ENSMUSG00000046782 | 31.31775638 | -0.144312248 | 0.393937475 | -0.366332876 | 0.7141167 | 0.94455746 | Ttc6 | tetratricopeptide repeat domain 6 [Source | 25.699 | 37.005 | 36.225 | 32.775 | 28.356 | 27.846 |
| 13802 | ENSMUSG00000046785 | 509.7697551 | 0.122790175 | 0.124537379 | 0.985970445 | 0.324147616 | 0.771003309 | Epm2aip1 | EPM2A (laforin) interacting protein 1 [Sou | 478.65 | 466.67 | 519.22 | 541.76 | 535.49 | 516.82 |
| 13803 | ENSMUSG00000046790 | 0 | NA | NA | NA | NA | NA | Olfr1341 | olfactory receptor 1341 [Source:MGI Symb | 0 | 0 | 0 | 0 | 0 | 0 |
| 13804 | ENSMUSG00000046791 | 102.0323901 | 0.175855809 | 0.233220038 | 0.754033875 | 0.450828881 | 0.855818389 | Riox1 | ribosomal oxygenase 1 [Source:MGI Symb | 104.4 | 92.513 | 90.562 | 125.32 | 103.61 | 95.791 |
| 13805 | ENSMUSG00000046792 | 479.7972803 | 0.112934885 | 0.133270447 | 0.84741132 | 0.396765897 | 0.824225434 | Zfp787 | zinc finger protein 787 [Source:MGI Symb | 460.18 | 461.54 | 460.86 | 454.03 | 488.6 | 553.58 |
| 13806 | ENSMUSG00000046793 | 0.17131964 | -0.796629176 | 4.080472857 | -0.195229623 | 0.84521319 | NA | Gpr61 | G protein-coupled receptor 61 [Source:M | 0 | 1.0279 | 0 | 0 | 0 | 0 |
| 13807 | ENSMUSG00000046794 | 7386.373723 | -1.258051214 | 0.473103146 | -2.659147852 | 0.007833858 | 0.090574674 | Ppp1r3b | protein phosphatase 1, regulatory subunit | 8202.1 | 11626 | 11424 | 3899.3 | 6890.5 | 2276.7 |
| 13808 | ENSMUSG00000046798 | 1460.04293 | 0.168646429 | 0.104164093 | 1.619045725 | 0.105437425 | 0.478521154 | Cldn12 | claudin 12 [Source:MGI Symbol;Acc:MGI:1 | 1345.2 | 1295.2 | 1484.2 | 1528.9 | 1607.6 | 1499.2 |
| 13809 | ENSMUSG00000046804 | 0 | NA | NA | NA | NA | NA | Phgr1 | proline/histidine/glycine-rich 1 [Source:M | 0 | 0 | 0 | 0 | 0 | 0 |
| 13810 | ENSMUSG00000046805 | 821.96055 | -0.264942526 | 0.193468537 | -1.369434688 | 0.170863437 | 0.597622477 | Mpeg1 | macrophage expressed gene 1 [Source:M | 949.27 | 1086.5 | 655.07 | 565.85 | 791.79 | 883.28 |
| 13811 | ENSMUSG00000046806 | 42.32397202 | -0.314640598 | 0.349529751 | -0.900182595 | 0.368023087 | 0.80627882 | Cyren | cell cycle regulator of NHEJ [Source:MGI S | 48.186 | 39.061 | 53.331 | 32.775 | 33.809 | 46.781 |
| 13812 | ENSMUSG00000046807 | 0.671009685 | -1.413087452 | 2.800223337 | -0.504633839 | 0.613816038 | NA | Lrrc75b | leucine rich repeat containing 75B [Source | 0 | 2.0558 | 1.0062 | 0.964 | 0 | 0 |
| 13813 | ENSMUSG00000046808 | 63.83791019 | -0.715468231 | 0.357565009 | -2.000945881 | 0.045398222 | 0.294697812 | Atp10d | ATPase, class V, type 10D [Source:MGI Sy | 88.341 | 72.982 | 76.475 | 36.631 | 78.525 | 30.074 |
| 13814 | ENSMUSG00000046811 | 1241.34287 | 0.139485079 | 0.102622572 | 1.359204672 | 0.174081741 | 0.604151393 | Gltpd2 | glycolipid transfer protein domain contair | 1163.7 | 1125.6 | 1254.8 | 1267.6 | 1312 | 1324.4 |
| 13815 | ENSMUSG00000046814 | 2085.587624 | 0.110085574 | 0.115787096 | 0.950758574 | 0.341726947 | 0.783459222 | Gchfr | GTP cyclohydrolase I feedback regulator [ | 1922.6 | 1903.7 | 2191.6 | 1998.3 | 2060.2 | 2437.1 |
| 13816 | ENSMUSG00000046818 | 9.179540063 | -0.259772623 | 0.717065615 | -0.362271761 | 0.717148958 | 0.9447719 | Ddit4l | DNA-damage-inducible transcript 4-like [S | 13.653 | 8.2233 | 8.05 | 8.6758 | 10.906 | 5.5692 |
| 13817 | ENSMUSG00000046822 | 212.8979625 | -0.062156037 | 0.178784536 | -0.347658911 | 0.728096358 | 0.946158959 | Slc39a3 | solute carrier family 39 (zinc transporter), | 199.97 | 244.64 | 208.29 | 207.26 | 231.21 | 186.01 |
| 13818 | ENSMUSG00000046826 | 63.5167654 | 0.036989523 | 0.329961747 | 0.112102459 | 0.910742168 | 0.98570598 | Fam187b | family with sequence similarity 187, meml | 81.113 | 37.005 | 69.431 | 56.875 | 67.618 | 69.058 |
| 13819 | ENSMUSG00000046828 | 0.160662974 | 1.126924175 | 4.080472857 | 0.276174898 | 0.782413736 | NA | Mettl21e | methyltransferase like 21E [Source:MGI Sy | 0 | 0 | 0 | 0.964 | 0 | 0 |
| 13820 | ENSMUSG00000046834 | 0 | NA | NA | NA | NA | NA | Krt1 | keratin 1 [Source:MGI Symbol;Acc:MGI:96 | 0 | 0 | 0 | 0 | 0 | 0 |
| 13821 | ENSMUSG00000046836 | 438.7289436 | 0.148388162 | 0.130605654 | 1.136154198 | 0.255892036 | 0.708918985 | Brox | BRO1 domain and CAAX motif containing | 414.4 | 429.67 | 404.51 | 453.07 | 491.87 | 438.85 |
| 13822 | ENSMUSG00000046840 | 193.9547803 | -0.135907925 | 0.241799803 | -0.562067971 | 0.574069709 | 0.9059841 | Hnf4aos | hepatic nuclear factor 4 alpha, opposite st | 169.45 | 182.97 | 257.6 | 221.71 | 206.13 | 125.86 |
| 13823 | ENSMUSG00000046841 | 28.20575924 | -1.06050953 | 0.428951009 | -2.472332526 | 0.013423459 | 0.132518282 | Ckap4 | cytoskeleton-associated protein 4 [Source | 44.974 | 38.033 | 31.194 | 15.424 | 22.903 | 16.708 |
| 13824 | ENSMUSG00000046844 | 0.167707431 | -0.796629176 | 4.080472857 | -0.195229623 | 0.84521319 | NA | Vat1l | vesicle amine transport protein 1 like [Sou | 0 | 0 | 1.0062 | 0 | 0 | 0 |
| 13825 | ENSMUSG00000046845 | 0 | NA | NA | NA | NA | NA | Il1f10 | interleukin 1 family, member 10 [Source:N | 0 | 0 | 0 | 0 | 0 | 0 |
| 13826 | ENSMUSG00000046846 | 0 | NA | NA | NA | NA | NA | Spesp1 | sperm equatorial segment protein 1 [Sour | 0 | 0 | 0 | 0 | 0 | 0 |
| 13827 | ENSMUSG00000046854 | 0.36353943 | 2.065989254 | 4.034137301 | 0.512126658 | 0.608562368 | NA | Pip5kl1 | phosphatidylinositol-4-phosphate 5-kinas | 0 | 0 | 0 | 0 | 2.1812 | 0 |
| 13828 | ENSMUSG00000046856 | 0 | NA | NA | NA | NA | NA | Gpr1 | G protein-coupled receptor 1 [Source:MG | 0 | 0 | 0 | 0 | 0 | 0 |

| | | | | | | | | | | | | | | | |
|---|---|---|---|---|---|---|---|---|---|---|---|---|---|---|---|
| 13829 | ENSMUSG00000046861 | 1331.934862 | 0.343043192 | 0.108284244 | 3.167988043 | 0.001534978 | 0.028984238 | Hectd3 | HECT domain E3 ubiquitin protein ligase 3 | 1263.3 | 1090.6 | 1168.2 | 1541.4 | 1528 | 1400.1 |
| 13830 | ENSMUSG00000046862 | 103.6895967 | 0.14506511 | 0.226885283 | 0.639376464 | 0.522578056 | 0.888334265 | Pramel12 | PRAME like 12 [Source:MGI Symbol;Acc:M | 93.16 | 101.76 | 100.62 | 107 | 97.065 | 122.52 |
| 13831 | ENSMUSG00000046865 | 483.3811406 | 0.115013844 | 0.144826088 | 0.794151419 | 0.427107287 | 0.842220356 | Fbl | fibrillarin [Source:MGI Symbol;Acc:MGI:9 | 426.45 | 497.51 | 468.91 | 477.17 | 578.03 | 452.22 |
| 13832 | ENSMUSG00000046873 | 153.9867554 | -0.16444204 | 0.2011943 | -0.817329516 | 0.413740144 | 0.835385466 | Mbtps2 | membrane-bound transcription factor pep | 171.86 | 166.52 | 149.93 | 161.95 | 121.06 | 152.6 |
| 13833 | ENSMUSG00000046876 | 551.7999427 | -1.015399347 | 0.474285425 | -2.140903542 | 0.032281817 | 0.240179886 | Atxn1 | ataxin 1 [Source:MGI Symbol;Acc:MGI:104 | 471.42 | 939.52 | 803.99 | 193.76 | 423.16 | 478.95 |
| 13834 | ENSMUSG00000046879 | 972.7921238 | 0.150399498 | 0.11524953 | 1.30499012 | 0.191896212 | 0.629099068 | Irgm1 | immunity-related GTPase family M memb | 925.17 | 905.6 | 934.8 | 911.92 | 1068.8 | 1090.5 |
| 13835 | ENSMUSG00000046881 | 0 | NA | NA | NA | NA | NA | Olfr374 | olfactory receptor 374 [Source:MGI Symb | 0 | 0 | 0 | 0 | 0 | 0 |
| 13836 | ENSMUSG00000046886 | 0 | NA | NA | NA | NA | NA | Zfp474 | zinc finger protein 474 [Source:MGI Symb | 0 | 0 | 0 | 0 | 0 | 0 |
| 13837 | ENSMUSG00000046897 | 477.6924769 | -0.101450665 | 0.135375778 | -0.749400418 | 0.453615899 | 0.857099982 | Zfp740 | zinc finger protein 740 [Source:MGI Symb | 489.09 | 539.66 | 454.82 | 453.07 | 496.23 | 433.29 |
| 13838 | ENSMUSG00000046899 | 0 | NA | NA | NA | NA | NA | Prl7a2 | prolactin family 7, subfamily a, member 2 | 0 | 0 | 0 | 0 | 0 | 0 |
| 13839 | ENSMUSG00000046908 | 20.90507488 | -0.581652239 | 0.589080742 | -0.987389669 | 0.323451647 | 0.770475661 | Ltb4r1 | leukotriene B4 receptor 1 [Source:MGI Sy | 23.29 | 40.089 | 12.075 | 27.955 | 11.997 | 10.025 |
| 13840 | ENSMUSG00000046909 | 239.523231 | 0.068032868 | 0.186195782 | 0.365383507 | 0.714825151 | 0.944595913 | Tefm | transcription elongation factor, mitocho | 263.42 | 218.95 | 218.36 | 209.18 | 295.56 | 231.68 |
| 13841 | ENSMUSG00000046913 | 0 | NA | NA | NA | NA | NA | Olfr1451 | olfactory receptor 1451 [Source:MGI Sym | 0 | 0 | 0 | 0 | 0 | 0 |
| 13842 | ENSMUSG00000046916 | 45.11751491 | -0.201991324 | 0.368032691 | -0.548840711 | 0.583114769 | 0.908645447 | Myct1 | myc target 1 [Source:MGI Symbol;Acc:MG | 40.155 | 44.2 | 60.375 | 29.883 | 38.172 | 57.92 |
| 13843 | ENSMUSG00000046922 | 0 | NA | NA | NA | NA | NA | Gpr6 | G protein-coupled receptor 6 [Source:MG | 0 | 0 | 0 | 0 | 0 | 0 |
| 13844 | ENSMUSG00000046924 | 0 | NA | NA | NA | NA | NA | Vmn1r179 | vomeronasal 1 receptor 179 [Source:MGI | 0 | 0 | 0 | 0 | 0 | 0 |
| 13845 | ENSMUSG00000046932 | 0 | NA | NA | NA | NA | NA | Vmn1r193 | vomeronasal 1 receptor 193 [Source:MGI | 0 | 0 | 0 | 0 | 0 | 0 |
| 13846 | ENSMUSG00000046934 | 0 | NA | NA | NA | NA | NA | Csl | citrate synthase like [Source:MGI Symbol; | 0 | 0 | 0 | 0 | 0 | 0 |
| 13847 | ENSMUSG00000046942 | 0 | NA | NA | NA | NA | NA | Mageb16 | MAGE family member B16 [Source:MGI Sy | 0 | 0 | 0 | 0 | 0 | 0 |
| 13848 | ENSMUSG00000046947 | 170.751191 | -0.216414578 | 0.187620314 | -1.153470931 | 0.248717142 | 0.70053822 | Adck2 | aarF domain containing kinase 2 [Source:I | 187.12 | 194.28 | 169.05 | 155.2 | 142.87 | 175.99 |
| 13849 | ENSMUSG00000046949 | 1509.69034 | 0.08496581 | 0.112817021 | 0.753129349 | 0.451372189 | 0.85613987 | Nqo2 | N-ribosyldihydronicotinamide quinone re | 1557.2 | 1365.1 | 1472.1 | 1467.2 | 1473.4 | 1723.1 |
| 13850 | ENSMUSG00000046952 | 0.181769715 | 1.126924175 | 4.080472857 | 0.276174898 | 0.782413736 | NA | Gm5815 | predicted pseudogene 5815 [Source:MGI | 0 | 0 | 0 | 0 | 1.0906 | 0 |
| 13851 | ENSMUSG00000046957 | 0 | NA | NA | NA | NA | NA | Spz1 | spermatogenic leucine zipper 1 [Source:M | 0 | 0 | 0 | 0 | 0 | 0 |
| 13852 | ENSMUSG00000046958 | 0 | NA | NA | NA | NA | NA | 4930432E11 | RIKEN cDNA 4930432E11 gene [Source:M | 0 | 0 | 0 | 0 | 0 | 0 |
| 13853 | ENSMUSG00000046959 | 3512.066595 | -0.015906825 | 0.122656975 | -0.129685454 | 0.896815292 | 0.982798755 | Slc26a1 | solute carrier family 26 (sulfate transporte | 3410.8 | 3642.9 | 3540 | 2929.5 | 4048.4 | 3500.8 |
| 13854 | ENSMUSG00000046961 | 0.17131964 | -0.796629176 | 4.080472857 | -0.195229623 | 0.84521319 | NA | Gpr156 | G protein-coupled receptor 156 [Source:M | 0 | 1.0279 | 0 | 0 | 0 | 0 |
| 13855 | ENSMUSG00000046962 | 158.8131193 | -0.165503007 | 0.219179541 | -0.755102445 | 0.450187516 | 0.855539645 | Zbtb21 | zinc finger and BTB domain containing 21 | 163.83 | 165.49 | 175.09 | 190.87 | 142.87 | 114.73 |
| 13856 | ENSMUSG00000046971 | 0.342639279 | -1.805976596 | 4.028547714 | -0.448294702 | 0.653940524 | NA | Pla2g4f | phospholipase A2, group IVF [Source:MGI | 0 | 2.0558 | 0 | 0 | 0 | 0 |
| 13857 | ENSMUSG00000046974 | 0 | NA | NA | NA | NA | NA | BC053393 | cDNA sequence BC053393 [Source:MGI Sy | 0 | 0 | 0 | 0 | 0 | 0 |
| 13858 | ENSMUSG00000046975 | 0 | NA | NA | NA | NA | NA | Olfr1020 | olfactory receptor 1020 [Source:MGI Sym | 0 | 0 | 0 | 0 | 0 | 0 |
| 13859 | ENSMUSG00000046982 | 389.4434958 | 0.034763295 | 0.144120571 | 0.241209807 | 0.80939251 | 0.966005805 | Tshz1 | teashirt zinc finger family member 1 [Sour | 371.84 | 360.8 | 421.62 | 382.7 | 367.54 | 432.17 |
| 13860 | ENSMUSG00000046985 | 1124.782888 | 0.034359077 | 0.116399772 | 0.295181649 | 0.767855119 | 0.95578799 | Tapt1 | transmembrane anterior posterior transfo | 1166.1 | 1174.9 | 993.16 | 1208.8 | 1137.5 | 1068.2 |
| 13861 | ENSMUSG00000046991 | 12.2521752 | 0.08066356 | 0.613310963 | 0.131521471 | 0.895362803 | 0.982798755 | Wdr27 | WD repeat domain 27 [Source:MGI Symbo | 10.44 | 9.2513 | 16.1 | 13.496 | 13.087 | 11.138 |
| 13862 | ENSMUSG00000046993 | 0 | NA | NA | NA | NA | NA | Gm5637 | predicted pseudogene 5637 [Source:MGI | 0 | 0 | 0 | 0 | 0 | 0 |
| 13863 | ENSMUSG00000046994 | 124.2359213 | -0.091501739 | 0.209944901 | -0.435836923 | 0.662955053 | 0.92906822 | Mars2 | methionine-tRNA synthetase 2 (mitochon | 141.35 | 114.1 | 128.8 | 125.32 | 117.79 | 118.07 |
| 13864 | ENSMUSG00000046995 | 0 | NA | NA | NA | NA | NA | 2610303G11 | RIKEN cDNA 2610303G11 gene [Source:M | 0 | 0 | 0 | 0 | 0 | 0 |
| 13865 | ENSMUSG00000046997 | 69.20301359 | 0.429210523 | 0.346240402 | 1.239631539 | 0.215111709 | 0.659728602 | Spsb4 | splA/ryanodine receptor domain and SOC | 44.171 | 53.452 | 79.493 | 65.55 | 56.712 | 115.84 |
| 13866 | ENSMUSG00000046999 | 0 | NA | NA | NA | NA | NA | 1110032F04 | RIKEN cDNA 1110032F04 gene [Source:M | 0 | 0 | 0 | 0 | 0 | 0 |
| 13867 | ENSMUSG00000047002 | 0 | NA | NA | NA | NA | NA | Msgn1 | mesogenin 1 [Source:MGI Symbol;Acc:MG | 0 | 0 | 0 | 0 | 0 | 0 |
| 13868 | ENSMUSG00000047003 | 35.49895848 | -0.068164163 | 0.393532478 | -0.173211023 | 0.862485559 | 0.976334255 | Zfp41 | zinc finger protein 41 [Source:MGI Symbo | 32.124 | 33.921 | 43.269 | 46.271 | 23.994 | 33.415 |
| 13869 | ENSMUSG00000047013 | 0.335414862 | -1.779047242 | 4.030804179 | -0.441362855 | 0.658950329 | NA | Fbxo41 | F-box protein 41 [Source:MGI Symbol;Acc | 0 | 0 | 2.0125 | 0 | 0 | 0 |
| 13870 | ENSMUSG00000047014 | 0 | NA | NA | NA | NA | NA | Catsperb | cation channel sperm associated auxiliary | 0 | 0 | 0 | 0 | 0 | 0 |
| 13871 | ENSMUSG00000047021 | 0 | NA | NA | NA | NA | NA | Cfap65 | cilia and flagella associated protein 65 [So | 0 | 0 | 0 | 0 | 0 | 0 |
| 13872 | ENSMUSG00000047022 | 52.42007637 | 0.686713164 | 0.341989343 | 2.007995799 | 0.044643738 | 0.291866583 | Mipol1 | mirror-image polydactyly 1 [Source:MGI S | 39.352 | 34.949 | 46.287 | 66.514 | 83.978 | 43.44 |
| 13873 | ENSMUSG00000047025 | 0 | NA | NA | NA | NA | NA | Ccer1 | coiled-coil glutamate-rich protein 1 [Sourc | 0 | 0 | 0 | 0 | 0 | 0 |
| 13874 | ENSMUSG00000047026 | 0 | NA | NA | NA | NA | NA | Acsm4 | acyl-CoA synthetase medium-chain family | 0 | 0 | 0 | 0 | 0 | 0 |
| 13875 | ENSMUSG00000047030 | 231.2897361 | 0.208615482 | 0.167922535 | 1.242331659 | 0.214114219 | 0.658874792 | Spata2 | spermatogenesis associated 2 [Source:MG | 224.87 | 211.75 | 207.29 | 275.7 | 246.48 | 221.65 |
| 13876 | ENSMUSG00000047031 | 0 | NA | NA | NA | NA | NA | Vmn1r68 | vomeronasal 1 receptor 68 [Source:MGI S | 0 | 0 | 0 | 0 | 0 | 0 |
| 13877 | ENSMUSG00000047033 | 0.813971537 | 1.950992709 | 2.892144212 | 0.674583481 | 0.499940431 | NA | Pcdhb15 | protocadherin beta 15 [Source:MGI Symb | 0 | 1.0279 | 0 | 3.8559 | 0 | 0 |
| 13878 | ENSMUSG00000047034 | 0 | NA | NA | NA | NA | NA | Ankrd33 | ankyrin repeat domain 33 [Source:MGI Sy | 0 | 0 | 0 | 0 | 0 | 0 |
| 13879 | ENSMUSG00000047036 | 2451.10627 | 0.09492474 | 0.160365181 | 0.591928618 | 0.55389839 | 0.896852827 | Zfp445 | zinc finger protein 445 [Source:MGI Symb | 2520.9 | 2375.5 | 2215.8 | 3238 | 2449.5 | 1906.9 |
| 13880 | ENSMUSG00000047037 | 17.09608012 | -0.375816622 | 0.536056161 | -0.701076956 | 0.483254991 | 0.871960882 | Nipa1 | non imprinted in Prader-Willi/Angelman s | 19.274 | 14.391 | 24.15 | 10.604 | 17.45 | 16.708 |
| 13881 | ENSMUSG00000047039 | 0 | NA | NA | NA | NA | NA | Olfr1151 | olfactory receptor 1151 [Source:MGI Sym | 0 | 0 | 0 | 0 | 0 | 0 |
| 13882 | ENSMUSG00000047040 | 10.50085734 | -0.043395693 | 0.773426083 | -0.056108391 | 0.95525546 | 0.991678478 | Prr15l | proline rich 15-like [Source:MGI Symbol;A | 5.6217 | 12.335 | 14.087 | 5.7839 | 20.722 | 4.4554 |
| 13883 | ENSMUSG00000047044 | 141.2814852 | 0.055697911 | 0.201316851 | 0.276667903 | 0.78203512 | 0.960298495 | D030056L22 | RIKEN cDNA D030056L22 gene [Source:M | 148.57 | 137.74 | 128.8 | 127.25 | 150.51 | 154.82 |
| 13884 | ENSMUSG00000047045 | 143.6750519 | -0.205475085 | 0.206839611 | -0.993402975 | 0.320513612 | 0.767391055 | Tmem164 | transmembrane protein 164 [Source:MGI | 155 | 167.55 | 138.86 | 121.46 | 123.24 | 155.94 |

| | | | | | | | | | | | | | | | |
|---|---|---|---|---|---|---|---|---|---|---|---|---|---|---|---|
| 13885 | ENSMUSG00000047046 | 0 | NA | NA | NA | NA | NA | 1700031F10 | RIKEN cDNA 1700031F10 gene [Source:M | 0 | 0 | 0 | 0 | 0 | 0 |
| 13886 | ENSMUSG00000047048 | 0 | NA | NA | NA | NA | NA | Olfr432 | olfactory receptor 432 [Source:MGI Symb | 0 | 0 | 0 | 0 | 0 | 0 |
| 13887 | ENSMUSG00000047050 | 0 | NA | NA | NA | NA | NA | Olfr914 | olfactory receptor 914 [Source:MGI Symb | 0 | 0 | 0 | 0 | 0 | 0 |
| 13888 | ENSMUSG00000047053 | 0.401551119 | -2.072982726 | 4.008251979 | -0.517178744 | 0.605031388 | NA | Teddm1a | transmembrane epididymal protein 1A [S | 2.4093 | 0 | 0 | 0 | 0 | 0 |
| 13889 | ENSMUSG00000047061 | 0 | NA | NA | NA | NA | NA | Gm9817 | predicted gene 9817 [Source:MGI Symbol | 0 | 0 | 0 | 0 | 0 | 0 |
| 13890 | ENSMUSG00000047067 | 263.73177 | -0.16240526 | 0.179624887 | -0.904135628 | 0.365923488 | 0.804235439 | Dusp28 | dual specificity phosphatase 28 [Source:M | 318.03 | 256.98 | 260.62 | 295.94 | 222.49 | 228.34 |
| 13891 | ENSMUSG00000047079 | 0 | NA | NA | NA | NA | NA | Ube2dnl2 | ubiquitin-conjugating enzyme E2D N-term | 0 | 0 | 0 | 0 | 0 | 0 |
| 13892 | ENSMUSG00000047084 | 397.4089568 | 0.137645854 | 0.134604305 | 1.022596226 | 0.306498795 | 0.755610724 | Ngrn | neugrin, neurite outgrowth associated [Sc | 370.23 | 358.74 | 406.52 | 414.51 | 410.07 | 424.37 |
| 13893 | ENSMUSG00000047085 | 0 | NA | NA | NA | NA | NA | Lrrc4b | leucine rich repeat containing 4B [Source: | 0 | 0 | 0 | 0 | 0 | 0 |
| 13894 | ENSMUSG00000047090 | 60.5459525 | -0.024301936 | 0.290854215 | -0.083553665 | 0.933411308 | 0.988420305 | Tmem198b | transmembrane protein 198b [Source:MG | 60.233 | 54.48 | 68.425 | 56.875 | 69.8 | 53.465 |
| 13895 | ENSMUSG00000047094 | 0 | NA | NA | NA | NA | NA | Ofcc1 | orofacial cleft 1 candidate 1 [Source:MGI | 0 | 0 | 0 | 0 | 0 | 0 |
| 13896 | ENSMUSG00000047098 | 319.5400648 | 0.494214106 | 0.166906033 | 2.961031999 | 0.003066101 | 0.047431104 | Rnf31 | ring finger protein 31 [Source:MGI Symbo | 258.6 | 247.73 | 289.8 | 383.66 | 309.74 | 427.72 |
| 13897 | ENSMUSG00000047102 | 0 | NA | NA | NA | NA | NA | Tas2r139 | taste receptor, type 2, member 139 [Sour | 0 | 0 | 0 | 0 | 0 | 0 |
| 13898 | ENSMUSG00000047104 | 0 | NA | NA | NA | NA | NA | Pbp2 | phosphatidylethanolamine binding protei | 0 | 0 | 0 | 0 | 0 | 0 |
| 13899 | ENSMUSG00000047108 | 2.471754952 | 1.511012174 | 1.504215105 | 1.004518682 | 0.315128671 | NA | Dnajb7 | DnaJ heat shock protein family (Hsp40) m | 0.8031 | 2.0558 | 1.0062 | 0.964 | 1.0906 | 8.9108 |
| 13900 | ENSMUSG00000047109 | 161.3389369 | 0.477846366 | 0.246263615 | 1.940385576 | 0.052332848 | 0.321555784 | Cldn14 | claudin 14 [Source:MGI Symbol;Acc:MGI: | 150.18 | 118.21 | 135.84 | 196.65 | 124.33 | 242.82 |
| 13901 | ENSMUSG00000047115 | 3.630129988 | 0.072037739 | 1.267736794 | 0.056823892 | 0.954685481 | NA | Fam221a | family with sequence similarity 221, meml | 1.6062 | 3.0838 | 6.0375 | 7.7118 | 0 | 3.3415 |
| 13902 | ENSMUSG00000047117 | 0 | NA | NA | NA | NA | NA | Ankdd1b | ankyrin repeat and death domain contain | 0 | 0 | 0 | 0 | 0 | 0 |
| 13903 | ENSMUSG00000047123 | 295.8815549 | 0.029280711 | 0.15353521 | 0.190710075 | 0.848752745 | 0.972888368 | Ticam1 | toll-like receptor adaptor molecule 1 [Sou | 313.21 | 293.98 | 270.68 | 276.66 | 297.74 | 323.01 |
| 13904 | ENSMUSG00000047126 | 2772.228864 | -0.041963727 | 0.10318921 | -0.406667785 | 0.684252004 | 0.935528319 | Cltc | clathrin, heavy polypeptide (Hc) [Source:N | 2753 | 3076.6 | 2608.2 | 2634.6 | 2894.5 | 2666.5 |
| 13905 | ENSMUSG00000047129 | 0.514011053 | 1.044022102 | 3.082659575 | 0.338675769 | 0.734853997 | NA | 1700113H08 | RIKEN cDNA 1700113H08 gene [Source:M | 0 | 0 | 1.0062 | 0.964 | 0 | 1.1138 |
| 13906 | ENSMUSG00000047139 | 23.94855103 | -0.383383917 | 0.461204527 | -0.831266596 | 0.405823041 | 0.829973745 | Cd24a | CD24a antigen [Source:MGI Symbol;Acc:N | 31.321 | 24.67 | 25.156 | 18.316 | 15.269 | 28.96 |
| 13907 | ENSMUSG00000047141 | 287.9667281 | 0.106007935 | 0.148180574 | 0.715396976 | 0.474363781 | 0.869056076 | Zfp654 | zinc finger protein 654 [Source:MGI Symb | 289.12 | 263.15 | 279.74 | 309.44 | 291.2 | 295.17 |
| 13908 | ENSMUSG00000047143 | 2.543499732 | 0.990671922 | 1.594217124 | 0.621415934 | 0.534325991 | NA | Dmrta2 | doublesex and mab-3 related transcriptio | 4.0155 | 0 | 1.0062 | 5.7839 | 0 | 4.4554 |
| 13909 | ENSMUSG00000047146 | 2.310735916 | 0.612205025 | 1.35459134 | 0.451948131 | 0.651306349 | NA | Tet1 | tet methylcytosine dioxygenase 1 [Source | 2.4093 | 1.0279 | 2.0125 | 2.8919 | 2.1812 | 3.3415 |
| 13910 | ENSMUSG00000047149 | 0 | NA | NA | NA | NA | NA | Olfr1052 | olfactory receptor 1052 [Source:MGI Syml | 0 | 0 | 0 | 0 | 0 | 0 |
| 13911 | ENSMUSG00000047150 | 159.24408 | 0.156114184 | 0.202017413 | 0.772775883 | 0.439655029 | 0.84844279 | 1700001C19 | RIKEN cDNA 1700001C19 gene [Source:M | 174.27 | 145.96 | 130.81 | 148.45 | 178.86 | 177.1 |
| 13912 | ENSMUSG00000047153 | 627.7175318 | 0.256021685 | 0.168160988 | 1.522479661 | 0.127888938 | 0.523750856 | Khnyn | KH and NYN domain containing [Source:M | 465.8 | 534.52 | 717.45 | 754.79 | 600.93 | 692.81 |
| 13913 | ENSMUSG00000047155 | 0 | NA | NA | NA | NA | NA | Cyp4x1 | cytochrome P450, family 4, subfamily x, p | 0 | 0 | 0 | 0 | 0 | 0 |
| 13914 | ENSMUSG00000047161 | 0.68145976 | 0.058694068 | 2.590797836 | 0.022654824 | 0.981925612 | NA | Chst9 | carbohydrate (N-acetylgalactosamine 4-0) | 0 | 1.0279 | 1.0062 | 0.964 | 1.0906 | 0 |
| 13915 | ENSMUSG00000047168 | 0 | NA | NA | NA | NA | NA | Gm6684 | predicted pseudogene 6684 [Source:MGI | 0 | 0 | 0 | 0 | 0 | 0 |
| 13916 | ENSMUSG00000047171 | 0 | NA | NA | NA | NA | NA | Helt | helt bHLH transcription factor [Source:MG | 0 | 0 | 0 | 0 | 0 | 0 |
| 13917 | ENSMUSG00000047180 | 56.97666197 | -0.426658151 | 0.344244167 | -1.239405605 | 0.215195327 | 0.659728602 | Neurl3 | neuralized E3 ubiquitin protein ligase 3 [S | 74.689 | 69.898 | 51.318 | 53.019 | 29.447 | 63.489 |
| 13918 | ENSMUSG00000047181 | 32.13775996 | -0.422346835 | 0.410234522 | -1.029525338 | 0.30323288 | 0.752484788 | Samd14 | sterile alpha motif domain containing 14 [ | 40.155 | 43.173 | 27.169 | 33.739 | 19.631 | 28.96 |
| 13919 | ENSMUSG00000047182 | 4.209548872 | -1.259371723 | 1.087909799 | -1.157606746 | 0.247024555 | NA | Irs3 | insulin receptor substrate 3 [Source:MGI | 5.6217 | 4.1117 | 8.05 | 1.928 | 1.0906 | 4.4554 |
| 13920 | ENSMUSG00000047187 | 2306.415925 | 0.083923011 | 0.106693071 | 0.786583514 | 0.431525713 | 0.844934876 | Rab2a | RAB2A, member RAS oncogene family [So | 2437.4 | 2212.1 | 2066.8 | 2199.8 | 2496.4 | 2426 |
| 13921 | ENSMUSG00000047189 | 0 | NA | NA | NA | NA | NA | Gm9818 | predicted gene 9818 [Source:MGI Symbol | 0 | 0 | 0 | 0 | 0 | 0 |
| 13922 | ENSMUSG00000047193 | 87.50893963 | -0.464571375 | 0.354447257 | -1.310692539 | 0.18996166 | 0.627369158 | Dync2h1 | dynein cytoplasmic 2 heavy chain 1 [Sourc | 79.507 | 99.708 | 125.78 | 98.326 | 89.431 | 32.301 |
| 13923 | ENSMUSG00000047197 | 0.160662974 | 1.126924175 | 4.080472857 | 0.276174898 | 0.782413736 | NA | Gjd3 | gap junction protein, delta 3 [Source:MGI | 0 | 0 | 0 | 0.964 | 0 | 0 |
| 13924 | ENSMUSG00000047203 | 0 | NA | NA | NA | NA | NA | Vmn1r54 | vomeronasal 1 receptor 54 [Source:MGI S | 0 | 0 | 0 | 0 | 0 | 0 |
| 13925 | ENSMUSG00000047205 | 0.624661033 | -1.234026074 | 2.854658177 | -0.432285058 | 0.665534246 | NA | Dusp18 | dual specificity phosphatase 18 [Source:M | 1.6062 | 1.0279 | 0 | 0 | 0 | 1.1138 |
| 13926 | ENSMUSG00000047207 | 0 | NA | NA | NA | NA | NA | Olfr1495 | olfactory receptor 1495 [Source:MGI Syml | 0 | 0 | 0 | 0 | 0 | 0 |
| 13927 | ENSMUSG00000047213 | 1469.531122 | 0.00827568 | 0.099391214 | 0.083263693 | 0.933641869 | 0.988420305 | Ythdf3 | YTH N6-methyladenosine RNA binding pro | 1458.4 | 1487.4 | 1450 | 1429.6 | 1569.4 | 1422.4 |
| 13928 | ENSMUSG00000047215 | 2924.66213 | -0.008357624 | 0.105520639 | -0.079203694 | 0.936870607 | 0.988420305 | Rpl9 | ribosomal protein L9 [Source:MGI Symbol | 2737 | 3109.5 | 2953.3 | 2757 | 2777.8 | 3213.4 |
| 13929 | ENSMUSG00000047216 | 0.335414862 | -1.779047242 | 4.030804179 | -0.441362855 | 0.658950329 | NA | Cdh19 | cadherin 19, type 2 [Source:MGI Symbol;A | 0 | 0 | 2.0125 | 0 | 0 | 0 |
| 13930 | ENSMUSG00000047220 | 1.566984201 | 4.18124924 | 2.053658707 | 2.036000055 | 0.04175035 | NA | Iho1 | interactor of HORMAD1 1 [Source:MGI Sy | 0 | 0 | 0 | 3.8559 | 1.0906 | 4.4554 |
| 13931 | ENSMUSG00000047221 | 184.2802067 | -0.216776716 | 0.188259339 | -1.151479215 | 0.249535142 | 0.701784229 | Fam185a | family with sequence similarity 185, meml | 199.97 | 209.7 | 184.14 | 143.63 | 177.77 | 190.47 |
| 13932 | ENSMUSG00000047222 | 3.769921776 | 1.417148027 | 1.41670647 | 1.000311679 | 0.317159697 | NA | Rnase2a | ribonuclease, RNase A family, 2A (liver, ec | 4.0155 | 0 | 2.0125 | 0 | 5.4531 | 11.138 |
| 13933 | ENSMUSG00000047225 | 0 | NA | NA | NA | NA | NA | Olfr684 | olfactory receptor 684 [Source:MGI Symb | 0 | 0 | 0 | 0 | 0 | 0 |
| 13934 | ENSMUSG00000047227 | 12.31245085 | -0.558439209 | 0.629778461 | -0.886723258 | 0.375227913 | 0.809332934 | Gm527 | predicted gene 527 [Source:MGI Symbol;A | 10.44 | 17.475 | 16.1 | 6.7478 | 13.087 | 10.025 |
| 13935 | ENSMUSG00000047228 | 174.9208397 | 0.367955201 | 0.259530943 | 1.417770064 | 0.156257907 | 0.574894692 | A2ml1 | alpha-2-macroglobulin like 1 [Source:MGI | 210.41 | 98.68 | 147.92 | 161.95 | 232.3 | 198.26 |
| 13936 | ENSMUSG00000047230 | 1591.996317 | 0.220433542 | 0.14574401 | 1.512470678 | 0.130414167 | 0.528822682 | Cldn2 | claudin 2 [Source:MGI Symbol;Acc:MGI:1 | 1816.6 | 1371.2 | 1222.6 | 1862.4 | 1622.8 | 1656.3 |
| 13937 | ENSMUSG00000047237 | 0 | NA | NA | NA | NA | NA | Fbxw21 | F-box and WD-40 domain protein 21 [Sou | 0 | 0 | 0 | 0 | 0 | 0 |
| 13938 | ENSMUSG00000047238 | 43.21165186 | -0.21772644 | 0.338307472 | -0.643575616 | 0.519850666 | 0.887802551 | Mageh1 | MAGE family member H1 [Source:MGI Syr | 40.958 | 48.312 | 50.312 | 42.415 | 32.719 | 44.554 |
| 13939 | ENSMUSG00000047242 | 11.07600638 | -0.796050773 | 0.654172963 | -1.216881189 | 0.223649424 | 0.669328629 | Taf9b | TATA-box binding protein associated facto | 13.653 | 15.419 | 13.081 | 7.7118 | 5.4531 | 11.138 |
| 13940 | ENSMUSG00000047246 | 11.10134237 | 0.334082082 | 0.683581869 | 0.488722855 | 0.625037921 | 0.919503423 | H2bc6 | H2B clustered histone 6 [Source:MGI Sym | 7.2279 | 7.1954 | 15.094 | 9.6398 | 18.541 | 8.9108 |

| | | | | | | | | | | | | | | | |
|---|---|---|---|---|---|---|---|---|---|---|---|---|---|---|---|
| 13941 | ENSMUSG00000047248 | 208.4009846 | 0.004687864 | 0.169601001 | 0.027640542 | 0.977948846 | 0.997746499 | C2cd3 | C2 calcium-dependent domain containing | 214.43 | 195.3 | 214.33 | 214.97 | 191.95 | 219.43 |
| 13942 | ENSMUSG00000047250 | 97.42160412 | -1.108621594 | 0.275769861 | -4.020097005 | 5.82E-05 | 0.002370229 | Ptgs1 | prostaglandin-endoperoxide synthase 1 [S | 126.89 | 160.36 | 111.69 | 44.343 | 62.165 | 79.083 |
| 13943 | ENSMUSG00000047253 | 0 | NA | NA | NA | NA | NA | Krtap1-5 | keratin associated protein 1-5 [Source:MG | 0 | 0 | 0 | 0 | 0 | 0 |
| 13944 | ENSMUSG00000047257 | 0 | NA | NA | NA | NA | NA | Prss45 | protease, serine 45 [Source:MGI Symbol;A | 0 | 0 | 0 | 0 | 0 | 0 |
| 13945 | ENSMUSG00000047259 | 0 | NA | NA | NA | NA | NA | Mc4r | melanocortin 4 receptor [Source:MGI Sym | 0 | 0 | 0 | 0 | 0 | 0 |
| 13946 | ENSMUSG00000047260 | 1485.80567 | 0.062563827 | 0.099257432 | 0.630318815 | 0.528486015 | 0.888334265 | Emc6 | ER membrane protein complex subunit 6 | 1525.1 | 1451.4 | 1383.6 | 1518.3 | 1562.9 | 1473.6 |
| 13947 | ENSMUSG00000047261 | 0 | NA | NA | NA | NA | NA | Gap43 | growth associated protein 43 [Source:MG | 0 | 0 | 0 | 0 | 0 | 0 |
| 13948 | ENSMUSG00000047264 | 152.5231126 | 0.317885978 | 0.224533039 | 1.415764826 | 0.156844371 | 0.575786484 | Zfp358 | zinc finger protein 358 [Source:MGI Symb | 133.31 | 140.82 | 133.83 | 220.75 | 149.41 | 137 |
| 13949 | ENSMUSG00000047281 | 14.5844637 | 0.162950924 | 0.593082453 | 0.274752563 | 0.783506345 | 0.960670212 | Sfn | stratifin [Source:MGI Symbol;Acc:MGI:18 | 8.031 | 12.335 | 21.131 | 17.352 | 14.178 | 14.48 |
| 13950 | ENSMUSG00000047284 | 463.089804 | 0.090373316 | 0.13153161 | 0.687084391 | 0.49202955 | 0.876368691 | Neurl4 | neuralized E3 ubiquitin protein ligase 4 [S | 458.57 | 424.53 | 462.87 | 516.69 | 456.97 | 458.9 |
| 13951 | ENSMUSG00000047286 | 0 | NA | NA | NA | NA | NA | Olfr370 | olfactory receptor 370 [Source:MGI Symb | 0 | 0 | 0 | 0 | 0 | 0 |
| 13952 | ENSMUSG00000047293 | 1.184762013 | -0.39060714 | 2.10457176 | -0.185599345 | 0.852758967 | NA | Gpr15 | G protein-coupled receptor 15 [Source:M | 0 | 3.0838 | 1.0062 | 1.928 | 1.0906 | 0 |
| 13953 | ENSMUSG00000047295 | 0 | NA | NA | NA | NA | NA | Smgc | submandibular gland protein C [Source:M | 0 | 0 | 0 | 0 | 0 | 0 |
| 13954 | ENSMUSG00000047298 | 0 | NA | NA | NA | NA | NA | Kcnv2 | potassium channel, subfamily V, member | 0 | 0 | 0 | 0 | 0 | 0 |
| 13955 | ENSMUSG00000047307 | 0.885292348 | 2.085359241 | 2.560462423 | 0.814446337 | 0.415389303 | NA | Pcdhb13 | protocadherin beta 13 [Source:MGI Symb | 0 | 0 | 1.0062 | 0.964 | 0 | 3.3415 |
| 13956 | ENSMUSG00000047324 | 0 | NA | NA | NA | NA | NA | 4931429P17 | RIKEN cDNA 4931429P17 gene [Source:M | 0 | 0 | 0 | 0 | 0 | 0 |
| 13957 | ENSMUSG00000047330 | 3.226531308 | -1.381952967 | 1.317084122 | -1.049251861 | 0.294062215 | NA | Kcne4 | potassium voltage-gated channel, Isk-rela | 4.8186 | 3.0838 | 6.0375 | 0.964 | 0 | 4.4554 |
| 13958 | ENSMUSG00000047342 | 0 | NA | NA | NA | NA | NA | Zfp286 | zinc finger protein 286 [Source:MGI Symb | 0 | 0 | 0 | 0 | 0 | 0 |
| 13959 | ENSMUSG00000047343 | 0.331982614 | 0.16514309 | 3.986751476 | 0.041422971 | 0.966958701 | NA | Mettl21c | methyltransferase like 21C [Source:MGI S | 0 | 1.0279 | 0 | 0.964 | 0 | 0 |
| 13960 | ENSMUSG00000047344 | 0.674621893 | -1.426050892 | 3.091195043 | -0.461326727 | 0.644564212 | NA | Lancl3 | LanC lantibiotic synthetase component C- | 0 | 3.0838 | 0 | 0.964 | 0 | 0 |
| 13961 | ENSMUSG00000047347 | 0 | NA | NA | NA | NA | NA | Tdg-ps | thymine DNA glycosylase, pseudogene [So | 0 | 0 | 0 | 0 | 0 | 0 |
| 13962 | ENSMUSG00000047352 | 0 | NA | NA | NA | NA | NA | Olfr976 | olfactory receptor 976 [Source:MGI Symb | 0 | 0 | 0 | 0 | 0 | 0 |
| 13963 | ENSMUSG00000047356 | 0 | NA | NA | NA | NA | NA | Lcn10 | lipocalin 10 [Source:MGI Symbol;Acc:MGI | 0 | 0 | 0 | 0 | 0 | 0 |
| 13964 | ENSMUSG00000047361 | 0.167707431 | -0.796629176 | 4.080472857 | -0.195229623 | 0.84521319 | NA | Gm973 | predicted gene 973 [Source:MGI Symbol;A | 0 | 0 | 1.0062 | 0 | 0 | 0 |
| 13965 | ENSMUSG00000047363 | 0.17131964 | -0.796629176 | 4.080472857 | -0.195229623 | 0.84521319 | NA | Cstad | CSA-conditional, T cell activation-depende | 0 | 1.0279 | 0 | 0 | 0 | 0 |
| 13966 | ENSMUSG00000047368 | 451.1001287 | 0.905496207 | 0.214650853 | 4.218460793 | 2.46E-05 | 0.001143985 | Abhd17b | abhydrolase domain containing 17B [Sour | 355.77 | 318.65 | 267.66 | 826.13 | 446.06 | 492.32 |
| 13967 | ENSMUSG00000047369 | 0.790591184 | -0.507346814 | 2.633409124 | -0.192657802 | 0.847226973 | NA | Dnah14 | dynein, axonemal, heavy chain 14 [Source | 0.8031 | 0 | 2.0125 | 1.928 | 0 | 0 |
| 13968 | ENSMUSG00000047370 | 2.244308791 | 2.75245528 | 1.717508418 | 1.602586195 | 0.109026044 | NA | Gm7367 | predicted pseudogene 7367 [Source:MGI | 1.6062 | 0 | 0 | 1.928 | 4.3625 | 5.5692 |
| 13969 | ENSMUSG00000047371 | 384.1791515 | 0.193607304 | 0.172374333 | 1.123179428 | 0.261361298 | 0.71553796 | Zfp768 | zinc finger protein 768 [Source:MGI Symb | 332.48 | 338.18 | 404.51 | 356.67 | 367.54 | 505.69 |
| 13970 | ENSMUSG00000047379 | 410.1583284 | -0.028310287 | 0.14478156 | -0.195537931 | 0.844971846 | 0.972803041 | B4gat1 | beta-1,4-glucuronyltransferase 1 [Source: | 424.84 | 357.72 | 459.85 | 418.37 | 398.08 | 402.1 |
| 13971 | ENSMUSG00000047383 | 0.622883753 | 0.113321311 | 2.884614835 | 0.039284729 | 0.968663382 | NA | | novel transcript | 0.8031 | 0 | 1.0062 | 1.928 | 0 | 0 |
| 13972 | ENSMUSG00000047384 | 7.482648414 | 2.101785291 | 0.914931261 | 2.297205682 | 0.021607039 | 0.182976706 | A730013G03 | RIKEN cDNA A730013G03 gene [Source:M | 2.4093 | 2.0558 | 4.025 | 8.6758 | 5.4531 | 22.277 |
| 13973 | ENSMUSG00000047388 | 193.5032722 | 0.113222049 | 0.187300719 | 0.604493403 | 0.545515654 | 0.89549457 | Atmin | ATM interactor [Source:MGI Symbol;Acc:I | 187.93 | 213.81 | 155.97 | 198.58 | 189.77 | 214.97 |
| 13974 | ENSMUSG00000047390 | 0 | NA | NA | NA | NA | NA | Defb9 | defensin beta 9 [Source:MGI Symbol;Acc: | 0 | 0 | 0 | 0 | 0 | 0 |
| 13975 | ENSMUSG00000047394 | 220.306565 | -0.487566434 | 0.185985664 | -2.621526977 | 0.008753683 | 0.098272117 | Odf3b | outer dense fiber of sperm tails 3B [Sourc | 244.14 | 230.25 | 296.84 | 156.16 | 202.86 | 191.58 |
| 13976 | ENSMUSG00000047407 | 210.9504498 | -0.138070159 | 0.242081435 | -0.570345921 | 0.568443101 | 0.903527882 | Tgif1 | TGFB-induced factor homeobox 1 [Source | 263.42 | 194.28 | 205.27 | 230.39 | 246.48 | 125.86 |
| 13977 | ENSMUSG00000047409 | 61.89265046 | 0.056428715 | 0.294389895 | 0.191680202 | 0.847992715 | 0.972803041 | Ctdspl | CTD (carboxy-terminal domain, RNA polyr | 69.067 | 54.48 | 58.362 | 71.334 | 50.168 | 67.944 |
| 13978 | ENSMUSG00000047412 | 1125.647154 | 0.106450503 | 0.112706491 | 0.944493099 | 0.344917727 | 0.786033695 | Zbtb44 | zinc finger and BTB domain containing 44 | 1032.8 | 1146.1 | 1074.7 | 1243.5 | 1197.5 | 1059.3 |
| 13979 | ENSMUSG00000047414 | 10.72463584 | 0.219325972 | 0.685335436 | 0.320027187 | 0.748947721 | 0.951074124 | Flrt2 | fibronectin leucine rich transmembrane p | 15.259 | 9.2513 | 5.0312 | 10.604 | 14.178 | 10.025 |
| 13980 | ENSMUSG00000047415 | 5.296191722 | -0.422933022 | 1.182481092 | -0.35766578 | 0.720593445 | 0.9447719 | Gpr68 | G protein-coupled receptor 68 [Source:M | 12.047 | 1.0279 | 5.0312 | 3.8559 | 9.8156 | 0 |
| 13981 | ENSMUSG00000047417 | 553.5907648 | 0.157720624 | 0.127552689 | 1.23651351 | 0.216267747 | 0.661699385 | Rexo1 | REX1, RNA exonuclease 1 [Source:MGI Sy | 473.03 | 543.77 | 554.44 | 607.31 | 566.03 | 576.97 |
| 13982 | ENSMUSG00000047419 | 0.804860058 | -1.761048096 | 2.420315956 | -0.727610828 | 0.46685185 | NA | Cmya5 | cardiomyopathy associated 5 [Source:MG | 0.8031 | 2.0558 | 1.0062 | 0.964 | 0 | 0 |
| 13983 | ENSMUSG00000047420 | 20.20639889 | -1.10130581 | 0.492533934 | -2.235999864 | 0.02535178 | 0.204853713 | Fam180a | family with sequence similarity 180, meml | 28.912 | 26.726 | 27.169 | 16.388 | 10.906 | 11.138 |
| 13984 | ENSMUSG00000047423 | 328.5587955 | -0.044639721 | 0.148853539 | -0.299890225 | 0.76426089 | 0.954980327 | AI837181 | expressed sequence AI837181 [Source:M | 346.14 | 313.51 | 341.12 | 337.39 | 289.01 | 344.18 |
| 13985 | ENSMUSG00000047428 | 0.513958919 | -2.394274821 | 3.756096013 | -0.637437066 | 0.523840191 | NA | Dlk2 | delta like non-canonical Notch ligand 2 [S | 0 | 3.0838 | 0 | 0 | 0 | 0 |
| 13986 | ENSMUSG00000047434 | 41.64966787 | -0.540280412 | 0.350182206 | -1.542855127 | 0.12286593 | 0.513267141 | Xxylt1 | xyloside xylosyltransferase 1 [Source:MGI | 55.414 | 46.256 | 46.287 | 35.667 | 26.175 | 40.098 |
| 13987 | ENSMUSG00000047441 | 0 | NA | NA | NA | NA | NA | Antxrl | anthrax toxin receptor-like [Source:MGI S | 0 | 0 | 0 | 0 | 0 | 0 |
| 13988 | ENSMUSG00000047443 | 0.649696354 | 1.537929172 | 3.118369374 | 0.493183773 | 0.621882743 | NA | Erfe | erythroferrone [Source:MGI Symbol;Acc:I | 0 | 0 | 1.0062 | 2.8919 | 0 | 0 |
| 13989 | ENSMUSG00000047444 | 0 | NA | NA | NA | NA | NA | Olfr139 | olfactory receptor 139 [Source:MGI Symb | 0 | 0 | 0 | 0 | 0 | 0 |
| 13990 | ENSMUSG00000047446 | 777.3661069 | -0.685462001 | 0.136715925 | -5.0137685 | 5.34E-07 | 4.68E-05 | Arl4a | ADP-ribosylation factor-like 4A [Source:M | 987.82 | 936.43 | 951.91 | 621.77 | 676.18 | 490.09 |
| 13991 | ENSMUSG00000047454 | 1394.287413 | 0.037465141 | 0.122657766 | 0.305444511 | 0.760027626 | 0.95402891 | Gphn | gephyrin [Source:MGI Symbol;Acc:MGI:1( | 1206.3 | 1420.6 | 1502.3 | 1273.4 | 1501.8 | 1461.4 |
| 13992 | ENSMUSG00000047459 | 1061.608475 | 0.060596109 | 0.128641603 | 0.471045972 | 0.637607906 | 0.923478764 | Dynlrb1 | dynein light chain roadblock-type 1 [Sourc | 1125.1 | 998.11 | 993.16 | 953.37 | 1056.8 | 1243 |
| 13993 | ENSMUSG00000047462 | 1.138517101 | -3.568011124 | 2.494578756 | -1.430306065 | 0.152629195 | NA | Gpr141b | G protein-coupled receptor 141B [Source: | 4.8186 | 0 | 2.0125 | 0 | 0 | 0 |
| 13994 | ENSMUSG00000047466 | 107.8111595 | 0.115794274 | 0.243489338 | 0.475561991 | 0.63438646 | 0.923478764 | 8030462N17 | RIKEN cDNA 8030462N17 gene [Source:M | 88.341 | 117.18 | 105.66 | 132.06 | 114.51 | 89.108 |
| 13995 | ENSMUSG00000047473 | 38.28373565 | 0.566068556 | 0.389624573 | 1.452856403 | 0.146263628 | 0.556855766 | Zfp30 | zinc finger protein 30 [Source:MGI Symbo | 21.684 | 40.089 | 31.194 | 47.235 | 33.809 | 55.692 |
| 13996 | ENSMUSG00000047485 | 0.185640647 | 1.126924175 | 4.080472857 | 0.276174898 | 0.782413736 | NA | Klhl34 | kelch-like 34 [Source:MGI Symbol;Acc:MG | 0 | 0 | 0 | 0 | 0 | 1.1138 |

| | | | | | | | | | | | | | | | |
|---|---|---|---|---|---|---|---|---|---|---|---|---|---|---|---|
| 13997 | ENSMUSG00000047492 | 1508.096081 | -0.103200907 | 0.17774488 | -0.580612544 | 0.561501616 | 0.900553224 | Inhbe | inhibin beta-E [Source:MGI Symbol;Acc:M | 1990.9 | 1174.9 | 1518.4 | 1199.2 | 1691.5 | 1473.6 |
| 13998 | ENSMUSG00000047495 | 0 | NA | NA | NA | NA | NA | Dlgap2 | DLG associated protein 2 [Source:MGI Syr | 0 | 0 | 0 | 0 | 0 | 0 |
| 13999 | ENSMUSG00000047496 | 782.5041159 | 0.681199055 | 0.158459288 | 4.298890033 | 1.72E-05 | 0.000833163 | Rnf152 | ring finger protein 152 [Source:MGI Symb | 651.32 | 471.81 | 679.22 | 887.82 | 892.13 | 1112.7 |
| 14000 | ENSMUSG00000047497 | 4.629373139 | -2.1559806 | 1.210517805 | -1.781039973 | 0.074905921 | NA | Adamts12 | a disintegrin-like and metallopeptidase (re | 6.4248 | 13.363 | 3.0187 | 3.8559 | 0 | 1.1138 |
| 14001 | ENSMUSG00000047501 | 0.638982041 | 0.404325377 | 3.14176733 | 0.128693609 | 0.897600092 | NA | Cldn4 | claudin 4 [Source:MGI Symbol;Acc:MGI:1: | 1.6062 | 0 | 0 | 0 | 0 | 2.2277 |
| 14002 | ENSMUSG00000047502 | 4.954312712 | -0.976706776 | 1.113445984 | -0.877192778 | 0.380381938 | NA | Mroh7 | maestro heat-like repeat family member 7 | 9.6372 | 4.1117 | 6.0375 | 7.7118 | 0 | 2.2277 |
| 14003 | ENSMUSG00000047507 | 2.261270859 | -1.300863991 | 1.661840928 | -0.782784904 | 0.433753434 | NA | Baiap3 | BAI1-associated protein 3 [Source:MGI Sy | 5.6217 | 3.0838 | 1.0062 | 3.8559 | 0 | 0 |
| 14004 | ENSMUSG00000047508 | 0 | NA | NA | NA | NA | NA | Mbd3l2 | methyl-CpG binding domain protein 3-like | 0 | 0 | 0 | 0 | 0 | 0 |
| 14005 | ENSMUSG00000047509 | 0 | NA | NA | NA | NA | NA | Gm6776 | predicted pseudogene 6776 [Source:MGI | 0 | 0 | 0 | 0 | 0 | 0 |
| 14006 | ENSMUSG00000047511 | 3.669324402 | -0.734732753 | 1.25442314 | -0.585713648 | 0.558067961 | NA | Olfr1396 | olfactory receptor 1396 [Source:MGI Syml | 1.6062 | 8.2233 | 4.025 | 4.8199 | 0 | 3.3415 |
| 14007 | ENSMUSG00000047514 | 451.3706325 | 0.150674097 | 0.132270179 | 1.139138831 | 0.254645254 | 0.708091239 | Tspyl1 | testis-specific protein, Y-encoded-like 1 [S | 444.12 | 415.28 | 423.63 | 473.31 | 439.52 | 512.37 |
| 14008 | ENSMUSG00000047515 | 6.260331721 | 0.421726167 | 0.967749033 | 0.43578051 | 0.662995986 | 0.92906822 | BC049715 | cDNA sequence BC049715 [Source:MGI Sy | 4.0155 | 3.0838 | 9.0562 | 11.568 | 8.7249 | 1.1138 |
| 14009 | ENSMUSG00000047517 | 0 | NA | NA | NA | NA | NA | Dmbt1 | deleted in malignant brain tumors 1 [Sour | 0 | 0 | 0 | 0 | 0 | 0 |
| 14010 | ENSMUSG00000047518 | 0 | NA | NA | NA | NA | NA | Slfnl1 | schlafen like 1 [Source:MGI Symbol;Acc:M | 0 | 0 | 0 | 0 | 0 | 0 |
| 14011 | ENSMUSG00000047528 | 12.24228239 | -1.095214436 | 0.705067383 | -1.553347187 | 0.120340213 | 0.507960396 | Flacc1 | flagellum associated containing coiled-coi | 22.487 | 9.2513 | 18.112 | 3.8559 | 14.178 | 5.5692 |
| 14012 | ENSMUSG00000047531 | 0 | NA | NA | NA | NA | NA | Rtp2 | receptor transporter protein 2 [Source:MG | 0 | 0 | 0 | 0 | 0 | 0 |
| 14013 | ENSMUSG00000047534 | 5.185334492 | 1.374573322 | 0.978623534 | 1.404598678 | 0.16014065 | 0.582563974 | Mis18bp1 | MIS18 binding protein 1 [Source:MGI Sym | 1.6062 | 4.1117 | 3.0187 | 5.7839 | 5.4531 | 11.138 |
| 14014 | ENSMUSG00000047535 | 0 | NA | NA | NA | NA | NA | Olfr67 | olfactory receptor 67 [Source:MGI Symbo | 0 | 0 | 0 | 0 | 0 | 0 |
| 14015 | ENSMUSG00000047539 | 197.4828356 | -0.13615585 | 0.174163876 | -0.7817686 | 0.434350578 | 0.845264432 | Fbxo28 | F-box protein 28 [Source:MGI Symbol;Acc | 191.94 | 208.67 | 220.37 | 202.44 | 181.04 | 180.44 |
| 14016 | ENSMUSG00000047544 | 0 | NA | NA | NA | NA | NA | Olfr616 | olfactory receptor 616 [Source:MGI Symb | 0 | 0 | 0 | 0 | 0 | 0 |
| 14017 | ENSMUSG00000047545 | 0 | NA | NA | NA | NA | NA | Olfr629 | olfactory receptor 629 [Source:MGI Symb | 0 | 0 | 0 | 0 | 0 | 0 |
| 14018 | ENSMUSG00000047547 | 996.6301161 | 0.051894733 | 0.121812943 | 0.426019866 | 0.67009337 | 0.931050001 | Cltb | clathrin, light polypeptide (Lcb) [Source:M | 1012.7 | 892.23 | 1031.4 | 1124 | 928.12 | 991.32 |
| 14019 | ENSMUSG00000047554 | 327.3762464 | 0.145665376 | 0.162194877 | 0.898088634 | 0.369138298 | 0.80653433 | Tmem41b | transmembrane protein 41B [Source:MGI | 353.36 | 300.15 | 278.73 | 387.52 | 333.73 | 310.76 |
| 14020 | ENSMUSG00000047557 | 11.72760837 | -0.208858874 | 0.651674286 | -0.320495803 | 0.748592512 | 0.951074124 | Lxn | latexin [Source:MGI Symbol;Acc:MGI:107( | 11.243 | 17.475 | 9.0562 | 11.568 | 6.5437 | 14.48 |
| 14021 | ENSMUSG00000047562 | 0 | NA | NA | NA | NA | NA | Mmp10 | matrix metallopeptidase 10 [Source:MGI S | 0 | 0 | 0 | 0 | 0 | 0 |
| 14022 | ENSMUSG00000047564 | 0 | NA | NA | NA | NA | NA | Krtap3-1 | keratin associated protein 3-1 [Source:MG | 0 | 0 | 0 | 0 | 0 | 0 |
| 14023 | ENSMUSG00000047565 | 0 | NA | NA | NA | NA | NA | Acot10 | acyl-CoA thioesterase 10 [Source:MGI Syn | 0 | 0 | 0 | 0 | 0 | 0 |
| 14024 | ENSMUSG00000047583 | 26.69104656 | -0.583522906 | 0.449831732 | -1.297202632 | 0.194561475 | 0.633957463 | Tyw3 | tRNA-yW synthesizing protein 3 homolog | 40.155 | 28.782 | 27.169 | 29.883 | 17.45 | 16.708 |
| 14025 | ENSMUSG00000047586 | 0.133850373 | -0.796629176 | 4.080472857 | -0.195229623 | 0.84521319 | NA | Nccrp1 | non-specific cytotoxic cell receptor protei | 0.8031 | 0 | 0 | 0 | 0 | 0 |
| 14026 | ENSMUSG00000047591 | 0.636972666 | -2.712703855 | 3.077964703 | -0.881330397 | 0.378139018 | NA | Mafa | v-maf musculoaponeurotic fibrosarcoma | 0.8031 | 0 | 3.0187 | 0 | 0 | 0 |
| 14027 | ENSMUSG00000047592 | 0.531246861 | 1.101376188 | 3.36099124 | 0.327693858 | 0.743143147 | NA | Nxpe5 | neurexophilin and PC-esterase domain fa | 0 | 0 | 1.0062 | 0 | 2.1812 | 0 |
| 14028 | ENSMUSG00000047594 | 0 | NA | NA | NA | NA | NA | Olfr1122 | olfactory receptor 1122 [Source:MGI Syml | 0 | 0 | 0 | 0 | 0 | 0 |
| 14029 | ENSMUSG00000047603 | 107.1685317 | -0.024039773 | 0.242491198 | -0.099136683 | 0.921029746 | 0.986685804 | Zfp235 | zinc finger protein 235 [Source:MGI Symb | 91.554 | 126.43 | 106.66 | 106.04 | 124.33 | 87.994 |
| 14030 | ENSMUSG00000047604 | 53.14950887 | -0.26845453 | 0.340282927 | -0.788915658 | 0.430161303 | 0.84347751 | Frat2 | frequently rearranged in advanced T cell l | 42.564 | 53.452 | 78.487 | 47.235 | 40.353 | 56.806 |
| 14031 | ENSMUSG00000047606 | 0 | NA | NA | NA | NA | NA | Ankrd34c | ankyrin repeat domain 34C [Source:MGI S | 0 | 0 | 0 | 0 | 0 | 0 |
| 14032 | ENSMUSG00000047613 | 634.0858042 | -0.086134206 | 0.115694065 | -0.744499781 | 0.456574174 | 0.858190757 | A430005L14 | RIKEN cDNA A430005L14 gene [Source:M | 659.35 | 661.98 | 637.96 | 643.94 | 600.93 | 600.36 |
| 14033 | ENSMUSG00000047617 | 221.9765253 | 0.171919474 | 0.220036224 | 0.781323504 | 0.43461225 | 0.845264432 | Paxx | non-homologous end joining factor [Sourc | 197.56 | 148.02 | 280.74 | 230.39 | 231.21 | 243.93 |
| 14034 | ENSMUSG00000047619 | 0 | NA | NA | NA | NA | NA | Ddi1 | DNA-damage inducible 1 [Source:MGI Syn | 0 | 0 | 0 | 0 | 0 | 0 |
| 14035 | ENSMUSG00000047626 | 0 | NA | NA | NA | NA | NA | Olfr791 | olfactory receptor 791 [Source:MGI Symb | 0 | 0 | 0 | 0 | 0 | 0 |
| 14036 | ENSMUSG00000047631 | 12793.95708 | 0.129664879 | 0.092763138 | 1.39780608 | 0.162171308 | 0.585529252 | Apof | apolipoprotein F [Source:MGI Symbol;Acc | 13084 | 11542 | 12030 | 12655 | 13416 | 14037 |
| 14037 | ENSMUSG00000047632 | 37.26235752 | -0.577297178 | 0.394111647 | -1.464806185 | 0.142973804 | 0.552624717 | Fgfbp3 | fibroblast growth factor binding protein 3 | 54.611 | 31.865 | 47.293 | 35.667 | 20.722 | 33.415 |
| 14038 | ENSMUSG00000047635 | 189.7536662 | 0.028122031 | 0.187200248 | 0.150224325 | 0.880587635 | 0.981448736 | 2810006K23 | RIKEN cDNA 2810006K23 gene [Source:M | 167.05 | 207.64 | 189.17 | 166.77 | 198.49 | 209.4 |
| 14039 | ENSMUSG00000047636 | 0 | NA | NA | NA | NA | NA | Cdcp2 | CUB domain containing protein 2 [Source: | 0 | 0 | 0 | 0 | 0 | 0 |
| 14040 | ENSMUSG00000047638 | 3262.067752 | 0.054354772 | 0.126362542 | 0.430149404 | 0.667086964 | 0.929709374 | Nr1h4 | nuclear receptor subfamily 1, group H, me | 3369 | 3293.4 | 2939.2 | 3393.2 | 3806.3 | 2771.2 |
| 14041 | ENSMUSG00000047641 | 0 | NA | NA | NA | NA | NA | Krt87 | keratin 87 [Source:MGI Symbol;Acc:MGI:3 | 0 | 0 | 0 | 0 | 0 | 0 |
| 14042 | ENSMUSG00000047642 | 0 | NA | NA | NA | NA | NA | D930020B18 | RIKEN cDNA D930020B18 gene [Source:M | 0 | 0 | 0 | 0 | 0 | 0 |
| 14043 | ENSMUSG00000047643 | 0 | NA | NA | NA | NA | NA | Gm5454 | predicted gene 5454 [Source:MGI Symbol | 0 | 0 | 0 | 0 | 0 | 0 |
| 14044 | ENSMUSG00000047648 | 188.9254903 | -0.034319484 | 0.186831796 | -0.183691882 | 0.854255178 | 0.973825032 | Fbxo30 | F-box protein 30 [Source:MGI Symbol;Acc | 184.71 | 184 | 205.27 | 212.08 | 158.14 | 189.35 |
| 14045 | ENSMUSG00000047649 | 114.6201193 | 0.097333261 | 0.221473466 | 0.439480461 | 0.660313437 | 0.927980329 | Cd3eap | CD3E antigen, epsilon polypeptide associa | 112.43 | 124.38 | 95.593 | 128.21 | 110.15 | 116.95 |
| 14046 | ENSMUSG00000047652 | 0 | NA | NA | NA | NA | NA | Pea15b-ps | phosphoprotein enriched in astrocytes 15 | 0 | 0 | 0 | 0 | 0 | 0 |
| 14047 | ENSMUSG00000047654 | 5.024716104 | 1.024809389 | 1.359013746 | 0.754083166 | 0.450799285 | NA | Tssk6 | testis-specific serine kinase 6 [Source:MG | 0 | 0 | 10.062 | 13.496 | 4.3625 | 2.2277 |
| 14048 | ENSMUSG00000047655 | 0 | NA | NA | NA | NA | NA | Vmn1r74 | vomeronasal 1 receptor 74 [Source:MGI S | 0 | 0 | 0 | 0 | 0 | 0 |
| 14049 | ENSMUSG00000047656 | 66.19893909 | 0.404684285 | 0.273163912 | 1.481470527 | 0.138481233 | 0.543828246 | Trpt1 | tRNA phosphotransferase 1 [Source:MGI | 61.839 | 56.535 | 52.325 | 72.298 | 81.796 | 72.4 |
| 14050 | ENSMUSG00000047657 | 0.181769715 | 1.126924175 | 4.080472857 | 0.276174898 | 0.782413736 | NA | Crry-ps | complement receptor related protein, pse | 0 | 0 | 0 | 0 | 1.0906 | 0 |
| 14051 | ENSMUSG00000047658 | 0 | NA | NA | NA | NA | NA | Gal3st3 | galactose-3-O-sulfotransferase 3 [Source: | 0 | 0 | 0 | 0 | 0 | 0 |
| 14052 | ENSMUSG00000047661 | 0 | NA | NA | NA | NA | NA | 1600012P17 | RIKEN cDNA 1600012P17 gene [Source:M | 0 | 0 | 0 | 0 | 0 | 0 |

| | | | | | | | | | | | | | | | |
|---|---|---|---|---|---|---|---|---|---|---|---|---|---|---|---|
| 14053 | ENSMUSG00000047667 | 0 | NA | NA | NA | NA | NA | Olfr26 | olfactory receptor 26 [Source:MGI Symbo | 0 | 0 | 0 | 0 | 0 | 0 |
| 14054 | ENSMUSG00000047669 | 19.22580206 | 0.362155485 | 0.54965189 | 0.658881541 | 0.50997184 | 0.882852507 | Msl3l2 | MSL3 like 2 [Source:MGI Symbol;Acc:MGI | 25.699 | 10.279 | 14.087 | 13.496 | 26.175 | 25.618 |
| 14055 | ENSMUSG00000047671 | 1.495431296 | 1.005864465 | 1.961955563 | 0.512684631 | 0.608171942 | NA | Dynlt4 | dynein light chain Tctex-type 4 [Source:M | 0 | 2.0558 | 1.0062 | 4.8199 | 1.0906 | 0 |
| 14056 | ENSMUSG00000047674 | 0 | NA | NA | NA | NA | NA | Pdha2 | pyruvate dehydrogenase E1 alpha 2 [Sour | 0 | 0 | 0 | 0 | 0 | 0 |
| 14057 | ENSMUSG00000047675 | 3211.023685 | 0.034159133 | 0.097024723 | 0.352066273 | 0.724788561 | 0.9448775 | Rps8 | ribosomal protein S8 [Source:MGI Symbol | 2942.6 | 3414.7 | 3162.6 | 3151.2 | 3247.9 | 3347.1 |
| 14058 | ENSMUSG00000047676 | 16.68408136 | -0.223320423 | 0.526393761 | -0.424245953 | 0.671386451 | 0.931191575 | Rpsa-ps10 | ribosomal protein SA, pseudogene 10 [So | 19.274 | 19.53 | 15.094 | 16.388 | 11.997 | 17.822 |
| 14059 | ENSMUSG00000047678 | 0 | NA | NA | NA | NA | NA | Gpr82 | G protein-coupled receptor 82 [Source:M | 0 | 0 | 0 | 0 | 0 | 0 |
| 14060 | ENSMUSG00000047686 | 0.670829724 | -2.782124174 | 3.401449285 | -0.817923168 | 0.41340106 | NA | Rtl3 | retrotransposon Gag like 3 [Source:MGI S | 0 | 0 | 4.025 | 0 | 0 | 0 |
| 14061 | ENSMUSG00000047692 | 0 | NA | NA | NA | NA | NA | 4930533K18 | RIKEN cDNA 4930533K18 gene [Source:M | 0 | 0 | 0 | 0 | 0 | 0 |
| 14062 | ENSMUSG00000047694 | 635.3175817 | -0.09841921 | 0.130295737 | -0.755352494 | 0.450037509 | 0.855479371 | Yipf6 | Yip1 domain family, member 6 [Source:M | 641.68 | 691.79 | 637.96 | 628.51 | 675.09 | 536.87 |
| 14063 | ENSMUSG00000047696 | 0 | NA | NA | NA | NA | NA | Ccdc144b | coiled-coil domain containing 144B [Sourc | 0 | 0 | 0 | 0 | 0 | 0 |
| 14064 | ENSMUSG00000047702 | 0 | NA | NA | NA | NA | NA | Olfr1388 | olfactory receptor 1388 [Source:MGI Syml | 0 | 0 | 0 | 0 | 0 | 0 |
| 14065 | ENSMUSG00000047710 | 125.8871506 | 0.273446328 | 0.21698297 | 1.260220229 | 0.207589928 | 0.652593069 | Champ1 | chromosome alignment maintaining phos | 126.09 | 94.568 | 120.75 | 127.25 | 137.42 | 149.26 |
| 14066 | ENSMUSG00000047712 | 8.791624395 | -2.017581354 | 0.809321743 | -2.492928643 | 0.01266943 | 0.127345825 | Ust | uronyl-2-sulfotransferase [Source:MGI Sy | 8.8341 | 18.503 | 15.094 | 4.8199 | 3.2719 | 2.2277 |
| 14067 | ENSMUSG00000047714 | 721.1693949 | -0.165337631 | 0.134884849 | -1.225768741 | 0.220285714 | 0.666571666 | Ppp1r2 | protein phosphatase 1, regulatory inhibito | 685.85 | 888.12 | 714.43 | 705.63 | 641.28 | 691.7 |
| 14068 | ENSMUSG00000047716 | 0 | NA | NA | NA | NA | NA | Olfr736 | olfactory receptor 736 [Source:MGI Symb | 0 | 0 | 0 | 0 | 0 | 0 |
| 14069 | ENSMUSG00000047719 | 202.6400631 | 0.046281582 | 0.197367774 | 0.234494117 | 0.814601399 | 0.967404825 | Ubiad1 | UbiA prenyltransferase domain containing | 218.44 | 166.52 | 212.32 | 167.73 | 222.49 | 228.34 |
| 14070 | ENSMUSG00000047720 | 0 | NA | NA | NA | NA | NA | Clec2m | C-type lectin domain family 2, member m | 0 | 0 | 0 | 0 | 0 | 0 |
| 14071 | ENSMUSG00000047721 | 376.9937898 | 0.039418403 | 0.152193958 | 0.259001101 | 0.795634391 | 0.962236316 | Bola2 | bolA-like 2 (E. coli) [Source:MGI Symbol;A | 376.65 | 322.77 | 415.58 | 348 | 402.44 | 396.53 |
| 14072 | ENSMUSG00000047728 | 3629.838378 | 0.120588955 | 0.112741447 | 1.069606235 | 0.284796588 | 0.737432482 | Ly6g2 | lymphocyte antigen 6 complex, locus G2 [ | 3863.7 | 3069.4 | 3500.7 | 4059.3 | 3753.9 | 3532 |
| 14073 | ENSMUSG00000047730 | 5.973889142 | -0.013902084 | 0.858408588 | -0.016195183 | 0.987078678 | 0.99859176 | Fcgbp | Fc fragment of IgG binding protein [Sourc | 4.8186 | 8.2233 | 5.0312 | 6.7478 | 5.4531 | 5.5692 |
| 14074 | ENSMUSG00000047731 | 6282.591384 | -0.234625569 | 0.118964312 | -1.972234905 | 0.048582797 | 0.306856582 | Wbp1l | WW domain binding protein 1 like [Sourc | 6132.5 | 7234.5 | 7011.5 | 6283.2 | 4911.1 | 6122.8 |
| 14075 | ENSMUSG00000047733 | 1.329604885 | -1.614769125 | 2.056107202 | -0.785352594 | 0.432246867 | NA | Tmem262 | transmembrane protein 262 [Source:MGI | 4.0155 | 1.0279 | 1.0062 | 1.928 | 0 | 0 |
| 14076 | ENSMUSG00000047735 | 404.1983798 | -0.317750987 | 0.167730307 | -1.894416057 | 0.058169807 | 0.342716124 | Samd9l | sterile alpha motif domain containing 9-lik | 517.2 | 384.44 | 442.75 | 312.33 | 422.07 | 346.41 |
| 14077 | ENSMUSG00000047746 | 19.83678237 | 0.591076063 | 0.56988171 | 1.037190793 | 0.299646948 | 0.750882442 | Fbxo40 | F-box protein 40 [Source:MGI Symbol;Acc | 25.699 | 9.2513 | 12.075 | 13.496 | 25.084 | 33.415 |
| 14078 | ENSMUSG00000047747 | 14.87397714 | -0.696497841 | 0.622686752 | -1.118536469 | 0.26333794 | 0.717851482 | Rnf150 | ring finger protein 150 [Source:MGI Symb | 13.653 | 25.698 | 16.1 | 18.316 | 5.4531 | 10.025 |
| 14079 | ENSMUSG00000047749 | 3.916579585 | -0.263079117 | 1.071800155 | -0.245455383 | 0.806103851 | NA | Zc3hav1l | zinc finger CCCH-type, antiviral 1-like [Sou | 5.6217 | 3.0838 | 4.025 | 1.928 | 3.2719 | 5.5692 |
| 14080 | ENSMUSG00000047751 | 0 | NA | NA | NA | NA | NA | Utf1 | undifferentiated embryonic cell transcript | 0 | 0 | 0 | 0 | 0 | 0 |
| 14081 | ENSMUSG00000047757 | 12.13508672 | 0.532484624 | 0.640503988 | 0.83135255 | 0.405774496 | 0.829973745 | Fancb | Fanconi anemia, complementation group | 15.259 | 8.2233 | 6.0375 | 13.496 | 13.087 | 16.708 |
| 14082 | ENSMUSG00000047759 | 9.2984815 | -0.748314197 | 0.705603927 | -1.060530091 | 0.288903508 | 0.739603195 | Hs3st3a1 | heparan sulfate (glucosamine) 3-O-sulfotr | 13.653 | 9.2513 | 12.075 | 8.6758 | 5.4531 | 6.6831 |
| 14083 | ENSMUSG00000047766 | 2.655752547 | -0.332122122 | 1.395554814 | -0.237985723 | 0.811892167 | NA | Lrrc49 | leucine rich repeat containing 49 [Source: | 4.8186 | 2.0558 | 2.0125 | 4.8199 | 0 | 2.2277 |
| 14084 | ENSMUSG00000047767 | 160.7270374 | 0.201473746 | 0.206477849 | 0.975764456 | 0.329181203 | 0.774065504 | Atg16l2 | autophagy related 16-like 2 (S. cerevisiae) | 139.74 | 149.05 | 159.99 | 175.44 | 137.42 | 202.72 |
| 14085 | ENSMUSG00000047773 | 0 | NA | NA | NA | NA | NA | Ankfn1 | ankyrin-repeat and fibronectin type III do | 0 | 0 | 0 | 0 | 0 | 0 |
| 14086 | ENSMUSG00000047777 | 116.171587 | 0.211129167 | 0.229004494 | 0.921943335 | 0.356558134 | 0.795995809 | Phf13 | PHD finger protein 13 [Source:MGI Symbc | 105.21 | 128.49 | 89.556 | 131.1 | 111.24 | 131.43 |
| 14087 | ENSMUSG00000047786 | 2.187103524 | -0.046751494 | 1.430004783 | -0.032693243 | 0.973919212 | NA | Lix1 | limb and CNS expressed 1 [Source:MGI Sy | 1.6062 | 1.0279 | 4.025 | 0.964 | 3.2719 | 2.2277 |
| 14088 | ENSMUSG00000047787 | 5.370722288 | -0.869961435 | 0.934966572 | -0.930473304 | 0.35212608 | 0.792945339 | Flrt1 | fibronectin leucine rich transmembrane p | 9.6372 | 5.1396 | 6.0375 | 4.8199 | 4.3625 | 2.2277 |
| 14089 | ENSMUSG00000047789 | 197.2979405 | 0.01271062 | 0.182271199 | 0.069734658 | 0.944404856 | 0.990077979 | Slc38a9 | solute carrier family 38, member 9 [Sourc | 205.59 | 206.61 | 177.1 | 214.97 | 171.23 | 208.29 |
| 14090 | ENSMUSG00000047793 | 235.0484101 | 0.205055901 | 0.193382955 | 1.060361812 | 0.288980028 | 0.739603195 | Sned1 | sushi, nidogen and EGF-like domains 1 [Sc | 224.87 | 188.11 | 242.5 | 309.44 | 217.03 | 228.34 |
| 14091 | ENSMUSG00000047794 | 0 | NA | NA | NA | NA | NA | Olfr685 | olfactory receptor 685 [Source:MGI Symb | 0 | 0 | 0 | 0 | 0 | 0 |
| 14092 | ENSMUSG00000047797 | 9849.183613 | 0.44563273 | 0.095154782 | 4.683240499 | 2.82E-06 | 0.000189087 | Gjb1 | gap junction protein, beta 1 [Source:MGI | 8080.8 | 7967.4 | 8971.7 | 10600 | 11562 | 11914 |
| 14093 | ENSMUSG00000047798 | 17.13083227 | -2.061125567 | 0.586182345 | -3.516184997 | 0.000437796 | 0.011270406 | Cd300lf | CD300 molecule like family member F [So | 24.093 | 27.754 | 31.194 | 8.6758 | 3.2719 | 7.7969 |
| 14094 | ENSMUSG00000047799 | 0 | NA | NA | NA | NA | NA | Oog4 | oogenesin 4 [Source:MGI Symbol;Acc:MG | 0 | 0 | 0 | 0 | 0 | 0 |
| 14095 | ENSMUSG00000047804 | 225.8960568 | -0.052346999 | 0.182102232 | -0.287459405 | 0.773760581 | 0.958261993 | Akap10 | A kinase (PRKA) anchor protein 10 [Sourc | 226.47 | 273.43 | 190.18 | 217.86 | 224.67 | 222.77 |
| 14096 | ENSMUSG00000047807 | 0.524408994 | -0.837196744 | 3.377823722 | -0.247850928 | 0.804249749 | NA | 9330154K18 | RIKEN cDNA 9330154K18 gene [Source:M | 0 | 2.0558 | 0 | 0 | 1.0906 | 0 |
| 14097 | ENSMUSG00000047810 | 26.50048477 | -0.526646483 | 0.479878712 | -1.097457483 | 0.272441457 | 0.727618887 | Ccdc88b | coiled-coil domain containing 88B [Source | 26.502 | 41.117 | 26.162 | 15.424 | 15.269 | 34.529 |
| 14098 | ENSMUSG00000047819 | 0 | NA | NA | NA | NA | NA | Tigd4 | tigger transposable element derived 4 [So | 0 | 0 | 0 | 0 | 0 | 0 |
| 14099 | ENSMUSG00000047821 | 3.133322071 | -1.383180876 | 1.24859524 | -1.107789644 | 0.267952674 | NA | Trim16 | tripartite motif-containing 16 [Source:MG | 6.4248 | 5.1396 | 2.0125 | 1.928 | 2.1812 | 1.1138 |
| 14100 | ENSMUSG00000047822 | 457.4897796 | -0.233837493 | 0.500930439 | -0.466806317 | 0.640638456 | 0.923932929 | Angptl8 | angiopoietin-like 8 [Source:MGI Symbol;A | 577.43 | 255.95 | 650.03 | 474.28 | 534.4 | 252.84 |
| 14101 | ENSMUSG00000047824 | 284.2231738 | 0.029666927 | 0.165845177 | 0.178883265 | 0.858029364 | 0.975287758 | Pygo2 | pygopus 2 [Source:MGI Symbol;Acc:MGI:1 | 292.33 | 255.95 | 294.83 | 245.81 | 292.29 | 324.13 |
| 14102 | ENSMUSG00000047828 | 0 | NA | NA | NA | NA | NA | A730018C14 | RIKEN cDNA A730018C14 gene [Source:M | 0 | 0 | 0 | 0 | 0 | 0 |
| 14103 | ENSMUSG00000047832 | 63.04512151 | 0.101887154 | 0.300414663 | 0.339155064 | 0.734492919 | 0.948109711 | Cdca4 | cell division cycle associated 4 [Source:M | 52.202 | 68.87 | 61.381 | 49.163 | 69.8 | 76.855 |
| 14104 | ENSMUSG00000047841 | 0.781711799 | -0.350673833 | 2.438876442 | -0.143784994 | 0.885670253 | NA | Fndc11 | fibronectin type III domain containing 11 [ | 1.6062 | 0 | 1.0062 | 0.964 | 0 | 1.1138 |
| 14105 | ENSMUSG00000047842 | 0 | NA | NA | NA | NA | NA | Diras2 | DIRAS family, GTP-binding RAS-like 2 [Sou | 0 | 0 | 0 | 0 | 0 | 0 |
| 14106 | ENSMUSG00000047843 | 2062.155338 | -0.007993908 | 0.102294635 | -0.078145916 | 0.937711983 | 0.988420305 | Bri3 | brain protein I3 [Source:MGI Symbol;Acc: | 2213.3 | 1953 | 2035.6 | 1948.2 | 2034 | 2188.7 |
| 14107 | ENSMUSG00000047844 | 3.723172097 | 0.171760005 | 1.119567171 | 0.153416436 | 0.878069884 | NA | Bex4 | brain expressed X-linked 4 [Source:MGI Sy | 2.4093 | 2.0558 | 6.0375 | 1.928 | 5.4531 | 4.4554 |
| 14108 | ENSMUSG00000047854 | 0.294513347 | 0.165143496 | 4.080472857 | 0.040471657 | 0.967717103 | NA | Stx19 | syntaxin 19 [Source:MGI Symbol;Acc:MGI | 0.8031 | 0 | 0 | 0.964 | 0 | 0 |

| | | | | | | | | | | | | | | | |
|---|---|---|---|---|---|---|---|---|---|---|---|---|---|---|---|
| 14109 | ENSMUSG00000047861 | 0 | NA | NA | NA | NA | NA | Foxi1 | forkhead box I1 [Source:MGI Symbol;Acc: | 0 | 0 | 0 | 0 | 0 | 0 |
| 14110 | ENSMUSG00000047866 | 6925.582567 | 0.050596005 | 0.099255912 | 0.509753057 | 0.610224477 | 0.916376001 | Lonp2 | lon peptidase 2, peroxisomal [Source:MGI | 7093.8 | 6652.7 | 6664.4 | 6320.8 | 7535.1 | 7286.8 |
| 14111 | ENSMUSG00000047867 | 110.3660149 | -0.498752256 | 0.261114847 | -1.910087698 | 0.056121923 | 0.335017536 | Gimap6 | GTPase, IMAP family member 6 [Source:M | 135.72 | 120.27 | 131.82 | 104.11 | 58.893 | 111.38 |
| 14112 | ENSMUSG00000047868 | 0 | NA | NA | NA | NA | NA | Olfr770 | olfactory receptor 770 [Source:MGI Symb | 0 | 0 | 0 | 0 | 0 | 0 |
| 14113 | ENSMUSG00000047875 | 85.61924834 | 0.144296534 | 0.253581954 | 0.569033133 | 0.569333655 | 0.903735951 | Gpr157 | G protein-coupled receptor 157 [Source:N | 83.523 | 70.926 | 89.556 | 92.542 | 101.43 | 75.741 |
| 14114 | ENSMUSG00000047878 | 3.374863188 | -0.216099432 | 1.309743555 | -0.164993697 | 0.868948943 | NA | A4galt | alpha 1,4-galactosyltransferase [Source:M | 0.8031 | 5.1396 | 5.0312 | 4.8199 | 0 | 4.4554 |
| 14115 | ENSMUSG00000047879 | 692.646996 | -0.003545668 | 0.117953636 | -0.030059841 | 0.976019328 | 0.996762237 | Usp14 | ubiquitin specific peptidase 14 [Source:M | 734.04 | 693.84 | 652.05 | 690.21 | 726.35 | 659.4 |
| 14116 | ENSMUSG00000047880 | 1.254330519 | -1.404465216 | 1.938986102 | -0.724329697 | 0.468863349 | NA | Cxcr5 | chemokine (C-X-C motif) receptor 5 [Sour | 2.4093 | 2.0558 | 1.0062 | 0.964 | 1.0906 | 0 |
| 14117 | ENSMUSG00000047881 | 162.4967325 | 0.338434095 | 0.222691318 | 1.51974535 | 0.128574989 | 0.524858587 | Rell1 | RELT-like 1 [Source:MGI Symbol;Acc:MGI: | 130.91 | 141.85 | 157.98 | 177.37 | 137.42 | 229.45 |
| 14118 | ENSMUSG00000047884 | 0.17131964 | -0.796629176 | 4.080472857 | -0.195229623 | 0.84521319 | NA | Klk9 | kallikrein related-peptidase 9 [Source:MG | 0 | 1.0279 | 0 | 0 | 0 | 0 |
| 14119 | ENSMUSG00000047888 | 630.1574272 | 0.068436535 | 0.123575634 | 0.55380282 | 0.579713798 | 0.908350165 | Tnrc6b | trinucleotide repeat containing 6b [Source | 579.04 | 655.81 | 611.8 | 687.32 | 599.84 | 647.14 |
| 14120 | ENSMUSG00000047889 | 0 | NA | NA | NA | NA | NA | Serpinb6d | serine (or cysteine) peptidase inhibitor, cl | 0 | 0 | 0 | 0 | 0 | 0 |
| 14121 | ENSMUSG00000047894 | 0 | NA | NA | NA | NA | NA | Ang2 | angiogenin, ribonuclease A family, membe | 0 | 0 | 0 | 0 | 0 | 0 |
| 14122 | ENSMUSG00000047897 | 0 | NA | NA | NA | NA | NA | Ripply2 | ripply transcriptional repressor 2 [Source: | 0 | 0 | 0 | 0 | 0 | 0 |
| 14123 | ENSMUSG00000047898 | 0.342639279 | -1.805976596 | 4.028547714 | -0.448294702 | 0.653940524 | NA | Ccr4 | chemokine (C-C motif) receptor 4 [Source | 0 | 2.0558 | 0 | 0 | 0 | 0 |
| 14124 | ENSMUSG00000047904 | 12.55040242 | 0.053643294 | 0.786449765 | 0.068209435 | 0.945618916 | 0.990309554 | Sstr2 | somatostatin receptor 2 [Source:MGI Sym | 2.4093 | 24.67 | 10.062 | 11.568 | 6.5437 | 20.049 |
| 14125 | ENSMUSG00000047905 | 5.698261057 | 0.013505312 | 0.90581265 | 0.014909608 | 0.988104294 | 0.99859176 | Gm8566 | predicted pseudogene 8566 [Source:MGI | 4.8186 | 4.1117 | 8.05 | 2.8919 | 7.6343 | 6.6831 |
| 14126 | ENSMUSG00000047907 | 31.10327361 | -0.367142938 | 0.402619295 | -0.911886099 | 0.36182868 | 0.800840096 | Tshz2 | teashirt zinc finger family member 2 [Sour | 37.746 | 37.005 | 30.187 | 23.135 | 22.903 | 35.643 |
| 14127 | ENSMUSG00000047909 | 304.6677706 | 0.239408028 | 0.166479668 | 1.438061661 | 0.150416559 | 0.564307976 | Ankrd16 | ankyrin repeat domain 16 [Source:MGI Sy | 260.21 | 311.46 | 267.66 | 381.74 | 318.46 | 288.49 |
| 14128 | ENSMUSG00000047910 | 2.864386468 | -1.748071285 | 1.493943069 | -1.170105689 | 0.241958439 | NA | Pcdhb16 | protocadherin beta 16 [Source:MGI Symb | 7.2279 | 3.0838 | 3.0187 | 3.8559 | 0 | 0 |
| 14129 | ENSMUSG00000047911 | 0.981621321 | -0.747528389 | 2.423078149 | -0.308503623 | 0.757699144 | NA | Npm2 | nucleophosmin/nucleoplasmin 2 [Source: | 1.6062 | 2.0558 | 0 | 0 | 0 | 2.2277 |
| 14130 | ENSMUSG00000047921 | 221.6130566 | -0.044294953 | 0.16530398 | -0.267960598 | 0.788729648 | 0.960932078 | Trappc9 | trafficking protein particle complex 9 [Sou | 228.88 | 206.61 | 239.49 | 223.64 | 209.4 | 221.65 |
| 14131 | ENSMUSG00000047935 | 0 | NA | NA | NA | NA | NA | Sox1ot | Sox1 overlapping transcript [Source:MGI S | 0 | 0 | 0 | 0 | 0 | 0 |
| 14132 | ENSMUSG00000047938 | 0.990320745 | -2.084832509 | 2.480930004 | -0.84034314 | 0.40071602 | NA | 4930483J18R | RIKEN cDNA 4930483J18 gene [Source:MG | 0.8031 | 0 | 4.025 | 0 | 0 | 1.1138 |
| 14133 | ENSMUSG00000047940 | 1.243823325 | 2.709383194 | 2.333818129 | 1.160923021 | 0.245673206 | NA | Stpg2 | sperm tail PG rich repeat containing 2 [So | 0.8031 | 0 | 0 | 0 | 1.0906 | 5.5692 |
| 14134 | ENSMUSG00000047945 | 109.0731695 | -1.090138395 | 0.242386316 | -4.497524508 | 6.87E-06 | 0.000402587 | Marcksl1 | MARCKS-like 1 [Source:MGI Symbol;Acc:M | 173.47 | 146.99 | 124.77 | 79.046 | 59.984 | 70.172 |
| 14135 | ENSMUSG00000047953 | 6.0029441 | 0.249641121 | 0.970799573 | 0.257150011 | 0.797062972 | 0.962236316 | Gp5 | glycoprotein 5 (platelet) [Source:MGI Sym | 11.243 | 1.0279 | 4.025 | 8.6758 | 4.3625 | 6.6831 |
| 14136 | ENSMUSG00000047959 | 2.285294054 | -2.423458937 | 1.689527462 | -1.434400441 | 0.151458022 | NA | Kcna3 | potassium voltage-gated channel, shaker- | 2.4093 | 3.0838 | 6.0375 | 0 | 2.1812 | 0 |
| 14137 | ENSMUSG00000047960 | 0 | NA | NA | NA | NA | NA | Olfr186 | olfactory receptor 186 [Source:MGI Symb | 0 | 0 | 0 | 0 | 0 | 0 |
| 14138 | ENSMUSG00000047963 | 1570.720575 | 0.430719664 | 0.141491369 | 3.044140894 | 0.002333458 | 0.039199931 | Stbd1 | starch binding domain 1 [Source:MGI Sym | 1296.2 | 1557.3 | 1160.2 | 1556.8 | 1783.2 | 2070.6 |
| 14139 | ENSMUSG00000047965 | 4.419624769 | -0.518403326 | 1.087356507 | -0.47675562 | 0.633536152 | NA | Rpl9-ps7 | ribosomal protein L9, pseudogene 7 [Sour | 2.4093 | 5.1396 | 8.05 | 0.964 | 3.2719 | 6.6831 |
| 14140 | ENSMUSG00000047969 | 0 | NA | NA | NA | NA | NA | Olfr1093 | olfactory receptor 1093 [Source:MGI Symb | 0 | 0 | 0 | 0 | 0 | 0 |
| 14141 | ENSMUSG00000047976 | 0 | NA | NA | NA | NA | NA | Kcna1 | potassium voltage-gated channel, shaker- | 0 | 0 | 0 | 0 | 0 | 0 |
| 14142 | ENSMUSG00000047977 | 0 | NA | NA | NA | NA | NA | Synb | syncytin b [Source:MGI Symbol;Acc:MGI:3 | 0 | 0 | 0 | 0 | 0 | 0 |
| 14143 | ENSMUSG00000047980 | 0 | NA | NA | NA | NA | NA | Pate5 | prostate and testis expressed 5 [Source:M | 0 | 0 | 0 | 0 | 0 | 0 |
| 14144 | ENSMUSG00000047986 | 1.155907894 | -0.174333386 | 1.958174908 | -0.089028506 | 0.929059256 | NA | Palm3 | paralemmin 3 [Source:MGI Symbol;Acc:M | 1.6062 | 1.0279 | 1.0062 | 0 | 2.1812 | 1.1138 |
| 14145 | ENSMUSG00000047988 | 1.401163304 | 0.51802129 | 1.968353158 | 0.263174973 | 0.792415719 | NA | 4933428G20 | RIKEN cDNA 4933428G20 gene [Source:M | 2.4093 | 1.0279 | 0 | 3.8559 | 0 | 1.1138 |
| 14146 | ENSMUSG00000047989 | 231.9896878 | -0.171130968 | 0.184673033 | -0.926670047 | 0.354097874 | 0.79449673 | Ino80c | INO80 complex subunit C [Source:MGI Sy | 294.74 | 232.31 | 209.3 | 214 | 237.75 | 203.83 |
| 14147 | ENSMUSG00000047990 | 0 | NA | NA | NA | NA | NA | C2cd4a | C2 calcium-dependent domain containing | 0 | 0 | 0 | 0 | 0 | 0 |
| 14148 | ENSMUSG00000047992 | 0 | NA | NA | NA | NA | NA | Dipk1c | divergent protein kinase domain 1C [Sour | 0 | 0 | 0 | 0 | 0 | 0 |
| 14149 | ENSMUSG00000047995 | 0 | NA | NA | NA | NA | NA | Cypt4 | cysteine-rich perinuclear theca 4 [Source: | 0 | 0 | 0 | 0 | 0 | 0 |
| 14150 | ENSMUSG00000047996 | 12.65119754 | -0.5867227 | 0.599446857 | -0.978773502 | 0.327691905 | 0.772937193 | Prrg1 | proline rich Gla (G-carboxyglutamic acid) 1 | 16.062 | 16.447 | 13.081 | 11.568 | 8.7249 | 10.025 |
| 14151 | ENSMUSG00000048000 | 651.1999281 | 0.148488172 | 0.128050441 | 1.159606874 | 0.246208901 | 0.696492381 | Gigyf2 | GRB10 interacting GYF protein 2 [Source:M | 605.54 | 632.17 | 615.82 | 693.1 | 757.98 | 602.59 |
| 14152 | ENSMUSG00000048001 | 0 | NA | NA | NA | NA | NA | Hes5 | hes family bHLH transcription factor 5 [So | 0 | 0 | 0 | 0 | 0 | 0 |
| 14153 | ENSMUSG00000048003 | 0.133850373 | -0.796629176 | 4.080472857 | -0.195229623 | 0.84521319 | NA | Catsper4 | cation channel, sperm associated 4 [Sourc | 0.8031 | 0 | 0 | 0 | 0 | 0 |
| 14154 | ENSMUSG00000048004 | 0 | NA | NA | NA | NA | NA | Tmem196 | transmembrane protein 196 [Source:MGI | 0 | 0 | 0 | 0 | 0 | 0 |
| 14155 | ENSMUSG00000048007 | 335.0525175 | -0.19550635 | 0.171436757 | -1.140399252 | 0.254120004 | 0.707252596 | Timm8a1 | translocase of inner mitochondrial membr | 362.2 | 339.21 | 371.3 | 270.88 | 388.26 | 278.46 |
| 14156 | ENSMUSG00000048012 | 2.261090899 | -1.840191451 | 1.673866275 | -1.099365869 | 0.271608512 | NA | Zfp473 | zinc finger protein 473 [Source:MGI Symb | 5.6217 | 1.0279 | 4.025 | 2.8919 | 0 | 0 |
| 14157 | ENSMUSG00000048013 | 0 | NA | NA | NA | NA | NA | Krt35 | keratin 35 [Source:MGI Symbol;Acc:MGI:1 | 0 | 0 | 0 | 0 | 0 | 0 |
| 14158 | ENSMUSG00000048015 | 0 | NA | NA | NA | NA | NA | Neurod4 | neurogenic differentiation 4 [Source:MGI | 0 | 0 | 0 | 0 | 0 | 0 |
| 14159 | ENSMUSG00000048022 | 14.68439997 | -1.03557195 | 0.595396904 | -1.739296834 | 0.081982564 | 0.421541345 | Tmem229a | transmembrane protein 229A [Source:MG | 14.456 | 20.558 | 24.15 | 5.7839 | 10.906 | 12.252 |
| 14160 | ENSMUSG00000048027 | 61.62405003 | 0.1360756 | 0.317337525 | 0.428804 | 0.668065868 | 0.930136128 | Rgmb | repulsive guidance molecule family memb | 64.248 | 55.508 | 56.35 | 74.226 | 41.443 | 77.969 |
| 14161 | ENSMUSG00000048029 | 1.080168213 | 2.425859534 | 2.660049518 | 0.911960292 | 0.361789621 | NA | Eno4 | enolase 4 [Source:MGI Symbol;Acc:MGI:2 | 0 | 1.0279 | 0 | 0 | 5.4531 | 0 |
| 14162 | ENSMUSG00000048031 | 0.167707431 | -0.796629176 | 4.080472857 | -0.195229623 | 0.84521319 | NA | Fcrl5 | Fc receptor-like 5 [Source:MGI Symbol;Ac | 0 | 0 | 1.0062 | 0 | 0 | 0 |
| 14163 | ENSMUSG00000048038 | 0.328370405 | 0.16514316 | 4.003149536 | 0.041253308 | 0.967093956 | NA | Ccdc187 | coiled-coil domain containing 187 [Source | 0 | 0 | 1.0062 | 0.964 | 0 | 0 |
| 14164 | ENSMUSG00000048039 | 269.9970572 | 0.109990208 | 0.165519494 | 0.66451513 | 0.506360668 | 0.881592135 | Isg20l2 | interferon stimulated exonuclease gene 2 | 238.52 | 289.87 | 251.56 | 308.47 | 249.75 | 281.8 |

| | | | | | | | | | | | | | | | |
|---|---|---|---|---|---|---|---|---|---|---|---|---|---|---|---|
| 14165 | ENSMUSG00000048040 | 0.904646783 | 0.28560953 | 2.491718272 | 0.114623524 | 0.908743533 | NA | Arxes2 | adipocyte-related X-chromosome express | 2.4093 | 0 | 0 | 1.928 | 1.0906 | 0 |
| 14166 | ENSMUSG00000048047 | 405.591677 | 0.260823648 | 0.133375419 | 1.955560101 | 0.050516997 | 0.315931181 | Zbtb33 | zinc finger and BTB domain containing 33 | 346.14 | 373.13 | 388.41 | 447.29 | 429.7 | 448.88 |
| 14167 | ENSMUSG00000048058 | 130.8526899 | 0.165781567 | 0.223453683 | 0.741905726 | 0.458144456 | 0.859322431 | Ldlrad3 | low density lipoprotein receptor class A d | 98.782 | 142.88 | 128.8 | 121.46 | 145.05 | 148.14 |
| 14168 | ENSMUSG00000048062 | 0 | NA | NA | NA | NA | NA | Fpr-rs4 | formyl peptide receptor, related sequence | 0 | 0 | 0 | 0 | 0 | 0 |
| 14169 | ENSMUSG00000048065 | 1.377679212 | 1.655421808 | 2.000381483 | 0.827553055 | 0.407923664 | NA | Cyb5r2 | cytochrome b5 reductase 2 [Source:MGI S | 0 | 2.0558 | 0 | 2.8919 | 1.0906 | 2.2277 |
| 14170 | ENSMUSG00000048067 | 0 | NA | NA | NA | NA | NA | Olfr1349 | olfactory receptor 1349 [Source:MGI Syml | 0 | 0 | 0 | 0 | 0 | 0 |
| 14171 | ENSMUSG00000048070 | 0 | NA | NA | NA | NA | NA | Pirt | phosphoinositide-interacting regulator of | 0 | 0 | 0 | 0 | 0 | 0 |
| 14172 | ENSMUSG00000048076 | 4346.497414 | 0.053941035 | 0.084757388 | 0.636416912 | 0.524504721 | 0.888334265 | Arf1 | ADP-ribosylation factor 1 [Source:MGI Syr | 4246 | 4370.7 | 4178.9 | 4330.2 | 4546.8 | 4406.4 |
| 14173 | ENSMUSG00000048077 | 0 | NA | NA | NA | NA | NA | H1f7 | H1.7 linker histone [Source:MGI Symbol;A | 0 | 0 | 0 | 0 | 0 | 0 |
| 14174 | ENSMUSG00000048078 | 0.880724006 | 0.715724162 | 2.808295084 | 0.25486074 | 0.79883066 | NA | Tenm4 | teneurin transmembrane protein 4 [Sourc | 0 | 0 | 2.0125 | 0 | 3.2719 | 0 |
| 14175 | ENSMUSG00000048080 | 0 | NA | NA | NA | NA | NA | Olfr731 | olfactory receptor 731 [Source:MGI Symb | 0 | 0 | 0 | 0 | 0 | 0 |
| 14176 | ENSMUSG00000048087 | 142.5748964 | -0.322612879 | 0.204410338 | -1.578261071 | 0.114505645 | 0.497223548 | Gm4737 | predicted gene 4737 [Source:MGI Symbol | 144.56 | 160.36 | 171.06 | 142.67 | 125.42 | 111.38 |
| 14177 | ENSMUSG00000048096 | 2.803535961 | -0.754184353 | 1.328735635 | -0.567595489 | 0.570309671 | NA | Lmod1 | leiomodin 1 (smooth muscle) [Source:MG | 2.4093 | 7.1954 | 1.0062 | 2.8919 | 1.0906 | 2.2277 |
| 14178 | ENSMUSG00000048100 | 339.9468716 | -0.01345322 | 0.143997637 | -0.093426673 | 0.925564601 | 0.987546253 | Taf13 | TATA-box binding protein associated facto | 343.73 | 317.63 | 363.25 | 343.18 | 354.45 | 317.45 |
| 14179 | ENSMUSG00000048101 | 0 | NA | NA | NA | NA | NA | Olfr19 | olfactory receptor 19 [Source:MGI Symbo | 0 | 0 | 0 | 0 | 0 | 0 |
| 14180 | ENSMUSG00000048106 | 172.2356699 | 0.037386422 | 0.198865037 | 0.187998968 | 0.85087746 | 0.973568625 | 4632415L05 | RIKEN cDNA 4632415L05 gene [Source:M( | 169.45 | 148.02 | 192.19 | 150.38 | 198.49 | 174.87 |
| 14181 | ENSMUSG00000048108 | 0.335414862 | -1.779047242 | 4.030804179 | -0.441362855 | 0.658950329 | NA | Tmem72 | transmembrane protein 72 [Source:MGI S | 0 | 0 | 2.0125 | 0 | 0 | 0 |
| 14182 | ENSMUSG00000048109 | 283.2221366 | 0.072716108 | 0.161775749 | 0.449487076 | 0.653080329 | 0.925756399 | Rbm15 | RNA binding motif protein 15 [Source:MG | 281.89 | 301.18 | 245.52 | 321.97 | 278.11 | 270.66 |
| 14183 | ENSMUSG00000048118 | 290.2194163 | 0.014409685 | 0.167629576 | 0.085961471 | 0.931497046 | 0.988420305 | Arid4a | AT rich interactive domain 4A (RBP1-like) | 244.95 | 311.46 | 310.93 | 303.65 | 311.92 | 258.41 |
| 14184 | ENSMUSG00000048120 | 41.92517427 | -0.632896228 | 0.372251383 | -1.700185027 | 0.089096128 | 0.439824707 | Entpd1 | ectonucleoside triphosphate diphosphohy | 49.792 | 65.787 | 37.231 | 26.991 | 30.537 | 41.212 |
| 14185 | ENSMUSG00000048126 | 99.21560072 | -0.68439847 | 0.290179505 | -2.358534832 | 0.018347238 | 0.16280641 | Col6a3 | collagen, type VI, alpha 3 [Source:MGI Syr | 174.27 | 99.708 | 92.575 | 87.722 | 73.071 | 67.944 |
| 14186 | ENSMUSG00000048138 | 2.210478363 | 0.962786249 | 1.64675276 | 0.584657437 | 0.558778078 | NA | Dmrt2 | doublesex and mab-3 related transcriptio | 2.4093 | 0 | 2.0125 | 0 | 3.2719 | 5.5692 |
| 14187 | ENSMUSG00000048142 | 1.407975666 | -2.72347108 | 1.994677936 | -1.36536883 | 0.172137156 | NA | Nat8l | N-acetyltransferase 8-like [Source:MGI Sy | 2.4093 | 2.0558 | 3.0187 | 0.964 | 0 | 0 |
| 14188 | ENSMUSG00000048148 | 0 | NA | NA | NA | NA | NA | Nwd1 | NACHT and WD repeat domain containing | 0 | 0 | 0 | 0 | 0 | 0 |
| 14189 | ENSMUSG00000048153 | 0.160662974 | 1.126924175 | 4.080472857 | 0.276174898 | 0.782413736 | NA | Olfr49 | olfactory receptor 49 [Source:MGI Symbo | 0 | 0 | 0 | 0.964 | 0 | 0 |
| 14190 | ENSMUSG00000048154 | 834.0151344 | -0.025767383 | 0.144660998 | -0.178122533 | 0.858626746 | 0.975439379 | Kmt2d | lysine (K)-specific methyltransferase 2D [S | 804.71 | 875.79 | 845.25 | 1010.2 | 729.62 | 738.48 |
| 14191 | ENSMUSG00000048155 | 0 | NA | NA | NA | NA | NA | H2bw2 | H2B.W histone 2 [Source:MGI Symbol;Acc | 0 | 0 | 0 | 0 | 0 | 0 |
| 14192 | ENSMUSG00000048163 | 22.6062332 | -0.976429344 | 0.494325648 | -1.975275506 | 0.048236891 | 0.305605956 | Selplg | selectin, platelet (p-selectin) ligand [Sourc | 21.684 | 41.117 | 27.169 | 12.532 | 13.087 | 20.049 |
| 14193 | ENSMUSG00000048164 | 0.181769715 | 1.126924175 | 4.080472857 | 0.276174898 | 0.782413736 | NA | Gm7204 | predicted pseudogene 7204 [Source:MGI | 0 | 0 | 0 | 0 | 1.0906 | 0 |
| 14194 | ENSMUSG00000048170 | 530.4937825 | -0.055692277 | 0.128519052 | -0.433338688 | 0.664768735 | 0.929301301 | Mcmbp | minichromosome maintenance complex b | 521.21 | 560.22 | 541.36 | 535.01 | 555.12 | 470.04 |
| 14195 | ENSMUSG00000048173 | 0 | NA | NA | NA | NA | NA | Olfr1324 | olfactory receptor 1324 [Source:MGI Syml | 0 | 0 | 0 | 0 | 0 | 0 |
| 14196 | ENSMUSG00000048174 | 57.12959737 | 0.219937822 | 0.321666151 | 0.68374562 | 0.494135811 | 0.877473722 | Tmem81 | transmembrane protein 81 [Source:MGI S | 45.777 | 63.731 | 49.306 | 76.154 | 63.256 | 44.554 |
| 14197 | ENSMUSG00000048175 | 403.2657189 | 0.051409586 | 0.139876681 | 0.367535071 | 0.713219937 | 0.944239215 | Asb8 | ankyrin repeat and SOCS box-containing 8 | 390.31 | 419.39 | 378.35 | 385.59 | 392.62 | 453.33 |
| 14198 | ENSMUSG00000048185 | 0 | NA | NA | NA | NA | NA | Gm7075 | predicted gene 7075 [Source:MGI Symbol | 0 | 0 | 0 | 0 | 0 | 0 |
| 14199 | ENSMUSG00000048186 | 201.1052698 | 0.316342137 | 0.19248668 | 1.643449495 | 0.100289967 | 0.466279035 | Bend7 | BEN domain containing 7 [Source:MGI Syr | 144.56 | 195.3 | 198.23 | 201.47 | 244.3 | 222.77 |
| 14200 | ENSMUSG00000048188 | 0.453341394 | -0.545391191 | 3.551113277 | -0.153583158 | 0.877938417 | NA | Gm8181 | predicted gene 8181 [Source:MGI Symbol | 1.6062 | 0 | 0 | 0 | 0 | 1.1138 |
| 14201 | ENSMUSG00000048191 | 1.380620642 | -1.484716045 | 2.178161988 | -0.681637111 | 0.495468443 | NA | NA | NA | 0 | 3.0838 | 3.0187 | 0 | 2.1812 | 0 |
| 14202 | ENSMUSG00000048197 | 0 | NA | NA | NA | NA | NA | Olfr1153 | olfactory receptor 1153 [Source:MGI Syml | 0 | 0 | 0 | 0 | 0 | 0 |
| 14203 | ENSMUSG00000048200 | 34.8766179 | -0.484594504 | 0.370843956 | -1.306734265 | 0.191302977 | 0.629099068 | Cracr2b | calcium release activated channel regulato | 46.58 | 40.089 | 35.219 | 28.919 | 27.265 | 31.188 |
| 14204 | ENSMUSG00000048206 | 0.133850373 | -0.796629176 | 4.080472857 | -0.195229623 | 0.84521319 | NA | Dnajb8 | DnaJ heat shock protein family (Hsp40) m | 0.8031 | 0 | 0 | 0 | 0 | 0 |
| 14205 | ENSMUSG00000048215 | 0 | NA | NA | NA | NA | NA | A630023P12 | RIKEN cDNA A630023P12 gene [Source:M | 0 | 0 | 0 | 0 | 0 | 0 |
| 14206 | ENSMUSG00000048216 | 0.321325949 | 1.911575114 | 4.048267893 | 0.472195804 | 0.636787034 | NA | Gpr85 | G protein-coupled receptor 85 [Source:M( | 0 | 0 | 0 | 1.928 | 0 | 0 |
| 14207 | ENSMUSG00000048217 | 1169.578124 | 0.00240269 | 0.122624453 | 0.01959389 | 0.984367338 | 0.998580735 | Nags | N-acetylglutamate synthase [Source:MGI | 1045.6 | 1140 | 1321.2 | 1184.7 | 1087.3 | 1238.6 |
| 14208 | ENSMUSG00000048218 | 181.361834 | -0.089266106 | 0.208505055 | -0.428124423 | 0.668560536 | 0.930350163 | Amigo2 | adhesion molecule with Ig like domain 2 [! | 224.07 | 168.58 | 168.04 | 209.18 | 169.05 | 149.26 |
| 14209 | ENSMUSG00000048222 | 133.1547303 | -0.214567551 | 0.215141518 | -0.997332139 | 0.318603319 | 0.7652277 | Mfap1b | microfibrillar-associated protein 1B [Sourc | 134.92 | 160.36 | 133.83 | 108.93 | 145.05 | 115.84 |
| 14210 | ENSMUSG00000048226 | 0 | NA | NA | NA | NA | NA | Olfr1212 | olfactory receptor 1212 [Source:MGI Syml | 0 | 0 | 0 | 0 | 0 | 0 |
| 14211 | ENSMUSG00000048230 | 0 | NA | NA | NA | NA | NA | Fbxo43 | F-box protein 43 [Source:MGI Symbol;Acc | 0 | 0 | 0 | 0 | 0 | 0 |
| 14212 | ENSMUSG00000048231 | 0 | NA | NA | NA | NA | NA | H2-M10.4 | histocompatibility 2, M region locus 10.4 [ | 0 | 0 | 0 | 0 | 0 | 0 |
| 14213 | ENSMUSG00000048232 | 12.32363267 | -0.163602958 | 0.657031249 | -0.249003314 | 0.803358218 | 0.963596811 | Fbxo10 | F-box protein 10 [Source:MGI Symbol;Acc | 17.668 | 13.363 | 8.05 | 18.316 | 7.6343 | 8.9108 |
| 14214 | ENSMUSG00000048234 | 1280.917714 | 0.119981853 | 0.132679364 | 0.904299273 | 0.365836732 | 0.804235439 | Rnf149 | ring finger protein 149 [Source:MGI Symb | 1074.6 | 1183.1 | 1425.8 | 1206.9 | 1339.3 | 1455.8 |
| 14215 | ENSMUSG00000048236 | 0 | NA | NA | NA | NA | NA | Ovch2 | ovochymase 2 [Source:MGI Symbol;Acc:M | 0 | 0 | 0 | 0 | 0 | 0 |
| 14216 | ENSMUSG00000048240 | 3.040269474 | -0.528580439 | 1.250858668 | -0.422574071 | 0.672606048 | NA | Gng7 | guanine nucleotide binding protein (G pro | 1.6062 | 7.1954 | 2.0125 | 1.928 | 3.2719 | 2.2277 |
| 14217 | ENSMUSG00000048249 | 610.6233247 | 0.321892673 | 0.121562006 | 2.647971053 | 0.008097646 | 0.092808518 | Crebrf | CREB3 regulatory factor [Source:MGI Sym | 535.67 | 551.99 | 541.36 | 734.55 | 665.28 | 634.89 |
| 14218 | ENSMUSG00000048251 | 2.797777966 | -2.05291442 | 1.636560862 | -1.254407622 | 0.209693883 | NA | Bcl11b | B cell leukemia/lymphoma 11B [Source:M | 3.2124 | 10.279 | 0 | 0 | 2.1812 | 1.1138 |
| 14219 | ENSMUSG00000048261 | 0 | NA | NA | NA | NA | NA | Gm4879 | predicted pseudogene 4879 [Source:MGI | 0 | 0 | 0 | 0 | 0 | 0 |
| 14220 | ENSMUSG00000048264 | 336.6456791 | -0.005268983 | 0.156595651 | -0.033647059 | 0.973158596 | 0.995820834 | Dip2c | disco interacting protein 2 homolog C [Sou | 304.38 | 355.66 | 352.19 | 315.22 | 383.9 | 308.53 |

| | | | | | | | | | | | | | | | |
|---|---|---|---|---|---|---|---|---|---|---|---|---|---|---|---|
| 14221 | ENSMUSG00000048271 | 909.6056553 | 0.343380065 | 0.157444721 | 2.180956352 | 0.029186645 | 0.224312239 | Rbm33 | RNA binding motif protein 33 [Source:MG | 806.31 | 871.67 | 728.52 | 1281.1 | 884.49 | 885.51 |
| 14222 | ENSMUSG00000048277 | 1171.895133 | 0.135410997 | 0.101648926 | 1.332143902 | 0.182812906 | 0.616796332 | Syngr2 | synaptogyrin 2 [Source:MGI Symbol;Acc:N | 1092.2 | 1107.1 | 1152.2 | 1263.8 | 1244.4 | 1171.8 |
| 14223 | ENSMUSG00000048279 | 50.62957869 | 0.064875806 | 0.362406605 | 0.179013863 | 0.857926816 | 0.975277277 | Sacs | sacsin [Source:MGI Symbol;Acc:MGI:1354 | 44.974 | 54.48 | 49.306 | 66.514 | 28.356 | 60.148 |
| 14224 | ENSMUSG00000048280 | 138.5806618 | 0.100722065 | 0.222403799 | 0.452879248 | 0.650635695 | 0.925643431 | Zfp738 | zinc finger protein 738 [Source:MGI Symb | 146.16 | 99.708 | 154.96 | 139.78 | 149.41 | 141.46 |
| 14225 | ENSMUSG00000048281 | 0.160662974 | 1.126924175 | 4.080472857 | 0.276174898 | 0.782413736 | NA | Dleu7 | deleted in lymphocytic leukemia, 7 [Sourc | 0 | 0 | 0 | 0.964 | 0 | 0 |
| 14226 | ENSMUSG00000048284 | 0 | NA | NA | NA | NA | NA | Tas2r126 | taste receptor, type 2, member 126 [Sour | 0 | 0 | 0 | 0 | 0 | 0 |
| 14227 | ENSMUSG00000048285 | 96.97979503 | -0.00570951 | 0.235819538 | -0.024211352 | 0.980684023 | 0.99820894 | Frmd6 | FERM domain containing 6 [Source:MGI S | 83.523 | 101.76 | 106.66 | 100.25 | 89.431 | 100.25 |
| 14228 | ENSMUSG00000048292 | 0 | NA | NA | NA | NA | NA | Olfr1417 | olfactory receptor 1417 [Source:MGI Symb | 0 | 0 | 0 | 0 | 0 | 0 |
| 14229 | ENSMUSG00000048294 | 0 | NA | NA | NA | NA | NA | Krtap4-13 | keratin associated protein 4-13 [Source:M | 0 | 0 | 0 | 0 | 0 | 0 |
| 14230 | ENSMUSG00000048299 | 0 | NA | NA | NA | NA | NA | Olfr148 | olfactory receptor 148 [Source:MGI Symb | 0 | 0 | 0 | 0 | 0 | 0 |
| 14231 | ENSMUSG00000048304 | 0 | NA | NA | NA | NA | NA | Slitrk3 | SLIT and NTRK-like family, member 3 [Sou | 0 | 0 | 0 | 0 | 0 | 0 |
| 14232 | ENSMUSG00000048307 | 1431.157478 | -0.016672109 | 0.122101141 | -0.13654343 | 0.891391694 | 0.982489666 | Ankrd46 | ankyrin repeat domain 46 [Source:MGI Sy | 1339.6 | 1370.2 | 1609 | 1373.7 | 1586.8 | 1307.7 |
| 14233 | ENSMUSG00000048310 | 727.8461941 | 0.071203527 | 0.117678819 | 0.605066636 | 0.545134722 | 0.895484962 | Pskh1 | protein serine kinase H1 [Source:MGI Sym | 689.86 | 750.38 | 689.28 | 690.21 | 774.34 | 773.01 |
| 14234 | ENSMUSG00000048312 | 0 | NA | NA | NA | NA | NA | Gm4884 | predicted gene 4884 [Source:MGI Symbol | 0 | 0 | 0 | 0 | 0 | 0 |
| 14235 | ENSMUSG00000048316 | 0 | NA | NA | NA | NA | NA | Gm9823 | predicted gene 9823 [Source:MGI Symbol | 0 | 0 | 0 | 0 | 0 | 0 |
| 14236 | ENSMUSG00000048327 | 2.938595449 | -0.864014256 | 1.30511498 | -0.662021561 | 0.507957404 | NA | Ckap2l | cytoskeleton associated protein 2-like [So | 7.2279 | 3.0838 | 1.0062 | 1.928 | 3.2719 | 1.1138 |
| 14237 | ENSMUSG00000048329 | 0.698954292 | 0.126053087 | 3.048972385 | 0.04134281 | 0.967022605 | NA | Mfsd6l | major facilitator superfamily domain cont | 0 | 0 | 2.0125 | 0 | 2.1812 | 0 |
| 14238 | ENSMUSG00000048330 | 9.323920929 | -0.367360473 | 0.777072135 | -0.472749512 | 0.636391898 | 0.923478764 | Ric3 | RIC3 acetylcholine receptor chaperone [S | 11.243 | 3.0838 | 17.106 | 5.7839 | 9.8156 | 8.9108 |
| 14239 | ENSMUSG00000048332 | 101.9433705 | -1.170617004 | 0.285433051 | -4.101196413 | 4.11E-05 | 0.001784488 | Lhfp | lipoma HMGIC fusion partner [Source:MG | 175.08 | 133.63 | 114.71 | 73.262 | 40.353 | 74.628 |
| 14240 | ENSMUSG00000048334 | 1.009830165 | 0.037117158 | 2.215730638 | 0.016751656 | 0.986634737 | NA | Gm8258 | predicted gene 8258 [Source:MGI Symbol | 0 | 1.0279 | 2.0125 | 1.928 | 1.0906 | 0 |
| 14241 | ENSMUSG00000048337 | 0 | NA | NA | NA | NA | NA | Npy4r | neuropeptide Y receptor Y4 [Source:MGI S | 0 | 0 | 0 | 0 | 0 | 0 |
| 14242 | ENSMUSG00000048347 | 1.190410227 | -1.156645952 | 2.318925491 | -0.498785302 | 0.617930643 | NA | Pcdhb18 | protocadherin beta 18 [Source:MGI Symb | 0.8031 | 4.1117 | 0 | 0 | 0 | 2.2277 |
| 14243 | ENSMUSG00000048349 | 0.965523032 | -0.944628791 | 2.267086424 | -0.416670834 | 0.67691919 | NA | Pou4f1 | POU domain, class 4, transcription factor 1 | 0.8031 | 2.0558 | 1.0062 | 1.928 | 0 | 0 |
| 14244 | ENSMUSG00000048351 | 430.0913424 | 0.241867454 | 0.144061762 | 1.678915004 | 0.09316861 | 0.449646571 | Coa7 | cytochrome c oxidase assembly factor 7 [S | 415.2 | 407.06 | 359.23 | 415.47 | 467.88 | 515.71 |
| 14245 | ENSMUSG00000048355 | 0 | NA | NA | NA | NA | NA | Arxes1 | adipocyte-related X-chromosome express | 0 | 0 | 0 | 0 | 0 | 0 |
| 14246 | ENSMUSG00000048356 | 0 | NA | NA | NA | NA | NA | Olfr1496 | olfactory receptor 1496 [Source:MGI Syml | 0 | 0 | 0 | 0 | 0 | 0 |
| 14247 | ENSMUSG00000048368 | 114.4006138 | -0.383331804 | 0.267442779 | -1.433322693 | 0.15176564 | 0.567340036 | Omd | osteomodulin [Source:MGI Symbol;Acc:M | 128.5 | 159.33 | 100.62 | 85.794 | 131.96 | 80.197 |
| 14248 | ENSMUSG00000048371 | 877.0950629 | 0.233709299 | 0.11987811 | 1.949557766 | 0.051228851 | 0.317878209 | Pdp2 | pyruvate dehyrogenase phosphatase cata | 876.99 | 744.21 | 795.94 | 875.29 | 1005.6 | 964.59 |
| 14249 | ENSMUSG00000048373 | 0.321325949 | 1.911575114 | 4.048267893 | 0.472195804 | 0.636787034 | NA | Fgfbp1 | fibroblast growth factor binding protein 1 | 0 | 0 | 0 | 1.928 | 0 | 0 |
| 14250 | ENSMUSG00000048376 | 667.3071405 | 0.10017529 | 0.206728916 | 0.484573188 | 0.627979136 | 0.920328819 | F2r | coagulation factor II (thrombin) receptor [ | 725.2 | 501.62 | 704.37 | 457.89 | 799.42 | 815.33 |
| 14251 | ENSMUSG00000048377 | 0 | NA | NA | NA | NA | NA | Foxi2 | forkhead box I2 [Source:MGI Symbol;Acc: | 0 | 0 | 0 | 0 | 0 | 0 |
| 14252 | ENSMUSG00000048378 | 0.185640647 | 1.126924175 | 4.080472857 | 0.276174898 | 0.782413736 | NA | Olfr1394 | olfactory receptor 1394 [Source:MGI Syml | 0 | 0 | 0 | 0 | 0 | 1.1138 |
| 14253 | ENSMUSG00000048379 | 399.6623121 | 0.06811199 | 0.141246151 | 0.482221923 | 0.629648305 | 0.92057725 | Socs4 | suppressor of cytokine signaling 4 [Source | 379.06 | 411.17 | 380.36 | 384.63 | 386.08 | 456.68 |
| 14254 | ENSMUSG00000048385 | 0 | NA | NA | NA | NA | NA | Scrt1 | scratch family zinc finger 1 [Source:MGI S | 0 | 0 | 0 | 0 | 0 | 0 |
| 14255 | ENSMUSG00000048387 | 0.569258551 | -2.569337407 | 3.18860282 | -0.805787849 | 0.420365185 | NA | Osr1 | odd-skipped related transcription factor 1 | 2.4093 | 0 | 1.0062 | 0 | 0 | 0 |
| 14256 | ENSMUSG00000048388 | 17.74577346 | -1.509938139 | 0.555968334 | -2.715870756 | 0.006610172 | 0.080919055 | Fam171b | family with sequence similarity 171, meml | 29.715 | 27.754 | 21.131 | 5.7839 | 8.7249 | 13.366 |
| 14257 | ENSMUSG00000048391 | 0 | NA | NA | NA | NA | NA | Olfr843 | olfactory receptor 843 [Source:MGI Symb | 0 | 0 | 0 | 0 | 0 | 0 |
| 14258 | ENSMUSG00000048398 | 0 | NA | NA | NA | NA | NA | Gm5065 | predicted gene 5065 [Source:MGI Symbol | 0 | 0 | 0 | 0 | 0 | 0 |
| 14259 | ENSMUSG00000048399 | 0 | NA | NA | NA | NA | NA | Tprg | transformation related protein 63 regulat | 0 | 0 | 0 | 0 | 0 | 0 |
| 14260 | ENSMUSG00000048400 | 0 | NA | NA | NA | NA | NA | Prss54 | protease, serine 54 [Source:MGI Symbol;A | 0 | 0 | 0 | 0 | 0 | 0 |
| 14261 | ENSMUSG00000048402 | 2.602774574 | 0.516269665 | 1.52992633 | 0.337447402 | 0.735779656 | NA | Gli2 | GLI-Kruppel family member GLI2 [Source:I | 2.4093 | 0 | 4.025 | 4.8199 | 4.3625 | 0 |
| 14262 | ENSMUSG00000048406 | 44.24561788 | 0.080663366 | 0.409333702 | 0.197060164 | 0.843780457 | 0.97250284 | B330016D1C | RIKEN cDNA B330016D10 gene [Source:M | 28.109 | 41.117 | 60.375 | 66.514 | 38.172 | 31.188 |
| 14263 | ENSMUSG00000048410 | 102.1135585 | -0.236886266 | 0.242332131 | -0.977527268 | 0.328308185 | 0.773161429 | Zfp407 | zinc finger protein 407 [Source:MGI Symb | 93.16 | 101.76 | 136.85 | 93.506 | 91.612 | 95.791 |
| 14264 | ENSMUSG00000048411 | 0 | NA | NA | NA | NA | NA | Gm597 | predicted gene 597 [Source:MGI Symbol;A | 0 | 0 | 0 | 0 | 0 | 0 |
| 14265 | ENSMUSG00000048416 | 0.847770947 | -0.376623742 | 2.576917887 | -0.146152791 | 0.883800772 | NA | Mlf1 | myeloid leukemia factor 1 [Source:MGI Sy | 0.8031 | 2.0558 | 0 | 0 | 0 | 2.2277 |
| 14266 | ENSMUSG00000048424 | 21.66427002 | 0.237831593 | 0.564570046 | 0.421261444 | 0.673564181 | 0.931915649 | Ranbp3l | RAN binding protein 3-like [Source:MGI Sy | 11.243 | 21.586 | 27.169 | 35.667 | 9.8156 | 24.505 |
| 14267 | ENSMUSG00000048425 | 0 | NA | NA | NA | NA | NA | Olfr686 | olfactory receptor 686 [Source:MGI Symb | 0 | 0 | 0 | 0 | 0 | 0 |
| 14268 | ENSMUSG00000048429 | 857.8169218 | -0.108418305 | 0.125758447 | -0.862115489 | 0.388623969 | 0.818441625 | Timm29 | translocase of inner mitochondrial membr | 837.64 | 900.46 | 932.79 | 842.52 | 912.85 | 720.66 |
| 14269 | ENSMUSG00000048439 | 80.79841356 | -0.164247002 | 0.2941071 | -0.558459832 | 0.576530426 | 0.9077105 | Nupl2 | nucleoporin like 2 [Source:MGI Symbol;Ac | 78.704 | 75.038 | 102.64 | 85.794 | 50.168 | 92.449 |
| 14270 | ENSMUSG00000048440 | 20.73027173 | -0.563574712 | 0.515466772 | -1.093328886 | 0.274249424 | 0.729449002 | Cyp4f16 | cytochrome P450, family 4, subfamily f, pc | 18.471 | 27.754 | 28.175 | 21.208 | 8.7249 | 20.049 |
| 14271 | ENSMUSG00000048442 | 4.63383507 | -0.732960501 | 1.017607247 | -0.720278382 | 0.471353612 | NA | Smim5 | small integral membrane protein 5 [Sourc | 7.2279 | 2.0558 | 8.05 | 3.8559 | 3.2719 | 3.3415 |
| 14272 | ENSMUSG00000048445 | 87.94742659 | -0.205742549 | 0.272252102 | -0.755706007 | 0.449825481 | 0.855321038 | Ccdc57 | coiled-coil domain containing 57 [Source:I | 94.766 | 94.568 | 93.581 | 96.398 | 93.793 | 54.578 |
| 14273 | ENSMUSG00000048450 | 7.972954967 | 0.766494951 | 0.823406591 | 0.930882701 | 0.351914244 | 0.792889192 | Msx1 | msh homeobox 1 [Source:MGI Symbol;Ac | 5.6217 | 2.0558 | 10.062 | 12.532 | 11.997 | 5.5692 |
| 14274 | ENSMUSG00000048455 | 0 | NA | NA | NA | NA | NA | Sprr1b | small proline-rich protein 1B [Source:MGI | 0 | 0 | 0 | 0 | 0 | 0 |
| 14275 | ENSMUSG00000048456 | 0 | NA | NA | NA | NA | NA | Olfr1448 | olfactory receptor 1448 [Source:MGI Syml | 0 | 0 | 0 | 0 | 0 | 0 |
| 14276 | ENSMUSG00000048458 | 1.540378191 | 1.16533341 | 1.910371369 | 0.610003599 | 0.541859424 | NA | Inka2 | inka box actin regulator 2 [Source:MGI Syı | 0.8031 | 2.0558 | 0 | 1.928 | 0 | 4.4554 |

| | | | | | | | | | | | | | | | |
|---|---|---|---|---|---|---|---|---|---|---|---|---|---|---|---|
| 14277 | ENSMUSG00000048469 | 0 | NA | NA | NA | NA | NA | Olfr564 | olfactory receptor 564 [Source:MGI Symb | 0 | 0 | 0 | 0 | 0 | 0 |
| 14278 | ENSMUSG00000048473 | 0.133850373 | -0.796629176 | 4.080472857 | -0.195229623 | 0.84521319 | NA | Sult6b2 | sulfotransferase family 6B, member 2 [Sou | 0.8031 | 0 | 0 | 0 | 0 | 0 |
| 14279 | ENSMUSG00000048478 | 5.857087449 | 0.07215164 | 0.904482061 | 0.079771223 | 0.936419212 | 0.988420305 | Spata33 | spermatogenesis associated 33 [Source:M | 8.8341 | 5.1396 | 3.0187 | 3.8559 | 8.7249 | 5.5692 |
| 14280 | ENSMUSG00000048480 | 1.445624893 | -1.749642993 | 2.160198385 | -0.809945515 | 0.417971491 | NA | Cxcr1 | chemokine (C-X-C motif) receptor 1 [Sourc | 1.6062 | 5.1396 | 0 | 1.928 | 0 | 0 |
| 14281 | ENSMUSG00000048481 | 22.21457494 | 0.14187781 | 0.493834687 | 0.287298186 | 0.773884012 | 0.958261993 | Mypop | Myb-related transcription factor, partner | 12.85 | 27.754 | 23.144 | 29.883 | 20.722 | 18.935 |
| 14282 | ENSMUSG00000048482 | 35.80127895 | 0.323091772 | 0.401874646 | 0.80396157 | 0.42141917 | 0.838928396 | Bdnf | brain derived neurotrophic factor [Source | 38.549 | 20.558 | 36.225 | 51.091 | 31.628 | 36.757 |
| 14283 | ENSMUSG00000048483 | 0.181769715 | 1.126924175 | 4.080472857 | 0.276174898 | 0.782413736 | NA | Zdhhc22 | zinc finger, DHHC-type containing 22 [Sou | 0 | 0 | 0 | 0 | 1.0906 | 0 |
| 14284 | ENSMUSG00000048484 | 0.17131964 | -0.796629176 | 4.080472857 | -0.195229623 | 0.84521319 | NA | Gm7461 | predicted gene 7461 [Source:MGI Symbol | 0 | 1.0279 | 0 | 0 | 0 | 0 |
| 14285 | ENSMUSG00000048485 | 0.167707431 | -0.796629176 | 4.080472857 | -0.195229623 | 0.84521319 | NA | Zbtb8b | zinc finger and BTB domain containing 8b | 0 | 0 | 1.0062 | 0 | 0 | 0 |
| 14286 | ENSMUSG00000048486 | 1973.92877 | -0.292828307 | 0.156873441 | -1.86665318 | 0.061950036 | 0.355884025 | Fitm2 | fat storage-inducing transmembrane prot | 2365.9 | 2403.3 | 1751.9 | 2057.1 | 1850.8 | 1414.6 |
| 14287 | ENSMUSG00000048489 | 832.7098235 | -0.1260998 | 0.555067881 | -0.22717906 | 0.82028451 | 0.969160361 | Depp1 | DEPP1 autophagy regulator [Source:MGI S | 1121.1 | 850.09 | 635.95 | 869.51 | 1188.8 | 330.81 |
| 14288 | ENSMUSG00000048490 | 215.3536248 | -0.087222122 | 0.194578578 | -0.448261686 | 0.653964349 | 0.925797231 | Nrip1 | nuclear receptor interacting protein 1 [So | 226.47 | 233.34 | 206.28 | 249.67 | 214.85 | 161.51 |
| 14289 | ENSMUSG00000048495 | 136.3373753 | 0.157739713 | 0.22079199 | 0.714426789 | 0.474963312 | 0.869368495 | Tyw5 | tRNA-yW synthesizing protein 5 [Source:N | 143.76 | 133.63 | 108.67 | 121.46 | 169.05 | 141.46 |
| 14290 | ENSMUSG00000048497 | 13.18344858 | -0.424586544 | 0.606866476 | -0.699637499 | 0.484153718 | 0.872431004 | Mmgt2 | membrane magnesium transporter 2 [Sou | 16.865 | 9.2513 | 19.119 | 9.6398 | 13.087 | 11.138 |
| 14291 | ENSMUSG00000048498 | 11.40875748 | -1.972272406 | 0.743034418 | -2.654348651 | 0.007946168 | 0.091542106 | Cd300e | CD300E molecule [Source:MGI Symbol;Ac | 17.668 | 25.698 | 11.069 | 1.928 | 7.6343 | 4.4554 |
| 14292 | ENSMUSG00000048500 | 0 | NA | NA | NA | NA | NA | Defb15 | defensin beta 15 [Source:MGI Symbol;Acc | 0 | 0 | 0 | 0 | 0 | 0 |
| 14293 | ENSMUSG00000048501 | 0 | NA | NA | NA | NA | NA | Olfr938 | olfactory receptor 938 [Source:MGI Symb | 0 | 0 | 0 | 0 | 0 | 0 |
| 14294 | ENSMUSG00000048502 | 0 | NA | NA | NA | NA | NA | Duxbl1 | double homeobox B-like 1 [Source:MGI Sy | 0 | 0 | 0 | 0 | 0 | 0 |
| 14295 | ENSMUSG00000048503 | 12.34979561 | -0.621448971 | 0.61815594 | -1.005327186 | 0.314739329 | 0.761014653 | Tlcd5 | TLC domain containing 5 [Source:MGI Sym | 18.471 | 11.307 | 15.094 | 11.568 | 7.6343 | 10.025 |
| 14296 | ENSMUSG00000048516 | 0 | NA | NA | NA | NA | NA | Adam26a | a disintegrin and metallopeptidase domai | 0 | 0 | 0 | 0 | 0 | 0 |
| 14297 | ENSMUSG00000048520 | 3.846261538 | -0.743450679 | 1.133604285 | -0.655829101 | 0.51193409 | NA | Fbxl13 | F-box and leucine-rich repeat protein 13 [ | 3.2124 | 7.1954 | 4.025 | 0.964 | 5.4531 | 2.2277 |
| 14298 | ENSMUSG00000048521 | 5.622941186 | -0.880747687 | 1.025432264 | -0.858903818 | 0.390393583 | 0.819763149 | Cxcr6 | chemokine (C-X-C motif) receptor 6 [Sourc | 5.6217 | 4.1117 | 12.075 | 1.928 | 1.0906 | 8.9108 |
| 14299 | ENSMUSG00000048528 | 0 | NA | NA | NA | NA | NA | Nkx1-2 | NK1 homeobox 2 [Source:MGI Symbol;Ac | 0 | 0 | 0 | 0 | 0 | 0 |
| 14300 | ENSMUSG00000048534 | 16.71544436 | -0.522118701 | 0.659049909 | -0.79222938 | 0.428226941 | 0.842753542 | Jaml | junction adhesion molecule like [Source:N | 16.062 | 32.893 | 10.062 | 4.8199 | 14.178 | 22.277 |
| 14301 | ENSMUSG00000048537 | 94.7513828 | -0.54517577 | 0.347281644 | -1.569837564 | 0.116452906 | 0.501750025 | Phldb1 | pleckstrin homology like domain, family B | 158.21 | 94.568 | 84.525 | 106.04 | 85.068 | 40.098 |
| 14302 | ENSMUSG00000048538 | 68.29635775 | 0.033232176 | 0.29974932 | 0.11086656 | 0.911722163 | 0.98589875 | Gm9826 | predicted gene 9826 [Source:MGI Symbol | 46.58 | 75.038 | 81.506 | 72.298 | 71.981 | 62.375 |
| 14303 | ENSMUSG00000048540 | 0 | NA | NA | NA | NA | NA | Nhlh2 | nescient helix loop helix 2 [Source:MGI Sy | 0 | 0 | 0 | 0 | 0 | 0 |
| 14304 | ENSMUSG00000048544 | 0.160662974 | 1.126924175 | 4.080472857 | 0.276174898 | 0.782413736 | NA | NA | NA | 0 | 0 | 0 | 0.964 | 0 | 0 |
| 14305 | ENSMUSG00000048546 | 1168.155484 | -0.735053137 | 0.366336956 | -2.006494637 | 0.0448035 | 0.292435844 | Tob2 | transducer of ERBB2, 2 [Source:MGI Symb | 1407.8 | 1530.6 | 1439.9 | 722.02 | 1333.8 | 574.74 |
| 14306 | ENSMUSG00000048550 | 89.67141432 | 0.050128036 | 0.245978365 | 0.203790425 | 0.838517276 | 0.971275236 | Thnsl1 | threonine synthase-like 1 (bacterial) [Sour | 97.175 | 86.345 | 80.5 | 85.794 | 105.79 | 82.424 |
| 14307 | ENSMUSG00000048559 | 0 | NA | NA | NA | NA | NA | 4930555K19 | RIKEN cDNA 4930555K19 gene [Source:M | 0 | 0 | 0 | 0 | 0 | 0 |
| 14308 | ENSMUSG00000048562 | 0 | NA | NA | NA | NA | NA | Sp8 | trans-acting transcription factor 8 [Source | 0 | 0 | 0 | 0 | 0 | 0 |
| 14309 | ENSMUSG00000048572 | 1.68997079 | -0.011258099 | 1.715256606 | -0.006563507 | 0.994763117 | NA | Tmem252 | transmembrane protein 252 [Source:MGI | 4.0155 | 1.0279 | 0 | 2.8919 | 1.0906 | 1.1138 |
| 14310 | ENSMUSG00000048573 | 0 | NA | NA | NA | NA | NA | Cypt3 | cysteine-rich perinuclear theca 3 [Source: | 0 | 0 | 0 | 0 | 0 | 0 |
| 14311 | ENSMUSG00000048574 | 0 | NA | NA | NA | NA | NA | Ccnb1-ps | cyclin B1, pseudogene [Source:MGI Symb | 0 | 0 | 0 | 0 | 0 | 0 |
| 14312 | ENSMUSG00000048578 | 2237.134614 | 0.323978532 | 0.130078095 | 2.490646346 | 0.012751097 | 0.127649303 | Mlec | malectin [Source:MGI Symbol;Acc:MGI:19 | 2285.6 | 1872.9 | 1801.2 | 2605.6 | 2143.1 | 2714.4 |
| 14313 | ENSMUSG00000048581 | 120.4681765 | -0.104373231 | 0.257220672 | -0.405773107 | 0.68490932 | 0.935737593 | E130311K13 | RIKEN cDNA E130311K13 gene [Source:M | 150.18 | 96.624 | 126.79 | 87.722 | 116.7 | 144.8 |
| 14314 | ENSMUSG00000048582 | 0 | NA | NA | NA | NA | NA | Gja3 | gap junction protein, alpha 3 [Source:MGI | 0 | 0 | 0 | 0 | 0 | 0 |
| 14315 | ENSMUSG00000048583 | 0.331982614 | 0.16514309 | 3.986751476 | 0.041422971 | 0.966958701 | NA | Igf2 | insulin-like growth factor 2 [Source:MGI S | 0 | 1.0279 | 0 | 0.964 | 0 | 0 |
| 14316 | ENSMUSG00000048592 | 0 | NA | NA | NA | NA | NA | Gm5946 | predicted gene 5946 [Source:MGI Symbol | 0 | 0 | 0 | 0 | 0 | 0 |
| 14317 | ENSMUSG00000048600 | 2.254433521 | 1.522468734 | 1.675972657 | 0.908409053 | 0.363662145 | NA | Gm5763 | predicted pseudogene 5763 [Source:MGI | 2.4093 | 1.0279 | 0 | 6.7478 | 0 | 3.3415 |
| 14318 | ENSMUSG00000048602 | 0 | NA | NA | NA | NA | NA | Morc2b | microrchidia 2B [Source:MGI Symbol;Acc: | 0 | 0 | 0 | 0 | 0 | 0 |
| 14319 | ENSMUSG00000048603 | 9.175743447 | 0.701941111 | 0.777910882 | 0.902341293 | 0.366875597 | 0.805239112 | Gm9828 | predicted gene 9828 [Source:MGI Symbol | 4.8186 | 4.1117 | 12.075 | 7.7118 | 18.541 | 7.7969 |
| 14320 | ENSMUSG00000048612 | 27.66168691 | -0.891632083 | 0.440443575 | -2.024395709 | 0.042929453 | 0.286011795 | Myof | myoferlin [Source:MGI Symbol;Acc:MGI:1 | 38.549 | 34.949 | 34.212 | 15.424 | 28.356 | 14.48 |
| 14321 | ENSMUSG00000048616 | 0 | NA | NA | NA | NA | NA | Nog | noggin [Source:MGI Symbol;Acc:MGI:104 | 0 | 0 | 0 | 0 | 0 | 0 |
| 14322 | ENSMUSG00000048617 | 2.247487829 | 1.824404243 | 1.576271912 | 1.157417213 | 0.247101945 | NA | Rtbdn | retbindin [Source:MGI Symbol;Acc:MGI:2 | 0 | 2.0558 | 1.0062 | 3.8559 | 5.4531 | 1.1138 |
| 14323 | ENSMUSG00000048620 | 0 | NA | NA | NA | NA | NA | Olfr1336 | olfactory receptor 1336 [Source:MGI Sym | 0 | 0 | 0 | 0 | 0 | 0 |
| 14324 | ENSMUSG00000048621 | 15.40220893 | -0.275993301 | 0.592808425 | -0.465569126 | 0.641523946 | 0.924019354 | Gm6377 | predicted gene 6377 [Source:MGI Symbol | 12.85 | 24.67 | 13.081 | 8.6758 | 13.087 | 20.049 |
| 14325 | ENSMUSG00000048626 | 0 | NA | NA | NA | NA | NA | Klf17 | Kruppel-like factor 17 [Source:MGI Symbo | 0 | 0 | 0 | 0 | 0 | 0 |
| 14326 | ENSMUSG00000048628 | 0 | NA | NA | NA | NA | NA | 4930542C16 | RIKEN cDNA 4930542C16 gene [Source:M | 0 | 0 | 0 | 0 | 0 | 0 |
| 14327 | ENSMUSG00000048636 | 0.17131964 | -0.796629176 | 4.080472857 | -0.195229623 | 0.84521319 | NA | A730049H05 | RIKEN cDNA A730049H05 gene [Source:M | 0 | 1.0279 | 0 | 0 | 0 | 0 |
| 14328 | ENSMUSG00000048643 | 0 | NA | NA | NA | NA | NA | Krtap19-9a | keratin associated protein 19-9A [Source: | 0 | 0 | 0 | 0 | 0 | 0 |
| 14329 | ENSMUSG00000048644 | 9.578087358 | 0.264976517 | 0.767293376 | 0.345339247 | 0.729839345 | 0.946534338 | Ctxn1 | cortexin 1 [Source:MGI Symbol;Acc:MGI:8 | 9.6372 | 11.307 | 5.0312 | 3.8559 | 9.8156 | 17.822 |
| 14330 | ENSMUSG00000048647 | 107.6019748 | 0.65890236 | 0.237979252 | 2.768738681 | 0.005627375 | 0.072286281 | Exd1 | exonuclease 3'-5' domain containing 1 [Sc | 90.751 | 69.898 | 89.556 | 145.56 | 140.69 | 109.16 |
| 14331 | ENSMUSG00000048652 | 0 | NA | NA | NA | NA | NA | Samd13 | sterile alpha motif domain containing 13 [ | 0 | 0 | 0 | 0 | 0 | 0 |
| 14332 | ENSMUSG00000048655 | 0.688762941 | -1.399623551 | 3.067774249 | -0.456234207 | 0.64822157 | NA | Ccdc169 | coiled-coil domain containing 169 [Source | 0 | 0 | 3.0187 | 0 | 0 | 1.1138 |

| | | | | | | | | | | | | | | | |
|---|---|---|---|---|---|---|---|---|---|---|---|---|---|---|---|
| 14333 | ENSMUSG00000048661 | 235.1621643 | 0.115372069 | 0.161875399 | 0.712721446 | 0.476018143 | 0.869483933 | Lemd3 | LEM domain containing 3 [Source:MGI Sy | 208 | 237.45 | 232.44 | 253.53 | 233.39 | 246.16 |
| 14334 | ENSMUSG00000048668 | 51.51907649 | 0.004711098 | 0.316585353 | 0.014880973 | 0.98812714 | 0.99859176 | Rhno1 | RAD9-HUS1-RAD1 interacting nuclear orp | 58.626 | 46.256 | 49.306 | 55.911 | 57.803 | 41.212 |
| 14335 | ENSMUSG00000048677 | 59.21586325 | -0.393524822 | 0.316825343 | -1.242087576 | 0.214204252 | 0.658874792 | Tpcn2 | two pore segment channel 2 [Source:MGI | 70.673 | 74.01 | 57.356 | 68.442 | 44.715 | 40.098 |
| 14336 | ENSMUSG00000048686 | 0 | NA | NA | NA | NA | NA | Hmgb4 | high-mobility group box 4 [Source:MGI Sy | 0 | 0 | 0 | 0 | 0 | 0 |
| 14337 | ENSMUSG00000048693 | 0 | NA | NA | NA | NA | NA | Olfr435 | olfactory receptor 435 [Source:MGI Symb | 0 | 0 | 0 | 0 | 0 | 0 |
| 14338 | ENSMUSG00000048696 | 61.90003662 | 0.043622891 | 0.333675732 | 0.130734383 | 0.895985432 | 0.982798755 | Mex3d | mex3 RNA binding family member D [Sour | 41.761 | 83.261 | 58.362 | 66.514 | 47.987 | 73.514 |
| 14339 | ENSMUSG00000048697 | 0 | NA | NA | NA | NA | NA | Vmn1r26 | vomeronasal 1 receptor 26 [Source:MGI S | 0 | 0 | 0 | 0 | 0 | 0 |
| 14340 | ENSMUSG00000048699 | 0 | NA | NA | NA | NA | NA | Krt90 | keratin 90 [Source:MGI Symbol;Acc:MGI:3 | 0 | 0 | 0 | 0 | 0 | 0 |
| 14341 | ENSMUSG00000048701 | 454.5937067 | 0.166980011 | 0.13298723 | 1.255609362 | 0.209257635 | 0.65350014 | Ccdc6 | coiled-coil domain containing 6 [Source:M | 428.05 | 438.92 | 417.59 | 448.25 | 466.78 | 527.96 |
| 14342 | ENSMUSG00000048703 | 0 | NA | NA | NA | NA | NA | Speer4b | spermatogenesis associated glutamate (E) | 0 | 0 | 0 | 0 | 0 | 0 |
| 14343 | ENSMUSG00000048706 | 1482.030603 | -0.209805699 | 0.136341744 | -1.53882217 | 0.123847716 | 0.515640001 | Lurap1l | leucine rich adaptor protein 1-like [Source | 1563.6 | 1896.5 | 1309.1 | 1419.9 | 1375.3 | 1327.7 |
| 14344 | ENSMUSG00000048707 | 59.17199107 | -0.163143538 | 0.341242913 | -0.47808623 | 0.632588833 | 0.922076476 | Tprn | taperin [Source:MGI Symbol;Acc:MGI:213 | 61.839 | 55.508 | 70.437 | 68.442 | 67.618 | 31.188 |
| 14345 | ENSMUSG00000048709 | 0 | NA | NA | NA | NA | NA | Gm8666 | predicted gene 8666 [Source:MGI Symbol | 0 | 0 | 0 | 0 | 0 | 0 |
| 14346 | ENSMUSG00000048720 | 190.474726 | 0.036194938 | 0.218806565 | 0.165419799 | 0.868613571 | 0.97847181 | Tbc1d12 | TBC1D12: TBC1 domain family, member 1 | 168.65 | 191.19 | 205.27 | 240.03 | 199.58 | 138.12 |
| 14347 | ENSMUSG00000048721 | 0 | NA | NA | NA | NA | NA | Fndc9 | fibronectin type III domain containing 9 [S | 0 | 0 | 0 | 0 | 0 | 0 |
| 14348 | ENSMUSG00000048728 | 14.92115146 | 0.221234894 | 0.585705301 | 0.377723907 | 0.705635706 | 0.94295036 | Zfp454 | zinc finger protein 454 [Source:MGI Symb | 8.8341 | 18.503 | 14.087 | 10.604 | 17.45 | 20.049 |
| 14349 | ENSMUSG00000048731 | 286.9832302 | 0.093701027 | 0.206465132 | 0.45383463 | 0.649947856 | 0.925643431 | Ggnbp1 | gametogenetin binding protein 1 [Source: | 229.69 | 276.51 | 328.04 | 377.88 | 278.11 | 231.68 |
| 14350 | ENSMUSG00000048732 | 43.20402145 | -0.272006812 | 0.352003793 | -0.772738299 | 0.439677276 | 0.84844279 | Klhl11 | kelch-like 11 [Source:MGI Symbol;Acc:MG | 47.383 | 52.424 | 42.262 | 51.091 | 35.99 | 30.074 |
| 14351 | ENSMUSG00000048745 | 0 | NA | NA | NA | NA | NA | Olfr820 | olfactory receptor 820 [Source:MGI Symb | 0 | 0 | 0 | 0 | 0 | 0 |
| 14352 | ENSMUSG00000048747 | 0 | NA | NA | NA | NA | NA | E130114P18 | RIKEN cDNA E130114P18 gene [Source:M | 0 | 0 | 0 | 0 | 0 | 0 |
| 14353 | ENSMUSG00000048752 | 14.97651983 | -0.895486333 | 0.565199834 | -1.584371187 | 0.113109275 | 0.493951959 | Prss50 | protease, serine 50 [Source:MGI Symbol;A | 16.865 | 20.558 | 21.131 | 12.532 | 7.6343 | 11.138 |
| 14354 | ENSMUSG00000048755 | 522.0737054 | 0.136820647 | 0.137893249 | 0.992221507 | 0.321089483 | 0.768277632 | Mcat | malonyl CoA:ACP acyltransferase (mitoch | 464.19 | 558.16 | 469.92 | 510.91 | 531.13 | 598.13 |
| 14355 | ENSMUSG00000048756 | 1138.340753 | 0.052584978 | 0.13711776 | 0.383502312 | 0.701347356 | 0.941991169 | Foxo3 | forkhead box O3 [Source:MGI Symbol;Acc | 1114.7 | 1161.5 | 1075.7 | 937.95 | 1331.6 | 1208.5 |
| 14356 | ENSMUSG00000048758 | 3013.705605 | 0.051649001 | 0.090907025 | 0.56815192 | 0.569931816 | 0.904090127 | Rpl29 | ribosomal protein L29 [Source:MGI Symbo | 2810.1 | 2970.7 | 3099.2 | 3016.3 | 3048.3 | 3137.7 |
| 14357 | ENSMUSG00000048763 | 2.55474598 | 1.669509611 | 1.411593688 | 1.182712579 | 0.236923077 | NA | Hoxb3 | homeobox B3 [Source:MGI Symbol;Acc:M | 1.6062 | 0 | 2.0125 | 2.8919 | 4.3625 | 4.4554 |
| 14358 | ENSMUSG00000048764 | 0 | NA | NA | NA | NA | NA | Tmprss11f | transmembrane protease, serine 11f [Sou | 0 | 0 | 0 | 0 | 0 | 0 |
| 14359 | ENSMUSG00000048766 | 0 | NA | NA | NA | NA | NA | Skint10 | selection and upkeep of intraepithelial T c | 0 | 0 | 0 | 0 | 0 | 0 |
| 14360 | ENSMUSG00000048772 | 349.1391424 | 0.165197632 | 0.168282376 | 0.981669237 | 0.326262826 | 0.772018857 | Tmem53 | transmembrane protein 53 [Source:MGI S | 375.85 | 312.49 | 298.85 | 427.04 | 311.92 | 368.68 |
| 14361 | ENSMUSG00000048775 | 0 | NA | NA | NA | NA | NA | Serpinb13 | serine (or cysteine) peptidase inhibitor, cl | 0 | 0 | 0 | 0 | 0 | 0 |
| 14362 | ENSMUSG00000048776 | 0 | NA | NA | NA | NA | NA | Pthlh | parathyroid hormone-like peptide [Source | 0 | 0 | 0 | 0 | 0 | 0 |
| 14363 | ENSMUSG00000048779 | 13.36535632 | -1.605943292 | 0.633535125 | -2.534892272 | 0.011248191 | 0.117808716 | P2ry6 | pyrimidinergic receptor P2Y, G-protein co | 17.668 | 26.726 | 16.1 | 8.6758 | 5.4531 | 5.5692 |
| 14364 | ENSMUSG00000048782 | 223.6953968 | 0.023468572 | 0.197120724 | 0.119056847 | 0.905230318 | 0.983972587 | Insc | INSC spindle orientation adaptor protein [ | 199.97 | 195.3 | 270.68 | 231.35 | 188.68 | 256.18 |
| 14365 | ENSMUSG00000048787 | 292.4569076 | -0.753474077 | 0.218815458 | -3.443422529 | 0.000574401 | 0.013709799 | Dcun1d3 | DCN1, defective in cullin neddylation 1, do | 346.94 | 451.26 | 303.89 | 252.56 | 250.84 | 149.26 |
| 14366 | ENSMUSG00000048794 | 15.04752177 | 0.004111866 | 0.666764436 | 0.006166895 | 0.995079561 | 0.998676967 | Cfap100 | cilia and flagella associated protein 100 [S | 5.6217 | 15.419 | 24.15 | 7.7118 | 22.903 | 14.48 |
| 14367 | ENSMUSG00000048796 | 149.3482442 | 0.23148474 | 0.191860863 | 1.20652402 | 0.227615503 | 0.673700979 | Cyb561d1 | cytochrome b-561 domain containing 1 [S | 148.57 | 127.46 | 135.84 | 166.77 | 157.05 | 160.39 |
| 14368 | ENSMUSG00000048799 | 231.5004527 | 0.030685077 | 0.174953481 | 0.175389918 | 0.860773262 | 0.975943719 | Cep120 | centrosomal protein 120 [Source:MGI Syn | 265.02 | 225.11 | 196.22 | 233.28 | 241.03 | 228.34 |
| 14369 | ENSMUSG00000048806 | 0 | NA | NA | NA | NA | NA | Ifnb1 | interferon beta 1, fibroblast [Source:MGI | 0 | 0 | 0 | 0 | 0 | 0 |
| 14370 | ENSMUSG00000048807 | 7.071667241 | -1.850547376 | 0.898644218 | -2.059265881 | 0.039468775 | 0.271845887 | Slc35e4 | solute carrier family 35, member E4 [Sour | 12.047 | 8.2233 | 13.081 | 5.7839 | 2.1812 | 1.1138 |
| 14371 | ENSMUSG00000048810 | 0 | NA | NA | NA | NA | NA | Olfr202 | olfactory receptor 202 [Source:MGI Symb | 0 | 0 | 0 | 0 | 0 | 0 |
| 14372 | ENSMUSG00000048814 | 0.335414862 | -1.779047242 | 4.030804179 | -0.441362855 | 0.658950329 | NA | Lonrf2 | LON peptidase N-terminal domain and rin | 0 | 0 | 2.0125 | 0 | 0 | 0 |
| 14373 | ENSMUSG00000048824 | 0.17131964 | -0.796629176 | 4.080472857 | -0.195229623 | 0.84521319 | NA | Gm568 | predicted gene 568 [Source:MGI Symbol;A | 0 | 1.0279 | 0 | 0 | 0 | 0 |
| 14374 | ENSMUSG00000048826 | 189.0055359 | 0.730995999 | 0.214716917 | 3.404463933 | 0.000662941 | 0.015293097 | Dact2 | dishevelled-binding antagonist of beta-ca | 144.56 | 133.63 | 147.92 | 221.71 | 182.13 | 304.08 |
| 14375 | ENSMUSG00000048827 | 11.86369667 | 0.222541738 | 0.681126811 | 0.326725853 | 0.743875245 | 0.949911274 | Pkd1l3 | polycystic kidney disease 1 like 3 [Source:I | 6.4248 | 18.503 | 8.05 | 9.6398 | 18.541 | 10.025 |
| 14376 | ENSMUSG00000048830 | 0 | NA | NA | NA | NA | NA | 2310057N15 | RIKEN cDNA 2310057N15 gene [Source:M | 0 | 0 | 0 | 0 | 0 | 0 |
| 14377 | ENSMUSG00000048832 | 205.9311502 | -0.117147005 | 0.170390125 | -0.687522268 | 0.491753673 | 0.876368691 | Vps37c | vacuolar protein sorting 37C [Source:MGI | 215.23 | 227.17 | 200.24 | 184.12 | 207.22 | 201.61 |
| 14378 | ENSMUSG00000048833 | 1163.253851 | -0.06269458 | 0.104952009 | -0.59736427 | 0.550264204 | 0.895703112 | Slc39a9 | solute carrier family 39 (zinc transporter), | 1249.6 | 1169.8 | 1145.1 | 1083.5 | 1192 | 1139.5 |
| 14379 | ENSMUSG00000048834 | 0 | NA | NA | NA | NA | NA | Vstm2a | V-set and transmembrane domain contair | 0 | 0 | 0 | 0 | 0 | 0 |
| 14380 | ENSMUSG00000048852 | 3.022801572 | -1.690141778 | 1.334985198 | -1.266037841 | 0.205499535 | NA | Gm12185 | predicted gene 12185 [Source:MGI Symbo | 1.6062 | 8.2233 | 4.025 | 0.964 | 1.0906 | 2.2277 |
| 14381 | ENSMUSG00000048856 | 19199.40191 | -0.730087766 | 0.140333908 | -5.202504342 | 1.97E-07 | 1.99E-05 | Slc25a47 | solute carrier family 25, member 47 [Sour | 28222 | 22444 | 21203 | 14004 | 17076 | 12249 |
| 14382 | ENSMUSG00000048865 | 39.62589883 | -1.417780525 | 0.413614699 | -3.427780803 | 0.000608537 | 0.014288755 | Arhgap30 | Rho GTPase activating protein 30 [Source: | 45.777 | 84.289 | 43.269 | 26.991 | 20.722 | 16.708 |
| 14383 | ENSMUSG00000048874 | 923.1903928 | -0.04147317 | 0.117906467 | -0.351746352 | 0.725028495 | 0.9448775 | Phf3 | PHD finger protein 3 [Source:MGI Symbol | 919.55 | 1036.1 | 854.3 | 951.45 | 888.85 | 888.85 |
| 14384 | ENSMUSG00000048875 | 0 | NA | NA | NA | NA | NA | Pudp | pseudouridine 5'-phosphatase [Source:M | 0 | 0 | 0 | 0 | 0 | 0 |
| 14385 | ENSMUSG00000048878 | 857.0040782 | -0.440159008 | 0.135762484 | -3.242125483 | 0.001186418 | 0.02363684 | Hexim1 | hexamethylene bis-acetamide inducible 1 | 887.43 | 1000.2 | 1073.7 | 757.69 | 803.79 | 619.3 |
| 14386 | ENSMUSG00000048895 | 9.706120046 | -0.08775452 | 0.723400442 | -0.121308358 | 0.9034468 | 0.983821605 | Cdk5r1 | cyclin-dependent kinase 5, regulatory sub | 13.653 | 7.1954 | 9.0562 | 10.604 | 4.3625 | 13.366 |
| 14387 | ENSMUSG00000048897 | 278.3768284 | 0.312919421 | 0.168306889 | 1.859219331 | 0.062996056 | 0.359154494 | Zfp710 | zinc finger protein 710 [Source:MGI Symb | 265.02 | 225.11 | 254.58 | 337.39 | 259.57 | 328.58 |
| 14388 | ENSMUSG00000048899 | 0.185640647 | 1.126924175 | 4.080472857 | 0.276174898 | 0.782413736 | NA | Rimkla | ribosomal modification protein rimK-like f | 0 | 0 | 0 | 0 | 0 | 1.1138 |

| | | | | | | | | | | | | | | | |
|---|---|---|---|---|---|---|---|---|---|---|---|---|---|---|---|
| 14389 | ENSMUSG00000048904 | 0 | NA | NA | NA | NA | NA | Neurog1 | neurogenin 1 [Source:MGI Symbol;Acc:M( | 0 | 0 | 0 | 0 | 0 | 0 |
| 14390 | ENSMUSG00000048905 | 5.594938704 | -1.781466024 | 1.043636009 | -1.706980219 | 0.087825729 | 0.43690328 | Bnip5 | BCL2 interacting protein 5 [Source:MGI Sy | 9.6372 | 7.1954 | 9.0562 | 0 | 5.4531 | 2.2277 |
| 14391 | ENSMUSG00000048911 | 83.82974003 | 0.466523645 | 0.344747638 | 1.353232316 | 0.175981424 | 0.606768849 | Rnf24 | ring finger protein 24 [Source:MGI Symbo | 97.978 | 61.675 | 51.318 | 145.56 | 79.615 | 66.831 |
| 14392 | ENSMUSG00000048915 | 115.296752 | -0.049622904 | 0.242517853 | -0.204615467 | 0.83787257 | 0.971073158 | Efna5 | ephrin A5 [Source:MGI Symbol;Acc:MGI:1 | 119.66 | 107.93 | 123.77 | 83.866 | 139.6 | 116.95 |
| 14393 | ENSMUSG00000048916 | 5.915803954 | 0.48731613 | 0.887762324 | 0.548926348 | 0.583055996 | 0.908645447 | Gm5540 | predicted pseudogene 5540 [Source:MGI | 5.6217 | 5.1396 | 4.025 | 9.6398 | 3.2719 | 7.7969 |
| 14394 | ENSMUSG00000048919 | 0 | NA | NA | NA | NA | NA | Olfr223 | olfactory receptor 223 [Source:MGI Symb | 0 | 0 | 0 | 0 | 0 | 0 |
| 14395 | ENSMUSG00000048920 | 267.6425164 | -0.093717552 | 0.164430427 | -0.569952615 | 0.568709837 | 0.903527882 | Fkrp | fukutin related protein [Source:MGI Symb | 253.78 | 271.37 | 303.89 | 230.39 | 283.56 | 262.87 |
| 14396 | ENSMUSG00000048921 | 32.52446311 | -0.737535446 | 0.416331695 | -1.771509243 | 0.076476057 | 0.404886416 | Zfp689 | zinc finger protein 689 [Source:MGI Symb | 29.715 | 50.368 | 42.262 | 30.847 | 17.45 | 24.505 |
| 14397 | ENSMUSG00000048922 | 4.87815345 | 1.326358247 | 0.995805776 | 1.331944721 | 0.182878354 | NA | Cdca2 | cell division cycle associated 2 [Source:M( | 3.2124 | 4.1117 | 1.0062 | 7.7118 | 6.5437 | 6.6831 |
| 14398 | ENSMUSG00000048924 | 123.439782 | 0.333552281 | 0.2359354 | 1.413744104 | 0.157437051 | 0.576703416 | Ccdc125 | coiled-coil domain containing 125 [Source | 110.83 | 100.74 | 115.72 | 110.86 | 126.51 | 175.99 |
| 14399 | ENSMUSG00000048930 | 207.9711168 | 0.06419069 | 0.195216728 | 0.328817571 | 0.742293581 | 0.949043964 | Tada3 | transcriptional adaptor 3 [Source:MGI Syr | 188.73 | 176.8 | 244.52 | 198.58 | 193.04 | 246.16 |
| 14400 | ENSMUSG00000048933 | 0 | NA | NA | NA | NA | NA | Olfr722 | olfactory receptor 722 [Source:MGI Symb | 0 | 0 | 0 | 0 | 0 | 0 |
| 14401 | ENSMUSG00000048938 | 33.62609283 | -0.934972773 | 0.383028178 | -2.441002585 | 0.01464655 | 0.140575779 | Nr1h5 | nuclear receptor subfamily 1, group H, me | 40.958 | 51.396 | 40.25 | 25.063 | 21.812 | 22.277 |
| 14402 | ENSMUSG00000048939 | 0.401551119 | -2.072982726 | 4.008251979 | -0.517178744 | 0.605031388 | NA | Atp13a5 | ATPase type 13A5 [Source:MGI Symbol;Ac | 2.4093 | 0 | 0 | 0 | 0 | 0 |
| 14403 | ENSMUSG00000048949 | 4.591000402 | -0.074158755 | 0.993861343 | -0.074616802 | 0.940519606 | NA | Gm6206 | predicted pseudogene 6206 [Source:MGI | 4.0155 | 6.1675 | 4.025 | 6.7478 | 4.3625 | 2.2277 |
| 14404 | ENSMUSG00000048960 | 159.9930966 | -0.12181207 | 0.202025859 | -0.60295286 | 0.546540049 | 0.89549457 | Prex2 | phosphatidylinositol-3,4,5-trisphosphate-( | 153.39 | 146.99 | 200.24 | 162.91 | 150.51 | 145.91 |
| 14405 | ENSMUSG00000048965 | 18.07228443 | 0.051678129 | 0.508508618 | 0.101626851 | 0.919052866 | 0.986374671 | Mrgpre | MAS-related GPR, member E [Source:MGI | 13.653 | 20.558 | 19.119 | 15.424 | 19.631 | 20.049 |
| 14406 | ENSMUSG00000048967 | 7.964410528 | 1.378854495 | 0.77789946 | 1.772535612 | 0.076305689 | 0.404235368 | Yjefn3 | YjeF N-terminal domain containing 3 [Sou | 3.2124 | 5.1396 | 5.0312 | 13.496 | 11.997 | 8.9108 |
| 14407 | ENSMUSG00000048970 | 375.1599078 | 0.073077152 | 0.1481372 | 0.493307229 | 0.621795521 | 0.918563009 | C1galt1c1 | C1GALT1-specific chaperone 1 [Source:M( | 385.49 | 313.51 | 397.47 | 378.84 | 399.17 | 376.48 |
| 14408 | ENSMUSG00000048978 | 0 | NA | NA | NA | NA | NA | Nrsn1 | neurensin 1 [Source:MGI Symbol;Acc:MG | 0 | 0 | 0 | 0 | 0 | 0 |
| 14409 | ENSMUSG00000048981 | 0 | NA | NA | NA | NA | NA | Krt31 | keratin 31 [Source:MGI Symbol;Acc:MGI:1 | 0 | 0 | 0 | 0 | 0 | 0 |
| 14410 | ENSMUSG00000048982 | 0.167707431 | -0.796629176 | 4.080472857 | -0.195229623 | 0.84521319 | NA | Gphb5 | glycoprotein hormone beta 5 [Source:MG | 0 | 0 | 1.0062 | 0 | 0 | 0 |
| 14411 | ENSMUSG00000048988 | 0.603115608 | -2.642822926 | 3.106511347 | -0.850736608 | 0.394915683 | NA | Elfn1 | leucine rich repeat and fibronectin type III | 1.6062 | 0 | 2.0125 | 0 | 0 | 0 |
| 14412 | ENSMUSG00000048992 | 0 | NA | NA | NA | NA | NA | Prss32 | protease, serine 32 [Source:MGI Symbol;A | 0 | 0 | 0 | 0 | 0 | 0 |
| 14413 | ENSMUSG00000048994 | 0 | NA | NA | NA | NA | NA | H2al3 | H2A histone family member L3 [Source:M | 0 | 0 | 0 | 0 | 0 | 0 |
| 14414 | ENSMUSG00000048996 | 0.342432689 | 1.990653086 | 3.900281263 | 0.51038706 | 0.609780321 | NA | Olfr1366 | olfactory receptor 1366 [Source:MGI Syml | 0 | 0 | 0 | 0.964 | 1.0906 | 0 |
| 14415 | ENSMUSG00000048997 | 26.70773458 | -0.375808143 | 0.496315518 | -0.757196037 | 0.448932424 | 0.854431786 | Atxn7l2 | ataxin 7-like 2 [Source:MGI Symbol;Acc:M | 28.912 | 23.642 | 38.237 | 37.595 | 20.722 | 11.138 |
| 14416 | ENSMUSG00000049001 | 0.531246861 | 1.101376188 | 3.36099124 | 0.327693858 | 0.743143147 | NA | Ndnf | neuron-derived neurotrophic factor [Sour | 0 | 0 | 1.0062 | 0 | 2.1812 | 0 |
| 14417 | ENSMUSG00000049008 | 0 | NA | NA | NA | NA | NA | BB014433 | expressed sequence BB014433 [Source:M | 0 | 0 | 0 | 0 | 0 | 0 |
| 14418 | ENSMUSG00000049010 | 0 | NA | NA | NA | NA | NA | Olfr982 | olfactory receptor 982 [Source:MGI Symb | 0 | 0 | 0 | 0 | 0 | 0 |
| 14419 | ENSMUSG00000049011 | 0 | NA | NA | NA | NA | NA | Olfr729 | olfactory receptor 729 [Source:MGI Symb | 0 | 0 | 0 | 0 | 0 | 0 |
| 14420 | ENSMUSG00000049013 | 0.531246861 | 1.101376188 | 3.36099124 | 0.327693858 | 0.743143147 | NA | Prss48 | protease, serine 48 [Source:MGI Symbol;A | 0 | 0 | 1.0062 | 0 | 2.1812 | 0 |
| 14421 | ENSMUSG00000049015 | 0 | NA | NA | NA | NA | NA | Olfr1467 | olfactory receptor 1467 [Source:MGI Syml | 0 | 0 | 0 | 0 | 0 | 0 |
| 14422 | ENSMUSG00000049018 | 0 | NA | NA | NA | NA | NA | Olfr368 | olfactory receptor 368 [Source:MGI Symb | 0 | 0 | 0 | 0 | 0 | 0 |
| 14423 | ENSMUSG00000049028 | 0 | NA | NA | NA | NA | NA | Olfr873 | olfactory receptor 873 [Source:MGI Symb | 0 | 0 | 0 | 0 | 0 | 0 |
| 14424 | ENSMUSG00000049036 | 0 | NA | NA | NA | NA | NA | Tmem121 | transmembrane protein 121 [Source:MGI | 0 | 0 | 0 | 0 | 0 | 0 |
| 14425 | ENSMUSG00000049037 | 20.70528743 | -2.338817039 | 0.679122464 | -3.443881132 | 0.000573428 | 0.013709799 | Clec4a1 | C-type lectin domain family 4, member a1 | 16.865 | 56.535 | 30.187 | 2.8919 | 3.2719 | 14.48 |
| 14426 | ENSMUSG00000049038 | 55.57644156 | 0.1445271 | 0.335063013 | 0.431343045 | 0.666218956 | 0.929596279 | Mterf2 | mitochondrial transcription termination fa | 63.445 | 40.089 | 54.337 | 42.415 | 75.253 | 57.92 |
| 14427 | ENSMUSG00000049041 | 0 | NA | NA | NA | NA | NA | Olfr398 | olfactory receptor 398 [Source:MGI Symb | 0 | 0 | 0 | 0 | 0 | 0 |
| 14428 | ENSMUSG00000049044 | 2924.89751 | 0.416145551 | 0.14861839 | 2.800094594 | 0.005108763 | 0.067544016 | Rapgef4 | Rap guanine nucleotide exchange factor (( | 3014 | 2078.4 | 2424 | 3521.4 | 3717.9 | 2793.5 |
| 14429 | ENSMUSG00000049047 | 90.77268022 | 0.298216 | 0.265252309 | 1.124272964 | 0.260897244 | 0.714835715 | Armcx3 | armadillo repeat containing, X-linked 3 [S( | 84.326 | 75.038 | 84.525 | 80.974 | 88.34 | 131.43 |
| 14430 | ENSMUSG00000049052 | 0 | NA | NA | NA | NA | NA | Olfr812 | olfactory receptor 812 [Source:MGI Symb | 0 | 0 | 0 | 0 | 0 | 0 |
| 14431 | ENSMUSG00000049057 | 0 | NA | NA | NA | NA | NA | Olfr1257 | olfactory receptor 1257 [Source:MGI Syml | 0 | 0 | 0 | 0 | 0 | 0 |
| 14432 | ENSMUSG00000049062 | 0 | NA | NA | NA | NA | NA | Gm9244 | predicted gene 9244 [Source:MGI Symbol | 0 | 0 | 0 | 0 | 0 | 0 |
| 14433 | ENSMUSG00000049073 | 0 | NA | NA | NA | NA | NA | Olfr985 | olfactory receptor 985 [Source:MGI Symb | 0 | 0 | 0 | 0 | 0 | 0 |
| 14434 | ENSMUSG00000049076 | 395.8190199 | -0.043682972 | 0.139221233 | -0.313766592 | 0.753698319 | 0.952989051 | Acap2 | ArfGAP with coiled-coil, ankyrin repeat an | 401.55 | 433.78 | 370.3 | 402.94 | 365.36 | 400.98 |
| 14435 | ENSMUSG00000049086 | 71.70592249 | -0.27811103 | 0.321979295 | -0.863754391 | 0.387722828 | 0.817734468 | Bmyc | brain expressed myelocytomatosis oncoge | 113.24 | 60.647 | 61.381 | 71.334 | 52.35 | 71.286 |
| 14436 | ENSMUSG00000049090 | 1964.813432 | 0.099935893 | 0.119839434 | 0.833914925 | 0.404328926 | 0.829271989 | Zadh2 | zinc binding alcohol dehydrogenase, dom | 1843.9 | 1855.4 | 1990.4 | 1721.7 | 2242.3 | 2135.2 |
| 14437 | ENSMUSG00000049091 | 14148.03539 | 0.102131536 | 0.090014284 | 1.134614772 | 0.256536761 | 0.709880947 | Sephs2 | selenophosphate synthetase 2 [Source:M | 13543 | 13355 | 14044 | 13552 | 14885 | 15510 |
| 14438 | ENSMUSG00000049092 | 16.01531278 | 0.874699188 | 0.606253279 | 1.442794981 | 0.149078233 | 0.561319344 | Gpr137c | G protein-coupled receptor 137C [Source: | 10.44 | 18.503 | 5.0312 | 21.208 | 14.178 | 26.732 |
| 14439 | ENSMUSG00000049093 | 0.133850373 | -0.796629176 | 4.080472857 | -0.195229623 | 0.84521319 | NA | Il23r | interleukin 23 receptor [Source:MGI Symb | 0.8031 | 0 | 0 | 0 | 0 | 0 |
| 14440 | ENSMUSG00000049094 | 0 | NA | NA | NA | NA | NA | Olfr367-ps | olfactory receptor 367, pseudogene [Sour | 0 | 0 | 0 | 0 | 0 | 0 |
| 14441 | ENSMUSG00000049097 | 0.267700746 | -1.497481649 | 4.057001436 | -0.369110456 | 0.712045396 | NA | Ankrd34a | ankyrin repeat domain 34A [Source:MGI S | 1.6062 | 0 | 0 | 0 | 0 | 0 |
| 14442 | ENSMUSG00000049098 | 0 | NA | NA | NA | NA | NA | Olfr147 | olfactory receptor 147 [Source:MGI Symb | 0 | 0 | 0 | 0 | 0 | 0 |
| 14443 | ENSMUSG00000049100 | 0 | NA | NA | NA | NA | NA | Pcdh10 | protocadherin 10 [Source:MGI Symbol;Ac | 0 | 0 | 0 | 0 | 0 | 0 |
| 14444 | ENSMUSG00000049103 | 16.48512283 | -1.159757747 | 0.543839199 | -2.13253798 | 0.032962646 | 0.242879117 | Ccr2 | chemokine (C-C motif) receptor 2 [Source | 26.502 | 21.586 | 20.125 | 8.6758 | 11.997 | 10.025 |

| | | | | | | | | | | | | | | | |
|---|---|---|---|---|---|---|---|---|---|---|---|---|---|---|---|
| 14445 | ENSMUSG00000049106 | 676.3865602 | 0.159532665 | 0.124349741 | 1.282935243 | 0.199514764 | 0.641427328 | Dcaf5 | DDB1 and CUL4 associated factor 5 [Sourc | 659.35 | 631.14 | 626.89 | 781.79 | 719.81 | 639.35 |
| 14446 | ENSMUSG00000049107 | 8.594083641 | -2.629659058 | 0.955686088 | -2.751592903 | 0.00593062 | 0.074656641 | Ntf3 | neurotrophin 3 [Source:MGI Symbol;Acc:I | 12.85 | 24.67 | 7.0437 | 4.8199 | 2.1812 | 0 |
| 14447 | ENSMUSG00000049109 | 117.2896683 | -0.597360889 | 1.21998868 | -0.489644616 | 0.624385399 | 0.919503423 | Themis | thymocyte selection associated [Source:M | 280.28 | 115.13 | 28.175 | 37.595 | 221.4 | 21.163 |
| 14448 | ENSMUSG00000049112 | 0.160662974 | 1.126924175 | 4.080472857 | 0.276174898 | 0.782413736 | NA | Oxtr | oxytocin receptor [Source:MGI Symbol;Ac | 0 | 0 | 0 | 0.964 | 0 | 0 |
| 14449 | ENSMUSG00000049115 | 782.5688108 | -0.086288601 | 0.154023684 | -0.560229434 | 0.575322953 | 0.906703762 | Agtr1a | angiotensin II receptor, type 1a [Source:M | 759.73 | 807.94 | 850.28 | 735.52 | 609.66 | 932.29 |
| 14450 | ENSMUSG00000049119 | 2.779949546 | 2.225543078 | 1.518373531 | 1.465741488 | 0.142718727 | NA | Fam110b | family with sequence similarity 110, meml | 0 | 0 | 3.0187 | 4.8199 | 3.2719 | 5.5692 |
| 14451 | ENSMUSG00000049122 | 0 | NA | NA | NA | NA | NA | Frmd3 | FERM domain containing 3 [Source:MGI S | 0 | 0 | 0 | 0 | 0 | 0 |
| 14452 | ENSMUSG00000049123 | 0 | NA | NA | NA | NA | NA | Catsperg2 | cation channel sperm associated auxiliary | 0 | 0 | 0 | 0 | 0 | 0 |
| 14453 | ENSMUSG00000049124 | 48.49622961 | 0.596154806 | 0.325443019 | 1.831825456 | 0.066977429 | 0.372457339 | Gm8186 | predicted gene 8186 [Source:MGI Symbol | 39.352 | 46.256 | 30.187 | 53.983 | 62.165 | 59.034 |
| 14454 | ENSMUSG00000049128 | 0 | NA | NA | NA | NA | NA | Ivl | involucrin [Source:MGI Symbol;Acc:MGI:9 | 0 | 0 | 0 | 0 | 0 | 0 |
| 14455 | ENSMUSG00000049130 | 11.77192816 | -1.213415133 | 0.749996641 | -1.61789409 | 0.105685424 | 0.479195603 | C5ar1 | complement component 5a receptor 1 [So | 29.715 | 12.335 | 7.0437 | 3.8559 | 6.5437 | 11.138 |
| 14456 | ENSMUSG00000049133 | 0 | NA | NA | NA | NA | NA | Flg2 | filaggrin family member 2 [Source:MGI Sy | 0 | 0 | 0 | 0 | 0 | 0 |
| 14457 | ENSMUSG00000049134 | 1.685790655 | -1.871880402 | 1.891152111 | -0.989809541 | 0.322267221 | NA | Nrap | nebulin-related anchoring protein [Source | 0.8031 | 4.1117 | 3.0187 | 0 | 2.1812 | 0 |
| 14458 | ENSMUSG00000049148 | 1.202488639 | 2.620099056 | 2.125692346 | 1.232586202 | 0.217730187 | NA | Plcxd3 | phosphatidylinositol-specific phospholipa | 0 | 1.0279 | 0 | 2.8919 | 2.1812 | 1.1138 |
| 14459 | ENSMUSG00000049149 | 0 | NA | NA | NA | NA | NA | Olfr1258 | olfactory receptor 1258 [Source:MGI Syml | 0 | 0 | 0 | 0 | 0 | 0 |
| 14460 | ENSMUSG00000049152 | 7811.898343 | 0.04444055 | 0.087258747 | 0.50929622 | 0.610544607 | 0.916376001 | Ugt3a2 | UDP glycosyltransferases 3 family, polype | 7301 | 7776.2 | 7998.6 | 7943.2 | 8204.7 | 7647.7 |
| 14461 | ENSMUSG00000049154 | 0.321325949 | 1.911575114 | 4.048267893 | 0.472195804 | 0.636787034 | NA | Fam183b | family with sequence similarity 183, meml | 0 | 0 | 0 | 1.928 | 0 | 0 |
| 14462 | ENSMUSG00000049160 | 0 | NA | NA | NA | NA | NA | Tex50 | testis expressed 50 [Source:MGI Symbol;A | 0 | 0 | 0 | 0 | 0 | 0 |
| 14463 | ENSMUSG00000049164 | 166.6134392 | -0.06418003 | 0.186138183 | -0.344797767 | 0.730246412 | 0.946534338 | Zfp518a | zinc finger protein 518A [Source:MGI Sym | 167.85 | 182.97 | 159.99 | 150.38 | 162.5 | 175.99 |
| 14464 | ENSMUSG00000049168 | 0 | NA | NA | NA | NA | NA | Olfr449 | olfactory receptor 449 [Source:MGI Symb | 0 | 0 | 0 | 0 | 0 | 0 |
| 14465 | ENSMUSG00000049173 | 0.185640647 | 1.126924175 | 4.080472857 | 0.276174898 | 0.782413736 | NA | Myoz3 | myozenin 3 [Source:MGI Symbol;Acc:MGI | 0 | 0 | 0 | 0 | 0 | 1.1138 |
| 14466 | ENSMUSG00000049176 | 0 | NA | NA | NA | NA | NA | Frmpd4 | FERM and PDZ domain containing 4 [Sour | 0 | 0 | 0 | 0 | 0 | 0 |
| 14467 | ENSMUSG00000049184 | 33.31020421 | 0.226815226 | 0.418058752 | 0.5425439 | 0.587443874 | 0.910765033 | Purg | purine-rich element binding protein G [So | 28.109 | 28.782 | 35.219 | 34.703 | 21.812 | 51.237 |
| 14468 | ENSMUSG00000049191 | 9.300775341 | -0.367309638 | 0.7317651 | -0.5019502 | 0.615702552 | 0.917509899 | Rtl5 | retrotransposon Gag like 5 [Source:MGI S | 8.031 | 14.391 | 9.0562 | 7.7118 | 4.3625 | 12.252 |
| 14469 | ENSMUSG00000049202 | 0 | NA | NA | NA | NA | NA | 4930515G13 | RIKEN cDNA 4930515G13 gene [Source:M | 0 | 0 | 0 | 0 | 0 | 0 |
| 14470 | ENSMUSG00000049214 | 0 | NA | NA | NA | NA | NA | Skint7 | selection and upkeep of intraepithelial T c | 0 | 0 | 0 | 0 | 0 | 0 |
| 14471 | ENSMUSG00000049217 | 0 | NA | NA | NA | NA | NA | Olfr788 | olfactory receptor 788 [Source:MGI Symb | 0 | 0 | 0 | 0 | 0 | 0 |
| 14472 | ENSMUSG00000049225 | 16.5755496 | -1.532001951 | 0.652545316 | -2.347732659 | 0.018888073 | 0.166685225 | Pdp1 | pyruvate dehyrogenase phosphatase cata | 20.078 | 20.558 | 33.206 | 4.8199 | 17.45 | 3.3415 |
| 14473 | ENSMUSG00000049229 | 0 | NA | NA | NA | NA | NA | Olfr844 | olfactory receptor 844 [Source:MGI Symb | 0 | 0 | 0 | 0 | 0 | 0 |
| 14474 | ENSMUSG00000049230 | 8.971675384 | -0.126865535 | 0.746620285 | -0.169919754 | 0.865073245 | 0.977227145 | Gm9833 | predicted gene 9833 [Source:MGI Symbol | 9.6372 | 12.335 | 6.0375 | 4.8199 | 7.6343 | 13.366 |
| 14475 | ENSMUSG00000049231 | 0 | NA | NA | NA | NA | NA | Gm12497 | predicted pseudogene 12497 [Source:MG | 0 | 0 | 0 | 0 | 0 | 0 |
| 14476 | ENSMUSG00000049232 | 237.7902419 | 0.164071669 | 0.167462235 | 0.979753251 | 0.327207933 | 0.772772049 | Tigd2 | tigger transposable element derived 2 [So | 240.13 | 221 | 211.31 | 254.49 | 223.58 | 276.23 |
| 14477 | ENSMUSG00000049233 | 2.423247454 | -0.825512089 | 1.343492214 | -0.614452455 | 0.538916382 | NA | Apoo-ps | apolipoprotein O, pseudogene [Source:M | 3.2124 | 2.0558 | 4.025 | 1.928 | 1.0906 | 2.2277 |
| 14478 | ENSMUSG00000049235 | 1.507478692 | 0.462710215 | 1.764905183 | 0.262172846 | 0.793188187 | NA | Gm7324 | predicted gene 7324 [Source:MGI Symbol | 0.8031 | 0 | 3.0187 | 1.928 | 2.1812 | 1.1138 |
| 14479 | ENSMUSG00000049241 | 1.69760483 | 0.835162576 | 1.854320462 | 0.450387403 | 0.652431127 | NA | Hcar1 | hydrocarboxylic acid receptor 1 [Source:M | 1.6062 | 0 | 2.0125 | 0 | 5.4531 | 1.1138 |
| 14480 | ENSMUSG00000049246 | 0.339027071 | -1.79174149 | 3.919585585 | -0.457125237 | 0.647581032 | NA | Hspe1-ps2 | heat shock protein 1 (chaperonin 10), pse | 0 | 1.0279 | 1.0062 | 0 | 0 | 0 |
| 14481 | ENSMUSG00000049249 | 0 | NA | NA | NA | NA | NA | Gm5316 | predicted gene 5316 [Source:MGI Symbol | 0 | 0 | 0 | 0 | 0 | 0 |
| 14482 | ENSMUSG00000049252 | 0 | NA | NA | NA | NA | NA | Lrp1b | low density lipoprotein-related protein 1B | 0 | 0 | 0 | 0 | 0 | 0 |
| 14483 | ENSMUSG00000049265 | 2.833130022 | -0.284709505 | 1.38392405 | -0.20572625 | 0.837004752 | NA | Kcnk3 | potassium channel, subfamily K, member | 3.2124 | 1.0279 | 5.0312 | 0 | 3.2719 | 4.4554 |
| 14484 | ENSMUSG00000049280 | 0 | NA | NA | NA | NA | NA | Olfr509 | olfactory receptor 509 [Source:MGI Symb | 0 | 0 | 0 | 0 | 0 | 0 |
| 14485 | ENSMUSG00000049281 | 1.179598576 | -3.615568919 | 2.240859027 | -1.61347451 | 0.106641457 | NA | Scn3b | sodium channel, voltage-gated, type III, b | 4.0155 | 2.0558 | 1.0062 | 0 | 0 | 0 |
| 14486 | ENSMUSG00000049285 | 31.9213911 | -0.229818364 | 0.405435402 | -0.566843357 | 0.570820611 | 0.904651508 | Mblac1 | metallo-beta-lactamase domain containin | 39.352 | 33.921 | 30.187 | 39.523 | 21.812 | 26.732 |
| 14487 | ENSMUSG00000049287 | 268.8889247 | 0.04112061 | 0.154970062 | 0.265345512 | 0.790743307 | 0.961019028 | Iba57 | IBA57 homolog, iron-sulfur cluster asseml | 261.81 | 277.54 | 255.59 | 249.67 | 282.47 | 286.26 |
| 14488 | ENSMUSG00000049288 | 100.4992716 | -0.074207784 | 0.260864357 | -0.284468851 | 0.776051102 | 0.95893551 | Lix1l | Lix1-like [Source:MGI Symbol;Acc:MGI:30 | 109.22 | 74.01 | 125.78 | 92.542 | 112.33 | 89.108 |
| 14489 | ENSMUSG00000049291 | 0 | NA | NA | NA | NA | NA | Prss38 | protease, serine 38 [Source:MGI Symbol;A | 0 | 0 | 0 | 0 | 0 | 0 |
| 14490 | ENSMUSG00000049295 | 447.5205223 | 0.015148675 | 0.151757386 | 0.099821666 | 0.920485907 | 0.986685804 | Zfp219 | zinc finger protein 219 [Source:MGI Symb | 409.58 | 432.75 | 494.07 | 500.3 | 381.72 | 466.7 |
| 14491 | ENSMUSG00000049299 | 406.0734377 | 0.105835392 | 0.160554779 | 0.659185562 | 0.509776617 | 0.882793133 | Trappc1 | trafficking protein particle complex 1 [Sou | 465.8 | 329.96 | 376.34 | 393.3 | 417.71 | 453.33 |
| 14492 | ENSMUSG00000049300 | 94.59294139 | 0.094739302 | 0.243800706 | 0.388593222 | 0.697577081 | 0.940422865 | Prmt6 | protein arginine N-methyltransferase 6 [S | 108.42 | 78.122 | 87.543 | 105.07 | 97.065 | 91.335 |
| 14493 | ENSMUSG00000049303 | 19.17311208 | -0.275572283 | 0.552933925 | -0.498381941 | 0.618214863 | 0.917509899 | Syt12 | synaptotagmin XII [Source:MGI Symbol;Ac | 26.502 | 9.2513 | 27.169 | 21.208 | 13.087 | 17.822 |
| 14494 | ENSMUSG00000049305 | 354.2718095 | 0.064568931 | 0.139123173 | 0.464113416 | 0.64256649 | 0.924876278 | Ccdc71 | coiled-coil domain containing 71 [Source:I | 365.41 | 337.16 | 336.09 | 369.2 | 344.64 | 373.14 |
| 14495 | ENSMUSG00000049307 | 2.728571924 | -3.746804255 | 1.690180186 | -2.216807584 | 0.026636243 | NA | Fut4 | fucosyltransferase 4 [Source:MGI Symbol; | 4.0155 | 8.2233 | 3.0187 | 0 | 0 | 1.1138 |
| 14496 | ENSMUSG00000049313 | 26.68700458 | -0.96212567 | 0.449672909 | -2.139612261 | 0.032386115 | 0.24029826 | Sorl1 | sortilin-related receptor, LDLR class A repe | 35.336 | 40.089 | 30.187 | 12.532 | 16.359 | 25.618 |
| 14497 | ENSMUSG00000049314 | 0 | NA | NA | NA | NA | NA | Fbxw13 | F-box and WD-40 domain protein 13 [Sou | 0 | 0 | 0 | 0 | 0 | 0 |
| 14498 | ENSMUSG00000049315 | 0 | NA | NA | NA | NA | NA | Olfr348 | olfactory receptor 348 [Source:MGI Symb | 0 | 0 | 0 | 0 | 0 | 0 |
| 14499 | ENSMUSG00000049321 | 101.2595676 | 0.303993899 | 0.22491261 | 1.35160896 | 0.176500443 | 0.607400048 | Zfp2 | zinc finger protein 2 [Source:MGI Symbol; | 90.751 | 88.401 | 92.575 | 102.18 | 116.7 | 116.95 |
| 14500 | ENSMUSG00000049323 | 562.7563686 | 0.086181192 | 0.143380269 | 0.601067304 | 0.547795159 | 0.895600915 | Smcr8 | Smith-Magenis syndrome chromosome re | 605.54 | 553.02 | 477.97 | 511.87 | 595.48 | 632.66 |

| | | | | | | | | | | | | | | | |
|---|---|---|---|---|---|---|---|---|---|---|---|---|---|---|---|
| 14501 | ENSMUSG00000049327 | 586.5643203 | 0.104444479 | 0.121261968 | 0.861312746 | 0.389065817 | 0.818776078 | Kmt5a | lysine methyltransferase 5A [Source:MGI | 596.7 | 551.99 | 546.39 | 574.53 | 626.01 | 623.75 |
| 14502 | ENSMUSG00000049334 | 0 | NA | NA | NA | NA | NA | Olfr902 | olfactory receptor 902 [Source:MGI Symb | 0 | 0 | 0 | 0 | 0 | 0 |
| 14503 | ENSMUSG00000049336 | 0 | NA | NA | NA | NA | NA | Tenm2 | teneurin transmembrane protein 2 [Sourc | 0 | 0 | 0 | 0 | 0 | 0 |
| 14504 | ENSMUSG00000049339 | 3648.095347 | 0.163400748 | 0.102471042 | 1.594604149 | 0.110800784 | 0.489424955 | Retreg2 | reticulophagy regulator family member 2 | 3089.5 | 3495.9 | 3741.2 | 3896.4 | 3713.6 | 3951.9 |
| 14505 | ENSMUSG00000049349 | 0 | NA | NA | NA | NA | NA | Gm5105 | predicted gene 5105 [Source:MGI Symbol | 0 | 0 | 0 | 0 | 0 | 0 |
| 14506 | ENSMUSG00000049350 | 0 | NA | NA | NA | NA | NA | Zg16 | zymogen granule protein 16 [Source:MGI | 0 | 0 | 0 | 0 | 0 | 0 |
| 14507 | ENSMUSG00000049353 | 33.32751886 | 0.972100781 | 0.43290097 | 2.245550018 | 0.024732845 | 0.201358738 | Rd3 | retinal degeneration 3 [Source:MGI Symb | 30.518 | 14.391 | 22.137 | 35.667 | 35.99 | 61.261 |
| 14508 | ENSMUSG00000049354 | 610.9623942 | 0.204379503 | 0.134956987 | 1.514404751 | 0.129923215 | 0.527655829 | Dcaf7 | DDB1 and CUL4 associated factor 7 [Sourc | 514.79 | 621.89 | 567.52 | 671.89 | 582.39 | 707.29 |
| 14509 | ENSMUSG00000049357 | 0.51034671 | -2.381743948 | 3.331081356 | -0.715006238 | 0.474605189 | NA | Brd8dc | BRD8 domain containing [Source:MGI Syn | 0 | 2.0558 | 1.0062 | 0 | 0 | 0 |
| 14510 | ENSMUSG00000049362 | 0 | NA | NA | NA | NA | NA | Olfr173 | olfactory receptor 173 [Source:MGI Symb | 0 | 0 | 0 | 0 | 0 | 0 |
| 14511 | ENSMUSG00000049372 | 0 | NA | NA | NA | NA | NA | Olfr1183 | olfactory receptor 1183 [Source:MGI Sym | 0 | 0 | 0 | 0 | 0 | 0 |
| 14512 | ENSMUSG00000049382 | 3612.16434 | 0.039686985 | 0.146004483 | 0.271820316 | 0.785760183 | 0.960775321 | Krt8 | keratin 8 [Source:MGI Symbol;Acc:MGI:96 | 4430.7 | 3384.9 | 2869.8 | 3373.9 | 3952.4 | 3661.2 |
| 14513 | ENSMUSG00000049387 | 0 | NA | NA | NA | NA | NA | Cox7b2 | cytochrome c oxidase subunit 7B2 [Source | 0 | 0 | 0 | 0 | 0 | 0 |
| 14514 | ENSMUSG00000049396 | 1.939810683 | 1.007264572 | 1.698777145 | 0.592935086 | 0.553224596 | NA | Gemin4 | gem nuclear organelle associated protein | 0.8031 | 2.0558 | 1.0062 | 0 | 1.0906 | 6.6831 |
| 14515 | ENSMUSG00000049401 | 845.0080001 | 0.134335496 | 0.133278936 | 1.007927433 | 0.313489301 | 0.760152431 | Ogfr | opioid growth factor receptor [Source:MG | 840.85 | 788.41 | 786.88 | 809.74 | 801.6 | 1042.6 |
| 14516 | ENSMUSG00000049404 | 1636.338394 | 0.168473416 | 0.338568013 | 0.497605827 | 0.618761896 | 0.917509899 | Rarres1 | retinoic acid receptor responder (tazarote | 1970.8 | 1423.7 | 1227.6 | 1166.4 | 2266.3 | 1763.2 |
| 14517 | ENSMUSG00000049409 | 0.631240176 | 0.380643921 | 3.15447101 | 0.120668068 | 0.903953952 | NA | Prokr1 | prokineticin receptor 1 [Source:MGI Symb | 1.6062 | 0 | 0 | 0 | 2.1812 | 0 |
| 14518 | ENSMUSG00000049410 | 3.028243195 | -1.204484202 | 1.375622571 | -0.875592061 | 0.381251843 | NA | Zfp683 | zinc finger protein 683 [Source:MGI Symb | 2.4093 | 7.1954 | 3.0187 | 0 | 1.0906 | 4.4554 |
| 14519 | ENSMUSG00000049411 | 115.0182531 | 0.031532902 | 0.245295068 | 0.128550902 | 0.897713019 | 0.982798755 | Tmem241 | transmembrane protein 241 [Source:MGI | 134.92 | 120.27 | 85.531 | 109.89 | 103.61 | 135.89 |
| 14520 | ENSMUSG00000049414 | 0.637152627 | -1.330052897 | 2.84179579 | -0.468032538 | 0.639761321 | NA | Gm5417 | predicted gene 5417 [Source:MGI Symbol | 0.8031 | 2.0558 | 0 | 0.964 | 0 | 0 |
| 14521 | ENSMUSG00000049420 | 0 | NA | NA | NA | NA | NA | Tmem200a | transmembrane protein 200A [Source:MG | 0 | 0 | 0 | 0 | 0 | 0 |
| 14522 | ENSMUSG00000049421 | 421.345454 | 0.121575056 | 0.138463148 | 0.878031867 | 0.379926425 | 0.812628154 | Zfp260 | zinc finger protein 260 [Source:MGI Symb | 405.57 | 374.16 | 430.67 | 407.76 | 475.51 | 434.4 |
| 14523 | ENSMUSG00000049422 | 4403.440278 | -0.245071645 | 0.087946186 | -2.786609119 | 0.005326268 | 0.069529913 | Chchd10 | coiled-coil-helix-coiled-coil-helix domain c | 4662.8 | 4814.8 | 4853.1 | 4199.1 | 4061.5 | 3829.4 |
| 14524 | ENSMUSG00000049427 | 2.533173397 | -0.514817222 | 1.443006963 | -0.35676697 | 0.721266265 | NA | Gm9837 | predicted gene 9837 [Source:MGI Symbol | 4.8186 | 3.0838 | 1.0062 | 1.928 | 4.3625 | 0 |
| 14525 | ENSMUSG00000049436 | 11.86108698 | 0.386293491 | 0.653289089 | 0.591305591 | 0.554315684 | 0.897428034 | Upk1b | uroplakin 1B [Source:MGI Symbol;Acc:MG | 10.44 | 11.307 | 9.0562 | 9.6398 | 21.812 | 8.9108 |
| 14526 | ENSMUSG00000049439 | 168.9918461 | 0.041536389 | 0.244570352 | 0.169834116 | 0.865140596 | 0.977227145 | Cyp20a1 | cytochrome P450, family 20, subfamily a, | 165.44 | 195.3 | 138.86 | 148.45 | 235.57 | 130.32 |
| 14527 | ENSMUSG00000049456 | 0 | NA | NA | NA | NA | NA | Olfr1404 | olfactory receptor 1404 [Source:MGI Sym | 0 | 0 | 0 | 0 | 0 | 0 |
| 14528 | ENSMUSG00000049460 | 0 | NA | NA | NA | NA | NA | Gm8396 | predicted gene 8396 [Source:MGI Symbol | 0 | 0 | 0 | 0 | 0 | 0 |
| 14529 | ENSMUSG00000049470 | 1530.687417 | 0.095716572 | 0.100091673 | 0.956289062 | 0.338926215 | 0.780858357 | Aff4 | AF4/FMR2 family, member 4 [Source:MGI | 1383.7 | 1520.3 | 1536.5 | 1550.1 | 1565 | 1628.4 |
| 14530 | ENSMUSG00000049476 | 0.572870759 | -2.577107685 | 3.182795569 | -0.809699407 | 0.418112959 | NA | 1700104B16 | RIKEN cDNA 1700104B16 gene [Source:M | 2.4093 | 1.0279 | 0 | 0 | 0 | 0 |
| 14531 | ENSMUSG00000049477 | 2.176394724 | -0.524315902 | 1.424468111 | -0.368078371 | 0.712814798 | NA | Gm8508 | predicted gene 8508 [Source:MGI Symbol | 1.6062 | 3.0838 | 3.0187 | 0.964 | 3.2719 | 1.1138 |
| 14532 | ENSMUSG00000049482 | 181.8695467 | 0.069988709 | 0.18068334 | 0.38735563 | 0.698492947 | 0.940696964 | Ctu2 | cytosolic thiouridylase subunit 2 [Source:I | 181.5 | 160.36 | 190.18 | 173.52 | 196.31 | 189.35 |
| 14533 | ENSMUSG00000049488 | 44.58996046 | -0.292860557 | 0.335291801 | -0.873449802 | 0.382417956 | 0.81414385 | Tmem67 | transmembrane protein 67 [Source:MGI S | 54.611 | 41.117 | 51.318 | 36.631 | 37.081 | 46.781 |
| 14534 | ENSMUSG00000049489 | 131.4024034 | -0.093323498 | 0.215817918 | -0.432417746 | 0.665437823 | 0.929426323 | Ccnq | cyclin Q [Source:MGI Symbol;Acc:MGI:19 | 132.51 | 136.71 | 137.86 | 133.99 | 101.43 | 145.91 |
| 14535 | ENSMUSG00000049491 | 0.167707431 | -0.796629176 | 4.080472857 | -0.195229623 | 0.84521319 | NA | Slc36a3 | solute carrier family 36 (proton/amino aci | 0 | 0 | 1.0062 | 0 | 0 | 0 |
| 14536 | ENSMUSG00000049493 | 19.44977713 | -0.627906679 | 0.599334602 | -1.047672999 | 0.294789292 | 0.745164683 | Pls1 | plastin 1 (I-isoform) [Source:MGI Symbol; | 36.943 | 26.726 | 7.0437 | 17.352 | 15.269 | 13.366 |
| 14537 | ENSMUSG00000049498 | 0 | NA | NA | NA | NA | NA | Olfr1449 | olfactory receptor 1449 [Source:MGI Sym | 0 | 0 | 0 | 0 | 0 | 0 |
| 14538 | ENSMUSG00000049502 | 1148.023259 | 0.205510124 | 0.108736004 | 1.889991505 | 0.058759096 | 0.344306009 | Dtx3l | deltex 3-like, E3 ubiquitin ligase [Source:M | 1158.9 | 1030 | 1009.3 | 1238.7 | 1267.3 | 1184 |
| 14539 | ENSMUSG00000049504 | 168.9238835 | 0.217206458 | 0.183885591 | 1.181204339 | 0.237521558 | 0.686377492 | Proser1 | proline and serine rich 1 [Source:MGI Sym | 156.6 | 156.24 | 155.97 | 187.98 | 194.13 | 162.62 |
| 14540 | ENSMUSG00000049506 | 0 | NA | NA | NA | NA | NA | Sppl2c | signal peptide peptidase 2C [Source:MGI | 0 | 0 | 0 | 0 | 0 | 0 |
| 14541 | ENSMUSG00000049511 | 0 | NA | NA | NA | NA | NA | Htr1b | 5-hydroxytryptamine (serotonin) receptor | 0 | 0 | 0 | 0 | 0 | 0 |
| 14542 | ENSMUSG00000049515 | 2.15347856 | 1.4212158 | 1.514370493 | 0.938486194 | 0.347994609 | NA | Espnl | espin-like [Source:MGI Symbol;Acc:MGI:2 | 2.4093 | 0 | 1.0062 | 2.8919 | 3.2719 | 3.3415 |
| 14543 | ENSMUSG00000049516 | 268.5609912 | -0.217916311 | 0.176263118 | -1.236312586 | 0.216342394 | 0.661699385 | Spty2d1 | SPT2 chromatin protein domain containin | 276.27 | 298.1 | 291.81 | 209.18 | 302.1 | 233.91 |
| 14544 | ENSMUSG00000049517 | 2706.432498 | 0.078062145 | 0.110360763 | 0.707336042 | 0.479357673 | 0.870236354 | Rps23 | ribosomal protein S23 [Source:MGI Symb | 2431 | 2872 | 2597.1 | 2589.2 | 2687.3 | 3062 |
| 14545 | ENSMUSG00000049521 | 564.0864393 | 0.029586806 | 0.154864518 | 0.191049613 | 0.848486724 | 0.972803041 | Cdc42ep1 | CDC42 effector protein (Rho GTPase bindi | 594.3 | 470.79 | 608.78 | 499.34 | 548.58 | 662.74 |
| 14546 | ENSMUSG00000049526 | 1.015219654 | -0.829334329 | 2.257901143 | -0.367303206 | 0.713392863 | NA | Tmem202 | transmembrane protein 202 [Source:MGI | 0.8031 | 3.0838 | 0 | 0 | 1.0906 | 1.1138 |
| 14547 | ENSMUSG00000049528 | 0 | NA | NA | NA | NA | NA | Olfr429 | olfactory receptor 429 [Source:MGI Symb | 0 | 0 | 0 | 0 | 0 | 0 |
| 14548 | ENSMUSG00000049530 | 0 | NA | NA | NA | NA | NA | Clrn2 | clarin 2 [Source:MGI Symbol;Acc:MGI:364 | 0 | 0 | 0 | 0 | 0 | 0 |
| 14549 | ENSMUSG00000049532 | 63.16906846 | 0.268101096 | 0.373914007 | 0.717012713 | 0.473366253 | 0.868952133 | Sall2 | spalt like transcription factor 2 [Source:M | 50.595 | 45.228 | 76.475 | 98.326 | 35.99 | 72.4 |
| 14550 | ENSMUSG00000049536 | 7.647652751 | -1.109901905 | 0.812754328 | -1.365605653 | 0.172062771 | 0.599979731 | Tceal1 | transcription elongation factor A (SII)-like | 16.062 | 7.1954 | 8.05 | 5.7839 | 5.4531 | 3.3415 |
| 14551 | ENSMUSG00000049537 | 0 | NA | NA | NA | NA | NA | Tecrl | trans-2,3-enoyl-CoA reductase-like [Sourc | 0 | 0 | 0 | 0 | 0 | 0 |
| 14552 | ENSMUSG00000049538 | 1.358401886 | 2.862656237 | 2.216651137 | 1.291432914 | 0.196553604 | NA | Adamts16 | a disintegrin-like and metallopeptidase (re | 0.8031 | 0 | 0 | 2.8919 | 0 | 4.4554 |
| 14553 | ENSMUSG00000049539 | 0 | NA | NA | NA | NA | NA | H1f1 | H1.1 linker histone, cluster member [Sour | 0 | 0 | 0 | 0 | 0 | 0 |
| 14554 | ENSMUSG00000049548 | 0 | NA | NA | NA | NA | NA | Krt82 | keratin 82 [Source:MGI Symbol;Acc:MGI:2 | 0 | 0 | 0 | 0 | 0 | 0 |
| 14555 | ENSMUSG00000049550 | 747.2129661 | -0.09174375 | 0.115372844 | -0.795193629 | 0.426500877 | 0.842053132 | Clip1 | CAP-GLY domain containing linker protein | 747.69 | 783.27 | 782.86 | 779.86 | 702.36 | 687.24 |
| 14556 | ENSMUSG00000049551 | 2.337659424 | -0.835309361 | 1.902325338 | -0.439099109 | 0.660589725 | NA | Fzd9 | frizzled class receptor 9 [Source:MGI Sym | 0 | 0 | 9.0562 | 3.8559 | 0 | 1.1138 |

| | | | | | | | | | | | | | | | |
|---|---|---|---|---|---|---|---|---|---|---|---|---|---|---|---|
| 14557 | ENSMUSG00000049553 | 389.6204327 | -0.282051348 | 0.172696914 | -1.633215916 | 0.102423594 | 0.470781813 | Polr1a | polymerase (RNA) I polypeptide A [Source | 429.66 | 449.2 | 403.5 | 300.76 | 446.06 | 308.53 |
| 14558 | ENSMUSG00000049555 | 644.6940048 | -0.452428997 | 0.166904739 | -2.710701925 | 0.006714095 | 0.08160729 | Tmie | transmembrane inner ear [Source:MGI Sy | 617.59 | 749.35 | 868.39 | 517.66 | 460.24 | 654.94 |
| 14559 | ENSMUSG00000049556 | 2.866087168 | 0.664556453 | 1.266817875 | 0.524587209 | 0.59987018 | NA | Lingo1 | leucine rich repeat and Ig domain contain | 1.6062 | 2.0558 | 3.0187 | 3.8559 | 1.0906 | 5.5692 |
| 14560 | ENSMUSG00000049560 | 0 | NA | NA | NA | NA | NA | Defb20 | defensin beta 20 [Source:MGI Symbol;Acc | 0 | 0 | 0 | 0 | 0 | 0 |
| 14561 | ENSMUSG00000049561 | 0 | NA | NA | NA | NA | NA | Olfr95 | olfactory receptor 95 [Source:MGI Symbo | 0 | 0 | 0 | 0 | 0 | 0 |
| 14562 | ENSMUSG00000049562 | 56.01410779 | -0.097945866 | 0.367182585 | -0.266749759 | 0.789661839 | 0.961015468 | Ap5b1 | adaptor-related protein complex 5, beta 1 | 52.202 | 63.731 | 58.362 | 85.794 | 40.353 | 35.643 |
| 14563 | ENSMUSG00000049565 | 0 | NA | NA | NA | NA | NA | Aknad1 | AKNA domain containing 1 [Source:MGI S | 0 | 0 | 0 | 0 | 0 | 0 |
| 14564 | ENSMUSG00000049571 | 0.503095663 | 2.549494177 | 3.348715477 | 0.761334964 | 0.446457022 | NA | Cfap46 | cilia and flagella associated protein 46 [So | 0 | 0 | 0 | 1.928 | 1.0906 | 0 |
| 14565 | ENSMUSG00000049573 | 0 | NA | NA | NA | NA | NA | Olfr780 | olfactory receptor 780 [Source:MGI Symb | 0 | 0 | 0 | 0 | 0 | 0 |
| 14566 | ENSMUSG00000049576 | 0 | NA | NA | NA | NA | NA | Zfa-ps | zinc finger protein, autosomal, pseudoger | 0 | 0 | 0 | 0 | 0 | 0 |
| 14567 | ENSMUSG00000049577 | 789.977604 | 0.320117713 | 0.133434246 | 2.399067123 | 0.016436901 | 0.152107064 | Zfpm1 | zinc finger protein, multitype 1 [Source:M | 648.1 | 650.67 | 810.03 | 907.1 | 797.24 | 926.72 |
| 14568 | ENSMUSG00000049580 | 3072.834502 | -0.80110986 | 0.339620853 | -2.358835902 | 0.01833236 | 0.162774254 | Tsku | tsukushi, small leucine rich proteoglycan [ | 3852.5 | 4200.1 | 3661.7 | 2299.1 | 3111.5 | 1312.1 |
| 14569 | ENSMUSG00000049583 | 0 | NA | NA | NA | NA | NA | Grm5 | glutamate receptor, metabotropic 5 [Sour | 0 | 0 | 0 | 0 | 0 | 0 |
| 14570 | ENSMUSG00000049588 | 2.387560982 | -1.340502459 | 1.456697302 | -0.920234051 | 0.357450464 | NA | Ccdc69 | coiled-coil domain containing 69 [Source:I | 3.2124 | 4.1117 | 3.0187 | 2.8919 | 1.0906 | 0 |
| 14571 | ENSMUSG00000049593 | 0 | NA | NA | NA | NA | NA | Lce1h | late cornified envelope 1H [Source:MGI Sy | 0 | 0 | 0 | 0 | 0 | 0 |
| 14572 | ENSMUSG00000049598 | 0.562214094 | -1.122836648 | 3.276281221 | -0.34271681 | 0.731811515 | NA | Vsig8 | V-set and immunoglobulin domain contai | 2.4093 | 0 | 0 | 0.964 | 0 | 0 |
| 14573 | ENSMUSG00000049600 | 154.1740753 | 0.372089647 | 0.270850466 | 1.37378256 | 0.169509197 | 0.596876778 | Zbtb45 | zinc finger and BTB domain containing 45 | 131.71 | 112.04 | 159.99 | 254.49 | 128.69 | 138.12 |
| 14574 | ENSMUSG00000049604 | 0 | NA | NA | NA | NA | NA | Hoxb13 | homeobox B13 [Source:MGI Symbol;Acc:I | 0 | 0 | 0 | 0 | 0 | 0 |
| 14575 | ENSMUSG00000049605 | 0 | NA | NA | NA | NA | NA | Olfr418 | olfactory receptor 418 [Source:MGI Symb | 0 | 0 | 0 | 0 | 0 | 0 |
| 14576 | ENSMUSG00000049606 | 304.9195736 | -0.001494312 | 0.144656635 | -0.010330059 | 0.991757952 | 0.99859176 | Zfp644 | zinc finger protein 644 [Source:MGI Symb | 301.97 | 318.65 | 294.83 | 313.29 | 294.47 | 306.31 |
| 14577 | ENSMUSG00000049608 | 2.422316826 | 0.810216642 | 1.576951551 | 0.513786642 | 0.607401168 | NA | Gpr55 | G protein-coupled receptor 55 [Source:M | 3.2124 | 0 | 2.0125 | 3.8559 | 5.4531 | 0 |
| 14578 | ENSMUSG00000049612 | 0.17131964 | -0.796629176 | 4.080472857 | -0.195229623 | 0.84521319 | NA | Omg | oligodendrocyte myelin glycoprotein [Sou | 0 | 1.0279 | 0 | 0 | 0 | 0 |
| 14579 | ENSMUSG00000049618 | 0 | NA | NA | NA | NA | NA | Olfr103 | olfactory receptor 103 [Source:MGI Symb | 0 | 0 | 0 | 0 | 0 | 0 |
| 14580 | ENSMUSG00000049620 | 0.133850373 | -0.796629176 | 4.080472857 | -0.195229623 | 0.84521319 | NA | Prss33 | protease, serine 33 [Source:MGI Symbol;/ | 0.8031 | 0 | 0 | 0 | 0 | 0 |
| 14581 | ENSMUSG00000049624 | 719.9301539 | 0.064558024 | 0.116570436 | 0.553811289 | 0.579708002 | 0.908350165 | Slc17a5 | solute carrier family 17 (anion/sugar trans | 681.03 | 673.29 | 757.7 | 758.65 | 741.62 | 707.29 |
| 14582 | ENSMUSG00000049625 | 22.18746882 | -1.183924037 | 0.500284503 | -2.366501523 | 0.0179571 | 0.16022918 | Tifab | TRAF-interacting protein with forkhead-as | 35.336 | 38.033 | 19.119 | 16.388 | 11.997 | 12.252 |
| 14583 | ENSMUSG00000049630 | 6.53587167 | 0.518243977 | 0.824859709 | 0.628281357 | 0.529819642 | 0.888334265 | C1ql3 | C1q-like 3 [Source:MGI Symbol;Acc:MGI:2 | 4.0155 | 5.1396 | 7.0437 | 8.6758 | 6.5437 | 7.7969 |
| 14584 | ENSMUSG00000049641 | 0 | NA | NA | NA | NA | NA | Vgll2 | vestigial like family member 2 [Source:MG | 0 | 0 | 0 | 0 | 0 | 0 |
| 14585 | ENSMUSG00000049643 | 97.67709554 | 0.008766474 | 0.240801832 | 0.036405347 | 0.970959151 | 0.994966572 | 2310022A10 | RIKEN cDNA 2310022A10 gene [Source:M | 108.42 | 97.652 | 85.531 | 82.902 | 107.97 | 103.59 |
| 14586 | ENSMUSG00000049648 | 0 | NA | NA | NA | NA | NA | Olfr273 | olfactory receptor 273 [Source:MGI Symb | 0 | 0 | 0 | 0 | 0 | 0 |
| 14587 | ENSMUSG00000049649 | 0 | NA | NA | NA | NA | NA | Gpr3 | G-protein coupled receptor 3 [Source:MG | 0 | 0 | 0 | 0 | 0 | 0 |
| 14588 | ENSMUSG00000049653 | 0.185640647 | 1.126924175 | 4.080472857 | 0.276174898 | 0.782413736 | NA | Spatc1 | spermatogenesis and centriole associated | 0 | 0 | 0 | 0 | 0 | 1.1138 |
| 14589 | ENSMUSG00000049657 | 246.6570985 | -0.01430405 | 0.170040182 | -0.084121589 | 0.93295976 | 0.988420305 | Zbtb5 | zinc finger and BTB domain containing 5 [! | 212.82 | 247.73 | 283.76 | 242.92 | 243.21 | 249.5 |
| 14590 | ENSMUSG00000049658 | 265.1064481 | 0.128996659 | 0.164222677 | 0.785498455 | 0.432161376 | 0.845264432 | Bdp1 | B double prime 1, subunit of RNA polyme | 218.44 | 276.51 | 265.65 | 273.77 | 292.29 | 263.98 |
| 14591 | ENSMUSG00000049659 | 659.9342915 | 0.047264893 | 0.121161836 | 0.3900972 | 0.696464673 | 0.939992732 | Aftph | aftiphilin [Source:MGI Symbol;Acc:MGI:1 | 607.15 | 709.26 | 631.92 | 693.1 | 656.55 | 661.62 |
| 14592 | ENSMUSG00000049670 | 1.772212504 | -0.637366019 | 1.609081053 | -0.396105602 | 0.692027126 | NA | Morn4 | MORN repeat containing 4 [Source:MGI S | 2.4093 | 2.0558 | 2.0125 | 1.928 | 0 | 2.2277 |
| 14593 | ENSMUSG00000049672 | 236.5509676 | 0.011395024 | 0.185530983 | 0.061418441 | 0.951025966 | 0.990883058 | Zbtb14 | zinc finger and BTB domain containing 14 | 201.58 | 237.45 | 268.67 | 251.6 | 261.75 | 198.26 |
| 14594 | ENSMUSG00000049674 | 0 | NA | NA | NA | NA | NA | Olfr714 | olfactory receptor 714 [Source:MGI Symb | 0 | 0 | 0 | 0 | 0 | 0 |
| 14595 | ENSMUSG00000049676 | 3.350810402 | -1.941155694 | 1.278322142 | -1.518518401 | 0.128883764 | NA | Catsperg1 | cation channel sperm associated auxiliary | 4.8186 | 5.1396 | 6.0375 | 1.928 | 2.1812 | 0 |
| 14596 | ENSMUSG00000049680 | 555.8415761 | -0.11860447 | 0.119675707 | -0.99104883 | 0.321661737 | 0.769010805 | Urgcp | upregulator of cell proliferation [Source:N | 561.37 | 607.5 | 567.52 | 534.04 | 533.31 | 531.3 |
| 14597 | ENSMUSG00000049685 | 10.61111897 | 1.904901866 | 0.754666572 | 2.524163568 | 0.011597395 | 0.119989074 | Cyp2g1 | cytochrome P450, family 2, subfamily g, p | 2.4093 | 3.0838 | 8.05 | 12.532 | 13.087 | 24.505 |
| 14598 | ENSMUSG00000049686 | 210.6855977 | 0.074514412 | 0.195792982 | 0.380577542 | 0.703516748 | 0.942540408 | Orai1 | ORAI calcium release-activated calcium m | 217.64 | 189.14 | 209.3 | 269.91 | 184.31 | 193.81 |
| 14599 | ENSMUSG00000049687 | 1.864954772 | 0.782515057 | 1.765860083 | 0.443135367 | 0.657667825 | NA | Pheta2 | PH domain containing endocytic traffickin | 4.0155 | 0 | 0 | 3.8559 | 1.0906 | 2.2277 |
| 14600 | ENSMUSG00000049690 | 4.174044015 | -1.542832011 | 1.201976801 | -1.283578859 | 0.199289351 | NA | Nckap5 | NCK-associated protein 5 [Source:MGI Syr | 2.4093 | 7.1954 | 9.0562 | 1.928 | 0 | 4.4554 |
| 14601 | ENSMUSG00000049691 | 0 | NA | NA | NA | NA | NA | Nkx3-2 | NK3 homeobox 2 [Source:MGI Symbol;Ac | 0 | 0 | 0 | 0 | 0 | 0 |
| 14602 | ENSMUSG00000049692 | 0 | NA | NA | NA | NA | NA | Tmem239 | transmembrane 239 [Source:MGI Symbol, | 0 | 0 | 0 | 0 | 0 | 0 |
| 14603 | ENSMUSG00000049694 | 14.13974629 | 0.046976656 | 0.606909884 | 0.077403017 | 0.938302941 | 0.988420305 | BC048671 | cDNA sequence BC048671 [Source:MGI S | 14.456 | 11.307 | 16.1 | 23.135 | 9.8156 | 10.025 |
| 14604 | ENSMUSG00000049699 | 0 | NA | NA | NA | NA | NA | Ucn2 | urocortin 2 [Source:MGI Symbol;Acc:MGI | 0 | 0 | 0 | 0 | 0 | 0 |
| 14605 | ENSMUSG00000049708 | 0 | NA | NA | NA | NA | NA | Olfr27 | olfactory receptor 27 [Source:MGI Symbo | 0 | 0 | 0 | 0 | 0 | 0 |
| 14606 | ENSMUSG00000049709 | 1.919634871 | 2.290278755 | 1.767588359 | 1.295708213 | 0.195076031 | NA | Nlrp10 | NLR family, pyrin domain containing 10 [S | 0 | 0 | 2.0125 | 2.8919 | 3.2719 | 3.3415 |
| 14607 | ENSMUSG00000049717 | 113.4147961 | 0.192636949 | 0.23395194 | 0.823403939 | 0.410278324 | 0.833241465 | Lig4 | ligase IV, DNA, ATP-dependent [Source:M | 105.21 | 116.15 | 96.599 | 147.49 | 101.43 | 113.61 |
| 14608 | ENSMUSG00000049719 | 0 | NA | NA | NA | NA | NA | Prss46 | protease, serine 46 [Source:MGI Symbol;/ | 0 | 0 | 0 | 0 | 0 | 0 |
| 14609 | ENSMUSG00000049721 | 42.77457765 | -0.90887164 | 0.557325675 | -1.630772957 | 0.102938236 | 0.47191716 | Gal3st1 | galactose-3-O-sulfotransferase 1 [Source:I | 82.72 | 66.815 | 18.112 | 55.911 | 15.269 | 17.822 |
| 14610 | ENSMUSG00000049723 | 12.0108141 | -1.70372845 | 0.686436924 | -2.481988352 | 0.013065155 | 0.129968636 | Mmp12 | matrix metallopeptidase 12 [Source:MGI S | 26.502 | 13.363 | 15.094 | 3.8559 | 5.4531 | 7.7969 |
| 14611 | ENSMUSG00000049728 | 39.17774649 | 0.005438863 | 0.343378682 | 0.015839258 | 0.987362629 | 0.99859176 | Zfp668 | zinc finger protein 668 [Source:MGI Symb | 40.958 | 39.061 | 37.231 | 39.523 | 37.081 | 41.212 |
| 14612 | ENSMUSG00000049734 | 0 | NA | NA | NA | NA | NA | Trex1 | three prime repair exonuclease 1 [Source: | 0 | 0 | 0 | 0 | 0 | 0 |

| | | | | | | | | | | | | | | | |
|---|---|---|---|---|---|---|---|---|---|---|---|---|---|---|---|
| 14613 | ENSMUSG00000049737 | 0 | NA | NA | NA | NA | NA | Olfr1361 | olfactory receptor 1361 [Source:MGI Syml | 0 | 0 | 0 | 0 | 0 | 0 |
| 14614 | ENSMUSG00000049739 | 272.3355469 | 0.0132722 | 0.164004932 | 0.080925616 | 0.935501108 | 0.988420305 | Zfp646 | zinc finger protein 646 [Source:MGI Symb | 241.73 | 283.71 | 288.79 | 298.83 | 277.02 | 243.93 |
| 14615 | ENSMUSG00000049740 | 0 | NA | NA | NA | NA | NA | 5330417H12 | RIKEN cDNA 5330417H12 gene [Source:M | 0 | 0 | 0 | 0 | 0 | 0 |
| 14616 | ENSMUSG00000049741 | 0 | NA | NA | NA | NA | NA | Rxfp4 | relaxin family peptide receptor 4 [Source: | 0 | 0 | 0 | 0 | 0 | 0 |
| 14617 | ENSMUSG00000049744 | 10.24532343 | -1.291051307 | 0.698137886 | -1.84927839 | 0.064417624 | 0.363533562 | Arhgap15 | Rho GTPase activating protein 15 [Source: | 11.243 | 13.363 | 19.119 | 6.7478 | 6.5437 | 4.4554 |
| 14618 | ENSMUSG00000049751 | 20.81286755 | 0.492595606 | 0.483540031 | 1.018727664 | 0.308332275 | 0.757131228 | Rpl36al | ribosomal protein L36A-like [Source:MGI ! | 14.456 | 19.53 | 18.112 | 30.847 | 18.541 | 23.391 |
| 14619 | ENSMUSG00000049755 | 254.7401531 | -0.034616556 | 0.154984058 | -0.223355595 | 0.823258762 | 0.96943211 | Zfp672 | zinc finger protein 672 [Source:MGI Symb | 259.4 | 263.15 | 250.55 | 236.17 | 256.3 | 262.87 |
| 14620 | ENSMUSG00000049758 | 0 | NA | NA | NA | NA | NA | Olfr1318 | olfactory receptor 1318 [Source:MGI Syml | 0 | 0 | 0 | 0 | 0 | 0 |
| 14621 | ENSMUSG00000049760 | 1251.084503 | 0.027266265 | 0.124709973 | 0.218637403 | 0.826932514 | 0.969748238 | Micos13 | mitochondrial contact site and cristae org | 1112.3 | 1265.4 | 1340.3 | 1106.6 | 1319.6 | 1362.2 |
| 14622 | ENSMUSG00000049761 | 0.506734502 | -2.372847156 | 3.338060886 | -0.710845978 | 0.477179685 | NA | Pmis2 | PMIS2 transmembrane protein [Source:M | 0 | 1.0279 | 2.0125 | 0 | 0 | 0 |
| 14623 | ENSMUSG00000049764 | 125.6090557 | 0.045374849 | 0.238737401 | 0.190061753 | 0.84926074 | 0.973071229 | Zfp280b | zinc finger protein 280B [Source:MGI Sym | 126.09 | 111.02 | 133.83 | 140.74 | 91.612 | 150.37 |
| 14624 | ENSMUSG00000049775 | 1031.729939 | -1.048062872 | 0.354263493 | -2.958427534 | 0.003092129 | 0.047680766 | Tmsb4x | thymosin, beta 4, X chromosome [Source: | 1301.8 | 1897.5 | 973.04 | 592.85 | 550.76 | 874.37 |
| 14625 | ENSMUSG00000049791 | 1203.631376 | 0.498449462 | 0.155383971 | 3.207856387 | 0.001337283 | 0.025963027 | Fzd4 | frizzled class receptor 4 [Source:MGI Syml | 1125.1 | 821.31 | 1046.5 | 1706.2 | 1267.3 | 1255.3 |
| 14626 | ENSMUSG00000049792 | 347.8012548 | 0.076713799 | 0.151154921 | 0.507517708 | 0.611791615 | 0.916420248 | Bag5 | BCL2-associated athanogene 5 [Source:M | 386.29 | 304.26 | 324.01 | 346.07 | 368.63 | 357.54 |
| 14627 | ENSMUSG00000049796 | 0 | NA | NA | NA | NA | NA | Crh | corticotropin releasing hormone [Source:I | 0 | 0 | 0 | 0 | 0 | 0 |
| 14628 | ENSMUSG00000049797 | 0 | NA | NA | NA | NA | NA | Olfr642 | olfactory receptor 642 [Source:MGI Symb | 0 | 0 | 0 | 0 | 0 | 0 |
| 14629 | ENSMUSG00000049799 | 1.838271653 | -0.64556191 | 1.658258399 | -0.389301155 | 0.697053382 | NA | Lrrc19 | leucine rich repeat containing 19 [Source: | 1.6062 | 4.1117 | 1.0062 | 0.964 | 0 | 3.3415 |
| 14630 | ENSMUSG00000049800 | 346.1542296 | 0.163334322 | 0.159565803 | 1.023617338 | 0.306016053 | 0.755069291 | Sertad2 | SERTA domain containing 2 [Source:MGI ! | 340.52 | 328.93 | 309.92 | 363.42 | 308.64 | 425.49 |
| 14631 | ENSMUSG00000049804 | 18.33306696 | -0.917995576 | 0.545359067 | -1.683286536 | 0.092319636 | 0.447351243 | Armcx4 | armadillo repeat containing, X-linked 4 [Sc | 28.912 | 28.782 | 14.087 | 10.604 | 10.906 | 16.708 |
| 14632 | ENSMUSG00000049806 | 0 | NA | NA | NA | NA | NA | Olfr466 | olfactory receptor 466 [Source:MGI Symb | 0 | 0 | 0 | 0 | 0 | 0 |
| 14633 | ENSMUSG00000049807 | 220.5532941 | -0.357456139 | 0.18362461 | -1.946667929 | 0.051574559 | 0.319065193 | Arhgap23 | Rho GTPase activating protein 23 [Source: | 242.54 | 252.87 | 248.54 | 234.25 | 164.68 | 180.44 |
| 14634 | ENSMUSG00000049809 | 0 | NA | NA | NA | NA | NA | Krtap9-3 | keratin associated protein 9-3 [Source:MC | 0 | 0 | 0 | 0 | 0 | 0 |
| 14635 | ENSMUSG00000049811 | 38.97347717 | 0.070081649 | 0.354021967 | 0.197958477 | 0.843077552 | 0.972273952 | Fam161a | family with sequence similarity 161, meml | 36.943 | 33.921 | 43.269 | 40.487 | 45.806 | 33.415 |
| 14636 | ENSMUSG00000049815 | 0 | NA | NA | NA | NA | NA | 4921511C20 | RIKEN cDNA 4921511C20 gene [Source:M | 0 | 0 | 0 | 0 | 0 | 0 |
| 14637 | ENSMUSG00000049823 | 67.5375232 | 0.884919667 | 0.288413262 | 3.068235005 | 0.002153272 | 0.036696139 | Zbtb12 | zinc finger and BTB domain containing 12 | 53.808 | 39.061 | 49.306 | 103.15 | 75.253 | 84.652 |
| 14638 | ENSMUSG00000049832 | 0.305170013 | -1.657836477 | 4.041480335 | -0.410205257 | 0.681655385 | NA | Rbx1-ps | ring-box 1, pseudogene [Source:MGI Sym | 0.8031 | 1.0279 | 0 | 0 | 0 | 0 |
| 14639 | ENSMUSG00000049843 | 0 | NA | NA | NA | NA | NA | Olfr1102 | olfactory receptor 1102 [Source:MGI Syml | 0 | 0 | 0 | 0 | 0 | 0 |
| 14640 | ENSMUSG00000049848 | 0.342639279 | -1.805976596 | 4.028547714 | -0.448294702 | 0.653940524 | NA | Ceacam19 | carcinoembryonic antigen-related cell adh | 0 | 2.0558 | 0 | 0 | 0 | 0 |
| 14641 | ENSMUSG00000049858 | 2346.155366 | -0.088726789 | 0.093815443 | -0.945758885 | 0.344271581 | 0.785808801 | Suox | sulfite oxidase [Source:MGI Symbol;Acc:N | 2413.3 | 2313.8 | 2527.7 | 2272.1 | 2356.8 | 2193.2 |
| 14642 | ENSMUSG00000049864 | 0 | NA | NA | NA | NA | NA | Olfr250 | olfactory receptor 250 [Source:MGI Symb | 0 | 0 | 0 | 0 | 0 | 0 |
| 14643 | ENSMUSG00000049866 | 27.79535091 | -0.576137765 | 0.431843782 | -1.334134678 | 0.182159722 | 0.615722956 | Arl4c | ADP-ribosylation factor-like 4C [Source:M | 38.549 | 27.754 | 33.206 | 15.424 | 23.994 | 27.846 |
| 14644 | ENSMUSG00000049871 | 4.663478194 | -0.476130694 | 1.011563845 | -0.470687734 | 0.637863745 | NA | Nlrc3 | NLR family, CARD domain containing 3 [Sc | 8.031 | 4.1117 | 4.025 | 1.928 | 6.5437 | 3.3415 |
| 14645 | ENSMUSG00000049872 | 5.276966768 | -0.528116948 | 0.980483472 | -0.538629118 | 0.590142792 | 0.911690353 | Calhm5 | calcium homeostasis modulator family me | 6.4248 | 5.1396 | 7.0437 | 0.964 | 7.6343 | 4.4554 |
| 14646 | ENSMUSG00000049878 | 284.1427013 | 0.309069786 | 0.242611138 | 1.273930738 | 0.202687964 | 0.647682949 | Rlf | rearranged L-myc fusion sequence [Sourc | 252.98 | 288.84 | 220.37 | 461.75 | 221.4 | 259.53 |
| 14647 | ENSMUSG00000049881 | 65.97823828 | -0.05760463 | 0.28132751 | -0.204760031 | 0.837759616 | 0.971073158 | 2810025M1! | RIKEN cDNA 2810025M15 gene [Source:N | 65.854 | 62.703 | 73.456 | 71.334 | 52.35 | 70.172 |
| 14648 | ENSMUSG00000049882 | 38.8379839 | -0.120848963 | 0.458864477 | -0.263365262 | 0.792269063 | 0.961434599 | Vcpkmt | valosin containing protein lysine (K) methy | 54.611 | 42.145 | 24.15 | 19.28 | 33.809 | 59.034 |
| 14649 | ENSMUSG00000049891 | 0.724629373 | 1.713042495 | 3.014019035 | 0.56835822 | 0.569791754 | NA | Gm7984 | predicted gene 7984 [Source:MGI Symbol | 0 | 0 | 1.0062 | 0 | 0 | 3.3415 |
| 14650 | ENSMUSG00000049892 | 7.177023654 | -2.258597583 | 0.926625278 | -2.437444386 | 0.014791489 | 0.141505508 | Rasd1 | RAS, dexamethasone-induced 1 [Source:N | 8.031 | 19.53 | 8.05 | 1.928 | 2.1812 | 3.3415 |
| 14651 | ENSMUSG00000049894 | 0 | NA | NA | NA | NA | NA | Olfr808 | olfactory receptor 808 [Source:MGI Symb | 0 | 0 | 0 | 0 | 0 | 0 |
| 14652 | ENSMUSG00000049897 | 19.33888351 | -0.236533627 | 0.527854012 | -0.448104252 | 0.654077959 | 0.925797231 | Stkld1 | serine/threonine kinase-like domain conta | 20.078 | 24.67 | 18.112 | 22.171 | 8.7249 | 22.277 |
| 14653 | ENSMUSG00000049902 | 0 | NA | NA | NA | NA | NA | 4921517D22 | RIKEN cDNA 4921517D22 gene [Source:M | 0 | 0 | 0 | 0 | 0 | 0 |
| 14654 | ENSMUSG00000049904 | 21.85878067 | -0.70047447 | 0.472802045 | -1.481538579 | 0.138463112 | 0.543828246 | Tmem17 | transmembrane protein 17 [Source:MGI S | 28.109 | 27.754 | 25.156 | 11.568 | 19.631 | 18.935 |
| 14655 | ENSMUSG00000049907 | 64.27537748 | -0.225101958 | 0.342075654 | -0.658047291 | 0.510507743 | 0.883146035 | Rasl11b | RAS-like, family 11, member B [Source:MC | 54.611 | 60.647 | 92.575 | 46.271 | 46.897 | 84.652 |
| 14656 | ENSMUSG00000049908 | 0.401551119 | -2.072982726 | 4.008251979 | -0.517178744 | 0.605031388 | NA | Gja8 | gap junction protein, alpha 8 [Source:MGI | 2.4093 | 0 | 0 | 0 | 0 | 0 |
| 14657 | ENSMUSG00000049916 | 0.181769715 | 1.126924175 | 4.080472857 | 0.276174898 | 0.782413736 | NA | 2610318N02 | RIKEN cDNA 2610318N02 gene [Source:M | 0 | 0 | 0 | 0 | 1.0906 | 0 |
| 14658 | ENSMUSG00000049922 | 303.1060529 | 0.114128119 | 0.155495889 | 0.733962287 | 0.462971723 | 0.861054778 | Slc35c1 | solute carrier family 35, member C1 [Sour | 313.21 | 253.9 | 305.9 | 331.61 | 299.92 | 314.1 |
| 14659 | ENSMUSG00000049926 | 0 | NA | NA | NA | NA | NA | Olfr921 | olfactory receptor 921 [Source:MGI Symb | 0 | 0 | 0 | 0 | 0 | 0 |
| 14660 | ENSMUSG00000049928 | 2.889928748 | -0.001656701 | 1.311289693 | -0.001263413 | 0.998991942 | NA | Glp2r | glucagon-like peptide 2 receptor [Source:I | 1.6062 | 1.0279 | 6.0375 | 0.964 | 4.3625 | 3.3415 |
| 14661 | ENSMUSG00000049929 | 1.784472004 | -3.094644138 | 1.879936648 | -1.646142779 | 0.099734367 | NA | Lpar4 | lysophosphatidic acid receptor 4 [Source:I | 1.6062 | 4.1117 | 4.025 | 0.964 | 0 | 0 |
| 14662 | ENSMUSG00000049932 | 153.0727095 | -0.048783327 | 0.224335767 | -0.217456751 | 0.827852408 | 0.969794868 | H2ax | H2A.X variant histone [Source:MGI Symbc | 165.44 | 182.97 | 118.74 | 174.48 | 129.78 | 147.03 |
| 14663 | ENSMUSG00000049939 | 0.305170013 | -1.657836477 | 4.041480335 | -0.410205257 | 0.681655385 | NA | Lrrc4 | leucine rich repeat containing 4 [Source:N | 0.8031 | 1.0279 | 0 | 0 | 0 | 0 |
| 14664 | ENSMUSG00000049940 | 1297.46642 | -0.082548993 | 0.119985576 | -0.687990972 | 0.491458466 | 0.876368691 | Pgrmc2 | progesterone receptor membrane compo | 1476.9 | 1357.9 | 1168.2 | 1326.4 | 1183.3 | 1272 |
| 14665 | ENSMUSG00000049946 | 0 | NA | NA | NA | NA | NA | BC030500 | cDNA sequence BC030500 [Source:MGI Sy | 0 | 0 | 0 | 0 | 0 | 0 |
| 14666 | ENSMUSG00000049950 | 48.98451084 | 0.204943518 | 0.325576772 | 0.629478316 | 0.52903596 | 0.888334265 | Rpp38 | ribonuclease P/MRP 38 subunit [Source:N | 53.808 | 38.033 | 44.275 | 44.343 | 59.984 | 53.465 |
| 14667 | ENSMUSG00000049957 | 186.0469761 | 0.048457558 | 0.190539092 | 0.254318195 | 0.799249744 | 0.962424862 | Ccdc137 | coiled-coil domain containing 137 [Source | 157.41 | 212.78 | 179.11 | 179.3 | 206.13 | 181.56 |
| 14668 | ENSMUSG00000049960 | 478.6815031 | 0.053480985 | 0.130296554 | 0.410455868 | 0.68147157 | 0.934522544 | Mrps16 | mitochondrial ribosomal protein S16 [Sou | 432.87 | 483.12 | 494.07 | 477.17 | 512.59 | 472.27 |

| | | | | | | | | | | | | | | | |
|---|---|---|---|---|---|---|---|---|---|---|---|---|---|---|---|
| 14669 | ENSMUSG00000049969 | 278.5671754 | 0.120922473 | 0.15013528 | 0.805423438 | 0.420575371 | 0.837906635 | Plekhf2 | pleckstrin homology domain containing, fa | 270.65 | 265.2 | 264.64 | 285.34 | 275.93 | 309.65 |
| 14670 | ENSMUSG00000049971 | 970.6619868 | -0.445966874 | 0.149042832 | -2.992206117 | 0.002769692 | 0.04430139 | Glt1d1 | glycosyltransferase 1 domain containing 1 | 1001.5 | 1150.2 | 1208.5 | 996.75 | 780.88 | 686.13 |
| 14671 | ENSMUSG00000049972 | 0 | NA | NA | NA | NA | NA | Skint9 | selection and upkeep of intraepithelial T c | 0 | 0 | 0 | 0 | 0 | 0 |
| 14672 | ENSMUSG00000049985 | 5.283303717 | 0.398781738 | 1.11790482 | 0.356722443 | 0.721299602 | 0.9447719 | Ankrd55 | ankyrin repeat domain 55 [Source:MGI Sy | 5.6217 | 2.0558 | 6.0375 | 12.532 | 5.4531 | 0 |
| 14673 | ENSMUSG00000049988 | 26.52220692 | -1.300925448 | 0.546745148 | -2.379400077 | 0.017340844 | 0.156761227 | Lrrc25 | leucine rich repeat containing 25 [Source: | 39.352 | 57.563 | 16.1 | 9.6398 | 13.087 | 23.391 |
| 14674 | ENSMUSG00000049999 | 6.579912611 | -0.065163853 | 0.935360195 | -0.069667123 | 0.944458611 | 0.990077979 | Ppp1r3d | protein phosphatase 1, regulatory subunit | 8.8341 | 8.2233 | 3.0187 | 3.8559 | 2.1812 | 13.366 |
| 14675 | ENSMUSG00000050002 | 532.2320516 | 0.06879106 | 0.132460189 | 0.519333847 | 0.603527954 | 0.916106578 | Idnk | idnK gluconokinase homolog (E. coli) [Sou | 524.43 | 491.34 | 543.37 | 608.27 | 516.95 | 509.03 |
| 14676 | ENSMUSG00000050010 | 3.211279672 | -0.388810377 | 1.190665899 | -0.32654868 | 0.744009265 | NA | Shisa3 | shisa family member 3 [Source:MGI Symb | 4.8186 | 2.0558 | 4.025 | 2.8919 | 4.3625 | 1.1138 |
| 14677 | ENSMUSG00000050014 | 3.467056991 | -0.810471567 | 1.17002329 | -0.692696952 | 0.488499752 | NA | Apol10b | apolipoprotein L 10B [Source:MGI Symbol | 4.0155 | 6.1675 | 3.0187 | 0.964 | 2.1812 | 4.4554 |
| 14678 | ENSMUSG00000050015 | 0 | NA | NA | NA | NA | NA | Olfr350 | olfactory receptor 350 [Source:MGI Symb | 0 | 0 | 0 | 0 | 0 | 0 |
| 14679 | ENSMUSG00000050017 | 730.3612369 | 0.074044651 | 0.120973101 | 0.612075333 | 0.540487917 | 0.893324212 | Pitpnb | phosphatidylinositol transfer protein, beta | 643.28 | 740.1 | 752.67 | 764.43 | 772.16 | 709.52 |
| 14680 | ENSMUSG00000050022 | 13.28352997 | -1.286944804 | 0.637999516 | -2.017156397 | 0.043679195 | 0.288482134 | Amz1 | archaelysin family metallopeptidase 1 [So | 12.85 | 24.67 | 19.119 | 8.6758 | 4.3625 | 10.025 |
| 14681 | ENSMUSG00000050023 | 0 | NA | NA | NA | NA | NA | Olfr1176 | olfactory receptor 1176 [Source:MGI Sym | 0 | 0 | 0 | 0 | 0 | 0 |
| 14682 | ENSMUSG00000050028 | 0 | NA | NA | NA | NA | NA | Olfr745 | olfactory receptor 745 [Source:MGI Symb | 0 | 0 | 0 | 0 | 0 | 0 |
| 14683 | ENSMUSG00000050029 | 756.5107223 | 0.010007102 | 0.116269073 | 0.086068477 | 0.931411982 | 0.988420305 | Rap2c | RAP2C, member of RAS oncogene family [ | 789.45 | 727.77 | 743.61 | 736.48 | 723.08 | 818.68 |
| 14684 | ENSMUSG00000050030 | 0 | NA | NA | NA | NA | NA | Olfr728 | olfactory receptor 728 [Source:MGI Symb | 0 | 0 | 0 | 0 | 0 | 0 |
| 14685 | ENSMUSG00000050035 | 0.371281295 | 2.09253362 | 4.031852165 | 0.519000582 | 0.603760335 | NA | Fhl4 | four and a half LIM domains 4 [Source:MC | 0 | 0 | 0 | 0 | 0 | 2.2277 |
| 14686 | ENSMUSG00000050043 | 736.1540461 | 0.054004957 | 0.141299868 | 0.382201046 | 0.702312246 | 0.942345172 | Tmx2 | thioredoxin-related transmembrane prote | 808.72 | 683.57 | 673.18 | 635.26 | 784.15 | 832.04 |
| 14687 | ENSMUSG00000050050 | 15.87336205 | 0.64613024 | 0.545880921 | 1.183646863 | 0.236552881 | 0.685340334 | Ccdc158 | coiled-coil domain containing 158 [Source | 8.8341 | 13.363 | 15.094 | 18.316 | 21.812 | 17.822 |
| 14688 | ENSMUSG00000050052 | 386.0938843 | -0.167394102 | 0.138598032 | -1.207766801 | 0.227136975 | 0.673495122 | Tdrp | testis development related protein [Sourc | 431.27 | 423.5 | 370.3 | 365.35 | 369.72 | 356.43 |
| 14689 | ENSMUSG00000050058 | 0 | NA | NA | NA | NA | NA | Prm3 | protamine 3 [Source:MGI Symbol;Acc:MG | 0 | 0 | 0 | 0 | 0 | 0 |
| 14690 | ENSMUSG00000050063 | 0 | NA | NA | NA | NA | NA | Klk6 | kallikrein related-peptidase 6 [Source:MG | 0 | 0 | 0 | 0 | 0 | 0 |
| 14691 | ENSMUSG00000050064 | 36.3209441 | -0.143087795 | 0.392747418 | -0.364325235 | 0.715615158 | 0.9447719 | Zfp697 | zinc finger protein 697 [Source:MGI Symb | 43.368 | 30.838 | 40.25 | 46.271 | 33.809 | 23.391 |
| 14692 | ENSMUSG00000050066 | 0 | NA | NA | NA | NA | NA | A130023I24I | RIKEN cDNA A130023I24 gene [Source:MC | 0 | 0 | 0 | 0 | 0 | 0 |
| 14693 | ENSMUSG00000050069 | 210.7335336 | 1.094777601 | 0.217997675 | 5.021969169 | 5.11E-07 | 4.54E-05 | Grem2 | gremlin 2, DAN family BMP antagonist [So | 154.2 | 153.16 | 95.593 | 339.32 | 273.75 | 248.39 |
| 14694 | ENSMUSG00000050071 | 3.117890702 | -0.24462165 | 1.320238452 | -0.185285961 | 0.853004749 | NA | Bex1 | brain expressed X-linked 1 [Source:MGI Sy | 4.0155 | 2.0558 | 4.025 | 1.928 | 0 | 6.6831 |
| 14695 | ENSMUSG00000050074 | 0 | NA | NA | NA | NA | NA | Spink8 | serine peptidase inhibitor, Kazal type 8 [Sc | 0 | 0 | 0 | 0 | 0 | 0 |
| 14696 | ENSMUSG00000050075 | 1.782171761 | -1.986261892 | 1.804224848 | -1.100894876 | 0.270942412 | NA | Gpr171 | G protein-coupled receptor 171 [Source:N | 2.4093 | 3.0838 | 3.0187 | 0 | 2.1812 | 0 |
| 14697 | ENSMUSG00000050079 | 385.6224989 | 0.186830304 | 0.14974102 | 1.247689545 | 0.212144772 | 0.656735971 | Rspry1 | ring finger and SPRY domain containing 1 | 340.52 | 385.47 | 356.21 | 388.48 | 371.9 | 471.16 |
| 14698 | ENSMUSG00000050085 | 0 | NA | NA | NA | NA | NA | Olfr622 | olfactory receptor 622 [Source:MGI Symb | 0 | 0 | 0 | 0 | 0 | 0 |
| 14699 | ENSMUSG00000050087 | 0.160662974 | 1.126924175 | 4.080472857 | 0.276174898 | 0.782413736 | NA | Cby3 | chibby family member 3 [Source:MGI Sym | 0 | 0 | 0 | 0.964 | 0 | 0 |
| 14700 | ENSMUSG00000050088 | 675.0104668 | -0.117426707 | 0.12253064 | -0.958345659 | 0.337888487 | 0.780207388 | 1600012H06 | RIKEN cDNA 1600012H06 gene [Source:M | 722.79 | 746.27 | 637.96 | 626.59 | 684.91 | 631.55 |
| 14701 | ENSMUSG00000050089 | 0 | NA | NA | NA | NA | NA | Akap4 | A kinase (PRKA) anchor protein 4 [Source: | 0 | 0 | 0 | 0 | 0 | 0 |
| 14702 | ENSMUSG00000050092 | 0 | NA | NA | NA | NA | NA | Sprr2b | small proline-rich protein 2B [Source:MGI | 0 | 0 | 0 | 0 | 0 | 0 |
| 14703 | ENSMUSG00000050097 | 27.71999865 | 0.916573092 | 0.568808122 | 1.611392415 | 0.10709422 | 0.483009289 | Ces2b | carboxyesterase 2B [Source:MGI Symbol;A | 4.0155 | 32.893 | 21.131 | 28.919 | 39.262 | 40.098 |
| 14704 | ENSMUSG00000050100 | 0 | NA | NA | NA | NA | NA | Hmx2 | H6 homeobox 2 [Source:MGI Symbol;Acc: | 0 | 0 | 0 | 0 | 0 | 0 |
| 14705 | ENSMUSG00000050102 | 0 | NA | NA | NA | NA | NA | Vmn1r235 | vomeronasal 1 receptor 235 [Source:MGI | 0 | 0 | 0 | 0 | 0 | 0 |
| 14706 | ENSMUSG00000050103 | 1449.618357 | -0.412125384 | 0.148416305 | -2.776820143 | 0.005489355 | 0.070827854 | Agmo | alkylglycerol monooxygenase [Source:MG | 1390.2 | 1706.3 | 1870.6 | 1310 | 1404.7 | 1015.8 |
| 14707 | ENSMUSG00000050105 | 1.470712347 | 0.977882427 | 2.157846864 | 0.453175081 | 0.650422675 | NA | Grrp1 | glycine/arginine rich protein 1 [Source:MC | 0 | 1.0279 | 2.0125 | 5.7839 | 0 | 0 |
| 14708 | ENSMUSG00000050106 | 22.41286182 | -1.178013928 | 0.508894948 | -2.314846969 | 0.020621308 | 0.176802967 | Tmc8 | transmembrane channel-like gene family | 38.549 | 24.67 | 30.187 | 21.208 | 8.7249 | 11.138 |
| 14709 | ENSMUSG00000050107 | 6.999382232 | -0.371696845 | 0.918192607 | -0.40481359 | 0.685614539 | 0.935893609 | Haspin | histone H3 associated protein kinase [Sou | 14.456 | 6.1675 | 3.0187 | 10.604 | 2.1812 | 5.5692 |
| 14710 | ENSMUSG00000050108 | 0 | NA | NA | NA | NA | NA | Bpifc | BPI fold containing family C [Source:MGI S | 0 | 0 | 0 | 0 | 0 | 0 |
| 14711 | ENSMUSG00000050114 | 0 | NA | NA | NA | NA | NA | Prdx6b | peroxiredoxin 6B [Source:MGI Symbol;Acc | 0 | 0 | 0 | 0 | 0 | 0 |
| 14712 | ENSMUSG00000050121 | 0 | NA | NA | NA | NA | NA | Opalin | oligodendrocytic myelin paranodal and in | 0 | 0 | 0 | 0 | 0 | 0 |
| 14713 | ENSMUSG00000050122 | 28.2520911 | -1.037705195 | 0.449022573 | -2.31103124 | 0.020831129 | 0.177968964 | Vwa3b | von Willebrand factor A domain containin | 34.533 | 53.452 | 26.162 | 21.208 | 17.45 | 16.708 |
| 14714 | ENSMUSG00000050128 | 0 | NA | NA | NA | NA | NA | Olfr1023 | olfactory receptor 1023 [Source:MGI Sym | 0 | 0 | 0 | 0 | 0 | 0 |
| 14715 | ENSMUSG00000050132 | 7.294707893 | -0.310256538 | 0.877993999 | -0.353369771 | 0.723811245 | 0.944786345 | Sarm1 | sterile alpha and HEAT/Armadillo motif co | 8.8341 | 8.2233 | 7.0437 | 0.964 | 10.906 | 7.7969 |
| 14716 | ENSMUSG00000050134 | 0 | NA | NA | NA | NA | NA | Olfr430 | olfactory receptor 430 [Source:MGI Symb | 0 | 0 | 0 | 0 | 0 | 0 |
| 14717 | ENSMUSG00000050138 | 0 | NA | NA | NA | NA | NA | Kcnk12 | potassium channel, subfamily K, member | 0 | 0 | 0 | 0 | 0 | 0 |
| 14718 | ENSMUSG00000050141 | 0.877291758 | 2.066311913 | 2.565974136 | 0.805273866 | 0.420661659 | NA | Fam205c | family with sequence similarity 205, meml | 0 | 1.0279 | 0 | 0.964 | 3.2719 | 0 |
| 14719 | ENSMUSG00000050144 | 1470.559045 | 0.099840314 | 0.122630317 | 0.814156866 | 0.415555092 | 0.83614452 | Slc25a44 | solute carrier family 25, member 44 [Sour | 1599 | 1376.4 | 1282 | 1415.1 | 1471.2 | 1679.7 |
| 14720 | ENSMUSG00000050147 | 0.465832987 | -0.698295221 | 3.18348759 | -0.219349126 | 0.826378096 | NA | F2rl3 | coagulation factor II (thrombin) receptor-l | 0.8031 | 1.0279 | 0 | 0.964 | 0 | 0 |
| 14721 | ENSMUSG00000050148 | 402.9699278 | -0.088428332 | 0.136901574 | -0.64592634 | 0.51832706 | 0.887543815 | Ubqln2 | ubiquilin 2 [Source:MGI Symbol;Acc:MGI: | 400.75 | 437.89 | 407.53 | 379.81 | 423.16 | 368.68 |
| 14722 | ENSMUSG00000050150 | 0.455176322 | 0.955156269 | 3.546674842 | 0.269310357 | 0.787690862 | NA | Slc9b1 | solute carrier family 9, subfamily B (NHA1 | 0.8031 | 0 | 0 | 1.928 | 0 | 0 |
| 14723 | ENSMUSG00000050157 | 3.109482446 | 0.92291409 | 1.345472625 | 0.685940444 | 0.492750665 | NA | Gm867 | predicted gene 867 [Source:MGI Symbol;A | 2.4093 | 1.0279 | 3.0187 | 6.7478 | 5.4531 | 0 |
| 14724 | ENSMUSG00000050158 | 0 | NA | NA | NA | NA | NA | Olfr165 | olfactory receptor 165 [Source:MGI Symb | 0 | 0 | 0 | 0 | 0 | 0 |

| | | | | | | | | | | | | | | | |
|---|---|---|---|---|---|---|---|---|---|---|---|---|---|---|---|
| 14725 | ENSMUSG00000050164 | 1.818735602 | -0.036198223 | 1.782016974 | -0.020313063 | 0.983793635 | NA | Mchr1 | melanin-concentrating hormone receptor | 2.4093 | 3.0838 | 0 | 0.964 | 0 | 4.4554 |
| 14726 | ENSMUSG00000050174 | 96.00779197 | -0.023439924 | 0.266745021 | -0.087873897 | 0.929976904 | 0.988405116 | Nudt6 | nudix (nucleoside diphosphate linked moi | 122.07 | 86.345 | 81.506 | 100.25 | 111.24 | 74.628 |
| 14727 | ENSMUSG00000050179 | 0 | NA | NA | NA | NA | NA | A930002I21I | RIKEN cDNA A930002I21 gene [Source:MG | 0 | 0 | 0 | 0 | 0 | 0 |
| 14728 | ENSMUSG00000050188 | 175.9588707 | 0.06325406 | 0.194570895 | 0.325095177 | 0.745109039 | 0.949997329 | Lsm10 | U7 snRNP-specific Sm-like protein LSM10 | 179.09 | 156.24 | 181.12 | 207.26 | 151.6 | 180.44 |
| 14729 | ENSMUSG00000050189 | 0 | NA | NA | NA | NA | NA | Etos1 | ectopic ossification 1 [Source:MGI Symbo | 0 | 0 | 0 | 0 | 0 | 0 |
| 14730 | ENSMUSG00000050190 | 0 | NA | NA | NA | NA | NA | Adam34l | a disintegrin and metallopeptidase domai | 0 | 0 | 0 | 0 | 0 | 0 |
| 14731 | ENSMUSG00000050192 | 6.507669466 | -0.879834417 | 0.872846284 | -1.008006144 | 0.313451513 | 0.760152431 | Eif5a2 | eukaryotic translation initiation factor 5A2 | 4.0155 | 11.307 | 10.062 | 4.8199 | 3.2719 | 5.5692 |
| 14732 | ENSMUSG00000050195 | 0 | NA | NA | NA | NA | NA | Scd4 | stearoyl-coenzyme A desaturase 4 [Source | 0 | 0 | 0 | 0 | 0 | 0 |
| 14733 | ENSMUSG00000050197 | 1.106102019 | 3.668747043 | 2.525738057 | 1.452544547 | 0.146350252 | NA | Rhox13 | reproductive homeobox 13 [Source:MGI S | 0 | 0 | 0 | 0 | 2.1812 | 4.4554 |
| 14734 | ENSMUSG00000050198 | 0 | NA | NA | NA | NA | NA | Olfr768 | olfactory receptor 768 [Source:MGI Symb | 0 | 0 | 0 | 0 | 0 | 0 |
| 14735 | ENSMUSG00000050199 | 1255.769367 | 0.286739054 | 0.11787094 | 2.432652658 | 0.014988672 | 0.142348094 | Lgr4 | leucine-rich repeat-containing G protein-c | 1021.5 | 1122.5 | 1250.8 | 1277.3 | 1465.8 | 1396.8 |
| 14736 | ENSMUSG00000050201 | 0.167707431 | -0.796629176 | 4.080472857 | -0.195229623 | 0.84521319 | NA | Otop2 | otopetrin 2 [Source:MGI Symbol;Acc:MGI | 0 | 0 | 1.0062 | 0 | 0 | 0 |
| 14737 | ENSMUSG00000050211 | 0 | NA | NA | NA | NA | NA | Pla2g4e | phospholipase A2, group IVE [Source:MGI | 0 | 0 | 0 | 0 | 0 | 0 |
| 14738 | ENSMUSG00000050212 | 69.68977736 | -0.454261522 | 0.301983738 | -1.504258225 | 0.132514891 | 0.533565316 | Eva1b | eva-1 homolog B (C. elegans) [Source:MG | 93.16 | 76.066 | 72.45 | 70.37 | 39.262 | 66.831 |
| 14739 | ENSMUSG00000050213 | 248.3580745 | 0.083328599 | 0.162032544 | 0.514270757 | 0.607062704 | 0.916376001 | Snip1 | Smad nuclear interacting protein 1 [Sourc | 244.95 | 263.15 | 215.34 | 247.74 | 265.02 | 253.96 |
| 14740 | ENSMUSG00000050215 | 0 | NA | NA | NA | NA | NA | Olfr70 | olfactory receptor 70 [Source:MGI Symbo | 0 | 0 | 0 | 0 | 0 | 0 |
| 14741 | ENSMUSG00000050217 | 0 | NA | NA | NA | NA | NA | Lgsn | lengsin, lens protein with glutamine synth | 0 | 0 | 0 | 0 | 0 | 0 |
| 14742 | ENSMUSG00000050222 | 0.906140364 | 2.113800522 | 2.518615882 | 0.839270703 | 0.401317419 | NA | Il17d | interleukin 17D [Source:MGI Symbol;Acc:I | 0 | 1.0279 | 0 | 0 | 2.1812 | 2.2277 |
| 14743 | ENSMUSG00000050224 | 0 | NA | NA | NA | NA | NA | Krtap13 | keratin associated protein 13 [Source:MG | 0 | 0 | 0 | 0 | 0 | 0 |
| 14744 | ENSMUSG00000050229 | 263.5829423 | -0.116147754 | 0.153285201 | -0.757723204 | 0.448616705 | 0.854238022 | Pigm | phosphatidylinositol glycan anchor biosyn | 282.69 | 262.12 | 277.72 | 265.09 | 241.03 | 252.84 |
| 14745 | ENSMUSG00000050232 | 3.906542691 | -1.711725354 | 1.333741971 | -1.283400681 | 0.199351735 | NA | Cxcr3 | chemokine (C-X-C motif) receptor 3 [Sourc | 5.6217 | 8.2233 | 4.025 | 0 | 0 | 5.5692 |
| 14746 | ENSMUSG00000050234 | 52.28196992 | -1.211290068 | 0.31170821 | -3.885974215 | 0.00010192 | 0.003694676 | Gja4 | gap junction protein, alpha 4 [Source:MGI | 70.673 | 74.01 | 74.462 | 32.775 | 28.356 | 33.415 |
| 14747 | ENSMUSG00000050239 | 0 | NA | NA | NA | NA | NA | Krtap24-1 | keratin associated protein 24-1 [Source:M | 0 | 0 | 0 | 0 | 0 | 0 |
| 14748 | ENSMUSG00000050240 | 37.71053264 | -0.10470881 | 0.384238399 | -0.27251001 | 0.785229896 | 0.960670212 | Hic2 | hypermethylated in cancer 2 [Source:MGI | 48.989 | 35.977 | 32.2 | 45.307 | 38.172 | 25.618 |
| 14749 | ENSMUSG00000050241 | 1.977234217 | -3.272776499 | 1.822488726 | -1.795773248 | 0.072530587 | NA | Klre1 | killer cell lectin-like receptor family E mem | 4.8186 | 2.0558 | 4.025 | 0.964 | 0 | 0 |
| 14750 | ENSMUSG00000050243 | 0 | NA | NA | NA | NA | NA | Gm5446 | predicted gene 5446 [Source:MGI Symbol | 0 | 0 | 0 | 0 | 0 | 0 |
| 14751 | ENSMUSG00000050244 | 409.4601235 | -0.00407291 | 0.168461431 | -0.024177105 | 0.98071134 | 0.99820894 | Heatr1 | HEAT repeat containing 1 [Source:MGI Sy | 387.9 | 457.42 | 384.39 | 323.9 | 426.43 | 476.73 |
| 14752 | ENSMUSG00000050248 | 12.16081712 | 0.000260794 | 0.74714522 | 0.000349054 | 0.999721496 | 0.999897338 | Evc2 | EvC ciliary complex subunit 2 [Source:MGI | 11.243 | 12.335 | 13.081 | 24.099 | 2.1812 | 10.025 |
| 14753 | ENSMUSG00000050251 | 0 | NA | NA | NA | NA | NA | Olfr809 | olfactory receptor 809 [Source:MGI Symb | 0 | 0 | 0 | 0 | 0 | 0 |
| 14754 | ENSMUSG00000050266 | 0 | NA | NA | NA | NA | NA | Olfr690 | olfactory receptor 690 [Source:MGI Symb | 0 | 0 | 0 | 0 | 0 | 0 |
| 14755 | ENSMUSG00000050270 | 441.6028715 | 0.010318738 | 0.13589536 | 0.075931498 | 0.939473597 | 0.988812319 | Tmem220 | transmembrane protein 220 [Source:MGI | 431.27 | 426.59 | 461.87 | 405.83 | 438.43 | 485.64 |
| 14756 | ENSMUSG00000050271 | 7.688230216 | -0.16572011 | 0.902612173 | -0.18360057 | 0.854326816 | 0.973825032 | Prag1 | PEAK1 related kinase activating pseudokir | 3.2124 | 10.279 | 11.069 | 11.568 | 1.0906 | 8.9108 |
| 14757 | ENSMUSG00000050272 | 0.335414862 | -1.779047242 | 4.030804179 | -0.441362855 | 0.658950329 | NA | Dscam | DS cell adhesion molecule [Source:MGI Sy | 0 | 0 | 2.0125 | 0 | 0 | 0 |
| 14758 | ENSMUSG00000050276 | 0 | NA | NA | NA | NA | NA | Mrgprg | MAS-related GPR, member G [Source:MG | 0 | 0 | 0 | 0 | 0 | 0 |
| 14759 | ENSMUSG00000050281 | 0 | NA | NA | NA | NA | NA | Olfr630 | olfactory receptor 630 [Source:MGI Symb | 0 | 0 | 0 | 0 | 0 | 0 |
| 14760 | ENSMUSG00000050288 | 3.340948783 | 1.064672102 | 1.210189017 | 0.87975687 | 0.378991034 | NA | Fzd2 | frizzled class receptor 2 [Source:MGI Syml | 2.4093 | 2.0558 | 2.0125 | 4.8199 | 7.6343 | 1.1138 |
| 14761 | ENSMUSG00000050295 | 3.75062446 | 0.223657702 | 1.239164479 | 0.180490731 | 0.856767331 | NA | Foxc1 | forkhead box C1 [Source:MGI Symbol;Acc | 3.2124 | 3.0838 | 4.025 | 0 | 3.2719 | 8.9108 |
| 14762 | ENSMUSG00000050296 | 0.167707431 | -0.796629176 | 4.080472857 | -0.195229623 | 0.84521319 | NA | Abca12 | ATP-binding cassette, sub-family A (ABC1) | 0 | 0 | 1.0062 | 0 | 0 | 0 |
| 14763 | ENSMUSG00000050299 | 26.95784401 | 0.196541717 | 0.440068969 | 0.446615714 | 0.655152548 | 0.92594552 | Gm9843 | predicted gene 9843 [Source:MGI Symbol | 21.684 | 29.81 | 24.15 | 36.631 | 30.537 | 18.935 |
| 14764 | ENSMUSG00000050304 | 2.297169964 | 1.400521658 | 1.58574082 | 0.883197078 | 0.3771298 | NA | Slc25a2 | solute carrier family 25 (mitochondrial car | 0.8031 | 1.0279 | 2.0125 | 7.7118 | 0 | 2.2277 |
| 14765 | ENSMUSG00000050310 | 373.3852178 | -0.037002199 | 0.157126508 | -0.235493041 | 0.813826079 | 0.966990915 | Rictor | RPTOR independent companion of MTOR, | 380.67 | 377.25 | 377.34 | 438.61 | 349 | 317.45 |
| 14766 | ENSMUSG00000050312 | 265.3525413 | 0.207386462 | 0.176978313 | 1.171818504 | 0.241269932 | 0.691388521 | Nsun3 | NOL1/NOP2/Sun domain family member 3 | 268.24 | 236.42 | 233.45 | 241.96 | 340.27 | 271.78 |
| 14767 | ENSMUSG00000050315 | 4.706859911 | -0.74169646 | 0.976830854 | -0.759288527 | 0.447679979 | NA | Synpo2 | synaptopodin 2 [Source:MGI Symbol;Acc: | 6.4248 | 7.1954 | 4.025 | 2.8919 | 4.3625 | 3.3415 |
| 14768 | ENSMUSG00000050321 | 0.267700746 | -1.497481649 | 4.057001436 | -0.369110456 | 0.712045396 | NA | Neto1 | neuropilin (NRP) and tolloid (TLL)-like 1 [S | 1.6062 | 0 | 0 | 0 | 0 | 0 |
| 14769 | ENSMUSG00000050323 | 299.1230999 | 0.247831854 | 0.147292836 | 1.682579149 | 0.092456591 | 0.447704095 | Ndufaf6 | NADH:ubiquinone oxidoreductase comple | 259.4 | 275.48 | 285.77 | 308.47 | 335.91 | 329.7 |
| 14770 | ENSMUSG00000050328 | 0 | NA | NA | NA | NA | NA | Hoxc12 | homeobox C12 [Source:MGI Symbol;Acc:I | 0 | 0 | 0 | 0 | 0 | 0 |
| 14771 | ENSMUSG00000050332 | 129.6582358 | 0.54048328 | 0.224380788 | 2.408776988 | 0.016006075 | 0.148963156 | Amer1 | APC membrane recruitment 1 [Source:MG | 117.25 | 100.74 | 98.612 | 160.98 | 122.15 | 178.22 |
| 14772 | ENSMUSG00000050334 | 0 | NA | NA | NA | NA | NA | C130071C03 | RIKEN cDNA C130071C03 gene [Source:M | 0 | 0 | 0 | 0 | 0 | 0 |
| 14773 | ENSMUSG00000050335 | 58.25918971 | -1.099468192 | 0.392534129 | -2.800949296 | 0.005095252 | 0.067488765 | Lgals3 | lectin, galactose binding, soluble 3 [Sourc | 93.963 | 95.596 | 48.3 | 22.171 | 32.719 | 56.806 |
| 14774 | ENSMUSG00000050343 | 0 | NA | NA | NA | NA | NA | Olfr1378 | olfactory receptor 1378 [Source:MGI Syml | 0 | 0 | 0 | 0 | 0 | 0 |
| 14775 | ENSMUSG00000050345 | 0 | NA | NA | NA | NA | NA | 4930486L24I | RIKEN cDNA 4930486L24 gene [Source:MG | 0 | 0 | 0 | 0 | 0 | 0 |
| 14776 | ENSMUSG00000050350 | 2.188084632 | -0.599341579 | 1.527906756 | -0.392263191 | 0.694863758 | NA | Gpr18 | G protein-coupled receptor 18 [Source:M | 4.8186 | 1.0279 | 2.0125 | 1.928 | 0 | 3.3415 |
| 14777 | ENSMUSG00000050357 | 5.154528116 | -0.129346453 | 0.919068434 | -0.140736477 | 0.888078127 | 0.982382091 | Carmil2 | capping protein regulator and myosin 1 lir | 4.0155 | 5.1396 | 7.0437 | 4.8199 | 5.4531 | 4.4554 |
| 14778 | ENSMUSG00000050359 | 1.896769784 | -3.201445209 | 1.915170015 | -1.671624547 | 0.094598381 | NA | Sprr1a | small proline-rich protein 1A [Source:MGI | 7.2279 | 2.0558 | 1.0062 | 0 | 1.0906 | 0 |
| 14779 | ENSMUSG00000050366 | 0 | NA | NA | NA | NA | NA | Olfr524 | olfactory receptor 524 [Source:MGI Symb | 0 | 0 | 0 | 0 | 0 | 0 |
| 14780 | ENSMUSG00000050368 | 0 | NA | NA | NA | NA | NA | Hoxd10 | homeobox D10 [Source:MGI Symbol;Acc:I | 0 | 0 | 0 | 0 | 0 | 0 |

| | | | | | | | | | | | | | | | |
|---|---|---|---|---|---|---|---|---|---|---|---|---|---|---|---|
| 14781 | ENSMUSG00000050370 | 0.803102239 | -3.069617558 | 3.178309912 | -0.965801839 | 0.334143348 | NA | Ch25h | cholesterol 25-hydroxylase [Source:MGI S | 4.8186 | 0 | 0 | 0 | 0 | 0 |
| 14782 | ENSMUSG00000050373 | 228.145341 | 0.109067399 | 0.164595818 | 0.662637728 | 0.507562601 | 0.882258527 | Snx21 | sorting nexin family member 21 [Source:N | 206.4 | 232.31 | 220.37 | 247.74 | 218.12 | 243.93 |
| 14783 | ENSMUSG00000050377 | 0 | NA | NA | NA | NA | NA | Il31ra | interleukin 31 receptor A [Source:MGI Syr | 0 | 0 | 0 | 0 | 0 | 0 |
| 14784 | ENSMUSG00000050379 | 14.93413892 | -1.321762401 | 0.612330599 | -2.158576433 | 0.030883043 | 0.233016344 | Septin6 | septin 6 [Source:MGI Symbol;Acc:MGI:18 | 20.078 | 25.698 | 18.112 | 5.7839 | 5.4531 | 14.48 |
| 14785 | ENSMUSG00000050382 | 7.564726538 | -1.110701585 | 0.854128305 | -1.300391965 | 0.193466662 | 0.631833452 | Kif7 | kinesin family member 7 [Source:MGI Sym | 8.8341 | 9.2513 | 13.081 | 8.6758 | 1.0906 | 4.4554 |
| 14786 | ENSMUSG00000050383 | 0 | NA | NA | NA | NA | NA | Svs3b | seminal vesicle secretory protein 3B [Sour | 0 | 0 | 0 | 0 | 0 | 0 |
| 14787 | ENSMUSG00000050390 | 2002.491787 | 0.593114378 | 0.133038604 | 4.458212576 | 8.26E-06 | 0.000455037 | C77080 | expressed sequence C77080 [Source:MGI | 1754.8 | 1586.1 | 1448 | 2524.7 | 1991.5 | 2710 |
| 14788 | ENSMUSG00000050394 | 6.981785615 | -0.042222474 | 0.890058462 | -0.047437866 | 0.96216425 | 0.992956886 | Armcx6 | armadillo repeat containing, X-linked 6 [S | 8.031 | 8.2233 | 5.0312 | 10.604 | 1.0906 | 8.9108 |
| 14789 | ENSMUSG00000050395 | 0.538988726 | 1.126003851 | 3.343747182 | 0.336749099 | 0.736306047 | NA | Tnfsf15 | tumor necrosis factor (ligand) superfamily | 0 | 0 | 1.0062 | 0 | 0 | 2.2277 |
| 14790 | ENSMUSG00000050397 | 0.185640647 | 1.126924175 | 4.080472857 | 0.276174898 | 0.782413736 | NA | Foxl2 | forkhead box L2 [Source:MGI Symbol;Acc: | 0 | 0 | 0 | 0 | 0 | 1.1138 |
| 14791 | ENSMUSG00000050410 | 46.04049409 | 0.246194857 | 0.332525196 | 0.740379557 | 0.45906972 | 0.859322431 | Tcf19 | transcription factor 19 [Source:MGI Symb | 44.974 | 40.089 | 41.256 | 54.947 | 38.172 | 56.806 |
| 14792 | ENSMUSG00000050423 | 22.82672446 | -0.901305028 | 0.596511058 | -1.51096114 | 0.130798352 | 0.529786436 | Ppp1r3g | protein phosphatase 1, regulatory subuni | 39.352 | 34.949 | 15.094 | 29.883 | 6.5437 | 11.138 |
| 14793 | ENSMUSG00000050424 | 0 | NA | NA | NA | NA | NA | Pnma5 | paraneoplastic antigen family 5 [Source:M | 0 | 0 | 0 | 0 | 0 | 0 |
| 14794 | ENSMUSG00000050425 | 0.167707431 | -0.796629176 | 4.080472857 | -0.195229623 | 0.84521319 | NA | Mrgprb2 | MAS-related GPR, member B2 [Source:M | 0 | 0 | 1.0062 | 0 | 0 | 0 |
| 14795 | ENSMUSG00000050428 | 119.0682871 | 0.174444099 | 0.228508733 | 0.76340233 | 0.445223491 | 0.852147532 | Fbxo46 | F-box protein 46 [Source:MGI Symbol;Acc | 97.175 | 107.93 | 130.81 | 122.43 | 110.15 | 145.91 |
| 14796 | ENSMUSG00000050435 | 0 | NA | NA | NA | NA | NA | Ube2dnl1 | ubiquitin-conjugating enzyme E2D N-term | 0 | 0 | 0 | 0 | 0 | 0 |
| 14797 | ENSMUSG00000050439 | 0 | NA | NA | NA | NA | NA | Enthd1 | ENTH domain containing 1 [Source:MGI S | 0 | 0 | 0 | 0 | 0 | 0 |
| 14798 | ENSMUSG00000050440 | 6006.785029 | -0.171616224 | 0.175257721 | -0.979222047 | 0.327470277 | 0.772937193 | Hamp | hepcidin antimicrobial peptide [Source:M | 4429.9 | 6885 | 7777.3 | 5654.7 | 4784.5 | 6509.3 |
| 14799 | ENSMUSG00000050445 | 6478.02031 | -0.547229506 | 0.145345073 | -3.765036499 | 0.000166525 | 0.005412616 | Cyp8b1 | cytochrome P450, family 8, subfamily b, p | 7122.7 | 8349.8 | 7603.2 | 4132.6 | 6478.3 | 5181.6 |
| 14800 | ENSMUSG00000050447 | 1.28116975 | -3.718814509 | 2.156698791 | -1.72430871 | 0.084652128 | NA | Lypd6 | LY6/PLAUR domain containing 6 [Source:N | 1.6062 | 2.0558 | 4.025 | 0 | 0 | 0 |
| 14801 | ENSMUSG00000050463 | 0 | NA | NA | NA | NA | NA | Krt78 | keratin 78 [Source:MGI Symbol;Acc:MGI:1 | 0 | 0 | 0 | 0 | 0 | 0 |
| 14802 | ENSMUSG00000050468 | 0 | NA | NA | NA | NA | NA | Astl | astacin-like metalloendopeptidase (M12 f | 0 | 0 | 0 | 0 | 0 | 0 |
| 14803 | ENSMUSG00000050471 | 101.8774557 | -0.144317888 | 0.22922428 | -0.629592503 | 0.52896123 | 0.888334265 | Fam118b | family with sequence similarity 118, meml | 114.04 | 114.1 | 92.575 | 94.47 | 102.52 | 93.563 |
| 14804 | ENSMUSG00000050473 | 0.185640647 | 1.126924175 | 4.080472857 | 0.276174898 | 0.782413736 | NA | Slc35d3 | solute carrier family 35, member D3 [Sour | 0 | 0 | 0 | 0 | 0 | 1.1138 |
| 14805 | ENSMUSG00000050478 | 0 | NA | NA | NA | NA | NA | Olfr807 | olfactory receptor 807 [Source:MGI Symb | 0 | 0 | 0 | 0 | 0 | 0 |
| 14806 | ENSMUSG00000050490 | 1.772775457 | 3.245735906 | 1.900899444 | 1.707473752 | 0.087734032 | NA | Psma5-ps | proteasome subunit alpha 5, pseudogene | 0 | 1.0279 | 0 | 1.928 | 5.4531 | 2.2277 |
| 14807 | ENSMUSG00000050493 | 55.81266634 | -0.706775717 | 0.343170722 | -2.059545501 | 0.039442011 | 0.271845887 | Fam167b | family with sequence similarity 167, meml | 60.233 | 80.178 | 67.418 | 46.271 | 25.084 | 55.692 |
| 14808 | ENSMUSG00000050503 | 6.592752481 | 0.209780588 | 1.107283165 | 0.189455231 | 0.84973604 | 0.973207825 | Fbxl22 | F-box and leucine-rich repeat protein 22 [ | 11.243 | 0 | 7.0437 | 13.496 | 1.0906 | 6.6831 |
| 14809 | ENSMUSG00000050504 | 0 | NA | NA | NA | NA | NA | Olfr1323 | olfactory receptor 1323 [Source:MGI Syml | 0 | 0 | 0 | 0 | 0 | 0 |
| 14810 | ENSMUSG00000050505 | 0.842329324 | -1.832359871 | 2.605893773 | -0.703159848 | 0.481956138 | NA | Pcdh20 | protocadherin 20 [Source:MGI Symbol;Ac | 0 | 3.0838 | 1.0062 | 0.964 | 0 | 0 |
| 14811 | ENSMUSG00000050511 | 0.167707431 | -0.796629176 | 4.080472857 | -0.195229623 | 0.84521319 | NA | Oprd1 | opioid receptor, delta 1 [Source:MGI Syml | 0 | 0 | 1.0062 | 0 | 0 | 0 |
| 14812 | ENSMUSG00000050520 | 3.867089553 | -0.813041046 | 1.093780953 | -0.743330777 | 0.457281442 | NA | Cldn8 | claudin 8 [Source:MGI Symbol;Acc:MGI:18 | 5.6217 | 4.1117 | 5.0312 | 2.8919 | 1.0906 | 4.4554 |
| 14813 | ENSMUSG00000050526 | 0 | NA | NA | NA | NA | NA | Mgat4f | MGAT4 family, member F [Source:MGI Sy | 0 | 0 | 0 | 0 | 0 | 0 |
| 14814 | ENSMUSG00000050530 | 42.71938677 | -0.299420821 | 0.365034412 | -0.820253682 | 0.412071505 | 0.834411626 | Fam171a1 | family with sequence similarity 171, meml | 49.792 | 44.2 | 47.293 | 37.595 | 25.084 | 52.351 |
| 14815 | ENSMUSG00000050533 | 0.915562172 | -0.740414916 | 2.315652983 | -0.319743468 | 0.749162805 | NA | Gm9845 | predicted pseudogene 9845 [Source:MGI | 2.4093 | 0 | 1.0062 | 0.964 | 0 | 1.1138 |
| 14816 | ENSMUSG00000050534 | 0 | NA | NA | NA | NA | NA | Htr5b | 5-hydroxytryptamine (serotonin) receptor | 0 | 0 | 0 | 0 | 0 | 0 |
| 14817 | ENSMUSG00000050538 | 2.518186226 | 1.096336636 | 1.396547909 | 0.785033316 | 0.432434035 | NA | B230217C12 | RIKEN cDNA B230217C12 gene [Source:M | 0.8031 | 3.0838 | 1.0062 | 5.7839 | 1.0906 | 3.3415 |
| 14818 | ENSMUSG00000050541 | 1274.331597 | -0.018711662 | 0.14490388 | -0.129131546 | 0.897253561 | 0.982798755 | Adra1b | adrenergic receptor, alpha 1b [Source:MG | 1081.8 | 1269.5 | 1498.3 | 1458.5 | 1204 | 1133.9 |
| 14819 | ENSMUSG00000050545 | 32.99123406 | -0.004830446 | 0.501845608 | -0.009625362 | 0.992320191 | 0.99859176 | Fam228b | family with sequence similarity 228, meml | 44.974 | 39.061 | 15.094 | 47.235 | 35.99 | 15.594 |
| 14820 | ENSMUSG00000050549 | 88.4277667 | 0.0898536 | 0.253254232 | 0.354796047 | 0.72274239 | 0.9447719 | Fam241a | family with sequence similarity 241, meml | 96.372 | 66.815 | 93.581 | 96.398 | 90.521 | 86.88 |
| 14821 | ENSMUSG00000050550 | 0.342432689 | 1.990653086 | 3.900281263 | 0.51038706 | 0.609780321 | NA | Gm11868 | predicted gene 11868 [Source:MGI Symbo | 0 | 0 | 0 | 0.964 | 1.0906 | 0 |
| 14822 | ENSMUSG00000050552 | 1046.135896 | -0.065765344 | 0.120988467 | -0.543567045 | 0.586739443 | 0.910448513 | Lamtor4 | late endosomal/lysosomal adaptor, MAPK | 1085 | 1040.3 | 1083.7 | 915.78 | 988.1 | 1164 |
| 14823 | ENSMUSG00000050553 | 0 | NA | NA | NA | NA | NA | Gk2 | glycerol kinase 2 [Source:MGI Symbol;Acc | 0 | 0 | 0 | 0 | 0 | 0 |
| 14824 | ENSMUSG00000050555 | 25.95165433 | 0.251454576 | 0.436876145 | 0.575574059 | 0.564903119 | 0.901542552 | Hyls1 | HYLS1, centriolar and ciliogenesis associat | 22.487 | 19.53 | 29.181 | 33.739 | 21.812 | 28.96 |
| 14825 | ENSMUSG00000050556 | 31.91770101 | -0.32392828 | 0.398789251 | -0.812279366 | 0.416631347 | 0.836878476 | Kcnb1 | potassium voltage gated channel, Shab-re | 30.518 | 29.81 | 46.287 | 30.847 | 25.084 | 28.96 |
| 14826 | ENSMUSG00000050558 | 0 | NA | NA | NA | NA | NA | Prokr2 | prokineticin receptor 2 [Source:MGI Symb | 0 | 0 | 0 | 0 | 0 | 0 |
| 14827 | ENSMUSG00000050565 | 2859.965733 | 0.278101679 | 0.104179413 | 2.669449472 | 0.00759757 | 0.088781756 | Tor1aip2 | torsin A interacting protein 2 [Source:MGI | 2719.3 | 2406.4 | 2628.3 | 2895.8 | 3103.9 | 3406.1 |
| 14828 | ENSMUSG00000050567 | 356.5734617 | 0.254656222 | 0.153929119 | 1.654373283 | 0.098051691 | 0.461810376 | Maml1 | mastermind like transcriptional coactivatc | 296.34 | 344.35 | 335.08 | 352.82 | 367.54 | 443.31 |
| 14829 | ENSMUSG00000050577 | 0 | NA | NA | NA | NA | NA | Olfr1038-ps | olfactory receptor 1038, pseudogene [Sou | 0 | 0 | 0 | 0 | 0 | 0 |
| 14830 | ENSMUSG00000050578 | 3.403064336 | -2.011858507 | 1.335756922 | -1.506156152 | 0.132027093 | NA | Mmp13 | matrix metallopeptidase 13 [Source:MGI S | 7.2279 | 7.1954 | 2.0125 | 2.8919 | 1.0906 | 0 |
| 14831 | ENSMUSG00000050587 | 11.60426608 | -0.9481438 | 0.652911099 | -1.452179021 | 0.146451834 | 0.556855766 | Lrrc4c | leucine rich repeat containing 4C [Source: | 18.471 | 13.363 | 14.087 | 11.568 | 5.4531 | 6.6831 |
| 14832 | ENSMUSG00000050592 | 12.27909622 | -1.239062881 | 0.652082544 | -1.900162628 | 0.057411781 | 0.340608695 | Fam78a | family with sequence similarity 78, memb | 15.259 | 17.475 | 19.119 | 9.6398 | 3.2719 | 8.9108 |
| 14833 | ENSMUSG00000050600 | 1.668296123 | -1.918873223 | 1.820971202 | -1.053763629 | 0.291991149 | NA | Zfp831 | zinc finger protein 831 [Source:MGI Symb | 0.8031 | 5.1396 | 2.0125 | 0.964 | 1.0906 | 0 |
| 14834 | ENSMUSG00000050603 | 0 | NA | NA | NA | NA | NA | Olfr1008 | olfactory receptor 1008 [Source:MGI Syml | 0 | 0 | 0 | 0 | 0 | 0 |
| 14835 | ENSMUSG00000050605 | 5.319836012 | -1.287717634 | 0.98167968 | -1.311749301 | 0.189604736 | 0.626663447 | Zfp61 | zinc finger protein 61 [Source:MGI Symbo | 10.44 | 7.1954 | 5.0312 | 4.8199 | 1.0906 | 3.3415 |
| 14836 | ENSMUSG00000050608 | 1055.747797 | 0.028446485 | 0.112733211 | 0.252334556 | 0.80078248 | 0.962424862 | Micos10 | mitochondrial contact site and cristae org | 1094.6 | 987.83 | 1052.5 | 1018 | 1016.5 | 1165.1 |

| | | | | | | | | | | | | | | | |
|---|---|---|---|---|---|---|---|---|---|---|---|---|---|---|---|
| 14837 | ENSMUSG00000050612 | 0 | NA | NA | NA | NA | NA | Txndc2 | thioredoxin domain containing 2 (spermat | 0 | 0 | 0 | 0 | 0 | 0 |
| 14838 | ENSMUSG00000050613 | 0 | NA | NA | NA | NA | NA | Olfr125 | olfactory receptor 125 [Source:MGI Symb | 0 | 0 | 0 | 0 | 0 | 0 |
| 14839 | ENSMUSG00000050619 | 290.1978865 | 0.091629831 | 0.153160028 | 0.598262041 | 0.5496651 | 0.895703112 | Zscan29 | zinc finger SCAN domains 29 [Source:MGI | 277.07 | 265.2 | 300.87 | 305.58 | 318.46 | 274.01 |
| 14840 | ENSMUSG00000050621 | 11.58556032 | 0.703636125 | 0.681793993 | 1.032036263 | 0.30205513 | 0.752248241 | Rps27rt | ribosomal protein S27, retrogene [Source: | 8.031 | 12.335 | 6.0375 | 6.7478 | 18.541 | 17.822 |
| 14841 | ENSMUSG00000050623 | 3.922957651 | 0.41202443 | 1.108548772 | 0.371679118 | 0.710131774 | NA | Catsperz | cation channel sperm associated auxiliary | 4.0155 | 2.0558 | 4.025 | 5.7839 | 6.5437 | 1.1138 |
| 14842 | ENSMUSG00000050625 | 0 | NA | NA | NA | NA | NA | Ccdc121 | coiled-coil domain containing 121 [Source | 0 | 0 | 0 | 0 | 0 | 0 |
| 14843 | ENSMUSG00000050627 | 924.5177878 | 0.304109791 | 0.110362813 | 2.755545853 | 0.005859429 | 0.074161384 | Gpd1l | glycerol-3-phosphate dehydrogenase 1-lik | 840.85 | 785.33 | 856.31 | 1084.5 | 1006.6 | 973.5 |
| 14844 | ENSMUSG00000050628 | 397.693026 | 0.193201451 | 0.145029629 | 1.332151591 | 0.18281038 | 0.616796332 | Ubald2 | UBA-like domain containing 2 [Source:MG | 361.4 | 371.08 | 381.37 | 463.67 | 366.45 | 442.2 |
| 14845 | ENSMUSG00000050635 | 0 | NA | NA | NA | NA | NA | Sprr2f | small proline-rich protein 2F [Source:MGI | 0 | 0 | 0 | 0 | 0 | 0 |
| 14846 | ENSMUSG00000050640 | 1.093926269 | -2.27368979 | 2.17768229 | -1.04408701 | 0.296445135 | NA | Tmem150c | transmembrane protein 150C [Source:MG | 2.4093 | 1.0279 | 2.0125 | 0 | 0 | 1.1138 |
| 14847 | ENSMUSG00000050641 | 0 | NA | NA | NA | NA | NA | BC048562 | cDNA sequence BC048562 [Source:MGI Sy | 0 | 0 | 0 | 0 | 0 | 0 |
| 14848 | ENSMUSG00000050645 | 0 | NA | NA | NA | NA | NA | Defb19 | defensin beta 19 [Source:MGI Symbol;Acc | 0 | 0 | 0 | 0 | 0 | 0 |
| 14849 | ENSMUSG00000050650 | 0 | NA | NA | NA | NA | NA | Mrgpra1 | MAS-related GPR, member A1 [Source:M( | 0 | 0 | 0 | 0 | 0 | 0 |
| 14850 | ENSMUSG00000050654 | 0 | NA | NA | NA | NA | NA | Olfr215 | olfactory receptor 215 [Source:MGI Symb | 0 | 0 | 0 | 0 | 0 | 0 |
| 14851 | ENSMUSG00000050663 | 66.67287186 | 0.600696037 | 0.300043015 | 2.002033065 | 0.045281175 | 0.294358167 | Trhde | TRH-degrading enzyme [Source:MGI Symb | 57.02 | 44.2 | 57.356 | 59.767 | 98.156 | 83.538 |
| 14852 | ENSMUSG00000050666 | 23.01414418 | -0.576637345 | 0.515432795 | -1.118743996 | 0.26324937 | 0.717837703 | Vstm4 | V-set and transmembrane domain contair | 34.533 | 25.698 | 22.137 | 12.532 | 11.997 | 31.188 |
| 14853 | ENSMUSG00000050668 | 147.8654741 | 0.311355462 | 0.208981367 | 1.489871878 | 0.136257927 | 0.538921153 | Gpatch11 | G patch domain containing 11 [Source:M( | 131.71 | 121.29 | 142.89 | 159.06 | 194.13 | 138.12 |
| 14854 | ENSMUSG00000050671 | 5.15353984 | 0.303678372 | 0.956144079 | 0.317607335 | 0.75078281 | 0.952074518 | Ism2 | isthmin 2 [Source:MGI Symbol;Acc:MGI:2 | 5.6217 | 4.1117 | 4.025 | 2.8919 | 9.8156 | 4.4554 |
| 14855 | ENSMUSG00000050675 | 9.630495131 | -0.236824993 | 0.730516201 | -0.324188557 | 0.745795285 | 0.949997329 | Gp1ba | glycoprotein 1b, alpha polypeptide [Sourc | 12.047 | 9.2513 | 10.062 | 15.424 | 6.5437 | 4.4554 |
| 14856 | ENSMUSG00000050677 | 5.200049578 | -1.183679969 | 0.983179112 | -1.20393116 | 0.228616187 | 0.675041782 | Ccdc96 | coiled-coil domain containing 96 [Source:I | 6.4248 | 6.1675 | 9.0562 | 2.8919 | 1.0906 | 5.5692 |
| 14857 | ENSMUSG00000050685 | 0 | NA | NA | NA | NA | NA | Ccdc54 | coiled-coil domain containing 54 [Source:I | 0 | 0 | 0 | 0 | 0 | 0 |
| 14858 | ENSMUSG00000050697 | 192.7085267 | 0.069133268 | 0.179242641 | 0.385696549 | 0.699721419 | 0.941362191 | Prkaa1 | protein kinase, AMP-activated, alpha 1 ca | 175.08 | 207.64 | 182.13 | 212.08 | 179.95 | 199.38 |
| 14859 | ENSMUSG00000050700 | 0 | NA | NA | NA | NA | NA | Emilin3 | elastin microfibril interfacer 3 [Source:MG | 0 | 0 | 0 | 0 | 0 | 0 |
| 14860 | ENSMUSG00000050702 | 0 | NA | NA | NA | NA | NA | 4930563M21 | RIKEN cDNA 4930563M21 gene [Source:M | 0 | 0 | 0 | 0 | 0 | 0 |
| 14861 | ENSMUSG00000050704 | 0 | NA | NA | NA | NA | NA | 2310061N02 | RIKEN cDNA 2310061N02 gene [Source:M | 0 | 0 | 0 | 0 | 0 | 0 |
| 14862 | ENSMUSG00000050705 | 551.5078012 | 0.085964548 | 0.145859504 | 0.58936542 | 0.555616169 | 0.897628185 | 2310061I04R | RIKEN cDNA 2310061I04 gene [Source:M( | 609.55 | 537.6 | 456.84 | 525.37 | 621.65 | 558.04 |
| 14863 | ENSMUSG00000050708 | 27106.13278 | -0.262738673 | 0.107345637 | -2.447595267 | 0.01438131 | 0.138952083 | Ftl1 | ferritin light polypeptide 1 [Source:MGI Sy | 33879 | 26340 | 28482 | 24741 | 24935 | 24260 |
| 14864 | ENSMUSG00000050711 | 0 | NA | NA | NA | NA | NA | Scg2 | secretogranin II [Source:MGI Symbol;Acc: | 0 | 0 | 0 | 0 | 0 | 0 |
| 14865 | ENSMUSG00000050714 | 94.39186732 | 0.321441803 | 0.250345696 | 1.283991732 | 0.199144848 | 0.641237767 | Zbtb26 | zinc finger and BTB domain containing 26 | 72.279 | 89.429 | 90.562 | 117.61 | 83.978 | 112.5 |
| 14866 | ENSMUSG00000050721 | 49.8792712 | -0.783565687 | 0.331208057 | -2.365780876 | 0.01799209 | 0.160343553 | Plekho2 | pleckstrin homology domain containing, fa | 57.823 | 79.15 | 52.325 | 32.775 | 35.99 | 41.212 |
| 14867 | ENSMUSG00000050730 | 1160.121338 | 0.15702049 | 0.135111743 | 1.162152797 | 0.245173405 | 0.695331005 | Arhgap42 | Rho GTPase activating protein 42 [Source: | 1001.5 | 1067 | 1222.6 | 1026.6 | 1312 | 1331 |
| 14868 | ENSMUSG00000050732 | 1731.701511 | 0.099160225 | 0.102937388 | 0.96330621 | 0.335393878 | 0.776679003 | Vamp8 | vesicle-associated membrane protein 8 [S | 1681.7 | 1594.3 | 1739.8 | 1656.1 | 1817 | 1901.3 |
| 14869 | ENSMUSG00000050737 | 6.20732629 | -0.824699169 | 0.908391816 | -0.907867237 | 0.363948369 | 0.802706498 | Ptges | prostaglandin E synthase [Source:MGI Syr | 5.6217 | 9.2513 | 9.0562 | 6.7478 | 5.4531 | 1.1138 |
| 14870 | ENSMUSG00000050742 | 0.401551119 | -2.072982726 | 4.008251979 | -0.517178744 | 0.605031388 | NA | Olfr164 | olfactory receptor 164 [Source:MGI Symb | 2.4093 | 0 | 0 | 0 | 0 | 0 |
| 14871 | ENSMUSG00000050747 | 3.821774683 | 0.030564381 | 1.057988087 | 0.028889154 | 0.976952996 | NA | Trim15 | tripartite motif-containing 15 [Source:MG | 3.2124 | 5.1396 | 3.0187 | 3.8559 | 4.3625 | 3.3415 |
| 14872 | ENSMUSG00000050751 | 0.185640647 | 1.126924175 | 4.080472857 | 0.276174898 | 0.782413736 | NA | Pgbd5 | piggyBac transposable element derived 5 | 0 | 0 | 0 | 0 | 0 | 1.1138 |
| 14873 | ENSMUSG00000050756 | 0 | NA | NA | NA | NA | NA | Defb6 | defensin beta 6 [Source:MGI Symbol;Acc: | 0 | 0 | 0 | 0 | 0 | 0 |
| 14874 | ENSMUSG00000050761 | 0 | NA | NA | NA | NA | NA | Gp1bb | glycoprotein Ib, beta polypeptide [Source | 0 | 0 | 0 | 0 | 0 | 0 |
| 14875 | ENSMUSG00000050762 | 0 | NA | NA | NA | NA | NA | Prss27 | protease, serine 27 [Source:MGI Symbol;A | 0 | 0 | 0 | 0 | 0 | 0 |
| 14876 | ENSMUSG00000050763 | 0 | NA | NA | NA | NA | NA | Olfr1395 | olfactory receptor 1395 [Source:MGI Sym | 0 | 0 | 0 | 0 | 0 | 0 |
| 14877 | ENSMUSG00000050765 | 0 | NA | NA | NA | NA | NA | Gm5084 | predicted gene 5084 [Source:MGI Symbol | 0 | 0 | 0 | 0 | 0 | 0 |
| 14878 | ENSMUSG00000050766 | 0 | NA | NA | NA | NA | NA | Ear14 | eosinophil-associated, ribonuclease A fam | 0 | 0 | 0 | 0 | 0 | 0 |
| 14879 | ENSMUSG00000050772 | 0 | NA | NA | NA | NA | NA | Olfr1124 | olfactory receptor 1124 [Source:MGI Sym | 0 | 0 | 0 | 0 | 0 | 0 |
| 14880 | ENSMUSG00000050776 | 0 | NA | NA | NA | NA | NA | Olfr1317 | olfactory receptor 1317 [Source:MGI Sym | 0 | 0 | 0 | 0 | 0 | 0 |
| 14881 | ENSMUSG00000050777 | 1285.215605 | -0.402823939 | 0.103328366 | -3.898483583 | 9.68E-05 | 0.003535533 | Tmem37 | transmembrane protein 37 [Source:MGI S | 1517.9 | 1467.9 | 1404.7 | 1145.2 | 1136.4 | 1039.2 |
| 14882 | ENSMUSG00000050783 | 0 | NA | NA | NA | NA | NA | Htr1f | 5-hydroxytryptamine (serotonin) receptor | 0 | 0 | 0 | 0 | 0 | 0 |
| 14883 | ENSMUSG00000050786 | 21.89756847 | -0.005780849 | 0.455241243 | -0.012698429 | 0.989868392 | 0.99859176 | Ccdc126 | coiled-coil domain containing 126 [Source | 20.078 | 22.614 | 23.144 | 19.28 | 23.994 | 22.277 |
| 14884 | ENSMUSG00000050788 | 0 | NA | NA | NA | NA | NA | Olfr419 | olfactory receptor 419 [Source:MGI Symb | 0 | 0 | 0 | 0 | 0 | 0 |
| 14885 | ENSMUSG00000050796 | 73.07444162 | -0.006608995 | 0.28688241 | -0.023037295 | 0.981620523 | 0.998323671 | B3galt6 | UDP-Gal:betaGal beta 1,3-galactosyltransf | 80.31 | 79.15 | 60.375 | 90.614 | 56.712 | 71.286 |
| 14886 | ENSMUSG00000050799 | 0 | NA | NA | NA | NA | NA | H2bc1 | H2B clustered histone 1 [Source:MGI Sym | 0 | 0 | 0 | 0 | 0 | 0 |
| 14887 | ENSMUSG00000050803 | 0 | NA | NA | NA | NA | NA | Olfr866 | olfactory receptor 866 [Source:MGI Symb | 0 | 0 | 0 | 0 | 0 | 0 |
| 14888 | ENSMUSG00000050808 | 0 | NA | NA | NA | NA | NA | Muc15 | mucin 15 [Source:MGI Symbol;Acc:MGI:2 | 0 | 0 | 0 | 0 | 0 | 0 |
| 14889 | ENSMUSG00000050810 | 0 | NA | NA | NA | NA | NA | Oog3 | oogenesin 3 [Source:MGI Symbol;Acc:MG | 0 | 0 | 0 | 0 | 0 | 0 |
| 14890 | ENSMUSG00000050812 | 1654.998454 | 0.131687085 | 0.103308322 | 1.274699681 | 0.20241556 | 0.647300167 | Ecpas | Ecm29 proteasome adaptor and scaffold [ | 1581.3 | 1630.3 | 1526.5 | 1611.8 | 1870.4 | 1709.8 |
| 14891 | ENSMUSG00000050813 | 0 | NA | NA | NA | NA | NA | Olfr332 | olfactory receptor 332 [Source:MGI Symb | 0 | 0 | 0 | 0 | 0 | 0 |
| 14892 | ENSMUSG00000050815 | 0 | NA | NA | NA | NA | NA | Olfr1441 | olfactory receptor 1441 [Source:MGI Sym | 0 | 0 | 0 | 0 | 0 | 0 |

| | | | | | | | | | | | | | | | |
|---|---|---|---|---|---|---|---|---|---|---|---|---|---|---|---|
| 14893 | ENSMUSG00000050818 | 0 | NA | NA | NA | NA | NA | Olfr330 | olfactory receptor 330 [Source:MGI Symb | 0 | 0 | 0 | 0 | 0 | 0 |
| 14894 | ENSMUSG00000050821 | 8.976712899 | -0.190559611 | 0.889635839 | -0.214199567 | 0.830391434 | 0.9702546 | Fam131a | family with sequence similarity 131, meml | 11.243 | 16.447 | 1.0062 | 9.6398 | 3.2719 | 12.252 |
| 14895 | ENSMUSG00000050822 | 0.185640647 | 1.126924175 | 4.080472857 | 0.276174898 | 0.782413736 | NA | Slc29a4 | solute carrier family 29 (nucleoside transp | 0 | 0 | 0 | 0 | 0 | 1.1138 |
| 14896 | ENSMUSG00000050824 | 0 | NA | NA | NA | NA | NA | Sstr5 | somatostatin receptor 5 [Source:MGI Sym | 0 | 0 | 0 | 0 | 0 | 0 |
| 14897 | ENSMUSG00000050830 | 1.297505646 | -0.703686153 | 2.034781063 | -0.345828928 | 0.729471284 | NA | Vwc2 | von Willebrand factor C domain containin | 0.8031 | 3.0838 | 1.0062 | 2.8919 | 0 | 0 |
| 14898 | ENSMUSG00000050840 | 0 | NA | NA | NA | NA | NA | Cdh20 | cadherin 20 [Source:MGI Symbol;Acc:MGI | 0 | 0 | 0 | 0 | 0 | 0 |
| 14899 | ENSMUSG00000050844 | 0.305170013 | -1.657836477 | 4.041480335 | -0.410205257 | 0.681655385 | NA | 1700020N01 | RIKEN cDNA 1700020N01 gene [Source:M | 0.8031 | 1.0279 | 0 | 0 | 0 | 0 |
| 14900 | ENSMUSG00000050846 | 232.4434345 | 0.085947776 | 0.17071383 | 0.503461119 | 0.61464011 | 0.917509899 | Zfp623 | zinc finger protein 623 [Source:MGI Symb | 225.67 | 217.92 | 233.45 | 274.73 | 229.03 | 213.86 |
| 14901 | ENSMUSG00000050853 | 0 | NA | NA | NA | NA | NA | Olfr958 | olfactory receptor 958 [Source:MGI Symb | 0 | 0 | 0 | 0 | 0 | 0 |
| 14902 | ENSMUSG00000050854 | 160.9392228 | 0.373673073 | 0.26084955 | 1.432523357 | 0.151994098 | 0.567340036 | Tmem125 | transmembrane protein 125 [Source:MGI | 126.09 | 100.74 | 194.21 | 217.86 | 138.51 | 188.24 |
| 14903 | ENSMUSG00000050855 | 20.80055515 | 0.150300038 | 0.586957701 | 0.256066217 | 0.797899707 | 0.962377049 | Zfp940 | zinc finger protein 940 [Source:MGI Symb | 5.6217 | 24.67 | 29.181 | 20.244 | 27.265 | 17.822 |
| 14904 | ENSMUSG00000050856 | 940.1649937 | 0.057567633 | 0.12938184 | 0.444943685 | 0.656360452 | 0.92672284 | Atp5k | ATP synthase, H+ transporting, mitochonc | 897.87 | 973.44 | 892.54 | 865.65 | 894.31 | 1117.2 |
| 14905 | ENSMUSG00000050860 | 8.155933735 | -0.342639053 | 0.807927137 | -0.424096478 | 0.671495454 | 0.931191575 | Phospho1 | phosphatase, orphan 1 [Source:MGI Symk | 4.0155 | 11.307 | 12.075 | 3.8559 | 6.5437 | 11.138 |
| 14906 | ENSMUSG00000050865 | 0 | NA | NA | NA | NA | NA | Olfr1494 | olfactory receptor 1494 [Source:MGI Syml | 0 | 0 | 0 | 0 | 0 | 0 |
| 14907 | ENSMUSG00000050866 | 0 | NA | NA | NA | NA | NA | Clrn3 | clarin 3 [Source:MGI Symbol;Acc:MGI:214 | 0 | 0 | 0 | 0 | 0 | 0 |
| 14908 | ENSMUSG00000050870 | 0 | NA | NA | NA | NA | NA | Mrgprb8 | MAS-related GPR, member B8 [Source:M( | 0 | 0 | 0 | 0 | 0 | 0 |
| 14909 | ENSMUSG00000050875 | 0.185640647 | 1.126924175 | 4.080472857 | 0.276174898 | 0.782413736 | NA | Minar2 | membrane integral NOTCH2 associated re | 0 | 0 | 0 | 0 | 0 | 1.1138 |
| 14910 | ENSMUSG00000050876 | 0 | NA | NA | NA | NA | NA | Spata31d1a | spermatogenesis associated 31 subfamily | 0 | 0 | 0 | 0 | 0 | 0 |
| 14911 | ENSMUSG00000050890 | 215.8046995 | 0.055648457 | 0.200248819 | 0.277896554 | 0.781091769 | 0.959993512 | Pdik1l | PDLIM1 interacting kinase 1 like [Source:N | 170.26 | 202.5 | 262.63 | 199.54 | 214.85 | 245.05 |
| 14912 | ENSMUSG00000050891 | 137.2483534 | 0.165835074 | 0.194766542 | 0.851455658 | 0.394516285 | 0.822827574 | Tatdn1 | TatD DNase domain containing 1 [Source: | 126.89 | 128.49 | 132.82 | 146.52 | 143.96 | 144.8 |
| 14913 | ENSMUSG00000050896 | 3.067262923 | -1.385714135 | 1.320194829 | -1.049628512 | 0.293888943 | NA | Rtn4rl2 | reticulon 4 receptor-like 2 [Source:MGI Sy | 7.2279 | 3.0838 | 3.0187 | 2.8919 | 2.1812 | 0 |
| 14914 | ENSMUSG00000050900 | 0.538730002 | 1.11098254 | 3.037664833 | 0.365735722 | 0.714562288 | NA | Gm7327 | predicted gene 7327 [Source:MGI Symbol | 0 | 1.0279 | 0 | 0 | 1.0906 | 1.1138 |
| 14915 | ENSMUSG00000050901 | 0 | NA | NA | NA | NA | NA | Mtnr1b | melatonin receptor 1B [Source:MGI Symb | 0 | 0 | 0 | 0 | 0 | 0 |
| 14916 | ENSMUSG00000050908 | 3.721294398 | 0.738365588 | 1.360510954 | 0.542711976 | 0.587328127 | NA | Tvp23a | trans-golgi network vesicle protein 23A [S | 7.2279 | 1.0279 | 0 | 9.6398 | 1.0906 | 3.3415 |
| 14917 | ENSMUSG00000050910 | 18.45740753 | -0.581720367 | 0.509001313 | -1.142866142 | 0.253094172 | 0.706295664 | Cdr2l | cerebellar degeneration-related protein 2 | 17.668 | 23.642 | 25.156 | 14.46 | 11.997 | 17.822 |
| 14918 | ENSMUSG00000050912 | 872.8905698 | 0.006527305 | 0.133690904 | 0.04882385 | 0.961059675 | 0.992557454 | Tmem123 | transmembrane protein 123 [Source:MGI | 784.63 | 901.48 | 926.75 | 752.87 | 977.19 | 894.42 |
| 14919 | ENSMUSG00000050914 | 10.50167592 | 1.24636178 | 0.727351624 | 1.713561554 | 0.086609279 | 0.433766142 | Ankrd37 | ankyrin repeat domain 37 [Source:MGI Sy | 11.243 | 3.0838 | 4.025 | 11.568 | 15.269 | 17.822 |
| 14920 | ENSMUSG00000050917 | 0 | NA | NA | NA | NA | NA | Fgf4 | fibroblast growth factor 4 [Source:MGI Sy | 0 | 0 | 0 | 0 | 0 | 0 |
| 14921 | ENSMUSG00000050919 | 11.69708315 | -0.169911014 | 0.811048841 | -0.209495416 | 0.834061514 | 0.971005543 | Zfp366 | zinc finger protein 366 [Source:MGI Symb | 17.668 | 16.447 | 3.0187 | 16.388 | 2.1812 | 14.48 |
| 14922 | ENSMUSG00000050921 | 2.524552736 | -0.463057095 | 1.370084129 | -0.337977125 | 0.735380426 | NA | P2ry10 | purinergic receptor P2Y, G-protein couple | 5.6217 | 2.0558 | 1.0062 | 0.964 | 3.2719 | 2.2277 |
| 14923 | ENSMUSG00000050926 | 0 | NA | NA | NA | NA | NA | Dcaf12l2 | DDB1 and CUL4 associated factor 12-like 2 | 0 | 0 | 0 | 0 | 0 | 0 |
| 14924 | ENSMUSG00000050930 | 7.522016128 | 0.546012944 | 0.867112682 | 0.629690875 | 0.528896854 | 0.888334265 | Map10 | microtubule-associated protein 10 [Sourc | 7.2279 | 2.0558 | 9.0562 | 13.496 | 3.2719 | 10.025 |
| 14925 | ENSMUSG00000050931 | 970.3250605 | 0.461054675 | 0.120295572 | 3.832682006 | 0.000126754 | 0.00434447 | Sgms2 | sphingomyelin synthase 2 [Source:MGI Sy | 787.84 | 776.08 | 886.5 | 1202.1 | 1008.8 | 1160.6 |
| 14926 | ENSMUSG00000050933 | 0 | NA | NA | NA | NA | NA | Vmn1r231 | vomeronasal 1 receptor 231 [Source:MGI | 0 | 0 | 0 | 0 | 0 | 0 |
| 14927 | ENSMUSG00000050936 | 0 | NA | NA | NA | NA | NA | Gm42743 | predicted gene 42743 [Source:MGI Symbo | 0 | 0 | 0 | 0 | 0 | 0 |
| 14928 | ENSMUSG00000050944 | 0.294513347 | 0.165143496 | 4.080472857 | 0.040471657 | 0.967717103 | NA | Efcab5 | EF-hand calcium binding domain 5 [Sourc | 0.8031 | 0 | 0 | 0.964 | 0 | 0 |
| 14929 | ENSMUSG00000050945 | 32.01536747 | 0.276119395 | 0.397739793 | 0.694221196 | 0.487543502 | 0.873910634 | Zfp438 | zinc finger protein 438 [Source:MGI Symb | 29.715 | 30.838 | 26.162 | 26.991 | 32.719 | 45.668 |
| 14930 | ENSMUSG00000050947 | 144.0526486 | 0.270831575 | 0.228911726 | 1.183126697 | 0.236758939 | 0.68558184 | Amigo1 | adhesion molecule with Ig like domain 1 [ | 97.978 | 137.74 | 156.97 | 178.34 | 140.69 | 152.6 |
| 14931 | ENSMUSG00000050953 | 58.04234452 | -1.026613022 | 0.299970117 | -3.422384311 | 0.000620745 | 0.014528247 | Gja1 | gap junction protein, alpha 1 [Source:MGI | 73.885 | 86.345 | 73.456 | 39.523 | 32.719 | 42.326 |
| 14932 | ENSMUSG00000050954 | 64.49474085 | 0.073350174 | 0.285157429 | 0.257226943 | 0.797003586 | 0.962236316 | Zfp169 | zinc finger protein 169 [Source:MGI Symb | 57.823 | 62.703 | 68.425 | 79.046 | 62.165 | 56.806 |
| 14933 | ENSMUSG00000050957 | 4.321852166 | -1.387419484 | 1.085059259 | -1.278657799 | 0.201017587 | NA | Insl6 | insulin-like 6 [Source:MGI Symbol;Acc:MG | 9.6372 | 4.1117 | 5.0312 | 3.8559 | 2.1812 | 1.1138 |
| 14934 | ENSMUSG00000050961 | 0 | NA | NA | NA | NA | NA | AY358078 | cDNA sequence AY358078 [Source:MGI Sy | 0 | 0 | 0 | 0 | 0 | 0 |
| 14935 | ENSMUSG00000050963 | 0 | NA | NA | NA | NA | NA | Kcns2 | K+ voltage-gated channel, subfamily S, 2 [ | 0 | 0 | 0 | 0 | 0 | 0 |
| 14936 | ENSMUSG00000050965 | 207.1648661 | 0.050899299 | 0.185073225 | 0.275022489 | 0.783298961 | 0.960670212 | Prkca | protein kinase C, alpha [Source:MGI Symk | 207.2 | 222.03 | 181.12 | 195.69 | 246.48 | 190.47 |
| 14937 | ENSMUSG00000050966 | 0 | NA | NA | NA | NA | NA | Lin28a | lin-28 homolog A (C. elegans) [Source:MG | 0 | 0 | 0 | 0 | 0 | 0 |
| 14938 | ENSMUSG00000050967 | 0.267700746 | -1.497481649 | 4.057001436 | -0.369110456 | 0.712045396 | NA | Creg2 | cellular repressor of E1A-stimulated gene: | 1.6062 | 0 | 0 | 0 | 0 | 0 |
| 14939 | ENSMUSG00000050973 | 70.1231308 | 0.157694249 | 0.298003749 | 0.529168674 | 0.596688446 | 0.914248059 | Gdpgp1 | GDP-D-glucose phosphorylase 1 [Source:N | 61.036 | 53.452 | 84.525 | 73.262 | 89.431 | 59.034 |
| 14940 | ENSMUSG00000050974 | 0.160662974 | 1.126924175 | 4.080472857 | 0.276174898 | 0.782413736 | NA | Gm9847 | predicted pseudogene 9847 [Source:MGI | 0 | 0 | 0 | 0.964 | 0 | 0 |
| 14941 | ENSMUSG00000050978 | 0 | NA | NA | NA | NA | NA | Gm5215 | predicted gene 5215 [Source:MGI Symbol | 0 | 0 | 0 | 0 | 0 | 0 |
| 14942 | ENSMUSG00000050982 | 0 | NA | NA | NA | NA | NA | Apol10a | apolipoprotein L 10A [Source:MGI Symbol | 0 | 0 | 0 | 0 | 0 | 0 |
| 14943 | ENSMUSG00000050989 | 29.94278461 | -1.259035983 | 0.430261603 | -2.926210415 | 0.003431188 | 0.051589086 | Selenon | selenoprotein N [Source:MGI Symbol;Acc | 40.155 | 44.2 | 42.262 | 15.424 | 11.997 | 25.618 |
| 14944 | ENSMUSG00000050994 | 20.40114381 | -2.077958689 | 0.639442282 | -3.249642301 | 0.001155503 | 0.023245046 | Adgb | androglobin [Source:MGI Symbol;Acc:MG | 19.274 | 56.535 | 23.144 | 4.8199 | 14.178 | 4.4554 |
| 14945 | ENSMUSG00000050996 | 0 | NA | NA | NA | NA | NA | Cetn1 | centrin 1 [Source:MGI Symbol;Acc:MGI:1 | 0 | 0 | 0 | 0 | 0 | 0 |
| 14946 | ENSMUSG00000051000 | 107.1450732 | -0.340531667 | 0.245713614 | -1.385888483 | 0.165780958 | 0.591770922 | Fam160a1 | family with sequence similarity 160, meml | 97.175 | 132.6 | 129.81 | 94.47 | 77.434 | 111.38 |
| 14947 | ENSMUSG00000051002 | 0 | NA | NA | NA | NA | NA | Gm9848 | predicted gene 9848 [Source:MGI Symbol | 0 | 0 | 0 | 0 | 0 | 0 |
| 14948 | ENSMUSG00000051003 | 0 | NA | NA | NA | NA | NA | Olfr161 | olfactory receptor 161 [Source:MGI Symb | 0 | 0 | 0 | 0 | 0 | 0 |

| | | | | | | | | | | | | | | | |
|---|---|---|---|---|---|---|---|---|---|---|---|---|---|---|---|
| 14949 | ENSMUSG00000051007 | 321.329323 | -0.142249839 | 0.159288858 | -0.893030689 | 0.371840738 | 0.807685567 | Gatd1 | glutamine amidotransferase like class 1 dc | 318.03 | 321.74 | 372.31 | 334.5 | 262.84 | 318.56 |
| 14950 | ENSMUSG00000051008 | 5.302654003 | -2.335882602 | 1.134324401 | -2.05927211 | 0.039468179 | 0.271845887 | 4930412M03 | RIKEN cDNA 4930412M03 gene [Source:M | 3.2124 | 12.335 | 11.069 | 1.928 | 3.2719 | 0 |
| 14951 | ENSMUSG00000051022 | 2.218018623 | -1.685169787 | 1.476016604 | -1.141701104 | 0.253578282 | NA | Hs3st1 | heparan sulfate (glucosamine) 3-O-sulfotr | 4.0155 | 4.1117 | 2.0125 | 0.964 | 1.0906 | 1.1138 |
| 14952 | ENSMUSG00000051029 | 0.513958919 | -2.394274821 | 3.756096013 | -0.637437066 | 0.523840191 | NA | Serpinb1b | serine (or cysteine) peptidase inhibitor, cl | 0 | 3.0838 | 0 | 0 | 0 | 0 |
| 14953 | ENSMUSG00000051034 | 29.50081784 | 0.478488064 | 0.426725477 | 1.121301845 | 0.2621594 | 0.716133499 | Zfp11 | zinc finger protein 11 [Source:MGI Symbo | 24.896 | 31.865 | 17.106 | 26.027 | 40.353 | 36.757 |
| 14954 | ENSMUSG00000051036 | 0 | NA | NA | NA | NA | NA | Ttc24 | tetratricopeptide repeat domain 24 [Sour | 0 | 0 | 0 | 0 | 0 | 0 |
| 14955 | ENSMUSG00000051037 | 27.21275279 | 0.169316254 | 0.461278213 | 0.367058857 | 0.713575116 | 0.94429556 | Zfp455 | zinc finger protein 455 [Source:MGI Symb | 32.124 | 22.614 | 22.137 | 43.379 | 19.631 | 23.391 |
| 14956 | ENSMUSG00000051038 | 0 | NA | NA | NA | NA | NA | Rhox11 | reproductive homeobox 11 [Source:MGI S | 0 | 0 | 0 | 0 | 0 | 0 |
| 14957 | ENSMUSG00000051041 | 390.5166138 | 0.246259684 | 0.162908572 | 1.511643499 | 0.130624579 | 0.529460957 | Olfml1 | olfactomedin-like 1 [Source:MGI Symbol;A | 400.75 | 376.22 | 293.82 | 377.88 | 470.06 | 424.37 |
| 14958 | ENSMUSG00000051043 | 1402.380989 | 0.334000748 | 0.137221538 | 2.434025689 | 0.014931935 | 0.142198341 | Gprc5c | G protein-coupled receptor, family C, grou | 1417.5 | 1132.8 | 1170.3 | 1413.2 | 1449.4 | 1831.2 |
| 14959 | ENSMUSG00000051046 | 0 | NA | NA | NA | NA | NA | Olfr214 | olfactory receptor 214 [Source:MGI Symb | 0 | 0 | 0 | 0 | 0 | 0 |
| 14960 | ENSMUSG00000051048 | 0.167707431 | -0.796629176 | 4.080472857 | -0.195229623 | 0.84521319 | NA | P4ha3 | procollagen-proline, 2-oxoglutarate 4-diox | 0 | 0 | 1.0062 | 0 | 0 | 0 |
| 14961 | ENSMUSG00000051050 | 0 | NA | NA | NA | NA | NA | Spink14 | serine peptidase inhibitor, Kazal type 14 [S | 0 | 0 | 0 | 0 | 0 | 0 |
| 14962 | ENSMUSG00000051051 | 0 | NA | NA | NA | NA | NA | Olfr523 | olfactory receptor 523 [Source:MGI Symb | 0 | 0 | 0 | 0 | 0 | 0 |
| 14963 | ENSMUSG00000051054 | 0.185640647 | 1.126924175 | 4.080472857 | 0.276174898 | 0.782413736 | NA | 1700014D04 | RIKEN cDNA 1700014D04 gene [Source:M | 0 | 0 | 0 | 0 | 0 | 1.1138 |
| 14964 | ENSMUSG00000051056 | 0 | NA | NA | NA | NA | NA | Gja10 | gap junction protein, alpha 10 [Source:MG | 0 | 0 | 0 | 0 | 0 | 0 |
| 14965 | ENSMUSG00000051062 | 0 | NA | NA | NA | NA | NA | Fbll1 | fibrillarin-like 1 [Source:MGI Symbol;Acc:M | 0 | 0 | 0 | 0 | 0 | 0 |
| 14966 | ENSMUSG00000051065 | 345.7067957 | 0.225170131 | 0.153638361 | 1.465585356 | 0.142761283 | 0.552480247 | Mb21d2 | Mab-21 domain containing 2 [Source:MGI | 333.29 | 299.12 | 324.01 | 417.4 | 322.82 | 377.59 |
| 14967 | ENSMUSG00000051067 | 0 | NA | NA | NA | NA | NA | Lingo3 | leucine rich repeat and Ig domain contain | 0 | 0 | 0 | 0 | 0 | 0 |
| 14968 | ENSMUSG00000051074 | 2.405494727 | 2.685831195 | 1.669283146 | 1.608972811 | 0.107622287 | NA | 4930579K19 | RIKEN cDNA 4930579K19 gene [Source:M | 0 | 0 | 2.0125 | 5.7839 | 2.1812 | 4.4554 |
| 14969 | ENSMUSG00000051076 | 2.002133655 | 1.793577454 | 1.59065019 | 1.127575042 | 0.259499432 | NA | Vtcn1 | V-set domain containing T cell activation i | 1.6062 | 1.0279 | 0 | 3.8559 | 2.1812 | 3.3415 |
| 14970 | ENSMUSG00000051079 | 0 | NA | NA | NA | NA | NA | Rgs13 | regulator of G-protein signaling 13 [Sourc | 0 | 0 | 0 | 0 | 0 | 0 |
| 14971 | ENSMUSG00000051081 | 0 | NA | NA | NA | NA | NA | Gm4847 | predicted gene 4847 [Source:MGI Symbol | 0 | 0 | 0 | 0 | 0 | 0 |
| 14972 | ENSMUSG00000051095 | 0 | NA | NA | NA | NA | NA | Olfr986 | olfactory receptor 986 [Source:MGI Symb | 0 | 0 | 0 | 0 | 0 | 0 |
| 14973 | ENSMUSG00000051098 | 141.8376765 | 0.401104007 | 0.195307756 | 2.0537024 | 0.040004504 | 0.273840581 | Mblac2 | metallo-beta-lactamase domain containin | 119.66 | 124.38 | 122.76 | 157.13 | 174.5 | 152.6 |
| 14974 | ENSMUSG00000051111 | 0.294513347 | 0.165143496 | 4.080472857 | 0.040471657 | 0.967717103 | NA | Sv2c | synaptic vesicle glycoprotein 2c [Source:M | 0.8031 | 0 | 0 | 0.964 | 0 | 0 |
| 14975 | ENSMUSG00000051113 | 18.64698369 | -0.317091703 | 0.545691415 | -0.581082447 | 0.561184888 | 0.900385826 | Fam71e1 | family with sequence similarity 71, membe | 17.668 | 14.391 | 30.187 | 23.135 | 10.906 | 15.594 |
| 14976 | ENSMUSG00000051116 | 1.088168803 | 2.43902648 | 2.378968265 | 1.025245488 | 0.30524737 | NA | Gm8121 | predicted pseudogene 8121 [Source:MGI | 0 | 0 | 1.0062 | 0 | 2.1812 | 3.3415 |
| 14977 | ENSMUSG00000051118 | 0 | NA | NA | NA | NA | NA | Olfr77 | olfactory receptor 77 [Source:MGI Symbo | 0 | 0 | 0 | 0 | 0 | 0 |
| 14978 | ENSMUSG00000051124 | 133.2450492 | 0.114353301 | 0.215859062 | 0.529759093 | 0.596278971 | 0.914007952 | Gimap9 | GTPase, IMAP family member 9 [Source:M | 106.01 | 126.43 | 151.94 | 133.03 | 145.05 | 137 |
| 14979 | ENSMUSG00000051133 | 0 | NA | NA | NA | NA | NA | Gm6293 | predicted pseudogene 6293 [Source:MGI | 0 | 0 | 0 | 0 | 0 | 0 |
| 14980 | ENSMUSG00000051136 | 0 | NA | NA | NA | NA | NA | Ghsr | growth hormone secretagogue receptor [S | 0 | 0 | 0 | 0 | 0 | 0 |
| 14981 | ENSMUSG00000051146 | 55.86712926 | 0.102377703 | 0.306246651 | 0.334298195 | 0.738154553 | 0.948137969 | Camk2n2 | calcium/calmodulin-dependent protein ki | 66.657 | 45.228 | 49.306 | 54.947 | 57.803 | 61.261 |
| 14982 | ENSMUSG00000051147 | 358.1730665 | 0.155185582 | 0.160467317 | 0.967085292 | 0.333501397 | 0.775366061 | Nat2 | N-acetyltransferase 2 (arylamine N-acetyl | 387.9 | 287.82 | 340.11 | 370.17 | 415.53 | 347.52 |
| 14983 | ENSMUSG00000051149 | 44.55828091 | -0.018299535 | 0.383573412 | -0.047708037 | 0.961948929 | 0.992908678 | Adnp | activity-dependent neuroprotective prote | 51.399 | 37.005 | 46.287 | 65.55 | 39.262 | 27.846 |
| 14984 | ENSMUSG00000051153 | 0 | NA | NA | NA | NA | NA | Tas2r105 | taste receptor, type 2, member 105 [Sour | 0 | 0 | 0 | 0 | 0 | 0 |
| 14985 | ENSMUSG00000051154 | 977.084725 | 0.128402097 | 0.125377083 | 1.024127332 | 0.305775136 | 0.755004385 | Commd3 | COMM domain containing 3 [Source:MGI | 950.07 | 925.13 | 924.74 | 899.39 | 989.19 | 1174 |
| 14986 | ENSMUSG00000051156 | 0 | NA | NA | NA | NA | NA | Olfr1491 | olfactory receptor 1491 [Source:MGI Sym | 0 | 0 | 0 | 0 | 0 | 0 |
| 14987 | ENSMUSG00000051159 | 0 | NA | NA | NA | NA | NA | Cited1 | Cbp/p300-interacting transactivator with | 0 | 0 | 0 | 0 | 0 | 0 |
| 14988 | ENSMUSG00000051160 | 0 | NA | NA | NA | NA | NA | Olfr853 | olfactory receptor 853 [Source:MGI Symb | 0 | 0 | 0 | 0 | 0 | 0 |
| 14989 | ENSMUSG00000051166 | 113.2160901 | 0.034584171 | 0.277524865 | 0.124616477 | 0.900827184 | 0.983000175 | Eml5 | echinoderm microtubule associated prote | 87.538 | 111.02 | 137.86 | 150.38 | 113.42 | 79.083 |
| 14990 | ENSMUSG00000051169 | 115.2200681 | 0.235547573 | 0.228313123 | 1.031686528 | 0.30221899 | 0.752374437 | Rpusd3 | RNA pseudouridylate synthase domain co | 96.372 | 114.1 | 107.67 | 149.42 | 110.15 | 113.61 |
| 14991 | ENSMUSG00000051172 | 0 | NA | NA | NA | NA | NA | Olfr672 | olfactory receptor 672 [Source:MGI Symb | 0 | 0 | 0 | 0 | 0 | 0 |
| 14992 | ENSMUSG00000051176 | 0 | NA | NA | NA | NA | NA | Zfp42 | zinc finger protein 42 [Source:MGI Symbo | 0 | 0 | 0 | 0 | 0 | 0 |
| 14993 | ENSMUSG00000051177 | 340.6641994 | -0.111981046 | 0.150985826 | -0.741665949 | 0.458289755 | 0.859322431 | Plcb1 | phospholipase C, beta 1 [Source:MGI Sym | 320.44 | 362.85 | 379.35 | 359.56 | 302.1 | 319.67 |
| 14994 | ENSMUSG00000051180 | 0 | NA | NA | NA | NA | NA | Olfr522 | olfactory receptor 522 [Source:MGI Symb | 0 | 0 | 0 | 0 | 0 | 0 |
| 14995 | ENSMUSG00000051182 | 0 | NA | NA | NA | NA | NA | Olfr655 | olfactory receptor 655 [Source:MGI Symb | 0 | 0 | 0 | 0 | 0 | 0 |
| 14996 | ENSMUSG00000051184 | 91.55053314 | 0.03276381 | 0.291911791 | 0.112238734 | 0.910634118 | 0.985662791 | Zfp524 | zinc finger protein 524 [Source:MGI Symb | 103.6 | 75.038 | 92.575 | 94.47 | 59.984 | 123.64 |
| 14997 | ENSMUSG00000051185 | 310.7017372 | 0.158677053 | 0.145844617 | 1.087987045 | 0.276600827 | 0.730038567 | Fam174a | family with sequence similarity 174, meml | 314.01 | 276.51 | 289.8 | 323.9 | 338.09 | 321.9 |
| 14998 | ENSMUSG00000051190 | 0 | NA | NA | NA | NA | NA | Olfr1356 | olfactory receptor 1356 [Source:MGI Sym | 0 | 0 | 0 | 0 | 0 | 0 |
| 14999 | ENSMUSG00000051198 | 0 | NA | NA | NA | NA | NA | 4930548G14 | RIKEN cDNA 4930548G14 gene [Source:M | 0 | 0 | 0 | 0 | 0 | 0 |
| 15000 | ENSMUSG00000051200 | 0 | NA | NA | NA | NA | NA | Olfr513 | olfactory receptor 513 [Source:MGI Symb | 0 | 0 | 0 | 0 | 0 | 0 |
| 15001 | ENSMUSG00000051207 | 0 | NA | NA | NA | NA | NA | Mrgprd | MAS-related GPR, member D [Source:MG | 0 | 0 | 0 | 0 | 0 | 0 |
| 15002 | ENSMUSG00000051209 | 0 | NA | NA | NA | NA | NA | Gpr119 | G-protein coupled receptor 119 [Source:M | 0 | 0 | 0 | 0 | 0 | 0 |
| 15003 | ENSMUSG00000051212 | 1.441806094 | -0.20332298 | 1.782286416 | -0.114079857 | 0.909174489 | NA | Gpr183 | G protein-coupled receptor 183 [Source:M | 1.6062 | 2.0558 | 1.0062 | 2.8919 | 1.0906 | 0 |
| 15004 | ENSMUSG00000051219 | 0 | NA | NA | NA | NA | NA | NA | NA | 0 | 0 | 0 | 0 | 0 | 0 |

| | | | | | | | | | | | | | | | |
|---|---|---|---|---|---|---|---|---|---|---|---|---|---|---|---|
| 15005 | ENSMUSG00000051220 | 3.285106189 | 1.132756558 | 1.210193335 | 0.936012888 | 0.349266552 | NA | Ercc6l | excision repair cross-complementing rode | 4.0155 | 1.0279 | 1.0062 | 4.8199 | 3.2719 | 5.5692 |
| 15006 | ENSMUSG00000051223 | 2959.966744 | 0.040901988 | 0.09959065 | 0.410701084 | 0.681291732 | 0.934452508 | Bzw1 | basic leucine zipper and W2 domains 1 [S | 2910.4 | 2982 | 2860.8 | 2711.7 | 3161.7 | 3133.2 |
| 15007 | ENSMUSG00000051224 | 46.48627048 | 0.32826849 | 0.341879353 | 0.96018811 | 0.33696055 | 0.779072142 | Tceanc | transcription elongation factor A (SII) N-te | 35.336 | 44.2 | 44.275 | 50.127 | 39.262 | 65.717 |
| 15008 | ENSMUSG00000051225 | 13.11379219 | -0.568244077 | 0.666164177 | -0.853009058 | 0.393654281 | 0.822567975 | Fam83a | family with sequence similarity 83, memb | 23.29 | 17.475 | 6.0375 | 8.6758 | 8.7249 | 14.48 |
| 15009 | ENSMUSG00000051228 | 1.05258607 | -2.221073771 | 2.388733928 | -0.929812125 | 0.352468367 | NA | Nyx | nyctalopin [Source:MGI Symbol;Acc:MGI: | 3.2124 | 0 | 2.0125 | 0 | 1.0906 | 0 |
| 15010 | ENSMUSG00000051232 | 181.5711921 | 0.218887494 | 0.196660214 | 1.113023774 | 0.265698219 | 0.721565724 | Tmem199 | transmembrane protein 199 [Source:MGI | 153.39 | 203.53 | 146.91 | 191.83 | 182.13 | 211.63 |
| 15011 | ENSMUSG00000051234 | 492.0470047 | -0.065429521 | 0.123351609 | -0.530431027 | 0.595813118 | 0.91381309 | Rnf7 | ring finger protein 7 [Source:MGI Symbol; | 486.68 | 513.96 | 509.16 | 469.46 | 492.96 | 480.07 |
| 15012 | ENSMUSG00000051235 | 29.9843033 | 0.197987795 | 0.454355811 | 0.435754953 | 0.663014531 | 0.92906822 | Gen1 | GEN1, Holliday junction 5' flap endonucle | 20.881 | 40.089 | 23.144 | 37.595 | 39.262 | 18.935 |
| 15013 | ENSMUSG00000051236 | 171.011013 | -0.032915994 | 0.207970176 | -0.158272664 | 0.874241948 | 0.980007404 | Msrb3 | methionine sulfoxide reductase B3 [Sourc | 192.74 | 163.44 | 162.01 | 152.31 | 145.05 | 210.52 |
| 15014 | ENSMUSG00000051237 | 0 | NA | NA | NA | NA | NA | 9230109A22 | RIKEN cDNA 9230109A22 gene [Source:M | 0 | 0 | 0 | 0 | 0 | 0 |
| 15015 | ENSMUSG00000051238 | 109.7756011 | -0.073721626 | 0.230892004 | -0.319290511 | 0.749506228 | 0.951356173 | Swsap1 | SWIM type zinc finger 7 associated protei | 126.89 | 108.96 | 101.63 | 109.89 | 121.06 | 90.221 |
| 15016 | ENSMUSG00000051242 | 0.844158739 | -0.371321695 | 2.388630557 | -0.155453799 | 0.876463574 | NA | Pcdhb9 | protocadherin beta 9 [Source:MGI Symbo | 0.8031 | 1.0279 | 1.0062 | 0 | 0 | 2.2277 |
| 15017 | ENSMUSG00000051243 | 0 | NA | NA | NA | NA | NA | Islr2 | immunoglobulin superfamily containing le | 0 | 0 | 0 | 0 | 0 | 0 |
| 15018 | ENSMUSG00000051246 | 8.50652509 | -0.205482293 | 0.750996744 | -0.273612761 | 0.784382226 | 0.960670212 | Msantd1 | Myb/SANT-like DNA-binding domain cont | 12.047 | 9.2513 | 6.0375 | 11.568 | 5.4531 | 6.6831 |
| 15019 | ENSMUSG00000051251 | 0 | NA | NA | NA | NA | NA | Nhlh1 | nescient helix loop helix 1 [Source:MGI Sy | 0 | 0 | 0 | 0 | 0 | 0 |
| 15020 | ENSMUSG00000051255 | 73.67724431 | -0.096022621 | 0.267922782 | -0.358396627 | 0.720046516 | 0.9447719 | Gm6563 | predicted pseudogene 6563 [Source:MGI | 66.657 | 88.401 | 73.456 | 63.623 | 73.071 | 76.855 |
| 15021 | ENSMUSG00000051256 | 441.4856045 | 0.026534265 | 0.127766103 | 0.207678442 | 0.835480051 | 0.971073158 | Jagn1 | jagunal homolog 1 [Source:MGI Symbol;A | 451.34 | 438.92 | 421.62 | 438.61 | 439.52 | 458.9 |
| 15022 | ENSMUSG00000051257 | 0 | NA | NA | NA | NA | NA | Trap1a | tumor rejection antigen P1A [Source:MGI | 0 | 0 | 0 | 0 | 0 | 0 |
| 15023 | ENSMUSG00000051258 | 0 | NA | NA | NA | NA | NA | Olfr1362 | olfactory receptor 1362 [Source:MGI Sym | 0 | 0 | 0 | 0 | 0 | 0 |
| 15024 | ENSMUSG00000051262 | 0.342639279 | -1.805976596 | 4.028547714 | -0.448294702 | 0.653940524 | NA | Nat8f3 | N-acetyltransferase 8 (GCN5-related) fam | 0 | 2.0558 | 0 | 0 | 0 | 0 |
| 15025 | ENSMUSG00000051276 | 0 | NA | NA | NA | NA | NA | Actrt2 | actin-related protein T2 [Source:MGI Sym | 0 | 0 | 0 | 0 | 0 | 0 |
| 15026 | ENSMUSG00000051278 | 11.05522386 | 0.160218174 | 0.646503624 | 0.247822546 | 0.80427171 | 0.963701061 | Zgrf1 | zinc finger, GRF-type containing 1 [Source | 12.85 | 10.279 | 8.05 | 8.6758 | 11.997 | 14.48 |
| 15027 | ENSMUSG00000051279 | 8.111890072 | -0.954731254 | 0.766139297 | -1.246158835 | 0.212706088 | 0.657529571 | Gdf6 | growth differentiation factor 6 [Source:M | 12.85 | 8.2233 | 11.069 | 7.7118 | 4.3625 | 4.4554 |
| 15028 | ENSMUSG00000051285 | 2389.490792 | 0.229424914 | 0.104986165 | 2.185287112 | 0.028867802 | 0.222927862 | Pcmtd1 | protein-L-isoaspartate (D-aspartate) O-me | 2209.3 | 2309.7 | 2079.9 | 2335.7 | 2766.9 | 2635.4 |
| 15029 | ENSMUSG00000051295 | 0.17131964 | -0.796629176 | 4.080472857 | -0.195229623 | 0.84521319 | NA | 9630028B13 | RIKEN cDNA 9630028B13 gene [Source:M | 0 | 1.0279 | 0 | 0 | 0 | 0 |
| 15030 | ENSMUSG00000051297 | 0 | NA | NA | NA | NA | NA | 2410124H12 | RIKEN cDNA 2410124H12 gene [Source:M | 0 | 0 | 0 | 0 | 0 | 0 |
| 15031 | ENSMUSG00000051306 | 150.7917946 | 0.023851954 | 0.190286188 | 0.125347793 | 0.900248217 | 0.982805511 | Usp42 | ubiquitin specific peptidase 42 [Source:M | 155.8 | 143.91 | 148.92 | 161.95 | 151.6 | 142.57 |
| 15032 | ENSMUSG00000051313 | 0 | NA | NA | NA | NA | NA | Olfr1262 | olfactory receptor 1262 [Source:MGI Sym | 0 | 0 | 0 | 0 | 0 | 0 |
| 15033 | ENSMUSG00000051314 | 0.353348078 | 0.165142701 | 3.895042568 | 0.042398176 | 0.966181282 | NA | Ffar2 | free fatty acid receptor 2 [Source:MGI Syr | 0 | 0 | 1.0062 | 0 | 0 | 1.1138 |
| 15034 | ENSMUSG00000051316 | 94.58353756 | 0.023747072 | 0.241561943 | 0.098306346 | 0.92168904 | 0.986685804 | Taf7 | TATA-box binding protein associated fact | 106.01 | 92.513 | 82.512 | 94.47 | 85.068 | 106.93 |
| 15035 | ENSMUSG00000051319 | 246.3159423 | 0.000178368 | 0.191635784 | 0.000930764 | 0.999257358 | 0.999764475 | Mtln | mitoregulin [Source:MGI Symbol;Acc:MGI | 274.66 | 205.58 | 257.6 | 197.62 | 261.75 | 280.69 |
| 15036 | ENSMUSG00000051323 | 0 | NA | NA | NA | NA | NA | Pcdh19 | protocadherin 19 [Source:MGI Symbol;Ac | 0 | 0 | 0 | 0 | 0 | 0 |
| 15037 | ENSMUSG00000051329 | 117.4074599 | -0.147834073 | 0.229462531 | -0.644262365 | 0.519405316 | 0.887731952 | Nup160 | nucleoporin 160 [Source:MGI Symbol;Acc | 127.69 | 129.52 | 112.7 | 93.506 | 136.33 | 104.7 |
| 15038 | ENSMUSG00000051331 | 1.759307202 | -3.080865191 | 1.942554244 | -1.585986698 | 0.112742327 | NA | Cacna1c | calcium channel, voltage-dependent, L typ | 6.4248 | 1.0279 | 2.0125 | 0 | 1.0906 | 0 |
| 15039 | ENSMUSG00000051335 | 47.91701718 | 0.136556635 | 0.363467502 | 0.375705214 | 0.707136061 | 0.943108897 | Gfod1 | glucose-fructose oxidoreductase domain | 51.399 | 32.893 | 52.325 | 41.451 | 39.262 | 70.172 |
| 15040 | ENSMUSG00000051339 | 1518.960236 | 0.39619441 | 0.148487066 | 2.668208219 | 0.007625698 | 0.088790029 | 2900026A02 | RIKEN cDNA 2900026A02 gene [Source:M | 1287.4 | 1265.4 | 1381.6 | 1366 | 1688.3 | 2125.2 |
| 15041 | ENSMUSG00000051340 | 0 | NA | NA | NA | NA | NA | Olfr645 | olfactory receptor 645 [Source:MGI Symb | 0 | 0 | 0 | 0 | 0 | 0 |
| 15042 | ENSMUSG00000051341 | 7.72042491 | -0.196256529 | 0.859650914 | -0.228297936 | 0.819414629 | 0.969160361 | Zfp52 | zinc finger protein 52 [Source:MGI Symbo | 10.44 | 11.307 | 3.0187 | 10.604 | 8.7249 | 2.2277 |
| 15043 | ENSMUSG00000051343 | 17.01517123 | -0.11926974 | 0.558972081 | -0.21337334 | 0.831035774 | 0.970505212 | Rab11fip5 | RAB11 family interacting protein 5 (class I | 13.653 | 22.614 | 17.106 | 22.171 | 8.7249 | 17.822 |
| 15044 | ENSMUSG00000051344 | 29.20103184 | -0.630544863 | 0.40969566 | -1.539056731 | 0.123790447 | 0.515549964 | Plekhm3 | pleckstrin homology domain containing, f | 40.155 | 30.838 | 35.219 | 18.316 | 26.175 | 24.505 |
| 15045 | ENSMUSG00000051346 | 615.9521603 | 0.103672173 | 0.13192984 | 0.785812922 | 0.431977095 | 0.845253882 | Spryd4 | SPRY domain containing 4 [Source:MGI Sy | 655.33 | 579.75 | 545.38 | 632.37 | 587.84 | 695.04 |
| 15046 | ENSMUSG00000051351 | 89.39960536 | -0.028802089 | 0.238087556 | -0.120972675 | 0.903712678 | 0.983821605 | Zfp46 | zinc finger protein 46 [Source:MGI Symbo | 94.766 | 94.568 | 81.506 | 92.542 | 87.249 | 85.766 |
| 15047 | ENSMUSG00000051354 | 1.333087851 | -3.772154312 | 2.446517662 | -1.541846344 | 0.123110937 | NA | Samd3 | sterile alpha motif domain containing 3 [S | 0.8031 | 7.1954 | 0 | 0 | 0 | 0 |
| 15048 | ENSMUSG00000051355 | 244.9456814 | -0.017911323 | 0.170051975 | -0.10532852 | 0.916115133 | 0.986374671 | Commd1 | COMM domain containing 1 [Source:MGI | 220.05 | 255.95 | 263.64 | 217.86 | 270.47 | 241.7 |
| 15049 | ENSMUSG00000051359 | 203.1441756 | 0.219462424 | 0.19375505 | 1.13267976 | 0.25734876 | 0.710872182 | Ncald | neurocalcin delta [Source:MGI Symbol;Ac | 167.85 | 182.97 | 212.32 | 178.34 | 230.12 | 247.27 |
| 15050 | ENSMUSG00000051361 | 1.138729733 | 1.305140952 | 2.579810336 | 0.505905777 | 0.612922797 | NA | 6030498E09 | RIKEN cDNA 6030498E09 gene [Source:M | 0 | 0 | 2.0125 | 4.8199 | 0 | 0 |
| 15051 | ENSMUSG00000051362 | 0 | NA | NA | NA | NA | NA | Olfr589 | olfactory receptor 589 [Source:MGI Symb | 0 | 0 | 0 | 0 | 0 | 0 |
| 15052 | ENSMUSG00000051367 | 0 | NA | NA | NA | NA | NA | Six1 | sine oculis-related homeobox 1 [Source:M | 0 | 0 | 0 | 0 | 0 | 0 |
| 15053 | ENSMUSG00000051373 | 13.90038514 | 0.019549626 | 0.610534287 | 0.032020521 | 0.974455686 | 0.996336532 | Plpp7 | phospholipid phosphatase 7 (inactive) [So | 20.881 | 8.2233 | 12.075 | 13.496 | 10.906 | 17.822 |
| 15054 | ENSMUSG00000051375 | 1568.288466 | 0.301501563 | 0.123878539 | 2.433848229 | 0.014939258 | 0.142198341 | Pcdh1 | protocadherin 1 [Source:MGI Symbol;Acc | 1285.8 | 1329.1 | 1600.9 | 1748.7 | 1561.8 | 1883.5 |
| 15055 | ENSMUSG00000051378 | 1.409494751 | 2.879175517 | 2.009903607 | 1.432494328 | 0.1520024 | NA | Kif18b | kinesin family member 18B [Source:MGI S | 0 | 0 | 1.0062 | 1.928 | 2.1812 | 3.3415 |
| 15056 | ENSMUSG00000051379 | 0.621048825 | -1.224891493 | 2.859518283 | -0.428355888 | 0.668392035 | NA | Flrt3 | fibronectin leucine rich transmembrane p | 1.6062 | 0 | 1.0062 | 0 | 0 | 1.1138 |
| 15057 | ENSMUSG00000051390 | 357.5727912 | 0.434932586 | 0.146787481 | 2.963008712 | 0.003046479 | 0.047316364 | Zbtb22 | zinc finger and BTB domain containing 22 | 315.62 | 281.65 | 314.95 | 449.21 | 372.99 | 411.01 |
| 15058 | ENSMUSG00000051391 | 1992.759674 | -0.143087143 | 0.120634933 | -1.186116982 | 0.235576105 | 0.683799474 | Ywhag | tyrosine 3-monooxygenase/tryptophan 5- | 2236.6 | 2254.2 | 1783.1 | 1872 | 2034 | 1776.6 |
| 15059 | ENSMUSG00000051392 | 0 | NA | NA | NA | NA | NA | Olfr1402 | olfactory receptor 1402 [Source:MGI Sym | 0 | 0 | 0 | 0 | 0 | 0 |
| 15060 | ENSMUSG00000051396 | 16.58304763 | -0.189736311 | 0.535905882 | -0.354047823 | 0.723303044 | 0.9447719 | Gm45902 | predicted gene 45902 [Source:MGI Symbo | 19.274 | 21.586 | 12.075 | 13.496 | 16.359 | 16.708 |

| | | | | | | | | | | | | | | | |
|---|---|---|---|---|---|---|---|---|---|---|---|---|---|---|---|
| 15061 | ENSMUSG00000051397 | 0.17131964 | -0.796629176 | 4.080472857 | -0.195229623 | 0.84521319 | NA | Tacstd2 | tumor-associated calcium signal transduc | 0 | 1.0279 | 0 | 0 | 0 | 0 |
| 15062 | ENSMUSG00000051401 | 0 | NA | NA | NA | NA | NA | Kctd16 | potassium channel tetramerisation domai | 0 | 0 | 0 | 0 | 0 | 0 |
| 15063 | ENSMUSG00000051403 | 1988.675482 | 0.230068403 | 0.106071375 | 2.168996144 | 0.030082977 | 0.229493763 | Ppp1r37 | protein phosphatase 1, regulatory subunit | 1902.5 | 1644.7 | 1943.1 | 2042.7 | 2223.8 | 2175.3 |
| 15064 | ENSMUSG00000051412 | 651.3637763 | 0.178782518 | 0.130193865 | 1.373202325 | 0.169689457 | 0.596876778 | NA | NA | 665.77 | 560.22 | 605.76 | 625.62 | 704.54 | 746.28 |
| 15065 | ENSMUSG00000051413 | 102.6043118 | -0.052151407 | 0.221202051 | -0.235763668 | 0.813616062 | 0.966842541 | Plagl2 | pleiomorphic adenoma gene-like 2 [Sourc | 106.81 | 101.76 | 104.65 | 97.362 | 100.34 | 104.7 |
| 15066 | ENSMUSG00000051414 | 0 | NA | NA | NA | NA | NA | Olfr829 | olfactory receptor 829 [Source:MGI Symb | 0 | 0 | 0 | 0 | 0 | 0 |
| 15067 | ENSMUSG00000051424 | 0 | NA | NA | NA | NA | NA | Olfr1184 | olfactory receptor 1184 [Source:MGI Sym | 0 | 0 | 0 | 0 | 0 | 0 |
| 15068 | ENSMUSG00000051427 | 394.9349632 | 0.22409417 | 0.152171455 | 1.472642619 | 0.140847426 | 0.548849969 | Ccdc157 | coiled-coil domain containing 157 [Source | 429.66 | 332.02 | 330.05 | 428.01 | 417.71 | 432.17 |
| 15069 | ENSMUSG00000051431 | 0 | NA | NA | NA | NA | NA | Gpr87 | G protein-coupled receptor 87 [Source:M | 0 | 0 | 0 | 0 | 0 | 0 |
| 15070 | ENSMUSG00000051435 | 94.2650649 | -0.336396178 | 0.293583836 | -1.145826633 | 0.251866893 | 0.704370213 | Fhad1 | forkhead-associated (FHA) phosphopeptic | 100.39 | 121.29 | 93.581 | 51.091 | 111.24 | 87.994 |
| 15071 | ENSMUSG00000051437 | 0 | NA | NA | NA | NA | NA | Ubqlnl | ubiquilin-like [Source:MGI Symbol;Acc:MC | 0 | 0 | 0 | 0 | 0 | 0 |
| 15072 | ENSMUSG00000051439 | 43.93519612 | -1.239822053 | 0.413935286 | -2.99520745 | 0.002742582 | 0.044003638 | Cd14 | CD14 antigen [Source:MGI Symbol;Acc:M | 43.368 | 97.652 | 44.275 | 23.135 | 23.994 | 31.188 |
| 15073 | ENSMUSG00000051444 | 5.321251718 | -1.764498276 | 1.019260295 | -1.731155706 | 0.083423997 | 0.425868616 | Bbs12 | Bardet-Biedl syndrome 12 (human) [Sourc | 14.456 | 5.1396 | 5.0312 | 2.8919 | 2.1812 | 2.2277 |
| 15074 | ENSMUSG00000051451 | 798.9126313 | 0.033888804 | 0.156906306 | 0.215981144 | 0.829002446 | 0.969955221 | Crebzf | CREB/ATF bZIP transcription factor [Sourc | 722.79 | 703.1 | 943.86 | 957.23 | 752.53 | 713.97 |
| 15075 | ENSMUSG00000051452 | 64.77123159 | -0.274309011 | 0.307260585 | -0.892756911 | 0.371987366 | 0.807750377 | Gm11437 | predicted gene 11437 [Source:MGI Symbo | 70.673 | 52.424 | 89.556 | 57.839 | 49.078 | 69.058 |
| 15076 | ENSMUSG00000051455 | 3.953790656 | 1.678338433 | 1.119167082 | 1.4996317 | 0.133709832 | NA | Meioc | meiosis specific with coiled-coil domain [S | 1.6062 | 1.0279 | 3.0187 | 4.8199 | 5.4531 | 7.7969 |
| 15077 | ENSMUSG00000051456 | 0 | NA | NA | NA | NA | NA | Hspb3 | heat shock protein 3 [Source:MGI Symbol | 0 | 0 | 0 | 0 | 0 | 0 |
| 15078 | ENSMUSG00000051457 | 6.597166086 | -1.519369176 | 0.982424545 | -1.546550504 | 0.121971673 | 0.511776306 | Spn | sialophorin [Source:MGI Symbol;Acc:MGI | 4.0155 | 17.475 | 8.05 | 6.7478 | 2.1812 | 1.1138 |
| 15079 | ENSMUSG00000051469 | 441.6796917 | 0.171605536 | 0.137796192 | 1.245357611 | 0.213000327 | 0.657724029 | Zfp24 | zinc finger protein 24 [Source:MGI Symbo | 381.47 | 425.56 | 439.73 | 439.57 | 510.41 | 453.33 |
| 15080 | ENSMUSG00000051481 | 0 | NA | NA | NA | NA | NA | Krtap31-2 | keratin associated protein 31-2 [Source:M | 0 | 0 | 0 | 0 | 0 | 0 |
| 15081 | ENSMUSG00000051483 | 1323.954956 | -0.546595822 | 0.150262549 | -3.637605157 | 0.000275185 | 0.008113038 | Cbr1 | carbonyl reductase 1 [Source:MGI Symbol | 1528.3 | 1566.5 | 1622.1 | 1366.9 | 952.11 | 907.78 |
| 15082 | ENSMUSG00000051486 | 1.14577419 | 0.394191739 | 2.55065941 | 0.154545031 | 0.877180008 | NA | Pcdhb11 | protocadherin beta 11 [Source:MGI Symb | 0 | 0 | 3.0187 | 3.8559 | 0 | 0 |
| 15083 | ENSMUSG00000051490 | 0 | NA | NA | NA | NA | NA | Foxd4 | forkhead box D4 [Source:MGI Symbol;Acc | 0 | 0 | 0 | 0 | 0 | 0 |
| 15084 | ENSMUSG00000051493 | 0 | NA | NA | NA | NA | NA | Olfr974 | olfactory receptor 974 [Source:MGI Symb | 0 | 0 | 0 | 0 | 0 | 0 |
| 15085 | ENSMUSG00000051495 | 819.1191334 | -0.848953809 | 0.156768975 | -5.415317715 | 6.12E-08 | 7.08E-06 | Irf2bp2 | interferon regulatory factor 2 binding prot | 847.27 | 1190.3 | 1124 | 630.44 | 630.38 | 492.32 |
| 15086 | ENSMUSG00000051497 | 12.52836874 | -1.121513718 | 0.72467638 | -1.547606283 | 0.121717118 | 0.511629292 | Kcnj16 | potassium inwardly-rectifying channel, su | 16.865 | 27.754 | 7.0437 | 12.532 | 7.6343 | 3.3415 |
| 15087 | ENSMUSG00000051498 | 10.48354081 | -0.441634516 | 0.730542154 | -0.604529818 | 0.545491451 | 0.89549457 | Tlr6 | toll-like receptor 6 [Source:MGI Symbol;A | 9.6372 | 20.558 | 6.0375 | 5.7839 | 13.087 | 7.7969 |
| 15088 | ENSMUSG00000051499 | 5.802436173 | -0.553138637 | 1.049385937 | -0.527106966 | 0.59811931 | 0.915029425 | Zfp786 | zinc finger protein 786 [Source:MGI Symb | 2.4093 | 9.2513 | 9.0562 | 0.964 | 10.906 | 2.2277 |
| 15089 | ENSMUSG00000051502 | 68.20972991 | -0.112251133 | 0.271976149 | -0.412724177 | 0.679808709 | 0.934260934 | Ufsp1 | UFM1-specific peptidase 1 [Source:MGI S | 67.461 | 72.982 | 72.45 | 75.19 | 63.256 | 57.92 |
| 15090 | ENSMUSG00000051503 | 0.749689669 | 0.81510874 | 2.956253303 | 0.275723579 | 0.782760384 | NA | Gm6583 | predicted gene 6583 [Source:MGI Symbol | 1.6062 | 0 | 0 | 2.8919 | 0 | 0 |
| 15091 | ENSMUSG00000051504 | 6.232406804 | 0.172098566 | 1.011079664 | 0.170212667 | 0.86484289 | 0.977227145 | Siglech | sialic acid binding Ig-like lectin H [Source:I | 6.4248 | 4.1117 | 7.0437 | 8.6758 | 0 | 11.138 |
| 15092 | ENSMUSG00000051506 | 115.7886688 | -0.984917564 | 0.270550586 | -3.640419266 | 0.000272194 | 0.008067665 | Wdfy4 | WD repeat and FYVE domain containing 4 | 184.71 | 134.66 | 141.88 | 74.226 | 105.79 | 53.465 |
| 15093 | ENSMUSG00000051509 | 0 | NA | NA | NA | NA | NA | Olfr414 | olfactory receptor 414 [Source:MGI Symb | 0 | 0 | 0 | 0 | 0 | 0 |
| 15094 | ENSMUSG00000051510 | 1982.81424 | 0.140810355 | 0.115651026 | 1.217545236 | 0.223396836 | 0.668933038 | Mafg | v-maf musculoaponeurotic fibrosarcoma | 1951.5 | 1626.2 | 2079.9 | 2023.4 | 2040.5 | 2175.3 |
| 15095 | ENSMUSG00000051515 | 0.160662974 | 1.126924175 | 4.080472857 | 0.276174898 | 0.782413736 | NA | Fam181b | family with sequence similarity 181, meml | 0 | 0 | 0 | 0.964 | 0 | 0 |
| 15096 | ENSMUSG00000051517 | 6.471433402 | -0.533049494 | 0.876236749 | -0.608339577 | 0.542962277 | 0.894344786 | Arhgef39 | Rho guanine nucleotide exchange factor ( | 5.6217 | 13.363 | 4.025 | 4.8199 | 6.5437 | 4.4554 |
| 15097 | ENSMUSG00000051518 | 217.3042699 | -0.129986961 | 0.175807281 | -0.739371889 | 0.459681207 | 0.859753653 | Rps19bp1 | ribosomal protein S19 binding protein 1 [S | 226.47 | 246.7 | 208.29 | 220.75 | 179.95 | 221.65 |
| 15098 | ENSMUSG00000051527 | 0 | NA | NA | NA | NA | NA | Usp29 | ubiquitin specific peptidase 29 [Source:M | 0 | 0 | 0 | 0 | 0 | 0 |
| 15099 | ENSMUSG00000051528 | 0 | NA | NA | NA | NA | NA | Olfr424 | olfactory receptor 424 [Source:MGI Symb | 0 | 0 | 0 | 0 | 0 | 0 |
| 15100 | ENSMUSG00000051537 | 0 | NA | NA | NA | NA | NA | Gm5124 | predicted pseudogene 5124 [Source:MGI | 0 | 0 | 0 | 0 | 0 | 0 |
| 15101 | ENSMUSG00000051548 | 0.133850373 | -0.796629176 | 4.080472857 | -0.195229623 | 0.84521319 | NA | Gm6365 | predicted pseudogene 6365 [Source:MGI | 0.8031 | 0 | 0 | 0 | 0 | 0 |
| 15102 | ENSMUSG00000051550 | 89.50663965 | 0.001304582 | 0.24305472 | 0.005367441 | 0.995717423 | 0.999003684 | Zfp579 | zinc finger protein 579 [Source:MGI Symb | 82.72 | 87.373 | 98.612 | 97.362 | 78.525 | 92.449 |
| 15103 | ENSMUSG00000051557 | 125.1237296 | 0.022546274 | 0.217715841 | 0.103558262 | 0.917519912 | 0.986374671 | Pusl1 | pseudouridylate synthase-like 1 [Source:N | 132.51 | 120.27 | 119.74 | 142.67 | 131.96 | 103.59 |
| 15104 | ENSMUSG00000051579 | 310.3644404 | -1.145710366 | 0.260997961 | -4.38972919 | 1.13E-05 | 0.000590485 | Tceal8 | transcription elongation factor A (SII)-like | 587.07 | 466.67 | 228.42 | 204.36 | 195.22 | 180.44 |
| 15105 | ENSMUSG00000051582 | 0.667397476 | -1.404797968 | 2.804497115 | -0.500909044 | 0.616435137 | NA | Otud6a | OTU domain containing 6A [Source:MGI S | 0 | 1.0279 | 2.0125 | 0.964 | 0 | 0 |
| 15106 | ENSMUSG00000051586 | 494.9165654 | -0.070287371 | 0.152843806 | -0.459864047 | 0.645613808 | 0.925643431 | Mical3 | microtubule associated monooxygenase, | 501.94 | 540.68 | 477.97 | 423.19 | 581.3 | 444.42 |
| 15107 | ENSMUSG00000051590 | 0.603115608 | -2.642822926 | 3.106511347 | -0.850736608 | 0.394915683 | NA | Map3k19 | mitogen-activated protein kinase kinase k | 1.6062 | 0 | 2.0125 | 0 | 0 | 0 |
| 15108 | ENSMUSG00000051591 | 0.17131964 | -0.796629176 | 4.080472857 | -0.195229623 | 0.84521319 | NA | Olfr697 | olfactory receptor 697 [Source:MGI Symb | 0 | 1.0279 | 0 | 0 | 0 | 0 |
| 15109 | ENSMUSG00000051592 | 0 | NA | NA | NA | NA | NA | Ccnb3 | cyclin B3 [Source:MGI Symbol;Acc:MGI:21 | 0 | 0 | 0 | 0 | 0 | 0 |
| 15110 | ENSMUSG00000051593 | 0.36353943 | 2.065989254 | 4.034137301 | 0.512126658 | 0.608562368 | NA | Olfr272 | olfactory receptor 272 [Source:MGI Symb | 0 | 0 | 0 | 0 | 2.1812 | 0 |
| 15111 | ENSMUSG00000051596 | 0.321325949 | 1.911575114 | 4.048267893 | 0.472195804 | 0.636787034 | NA | Otop1 | otopetrin 1 [Source:MGI Symbol;Acc:MGI | 0 | 0 | 0 | 1.928 | 0 | 0 |
| 15112 | ENSMUSG00000051599 | 0 | NA | NA | NA | NA | NA | Pcdhb2 | protocadherin beta 2 [Source:MGI Symbo | 0 | 0 | 0 | 0 | 0 | 0 |
| 15113 | ENSMUSG00000051606 | 0.888490847 | -1.926390864 | 2.548063777 | -0.756021447 | 0.449636337 | NA | 2010001K21 | RIKEN cDNA 2010001K21 gene [Source:M | 3.2124 | 1.0279 | 0 | 0 | 1.0906 | 0 |
| 15114 | ENSMUSG00000051611 | 0 | NA | NA | NA | NA | NA | Olfr112 | olfactory receptor 112 [Source:MGI Symb | 0 | 0 | 0 | 0 | 0 | 0 |
| 15115 | ENSMUSG00000051615 | 587.7279269 | 0.207133543 | 0.133150056 | 1.555639925 | 0.119793739 | 0.50648835 | Rap2a | RAS related protein 2a [Source:MGI Symb | 487.48 | 583.86 | 566.52 | 626.59 | 682.73 | 579.2 |
| 15116 | ENSMUSG00000051616 | 0 | NA | NA | NA | NA | NA | C230029F24 | RIKEN cDNA C230029F24 gene [Source:M | 0 | 0 | 0 | 0 | 0 | 0 |

| | | | | | | | | | | | | | | | |
|---|---|---|---|---|---|---|---|---|---|---|---|---|---|---|---|
| 15117 | ENSMUSG00000051617 | 0 | NA | NA | NA | NA | NA | Krt9 | keratin 9 [Source:MGI Symbol;Acc:MGI:96 | 0 | 0 | 0 | 0 | 0 | 0 |
| 15118 | ENSMUSG00000051618 | 0 | NA | NA | NA | NA | NA | Ubqln3 | ubiquilin 3 [Source:MGI Symbol;Acc:MGI:1 | 0 | 0 | 0 | 0 | 0 | 0 |
| 15119 | ENSMUSG00000051627 | 9.223728192 | -0.237989629 | 0.868397117 | -0.274056217 | 0.78404142 | 0.960670212 | H1f4 | H1.4 linker histone, cluster member [Sour | 4.8186 | 2.0558 | 23.144 | 7.7118 | 9.8156 | 7.7969 |
| 15120 | ENSMUSG00000051638 | 0 | NA | NA | NA | NA | NA | Gm9857 | predicted gene 9857 [Source:MGI Symbol | 0 | 0 | 0 | 0 | 0 | 0 |
| 15121 | ENSMUSG00000051639 | 0 | NA | NA | NA | NA | NA | Fbl-ps2 | fibrillarin, pseudogene 2 [Source:MGI Sym | 0 | 0 | 0 | 0 | 0 | 0 |
| 15122 | ENSMUSG00000051648 | 0 | NA | NA | NA | NA | NA | Kctd19 | potassium channel tetramerisation domai | 0 | 0 | 0 | 0 | 0 | 0 |
| 15123 | ENSMUSG00000051650 | 52.78691285 | -0.019676081 | 0.30191064 | -0.065171871 | 0.948037157 | 0.990682895 | B3gnt2 | UDP-GlcNAc:betaGal beta-1,3-N-acetylglu | 49.792 | 56.535 | 53.331 | 57.839 | 47.987 | 51.237 |
| 15124 | ENSMUSG00000051652 | 1982.937689 | 0.130344769 | 0.124222329 | 1.049286146 | 0.29404644 | 0.745164683 | Lrrc3 | leucine rich repeat containing 3 [Source:N | 1648.8 | 1799.9 | 2232.9 | 2084.1 | 2109.3 | 2022.7 |
| 15125 | ENSMUSG00000051663 | 0 | NA | NA | NA | NA | NA | Pcdhb1 | protocadherin beta 1 [Source:MGI Symbo | 0 | 0 | 0 | 0 | 0 | 0 |
| 15126 | ENSMUSG00000051669 | 28.9120611 | 0.138385805 | 0.418676504 | 0.330531578 | 0.740998337 | 0.948792385 | AU021092 | expressed sequence AU021092 [Source:M | 24.896 | 23.642 | 34.212 | 36.631 | 22.903 | 31.188 |
| 15127 | ENSMUSG00000051671 | 797.0104345 | -0.363674402 | 0.139520563 | -2.606600728 | 0.009144592 | 0.101502835 | Coa6 | cytochrome c oxidase assembly factor 6 [S | 908.31 | 893.26 | 888.51 | 600.56 | 837.59 | 653.83 |
| 15128 | ENSMUSG00000051674 | 658.9425344 | 0.432708541 | 0.135917265 | 3.183617199 | 0.001454472 | 0.027717373 | Dcun1d4 | DCN1, defective in cullin neddylation 1, do | 636.06 | 560.22 | 485.01 | 746.12 | 718.72 | 807.54 |
| 15129 | ENSMUSG00000051675 | 153.4098786 | -0.021780455 | 0.207385825 | -0.105023836 | 0.916356895 | 0.986374671 | Trim32 | tripartite motif-containing 32 [Source:MG | 142.15 | 132.6 | 189.17 | 151.34 | 157.05 | 148.14 |
| 15130 | ENSMUSG00000051678 | 0 | NA | NA | NA | NA | NA | Pcdhb6 | protocadherin beta 6 [Source:MGI Symbo | 0 | 0 | 0 | 0 | 0 | 0 |
| 15131 | ENSMUSG00000051680 | 0 | NA | NA | NA | NA | NA | Olfr693 | olfactory receptor 693 [Source:MGI Symb | 0 | 0 | 0 | 0 | 0 | 0 |
| 15132 | ENSMUSG00000051682 | 31.14535983 | -1.571795108 | 0.496954518 | -3.16285505 | 0.001562301 | 0.029385073 | Treml4 | triggering receptor expressed on myeloid | 42.564 | 70.926 | 26.162 | 8.6758 | 20.722 | 17.822 |
| 15133 | ENSMUSG00000051687 | 0 | NA | NA | NA | NA | NA | Vmn1r73 | vomeronasal 1 receptor 73 [Source:MGI S | 0 | 0 | 0 | 0 | 0 | 0 |
| 15134 | ENSMUSG00000051695 | 3200.90031 | -0.076076459 | 0.087539644 | -0.869051504 | 0.384818961 | 0.816437825 | Pcbp1 | poly(rC) binding protein 1 [Source:MGI Sy | 3407.6 | 3169.1 | 3278.3 | 3094.4 | 3105 | 3151.1 |
| 15135 | ENSMUSG00000051703 | 3.433670522 | -0.38972059 | 1.413441398 | -0.275724618 | 0.782759586 | NA | Tmem198 | transmembrane protein 198 [Source:MGI | 9.6372 | 0 | 2.0125 | 6.7478 | 1.0906 | 1.1138 |
| 15136 | ENSMUSG00000051705 | 50.58620011 | 0.220934458 | 0.323745646 | 0.682432214 | 0.494965692 | 0.877919732 | Senp8 | SUMO/sentrin specific peptidase 8 [Sourc | 51.399 | 40.089 | 48.3 | 42.415 | 56.712 | 64.603 |
| 15137 | ENSMUSG00000051706 | 0 | NA | NA | NA | NA | NA | Olfr1325 | olfactory receptor 1325 [Source:MGI Syml | 0 | 0 | 0 | 0 | 0 | 0 |
| 15138 | ENSMUSG00000051716 | 7062.994938 | 0.017346433 | 0.087712499 | 0.197764661 | 0.843229196 | 0.972273952 | Apon | apolipoprotein N [Source:MGI Symbol;Acc | 7407 | 6703.1 | 6950.1 | 6912.7 | 7334.4 | 7070.7 |
| 15139 | ENSMUSG00000051721 | 159.2578013 | 0.326054571 | 0.213937825 | 1.524062287 | 0.127493152 | 0.523285632 | Wdcp | WD repeat and coiled coil containing [Sou | 145.36 | 126.43 | 151.94 | 170.62 | 142.87 | 218.31 |
| 15140 | ENSMUSG00000051723 | 0 | NA | NA | NA | NA | NA | Rpl31-ps13 | ribosomal protein L31, pseudogene 13 [Sc | 0 | 0 | 0 | 0 | 0 | 0 |
| 15141 | ENSMUSG00000051726 | 0 | NA | NA | NA | NA | NA | Kcnf1 | potassium voltage-gated channel, subfam | 0 | 0 | 0 | 0 | 0 | 0 |
| 15142 | ENSMUSG00000051727 | 1.375223984 | 0.03098273 | 2.170995137 | 0.014271211 | 0.988613607 | NA | Kctd14 | potassium channel tetramerisation domai | 4.0155 | 0 | 0 | 0.964 | 3.2719 | 0 |
| 15143 | ENSMUSG00000051728 | 0.513958919 | -2.394274821 | 3.756096013 | -0.637437066 | 0.523840191 | NA | Fam243 | family with sequence similarity 243 [Sourc | 0 | 3.0838 | 0 | 0 | 0 | 0 |
| 15144 | ENSMUSG00000051729 | 0 | NA | NA | NA | NA | NA | Gm5087 | predicted gene 5087 [Source:MGI Symbol | 0 | 0 | 0 | 0 | 0 | 0 |
| 15145 | ENSMUSG00000051730 | 256.5738044 | -0.019432729 | 0.161071049 | -0.12064694 | 0.903970688 | 0.983821605 | Mettl5 | methyltransferase like 5 [Source:MGI Sym | 236.92 | 252.87 | 285.77 | 268.95 | 243.21 | 251.73 |
| 15146 | ENSMUSG00000051732 | 0 | NA | NA | NA | NA | NA | Pabpc2 | poly(A) binding protein, cytoplasmic 2 [So | 0 | 0 | 0 | 0 | 0 | 0 |
| 15147 | ENSMUSG00000051735 | 19.13439411 | -0.88670819 | 0.512603422 | -1.729813246 | 0.083663647 | 0.426388452 | Rinl | Ras and Rab interactor-like [Source:MGI S | 22.487 | 31.865 | 20.125 | 10.604 | 16.359 | 13.366 |
| 15148 | ENSMUSG00000051736 | 6.183913793 | -0.59011372 | 0.85061884 | -0.693746356 | 0.48784129 | 0.873910634 | Fam229b | family with sequence similarity 229, meml | 8.031 | 6.1675 | 8.05 | 3.8559 | 6.5437 | 4.4554 |
| 15149 | ENSMUSG00000051747 | 2.623189119 | -2.754942746 | 1.652964298 | -1.666668028 | 0.095580434 | NA | Ttn | titin [Source:MGI Symbol;Acc:MGI:98864] | 1.6062 | 6.1675 | 6.0375 | 1.928 | 0 | 0 |
| 15150 | ENSMUSG00000051748 | 6804.090147 | 0.318748686 | 0.095823079 | 3.326429193 | 0.000879664 | 0.019075645 | Wfdc21 | WAP four-disulfide core domain 21 [Sourc | 6149.4 | 6496.4 | 5520.3 | 7360.9 | 7478.4 | 7819.2 |
| 15151 | ENSMUSG00000051758 | 0 | NA | NA | NA | NA | NA | 4930544M13 | RIKEN cDNA 4930544M13 gene [Source:M | 0 | 0 | 0 | 0 | 0 | 0 |
| 15152 | ENSMUSG00000051768 | 231.9374197 | -0.020242843 | 0.158738511 | -0.127523203 | 0.89852631 | 0.982798755 | Xrcc1 | X-ray repair complementing defective rep | 240.93 | 224.09 | 235.46 | 230.39 | 226.85 | 233.91 |
| 15153 | ENSMUSG00000051769 | 0 | NA | NA | NA | NA | NA | Wfdc15a | WAP four-disulfide core domain 15A [Sou | 0 | 0 | 0 | 0 | 0 | 0 |
| 15154 | ENSMUSG00000051777 | 0 | NA | NA | NA | NA | NA | Iqcj | IQ motif containing J [Source:MGI Symbol | 0 | 0 | 0 | 0 | 0 | 0 |
| 15155 | ENSMUSG00000051786 | 211.8725508 | -0.170331557 | 0.173587588 | -0.981242716 | 0.326473064 | 0.772216908 | Tubgcp6 | tubulin, gamma complex associated prote | 223.26 | 208.67 | 241.5 | 220.75 | 182.13 | 194.92 |
| 15156 | ENSMUSG00000051788 | 0 | NA | NA | NA | NA | NA | 4930564D02 | RIKEN cDNA 4930564D02 gene [Source:M | 0 | 0 | 0 | 0 | 0 | 0 |
| 15157 | ENSMUSG00000051790 | 38.13943359 | -0.290112159 | 0.394581336 | -0.73524045 | 0.462193068 | 0.860650866 | Nlgn2 | neuroligin 2 [Source:MGI Symbol;Acc:MG | 37.746 | 42.145 | 46.287 | 46.271 | 19.631 | 36.757 |
| 15158 | ENSMUSG00000051793 | 0 | NA | NA | NA | NA | NA | Olfr284 | olfactory receptor 284 [Source:MGI Symb | 0 | 0 | 0 | 0 | 0 | 0 |
| 15159 | ENSMUSG00000051802 | 0 | NA | NA | NA | NA | NA | Krtap19-5 | keratin associated protein 19-5 [Source:M | 0 | 0 | 0 | 0 | 0 | 0 |
| 15160 | ENSMUSG00000051804 | 0 | NA | NA | NA | NA | NA | Adam6b | a disintegrin and metallopeptidase domai | 0 | 0 | 0 | 0 | 0 | 0 |
| 15161 | ENSMUSG00000051805 | 0.321325949 | 1.911575114 | 4.048267893 | 0.472195804 | 0.636787034 | NA | A530095I07I | RIKEN cDNA A530095I07 gene [Source:MG | 0 | 0 | 0 | 1.928 | 0 | 0 |
| 15162 | ENSMUSG00000051811 | 4.401720916 | 0.459995016 | 1.07768864 | 0.426834801 | 0.669499657 | NA | Cox6b2 | cytochrome c oxidase subunit 6B2 [Source | 4.0155 | 5.1396 | 2.0125 | 8.6758 | 5.4531 | 1.1138 |
| 15163 | ENSMUSG00000051817 | 127.1355854 | 0.230935884 | 0.315757723 | 0.7313705 | 0.464552879 | 0.862553639 | Sox12 | SRY (sex determining region Y)-box 12 [So | 65.854 | 101.76 | 184.14 | 161.95 | 123.24 | 125.86 |
| 15164 | ENSMUSG00000051827 | 0 | NA | NA | NA | NA | NA | Rhox2a | reproductive homeobox 2A [Source:MGI S | 0 | 0 | 0 | 0 | 0 | 0 |
| 15165 | ENSMUSG00000051832 | 0 | NA | NA | NA | NA | NA | E230016K23 | RIKEN cDNA E230016K23 gene [Source:MG | 0 | 0 | 0 | 0 | 0 | 0 |
| 15166 | ENSMUSG00000051839 | 2.197526969 | 1.966118561 | 1.555145955 | 1.264266261 | 0.206134475 | NA | Gypa | glycophorin A [Source:MGI Symbol;Acc:M | 1.6062 | 0 | 1.0062 | 2.8919 | 5.4531 | 2.2277 |
| 15167 | ENSMUSG00000051844 | 3.208979429 | -0.039029426 | 1.260275494 | -0.030968964 | 0.975294291 | NA | B230319C09 | RIKEN cDNA B230319C09 gene [Source:M | 5.6217 | 1.0279 | 3.0187 | 1.928 | 6.5437 | 1.1138 |
| 15168 | ENSMUSG00000051851 | 7.810067143 | -0.269224595 | 0.784730078 | -0.343079235 | 0.731538853 | 0.947352311 | Rtl8c | retrotransposon Gag like 8C [Source:MGI | 7.2279 | 13.363 | 5.0312 | 5.7839 | 6.5437 | 8.9108 |
| 15169 | ENSMUSG00000051853 | 1166.488174 | 0.097422881 | 0.108586996 | 0.897187368 | 0.369618945 | 0.80653433 | Arf3 | ADP-ribosylation factor 3 [Source:MGI Syr | 1087.4 | 1170.8 | 1123 | 1106.6 | 1231.3 | 1279.8 |
| 15170 | ENSMUSG00000051855 | 17.5548817 | 0.103702374 | 0.604378076 | 0.171585267 | 0.863763591 | 0.976954351 | Mest | mesoderm specific transcript [Source:MG | 24.093 | 9.2513 | 17.106 | 10.604 | 13.087 | 31.188 |
| 15171 | ENSMUSG00000051860 | 0 | NA | NA | NA | NA | NA | Samd7 | sterile alpha motif domain containing 7 [S | 0 | 0 | 0 | 0 | 0 | 0 |
| 15172 | ENSMUSG00000051864 | 280.8790001 | 0.101658252 | 0.16841473 | 0.603618533 | 0.546097289 | 0.89549457 | Tbc1d22a | TBC1 domain family, member 22a [Source | 280.28 | 275.48 | 256.59 | 259.31 | 267.2 | 346.41 |

| | | | | | | | | | | | | | | | |
|---|---|---|---|---|---|---|---|---|---|---|---|---|---|---|---|
| 15173 | ENSMUSG00000051877 | 0 | NA | NA | NA | NA | NA | Vmn2r72 | vomeronasal 2, receptor 72 [Source:MGI S | 0 | 0 | 0 | 0 | 0 | 0 |
| 15174 | ENSMUSG00000051879 | 0 | NA | NA | NA | NA | NA | Krt71 | keratin 71 [Source:MGI Symbol;Acc:MGI:1 | 0 | 0 | 0 | 0 | 0 | 0 |
| 15175 | ENSMUSG00000051885 | 0 | NA | NA | NA | NA | NA | Olfr664 | olfactory receptor 664 [Source:MGI Symb | 0 | 0 | 0 | 0 | 0 | 0 |
| 15176 | ENSMUSG00000051890 | 1.914942262 | -0.874285832 | 1.582094445 | -0.55261292 | 0.580528493 | NA | Klhdc1 | kelch domain containing 1 [Source:MGI Sy | 2.4093 | 2.0558 | 3.0187 | 2.8919 | 0 | 1.1138 |
| 15177 | ENSMUSG00000051896 | 0 | NA | NA | NA | NA | NA | Tex37 | testis expressed 37 [Source:MGI Symbol;A | 0 | 0 | 0 | 0 | 0 | 0 |
| 15178 | ENSMUSG00000051900 | 0 | NA | NA | NA | NA | NA | Abca16 | ATP-binding cassette, sub-family A (ABC1) | 0 | 0 | 0 | 0 | 0 | 0 |
| 15179 | ENSMUSG00000051906 | 0.640584875 | -2.718595279 | 2.766810251 | -0.982573806 | 0.325817242 | NA | Cd209f | CD209f antigen [Source:MGI Symbol;Acc:I | 0.8031 | 1.0279 | 2.0125 | 0 | 0 | 0 |
| 15180 | ENSMUSG00000051910 | 245.1897027 | 0.11782634 | 0.184290334 | 0.639351712 | 0.522594155 | 0.888334265 | Sox6 | SRY (sex determining region Y)-box 6 [Sou | 277.07 | 238.48 | 189.17 | 247.74 | 279.2 | 239.48 |
| 15181 | ENSMUSG00000051917 | 0 | NA | NA | NA | NA | NA | Tas2r144 | taste receptor, type 2, member 144 [Sour | 0 | 0 | 0 | 0 | 0 | 0 |
| 15182 | ENSMUSG00000051920 | 0 | NA | NA | NA | NA | NA | Rspo2 | R-spondin 2 [Source:MGI Symbol;Acc:MG | 0 | 0 | 0 | 0 | 0 | 0 |
| 15183 | ENSMUSG00000051934 | 83.56176715 | -0.173010437 | 0.251375793 | -0.688254168 | 0.491292738 | 0.876368691 | Spats2 | spermatogenesis associated, serine-rich 2 | 92.357 | 94.568 | 78.487 | 68.442 | 83.978 | 83.538 |
| 15184 | ENSMUSG00000051936 | 0 | NA | NA | NA | NA | NA | Prss58 | protease, serine 58 [Source:MGI Symbol;A | 0 | 0 | 0 | 0 | 0 | 0 |
| 15185 | ENSMUSG00000051940 | 0 | NA | NA | NA | NA | NA | 5031410I06F | RIKEN cDNA 5031410I06 gene [Source:MG | 0 | 0 | 0 | 0 | 0 | 0 |
| 15186 | ENSMUSG00000051950 | 35.35186088 | 0.015303689 | 0.417598655 | 0.036646882 | 0.970766562 | 0.994966572 | B3glct | beta-3-glucosyltransferase [Source:MGI S | 22.487 | 45.228 | 38.237 | 42.415 | 40.353 | 23.391 |
| 15187 | ENSMUSG00000051951 | 0.910352643 | 1.221794025 | 2.770215377 | 0.441046583 | 0.659179274 | NA | Xkr4 | X-linked Kx blood group related 4 [Source | 1.6062 | 0 | 0 | 3.8559 | 0 | 0 |
| 15188 | ENSMUSG00000051952 | 0 | NA | NA | NA | NA | NA | Olfr371 | olfactory receptor 371 [Source:MGI Symb | 0 | 0 | 0 | 0 | 0 | 0 |
| 15189 | ENSMUSG00000051956 | 0.17131964 | -0.796629176 | 4.080472857 | -0.195229623 | 0.84521319 | NA | Rnf133 | ring finger protein 133 [Source:MGI Symb | 0 | 1.0279 | 0 | 0 | 0 | 0 |
| 15190 | ENSMUSG00000051965 | 0 | NA | NA | NA | NA | NA | Nanos2 | nanos C2HC-type zinc finger 2 [Source:MG | 0 | 0 | 0 | 0 | 0 | 0 |
| 15191 | ENSMUSG00000051969 | 3.327764994 | -1.09388055 | 1.187998486 | -0.920776048 | 0.357167363 | NA | Tlr11 | toll-like receptor 11 [Source:MGI Symbol;A | 2.4093 | 7.1954 | 4.025 | 1.928 | 2.1812 | 2.2277 |
| 15192 | ENSMUSG00000051977 | 93.65152287 | 0.150505223 | 0.233321849 | 0.645054133 | 0.518892106 | 0.887592015 | Prdm9 | PR domain containing 9 [Source:MGI Sym | 84.326 | 97.652 | 84.525 | 100.25 | 93.793 | 101.36 |
| 15193 | ENSMUSG00000051978 | 53.2530257 | 0.214418911 | 0.319855255 | 0.670362321 | 0.502626848 | 0.881220881 | Erich1 | glutamate rich 1 [Source:MGI Symbol;Acc | 48.989 | 56.535 | 42.262 | 49.163 | 50.168 | 72.4 |
| 15194 | ENSMUSG00000051980 | 0 | NA | NA | NA | NA | NA | Casr | calcium-sensing receptor [Source:MGI Syr | 0 | 0 | 0 | 0 | 0 | 0 |
| 15195 | ENSMUSG00000051984 | 198.2668152 | 0.153608691 | 0.219188892 | 0.700805088 | 0.483424662 | 0.871960882 | Sec31b | Sec31 homolog B (S. cerevisiae) [Source:M | 187.93 | 163.44 | 212.32 | 266.06 | 205.04 | 154.82 |
| 15196 | ENSMUSG00000051985 | 0.160662974 | 1.126924175 | 4.080472857 | 0.276174898 | 0.782413736 | NA | Igfn1 | immunoglobulin-like and fibronectin type | 0 | 0 | 0 | 0.964 | 0 | 0 |
| 15197 | ENSMUSG00000051986 | 0.733533733 | -1.621434111 | 2.70314781 | -0.599831835 | 0.548618315 | NA | A530006G24 | RIKEN cDNA A530006G24 gene [Source:M | 2.4093 | 1.0279 | 0 | 0.964 | 0 | 0 |
| 15198 | ENSMUSG00000051989 | 194.9919374 | -0.063876969 | 0.183892694 | -0.347360016 | 0.728320868 | 0.946208446 | Smim11 | small integral membrane protein 11 [Sour | 196.76 | 202.5 | 198.23 | 158.09 | 208.31 | 206.06 |
| 15199 | ENSMUSG00000051998 | 2.570125443 | -0.024490512 | 1.647786004 | -0.014862678 | 0.988141735 | NA | Lax1 | lymphocyte transmembrane adaptor 1 [Sc | 1.6062 | 6.1675 | 0 | 0.964 | 0 | 6.6831 |
| 15200 | ENSMUSG00000052005 | 0 | NA | NA | NA | NA | NA | Gm9864 | predicted gene 9864 [Source:MGI Symbol | 0 | 0 | 0 | 0 | 0 | 0 |
| 15201 | ENSMUSG00000052012 | 0 | NA | NA | NA | NA | NA | Olfr796 | olfactory receptor 796 [Source:MGI Symb | 0 | 0 | 0 | 0 | 0 | 0 |
| 15202 | ENSMUSG00000052013 | 5.206060793 | -1.625873691 | 1.100488116 | -1.477411402 | 0.139565392 | 0.545861088 | Btla | B and T lymphocyte associated [Source:M | 3.2124 | 9.2513 | 11.069 | 0 | 4.3625 | 3.3415 |
| 15203 | ENSMUSG00000052014 | 0 | NA | NA | NA | NA | NA | A330070K13 | RIKEN cDNA A330070K13 gene [Source:M | 0 | 0 | 0 | 0 | 0 | 0 |
| 15204 | ENSMUSG00000052025 | 0 | NA | NA | NA | NA | NA | Ppp1r2-ps7 | protein phosphatase 1, regulatory (inhibit | 0 | 0 | 0 | 0 | 0 | 0 |
| 15205 | ENSMUSG00000052026 | 1.386662335 | -0.878148267 | 2.121038941 | -0.414017984 | 0.678860936 | NA | Slc6a7 | solute carrier family 6 (neurotransmitter t | 2.4093 | 0 | 3.0187 | 2.8919 | 0 | 0 |
| 15206 | ENSMUSG00000052031 | 152.7672236 | 0.458041862 | 0.222638385 | 2.05733554 | 0.039653961 | 0.272725843 | Tagap1 | T cell activation GTPase activating protein | 146.97 | 95.596 | 142.89 | 164.84 | 163.59 | 202.72 |
| 15207 | ENSMUSG00000052033 | 92.27831248 | -0.282806603 | 0.250829283 | -1.127486388 | 0.259536892 | 0.712728614 | Pfdn4 | prefoldin 4 [Source:MGI Symbol;Acc:MGI: | 93.16 | 123.35 | 87.543 | 81.938 | 76.343 | 91.335 |
| 15208 | ENSMUSG00000052040 | 1396.985197 | -0.083625217 | 0.113722768 | -0.735342785 | 0.462130758 | 0.860650866 | Klf13 | Kruppel-like factor 13 [Source:MGI Symbo | 1344.4 | 1557.3 | 1410.8 | 1232.9 | 1379.6 | 1456.9 |
| 15209 | ENSMUSG00000052056 | 160.8565797 | 0.1999032 | 0.203223201 | 0.983663276 | 0.325281102 | 0.771501263 | Zfp217 | zinc finger protein 217 [Source:MGI Symb | 133.31 | 146.99 | 169.05 | 157.13 | 157.05 | 201.61 |
| 15210 | ENSMUSG00000052058 | 0 | NA | NA | NA | NA | NA | Olfr901 | olfactory receptor 901 [Source:MGI Symb | 0 | 0 | 0 | 0 | 0 | 0 |
| 15211 | ENSMUSG00000052062 | 109.5718047 | 0.724765992 | 0.231735512 | 3.12755687 | 0.001762657 | 0.031868841 | Pard3b | par-3 family cell polarity regulator beta [S | 99.585 | 74.01 | 73.456 | 128.21 | 139.6 | 142.57 |
| 15212 | ENSMUSG00000052075 | 0 | NA | NA | NA | NA | NA | 1700029F12 | RIKEN cDNA 1700029F12 gene [Source:MG | 0 | 0 | 0 | 0 | 0 | 0 |
| 15213 | ENSMUSG00000052085 | 588.9822717 | 0.375286691 | 0.11932855 | 3.144986595 | 0.001660944 | 0.030548062 | Dock8 | dedicator of cytokinesis 8 [Source:MGI Sy | 508.36 | 497.51 | 532.3 | 633.33 | 672.91 | 689.47 |
| 15214 | ENSMUSG00000052087 | 2.016712858 | -1.501465314 | 1.622250494 | -0.92554468 | 0.354682654 | NA | Rgs14 | regulator of G-protein signaling 14 [Sourc | 4.8186 | 3.0838 | 1.0062 | 0.964 | 0 | 2.2277 |
| 15215 | ENSMUSG00000052099 | 0 | NA | NA | NA | NA | NA | Prss51 | protease, serine 51 [Source:MGI Symbol;A | 0 | 0 | 0 | 0 | 0 | 0 |
| 15216 | ENSMUSG00000052102 | 531.7801943 | 0.259693579 | 0.170172348 | 1.526062148 | 0.126994388 | 0.522128148 | Gnpda1 | glucosamine-6-phosphate deaminase 1 [S | 550.93 | 442 | 457.84 | 485.84 | 528.95 | 725.11 |
| 15217 | ENSMUSG00000052105 | 4.395187865 | -0.866637687 | 1.037921854 | -0.834973927 | 0.403732388 | NA | Mtcl1 | microtubule crosslinking factor 1 [Source: | 8.8341 | 5.1396 | 3.0187 | 3.8559 | 2.1812 | 3.3415 |
| 15218 | ENSMUSG00000052117 | 243.6445597 | 0.964357087 | 0.258529736 | 3.730159247 | 0.000191359 | 0.006096505 | D630039A03 | RIKEN cDNA D630039A03 gene [Source:M | 223.26 | 129.52 | 141.88 | 439.57 | 224.67 | 302.97 |
| 15219 | ENSMUSG00000052125 | 2.244624634 | -0.23538371 | 1.371958156 | -0.171567704 | 0.8637774 | NA | F730043M19 | RIKEN cDNA F730043M19 gene [Source:M | 3.2124 | 2.0558 | 2.0125 | 2.8919 | 2.1812 | 1.1138 |
| 15220 | ENSMUSG00000052131 | 0.842381458 | 0.63517692 | 2.579321552 | 0.246257362 | 0.805483014 | NA | Akr1b7 | aldo-keto reductase family 1, member B7 | 0 | 0 | 2.0125 | 1.928 | 0 | 1.1138 |
| 15221 | ENSMUSG00000052133 | 263.2880542 | -1.188804373 | 0.264636129 | -4.4922225 | 7.05E-06 | 0.000404554 | Sema5b | sema domain, seven thrombospondin rep | 433.68 | 269.31 | 395.45 | 211.11 | 179.95 | 90.221 |
| 15222 | ENSMUSG00000052135 | 1.416487074 | 0.12239615 | 1.960786118 | 0.062421979 | 0.950226792 | NA | Foxo6 | forkhead box O6 [Source:MGI Symbol;Acc | 0 | 3.0838 | 1.0062 | 0 | 2.1812 | 2.2277 |
| 15223 | ENSMUSG00000052137 | 56.04243619 | -0.150538206 | 0.299203806 | -0.503129315 | 0.614873357 | 0.917509899 | Rbm12b2 | RNA binding motif protein 12 B2 [Source:I | 57.823 | 60.647 | 58.362 | 50.127 | 45.806 | 63.489 |
| 15224 | ENSMUSG00000052139 | 324.2573898 | -0.129691195 | 0.147633246 | -0.878468761 | 0.379689383 | 0.812628154 | Babam2 | BRISC and BRCA1 A complex member 2 [S | 370.23 | 308.38 | 337.09 | 302.69 | 310.83 | 316.33 |
| 15225 | ENSMUSG00000052142 | 11.06027788 | -0.951491618 | 0.734798885 | -1.294900737 | 0.195354474 | 0.635641153 | Rasal3 | RAS protein activator like 3 [Source:MGI S | 20.078 | 13.363 | 10.062 | 1.928 | 10.906 | 10.025 |
| 15226 | ENSMUSG00000052143 | 2.141890965 | 0.976617732 | 1.71273127 | 0.570210721 | 0.568534785 | NA | Gm9869 | predicted gene 9869 [Source:MGI Symbol | 3.2124 | 1.0279 | 0 | 1.928 | 0 | 6.6831 |
| 15227 | ENSMUSG00000052144 | 600.1672399 | 0.012580159 | 0.127306226 | 0.098818101 | 0.921282697 | 0.986685804 | Ppp4r2 | protein phosphatase 4, regulatory subunit | 566.19 | 632.17 | 594.69 | 581.28 | 664.19 | 562.49 |
| 15228 | ENSMUSG00000052146 | 28.95485194 | -0.350827309 | 0.448921337 | -0.781489494 | 0.434514653 | 0.845264432 | Rps10 | ribosomal protein S10 [Source:MGI Symbo | 38.549 | 26.726 | 32.2 | 37.595 | 15.269 | 23.391 |

| | | | | | | | | | | | | | | | |
|---|---|---|---|---|---|---|---|---|---|---|---|---|---|---|---|
| 15229 | ENSMUSG00000052151 | 348.9803847 | 0.204606595 | 0.147871966 | 1.383674006 | 0.166458296 | 0.5925379 | Plpp2 | phospholipid phosphatase 2 [Source:MGI | 317.23 | 315.57 | 340.11 | 362.46 | 418.8 | 339.72 |
| 15230 | ENSMUSG00000052154 | 1.06319113 | 2.404631922 | 2.210715176 | 1.087716748 | 0.276720172 | NA | Gm9870 | predicted gene 9870 [Source:MGI Symbol | 0 | 0 | 1.0062 | 0.964 | 2.1812 | 2.2277 |
| 15231 | ENSMUSG00000052155 | 430.1647912 | -0.132758487 | 0.14841716 | -0.894495538 | 0.371056816 | 0.807214774 | Acvr2a | activin receptor IIA [Source:MGI Symbol;A | 430.46 | 503.68 | 416.59 | 455.96 | 410.07 | 364.23 |
| 15232 | ENSMUSG00000052157 | 0 | NA | NA | NA | NA | NA | Gm650 | predicted pseudogene 650 [Source:MGI S | 0 | 0 | 0 | 0 | 0 | 0 |
| 15233 | ENSMUSG00000052160 | 67.64867557 | -1.056996294 | 0.338372308 | -3.123767136 | 0.001785517 | 0.032166791 | Pld4 | phospholipase D family, member 4 [Sourc | 83.523 | 130.55 | 60.375 | 51.091 | 44.715 | 35.643 |
| 15234 | ENSMUSG00000052180 | 0 | NA | NA | NA | NA | NA | Serpinb6c | serine (or cysteine) peptidase inhibitor, cl | 0 | 0 | 0 | 0 | 0 | 0 |
| 15235 | ENSMUSG00000052182 | 0 | NA | NA | NA | NA | NA | Olfr849 | olfactory receptor 849 [Source:MGI Symb | 0 | 0 | 0 | 0 | 0 | 0 |
| 15236 | ENSMUSG00000052187 | 0 | NA | NA | NA | NA | NA | Hbb-y | hemoglobin Y, beta-like embryonic chain | | 0 | 0 | 0 | 0 | 0 | 0 |
| 15237 | ENSMUSG00000052188 | 7.759731288 | 0.738994826 | 0.8112031 | 0.910986195 | 0.362302647 | 0.80090868 | Gm14964 | predicted gene 14964 [Source:MGI Symbo | 2.4093 | 7.1954 | 8.05 | 13.496 | 8.7249 | 6.6831 |
| 15238 | ENSMUSG00000052192 | 1.568529388 | 0.516189757 | 1.786580691 | 0.288926081 | 0.772637945 | NA | Gm5963 | predicted pseudogene 5963 [Source:MGI | 0.8031 | 2.0558 | 1.0062 | 0 | 1.0906 | 4.4554 |
| 15239 | ENSMUSG00000052212 | 0 | NA | NA | NA | NA | NA | Cd177 | CD177 antigen [Source:MGI Symbol;Acc:N | 0 | 0 | 0 | 0 | 0 | 0 |
| 15240 | ENSMUSG00000052214 | 975.2173246 | 0.145481741 | 0.116579992 | 1.247913461 | 0.21206275 | 0.656735971 | Opa3 | optic atrophy 3 [Source:MGI Symbol;Acc:I | 970.95 | 896.34 | 910.65 | 1076.8 | 913.94 | 1082.7 |
| 15241 | ENSMUSG00000052217 | 0 | NA | NA | NA | NA | NA | Hbb-bh1 | hemoglobin Z, beta-like embryonic chain | | 0 | 0 | 0 | 0 | 0 | 0 |
| 15242 | ENSMUSG00000052221 | 0.331982614 | 0.16514309 | 3.986751476 | 0.041422971 | 0.966958701 | NA | Ppp1r36 | protein phosphatase 1, regulatory subunit | 0 | 1.0279 | 0 | 0.964 | 0 | 0 |
| 15243 | ENSMUSG00000052229 | 50.86081053 | -0.556768172 | 0.373374936 | -1.491177146 | 0.135914992 | 0.538690549 | Gpr17 | G protein-coupled receptor 17 [Source:M | 69.067 | 48.312 | 64.4 | 52.055 | 50.168 | 21.163 |
| 15244 | ENSMUSG00000052234 | 0 | NA | NA | NA | NA | NA | Epx | eosinophil peroxidase [Source:MGI Symbo | 0 | 0 | 0 | 0 | 0 | 0 |
| 15245 | ENSMUSG00000052236 | 0 | NA | NA | NA | NA | NA | Gm9871 | predicted gene 9871 [Source:MGI Symbol | 0 | 0 | 0 | 0 | 0 | 0 |
| 15246 | ENSMUSG00000052241 | 3.154222221 | -0.658181655 | 1.229899679 | -0.535150685 | 0.592545664 | NA | A930035D04 | RIKEN cDNA A930035D04 gene [Source:M | 6.4248 | 3.0838 | 2.0125 | 1.928 | 4.3625 | 1.1138 |
| 15247 | ENSMUSG00000052248 | 1.272470326 | -2.525347334 | 2.257707091 | -1.118545158 | 0.263334231 | NA | Zeb2os | zinc finger E-box binding homeobox 2, op | 2.4093 | 4.1117 | 0 | 0 | 0 | 1.1138 |
| 15248 | ENSMUSG00000052253 | 577.6479937 | 0.116903227 | 0.126496914 | 0.924158729 | 0.35540368 | 0.795424787 | Zfp622 | zinc finger protein 622 [Source:MGI Symb | 519.61 | 578.72 | 565.51 | 641.05 | 609.66 | 551.35 |
| 15249 | ENSMUSG00000052270 | 6.424500392 | -4.153361041 | 1.451854441 | -2.860728269 | 0.004226692 | 0.059702018 | Fpr2 | formyl peptide receptor 2 [Source:MGI Sy | 4.8186 | 28.782 | 3.0187 | 1.928 | 0 | 0 |
| 15250 | ENSMUSG00000052271 | 1.085201869 | 1.137592316 | 2.627090041 | 0.433023725 | 0.664997534 | NA | Bhlha15 | basic helix-loop-helix family, member a15 | 0 | 2.0558 | 0 | 0 | 0 | 4.4554 |
| 15251 | ENSMUSG00000052273 | 11.06556053 | -2.159033663 | 0.825718069 | -2.614734669 | 0.008929679 | 0.099736589 | Dnah3 | dynein, axonemal, heavy chain 3 [Source: | 24.896 | 17.475 | 12.075 | 8.6758 | 3.2719 | 0 |
| 15252 | ENSMUSG00000052276 | 0 | NA | NA | NA | NA | NA | Ostn | osteocrin [Source:MGI Symbol;Acc:MGI:2 | 0 | 0 | 0 | 0 | 0 | 0 |
| 15253 | ENSMUSG00000052281 | 0 | NA | NA | NA | NA | NA | Tspan32os | tetraspanin 32, opposite strand [Source:M | 0 | 0 | 0 | 0 | 0 | 0 |
| 15254 | ENSMUSG00000052291 | 27.15328662 | 0.261708577 | 0.454453432 | 0.575875454 | 0.564699368 | 0.901542552 | 5330438D12 | RIKEN cDNA 5330438D12 gene [Source:M | 19.274 | 37.005 | 18.112 | 35.667 | 28.356 | 24.505 |
| 15255 | ENSMUSG00000052293 | 2.490959916 | -0.555413893 | 1.445440504 | -0.384252338 | 0.70079143 | NA | Taf9 | TATA-box binding protein associated facto | 4.8186 | 3.0838 | 1.0062 | 3.8559 | 2.1812 | 0 |
| 15256 | ENSMUSG00000052295 | 0 | NA | NA | NA | NA | NA | 8030423F21 | RIKEN cDNA 8030423F21 gene [Source:M | 0 | 0 | 0 | 0 | 0 | 0 |
| 15257 | ENSMUSG00000052296 | 1805.727481 | 0.147308341 | 0.108248047 | 1.360840644 | 0.173564054 | 0.603759135 | Ppp6r1 | protein phosphatase 6, regulatory subunit | 1736.3 | 1685.8 | 1717.7 | 1684.1 | 1954.4 | 2056.2 |
| 15258 | ENSMUSG00000052298 | 206.5040019 | -0.042654544 | 0.17836433 | -0.239142795 | 0.810994859 | 0.966452061 | Cdc42se2 | CDC42 small effector 2 [Source:MGI Symb | 192.74 | 227.17 | 209.3 | 215.93 | 176.68 | 217.2 |
| 15259 | ENSMUSG00000052299 | 688.8768124 | -0.024747285 | 0.119187071 | -0.207633975 | 0.835514774 | 0.971073158 | Ltn1 | listerin E3 ubiquitin protein ligase 1 [Sourc | 681.03 | 744.21 | 659.09 | 649.72 | 721.99 | 677.22 |
| 15260 | ENSMUSG00000052301 | 0 | NA | NA | NA | NA | NA | Doc2a | double C2, alpha [Source:MGI Symbol;Acc | 0 | 0 | 0 | 0 | 0 | 0 |
| 15261 | ENSMUSG00000052302 | 257.2712746 | 1.045040946 | 0.215713973 | 4.844567691 | 1.27E-06 | 9.92E-05 | Tbc1d30 | TBC1 domain family, member 30 [Source: | 178.29 | 135.69 | 189.17 | 248.71 | 374.08 | 417.69 |
| 15262 | ENSMUSG00000052303 | 0 | NA | NA | NA | NA | NA | Mrgpra6 | MAS-related GPR, member A6 [Source:M | 0 | 0 | 0 | 0 | 0 | 0 |
| 15263 | ENSMUSG00000052305 | 2343.274728 | 0.290803637 | 0.167216689 | 1.739082611 | 0.082020233 | 0.421585163 | Hbb-bs | hemoglobin, beta adult s chain [Source:M | 1613.4 | 1966.4 | 2745 | 2540.1 | 2314.3 | 2880.4 |
| 15264 | ENSMUSG00000052310 | 1243.754726 | 0.275878114 | 0.100216517 | 2.752820804 | 0.005908423 | 0.07450361 | Slc39a1 | solute carrier family 39 (zinc transporter), | 1094.6 | 1171.8 | 1109.9 | 1404.5 | 1329.5 | 1352.2 |
| 15265 | ENSMUSG00000052316 | 45.15211593 | -1.200525328 | 0.400292901 | -2.999117206 | 0.002707631 | 0.043708904 | Lrrc15 | leucine rich repeat containing 15 [Source: | 68.264 | 57.563 | 63.393 | 45.307 | 17.45 | 18.935 |
| 15266 | ENSMUSG00000052331 | 76.71483311 | -0.167411794 | 0.296862706 | -0.563936766 | 0.572797168 | 0.905173631 | Ankrd44 | ankyrin repeat domain 44 [Source:MGI Sy | 76.295 | 56.535 | 110.69 | 72.298 | 67.618 | 76.855 |
| 15267 | ENSMUSG00000052334 | 0 | NA | NA | NA | NA | NA | 1700024B05 | RIKEN cDNA 1700024B05 gene [Source:M | 0 | 0 | 0 | 0 | 0 | 0 |
| 15268 | ENSMUSG00000052336 | 8.771876241 | -1.482592296 | 0.75979393 | -1.951308424 | 0.051020368 | 0.317128748 | Cx3cr1 | chemokine (C-X3-C motif) receptor 1 [Sou | 11.243 | 13.363 | 14.087 | 2.8919 | 4.3625 | 6.6831 |
| 15269 | ENSMUSG00000052337 | 2211.668201 | -0.155354251 | 0.10322505 | -1.50500533 | 0.132322706 | 0.533283479 | Immt | inner membrane protein, mitochondrial [S | 2397.3 | 2296.4 | 2297.3 | 1931.8 | 2302.3 | 2045 |
| 15270 | ENSMUSG00000052353 | 0.167707431 | -0.796629176 | 4.080472857 | -0.195229623 | 0.84521319 | NA | Cemip | cell migration inducing protein, hyalurona | 0 | 0 | 1.0062 | 0 | 0 | 0 |
| 15271 | ENSMUSG00000052363 | 0 | NA | NA | NA | NA | NA | Zdhhc19 | zinc finger, DHHC domain containing 19 [S | 0 | 0 | 0 | 0 | 0 | 0 |
| 15272 | ENSMUSG00000052364 | 23.47667795 | 0.300927357 | 0.466542791 | 0.645015555 | 0.518917106 | 0.887592015 | B630019K06 | RIKEN cDNA B630019K06 gene [Source:M | 14.456 | 23.642 | 25.156 | 20.244 | 26.175 | 31.188 |
| 15273 | ENSMUSG00000052368 | 0.167707431 | -0.796629176 | 4.080472857 | -0.195229623 | 0.84521319 | NA | Gm9873 | predicted gene 9873 [Source:MGI Symbol | 0 | 0 | 1.0062 | 0 | 0 | 0 |
| 15274 | ENSMUSG00000052369 | 290.1280294 | 0.141688338 | 0.150123502 | 0.943811833 | 0.345265813 | 0.786444837 | Tmem106c | transmembrane protein 106C [Source:MG | 281.09 | 284.73 | 261.62 | 292.09 | 327.19 | 294.05 |
| 15275 | ENSMUSG00000052371 | 0 | NA | NA | NA | NA | NA | Hoxd3os1 | homeobox D3, opposite strand 1 [Source: | 0 | 0 | 0 | 0 | 0 | 0 |
| 15276 | ENSMUSG00000052372 | 0 | NA | NA | NA | NA | NA | Il1rapl1 | interleukin 1 receptor accessory protein-li | 0 | 0 | 0 | 0 | 0 | 0 |
| 15277 | ENSMUSG00000052373 | 1.673737746 | -1.038170307 | 1.728080672 | -0.600764955 | 0.547996549 | NA | Mpp3 | membrane protein, palmitoylated 3 (MAG | 1.6062 | 4.1117 | 1.0062 | 0 | 1.0906 | 2.2277 |
| 15278 | ENSMUSG00000052374 | 0.133850373 | -0.796629176 | 4.080472857 | -0.195229623 | 0.84521319 | NA | Actn2 | actinin alpha 2 [Source:MGI Symbol;Acc:N | 0.8031 | 0 | 0 | 0 | 0 | 0 |
| 15279 | ENSMUSG00000052376 | 0 | NA | NA | NA | NA | NA | 4921508M14 | RIKEN cDNA 4921508M14 gene [Source:M | 0 | 0 | 0 | 0 | 0 | 0 |
| 15280 | ENSMUSG00000052382 | 0 | NA | NA | NA | NA | NA | Rnase9 | ribonuclease, RNase A family, 9 (non-activ | 0 | 0 | 0 | 0 | 0 | 0 |
| 15281 | ENSMUSG00000052384 | 54.47363798 | -0.515456194 | 0.342384774 | -1.505488075 | 0.132198641 | 0.533232592 | Nrros | negative regulator of reactive oxygen spec | 69.87 | 77.094 | 45.281 | 43.379 | 57.803 | 33.415 |
| 15282 | ENSMUSG00000052387 | 0.503095663 | 2.549494177 | 3.348715477 | 0.761334964 | 0.446457022 | NA | Trpm3 | transient receptor potential cation channe | 0 | 0 | 0 | 1.928 | 1.0906 | 0 |
| 15283 | ENSMUSG00000052392 | 2575.805321 | -0.588962716 | 0.418777262 | -1.406386568 | 0.159609367 | 0.581402021 | Acot4 | acyl-CoA thioesterase 4 [Source:MGI Syml | 3657.3 | 3157.8 | 2468.3 | 3075.1 | 1942.4 | 1153.9 |
| 15284 | ENSMUSG00000052395 | 360.8491941 | 0.245231503 | 0.149680387 | 1.638367648 | 0.101345024 | 0.469073415 | Rft1 | RFT1 homolog [Source:MGI Symbol;Acc:M | 315.62 | 339.21 | 336.09 | 367.28 | 448.24 | 358.66 |

| | | | | | | | | | | | | | | | |
|---|---|---|---|---|---|---|---|---|---|---|---|---|---|---|---|
| 15285 | ENSMUSG00000052396 | 9.523657978 | 1.841725301 | 0.871485111 | 2.113318149 | 0.034573546 | 0.249163814 | Mogat2 | monoacylglycerol O-acyltransferase 2 [Sou | 2.4093 | 2.0558 | 8.05 | 3.8559 | 20.722 | 20.049 |
| 15286 | ENSMUSG00000052397 | 108.9707801 | -0.189152876 | 0.247304902 | -0.764856959 | 0.444356726 | 0.851724159 | Ezr | ezrin [Source:MGI Symbol;Acc:MGI:98931 | 100.39 | 143.91 | 104.65 | 116.64 | 104.7 | 83.538 |
| 15287 | ENSMUSG00000052403 | 4.733988884 | -0.179737326 | 0.991978405 | -0.181190765 | 0.856217843 | NA | Fcnaos | ficolin A, opposite strand [Source:MGI Syr | 4.0155 | 5.1396 | 6.0375 | 7.7118 | 3.2719 | 2.2277 |
| 15288 | ENSMUSG00000052406 | 319.9781412 | 0.018455612 | 0.146823922 | 0.125698945 | 0.899970237 | 0.982798755 | Rexo4 | REX4, 3'-5' exonuclease [Source:MGI Syml | 333.29 | 314.54 | 305.9 | 338.36 | 333.73 | 294.05 |
| 15289 | ENSMUSG00000052407 | 110.4416627 | 0.1919082 | 0.249555045 | 0.769001484 | 0.441892429 | 0.850050074 | Ccdc171 | coiled-coil domain containing 171 [Source | 113.24 | 72.982 | 122.76 | 117.61 | 107.97 | 128.09 |
| 15290 | ENSMUSG00000052414 | 0 | NA | NA | NA | NA | NA | Gm28047 | predicted gene, 28047 [Source:MGI Symb | 0 | 0 | 0 | 0 | 0 | 0 |
| 15291 | ENSMUSG00000052415 | 35.48344899 | 0.036508065 | 0.430825151 | 0.084739864 | 0.932468203 | 0.988420305 | Tchh | trichohyalin [Source:MGI Symbol;Acc:MG | 45.777 | 22.614 | 36.225 | 22.171 | 35.99 | 50.123 |
| 15292 | ENSMUSG00000052417 | 0 | NA | NA | NA | NA | NA | Olfr720 | olfactory receptor 720 [Source:MGI Symb | 0 | 0 | 0 | 0 | 0 | 0 |
| 15293 | ENSMUSG00000052419 | 246.7658351 | -0.049333411 | 0.171843576 | -0.287083244 | 0.774048582 | 0.958301694 | 2610001J05I | RIKEN cDNA 2610001J05 gene [Source:MC | 277.87 | 214.83 | 259.61 | 242.92 | 222.49 | 262.87 |
| 15294 | ENSMUSG00000052423 | 98.69985994 | 0.165780472 | 0.244949255 | 0.676795167 | 0.498535925 | 0.879866965 | B4galt3 | UDP-Gal:betaGlcNAc beta 1,4-galactosyltr | 76.295 | 101.76 | 101.63 | 112.79 | 88.34 | 111.38 |
| 15295 | ENSMUSG00000052426 | 0 | NA | NA | NA | NA | NA | Gm9877 | predicted gene 9877 [Source:MGI Symbol | 0 | 0 | 0 | 0 | 0 | 0 |
| 15296 | ENSMUSG00000052428 | 1602.052559 | -0.061403108 | 0.094847147 | -0.647390147 | 0.517379471 | 0.887022094 | Tmco1 | transmembrane and coiled-coil domains 1 | 1641.5 | 1627.2 | 1639.2 | 1505.7 | 1582.5 | 1616.2 |
| 15297 | ENSMUSG00000052430 | 0.435408177 | -2.179616019 | 3.531869654 | -0.617128103 | 0.537150234 | NA | Bmpr1b | bone morphogenetic protein receptor, typ | 1.6062 | 0 | 1.0062 | 0 | 0 | 0 |
| 15298 | ENSMUSG00000052435 | 240.7121449 | 0.542877604 | 0.176296324 | 3.079347273 | 0.002074547 | 0.035806979 | Cebpe | CCAAT/enhancer binding protein (C/EBP), | 228.88 | 172.69 | 185.15 | 266.06 | 311.92 | 279.57 |
| 15299 | ENSMUSG00000052446 | 156.9928131 | -0.006777002 | 0.197739976 | -0.034272291 | 0.97266002 | 0.99573532 | Zfp961 | zinc finger protein 961 [Source:MGI Symb | 153.39 | 180.91 | 137.86 | 150.38 | 169.05 | 150.37 |
| 15300 | ENSMUSG00000052456 | 607.3796939 | 0.083443782 | 0.121823097 | 0.684958634 | 0.493370025 | 0.87710702 | Get3 | guided entry of tail-anchored proteins fac | 594.3 | 584.89 | 590.67 | 666.11 | 636.92 | 571.4 |
| 15301 | ENSMUSG00000052459 | 855.3611373 | -0.13070308 | 0.129154711 | -1.011988484 | 0.31154356 | 0.759830222 | Atp6v1a | ATPase, H+ transporting, lysosomal V1 sul | 893.85 | 995.02 | 792.92 | 749.01 | 812.51 | 888.85 |
| 15302 | ENSMUSG00000052468 | 0 | NA | NA | NA | NA | NA | Pmp2 | peripheral myelin protein 2 [Source:MGI S | 0 | 0 | 0 | 0 | 0 | 0 |
| 15303 | ENSMUSG00000052469 | 0 | NA | NA | NA | NA | NA | Tcp10c | t-complex protein 10c [Source:MGI Symbo | 0 | 0 | 0 | 0 | 0 | 0 |
| 15304 | ENSMUSG00000052477 | 0.160662974 | 1.126924175 | 4.080472857 | 0.276174898 | 0.782413736 | NA | C130026I21I | RIKEN cDNA C130026I21 gene [Source:MC | 0 | 0 | 0 | 0.964 | 0 | 0 |
| 15305 | ENSMUSG00000052479 | 0 | NA | NA | NA | NA | NA | A330008L17 | RIKEN cDNA A330008L17 gene [Source:M | 0 | 0 | 0 | 0 | 0 | 0 |
| 15306 | ENSMUSG00000052482 | 0 | NA | NA | NA | NA | NA | Gm8174 | predicted gene 8174 [Source:MGI Symbol | 0 | 0 | 0 | 0 | 0 | 0 |
| 15307 | ENSMUSG00000052485 | 1.659752571 | -1.932355961 | 1.807396807 | -1.069137642 | 0.285007652 | NA | Tmem171 | transmembrane protein 171 [Source:MGI | 4.8186 | 2.0558 | 1.0062 | 0.964 | 0 | 1.1138 |
| 15308 | ENSMUSG00000052488 | 417.1789241 | 0.09839717 | 0.133308325 | 0.738117212 | 0.460443225 | 0.859817816 | Cherp | calcium homeostasis endoplasmic reticulu | 427.25 | 388.55 | 392.44 | 427.04 | 413.34 | 454.45 |
| 15309 | ENSMUSG00000052496 | 40.34510959 | 0.182765211 | 0.466383136 | 0.39187783 | 0.695148485 | 0.939992732 | Pkdrej | polycystin (PKD) family receptor for egg je | 55.414 | 12.335 | 45.281 | 44.343 | 50.168 | 34.529 |
| 15310 | ENSMUSG00000052504 | 8.970885084 | -1.59200255 | 0.792236593 | -2.009503934 | 0.044483718 | 0.291685757 | Epha3 | Eph receptor A3 [Source:MGI Symbol;Acc | 20.078 | 11.307 | 9.0562 | 6.7478 | 2.1812 | 4.4554 |
| 15311 | ENSMUSG00000052508 | 0 | NA | NA | NA | NA | NA | Olfr536 | olfactory receptor 536 [Source:MGI Symb | 0 | 0 | 0 | 0 | 0 | 0 |
| 15312 | ENSMUSG00000052512 | 588.4533975 | -0.153764532 | 0.198286825 | -0.775465197 | 0.438064823 | 0.847534724 | Nav2 | neuron navigator 2 [Source:MGI Symbol;A | 630.44 | 683.57 | 545.38 | 597.67 | 369.72 | 703.95 |
| 15313 | ENSMUSG00000052516 | 26.96820258 | -0.645115917 | 0.495557712 | -1.301797755 | 0.192985531 | 0.631239875 | Robo2 | roundabout guidance receptor 2 [Source: | 41.761 | 19.53 | 37.231 | 18.316 | 32.719 | 12.252 |
| 15314 | ENSMUSG00000052520 | 11971.35803 | 0.054320043 | 0.085526297 | 0.635126796 | 0.525345722 | 0.888334265 | Cyp2j5 | cytochrome P450, family 2, subfamily j, po | 11219 | 12069 | 11951 | 11695 | 12645 | 12250 |
| 15315 | ENSMUSG00000052525 | 3.173990366 | -0.072203116 | 1.233342855 | -0.058542615 | 0.953316419 | NA | Spdya | speedy/RINGO cell cycle regulator family, | 5.6217 | 3.0838 | 1.0062 | 3.8559 | 4.3625 | 1.1138 |
| 15316 | ENSMUSG00000052533 | 157.9559582 | 0.147435014 | 0.195971797 | 0.752327716 | 0.451854004 | 0.856471272 | Nup188 | nucleoporin 188 [Source:MGI Symbol;Acc | 157.41 | 137.74 | 153.96 | 142.67 | 178.86 | 177.1 |
| 15317 | ENSMUSG00000052534 | 394.2793613 | -0.16532456 | 0.172200461 | -0.960070369 | 0.3370198 | 0.779072142 | Pbx1 | pre B cell leukemia homeobox 1 [Source:N | 374.25 | 502.65 | 374.32 | 374.99 | 424.25 | 315.22 |
| 15318 | ENSMUSG00000052537 | 0 | NA | NA | NA | NA | NA | Olfr198 | olfactory receptor 198 [Source:MGI Symb | 0 | 0 | 0 | 0 | 0 | 0 |
| 15319 | ENSMUSG00000052539 | 426.7996712 | 0.125580905 | 0.141088685 | 0.890084878 | 0.373420312 | 0.80865857 | Magi3 | membrane associated guanylate kinase, V | 357.38 | 419.39 | 448.79 | 434.75 | 447.15 | 453.33 |
| 15320 | ENSMUSG00000052544 | 3.420998081 | 0.604984535 | 1.165272119 | 0.519178761 | 0.603636089 | NA | St6galnac3 | ST6 (alpha-N-acetyl-neuraminyl-2,3-beta-( | 4.0155 | 3.0838 | 1.0062 | 5.7839 | 2.1812 | 4.4554 |
| 15321 | ENSMUSG00000052549 | 0 | NA | NA | NA | NA | NA | Arl13a | ADP-ribosylation factor-like 13A [Source:N | 0 | 0 | 0 | 0 | 0 | 0 |
| 15322 | ENSMUSG00000052551 | 0.506966596 | 2.558866737 | 3.339834942 | 0.766165628 | 0.443577757 | NA | Adarb2 | adenosine deaminase, RNA-specific, B2 [S | 0 | 0 | 0 | 1.928 | 0 | 1.1138 |
| 15323 | ENSMUSG00000052554 | 0 | NA | NA | NA | NA | NA | Defb34 | defensin beta 34 [Source:MGI Symbol;Acc | 0 | 0 | 0 | 0 | 0 | 0 |
| 15324 | ENSMUSG00000052557 | 175.0775367 | -0.178784872 | 0.223336129 | -0.800519256 | 0.423410012 | 0.84018348 | Gan | giant axonal neuropathy [Source:MGI Syn | 182.3 | 182.97 | 193.2 | 220.75 | 129.78 | 141.46 |
| 15325 | ENSMUSG00000052560 | 28.60675279 | -0.817411767 | 0.437941422 | -1.866486532 | 0.061973327 | 0.355884025 | Cpne8 | copine VIII [Source:MGI Symbol;Acc:MGI: | 36.14 | 43.173 | 30.187 | 21.208 | 13.087 | 27.846 |
| 15326 | ENSMUSG00000052562 | 2711.986251 | 0.547557421 | 0.100622562 | 5.441696248 | 5.28E-08 | 6.26E-06 | Slc22a30 | solute carrier family 22, member 30 [Sour | 2129.8 | 2230.6 | 2251 | 3402.8 | 3341.7 | 2916 |
| 15327 | ENSMUSG00000052563 | 140.4431581 | 0.012093788 | 0.277610507 | 0.043563869 | 0.965252053 | 0.993480943 | D930048N14 | RIKEN cDNA D930048N14 gene [Source:M | 130.1 | 88.401 | 201.25 | 158.09 | 117.79 | 147.03 |
| 15328 | ENSMUSG00000052565 | 8.996249587 | -0.352726988 | 0.873372107 | -0.403867933 | 0.686309838 | 0.936489198 | H1f3 | H1.3 linker histone, cluster member [Sour | 8.031 | 2.0558 | 20.125 | 3.8559 | 6.5437 | 13.366 |
| 15329 | ENSMUSG00000052566 | 547.6816802 | 0.595902708 | 0.162701903 | 3.662542947 | 0.000249724 | 0.007496527 | Hook2 | hook microtubule tethering protein 2 [Sou | 480.26 | 432.75 | 395.45 | 796.25 | 646.74 | 534.65 |
| 15330 | ENSMUSG00000052572 | 9.563498953 | 0.657375548 | 0.718465064 | 0.914972183 | 0.360206234 | 0.799598236 | Dlg2 | discs large MAGUK scaffold protein 2 [Sou | 8.031 | 9.2513 | 5.0312 | 16.388 | 11.997 | 6.6831 |
| 15331 | ENSMUSG00000052581 | 0 | NA | NA | NA | NA | NA | Lrrtm4 | leucine rich repeat transmembrane neuro | 0 | 0 | 0 | 0 | 0 | 0 |
| 15332 | ENSMUSG00000052584 | 2.335920179 | -0.218358273 | 1.387536244 | -0.157371221 | 0.874952293 | NA | Serp2 | stress-associated endoplasmic reticulum p | 2.4093 | 3.0838 | 2.0125 | 0.964 | 1.0906 | 4.4554 |
| 15333 | ENSMUSG00000052593 | 297.2833186 | -0.154155125 | 0.161046075 | -0.957211317 | 0.338460606 | 0.780780576 | Adam17 | a disintegrin and metallopeptidase domai | 273.86 | 355.66 | 310.93 | 297.87 | 280.29 | 265.09 |
| 15334 | ENSMUSG00000052595 | 2329.729724 | -0.013287572 | 0.168160237 | -0.079017327 | 0.937018841 | 0.988420305 | A1cf | APOBEC1 complementation factor [Source | 2104.1 | 2438.2 | 2480.4 | 2817.7 | 2510.6 | 1627.3 |
| 15335 | ENSMUSG00000052605 | 366.978554 | 0.040520974 | 0.175835502 | 0.230448196 | 0.817743514 | 0.968565186 | Isoc2b | isochorismatase domain containing 2b [Sc | 420.02 | 343.32 | 320.99 | 340.28 | 323.91 | 453.33 |
| 15336 | ENSMUSG00000052609 | 750.5653716 | -0.060562002 | 0.135737238 | -0.446170869 | 0.655473824 | 0.926075335 | Plekhg3 | pleckstrin homology domain containing, fa | 679.42 | 893.26 | 727.51 | 763.47 | 754.71 | 685.01 |
| 15337 | ENSMUSG00000052613 | 1.213558485 | -2.430141359 | 2.584716269 | -0.940196566 | 0.347116743 | NA | Pcdh15 | protocadherin 15 [Source:MGI Symbol;Ac | 0 | 6.1675 | 0 | 0 | 0 | 1.1138 |
| 15338 | ENSMUSG00000052616 | 0.160662974 | 1.126924175 | 4.080472857 | 0.276174898 | 0.782413736 | NA | Terb1 | telomere repeat binding bouquet formati | 0 | 0 | 0 | 0.964 | 0 | 0 |
| 15339 | ENSMUSG00000052625 | 0 | NA | NA | NA | NA | NA | Olfr851 | olfactory receptor 851 [Source:MGI Symb | 0 | 0 | 0 | 0 | 0 | 0 |
| 15340 | ENSMUSG00000052629 | 0 | NA | NA | NA | NA | NA | Gm9885 | predicted gene 9885 [Source:MGI Symbol | 0 | 0 | 0 | 0 | 0 | 0 |

| | | | | | | | | | | | | | | | |
|---|---|---|---|---|---|---|---|---|---|---|---|---|---|---|---|
| 15341 | ENSMUSG00000052631 | 0 | NA | NA | NA | NA | NA | Sh2d6 | SH2 domain containing 6 [Source:MGI Syr | 0 | 0 | 0 | 0 | 0 | 0 |
| 15342 | ENSMUSG00000052632 | 91.37389405 | -0.351088079 | 0.309877635 | -1.13298941 | 0.2572187 | 0.710872182 | Asap2 | ArfGAP with SH3 domain, ankyrin repeat a | 72.279 | 83.261 | 151.94 | 66.514 | 81.796 | 92.449 |
| 15343 | ENSMUSG00000052642 | 0 | NA | NA | NA | NA | NA | Lypd9 | LY6/PLAUR domain containing 9 [Source:N | 0 | 0 | 0 | 0 | 0 | 0 |
| 15344 | ENSMUSG00000052656 | 2178.359705 | -0.053728096 | 0.091722594 | -0.585767296 | 0.558031904 | 0.899217185 | Rnf103 | ring finger protein 103 [Source:MGI Symb | 2269.6 | 2267.6 | 2119.2 | 2115 | 2170.3 | 2128.6 |
| 15345 | ENSMUSG00000052658 | 15.09664203 | 0.208159106 | 0.581637853 | 0.357884385 | 0.720429837 | 0.9447719 | 5830454E08 | RIKEN cDNA 5830454E08 gene [Source:M | 13.653 | 16.447 | 12.075 | 23.135 | 16.359 | 8.9108 |
| 15346 | ENSMUSG00000052673 | 4.275556415 | 0.020298397 | 1.080777081 | 0.018781298 | 0.985015573 | NA | Gm9887 | predicted gene 9887 [Source:MGI Symbol | 1.6062 | 5.1396 | 6.0375 | 2.8919 | 2.1812 | 7.7969 |
| 15347 | ENSMUSG00000052675 | 32.37464038 | 0.858760162 | 0.387340642 | 2.217067017 | 0.026618512 | 0.21119592 | Zfp112 | zinc finger protein 112 [Source:MGI Symb | 18.471 | 24.67 | 26.162 | 43.379 | 40.353 | 41.212 |
| 15348 | ENSMUSG00000052676 | 113.7741908 | -0.184101412 | 0.24667396 | -0.74633501 | 0.455465068 | 0.857366983 | Zmat1 | zinc finger, matrin type 1 [Source:MGI Syr | 118.86 | 112.04 | 131.82 | 80.01 | 136.33 | 103.59 |
| 15349 | ENSMUSG00000052681 | 865.0993081 | -0.228813798 | 0.110108981 | -2.07806661 | 0.037703225 | 0.263449005 | Rap1b | RAS related protein 1b [Source:MGI Symb | 893.05 | 966.24 | 941.84 | 797.21 | 757.98 | 834.27 |
| 15350 | ENSMUSG00000052684 | 335.323495 | -1.30192275 | 0.626455724 | -2.078235859 | 0.037687641 | 0.263449005 | Jun | jun proto-oncogene [Source:MGI Symbol; | 723.6 | 495.46 | 212.32 | 228.46 | 255.2 | 96.904 |
| 15351 | ENSMUSG00000052688 | 15.79812345 | -1.002497322 | 0.577783987 | -1.735072874 | 0.082727898 | 0.42416743 | Rab7b | RAB7B, member RAS oncogene family [So | 18.471 | 25.698 | 19.119 | 11.568 | 5.4531 | 14.48 |
| 15352 | ENSMUSG00000052698 | 175.495491 | -0.76288091 | 0.229387861 | -3.325724845 | 0.000881889 | 0.019076633 | Tln2 | talin 2 [Source:MGI Symbol;Acc:MGI:1917 | 257.8 | 194.28 | 210.31 | 137.85 | 161.41 | 91.335 |
| 15353 | ENSMUSG00000052707 | 945.5901212 | 0.162634188 | 0.111362013 | 1.460409917 | 0.144177449 | 0.554380949 | Tnrc6a | trinucleotide repeat containing 6a [Source | 861.73 | 879.9 | 935.81 | 986.15 | 1071 | 938.97 |
| 15354 | ENSMUSG00000052712 | 1098.917611 | 0.138750789 | 0.135530299 | 1.023762136 | 0.305947639 | 0.755069291 | BC004004 | cDNA sequence BC004004 [Source:MGI Sy | 1166.9 | 1089.6 | 880.46 | 1055.6 | 1120.1 | 1280.9 |
| 15355 | ENSMUSG00000052713 | 61.24838478 | 0.068873701 | 0.354018991 | 0.19454804 | 0.845746784 | 0.972803041 | Zfp608 | zinc finger protein 608 [Source:MGI Symb | 73.885 | 58.591 | 46.287 | 35.667 | 82.887 | 70.172 |
| 15356 | ENSMUSG00000052724 | 1.142104334 | -1.08390987 | 2.30413889 | -0.470418634 | 0.638055953 | NA | Gm9888 | predicted gene 9888 [Source:MGI Symbol | 1.6062 | 0 | 3.0187 | 0 | 0 | 2.2277 |
| 15357 | ENSMUSG00000052726 | 29.49424269 | 0.023968615 | 0.485458922 | 0.049373107 | 0.96062196 | 0.992515024 | Kcnt2 | potassium channel, subfamily T, member | 38.549 | 33.921 | 15.094 | 27.955 | 43.625 | 17.822 |
| 15358 | ENSMUSG00000052727 | 3.874087918 | 0.196393112 | 1.142364325 | 0.171918106 | 0.863501912 | NA | Map1b | microtubule-associated protein 1B [Sourc | 0.8031 | 5.1396 | 5.0312 | 6.7478 | 2.1812 | 3.3415 |
| 15359 | ENSMUSG00000052730 | 0 | NA | NA | NA | NA | NA | Gm5111 | predicted gene 5111 [Source:MGI Symbol | 0 | 0 | 0 | 0 | 0 | 0 |
| 15360 | ENSMUSG00000052736 | 1.723829503 | -0.480059299 | 2.058165801 | -0.233246174 | 0.815570253 | NA | Klrc2 | killer cell lectin-like receptor subfamily C, | 0 | 0 | 6.0375 | 0.964 | 0 | 3.3415 |
| 15361 | ENSMUSG00000052738 | 4830.788703 | -0.070033815 | 0.086561454 | -0.809064673 | 0.418477947 | 0.837575303 | Suclg1 | succinate-CoA ligase, GDP-forming, alpha | 4996.9 | 5066.6 | 4780.7 | 4780.4 | 4838 | 4522.2 |
| 15362 | ENSMUSG00000052748 | 243.9243373 | 0.048643151 | 0.157755032 | 0.308346115 | 0.757818979 | 0.953466696 | Swt1 | SWT1 RNA endoribonuclease homolog (S. | 250.57 | 224.09 | 244.52 | 249.67 | 254.11 | 240.59 |
| 15363 | ENSMUSG00000052749 | 5.630088795 | 0.206776629 | 0.968539638 | 0.213493203 | 0.83094229 | 0.970505212 | Trim30b | tripartite motif-containing 30B [Source:M | 1.6062 | 4.1117 | 10.062 | 4.8199 | 8.7249 | 4.4554 |
| 15364 | ENSMUSG00000052751 | 802.05158 | 0.287168325 | 0.145648142 | 1.971658013 | 0.04864866 | 0.307138461 | Repin1 | replication initiator 1 [Source:MGI Symbol | 644.09 | 644.5 | 879.46 | 844.44 | 948.84 | 850.98 |
| 15365 | ENSMUSG00000052752 | 214.0224248 | 0.17012038 | 0.174861202 | 0.972888085 | 0.330608928 | 0.774747658 | Traf7 | TNF receptor-associated factor 7 [Source: | 203.18 | 179.89 | 221.37 | 248.71 | 212.67 | 218.31 |
| 15366 | ENSMUSG00000052759 | 0 | NA | NA | NA | NA | NA | Gpr25 | G protein-coupled receptor 25 [Source:M | 0 | 0 | 0 | 0 | 0 | 0 |
| 15367 | ENSMUSG00000052760 | 6.095771482 | -4.033890962 | 1.295740622 | -3.113193253 | 0.001850748 | 0.032845665 | A630001G21 | RIKEN cDNA A630001G21 gene [Source:M | 8.031 | 14.391 | 12.075 | 0.964 | 0 | 1.1138 |
| 15368 | ENSMUSG00000052763 | 141.1131693 | -0.082576039 | 0.200796556 | -0.411242306 | 0.680894869 | 0.934384556 | Zfp212 | Zinc finger protein 212 [Source:MGI Symb | 126.89 | 152.13 | 156.97 | 140.74 | 138.51 | 131.43 |
| 15369 | ENSMUSG00000052767 | 0 | NA | NA | NA | NA | NA | Gm12703 | predicted gene 12703 [Source:MGI Symbo | 0 | 0 | 0 | 0 | 0 | 0 |
| 15370 | ENSMUSG00000052769 | 1.6903845 | 2.172649989 | 1.916733125 | 1.133517213 | 0.256997117 | NA | Gm9889 | predicted gene 9889 [Source:MGI Symbol | 0.8031 | 1.0279 | 0 | 3.8559 | 0 | 4.4554 |
| 15371 | ENSMUSG00000052776 | 49.37693047 | -0.31389836 | 0.397584925 | -0.78951273 | 0.429812392 | 0.84307112 | Oas1a | 2'-5' oligoadenylate synthetase 1A [Sourc | 68.264 | 41.117 | 54.337 | 31.811 | 29.447 | 71.286 |
| 15372 | ENSMUSG00000052779 | 0.674621893 | -1.426050892 | 3.091195043 | -0.461326727 | 0.644564212 | NA | 4833418N17 | RIKEN cDNA 4833418N17 gene [Source:M | 0 | 3.0838 | 0 | 0.964 | 0 | 0 |
| 15373 | ENSMUSG00000052783 | 22.5624805 | 0.500468861 | 0.523294209 | 0.956381425 | 0.338879566 | 0.780858357 | Grk4 | G protein-coupled receptor kinase 4 [Sour | 9.6372 | 25.698 | 21.131 | 34.703 | 16.359 | 27.846 |
| 15374 | ENSMUSG00000052785 | 0 | NA | NA | NA | NA | NA | Olfr573-ps1 | olfactory receptor 573, pseudogene 1 [So | 0 | 0 | 0 | 0 | 0 | 0 |
| 15375 | ENSMUSG00000052790 | 0 | NA | NA | NA | NA | NA | Gm9890 | predicted gene 9890 [Source:MGI Symbol | 0 | 0 | 0 | 0 | 0 | 0 |
| 15376 | ENSMUSG00000052794 | 73.47278189 | -0.026742097 | 0.273353852 | -0.097829598 | 0.922067604 | 0.986685804 | 1700030K09 | RIKEN cDNA 1700030K09 gene [Source:M | 77.901 | 68.87 | 75.468 | 64.587 | 90.521 | 63.489 |
| 15377 | ENSMUSG00000052798 | 66.7829316 | -0.08317625 | 0.278663036 | -0.298483256 | 0.765334353 | 0.954980327 | Nup107 | nucleoporin 107 [Source:MGI Symbol;Acc | 71.476 | 75.038 | 59.368 | 55.911 | 67.618 | 71.286 |
| 15378 | ENSMUSG00000052812 | 376.1315165 | 0.061344379 | 0.141086628 | 0.434799388 | 0.663708049 | 0.929301301 | Atad2b | ATPase family, AAA domain containing 2B | 372.64 | 387.53 | 344.14 | 374.99 | 416.62 | 360.89 |
| 15379 | ENSMUSG00000052818 | 0 | NA | NA | NA | NA | NA | Olfr813 | olfactory receptor 813 [Source:MGI Symb | 0 | 0 | 0 | 0 | 0 | 0 |
| 15380 | ENSMUSG00000052819 | 0 | NA | NA | NA | NA | NA | Best2 | bestrophin 2 [Source:MGI Symbol;Acc:MG | 0 | 0 | 0 | 0 | 0 | 0 |
| 15381 | ENSMUSG00000052821 | 7.289761535 | -1.483516531 | 0.861792306 | -1.721431626 | 0.08517253 | 0.429994927 | Cysltr1 | cysteinyl leukotriene receptor 1 [Source:N | 16.865 | 8.2233 | 7.0437 | 3.8559 | 2.1812 | 5.5692 |
| 15382 | ENSMUSG00000052825 | 9.108312731 | -0.162345735 | 0.733513892 | -0.221326054 | 0.824838565 | 0.96943211 | Gm9892 | predicted gene 9892 [Source:MGI Symbol | 10.44 | 8.2233 | 10.062 | 3.8559 | 9.8156 | 12.252 |
| 15383 | ENSMUSG00000052833 | 328.8451684 | 0.0927923 | 0.156015628 | 0.594762852 | 0.552002002 | 0.895935901 | Sae1 | SUMO1 activating enzyme subunit 1 [Sour | 354.97 | 286.79 | 311.94 | 306.54 | 367.54 | 345.29 |
| 15384 | ENSMUSG00000052837 | 379.8828266 | 0.52980108 | 0.209943815 | 2.523537457 | 0.011618068 | 0.12011704 | Junb | jun B proto-oncogene [Source:MGI Symbo | 419.22 | 234.37 | 277.72 | 463.67 | 369.72 | 514.6 |
| 15385 | ENSMUSG00000052848 | 0 | NA | NA | NA | NA | NA | C130026L21 | RIKEN cDNA C130026L21 gene [Source:M | 0 | 0 | 0 | 0 | 0 | 0 |
| 15386 | ENSMUSG00000052850 | 0 | NA | NA | NA | NA | NA | Tas2r137 | taste receptor, type 2, member 137 [Sour | 0 | 0 | 0 | 0 | 0 | 0 |
| 15387 | ENSMUSG00000052852 | 8.970311812 | 0.321687631 | 0.745630409 | 0.431430408 | 0.666155444 | 0.929596279 | Reep1 | receptor accessory protein 1 [Source:MGI | 5.6217 | 11.307 | 7.0437 | 7.7118 | 6.5437 | 15.594 |
| 15388 | ENSMUSG00000052854 | 0 | NA | NA | NA | NA | NA | Nrk | Nik related kinase [Source:MGI Symbol;Ac | 0 | 0 | 0 | 0 | 0 | 0 |
| 15389 | ENSMUSG00000052861 | 0.160662974 | 1.126924175 | 4.080472857 | 0.276174898 | 0.782413736 | NA | Dnah6 | dynein, axonemal, heavy chain 6 [Source: | 0 | 0 | 0 | 0.964 | 0 | 0 |
| 15390 | ENSMUSG00000052865 | 0 | NA | NA | NA | NA | NA | Gm13619 | predicted gene 13619 [Source:MGI Symbo | 0 | 0 | 0 | 0 | 0 | 0 |
| 15391 | ENSMUSG00000052889 | 14.8315284 | -0.5950635 | 0.64084945 | -0.92855428 | 0.353120125 | 0.793464268 | Prkcb | protein kinase C, beta [Source:MGI Symbo | 13.653 | 28.782 | 11.069 | 5.7839 | 17.45 | 12.252 |
| 15392 | ENSMUSG00000052906 | 584.8609801 | 0.00832009 | 0.117112839 | 0.07104336 | 0.943363246 | 0.990077979 | Ubxn8 | UBX domain protein 8 [Source:MGI Symbo | 588.67 | 595.16 | 565.51 | 580.31 | 578.03 | 601.48 |
| 15393 | ENSMUSG00000052909 | 0 | NA | NA | NA | NA | NA | Gm9894 | predicted gene 9894 [Source:MGI Symbol | 0 | 0 | 0 | 0 | 0 | 0 |
| 15394 | ENSMUSG00000052911 | 59.28120098 | -0.92483373 | 0.33771815 | -2.738478022 | 0.006172429 | 0.076765268 | Lamb2 | laminin, beta 2 [Source:MGI Symbol;Acc:N | 75.492 | 89.429 | 68.425 | 52.055 | 46.897 | 23.391 |
| 15395 | ENSMUSG00000052912 | 0.133850373 | -0.796629176 | 4.080472857 | -0.195229623 | 0.84521319 | NA | Smarca5-ps | SWI/SNF related, matrix associated, actin | 0.8031 | 0 | 0 | 0 | 0 | 0 |
| 15396 | ENSMUSG00000052914 | 1500.534516 | 0.163721905 | 0.117265265 | 1.396167103 | 0.162664181 | 0.586288131 | Cyp2j6 | cytochrome P450, family 2, subfamily j, po | 1517.1 | 1375.4 | 1352.4 | 1386.2 | 1698.1 | 1674.1 |

| | | | | | | | | | | | | | | | |
|---|---|---|---|---|---|---|---|---|---|---|---|---|---|---|---|
| 15397 | ENSMUSG00000052915 | 889.4212841 | 0.26484737 | 0.139333495 | 1.900816232 | 0.057326087 | 0.340239855 | Msl1 | male specific lethal 1 [Source:MGI Symbol | 698.7 | 822.33 | 903.61 | 882.04 | 921.57 | 1108.3 |
| 15398 | ENSMUSG00000052917 | 137.2360931 | 0.165228639 | 0.200785986 | 0.822909221 | 0.410559619 | 0.833241465 | Senp7 | SUMO1/sentrin specific peptidase 7 [Sour | 122.07 | 121.29 | 144.9 | 139.78 | 147.23 | 148.14 |
| 15399 | ENSMUSG00000052920 | 5.414995861 | -0.606413065 | 1.12526769 | -0.538905604 | 0.58995199 | 0.911652306 | Prkg1 | protein kinase, cGMP-dependent, type I [! | 7.2279 | 12.335 | 0 | 1.928 | 6.5437 | 4.4554 |
| 15400 | ENSMUSG00000052921 | 95.61506679 | -0.64957386 | 0.252071918 | -2.576938611 | 0.009967965 | 0.108460061 | Arhgef15 | Rho guanine nucleotide exchange factor ( | 126.89 | 105.88 | 117.73 | 89.65 | 57.803 | 75.741 |
| 15401 | ENSMUSG00000052922 | 0 | NA | NA | NA | NA | NA | Bpi | bactericidal permeablility increasing prote | 0 | 0 | 0 | 0 | 0 | 0 |
| 15402 | ENSMUSG00000052926 | 360.8396032 | -0.06878649 | 0.159786145 | -0.430490954 | 0.666838546 | 0.929596279 | Rnaseh2a | ribonuclease H2, large subunit [Source:M | 358.99 | 335.1 | 414.57 | 366.31 | 390.44 | 299.62 |
| 15403 | ENSMUSG00000052928 | 645.4116569 | -0.095064991 | 0.120566315 | -0.788487152 | 0.430411811 | 0.843621458 | Ctif | CBP80/20-dependent translation initiation | 675.41 | 660.95 | 664.12 | 681.53 | 586.75 | 603.7 |
| 15404 | ENSMUSG00000052934 | 2306.439566 | -0.19557976 | 0.108381593 | -1.804547752 | 0.07114548 | 0.386047033 | Fbxo31 | F-box protein 31 [Source:MGI Symbol;Acc | 2320.2 | 2602.7 | 2465.3 | 2139.1 | 1947.8 | 2363.6 |
| 15405 | ENSMUSG00000052942 | 6.472855736 | -1.431851398 | 0.964655178 | -1.48431422 | 0.13772558 | 0.541918695 | Glis3 | GLIS family zinc finger 3 [Source:MGI Sym | 12.047 | 7.1954 | 9.0562 | 3.8559 | 0 | 6.6831 |
| 15406 | ENSMUSG00000052949 | 9.044783736 | -0.669475339 | 0.751332707 | -0.89105044 | 0.372902111 | 0.808400044 | Rnf157 | ring finger protein 157 [Source:MGI Symb | 16.865 | 11.307 | 5.0312 | 6.7478 | 7.6343 | 6.6831 |
| 15407 | ENSMUSG00000052951 | 0 | NA | NA | NA | NA | NA | C130021I20I | Riken cDNA C130021I20 gene [Source:M( | 0 | 0 | 0 | 0 | 0 | 0 |
| 15408 | ENSMUSG00000052955 | 0 | NA | NA | NA | NA | NA | Cpvl | carboxypeptidase, vitellogenic-like [Sourc | 0 | 0 | 0 | 0 | 0 | 0 |
| 15409 | ENSMUSG00000052957 | 450.9004018 | -0.025589453 | 0.15222517 | -0.168102643 | 0.86650253 | 0.977925452 | Gas1 | growth arrest specific 1 [Source:MGI Symb | 424.84 | 405 | 535.32 | 465.6 | 405.71 | 468.93 |
| 15410 | ENSMUSG00000052962 | 412.8893372 | 0.075913656 | 0.138681307 | 0.547396452 | 0.584106393 | 0.908955096 | Mrpl35 | mitochondrial ribosomal protein L35 [Sou | 434.48 | 374.16 | 396.46 | 395.23 | 451.52 | 425.49 |
| 15411 | ENSMUSG00000052974 | 14342.04131 | 0.14739406 | 0.133431524 | 1.104641957 | 0.269314763 | 0.724979259 | Cyp2f2 | cytochrome P450, family 2, subfamily f, pc | 11078 | 13976 | 15778 | 14170 | 16917 | 14135 |
| 15412 | ENSMUSG00000052981 | 1.353988001 | -1.451133495 | 2.240453263 | -0.647696392 | 0.517181338 | NA | Ube2ql1 | ubiquitin-conjugating enzyme E2Q family- | 0.8031 | 5.1396 | 0 | 0 | 2.1812 | 0 |
| 15413 | ENSMUSG00000052997 | 473.0334679 | 0.015400867 | 0.139049434 | 0.110758213 | 0.911808082 | 0.98589875 | Uba2 | ubiquitin-like modifier activating enzyme : | 408.78 | 493.4 | 510.17 | 455.96 | 483.14 | 486.75 |
| 15414 | ENSMUSG00000053004 | 0.658259367 | -1.329928433 | 2.812497761 | -0.472863819 | 0.636310339 | NA | Hrh1 | histamine receptor H1 [Source:MGI Symb | 0.8031 | 2.0558 | 0 | 0 | 1.0906 | 0 |
| 15415 | ENSMUSG00000053007 | 1.145199095 | -1.119908671 | 2.303879602 | -0.486096874 | 0.626898484 | NA | Creb5 | cAMP responsive element binding protein | 1.6062 | 3.0838 | 0 | 0 | 2.1812 | 0 |
| 15416 | ENSMUSG00000053012 | 1011.183226 | 0.119733011 | 0.132029819 | 0.906863405 | 0.364479035 | 0.802897466 | Krcc1 | lysine-rich coiled-coil 1 [Source:MGI Symb | 1027.2 | 981.66 | 897.57 | 883 | 1156.1 | 1121.6 |
| 15417 | ENSMUSG00000053024 | 11.18149044 | 0.156573758 | 0.698171754 | 0.224262522 | 0.822553037 | 0.969219124 | Cntn2 | contactin 2 [Source:MGI Symbol;Acc:MGI | 6.4248 | 9.2513 | 16.1 | 6.7478 | 18.541 | 10.025 |
| 15418 | ENSMUSG00000053025 | 0 | NA | NA | NA | NA | NA | Sv2b | synaptic vesicle glycoprotein 2 b [Source:I | 0 | 0 | 0 | 0 | 0 | 0 |
| 15419 | ENSMUSG00000053030 | 0 | NA | NA | NA | NA | NA | Spink2 | serine peptidase inhibitor, Kazal type 2 [S | 0 | 0 | 0 | 0 | 0 | 0 |
| 15420 | ENSMUSG00000053038 | 1.522444974 | 0.461067932 | 1.761655211 | 0.261724274 | 0.793534024 | NA | Gm6180 | predicted pseudogene 6180 [Source:MGI | 0.8031 | 2.0558 | 1.0062 | 1.928 | 0 | 3.3415 |
| 15421 | ENSMUSG00000053040 | 10.81292641 | 0.081152954 | 0.645427547 | 0.125735188 | 0.899941547 | 0.982798755 | Aph1c | aph1 homolog C, gamma secretase subun | 11.243 | 8.2233 | 12.075 | 13.496 | 9.8156 | 10.025 |
| 15422 | ENSMUSG00000053044 | 0.936410189 | -0.694018354 | 2.480552701 | -0.279783757 | 0.779643414 | NA | Cd8b1 | CD8 antigen, beta chain 1 [Source:MGI Sy | 2.4093 | 1.0279 | 0 | 0 | 2.1812 | 0 |
| 15423 | ENSMUSG00000053046 | 1.17776969 | 2.595769719 | 2.169671889 | 1.196388141 | 0.231545127 | NA | Brsk2 | BR serine/threonine kinase 2 [Source:MGI | 0 | 0 | 1.0062 | 3.8559 | 1.0906 | 1.1138 |
| 15424 | ENSMUSG00000053049 | 0 | NA | NA | NA | NA | NA | Gm15413 | predicted gene 15413 [Source:MGI Symbc | 0 | 0 | 0 | 0 | 0 | 0 |
| 15425 | ENSMUSG00000053054 | 1.269938518 | 2.700065798 | 2.266662151 | 1.191207872 | 0.23357199 | NA | Adh6a | alcohol dehydrogenase 6A (class V) [Sourc | 0 | 0 | 1.0062 | 0 | 3.2719 | 3.3415 |
| 15426 | ENSMUSG00000053062 | 106.86728 | -0.514370484 | 0.240087067 | -2.142433126 | 0.032158645 | 0.23951732 | Jam2 | junction adhesion molecule 2 [Source:MG | 118.86 | 135.69 | 122.76 | 91.578 | 67.618 | 104.7 |
| 15427 | ENSMUSG00000053063 | 42.7709086 | -1.455996611 | 0.381276379 | -3.818743278 | 0.000134133 | 0.004554234 | Clec12a | C-type lectin domain family 12, member a | 50.595 | 83.261 | 54.337 | 23.135 | 17.45 | 27.846 |
| 15428 | ENSMUSG00000053070 | 4.394692601 | 0.689172134 | 1.222428259 | 0.563773072 | 0.57290858 | NA | Cfap300 | cilia and flagella associated protein 300 [S | 0 | 6.1675 | 4.025 | 1.928 | 10.906 | 3.3415 |
| 15429 | ENSMUSG00000053080 | 55.95150566 | -0.460948948 | 0.350109609 | -1.316584679 | 0.187977877 | 0.624893591 | 2700081O15 | RIKEN cDNA 2700081O15 gene [Source:M | 56.217 | 88.401 | 50.312 | 63.623 | 38.172 | 38.985 |
| 15430 | ENSMUSG00000053081 | 0.895483699 | 3.367829889 | 2.453671272 | 1.372567682 | 0.169886785 | NA | 1700069B07 | RIKEN cDNA 1700069B07 gene [Source:M | 0 | 0 | 0 | 0.964 | 2.1812 | 2.2277 |
| 15431 | ENSMUSG00000053091 | 63.1417895 | 0.330578404 | 0.289282592 | 1.142752498 | 0.253141366 | 0.706295664 | Lins1 | lines homolog 1 [Source:MGI Symbol;Acc: | 48.186 | 59.619 | 60.375 | 74.226 | 78.525 | 57.92 |
| 15432 | ENSMUSG00000053093 | 0.331982614 | 0.16514309 | 3.986751476 | 0.041422971 | 0.966958701 | NA | Myh7 | myosin, heavy polypeptide 7, cardiac mus | 0 | 1.0279 | 0 | 0.964 | 0 | 0 |
| 15433 | ENSMUSG00000053094 | 978.8738473 | -0.103295976 | 0.108812637 | -0.949301284 | 0.342467401 | 0.784247367 | Tmem248 | transmembrane protein 248 [Source:MGI | 987.82 | 1083.4 | 971.03 | 967.83 | 904.12 | 959.02 |
| 15434 | ENSMUSG00000053101 | 0.806430748 | -0.300727144 | 2.416884806 | -0.124427587 | 0.900976732 | NA | Gpr141 | G protein-coupled receptor 141 [Source:N | 1.6062 | 1.0279 | 0 | 0 | 1.0906 | 1.1138 |
| 15435 | ENSMUSG00000053110 | 1331.228558 | 0.02959288 | 0.109532252 | 0.270175034 | 0.787025599 | 0.960870081 | Yap1 | yes-associated protein 1 [Source:MGI Sym | 1280.1 | 1285.9 | 1386.6 | 1301.4 | 1257.5 | 1475.8 |
| 15436 | ENSMUSG00000053111 | 0.455176322 | 0.955156269 | 3.546674842 | 0.269310357 | 0.787690862 | NA | Fank1 | fibronectin type 3 and ankyrin repeat dom | 0.8031 | 0 | 0 | 1.928 | 0 | 0 |
| 15437 | ENSMUSG00000053113 | 231.8867972 | 2.202464406 | 0.852054847 | 2.584885719 | 0.009741128 | 0.106497108 | Socs3 | suppressor of cytokine signaling 3 [Source | 48.989 | 105.88 | 93.581 | 235.21 | 110.15 | 797.51 |
| 15438 | ENSMUSG00000053117 | 0.353089354 | 0.165142706 | 3.896102244 | 0.042386646 | 0.966190474 | NA | E330013P04 | RIKEN cDNA E330013P04 gene [Source:M | 0 | 1.0279 | 0 | 0 | 1.0906 | 0 |
| 15439 | ENSMUSG00000053119 | 1065.272044 | 0.105194995 | 0.105891859 | 0.993419089 | 0.320505763 | 0.767391055 | Chmp3 | charged multivesicular body protein 3 [So | 1022.3 | 990.91 | 1065.6 | 1039.2 | 1139.7 | 1133.9 |
| 15440 | ENSMUSG00000053121 | 0.267700746 | -1.497481649 | 4.057001436 | -0.369110456 | 0.712045396 | NA | Gm5129 | predicted gene 5129 [Source:MGI Symbol | 1.6062 | 0 | 0 | 0 | 0 | 0 |
| 15441 | ENSMUSG00000053128 | 31.27314178 | 0.511238344 | 0.391563681 | 1.305632696 | 0.191677496 | 0.629099068 | Rnf26 | ring finger protein 26 [Source:MGI Symbo | 22.487 | 28.782 | 26.162 | 30.847 | 39.262 | 40.098 |
| 15442 | ENSMUSG00000053129 | 0 | NA | NA | NA | NA | NA | Gsx1 | GS homeobox 1 [Source:MGI Symbol;Acc: | 0 | 0 | 0 | 0 | 0 | 0 |
| 15443 | ENSMUSG00000053134 | 184.1897656 | 0.135990124 | 0.186191099 | 0.730379298 | 0.465158369 | 0.86312388 | Supt7l | SPT7-like, STAGA complex gamma subunit | 195.15 | 168.58 | 162.01 | 170.62 | 209.4 | 199.38 |
| 15444 | ENSMUSG00000053137 | 10.61835976 | -0.170087778 | 0.674474952 | -0.25217805 | 0.800903444 | 0.962424862 | Mapk11 | mitogen-activated protein kinase 11 [Sour | 10.44 | 13.363 | 10.062 | 14.46 | 9.8156 | 5.5692 |
| 15445 | ENSMUSG00000053141 | 6.987558456 | -1.224216064 | 0.862287381 | -1.419730929 | 0.15568603 | 0.574732247 | Ptprt | protein tyrosine phosphatase, receptor ty | 16.062 | 5.1396 | 8.05 | 3.8559 | 4.3625 | 4.4554 |
| 15446 | ENSMUSG00000053146 | 0 | NA | NA | NA | NA | NA | Olfr352 | olfactory receptor 352 [Source:MGI Symb | 0 | 0 | 0 | 0 | 0 | 0 |
| 15447 | ENSMUSG00000053153 | 0 | NA | NA | NA | NA | NA | Spag16 | sperm associated antigen 16 [Source:MGI | 0 | 0 | 0 | 0 | 0 | 0 |
| 15448 | ENSMUSG00000053158 | 112.6233462 | -0.21959991 | 0.292254948 | -0.751398434 | 0.452412905 | 0.856769845 | Fes | feline sarcoma oncogene [Source:MGI Syr | 86.735 | 150.08 | 126.79 | 70.37 | 101.43 | 140.34 |
| 15449 | ENSMUSG00000053161 | 0 | NA | NA | NA | NA | NA | Daw1 | dynein assembly factor with WDR repeat ( | 0 | 0 | 0 | 0 | 0 | 0 |
| 15450 | ENSMUSG00000053164 | 0.949367096 | -3.28842827 | 2.400393427 | -1.369953872 | 0.170701301 | NA | Gpr21 | G protein-coupled receptor 21 [Source:M | 1.6062 | 3.0838 | 1.0062 | 0 | 0 | 0 |
| 15451 | ENSMUSG00000053166 | 0 | NA | NA | NA | NA | NA | Cdh22 | cadherin 22 [Source:MGI Symbol;Acc:MGI | 0 | 0 | 0 | 0 | 0 | 0 |
| 15452 | ENSMUSG00000053168 | 332.8319359 | -0.721357791 | 0.23921389 | -3.015534725 | 0.002565266 | 0.04206803 | 9030619P08 | RIKEN cDNA 9030619P08 gene [Source:M | 580.64 | 284.73 | 377.34 | 309.44 | 242.12 | 202.72 |

| | | | | | | | | | | | | | | | |
|---|---|---|---|---|---|---|---|---|---|---|---|---|---|---|---|
| 15453 | ENSMUSG00000053173 | 5.935018119 | 0.073427391 | 0.92909595 | 0.07903101 | 0.937007958 | 0.988420305 | Rpl18-ps2 | ribosomal protein L18, pseudogene 2 [Sou | 8.031 | 6.1675 | 3.0187 | 2.8919 | 4.3625 | 11.138 |
| 15454 | ENSMUSG00000053175 | 335.9019636 | 0.375382307 | 0.234720125 | 1.599276187 | 0.109759248 | 0.486682332 | Bcl3 | B cell leukemia/lymphoma 3 [Source:MGI | 229.69 | 239.5 | 408.54 | 349.92 | 301.01 | 486.75 |
| 15455 | ENSMUSG00000053178 | 16.53059407 | 0.757979604 | 0.632528406 | 1.198332908 | 0.23078744 | 0.677910147 | Mterf1b | mitochondrial transcription termination fa | 11.243 | 20.558 | 5.0312 | 10.604 | 28.356 | 23.391 |
| 15456 | ENSMUSG00000053181 | 0.339027071 | -1.79174149 | 3.919585585 | -0.457125237 | 0.647581032 | NA | A830005F24 | RIKEN cDNA A830005F24 gene [Source:M | 0 | 1.0279 | 1.0062 | 0 | 0 | 0 |
| 15457 | ENSMUSG00000053182 | 1.195470812 | 0.486711373 | 1.942687574 | 0.250535073 | 0.802173585 | NA | Gm609 | predicted gene 609 [Source:MGI Symbol;A | 0 | 1.0279 | 2.0125 | 1.928 | 1.0906 | 1.1138 |
| 15458 | ENSMUSG00000053184 | 0 | NA | NA | NA | NA | NA | Spaca3 | sperm acrosome associated 3 [Source:MG | 0 | 0 | 0 | 0 | 0 | 0 |
| 15459 | ENSMUSG00000053185 | 0 | NA | NA | NA | NA | NA | Gm9898 | predicted gene 9898 [Source:MGI Symbol | 0 | 0 | 0 | 0 | 0 | 0 |
| 15460 | ENSMUSG00000053192 | 23.43196351 | -0.07170576 | 0.444352243 | -0.161371438 | 0.871800861 | 0.979369708 | Mllt11 | myeloid/lymphoid or mixed-lineage leuke | 23.29 | 20.558 | 28.175 | 21.208 | 25.084 | 22.277 |
| 15461 | ENSMUSG00000053194 | 0 | NA | NA | NA | NA | NA | Cib4 | calcium and integrin binding family memb | 0 | 0 | 0 | 0 | 0 | 0 |
| 15462 | ENSMUSG00000053198 | 3.995649201 | -0.836428659 | 1.178935965 | -0.7094776 | 0.478028147 | NA | Prx | periaxin [Source:MGI Symbol;Acc:MGI:10 | 3.2124 | 3.0838 | 9.0562 | 0.964 | 6.5437 | 1.1138 |
| 15463 | ENSMUSG00000053199 | 0.449470461 | -0.545136533 | 3.562105917 | -0.15303771 | 0.878368537 | NA | Arhgap20 | Rho GTPase activating protein 20 [Source: | 1.6062 | 0 | 0 | 0 | 1.0906 | 0 |
| 15464 | ENSMUSG00000053205 | 121.7212343 | 0.167285031 | 0.240553959 | 0.695415829 | 0.486794743 | 0.873681495 | Styx | serine/threonine/tyrosine interaction pro | 133.31 | 109.99 | 100.62 | 154.24 | 134.15 | 98.018 |
| 15465 | ENSMUSG00000053208 | 0.328370405 | 0.16514316 | 4.003149536 | 0.041253308 | 0.967093956 | NA | Kat2b-ps | K(lysine) acetyltransferase 2B, pseudogen | 0 | 0 | 1.0062 | 0.964 | 0 | 0 |
| 15466 | ENSMUSG00000053211 | 0.167707431 | -0.796629176 | 4.080472857 | -0.195229623 | 0.84521319 | NA | Zfy1 | zinc finger protein 1, Y-linked [Source:MG | 0 | 0 | 1.0062 | 0 | 0 | 0 |
| 15467 | ENSMUSG00000053214 | 0 | NA | NA | NA | NA | NA | Gm9899 | predicted gene 9899 [Source:MGI Symbol | 0 | 0 | 0 | 0 | 0 | 0 |
| 15468 | ENSMUSG00000053216 | 0 | NA | NA | NA | NA | NA | Btn2a2 | butyrophilin, subfamily 2, member A2 [So | 0 | 0 | 0 | 0 | 0 | 0 |
| 15469 | ENSMUSG00000053217 | 0 | NA | NA | NA | NA | NA | Tas2r136 | taste receptor, type 2, member 136 [Sour | 0 | 0 | 0 | 0 | 0 | 0 |
| 15470 | ENSMUSG00000053219 | 24.78801619 | -0.161660605 | 0.551616527 | -0.293067008 | 0.769470934 | 0.956153573 | Raet1e | retinoic acid early transcript 1E [Source:M | 43.368 | 24.67 | 10.062 | 15.424 | 22.903 | 32.301 |
| 15471 | ENSMUSG00000053226 | 69.02403781 | 0.322242782 | 0.290857279 | 1.107906886 | 0.267902032 | 0.723745795 | Dand5 | DAN domain family member 5, BMP antag | 61.839 | 64.759 | 57.356 | 67.478 | 100.34 | 62.375 |
| 15472 | ENSMUSG00000053228 | 0 | NA | NA | NA | NA | NA | Ceacam3 | carcinoembryonic antigen-related cell adh | 0 | 0 | 0 | 0 | 0 | 0 |
| 15473 | ENSMUSG00000053251 | 0 | NA | NA | NA | NA | NA | Olfr212 | olfactory receptor 212 [Source:MGI Symb | 0 | 0 | 0 | 0 | 0 | 0 |
| 15474 | ENSMUSG00000053253 | 692.298944 | 0.167800688 | 0.117238284 | 1.431278948 | 0.152350285 | 0.568229634 | Ndfip2 | Nedd4 family interacting protein 2 [Sourc | 669.79 | 681.51 | 604.75 | 733.59 | 757.98 | 706.18 |
| 15475 | ENSMUSG00000053263 | 0.160662974 | 1.126924175 | 4.080472857 | 0.276174898 | 0.782413736 | NA | Gm12592 | predicted gene 12592 [Source:MGI Symbo | 0 | 0 | 0 | 0.964 | 0 | 0 |
| 15476 | ENSMUSG00000053268 | 0 | NA | NA | NA | NA | NA | Dspp | dentin sialophosphoprotein [Source:MGI | 0 | 0 | 0 | 0 | 0 | 0 |
| 15477 | ENSMUSG00000053279 | 27783.10218 | -0.294668967 | 0.098315861 | -2.997166121 | 0.002725022 | 0.043871772 | Aldh1a1 | aldehyde dehydrogenase family 1, subfam | 30300 | 30140 | 31392 | 23836 | 27881 | 23150 |
| 15478 | ENSMUSG00000053286 | 378.3489929 | -0.108869809 | 0.149828576 | -0.726629142 | 0.467453171 | 0.864498487 | Trmt1l | tRNA methyltransferase 1 like [Source:MG | 410.39 | 342.3 | 424.64 | 346.07 | 398.08 | 348.63 |
| 15479 | ENSMUSG00000053287 | 0 | NA | NA | NA | NA | NA | Olfr1013 | olfactory receptor 1013 [Source:MGI Sym | 0 | 0 | 0 | 0 | 0 | 0 |
| 15480 | ENSMUSG00000053289 | 220.2708107 | 0.21477467 | 0.177108914 | 1.212670022 | 0.225256006 | 0.672301295 | Ddx10 | DEAD box helicase 10 [Source:MGI Symbo | 185.52 | 224.09 | 202.26 | 206.29 | 260.66 | 242.82 |
| 15481 | ENSMUSG00000053291 | 10.61324899 | 0.39757441 | 0.679237527 | 0.585324565 | 0.5583295 | 0.899286756 | Rab4b | RAB4B, member RAS oncogene family [So | 11.243 | 5.1396 | 11.069 | 16.388 | 9.8156 | 10.025 |
| 15482 | ENSMUSG00000053293 | 245.0869673 | -0.339230984 | 0.181851295 | -1.865430673 | 0.062121061 | 0.355990107 | Pom121 | nuclear pore membrane protein 121 [Sou | 260.21 | 251.84 | 309.92 | 252.56 | 182.13 | 213.86 |
| 15483 | ENSMUSG00000053297 | 0.631240176 | 0.380643921 | 3.15447101 | 0.120668068 | 0.903953952 | NA | AI854703 | expressed sequence AI854703 [Source:MG | 1.6062 | 0 | 0 | 0 | 2.1812 | 0 |
| 15484 | ENSMUSG00000053303 | 26.13154819 | -1.237994313 | 1.419519274 | -0.872122229 | 0.383141701 | 0.815324637 | Slc22a26 | solute carrier family 22 (organic cation tra | 22.487 | 56.535 | 31.194 | 35.667 | 10.906 | 0 |
| 15485 | ENSMUSG00000053310 | 8.51773046 | -0.197249667 | 0.831041152 | -0.237352466 | 0.812383363 | 0.966547089 | Nrgn | neurogranin [Source:MGI Symbol;Acc:MG | 4.0155 | 16.447 | 7.0437 | 12.532 | 3.2719 | 7.7969 |
| 15486 | ENSMUSG00000053317 | 1043.885746 | 0.197082924 | 0.121830294 | 1.617684051 | 0.105730705 | 0.479250679 | Sec61b | Sec61 beta subunit [Source:MGI Symbol;A | 1016.7 | 984.75 | 915.68 | 1046.9 | 1031.7 | 1267.6 |
| 15487 | ENSMUSG00000053318 | 7.284429403 | -3.725697638 | 1.15234465 | -3.233145257 | 0.001224353 | 0.02425896 | Slamf8 | SLAM family member 8 [Source:MGI Symb | 11.243 | 18.503 | 11.069 | 2.8919 | 0 | 0 |
| 15488 | ENSMUSG00000053329 | 2252.439339 | 0.000133338 | 0.09235061 | 0.001443825 | 0.998847995 | 0.999764475 | Gatd3a | glutamine amidotransferase like class 1 do | 2137.9 | 2301.5 | 2318.4 | 2221 | 2287 | 2248.9 |
| 15489 | ENSMUSG00000053332 | 788.7640278 | 0.134052626 | 0.120234786 | 1.11492381 | 0.264883072 | 0.720705184 | Gas5 | growth arrest specific 5 [Source:MGI Sym | 778.21 | 725.71 | 752.67 | 886.86 | 851.77 | 737.36 |
| 15490 | ENSMUSG00000053333 | 270.409684 | 0.051602202 | 0.197815708 | 0.260859982 | 0.794200487 | 0.962006183 | Dis3l2 | DIS3 like 3'-5' exoribonuclease 2 [Source:I | 267.43 | 312.49 | 216.34 | 215.93 | 319.55 | 290.71 |
| 15491 | ENSMUSG00000053334 | 150.228378 | -0.11823141 | 0.208189497 | -0.567902857 | 0.570100932 | 0.90421633 | Ficd | FIC domain containing [Source:MGI Symb | 141.35 | 179.89 | 147.92 | 121.46 | 157.05 | 153.71 |
| 15492 | ENSMUSG00000053338 | 0.342639279 | -1.805976596 | 4.028547714 | -0.448294702 | 0.653940524 | NA | Tarm1 | T cell-interacting, activating receptor on m | 0 | 2.0558 | 0 | 0 | 0 | 0 |
| 15493 | ENSMUSG00000053347 | 204.1543421 | 0.178967625 | 0.199772852 | 0.895855583 | 0.370329899 | 0.807048832 | Zfp943 | zinc finger prtoein 943 [Source:MGI Symb | 207.2 | 203.53 | 163.01 | 171.59 | 231.21 | 248.39 |
| 15494 | ENSMUSG00000053353 | 9.794585369 | 0.471781899 | 0.763095668 | 0.618247383 | 0.53641228 | 0.890560262 | 2310001K24 | RIKEN cDNA 2310001K24 gene [Source:M | 10.44 | 2.0558 | 12.075 | 15.424 | 7.6343 | 11.138 |
| 15495 | ENSMUSG00000053358 | 14.34499535 | 0.265113015 | 0.59496459 | 0.445594611 | 0.655890104 | 0.926444772 | Gm9905 | predicted gene 9905 [Source:MGI Symbol | 8.8341 | 13.363 | 17.106 | 20.244 | 9.8156 | 16.708 |
| 15496 | ENSMUSG00000053360 | 0 | NA | NA | NA | NA | NA | F730035P03 | RIKEN cDNA F730035P03 gene [Source:M | 0 | 0 | 0 | 0 | 0 | 0 |
| 15497 | ENSMUSG00000053367 | 0.160662974 | 1.126924175 | 4.080472857 | 0.276174898 | 0.782413736 | NA | Epp13 | epididymal protein 13 [Source:MGI Symbo | 0 | 0 | 0 | 0.964 | 0 | 0 |
| 15498 | ENSMUSG00000053368 | 0 | NA | NA | NA | NA | NA | Rxfp2 | relaxin/insulin-like family peptide recepto | 0 | 0 | 0 | 0 | 0 | 0 |
| 15499 | ENSMUSG00000053375 | 0 | NA | NA | NA | NA | NA | Atp6v1e2 | ATPase, H+ transporting, lysosomal V1 sul | 0 | 0 | 0 | 0 | 0 | 0 |
| 15500 | ENSMUSG00000053388 | 0 | NA | NA | NA | NA | NA | Trim50 | tripartite motif-containing 50 [Source:MG | 0 | 0 | 0 | 0 | 0 | 0 |
| 15501 | ENSMUSG00000053389 | 0 | NA | NA | NA | NA | NA | Tas2r107 | taste receptor, type 2, member 107 [Sour | 0 | 0 | 0 | 0 | 0 | 0 |
| 15502 | ENSMUSG00000053390 | 105.5252872 | 0.169984405 | 0.224117096 | 0.758462465 | 0.448174175 | 0.854156674 | Zfp952 | zinc finger protein 952 [Source:MGI Symb | 94.766 | 93.541 | 109.68 | 107.97 | 105.79 | 121.41 |
| 15503 | ENSMUSG00000053391 | 0 | NA | NA | NA | NA | NA | Olfr211 | olfactory receptor 211 [Source:MGI Symb | 0 | 0 | 0 | 0 | 0 | 0 |
| 15504 | ENSMUSG00000053395 | 0 | NA | NA | NA | NA | NA | Cacng8 | calcium channel, voltage-dependent, gam | 0 | 0 | 0 | 0 | 0 | 0 |
| 15505 | ENSMUSG00000053398 | 3.455417791 | 0.44971933 | 1.387248063 | 0.324180903 | 0.745801079 | NA | Phgdh | 3-phosphoglycerate dehydrogenase [Sour | 1.6062 | 6.1675 | 1.0062 | 1.928 | 0 | 10.025 |
| 15506 | ENSMUSG00000053399 | 0 | NA | NA | NA | NA | NA | Adamts18 | a disintegrin-like and metallopeptidase (re | 0 | 0 | 0 | 0 | 0 | 0 |
| 15507 | ENSMUSG00000053411 | 263.2885212 | 0.28808626 | 0.173732482 | 1.658217609 | 0.097273552 | 0.459192119 | Cbx7 | chromobox 7 [Source:MGI Symbol;Acc:MG | 207.2 | 230.25 | 274.7 | 310.4 | 302.1 | 255.07 |
| 15508 | ENSMUSG00000053414 | 49.04348021 | 0.491717506 | 0.357238014 | 1.376442278 | 0.168684746 | 0.595783862 | Hunk | hormonally upregulated Neu-associated k | 39.352 | 32.893 | 50.312 | 79.046 | 42.534 | 50.123 |

| | | | | | | | | | | | | | | | |
|---|---|---|---|---|---|---|---|---|---|---|---|---|---|---|---|
| 15509 | ENSMUSG00000053420 | 0 | NA | NA | NA | NA | NA | Gm4792 | predicted gene 4792 [Source:MGI Symbol | 0 | 0 | 0 | 0 | 0 | 0 |
| 15510 | ENSMUSG00000053436 | 1986.086902 | -0.033472865 | 0.109819311 | -0.30479945 | 0.7605189 | 0.95402891 | Mapk14 | mitogen-activated protein kinase 14 [Sour | 1852 | 1952 | 2224.8 | 2094.7 | 1865 | 1928.1 |
| 15511 | ENSMUSG00000053441 | 0 | NA | NA | NA | NA | NA | Adamts19 | a disintegrin-like and metallopeptidase (re | 0 | 0 | 0 | 0 | 0 | 0 |
| 15512 | ENSMUSG00000053442 | 2.852049859 | -0.463472606 | 1.33328479 | -0.34761711 | 0.728127755 | NA | 4930597O21 | RIKEN cDNA 4930597O21 gene [Source:M | 4.8186 | 3.0838 | 2.0125 | 3.8559 | 0 | 3.3415 |
| 15513 | ENSMUSG00000053453 | 450.6775534 | 0.099920813 | 0.146397013 | 0.682533143 | 0.494901893 | 0.877919732 | Thoc7 | THO complex 7 [Source:MGI Symbol;Acc:I | 444.92 | 403.97 | 455.83 | 425.11 | 540.95 | 433.29 |
| 15514 | ENSMUSG00000053460 | 841.8515727 | -0.18121822 | 0.130191063 | -1.391940548 | 0.163940402 | 0.588267729 | Ggcx | gamma-glutamyl carboxylase [Source:MG | 801.5 | 844.95 | 1038.4 | 810.71 | 756.89 | 798.63 |
| 15515 | ENSMUSG00000053461 | 14.30902652 | 0.638082151 | 0.670093174 | 0.952228997 | 0.340980858 | 0.783129922 | Hhipl2 | hedgehog interacting protein-like 2 [Sour | 20.078 | 5.1396 | 8.05 | 16.388 | 26.175 | 10.025 |
| 15516 | ENSMUSG00000053465 | 0 | NA | NA | NA | NA | NA | Hs6st3 | heparan sulfate 6-O-sulfotransferase 3 [S | 0 | 0 | 0 | 0 | 0 | 0 |
| 15517 | ENSMUSG00000053469 | 0.167707431 | -0.796629176 | 4.080472857 | -0.195229623 | 0.84521319 | NA | Tg | thyroglobulin [Source:MGI Symbol;Acc:M | 0 | 0 | 1.0062 | 0 | 0 | 0 |
| 15518 | ENSMUSG00000053470 | 227.0325575 | 0.254054358 | 0.22325139 | 1.137974363 | 0.255131188 | 0.708637501 | Kdm3a | lysine (K)-specific demethylase 3A [Source | 183.11 | 212.78 | 226.41 | 337.39 | 189.77 | 212.74 |
| 15519 | ENSMUSG00000053472 | 0 | NA | NA | NA | NA | NA | Gm13534 | predicted gene 13534 [Source:MGI Symbo | 0 | 0 | 0 | 0 | 0 | 0 |
| 15520 | ENSMUSG00000053475 | 0.663785267 | -1.400021131 | 3.106754789 | -0.450637796 | 0.652250621 | NA | Tnfaip6 | tumor necrosis factor alpha induced prote | 0 | 0 | 3.0187 | 0.964 | 0 | 0 |
| 15521 | ENSMUSG00000053477 | 204.6494019 | -0.193881744 | 0.189142236 | -1.025057904 | 0.305335867 | 0.754291713 | Tcf4 | transcription factor 4 [Source:MGI Symbo | 235.31 | 211.75 | 207.29 | 156.16 | 223.58 | 193.81 |
| 15522 | ENSMUSG00000053483 | 261.1997791 | -0.030154001 | 0.197439663 | -0.152725144 | 0.878615031 | 0.980976563 | Usp21 | ubiquitin specific peptidase 21 [Source:M | 237.72 | 244.64 | 309.92 | 283.41 | 195.22 | 296.28 |
| 15523 | ENSMUSG00000053490 | 0 | NA | NA | NA | NA | NA | Trim60 | tripartite motif-containing 60 [Source:MG | 0 | 0 | 0 | 0 | 0 | 0 |
| 15524 | ENSMUSG00000053499 | 0 | NA | NA | NA | NA | NA | Gm5444 | predicted gene 5444 [Source:MGI Symbol | 0 | 0 | 0 | 0 | 0 | 0 |
| 15525 | ENSMUSG00000053508 | 0 | NA | NA | NA | NA | NA | Gtsf2 | gametocyte specific factor 2 [Source:MGI | 0 | 0 | 0 | 0 | 0 | 0 |
| 15526 | ENSMUSG00000053510 | 1195.706076 | 0.176787998 | 0.102307658 | 1.72800356 | 0.083987586 | 0.427143617 | Nrd1 | nardilysin, N-arginine dibasic convertase, | 1101.1 | 1144.1 | 1123 | 1304.3 | 1194.2 | 1307.7 |
| 15527 | ENSMUSG00000053519 | 0 | NA | NA | NA | NA | NA | Kcnip1 | Kv channel-interacting protein 1 [Source:I | 0 | 0 | 0 | 0 | 0 | 0 |
| 15528 | ENSMUSG00000053522 | 0 | NA | NA | NA | NA | NA | Lgals7 | lectin, galactose binding, soluble 7 [Sourc | 0 | 0 | 0 | 0 | 0 | 0 |
| 15529 | ENSMUSG00000053528 | 0 | NA | NA | NA | NA | NA | A530021J07I | Riken cDNA A530021J07 gene [Source:MG | 0 | 0 | 0 | 0 | 0 | 0 |
| 15530 | ENSMUSG00000053536 | 476.1202458 | 0.348762778 | 0.150825072 | 2.312366059 | 0.020757519 | 0.177549827 | Cstf2t | cleavage stimulation factor, 3' pre-RNA su | 457.77 | 407.06 | 390.42 | 449.21 | 549.67 | 602.59 |
| 15531 | ENSMUSG00000053541 | 1.075993052 | -3.475052663 | 2.294985823 | -1.51419352 | 0.129976765 | NA | Gvin-ps6 | GTPase, very large interferon inducible, p | 2.4093 | 1.0279 | 3.0187 | 0 | 0 | 0 |
| 15532 | ENSMUSG00000053545 | 0 | NA | NA | NA | NA | NA | 6430503K07 | RIKEN cDNA 6430503K07 gene [Source:M | 0 | 0 | 0 | 0 | 0 | 0 |
| 15533 | ENSMUSG00000053550 | 0.698954292 | 0.126053087 | 3.048972385 | 0.04134281 | 0.967022605 | NA | Shisa7 | shisa family member 7 [Source:MGI Symb | 0 | 0 | 2.0125 | 0 | 2.1812 | 0 |
| 15534 | ENSMUSG00000053552 | 3.474243372 | -0.979393636 | 1.382256976 | -0.708546713 | 0.478605814 | NA | Ebf4 | early B cell factor 4 [Source:MGI Symbol;A | 0.8031 | 4.1117 | 9.0562 | 5.7839 | 1.0906 | 0 |
| 15535 | ENSMUSG00000053553 | 163.5693989 | 1.529826616 | 0.226033637 | 6.768136968 | 1.30E-11 | 4.19E-09 | 3110082I17I | RIKEN cDNA 3110082I17 gene [Source:MG | 87.538 | 68.87 | 95.593 | 198.58 | 231.21 | 299.62 |
| 15536 | ENSMUSG00000053559 | 227.3887869 | 0.176093765 | 0.191727409 | 0.918459003 | 0.358378613 | 0.798191951 | Smagp | small cell adhesion glycoprotein [Source:I | 212.02 | 214.83 | 213.32 | 206.29 | 212.67 | 305.19 |
| 15537 | ENSMUSG00000053560 | 475.5368594 | 0.279317602 | 0.166116574 | 1.681455351 | 0.092674502 | 0.448459015 | Ier2 | immediate early response 2 [Source:MGI | 403.16 | 366.97 | 519.22 | 555.25 | 442.79 | 565.83 |
| 15538 | ENSMUSG00000053565 | 806.2981284 | 0.081381353 | 0.11932388 | 0.682020675 | 0.495225877 | 0.877919732 | Eif3k | eukaryotic translation initiation factor 3, s | 836.03 | 716.46 | 796.95 | 772.15 | 847.41 | 868.8 |
| 15539 | ENSMUSG00000053574 | 1.458220754 | 1.095126875 | 1.97365769 | 0.554871739 | 0.578982393 | NA | 4930563E22 | RIKEN cDNA 4930563E22 gene [Source:M | 0.8031 | 0 | 2.0125 | 4.8199 | 0 | 1.1138 |
| 15540 | ENSMUSG00000053580 | 63.03195523 | -0.041818193 | 0.3138193 | -0.133255644 | 0.893991205 | 0.982575136 | Tanc2 | tetratricopeptide repeat, ankyrin repeat a | 61.839 | 64.759 | 65.406 | 72.298 | 39.262 | 74.628 |
| 15541 | ENSMUSG00000053581 | 204.8919153 | 0.030201792 | 0.177603385 | 0.170051895 | 0.864969325 | 0.977227145 | Zfand2a | zinc finger, AN1-type domain 2A [Source:I | 232.1 | 189.14 | 186.16 | 197.62 | 210.49 | 213.86 |
| 15542 | ENSMUSG00000053588 | 0.133850373 | -0.796629176 | 4.080472857 | -0.195229623 | 0.84521319 | NA | A730061H03 | RIKEN cDNA A730061H03 gene [Source:M | 0.8031 | 0 | 0 | 0 | 0 | 0 |
| 15543 | ENSMUSG00000053593 | 0 | NA | NA | NA | NA | NA | Fate1 | fetal and adult testis expressed 1 [Source: | 0 | 0 | 0 | 0 | 0 | 0 |
| 15544 | ENSMUSG00000053600 | 69.7734302 | 0.501216634 | 0.294763989 | 1.70039982 | 0.089055746 | 0.439824707 | Zfp472 | zinc finger protein 472 [Source:MGI Symb | 55.414 | 53.452 | 64.4 | 75.19 | 63.256 | 106.93 |
| 15545 | ENSMUSG00000053603 | 0.17131964 | -0.796629176 | 4.080472857 | -0.195229623 | 0.84521319 | NA | 4930442H23 | RIKEN cDNA 4930442H23 gene [Source:M | 0 | 1.0279 | 0 | 0 | 0 | 0 |
| 15546 | ENSMUSG00000053604 | 95.51115773 | -0.100987943 | 0.267596563 | -0.377388789 | 0.705884697 | 0.94295036 | Rpia | ribose 5-phosphate isomerase A [Source:I | 129.3 | 78.122 | 88.55 | 98.326 | 78.525 | 100.25 |
| 15547 | ENSMUSG00000053613 | 77.36658751 | -0.179581284 | 0.284983351 | -0.630146578 | 0.528598688 | 0.888334265 | Notumos | notum palmitoleoyl-protein carboxylester | 73.885 | 68.87 | 103.64 | 56.875 | 79.615 | 81.311 |
| 15548 | ENSMUSG00000053615 | 0.669251866 | -2.807208097 | 3.391018549 | -0.827836255 | 0.407763239 | NA | Gm9913 | predicted gene 9913 [Source:MGI Symbol | 4.0155 | 0 | 0 | 0 | 0 | 0 |
| 15549 | ENSMUSG00000053617 | 749.883882 | -0.022450602 | 0.214794446 | -0.104521333 | 0.916755639 | 0.986374671 | Sh3pxd2a | SH3 and PX domains 2A [Source:MGI Sym | 664.97 | 784.3 | 818.08 | 724.91 | 470.06 | 1037 |
| 15550 | ENSMUSG00000053624 | 0 | NA | NA | NA | NA | NA | Gykl1 | glycerol kinase-like 1 [Source:MGI Symbol | 0 | 0 | 0 | 0 | 0 | 0 |
| 15551 | ENSMUSG00000053626 | 0.342432689 | 1.990653086 | 3.900281263 | 0.51038706 | 0.609780321 | NA | Tll1 | tolloid-like [Source:MGI Symbol;Acc:MGI: | 0 | 0 | 0 | 0.964 | 1.0906 | 0 |
| 15552 | ENSMUSG00000053640 | 0 | NA | NA | NA | NA | NA | Gm9915 | predicted gene 9915 [Source:MGI Symbol | 0 | 0 | 0 | 0 | 0 | 0 |
| 15553 | ENSMUSG00000053641 | 697.9952566 | 0.036403131 | 0.114815545 | 0.31705751 | 0.751199965 | 0.952182657 | Dennd4a | DENN/MADD domain containing 4A [Sour | 674.61 | 714.4 | 679.22 | 740.33 | 709.99 | 669.42 |
| 15554 | ENSMUSG00000053644 | 10195.07725 | -0.041894354 | 0.091669757 | -0.457013914 | 0.647661044 | 0.925643431 | Aldh7a1 | aldehyde dehydrogenase family 7, memb | 10485 | 9749.8 | 10794 | 10070 | 10608 | 9463.2 |
| 15555 | ENSMUSG00000053646 | 1061.900499 | 0.752394517 | 0.135787179 | 5.540983501 | 3.01E-08 | 3.87E-06 | Plxnb1 | plexin B1 [Source:MGI Symbol;Acc:MGI:2 | 708.34 | 762.72 | 903.61 | 1473 | 1159.3 | 1364.5 |
| 15556 | ENSMUSG00000053647 | 0.36353943 | 2.065989254 | 4.034137301 | 0.512126658 | 0.608562368 | NA | Gper1 | G protein-coupled estrogen receptor 1 [S | 0 | 0 | 0 | 0 | 2.1812 | 0 |
| 15557 | ENSMUSG00000053654 | 0 | NA | NA | NA | NA | NA | Krt42 | keratin 42 [Source:MGI Symbol;Acc:MGI:1 | 0 | 0 | 0 | 0 | 0 | 0 |
| 15558 | ENSMUSG00000053675 | 0 | NA | NA | NA | NA | NA | Tgm5 | transglutaminase 5 [Source:MGI Symbol;A | 0 | 0 | 0 | 0 | 0 | 0 |
| 15559 | ENSMUSG00000053678 | 0.133850373 | -0.796629176 | 4.080472857 | -0.195229623 | 0.84521319 | NA | Defb40 | defensin beta 40 [Source:MGI Symbol;Acc | 0.8031 | 0 | 0 | 0 | 0 | 0 |
| 15560 | ENSMUSG00000053684 | 100.2807945 | -0.048215865 | 0.254127195 | -0.189731227 | 0.849519749 | 0.9731095 | BC048403 | cDNA sequence BC048403 [Source:MGI S | 79.507 | 113.07 | 113.71 | 94.47 | 83.978 | 116.95 |
| 15561 | ENSMUSG00000053687 | 0.36353943 | 2.065989254 | 4.034137301 | 0.512126658 | 0.608562368 | NA | Dpep2 | dipeptidase 2 [Source:MGI Symbol;Acc:M | 0 | 0 | 0 | 0 | 2.1812 | 0 |
| 15562 | ENSMUSG00000053693 | 10.16317588 | -0.058657959 | 0.890504354 | -0.065870491 | 0.947480934 | 0.990677354 | Mast1 | microtubule associated serine/threonine I | 12.047 | 3.0838 | 16.1 | 23.135 | 3.2719 | 3.3415 |
| 15563 | ENSMUSG00000053695 | 0 | NA | NA | NA | NA | NA | Defb37 | defensin beta 37 [Source:MGI Symbol;Acc | 0 | 0 | 0 | 0 | 0 | 0 |
| 15564 | ENSMUSG00000053702 | 5.876323129 | -0.950684096 | 1.095285575 | -0.867978286 | 0.38540622 | 0.816437825 | Nebl | nebulette [Source:MGI Symbol;Acc:MGI:1 | 12.047 | 9.2513 | 2.0125 | 8.6758 | 3.2719 | 0 |

| | | | | | | | | | | | | | | | |
|---|---|---|---|---|---|---|---|---|---|---|---|---|---|---|---|
| 15565 | ENSMUSG00000053706 | 0 | NA | NA | NA | NA | NA | B430305J03I | RIKEN cDNA B430305J03 gene [Source:MG | 0 | 0 | 0 | 0 | 0 | 0 |
| 15566 | ENSMUSG00000053714 | 20.3291577 | 0.027781555 | 0.524380836 | 0.05297973 | 0.957748058 | 0.991861398 | 4732471J01I | RIKEN cDNA 4732471J01 gene [Source:MG | 24.896 | 16.447 | 19.119 | 31.811 | 17.45 | 12.252 |
| 15567 | ENSMUSG00000053716 | 90.11699394 | -0.562747424 | 0.25316912 | -2.22281226 | 0.026228462 | 0.209595843 | Dusp7 | dual specificity phosphatase 7 [Source:MG | 106.01 | 93.541 | 122.76 | 66.514 | 86.159 | 65.717 |
| 15568 | ENSMUSG00000053719 | 0 | NA | NA | NA | NA | NA | Klk1b26 | kallikrein 1-related petidase b26 [Source:I | 0 | 0 | 0 | 0 | 0 | 0 |
| 15569 | ENSMUSG00000053720 | 0 | NA | NA | NA | NA | NA | Vmn2r43 | vomeronasal 2, receptor 43 [Source:MGI S | 0 | 0 | 0 | 0 | 0 | 0 |
| 15570 | ENSMUSG00000053729 | 0 | NA | NA | NA | NA | NA | Spinkl | serine protease inhibitor, Kazal type-like [ | 0 | 0 | 0 | 0 | 0 | 0 |
| 15571 | ENSMUSG00000053730 | 46.15134922 | -0.219218017 | 0.340994244 | -0.642878937 | 0.520302658 | 0.887802551 | Tmem39b | transmembrane protein 39b [Source:MGI | 46.58 | 51.396 | 51.318 | 55.911 | 32.719 | 38.985 |
| 15572 | ENSMUSG00000053740 | 1.149122161 | 0.670099062 | 1.962006334 | 0.341537665 | 0.732698855 | NA | Gm6457 | predicted pseudogene 6457 [Source:MGI | 1.6062 | 0 | 1.0062 | 0.964 | 1.0906 | 2.2277 |
| 15573 | ENSMUSG00000053742 | 0 | NA | NA | NA | NA | NA | Gm5114 | predicted gene 5114 [Source:MGI Symbol | 0 | 0 | 0 | 0 | 0 | 0 |
| 15574 | ENSMUSG00000053746 | 16.54238822 | 0.031175363 | 0.557084361 | 0.055961656 | 0.955372354 | 0.991678478 | Ptrh1 | peptidyl-tRNA hydrolase 1 homolog [Sour | 17.668 | 10.279 | 21.131 | 18.316 | 20.722 | 11.138 |
| 15575 | ENSMUSG00000053747 | 0 | NA | NA | NA | NA | NA | Sox14 | SRY (sex determining region Y)-box 14 [So | 0 | 0 | 0 | 0 | 0 | 0 |
| 15576 | ENSMUSG00000053749 | 9.694532752 | 0.409053928 | 0.725177382 | 0.564074306 | 0.572703564 | 0.905173631 | Gm9920 | predicted gene 9920 [Source:MGI Symbol | 13.653 | 4.1117 | 7.0437 | 13.496 | 8.7249 | 11.138 |
| 15577 | ENSMUSG00000053754 | 516.0982774 | -0.010802069 | 0.122929283 | -0.087872219 | 0.929978237 | 0.988405116 | Chd8 | chromodomain helicase DNA binding prot | 505.95 | 533.49 | 515.2 | 535.97 | 514.77 | 491.21 |
| 15578 | ENSMUSG00000053765 | 0 | NA | NA | NA | NA | NA | Oas1f | 2'-5' oligoadenylate synthetase 1F [Source | 0 | 0 | 0 | 0 | 0 | 0 |
| 15579 | ENSMUSG00000053768 | 2000.92447 | -0.008098398 | 0.102880754 | -0.078716353 | 0.937258238 | 0.988420305 | Chchd3 | coiled-coil-helix-coiled-coil-helix domain c | 1932.3 | 2142.2 | 1945.1 | 1845.1 | 2053.6 | 2087.3 |
| 15580 | ENSMUSG00000053769 | 95.86388319 | 0.449657376 | 0.26502644 | 1.696651006 | 0.089762661 | 0.442058946 | Lysmd1 | LysM, putative peptidoglycan-binding, do | 92.357 | 60.647 | 89.556 | 96.398 | 100.34 | 135.89 |
| 15581 | ENSMUSG00000053773 | 0 | NA | NA | NA | NA | NA | Rdh8 | retinol dehydrogenase 8 [Source:MGI Sym | 0 | 0 | 0 | 0 | 0 | 0 |
| 15582 | ENSMUSG00000053774 | 487.8095069 | -0.105749282 | 0.150117819 | -0.704441902 | 0.481157619 | 0.870585915 | Ubxn7 | UBX domain protein 7 [Source:MGI Symbo | 431.27 | 537.6 | 549.41 | 492.59 | 503.87 | 412.12 |
| 15583 | ENSMUSG00000053783 | 0 | NA | NA | NA | NA | NA | 1700016K19 | RIKEN cDNA 1700016K19 gene [Source:M | 0 | 0 | 0 | 0 | 0 | 0 |
| 15584 | ENSMUSG00000053790 | 0 | NA | NA | NA | NA | NA | Defb38 | defensin beta 38 [Source:MGI Symbol;Acc | 0 | 0 | 0 | 0 | 0 | 0 |
| 15585 | ENSMUSG00000053797 | 0 | NA | NA | NA | NA | NA | Krt16 | keratin 16 [Source:MGI Symbol;Acc:MGI:S | 0 | 0 | 0 | 0 | 0 | 0 |
| 15586 | ENSMUSG00000053799 | 366.224014 | -0.065529144 | 0.15636473 | -0.4190788 | 0.675158544 | 0.932446243 | Exoc6 | exocyst complex component 6 [Source:M | 347.74 | 429.67 | 347.15 | 394.27 | 357.72 | 320.79 |
| 15587 | ENSMUSG00000053801 | 154.9115064 | -0.056207317 | 0.200176236 | -0.28078916 | 0.778872118 | 0.959828458 | Grwd1 | glutamate-rich WD repeat containing 1 [S | 153.39 | 175.77 | 144.9 | 157.13 | 167.96 | 130.32 |
| 15588 | ENSMUSG00000053815 | 0 | NA | NA | NA | NA | NA | Olfr744 | olfactory receptor 744 [Source:MGI Symb | 0 | 0 | 0 | 0 | 0 | 0 |
| 15589 | ENSMUSG00000053819 | 144.4812091 | -0.494246285 | 0.214525982 | -2.30389942 | 0.02122829 | 0.180908987 | Camk2d | calcium/calmodulin-dependent protein ki | 147.77 | 201.47 | 157.98 | 114.71 | 109.06 | 135.89 |
| 15590 | ENSMUSG00000053820 | 0 | NA | NA | NA | NA | NA | Bcl2a1c | B cell leukemia/lymphoma 2 related prote | 0 | 0 | 0 | 0 | 0 | 0 |
| 15591 | ENSMUSG00000053821 | 1.728313339 | 0.041478476 | 1.851687519 | 0.022400365 | 0.982128589 | NA | Gm9922 | predicted gene 9922 [Source:MGI Symbol | 4.0155 | 1.0279 | 0 | 0.964 | 4.3625 | 0 |
| 15592 | ENSMUSG00000053825 | 2.592705536 | -0.141751344 | 1.302481036 | -0.108831791 | 0.913335908 | NA | Ppfia2 | protein tyrosine phosphatase, receptor ty | 4.0155 | 3.0838 | 1.0062 | 1.928 | 2.1812 | 3.3415 |
| 15593 | ENSMUSG00000053830 | 0.321325949 | 1.911575114 | 4.048267893 | 0.472195804 | 0.636787034 | NA | Gm9923 | predicted pseudogene 9923 [Source:MGI | 0 | 0 | 0 | 1.928 | 0 | 0 |
| 15594 | ENSMUSG00000053835 | 52.07149137 | -0.078124241 | 0.317621164 | -0.245966737 | 0.805707982 | 0.964425206 | H2-T24 | histocompatibility 2, T region locus 24 [So | 51.399 | 56.535 | 52.325 | 38.559 | 52.35 | 61.261 |
| 15595 | ENSMUSG00000053838 | 598.5957231 | -0.117925347 | 0.122878866 | -0.95968779 | 0.33721237 | 0.77926821 | Nudcd3 | NudC domain containing 3 [Source:MGI S | 599.92 | 640.39 | 628.9 | 542.72 | 623.83 | 555.81 |
| 15596 | ENSMUSG00000053841 | 434.3432235 | 0.196059041 | 0.139268077 | 1.407781641 | 0.159195738 | 0.580586777 | Txlna | taxilin alpha [Source:MGI Symbol;Acc:MG | 368.62 | 451.26 | 395.45 | 457.89 | 444.97 | 487.86 |
| 15597 | ENSMUSG00000053846 | 421.7271697 | -0.789230003 | 0.49092186 | -1.607648932 | 0.107912089 | 0.48458257 | Lipg | lipase, endothelial [Source:MGI Symbol;A | 653.73 | 569.47 | 379.35 | 156.16 | 482.05 | 289.6 |
| 15598 | ENSMUSG00000053852 | 0 | NA | NA | NA | NA | NA | Adgrg4 | adhesion G protein-coupled receptor G4 [ | 0 | 0 | 0 | 0 | 0 | 0 |
| 15599 | ENSMUSG00000053856 | 0 | NA | NA | NA | NA | NA | Dnajc5g | DnaJ heat shock protein family (Hsp40) m | 0 | 0 | 0 | 0 | 0 | 0 |
| 15600 | ENSMUSG00000053861 | 0 | NA | NA | NA | NA | NA | Gm9925 | predicted gene 9925 [Source:MGI Symbol | 0 | 0 | 0 | 0 | 0 | 0 |
| 15601 | ENSMUSG00000053862 | 4.55092169 | -0.411546295 | 1.032178621 | -0.398716158 | 0.69010236 | NA | Slc51b | solute carrier family 51, beta subunit [Sou | 2.4093 | 7.1954 | 6.0375 | 2.8919 | 6.5437 | 2.2277 |
| 15602 | ENSMUSG00000053863 | 0 | NA | NA | NA | NA | NA | Mepe | matrix extracellular phosphoglycoprotein | 0 | 0 | 0 | 0 | 0 | 0 |
| 15603 | ENSMUSG00000053868 | 0 | NA | NA | NA | NA | NA | Gm5142 | predicted gene 5142 [Source:MGI Symbol | 0 | 0 | 0 | 0 | 0 | 0 |
| 15604 | ENSMUSG00000053870 | 265.4789605 | 0.047144966 | 0.155810582 | 0.302578718 | 0.762210936 | 0.954414324 | Fpgt | fucose-1-phosphate guanylyltransferase [ | 267.43 | 266.23 | 249.55 | 261.24 | 294.47 | 253.96 |
| 15605 | ENSMUSG00000053873 | 0 | NA | NA | NA | NA | NA | Aym1 | activator of yeast meiotic promoters 1 [So | 0 | 0 | 0 | 0 | 0 | 0 |
| 15606 | ENSMUSG00000053877 | 41.75273082 | 0.072769738 | 0.374894469 | 0.194107259 | 0.846091898 | 0.972803041 | Srcap | Snf2-related CREBBP activator protein [So | 40.155 | 32.893 | 49.306 | 58.803 | 38.172 | 31.188 |
| 15607 | ENSMUSG00000053886 | 129.4920956 | -0.113542334 | 0.24149419 | -0.470165903 | 0.638236493 | 0.923699483 | Sh2d4a | SH2 domain containing 4A [Source:MGI S | 114.84 | 113.07 | 176.09 | 122.43 | 109.06 | 141.46 |
| 15608 | ENSMUSG00000053889 | 0.974421879 | -3.346724782 | 2.664166553 | -1.256199534 | 0.209043636 | NA | Kirrel3os | kirre like nephrin family adhesion molecul | 4.8186 | 1.0279 | 0 | 0 | 0 | 0 |
| 15609 | ENSMUSG00000053896 | 0 | NA | NA | NA | NA | NA | 4933409G03 | RIKEN cDNA 4933409G03 gene [Source:M | 0 | 0 | 0 | 0 | 0 | 0 |
| 15610 | ENSMUSG00000053897 | 499.5614043 | -0.020485345 | 0.148691759 | -0.137770544 | 0.890421764 | 0.982489666 | Slc39a8 | solute carrier family 39 (metal ion transpo | 458.57 | 533.49 | 517.21 | 428.01 | 484.23 | 575.86 |
| 15611 | ENSMUSG00000053898 | 8094.497998 | -0.297676863 | 0.091272632 | -3.261403294 | 0.001108622 | 0.022572786 | Ech1 | enoyl coenzyme A hydratase 1, peroxisom | 8537.8 | 9411.6 | 8831.8 | 7525.8 | 7452.2 | 6807.8 |
| 15612 | ENSMUSG00000053907 | 1658.525311 | -0.214055079 | 0.11947577 | -1.791619166 | 0.073193991 | 0.392921239 | Mat2a | methionine adenosyltransferase II, alpha | 1559.6 | 1852.3 | 1933 | 1457.5 | 1680.6 | 1468 |
| 15613 | ENSMUSG00000053909 | 0 | NA | NA | NA | NA | NA | Rhox10 | reproductive homeobox 10 [Source:MGI S | 0 | 0 | 0 | 0 | 0 | 0 |
| 15614 | ENSMUSG00000053914 | 0.367410362 | 2.078255025 | 3.800070157 | 0.54689912 | 0.58444804 | NA | Kdm4d | lysine (K)-specific demethylase 4D [Source | 0 | 0 | 0 | 0 | 1.0906 | 1.1138 |
| 15615 | ENSMUSG00000053916 | 5.083234464 | 0.669960087 | 1.059211647 | 0.632508233 | 0.527054831 | NA | Nanp | N-acetylneuraminic acid phosphatase [So | 0.8031 | 2.0558 | 9.0562 | 8.6758 | 5.4531 | 4.4554 |
| 15616 | ENSMUSG00000053925 | 17.19501735 | 1.072741867 | 0.536454235 | 1.999689437 | 0.04553381 | 0.295238834 | Gm9929 | predicted gene 9929 [Source:MGI Symbol | 12.85 | 9.2513 | 11.069 | 26.027 | 27.265 | 16.708 |
| 15617 | ENSMUSG00000053929 | 1578.41077 | 0.102465262 | 0.095086356 | 1.077602145 | 0.281211345 | 0.733710968 | Cyhr1 | cysteine and histidine rich 1 [Source:MGI | 1522.7 | 1514.1 | 1530.5 | 1669.6 | 1566.1 | 1667.4 |
| 15618 | ENSMUSG00000053930 | 0.610340026 | -2.657561537 | 3.094276798 | -0.858863544 | 0.390415805 | NA | Shisa6 | shisa family member 6 [Source:MGI Symb | 1.6062 | 2.0558 | 0 | 0 | 0 | 0 |
| 15619 | ENSMUSG00000053931 | 2063.23058 | 0.227329624 | 0.095787303 | 2.373275136 | 0.017631125 | 0.158493842 | Cnn3 | calponin 3, acidic [Source:MGI Symbol;Ac | 1794.9 | 1938.7 | 1970.2 | 2194 | 2170.3 | 2311.2 |
| 15620 | ENSMUSG00000053950 | 136.5086705 | -0.110775153 | 0.235228052 | -0.470926628 | 0.637693132 | 0.923478764 | Adnp2 | ADNP homeobox 2 [Source:MGI Symbol;A | 156.6 | 117.18 | 150.94 | 121.46 | 105.79 | 167.08 |

| | | | | | | | | | | | | | | | |
|---|---|---|---|---|---|---|---|---|---|---|---|---|---|---|---|
| 15621 | ENSMUSG00000053957 | 0.342432689 | 1.990653086 | 3.900281263 | 0.51038706 | 0.609780321 | NA | Gm12474 | predicted gene 12474 [Source:MGI Symbo | 0 | 0 | 0 | 0.964 | 1.0906 | 0 |
| 15622 | ENSMUSG00000053961 | 0 | NA | NA | NA | NA | NA | Ang5 | angiogenin, ribonuclease A family, membe | 0 | 0 | 0 | 0 | 0 | 0 |
| 15623 | ENSMUSG00000053963 | 1.749012112 | -1.207975817 | 1.685158079 | -0.716832345 | 0.473477551 | NA | Stum | mechanosensory transduction mediator [ | 3.2124 | 3.0838 | 1.0062 | 0.964 | 0 | 2.2277 |
| 15624 | ENSMUSG00000053964 | 358.6584026 | -0.324339647 | 0.154210981 | -2.103220175 | 0.035446527 | 0.252635616 | Lgals4 | lectin, galactose binding, soluble 4 [Sourc | 362.2 | 421.45 | 413.57 | 330.64 | 350.09 | 274.01 |
| 15625 | ENSMUSG00000053965 | 31.35393797 | 0.279385588 | 0.427885664 | 0.652944494 | 0.513792063 | 0.885004045 | Pde5a | phosphodiesterase 5A, cGMP-specific [So | 20.078 | 31.865 | 33.206 | 26.027 | 47.987 | 28.96 |
| 15626 | ENSMUSG00000053977 | 3.819252834 | -1.22903767 | 1.348930394 | -0.911120155 | 0.362232068 | NA | Cd8a | CD8 antigen, alpha chain [Source:MGI Syr | 8.031 | 4.1117 | 4.025 | 6.7478 | 0 | 0 |
| 15627 | ENSMUSG00000053979 | 0 | NA | NA | NA | NA | NA | A730045E13 | RIKEN cDNA A730045E13 gene [Source:M | 0 | 0 | 0 | 0 | 0 | 0 |
| 15628 | ENSMUSG00000053980 | 0 | NA | NA | NA | NA | NA | Gm9930 | predicted gene 9930 [Source:MGI Symbol | 0 | 0 | 0 | 0 | 0 | 0 |
| 15629 | ENSMUSG00000053985 | 23.04784469 | 0.190925777 | 0.463754665 | 0.411695647 | 0.680562516 | 0.934384556 | Zfp14 | zinc finger protein 14 [Source:MGI Symbo | 20.078 | 19.53 | 25.156 | 32.775 | 21.812 | 18.935 |
| 15630 | ENSMUSG00000054000 | 73.16698749 | 0.15472294 | 0.299513033 | 0.516581659 | 0.605448221 | 0.916376001 | Tusc1 | tumor suppressor candidate 1 [Source:MG | 68.264 | 80.178 | 59.368 | 81.938 | 52.35 | 96.904 |
| 15631 | ENSMUSG00000054003 | 0 | NA | NA | NA | NA | NA | Tdrd9 | tudor domain containing 9 [Source:MGI S | 0 | 0 | 0 | 0 | 0 | 0 |
| 15632 | ENSMUSG00000054005 | 0 | NA | NA | NA | NA | NA | Mill1 | MHC I like leukocyte 1 [Source:MGI Symbo | 0 | 0 | 0 | 0 | 0 | 0 |
| 15633 | ENSMUSG00000054006 | 0 | NA | NA | NA | NA | NA | D630008O14 | RIKEN cDNA D630008O14 gene [Source:M | 0 | 0 | 0 | 0 | 0 | 0 |
| 15634 | ENSMUSG00000054008 | 2263.686028 | 0.234814503 | 0.115573459 | 2.031733793 | 0.042180613 | 0.282846725 | Ndst1 | N-deacetylase/N-sulfotransferase (hepara | 1891.3 | 1995.2 | 2353.6 | 2276 | 2444.1 | 2622 |
| 15635 | ENSMUSG00000054013 | 0 | NA | NA | NA | NA | NA | Tmem179 | transmembrane protein 179 [Source:MGI | 0 | 0 | 0 | 0 | 0 | 0 |
| 15636 | ENSMUSG00000054021 | 289.483729 | -0.029832841 | 0.176183696 | -0.169328044 | 0.865538619 | 0.977447734 | Sirt5 | sirtuin 5 [Source:MGI Symbol;Acc:MGI:19 | 335.7 | 304.26 | 236.47 | 255.45 | 304.28 | 300.74 |
| 15637 | ENSMUSG00000054027 | 286.1260988 | -0.000363939 | 0.182676015 | -0.001992266 | 0.998410403 | 0.999764475 | Nt5dc3 | 5'-nucleotidase domain containing 3 [Sou | 247.36 | 263.15 | 348.16 | 252.56 | 280.29 | 325.24 |
| 15638 | ENSMUSG00000054031 | 7.789473763 | -0.502147109 | 0.787832997 | -0.637377606 | 0.523878912 | 0.888334265 | Rdh18-ps | retinol dehydrogenase 18, pseudogene [S | 8.031 | 8.2233 | 11.069 | 2.8919 | 8.7249 | 7.7969 |
| 15639 | ENSMUSG00000054033 | 0 | NA | NA | NA | NA | NA | Adam39 | a disintegrin and metallopeptidase domai | 0 | 0 | 0 | 0 | 0 | 0 |
| 15640 | ENSMUSG00000054034 | 0 | NA | NA | NA | NA | NA | Tceal5 | transcription elongation factor A (SII)-like | 0 | 0 | 0 | 0 | 0 | 0 |
| 15641 | ENSMUSG00000054036 | 0 | NA | NA | NA | NA | NA | Olfr279 | olfactory receptor 279 [Source:MGI Symb | 0 | 0 | 0 | 0 | 0 | 0 |
| 15642 | ENSMUSG00000054046 | 0 | NA | NA | NA | NA | NA | Klk13 | kallikrein related-peptidase 13 [Source:MG | 0 | 0 | 0 | 0 | 0 | 0 |
| 15643 | ENSMUSG00000054051 | 234.8325183 | 0.068757565 | 0.164239524 | 0.418642015 | 0.675477779 | 0.932529406 | Ercc6 | excision repair cross-complementing rode | 218.44 | 236.42 | 233.45 | 260.27 | 243.21 | 217.2 |
| 15644 | ENSMUSG00000054052 | 1.060224196 | 1.100879877 | 2.397023932 | 0.459269456 | 0.646040679 | NA | Rdh19 | retinol dehydrogenase 19 [Source:MGI Sy | 0 | 2.0558 | 0 | 0.964 | 0 | 3.3415 |
| 15645 | ENSMUSG00000054054 | 0 | NA | NA | NA | NA | NA | Olfr309 | olfactory receptor 309 [Source:MGI Symb | 0 | 0 | 0 | 0 | 0 | 0 |
| 15646 | ENSMUSG00000054057 | 34.47238941 | 0.500158641 | 0.458044768 | 1.091942702 | 0.274858287 | 0.729578574 | A930004D18 | RIKEN cDNA A930004D18 gene [Source:M | 20.881 | 19.53 | 45.281 | 24.099 | 45.806 | 51.237 |
| 15647 | ENSMUSG00000054061 | 0 | NA | NA | NA | NA | NA | Gm9934 | predicted gene 9934 [Source:MGI Symbol | 0 | 0 | 0 | 0 | 0 | 0 |
| 15648 | ENSMUSG00000054065 | 23.67874831 | -0.66245807 | 0.488985597 | -1.354759882 | 0.175494072 | 0.605810563 | Pkp3 | plakophilin 3 [Source:MGI Symbol;Acc:MG | 23.29 | 19.53 | 44.275 | 16.388 | 18.541 | 20.049 |
| 15649 | ENSMUSG00000054069 | 0 | NA | NA | NA | NA | NA | 6030452D12 | RIKEN cDNA 6030452D12 gene [Source:M | 0 | 0 | 0 | 0 | 0 | 0 |
| 15650 | ENSMUSG00000054072 | 7249.475359 | 0.350094981 | 0.104288697 | 3.356979134 | 0.000787991 | 0.017534609 | Iigp1 | interferon inducible GTPase 1 [Source:MG | 6789.4 | 5628.9 | 6702.6 | 7642.4 | 8408.7 | 8324.9 |
| 15651 | ENSMUSG00000054074 | 20.18999496 | 0.897285914 | 0.521714695 | 1.719878551 | 0.08545452 | 0.430973029 | Skida1 | SKI/DACH domain containing 1 [Source:M | 8.031 | 14.391 | 20.125 | 21.208 | 25.084 | 32.301 |
| 15652 | ENSMUSG00000054079 | 292.0537975 | -0.211645529 | 0.164720971 | -1.284873007 | 0.198836665 | 0.640976045 | Utp18 | UTP18 small subunit processome compon | 305.18 | 320.71 | 314.95 | 292.09 | 297.74 | 221.65 |
| 15653 | ENSMUSG00000054083 | 0.449470461 | -0.545136533 | 3.562105917 | -0.15303771 | 0.878368537 | NA | Capn12 | calpain 12 [Source:MGI Symbol;Acc:MGI: | 1.6062 | 0 | 0 | 0 | 1.0906 | 0 |
| 15654 | ENSMUSG00000054087 | 0 | NA | NA | NA | NA | NA | Fbxw28 | F-box and WD-40 domain protein 28 [Sou | 0 | 0 | 0 | 0 | 0 | 0 |
| 15655 | ENSMUSG00000054091 | 369.8832868 | 0.144066772 | 0.148108371 | 0.972711876 | 0.330696521 | 0.774747658 | 1810037I17F | RIKEN cDNA 1810037I17 gene [Source:MG | 372.64 | 361.83 | 318.98 | 349.92 | 390.44 | 425.49 |
| 15656 | ENSMUSG00000054099 | 84.12449627 | -0.213184065 | 0.258490986 | -0.824725334 | 0.409527545 | 0.833241465 | Slc25a40 | solute carrier family 25, member 40 [Sour | 107.62 | 88.401 | 74.462 | 72.298 | 82.887 | 79.083 |
| 15657 | ENSMUSG00000054102 | 0 | NA | NA | NA | NA | NA | Nlrp9a | NLR family, pyrin domain containing 9A [S | 0 | 0 | 0 | 0 | 0 | 0 |
| 15658 | ENSMUSG00000054106 | 0 | NA | NA | NA | NA | NA | Try4 | trypsin 4 [Source:MGI Symbol;Acc:MGI:10 | 0 | 0 | 0 | 0 | 0 | 0 |
| 15659 | ENSMUSG00000054115 | 48.36482932 | -0.578821694 | 0.368901086 | -1.569043073 | 0.116637903 | 0.501814922 | Skp2 | S-phase kinase-associated protein 2 [Sour | 47.383 | 42.145 | 84.525 | 42.415 | 42.534 | 31.188 |
| 15660 | ENSMUSG00000054116 | 0 | NA | NA | NA | NA | NA | E130116L18I | RIKEN cDNA E130116L18 gene [Source:MG | 0 | 0 | 0 | 0 | 0 | 0 |
| 15661 | ENSMUSG00000054117 | 0 | NA | NA | NA | NA | NA | Zdhhc25 | zinc finger, DHHC domain containing 25 [S | 0 | 0 | 0 | 0 | 0 | 0 |
| 15662 | ENSMUSG00000054128 | 0 | NA | NA | NA | NA | NA | H2-T3 | histocompatibility 2, T region locus 3 [Sou | 0 | 0 | 0 | 0 | 0 | 0 |
| 15663 | ENSMUSG00000054134 | 0 | NA | NA | NA | NA | NA | Umodl1 | uromodulin-like 1 [Source:MGI Symbol;Ac | 0 | 0 | 0 | 0 | 0 | 0 |
| 15664 | ENSMUSG00000054135 | 13.73325394 | -0.760941283 | 0.580115182 | -1.311707238 | 0.189618934 | 0.626663447 | A430110L20 | RIKEN cDNA A430110L20 gene [Source:M | 15.259 | 17.475 | 19.119 | 9.6398 | 11.997 | 8.9108 |
| 15665 | ENSMUSG00000054136 | 0 | NA | NA | NA | NA | NA | Adm2 | adrenomedullin 2 [Source:MGI Symbol;Ac | 0 | 0 | 0 | 0 | 0 | 0 |
| 15666 | ENSMUSG00000054141 | 0 | NA | NA | NA | NA | NA | Olfr24 | olfactory receptor 24 [Source:MGI Symbo | 0 | 0 | 0 | 0 | 0 | 0 |
| 15667 | ENSMUSG00000054142 | 0 | NA | NA | NA | NA | NA | Vmn1r236 | vomeronasal 1 receptor 236 [Source:MGI | 0 | 0 | 0 | 0 | 0 | 0 |
| 15668 | ENSMUSG00000054146 | 0 | NA | NA | NA | NA | NA | Krt15 | keratin 15 [Source:MGI Symbol;Acc:MGI:9 | 0 | 0 | 0 | 0 | 0 | 0 |
| 15669 | ENSMUSG00000054150 | 91.81209079 | 1.055428178 | 0.298435463 | 3.536537405 | 0.000405409 | 0.010719995 | Syne3 | spectrin repeat containing, nuclear envelo | 68.264 | 45.228 | 65.406 | 172.55 | 102.52 | 96.904 |
| 15670 | ENSMUSG00000054156 | 0 | NA | NA | NA | NA | NA | Gm4894 | predicted gene 4894 [Source:MGI Symbol | 0 | 0 | 0 | 0 | 0 | 0 |
| 15671 | ENSMUSG00000054160 | 0 | NA | NA | NA | NA | NA | Nkx2-4 | NK2 homeobox 4 [Source:MGI Symbol;Ac | 0 | 0 | 0 | 0 | 0 | 0 |
| 15672 | ENSMUSG00000054161 | 0 | NA | NA | NA | NA | NA | Fam83e | family with sequence similarity 83, membe | 0 | 0 | 0 | 0 | 0 | 0 |
| 15673 | ENSMUSG00000054162 | 0.74256259 | 3.09446134 | 3.273353887 | 0.945348852 | 0.344480805 | NA | Spock3 | sparc/osteonectin, cwcv and kazal-like do | 0 | 0 | 0 | 0 | 0 | 4.4554 |
| 15674 | ENSMUSG00000054169 | 0 | NA | NA | NA | NA | NA | Ceacam10 | carcinoembryonic antigen-related cell adh | 0 | 0 | 0 | 0 | 0 | 0 |
| 15675 | ENSMUSG00000054178 | 34.67580094 | -0.035667012 | 0.390572541 | -0.091319815 | 0.927238473 | 0.987712228 | NA | NA | 40.958 | 40.089 | 24.15 | 36.631 | 28.356 | 37.871 |
| 15676 | ENSMUSG00000054181 | 7.259181909 | 0.602921388 | 0.809320787 | 0.744972078 | 0.4562886 | 0.857895893 | A930012O16 | RIKEN cDNA A930012O16 gene [Source:M | 8.031 | 4.1117 | 5.0312 | 8.6758 | 5.4531 | 12.252 |

| | | | | | | | | | | | | | | | |
|---|---|---|---|---|---|---|---|---|---|---|---|---|---|---|---|
| 15677 | ENSMUSG00000054191 | 109.5888517 | 0.49377712 | 0.263957973 | 1.870665673 | 0.061391436 | 0.353912206 | Klf1 | Kruppel-like factor 1 (erythroid) [Source:N | 106.01 | 62.703 | 103.64 | 103.15 | 146.14 | 135.89 |
| 15678 | ENSMUSG00000054196 | 0 | NA | NA | NA | NA | NA | Cthrc1 | collagen triple helix repeat containing 1 [S | 0 | 0 | 0 | 0 | 0 | 0 |
| 15679 | ENSMUSG00000054199 | 185.3005007 | -0.001952349 | 0.188521552 | -0.010356105 | 0.991737172 | 0.99859176 | Gon4l | gon-4-like (C.elegans) [Source:MGI Symbo | 179.89 | 194.28 | 182.13 | 172.55 | 219.21 | 163.74 |
| 15680 | ENSMUSG00000054200 | 2.060218844 | 0.13497892 | 1.524436061 | 0.08854351 | 0.929444704 | NA | Ffar4 | free fatty acid receptor 4 [Source:MGI Syr | 0.8031 | 4.1117 | 1.0062 | 0.964 | 4.3625 | 1.1138 |
| 15681 | ENSMUSG00000054203 | 4.090198939 | -3.38104043 | 1.43089348 | -2.362887578 | 0.018133169 | NA | Ifi205 | interferon activated gene 205 [Source:MG | 8.031 | 9.2513 | 5.0312 | 0 | 0 | 2.2277 |
| 15682 | ENSMUSG00000054204 | 0 | NA | NA | NA | NA | NA | Alkal2 | ALK and LTK ligand 2 [Source:MGI Symbol | 0 | 0 | 0 | 0 | 0 | 0 |
| 15683 | ENSMUSG00000054206 | 3.242633984 | 1.111017933 | 1.291155868 | 0.8604832 | 0.38952274 | NA | Gzmm | granzyme M (lymphocyte met-ase 1) [Sou | 4.0155 | 2.0558 | 0 | 6.7478 | 2.1812 | 4.4554 |
| 15684 | ENSMUSG00000054215 | 0 | NA | NA | NA | NA | NA | Sprr2k | small proline-rich protein 2K [Source:MGI | 0 | 0 | 0 | 0 | 0 | 0 |
| 15685 | ENSMUSG00000054226 | 3045.048801 | 0.41670793 | 0.089661633 | 4.647561251 | 3.36E-06 | 0.000217655 | Tprkb | Tp53rk binding protein [Source:MGI Symb | 2535.4 | 2573.9 | 2715.9 | 3382.6 | 3468.2 | 3594.4 |
| 15686 | ENSMUSG00000054236 | 0 | NA | NA | NA | NA | NA | Olfr481 | olfactory receptor 481 [Source:MGI Symb | 0 | 0 | 0 | 0 | 0 | 0 |
| 15687 | ENSMUSG00000054237 | 307.2858323 | -0.015664548 | 0.159632639 | -0.098128731 | 0.921830073 | 0.986685804 | Fra10ac1 | FRA10AC1 homolog (human) [Source:MG | 343.73 | 312.49 | 269.67 | 279.55 | 311.92 | 326.36 |
| 15688 | ENSMUSG00000054247 | 1.515576391 | 0.562655084 | 1.95335792 | 0.288045052 | 0.773312253 | NA | Gm9939 | predicted gene 9939 [Source:MGI Symbol | 1.6062 | 1.0279 | 1.0062 | 0 | 5.4531 | 0 |
| 15689 | ENSMUSG00000054252 | 881.3261478 | 0.533966961 | 0.161022135 | 3.316109067 | 0.000912802 | 0.019559656 | Fgfr3 | fibroblast growth factor receptor 3 [Sourc | 753.31 | 629.09 | 777.83 | 1208.8 | 796.15 | 1122.8 |
| 15690 | ENSMUSG00000054256 | 2.402397783 | -0.516690636 | 1.654075462 | -0.312374283 | 0.754756093 | NA | Msi1 | musashi RNA-binding protein 1 [Source:M | 6.4248 | 2.0558 | 0 | 4.8199 | 0 | 1.1138 |
| 15691 | ENSMUSG00000054258 | 0 | NA | NA | NA | NA | NA | Gm5082 | predicted gene 5082 [Source:MGI Symbol | 0 | 0 | 0 | 0 | 0 | 0 |
| 15692 | ENSMUSG00000054263 | 4082.532972 | 0.882733245 | 0.151651729 | 5.820792499 | 5.86E-09 | 9.63E-07 | Lifr | LIF receptor alpha [Source:MGI Symbol;Ac | 2929.7 | 2321 | 3362.9 | 6373.8 | 4760.5 | 4747.2 |
| 15693 | ENSMUSG00000054266 | 0 | NA | NA | NA | NA | NA | Serpinb9d | serine (or cysteine) peptidase inhibitor, cl | 0 | 0 | 0 | 0 | 0 | 0 |
| 15694 | ENSMUSG00000054272 | 0 | NA | NA | NA | NA | NA | Zscan4c | zinc finger and SCAN domain containing 4 | 0 | 0 | 0 | 0 | 0 | 0 |
| 15695 | ENSMUSG00000054277 | 139.7365364 | 0.186877815 | 0.231367825 | 0.807708741 | 0.419258271 | 0.837591673 | Arfgap3 | ADP-ribosylation factor GTPase activating | 147.77 | 136.71 | 107.67 | 186.05 | 124.33 | 135.89 |
| 15696 | ENSMUSG00000054280 | 258.1692641 | 0.150986626 | 0.158690438 | 0.951453835 | 0.341374043 | 0.783129922 | Prr14l | proline rich 14-like [Source:MGI Symbol;A | 245.75 | 231.28 | 256.59 | 248.71 | 273.75 | 292.94 |
| 15697 | ENSMUSG00000054293 | 2.917025048 | -0.451468299 | 1.251630824 | -0.360704044 | 0.718320702 | NA | P2ry10b | purinergic receptor P2Y, G-protein couple | 4.0155 | 1.0279 | 5.0312 | 1.928 | 3.2719 | 2.2277 |
| 15698 | ENSMUSG00000054302 | 393.5511471 | 0.059751173 | 0.13665217 | 0.437250085 | 0.661929991 | 0.928760569 | Eapp | E2F-associated phosphoprotein [Source:N | 404.76 | 384.44 | 366.27 | 377.88 | 401.35 | 426.6 |
| 15699 | ENSMUSG00000054304 | 1.703776005 | 0.15434538 | 1.588566128 | 0.097160186 | 0.922599186 | NA | D130007C19 | RIKEN cDNA D130007C19 gene [Source:M | 0.8031 | 1.0279 | 3.0187 | 0.964 | 2.1812 | 2.2277 |
| 15700 | ENSMUSG00000054309 | 545.2500851 | 0.004771941 | 0.122425682 | 0.03897827 | 0.968907714 | 0.994555473 | Cpsf3 | cleavage and polyadenylation specificity fa | 551.73 | 528.35 | 552.43 | 512.84 | 572.57 | 553.58 |
| 15701 | ENSMUSG00000054310 | 0 | NA | NA | NA | NA | NA | Obox1 | oocyte specific homeobox 1 [Source:MGI | 0 | 0 | 0 | 0 | 0 | 0 |
| 15702 | ENSMUSG00000054312 | 653.4867042 | -0.176067354 | 0.118276072 | -1.488613472 | 0.136589182 | 0.539788505 | Mrps21 | mitochondrial ribosomal protein S21 [Sou | 656.94 | 697.96 | 725.5 | 603.45 | 646.74 | 590.34 |
| 15703 | ENSMUSG00000054320 | 0 | NA | NA | NA | NA | NA | Lrrc36 | leucine rich repeat containing 36 [Source: | 0 | 0 | 0 | 0 | 0 | 0 |
| 15704 | ENSMUSG00000054321 | 40.45425793 | 0.134569665 | 0.39665965 | 0.339257258 | 0.734415939 | 0.948109711 | Taf4b | TATA-box binding protein associated facto | 38.549 | 26.726 | 50.312 | 33.739 | 59.984 | 33.415 |
| 15705 | ENSMUSG00000054325 | 0 | NA | NA | NA | NA | NA | Lce3a | late cornified envelope 3A [Source:MGI Sy | 0 | 0 | 0 | 0 | 0 | 0 |
| 15706 | ENSMUSG00000054342 | 2.346893216 | 0.090768336 | 1.486033893 | 0.061080932 | 0.951294755 | NA | Kcnn4 | potassium intermediate/small conductanc | 0.8031 | 2.0558 | 4.025 | 3.8559 | 0 | 3.3415 |
| 15707 | ENSMUSG00000054351 | 0 | NA | NA | NA | NA | NA | 4930553M12 | RIKEN cDNA 4930553M12 gene [Source:M | 0 | 0 | 0 | 0 | 0 | 0 |
| 15708 | ENSMUSG00000054360 | 0 | NA | NA | NA | NA | NA | Bsx | brain specific homeobox [Source:MGI Syn | 0 | 0 | 0 | 0 | 0 | 0 |
| 15709 | ENSMUSG00000054362 | 0.513752329 | 1.031892132 | 3.082998058 | 0.334704114 | 0.737848298 | NA | Lexm | lymphocyte expansion molecule [Source:I | 0 | 1.0279 | 0 | 0.964 | 1.0906 | 0 |
| 15710 | ENSMUSG00000054364 | 692.0471327 | -0.544012905 | 0.132102052 | -4.118126087 | 3.82E-05 | 0.001689527 | Rhob | ras homolog family member B [Source:MG | 910.72 | 788.41 | 762.73 | 539.83 | 523.5 | 627.09 |
| 15711 | ENSMUSG00000054379 | 0 | NA | NA | NA | NA | NA | 2410024N13 | RIKEN cDNA 2410024N13 gene [Source:M | 0 | 0 | 0 | 0 | 0 | 0 |
| 15712 | ENSMUSG00000054381 | 81.12093796 | -0.066197067 | 0.277758217 | -0.238326226 | 0.811628082 | 0.96653525 | Zfp747 | zinc finger protein 747 [Source:MGI Symb | 66.657 | 88.401 | 94.587 | 99.29 | 67.618 | 70.172 |
| 15713 | ENSMUSG00000054383 | 2.260548475 | -0.720885109 | 1.461417254 | -0.493278088 | 0.621816109 | NA | Pnma1 | paraneoplastic antigen MA1 [Source:MGI | 2.4093 | 2.0558 | 4.025 | 2.8919 | 2.1812 | 0 |
| 15714 | ENSMUSG00000054385 | 3.208603605 | -1.934617936 | 1.422295771 | -1.360207894 | 0.173764145 | NA | Ceacam2 | carcinoembryonic antigen-related cell adh | 3.2124 | 5.1396 | 7.0437 | 3.8559 | 0 | 0 |
| 15715 | ENSMUSG00000054387 | 748.9766203 | 0.016527953 | 0.132731568 | 0.124521646 | 0.900902263 | 0.983000175 | Mdm4 | transformed mouse 3T3 cell double minut | 663.36 | 829.53 | 742.61 | 806.85 | 776.52 | 674.99 |
| 15716 | ENSMUSG00000054391 | 1.734200814 | 0.677350722 | 1.642746804 | 0.412328133 | 0.680098931 | NA | 4930517O19 | RIKEN cDNA 4930517O19 gene [Source:M | 0 | 2.0558 | 2.0125 | 1.928 | 2.1812 | 2.2277 |
| 15717 | ENSMUSG00000054400 | 11.05611837 | -1.155400901 | 0.760435887 | -1.519392918 | 0.128663623 | 0.524858587 | Cklf | chemokine-like factor [Source:MGI Symbo | 11.243 | 25.698 | 9.0562 | 11.568 | 6.5437 | 2.2277 |
| 15718 | ENSMUSG00000054404 | 140.7032019 | -0.629005629 | 0.216976445 | -2.898958121 | 0.003744049 | 0.054645736 | Slfn5 | schlafen 5 [Source:MGI Symbol;Acc:MGI:1 | 178.29 | 191.19 | 142.89 | 101.22 | 101.43 | 129.21 |
| 15719 | ENSMUSG00000054405 | 400.2940877 | -0.079812267 | 0.132366338 | -0.602964982 | 0.546531985 | 0.89549457 | Dnajc8 | DnaJ heat shock protein family (Hsp40) m | 395.13 | 430.7 | 408.54 | 381.74 | 399.17 | 386.5 |
| 15720 | ENSMUSG00000054406 | 0 | NA | NA | NA | NA | NA | Olfr411 | olfactory receptor 411 [Source:MGI Symb | 0 | 0 | 0 | 0 | 0 | 0 |
| 15721 | ENSMUSG00000054408 | 1061.856135 | 0.200956792 | 0.107638842 | 1.866954244 | 0.061907978 | 0.355884025 | Spcs3 | signal peptidase complex subunit 3 homo | 990.23 | 971.38 | 1002.2 | 1122.1 | 1063.4 | 1221.9 |
| 15722 | ENSMUSG00000054409 | 0.17131964 | -0.796629176 | 4.080472857 | -0.195229623 | 0.84521319 | NA | Tmem74 | transmembrane protein 74 [Source:MGI S | 0 | 1.0279 | 0 | 0 | 0 | 0 |
| 15723 | ENSMUSG00000054412 | 0 | NA | NA | NA | NA | NA | Gm4793 | predicted gene 4793 [Source:MGI Symbol | 0 | 0 | 0 | 0 | 0 | 0 |
| 15724 | ENSMUSG00000054414 | 373.7932917 | 0.059521132 | 0.144077244 | 0.413119588 | 0.679518999 | 0.93418523 | Slc30a7 | solute carrier family 30 (zinc transporter), | 339.71 | 366.97 | 392.44 | 406.8 | 390.44 | 346.41 |
| 15725 | ENSMUSG00000054417 | 0.667397476 | -1.404797968 | 2.804497115 | -0.500909044 | 0.616435137 | NA | Cyp3a44 | cytochrome P450, family 3, subfamily a, p | 0 | 1.0279 | 2.0125 | 0.964 | 0 | 0 |
| 15726 | ENSMUSG00000054418 | 0 | NA | NA | NA | NA | NA | 2900041M22 | RIKEN cDNA 2900041M22 gene [Source:M | 0 | 0 | 0 | 0 | 0 | 0 |
| 15727 | ENSMUSG00000054422 | 73615.8884 | 0.198825372 | 0.104880355 | 1.895735126 | 0.057995078 | 0.342343159 | Fabp1 | fatty acid binding protein 1, liver [Source:I | 76362 | 69428 | 59862 | 80819 | 77988 | 77236 |
| 15728 | ENSMUSG00000054423 | 0 | NA | NA | NA | NA | NA | Cadps | Ca2+-dependent secretion activator [Sour | 0 | 0 | 0 | 0 | 0 | 0 |
| 15729 | ENSMUSG00000054426 | 70.64545823 | -0.04801877 | 0.266867856 | -0.179934635 | 0.857203885 | 0.974809103 | A930005H10 | RIKEN cDNA A930005H10 gene [Source:M | 72.279 | 67.843 | 75.468 | 76.154 | 71.981 | 60.148 |
| 15730 | ENSMUSG00000054428 | 123.4213411 | -0.287449444 | 0.224224127 | -1.281973747 | 0.199851856 | 0.642000421 | Atpif1 | ATPase inhibitory factor 1 [Source:MGI Sy | 150.18 | 130.55 | 125.78 | 102.18 | 97.065 | 134.78 |
| 15731 | ENSMUSG00000054431 | 0 | NA | NA | NA | NA | NA | Olfr450 | olfactory receptor 450 [Source:MGI Symb | 0 | 0 | 0 | 0 | 0 | 0 |
| 15732 | ENSMUSG00000054434 | 54.41571655 | 0.846586544 | 0.326655248 | 2.591682053 | 0.009550799 | 0.104891999 | Tmem120b | transmembrane protein 120B [Source:MG | 48.989 | 41.117 | 26.162 | 70.37 | 75.253 | 64.603 |

| | | | | | | | | | | | | | | |
|---|---|---|---|---|---|---|---|---|---|---|---|---|---|---|
| 15733 | ENSMUSG00000054435 | 85.73452488 | -0.517293302 | 0.332022307 | -1.558007673 | 0.119231429 | 0.505144521 | Gimap4 | GTPase, IMAP family member 4 [Source:M | 85.932 | 86.345 | 130.81 | 82.902 | 37.081 | 91.335 |
| 15734 | ENSMUSG00000054446 | 0 | NA | NA | NA | NA | NA | Cpa1 | carboxypeptidase A1, pancreatic [Source: | 0 | 0 | 0 | 0 | 0 | 0 |
| 15735 | ENSMUSG00000054450 | 0 | NA | NA | NA | NA | NA | Gm9945 | predicted gene 9945 [Source:MGI Symbol | 0 | 0 | 0 | 0 | 0 | 0 |
| 15736 | ENSMUSG00000054452 | 11901.53801 | 0.046905658 | 0.09368693 | 0.500663833 | 0.61660773 | 0.917509899 | Tle5 | TLE family member 5, transcriptional mod | 12397 | 10665 | 12062 | 12225 | 11682 | 12379 |
| 15737 | ENSMUSG00000054453 | 0.885240214 | -0.466577448 | 2.807084628 | -0.166214244 | 0.867988351 | NA | Sytl5 | synaptotagmin-like 5 [Source:MGI Symbol | 0 | 3.0838 | 0 | 0 | 0 | 2.2277 |
| 15738 | ENSMUSG00000054455 | 1834.814874 | -0.00766295 | 0.103177135 | -0.074269847 | 0.94079567 | 0.989001277 | Vapb | vesicle-associated membrane protein, ass | 1752.4 | 1749.5 | 2017.5 | 1789.1 | 1823.5 | 1876.8 |
| 15739 | ENSMUSG00000054457 | 0 | NA | NA | NA | NA | NA | 9430021M05 | RIKEN cDNA 9430021M05 gene [Source:M | 0 | 0 | 0 | 0 | 0 | 0 |
| 15740 | ENSMUSG00000054459 | 0.160662974 | 1.126924175 | 4.080472857 | 0.276174898 | 0.782413736 | NA | Vsnl1 | visinin-like 1 [Source:MGI Symbol;Acc:MG | 0 | 0 | 0 | 0.964 | 0 | 0 |
| 15741 | ENSMUSG00000054469 | 810.4614671 | 0.070982633 | 0.121695635 | 0.583280026 | 0.559704806 | 0.899475387 | Lclat1 | lysocardiolipin acyltransferase 1 [Source:M | 832.82 | 762.72 | 774.81 | 739.37 | 895.4 | 857.66 |
| 15742 | ENSMUSG00000054474 | 825.9557756 | 0.117158418 | 0.142315909 | 0.823227837 | 0.410378442 | 0.833241465 | Thnsl2 | threonine synthase-like 2 (bacterial) [Sour | 926.78 | 750.38 | 698.33 | 766.36 | 863.77 | 950.11 |
| 15743 | ENSMUSG00000054477 | 563.9519143 | 0.163988173 | 0.160330412 | 1.02281389 | 0.306395849 | 0.755586718 | Kcnn2 | potassium intermediate/small conductanc | 425.64 | 612.64 | 558.47 | 551.4 | 560.58 | 674.99 |
| 15744 | ENSMUSG00000054484 | 230.3086832 | 0.179473353 | 0.199002984 | 0.901862623 | 0.367129848 | 0.805239112 | Tmem62 | transmembrane protein 62 [Source:MGI S | 216.84 | 186.05 | 244.52 | 192.8 | 246.48 | 295.17 |
| 15745 | ENSMUSG00000054488 | 3.013793234 | 1.593762732 | 1.30991769 | 1.216689219 | 0.223722483 | NA | Gm9946 | predicted gene 9946 [Source:MGI Symbol | 2.4093 | 0 | 2.0125 | 4.8199 | 3.2719 | 5.5692 |
| 15746 | ENSMUSG00000054493 | 0.167707431 | -0.796629176 | 4.080472857 | -0.195229623 | 0.84521319 | NA | Gm9947 | predicted gene 9947 [Source:MGI Symbol | 0 | 0 | 1.0062 | 0 | 0 | 0 |
| 15747 | ENSMUSG00000054497 | 0 | NA | NA | NA | NA | NA | Tas2r130 | taste receptor, type 2, member 130 [Sour | 0 | 0 | 0 | 0 | 0 | 0 |
| 15748 | ENSMUSG00000054498 | 0 | NA | NA | NA | NA | NA | Olfr308 | olfactory receptor 308 [Source:MGI Symb | 0 | 0 | 0 | 0 | 0 | 0 |
| 15749 | ENSMUSG00000054499 | 585.6480747 | 0.678206503 | 0.180384146 | 3.759789982 | 0.000170056 | 0.005502664 | Dedd2 | death effector domain-containing DNA bi | 517.2 | 369.02 | 463.88 | 610.2 | 638.01 | 915.58 |
| 15750 | ENSMUSG00000054509 | 246.797634 | -0.085186734 | 0.165817505 | -0.513737881 | 0.607435264 | 0.916376001 | Parp4 | poly (ADP-ribose) polymerase family, men | 255.39 | 246.7 | 260.62 | 270.88 | 235.57 | 211.63 |
| 15751 | ENSMUSG00000054510 | 0 | NA | NA | NA | NA | NA | Gm14461 | predicted gene 14461 [Source:MGI Symbo | 0 | 0 | 0 | 0 | 0 | 0 |
| 15752 | ENSMUSG00000054514 | 10.85982353 | 0.4855505 | 0.695885059 | 0.697745258 | 0.485336519 | 0.872797142 | Atad3aos | ATPase family, AAA domain containing 3A | 8.031 | 6.1675 | 13.081 | 20.244 | 8.7249 | 8.9108 |
| 15753 | ENSMUSG00000054517 | 168.2516458 | 0.089256285 | 0.228921402 | 0.389899261 | 0.696611041 | 0.939992732 | Trim65 | tripartite motif-containing 65 [Source:MG | 124.48 | 178.86 | 186.16 | 143.63 | 161.41 | 214.97 |
| 15754 | ENSMUSG00000054519 | 107.288185 | 0.583780971 | 0.230197234 | 2.536003414 | 0.011212564 | 0.11768024 | Zfp867 | zinc finger protein 867 [Source:MGI Symb | 86.735 | 90.457 | 80.5 | 138.81 | 106.88 | 140.34 |
| 15755 | ENSMUSG00000054520 | 415.9408153 | 0.180906694 | 0.154783832 | 1.168769969 | 0.242496323 | 0.692000002 | Sh3bp2 | SH3-domain binding protein 2 [Source:MG | 444.12 | 378.27 | 346.15 | 421.26 | 404.62 | 501.23 |
| 15756 | ENSMUSG00000054523 | 0 | NA | NA | NA | NA | NA | Ms4a5 | membrane-spanning 4-domains, subfamil | 0 | 0 | 0 | 0 | 0 | 0 |
| 15757 | ENSMUSG00000054526 | 0 | NA | NA | NA | NA | NA | Olfr1500 | olfactory receptor 1500 [Source:MGI Sym | 0 | 0 | 0 | 0 | 0 | 0 |
| 15758 | ENSMUSG00000054537 | 0.667397476 | -1.404797968 | 2.804497115 | -0.500909044 | 0.616435137 | NA | Tmprss11e | transmembrane protease, serine 11e [Sou | 0 | 1.0279 | 2.0125 | 0.964 | 0 | 0 |
| 15759 | ENSMUSG00000054545 | 154.6117911 | 0.397215132 | 0.2906707 | 1.366546857 | 0.171767384 | 0.599527859 | Ugt1a6a | UDP glucuronosyltransferase 1 family, pol | 181.5 | 98.68 | 119.74 | 216.9 | 205.04 | 105.82 |
| 15760 | ENSMUSG00000054555 | 3.834647313 | -0.190447099 | 1.236750417 | -0.153989921 | 0.877617683 | NA | Adam12 | a disintegrin and metallopeptidase domai | 4.0155 | 8.2233 | 0 | 1.928 | 3.2719 | 5.5692 |
| 15761 | ENSMUSG00000054556 | 3.828995541 | 1.569314988 | 1.298001231 | 1.209024268 | 0.226653522 | NA | Gm4876 | predicted gene 4876 [Source:MGI Symbol | 0.8031 | 0 | 5.0312 | 2.8919 | 10.906 | 3.3415 |
| 15762 | ENSMUSG00000054568 | 0 | NA | NA | NA | NA | NA | Usp17la | ubiquitin specific peptidase 17-like A [Sou | 0 | 0 | 0 | 0 | 0 | 0 |
| 15763 | ENSMUSG00000054580 | 43.26463737 | -0.680111807 | 0.341195756 | -1.993318486 | 0.046226583 | 0.297429405 | Pla2r1 | phospholipase A2 receptor 1 [Source:MGI | 44.171 | 58.591 | 57.356 | 36.631 | 30.537 | 32.301 |
| 15764 | ENSMUSG00000054582 | 0.677847551 | 0.063310748 | 2.809814301 | 0.022532004 | 0.982023583 | NA | Pabpc1l | poly(A) binding protein, cytoplasmic 1-like | 0 | 0 | 2.0125 | 0.964 | 1.0906 | 0 |
| 15765 | ENSMUSG00000054589 | 1.358194767 | -1.558749626 | 2.042024688 | -0.763335348 | 0.445263426 | NA | Gm9949 | predicted gene 9949 [Source:MGI Symbol | 4.0155 | 2.0558 | 0 | 0.964 | 0 | 1.1138 |
| 15766 | ENSMUSG00000054594 | 2.08367796 | 0.582180093 | 1.530312015 | 0.380432283 | 0.703624555 | NA | Oscar | osteoclast associated receptor [Source:M | 0 | 2.0558 | 3.0187 | 1.928 | 3.2719 | 2.2277 |
| 15767 | ENSMUSG00000054598 | 1.5524311 | 0.404206833 | 1.731706262 | 0.233415356 | 0.81543889 | NA | 9130230L23 | RIKEN cDNA 9130230L23 gene [Source:M | 0 | 2.0558 | 2.0125 | 1.928 | 1.0906 | 2.2277 |
| 15768 | ENSMUSG00000054604 | 838.6816703 | 0.020925254 | 0.11500406 | 0.181952308 | 0.855620153 | 0.974158771 | Cggbp1 | CGG triplet repeat binding protein 1 [Sour | 788.65 | 853.17 | 856.31 | 822.27 | 798.33 | 913.35 |
| 15769 | ENSMUSG00000054611 | 956.582155 | 0.259722958 | 0.113767949 | 2.282918524 | 0.022435168 | 0.187573567 | Kdm2a | lysine (K)-specific demethylase 2A [Source | 888.23 | 792.52 | 930.78 | 1011.2 | 1025.2 | 1091.6 |
| 15770 | ENSMUSG00000054612 | 494.4782469 | -0.145473934 | 0.141272949 | -1.029736656 | 0.303133643 | 0.752484788 | Mgmt | O-6-methylguanine-DNA methyltransferas | 551.73 | 472.84 | 533.31 | 481.02 | 516.95 | 411.01 |
| 15771 | ENSMUSG00000054618 | 0 | NA | NA | NA | NA | NA | Gm9951 | predicted gene 9951 [Source:MGI Symbol | 0 | 0 | 0 | 0 | 0 | 0 |
| 15772 | ENSMUSG00000054619 | 7736.268203 | 0.085007319 | 0.113704253 | 0.74761776 | 0.454690749 | 0.857337687 | Mettl7a1 | methyltransferase like 7A1 [Source:MGI S | 7000.6 | 6907.6 | 8617.5 | 7679 | 8643.1 | 7569.7 |
| 15773 | ENSMUSG00000054622 | 7.362373724 | 0.116634431 | 0.834299913 | 0.139799164 | 0.888818674 | 0.982382091 | D730045B01 | RIKEN cDNA D730045B01 gene [Source:M | 12.85 | 5.1396 | 3.0187 | 7.7118 | 6.5437 | 8.9108 |
| 15774 | ENSMUSG00000054626 | 3.339066935 | -1.872545742 | 1.400760994 | -1.336806029 | 0.181285961 | NA | Xlr | X-linked lymphocyte-regulated [Source:M | 2.4093 | 11.307 | 2.0125 | 0.964 | 0 | 3.3415 |
| 15775 | ENSMUSG00000054630 | 13176.02749 | 0.330800413 | 0.094979665 | 3.482855131 | 0.000496097 | 0.012307964 | Ugt2b5 | UDP glucuronosyltransferase 2 family, pol | 11943 | 10829 | 12244 | 15341 | 15059 | 13640 |
| 15776 | ENSMUSG00000054640 | 4.237362582 | -1.255228069 | 1.088554394 | -1.153114695 | 0.248863311 | NA | Slc8a1 | solute carrier family 8 (sodium/calcium ex | 5.6217 | 8.2233 | 4.025 | 0.964 | 4.3625 | 2.2277 |
| 15777 | ENSMUSG00000054641 | 3.023504255 | -0.120165791 | 1.571234187 | -0.0764786 | 0.939038339 | NA | Mmrn1 | multimerin 1 [Source:MGI Symbol;Acc:MG | 6.4248 | 1.0279 | 2.0125 | 8.6758 | 0 | 0 |
| 15778 | ENSMUSG00000054648 | 195.9593485 | 0.022397752 | 0.176235874 | 0.127089631 | 0.898869458 | 0.982798755 | Zfp869 | zinc finger protein 869 [Source:MGI Symb | 213.63 | 189.14 | 180.12 | 198.58 | 209.4 | 184.9 |
| 15779 | ENSMUSG00000054659 | 52.8081789 | -0.308353956 | 0.314769408 | -0.979618566 | 0.327274437 | 0.772772049 | Pm20d2 | peptidase M20 domain containing 2 [Sour | 58.626 | 47.284 | 69.431 | 50.127 | 50.168 | 41.212 |
| 15780 | ENSMUSG00000054662 | 2.839125095 | -0.881083075 | 1.362497071 | -0.646667867 | 0.517846924 | NA | Ano9 | anoctamin 9 [Source:MGI Symbol;Acc:MG | 4.0155 | 3.0838 | 4.025 | 4.8199 | 1.0906 | 0 |
| 15781 | ENSMUSG00000054666 | 0 | NA | NA | NA | NA | NA | Olfr63 | olfactory receptor 63 [Source:MGI Symbo | 0 | 0 | 0 | 0 | 0 | 0 |
| 15782 | ENSMUSG00000054667 | 0 | NA | NA | NA | NA | NA | Irs4 | insulin receptor substrate 4 [Source:MGI S | 0 | 0 | 0 | 0 | 0 | 0 |
| 15783 | ENSMUSG00000054672 | 0 | NA | NA | NA | NA | NA | Scart2 | scavenger receptor family member expres | 0 | 0 | 0 | 0 | 0 | 0 |
| 15784 | ENSMUSG00000054675 | 14.2639079 | -0.393397854 | 0.663400908 | -0.593001681 | 0.553180028 | 0.896245536 | Tmem119 | transmembrane protein 119 [Source:MGI | 28.109 | 11.307 | 9.0562 | 18.316 | 6.5437 | 12.252 |
| 15785 | ENSMUSG00000054676 | 3956.764459 | 0.183399395 | 0.085811639 | 2.137232159 | 0.032579114 | 0.241177094 | 1600014C10 | RIKEN cDNA 1600014C10 gene [Source:M | 3799.5 | 3587.4 | 3729.1 | 4136.4 | 4312.3 | 4175.8 |
| 15786 | ENSMUSG00000054679 | 0.367410362 | 2.078255025 | 3.800070157 | 0.54689912 | 0.58444804 | NA | Srsf12 | serine and arginine-rich splicing factor 12 | 0 | 0 | 0 | 0 | 1.0906 | 1.1138 |
| 15787 | ENSMUSG00000054690 | 28.15917105 | -0.932769252 | 0.438563651 | -2.126873146 | 0.033430625 | 0.244672745 | Emcn | endomucin [Source:MGI Symbol;Acc:MGI | 26.502 | 48.312 | 36.225 | 19.28 | 16.359 | 22.277 |
| 15788 | ENSMUSG00000054693 | 523.0074553 | -0.081102514 | 0.125342708 | -0.647046133 | 0.517602087 | 0.887022094 | Adam10 | a disintegrin and metallopeptidase domai | 528.44 | 556.1 | 528.28 | 470.42 | 523.5 | 531.3 |

| | | | | | | | | | | | | | | | |
|---|---|---|---|---|---|---|---|---|---|---|---|---|---|---|---|
| 15789 | ENSMUSG00000054702 | 9.720426577 | -0.555447418 | 0.732887505 | -0.757889054 | 0.448517403 | 0.854238022 | Ap1s3 | adaptor-related protein complex AP-1, sig | 10.44 | 13.363 | 11.069 | 13.496 | 3.2719 | 6.6831 |
| 15790 | ENSMUSG00000054708 | 87.95994226 | 0.107388971 | 0.236708399 | 0.453676217 | 0.650061887 | 0.925643431 | Ankrd24 | ankyrin repeat domain 24 [Source:MGI Sy | 80.31 | 86.345 | 87.543 | 90.614 | 91.612 | 91.335 |
| 15791 | ENSMUSG00000054715 | 78.74892704 | -0.072477764 | 0.275221492 | -0.263343401 | 0.79228591 | 0.961434599 | Zscan22 | zinc finger and SCAN domain containing 2 | 78.704 | 87.373 | 76.475 | 96.398 | 57.803 | 75.741 |
| 15792 | ENSMUSG00000054716 | 374.2284619 | 0.396635381 | 0.148622915 | 2.668736386 | 0.007613718 | 0.088790029 | Zfp771 | zinc finger protein 771 [Source:MGI Symb | 335.7 | 321.74 | 311.94 | 468.49 | 367.54 | 439.97 |
| 15793 | ENSMUSG00000054717 | 108.5242935 | -0.209278377 | 0.232752898 | -0.899144021 | 0.368575953 | 0.806506847 | Hmgb2 | high mobility group box 2 [Source:MGI Sy | 130.91 | 111.02 | 106.66 | 86.758 | 117.79 | 98.018 |
| 15794 | ENSMUSG00000054720 | 39.84365269 | -0.89847974 | 0.364922039 | -2.462114213 | 0.013812068 | 0.135350775 | Lrrc8c | leucine rich repeat containing 8 family, m | 53.808 | 47.284 | 54.337 | 25.063 | 21.812 | 36.757 |
| 15795 | ENSMUSG00000054723 | 271.2332182 | 0.193294234 | 0.159989537 | 1.208167973 | 0.226982658 | 0.67343689 | Vmac | vimentin-type intermediate filament asso | 228.88 | 274.45 | 256.59 | 288.23 | 311.92 | 267.32 |
| 15796 | ENSMUSG00000054727 | 0 | NA | NA | NA | NA | NA | 1700013H16 | RIKEN cDNA 1700013H16 gene [Source:M | 0 | 0 | 0 | 0 | 0 | 0 |
| 15797 | ENSMUSG00000054728 | 9.94326704 | 0.141413291 | 0.678319956 | 0.208475794 | 0.834857481 | 0.971073158 | Phactr1 | phosphatase and actin regulator 1 [Sourc | 12.047 | 7.1954 | 9.0562 | 11.568 | 11.997 | 7.7969 |
| 15798 | ENSMUSG00000054733 | 3139.478628 | -0.166383871 | 0.09619476 | -1.729656279 | 0.083691705 | 0.426388452 | Msra | methionine sulfoxide reductase A [Source | 3597.1 | 3206.1 | 3156.6 | 2968.1 | 3029.7 | 2879.3 |
| 15799 | ENSMUSG00000054737 | 81.72103905 | 0.241561917 | 0.258861633 | 0.933170026 | 0.350732188 | 0.791943999 | Zfp182 | zinc finger protein 182 [Source:MGI Symb | 77.098 | 64.759 | 82.512 | 75.19 | 90.521 | 100.25 |
| 15800 | ENSMUSG00000054745 | 0.733533733 | -1.621434111 | 2.70314781 | -0.599831835 | 0.548618315 | NA | Gm9955 | predicted gene 9955 [Source:MGI Symbol | 2.4093 | 1.0279 | 0 | 0.964 | 0 | 0 |
| 15801 | ENSMUSG00000054746 | 0 | NA | NA | NA | NA | NA | Abca15 | ATP-binding cassette, sub-family A (ABC1) | 0 | 0 | 0 | 0 | 0 | 0 |
| 15802 | ENSMUSG00000054752 | 7.977140316 | 0.332830006 | 0.805069527 | 0.413417717 | 0.679300596 | 0.934062537 | Fsd1l | fibronectin type III and SPRY domain cont | 8.031 | 5.1396 | 8.05 | 14.46 | 3.2719 | 8.9108 |
| 15803 | ENSMUSG00000054753 | 0 | NA | NA | NA | NA | NA | AU018091 | expressed sequence AU018091 [Source:M | 0 | 0 | 0 | 0 | 0 | 0 |
| 15804 | ENSMUSG00000054757 | 2081.672729 | -0.175871387 | 0.104549984 | -1.682175171 | 0.092534877 | 0.447933221 | Akr1c20 | aldo-keto reductase family 1, member C2 | 2037.5 | 2364.2 | 2224.8 | 1988.7 | 2049.3 | 1825.6 |
| 15805 | ENSMUSG00000054759 | 0 | NA | NA | NA | NA | NA | Krtap5-2 | keratin associated protein 5-2 [Source:MG | 0 | 0 | 0 | 0 | 0 | 0 |
| 15806 | ENSMUSG00000054763 | 0 | NA | NA | NA | NA | NA | Defb42 | defensin beta 42 [Source:MGI Symbol;Acc | 0 | 0 | 0 | 0 | 0 | 0 |
| 15807 | ENSMUSG00000054764 | 52.39503719 | -0.371051587 | 0.354975834 | -1.045286896 | 0.295890392 | 0.746382741 | Mtnr1a | melatonin receptor 1A [Source:MGI Symb | 48.989 | 63.731 | 64.4 | 27.955 | 45.806 | 63.489 |
| 15808 | ENSMUSG00000054766 | 1576.939667 | 0.184655264 | 0.108409958 | 1.703305379 | 0.088510935 | 0.438432246 | Set | SET nuclear oncogene [Source:MGI Symbo | 1414.3 | 1608.7 | 1405.7 | 1590.6 | 1644.7 | 1797.7 |
| 15809 | ENSMUSG00000054770 | 125.737142 | 0.015038273 | 0.207547165 | 0.072457135 | 0.942238117 | 0.989798252 | Kctd18 | potassium channel tetramerisation domai | 130.1 | 112.04 | 132.82 | 121.46 | 124.33 | 133.66 |
| 15810 | ENSMUSG00000054779 | 1.048817988 | -0.908682365 | 2.659333108 | -0.341695579 | 0.73258 | NA | Fgf2os | fibroblast growth factor 2, opposite stran | 0 | 4.1117 | 0 | 0 | 2.1812 | 0 |
| 15811 | ENSMUSG00000054792 | 147.8222886 | -0.074393249 | 0.237916372 | -0.312686549 | 0.754518816 | 0.953048681 | Klhl18 | kelch-like 18 [Source:MGI Symbol;Acc:MG | 105.21 | 157.27 | 193.2 | 143.63 | 145.05 | 142.57 |
| 15812 | ENSMUSG00000054793 | 179.0151121 | 0.771826585 | 0.256134651 | 3.013362627 | 0.0025837 | 0.042274474 | Cadm4 | cell adhesion molecule 4 [Source:MGI Syn | 114.84 | 100.74 | 181.12 | 162.91 | 267.2 | 247.27 |
| 15813 | ENSMUSG00000054808 | 2422.893878 | -0.2219687 | 0.103359039 | -2.147549957 | 0.031749525 | 0.237325645 | Actn4 | actinin alpha 4 [Source:MGI Symbol;Acc:N | 2369.2 | 2741.5 | 2717.9 | 2362.7 | 2146.3 | 2199.8 |
| 15814 | ENSMUSG00000054814 | 218.9151681 | 0.001630598 | 0.162529688 | 0.010032615 | 0.991995266 | 0.99859176 | Usp46 | ubiquitin specific peptidase 46 [Source:M | 219.25 | 218.95 | 218.36 | 230.39 | 211.58 | 214.97 |
| 15815 | ENSMUSG00000054822 | 0 | NA | NA | NA | NA | NA | 1700041G16 | RIKEN cDNA 1700041G16 gene [Source:M | 0 | 0 | 0 | 0 | 0 | 0 |
| 15816 | ENSMUSG00000054823 | 1183.353367 | 0.142331839 | 0.121736596 | 1.169178733 | 0.242331628 | 0.692000002 | Nsd3 | nuclear receptor binding SET domain prot | 1085.8 | 1190.3 | 1099.8 | 1309.1 | 1346.9 | 1068.2 |
| 15817 | ENSMUSG00000054827 | 11892.74301 | -0.179228181 | 0.124474949 | -1.439873501 | 0.149903191 | 0.563095827 | Cyp2c50 | cytochrome P450, family 2, subfamily c, p | 11040 | 11794 | 15058 | 11836 | 11288 | 10340 |
| 15818 | ENSMUSG00000054836 | 87.34084386 | 0.055324372 | 0.245239161 | 0.225593548 | 0.82151756 | 0.969160361 | Elp6 | elongator acetyltransferase complex subu | 97.978 | 77.094 | 81.506 | 87.722 | 85.068 | 94.677 |
| 15819 | ENSMUSG00000054843 | 461.1514189 | 0.249827306 | 0.14666951 | 1.703334968 | 0.0885054 | 0.438432246 | Atrnl1 | attractin like 1 [Source:MGI Symbol;Acc:M | 367.02 | 425.56 | 471.93 | 452.11 | 525.68 | 524.62 |
| 15820 | ENSMUSG00000054850 | 0.849605875 | 0.613471193 | 2.572643751 | 0.238459442 | 0.811524769 | NA | Smim10l2a | small integral membrane protein 10 like 2 | 0 | 2.0558 | 0 | 1.928 | 0 | 1.1138 |
| 15821 | ENSMUSG00000054855 | 187.4093617 | -0.335855321 | 0.280891424 | -1.195676662 | 0.231822762 | 0.67889946 | Rnd1 | Rho family GTPase 1 [Source:MGI Symbol | 134.92 | 185.03 | 307.91 | 162.91 | 126.51 | 207.17 |
| 15822 | ENSMUSG00000054863 | 55.28604243 | 0.123129394 | 0.333805244 | 0.368865966 | 0.712227633 | 0.944239215 | Tafa5 | TAFA chemokine like family member 5 [So | 70.673 | 47.284 | 40.25 | 52.055 | 50.168 | 71.286 |
| 15823 | ENSMUSG00000054871 | 12.10147118 | -0.602156628 | 0.642122861 | -0.937759211 | 0.348368167 | 0.789711685 | Tmem158 | transmembrane protein 158 [Source:MGI | 11.243 | 14.391 | 18.112 | 6.7478 | 7.6343 | 14.48 |
| 15824 | ENSMUSG00000054874 | 741.4581164 | 0.095844172 | 0.110754565 | 0.865374458 | 0.386833295 | 0.817407857 | Pcnx3 | pecanex homolog 3 [Source:MGI Symbol; | 721.99 | 706.18 | 722.48 | 793.35 | 735.08 | 769.67 |
| 15825 | ENSMUSG00000054885 | 0 | NA | NA | NA | NA | NA | 4930578G10 | RIKEN cDNA 4930578G10 gene [Source:M | 0 | 0 | 0 | 0 | 0 | 0 |
| 15826 | ENSMUSG00000054889 | 1034.850239 | 0.150272125 | 0.144721053 | 1.038357044 | 0.29910386 | 0.750301462 | Dsp | desmoplakin [Source:MGI Symbol;Acc:MG | 832.01 | 1133.8 | 979.08 | 1251.2 | 1033.9 | 979.07 |
| 15827 | ENSMUSG00000054890 | 0.74256259 | 3.09446134 | 3.273353887 | 0.945348852 | 0.344480805 | NA | Olfr1535 | olfactory receptor 1535 [Source:MGI Sym | 0 | 0 | 0 | 0 | 0 | 4.4554 |
| 15828 | ENSMUSG00000054892 | 2.991704857 | -0.503497139 | 1.229019845 | -0.409673726 | 0.682045306 | NA | Txk | TXK tyrosine kinase [Source:MGI Symbol; | 2.4093 | 4.1117 | 4.025 | 1.928 | 4.3625 | 1.1138 |
| 15829 | ENSMUSG00000054893 | 54.15844526 | 0.189522415 | 0.340916564 | 0.555920231 | 0.578265387 | 0.908137944 | Zfp667 | zinc finger protein 667 [Source:MGI Symb | 31.321 | 60.647 | 60.375 | 55.911 | 64.346 | 52.351 |
| 15830 | ENSMUSG00000054894 | 118.3356952 | -0.053111977 | 0.242342911 | -0.219160432 | 0.826525077 | 0.969648181 | Dmac2l | distal membrane arm assembly complex 2 | 130.1 | 113.07 | 117.73 | 84.83 | 142.87 | 121.41 |
| 15831 | ENSMUSG00000054901 | 0.17131964 | -0.796629176 | 4.080472857 | -0.195229623 | 0.84521319 | NA | Arhgef33 | Rho guanine nucleotide exchange factor ( | 0 | 1.0279 | 0 | 0 | 0 | 0 |
| 15832 | ENSMUSG00000054905 | 0.17131964 | -0.796629176 | 4.080472857 | -0.195229623 | 0.84521319 | NA | Stfa3 | stefin A3 [Source:MGI Symbol;Acc:MGI:1 | 0 | 1.0279 | 0 | 0 | 0 | 0 |
| 15833 | ENSMUSG00000054909 | 0.818715751 | 2.030181432 | 2.59128962 | 0.78346373 | 0.433354844 | NA | Wbscr25 | Williams Beuren syndrome chromosome r | 0.8031 | 0 | 0 | 1.928 | 2.1812 | 0 |
| 15834 | ENSMUSG00000054910 | 0 | NA | NA | NA | NA | NA | 4931415C17 | RIKEN cDNA 4931415C17 gene [Source:M | 0 | 0 | 0 | 0 | 0 | 0 |
| 15835 | ENSMUSG00000054920 | 785.3783507 | 0.002455115 | 0.133842938 | 0.018343252 | 0.985365023 | 0.99859176 | Klhl5 | kelch-like 5 [Source:MGI Symbol;Acc:MGI | 804.71 | 801.78 | 746.63 | 660.32 | 879.04 | 819.79 |
| 15836 | ENSMUSG00000054931 | 21.92175436 | 0.319488096 | 0.460118565 | 0.694360369 | 0.487456241 | 0.873910634 | Zkscan4 | zinc finger with KRAB and SCAN domains | 20.881 | 20.558 | 17.106 | 28.919 | 22.903 | 21.163 |
| 15837 | ENSMUSG00000054932 | 47.52326353 | 0.246804244 | 0.372995435 | 0.661681674 | 0.508175252 | 0.882700473 | Afp | alpha fetoprotein [Source:MGI Symbol;Ac | 57.02 | 28.782 | 44.275 | 53.983 | 65.437 | 35.643 |
| 15838 | ENSMUSG00000054934 | 0 | NA | NA | NA | NA | NA | Kcnmb4 | potassium large conductance calcium-acti | 0 | 0 | 0 | 0 | 0 | 0 |
| 15839 | ENSMUSG00000054938 | 0 | NA | NA | NA | NA | NA | Olfr1346 | olfactory receptor 1346 [Source:MGI Sym | 0 | 0 | 0 | 0 | 0 | 0 |
| 15840 | ENSMUSG00000054939 | 56.08145203 | 0.04135864 | 0.305348067 | 0.135447524 | 0.892258052 | 0.982489666 | Zfp174 | zinc finger protein 174 [Source:MGI Symb | 61.839 | 60.647 | 43.269 | 61.695 | 57.803 | 51.237 |
| 15841 | ENSMUSG00000054940 | 0 | NA | NA | NA | NA | NA | Olfr137 | olfactory receptor 137 [Source:MGI Symb | 0 | 0 | 0 | 0 | 0 | 0 |
| 15842 | ENSMUSG00000054942 | 21.3088864 | -0.809247121 | 0.515107899 | -1.571024483 | 0.116176962 | 0.500730933 | Miga1 | mitoguardin 1 [Source:MGI Symbol;Acc:M | 19.274 | 42.145 | 20.125 | 15.424 | 14.178 | 16.708 |
| 15843 | ENSMUSG00000054944 | 0 | NA | NA | NA | NA | NA | 5330416C01 | RIKEN cDNA 5330416C01 gene [Source:M | 0 | 0 | 0 | 0 | 0 | 0 |
| 15844 | ENSMUSG00000054945 | 26.25185408 | 0.354910954 | 0.45982518 | 0.771838885 | 0.440209858 | 0.84892746 | Gm9958 | predicted gene 9958 [Source:MGI Symbol | 17.668 | 20.558 | 31.194 | 39.523 | 20.722 | 27.846 |

| | | | | | | | | | | | | | | | |
|---|---|---|---|---|---|---|---|---|---|---|---|---|---|---|---|
| 15845 | ENSMUSG00000054951 | 3.856587873 | 0.26736283 | 1.110674265 | 0.240721189 | 0.809771215 | NA | 9130008F23 | RIKEN cDNA 9130008F23 gene [Source:M | 2.4093 | 4.1117 | 4.025 | 4.8199 | 1.0906 | 6.6831 |
| 15846 | ENSMUSG00000054957 | 0.342432689 | 1.990653086 | 3.900281263 | 0.51038706 | 0.609780321 | NA | Gm9959 | predicted gene 9959 [Source:MGI Symbol | 0 | 0 | 0 | 0.964 | 1.0906 | 0 |
| 15847 | ENSMUSG00000054958 | 0 | NA | NA | NA | NA | NA | Nt5c1a | 5'-nucleotidase, cytosolic IA [Source:MGI | 0 | 0 | 0 | 0 | 0 | 0 |
| 15848 | ENSMUSG00000054966 | 0 | NA | NA | NA | NA | NA | Lmntd1 | lamin tail domain containing 1 [Source:M | 0 | 0 | 0 | 0 | 0 | 0 |
| 15849 | ENSMUSG00000054967 | 8.462773591 | 0.097979611 | 0.846931536 | 0.115687758 | 0.907900009 | 0.984639154 | Zfp647 | zinc finger protein 647 [Source:MGI Symb | 6.4248 | 8.2233 | 10.062 | 18.316 | 2.1812 | 5.5692 |
| 15850 | ENSMUSG00000054976 | 0 | NA | NA | NA | NA | NA | Nyap2 | neuronal tyrosine-phophorylated phosph | 0 | 0 | 0 | 0 | 0 | 0 |
| 15851 | ENSMUSG00000054978 | 2.906342879 | -2.131596735 | 1.41737034 | -1.503909512 | 0.132604668 | NA | Kbtbd13 | kelch repeat and BTB (POZ) domain conta | 4.0155 | 3.0838 | 7.0437 | 0 | 2.1812 | 1.1138 |
| 15852 | ENSMUSG00000054986 | 0 | NA | NA | NA | NA | NA | Sec14l3 | SEC14-like lipid binding 3 [Source:MGI Syr | 0 | 0 | 0 | 0 | 0 | 0 |
| 15853 | ENSMUSG00000054988 | 0 | NA | NA | NA | NA | NA | Agtr1b | angiotensin II receptor, type 1b [Source:N | 0 | 0 | 0 | 0 | 0 | 0 |
| 15854 | ENSMUSG00000054994 | 0 | NA | NA | NA | NA | NA | AV320801 | expressed sequence AV320801 [Source:M | 0 | 0 | 0 | 0 | 0 | 0 |
| 15855 | ENSMUSG00000054999 | 0.956437057 | 1.282470111 | 2.271719443 | 0.56453719 | 0.572388597 | NA | Naaladl1 | N-acetylated alpha-linked acidic dipeptida | 1.6062 | 0 | 0 | 1.928 | 1.0906 | 1.1138 |
| 15856 | ENSMUSG00000055003 | 0.706178709 | 0.10198885 | 3.03884937 | 0.033561667 | 0.97322669 | NA | Lrtm2 | leucine-rich repeats and transmembrane | 0 | 2.0558 | 0 | 0 | 2.1812 | 0 |
| 15857 | ENSMUSG00000055010 | 0 | NA | NA | NA | NA | NA | A830031A19 | RIKEN cDNA A830031A19 gene [Source:M | 0 | 0 | 0 | 0 | 0 | 0 |
| 15858 | ENSMUSG00000055013 | 224.8450501 | -0.114094974 | 0.180614379 | -0.631704823 | 0.527579776 | 0.888334265 | Agap1 | ArfGAP with GTPase domain, ankyrin repe | 224.87 | 241.56 | 235.46 | 248.71 | 222.49 | 175.99 |
| 15859 | ENSMUSG00000055015 | 1.517229932 | 2.125958547 | 2.131593866 | 0.997356288 | 0.318591601 | NA | Gm9961 | predicted gene 9961 [Source:MGI Symbol | 1.6062 | 0 | 0 | 1.928 | 0 | 5.5692 |
| 15860 | ENSMUSG00000055022 | 0 | NA | NA | NA | NA | NA | Cntn1 | contactin 1 [Source:MGI Symbol;Acc:MGI | 0 | 0 | 0 | 0 | 0 | 0 |
| 15861 | ENSMUSG00000055024 | 550.2117069 | -0.262000561 | 0.141913156 | -1.846203473 | 0.064862667 | 0.365332404 | Ep300 | E1A binding protein p300 [Source:MGI Sy | 518 | 648.62 | 634.94 | 544.65 | 499.5 | 455.56 |
| 15862 | ENSMUSG00000055026 | 0 | NA | NA | NA | NA | NA | Gabrg3 | gamma-aminobutyric acid (GABA) A recep | 0 | 0 | 0 | 0 | 0 | 0 |
| 15863 | ENSMUSG00000055027 | 94.92822427 | -0.233707434 | 0.245880507 | -0.950491916 | 0.34186236 | 0.783504543 | Smyd1 | SET and MYND domain containing 1 [Sour | 102.8 | 90.457 | 114.71 | 95.434 | 94.884 | 71.286 |
| 15864 | ENSMUSG00000055030 | 0 | NA | NA | NA | NA | NA | Sprr2e | small proline-rich protein 2E [Source:MGI | 0 | 0 | 0 | 0 | 0 | 0 |
| 15865 | ENSMUSG00000055033 | 0 | NA | NA | NA | NA | NA | Olfr420 | olfactory receptor 420 [Source:MGI Symb | 0 | 0 | 0 | 0 | 0 | 0 |
| 15866 | ENSMUSG00000055041 | 5.750335559 | 1.073862857 | 1.010054651 | 1.063173023 | 0.287703494 | 0.73951397 | Commd5 | COMM domain containing 5 [Source:MGI | 4.0155 | 0 | 7.0437 | 5.7839 | 7.6343 | 10.025 |
| 15867 | ENSMUSG00000055044 | 388.0751239 | -0.264621599 | 0.14919944 | -1.773609864 | 0.076127705 | 0.403877794 | Pdlim1 | PDZ and LIM domain 1 (elfin) [Source:MG | 420.83 | 367.99 | 481.99 | 363.42 | 352.27 | 341.95 |
| 15868 | ENSMUSG00000055045 | 0 | NA | NA | NA | NA | NA | Gm11190 | predicted gene 11190 [Source:MGI Symbo | 0 | 0 | 0 | 0 | 0 | 0 |
| 15869 | ENSMUSG00000055048 | 0 | NA | NA | NA | NA | NA | Gm9962 | predicted gene 9962 [Source:MGI Symbol | 0 | 0 | 0 | 0 | 0 | 0 |
| 15870 | ENSMUSG00000055053 | 3008.640411 | 0.058244849 | 0.105053201 | 0.554431934 | 0.579283276 | 0.908337213 | Nfic | nuclear factor I/C [Source:MGI Symbol;Ac | 2903.2 | 2744.5 | 3196.8 | 3334.4 | 2950.1 | 2922.7 |
| 15871 | ENSMUSG00000055061 | 0.328370405 | 0.16514316 | 4.003149536 | 0.041253308 | 0.967093956 | NA | 4931431C16 | RIKEN cDNA 4931431C16 gene [Source:M | 0 | 0 | 1.0062 | 0.964 | 0 | 0 |
| 15872 | ENSMUSG00000055065 | 3651.968615 | 0.101935662 | 0.11548284 | 0.882690988 | 0.377403253 | 0.81030298 | Ddx17 | DEAD box helicase 17 [Source:MGI Symbo | 3470.2 | 3387 | 3713 | 4358.1 | 3626.3 | 3357.1 |
| 15873 | ENSMUSG00000055067 | 83.52974205 | 0.043502263 | 0.273833468 | 0.158863938 | 0.873776073 | 0.979925081 | Smyd3 | SET and MYND domain containing 3 [Sour | 73.082 | 79.15 | 94.587 | 68.442 | 109.06 | 76.855 |
| 15874 | ENSMUSG00000055069 | 7.777736697 | -1.101250317 | 0.772536555 | -1.425499299 | 0.154012923 | 0.57119049 | Rab39 | RAB39, member RAS oncogene family [So | 10.44 | 10.279 | 11.069 | 3.8559 | 5.4531 | 5.5692 |
| 15875 | ENSMUSG00000055072 | 0 | NA | NA | NA | NA | NA | Gm9964 | predicted gene 9964 [Source:MGI Symbol | 0 | 0 | 0 | 0 | 0 | 0 |
| 15876 | ENSMUSG00000055078 | 0.342639279 | -1.805976596 | 4.028547714 | -0.448294702 | 0.653940524 | NA | Gabra5 | gamma-aminobutyric acid (GABA) A recep | 0 | 2.0558 | 0 | 0 | 0 | 0 |
| 15877 | ENSMUSG00000055088 | 0 | NA | NA | NA | NA | NA | Olfr354 | olfactory receptor 354 [Source:MGI Symb | 0 | 0 | 0 | 0 | 0 | 0 |
| 15878 | ENSMUSG00000055093 | 7.402495736 | 0.905023817 | 0.882124485 | 1.025959298 | 0.30491077 | 0.753627713 | Rps27a-ps3 | ribosomal protein S27A, pseudogene 3 [S | 7.2279 | 1.0279 | 7.0437 | 4.8199 | 9.8156 | 14.48 |
| 15879 | ENSMUSG00000055095 | 0 | NA | NA | NA | NA | NA | Spink6 | serine peptidase inhibitor, Kazal type 6 [S | 0 | 0 | 0 | 0 | 0 | 0 |
| 15880 | ENSMUSG00000055102 | 0 | NA | NA | NA | NA | NA | Zfp819 | zinc finger protein 819 [Source:MGI Symb | 0 | 0 | 0 | 0 | 0 | 0 |
| 15881 | ENSMUSG00000055108 | 0.167707431 | -0.796629176 | 4.080472857 | -0.195229623 | 0.84521319 | NA | Phxr2 | per-hexamer repeat gene 2 [Source:MGI S | 0 | 0 | 1.0062 | 0 | 0 | 0 |
| 15882 | ENSMUSG00000055109 | 0 | NA | NA | NA | NA | NA | Gm15155 | predicted gene 15155 [Source:MGI Symbo | 0 | 0 | 0 | 0 | 0 | 0 |
| 15883 | ENSMUSG00000055110 | 0 | NA | NA | NA | NA | NA | A630012P03 | RIKEN cDNA A630012P03 gene [Source:M | 0 | 0 | 0 | 0 | 0 | 0 |
| 15884 | ENSMUSG00000055114 | 7.069901895 | -0.048345991 | 0.831882267 | -0.058116387 | 0.953655922 | 0.9915542 | Anxa13 | annexin A13 [Source:MGI Symbol;Acc:MG | 11.243 | 7.1954 | 3.0187 | 7.7118 | 5.4531 | 7.7969 |
| 15885 | ENSMUSG00000055116 | 69.0473488 | -2.50174744 | 0.466734589 | -5.360107217 | 8.32E-08 | 9.25E-06 | Arntl | aryl hydrocarbon receptor nuclear translo | 37.746 | 123.35 | 191.19 | 21.208 | 19.631 | 21.163 |
| 15886 | ENSMUSG00000055125 | 0 | NA | NA | NA | NA | NA | M5C1000I18 | RIKEN cDNA M5C1000I18 gene [Source:M | 0 | 0 | 0 | 0 | 0 | 0 |
| 15887 | ENSMUSG00000055128 | 669.5495804 | 0.04828228 | 0.113556362 | 0.425183401 | 0.670702983 | 0.93116767 | Cgrrf1 | cell growth regulator with ring finger dom | 640.07 | 677.4 | 658.08 | 693.1 | 690.36 | 658.28 |
| 15888 | ENSMUSG00000055134 | 0 | NA | NA | NA | NA | NA | 9130017K11 | RIKEN cDNA 9130017K11 gene [Source:M | 0 | 0 | 0 | 0 | 0 | 0 |
| 15889 | ENSMUSG00000055137 | 961.2238668 | -0.225302062 | 0.112964597 | -1.994448422 | 0.046103071 | 0.297163469 | Sugct | succinyl-CoA glutarate-CoA transferase [S | 1016.7 | 1118.4 | 974.04 | 937.95 | 867.04 | 853.2 |
| 15890 | ENSMUSG00000055138 | 0 | NA | NA | NA | NA | NA | Gm4861 | predicted gene 4861 [Source:MGI Symbol | 0 | 0 | 0 | 0 | 0 | 0 |
| 15891 | ENSMUSG00000055148 | 125.9195751 | -1.015837968 | 0.252836565 | -4.017765257 | 5.88E-05 | 0.00237374 | Klf2 | Kruppel-like factor 2 (lung) [Source:MGI S | 156.6 | 210.72 | 138.86 | 106.04 | 70.89 | 72.4 |
| 15892 | ENSMUSG00000055150 | 19.64458884 | -0.092884059 | 0.507107044 | -0.1831646 | 0.85466887 | 0.97395562 | Zfp78 | zinc finger protein 78 [Source:MGI Symbo | 15.259 | 19.53 | 26.162 | 17.352 | 25.084 | 14.48 |
| 15893 | ENSMUSG00000055159 | 0.709843051 | 3.034749291 | 2.67440342 | 1.134738786 | 0.256484781 | NA | 4930583K01 | RIKEN cDNA 4930583K01 gene [Source:M | 0 | 0 | 0 | 0.964 | 2.1812 | 1.1138 |
| 15894 | ENSMUSG00000055170 | 0 | NA | NA | NA | NA | NA | Ifng | interferon gamma [Source:MGI Symbol;A | 0 | 0 | 0 | 0 | 0 | 0 |
| 15895 | ENSMUSG00000055172 | 4366.71829 | 0.223290807 | 0.106215777 | 2.102237674 | 0.035532461 | 0.253048507 | C1ra | complement component 1, r subcompone | 3601.1 | 4340.9 | 4147.7 | 4541.3 | 5062.7 | 4506.6 |
| 15896 | ENSMUSG00000055177 | 0 | NA | NA | NA | NA | NA | Cstl1 | cystatin-like 1 [Source:MGI Symbol;Acc:M | 0 | 0 | 0 | 0 | 0 | 0 |
| 15897 | ENSMUSG00000055184 | 19.05822177 | -0.179686071 | 0.484463645 | -0.370896913 | 0.710714315 | 0.944239215 | Fam72a | family with sequence similarity 72, memb | 20.078 | 19.53 | 21.131 | 18.316 | 16.359 | 18.935 |
| 15898 | ENSMUSG00000055188 | 0.824421612 | 3.264491806 | 2.808832384 | 1.162223786 | 0.245144575 | NA | Rbm3os | RNA binding motif protein 3, opposite stra | 0 | 0 | 0 | 3.8559 | 1.0906 | 0 |
| 15899 | ENSMUSG00000055193 | 0 | NA | NA | NA | NA | NA | Klk15 | kallikrein related-peptidase 15 [Source:M | 0 | 0 | 0 | 0 | 0 | 0 |
| 15900 | ENSMUSG00000055194 | 0 | NA | NA | NA | NA | NA | Actbl2 | actin, beta-like 2 [Source:MGI Symbol;Acc | 0 | 0 | 0 | 0 | 0 | 0 |

| | | | | | | | | | | | | | | | |
|---|---|---|---|---|---|---|---|---|---|---|---|---|---|---|---|
| 15901 | ENSMUSG00000055197 | 0.167707431 | -0.796629176 | 4.080472857 | -0.195229623 | 0.84521319 | NA | Fev | FEV transcription factor, ETS family memb | 0 | 0 | 1.0062 | 0 | 0 | 0 |
| 15902 | ENSMUSG00000055198 | 0 | NA | NA | NA | NA | NA | Gm12830 | predicted gene 12830 [Source:MGI Symbo | 0 | 0 | 0 | 0 | 0 | 0 |
| 15903 | ENSMUSG00000055200 | 63.50666612 | -0.084041617 | 0.303319876 | -0.277072567 | 0.781724386 | 0.960079945 | Sertad3 | SERTA domain containing 3 [Source:MGI S | 70.673 | 55.508 | 69.431 | 44.343 | 69.8 | 71.286 |
| 15904 | ENSMUSG00000055202 | 1.544895825 | -0.936377367 | 1.917959155 | -0.488215489 | 0.625397215 | NA | Zfp811 | zinc finger protein 811 [Source:MGI Symb | 0 | 4.1117 | 2.0125 | 0.964 | 2.1812 | 0 |
| 15905 | ENSMUSG00000055204 | 1012.944416 | 0.131970204 | 0.124154467 | 1.06295172 | 0.287803847 | 0.73951397 | Ankrd17 | ankyrin repeat domain 17 [Source:MGI Sy | 911.52 | 1074.2 | 914.68 | 1002.5 | 1177.9 | 996.89 |
| 15906 | ENSMUSG00000055210 | 0.335414862 | -1.779047242 | 4.030804179 | -0.441362855 | 0.658950329 | NA | Foxd2 | forkhead box D2 [Source:MGI Symbol;Acc | 0 | 0 | 2.0125 | 0 | 0 | 0 |
| 15907 | ENSMUSG00000055214 | 0 | NA | NA | NA | NA | NA | Pld5 | phospholipase D family, member 5 [Sourc | 0 | 0 | 0 | 0 | 0 | 0 |
| 15908 | ENSMUSG00000055216 | 2.564006402 | 1.256373292 | 1.459067795 | 0.861079448 | 0.389194287 | NA | 9430025C20 | RIKEN cDNA 9430025C20 gene [Source:M | 2.4093 | 2.0558 | 0 | 0.964 | 3.2719 | 6.6831 |
| 15909 | ENSMUSG00000055228 | 0 | NA | NA | NA | NA | NA | Zfp935 | zinc finger protein 935 [Source:MGI Symb | 0 | 0 | 0 | 0 | 0 | 0 |
| 15910 | ENSMUSG00000055235 | 9.881000299 | -0.113638196 | 0.717494683 | -0.158381935 | 0.874155848 | 0.979986838 | Wdr86 | WD repeat domain 86 [Source:MGI Symbo | 6.4248 | 17.475 | 7.0437 | 9.6398 | 10.906 | 7.7969 |
| 15911 | ENSMUSG00000055239 | 1127.208365 | 0.108684774 | 0.126358517 | 0.860130182 | 0.389717286 | 0.819052248 | Kcmf1 | potassium channel modulatory factor 1 [S | 1096.2 | 1168.7 | 990.14 | 1335.1 | 1102.6 | 1070.4 |
| 15912 | ENSMUSG00000055240 | 115.5641685 | -0.108780829 | 0.241792744 | -0.44989286 | 0.652787696 | 0.925756399 | Zfp101 | zinc finger protein 101 [Source:MGI Symb | 109.22 | 102.79 | 147.92 | 113.75 | 91.612 | 128.09 |
| 15913 | ENSMUSG00000055254 | 105.8346164 | -0.153181904 | 0.335119127 | -0.457096869 | 0.647601421 | 0.925643431 | Ntrk2 | neurotrophic tyrosine kinase, receptor, ty | 150.98 | 115.13 | 67.418 | 58.803 | 111.24 | 131.43 |
| 15914 | ENSMUSG00000055271 | 1.060017606 | 3.614174932 | 2.319997292 | 1.557835841 | 0.119272167 | NA | 9330161L09 | RIKEN cDNA 9330161L09 gene [Source:M | 0 | 0 | 0 | 1.928 | 1.0906 | 3.3415 |
| 15915 | ENSMUSG00000055296 | 868.9393925 | -0.041529302 | 0.149685857 | -0.277443061 | 0.781439921 | 0.960079945 | Tmem245 | transmembrane protein 245 [Source:MGI | 871.37 | 952.88 | 821.1 | 1014.1 | 871.4 | 682.79 |
| 15916 | ENSMUSG00000055298 | 0.556921942 | 2.679043159 | 3.638243063 | 0.736356289 | 0.461513899 | NA | Ctsj | cathepsin J [Source:MGI Symbol;Acc:MGI: | 0 | 0 | 0 | 0 | 0 | 3.3415 |
| 15917 | ENSMUSG00000055301 | 570.6891832 | 0.274278804 | 0.162216233 | 1.69082217 | 0.090870772 | 0.444639662 | Adh7 | alcohol dehydrogenase 7 (class IV), mu or | 491.5 | 520.13 | 539.35 | 787.57 | 542.04 | 543.56 |
| 15918 | ENSMUSG00000055302 | 2702.304475 | 0.004999015 | 0.095547304 | 0.05231979 | 0.958273885 | 0.991891687 | Mrfap1 | Morf4 family associated protein 1 [Source | 2500.1 | 2779.5 | 2814.5 | 2705.9 | 2637.1 | 2776.8 |
| 15919 | ENSMUSG00000055305 | 82.13306006 | 0.031724878 | 0.249252235 | 0.127280216 | 0.898718619 | 0.982798755 | Zfp93 | zinc finger protein 93 [Source:MGI Symbo | 73.885 | 90.457 | 79.493 | 78.082 | 82.887 | 87.994 |
| 15920 | ENSMUSG00000055312 | 461.2192358 | -0.234530463 | 0.180390489 | -1.300126542 | 0.193557602 | 0.631833452 | Them7 | thioesterase superfamily member 7 [Sour | 610.36 | 471.81 | 412.56 | 384.63 | 512.59 | 375.37 |
| 15921 | ENSMUSG00000055313 | 11.99489238 | 0.505015217 | 0.6266097 | 0.80594861 | 0.420272481 | 0.837906635 | Pgbd1 | piggyBac transposable element derived 1 | 10.44 | 6.1675 | 13.081 | 12.532 | 15.269 | 14.48 |
| 15922 | ENSMUSG00000055319 | 342.5288046 | 0.100441258 | 0.156894003 | 0.640185451 | 0.52205204 | 0.888334265 | Sec23ip | Sec23 interacting protein [Source:MGI Syr | 300.36 | 347.44 | 344.14 | 314.26 | 343.54 | 405.44 |
| 15923 | ENSMUSG00000055320 | 166.7165852 | -0.86178546 | 0.242269339 | -3.557137956 | 0.000374917 | 0.010261584 | Tead1 | TEA domain family member 1 [Source:MG | 244.14 | 239.5 | 162.01 | 155.2 | 101.43 | 98.018 |
| 15924 | ENSMUSG00000055322 | 3273.075039 | -0.626964046 | 0.12089873 | -5.185861306 | 2.15E-07 | 2.14E-05 | Tns1 | tensin 1 [Source:MGI Symbol;Acc:MGI:10 | 3576.2 | 4605.1 | 3740.2 | 2809 | 2443 | 2464.9 |
| 15925 | ENSMUSG00000055323 | 4.849234136 | 0.333590297 | 1.126746612 | 0.296065054 | 0.767180401 | NA | Gm9967 | predicted gene 9967 [Source:MGI Symbol | 4.8186 | 2.0558 | 6.0375 | 9.6398 | 6.5437 | 0 |
| 15926 | ENSMUSG00000055333 | 0 | NA | NA | NA | NA | NA | Fat2 | FAT atypical cadherin 2 [Source:MGI Symb | 0 | 0 | 0 | 0 | 0 | 0 |
| 15927 | ENSMUSG00000055334 | 104.1191808 | 0.236271086 | 0.224755667 | 1.051235282 | 0.293150543 | 0.744143463 | Snupn | snurportin 1 [Source:MGI Symbol;Acc:MG | 98.782 | 88.401 | 99.618 | 119.53 | 101.43 | 116.95 |
| 15928 | ENSMUSG00000055341 | 6.872190531 | 0.203890564 | 0.841723622 | 0.242229823 | 0.808602088 | 0.965682062 | Zfp457 | zinc finger protein 457 [Source:MGI Symb | 4.0155 | 7.1954 | 8.05 | 8.6758 | 3.2719 | 10.025 |
| 15929 | ENSMUSG00000055357 | 0 | NA | NA | NA | NA | NA | 4933400A11 | RIKEN cDNA 4933400A11 gene [Source:M | 0 | 0 | 0 | 0 | 0 | 0 |
| 15930 | ENSMUSG00000055360 | 0 | NA | NA | NA | NA | NA | Prl2c5 | prolactin family 2, subfamily c, member 5 | 0 | 0 | 0 | 0 | 0 | 0 |
| 15931 | ENSMUSG00000055368 | 0 | NA | NA | NA | NA | NA | Slc6a2 | solute carrier family 6 (neurotransmitter t | 0 | 0 | 0 | 0 | 0 | 0 |
| 15932 | ENSMUSG00000055370 | 3.189810468 | 0.57651083 | 1.254287331 | 0.459632188 | 0.645780251 | NA | Gm9968 | predicted gene 9968 [Source:MGI Symbol | 1.6062 | 5.1396 | 1.0062 | 4.8199 | 5.4531 | 1.1138 |
| 15933 | ENSMUSG00000055371 | 270.8335689 | 0.073861746 | 0.171146259 | 0.431570904 | 0.666053309 | 0.929596279 | Stam2 | signal transducing adaptor molecule (SH3 | 256.99 | 311.46 | 223.39 | 267.99 | 292.29 | 272.89 |
| 15934 | ENSMUSG00000055373 | 0 | NA | NA | NA | NA | NA | Fut9 | fucosyltransferase 9 [Source:MGI Symbol; | 0 | 0 | 0 | 0 | 0 | 0 |
| 15935 | ENSMUSG00000055385 | 0.294513347 | 0.165143496 | 4.080472857 | 0.040471657 | 0.967717103 | NA | Rnf212 | ring finger protein 212 [Source:MGI Symb | 0.8031 | 0 | 0 | 0.964 | 0 | 0 |
| 15936 | ENSMUSG00000055401 | 557.1517082 | 0.071937249 | 0.144318926 | 0.498460259 | 0.618159673 | 0.917509899 | Fbxo6 | F-box protein 6 [Source:MGI Symbol;Acc:I | 603.13 | 522.18 | 503.12 | 510.91 | 546.4 | 657.17 |
| 15937 | ENSMUSG00000055403 | 0 | NA | NA | NA | NA | NA | 4933427D06 | RIKEN cDNA 4933427D06 gene [Source:M | 0 | 0 | 0 | 0 | 0 | 0 |
| 15938 | ENSMUSG00000055407 | 0.267700746 | -1.497481649 | 4.057001436 | -0.369110456 | 0.712045396 | NA | Map6 | microtubule-associated protein 6 [Source | 1.6062 | 0 | 0 | 0 | 0 | 0 |
| 15939 | ENSMUSG00000055408 | 0 | NA | NA | NA | NA | NA | Hottip | Hoxa distal transcript antisense RNA [Sour | 0 | 0 | 0 | 0 | 0 | 0 |
| 15940 | ENSMUSG00000055409 | 0 | NA | NA | NA | NA | NA | Nell1 | NEL-like 1 [Source:MGI Symbol;Acc:MGI:2 | 0 | 0 | 0 | 0 | 0 | 0 |
| 15941 | ENSMUSG00000055413 | 126.001907 | 0.268714456 | 0.268895832 | 0.999325479 | 0.317637047 | 0.764257918 | H2-Q5 | histocompatibility 2, Q region locus 5 [Sou | 90.751 | 108.96 | 143.89 | 171.59 | 93.793 | 147.03 |
| 15942 | ENSMUSG00000055415 | 0.133850373 | -0.796629176 | 4.080472857 | -0.195229623 | 0.84521319 | NA | Atp10b | ATPase, class V, type 10B [Source:MGI Syr | 0.8031 | 0 | 0 | 0 | 0 | 0 |
| 15943 | ENSMUSG00000055416 | 0.267700746 | -1.497481649 | 4.057001436 | -0.369110456 | 0.712045396 | NA | Fam136b-ps | family with sequence similarity 136, meml | 1.6062 | 0 | 0 | 0 | 0 | 0 |
| 15944 | ENSMUSG00000055421 | 1.166907561 | -2.40325111 | 2.613065148 | -0.919705776 | 0.357726533 | NA | Pcdh9 | protocadherin 9 [Source:MGI Symbol;Acc | 0 | 0 | 6.0375 | 0.964 | 0 | 0 |
| 15945 | ENSMUSG00000055424 | 0.534859069 | 1.101323472 | 3.351640271 | 0.328592386 | 0.742463804 | NA | Gm9970 | predicted gene 9970 [Source:MGI Symbol | 0 | 1.0279 | 0 | 0 | 2.1812 | 0 |
| 15946 | ENSMUSG00000055430 | 2.584092044 | -0.499921161 | 1.584889639 | -0.315429636 | 0.752435468 | NA | Nap1l5 | nucleosome assembly protein 1-like 5 [So | 0 | 3.0838 | 6.0375 | 1.928 | 0 | 4.4554 |
| 15947 | ENSMUSG00000055435 | 475.5282753 | -0.702601942 | 0.145470796 | -4.82984874 | 1.37E-06 | 0.000104567 | Maf | avian musculoaponeurotic fibrosarcoma c | 554.14 | 609.56 | 603.75 | 360.53 | 309.74 | 415.46 |
| 15948 | ENSMUSG00000055436 | 1262.803526 | 0.077139452 | 0.122186602 | 0.631324962 | 0.527828069 | 0.888334265 | Srsf11 | serine and arginine-rich splicing factor 11 | 1317.9 | 1175.9 | 1193.4 | 1455.6 | 1296.7 | 1137.2 |
| 15949 | ENSMUSG00000055446 | 0 | NA | NA | NA | NA | NA | A630091E08 | RIKEN cDNA A630091E08 gene [Source:M | 0 | 0 | 0 | 0 | 0 | 0 |
| 15950 | ENSMUSG00000055447 | 1235.214041 | 0.056678934 | 0.121711184 | 0.465683865 | 0.641441804 | 0.924019354 | Cd47 | CD47 antigen (Rh-related antigen, integrir | 1113.1 | 1364 | 1156.2 | 1180.9 | 1221.5 | 1375.6 |
| 15951 | ENSMUSG00000055452 | 3.991963204 | -1.255444856 | 1.171379857 | -1.071765789 | 0.283825243 | NA | Gm7353 | predicted pseudogene 7353 [Source:MGI | 8.8341 | 2.0558 | 6.0375 | 4.8199 | 1.0906 | 1.1138 |
| 15952 | ENSMUSG00000055456 | 0 | NA | NA | NA | NA | NA | Gm9972 | predicted gene 9972 [Source:MGI Symbol | 0 | 0 | 0 | 0 | 0 | 0 |
| 15953 | ENSMUSG00000055471 | 0.17131964 | -0.796629176 | 4.080472857 | -0.195229623 | 0.84521319 | NA | Alk | anaplastic lymphoma kinase [Source:MGI | 0 | 1.0279 | 0 | 0 | 0 | 0 |
| 15954 | ENSMUSG00000055480 | 24.95306648 | -0.715043783 | 0.506832457 | -1.410808981 | 0.158300946 | 0.578929172 | Zfp458 | zinc finger protein 458 [Source:MGI Symb | 31.321 | 26.726 | 35.219 | 29.883 | 7.6343 | 18.935 |
| 15955 | ENSMUSG00000055485 | 20.32037648 | -0.562606894 | 0.519005973 | -1.084008514 | 0.278361024 | 0.73110838 | Soga1 | suppressor of glucose, autophagy associa | 24.896 | 25.698 | 22.137 | 19.28 | 7.6343 | 22.277 |
| 15956 | ENSMUSG00000055489 | 1.058182678 | 2.444751467 | 2.435756566 | 1.003692857 | 0.315526681 | NA | Ano5 | anoctamin 5 [Source:MGI Symbol;Acc:MG | 0.8031 | 0 | 0 | 0 | 1.0906 | 4.4554 |

| | | | | | | | | | | | | | | | |
|---|---|---|---|---|---|---|---|---|---|---|---|---|---|---|---|
| 15957 | ENSMUSG00000055491 | 199.7274926 | 0.115332907 | 0.174887205 | 0.659470235 | 0.509593853 | 0.882793133 | Pprc1 | peroxisome proliferative activated recept | 199.97 | 186.05 | 189.17 | 216.9 | 222.49 | 183.78 |
| 15958 | ENSMUSG00000055493 | 166.3257989 | 0.183003642 | 0.189424801 | 0.966101806 | 0.333993241 | 0.775900437 | Epm2a | epilepsy, progressive myoclonic epilepsy, | 172.67 | 158.3 | 135.84 | 169.66 | 181.04 | 180.44 |
| 15959 | ENSMUSG00000055494 | 6.112068954 | -2.569940544 | 1.084892341 | -2.368843845 | 0.017843785 | 0.159316366 | Gm14168 | predicted gene 14168 [Source:MGI Symbo | 20.078 | 6.1675 | 5.0312 | 0.964 | 1.0906 | 3.3415 |
| 15960 | ENSMUSG00000055497 | 1.506289039 | -0.791643788 | 1.91304026 | -0.413814494 | 0.679009968 | NA | Gm9974 | predicted gene 9974 [Source:MGI Symbol | 1.6062 | 3.0838 | 1.0062 | 0 | 0 | 3.3415 |
| 15961 | ENSMUSG00000055515 | 0 | NA | NA | NA | NA | NA | Vmn2r81 | vomeronasal 2, receptor 81 [Source:MGI S | 0 | 0 | 0 | 0 | 0 | 0 |
| 15962 | ENSMUSG00000055523 | 0.353089354 | 0.165142706 | 3.896102244 | 0.042386646 | 0.966190474 | NA | Gucy2g | guanylate cyclase 2g [Source:MGI Symbol | 0 | 1.0279 | 0 | 0 | 1.0906 | 0 |
| 15963 | ENSMUSG00000055531 | 633.9664606 | 0.047361905 | 0.146170262 | 0.324018746 | 0.745923842 | 0.949997329 | Cpsf6 | cleavage and polyadenylation specific fact | 598.31 | 690.76 | 582.62 | 727.8 | 668.55 | 535.76 |
| 15964 | ENSMUSG00000055538 | 2835.092557 | 0.168242022 | 0.092150933 | 1.825722403 | 0.067892087 | 0.375219813 | Zcchc24 | zinc finger, CCHC domain containing 24 [S | 2575.5 | 2677.7 | 2757.1 | 2978.7 | 2874.9 | 3146.6 |
| 15965 | ENSMUSG00000055540 | 0 | NA | NA | NA | NA | NA | Epha6 | Eph receptor A6 [Source:MGI Symbol;Acc | 0 | 0 | 0 | 0 | 0 | 0 |
| 15966 | ENSMUSG00000055541 | 25.34886628 | -0.776270846 | 0.46749785 | -1.660480035 | 0.096817921 | 0.458530879 | Lair1 | leukocyte-associated Ig-like receptor 1 [Sc | 25.699 | 46.256 | 24.15 | 17.352 | 16.359 | 22.277 |
| 15967 | ENSMUSG00000055546 | 38.49962961 | -2.050581432 | 0.490631516 | -4.179473527 | 2.92E-05 | 0.001345911 | Timd4 | T cell immunoglobulin and mucin domain | 40.958 | 99.708 | 45.281 | 9.6398 | 10.906 | 24.505 |
| 15968 | ENSMUSG00000055547 | 0 | NA | NA | NA | NA | NA | Apobec4 | apolipoprotein B mRNA editing enzyme, c | 0 | 0 | 0 | 0 | 0 | 0 |
| 15969 | ENSMUSG00000055553 | 730.6566667 | 0.067939756 | 0.110711752 | 0.613663453 | 0.539437744 | 0.89221584 | Kxd1 | KxDL motif containing 1 [Source:MGI Sym | 707.53 | 723.65 | 709.4 | 753.83 | 769.98 | 719.54 |
| 15970 | ENSMUSG00000055555 | 0 | NA | NA | NA | NA | NA | Ct55 | cancer/testis antigen 55 [Source:MGI Sym | 0 | 0 | 0 | 0 | 0 | 0 |
| 15971 | ENSMUSG00000055560 | 1.887148553 | -0.725322522 | 1.793590459 | -0.404396956 | 0.685920838 | NA | Zfp459 | zinc finger protein 459 [Source:MGI Symb | 0 | 2.0558 | 5.0312 | 0.964 | 3.2719 | 0 |
| 15972 | ENSMUSG00000055561 | 10.40303146 | 0.656844111 | 0.806024319 | 0.814918478 | 0.415118976 | 0.836001897 | Spink5 | serine peptidase inhibitor, Kazal type 5 [S | 16.865 | 2.0558 | 5.0312 | 15.424 | 16.359 | 6.6831 |
| 15973 | ENSMUSG00000055567 | 0.480153995 | 1.043847617 | 3.151673025 | 0.331204287 | 0.740490182 | NA | Unc80 | unc-80, NALCN activator [Source:MGI Sym | 0.8031 | 0 | 0 | 0.964 | 0 | 1.1138 |
| 15974 | ENSMUSG00000055571 | 0 | NA | NA | NA | NA | NA | Olfr305 | olfactory receptor 305 [Source:MGI Symb | 0 | 0 | 0 | 0 | 0 | 0 |
| 15975 | ENSMUSG00000055594 | 0 | NA | NA | NA | NA | NA | 5530400C23 | RIKEN cDNA 5530400C23 gene [Source:M | 0 | 0 | 0 | 0 | 0 | 0 |
| 15976 | ENSMUSG00000055602 | 3.067295595 | -0.060604025 | 1.379246537 | -0.043939951 | 0.96495227 | NA | Tcp10b | t-complex protein 10b [Source:MGI Symb | 2.4093 | 1.0279 | 6.0375 | 6.7478 | 2.1812 | 0 |
| 15977 | ENSMUSG00000055609 | 0 | NA | NA | NA | NA | NA | Hba-x | hemoglobin X, alpha-like embryonic chain | 0 | 0 | 0 | 0 | 0 | 0 |
| 15978 | ENSMUSG00000055610 | 0 | NA | NA | NA | NA | NA | Olfr307 | olfactory receptor 307 [Source:MGI Symb | 0 | 0 | 0 | 0 | 0 | 0 |
| 15979 | ENSMUSG00000055612 | 55.80126506 | -0.009955936 | 0.328305077 | -0.030325257 | 0.975807654 | 0.996622832 | Cdca7 | cell division cycle associated 7 [Source:M | 51.399 | 41.117 | 75.468 | 50.127 | 64.346 | 52.351 |
| 15980 | ENSMUSG00000055621 | 0 | NA | NA | NA | NA | NA | Olfr636-ps1 | olfactory receptor 636, pseudogene 1 [So | 0 | 0 | 0 | 0 | 0 | 0 |
| 15981 | ENSMUSG00000055629 | 4.154237088 | 0.919700105 | 1.303052282 | 0.705804454 | 0.480309752 | NA | B4galnt4 | beta-1,4-N-acetyl-galactosaminyl transfer | 6.4248 | 0 | 2.0125 | 0.964 | 3.2719 | 12.252 |
| 15982 | ENSMUSG00000055632 | 84.74753115 | -0.17571444 | 0.251878961 | -0.697614599 | 0.485418249 | 0.872835599 | Hmcn2 | hemicentin 2 [Source:MGI Symbol;Acc:M | 85.129 | 82.233 | 102.64 | 89.65 | 71.981 | 76.855 |
| 15983 | ENSMUSG00000055633 | 12.45792503 | -0.232878206 | 0.652702208 | -0.356790897 | 0.721248351 | 0.9447719 | Zfp580 | zinc finger protein 580 [Source:MGI Symb | 12.047 | 12.335 | 16.1 | 15.424 | 4.3625 | 14.48 |
| 15984 | ENSMUSG00000055639 | 0.74057819 | -2.943622841 | 2.624979752 | -1.12138878 | 0.26212241 | NA | Dach1 | dachshund family transcription factor 1 [S | 2.4093 | 1.0279 | 1.0062 | 0 | 0 | 0 |
| 15985 | ENSMUSG00000055643 | 0 | NA | NA | NA | NA | NA | Ubqln5 | ubiquilin 5 [Source:MGI Symbol;Acc:MGI: | 0 | 0 | 0 | 0 | 0 | 0 |
| 15986 | ENSMUSG00000055652 | 387.2006945 | -0.505935557 | 0.165827019 | -3.050983851 | 0.002280928 | 0.038406692 | Klhl25 | kelch-like 25 [Source:MGI Symbol;Acc:MG | 403.96 | 440.98 | 518.22 | 271.84 | 372.99 | 315.22 |
| 15987 | ENSMUSG00000055653 | 54.54785139 | -0.069409746 | 0.323953069 | -0.214258644 | 0.830345366 | 0.9702546 | Gpc3 | glypican 3 [Source:MGI Symbol;Acc:MGI:1 | 62.642 | 53.452 | 51.318 | 51.091 | 69.8 | 38.985 |
| 15988 | ENSMUSG00000055660 | 101.3939236 | 0.18611599 | 0.240105792 | 0.77514161 | 0.438255986 | 0.847566655 | Mettl4 | methyltransferase like 4 [Source:MGI Sym | 80.31 | 116.15 | 88.55 | 104.11 | 113.42 | 105.82 |
| 15989 | ENSMUSG00000055670 | 759.2411797 | -0.11229452 | 0.148313256 | -0.75714419 | 0.448963482 | 0.854431786 | Zzef1 | zinc finger, ZZ-type with EF hand domain : | 672.2 | 870.65 | 825.12 | 822.27 | 749.25 | 615.96 |
| 15990 | ENSMUSG00000055675 | 40.37019683 | -1.931466648 | 0.578490494 | -3.33880447 | 0.000841398 | 0.018439357 | Kbtbd11 | kelch repeat and BTB (POZ) domain conta | 110.03 | 59.619 | 22.137 | 15.424 | 29.447 | 5.5692 |
| 15991 | ENSMUSG00000055676 | 0.335414862 | -1.779047242 | 4.030804179 | -0.441362855 | 0.658950329 | NA | Gm5069 | predicted pseudogene 5069 [Source:MGI | 0 | 0 | 2.0125 | 0 | 0 | 0 |
| 15992 | ENSMUSG00000055679 | 0 | NA | NA | NA | NA | NA | Ctsr | cathepsin R [Source:MGI Symbol;Acc:MGI | 0 | 0 | 0 | 0 | 0 | 0 |
| 15993 | ENSMUSG00000055681 | 1823.75393 | 0.113300145 | 0.092513208 | 1.224691555 | 0.220691457 | 0.667237604 | Cope | coatomer protein complex, subunit epsilo | 1775.7 | 1750.5 | 1729.7 | 1835.4 | 1906.4 | 1944.8 |
| 15994 | ENSMUSG00000055691 | 1.878707853 | -0.124098722 | 1.51437336 | -0.081947243 | 0.934688666 | NA | Gja6 | gap junction protein, alpha 6 [Source:MG | 0.8031 | 3.0838 | 2.0125 | 0.964 | 2.1812 | 2.2277 |
| 15995 | ENSMUSG00000055692 | 13.76223143 | 0.217413904 | 0.581801704 | 0.373690731 | 0.708634422 | 0.943888782 | Tmem191c | transmembrane protein 191C [Source:MG | 8.8341 | 14.391 | 15.094 | 14.46 | 13.087 | 16.708 |
| 15996 | ENSMUSG00000055700 | 0 | NA | NA | NA | NA | NA | 4933401B06 | RIKEN cDNA 4933401B06 gene [Source:M | 0 | 0 | 0 | 0 | 0 | 0 |
| 15997 | ENSMUSG00000055704 | 0 | NA | NA | NA | NA | NA | Gm9978 | predicted gene 9978 [Source:MGI Symbol | 0 | 0 | 0 | 0 | 0 | 0 |
| 15998 | ENSMUSG00000055707 | 111.1122897 | -0.012091378 | 0.309444272 | -0.039074494 | 0.968830996 | 0.994547305 | Klhl26 | kelch-like 26 [Source:MGI Symbol;Acc:MG | 72.279 | 116.15 | 146.91 | 112.79 | 146.14 | 72.4 |
| 15999 | ENSMUSG00000055717 | 2.163003219 | 0.829050955 | 1.520707148 | 0.545174629 | 0.585633419 | NA | Slain1 | SLAIN motif family, member 1 [Source:MG | 1.6062 | 3.0838 | 0 | 3.8559 | 1.0906 | 3.3415 |
| 16000 | ENSMUSG00000055720 | 638.5768215 | -0.110998804 | 0.124850615 | -0.889052924 | 0.373974637 | 0.808908203 | Ubl7 | ubiquitin-like 7 (bone marrow stromal cell | 725.2 | 609.56 | 654.06 | 646.83 | 596.57 | 599.25 |
| 16001 | ENSMUSG00000055723 | 320.1908248 | -0.01230538 | 0.154402158 | -0.079696945 | 0.936478289 | 0.988420305 | Rras2 | related RAS viral (r-ras) oncogene 2 [Sourc | 323.65 | 310.43 | 330.05 | 286.3 | 305.37 | 365.34 |
| 16002 | ENSMUSG00000055725 | 97.43893129 | 0.047136901 | 0.25623284 | 0.183961199 | 0.854043894 | 0.973825032 | Paqr3 | progestin and adipoQ receptor family mer | 98.782 | 79.15 | 109.68 | 113.75 | 76.343 | 106.93 |
| 16003 | ENSMUSG00000055730 | 3114.414816 | 0.149922471 | 0.126390808 | 1.186181763 | 0.235550527 | 0.683799474 | Ces2a | carboxylesterase 2A [Source:MGI Symbol; | 2496.8 | 3041.6 | 3321.6 | 3452 | 3457.3 | 2917.2 |
| 16004 | ENSMUSG00000055732 | 0 | NA | NA | NA | NA | NA | Gm9979 | predicted gene 9979 [Source:MGI Symbol | 0 | 0 | 0 | 0 | 0 | 0 |
| 16005 | ENSMUSG00000055733 | 13.79961807 | 0.250485888 | 0.625427583 | 0.400503424 | 0.688785762 | 0.936908509 | Nap1l3 | nucleosome assembly protein 1-like 3 [So | 6.4248 | 15.419 | 16.1 | 10.604 | 13.087 | 21.163 |
| 16006 | ENSMUSG00000055737 | 11555.63319 | 0.111703669 | 0.104609223 | 1.067818554 | 0.285602367 | 0.738065506 | Ghr | growth hormone receptor [Source:MGI Sy | 10830 | 11310 | 11185 | 11189 | 13665 | 11154 |
| 16007 | ENSMUSG00000055745 | 4.194808055 | 0.281348493 | 1.204559032 | 0.233569701 | 0.815319052 | NA | Rtl6 | retrotransposon Gag like 6 [Source:MGI S | 3.2124 | 5.1396 | 3.0187 | 2.8919 | 10.906 | 0 |
| 16008 | ENSMUSG00000055746 | 0 | NA | NA | NA | NA | NA | Magea2 | MAGE family member A2 [Source:MGI Syr | 0 | 0 | 0 | 0 | 0 | 0 |
| 16009 | ENSMUSG00000055748 | 0.160662974 | 1.126924175 | 4.080472857 | 0.276174898 | 0.782413736 | NA | Gsdmc4 | gasdermin C4 [Source:MGI Symbol;Acc:M | 0 | 0 | 0 | 0.964 | 0 | 0 |
| 16010 | ENSMUSG00000055760 | 74.47709783 | 0.179419348 | 0.271360721 | 0.66118393 | 0.508494366 | 0.882793133 | Gemin6 | gem nuclear organelle associated protein | 69.067 | 83.261 | 57.356 | 81.938 | 85.068 | 70.172 |
| 16011 | ENSMUSG00000055761 | 0.371281295 | 2.09253362 | 4.031852165 | 0.519000582 | 0.603760335 | NA | Nkain3 | Na+/K+ transporting ATPase interacting 3 | 0 | 0 | 0 | 0 | 0 | 2.2277 |
| 16012 | ENSMUSG00000055762 | 1346.806325 | -0.149524176 | 0.111164722 | -1.345068592 | 0.178603085 | 0.610813333 | Eef1d | eukaryotic translation elongation factor 1 | 1272.9 | 1518.2 | 1460.1 | 1339.9 | 1243.3 | 1246.4 |

| | | | | | | | | | | | | | | | |
|---|---|---|---|---|---|---|---|---|---|---|---|---|---|---|---|
| 16013 | ENSMUSG00000055763 | 0.503302254 | -0.837877584 | 3.426698643 | -0.244514523 | 0.806832356 | NA | Rybp-ps | RING1 and YY1 binding protein, pseudoge | 0 | 2.0558 | 0 | 0.964 | 0 | 0 |
| 16014 | ENSMUSG00000055771 | 0 | NA | NA | NA | NA | NA | Gm7936 | predicted pseudogene 7936 [Source:MGI | 0 | 0 | 0 | 0 | 0 | 0 |
| 16015 | ENSMUSG00000055775 | 0 | NA | NA | NA | NA | NA | Myh8 | myosin, heavy polypeptide 8, skeletal mus | 0 | 0 | 0 | 0 | 0 | 0 |
| 16016 | ENSMUSG00000055780 | 0 | NA | NA | NA | NA | NA | Usp26 | ubiquitin specific peptidase 26 [Source:M | 0 | 0 | 0 | 0 | 0 | 0 |
| 16017 | ENSMUSG00000055782 | 468.6195694 | -0.601301171 | 0.244672443 | -2.457576197 | 0.013987813 | 0.136282395 | Abcd2 | ATP-binding cassette, sub-family D (ALD), | 668.18 | 685.62 | 341.12 | 486.81 | 282.47 | 347.52 |
| 16018 | ENSMUSG00000055789 | 0 | NA | NA | NA | NA | NA | Ftdc2 | ferritin domain containing 2 [Source:MGI | 0 | 0 | 0 | 0 | 0 | 0 |
| 16019 | ENSMUSG00000055795 | 0.17131964 | -0.796629176 | 4.080472857 | -0.195229623 | 0.84521319 | NA | Gm5160 | predicted gene 5160 [Source:MGI Symbol | 0 | 1.0279 | 0 | 0 | 0 | 0 |
| 16020 | ENSMUSG00000055799 | 91.90270473 | -0.357144932 | 0.26970434 | -1.324209064 | 0.185433621 | 0.621059253 | Tcf7l1 | transcription factor 7 like 1 (T cell specific, | 80.31 | 116.15 | 113.71 | 87.722 | 61.075 | 92.449 |
| 16021 | ENSMUSG00000055805 | 17.86469627 | -0.824596561 | 0.514975514 | -1.601234502 | 0.10932499 | 0.486233674 | Fmnl1 | formin-like 1 [Source:MGI Symbol;Acc:MG | 20.881 | 21.586 | 26.162 | 15.424 | 11.997 | 11.138 |
| 16022 | ENSMUSG00000055809 | 2.196189499 | -1.05483598 | 1.479316341 | -0.713056396 | 0.47581086 | NA | Dnaaf3 | dynein, axonemal assembly factor 3 [Sour | 0.8031 | 3.0838 | 5.0312 | 0.964 | 2.1812 | 1.1138 |
| 16023 | ENSMUSG00000055811 | 0 | NA | NA | NA | NA | NA | Cldn17 | claudin 17 [Source:MGI Symbol;Acc:MGI:2 | 0 | 0 | 0 | 0 | 0 | 0 |
| 16024 | ENSMUSG00000055817 | 188.1239051 | 0.055173888 | 0.18458391 | 0.298909523 | 0.76500908 | 0.954980327 | Mta3 | metastasis associated 3 [Source:MGI Syml | 179.09 | 203.53 | 171.06 | 183.16 | 217.03 | 174.87 |
| 16025 | ENSMUSG00000055818 | 48.4775042 | -0.474069565 | 0.435767711 | -1.087895117 | 0.276641412 | 0.730038567 | A230083G16 | RIKEN cDNA A230083G16 gene [Source:M | 65.051 | 25.698 | 78.487 | 57.839 | 38.172 | 25.618 |
| 16026 | ENSMUSG00000055820 | 0 | NA | NA | NA | NA | NA | Olfr967 | olfactory receptor 967 [Source:MGI Symb | 0 | 0 | 0 | 0 | 0 | 0 |
| 16027 | ENSMUSG00000055826 | 0 | NA | NA | NA | NA | NA | Tescl | tescalcin-like [Source:MGI Symbol;Acc:MG | 0 | 0 | 0 | 0 | 0 | 0 |
| 16028 | ENSMUSG00000055827 | 0 | NA | NA | NA | NA | NA | Gsdmc3 | gasdermin C3 [Source:MGI Symbol;Acc:M | 0 | 0 | 0 | 0 | 0 | 0 |
| 16029 | ENSMUSG00000055833 | 4.271735962 | 0.648475872 | 1.01958064 | 0.636022151 | 0.524761984 | NA | 1700034H15 | RIKEN cDNA 1700034H15 gene [Source:M | 4.8186 | 3.0838 | 2.0125 | 5.7839 | 4.3625 | 5.5692 |
| 16030 | ENSMUSG00000055835 | 132.6896861 | 0.183569738 | 0.220611219 | 0.832096117 | 0.405354693 | 0.829973745 | Zfp1 | zinc finger protein 1 [Source:MGI Symbol; | 119.66 | 143.91 | 109.68 | 161.95 | 142.87 | 118.07 |
| 16031 | ENSMUSG00000055838 | 0 | NA | NA | NA | NA | NA | Olfr357 | olfactory receptor 357 [Source:MGI Symb | 0 | 0 | 0 | 0 | 0 | 0 |
| 16032 | ENSMUSG00000055839 | 1156.075264 | -0.054768953 | 0.104289332 | -0.525163521 | 0.599469521 | 0.915117178 | Elob | elongin B [Source:MGI Symbol;Acc:MGI:1 | 1214.3 | 1137.9 | 1181.3 | 1134.6 | 1073.2 | 1195.2 |
| 16033 | ENSMUSG00000055850 | 1515.223432 | 0.202517055 | 0.113606775 | 1.782614239 | 0.07464912 | 0.399158919 | Rnf181 | ring finger protein 181 [Source:MGI Symb | 1475.3 | 1232.5 | 1518.4 | 1571.3 | 1610.8 | 1683 |
| 16034 | ENSMUSG00000055862 | 179.1060888 | 0.407038063 | 0.20977116 | 1.940390963 | 0.052332194 | 0.321555784 | Izumo4 | IZUMO family member 4 [Source:MGI Syn | 145.36 | 126.43 | 190.18 | 188.94 | 187.59 | 236.13 |
| 16035 | ENSMUSG00000055865 | 1.486398581 | -1.702287155 | 2.053289899 | -0.829053489 | 0.407074141 | NA | Tafa3 | TAFA chemokine like family member 3 [So | 0.8031 | 0 | 6.0375 | 0.964 | 0 | 1.1138 |
| 16036 | ENSMUSG00000055866 | 836.7870443 | -0.051190563 | 0.129801336 | -0.394376244 | 0.69330328 | 0.938857658 | Per2 | period circadian clock 2 [Source:MGI Syml | 823.98 | 910.74 | 821.1 | 890.72 | 873.59 | 700.61 |
| 16037 | ENSMUSG00000055872 | 0 | NA | NA | NA | NA | NA | Gm9985 | predicted gene 9985 [Source:MGI Symbol | 0 | 0 | 0 | 0 | 0 | 0 |
| 16038 | ENSMUSG00000055874 | 0 | NA | NA | NA | NA | NA | Foxi3 | forkhead box I3 [Source:MGI Symbol;Acc: | 0 | 0 | 0 | 0 | 0 | 0 |
| 16039 | ENSMUSG00000055882 | 0.480153995 | 1.043847617 | 3.151673025 | 0.331204287 | 0.740490182 | NA | Abhd16b | abhydrolase domain containing 16B [Sour | 0.8031 | 0 | 0 | 0.964 | 0 | 1.1138 |
| 16040 | ENSMUSG00000055884 | 80.98330124 | -0.068571963 | 0.265888894 | -0.257897057 | 0.796486356 | 0.962236316 | Fancm | Fanconi anemia, complementation group | 77.901 | 96.624 | 74.462 | 82.902 | 90.521 | 63.489 |
| 16041 | ENSMUSG00000055891 | 0 | NA | NA | NA | NA | NA | Ubl4b | ubiquitin-like 4B [Source:MGI Symbol;Acc | 0 | 0 | 0 | 0 | 0 | 0 |
| 16042 | ENSMUSG00000055895 | 0 | NA | NA | NA | NA | NA | Oosp2 | oocyte secreted protein 2 [Source:MGI Sy | 0 | 0 | 0 | 0 | 0 | 0 |
| 16043 | ENSMUSG00000055897 | 219.5326044 | 0.117015509 | 0.178889564 | 0.654121496 | 0.513033533 | 0.884860213 | Ppp4r1l-ps | protein phosphatase 4, regulatory subunit | 220.05 | 217.92 | 194.21 | 266.06 | 201.76 | 217.2 |
| 16044 | ENSMUSG00000055900 | 241.9935061 | 0.061469807 | 0.164378539 | 0.373952754 | 0.708439467 | 0.943888782 | Tmem69 | transmembrane protein 69 [Source:MGI S | 248.16 | 237.45 | 224.39 | 226.53 | 274.84 | 240.59 |
| 16045 | ENSMUSG00000055912 | 1769.057203 | 0.193071694 | 0.109931113 | 1.756297082 | 0.079037695 | 0.412857069 | Tmem150a | transmembrane protein 150A [Source:MG | 1517.1 | 1628.2 | 1808.2 | 1848.9 | 2023.1 | 1788.8 |
| 16046 | ENSMUSG00000055917 | 221.6336528 | 0.416361306 | 0.17498507 | 2.379410461 | 0.017340355 | 0.156761227 | Zfp277 | zinc finger protein 277 [Source:MGI Symb | 210.41 | 166.52 | 192.19 | 269.91 | 230.12 | 260.64 |
| 16047 | ENSMUSG00000055923 | 176.3863 | 0.175348284 | 0.192773616 | 0.909607275 | 0.363029661 | 0.80202551 | Aasdh | aminoadipate-semialdehyde dehydrogena | 187.93 | 162.41 | 145.91 | 167.73 | 207.22 | 187.13 |
| 16048 | ENSMUSG00000055926 | 5.531049429 | -0.718892078 | 0.910522832 | -0.789537673 | 0.42979782 | 0.84307112 | Gm14137 | predicted gene 14137 [Source:MGI Symbo | 6.4248 | 8.2233 | 6.0375 | 4.8199 | 5.4531 | 2.2277 |
| 16049 | ENSMUSG00000055932 | 522.6809418 | -0.052718397 | 0.129621219 | -0.406711164 | 0.68422014 | 0.935528319 | Fto | fat mass and obesity associated [Source:N | 550.93 | 561.24 | 484 | 493.56 | 501.68 | 544.67 |
| 16050 | ENSMUSG00000055933 | 0 | NA | NA | NA | NA | NA | Oosp3 | oocyte secreted protein 3 [Source:MGI Sy | 0 | 0 | 0 | 0 | 0 | 0 |
| 16051 | ENSMUSG00000055937 | 0 | NA | NA | NA | NA | NA | Krt28 | keratin 28 [Source:MGI Symbol;Acc:MGI:1 | 0 | 0 | 0 | 0 | 0 | 0 |
| 16052 | ENSMUSG00000055942 | 0 | NA | NA | NA | NA | NA | Obox7 | oocyte specific homeobox 7 [Source:MGI | 0 | 0 | 0 | 0 | 0 | 0 |
| 16053 | ENSMUSG00000055943 | 1297.678504 | 0.025602694 | 0.112237683 | 0.22811139 | 0.819559646 | 0.969160361 | Emc7 | ER membrane protein complex subunit 7 [ | 1173.3 | 1336.3 | 1350.4 | 1396.8 | 1216 | 1313.2 |
| 16054 | ENSMUSG00000055944 | 0 | NA | NA | NA | NA | NA | E130018O15 | RIKEN cDNA E130018O15 gene [Source:M | 0 | 0 | 0 | 0 | 0 | 0 |
| 16055 | ENSMUSG00000055945 | 4.908515855 | 5.822209095 | 1.688240737 | 3.448684163 | 0.000563325 | NA | Prr18 | proline rich 18 [Source:MGI Symbol;Acc:N | 0 | 0 | 0 | 3.8559 | 1.0906 | 24.505 |
| 16056 | ENSMUSG00000055958 | 0 | NA | NA | NA | NA | NA | Gm9987 | predicted gene 9987 [Source:MGI Symbol | 0 | 0 | 0 | 0 | 0 | 0 |
| 16057 | ENSMUSG00000055960 | 0 | NA | NA | NA | NA | NA | Skint4 | selection and upkeep of intraepithelial T c | 0 | 0 | 0 | 0 | 0 | 0 |
| 16058 | ENSMUSG00000055961 | 0 | NA | NA | NA | NA | NA | BC051076 | cDNA sequence BC051076 [Source:MGI Sy | 0 | 0 | 0 | 0 | 0 | 0 |
| 16059 | ENSMUSG00000055963 | 1.723563611 | 0.139935013 | 1.996457141 | 0.070091669 | 0.944120698 | NA | Triqk | triple QxxK/R motif containing [Source:MG | 4.8186 | 0 | 0 | 0 | 2.1812 | 3.3415 |
| 16060 | ENSMUSG00000055968 | 0 | NA | NA | NA | NA | NA | Gm9988 | predicted gene 9988 [Source:MGI Symbol | 0 | 0 | 0 | 0 | 0 | 0 |
| 16061 | ENSMUSG00000055971 | 0 | NA | NA | NA | NA | NA | Olfr378 | olfactory receptor 378 [Source:MGI Symb | 0 | 0 | 0 | 0 | 0 | 0 |
| 16062 | ENSMUSG00000055972 | 0 | NA | NA | NA | NA | NA | 2810407A14 | RIKEN cDNA 2810407A14 gene [Source:M | 0 | 0 | 0 | 0 | 0 | 0 |
| 16063 | ENSMUSG00000055976 | 1.477699157 | -0.83265173 | 1.762683391 | -0.472377362 | 0.63665746 | NA | Cldn23 | claudin 23 [Source:MGI Symbol;Acc:MGI:1 | 1.6062 | 2.0558 | 2.0125 | 0.964 | 0 | 2.2277 |
| 16064 | ENSMUSG00000055978 | 2.125850324 | 1.835147373 | 1.759252684 | 1.043140301 | 0.296883318 | NA | Fut2 | fucosyltransferase 2 [Source:MGI Symbol; | 0.8031 | 0 | 2.0125 | 7.7118 | 0 | 2.2277 |
| 16065 | ENSMUSG00000055980 | 159.1456167 | -0.191153556 | 0.210279557 | -0.909044883 | 0.363326434 | 0.802199995 | Irs1 | insulin receptor substrate 1 [Source:MGI S | 142.95 | 159.33 | 207.29 | 155.2 | 133.06 | 157.05 |
| 16066 | ENSMUSG00000055991 | 315.7625218 | 0.487454526 | 0.16041689 | 3.038673335 | 0.002376224 | 0.039642097 | Zkscan5 | zinc finger with KRAB and SCAN domains 5 | 267.43 | 247.73 | 273.7 | 403.91 | 309.74 | 392.07 |
| 16067 | ENSMUSG00000055994 | 2.094023768 | -0.906000804 | 1.954214766 | -0.463613733 | 0.642924513 | NA | Nod2 | nucleotide-binding oligomerization domai | 0 | 7.1954 | 1.0062 | 0 | 4.3625 | 0 |
| 16068 | ENSMUSG00000056004 | 4.940142841 | -1.845540238 | 1.197461288 | -1.541210774 | 0.123265496 | NA | Elapor2 | endosome-lysosome associated apoptosis | 0.8031 | 12.335 | 10.062 | 0.964 | 4.3625 | 1.1138 |

| | | | | | | | | | | | | | | | |
|---|---|---|---|---|---|---|---|---|---|---|---|---|---|---|---|
| 16069 | ENSMUSG00000056008 | 0.349477146 | 0.165142768 | 3.911021834 | 0.042224967 | 0.966319359 | NA | Gm11541 | predicted gene 11541 [Source:MGI Symbo | 0 | 0 | 1.0062 | 0 | 1.0906 | 0 |
| 16070 | ENSMUSG00000056014 | 101.2267461 | 0.249145962 | 0.23272021 | 1.070581546 | 0.284357625 | 0.736709833 | A430033K04 | RIKEN cDNA A430033K04 gene [Source:M | 100.39 | 78.122 | 98.612 | 102.18 | 117.79 | 110.27 |
| 16071 | ENSMUSG00000056018 | 0.36353943 | 2.065989254 | 4.034137301 | 0.512126658 | 0.608562368 | NA | Ccdc7b | coiled-coil domain containing 7B [Source: | 0 | 0 | 0 | 0 | 2.1812 | 0 |
| 16072 | ENSMUSG00000056019 | 61.11315386 | 0.006174516 | 0.310586542 | 0.019880179 | 0.984138957 | 0.998580735 | Zfp709 | zinc finger protein 709 [Source:MGI Symb | 77.901 | 51.396 | 53.331 | 70.37 | 49.078 | 64.603 |
| 16073 | ENSMUSG00000056023 | 0.781659665 | -3.010437285 | 2.830777252 | -1.063466679 | 0.287570368 | NA | Gm9989 | predicted gene 9989 [Source:MGI Symbol | 1.6062 | 3.0838 | 0 | 0 | 0 | 0 |
| 16074 | ENSMUSG00000056025 | 56.13081798 | -1.034061559 | 0.329928242 | -3.134201407 | 0.001723225 | 0.031430935 | Clca3a1 | chloride channel accessory 3A1 [Source:M | 91.554 | 77.094 | 57.356 | 35.667 | 29.447 | 45.668 |
| 16075 | ENSMUSG00000056031 | 0 | NA | NA | NA | NA | NA | 9330154J02I | RIKEN cDNA 9330154J02 gene [Source:MC | 0 | 0 | 0 | 0 | 0 | 0 |
| 16076 | ENSMUSG00000056032 | 0 | NA | NA | NA | NA | NA | BC018473 | cDNA sequence BC018473 [Source:MGI Sy | 0 | 0 | 0 | 0 | 0 | 0 |
| 16077 | ENSMUSG00000056035 | 12158.12014 | -1.380858002 | 0.294413127 | -4.69020528 | 2.73E-06 | 0.000184471 | Cyp3a11 | cytochrome P450, family 3, subfamily a, p | 17423 | 21267 | 14019 | 5980.5 | 8949.6 | 5309.7 |
| 16078 | ENSMUSG00000056043 | 3.703195936 | 0.784495597 | 1.242688332 | 0.631289099 | 0.527851513 | NA | Rgs9bp | regulator of G-protein signalling 9 binding | 4.0155 | 1.0279 | 3.0187 | 1.928 | 1.0906 | 11.138 |
| 16079 | ENSMUSG00000056050 | 2629.558287 | 0.039406107 | 0.123119532 | 0.320063816 | 0.748919955 | 0.951074124 | Mia3 | melanoma inhibitory activity 3 [Source:M( | 2519.3 | 2635.6 | 2627.3 | 3114.6 | 2265.2 | 2615.3 |
| 16080 | ENSMUSG00000056054 | 3.377416412 | -0.86399881 | 1.236526474 | -0.698730539 | 0.484720445 | NA | S100a8 | S100 calcium binding protein A8 (calgranu | 4.0155 | 3.0838 | 6.0375 | 3.8559 | 3.2719 | 0 |
| 16081 | ENSMUSG00000056055 | 1.17227042 | -0.274240045 | 1.851457481 | -0.148121168 | 0.882247144 | NA | Sag | S-antigen, retina and pineal gland (arresti | 0.8031 | 2.0558 | 1.0062 | 0.964 | 1.0906 | 1.1138 |
| 16082 | ENSMUSG00000056061 | 0 | NA | NA | NA | NA | NA | Gata5os | GATA binding protein 5, opposite strand [ | 0 | 0 | 0 | 0 | 0 | 0 |
| 16083 | ENSMUSG00000056069 | 23.6293798 | -0.90550369 | 0.578559352 | -1.565100773 | 0.117559282 | 0.502806291 | Otulinl | OTU deubiquitinase with linear linkage sp | 36.943 | 43.173 | 12.075 | 8.6758 | 14.178 | 26.732 |
| 16084 | ENSMUSG00000056071 | 5.915745778 | -1.028449615 | 0.952853041 | -1.079337076 | 0.280437491 | 0.733413074 | S100a9 | S100 calcium binding protein A9 (calgranu | 10.44 | 10.279 | 3.0187 | 2.8919 | 2.1812 | 6.6831 |
| 16085 | ENSMUSG00000056073 | 0 | NA | NA | NA | NA | NA | Grik2 | glutamate receptor, ionotropic, kainate 2 | 0 | 0 | 0 | 0 | 0 | 0 |
| 16086 | ENSMUSG00000056076 | 1815.869579 | -0.037221115 | 0.097474386 | -0.38185534 | 0.702568668 | 0.942405009 | Eif3b | eukaryotic translation initiation factor 3, s | 1750 | 1844.1 | 1924.9 | 1864.3 | 1728.6 | 1783.3 |
| 16087 | ENSMUSG00000056078 | 0 | NA | NA | NA | NA | NA | Lipm | lipase, family member M [Source:MGI Syn | 0 | 0 | 0 | 0 | 0 | 0 |
| 16088 | ENSMUSG00000056089 | 0.267700746 | -1.497481649 | 4.057001436 | -0.369110456 | 0.712045396 | NA | Gm5468 | predicted gene 5468 [Source:MGI Symbol | 1.6062 | 0 | 0 | 0 | 0 | 0 |
| 16089 | ENSMUSG00000056091 | 5833.553754 | -0.813060284 | 0.090262658 | -9.007714794 | 2.10E-19 | 2.34E-16 | St3gal5 | ST3 beta-galactoside alpha-2,3-sialyltrans | 7190.2 | 7964.3 | 7151.4 | 4179.8 | 4169.4 | 4346.2 |
| 16090 | ENSMUSG00000056115 | 0 | NA | NA | NA | NA | NA | Tas2r134 | taste receptor, type 2, member 134 [Sour | 0 | 0 | 0 | 0 | 0 | 0 |
| 16091 | ENSMUSG00000056116 | 2123.809394 | 0.029780453 | 0.096315467 | 0.309196999 | 0.757171676 | 0.953260973 | H2-T22 | histocompatibility 2, T region locus 22 [So | 2232.6 | 2026 | 2045.7 | 2092.8 | 2115.8 | 2229.9 |
| 16092 | ENSMUSG00000056121 | 706.4627713 | -0.027025943 | 0.129955303 | -0.207963373 | 0.835257566 | 0.971073158 | Fez2 | fasciculation and elongation protein zeta | 787.04 | 642.45 | 708.4 | 646.83 | 761.25 | 692.81 |
| 16093 | ENSMUSG00000056124 | 22.17183052 | -0.611264692 | 0.520284263 | -1.17486677 | 0.240048022 | 0.689857856 | B4galt6 | UDP-Gal:betaGlcNAc beta 1,4-galactosyltr | 19.274 | 44.2 | 17.106 | 18.316 | 18.541 | 15.594 |
| 16094 | ENSMUSG00000056128 | 0 | NA | NA | NA | NA | NA | Gm9991 | predicted gene 9991 [Source:MGI Symbol | 0 | 0 | 0 | 0 | 0 | 0 |
| 16095 | ENSMUSG00000056130 | 1.803251872 | -0.054540675 | 1.784784218 | -0.030558694 | 0.975621484 | NA | Ticam2 | toll-like receptor adaptor molecule 2 [Sou | 2.4093 | 3.0838 | 0 | 0.964 | 4.3625 | 0 |
| 16096 | ENSMUSG00000056131 | 420.935357 | -0.294402972 | 0.171559446 | -1.716040581 | 0.086154615 | 0.432482144 | Pgm3 | phosphoglucomutase 3 [Source:MGI Syml | 548.52 | 479.01 | 363.25 | 419.33 | 346.82 | 368.68 |
| 16097 | ENSMUSG00000056133 | 34.45355048 | 0.376329559 | 0.411151129 | 0.915307127 | 0.360030419 | 0.799598236 | Unc93a2 | unc-93 homolog A2 [Source:MGI Symbol;/ | 37.746 | 22.614 | 29.181 | 29.883 | 32.719 | 54.578 |
| 16098 | ENSMUSG00000056144 | 80.25323068 | 0.710931801 | 0.293052091 | 2.425957102 | 0.015268075 | 0.144358963 | Trim34a | tripartite motif-containing 34A [Source:M | 80.31 | 47.284 | 54.337 | 104.11 | 78.525 | 116.95 |
| 16099 | ENSMUSG00000056145 | 2.334194504 | -2.515471611 | 1.596774557 | -1.575345499 | 0.115176717 | NA | AI504432 | expressed sequence AI504432 [Source:M( | 4.8186 | 3.0838 | 4.025 | 0.964 | 0 | 1.1138 |
| 16100 | ENSMUSG00000056148 | 237.4492525 | -0.039199758 | 0.202452315 | -0.193624649 | 0.846469796 | 0.972803041 | Rdh9 | retinol dehydrogenase 9 [Source:MGI Sym | 187.12 | 307.35 | 228.42 | 229.43 | 256.3 | 216.09 |
| 16101 | ENSMUSG00000056153 | 251.7269182 | -0.084525976 | 0.181015357 | -0.466954724 | 0.640532272 | 0.923932929 | Socs6 | suppressor of cytokine signaling 6 [Source | 229.69 | 317.63 | 230.43 | 234.25 | 238.85 | 259.53 |
| 16102 | ENSMUSG00000056155 | 0.331982614 | 0.16514309 | 3.986751476 | 0.041422971 | 0.966958701 | NA | Nanos3 | nanos C2HC-type zinc finger 3 [Source:MC | 0 | 1.0279 | 0 | 0.964 | 0 | 0 |
| 16103 | ENSMUSG00000056158 | 0 | NA | NA | NA | NA | NA | Car10 | carbonic anhydrase 10 [Source:MGI Symb | 0 | 0 | 0 | 0 | 0 | 0 |
| 16104 | ENSMUSG00000056162 | 10.88849754 | 0.096233835 | 0.69373405 | 0.138718627 | 0.889672498 | 0.982382091 | Cndp1 | carnosine dipeptidase 1 (metallopeptidas | 16.062 | 10.279 | 5.0312 | 9.6398 | 8.7249 | 15.594 |
| 16105 | ENSMUSG00000056167 | 382.3715678 | 0.056591171 | 0.1415093 | 0.399911323 | 0.689221832 | 0.937015187 | Cnot10 | CCR4-NOT transcription complex, subunit | 392.72 | 355.66 | 376.34 | 428.01 | 380.63 | 360.89 |
| 16106 | ENSMUSG00000056174 | 6.579220716 | -0.263687679 | 0.843754908 | -0.312516913 | 0.754647712 | 0.953048681 | Col8a2 | collagen, type VIII, alpha 2 [Source:MGI Sy | 7.2279 | 10.279 | 4.025 | 5.7839 | 4.3625 | 7.7969 |
| 16107 | ENSMUSG00000056184 | 0 | NA | NA | NA | NA | NA | Olfr283 | olfactory receptor 283 [Source:MGI Symb | 0 | 0 | 0 | 0 | 0 | 0 |
| 16108 | ENSMUSG00000056185 | 86.86905753 | 0.127257334 | 0.270639272 | 0.470210154 | 0.63820488 | 0.923699483 | Snx32 | sorting nexin 32 [Source:MGI Symbol;Acc: | 86.735 | 72.982 | 89.556 | 107 | 100.34 | 64.603 |
| 16109 | ENSMUSG00000056197 | 0 | NA | NA | NA | NA | NA | 4931417E11 | RIKEN cDNA 4931417E11 gene [Source:M( | 0 | 0 | 0 | 0 | 0 | 0 |
| 16110 | ENSMUSG00000056201 | 3480.586113 | -0.119795501 | 0.090233222 | -1.327620779 | 0.184303416 | 0.619137675 | Cfl1 | cofilin 1, non-muscle [Source:MGI Symbol | 3687 | 3676.9 | 3510.8 | 3342.1 | 3173.7 | 3493 |
| 16111 | ENSMUSG00000056203 | 0.185640647 | 1.126924175 | 4.080472857 | 0.276174898 | 0.782413736 | NA | Tas2r135 | taste receptor, type 2, member 135 [Sour | 0 | 0 | 0 | 0 | 0 | 1.1138 |
| 16112 | ENSMUSG00000056204 | 1600.69843 | 0.272333691 | 0.140626892 | 1.936569086 | 0.05279804 | 0.32296796 | Pgpep1 | pyroglutamyl-peptidase I [Source:MGI Syr | 1531.5 | 1201.6 | 1616 | 1613.7 | 1614.1 | 2027.2 |
| 16113 | ENSMUSG00000056209 | 262.7696404 | -0.225853502 | 0.172196245 | -1.311605268 | 0.189653354 | 0.626663447 | Npm3 | nucleoplasmin 3 [Source:MGI Symbol;Acc | 252.98 | 326.88 | 270.68 | 243.89 | 266.11 | 216.09 |
| 16114 | ENSMUSG00000056211 | 365.6095128 | -0.152825092 | 0.14629963 | -1.044603402 | 0.296206306 | 0.74691911 | R3hdm1 | R3H domain containing 1 [Source:MGI Syr | 388.7 | 378.27 | 387.4 | 311.36 | 391.53 | 336.38 |
| 16115 | ENSMUSG00000056214 | 13.08432478 | -0.95503691 | 0.691681452 | -1.380746741 | 0.167356844 | 0.593441872 | Pard6g | par-6 family cell polarity regulator gamma | 14.456 | 14.391 | 23.144 | 15.424 | 2.1812 | 8.9108 |
| 16116 | ENSMUSG00000056215 | 0 | NA | NA | NA | NA | NA | Lrguk | leucine-rich repeats and guanylate kinase | 0 | 0 | 0 | 0 | 0 | 0 |
| 16117 | ENSMUSG00000056216 | 1218.650023 | 0.092089603 | 0.114494045 | 0.804317844 | 0.421213435 | 0.838715738 | Cebpg | CCAAT/enhancer binding protein (C/EBP), | 1174.9 | 1261.3 | 1103.9 | 1372.7 | 1153.9 | 1245.3 |
| 16118 | ENSMUSG00000056219 | 0 | NA | NA | NA | NA | NA | Tmem229b-| | transmembrane protein 229B, pseudogen | 0 | 0 | 0 | 0 | 0 | 0 |
| 16119 | ENSMUSG00000056220 | 12.60801993 | -1.794543478 | 0.731837805 | -2.452105461 | 0.014202303 | 0.137774723 | Pla2g4a | phospholipase A2, group IVA (cytosolic, ca | 24.896 | 24.67 | 9.0562 | 4.8199 | 2.1812 | 10.025 |
| 16120 | ENSMUSG00000056222 | 0 | NA | NA | NA | NA | NA | Spock1 | sparc/osteonectin, cwcv and kazal-like do | 0 | 0 | 0 | 0 | 0 | 0 |
| 16121 | ENSMUSG00000056223 | 0 | NA | NA | NA | NA | NA | Spata31 | spermatogenesis associated 31 [Source:M | 0 | 0 | 0 | 0 | 0 | 0 |
| 16122 | ENSMUSG00000056228 | 236.589001 | -0.079084096 | 0.172297703 | -0.458996812 | 0.646236455 | 0.925643431 | Cars2 | cysteinyl-tRNA synthetase 2 (mitochondri | 245.75 | 237.45 | 246.53 | 269.91 | 210.49 | 209.4 |
| 16123 | ENSMUSG00000056234 | 2155.097476 | -0.321503055 | 0.11516184 | -2.791749901 | 0.005242386 | 0.068745122 | Ncoa4 | nuclear receptor coactivator 4 [Source:M( | 2102.5 | 2618.1 | 2463.3 | 1878.8 | 2063.4 | 1804.4 |
| 16124 | ENSMUSG00000056257 | 0.706696157 | 0.149852971 | 3.037999338 | 0.049326203 | 0.960659338 | NA | Gm5447 | predicted gene 5447 [Source:MGI Symbol | 0 | 0 | 2.0125 | 0 | 0 | 2.2277 |

| | | | | | | | | | | | | | | | |
|---|---|---|---|---|---|---|---|---|---|---|---|---|---|---|---|
| 16125 | ENSMUSG00000056258 | 0 | NA | NA | NA | NA | NA | Kcnq3 | potassium voltage-gated channel, subfam | 0 | 0 | 0 | 0 | 0 | 0 |
| 16126 | ENSMUSG00000056260 | 153.5645689 | -0.086309147 | 0.189091237 | -0.456441812 | 0.648072305 | 0.925643431 | Lrif1 | ligand dependent nuclear receptor intera | 162.23 | 161.38 | 150.94 | 157.13 | 151.6 | 138.12 |
| 16127 | ENSMUSG00000056267 | 88.44010552 | 0.010373591 | 0.259747269 | 0.039937247 | 0.968143156 | 0.994231936 | Cep70 | centrosomal protein 70 [Source:MGI Syml | 99.585 | 80.178 | 84.525 | 100.25 | 98.156 | 67.944 |
| 16128 | ENSMUSG00000056268 | 346.8456007 | 0.079838049 | 0.174308192 | 0.458028091 | 0.64693226 | 0.925643431 | Dennd1b | DENN/MADD domain containing 1B [Sour | 388.7 | 352.58 | 269.67 | 340.28 | 405.71 | 324.13 |
| 16129 | ENSMUSG00000056270 | 0 | NA | NA | NA | NA | NA | Prr9 | proline rich 9 [Source:MGI Symbol;Acc:M( | 0 | 0 | 0 | 0 | 0 | 0 |
| 16130 | ENSMUSG00000056271 | 0 | NA | NA | NA | NA | NA | Lman1l | lectin, mannose-binding 1 like [Source:MC | 0 | 0 | 0 | 0 | 0 | 0 |
| 16131 | ENSMUSG00000056281 | 0 | NA | NA | NA | NA | NA | Olfr1322 | olfactory receptor 1322 [Source:MGI Syml | 0 | 0 | 0 | 0 | 0 | 0 |
| 16132 | ENSMUSG00000056288 | 0.17131964 | -0.796629176 | 4.080472857 | -0.195229623 | 0.84521319 | NA | Gm11961 | predicted gene 11961 [Source:MGI Symbc | 0 | 1.0279 | 0 | 0 | 0 | 0 |
| 16133 | ENSMUSG00000056290 | 6.973914866 | -2.277852845 | 0.941095589 | -2.420426652 | 0.015502306 | 0.145412036 | Ms4a4b | membrane-spanning 4-domains, subfamil | 14.456 | 11.307 | 9.0562 | 4.8199 | 1.0906 | 1.1138 |
| 16134 | ENSMUSG00000056293 | 0 | NA | NA | NA | NA | NA | Gsdmc2 | gasdermin C2 [Source:MGI Symbol;Acc:M | 0 | 0 | 0 | 0 | 0 | 0 |
| 16135 | ENSMUSG00000056296 | 0.160662974 | 1.126924175 | 4.080472857 | 0.276174898 | 0.782413736 | NA | Synpr | synaptoporin [Source:MGI Symbol;Acc:M( | 0 | 0 | 0 | 0.964 | 0 | 0 |
| 16136 | ENSMUSG00000056300 | 3.222815361 | -0.317745747 | 1.224623684 | -0.25946399 | 0.795277263 | NA | Zfp981 | zinc finger protein 981 [Source:MGI Symb | 1.6062 | 4.1117 | 5.0312 | 1.928 | 1.0906 | 5.5692 |
| 16137 | ENSMUSG00000056305 | 408.2939421 | 0.037664572 | 0.134892658 | 0.279218843 | 0.780076884 | 0.959870443 | Usp39 | ubiquitin specific peptidase 39 [Source:M( | 379.06 | 397.8 | 432.69 | 427.04 | 415.53 | 397.64 |
| 16138 | ENSMUSG00000056306 | 0 | NA | NA | NA | NA | NA | Sertm1 | serine rich and transmembrane domain cc | 0 | 0 | 0 | 0 | 0 | 0 |
| 16139 | ENSMUSG00000056310 | 345.5376654 | -0.02657758 | 0.153066974 | -0.173633667 | 0.862153371 | 0.976334255 | Tyw1 | tRNA-yW synthesizing protein 1 homolog | 314.01 | 355.66 | 377.34 | 355.71 | 367.54 | 302.97 |
| 16140 | ENSMUSG00000056313 | 923.1495839 | -1.54639908 | 0.53249057 | -2.904087262 | 0.003683255 | 0.054140852 | Tcim | transcriptional and immune response regu | 1274.5 | 1625.1 | 1226.6 | 445.36 | 763.43 | 203.83 |
| 16141 | ENSMUSG00000056316 | 0 | NA | NA | NA | NA | NA | Gm9996 | predicted gene 9996 [Source:MGI Symbol | 0 | 0 | 0 | 0 | 0 | 0 |
| 16142 | ENSMUSG00000056328 | 0 | NA | NA | NA | NA | NA | Myh1 | myosin, heavy polypeptide 1, skeletal mus | 0 | 0 | 0 | 0 | 0 | 0 |
| 16143 | ENSMUSG00000056329 | 0 | NA | NA | NA | NA | NA | 1810010K12 | RIKEN cDNA 1810010K12 gene [Source:M( | 0 | 0 | 0 | 0 | 0 | 0 |
| 16144 | ENSMUSG00000056332 | 0 | NA | NA | NA | NA | NA | Gm16294 | predicted gene 16294 [Source:MGI Symbc | 0 | 0 | 0 | 0 | 0 | 0 |
| 16145 | ENSMUSG00000056342 | 614.6692698 | -0.037440827 | 0.147606147 | -0.253653577 | 0.799763202 | 0.962424862 | Usp34 | ubiquitin specific peptidase 34 [Source:M( | 540.49 | 640.39 | 688.27 | 640.08 | 664.19 | 514.6 |
| 16146 | ENSMUSG00000056350 | 0.181769715 | 1.126924175 | 4.080472857 | 0.276174898 | 0.782413736 | NA | Krtap13-1 | keratin associated protein 13-1 [Source:M | 0 | 0 | 0 | 0 | 1.0906 | 0 |
| 16147 | ENSMUSG00000056359 | 0.339027071 | -1.79174149 | 3.919585585 | -0.457125237 | 0.647581032 | NA | D830013O2( | RIKEN cDNA D830013O20 gene [Source:M | 0 | 1.0279 | 1.0062 | 0 | 0 | 0 |
| 16148 | ENSMUSG00000056366 | 0.17131964 | -0.796629176 | 4.080472857 | -0.195229623 | 0.84521319 | NA | Fabp3-ps1 | fatty acid binding protein 3, muscle and h | 0 | 1.0279 | 0 | 0 | 0 | 0 |
| 16149 | ENSMUSG00000056367 | 3.433903383 | 1.647614874 | 1.370449271 | 1.202244336 | 0.229268876 | NA | Actr3b | ARP3 actin-related protein 3B [Source:MG | 0 | 4.1117 | 1.0062 | 7.7118 | 1.0906 | 6.6831 |
| 16150 | ENSMUSG00000056370 | 0 | NA | NA | NA | NA | NA | Sftpb | surfactant associated protein B [Source:M | 0 | 0 | 0 | 0 | 0 | 0 |
| 16151 | ENSMUSG00000056379 | 0 | NA | NA | NA | NA | NA | Taar1 | trace amine-associated receptor 1 [Source | 0 | 0 | 0 | 0 | 0 | 0 |
| 16152 | ENSMUSG00000056380 | 0 | NA | NA | NA | NA | NA | Gpr50 | G-protein-coupled receptor 50 [Source:M | 0 | 0 | 0 | 0 | 0 | 0 |
| 16153 | ENSMUSG00000056383 | 227.2566689 | 0.061853846 | 0.169512836 | 0.364891816 | 0.715192164 | 0.944595913 | AI987944 | expressed sequence AI987944 [Source:M( | 235.31 | 211.75 | 219.36 | 204.36 | 241.03 | 251.73 |
| 16154 | ENSMUSG00000056394 | 38.77919391 | -0.077227213 | 0.354861184 | -0.217626545 | 0.8277201 | 0.969794868 | Lig1 | ligase I, DNA, ATP-dependent [Source:MG | 44.171 | 33.921 | 41.256 | 40.487 | 31.628 | 41.212 |
| 16155 | ENSMUSG00000056399 | 0.133850373 | -0.796629176 | 4.080472857 | -0.195229623 | 0.84521319 | NA | Prss34 | protease, serine 34 [Source:MGI Symbol;/ | 0.8031 | 0 | 0 | 0 | 0 | 0 |
| 16156 | ENSMUSG00000056411 | 13.20001604 | -0.529590551 | 0.592217174 | -0.89425058 | 0.371187835 | 0.807214774 | Gm12500 | predicted gene 12500 [Source:MGI Symbc | 19.274 | 13.363 | 14.087 | 11.568 | 11.997 | 8.9108 |
| 16157 | ENSMUSG00000056412 | 3.490541611 | 0.899284581 | 1.161958595 | 0.773938577 | 0.438967116 | NA | Psenen-ps | presenilin enhancer gamma secretase sub | 3.2124 | 1.0279 | 3.0187 | 4.8199 | 2.1812 | 6.6831 |
| 16158 | ENSMUSG00000056413 | 14.00012607 | -1.030928437 | 0.680625379 | -1.51467822 | 0.129853913 | 0.527586234 | Adap1 | ArfGAP with dual PH domains 1 [Source:M | 21.684 | 18.503 | 16.1 | 7.7118 | 2.1812 | 17.822 |
| 16159 | ENSMUSG00000056418 | 1.484278299 | -0.134086198 | 1.659844494 | -0.080782386 | 0.935615016 | NA | BC043934 | cDNA sequence BC043934 [Source:MGI Sy | 1.6062 | 1.0279 | 2.0125 | 0.964 | 2.1812 | 1.1138 |
| 16160 | ENSMUSG00000056423 | 0 | NA | NA | NA | NA | NA | Uts2b | urotensin 2B [Source:MGI Symbol;Acc:MC | 0 | 0 | 0 | 0 | 0 | 0 |
| 16161 | ENSMUSG00000056426 | 0 | NA | NA | NA | NA | NA | Nat3 | N-acetyltransferase 3 [Source:MGI Symbo | 0 | 0 | 0 | 0 | 0 | 0 |
| 16162 | ENSMUSG00000056427 | 1.731368637 | 0.904355405 | 1.882585287 | 0.480379514 | 0.630957559 | NA | Slit3 | slit guidance ligand 3 [Source:MGI Symbol | 1.6062 | 1.0279 | 1.0062 | 6.7478 | 0 | 0 |
| 16163 | ENSMUSG00000056429 | 4860.643944 | -0.297786692 | 0.101638204 | -2.92986967 | 0.003391042 | 0.051091699 | Tgoln1 | trans-golgi network protein [Source:MGI S | 5001.7 | 5382.2 | 5699.4 | 4715.8 | 4348.3 | 4016.5 |
| 16164 | ENSMUSG00000056436 | 0 | NA | NA | NA | NA | NA | Cyct | cytochrome c, testis [Source:MGI Symbol; | 0 | 0 | 0 | 0 | 0 | 0 |
| 16165 | ENSMUSG00000056445 | 2.03970666 | 0.018547965 | 1.55243814 | 0.011947635 | 0.990467393 | NA | Hoxaas2 | Hoxa cluster antisense RNA 2 [Source:MG | 4.0155 | 0 | 2.0125 | 2.8919 | 1.0906 | 2.2277 |
| 16166 | ENSMUSG00000056457 | 0 | NA | NA | NA | NA | NA | Prl2c3 | prolactin family 2, subfamily c, member 3 | 0 | 0 | 0 | 0 | 0 | 0 |
| 16167 | ENSMUSG00000056458 | 2.04262146 | -0.454237638 | 1.554757094 | -0.292159875 | 0.770164389 | NA | Mok | MOK protein kinase [Source:MGI Symbol; | 4.0155 | 1.0279 | 2.0125 | 1.928 | 3.2719 | 0 |
| 16168 | ENSMUSG00000056459 | 38.26569647 | -0.237874235 | 0.37373301 | -0.636481735 | 0.524462482 | 0.888334265 | Zbtb25 | zinc finger and BTB domain containing 25 | 53.808 | 38.033 | 32.2 | 40.487 | 30.537 | 34.529 |
| 16169 | ENSMUSG00000056468 | 0.305170013 | -1.657836477 | 4.041480335 | -0.410205257 | 0.681655385 | NA | 5730596B20 | RIKEN cDNA 5730596B20 gene [Source:M | 0.8031 | 1.0279 | 0 | 0 | 0 | 0 |
| 16170 | ENSMUSG00000056476 | 18.04447534 | 0.547661955 | 0.555334327 | 0.986184228 | 0.324042717 | 0.77088057 | Med12l | mediator complex subunit 12-like [Source | 9.6372 | 15.419 | 19.119 | 23.135 | 11.997 | 28.96 |
| 16171 | ENSMUSG00000056481 | 6.839010125 | -0.942808122 | 0.989931983 | -0.95239687 | 0.340895746 | 0.783129922 | Cd248 | CD248 antigen, endosialin [Source:MGI Sy | 12.85 | 7.1954 | 7.0437 | 10.604 | 0 | 3.3415 |
| 16172 | ENSMUSG00000056486 | 2.234253322 | -1.151215378 | 1.650810663 | -0.697363668 | 0.485575233 | NA | Chn1 | chimerin 1 [Source:MGI Symbol;Acc:MGI: | 3.2124 | 0 | 6.0375 | 1.928 | 0 | 2.2277 |
| 16173 | ENSMUSG00000056487 | 0.524408994 | -0.837196744 | 3.377823722 | -0.247850928 | 0.804249749 | NA | Mettl7a2 | methyltransferase like 7A2 [Source:MGI S | 0 | 2.0558 | 0 | 0 | 1.0906 | 0 |
| 16174 | ENSMUSG00000056492 | 315.5277688 | -0.766333898 | 0.191713935 | -3.997278016 | 6.41E-05 | 0.002539127 | Adgrf5 | adhesion G protein-coupled receptor F5 [! | 402.35 | 410.14 | 380.36 | 287.27 | 165.77 | 247.27 |
| 16175 | ENSMUSG00000056493 | 515.5461062 | 0.40313427 | 0.141598604 | 2.8470215 | 0.004413038 | 0.061493434 | Foxk1 | forkhead box K1 [Source:MGI Symbol;Acc | 468.21 | 378.27 | 485.01 | 608.27 | 543.13 | 610.39 |
| 16176 | ENSMUSG00000056494 | 0.842149364 | -3.107408315 | 2.805524882 | -1.107603192 | 0.268033224 | NA | Cngb3 | cyclic nucleotide gated channel beta 3 [So | 0 | 1.0279 | 4.025 | 0 | 0 | 0 |
| 16177 | ENSMUSG00000056498 | 2.066461614 | -0.35359172 | 1.629834977 | -0.2169494 | 0.828247779 | NA | Tmem154 | transmembrane protein 154 [Source:MGI | 4.8186 | 1.0279 | 1.0062 | 0 | 1.0906 | 4.4554 |
| 16178 | ENSMUSG00000056501 | 4270.3532 | -0.61110286 | 0.122551745 | -4.986488431 | 6.15E-07 | 5.33E-05 | Cebpb | CCAAT/enhancer binding protein (C/EBP), | 5355.1 | 5071.7 | 5057.4 | 3297.8 | 3974.2 | 2865.9 |
| 16179 | ENSMUSG00000056508 | 0.849373781 | -3.117833363 | 2.761285217 | -1.129123983 | 0.258845535 | NA | 1700001K19 | RIKEN cDNA 1700001K19 gene [Source:M( | 0 | 3.0838 | 2.0125 | 0 | 0 | 0 |
| 16180 | ENSMUSG00000056509 | 0 | NA | NA | NA | NA | NA | Gm9999 | predicted gene 9999 [Source:MGI Symbol | 0 | 0 | 0 | 0 | 0 | 0 |

| | | | | | | | | | | | | | | | |
|---|---|---|---|---|---|---|---|---|---|---|---|---|---|---|---|
| 16181 | ENSMUSG00000056515 | 47.67046033 | -0.85810636 | 0.356521433 | -2.406885759 | 0.016089202 | 0.149271467 | Rab31 | RAB31, member RAS oncogene family [So | 61.839 | 75.038 | 47.293 | 28.919 | 27.265 | 45.668 |
| 16182 | ENSMUSG00000056529 | 14.05018427 | -0.931010805 | 0.720674166 | -1.291860939 | 0.196405307 | 0.637356187 | Ptafr | platelet-activating factor receptor [Source | 18.471 | 22.614 | 14.087 | 5.7839 | 2.1812 | 21.163 |
| 16183 | ENSMUSG00000056531 | 6.556359766 | 0.228899107 | 0.900079345 | 0.254309921 | 0.799256136 | 0.962424862 | Ccdc18 | coiled-coil domain containing 18 [Source:I | 4.0155 | 2.0558 | 12.075 | 5.7839 | 8.7249 | 6.6831 |
| 16184 | ENSMUSG00000056536 | 203.0579334 | 0.063973215 | 0.193446031 | 0.330703169 | 0.740868708 | 0.948792385 | Pign | phosphatidylinositol glycan anchor biosyn | 183.11 | 208.67 | 204.27 | 204.36 | 249.75 | 168.19 |
| 16185 | ENSMUSG00000056537 | 669.7450634 | 0.077950307 | 0.117200979 | 0.665099454 | 0.505986884 | 0.881592135 | Rlim | ring finger protein, LIM domain interacting | 673 | 659.92 | 621.86 | 708.52 | 645.65 | 709.52 |
| 16186 | ENSMUSG00000056544 | 0 | NA | NA | NA | NA | NA | Defb21 | defensin beta 21 [Source:MGI Symbol;Acc | 0 | 0 | 0 | 0 | 0 | 0 |
| 16187 | ENSMUSG00000056553 | 0.331982614 | 0.16514309 | 3.986751476 | 0.041422971 | 0.966958701 | NA | Ptprn2 | protein tyrosine phosphatase, receptor ty | 0 | 1.0279 | 0 | 0.964 | 0 | 0 |
| 16188 | ENSMUSG00000056564 | 0 | NA | NA | NA | NA | NA | Olfr324 | olfactory receptor 324 [Source:MGI Symb | 0 | 0 | 0 | 0 | 0 | 0 |
| 16189 | ENSMUSG00000056569 | 2.200524091 | -1.718360931 | 1.580401836 | -1.087293682 | 0.27690704 | NA | Mpz | myelin protein zero [Source:MGI Symbol;A | 4.0155 | 5.1396 | 1.0062 | 1.928 | 0 | 1.1138 |
| 16190 | ENSMUSG00000056579 | 649.6532047 | 0.143755361 | 0.129632479 | 1.108945555 | 0.26745367 | 0.72293778 | Tug1 | taurine upregulated gene 1 [Source:MGI S | 653.73 | 604.42 | 592.68 | 619.84 | 659.82 | 767.44 |
| 16191 | ENSMUSG00000056586 | 0 | NA | NA | NA | NA | NA | Zar1l | zygote arrest 1-like [Source:MGI Symbol;A | 0 | 0 | 0 | 0 | 0 | 0 |
| 16192 | ENSMUSG00000056592 | 17.6826326 | -0.506242954 | 0.55282779 | -0.915733549 | 0.359806664 | 0.799452417 | Zfp658 | zinc finger protein 658 [Source:MGI Symb | 26.502 | 22.614 | 13.081 | 17.352 | 8.7249 | 17.822 |
| 16193 | ENSMUSG00000056596 | 1.557613999 | 1.184167365 | 1.764360259 | 0.671159623 | 0.502118848 | NA | Trnp1 | TMF1-regulated nuclear protein 1 [Source | 0.8031 | 2.0558 | 0 | 0.964 | 2.1812 | 3.3415 |
| 16194 | ENSMUSG00000056598 | 14.21611667 | -0.108302983 | 0.573385493 | -0.188883367 | 0.85018423 | 0.973301217 | Drc3 | dynein regulatory complex subunit 3 [Sou | 12.85 | 12.335 | 19.119 | 13.496 | 16.359 | 11.138 |
| 16195 | ENSMUSG00000056600 | 0 | NA | NA | NA | NA | NA | Olfr90 | olfactory receptor 90 [Source:MGI Symbo | 0 | 0 | 0 | 0 | 0 | 0 |
| 16196 | ENSMUSG00000056602 | 39.23572379 | -0.155206383 | 0.386936744 | -0.401115649 | 0.68833498 | 0.936908509 | Fry | FRY microtubule binding protein [Source:I | 51.399 | 45.228 | 27.169 | 37.595 | 28.356 | 45.668 |
| 16197 | ENSMUSG00000056605 | 0 | NA | NA | NA | NA | NA | Krt72 | keratin 72 [Source:MGI Symbol;Acc:MGI:2 | 0 | 0 | 0 | 0 | 0 | 0 |
| 16198 | ENSMUSG00000056608 | 545.9789122 | 0.073199338 | 0.163113811 | 0.448762353 | 0.653603099 | 0.925797231 | Chd9 | chromodomain helicase DNA binding prot | 511.58 | 614.69 | 470.92 | 625.62 | 605.29 | 447.77 |
| 16199 | ENSMUSG00000056612 | 347.7596424 | -0.055586306 | 0.137030454 | -0.405649286 | 0.68500031 | 0.935761662 | Ppp1r14b | protein phosphatase 1, regulatory inhibito | 344.53 | 359.77 | 359.23 | 337.39 | 337 | 348.63 |
| 16200 | ENSMUSG00000056617 | 0.335414862 | -1.779047242 | 4.030804179 | -0.441362855 | 0.658950329 | NA | 4931429L15I | RIKEN cDNA 4931429L15 gene [Source:MG | 0 | 0 | 2.0125 | 0 | 0 | 0 |
| 16201 | ENSMUSG00000056629 | 1085.606491 | 0.08483243 | 0.110201736 | 0.769792139 | 0.441423204 | 0.849828766 | Fkbp2 | FK506 binding protein 2 [Source:MGI Sym | 1024.8 | 1039.2 | 1097.8 | 1183.8 | 1020.8 | 1147.3 |
| 16202 | ENSMUSG00000056632 | 0 | NA | NA | NA | NA | NA | Dsg3 | desmoglein 3 [Source:MGI Symbol;Acc:MG | 0 | 0 | 0 | 0 | 0 | 0 |
| 16203 | ENSMUSG00000056643 | 161.9682209 | -0.201293765 | 0.197355217 | -1.019956644 | 0.307749023 | 0.756343859 | Chst13 | carbohydrate sulfotransferase 13 [Source | 153.39 | 162.41 | 204.27 | 146.52 | 155.96 | 149.26 |
| 16204 | ENSMUSG00000056648 | 1.370609251 | -1.535950181 | 2.095342155 | -0.733030726 | 0.463539689 | NA | Hoxb8 | homeobox B8 [Source:MGI Symbol;Acc:M | 0 | 5.1396 | 1.0062 | 0.964 | 0 | 1.1138 |
| 16205 | ENSMUSG00000056656 | 0.160662974 | 1.126924175 | 4.080472857 | 0.276174898 | 0.782413736 | NA | Apol8 | apolipoprotein L 8 [Source:MGI Symbol;Ac | 0 | 0 | 0 | 0.964 | 0 | 0 |
| 16206 | ENSMUSG00000056665 | 97.8074907 | -0.356202224 | 0.259495578 | -1.372671655 | 0.169854445 | 0.596876778 | Them6 | thioesterase superfamily member 6 [Sour | 117.25 | 109.99 | 101.63 | 62.659 | 86.159 | 109.16 |
| 16207 | ENSMUSG00000056666 | 9603.741964 | -0.893688561 | 0.317093162 | -2.818378533 | 0.004826687 | 0.065256728 | Retsat | retinol saturase (all trans retinol 13,14 red | 8705.6 | 14351 | 14404 | 8752.9 | 6292.9 | 5115.9 |
| 16208 | ENSMUSG00000056671 | 50.41062723 | 2.702817439 | 0.369731646 | 7.310213953 | 2.67E-13 | 1.24E-10 | Prelid2 | PRELI domain containing 2 [Source:MGI Sy | 12.85 | 18.503 | 9.0562 | 94.47 | 80.706 | 86.88 |
| 16209 | ENSMUSG00000056673 | 363.5981866 | -0.033558205 | 0.14571511 | -0.230300103 | 0.81785858 | 0.968599566 | Kdm5d | lysine (K)-specific demethylase 5D [Source | 336.5 | 387.53 | 380.36 | 377.88 | 374.08 | 325.24 |
| 16210 | ENSMUSG00000056679 | 0.589026695 | 0.241472922 | 3.228838524 | 0.074786311 | 0.940384733 | NA | Gpr173 | G-protein coupled receptor 173 [Source:N | 1.6062 | 0 | 0 | 1.928 | 0 | 0 |
| 16211 | ENSMUSG00000056682 | 0 | NA | NA | NA | NA | NA | 4930465K10 | RIKEN cDNA 4930465K10 gene [Source:MG | 0 | 0 | 0 | 0 | 0 | 0 |
| 16212 | ENSMUSG00000056687 | 10.12803441 | 0.175352546 | 0.666701709 | 0.263014994 | 0.792539022 | 0.961523605 | Gm11696 | predicted gene 11696 [Source:MGI Symbo | 11.243 | 8.2233 | 9.0562 | 13.496 | 8.7249 | 10.025 |
| 16213 | ENSMUSG00000056692 | 6003.65307 | 0.02372905 | 0.084062104 | 0.282279987 | 0.77772883 | 0.959559604 | Ilrun | inflammation and lipid regulator with UBA | 5784.7 | 5990.7 | 6087.8 | 5991.1 | 5913.3 | 6254.2 |
| 16214 | ENSMUSG00000056696 | 0 | NA | NA | NA | NA | NA | Olfr1350 | olfactory receptor 1350 [Source:MGI Sym | 0 | 0 | 0 | 0 | 0 | 0 |
| 16215 | ENSMUSG00000056698 | 551.7877585 | -0.419863646 | 0.166195593 | -2.526322377 | 0.011526366 | 0.119817569 | Elmod3 | ELMO/CED-12 domain containing 3 [Sourc | 615.18 | 598.25 | 682.23 | 586.1 | 458.06 | 370.91 |
| 16216 | ENSMUSG00000056699 | 0 | NA | NA | NA | NA | NA | Gm5533 | predicted gene 5533 [Source:MGI Symbol | 0 | 0 | 0 | 0 | 0 | 0 |
| 16217 | ENSMUSG00000056706 | 0 | NA | NA | NA | NA | NA | Krtap7-1 | keratin associated protein 7-1 [Source:MG | 0 | 0 | 0 | 0 | 0 | 0 |
| 16218 | ENSMUSG00000056708 | 116.4872849 | -0.712873484 | 0.299898654 | -2.377047965 | 0.017451819 | 0.157371013 | Ier5 | immediate early response 5 [Source:MGI S | 95.569 | 144.94 | 194.21 | 105.07 | 58.893 | 100.25 |
| 16219 | ENSMUSG00000056716 | 0.936952612 | -3.291460614 | 3.015666682 | -1.091453719 | 0.275073286 | NA | Gm5420 | predicted gene 5420 [Source:MGI Symbol | 5.6217 | 0 | 0 | 0 | 0 | 0 |
| 16220 | ENSMUSG00000056718 | 2.9820293 | -0.581573511 | 1.242376251 | -0.468113835 | 0.639703186 | NA | Gm13199 | predicted gene 13199 [Source:MGI Symbo | 5.6217 | 3.0838 | 2.0125 | 3.8559 | 1.0906 | 2.2277 |
| 16221 | ENSMUSG00000056724 | 147.377155 | -0.411208926 | 0.21941208 | -1.874139863 | 0.06091115 | 0.352269825 | Nbeal2 | neurobeachin-like 2 [Source:MGI Symbol; | 135.72 | 165.49 | 204.27 | 140.74 | 119.97 | 118.07 |
| 16222 | ENSMUSG00000056728 | 0 | NA | NA | NA | NA | NA | Ctsll3 | cathepsin L-like 3 [Source:MGI Symbol;Ac | 0 | 0 | 0 | 0 | 0 | 0 |
| 16223 | ENSMUSG00000056735 | 9.554408591 | 0.088033774 | 0.677765949 | 0.129888162 | 0.89665491 | 0.982798755 | A930024E05 | RIKEN cDNA A930024E05 gene [Source:M | 10.44 | 8.2233 | 9.0562 | 8.6758 | 10.906 | 10.025 |
| 16224 | ENSMUSG00000056737 | 24.44883815 | -1.521493808 | 0.490769596 | -3.100220186 | 0.001933768 | 0.033985449 | Capg | capping protein (actin filament), gelsolin-l | 39.352 | 46.256 | 23.144 | 12.532 | 9.8156 | 15.594 |
| 16225 | ENSMUSG00000056738 | 4.872118157 | 0.173390723 | 1.012325569 | 0.171279605 | 0.864003917 | NA | A730036I17I | RIKEN cDNA A730036I17 gene [Source:MG | 5.6217 | 3.0838 | 5.0312 | 6.7478 | 7.6343 | 1.1138 |
| 16226 | ENSMUSG00000056749 | 331.2432021 | -0.65189815 | 0.245263431 | -2.657950867 | 0.007861735 | 0.090824393 | Nfil3 | nuclear factor, interleukin 3, regulated [Sc | 260.21 | 409.11 | 546.39 | 325.82 | 243.21 | 202.72 |
| 16227 | ENSMUSG00000056752 | 0.167707431 | -0.796629176 | 4.080472857 | -0.195229623 | 0.84521319 | NA | Dnah9 | dynein, axonemal, heavy chain 9 [Source:I | 0 | 0 | 1.0062 | 0 | 0 | 0 |
| 16228 | ENSMUSG00000056753 | 7.880029129 | -0.230774996 | 0.765146924 | -0.301608735 | 0.762950348 | 0.95469451 | C330011M1 | RIKEN cDNA C330011M18 gene [Source:M | 11.243 | 8.2233 | 6.0375 | 9.6398 | 5.4531 | 6.6831 |
| 16229 | ENSMUSG00000056755 | 0 | NA | NA | NA | NA | NA | Grm7 | glutamate receptor, metabotropic 7 [Sour | 0 | 0 | 0 | 0 | 0 | 0 |
| 16230 | ENSMUSG00000056758 | 0 | NA | NA | NA | NA | NA | Hmga2 | high mobility group AT-hook 2 [Source:MG | 0 | 0 | 0 | 0 | 0 | 0 |
| 16231 | ENSMUSG00000056763 | 251.7993099 | 0.215670749 | 0.175277235 | 1.230454987 | 0.218526773 | 0.663735712 | Cspp1 | centrosome and spindle pole associated p | 212.02 | 237.45 | 250.55 | 316.18 | 259.57 | 235.02 |
| 16232 | ENSMUSG00000056770 | 1948.730435 | 0.063607171 | 0.098337423 | 0.646825678 | 0.517744771 | 0.887022094 | Setd3 | SET domain containing 3 [Source:MGI Sym | 1822.2 | 1885.2 | 2010.5 | 1963.6 | 2097.3 | 1913.6 |
| 16233 | ENSMUSG00000056771 | 1.666254604 | -1.041201082 | 1.69569715 | -0.614025377 | 0.539198558 | NA | Gm10010 | predicted gene 10010 [Source:MGI Symbo | 1.6062 | 3.0838 | 2.0125 | 0 | 2.1812 | 1.1138 |
| 16234 | ENSMUSG00000056772 | 0.606727817 | -2.651277273 | 2.811524265 | -0.943003518 | 0.345679103 | NA | Rps6-ps2 | ribosomal protein S6, pseudogene 2 [Sour | 1.6062 | 1.0279 | 1.0062 | 0 | 0 | 0 |
| 16235 | ENSMUSG00000056777 | 0 | NA | NA | NA | NA | NA | Rpl21-ps13 | ribosomal protein L21-ps13 [Source:MGI S | 0 | 0 | 0 | 0 | 0 | 0 |
| 16236 | ENSMUSG00000056782 | 0 | NA | NA | NA | NA | NA | Olfr667 | olfactory receptor 667 [Source:MGI Symb | 0 | 0 | 0 | 0 | 0 | 0 |

| | | | | | | | | | | | | | | | |
|---|---|---|---|---|---|---|---|---|---|---|---|---|---|---|---|
| 16237 | ENSMUSG00000056797 | 0 | NA | NA | NA | NA | NA | Gm4889 | predicted gene 4889 [Source:MGI Symbol | 0 | 0 | 0 | 0 | 0 | 0 |
| 16238 | ENSMUSG00000056812 | 10.97365382 | 0.229010335 | 0.759188256 | 0.301651577 | 0.762917686 | 0.95469451 | St8sia3 | ST8 alpha-N-acetyl-neuraminide alpha-2,8 | 20.881 | 3.0838 | 6.0375 | 11.568 | 10.906 | 13.366 |
| 16239 | ENSMUSG00000056815 | 0 | NA | NA | NA | NA | NA | Gm6812 | predicted gene 6812 [Source:MGI Symbol | 0 | 0 | 0 | 0 | 0 | 0 |
| 16240 | ENSMUSG00000056820 | 711.2958893 | -0.134963817 | 0.119197735 | -1.132268299 | 0.257521653 | 0.710872182 | Tsnax | translin-associated factor X [Source:MGI S | 729.22 | 767.85 | 736.57 | 648.76 | 743.8 | 641.57 |
| 16241 | ENSMUSG00000056821 | 0 | NA | NA | NA | NA | NA | 1700028I16Rik | RIKEN cDNA 1700028I16 gene [Source:MG | 0 | 0 | 0 | 0 | 0 | 0 |
| 16242 | ENSMUSG00000056822 | 0 | NA | NA | NA | NA | NA | Olfr166 | olfactory receptor 166 [Source:MGI Symb | 0 | 0 | 0 | 0 | 0 | 0 |
| 16243 | ENSMUSG00000056824 | 0 | NA | NA | NA | NA | NA | Zfp663 | zinc finger protein 663 [Source:MGI Symb | 0 | 0 | 0 | 0 | 0 | 0 |
| 16244 | ENSMUSG00000056829 | 0 | NA | NA | NA | NA | NA | Foxb2 | forkhead box B2 [Source:MGI Symbol;Acc | 0 | 0 | 0 | 0 | 0 | 0 |
| 16245 | ENSMUSG00000056832 | 7.178283777 | -0.519923542 | 0.822579454 | -0.632064829 | 0.527344516 | 0.888334265 | Ttc26 | tetratricopeptide repeat domain 26 [Sour | 12.047 | 9.2513 | 4.025 | 6.7478 | 6.5437 | 4.4554 |
| 16246 | ENSMUSG00000056836 | 2.137581348 | 0.499385096 | 1.535155167 | 0.325299427 | 0.744954464 | NA | Gm6851 | predicted pseudogene 6851 [Source:MGI | 3.2124 | 0 | 2.0125 | 0.964 | 2.1812 | 4.4554 |
| 16247 | ENSMUSG00000056849 | 0 | NA | NA | NA | NA | NA | Gm16415 | predicted pseudogene 16415 [Source:MG | 0 | 0 | 0 | 0 | 0 | 0 |
| 16248 | ENSMUSG00000056851 | 1851.050847 | -0.080153744 | 0.118210096 | -0.678061746 | 0.497732544 | 0.879037876 | Pcbp2 | poly(rC) binding protein 2 [Source:MGI Sy | 1704.2 | 2064.1 | 1941 | 1990.6 | 1765.7 | 1640.7 |
| 16249 | ENSMUSG00000056853 | 0 | NA | NA | NA | NA | NA | Olfr765 | olfactory receptor 765 [Source:MGI Symb | 0 | 0 | 0 | 0 | 0 | 0 |
| 16250 | ENSMUSG00000056854 | 0 | NA | NA | NA | NA | NA | Pou3f4 | POU domain, class 3, transcription factor 4 | 0 | 0 | 0 | 0 | 0 | 0 |
| 16251 | ENSMUSG00000056856 | 3.299351514 | 0.874598998 | 1.412887924 | 0.619015127 | 0.535906393 | NA | Jakmip3 | janus kinase and microtubule interacting p | 0 | 1.0279 | 6.0375 | 2.8919 | 8.7249 | 1.1138 |
| 16252 | ENSMUSG00000056858 | 0 | NA | NA | NA | NA | NA | Olfr1502 | olfactory receptor 1502 [Source:MGI Symb | 0 | 0 | 0 | 0 | 0 | 0 |
| 16253 | ENSMUSG00000056863 | 0.267700746 | -1.497481649 | 4.057001436 | -0.369110456 | 0.712045396 | NA | Olfr702 | olfactory receptor 702 [Source:MGI Symb | 1.6062 | 0 | 0 | 0 | 0 | 0 |
| 16254 | ENSMUSG00000056870 | 16.88080844 | 1.824231294 | 0.744194132 | 2.451284167 | 0.014234753 | 0.137996958 | Gulp1 | GULP, engulfment adaptor PTB domain co | 3.2124 | 6.1675 | 13.081 | 8.6758 | 53.44 | 16.708 |
| 16255 | ENSMUSG00000056877 | 0 | NA | NA | NA | NA | NA | Rps15a-ps1 | ribosomal protein S15A, pseudogene 1 [So | 0 | 0 | 0 | 0 | 0 | 0 |
| 16256 | ENSMUSG00000056880 | 0 | NA | NA | NA | NA | NA | Gadl1 | glutamate decarboxylase-like 1 [Source:M | 0 | 0 | 0 | 0 | 0 | 0 |
| 16257 | ENSMUSG00000056883 | 0 | NA | NA | NA | NA | NA | Olfr533 | olfactory receptor 533 [Source:MGI Symb | 0 | 0 | 0 | 0 | 0 | 0 |
| 16258 | ENSMUSG00000056885 | 0 | NA | NA | NA | NA | NA | Gm4559 | predicted gene 4559 [Source:MGI Symbol | 0 | 0 | 0 | 0 | 0 | 0 |
| 16259 | ENSMUSG00000056888 | 2.240960292 | -1.169648768 | 1.479593417 | -0.790520392 | 0.429223918 | NA | Glipr1 | GLI pathogenesis-related 1 (glioma) [Sour | 3.2124 | 4.1117 | 2.0125 | 1.928 | 2.1812 | 0 |
| 16260 | ENSMUSG00000056895 | 1.15294096 | -1.100079245 | 2.299017066 | -0.478499817 | 0.632294504 | NA | H2bu2 | H2B.U histone 2 [Source:MGI Symbol;Acc | 1.6062 | 3.0838 | 0 | 0 | 0 | 2.2277 |
| 16261 | ENSMUSG00000056899 | 587.0460723 | -0.233424237 | 0.134689392 | -1.733055834 | 0.083085745 | 0.425083636 | Immp2l | IMP2 inner mitochondrial membrane pep | 625.62 | 654.78 | 622.87 | 512.84 | 621.65 | 484.52 |
| 16262 | ENSMUSG00000056900 | 0.65103495 | -1.312578592 | 2.821398903 | -0.465222621 | 0.641772041 | NA | Usp13 | ubiquitin specific peptidase 13 (isopeptida | 0.8031 | 0 | 2.0125 | 0 | 1.0906 | 0 |
| 16263 | ENSMUSG00000056901 | 0 | NA | NA | NA | NA | NA | Tas2r102 | taste receptor, type 2, member 102 [Sour | 0 | 0 | 0 | 0 | 0 | 0 |
| 16264 | ENSMUSG00000056904 | 0.185640647 | 1.126924175 | 4.080472857 | 0.276174898 | 0.782413736 | NA | Gm5620 | predicted gene 5620 [Source:MGI Symbol | 0 | 0 | 0 | 0 | 0 | 1.1138 |
| 16265 | ENSMUSG00000056910 | 0 | NA | NA | NA | NA | NA | Vmn2r107 | vomeronasal 2, receptor 107 [Source:MGI | 0 | 0 | 0 | 0 | 0 | 0 |
| 16266 | ENSMUSG00000056912 | 0 | NA | NA | NA | NA | NA | 1700017N19Rik | RIKEN cDNA 1700017N19 gene [Source:M | 0 | 0 | 0 | 0 | 0 | 0 |
| 16267 | ENSMUSG00000056917 | 168.3403859 | -0.256574392 | 0.213701624 | -1.20061976 | 0.229898732 | 0.676552445 | Sipa1 | signal-induced proliferation associated ge | 170.26 | 164.47 | 215.34 | 162.91 | 119.97 | 177.1 |
| 16268 | ENSMUSG00000056919 | 69.07569298 | -0.221152764 | 0.305079593 | -0.724901857 | 0.468512242 | 0.865461183 | Cep162 | centrosomal protein 162 [Source:MGI Syn | 75.492 | 87.373 | 60.375 | 75.19 | 43.625 | 72.4 |
| 16269 | ENSMUSG00000056921 | 0 | NA | NA | NA | NA | NA | Olfr394 | olfactory receptor 394 [Source:MGI Symb | 0 | 0 | 0 | 0 | 0 | 0 |
| 16270 | ENSMUSG00000056926 | 0 | NA | NA | NA | NA | NA | Tas2r113 | taste receptor, type 2, member 113 [Sour | 0 | 0 | 0 | 0 | 0 | 0 |
| 16271 | ENSMUSG00000056938 | 1110.350401 | 0.138232857 | 0.103589853 | 1.334424678 | 0.182064717 | 0.615708221 | Acbd4 | acyl-Coenzyme A binding domain containi | 1076.2 | 999.14 | 1095.8 | 1145.2 | 1148.4 | 1197.4 |
| 16272 | ENSMUSG00000056941 | 282.7603267 | 0.0748335 | 0.155196675 | 0.482184942 | 0.629674573 | 0.92057725 | Commd7 | COMM domain containing 7 [Source:MGI | 268.24 | 266.23 | 291.81 | 283.41 | 267.2 | 319.67 |
| 16273 | ENSMUSG00000056943 | 0.17131964 | -0.796629176 | 4.080472857 | -0.195229623 | 0.84521319 | NA | Gm10015 | predicted gene 10015 [Source:MGI Symbo | 0 | 1.0279 | 0 | 0 | 0 | 0 |
| 16274 | ENSMUSG00000056946 | 0 | NA | NA | NA | NA | NA | Olfr512 | olfactory receptor 512 [Source:MGI Symb | 0 | 0 | 0 | 0 | 0 | 0 |
| 16275 | ENSMUSG00000056947 | 0 | NA | NA | NA | NA | NA | Mab21l1 | mab-21-like 1 [Source:MGI Symbol;Acc:M | 0 | 0 | 0 | 0 | 0 | 0 |
| 16276 | ENSMUSG00000056952 | 578.2522091 | -0.18568485 | 0.142281544 | -1.305052261 | 0.191875053 | 0.629099068 | Tatdn2 | TatD DNase domain containing 2 [Source: | 531.65 | 667.12 | 649.03 | 602.49 | 520.22 | 499 |
| 16277 | ENSMUSG00000056959 | 0 | NA | NA | NA | NA | NA | Olfr315 | olfactory receptor 315 [Source:MGI Symb | 0 | 0 | 0 | 0 | 0 | 0 |
| 16278 | ENSMUSG00000056961 | 0 | NA | NA | NA | NA | NA | Olfr919 | olfactory receptor 919 [Source:MGI Symb | 0 | 0 | 0 | 0 | 0 | 0 |
| 16279 | ENSMUSG00000056962 | 321.1602347 | 0.180387101 | 0.144866678 | 1.245193884 | 0.21306049 | 0.657724029 | Jmjd6 | jumonji domain containing 6 [Source:MGI | 289.92 | 314.54 | 298.85 | 317.15 | 351.18 | 355.32 |
| 16280 | ENSMUSG00000056966 | 29.26904838 | -0.519249541 | 0.411568805 | -1.261634834 | 0.207080216 | 0.652364421 | Gjc3 | gap junction protein, gamma 3 [Source:M | 30.518 | 31.865 | 41.256 | 27.955 | 25.084 | 18.935 |
| 16281 | ENSMUSG00000056972 | 0 | NA | NA | NA | NA | NA | Magel2 | MAGE family member L2 [Source:MGI Syn | 0 | 0 | 0 | 0 | 0 | 0 |
| 16282 | ENSMUSG00000056973 | 18816.19831 | -0.730719544 | 0.433243028 | -1.686627358 | 0.091675028 | 0.446623824 | Ces1d | carboxylesterase 1D [Source:MGI Symbol; | 15706 | 26894 | 27846 | 18207 | 17151 | 7093 |
| 16283 | ENSMUSG00000056978 | 428.5569617 | -1.314117208 | 0.249404005 | -5.269030101 | 1.37E-07 | 1.46E-05 | Hamp2 | hepcidin antimicrobial peptide 2 [Source:I | 414.4 | 616.75 | 803.99 | 347.03 | 188.68 | 200.49 |
| 16284 | ENSMUSG00000056987 | 0 | NA | NA | NA | NA | NA | Fam71d | family with sequence similarity 71, memb | 0 | 0 | 0 | 0 | 0 | 0 |
| 16285 | ENSMUSG00000056995 | 0 | NA | NA | NA | NA | NA | Olfr1178 | olfactory receptor 1178 [Source:MGI Symb | 0 | 0 | 0 | 0 | 0 | 0 |
| 16286 | ENSMUSG00000056999 | 2647.157685 | -0.233058412 | 0.138951881 | -1.677259858 | 0.093491679 | 0.450459278 | Ide | insulin degrading enzyme [Source:MGI Sy | 2949 | 2896.7 | 2737 | 3016.3 | 2241.2 | 2042.8 |
| 16287 | ENSMUSG00000057000 | 0 | NA | NA | NA | NA | NA | Nxf3 | nuclear RNA export factor 3 [Source:MGI | 0 | 0 | 0 | 0 | 0 | 0 |
| 16288 | ENSMUSG00000057003 | 0 | NA | NA | NA | NA | NA | Myh4 | myosin, heavy polypeptide 4, skeletal mus | 0 | 0 | 0 | 0 | 0 | 0 |
| 16289 | ENSMUSG00000057021 | 0 | NA | NA | NA | NA | NA | Vmn2r129 | vomeronasal 2, receptor 129 [Source:MGI | 0 | 0 | 0 | 0 | 0 | 0 |
| 16290 | ENSMUSG00000057036 | 26.84091208 | 0.757278273 | 0.437662892 | 1.730277544 | 0.0835807 | 0.426273357 | Gm7536 | predicted gene 7536 [Source:MGI Symbol | 20.078 | 19.53 | 20.125 | 23.135 | 42.534 | 35.643 |
| 16291 | ENSMUSG00000057037 | 3015.771094 | 0.12339249 | 0.109489032 | 1.126984935 | 0.259748851 | 0.713039927 | Cfhr1 | complement factor H-related 1 [Source:M | 2621.3 | 2899.8 | 3140.5 | 3018.2 | 3442 | 2972.8 |
| 16292 | ENSMUSG00000057047 | 0 | NA | NA | NA | NA | NA | 1700010B08 | RIKEN cDNA 1700010B08 gene [Source:M | 0 | 0 | 0 | 0 | 0 | 0 |

| | | | | | | | | | | | | | | | |
|---|---|---|---|---|---|---|---|---|---|---|---|---|---|---|---|
| 16293 | ENSMUSG00000057050 | 0 | NA | NA | NA | NA | NA | Olfr397 | olfactory receptor 397 [Source:MGI Symb | 0 | 0 | 0 | 0 | 0 | 0 |
| 16294 | ENSMUSG00000057054 | 267.3109185 | 0.215618529 | 0.187380579 | 1.150698379 | 0.249856344 | 0.702104867 | Inca1 | inhibitor of CDK, cyclin A1 interacting prot | 212.82 | 248.76 | 280.74 | 236.17 | 342.45 | 282.92 |
| 16295 | ENSMUSG00000057058 | 1.176141353 | -0.268928376 | 1.952237713 | -0.137753909 | 0.890434912 | NA | Skap1 | src family associated phosphoprotein 1 [S | 0.8031 | 2.0558 | 1.0062 | 0.964 | 0 | 2.2277 |
| 16296 | ENSMUSG00000057060 | 0.305170013 | -1.657836477 | 4.041480335 | -0.410205257 | 0.681655385 | NA | Slc35f3 | solute carrier family 35, member F3 [Sour | 0.8031 | 1.0279 | 0 | 0 | 0 | 0 |
| 16297 | ENSMUSG00000057067 | 0 | NA | NA | NA | NA | NA | Olfr297 | olfactory receptor 297 [Source:MGI Symb | 0 | 0 | 0 | 0 | 0 | 0 |
| 16298 | ENSMUSG00000057068 | 466.1943962 | 0.262772679 | 0.138238106 | 1.900870069 | 0.057319033 | 0.340239855 | Fam47e | family with sequence similarity 47, memb | 387.9 | 444.06 | 440.74 | 555.25 | 462.42 | 506.8 |
| 16299 | ENSMUSG00000057069 | 1026.049357 | 0.402522263 | 0.137789521 | 2.921283563 | 0.003485924 | 0.052141058 | Ero1b | endoplasmic reticulum oxidoreductase 1 b | 855.3 | 804.86 | 992.16 | 1345.7 | 1011 | 1147.3 |
| 16300 | ENSMUSG00000057072 | 1.008278937 | -3.39371293 | 2.386458974 | -1.42207051 | 0.155005783 | NA | Spata45 | spermatogenesis associated 45 [Source:M | 4.0155 | 1.0279 | 1.0062 | 0 | 0 | 0 |
| 16301 | ENSMUSG00000057074 | 3987.086826 | -0.413230163 | 0.185087717 | -2.232617973 | 0.025574148 | 0.206060398 | Ces1g | carboxylesterase 1G [Source:MGI Symbol; | 3746.5 | 5114.9 | 4801.8 | 3076.1 | 4650.4 | 2532.9 |
| 16302 | ENSMUSG00000057092 | 6.390509396 | -0.02511892 | 0.991264486 | -0.02534028 | 0.979783546 | 0.99820894 | Fxyd3 | FXYD domain-containing ion transport reg | 4.0155 | 14.391 | 1.0062 | 6.7478 | 3.2719 | 8.9108 |
| 16303 | ENSMUSG00000057093 | 4.406330374 | 0.173930034 | 1.062258662 | 0.163736047 | 0.869938939 | NA | Zfp607b | zinc finger protein 607B [Source:MGI Sym | 3.2124 | 7.1954 | 2.0125 | 1.928 | 7.6343 | 4.4554 |
| 16304 | ENSMUSG00000057098 | 1.568638641 | -1.893779825 | 1.886599435 | -1.003805996 | 0.315472134 | NA | Ebf1 | early B cell factor 1 [Source:MGI Symbol;A | 2.4093 | 2.0558 | 3.0187 | 1.928 | 0 | 0 |
| 16305 | ENSMUSG00000057101 | 272.651633 | 0.212206466 | 0.15333477 | 1.383942244 | 0.16637614 | 0.5925379 | Zfp180 | zinc finger protein 180 [Source:MGI Symb | 238.52 | 256.98 | 262.63 | 283.41 | 281.38 | 312.99 |
| 16306 | ENSMUSG00000057103 | 3845.092923 | 0.504835672 | 0.135709306 | 3.719978291 | 0.00019924 | 0.006292153 | Nat8f1 | N-acetyltransferase 8 (GCN5-related) fam | 2727.3 | 2860.7 | 3950.5 | 4504.7 | 4462.8 | 4564.5 |
| 16307 | ENSMUSG00000057110 | 676.1887674 | -0.439190972 | 0.133392236 | -3.292477767 | 0.000993087 | 0.020817417 | Cntrl | centriolin [Source:MGI Symbol;Acc:MGI:1 | 786.24 | 856.26 | 692.3 | 548.5 | 634.74 | 539.1 |
| 16308 | ENSMUSG00000057113 | 2040.312317 | 0.015997753 | 0.091252123 | 0.175313759 | 0.860833101 | 0.975943719 | Npm1 | nucleophosmin 1 [Source:MGI Symbol;Ac | 1982.1 | 2057.9 | 2047.7 | 2085.1 | 2079.8 | 1989.3 |
| 16309 | ENSMUSG00000057116 | 0 | NA | NA | NA | NA | NA | AF366264 | cDNA sequence AF366264 [Source:MGI Sy | 0 | 0 | 0 | 0 | 0 | 0 |
| 16310 | ENSMUSG00000057123 | 6.273476495 | -1.13005707 | 0.95182034 | -1.187258796 | 0.235125556 | 0.683240713 | Gja5 | gap junction protein, alpha 5 [Source:MGI | 14.456 | 6.1675 | 5.0312 | 0.964 | 5.4531 | 5.5692 |
| 16311 | ENSMUSG00000057130 | 206.7149599 | 0.058621448 | 0.175016284 | 0.334948535 | 0.737663909 | 0.948137969 | Txnl4a | thioredoxin-like 4A [Source:MGI Symbol;A | 223.26 | 204.56 | 179.11 | 201.47 | 222.49 | 209.4 |
| 16312 | ENSMUSG00000057132 | 14.95766436 | 0.382508434 | 0.557824055 | 0.6857152 | 0.49289272 | 0.876684739 | Rpgrip1 | retinitis pigmentosa GTPase regulator inte | 14.456 | 11.307 | 13.081 | 13.496 | 21.812 | 15.594 |
| 16313 | ENSMUSG00000057133 | 269.1572825 | -0.056807747 | 0.18290117 | -0.310592582 | 0.756110367 | 0.953085143 | Chd6 | chromodomain helicase DNA binding prot | 245.75 | 292.96 | 285.77 | 322.93 | 222.49 | 245.05 |
| 16314 | ENSMUSG00000057134 | 349.9024821 | -0.153392536 | 0.138893481 | -1.104389745 | 0.269424107 | 0.724979259 | Ado | 2-aminoethanethiol (cysteamine) dioxyge | 349.35 | 367.99 | 388.41 | 325.82 | 335.91 | 331.93 |
| 16315 | ENSMUSG00000057135 | 7.493904103 | -0.997531432 | 0.865630235 | -1.152375913 | 0.249166634 | 0.701293287 | Scimp | SLP adaptor and CSK interacting membran | 6.4248 | 18.503 | 5.0312 | 2.8919 | 6.5437 | 5.5692 |
| 16316 | ENSMUSG00000057137 | 89.90950159 | -0.08741439 | 0.243059008 | -0.359642665 | 0.719114373 | 0.9447719 | Tmem140 | transmembrane protein 140 [Source:MGI | 107.62 | 86.345 | 83.518 | 86.758 | 88.34 | 86.88 |
| 16317 | ENSMUSG00000057143 | 284.766386 | 0.247601871 | 0.175641701 | 1.40969866 | 0.158628681 | 0.579569253 | Trim12c | tripartite motif-containing 12C [Source:M | 289.92 | 248.76 | 241.5 | 261.24 | 366.45 | 300.74 |
| 16318 | ENSMUSG00000057147 | 64.36002512 | -0.622653703 | 0.293578566 | -2.120909951 | 0.033929381 | 0.246610335 | Dph6 | diphthamine biosynthesis 6 [Source:MGI S | 64.248 | 96.624 | 73.456 | 48.199 | 50.168 | 53.465 |
| 16319 | ENSMUSG00000057149 | 0 | NA | NA | NA | NA | NA | Olfr1301 | olfactory receptor 1301 [Source:MGI Symb | 0 | 0 | 0 | 0 | 0 | 0 |
| 16320 | ENSMUSG00000057156 | 95.14905879 | 0.108547435 | 0.263896888 | 0.411325178 | 0.68083411 | 0.934384556 | Homez | homeodomain leucine zipper-encoding ge | 76.295 | 82.233 | 116.72 | 114.71 | 81.796 | 99.132 |
| 16321 | ENSMUSG00000057157 | 0 | NA | NA | NA | NA | NA | Gm6054 | predicted gene 6054 [Source:MGI Symbol | 0 | 0 | 0 | 0 | 0 | 0 |
| 16322 | ENSMUSG00000057160 | 0 | NA | NA | NA | NA | NA | Gm16372 | predicted pseudogene 16372 [Source:MG | 0 | 0 | 0 | 0 | 0 | 0 |
| 16323 | ENSMUSG00000057163 | 0 | NA | NA | NA | NA | NA | Prss2 | protease, serine 2 [Source:MGI Symbol;Ac | 0 | 0 | 0 | 0 | 0 | 0 |
| 16324 | ENSMUSG00000057170 | 0 | NA | NA | NA | NA | NA | Prl3d1 | prolactin family 3, subfamily d, member 1 | 0 | 0 | 0 | 0 | 0 | 0 |
| 16325 | ENSMUSG00000057173 | 0 | NA | NA | NA | NA | NA | Rfx8 | regulatory factor X 8 [Source:MGI Symbol | 0 | 0 | 0 | 0 | 0 | 0 |
| 16326 | ENSMUSG00000057174 | 0 | NA | NA | NA | NA | NA | Krtap19-9b | keratin associated protein 19-9B [Source:I | 0 | 0 | 0 | 0 | 0 | 0 |
| 16327 | ENSMUSG00000057176 | 18.33494534 | -0.478476615 | 0.571362649 | -0.837430687 | 0.402350523 | 0.828953151 | Ccdc189 | coiled-coil domain containing 189 [Source | 26.502 | 21.586 | 16.1 | 26.027 | 11.997 | 7.7969 |
| 16328 | ENSMUSG00000057177 | 669.3483908 | -0.025660498 | 0.115993547 | -0.221223497 | 0.824918415 | 0.96943211 | Gsk3a | glycogen synthase kinase 3 alpha [Source: | 707.53 | 643.48 | 674.18 | 653.58 | 646.74 | 690.58 |
| 16329 | ENSMUSG00000057179 | 0 | NA | NA | NA | NA | NA | Olfr747 | olfactory receptor 747 [Source:MGI Symb | 0 | 0 | 0 | 0 | 0 | 0 |
| 16330 | ENSMUSG00000057181 | 389.1475266 | -0.022462118 | 0.148327848 | -0.151435607 | 0.879632103 | 0.98128671 | 5730455P16 | RIKEN cDNA 5730455P16 gene [Source:M | 439.3 | 354.63 | 381.37 | 351.85 | 408.98 | 398.76 |
| 16331 | ENSMUSG00000057182 | 1.766920353 | 0.704500416 | 1.840507134 | 0.382775162 | 0.70188648 | NA | Scn3a | sodium channel, voltage-gated, type III, al | 0 | 2.0558 | 2.0125 | 0.964 | 0 | 5.5692 |
| 16332 | ENSMUSG00000057191 | 32.76015969 | -1.834171226 | 0.518770338 | -3.535613144 | 0.00040683 | 0.010737936 | AB124611 | cDNA sequence AB124611 [Source:MGI Sy | 48.989 | 75.038 | 29.181 | 5.7839 | 14.178 | 23.391 |
| 16333 | ENSMUSG00000057193 | 81.35595677 | -0.783612289 | 0.258736656 | -3.028609474 | 0.00245682 | 0.040782715 | Slc44a2 | solute carrier family 44, member 2 [Sourc | 117.25 | 102.79 | 88.55 | 62.659 | 55.622 | 61.261 |
| 16334 | ENSMUSG00000057195 | 0 | NA | NA | NA | NA | NA | Ceacam13 | carcinoembryonic antigen-related cell adh | 0 | 0 | 0 | 0 | 0 | 0 |
| 16335 | ENSMUSG00000057203 | 0 | NA | NA | NA | NA | NA | Vmn1r234 | vomeronasal 1 receptor 234 [Source:MGI | 0 | 0 | 0 | 0 | 0 | 0 |
| 16336 | ENSMUSG00000057207 | 0 | NA | NA | NA | NA | NA | Olfr1028 | olfactory receptor 1028 [Source:MGI Symb | 0 | 0 | 0 | 0 | 0 | 0 |
| 16337 | ENSMUSG00000057215 | 0 | NA | NA | NA | NA | NA | Platr28 | pluripotency associated transcript 28 [Sou | 0 | 0 | 0 | 0 | 0 | 0 |
| 16338 | ENSMUSG00000057216 | 0 | NA | NA | NA | NA | NA | Gm10807 | predicted gene 10807 [Source:MGI Symbo | 0 | 0 | 0 | 0 | 0 | 0 |
| 16339 | ENSMUSG00000057219 | 54.05043404 | -0.639623159 | 0.299883052 | -2.132908663 | 0.03293222 | 0.242815663 | Armc7 | armadillo repeat containing 7 [Source:MG | 67.461 | 63.731 | 66.412 | 46.271 | 41.443 | 38.985 |
| 16340 | ENSMUSG00000057223 | 0 | NA | NA | NA | NA | NA | Gm6578 | predicted gene 6578 [Source:MGI Symbol | 0 | 0 | 0 | 0 | 0 | 0 |
| 16341 | ENSMUSG00000057228 | 1906.643057 | 0.284637939 | 0.122367106 | 2.326098483 | 0.020013297 | 0.17340647 | Aadat | aminoadipate aminotransferase [Source:M | 1627.9 | 1739.2 | 1790.1 | 1778.5 | 2367.7 | 2136.4 |
| 16342 | ENSMUSG00000057229 | 825.6946363 | 0.350482305 | 0.106535225 | 3.289825551 | 0.001002495 | 0.02092365 | Dmac2 | distal membrane arm assembly complex 2 | 718.78 | 715.43 | 743.61 | 928.31 | 933.57 | 914.47 |
| 16343 | ENSMUSG00000057230 | 299.0232618 | 0.001549674 | 0.148627375 | 0.010426575 | 0.991680948 | 0.99859176 | Aak1 | AP2 associated kinase 1 [Source:MGI Sym | 293.13 | 295.01 | 308.92 | 321.97 | 296.65 | 278.46 |
| 16344 | ENSMUSG00000057234 | 79.6951091 | -0.112907943 | 0.271576841 | -0.415749527 | 0.677593296 | 0.93325821 | Mettl15 | methyltransferase like 15 [Source:MGI Sy | 76.295 | 90.457 | 81.506 | 56.875 | 86.159 | 86.88 |
| 16345 | ENSMUSG00000057236 | 1736.384451 | 0.229277322 | 0.103678821 | 2.211419061 | 0.027006831 | 0.212875643 | Rbbp4 | retinoblastoma binding protein 4, chroma | 1460 | 1652.9 | 1684.5 | 1885.5 | 1799.5 | 1935.9 |
| 16346 | ENSMUSG00000057240 | 0 | NA | NA | NA | NA | NA | Ms4a13 | membrane-spanning 4-domains, subfamil | 0 | 0 | 0 | 0 | 0 | 0 |
| 16347 | ENSMUSG00000057244 | 0.319491021 | 0.16514334 | 4.044653141 | 0.040830038 | 0.967431393 | NA | Gm6139 | predicted gene 6139 [Source:MGI Symbol | 0.8031 | 0 | 0 | 0 | 0 | 1.1138 |
| 16348 | ENSMUSG00000057246 | 0.339027071 | -1.79174149 | 3.919585585 | -0.457125237 | 0.647581032 | NA | BC051142 | cDNA sequence BC051142 [Source:MGI Sy | 0 | 1.0279 | 1.0062 | 0 | 0 | 0 |

| | | | | | | | | | | | | | | | |
|---|---|---|---|---|---|---|---|---|---|---|---|---|---|---|---|
| 16349 | ENSMUSG00000057262 | 1.35097035 | -1.550751123 | 2.044013871 | -0.758679354 | 0.44804439 | NA | Rpl15-ps6 | ribosomal protein L15, pseudogene 6 [Sou | 4.0155 | 0 | 2.0125 | 0.964 | 0 | 1.1138 |
| 16350 | ENSMUSG00000057265 | 11.36221598 | -0.40037809 | 0.670927839 | -0.596752834 | 0.550672412 | 0.895703112 | Bbof1 | basal body orientation factor 1 [Source:M | 6.4248 | 17.475 | 15.094 | 11.568 | 9.8156 | 7.7969 |
| 16351 | ENSMUSG00000057270 | 0 | NA | NA | NA | NA | NA | Olfr1501 | olfactory receptor 1501 [Source:MGI Symb | 0 | 0 | 0 | 0 | 0 | 0 |
| 16352 | ENSMUSG00000057278 | 175.9380285 | 0.108495852 | 0.184125632 | 0.589249043 | 0.555694222 | 0.897628185 | Snrpg | small nuclear ribonucleoprotein polypepti | 157.41 | 182.97 | 168.04 | 186.05 | 196.31 | 164.85 |
| 16353 | ENSMUSG00000057280 | 0 | NA | NA | NA | NA | NA | Musk | muscle, skeletal, receptor tyrosine kinase | 0 | 0 | 0 | 0 | 0 | 0 |
| 16354 | ENSMUSG00000057286 | 15.68240147 | -1.311687694 | 0.649557831 | -2.019354753 | 0.043450361 | 0.287364435 | St6galnac2 | ST6 (alpha-N-acetyl-neuraminyl-2,3-beta-g | 14.456 | 23.642 | 29.181 | 13.496 | 2.1812 | 11.138 |
| 16355 | ENSMUSG00000057290 | 0 | NA | NA | NA | NA | NA | Gm15732 | predicted gene 15732 [Source:MGI Symbo | 0 | 0 | 0 | 0 | 0 | 0 |
| 16356 | ENSMUSG00000057315 | 136.9649196 | -0.530730881 | 0.209437848 | -2.534073405 | 0.011274511 | 0.117998936 | Arhgap24 | Rho GTPase activating protein 24 [Source: | 159.01 | 158.3 | 168.04 | 100.25 | 102.52 | 133.66 |
| 16357 | ENSMUSG00000057321 | 0 | NA | NA | NA | NA | NA | Usp17ld | ubiquitin specific peptidase 17-like D [Sou | 0 | 0 | 0 | 0 | 0 | 0 |
| 16358 | ENSMUSG00000057322 | 1217.871983 | 0.091030891 | 0.117899274 | 0.772107309 | 0.440050873 | 0.848765947 | Rpl38 | ribosomal protein L38 [Source:MGI Symbo | 1059.3 | 1315.7 | 1164.2 | 1200.2 | 1235.7 | 1332.2 |
| 16359 | ENSMUSG00000057329 | 14.86474017 | -0.191611069 | 0.571033467 | -0.335551381 | 0.737209192 | 0.948137969 | Bcl2 | B cell leukemia/lymphoma 2 [Source:MGI | 16.865 | 16.447 | 14.087 | 8.6758 | 14.178 | 18.935 |
| 16360 | ENSMUSG00000057335 | 108.8506552 | 0.286283137 | 0.231113228 | 1.238713767 | 0.215451518 | 0.660231093 | Cep170 | centrosomal protein 170 [Source:MGI Syn | 101.19 | 100.74 | 92.575 | 143.63 | 104.7 | 110.27 |
| 16361 | ENSMUSG00000057337 | 3.372639526 | -2.482447251 | 1.363526572 | -1.820607901 | 0.068666481 | NA | Chst3 | carbohydrate sulfotransferase 3 [Source:M | 8.031 | 6.1675 | 3.0187 | 1.928 | 1.0906 | 0 |
| 16362 | ENSMUSG00000057342 | 1056.594652 | -0.18832055 | 0.134578191 | -1.399339283 | 0.161711265 | 0.584894158 | Sphk2 | sphingosine kinase 2 [Source:MGI Symbol | 1096.2 | 950.82 | 1329.2 | 962.05 | 1011 | 990.21 |
| 16363 | ENSMUSG00000057346 | 1134.041626 | 0.459949713 | 0.155289282 | 2.961889626 | 0.003057574 | 0.047349835 | Apol9a | apolipoprotein L 9a [Source:MGI Symbol;A | 1044.8 | 758.6 | 1059.6 | 1094.1 | 1458.2 | 1389 |
| 16364 | ENSMUSG00000057349 | 0 | NA | NA | NA | NA | NA | Olfr948 | olfactory receptor 948 [Source:MGI Symb | 0 | 0 | 0 | 0 | 0 | 0 |
| 16365 | ENSMUSG00000057359 | 5.929490714 | 0.723265836 | 1.014983259 | 0.712588932 | 0.476100163 | 0.869483933 | Gm17494 | predicted gene, 17494 [Source:MGI Symb | 2.4093 | 5.1396 | 6.0375 | 15.424 | 5.4531 | 1.1138 |
| 16366 | ENSMUSG00000057363 | 183.6540228 | 0.001760414 | 0.197643445 | 0.00890702 | 0.99289332 | 0.99859176 | Uxs1 | UDP-glucuronate decarboxylase 1 [Source | 150.18 | 210.72 | 190.18 | 163.88 | 187.59 | 199.38 |
| 16367 | ENSMUSG00000057367 | 448.5706173 | 0.104323194 | 0.137282867 | 0.759914156 | 0.447305899 | 0.85365253 | Birc2 | baculoviral IAP repeat-containing 2 [Sourc | 477.04 | 429.67 | 389.42 | 455.96 | 453.7 | 485.64 |
| 16368 | ENSMUSG00000057370 | 0.401551119 | -2.072982726 | 4.008251979 | -0.517178744 | 0.605031388 | NA | Gm8724 | predicted pseudogene 8724 [Source:MGI | 2.4093 | 0 | 0 | 0 | 0 | 0 |
| 16369 | ENSMUSG00000057375 | 606.5985854 | 0.102101017 | 0.130639377 | 0.781548561 | 0.434479927 | 0.845264432 | Yipf1 | Yip1 domain family, member 1 [Source:M | 623.21 | 534.52 | 596.7 | 595.74 | 594.39 | 695.04 |
| 16370 | ENSMUSG00000057378 | 1.470395976 | -0.172330405 | 1.775199961 | -0.097076616 | 0.922665551 | NA | Ryr3 | ryanodine receptor 3 [Source:MGI Symbo | 1.6062 | 3.0838 | 0 | 1.928 | 1.0906 | 1.1138 |
| 16371 | ENSMUSG00000057381 | 0 | NA | NA | NA | NA | NA | Tas2r123 | taste receptor, type 2, member 123 [Sour | 0 | 0 | 0 | 0 | 0 | 0 |
| 16372 | ENSMUSG00000057388 | 665.6014938 | 0.072047853 | 0.11967829 | 0.602012724 | 0.547165668 | 0.895518681 | Mrpl18 | mitochondrial ribosomal protein L18 [Sou | 613.57 | 700.01 | 633.93 | 669 | 714.35 | 662.74 |
| 16373 | ENSMUSG00000057396 | 76.37764959 | 0.011531187 | 0.271619018 | 0.042453534 | 0.966137153 | 0.993855397 | Zfp759 | zinc finger protein 759 [Source:MGI Symb | 81.916 | 57.563 | 88.55 | 78.082 | 73.071 | 79.083 |
| 16374 | ENSMUSG00000057400 | 45196.7764 | -0.093136255 | 0.130042781 | -0.716197041 | 0.47386969 | 0.868952133 | Ces1c | carboxylesterase 1C [Source:MGI Symbol; | 38626 | 47279 | 54063 | 49128 | 41154 | 40930 |
| 16375 | ENSMUSG00000057402 | 0 | NA | NA | NA | NA | NA | Cldn34b2 | claudin 34B2 [Source:MGI Symbol;Acc:MG | 0 | 0 | 0 | 0 | 0 | 0 |
| 16376 | ENSMUSG00000057406 | 305.6968981 | 0.04189479 | 0.155572465 | 0.269294381 | 0.787703156 | 0.960870081 | Nsd2 | nuclear receptor binding SET domain prot | 276.27 | 313.51 | 314.95 | 341.25 | 309.74 | 278.46 |
| 16377 | ENSMUSG00000057409 | 84.87520397 | 0.068684342 | 0.245794879 | 0.279437646 | 0.779908984 | 0.959870443 | Zfp53 | zinc finger protein 53 [Source:MGI Symbo | 78.704 | 87.373 | 82.512 | 81.938 | 80.706 | 98.018 |
| 16378 | ENSMUSG00000057411 | 657.7159005 | -0.005179441 | 0.117056924 | -0.044247195 | 0.964707363 | 0.993480943 | Antkmt | adenine nucleotide translocase lysine met | 676.21 | 615.72 | 684.25 | 640.08 | 675.09 | 654.94 |
| 16379 | ENSMUSG00000057417 | 0 | NA | NA | NA | NA | NA | Dcpp3 | demilune cell and parotid protein 3 [Sourc | 0 | 0 | 0 | 0 | 0 | 0 |
| 16380 | ENSMUSG00000057421 | 580.0766205 | 0.098360561 | 0.124572966 | 0.789581914 | 0.429771974 | 0.84307112 | Las1l | LAS1-like (S. cerevisiae) [Source:MGI Syml | 520.41 | 560.22 | 600.73 | 571.64 | 627.11 | 600.36 |
| 16381 | ENSMUSG00000057424 | 0 | NA | NA | NA | NA | NA | Olfr934 | olfactory receptor 934 [Source:MGI Symb | 0 | 0 | 0 | 0 | 0 | 0 |
| 16382 | ENSMUSG00000057425 | 43.78620298 | -0.461102059 | 0.375206071 | -1.228930165 | 0.219097991 | 0.664506885 | Ugt2b37 | UDP glucuronosyltransferase 2 family, pol | 41.761 | 42.145 | 68.425 | 35.667 | 47.987 | 26.732 |
| 16383 | ENSMUSG00000057439 | 0 | NA | NA | NA | NA | NA | Kir3dl2 | killer cell immunoglobulin-like receptor, th | 0 | 0 | 0 | 0 | 0 | 0 |
| 16384 | ENSMUSG00000057440 | 93.53081019 | 0.27795047 | 0.269057615 | 1.033051863 | 0.301579629 | 0.752246851 | Mpp7 | membrane protein, palmitoylated 7 (MAG | 97.175 | 97.652 | 58.362 | 88.686 | 110.15 | 109.16 |
| 16385 | ENSMUSG00000057443 | 0 | NA | NA | NA | NA | NA | Olfr138 | olfactory receptor 138 [Source:MGI Symb | 0 | 0 | 0 | 0 | 0 | 0 |
| 16386 | ENSMUSG00000057444 | 0 | NA | NA | NA | NA | NA | Olfr915 | olfactory receptor 915 [Source:MGI Symb | 0 | 0 | 0 | 0 | 0 | 0 |
| 16387 | ENSMUSG00000057446 | 0 | NA | NA | NA | NA | NA | Cts8 | cathepsin 8 [Source:MGI Symbol;Acc:MGI | 0 | 0 | 0 | 0 | 0 | 0 |
| 16388 | ENSMUSG00000057447 | 0 | NA | NA | NA | NA | NA | Olfr1205 | olfactory receptor 1205 [Source:MGI Syml | 0 | 0 | 0 | 0 | 0 | 0 |
| 16389 | ENSMUSG00000057454 | 0 | NA | NA | NA | NA | NA | Lypd3 | Ly6/Plaur domain containing 3 [Source:MG | 0 | 0 | 0 | 0 | 0 | 0 |
| 16390 | ENSMUSG00000057455 | 0 | NA | NA | NA | NA | NA | Rit2 | Ras-like without CAAX 2 [Source:MGI Sym | 0 | 0 | 0 | 0 | 0 | 0 |
| 16391 | ENSMUSG00000057457 | 0 | NA | NA | NA | NA | NA | Phex | phosphate regulating endopeptidase hom | 0 | 0 | 0 | 0 | 0 | 0 |
| 16392 | ENSMUSG00000057461 | 0 | NA | NA | NA | NA | NA | Olfr591 | olfactory receptor 591 [Source:MGI Symb | 0 | 0 | 0 | 0 | 0 | 0 |
| 16393 | ENSMUSG00000057464 | 0 | NA | NA | NA | NA | NA | Olfr1415 | olfactory receptor 1415 [Source:MGI Syml | 0 | 0 | 0 | 0 | 0 | 0 |
| 16394 | ENSMUSG00000057465 | 443.8742547 | -1.448876622 | 0.547833891 | -2.644737108 | 0.008175441 | 0.093256761 | Saa2 | serum amyloid A 2 [Source:MGI Symbol;A | 836.03 | 814.11 | 298.85 | 163.88 | 202.86 | 347.52 |
| 16395 | ENSMUSG00000057469 | 289.7363705 | 0.220986737 | 0.174621601 | 1.265517759 | 0.205685786 | 0.650907725 | E2f6 | E2F transcription factor 6 [Source:MGI Syr | 270.65 | 244.64 | 286.78 | 249.67 | 340.27 | 346.41 |
| 16396 | ENSMUSG00000057497 | 1039.065857 | 0.208462463 | 0.110238244 | 1.891017642 | 0.058621989 | 0.343700224 | Fam136a | family with sequence similarity 136, meml | 1003.1 | 931.29 | 956.94 | 1030.5 | 1191 | 1121.6 |
| 16397 | ENSMUSG00000057499 | 1.19046236 | 0.59832885 | 1.969476722 | 0.303800925 | 0.761279562 | NA | Gm6177 | predicted gene 6177 [Source:MGI Symbol | 0.8031 | 1.0279 | 1.0062 | 0.964 | 0 | 3.3415 |
| 16398 | ENSMUSG00000057503 | 0 | NA | NA | NA | NA | NA | Olfr1459 | olfactory receptor 1459 [Source:MGI Syml | 0 | 0 | 0 | 0 | 0 | 0 |
| 16399 | ENSMUSG00000057506 | 235.2263377 | 0.139981017 | 0.168051044 | 0.832967257 | 0.404863196 | 0.829488307 | Bloc1s2 | biogenesis of lysosomal organelles comple | 217.64 | 247.73 | 206.28 | 247.74 | 225.76 | 266.21 |
| 16400 | ENSMUSG00000057513 | 0 | NA | NA | NA | NA | NA | Vmn1r177 | vomeronasal 1 receptor 177 [Source:MGI | 0 | 0 | 0 | 0 | 0 | 0 |
| 16401 | ENSMUSG00000057522 | 2163.887036 | 0.159529084 | 0.102482163 | 1.556652193 | 0.119553085 | 0.505914517 | Spop | speckle-type BTB/POZ protein [Source:MG | 1980.5 | 2089.8 | 2062.8 | 2157.4 | 2182.3 | 2510.6 |
| 16402 | ENSMUSG00000057530 | 4413.353812 | -0.075932202 | 0.099948131 | -0.759716077 | 0.447424317 | 0.85368993 | Ece1 | endothelin converting enzyme 1 [Source:I | 4299 | 4927.8 | 4363.1 | 4578.9 | 4245.8 | 4065.5 |
| 16403 | ENSMUSG00000057531 | 613.2555545 | -0.012323805 | 0.124020152 | -0.099369375 | 0.920844997 | 0.986685804 | Dtnbp1 | dystrobrevin binding protein 1 [Source:MG | 583.05 | 670.2 | 594.69 | 588.99 | 596.57 | 646.03 |
| 16404 | ENSMUSG00000057534 | 0 | NA | NA | NA | NA | NA | Elobl | elongin B-like [Source:MGI Symbol;Acc:M | 0 | 0 | 0 | 0 | 0 | 0 |

| | | | | | | | | | | | | | | | |
|---|---|---|---|---|---|---|---|---|---|---|---|---|---|---|---|
| 16405 | ENSMUSG00000057540 | 0 | NA | NA | NA | NA | NA | Olfr310 | olfactory receptor 310 [Source:MGI Symb | 0 | 0 | 0 | 0 | 0 | 0 |
| 16406 | ENSMUSG00000057541 | 246.7973482 | -0.206299779 | 0.194435875 | -1.061017053 | 0.288682151 | 0.73951397 | Pus7 | pseudouridylate synthase 7 [Source:MGI S | 273.05 | 281.65 | 238.48 | 223.64 | 285.74 | 178.22 |
| 16407 | ENSMUSG00000057551 | 216.164117 | 0.109291754 | 0.177710091 | 0.615000269 | 0.538554543 | 0.891577636 | Zfp317 | zinc finger protein 317 [Source:MGI Symb | 211.22 | 211.75 | 201.25 | 253.53 | 188.68 | 230.57 |
| 16408 | ENSMUSG00000057554 | 3043.770011 | 0.156639727 | 0.122790228 | 1.275669319 | 0.202072438 | 0.646631803 | Lgals8 | lectin, galactose binding, soluble 8 [Sourc | 3178.7 | 2650 | 2805.4 | 2730 | 3426.7 | 3471.9 |
| 16409 | ENSMUSG00000057561 | 707.0302894 | -0.311685566 | 0.142516103 | -2.187019991 | 0.028741065 | 0.222305219 | Eif1a | eukaryotic translation initiation factor 1A | 713.96 | 934.38 | 701.35 | 593.81 | 630.38 | 668.31 |
| 16410 | ENSMUSG00000057564 | 0 | NA | NA | NA | NA | NA | Olfr1508 | olfactory receptor 1508 [Source:MGI Syml | 0 | 0 | 0 | 0 | 0 | 0 |
| 16411 | ENSMUSG00000057572 | 462.0025924 | 0.362039103 | 0.128894069 | 2.808811197 | 0.00497248 | 0.066532792 | Zbtb8os | zinc finger and BTB domain containing 8 o | 382.28 | 426.59 | 404.51 | 499.34 | 521.32 | 537.99 |
| 16412 | ENSMUSG00000057577 | 0 | NA | NA | NA | NA | NA | Gm10025 | predicted gene 10025 [Source:MGI Symbo | 0 | 0 | 0 | 0 | 0 | 0 |
| 16413 | ENSMUSG00000057580 | 3.047939505 | 0.603126225 | 1.262756689 | 0.477626632 | 0.632915972 | NA | Cox7c-ps1 | cytochrome c oxidase subunit 7C, pseudo | 3.2124 | 2.0558 | 2.0125 | 7.7118 | 2.1812 | 1.1138 |
| 16414 | ENSMUSG00000057592 | 0 | NA | NA | NA | NA | NA | Vmn1r48 | vomeronasal 1 receptor 48 [Source:MGI S | 0 | 0 | 0 | 0 | 0 | 0 |
| 16415 | ENSMUSG00000057594 | 113.6608318 | 0.113960482 | 0.221918562 | 0.513523883 | 0.607584909 | 0.916376001 | Arl16 | ADP-ribosylation factor-like 16 [Source:M | 114.04 | 93.541 | 119.74 | 115.68 | 128.69 | 110.27 |
| 16416 | ENSMUSG00000057596 | 142.7945341 | 0.441512888 | 0.236926023 | 1.86350525 | 0.062391215 | 0.356831368 | Trim30d | tripartite motif-containing 30D [Source:M | 128.5 | 105.88 | 128.8 | 176.41 | 116.7 | 200.49 |
| 16417 | ENSMUSG00000057604 | 5.860661321 | -1.03476104 | 1.002394039 | -1.032289698 | 0.301936426 | 0.752248241 | Lmcd1 | LIM and cysteine-rich domains 1 [Source:I | 6.4248 | 6.1675 | 11.069 | 4.8199 | 0 | 6.6831 |
| 16418 | ENSMUSG00000057605 | 3.169092821 | -1.431331411 | 1.262290799 | -1.133915744 | 0.256829892 | NA | Gm6807 | predicted gene 6807 [Source:MGI Symbol | 4.8186 | 2.0558 | 7.0437 | 2.8919 | 1.0906 | 1.1138 |
| 16419 | ENSMUSG00000057606 | 0 | NA | NA | NA | NA | NA | Colq | collagen-like tail subunit (single strand of | 0 | 0 | 0 | 0 | 0 | 0 |
| 16420 | ENSMUSG00000057609 | 0 | NA | NA | NA | NA | NA | Lce1a1 | late cornified envelope 1A1 [Source:MGI S | 0 | 0 | 0 | 0 | 0 | 0 |
| 16421 | ENSMUSG00000057614 | 26.16206024 | 0.189933001 | 0.549331922 | 0.345752709 | 0.729528568 | 0.946534338 | Gnai1 | guanine nucleotide binding protein (G pro | 18.471 | 24.67 | 30.187 | 13.496 | 53.44 | 16.708 |
| 16422 | ENSMUSG00000057615 | 0 | NA | NA | NA | NA | NA | Ldoc1 | regulator of NFKB signaling [Source:MGI S | 0 | 0 | 0 | 0 | 0 | 0 |
| 16423 | ENSMUSG00000057637 | 634.0751896 | 0.049005384 | 0.12453446 | 0.393508628 | 0.693943853 | 0.939133502 | Prdm2 | PR domain containing 2, with ZNF domain | 590.28 | 691.79 | 588.65 | 663.22 | 645.65 | 624.87 |
| 16424 | ENSMUSG00000057649 | 159.6045585 | 0.143180643 | 0.19168112 | 0.746973115 | 0.45507979 | 0.857337687 | Brd9 | bromodomain containing 9 [Source:MGI S | 150.18 | 160.36 | 144.9 | 187.98 | 151.6 | 162.62 |
| 16425 | ENSMUSG00000057650 | 0 | NA | NA | NA | NA | NA | Krtap19-2 | keratin associated protein 19-2 [Source:M | 0 | 0 | 0 | 0 | 0 | 0 |
| 16426 | ENSMUSG00000057654 | 0 | NA | NA | NA | NA | NA | Olfr328 | olfactory receptor 328 [Source:MGI Symb | 0 | 0 | 0 | 0 | 0 | 0 |
| 16427 | ENSMUSG00000057657 | 2.782217116 | 0.060641526 | 1.308068684 | 0.046359588 | 0.963023646 | NA | Rps18-ps3 | ribosomal protein S18, pseudogene 3 [Sou | 4.0155 | 3.0838 | 1.0062 | 1.928 | 1.0906 | 5.5692 |
| 16428 | ENSMUSG00000057666 | 29865.15807 | -0.170825651 | 0.106110091 | -1.609890719 | 0.107421715 | 0.48315751 | Gapdh | glyceraldehyde-3-phosphate dehydrogena | 36279 | 28907 | 29706 | 28567 | 28290 | 27443 |
| 16429 | ENSMUSG00000057667 | 114.1851599 | -0.046472753 | 0.243717461 | -0.190682905 | 0.848774033 | 0.972888368 | Bloc1s3 | biogenesis of lysosomal organelles comple | 150.18 | 96.624 | 100.62 | 119.53 | 112.33 | 105.82 |
| 16430 | ENSMUSG00000057672 | 256.9609289 | -0.084029743 | 0.187141121 | -0.449018057 | 0.653418631 | 0.925756399 | Pkn1 | protein kinase N1 [Source:MGI Symbol;Ac | 248.16 | 275.48 | 270.68 | 314.26 | 213.76 | 219.43 |
| 16431 | ENSMUSG00000057674 | 0 | NA | NA | NA | NA | NA | Gm11938 | predicted gene 11938 [Source:MGI Symbo | 0 | 0 | 0 | 0 | 0 | 0 |
| 16432 | ENSMUSG00000057685 | 0 | NA | NA | NA | NA | NA | Gm8526 | predicted gene 8526 [Source:MGI Symbol | 0 | 0 | 0 | 0 | 0 | 0 |
| 16433 | ENSMUSG00000057691 | 458.6909245 | -0.239212437 | 0.128177018 | -1.866266205 | 0.062004131 | 0.355884025 | Zfp746 | zinc finger protein 746 [Source:MGI Symb | 520.41 | 490.32 | 478.97 | 432.83 | 427.52 | 402.1 |
| 16434 | ENSMUSG00000057696 | 1.111917132 | 0.481281919 | 2.335886133 | 0.206038262 | 0.836761024 | NA | Rpl30-ps1 | ribosomal protein L30, pseudogene 1 [Sou | 0.8031 | 0 | 2.0125 | 3.8559 | 0 | 0 |
| 16435 | ENSMUSG00000057697 | 1.50378772 | 1.125726338 | 1.907977681 | 0.590010223 | 0.555183796 | NA | Vmn1r53 | vomeronasal 1 receptor 53 [Source:MGI S | 0.8031 | 2.0558 | 0 | 2.8919 | 3.2719 | 0 |
| 16436 | ENSMUSG00000057699 | 0 | NA | NA | NA | NA | NA | Tas2r131 | taste receptor, type 2, member 131 [Sour | 0 | 0 | 0 | 0 | 0 | 0 |
| 16437 | ENSMUSG00000057706 | 11.08930825 | -0.634379389 | 0.707813466 | -0.896252218 | 0.370118072 | 0.806719078 | Mex3b | mex3 RNA binding family member B [Sour | 7.2279 | 10.279 | 23.144 | 11.568 | 7.6343 | 6.6831 |
| 16438 | ENSMUSG00000057710 | 0 | NA | NA | NA | NA | NA | 9630041A04 | RIKEN cDNA 9630041A04 gene [Source:M | 0 | 0 | 0 | 0 | 0 | 0 |
| 16439 | ENSMUSG00000057715 | 0 | NA | NA | NA | NA | NA | A830018L16 | RIKEN cDNA A830018L16 gene [Source:M | 0 | 0 | 0 | 0 | 0 | 0 |
| 16440 | ENSMUSG00000057716 | 0.133850373 | -0.796629176 | 4.080472857 | -0.195229623 | 0.84521319 | NA | Tmem178b | transmembrane protein 178B [Source:MG | 0.8031 | 0 | 0 | 0 | 0 | 0 |
| 16441 | ENSMUSG00000057719 | 2.811247109 | -0.320154177 | 1.302781222 | -0.245746693 | 0.805878325 | NA | Sh3rf2 | SH3 domain containing ring finger 2 [Sour | 3.2124 | 4.1117 | 2.0125 | 0.964 | 5.4531 | 1.1138 |
| 16442 | ENSMUSG00000057722 | 117.0623001 | -1.13926976 | 0.372587232 | -3.057726251 | 0.002230232 | 0.037728745 | Lepr | leptin receptor [Source:MGI Symbol;Acc:M | 136.53 | 195.3 | 151.94 | 123.39 | 27.265 | 67.944 |
| 16443 | ENSMUSG00000057723 | 0 | NA | NA | NA | NA | NA | Krt33b | keratin 33B [Source:MGI Symbol;Acc:MGI | 0 | 0 | 0 | 0 | 0 | 0 |
| 16444 | ENSMUSG00000057726 | 0 | NA | NA | NA | NA | NA | Serpinb9g | serine (or cysteine) peptidase inhibitor, cl | 0 | 0 | 0 | 0 | 0 | 0 |
| 16445 | ENSMUSG00000057729 | 4.897884835 | 0.47636585 | 1.109504463 | 0.429350098 | 0.667668464 | NA | Prtn3 | proteinase 3 [Source:MGI Symbol;Acc:MG | 8.031 | 3.0838 | 1.0062 | 1.928 | 11.997 | 3.3415 |
| 16446 | ENSMUSG00000057735 | 0 | NA | NA | NA | NA | NA | Olfr1185-ps1 | olfactory receptor 1185, pseudogene 1 [S | 0 | 0 | 0 | 0 | 0 | 0 |
| 16447 | ENSMUSG00000057738 | 1677.691315 | -0.209462156 | 0.118104641 | -1.773530273 | 0.07614088 | 0.403877794 | Sptan1 | spectrin alpha, non-erythrocytic 1 [Source | 1763.6 | 1902.7 | 1732.8 | 1748.7 | 1560.7 | 1357.8 |
| 16448 | ENSMUSG00000057751 | 13.200098 | -0.141038253 | 0.629247802 | -0.224137856 | 0.822650037 | 0.969219124 | Megf6 | multiple EGF-like-domains 6 [Source:MGI | 12.047 | 19.53 | 10.062 | 15.424 | 6.5437 | 15.594 |
| 16449 | ENSMUSG00000057754 | 0 | NA | NA | NA | NA | NA | Tas2r106 | taste receptor, type 2, member 106 [Sour | 0 | 0 | 0 | 0 | 0 | 0 |
| 16450 | ENSMUSG00000057761 | 0 | NA | NA | NA | NA | NA | Olfr1022 | olfactory receptor 1022 [Source:MGI Syml | 0 | 0 | 0 | 0 | 0 | 0 |
| 16451 | ENSMUSG00000057762 | 0 | NA | NA | NA | NA | NA | Gm6169 | predicted gene 6169 [Source:MGI Symbol | 0 | 0 | 0 | 0 | 0 | 0 |
| 16452 | ENSMUSG00000057766 | 2.853775006 | -1.589676646 | 1.56520011 | -1.01563796 | 0.309801821 | NA | Ankrd29 | ankyrin repeat domain 29 [Source:MGI Sy | 5.6217 | 7.1954 | 0 | 0.964 | 0 | 3.3415 |
| 16453 | ENSMUSG00000057770 | 0 | NA | NA | NA | NA | NA | Olfr668 | olfactory receptor 668 [Source:MGI Symb | 0 | 0 | 0 | 0 | 0 | 0 |
| 16454 | ENSMUSG00000057777 | 9.452022142 | -0.942813405 | 0.781256565 | -1.206791018 | 0.227512636 | 0.673700979 | Mab21l2 | mab-21-like 2 [Source:MGI Symbol;Acc:M | 20.078 | 6.1675 | 11.069 | 10.604 | 5.4531 | 3.3415 |
| 16455 | ENSMUSG00000057778 | 300.2976101 | 0.144803896 | 0.150604175 | 0.961486596 | 0.336307562 | 0.778173248 | Cyb5d2 | cytochrome b5 domain containing 2 [Sour | 278.68 | 306.32 | 270.68 | 298.83 | 307.55 | 339.72 |
| 16456 | ENSMUSG00000057788 | 255.1976329 | 0.144789892 | 0.15442154 | 0.937627558 | 0.348435844 | 0.789711685 | Ddx49 | DEAD box helicase 49 [Source:MGI Symbo | 256.99 | 235.39 | 234.45 | 269.91 | 271.56 | 262.87 |
| 16457 | ENSMUSG00000057789 | 256.8539849 | 0.046407084 | 0.176263345 | 0.263282668 | 0.792332717 | 0.961434599 | Bak1 | BCL2-antagonist/killer 1 [Source:MGI Sym | 273.05 | 245.67 | 238.48 | 215.93 | 265.02 | 302.97 |
| 16458 | ENSMUSG00000057801 | 0 | NA | NA | NA | NA | NA | Olfr135 | olfactory receptor 135 [Source:MGI Symb | 0 | 0 | 0 | 0 | 0 | 0 |
| 16459 | ENSMUSG00000057802 | 24.23732654 | -0.353873115 | 0.453365986 | -0.780546239 | 0.435069422 | 0.845325456 | Gm10030 | predicted gene 10030 [Source:MGI Symbo | 20.881 | 27.754 | 33.206 | 25.063 | 21.812 | 16.708 |
| 16460 | ENSMUSG00000057805 | 0 | NA | NA | NA | NA | NA | 1700084M14 | RIKEN cDNA 1700084M14 gene [Source:M | 0 | 0 | 0 | 0 | 0 | 0 |

| | | | | | | | | | | | | | | | |
|---|---|---|---|---|---|---|---|---|---|---|---|---|---|---|---|
| 16461 | ENSMUSG00000057816 | 0.160662974 | 1.126924175 | 4.080472857 | 0.276174898 | 0.782413736 | NA | Cfap299 | cilia and flagella associated protein 299 [S | 0 | 0 | 0 | 0.964 | 0 | 0 |
| 16462 | ENSMUSG00000057817 | 0 | NA | NA | NA | NA | NA | Olfr1446 | olfactory receptor 1446 [Source:MGI Symb | 0 | 0 | 0 | 0 | 0 | 0 |
| 16463 | ENSMUSG00000057818 | 0 | NA | NA | NA | NA | NA | 1600029O15 | RIKEN cDNA 1600029O15 gene [Source:M | 0 | 0 | 0 | 0 | 0 | 0 |
| 16464 | ENSMUSG00000057829 | 0 | NA | NA | NA | NA | NA | Gm5278 | predicted pseudogene 5278 [Source:MGI | 0 | 0 | 0 | 0 | 0 | 0 |
| 16465 | ENSMUSG00000057835 | 59.42687054 | -0.282901552 | 0.307049068 | -0.921356168 | 0.356864505 | 0.796166767 | Zfp119a | zinc finger protein 119a [Source:MGI Sym | 62.642 | 52.424 | 80.5 | 46.271 | 52.35 | 62.375 |
| 16466 | ENSMUSG00000057836 | 124.7342073 | -0.890213076 | 1.065585497 | -0.835421539 | 0.403480407 | 0.828953151 | Xlr3a | X-linked lymphocyte-regulated 3A [Source | 226.47 | 236.42 | 23.144 | 55.911 | 33.809 | 172.65 |
| 16467 | ENSMUSG00000057841 | 2351.460856 | -0.018471498 | 0.100387849 | -0.184001337 | 0.854012406 | 0.973825032 | Rpl32 | ribosomal protein L32 [Source:MGI Symbo | 2198.1 | 2515.3 | 2386.8 | 2249.9 | 2288.1 | 2470.5 |
| 16468 | ENSMUSG00000057842 | 98.92975811 | 0.661036968 | 0.282152407 | 2.342836535 | 0.019137766 | 0.168170501 | Zfp595 | zinc finger protein 595 [Source:MGI Symb | 53.005 | 86.345 | 91.568 | 152.31 | 112.33 | 98.018 |
| 16469 | ENSMUSG00000057855 | 0 | NA | NA | NA | NA | NA | Krtap6-3 | keratin associated protein 6-3 [Source:MG | 0 | 0 | 0 | 0 | 0 | 0 |
| 16470 | ENSMUSG00000057858 | 94.34206751 | -0.352869452 | 0.242870322 | -1.452913018 | 0.146247906 | 0.556855766 | Fam204a | family with sequence similarity 204, meml | 114.84 | 108.96 | 93.581 | 85.794 | 92.703 | 70.172 |
| 16471 | ENSMUSG00000057863 | 2473.948504 | 0.005364284 | 0.100969452 | 0.053127791 | 0.957630088 | 0.991861398 | Rpl36 | ribosomal protein L36 [Source:MGI Symbo | 2241.5 | 2530.7 | 2637.4 | 2533.3 | 2411.4 | 2489.4 |
| 16472 | ENSMUSG00000057880 | 6999.289873 | 0.05766317 | 0.106967747 | 0.539070618 | 0.589838128 | 0.911573746 | Abat | 4-aminobutyrate aminotransferase [Sourc | 6561.3 | 7113.2 | 6903.8 | 6580.1 | 8119.7 | 6717.6 |
| 16473 | ENSMUSG00000057886 | 0 | NA | NA | NA | NA | NA | Cbx3-ps6 | chromobox 3, pseudogene 6 [Source:MGI | 0 | 0 | 0 | 0 | 0 | 0 |
| 16474 | ENSMUSG00000057890 | 0 | NA | NA | NA | NA | NA | Olfr1375 | olfactory receptor 1375 [Source:MGI Symb | 0 | 0 | 0 | 0 | 0 | 0 |
| 16475 | ENSMUSG00000057894 | 95.81943861 | 0.182226781 | 0.242221584 | 0.752314381 | 0.451862021 | 0.856471272 | Zfp329 | zinc finger protein 329 [Source:MGI Symb | 74.689 | 88.401 | 106.66 | 101.22 | 99.246 | 104.7 |
| 16476 | ENSMUSG00000057895 | 14.1347838 | 0.626570847 | 0.706098161 | 0.88737074 | 0.374879331 | 0.809332934 | Zfp105 | zinc finger protein 105 [Source:MGI Symb | 8.031 | 12.335 | 13.081 | 20.244 | 3.2719 | 27.846 |
| 16477 | ENSMUSG00000057897 | 477.0099946 | 0.844806208 | 0.413008717 | 2.04549244 | 0.040806334 | 0.276580512 | Camk2b | calcium/calmodulin-dependent protein ki | 365.41 | 328.93 | 329.04 | 430.9 | 424.25 | 983.52 |
| 16478 | ENSMUSG00000057899 | 1.428849187 | 1.291261643 | 2.100866506 | 0.614632886 | 0.538797191 | NA | Adgrf2 | adhesion G protein-coupled receptor F2 [ | 2.4093 | 0 | 0 | 2.8919 | 3.2719 | 0 |
| 16479 | ENSMUSG00000057903 | 0 | NA | NA | NA | NA | NA | Olfr739 | olfactory receptor 739 [Source:MGI Symb | 0 | 0 | 0 | 0 | 0 | 0 |
| 16480 | ENSMUSG00000057913 | 34.61604444 | -1.654520677 | 0.490634991 | -3.372202773 | 0.000745695 | 0.016826421 | Gm10032 | predicted gene 10032 [Source:MGI Symbo | 65.051 | 49.34 | 43.269 | 18.316 | 27.265 | 4.4554 |
| 16481 | ENSMUSG00000057914 | 3.213761529 | -0.685420001 | 1.261587619 | -0.543299562 | 0.586923566 | NA | Cacnb2 | calcium channel, voltage-dependent, beta | 0.8031 | 4.1117 | 7.0437 | 2.8919 | 1.0906 | 3.3415 |
| 16482 | ENSMUSG00000057933 | 226.2790983 | -0.618010834 | 0.195580364 | -3.159881801 | 0.001578332 | 0.029454792 | Gsta2 | glutathione S-transferase, alpha 2 (Yc2) [S | 225.67 | 264.17 | 333.07 | 199.54 | 159.23 | 175.99 |
| 16483 | ENSMUSG00000057945 | 0 | NA | NA | NA | NA | NA | Gm17136 | predicted gene 17136 [Source:MGI Symbo | 0 | 0 | 0 | 0 | 0 | 0 |
| 16484 | ENSMUSG00000057948 | 9.18640095 | -0.952717889 | 0.811999133 | -1.173299145 | 0.240675865 | 0.690975727 | Unc13d | unc-13 homolog D [Source:MGI Symbol;A | 20.881 | 12.335 | 3.0187 | 6.7478 | 5.4531 | 6.6831 |
| 16485 | ENSMUSG00000057963 | 367.4875603 | 0.130580545 | 0.145171964 | 0.899488726 | 0.368392398 | 0.80627882 | Itpk1 | inositol 1,3,4-triphosphate 5/6 kinase [Sou | 316.42 | 344.35 | 392.44 | 377.88 | 379.54 | 394.3 |
| 16486 | ENSMUSG00000057967 | 1.204524644 | 1.538179148 | 2.000264724 | 0.768987789 | 0.441900559 | NA | Fgf18 | fibroblast growth factor 18 [Source:MGI S | 0.8031 | 1.0279 | 0 | 0.964 | 1.0906 | 3.3415 |
| 16487 | ENSMUSG00000057969 | 18.03752491 | 0.768470831 | 0.574809885 | 1.336913041 | 0.181251023 | 0.614562765 | Sema3b | sema domain, immunoglobulin domain (Ig | 14.456 | 19.53 | 6.0375 | 25.063 | 14.178 | 28.96 |
| 16488 | ENSMUSG00000057977 | 0 | NA | NA | NA | NA | NA | Skint11 | selection and upkeep of intraepithelial T c | 0 | 0 | 0 | 0 | 0 | 0 |
| 16489 | ENSMUSG00000057981 | 0 | NA | NA | NA | NA | NA | Vmn1r12 | vomeronasal 1 receptor 12 [Source:MGI S | 0 | 0 | 0 | 0 | 0 | 0 |
| 16490 | ENSMUSG00000057982 | 452.8937202 | 0.640524222 | 0.467233584 | 1.370886519 | 0.170410334 | 0.597556794 | Zfp809 | zinc finger protein 809 [Source:MGI Symb | 363 | 321.74 | 377.34 | 864.69 | 272.65 | 517.94 |
| 16491 | ENSMUSG00000057990 | 1.329269052 | -1.494954196 | 1.953578141 | -0.765239007 | 0.444129236 | NA | Gm53055 | predicted gene 53055 [Source:MGI Symbo | 0.8031 | 4.1117 | 1.0062 | 0.964 | 1.0906 | 0 |
| 16492 | ENSMUSG00000057997 | 4.503699228 | -2.423978619 | 1.155939656 | -2.096976781 | 0.035995631 | NA | Olfr541 | olfactory receptor 541 [Source:MGI Symb | 6.4248 | 11.307 | 5.0312 | 0.964 | 2.1812 | 1.1138 |
| 16493 | ENSMUSG00000058005 | 0 | NA | NA | NA | NA | NA | Tdpoz3 | TD and POZ domain containing 3 [Source: | 0 | 0 | 0 | 0 | 0 | 0 |
| 16494 | ENSMUSG00000058006 | 200.9769824 | -0.042232946 | 0.190053558 | -0.222216026 | 0.824145719 | 0.96943211 | Mdn1 | midasin AAA ATPase 1 [Source:MGI Symb | 191.94 | 222.03 | 198.23 | 205.33 | 227.94 | 160.39 |
| 16495 | ENSMUSG00000058013 | 122.9670527 | 0.201154641 | 0.286256467 | 0.702707763 | 0.482237889 | 0.871593699 | Septin11 | septin 11 [Source:MGI Symbol;Acc:MGI:1 | 122.87 | 115.13 | 104.65 | 86.758 | 111.24 | 197.15 |
| 16496 | ENSMUSG00000058014 | 0 | NA | NA | NA | NA | NA | Olfr502 | olfactory receptor 502 [Source:MGI Symb | 0 | 0 | 0 | 0 | 0 | 0 |
| 16497 | ENSMUSG00000058017 | 0 | NA | NA | NA | NA | NA | Serpinb3d | serine (or cysteine) peptidase inhibitor, cl | 0 | 0 | 0 | 0 | 0 | 0 |
| 16498 | ENSMUSG00000058019 | 0 | NA | NA | NA | NA | NA | Ces5a | carboxylesterase 5A [Source:MGI Symbol; | 0 | 0 | 0 | 0 | 0 | 0 |
| 16499 | ENSMUSG00000058022 | 2594.502711 | 0.138391336 | 0.092886824 | 1.489891992 | 0.136252637 | 0.538921153 | Adtrp | androgen dependent TFPI regulating prot | 2458.3 | 2403.3 | 2548.8 | 2666.4 | 2870.5 | 2619.8 |
| 16500 | ENSMUSG00000058028 | 0 | NA | NA | NA | NA | NA | Zscan5b | zinc finger and SCAN domain containing 5 | 0 | 0 | 0 | 0 | 0 | 0 |
| 16501 | ENSMUSG00000058030 | 0 | NA | NA | NA | NA | NA | Vmn1r237 | vomeronasal 1 receptor 237 [Source:MGI | 0 | 0 | 0 | 0 | 0 | 0 |
| 16502 | ENSMUSG00000058046 | 3.618511148 | -2.082776367 | 1.247332523 | -1.669784382 | 0.094962031 | NA | 4933430I17F | RIKEN cDNA 4933430I17 gene [Source:MG | 6.4248 | 5.1396 | 6.0375 | 1.928 | 2.1812 | 0 |
| 16503 | ENSMUSG00000058050 | 0.614004368 | 0.315622926 | 2.898299649 | 0.108899342 | 0.913282328 | NA | Gm9234 | predicted pseudogene 9234 [Source:MGI | 1.6062 | 0 | 0 | 0.964 | 0 | 1.1138 |
| 16504 | ENSMUSG00000058052 | 0 | NA | NA | NA | NA | NA | Defb35 | defensin beta 35 [Source:MGI Symbol;Acc | 0 | 0 | 0 | 0 | 0 | 0 |
| 16505 | ENSMUSG00000058056 | 934.6463269 | -0.094983116 | 0.122843077 | -0.773206906 | 0.43939994 | 0.848415529 | Palld | palladin, cytoskeletal associated protein [ | 958.1 | 910.74 | 1027.4 | 918.67 | 798.33 | 994.66 |
| 16506 | ENSMUSG00000058057 | 0.342432689 | 1.990653086 | 3.900281263 | 0.51038706 | 0.609780321 | NA | Mettl7a3 | methyltransferase like 7A3 [Source:MGI S | 0 | 0 | 0 | 0.964 | 1.0906 | 0 |
| 16507 | ENSMUSG00000058063 | 0 | NA | NA | NA | NA | NA | Trim31 | tripartite motif-containing 31 [Source:MG | 0 | 0 | 0 | 0 | 0 | 0 |
| 16508 | ENSMUSG00000058064 | 2.967038042 | 0.153061799 | 1.330996898 | 0.114997863 | 0.908446816 | NA | Gm10036 | predicted gene 10036 [Source:MGI Symbo | 2.4093 | 0 | 6.0375 | 3.8559 | 3.2719 | 2.2277 |
| 16509 | ENSMUSG00000058070 | 28.49824504 | -0.481120288 | 0.465895861 | -1.032677746 | 0.301754731 | 0.752246851 | Eml1 | echinoderm microtubule associated prote | 46.58 | 30.838 | 22.137 | 32.775 | 15.269 | 23.391 |
| 16510 | ENSMUSG00000058071 | 0 | NA | NA | NA | NA | NA | Olfr818 | olfactory receptor 818 [Source:MGI Symb | 0 | 0 | 0 | 0 | 0 | 0 |
| 16511 | ENSMUSG00000058073 | 0.133850373 | -0.796629176 | 4.080472857 | -0.195229623 | 0.84521319 | NA | Gm11189 | predicted gene 11189 [Source:MGI Symbo | 0.8031 | 0 | 0 | 0 | 0 | 0 |
| 16512 | ENSMUSG00000058076 | 3865.331736 | -0.120067826 | 0.092181053 | -1.302521746 | 0.192738089 | 0.630757769 | Sdhc | succinate dehydrogenase complex, subun | 4147.2 | 3941 | 3988.8 | 3463.6 | 3903.3 | 3748.1 |
| 16513 | ENSMUSG00000058078 | 0 | NA | NA | NA | NA | NA | Olfr1503-ps1 | olfactory receptor 1503, pseudogene 1 [S | 0 | 0 | 0 | 0 | 0 | 0 |
| 16514 | ENSMUSG00000058084 | 0 | NA | NA | NA | NA | NA | Olfr825 | olfactory receptor 825 [Source:MGI Symb | 0 | 0 | 0 | 0 | 0 | 0 |
| 16515 | ENSMUSG00000058093 | 124.7455715 | 0.151059903 | 0.227072214 | 0.665250495 | 0.505890289 | 0.881592135 | Zfp729b | zinc finger protein 729b [Source:MGI Sym | 129.3 | 125.41 | 99.618 | 129.17 | 110.15 | 154.82 |
| 16516 | ENSMUSG00000058099 | 42.32150633 | -1.224933918 | 0.408622682 | -2.997713955 | 0.002720128 | 0.043861693 | Nfam1 | Nfat activating molecule with ITAM motif | 60.233 | 71.954 | 45.281 | 12.532 | 31.628 | 32.301 |

| | | | | | | | | | | | | | | | |
|---|---|---|---|---|---|---|---|---|---|---|---|---|---|---|---|
| 16517 | ENSMUSG00000058101 | 0.160662974 | 1.126924175 | 4.080472857 | 0.276174898 | 0.782413736 | NA | 1700084J12I | RIKEN cDNA 1700084J12 gene [Source:MG | 0 | 0 | 0 | 0.964 | 0 | 0 |
| 16518 | ENSMUSG00000058114 | 0 | NA | NA | NA | NA | NA | Olfr127 | olfactory receptor 127 [Source:MGI Symb | 0 | 0 | 0 | 0 | 0 | 0 |
| 16519 | ENSMUSG00000058119 | 0 | NA | NA | NA | NA | NA | Gm5771 | predicted gene 5771 [Source:MGI Symbol | 0 | 0 | 0 | 0 | 0 | 0 |
| 16520 | ENSMUSG00000058124 | 0 | NA | NA | NA | NA | NA | H2-M10.3 | histocompatibility 2, M region locus 10.3 [ | 0 | 0 | 0 | 0 | 0 | 0 |
| 16521 | ENSMUSG00000058126 | 4.018459434 | -1.505576895 | 1.115659164 | -1.349495387 | 0.177177901 | NA | Tpm3-rs7 | tropomyosin 3, related sequence 7 [Sourc | 9.6372 | 4.1117 | 4.025 | 1.928 | 2.1812 | 2.2277 |
| 16522 | ENSMUSG00000058132 | 0 | NA | NA | NA | NA | NA | Vmn1r82 | vomeronasal 1 receptor 82 [Source:MGI S | 0 | 0 | 0 | 0 | 0 | 0 |
| 16523 | ENSMUSG00000058135 | 13746.18318 | -0.418430222 | 0.144482075 | -2.896070137 | 0.003778679 | 0.054984727 | Gstm1 | glutathione S-transferase, mu 1 [Source:M | 18851 | 12572 | 15753 | 12958 | 10134 | 12209 |
| 16524 | ENSMUSG00000058145 | 0 | NA | NA | NA | NA | NA | Adamts17 | a disintegrin-like and metallopeptidase (re | 0 | 0 | 0 | 0 | 0 | 0 |
| 16525 | ENSMUSG00000058147 | 0 | NA | NA | NA | NA | NA | Xlr3c | X-linked lymphocyte-regulated 3C [Source | 0 | 0 | 0 | 0 | 0 | 0 |
| 16526 | ENSMUSG00000058152 | 64.60311644 | 0.72139549 | 0.301176332 | 2.395259563 | 0.016608603 | 0.152913326 | Chsy3 | chondroitin sulfate synthase 3 [Source:MG | 49.792 | 55.508 | 41.256 | 102.18 | 68.709 | 70.172 |
| 16527 | ENSMUSG00000058153 | 0.181769715 | 1.126924175 | 4.080472857 | 0.276174898 | 0.782413736 | NA | Sez6l | seizure related 6 homolog like [Source:MG | 0 | 0 | 0 | 0 | 1.0906 | 0 |
| 16528 | ENSMUSG00000058159 | 0.167707431 | -0.796629176 | 4.080472857 | -0.195229623 | 0.84521319 | NA | T2 | brachyury 2 [Source:MGI Symbol;Acc:MG | 0 | 0 | 1.0062 | 0 | 0 | 0 |
| 16529 | ENSMUSG00000058163 | 6.96715375 | -0.476155477 | 0.900213978 | -0.528935885 | 0.596849928 | 0.91428215 | Gm5431 | predicted gene 5431 [Source:MGI Symbol | 8.031 | 13.363 | 3.0187 | 9.6398 | 2.1812 | 5.5692 |
| 16530 | ENSMUSG00000058172 | 0 | NA | NA | NA | NA | NA | Krtap6-1 | keratin associated protein 6-1 [Source:MG | 0 | 0 | 0 | 0 | 0 | 0 |
| 16531 | ENSMUSG00000058173 | 143.265226 | -0.199948369 | 0.227761435 | -0.877885097 | 0.380006078 | 0.812628154 | Smco4 | single-pass membrane protein with coiled | 172.67 | 157.27 | 128.8 | 104.11 | 135.24 | 161.51 |
| 16532 | ENSMUSG00000058174 | 1.641819355 | -2.983537005 | 1.927417618 | -1.54794528 | 0.121635472 | NA | Gm5148 | predicted gene 5148 [Source:MGI Symbol | 4.8186 | 2.0558 | 2.0125 | 0.964 | 0 | 0 |
| 16533 | ENSMUSG00000058176 | 0 | NA | NA | NA | NA | NA | Gm5407 | predicted gene 5407 [Source:MGI Symbol | 0 | 0 | 0 | 0 | 0 | 0 |
| 16534 | ENSMUSG00000058183 | 0 | NA | NA | NA | NA | NA | Mmel1 | membrane metallo-endopeptidase-like 1 | 0 | 0 | 0 | 0 | 0 | 0 |
| 16535 | ENSMUSG00000058186 | 6.649838007 | 2.248255727 | 1.035416465 | 2.171354042 | 0.02990442 | 0.228601666 | Zfp980 | zinc finger protein 980 [Source:MGI Symb | 5.6217 | 0 | 1.0062 | 6.7478 | 9.8156 | 16.708 |
| 16536 | ENSMUSG00000058188 | 0 | NA | NA | NA | NA | NA | Olfr746 | olfactory receptor 746 [Source:MGI Symb | 0 | 0 | 0 | 0 | 0 | 0 |
| 16537 | ENSMUSG00000058192 | 56.67719502 | 0.231194765 | 0.308323455 | 0.749844883 | 0.453348133 | 0.857073339 | Zfp846 | zinc finger protein 846 [Source:MGI Symb | 40.958 | 51.396 | 64.4 | 59.767 | 56.712 | 66.831 |
| 16538 | ENSMUSG00000058194 | 0 | NA | NA | NA | NA | NA | Olfr1126 | olfactory receptor 1126 [Source:MGI Sym | 0 | 0 | 0 | 0 | 0 | 0 |
| 16539 | ENSMUSG00000058200 | 0 | NA | NA | NA | NA | NA | Olfr66 | olfactory receptor 66 [Source:MGI Symbo | 0 | 0 | 0 | 0 | 0 | 0 |
| 16540 | ENSMUSG00000058207 | 188792.099 | 0.463365827 | 0.104841556 | 4.419677117 | 9.88E-06 | 0.000527581 | Serpina3k | serine (or cysteine) peptidase inhibitor, cl | 142622 | 160604 | 172970 | 200683 | 221488 | 234385 |
| 16541 | ENSMUSG00000058216 | 312.4590738 | -0.068105047 | 0.165830836 | -0.410689886 | 0.681299944 | 0.934452508 | Gstp3 | glutathione S-transferase pi 3 [Source:MG | 345.33 | 299.12 | 313.95 | 251.6 | 323.91 | 340.84 |
| 16542 | ENSMUSG00000058230 | 1767.362253 | 0.02009501 | 0.114853458 | 0.174962166 | 0.861109363 | 0.976013118 | Arhgap35 | Rho GTPase activating protein 35 [Source: | 1519.5 | 1874.9 | 1872.6 | 1852.8 | 1764.6 | 1719.8 |
| 16543 | ENSMUSG00000058239 | 941.0328708 | 0.117036762 | 0.134888808 | 0.867653614 | 0.385583987 | 0.816437825 | Usf2 | upstream transcription factor 2 [Source:M | 776.6 | 987.83 | 945.87 | 1071.9 | 865.95 | 998 |
| 16544 | ENSMUSG00000058240 | 197.1758918 | 0.13099096 | 0.179447273 | 0.729969075 | 0.465409088 | 0.863256867 | Cryzl1 | crystallin, zeta (quinone reductase)-like 1 | 196.76 | 172.69 | 195.21 | 224.61 | 181.04 | 212.74 |
| 16545 | ENSMUSG00000058244 | 0 | NA | NA | NA | NA | NA | Olfr506 | olfactory receptor 506 [Source:MGI Symb | 0 | 0 | 0 | 0 | 0 | 0 |
| 16546 | ENSMUSG00000058246 | 11.9946403 | -0.175688466 | 0.631524794 | -0.278197258 | 0.78086094 | 0.959993512 | Gm10037 | predicted gene 10037 [Source:MGI Symbo | 8.8341 | 12.335 | 17.106 | 10.604 | 14.178 | 8.9108 |
| 16547 | ENSMUSG00000058248 | 0 | NA | NA | NA | NA | NA | Kcnh1 | potassium voltage-gated channel, subfam | 0 | 0 | 0 | 0 | 0 | 0 |
| 16548 | ENSMUSG00000058250 | 0.367410362 | 2.078255025 | 3.800070157 | 0.54689912 | 0.58444804 | NA | Tas2r138 | taste receptor, type 2, member 138 [Sour | 0 | 0 | 0 | 0 | 1.0906 | 1.1138 |
| 16549 | ENSMUSG00000058251 | 0 | NA | NA | NA | NA | NA | Olfr822 | olfactory receptor 822 [Source:MGI Symb | 0 | 0 | 0 | 0 | 0 | 0 |
| 16550 | ENSMUSG00000058252 | 0 | NA | NA | NA | NA | NA | Tcp11x2 | t-complex 11 family, X-linked 2 [Source:M | 0 | 0 | 0 | 0 | 0 | 0 |
| 16551 | ENSMUSG00000058254 | 141.5829636 | -0.482224152 | 0.236265673 | -2.041025027 | 0.04124834 | 0.278402232 | Tspan7 | tetraspanin 7 [Source:MGI Symbol;Acc:MG | 180.7 | 143.91 | 170.06 | 112.79 | 88.34 | 153.71 |
| 16552 | ENSMUSG00000058258 | 331.5984283 | 1.596844056 | 0.446766411 | 3.574225852 | 0.000351266 | 0.009865449 | Idi1 | isopentenyl-diphosphate delta isomerase | 203.99 | 202.5 | 87.543 | 359.56 | 476.6 | 659.4 |
| 16553 | ENSMUSG00000058260 | 23.49944753 | 1.428026527 | 0.48364844 | 2.952612699 | 0.00315097 | 0.048381777 | Serpina9 | serine (or cysteine) peptidase inhibitor, cl | 8.031 | 16.447 | 14.087 | 39.523 | 27.265 | 35.643 |
| 16554 | ENSMUSG00000058267 | 553.2494706 | -0.021993641 | 0.131023216 | -0.167860639 | 0.866692916 | 0.977930657 | Mrps14 | mitochondrial ribosomal protein S14 [Sou | 604.74 | 519.1 | 547.4 | 515.73 | 534.4 | 598.13 |
| 16555 | ENSMUSG00000058270 | 0 | NA | NA | NA | NA | NA | Olfr25 | olfactory receptor 25 [Source:MGI Symbo | 0 | 0 | 0 | 0 | 0 | 0 |
| 16556 | ENSMUSG00000058275 | 0 | NA | NA | NA | NA | NA | Olfr412 | olfactory receptor 412 [Source:MGI Symb | 0 | 0 | 0 | 0 | 0 | 0 |
| 16557 | ENSMUSG00000058281 | 0.887948424 | 0.692526802 | 2.801019149 | 0.247241009 | 0.804721709 | NA | Gm10038 | predicted gene 10038 [Source:MGI Symbo | 0 | 2.0558 | 0 | 0 | 3.2719 | 0 |
| 16558 | ENSMUSG00000058287 | 0.160662974 | 1.126924175 | 4.080472857 | 0.276174898 | 0.782413736 | NA | Gm12253 | predicted gene 12253 [Source:MGI Symbo | 0 | 0 | 0 | 0.964 | 0 | 0 |
| 16559 | ENSMUSG00000058290 | 3.724870614 | 0.196111715 | 1.185729566 | 0.165393291 | 0.868634434 | NA | Espl1 | extra spindle pole bodies 1, separase [Sou | 3.2124 | 6.1675 | 1.0062 | 0.964 | 6.5437 | 4.4554 |
| 16560 | ENSMUSG00000058291 | 225.3152767 | 0.180407908 | 0.165874455 | 1.087617185 | 0.276764141 | 0.730041765 | Zfp68 | zinc finger protein 68 [Source:MGI Symbo | 195.15 | 207.64 | 231.44 | 245.81 | 229.03 | 242.82 |
| 16561 | ENSMUSG00000058295 | 0 | NA | NA | NA | NA | NA | Prp2 | proline rich protein 2 [Source:MGI Symbo | 0 | 0 | 0 | 0 | 0 | 0 |
| 16562 | ENSMUSG00000058297 | 28.10958119 | -0.345482332 | 0.441013678 | -0.78338235 | 0.433402618 | 0.845264432 | Spock2 | sparc/osteonectin, cwcv and kazal-like do | 34.533 | 31.865 | 28.175 | 36.631 | 19.631 | 17.822 |
| 16563 | ENSMUSG00000058298 | 109.9022675 | 0.096293928 | 0.280232665 | 0.343621353 | 0.731131064 | 0.94705581 | Mcm9 | minichromosome maintenance 9 homolog | 101.99 | 84.289 | 132.82 | 144.6 | 119.97 | 75.741 |
| 16564 | ENSMUSG00000058301 | 1501.201058 | -0.256724983 | 0.111948752 | -2.293236678 | 0.021834383 | 0.183932737 | Upf1 | UPF1 regulator of nonsense transcripts hc | 1609.4 | 1576.8 | 1717.7 | 1434.4 | 1458.2 | 1210.7 |
| 16565 | ENSMUSG00000058317 | 347.2225715 | -0.08847501 | 0.147799287 | -0.598615948 | 0.549429019 | 0.895703112 | Ube2e2 | ubiquitin-conjugating enzyme E2E 2 [Sour | 391.11 | 356.69 | 325.02 | 326.79 | 321.73 | 362 |
| 16566 | ENSMUSG00000058318 | 268.9081376 | 0.101899394 | 0.175751713 | 0.579791753 | 0.56205506 | 0.900565925 | Phf21a | PHD finger protein 21A [Source:MGI Symb | 276.27 | 239.5 | 262.63 | 328.72 | 231.21 | 275.12 |
| 16567 | ENSMUSG00000058325 | 282.2049505 | 0.151496991 | 0.174425897 | 0.868546436 | 0.385095263 | 0.816437825 | Dock1 | dedicator of cytokinesis 1 [Source:MGI Sy | 267.43 | 236.42 | 297.85 | 259.31 | 282.47 | 349.75 |
| 16568 | ENSMUSG00000058328 | 0.17131964 | -0.796629176 | 4.080472857 | -0.195229623 | 0.84521319 | NA | Xlr5a | X-linked lymphocyte-regulated 5A [Source | 0 | 1.0279 | 0 | 0 | 0 | 0 |
| 16569 | ENSMUSG00000058331 | 39.83345328 | 0.019216922 | 0.370445755 | 0.051875132 | 0.95862819 | 0.992027194 | Zfp85 | zinc finger protein 85 [Source:MGI Symbo | 51.399 | 27.754 | 39.244 | 43.379 | 34.9 | 42.326 |
| 16570 | ENSMUSG00000058349 | 0 | NA | NA | NA | NA | NA | Tas2r117 | taste receptor, type 2, member 117 [Sour | 0 | 0 | 0 | 0 | 0 | 0 |
| 16571 | ENSMUSG00000058351 | 233.9526149 | 0.15260865 | 0.167250522 | 0.912455447 | 0.361529013 | 0.800421803 | Smim4 | small integral membrane protein 4 [Sourc | 247.36 | 196.33 | 220.37 | 245.81 | 242.12 | 251.73 |
| 16572 | ENSMUSG00000058354 | 0 | NA | NA | NA | NA | NA | Krt6a | keratin 6A [Source:MGI Symbol;Acc:MGI: | 0 | 0 | 0 | 0 | 0 | 0 |

| | | | | | | | | | | | | | | | |
|---|---|---|---|---|---|---|---|---|---|---|---|---|---|---|---|
| 16573 | ENSMUSG00000058355 | 1396.973361 | -0.103690985 | 0.107565793 | -0.963977324 | 0.335057295 | 0.776159726 | Abce1 | ATP-binding cassette, sub-family E (OABP) | 1325.9 | 1514.1 | 1502.3 | 1312 | 1426.5 | 1301 |
| 16574 | ENSMUSG00000058357 | 0.167707431 | -0.796629176 | 4.080472857 | -0.195229623 | 0.84521319 | NA | Gm6612 | predicted gene 6612 [Source:MGI Symbol | 0 | 0 | 1.0062 | 0 | 0 | 0 |
| 16575 | ENSMUSG00000058360 | 0.133850373 | -0.796629176 | 4.080472857 | -0.195229623 | 0.84521319 | NA | Gm10040 | predicted gene 10040 [Source:MGI Symbo | 0.8031 | 0 | 0 | 0 | 0 | 0 |
| 16576 | ENSMUSG00000058368 | 0 | NA | NA | NA | NA | NA | Krtap21-1 | keratin associated protein 21-1 [Source:M | 0 | 0 | 0 | 0 | 0 | 0 |
| 16577 | ENSMUSG00000058385 | 4.538894285 | 0.938390945 | 0.992011708 | 0.94594745 | 0.34417539 | NA | H2bc8 | H2B clustered histone 8 [Source:MGI Sym | 3.2124 | 4.1117 | 2.0125 | 5.7839 | 6.5437 | 5.5692 |
| 16578 | ENSMUSG00000058388 | 114.3694285 | 0.059404082 | 0.216210862 | 0.274750683 | 0.78350779 | 0.960670212 | Phtf1 | putative homeodomain transcription fact | 114.04 | 116.15 | 105.66 | 109.89 | 110.15 | 130.32 |
| 16579 | ENSMUSG00000058392 | 117.3865773 | 0.113202526 | 0.22698706 | 0.498717971 | 0.617978083 | 0.917509899 | Rrp1b | ribosomal RNA processing 1B [Source:MG | 132.51 | 92.513 | 112.7 | 119.53 | 114.51 | 132.55 |
| 16580 | ENSMUSG00000058396 | 262.96587 | -0.839699 | 0.180589244 | -4.649773042 | 3.32E-06 | 0.000216504 | Gpr182 | G protein-coupled receptor 182 [Source:N | 302.77 | 323.79 | 385.39 | 160.98 | 184.31 | 220.54 |
| 16581 | ENSMUSG00000058398 | 0 | NA | NA | NA | NA | NA | Prss43 | protease, serine 43 [Source:MGI Symbol;A | 0 | 0 | 0 | 0 | 0 | 0 |
| 16582 | ENSMUSG00000058399 | 0 | NA | NA | NA | NA | NA | NA | NA | 0 | 0 | 0 | 0 | 0 | 0 |
| 16583 | ENSMUSG00000058400 | 0 | NA | NA | NA | NA | NA | Qrfpr | pyroglutamylated RFamide peptide recep | 0 | 0 | 0 | 0 | 0 | 0 |
| 16584 | ENSMUSG00000058402 | 43.86207844 | -0.020998983 | 0.329088232 | -0.063809584 | 0.949121847 | 0.990731019 | Zfp420 | zinc finger protein 420 [Source:MGI Symb | 44.974 | 46.256 | 41.256 | 42.415 | 39.262 | 49.009 |
| 16585 | ENSMUSG00000058407 | 635.8517275 | 0.039219478 | 0.14576007 | 0.269068734 | 0.78787679 | 0.960870081 | Txndc9 | thioredoxin domain containing 9 [Source: | 700.31 | 667.12 | 513.18 | 601.52 | 693.63 | 639.35 |
| 16586 | ENSMUSG00000058420 | 0.945986982 | 0.197573854 | 2.252218024 | 0.087724124 | 0.930095945 | NA | Syt17 | synaptotagmin XVII [Source:MGI Symbol;A | 1.6062 | 1.0279 | 0 | 1.928 | 0 | 1.1138 |
| 16587 | ENSMUSG00000058427 | 1.111859485 | -1.041840268 | 2.15740327 | -0.482914012 | 0.629156792 | NA | Cxcl2 | chemokine (C-X-C motif) ligand 2 [Source: | 2.4093 | 1.0279 | 1.0062 | 0 | 0 | 2.2277 |
| 16588 | ENSMUSG00000058430 | 0 | NA | NA | NA | NA | NA | Rpl27a-ps2 | ribosomal protein L27A, pseudogene 2 [Sc | 0 | 0 | 0 | 0 | 0 | 0 |
| 16589 | ENSMUSG00000058435 | 0 | NA | NA | NA | NA | NA | Btnl4 | butyrophilin-like 4 [Source:MGI Symbol;Ac | 0 | 0 | 0 | 0 | 0 | 0 |
| 16590 | ENSMUSG00000058440 | 170.3755672 | 0.206405647 | 0.181632263 | 1.136393085 | 0.255792089 | 0.708918985 | Nrf1 | nuclear respiratory factor 1 [Source:MGI S | 158.21 | 168.58 | 147.92 | 187.01 | 173.41 | 187.13 |
| 16591 | ENSMUSG00000058441 | 2.332106894 | 0.501235031 | 1.401080623 | 0.357748885 | 0.720531246 | NA | Panx2 | pannexin 2 [Source:MGI Symbol;Acc:MGI: | 0.8031 | 2.0558 | 3.0187 | 4.8199 | 2.1812 | 1.1138 |
| 16592 | ENSMUSG00000058443 | 2.580421062 | 2.243631037 | 1.503487096 | 1.492284864 | 0.135624483 | NA | Rpl10-ps3 | ribosomal protein L10, pseudogene 3 [Sou | 1.6062 | 0 | 1.0062 | 2.8919 | 2.1812 | 7.7969 |
| 16593 | ENSMUSG00000058444 | 372.9460478 | 0.199806798 | 0.151346562 | 1.320193836 | 0.18677031 | 0.623054278 | Map2k5 | mitogen-activated protein kinase kinase 5 | 358.99 | 367.99 | 313.95 | 368.24 | 375.17 | 453.33 |
| 16594 | ENSMUSG00000058446 | 1336.453024 | 0.179718191 | 0.104330953 | 1.722577869 | 0.08496489 | 0.429396285 | Znrf2 | zinc and ring finger 2 [Source:MGI Symbol | 1195.8 | 1316.8 | 1247.7 | 1372.7 | 1373.1 | 1512.6 |
| 16595 | ENSMUSG00000058447 | 0 | NA | NA | NA | NA | NA | Gm26920 | predicted gene, 26920 [Source:MGI Symb | 0 | 0 | 0 | 0 | 0 | 0 |
| 16596 | ENSMUSG00000058454 | 706.4728353 | 0.520894149 | 0.212352139 | 2.452973405 | 0.014168081 | 0.137534985 | Dhcr7 | 7-dehydrocholesterol reductase [Source:N | 747.69 | 544.8 | 446.77 | 581.28 | 876.86 | 1041.4 |
| 16597 | ENSMUSG00000058463 | 0 | NA | NA | NA | NA | NA | Gm5471 | predicted pseudogene 5471 [Source:MGI | 0 | 0 | 0 | 0 | 0 | 0 |
| 16598 | ENSMUSG00000058470 | 1.545670342 | -0.393581507 | 1.873957311 | -0.210026933 | 0.833646652 | NA | Gm8369 | predicted gene 8369 [Source:MGI Symbol | 3.2124 | 2.0558 | 0 | 2.8919 | 0 | 1.1138 |
| 16599 | ENSMUSG00000058486 | 343.9894694 | -0.021511159 | 0.185432059 | -0.116005609 | 0.907648097 | 0.984592636 | Wdr91 | WD repeat domain 91 [Source:MGI Symbo | 335.7 | 374.16 | 330.05 | 364.38 | 249.75 | 409.89 |
| 16600 | ENSMUSG00000058488 | 0.367410362 | 2.078255025 | 3.800070157 | 0.54689912 | 0.58444804 | NA | Kl | klotho [Source:MGI Symbol;Acc:MGI:1101 | 0 | 0 | 0 | 0 | 1.0906 | 1.1138 |
| 16601 | ENSMUSG00000058491 | 0 | NA | NA | NA | NA | NA | Olfr869 | olfactory receptor 869 [Source:MGI Symb | 0 | 0 | 0 | 0 | 0 | 0 |
| 16602 | ENSMUSG00000058492 | 189.3234339 | 0.092594348 | 0.18776534 | 0.493138662 | 0.621914615 | 0.918578091 | Scp2-ps2 | sterol carrier protein 2, pseudogene 2 [So | 174.27 | 198.39 | 177.1 | 174.48 | 229.03 | 182.67 |
| 16603 | ENSMUSG00000058498 | 0.181769715 | 1.126924175 | 4.080472857 | 0.276174898 | 0.782413736 | NA | Rnf207 | ring finger protein 207 [Source:MGI Symb | 0 | 0 | 0 | 0 | 1.0906 | 0 |
| 16604 | ENSMUSG00000058499 | 0 | NA | NA | NA | NA | NA | Pip | prolactin induced protein [Source:MGI Sy | 0 | 0 | 0 | 0 | 0 | 0 |
| 16605 | ENSMUSG00000058503 | 157.7957832 | 0.018125661 | 0.183848765 | 0.09859006 | 0.921463763 | 0.986685804 | Fam133b | family with sequence similarity 133, meml | 158.21 | 154.19 | 157.98 | 160.02 | 154.87 | 161.51 |
| 16606 | ENSMUSG00000058508 | 0 | NA | NA | NA | NA | NA | Gm10042 | predicted gene 10042 [Source:MGI Symbo | 0 | 0 | 0 | 0 | 0 | 0 |
| 16607 | ENSMUSG00000058513 | 0 | NA | NA | NA | NA | NA | Olfr801 | olfactory receptor 801 [Source:MGI Symb | 0 | 0 | 0 | 0 | 0 | 0 |
| 16608 | ENSMUSG00000058515 | 0 | NA | NA | NA | NA | NA | Olfr933 | olfactory receptor 933 [Source:MGI Symb | 0 | 0 | 0 | 0 | 0 | 0 |
| 16609 | ENSMUSG00000058523 | 0.181769715 | 1.126924175 | 4.080472857 | 0.276174898 | 0.782413736 | NA | Mup5 | major urinary protein 5 [Source:MGI Syml | 0 | 0 | 0 | 0 | 1.0906 | 0 |
| 16610 | ENSMUSG00000058537 | 0 | NA | NA | NA | NA | NA | AW822073 | expressed sequence AW822073 [Source:N | 0 | 0 | 0 | 0 | 0 | 0 |
| 16611 | ENSMUSG00000058542 | 23.20844918 | -0.414188317 | 0.461077428 | -0.898305343 | 0.369022785 | 0.80653433 | Gm15590 | predicted gene 15590 [Source:MGI Symbo | 25.699 | 28.782 | 25.156 | 23.135 | 13.087 | 23.391 |
| 16612 | ENSMUSG00000058546 | 1506.977044 | 0.022655725 | 0.108832107 | 0.208171338 | 0.835095188 | 0.971073158 | Rpl23a | ribosomal protein L23A [Source:MGI Syml | 1440.8 | 1654.9 | 1390.6 | 1580 | 1494.1 | 1481.4 |
| 16613 | ENSMUSG00000058550 | 0 | NA | NA | NA | NA | NA | Dppa4 | developmental pluripotency associated 4 | 0 | 0 | 0 | 0 | 0 | 0 |
| 16614 | ENSMUSG00000058558 | 3715.527782 | 0.060457021 | 0.107211638 | 0.56390353 | 0.572819787 | 0.905173631 | Rpl5 | ribosomal protein L5 [Source:MGI Symbol | 3299.9 | 3833.1 | 3780.5 | 3436.6 | 3911 | 4032.1 |
| 16615 | ENSMUSG00000058567 | 0 | NA | NA | NA | NA | NA | Gm2531 | predicted gene 2531 [Source:MGI Symbol | 0 | 0 | 0 | 0 | 0 | 0 |
| 16616 | ENSMUSG00000058568 | 0 | NA | NA | NA | NA | NA | Defb50 | defensin beta 50 [Source:MGI Symbol;Acc | 0 | 0 | 0 | 0 | 0 | 0 |
| 16617 | ENSMUSG00000058569 | 958.762332 | 0.331774848 | 0.124083647 | 2.673799935 | 0.007499718 | 0.087977225 | Tmed9 | transmembrane p24 trafficking protein 9 [ | 913.13 | 793.55 | 839.21 | 1035.3 | 970.65 | 1200.7 |
| 16618 | ENSMUSG00000058571 | 5.5079278 | -1.087293046 | 0.915759275 | -1.187313169 | 0.235104116 | 0.683240713 | Gpc6 | glypican 6 [Source:MGI Symbol;Acc:MGI:1 | 7.2279 | 6.1675 | 9.0562 | 2.8919 | 4.3625 | 3.3415 |
| 16619 | ENSMUSG00000058579 | 5.761817988 | 2.639584501 | 1.038173926 | 2.542526291 | 0.011005432 | 0.116191657 | Cela2a | chymotrypsin-like elastase family, membe | 0.8031 | 2.0558 | 2.0125 | 8.6758 | 6.5437 | 14.48 |
| 16620 | ENSMUSG00000058581 | 0.435408177 | -2.179616019 | 3.531869654 | -0.617128103 | 0.537150234 | NA | Gm5801 | predicted pseudogene 5801 [Source:MGI | 1.6062 | 0 | 1.0062 | 0 | 0 | 0 |
| 16621 | ENSMUSG00000058586 | 278.8484028 | 0.052729494 | 0.14987363 | 0.351826361 | 0.724968488 | 0.9448775 | Serhl | serine hydrolase-like [Source:MGI Symbol | 277.07 | 274.45 | 269.67 | 282.45 | 301.01 | 268.44 |
| 16622 | ENSMUSG00000058587 | 392.7478191 | -0.16185912 | 0.139611232 | -1.159356004 | 0.246311102 | 0.696645245 | Tmod3 | tropomodulin 3 [Source:MGI Symbol;Acc: | 399.14 | 413.22 | 431.68 | 334.5 | 395.89 | 382.05 |
| 16623 | ENSMUSG00000058589 | 0.335414862 | -1.779047242 | 4.030804179 | -0.441362855 | 0.658950329 | NA | Anks1b | ankyrin repeat and sterile alpha motif dor | 0 | 0 | 2.0125 | 0 | 0 | 0 |
| 16624 | ENSMUSG00000058590 | 0 | NA | NA | NA | NA | NA | Gm6462 | predicted gene 6462 [Source:MGI Symbol | 0 | 0 | 0 | 0 | 0 | 0 |
| 16625 | ENSMUSG00000058594 | 510.4335796 | 0.198164712 | 0.123647131 | 1.602663246 | 0.109009022 | 0.485738292 | Fbh1 | F-box DNA helicase 1 [Source:MGI Symbo | 496.32 | 455.37 | 473.94 | 527.3 | 561.67 | 548.01 |
| 16626 | ENSMUSG00000058600 | 2160.502181 | 0.021414044 | 0.106160087 | 0.201714647 | 0.840139812 | 0.971838962 | Rpl30 | ribosomal protein L30 [Source:MGI Symbo | 2014.2 | 2198.7 | 2220.8 | 2067.7 | 2061.3 | 2400.3 |
| 16627 | ENSMUSG00000058603 | 12.23451406 | 0.701453495 | 0.625640374 | 1.121176837 | 0.262212597 | 0.716133499 | Rpl28-ps1 | ribosomal protein L28, pseudogene 1 [Sou | 9.6372 | 7.1954 | 11.069 | 12.532 | 20.722 | 12.252 |
| 16628 | ENSMUSG00000058607 | 0.181769715 | 1.126924175 | 4.080472857 | 0.276174898 | 0.782413736 | NA | Rpl31-ps20 | ribosomal protein L31, pseudogene 20 [Sc | 0 | 0 | 0 | 0 | 1.0906 | 0 |

| | | | | | | | | | | | | | | | |
|---|---|---|---|---|---|---|---|---|---|---|---|---|---|---|---|
| 16629 | ENSMUSG00000058613 | 40.08499809 | 0.502019326 | 0.402648158 | 1.246794046 | 0.212473024 | 0.657146732 | Cyp2d41-ps | cytochrome P450, family 2, subfamily d, m | 29.715 | 22.614 | 47.293 | 45.307 | 62.165 | 33.415 |
| 16630 | ENSMUSG00000058618 | 0 | NA | NA | NA | NA | NA | Defa39 | defensin, alpha, 39 [Source:MGI Symbol;A | 0 | 0 | 0 | 0 | 0 | 0 |
| 16631 | ENSMUSG00000058620 | 55.94320926 | -0.79109035 | 0.316949363 | -2.495951853 | 0.012561965 | 0.126529431 | Adra2b | adrenergic receptor, alpha 2b [Source:MG | 74.689 | 52.424 | 85.531 | 41.451 | 40.353 | 41.212 |
| 16632 | ENSMUSG00000058622 | 0 | NA | NA | NA | NA | NA | Gm5265 | predicted pseudogene 5265 [Source:MGI | 0 | 0 | 0 | 0 | 0 | 0 |
| 16633 | ENSMUSG00000058624 | 871.5008105 | -0.450584137 | 0.170585417 | -2.641398924 | 0.008256444 | 0.094106544 | Gda | guanine deaminase [Source:MGI Symbol;A | 864.94 | 1107.1 | 1048.5 | 783.71 | 881.22 | 543.56 |
| 16634 | ENSMUSG00000058625 | 0 | NA | NA | NA | NA | NA | Gm17383 | predicted gene, 17383 [Source:MGI Symb | 0 | 0 | 0 | 0 | 0 | 0 |
| 16635 | ENSMUSG00000058626 | 14.60464939 | 5.655650208 | 2.159702045 | 2.618717809 | 0.008826093 | 0.098938285 | Capn11 | calpain 11 [Source:MGI Symbol;Acc:MGI:1 | 1.6062 | 0 | 0 | 1.928 | 78.525 | 5.5692 |
| 16636 | ENSMUSG00000058628 | 0 | NA | NA | NA | NA | NA | Olfr875 | olfactory receptor 875 [Source:MGI Symb | 0 | 0 | 0 | 0 | 0 | 0 |
| 16637 | ENSMUSG00000058631 | 0 | NA | NA | NA | NA | NA | Vmn1r63 | vomeronasal 1 receptor 63 [Source:MGI S | 0 | 0 | 0 | 0 | 0 | 0 |
| 16638 | ENSMUSG00000058638 | 515.0851968 | -0.067303972 | 0.149262428 | -0.450910339 | 0.652054171 | 0.925643431 | Zfp110 | zinc finger protein 110 [Source:MGI Symb | 495.51 | 516.01 | 570.54 | 537.9 | 557.31 | 413.24 |
| 16639 | ENSMUSG00000058643 | 0 | NA | NA | NA | NA | NA | Speer4f1 | spermatogenesis associated glutamate (E) | 0 | 0 | 0 | 0 | 0 | 0 |
| 16640 | ENSMUSG00000058655 | 5871.85721 | 0.283014383 | 0.100094264 | 2.827478542 | 0.004691616 | 0.064018427 | Eif4b | eukaryotic translation initiation factor 4B | 4882.1 | 5820.1 | 5192.2 | 6197.4 | 6777.1 | 6362.3 |
| 16641 | ENSMUSG00000058656 | 5.276461463 | -1.265462637 | 0.966434332 | -1.309413992 | 0.190394154 | 0.627779796 | Samd12 | sterile alpha motif domain containing 12 [ | 7.2279 | 5.1396 | 10.062 | 4.8199 | 2.1812 | 2.2277 |
| 16642 | ENSMUSG00000058659 | 0 | NA | NA | NA | NA | NA | Olfr58 | olfactory receptor 58 [Source:MGI Symbo | 0 | 0 | 0 | 0 | 0 | 0 |
| 16643 | ENSMUSG00000058662 | 0 | NA | NA | NA | NA | NA | Olfr69 | olfactory receptor 69 [Source:MGI Symbo | 0 | 0 | 0 | 0 | 0 | 0 |
| 16644 | ENSMUSG00000058665 | 0 | NA | NA | NA | NA | NA | En1 | engrailed 1 [Source:MGI Symbol;Acc:MGI: | 0 | 0 | 0 | 0 | 0 | 0 |
| 16645 | ENSMUSG00000058669 | 0 | NA | NA | NA | NA | NA | Nkx2-9 | NK2 homeobox 9 [Source:MGI Symbol;Ac | 0 | 0 | 0 | 0 | 0 | 0 |
| 16646 | ENSMUSG00000058670 | 0 | NA | NA | NA | NA | NA | 4932411N23 | RIKEN cDNA 4932411N23 gene [Source:M | 0 | 0 | 0 | 0 | 0 | 0 |
| 16647 | ENSMUSG00000058672 | 426.37237 | -1.323990422 | 0.210982359 | -6.275360794 | 3.49E-10 | 8.14E-08 | Tubb2a | tubulin, beta 2A class IIA [Source:MGI Syn | 836.83 | 546.85 | 443.75 | 256.42 | 214.85 | 259.53 |
| 16648 | ENSMUSG00000058685 | 0 | NA | NA | NA | NA | NA | Vmn2r51 | vomeronasal 2, receptor 51 [Source:MGI S | 0 | 0 | 0 | 0 | 0 | 0 |
| 16649 | ENSMUSG00000058690 | 739.5651974 | 0.182612348 | 0.12444402 | 1.467425652 | 0.142260301 | 0.551945544 | Ccser2 | coiled-coil serine rich 2 [Source:MGI Symb | 769.37 | 652.73 | 655.07 | 758.65 | 843.05 | 758.53 |
| 16650 | ENSMUSG00000058692 | 0 | NA | NA | NA | NA | NA | Olfr846 | olfactory receptor 846 [Source:MGI Symb | 0 | 0 | 0 | 0 | 0 | 0 |
| 16651 | ENSMUSG00000058700 | 0 | NA | NA | NA | NA | NA | Rpl21-ps12 | ribosomal protein L21, pseudogene 12 [Sc | 0 | 0 | 0 | 0 | 0 | 0 |
| 16652 | ENSMUSG00000058704 | 459.6813244 | 0.197910983 | 0.131581658 | 1.504092483 | 0.132557555 | 0.533565316 | Memo1 | mediator of cell motility 1 [Source:MGI Sy | 424.84 | 433.78 | 425.64 | 449.21 | 528.95 | 495.66 |
| 16653 | ENSMUSG00000058706 | 682.8232604 | 0.429636964 | 0.172754583 | 2.48697868 | 0.012883312 | 0.128513258 | 0610030E20 | RIKEN cDNA 0610030E20 gene [Source:M | 575.82 | 530.41 | 638.97 | 673.82 | 657.64 | 1020.3 |
| 16654 | ENSMUSG00000058709 | 1302.549728 | 0.090750047 | 0.113226844 | 0.801488791 | 0.42284873 | 0.84005258 | Egln2 | egl-9 family hypoxia-inducible factor 2 [So | 1179.8 | 1333.2 | 1271.9 | 1211.7 | 1379.6 | 1439.1 |
| 16655 | ENSMUSG00000058715 | 83.07512816 | -1.740596975 | 0.346291273 | -5.026395729 | 5.00E-07 | 4.47E-05 | Fcer1g | Fc receptor, IgE, high affinity I, gamma po | 117.25 | 180.91 | 85.531 | 39.523 | 23.994 | 51.237 |
| 16656 | ENSMUSG00000058717 | 0 | NA | NA | NA | NA | NA | Lypd11 | Ly6/PLAUR domain containing 11 [Source: | 0 | 0 | 0 | 0 | 0 | 0 |
| 16657 | ENSMUSG00000058725 | 0 | NA | NA | NA | NA | NA | Gm11937 | predicted gene 11937 [Source:MGI Symbo | 0 | 0 | 0 | 0 | 0 | 0 |
| 16658 | ENSMUSG00000058728 | 0.806689472 | -0.274753491 | 2.612595279 | -0.105164965 | 0.91624491 | NA | Cd300c | CD300C molecule [Source:MGI Symbol;Ac | 1.6062 | 0 | 1.0062 | 0 | 0 | 2.2277 |
| 16659 | ENSMUSG00000058729 | 78.02325482 | 0.566092773 | 0.279019076 | 2.028867631 | 0.042471774 | 0.283892783 | Lin9 | lin-9 homolog (C. elegans) [Source:MGI Sy | 71.476 | 58.591 | 58.362 | 90.614 | 116.7 | 72.4 |
| 16660 | ENSMUSG00000058740 | 0.401551119 | -2.072982726 | 4.008251979 | -0.517178744 | 0.605031388 | NA | Kcnt1 | potassium channel, subfamily T, member | 2.4093 | 0 | 0 | 0 | 0 | 0 |
| 16661 | ENSMUSG00000058741 | 0 | NA | NA | NA | NA | NA | Prr19 | proline rich 19 [Source:MGI Symbol;Acc:M | 0 | 0 | 0 | 0 | 0 | 0 |
| 16662 | ENSMUSG00000058743 | 0.185640647 | 1.126924175 | 4.080472857 | 0.276174898 | 0.782413736 | NA | Kcnj14 | potassium inwardly-rectifying channel, su | 0 | 0 | 0 | 0 | 0 | 1.1138 |
| 16663 | ENSMUSG00000058745 | 0.349477146 | 0.165142768 | 3.911021834 | 0.042224967 | 0.966319359 | NA | Gm9174 | predicted pseudogene 9174 [Source:MGI | 0 | 0 | 1.0062 | 0 | 1.0906 | 0 |
| 16664 | ENSMUSG00000058748 | 91.9645592 | 0.121978936 | 0.252843975 | 0.482427693 | 0.629502152 | 0.92057725 | Zfp958 | zinc finger protein 958 [Source:MGI Symb | 102.8 | 86.345 | 74.462 | 81.938 | 95.974 | 110.27 |
| 16665 | ENSMUSG00000058755 | 0 | NA | NA | NA | NA | NA | Osm | oncostatin M [Source:MGI Symbol;Acc:MG | 0 | 0 | 0 | 0 | 0 | 0 |
| 16666 | ENSMUSG00000058756 | 198.3994897 | 0.124826134 | 0.17580328 | 0.710033024 | 0.477683658 | 0.869794311 | Thra | thyroid hormone receptor alpha [Source:I | 187.12 | 191.19 | 191.19 | 208.22 | 183.22 | 229.45 |
| 16667 | ENSMUSG00000058761 | 1093.092079 | -0.039635634 | 0.108433044 | -0.36553095 | 0.714715108 | 0.944595913 | Rnf169 | ring finger protein 169 [Source:MGI Symb | 1050.5 | 1185.2 | 1089.8 | 1121.1 | 1089.5 | 1022.5 |
| 16668 | ENSMUSG00000058773 | 1.035066562 | 0.098376889 | 2.401477279 | 0.040965155 | 0.967323675 | NA | H1f5 | H1.5 linker histone, cluster member [Sour | 0 | 0 | 3.0187 | 0.964 | 0 | 2.2277 |
| 16669 | ENSMUSG00000058793 | 1785.824338 | 0.114366737 | 0.109454512 | 1.044879144 | 0.29607883 | 0.746727847 | Cds2 | CDP-diacylglycerol synthase (phosphatida | 1766 | 1672.4 | 1706.6 | 1934.7 | 1639.2 | 1996 |
| 16670 | ENSMUSG00000058794 | 4.407447893 | 0.063667254 | 0.992858146 | 0.064125227 | 0.948870515 | NA | Nfe2 | nuclear factor, erythroid derived 2 [Sourc | 4.8186 | 3.0838 | 5.0312 | 5.7839 | 3.2719 | 4.4554 |
| 16671 | ENSMUSG00000058799 | 1042.800816 | -0.250179628 | 0.106110245 | -2.357733021 | 0.018386911 | 0.163058422 | Nap1l1 | nucleosome assembly protein 1-like 1 [So | 1112.3 | 1183.1 | 1103.9 | 948.55 | 1001.2 | 907.78 |
| 16672 | ENSMUSG00000058802 | 0 | NA | NA | NA | NA | NA | Olfr97 | olfactory receptor 97 [Source:MGI Symbo | 0 | 0 | 0 | 0 | 0 | 0 |
| 16673 | ENSMUSG00000058806 | 59.38529376 | -0.433532124 | 0.42408794 | -1.0222694 | 0.306653412 | 0.755610724 | Col13a1 | collagen, type XIII, alpha 1 [Source:MGI Sy | 51.399 | 68.87 | 84.525 | 47.235 | 19.631 | 84.652 |
| 16674 | ENSMUSG00000058807 | 0 | NA | NA | NA | NA | NA | Olfr331 | olfactory receptor 331 [Source:MGI Symb | 0 | 0 | 0 | 0 | 0 | 0 |
| 16675 | ENSMUSG00000058809 | 13.84645037 | 0.525517237 | 0.593559635 | 0.885365524 | 0.37595952 | 0.809690069 | Hspd1-ps3 | heat shock protein 1 (chaperonin), pseudo | 9.6372 | 13.363 | 11.069 | 12.532 | 13.087 | 23.391 |
| 16676 | ENSMUSG00000058812 | 7.430578668 | -0.163416604 | 0.83625552 | -0.195414679 | 0.845068326 | 0.972803041 | 0610039K10 | RIKEN cDNA 0610039K10 gene [Source:M | 7.2279 | 12.335 | 4.025 | 6.7478 | 10.906 | 3.3415 |
| 16677 | ENSMUSG00000058816 | 0 | NA | NA | NA | NA | NA | Ppp1r2-ps3 | protein phosphatase 1, regulatory (inhibit | 0 | 0 | 0 | 0 | 0 | 0 |
| 16678 | ENSMUSG00000058818 | 43.28975815 | -1.43765159 | 0.446258702 | -3.221565394 | 0.001274924 | 0.02508911 | Pirb | paired Ig-like receptor B [Source:MGI Sym | 65.051 | 95.596 | 29.181 | 27.955 | 17.45 | 24.505 |
| 16679 | ENSMUSG00000058820 | 0 | NA | NA | NA | NA | NA | Olfr146 | olfactory receptor 146 [Source:MGI Symb | 0 | 0 | 0 | 0 | 0 | 0 |
| 16680 | ENSMUSG00000058831 | 2.992810233 | 0.535619324 | 1.274610455 | 0.420221976 | 0.674323302 | NA | Opn1sw | opsin 1 (cone pigments), short-wave-sens | 3.2124 | 2.0558 | 2.0125 | 1.928 | 7.6343 | 1.1138 |
| 16681 | ENSMUSG00000058833 | 693.4213138 | 0.036279703 | 0.128872007 | 0.281517326 | 0.778313641 | 0.959559604 | Rex1bd | required for excision 1-B domain containi | 766.16 | 613.67 | 673.18 | 699.85 | 742.71 | 664.96 |
| 16682 | ENSMUSG00000058835 | 697.1294119 | 0.064319345 | 0.12820526 | 0.501690374 | 0.615885337 | 0.917509899 | Abi1 | abl interactor 1 [Source:MGI Symbol;Acc:I | 686.65 | 674.31 | 683.24 | 641.05 | 812.51 | 685.01 |
| 16683 | ENSMUSG00000058838 | 7.82268412 | -0.108817405 | 0.803824396 | -0.135374599 | 0.892315707 | 0.982489666 | Rps27a-ps2 | ribosomal protein S27A, pseudogene 2 [Sc | 8.031 | 6.1675 | 10.062 | 3.8559 | 5.4531 | 13.366 |
| 16684 | ENSMUSG00000058856 | 0 | NA | NA | NA | NA | NA | Olfr952 | olfactory receptor 952 [Source:MGI Symb | 0 | 0 | 0 | 0 | 0 | 0 |

| | | | | | | | | | | | | | | | |
|---|---|---|---|---|---|---|---|---|---|---|---|---|---|---|---|
| 16685 | ENSMUSG00000058869 | 0 | NA | NA | NA | NA | NA | Gm4832 | predicted gene 4832 [Source:MGI Symbol | 0 | 0 | 0 | 0 | 0 | 0 |
| 16686 | ENSMUSG00000058873 | 0.863436065 | -1.832005063 | 2.586421136 | -0.708316615 | 0.478748662 | NA | Gm5582 | predicted gene 5582 [Source:MGI Symbol | 0 | 3.0838 | 1.0062 | 0 | 1.0906 | 0 |
| 16687 | ENSMUSG00000058881 | 92.47562024 | -0.135439475 | 0.287208332 | -0.471572237 | 0.637232147 | 0.923478764 | Zfp516 | zinc finger protein 516 [Source:MGI Symb | 87.538 | 101.76 | 101.63 | 114.71 | 54.531 | 94.677 |
| 16688 | ENSMUSG00000058883 | 0.133850373 | -0.796629176 | 4.080472857 | -0.195229623 | 0.84521319 | NA | Zfp708 | zinc finger protein 708 [Source:MGI Symb | 0.8031 | 0 | 0 | 0 | 0 | 0 |
| 16689 | ENSMUSG00000058884 | 0 | NA | NA | NA | NA | NA | Olfr1025-ps1 | olfactory receptor 1025, pseudogene 1 [S | 0 | 0 | 0 | 0 | 0 | 0 |
| 16690 | ENSMUSG00000058886 | 209.7980114 | -0.006276512 | 0.165302931 | -0.037969755 | 0.969711797 | 0.994674947 | Deaf1 | DEAF1, transcription factor [Source:MGI S | 212.82 | 210.72 | 207.29 | 215.93 | 213.76 | 198.26 |
| 16691 | ENSMUSG00000058897 | 10.35634172 | -0.08487594 | 0.855513397 | -0.099210532 | 0.920971113 | 0.986685804 | Col25a1 | collagen, type XXV, alpha 1 [Source:MGI S | 6.4248 | 20.558 | 5.0312 | 3.8559 | 21.812 | 4.4554 |
| 16692 | ENSMUSG00000058900 | 47.1060499 | 0.18163441 | 0.323652123 | 0.56120259 | 0.574659438 | 0.906163061 | Rsl1 | regulator of sex limited protein 1 [Source: | 49.792 | 45.228 | 37.231 | 51.091 | 44.715 | 54.578 |
| 16693 | ENSMUSG00000058904 | 0 | NA | NA | NA | NA | NA | Olfr1413 | olfactory receptor 1413 [Source:MGI Sym | 0 | 0 | 0 | 0 | 0 | 0 |
| 16694 | ENSMUSG00000058905 | 1.147551471 | -0.308736011 | 1.963633334 | -0.157226915 | 0.875066017 | NA | Gm10051 | predicted pseudogene 10051 [Source:MG | 0.8031 | 1.0279 | 2.0125 | 1.928 | 0 | 1.1138 |
| 16695 | ENSMUSG00000058908 | 0 | NA | NA | NA | NA | NA | Pla2g2a | phospholipase A2, group IIA (platelets, sy | 0 | 0 | 0 | 0 | 0 | 0 |
| 16696 | ENSMUSG00000058914 | 9.650117793 | 0.459257631 | 0.715853222 | 0.641552789 | 0.521163595 | 0.887887809 | C1qtnf3 | C1q and tumor necrosis factor related pro | 12.047 | 4.1117 | 8.05 | 10.604 | 14.178 | 8.9108 |
| 16697 | ENSMUSG00000058921 | 852.3144929 | 0.219178611 | 0.157669705 | 1.3901124 | 0.164494749 | 0.589361417 | Slc10a5 | solute carrier family 10 (sodium/bile acid | 615.98 | 780.19 | 968.01 | 852.16 | 961.93 | 935.63 |
| 16698 | ENSMUSG00000058922 | 6.732375574 | 0.142265932 | 0.8220059 | 0.173071668 | 0.862595095 | 0.976334255 | Gm10052 | predicted pseudogene 10052 [Source:MG | 4.0155 | 8.2233 | 7.0437 | 6.7478 | 5.4531 | 8.9108 |
| 16699 | ENSMUSG00000058925 | 0 | NA | NA | NA | NA | NA | Ccdc192 | coiled-coil domain containing 192 [Source | 0 | 0 | 0 | 0 | 0 | 0 |
| 16700 | ENSMUSG00000058927 | 11.7581714 | 0.100090308 | 0.684514808 | 0.146220807 | 0.88374708 | 0.981769403 | Gm10053 | predicted gene 10053 [Source:MGI Symbo | 14.456 | 13.363 | 6.0375 | 5.7839 | 13.087 | 17.822 |
| 16701 | ENSMUSG00000058932 | 0 | NA | NA | NA | NA | NA | Gm2174 | predicted gene 2174 [Source:MGI Symbol | 0 | 0 | 0 | 0 | 0 | 0 |
| 16702 | ENSMUSG00000058933 | 0 | NA | NA | NA | NA | NA | Gm5185 | predicted gene 5185 [Source:MGI Symbol | 0 | 0 | 0 | 0 | 0 | 0 |
| 16703 | ENSMUSG00000058934 | 25.36417849 | 0.196669801 | 0.43782464 | 0.449197652 | 0.653289081 | 0.925756399 | Igf1os | insulin-like growth factor 1, opposite stra | 28.912 | 23.642 | 18.112 | 23.135 | 30.537 | 27.846 |
| 16704 | ENSMUSG00000058935 | 0 | NA | NA | NA | NA | NA | Tex48 | testis expressed 48 [Source:MGI Symbol;A | 0 | 0 | 0 | 0 | 0 | 0 |
| 16705 | ENSMUSG00000058952 | 10487.79917 | 0.055433627 | 0.094033902 | 0.589506823 | 0.555521336 | 0.897628185 | Cfi | complement component factor i [Source:I | 9968.9 | 10354 | 10537 | 11699 | 10161 | 10206 |
| 16706 | ENSMUSG00000058953 | 0 | NA | NA | NA | NA | NA | Gm4822 | predicted pseudogene 4822 [Source:MGI | 0 | 0 | 0 | 0 | 0 | 0 |
| 16707 | ENSMUSG00000058966 | 3.091981872 | -1.366535096 | 1.330272834 | -1.027259267 | 0.304298397 | NA | Tlcd3b | TLC domain containing 3B [Source:MGI Sy | 7.2279 | 4.1117 | 2.0125 | 1.928 | 3.2719 | 0 |
| 16708 | ENSMUSG00000058975 | 0 | NA | NA | NA | NA | NA | Kcnc1 | potassium voltage gated channel, Shaw-re | 0 | 0 | 0 | 0 | 0 | 0 |
| 16709 | ENSMUSG00000058976 | 0 | NA | NA | NA | NA | NA | Usp17lc | ubiquitin specific peptidase 17-like C [Sou | 0 | 0 | 0 | 0 | 0 | 0 |
| 16710 | ENSMUSG00000058979 | 516.6897186 | -0.026892273 | 0.136929116 | -0.196395576 | 0.844300558 | 0.972803041 | Hdhd5 | haloacid dehalogenase like hydrolase dom | 454.56 | 536.57 | 574.57 | 526.33 | 519.13 | 488.98 |
| 16711 | ENSMUSG00000058981 | 0 | NA | NA | NA | NA | NA | Olfr1406 | olfactory receptor 1406 [Source:MGI Sym | 0 | 0 | 0 | 0 | 0 | 0 |
| 16712 | ENSMUSG00000058997 | 5369.433526 | -0.158402004 | 0.107816918 | -1.469175773 | 0.14178512 | 0.551284939 | Vwa8 | von Willebrand factor A domain containin | 5767.1 | 5939.3 | 5285.8 | 5352 | 5420.4 | 4452 |
| 16713 | ENSMUSG00000059003 | 0 | NA | NA | NA | NA | NA | Grin2a | glutamate receptor, ionotropic, NMDA2A | 0 | 0 | 0 | 0 | 0 | 0 |
| 16714 | ENSMUSG00000059005 | 2597.664665 | 0.201038493 | 0.094452944 | 2.128451321 | 0.033299682 | 0.243995238 | Hnrnpa3 | heterogeneous nuclear ribonucleoprotein | 2295.3 | 2402.2 | 2553.8 | 2679.9 | 2782.2 | 2872.6 |
| 16715 | ENSMUSG00000059013 | 128.7513548 | -0.258665515 | 0.225065944 | -1.149287676 | 0.250437378 | 0.702302228 | Sh2d3c | SH2 domain containing 3C [Source:MGI Sy | 119.66 | 132.6 | 169.05 | 127.25 | 101.43 | 122.52 |
| 16716 | ENSMUSG00000059014 | 0 | NA | NA | NA | NA | NA | Gm5409 | predicted pseudogene 5409 [Source:MGI | 0 | 0 | 0 | 0 | 0 | 0 |
| 16717 | ENSMUSG00000059022 | 96.33339352 | 0.688488011 | 0.247927271 | 2.776975716 | 0.005486728 | 0.070827854 | Kcp | kielin/chordin-like protein [Source:MGI Sy | 73.082 | 61.675 | 86.537 | 115.68 | 136.33 | 104.7 |
| 16718 | ENSMUSG00000059023 | 0 | NA | NA | NA | NA | NA | Olfr1201 | olfactory receptor 1201 [Source:MGI Sym | 0 | 0 | 0 | 0 | 0 | 0 |
| 16719 | ENSMUSG00000059027 | 187.9508512 | 0.048932312 | 0.216850922 | 0.225649545 | 0.821474003 | 0.969160361 | 9630013D21 | RIKEN cDNA 9630013D21 gene [Source:M | 206.4 | 186.05 | 161 | 142.67 | 236.66 | 194.92 |
| 16720 | ENSMUSG00000059030 | 0 | NA | NA | NA | NA | NA | Olfr128 | olfactory receptor 128 [Source:MGI Symb | 0 | 0 | 0 | 0 | 0 | 0 |
| 16721 | ENSMUSG00000059031 | 0 | NA | NA | NA | NA | NA | Olfr482 | olfactory receptor 482 [Source:MGI Symb | 0 | 0 | 0 | 0 | 0 | 0 |
| 16722 | ENSMUSG00000059033 | 0.185640647 | 1.126924175 | 4.080472857 | 0.276174898 | 0.782413736 | NA | Rpl18a-ps1 | ribosomal protein L18A, pseudogene 1 [S | 0 | 0 | 0 | 0 | 0 | 1.1138 |
| 16723 | ENSMUSG00000059040 | 5.210856015 | 0.464150994 | 0.958806116 | 0.48409265 | 0.628320117 | 0.920328819 | Eno1b | enolase 1B, retrotransposed [Source:MGI | 4.0155 | 2.0558 | 7.0437 | 3.8559 | 8.7249 | 5.5692 |
| 16724 | ENSMUSG00000059042 | 0 | NA | NA | NA | NA | NA | Klk1b9 | kallikrein 1-related peptidase b9 [Source: | 0 | 0 | 0 | 0 | 0 | 0 |
| 16725 | ENSMUSG00000059043 | 0 | NA | NA | NA | NA | NA | Olfr15 | olfactory receptor 15 [Source:MGI Symbo | 0 | 0 | 0 | 0 | 0 | 0 |
| 16726 | ENSMUSG00000059049 | 5.591693584 | -1.575715436 | 0.978658823 | -1.61007636 | 0.107381187 | 0.48315751 | Frem1 | Fras1 related extracellular matrix protein | 4.8186 | 11.307 | 9.0562 | 2.8919 | 4.3625 | 1.1138 |
| 16727 | ENSMUSG00000059058 | 2.404739671 | 1.059227085 | 1.391450862 | 0.761239303 | 0.446514147 | NA | Tma7-ps | translational machinery associated 7 hom | 1.6062 | 1.0279 | 2.0125 | 0.964 | 4.3625 | 4.4554 |
| 16728 | ENSMUSG00000059060 | 47.03893096 | -2.273650504 | 0.474760311 | -4.789049229 | 1.68E-06 | 0.000124296 | Rad51b | RAD51 paralog B [Source:MGI Symbol;Acc | 89.144 | 104.85 | 40.25 | 26.991 | 7.6343 | 13.366 |
| 16729 | ENSMUSG00000059064 | 0.331982614 | 0.16514309 | 3.986751476 | 0.041422971 | 0.966958701 | NA | Gm10059 | predicted pseudogene 10059 [Source:MG | 0 | 1.0279 | 0 | 0.964 | 0 | 0 |
| 16730 | ENSMUSG00000059069 | 0 | NA | NA | NA | NA | NA | Olfr749 | olfactory receptor 749 [Source:MGI Symb | 0 | 0 | 0 | 0 | 0 | 0 |
| 16731 | ENSMUSG00000059070 | 3253.193956 | 0.134947501 | 0.093647376 | 1.441017421 | 0.149579761 | 0.562391907 | Rpl18 | ribosomal protein L18 [Source:MGI Symbo | 3018.1 | 3072.4 | 3212.9 | 3285.2 | 3288.2 | 3642.3 |
| 16732 | ENSMUSG00000059077 | 0 | NA | NA | NA | NA | NA | Pth | parathyroid hormone [Source:MGI Symbo | 0 | 0 | 0 | 0 | 0 | 0 |
| 16733 | ENSMUSG00000059087 | 0.331982614 | 0.16514309 | 3.986751476 | 0.041422971 | 0.966958701 | NA | Olfr698 | olfactory receptor 698 [Source:MGI Symb | 0 | 1.0279 | 0 | 0.964 | 0 | 0 |
| 16734 | ENSMUSG00000059089 | 42.52924544 | -1.204342757 | 0.456952461 | -2.635597485 | 0.00839893 | 0.095354881 | Fcgr4 | Fc receptor, IgG, low affinity IV [Source:M | 60.233 | 91.485 | 26.162 | 23.135 | 19.631 | 34.529 |
| 16735 | ENSMUSG00000059105 | 0 | NA | NA | NA | NA | NA | Olfr1504 | olfactory receptor 1504 [Source:MGI Sym | 0 | 0 | 0 | 0 | 0 | 0 |
| 16736 | ENSMUSG00000059106 | 0 | NA | NA | NA | NA | NA | Olfr961 | olfactory receptor 961 [Source:MGI Symb | 0 | 0 | 0 | 0 | 0 | 0 |
| 16737 | ENSMUSG00000059108 | 1.210023386 | -2.482020224 | 2.285496674 | -1.085987239 | 0.277484638 | NA | Ifitm6 | interferon induced transmembrane prote | 3.2124 | 3.0838 | 0 | 0.964 | 0 | 0 |
| 16738 | ENSMUSG00000059112 | 0 | NA | NA | NA | NA | NA | Olfr1263 | olfactory receptor 1263 [Source:MGI Sym | 0 | 0 | 0 | 0 | 0 | 0 |
| 16739 | ENSMUSG00000059113 | 0 | NA | NA | NA | NA | NA | Gm10061 | predicted gene 10061 [Source:MGI Symbo | 0 | 0 | 0 | 0 | 0 | 0 |
| 16740 | ENSMUSG00000059114 | 0 | NA | NA | NA | NA | NA | Lrrc74a | leucine rich repeat containing 74A [Sourc | 0 | 0 | 0 | 0 | 0 | 0 |

| | | | | | | | | | | | | | | | |
|---|---|---|---|---|---|---|---|---|---|---|---|---|---|---|---|
| 16741 | ENSMUSG00000059119 | 917.3406398 | -0.029084126 | 0.112829799 | -0.257769903 | 0.796584494 | 0.962236316 | Nap1l4 | nucleosome assembly protein 1-like 4 [Sou | 902.69 | 972.41 | 904.61 | 870.47 | 869.22 | 984.64 |
| 16742 | ENSMUSG00000059128 | 0 | NA | NA | NA | NA | NA | Ifnl2 | interferon lambda 2 [Source:MGI Symbol; | 0 | 0 | 0 | 0 | 0 | 0 |
| 16743 | ENSMUSG00000059134 | 0 | NA | NA | NA | NA | NA | Olfr814 | olfactory receptor 814 [Source:MGI Symb | 0 | 0 | 0 | 0 | 0 | 0 |
| 16744 | ENSMUSG00000059136 | 3.886775375 | 1.382200282 | 1.193130229 | 1.158465563 | 0.246674097 | NA | Olfr539 | olfactory receptor 539 [Source:MGI Symb | 2.4093 | 0 | 4.025 | 4.8199 | 8.7249 | 3.3415 |
| 16745 | ENSMUSG00000059142 | 189.9639725 | 0.288519214 | 0.181197196 | 1.592294036 | 0.111318652 | 0.489450944 | Zfp945 | zinc finger protein 945 [Source:MGI Symb | 167.05 | 191.19 | 154.96 | 201.47 | 226.85 | 198.26 |
| 16746 | ENSMUSG00000059146 | 1.741735561 | -3.042880744 | 1.882697693 | -1.616234383 | 0.106043647 | NA | Ntrk3 | neurotrophic tyrosine kinase, receptor, ty | 3.2124 | 4.1117 | 2.0125 | 0 | 0 | 1.1138 |
| 16747 | ENSMUSG00000059149 | 271.5071158 | 0.265562567 | 0.159909534 | 1.660705032 | 0.096772702 | 0.458530879 | Mfsd4a | major facilitator superfamily domain cont | 244.95 | 245.67 | 249.55 | 322.93 | 308.64 | 257.3 |
| 16748 | ENSMUSG00000059150 | 0.167707431 | -0.796629176 | 4.080472857 | -0.195229623 | 0.84521319 | NA | Rpl26-ps4 | ribosomal protein L26, pseudogene 4 [Sou | 0 | 0 | 1.0062 | 0 | 0 | 0 |
| 16749 | ENSMUSG00000059159 | 1.891612627 | -3.160701012 | 2.083290426 | -1.517167733 | 0.129224341 | NA | Gm8129 | predicted pseudogene 8129 [Source:MGI | 0 | 8.2233 | 2.0125 | 0 | 0 | 1.1138 |
| 16750 | ENSMUSG00000059169 | 0 | NA | NA | NA | NA | NA | Krt40 | keratin 40 [Source:MGI Symbol;Acc:MGI:3 | 0 | 0 | 0 | 0 | 0 | 0 |
| 16751 | ENSMUSG00000059173 | 34.0053876 | -0.547344094 | 0.402821182 | -1.358776844 | 0.174217313 | 0.604151393 | Pde1a | phosphodiesterase 1A, calmodulin-depen | 33.73 | 40.089 | 47.293 | 20.244 | 38.172 | 24.505 |
| 16752 | ENSMUSG00000059179 | 0.685330692 | 0.069534182 | 2.586556116 | 0.02688292 | 0.978553116 | NA | Gm8991 | predicted pseudogene 8991 [Source:MGI | 0 | 1.0279 | 1.0062 | 0.964 | 0 | 1.1138 |
| 16753 | ENSMUSG00000059182 | 433.6547834 | -0.049759013 | 0.139869304 | -0.355753631 | 0.722025077 | 0.9447719 | Skap2 | src family associated phosphoprotein 2 [S | 409.58 | 468.73 | 445.77 | 428.97 | 464.6 | 384.28 |
| 16754 | ENSMUSG00000059183 | 170.9099972 | 0.054204466 | 0.197995663 | 0.273765923 | 0.784264513 | 0.960670212 | Mtfmt | mitochondrial methionyl-tRNA formyltran | 142.95 | 164.47 | 196.22 | 175.44 | 158.14 | 188.24 |
| 16755 | ENSMUSG00000059187 | 0.449470461 | -0.545136533 | 3.562105917 | -0.15303771 | 0.878368537 | NA | Tafa1 | TAFA chemokine like family member 1 [So | 1.6062 | 0 | 0 | 0 | 1.0906 | 0 |
| 16756 | ENSMUSG00000059189 | 0 | NA | NA | NA | NA | NA | Olfr913 | olfactory receptor 913 [Source:MGI Symb | 0 | 0 | 0 | 0 | 0 | 0 |
| 16757 | ENSMUSG00000059195 | 2.851352451 | -0.817516563 | 1.247742358 | -0.655196611 | 0.512341176 | NA | Gm12715 | predicted gene 12715 [Source:MGI Symbo | 4.8186 | 3.0838 | 3.0187 | 2.8919 | 2.1812 | 1.1138 |
| 16758 | ENSMUSG00000059201 | 0 | NA | NA | NA | NA | NA | Lep | leptin [Source:MGI Symbol;Acc:MGI:1046 | 0 | 0 | 0 | 0 | 0 | 0 |
| 16759 | ENSMUSG00000059203 | 0 | NA | NA | NA | NA | NA | Il1rapl2 | interleukin 1 receptor accessory protein-li | 0 | 0 | 0 | 0 | 0 | 0 |
| 16760 | ENSMUSG00000059205 | 0 | NA | NA | NA | NA | NA | Olfr1130 | olfactory receptor 1130 [Source:MGI Sym | 0 | 0 | 0 | 0 | 0 | 0 |
| 16761 | ENSMUSG00000059206 | 0 | NA | NA | NA | NA | NA | Vmn1r71 | vomeronasal 1 receptor 71 [Source:MGI S | 0 | 0 | 0 | 0 | 0 | 0 |
| 16762 | ENSMUSG00000059208 | 1548.119054 | 0.049027308 | 0.10497709 | 0.467028643 | 0.640479385 | 0.923932929 | Hnrnpm | heterogeneous nuclear ribonucleoprotein | 1426.3 | 1569.6 | 1569.7 | 1462.4 | 1599.9 | 1660.7 |
| 16763 | ENSMUSG00000059213 | 0 | NA | NA | NA | NA | NA | Ddn | dendrin [Source:MGI Symbol;Acc:MGI:10 | 0 | 0 | 0 | 0 | 0 | 0 |
| 16764 | ENSMUSG00000059218 | 0 | NA | NA | NA | NA | NA | Pramel26 | PRAME like 26 [Source:MGI Symbol;Acc:M | 0 | 0 | 0 | 0 | 0 | 0 |
| 16765 | ENSMUSG00000059229 | 0 | NA | NA | NA | NA | NA | Gm6802 | predicted gene 6802 [Source:MGI Symbol | 0 | 0 | 0 | 0 | 0 | 0 |
| 16766 | ENSMUSG00000059230 | 0 | NA | NA | NA | NA | NA | Defb4 | defensin beta 4 [Source:MGI Symbol;Acc: | 0 | 0 | 0 | 0 | 0 | 0 |
| 16767 | ENSMUSG00000059237 | 0.898940922 | -0.606261109 | 2.77650979 | -0.218353672 | 0.827153559 | NA | Ppp1r2-ps4 | protein phosphatase 1, regulatory (inhibit | 3.2124 | 0 | 0 | 0 | 2.1812 | 0 |
| 16768 | ENSMUSG00000059244 | 0 | NA | NA | NA | NA | NA | Gm10062 | predicted gene 10062 [Source:MGI Symbo | 0 | 0 | 0 | 0 | 0 | 0 |
| 16769 | ENSMUSG00000059246 | 0 | NA | NA | NA | NA | NA | Foxb1 | forkhead box B1 [Source:MGI Symbol;Acc | 0 | 0 | 0 | 0 | 0 | 0 |
| 16770 | ENSMUSG00000059248 | 1251.497431 | -0.132001188 | 0.107305063 | -1.230148734 | 0.218641413 | 0.663735712 | Septin9 | septin 9 [Source:MGI Symbol;Acc:MGI:18 | 1311.5 | 1298.3 | 1316.2 | 1182.8 | 1100.4 | 1299.9 |
| 16771 | ENSMUSG00000059251 | 0 | NA | NA | NA | NA | NA | Olfr345 | olfactory receptor 345 [Source:MGI Symb | 0 | 0 | 0 | 0 | 0 | 0 |
| 16772 | ENSMUSG00000059252 | 4.21511885 | 0.52847549 | 1.056691187 | 0.500122928 | 0.616988523 | NA | Tpt1-ps5 | tumor protein, translationally-controlled, | 3.2124 | 4.1117 | 3.0187 | 3.8559 | 2.1812 | 8.9108 |
| 16773 | ENSMUSG00000059256 | 0 | NA | NA | NA | NA | NA | Gzmd | granzyme D [Source:MGI Symbol;Acc:MGI | 0 | 0 | 0 | 0 | 0 | 0 |
| 16774 | ENSMUSG00000059263 | 1770.268673 | 0.129269358 | 0.096308526 | 1.342242104 | 0.179517509 | 0.611676727 | Usp47 | ubiquitin specific peptidase 47 [Source:M | 1621.5 | 1692 | 1759.9 | 1779.5 | 1905.3 | 1863.5 |
| 16775 | ENSMUSG00000059273 | 484.5046594 | -0.067674626 | 0.134400364 | -0.503530076 | 0.61459164 | 0.917509899 | Zc3h4 | zinc finger CCCH-type containing 4 [Sourc | 478.65 | 544.8 | 464.88 | 503.2 | 474.42 | 441.08 |
| 16776 | ENSMUSG00000059277 | 14.06374297 | -0.84171527 | 0.582705006 | -1.444496377 | 0.148599398 | 0.560454158 | R74862 | expressed sequence R74862 [Source:MGI | 17.668 | 17.475 | 19.119 | 12.532 | 10.906 | 6.6831 |
| 16777 | ENSMUSG00000059278 | 351.1052575 | 0.164132059 | 0.149184237 | 1.100197057 | 0.271246273 | 0.726942021 | Naa38 | N(alpha)-acetyltransferase 38, NatC auxili | 359.79 | 318.65 | 313.95 | 331.61 | 384.99 | 397.64 |
| 16778 | ENSMUSG00000059279 | 0 | NA | NA | NA | NA | NA | Olfr224 | olfactory receptor 224 [Source:MGI Symb | 0 | 0 | 0 | 0 | 0 | 0 |
| 16779 | ENSMUSG00000059280 | 0 | NA | NA | NA | NA | NA | Vpreb2 | pre-B lymphocyte gene 2 [Source:MGI Syr | 0 | 0 | 0 | 0 | 0 | 0 |
| 16780 | ENSMUSG00000059288 | 102.3940909 | 0.216700204 | 0.232725939 | 0.931139028 | 0.351781652 | 0.792889192 | Cdyl | chromodomain protein, Y chromosome-lil | 81.113 | 109.99 | 93.581 | 114.71 | 104.7 | 110.27 |
| 16781 | ENSMUSG00000059291 | 3956.198993 | 0.06379456 | 0.093360561 | 0.683313803 | 0.494408574 | 0.877473722 | Rpl11 | ribosomal protein L11 [Source:MGI Symbo | 3937.6 | 3914.3 | 3753.3 | 3792.3 | 4011.3 | 4328.4 |
| 16782 | ENSMUSG00000059301 | 0 | NA | NA | NA | NA | NA | Gm5434 | predicted gene 5434 [Source:MGI Symbol | 0 | 0 | 0 | 0 | 0 | 0 |
| 16783 | ENSMUSG00000059303 | 0 | NA | NA | NA | NA | NA | Olfr836 | olfactory receptor 836 [Source:MGI Symb | 0 | 0 | 0 | 0 | 0 | 0 |
| 16784 | ENSMUSG00000059305 | 1.798502144 | 0.030273587 | 1.759132864 | 0.017209381 | 0.986269578 | NA | Vpreb1 | pre-B lymphocyte gene 1 [Source:MGI Syr | 3.2124 | 2.0558 | 0 | 0 | 2.1812 | 3.3415 |
| 16785 | ENSMUSG00000059316 | 671.8065978 | -0.052321589 | 0.115989695 | -0.451088251 | 0.651925945 | 0.925643431 | Slc27a4 | solute carrier family 27 (fatty acid transpo | 645.69 | 712.35 | 694.31 | 638.15 | 662.01 | 678.33 |
| 16786 | ENSMUSG00000059319 | 0 | NA | NA | NA | NA | NA | Olfr295 | olfactory receptor 295 [Source:MGI Symb | 0 | 0 | 0 | 0 | 0 | 0 |
| 16787 | ENSMUSG00000059323 | 24.21947279 | 0.60794749 | 0.467539012 | 1.300313929 | 0.193493396 | 0.631833452 | Tonsl | tonsoku-like, DNA repair protein [Source:I | 20.078 | 18.503 | 19.119 | 42.415 | 21.812 | 23.391 |
| 16788 | ENSMUSG00000059325 | 1056.835683 | 0.22788354 | 0.122689378 | 1.857402364 | 0.06325393 | 0.359877642 | Hopx | HOP homeobox [Source:MGI Symbol;Acc: | 1095.4 | 981.66 | 842.23 | 1132.7 | 1147.3 | 1141.7 |
| 16789 | ENSMUSG00000059326 | 57.32401763 | -1.183816896 | 0.337381932 | -3.50883312 | 0.000450077 | 0.011481335 | Csf2ra | colony stimulating factor 2 receptor, alpha | 86.735 | 99.708 | 52.325 | 34.703 | 38.172 | 32.301 |
| 16790 | ENSMUSG00000059327 | 42.53914576 | -0.767582419 | 0.359886234 | -2.132847402 | 0.032937246 | 0.242815663 | Eda | ectodysplasin-A [Source:MGI Symbol;Acc: | 56.217 | 43.173 | 61.381 | 31.811 | 39.262 | 23.391 |
| 16791 | ENSMUSG00000059334 | 0 | NA | NA | NA | NA | NA | Zfp36l3 | zinc finger protein 36, C3H type-like 3 [Sou | 0 | 0 | 0 | 0 | 0 | 0 |
| 16792 | ENSMUSG00000059336 | 1.499064621 | -0.783124799 | 1.914747145 | -0.408996457 | 0.682542261 | NA | Slc14a1 | solute carrier family 14 (urea transporter) | 1.6062 | 1.0279 | 3.0187 | 0 | 0 | 3.3415 |
| 16793 | ENSMUSG00000059343 | 0 | NA | NA | NA | NA | NA | Aldoart1 | aldolase 1 A, retrogene 1 [Source:MGI Syr | 0 | 0 | 0 | 0 | 0 | 0 |
| 16794 | ENSMUSG00000059352 | 0 | NA | NA | NA | NA | NA | Gm10064 | predicted gene 10064 [Source:MGI Symbo | 0 | 0 | 0 | 0 | 0 | 0 |
| 16795 | ENSMUSG00000059355 | 361.1840043 | -0.206109816 | 0.141911298 | -1.452384829 | 0.146394632 | 0.556855766 | Wdr83os | WD repeat domain 83 opposite strand [Sc | 390.31 | 373.13 | 397.47 | 342.21 | 359.9 | 304.08 |
| 16796 | ENSMUSG00000059361 | 0.371281295 | 2.09253362 | 4.031852165 | 0.519000582 | 0.603760335 | NA | Nrsn2 | neurensin 2 [Source:MGI Symbol;Acc:MG | 0 | 0 | 0 | 0 | 0 | 2.2277 |

| | | | | | | | | | | | | | | | |
|---|---|---|---|---|---|---|---|---|---|---|---|---|---|---|---|
| 16797 | ENSMUSG00000059363 | 296.6125897 | 0.134245906 | 0.152613774 | 0.879644758 | 0.379051784 | 0.812043903 | Fxn | frataxin [Source:MGI Symbol;Acc:MGI:10! | 299.56 | 277.54 | 270.68 | 291.12 | 298.83 | 341.95 |
| 16798 | ENSMUSG00000059366 | 0 | NA | NA | NA | NA | NA | Olfr959 | olfactory receptor 959 [Source:MGI Symb | 0 | 0 | 0 | 0 | 0 | 0 |
| 16799 | ENSMUSG00000059371 | 0 | NA | NA | NA | NA | NA | Olfr427 | olfactory receptor 427 [Source:MGI Symb | 0 | 0 | 0 | 0 | 0 | 0 |
| 16800 | ENSMUSG00000059375 | 0 | NA | NA | NA | NA | NA | Vmn1r33 | vomeronasal 1 receptor 33 [Source:MGI S | 0 | 0 | 0 | 0 | 0 | 0 |
| 16801 | ENSMUSG00000059379 | 0 | NA | NA | NA | NA | NA | Olfr1014 | olfactory receptor 1014 [Source:MGI Sym | 0 | 0 | 0 | 0 | 0 | 0 |
| 16802 | ENSMUSG00000059382 | 0 | NA | NA | NA | NA | NA | Tas2r120 | taste receptor, type 2, member 120 [Sour | 0 | 0 | 0 | 0 | 0 | 0 |
| 16803 | ENSMUSG00000059383 | 0 | NA | NA | NA | NA | NA | Gfral | GDNF family receptor alpha like [Source:N | 0 | 0 | 0 | 0 | 0 | 0 |
| 16804 | ENSMUSG00000059395 | 8.995988321 | 0.203923821 | 0.729776904 | 0.279433098 | 0.779912474 | 0.959870443 | Nkapl | NFKB activating protein-like [Source:MGI ! | 4.8186 | 10.279 | 10.062 | 6.7478 | 9.8156 | 12.252 |
| 16805 | ENSMUSG00000059397 | 0 | NA | NA | NA | NA | NA | Olfr54 | olfactory receptor 54 [Source:MGI Symbo | 0 | 0 | 0 | 0 | 0 | 0 |
| 16806 | ENSMUSG00000059401 | 9.913255411 | -0.341159635 | 0.669159366 | -0.509833161 | 0.610168351 | 0.916376001 | Mamld1 | mastermind-like domain containing 1 [Sou | 12.85 | 10.279 | 10.062 | 8.6758 | 9.8156 | 7.7969 |
| 16807 | ENSMUSG00000059406 | 3.316973334 | -0.122312747 | 1.294915715 | -0.094456145 | 0.924746818 | NA | Tmprss9 | transmembrane protease, serine 9 [Sourc | 7.2279 | 0 | 3.0187 | 1.928 | 3.2719 | 4.4554 |
| 16808 | ENSMUSG00000059408 | 0 | NA | NA | NA | NA | NA | Mrgprh | MAS-related GPR, member H [Source:MG | 0 | 0 | 0 | 0 | 0 | 0 |
| 16809 | ENSMUSG00000059409 | 547.838683 | 0.074076778 | 0.143367948 | 0.516689948 | 0.605372614 | 0.916376001 | Ppp2r5d | protein phosphatase 2, regulatory subuni | 469.81 | 614.69 | 518.22 | 612.13 | 544.22 | 527.96 |
| 16810 | ENSMUSG00000059410 | 0 | NA | NA | NA | NA | NA | Tas2r125 | taste receptor, type 2, member 125 [Sour | 0 | 0 | 0 | 0 | 0 | 0 |
| 16811 | ENSMUSG00000059411 | 0 | NA | NA | NA | NA | NA | Olfr434 | olfactory receptor 434 [Source:MGI Symb | 0 | 0 | 0 | 0 | 0 | 0 |
| 16812 | ENSMUSG00000059412 | 0.717584917 | 3.048786435 | 2.951298099 | 1.033032358 | 0.301588756 | NA | Fxyd2 | FXYD domain-containing ion transport reg | 0 | 0 | 0 | 0.964 | 0 | 3.3415 |
| 16813 | ENSMUSG00000059422 | 2.734247596 | 0.868235569 | 1.364008727 | 0.636532268 | 0.524429556 | NA | Gm8116 | predicted gene 8116 [Source:MGI Symbol | 0.8031 | 1.0279 | 4.025 | 2.8919 | 6.5437 | 1.1138 |
| 16814 | ENSMUSG00000059423 | 96.41740467 | 0.015538577 | 0.233535988 | 0.066536112 | 0.946951008 | 0.990677354 | Zfp933 | zinc finger protein 933 [Source:MGI Symb | 96.372 | 97.652 | 93.581 | 94.47 | 86.159 | 110.27 |
| 16815 | ENSMUSG00000059429 | 0 | NA | NA | NA | NA | NA | Olfr365 | olfactory receptor 365 [Source:MGI Symb | 0 | 0 | 0 | 0 | 0 | 0 |
| 16816 | ENSMUSG00000059430 | 1.367074152 | -1.655813959 | 2.003731625 | -0.826365137 | 0.408596994 | NA | Actg2 | actin, gamma 2, smooth muscle, enteric [! | 3.2124 | 2.0558 | 1.0062 | 1.928 | 0 | 0 |
| 16817 | ENSMUSG00000059434 | 4240.993775 | 0.037000506 | 0.093582427 | 0.395378782 | 0.692563366 | 0.938470725 | Gckr | glucokinase regulatory protein [Source:M | 4379.3 | 4084.9 | 4095.4 | 4568.3 | 4036.4 | 4281.6 |
| 16818 | ENSMUSG00000059436 | 351.870677 | -0.025557373 | 0.18212245 | -0.140330711 | 0.8883987 | 0.982382091 | Max | Max protein [Source:MGI Symbol;Acc:MG | 405.57 | 342.3 | 315.96 | 298.83 | 310.83 | 437.74 |
| 16819 | ENSMUSG00000059439 | 409.0247168 | 0.040126568 | 0.142331835 | 0.281922645 | 0.778002825 | 0.959559604 | Bcas3 | breast carcinoma amplified sequence 3 [S | 404.76 | 397.8 | 407.53 | 417.4 | 462.42 | 364.23 |
| 16820 | ENSMUSG00000059447 | 4444.892345 | -0.100686562 | 0.091304073 | -1.102760901 | 0.270131012 | 0.725566379 | Hadhb | hydroxyacyl-CoA dehydrogenase trifuncti | 4517.5 | 4743.8 | 4539.2 | 4358.1 | 4504.3 | 4006.5 |
| 16821 | ENSMUSG00000059455 | 0 | NA | NA | NA | NA | NA | Plac8l1 | PLAC8-like 1 [Source:MGI Symbol;Acc:MG | 0 | 0 | 0 | 0 | 0 | 0 |
| 16822 | ENSMUSG00000059456 | 928.3869607 | 0.164395767 | 0.107406959 | 1.530587667 | 0.125871332 | 0.520320932 | Ptk2b | PTK2 protein tyrosine kinase 2 beta [Sour | 878.59 | 850.09 | 897.57 | 949.52 | 957.56 | 1037 |
| 16823 | ENSMUSG00000059459 | 0 | NA | NA | NA | NA | NA | NA | NA | 0 | 0 | 0 | 0 | 0 | 0 |
| 16824 | ENSMUSG00000059460 | 0 | NA | NA | NA | NA | NA | Olfr286 | olfactory receptor 286 [Source:MGI Symb | 0 | 0 | 0 | 0 | 0 | 0 |
| 16825 | ENSMUSG00000059461 | 6.997490724 | 0.59920493 | 0.834733259 | 0.717840009 | 0.472855941 | 0.86857093 | Gm7331 | predicted gene 7331 [Source:MGI Symbol | 6.4248 | 8.2233 | 2.0125 | 7.7118 | 9.8156 | 7.7969 |
| 16826 | ENSMUSG00000059463 | 0 | NA | NA | NA | NA | NA | Spag11b | sperm associated antigen 11B [Source:MC | 0 | 0 | 0 | 0 | 0 | 0 |
| 16827 | ENSMUSG00000059473 | 0 | NA | NA | NA | NA | NA | Olfr979 | olfactory receptor 979 [Source:MGI Symb | 0 | 0 | 0 | 0 | 0 | 0 |
| 16828 | ENSMUSG00000059474 | 342.717699 | 0.259731153 | 0.143509773 | 1.809849931 | 0.070319062 | 0.383450475 | Mbtd1 | mbt domain containing 1 [Source:MGI Syr | 289.92 | 319.68 | 327.03 | 394.27 | 352.27 | 373.14 |
| 16829 | ENSMUSG00000059475 | 153.2208252 | 0.16868543 | 0.227107606 | 0.742755527 | 0.457629705 | 0.859184328 | Zfp426 | zinc finger protein 426 [Source:MGI Symb | 174.27 | 140.82 | 116.72 | 129.17 | 172.32 | 186.01 |
| 16830 | ENSMUSG00000059479 | 47.15984762 | 0.017150316 | 0.336872203 | 0.050910451 | 0.959396877 | 0.992148164 | B3gnt8 | UDP-GlcNAc:betaGal beta-1,3-N-acetylglu | 47.383 | 34.949 | 58.362 | 53.019 | 45.806 | 43.44 |
| 16831 | ENSMUSG00000059481 | 32314.87536 | 0.162125753 | 0.097621228 | 1.660763311 | 0.096760992 | 0.458530879 | Plg | plasminogen [Source:MGI Symbol;Acc:MC | 27132 | 31423 | 32949 | 33993 | 34604 | 33787 |
| 16832 | ENSMUSG00000059482 | 139.6592744 | -0.005552029 | 0.22024133 | -0.025208841 | 0.979888385 | 0.99820894 | 2610301B20 | RIKEN cDNA 2610301B20 gene [Source:M | 155 | 115.13 | 148.92 | 114.71 | 151.6 | 152.6 |
| 16833 | ENSMUSG00000059483 | 0 | NA | NA | NA | NA | NA | Gm5528 | predicted gene 5528 [Source:MGI Symbol | 0 | 0 | 0 | 0 | 0 | 0 |
| 16834 | ENSMUSG00000059486 | 263.0045804 | -0.038808077 | 0.160378253 | -0.241978425 | 0.808796881 | 0.965682062 | Kbtbd2 | kelch repeat and BTB (POZ) domain conta | 238.52 | 297.07 | 264.64 | 255.45 | 263.93 | 258.41 |
| 16835 | ENSMUSG00000059488 | 0 | NA | NA | NA | NA | NA | Olfr727 | olfactory receptor 727 [Source:MGI Symb | 0 | 0 | 0 | 0 | 0 | 0 |
| 16836 | ENSMUSG00000059493 | 0 | NA | NA | NA | NA | NA | Nhs | NHS actin remodeling regulator [Source:N | 0 | 0 | 0 | 0 | 0 | 0 |
| 16837 | ENSMUSG00000059495 | 2373.266555 | 0.081681206 | 0.091061896 | 0.89698557 | 0.369726617 | 0.80653433 | Arhgef12 | Rho guanine nucleotide exchange factor ( | 2263.1 | 2288.1 | 2367.7 | 2526.6 | 2351.4 | 2442.7 |
| 16838 | ENSMUSG00000059498 | 60.36082852 | -1.489876239 | 0.35067319 | -4.248617465 | 2.15E-05 | 0.001026771 | Fcgr3 | Fc receptor, IgG, low affinity III [Source:M | 92.357 | 119.24 | 55.343 | 27.955 | 31.628 | 35.643 |
| 16839 | ENSMUSG00000059503 | 0 | NA | NA | NA | NA | NA | Olfr248 | olfactory receptor 248 [Source:MGI Symb | 0 | 0 | 0 | 0 | 0 | 0 |
| 16840 | ENSMUSG00000059504 | 0 | NA | NA | NA | NA | NA | Olfr314 | olfactory receptor 314 [Source:MGI Symb | 0 | 0 | 0 | 0 | 0 | 0 |
| 16841 | ENSMUSG00000059518 | 597.6208631 | 0.11835541 | 0.133288213 | 0.887966064 | 0.374559006 | 0.809332934 | Znhit1 | zinc finger, HIT domain containing 1 [Sour | 640.88 | 543.77 | 533.31 | 584.17 | 606.38 | 677.22 |
| 16842 | ENSMUSG00000059534 | 2103.360078 | -0.059245869 | 0.093829908 | -0.631417747 | 0.527767415 | 0.888334265 | Uqcr10 | ubiquinol-cytochrome c reductase, compl | 2222.2 | 2078.4 | 2138.3 | 2053.3 | 1983.8 | 2144.1 |
| 16843 | ENSMUSG00000059540 | 10.96827173 | -0.109712888 | 0.643828785 | -0.170406931 | 0.864690122 | 0.977227145 | Tcea2 | transcription elongation factor A (SII), 2 [S | 8.8341 | 11.307 | 14.087 | 10.604 | 8.7249 | 12.252 |
| 16844 | ENSMUSG00000059547 | 0 | NA | NA | NA | NA | NA | Fbxw26 | F-box and WD-40 domain protein 26 [Sou | 0 | 0 | 0 | 0 | 0 | 0 |
| 16845 | ENSMUSG00000059552 | 328.0000636 | 0.067576887 | 0.153891315 | 0.439120864 | 0.660573962 | 0.928166095 | Trp53 | transformation related protein 53 [Source | 327.67 | 280.62 | 352.19 | 316.18 | 330.46 | 360.89 |
| 16846 | ENSMUSG00000059554 | 265.6801064 | 0.145595594 | 0.186001237 | 0.782766808 | 0.433764062 | 0.845264432 | Ccdc28a | coiled-coil domain containing 28A [Source | 278.68 | 202.5 | 274.7 | 234.25 | 302.1 | 301.85 |
| 16847 | ENSMUSG00000059555 | 77.80354467 | -0.29978019 | 0.264748273 | -1.132321605 | 0.25749925 | 0.710872182 | Tor4a | torsin family 4, member A [Source:MGI Sy | 91.554 | 78.122 | 87.543 | 61.695 | 63.256 | 84.652 |
| 16848 | ENSMUSG00000059562 | 0.439020386 | -2.189747911 | 3.52148864 | -0.621824499 | 0.534057275 | NA | Ccdc154 | coiled-coil domain containing 154 [Source | 1.6062 | 1.0279 | 0 | 0 | 0 | 0 |
| 16849 | ENSMUSG00000059565 | 0 | NA | NA | NA | NA | NA | Gm5292 | predicted gene 5292 [Source:MGI Symbol | 0 | 0 | 0 | 0 | 0 | 0 |
| 16850 | ENSMUSG00000059571 | 0.349477146 | 0.165142768 | 3.911021834 | 0.042224967 | 0.966319359 | NA | Gm16218 | predicted gene 16218 [Source:MGI Symbo | 0 | 0 | 1.0062 | 0 | 1.0906 | 0 |
| 16851 | ENSMUSG00000059585 | 0 | NA | NA | NA | NA | NA | Ube2nl | ubiquitin-conjugating enzyme E2N-like [Sc | 0 | 0 | 0 | 0 | 0 | 0 |
| 16852 | ENSMUSG00000059586 | 142.3634175 | -0.195856158 | 0.197074281 | -0.993818966 | 0.320311011 | 0.767391055 | Nsmce2 | NSE2/MMS21 homolog, SMC5-SMC6 com | 156.6 | 161.38 | 137.86 | 128.21 | 129.78 | 140.34 |

| | | | | | | | | | | | | | | | |
|---|---|---|---|---|---|---|---|---|---|---|---|---|---|---|---|
| 16853 | ENSMUSG00000059588 | 132.2351389 | -1.257621265 | 0.221513249 | -5.677408795 | 1.37E-08 | 2.00E-06 | Calcrl | calcitonin receptor-like [Source:MGI Symb | 173.47 | 205.58 | 180.12 | 66.514 | 74.162 | 93.563 |
| 16854 | ENSMUSG00000059595 | 0 | NA | NA | NA | NA | NA | Olfr935 | olfactory receptor 935 [Source:MGI Symb | 0 | 0 | 0 | 0 | 0 | 0 |
| 16855 | ENSMUSG00000059602 | 14.72485837 | -0.496090213 | 0.579149607 | -0.856583873 | 0.391674902 | 0.820404559 | Syn3 | synapsin III [Source:MGI Symbol;Acc:MGI | 20.078 | 18.503 | 13.081 | 12.532 | 16.359 | 7.7969 |
| 16856 | ENSMUSG00000059606 | 0 | NA | NA | NA | NA | NA | Rnase2b | ribonuclease, RNase A family, 2B (liver, ec | 0 | 0 | 0 | 0 | 0 | 0 |
| 16857 | ENSMUSG00000059610 | 0 | NA | NA | NA | NA | NA | Olfr222 | olfactory receptor 222 [Source:MGI Symb | 0 | 0 | 0 | 0 | 0 | 0 |
| 16858 | ENSMUSG00000059623 | 0 | NA | NA | NA | NA | NA | Olfr39 | olfactory receptor 39 [Source:MGI Symbo | 0 | 0 | 0 | 0 | 0 | 0 |
| 16859 | ENSMUSG00000059625 | 0 | NA | NA | NA | NA | NA | Sohlh1 | spermatogenesis and oogenesis specific b | 0 | 0 | 0 | 0 | 0 | 0 |
| 16860 | ENSMUSG00000059626 | 0 | NA | NA | NA | NA | NA | Rhox3e | reproductive homeobox 3E [Source:MGI S | 0 | 0 | 0 | 0 | 0 | 0 |
| 16861 | ENSMUSG00000059631 | 0.481988923 | 2.492401104 | 3.837077151 | 0.649557203 | 0.515978285 | NA | 1500035N22 | RIKEN cDNA 1500035N22 gene [Source:M | 0 | 0 | 0 | 2.8919 | 0 | 0 |
| 16862 | ENSMUSG00000059632 | 0 | NA | NA | NA | NA | NA | Krtap8-1 | keratin associated protein 8-1 [Source:MG | 0 | 0 | 0 | 0 | 0 | 0 |
| 16863 | ENSMUSG00000059639 | 0 | NA | NA | NA | NA | NA | Clec4a4 | C-type lectin domain family 4, member a4 | 0 | 0 | 0 | 0 | 0 | 0 |
| 16864 | ENSMUSG00000059645 | 0 | NA | NA | NA | NA | NA | Gm7361 | predicted gene 7361 [Source:MGI Symbol | 0 | 0 | 0 | 0 | 0 | 0 |
| 16865 | ENSMUSG00000059647 | 0.353089354 | 0.165142706 | 3.896102244 | 0.042386646 | 0.966190474 | NA | Cbx3-ps7 | chromobox 3, pseudogene 7 [Source:MGI | 0 | 1.0279 | 0 | 0 | 1.0906 | 0 |
| 16866 | ENSMUSG00000059648 | 0 | NA | NA | NA | NA | NA | Rnase11 | ribonuclease, RNase A family, 11 (non-act | 0 | 0 | 0 | 0 | 0 | 0 |
| 16867 | ENSMUSG00000059654 | 0 | NA | NA | NA | NA | NA | Reg1 | regenerating islet-derived 1 [Source:MGI S | 0 | 0 | 0 | 0 | 0 | 0 |
| 16868 | ENSMUSG00000059657 | 0 | NA | NA | NA | NA | NA | Stfa2l1 | stefin A2 like 1 [Source:MGI Symbol;Acc:N | 0 | 0 | 0 | 0 | 0 | 0 |
| 16869 | ENSMUSG00000059658 | 0 | NA | NA | NA | NA | NA | Mif-ps6 | macrophage migration inhibitory factor, p | 0 | 0 | 0 | 0 | 0 | 0 |
| 16870 | ENSMUSG00000059659 | 13.79652513 | 0.381676468 | 0.714094051 | 0.534490475 | 0.593002238 | 0.913011006 | Gm10069 | predicted gene 10069 [Source:MGI Symbo | 8.8341 | 7.1954 | 20.125 | 28.919 | 5.4531 | 12.252 |
| 16871 | ENSMUSG00000059663 | 0 | NA | NA | NA | NA | NA | Samt2b | spermatogenesis associated multipass tra | 0 | 0 | 0 | 0 | 0 | 0 |
| 16872 | ENSMUSG00000059668 | 0.556921942 | 2.679043159 | 3.638243063 | 0.736356289 | 0.461513899 | NA | Krt4 | keratin 4 [Source:MGI Symbol;Acc:MGI:96 | 0 | 0 | 0 | 0 | 0 | 3.3415 |
| 16873 | ENSMUSG00000059669 | 98.24457067 | 0.179396362 | 0.27171589 | 0.66023508 | 0.509102984 | 0.882793133 | Taf1b | TATA-box binding protein associated facto | 85.932 | 113.07 | 77.481 | 88.686 | 137.42 | 86.88 |
| 16874 | ENSMUSG00000059674 | 3.601658332 | -0.630230883 | 1.174745573 | -0.536482875 | 0.591624868 | NA | Cdh24 | cadherin-like 24 [Source:MGI Symbol;Acc: | 4.0155 | 2.0558 | 7.0437 | 1.928 | 5.4531 | 1.1138 |
| 16875 | ENSMUSG00000059687 | 0 | NA | NA | NA | NA | NA | Olfr108 | olfactory receptor 108 [Source:MGI Symb | 0 | 0 | 0 | 0 | 0 | 0 |
| 16876 | ENSMUSG00000059689 | 243.9906977 | -0.029818354 | 0.165133156 | -0.180571573 | 0.856703871 | 0.97470029 | Zfp637 | zinc finger protein 637 [Source:MGI Symb | 269.04 | 248.76 | 221.37 | 246.78 | 254.11 | 223.88 |
| 16877 | ENSMUSG00000059690 | 0 | NA | NA | NA | NA | NA | 1700020N15 | RIKEN cDNA 1700020N15 gene [Source:M | 0 | 0 | 0 | 0 | 0 | 0 |
| 16878 | ENSMUSG00000059695 | 0 | NA | NA | NA | NA | NA | 1700019M2: | RIKEN cDNA 1700019M22 gene [Source:M | 0 | 0 | 0 | 0 | 0 | 0 |
| 16879 | ENSMUSG00000059706 | 0.481988923 | 2.492401104 | 3.837077151 | 0.649557203 | 0.515978285 | NA | A830035A12 | RIKEN cDNA A830035A12 gene [Source:M | 0 | 0 | 0 | 2.8919 | 0 | 0 |
| 16880 | ENSMUSG00000059708 | 82.10861325 | 0.562161673 | 0.269034618 | 2.089551437 | 0.036658113 | 0.258645339 | Akap17b | A kinase (PRKA) anchor protein 17B [Sour | 65.051 | 53.452 | 80.5 | 106.04 | 81.796 | 105.82 |
| 16881 | ENSMUSG00000059713 | 15.2027327 | -0.682848382 | 0.626953639 | -1.089152912 | 0.27608646 | 0.730038567 | Rcan3 | regulator of calcineurin 3 [Source:MGI Syr | 30.518 | 10.279 | 15.094 | 7.7118 | 10.906 | 16.708 |
| 16882 | ENSMUSG00000059714 | 405.8910733 | 0.074950615 | 0.136473682 | 0.549194643 | 0.58287188 | 0.908645447 | Flot1 | flotillin 1 [Source:MGI Symbol;Acc:MGI:11 | 429.66 | 373.13 | 382.37 | 406.8 | 408.98 | 434.4 |
| 16883 | ENSMUSG00000059729 | 0 | NA | NA | NA | NA | NA | Olfr1385 | olfactory receptor 1385 [Source:MGI Syml | 0 | 0 | 0 | 0 | 0 | 0 |
| 16884 | ENSMUSG00000059734 | 1459.955401 | -0.020422004 | 0.106694003 | -0.191407233 | 0.848206554 | 0.972803041 | Ndufs8 | NADH:ubiquinone oxidoreductase core su | 1600.6 | 1368.2 | 1440.9 | 1461.4 | 1390.5 | 1498.1 |
| 16885 | ENSMUSG00000059741 | 0 | NA | NA | NA | NA | NA | Myl3 | myosin, light polypeptide 3 [Source:MGI S | 0 | 0 | 0 | 0 | 0 | 0 |
| 16886 | ENSMUSG00000059742 | 14.08274415 | 1.437008965 | 0.665818641 | 2.158258835 | 0.030907713 | 0.2330809 | Kcnh7 | potassium voltage-gated channel, subfam | 10.44 | 5.1396 | 7.0437 | 20.244 | 32.719 | 8.9108 |
| 16887 | ENSMUSG00000059743 | 32.90413842 | 1.409755345 | 0.466188152 | 3.024005088 | 0.002494522 | 0.041235169 | Fdps | farnesyl diphosphate synthetase [Source:I | 20.078 | 24.67 | 9.0562 | 28.919 | 53.44 | 61.261 |
| 16888 | ENSMUSG00000059751 | 2.723046552 | 0.147488273 | 1.412071032 | 0.104448196 | 0.916813677 | NA | Rps3a3 | ribosomal protein S3A3 [Source:MGI Syml | 1.6062 | 6.1675 | 0 | 1.928 | 2.1812 | 4.4554 |
| 16889 | ENSMUSG00000059762 | 0 | NA | NA | NA | NA | NA | Olfr767 | olfactory receptor 767 [Source:MGI Symb | 0 | 0 | 0 | 0 | 0 | 0 |
| 16890 | ENSMUSG00000059763 | 0 | NA | NA | NA | NA | NA | Taar2 | trace amine-associated receptor 2 [Source | 0 | 0 | 0 | 0 | 0 | 0 |
| 16891 | ENSMUSG00000059768 | 0 | NA | NA | NA | NA | NA | Olfr682-ps1 | olfactory receptor 682, pseudogene 1 [Sou | 0 | 0 | 0 | 0 | 0 | 0 |
| 16892 | ENSMUSG00000059772 | 144.0351854 | 0.269922333 | 0.193141018 | 1.397540182 | 0.162251192 | 0.585529252 | Slx1b | SLX1 structure-specific endonuclease subu | 124.48 | 130.55 | 136.85 | 149.42 | 161.41 | 161.51 |
| 16893 | ENSMUSG00000059775 | 7.31744653 | -0.660529599 | 0.84888619 | -0.778113258 | 0.436502248 | 0.846663338 | Rps26-ps1 | ribosomal protein S26, pseudogene 1 [Sou | 10.44 | 13.363 | 3.0187 | 3.8559 | 6.5437 | 6.6831 |
| 16894 | ENSMUSG00000059776 | 0.133850373 | -0.796629176 | 4.080472857 | -0.195229623 | 0.84521319 | NA | Rpl13-ps6 | ribosomal protein L13, pseudogene 6 [Sou | 0.8031 | 0 | 0 | 0 | 0 | 0 |
| 16895 | ENSMUSG00000059791 | 10.27924545 | -0.921294417 | 0.742280416 | -1.241167619 | 0.214543834 | 0.659265353 | Nrm | nurim (nuclear envelope membrane prote | 12.85 | 13.363 | 14.087 | 3.8559 | 14.178 | 3.3415 |
| 16896 | ENSMUSG00000059796 | 5079.553102 | 0.041089486 | 0.094511169 | 0.434757989 | 0.663738102 | 0.929301301 | Eif4a1 | eukaryotic translation initiation factor 4A1 | 5115 | 5171.5 | 4734.4 | 4837.2 | 5496.7 | 5122.6 |
| 16897 | ENSMUSG00000059810 | 209.8287115 | 1.637834345 | 0.656039766 | 2.496547358 | 0.012540892 | 0.126529431 | Rgs3 | regulator of G-protein signaling 3 [Source: | 110.03 | 93.541 | 102.64 | 549.47 | 100.34 | 302.97 |
| 16898 | ENSMUSG00000059811 | 2894.586466 | -0.312007858 | 0.09018379 | -3.459688894 | 0.0005408 | 0.013124377 | Atl2 | atlastin GTPase 2 [Source:MGI Symbol;Acc | 3212.4 | 3308.9 | 3098.2 | 2641.3 | 2627.3 | 2479.4 |
| 16899 | ENSMUSG00000059816 | 0.160662974 | 1.126924175 | 4.080472857 | 0.276174898 | 0.782413736 | NA | Shisal2a | shisa like 2A [Source:MGI Symbol;Acc:MG | 0 | 0 | 0 | 0.964 | 0 | 0 |
| 16900 | ENSMUSG00000059820 | 208.5307907 | 0.017592423 | 0.166140061 | 0.105889107 | 0.915670337 | 0.98625062 | Nkapd1 | NKAP domain containing 1 [Source:MGI S | 199.17 | 205.58 | 217.35 | 216.9 | 206.13 | 206.06 |
| 16901 | ENSMUSG00000059821 | 0 | NA | NA | NA | NA | NA | Olfr847 | olfactory receptor 847 [Source:MGI Symb | 0 | 0 | 0 | 0 | 0 | 0 |
| 16902 | ENSMUSG00000059824 | 1835.48124 | 1.820785042 | 0.62109506 | 2.931572248 | 0.003372509 | 0.050865455 | Dbp | D site albumin promoter binding protein [ | 1554.8 | 469.76 | 404.51 | 2708.8 | 2690.6 | 3184.5 |
| 16903 | ENSMUSG00000059827 | 0 | NA | NA | NA | NA | NA | Gm5670 | predicted gene 5670 [Source:MGI Symbol | 0 | 0 | 0 | 0 | 0 | 0 |
| 16904 | ENSMUSG00000059830 | 0 | NA | NA | NA | NA | NA | Gm16380 | predicted pseudogene 16380 [Source:MG | 0 | 0 | 0 | 0 | 0 | 0 |
| 16905 | ENSMUSG00000059832 | 0 | NA | NA | NA | NA | NA | Kprp | keratinocyte expressed, proline-rich [Sour | 0 | 0 | 0 | 0 | 0 | 0 |
| 16906 | ENSMUSG00000059834 | 120.4117045 | 0.128170724 | 0.207963396 | 0.616313863 | 0.537687391 | 0.891052982 | Sclt1 | sodium channel and clathrin linker 1 [Sou | 113.24 | 119.24 | 112.7 | 121.46 | 121.06 | 134.78 |
| 16907 | ENSMUSG00000059835 | 0.181769715 | 1.126924175 | 4.080472857 | 0.276174898 | 0.782413736 | NA | Rpl13-ps3 | ribosomal protein L13, pseudogene 3 [Sou | 0 | 0 | 0 | 0 | 1.0906 | 0 |
| 16908 | ENSMUSG00000059839 | 98.61094209 | 0.144279474 | 0.239240873 | 0.603072011 | 0.546460785 | 0.89549457 | Zfp874b | zinc finger protein 874b [Source:MGI Sym | 106.01 | 84.289 | 90.562 | 113.75 | 109.06 | 87.994 |

| | | | | | | | | | | | | | | | |
|---|---|---|---|---|---|---|---|---|---|---|---|---|---|---|---|
| 16909 | ENSMUSG00000059842 | 36.43899395 | -0.143184666 | 0.373283963 | -0.383581079 | 0.701288966 | 0.941991169 | Zfp341 | zinc finger protein 341 [Source:MGI Symb | 30.518 | 48.312 | 36.225 | 38.559 | 32.719 | 32.301 |
| 16910 | ENSMUSG00000059845 | 0.17131964 | -0.796629176 | 4.080472857 | -0.195229623 | 0.84521319 | NA | Gm11567 | predicted gene 11567 [Source:MGI Symbo | 0 | 1.0279 | 0 | 0 | 0 | 0 |
| 16911 | ENSMUSG00000059851 | 196.6613252 | 0.540872006 | 0.179201873 | 3.018227417 | 0.00254258 | 0.041884839 | Kmt5c | lysine methyltransferase 5C [Source:MGI S | 168.65 | 143.91 | 168.04 | 257.38 | 219.21 | 222.77 |
| 16912 | ENSMUSG00000059852 | 1.017081212 | -3.37846085 | 2.590925565 | -1.303959054 | 0.192247543 | NA | Kcng2 | potassium voltage-gated channel, subfam | 0 | 3.0838 | 3.0187 | 0 | 0 | 0 |
| 16913 | ENSMUSG00000059854 | 0.160662974 | 1.126924175 | 4.080472857 | 0.276174898 | 0.782413736 | NA | Hydin | HYDIN, axonemal central pair apparatus p | 0 | 0 | 0 | 0.964 | 0 | 0 |
| 16914 | ENSMUSG00000059857 | 0 | NA | NA | NA | NA | NA | Ntng1 | netrin G1 [Source:MGI Symbol;Acc:MGI:1 | 0 | 0 | 0 | 0 | 0 | 0 |
| 16915 | ENSMUSG00000059862 | 0 | NA | NA | NA | NA | NA | Olfr826 | olfactory receptor 826 [Source:MGI Symb | 0 | 0 | 0 | 0 | 0 | 0 |
| 16916 | ENSMUSG00000059864 | 4.25429469 | -0.744931882 | 1.062681176 | -0.700992827 | 0.483307492 | NA | Olfr1393 | olfactory receptor 1393 [Source:MGI Symb | 4.8186 | 4.1117 | 7.0437 | 2.8919 | 1.0906 | 5.5692 |
| 16917 | ENSMUSG00000059866 | 102.2477818 | 0.160585575 | 0.266145314 | 0.60337555 | 0.546258884 | 0.89549457 | Tnip2 | TNFAIP3 interacting protein 2 [Source:MG | 111.63 | 95.596 | 82.512 | 141.7 | 81.796 | 100.25 |
| 16918 | ENSMUSG00000059867 | 0 | NA | NA | NA | NA | NA | Olfr960 | olfactory receptor 960 [Source:MGI Symb | 0 | 0 | 0 | 0 | 0 | 0 |
| 16919 | ENSMUSG00000059873 | 1.831978164 | 0.370978154 | 1.77833578 | 0.208609734 | 0.834752911 | NA | Olfr1029 | olfactory receptor 1029 [Source:MGI Symb | 0.8031 | 0 | 4.025 | 2.8919 | 3.2719 | 0 |
| 16920 | ENSMUSG00000059874 | 0 | NA | NA | NA | NA | NA | Olfr603 | olfactory receptor 603 [Source:MGI Symb | 0 | 0 | 0 | 0 | 0 | 0 |
| 16921 | ENSMUSG00000059877 | 0 | NA | NA | NA | NA | NA | Dph3b-ps | diphthamide biosynthesis 3, pseudogene | 0 | 0 | 0 | 0 | 0 | 0 |
| 16922 | ENSMUSG00000059878 | 323.1370324 | 0.050217058 | 0.147461692 | 0.340543077 | 0.733447588 | 0.947915087 | Zfp422 | zinc finger protein 422 [Source:MGI Symb | 332.48 | 295.01 | 325.02 | 352.82 | 327.19 | 306.31 |
| 16923 | ENSMUSG00000059883 | 218.5699585 | -0.17478121 | 0.163598369 | -1.068355459 | 0.285360201 | 0.737967092 | Irak4 | interleukin-1 receptor-associated kinase 4 | 229.69 | 226.14 | 239.49 | 198.58 | 217.03 | 200.49 |
| 16924 | ENSMUSG00000059887 | 0 | NA | NA | NA | NA | NA | Olfr1507 | olfactory receptor 1507 [Source:MGI Symb | 0 | 0 | 0 | 0 | 0 | 0 |
| 16925 | ENSMUSG00000059890 | 1679.884609 | 0.105736215 | 0.092633566 | 1.141446016 | 0.253684365 | 0.706909089 | Ube4a | ubiquitination factor E4A [Source:MGI Syr | 1639.9 | 1612.8 | 1601.9 | 1714.9 | 1778.8 | 1730.9 |
| 16926 | ENSMUSG00000059891 | 2.794063435 | 0.373402714 | 1.242919115 | 0.300423985 | 0.763853771 | NA | Tsks | testis-specific serine kinase substrate [Sou | 3.2124 | 1.0279 | 3.0187 | 2.8919 | 3.2719 | 3.3415 |
| 16927 | ENSMUSG00000059895 | 91.90394847 | -0.721421407 | 0.27439885 | -2.629097781 | 0.008561174 | 0.096816905 | Ptp4a3 | protein tyrosine phosphatase 4a3 [Source | 126.09 | 127.46 | 89.556 | 66.514 | 88.34 | 53.465 |
| 16928 | ENSMUSG00000059897 | 77.76278032 | 0.11911126 | 0.274743674 | 0.433535951 | 0.664625454 | 0.929301301 | Zfp930 | zinc finger protein 930 [Source:MGI Symb | 78.704 | 88.401 | 56.35 | 73.262 | 78.525 | 91.335 |
| 16929 | ENSMUSG00000059898 | 0 | NA | NA | NA | NA | NA | Dsc3 | desmocollin 3 [Source:MGI Symbol;Acc:M | 0 | 0 | 0 | 0 | 0 | 0 |
| 16930 | ENSMUSG00000059900 | 0.678286236 | 1.600482029 | 2.782216593 | 0.575254289 | 0.565119331 | NA | Tmem40 | transmembrane protein 40 [Source:MGI S | 0 | 1.0279 | 0 | 1.928 | 0 | 1.1138 |
| 16931 | ENSMUSG00000059901 | 18.08920898 | -0.901511165 | 0.577594714 | -1.560802313 | 0.118570402 | 0.504308497 | Adamts14 | a disintegrin-like and metallopeptidase (re | 32.927 | 17.475 | 20.125 | 10.604 | 20.722 | 6.6831 |
| 16932 | ENSMUSG00000059908 | 54002.58973 | 0.036549673 | 0.134184279 | 0.272384169 | 0.785326644 | 0.960670212 | Mug1 | murinoglobulin 1 [Source:MGI Symbol;Acc | 41261 | 60077 | 58620 | 52652 | 57432 | 53975 |
| 16933 | ENSMUSG00000059910 | 0 | NA | NA | NA | NA | NA | Olfr1265 | olfactory receptor 1265 [Source:MGI Symb | 0 | 0 | 0 | 0 | 0 | 0 |
| 16934 | ENSMUSG00000059912 | 0.531246861 | 1.101376188 | 3.36099124 | 0.327693858 | 0.743143147 | NA | Rpl21-ps6 | ribosomal protein L21, pseudogene 6 [Sou | 0 | 0 | 1.0062 | 0 | 2.1812 | 0 |
| 16935 | ENSMUSG00000059920 | 479.1381351 | -0.024508575 | 0.131241975 | -0.186743418 | 0.851861812 | 0.973633287 | 4930453N24 | RIKEN cDNA 4930453N24 gene [Source:M | 492.3 | 508.82 | 448.79 | 508.98 | 453.7 | 462.25 |
| 16936 | ENSMUSG00000059921 | 0 | NA | NA | NA | NA | NA | Unc5c | unc-5 netrin receptor C [Source:MGI Symb | 0 | 0 | 0 | 0 | 0 | 0 |
| 16937 | ENSMUSG00000059923 | 896.7961457 | -0.204504849 | 0.113905588 | -1.795389072 | 0.072591732 | 0.390758026 | Grb2 | growth factor receptor bound protein 2 [S | 913.13 | 1022.8 | 944.86 | 776 | 847.41 | 876.6 |
| 16938 | ENSMUSG00000059934 | 0 | NA | NA | NA | NA | NA | Prh1 | proline rich protein HaeIII subfamily 1 [So | 0 | 0 | 0 | 0 | 0 | 0 |
| 16939 | ENSMUSG00000059939 | 82.85159235 | 0.203121345 | 0.266424116 | 0.762398495 | 0.445822204 | 0.852395554 | 9430015G10 | RIKEN cDNA 9430015G10 gene [Source:M | 69.067 | 71.954 | 90.562 | 107.97 | 80.706 | 76.855 |
| 16940 | ENSMUSG00000059956 | 0 | NA | NA | NA | NA | NA | Serpinb12 | serine (or cysteine) peptidase inhibitor, cl | 0 | 0 | 0 | 0 | 0 | 0 |
| 16941 | ENSMUSG00000059964 | 0 | NA | NA | NA | NA | NA | Olfr119 | olfactory receptor 119 [Source:MGI Symb | 0 | 0 | 0 | 0 | 0 | 0 |
| 16942 | ENSMUSG00000059970 | 40.95632366 | -0.947511573 | 0.407365841 | -2.325947532 | 0.02002135 | 0.17340647 | Hspa2 | heat shock protein 2 [Source:MGI Symbol | 67.461 | 41.117 | 53.331 | 36.631 | 32.719 | 14.48 |
| 16943 | ENSMUSG00000059972 | 0 | NA | NA | NA | NA | NA | Gm14773 | predicted gene 14773 [Source:MGI Symbo | 0 | 0 | 0 | 0 | 0 | 0 |
| 16944 | ENSMUSG00000059974 | 22.65465527 | -1.019731265 | 0.478529084 | -2.130970299 | 0.033091589 | 0.243457144 | Ntm | neurotrimin [Source:MGI Symbol;Acc:MG | 24.093 | 37.005 | 30.187 | 19.28 | 11.997 | 13.366 |
| 16945 | ENSMUSG00000059975 | 70.8957637 | 0.118934655 | 0.278760248 | 0.426655723 | 0.669630105 | 0.930837335 | Zfp74 | zinc finger protein 74 [Source:MGI Symbo | 59.43 | 81.206 | 63.393 | 63.623 | 73.071 | 84.652 |
| 16946 | ENSMUSG00000059981 | 939.8228422 | -0.256799703 | 0.121501279 | -2.113555555 | 0.034553245 | 0.249141645 | Taok2 | TAO kinase 2 [Source:MGI Symbol;Acc:MG | 979.78 | 1049.5 | 1041.5 | 916.74 | 907.39 | 744.05 |
| 16947 | ENSMUSG00000059991 | 0 | NA | NA | NA | NA | NA | Nptx2 | neuronal pentraxin 2 [Source:MGI Symbo | 0 | 0 | 0 | 0 | 0 | 0 |
| 16948 | ENSMUSG00000059994 | 4.100888276 | -1.899800154 | 1.197284403 | -1.586757624 | 0.11256755 | NA | Fcrl1 | Fc receptor-like 1 [Source:MGI Symbol;Ac | 3.2124 | 8.2233 | 8.05 | 2.8919 | 0 | 2.2277 |
| 16949 | ENSMUSG00000059995 | 684.1879706 | 0.054439911 | 0.151263705 | 0.359900687 | 0.718921403 | 0.9447719 | Atxn7l3 | ataxin 7-like 3 [Source:MGI Symbol;Acc:M | 618.39 | 630.11 | 765.75 | 707.56 | 579.12 | 804.2 |
| 16950 | ENSMUSG00000060002 | 5761.316985 | -0.342956743 | 0.108782317 | -3.152688349 | 0.001617744 | 0.029998785 | Chpt1 | choline phosphotransferase 1 [Source:MG | 6772.6 | 6755.5 | 5801 | 5493.7 | 5171.7 | 4573.4 |
| 16951 | ENSMUSG00000060012 | 887.3497204 | -0.014307682 | 0.111462444 | -0.128363252 | 0.897861511 | 0.982798755 | Kif13b | kinesin family member 13B [Source:MGI S | 828.8 | 910.74 | 936.81 | 909.03 | 888.85 | 849.86 |
| 16952 | ENSMUSG00000060017 | 0 | NA | NA | NA | NA | NA | Olfr121 | olfactory receptor 121 [Source:MGI Symb | 0 | 0 | 0 | 0 | 0 | 0 |
| 16953 | ENSMUSG00000060019 | 3.637090168 | 0.156428532 | 1.12443826 | 0.139117049 | 0.889357656 | NA | Gm10073 | predicted pseudogene 10073 [Source:MG | 3.2124 | 4.1117 | 3.0187 | 4.8199 | 1.0906 | 5.5692 |
| 16954 | ENSMUSG00000060024 | 0 | NA | NA | NA | NA | NA | Vmn1r213 | vomeronasal 1 receptor 213 [Source:MGI | 0 | 0 | 0 | 0 | 0 | 0 |
| 16955 | ENSMUSG00000060029 | 3.535008714 | -0.075672064 | 1.207658201 | -0.062660167 | 0.950037118 | NA | 4930473A02 | RIKEN cDNA 4930473A02 gene [Source:M | 0.8031 | 7.1954 | 3.0187 | 5.7839 | 2.1812 | 2.2277 |
| 16956 | ENSMUSG00000060030 | 0 | NA | NA | NA | NA | NA | Olfr317 | olfactory receptor 317 [Source:MGI Symb | 0 | 0 | 0 | 0 | 0 | 0 |
| 16957 | ENSMUSG00000060032 | 929.1985126 | -0.003713035 | 0.117732997 | -0.03153776 | 0.974840679 | 0.996336532 | H2aj | H2J.A histone [Source:MGI Symbol;Acc:M | 1014.3 | 838.78 | 936.81 | 903.25 | 906.3 | 975.73 |
| 16958 | ENSMUSG00000060034 | 3.989328783 | -1.117876912 | 1.141873905 | -0.978984551 | 0.327587613 | NA | Ctf2 | cardiotrophin 2 [Source:MGI Symbol;Acc: | 3.2124 | 3.0838 | 10.062 | 0.964 | 3.2719 | 3.3415 |
| 16959 | ENSMUSG00000060036 | 3877.702351 | 0.010104152 | 0.148394081 | 0.068089991 | 0.945713998 | 0.990309554 | Rpl3 | ribosomal protein L3 [Source:MGI Symbol | 3317.6 | 4610.2 | 3664.7 | 3147.4 | 4194.5 | 4331.7 |
| 16960 | ENSMUSG00000060038 | 220.2088729 | 0.112564478 | 0.166851079 | 0.674640394 | 0.499904263 | 0.880716237 | Dhps | deoxyhypusine synthase [Source:MGI Syn | 212.82 | 212.78 | 209.3 | 229.43 | 249.75 | 207.17 |
| 16961 | ENSMUSG00000060044 | 9.217270794 | -5.577134084 | 1.414367787 | -3.94319931 | 8.04E-05 | 0.003044323 | Tmem26 | transmembrane protein 26 [Source:MGI S | 15.259 | 32.893 | 6.0375 | 0 | 0 | 1.1138 |
| 16962 | ENSMUSG00000060049 | 0 | NA | NA | NA | NA | NA | Olfr235 | olfactory receptor 235 [Source:MGI Symb | 0 | 0 | 0 | 0 | 0 | 0 |
| 16963 | ENSMUSG00000060057 | 0 | NA | NA | NA | NA | NA | Olfr193 | olfactory receptor 193 [Source:MGI Symb | 0 | 0 | 0 | 0 | 0 | 0 |
| 16964 | ENSMUSG00000060063 | 13.13129389 | -0.650461099 | 0.602824267 | -1.079022751 | 0.280577586 | 0.733467233 | Alox5ap | arachidonate 5-lipoxygenase activating pr | 18.471 | 17.475 | 12.075 | 8.6758 | 8.7249 | 13.366 |

| | | | | | | | | | | | | | | | |
|---|---|---|---|---|---|---|---|---|---|---|---|---|---|---|---|
| 16965 | ENSMUSG00000060068 | 0 | NA | NA | NA | NA | NA | Gm12222 | predicted gene 12222 [Source:MGI Symbo | 0 | 0 | 0 | 0 | 0 | 0 |
| 16966 | ENSMUSG00000060070 | 0 | NA | NA | NA | NA | NA | Defa26 | defensin, alpha, 26 [Source:MGI Symbol;A | 0 | 0 | 0 | 0 | 0 | 0 |
| 16967 | ENSMUSG00000060073 | 2273.072606 | 0.120658349 | 0.09539652 | 1.264808712 | 0.205939907 | 0.651085207 | Psma3 | proteasome subunit alpha 3 [Source:MGI | 2332.2 | 2089.8 | 2111.1 | 2387.8 | 2331.7 | 2385.9 |
| 16968 | ENSMUSG00000060081 | 0 | NA | NA | NA | NA | NA | H2ac1 | H2A clustered histone 1 [Source:MGI Sym | 0 | 0 | 0 | 0 | 0 | 0 |
| 16969 | ENSMUSG00000060084 | 0 | NA | NA | NA | NA | NA | Olfr748 | olfactory receptor 748 [Source:MGI Symb | 0 | 0 | 0 | 0 | 0 | 0 |
| 16970 | ENSMUSG00000060090 | 190.8623285 | 0.065401757 | 0.184327968 | 0.354811904 | 0.72273051 | 0.9447719 | Rp2 | retinitis pigmentosa 2 homolog [Source:M | 161.42 | 213.81 | 185.15 | 203.4 | 187.59 | 193.81 |
| 16971 | ENSMUSG00000060093 | 1.552689824 | 0.408349768 | 1.891165486 | 0.215924926 | 0.829046267 | NA | H4c1 | H4 clustered histone 1 [Source:MGI Symb | 0 | 1.0279 | 3.0187 | 1.928 | 0 | 3.3415 |
| 16972 | ENSMUSG00000060096 | 1.192065194 | -1.259467627 | 2.132120447 | -0.590711293 | 0.55471388 | NA | Amd-ps3 | S-adenosylmethionine decarboxylase, pse | 0 | 2.0558 | 3.0187 | 0.964 | 0 | 1.1138 |
| 16973 | ENSMUSG00000060098 | 263.5794692 | -0.257911753 | 0.174221708 | -1.480365199 | 0.138775812 | 0.544447711 | Prmt7 | protein arginine N-methyltransferase 7 [S | 299.56 | 268.29 | 292.82 | 210.15 | 289.01 | 221.65 |
| 16974 | ENSMUSG00000060105 | 0 | NA | NA | NA | NA | NA | Olfr504 | olfactory receptor 504 [Source:MGI Symb | 0 | 0 | 0 | 0 | 0 | 0 |
| 16975 | ENSMUSG00000060112 | 0 | NA | NA | NA | NA | NA | Olfr60 | olfactory receptor 60 [Source:MGI Symbo | 0 | 0 | 0 | 0 | 0 | 0 |
| 16976 | ENSMUSG00000060114 | 0 | NA | NA | NA | NA | NA | Olfr910 | olfactory receptor 910 [Source:MGI Symb | 0 | 0 | 0 | 0 | 0 | 0 |
| 16977 | ENSMUSG00000060121 | 308.9395062 | -0.078935283 | 0.14212151 | -0.555407012 | 0.578616298 | 0.908137944 | Gemin2 | gem nuclear organelle associated protein | 311.6 | 321.74 | 318.98 | 301.73 | 297.74 | 301.85 |
| 16978 | ENSMUSG00000060126 | 14762.25104 | -0.010107181 | 0.089118713 | -0.113412551 | 0.90970349 | 0.985435587 | Tpt1 | tumor protein, translationally-controlled 1 | 14554 | 15472 | 14415 | 13837 | 14715 | 15580 |
| 16979 | ENSMUSG00000060128 | 0.528279927 | -0.837071592 | 3.369085264 | -0.248456636 | 0.803781118 | NA | Gm2710 | predicted gene 2710 [Source:MGI Symbol | 0 | 2.0558 | 0 | 0 | 0 | 1.1138 |
| 16980 | ENSMUSG00000060131 | 23.39937898 | 0.503686509 | 0.493961851 | 1.019687061 | 0.3078769 | 0.756343859 | Atp8b4 | ATPase, class I, type 8B, member 4 [Sourc | 15.259 | 28.782 | 14.087 | 18.316 | 30.537 | 33.415 |
| 16981 | ENSMUSG00000060143 | 763.2856391 | 0.006591374 | 0.113675024 | 0.057984361 | 0.953761086 | 0.9915542 | Gm10076 | predicted gene 10076 [Source:MGI Symbo | 749.29 | 724.68 | 811.03 | 789.5 | 765.61 | 739.59 |
| 16982 | ENSMUSG00000060147 | 151.682511 | -0.586500712 | 0.240986659 | -2.433747636 | 0.01494341 | 0.142198341 | Serpinb6a | serine (or cysteine) peptidase inhibitor, cl | 155 | 246.7 | 144.9 | 118.57 | 110.15 | 134.78 |
| 16983 | ENSMUSG00000060149 | 41.72047975 | 0.036739653 | 0.376815544 | 0.097500364 | 0.922329045 | 0.986685804 | BC002059 | cDNA sequence BC002059 [Source:MGI Sy | 47.383 | 26.726 | 49.306 | 46.271 | 31.628 | 49.009 |
| 16984 | ENSMUSG00000060152 | 499.9874024 | 0.090754085 | 0.133274059 | 0.680958359 | 0.495897839 | 0.878034808 | Pop5 | processing of precursor 5, ribonuclease P/ | 456.97 | 463.59 | 532.3 | 478.13 | 539.86 | 529.08 |
| 16985 | ENSMUSG00000060161 | 0 | NA | NA | NA | NA | NA | Klk1b7-ps | kallikrein 1-related peptidase b7, pseudog | 0 | 0 | 0 | 0 | 0 | 0 |
| 16986 | ENSMUSG00000060166 | 167.9259037 | -0.296122756 | 0.210377063 | -1.407580998 | 0.159255178 | 0.580657143 | Zdhhc8 | zinc finger, DHHC domain containing 8 [Sc | 167.05 | 162.41 | 226.41 | 171.59 | 131.96 | 148.14 |
| 16987 | ENSMUSG00000060170 | 0.331982614 | 0.16514309 | 3.986751476 | 0.041422971 | 0.966958701 | NA | Olfr1371 | olfactory receptor 1371 [Source:MGI Syml | 0 | 1.0279 | 0 | 0.964 | 0 | 0 |
| 16988 | ENSMUSG00000060176 | 0.936410189 | -0.694018354 | 2.480552701 | -0.279783757 | 0.779643414 | NA | Kif27 | kinesin family member 27 [Source:MGI Sy | 2.4093 | 1.0279 | 0 | 0 | 2.1812 | 0 |
| 16989 | ENSMUSG00000060177 | 0 | NA | NA | NA | NA | NA | Klk1b22 | kallikrein 1-related peptidase b22 [Source | 0 | 0 | 0 | 0 | 0 | 0 |
| 16990 | ENSMUSG00000060180 | 0 | NA | NA | NA | NA | NA | Myh13 | myosin, heavy polypeptide 13, skeletal mu | 0 | 0 | 0 | 0 | 0 | 0 |
| 16991 | ENSMUSG00000060181 | 308.7602234 | -0.140736428 | 0.209583127 | -0.671506479 | 0.501897934 | 0.881001421 | Slc35e3 | solute carrier family 35, member E3 [Sour | 324.45 | 223.06 | 423.63 | 270.88 | 306.46 | 304.08 |
| 16992 | ENSMUSG00000060183 | 35.30423728 | 0.154824389 | 0.382422469 | 0.404851707 | 0.685586519 | 0.935893609 | Cxcl11 | chemokine (C-X-C motif) ligand 11 [Source | 31.321 | 24.67 | 44.275 | 36.631 | 38.172 | 36.757 |
| 16993 | ENSMUSG00000060187 | 0 | NA | NA | NA | NA | NA | Lrrc10 | leucine rich repeat containing 10 [Source: | 0 | 0 | 0 | 0 | 0 | 0 |
| 16994 | ENSMUSG00000060188 | 0.353348078 | 0.165142701 | 3.895042568 | 0.042398176 | 0.966181282 | NA | Cxcl17 | chemokine (C-X-C motif) ligand 17 [Source | 0 | 0 | 1.0062 | 0 | 0 | 1.1138 |
| 16995 | ENSMUSG00000060198 | 3.123720303 | -0.904521752 | 1.287689835 | -0.702437596 | 0.482406306 | NA | Gm11353 | predicted gene 11353 [Source:MGI Symbo | 4.0155 | 4.1117 | 4.025 | 0 | 4.3625 | 2.2277 |
| 16996 | ENSMUSG00000060201 | 0 | NA | NA | NA | NA | NA | Spink7 | serine peptidase inhibitor, Kazal type 7 (p | 0 | 0 | 0 | 0 | 0 | 0 |
| 16997 | ENSMUSG00000060204 | 0 | NA | NA | NA | NA | NA | Slc10a4-ps | solute carrier family 10 (sodium/bile acid | 0 | 0 | 0 | 0 | 0 | 0 |
| 16998 | ENSMUSG00000060205 | 0 | NA | NA | NA | NA | NA | Olfr57 | olfactory receptor 57 [Source:MGI Symbo | 0 | 0 | 0 | 0 | 0 | 0 |
| 16999 | ENSMUSG00000060206 | 6.744717695 | -0.393676799 | 0.868808653 | -0.453122558 | 0.650460493 | 0.925643431 | Zfp462 | zinc finger protein 462 [Source:MGI Symb | 4.8186 | 10.279 | 8.05 | 9.6398 | 5.4531 | 2.2277 |
| 17000 | ENSMUSG00000060208 | 0 | NA | NA | NA | NA | NA | Defa17 | defensin, alpha, 17 [Source:MGI Symbol;A | 0 | 0 | 0 | 0 | 0 | 0 |
| 17001 | ENSMUSG00000060212 | 5.823145963 | -0.167390834 | 0.99511329 | -0.168212842 | 0.866415839 | 0.977925452 | Pcnx2 | pecanex homolog 2 [Source:MGI Symbol;A | 2.4093 | 6.1675 | 10.062 | 9.6398 | 1.0906 | 5.5692 |
| 17002 | ENSMUSG00000060216 | 198.9936346 | -0.383235339 | 0.189660598 | -2.020637612 | 0.043317292 | 0.287075127 | Arrb2 | arrestin, beta 2 [Source:MGI Symbol;Acc:I | 253.78 | 229.23 | 192.19 | 166.77 | 155.96 | 196.04 |
| 17003 | ENSMUSG00000060224 | 465.8349337 | -0.133347439 | 0.150665515 | -0.885056138 | 0.376126355 | 0.809731241 | Pyroxd2 | pyridine nucleotide-disulphide oxidoredu | 549.32 | 406.03 | 506.14 | 475.24 | 441.7 | 416.58 |
| 17004 | ENSMUSG00000060227 | 94.59010043 | -0.317534389 | 0.288899344 | -1.099117724 | 0.27171672 | 0.727526428 | Golm2 | golgi membrane protein 2 [Source:MGI Sy | 126.09 | 120.27 | 68.425 | 95.434 | 91.612 | 65.717 |
| 17005 | ENSMUSG00000060240 | 1.042111019 | -0.860289347 | 2.663009232 | -0.323051583 | 0.746656177 | NA | Cend1 | cell cycle exit and neuronal differentiatior | 0 | 0 | 4.025 | 0 | 0 | 2.2277 |
| 17006 | ENSMUSG00000060244 | 30.95551268 | 0.071692394 | 0.390640843 | 0.183525084 | 0.854386038 | 0.973825032 | Alyref2 | Aly/REF export factor 2 [Source:MGI Symb | 30.518 | 26.726 | 33.206 | 26.991 | 35.99 | 32.301 |
| 17007 | ENSMUSG00000060245 | 0 | NA | NA | NA | NA | NA | Vmn1r228 | vomeronasal 1 receptor 228 [Source:MGI | 0 | 0 | 0 | 0 | 0 | 0 |
| 17008 | ENSMUSG00000060254 | 0 | NA | NA | NA | NA | NA | Olfr980 | olfactory receptor 980 [Source:MGI Symb | 0 | 0 | 0 | 0 | 0 | 0 |
| 17009 | ENSMUSG00000060256 | 0 | NA | NA | NA | NA | NA | Tdpoz4 | TD and POZ domain containing 4 [Source: | 0 | 0 | 0 | 0 | 0 | 0 |
| 17010 | ENSMUSG00000060257 | 0 | NA | NA | NA | NA | NA | Scrt2 | scratch family zinc finger 2 [Source:MGI S | 0 | 0 | 0 | 0 | 0 | 0 |
| 17011 | ENSMUSG00000060260 | 46.22101402 | 0.395974177 | 0.334551438 | 1.183597296 | 0.236572511 | 0.685340334 | Pwwp2b | PWWP domain containing 2B [Source:MG | 30.518 | 47.284 | 42.262 | 48.199 | 56.712 | 52.351 |
| 17012 | ENSMUSG00000060261 | 2004.082015 | -0.0113381 | 0.098171558 | -0.115492714 | 0.908054595 | 0.984639154 | Gtf2i | general transcription factor II I [Source:M | 1995.7 | 2040.4 | 1999.4 | 1884.6 | 2150.7 | 1953.7 |
| 17013 | ENSMUSG00000060268 | 0 | NA | NA | NA | NA | NA | Armh1 | armadillo-like helical domain containing 1 | 0 | 0 | 0 | 0 | 0 | 0 |
| 17014 | ENSMUSG00000060275 | 1.583470467 | 1.089112889 | 2.103508062 | 0.517760264 | 0.604625546 | NA | Nrg2 | neuregulin 2 [Source:MGI Symbol;Acc:MG | 0 | 3.0838 | 0 | 0.964 | 5.4531 | 0 |
| 17015 | ENSMUSG00000060279 | 498.2506909 | 0.095537006 | 0.126194737 | 0.757060149 | 0.449013828 | 0.854431786 | Ap2a1 | adaptor-related protein complex 2, alpha | 466.6 | 492.37 | 487.02 | 550.43 | 492.96 | 500.12 |
| 17016 | ENSMUSG00000060284 | 0 | NA | NA | NA | NA | NA | Sp7 | Sp7 transcription factor 7 [Source:MGI Sy | 0 | 0 | 0 | 0 | 0 | 0 |
| 17017 | ENSMUSG00000060288 | 82.67601612 | 0.108227192 | 0.254381517 | 0.425452264 | 0.670507013 | 0.93116767 | Ppih | peptidyl prolyl isomerase H [Source:MGI S | 89.947 | 74.01 | 74.462 | 84.83 | 97.065 | 75.741 |
| 17018 | ENSMUSG00000060301 | 169.6466351 | 0.214435031 | 0.196731303 | 1.089989381 | 0.275717822 | 0.730038567 | 2610008E11 | RIKEN cDNA 2610008E11 gene [Source:M | 165.44 | 158.3 | 146.91 | 160.98 | 169.05 | 217.2 |
| 17019 | ENSMUSG00000060303 | 0 | NA | NA | NA | NA | NA | Olfr1447 | olfactory receptor 1447 [Source:MGI Syml | 0 | 0 | 0 | 0 | 0 | 0 |
| 17020 | ENSMUSG00000060311 | 0 | NA | NA | NA | NA | NA | Mucl1 | mucin-like 1 [Source:MGI Symbol;Acc:MG | 0 | 0 | 0 | 0 | 0 | 0 |

| | | | | | | | | | | | | | | | |
|---|---|---|---|---|---|---|---|---|---|---|---|---|---|---|---|
| 17021 | ENSMUSG00000060314 | 11.08394737 | 1.024409096 | 0.74356139 | 1.377706145 | 0.168294033 | 0.595160122 | Zfp941 | zinc finger protein 941 [Source:MGI Symb | 14.456 | 5.1396 | 2.0125 | 17.352 | 14.178 | 13.366 |
| 17022 | ENSMUSG00000060317 | 613.5119735 | -1.79140866 | 0.418898849 | -4.276470715 | 1.90E-05 | 0.000915472 | Acnat2 | acyl-coenzyme A amino acid N-acyltransfe | 740.46 | 850.09 | 1265.9 | 390.41 | 163.59 | 270.66 |
| 17023 | ENSMUSG00000060332 | 0 | NA | NA | NA | NA | NA | Tmc2 | transmembrane channel-like gene family | 0 | 0 | 0 | 0 | 0 | 0 |
| 17024 | ENSMUSG00000060335 | 0 | NA | NA | NA | NA | NA | Olfr384 | olfactory receptor 384 [Source:MGI Symb | 0 | 0 | 0 | 0 | 0 | 0 |
| 17025 | ENSMUSG00000060336 | 14.88009994 | -0.092629164 | 0.563395373 | -0.164412362 | 0.869406533 | 0.978621761 | Zfp937 | zinc finger protein 937 [Source:MGI Symb | 18.471 | 14.391 | 13.081 | 13.496 | 10.906 | 18.935 |
| 17026 | ENSMUSG00000060371 | 132.9371411 | -1.174826252 | 0.333829226 | -3.519243252 | 0.00043278 | 0.011212784 | Caln1 | calneuron 1 [Source:MGI Symbol;Acc:MG | 107.62 | 242.59 | 203.26 | 99.29 | 101.43 | 43.44 |
| 17027 | ENSMUSG00000060373 | 1890.435119 | -0.010794441 | 0.104436358 | -0.103359038 | 0.917678021 | 0.986374671 | Hnrnpc | heterogeneous nuclear ribonucleoprotein | 1867.2 | 1855.4 | 1969.2 | 1702.4 | 1943.5 | 2004.9 |
| 17028 | ENSMUSG00000060375 | 0 | NA | NA | NA | NA | NA | 3110018I06R | RIKEN cDNA 3110018I06 gene [Source:MG | 0 | 0 | 0 | 0 | 0 | 0 |
| 17029 | ENSMUSG00000060376 | 1588.697318 | 0.060441919 | 0.096033266 | 0.629385228 | 0.529096886 | 0.888334265 | Bckdha | branched chain ketoacid dehydrogenase E | 1581.3 | 1563.5 | 1521.4 | 1643.6 | 1671.9 | 1550.5 |
| 17030 | ENSMUSG00000060377 | 1.359539405 | 2.831315047 | 2.05728984 | 1.376235371 | 0.168748774 | NA | Rpl36a-ps1 | ribosomal protein L36A, pseudogene 1 [So | 0 | 0 | 1.0062 | 3.8559 | 2.1812 | 1.1138 |
| 17031 | ENSMUSG00000060380 | 0.339027071 | -1.79174149 | 3.919585585 | -0.457125237 | 0.647581032 | NA | C030014I23R | RIKEN cDNA C030014I23 gene [Source:MG | 0 | 1.0279 | 1.0062 | 0 | 0 | 0 |
| 17032 | ENSMUSG00000060397 | 43.37678794 | -0.03941545 | 0.328140331 | -0.120117664 | 0.90438994 | 0.98383695 | Zfp128 | zinc finger protein 128 [Source:MGI Symb | 47.383 | 41.117 | 43.269 | 41.451 | 44.715 | 42.326 |
| 17033 | ENSMUSG00000060402 | 0.922606629 | -1.977583741 | 2.486779435 | -0.795238899 | 0.426474548 | NA | Chst8 | carbohydrate sulfotransferase 8 [Source:M | 2.4093 | 0 | 2.0125 | 0 | 0 | 1.1138 |
| 17034 | ENSMUSG00000060404 | 0 | NA | NA | NA | NA | NA | Olfr1369-ps1 | olfactory receptor 1369, pseudogene 1 [So | 0 | 0 | 0 | 0 | 0 | 0 |
| 17035 | ENSMUSG00000060407 | 11451.53047 | -0.101194317 | 0.098364159 | -1.028772255 | 0.303586707 | 0.752690022 | Cyp2a12 | cytochrome P450, family 2, subfamily a, p | 11409 | 11547 | 12603 | 10147 | 12003 | 11000 |
| 17036 | ENSMUSG00000060411 | 0 | NA | NA | NA | NA | NA | Npn2 | neoplastic progression 2 [Source:MGI Sym | 0 | 0 | 0 | 0 | 0 | 0 |
| 17037 | ENSMUSG00000060412 | 0 | NA | NA | NA | NA | NA | Tas2r124 | taste receptor, type 2, member 124 [Sour | 0 | 0 | 0 | 0 | 0 | 0 |
| 17038 | ENSMUSG00000060416 | 1.744392165 | 1.320498094 | 1.708030964 | 0.773111332 | 0.439456495 | NA | Gm839 | predicted gene 839 [Source:MGI Symbol;A | 0 | 2.0558 | 1.0062 | 1.928 | 4.3625 | 1.1138 |
| 17039 | ENSMUSG00000060417 | 0 | NA | NA | NA | NA | NA | Gm8603 | predicted gene 8603 [Source:MGI Symbol | 0 | 0 | 0 | 0 | 0 | 0 |
| 17040 | ENSMUSG00000060419 | 27.95086241 | 0.107869326 | 0.439538668 | 0.245414872 | 0.806135215 | 0.964428432 | Rps16-ps2 | ribosomal protein S16, pseudogene 2 [Sou | 16.865 | 33.921 | 30.187 | 26.027 | 26.175 | 34.529 |
| 17041 | ENSMUSG00000060424 | 0 | NA | NA | NA | NA | NA | Pantr1 | POU domain, class 3, transcription factor | 0 | 0 | 0 | 0 | 0 | 0 |
| 17042 | ENSMUSG00000060427 | 119.0934771 | 0.052127606 | 0.222928708 | 0.233830834 | 0.815116311 | 0.967689186 | Zfp868 | zinc finger protein 868 [Source:MGI Symb | 107.62 | 139.8 | 103.64 | 115.68 | 130.87 | 116.95 |
| 17043 | ENSMUSG00000060429 | 1244.233262 | 0.109409507 | 0.12103737 | 0.903931627 | 0.366031657 | 0.804235439 | Sntb1 | syntrophin, basic 1 [Source:MGI Symbol;A | 1214.3 | 1318.8 | 1057.6 | 1193.4 | 1293.5 | 1387.8 |
| 17044 | ENSMUSG00000060438 | 1.868516502 | -0.676272514 | 1.644036753 | -0.411348781 | 0.680816805 | NA | Rps10-ps1 | ribosomal protein S10, pseudogene 1 [Sou | 0.8031 | 3.0838 | 3.0187 | 0.964 | 0 | 3.3415 |
| 17045 | ENSMUSG00000060441 | 46.55704506 | 0.125123005 | 0.355189529 | 0.352271097 | 0.724634962 | 0.9448775 | Trim5 | tripartite motif-containing 5 [Source:MGI | 53.808 | 39.061 | 40.25 | 32.775 | 59.984 | 53.465 |
| 17046 | ENSMUSG00000060445 | 0.17131964 | -0.796629176 | 4.080472857 | -0.195229623 | 0.84521319 | NA | Sycp2 | synaptonemal complex protein 2 [Source: | 0 | 1.0279 | 0 | 0 | 0 | 0 |
| 17047 | ENSMUSG00000060450 | 1213.099002 | 0.184377388 | 0.117583912 | 1.568049447 | 0.116869593 | 0.50225288 | Rnf14 | ring finger protein 14 [Source:MGI Symbo | 1119.5 | 1239.7 | 1047.5 | 1174.1 | 1305.5 | 1392.3 |
| 17048 | ENSMUSG00000060459 | 4997.803048 | 0.094052933 | 0.110344659 | 0.852356008 | 0.394016529 | 0.822827574 | Kng2 | kininogen 2 [Source:MGI Symbol;Acc:MGI | 4567.2 | 4995.7 | 4943.7 | 5822.4 | 5089.9 | 4567.9 |
| 17049 | ENSMUSG00000060461 | 0 | NA | NA | NA | NA | NA | Dppa5a | developmental pluripotency associated 5A | 0 | 0 | 0 | 0 | 0 | 0 |
| 17050 | ENSMUSG00000060467 | 2.028708648 | 0.772851659 | 1.705150091 | 0.453245532 | 0.65037195 | NA | Gm10080 | predicted gene 10080 [Source:MGI Symbo | 2.4093 | 0 | 2.0125 | 0 | 2.1812 | 5.5692 |
| 17051 | ENSMUSG00000060469 | 0 | NA | NA | NA | NA | NA | Krtap19-3 | keratin associated protein 19-3 [Source:M | 0 | 0 | 0 | 0 | 0 | 0 |
| 17052 | ENSMUSG00000060470 | 46.4194468 | -0.016294486 | 0.358344311 | -0.045471592 | 0.963731418 | 0.993245949 | Adgrg3 | adhesion G protein-coupled receptor G3 [ | 51.399 | 56.535 | 32.2 | 57.839 | 35.99 | 44.554 |
| 17053 | ENSMUSG00000060475 | 469.0303502 | 0.142037185 | 0.128008052 | 1.109595704 | 0.267173284 | 0.722778036 | Wtap | Wilms tumour 1-associating protein [Sour | 455.36 | 459.48 | 422.62 | 468.49 | 513.68 | 494.55 |
| 17054 | ENSMUSG00000060477 | 371.1299347 | -0.16895234 | 0.157956965 | -1.069609941 | 0.284794919 | 0.737432482 | Irak2 | interleukin-1 receptor-associated kinase 2 | 381.47 | 340.24 | 456.84 | 341.25 | 380.63 | 326.36 |
| 17055 | ENSMUSG00000060480 | 0 | NA | NA | NA | NA | NA | Olfr171 | olfactory receptor 171 [Source:MGI Symb | 0 | 0 | 0 | 0 | 0 | 0 |
| 17056 | ENSMUSG00000060487 | 12.20117696 | -0.92456616 | 0.655192641 | -1.411136362 | 0.15820441 | 0.578722455 | Samd5 | sterile alpha motif domain containing 5 [S | 19.274 | 20.558 | 8.05 | 7.7118 | 9.8156 | 7.7969 |
| 17057 | ENSMUSG00000060491 | 0 | NA | NA | NA | NA | NA | 4930522H14 | RIKEN cDNA 4930522H14 gene [Source:M | 0 | 0 | 0 | 0 | 0 | 0 |
| 17058 | ENSMUSG00000060499 | 0 | NA | NA | NA | NA | NA | Rpl10l | ribosomal protein L10-like [Source:MGI Sy | 0 | 0 | 0 | 0 | 0 | 0 |
| 17059 | ENSMUSG00000060503 | 0 | NA | NA | NA | NA | NA | Olfr715 | olfactory receptor 715 [Source:MGI Symb | 0 | 0 | 0 | 0 | 0 | 0 |
| 17060 | ENSMUSG00000060508 | 0 | NA | NA | NA | NA | NA | Nlrp9b | NLR family, pyrin domain containing 9B [S | 0 | 0 | 0 | 0 | 0 | 0 |
| 17061 | ENSMUSG00000060509 | 4.661267442 | -2.105442976 | 1.122689897 | -1.87535577 | 0.060743796 | NA | Xcr1 | chemokine (C motif) receptor 1 [Source:M | 6.4248 | 8.2233 | 8.05 | 1.928 | 0 | 3.3415 |
| 17062 | ENSMUSG00000060510 | 393.6414322 | 0.262365824 | 0.142651251 | 1.839211523 | 0.065884081 | 0.368645006 | Zfp266 | zinc finger protein 266 [Source:MGI Symb | 383.08 | 358.74 | 331.05 | 388.48 | 447.15 | 453.33 |
| 17063 | ENSMUSG00000060512 | 617.0508324 | 0.272877347 | 0.127679731 | 2.137201763 | 0.032581585 | 0.241177094 | 0610040J01R | RIKEN cDNA 0610040J01 gene [Source:MG | 529.24 | 591.05 | 556.45 | 631.41 | 645.65 | 748.5 |
| 17064 | ENSMUSG00000060519 | 170.7003149 | -1.233953932 | 0.242493873 | -5.088598387 | 3.61E-07 | 3.28E-05 | Tor3a | torsin family 3, member A [Source:MGI Sy | 200.78 | 211.75 | 306.9 | 124.35 | 105.79 | 74.628 |
| 17065 | ENSMUSG00000060523 | 0 | NA | NA | NA | NA | NA | Olfr726 | olfactory receptor 726 [Source:MGI Symb | 0 | 0 | 0 | 0 | 0 | 0 |
| 17066 | ENSMUSG00000060530 | 0.788497532 | -1.691662715 | 2.429158853 | -0.696398555 | 0.486179268 | NA | A930017M0 | RIKEN cDNA A930017M01 gene [Source:M | 1.6062 | 1.0279 | 1.0062 | 0 | 1.0906 | 0 |
| 17067 | ENSMUSG00000060534 | 0 | NA | NA | NA | NA | NA | Dcc | deleted in colorectal carcinoma [Source:M | 0 | 0 | 0 | 0 | 0 | 0 |
| 17068 | ENSMUSG00000060538 | 569.9563009 | 0.084334956 | 0.135385689 | 0.622923713 | 0.533334655 | 0.889143097 | Tmem219 | transmembrane protein 219 [Source:MGI | 492.3 | 553.02 | 615.82 | 621.77 | 594.39 | 542.44 |
| 17069 | ENSMUSG00000060548 | 117.3307126 | -0.071986197 | 0.319519342 | -0.225295272 | 0.821749578 | 0.969160361 | Tnfrsf19 | tumor necrosis factor receptor superfamil | 176.68 | 120.27 | 63.393 | 138.81 | 110.15 | 94.677 |
| 17070 | ENSMUSG00000060550 | 1473.977921 | -0.00183409 | 0.117326895 | -0.015632304 | 0.987527734 | 0.99859176 | H2-Q7 | histocompatibility 2, Q region locus 7 [Sou | 1328.3 | 1574.8 | 1523.5 | 1626.2 | 1369.8 | 1421.3 |
| 17071 | ENSMUSG00000060556 | 0 | NA | NA | NA | NA | NA | Olfr1418 | olfactory receptor 1418 [Source:MGI Sym | 0 | 0 | 0 | 0 | 0 | 0 |
| 17072 | ENSMUSG00000060560 | 105.6611189 | 1.332907124 | 0.314707039 | 4.235390248 | 2.28E-05 | 0.001078443 | Ces4a | carboxylesterase 4A [Source:MGI Symbol; | 55.414 | 58.591 | 66.412 | 171.59 | 201.76 | 80.197 |
| 17073 | ENSMUSG00000060565 | 0 | NA | NA | NA | NA | NA | Gm5591 | predicted gene 5591 [Source:MGI Symbol | 0 | 0 | 0 | 0 | 0 | 0 |
| 17074 | ENSMUSG00000060566 | 0 | NA | NA | NA | NA | NA | Rpl21-ps3 | ribosomal protein L21, pseudogene 3 [Sou | 0 | 0 | 0 | 0 | 0 | 0 |
| 17075 | ENSMUSG00000060568 | 1.215440034 | 0.637618514 | 2.123998586 | 0.30019724 | 0.76402671 | NA | Fam78b | family with sequence similarity 78, membe | 0.8031 | 1.0279 | 1.0062 | 0 | 0 | 4.4554 |
| 17076 | ENSMUSG00000060572 | 0.978009112 | -0.745988154 | 2.250857147 | -0.331424033 | 0.740324214 | NA | Mfap2 | microfibrillar-associated protein 2 [Source | 1.6062 | 1.0279 | 1.0062 | 0 | 0 | 2.2277 |

| | | | | | | | | | | | | | | | |
|---|---|---|---|---|---|---|---|---|---|---|---|---|---|---|---|
| 17077 | ENSMUSG00000060579 | 43.70687128 | 0.482797193 | 0.333644376 | 1.447041305 | 0.147885355 | 0.559219289 | Fhit | fragile histidine triad gene [Source:MGI Sy | 40.958 | 35.977 | 32.2 | 46.271 | 56.712 | 50.123 |
| 17078 | ENSMUSG00000060583 | 0 | NA | NA | NA | NA | NA | Olfr881 | olfactory receptor 881 [Source:MGI Symb | 0 | 0 | 0 | 0 | 0 | 0 |
| 17079 | ENSMUSG00000060586 | 257.293838 | -1.031931101 | 0.249255398 | -4.14005517 | 3.47E-05 | 0.001569445 | H2-Eb1 | histocompatibility 2, class II antigen E beta | 358.18 | 439.95 | 238.48 | 167.73 | 118.88 | 220.54 |
| 17080 | ENSMUSG00000060591 | 980.1297036 | -0.257998321 | 0.160629906 | -1.606166171 | 0.108237404 | 0.4851397 | Ifitm2 | interferon induced transmembrane prote | 901.88 | 1052.6 | 1249.8 | 1085.4 | 759.07 | 832.04 |
| 17081 | ENSMUSG00000060594 | 12.1641807 | -0.563454307 | 0.616788293 | -0.913529509 | 0.360964121 | 0.800074801 | Layn | layilin [Source:MGI Symbol;Acc:MGI:2685 | 12.047 | 17.475 | 14.087 | 10.604 | 7.6343 | 11.138 |
| 17082 | ENSMUSG00000060600 | 81.49631968 | 0.04752346 | 0.261667518 | 0.181617727 | 0.855882736 | 0.974266258 | Eno3 | enolase 3, beta muscle [Source:MGI Symb | 90.751 | 74.01 | 75.468 | 84.83 | 95.974 | 67.944 |
| 17083 | ENSMUSG00000060601 | 732.8094319 | 0.134299347 | 0.118710974 | 1.131313667 | 0.257923094 | 0.711453498 | Nr1h2 | nuclear receptor subfamily 1, group H, me | 746.89 | 689.73 | 659.09 | 804.92 | 716.54 | 779.69 |
| 17084 | ENSMUSG00000060613 | 7036.376505 | 0.324456573 | 0.16691436 | 1.943850568 | 0.051913474 | 0.320245495 | Cyp2c70 | cytochrome P450, family 2, subfamily c, p | 5457.9 | 5055.3 | 8233.1 | 8369.3 | 8519.9 | 6582.8 |
| 17085 | ENSMUSG00000060615 | 0 | NA | NA | NA | NA | NA | Ang4 | angiogenin, ribonuclease A family, membe | 0 | 0 | 0 | 0 | 0 | 0 |
| 17086 | ENSMUSG00000060621 | 0.335414862 | -1.779047242 | 4.030804179 | -0.441362855 | 0.658950329 | NA | Nkpd1 | NTPase, KAP family P-loop domain contair | 0 | 0 | 2.0125 | 0 | 0 | 0 |
| 17087 | ENSMUSG00000060630 | 0 | NA | NA | NA | NA | NA | Zfp735 | zinc finger protein 735 [Source:MGI Symb | 0 | 0 | 0 | 0 | 0 | 0 |
| 17088 | ENSMUSG00000060636 | 2593.891324 | 0.166152437 | 0.114349376 | 1.453024429 | 0.146216971 | 0.556855766 | Rpl35a | ribosomal protein L35A [Source:MGI Syml | 2124.2 | 2733.2 | 2478.4 | 2764.7 | 2850.9 | 2612 |
| 17089 | ENSMUSG00000060639 | 56.57622374 | -0.013589386 | 0.316615596 | -0.042920772 | 0.965764691 | 0.993727003 | H4c9 | H4 clustered histone 9 [Source:MGI Symb | 61.839 | 42.145 | 66.412 | 59.767 | 45.806 | 63.489 |
| 17090 | ENSMUSG00000060647 | 0 | NA | NA | NA | NA | NA | Gm7099 | predicted gene 7099 [Source:MGI Symbol | 0 | 0 | 0 | 0 | 0 | 0 |
| 17091 | ENSMUSG00000060657 | 1181.355884 | 0.296980696 | 0.121881725 | 2.43663023 | 0.01482483 | 0.141629025 | Marf1 | meiosis regulator and mRNA stability 1 [Sc | 955.69 | 1040.3 | 1185.4 | 1253.2 | 1232.4 | 1421.3 |
| 17092 | ENSMUSG00000060663 | 0 | NA | NA | NA | NA | NA | Olfr175 | olfactory receptor 175 [Source:MGI Symb | 0 | 0 | 0 | 0 | 0 | 0 |
| 17093 | ENSMUSG00000060671 | 31.14042378 | -0.70416358 | 0.463601588 | -1.518898121 | 0.128788142 | 0.525173862 | Atp8b2 | ATPase, class I, type 8B, member 2 [Sourc | 52.202 | 40.089 | 23.144 | 18.316 | 17.45 | 35.643 |
| 17094 | ENSMUSG00000060673 | 0 | NA | NA | NA | NA | NA | 4930595M1ε | RIKEN cDNA 4930595M18 gene [Source:M | 0 | 0 | 0 | 0 | 0 | 0 |
| 17095 | ENSMUSG00000060675 | 90.81956444 | 0.241988167 | 0.283031763 | 0.854985902 | 0.392558953 | 0.821375223 | Plaat3 | phospholipase A and acyltransferase 3 [Sc | 77.901 | 104.85 | 67.418 | 128.21 | 77.434 | 89.108 |
| 17096 | ENSMUSG00000060678 | 3.199027341 | 0.244049508 | 1.275414727 | 0.191349138 | 0.848252066 | NA | H4c3 | H4 clustered histone 3 [Source:MGI Symb | 0.8031 | 1.0279 | 7.0437 | 4.8199 | 3.2719 | 2.2277 |
| 17097 | ENSMUSG00000060679 | 446.4517253 | 0.121754214 | 0.136131851 | 0.894384478 | 0.371116215 | 0.807214774 | Mrps9 | mitochondrial ribosomal protein S9 [Sourc | 394.32 | 444.06 | 444.76 | 427.04 | 495.14 | 473.38 |
| 17098 | ENSMUSG00000060680 | 0.342432689 | 1.990653086 | 3.900281263 | 0.51038706 | 0.609780321 | NA | Gm8894 | predicted gene 8894 [Source:MGI Symbol | 0 | 0 | 0 | 0.964 | 1.0906 | 0 |
| 17099 | ENSMUSG00000060681 | 241.8750385 | 0.027559756 | 0.180480605 | 0.152702039 | 0.878633253 | 0.980976563 | Slc9a6 | solute carrier family 9 (sodium/hydrogen | 220.05 | 291.93 | 207.29 | 251.6 | 246.48 | 233.91 |
| 17100 | ENSMUSG00000060685 | 0 | NA | NA | NA | NA | NA | Fthl17c | ferritin, heavy polypeptide-like 17, membe | 0 | 0 | 0 | 0 | 0 | 0 |
| 17101 | ENSMUSG00000060688 | 0 | NA | NA | NA | NA | NA | Olfr292 | olfactory receptor 292 [Source:MGI Symb | 0 | 0 | 0 | 0 | 0 | 0 |
| 17102 | ENSMUSG00000060691 | 0 | NA | NA | NA | NA | NA | Krtap19-1 | keratin associated protein 19-1 [Source:M | 0 | 0 | 0 | 0 | 0 | 0 |
| 17103 | ENSMUSG00000060699 | 0 | NA | NA | NA | NA | NA | Vmn1r35 | vomeronasal 1 receptor 35 [Source:MGI S | 0 | 0 | 0 | 0 | 0 | 0 |
| 17104 | ENSMUSG00000060701 | 0 | NA | NA | NA | NA | NA | Fpr-rs3 | formyl peptide receptor, related sequence | 0 | 0 | 0 | 0 | 0 | 0 |
| 17105 | ENSMUSG00000060703 | 4842.57424 | -0.063261824 | 0.095545231 | -0.662113887 | 0.507898237 | 0.882537253 | Cd302 | CD302 antigen [Source:MGI Symbol;Acc:N | 4769.6 | 4986.4 | 5091.6 | 5145.7 | 4453 | 4609.1 |
| 17106 | ENSMUSG00000060708 | 254.2005026 | 0.130820943 | 0.163489587 | 0.800179052 | 0.423607065 | 0.84018348 | Bloc1s4 | biogenesis of lysosomal organelles comple | 244.14 | 231.28 | 252.57 | 270.88 | 233.39 | 292.94 |
| 17107 | ENSMUSG00000060715 | 1.032248872 | 1.23506196 | 2.454802548 | 0.503120693 | 0.614879419 | NA | 1700019A02 | RIKEN cDNA 1700019A02 gene [Source:M | 0.8031 | 1.0279 | 0 | 0 | 4.3625 | 0 |
| 17108 | ENSMUSG00000060716 | 3.685649042 | -0.147372943 | 1.277551774 | -0.11535575 | 0.908163151 | NA | Plekhh1 | pleckstrin homology domain containing, fa | 6.4248 | 5.1396 | 0 | 2.8919 | 6.5437 | 1.1138 |
| 17109 | ENSMUSG00000060724 | 0 | NA | NA | NA | NA | NA | Vmn1r47 | vomeronasal 1 receptor 47 [Source:MGI S | 0 | 0 | 0 | 0 | 0 | 0 |
| 17110 | ENSMUSG00000060726 | 0 | NA | NA | NA | NA | NA | Tmsb15a | thymosin beta 15a [Source:MGI Symbol;A | 0 | 0 | 0 | 0 | 0 | 0 |
| 17111 | ENSMUSG00000060727 | 1.238665401 | -0.445718538 | 2.061328571 | -0.216228768 | 0.82880943 | NA | Gm12571 | predicted gene 12571 [Source:MGI Symbo | 3.2124 | 1.0279 | 0 | 0.964 | 0 | 2.2277 |
| 17112 | ENSMUSG00000060730 | 0 | NA | NA | NA | NA | NA | Gm10086 | predicted pseudogene 10086 [Source:MG | 0 | 0 | 0 | 0 | 0 | 0 |
| 17113 | ENSMUSG00000060733 | 1678.78084 | -0.055925584 | 0.124862082 | -0.44789886 | 0.654226191 | 0.925797231 | Ipmk | inositol polyphosphate multikinase [Sourc | 1491.4 | 1633.4 | 2010.5 | 1686 | 1613 | 1638.5 |
| 17114 | ENSMUSG00000060735 | 0 | NA | NA | NA | NA | NA | Rxfp3 | relaxin family peptide receptor 3 [Source: | 0 | 0 | 0 | 0 | 0 | 0 |
| 17115 | ENSMUSG00000060738 | 0 | NA | NA | NA | NA | NA | Prl7c1 | prolactin family 7, subfamily c, member 1 | 0 | 0 | 0 | 0 | 0 | 0 |
| 17116 | ENSMUSG00000060739 | 721.5645037 | -0.049509037 | 0.116754783 | -0.424042905 | 0.671534523 | 0.931191575 | Nsa2 | NSA2 ribosome biogenesis homolog [Sour | 717.17 | 726.74 | 757.7 | 660.32 | 760.16 | 707.29 |
| 17117 | ENSMUSG00000060742 | 0 | NA | NA | NA | NA | NA | Olfr1076 | olfactory receptor 1076 [Source:MGI Syml | 0 | 0 | 0 | 0 | 0 | 0 |
| 17118 | ENSMUSG00000060743 | 1299.90395 | 0.024953279 | 0.116428853 | 0.214322125 | 0.830295866 | 0.9702546 | H3f3a | H3.3 histone A [Source:MGI Symbol;Acc:N | 1162.1 | 1252 | 1453 | 1324.5 | 1345.8 | 1262 |
| 17119 | ENSMUSG00000060747 | 0 | NA | NA | NA | NA | NA | Ifnl3 | interferon lambda 3 [Source:MGI Symbol; | 0 | 0 | 0 | 0 | 0 | 0 |
| 17120 | ENSMUSG00000060756 | 0 | NA | NA | NA | NA | NA | Krtap2-4 | keratin associated protein 2-4 [Source:MG | 0 | 0 | 0 | 0 | 0 | 0 |
| 17121 | ENSMUSG00000060759 | 0 | NA | NA | NA | NA | NA | Olfr503 | olfactory receptor 503 [Source:MGI Symb | 0 | 0 | 0 | 0 | 0 | 0 |
| 17122 | ENSMUSG00000060765 | 0 | NA | NA | NA | NA | NA | Olfr325 | olfactory receptor 325 [Source:MGI Symb | 0 | 0 | 0 | 0 | 0 | 0 |
| 17123 | ENSMUSG00000060771 | 34.8145437 | -0.241482049 | 0.429792226 | -0.561857646 | 0.574213012 | 0.906012546 | Tsga10 | testis specific 10 [Source:MGI Symbol;Acc | 35.336 | 32.893 | 45.281 | 50.127 | 19.631 | 25.618 |
| 17124 | ENSMUSG00000060777 | 0 | NA | NA | NA | NA | NA | Gm10087 | predicted gene 10087 [Source:MGI Symbo | 0 | 0 | 0 | 0 | 0 | 0 |
| 17125 | ENSMUSG00000060780 | 0 | NA | NA | NA | NA | NA | Lrrtm1 | leucine rich repeat transmembrane neuro | 0 | 0 | 0 | 0 | 0 | 0 |
| 17126 | ENSMUSG00000060787 | 0 | NA | NA | NA | NA | NA | Olfr464 | olfactory receptor 464 [Source:MGI Symb | 0 | 0 | 0 | 0 | 0 | 0 |
| 17127 | ENSMUSG00000060791 | 85.61774188 | -0.316325169 | 0.268274456 | -1.179110282 | 0.238354265 | 0.687311819 | Gmfg | glia maturation factor, gamma [Source:MG | 98.782 | 75.038 | 110.69 | 62.659 | 77.434 | 89.108 |
| 17128 | ENSMUSG00000060794 | 0 | NA | NA | NA | NA | NA | Tssk5 | testis-specific serine kinase 5 [Source:MG | 0 | 0 | 0 | 0 | 0 | 0 |
| 17129 | ENSMUSG00000060795 | 1.037954732 | 2.366557412 | 2.405105848 | 0.98397225 | 0.325129157 | NA | Gm13363 | predicted gene 13363 [Source:MGI Symbo | 0 | 1.0279 | 0 | 1.928 | 3.2719 | 0 |
| 17130 | ENSMUSG00000060798 | 69.42786993 | 0.404362329 | 0.265937699 | 1.520515256 | 0.128381528 | 0.524402828 | Intu | inturned planar cell polarity protein [Sour | 60.233 | 57.563 | 61.381 | 73.262 | 86.159 | 77.969 |
| 17131 | ENSMUSG00000060802 | 26981.85991 | -0.147561114 | 0.085938263 | -1.71705954 | 0.085968294 | 0.432051911 | B2m | beta-2 microglobulin [Source:MGI Symbol | 29940 | 27503 | 27639 | 26713 | 25321 | 24776 |
| 17132 | ENSMUSG00000060803 | 17312.32058 | 0.995907513 | 0.318208101 | 3.129736511 | 0.001749632 | 0.031752411 | Gstp1 | glutathione S-transferase, pi 1 [Source:MG | 11537 | 9097.1 | 14056 | 14687 | 26473 | 28024 |

| | | | | | | | | | | | | | | | |
|---|---|---|---|---|---|---|---|---|---|---|---|---|---|---|---|
| 17133 | ENSMUSG00000060807 | 5593.531101 | -0.171249634 | 0.125473353 | -1.364828706 | 0.172306894 | 0.600396752 | Serpina6 | serine (or cysteine) peptidase inhibitor, cl | 5725.3 | 5464.4 | 6586.9 | 6031.6 | 5275.3 | 4477.7 |
| 17134 | ENSMUSG00000060808 | 0 | NA | NA | NA | NA | NA | B9d1os | B9 protein domain 1, opposite strand [Sou | 0 | 0 | 0 | 0 | 0 | 0 |
| 17135 | ENSMUSG00000060811 | 0 | NA | NA | NA | NA | NA | Gm10088 | predicted gene 10088 [Source:MGI Symbo | 0 | 0 | 0 | 0 | 0 | 0 |
| 17136 | ENSMUSG00000060816 | 0 | NA | NA | NA | NA | NA | Vmn1r52 | vomeronasal 1 receptor 52 [Source:MGI S | 0 | 0 | 0 | 0 | 0 | 0 |
| 17137 | ENSMUSG00000060827 | 0 | NA | NA | NA | NA | NA | Olfr1193 | olfactory receptor 1193 [Source:MGI Syml | 0 | 0 | 0 | 0 | 0 | 0 |
| 17138 | ENSMUSG00000060843 | 0 | NA | NA | NA | NA | NA | Ctnna3 | catenin (cadherin associated protein), alp | 0 | 0 | 0 | 0 | 0 | 0 |
| 17139 | ENSMUSG00000060860 | 269.268894 | 0.106735446 | 0.155629031 | 0.685832494 | 0.492818743 | 0.876660964 | Ube2s | ubiquitin-conjugating enzyme E2S [Source | 281.89 | 260.06 | 235.46 | 280.52 | 278.11 | 279.57 |
| 17140 | ENSMUSG00000060862 | 155.101503 | -0.217952282 | 0.197514422 | -1.103475283 | 0.26982082 | 0.725011151 | Zbtb40 | zinc finger and BTB domain containing 40 | 175.08 | 162.41 | 163.01 | 157.13 | 152.69 | 120.3 |
| 17141 | ENSMUSG00000060863 | 0 | NA | NA | NA | NA | NA | Tcl1b2 | T cell leukemia/lymphoma 1B, 2 [Source:N | 0 | 0 | 0 | 0 | 0 | 0 |
| 17142 | ENSMUSG00000060878 | 0.808265676 | 0.728755068 | 2.41725015 | 0.301481031 | 0.763047712 | NA | Olfr1420 | olfactory receptor 1420 [Source:MGI Syml | 0.8031 | 1.0279 | 0 | 1.928 | 1.0906 | 0 |
| 17143 | ENSMUSG00000060879 | 0 | NA | NA | NA | NA | NA | Gm6564 | predicted pseudogene 6564 [Source:MGI | 0 | 0 | 0 | 0 | 0 | 0 |
| 17144 | ENSMUSG00000060882 | 0.642651897 | 2.906894826 | 3.440535777 | 0.844895974 | 0.398168923 | NA | Kcnd2 | potassium voltage-gated channel, Shal-rel | 0 | 0 | 0 | 3.8559 | 0 | 0 |
| 17145 | ENSMUSG00000060888 | 0 | NA | NA | NA | NA | NA | Olfr566 | olfactory receptor 566 [Source:MGI Symb | 0 | 0 | 0 | 0 | 0 | 0 |
| 17146 | ENSMUSG00000060890 | 0 | NA | NA | NA | NA | NA | Arr3 | arrestin 3, retinal [Source:MGI Symbol;Ac | 0 | 0 | 0 | 0 | 0 | 0 |
| 17147 | ENSMUSG00000060904 | 2371.575652 | 0.044807614 | 0.09744522 | 0.45982362 | 0.645642827 | 0.925643431 | Arl1 | ADP-ribosylation factor-like 1 [Source:MG | 2264.7 | 2425.9 | 2314.4 | 2437.9 | 2545.5 | 2241.1 |
| 17148 | ENSMUSG00000060913 | 0 | NA | NA | NA | NA | NA | Trim55 | tripartite motif-containing 55 [Source:MG | 0 | 0 | 0 | 0 | 0 | 0 |
| 17149 | ENSMUSG00000060918 | 0 | NA | NA | NA | NA | NA | Olfr51 | olfactory receptor 51 [Source:MGI Symbo | 0 | 0 | 0 | 0 | 0 | 0 |
| 17150 | ENSMUSG00000060923 | 24.90589156 | -0.651763876 | 0.47568388 | -1.370161789 | 0.170636403 | 0.597556794 | Acyp2 | acylphosphatase 2, muscle type [Source:N | 36.14 | 30.838 | 24.15 | 17.352 | 11.997 | 28.96 |
| 17151 | ENSMUSG00000060924 | 0.319491021 | 0.16514334 | 4.044653141 | 0.040830038 | 0.967431393 | NA | Csmd1 | CUB and Sushi multiple domains 1 [Source | 0.8031 | 0 | 0 | 0 | 0 | 1.1138 |
| 17152 | ENSMUSG00000060935 | 137.6461038 | 0.128634744 | 0.208114273 | 0.618096693 | 0.536511602 | 0.890560262 | Tmem263 | transmembrane protein 263 [Source:MGI | 146.97 | 114.1 | 132.82 | 130.14 | 157.05 | 144.8 |
| 17153 | ENSMUSG00000060938 | 2789.734558 | 0.002910785 | 0.086329396 | 0.033717194 | 0.973102668 | 0.995820834 | Rpl26 | ribosomal protein L26 [Source:MGI Symbo | 2741 | 2819.6 | 2800.4 | 2757.9 | 2816 | 2803.5 |
| 17154 | ENSMUSG00000060950 | 99.25048593 | -0.023206373 | 0.260987001 | -0.088917734 | 0.92914729 | 0.988405116 | Trmt61a | tRNA methyltransferase 61A [Source:MGI | 93.16 | 129.52 | 77.481 | 84.83 | 104.7 | 105.82 |
| 17155 | ENSMUSG00000060956 | 0 | NA | NA | NA | NA | NA | Gm10092 | predicted gene 10092 [Source:MGI Symbo | 0 | 0 | 0 | 0 | 0 | 0 |
| 17156 | ENSMUSG00000060961 | 1299.559866 | -0.279609627 | 0.12454085 | -2.2451238 | 0.024760186 | 0.201358738 | Slc4a4 | solute carrier family 4 (anion exchanger), | 1337.2 | 1615.9 | 1322.2 | 1052.7 | 1253.1 | 1216.3 |
| 17157 | ENSMUSG00000060962 | 0.531944269 | 2.618916564 | 3.285123409 | 0.797204926 | 0.425332026 | NA | Dmkn | dermokine [Source:MGI Symbol;Acc:MGI: | 0 | 0 | 0 | 0.964 | 0 | 2.2277 |
| 17158 | ENSMUSG00000060967 | 0 | NA | NA | NA | NA | NA | Etd | embryonic testis differentiation [Source:N | 0 | 0 | 0 | 0 | 0 | 0 |
| 17159 | ENSMUSG00000060969 | 29.24153575 | 1.304205566 | 0.502972473 | 2.592995911 | 0.00951439 | 0.104661272 | Irx1 | Iroquois homeobox 1 [Source:MGI Symbo | 11.243 | 11.307 | 28.175 | 31.811 | 30.537 | 62.375 |
| 17160 | ENSMUSG00000060974 | 0 | NA | NA | NA | NA | NA | Olfr530 | olfactory receptor 530 [Source:MGI Symb | 0 | 0 | 0 | 0 | 0 | 0 |
| 17161 | ENSMUSG00000060981 | 38.77637238 | 1.055879775 | 0.384848654 | 2.743623409 | 0.00607652 | 0.076030095 | H4c8 | H4 clustered histone 8 [Source:MGI Symb | 35.336 | 18.503 | 21.131 | 45.307 | 57.803 | 54.578 |
| 17162 | ENSMUSG00000060985 | 0 | NA | NA | NA | NA | NA | Tdrd5 | tudor domain containing 5 [Source:MGI S | 0 | 0 | 0 | 0 | 0 | 0 |
| 17163 | ENSMUSG00000060988 | 0 | NA | NA | NA | NA | NA | Galnt13 | polypeptide N-acetylgalactosaminyltransf | 0 | 0 | 0 | 0 | 0 | 0 |
| 17164 | ENSMUSG00000060989 | 3.322839693 | 1.006313023 | 1.192321813 | 0.843994475 | 0.398672493 | NA | Gm11847 | predicted gene 11847 [Source:MGI Symbo | 1.6062 | 3.0838 | 2.0125 | 7.7118 | 2.1812 | 3.3415 |
| 17165 | ENSMUSG00000060992 | 2204.557238 | 0.195539215 | 0.104569334 | 1.869947974 | 0.061491043 | 0.354345197 | Copz1 | coatomer protein complex, subunit zeta 1 | 2233.4 | 1996.2 | 1935 | 2302 | 2242.3 | 2518.4 |
| 17166 | ENSMUSG00000061000 | 0 | NA | NA | NA | NA | NA | Olfr507 | olfactory receptor 507 [Source:MGI Symb | 0 | 0 | 0 | 0 | 0 | 0 |
| 17167 | ENSMUSG00000061013 | 21.71661594 | 0.211316847 | 0.531345051 | 0.397701732 | 0.690850061 | 0.937606823 | Mkx | mohawk homeobox [Source:MGI Symbol; | 18.471 | 32.893 | 9.0562 | 20.244 | 22.903 | 26.732 |
| 17168 | ENSMUSG00000061022 | 0 | NA | NA | NA | NA | NA | Vmn1r222 | vomeronasal 1 receptor 222 [Source:MGI | 0 | 0 | 0 | 0 | 0 | 0 |
| 17169 | ENSMUSG00000061024 | 444.2028262 | -0.072734974 | 0.155520045 | -0.467688741 | 0.640007194 | 0.923932929 | Rrs1 | ribosome biogenesis regulator 1 [Source:I | 493.1 | 443.03 | 429.67 | 455.96 | 353.36 | 490.09 |
| 17170 | ENSMUSG00000061028 | 209.334393 | 0.146137929 | 0.1771932 | 0.824737794 | 0.40952047 | 0.833241465 | Clasrp | CLK4-associating serine/arginine rich prot | 216.84 | 188.11 | 191.19 | 248.71 | 201.76 | 209.4 |
| 17171 | ENSMUSG00000061032 | 1236.697253 | -0.05948151 | 0.106620321 | -0.557881546 | 0.576925273 | 0.907752116 | Rrp1 | ribosomal RNA processing 1 [Source:MGI | 1174.9 | 1296.2 | 1316.2 | 1181.8 | 1280.4 | 1170.6 |
| 17172 | ENSMUSG00000061039 | 2.190773379 | 0.717842228 | 1.686653326 | 0.425601525 | 0.670398229 | NA | Olfr920 | olfactory receptor 920 [Source:MGI Symb | 0 | 1.0279 | 4.025 | 4.8199 | 3.2719 | 0 |
| 17173 | ENSMUSG00000061046 | 61.5515288 | 0.327498005 | 0.295023772 | 1.110073276 | 0.266967452 | 0.722778036 | Haghl | hydroxyacylglutathione hydrolase-like [So | 48.989 | 67.843 | 47.293 | 75.19 | 67.618 | 62.375 |
| 17174 | ENSMUSG00000061048 | 5.826983336 | -0.100482086 | 1.113539459 | -0.090236664 | 0.92809915 | 0.988077882 | Cdh3 | cadherin 3 [Source:MGI Symbol;Acc:MGI: | 0.8031 | 14.391 | 3.0187 | 6.7478 | 1.0906 | 8.9108 |
| 17175 | ENSMUSG00000061062 | 2.544088786 | -0.911591461 | 1.349257802 | -0.675624376 | 0.499279161 | NA | Hdac1-ps | histone deacetylase 1, pseudogene [Sourc | 4.8186 | 3.0838 | 2.0125 | 0.964 | 3.2719 | 1.1138 |
| 17176 | ENSMUSG00000061065 | 0 | NA | NA | NA | NA | NA | H2al1o | H2A histone family member L1O [Source:I | 0 | 0 | 0 | 0 | 0 | 0 |
| 17177 | ENSMUSG00000061068 | 0.82991484 | -1.853997749 | 2.875131703 | -0.644839242 | 0.519031369 | NA | Mcpt4 | mast cell protease 4 [Source:MGI Symbol; | 4.0155 | 0 | 0 | 0.964 | 0 | 0 |
| 17178 | ENSMUSG00000061079 | 81.65238385 | -0.0622928 | 0.25033206 | -0.248840678 | 0.803484024 | 0.963596811 | Zfp143 | zinc finger protein 143 [Source:MGI Symb | 87.538 | 84.289 | 78.487 | 89.65 | 71.981 | 77.969 |
| 17179 | ENSMUSG00000061080 | 2.207516943 | 0.332563764 | 1.447698702 | 0.229718907 | 0.818310202 | NA | Lsamp | limbic system-associated membrane prote | 0.8031 | 4.1117 | 1.0062 | 2.8919 | 1.0906 | 3.3415 |
| 17180 | ENSMUSG00000061082 | 0 | NA | NA | NA | NA | NA | Plac1 | placental specific protein 1 [Source:MGI S | 0 | 0 | 0 | 0 | 0 | 0 |
| 17181 | ENSMUSG00000061086 | 9.120861084 | 0.482655308 | 0.807008816 | 0.59807935 | 0.549786988 | 0.895703112 | Myl4 | myosin, light polypeptide 4 [Source:MGI S | 1.6062 | 14.391 | 7.0437 | 9.6398 | 10.906 | 11.138 |
| 17182 | ENSMUSG00000061099 | 44.55829576 | 0.097666363 | 0.328513699 | 0.297297688 | 0.766239247 | 0.955091733 | Gapdhs | glyceraldehyde-3-phosphate dehydrogena | 41.761 | 39.061 | 48.3 | 42.415 | 51.259 | 44.554 |
| 17183 | ENSMUSG00000061100 | 1.486578542 | -0.91453072 | 1.787235692 | -0.51170124 | 0.608860118 | NA | Retnla | resistin like alpha [Source:MGI Symbol;Ac | 0.8031 | 2.0558 | 3.0187 | 1.928 | 0 | 1.1138 |
| 17184 | ENSMUSG00000061104 | 16.72478985 | -0.505304821 | 0.554644813 | -0.911042181 | 0.362273149 | 0.80090868 | Sap18b | Sin3-associated polypeptide 18B [Source:I | 18.471 | 14.391 | 26.162 | 19.28 | 10.906 | 11.138 |
| 17185 | ENSMUSG00000061111 | 234.1249425 | 0.037790054 | 0.174043262 | 0.217130235 | 0.828106852 | 0.969935825 | Mcrip1 | MAPK regulated corepressor interacting p | 233.7 | 242.59 | 216.34 | 199.54 | 251.93 | 260.64 |
| 17186 | ENSMUSG00000061115 | 0 | NA | NA | NA | NA | NA | Gm17113 | predicted gene 17113 [Source:MGI Symbo | 0 | 0 | 0 | 0 | 0 | 0 |
| 17187 | ENSMUSG00000061118 | 291.6331285 | -0.184804825 | 0.170781022 | -1.082115699 | 0.279201121 | 0.731853029 | Dnajc30 | DnaJ heat shock protein family (Hsp40) m | 330.08 | 255.95 | 344.14 | 246.78 | 297.74 | 275.12 |
| 17188 | ENSMUSG00000061119 | 117.5593411 | -0.812168138 | 0.228987968 | -3.5467721 | 0.000389982 | 0.010484566 | Prcp | prolylcarboxypeptidase (angiotensinase C | 138.94 | 150.08 | 161 | 101.22 | 85.068 | 69.058 |

| | | | | | | | | | | | | | | | |
|---|---|---|---|---|---|---|---|---|---|---|---|---|---|---|---|
| 17189 | ENSMUSG00000061126 | 1.705844153 | -2.995191587 | 2.106062321 | -1.422176142 | 0.154975123 | NA | Cyp4f39 | cytochrome P450, family 4, subfamily f, po | 0 | 3.0838 | 6.0375 | 0 | 0 | 1.1138 |
| 17190 | ENSMUSG00000061130 | 2646.690397 | -0.019014555 | 0.103967818 | -0.182888855 | 0.854885228 | 0.974086965 | Ppm1b | protein phosphatase 1B, magnesium depe | 2537.8 | 2719.9 | 2735 | 2551.6 | 2433.2 | 2902.7 |
| 17191 | ENSMUSG00000061132 | 45.57972446 | 0.237747702 | 0.363231898 | 0.654534207 | 0.512767696 | 0.884860213 | Blnk | B cell linker [Source:MGI Symbol;Acc:MGI | 53.005 | 33.921 | 38.237 | 47.235 | 65.437 | 35.643 |
| 17192 | ENSMUSG00000061136 | 888.3604318 | -0.014165943 | 0.11883929 | -0.119202525 | 0.905114906 | 0.983972587 | Prpf40a | pre-mRNA processing factor 40A [Source: | 810.33 | 960.08 | 908.64 | 857.94 | 946.66 | 846.52 |
| 17193 | ENSMUSG00000061143 | 48.82175279 | 0.130935184 | 0.365788359 | 0.357953391 | 0.720378194 | 0.9447719 | Maml3 | mastermind like transcriptional coactivato | 36.14 | 49.34 | 54.337 | 30.847 | 64.346 | 57.92 |
| 17194 | ENSMUSG00000061144 | 0 | NA | NA | NA | NA | NA | Spink12 | serine peptidase inhibitor, Kazal type 12 [S | 0 | 0 | 0 | 0 | 0 | 0 |
| 17195 | ENSMUSG00000061165 | 0 | NA | NA | NA | NA | NA | Olfr160 | olfactory receptor 160 [Source:MGI Symb | 0 | 0 | 0 | 0 | 0 | 0 |
| 17196 | ENSMUSG00000061167 | 1.227672903 | 0.505246938 | 2.08258656 | 0.242605492 | 0.808311026 | NA | Rpl15-ps3 | ribosomal protein L15, pseudogene 3 [Sou | 0 | 3.0838 | 0 | 0.964 | 1.0906 | 2.2277 |
| 17197 | ENSMUSG00000061171 | 14.65691291 | -0.109760297 | 0.563161457 | -0.194900228 | 0.845471056 | 0.972803041 | Slc38a11 | solute carrier family 38, member 11 [Sour | 16.062 | 15.419 | 14.087 | 12.532 | 10.906 | 18.935 |
| 17198 | ENSMUSG00000061175 | 979.1326292 | 0.04056833 | 0.125009727 | 0.324521389 | 0.745543332 | 0.949997329 | Fnip2 | folliculin interacting protein 2 [Source:MG | 1060.9 | 912.79 | 920.71 | 877.22 | 1035 | 1068.2 |
| 17199 | ENSMUSG00000061184 | 0 | NA | NA | NA | NA | NA | Tmprss11c | transmembrane protease, serine 11c [Sou | 0 | 0 | 0 | 0 | 0 | 0 |
| 17200 | ENSMUSG00000061186 | 8.54715828 | -0.496737965 | 0.790291044 | -0.62855067 | 0.529643264 | 0.888334265 | Sfmbt2 | Scm-like with four mbt domains 2 [Source | 13.653 | 5.1396 | 11.069 | 3.8559 | 11.997 | 5.5692 |
| 17201 | ENSMUSG00000061195 | 0 | NA | NA | NA | NA | NA | Olfr1289 | olfactory receptor 1289 [Source:MGI Syml | 0 | 0 | 0 | 0 | 0 | 0 |
| 17202 | ENSMUSG00000061207 | 277.1005545 | 0.1575418 | 0.178072233 | 0.884707272 | 0.376314533 | 0.809731241 | Stk19 | serine/threonine kinase 19 [Source:MGI S | 255.39 | 233.34 | 297.85 | 345.1 | 279.2 | 251.73 |
| 17203 | ENSMUSG00000061208 | 0.503122293 | -2.366675966 | 3.785596405 | -0.625179156 | 0.531853481 | NA | Vmn1r8 | vomeronasal 1 receptor 8 [Source:MGI Sy | 0 | 0 | 3.0187 | 0 | 0 | 0 |
| 17204 | ENSMUSG00000061210 | 0 | NA | NA | NA | NA | NA | Olfr47 | olfactory receptor 47 [Source:MGI Symbo | 0 | 0 | 0 | 0 | 0 | 0 |
| 17205 | ENSMUSG00000061232 | 30156.35438 | -0.092946605 | 0.087724966 | -1.059522844 | 0.289361731 | 0.739770202 | H2-K1 | histocompatibility 2, K1, K region [Source: | 30966 | 32732 | 29685 | 30747 | 28786 | 28022 |
| 17206 | ENSMUSG00000061242 | 0 | NA | NA | NA | NA | NA | Gm9104 | predicted gene 9104 [Source:MGI Symbol | 0 | 0 | 0 | 0 | 0 | 0 |
| 17207 | ENSMUSG00000061244 | 448.6371384 | 0.052288356 | 0.136081868 | 0.384241906 | 0.700799161 | 0.941778228 | Exoc5 | exocyst complex component 5 [Source:M | 415.2 | 492.37 | 414.57 | 464.64 | 442.79 | 462.25 |
| 17208 | ENSMUSG00000061246 | 0 | NA | NA | NA | NA | NA | Olfr387-ps1 | olfactory receptor 387, pseudogene 1 [Sou | 0 | 0 | 0 | 0 | 0 | 0 |
| 17209 | ENSMUSG00000061259 | 0 | NA | NA | NA | NA | NA | Tmprss11d | transmembrane protease, serine 11d [Sou | 0 | 0 | 0 | 0 | 0 | 0 |
| 17210 | ENSMUSG00000061272 | 0 | NA | NA | NA | NA | NA | Gm14173 | predicted gene 14173 [Source:MGI Symbo | 0 | 0 | 0 | 0 | 0 | 0 |
| 17211 | ENSMUSG00000061273 | 467.7993998 | -0.110122213 | 0.133217503 | -0.826634718 | 0.408444133 | 0.832151413 | Mmgt1 | membrane magnesium transporter 1 [Sou | 465 | 524.24 | 467.9 | 429.93 | 485.33 | 434.4 |
| 17212 | ENSMUSG00000061286 | 647.6032673 | 0.154935815 | 0.13338374 | 1.161579475 | 0.245406323 | 0.695718749 | Exosc5 | exosome component 5 [Source:MGI Symb | 603.13 | 582.83 | 652.05 | 589.95 | 749.25 | 708.4 |
| 17213 | ENSMUSG00000061288 | 1016.161186 | -0.16583625 | 0.168982432 | -0.981381601 | 0.326404596 | 0.772181236 | Taok3 | TAO kinase 3 [Source:MGI Symbol;Acc:MG | 1017.5 | 1162.6 | 1043.5 | 943.73 | 709.99 | 1219.7 |
| 17214 | ENSMUSG00000061292 | 402.2556564 | -1.174873165 | 0.195001339 | -6.024949206 | 1.69E-09 | 3.26E-07 | Cyp3a59 | cytochrome P450, family 3, subfamily a, p | 685.05 | 408.08 | 578.59 | 218.82 | 285.74 | 237.25 |
| 17215 | ENSMUSG00000061295 | 0 | NA | NA | NA | NA | NA | Olfr1261 | olfactory receptor 1261 [Source:MGI Syml | 0 | 0 | 0 | 0 | 0 | 0 |
| 17216 | ENSMUSG00000061296 | 0 | NA | NA | NA | NA | NA | Vmn1r210 | vomeronasal 1 receptor 210 [Source:MGI | 0 | 0 | 0 | 0 | 0 | 0 |
| 17217 | ENSMUSG00000061298 | 0 | NA | NA | NA | NA | NA | Agbl4 | ATP/GTP binding protein-like 4 [Source:M | 0 | 0 | 0 | 0 | 0 | 0 |
| 17218 | ENSMUSG00000061306 | 1923.260904 | 0.065549846 | 0.092364779 | 0.709684436 | 0.477899846 | 0.869794311 | Slc38a10 | solute carrier family 38, member 10 [Sour | 1869.6 | 1935.6 | 1833.4 | 1913.5 | 1990.4 | 1997.1 |
| 17219 | ENSMUSG00000061311 | 0 | NA | NA | NA | NA | NA | Rag1 | recombination activating 1 [Source:MGI S | 0 | 0 | 0 | 0 | 0 | 0 |
| 17220 | ENSMUSG00000061313 | 660.2516352 | -0.377287171 | 0.169090781 | -2.231269907 | 0.025663256 | 0.206447904 | Ddhd2 | DDHD domain containing 2 [Source:MGI S | 778.21 | 835.7 | 624.88 | 654.54 | 627.11 | 441.08 |
| 17221 | ENSMUSG00000061315 | 3625.364193 | 0.05204333 | 0.088907699 | 0.585363592 | 0.558303264 | 0.899286756 | Naca | nascent polypeptide-associated complex a | 3449.3 | 3730.3 | 3500.7 | 3604.3 | 3804.1 | 3663.4 |
| 17222 | ENSMUSG00000061322 | 1.448120999 | 0.705574662 | 2.138746123 | 0.329901083 | 0.741474705 | NA | Dnai1 | dynein axonemal intermediate chain 1 [So | 3.2124 | 0 | 0 | 0 | 4.3625 | 1.1138 |
| 17223 | ENSMUSG00000061330 | 0 | NA | NA | NA | NA | NA | Gm6436 | predicted gene 6436 [Source:MGI Symbol | 0 | 0 | 0 | 0 | 0 | 0 |
| 17224 | ENSMUSG00000061331 | 0 | NA | NA | NA | NA | NA | Gm17132 | predicted gene 17132 [Source:MGI Symbo | 0 | 0 | 0 | 0 | 0 | 0 |
| 17225 | ENSMUSG00000061336 | 0 | NA | NA | NA | NA | NA | Olfr116 | olfactory receptor 116 [Source:MGI Symb | 0 | 0 | 0 | 0 | 0 | 0 |
| 17226 | ENSMUSG00000061342 | 0 | NA | NA | NA | NA | NA | Olfr1168 | olfactory receptor 1168 [Source:MGI Syml | 0 | 0 | 0 | 0 | 0 | 0 |
| 17227 | ENSMUSG00000061353 | 6051.022629 | -0.095183506 | 0.112412042 | -0.846737624 | 0.397141383 | 0.824612828 | Cxcl12 | chemokine (C-X-C motif) ligand 12 [Source | 5976.7 | 6223 | 6553.7 | 6731.5 | 5581.8 | 5239.5 |
| 17228 | ENSMUSG00000061356 | 2.836747744 | -0.399282416 | 1.257107688 | -0.317619899 | 0.750773278 | NA | Nuggc | nuclear GTPase, germinal center associate | 1.6062 | 4.1117 | 4.025 | 2.8919 | 3.2719 | 1.1138 |
| 17229 | ENSMUSG00000061360 | 401.2771668 | 0.117919655 | 0.134563673 | 0.87631121 | 0.380860873 | 0.812628154 | Phf5a | PHD finger protein 5A [Source:MGI Symbo | 394.32 | 390.61 | 369.29 | 395.23 | 444.97 | 413.24 |
| 17230 | ENSMUSG00000061361 | 0 | NA | NA | NA | NA | NA | Olfr168 | olfactory receptor 168 [Source:MGI Symb | 0 | 0 | 0 | 0 | 0 | 0 |
| 17231 | ENSMUSG00000061367 | 0 | NA | NA | NA | NA | NA | Olfr771 | olfactory receptor 771 [Source:MGI Symb | 0 | 0 | 0 | 0 | 0 | 0 |
| 17232 | ENSMUSG00000061371 | 54.30120345 | 0.334955993 | 0.312114007 | 1.073184751 | 0.283188231 | 0.735802616 | Zfp873 | zinc finger protein 873 [Source:MGI Symb | 60.233 | 41.117 | 42.262 | 59.767 | 56.712 | 65.717 |
| 17233 | ENSMUSG00000061374 | 297.3119624 | 0.191187464 | 0.150571446 | 1.269745816 | 0.204175187 | 0.649416772 | Fiz1 | Flt3 interacting zinc finger protein 1 [Sour | 256.99 | 293.98 | 282.75 | 337.39 | 307.55 | 305.19 |
| 17234 | ENSMUSG00000061376 | 0 | NA | NA | NA | NA | NA | Vmn1r219 | vomeronasal 1 receptor 219 [Source:MGI | 0 | 0 | 0 | 0 | 0 | 0 |
| 17235 | ENSMUSG00000061387 | 0 | NA | NA | NA | NA | NA | Olfr1490 | olfactory receptor 1490 [Source:MGI Syml | 0 | 0 | 0 | 0 | 0 | 0 |
| 17236 | ENSMUSG00000061388 | 0.133850373 | -0.796629176 | 4.080472857 | -0.195229623 | 0.84521319 | NA | Csn1s2b | casein alpha s2-like B [Source:MGI Symbo | 0.8031 | 0 | 0 | 0 | 0 | 0 |
| 17237 | ENSMUSG00000061390 | 0.476489653 | -2.293616034 | 3.421506946 | -0.670352587 | 0.502633051 | NA | Uba52-ps | ubitquitin A-52 residue ribosomal protein | 0.8031 | 2.0558 | 0 | 0 | 0 | 0 |
| 17238 | ENSMUSG00000061392 | 0 | NA | NA | NA | NA | NA | Mageb5 | MAGE family member B5 [Source:MGI Syr | 0 | 0 | 0 | 0 | 0 | 0 |
| 17239 | ENSMUSG00000061393 | 61.18278676 | 0.925653517 | 0.322949104 | 2.866252007 | 0.004153635 | 0.058842478 | Acvr2b | activin receptor IIB [Source:MGI Symbol;A | 33.73 | 34.949 | 58.362 | 92.542 | 81.796 | 65.717 |
| 17240 | ENSMUSG00000061397 | 1.360619277 | 0.910684626 | 1.897680624 | 0.479893516 | 0.631303112 | NA | Krt79 | keratin 79 [Source:MGI Symbol;Acc:MGI:2 | 0.8031 | 1.0279 | 1.0062 | 0.964 | 4.3625 | 0 |
| 17241 | ENSMUSG00000061410 | 892.308739 | 0.037186883 | 0.118353078 | 0.314202924 | 0.753366922 | 0.952989051 | Zcchc14 | zinc finger, CCHC domain containing 14 [S | 899.47 | 933.35 | 809.02 | 856.01 | 873.59 | 982.41 |
| 17242 | ENSMUSG00000061411 | 10.67775691 | 0.109764364 | 0.665812156 | 0.164857855 | 0.869055866 | 0.978605211 | Nol4l | nucleolar protein 4-like [Source:MGI Symb | 14.456 | 7.1954 | 9.0562 | 13.496 | 8.7249 | 11.138 |
| 17243 | ENSMUSG00000061414 | 2.653401825 | -0.97099755 | 1.326154362 | -0.732190443 | 0.464052337 | NA | Cracr2a | calcium release activated channel regulat | 2.4093 | 3.0838 | 5.0312 | 0.964 | 1.0906 | 3.3415 |
| 17244 | ENSMUSG00000061436 | 389.1865793 | -0.118730243 | 0.187558262 | -0.633031263 | 0.526713228 | 0.888334265 | Hipk2 | homeodomain interacting protein kinase 2 | 382.28 | 513.96 | 319.99 | 407.76 | 394.8 | 316.33 |

| | | | | | | | | | | | | | | | |
|---|---|---|---|---|---|---|---|---|---|---|---|---|---|---|---|
| 17245 | ENSMUSG00000061451 | 11.14092423 | -0.408102872 | 0.680575979 | -0.599643368 | 0.548743938 | 0.895703112 | Tmem151a | transmembrane protein 151A [Source:MG | 13.653 | 10.279 | 14.087 | 5.7839 | 16.359 | 6.6831 |
| 17246 | ENSMUSG00000061455 | 647.1680834 | 0.049900722 | 0.115160807 | 0.433313412 | 0.664787096 | 0.929301301 | Stx17 | syntaxin 17 [Source:MGI Symbol;Acc:MGI | 644.09 | 650.67 | 612.8 | 634.3 | 676.18 | 664.96 |
| 17247 | ENSMUSG00000061457 | 0 | NA | NA | NA | NA | NA | Olfr871 | olfactory receptor 871 [Source:MGI Symb | 0 | 0 | 0 | 0 | 0 | 0 |
| 17248 | ENSMUSG00000061458 | 110.882802 | -0.470287329 | 0.24694802 | -1.90439806 | 0.056858362 | 0.338327228 | Nol10 | nucleolar protein 10 [Source:MGI Symbol; | 132.51 | 146.99 | 106.66 | 80.01 | 116.7 | 82.424 |
| 17249 | ENSMUSG00000061461 | 488.0882618 | 0.075914862 | 0.131208401 | 0.57858233 | 0.562871031 | 0.900693283 | Smim20 | small integral membrane protein 20 [Sour | 516.39 | 473.87 | 434.7 | 500.3 | 485.33 | 517.94 |
| 17250 | ENSMUSG00000061462 | 1.956581176 | 2.412019647 | 1.857060974 | 1.298837077 | 0.193999847 | NA | Obscn | obscurin, cytoskeletal calmodulin and titir | 0.8031 | 1.0279 | 0 | 0 | 5.4531 | 4.4554 |
| 17251 | ENSMUSG00000061469 | 0 | NA | NA | NA | NA | NA | Msantd5 | Myb/SANT DNA binding domain containir | 0 | 0 | 0 | 0 | 0 | 0 |
| 17252 | ENSMUSG00000061474 | 153.3495478 | -0.042256877 | 0.219288212 | -0.192700173 | 0.847193788 | 0.972803041 | Mrps36 | mitochondrial ribosomal protein S36 [Sou | 154.2 | 171.66 | 140.87 | 148.45 | 117.79 | 187.13 |
| 17253 | ENSMUSG00000061477 | 3855.578091 | 0.010644865 | 0.086585159 | 0.122940986 | 0.90215383 | 0.983821605 | Rps7 | ribosomal protein S7 [Source:MGI Symbol | 3672.6 | 3947.2 | 3905.2 | 3896.4 | 3828.1 | 3884 |
| 17254 | ENSMUSG00000061479 | 449.1231379 | 0.17870053 | 0.127465424 | 1.401952972 | 0.160929291 | 0.583964189 | Snrpa | small nuclear ribonucleoprotein polypepti | 436.08 | 399.86 | 427.65 | 473.31 | 474.42 | 483.41 |
| 17255 | ENSMUSG00000061482 | 2.469098648 | 1.269995629 | 1.435889339 | 0.884466229 | 0.376444585 | NA | H4c4 | H4 clustered histone 4 [Source:MGI Symb | 3.2124 | 0 | 1.0062 | 2.8919 | 4.3625 | 3.3415 |
| 17256 | ENSMUSG00000061486 | 0 | NA | NA | NA | NA | NA | Gm5161 | predicted pseudogene 5161 [Source:MGI | 0 | 0 | 0 | 0 | 0 | 0 |
| 17257 | ENSMUSG00000061488 | 1.348677275 | -1.529254853 | 2.290768786 | -0.667572765 | 0.504406347 | NA | Rpl27a-ps1 | ribosomal protein L27A, pseudogene 1 [So | 0 | 0 | 6.0375 | 0.964 | 1.0906 | 0 |
| 17258 | ENSMUSG00000061489 | 0 | NA | NA | NA | NA | NA | Olfr525 | olfactory receptor 525 [Source:MGI Symb | 0 | 0 | 0 | 0 | 0 | 0 |
| 17259 | ENSMUSG00000061490 | 0 | NA | NA | NA | NA | NA | Ube2d4 | ubiquitin-conjugating enzyme E2D 4 [Sour | 0 | 0 | 0 | 0 | 0 | 0 |
| 17260 | ENSMUSG00000061510 | 5.923178143 | 2.478028582 | 1.027259674 | 2.412270864 | 0.015853499 | 0.147843332 | Gm10101 | predicted gene 10101 [Source:MGI Symbo | 3.2124 | 2.0558 | 0 | 11.568 | 10.906 | 7.7969 |
| 17261 | ENSMUSG00000061517 | 0 | NA | NA | NA | NA | NA | Sox21 | SRY (sex determining region Y)-box 21 [So | 0 | 0 | 0 | 0 | 0 | 0 |
| 17262 | ENSMUSG00000061518 | 3025.099548 | 0.035535586 | 0.106463896 | 0.333780625 | 0.738545105 | 0.948137969 | Cox5b | cytochrome c oxidase subunit 5B [Source: | 3047 | 3038.5 | 2876.9 | 2740.6 | 3051.5 | 3396.1 |
| 17263 | ENSMUSG00000061520 | 0 | NA | NA | NA | NA | NA | Olfr153 | olfactory receptor 153 [Source:MGI Symb | 0 | 0 | 0 | 0 | 0 | 0 |
| 17264 | ENSMUSG00000061524 | 0 | NA | NA | NA | NA | NA | Zic2 | zinc finger protein of the cerebellum 2 [Sc | 0 | 0 | 0 | 0 | 0 | 0 |
| 17265 | ENSMUSG00000061525 | 0 | NA | NA | NA | NA | NA | 4921509C19 | RIKEN cDNA 4921509C19 gene [Source:M | 0 | 0 | 0 | 0 | 0 | 0 |
| 17266 | ENSMUSG00000061527 | 0 | NA | NA | NA | NA | NA | Krt5 | keratin 5 [Source:MGI Symbol;Acc:MGI:96 | 0 | 0 | 0 | 0 | 0 | 0 |
| 17267 | ENSMUSG00000061531 | 0.185640647 | 1.126924175 | 4.080472857 | 0.276174898 | 0.782413736 | NA | Tmem236 | transmembrane protein 236 [Source:MGI | 0 | 0 | 0 | 0 | 0 | 1.1138 |
| 17268 | ENSMUSG00000061533 | 16.63235801 | 0.221511981 | 0.551059403 | 0.401974777 | 0.687702589 | 0.936885029 | Cep128 | centrosomal protein 128 [Source:MGI Syn | 18.471 | 14.391 | 13.081 | 15.424 | 26.175 | 12.252 |
| 17269 | ENSMUSG00000061535 | 14.28320108 | -1.309273325 | 0.636819916 | -2.05595537 | 0.039786818 | 0.273126027 | C1qtnf7 | C1q and tumor necrosis factor related pro | 20.078 | 27.754 | 13.081 | 3.8559 | 10.906 | 10.025 |
| 17270 | ENSMUSG00000061536 | 576.3751392 | 0.217258454 | 0.141627451 | 1.534013732 | 0.125026275 | 0.518510965 | Sec22c | SEC22 homolog C, vesicle trafficking prote | 618.39 | 484.15 | 495.07 | 578.39 | 669.64 | 612.61 |
| 17271 | ENSMUSG00000061540 | 120.0437741 | -0.806327048 | 0.251043553 | -3.211901039 | 0.001318598 | 0.025669179 | Orm2 | orosomucoid 2 [Source:MGI Symbol;Acc:I | 168.65 | 179.89 | 109.68 | 95.434 | 74.162 | 92.449 |
| 17272 | ENSMUSG00000061544 | 103.0323173 | 0.009582675 | 0.234047543 | 0.040943286 | 0.967341109 | 0.994231936 | Zfp229 | zinc finger protein 229 [Source:MGI Symb | 110.83 | 83.261 | 113.71 | 102.18 | 107.97 | 100.25 |
| 17273 | ENSMUSG00000061549 | 0 | NA | NA | NA | NA | NA | Olfr301 | olfactory receptor 301 [Source:MGI Symb | 0 | 0 | 0 | 0 | 0 | 0 |
| 17274 | ENSMUSG00000061559 | 897.7319665 | 0.035714934 | 0.111882633 | 0.319217851 | 0.749561321 | 0.951356173 | Wdr61 | WD repeat domain 61 [Source:MGI Symbo | 929.99 | 859.34 | 869.4 | 850.23 | 912.85 | 964.59 |
| 17275 | ENSMUSG00000061561 | 0 | NA | NA | NA | NA | NA | Olfr373 | olfactory receptor 373 [Source:MGI Symb | 0 | 0 | 0 | 0 | 0 | 0 |
| 17276 | ENSMUSG00000061576 | 0.79210974 | -1.697240781 | 2.618300099 | -0.648222403 | 0.516841114 | NA | Dpp6 | dipeptidylpeptidase 6 [Source:MGI Symbo | 1.6062 | 2.0558 | 0 | 0 | 1.0906 | 0 |
| 17277 | ENSMUSG00000061577 | 1.566338398 | -1.003637873 | 1.747785475 | -0.574234016 | 0.565809454 | NA | Adgrg5 | adhesion G protein-coupled receptor G5 [ | 3.2124 | 1.0279 | 2.0125 | 0.964 | 2.1812 | 0 |
| 17278 | ENSMUSG00000061578 | 2.237761264 | -4.523669626 | 1.819095622 | -2.486768463 | 0.012890927 | NA | Ksr2 | kinase suppressor of ras 2 [Source:MGI Sy | 3.2124 | 7.1954 | 3.0187 | 0 | 0 | 0 |
| 17279 | ENSMUSG00000061584 | 0 | NA | NA | NA | NA | NA | Lyg2 | lysozyme G-like 2 [Source:MGI Symbol;Ac | 0 | 0 | 0 | 0 | 0 | 0 |
| 17280 | ENSMUSG00000061589 | 445.6711265 | -0.265932865 | 0.151547969 | -1.754776835 | 0.079297487 | 0.413467505 | Dot1l | DOT1-like, histone H3 methyltransferase ( | 455.36 | 502.65 | 503.12 | 480.06 | 369.72 | 363.11 |
| 17281 | ENSMUSG00000061601 | 13.04033487 | 1.390578597 | 0.619071693 | 2.246231918 | 0.024689157 | 0.20129874 | Pclo | piccolo (presynaptic cytomatrix protein) [! | 6.4248 | 7.1954 | 8.05 | 21.208 | 13.087 | 22.277 |
| 17282 | ENSMUSG00000061602 | 0 | NA | NA | NA | NA | NA | Vmn1r78 | vomeronasal 1 receptor 78 [Source:MGI S | 0 | 0 | 0 | 0 | 0 | 0 |
| 17283 | ENSMUSG00000061603 | 0.706721132 | -2.881789029 | 2.959064625 | -0.973885127 | 0.330113579 | NA | Akap6 | A kinase (PRKA) anchor protein 6 [Source: | 3.2124 | 1.0279 | 0 | 0 | 0 | 0 |
| 17284 | ENSMUSG00000061607 | 173.3760163 | 0.360876694 | 0.220837921 | 1.63412467 | 0.102232676 | 0.470577139 | Mdc1 | mediator of DNA damage checkpoint 1 [S | 141.35 | 144.94 | 169.05 | 170.62 | 159.23 | 255.07 |
| 17285 | ENSMUSG00000061613 | 657.7455035 | -0.084126905 | 0.123472794 | -0.681339604 | 0.495656629 | 0.877965876 | U2af1 | U2 small nuclear ribonucleoprotein auxilia | 611.16 | 691.79 | 728.52 | 616.95 | 659.82 | 638.23 |
| 17286 | ENSMUSG00000061614 | 0 | NA | NA | NA | NA | NA | Olfr845 | olfactory receptor 845 [Source:MGI Symb | 0 | 0 | 0 | 0 | 0 | 0 |
| 17287 | ENSMUSG00000061615 | 0.349477146 | 0.165142768 | 3.911021834 | 0.042224967 | 0.966319359 | NA | H2ac4 | H2A clustered histone 4 [Source:MGI Sym | 0 | 0 | 1.0062 | 0 | 1.0906 | 0 |
| 17288 | ENSMUSG00000061616 | 0 | NA | NA | NA | NA | NA | Olfr12 | olfactory receptor 12 [Source:MGI Symbo | 0 | 0 | 0 | 0 | 0 | 0 |
| 17289 | ENSMUSG00000061619 | 0 | NA | NA | NA | NA | NA | Vma21-ps | VMA21 vacuolar H+-ATPase homolog (S. c | 0 | 0 | 0 | 0 | 0 | 0 |
| 17290 | ENSMUSG00000061626 | 0 | NA | NA | NA | NA | NA | Olfr68 | olfactory receptor 68 [Source:MGI Symbo | 0 | 0 | 0 | 0 | 0 | 0 |
| 17291 | ENSMUSG00000061633 | 0 | NA | NA | NA | NA | NA | Ndufb11b | NADH:ubiquinone oxidoreductase subuni | 0 | 0 | 0 | 0 | 0 | 0 |
| 17292 | ENSMUSG00000061637 | 0 | NA | NA | NA | NA | NA | Olfr1457 | olfactory receptor 1457 [Source:MGI Syml | 0 | 0 | 0 | 0 | 0 | 0 |
| 17293 | ENSMUSG00000061650 | 197.8735532 | 0.182400112 | 0.170854984 | 1.067572672 | 0.285713316 | 0.73822033 | Med9 | mediator complex subunit 9 [Source:MGI | 187.12 | 173.72 | 195.21 | 211.11 | 201.76 | 218.31 |
| 17294 | ENSMUSG00000061653 | 0 | NA | NA | NA | NA | NA | Vmn1r46 | vomeronasal 1 receptor 46 [Source:MGI S | 0 | 0 | 0 | 0 | 0 | 0 |
| 17295 | ENSMUSG00000061654 | 0.181769715 | 1.126924175 | 4.080472857 | 0.276174898 | 0.782413736 | NA | NA | NA | 0 | 0 | 0 | 0 | 1.0906 | 0 |
| 17296 | ENSMUSG00000061665 | 699.6461209 | -0.042848244 | 0.129554475 | -0.330735346 | 0.740844401 | 0.948792385 | Cd2ap | CD2-associated protein [Source:MGI Syml | 637.66 | 805.89 | 687.27 | 659.36 | 689.27 | 718.43 |
| 17297 | ENSMUSG00000061666 | 4.772124541 | -0.993670491 | 1.089865352 | -0.911736931 | 0.361907218 | NA | Gdpd1 | glycerophosphodiester phosphodiesteras | 4.8186 | 7.1954 | 7.0437 | 2.8919 | 0 | 6.6831 |
| 17298 | ENSMUSG00000061669 | 0 | NA | NA | NA | NA | NA | Rpl34-ps2 | ribosomal protein L34, pseudogene 2 [Sou | 0 | 0 | 0 | 0 | 0 | 0 |
| 17299 | ENSMUSG00000061684 | 0.185640647 | 1.126924175 | 4.080472857 | 0.276174898 | 0.782413736 | NA | Rpl21-ps8 | ribosomal protein L21, pseudogene 8 [Sou | 0 | 0 | 0 | 0 | 0 | 1.1138 |
| 17300 | ENSMUSG00000061689 | 294.6381868 | 0.144464495 | 0.166314083 | 0.868624545 | 0.385052525 | 0.816437825 | Dlgap4 | DLG associated protein 4 [Source:MGI Syr | 273.05 | 279.59 | 287.79 | 367.28 | 268.29 | 291.83 |

| | | | | | | | | | | | | | | | |
|---|---|---|---|---|---|---|---|---|---|---|---|---|---|---|---|
| 17301 | ENSMUSG00000061701 | 0 | NA | NA | NA | NA | NA | Fbxw20 | F-box and WD-40 domain protein 20 [Sou | 0 | 0 | 0 | 0 | 0 | 0 |
| 17302 | ENSMUSG00000061702 | 1.580400682 | -0.334602804 | 1.754608627 | -0.190699395 | 0.848761112 | NA | Tmem91 | transmembrane protein 91 [Source:MGI S | 3.2124 | 1.0279 | 1.0062 | 0.964 | 3.2719 | 0 |
| 17303 | ENSMUSG00000061707 | 0 | NA | NA | NA | NA | NA | Gm4871 | predicted gene 4871 [Source:MGI Symbol | 0 | 0 | 0 | 0 | 0 | 0 |
| 17304 | ENSMUSG00000061718 | 257.2701494 | -0.801171743 | 0.267327691 | -2.996965032 | 0.00272682 | 0.043871772 | Ppp1r1b | protein phosphatase 1, regulatory inhibitc | 224.07 | 370.05 | 387.4 | 205.33 | 245.39 | 111.38 |
| 17305 | ENSMUSG00000061723 | 0 | NA | NA | NA | NA | NA | Tnnt3 | troponin T3, skeletal, fast [Source:MGI Sy | 0 | 0 | 0 | 0 | 0 | 0 |
| 17306 | ENSMUSG00000061724 | 0.66192371 | 1.647602865 | 2.590567345 | 0.636000785 | 0.52477591 | NA | Gm2423 | predicted gene 2423 [Source:MGI Symbol | 0.8031 | 0 | 0 | 0.964 | 1.0906 | 1.1138 |
| 17307 | ENSMUSG00000061728 | 0 | NA | NA | NA | NA | NA | Btnl7-ps | butyrophilin-like 7, pseudogene [Source:N | 0 | 0 | 0 | 0 | 0 | 0 |
| 17308 | ENSMUSG00000061731 | 1029.575068 | -0.152121265 | 0.113433417 | -1.341062182 | 0.179900265 | 0.612025777 | Ext1 | exostosin glycosyltransferase 1 [Source:M | 1104.3 | 1043.3 | 1102.8 | 893.61 | 964.11 | 1069.3 |
| 17309 | ENSMUSG00000061740 | 5596.454314 | -0.145082018 | 0.091992797 | -1.577101934 | 0.114772072 | 0.497545211 | Cyp2d22 | cytochrome P450, family 2, subfamily d, p | 5498.8 | 5916.7 | 6218.6 | 5392.5 | 5460.7 | 5091.4 |
| 17310 | ENSMUSG00000061742 | 0 | NA | NA | NA | NA | NA | Slc22a12 | solute carrier family 22 (organic anion/cat | 0 | 0 | 0 | 0 | 0 | 0 |
| 17311 | ENSMUSG00000061751 | 778.8018308 | 0.26359006 | 0.12227843 | 2.155654608 | 0.031110645 | 0.234122981 | Kalrn | kalirin, RhoGEF kinase [Source:MGI Symbo | 653.73 | 719.54 | 750.66 | 794.32 | 824.51 | 930.06 |
| 17312 | ENSMUSG00000061755 | 354.2525097 | -0.055023514 | 0.144526158 | -0.380716642 | 0.703413519 | 0.942540408 | Bod1l | biorientation of chromosomes in cell divis | 376.65 | 364.91 | 341.12 | 348.96 | 316.28 | 377.59 |
| 17313 | ENSMUSG00000061758 | 18.07077276 | -0.297056098 | 0.584050494 | -0.508613726 | 0.611023006 | 0.916376001 | Akr1b10 | aldo-keto reductase family 1, member B1( | 24.093 | 23.642 | 12.075 | 23.135 | 6.5437 | 18.935 |
| 17314 | ENSMUSG00000061759 | 270.2938708 | -0.033211204 | 0.162105983 | -0.204873398 | 0.83767104 | 0.971073158 | Armt1 | acidic residue methyltransferase 1 [Source | 290.72 | 249.78 | 279.74 | 298.83 | 245.39 | 257.3 |
| 17315 | ENSMUSG00000061762 | 0 | NA | NA | NA | NA | NA | Tac1 | tachykinin 1 [Source:MGI Symbol;Acc:MG | 0 | 0 | 0 | 0 | 0 | 0 |
| 17316 | ENSMUSG00000061769 | 0 | NA | NA | NA | NA | NA | Klra6 | killer cell lectin-like receptor, subfamily A, | 0 | 0 | 0 | 0 | 0 | 0 |
| 17317 | ENSMUSG00000061778 | 691.504355 | 0.016523805 | 0.130272398 | 0.126840418 | 0.899066705 | 0.982798755 | Mospd2 | motile sperm domain containing 2 [Source | 769.37 | 649.64 | 642.99 | 743.23 | 708.9 | 634.89 |
| 17318 | ENSMUSG00000061780 | 153.6501406 | -1.633049843 | 1.746028187 | -0.935294089 | 0.349636761 | 0.790920567 | Cfd | complement factor D (adipsin) [Source:M | 469.01 | 224.09 | 4.025 | 74.226 | 148.32 | 2.2277 |
| 17319 | ENSMUSG00000061787 | 3104.499537 | 0.046863294 | 0.107851441 | 0.43451709 | 0.663912988 | 0.929301301 | Rps17 | ribosomal protein S17 [Source:MGI Symbo | 2921.7 | 3175.2 | 3065 | 2846.6 | 3087.5 | 3530.9 |
| 17320 | ENSMUSG00000061798 | 0 | NA | NA | NA | NA | NA | Olfr1204 | olfactory receptor 1204 [Source:MGI Sym | 0 | 0 | 0 | 0 | 0 | 0 |
| 17321 | ENSMUSG00000061802 | 0 | NA | NA | NA | NA | NA | Odad2 | outer dynein arm docking complex subuni | 0 | 0 | 0 | 0 | 0 | 0 |
| 17322 | ENSMUSG00000061808 | 112447.6408 | -0.081148022 | 0.089690582 | -0.904755214 | 0.365595083 | 0.804235439 | Ttr | transthyretin [Source:MGI Symbol;Acc:M( | 119174 | 108091 | 119563 | 114444 | 103238 | 110176 |
| 17323 | ENSMUSG00000061809 | 0 | NA | NA | NA | NA | NA | Tbpl2 | TATA box binding protein like 2 [Source:M | 0 | 0 | 0 | 0 | 0 | 0 |
| 17324 | ENSMUSG00000061815 | 32.90679449 | -1.76762432 | 0.495293617 | -3.568841306 | 0.000358564 | 0.009973582 | Rufy4 | RUN and FYVE domain containing 4 [Sour | 58.626 | 71.954 | 22.137 | 18.316 | 15.269 | 11.138 |
| 17325 | ENSMUSG00000061816 | 0.160662974 | 1.126924175 | 4.080472857 | 0.276174898 | 0.782413736 | NA | Myl1 | myosin, light polypeptide 1 [Source:MGI S | 0 | 0 | 0 | 0.964 | 0 | 0 |
| 17326 | ENSMUSG00000061825 | 926.2336819 | 0.591790681 | 0.371595822 | 1.592565489 | 0.111257701 | 0.489450944 | Ces2c | carboxylesterase 2C [Source:MGI Symbol; | 811.94 | 603.39 | 800.97 | 658.4 | 1542.1 | 1140.6 |
| 17327 | ENSMUSG00000061829 | 0 | NA | NA | NA | NA | NA | Vmn1r214 | vomeronasal 1 receptor 214 [Source:MGI | 0 | 0 | 0 | 0 | 0 | 0 |
| 17328 | ENSMUSG00000061833 | 0.305170013 | -1.657836477 | 4.041480335 | -0.410205257 | 0.681655385 | NA | Gm6311 | predicted gene 6311 [Source:MGI Symbol | 0.8031 | 1.0279 | 0 | 0 | 0 | 0 |
| 17329 | ENSMUSG00000061838 | 6034.609475 | -0.258125757 | 0.094977123 | -2.717767696 | 0.006572396 | 0.080561986 | Suclg2 | succinate-Coenzyme A ligase, GDP-formin | 6522 | 6905.6 | 6292 | 5587.2 | 5821.7 | 5079.1 |
| 17330 | ENSMUSG00000061843 | 3.953154454 | 0.330609011 | 1.193311793 | 0.277051658 | 0.781740441 | NA | Vmn1r-ps32 | vomeronasal 1 receptor, pseudogene 32 [ | 2.4093 | 7.1954 | 1.0062 | 8.6758 | 1.0906 | 3.3415 |
| 17331 | ENSMUSG00000061845 | 0 | NA | NA | NA | NA | NA | Defa35 | defensin, alpha, 35 [Source:MGI Symbol;A | 0 | 0 | 0 | 0 | 0 | 0 |
| 17332 | ENSMUSG00000061847 | 0 | NA | NA | NA | NA | NA | Defb39 | defensin beta 39 [Source:MGI Symbol;Acc | 0 | 0 | 0 | 0 | 0 | 0 |
| 17333 | ENSMUSG00000061848 | 0.353089354 | 0.165142706 | 3.896102244 | 0.042386646 | 0.966190474 | NA | Gm5805 | predicted gene 5805 [Source:MGI Symbol | 0 | 1.0279 | 0 | 0 | 1.0906 | 0 |
| 17334 | ENSMUSG00000061852 | 0.469265235 | -2.274672822 | 3.439794491 | -0.661281605 | 0.508431736 | NA | Gm53056 | predicted gene 53056 [Source:MGI Symbo | 0.8031 | 0 | 2.0125 | 0 | 0 | 0 |
| 17335 | ENSMUSG00000061859 | 207.9350669 | -0.120629786 | 0.172944233 | -0.697506847 | 0.485485656 | 0.872848294 | Patj | PATJ, crumbs cell polarity complex compo | 218.44 | 215.86 | 215.34 | 185.08 | 225.76 | 187.13 |
| 17336 | ENSMUSG00000061863 | 0.160662974 | 1.126924175 | 4.080472857 | 0.276174898 | 0.782413736 | NA | Gm6822 | predicted pseudogene 6822 [Source:MGI | 0 | 0 | 0 | 0.964 | 0 | 0 |
| 17337 | ENSMUSG00000061875 | 0 | NA | NA | NA | NA | NA | Olfr1148 | olfactory receptor 1148 [Source:MGI Sym | 0 | 0 | 0 | 0 | 0 | 0 |
| 17338 | ENSMUSG00000061877 | 0 | NA | NA | NA | NA | NA | BC048679 | cDNA sequence BC048679 [Source:MGI Sy | 0 | 0 | 0 | 0 | 0 | 0 |
| 17339 | ENSMUSG00000061878 | 6.4602352 | -0.714882185 | 0.858163417 | -0.833037357 | 0.404823661 | 0.829488307 | Sphk1 | sphingosine kinase 1 [Source:MGI Symbol | 8.8341 | 7.1954 | 8.05 | 4.8199 | 7.6343 | 2.2277 |
| 17340 | ENSMUSG00000061882 | 32.41949749 | 0.247284246 | 0.431169118 | 0.573520308 | 0.566292453 | 0.902076435 | Ccdc62 | coiled-coil domain containing 62 [Source:I | 36.943 | 30.838 | 21.131 | 46.271 | 38.172 | 21.163 |
| 17341 | ENSMUSG00000061887 | 948.0306427 | 0.025007598 | 0.117169281 | 0.213431349 | 0.830990531 | 0.970505212 | Ssbp3 | single-stranded DNA binding protein 3 [Sc | 849.68 | 964.19 | 1006.2 | 901.32 | 1007.7 | 959.02 |
| 17342 | ENSMUSG00000061894 | 44.81867099 | 0.454856803 | 0.401110185 | 1.133994647 | 0.256796793 | 0.710192891 | Zscan20 | zinc finger and SCAN domains 20 [Source: | 48.186 | 24.67 | 40.25 | 53.983 | 30.537 | 71.286 |
| 17343 | ENSMUSG00000061897 | 0.133850373 | -0.796629176 | 4.080472857 | -0.195229623 | 0.84521319 | NA | Gm14292 | predicted gene 14292 [Source:MGI Symbo | 0.8031 | 0 | 0 | 0 | 0 | 0 |
| 17344 | ENSMUSG00000061898 | 69.14976712 | 0.197234729 | 0.302800547 | 0.65136847 | 0.514808659 | 0.885502728 | Rbak | RB-associated KRAB zinc finger [Source:M | 46.58 | 70.926 | 76.475 | 87.722 | 74.162 | 59.034 |
| 17345 | ENSMUSG00000061904 | 5405.658331 | -0.022236545 | 0.095891222 | -0.231893438 | 0.816620781 | 0.967767553 | Slc25a3 | solute carrier family 25 (mitochondrial car | 5100.5 | 5831.4 | 5410.6 | 5095.6 | 5333.1 | 5662.8 |
| 17346 | ENSMUSG00000061906 | 192.5357543 | 1.46427717 | 0.528215349 | 2.772121586 | 0.005569223 | 0.071602882 | Ugt2b38 | UDP glucuronosyltransferase 2 family, pol | 48.186 | 137.74 | 121.76 | 216.9 | 408.98 | 221.65 |
| 17347 | ENSMUSG00000061911 | 0 | NA | NA | NA | NA | NA | Myt1l | myelin transcription factor 1-like [Source:I | 0 | 0 | 0 | 0 | 0 | 0 |
| 17348 | ENSMUSG00000061923 | 0 | NA | NA | NA | NA | NA | Odf1 | outer dense fiber of sperm tails 1 [Source: | 0 | 0 | 0 | 0 | 0 | 0 |
| 17349 | ENSMUSG00000061928 | 0 | NA | NA | NA | NA | NA | Dsg1b | desmoglein 1 beta [Source:MGI Symbol;A | 0 | 0 | 0 | 0 | 0 | 0 |
| 17350 | ENSMUSG00000061937 | 0 | NA | NA | NA | NA | NA | Csn1s2a | casein alpha s2-like A [Source:MGI Symbo | 0 | 0 | 0 | 0 | 0 | 0 |
| 17351 | ENSMUSG00000061947 | 3283.2825 | 0.464909912 | 0.087954797 | 5.285782327 | 1.25E-07 | 1.34E-05 | Serpina10 | serine (or cysteine) peptidase inhibitor, cl | 2816.5 | 2749.7 | 2709.8 | 3847.2 | 3655.8 | 3920.7 |
| 17352 | ENSMUSG00000061950 | 359.2427897 | 0.008221639 | 0.154330457 | 0.05327295 | 0.957514432 | 0.991861398 | Ppp4r1 | protein phosphatase 4, regulatory subunit | 317.23 | 364.91 | 393.44 | 373.06 | 388.26 | 318.56 |
| 17353 | ENSMUSG00000061952 | 0 | NA | NA | NA | NA | NA | Olfr1377 | olfactory receptor 1377 [Source:MGI Sym | 0 | 0 | 0 | 0 | 0 | 0 |
| 17354 | ENSMUSG00000061958 | 0 | NA | NA | NA | NA | NA | Defa38 | defensin, alpha, 38 [Source:MGI Symbol;A | 0 | 0 | 0 | 0 | 0 | 0 |
| 17355 | ENSMUSG00000061959 | 2824.440234 | -0.098767094 | 0.15743571 | -0.627348734 | 0.530430665 | 0.888700236 | Ces1e | carboxylesterase 1E [Source:MGI Symbol; | 2420.6 | 3610 | 2734 | 2811.9 | 3123.5 | 2246.6 |
| 17356 | ENSMUSG00000061961 | 0 | NA | NA | NA | NA | NA | Olfr815 | olfactory receptor 815 [Source:MGI Symb | 0 | 0 | 0 | 0 | 0 | 0 |

| | | | | | | | | | | | | | | | |
|---|---|---|---|---|---|---|---|---|---|---|---|---|---|---|---|
| 17357 | ENSMUSG00000061969 | 0.181769715 | 1.126924175 | 4.080472857 | 0.276174898 | 0.782413736 | NA | Gm5581 | predicted gene 5581 [Source:MGI Symbol | 0 | 0 | 0 | 0 | 1.0906 | 0 |
| 17358 | ENSMUSG00000061972 | 2.247953143 | 0.760280361 | 1.480168158 | 0.513644586 | 0.607500501 | NA | Olfr99 | olfactory receptor 99 [Source:MGI Symbo | 0 | 3.0838 | 2.0125 | 2.8919 | 3.2719 | 2.2277 |
| 17359 | ENSMUSG00000061974 | 0 | NA | NA | NA | NA | NA | Cldn24 | claudin 24 [Source:MGI Symbol;Acc:MGI:3 | 0 | 0 | 0 | 0 | 0 | 0 |
| 17360 | ENSMUSG00000061977 | 0 | NA | NA | NA | NA | NA | Tas2r104 | taste receptor, type 2, member 104 [Sour | 0 | 0 | 0 | 0 | 0 | 0 |
| 17361 | ENSMUSG00000061979 | 355.6269213 | 0.040266724 | 0.158938305 | 0.253348139 | 0.7999992 | 0.962424862 | Rcc1l | reculator of chromosome condensation 1 | 322.04 | 347.44 | 382.37 | 302.69 | 388.26 | 390.96 |
| 17362 | ENSMUSG00000061981 | 954.0337169 | -0.14856847 | 0.143839318 | -1.032878025 | 0.301660984 | 0.752246851 | Flot2 | flotillin 2 [Source:MGI Symbol;Acc:MGI:1( | 879.4 | 1041.3 | 1089.8 | 930.24 | 1033.9 | 749.62 |
| 17363 | ENSMUSG00000061983 | 2344.843876 | -0.024729097 | 0.113128752 | -0.218592499 | 0.826967497 | 0.969748238 | Rps12 | ribosomal protein S12 [Source:MGI Symbo | 2368.3 | 2626.3 | 2100 | 2280.8 | 2204.1 | 2489.4 |
| 17364 | ENSMUSG00000061984 | 0 | NA | NA | NA | NA | NA | Olfr392 | olfactory receptor 392 [Source:MGI Symb | 0 | 0 | 0 | 0 | 0 | 0 |
| 17365 | ENSMUSG00000061988 | 0 | NA | NA | NA | NA | NA | Rpl10a-ps2 | ribosomal protein L10A, pseudogene 2 [Sc | 0 | 0 | 0 | 0 | 0 | 0 |
| 17366 | ENSMUSG00000061991 | 0 | NA | NA | NA | NA | NA | H2ac10 | H2A clustered histone 10 [Source:MGI Syr | 0 | 0 | 0 | 0 | 0 | 0 |
| 17367 | ENSMUSG00000061992 | 0 | NA | NA | NA | NA | NA | Gm4953 | predicted pseudogene 4953 [Source:MGI | 0 | 0 | 0 | 0 | 0 | 0 |
| 17368 | ENSMUSG00000062006 | 2355.935019 | 0.075452539 | 0.093017218 | 0.811167449 | 0.417269516 | 0.836969392 | Rpl34 | ribosomal protein L34 [Source:MGI Symbo | 2264.7 | 2357 | 2261 | 2320.3 | 2386.3 | 2546.2 |
| 17369 | ENSMUSG00000062007 | 0.722877068 | -0.331317305 | 2.989821669 | -0.110815073 | 0.911762993 | NA | Hsh2d | hematopoietic SH2 domain containing [Sc | 2.4093 | 0 | 0 | 1.928 | 0 | 0 |
| 17370 | ENSMUSG00000062012 | 11.08951961 | 0.23963432 | 0.64910128 | 0.369178628 | 0.711994585 | 0.944239215 | Zfp13 | zinc finger protein 13 [Source:MGI Symbo | 12.047 | 11.307 | 7.0437 | 9.6398 | 10.906 | 15.594 |
| 17371 | ENSMUSG00000062014 | 708.6947128 | 0.119344261 | 0.140254524 | 0.850912028 | 0.394818223 | 0.822933206 | Gmfb | glia maturation factor, beta [Source:MGI S | 793.47 | 644.5 | 598.72 | 698.88 | 805.97 | 710.63 |
| 17372 | ENSMUSG00000062017 | 5.68939216 | -0.461362088 | 0.972417543 | -0.474448545 | 0.635180083 | 0.923478764 | Abca14 | ATP-binding cassette, sub-family A (ABC1) | 9.6372 | 3.0838 | 7.0437 | 7.7118 | 1.0906 | 5.5692 |
| 17373 | ENSMUSG00000062028 | 0.453341394 | -0.545391191 | 3.551113277 | -0.153583158 | 0.877938417 | NA | Irgc1 | immunity-related GTPase family, cinema 1 | 1.6062 | 0 | 0 | 0 | 0 | 1.1138 |
| 17374 | ENSMUSG00000062031 | 829.1327932 | 0.118105554 | 0.123494593 | 0.956362151 | 0.3388893 | 0.780858357 | Pgghg | protein glucosylgalactosylhydroxylysine gl | 883.41 | 730.85 | 769.78 | 805.89 | 865.95 | 918.92 |
| 17375 | ENSMUSG00000062036 | 0 | NA | NA | NA | NA | NA | 4932415M13 | RIKEN cDNA 4932415M13 gene [Source:M | 0 | 0 | 0 | 0 | 0 | 0 |
| 17376 | ENSMUSG00000062037 | 0 | NA | NA | NA | NA | NA | Olfr285 | olfactory receptor 285 [Source:MGI Symb | 0 | 0 | 0 | 0 | 0 | 0 |
| 17377 | ENSMUSG00000062038 | 1.215522656 | 0.167846246 | 2.251768889 | 0.074539731 | 0.940580929 | NA | Gm10108 | predicted pseudogene 10108 [Source:MG | 2.4093 | 1.0279 | 0 | 3.8559 | 0 | 0 |
| 17378 | ENSMUSG00000062040 | 88.68399627 | 0.20777971 | 0.251104273 | 0.827463855 | 0.407974201 | 0.832051384 | Zfp27 | zinc finger protein 27 [Source:MGI Symbo | 76.295 | 91.485 | 79.493 | 106.04 | 77.434 | 101.36 |
| 17379 | ENSMUSG00000062042 | 0 | NA | NA | NA | NA | NA | Olfr294 | olfactory receptor 294 [Source:MGI Symb | 0 | 0 | 0 | 0 | 0 | 0 |
| 17380 | ENSMUSG00000062044 | 1.952179435 | -3.236084722 | 1.840920223 | -1.757862552 | 0.078770899 | NA | Lmtk3 | lemur tyrosine kinase 3 [Source:MGI Symb | 1.6062 | 4.1117 | 5.0312 | 0.964 | 0 | 0 |
| 17381 | ENSMUSG00000062046 | 0 | NA | NA | NA | NA | NA | 5730460C07 | RIKEN cDNA 5730460C07 gene [Source:M | 0 | 0 | 0 | 0 | 0 | 0 |
| 17382 | ENSMUSG00000062054 | 2091.588488 | 0.101501036 | 0.08960176 | 1.132801805 | 0.257297493 | 0.710872182 | Iah1 | isoamyl acetate-hydrolyzing esterase 1 ho | 2050.3 | 1979.8 | 2023.6 | 2138.1 | 2176.9 | 2180.9 |
| 17383 | ENSMUSG00000062061 | 73.63987262 | 0.270289651 | 0.45276457 | 0.596976153 | 0.550523302 | 0.895703112 | Obp2a | odorant binding protein 2A [Source:MGI S | 60.233 | 47.284 | 92.575 | 44.343 | 40.353 | 157.05 |
| 17384 | ENSMUSG00000062064 | 0 | NA | NA | NA | NA | NA | Slc2a7 | solute carrier family 2 (facilitated glucose | 0 | 0 | 0 | 0 | 0 | 0 |
| 17385 | ENSMUSG00000062070 | 5347.673755 | -0.091413997 | 0.093391511 | -0.978825547 | 0.327666185 | 0.772937193 | Pgk1 | phosphoglycerate kinase 1 [Source:MGI S | 5167.2 | 5672.1 | 5712.5 | 4921.1 | 5465.1 | 5148.2 |
| 17386 | ENSMUSG00000062073 | 0.160662974 | 1.126924175 | 4.080472857 | 0.276174898 | 0.782413736 | NA | Gm10109 | predicted gene 10109 [Source:MGI Symbo | 0 | 0 | 0 | 0.964 | 0 | 0 |
| 17387 | ENSMUSG00000062074 | 1.910864739 | 0.157924021 | 1.705545517 | 0.092594434 | 0.926225766 | NA | Ccn6 | cellular communication network factor 6 [ | 2.4093 | 0 | 3.0187 | 3.8559 | 2.1812 | 0 |
| 17388 | ENSMUSG00000062075 | 226.6198944 | 0.152360098 | 0.177832191 | 0.856763316 | 0.391575703 | 0.820404559 | Lmnb2 | lamin B2 [Source:MGI Symbol;Acc:MGI:96 | 236.11 | 203.53 | 204.27 | 274.73 | 209.4 | 231.68 |
| 17389 | ENSMUSG00000062077 | 0 | NA | NA | NA | NA | NA | Trim54 | tripartite motif-containing 54 [Source:MG | 0 | 0 | 0 | 0 | 0 | 0 |
| 17390 | ENSMUSG00000062078 | 1180.92055 | -0.066414394 | 0.104330499 | -0.636576984 | 0.524400421 | 0.888334265 | Qk | quaking, KH domain containing RNA bindi | 1181.4 | 1295.2 | 1148.1 | 1150 | 1169.1 | 1141.7 |
| 17391 | ENSMUSG00000062081 | 3.898883977 | 0.146905045 | 1.070872029 | 0.137182634 | 0.890886437 | NA | Gm6055 | predicted gene 6055 [Source:MGI Symbol | 4.0155 | 4.1117 | 3.0187 | 6.7478 | 3.2719 | 2.2277 |
| 17392 | ENSMUSG00000062082 | 1.884645279 | -2.224255567 | 1.983889701 | -1.121158886 | 0.262220237 | NA | Cd200r4 | CD200 receptor 4 [Source:MGI Symbol;Ac | 3.2124 | 6.1675 | 0 | 1.928 | 0 | 0 |
| 17393 | ENSMUSG00000062083 | 0 | NA | NA | NA | NA | NA | Rpl13a-ps1 | ribosomal protein 13A, pseudogene 1 [Sou | 0 | 0 | 0 | 0 | 0 | 0 |
| 17394 | ENSMUSG00000062093 | 6.50311946 | -0.090585406 | 0.843589915 | -0.107380854 | 0.914486843 | 0.98625062 | Gm10110 | predicted gene 10110 [Source:MGI Symbo | 8.8341 | 7.1954 | 4.025 | 5.7839 | 8.7249 | 4.4554 |
| 17395 | ENSMUSG00000062098 | 104.4825029 | -0.25972382 | 0.224460872 | -1.157100644 | 0.247231244 | 0.697450133 | Btbd3 | BTB (POZ) domain containing 3 [Source:M | 110.83 | 123.35 | 107.67 | 103.15 | 88.34 | 93.563 |
| 17396 | ENSMUSG00000062101 | 63.15836414 | -0.066880595 | 0.295780847 | -0.226115367 | 0.821111693 | 0.969160361 | Zfp119b | zinc finger protein 119b [Source:MGI Sym | 70.673 | 50.368 | 72.45 | 52.055 | 64.346 | 69.058 |
| 17397 | ENSMUSG00000062103 | 0 | NA | NA | NA | NA | NA | Olfr924 | olfactory receptor 924 [Source:MGI Symb | 0 | 0 | 0 | 0 | 0 | 0 |
| 17398 | ENSMUSG00000062105 | 0 | NA | NA | NA | NA | NA | Olfr183 | olfactory receptor 183 [Source:MGI Symb | 0 | 0 | 0 | 0 | 0 | 0 |
| 17399 | ENSMUSG00000062110 | 159.148529 | 0.028234348 | 0.198836838 | 0.141997572 | 0.887081923 | 0.982382091 | Scfd2 | Sec1 family domain containing 2 [Source:I | 154.2 | 142.88 | 176.09 | 184.12 | 147.23 | 150.37 |
| 17400 | ENSMUSG00000062115 | 320.3317653 | 0.224885268 | 0.173100991 | 1.299156444 | 0.193890246 | 0.632197591 | Rai1 | retinoic acid induced 1 [Source:MGI Symb | 310.8 | 280.62 | 294.83 | 369.2 | 396.99 | 269.55 |
| 17401 | ENSMUSG00000062116 | 95.11095852 | -0.166449341 | 0.240511226 | -0.692064749 | 0.488896668 | 0.874706988 | Zfp954 | zinc finger protein 954 [Source:MGI Symb | 98.782 | 95.596 | 107.67 | 103.15 | 75.253 | 90.221 |
| 17402 | ENSMUSG00000062121 | 0 | NA | NA | NA | NA | NA | Olfr149 | olfactory receptor 149 [Source:MGI Symb | 0 | 0 | 0 | 0 | 0 | 0 |
| 17403 | ENSMUSG00000062124 | 0 | NA | NA | NA | NA | NA | Defb45 | defensin beta 45 [Source:MGI Symbol;Acc | 0 | 0 | 0 | 0 | 0 | 0 |
| 17404 | ENSMUSG00000062127 | 50.70345295 | -1.006684104 | 0.385093199 | -2.614131091 | 0.008945471 | 0.099758896 | Cttnbp2nl | CTTNBP2 N-terminal like [Source:MGI Syn | 64.248 | 87.373 | 51.318 | 26.027 | 22.903 | 52.351 |
| 17405 | ENSMUSG00000062128 | 0 | NA | NA | NA | NA | NA | Olfr20 | olfactory receptor 20 [Source:MGI Symbo | 0 | 0 | 0 | 0 | 0 | 0 |
| 17406 | ENSMUSG00000062132 | 0.647602702 | 0.168575675 | 2.628959883 | 0.064122574 | 0.948872627 | NA | Arhgap33os | Rho GTPase activating protein 33, opposit | 0.8031 | 1.0279 | 0 | 0.964 | 1.0906 | 0 |
| 17407 | ENSMUSG00000062142 | 0 | NA | NA | NA | NA | NA | Olfr569 | olfactory receptor 569 [Source:MGI Symb | 0 | 0 | 0 | 0 | 0 | 0 |
| 17408 | ENSMUSG00000062148 | 0 | NA | NA | NA | NA | NA | Ear6 | eosinophil-associated, ribonuclease A fan | 0 | 0 | 0 | 0 | 0 | 0 |
| 17409 | ENSMUSG00000062151 | 0 | NA | NA | NA | NA | NA | Unc13c | unc-13 homolog C [Source:MGI Symbol;Ac | 0 | 0 | 0 | 0 | 0 | 0 |
| 17410 | ENSMUSG00000062152 | 0.371281295 | 2.09253362 | 4.031852165 | 0.519000582 | 0.603760335 | NA | Rpl21-ps14 | ribosomal protein L21, pseudogene 14 [Sc | 0 | 0 | 0 | 0 | 0 | 2.2277 |
| 17411 | ENSMUSG00000062154 | 0 | NA | NA | NA | NA | NA | Tex33 | testis expressed 33 [Source:MGI Symbol;A | 0 | 0 | 0 | 0 | 0 | 0 |
| 17412 | ENSMUSG00000062157 | 2.60385302 | 0.620014032 | 1.430689991 | 0.433367142 | 0.664748068 | NA | Ifnlr1 | interferon lambda receptor 1 [Source:MG | 4.0155 | 2.0558 | 0 | 2.8919 | 1.0906 | 5.5692 |

| | | | | | | | | | | | | | | | |
|---|---|---|---|---|---|---|---|---|---|---|---|---|---|---|---|
| 17413 | ENSMUSG00000062162 | 0 | NA | NA | NA | NA | NA | Mageb1 | MAGE family member B1 [Source:MGI Syr | 0 | 0 | 0 | 0 | 0 | 0 |
| 17414 | ENSMUSG00000062165 | 0 | NA | NA | NA | NA | NA | Vmn1r232 | vomeronasal 1 receptor 232 [Source:MGI | 0 | 0 | 0 | 0 | 0 | 0 |
| 17415 | ENSMUSG00000062168 | 0 | NA | NA | NA | NA | NA | Ppef1 | protein phosphatase with EF hand calciun | 0 | 0 | 0 | 0 | 0 | 0 |
| 17416 | ENSMUSG00000062169 | 225.574037 | 0.018546906 | 0.169579782 | 0.109369794 | 0.912909191 | 0.98625062 | Cnih4 | cornichon family AMPA receptor auxiliary | 247.36 | 218.95 | 205.27 | 211.11 | 227.94 | 242.82 |
| 17417 | ENSMUSG00000062170 | 16.30994136 | -0.807648018 | 0.83234304 | -0.970330716 | 0.331881671 | 0.774747658 | Fmr1nb | Fmr1 neighbor [Source:MGI Symbol;Acc:N | 20.078 | 41.117 | 1.0062 | 4.8199 | 16.359 | 14.48 |
| 17418 | ENSMUSG00000062175 | 3.237569311 | -0.025872893 | 1.24106997 | -0.020847248 | 0.983367508 | NA | Tgif2 | TGFB-induced factor homeobox 2 [Source | 5.6217 | 2.0558 | 2.0125 | 0.964 | 6.5437 | 2.2277 |
| 17419 | ENSMUSG00000062181 | 10940.22553 | 1.097846085 | 0.154607241 | 7.100871088 | 1.24E-12 | 4.98E-10 | Ces3b | carboxylesterase 3B [Source:MGI Symbol; | 5869.1 | 6730.8 | 8304.5 | 18499 | 13000 | 13238 |
| 17420 | ENSMUSG00000062184 | 0.267700746 | -1.497481649 | 4.057001436 | -0.369110456 | 0.712045396 | NA | Hs6st2 | heparan sulfate 6-O-sulfotransferase 2 [Sc | 1.6062 | 0 | 0 | 0 | 0 | 0 |
| 17421 | ENSMUSG00000062186 | 0 | NA | NA | NA | NA | NA | Olfr395 | olfactory receptor 395 [Source:MGI Symb | 0 | 0 | 0 | 0 | 0 | 0 |
| 17422 | ENSMUSG00000062190 | 169.9281455 | 0.185794275 | 0.185930741 | 0.999266039 | 0.317665833 | 0.764257918 | Lancl2 | LanC (bacterial lantibiotic synthetase com | 164.64 | 156.24 | 155.97 | 178.34 | 201.76 | 162.62 |
| 17423 | ENSMUSG00000062198 | 100.9158855 | -0.021921489 | 0.227415695 | -0.096393912 | 0.923207727 | 0.98705827 | 2700097O09 | RIKEN cDNA 2700097O09 gene [Source:M | 102.8 | 104.85 | 97.606 | 111.82 | 95.974 | 92.449 |
| 17424 | ENSMUSG00000062199 | 0 | NA | NA | NA | NA | NA | Olfr1465 | olfactory receptor 1465 [Source:MGI Syml | 0 | 0 | 0 | 0 | 0 | 0 |
| 17425 | ENSMUSG00000062200 | 0 | NA | NA | NA | NA | NA | Vmn2r7 | vomeronasal 2, receptor 7 [Source:MGI Sy | 0 | 0 | 0 | 0 | 0 | 0 |
| 17426 | ENSMUSG00000062201 | 0 | NA | NA | NA | NA | NA | Prl3d3 | prolactin family 3, subfamily d, member 3 | 0 | 0 | 0 | 0 | 0 | 0 |
| 17427 | ENSMUSG00000062202 | 283.8064094 | 0.123462586 | 0.154786204 | 0.797633008 | 0.42508349 | 0.840865372 | Btbd9 | BTB (POZ) domain containing 9 [Source:M | 251.37 | 269.31 | 294.83 | 301.73 | 274.84 | 310.76 |
| 17428 | ENSMUSG00000062203 | 1445.49728 | 0.112168448 | 0.115650956 | 0.969887773 | 0.332102435 | 0.775012846 | Gspt1 | G1 to S phase transition 1 [Source:MGI Sy | 1254.4 | 1365.1 | 1549.6 | 1505.7 | 1585.8 | 1412.4 |
| 17429 | ENSMUSG00000062204 | 0 | NA | NA | NA | NA | NA | Olfr1373 | olfactory receptor 1373 [Source:MGI Syml | 0 | 0 | 0 | 0 | 0 | 0 |
| 17430 | ENSMUSG00000062209 | 70.15708365 | 0.690804841 | 0.344358692 | 2.00606187 | 0.044849647 | 0.292602477 | Erbb4 | erb-b2 receptor tyrosine kinase 4 [Source: | 66.657 | 55.508 | 38.237 | 53.019 | 88.34 | 119.18 |
| 17431 | ENSMUSG00000062210 | 26.82530081 | -1.741379066 | 0.46264026 | -3.764002437 | 0.000167215 | 0.005422873 | Tnfaip8 | tumor necrosis factor, alpha-induced prot | 47.383 | 39.061 | 37.231 | 9.6398 | 9.8156 | 17.822 |
| 17432 | ENSMUSG00000062224 | 0 | NA | NA | NA | NA | NA | 4933411G06 | RIKEN cDNA 4933411G06 gene [Source:M | 0 | 0 | 0 | 0 | 0 | 0 |
| 17433 | ENSMUSG00000062226 | 0 | NA | NA | NA | NA | NA | Gm10112 | predicted pseudogene 10112 [Source:MG | 0 | 0 | 0 | 0 | 0 | 0 |
| 17434 | ENSMUSG00000062232 | 248.2255305 | -0.074495841 | 0.163317536 | -0.456141103 | 0.648288515 | 0.925643431 | Rapgef2 | Rap guanine nucleotide exchange factor ( | 238.52 | 265.2 | 260.62 | 250.63 | 214.85 | 259.53 |
| 17435 | ENSMUSG00000062234 | 1228.215098 | -0.095431229 | 0.104992689 | -0.908932137 | 0.363385949 | 0.802199995 | Gak | cyclin G associated kinase [Source:MGI Sy | 1198.2 | 1303.4 | 1306.1 | 1254.1 | 1172.4 | 1135 |
| 17436 | ENSMUSG00000062235 | 0 | NA | NA | NA | NA | NA | Btf3-ps5 | basic transcription factor 3, pseudogene 5 | 0 | 0 | 0 | 0 | 0 | 0 |
| 17437 | ENSMUSG00000062245 | 0 | NA | NA | NA | NA | NA | Olfr170 | olfactory receptor 170 [Source:MGI Symb | 0 | 0 | 0 | 0 | 0 | 0 |
| 17438 | ENSMUSG00000062248 | 264.8447379 | 0.065443211 | 0.174017927 | 0.376071661 | 0.706863622 | 0.943006403 | Cks2 | CDC28 protein kinase regulatory subunit 2 | 225.67 | 272.4 | 278.73 | 235.21 | 309.74 | 267.32 |
| 17439 | ENSMUSG00000062252 | 0.545309144 | 2.651974629 | 3.666219369 | 0.72335405 | 0.469462394 | NA | Lhfpl5 | lipoma HMGIC fusion partner-like 5 [Sourc | 0 | 0 | 0 | 0 | 3.2719 | 0 |
| 17440 | ENSMUSG00000062257 | 0.133850373 | -0.796629176 | 4.080472857 | -0.195229623 | 0.84521319 | NA | Opcml | opioid binding protein/cell adhesion mole | 0.8031 | 0 | 0 | 0 | 0 | 0 |
| 17441 | ENSMUSG00000062270 | 2155.821545 | 0.013547583 | 0.100863426 | 0.134316109 | 0.893152615 | 0.982562372 | Morf4l1 | mortality factor 4 like 1 [Source:MGI Symb | 1978.8 | 2205.9 | 2253 | 2055.2 | 2197.6 | 2244.4 |
| 17442 | ENSMUSG00000062272 | 0 | NA | NA | NA | NA | NA | Olfr1129 | olfactory receptor 1129 [Source:MGI Syml | 0 | 0 | 0 | 0 | 0 | 0 |
| 17443 | ENSMUSG00000062275 | 0 | NA | NA | NA | NA | NA | Fbxw24 | F-box and WD-40 domain protein 24 [Sou | 0 | 0 | 0 | 0 | 0 | 0 |
| 17444 | ENSMUSG00000062278 | 0 | NA | NA | NA | NA | NA | Gm11562 | predicted gene 11562 [Source:MGI Symbc | 0 | 0 | 0 | 0 | 0 | 0 |
| 17445 | ENSMUSG00000062279 | 0 | NA | NA | NA | NA | NA | Gm10237 | predicted gene 10237 [Source:MGI Symbc | 0 | 0 | 0 | 0 | 0 | 0 |
| 17446 | ENSMUSG00000062282 | 0 | NA | NA | NA | NA | NA | Gm10113 | predicted gene 10113 [Source:MGI Symbc | 0 | 0 | 0 | 0 | 0 | 0 |
| 17447 | ENSMUSG00000062284 | 0.870273931 | -0.470854203 | 2.55968665 | -0.183949939 | 0.854052728 | NA | Gm6030 | predicted gene 6030 [Source:MGI Symbol | 0 | 1.0279 | 2.0125 | 0 | 2.1812 | 0 |
| 17448 | ENSMUSG00000062296 | 0 | NA | NA | NA | NA | NA | Trank1 | tetratricopeptide repeat and ankyrin repe | 0 | 0 | 0 | 0 | 0 | 0 |
| 17449 | ENSMUSG00000062300 | 1322.467384 | 0.185351665 | 0.104958034 | 1.765959763 | 0.0774026 | 0.40712138 | Nectin2 | nectin cell adhesion molecule 2 [Source:N | 1289 | 1185.2 | 1238.7 | 1510.6 | 1345.8 | 1365.6 |
| 17450 | ENSMUSG00000062309 | 1.927583327 | 0.910635701 | 1.754255277 | 0.519101019 | 0.603690298 | NA | Rpp25 | ribonuclease P/MRP 25 subunit [Source:N | 0 | 2.0558 | 2.0125 | 1.928 | 0 | 5.5692 |
| 17451 | ENSMUSG00000062310 | 1.157742822 | 0.550146268 | 1.956961148 | 0.281122734 | 0.778616265 | NA | Glrp1 | glutamine repeat protein 1 [Source:MGI S | 0.8031 | 1.0279 | 1.0062 | 1.928 | 2.1812 | 0 |
| 17452 | ENSMUSG00000062312 | 13.7310044 | -0.369494873 | 0.586386193 | -0.630122055 | 0.528614731 | 0.888334265 | Erbb2 | erb-b2 receptor tyrosine kinase 2 [Source: | 16.865 | 14.391 | 15.094 | 10.604 | 8.7249 | 16.708 |
| 17453 | ENSMUSG00000062314 | 0 | NA | NA | NA | NA | NA | Olfr1505 | olfactory receptor 1505 [Source:MGI Syml | 0 | 0 | 0 | 0 | 0 | 0 |
| 17454 | ENSMUSG00000062319 | 0 | NA | NA | NA | NA | NA | Gm10115 | predicted gene 10115 [Source:MGI Symbc | 0 | 0 | 0 | 0 | 0 | 0 |
| 17455 | ENSMUSG00000062327 | 0 | NA | NA | NA | NA | NA | T | brachyury, T-box transcription factor T [Sc | 0 | 0 | 0 | 0 | 0 | 0 |
| 17456 | ENSMUSG00000062328 | 4567.753474 | 0.053782251 | 0.096274558 | 0.558634095 | 0.576411466 | 0.907699014 | Rpl17 | ribosomal protein L17 [Source:MGI Symbc | 4482.9 | 4653.4 | 4310.8 | 4254 | 4840.2 | 4865.3 |
| 17457 | ENSMUSG00000062329 | 0 | NA | NA | NA | NA | NA | Cytl1 | cytokine-like 1 [Source:MGI Symbol;Acc:N | 0 | 0 | 0 | 0 | 0 | 0 |
| 17458 | ENSMUSG00000062342 | 0 | NA | NA | NA | NA | NA | Serpinb9e | serine (or cysteine) peptidase inhibitor, cl | 0 | 0 | 0 | 0 | 0 | 0 |
| 17459 | ENSMUSG00000062345 | 1.489752066 | -1.713049692 | 1.866114832 | -0.91797657 | 0.358631133 | NA | Serpinb2 | serine (or cysteine) peptidase inhibitor, cl | 0.8031 | 2.0558 | 4.025 | 0.964 | 1.0906 | 0 |
| 17460 | ENSMUSG00000062352 | 167.0556758 | -0.16046647 | 0.182282566 | -0.880317155 | 0.378687522 | 0.811969331 | Itgb1bp1 | integrin beta 1 binding protein 1 [Source:I | 187.93 | 173.72 | 167.04 | 156.16 | 154.87 | 162.62 |
| 17461 | ENSMUSG00000062353 | 4.256174583 | 0.17536963 | 1.196683723 | 0.146546348 | 0.883490104 | NA | Gm15772 | predicted gene 15772 [Source:MGI Symbc | 8.8341 | 0 | 3.0187 | 4.8199 | 2.1812 | 6.6831 |
| 17462 | ENSMUSG00000062365 | 0.181769715 | 1.126924175 | 4.080472857 | 0.276174898 | 0.782413736 | NA | Gm4968 | predicted gene 4968 [Source:MGI Symbol | 0 | 0 | 0 | 0 | 1.0906 | 0 |
| 17463 | ENSMUSG00000062369 | 0 | NA | NA | NA | NA | NA | Usp17lb | ubiquitin specific peptidase 17-like B [Sou | 0 | 0 | 0 | 0 | 0 | 0 |
| 17464 | ENSMUSG00000062372 | 0.342639279 | -1.805976596 | 4.028547714 | -0.448294702 | 0.653940524 | NA | Otof | otoferlin [Source:MGI Symbol;Acc:MGI:18 | 0 | 2.0558 | 0 | 0 | 0 | 0 |
| 17465 | ENSMUSG00000062373 | 101.9522046 | -0.152826982 | 0.248366515 | -0.615328448 | 0.538337834 | 0.891577636 | Tmem65 | transmembrane protein 65 [Source:MGI S | 101.19 | 124.38 | 96.599 | 97.362 | 76.343 | 115.84 |
| 17466 | ENSMUSG00000062376 | 162.9258637 | -0.146078149 | 0.219262733 | -0.666224246 | 0.505267778 | 0.881592135 | Borcs7 | BLOC-1 related complex subunit 7 [Source | 214.43 | 155.22 | 142.89 | 136.88 | 177.77 | 150.37 |
| 17467 | ENSMUSG00000062380 | 0.342639279 | -1.805976596 | 4.028547714 | -0.448294702 | 0.653940524 | NA | Tubb3 | tubulin, beta 3 class III [Source:MGI Symb | 0 | 2.0558 | 0 | 0 | 0 | 0 |
| 17468 | ENSMUSG00000062382 | 95.70807361 | -0.344700555 | 0.259206377 | -1.329830534 | 0.183574113 | 0.617851749 | Ftl1-ps1 | ferritin light polypeptide 1, pseudogene 1 | 126.89 | 93.541 | 100.62 | 96.398 | 64.346 | 92.449 |

| | | | | | | | | | | | | | | | |
|---|---|---|---|---|---|---|---|---|---|---|---|---|---|---|---|
| 17469 | ENSMUSG00000062391 | 0 | NA | NA | NA | NA | NA | 4932435O22 | RIKEN cDNA 4932435O22 gene [Source:M | 0 | 0 | 0 | 0 | 0 | 0 |
| 17470 | ENSMUSG00000062393 | 0 | NA | NA | NA | NA | NA | Dgkk | diacylglycerol kinase kappa [Source:MGI S | 0 | 0 | 0 | 0 | 0 | 0 |
| 17471 | ENSMUSG00000062397 | 1961.039282 | 0.027844165 | 0.098420643 | 0.28290981 | 0.777245974 | 0.959467933 | Zfp706 | zinc finger protein 706 [Source:MGI Symb | 1916.2 | 1882.1 | 2027.6 | 1871.1 | 1969.7 | 2099.6 |
| 17472 | ENSMUSG00000062400 | 0 | NA | NA | NA | NA | NA | Krtap6-5 | keratin associated protein 6-5 [Source:MG | 0 | 0 | 0 | 0 | 0 | 0 |
| 17473 | ENSMUSG00000062410 | 3782.796551 | 0.469756364 | 0.121214858 | 3.875402492 | 0.000106449 | 0.003820527 | Hsd3b3 | hydroxy-delta-5-steroid dehydrogenase, 3 | 3205.2 | 3200.9 | 3110.3 | 3939.8 | 5224.1 | 4016.5 |
| 17474 | ENSMUSG00000062421 | 164.6358528 | 0.045571946 | 0.181588083 | 0.250963309 | 0.801842476 | 0.962796744 | Arf2 | ADP-ribosylation factor 2 [Source:MGI Syr | 160.62 | 160.36 | 165.02 | 159.06 | 171.23 | 171.53 |
| 17475 | ENSMUSG00000062426 | 0 | NA | NA | NA | NA | NA | Olfr304 | olfactory receptor 304 [Source:MGI Symb | 0 | 0 | 0 | 0 | 0 | 0 |
| 17476 | ENSMUSG00000062432 | 3.371333602 | -0.377133144 | 1.439320556 | -0.262021648 | 0.793304752 | NA | Cyp26c1 | cytochrome P450, family 26, subfamily c, | 2.4093 | 4.1117 | 5.0312 | 8.6758 | 0 | 0 |
| 17477 | ENSMUSG00000062433 | 0 | NA | NA | NA | NA | NA | Krtap6-2 | keratin associated protein 6-2 [Source:MG | 0 | 0 | 0 | 0 | 0 | 0 |
| 17478 | ENSMUSG00000062434 | 0 | NA | NA | NA | NA | NA | Olfr516 | olfactory receptor 516 [Source:MGI Symb | 0 | 0 | 0 | 0 | 0 | 0 |
| 17479 | ENSMUSG00000062438 | 0 | NA | NA | NA | NA | NA | Adam1b | a disintegrin and metallopeptidase domai | 0 | 0 | 0 | 0 | 0 | 0 |
| 17480 | ENSMUSG00000062444 | 0 | NA | NA | NA | NA | NA | Ap3b2 | adaptor-related protein complex 3, beta 2 | 0 | 0 | 0 | 0 | 0 | 0 |
| 17481 | ENSMUSG00000062456 | 3.784041178 | 0.130222735 | 1.140020967 | 0.114228368 | 0.909056765 | NA | Rpl9-ps6 | ribosomal protein L9, pseudogene 6 [Sour | 5.6217 | 3.0838 | 2.0125 | 0.964 | 5.4531 | 5.5692 |
| 17482 | ENSMUSG00000062458 | 1.674954029 | -1.861921599 | 1.953892182 | -0.952929551 | 0.340625766 | NA | Gm8623 | predicted gene 8623 [Source:MGI Symbol | 0.8031 | 1.0279 | 6.0375 | 0 | 2.1812 | 0 |
| 17483 | ENSMUSG00000062461 | 0.503095663 | 2.549494177 | 3.348715477 | 0.761334964 | 0.446457022 | NA | Rpl27a-ps4 | ribosomal protein L27A, pseudogene 4 [Sc | 0 | 0 | 0 | 1.928 | 1.0906 | 0 |
| 17484 | ENSMUSG00000062464 | 0 | NA | NA | NA | NA | NA | Cyp4f37 | cytochrome P450, family 4, subfamily f, pc | 0 | 0 | 0 | 0 | 0 | 0 |
| 17485 | ENSMUSG00000062470 | 3.419881449 | -0.497692211 | 1.276835603 | -0.389785662 | 0.696695047 | NA | Fbxl12os | F-box and leucine-rich repeat protein 12, | 8.8341 | 1.0279 | 2.0125 | 0.964 | 5.4531 | 2.2277 |
| 17486 | ENSMUSG00000062472 | 0 | NA | NA | NA | NA | NA | Gm4945 | predicted gene 4945 [Source:MGI Symbol | 0 | 0 | 0 | 0 | 0 | 0 |
| 17487 | ENSMUSG00000062477 | 0 | NA | NA | NA | NA | NA | Hmgb1l | high mobility group box 1-like [Source:MG | 0 | 0 | 0 | 0 | 0 | 0 |
| 17488 | ENSMUSG00000062478 | 0 | NA | NA | NA | NA | NA | Ctrc | chymotrypsin C (caldecrin) [Source:MGI S | 0 | 0 | 0 | 0 | 0 | 0 |
| 17489 | ENSMUSG00000062480 | 2656.939272 | 0.217618564 | 0.109533519 | 1.986775979 | 0.046947228 | 0.300126868 | Acat3 | acetyl-Coenzyme A acetyltransferase 3 [Sc | 2352.3 | 2299.5 | 2718.9 | 2628.8 | 2881.4 | 3060.8 |
| 17490 | ENSMUSG00000062483 | 0 | NA | NA | NA | NA | NA | Vmn1r171 | vomeronasal 1 receptor 171 [Source:MGI | 0 | 0 | 0 | 0 | 0 | 0 |
| 17491 | ENSMUSG00000062488 | 75.41665228 | 0.052703125 | 0.317620113 | 0.16593132 | 0.868210999 | 0.978326885 | Ifit3b | interferon-induced protein with tetratrico | 93.16 | 67.843 | 60.375 | 48.199 | 92.703 | 90.221 |
| 17492 | ENSMUSG00000062496 | 0 | NA | NA | NA | NA | NA | 4930431F12 | RIKEN cDNA 4930431F12 gene [Source:M | 0 | 0 | 0 | 0 | 0 | 0 |
| 17493 | ENSMUSG00000062497 | 0 | NA | NA | NA | NA | NA | Olfr1411 | olfactory receptor 1411 [Source:MGI Sym | 0 | 0 | 0 | 0 | 0 | 0 |
| 17494 | ENSMUSG00000062510 | 6.351078801 | -0.303203393 | 0.858789692 | -0.353058956 | 0.724044242 | 0.944786345 | Nsl1 | NSL1, MIS12 kinetochore complex compo | 4.8186 | 7.1954 | 9.0562 | 3.8559 | 8.7249 | 4.4554 |
| 17495 | ENSMUSG00000062511 | 0 | NA | NA | NA | NA | NA | 4930512M0 | RIKEN cDNA 4930512M02 gene [Source:M | 0 | 0 | 0 | 0 | 0 | 0 |
| 17496 | ENSMUSG00000062515 | 106.0830656 | -0.711105348 | 0.300854333 | -2.363620099 | 0.01809736 | 0.161083208 | Fabp4 | fatty acid binding protein 4, adipocyte [So | 177.49 | 130.55 | 86.537 | 76.154 | 61.075 | 104.7 |
| 17497 | ENSMUSG00000062518 | 0 | NA | NA | NA | NA | NA | Zfp534 | zinc finger protein 534 [Source:MGI Symb | 0 | 0 | 0 | 0 | 0 | 0 |
| 17498 | ENSMUSG00000062519 | 155.0829705 | 0.11297768 | 0.203080694 | 0.556319157 | 0.577992693 | 0.908137944 | Zfp398 | zinc finger protein 398 [Source:MGI Symb | 131.71 | 171.66 | 143.89 | 141.7 | 176.68 | 164.85 |
| 17499 | ENSMUSG00000062524 | 0.305170013 | -1.657836477 | 4.041480335 | -0.410205257 | 0.681655385 | NA | Ncr1 | natural cytotoxicity triggering receptor 1 [ | 0.8031 | 1.0279 | 0 | 0 | 0 | 0 |
| 17500 | ENSMUSG00000062526 | 227.9547582 | 0.047847182 | 0.202611983 | 0.236151785 | 0.813314891 | 0.966802988 | Mppe1 | metallophosphoesterase 1 [Source:MGI S | 221.66 | 211.75 | 238.48 | 168.7 | 247.57 | 279.57 |
| 17501 | ENSMUSG00000062527 | 0 | NA | NA | NA | NA | NA | Olfr1408 | olfactory receptor 1408 [Source:MGI Sym | 0 | 0 | 0 | 0 | 0 | 0 |
| 17502 | ENSMUSG00000062528 | 0 | NA | NA | NA | NA | NA | Tas2r109 | taste receptor, type 2, member 109 [Sour | 0 | 0 | 0 | 0 | 0 | 0 |
| 17503 | ENSMUSG00000062542 | 10.73393504 | -0.942733858 | 0.680517492 | -1.385319068 | 0.165954926 | 0.592099665 | Syt9 | synaptotagmin IX [Source:MGI Symbol;Ac | 12.85 | 18.503 | 11.069 | 7.7118 | 9.8156 | 4.4554 |
| 17504 | ENSMUSG00000062543 | 0 | NA | NA | NA | NA | NA | Gsdmcl1 | gasdermin C-like 1 [Source:MGI Symbol;A | 0 | 0 | 0 | 0 | 0 | 0 |
| 17505 | ENSMUSG00000062545 | 415.1043673 | -0.427617445 | 0.184486786 | -2.317875732 | 0.020456076 | 0.176221965 | Tlr12 | toll-like receptor 12 [Source:MGI Symbol; | 607.95 | 453.31 | 366.27 | 352.82 | 382.81 | 327.47 |
| 17506 | ENSMUSG00000062546 | 0.371281295 | 2.09253362 | 4.031852165 | 0.519000582 | 0.603760335 | NA | V1ra8 | vomeronasal 1 receptor, A8 [Source:MGI | 0 | 0 | 0 | 0 | 0 | 2.2277 |
| 17507 | ENSMUSG00000062551 | 0 | NA | NA | NA | NA | NA | Prl2c1 | Prolactin family 2, subfamily c, member 1 | 0 | 0 | 0 | 0 | 0 | 0 |
| 17508 | ENSMUSG00000062553 | 0 | NA | NA | NA | NA | NA | Olfr709-ps1 | olfactory receptor 709, pseudogene 1 [So | 0 | 0 | 0 | 0 | 0 | 0 |
| 17509 | ENSMUSG00000062554 | 1.004769579 | -2.106122861 | 2.47240066 | -0.851853381 | 0.394295474 | NA | Gm12751 | predicted gene 12751 [Source:MGI Symbo | 0.8031 | 4.1117 | 0 | 0 | 0 | 1.1138 |
| 17510 | ENSMUSG00000062556 | 0 | NA | NA | NA | NA | NA | Scgb1b2 | secretoglobin, family 1B, member 2 [Sour | 0 | 0 | 0 | 0 | 0 | 0 |
| 17511 | ENSMUSG00000062561 | 0.531246861 | 1.101376188 | 3.36099124 | 0.327693858 | 0.743143147 | NA | Gm10118 | predicted gene 10118 [Source:MGI Symbo | 0 | 0 | 1.0062 | 0 | 2.1812 | 0 |
| 17512 | ENSMUSG00000062563 | 106.6011024 | 0.298745416 | 0.268400018 | 1.113060346 | 0.265682513 | 0.721565724 | Cys1 | cystin 1 [Source:MGI Symbol;Acc:MGI:217 | 106.01 | 85.317 | 95.593 | 156.16 | 81.796 | 114.73 |
| 17513 | ENSMUSG00000062564 | 0 | NA | NA | NA | NA | NA | Tex28 | testis expressed 28 [Source:MGI Symbol;A | 0 | 0 | 0 | 0 | 0 | 0 |
| 17514 | ENSMUSG00000062580 | 515.1587836 | 0.208126189 | 0.146018744 | 1.425338857 | 0.154059273 | 0.571215925 | Timm17a | translocase of inner mitochondrial memb | 468.21 | 497.51 | 467.9 | 459.82 | 569.3 | 628.21 |
| 17515 | ENSMUSG00000062582 | 0.181769715 | 1.126924175 | 4.080472857 | 0.276174898 | 0.782413736 | NA | Rpl30-ps8 | ribosomal protein L30, pseudogene 8 [Sou | 0 | 0 | 0 | 0 | 1.0906 | 0 |
| 17516 | ENSMUSG00000062585 | 27.32110871 | -0.449365084 | 0.508569271 | -0.883586778 | 0.376919319 | 0.81030298 | Cnr2 | cannabinoid receptor 2 (macrophage) [So | 43.368 | 35.977 | 15.094 | 24.099 | 13.087 | 32.301 |
| 17517 | ENSMUSG00000062588 | 0.812136609 | 0.736604728 | 2.414276032 | 0.305103774 | 0.760287117 | NA | Gm6104 | predicted gene 6104 [Source:MGI Symbol | 0.8031 | 1.0279 | 0 | 1.928 | 0 | 1.1138 |
| 17518 | ENSMUSG00000062590 | 74.94002905 | -0.044998917 | 0.298364011 | -0.150818851 | 0.880118615 | 0.981359269 | Armc9 | armadillo repeat containing 9 [Source:MG | 53.808 | 93.541 | 81.506 | 72.298 | 88.34 | 60.148 |
| 17519 | ENSMUSG00000062591 | 1.886268103 | -0.236130036 | 1.606040561 | -0.147026197 | 0.883111343 | NA | Tubb4a | tubulin, beta 4A class IVA [Source:MGI Sy | 4.0155 | 2.0558 | 0 | 1.928 | 1.0906 | 2.2277 |
| 17520 | ENSMUSG00000062593 | 0 | NA | NA | NA | NA | NA | Gm49339 | predicted gene, 49339 [Source:MGI Symb | 0 | 0 | 0 | 0 | 0 | 0 |
| 17521 | ENSMUSG00000062598 | 0 | NA | NA | NA | NA | NA | Vmn1r178 | vomeronasal 1 receptor 178 [Source:MGI | 0 | 0 | 0 | 0 | 0 | 0 |
| 17522 | ENSMUSG00000062604 | 320.8639732 | 0.14028366 | 0.163105198 | 0.860080867 | 0.389744467 | 0.819052248 | Srpk2 | serine/arginine-rich protein specific kinas | 360.59 | 265.2 | 288.79 | 321.97 | 353.36 | 335.27 |
| 17523 | ENSMUSG00000062608 | 0 | NA | NA | NA | NA | NA | Olfr195 | olfactory receptor 195 [Source:MGI Symb | 0 | 0 | 0 | 0 | 0 | 0 |
| 17524 | ENSMUSG00000062609 | 1.982656378 | -0.937197253 | 1.62709891 | -0.575992798 | 0.56462005 | NA | Kcnj15 | potassium inwardly-rectifying channel, su | 0.8031 | 2.0558 | 5.0312 | 2.8919 | 0 | 1.1138 |

| | | | | | | | | | | | | | | | |
|---|---|---|---|---|---|---|---|---|---|---|---|---|---|---|---|
| 17525 | ENSMUSG00000062611 | 6.816471677 | -0.178516354 | 0.822087657 | -0.217150024 | 0.82809143 | 0.969935825 | Rps3a2 | ribosomal protein S3A2 [Source:MGI Symb | 6.4248 | 8.2233 | 7.0437 | 4.8199 | 4.3625 | 10.025 |
| 17526 | ENSMUSG00000062619 | 781.5097734 | -0.003181739 | 0.113038951 | -0.028147282 | 0.977544683 | 0.997564159 | 2310039H08 | RIKEN cDNA 2310039H08 gene [Source:M | 774.99 | 820.28 | 751.66 | 742.26 | 819.05 | 780.8 |
| 17527 | ENSMUSG00000062621 | 0 | NA | NA | NA | NA | NA | Olfr911-ps1 | olfactory receptor 911, pseudogene 1 [So | 0 | 0 | 0 | 0 | 0 | 0 |
| 17528 | ENSMUSG00000062624 | 3766.037599 | 0.33202038 | 0.146298492 | 2.269472331 | 0.023239618 | 0.192524645 | Cyp2c67 | cytochrome P450, family 2, subfamily c, p | 2653.4 | 3375.7 | 3976.7 | 4790 | 3810.6 | 3989.8 |
| 17529 | ENSMUSG00000062627 | 280.3658175 | 0.120195401 | 0.162615756 | 0.739137486 | 0.459823516 | 0.859753653 | Mysm1 | myb-like, SWIRM and MPN domains 1 [So | 281.89 | 237.45 | 286.78 | 321.97 | 289.01 | 265.09 |
| 17530 | ENSMUSG00000062629 | 0 | NA | NA | NA | NA | NA | Olfr114 | olfactory receptor 114 [Source:MGI Symb | 0 | 0 | 0 | 0 | 0 | 0 |
| 17531 | ENSMUSG00000062638 | 0 | NA | NA | NA | NA | NA | Btnl1 | butyrophilin-like 1 [Source:MGI Symbol;A | 0 | 0 | 0 | 0 | 0 | 0 |
| 17532 | ENSMUSG00000062646 | 447.4878344 | -0.051926794 | 0.148581573 | -0.349483402 | 0.72672643 | 0.945777495 | Ganc | glucosidase, alpha; neutral C [Source:MGI | 492.3 | 430.7 | 443.75 | 503.2 | 435.16 | 379.82 |
| 17533 | ENSMUSG00000062647 | 5454.299396 | -0.013488304 | 0.084111045 | -0.160363057 | 0.872595089 | 0.979419285 | Rpl7a | ribosomal protein L7A [Source:MGI Symb | 5356.7 | 5687.5 | 5395.5 | 5321.2 | 5477.1 | 5487.9 |
| 17534 | ENSMUSG00000062649 | 0 | NA | NA | NA | NA | NA | Olfr44 | olfactory receptor 44 [Source:MGI Symbo | 0 | 0 | 0 | 0 | 0 | 0 |
| 17535 | ENSMUSG00000062651 | 0 | NA | NA | NA | NA | NA | H2al2a | H2A histone family member L2A [Source:I | 0 | 0 | 0 | 0 | 0 | 0 |
| 17536 | ENSMUSG00000062661 | 1.905333325 | 0.399188399 | 1.923036261 | 0.207582357 | 0.835555082 | NA | Ncs1 | neuronal calcium sensor 1 [Source:MGI Sy | 4.8186 | 0 | 0 | 0 | 3.2719 | 3.3415 |
| 17537 | ENSMUSG00000062683 | 1484.633051 | 0.067198156 | 0.103488514 | 0.64932961 | 0.516125351 | 0.886609349 | Atp5g2 | ATP synthase, H+ transporting, mitochonc | 1501.8 | 1461.7 | 1385.6 | 1409.3 | 1596.7 | 1552.7 |
| 17538 | ENSMUSG00000062687 | 0 | NA | NA | NA | NA | NA | Cks1brt | CDC28 protein kinase 1b, retrogene [Sour | 0 | 0 | 0 | 0 | 0 | 0 |
| 17539 | ENSMUSG00000062691 | 100.7526667 | 0.080364213 | 0.236091851 | 0.340393845 | 0.733559954 | 0.947915087 | Cebpzos | CCAAT/enhancer binding protein (C/EBP), | 116.45 | 84.289 | 92.575 | 105.07 | 101.43 | 104.7 |
| 17540 | ENSMUSG00000062694 | 0 | NA | NA | NA | NA | NA | Cav3 | caveolin 3 [Source:MGI Symbol;Acc:MGI:1 | 0 | 0 | 0 | 0 | 0 | 0 |
| 17541 | ENSMUSG00000062704 | 3.825754868 | -0.883759319 | 1.185960103 | -0.745184696 | 0.456160073 | NA | 9430002A10 | RIKEN cDNA 9430002A10 gene [Source:M | 4.8186 | 5.1396 | 5.0312 | 5.7839 | 2.1812 | 0 |
| 17542 | ENSMUSG00000062705 | 0 | NA | NA | NA | NA | NA | Tpbpb | trophoblast specific protein beta [Source: | 0 | 0 | 0 | 0 | 0 | 0 |
| 17543 | ENSMUSG00000062712 | 0.17131964 | -0.796629176 | 4.080472857 | -0.195229623 | 0.84521319 | NA | Olfr531 | olfactory receptor 531 [Source:MGI Symb | 0 | 1.0279 | 0 | 0 | 0 | 0 |
| 17544 | ENSMUSG00000062713 | 0 | NA | NA | NA | NA | NA | Sim2 | single-minded family bHLH transcription f | 0 | 0 | 0 | 0 | 0 | 0 |
| 17545 | ENSMUSG00000062727 | 1.099193445 | -2.312770509 | 2.368100974 | -0.976635091 | 0.32874984 | NA | H2bc12 | H2B clustered histone 12 [Source:MGI Syr | 1.6062 | 0 | 4.025 | 0.964 | 0 | 0 |
| 17546 | ENSMUSG00000062729 | 343.9764208 | 0.494431442 | 0.146417084 | 3.376869892 | 0.000733157 | 0.016647388 | Ppox | protoporphyrinogen oxidase [Source:MGI | 292.33 | 280.62 | 283.76 | 418.37 | 357.72 | 431.06 |
| 17547 | ENSMUSG00000062732 | 0 | NA | NA | NA | NA | NA | Lypd4 | Ly6/Plaur domain containing 4 [Source:M | 0 | 0 | 0 | 0 | 0 | 0 |
| 17548 | ENSMUSG00000062737 | 0 | NA | NA | NA | NA | NA | Prl3d2 | prolactin family 3, subfamily d, member 1 | 0 | 0 | 0 | 0 | 0 | 0 |
| 17549 | ENSMUSG00000062742 | 0.133850373 | -0.796629176 | 4.080472857 | -0.195229623 | 0.84521319 | NA | Gm5239 | predicted pseudogene 5239 [Source:MGI | 0.8031 | 0 | 0 | 0 | 0 | 0 |
| 17550 | ENSMUSG00000062743 | 8.483609164 | -0.149766811 | 0.788368144 | -0.189970653 | 0.849332127 | 0.973071229 | Zfp677 | zinc finger protein 677 [Source:MGI Symb | 6.4248 | 15.419 | 5.0312 | 9.6398 | 4.3625 | 10.025 |
| 17551 | ENSMUSG00000062751 | 0 | NA | NA | NA | NA | NA | Prss1 | protease, serine 1 (trypsin 1) [Source:MGI | 0 | 0 | 0 | 0 | 0 | 0 |
| 17552 | ENSMUSG00000062753 | 55.33191555 | -0.356690875 | 0.332059264 | -1.074178358 | 0.28274275 | 0.735558641 | AI413582 | expressed sequence AI413582 [Source:M | 76.295 | 48.312 | 61.381 | 44.343 | 38.172 | 63.489 |
| 17553 | ENSMUSG00000062757 | 0 | NA | NA | NA | NA | NA | Olfr1206 | olfactory receptor 1206 [Source:MGI Sym | 0 | 0 | 0 | 0 | 0 | 0 |
| 17554 | ENSMUSG00000062758 | 3.038538813 | 0.518987192 | 1.229989285 | 0.421944483 | 0.673065541 | NA | Gm16477 | predicted pseudogene 16477 [Source:MG | 2.4093 | 1.0279 | 4.025 | 1.928 | 3.2719 | 5.5692 |
| 17555 | ENSMUSG00000062760 | 0.472877444 | -2.283655516 | 3.093270275 | -0.738265755 | 0.460352972 | NA | Shisal1 | shisa like 1 [Source:MGI Symbol;Acc:MGI: | 0.8031 | 1.0279 | 1.0062 | 0 | 0 | 0 |
| 17556 | ENSMUSG00000062761 | 173.4421037 | 0.271039492 | 0.202910793 | 1.335756901 | 0.181628744 | 0.615095921 | Zfp512 | zinc finger protein 512 [Source:MGI Symb | 129.3 | 169.61 | 173.07 | 162.91 | 194.13 | 211.63 |
| 17557 | ENSMUSG00000062762 | 4385.050415 | -0.28027612 | 0.102432382 | -2.736206205 | 0.006215207 | 0.077098412 | Ei24 | etoposide induced 2.4 mRNA [Source:MG | 4548.8 | 5144.7 | 4735.4 | 3655.4 | 3909.9 | 4316.1 |
| 17558 | ENSMUSG00000062773 | 0 | NA | NA | NA | NA | NA | Tex101 | testis expressed gene 101 [Source:MGI Sy | 0 | 0 | 0 | 0 | 0 | 0 |
| 17559 | ENSMUSG00000062778 | 0 | NA | NA | NA | NA | NA | Chia1 | chitinase, acidic 1 [Source:MGI Symbol;Ac | 0 | 0 | 0 | 0 | 0 | 0 |
| 17560 | ENSMUSG00000062782 | 0 | NA | NA | NA | NA | NA | Olfr527 | olfactory receptor 527 [Source:MGI Symb | 0 | 0 | 0 | 0 | 0 | 0 |
| 17561 | ENSMUSG00000062783 | 1.657459497 | -1.909542199 | 1.822533891 | -1.047740297 | 0.294758276 | NA | | component of Sp100-rs [Source:NCBI gen | 0.8031 | 2.0558 | 5.0312 | 0.964 | 1.0906 | 0 |
| 17562 | ENSMUSG00000062785 | 112.3317799 | -0.384691923 | 0.237476805 | -1.619913668 | 0.105250823 | 0.478186913 | Kcnc3 | potassium voltage gated channel, Shaw-re | 106.01 | 122.32 | 153.96 | 109.89 | 92.703 | 89.108 |
| 17563 | ENSMUSG00000062791 | 0 | NA | NA | NA | NA | NA | Gm382 | predicted gene 382 [Source:MGI Symbol;A | 0 | 0 | 0 | 0 | 0 | 0 |
| 17564 | ENSMUSG00000062793 | 0 | NA | NA | NA | NA | NA | Olfr1158 | olfactory receptor 1158 [Source:MGI Sym | 0 | 0 | 0 | 0 | 0 | 0 |
| 17565 | ENSMUSG00000062794 | 3.846810003 | 0.481709638 | 1.098107215 | 0.438672683 | 0.660898724 | NA | Zfp599 | zinc finger protein 599 [Source:MGI Symb | 1.6062 | 4.1117 | 4.025 | 6.7478 | 4.3625 | 2.2277 |
| 17566 | ENSMUSG00000062797 | 323.821081 | 0.016556253 | 0.16815189 | 0.098460103 | 0.921566951 | 0.986685804 | Hikeshi | heat shock protein nuclear import factor [ | 327.67 | 325.85 | 311.94 | 301.73 | 278.11 | 397.64 |
| 17567 | ENSMUSG00000062814 | 0 | NA | NA | NA | NA | NA | Ssxa1 | synovial sarcoma, X member A1 [Source:N | 0 | 0 | 0 | 0 | 0 | 0 |
| 17568 | ENSMUSG00000062818 | 11.18116723 | -0.841867647 | 0.758713285 | -1.109599191 | 0.26717178 | 0.722778036 | Vmn1r51 | vomeronasal 1 receptor 51 [Source:MGI S | 6.4248 | 28.782 | 8.05 | 10.604 | 6.5437 | 6.6831 |
| 17569 | ENSMUSG00000062822 | 548.1107175 | -0.014246441 | 0.14126963 | -0.100845746 | 0.919672912 | 0.986685804 | 4833420G17 | RIKEN cDNA 4833420G17 gene [Source:M | 554.14 | 530.41 | 568.53 | 615.02 | 561.67 | 458.9 |
| 17570 | ENSMUSG00000062825 | 1406.401899 | -0.739753344 | 0.133056456 | -5.559695245 | 2.70E-08 | 3.62E-06 | Actg1 | actin, gamma, cytoplasmic 1 [Source:MGI | 2033.5 | 1547 | 1695.5 | 929.27 | 1103.7 | 1129.4 |
| 17571 | ENSMUSG00000062826 | 0.133850373 | -0.796629176 | 4.080472857 | -0.195229623 | 0.84521319 | NA | Ces2f | carboxylesterase 2F [Source:MGI Symbol; | 0.8031 | 0 | 0 | 0 | 0 | 0 |
| 17572 | ENSMUSG00000062833 | 0 | NA | NA | NA | NA | NA | Sval3 | seminal vesicle antigen-like 3 [Source:MG | 0 | 0 | 0 | 0 | 0 | 0 |
| 17573 | ENSMUSG00000062844 | 0 | NA | NA | NA | NA | NA | Olfr1458 | olfactory receptor 1458 [Source:MGI Sym | 0 | 0 | 0 | 0 | 0 | 0 |
| 17574 | ENSMUSG00000062846 | 0.760346335 | -0.431432104 | 2.662343224 | -0.162049769 | 0.871266662 | NA | Gm14176 | predicted gene 14176 [Source:MGI Symbo | 1.6062 | 1.0279 | 0 | 1.928 | 0 | 0 |
| 17575 | ENSMUSG00000062859 | 0 | NA | NA | NA | NA | NA | Tcp11 | t-complex protein 11 [Source:MGI Symbol | 0 | 0 | 0 | 0 | 0 | 0 |
| 17576 | ENSMUSG00000062861 | 32.54070054 | -0.445032825 | 0.430055257 | -1.03482708 | 0.300749674 | 0.751875664 | Zfp28 | zinc finger protein 28 [Source:MGI Symbo | 30.518 | 46.256 | 36.225 | 41.451 | 19.631 | 21.163 |
| 17577 | ENSMUSG00000062866 | 66.26893574 | -0.759866353 | 0.30119726 | -2.522819605 | 0.011641811 | 0.120276534 | Phactr2 | phosphatase and actin regulator 2 [Sourc | 80.31 | 75.038 | 94.587 | 48.199 | 35.99 | 63.489 |
| 17578 | ENSMUSG00000062867 | 636.8045592 | 0.004999943 | 0.123737619 | 0.040407619 | 0.967768156 | 0.994231936 | Impdh2 | inosine monophosphate dehydrogenase 2 | 576.63 | 660.95 | 670.16 | 607.31 | 663.1 | 642.69 |
| 17579 | ENSMUSG00000062868 | 0 | NA | NA | NA | NA | NA | Olfr830 | olfactory receptor 830 [Source:MGI Symb | 0 | 0 | 0 | 0 | 0 | 0 |
| 17580 | ENSMUSG00000062873 | 0 | NA | NA | NA | NA | NA | Olfr1355 | olfactory receptor 1355 [Source:MGI Sym | 0 | 0 | 0 | 0 | 0 | 0 |

| | | | | | | | | | | | | | | | |
|---|---|---|---|---|---|---|---|---|---|---|---|---|---|---|---|
| 17581 | ENSMUSG00000062874 | 0 | NA | NA | NA | NA | NA | Gm4978 | predicted gene 4978 [Source:MGI Symbol | 0 | 0 | 0 | 0 | 0 | 0 |
| 17582 | ENSMUSG00000062878 | 0 | NA | NA | NA | NA | NA | Olfr298 | olfactory receptor 298 [Source:MGI Symb | 0 | 0 | 0 | 0 | 0 | 0 |
| 17583 | ENSMUSG00000062892 | 0 | NA | NA | NA | NA | NA | Olfr1450 | olfactory receptor 1450 [Source:MGI Sym | 0 | 0 | 0 | 0 | 0 | 0 |
| 17584 | ENSMUSG00000062896 | 1.464742249 | 0.602336868 | 1.775288764 | 0.339289517 | 0.734391639 | NA | Rpl31-ps11 | ribosomal protein L31, pseudogene 11 [Sc | 2.4093 | 0 | 1.0062 | 0.964 | 2.1812 | 2.2277 |
| 17585 | ENSMUSG00000062901 | 1573.081695 | 0.414775178 | 0.109916425 | 3.773550478 | 0.000160941 | 0.005290558 | Klhl24 | kelch-like 24 [Source:MGI Symbol;Acc:MG | 1486.5 | 1300.3 | 1256.8 | 1698.5 | 1803.9 | 1892.4 |
| 17586 | ENSMUSG00000062905 | 0 | NA | NA | NA | NA | NA | Vmn1r32 | vomeronasal 1 receptor 32 [Source:MGI S | 0 | 0 | 0 | 0 | 0 | 0 |
| 17587 | ENSMUSG00000062906 | 214.6347925 | -0.055514392 | 0.172692896 | -0.32146309 | 0.747859479 | 0.951032135 | Hdac10 | histone deacetylase 10 [Source:MGI Symb | 202.38 | 218.95 | 235.46 | 215.93 | 226.85 | 188.24 |
| 17588 | ENSMUSG00000062908 | 9555.787545 | -0.423132029 | 0.099495368 | -4.252781186 | 2.11E-05 | 0.001011196 | Acadm | acyl-Coenzyme A dehydrogenase, mediun | 10218 | 11949 | 10676 | 8271.9 | 8611.5 | 7608.7 |
| 17589 | ENSMUSG00000062914 | 0 | NA | NA | NA | NA | NA | Olfr777 | olfactory receptor 777 [Source:MGI Symb | 0 | 0 | 0 | 0 | 0 | 0 |
| 17590 | ENSMUSG00000062929 | 1080.05039 | -0.039631397 | 0.10208947 | -0.388202593 | 0.697866114 | 0.940502785 | Cfl2 | cofilin 2, muscle [Source:MGI Symbol;Acc | 1077.8 | 1100.9 | 1105.9 | 1022.8 | 1103.7 | 1069.3 |
| 17591 | ENSMUSG00000062931 | 78.72323943 | -0.042253957 | 0.274804001 | -0.153760341 | 0.877798705 | 0.98072613 | Zfp938 | zinc finger protein 938 [Source:MGI Symb | 94.766 | 63.731 | 80.5 | 63.623 | 85.068 | 84.652 |
| 17592 | ENSMUSG00000062933 | 157.5849952 | 0.253891031 | 0.193975174 | 1.308884151 | 0.190573596 | 0.627894417 | Gm10123 | predicted pseudogene 10123 [Source:MG | 140.54 | 158.3 | 132.82 | 189.9 | 165.77 | 158.17 |
| 17593 | ENSMUSG00000062937 | 358.4786173 | -0.117100316 | 0.140766885 | -0.831874033 | 0.405480051 | 0.829973745 | Mtap | methylthioadenosine phosphorylase [Sou | 386.29 | 359.77 | 372.31 | 318.11 | 346.82 | 367.57 |
| 17594 | ENSMUSG00000062939 | 1.798559791 | 0.971057846 | 1.557165108 | 0.62360622 | 0.532886225 | NA | Stat4 | signal transducer and activator of transcri | 1.6062 | 1.0279 | 1.0062 | 3.8559 | 2.1812 | 1.1138 |
| 17595 | ENSMUSG00000062944 | 73.99280205 | 0.207625543 | 0.289297688 | 0.717688222 | 0.472949547 | 0.86857093 | 9130023H24 | RIKEN cDNA 9130023H24 gene [Source:M | 65.051 | 61.675 | 79.493 | 83.866 | 97.065 | 56.806 |
| 17596 | ENSMUSG00000062949 | 4108.12751 | 0.273200486 | 0.100222225 | 2.725947125 | 0.006411727 | 0.079061566 | Atp11c | ATPase, class VI, type 11C [Source:MGI Sy | 3395.5 | 3934.9 | 3831.8 | 4350.4 | 4792.2 | 4344 |
| 17597 | ENSMUSG00000062952 | 0 | NA | NA | NA | NA | NA | Tas2r110 | taste receptor, type 2, member 110 [Sour | 0 | 0 | 0 | 0 | 0 | 0 |
| 17598 | ENSMUSG00000062960 | 372.3591168 | -0.547928376 | 0.185742897 | -2.949929099 | 0.003178469 | 0.048599236 | Kdr | kinase insert domain protein receptor [So | 469.81 | 391.64 | 464.88 | 305.58 | 225.76 | 376.48 |
| 17599 | ENSMUSG00000062961 | 0 | NA | NA | NA | NA | NA | Ccdc177 | coiled-coil domain containing 177 [Source | 0 | 0 | 0 | 0 | 0 | 0 |
| 17600 | ENSMUSG00000062962 | 0 | NA | NA | NA | NA | NA | Gm6378 | predicted pseudogene 6378 [Source:MGI | 0 | 0 | 0 | 0 | 0 | 0 |
| 17601 | ENSMUSG00000062963 | 1336.775336 | 0.061779087 | 0.106413258 | 0.580558173 | 0.561538269 | 0.900553224 | Ufc1 | ubiquitin-fold modifier conjugating enzym | 1301 | 1366.1 | 1257.8 | 1432.5 | 1253.1 | 1410.1 |
| 17602 | ENSMUSG00000062980 | 943.057254 | 0.014867282 | 0.115787454 | 0.128401494 | 0.897831249 | 0.982798755 | Cped1 | cadherin-like and PC-esterase domain con | 934.01 | 962.13 | 918.7 | 955.3 | 1035 | 853.2 |
| 17603 | ENSMUSG00000062981 | 662.9806991 | 0.043837783 | 0.124974987 | 0.350772456 | 0.725759063 | 0.945362372 | Mrpl42 | mitochondrial ribosomal protein L42 [Sou | 638.47 | 647.59 | 672.17 | 595.74 | 712.17 | 711.75 |
| 17604 | ENSMUSG00000062987 | 0 | NA | NA | NA | NA | NA | Olfr715b | olfactory receptor 715B [Source:MGI Sym | 0 | 0 | 0 | 0 | 0 | 0 |
| 17605 | ENSMUSG00000062991 | 2.544527471 | -0.510849431 | 1.419817935 | -0.359799252 | 0.718997262 | NA | Nrg1 | neuregulin 1 [Source:MGI Symbol;Acc:MG | 4.8186 | 4.1117 | 0 | 1.928 | 2.1812 | 2.2277 |
| 17606 | ENSMUSG00000062995 | 17.50961604 | 0.451178942 | 0.519601023 | 0.868318041 | 0.385220248 | 0.816437825 | Ica1 | islet cell autoantigen 1 [Source:MGI Symb | 12.047 | 14.391 | 18.112 | 24.099 | 16.359 | 20.049 |
| 17607 | ENSMUSG00000062997 | 1368.644698 | 0.087295647 | 0.11479527 | 0.760446375 | 0.446987812 | 0.853398186 | Rpl35 | ribosomal protein L35 [Source:MGI Symbo | 1291.4 | 1395.9 | 1294 | 1270.5 | 1387.3 | 1572.7 |
| 17608 | ENSMUSG00000063001 | 0.835284868 | -0.623834031 | 2.861223713 | -0.218030498 | 0.827405349 | NA | Gm9701 | predicted gene 9701 [Source:MGI Symbol | 0 | 3.0838 | 0 | 1.928 | 0 | 0 |
| 17609 | ENSMUSG00000063011 | 62.52581817 | 0.505504951 | 0.280772849 | 1.800405388 | 0.071796651 | 0.387921836 | Msln | mesothelin [Source:MGI Symbol;Acc:MGI | 51.399 | 53.452 | 50.312 | 79.046 | 76.343 | 64.603 |
| 17610 | ENSMUSG00000063015 | 2725.482143 | -0.014034327 | 0.094527694 | -0.148467882 | 0.881973532 | 0.981738667 | Ccni | cyclin I [Source:MGI Symbol;Acc:MGI:134 | 2589.2 | 2925.5 | 2702.8 | 2774.3 | 2657.8 | 2703.3 |
| 17611 | ENSMUSG00000063018 | 2.824359889 | -1.591317279 | 1.366398164 | -1.164607302 | 0.244177987 | NA | 2010204K13 | RIKEN cDNA 2010204K13 gene [Source:M | 5.6217 | 1.0279 | 6.0375 | 0.964 | 2.1812 | 1.1138 |
| 17612 | ENSMUSG00000063019 | 153.9975054 | 0.056485513 | 0.20878422 | 0.270544933 | 0.786741054 | 0.960870081 | Manbal | mannosidase, beta A, lysosomal-like [Sou | 166.24 | 166.52 | 119.74 | 149.42 | 149.41 | 172.65 |
| 17613 | ENSMUSG00000063020 | 0 | NA | NA | NA | NA | NA | Olfr194 | olfactory receptor 194 [Source:MGI Symb | 0 | 0 | 0 | 0 | 0 | 0 |
| 17614 | ENSMUSG00000063021 | 0 | NA | NA | NA | NA | NA | H2ac15 | H2A clustered histone 15 [Source:MGI Syr | 0 | 0 | 0 | 0 | 0 | 0 |
| 17615 | ENSMUSG00000063047 | 114.547797 | -0.029101178 | 0.233148331 | -0.124818297 | 0.900667402 | 0.98296637 | Zfp780b | zinc finger protein 780B [Source:MGI Sym | 116.45 | 93.541 | 136.85 | 100.25 | 123.24 | 116.95 |
| 17616 | ENSMUSG00000063049 | 159.0303793 | 0.346489885 | 0.204874713 | 1.691228167 | 0.090793233 | 0.444602826 | Ing2 | inhibitor of growth family, member 2 [Sou | 122.07 | 132.6 | 166.03 | 193.76 | 157.05 | 182.67 |
| 17617 | ENSMUSG00000063052 | 231.2530145 | 0.040262613 | 0.17511856 | 0.229916308 | 0.818156804 | 0.968873425 | Lrrc40 | leucine rich repeat containing 40 [Source: | 221.66 | 254.92 | 207.29 | 205.33 | 241.03 | 257.3 |
| 17618 | ENSMUSG00000063058 | 0 | NA | NA | NA | NA | NA | Prr23a2 | proline rich 23A, member 2 [Source:MGI S | 0 | 0 | 0 | 0 | 0 | 0 |
| 17619 | ENSMUSG00000063060 | 29.81891008 | -0.104386442 | 0.462208189 | -0.225842909 | 0.821323602 | 0.969160361 | Sox7 | SRY (sex determining region Y)-box 7 [Sou | 29.715 | 21.586 | 41.256 | 22.171 | 19.631 | 44.554 |
| 17620 | ENSMUSG00000063063 | 0.160662974 | 1.126924175 | 4.080472857 | 0.276174898 | 0.782413736 | NA | Ctnna2 | catenin (cadherin associated protein), alp | 0 | 0 | 0 | 0.964 | 0 | 0 |
| 17621 | ENSMUSG00000063065 | 595.9033212 | -0.056472202 | 0.120992959 | -0.466739572 | 0.640686214 | 0.923932929 | Mapk3 | mitogen-activated protein kinase 3 [Sourc | 638.47 | 575.63 | 607.77 | 563.93 | 573.67 | 615.96 |
| 17622 | ENSMUSG00000063077 | 3237.534409 | -0.015113753 | 0.096101867 | -0.157268044 | 0.875033604 | 0.980503106 | Kif1b | kinesin family member 1B [Source:MGI Sy | 3109.6 | 3541.2 | 3113.3 | 3159 | 3265.3 | 3236.8 |
| 17623 | ENSMUSG00000063087 | 4.083744593 | 0.166393797 | 1.058076209 | 0.157260693 | 0.875039397 | NA | Gm10125 | predicted gene 10125 [Source:MGI Symbo | 2.4093 | 4.1117 | 5.0312 | 1.928 | 5.4531 | 5.5692 |
| 17624 | ENSMUSG00000063089 | 0 | NA | NA | NA | NA | NA | Klk1b8 | kallikrein 1-related peptidase b8 [Source:I | 0 | 0 | 0 | 0 | 0 | 0 |
| 17625 | ENSMUSG00000063106 | 0 | NA | NA | NA | NA | NA | Olfr1510 | olfactory receptor 1510 [Source:MGI Sym | 0 | 0 | 0 | 0 | 0 | 0 |
| 17626 | ENSMUSG00000063108 | 140.5942929 | -0.148764831 | 0.20100925 | -0.740089481 | 0.459245701 | 0.859442977 | Zfp26 | zinc finger protein 26 [Source:MGI Symbo | 155.8 | 132.6 | 154.96 | 135.92 | 142.87 | 121.41 |
| 17627 | ENSMUSG00000063109 | 0.428363721 | -0.543730382 | 3.624677344 | -0.15000794 | 0.880758351 | NA | Dgkeos | diacylglycerol kinase, epsilon, opposite str | 1.6062 | 0 | 0 | 0.964 | 0 | 0 |
| 17628 | ENSMUSG00000063116 | 0 | NA | NA | NA | NA | NA | Olfr410 | olfactory receptor 410 [Source:MGI Symb | 0 | 0 | 0 | 0 | 0 | 0 |
| 17629 | ENSMUSG00000063120 | 0 | NA | NA | NA | NA | NA | Olfr480 | olfactory receptor 480 [Source:MGI Symb | 0 | 0 | 0 | 0 | 0 | 0 |
| 17630 | ENSMUSG00000063129 | 0 | NA | NA | NA | NA | NA | Aldoart2 | aldolase 1 A, retrogene 2 [Source:MGI Syr | 0 | 0 | 0 | 0 | 0 | 0 |
| 17631 | ENSMUSG00000063130 | 0 | NA | NA | NA | NA | NA | Calml3 | calmodulin-like 3 [Source:MGI Symbol;Acc | 0 | 0 | 0 | 0 | 0 | 0 |
| 17632 | ENSMUSG00000063133 | 0 | NA | NA | NA | NA | NA | Klk1b1 | kallikrein 1-related peptidase b1 [Source:I | 0 | 0 | 0 | 0 | 0 | 0 |
| 17633 | ENSMUSG00000063137 | 0 | NA | NA | NA | NA | NA | Olfr177 | olfactory receptor 177 [Source:MGI Symb | 0 | 0 | 0 | 0 | 0 | 0 |
| 17634 | ENSMUSG00000063142 | 4.089527563 | 0.311586484 | 1.031579983 | 0.302047819 | 0.76261561 | NA | Kcnma1 | potassium large conductance calcium-acti | 4.8186 | 2.0558 | 4.025 | 4.8199 | 4.3625 | 4.4554 |
| 17635 | ENSMUSG00000063145 | 69.22398674 | -0.099151896 | 0.285565093 | -0.34721294 | 0.72843135 | 0.946208446 | Bbs5 | Bardet-Biedl syndrome 5 (human) [Source | 84.326 | 55.508 | 74.462 | 58.803 | 67.618 | 74.628 |
| 17636 | ENSMUSG00000063146 | 35.49710176 | -0.302412599 | 0.383561106 | -0.788433952 | 0.430442918 | 0.843621458 | Clip2 | CAP-GLY domain containing linker protein | 44.171 | 44.2 | 29.181 | 34.703 | 25.084 | 35.643 |

| | | | | | | | | | | | | | | | |
|---|---|---|---|---|---|---|---|---|---|---|---|---|---|---|---|
| 17637 | ENSMUSG00000063157 | 0.640584875 | -2.718595279 | 2.766810251 | -0.982573806 | 0.325817242 | NA | Csn2 | casein beta [Source:MGI Symbol;Acc:MGI | 0.8031 | 1.0279 | 2.0125 | 0 | 0 | 0 |
| 17638 | ENSMUSG00000063160 | 10.22776438 | -0.729919856 | 0.752055225 | -0.97056683 | 0.33176403 | 0.774747658 | Numbl | numb-like [Source:MGI Symbol;Acc:MGI:8 | 16.865 | 10.279 | 11.069 | 6.7478 | 14.178 | 2.2277 |
| 17639 | ENSMUSG00000063163 | 0 | NA | NA | NA | NA | NA | Speer2 | spermatogenesis associated glutamate (E) | 0 | 0 | 0 | 0 | 0 | 0 |
| 17640 | ENSMUSG00000063166 | 0 | NA | NA | NA | NA | NA | Gm5449 | predicted pseudogene 5449 [Source:MGI | 0 | 0 | 0 | 0 | 0 | 0 |
| 17641 | ENSMUSG00000063171 | 391.0210264 | 0.365537545 | 0.252889016 | 1.445446506 | 0.148332509 | 0.56003169 | Rps4l | ribosomal protein S4-like [Source:MGI Syr | 311.6 | 234.37 | 478.97 | 291.12 | 534.4 | 495.66 |
| 17642 | ENSMUSG00000063172 | 52.82742956 | 0.342658562 | 0.302831669 | 1.131514954 | 0.257838413 | 0.711441206 | Hspb11 | heat shock protein family B (small), memb | 44.171 | 49.34 | 46.287 | 54.947 | 66.528 | 55.692 |
| 17643 | ENSMUSG00000063173 | 0 | NA | NA | NA | NA | NA | Olfr811 | olfactory receptor 811 [Source:MGI Symb | 0 | 0 | 0 | 0 | 0 | 0 |
| 17644 | ENSMUSG00000063176 | 0 | NA | NA | NA | NA | NA | Olfr955 | olfactory receptor 955 [Source:MGI Symb | 0 | 0 | 0 | 0 | 0 | 0 |
| 17645 | ENSMUSG00000063177 | 0.401551119 | -2.072982726 | 4.008251979 | -0.517178744 | 0.605031388 | NA | Klk1b27 | kallikrein 1-related peptidase b27 [Source | 2.4093 | 0 | 0 | 0 | 0 | 0 |
| 17646 | ENSMUSG00000063179 | 166.6240984 | -0.192516078 | 0.18067542 | -1.065535517 | 0.286633662 | 0.738882425 | Pstk | phosphoseryl-tRNA kinase [Source:MGI Sy | 174.27 | 174.75 | 184.14 | 151.34 | 155.96 | 159.28 |
| 17647 | ENSMUSG00000063188 | 0 | NA | NA | NA | NA | NA | Olfr107 | olfactory receptor 107 [Source:MGI Symb | 0 | 0 | 0 | 0 | 0 | 0 |
| 17648 | ENSMUSG00000063193 | 10.55669744 | -0.761358312 | 0.692472727 | -1.099477687 | 0.271559761 | 0.727377849 | Cd300lb | CD300 molecule like family member B [So | 15.259 | 13.363 | 11.069 | 3.8559 | 11.997 | 7.7969 |
| 17649 | ENSMUSG00000063200 | 855.6658276 | -0.045685927 | 0.121665617 | -0.375504011 | 0.707285664 | 0.943221449 | Nol7 | nucleolar protein 7 [Source:MGI Symbol;A | 805.51 | 905.6 | 896.56 | 754.79 | 864.86 | 906.67 |
| 17650 | ENSMUSG00000063206 | 0 | NA | NA | NA | NA | NA | Defa34 | defensin, alpha, 34 [Source:MGI Symbol;A | 0 | 0 | 0 | 0 | 0 | 0 |
| 17651 | ENSMUSG00000063216 | 0 | NA | NA | NA | NA | NA | Olfr1351 | olfactory receptor 1351 [Source:MGI Syml | 0 | 0 | 0 | 0 | 0 | 0 |
| 17652 | ENSMUSG00000063221 | 0 | NA | NA | NA | NA | NA | Olfr930 | olfactory receptor 930 [Source:MGI Symb | 0 | 0 | 0 | 0 | 0 | 0 |
| 17653 | ENSMUSG00000063225 | 0 | NA | NA | NA | NA | NA | Olfr917 | olfactory receptor 917 [Source:MGI Symb | 0 | 0 | 0 | 0 | 0 | 0 |
| 17654 | ENSMUSG00000063229 | 16598.39914 | -0.013154443 | 0.084423946 | -0.15581412 | 0.876179541 | 0.980632845 | Ldha | lactate dehydrogenase A [Source:MGI Syr | 17626 | 16055 | 16339 | 16492 | 16219 | 16859 |
| 17655 | ENSMUSG00000063230 | 0 | NA | NA | NA | NA | NA | Olfr535 | olfactory receptor 535 [Source:MGI Symb | 0 | 0 | 0 | 0 | 0 | 0 |
| 17656 | ENSMUSG00000063232 | 650.0944158 | 0.423394596 | 0.121264473 | 3.491497422 | 0.000480321 | 0.012061397 | Serpina11 | serine (or cysteine) peptidase inhibitor, cl | 535.67 | 579.75 | 551.42 | 784.68 | 682.73 | 766.32 |
| 17657 | ENSMUSG00000063234 | 3.661112237 | 2.200410738 | 1.271578807 | 1.730455656 | 0.083548897 | NA | Gpr84 | G protein-coupled receptor 84 [Source:M | 0 | 1.0279 | 3.0187 | 5.7839 | 5.4531 | 6.6831 |
| 17658 | ENSMUSG00000063235 | 855.6453533 | -0.227656829 | 0.116516543 | -1.953858427 | 0.050717962 | 0.316199857 | Ptpmt1 | protein tyrosine phosphatase, mitochond | 973.36 | 930.27 | 864.36 | 729.73 | 840.87 | 795.28 |
| 17659 | ENSMUSG00000063236 | 109.3189342 | -0.10880682 | 0.224303698 | -0.485087052 | 0.627614597 | 0.920328819 | 1110038F14 | RIKEN cDNA 1110038F14 gene [Source:M | 111.63 | 100.74 | 127.79 | 96.398 | 107.97 | 111.38 |
| 17660 | ENSMUSG00000063239 | 0 | NA | NA | NA | NA | NA | Grm4 | glutamate receptor, metabotropic 4 [Sour | 0 | 0 | 0 | 0 | 0 | 0 |
| 17661 | ENSMUSG00000063240 | 0 | NA | NA | NA | NA | NA | Olfr133 | olfactory receptor 133 [Source:MGI Symb | 0 | 0 | 0 | 0 | 0 | 0 |
| 17662 | ENSMUSG00000063245 | 4.394956299 | 0.092506229 | 1.010952495 | 0.091504031 | 0.927092103 | NA | Zfp993 | zinc finger protein 993 [Source:MGI Symb | 5.6217 | 2.0558 | 5.0312 | 4.8199 | 3.2719 | 5.5692 |
| 17663 | ENSMUSG00000063251 | 0 | NA | NA | NA | NA | NA | Krtap4-1 | keratin associated protein 4-1 [Source:MG | 0 | 0 | 0 | 0 | 0 | 0 |
| 17664 | ENSMUSG00000063252 | 0 | NA | NA | NA | NA | NA | Gm4744 | predicted gene 4744 [Source:MGI Symbol | 0 | 0 | 0 | 0 | 0 | 0 |
| 17665 | ENSMUSG00000063253 | 176.1173578 | -0.057928239 | 0.197410527 | -0.293440474 | 0.769185493 | 0.956045284 | Scoc | short coiled-coil protein [Source:MGI Sym | 183.91 | 157.27 | 197.22 | 143.63 | 189.77 | 184.9 |
| 17666 | ENSMUSG00000063260 | 0 | NA | NA | NA | NA | NA | Syt10 | synaptotagmin X [Source:MGI Symbol;Acc | 0 | 0 | 0 | 0 | 0 | 0 |
| 17667 | ENSMUSG00000063268 | 520.0406448 | 0.142358868 | 0.1302542 | 1.092931113 | 0.274424047 | 0.729487777 | Parp10 | poly (ADP-ribose) polymerase family, men | 531.65 | 466.67 | 484 | 524.4 | 591.12 | 522.39 |
| 17668 | ENSMUSG00000063273 | 649.0839363 | 0.067518825 | 0.126217317 | 0.534941058 | 0.592690616 | 0.912925376 | Naa15 | N(alpha)-acetyltransferase 15, NatA auxili | 671.39 | 673.29 | 556.45 | 656.47 | 666.37 | 670.53 |
| 17669 | ENSMUSG00000063275 | 102.9342055 | 0.359846825 | 0.229832333 | 1.565692783 | 0.117420557 | 0.502643331 | Hacd1 | 3-hydroxyacyl-CoA dehydratase 1 [Source | 96.372 | 78.122 | 95.593 | 105.07 | 122.15 | 120.3 |
| 17670 | ENSMUSG00000063277 | 0 | NA | NA | NA | NA | NA | Gm10128 | predicted gene 10128 [Source:MGI Symbo | 0 | 0 | 0 | 0 | 0 | 0 |
| 17671 | ENSMUSG00000063281 | 175.8075201 | -0.035892045 | 0.186559009 | -0.192389771 | 0.847436904 | 0.972803041 | Zfp35 | zinc finger protein 35 [Source:MGI Symbo | 195.96 | 161.38 | 176.09 | 170.62 | 160.32 | 190.47 |
| 17672 | ENSMUSG00000063286 | 38.40634677 | -1.300010564 | 0.455662575 | -2.853011495 | 0.004330706 | 0.060746309 | Gvin-ps7 | GTPase, very large interferon inducible, ps | 55.414 | 69.898 | 38.237 | 11.568 | 17.45 | 37.871 |
| 17673 | ENSMUSG00000063296 | 0.401551119 | -2.072982726 | 4.008251979 | -0.517178744 | 0.605031388 | NA | Tmem117 | transmembrane protein 117 [Source:MGI | 2.4093 | 0 | 0 | 0 | 0 | 0 |
| 17674 | ENSMUSG00000063297 | 0 | NA | NA | NA | NA | NA | Luzp2 | leucine zipper protein 2 [Source:MGI Sym | 0 | 0 | 0 | 0 | 0 | 0 |
| 17675 | ENSMUSG00000063305 | 0 | NA | NA | NA | NA | NA | Psg20 | pregnancy-specific glycoprotein 20 [Sourc | 0 | 0 | 0 | 0 | 0 | 0 |
| 17676 | ENSMUSG00000063314 | 1.045723228 | -0.867386806 | 2.411862378 | -0.359633623 | 0.719121136 | NA | Gm12657 | predicted gene 12657 [Source:MGI Symbo | 0 | 1.0279 | 3.0187 | 0 | 0 | 2.2277 |
| 17677 | ENSMUSG00000063316 | 2496.849217 | 0.013567582 | 0.096909004 | 0.140003313 | 0.888657373 | 0.982382091 | Rpl27 | ribosomal protein L27 [Source:MGI Symbo | 2347.5 | 2429 | 2679.6 | 2501.5 | 2455 | 2568.5 |
| 17678 | ENSMUSG00000063317 | 129.6794619 | -0.145577383 | 0.235817094 | -0.617331765 | 0.537015919 | 0.890705151 | Usp31 | ubiquitin specific peptidase 31 [Source:M | 139.74 | 151.1 | 117.73 | 125.32 | 92.703 | 151.48 |
| 17679 | ENSMUSG00000063320 | 197.0444026 | 0.006899503 | 0.173333871 | 0.039804701 | 0.968248828 | 0.994231936 | 1190007I07F | RIKEN cDNA 1190007I07 gene [Source:MG | 212.02 | 191.19 | 186.16 | 198.58 | 208.31 | 186.01 |
| 17680 | ENSMUSG00000063328 | 0.167707431 | -0.796629176 | 4.080472857 | -0.195229623 | 0.84521319 | NA | Gm9396 | predicted gene 9396 [Source:MGI Symbol | 0 | 0 | 1.0062 | 0 | 0 | 0 |
| 17681 | ENSMUSG00000063334 | 321.5038534 | 0.213496363 | 0.148024612 | 1.44230314 | 0.149216875 | 0.561319344 | Krr1 | KRR1, small subunit (SSU) processome co | 281.89 | 324.82 | 286.78 | 324.86 | 364.27 | 346.41 |
| 17682 | ENSMUSG00000063350 | 0 | NA | NA | NA | NA | NA | Olfr874 | olfactory receptor 874 [Source:MGI Symb | 0 | 0 | 0 | 0 | 0 | 0 |
| 17683 | ENSMUSG00000063354 | 336.0343079 | -0.867717634 | 0.243022465 | -3.570524369 | 0.000356267 | 0.009958034 | Slc39a4 | solute carrier family 39 (zinc transporter), | 363 | 514.99 | 424.64 | 205.33 | 350.09 | 158.17 |
| 17684 | ENSMUSG00000063358 | 1629.429587 | 0.122549016 | 0.097144367 | 1.261514386 | 0.20712358 | 0.652364421 | Mapk1 | mitogen-activated protein kinase 1 [Sourc | 1501 | 1545 | 1635.1 | 1647.4 | 1746.1 | 1702 |
| 17685 | ENSMUSG00000063362 | 580.4648205 | 0.187605052 | 0.130298972 | 1.439804539 | 0.149922707 | 0.563095827 | Alg11 | asparagine-linked glycosylation 11 (alpha- | 516.39 | 578.72 | 534.32 | 687.32 | 582.39 | 583.65 |
| 17686 | ENSMUSG00000063364 | 10.2265909 | 0.594975804 | 0.727836768 | 0.817457745 | 0.413666887 | 0.835385466 | 3300002I08F | RIKEN cDNA 3300002I08 gene [Source:MG | 7.2279 | 12.335 | 5.0312 | 20.244 | 8.7249 | 7.7969 |
| 17687 | ENSMUSG00000063374 | 0 | NA | NA | NA | NA | NA | Olfr763 | olfactory receptor 763 [Source:MGI Symb | 0 | 0 | 0 | 0 | 0 | 0 |
| 17688 | ENSMUSG00000063376 | 0 | NA | NA | NA | NA | NA | Ifna13 | interferon alpha 13 [Source:MGI Symbol;A | 0 | 0 | 0 | 0 | 0 | 0 |
| 17689 | ENSMUSG00000063380 | 0 | NA | NA | NA | NA | NA | Olfr945 | olfactory receptor 945 [Source:MGI Symb | 0 | 0 | 0 | 0 | 0 | 0 |
| 17690 | ENSMUSG00000063382 | 1023.031492 | -0.026278556 | 0.128348599 | -0.204743615 | 0.837772442 | 0.971073158 | Bcl9l | B cell CLL/lymphoma 9-like [Source:MGI S | 912.32 | 1114.3 | 1071.7 | 1050.7 | 894.31 | 1094.9 |
| 17691 | ENSMUSG00000063383 | 19.66096326 | -0.912253325 | 0.625413124 | -1.45864116 | 0.144663899 | 0.554723921 | Zfp947 | zinc finger protein 947 [Source:MGI Symb | 43.368 | 11.307 | 22.137 | 11.568 | 22.903 | 6.6831 |
| 17692 | ENSMUSG00000063386 | 0 | NA | NA | NA | NA | NA | Olfr1387 | olfactory receptor 1387 [Source:MGI Syml | 0 | 0 | 0 | 0 | 0 | 0 |

| | | | | | | | | | | | | | | | |
|---|---|---|---|---|---|---|---|---|---|---|---|---|---|---|---|
| 17693 | ENSMUSG00000063388 | 91.44770957 | 0.415621151 | 0.314526107 | 1.321420201 | 0.186361294 | 0.622503355 | BC023105 | cDNA sequence BC023105 [Source:MGI Sy | 89.947 | 103.82 | 41.256 | 117.61 | 103.61 | 92.449 |
| 17694 | ENSMUSG00000063394 | 0 | NA | NA | NA | NA | NA | Olfr293 | olfactory receptor 293 [Source:MGI Symb | 0 | 0 | 0 | 0 | 0 | 0 |
| 17695 | ENSMUSG00000063406 | 2343.730074 | -0.134991263 | 0.114329613 | -1.180720021 | 0.237713965 | 0.686377492 | Tmed5 | transmembrane p24 trafficking protein 5 [ | 2619.7 | 2490.6 | 2249 | 2225.8 | 2477.9 | 1999.3 |
| 17696 | ENSMUSG00000063409 | 0 | NA | NA | NA | NA | NA | Lrrc43 | leucine rich repeat containing 43 [Source: | 0 | 0 | 0 | 0 | 0 | 0 |
| 17697 | ENSMUSG00000063410 | 383.9097851 | -0.068013116 | 0.146900672 | -0.462987097 | 0.643373616 | 0.925477092 | Stk24 | serine/threonine kinase 24 [Source:MGI S | 346.94 | 422.47 | 410.55 | 399.09 | 346.82 | 377.59 |
| 17698 | ENSMUSG00000063412 | 1.948818782 | -2.144881517 | 1.831019099 | -1.171414061 | 0.241432383 | NA | Gm10131 | predicted pseudogene 10131 [Source:MG | 6.4248 | 1.0279 | 2.0125 | 0 | 0 | 2.2277 |
| 17699 | ENSMUSG00000063415 | 74.5993125 | 0.062619811 | 1.112232319 | 0.056301017 | 0.955102008 | 0.991678478 | Cyp26b1 | cytochrome P450, family 26, subfamily b, | 66.657 | 63.731 | 88.55 | 76.154 | 4.3625 | 148.14 |
| 17700 | ENSMUSG00000063428 | 619.086479 | -0.069412468 | 0.119892155 | -0.578957551 | 0.562617816 | 0.900565925 | Ddo | D-aspartate oxidase [Source:MGI Symbol; | 648.1 | 650.67 | 602.74 | 580.31 | 643.46 | 589.22 |
| 17701 | ENSMUSG00000063430 | 0.696064467 | -1.534444742 | 3.043667477 | -0.504143358 | 0.61416064 | NA | Wscd2 | WSC domain containing 2 [Source:MGI Sy | 3.2124 | 0 | 0 | 0.964 | 0 | 0 |
| 17702 | ENSMUSG00000063434 | 0.167707431 | -0.796629176 | 4.080472857 | -0.195229623 | 0.84521319 | NA | Sorcs3 | sortilin-related VPS10 domain containing | 0 | 0 | 1.0062 | 0 | 0 | 0 |
| 17703 | ENSMUSG00000063439 | 102.3968369 | 0.194074926 | 0.242125487 | 0.801546871 | 0.422815121 | 0.84005258 | B9d2 | B9 protein domain 2 [Source:MGI Symbol | 109.22 | 87.373 | 89.556 | 111.82 | 89.431 | 126.98 |
| 17704 | ENSMUSG00000063442 | 0 | NA | NA | NA | NA | NA | Gm7117 | predicted pseudogene 7117 [Source:MGI | 0 | 0 | 0 | 0 | 0 | 0 |
| 17705 | ENSMUSG00000063445 | 122.9066933 | -0.247201842 | 0.215106837 | -1.149204952 | 0.25047148 | 0.702302228 | Nmral1 | NmrA-like family domain containing 1 [Sou | 128.5 | 138.77 | 132.82 | 101.22 | 104.7 | 131.43 |
| 17706 | ENSMUSG00000063446 | 2.682151677 | -3.725827709 | 1.802401108 | -2.067146814 | 0.038720322 | NA | Plppr1 | phospholipid phosphatase related 1 [Sour | 0.8031 | 11.307 | 3.0187 | 0.964 | 0 | 0 |
| 17707 | ENSMUSG00000063447 | 2.607141838 | 1.653260115 | 1.65191244 | 1.000815827 | 0.316915857 | NA | Ube2d2b | ubiquitin-conjugating enzyme E2D 2B [Sou | 0.8031 | 0 | 3.0187 | 9.6398 | 2.1812 | 0 |
| 17708 | ENSMUSG00000063450 | 698.8056974 | 0.287374642 | 0.167762295 | 1.71298707 | 0.086714918 | 0.433994664 | Syne2 | spectrin repeat containing, nuclear envelo | 522.02 | 738.05 | 629.91 | 908.07 | 772.16 | 622.64 |
| 17709 | ENSMUSG00000063455 | 114.736093 | 0.318112005 | 0.261103672 | 1.218336008 | 0.223096312 | 0.668407528 | D630045J12 | RIKEN cDNA D630045J12 gene [Source:M | 110.83 | 102.79 | 92.575 | 118.57 | 172.32 | 91.335 |
| 17710 | ENSMUSG00000063457 | 3077.724454 | 0.011417126 | 0.089803917 | 0.127133947 | 0.898834384 | 0.982798755 | Rps15 | ribosomal protein S15 [Source:MGI Symbo | 2988.3 | 3190.7 | 3017.7 | 2985.4 | 3091.9 | 3192.3 |
| 17711 | ENSMUSG00000063458 | 1.943352184 | -1.413309866 | 1.644475079 | -0.859429178 | 0.390103778 | NA | Lrmda | leucine rich melanocyte differentiation as | 2.4093 | 1.0279 | 5.0312 | 0.964 | 0 | 2.2277 |
| 17712 | ENSMUSG00000063478 | 0 | NA | NA | NA | NA | NA | Tas2r114 | taste receptor, type 2, member 114 [Sour | 0 | 0 | 0 | 0 | 0 | 0 |
| 17713 | ENSMUSG00000063480 | 529.3207718 | 0.056082438 | 0.120734368 | 0.464510966 | 0.642281705 | 0.924834503 | Snu13 | SNU13 homolog, small nuclear ribonucleo | 506.76 | 535.55 | 515.2 | 551.4 | 524.59 | 542.44 |
| 17714 | ENSMUSG00000063488 | 418.3651313 | 0.089934596 | 0.158741625 | 0.566547029 | 0.571021972 | 0.904651508 | Zkscan7 | zinc finger with KRAB and SCAN domains 7 | 426.45 | 376.22 | 413.57 | 491.63 | 453.7 | 348.63 |
| 17715 | ENSMUSG00000063506 | 3.740830448 | 0.438115303 | 1.242680971 | 0.35255654 | 0.724420924 | NA | Arhgap22 | Rho GTPase activating protein 22 [Source: | 6.4248 | 2.0558 | 1.0062 | 9.6398 | 1.0906 | 2.2277 |
| 17716 | ENSMUSG00000063507 | 0.48903338 | 0.95637552 | 3.456690196 | 0.276673773 | 0.782030612 | NA | Gm1330 | predicted gene 1330 [Source:MGI Symbol | 0 | 0 | 1.0062 | 1.928 | 0 | 0 |
| 17717 | ENSMUSG00000063511 | 1485.682339 | 0.153623158 | 0.122576001 | 1.253289037 | 0.210100537 | 0.654024485 | Snrnp70 | small nuclear ribonucleoprotein 70 (U1) [S | 1313.9 | 1358.9 | 1548.6 | 1759.3 | 1536.7 | 1396.8 |
| 17718 | ENSMUSG00000063514 | 0 | NA | NA | NA | NA | NA | Gm6756 | predicted gene 6756 [Source:MGI Symbol | 0 | 0 | 0 | 0 | 0 | 0 |
| 17719 | ENSMUSG00000063522 | 0.439020386 | -2.189747911 | 3.52148864 | -0.621824499 | 0.534057275 | NA | Ly6m | lymphocyte antigen 6 complex, locus M [S | 1.6062 | 1.0279 | 0 | 0 | 0 | 0 |
| 17720 | ENSMUSG00000063524 | 10344.45114 | -0.334896764 | 0.100414126 | -3.335155895 | 0.000852516 | 0.018598477 | Eno1 | enolase 1, alpha non-neuron [Source:MGI | 11756 | 11758 | 11106 | 10006 | 9259.3 | 8181.2 |
| 17721 | ENSMUSG00000063529 | 0.133850373 | -0.796629176 | 4.080472857 | -0.195229623 | 0.84521319 | NA | Stmnd1 | stathmin domain containing 1 [Source:MG | 0.8031 | 0 | 0 | 0 | 0 | 0 |
| 17722 | ENSMUSG00000063531 | 2.53949547 | -0.413882259 | 1.604906658 | -0.257885564 | 0.796495226 | NA | Sema3e | sema domain, immunoglobulin domain (Ig | 1.6062 | 2.0558 | 5.0312 | 0 | 6.5437 | 0 |
| 17723 | ENSMUSG00000063535 | 7.848242402 | 0.073079097 | 0.828163252 | 0.088242381 | 0.929684034 | 0.988405116 | Zfp773 | zinc finger protein 773 [Source:MGI Symb | 4.8186 | 5.1396 | 13.081 | 9.6398 | 3.2719 | 11.138 |
| 17724 | ENSMUSG00000063543 | 0 | NA | NA | NA | NA | NA | Gm5616 | predicted gene 5616 [Source:MGI Symbol | 0 | 0 | 0 | 0 | 0 | 0 |
| 17725 | ENSMUSG00000063548 | 0 | NA | NA | NA | NA | NA | Gm4825 | predicted pseudogene 4825 [Source:MGI | 0 | 0 | 0 | 0 | 0 | 0 |
| 17726 | ENSMUSG00000063549 | 0 | NA | NA | NA | NA | NA | Olfr322 | olfactory receptor 322 [Source:MGI Symb | 0 | 0 | 0 | 0 | 0 | 0 |
| 17727 | ENSMUSG00000063550 | 323.817303 | -0.150952133 | 0.166321849 | -0.907590518 | 0.364094606 | 0.802784204 | Nup98 | nucleoporin 98 [Source:MGI Symbol;Acc:M | 334.89 | 392.66 | 294.83 | 300.76 | 344.64 | 275.12 |
| 17728 | ENSMUSG00000063556 | 1.354298859 | 0.160356331 | 1.740743988 | 0.092119423 | 0.926603157 | NA | Gm10132 | predicted gene 10132 [Source:MGI Symbo | 0.8031 | 1.0279 | 2.0125 | 0.964 | 1.0906 | 2.2277 |
| 17729 | ENSMUSG00000063558 | 1627.221536 | -0.083871149 | 0.166935071 | -0.502417788 | 0.615373669 | 0.917509899 | Aox1 | aldehyde oxidase 1 [Source:MGI Symbol;A | 1836.7 | 1606.6 | 1580.8 | 2055.2 | 1482.2 | 1201.8 |
| 17730 | ENSMUSG00000063564 | 6.19754148 | -0.291868522 | 0.855954563 | -0.340985999 | 0.733114122 | 0.947915087 | Col23a1 | collagen, type XXIII, alpha 1 [Source:MGI S | 7.2279 | 8.2233 | 5.0312 | 6.7478 | 3.2719 | 6.6831 |
| 17731 | ENSMUSG00000063568 | 4.854773097 | 0.899966777 | 0.951884723 | 0.945457738 | 0.344425237 | NA | Jazf1 | JAZF zinc finger 1 [Source:MGI Symbol;Ac | 4.0155 | 3.0838 | 3.0187 | 5.7839 | 6.5437 | 6.6831 |
| 17732 | ENSMUSG00000063576 | 1242.204046 | 0.152750291 | 0.116938416 | 1.306245596 | 0.191469052 | 0.629099068 | Klhdc3 | kelch domain containing 3 [Source:MGI Sy | 1272.1 | 1024.8 | 1231.6 | 1324.5 | 1340.4 | 1259.8 |
| 17733 | ENSMUSG00000063582 | 0 | NA | NA | NA | NA | NA | Olfr666 | olfactory receptor 666 [Source:MGI Symb | 0 | 0 | 0 | 0 | 0 | 0 |
| 17734 | ENSMUSG00000063583 | 0 | NA | NA | NA | NA | NA | Olfr1410 | olfactory receptor 1410 [Source:MGI Sym | 0 | 0 | 0 | 0 | 0 | 0 |
| 17735 | ENSMUSG00000063586 | 0 | NA | NA | NA | NA | NA | Gm5513 | predicted pseudogene 5513 [Source:MGI | 0 | 0 | 0 | 0 | 0 | 0 |
| 17736 | ENSMUSG00000063590 | 228.4153649 | 1.693746106 | 0.183256627 | 9.242482157 | 2.41E-20 | 3.87E-17 | Slc22a28 | solute carrier family 22, member 28 [Sour | 116.45 | 101.76 | 104.65 | 296.91 | 366.45 | 384.28 |
| 17737 | ENSMUSG00000063594 | 1.024357763 | -0.938111495 | 2.232331503 | -0.420238434 | 0.674311281 | NA | Gng8 | guanine nucleotide binding protein (G pro | 0 | 2.0558 | 2.0125 | 0.964 | 0 | 1.1138 |
| 17738 | ENSMUSG00000063600 | 8.069702095 | -0.337153776 | 0.815027767 | -0.413671521 | 0.679114687 | 0.934062537 | Egfem1 | EGF-like and EMI domain containing 1 [So | 12.85 | 5.1396 | 9.0562 | 12.532 | 3.2719 | 5.5692 |
| 17739 | ENSMUSG00000063605 | 17.64787855 | -1.705325706 | 0.5680554 | -3.002041183 | 0.002681759 | 0.043485382 | Ccdc102a | coiled-coil domain containing 102A [Sourc | 35.336 | 23.642 | 22.137 | 11.568 | 7.6343 | 5.5692 |
| 17740 | ENSMUSG00000063611 | 2.843740067 | -1.588511802 | 1.424143972 | -1.115415178 | 0.264672548 | NA | Gm10134 | predicted gene 10134 [Source:MGI Symbo | 1.6062 | 7.1954 | 4.025 | 0.964 | 3.2719 | 0 |
| 17741 | ENSMUSG00000063615 | 0 | NA | NA | NA | NA | NA | Olfr64 | olfactory receptor 64 [Source:MGI Symbo | 0 | 0 | 0 | 0 | 0 | 0 |
| 17742 | ENSMUSG00000063623 | 5.401438493 | 0.028516504 | 0.968772142 | 0.029435718 | 0.976517086 | 0.997059377 | C230062I16R | RIKEN cDNA C230062I16 gene [Source:M | 4.0155 | 3.0838 | 9.0562 | 9.6398 | 3.2719 | 3.3415 |
| 17743 | ENSMUSG00000063626 | 0 | NA | NA | NA | NA | NA | Unc5d | unc-5 netrin receptor D [Source:MGI Sym | 0 | 0 | 0 | 0 | 0 | 0 |
| 17744 | ENSMUSG00000063628 | 0.371281295 | 2.09253362 | 4.031852165 | 0.519000582 | 0.603760335 | NA | Gm7665 | predicted pseudogene 7665 [Source:MGI | 0 | 0 | 0 | 0 | 0 | 2.2277 |
| 17745 | ENSMUSG00000063632 | 0.160662974 | 1.126924175 | 4.080472857 | 0.276174898 | 0.782413736 | NA | Sox11 | SRY (sex determining region Y)-box 11 [So | 0 | 0 | 0 | 0.964 | 0 | 0 |
| 17746 | ENSMUSG00000063646 | 0.486939728 | -0.698472448 | 3.137608162 | -0.222613026 | 0.823836697 | NA | Jakmip1 | janus kinase and microtubule interacting p | 0.8031 | 1.0279 | 0 | 0 | 1.0906 | 0 |
| 17747 | ENSMUSG00000063651 | 0 | NA | NA | NA | NA | NA | Cnfn | cornifelin [Source:MGI Symbol;Acc:MGI:1 | 0 | 0 | 0 | 0 | 0 | 0 |
| 17748 | ENSMUSG00000063652 | 15.85900544 | 0.702661506 | 0.581302844 | 1.208770116 | 0.226751176 | 0.67343689 | Slc22a21 | solute carrier family 22 (organic cation tra | 8.031 | 14.391 | 14.087 | 28.919 | 16.359 | 13.366 |

| | | | | | | | | | | | | | | | |
|---|---|---|---|---|---|---|---|---|---|---|---|---|---|---|---|
| 17749 | ENSMUSG00000063656 | 0 | NA | NA | NA | NA | NA | Gm10135 | predicted gene 10135 [Source:MGI Symbo | 0 | 0 | 0 | 0 | 0 | 0 |
| 17750 | ENSMUSG00000063659 | 483.1374147 | 0.113884413 | 0.145478113 | 0.782828502 | 0.433727828 | 0.845264432 | Zbtb18 | zinc finger and BTB domain containing 18 | 411.99 | 495.46 | 486.02 | 552.36 | 439.52 | 513.48 |
| 17751 | ENSMUSG00000063660 | 0.185640647 | 1.126924175 | 4.080472857 | 0.276174898 | 0.782413736 | NA | Olfr98 | olfactory receptor 98 [Source:MGI Symbo | 0 | 0 | 0 | 0 | 0 | 1.1138 |
| 17752 | ENSMUSG00000063661 | 0 | NA | NA | NA | NA | NA | Krt73 | keratin 73 [Source:MGI Symbol;Acc:MGI:3 | 0 | 0 | 0 | 0 | 0 | 0 |
| 17753 | ENSMUSG00000063663 | 180.6959137 | 0.393117063 | 0.196613606 | 1.99943977 | 0.045560792 | 0.295246999 | Brwd3 | bromodomain and WD repeat domain cor | 146.97 | 185.03 | 136.85 | 192.8 | 190.86 | 231.68 |
| 17754 | ENSMUSG00000063672 | 0 | NA | NA | NA | NA | NA | Nkx6-3 | NK6 homeobox 3 [Source:MGI Symbol;Ac | 0 | 0 | 0 | 0 | 0 | 0 |
| 17755 | ENSMUSG00000063681 | 0 | NA | NA | NA | NA | NA | Crb1 | crumbs family member 1, photoreceptor r | 0 | 0 | 0 | 0 | 0 | 0 |
| 17756 | ENSMUSG00000063683 | 6314.916756 | 0.326197653 | 0.11442143 | 2.850844063 | 0.004360335 | 0.060970118 | Glyat | glycine-N-acyltransferase [Source:MGI Syı | 5855.4 | 5130.3 | 5826.2 | 7390.8 | 7643.1 | 6043.7 |
| 17757 | ENSMUSG00000063684 | 20.01780188 | -0.279132275 | 0.502984004 | -0.554952589 | 0.57892709 | 0.908337213 | Hadhb-ps | hydroxyacyl-CoA dehydrogenase trifuncti | 16.062 | 25.698 | 24.15 | 13.496 | 23.994 | 16.708 |
| 17758 | ENSMUSG00000063687 | 0 | NA | NA | NA | NA | NA | Pcdhb5 | protocadherin beta 5 [Source:MGI Symbo | 0 | 0 | 0 | 0 | 0 | 0 |
| 17759 | ENSMUSG00000063689 | 0 | NA | NA | NA | NA | NA | H2ac21 | H2A clustered histone 21 [Source:MGI Syr | 0 | 0 | 0 | 0 | 0 | 0 |
| 17760 | ENSMUSG00000063694 | 2170.18097 | -0.130385841 | 0.096090303 | -1.356909465 | 0.174809979 | 0.604813215 | Cycs | cytochrome c, somatic [Source:MGI Symb | 2268 | 2273.8 | 2262 | 1921.2 | 2143.1 | 2153.1 |
| 17761 | ENSMUSG00000063696 | 6.014091822 | -0.003895446 | 0.919341295 | -0.004237214 | 0.996619202 | 0.999154389 | Gm8730 | predicted pseudogene 8730 [Source:MGI | 4.8186 | 9.2513 | 4.025 | 5.7839 | 2.1812 | 10.025 |
| 17762 | ENSMUSG00000063698 | 5.15134855 | -1.35054646 | 1.059845555 | -1.274286102 | 0.20256204 | 0.647482287 | Sfxn4 | sideroflexin 4 [Source:MGI Symbol;Acc:M | 8.8341 | 7.1954 | 6.0375 | 0 | 3.2719 | 5.5692 |
| 17763 | ENSMUSG00000063704 | 224.4957774 | 0.231166694 | 0.224237755 | 1.030899967 | 0.302587732 | 0.752385931 | Mapk15 | mitogen-activated protein kinase 15 [Sour | 237.72 | 173.72 | 208.29 | 325.82 | 188.68 | 212.74 |
| 17764 | ENSMUSG00000063713 | 0 | NA | NA | NA | NA | NA | Klk1b24 | kallikrein 1-related peptidase b24 [Source | 0 | 0 | 0 | 0 | 0 | 0 |
| 17765 | ENSMUSG00000063714 | 10.97563792 | -0.34871542 | 0.715036212 | -0.487689175 | 0.625770021 | 0.919638611 | Sp3os | trans-acting transcription factor 3, opposi | 15.259 | 15.419 | 6.0375 | 4.8199 | 8.7249 | 15.594 |
| 17766 | ENSMUSG00000063715 | 0 | NA | NA | NA | NA | NA | Olfr816 | olfactory receptor 816 [Source:MGI Symb | 0 | 0 | 0 | 0 | 0 | 0 |
| 17767 | ENSMUSG00000063727 | 34.55850153 | -1.169157615 | 0.395243104 | -2.958072143 | 0.003095697 | 0.047684937 | Tnfrsf11b | tumor necrosis factor receptor superfamil | 41.761 | 47.284 | 54.337 | 15.424 | 21.812 | 26.732 |
| 17768 | ENSMUSG00000063728 | 0 | NA | NA | NA | NA | NA | Magea6 | MAGE family member A6 [Source:MGI Syr | 0 | 0 | 0 | 0 | 0 | 0 |
| 17769 | ENSMUSG00000063730 | 1403.643657 | 0.483314923 | 0.170514865 | 2.834444517 | 0.004590543 | 0.063175654 | Hsd3b2 | hydroxy-delta-5-steroid dehydrogenase, 3 | 1001.5 | 1111.2 | 1400.7 | 1847.9 | 1843.1 | 1217.4 |
| 17770 | ENSMUSG00000063739 | 37.73594633 | 0.390020532 | 0.410080086 | 0.951083815 | 0.34156183 | 0.783312242 | Gm4963 | predicted gene 4963 [Source:MGI Symbol | 33.73 | 46.256 | 18.112 | 51.091 | 34.9 | 42.326 |
| 17771 | ENSMUSG00000063751 | 0.833243349 | 0.77898031 | 2.246377207 | 0.346771819 | 0.72876275 | NA | Platr8 | pluripotency associated transcript 8 [Sour | 0.8031 | 1.0279 | 0 | 0.964 | 1.0906 | 1.1138 |
| 17772 | ENSMUSG00000063754 | 1.130930221 | -0.219351681 | 1.968366848 | -0.111438415 | 0.911268699 | NA | Gm10136 | predicted pseudogene 10136 [Source:MG | 1.6062 | 1.0279 | 1.0062 | 0.964 | 2.1812 | 0 |
| 17773 | ENSMUSG00000063760 | 590.2763824 | 0.071221788 | 0.135089709 | 0.527218457 | 0.598041893 | 0.915029425 | Rnf217 | ring finger protein 217 [Source:MGI Symb | 556.55 | 624.97 | 546.39 | 647.79 | 639.1 | 526.85 |
| 17774 | ENSMUSG00000063762 | 0 | NA | NA | NA | NA | NA | Tas2r129 | taste receptor, type 2, member 129 [Sour | 0 | 0 | 0 | 0 | 0 | 0 |
| 17775 | ENSMUSG00000063764 | 0 | NA | NA | NA | NA | NA | Olfr508 | olfactory receptor 508 [Source:MGI Symb | 0 | 0 | 0 | 0 | 0 | 0 |
| 17776 | ENSMUSG00000063765 | 29.94906196 | -0.363635825 | 0.4238518 | -0.857931534 | 0.39093027 | 0.820195366 | Chadl | chondroadherin-like [Source:MGI Symbol, | 40.155 | 31.865 | 29.181 | 36.631 | 21.812 | 20.049 |
| 17777 | ENSMUSG00000063767 | 0.185640647 | 1.126924175 | 4.080472857 | 0.276174898 | 0.782413736 | NA | S100a7a | S100 calcium binding protein A7A [Source | 0 | 0 | 0 | 0 | 0 | 1.1138 |
| 17778 | ENSMUSG00000063777 | 0 | NA | NA | NA | NA | NA | Olfr1469 | olfactory receptor 1469 [Source:MGI Syml | 0 | 0 | 0 | 0 | 0 | 0 |
| 17779 | ENSMUSG00000063779 | 0 | NA | NA | NA | NA | NA | Chil4 | chitinase-like 4 [Source:MGI Symbol;Acc:N | 0 | 0 | 0 | 0 | 0 | 0 |
| 17780 | ENSMUSG00000063780 | 0 | NA | NA | NA | NA | NA | Olfr282 | olfactory receptor 282 [Source:MGI Symb | 0 | 0 | 0 | 0 | 0 | 0 |
| 17781 | ENSMUSG00000063785 | 175.265814 | 0.368535171 | 0.207490095 | 1.776157902 | 0.075706895 | 0.402878781 | Utp14a | UTP14A small subunit processome compo | 179.09 | 153.16 | 125.78 | 165.8 | 207.22 | 220.54 |
| 17782 | ENSMUSG00000063787 | 502.5365815 | 0.065582383 | 0.13253604 | 0.494826787 | 0.620722394 | 0.918069106 | Chchd1 | coiled-coil-helix-coiled-coil-helix domain c | 477.04 | 462.56 | 534.32 | 546.58 | 520.22 | 474.5 |
| 17783 | ENSMUSG00000063796 | 0.339027071 | -1.79174149 | 3.919585585 | -0.457125237 | 0.647581032 | NA | Slc22a8 | solute carrier family 22 (organic anion trai | 0 | 1.0279 | 1.0062 | 0 | 0 | 0 |
| 17784 | ENSMUSG00000063800 | 163.0426775 | 0.123424126 | 0.209018285 | 0.590494396 | 0.554859241 | 0.897628185 | Prpf38a | PRP38 pre-mRNA processing factor 38 (ye | 136.53 | 178.86 | 152.95 | 143.63 | 164.68 | 201.61 |
| 17785 | ENSMUSG00000063801 | 527.6365858 | 0.072093546 | 0.128867009 | 0.559441446 | 0.575860482 | 0.907226448 | Ap3s2 | adaptor-related protein complex 3, sigma | 527.64 | 510.88 | 504.13 | 490.66 | 587.84 | 544.67 |
| 17786 | ENSMUSG00000063802 | 378.3460427 | -0.188613401 | 0.144537752 | -1.304942124 | 0.191912556 | 0.629099068 | Hspbp1 | HSPA (heat shock 70kDa) binding protein, | 396.73 | 400.89 | 411.55 | 343.18 | 399.17 | 318.56 |
| 17787 | ENSMUSG00000063804 | 0 | NA | NA | NA | NA | NA | Lin28b | lin-28 homolog B (C. elegans) [Source:MG | 0 | 0 | 0 | 0 | 0 | 0 |
| 17788 | ENSMUSG00000063808 | 181.7796007 | -0.107240614 | 0.185093494 | -0.579386185 | 0.562328624 | 0.900565925 | Gpatch1 | G patch domain containing 1 [Source:MGI | 191.94 | 188.11 | 185.15 | 153.27 | 200.67 | 171.53 |
| 17789 | ENSMUSG00000063810 | 39.68329229 | -0.064567212 | 0.383452685 | -0.168383779 | 0.866281369 | 0.977925452 | Alms1 | ALMS1, centrosome and basal body assoc | 34.533 | 32.893 | 54.337 | 32.775 | 51.259 | 32.301 |
| 17790 | ENSMUSG00000063816 | 0 | NA | NA | NA | NA | NA | Gm4987 | predicted gene 4987 [Source:MGI Symbol | 0 | 0 | 0 | 0 | 0 | 0 |
| 17791 | ENSMUSG00000063820 | 0.506966596 | 2.558866737 | 3.339834942 | 0.766165628 | 0.443577757 | NA | Arl9 | ADP-ribosylation factor-like 9 [Source:MG | 0 | 0 | 0 | 1.928 | 0 | 1.1138 |
| 17792 | ENSMUSG00000063821 | 0 | NA | NA | NA | NA | NA | Dupd1 | dual specificity phosphatase and pro isom | 0 | 0 | 0 | 0 | 0 | 0 |
| 17793 | ENSMUSG00000063823 | 0 | NA | NA | NA | NA | NA | Olfr532 | olfactory receptor 532 [Source:MGI Symb | 0 | 0 | 0 | 0 | 0 | 0 |
| 17794 | ENSMUSG00000063827 | 0 | NA | NA | NA | NA | NA | Olfr1382 | olfactory receptor 1382 [Source:MGI Syml | 0 | 0 | 0 | 0 | 0 | 0 |
| 17795 | ENSMUSG00000063830 | 0 | NA | NA | NA | NA | NA | Gm6065 | predicted gene 6065 [Source:MGI Symbol | 0 | 0 | 0 | 0 | 0 | 0 |
| 17796 | ENSMUSG00000063838 | 34.39703003 | -0.486645943 | 0.403475797 | -1.206134162 | 0.227765765 | 0.673977909 | Cdc42ep5 | CDC42 effector protein (Rho GTPase bindi | 43.368 | 37.005 | 40.25 | 40.487 | 18.541 | 26.732 |
| 17797 | ENSMUSG00000063842 | 0 | NA | NA | NA | NA | NA | Olfr862 | olfactory receptor 862 [Source:MGI Symb | 0 | 0 | 0 | 0 | 0 | 0 |
| 17798 | ENSMUSG00000063844 | 0 | NA | NA | NA | NA | NA | Olfr1276 | olfactory receptor 1276 [Source:MGI Syml | 0 | 0 | 0 | 0 | 0 | 0 |
| 17799 | ENSMUSG00000063846 | 0 | NA | NA | NA | NA | NA | Gm21731 | predicted gene, 21731 [Source:MGI Symb | 0 | 0 | 0 | 0 | 0 | 0 |
| 17800 | ENSMUSG00000063849 | 299.0734522 | -0.076800649 | 0.154462207 | -0.497213202 | 0.619038713 | 0.917509899 | Ppcdc | phosphopantothenoylcysteine decarboxy | 331.68 | 307.35 | 281.75 | 298.83 | 309.74 | 265.09 |
| 17801 | ENSMUSG00000063851 | 0 | NA | NA | NA | NA | NA | Rnf183 | ring finger protein 183 [Source:MGI Symb | 0 | 0 | 0 | 0 | 0 | 0 |
| 17802 | ENSMUSG00000063856 | 36121.49297 | -0.098298938 | 0.092884012 | -1.058297712 | 0.289919736 | 0.740058153 | Gpx1 | glutathione peroxidase 1 [Source:MGI Syr | 34842 | 36208 | 41005 | 34467 | 35220 | 34986 |
| 17803 | ENSMUSG00000063867 | 0 | NA | NA | NA | NA | NA | Olfr1511 | olfactory receptor 1511 [Source:MGI Syml | 0 | 0 | 0 | 0 | 0 | 0 |
| 17804 | ENSMUSG00000063870 | 1966.558655 | -0.308650914 | 0.153070513 | -2.016396938 | 0.043758485 | 0.288610458 | Chd4 | chromodomain helicase DNA binding prot | 1676.1 | 2466 | 2387.8 | 1788.2 | 1962 | 1519.3 |

| | | | | | | | | | | | | | | | |
|---|---|---|---|---|---|---|---|---|---|---|---|---|---|---|---|
| 17805 | ENSMUSG00000063873 | 6.180120208 | -1.28025355 | 0.958398674 | -1.33582567 | 0.18160626 | 0.615095921 | Slc24a3 | solute carrier family 24 (sodium/potassiur | 4.8186 | 13.363 | 8.05 | 0.964 | 6.5437 | 3.3415 |
| 17806 | ENSMUSG00000063875 | 1.143912632 | 2.595556111 | 2.186080874 | 1.187310196 | 0.235105288 | NA | Rps6-ps1 | ribosomal protein S6, pseudogene 1 [Sour | 0.8031 | 0 | 0 | 3.8559 | 1.0906 | 1.1138 |
| 17807 | ENSMUSG00000063881 | 0 | NA | NA | NA | NA | NA | Olfr376 | olfactory receptor 376 [Source:MGI Symb | 0 | 0 | 0 | 0 | 0 | 0 |
| 17808 | ENSMUSG00000063882 | 2578.34433 | 0.068190383 | 0.102003275 | 0.668511708 | 0.503807011 | 0.881510796 | Uqcrh | ubiquinol-cytochrome c reductase hinge p | 2326.6 | 2613 | 2613.2 | 2477.4 | 2621.8 | 2818 |
| 17809 | ENSMUSG00000063884 | 710.7605936 | -0.014352406 | 0.121904997 | -0.117734354 | 0.906278145 | 0.984171996 | Ptcd3 | pentatricopeptide repeat domain 3 [Sourc | 682.64 | 733.93 | 726.51 | 669 | 783.06 | 669.42 |
| 17810 | ENSMUSG00000063885 | 0 | NA | NA | NA | NA | NA | Gm6498 | predicted gene 6498 [Source:MGI Symbol | 0 | 0 | 0 | 0 | 0 | 0 |
| 17811 | ENSMUSG00000063887 | 0 | NA | NA | NA | NA | NA | Nlgn1 | neuroligin 1 [Source:MGI Symbol;Acc:MG | 0 | 0 | 0 | 0 | 0 | 0 |
| 17812 | ENSMUSG00000063888 | 1117.777592 | 0.047877628 | 0.1146386 | 0.417639673 | 0.676210586 | 0.932739835 | Rpl7l1 | ribosomal protein L7-like 1 [Source:MGI S | 993.44 | 1088.6 | 1216.5 | 1103.8 | 1159.3 | 1145 |
| 17813 | ENSMUSG00000063889 | 256.8335689 | 0.143977569 | 0.170254757 | 0.845659597 | 0.397742669 | 0.825387369 | Crem | cAMP responsive element modulator [Sou | 254.58 | 226.14 | 251.56 | 311.36 | 234.48 | 262.87 |
| 17814 | ENSMUSG00000063894 | 227.2483675 | 0.379648304 | 0.179285573 | 2.117561929 | 0.034212185 | 0.247603883 | Zkscan8 | zinc finger with KRAB and SCAN domains { | 186.32 | 210.72 | 196.22 | 289.19 | 268.29 | 212.74 |
| 17815 | ENSMUSG00000063895 | 0 | NA | NA | NA | NA | NA | Nupl1 | nucleoporin like 1 [Source:MGI Symbol;Ac | 0 | 0 | 0 | 0 | 0 | 0 |
| 17816 | ENSMUSG00000063897 | 534.604119 | 0.117448396 | 0.12422555 | 0.945444765 | 0.344431857 | 0.785874854 | NA | NA | 513.18 | 489.29 | 536.33 | 585.13 | 526.77 | 556.92 |
| 17817 | ENSMUSG00000063900 | 0 | NA | NA | NA | NA | NA | Adam26b | a disintegrin and metallopeptidase domai | 0 | 0 | 0 | 0 | 0 | 0 |
| 17818 | ENSMUSG00000063902 | 148.1206652 | 0.061606485 | 0.194630816 | 0.316529962 | 0.751600287 | 0.952272823 | Gm7964 | predicted gene 7964 [Source:MGI Symbol | 142.95 | 149.05 | 142.89 | 145.56 | 169.05 | 139.23 |
| 17819 | ENSMUSG00000063903 | 1.138206772 | 1.608612704 | 2.129587282 | 0.755363595 | 0.450030851 | NA | Klk1 | kallikrein 1 [Source:MGI Symbol;Acc:MGI: | 1.6062 | 0 | 0 | 1.928 | 2.1812 | 1.1138 |
| 17820 | ENSMUSG00000063904 | 1068.794661 | 0.27087278 | 0.143577834 | 1.886591914 | 0.059215235 | 0.346085052 | Dpp3 | dipeptidylpeptidase 3 [Source:MGI Symbc | 925.98 | 882.98 | 1096.8 | 957.23 | 1298.9 | 1250.8 |
| 17821 | ENSMUSG00000063905 | 0 | NA | NA | NA | NA | NA | Gm10139 | predicted gene 10139 [Source:MGI Symbc | 0 | 0 | 0 | 0 | 0 | 0 |
| 17822 | ENSMUSG00000063911 | 0 | NA | NA | NA | NA | NA | Tdgf1-ps2 | teratocarcinoma-derived growth factor, p | 0 | 0 | 0 | 0 | 0 | 0 |
| 17823 | ENSMUSG00000063919 | 66.59707165 | -0.196317938 | 0.357407245 | -0.549283599 | 0.582810841 | 0.908645447 | Srrm4 | serine/arginine repetitive matrix 4 [Source | 103.6 | 62.703 | 46.287 | 41.451 | 69.8 | 75.741 |
| 17824 | ENSMUSG00000063929 | 761.0231875 | -0.117992093 | 0.17287517 | -0.682527706 | 0.49490533 | 0.877919732 | Cyp4a32 | cytochrome P450, family 4, subfamily a, p | 821.57 | 715.43 | 840.21 | 957.23 | 638.01 | 593.68 |
| 17825 | ENSMUSG00000063931 | 971.0866542 | -0.189730082 | 0.111123101 | -1.707386502 | 0.087750237 | 0.43690328 | Pepd | peptidase D [Source:MGI Symbol;Acc:MG | 1087.4 | 1007.4 | 1009.3 | 904.21 | 970.65 | 847.64 |
| 17826 | ENSMUSG00000063932 | 0 | NA | NA | NA | NA | NA | Poteg | POTE ankyrin domain family, member G [S | 0 | 0 | 0 | 0 | 0 | 0 |
| 17827 | ENSMUSG00000063935 | 0 | NA | NA | NA | NA | NA | Zar1 | zygote arrest 1 [Source:MGI Symbol;Acc:N | 0 | 0 | 0 | 0 | 0 | 0 |
| 17828 | ENSMUSG00000063952 | 633.7158423 | 0.122031716 | 0.141212161 | 0.864172847 | 0.387492946 | 0.817618864 | Brpf3 | bromodomain and PHD finger containing, | 545.31 | 672.26 | 604.75 | 755.76 | 622.74 | 601.48 |
| 17829 | ENSMUSG00000063953 | 4.330338599 | -0.042107353 | 0.999217887 | -0.042140311 | 0.966386845 | NA | Amd2 | S-adenosylmethionine decarboxylase 2 [S | 4.0155 | 4.1117 | 5.0312 | 2.8919 | 4.3625 | 5.5692 |
| 17830 | ENSMUSG00000063954 | 0 | NA | NA | NA | NA | NA | H2ac19 | H2A clustered histone 19 [Source:MGI Syr | 0 | 0 | 0 | 0 | 0 | 0 |
| 17831 | ENSMUSG00000063971 | 0 | NA | NA | NA | NA | NA | Krt88 | keratin 88 [Source:MGI Symbol;Acc:MGI:1 | 0 | 0 | 0 | 0 | 0 | 0 |
| 17832 | ENSMUSG00000063972 | 146.8051753 | 0.075329161 | 0.224051023 | 0.336214315 | 0.736709259 | 0.948137969 | Nr6a1 | nuclear receptor subfamily 6, group A, me | 124.48 | 155.22 | 149.93 | 169.66 | 169.05 | 112.5 |
| 17833 | ENSMUSG00000063975 | 0 | NA | NA | NA | NA | NA | Slco1a5 | solute carrier organic anion transporter fa | 0 | 0 | 0 | 0 | 0 | 0 |
| 17834 | ENSMUSG00000063994 | 0 | NA | NA | NA | NA | NA | Olfr120 | olfactory receptor 120 [Source:MGI Symb | 0 | 0 | 0 | 0 | 0 | 0 |
| 17835 | ENSMUSG00000063998 | 0 | NA | NA | NA | NA | NA | Vmn1r211 | vomeronasal 1 receptor 211 [Source:MGI | 0 | 0 | 0 | 0 | 0 | 0 |
| 17836 | ENSMUSG00000064006 | 0 | NA | NA | NA | NA | NA | Olfr190 | olfactory receptor 190 [Source:MGI Symb | 0 | 0 | 0 | 0 | 0 | 0 |
| 17837 | ENSMUSG00000064010 | 0 | NA | NA | NA | NA | NA | Dppa1 | developmental pluripotency associated 1 | 0 | 0 | 0 | 0 | 0 | 0 |
| 17838 | ENSMUSG00000064016 | 0 | NA | NA | NA | NA | NA | Gm648 | predicted gene 648 [Source:MGI Symbol;/ | 0 | 0 | 0 | 0 | 0 | 0 |
| 17839 | ENSMUSG00000064023 | 4.544626247 | -1.164262425 | 1.01057096 | -1.152083793 | 0.249286642 | NA | Klk8 | kallikrein related-peptidase 8 [Source:MG | 5.6217 | 8.2233 | 5.0312 | 2.8919 | 3.2719 | 2.2277 |
| 17840 | ENSMUSG00000064030 | 220.5107024 | -0.161320917 | 0.17899133 | -0.901277824 | 0.367440621 | 0.805495779 | Pym1 | PYM homolog 1, exon junction complex a: | 238.52 | 200.44 | 259.61 | 227.5 | 187.59 | 209.4 |
| 17841 | ENSMUSG00000064032 | 0.826457617 | 2.043192078 | 2.584773689 | 0.790472329 | 0.429251976 | NA | Gm10143 | predicted gene 10143 [Source:MGI Symbc | 0.8031 | 0 | 0 | 1.928 | 0 | 2.2277 |
| 17842 | ENSMUSG00000064036 | 0 | NA | NA | NA | NA | NA | Mro | maestro [Source:MGI Symbol;Acc:MGI:21 | 0 | 0 | 0 | 0 | 0 | 0 |
| 17843 | ENSMUSG00000064037 | 208.1541492 | 0.121662655 | 0.18649965 | 0.6523479 | 0.514176766 | 0.885004045 | Gpn1 | GPN-loop GTPase 1 [Source:MGI Symbol;/ | 187.12 | 204.56 | 206.28 | 183.16 | 209.4 | 258.41 |
| 17844 | ENSMUSG00000064039 | 0.133850373 | -0.796629176 | 4.080472857 | -0.195229623 | 0.84521319 | NA | Ccr1l1 | chemokine (C-C motif) receptor 1-like 1 [S | 0.8031 | 0 | 0 | 0 | 0 | 0 |
| 17845 | ENSMUSG00000064043 | 11.66273835 | -0.663753205 | 0.631149887 | -1.051657014 | 0.29295694 | 0.743850082 | Trerf1 | transcriptional regulating factor 1 [Source | 14.456 | 16.447 | 12.075 | 11.568 | 7.6343 | 7.7969 |
| 17846 | ENSMUSG00000064044 | 0 | NA | NA | NA | NA | NA | Olfr319 | olfactory receptor 319 [Source:MGI Symb | 0 | 0 | 0 | 0 | 0 | 0 |
| 17847 | ENSMUSG00000064052 | 0 | NA | NA | NA | NA | NA | Gm5089 | predicted gene 5089 [Source:MGI Symbol | 0 | 0 | 0 | 0 | 0 | 0 |
| 17848 | ENSMUSG00000064057 | 1.697273971 | -0.555809743 | 1.798872636 | -0.308976707 | 0.757339246 | NA | Scgb3a1 | secretoglobin, family 3A, member 1 [Sour | 4.0155 | 0 | 2.0125 | 1.928 | 0 | 2.2277 |
| 17849 | ENSMUSG00000064061 | 39.35370585 | -0.137202648 | 0.360167641 | -0.380941075 | 0.703246973 | 0.942506832 | Dzip3 | DAZ interacting protein 3, zinc finger [Sou | 44.171 | 45.228 | 34.212 | 36.631 | 45.806 | 30.074 |
| 17850 | ENSMUSG00000064063 | 0 | NA | NA | NA | NA | NA | BC048507 | cDNA sequence BC048507 [Source:MGI Sy | 0 | 0 | 0 | 0 | 0 | 0 |
| 17851 | ENSMUSG00000064065 | 2.505564861 | -3.607905398 | 1.843712118 | -1.956870252 | 0.050362725 | NA | Ipcef1 | interaction protein for cytohesin exchang | 1.6062 | 11.307 | 1.0062 | 0 | 0 | 1.1138 |
| 17852 | ENSMUSG00000064068 | 428.8336074 | -0.114418031 | 0.137285819 | -0.833429348 | 0.404602629 | 0.829271989 | Mtx1 | metaxin 1 [Source:MGI Symbol;Acc:MGI:1 | 441.71 | 487.23 | 408.54 | 397.16 | 435.16 | 403.21 |
| 17853 | ENSMUSG00000064080 | 20.31585898 | -1.06614696 | 0.518264495 | -2.05714837 | 0.039671956 | 0.272725843 | Fbln2 | fibulin 2 [Source:MGI Symbol;Acc:MGI:95 | 37.746 | 21.586 | 23.144 | 17.352 | 9.8156 | 12.252 |
| 17854 | ENSMUSG00000064081 | 0 | NA | NA | NA | NA | NA | Gm8973 | predicted gene 8973 [Source:MGI Symbol | 0 | 0 | 0 | 0 | 0 | 0 |
| 17855 | ENSMUSG00000064084 | 0 | NA | NA | NA | NA | NA | Olfr1202 | olfactory receptor 1202 [Source:MGI Syml | 0 | 0 | 0 | 0 | 0 | 0 |
| 17856 | ENSMUSG00000064090 | 71.69659077 | -0.024046876 | 0.282630002 | -0.085082532 | 0.932195777 | 0.988420305 | Vrk2 | vaccinia related kinase 2 [Source:MGI Syn | 68.264 | 90.457 | 58.362 | 75.19 | 62.165 | 75.741 |
| 17857 | ENSMUSG00000064105 | 160.8844746 | -0.233956988 | 0.216504626 | -1.080609649 | 0.279870788 | 0.732834426 | Cnnm2 | cyclin M2 [Source:MGI Symbol;Acc:MGI:2 | 137.33 | 215.86 | 169.05 | 139.78 | 141.78 | 161.51 |
| 17858 | ENSMUSG00000064109 | 3.909353752 | -1.034963363 | 1.274707779 | -0.811922058 | 0.416836357 | NA | Hcst | hematopoietic cell signal transducer [Sour | 2.4093 | 11.307 | 2.0125 | 0 | 3.2719 | 4.4554 |
| 17859 | ENSMUSG00000064110 | 0 | NA | NA | NA | NA | NA | Olfr963 | olfactory receptor 963 [Source:MGI Symb | 0 | 0 | 0 | 0 | 0 | 0 |
| 17860 | ENSMUSG00000064115 | 0.167707431 | -0.796629176 | 4.080472857 | -0.195229623 | 0.84521319 | NA | Cadm2 | cell adhesion molecule 2 [Source:MGI Syn | 0 | 0 | 1.0062 | 0 | 0 | 0 |

| | | | | | | | | | | | | | | | |
|---|---|---|---|---|---|---|---|---|---|---|---|---|---|---|---|
| 17861 | ENSMUSG00000064120 | 2299.049075 | 0.047053068 | 0.094890067 | 0.495869272 | 0.619986647 | 0.917977899 | Mocs1 | molybdenum cofactor synthesis 1 [Source | 2153.1 | 2290.2 | 2342.5 | 2430.2 | 2232.5 | 2345.8 |
| 17862 | ENSMUSG00000064121 | 0 | NA | NA | NA | NA | NA | Olfr96 | olfactory receptor 96 [Source:MGI Symbo | 0 | 0 | 0 | 0 | 0 | 0 |
| 17863 | ENSMUSG00000064125 | 6.367860311 | 0.67941088 | 0.946253177 | 0.718001161 | 0.472756571 | 0.86857093 | Prr36 | proline rich 36 [Source:MGI Symbol;Acc:M | 5.6217 | 1.0279 | 8.05 | 12.532 | 7.6343 | 3.3415 |
| 17864 | ENSMUSG00000064127 | 301.2966083 | -0.220667599 | 0.179224406 | -1.231236324 | 0.218234489 | 0.663697152 | Med14 | mediator complex subunit 14 [Source:MG | 259.4 | 390.61 | 324.01 | 294.01 | 283.56 | 256.18 |
| 17865 | ENSMUSG00000064128 | 74.54410048 | 0.340736541 | 0.32998274 | 1.032588981 | 0.301796287 | 0.752246851 | Cenpj | centromere protein J [Source:MGI Symbo | 83.523 | 71.954 | 41.256 | 57.839 | 104.7 | 87.994 |
| 17866 | ENSMUSG00000064129 | 0 | NA | NA | NA | NA | NA | Mageb5b | MAGE family member B5B [Source:MGI Sy | 0 | 0 | 0 | 0 | 0 | 0 |
| 17867 | ENSMUSG00000064137 | 0 | NA | NA | NA | NA | NA | Rhox8 | reproductive homeobox 8 [Source:MGI Sy | 0 | 0 | 0 | 0 | 0 | 0 |
| 17868 | ENSMUSG00000064138 | 257.1628734 | 0.164252786 | 0.196603347 | 0.835452646 | 0.403462898 | 0.828953151 | Fam172a | family with sequence similarity 172, meml | 248.96 | 234.37 | 243.51 | 195.69 | 311.92 | 308.53 |
| 17869 | ENSMUSG00000064140 | 0.294513347 | 0.165143496 | 4.080472857 | 0.040471657 | 0.967717103 | NA | Trim38 | tripartite motif-containing 38 [Source:MG | 0.8031 | 0 | 0 | 0.964 | 0 | 0 |
| 17870 | ENSMUSG00000064141 | 20.66302837 | 0.4014917 | 0.543611151 | 0.738564136 | 0.460171709 | 0.859817816 | Zfp69 | zinc finger protein 69 [Source:MGI Symbo | 12.047 | 19.53 | 22.137 | 33.739 | 10.906 | 25.618 |
| 17871 | ENSMUSG00000064145 | 885.8209682 | 0.113279461 | 0.118488306 | 0.956039159 | 0.339052451 | 0.780887145 | Arih2 | ariadne RBR E3 ubiquitin protein ligase 2 [ | 808.72 | 852.14 | 893.55 | 1013.1 | 900.85 | 846.52 |
| 17872 | ENSMUSG00000064147 | 0.569258551 | -2.569337407 | 3.18860282 | -0.805787849 | 0.420365185 | NA | Rab44 | RAB44, member RAS oncogene family [So | 2.4093 | 0 | 1.0062 | 0 | 0 | 0 |
| 17873 | ENSMUSG00000064156 | 0 | NA | NA | NA | NA | NA | Prol1 | proline rich, lacrimal 1 [Source:MGI Symb | 0 | 0 | 0 | 0 | 0 | 0 |
| 17874 | ENSMUSG00000064158 | 1.138729733 | 1.305140952 | 2.579810336 | 0.505905777 | 0.612922797 | NA | Izumo1 | izumo sperm-egg fusion 1 [Source:MGI Sy | 0 | 0 | 2.0125 | 4.8199 | 0 | 0 |
| 17875 | ENSMUSG00000064165 | 0 | NA | NA | NA | NA | NA | Krt39 | keratin 39 [Source:MGI Symbol;Acc:MGI:3 | 0 | 0 | 0 | 0 | 0 | 0 |
| 17876 | ENSMUSG00000064177 | 1.611084216 | 0.213116484 | 1.602158528 | 0.13301835 | 0.894178868 | NA | Ghrl | ghrelin [Source:MGI Symbol;Acc:MGI:193 | 2.4093 | 1.0279 | 1.0062 | 1.928 | 2.1812 | 1.1138 |
| 17877 | ENSMUSG00000064179 | 14.09223516 | -0.11671822 | 0.599479426 | -0.194699292 | 0.845628366 | 0.972803041 | Tnnt1 | troponin T1, skeletal, slow [Source:MGI Sy | 9.6372 | 18.503 | 16.1 | 18.316 | 13.087 | 8.9108 |
| 17878 | ENSMUSG00000064181 | 585.1489676 | 0.238005599 | 0.119856159 | 1.985760276 | 0.047059949 | 0.300386185 | Rab3ip | RAB3A interacting protein [Source:MGI Sy | 567.79 | 529.38 | 513.18 | 632.37 | 651.1 | 617.07 |
| 17879 | ENSMUSG00000064193 | 4.085042072 | 0.275670252 | 1.094908536 | 0.251774685 | 0.801215225 | NA | Gm4735 | predicted gene 4735 [Source:MGI Symbol | 4.0155 | 2.0558 | 5.0312 | 6.7478 | 1.0906 | 5.5692 |
| 17880 | ENSMUSG00000064194 | 0 | NA | NA | NA | NA | NA | Zfp936 | zinc finger protein 936 [Source:MGI Symb | 0 | 0 | 0 | 0 | 0 | 0 |
| 17881 | ENSMUSG00000064201 | 0 | NA | NA | NA | NA | NA | Krt2 | keratin 2 [Source:MGI Symbol;Acc:MGI:96 | 0 | 0 | 0 | 0 | 0 | 0 |
| 17882 | ENSMUSG00000064202 | 30.92226387 | 0.170230977 | 0.405287269 | 0.420025473 | 0.674466845 | 0.932114677 | Spata6l | spermatogenesis associated 6 like [Source | 31.321 | 30.838 | 25.156 | 38.559 | 22.903 | 36.757 |
| 17883 | ENSMUSG00000064208 | 1.704214689 | 0.73303919 | 1.590512199 | 0.460882469 | 0.64488293 | NA | Gm10145 | predicted gene 10145 [Source:MGI Symbo | 0.8031 | 2.0558 | 1.0062 | 1.928 | 1.0906 | 3.3415 |
| 17884 | ENSMUSG00000064210 | 78.66876064 | -0.735353713 | 0.307728265 | -2.389620312 | 0.016865799 | 0.154222752 | Ano6 | anoctamin 6 [Source:MGI Symbol;Acc:MG | 112.43 | 116.15 | 66.412 | 73.262 | 44.715 | 59.034 |
| 17885 | ENSMUSG00000064213 | 0 | NA | NA | NA | NA | NA | Defa24 | defensin, alpha, 24 [Source:MGI Symbol;A | 0 | 0 | 0 | 0 | 0 | 0 |
| 17886 | ENSMUSG00000064215 | 2216.663719 | 0.166317658 | 0.152285002 | 1.092147328 | 0.27476835 | 0.729487777 | Ifi27 | interferon, alpha-inducible protein 27 [So | 2207.7 | 1716.6 | 2341.5 | 1953 | 2261.9 | 2819.1 |
| 17887 | ENSMUSG00000064220 | 0 | NA | NA | NA | NA | NA | H2ac18 | H2A clustered histone 18 [Source:MGI Syr | 0 | 0 | 0 | 0 | 0 | 0 |
| 17888 | ENSMUSG00000064223 | 0 | NA | NA | NA | NA | NA | Olfr694 | olfactory receptor 694 [Source:MGI Symb | 0 | 0 | 0 | 0 | 0 | 0 |
| 17889 | ENSMUSG00000064224 | 0 | NA | NA | NA | NA | NA | Gsdma3 | gasdermin A3 [Source:MGI Symbol;Acc:M | 0 | 0 | 0 | 0 | 0 | 0 |
| 17890 | ENSMUSG00000064225 | 7436.484527 | 0.112286859 | 0.167506863 | 0.670341841 | 0.5026399 | 0.881220881 | Paqr9 | progestin and adipoQ receptor family mer | 8404.5 | 7314.7 | 5722.5 | 9566.5 | 7656.1 | 5954.6 |
| 17891 | ENSMUSG00000064231 | 0 | NA | NA | NA | NA | NA | Gm5321 | predicted gene 5321 [Source:MGI Symbol | 0 | 0 | 0 | 0 | 0 | 0 |
| 17892 | ENSMUSG00000064232 | 0 | NA | NA | NA | NA | NA | Gm5414 | predicted gene 5414 [Source:MGI Symbol | 0 | 0 | 0 | 0 | 0 | 0 |
| 17893 | ENSMUSG00000064246 | 3.071004613 | -2.265170367 | 1.358021898 | -1.667992519 | 0.095317212 | NA | Chil1 | chitinase-like 1 [Source:MGI Symbol;Acc:M | 4.0155 | 8.2233 | 3.0187 | 0.964 | 1.0906 | 1.1138 |
| 17894 | ENSMUSG00000064247 | 32.2003286 | -0.462028642 | 0.432289408 | -1.068794732 | 0.285162173 | 0.737769696 | Plcxd1 | phosphatidylinositol-specific phospholipa | 40.958 | 38.033 | 33.206 | 42.415 | 18.541 | 20.049 |
| 17895 | ENSMUSG00000064252 | 0 | NA | NA | NA | NA | NA | Olfr329 | olfactory receptor 329 [Source:MGI Symb | 0 | 0 | 0 | 0 | 0 | 0 |
| 17896 | ENSMUSG00000064254 | 573.6301112 | 0.269720027 | 0.165805838 | 1.626722139 | 0.103796125 | 0.474047094 | Ethe1 | ethylmalonic encephalopathy 1 [Source:M | 570.2 | 427.61 | 561.48 | 534.04 | 595.48 | 752.96 |
| 17897 | ENSMUSG00000064259 | 0 | NA | NA | NA | NA | NA | Vmn1r13 | vomeronasal 1 receptor 13 [Source:MGI S | 0 | 0 | 0 | 0 | 0 | 0 |
| 17898 | ENSMUSG00000064262 | 41.94333022 | -0.538893639 | 0.404063181 | -1.333686572 | 0.182306597 | 0.61573286 | Gimap8 | GTPase, IMAP family member 8 [Source:M | 46.58 | 55.508 | 47.293 | 49.163 | 16.359 | 36.757 |
| 17899 | ENSMUSG00000064263 | 0 | NA | NA | NA | NA | NA | Platr26 | pluripotency associated transcript 26 [Sou | 0 | 0 | 0 | 0 | 0 | 0 |
| 17900 | ENSMUSG00000064264 | 28.81277712 | 0.553916936 | 0.417364389 | 1.327178241 | 0.184449728 | 0.619137675 | Zfp428 | zinc finger protein 428 [Source:MGI Symb | 23.29 | 23.642 | 23.144 | 37.595 | 23.994 | 41.212 |
| 17901 | ENSMUSG00000064267 | 6.80827664 | -0.416706129 | 0.899463307 | -0.463283078 | 0.643161474 | 0.925364325 | Hvcn1 | hydrogen voltage-gated channel 1 [Source | 7.2279 | 5.1396 | 11.069 | 9.6398 | 1.0906 | 6.6831 |
| 17902 | ENSMUSG00000064272 | 0 | NA | NA | NA | NA | NA | Gpbar1 | G protein-coupled bile acid receptor 1 [So | 0 | 0 | 0 | 0 | 0 | 0 |
| 17903 | ENSMUSG00000064280 | 6.022812663 | 0.342713156 | 0.947038589 | 0.361878766 | 0.717442628 | 0.9447719 | Ccdc146 | coiled-coil domain containing 146 [Source | 4.8186 | 1.0279 | 10.062 | 4.8199 | 8.7249 | 6.6831 |
| 17904 | ENSMUSG00000064281 | 0.709843051 | 3.034749291 | 2.67440342 | 1.134738786 | 0.256484781 | NA | Rpl19-ps1 | ribosomal protein L19, pseudogene 1 [Sou | 0 | 0 | 0 | 0.964 | 2.1812 | 1.1138 |
| 17905 | ENSMUSG00000064284 | 75.88583206 | 0.376337194 | 0.285067334 | 1.320169479 | 0.18677844 | 0.623054278 | Cdpf1 | cysteine rich, DPF motif domain containin | 63.445 | 55.508 | 79.493 | 106.04 | 82.887 | 67.944 |
| 17906 | ENSMUSG00000064288 | 1.241373611 | 0.346687691 | 2.239815274 | 0.154784055 | 0.876991562 | NA | H4c12 | H4 clustered histone 12 [Source:MGI Sym | 3.2124 | 0 | 0 | 0.964 | 3.2719 | 0 |
| 17907 | ENSMUSG00000064289 | 368.6596956 | -0.032251638 | 0.161755019 | -0.1993857 | 0.841961046 | 0.971854408 | Tank | TRAF family member-associated Nf-kappa | 425.64 | 336.13 | 356.21 | 405.83 | 376.26 | 311.88 |
| 17908 | ENSMUSG00000064293 | 0 | NA | NA | NA | NA | NA | Cntn4 | contactin 4 [Source:MGI Symbol;Acc:MGI | 0 | 0 | 0 | 0 | 0 | 0 |
| 17909 | ENSMUSG00000064294 | 10080.00744 | 0.653395805 | 0.110544672 | 5.910694708 | 3.41E-09 | 6.01E-07 | Aox3 | aldehyde oxidase 3 [Source:MGI Symbol;A | 8205.3 | 7678.5 | 7622.3 | 11984 | 14092 | 10898 |
| 17910 | ENSMUSG00000064299 | 0 | NA | NA | NA | NA | NA | 4921528I07R | RIKEN cDNA 4921528I07 gene [Source:MG | 0 | 0 | 0 | 0 | 0 | 0 |
| 17911 | ENSMUSG00000064302 | 256.8503237 | 0.032827628 | 0.157413769 | 0.208543563 | 0.834804572 | 0.971073158 | Clasp1 | CLIP associating protein 1 [Source:MGI Sy | 255.39 | 263.15 | 243.51 | 275.7 | 266.11 | 237.25 |
| 17912 | ENSMUSG00000064307 | 11.51118827 | -0.217115676 | 0.690900949 | -0.314250076 | 0.753331112 | 0.952989051 | Lrrc51 | leucine rich repeat containing 51 [Source: | 16.865 | 5.1396 | 15.094 | 15.424 | 7.6343 | 8.9108 |
| 17913 | ENSMUSG00000064308 | 0 | NA | NA | NA | NA | NA | 2410137M14 | RIKEN cDNA 2410137M14 gene [Source:M | 0 | 0 | 0 | 0 | 0 | 0 |
| 17914 | ENSMUSG00000064310 | 0 | NA | NA | NA | NA | NA | Zpld1 | zona pellucida like domain containing 1 [S | 0 | 0 | 0 | 0 | 0 | 0 |
| 17915 | ENSMUSG00000064317 | 2.026879234 | 0.186052606 | 1.443696135 | 0.128872414 | 0.897458605 | NA | Gm10146 | predicted gene 10146 [Source:MGI Symbo | 1.6062 | 2.0558 | 2.0125 | 0.964 | 2.1812 | 3.3415 |
| 17916 | ENSMUSG00000064325 | 37.1753287 | -1.912244204 | 0.424745499 | -4.502094094 | 6.73E-06 | 0.000398765 | Hhip | Hedgehog-interacting protein [Source:MG | 44.171 | 53.452 | 78.487 | 10.604 | 19.631 | 16.708 |

| | | | | | | | | | | | | | | | |
|---|---|---|---|---|---|---|---|---|---|---|---|---|---|---|---|
| 17917 | ENSMUSG00000064326 | 83.81616291 | 0.288574219 | 0.306181664 | 0.942493471 | 0.345940052 | 0.786899236 | Siva1 | SIVA1, apoptosis-inducing factor [Source:I | 70.673 | 72.982 | 82.512 | 57.839 | 129.78 | 89.108 |
| 17918 | ENSMUSG00000064329 | 0 | NA | NA | NA | NA | NA | Scn1a | sodium channel, voltage-gated, type I, alp | 0 | 0 | 0 | 0 | 0 | 0 |
| 17919 | ENSMUSG00000064330 | 0.994779605 | 1.370136611 | 2.701221166 | 0.507228593 | 0.611994435 | NA | Pde6h | phosphodiesterase 6H, cGMP-specific, coı | 1.6062 | 0 | 0 | 0 | 4.3625 | 0 |
| 17920 | ENSMUSG00000064333 | 0 | NA | NA | NA | NA | NA | Olfr926 | olfactory receptor 926 [Source:MGI Symb | 0 | 0 | 0 | 0 | 0 | 0 |
| 17921 | ENSMUSG00000064336 | 0 | NA | NA | NA | NA | NA | mt-Tf | mitochondrially encoded tRNA phenylalar | 0 | 0 | 0 | 0 | 0 | 0 |
| 17922 | ENSMUSG00000064337 | 8353.026297 | 0.534281679 | 0.166435991 | 3.210133067 | 0.001326735 | 0.025792877 | mt-Rnr1 | mitochondrially encoded 12S rRNA [Sourc | 7276.9 | 6746.2 | 6447 | 7144 | 12925 | 9579.1 |
| 17923 | ENSMUSG00000064338 | 0.765090549 | -0.195763385 | 2.933488969 | -0.066733977 | 0.946793485 | NA | mt-Tv | mitochondrially encoded tRNA valine [Sou | 2.4093 | 0 | 0 | 0 | 2.1812 | 0 |
| 17924 | ENSMUSG00000064339 | 12009.25417 | 0.445422022 | 0.156884989 | 2.839162775 | 0.004523208 | 0.062486797 | mt-Rnr2 | mitochondrially encoded 16S rRNA [Sourc | 10583 | 11281 | 8644.6 | 11907 | 17795 | 11845 |
| 17925 | ENSMUSG00000064340 | 86.32156951 | 0.734828705 | 0.291582738 | 2.520137885 | 0.011730887 | 0.120765517 | mt-Tl1 | mitochondrially encoded tRNA leucine 1 [ | 71.476 | 54.48 | 68.425 | 130.14 | 123.24 | 70.172 |
| 17926 | ENSMUSG00000064341 | 91262.62369 | -0.188818711 | 0.082563226 | -2.286958984 | 0.022198217 | 0.186216956 | mt-Nd1 | mitochondrially encoded NADH dehydrog | 98645 | 99425 | 93608 | 82888 | 89136 | 83874 |
| 17927 | ENSMUSG00000064342 | 10.04506479 | 0.441497819 | 0.692873594 | 0.637198218 | 0.523995739 | 0.888334265 | mt-Ti | mitochondrially encoded tRNA isoleucine | 11.243 | 4.1117 | 10.062 | 10.604 | 11.997 | 12.252 |
| 17928 | ENSMUSG00000064343 | 11.6337239 | 0.723063551 | 0.630603335 | 1.146621831 | 0.25153795 | 0.703994757 | mt-Tq | mitochondrially encoded tRNA glutamine | 8.031 | 9.2513 | 9.0562 | 12.532 | 11.997 | 18.935 |
| 17929 | ENSMUSG00000064344 | 18.01391176 | 0.565883147 | 0.564290039 | 1.002823207 | 0.315946169 | 0.762646719 | mt-Tm | mitochondrially encoded tRNA methionin | 11.243 | 10.279 | 22.137 | 18.316 | 31.628 | 14.48 |
| 17930 | ENSMUSG00000064345 | 108691.2442 | 0.092141413 | 0.086285215 | 1.067870236 | 0.28557905 | 0.738065506 | mt-Nd2 | mitochondrially encoded NADH dehydrog | 106967 | 103066 | 105631 | 106199 | 119967 | 110318 |
| 17931 | ENSMUSG00000064346 | 63.76970318 | 0.176169686 | 0.312410626 | 0.563904269 | 0.572819285 | 0.905173631 | mt-Tw | mitochondrially encoded tRNA tryptophai | 78.704 | 46.256 | 54.337 | 80.01 | 67.618 | 55.692 |
| 17932 | ENSMUSG00000064347 | 6.13264008 | -0.036410695 | 0.898882146 | -0.040506639 | 0.967689214 | 0.994231936 | mt-Ta | mitochondrially encoded tRNA alanine [Sc | 2.4093 | 10.279 | 6.0375 | 4.8199 | 5.4531 | 7.7969 |
| 17933 | ENSMUSG00000064348 | 400.2130756 | 0.669216394 | 0.571737329 | 1.170496241 | 0.241801326 | 0.691791181 | mt-Tn | mitochondrially encoded tRNA asparagine | 454.56 | 296.04 | 176.09 | 262.2 | 446.06 | 766.32 |
| 17934 | ENSMUSG00000064349 | 525.7509926 | 0.61913916 | 0.487521192 | 1.269973838 | 0.20409395 | 0.649416772 | mt-Tc | mitochondrially encoded tRNA cysteine [S | 647.3 | 352.58 | 243.51 | 434.75 | 655.46 | 820.9 |
| 17935 | ENSMUSG00000064350 | 145.3216238 | 0.511852394 | 0.220974297 | 2.316343574 | 0.020539517 | 0.176557791 | mt-Ty | mitochondrially encoded tRNA tyrosine [S | 146.97 | 117.18 | 94.587 | 177.37 | 183.22 | 152.6 |
| 17936 | ENSMUSG00000064351 | 398809.8332 | -0.024511684 | 0.094344365 | -0.259810791 | 0.795009727 | 0.962189806 | mt-Co1 | mitochondrially encoded cytochrome c ox | 423662 | 399228 | 383701 | 360583 | 408834 | 416852 |
| 17937 | ENSMUSG00000064352 | 0.353348078 | 0.165142701 | 3.895042568 | 0.042398176 | 0.966181282 | NA | mt-Ts1 | mitochondrially encoded tRNA serine 1 [S | 0 | 0 | 1.0062 | 0 | 0 | 1.1138 |
| 17938 | ENSMUSG00000064353 | 0 | NA | NA | NA | NA | NA | mt-Td | mitochondrially encoded tRNA aspartic ac | 0 | 0 | 0 | 0 | 0 | 0 |
| 17939 | ENSMUSG00000064354 | 480.7067975 | 0.319599618 | 0.158071771 | 2.021863965 | 0.043190407 | 0.286918392 | mt-Co2 | mitochondrially encoded cytochrome c ox | 426.45 | 411.17 | 444.76 | 445.36 | 507.14 | 649.37 |
| 17940 | ENSMUSG00000064355 | 0 | NA | NA | NA | NA | NA | mt-Tk | mitochondrially encoded tRNA lysine [Sou | 0 | 0 | 0 | 0 | 0 | 0 |
| 17941 | ENSMUSG00000064356 | 16.51360362 | 0.226451454 | 0.555411288 | 0.407718494 | 0.683480358 | 0.9354523 | mt-Atp8 | mitochondrially encoded ATP synthase 8 [ | 12.047 | 20.558 | 13.081 | 12.532 | 16.359 | 24.505 |
| 17942 | ENSMUSG00000064357 | 500.8439935 | 0.075774641 | 0.137211286 | 0.552247876 | 0.580778537 | 0.908645447 | mt-Atp6 | mitochondrially encoded ATP synthase 6 [ | 507.56 | 460.51 | 494.07 | 451.14 | 564.94 | 526.85 |
| 17943 | ENSMUSG00000064358 | 817.4976094 | -1.01894912 | 0.717838724 | -1.41946803 | 0.155762611 | 0.574732247 | mt-Co3 | mitochondrially encoded cytochrome c ox | 2258.3 | 541.71 | 484 | 474.28 | 550.76 | 595.91 |
| 17944 | ENSMUSG00000064359 | 44.79466899 | -0.633450277 | 0.378786128 | -1.672316461 | 0.094461935 | 0.452866235 | mt-Tg | mitochondrially encoded tRNA glycine [Sc | 69.87 | 50.368 | 43.269 | 50.127 | 26.175 | 28.96 |
| 17945 | ENSMUSG00000064360 | 716.4691685 | -0.195575295 | 0.159794908 | -1.223914438 | 0.220984507 | 0.667400313 | mt-Nd3 | mitochondrially encoded NADH dehydrog | 897.07 | 805.89 | 590.67 | 602.49 | 714.35 | 688.36 |
| 17946 | ENSMUSG00000064361 | 15.48194577 | 0.155404003 | 0.666342659 | 0.233219352 | 0.815591079 | 0.967767553 | mt-Tr | mitochondrially encoded tRNA arginine [S | 23.29 | 17.475 | 3.0187 | 18.316 | 18.541 | 12.252 |
| 17947 | ENSMUSG00000064363 | 75210.53012 | -0.000711773 | 0.085271304 | -0.008347162 | 0.993340005 | 0.99859176 | mt-Nd4 | mitochondrially encoded NADH dehydrog | 77499 | 74512 | 73676 | 73687 | 79923 | 71967 |
| 17948 | ENSMUSG00000064364 | 53.35196812 | -0.121670935 | 0.334430103 | -0.363815739 | 0.715995609 | 0.9447719 | mt-Th | mitochondrially encoded tRNA histidine [! | 62.642 | 66.815 | 37.231 | 51.091 | 58.893 | 43.44 |
| 17949 | ENSMUSG00000064365 | 71.3120262 | -0.330502599 | 0.282776584 | -1.168776402 | 0.24249373 | 0.692000002 | mt-Ts2 | mitochondrially encoded tRNA serine 2 [S | 88.341 | 87.373 | 62.387 | 58.803 | 74.162 | 56.806 |
| 17950 | ENSMUSG00000064366 | 355.0496702 | -0.051792104 | 0.203632857 | -0.254340607 | 0.799232431 | 0.962424862 | mt-Tl2 | mitochondrially encoded tRNA leucine 2 [ | 436.08 | 397.8 | 249.55 | 318.11 | 404.62 | 324.13 |
| 17951 | ENSMUSG00000064367 | 82777.54614 | -0.094663423 | 0.134074561 | -0.706050592 | 0.480156676 | 0.870278888 | mt-Nd5 | mitochondrially encoded NADH dehydrog | 92170 | 95457 | 68849 | 75756 | 90825 | 73608 |
| 17952 | ENSMUSG00000064368 | 20865.01957 | 0.074665023 | 0.146985219 | 0.507976406 | 0.61146989 | 0.916376001 | mt-Nd6 | mitochondrially encoded NADH dehydrog | 22225 | 22874 | 15876 | 19529 | 25117 | 19569 |
| 17953 | ENSMUSG00000064369 | 518.9866676 | -0.02137264 | 0.224077717 | -0.095380481 | 0.92401262 | 0.987408571 | mt-Te | mitochondrially encoded tRNA glutamic a | 624.81 | 546.85 | 396.46 | 532.12 | 676.18 | 337.49 |
| 17954 | ENSMUSG00000064370 | 187819.9239 | -0.054714129 | 0.098752893 | -0.554050899 | 0.579544012 | 0.908350165 | mt-Cytb | mitochondrially encoded cytochrome b [S | 204883 | 189322 | 179937 | 167617 | 197584 | 187577 |
| 17955 | ENSMUSG00000064371 | 1.209480962 | -0.360464025 | 2.290296351 | -0.157387504 | 0.874939461 | NA | mt-Tt | mitochondrially encoded tRNA threonine | 0 | 4.1117 | 0 | 0.964 | 2.1812 | 0 |
| 17956 | ENSMUSG00000064372 | 1970.939142 | 0.320210043 | 0.108290922 | 2.956942625 | 0.003107059 | 0.047809048 | mt-Tp | mitochondrially encoded tRNA proline [Sc | 1731.5 | 1785.5 | 1741.8 | 1956.9 | 2416.8 | 2193.2 |
| 17957 | ENSMUSG00000064373 | 106860.5371 | -0.216147121 | 0.087440144 | -2.471943783 | 0.013438065 | 0.132518282 | Selenop | selenoprotein P [Source:MGI Symbol;Acc: | 108802 | 115137 | 120615 | 104076 | 97318 | 95216 |
| 17958 | ENSMUSG00000064377 | 0 | NA | NA | NA | NA | NA | Gm24966 | predicted gene, 24966 [Source:MGI Symb | 0 | 0 | 0 | 0 | 0 | 0 |
| 17959 | ENSMUSG00000064380 | 1.645147864 | -1.074581611 | 1.622213076 | -0.662417056 | 0.507703976 | NA | Gm26448 | predicted gene, 26448 [Source:MGI Symb | 1.6062 | 3.0838 | 2.0125 | 0.964 | 1.0906 | 1.1138 |
| 17960 | ENSMUSG00000064382 | 0 | NA | NA | NA | NA | NA | Gm26447 | predicted gene, 26447 [Source:MGI Symb | 0 | 0 | 0 | 0 | 0 | 0 |
| 17961 | ENSMUSG00000064386 | 0 | NA | NA | NA | NA | NA | Gm26449 | predicted gene, 26449 [Source:MGI Symb | 0 | 0 | 0 | 0 | 0 | 0 |
| 17962 | ENSMUSG00000064387 | 0 | NA | NA | NA | NA | NA | Snora73a | small nucleolar RNA, H/ACA box 73a [Sour | 0 | 0 | 0 | 0 | 0 | 0 |
| 17963 | ENSMUSG00000064389 | 0.328370405 | 0.16514316 | 4.003149536 | 0.041253308 | 0.967093956 | NA | Gm26446 | predicted gene, 26446 [Source:MGI Symb | 0 | 0 | 1.0062 | 0.964 | 0 | 0 |
| 17964 | ENSMUSG00000064390 | 3.233373562 | 1.413382041 | 1.315142727 | 1.074698596 | 0.282509693 | NA | Snord73b | small nucleolar RNA, C/D box U73B [Sourc | 3.2124 | 0 | 2.0125 | 8.6758 | 3.2719 | 2.2277 |
| 17965 | ENSMUSG00000064391 | 0 | NA | NA | NA | NA | NA | Gm23625 | predicted gene, 23625 [Source:MGI Symb | 0 | 0 | 0 | 0 | 0 | 0 |
| 17966 | ENSMUSG00000064392 | 0 | NA | NA | NA | NA | NA | Gm23626 | predicted gene, 23626 [Source:MGI Symb | 0 | 0 | 0 | 0 | 0 | 0 |
| 17967 | ENSMUSG00000064393 | 0 | NA | NA | NA | NA | NA | NA | NA | 0 | 0 | 0 | 0 | 0 | 0 |
| 17968 | ENSMUSG00000064394 | 0 | NA | NA | NA | NA | NA | Gm23622 | predicted gene, 23622 [Source:MGI Symb | 0 | 0 | 0 | 0 | 0 | 0 |
| 17969 | ENSMUSG00000064395 | 0 | NA | NA | NA | NA | NA | Gm23623 | predicted gene, 23623 [Source:MGI Symb | 0 | 0 | 0 | 0 | 0 | 0 |
| 17970 | ENSMUSG00000064397 | 0 | NA | NA | NA | NA | NA | NA | NA | 0 | 0 | 0 | 0 | 0 | 0 |
| 17971 | ENSMUSG00000064398 | 1.017133346 | -0.93167857 | 2.468499519 | -0.377427082 | 0.705856244 | NA | Gm23628 | predicted gene, 23628 [Source:MGI Symb | 0 | 0 | 4.025 | 0.964 | 0 | 1.1138 |
| 17972 | ENSMUSG00000064399 | 0 | NA | NA | NA | NA | NA | Gm23629 | predicted gene, 23629 [Source:MGI Symb | 0 | 0 | 0 | 0 | 0 | 0 |

| | | | | | | | | | | | | | | | |
|---|---|---|---|---|---|---|---|---|---|---|---|---|---|---|---|
| 17973 | ENSMUSG00000064400 | 0 | NA | NA | NA | NA | NA | Snord3b-ps1 | small nucleolar RNA, C/D box 3B, pseudog | 0 | 0 | 0 | 0 | 0 | 0 |
| 17974 | ENSMUSG00000064403 | 0 | NA | NA | NA | NA | NA | Gm23928 | predicted gene, 23928 [Source:MGI Symb | 0 | 0 | 0 | 0 | 0 | 0 |
| 17975 | ENSMUSG00000064405 | 0 | NA | NA | NA | NA | NA | Gm23925 | predicted gene, 23925 [Source:MGI Symb | 0 | 0 | 0 | 0 | 0 | 0 |
| 17976 | ENSMUSG00000064407 | 0 | NA | NA | NA | NA | NA | Gm23926 | predicted gene, 23926 [Source:MGI Symb | 0 | 0 | 0 | 0 | 0 | 0 |
| 17977 | ENSMUSG00000064408 | 0 | NA | NA | NA | NA | NA | Gm23924 | predicted gene, 23924 [Source:MGI Symb | 0 | 0 | 0 | 0 | 0 | 0 |
| 17978 | ENSMUSG00000064409 | 0.328370405 | 0.16514316 | 4.003149536 | 0.041253308 | 0.967093956 | NA | NA | NA | 0 | 0 | 1.0062 | 0.964 | 0 | 0 |
| 17979 | ENSMUSG00000064410 | 0 | NA | NA | NA | NA | NA | Gm22247 | predicted gene, 22247 [Source:MGI Symb | 0 | 0 | 0 | 0 | 0 | 0 |
| 17980 | ENSMUSG00000064412 | 0 | NA | NA | NA | NA | NA | NA | NA | 0 | 0 | 0 | 0 | 0 | 0 |
| 17981 | ENSMUSG00000064413 | 0.690772315 | 1.713484714 | 3.063368077 | 0.559346664 | 0.575925154 | NA | Gm22245 | predicted gene, 22245 [Source:MGI Symb | 0.8031 | 0 | 0 | 0 | 0 | 3.3415 |
| 17982 | ENSMUSG00000064414 | 0 | NA | NA | NA | NA | NA | NA | NA | 0 | 0 | 0 | 0 | 0 | 0 |
| 17983 | ENSMUSG00000064416 | 0 | NA | NA | NA | NA | NA | Gm22248 | predicted gene, 22248 [Source:MGI Symb | 0 | 0 | 0 | 0 | 0 | 0 |
| 17984 | ENSMUSG00000064417 | 0 | NA | NA | NA | NA | NA | DQ267102 | snoRNA DQ267102 [Source:MGI Symbol;A | 0 | 0 | 0 | 0 | 0 | 0 |
| 17985 | ENSMUSG00000064419 | 0 | NA | NA | NA | NA | NA | Gm22249 | predicted gene, 22249 [Source:MGI Symb | 0 | 0 | 0 | 0 | 0 | 0 |
| 17986 | ENSMUSG00000064422 | 0.181769715 | 1.126924175 | 4.080472857 | 0.276174898 | 0.782413736 | NA | NA | NA | 0 | 0 | 0 | 0 | 1.0906 | 0 |
| 17987 | ENSMUSG00000064424 | 0 | NA | NA | NA | NA | NA | NA | NA | 0 | 0 | 0 | 0 | 0 | 0 |
| 17988 | ENSMUSG00000064427 | 3.61313502 | 0.782444907 | 1.337162792 | 0.585153065 | 0.5584448 | NA | Gm22748 | predicted gene, 22748 [Source:MGI Symb | 4.8186 | 2.0558 | 1.0062 | 2.8919 | 10.906 | 0 |
| 17989 | ENSMUSG00000064428 | 0 | NA | NA | NA | NA | NA | NA | NA | 0 | 0 | 0 | 0 | 0 | 0 |
| 17990 | ENSMUSG00000064429 | 0 | NA | NA | NA | NA | NA | NA | NA | 0 | 0 | 0 | 0 | 0 | 0 |
| 17991 | ENSMUSG00000064432 | 0 | NA | NA | NA | NA | NA | Gm25587 | predicted gene, 25587 [Source:MGI Symb | 0 | 0 | 0 | 0 | 0 | 0 |
| 17992 | ENSMUSG00000064433 | 0 | NA | NA | NA | NA | NA | NA | NA | 0 | 0 | 0 | 0 | 0 | 0 |
| 17993 | ENSMUSG00000064435 | 0 | NA | NA | NA | NA | NA | Gm25588 | predicted gene, 25588 [Source:MGI Symb | 0 | 0 | 0 | 0 | 0 | 0 |
| 17994 | ENSMUSG00000064436 | 0 | NA | NA | NA | NA | NA | Gm25589 | predicted gene, 25589 [Source:MGI Symb | 0 | 0 | 0 | 0 | 0 | 0 |
| 17995 | ENSMUSG00000064437 | 0.339027071 | -1.79174149 | 3.919585585 | -0.457125237 | 0.647581032 | NA | Snord49b | small nucleolar RNA, C/D box 49B [Source | 0 | 1.0279 | 1.0062 | 0 | 0 | 0 |
| 17996 | ENSMUSG00000064440 | 0 | NA | NA | NA | NA | NA | Gm26224 | predicted gene, 26224 [Source:MGI Symb | 0 | 0 | 0 | 0 | 0 | 0 |
| 17997 | ENSMUSG00000064441 | 0.496077836 | -0.812642266 | 3.444120572 | -0.235950586 | 0.813471014 | NA | Snord37 | small nucleolar RNA, C/D box 37 [Source:I | 0 | 0 | 2.0125 | 0.964 | 0 | 0 |
| 17998 | ENSMUSG00000064442 | 0 | NA | NA | NA | NA | NA | Gm26225 | predicted gene, 26225 [Source:MGI Symb | 0 | 0 | 0 | 0 | 0 | 0 |
| 17999 | ENSMUSG00000064443 | 0 | NA | NA | NA | NA | NA | Gm26226 | predicted gene, 26226 [Source:MGI Symb | 0 | 0 | 0 | 0 | 0 | 0 |
| 18000 | ENSMUSG00000064444 | 0 | NA | NA | NA | NA | NA | Gm26227 | predicted gene, 26227 [Source:MGI Symb | 0 | 0 | 0 | 0 | 0 | 0 |
| 18001 | ENSMUSG00000064446 | 0 | NA | NA | NA | NA | NA | Gm26228 | predicted gene, 26228 [Source:MGI Symb | 0 | 0 | 0 | 0 | 0 | 0 |
| 18002 | ENSMUSG00000064450 | 0 | NA | NA | NA | NA | NA | Snord68 | small nucleolar RNA, C/D box 68 [Source:I | 0 | 0 | 0 | 0 | 0 | 0 |
| 18003 | ENSMUSG00000064451 | 0.986888497 | -0.8704229 | 2.113455788 | -0.411848171 | 0.680450712 | NA | Snora23 | small nucleolar RNA, H/ACA box 23 [Sourc | 0.8031 | 1.0279 | 2.0125 | 0.964 | 0 | 1.1138 |
| 18004 | ENSMUSG00000064452 | 0 | NA | NA | NA | NA | NA | Gm24564 | predicted gene, 24564 [Source:MGI Symb | 0 | 0 | 0 | 0 | 0 | 0 |
| 18005 | ENSMUSG00000064453 | 0 | NA | NA | NA | NA | NA | Snord21 | small nucleolar RNA, C/D box 21 [Source:I | 0 | 0 | 0 | 0 | 0 | 0 |
| 18006 | ENSMUSG00000064455 | 0 | NA | NA | NA | NA | NA | NA | NA | 0 | 0 | 0 | 0 | 0 | 0 |
| 18007 | ENSMUSG00000064460 | 0 | NA | NA | NA | NA | NA | Gm25080 | predicted gene, 25080 [Source:MGI Symb | 0 | 0 | 0 | 0 | 0 | 0 |
| 18008 | ENSMUSG00000064463 | 0 | NA | NA | NA | NA | NA | Gm25079 | predicted gene, 25079 [Source:MGI Symb | 0 | 0 | 0 | 0 | 0 | 0 |
| 18009 | ENSMUSG00000064464 | 0 | NA | NA | NA | NA | NA | NA | NA | 0 | 0 | 0 | 0 | 0 | 0 |
| 18010 | ENSMUSG00000064467 | 0 | NA | NA | NA | NA | NA | NA | NA | 0 | 0 | 0 | 0 | 0 | 0 |
| 18011 | ENSMUSG00000064468 | 0 | NA | NA | NA | NA | NA | Gm25081 | predicted gene, 25081 [Source:MGI Symb | 0 | 0 | 0 | 0 | 0 | 0 |
| 18012 | ENSMUSG00000064471 | 0 | NA | NA | NA | NA | NA | Gm23406 | predicted gene, 23406 [Source:MGI Symb | 0 | 0 | 0 | 0 | 0 | 0 |
| 18013 | ENSMUSG00000064472 | 0.167707431 | -0.796629176 | 4.080472857 | -0.195229623 | 0.84521319 | NA | Gm23407 | predicted gene, 23407 [Source:MGI Symb | 0 | 0 | 1.0062 | 0 | 0 | 0 |
| 18014 | ENSMUSG00000064476 | 0 | NA | NA | NA | NA | NA | Gm23405 | predicted gene, 23405 [Source:MGI Symb | 0 | 0 | 0 | 0 | 0 | 0 |
| 18015 | ENSMUSG00000064477 | 0 | NA | NA | NA | NA | NA | Gm23404 | predicted gene, 23404 [Source:MGI Symb | 0 | 0 | 0 | 0 | 0 | 0 |
| 18016 | ENSMUSG00000064481 | 0 | NA | NA | NA | NA | NA | Gm23737 | predicted gene, 23737 [Source:MGI Symb | 0 | 0 | 0 | 0 | 0 | 0 |
| 18017 | ENSMUSG00000064485 | 0 | NA | NA | NA | NA | NA | Gm23735 | predicted gene, 23735 [Source:MGI Symb | 0 | 0 | 0 | 0 | 0 | 0 |
| 18018 | ENSMUSG00000064486 | 0 | NA | NA | NA | NA | NA | Gm23293 | predicted gene, 23293 [Source:MGI Symb | 0 | 0 | 0 | 0 | 0 | 0 |
| 18019 | ENSMUSG00000064487 | 0 | NA | NA | NA | NA | NA | Gm23736 | predicted gene, 23736 [Source:MGI Symb | 0 | 0 | 0 | 0 | 0 | 0 |
| 18020 | ENSMUSG00000064489 | 0 | NA | NA | NA | NA | NA | NA | NA | 0 | 0 | 0 | 0 | 0 | 0 |
| 18021 | ENSMUSG00000064490 | 0 | NA | NA | NA | NA | NA | Gm22283 | predicted gene, 22283 [Source:MGI Symb | 0 | 0 | 0 | 0 | 0 | 0 |
| 18022 | ENSMUSG00000064491 | 0 | NA | NA | NA | NA | NA | NA | NA | 0 | 0 | 0 | 0 | 0 | 0 |
| 18023 | ENSMUSG00000064492 | 0 | NA | NA | NA | NA | NA | Gm22060 | predicted gene, 22060 [Source:MGI Symb | 0 | 0 | 0 | 0 | 0 | 0 |
| 18024 | ENSMUSG00000064493 | 0 | NA | NA | NA | NA | NA | Snora28 | small nucleolar RNA, H/ACA box 28 [Sourc | 0 | 0 | 0 | 0 | 0 | 0 |
| 18025 | ENSMUSG00000064494 | 0.371281295 | 2.09253362 | 4.031852165 | 0.519000582 | 0.603760335 | NA | NA | NA | 0 | 0 | 0 | 0 | 0 | 2.2277 |
| 18026 | ENSMUSG00000064495 | 0 | NA | NA | NA | NA | NA | Gm24313 | predicted gene, 24313 [Source:MGI Symb | 0 | 0 | 0 | 0 | 0 | 0 |
| 18027 | ENSMUSG00000064496 | 0 | NA | NA | NA | NA | NA | DQ267101 | snoRNA DQ267101 [Source:MGI Symbol;A | 0 | 0 | 0 | 0 | 0 | 0 |
| 18028 | ENSMUSG00000064497 | 0 | NA | NA | NA | NA | NA | NA | NA | 0 | 0 | 0 | 0 | 0 | 0 |

| | | | | | | | | | | | | | | | |
|---|---|---|---|---|---|---|---|---|---|---|---|---|---|---|---|
| 18029 | ENSMUSG00000064500 | 0 | NA | NA | NA | NA | NA | Gm25296 | predicted gene, 25296 [Source:MGI Symb | 0 | 0 | 0 | 0 | 0 | 0 |
| 18030 | ENSMUSG00000064507 | 0 | NA | NA | NA | NA | NA | NA | NA | 0 | 0 | 0 | 0 | 0 | 0 |
| 18031 | ENSMUSG00000064508 | 1.649095906 | -1.176527232 | 1.762574148 | -0.6675051 | 0.504449553 | NA | NA | NA | 4.8186 | 1.0279 | 1.0062 | 1.928 | 0 | 1.1138 |
| 18032 | ENSMUSG00000064510 | 0 | NA | NA | NA | NA | NA | NA | NA | 0 | 0 | 0 | 0 | 0 | 0 |
| 18033 | ENSMUSG00000064511 | 0 | NA | NA | NA | NA | NA | Gm22459 | predicted gene, 22459 [Source:MGI Symb | 0 | 0 | 0 | 0 | 0 | 0 |
| 18034 | ENSMUSG00000064512 | 0 | NA | NA | NA | NA | NA | Gm22456 | predicted gene, 22456 [Source:MGI Symb | 0 | 0 | 0 | 0 | 0 | 0 |
| 18035 | ENSMUSG00000064513 | 0 | NA | NA | NA | NA | NA | Snora9 | small nucleolar RNA, H/ACA box 9 [Source | 0 | 0 | 0 | 0 | 0 | 0 |
| 18036 | ENSMUSG00000064514 | 0 | NA | NA | NA | NA | NA | Gm22455 | predicted gene, 22455 [Source:MGI Symb | 0 | 0 | 0 | 0 | 0 | 0 |
| 18037 | ENSMUSG00000064515 | 0 | NA | NA | NA | NA | NA | n-R5s162 | nuclear encoded rRNA 5S 162 [Source:MG | 0 | 0 | 0 | 0 | 0 | 0 |
| 18038 | ENSMUSG00000064517 | 0 | NA | NA | NA | NA | NA | Gm22454 | predicted gene, 22454 [Source:MGI Symb | 0 | 0 | 0 | 0 | 0 | 0 |
| 18039 | ENSMUSG00000064519 | 0 | NA | NA | NA | NA | NA | NA | NA | 0 | 0 | 0 | 0 | 0 | 0 |
| 18040 | ENSMUSG00000064522 | 0 | NA | NA | NA | NA | NA | Gm26442 | predicted gene, 26442 [Source:MGI Symb | 0 | 0 | 0 | 0 | 0 | 0 |
| 18041 | ENSMUSG00000064524 | 0 | NA | NA | NA | NA | NA | Gm26441 | predicted gene, 26441 [Source:MGI Symb | 0 | 0 | 0 | 0 | 0 | 0 |
| 18042 | ENSMUSG00000064525 | 0.267700746 | -1.497481649 | 4.057001436 | -0.369110456 | 0.712045396 | NA | Gm26440 | predicted gene, 26440 [Source:MGI Symb | 1.6062 | 0 | 0 | 0 | 0 | 0 |
| 18043 | ENSMUSG00000064526 | 0 | NA | NA | NA | NA | NA | Gm26439 | predicted gene, 26439 [Source:MGI Symb | 0 | 0 | 0 | 0 | 0 | 0 |
| 18044 | ENSMUSG00000064527 | 0 | NA | NA | NA | NA | NA | Gm26438 | predicted gene, 26438 [Source:MGI Symb | 0 | 0 | 0 | 0 | 0 | 0 |
| 18045 | ENSMUSG00000064529 | 0 | NA | NA | NA | NA | NA | Gm26437 | predicted gene, 26437 [Source:MGI Symb | 0 | 0 | 0 | 0 | 0 | 0 |
| 18046 | ENSMUSG00000064530 | 0 | NA | NA | NA | NA | NA | Gm23613 | predicted gene, 23613 [Source:MGI Symb | 0 | 0 | 0 | 0 | 0 | 0 |
| 18047 | ENSMUSG00000064532 | 0 | NA | NA | NA | NA | NA | Gm23614 | predicted gene, 23614 [Source:MGI Symb | 0 | 0 | 0 | 0 | 0 | 0 |
| 18048 | ENSMUSG00000064536 | 0 | NA | NA | NA | NA | NA | Gm23615 | predicted gene, 23615 [Source:MGI Symb | 0 | 0 | 0 | 0 | 0 | 0 |
| 18049 | ENSMUSG00000064540 | 0 | NA | NA | NA | NA | NA | Snord42a | small nucleolar RNA, C/D box 42A [Source | 0 | 0 | 0 | 0 | 0 | 0 |
| 18050 | ENSMUSG00000064541 | 0 | NA | NA | NA | NA | NA | Gm22978 | predicted gene, 22978 [Source:MGI Symb | 0 | 0 | 0 | 0 | 0 | 0 |
| 18051 | ENSMUSG00000064542 | 0.481988923 | 2.492401104 | 3.837077151 | 0.649557203 | 0.515978285 | NA | Gm22980 | predicted gene, 22980 [Source:MGI Symb | 0 | 0 | 0 | 2.8919 | 0 | 0 |
| 18052 | ENSMUSG00000064543 | 0 | NA | NA | NA | NA | NA | Gm22979 | predicted gene, 22979 [Source:MGI Symb | 0 | 0 | 0 | 0 | 0 | 0 |
| 18053 | ENSMUSG00000064545 | 0 | NA | NA | NA | NA | NA | Gm22981 | predicted gene, 22981 [Source:MGI Symb | 0 | 0 | 0 | 0 | 0 | 0 |
| 18054 | ENSMUSG00000064547 | 0 | NA | NA | NA | NA | NA | NA | NA | 0 | 0 | 0 | 0 | 0 | 0 |
| 18055 | ENSMUSG00000064548 | 0 | NA | NA | NA | NA | NA | n-R5s65 | nuclear encoded rRNA 5S 65 [Source:MGI | 0 | 0 | 0 | 0 | 0 | 0 |
| 18056 | ENSMUSG00000064549 | 0 | NA | NA | NA | NA | NA | Gm22977 | predicted gene, 22977 [Source:MGI Symb | 0 | 0 | 0 | 0 | 0 | 0 |
| 18057 | ENSMUSG00000064550 | 0 | NA | NA | NA | NA | NA | Gm24623 | predicted gene, 24623 [Source:MGI Symb | 0 | 0 | 0 | 0 | 0 | 0 |
| 18058 | ENSMUSG00000064551 | 0 | NA | NA | NA | NA | NA | Gm24624 | predicted gene, 24624 [Source:MGI Symb | 0 | 0 | 0 | 0 | 0 | 0 |
| 18059 | ENSMUSG00000064553 | 0 | NA | NA | NA | NA | NA | Gm24622 | predicted gene, 24622 [Source:MGI Symb | 0 | 0 | 0 | 0 | 0 | 0 |
| 18060 | ENSMUSG00000064554 | 0 | NA | NA | NA | NA | NA | n-R5s173 | nuclear encoded rRNA 5S 173 [Source:MG | 0 | 0 | 0 | 0 | 0 | 0 |
| 18061 | ENSMUSG00000064555 | 0 | NA | NA | NA | NA | NA | Gm24626 | predicted gene, 24626 [Source:MGI Symb | 0 | 0 | 0 | 0 | 0 | 0 |
| 18062 | ENSMUSG00000064556 | 0 | NA | NA | NA | NA | NA | Gm24625 | predicted gene, 24625 [Source:MGI Symb | 0 | 0 | 0 | 0 | 0 | 0 |
| 18063 | ENSMUSG00000064558 | 0.133850373 | -0.796629176 | 4.080472857 | -0.195229623 | 0.84521319 | NA | Gm24620 | predicted gene, 24620 [Source:MGI Symb | 0.8031 | 0 | 0 | 0 | 0 | 0 |
| 18064 | ENSMUSG00000064563 | 0 | NA | NA | NA | NA | NA | Snora7a | small nucleolar RNA, H/ACA box 7A [Sourc | 0 | 0 | 0 | 0 | 0 | 0 |
| 18065 | ENSMUSG00000064565 | 0 | NA | NA | NA | NA | NA | Gm24145 | predicted gene, 24145 [Source:MGI Symb | 0 | 0 | 0 | 0 | 0 | 0 |
| 18066 | ENSMUSG00000064566 | 0 | NA | NA | NA | NA | NA | Gm24144 | predicted gene, 24144 [Source:MGI Symb | 0 | 0 | 0 | 0 | 0 | 0 |
| 18067 | ENSMUSG00000064567 | 0.771003 | -1.69715066 | 2.640552792 | -0.642725518 | 0.52040222 | NA | Gm24143 | predicted gene, 24143 [Source:MGI Symb | 1.6062 | 2.0558 | 0 | 0.964 | 0 | 0 |
| 18068 | ENSMUSG00000064569 | 0 | NA | NA | NA | NA | NA | Gm25592 | predicted gene, 25592 [Source:MGI Symb | 0 | 0 | 0 | 0 | 0 | 0 |
| 18069 | ENSMUSG00000064570 | 0 | NA | NA | NA | NA | NA | Gm25800 | predicted gene, 25800 [Source:MGI Symb | 0 | 0 | 0 | 0 | 0 | 0 |
| 18070 | ENSMUSG00000064572 | 0 | NA | NA | NA | NA | NA | Gm25801 | predicted gene, 25801 [Source:MGI Symb | 0 | 0 | 0 | 0 | 0 | 0 |
| 18071 | ENSMUSG00000064573 | 0 | NA | NA | NA | NA | NA | Gm25802 | predicted gene, 25802 [Source:MGI Symb | 0 | 0 | 0 | 0 | 0 | 0 |
| 18072 | ENSMUSG00000064574 | 0 | NA | NA | NA | NA | NA | NA | NA | 0 | 0 | 0 | 0 | 0 | 0 |
| 18073 | ENSMUSG00000064575 | 0 | NA | NA | NA | NA | NA | NA | NA | 0 | 0 | 0 | 0 | 0 | 0 |
| 18074 | ENSMUSG00000064577 | 0 | NA | NA | NA | NA | NA | NA | NA | 0 | 0 | 0 | 0 | 0 | 0 |
| 18075 | ENSMUSG00000064579 | 0 | NA | NA | NA | NA | NA | Gm25796 | predicted gene, 25796 [Source:MGI Symb | 0 | 0 | 0 | 0 | 0 | 0 |
| 18076 | ENSMUSG00000064580 | 0.517184577 | -0.812115922 | 3.393570547 | -0.239310163 | 0.810865085 | NA | Gm25134 | predicted gene, 25134 [Source:MGI Symb | 0 | 0 | 2.0125 | 0 | 1.0906 | 0 |
| 18077 | ENSMUSG00000064581 | 0 | NA | NA | NA | NA | NA | Gm25133 | predicted gene, 25133 [Source:MGI Symb | 0 | 0 | 0 | 0 | 0 | 0 |
| 18078 | ENSMUSG00000064582 | 0.167707431 | -0.796629176 | 4.080472857 | -0.195229623 | 0.84521319 | NA | Gm25135 | predicted gene, 25135 [Source:MGI Symb | 0 | 0 | 1.0062 | 0 | 0 | 0 |
| 18079 | ENSMUSG00000064584 | 0 | NA | NA | NA | NA | NA | Gm25130 | predicted gene, 25130 [Source:MGI Symb | 0 | 0 | 0 | 0 | 0 | 0 |
| 18080 | ENSMUSG00000064585 | 0.181769715 | 1.126924175 | 4.080472857 | 0.276174898 | 0.782413736 | NA | Gm25129 | predicted gene, 25129 [Source:MGI Symb | 0 | 0 | 0 | 0 | 1.0906 | 0 |
| 18081 | ENSMUSG00000064586 | 0 | NA | NA | NA | NA | NA | Gm25132 | predicted gene, 25132 [Source:MGI Symb | 0 | 0 | 0 | 0 | 0 | 0 |
| 18082 | ENSMUSG00000064587 | 0 | NA | NA | NA | NA | NA | Gm25131 | predicted gene, 25131 [Source:MGI Symb | 0 | 0 | 0 | 0 | 0 | 0 |
| 18083 | ENSMUSG00000064588 | 0 | NA | NA | NA | NA | NA | NA | NA | 0 | 0 | 0 | 0 | 0 | 0 |
| 18084 | ENSMUSG00000064590 | 0 | NA | NA | NA | NA | NA | Gm22301 | predicted gene, 22301 [Source:MGI Symb | 0 | 0 | 0 | 0 | 0 | 0 |

| | | | | | | | | | | | | | | | |
|---|---|---|---|---|---|---|---|---|---|---|---|---|---|---|---|
| 18085 | ENSMUSG00000064591 | 0 | NA | NA | NA | NA | NA | Gm22302 | predicted gene, 22302 [Source:MGI Symb | 0 | 0 | 0 | 0 | 0 | 0 |
| 18086 | ENSMUSG00000064595 | 0.301557804 | -1.64335916 | 4.042814367 | -0.406488899 | 0.684383412 | NA | Gm22300 | predicted gene, 22300 [Source:MGI Symb | 0.8031 | 0 | 1.0062 | 0 | 0 | 0 |
| 18087 | ENSMUSG00000064596 | 0 | NA | NA | NA | NA | NA | Gm22298 | predicted gene, 22298 [Source:MGI Symb | 0 | 0 | 0 | 0 | 0 | 0 |
| 18088 | ENSMUSG00000064597 | 0.667397476 | -1.404797968 | 2.804497115 | -0.500909044 | 0.616435137 | NA | Gm22299 | predicted gene, 22299 [Source:MGI Symb | 0 | 1.0279 | 2.0125 | 0.964 | 0 | 0 |
| 18089 | ENSMUSG00000064600 | 0 | NA | NA | NA | NA | NA | Gm25636 | predicted gene, 25636 [Source:MGI Symb | 0 | 0 | 0 | 0 | 0 | 0 |
| 18090 | ENSMUSG00000064601 | 0 | NA | NA | NA | NA | NA | Gm25634 | predicted gene, 25634 [Source:MGI Symb | 0 | 0 | 0 | 0 | 0 | 0 |
| 18091 | ENSMUSG00000064602 | 1.586803951 | 0.79939641 | 1.860506363 | 0.429666045 | 0.667438587 | NA | Snora41 | small nucleolar RNA, H/ACA box 41 [Sourc | 2.4093 | 1.0279 | 0 | 3.8559 | 0 | 2.2277 |
| 18092 | ENSMUSG00000064604 | 0 | NA | NA | NA | NA | NA | Snora44 | small nucleolar RNA, H/ACA box 44 [Sourc | 0 | 0 | 0 | 0 | 0 | 0 |
| 18093 | ENSMUSG00000064605 | 0 | NA | NA | NA | NA | NA | Gm22220 | predicted gene, 22220 [Source:MGI Symb | 0 | 0 | 0 | 0 | 0 | 0 |
| 18094 | ENSMUSG00000064606 | 0.695728634 | -1.425610641 | 3.05889997 | -0.46605337 | 0.641177299 | NA | Gm25899 | predicted gene, 25899 [Source:MGI Symb | 0 | 3.0838 | 0 | 0 | 1.0906 | 0 |
| 18095 | ENSMUSG00000064609 | 0 | NA | NA | NA | NA | NA | Gm25635 | predicted gene, 25635 [Source:MGI Symb | 0 | 0 | 0 | 0 | 0 | 0 |
| 18096 | ENSMUSG00000064612 | 0 | NA | NA | NA | NA | NA | Gm26457 | predicted gene, 26457 [Source:MGI Symb | 0 | 0 | 0 | 0 | 0 | 0 |
| 18097 | ENSMUSG00000064613 | 0 | NA | NA | NA | NA | NA | Gm24621 | predicted gene, 24621 [Source:MGI Symb | 0 | 0 | 0 | 0 | 0 | 0 |
| 18098 | ENSMUSG00000064616 | 0 | NA | NA | NA | NA | NA | NA | NA | 0 | 0 | 0 | 0 | 0 | 0 |
| 18099 | ENSMUSG00000064617 | 0 | NA | NA | NA | NA | NA | NA | NA | 0 | 0 | 0 | 0 | 0 | 0 |
| 18100 | ENSMUSG00000064618 | 0 | NA | NA | NA | NA | NA | NA | NA | 0 | 0 | 0 | 0 | 0 | 0 |
| 18101 | ENSMUSG00000064619 | 0 | NA | NA | NA | NA | NA | Gm23975 | predicted gene, 23975 [Source:MGI Symb | 0 | 0 | 0 | 0 | 0 | 0 |
| 18102 | ENSMUSG00000064620 | 0 | NA | NA | NA | NA | NA | Gm22303 | predicted gene, 22303 [Source:MGI Symb | 0 | 0 | 0 | 0 | 0 | 0 |
| 18103 | ENSMUSG00000064621 | 0 | NA | NA | NA | NA | NA | Gm26922 | predicted gene, 26922 [Source:MGI Symb | 0 | 0 | 0 | 0 | 0 | 0 |
| 18104 | ENSMUSG00000064622 | 0 | NA | NA | NA | NA | NA | Gm22304 | predicted gene, 22304 [Source:MGI Symb | 0 | 0 | 0 | 0 | 0 | 0 |
| 18105 | ENSMUSG00000064624 | 0 | NA | NA | NA | NA | NA | Gm22305 | predicted gene, 22305 [Source:MGI Symb | 0 | 0 | 0 | 0 | 0 | 0 |
| 18106 | ENSMUSG00000064626 | 0 | NA | NA | NA | NA | NA | NA | NA | 0 | 0 | 0 | 0 | 0 | 0 |
| 18107 | ENSMUSG00000064630 | 0 | NA | NA | NA | NA | NA | Gm25143 | predicted gene, 25143 [Source:MGI Symb | 0 | 0 | 0 | 0 | 0 | 0 |
| 18108 | ENSMUSG00000064631 | 0 | NA | NA | NA | NA | NA | NA | NA | 0 | 0 | 0 | 0 | 0 | 0 |
| 18109 | ENSMUSG00000064632 | 0 | NA | NA | NA | NA | NA | Gm25142 | predicted gene, 25142 [Source:MGI Symb | 0 | 0 | 0 | 0 | 0 | 0 |
| 18110 | ENSMUSG00000064634 | 0 | NA | NA | NA | NA | NA | Gm22620 | predicted gene, 22620 [Source:MGI Symb | 0 | 0 | 0 | 0 | 0 | 0 |
| 18111 | ENSMUSG00000064637 | 44.87716775 | 0.074690787 | 0.380020189 | 0.196544262 | 0.844184192 | 0.972803041 | Snora20 | small nucleolar RNA, H/ACA box 20 [Sourc | 40.155 | 26.726 | 64.4 | 52.055 | 44.715 | 41.212 |
| 18112 | ENSMUSG00000064642 | 0 | NA | NA | NA | NA | NA | NA | NA | 0 | 0 | 0 | 0 | 0 | 0 |
| 18113 | ENSMUSG00000064644 | 0 | NA | NA | NA | NA | NA | Gm23302 | predicted gene, 23302 [Source:MGI Symb | 0 | 0 | 0 | 0 | 0 | 0 |
| 18114 | ENSMUSG00000064646 | 0 | NA | NA | NA | NA | NA | Gm23300 | predicted gene, 23300 [Source:MGI Symb | 0 | 0 | 0 | 0 | 0 | 0 |
| 18115 | ENSMUSG00000064647 | 3.231049471 | -0.039091517 | 1.276368704 | -0.030627136 | 0.975566901 | NA | Gm23301 | predicted gene, 23301 [Source:MGI Symb | 0.8031 | 1.0279 | 8.05 | 2.8919 | 3.2719 | 3.3415 |
| 18116 | ENSMUSG00000064648 | 0.531944269 | 2.618916564 | 3.285123409 | 0.797204926 | 0.425332026 | NA | Gm23303 | predicted gene, 23303 [Source:MGI Symb | 0 | 0 | 0 | 0.964 | 0 | 2.2277 |
| 18117 | ENSMUSG00000064652 | 0 | NA | NA | NA | NA | NA | Gm24165 | predicted gene, 24165 [Source:MGI Symb | 0 | 0 | 0 | 0 | 0 | 0 |
| 18118 | ENSMUSG00000064653 | 0 | NA | NA | NA | NA | NA | Gm26129 | predicted gene, 26129 [Source:MGI Symb | 0 | 0 | 0 | 0 | 0 | 0 |
| 18119 | ENSMUSG00000064655 | 1.475869743 | -1.772536753 | 1.977339671 | -0.896425019 | 0.37002581 | NA | Gm25788 | predicted gene, 25788 [Source:MGI Symb | 0.8031 | 4.1117 | 2.0125 | 1.928 | 0 | 0 |
| 18120 | ENSMUSG00000064658 | 0 | NA | NA | NA | NA | NA | Gm24166 | predicted gene, 24166 [Source:MGI Symb | 0 | 0 | 0 | 0 | 0 | 0 |
| 18121 | ENSMUSG00000064659 | 0 | NA | NA | NA | NA | NA | Gm26131 | predicted gene, 26131 [Source:MGI Symb | 0 | 0 | 0 | 0 | 0 | 0 |
| 18122 | ENSMUSG00000064663 | 0 | NA | NA | NA | NA | NA | NA | NA | 0 | 0 | 0 | 0 | 0 | 0 |
| 18123 | ENSMUSG00000064664 | 0 | NA | NA | NA | NA | NA | Gm24459 | predicted gene, 24459 [Source:MGI Symb | 0 | 0 | 0 | 0 | 0 | 0 |
| 18124 | ENSMUSG00000064665 | 0 | NA | NA | NA | NA | NA | Gm24460 | predicted gene, 24460 [Source:MGI Symb | 0 | 0 | 0 | 0 | 0 | 0 |
| 18125 | ENSMUSG00000064666 | 0 | NA | NA | NA | NA | NA | Snora52 | small nucleolar RNA, H/ACA box 52 [Sourc | 0 | 0 | 0 | 0 | 0 | 0 |
| 18126 | ENSMUSG00000064669 | 0 | NA | NA | NA | NA | NA | Gm24461 | predicted gene, 24461 [Source:MGI Symb | 0 | 0 | 0 | 0 | 0 | 0 |
| 18127 | ENSMUSG00000064670 | 0 | NA | NA | NA | NA | NA | Gm22808 | predicted gene, 22808 [Source:MGI Symb | 0 | 0 | 0 | 0 | 0 | 0 |
| 18128 | ENSMUSG00000064671 | 0 | NA | NA | NA | NA | NA | Gm22807 | predicted gene, 22807 [Source:MGI Symb | 0 | 0 | 0 | 0 | 0 | 0 |
| 18129 | ENSMUSG00000064672 | 5.848241862 | -0.589197008 | 0.915349146 | -0.643685539 | 0.519779369 | 0.887802551 | Gm22806 | predicted gene, 22806 [Source:MGI Symb | 4.8186 | 6.1675 | 10.062 | 1.928 | 6.5437 | 5.5692 |
| 18130 | ENSMUSG00000064673 | 0 | NA | NA | NA | NA | NA | Gm22805 | predicted gene, 22805 [Source:MGI Symb | 0 | 0 | 0 | 0 | 0 | 0 |
| 18131 | ENSMUSG00000064675 | 0 | NA | NA | NA | NA | NA | Gm22810 | predicted gene, 22810 [Source:MGI Symb | 0 | 0 | 0 | 0 | 0 | 0 |
| 18132 | ENSMUSG00000064677 | 0 | NA | NA | NA | NA | NA | Gm22809 | predicted gene, 22809 [Source:MGI Symb | 0 | 0 | 0 | 0 | 0 | 0 |
| 18133 | ENSMUSG00000064679 | 0 | NA | NA | NA | NA | NA | Gm25357 | predicted gene, 25357 [Source:MGI Symb | 0 | 0 | 0 | 0 | 0 | 0 |
| 18134 | ENSMUSG00000064682 | 0 | NA | NA | NA | NA | NA | Gm25813 | predicted gene, 25813 [Source:MGI Symb | 0 | 0 | 0 | 0 | 0 | 0 |
| 18135 | ENSMUSG00000064685 | 0 | NA | NA | NA | NA | NA | Gm25812 | predicted gene, 25812 [Source:MGI Symb | 0 | 0 | 0 | 0 | 0 | 0 |
| 18136 | ENSMUSG00000064686 | 0 | NA | NA | NA | NA | NA | Gm25810 | predicted gene, 25810 [Source:MGI Symb | 0 | 0 | 0 | 0 | 0 | 0 |
| 18137 | ENSMUSG00000064687 | 0 | NA | NA | NA | NA | NA | Gm25811 | predicted gene, 25811 [Source:MGI Symb | 0 | 0 | 0 | 0 | 0 | 0 |
| 18138 | ENSMUSG00000064688 | 0 | NA | NA | NA | NA | NA | Gm25814 | predicted gene, 25814 [Source:MGI Symb | 0 | 0 | 0 | 0 | 0 | 0 |
| 18139 | ENSMUSG00000064689 | 0 | NA | NA | NA | NA | NA | Gm25815 | predicted gene, 25815 [Source:MGI Symb | 0 | 0 | 0 | 0 | 0 | 0 |
| 18140 | ENSMUSG00000064691 | 0 | NA | NA | NA | NA | NA | NA | NA | 0 | 0 | 0 | 0 | 0 | 0 |

| | | | | | | | | | | | | | | | |
|---|---|---|---|---|---|---|---|---|---|---|---|---|---|---|---|
| 18141 | ENSMUSG00000064692 | 0 | NA | NA | NA | NA | NA | Gm24151 | predicted gene, 24151 [Source:MGI Symb | 0 | 0 | 0 | 0 | 0 | 0 |
| 18142 | ENSMUSG00000064693 | 0 | NA | NA | NA | NA | NA | Gm24150 | predicted gene, 24150 [Source:MGI Symb | 0 | 0 | 0 | 0 | 0 | 0 |
| 18143 | ENSMUSG00000064694 | 0 | NA | NA | NA | NA | NA | NA | NA | 0 | 0 | 0 | 0 | 0 | 0 |
| 18144 | ENSMUSG00000064696 | 0 | NA | NA | NA | NA | NA | NA | NA | 0 | 0 | 0 | 0 | 0 | 0 |
| 18145 | ENSMUSG00000064697 | 0 | NA | NA | NA | NA | NA | Gm24147 | predicted gene, 24147 [Source:MGI Symb | 0 | 0 | 0 | 0 | 0 | 0 |
| 18146 | ENSMUSG00000064698 | 0 | NA | NA | NA | NA | NA | Gm24152 | predicted gene, 24152 [Source:MGI Symb | 0 | 0 | 0 | 0 | 0 | 0 |
| 18147 | ENSMUSG00000064702 | 0 | NA | NA | NA | NA | NA | NA | NA | 0 | 0 | 0 | 0 | 0 | 0 |
| 18148 | ENSMUSG00000064704 | 0 | NA | NA | NA | NA | NA | NA | NA | 0 | 0 | 0 | 0 | 0 | 0 |
| 18149 | ENSMUSG00000064706 | 0 | NA | NA | NA | NA | NA | NA | NA | 0 | 0 | 0 | 0 | 0 | 0 |
| 18150 | ENSMUSG00000064707 | 0 | NA | NA | NA | NA | NA | Gm22507 | predicted gene, 22507 [Source:MGI Symb | 0 | 0 | 0 | 0 | 0 | 0 |
| 18151 | ENSMUSG00000064709 | 0 | NA | NA | NA | NA | NA | NA | NA | 0 | 0 | 0 | 0 | 0 | 0 |
| 18152 | ENSMUSG00000064714 | 0 | NA | NA | NA | NA | NA | NA | NA | 0 | 0 | 0 | 0 | 0 | 0 |
| 18153 | ENSMUSG00000064715 | 0 | NA | NA | NA | NA | NA | Gm24207 | predicted gene, 24207 [Source:MGI Symb | 0 | 0 | 0 | 0 | 0 | 0 |
| 18154 | ENSMUSG00000064717 | 0 | NA | NA | NA | NA | NA | Gm24208 | predicted gene, 24208 [Source:MGI Symb | 0 | 0 | 0 | 0 | 0 | 0 |
| 18155 | ENSMUSG00000064718 | 0 | NA | NA | NA | NA | NA | NA | NA | 0 | 0 | 0 | 0 | 0 | 0 |
| 18156 | ENSMUSG00000064719 | 0.133850373 | -0.796629176 | 4.080472857 | -0.195229623 | 0.84521319 | NA | Gm25930 | predicted gene, 25930 [Source:MGI Symb | 0.8031 | 0 | 0 | 0 | 0 | 0 |
| 18157 | ENSMUSG00000064720 | 0 | NA | NA | NA | NA | NA | Gm25856 | predicted gene, 25856 [Source:MGI Symb | 0 | 0 | 0 | 0 | 0 | 0 |
| 18158 | ENSMUSG00000064721 | 0 | NA | NA | NA | NA | NA | Gm25855 | predicted gene, 25855 [Source:MGI Symb | 0 | 0 | 0 | 0 | 0 | 0 |
| 18159 | ENSMUSG00000064722 | 0 | NA | NA | NA | NA | NA | Gm25857 | predicted gene, 25857 [Source:MGI Symb | 0 | 0 | 0 | 0 | 0 | 0 |
| 18160 | ENSMUSG00000064724 | 0 | NA | NA | NA | NA | NA | Gm25852 | predicted gene, 25852 [Source:MGI Symb | 0 | 0 | 0 | 0 | 0 | 0 |
| 18161 | ENSMUSG00000064725 | 0 | NA | NA | NA | NA | NA | Snord96a | small nucleolar RNA, C/D box 96A [Source | 0 | 0 | 0 | 0 | 0 | 0 |
| 18162 | ENSMUSG00000064726 | 0 | NA | NA | NA | NA | NA | Gm25854 | predicted gene, 25854 [Source:MGI Symb | 0 | 0 | 0 | 0 | 0 | 0 |
| 18163 | ENSMUSG00000064727 | 0 | NA | NA | NA | NA | NA | Gm25853 | predicted gene, 25853 [Source:MGI Symb | 0 | 0 | 0 | 0 | 0 | 0 |
| 18164 | ENSMUSG00000064728 | 0 | NA | NA | NA | NA | NA | NA | NA | 0 | 0 | 0 | 0 | 0 | 0 |
| 18165 | ENSMUSG00000064731 | 0 | NA | NA | NA | NA | NA | Snord45b | small nucleolar RNA, C/D box 45B [Source | 0 | 0 | 0 | 0 | 0 | 0 |
| 18166 | ENSMUSG00000064732 | 0 | NA | NA | NA | NA | NA | Gm23447 | predicted gene, 23447 [Source:MGI Symb | 0 | 0 | 0 | 0 | 0 | 0 |
| 18167 | ENSMUSG00000064734 | 0.685278559 | -2.809870862 | 3.376578027 | -0.832165239 | 0.405315682 | NA | Gm23445 | predicted gene, 23445 [Source:MGI Symb | 0 | 4.1117 | 0 | 0 | 0 | 0 |
| 18168 | ENSMUSG00000064735 | 0 | NA | NA | NA | NA | NA | Gm23057 | predicted gene, 23057 [Source:MGI Symb | 0 | 0 | 0 | 0 | 0 | 0 |
| 18169 | ENSMUSG00000064737 | 0 | NA | NA | NA | NA | NA | Gm23056 | predicted gene, 23056 [Source:MGI Symb | 0 | 0 | 0 | 0 | 0 | 0 |
| 18170 | ENSMUSG00000064738 | 0 | NA | NA | NA | NA | NA | Gm23054 | predicted gene, 23054 [Source:MGI Symb | 0 | 0 | 0 | 0 | 0 | 0 |
| 18171 | ENSMUSG00000064739 | 0 | NA | NA | NA | NA | NA | Gm23055 | predicted gene, 23055 [Source:MGI Symb | 0 | 0 | 0 | 0 | 0 | 0 |
| 18172 | ENSMUSG00000064740 | 0 | NA | NA | NA | NA | NA | NA | NA | 0 | 0 | 0 | 0 | 0 | 0 |
| 18173 | ENSMUSG00000064741 | 0 | NA | NA | NA | NA | NA | Snord14a | small nucleolar RNA, C/D box 14A [Source | 0 | 0 | 0 | 0 | 0 | 0 |
| 18174 | ENSMUSG00000064742 | 0 | NA | NA | NA | NA | NA | Rnu6-ps1 | U6 small nuclear RNA, pseudogene 1 [Sou | 0 | 0 | 0 | 0 | 0 | 0 |
| 18175 | ENSMUSG00000064743 | 0 | NA | NA | NA | NA | NA | NA | NA | 0 | 0 | 0 | 0 | 0 | 0 |
| 18176 | ENSMUSG00000064745 | 0.36353943 | 2.065989254 | 4.034137301 | 0.512126658 | 0.608562368 | NA | Gm24708 | predicted gene, 24708 [Source:MGI Symb | 0 | 0 | 0 | 0 | 2.1812 | 0 |
| 18177 | ENSMUSG00000064746 | 0 | NA | NA | NA | NA | NA | NA | NA | 0 | 0 | 0 | 0 | 0 | 0 |
| 18178 | ENSMUSG00000064748 | 0 | NA | NA | NA | NA | NA | n-R5s28 | nuclear encoded rRNA 5S 28 [Source:MGI | 0 | 0 | 0 | 0 | 0 | 0 |
| 18179 | ENSMUSG00000064751 | 0 | NA | NA | NA | NA | NA | Gm26330 | predicted gene, 26330 [Source:MGI Symb | 0 | 0 | 0 | 0 | 0 | 0 |
| 18180 | ENSMUSG00000064755 | 0 | NA | NA | NA | NA | NA | Gm26328 | predicted gene, 26328 [Source:MGI Symb | 0 | 0 | 0 | 0 | 0 | 0 |
| 18181 | ENSMUSG00000064758 | 0 | NA | NA | NA | NA | NA | NA | NA | 0 | 0 | 0 | 0 | 0 | 0 |
| 18182 | ENSMUSG00000064759 | 0 | NA | NA | NA | NA | NA | Gm25475 | predicted gene, 25475 [Source:MGI Symb | 0 | 0 | 0 | 0 | 0 | 0 |
| 18183 | ENSMUSG00000064761 | 0 | NA | NA | NA | NA | NA | Gm23514 | predicted gene, 23514 [Source:MGI Symb | 0 | 0 | 0 | 0 | 0 | 0 |
| 18184 | ENSMUSG00000064762 | 0 | NA | NA | NA | NA | NA | Gm23738 | predicted gene, 23738 [Source:MGI Symb | 0 | 0 | 0 | 0 | 0 | 0 |
| 18185 | ENSMUSG00000064763 | 0 | NA | NA | NA | NA | NA | Gm23515 | predicted gene, 23515 [Source:MGI Symb | 0 | 0 | 0 | 0 | 0 | 0 |
| 18186 | ENSMUSG00000064764 | 0 | NA | NA | NA | NA | NA | Gm23517 | predicted gene, 23517 [Source:MGI Symb | 0 | 0 | 0 | 0 | 0 | 0 |
| 18187 | ENSMUSG00000064765 | 0 | NA | NA | NA | NA | NA | Gm26456 | predicted gene, 26456 [Source:MGI Symb | 0 | 0 | 0 | 0 | 0 | 0 |
| 18188 | ENSMUSG00000064766 | 0 | NA | NA | NA | NA | NA | Gm26458 | predicted gene, 26458 [Source:MGI Symb | 0 | 0 | 0 | 0 | 0 | 0 |
| 18189 | ENSMUSG00000064767 | 0 | NA | NA | NA | NA | NA | Snord35b | small nucleolar RNA, C/D box 35B [Source | 0 | 0 | 0 | 0 | 0 | 0 |
| 18190 | ENSMUSG00000064768 | 0 | NA | NA | NA | NA | NA | Snord60 | small nucleolar RNA, C/D box 60 [Source:I | 0 | 0 | 0 | 0 | 0 | 0 |
| 18191 | ENSMUSG00000064769 | 0 | NA | NA | NA | NA | NA | Gm23518 | predicted gene, 23518 [Source:MGI Symb | 0 | 0 | 0 | 0 | 0 | 0 |
| 18192 | ENSMUSG00000064770 | 0 | NA | NA | NA | NA | NA | NA | NA | 0 | 0 | 0 | 0 | 0 | 0 |
| 18193 | ENSMUSG00000064772 | 0 | NA | NA | NA | NA | NA | Gm25202 | predicted gene, 25202 [Source:MGI Symb | 0 | 0 | 0 | 0 | 0 | 0 |
| 18194 | ENSMUSG00000064775 | 0 | NA | NA | NA | NA | NA | Gm25205 | predicted gene, 25205 [Source:MGI Symb | 0 | 0 | 0 | 0 | 0 | 0 |
| 18195 | ENSMUSG00000064776 | 0 | NA | NA | NA | NA | NA | Gm25203 | predicted gene, 25203 [Source:MGI Symb | 0 | 0 | 0 | 0 | 0 | 0 |
| 18196 | ENSMUSG00000064777 | 0 | NA | NA | NA | NA | NA | Gm25204 | predicted gene, 25204 [Source:MGI Symb | 0 | 0 | 0 | 0 | 0 | 0 |

| | | | | | | | | | | | | | | | |
|---|---|---|---|---|---|---|---|---|---|---|---|---|---|---|---|
| 18197 | ENSMUSG00000064778 | 0.167707431 | -0.796629176 | 4.080472857 | -0.195229623 | 0.84521319 | NA | Snord59a | small nucleolar RNA, C/D box 59A [Source | 0 | 0 | 1.0062 | 0 | 0 | 0 |
| 18198 | ENSMUSG00000064779 | 0 | NA | NA | NA | NA | NA | NA | NA | 0 | 0 | 0 | 0 | 0 | 0 |
| 18199 | ENSMUSG00000064780 | 0 | NA | NA | NA | NA | NA | Snord95 | small nucleolar RNA, C/D box 95 [Source:I | 0 | 0 | 0 | 0 | 0 | 0 |
| 18200 | ENSMUSG00000064781 | 0 | NA | NA | NA | NA | NA | Gm22686 | predicted gene, 22686 [Source:MGI Symb | 0 | 0 | 0 | 0 | 0 | 0 |
| 18201 | ENSMUSG00000064782 | 0 | NA | NA | NA | NA | NA | Gm22685 | predicted gene, 22685 [Source:MGI Symb | 0 | 0 | 0 | 0 | 0 | 0 |
| 18202 | ENSMUSG00000064783 | 0 | NA | NA | NA | NA | NA | Gm22684 | predicted gene, 22684 [Source:MGI Symb | 0 | 0 | 0 | 0 | 0 | 0 |
| 18203 | ENSMUSG00000064788 | 0 | NA | NA | NA | NA | NA | Gm22683 | predicted gene, 22683 [Source:MGI Symb | 0 | 0 | 0 | 0 | 0 | 0 |
| 18204 | ENSMUSG00000064791 | 0 | NA | NA | NA | NA | NA | NA | NA | 0 | 0 | 0 | 0 | 0 | 0 |
| 18205 | ENSMUSG00000064792 | 0 | NA | NA | NA | NA | NA | Gm24355 | predicted gene, 24355 [Source:MGI Symb | 0 | 0 | 0 | 0 | 0 | 0 |
| 18206 | ENSMUSG00000064793 | 0 | NA | NA | NA | NA | NA | Gm24356 | predicted gene, 24356 [Source:MGI Symb | 0 | 0 | 0 | 0 | 0 | 0 |
| 18207 | ENSMUSG00000064795 | 0 | NA | NA | NA | NA | NA | Gm22423 | predicted gene, 22423 [Source:MGI Symb | 0 | 0 | 0 | 0 | 0 | 0 |
| 18208 | ENSMUSG00000064796 | 8.175592388 | 0.751682745 | 0.797996055 | 0.941962983 | 0.346211592 | 0.786987972 | NA | NA | 8.031 | 2.0558 | 8.05 | 7.7118 | 8.7249 | 14.48 |
| 18209 | ENSMUSG00000064797 | 0 | NA | NA | NA | NA | NA | Gm24357 | predicted gene, 24357 [Source:MGI Symb | 0 | 0 | 0 | 0 | 0 | 0 |
| 18210 | ENSMUSG00000064798 | 0 | NA | NA | NA | NA | NA | NA | NA | 0 | 0 | 0 | 0 | 0 | 0 |
| 18211 | ENSMUSG00000064800 | 0 | NA | NA | NA | NA | NA | Gm24948 | predicted gene, 24948 [Source:MGI Symb | 0 | 0 | 0 | 0 | 0 | 0 |
| 18212 | ENSMUSG00000064801 | 0 | NA | NA | NA | NA | NA | NA | NA | 0 | 0 | 0 | 0 | 0 | 0 |
| 18213 | ENSMUSG00000064802 | 0 | NA | NA | NA | NA | NA | Gm24036 | predicted gene, 24036 [Source:MGI Symb | 0 | 0 | 0 | 0 | 0 | 0 |
| 18214 | ENSMUSG00000064805 | 0 | NA | NA | NA | NA | NA | Gm24037 | predicted gene, 24037 [Source:MGI Symb | 0 | 0 | 0 | 0 | 0 | 0 |
| 18215 | ENSMUSG00000064806 | 0 | NA | NA | NA | NA | NA | Gm24038 | predicted gene, 24038 [Source:MGI Symb | 0 | 0 | 0 | 0 | 0 | 0 |
| 18216 | ENSMUSG00000064811 | 0.777840866 | -0.360978154 | 2.442491672 | -0.147790945 | 0.882507755 | NA | Gm22358 | predicted gene, 22358 [Source:MGI Symb | 1.6062 | 0 | 1.0062 | 0.964 | 1.0906 | 0 |
| 18217 | ENSMUSG00000064814 | 0 | NA | NA | NA | NA | NA | Gm23939 | predicted gene, 23939 [Source:MGI Symb | 0 | 0 | 0 | 0 | 0 | 0 |
| 18218 | ENSMUSG00000064816 | 0 | NA | NA | NA | NA | NA | Gm22357 | predicted gene, 22357 [Source:MGI Symb | 0 | 0 | 0 | 0 | 0 | 0 |
| 18219 | ENSMUSG00000064818 | 0 | NA | NA | NA | NA | NA | Gm22359 | predicted gene, 22359 [Source:MGI Symb | 0 | 0 | 0 | 0 | 0 | 0 |
| 18220 | ENSMUSG00000064822 | 0 | NA | NA | NA | NA | NA | Gm24292 | predicted gene, 24292 [Source:MGI Symb | 0 | 0 | 0 | 0 | 0 | 0 |
| 18221 | ENSMUSG00000064823 | 0 | NA | NA | NA | NA | NA | Snord82 | small nucleolar RNA, C/D box 82 [Source:I | 0 | 0 | 0 | 0 | 0 | 0 |
| 18222 | ENSMUSG00000064824 | 0 | NA | NA | NA | NA | NA | NA | NA | 0 | 0 | 0 | 0 | 0 | 0 |
| 18223 | ENSMUSG00000064829 | 0 | NA | NA | NA | NA | NA | Gm22879 | predicted gene, 22879 [Source:MGI Symb | 0 | 0 | 0 | 0 | 0 | 0 |
| 18224 | ENSMUSG00000064830 | 0 | NA | NA | NA | NA | NA | Gm25699 | predicted gene, 25699 [Source:MGI Symb | 0 | 0 | 0 | 0 | 0 | 0 |
| 18225 | ENSMUSG00000064831 | 0.185640647 | 1.126924175 | 4.080472857 | 0.276174898 | 0.782413736 | NA | NA | NA | 0 | 0 | 0 | 0 | 0 | 1.1138 |
| 18226 | ENSMUSG00000064833 | 0 | NA | NA | NA | NA | NA | Gm25926 | predicted gene, 25926 [Source:MGI Symb | 0 | 0 | 0 | 0 | 0 | 0 |
| 18227 | ENSMUSG00000064834 | 0 | NA | NA | NA | NA | NA | NA | NA | 0 | 0 | 0 | 0 | 0 | 0 |
| 18228 | ENSMUSG00000064836 | 0 | NA | NA | NA | NA | NA | Gm25697 | predicted gene, 25697 [Source:MGI Symb | 0 | 0 | 0 | 0 | 0 | 0 |
| 18229 | ENSMUSG00000064837 | 0 | NA | NA | NA | NA | NA | Snora75 | small nucleolar RNA, H/ACA box 75 [Sourc | 0 | 0 | 0 | 0 | 0 | 0 |
| 18230 | ENSMUSG00000064840 | 0 | NA | NA | NA | NA | NA | NA | NA | 0 | 0 | 0 | 0 | 0 | 0 |
| 18231 | ENSMUSG00000064841 | 0 | NA | NA | NA | NA | NA | Gm26205 | predicted gene, 26205 [Source:MGI Symb | 0 | 0 | 0 | 0 | 0 | 0 |
| 18232 | ENSMUSG00000064842 | 0 | NA | NA | NA | NA | NA | Gm26206 | predicted gene, 26206 [Source:MGI Symb | 0 | 0 | 0 | 0 | 0 | 0 |
| 18233 | ENSMUSG00000064843 | 0 | NA | NA | NA | NA | NA | Gm26207 | predicted gene, 26207 [Source:MGI Symb | 0 | 0 | 0 | 0 | 0 | 0 |
| 18234 | ENSMUSG00000064844 | 1.198617707 | 2.614666659 | 2.294845084 | 1.139365214 | 0.254550859 | NA | Gm26202 | predicted gene, 26202 [Source:MGI Symb | 0 | 1.0279 | 0 | 2.8919 | 3.2719 | 0 |
| 18235 | ENSMUSG00000064845 | 4.537821052 | -0.491182951 | 1.119845748 | -0.438616615 | 0.660939356 | NA | Gm26203 | predicted gene, 26203 [Source:MGI Symb | 0.8031 | 8.2233 | 7.0437 | 6.7478 | 2.1812 | 2.2277 |
| 18236 | ENSMUSG00000064850 | 0 | NA | NA | NA | NA | NA | Gm23443 | predicted gene, 23443 [Source:MGI Symb | 0 | 0 | 0 | 0 | 0 | 0 |
| 18237 | ENSMUSG00000064851 | 0 | NA | NA | NA | NA | NA | Gm24525 | predicted gene, 24525 [Source:MGI Symb | 0 | 0 | 0 | 0 | 0 | 0 |
| 18238 | ENSMUSG00000064853 | 0.342639279 | -1.805976596 | 4.028547714 | -0.448294702 | 0.653940524 | NA | Gm23442 | predicted gene, 23442 [Source:MGI Symb | 0 | 2.0558 | 0 | 0 | 0 | 0 |
| 18239 | ENSMUSG00000064854 | 0 | NA | NA | NA | NA | NA | Gm24527 | predicted gene, 24527 [Source:MGI Symb | 0 | 0 | 0 | 0 | 0 | 0 |
| 18240 | ENSMUSG00000064855 | 0 | NA | NA | NA | NA | NA | Gm24526 | predicted gene, 24526 [Source:MGI Symb | 0 | 0 | 0 | 0 | 0 | 0 |
| 18241 | ENSMUSG00000064856 | 0 | NA | NA | NA | NA | NA | NA | NA | 0 | 0 | 0 | 0 | 0 | 0 |
| 18242 | ENSMUSG00000064857 | 0 | NA | NA | NA | NA | NA | n-R5s25 | nuclear encoded rRNA 5S 25 [Source:MGI | 0 | 0 | 0 | 0 | 0 | 0 |
| 18243 | ENSMUSG00000064858 | 0 | NA | NA | NA | NA | NA | Snora43 | small nucleolar RNA, H/ACA box 43 [Sourc | 0 | 0 | 0 | 0 | 0 | 0 |
| 18244 | ENSMUSG00000064859 | 0 | NA | NA | NA | NA | NA | Gm24524 | predicted gene, 24524 [Source:MGI Symb | 0 | 0 | 0 | 0 | 0 | 0 |
| 18245 | ENSMUSG00000064860 | 0 | NA | NA | NA | NA | NA | NA | NA | 0 | 0 | 0 | 0 | 0 | 0 |
| 18246 | ENSMUSG00000064862 | 0 | NA | NA | NA | NA | NA | Gm25038 | predicted gene, 25038 [Source:MGI Symb | 0 | 0 | 0 | 0 | 0 | 0 |
| 18247 | ENSMUSG00000064867 | 0.506966596 | 2.558866737 | 3.339834942 | 0.766165628 | 0.443577757 | NA | Gm25039 | predicted gene, 25039 [Source:MGI Symb | 0 | 0 | 0 | 1.928 | 0 | 1.1138 |
| 18248 | ENSMUSG00000064871 | 1.727207963 | -2.000414471 | 1.741600658 | -1.148606865 | 0.250718125 | NA | Snord58b | small nucleolar RNA, C/D box 58B [Source | 3.2124 | 3.0838 | 2.0125 | 0.964 | 1.0906 | 0 |
| 18249 | ENSMUSG00000064872 | 0 | NA | NA | NA | NA | NA | Gm23376 | predicted gene, 23376 [Source:MGI Symb | 0 | 0 | 0 | 0 | 0 | 0 |
| 18250 | ENSMUSG00000064876 | 0 | NA | NA | NA | NA | NA | Gm23378 | predicted gene, 23378 [Source:MGI Symb | 0 | 0 | 0 | 0 | 0 | 0 |
| 18251 | ENSMUSG00000064877 | 0 | NA | NA | NA | NA | NA | Gm23377 | predicted gene, 23377 [Source:MGI Symb | 0 | 0 | 0 | 0 | 0 | 0 |
| 18252 | ENSMUSG00000064880 | 0 | NA | NA | NA | NA | NA | Gm24201 | predicted gene, 24201 [Source:MGI Symb | 0 | 0 | 0 | 0 | 0 | 0 |

| | | | | | | | | | | | | | | | |
|---|---|---|---|---|---|---|---|---|---|---|---|---|---|---|---|
| 18253 | ENSMUSG00000064882 | 0 | NA | NA | NA | NA | NA | Gm24202 | predicted gene, 24202 [Source:MGI Symb | 0 | 0 | 0 | 0 | 0 | 0 |
| 18254 | ENSMUSG00000064883 | 0 | NA | NA | NA | NA | NA | Gm24203 | predicted gene, 24203 [Source:MGI Symb | 0 | 0 | 0 | 0 | 0 | 0 |
| 18255 | ENSMUSG00000064884 | 0 | NA | NA | NA | NA | NA | Gm24204 | predicted gene, 24204 [Source:MGI Symb | 0 | 0 | 0 | 0 | 0 | 0 |
| 18256 | ENSMUSG00000064887 | 0 | NA | NA | NA | NA | NA | NA | NA | 0 | 0 | 0 | 0 | 0 | 0 |
| 18257 | ENSMUSG00000064889 | 0 | NA | NA | NA | NA | NA | Gm24200 | predicted gene, 24200 [Source:MGI Symb | 0 | 0 | 0 | 0 | 0 | 0 |
| 18258 | ENSMUSG00000064890 | 0 | NA | NA | NA | NA | NA | Gm22505 | predicted gene, 22505 [Source:MGI Symb | 0 | 0 | 0 | 0 | 0 | 0 |
| 18259 | ENSMUSG00000064897 | 0 | NA | NA | NA | NA | NA | Gm22504 | predicted gene, 22504 [Source:MGI Symb | 0 | 0 | 0 | 0 | 0 | 0 |
| 18260 | ENSMUSG00000064899 | 0 | NA | NA | NA | NA | NA | Snord118 | small nucleolar RNA, C/D box 118 [Source | 0 | 0 | 0 | 0 | 0 | 0 |
| 18261 | ENSMUSG00000064900 | 0 | NA | NA | NA | NA | NA | Gm25402 | predicted gene, 25402 [Source:MGI Symb | 0 | 0 | 0 | 0 | 0 | 0 |
| 18262 | ENSMUSG00000064901 | 1.58754799 | -0.218344156 | 2.10395347 | -0.103778035 | 0.917345498 | NA | Snora21 | small nucleolar RNA, H/ACA box 21 [Sourc | 0 | 5.1396 | 0 | 0 | 3.2719 | 1.1138 |
| 18263 | ENSMUSG00000064904 | 0 | NA | NA | NA | NA | NA | NA | NA | 0 | 0 | 0 | 0 | 0 | 0 |
| 18264 | ENSMUSG00000064905 | 0 | NA | NA | NA | NA | NA | NA | NA | 0 | 0 | 0 | 0 | 0 | 0 |
| 18265 | ENSMUSG00000064906 | 0 | NA | NA | NA | NA | NA | n-R5s204 | nuclear encoded rRNA 5S 204 [Source:MG | 0 | 0 | 0 | 0 | 0 | 0 |
| 18266 | ENSMUSG00000064907 | 0 | NA | NA | NA | NA | NA | NA | NA | 0 | 0 | 0 | 0 | 0 | 0 |
| 18267 | ENSMUSG00000064909 | 0 | NA | NA | NA | NA | NA | Gm25401 | predicted gene, 25401 [Source:MGI Symb | 0 | 0 | 0 | 0 | 0 | 0 |
| 18268 | ENSMUSG00000064915 | 0 | NA | NA | NA | NA | NA | Gm22575 | predicted gene, 22575 [Source:MGI Symb | 0 | 0 | 0 | 0 | 0 | 0 |
| 18269 | ENSMUSG00000064916 | 2.086618264 | 3.511705996 | 1.84646263 | 1.901855981 | 0.057189985 | NA | Gm22573 | predicted gene, 22573 [Source:MGI Symb | 0 | 0 | 1.0062 | 3.8559 | 6.5437 | 1.1138 |
| 18270 | ENSMUSG00000064917 | 3.622801303 | -1.060928566 | 1.304893087 | -0.81303869 | 0.416195876 | NA | Gm22574 | predicted gene, 22574 [Source:MGI Symb | 1.6062 | 7.1954 | 6.0375 | 5.7839 | 0 | 1.1138 |
| 18271 | ENSMUSG00000064918 | 1.241373611 | 0.346687691 | 2.239815274 | 0.154784055 | 0.876991562 | NA | Gm22571 | predicted gene, 22571 [Source:MGI Symb | 3.2124 | 0 | 0 | 0.964 | 3.2719 | 0 |
| 18272 | ENSMUSG00000064919 | 3.032139631 | 0.500894572 | 1.27014043 | 0.394361568 | 0.693314113 | NA | Gm22572 | predicted gene, 22572 [Source:MGI Symb | 2.4093 | 4.1117 | 1.0062 | 2.8919 | 1.0906 | 6.6831 |
| 18273 | ENSMUSG00000064921 | 0 | NA | NA | NA | NA | NA | Gm22229 | predicted gene, 22229 [Source:MGI Symb | 0 | 0 | 0 | 0 | 0 | 0 |
| 18274 | ENSMUSG00000064923 | 0 | NA | NA | NA | NA | NA | Gm22042 | predicted gene, 22042 [Source:MGI Symb | 0 | 0 | 0 | 0 | 0 | 0 |
| 18275 | ENSMUSG00000064925 | 1.198617707 | 2.614666659 | 2.294845084 | 1.139365214 | 0.254550859 | NA | Snora62 | small nucleolar RNA, H/ACA box 62 [Sourc | 0 | 1.0279 | 0 | 2.8919 | 3.2719 | 0 |
| 18276 | ENSMUSG00000064926 | 0 | NA | NA | NA | NA | NA | NA | NA | 0 | 0 | 0 | 0 | 0 | 0 |
| 18277 | ENSMUSG00000064927 | 0.660353019 | -0.01500701 | 2.83315355 | -0.005296928 | 0.995773683 | NA | Gm22043 | predicted gene, 22043 [Source:MGI Symb | 0 | 1.0279 | 1.0062 | 1.928 | 0 | 0 |
| 18278 | ENSMUSG00000064928 | 0 | NA | NA | NA | NA | NA | Gm22045 | predicted gene, 22045 [Source:MGI Symb | 0 | 0 | 0 | 0 | 0 | 0 |
| 18279 | ENSMUSG00000064930 | 0 | NA | NA | NA | NA | NA | Gm23723 | predicted gene, 23723 [Source:MGI Symb | 0 | 0 | 0 | 0 | 0 | 0 |
| 18280 | ENSMUSG00000064932 | 0 | NA | NA | NA | NA | NA | NA | NA | 0 | 0 | 0 | 0 | 0 | 0 |
| 18281 | ENSMUSG00000064936 | 0.822613314 | -1.749415261 | 2.618121802 | -0.668194757 | 0.504009282 | NA | Gm23722 | predicted gene, 23722 [Source:MGI Symb | 0.8031 | 0 | 3.0187 | 0 | 0 | 1.1138 |
| 18282 | ENSMUSG00000064938 | 0 | NA | NA | NA | NA | NA | NA | NA | 0 | 0 | 0 | 0 | 0 | 0 |
| 18283 | ENSMUSG00000064939 | 0 | NA | NA | NA | NA | NA | NA | NA | 0 | 0 | 0 | 0 | 0 | 0 |
| 18284 | ENSMUSG00000064940 | 0 | NA | NA | NA | NA | NA | Gm23239 | predicted gene, 23239 [Source:MGI Symb | 0 | 0 | 0 | 0 | 0 | 0 |
| 18285 | ENSMUSG00000064941 | 0 | NA | NA | NA | NA | NA | Gm23238 | predicted gene, 23238 [Source:MGI Symb | 0 | 0 | 0 | 0 | 0 | 0 |
| 18286 | ENSMUSG00000064943 | 0 | NA | NA | NA | NA | NA | Gm23240 | predicted gene, 23240 [Source:MGI Symb | 0 | 0 | 0 | 0 | 0 | 0 |
| 18287 | ENSMUSG00000064945 | 0 | NA | NA | NA | NA | NA | Rny3 | RNA, Y3 small cytoplasmic (associated wit | 0 | 0 | 0 | 0 | 0 | 0 |
| 18288 | ENSMUSG00000064948 | 0 | NA | NA | NA | NA | NA | Gm23241 | predicted gene, 23241 [Source:MGI Symb | 0 | 0 | 0 | 0 | 0 | 0 |
| 18289 | ENSMUSG00000064949 | 0 | NA | NA | NA | NA | NA | Snora61 | small nucleolar RNA, H/ACA box 61 [Sourc | 0 | 0 | 0 | 0 | 0 | 0 |
| 18290 | ENSMUSG00000064952 | 0 | NA | NA | NA | NA | NA | Gm24920 | predicted gene, 24920 [Source:MGI Symb | 0 | 0 | 0 | 0 | 0 | 0 |
| 18291 | ENSMUSG00000064954 | 0 | NA | NA | NA | NA | NA | Gm24919 | predicted gene, 24919 [Source:MGI Symb | 0 | 0 | 0 | 0 | 0 | 0 |
| 18292 | ENSMUSG00000064956 | 0 | NA | NA | NA | NA | NA | Gm24918 | predicted gene, 24918 [Source:MGI Symb | 0 | 0 | 0 | 0 | 0 | 0 |
| 18293 | ENSMUSG00000064958 | 0 | NA | NA | NA | NA | NA | NA | NA | 0 | 0 | 0 | 0 | 0 | 0 |
| 18294 | ENSMUSG00000064959 | 0 | NA | NA | NA | NA | NA | n-R5s41 | nuclear encoded rRNA 5S 41 [Source:MGI | 0 | 0 | 0 | 0 | 0 | 0 |
| 18295 | ENSMUSG00000064960 | 0 | NA | NA | NA | NA | NA | NA | NA | 0 | 0 | 0 | 0 | 0 | 0 |
| 18296 | ENSMUSG00000064961 | 0 | NA | NA | NA | NA | NA | Gm22704 | predicted gene, 22704 [Source:MGI Symb | 0 | 0 | 0 | 0 | 0 | 0 |
| 18297 | ENSMUSG00000064964 | 0.367410362 | 2.078255025 | 3.800070157 | 0.54689912 | 0.58444804 | NA | Gm25147 | predicted gene, 25147 [Source:MGI Symb | 0 | 0 | 0 | 0 | 1.0906 | 1.1138 |
| 18298 | ENSMUSG00000064966 | 0 | NA | NA | NA | NA | NA | Snord15b | small nucleolar RNA, C/D box 14B [Source | 0 | 0 | 0 | 0 | 0 | 0 |
| 18299 | ENSMUSG00000064967 | 0 | NA | NA | NA | NA | NA | Gm24405 | predicted gene, 24405 [Source:MGI Symb | 0 | 0 | 0 | 0 | 0 | 0 |
| 18300 | ENSMUSG00000064968 | 0 | NA | NA | NA | NA | NA | Snord47 | small nucleolar RNA, C/D box 47 [Source:I | 0 | 0 | 0 | 0 | 0 | 0 |
| 18301 | ENSMUSG00000064970 | 0 | NA | NA | NA | NA | NA | Gm23690 | predicted gene, 23690 [Source:MGI Symb | 0 | 0 | 0 | 0 | 0 | 0 |
| 18302 | ENSMUSG00000064971 | 0 | NA | NA | NA | NA | NA | NA | NA | 0 | 0 | 0 | 0 | 0 | 0 |
| 18303 | ENSMUSG00000064972 | 0 | NA | NA | NA | NA | NA | NA | NA | 0 | 0 | 0 | 0 | 0 | 0 |
| 18304 | ENSMUSG00000064977 | 0 | NA | NA | NA | NA | NA | Gm26056 | predicted gene, 26056 [Source:MGI Symb | 0 | 0 | 0 | 0 | 0 | 0 |
| 18305 | ENSMUSG00000064981 | 0 | NA | NA | NA | NA | NA | NA | NA | 0 | 0 | 0 | 0 | 0 | 0 |
| 18306 | ENSMUSG00000064982 | 1.347202268 | -0.618342174 | 1.905496975 | -0.324504411 | 0.745556183 | NA | Gm25259 | predicted gene, 25259 [Source:MGI Symb | 0.8031 | 4.1117 | 0 | 0.964 | 1.0906 | 1.1138 |
| 18307 | ENSMUSG00000064983 | 0 | NA | NA | NA | NA | NA | NA | NA | 0 | 0 | 0 | 0 | 0 | 0 |
| 18308 | ENSMUSG00000064984 | 2.175134892 | -0.119623437 | 1.585778988 | -0.075435126 | 0.939868513 | NA | Snord73a | small nucleolar RNA, C/D box U73A [Sourc | 0.8031 | 0 | 6.0375 | 2.8919 | 1.0906 | 2.2277 |

| | | | | | | | | | | | | | | | |
|---|---|---|---|---|---|---|---|---|---|---|---|---|---|---|---|
| 18309 | ENSMUSG00000064987 | 0 | NA | NA | NA | NA | NA | Gm25260 | predicted gene, 25260 [Source:MGI Symb | 0 | 0 | 0 | 0 | 0 | 0 |
| 18310 | ENSMUSG00000064992 | 0 | NA | NA | NA | NA | NA | NA | NA | 0 | 0 | 0 | 0 | 0 | 0 |
| 18311 | ENSMUSG00000064994 | 0.133850373 | -0.796629176 | 4.080472857 | -0.195229623 | 0.84521319 | NA | Gm22422 | predicted gene, 22422 [Source:MGI Symb | 0.8031 | 0 | 0 | 0 | 0 | 0 |
| 18312 | ENSMUSG00000064995 | 0 | NA | NA | NA | NA | NA | Gm26003 | predicted gene, 26003 [Source:MGI Symb | 0 | 0 | 0 | 0 | 0 | 0 |
| 18313 | ENSMUSG00000064999 | 0 | NA | NA | NA | NA | NA | NA | NA | 0 | 0 | 0 | 0 | 0 | 0 |
| 18314 | ENSMUSG00000065004 | 0 | NA | NA | NA | NA | NA | Gm26326 | predicted gene, 26326 [Source:MGI Symb | 0 | 0 | 0 | 0 | 0 | 0 |
| 18315 | ENSMUSG00000065005 | 0 | NA | NA | NA | NA | NA | NA | NA | 0 | 0 | 0 | 0 | 0 | 0 |
| 18316 | ENSMUSG00000065006 | 0 | NA | NA | NA | NA | NA | Gm26325 | predicted gene, 26325 [Source:MGI Symb | 0 | 0 | 0 | 0 | 0 | 0 |
| 18317 | ENSMUSG00000065007 | 0 | NA | NA | NA | NA | NA | NA | NA | 0 | 0 | 0 | 0 | 0 | 0 |
| 18318 | ENSMUSG00000065011 | 0 | NA | NA | NA | NA | NA | NA | NA | 0 | 0 | 0 | 0 | 0 | 0 |
| 18319 | ENSMUSG00000065012 | 0 | NA | NA | NA | NA | NA | Gm23507 | predicted gene, 23507 [Source:MGI Symb | 0 | 0 | 0 | 0 | 0 | 0 |
| 18320 | ENSMUSG00000065013 | 0 | NA | NA | NA | NA | NA | Gm23508 | predicted gene, 23508 [Source:MGI Symb | 0 | 0 | 0 | 0 | 0 | 0 |
| 18321 | ENSMUSG00000065014 | 0 | NA | NA | NA | NA | NA | Gm23509 | predicted gene, 23509 [Source:MGI Symb | 0 | 0 | 0 | 0 | 0 | 0 |
| 18322 | ENSMUSG00000065016 | 0 | NA | NA | NA | NA | NA | Snora3 | small nucleolar RNA, H/ACA box 3 [Source | 0 | 0 | 0 | 0 | 0 | 0 |
| 18323 | ENSMUSG00000065017 | 0 | NA | NA | NA | NA | NA | NA | NA | 0 | 0 | 0 | 0 | 0 | 0 |
| 18324 | ENSMUSG00000065018 | 0 | NA | NA | NA | NA | NA | NA | NA | 0 | 0 | 0 | 0 | 0 | 0 |
| 18325 | ENSMUSG00000065022 | 0 | NA | NA | NA | NA | NA | AF357341 | snoRNA AF357341 [Source:MGI Symbol;A | 0 | 0 | 0 | 0 | 0 | 0 |
| 18326 | ENSMUSG00000065027 | 0 | NA | NA | NA | NA | NA | Gm25201 | predicted gene, 25201 [Source:MGI Symb | 0 | 0 | 0 | 0 | 0 | 0 |
| 18327 | ENSMUSG00000065028 | 0 | NA | NA | NA | NA | NA | Gm25200 | predicted gene, 25200 [Source:MGI Symb | 0 | 0 | 0 | 0 | 0 | 0 |
| 18328 | ENSMUSG00000065029 | 0 | NA | NA | NA | NA | NA | Gm25199 | predicted gene, 25199 [Source:MGI Symb | 0 | 0 | 0 | 0 | 0 | 0 |
| 18329 | ENSMUSG00000065030 | 0 | NA | NA | NA | NA | NA | NA | NA | 0 | 0 | 0 | 0 | 0 | 0 |
| 18330 | ENSMUSG00000065031 | 0 | NA | NA | NA | NA | NA | Gm22364 | predicted gene, 22364 [Source:MGI Symb | 0 | 0 | 0 | 0 | 0 | 0 |
| 18331 | ENSMUSG00000065034 | 0 | NA | NA | NA | NA | NA | Gm22363 | predicted gene, 22363 [Source:MGI Symb | 0 | 0 | 0 | 0 | 0 | 0 |
| 18332 | ENSMUSG00000065036 | 0.267700746 | -1.497481649 | 4.057001436 | -0.369110456 | 0.712045396 | NA | Gm22362 | predicted gene, 22362 [Source:MGI Symb | 1.6062 | 0 | 0 | 0 | 0 | 0 |
| 18333 | ENSMUSG00000065037 | 38.88359478 | 0.762249404 | 1.28456066 | 0.593393078 | 0.55291812 | 0.896226672 | NA | NA | 11.243 | 41.117 | 34.212 | 5.7839 | 128.69 | 12.252 |
| 18334 | ENSMUSG00000065040 | 0 | NA | NA | NA | NA | NA | Gm24977 | predicted gene, 24977 [Source:MGI Symb | 0 | 0 | 0 | 0 | 0 | 0 |
| 18335 | ENSMUSG00000065041 | 0 | NA | NA | NA | NA | NA | Gm24044 | predicted gene, 24044 [Source:MGI Symb | 0 | 0 | 0 | 0 | 0 | 0 |
| 18336 | ENSMUSG00000065042 | 0 | NA | NA | NA | NA | NA | Gm24043 | predicted gene, 24043 [Source:MGI Symb | 0 | 0 | 0 | 0 | 0 | 0 |
| 18337 | ENSMUSG00000065044 | 0 | NA | NA | NA | NA | NA | NA | NA | 0 | 0 | 0 | 0 | 0 | 0 |
| 18338 | ENSMUSG00000065045 | 0 | NA | NA | NA | NA | NA | Gm24046 | predicted gene, 24046 [Source:MGI Symb | 0 | 0 | 0 | 0 | 0 | 0 |
| 18339 | ENSMUSG00000065047 | 0.587191767 | -1.12394062 | 3.226482889 | -0.348348545 | 0.72757844 | NA | Gm24045 | predicted gene, 24045 [Source:MGI Symb | 2.4093 | 0 | 0 | 0 | 0 | 1.1138 |
| 18340 | ENSMUSG00000065048 | 0 | NA | NA | NA | NA | NA | Gm24042 | predicted gene, 24042 [Source:MGI Symb | 0 | 0 | 0 | 0 | 0 | 0 |
| 18341 | ENSMUSG00000065049 | 0 | NA | NA | NA | NA | NA | Gm24041 | predicted gene, 24041 [Source:MGI Symb | 0 | 0 | 0 | 0 | 0 | 0 |
| 18342 | ENSMUSG00000065050 | 0 | NA | NA | NA | NA | NA | Gm25704 | predicted gene, 25704 [Source:MGI Symb | 0 | 0 | 0 | 0 | 0 | 0 |
| 18343 | ENSMUSG00000065053 | 0 | NA | NA | NA | NA | NA | Gm25705 | predicted gene, 25705 [Source:MGI Symb | 0 | 0 | 0 | 0 | 0 | 0 |
| 18344 | ENSMUSG00000065055 | 0 | NA | NA | NA | NA | NA | NA | NA | 0 | 0 | 0 | 0 | 0 | 0 |
| 18345 | ENSMUSG00000065057 | 0.986836363 | -3.337272459 | 2.415410927 | -1.38165826 | 0.167076656 | NA | Gm25703 | predicted gene, 25703 [Source:MGI Symb | 0.8031 | 4.1117 | 1.0062 | 0 | 0 | 0 |
| 18346 | ENSMUSG00000065061 | 0 | NA | NA | NA | NA | NA | Gm22884 | predicted gene, 22884 [Source:MGI Symb | 0 | 0 | 0 | 0 | 0 | 0 |
| 18347 | ENSMUSG00000065068 | 0 | NA | NA | NA | NA | NA | NA | NA | 0 | 0 | 0 | 0 | 0 | 0 |
| 18348 | ENSMUSG00000065070 | 0 | NA | NA | NA | NA | NA | NA | NA | 0 | 0 | 0 | 0 | 0 | 0 |
| 18349 | ENSMUSG00000065072 | 0 | NA | NA | NA | NA | NA | Gm24533 | predicted gene, 24533 [Source:MGI Symb | 0 | 0 | 0 | 0 | 0 | 0 |
| 18350 | ENSMUSG00000065074 | 0 | NA | NA | NA | NA | NA | Gm24535 | predicted gene, 24535 [Source:MGI Symb | 0 | 0 | 0 | 0 | 0 | 0 |
| 18351 | ENSMUSG00000065078 | 0 | NA | NA | NA | NA | NA | Gm24536 | predicted gene, 24536 [Source:MGI Symb | 0 | 0 | 0 | 0 | 0 | 0 |
| 18352 | ENSMUSG00000065079 | 0 | NA | NA | NA | NA | NA | Gm23464 | predicted gene, 23464 [Source:MGI Symb | 0 | 0 | 0 | 0 | 0 | 0 |
| 18353 | ENSMUSG00000065080 | 0 | NA | NA | NA | NA | NA | Gm26497 | predicted gene, 26497 [Source:MGI Symb | 0 | 0 | 0 | 0 | 0 | 0 |
| 18354 | ENSMUSG00000065083 | 0 | NA | NA | NA | NA | NA | NA | NA | 0 | 0 | 0 | 0 | 0 | 0 |
| 18355 | ENSMUSG00000065084 | 0 | NA | NA | NA | NA | NA | NA | NA | 0 | 0 | 0 | 0 | 0 | 0 |
| 18356 | ENSMUSG00000065086 | 0 | NA | NA | NA | NA | NA | Gm26495 | predicted gene, 26495 [Source:MGI Symb | 0 | 0 | 0 | 0 | 0 | 0 |
| 18357 | ENSMUSG00000065087 | 0 | NA | NA | NA | NA | NA | Snord22 | small nucleolar RNA, C/D box 22 [Source:I | 0 | 0 | 0 | 0 | 0 | 0 |
| 18358 | ENSMUSG00000065088 | 0 | NA | NA | NA | NA | NA | NA | NA | 0 | 0 | 0 | 0 | 0 | 0 |
| 18359 | ENSMUSG00000065089 | 0.453341394 | -0.545391191 | 3.551113277 | -0.153583158 | 0.877938417 | NA | Gm26493 | predicted gene, 26493 [Source:MGI Symb | 1.6062 | 0 | 0 | 0 | 0 | 1.1138 |
| 18360 | ENSMUSG00000065090 | 0 | NA | NA | NA | NA | NA | Gm23677 | predicted gene, 23677 [Source:MGI Symb | 0 | 0 | 0 | 0 | 0 | 0 |
| 18361 | ENSMUSG00000065091 | 0 | NA | NA | NA | NA | NA | NA | NA | 0 | 0 | 0 | 0 | 0 | 0 |
| 18362 | ENSMUSG00000065094 | 0 | NA | NA | NA | NA | NA | Snord1a | small nucleolar RNA, C/D box 1A [Source:I | 0 | 0 | 0 | 0 | 0 | 0 |
| 18363 | ENSMUSG00000065095 | 0 | NA | NA | NA | NA | NA | NA | NA | 0 | 0 | 0 | 0 | 0 | 0 |
| 18364 | ENSMUSG00000065096 | 0 | NA | NA | NA | NA | NA | Gm23680 | predicted gene, 23680 [Source:MGI Symb | 0 | 0 | 0 | 0 | 0 | 0 |

| | | | | | | | | | | | | | | | |
|---|---|---|---|---|---|---|---|---|---|---|---|---|---|---|---|
| 18365 | ENSMUSG00000065097 | 0 | NA | NA | NA | NA | NA | Snora16a | small nucleolar RNA, H/ACA box 16A [Sou | 0 | 0 | 0 | 0 | 0 | 0 |
| 18366 | ENSMUSG00000065100 | 0 | NA | NA | NA | NA | NA | Gm26132 | predicted gene, 26132 [Source:MGI Symb | 0 | 0 | 0 | 0 | 0 | 0 |
| 18367 | ENSMUSG00000065102 | 0 | NA | NA | NA | NA | NA | NA | NA | 0 | 0 | 0 | 0 | 0 | 0 |
| 18368 | ENSMUSG00000065104 | 0 | NA | NA | NA | NA | NA | Gm26128 | predicted gene, 26128 [Source:MGI Symb | 0 | 0 | 0 | 0 | 0 | 0 |
| 18369 | ENSMUSG00000065105 | 0 | NA | NA | NA | NA | NA | Gm26130 | predicted gene, 26130 [Source:MGI Symb | 0 | 0 | 0 | 0 | 0 | 0 |
| 18370 | ENSMUSG00000065107 | 0 | NA | NA | NA | NA | NA | n-R5s88 | nuclear encoded rRNA 5S 88 [Source:MGI | 0 | 0 | 0 | 0 | 0 | 0 |
| 18371 | ENSMUSG00000065110 | 0 | NA | NA | NA | NA | NA | Snord61 | small nucleolar RNA, C/D box 61 [Source:I | 0 | 0 | 0 | 0 | 0 | 0 |
| 18372 | ENSMUSG00000065111 | 0 | NA | NA | NA | NA | NA | Gm25141 | predicted gene, 25141 [Source:MGI Symb | 0 | 0 | 0 | 0 | 0 | 0 |
| 18373 | ENSMUSG00000065115 | 0 | NA | NA | NA | NA | NA | NA | NA | 0 | 0 | 0 | 0 | 0 | 0 |
| 18374 | ENSMUSG00000065118 | 0 | NA | NA | NA | NA | NA | Gm23297 | predicted gene, 23297 [Source:MGI Symb | 0 | 0 | 0 | 0 | 0 | 0 |
| 18375 | ENSMUSG00000065122 | 0 | NA | NA | NA | NA | NA | Gm26126 | predicted gene, 26126 [Source:MGI Symb | 0 | 0 | 0 | 0 | 0 | 0 |
| 18376 | ENSMUSG00000065124 | 0 | NA | NA | NA | NA | NA | Snora65 | small nucleolar RNA, H/ACA box 65 [Sourc | 0 | 0 | 0 | 0 | 0 | 0 |
| 18377 | ENSMUSG00000065126 | 48.37257514 | 0.189146844 | 0.366974081 | 0.515422897 | 0.606257536 | 0.916376001 | Snord104 | small nucleolar RNA, C/D box 104 [Source | 28.109 | 65.787 | 42.262 | 54.947 | 52.35 | 46.781 |
| 18378 | ENSMUSG00000065127 | 0 | NA | NA | NA | NA | NA | NA | NA | 0 | 0 | 0 | 0 | 0 | 0 |
| 18379 | ENSMUSG00000065130 | 0.492645588 | 0.9565034 | 3.447648281 | 0.277436479 | 0.781444974 | NA | Gm24457 | predicted gene, 24457 [Source:MGI Symb | 0 | 1.0279 | 0 | 1.928 | 0 | 0 |
| 18380 | ENSMUSG00000065132 | 0 | NA | NA | NA | NA | NA | NA | NA | 0 | 0 | 0 | 0 | 0 | 0 |
| 18381 | ENSMUSG00000065138 | 0 | NA | NA | NA | NA | NA | NA | NA | 0 | 0 | 0 | 0 | 0 | 0 |
| 18382 | ENSMUSG00000065141 | 0 | NA | NA | NA | NA | NA | Gm22652 | predicted gene, 22652 [Source:MGI Symb | 0 | 0 | 0 | 0 | 0 | 0 |
| 18383 | ENSMUSG00000065143 | 0 | NA | NA | NA | NA | NA | Gm25644 | predicted gene, 25644 [Source:MGI Symb | 0 | 0 | 0 | 0 | 0 | 0 |
| 18384 | ENSMUSG00000065145 | 0.339027071 | -1.79174149 | 3.919585585 | -0.457125237 | 0.647581032 | NA | NA | NA | 0 | 1.0279 | 1.0062 | 0 | 0 | 0 |
| 18385 | ENSMUSG00000065147 | 7.670755447 | 0.332540042 | 0.789854414 | 0.42101435 | 0.673744603 | 0.931915649 | Snora31 | small nucleolar RNA, H/ACA box 31 [Sourc | 7.2279 | 6.1675 | 7.0437 | 13.496 | 7.6343 | 4.4554 |
| 18386 | ENSMUSG00000065151 | 0 | NA | NA | NA | NA | NA | Gm25480 | predicted gene, 25480 [Source:MGI Symb | 0 | 0 | 0 | 0 | 0 | 0 |
| 18387 | ENSMUSG00000065155 | 0 | NA | NA | NA | NA | NA | Gm25481 | predicted gene, 25481 [Source:MGI Symb | 0 | 0 | 0 | 0 | 0 | 0 |
| 18388 | ENSMUSG00000065158 | 0 | NA | NA | NA | NA | NA | NA | NA | 0 | 0 | 0 | 0 | 0 | 0 |
| 18389 | ENSMUSG00000065159 | 0 | NA | NA | NA | NA | NA | Gm25482 | predicted gene, 25482 [Source:MGI Symb | 0 | 0 | 0 | 0 | 0 | 0 |
| 18390 | ENSMUSG00000065160 | 0 | NA | NA | NA | NA | NA | Gm50452 | predicted gene, 50452 [Source:MGI Symb | 0 | 0 | 0 | 0 | 0 | 0 |
| 18391 | ENSMUSG00000065162 | 0.935098222 | -2.037664976 | 2.285197225 | -0.891680138 | 0.372564403 | NA | Gm23645 | predicted gene, 23645 [Source:MGI Symb | 1.6062 | 1.0279 | 2.0125 | 0.964 | 0 | 0 |
| 18392 | ENSMUSG00000065164 | 0.185640647 | 1.126924175 | 4.080472857 | 0.276174898 | 0.782413736 | NA | Gm23814 | predicted gene, 23814 [Source:MGI Symb | 0 | 0 | 0 | 0 | 0 | 1.1138 |
| 18393 | ENSMUSG00000065165 | 0 | NA | NA | NA | NA | NA | NA | NA | 0 | 0 | 0 | 0 | 0 | 0 |
| 18394 | ENSMUSG00000065173 | 0 | NA | NA | NA | NA | NA | Gm22127 | predicted gene, 22127 [Source:MGI Symb | 0 | 0 | 0 | 0 | 0 | 0 |
| 18395 | ENSMUSG00000065176 | 0 | NA | NA | NA | NA | NA | Rnu12 | RNA U12, small nuclear [Source:MGI Syml | 0 | 0 | 0 | 0 | 0 | 0 |
| 18396 | ENSMUSG00000065178 | 0 | NA | NA | NA | NA | NA | Gm22126 | predicted gene, 22126 [Source:MGI Symb | 0 | 0 | 0 | 0 | 0 | 0 |
| 18397 | ENSMUSG00000065181 | 0 | NA | NA | NA | NA | NA | Gm25139 | predicted gene, 25139 [Source:MGI Symb | 0 | 0 | 0 | 0 | 0 | 0 |
| 18398 | ENSMUSG00000065182 | 0 | NA | NA | NA | NA | NA | Gm25138 | predicted gene, 25138 [Source:MGI Symb | 0 | 0 | 0 | 0 | 0 | 0 |
| 18399 | ENSMUSG00000065185 | 0 | NA | NA | NA | NA | NA | Gm25137 | predicted gene, 25137 [Source:MGI Symb | 0 | 0 | 0 | 0 | 0 | 0 |
| 18400 | ENSMUSG00000065187 | 0 | NA | NA | NA | NA | NA | Gm25136 | predicted gene, 25136 [Source:MGI Symb | 0 | 0 | 0 | 0 | 0 | 0 |
| 18401 | ENSMUSG00000065188 | 0 | NA | NA | NA | NA | NA | Gm25140 | predicted gene, 25140 [Source:MGI Symb | 0 | 0 | 0 | 0 | 0 | 0 |
| 18402 | ENSMUSG00000065191 | 3.101378705 | -1.397547636 | 1.267672501 | -1.102451646 | 0.270265369 | NA | Gm23458 | predicted gene, 23458 [Source:MGI Symb | 6.4248 | 2.0558 | 5.0312 | 2.8919 | 1.0906 | 1.1138 |
| 18403 | ENSMUSG00000065192 | 0 | NA | NA | NA | NA | NA | Gm23460 | predicted gene, 23460 [Source:MGI Symb | 0 | 0 | 0 | 0 | 0 | 0 |
| 18404 | ENSMUSG00000065193 | 0 | NA | NA | NA | NA | NA | Gm23459 | predicted gene, 23459 [Source:MGI Symb | 0 | 0 | 0 | 0 | 0 | 0 |
| 18405 | ENSMUSG00000065195 | 0 | NA | NA | NA | NA | NA | NA | NA | 0 | 0 | 0 | 0 | 0 | 0 |
| 18406 | ENSMUSG00000065196 | 0 | NA | NA | NA | NA | NA | Snord85 | small nucleolar RNA, C/D box 85 [Source:I | 0 | 0 | 0 | 0 | 0 | 0 |
| 18407 | ENSMUSG00000065198 | 0.181769715 | 1.126924175 | 4.080472857 | 0.276174898 | 0.782413736 | NA | Gm23462 | predicted gene, 23462 [Source:MGI Symb | 0 | 0 | 0 | 0 | 1.0906 | 0 |
| 18408 | ENSMUSG00000065199 | 0 | NA | NA | NA | NA | NA | NA | NA | 0 | 0 | 0 | 0 | 0 | 0 |
| 18409 | ENSMUSG00000065200 | 0 | NA | NA | NA | NA | NA | Gm24613 | predicted gene, 24613 [Source:MGI Symb | 0 | 0 | 0 | 0 | 0 | 0 |
| 18410 | ENSMUSG00000065201 | 0 | NA | NA | NA | NA | NA | NA | NA | 0 | 0 | 0 | 0 | 0 | 0 |
| 18411 | ENSMUSG00000065202 | 0 | NA | NA | NA | NA | NA | Gm24615 | predicted gene, 24615 [Source:MGI Symb | 0 | 0 | 0 | 0 | 0 | 0 |
| 18412 | ENSMUSG00000065203 | 0 | NA | NA | NA | NA | NA | Gm24614 | predicted gene, 24614 [Source:MGI Symb | 0 | 0 | 0 | 0 | 0 | 0 |
| 18413 | ENSMUSG00000065204 | 0 | NA | NA | NA | NA | NA | Gm24610 | predicted gene, 24610 [Source:MGI Symb | 0 | 0 | 0 | 0 | 0 | 0 |
| 18414 | ENSMUSG00000065206 | 0 | NA | NA | NA | NA | NA | NA | NA | 0 | 0 | 0 | 0 | 0 | 0 |
| 18415 | ENSMUSG00000065208 | 0.849167191 | -0.527085716 | 2.389288686 | -0.220603613 | 0.825401088 | NA | Gm24616 | predicted gene, 24616 [Source:MGI Symb | 0 | 1.0279 | 2.0125 | 0.964 | 1.0906 | 0 |
| 18416 | ENSMUSG00000065211 | 0 | NA | NA | NA | NA | NA | Gm26265 | predicted gene, 26265 [Source:MGI Symb | 0 | 0 | 0 | 0 | 0 | 0 |
| 18417 | ENSMUSG00000065212 | 0 | NA | NA | NA | NA | NA | Gm26264 | predicted gene, 26264 [Source:MGI Symb | 0 | 0 | 0 | 0 | 0 | 0 |
| 18418 | ENSMUSG00000065215 | 0 | NA | NA | NA | NA | NA | Gm26263 | predicted gene, 26263 [Source:MGI Symb | 0 | 0 | 0 | 0 | 0 | 0 |
| 18419 | ENSMUSG00000065219 | 0.663965228 | -0.037580459 | 3.10181903 | -0.012115619 | 0.990333371 | NA | Snord32a | small nucleolar RNA, C/D box 32A [Source | 0 | 2.0558 | 0 | 1.928 | 0 | 0 |
| 18420 | ENSMUSG00000065220 | 0 | NA | NA | NA | NA | NA | Gm25794 | predicted gene, 25794 [Source:MGI Symb | 0 | 0 | 0 | 0 | 0 | 0 |

| | | | | | | | | | | | | | | | |
|---|---|---|---|---|---|---|---|---|---|---|---|---|---|---|---|
| 18421 | ENSMUSG00000065221 | 0 | NA | NA | NA | NA | NA | NA | NA | 0 | 0 | 0 | 0 | 0 | 0 |
| 18422 | ENSMUSG00000065223 | 0 | NA | NA | NA | NA | NA | Gm25792 | predicted gene, 25792 [Source:MGI Symb | 0 | 0 | 0 | 0 | 0 | 0 |
| 18423 | ENSMUSG00000065226 | 0 | NA | NA | NA | NA | NA | Gm25791 | predicted gene, 25791 [Source:MGI Symb | 0 | 0 | 0 | 0 | 0 | 0 |
| 18424 | ENSMUSG00000065227 | 0 | NA | NA | NA | NA | NA | NA | NA | 0 | 0 | 0 | 0 | 0 | 0 |
| 18425 | ENSMUSG00000065228 | 0 | NA | NA | NA | NA | NA | Gm25789 | predicted gene, 25789 [Source:MGI Symb | 0 | 0 | 0 | 0 | 0 | 0 |
| 18426 | ENSMUSG00000065231 | 0 | NA | NA | NA | NA | NA | Gm22972 | predicted gene, 22972 [Source:MGI Symb | 0 | 0 | 0 | 0 | 0 | 0 |
| 18427 | ENSMUSG00000065232 | 0 | NA | NA | NA | NA | NA | Gm22973 | predicted gene, 22973 [Source:MGI Symb | 0 | 0 | 0 | 0 | 0 | 0 |
| 18428 | ENSMUSG00000065233 | 0 | NA | NA | NA | NA | NA | NA | NA | 0 | 0 | 0 | 0 | 0 | 0 |
| 18429 | ENSMUSG00000065235 | 0 | NA | NA | NA | NA | NA | Gm22974 | predicted gene, 22974 [Source:MGI Symb | 0 | 0 | 0 | 0 | 0 | 0 |
| 18430 | ENSMUSG00000065236 | 0.852599439 | -1.816262765 | 2.863514483 | -0.634277485 | 0.525899747 | NA | NA | NA | 0 | 0 | 4.025 | 0 | 1.0906 | 0 |
| 18431 | ENSMUSG00000065237 | 0 | NA | NA | NA | NA | NA | NA | NA | 0 | 0 | 0 | 0 | 0 | 0 |
| 18432 | ENSMUSG00000065238 | 0 | NA | NA | NA | NA | NA | Gm22970 | predicted gene, 22970 [Source:MGI Symb | 0 | 0 | 0 | 0 | 0 | 0 |
| 18433 | ENSMUSG00000065239 | 0 | NA | NA | NA | NA | NA | Gm22971 | predicted gene, 22971 [Source:MGI Symb | 0 | 0 | 0 | 0 | 0 | 0 |
| 18434 | ENSMUSG00000065242 | 0 | NA | NA | NA | NA | NA | NA | NA | 0 | 0 | 0 | 0 | 0 | 0 |
| 18435 | ENSMUSG00000065244 | 0 | NA | NA | NA | NA | NA | NA | NA | 0 | 0 | 0 | 0 | 0 | 0 |
| 18436 | ENSMUSG00000065245 | 0.321325949 | 1.911575114 | 4.048267893 | 0.472195804 | 0.636787034 | NA | NA | NA | 0 | 0 | 0 | 1.928 | 0 | 0 |
| 18437 | ENSMUSG00000065246 | 0 | NA | NA | NA | NA | NA | Gm22297 | predicted gene, 22297 [Source:MGI Symb | 0 | 0 | 0 | 0 | 0 | 0 |
| 18438 | ENSMUSG00000065248 | 0 | NA | NA | NA | NA | NA | Gm22293 | predicted gene, 22293 [Source:MGI Symb | 0 | 0 | 0 | 0 | 0 | 0 |
| 18439 | ENSMUSG00000065251 | 0.17131964 | -0.796629176 | 4.080472857 | -0.195229623 | 0.84521319 | NA | NA | NA | 0 | 1.0279 | 0 | 0 | 0 | 0 |
| 18440 | ENSMUSG00000065253 | 0 | NA | NA | NA | NA | NA | NA | NA | 0 | 0 | 0 | 0 | 0 | 0 |
| 18441 | ENSMUSG00000065254 | 0 | NA | NA | NA | NA | NA | NA | NA | 0 | 0 | 0 | 0 | 0 | 0 |
| 18442 | ENSMUSG00000065255 | 0 | NA | NA | NA | NA | NA | Gm23974 | predicted gene, 23974 [Source:MGI Symb | 0 | 0 | 0 | 0 | 0 | 0 |
| 18443 | ENSMUSG00000065256 | 0 | NA | NA | NA | NA | NA | Gm23972 | predicted gene, 23972 [Source:MGI Symb | 0 | 0 | 0 | 0 | 0 | 0 |
| 18444 | ENSMUSG00000065258 | 0.831672659 | -0.604339699 | 2.59720557 | -0.232688435 | 0.81600335 | NA | Gm23969 | predicted gene, 23969 [Source:MGI Symb | 0 | 2.0558 | 1.0062 | 1.928 | 0 | 0 |
| 18445 | ENSMUSG00000065259 | 0.82806045 | -0.59635427 | 2.600266913 | -0.229343483 | 0.81860196 | NA | Snora30 | small nucleolar RNA, H/ACA box 30 [Sourc | 0 | 1.0279 | 2.0125 | 1.928 | 0 | 0 |
| 18446 | ENSMUSG00000065260 | 0 | NA | NA | NA | NA | NA | Gm23453 | predicted gene, 23453 [Source:MGI Symb | 0 | 0 | 0 | 0 | 0 | 0 |
| 18447 | ENSMUSG00000065261 | 0 | NA | NA | NA | NA | NA | NA | NA | 0 | 0 | 0 | 0 | 0 | 0 |
| 18448 | ENSMUSG00000065262 | 0 | NA | NA | NA | NA | NA | Gm23451 | predicted gene, 23451 [Source:MGI Symb | 0 | 0 | 0 | 0 | 0 | 0 |
| 18449 | ENSMUSG00000065265 | 0 | NA | NA | NA | NA | NA | Gm23455 | predicted gene, 23455 [Source:MGI Symb | 0 | 0 | 0 | 0 | 0 | 0 |
| 18450 | ENSMUSG00000065266 | 0 | NA | NA | NA | NA | NA | Gm23454 | predicted gene, 23454 [Source:MGI Symb | 0 | 0 | 0 | 0 | 0 | 0 |
| 18451 | ENSMUSG00000065269 | 0 | NA | NA | NA | NA | NA | NA | NA | 0 | 0 | 0 | 0 | 0 | 0 |
| 18452 | ENSMUSG00000065270 | 0 | NA | NA | NA | NA | NA | NA | NA | 0 | 0 | 0 | 0 | 0 | 0 |
| 18453 | ENSMUSG00000065272 | 0 | NA | NA | NA | NA | NA | Snord57 | small nucleolar RNA, C/D box 57 [Source:I | 0 | 0 | 0 | 0 | 0 | 0 |
| 18454 | ENSMUSG00000065273 | 1.198850927 | -2.412458447 | 2.309606273 | -1.044532341 | 0.296239164 | NA | Gm25128 | predicted gene, 25128 [Source:MGI Symb | 0 | 3.0838 | 3.0187 | 0 | 1.0906 | 0 |
| 18455 | ENSMUSG00000065274 | 0 | NA | NA | NA | NA | NA | Gm25125 | predicted gene, 25125 [Source:MGI Symb | 0 | 0 | 0 | 0 | 0 | 0 |
| 18456 | ENSMUSG00000065275 | 0 | NA | NA | NA | NA | NA | NA | NA | 0 | 0 | 0 | 0 | 0 | 0 |
| 18457 | ENSMUSG00000065277 | 0 | NA | NA | NA | NA | NA | Gm25126 | predicted gene, 25126 [Source:MGI Symb | 0 | 0 | 0 | 0 | 0 | 0 |
| 18458 | ENSMUSG00000065279 | 0 | NA | NA | NA | NA | NA | Gm25124 | predicted gene, 25124 [Source:MGI Symb | 0 | 0 | 0 | 0 | 0 | 0 |
| 18459 | ENSMUSG00000065280 | 0 | NA | NA | NA | NA | NA | Gm24453 | predicted gene, 24453 [Source:MGI Symb | 0 | 0 | 0 | 0 | 0 | 0 |
| 18460 | ENSMUSG00000065281 | 0 | NA | NA | NA | NA | NA | Gm24452 | predicted gene, 24452 [Source:MGI Symb | 0 | 0 | 0 | 0 | 0 | 0 |
| 18461 | ENSMUSG00000065282 | 0 | NA | NA | NA | NA | NA | Gm24455 | predicted gene, 24455 [Source:MGI Symb | 0 | 0 | 0 | 0 | 0 | 0 |
| 18462 | ENSMUSG00000065283 | 0 | NA | NA | NA | NA | NA | Gm24454 | predicted gene, 24454 [Source:MGI Symb | 0 | 0 | 0 | 0 | 0 | 0 |
| 18463 | ENSMUSG00000065287 | 0 | NA | NA | NA | NA | NA | Gm24451 | predicted gene, 24451 [Source:MGI Symb | 0 | 0 | 0 | 0 | 0 | 0 |
| 18464 | ENSMUSG00000065289 | 0 | NA | NA | NA | NA | NA | Gm23650 | predicted gene, 23650 [Source:MGI Symb | 0 | 0 | 0 | 0 | 0 | 0 |
| 18465 | ENSMUSG00000065291 | 0 | NA | NA | NA | NA | NA | Gm26125 | predicted gene, 26125 [Source:MGI Symb | 0 | 0 | 0 | 0 | 0 | 0 |
| 18466 | ENSMUSG00000065293 | 0 | NA | NA | NA | NA | NA | Gm26124 | predicted gene, 26124 [Source:MGI Symb | 0 | 0 | 0 | 0 | 0 | 0 |
| 18467 | ENSMUSG00000065295 | 0 | NA | NA | NA | NA | NA | Gm24134 | predicted gene, 24134 [Source:MGI Symb | 0 | 0 | 0 | 0 | 0 | 0 |
| 18468 | ENSMUSG00000065296 | 0.335414862 | -1.779047242 | 4.030804179 | -0.441362855 | 0.658950329 | NA | Gm26122 | predicted gene, 26122 [Source:MGI Symb | 0 | 0 | 2.0125 | 0 | 0 | 0 |
| 18469 | ENSMUSG00000065297 | 0 | NA | NA | NA | NA | NA | Gm26123 | predicted gene, 26123 [Source:MGI Symb | 0 | 0 | 0 | 0 | 0 | 0 |
| 18470 | ENSMUSG00000065298 | 0 | NA | NA | NA | NA | NA | Gm26121 | predicted gene, 26121 [Source:MGI Symb | 0 | 0 | 0 | 0 | 0 | 0 |
| 18471 | ENSMUSG00000065299 | 0 | NA | NA | NA | NA | NA | Gm23138 | predicted gene, 23138 [Source:MGI Symb | 0 | 0 | 0 | 0 | 0 | 0 |
| 18472 | ENSMUSG00000065301 | 0 | NA | NA | NA | NA | NA | Gm23248 | predicted gene, 23248 [Source:MGI Symb | 0 | 0 | 0 | 0 | 0 | 0 |
| 18473 | ENSMUSG00000065303 | 0 | NA | NA | NA | NA | NA | Gm23249 | predicted gene, 23249 [Source:MGI Symb | 0 | 0 | 0 | 0 | 0 | 0 |
| 18474 | ENSMUSG00000065304 | 0 | NA | NA | NA | NA | NA | Gm23245 | predicted gene, 23245 [Source:MGI Symb | 0 | 0 | 0 | 0 | 0 | 0 |
| 18475 | ENSMUSG00000065305 | 0 | NA | NA | NA | NA | NA | Gm23246 | predicted gene, 23246 [Source:MGI Symb | 0 | 0 | 0 | 0 | 0 | 0 |
| 18476 | ENSMUSG00000065307 | 0 | NA | NA | NA | NA | NA | NA | NA | 0 | 0 | 0 | 0 | 0 | 0 |

| | | | | | | | | | | | | | | | |
|---|---|---|---|---|---|---|---|---|---|---|---|---|---|---|---|
| 18477 | ENSMUSG00000065309 | 0 | NA | NA | NA | NA | NA | Gm23244 | predicted gene, 23244 [Source:MGI Symb | 0 | 0 | 0 | 0 | 0 | 0 |
| 18478 | ENSMUSG00000065311 | 0 | NA | NA | NA | NA | NA | NA | NA | 0 | 0 | 0 | 0 | 0 | 0 |
| 18479 | ENSMUSG00000065312 | 0 | NA | NA | NA | NA | NA | Gm26063 | predicted gene, 26063 [Source:MGI Symb | 0 | 0 | 0 | 0 | 0 | 0 |
| 18480 | ENSMUSG00000065313 | 0 | NA | NA | NA | NA | NA | NA | NA | 0 | 0 | 0 | 0 | 0 | 0 |
| 18481 | ENSMUSG00000065314 | 0 | NA | NA | NA | NA | NA | NA | NA | 0 | 0 | 0 | 0 | 0 | 0 |
| 18482 | ENSMUSG00000065315 | 0 | NA | NA | NA | NA | NA | Gm26064 | predicted gene, 26064 [Source:MGI Symb | 0 | 0 | 0 | 0 | 0 | 0 |
| 18483 | ENSMUSG00000065319 | 0 | NA | NA | NA | NA | NA | NA | NA | 0 | 0 | 0 | 0 | 0 | 0 |
| 18484 | ENSMUSG00000065320 | 0.353089354 | 0.165142706 | 3.896102244 | 0.042386646 | 0.966190474 | NA | Gm22056 | predicted gene, 22056 [Source:MGI Symb | 0 | 1.0279 | 0 | 0 | 1.0906 | 0 |
| 18485 | ENSMUSG00000065321 | 0 | NA | NA | NA | NA | NA | NA | NA | 0 | 0 | 0 | 0 | 0 | 0 |
| 18486 | ENSMUSG00000065322 | 0 | NA | NA | NA | NA | NA | Gm22055 | predicted gene, 22055 [Source:MGI Symb | 0 | 0 | 0 | 0 | 0 | 0 |
| 18487 | ENSMUSG00000065327 | 0 | NA | NA | NA | NA | NA | NA | NA | 0 | 0 | 0 | 0 | 0 | 0 |
| 18488 | ENSMUSG00000065328 | 0 | NA | NA | NA | NA | NA | Gm22053 | predicted gene, 22053 [Source:MGI Symb | 0 | 0 | 0 | 0 | 0 | 0 |
| 18489 | ENSMUSG00000065329 | 0 | NA | NA | NA | NA | NA | Gm22054 | predicted gene, 22054 [Source:MGI Symb | 0 | 0 | 0 | 0 | 0 | 0 |
| 18490 | ENSMUSG00000065330 | 0 | NA | NA | NA | NA | NA | NA | NA | 0 | 0 | 0 | 0 | 0 | 0 |
| 18491 | ENSMUSG00000065331 | 6.339108515 | -0.212981053 | 0.863652124 | -0.24660514 | 0.805213827 | 0.96404377 | Gm24927 | predicted gene, 24927 [Source:MGI Symb | 7.2279 | 7.1954 | 6.0375 | 7.7118 | 7.6343 | 2.2277 |
| 18492 | ENSMUSG00000065333 | 0 | NA | NA | NA | NA | NA | Gm24928 | predicted gene, 24928 [Source:MGI Symb | 0 | 0 | 0 | 0 | 0 | 0 |
| 18493 | ENSMUSG00000065336 | 1.038213457 | 2.372140906 | 2.221591093 | 1.067766662 | 0.28562578 | NA | NA | NA | 0 | 0 | 1.0062 | 1.928 | 2.1812 | 1.1138 |
| 18494 | ENSMUSG00000065337 | 0 | NA | NA | NA | NA | NA | Gm24929 | predicted gene, 24929 [Source:MGI Symb | 0 | 0 | 0 | 0 | 0 | 0 |
| 18495 | ENSMUSG00000065340 | 0 | NA | NA | NA | NA | NA | Gm25578 | predicted gene, 25578 [Source:MGI Symb | 0 | 0 | 0 | 0 | 0 | 0 |
| 18496 | ENSMUSG00000065341 | 0 | NA | NA | NA | NA | NA | Gm25579 | predicted gene, 25579 [Source:MGI Symb | 0 | 0 | 0 | 0 | 0 | 0 |
| 18497 | ENSMUSG00000065343 | 0.160662974 | 1.126924175 | 4.080472857 | 0.276174898 | 0.782413736 | NA | NA | NA | 0 | 0 | 0 | 0.964 | 0 | 0 |
| 18498 | ENSMUSG00000065344 | 0 | NA | NA | NA | NA | NA | Gm25581 | predicted gene, 25581 [Source:MGI Symb | 0 | 0 | 0 | 0 | 0 | 0 |
| 18499 | ENSMUSG00000065346 | 0 | NA | NA | NA | NA | NA | Gm25582 | predicted gene, 25582 [Source:MGI Symb | 0 | 0 | 0 | 0 | 0 | 0 |
| 18500 | ENSMUSG00000065347 | 0 | NA | NA | NA | NA | NA | NA | NA | 0 | 0 | 0 | 0 | 0 | 0 |
| 18501 | ENSMUSG00000065348 | 0 | NA | NA | NA | NA | NA | NA | NA | 0 | 0 | 0 | 0 | 0 | 0 |
| 18502 | ENSMUSG00000065353 | 0 | NA | NA | NA | NA | NA | Snora73b | small nucleolar RNA, H/ACA box 73b [Sou | 0 | 0 | 0 | 0 | 0 | 0 |
| 18503 | ENSMUSG00000065360 | 0.449470461 | -0.545136533 | 3.562105917 | -0.15303771 | 0.878368537 | NA | Gm24412 | predicted gene, 24412 [Source:MGI Symb | 1.6062 | 0 | 0 | 0 | 1.0906 | 0 |
| 18504 | ENSMUSG00000065361 | 0 | NA | NA | NA | NA | NA | NA | NA | 0 | 0 | 0 | 0 | 0 | 0 |
| 18505 | ENSMUSG00000065362 | 0 | NA | NA | NA | NA | NA | Gm24411 | predicted gene, 24411 [Source:MGI Symb | 0 | 0 | 0 | 0 | 0 | 0 |
| 18506 | ENSMUSG00000065367 | 0 | NA | NA | NA | NA | NA | NA | NA | 0 | 0 | 0 | 0 | 0 | 0 |
| 18507 | ENSMUSG00000065368 | 0 | NA | NA | NA | NA | NA | NA | NA | 0 | 0 | 0 | 0 | 0 | 0 |
| 18508 | ENSMUSG00000065371 | 0 | NA | NA | NA | NA | NA | Gm22739 | predicted gene, 22739 [Source:MGI Symb | 0 | 0 | 0 | 0 | 0 | 0 |
| 18509 | ENSMUSG00000065372 | 0.17131964 | -0.796629176 | 4.080472857 | -0.195229623 | 0.84521319 | NA | Gm22741 | predicted gene, 22741 [Source:MGI Symb | 0 | 1.0279 | 0 | 0 | 0 | 0 |
| 18510 | ENSMUSG00000065373 | 0 | NA | NA | NA | NA | NA | Gm22740 | predicted gene, 22740 [Source:MGI Symb | 0 | 0 | 0 | 0 | 0 | 0 |
| 18511 | ENSMUSG00000065374 | 0.133850373 | -0.796629176 | 4.080472857 | -0.195229623 | 0.84521319 | NA | Gm22738 | predicted gene, 22738 [Source:MGI Symb | 0.8031 | 0 | 0 | 0 | 0 | 0 |
| 18512 | ENSMUSG00000065378 | 0 | NA | NA | NA | NA | NA | Gm22744 | predicted gene, 22744 [Source:MGI Symb | 0 | 0 | 0 | 0 | 0 | 0 |
| 18513 | ENSMUSG00000065379 | 0 | NA | NA | NA | NA | NA | Gm22743 | predicted gene, 22743 [Source:MGI Symb | 0 | 0 | 0 | 0 | 0 | 0 |
| 18514 | ENSMUSG00000065380 | 0 | NA | NA | NA | NA | NA | Gm23095 | predicted gene, 23095 [Source:MGI Symb | 0 | 0 | 0 | 0 | 0 | 0 |
| 18515 | ENSMUSG00000065381 | 0 | NA | NA | NA | NA | NA | NA | NA | 0 | 0 | 0 | 0 | 0 | 0 |
| 18516 | ENSMUSG00000065382 | 0 | NA | NA | NA | NA | NA | Gm23097 | predicted gene, 23097 [Source:MGI Symb | 0 | 0 | 0 | 0 | 0 | 0 |
| 18517 | ENSMUSG00000065384 | 0 | NA | NA | NA | NA | NA | Gm23093 | predicted gene, 23093 [Source:MGI Symb | 0 | 0 | 0 | 0 | 0 | 0 |
| 18518 | ENSMUSG00000065385 | 0 | NA | NA | NA | NA | NA | NA | NA | 0 | 0 | 0 | 0 | 0 | 0 |
| 18519 | ENSMUSG00000065387 | 0.17131964 | -0.796629176 | 4.080472857 | -0.195229623 | 0.84521319 | NA | n-R5s85 | nuclear encoded rRNA 5S 85 [Source:MGI | 0 | 1.0279 | 0 | 0 | 0 | 0 |
| 18520 | ENSMUSG00000065388 | 0 | NA | NA | NA | NA | NA | Gm23099 | predicted gene, 23099 [Source:MGI Symb | 0 | 0 | 0 | 0 | 0 | 0 |
| 18521 | ENSMUSG00000065389 | 0 | NA | NA | NA | NA | NA | Gm23100 | predicted gene, 23100 [Source:MGI Symb | 0 | 0 | 0 | 0 | 0 | 0 |
| 18522 | ENSMUSG00000065392 | 0 | NA | NA | NA | NA | NA | Gm25894 | predicted gene, 25894 [Source:MGI Symb | 0 | 0 | 0 | 0 | 0 | 0 |
| 18523 | ENSMUSG00000065394 | 0 | NA | NA | NA | NA | NA | Mir25 | microRNA 25 [Source:MGI Symbol;Acc:M( | 0 | 0 | 0 | 0 | 0 | 0 |
| 18524 | ENSMUSG00000065395 | 0 | NA | NA | NA | NA | NA | Mir193a | microRNA 193a [Source:MGI Symbol;Acc: | 0 | 0 | 0 | 0 | 0 | 0 |
| 18525 | ENSMUSG00000065396 | 0 | NA | NA | NA | NA | NA | Mir99b | microRNA 99b [Source:MGI Symbol;Acc:N | 0 | 0 | 0 | 0 | 0 | 0 |
| 18526 | ENSMUSG00000065397 | 0 | NA | NA | NA | NA | NA | Mir155 | microRNA 155 [Source:MGI Symbol;Acc:N | 0 | 0 | 0 | 0 | 0 | 0 |
| 18527 | ENSMUSG00000065398 | 0 | NA | NA | NA | NA | NA | Mir188 | microRNA 188 [Source:MGI Symbol;Acc:N | 0 | 0 | 0 | 0 | 0 | 0 |
| 18528 | ENSMUSG00000065399 | 0 | NA | NA | NA | NA | NA | Mir133a-1 | microRNA 133a-1 [Source:MGI Symbol;Ac | 0 | 0 | 0 | 0 | 0 | 0 |
| 18529 | ENSMUSG00000065400 | 0 | NA | NA | NA | NA | NA | Mir200a | microRNA 200a [Source:MGI Symbol;Acc: | 0 | 0 | 0 | 0 | 0 | 0 |
| 18530 | ENSMUSG00000065401 | 0 | NA | NA | NA | NA | NA | Mir144 | microRNA 144 [Source:MGI Symbol;Acc:N | 0 | 0 | 0 | 0 | 0 | 0 |
| 18531 | ENSMUSG00000065402 | 0.831492699 | -1.816502146 | 2.887260166 | -0.629143909 | 0.529254847 | NA | Mir122 | microRNA 122 [Source:MGI Symbol;Acc:N | 0 | 0 | 4.025 | 0.964 | 0 | 0 |
| 18532 | ENSMUSG00000065403 | 0 | NA | NA | NA | NA | NA | Mir18 | microRNA 18 [Source:MGI Symbol;Acc:M( | 0 | 0 | 0 | 0 | 0 | 0 |

| | | | | | | | | | | | | | | | |
|---|---|---|---|---|---|---|---|---|---|---|---|---|---|---|---|
| 18533 | ENSMUSG00000065404 | 0 | NA | NA | NA | NA | NA | Mir201 | microRNA 201 [Source:MGI Symbol;Acc:M | 0 | 0 | 0 | 0 | 0 | 0 |
| 18534 | ENSMUSG00000065405 | 0 | NA | NA | NA | NA | NA | Mir30a | microRNA 30a [Source:MGI Symbol;Acc:M | 0 | 0 | 0 | 0 | 0 | 0 |
| 18535 | ENSMUSG00000065406 | 0 | NA | NA | NA | NA | NA | Mirlet7i | microRNA let7i [Source:MGI Symbol;Acc:M | 0 | 0 | 0 | 0 | 0 | 0 |
| 18536 | ENSMUSG00000065407 | 0 | NA | NA | NA | NA | NA | Mir135a-1 | microRNA 135a-1 [Source:MGI Symbol;Ac | 0 | 0 | 0 | 0 | 0 | 0 |
| 18537 | ENSMUSG00000065408 | 0 | NA | NA | NA | NA | NA | Mir31 | microRNA 31 [Source:MGI Symbol;Acc:MG | 0 | 0 | 0 | 0 | 0 | 0 |
| 18538 | ENSMUSG00000065409 | 0 | NA | NA | NA | NA | NA | NA | NA | 0 | 0 | 0 | 0 | 0 | 0 |
| 18539 | ENSMUSG00000065410 | 0 | NA | NA | NA | NA | NA | Mir298 | microRNA 298 [Source:MGI Symbol;Acc:M | 0 | 0 | 0 | 0 | 0 | 0 |
| 18540 | ENSMUSG00000065411 | 0 | NA | NA | NA | NA | NA | Mir195a | microRNA 195a [Source:MGI Symbol;Acc: | 0 | 0 | 0 | 0 | 0 | 0 |
| 18541 | ENSMUSG00000065412 | 0 | NA | NA | NA | NA | NA | Mir29b-2 | microRNA 29b-2 [Source:MGI Symbol;Acc | 0 | 0 | 0 | 0 | 0 | 0 |
| 18542 | ENSMUSG00000065413 | 0 | NA | NA | NA | NA | NA | Mir367 | microRNA 367 [Source:MGI Symbol;Acc:M | 0 | 0 | 0 | 0 | 0 | 0 |
| 18543 | ENSMUSG00000065414 | 0 | NA | NA | NA | NA | NA | Mir138-1 | microRNA 138-1 [Source:MGI Symbol;Acc | 0 | 0 | 0 | 0 | 0 | 0 |
| 18544 | ENSMUSG00000065415 | 0 | NA | NA | NA | NA | NA | Mir217 | microRNA 217 [Source:MGI Symbol;Acc:M | 0 | 0 | 0 | 0 | 0 | 0 |
| 18545 | ENSMUSG00000065416 | 0 | NA | NA | NA | NA | NA | Mir19a | microRNA 19a [Source:MGI Symbol;Acc:M | 0 | 0 | 0 | 0 | 0 | 0 |
| 18546 | ENSMUSG00000065417 | 0 | NA | NA | NA | NA | NA | Mir340 | microRNA 340 [Source:MGI Symbol;Acc:M | 0 | 0 | 0 | 0 | 0 | 0 |
| 18547 | ENSMUSG00000065418 | 0 | NA | NA | NA | NA | NA | Mir322 | microRNA 322 [Source:MGI Symbol;Acc:M | 0 | 0 | 0 | 0 | 0 | 0 |
| 18548 | ENSMUSG00000065420 | 0.335414862 | -1.779047242 | 4.030804179 | -0.441362855 | 0.658950329 | NA | Mir142b | microRNA 142b [Source:MGI Symbol;Acc: | 0 | 0 | 2.0125 | 0 | 0 | 0 |
| 18549 | ENSMUSG00000065421 | 0.342432689 | 1.990653086 | 3.900281263 | 0.51038706 | 0.609780321 | NA | Mirlet7a-1 | microRNA let7a-1 [Source:MGI Symbol;Ac | 0 | 0 | 0 | 0.964 | 1.0906 | 0 |
| 18550 | ENSMUSG00000065422 | 0.181769715 | 1.126924175 | 4.080472857 | 0.276174898 | 0.782413736 | NA | Mir221 | microRNA 221 [Source:MGI Symbol;Acc:M | 0 | 0 | 0 | 0 | 1.0906 | 0 |
| 18551 | ENSMUSG00000065423 | 0 | NA | NA | NA | NA | NA | Mir181a-2 | microRNA 181a-2 [Source:MGI Symbol;Ac | 0 | 0 | 0 | 0 | 0 | 0 |
| 18552 | ENSMUSG00000065424 | 0 | NA | NA | NA | NA | NA | Mir9-2 | microRNA 9-2 [Source:MGI Symbol;Acc:M | 0 | 0 | 0 | 0 | 0 | 0 |
| 18553 | ENSMUSG00000065425 | 0 | NA | NA | NA | NA | NA | Mir325 | microRNA 325 [Source:MGI Symbol;Acc:M | 0 | 0 | 0 | 0 | 0 | 0 |
| 18554 | ENSMUSG00000065426 | 0 | NA | NA | NA | NA | NA | Mir134 | microRNA 134 [Source:MGI Symbol;Acc:M | 0 | 0 | 0 | 0 | 0 | 0 |
| 18555 | ENSMUSG00000065427 | 0 | NA | NA | NA | NA | NA | Mir429 | microRNA 429 [Source:MGI Symbol;Acc:M | 0 | 0 | 0 | 0 | 0 | 0 |
| 18556 | ENSMUSG00000065428 | 0 | NA | NA | NA | NA | NA | Mir382 | microRNA 382 [Source:MGI Symbol;Acc:M | 0 | 0 | 0 | 0 | 0 | 0 |
| 18557 | ENSMUSG00000065429 | 0 | NA | NA | NA | NA | NA | Mir345 | microRNA 345 [Source:MGI Symbol;Acc:M | 0 | 0 | 0 | 0 | 0 | 0 |
| 18558 | ENSMUSG00000065430 | 0.160662974 | 1.126924175 | 4.080472857 | 0.276174898 | 0.782413736 | NA | Mir26a-2 | microRNA 26a-2 [Source:MGI Symbol;Acc | 0 | 0 | 0 | 0.964 | 0 | 0 |
| 18559 | ENSMUSG00000065431 | 5.80673915 | 1.074748052 | 0.927277716 | 1.159035781 | 0.2464416 | 0.696725814 | Mir186 | microRNA 186 [Source:MGI Symbol;Acc:M | 4.0155 | 5.1396 | 2.0125 | 3.8559 | 10.906 | 8.9108 |
| 18560 | ENSMUSG00000065432 | 0 | NA | NA | NA | NA | NA | Mir208a | microRNA 208a [Source:MGI Symbol;Acc: | 0 | 0 | 0 | 0 | 0 | 0 |
| 18561 | ENSMUSG00000065433 | 0 | NA | NA | NA | NA | NA | Mir370 | microRNA 370 [Source:MGI Symbol;Acc:M | 0 | 0 | 0 | 0 | 0 | 0 |
| 18562 | ENSMUSG00000065434 | 5.856809263 | 0.395730029 | 0.898938636 | 0.44021918 | 0.65977837 | 0.927787797 | Mir7-1 | microRNA 7-1 [Source:MGI Symbol;Acc:M | 4.0155 | 7.1954 | 4.025 | 6.7478 | 9.8156 | 3.3415 |
| 18563 | ENSMUSG00000065435 | 0 | NA | NA | NA | NA | NA | Mir135b | microRNA 135b [Source:MGI Symbol;Acc: | 0 | 0 | 0 | 0 | 0 | 0 |
| 18564 | ENSMUSG00000065436 | 0 | NA | NA | NA | NA | NA | Mir342 | microRNA 342 [Source:MGI Symbol;Acc:M | 0 | 0 | 0 | 0 | 0 | 0 |
| 18565 | ENSMUSG00000065437 | 0 | NA | NA | NA | NA | NA | Mir30d | microRNA 30d [Source:MGI Symbol;Acc:M | 0 | 0 | 0 | 0 | 0 | 0 |
| 18566 | ENSMUSG00000065438 | 0 | NA | NA | NA | NA | NA | Mir377 | microRNA 377 [Source:MGI Symbol;Acc:M | 0 | 0 | 0 | 0 | 0 | 0 |
| 18567 | ENSMUSG00000065439 | 0 | NA | NA | NA | NA | NA | Mir140 | microRNA 140 [Source:MGI Symbol;Acc:M | 0 | 0 | 0 | 0 | 0 | 0 |
| 18568 | ENSMUSG00000065440 | 0 | NA | NA | NA | NA | NA | Mirlet7g | microRNA let7g [Source:MGI Symbol;Acc: | 0 | 0 | 0 | 0 | 0 | 0 |
| 18569 | ENSMUSG00000065441 | 0 | NA | NA | NA | NA | NA | Mir128-2 | microRNA 128-2 [Source:MGI Symbol;Acc | 0 | 0 | 0 | 0 | 0 | 0 |
| 18570 | ENSMUSG00000065442 | 0 | NA | NA | NA | NA | NA | Mir20a | microRNA 20a [Source:MGI Symbol;Acc:M | 0 | 0 | 0 | 0 | 0 | 0 |
| 18571 | ENSMUSG00000065443 | 0 | NA | NA | NA | NA | NA | Mir196b | microRNA 196b [Source:MGI Symbol;Acc: | 0 | 0 | 0 | 0 | 0 | 0 |
| 18572 | ENSMUSG00000065444 | 0 | NA | NA | NA | NA | NA | Mir27a | microRNA 27a [Source:MGI Symbol;Acc:M | 0 | 0 | 0 | 0 | 0 | 0 |
| 18573 | ENSMUSG00000065445 | 0 | NA | NA | NA | NA | NA | Mir143 | microRNA 143 [Source:MGI Symbol;Acc:M | 0 | 0 | 0 | 0 | 0 | 0 |
| 18574 | ENSMUSG00000065446 | 0 | NA | NA | NA | NA | NA | Mir139 | microRNA 139 [Source:MGI Symbol;Acc:M | 0 | 0 | 0 | 0 | 0 | 0 |
| 18575 | ENSMUSG00000065447 | 0 | NA | NA | NA | NA | NA | Mir302d | microRNA 302d [Source:MGI Symbol;Acc: | 0 | 0 | 0 | 0 | 0 | 0 |
| 18576 | ENSMUSG00000065448 | 0 | NA | NA | NA | NA | NA | Mir154 | microRNA 154 [Source:MGI Symbol;Acc:M | 0 | 0 | 0 | 0 | 0 | 0 |
| 18577 | ENSMUSG00000065449 | 0 | NA | NA | NA | NA | NA | Mir365-1 | microRNA 365-1 [Source:MGI Symbol;Acc | 0 | 0 | 0 | 0 | 0 | 0 |
| 18578 | ENSMUSG00000065450 | 0 | NA | NA | NA | NA | NA | Mir448 | microRNA 448 [Source:MGI Symbol;Acc:M | 0 | 0 | 0 | 0 | 0 | 0 |
| 18579 | ENSMUSG00000065451 | 0 | NA | NA | NA | NA | NA | Mir101a | microRNA 101a [Source:MGI Symbol;Acc: | 0 | 0 | 0 | 0 | 0 | 0 |
| 18580 | ENSMUSG00000065452 | 0 | NA | NA | NA | NA | NA | Mir207 | microRNA 207 [Source:MGI Symbol;Acc:M | 0 | 0 | 0 | 0 | 0 | 0 |
| 18581 | ENSMUSG00000065453 | 0 | NA | NA | NA | NA | NA | Mirlet7d | microRNA let7d [Source:MGI Symbol;Acc: | 0 | 0 | 0 | 0 | 0 | 0 |
| 18582 | ENSMUSG00000065454 | 0 | NA | NA | NA | NA | NA | Mir124a-3 | microRNA 124a-3 [Source:MGI Symbol;Ac | 0 | 0 | 0 | 0 | 0 | 0 |
| 18583 | ENSMUSG00000065455 | 0 | NA | NA | NA | NA | NA | Mir21a | microRNA 21a [Source:MGI Symbol;Acc:M | 0 | 0 | 0 | 0 | 0 | 0 |
| 18584 | ENSMUSG00000065456 | 0 | NA | NA | NA | NA | NA | Mir106a | microRNA 106a [Source:MGI Symbol;Acc: | 0 | 0 | 0 | 0 | 0 | 0 |
| 18585 | ENSMUSG00000065457 | 0 | NA | NA | NA | NA | NA | Mir383 | microRNA 383 [Source:MGI Symbol;Acc:M | 0 | 0 | 0 | 0 | 0 | 0 |
| 18586 | ENSMUSG00000065458 | 0 | NA | NA | NA | NA | NA | Mir181b-1 | microRNA 181b-1 [Source:MGI Symbol;Ac | 0 | 0 | 0 | 0 | 0 | 0 |
| 18587 | ENSMUSG00000065460 | 0 | NA | NA | NA | NA | NA | Mir133a-2 | microRNA 133a-2 [Source:MGI Symbol;Ac | 0 | 0 | 0 | 0 | 0 | 0 |
| 18588 | ENSMUSG00000065462 | 0 | NA | NA | NA | NA | NA | Mir200c | microRNA 200c [Source:MGI Symbol;Acc: | 0 | 0 | 0 | 0 | 0 | 0 |

| | | | | | | | | | | | | | | | |
|---|---|---|---|---|---|---|---|---|---|---|---|---|---|---|---|
| 18589 | ENSMUSG00000065463 | 0 | NA | NA | NA | NA | NA | Snord45c | small nucleolar RNA, C/D box 45C [Source | 0 | 0 | 0 | 0 | 0 | 0 |
| 18590 | ENSMUSG00000065464 | 0 | NA | NA | NA | NA | NA | Mir185 | microRNA 185 [Source:MGI Symbol;Acc:M | 0 | 0 | 0 | 0 | 0 | 0 |
| 18591 | ENSMUSG00000065465 | 0 | NA | NA | NA | NA | NA | Mir33 | microRNA 33 [Source:MGI Symbol;Acc:M( | 0 | 0 | 0 | 0 | 0 | 0 |
| 18592 | ENSMUSG00000065466 | 0 | NA | NA | NA | NA | NA | NA | NA | 0 | 0 | 0 | 0 | 0 | 0 |
| 18593 | ENSMUSG00000065468 | 0.835104907 | -1.8196403 | 2.6116807 | -0.696731534 | 0.48597082 | NA | Mir26b | microRNA 26b [Source:MGI Symbol;Acc:M | 0 | 1.0279 | 3.0187 | 0.964 | 0 | 0 |
| 18594 | ENSMUSG00000065469 | 0 | NA | NA | NA | NA | NA | Mir129-1 | microRNA 129-1 [Source:MGI Symbol;Acc | 0 | 0 | 0 | 0 | 0 | 0 |
| 18595 | ENSMUSG00000065470 | 0 | NA | NA | NA | NA | NA | Mir149 | microRNA 149 [Source:MGI Symbol;Acc:M | 0 | 0 | 0 | 0 | 0 | 0 |
| 18596 | ENSMUSG00000065471 | 0 | NA | NA | NA | NA | NA | Mir222 | microRNA 222 [Source:MGI Symbol;Acc:M | 0 | 0 | 0 | 0 | 0 | 0 |
| 18597 | ENSMUSG00000065472 | 0 | NA | NA | NA | NA | NA | Mir125b-2 | microRNA 125b-2 [Source:MGI Symbol;Ac | 0 | 0 | 0 | 0 | 0 | 0 |
| 18598 | ENSMUSG00000065473 | 0 | NA | NA | NA | NA | NA | Mir19b-2 | microRNA 19b-2 [Source:MGI Symbol;Acc | 0 | 0 | 0 | 0 | 0 | 0 |
| 18599 | ENSMUSG00000065474 | 0 | NA | NA | NA | NA | NA | Mir141 | microRNA 141 [Source:MGI Symbol;Acc:M | 0 | 0 | 0 | 0 | 0 | 0 |
| 18600 | ENSMUSG00000065475 | 0 | NA | NA | NA | NA | NA | Mir27b | microRNA 27b [Source:MGI Symbol;Acc:M | 0 | 0 | 0 | 0 | 0 | 0 |
| 18601 | ENSMUSG00000065476 | 0 | NA | NA | NA | NA | NA | Mir30b | microRNA 30b [Source:MGI Symbol;Acc:M | 0 | 0 | 0 | 0 | 0 | 0 |
| 18602 | ENSMUSG00000065477 | 0 | NA | NA | NA | NA | NA | Mir411 | microRNA 411 [Source:MGI Symbol;Acc:M | 0 | 0 | 0 | 0 | 0 | 0 |
| 18603 | ENSMUSG00000065479 | 0 | NA | NA | NA | NA | NA | Mir125a | microRNA 125a [Source:MGI Symbol;Acc: | 0 | 0 | 0 | 0 | 0 | 0 |
| 18604 | ENSMUSG00000065480 | 0 | NA | NA | NA | NA | NA | Mir133b | microRNA 133b [Source:MGI Symbol;Acc: | 0 | 0 | 0 | 0 | 0 | 0 |
| 18605 | ENSMUSG00000065481 | 0 | NA | NA | NA | NA | NA | Mir346 | microRNA 346 [Source:MGI Symbol;Acc:M | 0 | 0 | 0 | 0 | 0 | 0 |
| 18606 | ENSMUSG00000065482 | 0 | NA | NA | NA | NA | NA | Mir302c | microRNA 302c [Source:MGI Symbol;Acc:I | 0 | 0 | 0 | 0 | 0 | 0 |
| 18607 | ENSMUSG00000065483 | 0 | NA | NA | NA | NA | NA | Mir181c | microRNA 181c [Source:MGI Symbol;Acc:I | 0 | 0 | 0 | 0 | 0 | 0 |
| 18608 | ENSMUSG00000065484 | 0 | NA | NA | NA | NA | NA | Mir130a | microRNA 130a [Source:MGI Symbol;Acc: | 0 | 0 | 0 | 0 | 0 | 0 |
| 18609 | ENSMUSG00000065485 | 0 | NA | NA | NA | NA | NA | Mir219a-2 | microRNA 219a-2 [Source:MGI Symbol;Ac | 0 | 0 | 0 | 0 | 0 | 0 |
| 18610 | ENSMUSG00000065486 | 0 | NA | NA | NA | NA | NA | Mir450-1 | microRNA 450-1 [Source:MGI Symbol;Acc | 0 | 0 | 0 | 0 | 0 | 0 |
| 18611 | ENSMUSG00000065488 | 0 | NA | NA | NA | NA | NA | Mir196a-2 | microRNA 196a-2 [Source:MGI Symbol;Ac | 0 | 0 | 0 | 0 | 0 | 0 |
| 18612 | ENSMUSG00000065489 | 1.354323834 | -1.567540608 | 2.043960996 | -0.766913171 | 0.443133142 | NA | Mir365-2 | microRNA 365-2 [Source:MGI Symbol;Acc | 4.0155 | 2.0558 | 0 | 0.964 | 1.0906 | 0 |
| 18613 | ENSMUSG00000065490 | 0.185640647 | 1.126924175 | 4.080472857 | 0.276174898 | 0.782413736 | NA | Mir30c-1 | microRNA 30c-1 [Source:MGI Symbol;Acc | 0 | 0 | 0 | 0 | 0 | 1.1138 |
| 18614 | ENSMUSG00000065491 | 0 | NA | NA | NA | NA | NA | Mir7b | microRNA 7b [Source:MGI Symbol;Acc:M( | 0 | 0 | 0 | 0 | 0 | 0 |
| 18615 | ENSMUSG00000065492 | 0 | NA | NA | NA | NA | NA | Mir34b | microRNA 34b [Source:MGI Symbol;Acc:M | 0 | 0 | 0 | 0 | 0 | 0 |
| 18616 | ENSMUSG00000065493 | 0 | NA | NA | NA | NA | NA | Mir34a | microRNA 34a [Source:MGI Symbol;Acc:M | 0 | 0 | 0 | 0 | 0 | 0 |
| 18617 | ENSMUSG00000065494 | 0 | NA | NA | NA | NA | NA | Mir28a | microRNA 28a [Source:MGI Symbol;Acc:M | 0 | 0 | 0 | 0 | 0 | 0 |
| 18618 | ENSMUSG00000065495 | 0 | NA | NA | NA | NA | NA | Mir150 | microRNA 150 [Source:MGI Symbol;Acc:M | 0 | 0 | 0 | 0 | 0 | 0 |
| 18619 | ENSMUSG00000065497 | 0 | NA | NA | NA | NA | NA | Mir410 | microRNA 410 [Source:MGI Symbol;Acc:M | 0 | 0 | 0 | 0 | 0 | 0 |
| 18620 | ENSMUSG00000065498 | 0 | NA | NA | NA | NA | NA | Mir379 | microRNA 379 [Source:MGI Symbol;Acc:M | 0 | 0 | 0 | 0 | 0 | 0 |
| 18621 | ENSMUSG00000065499 | 0 | NA | NA | NA | NA | NA | Mir384 | microRNA 384 [Source:MGI Symbol;Acc:M | 0 | 0 | 0 | 0 | 0 | 0 |
| 18622 | ENSMUSG00000065500 | 0 | NA | NA | NA | NA | NA | Mir10b | microRNA 10b [Source:MGI Symbol;Acc:M | 0 | 0 | 0 | 0 | 0 | 0 |
| 18623 | ENSMUSG00000065501 | 0 | NA | NA | NA | NA | NA | Mir335 | microRNA 335 [Source:MGI Symbol;Acc:M | 0 | 0 | 0 | 0 | 0 | 0 |
| 18624 | ENSMUSG00000065502 | 0 | NA | NA | NA | NA | NA | Mir202 | microRNA 202 [Source:MGI Symbol;Acc:M | 0 | 0 | 0 | 0 | 0 | 0 |
| 18625 | ENSMUSG00000065503 | 0 | NA | NA | NA | NA | NA | Mir351 | microRNA 351 [Source:MGI Symbol;Acc:M | 0 | 0 | 0 | 0 | 0 | 0 |
| 18626 | ENSMUSG00000065505 | 1.472050944 | -0.245844811 | 1.799807761 | -0.136595039 | 0.891350898 | NA | Mir148a | microRNA 148a [Source:MGI Symbol;Acc: | 0.8031 | 1.0279 | 3.0187 | 2.8919 | 1.0906 | 0 |
| 18627 | ENSMUSG00000065507 | 0 | NA | NA | NA | NA | NA | Mir204 | microRNA 204 [Source:MGI Symbol;Acc:M | 0 | 0 | 0 | 0 | 0 | 0 |
| 18628 | ENSMUSG00000065508 | 0 | NA | NA | NA | NA | NA | Mir17 | microRNA 17 [Source:MGI Symbol;Acc:M( | 0 | 0 | 0 | 0 | 0 | 0 |
| 18629 | ENSMUSG00000065509 | 0.321325949 | 1.911575114 | 4.048267893 | 0.472195804 | 0.636787034 | NA | Mir421 | microRNA 421 [Source:MGI Symbol;Acc:M | 0 | 0 | 0 | 1.928 | 0 | 0 |
| 18630 | ENSMUSG00000065510 | 0.706721132 | -2.881789029 | 2.959064625 | -0.973885127 | 0.330113579 | NA | Mir361 | microRNA 361 [Source:MGI Symbol;Acc:M | 3.2124 | 1.0279 | 0 | 0 | 0 | 0 |
| 18631 | ENSMUSG00000065511 | 0 | NA | NA | NA | NA | NA | Mir129-2 | microRNA 129-2 [Source:MGI Symbol;Acc | 0 | 0 | 0 | 0 | 0 | 0 |
| 18632 | ENSMUSG00000065512 | 0 | NA | NA | NA | NA | NA | Mir138-2 | microRNA 138-2 [Source:MGI Symbol;Acc | 0 | 0 | 0 | 0 | 0 | 0 |
| 18633 | ENSMUSG00000065513 | 0 | NA | NA | NA | NA | NA | Mir26a-1 | microRNA 26a-1 [Source:MGI Symbol;Acc | 0 | 0 | 0 | 0 | 0 | 0 |
| 18634 | ENSMUSG00000065514 | 0 | NA | NA | NA | NA | NA | Mir106b | microRNA 106b [Source:MGI Symbol;Acc: | 0 | 0 | 0 | 0 | 0 | 0 |
| 18635 | ENSMUSG00000065515 | 0 | NA | NA | NA | NA | NA | Mir152 | microRNA 152 [Source:MGI Symbol;Acc:M | 0 | 0 | 0 | 0 | 0 | 0 |
| 18636 | ENSMUSG00000065516 | 0 | NA | NA | NA | NA | NA | Mir214 | microRNA 214 [Source:MGI Symbol;Acc:M | 0 | 0 | 0 | 0 | 0 | 0 |
| 18637 | ENSMUSG00000065518 | 0.185640647 | 1.126924175 | 4.080472857 | 0.276174898 | 0.782413736 | NA | Mir423 | microRNA 423 [Source:MGI Symbol;Acc:M | 0 | 0 | 0 | 0 | 0 | 1.1138 |
| 18638 | ENSMUSG00000065519 | 0.663965228 | -0.037580459 | 3.10181903 | -0.012115619 | 0.990333371 | NA | Mir10a | microRNA 10a [Source:MGI Symbol;Acc:M | 0 | 2.0558 | 0 | 1.928 | 0 | 0 |
| 18639 | ENSMUSG00000065520 | 0 | NA | NA | NA | NA | NA | Mir128-1 | microRNA 128-1 [Source:MGI Symbol;Acc | 0 | 0 | 0 | 0 | 0 | 0 |
| 18640 | ENSMUSG00000065521 | 0 | NA | NA | NA | NA | NA | Mir296 | microRNA 296 [Source:MGI Symbol;Acc:M | 0 | 0 | 0 | 0 | 0 | 0 |
| 18641 | ENSMUSG00000065523 | 0.980717322 | 0.289172824 | 2.436251824 | 0.118695786 | 0.905516375 | NA | Mir192 | microRNA 192 [Source:MGI Symbol;Acc:M | 1.6062 | 0 | 1.0062 | 0 | 3.2719 | 0 |
| 18642 | ENSMUSG00000065524 | 0 | NA | NA | NA | NA | NA | Mir135a-2 | microRNA 135a-2 [Source:MGI Symbol;Ac | 0 | 0 | 0 | 0 | 0 | 0 |
| 18643 | ENSMUSG00000065526 | 0 | NA | NA | NA | NA | NA | Mir337 | microRNA 337 [Source:MGI Symbol;Acc:M | 0 | 0 | 0 | 0 | 0 | 0 |
| 18644 | ENSMUSG00000065527 | 0 | NA | NA | NA | NA | NA | Mir93 | microRNA 93 [Source:MGI Symbol;Acc:M( | 0 | 0 | 0 | 0 | 0 | 0 |

| | | | | | | | | | | | | | | | |
|---|---|---|---|---|---|---|---|---|---|---|---|---|---|---|---|
| 18645 | ENSMUSG00000065528 | 0.181769715 | 1.126924175 | 4.080472857 | 0.276174898 | 0.782413736 | NA | Mir320 | microRNA 320 [Source:MGI Symbol;Acc:M | 0 | 0 | 0 | 0 | 1.0906 | 0 |
| 18646 | ENSMUSG00000065529 | 0 | NA | NA | NA | NA | NA | Mir22 | microRNA 22 [Source:MGI Symbol;Acc:M( | 0 | 0 | 0 | 0 | 0 | 0 |
| 18647 | ENSMUSG00000065530 | 0.181769715 | 1.126924175 | 4.080472857 | 0.276174898 | 0.782413736 | NA | Mir99a | microRNA 99a [Source:MGI Symbol;Acc:M | 0 | 0 | 0 | 0 | 1.0906 | 0 |
| 18648 | ENSMUSG00000065532 | 0 | NA | NA | NA | NA | NA | Mir187 | microRNA 187 [Source:MGI Symbol;Acc:M | 0 | 0 | 0 | 0 | 0 | 0 |
| 18649 | ENSMUSG00000065533 | 0 | NA | NA | NA | NA | NA | Mir205 | microRNA 205 [Source:MGI Symbol;Acc:M | 0 | 0 | 0 | 0 | 0 | 0 |
| 18650 | ENSMUSG00000065534 | 0 | NA | NA | NA | NA | NA | Mir324 | microRNA 324 [Source:MGI Symbol;Acc:M | 0 | 0 | 0 | 0 | 0 | 0 |
| 18651 | ENSMUSG00000065536 | 0 | NA | NA | NA | NA | NA | Mir98 | microRNA 98 [Source:MGI Symbol;Acc:M( | 0 | 0 | 0 | 0 | 0 | 0 |
| 18652 | ENSMUSG00000065537 | 0 | NA | NA | NA | NA | NA | Mir132 | microRNA 132 [Source:MGI Symbol;Acc:M | 0 | 0 | 0 | 0 | 0 | 0 |
| 18653 | ENSMUSG00000065538 | 0 | NA | NA | NA | NA | NA | Mir153 | microRNA 153 [Source:MGI Symbol;Acc:M | 0 | 0 | 0 | 0 | 0 | 0 |
| 18654 | ENSMUSG00000065539 | 0 | NA | NA | NA | NA | NA | Mir216a | microRNA 216a [Source:MGI Symbol;Acc: | 0 | 0 | 0 | 0 | 0 | 0 |
| 18655 | ENSMUSG00000065540 | 0 | NA | NA | NA | NA | NA | Mir126a | microRNA 126a [Source:MGI Symbol;Acc: | 0 | 0 | 0 | 0 | 0 | 0 |
| 18656 | ENSMUSG00000065541 | 0 | NA | NA | NA | NA | NA | Mir24-2 | microRNA 24-2 [Source:MGI Symbol;Acc:I | 0 | 0 | 0 | 0 | 0 | 0 |
| 18657 | ENSMUSG00000065542 | 0 | NA | NA | NA | NA | NA | Mir224 | microRNA 224 [Source:MGI Symbol;Acc:M | 0 | 0 | 0 | 0 | 0 | 0 |
| 18658 | ENSMUSG00000065543 | 0 | NA | NA | NA | NA | NA | Mir330 | microRNA 330 [Source:MGI Symbol;Acc:M | 0 | 0 | 0 | 0 | 0 | 0 |
| 18659 | ENSMUSG00000065544 | 0.545309144 | 2.651974629 | 3.666219369 | 0.72335405 | 0.469462394 | NA | Mir32 | microRNA 32 [Source:MGI Symbol;Acc:M( | 0 | 0 | 0 | 0 | 3.2719 | 0 |
| 18660 | ENSMUSG00000065546 | 0 | NA | NA | NA | NA | NA | Mir196a-1 | microRNA 196a-1 [Source:MGI Symbol;Ac | 0 | 0 | 0 | 0 | 0 | 0 |
| 18661 | ENSMUSG00000065547 | 0 | NA | NA | NA | NA | NA | Mir199a-1 | microRNA 199a-1 [Source:MGI Symbol;Ac | 0 | 0 | 0 | 0 | 0 | 0 |
| 18662 | ENSMUSG00000065548 | 0 | NA | NA | NA | NA | NA | Mir29c | microRNA 29c [Source:MGI Symbol;Acc:M | 0 | 0 | 0 | 0 | 0 | 0 |
| 18663 | ENSMUSG00000065549 | 0 | NA | NA | NA | NA | NA | Mir200b | microRNA 200b [Source:MGI Symbol;Acc: | 0 | 0 | 0 | 0 | 0 | 0 |
| 18664 | ENSMUSG00000065551 | 0 | NA | NA | NA | NA | NA | Mir210 | microRNA 210 [Source:MGI Symbol;Acc:M | 0 | 0 | 0 | 0 | 0 | 0 |
| 18665 | ENSMUSG00000065552 | 0 | NA | NA | NA | NA | NA | Mir302a | microRNA 302a [Source:MGI Symbol;Acc: | 0 | 0 | 0 | 0 | 0 | 0 |
| 18666 | ENSMUSG00000065553 | 0 | NA | NA | NA | NA | NA | Mir103-1 | microRNA 103-1 [Source:MGI Symbol;Acc | 0 | 0 | 0 | 0 | 0 | 0 |
| 18667 | ENSMUSG00000065555 | 0 | NA | NA | NA | NA | NA | Mir219a-1 | microRNA 219a-1 [Source:MGI Symbol;Ac | 0 | 0 | 0 | 0 | 0 | 0 |
| 18668 | ENSMUSG00000065556 | 0.713920574 | 0.125792037 | 3.028037396 | 0.041542432 | 0.966863466 | NA | Mir101b | microRNA 101b [Source:MGI Symbol;Acc: | 0 | 2.0558 | 0 | 0 | 0 | 2.2277 |
| 18669 | ENSMUSG00000065557 | 0.181769715 | 1.126924175 | 4.080472857 | 0.276174898 | 0.782413736 | NA | Mirlet7c-1 | microRNA let7c-1 [Source:MGI Symbol;Ac | 0 | 0 | 0 | 0 | 1.0906 | 0 |
| 18670 | ENSMUSG00000065559 | 0 | NA | NA | NA | NA | NA | Mir206 | microRNA 206 [Source:MGI Symbol;Acc:M | 0 | 0 | 0 | 0 | 0 | 0 |
| 18671 | ENSMUSG00000065560 | 0 | NA | NA | NA | NA | NA | Mir148b | microRNA 148b [Source:MGI Symbol;Acc: | 0 | 0 | 0 | 0 | 0 | 0 |
| 18672 | ENSMUSG00000065561 | 0 | NA | NA | NA | NA | NA | Mir369 | microRNA 369 [Source:MGI Symbol;Acc:M | 0 | 0 | 0 | 0 | 0 | 0 |
| 18673 | ENSMUSG00000065562 | 0 | NA | NA | NA | NA | NA | Mir215 | microRNA 215 [Source:MGI Symbol;Acc:M | 0 | 0 | 0 | 0 | 0 | 0 |
| 18674 | ENSMUSG00000065563 | 0.167707431 | -0.796629176 | 4.080472857 | -0.195229623 | 0.84521319 | NA | Mir103-2 | microRNA 103-2 [Source:MGI Symbol;Acc | 0 | 0 | 1.0062 | 0 | 0 | 0 |
| 18675 | ENSMUSG00000065564 | 0 | NA | NA | NA | NA | NA | Mirlet7b | microRNA let7b [Source:MGI Symbol;Acc: | 0 | 0 | 0 | 0 | 0 | 0 |
| 18676 | ENSMUSG00000065565 | 0 | NA | NA | NA | NA | NA | Mir181a-1 | microRNA 181a-1 [Source:MGI Symbol;Ac | 0 | 0 | 0 | 0 | 0 | 0 |
| 18677 | ENSMUSG00000065567 | 0 | NA | NA | NA | NA | NA | Mir30c-2 | microRNA 30c-2 [Source:MGI Symbol;Acc | 0 | 0 | 0 | 0 | 0 | 0 |
| 18678 | ENSMUSG00000065568 | 0 | NA | NA | NA | NA | NA | Mir344 | microRNA 344 [Source:MGI Symbol;Acc:M | 0 | 0 | 0 | 0 | 0 | 0 |
| 18679 | ENSMUSG00000065569 | 0 | NA | NA | NA | NA | NA | Mir137 | microRNA 137 [Source:MGI Symbol;Acc:M | 0 | 0 | 0 | 0 | 0 | 0 |
| 18680 | ENSMUSG00000065570 | 0 | NA | NA | NA | NA | NA | Mir412 | microRNA 412 [Source:MGI Symbol;Acc:M | 0 | 0 | 0 | 0 | 0 | 0 |
| 18681 | ENSMUSG00000065571 | 0 | NA | NA | NA | NA | NA | Mir326 | microRNA 326 [Source:MGI Symbol;Acc:M | 0 | 0 | 0 | 0 | 0 | 0 |
| 18682 | ENSMUSG00000065572 | 0 | NA | NA | NA | NA | NA | Mir130b | microRNA 130b [Source:MGI Symbol;Acc: | 0 | 0 | 0 | 0 | 0 | 0 |
| 18683 | ENSMUSG00000065573 | 0 | NA | NA | NA | NA | NA | Mir350 | microRNA 350 [Source:MGI Symbol;Acc:M | 0 | 0 | 0 | 0 | 0 | 0 |
| 18684 | ENSMUSG00000065574 | 0 | NA | NA | NA | NA | NA | Mir203 | microRNA 203 [Source:MGI Symbol;Acc:M | 0 | 0 | 0 | 0 | 0 | 0 |
| 18685 | ENSMUSG00000065575 | 0 | NA | NA | NA | NA | NA | Mir449a | microRNA 449a [Source:MGI Symbol;Acc: | 0 | 0 | 0 | 0 | 0 | 0 |
| 18686 | ENSMUSG00000065577 | 0 | NA | NA | NA | NA | NA | Mir329 | microRNA 329 [Source:MGI Symbol;Acc:M | 0 | 0 | 0 | 0 | 0 | 0 |
| 18687 | ENSMUSG00000065578 | 0 | NA | NA | NA | NA | NA | Mir181b-2 | microRNA 181b-2 [Source:MGI Symbol;Ac | 0 | 0 | 0 | 0 | 0 | 0 |
| 18688 | ENSMUSG00000065579 | 0 | NA | NA | NA | NA | NA | Mir425 | microRNA 425 [Source:MGI Symbol;Acc:M | 0 | 0 | 0 | 0 | 0 | 0 |
| 18689 | ENSMUSG00000065580 | 0 | NA | NA | NA | NA | NA | Mir15b | microRNA 15b [Source:MGI Symbol;Acc:M | 0 | 0 | 0 | 0 | 0 | 0 |
| 18690 | ENSMUSG00000065581 | 0.353089354 | 0.165142706 | 3.896102244 | 0.042386646 | 0.966190474 | NA | Mir194-1 | microRNA 194-1 [Source:MGI Symbol;Acc | 0 | 1.0279 | 0 | 0 | 1.0906 | 0 |
| 18691 | ENSMUSG00000065582 | 0.401551119 | -2.072982726 | 4.008251979 | -0.517178744 | 0.605031388 | NA | Mir194-2 | microRNA 194-2 [Source:MGI Symbol;Acc | 2.4093 | 0 | 0 | 0 | 0 | 0 |
| 18692 | ENSMUSG00000065583 | 0 | NA | NA | NA | NA | NA | Mir218-2 | microRNA 218-2 [Source:MGI Symbol;Acc | 0 | 0 | 0 | 0 | 0 | 0 |
| 18693 | ENSMUSG00000065585 | 0 | NA | NA | NA | NA | NA | Mir211 | microRNA 211 [Source:MGI Symbol;Acc:M | 0 | 0 | 0 | 0 | 0 | 0 |
| 18694 | ENSMUSG00000065586 | 0 | NA | NA | NA | NA | NA | Mir96 | microRNA 96 [Source:MGI Symbol;Acc:M( | 0 | 0 | 0 | 0 | 0 | 0 |
| 18695 | ENSMUSG00000065587 | 0 | NA | NA | NA | NA | NA | Mir34c | microRNA 34c [Source:MGI Symbol;Acc:M | 0 | 0 | 0 | 0 | 0 | 0 |
| 18696 | ENSMUSG00000065589 | 0.439020386 | -2.189747911 | 3.52148864 | -0.621824499 | 0.534057275 | NA | Mir301 | microRNA 301 [Source:MGI Symbol;Acc:M | 1.6062 | 1.0279 | 0 | 0 | 0 | 0 |
| 18697 | ENSMUSG00000065590 | 0 | NA | NA | NA | NA | NA | Mir212 | microRNA 212 [Source:MGI Symbol;Acc:M | 0 | 0 | 0 | 0 | 0 | 0 |
| 18698 | ENSMUSG00000065591 | 4.722965368 | -0.947523812 | 1.030718811 | -0.919284486 | 0.35794679 | NA | Gm22753 | predicted gene, 22753 [Source:MGI Symb | 2.4093 | 8.2233 | 8.05 | 1.928 | 3.2719 | 4.4554 |
| 18699 | ENSMUSG00000065592 | 0.294513347 | 0.165143496 | 4.080472857 | 0.040471657 | 0.967717103 | NA | Mir145a | microRNA 145a [Source:MGI Symbol;Acc: | 0.8031 | 0 | 0 | 0.964 | 0 | 0 |
| 18700 | ENSMUSG00000065593 | 2.944508428 | 1.211497618 | 1.275442625 | 0.949864458 | 0.342181128 | NA | Mir339 | microRNA 339 [Source:MGI Symbol;Acc:M | 3.2124 | 1.0279 | 1.0062 | 5.7839 | 2.1812 | 4.4554 |

| | | | | | | | | | | | | | | | |
|---|---|---|---|---|---|---|---|---|---|---|---|---|---|---|---|
| 18701 | ENSMUSG00000065594 | 0.294513347 | 0.165143496 | 4.080472857 | 0.040471657 | 0.967717103 | NA | Mir107 | microRNA 107 [Source:MGI Symbol;Acc:M | 0.8031 | 0 | 0 | 0.964 | 0 | 0 |
| 18702 | ENSMUSG00000065596 | 0 | NA | NA | NA | NA | NA | Mir184 | microRNA 184 [Source:MGI Symbol;Acc:M | 0 | 0 | 0 | 0 | 0 | 0 |
| 18703 | ENSMUSG00000065597 | 0 | NA | NA | NA | NA | NA | Mir124a-1 | microRNA 124a-1 [Source:MGI Symbol;Ac | 0 | 0 | 0 | 0 | 0 | 0 |
| 18704 | ENSMUSG00000065599 | 0.321325949 | 1.911575114 | 4.048267893 | 0.472195804 | 0.636787034 | NA | Mir23b | microRNA 23b [Source:MGI Symbol;Acc:M | 0 | 0 | 0 | 1.928 | 0 | 0 |
| 18705 | ENSMUSG00000065600 | 0 | NA | NA | NA | NA | NA | Mir338 | microRNA 338 [Source:MGI Symbol;Acc:M | 0 | 0 | 0 | 0 | 0 | 0 |
| 18706 | ENSMUSG00000065601 | 0.160662974 | 1.126924175 | 4.080472857 | 0.276174898 | 0.782413736 | NA | Mir146 | microRNA 146 [Source:MGI Symbol;Acc:M | 0 | 0 | 0 | 0.964 | 0 | 0 |
| 18707 | ENSMUSG00000065602 | 0 | NA | NA | NA | NA | NA | Mirlet7f-2 | microRNA let7f-2 [Source:MGI Symbol;Ac | 0 | 0 | 0 | 0 | 0 | 0 |
| 18708 | ENSMUSG00000065603 | 0 | NA | NA | NA | NA | NA | Mir218-1 | microRNA 218-1 [Source:MGI Symbol;Acc | 0 | 0 | 0 | 0 | 0 | 0 |
| 18709 | ENSMUSG00000065604 | 0 | NA | NA | NA | NA | NA | Mir29b-1 | microRNA 29b-1 [Source:MGI Symbol;Acc | 0 | 0 | 0 | 0 | 0 | 0 |
| 18710 | ENSMUSG00000065606 | 0 | NA | NA | NA | NA | NA | Mir16-2 | microRNA 16-2 [Source:MGI Symbol;Acc:I | 0 | 0 | 0 | 0 | 0 | 0 |
| 18711 | ENSMUSG00000065607 | 0 | NA | NA | NA | NA | NA | Mir331 | microRNA 331 [Source:MGI Symbol;Acc:M | 0 | 0 | 0 | 0 | 0 | 0 |
| 18712 | ENSMUSG00000065608 | 0 | NA | NA | NA | NA | NA | Mirlet7c-2 | microRNA let7c-2 [Source:MGI Symbol;Ac | 0 | 0 | 0 | 0 | 0 | 0 |
| 18713 | ENSMUSG00000065609 | 0 | NA | NA | NA | NA | NA | Mir7-2 | microRNA 7-2 [Source:MGI Symbol;Acc:M | 0 | 0 | 0 | 0 | 0 | 0 |
| 18714 | ENSMUSG00000065610 | 0 | NA | NA | NA | NA | NA | Mir29a | microRNA 29a [Source:MGI Symbol;Acc:M | 0 | 0 | 0 | 0 | 0 | 0 |
| 18715 | ENSMUSG00000065611 | 0.17131964 | -0.796629176 | 4.080472857 | -0.195229623 | 0.84521319 | NA | Mir23a | microRNA 23a [Source:MGI Symbol;Acc:M | 0 | 1.0279 | 0 | 0 | 0 | 0 |
| 18716 | ENSMUSG00000065612 | 0 | NA | NA | NA | NA | NA | Mir151 | microRNA 151 [Source:MGI Symbol;Acc:M | 0 | 0 | 0 | 0 | 0 | 0 |
| 18717 | ENSMUSG00000065613 | 0 | NA | NA | NA | NA | NA | Mir92-2 | microRNA 92-2 [Source:MGI Symbol;Acc:I | 0 | 0 | 0 | 0 | 0 | 0 |
| 18718 | ENSMUSG00000065616 | 0 | NA | NA | NA | NA | NA | Mir375 | microRNA 375 [Source:MGI Symbol;Acc:M | 0 | 0 | 0 | 0 | 0 | 0 |
| 18719 | ENSMUSG00000065617 | 0 | NA | NA | NA | NA | NA | Mir323 | microRNA 323 [Source:MGI Symbol;Acc:M | 0 | 0 | 0 | 0 | 0 | 0 |
| 18720 | ENSMUSG00000065619 | 0 | NA | NA | NA | NA | NA | Mir183 | microRNA 183 [Source:MGI Symbol;Acc:M | 0 | 0 | 0 | 0 | 0 | 0 |
| 18721 | ENSMUSG00000065620 | 0 | NA | NA | NA | NA | NA | Gm24764 | predicted gene, 24764 [Source:MGI Symb | 0 | 0 | 0 | 0 | 0 | 0 |
| 18722 | ENSMUSG00000065622 | 0 | NA | NA | NA | NA | NA | Gm24763 | predicted gene, 24763 [Source:MGI Symb | 0 | 0 | 0 | 0 | 0 | 0 |
| 18723 | ENSMUSG00000065623 | 0.986449812 | -2.084851164 | 2.483799425 | -0.83937984 | 0.401256193 | NA | Gm24762 | predicted gene, 24762 [Source:MGI Symb | 0.8031 | 0 | 4.025 | 0 | 1.0906 | 0 |
| 18724 | ENSMUSG00000065625 | 0 | NA | NA | NA | NA | NA | Gm24765 | predicted gene, 24765 [Source:MGI Symb | 0 | 0 | 0 | 0 | 0 | 0 |
| 18725 | ENSMUSG00000065627 | 0 | NA | NA | NA | NA | NA | NA | NA | 0 | 0 | 0 | 0 | 0 | 0 |
| 18726 | ENSMUSG00000065628 | 0 | NA | NA | NA | NA | NA | NA | NA | 0 | 0 | 0 | 0 | 0 | 0 |
| 18727 | ENSMUSG00000065629 | 0 | NA | NA | NA | NA | NA | Gm24826 | predicted gene, 24826 [Source:MGI Symb | 0 | 0 | 0 | 0 | 0 | 0 |
| 18728 | ENSMUSG00000065630 | 0 | NA | NA | NA | NA | NA | Gm24256 | predicted gene, 24256 [Source:MGI Symb | 0 | 0 | 0 | 0 | 0 | 0 |
| 18729 | ENSMUSG00000065634 | 0 | NA | NA | NA | NA | NA | Gm24252 | predicted gene, 24252 [Source:MGI Symb | 0 | 0 | 0 | 0 | 0 | 0 |
| 18730 | ENSMUSG00000065635 | 0 | NA | NA | NA | NA | NA | Gm24253 | predicted gene, 24253 [Source:MGI Symb | 0 | 0 | 0 | 0 | 0 | 0 |
| 18731 | ENSMUSG00000065636 | 0 | NA | NA | NA | NA | NA | Gm24254 | predicted gene, 24254 [Source:MGI Symb | 0 | 0 | 0 | 0 | 0 | 0 |
| 18732 | ENSMUSG00000065637 | 3.456245868 | 0.637608173 | 1.222181992 | 0.521696586 | 0.601881602 | NA | Gm26397 | predicted gene, 26397 [Source:MGI Symb | 4.0155 | 0 | 4.025 | 3.8559 | 3.2719 | 5.5692 |
| 18733 | ENSMUSG00000065640 | 0 | NA | NA | NA | NA | NA | Snord1c | small nucleolar RNA, C/D box 1C [Source:I | 0 | 0 | 0 | 0 | 0 | 0 |
| 18734 | ENSMUSG00000065642 | 1.518135357 | -0.178188922 | 1.675773527 | -0.106332341 | 0.915318673 | NA | Snora69 | small nucleolar RNA, H/ACA box 69 [Sourc | 0.8031 | 1.0279 | 3.0187 | 0.964 | 2.1812 | 1.1138 |
| 18735 | ENSMUSG00000065643 | 0 | NA | NA | NA | NA | NA | Gm25897 | predicted gene, 25897 [Source:MGI Symb | 0 | 0 | 0 | 0 | 0 | 0 |
| 18736 | ENSMUSG00000065644 | 0 | NA | NA | NA | NA | NA | NA | NA | 0 | 0 | 0 | 0 | 0 | 0 |
| 18737 | ENSMUSG00000065645 | 0 | NA | NA | NA | NA | NA | NA | NA | 0 | 0 | 0 | 0 | 0 | 0 |
| 18738 | ENSMUSG00000065647 | 0.17131964 | -0.796629176 | 4.080472857 | -0.195229623 | 0.84521319 | NA | Gm25896 | predicted gene, 25896 [Source:MGI Symb | 0 | 1.0279 | 0 | 0 | 0 | 0 |
| 18739 | ENSMUSG00000065648 | 0 | NA | NA | NA | NA | NA | Gm25895 | predicted gene, 25895 [Source:MGI Symb | 0 | 0 | 0 | 0 | 0 | 0 |
| 18740 | ENSMUSG00000065649 | 0 | NA | NA | NA | NA | NA | Snora74a | small nucleolar RNA, H/ACA box 74A [Sou | 0 | 0 | 0 | 0 | 0 | 0 |
| 18741 | ENSMUSG00000065651 | 0.342432689 | 1.990653086 | 3.900281263 | 0.51038706 | 0.609780321 | NA | Gm25973 | predicted gene, 25973 [Source:MGI Symb | 0 | 0 | 0 | 0.964 | 1.0906 | 0 |
| 18742 | ENSMUSG00000065653 | 0 | NA | NA | NA | NA | NA | Gm23105 | predicted gene, 23105 [Source:MGI Symb | 0 | 0 | 0 | 0 | 0 | 0 |
| 18743 | ENSMUSG00000065656 | 0 | NA | NA | NA | NA | NA | Gm23104 | predicted gene, 23104 [Source:MGI Symb | 0 | 0 | 0 | 0 | 0 | 0 |
| 18744 | ENSMUSG00000065657 | 0 | NA | NA | NA | NA | NA | Gm25337 | predicted gene, 25337 [Source:MGI Symb | 0 | 0 | 0 | 0 | 0 | 0 |
| 18745 | ENSMUSG00000065658 | 0 | NA | NA | NA | NA | NA | Gm23102 | predicted gene, 23102 [Source:MGI Symb | 0 | 0 | 0 | 0 | 0 | 0 |
| 18746 | ENSMUSG00000065659 | 0 | NA | NA | NA | NA | NA | NA | NA | 0 | 0 | 0 | 0 | 0 | 0 |
| 18747 | ENSMUSG00000065660 | 0 | NA | NA | NA | NA | NA | NA | NA | 0 | 0 | 0 | 0 | 0 | 0 |
| 18748 | ENSMUSG00000065661 | 0 | NA | NA | NA | NA | NA | NA | NA | 0 | 0 | 0 | 0 | 0 | 0 |
| 18749 | ENSMUSG00000065662 | 0 | NA | NA | NA | NA | NA | NA | NA | 0 | 0 | 0 | 0 | 0 | 0 |
| 18750 | ENSMUSG00000065663 | 0 | NA | NA | NA | NA | NA | Gm22579 | predicted gene, 22579 [Source:MGI Symb | 0 | 0 | 0 | 0 | 0 | 0 |
| 18751 | ENSMUSG00000065664 | 0 | NA | NA | NA | NA | NA | Gm22578 | predicted gene, 22578 [Source:MGI Symb | 0 | 0 | 0 | 0 | 0 | 0 |
| 18752 | ENSMUSG00000065665 | 0 | NA | NA | NA | NA | NA | Gm22577 | predicted gene, 22577 [Source:MGI Symb | 0 | 0 | 0 | 0 | 0 | 0 |
| 18753 | ENSMUSG00000065667 | 0 | NA | NA | NA | NA | NA | Gm22576 | predicted gene, 22576 [Source:MGI Symb | 0 | 0 | 0 | 0 | 0 | 0 |
| 18754 | ENSMUSG00000065669 | 1.377673698 | 1.95369361 | 2.234803055 | 0.874212878 | 0.382002334 | NA | Gm22581 | predicted gene, 22581 [Source:MGI Symb | 1.6062 | 0 | 0 | 0 | 1.0906 | 5.5692 |
| 18755 | ENSMUSG00000065673 | 0.167707431 | -0.796629176 | 4.080472857 | -0.195229623 | 0.84521319 | NA | Gm24258 | predicted gene, 24258 [Source:MGI Symb | 0 | 0 | 1.0062 | 0 | 0 | 0 |
| 18756 | ENSMUSG00000065674 | 0.401551119 | -2.072982726 | 4.008251979 | -0.517178744 | 0.605031388 | NA | Gm24259 | predicted gene, 24259 [Source:MGI Symb | 2.4093 | 0 | 0 | 0 | 0 | 0 |

| | | | | | | | | | | | | | | | |
|---|---|---|---|---|---|---|---|---|---|---|---|---|---|---|---|
| 18757 | ENSMUSG00000065675 | 0 | NA | NA | NA | NA | NA | NA | NA | 0 | 0 | 0 | 0 | 0 | 0 |
| 18758 | ENSMUSG00000065676 | 1.007736513 | -0.832948274 | 2.237007798 | -0.372349294 | 0.709632802 | NA | Snord42b | small nucleolar RNA, C/D box 42B [Source | 0.8031 | 2.0558 | 1.0062 | 0 | 2.1812 | 0 |
| 18759 | ENSMUSG00000065678 | 0 | NA | NA | NA | NA | NA | Gm24261 | predicted gene, 24261 [Source:MGI Symb | 0 | 0 | 0 | 0 | 0 | 0 |
| 18760 | ENSMUSG00000065680 | 0 | NA | NA | NA | NA | NA | Snord38a | small nucleolar RNA, C/D box 38A [Source | 0 | 0 | 0 | 0 | 0 | 0 |
| 18761 | ENSMUSG00000065683 | 0 | NA | NA | NA | NA | NA | NA | NA | 0 | 0 | 0 | 0 | 0 | 0 |
| 18762 | ENSMUSG00000065684 | 0 | NA | NA | NA | NA | NA | Gm23733 | predicted gene, 23733 [Source:MGI Symb | 0 | 0 | 0 | 0 | 0 | 0 |
| 18763 | ENSMUSG00000065685 | 0 | NA | NA | NA | NA | NA | NA | NA | 0 | 0 | 0 | 0 | 0 | 0 |
| 18764 | ENSMUSG00000065686 | 0 | NA | NA | NA | NA | NA | Snora5c | small nucleolar RNA, H/ACA box 5C [Sourc | 0 | 0 | 0 | 0 | 0 | 0 |
| 18765 | ENSMUSG00000065687 | 0 | NA | NA | NA | NA | NA | Gm23734 | predicted gene, 23734 [Source:MGI Symb | 0 | 0 | 0 | 0 | 0 | 0 |
| 18766 | ENSMUSG00000065689 | 0 | NA | NA | NA | NA | NA | n-R5s7 | nuclear encoded rRNA 5S 7 [Source:MGI S | 0 | 0 | 0 | 0 | 0 | 0 |
| 18767 | ENSMUSG00000065691 | 0 | NA | NA | NA | NA | NA | NA | NA | 0 | 0 | 0 | 0 | 0 | 0 |
| 18768 | ENSMUSG00000065692 | 0 | NA | NA | NA | NA | NA | Gm25410 | predicted gene, 25410 [Source:MGI Symb | 0 | 0 | 0 | 0 | 0 | 0 |
| 18769 | ENSMUSG00000065694 | 0 | NA | NA | NA | NA | NA | Gm25411 | predicted gene, 25411 [Source:MGI Symb | 0 | 0 | 0 | 0 | 0 | 0 |
| 18770 | ENSMUSG00000065698 | 0 | NA | NA | NA | NA | NA | Gm25409 | predicted gene, 25409 [Source:MGI Symb | 0 | 0 | 0 | 0 | 0 | 0 |
| 18771 | ENSMUSG00000065700 | 0 | NA | NA | NA | NA | NA | Gm22827 | predicted gene, 22827 [Source:MGI Symb | 0 | 0 | 0 | 0 | 0 | 0 |
| 18772 | ENSMUSG00000065701 | 0 | NA | NA | NA | NA | NA | Rny1 | RNA, Y1 small cytoplasmic, Ro-associated | 0 | 0 | 0 | 0 | 0 | 0 |
| 18773 | ENSMUSG00000065702 | 0 | NA | NA | NA | NA | NA | Gm22828 | predicted gene, 22828 [Source:MGI Symb | 0 | 0 | 0 | 0 | 0 | 0 |
| 18774 | ENSMUSG00000065705 | 0 | NA | NA | NA | NA | NA | Gm22829 | predicted gene, 22829 [Source:MGI Symb | 0 | 0 | 0 | 0 | 0 | 0 |
| 18775 | ENSMUSG00000065706 | 0 | NA | NA | NA | NA | NA | Gm22830 | predicted gene, 22830 [Source:MGI Symb | 0 | 0 | 0 | 0 | 0 | 0 |
| 18776 | ENSMUSG00000065708 | 0 | NA | NA | NA | NA | NA | Gm22826 | predicted gene, 22826 [Source:MGI Symb | 0 | 0 | 0 | 0 | 0 | 0 |
| 18777 | ENSMUSG00000065709 | 0 | NA | NA | NA | NA | NA | Gm22341 | predicted gene, 22341 [Source:MGI Symb | 0 | 0 | 0 | 0 | 0 | 0 |
| 18778 | ENSMUSG00000065714 | 0 | NA | NA | NA | NA | NA | Gm25652 | predicted gene, 25652 [Source:MGI Symb | 0 | 0 | 0 | 0 | 0 | 0 |
| 18779 | ENSMUSG00000065715 | 1.338633741 | 1.010616246 | 2.026399667 | 0.498725036 | 0.617973105 | NA | Snord7 | small nucleolar RNA, C/D box 7 [Source:M | 1.6062 | 0 | 1.0062 | 0.964 | 0 | 4.4554 |
| 18780 | ENSMUSG00000065716 | 0 | NA | NA | NA | NA | NA | Gm25651 | predicted gene, 25651 [Source:MGI Symb | 0 | 0 | 0 | 0 | 0 | 0 |
| 18781 | ENSMUSG00000065718 | 0 | NA | NA | NA | NA | NA | Gm25650 | predicted gene, 25650 [Source:MGI Symb | 0 | 0 | 0 | 0 | 0 | 0 |
| 18782 | ENSMUSG00000065719 | 0 | NA | NA | NA | NA | NA | NA | NA | 0 | 0 | 0 | 0 | 0 | 0 |
| 18783 | ENSMUSG00000065721 | 0 | NA | NA | NA | NA | NA | Gm26168 | predicted gene, 26168 [Source:MGI Symb | 0 | 0 | 0 | 0 | 0 | 0 |
| 18784 | ENSMUSG00000065722 | 0 | NA | NA | NA | NA | NA | Gm26166 | predicted gene, 26166 [Source:MGI Symb | 0 | 0 | 0 | 0 | 0 | 0 |
| 18785 | ENSMUSG00000065725 | 0 | NA | NA | NA | NA | NA | Gm26165 | predicted gene, 26165 [Source:MGI Symb | 0 | 0 | 0 | 0 | 0 | 0 |
| 18786 | ENSMUSG00000065726 | 0 | NA | NA | NA | NA | NA | Gm26163 | predicted gene, 26163 [Source:MGI Symb | 0 | 0 | 0 | 0 | 0 | 0 |
| 18787 | ENSMUSG00000065727 | 0 | NA | NA | NA | NA | NA | Gm26164 | predicted gene, 26164 [Source:MGI Symb | 0 | 0 | 0 | 0 | 0 | 0 |
| 18788 | ENSMUSG00000065728 | 0.301557804 | -1.64335916 | 4.042814367 | -0.406488899 | 0.684383412 | NA | Gm26175 | predicted gene, 26175 [Source:MGI Symb | 0.8031 | 0 | 1.0062 | 0 | 0 | 0 |
| 18789 | ENSMUSG00000065729 | 0 | NA | NA | NA | NA | NA | Gm26176 | predicted gene, 26176 [Source:MGI Symb | 0 | 0 | 0 | 0 | 0 | 0 |
| 18790 | ENSMUSG00000065732 | 0 | NA | NA | NA | NA | NA | Gm24492 | predicted gene, 24492 [Source:MGI Symb | 0 | 0 | 0 | 0 | 0 | 0 |
| 18791 | ENSMUSG00000065733 | 0 | NA | NA | NA | NA | NA | Gm24491 | predicted gene, 24491 [Source:MGI Symb | 0 | 0 | 0 | 0 | 0 | 0 |
| 18792 | ENSMUSG00000065734 | 0.17131964 | -0.796629176 | 4.080472857 | -0.195229623 | 0.84521319 | NA | Snord49a | small nucleolar RNA, C/D box 49A [Source | 0 | 1.0279 | 0 | 0 | 0 | 0 |
| 18793 | ENSMUSG00000065735 | 0 | NA | NA | NA | NA | NA | Snord80 | small nucleolar RNA, C/D box 80 [Source:I | 0 | 0 | 0 | 0 | 0 | 0 |
| 18794 | ENSMUSG00000065737 | 0 | NA | NA | NA | NA | NA | Gm24490 | predicted gene, 24490 [Source:MGI Symb | 0 | 0 | 0 | 0 | 0 | 0 |
| 18795 | ENSMUSG00000065738 | 5.900061199 | -0.494364253 | 0.948327331 | -0.521301282 | 0.602156908 | 0.915461118 | Gm24494 | predicted gene, 24494 [Source:MGI Symb | 6.4248 | 10.279 | 4.025 | 5.7839 | 1.0906 | 7.7969 |
| 18796 | ENSMUSG00000065739 | 0 | NA | NA | NA | NA | NA | NA | NA | 0 | 0 | 0 | 0 | 0 | 0 |
| 18797 | ENSMUSG00000065742 | 0 | NA | NA | NA | NA | NA | Gm25007 | predicted gene, 25007 [Source:MGI Symb | 0 | 0 | 0 | 0 | 0 | 0 |
| 18798 | ENSMUSG00000065743 | 0 | NA | NA | NA | NA | NA | Gm25008 | predicted gene, 25008 [Source:MGI Symb | 0 | 0 | 0 | 0 | 0 | 0 |
| 18799 | ENSMUSG00000065745 | 0 | NA | NA | NA | NA | NA | Gm25004 | predicted gene, 25004 [Source:MGI Symb | 0 | 0 | 0 | 0 | 0 | 0 |
| 18800 | ENSMUSG00000065746 | 0 | NA | NA | NA | NA | NA | Gm25005 | predicted gene, 25005 [Source:MGI Symb | 0 | 0 | 0 | 0 | 0 | 0 |
| 18801 | ENSMUSG00000065747 | 0 | NA | NA | NA | NA | NA | Gm25006 | predicted gene, 25006 [Source:MGI Symb | 0 | 0 | 0 | 0 | 0 | 0 |
| 18802 | ENSMUSG00000065749 | 0 | NA | NA | NA | NA | NA | AF357428 | snoRNA AF357428 [Source:MGI Symbol;A | 0 | 0 | 0 | 0 | 0 | 0 |
| 18803 | ENSMUSG00000065750 | 0.401551119 | -2.072982726 | 4.008251979 | -0.517178744 | 0.605031388 | NA | Gm23346 | predicted gene, 23346 [Source:MGI Symb | 2.4093 | 0 | 0 | 0 | 0 | 0 |
| 18804 | ENSMUSG00000065751 | 0 | NA | NA | NA | NA | NA | Gm23345 | predicted gene, 23345 [Source:MGI Symb | 0 | 0 | 0 | 0 | 0 | 0 |
| 18805 | ENSMUSG00000065752 | 0 | NA | NA | NA | NA | NA | Gm23344 | predicted gene, 23344 [Source:MGI Symb | 0 | 0 | 0 | 0 | 0 | 0 |
| 18806 | ENSMUSG00000065757 | 0 | NA | NA | NA | NA | NA | Gm23347 | predicted gene, 23347 [Source:MGI Symb | 0 | 0 | 0 | 0 | 0 | 0 |
| 18807 | ENSMUSG00000065758 | 0 | NA | NA | NA | NA | NA | NA | NA | 0 | 0 | 0 | 0 | 0 | 0 |
| 18808 | ENSMUSG00000065760 | 0 | NA | NA | NA | NA | NA | Gm23848 | predicted gene, 23848 [Source:MGI Symb | 0 | 0 | 0 | 0 | 0 | 0 |
| 18809 | ENSMUSG00000065763 | 0 | NA | NA | NA | NA | NA | Gm23847 | predicted gene, 23847 [Source:MGI Symb | 0 | 0 | 0 | 0 | 0 | 0 |
| 18810 | ENSMUSG00000065764 | 0 | NA | NA | NA | NA | NA | Gm23850 | predicted gene, 23850 [Source:MGI Symb | 0 | 0 | 0 | 0 | 0 | 0 |
| 18811 | ENSMUSG00000065765 | 1.451124163 | 0.347340109 | 2.143418406 | 0.162049606 | 0.87126679 | NA | NA | NA | 0.8031 | 3.0838 | 0 | 4.8199 | 0 | 0 |
| 18812 | ENSMUSG00000065767 | 3.551159136 | 0.544382228 | 1.129469235 | 0.481980572 | 0.629819748 | NA | NA | NA | 1.6062 | 4.1117 | 3.0187 | 4.8199 | 2.1812 | 5.5692 |

| | | | | | | | | | | | | | | | |
|---|---|---|---|---|---|---|---|---|---|---|---|---|---|---|---|
| 18813 | ENSMUSG00000065768 | 0 | NA | NA | NA | NA | NA | Gm23845 | predicted gene, 23845 [Source:MGI Symb | 0 | 0 | 0 | 0 | 0 | 0 |
| 18814 | ENSMUSG00000065769 | 0 | NA | NA | NA | NA | NA | NA | NA | 0 | 0 | 0 | 0 | 0 | 0 |
| 18815 | ENSMUSG00000065770 | 0 | NA | NA | NA | NA | NA | NA | NA | 0 | 0 | 0 | 0 | 0 | 0 |
| 18816 | ENSMUSG00000065771 | 0 | NA | NA | NA | NA | NA | NA | NA | 0 | 0 | 0 | 0 | 0 | 0 |
| 18817 | ENSMUSG00000065773 | 0 | NA | NA | NA | NA | NA | NA | NA | 0 | 0 | 0 | 0 | 0 | 0 |
| 18818 | ENSMUSG00000065778 | 0.305170013 | -1.657836477 | 4.041480335 | -0.410205257 | 0.681655385 | NA | Gm22154 | predicted gene, 22154 [Source:MGI Symb | 0.8031 | 1.0279 | 0 | 0 | 0 | 0 |
| 18819 | ENSMUSG00000065782 | 0 | NA | NA | NA | NA | NA | Gm22680 | predicted gene, 22680 [Source:MGI Symb | 0 | 0 | 0 | 0 | 0 | 0 |
| 18820 | ENSMUSG00000065787 | 0 | NA | NA | NA | NA | NA | Gm22681 | predicted gene, 22681 [Source:MGI Symb | 0 | 0 | 0 | 0 | 0 | 0 |
| 18821 | ENSMUSG00000065788 | 0 | NA | NA | NA | NA | NA | NA | NA | 0 | 0 | 0 | 0 | 0 | 0 |
| 18822 | ENSMUSG00000065789 | 0 | NA | NA | NA | NA | NA | NA | NA | 0 | 0 | 0 | 0 | 0 | 0 |
| 18823 | ENSMUSG00000065791 | 0 | NA | NA | NA | NA | NA | NA | NA | 0 | 0 | 0 | 0 | 0 | 0 |
| 18824 | ENSMUSG00000065792 | 0 | NA | NA | NA | NA | NA | Gm25515 | predicted gene, 25515 [Source:MGI Symb | 0 | 0 | 0 | 0 | 0 | 0 |
| 18825 | ENSMUSG00000065794 | 0 | NA | NA | NA | NA | NA | Gm25514 | predicted gene, 25514 [Source:MGI Symb | 0 | 0 | 0 | 0 | 0 | 0 |
| 18826 | ENSMUSG00000065795 | 0 | NA | NA | NA | NA | NA | Gm25513 | predicted gene, 25513 [Source:MGI Symb | 0 | 0 | 0 | 0 | 0 | 0 |
| 18827 | ENSMUSG00000065797 | 0 | NA | NA | NA | NA | NA | Gm25512 | predicted gene, 25512 [Source:MGI Symb | 0 | 0 | 0 | 0 | 0 | 0 |
| 18828 | ENSMUSG00000065798 | 0.349477146 | 0.165142768 | 3.911021834 | 0.042224967 | 0.966319359 | NA | NA | NA | 0 | 0 | 1.0062 | 0 | 1.0906 | 0 |
| 18829 | ENSMUSG00000065799 | 0 | NA | NA | NA | NA | NA | Gm25517 | predicted gene, 25517 [Source:MGI Symb | 0 | 0 | 0 | 0 | 0 | 0 |
| 18830 | ENSMUSG00000065800 | 0 | NA | NA | NA | NA | NA | Gm23042 | predicted gene, 23042 [Source:MGI Symb | 0 | 0 | 0 | 0 | 0 | 0 |
| 18831 | ENSMUSG00000065802 | 0 | NA | NA | NA | NA | NA | Gm23041 | predicted gene, 23041 [Source:MGI Symb | 0 | 0 | 0 | 0 | 0 | 0 |
| 18832 | ENSMUSG00000065803 | 0 | NA | NA | NA | NA | NA | Gm23040 | predicted gene, 23040 [Source:MGI Symb | 0 | 0 | 0 | 0 | 0 | 0 |
| 18833 | ENSMUSG00000065804 | 0 | NA | NA | NA | NA | NA | Gm23039 | predicted gene, 23039 [Source:MGI Symb | 0 | 0 | 0 | 0 | 0 | 0 |
| 18834 | ENSMUSG00000065806 | 0 | NA | NA | NA | NA | NA | NA | NA | 0 | 0 | 0 | 0 | 0 | 0 |
| 18835 | ENSMUSG00000065807 | 0 | NA | NA | NA | NA | NA | Gm23038 | predicted gene, 23038 [Source:MGI Symb | 0 | 0 | 0 | 0 | 0 | 0 |
| 18836 | ENSMUSG00000065808 | 1.414786903 | 0.546021487 | 1.927219075 | 0.283320923 | 0.77693084 | NA | NA | NA | 2.4093 | 0 | 1.0062 | 2.8919 | 2.1812 | 0 |
| 18837 | ENSMUSG00000065811 | 0 | NA | NA | NA | NA | NA | Gm24695 | predicted gene, 24695 [Source:MGI Symb | 0 | 0 | 0 | 0 | 0 | 0 |
| 18838 | ENSMUSG00000065812 | 0 | NA | NA | NA | NA | NA | Gm24696 | predicted gene, 24696 [Source:MGI Symb | 0 | 0 | 0 | 0 | 0 | 0 |
| 18839 | ENSMUSG00000065813 | 0 | NA | NA | NA | NA | NA | NA | NA | 0 | 0 | 0 | 0 | 0 | 0 |
| 18840 | ENSMUSG00000065815 | 0 | NA | NA | NA | NA | NA | Gm24697 | predicted gene, 24697 [Source:MGI Symb | 0 | 0 | 0 | 0 | 0 | 0 |
| 18841 | ENSMUSG00000065817 | 0.503095663 | 2.549494177 | 3.348715477 | 0.761334964 | 0.446457022 | NA | Gm24698 | predicted gene, 24698 [Source:MGI Symb | 0 | 0 | 0 | 1.928 | 1.0906 | 0 |
| 18842 | ENSMUSG00000065818 | 0.36353943 | 2.065989254 | 4.034137301 | 0.512126658 | 0.608562368 | NA | Snord35a | small nucleolar RNA, C/D box 35A [Source | 0 | 0 | 0 | 0 | 2.1812 | 0 |
| 18843 | ENSMUSG00000065819 | 0 | NA | NA | NA | NA | NA | NA | NA | 0 | 0 | 0 | 0 | 0 | 0 |
| 18844 | ENSMUSG00000065820 | 0 | NA | NA | NA | NA | NA | NA | NA | 0 | 0 | 0 | 0 | 0 | 0 |
| 18845 | ENSMUSG00000065822 | 0 | NA | NA | NA | NA | NA | Snord15a | small nucleolar RNA, C/D box 15A [Source | 0 | 0 | 0 | 0 | 0 | 0 |
| 18846 | ENSMUSG00000065823 | 0 | NA | NA | NA | NA | NA | NA | NA | 0 | 0 | 0 | 0 | 0 | 0 |
| 18847 | ENSMUSG00000065824 | 0 | NA | NA | NA | NA | NA | NA | NA | 0 | 0 | 0 | 0 | 0 | 0 |
| 18848 | ENSMUSG00000065825 | 0 | NA | NA | NA | NA | NA | Gm26314 | predicted gene, 26314 [Source:MGI Symb | 0 | 0 | 0 | 0 | 0 | 0 |
| 18849 | ENSMUSG00000065828 | 0 | NA | NA | NA | NA | NA | NA | NA | 0 | 0 | 0 | 0 | 0 | 0 |
| 18850 | ENSMUSG00000065836 | 0 | NA | NA | NA | NA | NA | Gm23502 | predicted gene, 23502 [Source:MGI Symb | 0 | 0 | 0 | 0 | 0 | 0 |
| 18851 | ENSMUSG00000065837 | 0 | NA | NA | NA | NA | NA | NA | NA | 0 | 0 | 0 | 0 | 0 | 0 |
| 18852 | ENSMUSG00000065840 | 0 | NA | NA | NA | NA | NA | Gm25187 | predicted gene, 25187 [Source:MGI Symb | 0 | 0 | 0 | 0 | 0 | 0 |
| 18853 | ENSMUSG00000065843 | 0 | NA | NA | NA | NA | NA | Gm25186 | predicted gene, 25186 [Source:MGI Symb | 0 | 0 | 0 | 0 | 0 | 0 |
| 18854 | ENSMUSG00000065845 | 0 | NA | NA | NA | NA | NA | Gm25189 | predicted gene, 25189 [Source:MGI Symb | 0 | 0 | 0 | 0 | 0 | 0 |
| 18855 | ENSMUSG00000065847 | 0.353089354 | 0.165142706 | 3.896102244 | 0.042386646 | 0.966190474 | NA | Gm25188 | predicted gene, 25188 [Source:MGI Symb | 0 | 1.0279 | 0 | 0 | 1.0906 | 0 |
| 18856 | ENSMUSG00000065851 | 0 | NA | NA | NA | NA | NA | NA | NA | 0 | 0 | 0 | 0 | 0 | 0 |
| 18857 | ENSMUSG00000065852 | 0 | NA | NA | NA | NA | NA | Gm22353 | predicted gene, 22353 [Source:MGI Symb | 0 | 0 | 0 | 0 | 0 | 0 |
| 18858 | ENSMUSG00000065853 | 0 | NA | NA | NA | NA | NA | Gm22354 | predicted gene, 22354 [Source:MGI Symb | 0 | 0 | 0 | 0 | 0 | 0 |
| 18859 | ENSMUSG00000065858 | 0 | NA | NA | NA | NA | NA | Gm22350 | predicted gene, 22350 [Source:MGI Symb | 0 | 0 | 0 | 0 | 0 | 0 |
| 18860 | ENSMUSG00000065859 | 0 | NA | NA | NA | NA | NA | Gm22351 | predicted gene, 22351 [Source:MGI Symb | 0 | 0 | 0 | 0 | 0 | 0 |
| 18861 | ENSMUSG00000065862 | 0 | NA | NA | NA | NA | NA | Gm24029 | predicted gene, 24029 [Source:MGI Symb | 0 | 0 | 0 | 0 | 0 | 0 |
| 18862 | ENSMUSG00000065864 | 0 | NA | NA | NA | NA | NA | Gm24031 | predicted gene, 24031 [Source:MGI Symb | 0 | 0 | 0 | 0 | 0 | 0 |
| 18863 | ENSMUSG00000065865 | 0 | NA | NA | NA | NA | NA | Gm24030 | predicted gene, 24030 [Source:MGI Symb | 0 | 0 | 0 | 0 | 0 | 0 |
| 18864 | ENSMUSG00000065866 | 0 | NA | NA | NA | NA | NA | Gm24032 | predicted gene, 24032 [Source:MGI Symb | 0 | 0 | 0 | 0 | 0 | 0 |
| 18865 | ENSMUSG00000065869 | 0.17131964 | -0.796629176 | 4.080472857 | -0.195229623 | 0.84521319 | NA | Gm24917 | predicted gene, 24917 [Source:MGI Symb | 0 | 1.0279 | 0 | 0 | 0 | 0 |
| 18866 | ENSMUSG00000065870 | 0 | NA | NA | NA | NA | NA | NA | NA | 0 | 0 | 0 | 0 | 0 | 0 |
| 18867 | ENSMUSG00000065872 | 0 | NA | NA | NA | NA | NA | Gm25681 | predicted gene, 25681 [Source:MGI Symb | 0 | 0 | 0 | 0 | 0 | 0 |
| 18868 | ENSMUSG00000065873 | 0.305170013 | -1.657836477 | 4.041480335 | -0.410205257 | 0.681655385 | NA | NA | NA | 0.8031 | 1.0279 | 0 | 0 | 0 | 0 |

| | | | | | | | | | | | | | | | |
|---|---|---|---|---|---|---|---|---|---|---|---|---|---|---|---|
| 18869 | ENSMUSG00000065874 | 0 | NA | NA | NA | NA | NA | Gm25683 | predicted gene, 25683 [Source:MGI Symb | 0 | 0 | 0 | 0 | 0 | 0 |
| 18870 | ENSMUSG00000065875 | 0 | NA | NA | NA | NA | NA | NA | NA | 0 | 0 | 0 | 0 | 0 | 0 |
| 18871 | ENSMUSG00000065876 | 0 | NA | NA | NA | NA | NA | Gm25682 | predicted gene, 25682 [Source:MGI Symb | 0 | 0 | 0 | 0 | 0 | 0 |
| 18872 | ENSMUSG00000065878 | 0 | NA | NA | NA | NA | NA | Snord34 | small nucleolar RNA, C/D box 34 [Source:I | 0 | 0 | 0 | 0 | 0 | 0 |
| 18873 | ENSMUSG00000065881 | 1.446756899 | -1.598585954 | 1.96261736 | -0.814517382 | 0.415348619 | NA | Gm22866 | predicted gene, 22866 [Source:MGI Symb | 2.4093 | 3.0838 | 1.0062 | 0 | 2.1812 | 0 |
| 18874 | ENSMUSG00000065882 | 0 | NA | NA | NA | NA | NA | NA | NA | 0 | 0 | 0 | 0 | 0 | 0 |
| 18875 | ENSMUSG00000065883 | 0.133850373 | -0.796629176 | 4.080472857 | -0.195229623 | 0.84521319 | NA | Gm22865 | predicted gene, 22865 [Source:MGI Symb | 0.8031 | 0 | 0 | 0 | 0 | 0 |
| 18876 | ENSMUSG00000065884 | 0 | NA | NA | NA | NA | NA | NA | NA | 0 | 0 | 0 | 0 | 0 | 0 |
| 18877 | ENSMUSG00000065887 | 0 | NA | NA | NA | NA | NA | n-R5s185 | nuclear encoded rRNA 5S 185 [Source:MG | 0 | 0 | 0 | 0 | 0 | 0 |
| 18878 | ENSMUSG00000065889 | 0 | NA | NA | NA | NA | NA | NA | NA | 0 | 0 | 0 | 0 | 0 | 0 |
| 18879 | ENSMUSG00000065892 | 0 | NA | NA | NA | NA | NA | Gm24521 | predicted gene, 24521 [Source:MGI Symb | 0 | 0 | 0 | 0 | 0 | 0 |
| 18880 | ENSMUSG00000065893 | 0 | NA | NA | NA | NA | NA | NA | NA | 0 | 0 | 0 | 0 | 0 | 0 |
| 18881 | ENSMUSG00000065894 | 0 | NA | NA | NA | NA | NA | NA | NA | 0 | 0 | 0 | 0 | 0 | 0 |
| 18882 | ENSMUSG00000065898 | 0 | NA | NA | NA | NA | NA | Gm24522 | predicted gene, 24522 [Source:MGI Symb | 0 | 0 | 0 | 0 | 0 | 0 |
| 18883 | ENSMUSG00000065899 | 0 | NA | NA | NA | NA | NA | Gm24523 | predicted gene, 24523 [Source:MGI Symb | 0 | 0 | 0 | 0 | 0 | 0 |
| 18884 | ENSMUSG00000065903 | 0 | NA | NA | NA | NA | NA | Gm26111 | predicted gene, 26111 [Source:MGI Symb | 0 | 0 | 0 | 0 | 0 | 0 |
| 18885 | ENSMUSG00000065904 | 1.160275158 | 2.568144844 | 2.365095751 | 1.085852377 | 0.277544309 | NA | Gm26109 | predicted gene, 26109 [Source:MGI Symb | 0 | 1.0279 | 0 | 4.8199 | 0 | 1.1138 |
| 18886 | ENSMUSG00000065905 | 0.17131964 | -0.796629176 | 4.080472857 | -0.195229623 | 0.84521319 | NA | Gm26110 | predicted gene, 26110 [Source:MGI Symb | 0 | 1.0279 | 0 | 0 | 0 | 0 |
| 18887 | ENSMUSG00000065906 | 0 | NA | NA | NA | NA | NA | NA | NA | 0 | 0 | 0 | 0 | 0 | 0 |
| 18888 | ENSMUSG00000065907 | 0 | NA | NA | NA | NA | NA | Gm26108 | predicted gene, 26108 [Source:MGI Symb | 0 | 0 | 0 | 0 | 0 | 0 |
| 18889 | ENSMUSG00000065911 | 0 | NA | NA | NA | NA | NA | NA | NA | 0 | 0 | 0 | 0 | 0 | 0 |
| 18890 | ENSMUSG00000065913 | 0 | NA | NA | NA | NA | NA | NA | NA | 0 | 0 | 0 | 0 | 0 | 0 |
| 18891 | ENSMUSG00000065915 | 0 | NA | NA | NA | NA | NA | NA | NA | 0 | 0 | 0 | 0 | 0 | 0 |
| 18892 | ENSMUSG00000065916 | 0 | NA | NA | NA | NA | NA | Gm24446 | predicted gene, 24446 [Source:MGI Symb | 0 | 0 | 0 | 0 | 0 | 0 |
| 18893 | ENSMUSG00000065919 | 0.17131964 | -0.796629176 | 4.080472857 | -0.195229623 | 0.84521319 | NA | Gm24449 | predicted gene, 24449 [Source:MGI Symb | 0 | 1.0279 | 0 | 0 | 0 | 0 |
| 18894 | ENSMUSG00000065922 | 0 | NA | NA | NA | NA | NA | NA | NA | 0 | 0 | 0 | 0 | 0 | 0 |
| 18895 | ENSMUSG00000065924 | 0 | NA | NA | NA | NA | NA | Gm22797 | predicted gene, 22797 [Source:MGI Symb | 0 | 0 | 0 | 0 | 0 | 0 |
| 18896 | ENSMUSG00000065926 | 0 | NA | NA | NA | NA | NA | NA | NA | 0 | 0 | 0 | 0 | 0 | 0 |
| 18897 | ENSMUSG00000065934 | 0 | NA | NA | NA | NA | NA | Gm25630 | predicted gene, 25630 [Source:MGI Symb | 0 | 0 | 0 | 0 | 0 | 0 |
| 18898 | ENSMUSG00000065935 | 0 | NA | NA | NA | NA | NA | NA | NA | 0 | 0 | 0 | 0 | 0 | 0 |
| 18899 | ENSMUSG00000065936 | 0 | NA | NA | NA | NA | NA | Gm25628 | predicted gene, 25628 [Source:MGI Symb | 0 | 0 | 0 | 0 | 0 | 0 |
| 18900 | ENSMUSG00000065937 | 0 | NA | NA | NA | NA | NA | Gm25627 | predicted gene, 25627 [Source:MGI Symb | 0 | 0 | 0 | 0 | 0 | 0 |
| 18901 | ENSMUSG00000065939 | 0.294513347 | 0.165143496 | 4.080472857 | 0.040471657 | 0.967717103 | NA | Snora2b | small nucleolar RNA, H/ACA box 2B [Sourc | 0.8031 | 0 | 0 | 0.964 | 0 | 0 |
| 18902 | ENSMUSG00000065940 | 0 | NA | NA | NA | NA | NA | Gm23795 | predicted gene, 23795 [Source:MGI Symb | 0 | 0 | 0 | 0 | 0 | 0 |
| 18903 | ENSMUSG00000065943 | 0 | NA | NA | NA | NA | NA | NA | NA | 0 | 0 | 0 | 0 | 0 | 0 |
| 18904 | ENSMUSG00000065944 | 0.160662974 | 1.126924175 | 4.080472857 | 0.276174898 | 0.782413736 | NA | NA | NA | 0 | 0 | 0 | 0.964 | 0 | 0 |
| 18905 | ENSMUSG00000065945 | 0 | NA | NA | NA | NA | NA | Gm23796 | predicted gene, 23796 [Source:MGI Symb | 0 | 0 | 0 | 0 | 0 | 0 |
| 18906 | ENSMUSG00000065947 | 138.6438205 | 0.01259597 | 0.206452294 | 0.061011529 | 0.951350027 | 0.990883058 | mt-Nd4l | mitochondrially encoded NADH dehydrog | 149.38 | 145.96 | 118.74 | 153.27 | 131.96 | 132.55 |
| 18907 | ENSMUSG00000065952 | 30.40815531 | 1.091543964 | 0.455657321 | 2.395536982 | 0.01659604 | 0.152894982 | Rps23rg1 | ribosomal protein S23, retrogene 1 [Sourc | 26.502 | 8.2233 | 23.144 | 38.559 | 40.353 | 45.668 |
| 18908 | ENSMUSG00000065954 | 892.5195714 | 0.018675336 | 0.148269757 | 0.125955127 | 0.899767445 | 0.982798755 | Tacc1 | transforming, acidic coiled-coil containing | 787.84 | 857.28 | 1016.3 | 981.33 | 976.1 | 736.25 |
| 18909 | ENSMUSG00000065956 | 0 | NA | NA | NA | NA | NA | Defa37 | defensin, alpha, 37 [Source:MGI Symbol;A | 0 | 0 | 0 | 0 | 0 | 0 |
| 18910 | ENSMUSG00000065968 | 0 | NA | NA | NA | NA | NA | Ifitm7 | interferon induced transmembrane prote | 0 | 0 | 0 | 0 | 0 | 0 |
| 18911 | ENSMUSG00000065979 | 1552.969994 | -0.123796256 | 0.116846403 | -1.059478534 | 0.2893819 | 0.739770202 | Cpped1 | calcineurin-like phosphoesterase domain | 1617.4 | 1631.3 | 1610 | 1464.3 | 1689.4 | 1305.4 |
| 18912 | ENSMUSG00000065987 | 3.190586112 | -1.845975639 | 1.258428072 | -1.466890067 | 0.142405963 | NA | Cd209b | CD209b antigen [Source:MGI Symbol;Acc: | 4.8186 | 6.1675 | 4.025 | 1.928 | 1.0906 | 1.1138 |
| 18913 | ENSMUSG00000065990 | 1242.278799 | 0.064517033 | 0.117616041 | 0.548539408 | 0.583321578 | 0.908645447 | Aurkaip1 | aurora kinase A interacting protein 1 [Sou | 1225.5 | 1145.1 | 1271.9 | 1113.4 | 1358.9 | 1338.8 |
| 18914 | ENSMUSG00000065999 | 0.535401493 | -2.48596965 | 3.684117979 | -0.674780141 | 0.499815459 | NA | Zfp985 | zinc finger protein 985 [Source:MGI Symb | 3.2124 | 0 | 0 | 0 | 0 | 0 |
| 18915 | ENSMUSG00000066000 | 39.94974134 | -0.207755131 | 0.52032504 | -0.399279517 | 0.689687258 | 0.937261276 | Zfp979 | zinc finger protein 979 [Source:MGI Symb | 53.005 | 33.921 | 41.256 | 17.352 | 76.343 | 17.822 |
| 18916 | ENSMUSG00000066007 | 3.229678203 | -0.692174119 | 1.162281391 | -0.595530587 | 0.55148886 | NA | Zfp600 | zinc finger protein 600 [Source:MGI Symb | 4.8186 | 3.0838 | 4.025 | 1.928 | 2.1812 | 3.3415 |
| 18917 | ENSMUSG00000066009 | 1.419686102 | 4.03484167 | 2.091603474 | 1.92906625 | 0.053722638 | NA | Zfp987 | zinc finger protein 987 [Source:MGI Symb | 0 | 0 | 0 | 1.928 | 4.3625 | 2.2277 |
| 18918 | ENSMUSG00000066026 | 2101.946254 | 0.123206163 | 0.135673457 | 0.908108088 | 0.363821118 | 0.802548215 | Dhrs3 | dehydrogenase/reductase (SDR family) m | 1738.7 | 1869.8 | 2429.1 | 2132.3 | 2050.4 | 2391.4 |
| 18919 | ENSMUSG00000066027 | 0 | NA | NA | NA | NA | NA | Pramel51 | PRAME like 51 [Source:MGI Symbol;Acc:N | 0 | 0 | 0 | 0 | 0 | 0 |
| 18920 | ENSMUSG00000066030 | 0 | NA | NA | NA | NA | NA | Oog2 | oogenesin 2 [Source:MGI Symbol;Acc:MG | 0 | 0 | 0 | 0 | 0 | 0 |
| 18921 | ENSMUSG00000066031 | 0 | NA | NA | NA | NA | NA | Pramel25 | PRAME like 25 [Source:MGI Symbol;Acc:N | 0 | 0 | 0 | 0 | 0 | 0 |
| 18922 | ENSMUSG00000066036 | 2360.107881 | -0.176656162 | 0.142479413 | -1.239871491 | 0.21502293 | 0.659728602 | Ubr4 | ubiquitin protein ligase E3 component n-r | 2315.3 | 2739.4 | 2460.3 | 2743.5 | 1918.4 | 1983.8 |
| 18923 | ENSMUSG00000066037 | 553.366849 | 0.190159555 | 0.141002013 | 1.348630076 | 0.177455815 | 0.608660401 | Hnrnpr | heterogeneous nuclear ribonucleoprotein | 482.66 | 524.24 | 544.38 | 564.89 | 681.64 | 522.39 |
| 18924 | ENSMUSG00000066042 | 34.49768242 | -0.021774698 | 0.383089811 | -0.056839669 | 0.954672914 | 0.991678478 | Med18 | mediator complex subunit 18 [Source:MG | 36.943 | 34.949 | 32.2 | 26.991 | 44.715 | 31.188 |

| | | | | | | | | | | | | | | |
|---|---|---|---|---|---|---|---|---|---|---|---|---|---|---|
| 18925 | ENSMUSG00000066043 | 609.618405 | -0.045834383 | 0.121607708 | -0.376903598 | 0.706245248 | 0.94295036 | Phactr4 | phosphatase and actin regulator 4 [Source | 632.84 | 570.49 | 654.06 | 582.24 | 597.66 | 620.41 |
| 18926 | ENSMUSG00000066057 | 11.13774376 | -0.103715402 | 0.701445449 | -0.147859542 | 0.882453618 | 0.981769403 | Gm1976 | predicted gene 1976 [Source:MGI Symbol | 16.062 | 9.2513 | 9.0562 | 4.8199 | 9.8156 | 17.822 |
| 18927 | ENSMUSG00000066058 | 0 | NA | NA | NA | NA | NA | Cldn19 | claudin 19 [Source:MGI Symbol;Acc:MGI:3 | 0 | 0 | 0 | 0 | 0 | 0 |
| 18928 | ENSMUSG00000066060 | 0 | NA | NA | NA | NA | NA | Gm12866 | predicted gene 12866 [Source:MGI Symbo | 0 | 0 | 0 | 0 | 0 | 0 |
| 18929 | ENSMUSG00000066061 | 0 | NA | NA | NA | NA | NA | Olfr1335 | olfactory receptor 1335 [Source:MGI Sym | 0 | 0 | 0 | 0 | 0 | 0 |
| 18930 | ENSMUSG00000066068 | 1.027504657 | 1.047266995 | 2.389110177 | 0.43835023 | 0.661132419 | NA | Gm13611 | predicted gene 13611 [Source:MGI Symbo | 0 | 2.0558 | 0 | 1.928 | 2.1812 | 0 |
| 18931 | ENSMUSG00000066071 | 2394.37511 | 0.077695497 | 0.368507933 | 0.210838058 | 0.83301364 | 0.970611245 | Cyp4a12a | cytochrome P450, family 4, subfamily a, p | 2060 | 3212.2 | 1717.7 | 2139.1 | 3346 | 1891.3 |
| 18932 | ENSMUSG00000066072 | 9423.35959 | -1.764867558 | 0.47826811 | -3.690121759 | 0.000224147 | 0.006883351 | Cyp4a10 | cytochrome P450, family 4, subfamily a, p | 12940 | 17027 | 13718 | 7004.3 | 3242.4 | 2607.5 |
| 18933 | ENSMUSG00000066090 | 0.17131964 | -0.796629176 | 4.080472857 | -0.195229623 | 0.84521319 | NA | Insl5 | insulin-like 5 [Source:MGI Symbol;Acc:MG | 0 | 1.0279 | 0 | 0 | 0 | 0 |
| 18934 | ENSMUSG00000066097 | 0 | NA | NA | NA | NA | NA | Cyp2j11 | cytochrome P450, family 2, subfamily j, pc | 0 | 0 | 0 | 0 | 0 | 0 |
| 18935 | ENSMUSG00000066100 | 0 | NA | NA | NA | NA | NA | Krtap5-1 | keratin associated protein 5-1 [Source:MG | 0 | 0 | 0 | 0 | 0 | 0 |
| 18936 | ENSMUSG00000066101 | 0 | NA | NA | NA | NA | NA | Gm10153 | predicted gene 10153 [Source:MGI Symbo | 0 | 0 | 0 | 0 | 0 | 0 |
| 18937 | ENSMUSG00000066107 | 0 | NA | NA | NA | NA | NA | Gm12666 | predicted gene 12666 [Source:MGI Symbo | 0 | 0 | 0 | 0 | 0 | 0 |
| 18938 | ENSMUSG00000066108 | 0.181769715 | 1.126924175 | 4.080472857 | 0.276174898 | 0.782413736 | NA | Muc5b | mucin 5, subtype B, tracheobronchial [Sou | 0 | 0 | 0 | 0 | 1.0906 | 0 |
| 18939 | ENSMUSG00000066113 | 9.873223359 | 0.809479051 | 0.828015085 | 0.977613893 | 0.328265323 | 0.773161429 | Adamtsl1 | ADAMTS-like 1 [Source:MGI Symbol;Acc:M | 2.4093 | 13.363 | 6.0375 | 23.135 | 8.7249 | 5.5692 |
| 18940 | ENSMUSG00000066116 | 1.163184445 | 1.640230721 | 2.119996406 | 0.773695048 | 0.439111149 | NA | Gm10154 | predicted gene 10154 [Source:MGI Symbo | 1.6062 | 0 | 0 | 0.964 | 2.1812 | 2.2277 |
| 18941 | ENSMUSG00000066122 | 4.343042057 | 0.207888839 | 1.335155806 | 0.155703805 | 0.876266498 | NA | Olfr45 | olfactory receptor 45 [Source:MGI Symbo | 8.031 | 0 | 4.025 | 9.6398 | 4.3625 | 0 |
| 18942 | ENSMUSG00000066129 | 0 | NA | NA | NA | NA | NA | Kndc1 | kinase non-catalytic C-lobe domain (KIND | 0 | 0 | 0 | 0 | 0 | 0 |
| 18943 | ENSMUSG00000066137 | 0 | NA | NA | NA | NA | NA | Gm11487 | predicted gene 11487 [Source:MGI Symbo | 0 | 0 | 0 | 0 | 0 | 0 |
| 18944 | ENSMUSG00000066141 | 0 | NA | NA | NA | NA | NA | Gm11232 | predicted gene 11232 [Source:MGI Symbo | 0 | 0 | 0 | 0 | 0 | 0 |
| 18945 | ENSMUSG00000066148 | 249.8673982 | -0.022221097 | 0.164047489 | -0.135455268 | 0.892251929 | 0.982489666 | Prpf4 | pre-mRNA processing factor 4 [Source:MG | 241.73 | 239.5 | 274.7 | 272.81 | 242.12 | 228.34 |
| 18946 | ENSMUSG00000066149 | 337.9007316 | 0.064077842 | 0.15655297 | 0.409304542 | 0.682316183 | 0.93502957 | Cdc26 | cell division cycle 26 [Source:MGI Symbol; | 359.79 | 348.46 | 282.75 | 374.99 | 323.91 | 337.49 |
| 18947 | ENSMUSG00000066150 | 4032.929336 | 0.077619554 | 0.088513577 | 0.876922583 | 0.380528689 | 0.812628154 | Slc31a1 | solute carrier family 31, member 1 [Sourc | 3746.5 | 3899.9 | 4127.6 | 4066.1 | 4168.3 | 4189.2 |
| 18948 | ENSMUSG00000066151 | 666.7366705 | 0.086315197 | 0.126299402 | 0.683417304 | 0.494343189 | 0.877473722 | Fkbp15 | FK506 binding protein 15 [Source:MGI Syr | 663.36 | 690.76 | 585.63 | 631.41 | 723.08 | 706.18 |
| 18949 | ENSMUSG00000066152 | 141.6566244 | -0.051417973 | 0.213662845 | -0.240650046 | 0.809826359 | 0.966157155 | Slc31a2 | solute carrier family 31, member 2 [Sourc | 151.79 | 127.46 | 152.95 | 133.99 | 117.79 | 165.96 |
| 18950 | ENSMUSG00000066153 | 4068.236252 | 1.580205888 | 0.435668056 | 3.627086878 | 0.000286637 | 0.008375587 | Mup21 | major urinary protein 21 [Source:MGI Sym | 2329 | 2131.9 | 1656.3 | 3490.6 | 5396.4 | 9405.3 |
| 18951 | ENSMUSG00000066154 | 182899.5748 | 0.431889884 | 0.140703245 | 3.069509049 | 0.002144109 | 0.036657676 | Mup3 | major urinary protein 3 [Source:MGI Symt | 164090 | 121831 | 181255 | 198573 | 194980 | 236670 |
| 18952 | ENSMUSG00000066158 | 0.328370405 | 0.16514316 | 4.003149536 | 0.041253308 | 0.967093956 | NA | AY512931 | cDNA sequence AY512931 [Source:MGI Sy | 0 | 0 | 1.0062 | 0.964 | 0 | 0 |
| 18953 | ENSMUSG00000066170 | 22.90516311 | -1.935964365 | 0.574370697 | -3.370583448 | 0.000750092 | 0.016872984 | E230001N04 | RIKEN cDNA E230001N04 gene [Source:M | 33.73 | 35.977 | 39.244 | 7.7118 | 18.541 | 2.2277 |
| 18954 | ENSMUSG00000066175 | 13.80678385 | -0.229655004 | 0.614962165 | -0.373445745 | 0.708816718 | 0.94395774 | 2510046G10 | RIKEN cDNA 2510046G10 gene [Source:M | 20.078 | 13.363 | 11.069 | 8.6758 | 19.631 | 10.025 |
| 18955 | ENSMUSG00000066176 | 0 | NA | NA | NA | NA | NA | Gm12511 | predicted gene 12511 [Source:MGI Symbo | 0 | 0 | 0 | 0 | 0 | 0 |
| 18956 | ENSMUSG00000066178 | 2.526103964 | 0.74670049 | 1.300461328 | 0.574181234 | 0.565845167 | NA | 6030445D17 | RIKEN cDNA 6030445D17 gene [Source:M | 1.6062 | 2.0558 | 2.0125 | 2.8919 | 4.3625 | 2.2277 |
| 18957 | ENSMUSG00000066180 | 0.181769715 | 1.126924175 | 4.080472857 | 0.276174898 | 0.782413736 | NA | Gm10155 | predicted gene 10155 [Source:MGI Symbo | 0 | 0 | 0 | 0 | 1.0906 | 0 |
| 18958 | ENSMUSG00000066189 | 0 | NA | NA | NA | NA | NA | Cacng3 | calcium channel, voltage-dependent, gam | 0 | 0 | 0 | 0 | 0 | 0 |
| 18959 | ENSMUSG00000066191 | 5.914505946 | -0.813225166 | 0.927184007 | -0.877091452 | 0.380436967 | 0.812628154 | Anks6 | ankyrin repeat and sterile alpha motif dor | 7.2279 | 11.307 | 4.025 | 1.928 | 6.5437 | 4.4554 |
| 18960 | ENSMUSG00000066196 | 4.736853205 | -1.008119138 | 1.127887183 | -0.893812034 | 0.371422469 | NA | Spag8 | sperm associated antigen 8 [Source:MGI S | 0.8031 | 8.2233 | 10.062 | 3.8559 | 4.3625 | 1.1138 |
| 18961 | ENSMUSG00000066197 | 0 | NA | NA | NA | NA | NA | Gpr139 | G protein-coupled receptor 139 [Source:N | 0 | 0 | 0 | 0 | 0 | 0 |
| 18962 | ENSMUSG00000066224 | 0 | NA | NA | NA | NA | NA | Arid3c | AT rich interactive domain 3C (BRIGHT-like | 0 | 0 | 0 | 0 | 0 | 0 |
| 18963 | ENSMUSG00000066232 | 1936.910223 | 0.050962129 | 0.111350215 | 0.457674275 | 0.647186472 | 0.925643431 | Ipo7 | importin 7 [Source:MGI Symbol;Acc:MGI:1 | 1706.6 | 2144.2 | 1858.5 | 1946.3 | 1962 | 2003.8 |
| 18964 | ENSMUSG00000066233 | 74.4813258 | 0.058452717 | 0.272451267 | 0.214543752 | 0.830123052 | 0.9702546 | Tmem42 | transmembrane protein 42 [Source:MGI S | 58.626 | 87.373 | 73.456 | 81.938 | 71.981 | 73.514 |
| 18965 | ENSMUSG00000066235 | 72.9296609 | -0.334218517 | 0.291535413 | -1.146407956 | 0.251626392 | 0.704106043 | Pomgnt2 | protein O-linked mannose beta 1,4-N-acet | 79.507 | 62.703 | 101.63 | 56.875 | 58.893 | 77.969 |
| 18966 | ENSMUSG00000066239 | 0 | NA | NA | NA | NA | NA | Olfr519 | olfactory receptor 519 [Source:MGI Symb | 0 | 0 | 0 | 0 | 0 | 0 |
| 18967 | ENSMUSG00000066240 | 0 | NA | NA | NA | NA | NA | Olfr517 | olfactory receptor 517 [Source:MGI Symb | 0 | 0 | 0 | 0 | 0 | 0 |
| 18968 | ENSMUSG00000066241 | 0 | NA | NA | NA | NA | NA | Olfr514 | olfactory receptor 514 [Source:MGI Symb | 0 | 0 | 0 | 0 | 0 | 0 |
| 18969 | ENSMUSG00000066242 | 0 | NA | NA | NA | NA | NA | Olfr467 | olfactory receptor 467 [Source:MGI Symb | 0 | 0 | 0 | 0 | 0 | 0 |
| 18970 | ENSMUSG00000066245 | 0.587191767 | -1.12394062 | 3.226482889 | -0.348348545 | 0.72757844 | NA | Gm10156 | predicted gene 10156 [Source:MGI Symbo | 2.4093 | 0 | 0 | 0 | 0 | 1.1138 |
| 18971 | ENSMUSG00000066257 | 0 | NA | NA | NA | NA | NA | Olfr206 | olfactory receptor 206 [Source:MGI Symb | 0 | 0 | 0 | 0 | 0 | 0 |
| 18972 | ENSMUSG00000066258 | 181.2840951 | 0.298813517 | 0.199873861 | 1.49501048 | 0.134911707 | 0.537616739 | Trim12a | tripartite motif-containing 12A [Source:M | 194.35 | 128.49 | 164.02 | 197.62 | 208.31 | 194.92 |
| 18973 | ENSMUSG00000066262 | 0 | NA | NA | NA | NA | NA | Olfr640 | olfactory receptor 640 [Source:MGI Symb | 0 | 0 | 0 | 0 | 0 | 0 |
| 18974 | ENSMUSG00000066263 | 0 | NA | NA | NA | NA | NA | Olfr639 | olfactory receptor 639 [Source:MGI Symb | 0 | 0 | 0 | 0 | 0 | 0 |
| 18975 | ENSMUSG00000066268 | 0 | NA | NA | NA | NA | NA | Olfr586 | olfactory receptor 586 [Source:MGI Symb | 0 | 0 | 0 | 0 | 0 | 0 |
| 18976 | ENSMUSG00000066269 | 0 | NA | NA | NA | NA | NA | Olfr575 | olfactory receptor 575 [Source:MGI Symb | 0 | 0 | 0 | 0 | 0 | 0 |
| 18977 | ENSMUSG00000066270 | 2.513301513 | -0.007583087 | 1.498263575 | -0.00506125 | 0.995961724 | NA | Gm10157 | predicted gene 10157 [Source:MGI Symbo | 2.4093 | 5.1396 | 0 | 0.964 | 5.4531 | 1.1138 |
| 18978 | ENSMUSG00000066272 | 0 | NA | NA | NA | NA | NA | Olfr559 | olfactory receptor 559 [Source:MGI Symb | 0 | 0 | 0 | 0 | 0 | 0 |
| 18979 | ENSMUSG00000066273 | 0 | NA | NA | NA | NA | NA | Olfr33 | olfactory receptor 33 [Source:MGI Symbo | 0 | 0 | 0 | 0 | 0 | 0 |
| 18980 | ENSMUSG00000066278 | 176.460887 | -0.496910299 | 0.224042674 | -2.217927019 | 0.026559809 | 0.210915839 | Vps37b | vacuolar protein sorting 37B [Source:MGI | 180.7 | 178.86 | 260.62 | 158.09 | 165.77 | 114.73 |

| | | | | | | | | | | | | | | | |
|---|---|---|---|---|---|---|---|---|---|---|---|---|---|---|---|
| 18981 | ENSMUSG00000066279 | 0.160662974 | 1.126924175 | 4.080472857 | 0.276174898 | 0.782413736 | NA | Chrna10 | cholinergic receptor, nicotinic, alpha poly | 0 | 0 | 0 | 0.964 | 0 | 0 |
| 18982 | ENSMUSG00000066306 | 1064.471183 | 0.011976915 | 0.130979009 | 0.091441481 | 0.927141803 | 0.987712228 | Numa1 | nuclear mitotic apparatus protein 1 [Sourc | 934.01 | 1078.3 | 1169.3 | 1156.8 | 934.66 | 1113.8 |
| 18983 | ENSMUSG00000066315 | 5.674859587 | 0.14901647 | 0.899703407 | 0.165628438 | 0.868449366 | 0.978366823 | Gm12918 | predicted gene 12918 [Source:MGI Symbo | 4.0155 | 5.1396 | 7.0437 | 5.7839 | 8.7249 | 3.3415 |
| 18984 | ENSMUSG00000066319 | 439.3625135 | 0.019674632 | 0.131854471 | 0.149214751 | 0.881384181 | 0.981623222 | Rtp3 | receptor transporter protein 3 [Source:M | 421.63 | 451.26 | 436.71 | 452.11 | 465.69 | 408.78 |
| 18985 | ENSMUSG00000066324 | 1093.280959 | -0.207503658 | 0.115393706 | -1.798223366 | 0.072141618 | 0.389315961 | Bpnt2 | 3'(2'), 5'-bisphosphate nucleotidase 2 [Sou | 1176.5 | 1214 | 1124 | 903.25 | 1097.2 | 1044.8 |
| 18986 | ENSMUSG00000066357 | 212.1277543 | 0.265163087 | 0.16828924 | 1.575638979 | 0.115109028 | 0.49818581 | Wdr6 | WD repeat domain 6 [Source:MGI Symbol | 206.4 | 190.16 | 181.12 | 231.35 | 244.3 | 219.43 |
| 18987 | ENSMUSG00000066359 | 0 | NA | NA | NA | NA | NA | Tcl1b1 | T cell leukemia/lymphoma 1B, 1 [Source:N | 0 | 0 | 0 | 0 | 0 | 0 |
| 18988 | ENSMUSG00000066361 | 6.062555014 | -0.555421769 | 0.859315209 | -0.646353938 | 0.518050164 | 0.887376101 | Serpina3c | serine (or cysteine) peptidase inhibitor, cl | 6.4248 | 7.1954 | 8.05 | 4.8199 | 6.5437 | 3.3415 |
| 18989 | ENSMUSG00000066362 | 6.172331712 | -0.790626822 | 0.857195618 | -0.922341185 | 0.356350638 | 0.795787977 | Rps13-ps1 | ribosomal protein S13, pseudogene 1 [Sou | 7.2279 | 9.2513 | 7.0437 | 5.7839 | 3.2719 | 4.4554 |
| 18990 | ENSMUSG00000066363 | 28.89307777 | -0.780800413 | 0.474051437 | -1.647079521 | 0.099541702 | 0.464748317 | Serpina3f | serine (or cysteine) peptidase inhibitor, cl | 28.912 | 56.535 | 24.15 | 17.352 | 17.45 | 28.96 |
| 18991 | ENSMUSG00000066364 | 1.08158966 | 1.151574547 | 2.421847399 | 0.475494264 | 0.634434721 | NA | Serpina3b | serine (or cysteine) peptidase inhibitor, cl | 0 | 1.0279 | 1.0062 | 0 | 0 | 4.4554 |
| 18992 | ENSMUSG00000066366 | 28553.68329 | 0.339897786 | 0.11723718 | 2.899232023 | 0.00374078 | 0.054645736 | Serpina1a | serine (or cysteine) peptidase inhibitor, cl | 21747 | 26644 | 27227 | 29964 | 29978 | 35762 |
| 18993 | ENSMUSG00000066368 | 0 | NA | NA | NA | NA | NA | Actl11 | actin-like 11 [Source:MGI Symbol;Acc:MG | 0 | 0 | 0 | 0 | 0 | 0 |
| 18994 | ENSMUSG00000066372 | 0 | NA | NA | NA | NA | NA | Vmn2r65 | vomeronasal 2, receptor 65 [Source:MGI S | 0 | 0 | 0 | 0 | 0 | 0 |
| 18995 | ENSMUSG00000066378 | 0 | NA | NA | NA | NA | NA | Gm10160 | predicted gene 10160 [Source:MGI Symbo | 0 | 0 | 0 | 0 | 0 | 0 |
| 18996 | ENSMUSG00000066382 | 0 | NA | NA | NA | NA | NA | Iqcf5 | IQ motif containing F5 [Source:MGI Symb | 0 | 0 | 0 | 0 | 0 | 0 |
| 18997 | ENSMUSG00000066383 | 0 | NA | NA | NA | NA | NA | Iqcf1 | IQ motif containing F1 [Source:MGI Symb | 0 | 0 | 0 | 0 | 0 | 0 |
| 18998 | ENSMUSG00000066389 | 0 | NA | NA | NA | NA | NA | Rpl31-ps1 | ribosomal protein L31, pseudogene 1 [Sou | 0 | 0 | 0 | 0 | 0 | 0 |
| 18999 | ENSMUSG00000066392 | 1.633094955 | -0.366694862 | 1.842455547 | -0.19902508 | 0.842243126 | NA | Nrxn3 | neurexin III [Source:MGI Symbol;Acc:MGI | 2.4093 | 3.0838 | 0 | 0.964 | 0 | 3.3415 |
| 19000 | ENSMUSG00000066406 | 920.5833141 | -0.090958416 | 0.135788236 | -0.669854907 | 0.502950288 | 0.881220881 | Akap13 | A kinase (PRKA) anchor protein 13 [Source | 899.47 | 1095.8 | 854.3 | 915.78 | 968.47 | 789.72 |
| 19001 | ENSMUSG00000066407 | 1.563087765 | -0.228300964 | 1.75634412 | -0.129986465 | 0.896577135 | NA | Gm10263 | predicted gene 10263 [Source:MGI Symbo | 0 | 3.0838 | 2.0125 | 0.964 | 1.0906 | 2.2277 |
| 19002 | ENSMUSG00000066415 | 507.937976 | 0.205960254 | 0.138541012 | 1.486637424 | 0.137110598 | 0.541092888 | Msl2 | MSL complex subunit 2 [Source:MGI Symb | 420.02 | 464.62 | 531.3 | 531.15 | 520.22 | 580.31 |
| 19003 | ENSMUSG00000066438 | 0.535401493 | -2.48596965 | 3.684117979 | -0.674780141 | 0.499815459 | NA | Plekhd1 | pleckstrin homology domain containing, f | 3.2124 | 0 | 0 | 0 | 0 | 0 |
| 19004 | ENSMUSG00000066440 | 612.2274011 | 0.261283913 | 0.163880033 | 1.594360877 | 0.11085523 | 0.489424955 | Zfyve26 | zinc finger, FYVE domain containing 26 [Sc | 692.27 | 510.88 | 465.89 | 610.2 | 646.74 | 747.39 |
| 19005 | ENSMUSG00000066441 | 401.3252253 | 0.936344626 | 0.196064003 | 4.775709013 | 1.79E-06 | 0.000129507 | Rdh11 | retinol dehydrogenase 11 [Source:MGI Sy | 322.04 | 298.1 | 205.27 | 468.49 | 461.33 | 652.71 |
| 19006 | ENSMUSG00000066442 | 153.2359405 | -0.072797118 | 0.271502563 | -0.268126818 | 0.788601703 | 0.960872359 | Mthfs | 5, 10-methenyltetrahydrofolate synthetas | 208.81 | 101.76 | 159.99 | 138.81 | 190.86 | 119.18 |
| 19007 | ENSMUSG00000066443 | 0 | NA | NA | NA | NA | NA | Gm10163 | predicted pseudogene 10163 [Source:MG | 0 | 0 | 0 | 0 | 0 | 0 |
| 19008 | ENSMUSG00000066456 | 7.084017967 | 0.271234847 | 0.825427125 | 0.328599386 | 0.742458512 | 0.949087045 | Hmgn3 | high mobility group nucleosomal binding | 8.031 | 3.0838 | 8.05 | 6.7478 | 5.4531 | 11.138 |
| 19009 | ENSMUSG00000066457 | 0 | NA | NA | NA | NA | NA | S100a11-ps | S100 calcium binding protein A11, pseudo | 0 | 0 | 0 | 0 | 0 | 0 |
| 19010 | ENSMUSG00000066463 | 0 | NA | NA | NA | NA | NA | Omt2a | oocyte maturation, alpha [Source:MGI Sy | 0 | 0 | 0 | 0 | 0 | 0 |
| 19011 | ENSMUSG00000066475 | 2.382470208 | 1.538610751 | 1.533328321 | 1.003445075 | 0.315646165 | NA | Gm10268 | predicted gene 10268 [Source:MGI Symbo | 1.6062 | 0 | 2.0125 | 1.928 | 7.6343 | 1.1138 |
| 19012 | ENSMUSG00000066477 | 97.96776801 | 1.393359919 | 0.265155373 | 5.254880957 | 1.48E-07 | 1.56E-05 | Gm16551 | predicted gene 16551 [Source:MGI Symbo | 36.943 | 62.703 | 63.393 | 151.34 | 131.96 | 141.46 |
| 19013 | ENSMUSG00000066478 | 0 | NA | NA | NA | NA | NA | Gm5745 | predicted gene 5745 [Source:MGI Symbol | 0 | 0 | 0 | 0 | 0 | 0 |
| 19014 | ENSMUSG00000066487 | 1.725786744 | -0.413205325 | 1.835673997 | -0.225097335 | 0.821903554 | NA | Gm5786 | predicted pseudogene 5786 [Source:MGI | 0.8031 | 3.0838 | 2.0125 | 0 | 0 | 4.4554 |
| 19015 | ENSMUSG00000066489 | 0 | NA | NA | NA | NA | NA | Rpl26-ps2 | ribosomal protein L26, pseudogene 2 [Sou | 0 | 0 | 0 | 0 | 0 | 0 |
| 19016 | ENSMUSG00000066491 | 2.492738851 | -1.271204853 | 1.454718219 | -0.873849545 | 0.382200195 | NA | Cox6c2 | cytochrome c oxidase subunit 6C2 [Source | 2.4093 | 3.0838 | 5.0312 | 0 | 1.0906 | 3.3415 |
| 19017 | ENSMUSG00000066500 | 0 | NA | NA | NA | NA | NA | Izumo2 | IZUMO family member 2 [Source:MGI Sym | 0 | 0 | 0 | 0 | 0 | 0 |
| 19018 | ENSMUSG00000066510 | 0 | NA | NA | NA | NA | NA | Ankdd1a | ankyrin repeat and death domain contain | 0 | 0 | 0 | 0 | 0 | 0 |
| 19019 | ENSMUSG00000066512 | 0.267700746 | -1.497481649 | 4.057001436 | -0.369110456 | 0.712045396 | NA | Klk1b5 | kallikrein 1-related peptidase b5 [Source:I | 1.6062 | 0 | 0 | 0 | 0 | 0 |
| 19020 | ENSMUSG00000066513 | 45.04083933 | 0.552184845 | 0.352548776 | 1.566265106 | 0.117286566 | 0.502643331 | Klk1b4 | kallikrein 1-related pepidase b4 [Source:M | 48.186 | 24.67 | 36.225 | 52.055 | 54.531 | 54.578 |
| 19021 | ENSMUSG00000066515 | 0 | NA | NA | NA | NA | NA | Klk1b3 | kallikrein 1-related peptidase b3 [Source:I | 0 | 0 | 0 | 0 | 0 | 0 |
| 19022 | ENSMUSG00000066516 | 0 | NA | NA | NA | NA | NA | Klk1b21 | kallikrein 1-related peptidase b21 [Source | 0 | 0 | 0 | 0 | 0 | 0 |
| 19023 | ENSMUSG00000066537 | 17.15618571 | -0.465316119 | 0.535892853 | -0.868300661 | 0.385229761 | 0.816437825 | Vmn2r57 | vomeronasal 2, receptor 57 [Source:MGI S | 16.062 | 21.586 | 22.137 | 13.496 | 19.631 | 10.025 |
| 19024 | ENSMUSG00000066538 | 3.815551374 | -0.197773822 | 1.204135859 | -0.164245438 | 0.869537932 | NA | Gm6254 | predicted gene 6254 [Source:MGI Symbol | 4.0155 | 7.1954 | 1.0062 | 1.928 | 7.6343 | 1.1138 |
| 19025 | ENSMUSG00000066543 | 1.218845652 | 2.679623599 | 2.159461417 | 1.240875886 | 0.214651602 | NA | Rpl17-ps9 | ribosomal protein L17, pseudogene 9 [Sou | 0.8031 | 0 | 0 | 0.964 | 1.0906 | 4.4554 |
| 19026 | ENSMUSG00000066551 | 2746.198212 | 0.014613103 | 0.086906191 | 0.16814801 | 0.86646684 | 0.977925452 | Hmgb1 | high mobility group box 1 [Source:MGI Sy | 2728.9 | 2772.3 | 2695.7 | 2760.8 | 2810.5 | 2708.9 |
| 19027 | ENSMUSG00000066553 | 44.9001737 | -0.054678126 | 0.348845115 | -0.156740407 | 0.87544944 | 0.980525066 | Gm6969 | predicted pseudogene 6969 [Source:MGI | 54.611 | 34.949 | 47.293 | 35.667 | 53.44 | 43.44 |
| 19028 | ENSMUSG00000066554 | 3.746318701 | 0.064830978 | 1.164992075 | 0.055649287 | 0.9556212 | NA | Gm10167 | predicted pseudogene 10167 [Source:MG | 4.8186 | 5.1396 | 1.0062 | 3.8559 | 6.5437 | 1.1138 |
| 19029 | ENSMUSG00000066568 | 619.8786882 | -0.118719816 | 0.11773898 | -1.008330601 | 0.313295777 | 0.760152431 | Lsm14a | LSM14A mRNA processing body assembly | 611.16 | 674.31 | 651.04 | 581.28 | 592.21 | 609.27 |
| 19030 | ENSMUSG00000066571 | 114.3216866 | 0.015316279 | 0.241090204 | 0.063529245 | 0.949345072 | 0.990731019 | Garre1 | granule associated Rac and RHOG effecto | 100.39 | 139.8 | 101.63 | 135.92 | 109.06 | 99.132 |
| 19031 | ENSMUSG00000066583 | 0 | NA | NA | NA | NA | NA | Scgb1b27 | secretoglobin, family 1B, member 27 [Sou | 0 | 0 | 0 | 0 | 0 | 0 |
| 19032 | ENSMUSG00000066584 | 0 | NA | NA | NA | NA | NA | Scgb2b27 | secretoglobin, family 2B, member 27 [Sou | 0 | 0 | 0 | 0 | 0 | 0 |
| 19033 | ENSMUSG00000066586 | 0 | NA | NA | NA | NA | NA | Scgb2b26 | secretoglobin, family 2B, member 26 [Sou | 0 | 0 | 0 | 0 | 0 | 0 |
| 19034 | ENSMUSG00000066592 | 0.531944269 | 2.618916564 | 3.285123409 | 0.797204926 | 0.425332026 | NA | Gm7589 | predicted gene 7589 [Source:MGI Symbol | 0 | 0 | 0 | 0.964 | 0 | 2.2277 |
| 19035 | ENSMUSG00000066595 | 52.65401731 | -0.195206921 | 0.307314722 | -0.635201985 | 0.525296689 | 0.888334265 | Flvcr1 | feline leukemia virus subgroup C cellular r | 61.839 | 54.48 | 52.325 | 56.875 | 43.625 | 46.781 |
| 19036 | ENSMUSG00000066607 | 1.392128934 | -1.720765878 | 2.439558825 | -0.705359453 | 0.480586571 | NA | Insyn1 | inhibitory synaptic factor 1 [Source:MGI S | 6.4248 | 0 | 0 | 1.928 | 0 | 0 |

| | | | | | | | | | | | | | | | |
|---|---|---|---|---|---|---|---|---|---|---|---|---|---|---|---|
| 19037 | ENSMUSG00000066613 | 100.8495457 | 0.075124658 | 0.256264238 | 0.293153108 | 0.769405125 | 0.956153573 | Zfp932 | zinc finger protein 932 [Source:MGI Symb | 92.357 | 86.345 | 115.72 | 79.046 | 106.88 | 124.75 |
| 19038 | ENSMUSG00000066620 | 0 | NA | NA | NA | NA | NA | Gm4883 | predicted gene 4883 [Source:MGI Symbol | 0 | 0 | 0 | 0 | 0 | 0 |
| 19039 | ENSMUSG00000066621 | 233.1352671 | 0.236105709 | 0.164814714 | 1.432552374 | 0.151985801 | 0.567340036 | Tecpr1 | tectonin beta-propeller repeat containing | 199.17 | 218.95 | 224.39 | 239.07 | 242.12 | 275.12 |
| 19040 | ENSMUSG00000066626 | 0 | NA | NA | NA | NA | NA | Gm5121 | predicted gene 5121 [Source:MGI Symbol | 0 | 0 | 0 | 0 | 0 | 0 |
| 19041 | ENSMUSG00000066629 | 0 | NA | NA | NA | NA | NA | Rpl36-ps3 | ribosomal protein L36, pseudogene 3 [Sou | 0 | 0 | 0 | 0 | 0 | 0 |
| 19042 | ENSMUSG00000066632 | 1.407742446 | 1.267490155 | 2.119347463 | 0.598056797 | 0.549802036 | NA | Pgk1-rs7 | phosphoglycerate kinase-1, related seque | 2.4093 | 0 | 0 | 3.8559 | 2.1812 | 0 |
| 19043 | ENSMUSG00000066637 | 260.3325313 | -0.015025263 | 0.178669299 | -0.084095381 | 0.932980597 | 0.988420305 | Ttc32 | tetratricopeptide repeat domain 32 [Sour | 305.98 | 232.31 | 245.52 | 222.68 | 275.93 | 279.57 |
| 19044 | ENSMUSG00000066640 | 206.9167103 | 0.196966795 | 0.178336018 | 1.104470075 | 0.269389278 | 0.724979259 | Fbxl18 | F-box and leucine-rich repeat protein 18 [ | 211.22 | 163.44 | 203.26 | 223.64 | 210.49 | 229.45 |
| 19045 | ENSMUSG00000066643 | 76.86811143 | -0.376533499 | 0.263951556 | -1.426525022 | 0.15371685 | 0.570987433 | Wdr35 | WD repeat domain 35 [Source:MGI Symb | 81.916 | 79.15 | 99.618 | 68.442 | 74.162 | 57.92 |
| 19046 | ENSMUSG00000066647 | 103.0148602 | 0.537713999 | 0.231576898 | 2.321967361 | 0.020234693 | 0.17452272 | Gm5113 | predicted gene 5113 [Source:MGI Symbol | 85.129 | 92.513 | 74.462 | 124.35 | 107.97 | 133.66 |
| 19047 | ENSMUSG00000066652 | 0 | NA | NA | NA | NA | NA | Lefty2 | left-right determination factor 2 [Source:N | 0 | 0 | 0 | 0 | 0 | 0 |
| 19048 | ENSMUSG00000066671 | 0 | NA | NA | NA | NA | NA | Olfr220 | olfactory receptor 220 [Source:MGI Symb | 0 | 0 | 0 | 0 | 0 | 0 |
| 19049 | ENSMUSG00000066672 | 0 | NA | NA | NA | NA | NA | Olfr417 | olfactory receptor 417 [Source:MGI Symb | 0 | 0 | 0 | 0 | 0 | 0 |
| 19050 | ENSMUSG00000066677 | 8.893833078 | -0.070441684 | 0.738967773 | -0.095324433 | 0.924057137 | 0.987408571 | Ifi208 | interferon activated gene 208 [Source:MG | 8.031 | 13.363 | 6.0375 | 11.568 | 5.4531 | 8.9108 |
| 19051 | ENSMUSG00000066680 | 0 | NA | NA | NA | NA | NA | Olfr1407-ps1 | olfactory receptor 1407, pseudogene 1 [So | 0 | 0 | 0 | 0 | 0 | 0 |
| 19052 | ENSMUSG00000066682 | 10.36084545 | -1.540917602 | 0.718672148 | -2.144117602 | 0.032023465 | 0.238756391 | Pilrb2 | paired immunoglobin-like type 2 receptor | 13.653 | 18.503 | 14.087 | 3.8559 | 8.7249 | 3.3415 |
| 19053 | ENSMUSG00000066684 | 14.78278904 | -1.113046733 | 0.640654079 | -1.73735994 | 0.082323658 | 0.422844241 | Pilrb1 | paired immunoglobin-like type 2 receptor | 13.653 | 29.81 | 17.106 | 3.8559 | 10.906 | 13.366 |
| 19054 | ENSMUSG00000066687 | 725.170752 | -1.827632564 | 0.50350413 | -3.629826359 | 0.000283612 | 0.008319729 | Zbtb16 | zinc finger and BTB domain containing 16 | 944.45 | 1376.4 | 1073.7 | 178.34 | 540.95 | 237.25 |
| 19055 | ENSMUSG00000066688 | 0 | NA | NA | NA | NA | NA | Pramel21 | PRAME like 21 [Source:MGI Symbol;Acc:M | 0 | 0 | 0 | 0 | 0 | 0 |
| 19056 | ENSMUSG00000066693 | 0 | NA | NA | NA | NA | NA | Gm7493 | predicted gene 7493 [Source:MGI Symbol | 0 | 0 | 0 | 0 | 0 | 0 |
| 19057 | ENSMUSG00000066697 | 0 | NA | NA | NA | NA | NA | Gm10172 | predicted gene 10172 [Source:MGI Symbo | 0 | 0 | 0 | 0 | 0 | 0 |
| 19058 | ENSMUSG00000066704 | 0 | NA | NA | NA | NA | NA | Cyp2b19 | cytochrome P450, family 2, subfamily b, p | 0 | 0 | 0 | 0 | 0 | 0 |
| 19059 | ENSMUSG00000066705 | 11.59922625 | -0.33069214 | 0.66785789 | -0.495153453 | 0.620491805 | 0.917977899 | Fxyd6 | FXYD domain-containing ion transport reg | 11.243 | 16.447 | 11.069 | 8.6758 | 5.4531 | 16.708 |
| 19060 | ENSMUSG00000066714 | 0 | NA | NA | NA | NA | NA | Gm8062 | predicted pseudogene 8062 [Source:MGI | 0 | 0 | 0 | 0 | 0 | 0 |
| 19061 | ENSMUSG00000066720 | 0 | NA | NA | NA | NA | NA | Cldn9 | claudin 9 [Source:MGI Symbol;Acc:MGI:1 | 0 | 0 | 0 | 0 | 0 | 0 |
| 19062 | ENSMUSG00000066721 | 0.699599566 | -1.425530384 | 3.053137229 | -0.466906751 | 0.640566595 | NA | Zfp575 | zinc finger protein 575 [Source:MGI Symb | 0 | 3.0838 | 0 | 0 | 0 | 1.1138 |
| 19063 | ENSMUSG00000066723 | 0 | NA | NA | NA | NA | NA | NA | NA | 0 | 0 | 0 | 0 | 0 | 0 |
| 19064 | ENSMUSG00000066724 | 5.682679688 | -1.353542463 | 0.932856429 | -1.450965467 | 0.146789477 | 0.55769976 | Gm10175 | predicted gene 10175 [Source:MGI Symbo | 7.2279 | 6.1675 | 11.069 | 1.928 | 4.3625 | 3.3415 |
| 19065 | ENSMUSG00000066735 | 1048.065198 | 0.090423162 | 0.109633147 | 0.824779401 | 0.409496843 | 0.833241465 | Vkorc1l1 | vitamin K epoxide reductase complex, sub | 1078.6 | 1005.3 | 960.96 | 1060.4 | 1148.4 | 1034.8 |
| 19066 | ENSMUSG00000066747 | 0 | NA | NA | NA | NA | NA | Olfr878 | olfactory receptor 878 [Source:MGI Symb | 0 | 0 | 0 | 0 | 0 | 0 |
| 19067 | ENSMUSG00000066748 | 0 | NA | NA | NA | NA | NA | Olfr145 | olfactory receptor 145 [Source:MGI Symb | 0 | 0 | 0 | 0 | 0 | 0 |
| 19068 | ENSMUSG00000066749 | 0 | NA | NA | NA | NA | NA | Olfr877 | olfactory receptor 877 [Source:MGI Symb | 0 | 0 | 0 | 0 | 0 | 0 |
| 19069 | ENSMUSG00000066750 | 0 | NA | NA | NA | NA | NA | Olfr876 | olfactory receptor 876 [Source:MGI Symb | 0 | 0 | 0 | 0 | 0 | 0 |
| 19070 | ENSMUSG00000066752 | 0 | NA | NA | NA | NA | NA | Gm10176 | predicted gene 10176 [Source:MGI Symbo | 0 | 0 | 0 | 0 | 0 | 0 |
| 19071 | ENSMUSG00000066755 | 0 | NA | NA | NA | NA | NA | Tnfsf18 | tumor necrosis factor (ligand) superfamily | 0 | 0 | 0 | 0 | 0 | 0 |
| 19072 | ENSMUSG00000066756 | 0 | NA | NA | NA | NA | NA | Igfl3 | IGF-like family member 3 [Source:MGI Syr | 0 | 0 | 0 | 0 | 0 | 0 |
| 19073 | ENSMUSG00000066760 | 0 | NA | NA | NA | NA | NA | Psg16 | pregnancy specific glycoprotein 16 [Sourc | 0 | 0 | 0 | 0 | 0 | 0 |
| 19074 | ENSMUSG00000066767 | 0 | NA | NA | NA | NA | NA | Gm3867 | predicted gene 3867 [Source:MGI Symbol | 0 | 0 | 0 | 0 | 0 | 0 |
| 19075 | ENSMUSG00000066771 | 0 | NA | NA | NA | NA | NA | Gm5614 | predicted gene 5614 [Source:MGI Symbol | 0 | 0 | 0 | 0 | 0 | 0 |
| 19076 | ENSMUSG00000066772 | 0 | NA | NA | NA | NA | NA | Obox3 | oocyte specific homeobox 3 [Source:MGI | 0 | 0 | 0 | 0 | 0 | 0 |
| 19077 | ENSMUSG00000066797 | 0 | NA | NA | NA | NA | NA | Zfp648 | zinc finger protein 648 [Source:MGI Symb | 0 | 0 | 0 | 0 | 0 | 0 |
| 19078 | ENSMUSG00000066798 | 154.1564391 | 0.18698423 | 0.191461525 | 0.976615172 | 0.328759705 | 0.77362974 | Zbtb6 | zinc finger and BTB domain containing 6 [ | 148.57 | 153.16 | 130.81 | 169.66 | 170.14 | 152.6 |
| 19079 | ENSMUSG00000066800 | 33.44205193 | -0.773320443 | 0.434811301 | -1.778519653 | 0.075318547 | 0.401550854 | Rnasel | ribonuclease L (2', 5'-oligoisoadenylate sy | 48.186 | 45.228 | 33.206 | 30.847 | 11.997 | 31.188 |
| 19080 | ENSMUSG00000066803 | 0 | NA | NA | NA | NA | NA | Vmn1r84 | vomeronasal 1 receptor 84 [Source:MGI S | 0 | 0 | 0 | 0 | 0 | 0 |
| 19081 | ENSMUSG00000066804 | 0 | NA | NA | NA | NA | NA | Vmn1r83 | vomeronasal 1 receptor 83 [Source:MGI S | 0 | 0 | 0 | 0 | 0 | 0 |
| 19082 | ENSMUSG00000066807 | 0.167707431 | -0.796629176 | 4.080472857 | -0.195229623 | 0.84521319 | NA | Gm10179 | predicted gene 10179 [Source:MGI Symbo | 0 | 0 | 1.0062 | 0 | 0 | 0 |
| 19083 | ENSMUSG00000066809 | 25.35118898 | 0.074672712 | 0.447197011 | 0.166979453 | 0.867386216 | 0.97808328 | Gm10180 | predicted gene 10180 [Source:MGI Symbo | 23.29 | 23.642 | 27.169 | 26.027 | 17.45 | 34.529 |
| 19084 | ENSMUSG00000066810 | 0 | NA | NA | NA | NA | NA | Gm10181 | predicted gene 10181 [Source:MGI Symbo | 0 | 0 | 0 | 0 | 0 | 0 |
| 19085 | ENSMUSG00000066820 | 0 | NA | NA | NA | NA | NA | Vmn2r28 | vomeronasal 2, receptor 28 [Source:MGI S | 0 | 0 | 0 | 0 | 0 | 0 |
| 19086 | ENSMUSG00000066828 | 0 | NA | NA | NA | NA | NA | Vmn2r37 | vomeronasal 2, receptor 37 [Source:MGI S | 0 | 0 | 0 | 0 | 0 | 0 |
| 19087 | ENSMUSG00000066829 | 71.17829791 | 0.003045531 | 0.265784589 | 0.011458646 | 0.990857523 | 0.99859176 | Zfp810 | zinc finger protein 810 [Source:MGI Symb | 77.901 | 69.898 | 65.406 | 77.118 | 64.346 | 72.4 |
| 19088 | ENSMUSG00000066838 | 32.35959946 | -0.088706031 | 0.417834482 | -0.21229945 | 0.831873424 | 0.970533397 | Zfp772 | zinc finger protein 772 [Source:MGI Symb | 31.321 | 28.782 | 40.25 | 45.307 | 23.994 | 24.505 |
| 19089 | ENSMUSG00000066839 | 817.3729487 | -0.040783754 | 0.111715611 | -0.365067636 | 0.715060919 | 0.944595913 | Ecsit | ECSIT signalling integrator [Source:MGI Sy | 849.68 | 785.33 | 851.28 | 798.17 | 773.25 | 846.52 |
| 19090 | ENSMUSG00000066842 | 18.45330066 | -0.786553397 | 0.536278151 | -1.466689245 | 0.14246061 | 0.55198239 | Hmcn1 | hemicentin 1 [Source:MGI Symbol;Acc:MG | 31.321 | 21.586 | 17.106 | 16.388 | 8.7249 | 15.594 |
| 19091 | ENSMUSG00000066850 | 0 | NA | NA | NA | NA | NA | Vmn1r65 | vomeronasal 1 receptor 65 [Source:MGI S | 0 | 0 | 0 | 0 | 0 | 0 |
| 19092 | ENSMUSG00000066861 | 14.22010388 | -0.645661646 | 0.610155365 | -1.058192197 | 0.289967828 | 0.740058153 | Oas1g | 2'-5' oligoadenylate synthetase 1G [Sourc | 19.274 | 17.475 | 15.094 | 4.8199 | 14.178 | 14.48 |

| | | | | | | | | | | | | | | | |
|---|---|---|---|---|---|---|---|---|---|---|---|---|---|---|---|
| 19093 | ENSMUSG00000066867 | 0 | NA | NA | NA | NA | NA | Oas1e | 2'-5' oligoadenylate synthetase 1E [Source | 0 | 0 | 0 | 0 | 0 | 0 |
| 19094 | ENSMUSG00000066877 | 46.3901508 | -0.853795502 | 0.403155503 | -2.117782086 | 0.034193527 | 0.247603883 | Nck2 | non-catalytic region of tyrosine kinase ada | 42.564 | 64.759 | 72.45 | 51.091 | 19.631 | 27.846 |
| 19095 | ENSMUSG00000066878 | 1.039990737 | 1.242706646 | 2.220478094 | 0.559657242 | 0.575713253 | NA | Gm10184 | predicted pseudogene 10184 [Source:MG | 0.8031 | 1.0279 | 0 | 0 | 2.1812 | 2.2277 |
| 19096 | ENSMUSG00000066880 | 301.5752929 | 0.017299659 | 0.16014723 | 0.108023467 | 0.913977077 | 0.98625062 | Zfp617 | zinc finger protein 617 [Source:MGI Symb | 305.98 | 282.68 | 309.92 | 260.27 | 308.64 | 341.95 |
| 19097 | ENSMUSG00000066892 | 100.6334414 | 0.023783007 | 0.238597173 | 0.099678496 | 0.920599573 | 0.986685804 | Fbxl12 | F-box and leucine-rich repeat protein 12 [ | 114.04 | 96.624 | 88.55 | 113.75 | 87.249 | 103.59 |
| 19098 | ENSMUSG00000066894 | 15.46161686 | 0.106854043 | 0.556595055 | 0.191978067 | 0.847759386 | 0.972803041 | Vsig10 | V-set and immunoglobulin domain contai | 20.078 | 13.363 | 11.069 | 17.352 | 13.087 | 17.822 |
| 19099 | ENSMUSG00000066896 | 0 | NA | NA | NA | NA | NA | Olfr18 | olfactory receptor 18 [Source:MGI Symbo | 0 | 0 | 0 | 0 | 0 | 0 |
| 19100 | ENSMUSG00000066897 | 0 | NA | NA | NA | NA | NA | Olfr872 | olfactory receptor 872 [Source:MGI Symb | 0 | 0 | 0 | 0 | 0 | 0 |
| 19101 | ENSMUSG00000066899 | 0 | NA | NA | NA | NA | NA | Olfr870 | olfactory receptor 870 [Source:MGI Symb | 0 | 0 | 0 | 0 | 0 | 0 |
| 19102 | ENSMUSG00000066900 | 2212.474524 | 0.095024914 | 0.098299429 | 0.966688359 | 0.333699848 | 0.775487565 | Suds3 | suppressor of defective silencing 3 homol | 2053.5 | 2115.5 | 2251 | 2405.1 | 2146.3 | 2303.4 |
| 19103 | ENSMUSG00000066902 | 0 | NA | NA | NA | NA | NA | Rps23-ps2 | ribosomal protein S23, pseudogene 2 [Sou | 0 | 0 | 0 | 0 | 0 | 0 |
| 19104 | ENSMUSG00000066905 | 0 | NA | NA | NA | NA | NA | Olfr860 | olfactory receptor 860 [Source:MGI Symb | 0 | 0 | 0 | 0 | 0 | 0 |
| 19105 | ENSMUSG00000066912 | 0 | NA | NA | NA | NA | NA | Olfr833-ps1 | olfactory receptor 833, pseudogene 1 [So | 0 | 0 | 0 | 0 | 0 | 0 |
| 19106 | ENSMUSG00000066914 | 0 | NA | NA | NA | NA | NA | Olfr831-ps1 | olfactory receptor 831, pseudogene 1 [So | 0 | 0 | 0 | 0 | 0 | 0 |
| 19107 | ENSMUSG00000066936 | 0 | NA | NA | NA | NA | NA | Gm10188 | predicted gene 10188 [Source:MGI Symbo | 0 | 0 | 0 | 0 | 0 | 0 |
| 19108 | ENSMUSG00000066938 | 0 | NA | NA | NA | NA | NA | Gm10190 | predicted gene 10190 [Source:MGI Symbo | 0 | 0 | 0 | 0 | 0 | 0 |
| 19109 | ENSMUSG00000066952 | 0.315620088 | 0.165143421 | 4.063300985 | 0.040642675 | 0.967580763 | NA | Myo1h | myosin 1H [Source:MGI Symbol;Acc:MGI: | 0.8031 | 0 | 0 | 0 | 1.0906 | 0 |
| 19110 | ENSMUSG00000066958 | 0 | NA | NA | NA | NA | NA | Gm16391 | predicted pseudogene 16391 [Source:MG | 0 | 0 | 0 | 0 | 0 | 0 |
| 19111 | ENSMUSG00000066964 | 0 | NA | NA | NA | NA | NA | Tmem211 | transmembrane protein 211 [Source:MGI | 0 | 0 | 0 | 0 | 0 | 0 |
| 19112 | ENSMUSG00000066975 | 1.395586924 | -1.47417701 | 2.225058157 | -0.662534148 | 0.507628958 | NA | Cryba4 | crystallin, beta A4 [Source:MGI Symbol;Ac | 0 | 5.1396 | 1.0062 | 0 | 0 | 2.2277 |
| 19113 | ENSMUSG00000066979 | 421.5154867 | -0.008293699 | 0.142413107 | -0.058236907 | 0.953559923 | 0.9915542 | Bub3 | BUB3 mitotic checkpoint protein [Source:I | 422.43 | 458.45 | 387.4 | 417.4 | 460.24 | 383.16 |
| 19114 | ENSMUSG00000066983 | 0 | NA | NA | NA | NA | NA | Gm16519 | predicted gene, 16519 [Source:MGI Symb | 0 | 0 | 0 | 0 | 0 | 0 |
| 19115 | ENSMUSG00000066990 | 0 | NA | NA | NA | NA | NA | NA | NA | 0 | 0 | 0 | 0 | 0 | 0 |
| 19116 | ENSMUSG00000067001 | 0.535401493 | -2.48596965 | 3.684117979 | -0.674780141 | 0.499815459 | NA | Serpinb7 | serine (or cysteine) peptidase inhibitor, cl | 3.2124 | 0 | 0 | 0 | 0 | 0 |
| 19117 | ENSMUSG00000067006 | 0 | NA | NA | NA | NA | NA | Serpinb5 | serine (or cysteine) peptidase inhibitor, cl | 0 | 0 | 0 | 0 | 0 | 0 |
| 19118 | ENSMUSG00000067010 | 0 | NA | NA | NA | NA | NA | Vmn2r10 | vomeronasal 2, receptor 10 [Source:MGI S | 0 | 0 | 0 | 0 | 0 | 0 |
| 19119 | ENSMUSG00000067017 | 7.344131305 | -0.388964713 | 0.779011685 | -0.499305365 | 0.617564276 | 0.917509899 | Capza1-ps1 | capping protein (actin filament) muscle Z- | 9.6372 | 8.2233 | 7.0437 | 4.8199 | 6.5437 | 7.7969 |
| 19120 | ENSMUSG00000067028 | 0 | NA | NA | NA | NA | NA | Cntnap5b | contactin associated protein-like 5B [Sour | 0 | 0 | 0 | 0 | 0 | 0 |
| 19121 | ENSMUSG00000067038 | 2.193273044 | -0.228215424 | 1.538705042 | -0.14831655 | 0.882092955 | NA | Rps12-ps3 | ribosomal protein S12, pseudogene 3 [Sou | 4.0155 | 3.0838 | 0 | 3.8559 | 1.0906 | 1.1138 |
| 19122 | ENSMUSG00000067047 | 0 | NA | NA | NA | NA | NA | Olfr719-ps | olfactory receptor 719, pseudogene [Sour | 0 | 0 | 0 | 0 | 0 | 0 |
| 19123 | ENSMUSG00000067049 | 5.661514174 | -0.189931599 | 0.896900916 | -0.211764305 | 0.832290918 | 0.970533397 | Unc93a | unc-93 homolog A [Source:MGI Symbol;A | 8.8341 | 4.1117 | 5.0312 | 3.8559 | 5.4531 | 6.6831 |
| 19124 | ENSMUSG00000067058 | 0.349477146 | 0.165142768 | 3.911021834 | 0.042224967 | 0.966319359 | NA | Rps15a-ps5 | ribosomal protein S15A, pseudogene 5 [So | 0 | 0 | 1.0062 | 0 | 1.0906 | 0 |
| 19125 | ENSMUSG00000067063 | 0.167707431 | -0.796629176 | 4.080472857 | -0.195229623 | 0.84521319 | NA | Mif-ps9 | macrophage migration inhibitory factor, p | 0 | 0 | 1.0062 | 0 | 0 | 0 |
| 19126 | ENSMUSG00000067064 | 0 | NA | NA | NA | NA | NA | Olfr1416 | olfactory receptor 1416 [Source:MGI Sym | 0 | 0 | 0 | 0 | 0 | 0 |
| 19127 | ENSMUSG00000067071 | 1453.195927 | 0.813899947 | 0.128783946 | 6.319886711 | 2.62E-10 | 6.53E-08 | Hes6 | hairy and enhancer of split 6 [Source:MGI | 1125.1 | 889.15 | 1146.1 | 1764.1 | 2048.2 | 1746.5 |
| 19128 | ENSMUSG00000067081 | 0 | NA | NA | NA | NA | NA | Asb18 | ankyrin repeat and SOCS box-containing 1 | 0 | 0 | 0 | 0 | 0 | 0 |
| 19129 | ENSMUSG00000067085 | 0 | NA | NA | NA | NA | NA | Gm10197 | predicted gene 10197 [Source:MGI Symbo | 0 | 0 | 0 | 0 | 0 | 0 |
| 19130 | ENSMUSG00000067101 | 0 | NA | NA | NA | NA | NA | 1700010H22 | RIKEN cDNA 1700010H22 gene [Source:M | 0 | 0 | 0 | 0 | 0 | 0 |
| 19131 | ENSMUSG00000067103 | 0 | NA | NA | NA | NA | NA | AY702103 | cDNA sequence AY702103 [Source:MGI Sy | 0 | 0 | 0 | 0 | 0 | 0 |
| 19132 | ENSMUSG00000067106 | 2.709063032 | -0.577324224 | 1.432197636 | -0.40310374 | 0.686871909 | NA | Gm5529 | predicted pseudogene 5529 [Source:MGI | 1.6062 | 3.0838 | 5.0312 | 0.964 | 0 | 5.5692 |
| 19133 | ENSMUSG00000067121 | 1.282972534 | 0.279055159 | 1.879520567 | 0.148471458 | 0.881970709 | NA | Gm7027 | predicted gene 7027 [Source:MGI Symbol | 2.4093 | 0 | 1.0062 | 0.964 | 1.0906 | 2.2277 |
| 19134 | ENSMUSG00000067122 | 0 | NA | NA | NA | NA | NA | Gm10198 | predicted gene 10198 [Source:MGI Symbo | 0 | 0 | 0 | 0 | 0 | 0 |
| 19135 | ENSMUSG00000067144 | 1271.247202 | 0.688078744 | 0.165433762 | 4.159240152 | 3.19E-05 | 0.001456931 | Slc22a7 | solute carrier family 22 (organic anion tra | 1024 | 826.45 | 1069.6 | 1171.2 | 1842.1 | 1694.2 |
| 19136 | ENSMUSG00000067147 | 0 | NA | NA | NA | NA | NA | Rpl7a-ps11 | ribosomal protein L7A, pseudogene 11 [So | 0 | 0 | 0 | 0 | 0 | 0 |
| 19137 | ENSMUSG00000067148 | 300.433037 | 0.299958528 | 0.181940232 | 1.648665193 | 0.099216245 | 0.464258353 | Polr1c | polymerase (RNA) I polypeptide C [Source | 236.92 | 279.59 | 291.81 | 316.18 | 272.65 | 405.44 |
| 19138 | ENSMUSG00000067149 | 18.2720118 | -0.107324946 | 0.518290496 | -0.207074887 | 0.835951374 | 0.971073158 | Jchain | immunoglobulin joining chain [Source:MG | 24.093 | 16.447 | 16.1 | 15.424 | 14.178 | 23.391 |
| 19139 | ENSMUSG00000067150 | 316.7303563 | 0.024941555 | 0.151014632 | 0.165159856 | 0.868818162 | 0.97847745 | Xpo5 | exportin 5 [Source:MGI Symbol;Acc:MGI:1 | 291.53 | 325.85 | 325.02 | 309.44 | 352.27 | 296.28 |
| 19140 | ENSMUSG00000067156 | 0 | NA | NA | NA | NA | NA | Gm7337 | predicted gene 7337 [Source:MGI Symbol | 0 | 0 | 0 | 0 | 0 | 0 |
| 19141 | ENSMUSG00000067158 | 11.29044932 | -0.407956993 | 0.669082644 | -0.609725863 | 0.542043419 | 0.894071846 | Col4a4 | collagen, type IV, alpha 4 [Source:MGI Syr | 11.243 | 16.447 | 11.069 | 12.532 | 11.997 | 4.4554 |
| 19142 | ENSMUSG00000067161 | 2.762145282 | -2.011558086 | 1.63676496 | -1.228984085 | 0.219077773 | NA | Gm5560 | predicted pseudogene 5560 [Source:MGI | 0 | 8.2233 | 5.0312 | 0 | 1.0906 | 2.2277 |
| 19143 | ENSMUSG00000067173 | 0 | NA | NA | NA | NA | NA | Mrgpra4 | MAS-related GPR, member A4 [Source:MG | 0 | 0 | 0 | 0 | 0 | 0 |
| 19144 | ENSMUSG00000067186 | 0 | NA | NA | NA | NA | NA | Olfr132 | olfactory receptor 132 [Source:MGI Symb | 0 | 0 | 0 | 0 | 0 | 0 |
| 19145 | ENSMUSG00000067189 | 0.305170013 | -1.657836477 | 4.041480335 | -0.410205257 | 0.681655385 | NA | Gm7335 | predicted gene 7335 [Source:MGI Symbol | 0.8031 | 1.0279 | 0 | 0 | 0 | 0 |
| 19146 | ENSMUSG00000067194 | 699.8381846 | -0.132903788 | 0.122152152 | -1.088018395 | 0.276586987 | 0.730038567 | Eif1ax | eukaryotic translation initiation factor 1A, | 725.2 | 671.23 | 799.96 | 694.06 | 636.92 | 671.65 |
| 19147 | ENSMUSG00000067199 | 169.1224767 | -0.209663728 | 0.206497669 | -1.015332176 | 0.309947511 | 0.758686885 | Frat1 | frequently rearranged in advanced T cell l | 155.8 | 174.75 | 214.33 | 176.41 | 133.06 | 160.39 |
| 19148 | ENSMUSG00000067201 | 0 | NA | NA | NA | NA | NA | H2-M9 | histocompatibility 2, M region locus 9 [Sou | 0 | 0 | 0 | 0 | 0 | 0 |

| | | | | | | | | | | | | | | | |
|---|---|---|---|---|---|---|---|---|---|---|---|---|---|---|---|
| 19149 | ENSMUSG00000067203 | 117.4827981 | 0.089383018 | 0.24252116 | 0.368557608 | 0.712457498 | 0.944239215 | H2-K2 | histocompatibility 2, K region locus 2 [Sou | 121.27 | 123.35 | 96.599 | 104.11 | 154.87 | 104.7 |
| 19150 | ENSMUSG00000067206 | 0.48903338 | 0.95637552 | 3.456690196 | 0.276673773 | 0.782030612 | NA | Lrrc66 | leucine rich repeat containing 66 [Source: | 0 | 0 | 1.0062 | 1.928 | 0 | 0 |
| 19151 | ENSMUSG00000067212 | 4917.855912 | 0.110029646 | 0.099571251 | 1.105034277 | 0.269144737 | 0.724979259 | H2-T23 | histocompatibility 2, T region locus 23 [So | 4568.8 | 4641 | 4980.9 | 4630.9 | 5217.5 | 5467.9 |
| 19152 | ENSMUSG00000067215 | 0.342639279 | -1.805976596 | 4.028547714 | -0.448294702 | 0.653940524 | NA | Usp51 | ubiquitin specific protease 51 [Source:MG | 0 | 2.0558 | 0 | 0 | 0 | 0 |
| 19153 | ENSMUSG00000067219 | 72.74408966 | -1.363618444 | 1.109328596 | -1.229228607 | 0.218986106 | 0.664306846 | Nipal1 | NIPA-like domain containing 1 [Source:M( | 90.751 | 154.19 | 69.431 | 59.767 | 2.1812 | 60.148 |
| 19154 | ENSMUSG00000067220 | 0 | NA | NA | NA | NA | NA | Cnga1 | cyclic nucleotide gated channel alpha 1 [S | 0 | 0 | 0 | 0 | 0 | 0 |
| 19155 | ENSMUSG00000067224 | 0.631240176 | 0.380643921 | 3.15447101 | 0.120668068 | 0.903953952 | NA | Gm3695 | predicted gene 3695 [Source:MGI Symbol | 1.6062 | 0 | 0 | 0 | 2.1812 | 0 |
| 19156 | ENSMUSG00000067225 | 4237.620002 | -0.084903717 | 0.314463814 | -0.26999519 | 0.787163954 | 0.960870081 | Cyp2c54 | cytochrome P450, family 2, subfamily c, p | 2730.5 | 4562.9 | 5794 | 4441 | 3866.2 | 4031 |
| 19157 | ENSMUSG00000067229 | 0 | NA | NA | NA | NA | NA | Cyp2c66 | cytochrome P450, family 2, subfamily c, p | 0 | 0 | 0 | 0 | 0 | 0 |
| 19158 | ENSMUSG00000067231 | 0.133850373 | -0.796629176 | 4.080472857 | -0.195229623 | 0.84521319 | NA | Cyp2c65 | cytochrome P450, family 2, subfamily c, p | 0.8031 | 0 | 0 | 0 | 0 | 0 |
| 19159 | ENSMUSG00000067235 | 44556.03711 | 0.168317221 | 0.113080719 | 1.488469665 | 0.136627076 | 0.539790776 | H2-Q10 | histocompatibility 2, Q region locus 10 [So | 47478 | 42979 | 35421 | 46667 | 46929 | 47862 |
| 19160 | ENSMUSG00000067242 | 0 | NA | NA | NA | NA | NA | Lgi1 | leucine-rich repeat LGI family, member 1 | | 0 | 0 | 0 | 0 | 0 | 0 |
| 19161 | ENSMUSG00000067261 | 0 | NA | NA | NA | NA | NA | Foxd3 | forkhead box D3 [Source:MGI Symbol;Acc | 0 | 0 | 0 | 0 | 0 | 0 |
| 19162 | ENSMUSG00000067262 | 0 | NA | NA | NA | NA | NA | Vmn1r239-p | vomeronasal 1 receptor 239, pseudogene | 0 | 0 | 0 | 0 | 0 | 0 |
| 19163 | ENSMUSG00000067274 | 6742.715026 | 0.081914816 | 0.091858458 | 0.891750391 | 0.372526738 | 0.808133324 | Rplp0 | ribosomal protein, large, P0 [Source:MGI ! | 6198.3 | 6706.1 | 6749.9 | 6577.2 | 6861.1 | 7363.6 |
| 19164 | ENSMUSG00000067276 | 14.52405163 | -0.427142603 | 0.698556352 | -0.611464775 | 0.540891931 | 0.893466372 | Capn6 | calpain 6 [Source:MGI Symbol;Acc:MGI:1 | 31.321 | 10.279 | 8.05 | 6.7478 | 20.722 | 10.025 |
| 19165 | ENSMUSG00000067279 | 1287.05679 | -1.997473155 | 0.515997071 | -3.871093984 | 0.000108348 | 0.003859964 | Ppp1r3c | protein phosphatase 1, regulatory subuni | 1219.1 | 2895.6 | 2060.8 | 272.81 | 691.45 | 582.54 |
| 19166 | ENSMUSG00000067285 | 0 | NA | NA | NA | NA | NA | Gm16223 | predicted gene 16223 [Source:MGI Symbo | 0 | 0 | 0 | 0 | 0 | 0 |
| 19167 | ENSMUSG00000067288 | 861.7289201 | -0.146009552 | 0.15349655 | -0.951223671 | 0.341490845 | 0.78327364 | Rps28 | ribosomal protein S28 [Source:MGI Symbo | 750.9 | 1077.3 | 888.51 | 773.11 | 741.62 | 938.97 |
| 19168 | ENSMUSG00000067292 | 0 | NA | NA | NA | NA | NA | Gm10203 | predicted gene 10203 [Source:MGI Symbo | 0 | 0 | 0 | 0 | 0 | 0 |
| 19169 | ENSMUSG00000067297 | 1.88219058 | 0.798301626 | 1.751732767 | 0.455721124 | 0.64859053 | NA | Ifit1bl2 | interferon induced protein with tetratrico | 4.0155 | 0 | 0 | 2.8919 | 3.2719 | 1.1138 |
| 19170 | ENSMUSG00000067299 | 0 | NA | NA | NA | NA | NA | Crygd | crystallin, gamma D [Source:MGI Symbol;/ | 0 | 0 | 0 | 0 | 0 | 0 |
| 19171 | ENSMUSG00000067321 | 0 | NA | NA | NA | NA | NA | Gm7931 | predicted pseudogene 7931 [Source:MGI | 0 | 0 | 0 | 0 | 0 | 0 |
| 19172 | ENSMUSG00000067331 | 0 | NA | NA | NA | NA | NA | Gm10205 | predicted gene 10205 [Source:MGI Symbo | 0 | 0 | 0 | 0 | 0 | 0 |
| 19173 | ENSMUSG00000067336 | 649.1381504 | -0.175925902 | 0.1253537 | -1.403436056 | 0.160486848 | 0.583090119 | Bmpr2 | bone morphogenetic protein receptor, typ | 651.32 | 758.6 | 657.08 | 649.72 | 591.12 | 587 |
| 19174 | ENSMUSG00000067338 | 0 | NA | NA | NA | NA | NA | Tuba3b | tubulin, alpha 3B [Source:MGI Symbol;Acc | 0 | 0 | 0 | 0 | 0 | 0 |
| 19175 | ENSMUSG00000067341 | 0 | NA | NA | NA | NA | NA | H2-Eb2 | histocompatibility 2, class II antigen E beta | 0 | 0 | 0 | 0 | 0 | 0 |
| 19176 | ENSMUSG00000067344 | 1.737813023 | 0.671249826 | 1.665698391 | 0.402984016 | 0.686959984 | NA | Rps25-ps1 | ribosomal protein S25, pseudogene 1 [Sou | 0 | 3.0838 | 1.0062 | 1.928 | 2.1812 | 2.2277 |
| 19177 | ENSMUSG00000067351 | 0 | NA | NA | NA | NA | NA | Rps15a-ps2 | ribosomal protein S15A, pseudogene 2 [S | 0 | 0 | 0 | 0 | 0 | 0 |
| 19178 | ENSMUSG00000067352 | 0 | NA | NA | NA | NA | NA | Gm14149 | predicted gene 14149 [Source:MGI Symbo | 0 | 0 | 0 | 0 | 0 | 0 |
| 19179 | ENSMUSG00000067356 | 2.994443784 | -0.559299317 | 1.25679612 | -0.445019927 | 0.656305354 | NA | B430203G13 | RIKEN cDNA B430203G13 gene [Source:M | 1.6062 | 6.1675 | 3.0187 | 3.8559 | 1.0906 | 2.2277 |
| 19180 | ENSMUSG00000067360 | 0 | NA | NA | NA | NA | NA | Pramel3 | PRAME like 3 [Source:MGI Symbol;Acc:M( | 0 | 0 | 0 | 0 | 0 | 0 |
| 19181 | ENSMUSG00000067365 | 247.2627451 | 0.0606518 | 0.163550676 | 0.37084408 | 0.710753667 | 0.944239215 | Tmem128 | transmembrane protein 128 [Source:MGI | 220.05 | 255.95 | 250.55 | 237.14 | 274.84 | 245.05 |
| 19182 | ENSMUSG00000067367 | 217.1088015 | -0.133022736 | 0.178239025 | -0.74631656 | 0.45547621 | 0.857366983 | Lyar | Ly1 antibody reactive clone [Source:MGI S | 216.03 | 263.15 | 202.26 | 193.76 | 221.4 | 206.06 |
| 19183 | ENSMUSG00000067369 | 676.6278676 | 0.060651631 | 0.11957899 | 0.507209757 | 0.61200765 | 0.916441371 | Trmt2b | TRM2 tRNA methyltransferase 2B [Source | 638.47 | 683.57 | 665.13 | 638.15 | 740.53 | 693.92 |
| 19184 | ENSMUSG00000067370 | 3.276549815 | -0.845868551 | 1.347551223 | -0.627707902 | 0.530195308 | NA | B3galt4 | UDP-Gal:betaGlcNAc beta 1,3-galactosyltr | 1.6062 | 4.1117 | 7.0437 | 5.7839 | 0 | 1.1138 |
| 19185 | ENSMUSG00000067377 | 4.244639133 | -0.727277489 | 1.044154009 | -0.696523198 | 0.486101235 | NA | Tspan6 | tetraspanin 6 [Source:MGI Symbol;Acc:M( | 5.6217 | 7.1954 | 3.0187 | 1.928 | 4.3625 | 3.3415 |
| 19186 | ENSMUSG00000067389 | 35.42475672 | -0.61639201 | 0.51280014 | -1.202012172 | 0.229358812 | 0.675988467 | Gm17080 | predicted gene 17080 [Source:MGI Symbo | 31.321 | 42.145 | 55.343 | 27.955 | 47.987 | 7.7969 |
| 19187 | ENSMUSG00000067399 | 3.754480677 | -0.309562175 | 1.191039338 | -0.259909278 | 0.794933754 | NA | Trim43c | tripartite motif-containing 43C [Source:M | 6.4248 | 1.0279 | 5.0312 | 6.7478 | 2.1812 | 1.1138 |
| 19188 | ENSMUSG00000067424 | 48.5187629 | -0.28303623 | 0.360704705 | -0.784675737 | 0.432643712 | 0.845264432 | Zfp563 | zinc finger protein 563 [Source:MGI Symb | 43.368 | 52.424 | 64.4 | 56.875 | 27.265 | 46.781 |
| 19189 | ENSMUSG00000067430 | 73.41071622 | 0.323707997 | 0.293133065 | 1.104303933 | 0.269461317 | 0.724979259 | Zfp763 | zinc finger protein 763 [Source:MGI Symb | 79.507 | 49.34 | 66.412 | 90.614 | 63.256 | 91.335 |
| 19190 | ENSMUSG00000067438 | 0.371281295 | 2.09253362 | 4.031852165 | 0.519000582 | 0.603760335 | NA | Hmx1 | H6 homeobox 1 [Source:MGI Symbol;Acc: | 0 | 0 | 0 | 0 | 0 | 2.2277 |
| 19191 | ENSMUSG00000067441 | 0 | NA | NA | NA | NA | NA | H2ab1 | H2A.B variant histone 1 [Source:MGI Sym | 0 | 0 | 0 | 0 | 0 | 0 |
| 19192 | ENSMUSG00000067455 | 1.409805081 | -1.525348824 | 2.144187023 | -0.711387956 | 0.47684386 | NA | H4c11 | H4 clustered histone 11 [Source:MGI Sym | 3.2124 | 0 | 3.0187 | 0 | 0 | 2.2277 |
| 19193 | ENSMUSG00000067482 | 0 | NA | NA | NA | NA | NA | Olfr1456-ps1 | olfactory receptor 1456, pseudogene 1 [S | 0 | 0 | 0 | 0 | 0 | 0 |
| 19194 | ENSMUSG00000067513 | 0 | NA | NA | NA | NA | NA | Olfr1436 | olfactory receptor 1436 [Source:MGI Sym | 0 | 0 | 0 | 0 | 0 | 0 |
| 19195 | ENSMUSG00000067519 | 0 | NA | NA | NA | NA | NA | Olfr262 | olfactory receptor 262 [Source:MGI Symb | 0 | 0 | 0 | 0 | 0 | 0 |
| 19196 | ENSMUSG00000067522 | 0 | NA | NA | NA | NA | NA | Olfr76 | olfactory receptor 76 [Source:MGI Symbo | 0 | 0 | 0 | 0 | 0 | 0 |
| 19197 | ENSMUSG00000067524 | 0 | NA | NA | NA | NA | NA | Olfr1428 | olfactory receptor 1428 [Source:MGI Sym | 0 | 0 | 0 | 0 | 0 | 0 |
| 19198 | ENSMUSG00000067525 | 0 | NA | NA | NA | NA | NA | Olfr1427 | olfactory receptor 1427 [Source:MGI Sym | 0 | 0 | 0 | 0 | 0 | 0 |
| 19199 | ENSMUSG00000067526 | 0 | NA | NA | NA | NA | NA | Olfr1425 | olfactory receptor 1425 [Source:MGI Sym | 0 | 0 | 0 | 0 | 0 | 0 |
| 19200 | ENSMUSG00000067528 | 0 | NA | NA | NA | NA | NA | Olfr1424 | olfactory receptor 1424 [Source:MGI Sym | 0 | 0 | 0 | 0 | 0 | 0 |
| 19201 | ENSMUSG00000067529 | 0 | NA | NA | NA | NA | NA | Olfr1423 | olfactory receptor 1423 [Source:MGI Sym | 0 | 0 | 0 | 0 | 0 | 0 |
| 19202 | ENSMUSG00000067541 | 0 | NA | NA | NA | NA | NA | A630073D07 | RIKEN cDNA A630073D07 gene [Source:M | 0 | 0 | 0 | 0 | 0 | 0 |
| 19203 | ENSMUSG00000067543 | 0 | NA | NA | NA | NA | NA | Prb1 | proline-rich protein BstNI subfamily 1 [Sou | 0 | 0 | 0 | 0 | 0 | 0 |
| 19204 | ENSMUSG00000067545 | 0 | NA | NA | NA | NA | NA | Olfr1419 | olfactory receptor 1419 [Source:MGI Sym | 0 | 0 | 0 | 0 | 0 | 0 |

| | | | | | | | | | | | | | | | |
|---|---|---|---|---|---|---|---|---|---|---|---|---|---|---|---|
| 19205 | ENSMUSG00000067547 | 5.748944301 | -0.238080181 | 0.867528209 | -0.274435088 | 0.783750281 | 0.960670212 | Gm7666 | predicted pseudogene 7666 [Source:MGI | 6.4248 | 7.1954 | 5.0312 | 4.8199 | 5.4531 | 5.5692 |
| 19206 | ENSMUSG00000067555 | 0.267700746 | -1.497481649 | 4.057001436 | -0.369110456 | 0.712045396 | NA | Gm10212 | predicted pseudogene 10212 [Source:MG | 1.6062 | 0 | 0 | 0 | 0 | 0 |
| 19207 | ENSMUSG00000067561 | 0 | NA | NA | NA | NA | NA | Dmrtc1c2 | DMRT-like family C1c2 [Source:MGI Symb | 0 | 0 | 0 | 0 | 0 | 0 |
| 19208 | ENSMUSG00000067562 | 0 | NA | NA | NA | NA | NA | Dmrtc1c1 | DMRT-like family C1c1 [Source:MGI Symb | 0 | 0 | 0 | 0 | 0 | 0 |
| 19209 | ENSMUSG00000067567 | 75.20834171 | 0.151067177 | 0.272737091 | 0.553893042 | 0.579652047 | 0.908350165 | Hdac8 | histone deacetylase 8 [Source:MGI Symbo | 63.445 | 77.094 | 73.456 | 77.118 | 64.346 | 95.791 |
| 19210 | ENSMUSG00000067571 | 0 | NA | NA | NA | NA | NA | Ms4a15 | membrane-spanning 4-domains, subfamil | 0 | 0 | 0 | 0 | 0 | 0 |
| 19211 | ENSMUSG00000067575 | 0.538730002 | 1.11098254 | 3.037664833 | 0.365735722 | 0.714562288 | NA | Rpl35a-ps3 | ribosomal protein L35A, pseudogene 3 [So | 0 | 1.0279 | 0 | 0 | 1.0906 | 1.1138 |
| 19212 | ENSMUSG00000067577 | 0.356960287 | 0.16514264 | 3.880370363 | 0.042558474 | 0.966053499 | NA | A430093F15 | RIKEN cDNA A430093F15 gene [Source:M | 0 | 1.0279 | 0 | 0 | 0 | 1.1138 |
| 19213 | ENSMUSG00000067578 | 0 | NA | NA | NA | NA | NA | Cbln4 | cerebellin 4 precursor protein [Source:MC | 0 | 0 | 0 | 0 | 0 | 0 |
| 19214 | ENSMUSG00000067586 | 41.21113487 | -1.34165393 | 0.388103404 | -3.456949658 | 0.000546327 | 0.013192107 | S1pr3 | sphingosine-1-phosphate receptor 3 [Sou | 68.264 | 49.34 | 59.368 | 18.316 | 17.45 | 34.529 |
| 19215 | ENSMUSG00000067591 | 0.877498349 | -0.489540585 | 2.815048567 | -0.173901293 | 0.861943037 | NA | Klra3 | killer cell lectin-like receptor, subfamily A, | 0 | 3.0838 | 0 | 0 | 2.1812 | 0 |
| 19216 | ENSMUSG00000067594 | 0 | NA | NA | NA | NA | NA | Krt77 | keratin 77 [Source:MGI Symbol;Acc:MGI:3 | 0 | 0 | 0 | 0 | 0 | 0 |
| 19217 | ENSMUSG00000067596 | 0 | NA | NA | NA | NA | NA | Krt74 | keratin 74 [Source:MGI Symbol;Acc:MGI:3 | 0 | 0 | 0 | 0 | 0 | 0 |
| 19218 | ENSMUSG00000067597 | 0.160662974 | 1.126924175 | 4.080472857 | 0.276174898 | 0.782413736 | NA | Dgat2l6 | diacylglycerol O-acyltransferase 2-like 6 [S | 0 | 0 | 0 | 0.964 | 0 | 0 |
| 19219 | ENSMUSG00000067599 | 0.185640647 | 1.126924175 | 4.080472857 | 0.276174898 | 0.782413736 | NA | Klra7 | killer cell lectin-like receptor, subfamily A, | 0 | 0 | 0 | 0 | 0 | 1.1138 |
| 19220 | ENSMUSG00000067604 | 0 | NA | NA | NA | NA | NA | Nms | neuromedin S [Source:MGI Symbol;Acc:M | 0 | 0 | 0 | 0 | 0 | 0 |
| 19221 | ENSMUSG00000067608 | 0.815310133 | -0.44746426 | 2.258708535 | -0.198106242 | 0.842961941 | NA | Pcna-ps2 | proliferating cell nuclear antigen pseudog | 0.8031 | 1.0279 | 1.0062 | 0.964 | 1.0906 | 0 |
| 19222 | ENSMUSG00000067610 | 0.439020386 | -2.189747911 | 3.52148864 | -0.621824499 | 0.534057275 | NA | Klri1 | killer cell lectin-like receptor family I mem | 1.6062 | 1.0279 | 0 | 0 | 0 | 0 |
| 19223 | ENSMUSG00000067613 | 0 | NA | NA | NA | NA | NA | Krt83 | keratin 83 [Source:MGI Symbol;Acc:MGI:3 | 0 | 0 | 0 | 0 | 0 | 0 |
| 19224 | ENSMUSG00000067614 | 0 | NA | NA | NA | NA | NA | Krt86 | keratin 86 [Source:MGI Symbol;Acc:MGI:1 | 0 | 0 | 0 | 0 | 0 | 0 |
| 19225 | ENSMUSG00000067615 | 0 | NA | NA | NA | NA | NA | Krt81 | keratin 81 [Source:MGI Symbol;Acc:MGI:1 | 0 | 0 | 0 | 0 | 0 | 0 |
| 19226 | ENSMUSG00000067616 | 0 | NA | NA | NA | NA | NA | Klk11 | kallikrein related-peptidase 11 [Source:M | 0 | 0 | 0 | 0 | 0 | 0 |
| 19227 | ENSMUSG00000067627 | 0 | NA | NA | NA | NA | NA | Gm10217 | predicted gene 10217 [Source:MGI Symbo | 0 | 0 | 0 | 0 | 0 | 0 |
| 19228 | ENSMUSG00000067629 | 26.78631474 | -0.047309134 | 0.464724395 | -0.10180041 | 0.918915101 | 0.986374671 | Syngap1 | synaptic Ras GTPase activating protein 1 h | 24.896 | 21.586 | 35.219 | 26.991 | 15.269 | 36.757 |
| 19229 | ENSMUSG00000067642 | 0.331982614 | 0.16514309 | 3.986751476 | 0.041422971 | 0.966958701 | NA | Adgrf3 | adhesion G protein-coupled receptor F3 [ | 0 | 1.0279 | 0 | 0.964 | 0 | 0 |
| 19230 | ENSMUSG00000067649 | 0 | NA | NA | NA | NA | NA | Mageb18 | MAGE family member B18 [Source:MGI Sy | 0 | 0 | 0 | 0 | 0 | 0 |
| 19231 | ENSMUSG00000067653 | 4.683170117 | -0.963932564 | 1.190250894 | -0.809856619 | 0.418022587 | NA | Ankrd23 | ankyrin repeat domain 23 [Source:MGI Sy | 10.44 | 0 | 8.05 | 1.928 | 5.4531 | 2.2277 |
| 19232 | ENSMUSG00000067656 | 67.76090142 | -2.376357626 | 0.922622144 | -2.575656396 | 0.010005002 | 0.108724524 | Slc22a27 | solute carrier family 22, member 27 [Sour | 111.63 | 148.02 | 81.506 | 51.091 | 7.6343 | 6.6831 |
| 19233 | ENSMUSG00000067679 | 0 | NA | NA | NA | NA | NA | Obp1b | odorant binding protein IB [Source:MGI S | 0 | 0 | 0 | 0 | 0 | 0 |
| 19234 | ENSMUSG00000067684 | 0 | NA | NA | NA | NA | NA | Obp1a | odorant binding protein IA [Source:MGI S | 0 | 0 | 0 | 0 | 0 | 0 |
| 19235 | ENSMUSG00000067698 | 0 | NA | NA | NA | NA | NA | Gm10220 | predicted gene 10220 [Source:MGI Symbo | 0 | 0 | 0 | 0 | 0 | 0 |
| 19236 | ENSMUSG00000067700 | 0 | NA | NA | NA | NA | NA | Gm5862 | predicted gene 5862 [Source:MGI Symbol | 0 | 0 | 0 | 0 | 0 | 0 |
| 19237 | ENSMUSG00000067702 | 0 | NA | NA | NA | NA | NA | Tuba3a | tubulin, alpha 3A [Source:MGI Symbol;Acc | 0 | 0 | 0 | 0 | 0 | 0 |
| 19238 | ENSMUSG00000067704 | 0 | NA | NA | NA | NA | NA | Wfdc13 | WAP four-disulfide core domain 13 [Sourc | 0 | 0 | 0 | 0 | 0 | 0 |
| 19239 | ENSMUSG00000067713 | 241.4024981 | -0.059149649 | 0.160502712 | -0.368527412 | 0.712480009 | 0.944239215 | Prkag1 | protein kinase, AMP-activated, gamma 1 r | 265.83 | 238.48 | 234.45 | 245.81 | 238.85 | 225 |
| 19240 | ENSMUSG00000067714 | 2.969234017 | -1.746694063 | 1.401554462 | -1.246254862 | 0.212670843 | NA | Lpar5 | lysophosphatidic acid receptor 5 [Source:I | 1.6062 | 7.1954 | 5.0312 | 2.8919 | 1.0906 | 0 |
| 19241 | ENSMUSG00000067716 | 0 | NA | NA | NA | NA | NA | Gm7275 | predicted gene 7275 [Source:MGI Symbol | 0 | 0 | 0 | 0 | 0 | 0 |
| 19242 | ENSMUSG00000067719 | 17.64458628 | 0.643511599 | 0.512512242 | 1.255602395 | 0.209260163 | 0.65350014 | Gm10221 | predicted gene 10221 [Source:MGI Symbo | 12.85 | 14.391 | 14.087 | 18.316 | 26.175 | 20.049 |
| 19243 | ENSMUSG00000067722 | 568.3614994 | 0.146973256 | 0.138659005 | 1.059961854 | 0.289161954 | 0.7396939 | BC003965 | cDNA sequence BC003965 [Source:MGI Sy | 592.69 | 472.84 | 551.42 | 539.83 | 612.93 | 640.46 |
| 19244 | ENSMUSG00000067724 | 0 | NA | NA | NA | NA | NA | Gbx1 | gastrulation brain homeobox 1 [Source:M | 0 | 0 | 0 | 0 | 0 | 0 |
| 19245 | ENSMUSG00000067736 | 117.9776732 | -0.104618349 | 0.230741063 | -0.45340152 | 0.650259642 | 0.925643431 | Gm10222 | predicted gene 10222 [Source:MGI Symbo | 105.21 | 116.15 | 145.91 | 124.35 | 97.065 | 119.18 |
| 19246 | ENSMUSG00000067745 | 0 | NA | NA | NA | NA | NA | Gm14243 | predicted gene 14243 [Source:MGI Symbo | 0 | 0 | 0 | 0 | 0 | 0 |
| 19247 | ENSMUSG00000067750 | 0 | NA | NA | NA | NA | NA | Khdc1a | KH domain containing 1A [Source:MGI Syr | 0 | 0 | 0 | 0 | 0 | 0 |
| 19248 | ENSMUSG00000067764 | 0 | NA | NA | NA | NA | NA | Xlr5c | X-linked lymphocyte-regulated 5C [Source | 0 | 0 | 0 | 0 | 0 | 0 |
| 19249 | ENSMUSG00000067767 | 0 | NA | NA | NA | NA | NA | Clec4b2 | C-type lectin domain family 4, member b2 | 0 | 0 | 0 | 0 | 0 | 0 |
| 19250 | ENSMUSG00000067768 | 6.454611134 | -6.059464094 | 1.434996026 | -4.222634756 | 2.41E-05 | 0.001126619 | Xlr4b | X-linked lymphocyte-regulated 4B [Source | 15.259 | 15.419 | 8.05 | 0 | 0 | 0 |
| 19251 | ENSMUSG00000067771 | 0 | NA | NA | NA | NA | NA | Pwwp4a | PWWP domain containing 4A [Source:MG | 0 | 0 | 0 | 0 | 0 | 0 |
| 19252 | ENSMUSG00000067773 | 0 | NA | NA | NA | NA | NA | Defb41 | defensin beta 41 [Source:MGI Symbol;Acc | 0 | 0 | 0 | 0 | 0 | 0 |
| 19253 | ENSMUSG00000067780 | 1.459713807 | -3.904553905 | 2.082760685 | -1.874701176 | 0.060833845 | NA | Pi15 | peptidase inhibitor 15 [Source:MGI Symb | 1.6062 | 5.1396 | 2.0125 | 0 | 0 | 0 |
| 19254 | ENSMUSG00000067786 | 21.40265938 | -0.439180125 | 0.549865828 | -0.798704161 | 0.424461973 | 0.840450984 | Nnat | neuronatin [Source:MGI Symbol;Acc:MGI | 18.471 | 19.53 | 36.225 | 28.919 | 16.359 | 8.9108 |
| 19255 | ENSMUSG00000067787 | 464.1726901 | -0.004393643 | 0.135823491 | -0.032348182 | 0.974194386 | 0.996336532 | Blcap | bladder cancer associated protein [Source | 432.07 | 464.62 | 498.09 | 430.9 | 454.79 | 504.57 |
| 19256 | ENSMUSG00000067795 | 0 | NA | NA | NA | NA | NA | 4930444P10 | RIKEN cDNA 4930444P10 gene [Source:M | 0 | 0 | 0 | 0 | 0 | 0 |
| 19257 | ENSMUSG00000067813 | 247.2713141 | -0.007533342 | 0.184803604 | -0.040764044 | 0.967484004 | 0.994231936 | Xkr9 | X-linked Kx blood group related 9 [Source | 297.95 | 241.56 | 203.26 | 234.25 | 270.47 | 236.13 |
| 19258 | ENSMUSG00000067818 | 21.21981262 | -0.300126234 | 0.504277463 | -0.59516091 | 0.551735917 | 0.895935901 | Myl9 | myosin, light polypeptide 9, regulatory [Sc | 22.487 | 31.865 | 16.1 | 26.027 | 16.359 | 14.48 |
| 19259 | ENSMUSG00000067825 | 985.6214825 | 0.781037132 | 0.199287804 | 3.919141638 | 8.89E-05 | 0.003295748 | Pex26 | peroxisomal biogenesis factor 26 [Source: | 818.36 | 657.87 | 699.34 | 1717.8 | 1158.2 | 862.12 |
| 19260 | ENSMUSG00000067835 | 0.181769715 | 1.126924175 | 4.080472857 | 0.276174898 | 0.782413736 | NA | Gm14661 | predicted gene 14661 [Source:MGI Symbo | 0 | 0 | 0 | 0 | 1.0906 | 0 |

| | | | | | | | | | | | | | | | |
|---|---|---|---|---|---|---|---|---|---|---|---|---|---|---|---|
| 19261 | ENSMUSG00000067847 | 696.2814421 | 0.010470075 | 0.135325585 | 0.077369515 | 0.938329592 | 0.988420305 | Romo1 | reactive oxygen species modulator 1 [Sou | 640.88 | 705.15 | 735.56 | 675.75 | 618.38 | 801.97 |
| 19262 | ENSMUSG00000067848 | 0 | NA | NA | NA | NA | NA | 4933402N22 | RIKEN cDNA 4933402N22 gene [Source:M | 0 | 0 | 0 | 0 | 0 | 0 |
| 19263 | ENSMUSG00000067851 | 1676.481506 | 0.002022527 | 0.107201948 | 0.018866513 | 0.984947594 | 0.99859176 | Arfgef1 | ADP-ribosylation factor guanine nucleotid | 1540.4 | 1764.9 | 1721.7 | 1655.1 | 1797.3 | 1579.4 |
| 19264 | ENSMUSG00000067855 | 0 | NA | NA | NA | NA | NA | Speer3 | spermatogenesis associated glutamate (E) | 0 | 0 | 0 | 0 | 0 | 0 |
| 19265 | ENSMUSG00000067860 | 0 | NA | NA | NA | NA | NA | Zic3 | zinc finger protein of the cerebellum 3 [Sc | 0 | 0 | 0 | 0 | 0 | 0 |
| 19266 | ENSMUSG00000067869 | 3.37041716 | -0.740308872 | 1.221283512 | -0.606172821 | 0.544399999 | NA | Tcea1-ps1 | transcription elongation factor A (SII) 1, ps | 2.4093 | 8.2233 | 2.0125 | 0.964 | 3.2719 | 3.3415 |
| 19267 | ENSMUSG00000067870 | 20.15084742 | 0.700171665 | 0.508810516 | 1.376095113 | 0.168792188 | 0.595822959 | Rpl31-ps8 | ribosomal protein L31, pseudogene 8 [Sou | 14.456 | 22.614 | 9.0562 | 24.099 | 26.175 | 24.505 |
| 19268 | ENSMUSG00000067872 | 1.811568832 | 0.904093907 | 1.798169257 | 0.502785766 | 0.615114902 | NA | Ccdc87 | coiled-coil domain containing 87 [Source:I | 0.8031 | 0 | 3.0187 | 4.8199 | 0 | 2.2277 |
| 19269 | ENSMUSG00000067873 | 546.8685165 | 0.050977697 | 0.127914102 | 0.398530703 | 0.690239031 | 0.937454779 | Htatsf1 | HIV TAT specific factor 1 [Source:MGI Sym | 495.51 | 548.91 | 567.52 | 517.66 | 580.21 | 571.4 |
| 19270 | ENSMUSG00000067878 | 0.517184577 | -0.812115922 | 3.393570547 | -0.239310163 | 0.810865085 | NA | Map7d3 | MAP7 domain containing 3 [Source:MGI S | 0 | 0 | 2.0125 | 0 | 1.0906 | 0 |
| 19271 | ENSMUSG00000067879 | 0.52105551 | -0.812028029 | 3.385364865 | -0.239864257 | 0.810435491 | NA | Vxn | vexin [Source:MGI Symbol;Acc:MGI:19242 | 0 | 0 | 2.0125 | 0 | 0 | 1.1138 |
| 19272 | ENSMUSG00000067882 | 15.45161934 | 0.109918055 | 0.629832798 | 0.174519421 | 0.861457269 | 0.976192255 | Casp16 | caspase 16, apoptosis-related cysteine pe | 19.274 | 8.2233 | 17.106 | 25.063 | 16.359 | 6.6831 |
| 19273 | ENSMUSG00000067889 | 1789.024456 | 0.42057844 | 0.148536031 | 2.831491039 | 0.004633153 | 0.063460159 | Sptbn2 | spectrin beta, non-erythrocytic 2 [Source: | 1313.9 | 1331.2 | 1946.1 | 1989.7 | 2150.7 | 2002.7 |
| 19274 | ENSMUSG00000067909 | 0 | NA | NA | NA | NA | NA | Slxl1 | Slx-like 1 [Source:MGI Symbol;Acc:MGI:19 | 0 | 0 | 0 | 0 | 0 | 0 |
| 19275 | ENSMUSG00000067916 | 10.60224992 | -0.813019902 | 0.660270381 | -1.231343895 | 0.218194271 | 0.663697152 | Zfp991 | zinc finger protein 991 [Source:MGI Symb | 16.062 | 13.363 | 11.069 | 7.7118 | 8.7249 | 6.6831 |
| 19276 | ENSMUSG00000067919 | 0.524202404 | 2.600864256 | 3.301637836 | 0.787749713 | 0.430843121 | NA | Rex2 | reduced expression 2 [Source:MGI Symbo | 0 | 0 | 0 | 0.964 | 2.1812 | 0 |
| 19277 | ENSMUSG00000067924 | 30.62963689 | -0.742016872 | 0.413642307 | -1.793861169 | 0.072835331 | 0.391632053 | Rtl8b | retrotransposon Gag like 8B [Source:MGI | 41.761 | 43.173 | 30.187 | 27.955 | 23.994 | 16.708 |
| 19278 | ENSMUSG00000067925 | 48.20023786 | -0.667831699 | 0.334458933 | -1.996752464 | 0.04585208 | 0.296229283 | Rtl8a | retrotransposon Gag like 8A [Source:MGI | 71.476 | 47.284 | 58.362 | 33.739 | 34.9 | 43.44 |
| 19279 | ENSMUSG00000067928 | 110.9797031 | 0.550970031 | 0.290785021 | 1.89476758 | 0.0581232 | 0.342581078 | Zfp760 | zinc finger protein 760 [Source:MGI Symb | 122.07 | 65.787 | 81.506 | 137.85 | 92.703 | 165.96 |
| 19280 | ENSMUSG00000067929 | 12.41626325 | 0.122344553 | 0.616732874 | 0.198375275 | 0.842751461 | 0.97213152 | Gm10226 | predicted gene 10226 [Source:MGI Symbo | 15.259 | 7.1954 | 13.081 | 12.532 | 14.178 | 12.252 |
| 19281 | ENSMUSG00000067931 | 58.85050711 | -0.123933285 | 0.295451411 | -0.419470954 | 0.674871978 | 0.932315978 | Zfp948 | zinc finger protein 948 [Source:MGI Symb | 65.051 | 51.396 | 67.418 | 50.127 | 55.622 | 63.489 |
| 19282 | ENSMUSG00000067941 | 0 | NA | NA | NA | NA | NA | Gm7168 | predicted gene 7168 [Source:MGI Symbol | 0 | 0 | 0 | 0 | 0 | 0 |
| 19283 | ENSMUSG00000067942 | 145.1904045 | -0.097224839 | 0.193236234 | -0.503139796 | 0.614865989 | 0.917509899 | Zfp160 | zinc finger protein 160 [Source:MGI Symb | 146.97 | 158.3 | 144.9 | 131.1 | 143.96 | 145.91 |
| 19284 | ENSMUSG00000067949 | 0 | NA | NA | NA | NA | NA | Vmn1r-ps14 | vomeronasal 1 receptor, pseudogene 144 | 0 | 0 | 0 | 0 | 0 | 0 |
| 19285 | ENSMUSG00000067951 | 0 | NA | NA | NA | NA | NA | Vmn1r227 | vomeronasal 1 receptor 227 [Source:MGI | 0 | 0 | 0 | 0 | 0 | 0 |
| 19286 | ENSMUSG00000067971 | 0 | NA | NA | NA | NA | NA | Olfr1321 | olfactory receptor 1321 [Source:MGI Sym | 0 | 0 | 0 | 0 | 0 | 0 |
| 19287 | ENSMUSG00000067973 | 0 | NA | NA | NA | NA | NA | Gm6994 | predicted gene 6994 [Source:MGI Symbol | 0 | 0 | 0 | 0 | 0 | 0 |
| 19288 | ENSMUSG00000067995 | 157.3222856 | -0.022878752 | 0.189267856 | -0.120880281 | 0.903785861 | 0.983821605 | Gtf2f2 | general transcription factor IIF, polypeptic | 160.62 | 160.36 | 154.96 | 172.55 | 143.96 | 151.48 |
| 19289 | ENSMUSG00000067996 | 0 | NA | NA | NA | NA | NA | Bpifb9b | BPI fold containing family B, member 9B [ | 0 | 0 | 0 | 0 | 0 | 0 |
| 19290 | ENSMUSG00000067998 | 0 | NA | NA | NA | NA | NA | Bpifb9a | BPI fold containing family B, member 9A [ | 0 | 0 | 0 | 0 | 0 | 0 |
| 19291 | ENSMUSG00000068008 | 0 | NA | NA | NA | NA | NA | Bpifb3 | BPI fold containing family B, member 3 [Sc | 0 | 0 | 0 | 0 | 0 | 0 |
| 19292 | ENSMUSG00000068009 | 0 | NA | NA | NA | NA | NA | Bpifb6 | BPI fold containing family B, member 6 [Sc | 0 | 0 | 0 | 0 | 0 | 0 |
| 19293 | ENSMUSG00000068011 | 319.6567428 | 0.00943868 | 0.151211435 | 0.062420411 | 0.950228041 | 0.990815588 | Mkrn2os | makorin, ring finger protein 2, opposite st | 337.3 | 287.82 | 330.05 | 300.76 | 347.91 | 314.1 |
| 19294 | ENSMUSG00000068014 | 0 | NA | NA | NA | NA | NA | Gm14648 | predicted gene 14648 [Source:MGI Symbo | 0 | 0 | 0 | 0 | 0 | 0 |
| 19295 | ENSMUSG00000068015 | 160.0865937 | -0.024774942 | 0.196457413 | -0.126108461 | 0.89964607 | 0.982798755 | Lrch1 | leucine-rich repeats and calponin homolo | 168.65 | 174.75 | 140.87 | 162.91 | 140.69 | 172.65 |
| 19296 | ENSMUSG00000068036 | 1027.481782 | 0.165088799 | 0.102110377 | 1.616768083 | 0.105928351 | 0.479845811 | Afdn | afadin, adherens junction formation facto | 991.83 | 970.35 | 943.86 | 1102.8 | 1062.3 | 1093.8 |
| 19297 | ENSMUSG00000068037 | 7.13841663 | -1.455732031 | 0.892020237 | -1.631949557 | 0.102690113 | 0.471078273 | Mas1 | MAS1 oncogene [Source:MGI Symbol;Acc | 16.062 | 9.2513 | 6.0375 | 4.8199 | 1.0906 | 5.5692 |
| 19298 | ENSMUSG00000068039 | 2318.181111 | 0.054368918 | 0.117026001 | 0.464588359 | 0.64222627 | 0.924834503 | Tcp1 | t-complex protein 1 [Source:MGI Symbol;A | 2079.2 | 2421.8 | 2324.4 | 2671.2 | 2276.1 | 2136.4 |
| 19299 | ENSMUSG00000068040 | 783.629362 | -0.141389898 | 0.110160221 | -1.283493226 | 0.199319331 | 0.641417519 | Tm9sf4 | transmembrane 9 superfamily member 4 | 819.97 | 856.26 | 789.9 | 747.08 | 762.34 | 726.23 |
| 19300 | ENSMUSG00000068048 | 0 | NA | NA | NA | NA | NA | Rhox9 | reproductive homeobox 9 [Source:MGI Sy | 0 | 0 | 0 | 0 | 0 | 0 |
| 19301 | ENSMUSG00000068062 | 1.060069211 | -2.228626096 | 2.226826576 | -1.00080811 | 0.316919588 | NA | Gm14164 | predicted gene 14164 [Source:MGI Symbo | 3.2124 | 1.0279 | 1.0062 | 0 | 0 | 1.1138 |
| 19302 | ENSMUSG00000068067 | 0 | NA | NA | NA | NA | NA | 1110057P08 | RIKEN cDNA 1110057P08 gene [Source:M | 0 | 0 | 0 | 0 | 0 | 0 |
| 19303 | ENSMUSG00000068068 | 0 | NA | NA | NA | NA | NA | Gm7735 | predicted gene 7735 [Source:MGI Symbol | 0 | 0 | 0 | 0 | 0 | 0 |
| 19304 | ENSMUSG00000068071 | 0 | NA | NA | NA | NA | NA | Gm6358 | predicted gene 6358 [Source:MGI Symbol | 0 | 0 | 0 | 0 | 0 | 0 |
| 19305 | ENSMUSG00000068073 | 0 | NA | NA | NA | NA | NA | 1110025L11 | RIKEN cDNA 1110025L11 gene [Source:M | 0 | 0 | 0 | 0 | 0 | 0 |
| 19306 | ENSMUSG00000068074 | 0 | NA | NA | NA | NA | NA | Gm10228 | predicted gene 10228 [Source:MGI Symbo | 0 | 0 | 0 | 0 | 0 | 0 |
| 19307 | ENSMUSG00000068075 | 0 | NA | NA | NA | NA | NA | Gm10229 | predicted gene 10229 [Source:MGI Symbo | 0 | 0 | 0 | 0 | 0 | 0 |
| 19308 | ENSMUSG00000068078 | 0 | NA | NA | NA | NA | NA | 2310034C09 | RIKEN cDNA 2310034C09 gene [Source:M | 0 | 0 | 0 | 0 | 0 | 0 |
| 19309 | ENSMUSG00000068079 | 0.181769715 | 1.126924175 | 4.080472857 | 0.276174898 | 0.782413736 | NA | Tcf15 | transcription factor 15 [Source:MGI Symb | 0 | 0 | 0 | 0 | 1.0906 | 0 |
| 19310 | ENSMUSG00000068082 | 0 | NA | NA | NA | NA | NA | Grxcr1 | glutaredoxin, cysteine rich 1 [Source:MGI | 0 | 0 | 0 | 0 | 0 | 0 |
| 19311 | ENSMUSG00000068083 | 2381.754476 | 0.068048243 | 0.156586667 | 0.434572394 | 0.663872836 | 0.929301301 | Cyp2d40 | cytochrome P450, family 2, subfamily d, p | 2724.9 | 1692 | 2558.9 | 2516 | 2546.6 | 2252.2 |
| 19312 | ENSMUSG00000068085 | 14.02676732 | -0.114388602 | 0.713529257 | -0.160313821 | 0.872633872 | 0.979419285 | Cyp2d11 | cytochrome P450, family 2, subfamily d, p | 12.047 | 18.503 | 13.081 | 2.8919 | 10.906 | 26.732 |
| 19313 | ENSMUSG00000068086 | 30816.4291 | 0.046291311 | 0.122716961 | 0.377220158 | 0.706010001 | 0.94295036 | Cyp2d9 | cytochrome P450, family 2, subfamily d, p | 31677 | 28398 | 30890 | 26011 | 36322 | 31601 |
| 19314 | ENSMUSG00000068099 | 0 | NA | NA | NA | NA | NA | 1500009C09 | RIKEN cDNA 1500009C09 gene [Source:M | 0 | 0 | 0 | 0 | 0 | 0 |
| 19315 | ENSMUSG00000068101 | 28.81110718 | -0.095210892 | 0.432397549 | -0.22019295 | 0.825720887 | 0.96943211 | Cenpm | centromere protein M [Source:MGI Symb | 36.14 | 25.698 | 27.169 | 21.208 | 39.262 | 23.391 |
| 19316 | ENSMUSG00000068105 | 0.401551119 | -2.072982726 | 4.008251979 | -0.517178744 | 0.605031388 | NA | Tnfrsf13c | tumor necrosis factor receptor superfamil | 2.4093 | 0 | 0 | 0 | 0 | 0 |

| | | | | | | | | | | | | | | | |
|---|---|---|---|---|---|---|---|---|---|---|---|---|---|---|---|
| 19317 | ENSMUSG00000068113 | 0 | NA | NA | NA | NA | NA | Tesl1 | testin LIM domain protein like 1 [Source:N | 0 | 0 | 0 | 0 | 0 | 0 |
| 19318 | ENSMUSG00000068114 | 79.69692312 | -0.247462108 | 0.255718298 | -0.967713731 | 0.333187359 | 0.77523112 | Ccdc134 | coiled-coil domain containing 134 [Source | 74.689 | 90.457 | 94.587 | 67.478 | 76.343 | 74.628 |
| 19319 | ENSMUSG00000068115 | 17.48809168 | -0.307276439 | 0.659146507 | -0.466173204 | 0.641091528 | 0.924019354 | Ninl | ninein-like [Source:MGI Symbol;Acc:MGI: | 23.29 | 24.67 | 10.062 | 20.244 | 3.2719 | 23.391 |
| 19320 | ENSMUSG00000068117 | 0 | NA | NA | NA | NA | NA | Mei1 | meiotic double-stranded break formation | 0 | 0 | 0 | 0 | 0 | 0 |
| 19321 | ENSMUSG00000068120 | 0.784885323 | -1.683590003 | 2.624742731 | -0.641430485 | 0.521243032 | NA | Gm10231 | predicted pseudogene 10231 [Source:MG | 1.6062 | 0 | 2.0125 | 0 | 1.0906 | 0 |
| 19322 | ENSMUSG00000068122 | 0 | NA | NA | NA | NA | NA | Agtr2 | angiotensin II receptor, type 2 [Source:MC | 0 | 0 | 0 | 0 | 0 | 0 |
| 19323 | ENSMUSG00000068129 | 0.160662974 | 1.126924175 | 4.080472857 | 0.276174898 | 0.782413736 | NA | Cst7 | cystatin F (leukocystatin) [Source:MGI Syr | 0 | 0 | 0 | 0.964 | 0 | 0 |
| 19324 | ENSMUSG00000068130 | 14.91990354 | -0.051974179 | 0.569452283 | -0.091270473 | 0.927277679 | 0.987712228 | Zfp442 | zinc finger protein 442 [Source:MGI Symb | 11.243 | 13.363 | 21.131 | 17.352 | 14.178 | 12.252 |
| 19325 | ENSMUSG00000068134 | 164.0641307 | 0.459127426 | 0.21429632 | 2.142488617 | 0.032154184 | 0.23951732 | Zfp120 | zinc finger protein 120 [Source:MGI Symb | 127.69 | 137.74 | 148.92 | 147.49 | 190.86 | 231.68 |
| 19326 | ENSMUSG00000068141 | 2.633639194 | -2.036624591 | 1.483728284 | -1.372639865 | 0.169864332 | NA | Gm10232 | predicted pseudogene 10232 [Source:MG | 1.6062 | 5.1396 | 6.0375 | 1.928 | 1.0906 | 0 |
| 19327 | ENSMUSG00000068149 | 0 | NA | NA | NA | NA | NA | Tex13c3 | TEX13 family member C3 [Source:MGI Syr | 0 | 0 | 0 | 0 | 0 | 0 |
| 19328 | ENSMUSG00000068151 | 0 | NA | NA | NA | NA | NA | A230006K03 | RIKEN cDNA A230006K03 gene [Source:M | 0 | 0 | 0 | 0 | 0 | 0 |
| 19329 | ENSMUSG00000068154 | 0.160662974 | 1.126924175 | 4.080472857 | 0.276174898 | 0.782413736 | NA | Insm1 | insulinoma-associated 1 [Source:MGI Sym | 0 | 0 | 0 | 0.964 | 0 | 0 |
| 19330 | ENSMUSG00000068165 | 0 | NA | NA | NA | NA | NA | Gm10233 | predicted pseudogene 10233 [Source:MG | 0 | 0 | 0 | 0 | 0 | 0 |
| 19331 | ENSMUSG00000068167 | 0 | NA | NA | NA | NA | NA | Csnka2ip | casein kinase 2, alpha prime interacting pı | 0 | 0 | 0 | 0 | 0 | 0 |
| 19332 | ENSMUSG00000068173 | 0 | NA | NA | NA | NA | NA | Btg1b | BTG anti-proliferation factor 1B [Source:N | 0 | 0 | 0 | 0 | 0 | 0 |
| 19333 | ENSMUSG00000068180 | 0 | NA | NA | NA | NA | NA | AY512915 | cDNA sequence AY512915 [Source:MGI Sy | 0 | 0 | 0 | 0 | 0 | 0 |
| 19334 | ENSMUSG00000068181 | 0 | NA | NA | NA | NA | NA | Gm10234 | predicted gene 10234 [Source:MGI Symbc | 0 | 0 | 0 | 0 | 0 | 0 |
| 19335 | ENSMUSG00000068182 | 0 | NA | NA | NA | NA | NA | Olfr203 | olfactory receptor 203 [Source:MGI Symb | 0 | 0 | 0 | 0 | 0 | 0 |
| 19336 | ENSMUSG00000068184 | 210.6357721 | -0.049328205 | 0.190122075 | -0.25945543 | 0.795283867 | 0.962236316 | Ndufaf2 | NADH:ubiquinone oxidoreductase comple | 208.81 | 226.14 | 207.29 | 162.91 | 220.3 | 238.36 |
| 19337 | ENSMUSG00000068196 | 2.921050438 | -2.132655875 | 1.402927695 | -1.520146677 | 0.128474115 | NA | Col8a1 | collagen, type VIII, alpha 1 [Source:MGI Sy | 4.0155 | 6.1675 | 4.025 | 0 | 1.0906 | 2.2277 |
| 19338 | ENSMUSG00000068200 | 0.499690045 | -0.82080011 | 3.113553401 | -0.263621658 | 0.792071468 | NA | Gm5931 | predicted pseudogene 5931 [Source:MGI | 0 | 1.0279 | 1.0062 | 0.964 | 0 | 0 |
| 19339 | ENSMUSG00000068205 | 69.17217736 | -0.304243822 | 0.26951733 | -1.128846974 | 0.258962393 | 0.712239465 | Macrod2 | mono-ADP ribosylhydrolase 2 [Source:MG | 69.87 | 84.289 | 75.468 | 67.478 | 58.893 | 59.034 |
| 19340 | ENSMUSG00000068206 | 115.8222767 | 0.109948093 | 0.220298221 | 0.499087523 | 0.617717727 | 0.917509899 | Pick1 | protein interacting with C kinase 1 [Source | 124.48 | 95.596 | 113.71 | 116.64 | 129.78 | 114.73 |
| 19341 | ENSMUSG00000068218 | 0 | NA | NA | NA | NA | NA | Ssxb9 | synovial sarcoma, X member B9 [Source:N | 0 | 0 | 0 | 0 | 0 | 0 |
| 19342 | ENSMUSG00000068219 | 0 | NA | NA | NA | NA | NA | Ssxb10 | synovial sarcoma, X member B10 [Source: | 0 | 0 | 0 | 0 | 0 | 0 |
| 19343 | ENSMUSG00000068220 | 483.0515234 | -1.121401083 | 0.579385859 | -1.935499575 | 0.052929021 | 0.323119557 | Lgals1 | lectin, galactose binding, soluble 1 [Sourc | 993.44 | 715.43 | 276.72 | 363.42 | 357.72 | 191.58 |
| 19344 | ENSMUSG00000068226 | 2.244444673 | -0.661397581 | 1.513566938 | -0.436979405 | 0.662126284 | NA | Gm6723 | predicted pseudogene 6723 [Source:MGI | 3.2124 | 0 | 5.0312 | 1.928 | 2.1812 | 1.1138 |
| 19345 | ENSMUSG00000068227 | 22.42626391 | -2.257843405 | 0.542446159 | -4.162336419 | 3.15E-05 | 0.001446436 | Il2rb | interleukin 2 receptor, beta chain [Source | 41.761 | 46.256 | 23.144 | 5.7839 | 9.8156 | 7.7969 |
| 19346 | ENSMUSG00000068231 | 0 | NA | NA | NA | NA | NA | Vmn1r43 | vomeronasal 1 receptor 43 [Source:MGI S | 0 | 0 | 0 | 0 | 0 | 0 |
| 19347 | ENSMUSG00000068232 | 0 | NA | NA | NA | NA | NA | Vmn1r42 | vomeronasal 1 receptor 42 [Source:MGI S | 0 | 0 | 0 | 0 | 0 | 0 |
| 19348 | ENSMUSG00000068234 | 0 | NA | NA | NA | NA | NA | Vmn1r44 | vomeronasal 1 receptor 44 [Source:MGI S | 0 | 0 | 0 | 0 | 0 | 0 |
| 19349 | ENSMUSG00000068240 | 19.13207522 | 0.221745624 | 0.490137161 | 0.452415449 | 0.650969719 | 0.925643431 | Gm11808 | predicted gene 11808 [Source:MGI Symbc | 18.471 | 14.391 | 20.125 | 22.171 | 21.812 | 17.822 |
| 19350 | ENSMUSG00000068243 | 0 | NA | NA | NA | NA | NA | Gm7079 | predicted pseudogene 7079 [Source:MGI | 0 | 0 | 0 | 0 | 0 | 0 |
| 19351 | ENSMUSG00000068245 | 35.85213068 | -0.623869328 | 0.37515092 | -1.662982268 | 0.096315985 | 0.458386897 | Phf11d | PHD finger protein 11D [Source:MGI Symt | 48.989 | 46.256 | 35.219 | 31.811 | 29.447 | 23.391 |
| 19352 | ENSMUSG00000068246 | 2045.484559 | 0.303794363 | 0.114640846 | 2.649966159 | 0.008049983 | 0.092428861 | Apol9b | apolipoprotein L 9b [Source:MGI Symbol;/ | 1990.1 | 1591.2 | 1909.9 | 2122.7 | 2360.1 | 2299 |
| 19353 | ENSMUSG00000068250 | 296.9490473 | 0.465258413 | 0.245793077 | 1.892886565 | 0.058372956 | 0.343085466 | Amn1 | antagonist of mitotic exit network 1 [Sour | 247.36 | 271.37 | 230.43 | 518.62 | 241.03 | 272.89 |
| 19354 | ENSMUSG00000068252 | 0 | NA | NA | NA | NA | NA | Apol7b | apolipoprotein L 7b [Source:MGI Symbol;/ | 0 | 0 | 0 | 0 | 0 | 0 |
| 19355 | ENSMUSG00000068259 | 0.181769715 | 1.126924175 | 4.080472857 | 0.276174898 | 0.782413736 | NA | Olfr461 | olfactory receptor 461 [Source:MGI Symb | 0 | 0 | 0 | 0 | 1.0906 | 0 |
| 19356 | ENSMUSG00000068262 | 0.870532655 | -0.452120295 | 2.585085873 | -0.174895658 | 0.861161622 | NA | Gm5879 | predicted gene 5879 [Source:MGI Symbol | 0 | 0 | 3.0187 | 0 | 1.0906 | 1.1138 |
| 19357 | ENSMUSG00000068263 | 7.415715227 | 0.65225488 | 0.784696045 | 0.831219788 | 0.405849478 | 0.829973745 | Efcc1 | EF hand and coiled-coil domain containing | 8.031 | 5.1396 | 4.025 | 9.6398 | 7.6343 | 10.025 |
| 19358 | ENSMUSG00000068264 | 262.8137207 | 0.240620162 | 0.210663069 | 1.142203821 | 0.253369308 | 0.706566564 | Ap5s1 | adaptor-related protein 5 complex, sigma | 310 | 213.81 | 198.23 | 323.9 | 225.76 | 305.19 |
| 19359 | ENSMUSG00000068267 | 2681.988783 | -0.020216396 | 0.117858248 | -0.17153145 | 0.863805903 | 0.976954351 | Cenpb | centromere protein B [Source:MGI Symbc | 2729.7 | 2577 | 2794.3 | 2251.9 | 2801.8 | 2937.2 |
| 19360 | ENSMUSG00000068270 | 11.51236548 | -0.248302839 | 0.637805361 | -0.389308171 | 0.697048192 | 0.94023175 | Shroom4 | shroom family member 4 [Source:MGI Syr | 16.062 | 8.2233 | 13.081 | 9.6398 | 9.8156 | 12.252 |
| 19361 | ENSMUSG00000068284 | 895.9954679 | 0.115206301 | 0.121568909 | 0.947662546 | 0.343301274 | 0.78518495 | Usf3 | upstream transcription factor family mem | 850.49 | 899.43 | 831.16 | 975.55 | 815.78 | 1003.6 |
| 19362 | ENSMUSG00000068289 | 0 | NA | NA | NA | NA | NA | Cma2 | chymase 2, mast cell [Source:MGI Symbol | 0 | 0 | 0 | 0 | 0 | 0 |
| 19363 | ENSMUSG00000068290 | 2801.218222 | -0.066347006 | 0.103414622 | -0.641563099 | 0.521156899 | 0.887887809 | Ddrgk1 | DDRGK domain containing 1 [Source:MGI | 2662.3 | 2931.6 | 3004.6 | 2969.1 | 2554.2 | 2685.5 |
| 19364 | ENSMUSG00000068299 | 144.0102521 | 0.53928727 | 0.275676931 | 1.956229228 | 0.050438157 | 0.315680442 | Nat8f4 | N-acetyltransferase 8 (GCN5-related) fam | 113.24 | 100.74 | 137.86 | 100.25 | 215.94 | 196.04 |
| 19365 | ENSMUSG00000068302 | 0 | NA | NA | NA | NA | NA | Noto | notochord homeobox [Source:MGI Symbc | 0 | 0 | 0 | 0 | 0 | 0 |
| 19366 | ENSMUSG00000068303 | 3.887106234 | 2.297657958 | 1.446890793 | 1.587996806 | 0.112287061 | NA | Spr-ps1 | sepiapterin reductase pseudogene 1 [Sou | 0 | 0 | 4.025 | 2.8919 | 14.178 | 2.2277 |
| 19367 | ENSMUSG00000068314 | 0 | NA | NA | NA | NA | NA | Gm6899 | predicted gene 6899 [Source:MGI Symbol | 0 | 0 | 0 | 0 | 0 | 0 |
| 19368 | ENSMUSG00000068323 | 2.402315161 | -0.303978789 | 1.526194027 | -0.199174406 | 0.842126319 | NA | Slc4a5 | solute carrier family 4, sodium bicarbonat | 4.8186 | 2.0558 | 1.0062 | 0.964 | 0 | 5.5692 |
| 19369 | ENSMUSG00000068324 | 0.185640647 | 1.126924175 | 4.080472857 | 0.276174898 | 0.782413736 | NA | Gm10029 | predicted gene 10029 [Source:MGI Symb | 0 | 0 | 0 | 0 | 0 | 1.1138 |
| 19370 | ENSMUSG00000068327 | 1.569873498 | -0.891123148 | 1.910581038 | -0.466414735 | 0.640918667 | NA | Tlx2 | T cell leukemia, homeobox 2 [Source:MGI | 0 | 4.1117 | 2.0125 | 0 | 2.1812 | 1.1138 |
| 19371 | ENSMUSG00000068328 | 2017.956274 | 0.023085193 | 0.094393502 | 0.244563366 | 0.806794532 | 0.964718338 | Aup1 | ancient ubiquitous protein 1 [Source:MGI | 1940.3 | 1968.5 | 2097 | 2019.5 | 1978.4 | 2104.1 |
| 19372 | ENSMUSG00000068329 | 189.1200449 | -0.085304193 | 0.185207193 | -0.460587908 | 0.645094288 | 0.925643431 | Htra2 | HtrA serine peptidase 2 [Source:MGI Syml | 164.64 | 212.78 | 207.29 | 177.37 | 179.95 | 192.69 |

| | | | | | | | | | | | | | | | |
|---|---|---|---|---|---|---|---|---|---|---|---|---|---|---|---|
| 19373 | ENSMUSG00000068335 | 18.57578647 | -0.647565382 | 0.516241119 | -1.254385514 | 0.209701915 | 0.653550636 | Dok1 | docking protein 1 [Source:MGI Symbol;Ac | 28.109 | 21.586 | 18.112 | 10.604 | 17.45 | 15.594 |
| 19374 | ENSMUSG00000068341 | 0 | NA | NA | NA | NA | NA | Reg3d | regenerating islet-derived 3 delta [Source: | 0 | 0 | 0 | 0 | 0 | 0 |
| 19375 | ENSMUSG00000068348 | 0 | NA | NA | NA | NA | NA | Gm10238 | predicted pseudogene 10238 [Source:MG | 0 | 0 | 0 | 0 | 0 | 0 |
| 19376 | ENSMUSG00000068349 | 0 | NA | NA | NA | NA | NA | Gml | glycosylphosphatidylinositol anchored mc | 0 | 0 | 0 | 0 | 0 | 0 |
| 19377 | ENSMUSG00000068373 | 4.113058813 | 1.159300641 | 1.129646886 | 1.026250464 | 0.30477354 | NA | D430041D05 | RIKEN cDNA D430041D05 gene [Source:M | 1.6062 | 1.0279 | 5.0312 | 3.8559 | 9.8156 | 3.3415 |
| 19378 | ENSMUSG00000068391 | 241.8475911 | -0.050449663 | 0.168386373 | -0.299606565 | 0.764477276 | 0.954980327 | Chrac1 | chromatin accessibility complex 1 [Source | 247.36 | 269.31 | 221.37 | 220.75 | 262.84 | 229.45 |
| 19379 | ENSMUSG00000068392 | 15.16230634 | 0.868778055 | 0.639431431 | 1.358672741 | 0.174250314 | 0.604151393 | Rnase13 | ribonuclease, RNase A family, 13 (non-act | 17.668 | 10.279 | 4.025 | 26.027 | 20.722 | 12.252 |
| 19380 | ENSMUSG00000068394 | 73.27682943 | 0.114155555 | 0.286131432 | 0.398961953 | 0.689921239 | 0.937261276 | Cep152 | centrosomal protein 152 [Source:MGI Syn | 77.901 | 66.815 | 66.412 | 91.578 | 54.531 | 82.424 |
| 19381 | ENSMUSG00000068396 | 6.585444025 | -0.131552916 | 0.899080132 | -0.146319456 | 0.883669208 | 0.981769403 | Rpl34-ps1 | ribosomal protein L34, pseudogene 1 [Sou | 6.4248 | 8.2233 | 6.0375 | 7.7118 | 1.0906 | 10.025 |
| 19382 | ENSMUSG00000068397 | 0 | NA | NA | NA | NA | NA | Gm10240 | predicted gene 10240 [Source:MGI Symbo | 0 | 0 | 0 | 0 | 0 | 0 |
| 19383 | ENSMUSG00000068399 | 0 | NA | NA | NA | NA | NA | Gm7247 | predicted gene 7247 [Source:MGI Symbol | 0 | 0 | 0 | 0 | 0 | 0 |
| 19384 | ENSMUSG00000068407 | 0 | NA | NA | NA | NA | NA | Rnase12 | ribonuclease, RNase A family, 12 (non-act | 0 | 0 | 0 | 0 | 0 | 0 |
| 19385 | ENSMUSG00000068417 | 4.122455118 | -0.662043472 | 1.115967174 | -0.593246367 | 0.553016286 | NA | Pnp2 | purine-nucleoside phosphorylase 2 [Sourc | 4.0155 | 3.0838 | 8.05 | 1.928 | 6.5437 | 1.1138 |
| 19386 | ENSMUSG00000068428 | 0 | NA | NA | NA | NA | NA | Gmnc | geminin coiled-coil domain containing [So | 0 | 0 | 0 | 0 | 0 | 0 |
| 19387 | ENSMUSG00000068431 | 0 | NA | NA | NA | NA | NA | Olfr742 | olfactory receptor 742 [Source:MGI Symb | 0 | 0 | 0 | 0 | 0 | 0 |
| 19388 | ENSMUSG00000068437 | 0 | NA | NA | NA | NA | NA | Olfr725 | olfactory receptor 725 [Source:MGI Symb | 0 | 0 | 0 | 0 | 0 | 0 |
| 19389 | ENSMUSG00000068452 | 0.181769715 | 1.126924175 | 4.080472857 | 0.276174898 | 0.782413736 | NA | Duox2 | dual oxidase 2 [Source:MGI Symbol;Acc:M | 0 | 0 | 0 | 0 | 1.0906 | 0 |
| 19390 | ENSMUSG00000068457 | 272.79439 | -0.062163351 | 0.155791103 | -0.399017338 | 0.689880429 | 0.937261276 | Uty | ubiquitously transcribed tetratricopeptide | 280.28 | 284.73 | 270.68 | 244.85 | 294.47 | 261.75 |
| 19391 | ENSMUSG00000068463 | 1313.828912 | 0.124660034 | 0.103679401 | 1.202360671 | 0.22922382 | 0.675803777 | B630019A10 | RIKEN cDNA B630019A10 gene [Source:M | 1236 | 1260.2 | 1275.9 | 1452.7 | 1388.4 | 1269.8 |
| 19392 | ENSMUSG00000068466 | 1.287980986 | 0.180143824 | 1.974544931 | 0.091233084 | 0.927307387 | NA | Gm5518 | predicted gene 5518 [Source:MGI Symbol | 1.6062 | 0 | 2.0125 | 1.928 | 2.1812 | 0 |
| 19393 | ENSMUSG00000068479 | 133.5309821 | -0.146755954 | 0.220705703 | -0.664939564 | 0.506089149 | 0.881592135 | Mfap1a | microfibrillar-associated protein 1A [Sourc | 130.91 | 141.85 | 147.92 | 99.29 | 131.96 | 149.26 |
| 19394 | ENSMUSG00000068480 | 0 | NA | NA | NA | NA | NA | Gm7551 | predicted gene 7551 [Source:MGI Symbol | 0 | 0 | 0 | 0 | 0 | 0 |
| 19395 | ENSMUSG00000068487 | 0 | NA | NA | NA | NA | NA | Ndufa11b | NADH:ubiquinone oxidoreductase subuni | 0 | 0 | 0 | 0 | 0 | 0 |
| 19396 | ENSMUSG00000068506 | 0 | NA | NA | NA | NA | NA | Gm5800 | predicted gene 5800 [Source:MGI Symbol | 0 | 0 | 0 | 0 | 0 | 0 |
| 19397 | ENSMUSG00000068522 | 24.21636217 | -1.13186463 | 0.451404161 | -2.507430651 | 0.012161244 | 0.123873406 | Aard | alanine and arginine rich domain containi | 30.518 | 37.005 | 32.2 | 12.532 | 17.45 | 15.594 |
| 19398 | ENSMUSG00000068523 | 1365.626062 | 0.044572053 | 0.102730925 | 0.433871821 | 0.664381524 | 0.929301301 | Gng5 | guanine nucleotide binding protein (G prc | 1430.3 | 1279.8 | 1322.2 | 1330.3 | 1422.2 | 1409 |
| 19399 | ENSMUSG00000068535 | 0 | NA | NA | NA | NA | NA | Olfr169 | olfactory receptor 169 [Source:MGI Symb | 0 | 0 | 0 | 0 | 0 | 0 |
| 19400 | ENSMUSG00000068536 | 0 | NA | NA | NA | NA | NA | Doxl2 | diamine oxidase-like protein 2 [Source:M( | 0 | 0 | 0 | 0 | 0 | 0 |
| 19401 | ENSMUSG00000068537 | 0 | NA | NA | NA | NA | NA | Gm10242 | predicted gene 10242 [Source:MGI Symbo | 0 | 0 | 0 | 0 | 0 | 0 |
| 19402 | ENSMUSG00000068547 | 0 | NA | NA | NA | NA | NA | Clca4a | chloride channel accessory 4A [Source:M( | 0 | 0 | 0 | 0 | 0 | 0 |
| 19403 | ENSMUSG00000068551 | 421.7164006 | 0.44457231 | 0.147988078 | 3.004108948 | 0.002663599 | 0.04333667 | Zfp467 | zinc finger protein 467 [Source:MGI Symb | 315.62 | 344.35 | 412.56 | 482.95 | 461.33 | 513.48 |
| 19404 | ENSMUSG00000068566 | 442.2148133 | 0.019181975 | 0.139914369 | 0.137097963 | 0.890953362 | 0.982489666 | Myadm | myeloid-associated differentiation marker | 465 | 453.31 | 399.48 | 474.28 | 460.24 | 400.98 |
| 19405 | ENSMUSG00000068574 | 0 | NA | NA | NA | NA | NA | Olfr458 | olfactory receptor 458 [Source:MGI Symb | 0 | 0 | 0 | 0 | 0 | 0 |
| 19406 | ENSMUSG00000068579 | 0.492645588 | 0.9565034 | 3.447648281 | 0.277436479 | 0.781444974 | NA | Rpl7a-ps3 | ribosomal protein L7A, pseudogene 3 [Sou | 0 | 1.0279 | 0 | 1.928 | 0 | 0 |
| 19407 | ENSMUSG00000068580 | 350.9599422 | 0.029378857 | 0.147893145 | 0.198649214 | 0.842537154 | 0.971961827 | Zfyve19 | zinc finger, FYVE domain containing 19 [Sc | 320.44 | 338.18 | 384.39 | 381.74 | 344.64 | 336.38 |
| 19408 | ENSMUSG00000068587 | 2157.233841 | 0.1214124 | 0.11064571 | 1.097307794 | 0.272506865 | 0.727618887 | Mgam | maltase-glucoamylase [Source:MGI Symb | 1911.4 | 2251.1 | 2037.6 | 2105.3 | 2464.8 | 2173.1 |
| 19409 | ENSMUSG00000068600 | 0 | NA | NA | NA | NA | NA | Gml2 | glycosylphosphatidylinositol anchored mc | 0 | 0 | 0 | 0 | 0 | 0 |
| 19410 | ENSMUSG00000068601 | 1.160037551 | -0.144733627 | 2.274505547 | -0.063633007 | 0.949262449 | NA | Gm10244 | predicted gene 10244 [Source:MGI Symbo | 1.6062 | 0 | 2.0125 | 0 | 0 | 3.3415 |
| 19411 | ENSMUSG00000068604 | 0 | NA | NA | NA | NA | NA | Gm8225 | predicted gene 8225 [Source:MGI Symbol | 0 | 0 | 0 | 0 | 0 | 0 |
| 19412 | ENSMUSG00000068606 | 14.82852319 | -0.450241044 | 0.566133842 | -0.795290815 | 0.426444355 | 0.842053132 | Gm4841 | predicted gene 4841 [Source:MGI Symbol | 16.865 | 17.475 | 17.106 | 14.46 | 15.269 | 7.7969 |
| 19413 | ENSMUSG00000068614 | 0.17131964 | -0.796629176 | 4.080472857 | -0.195229623 | 0.84521319 | NA | Actc1 | actin, alpha, cardiac muscle 1 [Source:MG | 0 | 1.0279 | 0 | 0 | 0 | 0 |
| 19414 | ENSMUSG00000068615 | 0 | NA | NA | NA | NA | NA | Gjd2 | gap junction protein, delta 2 [Source:MGI | 0 | 0 | 0 | 0 | 0 | 0 |
| 19415 | ENSMUSG00000068617 | 0 | NA | NA | NA | NA | NA | Efcab1 | EF-hand calcium binding domain 1 [Sourc | 0 | 0 | 0 | 0 | 0 | 0 |
| 19416 | ENSMUSG00000068629 | 0 | NA | NA | NA | NA | NA | Gm6155 | predicted pseudogene 6155 [Source:MGI | 0 | 0 | 0 | 0 | 0 | 0 |
| 19417 | ENSMUSG00000068631 | 0 | NA | NA | NA | NA | NA | Gm7676 | predicted gene 7676 [Source:MGI Symbol | 0 | 0 | 0 | 0 | 0 | 0 |
| 19418 | ENSMUSG00000068647 | 0 | NA | NA | NA | NA | NA | Olfr1278 | olfactory receptor 1278 [Source:MGI Syml | 0 | 0 | 0 | 0 | 0 | 0 |
| 19419 | ENSMUSG00000068663 | 364.6975201 | -0.10606715 | 0.156730464 | -0.676748777 | 0.498565363 | 0.879866965 | Clec16a | C-type lectin domain family 16, member A | 335.7 | 367.99 | 431.68 | 388.48 | 343.54 | 320.79 |
| 19420 | ENSMUSG00000068674 | 0.677847551 | 0.063310748 | 2.809814301 | 0.022532004 | 0.982023583 | NA | Gm10247 | predicted gene 10247 [Source:MGI Symbo | 0 | 0 | 2.0125 | 0.964 | 1.0906 | 0 |
| 19421 | ENSMUSG00000068680 | 0 | NA | NA | NA | NA | NA | Gm10248 | predicted gene 10248 [Source:MGI Symbo | 0 | 0 | 0 | 0 | 0 | 0 |
| 19422 | ENSMUSG00000068686 | 95.89416887 | 0.094814297 | 0.234009965 | 0.405172049 | 0.685351049 | 0.93583845 | Cd59b | CD59b antigen [Source:MGI Symbol;Acc:N | 101.99 | 82.233 | 93.581 | 92.542 | 101.43 | 103.59 |
| 19423 | ENSMUSG00000068689 | 0 | NA | NA | NA | NA | NA | Rps19-ps9 | ribosomal protein S19, pseudogene 9 [Sou | 0 | 0 | 0 | 0 | 0 | 0 |
| 19424 | ENSMUSG00000068696 | 0.660353019 | -0.01500701 | 2.83315355 | -0.005296928 | 0.995773683 | NA | Gpr88 | G-protein coupled receptor 88 [Source:M | 0 | 1.0279 | 1.0062 | 1.928 | 0 | 0 |
| 19425 | ENSMUSG00000068697 | 0.749689669 | 0.81510874 | 2.956253303 | 0.275723579 | 0.782760384 | NA | Myoz1 | myozenin 1 [Source:MGI Symbol;Acc:MGI | 1.6062 | 0 | 0 | 2.8919 | 0 | 0 |
| 19426 | ENSMUSG00000068699 | 2.112835241 | -1.607579035 | 1.649259072 | -0.974728023 | 0.329695189 | NA | Flnc | filamin C, gamma [Source:MGI Symbol;Ac | 6.4248 | 3.0838 | 0 | 0.964 | 1.0906 | 1.1138 |
| 19427 | ENSMUSG00000068706 | 12.94942683 | -0.818767165 | 0.604868122 | -1.353629221 | 0.175854699 | 0.606476483 | Gm10250 | predicted pseudogene 10250 [Source:MG | 20.078 | 15.419 | 14.087 | 11.568 | 7.6343 | 8.9108 |
| 19428 | ENSMUSG00000068732 | 500.1125724 | 0.16412849 | 0.127295806 | 1.289347188 | 0.197277413 | 0.638920846 | Tmem167b | transmembrane protein 167B [Source:MG | 466.6 | 446.12 | 502.12 | 501.27 | 537.67 | 546.9 |

| | | | | | | | | | | | | | | | |
|---|---|---|---|---|---|---|---|---|---|---|---|---|---|---|---|
| 19429 | ENSMUSG00000068735 | 77.04189883 | -0.639168034 | 0.331285348 | -1.929357991 | 0.053686435 | 0.325995213 | Trp53i11 | transformation related protein 53 inducib | 124.48 | 86.345 | 70.437 | 70.37 | 35.99 | 74.628 |
| 19430 | ENSMUSG00000068739 | 1014.813289 | 0.093332636 | 0.114149905 | 0.817632183 | 0.413567245 | 0.835385466 | Sars | seryl-aminoacyl-tRNA synthetase [Source: | 1081 | 971.38 | 892.54 | 1056.5 | 1040.4 | 1047 |
| 19431 | ENSMUSG00000068740 | 7.380910331 | -0.683345862 | 0.830089077 | -0.823219918 | 0.410382944 | 0.833241465 | Celsr2 | cadherin, EGF LAG seven-pass G-type rece | 4.0155 | 10.279 | 13.081 | 4.8199 | 7.6343 | 4.4554 |
| 19432 | ENSMUSG00000068742 | 843.9479696 | 0.307056044 | 0.121096086 | 2.535639697 | 0.011224215 | 0.11768024 | Cry2 | cryptochrome 2 (photolyase-like) [Source | 730.82 | 755.52 | 777.83 | 1009.3 | 823.42 | 966.82 |
| 19433 | ENSMUSG00000068744 | 5.833556198 | -0.473388136 | 0.951550981 | -0.497491091 | 0.618842784 | 0.917509899 | Psrc1 | proline/serine-rich coiled-coil 1 [Source:M | 12.047 | 6.1675 | 2.0125 | 4.8199 | 3.2719 | 6.6831 |
| 19434 | ENSMUSG00000068745 | 0 | NA | NA | NA | NA | NA | Mybphl | myosin binding protein H-like [Source:MG | 0 | 0 | 0 | 0 | 0 | 0 |
| 19435 | ENSMUSG00000068747 | 205.1661506 | 1.327430869 | 0.234343819 | 5.664458637 | 1.47E-08 | 2.11E-06 | Sort1 | sortilin 1 [Source:MGI Symbol;Acc:MGI:13 | 134.12 | 115.13 | 100.62 | 198.58 | 325 | 357.54 |
| 19436 | ENSMUSG00000068748 | 0.596071152 | -1.224343926 | 2.897583865 | -0.4225396 | 0.672631203 | NA | Ptprz1 | protein tyrosine phosphatase, receptor ty | 1.6062 | 0 | 1.0062 | 0.964 | 0 | 0 |
| 19437 | ENSMUSG00000068749 | 943.5753113 | 0.127539157 | 0.120702873 | 1.056637298 | 0.290677151 | 0.740463951 | Psma5 | proteasome subunit alpha 5 [Source:MGI | 985.41 | 930.27 | 788.9 | 955.3 | 993.55 | 1008 |
| 19438 | ENSMUSG00000068758 | 43.02295396 | -0.384521959 | 0.344191176 | -1.11717553 | 0.263919281 | 0.71906153 | Il3ra | interleukin 3 receptor, alpha chain [Sourc | 47.383 | 53.452 | 45.281 | 34.703 | 30.537 | 46.781 |
| 19439 | ENSMUSG00000068762 | 1285.559087 | 0.405599385 | 0.125972294 | 3.219750725 | 0.001283021 | 0.025183934 | Gstm6 | glutathione S-transferase, mu 6 [Source:M | 1229.5 | 997.08 | 1089.8 | 1361.1 | 1384 | 1651.8 |
| 19440 | ENSMUSG00000068776 | 0 | NA | NA | NA | NA | NA | 4930469B13 | RIKEN cDNA 4930469B13 gene [Source:M | 0 | 0 | 0 | 0 | 0 | 0 |
| 19441 | ENSMUSG00000068794 | 1.209687553 | -2.430175621 | 2.587233741 | -0.939294963 | 0.347579324 | NA | Col28a1 | collagen, type XXVIII, alpha 1 [Source:MG | 0 | 6.1675 | 0 | 0 | 1.0906 | 0 |
| 19442 | ENSMUSG00000068798 | 1195.61986 | -0.069113415 | 0.107435008 | -0.643304416 | 0.520026592 | 0.887802551 | Rap1a | RAS-related protein 1a [Source:MGI Symb | 1166.9 | 1197.5 | 1309.1 | 1192.4 | 1210.6 | 1097.1 |
| 19443 | ENSMUSG00000068806 | 0 | NA | NA | NA | NA | NA | Olfr1259 | olfactory receptor 1259 [Source:MGI Sym | 0 | 0 | 0 | 0 | 0 | 0 |
| 19444 | ENSMUSG00000068808 | 0 | NA | NA | NA | NA | NA | Olfr1189 | olfactory receptor 1189 [Source:MGI Sym | 0 | 0 | 0 | 0 | 0 | 0 |
| 19445 | ENSMUSG00000068809 | 0 | NA | NA | NA | NA | NA | Olfr1188 | olfactory receptor 1188 [Source:MGI Sym | 0 | 0 | 0 | 0 | 0 | 0 |
| 19446 | ENSMUSG00000068810 | 0 | NA | NA | NA | NA | NA | Olfr1187-ps1 | olfactory receptor 1187, pseudogene 1 [S | 0 | 0 | 0 | 0 | 0 | 0 |
| 19447 | ENSMUSG00000068814 | 0 | NA | NA | NA | NA | NA | Olfr1145 | olfactory receptor 1145 [Source:MGI Sym | 0 | 0 | 0 | 0 | 0 | 0 |
| 19448 | ENSMUSG00000068815 | 0 | NA | NA | NA | NA | NA | Olfr1143 | olfactory receptor 1143 [Source:MGI Sym | 0 | 0 | 0 | 0 | 0 | 0 |
| 19449 | ENSMUSG00000068816 | 0 | NA | NA | NA | NA | NA | Olfr152 | olfactory receptor 152 [Source:MGI Symb | 0 | 0 | 0 | 0 | 0 | 0 |
| 19450 | ENSMUSG00000068817 | 0 | NA | NA | NA | NA | NA | Olfr1140 | olfactory receptor 1140 [Source:MGI Sym | 0 | 0 | 0 | 0 | 0 | 0 |
| 19451 | ENSMUSG00000068818 | 0 | NA | NA | NA | NA | NA | Olfr1132 | olfactory receptor 1132 [Source:MGI Sym | 0 | 0 | 0 | 0 | 0 | 0 |
| 19452 | ENSMUSG00000068819 | 0 | NA | NA | NA | NA | NA | Olfr1131 | olfactory receptor 1131 [Source:MGI Sym | 0 | 0 | 0 | 0 | 0 | 0 |
| 19453 | ENSMUSG00000068823 | 3544.023493 | 0.045202061 | 0.094146207 | 0.480126203 | 0.631137657 | 0.921353964 | Csde1 | cold shock domain containing E1, RNA bin | 3371.4 | 3683 | 3411.2 | 3401.9 | 3784.4 | 3612.2 |
| 19454 | ENSMUSG00000068824 | 0 | NA | NA | NA | NA | NA | Olfr1083-ps | olfactory receptor 1083, pseudogene [Sou | 0 | 0 | 0 | 0 | 0 | 0 |
| 19455 | ENSMUSG00000068854 | 43.54403442 | 0.288520093 | 0.355352555 | 0.811926322 | 0.41683391 | 0.836878476 | H2bc21 | H2B clustered histone 21 [Source:MGI Syr | 34.533 | 29.81 | 53.331 | 44.343 | 46.897 | 52.351 |
| 19456 | ENSMUSG00000068855 | 0.17131964 | -0.796629176 | 4.080472857 | -0.195229623 | 0.84521319 | NA | H2ac20 | H2A clustered histone 20 [Source:MGI Syr | 0 | 1.0279 | 0 | 0 | 0 | 0 |
| 19457 | ENSMUSG00000068856 | 261.0302586 | -0.310271998 | 0.180452784 | -1.719408212 | 0.085540068 | 0.430973029 | Sf3b4 | splicing factor 3b, subunit 4 [Source:MGI | 276.27 | 266.23 | 325.02 | 266.06 | 188.68 | 243.93 |
| 19458 | ENSMUSG00000068859 | 0 | NA | NA | NA | NA | NA | Sp9 | trans-acting transcription factor 9 [Source | 0 | 0 | 0 | 0 | 0 | 0 |
| 19459 | ENSMUSG00000068860 | 0 | NA | NA | NA | NA | NA | Gm128 | predicted gene 128 [Source:MGI Symbol;/ | 0 | 0 | 0 | 0 | 0 | 0 |
| 19460 | ENSMUSG00000068874 | 6381.482987 | -0.030632101 | 0.089218978 | -0.343336154 | 0.731345585 | 0.947186832 | Selenbp1 | selenium binding protein 1 [Source:MGI S | 6458.5 | 6459.4 | 6428.9 | 5885.1 | 6673.5 | 6383.4 |
| 19461 | ENSMUSG00000068876 | 584.3764084 | 0.183446297 | 0.20263268 | 0.905314468 | 0.365298814 | 0.803870323 | Cgn | cingulin [Source:MGI Symbol;Acc:MGI:192 | 445.72 | 460.51 | 736.57 | 735.52 | 488.6 | 639.35 |
| 19462 | ENSMUSG00000068877 | 10881.95914 | 2.385322186 | 0.700124117 | 3.406999028 | 0.000656814 | 0.01519651 | Selenbp2 | selenium binding protein 2 [Source:MGI S | 3149 | 1466.8 | 5873.4 | 7068.8 | 26471 | 21262 |
| 19463 | ENSMUSG00000068879 | 0 | NA | NA | NA | NA | NA | Gm5773 | predicted pseudogene 5773 [Source:MGI | 0 | 0 | 0 | 0 | 0 | 0 |
| 19464 | ENSMUSG00000068882 | 784.3127566 | 0.044202569 | 0.120055341 | 0.368184948 | 0.712735333 | 0.944239215 | Ssb | Sjogren syndrome antigen B [Source:MGI | 705.12 | 825.42 | 786.88 | 743.23 | 833.23 | 811.99 |
| 19465 | ENSMUSG00000068885 | 0 | NA | NA | NA | NA | NA | Lce3f | late cornified envelope 3F [Source:MGI Sy | 0 | 0 | 0 | 0 | 0 | 0 |
| 19466 | ENSMUSG00000068887 | 0 | NA | NA | NA | NA | NA | Lce1j | late cornified envelope 1J [Source:MGI Sy | 0 | 0 | 0 | 0 | 0 | 0 |
| 19467 | ENSMUSG00000068888 | 0 | NA | NA | NA | NA | NA | Lce1i | late cornified envelope 1I [Source:MGI Syı | 0 | 0 | 0 | 0 | 0 | 0 |
| 19468 | ENSMUSG00000068889 | 0 | NA | NA | NA | NA | NA | Lce1e | late cornified envelope 1E [Source:MGI Sy | 0 | 0 | 0 | 0 | 0 | 0 |
| 19469 | ENSMUSG00000068890 | 0 | NA | NA | NA | NA | NA | Lce1a2 | late cornified envelope 1A2 [Source:MGI S | 0 | 0 | 0 | 0 | 0 | 0 |
| 19470 | ENSMUSG00000068893 | 0 | NA | NA | NA | NA | NA | Sprr2a2 | small proline-rich protein 2A2 [Source:MG | 0 | 0 | 0 | 0 | 0 | 0 |
| 19471 | ENSMUSG00000068917 | 271.8210854 | 0.145146547 | 0.160940901 | 0.901862401 | 0.367129967 | 0.805239112 | Clk2 | CDC-like kinase 2 [Source:MGI Symbol;Acc | 268.24 | 254.92 | 251.56 | 324.86 | 259.57 | 271.78 |
| 19472 | ENSMUSG00000068921 | 716.3341757 | 0.010940019 | 0.124947636 | 0.087556834 | 0.930228912 | 0.988420305 | Dap3 | death associated protein 3 [Source:MGI S | 778.21 | 735.99 | 625.88 | 722.98 | 717.63 | 717.32 |
| 19473 | ENSMUSG00000068922 | 127.4360198 | -0.014689359 | 0.216688363 | -0.067790253 | 0.945952603 | 0.990319806 | Msto1 | misato 1, mitochondrial distribution and r | 147.77 | 127.46 | 108.67 | 135.92 | 116.7 | 128.09 |
| 19474 | ENSMUSG00000068923 | 80.395038 | 0.072429866 | 0.298868026 | 0.242347323 | 0.808511048 | 0.965682062 | Syt11 | synaptotagmin XI [Source:MGI Symbol;Ac | 93.963 | 76.066 | 64.4 | 54.947 | 90.521 | 102.47 |
| 19475 | ENSMUSG00000068940 | 0 | NA | NA | NA | NA | NA | Tcl1b3 | T cell leukemia/lymphoma 1B, 3 [Source:N | 0 | 0 | 0 | 0 | 0 | 0 |
| 19476 | ENSMUSG00000068947 | 0 | NA | NA | NA | NA | NA | Olfr366 | olfactory receptor 366 [Source:MGI Symb | 0 | 0 | 0 | 0 | 0 | 0 |
| 19477 | ENSMUSG00000068948 | 0 | NA | NA | NA | NA | NA | Olfr363-ps | olfactory receptor 363, pseudogene [Sour | 0 | 0 | 0 | 0 | 0 | 0 |
| 19478 | ENSMUSG00000068950 | 0 | NA | NA | NA | NA | NA | Olfr338 | olfactory receptor 338 [Source:MGI Symb | 0 | 0 | 0 | 0 | 0 | 0 |
| 19479 | ENSMUSG00000068957 | 0.133850373 | -0.796629176 | 4.080472857 | -0.195229623 | 0.84521319 | NA | 4930589L23 | RIKEN cDNA 4930589L23 gene [Source:M( | 0.8031 | 0 | 0 | 0 | 0 | 0 |
| 19480 | ENSMUSG00000068959 | 28.23851584 | -0.620186174 | 0.4163924 | -1.489427219 | 0.136374906 | 0.539236368 | Zfp619 | zinc finger protein 619 [Source:MGI Symb | 38.549 | 34.949 | 29.181 | 26.027 | 22.903 | 17.822 |
| 19481 | ENSMUSG00000068962 | 1.352496074 | -3.790736611 | 2.368332372 | -1.600593167 | 0.109467057 | NA | Zfp114 | zinc finger protein 114 [Source:MGI Symb | 0 | 3.0838 | 5.0312 | 0 | 0 | 0 |
| 19482 | ENSMUSG00000068964 | 0 | NA | NA | NA | NA | NA | Gm6361 | predicted gene 6361 [Source:MGI Symbol | 0 | 0 | 0 | 0 | 0 | 0 |
| 19483 | ENSMUSG00000068966 | 82.14414452 | -0.498230138 | 0.271810284 | -1.833006946 | 0.066801538 | 0.37192178 | Zbtb34 | zinc finger and BTB domain containing 34 | 71.476 | 106.9 | 110.69 | 70.37 | 63.256 | 70.172 |
| 19484 | ENSMUSG00000068969 | 0 | NA | NA | NA | NA | NA | Rpl32-ps | ribosomal protein L32, pseudogene [Sourc | 0 | 0 | 0 | 0 | 0 | 0 |

| | | | | | | | | | | | | | | | |
|---|---|---|---|---|---|---|---|---|---|---|---|---|---|---|---|
| 19485 | ENSMUSG00000068999 | 0 | NA | NA | NA | NA | NA | Vmn2r5 | vomeronasal 2, receptor 5 [Source:MGI Sy | 0 | 0 | 0 | 0 | 0 | 0 |
| 19486 | ENSMUSG00000069008 | 0 | NA | NA | NA | NA | NA | Gm5537 | predicted gene 5537 [Source:MGI Symbol | 0 | 0 | 0 | 0 | 0 | 0 |
| 19487 | ENSMUSG00000069011 | 0.531944269 | 2.618916564 | 3.285123409 | 0.797204926 | 0.425332026 | NA | Gm10254 | predicted gene 10254 [Source:MGI Symbo | 0 | 0 | 0 | 0.964 | 0 | 2.2277 |
| 19488 | ENSMUSG00000069014 | 0.528073336 | 2.60973587 | 2.979250867 | 0.875970499 | 0.381046072 | NA | Gm5641 | predicted gene 5641 [Source:MGI Symbol | 0 | 0 | 0 | 0.964 | 1.0906 | 1.1138 |
| 19489 | ENSMUSG00000069020 | 278.691601 | 0.143206731 | 0.170814531 | 0.838375579 | 0.4018198 | 0.828853621 | Urm1 | ubiquitin related modifier 1 [Source:MGI S | 301.16 | 224.09 | 268.67 | 296.91 | 318.46 | 262.87 |
| 19490 | ENSMUSG00000069036 | 0 | NA | NA | NA | NA | NA | Sry | sex determining region of Chr Y [Source:M | 0 | 0 | 0 | 0 | 0 | 0 |
| 19491 | ENSMUSG00000069038 | 0 | NA | NA | NA | NA | NA | H2al1j | H2A histone family member L1J [Source:M | 0 | 0 | 0 | 0 | 0 | 0 |
| 19492 | ENSMUSG00000069041 | 1.027969972 | -0.944837183 | 2.255793195 | -0.418849204 | 0.675326343 | NA | Slc25a31 | solute carrier family 25 (mitochondrial car | 0 | 3.0838 | 1.0062 | 0.964 | 0 | 1.1138 |
| 19493 | ENSMUSG00000069044 | 0 | NA | NA | NA | NA | NA | Usp9y | ubiquitin specific peptidase 9, Y chromoso | 0 | 0 | 0 | 0 | 0 | 0 |
| 19494 | ENSMUSG00000069045 | 1464.126196 | -0.058756148 | 0.132333271 | -0.444001325 | 0.657041626 | 0.926984986 | Ddx3y | DEAD box helicase 3, Y-linked [Source:MG | 1419.9 | 1670.4 | 1392.6 | 1556.8 | 1544.3 | 1200.7 |
| 19495 | ENSMUSG00000069049 | 990.1894604 | -0.055947799 | 0.138401751 | -0.404241985 | 0.686034783 | 0.93637889 | Eif2s3y | eukaryotic translation initiation factor 2, s | 999.86 | 992.97 | 1036.4 | 1158.7 | 942.29 | 810.88 |
| 19496 | ENSMUSG00000069053 | 0 | NA | NA | NA | NA | NA | Uba1y | ubiquitin-activating enzyme, Chr Y [Source | 0 | 0 | 0 | 0 | 0 | 0 |
| 19497 | ENSMUSG00000069072 | 0 | NA | NA | NA | NA | NA | Slc7a14 | solute carrier family 7 (cationic amino acid | 0 | 0 | 0 | 0 | 0 | 0 |
| 19498 | ENSMUSG00000069074 | 0 | NA | NA | NA | NA | NA | Gm10258 | predicted gene 10258 [Source:MGI Symbo | 0 | 0 | 0 | 0 | 0 | 0 |
| 19499 | ENSMUSG00000069080 | 0 | NA | NA | NA | NA | NA | Lcn11 | lipocalin 11 [Source:MGI Symbol;Acc:MGI | 0 | 0 | 0 | 0 | 0 | 0 |
| 19500 | ENSMUSG00000069083 | 0.133850373 | -0.796629176 | 4.080472857 | -0.195229623 | 0.84521319 | NA | Gm10259 | predicted pseudogene 10259 [Source:MG | 0.8031 | 0 | 0 | 0 | 0 | 0 |
| 19501 | ENSMUSG00000069085 | 0 | NA | NA | NA | NA | NA | Dytn | dystrotelin [Source:MGI Symbol;Acc:MGI: | 0 | 0 | 0 | 0 | 0 | 0 |
| 19502 | ENSMUSG00000069089 | 200.0062902 | -0.140329166 | 0.18633359 | -0.753107187 | 0.451385506 | 0.85613987 | Cdk7 | cyclin-dependent kinase 7 [Source:MGI Sy | 230.49 | 217.92 | 180.12 | 170.62 | 213.76 | 187.13 |
| 19503 | ENSMUSG00000069094 | 121.061799 | -0.032152048 | 0.233120432 | -0.137920334 | 0.89030338 | 0.982489666 | Pde7a | phosphodiesterase 7A [Source:MGI Symb | 117.25 | 139.8 | 110.69 | 140.74 | 124.33 | 93.563 |
| 19504 | ENSMUSG00000069114 | 92.2162241 | 0.423346225 | 0.255621558 | 1.656144452 | 0.097692569 | 0.460719048 | Zbtb10 | zinc finger and BTB domain containing 10 | 81.916 | 78.122 | 76.475 | 129.17 | 81.796 | 105.82 |
| 19505 | ENSMUSG00000069117 | 0 | NA | NA | NA | NA | NA | Rps18-ps6 | ribosomal protein S18, pseudogene 6 [Sou | 0 | 0 | 0 | 0 | 0 | 0 |
| 19506 | ENSMUSG00000069118 | 0 | NA | NA | NA | NA | NA | 1700008P02 | RIKEN cDNA 1700008P02 gene [Source:M | 0 | 0 | 0 | 0 | 0 | 0 |
| 19507 | ENSMUSG00000069125 | 0 | NA | NA | NA | NA | NA | Rps24-ps2 | ribosomal protein S24, pseudogene 2 [Sou | 0 | 0 | 0 | 0 | 0 | 0 |
| 19508 | ENSMUSG00000069132 | 0 | NA | NA | NA | NA | NA | Nxph2 | neurexophilin 2 [Source:MGI Symbol;Acc: | 0 | 0 | 0 | 0 | 0 | 0 |
| 19509 | ENSMUSG00000069135 | 299.7212346 | 0.319371361 | 0.185079841 | 1.725586961 | 0.084421747 | 0.42844777 | Cep43 | centrosomal protein 43 [Source:MGI Syml | 281.89 | 279.59 | 238.48 | 364.38 | 251.93 | 382.05 |
| 19510 | ENSMUSG00000069136 | 2.196964016 | -0.655654238 | 1.506048235 | -0.435347436 | 0.663310258 | NA | A130006I12I | RIKEN cDNA A130006I12 gene [Source:M | 4.0155 | 1.0279 | 3.0187 | 2.8919 | 0 | 2.2277 |
| 19511 | ENSMUSG00000069170 | 298.5818279 | 0.112618834 | 0.641019662 | 0.17568702 | 0.860539833 | 0.975943719 | Adgrv1 | adhesion G protein-coupled receptor V1 [ | 486.68 | 215.86 | 157.98 | 255.45 | 498.41 | 177.1 |
| 19512 | ENSMUSG00000069171 | 7.439259218 | -1.523619213 | 0.833063727 | -1.828934768 | 0.067409381 | 0.373457298 | Nr2f1 | nuclear receptor subfamily 2, group F, me | 8.8341 | 9.2513 | 15.094 | 4.8199 | 2.1812 | 4.4554 |
| 19513 | ENSMUSG00000069184 | 48.76546764 | -0.218129209 | 0.327238947 | -0.66657472 | 0.505043821 | 0.881592135 | Zfp72 | zinc finger protein 72 [Source:MGI Symbo | 54.611 | 42.145 | 60.375 | 38.559 | 52.35 | 44.554 |
| 19514 | ENSMUSG00000069188 | 3.274399045 | 0.592561472 | 1.229591325 | 0.481917415 | 0.629864615 | NA | Gm13192 | predicted gene 13192 [Source:MGI Symbo | 0.8031 | 2.0558 | 5.0312 | 2.8919 | 2.1812 | 6.6831 |
| 19515 | ENSMUSG00000069189 | 0 | NA | NA | NA | NA | NA | Zdhhc11 | zinc finger, DHHC domain containing 11 [S | 0 | 0 | 0 | 0 | 0 | 0 |
| 19516 | ENSMUSG00000069196 | 0 | NA | NA | NA | NA | NA | Gm10161 | predicted pseudogene 10161 [Source:MG | 0 | 0 | 0 | 0 | 0 | 0 |
| 19517 | ENSMUSG00000069206 | 425.7687653 | 0.03568717 | 0.131893638 | 0.270575368 | 0.786717643 | 0.960870081 | Zfp874a | zinc finger protein 874a [Source:MGI Sym | 431.27 | 409.11 | 420.61 | 412.58 | 418.8 | 462.25 |
| 19518 | ENSMUSG00000069208 | 238.3052117 | 0.598056406 | 0.160176841 | 3.733725805 | 0.000188668 | 0.006024045 | Zfp825 | zinc finger protein 825 [Source:MGI Symb | 203.18 | 179.89 | 185.15 | 278.59 | 291.2 | 291.83 |
| 19519 | ENSMUSG00000069223 | 5.029193598 | -1.394959224 | 1.061001218 | -1.314757421 | 0.188591442 | NA | 4930451E10 | RIKEN cDNA 4930451E10 gene [Source:M | 9.6372 | 7.1954 | 5.0312 | 3.8559 | 0 | 4.4554 |
| 19520 | ENSMUSG00000069227 | 0 | NA | NA | NA | NA | NA | Gprin1 | G protein-regulated inducer of neurite ou | 0 | 0 | 0 | 0 | 0 | 0 |
| 19521 | ENSMUSG00000069236 | 0 | NA | NA | NA | NA | NA | Gm7251 | predicted pseudogene 7251 [Source:MGI | 0 | 0 | 0 | 0 | 0 | 0 |
| 19522 | ENSMUSG00000069237 | 1708.896573 | 0.201609236 | 0.095246229 | 2.116716207 | 0.03428394 | 0.247817549 | Fam8a1 | family with sequence similarity 8, membe | 1558 | 1626.2 | 1584.8 | 1748.7 | 1889 | 1846.8 |
| 19523 | ENSMUSG00000069248 | 0 | NA | NA | NA | NA | NA | Serpinb6e | serine (or cysteine) peptidase inhibitor, cl | 0 | 0 | 0 | 0 | 0 | 0 |
| 19524 | ENSMUSG00000069255 | 375.9332621 | 0.074376167 | 0.144772627 | 0.51374468 | 0.607430509 | 0.916376001 | Dusp22 | dual specificity phosphatase 22 [Source:M | 404.76 | 323.79 | 369.29 | 379.81 | 395.89 | 382.05 |
| 19525 | ENSMUSG00000069258 | 0 | NA | NA | NA | NA | NA | Prl2b1 | prolactin family 2, subfamily b, member 1 | 0 | 0 | 0 | 0 | 0 | 0 |
| 19526 | ENSMUSG00000069259 | 0 | NA | NA | NA | NA | NA | Prl6a1 | prolactin family 6, subfamily a, member 1 | 0 | 0 | 0 | 0 | 0 | 0 |
| 19527 | ENSMUSG00000069265 | 2.196266608 | -1.137207902 | 1.522807659 | -0.746783676 | 0.455194151 | NA | H3c1 | H3 clustered histone 1 [Source:MGI Symb | 4.0155 | 1.0279 | 4.025 | 1.928 | 2.1812 | 0 |
| 19528 | ENSMUSG00000069266 | 0 | NA | NA | NA | NA | NA | H4c2 | H4 clustered histone 2 [Source:MGI Symb | 0 | 0 | 0 | 0 | 0 | 0 |
| 19529 | ENSMUSG00000069267 | 0.756734126 | -0.422689986 | 2.665959267 | -0.158550804 | 0.874022791 | NA | H3c2 | H3 clustered histone 2 [Source:MGI Symb | 1.6062 | 0 | 1.0062 | 1.928 | 0 | 0 |
| 19530 | ENSMUSG00000069268 | 0.993467639 | 0.162634227 | 2.224152761 | 0.073121878 | 0.941709132 | NA | H2bc7 | H2B clustered histone 7 [Source:MGI Sym | 0.8031 | 0 | 2.0125 | 0.964 | 2.1812 | 0 |
| 19531 | ENSMUSG00000069270 | 5.007100764 | 0.39960055 | 1.029564435 | 0.388125829 | 0.697922919 | NA | H2ac6 | H2A clustered histone 6 [Source:MGI Sym | 4.8186 | 1.0279 | 7.0437 | 3.8559 | 3.2719 | 10.025 |
| 19532 | ENSMUSG00000069272 | 4.519028803 | 0.178406715 | 1.055770948 | 0.168982406 | 0.86581048 | NA | H2ac8 | H2A clustered histone 8 [Source:MGI Sym | 5.6217 | 2.0558 | 5.0312 | 6.7478 | 6.5437 | 1.1138 |
| 19533 | ENSMUSG00000069273 | 2.703182725 | -1.473490017 | 1.420965188 | -1.036964191 | 0.299752546 | NA | H3c6 | H3 clustered histone 6 [Source:MGI Symb | 0.8031 | 3.0838 | 8.05 | 0.964 | 1.0906 | 2.2277 |
| 19534 | ENSMUSG00000069274 | 0.401551119 | -2.072982726 | 4.008251979 | -0.517178744 | 0.605031388 | NA | H4c6 | H4 clustered histone 6 [Source:MGI Symb | 2.4093 | 0 | 0 | 0 | 0 | 0 |
| 19535 | ENSMUSG00000069280 | 0 | NA | NA | NA | NA | NA | Vmn1r223 | vomeronasal 1 receptor 223 [Source:MGI | 0 | 0 | 0 | 0 | 0 | 0 |
| 19536 | ENSMUSG00000069289 | 0 | NA | NA | NA | NA | NA | Vmn1r203 | vomeronasal 1 receptor 203 [Source:MGI | 0 | 0 | 0 | 0 | 0 | 0 |
| 19537 | ENSMUSG00000069292 | 0 | NA | NA | NA | NA | NA | Vmn1r199 | vomeronasal 1 receptor 199 [Source:MGI | 0 | 0 | 0 | 0 | 0 | 0 |
| 19538 | ENSMUSG00000069294 | 0 | NA | NA | NA | NA | NA | Vmn1r197 | vomeronasal 1 receptor 197 [Source:MGI | 0 | 0 | 0 | 0 | 0 | 0 |
| 19539 | ENSMUSG00000069295 | 0 | NA | NA | NA | NA | NA | Vmn1r196 | vomeronasal 1 receptor 196 [Source:MGI | 0 | 0 | 0 | 0 | 0 | 0 |
| 19540 | ENSMUSG00000069296 | 0 | NA | NA | NA | NA | NA | Vmn1r195 | vomeronasal 1 receptor 195 [Source:MGI | 0 | 0 | 0 | 0 | 0 | 0 |

| | | | | | | | | | | | | | | | |
|---|---|---|---|---|---|---|---|---|---|---|---|---|---|---|---|
| 19541 | ENSMUSG00000069297 | 0 | NA | NA | NA | NA | NA | Vmn1r194 | vomeronasal 1 receptor 194 [Source:MGI | 0 | 0 | 0 | 0 | 0 | 0 |
| 19542 | ENSMUSG00000069299 | 0 | NA | NA | NA | NA | NA | Vmn1r188 | vomeronasal 1 receptor 188 [Source:MGI | 0 | 0 | 0 | 0 | 0 | 0 |
| 19543 | ENSMUSG00000069300 | 0.706178709 | 0.10198885 | 3.03884937 | 0.033561667 | 0.97322669 | NA | H2bc11 | H2B clustered histone 11 [Source:MGI Syr | 0 | 2.0558 | 0 | 0 | 2.1812 | 0 |
| 19544 | ENSMUSG00000069301 | 1.04956753 | 2.382147202 | 2.398712306 | 0.993094168 | 0.320664066 | NA | H2ac11 | H2A clustered histone 11 [Source:MGI Syr | 0 | 1.0279 | 0 | 1.928 | 0 | 3.3415 |
| 19545 | ENSMUSG00000069302 | 0.267700746 | -1.497481649 | 4.057001436 | -0.369110456 | 0.712045396 | NA | H2ac12 | H2A clustered histone 12 [Source:MGI Syr | 1.6062 | 0 | 0 | 0 | 0 | 0 |
| 19546 | ENSMUSG00000069303 | 0 | NA | NA | NA | NA | NA | H2bc24 | H2B clustered histone 24 [Source:MGI Syr | 0 | 0 | 0 | 0 | 0 | 0 |
| 19547 | ENSMUSG00000069305 | 0.888465872 | 0.734664603 | 2.541756127 | 0.289038195 | 0.772552149 | NA | H4c18 | H4 clustered histone 18 [Source:MGI Sym | 0 | 0 | 2.0125 | 0 | 1.0906 | 2.2277 |
| 19548 | ENSMUSG00000069306 | 1.40469289 | 0.955825871 | 2.192946384 | 0.435863767 | 0.662935575 | NA | H4c17 | H4 clustered histone 17 [Source:MGI Sym | 0.8031 | 2.0558 | 0 | 0 | 0 | 5.5692 |
| 19549 | ENSMUSG00000069307 | 0.331982614 | 0.16514309 | 3.986751476 | 0.041422971 | 0.966958701 | NA | H2bc23 | H2B clustered histone 23 [Source:MGI Syr | 0 | 1.0279 | 0 | 0.964 | 0 | 0 |
| 19550 | ENSMUSG00000069308 | 0.181769715 | 1.126924175 | 4.080472857 | 0.276174898 | 0.782413736 | NA | H2bc22 | H2B clustered histone 22 [Source:MGI Syr | 0 | 0 | 0 | 0 | 1.0906 | 0 |
| 19551 | ENSMUSG00000069309 | 0 | NA | NA | NA | NA | NA | H2ac22 | H2A clustered histone 22 [Source:MGI Syr | 0 | 0 | 0 | 0 | 0 | 0 |
| 19552 | ENSMUSG00000069310 | 0.669251866 | -2.807208097 | 3.391018549 | -0.827836255 | 0.407763239 | NA | H3c3 | H3 clustered histone 3 [Source:MGI Symb | 4.0155 | 0 | 0 | 0 | 0 | 0 |
| 19553 | ENSMUSG00000069315 | 0 | NA | NA | NA | NA | NA | Gm7731 | predicted gene 7731 [Source:MGI Symbol | 0 | 0 | 0 | 0 | 0 | 0 |
| 19554 | ENSMUSG00000069324 | 229.2022327 | -0.270137149 | 0.214843973 | -1.257364335 | 0.208621738 | 0.653455953 | Gm5096 | predicted gene 5096 [Source:MGI Symbol | 231.29 | 279.59 | 240.49 | 143.63 | 255.2 | 225 |
| 19555 | ENSMUSG00000069355 | 0 | NA | NA | NA | NA | NA | Gm5152 | predicted gene 5152 [Source:MGI Symbol | 0 | 0 | 0 | 0 | 0 | 0 |
| 19556 | ENSMUSG00000069372 | 0 | NA | NA | NA | NA | NA | Ctxn3 | cortexin 3 [Source:MGI Symbol;Acc:MGI:3 | 0 | 0 | 0 | 0 | 0 | 0 |
| 19557 | ENSMUSG00000069376 | 0 | NA | NA | NA | NA | NA | Gm5507 | predicted gene 5507 [Source:MGI Symbol | 0 | 0 | 0 | 0 | 0 | 0 |
| 19558 | ENSMUSG00000069378 | 1.494676241 | -0.811671562 | 1.916581503 | -0.423499632 | 0.671930767 | NA | Prdm6 | PR domain containing 6 [Source:MGI Sym | 1.6062 | 3.0838 | 1.0062 | 0 | 3.2719 | 0 |
| 19559 | ENSMUSG00000069379 | 0.503122293 | -2.366675966 | 3.785596405 | -0.625179156 | 0.531853481 | NA | Gm4950 | predicted pseudogene 4950 [Source:MGI | 0 | 0 | 3.0187 | 0 | 0 | 0 |
| 19560 | ENSMUSG00000069385 | 0 | NA | NA | NA | NA | NA | Gm10267 | predicted gene 10267 [Source:MGI Symbo | 0 | 0 | 0 | 0 | 0 | 0 |
| 19561 | ENSMUSG00000069390 | 0 | NA | NA | NA | NA | NA | Olfr707 | olfactory receptor 707 [Source:MGI Symb | 0 | 0 | 0 | 0 | 0 | 0 |
| 19562 | ENSMUSG00000069421 | 0 | NA | NA | NA | NA | NA | Olfr810 | olfactory receptor 810 [Source:MGI Symb | 0 | 0 | 0 | 0 | 0 | 0 |
| 19563 | ENSMUSG00000069430 | 0 | NA | NA | NA | NA | NA | Olfr9 | olfactory receptor 9 [Source:MGI Symbol; | 0 | 0 | 0 | 0 | 0 | 0 |
| 19564 | ENSMUSG00000069439 | 0 | NA | NA | NA | NA | NA | Gm8444 | predicted gene 8444 [Source:MGI Symbol | 0 | 0 | 0 | 0 | 0 | 0 |
| 19565 | ENSMUSG00000069441 | 0 | NA | NA | NA | NA | NA | Dsg1a | desmoglein 1 alpha [Source:MGI Symbol;/ | 0 | 0 | 0 | 0 | 0 | 0 |
| 19566 | ENSMUSG00000069456 | 2002.98858 | -0.266424607 | 0.126595523 | -2.104534186 | 0.035331876 | 0.25220602 | Rdh16 | retinol dehydrogenase 16 [Source:MGI Sy | 2215 | 2441.3 | 1906.8 | 2052.3 | 1738.4 | 1664.1 |
| 19567 | ENSMUSG00000069475 | 0 | NA | NA | NA | NA | NA | Gm6020 | predicted gene 6020 [Source:MGI Symbol | 0 | 0 | 0 | 0 | 0 | 0 |
| 19568 | ENSMUSG00000069476 | 0 | NA | NA | NA | NA | NA | Zfp616 | zinc finger protein 616 [Source:MGI Symb | 0 | 0 | 0 | 0 | 0 | 0 |
| 19569 | ENSMUSG00000069495 | 212.2083466 | -0.039465853 | 0.16605914 | -0.237661434 | 0.812143698 | 0.96653525 | Epc2 | enhancer of polycomb homolog 2 [Source | 203.18 | 219.97 | 222.38 | 202.44 | 219.21 | 206.06 |
| 19570 | ENSMUSG00000069515 | 3.486729452 | -0.91059562 | 1.222262885 | -0.745007994 | 0.456266887 | NA | Lyz1 | lysozyme 1 [Source:MGI Symbol;Acc:MGI: | 1.6062 | 3.0838 | 9.0562 | 3.8559 | 1.0906 | 2.2277 |
| 19571 | ENSMUSG00000069516 | 556.5875405 | -1.233886654 | 0.405996383 | -3.039156762 | 0.002372414 | 0.039642097 | Lyz2 | lysozyme 2 [Source:MGI Symbol;Acc:MGI: | 765.36 | 1092.7 | 485.01 | 246.78 | 309.74 | 439.97 |
| 19572 | ENSMUSG00000069518 | 0 | NA | NA | NA | NA | NA | Gm10271 | predicted gene 10271 [Source:MGI Symbo | 0 | 0 | 0 | 0 | 0 | 0 |
| 19573 | ENSMUSG00000069520 | 2671.058892 | 0.213081092 | 0.110501396 | 1.928311319 | 0.053816415 | 0.326281119 | Tmem19 | transmembrane protein 19 [Source:MGI S | 2439.8 | 2433.1 | 2548.8 | 2540.1 | 2840 | 3224.6 |
| 19574 | ENSMUSG00000069539 | 803.8778589 | -0.182844364 | 0.115004609 | -1.589887276 | 0.111860216 | 0.490732838 | Scyl2 | SCY1-like 2 (S. cerevisiae) [Source:MGI Syr | 822.38 | 928.21 | 814.05 | 740.33 | 778.7 | 739.59 |
| 19575 | ENSMUSG00000069540 | 4.403055653 | -0.032100987 | 1.098256876 | -0.029229033 | 0.976681926 | NA | Gm4925 | predicted gene 4925 [Source:MGI Symbol | 3.2124 | 2.0558 | 8.05 | 0.964 | 5.4531 | 6.6831 |
| 19576 | ENSMUSG00000069554 | 0.133850373 | -0.796629176 | 4.080472857 | -0.195229623 | 0.84521319 | NA | I830134H01I | RIKEN cDNA I830134H01 gene [Source:M( | 0.8031 | 0 | 0 | 0 | 0 | 0 |
| 19577 | ENSMUSG00000069565 | 914.8423837 | 0.217315741 | 0.129744112 | 1.674956475 | 0.093942772 | 0.451274743 | Dazap1 | DAZ associated protein 1 [Source:MGI Syr | 938.02 | 844.95 | 753.68 | 875.29 | 1055.7 | 1021.4 |
| 19578 | ENSMUSG00000069581 | 3.809696798 | 0.936252901 | 1.135245961 | 0.824713704 | 0.40953415 | NA | Tspear | thrombospondin type laminin G domain a | 0.8031 | 4.1117 | 3.0187 | 3.8559 | 3.2719 | 7.7969 |
| 19579 | ENSMUSG00000069582 | 0 | NA | NA | NA | NA | NA | Krtap10-4 | keratin associated protein 10-4 [Source:M | 0 | 0 | 0 | 0 | 0 | 0 |
| 19580 | ENSMUSG00000069583 | 0 | NA | NA | NA | NA | NA | Krtap12-1 | keratin associated protein 12-1 [Source:M | 0 | 0 | 0 | 0 | 0 | 0 |
| 19581 | ENSMUSG00000069584 | 0 | NA | NA | NA | NA | NA | Gm10272 | predicted gene 10272 [Source:MGI Symbo | 0 | 0 | 0 | 0 | 0 | 0 |
| 19582 | ENSMUSG00000069588 | 0 | NA | NA | NA | NA | NA | Gm11733 | predicted gene 11733 [Source:MGI Symbo | 0 | 0 | 0 | 0 | 0 | 0 |
| 19583 | ENSMUSG00000069601 | 276.2499101 | 0.34012052 | 0.205946398 | 1.651500211 | 0.098636478 | 0.463186297 | Ank3 | ankyrin 3, epithelial [Source:MGI Symbol;/ | 256.19 | 270.34 | 204.27 | 230.39 | 306.46 | 389.85 |
| 19584 | ENSMUSG00000069607 | 0.331982614 | 0.16514309 | 3.986751476 | 0.041422971 | 0.966958701 | NA | Cd300ld3 | CD300 molecule like family member D3 [S | 0 | 1.0279 | 0 | 0.964 | 0 | 0 |
| 19585 | ENSMUSG00000069609 | 0 | NA | NA | NA | NA | NA | Cd300ld4 | CD300 molecule like family member D4 [S | 0 | 0 | 0 | 0 | 0 | 0 |
| 19586 | ENSMUSG00000069620 | 0 | NA | NA | NA | NA | NA | Gm5250 | predicted gene 5250 [Source:MGI Symbol | 0 | 0 | 0 | 0 | 0 | 0 |
| 19587 | ENSMUSG00000069622 | 0.788756256 | -1.683618556 | 2.620704039 | -0.64242987 | 0.52059411 | NA | Gm10273 | predicted pseudogene 10273 [Source:MG | 1.6062 | 0 | 2.0125 | 0 | 0 | 1.1138 |
| 19588 | ENSMUSG00000069631 | 55.00302573 | 0.034301264 | 0.317035976 | 0.108193601 | 0.913842121 | 0.98625062 | Strada | STE20-related kinase adaptor alpha [Sour | 60.233 | 40.089 | 62.387 | 48.199 | 55.622 | 63.489 |
| 19589 | ENSMUSG00000069633 | 579.4142559 | 0.116045618 | 0.127532056 | 0.909932936 | 0.362857879 | 0.801768464 | Pex11g | peroxisomal biogenesis factor 11 gamma | | 591.89 | 510.88 | 564.5 | 557.18 | 623.83 | 628.21 |
| 19590 | ENSMUSG00000069662 | 213.0339152 | -1.44367279 | 0.219169517 | -6.587014522 | 4.49E-11 | 1.35E-08 | Marcks | myristoylated alanine rich protein kinase ( | 281.09 | 388.55 | 264.64 | 94.47 | 106.88 | 142.57 |
| 19591 | ENSMUSG00000069668 | 1.54910259 | -1.032840645 | 1.75256384 | -0.589331254 | 0.555639083 | NA | Sult3a1 | sulfotransferase family 3A, member 1 [Sou | 3.2124 | 1.0279 | 2.0125 | 1.928 | 0 | 1.1138 |
| 19592 | ENSMUSG00000069670 | 3.797953991 | -0.512838343 | 1.227389006 | -0.417828692 | 0.676072371 | NA | Nkain2 | Na+/K+ transporting ATPase interacting 2 | 7.2279 | 4.1117 | 2.0125 | 2.8919 | 6.5437 | 0 |
| 19593 | ENSMUSG00000069678 | 97.29082472 | 0.259443697 | 0.267348788 | 0.970431542 | 0.331831433 | 0.774747658 | Pcgf1 | polycomb group ring finger 1 [Source:MG | 95.569 | 77.094 | 92.575 | 88.686 | 87.249 | 142.57 |
| 19594 | ENSMUSG00000069682 | 18.42857779 | 0.126694368 | 0.496358658 | 0.255247623 | 0.798531851 | 0.962424862 | Gm10275 | predicted pseudogene 10275 [Source:MG | 19.274 | 15.419 | 18.112 | 20.244 | 16.359 | 21.163 |
| 19595 | ENSMUSG00000069706 | 0 | NA | NA | NA | NA | NA | Taar5 | trace amine-associated receptor 5 [Source | 0 | 0 | 0 | 0 | 0 | 0 |
| 19596 | ENSMUSG00000069707 | 0 | NA | NA | NA | NA | NA | Taar4 | trace amine-associated receptor 4 [Source | 0 | 0 | 0 | 0 | 0 | 0 |

| | | | | | | | | | | | | | | | |
|---|---|---|---|---|---|---|---|---|---|---|---|---|---|---|---|
| 19597 | ENSMUSG00000069708 | 0 | NA | NA | NA | NA | NA | Taar3 | trace amine-associated receptor 3 [Source | 0 | 0 | 0 | 0 | 0 | 0 |
| 19598 | ENSMUSG00000069712 | 0 | NA | NA | NA | NA | NA | 4930444G20 | RIKEN cDNA 4930444G20 gene [Source:M | 0 | 0 | 0 | 0 | 0 | 0 |
| 19599 | ENSMUSG00000069713 | 4.641421062 | -0.478277312 | 0.989673757 | -0.483267651 | 0.628905706 | NA | 4933406P04 | RIKEN cDNA 4933406P04 gene [Source:M | 4.0155 | 7.1954 | 5.0312 | 3.8559 | 2.1812 | 5.5692 |
| 19600 | ENSMUSG00000069717 | 0 | NA | NA | NA | NA | NA | Gm11568 | predicted gene 11568 [Source:MGI Symbo | 0 | 0 | 0 | 0 | 0 | 0 |
| 19601 | ENSMUSG00000069718 | 0 | NA | NA | NA | NA | NA | Gm11563 | predicted gene 11563 [Source:MGI Symbo | 0 | 0 | 0 | 0 | 0 | 0 |
| 19602 | ENSMUSG00000069720 | 0 | NA | NA | NA | NA | NA | 4930572O03 | RIKEN cDNA 4930572O03 gene [Source:M | 0 | 0 | 0 | 0 | 0 | 0 |
| 19603 | ENSMUSG00000069721 | 0 | NA | NA | NA | NA | NA | Krtap3-2 | keratin associated protein 3-2 [Source:MG | 0 | 0 | 0 | 0 | 0 | 0 |
| 19604 | ENSMUSG00000069722 | 0 | NA | NA | NA | NA | NA | Krtap3-3 | keratin associated protein 3-3 [Source:MG | 0 | 0 | 0 | 0 | 0 | 0 |
| 19605 | ENSMUSG00000069727 | 65.62486931 | 0.331211731 | 0.282716726 | 1.171532139 | 0.241384947 | 0.691388521 | Zfp975 | zinc finger protein 975 [Source:MGI Symb | 56.217 | 62.703 | 55.343 | 59.767 | 83.978 | 75.741 |
| 19606 | ENSMUSG00000069729 | 527.1646173 | 0.018483188 | 0.136928412 | 0.134984314 | 0.892624276 | 0.982512308 | Arid1b | AT rich interactive domain 1B (SWI-like) [S | 497.92 | 548.91 | 524.25 | 460.78 | 548.58 | 582.54 |
| 19607 | ENSMUSG00000069733 | 55.07615755 | -0.507892654 | 0.312599779 | -1.624737723 | 0.104218456 | 0.475225645 | Ube2u | ubiquitin-conjugating enzyme E2U (putati | 69.067 | 52.424 | 72.45 | 48.199 | 37.081 | 51.237 |
| 19608 | ENSMUSG00000069743 | 50.48282379 | -0.340738354 | 0.330979399 | -1.029485084 | 0.303251786 | 0.752484788 | Zfp820 | zinc finger protein 820 [Source:MGI Symb | 53.808 | 45.228 | 70.437 | 53.019 | 42.534 | 37.871 |
| 19609 | ENSMUSG00000069744 | 1599.873069 | 0.133089139 | 0.100572821 | 1.323311181 | 0.185731915 | 0.621570203 | Psmb3 | proteasome (prosome, macropain) subun | 1623.1 | 1422.6 | 1531.5 | 1645.5 | 1704.6 | 1671.9 |
| 19610 | ENSMUSG00000069755 | 96.4692789 | -0.147028695 | 0.239193286 | -0.614685709 | 0.5387623 | 0.8916993 | Zfp125 | zinc finger protein 125 [Source:MGI Symb | 110.83 | 94.568 | 98.612 | 95.434 | 102.52 | 76.855 |
| 19611 | ENSMUSG00000069763 | 9.54549391 | -3.337839544 | 0.979191816 | -3.408769855 | 0.000652565 | 0.015126124 | Tmem100 | transmembrane protein 100 [Source:MGI | 10.44 | 28.782 | 13.081 | 3.8559 | 0 | 1.1138 |
| 19612 | ENSMUSG00000069769 | 355.424259 | 0.097831578 | 0.168575392 | 0.580343172 | 0.561683219 | 0.900565925 | Msi2 | musashi RNA-binding protein 2 [Source:M | 297.15 | 363.88 | 369.29 | 312.33 | 357.72 | 432.17 |
| 19613 | ENSMUSG00000069785 | 0 | NA | NA | NA | NA | NA | Gm11444 | predicted gene 11444 [Source:MGI Symbo | 0 | 0 | 0 | 0 | 0 | 0 |
| 19614 | ENSMUSG00000069792 | 83.08831666 | -1.202574425 | 0.338583891 | -3.551776852 | 0.000382639 | 0.010379287 | Wfdc17 | WAP four-disulfide core domain 17 [Sourc | 112.43 | 152.13 | 82.512 | 27.955 | 58.893 | 64.603 |
| 19615 | ENSMUSG00000069793 | 88.35159851 | -0.222816842 | 0.259869935 | -0.857416778 | 0.39121459 | 0.820195366 | Slfn9 | schlafen 9 [Source:MGI Symbol;Acc:MGI:2 | 117.25 | 80.178 | 87.543 | 80.01 | 91.612 | 73.514 |
| 19616 | ENSMUSG00000069796 | 0 | NA | NA | NA | NA | NA | Gm11426 | predicted gene 11426 [Source:MGI Symbo | 0 | 0 | 0 | 0 | 0 | 0 |
| 19617 | ENSMUSG00000069804 | 3.404669353 | -0.525400148 | 1.354622283 | -0.387857305 | 0.698121635 | NA | Gm10277 | predicted gene 10277 [Source:MGI Symbo | 0 | 3.0838 | 9.0562 | 3.8559 | 1.0906 | 3.3415 |
| 19618 | ENSMUSG00000069805 | 20510.26872 | -0.000747263 | 0.087521529 | -0.008538051 | 0.993187703 | 0.99859176 | Fbp1 | fructose bisphosphatase 1 [Source:MGI Sy | 19604 | 19871 | 22073 | 20361 | 20351 | 20802 |
| 19619 | ENSMUSG00000069806 | 18.94636359 | -0.616443918 | 0.633364239 | -0.973285008 | 0.330411671 | 0.774747658 | Cacng7 | calcium channel, voltage-dependent, gam | 32.124 | 25.698 | 11.069 | 25.063 | 15.269 | 4.4554 |
| 19620 | ENSMUSG00000069808 | 59.63676548 | -0.333387792 | 0.31805112 | -1.048220777 | 0.2945369 | 0.745164683 | Tlcd3a | TLC domain containing 3A [Source:MGI Sy | 71.476 | 46.256 | 81.506 | 46.271 | 61.075 | 51.237 |
| 19621 | ENSMUSG00000069814 | 3.146304483 | -0.399198035 | 1.250305582 | -0.319280375 | 0.749513913 | NA | Ccdc92b | coiled-coil domain containing 92B [Source | 5.6217 | 4.1117 | 1.0062 | 4.8199 | 1.0906 | 2.2277 |
| 19622 | ENSMUSG00000069816 | 0 | NA | NA | NA | NA | NA | Olfr23 | olfactory receptor 23 [Source:MGI Symbo | 0 | 0 | 0 | 0 | 0 | 0 |
| 19623 | ENSMUSG00000069818 | 0 | NA | NA | NA | NA | NA | Olfr390 | olfactory receptor 390 [Source:MGI Symb | 0 | 0 | 0 | 0 | 0 | 0 |
| 19624 | ENSMUSG00000069823 | 0 | NA | NA | NA | NA | NA | Olfr1 | olfactory receptor 1 [Source:MGI Symbol; | 0 | 0 | 0 | 0 | 0 | 0 |
| 19625 | ENSMUSG00000069824 | 0 | NA | NA | NA | NA | NA | Olfr377-ps1 | olfactory receptor 377, pseudogene 1 [Sou | 0 | 0 | 0 | 0 | 0 | 0 |
| 19626 | ENSMUSG00000069825 | 0 | NA | NA | NA | NA | NA | Gm49340 | predicted gene, 49340 [Source:MGI Symb | 0 | 0 | 0 | 0 | 0 | 0 |
| 19627 | ENSMUSG00000069830 | 1.738175486 | -1.1958362 | 1.82076097 | -0.656778248 | 0.511323513 | NA | Nlrp1a | NLR family, pyrin domain containing 1A [S | 3.2124 | 0 | 4.025 | 0.964 | 0 | 2.2277 |
| 19628 | ENSMUSG00000069833 | 309.1708188 | -0.234234597 | 0.15999718 | -1.463992032 | 0.143196125 | 0.552906769 | Ahnak | AHNAK nucleoprotein (desmoyokin) [Sour | 297.15 | 384.44 | 322 | 302.69 | 279.2 | 269.55 |
| 19629 | ENSMUSG00000069835 | 238.3239467 | 0.012318872 | 0.170535674 | 0.072236336 | 0.942413829 | 0.989839054 | Sat2 | spermidine/spermine N1-acetyl transfera | 251.37 | 200.44 | 259.61 | 231.35 | 242.12 | 245.05 |
| 19630 | ENSMUSG00000069844 | 277.7959644 | -0.050069935 | 0.151307787 | -0.33091446 | 0.740709099 | 0.948792385 | Sco1 | SCO1 cytochrome c oxidase assembly prot | 301.16 | 273.43 | 272.69 | 266.06 | 268.29 | 285.14 |
| 19631 | ENSMUSG00000069855 | 3.118620555 | 0.879016177 | 1.481008982 | 0.593525216 | 0.552829712 | NA | Slc47a2 | solute carrier family 47, member 2 [Sourc | 1.6062 | 0 | 5.0312 | 7.7118 | 4.3625 | 0 |
| 19632 | ENSMUSG00000069862 | 4.868019217 | -0.066287548 | 1.003377744 | -0.0660644 | 0.947326554 | NA | Rps12-ps9 | ribosomal protein S12, pseudogene 9 [Sou | 4.8186 | 7.1954 | 3.0187 | 8.6758 | 3.2719 | 2.2277 |
| 19633 | ENSMUSG00000069867 | 0 | NA | NA | NA | NA | NA | Pabpn1l | poly(A)binding protein nuclear 1-like [Sou | 0 | 0 | 0 | 0 | 0 | 0 |
| 19634 | ENSMUSG00000069873 | 0 | NA | NA | NA | NA | NA | 4930438A08 | RIKEN cDNA 4930438A08 gene [Source:M | 0 | 0 | 0 | 0 | 0 | 0 |
| 19635 | ENSMUSG00000069874 | 859.3936657 | 0.466283286 | 0.150769298 | 3.092693892 | 0.001983486 | 0.034648724 | Irgm2 | immunity-related GTPase family M memb | 823.18 | 713.37 | 627.9 | 1112.4 | 816.87 | 1062.6 |
| 19636 | ENSMUSG00000069890 | 0 | NA | NA | NA | NA | NA | 1700024J04R | RIKEN cDNA 1700024J04 gene [Source:MG | 0 | 0 | 0 | 0 | 0 | 0 |
| 19637 | ENSMUSG00000069892 | 15.2301242 | -1.547020728 | 0.609791145 | -2.536968172 | 0.011181712 | 0.117452637 | 9930111J21R | RIKEN cDNA 9930111J21 gene 2 [Source:N | 20.078 | 32.893 | 15.094 | 6.7478 | 6.5437 | 10.025 |
| 19638 | ENSMUSG00000069893 | 4.641905839 | -1.392590351 | 1.276996878 | -1.090519777 | 0.275484246 | NA | 9930111J21R | RIKEN cDNA 9930111J21 gene 1 [Source:N | 8.8341 | 7.1954 | 4.025 | 0 | 0 | 7.7969 |
| 19639 | ENSMUSG00000069895 | 411.0804331 | 0.076816459 | 0.141787198 | 0.541772885 | 0.587974973 | 0.910798643 | Atxn1l | ataxin 1-like [Source:MGI Symbol;Acc:MG | 369.43 | 424.53 | 407.53 | 464.64 | 389.35 | 411.01 |
| 19640 | ENSMUSG00000069899 | 67.87721663 | -0.125638931 | 0.274031249 | -0.458483956 | 0.646604787 | 0.925643431 | Gm12166 | predicted gene 12166 [Source:MGI Symbo | 81.113 | 65.787 | 65.406 | 70.37 | 59.984 | 64.603 |
| 19641 | ENSMUSG00000069910 | 5.490607715 | -0.846250486 | 0.941547974 | -0.89878637 | 0.368766461 | 0.806506847 | Spdl1 | spindle apparatus coiled-coil protein 1 [Sc | 8.8341 | 7.1954 | 5.0312 | 1.928 | 3.2719 | 6.6831 |
| 19642 | ENSMUSG00000069911 | 0 | NA | NA | NA | NA | NA | Insyn2b | inhibitory synaptic factor family member 2 | 0 | 0 | 0 | 0 | 0 | 0 |
| 19643 | ENSMUSG00000069913 | 0 | NA | NA | NA | NA | NA | Anp32-ps | acidic (leucine-rich) nuclear phosphoprote | 0 | 0 | 0 | 0 | 0 | 0 |
| 19644 | ENSMUSG00000069917 | 180.5610118 | 0.279931175 | 0.232349402 | 1.204785435 | 0.228286144 | 0.674708584 | Hba-a2 | hemoglobin alpha, adult chain 2 [Source:N | 126.89 | 164.47 | 198.23 | 160.98 | 179.95 | 252.84 |
| 19645 | ENSMUSG00000069919 | 1338.33003 | 0.005720056 | 0.184224411 | 0.031049392 | 0.97523015 | 0.996378391 | Hba-a1 | hemoglobin alpha, adult chain 1 [Source:N | 1036 | 1181.1 | 1791.1 | 1498 | 1108.1 | 1415.7 |
| 19646 | ENSMUSG00000069920 | 9.452229499 | -0.775318293 | 0.704446475 | -1.100606392 | 0.271068002 | 0.726733566 | B3gnt9 | UDP-GlcNAc:betaGal beta-1,3-N-acetylglu | 10.44 | 13.363 | 12.075 | 8.6758 | 4.3625 | 7.7969 |
| 19647 | ENSMUSG00000069922 | 32607.44766 | 0.438159212 | 0.084806538 | 5.1665735 | 2.38E-07 | 2.33E-05 | Ces3a | carboxylesterase 3A [Source:MGI Symbol; | 26461 | 29342 | 27278 | 36613 | 38312 | 37638 |
| 19648 | ENSMUSG00000069925 | 0 | NA | NA | NA | NA | NA | 4932416K20 | RIKEN cDNA 4932416K20 gene [Source:M | 0 | 0 | 0 | 0 | 0 | 0 |
| 19649 | ENSMUSG00000069939 | 1.355869549 | 1.030873258 | 1.984610048 | 0.519433659 | 0.603458365 | NA | Gm12070 | predicted gene 12070 [Source:MGI Symbo | 1.6062 | 0 | 1.0062 | 0 | 2.1812 | 3.3415 |
| 19650 | ENSMUSG00000069962 | 0 | NA | NA | NA | NA | NA | Gm5356 | predicted pseudogene 5356 [Source:MGI | 0 | 0 | 0 | 0 | 0 | 0 |
| 19651 | ENSMUSG00000069971 | 0 | NA | NA | NA | NA | NA | 4933402J07R | RIKEN cDNA 4933402J07 gene [Source:MG | 0 | 0 | 0 | 0 | 0 | 0 |
| 19652 | ENSMUSG00000069972 | 7.376204681 | 0.347553589 | 0.843182755 | 0.412192478 | 0.68019835 | 0.934384556 | Rps13-ps2 | ribosomal protein S13, pseudogene 2 [Sou | 3.2124 | 11.307 | 5.0312 | 4.8199 | 7.6343 | 12.252 |

| | | | | | | | | | | | | | | | |
|---|---|---|---|---|---|---|---|---|---|---|---|---|---|---|---|
| 19653 | ENSMUSG00000069986 | 0 | NA | NA | NA | NA | NA | Gm8054 | predicted pseudogene 8054 [Source:MGI | 0 | 0 | 0 | 0 | 0 | 0 |
| 19654 | ENSMUSG00000069996 | 0 | NA | NA | NA | NA | NA | Gm7125 | predicted pseudogene 7125 [Source:MGI | 0 | 0 | 0 | 0 | 0 | 0 |
| 19655 | ENSMUSG00000069998 | 0 | NA | NA | NA | NA | NA | Olfr372 | olfactory receptor 372 [Source:MGI Symb | 0 | 0 | 0 | 0 | 0 | 0 |
| 19656 | ENSMUSG00000070000 | 3.818214079 | -1.398423261 | 1.121377141 | -1.247058826 | 0.212375929 | NA | Fcho1 | FCH domain only 1 [Source:MGI Symbol;A | 6.4248 | 5.1396 | 5.0312 | 1.928 | 3.2719 | 1.1138 |
| 19657 | ENSMUSG00000070002 | 674.1908197 | 0.299696465 | 0.13284074 | 2.256058389 | 0.024066975 | 0.197338277 | Ell | elongation factor RNA polymerase II [Sour | 665.77 | 567.41 | 579.6 | 824.2 | 667.46 | 740.71 |
| 19658 | ENSMUSG00000070003 | 79.26676145 | -1.064460243 | 0.265432344 | -4.010288378 | 6.06E-05 | 0.002436567 | Ssbp4 | single stranded DNA binding protein 4 [So | 102.8 | 125.41 | 93.581 | 49.163 | 54.531 | 50.123 |
| 19659 | ENSMUSG00000070023 | 0 | NA | NA | NA | NA | NA | Gm10283 | predicted gene 10283 [Source:MGI Symbo | 0 | 0 | 0 | 0 | 0 | 0 |
| 19660 | ENSMUSG00000070025 | 0.294513347 | 0.165143496 | 4.080472857 | 0.040471657 | 0.967717103 | NA | Gm10284 | predicted pseudogene 10284 [Source:MG | 0.8031 | 0 | 0 | 0.964 | 0 | 0 |
| 19661 | ENSMUSG00000070031 | 41.09374628 | -0.82578743 | 0.365854666 | -2.257146094 | 0.023998947 | 0.197242243 | Sp140 | Sp140 nuclear body protein [Source:MGI S | 61.839 | 54.48 | 41.256 | 31.811 | 34.9 | 22.277 |
| 19662 | ENSMUSG00000070034 | 19.1935471 | -0.768393525 | 0.596712075 | -1.287712378 | 0.197846103 | 0.639811551 | Sp110 | Sp110 nuclear body protein [Source:MGI S | 29.715 | 28.782 | 14.087 | 19.28 | 4.3625 | 18.935 |
| 19663 | ENSMUSG00000070044 | 479.4447296 | 0.148102022 | 0.147429229 | 1.004563497 | 0.315107081 | 0.761215823 | Fam149a | family with sequence similarity 149, meml | 429.66 | 396.78 | 538.34 | 494.52 | 502.78 | 514.6 |
| 19664 | ENSMUSG00000070047 | 550.8933655 | 0.073283748 | 0.147906615 | 0.49547309 | 0.620266213 | 0.917977899 | Fat1 | FAT atypical cadherin 1 [Source:MGI Symt | 474.63 | 547.88 | 588.65 | 532.12 | 507.14 | 654.94 |
| 19665 | ENSMUSG00000070056 | 532.9221091 | 0.03222543 | 0.15034112 | 0.214348743 | 0.83027511 | 0.9702546 | Mfhas1 | malignant fibrous histiocytoma amplified | 458.57 | 487.23 | 635.95 | 542.72 | 555.12 | 517.94 |
| 19666 | ENSMUSG00000070063 | 0 | NA | NA | NA | NA | NA | Snora33 | small nucleolar RNA, H/ACA box 33 [Sourc | 0 | 0 | 0 | 0 | 0 | 0 |
| 19667 | ENSMUSG00000070065 | 0 | NA | NA | NA | NA | NA | NA | NA | 0 | 0 | 0 | 0 | 0 | 0 |
| 19668 | ENSMUSG00000070071 | 0 | NA | NA | NA | NA | NA | Mir465 | microRNA 465 [Source:MGI Symbol;Acc:M | 0 | 0 | 0 | 0 | 0 | 0 |
| 19669 | ENSMUSG00000070072 | 0 | NA | NA | NA | NA | NA | Mir433 | microRNA 433 [Source:MGI Symbol;Acc:M | 0 | 0 | 0 | 0 | 0 | 0 |
| 19670 | ENSMUSG00000070073 | 0 | NA | NA | NA | NA | NA | Mir470 | microRNA 470 [Source:MGI Symbol;Acc:M | 0 | 0 | 0 | 0 | 0 | 0 |
| 19671 | ENSMUSG00000070074 | 0 | NA | NA | NA | NA | NA | Mir484 | microRNA 484 [Source:MGI Symbol;Acc:M | 0 | 0 | 0 | 0 | 0 | 0 |
| 19672 | ENSMUSG00000070075 | 0 | NA | NA | NA | NA | NA | Mir490 | microRNA 490 [Source:MGI Symbol;Acc:M | 0 | 0 | 0 | 0 | 0 | 0 |
| 19673 | ENSMUSG00000070076 | 0 | NA | NA | NA | NA | NA | Mir127 | microRNA 127 [Source:MGI Symbol;Acc:M | 0 | 0 | 0 | 0 | 0 | 0 |
| 19674 | ENSMUSG00000070077 | 0 | NA | NA | NA | NA | NA | Mir491 | microRNA 491 [Source:MGI Symbol;Acc:M | 0 | 0 | 0 | 0 | 0 | 0 |
| 19675 | ENSMUSG00000070078 | 0 | NA | NA | NA | NA | NA | Mir463 | microRNA 463 [Source:MGI Symbol;Acc:M | 0 | 0 | 0 | 0 | 0 | 0 |
| 19676 | ENSMUSG00000070080 | 0 | NA | NA | NA | NA | NA | Mir431 | microRNA 431 [Source:MGI Symbol;Acc:M | 0 | 0 | 0 | 0 | 0 | 0 |
| 19677 | ENSMUSG00000070081 | 0.160662974 | 1.126924175 | 4.080472857 | 0.276174898 | 0.782413736 | NA | Mir505 | microRNA 505 [Source:MGI Symbol;Acc:M | 0 | 0 | 0 | 0.964 | 0 | 0 |
| 19678 | ENSMUSG00000070084 | 0.36353943 | 2.065989254 | 4.034137301 | 0.512126658 | 0.608562368 | NA | Mir486 | microRNA 486 [Source:MGI Symbol;Acc:M | 0 | 0 | 0 | 0 | 2.1812 | 0 |
| 19679 | ENSMUSG00000070099 | 0 | NA | NA | NA | NA | NA | Mir466 | microRNA 466 [Source:MGI Symbol;Acc:M | 0 | 0 | 0 | 0 | 0 | 0 |
| 19680 | ENSMUSG00000070100 | 0 | NA | NA | NA | NA | NA | Mir362 | microRNA 362 [Source:MGI Symbol;Acc:M | 0 | 0 | 0 | 0 | 0 | 0 |
| 19681 | ENSMUSG00000070101 | 0 | NA | NA | NA | NA | NA | Mir341 | microRNA 341 [Source:MGI Symbol;Acc:M | 0 | 0 | 0 | 0 | 0 | 0 |
| 19682 | ENSMUSG00000070102 | 0 | NA | NA | NA | NA | NA | Mir455 | microRNA 455 [Source:MGI Symbol;Acc:M | 0 | 0 | 0 | 0 | 0 | 0 |
| 19683 | ENSMUSG00000070103 | 0 | NA | NA | NA | NA | NA | Mir488 | microRNA 488 [Source:MGI Symbol;Acc:M | 0 | 0 | 0 | 0 | 0 | 0 |
| 19684 | ENSMUSG00000070104 | 0 | NA | NA | NA | NA | NA | Mir471 | microRNA 471 [Source:MGI Symbol;Acc:M | 0 | 0 | 0 | 0 | 0 | 0 |
| 19685 | ENSMUSG00000070105 | 0 | NA | NA | NA | NA | NA | Mir495 | microRNA 495 [Source:MGI Symbol;Acc:M | 0 | 0 | 0 | 0 | 0 | 0 |
| 19686 | ENSMUSG00000070106 | 0 | NA | NA | NA | NA | NA | Mir363 | microRNA 363 [Source:MGI Symbol;Acc:M | 0 | 0 | 0 | 0 | 0 | 0 |
| 19687 | ENSMUSG00000070107 | 0 | NA | NA | NA | NA | NA | Mir489 | microRNA 489 [Source:MGI Symbol;Acc:M | 0 | 0 | 0 | 0 | 0 | 0 |
| 19688 | ENSMUSG00000070108 | 0 | NA | NA | NA | NA | NA | Mir500 | microRNA 500 [Source:MGI Symbol;Acc:M | 0 | 0 | 0 | 0 | 0 | 0 |
| 19689 | ENSMUSG00000070109 | 0 | NA | NA | NA | NA | NA | Mir497b | microRNA 497b [Source:MGI Symbol;Acc: | 0 | 0 | 0 | 0 | 0 | 0 |
| 19690 | ENSMUSG00000070110 | 0 | NA | NA | NA | NA | NA | Mir504 | microRNA 504 [Source:MGI Symbol;Acc:M | 0 | 0 | 0 | 0 | 0 | 0 |
| 19691 | ENSMUSG00000070111 | 0 | NA | NA | NA | NA | NA | Gm10286 | predicted gene 10286 [Source:MGI Symbo | 0 | 0 | 0 | 0 | 0 | 0 |
| 19692 | ENSMUSG00000070126 | 0 | NA | NA | NA | NA | NA | Mir199a-2 | microRNA 199a-2 [Source:MGI Symbol;Ac | 0 | 0 | 0 | 0 | 0 | 0 |
| 19693 | ENSMUSG00000070127 | 0 | NA | NA | NA | NA | NA | Mir146b | microRNA 146b [Source:MGI Symbol;Acc: | 0 | 0 | 0 | 0 | 0 | 0 |
| 19694 | ENSMUSG00000070128 | 0 | NA | NA | NA | NA | NA | Mir485 | microRNA 485 [Source:MGI Symbol;Acc:M | 0 | 0 | 0 | 0 | 0 | 0 |
| 19695 | ENSMUSG00000070129 | 0 | NA | NA | NA | NA | NA | Mir3071 | microRNA 3071 [Source:MGI Symbol;Acc: | 0 | 0 | 0 | 0 | 0 | 0 |
| 19696 | ENSMUSG00000070130 | 0 | NA | NA | NA | NA | NA | Mir328 | microRNA 328 [Source:MGI Symbol;Acc:M | 0 | 0 | 0 | 0 | 0 | 0 |
| 19697 | ENSMUSG00000070133 | 0 | NA | NA | NA | NA | NA | Mir434 | microRNA 434 [Source:MGI Symbol;Acc:M | 0 | 0 | 0 | 0 | 0 | 0 |
| 19698 | ENSMUSG00000070136 | 0 | NA | NA | NA | NA | NA | Mir496a | microRNA 496a [Source:MGI Symbol;Acc: | 0 | 0 | 0 | 0 | 0 | 0 |
| 19699 | ENSMUSG00000070138 | 0 | NA | NA | NA | NA | NA | Mir452 | microRNA 452 [Source:MGI Symbol;Acc:M | 0 | 0 | 0 | 0 | 0 | 0 |
| 19700 | ENSMUSG00000070139 | 0 | NA | NA | NA | NA | NA | Mir532 | microRNA 532 [Source:MGI Symbol;Acc:M | 0 | 0 | 0 | 0 | 0 | 0 |
| 19701 | ENSMUSG00000070140 | 0 | NA | NA | NA | NA | NA | Mir483 | microRNA 483 [Source:MGI Symbol;Acc:M | 0 | 0 | 0 | 0 | 0 | 0 |
| 19702 | ENSMUSG00000070141 | 0 | NA | NA | NA | NA | NA | Mir494 | microRNA 494 [Source:MGI Symbol;Acc:M | 0 | 0 | 0 | 0 | 0 | 0 |
| 19703 | ENSMUSG00000070144 | 0 | NA | NA | NA | NA | NA | Mir1a-1 | microRNA 1a-1 [Source:MGI Symbol;Acc:N | 0 | 0 | 0 | 0 | 0 | 0 |
| 19704 | ENSMUSG00000070150 | 0 | NA | NA | NA | NA | NA | NA | NA | 0 | 0 | 0 | 0 | 0 | 0 |
| 19705 | ENSMUSG00000070166 | 0.321325949 | 1.911575114 | 4.048267893 | 0.472195804 | 0.636787034 | NA | Gm25821 | predicted gene, 25821 [Source:MGI Symb | 0 | 0 | 0 | 1.928 | 0 | 0 |
| 19706 | ENSMUSG00000070167 | 10.64449494 | 0.307739894 | 0.659120944 | 0.466894425 | 0.640575414 | 0.923932929 | Snora57 | small nucleolar RNA, H/ACA box 57 [Sourc | 11.243 | 8.2233 | 9.0562 | 14.46 | 13.087 | 7.7969 |
| 19707 | ENSMUSG00000070170 | 0 | NA | NA | NA | NA | NA | Gm24163 | predicted gene, 24163 [Source:MGI Symb | 0 | 0 | 0 | 0 | 0 | 0 |
| 19708 | ENSMUSG00000070173 | 0 | NA | NA | NA | NA | NA | Gm24164 | predicted gene, 24164 [Source:MGI Symb | 0 | 0 | 0 | 0 | 0 | 0 |

| | | | | | | | | | | | | | | | |
|---|---|---|---|---|---|---|---|---|---|---|---|---|---|---|---|
| 19709 | ENSMUSG00000070175 | 0 | NA | NA | NA | NA | NA | Gm24162 | predicted gene, 24162 [Source:MGI Symb | 0 | 0 | 0 | 0 | 0 | 0 |
| 19710 | ENSMUSG00000070178 | 0 | NA | NA | NA | NA | NA | NA | NA | 0 | 0 | 0 | 0 | 0 | 0 |
| 19711 | ENSMUSG00000070182 | 0 | NA | NA | NA | NA | NA | Gm24817 | predicted gene, 24817 [Source:MGI Symb | 0 | 0 | 0 | 0 | 0 | 0 |
| 19712 | ENSMUSG00000070192 | 0 | NA | NA | NA | NA | NA | NA | NA | 0 | 0 | 0 | 0 | 0 | 0 |
| 19713 | ENSMUSG00000070203 | 0 | NA | NA | NA | NA | NA | Gm22444 | predicted gene, 22444 [Source:MGI Symb | 0 | 0 | 0 | 0 | 0 | 0 |
| 19714 | ENSMUSG00000070209 | 0 | NA | NA | NA | NA | NA | NA | NA | 0 | 0 | 0 | 0 | 0 | 0 |
| 19715 | ENSMUSG00000070214 | 0 | NA | NA | NA | NA | NA | Gm23058 | predicted gene, 23058 [Source:MGI Symb | 0 | 0 | 0 | 0 | 0 | 0 |
| 19716 | ENSMUSG00000070219 | 0 | NA | NA | NA | NA | NA | NA | NA | 0 | 0 | 0 | 0 | 0 | 0 |
| 19717 | ENSMUSG00000070225 | 0 | NA | NA | NA | NA | NA | n-R5s33 | nuclear encoded rRNA 5S 33 [Source:MGI | 0 | 0 | 0 | 0 | 0 | 0 |
| 19718 | ENSMUSG00000070227 | 0 | NA | NA | NA | NA | NA | Gm24704 | predicted gene, 24704 [Source:MGI Symb | 0 | 0 | 0 | 0 | 0 | 0 |
| 19719 | ENSMUSG00000070235 | 0 | NA | NA | NA | NA | NA | Gm26329 | predicted gene, 26329 [Source:MGI Symb | 0 | 0 | 0 | 0 | 0 | 0 |
| 19720 | ENSMUSG00000070236 | 0 | NA | NA | NA | NA | NA | Gm26327 | predicted gene, 26327 [Source:MGI Symb | 0 | 0 | 0 | 0 | 0 | 0 |
| 19721 | ENSMUSG00000070240 | 0 | NA | NA | NA | NA | NA | NA | NA | 0 | 0 | 0 | 0 | 0 | 0 |
| 19722 | ENSMUSG00000070249 | 0 | NA | NA | NA | NA | NA | NA | NA | 0 | 0 | 0 | 0 | 0 | 0 |
| 19723 | ENSMUSG00000070258 | 0 | NA | NA | NA | NA | NA | NA | NA | 0 | 0 | 0 | 0 | 0 | 0 |
| 19724 | ENSMUSG00000070263 | 0 | NA | NA | NA | NA | NA | Gm22365 | predicted gene, 22365 [Source:MGI Symb | 0 | 0 | 0 | 0 | 0 | 0 |
| 19725 | ENSMUSG00000070271 | 0.167707431 | -0.796629176 | 4.080472857 | -0.195229623 | 0.84521319 | NA | Gm13268 | predicted gene 13268 [Source:MGI Symbo | 0 | 0 | 1.0062 | 0 | 0 | 0 |
| 19726 | ENSMUSG00000070280 | 2.092634454 | 0.391917309 | 1.644275662 | 0.238352557 | 0.811607661 | NA | Slc22a14 | solute carrier family 22 (organic cation tra | 2.4093 | 0 | 3.0187 | 3.8559 | 3.2719 | 0 |
| 19727 | ENSMUSG00000070282 | 16.35894378 | -0.228185891 | 0.533384323 | -0.427807645 | 0.66879117 | 0.930581576 | 3000002C10 | RIKEN cDNA 3000002C10 gene [Source:M | 14.456 | 17.475 | 21.131 | 16.388 | 11.997 | 16.708 |
| 19728 | ENSMUSG00000070283 | 235.4189721 | -0.057133882 | 0.165676941 | -0.344851143 | 0.730206282 | 0.946534338 | Ndufaf3 | NADH:ubiquinone oxidoreductase comple | 249.76 | 247.73 | 222.38 | 216.9 | 255.2 | 220.54 |
| 19729 | ENSMUSG00000070284 | 226.2175835 | 0.281097372 | 0.165729342 | 1.696123139 | 0.089862563 | 0.442400308 | Gmppb | GDP-mannose pyrophosphorylase B [Sour | 225.67 | 196.33 | 190.18 | 253.53 | 243.21 | 248.39 |
| 19730 | ENSMUSG00000070287 | 9.705814463 | 0.036545106 | 0.797947699 | 0.045798874 | 0.963470556 | 0.993245949 | Slc35g2 | solute carrier family 35, member G2 [Sour | 18.471 | 4.1117 | 6.0375 | 16.388 | 6.5437 | 6.6831 |
| 19731 | ENSMUSG00000070291 | 0.17131964 | -0.796629176 | 4.080472857 | -0.195229623 | 0.84521319 | NA | Ddx43 | DEAD box helicase 43 [Source:MGI Symbo | 0 | 1.0279 | 0 | 0 | 0 | 0 |
| 19732 | ENSMUSG00000070298 | 0.294513347 | 0.165143496 | 4.080472857 | 0.040471657 | 0.967717103 | NA | Trcg1 | taste receptor cell gene 1 [Source:MGI Sy | 0.8031 | 0 | 0 | 0.964 | 0 | 0 |
| 19733 | ENSMUSG00000070304 | 0.181769715 | 1.126924175 | 4.080472857 | 0.276174898 | 0.782413736 | NA | Scn2b | sodium channel, voltage-gated, type II, be | 0 | 0 | 0 | 0 | 1.0906 | 0 |
| 19734 | ENSMUSG00000070305 | 175.5099603 | 0.046573515 | 0.215269094 | 0.216350217 | 0.828714768 | 0.969955221 | Mpzl3 | myelin protein zero-like 3 [Source:MGI Sy | 182.3 | 193.25 | 141.88 | 137.85 | 202.86 | 194.92 |
| 19735 | ENSMUSG00000070306 | 0 | NA | NA | NA | NA | NA | Ccdc153 | coiled-coil domain containing 153 [Source | 0 | 0 | 0 | 0 | 0 | 0 |
| 19736 | ENSMUSG00000070311 | 0 | NA | NA | NA | NA | NA | Olfr894 | olfactory receptor 894 [Source:MGI Symb | 0 | 0 | 0 | 0 | 0 | 0 |
| 19737 | ENSMUSG00000070313 | 0 | NA | NA | NA | NA | NA | Pate14 | prostate and testis expressed 14 [Source:I | 0 | 0 | 0 | 0 | 0 | 0 |
| 19738 | ENSMUSG00000070315 | 599.0041145 | 0.135122166 | 0.121607455 | 1.111133908 | 0.266510715 | 0.722551261 | 4930581F22 | RIKEN cDNA 4930581F22 gene [Source:M | 597.51 | 522.18 | 592.68 | 627.55 | 631.47 | 622.64 |
| 19739 | ENSMUSG00000070319 | 971.5413337 | 0.107216588 | 0.108120742 | 0.991637551 | 0.321374364 | 0.76866499 | Eif3g | eukaryotic translation initiation factor 3, s | 877.79 | 975.49 | 953.92 | 1012.2 | 970.65 | 1039.2 |
| 19740 | ENSMUSG00000070323 | 7.313400083 | -2.593556175 | 0.995200859 | -2.606063038 | 0.00915896 | 0.101513565 | Mmp27 | matrix metallopeptidase 27 [Source:MGI S | 19.274 | 12.335 | 6.0375 | 2.8919 | 0 | 3.3415 |
| 19741 | ENSMUSG00000070324 | 0.181769715 | 1.126924175 | 4.080472857 | 0.276174898 | 0.782413736 | NA | Fbxw22 | F-box and WD-40 domain protein 22 [Sou | 0 | 0 | 0 | 0 | 1.0906 | 0 |
| 19742 | ENSMUSG00000070327 | 1042.839497 | 0.118264437 | 0.105931568 | 1.116422987 | 0.264241118 | 0.719492304 | Rnf213 | ring finger protein 213 [Source:MGI Symb | 1047.2 | 965.21 | 987.13 | 1065.2 | 1140.8 | 1051.5 |
| 19743 | ENSMUSG00000070330 | 0 | NA | NA | NA | NA | NA | Tmem235 | transmembrane protein 235 [Source:MGI | 0 | 0 | 0 | 0 | 0 | 0 |
| 19744 | ENSMUSG00000070331 | 0.428363721 | -0.543730382 | 3.624677344 | -0.15000794 | 0.880758351 | NA | Qrich2 | glutamine rich 2 [Source:MGI Symbol;Acc | 1.6062 | 0 | 0 | 0.964 | 0 | 0 |
| 19745 | ENSMUSG00000070332 | 54.30196417 | -0.348205257 | 0.44548423 | -0.781633183 | 0.434430179 | 0.845264432 | Trim80 | tripartite motif-containing 80 [Source:MG | 85.932 | 51.396 | 45.281 | 85.794 | 23.994 | 33.415 |
| 19746 | ENSMUSG00000070334 | 0 | NA | NA | NA | NA | NA | Krtap31-1 | keratin associated protein 31-1 [Source:M | 0 | 0 | 0 | 0 | 0 | 0 |
| 19747 | ENSMUSG00000070335 | 0 | NA | NA | NA | NA | NA | Krtap9-1 | keratin associated protein 9-1 [Source:MG | 0 | 0 | 0 | 0 | 0 | 0 |
| 19748 | ENSMUSG00000070336 | 10.3195875 | 0.114827672 | 0.671822634 | 0.170919624 | 0.864286966 | 0.977227145 | Fbxo47 | F-box protein 47 [Source:MGI Symbol;Acc | 10.44 | 10.279 | 9.0562 | 14.46 | 6.5437 | 11.138 |
| 19749 | ENSMUSG00000070337 | 2.10058291 | -0.53404381 | 1.498062427 | -0.35648969 | 0.721473872 | NA | Gpr179 | G protein-coupled receptor 179 [Source:M | 2.4093 | 2.0558 | 3.0187 | 2.8919 | 0 | 2.2277 |
| 19750 | ENSMUSG00000070342 | 0 | NA | NA | NA | NA | NA | Gm10287 | predicted gene 10287 [Source:MGI Symbo | 0 | 0 | 0 | 0 | 0 | 0 |
| 19751 | ENSMUSG00000070343 | 18.29967414 | -0.043206911 | 0.518509597 | -0.083329049 | 0.933589903 | 0.988420305 | Gm10288 | predicted gene 10288 [Source:MGI Symbo | 20.078 | 13.363 | 22.137 | 13.496 | 22.903 | 17.822 |
| 19752 | ENSMUSG00000070345 | 0.863926883 | 2.043289473 | 2.549701854 | 0.801383687 | 0.422909556 | NA | Hsf5 | heat shock transcription factor family mer | 0 | 1.0279 | 0 | 1.928 | 0 | 2.2277 |
| 19753 | ENSMUSG00000070348 | 288.9355219 | 0.625376855 | 0.257945867 | 2.424449991 | 0.015331595 | 0.14470442 | Ccnd1 | cyclin D1 [Source:MGI Symbol;Acc:MGI:88 | 352.56 | 173.72 | 153.96 | 314.26 | 388.26 | 350.86 |
| 19754 | ENSMUSG00000070354 | 0.528279927 | -0.837071592 | 3.369085264 | -0.248456636 | 0.803781118 | NA | Evi2 | ecotropic viral integration site 2 [Source:M | 0 | 2.0558 | 0 | 0 | 0 | 1.1138 |
| 19755 | ENSMUSG00000070360 | 0 | NA | NA | NA | NA | NA | Amy2a1 | amylase 2a1 [Source:MGI Symbol;Acc:MG | 0 | 0 | 0 | 0 | 0 | 0 |
| 19756 | ENSMUSG00000070366 | 0.513958919 | -2.394274821 | 3.756096013 | -0.637437066 | 0.523840191 | NA | Plpp4 | phospholipid phosphatase 4 [Source:MGI | 0 | 3.0838 | 0 | 0 | 0 | 0 |
| 19757 | ENSMUSG00000070368 | 90.37289657 | 0.462485908 | 0.271313167 | 1.704620212 | 0.088265279 | 0.438115648 | Prok1 | prokineticin 1 [Source:MGI Symbol;Acc:M | 70.673 | 90.457 | 67.418 | 132.06 | 101.43 | 80.197 |
| 19758 | ENSMUSG00000070369 | 0.267700746 | -1.497481649 | 4.057001436 | -0.369110456 | 0.712045396 | NA | Itgad | integrin, alpha D [Source:MGI Symbol;Acc | 1.6062 | 0 | 0 | 0 | 0 | 0 |
| 19759 | ENSMUSG00000070371 | 148.4388993 | 0.069395827 | 0.233863629 | 0.296736296 | 0.766667844 | 0.955295274 | Prss36 | protease, serine 36 [Source:MGI Symbol;A | 161.42 | 143.91 | 128.8 | 131.1 | 201.76 | 123.64 |
| 19760 | ENSMUSG00000070372 | 364.5670274 | -0.042091032 | 0.142760108 | -0.294837493 | 0.768118024 | 0.955856567 | Capza1 | capping protein (actin filament) muscle Z- | 356.58 | 403.97 | 349.17 | 338.36 | 378.44 | 360.89 |
| 19761 | ENSMUSG00000070374 | 0 | NA | NA | NA | NA | NA | Olfr59 | olfactory receptor 59 [Source:MGI Symbo | 0 | 0 | 0 | 0 | 0 | 0 |
| 19762 | ENSMUSG00000070375 | 0 | NA | NA | NA | NA | NA | Olfr406 | olfactory receptor 406 [Source:MGI Symb | 0 | 0 | 0 | 0 | 0 | 0 |
| 19763 | ENSMUSG00000070377 | 0 | NA | NA | NA | NA | NA | Olfr43 | olfactory receptor 43 [Source:MGI Symbo | 0 | 0 | 0 | 0 | 0 | 0 |
| 19764 | ENSMUSG00000070378 | 0 | NA | NA | NA | NA | NA | Olfr403 | olfactory receptor 403 [Source:MGI Symb | 0 | 0 | 0 | 0 | 0 | 0 |

| | | | | | | | | | | | | | | |
|---|---|---|---|---|---|---|---|---|---|---|---|---|---|---|
| 19765 | ENSMUSG00000070379 | 0 | NA | NA | NA | NA | NA | Olfr402 | olfactory receptor 402 [Source:MGI Symb | 0 | 0 | 0 | 0 | 0 | 0 |
| 19766 | ENSMUSG00000070380 | 0 | NA | NA | NA | NA | NA | Olfr401 | olfactory receptor 401 [Source:MGI Symb | 0 | 0 | 0 | 0 | 0 | 0 |
| 19767 | ENSMUSG00000070382 | 0 | NA | NA | NA | NA | NA | Olfr391 | olfactory receptor 391 [Source:MGI Symb | 0 | 0 | 0 | 0 | 0 | 0 |
| 19768 | ENSMUSG00000070383 | 0 | NA | NA | NA | NA | NA | Olfr389 | olfactory receptor 389 [Source:MGI Symb | 0 | 0 | 0 | 0 | 0 | 0 |
| 19769 | ENSMUSG00000070385 | 0 | NA | NA | NA | NA | NA | Ampd1 | adenosine monophosphate deaminase 1 \| | 0 | 0 | 0 | 0 | 0 | 0 |
| 19770 | ENSMUSG00000070388 | 0.328370405 | 0.16514316 | 4.003149536 | 0.041253308 | 0.967093956 | NA | Fbxo39 | F-box protein 39 [Source:MGI Symbol;Acc | 0 | 0 | 1.0062 | 0.964 | 0 | 0 |
| 19771 | ENSMUSG00000070390 | 6.977139108 | -2.825207322 | 1.024753188 | -2.756963682 | 0.005834083 | 0.074021209 | Nlrp1b | NLR family, pyrin domain containing 1B [S | 11.243 | 18.503 | 7.0437 | 2.8919 | 2.1812 | 0 |
| 19772 | ENSMUSG00000070392 | 0.342432689 | 1.990653086 | 3.900281263 | 0.51038706 | 0.609780321 | NA | Gm20634 | predicted gene 20634 [Source:MGI Symbo | 0 | 0 | 0 | 0.964 | 1.0906 | 0 |
| 19773 | ENSMUSG00000070394 | 1343.325656 | 0.144909179 | 0.122289457 | 1.184968706 | 0.23602982 | 0.684566938 | Tmem256 | transmembrane protein 256 [Source:MGI | 1309.9 | 1195.5 | 1321.2 | 1214.6 | 1450.5 | 1568.3 |
| 19774 | ENSMUSG00000070407 | 1633.577977 | 0.163855926 | 0.121790571 | 1.345390902 | 0.178499032 | 0.610813333 | Hs3st3b1 | heparan sulfate (glucosamine) 3-O-sulfotr | 1533.1 | 1564.5 | 1525.5 | 1843.1 | 1445.1 | 1890.2 |
| 19775 | ENSMUSG00000070417 | 0 | NA | NA | NA | NA | NA | Olfr2 | olfactory receptor 2 [Source:MGI Symbol; | 0 | 0 | 0 | 0 | 0 | 0 |
| 19776 | ENSMUSG00000070419 | 0.572870759 | -2.577107685 | 3.182795569 | -0.809699407 | 0.418112959 | NA | Cyp3a57 | cytochrome P450, family 3, subfamily a, p | 2.4093 | 1.0279 | 0 | 0 | 0 | 0 |
| 19777 | ENSMUSG00000070420 | 113.3390189 | 0.265020131 | 0.229057246 | 1.157003916 | 0.24727076 | 0.697450133 | Zscan25 | zinc finger and SCAN domain containing 2 | 97.978 | 87.373 | 123.77 | 133.99 | 119.97 | 116.95 |
| 19778 | ENSMUSG00000070421 | 0 | NA | NA | NA | NA | NA | Olfr658 | olfactory receptor 658 [Source:MGI Symb | 0 | 0 | 0 | 0 | 0 | 0 |
| 19779 | ENSMUSG00000070423 | 0 | NA | NA | NA | NA | NA | Olfr558 | olfactory receptor 558 [Source:MGI Symb | 0 | 0 | 0 | 0 | 0 | 0 |
| 19780 | ENSMUSG00000070424 | 0 | NA | NA | NA | NA | NA | Art5 | ADP-ribosyltransferase 5 [Source:MGI Syn | 0 | 0 | 0 | 0 | 0 | 0 |
| 19781 | ENSMUSG00000070425 | 2.294527846 | -1.110370286 | 1.514121878 | -0.73334274 | 0.463349414 | NA | Xntrpc | Xndc1-transient receptor potential cation | 3.2124 | 5.1396 | 1.0062 | 0 | 2.1812 | 2.2277 |
| 19782 | ENSMUSG00000070426 | 340.2984967 | 0.125380715 | 0.139382595 | 0.899543556 | 0.368363207 | 0.80627882 | Rnf121 | ring finger protein 121 [Source:MGI Symb | 326.06 | 319.68 | 331.05 | 374.02 | 347.91 | 343.06 |
| 19783 | ENSMUSG00000070427 | 111.5342969 | -0.909541544 | 0.268347036 | -3.389422736 | 0.000700399 | 0.015953665 | Il18bp | interleukin 18 binding protein [Source:MG | 138.13 | 184 | 114.71 | 80.974 | 56.712 | 94.677 |
| 19784 | ENSMUSG00000070436 | 175.0633743 | -0.795899671 | 0.273430376 | -2.910794635 | 0.003605109 | 0.053481328 | Serpinh1 | serine (or cysteine) peptidase inhibitor, cl | 310 | 182.97 | 173.07 | 145.56 | 85.068 | 153.71 |
| 19785 | ENSMUSG00000070438 | 0 | NA | NA | NA | NA | NA | Olfr313 | olfactory receptor 313 [Source:MGI Symb | 0 | 0 | 0 | 0 | 0 | 0 |
| 19786 | ENSMUSG00000070443 | 0.428363721 | -0.543730382 | 3.624677344 | -0.15000794 | 0.880758351 | NA | Gm10291 | predicted pseudogene 10291 [Source:MG | 1.6062 | 0 | 0 | 0.964 | 0 | 0 |
| 19787 | ENSMUSG00000070448 | 0 | NA | NA | NA | NA | NA | Vmn2r89 | vomeronasal 2, receptor 89 [Source:MGI S | 0 | 0 | 0 | 0 | 0 | 0 |
| 19788 | ENSMUSG00000070458 | 0 | NA | NA | NA | NA | NA | Vmn2r73 | vomeronasal 2, receptor 73 [Source:MGI S | 0 | 0 | 0 | 0 | 0 | 0 |
| 19789 | ENSMUSG00000070459 | 0 | NA | NA | NA | NA | NA | Olfr290 | olfactory receptor 290 [Source:MGI Symb | 0 | 0 | 0 | 0 | 0 | 0 |
| 19790 | ENSMUSG00000070460 | 0 | NA | NA | NA | NA | NA | Olfr291 | olfactory receptor 291 [Source:MGI Symb | 0 | 0 | 0 | 0 | 0 | 0 |
| 19791 | ENSMUSG00000070461 | 35.93165534 | 0.251933219 | 0.380921011 | 0.661379161 | 0.508369187 | 0.882793133 | 9230112E08 | RIKEN cDNA 9230112E08 gene [Source:M( | 39.352 | 23.642 | 35.219 | 41.451 | 43.625 | 32.301 |
| 19792 | ENSMUSG00000070462 | 123.6018361 | -0.053341723 | 0.237016038 | -0.225055332 | 0.82193623 | 0.969160361 | Tlnrd1 | talin rod domain containing 1 [Source:MG | 126.89 | 121.29 | 129.81 | 149.42 | 88.34 | 125.86 |
| 19793 | ENSMUSG00000070464 | 0 | NA | NA | NA | NA | NA | Ccl26 | chemokine (C-C motif) ligand 26 [Source:M | 0 | 0 | 0 | 0 | 0 | 0 |
| 19794 | ENSMUSG00000070465 | 0 | NA | NA | NA | NA | NA | Gm9696 | predicted gene 9696 [Source:MGI Symbol | 0 | 0 | 0 | 0 | 0 | 0 |
| 19795 | ENSMUSG00000070469 | 3.127907607 | -0.32892421 | 1.200175154 | -0.274063505 | 0.784035819 | NA | Adamtsl3 | ADAMTS-like 3 [Source:MGI Symbol;Acc:M | 2.4093 | 2.0558 | 6.0375 | 3.8559 | 2.1812 | 2.2277 |
| 19796 | ENSMUSG00000070471 | 0.353348078 | 0.165142701 | 3.895042568 | 0.042398176 | 0.966181282 | NA | Erich6 | glutamate rich 6 [Source:MGI Symbol;Acc | 0 | 0 | 1.0062 | 0 | 0 | 1.1138 |
| 19797 | ENSMUSG00000070473 | 1528.078083 | 0.346108882 | 0.128385119 | 2.695864474 | 0.007020622 | 0.08425311 | Cldn3 | claudin 3 [Source:MGI Symbol;Acc:MGI:1: | 1296.2 | 1188.3 | 1552.6 | 1708.2 | 1553 | 1870.1 |
| 19798 | ENSMUSG00000070476 | 6.404520433 | -0.863160338 | 0.893865546 | -0.965648964 | 0.334219866 | 0.775945291 | Fam217b | family with sequence similarity 217, meml | 10.44 | 11.307 | 3.0187 | 4.8199 | 3.2719 | 5.5692 |
| 19799 | ENSMUSG00000070489 | 0 | NA | NA | NA | NA | NA | 4930527J03I | RIKEN cDNA 4930527J03 gene [Source:M( | 0 | 0 | 0 | 0 | 0 | 0 |
| 19800 | ENSMUSG00000070490 | 1.147319377 | -3.555516624 | 2.272519517 | -1.564570336 | 0.117683689 | NA | Gm10293 | predicted pseudogene 10293 [Source:MG | 0.8031 | 2.0558 | 4.025 | 0 | 0 | 0 |
| 19801 | ENSMUSG00000070493 | 9962.113869 | -0.044491422 | 0.080557387 | -0.552294755 | 0.580746423 | 0.908645447 | Chchd2 | coiled-coil-helix-coiled-coil-helix domain c | 10227 | 9799.1 | 10321 | 9836.4 | 9876.6 | 9712.7 |
| 19802 | ENSMUSG00000070495 | 77.53584925 | -1.033226165 | 0.337832395 | -3.05839872 | 0.002225233 | 0.037688253 | Ctcfl | CCCTC-binding factor (zinc finger protein)- | 118.86 | 87.373 | 106.66 | 73.262 | 53.44 | 25.618 |
| 19803 | ENSMUSG00000070498 | 0.133850373 | -0.796629176 | 4.080472857 | -0.195229623 | 0.84521319 | NA | Tmem132b | transmembrane protein 132B [Source:MG | 0.8031 | 0 | 0 | 0 | 0 | 0 |
| 19804 | ENSMUSG00000070501 | 0.692375149 | -1.404276945 | 2.772265464 | -0.506544905 | 0.612474175 | NA | Ifi214 | interferon activated gene 214 [Source:MG | 0 | 1.0279 | 2.0125 | 0 | 0 | 1.1138 |
| 19805 | ENSMUSG00000070504 | 0 | NA | NA | NA | NA | NA | Fcrl6 | Fc receptor-like 6 [Source:MGI Symbol;Ac | 0 | 0 | 0 | 0 | 0 | 0 |
| 19806 | ENSMUSG00000070509 | 7.242998815 | -0.226696937 | 0.86014099 | -0.263557881 | 0.792120617 | 0.961434599 | Rgma | repulsive guidance molecule family memb | 12.047 | 9.2513 | 2.0125 | 5.7839 | 5.4531 | 8.9108 |
| 19807 | ENSMUSG00000070511 | 0 | NA | NA | NA | NA | NA | NA | NA | 0 | 0 | 0 | 0 | 0 | 0 |
| 19808 | ENSMUSG00000070520 | 413.9923975 | -0.016987448 | 0.132427086 | -0.128277742 | 0.897929179 | 0.982798755 | Nsmce3 | NSE3 homolog, SMC5-SMC6 complex com | 398.34 | 442 | 409.54 | 424.15 | 411.16 | 398.76 |
| 19809 | ENSMUSG00000070522 | 0.480153995 | 1.043847617 | 3.151673025 | 0.331204287 | 0.740490182 | NA | Gm6505 | predicted pseudogene 6505 [Source:MGI | 0.8031 | 0 | 0 | 0.964 | 0 | 1.1138 |
| 19810 | ENSMUSG00000070524 | 3.43302436 | 1.174331247 | 1.258931245 | 0.932800144 | 0.350923167 | NA | Fcrlb | Fc receptor-like B [Source:MGI Symbol;Ac | 3.2124 | 3.0838 | 0 | 7.7118 | 4.3625 | 2.2277 |
| 19811 | ENSMUSG00000070526 | 1.06339772 | 0.128965766 | 2.187061608 | 0.058967597 | 0.952977917 | NA | Peg12 | paternally expressed 12 [Source:MGI Sym | 0 | 2.0558 | 1.0062 | 0 | 1.0906 | 2.2277 |
| 19812 | ENSMUSG00000070527 | 7.390197144 | -0.704623359 | 0.838088697 | -0.840750341 | 0.400487812 | 0.827266295 | Mkrn3 | makorin, ring finger protein, 3 [Source:M( | 11.243 | 3.0838 | 13.081 | 4.8199 | 6.5437 | 5.5692 |
| 19813 | ENSMUSG00000070529 | 0 | NA | NA | NA | NA | NA | Wfdc10 | WAP four-disulfide core domain 10 [Sourc | 0 | 0 | 0 | 0 | 0 | 0 |
| 19814 | ENSMUSG00000070530 | 0.439020386 | -2.189747911 | 3.52148864 | -0.621824499 | 0.534057275 | NA | Wfdc16 | WAP four-disulfide core domain 16 [Sourc | 1.6062 | 1.0279 | 0 | 0 | 0 | 0 |
| 19815 | ENSMUSG00000070531 | 0 | NA | NA | NA | NA | NA | Wfdc6b | WAP four-disulfide core domain 6B [Sourc | 0 | 0 | 0 | 0 | 0 | 0 |
| 19816 | ENSMUSG00000070532 | 0.160662974 | 1.126924175 | 4.080472857 | 0.276174898 | 0.782413736 | NA | Ccdc190 | coiled-coil domain containing 190 [Source | 0 | 0 | 0 | 0.964 | 0 | 0 |
| 19817 | ENSMUSG00000070533 | 0 | NA | NA | NA | NA | NA | Wfdc8 | WAP four-disulfide core domain 8 [Source | 0 | 0 | 0 | 0 | 0 | 0 |
| 19818 | ENSMUSG00000070544 | 1011.199173 | -0.159799665 | 0.12017664 | -1.32970655 | 0.183614976 | 0.617851749 | Top1 | topoisomerase (DNA) I [Source:MGI Symb | 997.45 | 1107.1 | 1096.8 | 846.37 | 1041.5 | 977.95 |
| 19819 | ENSMUSG00000070546 | 0 | NA | NA | NA | NA | NA | Mrgprb3 | MAS-related GPR, member B3 [Source:M( | 0 | 0 | 0 | 0 | 0 | 0 |
| 19820 | ENSMUSG00000070547 | 0.315620088 | 0.165143421 | 4.063300985 | 0.040642675 | 0.967580763 | NA | Mrgprb1 | MAS-related GPR, member B1 [Source:M( | 0.8031 | 0 | 0 | 0 | 1.0906 | 0 |

| | | | | | | | | | | | | | | | |
|---|---|---|---|---|---|---|---|---|---|---|---|---|---|---|---|
| 19821 | ENSMUSG00000070550 | 0 | NA | NA | NA | NA | NA | Mrgprb4 | MAS-related GPR, member B4 [Source:M( | 0 | 0 | 0 | 0 | 0 | 0 |
| 19822 | ENSMUSG00000070551 | 0 | NA | NA | NA | NA | NA | Mrgprb5 | MAS-related GPR, member B5 [Source:M( | 0 | 0 | 0 | 0 | 0 | 0 |
| 19823 | ENSMUSG00000070552 | 0 | NA | NA | NA | NA | NA | Mrgprx1 | MAS-related GPR, member X1 [Source:M( | 0 | 0 | 0 | 0 | 0 | 0 |
| 19824 | ENSMUSG00000070563 | 0 | NA | NA | NA | NA | NA | Spaca4 | sperm acrosome associated 4 [Source:MG | 0 | 0 | 0 | 0 | 0 | 0 |
| 19825 | ENSMUSG00000070564 | 0 | NA | NA | NA | NA | NA | Ntn5 | netrin 5 [Source:MGI Symbol;Acc:MGI:26 | 0 | 0 | 0 | 0 | 0 | 0 |
| 19826 | ENSMUSG00000070565 | 247.1756984 | -0.180561212 | 0.244388686 | -0.738828033 | 0.460011427 | 0.859753653 | Rasal2 | RAS protein activator like 2 [Source:MGI S | 297.95 | 269.31 | 220.37 | 225.57 | 322.82 | 147.03 |
| 19827 | ENSMUSG00000070568 | 3.734067496 | -1.666359883 | 1.278504567 | -1.303366391 | 0.192449705 | NA | Slc6a21 | solute carrier family 6 member 21 [Source | 0.8031 | 7.1954 | 9.0562 | 0.964 | 3.2719 | 1.1138 |
| 19828 | ENSMUSG00000070570 | 0.321325949 | 1.911575114 | 4.048267893 | 0.472195804 | 0.636787034 | NA | Slc17a7 | solute carrier family 17 (sodium-depende | 0 | 0 | 0 | 1.928 | 0 | 0 |
| 19829 | ENSMUSG00000070572 | 2.9940051 | -0.901238702 | 1.253582023 | -0.71893078 | 0.47218357 | NA | Trmt112-ps2 | tRNA methyltransferase 11-2, pseudogen | 1.6062 | 5.1396 | 5.0312 | 2.8919 | 2.1812 | 1.1138 |
| 19830 | ENSMUSG00000070574 | 7.286500942 | 0.145648899 | 0.896809916 | 0.162407771 | 0.870984752 | 0.978935695 | 2310016G11 | RIKEN cDNA 2310016G11 gene [Source:M | 6.4248 | 7.1954 | 7.0437 | 0.964 | 8.7249 | 13.366 |
| 19831 | ENSMUSG00000070576 | 391.4540805 | 0.318426923 | 0.200487029 | 1.588266958 | 0.112225986 | 0.491740884 | Mn1 | meningioma 1 [Source:MGI Symbol;Acc:M | 371.03 | 343.32 | 330.05 | 326.79 | 386.08 | 591.45 |
| 19832 | ENSMUSG00000070577 | 0.167707431 | -0.796629176 | 4.080472857 | -0.195229623 | 0.84521319 | NA | Gm572 | predicted gene 572 [Source:MGI Symbol;/ | 0 | 0 | 1.0062 | 0 | 0 | 0 |
| 19833 | ENSMUSG00000070583 | 34.93999471 | 0.660795745 | 0.422436655 | 1.564248124 | 0.11775931 | 0.502838264 | Fv1 | Friend virus susceptibility 1 [Source:MGI S | 36.14 | 15.419 | 29.181 | 31.811 | 43.625 | 53.465 |
| 19834 | ENSMUSG00000070594 | 5536.838802 | 0.220502034 | 0.117184546 | 1.881664789 | 0.059881543 | 0.348261618 | Cfhr4 | complement factor H-related 4 [Source:M | 4392.2 | 5104.6 | 5848.3 | 5847.5 | 6326.7 | 5701.8 |
| 19835 | ENSMUSG00000070601 | 0.342432689 | 1.990653086 | 3.900281263 | 0.51038706 | 0.609780321 | NA | Vmn2r84 | vomeronasal 2, receptor 84 [Source:MGI | 0 | 0 | 0 | 0.964 | 1.0906 | 0 |
| 19836 | ENSMUSG00000070604 | 25.5129246 | 0.747513339 | 0.427419735 | 1.748897576 | 0.080308726 | 0.41597095 | Vsig10l | V-set and immunoglobulin domain contai | 18.471 | 21.586 | 17.106 | 30.847 | 30.537 | 34.529 |
| 19837 | ENSMUSG00000070605 | 81.8503895 | 0.362587848 | 0.346738285 | 1.04571045 | 0.295694736 | 0.746149454 | Zfp992 | zinc finger protein 992 [Source:MGI Symb | 80.31 | 58.591 | 75.468 | 49.163 | 140.69 | 86.88 |
| 19838 | ENSMUSG00000070606 | 0 | NA | NA | NA | NA | NA | Gm10298 | predicted gene 10298 [Source:MGI Symbo | 0 | 0 | 0 | 0 | 0 | 0 |
| 19839 | ENSMUSG00000070609 | 0 | NA | NA | NA | NA | NA | Aadacl4 | arylacetamide deacetylase like 4 [Source:I | 0 | 0 | 0 | 0 | 0 | 0 |
| 19840 | ENSMUSG00000070610 | 8.759035535 | 0.533567407 | 0.801162356 | 0.665991609 | 0.505416464 | 0.881592135 | Gm13127 | predicted gene 13127 [Source:MGI Symbo | 12.047 | 5.1396 | 4.025 | 4.8199 | 9.8156 | 16.708 |
| 19841 | ENSMUSG00000070616 | 0 | NA | NA | NA | NA | NA | Gm13040 | predicted gene 13040 [Source:MGI Symbo | 0 | 0 | 0 | 0 | 0 | 0 |
| 19842 | ENSMUSG00000070617 | 0 | NA | NA | NA | NA | NA | Pramel23 | PRAME like 23 [Source:MGI Symbol;Acc:M | 0 | 0 | 0 | 0 | 0 | 0 |
| 19843 | ENSMUSG00000070618 | 0 | NA | NA | NA | NA | NA | Pramel20 | PRAME like 20 [Source:MGI Symbol;Acc:M | 0 | 0 | 0 | 0 | 0 | 0 |
| 19844 | ENSMUSG00000070619 | 0 | NA | NA | NA | NA | NA | Pramel31 | PRAME like 31 [Source:MGI Symbol;Acc:M | 0 | 0 | 0 | 0 | 0 | 0 |
| 19845 | ENSMUSG00000070621 | 0 | NA | NA | NA | NA | NA | Vmn2r-ps60 | vomeronasal 2, receptor, pseudogene 60 | 0 | 0 | 0 | 0 | 0 | 0 |
| 19846 | ENSMUSG00000070637 | 0 | NA | NA | NA | NA | NA | Srarp | steroid receptor associated and regulated | 0 | 0 | 0 | 0 | 0 | 0 |
| 19847 | ENSMUSG00000070639 | 23.33803541 | 0.190996923 | 0.575223567 | 0.33203946 | 0.739859463 | 0.948393197 | Lrrc8b | leucine rich repeat containing 8 family, m | 16.865 | 10.279 | 38.237 | 19.28 | 15.269 | 40.098 |
| 19848 | ENSMUSG00000070641 | 0 | NA | NA | NA | NA | NA | Gm16410 | predicted gene 16410 [Source:MGI Symbo | 0 | 0 | 0 | 0 | 0 | 0 |
| 19849 | ENSMUSG00000070643 | 180.4239629 | 0.130464214 | 0.188934253 | 0.690527058 | 0.4898628 | 0.875278016 | Sox13 | SRY (sex determining region Y)-box 13 [So | 153.39 | 164.47 | 199.24 | 175.44 | 201.76 | 188.24 |
| 19850 | ENSMUSG00000070644 | 7895.964527 | 0.343765376 | 0.140771952 | 2.442001915 | 0.014606069 | 0.140429803 | Etnk2 | ethanolamine kinase 2 [Source:MGI Symb | 7100.2 | 6636.2 | 7141.3 | 6904 | 8822 | 10772 |
| 19851 | ENSMUSG00000070645 | 1.359740482 | 1.037305902 | 2.019972982 | 0.513524642 | 0.607584378 | NA | Ren1 | renin 1 structural [Source:MGI Symbol;Ac | 1.6062 | 0 | 1.0062 | 0 | 1.0906 | 4.4554 |
| 19852 | ENSMUSG00000070661 | 254.6592658 | 1.353791165 | 0.232370198 | 5.8260103 | 5.68E-09 | 9.55E-07 | Rnf186 | ring finger protein 186 [Source:MGI Symb | 175.08 | 108.96 | 144.9 | 426.08 | 253.02 | 419.92 |
| 19853 | ENSMUSG00000070667 | 0 | NA | NA | NA | NA | NA | Rpl31-ps10 | ribosomal protein L31, pseudogene 10 [Sc | 0 | 0 | 0 | 0 | 0 | 0 |
| 19854 | ENSMUSG00000070677 | 0 | NA | NA | NA | NA | NA | Pramel48 | PRAME like 48 [Source:MGI Symbol;Acc:M | 0 | 0 | 0 | 0 | 0 | 0 |
| 19855 | ENSMUSG00000070683 | 0 | NA | NA | NA | NA | NA | Lactbl1 | lactamase, beta-like 1 [Source:MGI Symbo | 0 | 0 | 0 | 0 | 0 | 0 |
| 19856 | ENSMUSG00000070686 | 0 | NA | NA | NA | NA | NA | Pramel34 | PRAME like 34 [Source:MGI Symbol;Acc:M | 0 | 0 | 0 | 0 | 0 | 0 |
| 19857 | ENSMUSG00000070687 | 0.185640647 | 1.126924175 | 4.080472857 | 0.276174898 | 0.782413736 | NA | Htr1d | 5-hydroxytryptamine (serotonin) receptor | 0 | 0 | 0 | 0 | 0 | 1.1138 |
| 19858 | ENSMUSG00000070690 | 581.4755692 | 0.12236863 | 0.15110777 | 0.809810308 | 0.418049208 | 0.837488052 | Albfm1 | albumin superfamily member 1 [Source:M | 545.31 | 584.89 | 541.36 | 735.52 | 563.85 | 517.94 |
| 19859 | ENSMUSG00000070691 | 4.140084106 | -1.895301199 | 1.20360486 | -1.574687227 | 0.115328658 | NA | Runx3 | runt related transcription factor 3 [Source | 6.4248 | 3.0838 | 10.062 | 1.928 | 0 | 3.3415 |
| 19860 | ENSMUSG00000070692 | 0.685278559 | -2.809870862 | 3.376578027 | -0.832165239 | 0.405315682 | NA | Rps12-ps10 | ribosomal protein S12, pseudogene 10 [Sc | 0 | 4.1117 | 0 | 0 | 0 | 0 |
| 19861 | ENSMUSG00000070695 | 0 | NA | NA | NA | NA | NA | Cntnap5a | contactin associated protein-like 5A [Sour | 0 | 0 | 0 | 0 | 0 | 0 |
| 19862 | ENSMUSG00000070697 | 619.1755493 | 0.235146774 | 0.124272623 | 1.892184852 | 0.058466356 | 0.343085466 | Utp3 | UTP3 small subunit processome compone | 572.61 | 539.66 | 593.68 | 629.48 | 647.83 | 731.8 |
| 19863 | ENSMUSG00000070699 | 175.1769586 | -0.100886559 | 0.184721151 | -0.546155965 | 0.584958733 | 0.90908382 | Sars2 | seryl-aminoacyl-tRNA synthetase 2 [Sourc | 159.01 | 195.3 | 190.18 | 172.55 | 163.59 | 170.42 |
| 19864 | ENSMUSG00000070702 | 0.17131964 | -0.796629176 | 4.080472857 | -0.195229623 | 0.84521319 | NA | Csn1s1 | casein alpha s1 [Source:MGI Symbol;Acc:I | 0 | 1.0279 | 0 | 0 | 0 | 0 |
| 19865 | ENSMUSG00000070704 | 12162.29757 | 0.022282726 | 0.093352575 | 0.238694282 | 0.811342651 | 0.966527993 | Ugt2b36 | UDP glucuronosyltransferase 2 family, pol | 12341 | 11038 | 12826 | 12235 | 12672 | 11862 |
| 19866 | ENSMUSG00000070705 | 129.3548722 | -0.119254592 | 0.212518434 | -0.561149404 | 0.574695692 | 0.906163061 | Eid2b | EP300 interacting inhibitor of differentiati | 130.91 | 141.85 | 131.82 | 145.56 | 110.15 | 115.84 |
| 19867 | ENSMUSG00000070708 | 0 | NA | NA | NA | NA | NA | Gtsf1l | gametocyte specific factor 1-like [Source:I | 0 | 0 | 0 | 0 | 0 | 0 |
| 19868 | ENSMUSG00000070709 | 61.74849242 | -0.060627448 | 0.303318532 | -0.19988046 | 0.841574072 | 0.971854408 | Zfp974 | zinc finger protein 974 [Source:MGI Symb | 52.202 | 71.954 | 65.406 | 69.406 | 45.806 | 65.717 |
| 19869 | ENSMUSG00000070713 | 49.89612235 | -0.203128821 | 0.406091024 | -0.500205149 | 0.616930633 | 0.917509899 | Gm10282 | predicted pseudogene 10282 [Source:MG | 65.854 | 62.703 | 31.194 | 25.063 | 59.984 | 54.578 |
| 19870 | ENSMUSG00000070717 | 13.96695307 | 0.008935028 | 0.643285707 | 0.013889673 | 0.988918001 | 0.99859176 | Gm10300 | predicted gene 10300 [Source:MGI Symbo | 14.456 | 13.363 | 14.087 | 22.171 | 15.269 | 4.4554 |
| 19871 | ENSMUSG00000070719 | 0 | NA | NA | NA | NA | NA | Pla2g4d | phospholipase A2, group IVD [Source:MG | 0 | 0 | 0 | 0 | 0 | 0 |
| 19872 | ENSMUSG00000070720 | 36.11088274 | -0.676070705 | 0.386589213 | -1.748809024 | 0.080324036 | 0.41597095 | Tmem200b | transmembrane protein 200B [Source:MG | 40.958 | 33.921 | 58.362 | 25.063 | 31.628 | 26.732 |
| 19873 | ENSMUSG00000070729 | 2.588189329 | 0.246405066 | 1.434201334 | 0.171806468 | 0.86358968 | NA | Gm12966 | predicted gene 12966 [Source:MGI Symbo | 4.0155 | 0 | 3.0187 | 1.928 | 5.4531 | 1.1138 |
| 19874 | ENSMUSG00000070730 | 1713.688867 | -0.15172367 | 0.102611611 | -1.478620885 | 0.139241668 | 0.545353772 | Rmdn3 | regulator of microtubule dynamics 3 [Sou | 1852 | 1764.9 | 1794.1 | 1644.5 | 1737.4 | 1489.2 |
| 19875 | ENSMUSG00000070732 | 0 | NA | NA | NA | NA | NA | Rbm44 | RNA binding motif protein 44 [Source:MG | 0 | 0 | 0 | 0 | 0 | 0 |
| 19876 | ENSMUSG00000070733 | 290.7777603 | 0.131504999 | 0.158157329 | 0.831482175 | 0.405701294 | 0.829973745 | Fryl | FRY like transcription coactivator [Source: | 255.39 | 296.04 | 281.75 | 302.69 | 335.91 | 272.89 |

| | | | | | | | | | | | | | | | |
|---|---|---|---|---|---|---|---|---|---|---|---|---|---|---|---|
| 19877 | ENSMUSG00000070737 | 114.0075392 | -0.216238323 | 0.219234835 | -0.986331955 | 0.323970244 | 0.77088057 | Tmem35b | transmembrane protein 35B [Source:MGI | 136.53 | 117.18 | 113.71 | 113.75 | 97.065 | 105.82 |
| 19878 | ENSMUSG00000070738 | 472.8486223 | 0.303170868 | 0.128421706 | 2.360744752 | 0.018238279 | 0.162137963 | Dgkd | diacylglycerol kinase, delta [Source:MGI S | 423.23 | 407.06 | 439.73 | 528.26 | 489.69 | 549.13 |
| 19879 | ENSMUSG00000070777 | 0 | NA | NA | NA | NA | NA | Ceacam20 | carcinoembryonic antigen-related cell adh | 0 | 0 | 0 | 0 | 0 | 0 |
| 19880 | ENSMUSG00000070778 | 0 | NA | NA | NA | NA | NA | Rpl7a-ps8 | ribosomal protein L7A, pseudogene 8 [Sou | 0 | 0 | 0 | 0 | 0 | 0 |
| 19881 | ENSMUSG00000070780 | 1258.67526 | 0.418642551 | 0.159371432 | 2.626835607 | 0.008618296 | 0.097386744 | Rbm47 | RNA binding motif protein 47 [Source:MG | 1105.9 | 1174.9 | 951.91 | 1818.1 | 1169.1 | 1332.2 |
| 19882 | ENSMUSG00000070794 | 0 | NA | NA | NA | NA | NA | BC016548 | cDNA sequence BC016548 [Source:MGI Sy | 0 | 0 | 0 | 0 | 0 | 0 |
| 19883 | ENSMUSG00000070796 | 0 | NA | NA | NA | NA | NA | Psg21 | pregnancy-specific glycoprotein 21 [Sourc | 0 | 0 | 0 | 0 | 0 | 0 |
| 19884 | ENSMUSG00000070797 | 0 | NA | NA | NA | NA | NA | Psg27 | pregnancy-specific glycoprotein 27 [Sourc | 0 | 0 | 0 | 0 | 0 | 0 |
| 19885 | ENSMUSG00000070798 | 0 | NA | NA | NA | NA | NA | Psg25 | pregnancy-specific glycoprotein 25 [Sourc | 0 | 0 | 0 | 0 | 0 | 0 |
| 19886 | ENSMUSG00000070799 | 0 | NA | NA | NA | NA | NA | Psg26 | pregnancy-specific glycoprotein 26 [Sourc | 0 | 0 | 0 | 0 | 0 | 0 |
| 19887 | ENSMUSG00000070802 | 1.639494137 | -0.329289297 | 1.814570189 | -0.181469584 | 0.855999005 | NA | Pnmal2 | PNMA-like 2 [Source:MGI Symbol;Acc:MG | 2.4093 | 0 | 3.0187 | 0 | 2.1812 | 2.2277 |
| 19888 | ENSMUSG00000070803 | 9.740960705 | 0.528619963 | 0.722115328 | 0.732043681 | 0.464141907 | 0.862122582 | Cited4 | Cbp/p300-interacting transactivator, with | 4.8186 | 7.1954 | 12.075 | 13.496 | 14.178 | 6.6831 |
| 19889 | ENSMUSG00000070806 | 3.742685076 | -0.793304541 | 1.399418075 | -0.566881731 | 0.570794537 | NA | Zmynd12 | zinc finger, MYND domain containing 12 [! | 4.0155 | 10.279 | 0 | 4.8199 | 0 | 3.3415 |
| 19890 | ENSMUSG00000070808 | 110.5282334 | -0.036347167 | 0.234602186 | -0.154931066 | 0.876875662 | 0.980632845 | Bicra | BRD4 interacting chromatin remodeling co | 97.978 | 135.69 | 102.64 | 119.53 | 97.065 | 110.27 |
| 19891 | ENSMUSG00000070810 | 0 | NA | NA | NA | NA | NA | Sult2a6 | sulfotransferase family 2A, dehydroepiand | 0 | 0 | 0 | 0 | 0 | 0 |
| 19892 | ENSMUSG00000070811 | 0.160662974 | 1.126924175 | 4.080472857 | 0.276174898 | 0.782413736 | NA | Sult2a2 | sulfotransferase family 2A, dehydroepiand | 0 | 0 | 0 | 0.964 | 0 | 0 |
| 19893 | ENSMUSG00000070814 | 48.75640984 | -0.096997293 | 0.310460429 | -0.312430455 | 0.754713408 | 0.953048681 | Zswim9 | zinc finger SWIM-type containing 9 [Sourc | 51.399 | 50.368 | 49.306 | 43.379 | 49.078 | 49.009 |
| 19894 | ENSMUSG00000070815 | 0 | NA | NA | NA | NA | NA | Vmn1r87 | vomeronasal 1 receptor 87 [Source:MGI S | 0 | 0 | 0 | 0 | 0 | 0 |
| 19895 | ENSMUSG00000070816 | 0 | NA | NA | NA | NA | NA | Vmn1r86 | vomeronasal 1 receptor 86 [Source:MGI S | 0 | 0 | 0 | 0 | 0 | 0 |
| 19896 | ENSMUSG00000070817 | 0 | NA | NA | NA | NA | NA | Vmn1r85 | vomeronasal 1 receptor 85 [Source:MGI S | 0 | 0 | 0 | 0 | 0 | 0 |
| 19897 | ENSMUSG00000070820 | 0 | NA | NA | NA | NA | NA | Olfr1339 | olfactory receptor 1339 [Source:MGI Syml | 0 | 0 | 0 | 0 | 0 | 0 |
| 19898 | ENSMUSG00000070821 | 0 | NA | NA | NA | NA | NA | Olfr1340 | olfactory receptor 1340 [Source:MGI Syml | 0 | 0 | 0 | 0 | 0 | 0 |
| 19899 | ENSMUSG00000070822 | 2.833135535 | -0.397225876 | 1.263550589 | -0.314372752 | 0.753237948 | NA | Zscan18 | zinc finger and SCAN domain containing 1 | 1.6062 | 3.0838 | 5.0312 | 2.8919 | 3.2719 | 1.1138 |
| 19900 | ENSMUSG00000070828 | 0 | NA | NA | NA | NA | NA | Zscan4f | zinc finger and SCAN domain containing 4 | 0 | 0 | 0 | 0 | 0 | 0 |
| 19901 | ENSMUSG00000070834 | 0 | NA | NA | NA | NA | NA | Gm13015 | predicted gene 13015 [Source:MGI Symbo | 0 | 0 | 0 | 0 | 0 | 0 |
| 19902 | ENSMUSG00000070837 | 0.305170013 | -1.657836477 | 4.041480335 | -0.410205257 | 0.681655385 | NA | Aurkc | aurora kinase C [Source:MGI Symbol;Acc:I | 0.8031 | 1.0279 | 0 | 0 | 0 | 0 |
| 19903 | ENSMUSG00000070841 | 0 | NA | NA | NA | NA | NA | Vmn2r34 | vomeronasal 2, receptor 34 [Source:MGI S | 0 | 0 | 0 | 0 | 0 | 0 |
| 19904 | ENSMUSG00000070844 | 0 | NA | NA | NA | NA | NA | Vmn2r42 | vomeronasal 2, receptor 42 [Source:MGI S | 0 | 0 | 0 | 0 | 0 | 0 |
| 19905 | ENSMUSG00000070847 | 0.160662974 | 1.126924175 | 4.080472857 | 0.276174898 | 0.782413736 | NA | Vmn2r30 | vomeronasal 2, receptor 30 [Source:MGI S | 0 | 0 | 0 | 0.964 | 0 | 0 |
| 19906 | ENSMUSG00000070852 | 0 | NA | NA | NA | NA | NA | Olfr1121 | olfactory receptor 1121 [Source:MGI Syml | 0 | 0 | 0 | 0 | 0 | 0 |
| 19907 | ENSMUSG00000070853 | 0 | NA | NA | NA | NA | NA | Olfr1150-ps1 | olfactory receptor 1150, pseudogene 1 [So | 0 | 0 | 0 | 0 | 0 | 0 |
| 19908 | ENSMUSG00000070855 | 0 | NA | NA | NA | NA | NA | Olfr1116 | olfactory receptor 1116 [Source:MGI Syml | 0 | 0 | 0 | 0 | 0 | 0 |
| 19909 | ENSMUSG00000070856 | 0 | NA | NA | NA | NA | NA | Olfr1115 | olfactory receptor 1115 [Source:MGI Syml | 0 | 0 | 0 | 0 | 0 | 0 |
| 19910 | ENSMUSG00000070857 | 0 | NA | NA | NA | NA | NA | Olfr1113 | olfactory receptor 1113 [Source:MGI Syml | 0 | 0 | 0 | 0 | 0 | 0 |
| 19911 | ENSMUSG00000070858 | 5.599759896 | 0.70746777 | 1.052141873 | 0.672407199 | 0.501324501 | 0.880974509 | Gm1673 | predicted gene 1673 [Source:MGI Symbol | 1.6062 | 6.1675 | 5.0312 | 1.928 | 3.2719 | 15.594 |
| 19912 | ENSMUSG00000070866 | 0 | NA | NA | NA | NA | NA | Zfp804a | zinc finger protein 804A [Source:MGI Sym | 0 | 0 | 0 | 0 | 0 | 0 |
| 19913 | ENSMUSG00000070867 | 4.724029627 | 0.223483769 | 1.036591667 | 0.215594796 | 0.829303612 | NA | Trabd2b | TraB domain containing 2B [Source:MGI S | 4.0155 | 4.1117 | 5.0312 | 9.6398 | 1.0906 | 4.4554 |
| 19914 | ENSMUSG00000070868 | 0.517184577 | -0.812115922 | 3.393570547 | -0.239310163 | 0.810865085 | NA | Skint3 | selection and upkeep of intraepithelial T c | 0 | 0 | 2.0125 | 0 | 1.0906 | 0 |
| 19915 | ENSMUSG00000070870 | 0 | NA | NA | NA | NA | NA | Cryge | crystallin, gamma E [Source:MGI Symbol;A | 0 | 0 | 0 | 0 | 0 | 0 |
| 19916 | ENSMUSG00000070871 | 113.9564041 | 0.048390803 | 0.213721155 | 0.226420277 | 0.820874559 | 0.969160361 | Ccnyl1 | cyclin Y-like 1 [Source:MGI Symbol;Acc:M | 121.27 | 109.99 | 104.65 | 118.57 | 113.42 | 115.84 |
| 19917 | ENSMUSG00000070873 | 30.94085775 | -1.180056453 | 0.479026489 | -2.463447179 | 0.013760817 | 0.134939971 | Lilra5 | leukocyte immunoglobulin-like receptor, s | 33.73 | 63.731 | 31.194 | 10.604 | 18.541 | 27.846 |
| 19918 | ENSMUSG00000070875 | 0 | NA | NA | NA | NA | NA | Olfr1100 | olfactory receptor 1100 [Source:MGI Syml | 0 | 0 | 0 | 0 | 0 | 0 |
| 19919 | ENSMUSG00000070877 | 0 | NA | NA | NA | NA | NA | Ldlrad1 | low density lipoprotein receptor class A de | 0 | 0 | 0 | 0 | 0 | 0 |
| 19920 | ENSMUSG00000070880 | 0 | NA | NA | NA | NA | NA | Gad1 | glutamate decarboxylase 1 [Source:MGI S | 0 | 0 | 0 | 0 | 0 | 0 |
| 19921 | ENSMUSG00000070883 | 4.524826487 | 0.775047115 | 1.026209471 | 0.75525235 | 0.450097583 | NA | Ccdc173 | coiled-coil domain containing 173 [Source | 4.8186 | 2.0558 | 3.0187 | 2.8919 | 5.4531 | 8.9108 |
| 19922 | ENSMUSG00000070890 | 0 | NA | NA | NA | NA | NA | Pramel19 | PRAME like 19 [Source:MGI Symbol;Acc:M | 0 | 0 | 0 | 0 | 0 | 0 |
| 19923 | ENSMUSG00000070891 | 5.915384731 | 0.006965077 | 0.90991857 | 0.007654615 | 0.99389256 | 0.998618081 | Gm12689 | predicted gene 12689 [Source:MGI Symbo | 10.44 | 3.0838 | 4.025 | 5.7839 | 4.3625 | 7.7969 |
| 19924 | ENSMUSG00000070896 | 0 | NA | NA | NA | NA | NA | Rpl23a-ps1 | ribosomal protein 23A, pseudogene 1 [Sou | 0 | 0 | 0 | 0 | 0 | 0 |
| 19925 | ENSMUSG00000070900 | 0 | NA | NA | NA | NA | NA | NA | NA | 0 | 0 | 0 | 0 | 0 | 0 |
| 19926 | ENSMUSG00000070902 | 0 | NA | NA | NA | NA | NA | Zfp352 | zinc finger protein 352 [Source:MGI Symb | 0 | 0 | 0 | 0 | 0 | 0 |
| 19927 | ENSMUSG00000070904 | 0 | NA | NA | NA | NA | NA | Ifna4 | interferon alpha 4 [Source:MGI Symbol;Ac | 0 | 0 | 0 | 0 | 0 | 0 |
| 19928 | ENSMUSG00000070908 | 0 | NA | NA | NA | NA | NA | Gm13288 | predicted gene 13288 [Source:MGI Symbo | 0 | 0 | 0 | 0 | 0 | 0 |
| 19929 | ENSMUSG00000070923 | 1013.415002 | 0.026629184 | 0.111417107 | 0.239004446 | 0.811102135 | 0.966452061 | Klhl9 | kelch-like 9 [Source:MGI Symbol;Acc:MGI | 1068.9 | 972.41 | 970.02 | 1007.4 | 1096.1 | 965.7 |
| 19930 | ENSMUSG00000070933 | 0 | NA | NA | NA | NA | NA | Speer4d | spermatogenesis associated glutamate (E) | 0 | 0 | 0 | 0 | 0 | 0 |
| 19931 | ENSMUSG00000070934 | 729.6892783 | 0.020852008 | 0.111263764 | 0.187410595 | 0.851338716 | 0.973568625 | Rraga | Ras-related GTP binding A [Source:MGI Sy | 693.88 | 747.3 | 732.55 | 730.7 | 727.44 | 746.28 |
| 19932 | ENSMUSG00000070939 | 854.6862748 | -0.087893582 | 0.115003953 | -0.764265742 | 0.444708896 | 0.851914525 | Tgfbrap1 | transforming growth factor, beta receptor | 881.81 | 888.12 | 872.41 | 839.62 | 894.31 | 751.84 |

| | | | | | | | | | | | | | | | |
|---|---|---|---|---|---|---|---|---|---|---|---|---|---|---|---|
| 19933 | ENSMUSG00000070942 | 3.634563345 | -0.426260118 | 1.136995802 | -0.374900345 | 0.707734583 | NA | Il1rl2 | interleukin 1 receptor-like 2 [Source:MGI | 2.4093 | 6.1675 | 4.025 | 4.8199 | 3.2719 | 1.1138 |
| 19934 | ENSMUSG00000070943 | 0 | NA | NA | NA | NA | NA | Olfr356 | olfactory receptor 356 [Source:MGI Symb | 0 | 0 | 0 | 0 | 0 | 0 |
| 19935 | ENSMUSG00000070953 | 174.5822084 | 0.077411711 | 0.18296232 | 0.42310193 | 0.672220894 | 0.931191575 | Rabepk | Rab9 effector protein with kelch motifs [S | 178.29 | 150.08 | 181.12 | 183.16 | 178.86 | 175.99 |
| 19936 | ENSMUSG00000070960 | 4.810259373 | 1.140601576 | 0.985384852 | 1.157518885 | 0.247060428 | NA | Gm19680 | predicted gene, 19680 [Source:MGI Symb | 4.8186 | 2.0558 | 2.0125 | 6.7478 | 6.5437 | 6.6831 |
| 19937 | ENSMUSG00000070972 | 425.7397189 | -0.226478603 | 0.141426193 | -1.601390784 | 0.109290393 | 0.486233674 | Dnajc25 | DnaJ heat shock protein family (Hsp40) m | 437.69 | 505.74 | 434.7 | 426.08 | 389.35 | 360.89 |
| 19938 | ENSMUSG00000070977 | 0 | NA | NA | NA | NA | NA | Defb44-ps | defensin beta 44, pseudogene [Source:M | 0 | 0 | 0 | 0 | 0 | 0 |
| 19939 | ENSMUSG00000070979 | 0 | NA | NA | NA | NA | NA | Actl7a | actin-like 7a [Source:MGI Symbol;Acc:MG | 0 | 0 | 0 | 0 | 0 | 0 |
| 19940 | ENSMUSG00000070980 | 0 | NA | NA | NA | NA | NA | Actl7b | actin-like 7b [Source:MGI Symbol;Acc:MG | 0 | 0 | 0 | 0 | 0 | 0 |
| 19941 | ENSMUSG00000070983 | 0 | NA | NA | NA | NA | NA | Olfr270 | olfactory receptor 270 [Source:MGI Symb | 0 | 0 | 0 | 0 | 0 | 0 |
| 19942 | ENSMUSG00000070985 | 686.7475706 | -0.276865527 | 0.176456727 | -1.569027898 | 0.116641439 | 0.501814922 | Acnat1 | acyl-coenzyme A amino acid N-acyltransfe | 585.46 | 771.97 | 900.59 | 593.81 | 523.5 | 745.16 |
| 19943 | ENSMUSG00000070990 | 0 | NA | NA | NA | NA | NA | Foxe1 | forkhead box E1 [Source:MGI Symbol;Acc | 0 | 0 | 0 | 0 | 0 | 0 |
| 19944 | ENSMUSG00000070995 | 0 | NA | NA | NA | NA | NA | Gm6313 | predicted gene 6313 [Source:MGI Symbol | 0 | 0 | 0 | 0 | 0 | 0 |
| 19945 | ENSMUSG00000070997 | 22.72888019 | -0.060902326 | 0.511052949 | -0.119170286 | 0.905140447 | 0.983972587 | 1700055D18 | RIKEN cDNA 1700055D18 gene [Source:M | 20.078 | 26.726 | 23.144 | 36.631 | 13.087 | 16.708 |
| 19946 | ENSMUSG00000070999 | 0 | NA | NA | NA | NA | NA | Ccin | calicin [Source:MGI Symbol;Acc:MGI:3045 | 0 | 0 | 0 | 0 | 0 | 0 |
| 19947 | ENSMUSG00000071000 | 0 | NA | NA | NA | NA | NA | Olfr155 | olfactory receptor 155 [Source:MGI Symb | 0 | 0 | 0 | 0 | 0 | 0 |
| 19948 | ENSMUSG00000071001 | 3.827500005 | -0.242110508 | 1.124851382 | -0.215237774 | 0.829581942 | NA | Hrct1 | histidine rich carboxyl terminus 1 [Source: | 7.2279 | 4.1117 | 1.0062 | 2.8919 | 3.2719 | 4.4554 |
| 19949 | ENSMUSG00000071005 | 0 | NA | NA | NA | NA | NA | Ccl19 | chemokine (C-C motif) ligand 19 [Source:N | 0 | 0 | 0 | 0 | 0 | 0 |
| 19950 | ENSMUSG00000071014 | 1034.027919 | 0.053476757 | 0.120626099 | 0.443326591 | 0.657529525 | 0.927303282 | Ndufb6 | NADH:ubiquinone oxidoreductase subuni | 1005.5 | 909.71 | 1129 | 987.11 | 1110.2 | 1062.6 |
| 19951 | ENSMUSG00000071015 | 0 | NA | NA | NA | NA | NA | Gm136 | predicted gene 136 [Source:MGI Symbol;A | 0 | 0 | 0 | 0 | 0 | 0 |
| 19952 | ENSMUSG00000071019 | 0 | NA | NA | NA | NA | NA | Sdr16c6 | short chain dehydrogenase/reductase fan | 0 | 0 | 0 | 0 | 0 | 0 |
| 19953 | ENSMUSG00000071033 | 0 | NA | NA | NA | NA | NA | Gm10308 | predicted gene 10308 [Source:MGI Symbo | 0 | 0 | 0 | 0 | 0 | 0 |
| 19954 | ENSMUSG00000071035 | 1.002992299 | -1.021045543 | 2.445409081 | -0.41753568 | 0.676286632 | NA | Gm5499 | predicted pseudogene 5499 [Source:MGI | 0 | 3.0838 | 1.0062 | 1.928 | 0 | 0 |
| 19955 | ENSMUSG00000071036 | 3.603822692 | 0.861562474 | 1.116349333 | 0.771767804 | 0.440251964 | NA | Gm10309 | predicted gene 10309 [Source:MGI Symbo | 1.6062 | 3.0838 | 3.0187 | 2.8919 | 5.4531 | 5.5692 |
| 19956 | ENSMUSG00000071037 | 13.13239428 | 0.157127324 | 0.628978377 | 0.249813554 | 0.802731537 | 0.963464356 | Camkmt | calmodulin-lysine N-methyltransferase [S | 17.668 | 9.2513 | 10.062 | 8.6758 | 13.087 | 20.049 |
| 19957 | ENSMUSG00000071040 | 0.181769715 | 1.126924175 | 4.080472857 | 0.276174898 | 0.782413736 | NA | Gm9386 | predicted pseudogene 9386 [Source:MGI | 0 | 0 | 0 | 0 | 1.0906 | 0 |
| 19958 | ENSMUSG00000071041 | 3.638047727 | -0.13916506 | 1.133271405 | -0.12279941 | 0.902265941 | NA | Impdh2-ps | inosine monophosphate dehydrogenase 2 | 2.4093 | 2.0558 | 7.0437 | 4.8199 | 3.2719 | 2.2277 |
| 19959 | ENSMUSG00000071042 | 53.59510736 | -0.161075986 | 0.335994475 | -0.479400699 | 0.631653596 | 0.921549084 | Rasgrp3 | RAS, guanyl releasing protein 3 [Source:M | 66.657 | 54.48 | 48.3 | 48.199 | 35.99 | 67.944 |
| 19960 | ENSMUSG00000071047 | 2.458680716 | -0.496918048 | 1.422761982 | -0.34926295 | 0.72689191 | NA | Ces1a | carboxylesterase 1A [Source:MGI Symbol; | 1.6062 | 3.0838 | 4.025 | 3.8559 | 2.1812 | 0 |
| 19961 | ENSMUSG00000071052 | 1.420358007 | 0.127664639 | 1.985043573 | 0.064313268 | 0.948720789 | NA | Rpl7a-ps5 | ribosomal protein L7A, pseudogene 5 [Sou | 0 | 3.0838 | 1.0062 | 0 | 1.0906 | 3.3415 |
| 19962 | ENSMUSG00000071054 | 896.7060458 | -0.178194754 | 0.119135137 | -1.495736341 | 0.134722376 | 0.537616739 | Safb | scaffold attachment factor B [Source:MGI | 876.18 | 948.77 | 1032.4 | 903.25 | 832.14 | 787.49 |
| 19963 | ENSMUSG00000071064 | 2.330582295 | -2.513441086 | 1.612889905 | -1.558346344 | 0.119151169 | NA | Zfp827 | zinc finger protein 827 [Source:MGI Symb | 4.8186 | 2.0558 | 5.0312 | 0.964 | 0 | 1.1138 |
| 19964 | ENSMUSG00000071065 | 0 | NA | NA | NA | NA | NA | Olfr806 | olfactory receptor 806 [Source:MGI Symb | 0 | 0 | 0 | 0 | 0 | 0 |
| 19965 | ENSMUSG00000071068 | 5.683640806 | -3.344545496 | 1.213538982 | -2.756026419 | 0.005850827 | 0.074161384 | Treml2 | triggering receptor expressed on myeloid | 8.8341 | 11.307 | 11.069 | 2.8919 | 0 | 0 |
| 19966 | ENSMUSG00000071072 | 1869.207342 | 0.08116288 | 0.095353355 | 0.851180122 | 0.394669303 | 0.822855164 | Ptges3 | prostaglandin E synthase 3 [Source:MGI S | 1794.1 | 1849.2 | 1806.2 | 1813.2 | 1963.1 | 1989.3 |
| 19967 | ENSMUSG00000071073 | 1.06918069 | 0.688566554 | 2.350891365 | 0.292895947 | 0.769601687 | NA | Lrrc73 | leucine rich repeat containing 73 [Source: | 2.4093 | 0 | 0 | 2.8919 | 0 | 1.1138 |
| 19968 | ENSMUSG00000071074 | 1127.904521 | 0.154536941 | 0.106084886 | 1.456729103 | 0.145191173 | 0.555361204 | Yipf3 | Yip1 domain family, member 3 [Source:M | 1024.8 | 1048.5 | 1130 | 1207.9 | 1125.5 | 1230.8 |
| 19969 | ENSMUSG00000071076 | 618.646461 | -0.117454853 | 0.123623974 | -0.950097692 | 0.342062616 | 0.783839302 | Jund | jun D proto-oncogene [Source:MGI Symbo | 636.86 | 631.14 | 664.12 | 642.97 | 596.57 | 540.21 |
| 19970 | ENSMUSG00000071078 | 15.31084433 | -0.622021569 | 0.571648289 | -1.088119358 | 0.276542419 | 0.730038567 | Nr2c2ap | nuclear receptor 2C2-associated protein [ | 12.047 | 20.558 | 23.144 | 9.6398 | 11.997 | 14.48 |
| 19971 | ENSMUSG00000071089 | 0 | NA | NA | NA | NA | NA | Trim75 | tripartite motif-containing 75 [Source:MG | 0 | 0 | 0 | 0 | 0 | 0 |
| 19972 | ENSMUSG00000071102 | 0 | NA | NA | NA | NA | NA | Gm10313 | predicted pseudogene 10313 [Source:MG | 0 | 0 | 0 | 0 | 0 | 0 |
| 19973 | ENSMUSG00000071103 | 12.24948035 | 0.70091184 | 0.609942055 | 1.149144963 | 0.250496211 | 0.702302228 | 1700029J07I | RIKEN cDNA 1700029J07 gene [Source:M | 9.6372 | 9.2513 | 9.0562 | 12.532 | 18.541 | 14.48 |
| 19974 | ENSMUSG00000071104 | 0 | NA | NA | NA | NA | NA | Ccdc110 | coiled-coil domain containing 110 [Source | 0 | 0 | 0 | 0 | 0 | 0 |
| 19975 | ENSMUSG00000071112 | 1.002521471 | 1.366653481 | 2.416683574 | 0.565507829 | 0.571728401 | NA | Spx | spexin hormone [Source:MGI Symbol;Acc | 1.6062 | 0 | 0 | 0 | 2.1812 | 2.2277 |
| 19976 | ENSMUSG00000071113 | 0 | NA | NA | NA | NA | NA | Mboat4 | membrane bound O-acyltransferase doma | 0 | 0 | 0 | 0 | 0 | 0 |
| 19977 | ENSMUSG00000071138 | 0 | NA | NA | NA | NA | NA | Tex24 | testis expressed gene 24 [Source:MGI Sym | 0 | 0 | 0 | 0 | 0 | 0 |
| 19978 | ENSMUSG00000071141 | 9.621170661 | 0.153120403 | 0.687177625 | 0.222825071 | 0.823671654 | 0.96943211 | Rpl36a-ps3 | ribosomal protein L36A, pseudogene 3 [S | 8.031 | 12.335 | 7.0437 | 11.568 | 8.7249 | 10.025 |
| 19979 | ENSMUSG00000071147 | 0 | NA | NA | NA | NA | NA | Tas2r140 | taste receptor, type 2, member 140 [Sour | 0 | 0 | 0 | 0 | 0 | 0 |
| 19980 | ENSMUSG00000071149 | 0 | NA | NA | NA | NA | NA | Tas2r115 | taste receptor, type 2, member 115 [Sour | 0 | 0 | 0 | 0 | 0 | 0 |
| 19981 | ENSMUSG00000071150 | 0 | NA | NA | NA | NA | NA | Tas2r121 | taste receptor, type 2, member 121 [Sour | 0 | 0 | 0 | 0 | 0 | 0 |
| 19982 | ENSMUSG00000071151 | 1.150957089 | 1.484380014 | 2.018008976 | 0.735566606 | 0.461994491 | NA | Gm4799 | predicted gene 4799 [Source:MGI Symbol | 0.8031 | 0 | 1.0062 | 2.8919 | 1.0906 | 1.1138 |
| 19983 | ENSMUSG00000071158 | 0 | NA | NA | NA | NA | NA | Klrh1 | killer cell lectin-like receptor subfamily H, | 0 | 0 | 0 | 0 | 0 | 0 |
| 19984 | ENSMUSG00000071165 | 0 | NA | NA | NA | NA | NA | Gm6040 | predicted gene 6040 [Source:MGI Symbol | 0 | 0 | 0 | 0 | 0 | 0 |
| 19985 | ENSMUSG00000071169 | 0 | NA | NA | NA | NA | NA | Defb46 | defensin beta 46 [Source:MGI Symbol;Acc | 0 | 0 | 0 | 0 | 0 | 0 |
| 19986 | ENSMUSG00000071172 | 1071.506035 | -0.256329289 | 0.129012956 | -1.986849201 | 0.04693911 | 0.300126868 | Srsf3 | serine and arginine-rich splicing factor 3 [ | 1044 | 1115.3 | 1340.3 | 895.54 | 1047 | 986.87 |
| 19987 | ENSMUSG00000071176 | 34.72856293 | -0.491333423 | 0.393802094 | -1.247665846 | 0.212153454 | 0.656735971 | Arhgef10 | Rho guanine nucleotide exchange factor ( | 46.58 | 32.893 | 42.262 | 34.703 | 19.631 | 32.301 |
| 19988 | ENSMUSG00000071177 | 42547.86567 | 0.28277974 | 0.103551508 | 2.730812374 | 0.006317843 | 0.078036959 | Serpina1d | serine (or cysteine) peptidase inhibitor, cl | 33724 | 39328 | 42123 | 45298 | 45986 | 48828 |

| | | | | | | | | | | | | | | | |
|---|---|---|---|---|---|---|---|---|---|---|---|---|---|---|---|
| 19989 | ENSMUSG00000071178 | 66920.00225 | 0.027659814 | 0.092523726 | 0.298948334 | 0.764979467 | 0.954980327 | Serpina1b | serine (or cysteine) preptidase inhibitor, c | 64998 | 70740 | 63098 | 69097 | 62856 | 70731 |
| 19990 | ENSMUSG00000071179 | 0 | NA | NA | NA | NA | NA | Serpina16 | serine (or cysteine) peptidase inhibitor, cl | 0 | 0 | 0 | 0 | 0 | 0 |
| 19991 | ENSMUSG00000071180 | 314.4503759 | 0.076347968 | 0.159944829 | 0.477339393 | 0.633120464 | 0.922107984 | Smim15 | small integral membrane protein 15 [Sour | 334.89 | 276.51 | 305.9 | 280.52 | 341.36 | 347.52 |
| 19992 | ENSMUSG00000071181 | 9.75564969 | 0.271842846 | 0.762808931 | 0.356370823 | 0.721562877 | 0.9447719 | 3830408C21 | RIKEN cDNA 3830408C21 gene [Source:M | 6.4248 | 5.1396 | 15.094 | 15.424 | 11.997 | 4.4554 |
| 19993 | ENSMUSG00000071185 | 0 | NA | NA | NA | NA | NA | Olfr1357 | olfactory receptor 1357 [Source:MGI Syml | 0 | 0 | 0 | 0 | 0 | 0 |
| 19994 | ENSMUSG00000071192 | 7.221382682 | -0.998904844 | 0.826689289 | -1.208319567 | 0.226924365 | 0.67343689 | Wfikkn1 | WAP, FS, Ig, KU, and NTR-containing prote | 13.653 | 6.1675 | 9.0562 | 6.7478 | 4.3625 | 3.3415 |
| 19995 | ENSMUSG00000071195 | 0 | NA | NA | NA | NA | NA | Gm10318 | predicted gene 10318 [Source:MGI Symbo | 0 | 0 | 0 | 0 | 0 | 0 |
| 19996 | ENSMUSG00000071202 | 1.695392952 | 2.069057133 | 1.795867266 | 1.152121413 | 0.249271185 | NA | Ccdc78 | coiled-coil domain containing 78 [Source:I | 0 | 1.0279 | 1.0062 | 4.8199 | 1.0906 | 2.2277 |
| 19997 | ENSMUSG00000071203 | 31.59936141 | -0.866291122 | 0.4002425 | -2.164415626 | 0.030432466 | 0.230941859 | Naip5 | NLR family, apoptosis inhibitory protein 5 | 40.958 | 49.34 | 32.2 | 24.099 | 20.722 | 22.277 |
| 19998 | ENSMUSG00000071204 | 568.2525785 | 0.5783031 | 0.206506103 | 2.800416504 | 0.005103671 | 0.067538422 | Gm10319 | predicted pseudogene 10319 [Source:MG | 604.74 | 382.39 | 379.35 | 749.97 | 789.61 | 503.46 |
| 19999 | ENSMUSG00000071207 | 0 | NA | NA | NA | NA | NA | Gm8587 | predicted pseudogene 8587 [Source:MGI | 0 | 0 | 0 | 0 | 0 | 0 |
| 20000 | ENSMUSG00000071226 | 343.7459516 | 0.270060005 | 0.182506228 | 1.479730352 | 0.138945222 | 0.544782785 | Cecr2 | CECR2, histone acetyl-lysine reader [Sourc | 236.11 | 319.68 | 380.36 | 380.77 | 347.91 | 397.64 |
| 20001 | ENSMUSG00000071229 | 0 | NA | NA | NA | NA | NA | Timm8a2 | translocase of inner mitochondrial membr | 0 | 0 | 0 | 0 | 0 | 0 |
| 20002 | ENSMUSG00000071230 | 0 | NA | NA | NA | NA | NA | Npw | neuropeptide W [Source:MGI Symbol;Acc | 0 | 0 | 0 | 0 | 0 | 0 |
| 20003 | ENSMUSG00000071234 | 0 | NA | NA | NA | NA | NA | Syndig1l | synapse differentiation inducing 1 like [So | 0 | 0 | 0 | 0 | 0 | 0 |
| 20004 | ENSMUSG00000071235 | 0 | NA | NA | NA | NA | NA | Vrtn | vertebrae development associated [Sourc | 0 | 0 | 0 | 0 | 0 | 0 |
| 20005 | ENSMUSG00000071237 | 0 | NA | NA | NA | NA | NA | Gemin6-ps | gem nuclear organelle associated protein | 0 | 0 | 0 | 0 | 0 | 0 |
| 20006 | ENSMUSG00000071252 | 50.58343783 | -0.130008382 | 0.347016388 | -0.374646231 | 0.707923586 | 0.943550198 | 2210408I21F | RIKEN cDNA 2210408I21 gene [Source:MC | 47.383 | 67.843 | 43.269 | 33.739 | 57.803 | 53.465 |
| 20007 | ENSMUSG00000071253 | 1829.691034 | -0.068584488 | 0.107290199 | -0.639242812 | 0.522664985 | 0.888334265 | Slc25a16 | solute carrier family 25 (mitochondrial car | 1790.9 | 1830.7 | 1998.4 | 1765 | 1944.6 | 1648.5 |
| 20008 | ENSMUSG00000071256 | 64.44545998 | 0.098004014 | 0.278317861 | 0.352129804 | 0.724740918 | 0.9448775 | Zfp213 | zinc finger protein 213 [Source:MGI Symb | 53.808 | 69.898 | 63.393 | 68.442 | 66.528 | 64.603 |
| 20009 | ENSMUSG00000071262 | 1.150957089 | 1.484380014 | 2.018008976 | 0.735566606 | 0.461994491 | NA | Zfp957 | zinc finger protein 957 [Source:MGI Symb | 0.8031 | 0 | 1.0062 | 2.8919 | 1.0906 | 1.1138 |
| 20010 | ENSMUSG00000071265 | 0 | NA | NA | NA | NA | NA | 1700086L19I | RIKEN cDNA 1700086L19 gene [Source:MC | 0 | 0 | 0 | 0 | 0 | 0 |
| 20011 | ENSMUSG00000071266 | 281.311811 | 0.21518183 | 0.15708955 | 1.369803599 | 0.170748218 | 0.597556794 | Zfp946 | zinc finger protein 946 [Source:MGI Symb | 248.96 | 243.62 | 288.79 | 300.76 | 322.82 | 282.92 |
| 20012 | ENSMUSG00000071267 | 212.3122732 | -0.002048256 | 0.182553292 | -0.011220043 | 0.991047889 | 0.99859176 | Zfp942 | zinc finger protein 942 [Source:MGI Symb | 188.73 | 197.36 | 251.56 | 198.58 | 213.76 | 223.88 |
| 20013 | ENSMUSG00000071273 | 0 | NA | NA | NA | NA | NA | Gm5145 | predicted pseudogene 5145 [Source:MGI | 0 | 0 | 0 | 0 | 0 | 0 |
| 20014 | ENSMUSG00000071275 | 0 | NA | NA | NA | NA | NA | Fpr-rs6 | formyl peptide receptor, related sequence | 0 | 0 | 0 | 0 | 0 | 0 |
| 20015 | ENSMUSG00000071276 | 0 | NA | NA | NA | NA | NA | Fpr-rs7 | formyl peptide receptor, related sequence | 0 | 0 | 0 | 0 | 0 | 0 |
| 20016 | ENSMUSG00000071280 | 0 | NA | NA | NA | NA | NA | Gm10322 | predicted gene 10322 [Source:MGI Symbo | 0 | 0 | 0 | 0 | 0 | 0 |
| 20017 | ENSMUSG00000071281 | 127.1465289 | 0.135874249 | 0.216669192 | 0.627104611 | 0.530590665 | 0.888794797 | Zfp65 | zinc finger protein 65 [Source:MGI Symbo | 119.66 | 113.07 | 130.81 | 138.81 | 110.15 | 150.37 |
| 20018 | ENSMUSG00000071291 | 42.24667693 | 0.175894118 | 0.363204159 | 0.484284428 | 0.628184026 | 0.920328819 | Zfp58 | zinc finger protein 58 [Source:MGI Symbo | 28.912 | 45.228 | 45.281 | 50.127 | 33.809 | 50.123 |
| 20019 | ENSMUSG00000071294 | 0 | NA | NA | NA | NA | NA | Cbx3-ps4 | chromobox 3, pseudogene 4 [Source:MGI | 0 | 0 | 0 | 0 | 0 | 0 |
| 20020 | ENSMUSG00000071302 | 97.55333173 | 0.397035683 | 0.239346336 | 1.658833342 | 0.09714938 | 0.45892125 | 2610044O15 | RIKEN cDNA 2610044O15 gene [Source:M | 73.082 | 81.206 | 98.612 | 102.18 | 119.97 | 110.27 |
| 20021 | ENSMUSG00000071303 | 0.663965228 | -0.037580459 | 3.10181903 | -0.012115619 | 0.990333371 | NA | Rps8-ps1 | ribosomal protein S8, pseudogene 1 [Sour | 0 | 2.0558 | 0 | 1.928 | 0 | 0 |
| 20022 | ENSMUSG00000071311 | 0.483327519 | -0.685704547 | 3.145069396 | -0.218025252 | 0.827409436 | NA | Gpr31b | G protein-coupled receptor 31, D17Leh66 | 0.8031 | 0 | 1.0062 | 0 | 1.0906 | 0 |
| 20023 | ENSMUSG00000071317 | 0 | NA | NA | NA | NA | NA | Bves | blood vessel epicardial substance [Source | 0 | 0 | 0 | 0 | 0 | 0 |
| 20024 | ENSMUSG00000071322 | 0.503122293 | -2.366675966 | 3.785596405 | -0.625179156 | 0.531853481 | NA | Tcp10a | t-complex protein 10a [Source:MGI Symbo | 0 | 0 | 3.0187 | 0 | 0 | 0 |
| 20025 | ENSMUSG00000071324 | 1.577640338 | -2.883093086 | 2.124940614 | -1.356787605 | 0.174848707 | NA | Armc2 | armadillo repeat containing 2 [Source:MG | 3.2124 | 5.1396 | 0 | 0 | 0 | 1.1138 |
| 20026 | ENSMUSG00000071332 | 2.148347266 | -1.690538127 | 1.588768321 | -1.064055788 | 0.287303428 | NA | E230015B07 | RIKEN cDNA E230015B07 gene [Source:M | 4.8186 | 1.0279 | 4.025 | 1.928 | 1.0906 | 0 |
| 20027 | ENSMUSG00000071335 | 208.5283989 | 0.038362641 | 0.176122103 | 0.217818437 | 0.827570579 | 0.969794868 | Mfsd4b3-ps | major facilitator superfamily domain cont | 193.55 | 199.42 | 224.39 | 202.44 | 194.13 | 237.25 |
| 20028 | ENSMUSG00000071337 | 186.4652098 | 0.063464568 | 0.187311759 | 0.338817852 | 0.734746952 | 0.948109711 | Tia1 | cytotoxic granule-associated RNA binding | 154.2 | 205.58 | 188.17 | 199.54 | 190.86 | 180.44 |
| 20029 | ENSMUSG00000071340 | 0 | NA | NA | NA | NA | NA | Trappc3l | trafficking protein particle complex 3 like [ | 0 | 0 | 0 | 0 | 0 | 0 |
| 20030 | ENSMUSG00000071341 | 0 | NA | NA | NA | NA | NA | Egr4 | early growth response 4 [Source:MGI Sym | 0 | 0 | 0 | 0 | 0 | 0 |
| 20031 | ENSMUSG00000071342 | 0 | NA | NA | NA | NA | NA | Lsmem1 | leucine-rich single-pass membrane protei | 0 | 0 | 0 | 0 | 0 | 0 |
| 20032 | ENSMUSG00000071343 | 0.950731196 | 0.382757398 | 2.457599045 | 0.155744444 | 0.876234463 | NA | Gm10327 | predicted pseudogene 10327 [Source:MG | 2.4093 | 0 | 0 | 0 | 2.1812 | 1.1138 |
| 20033 | ENSMUSG00000071347 | 5.321825925 | -0.480112622 | 0.979391785 | -0.49021508 | 0.623981711 | 0.919503423 | C1qtnf9 | C1q and tumor necrosis factor related pro | 6.4248 | 8.2233 | 4.025 | 7.7118 | 1.0906 | 4.4554 |
| 20034 | ENSMUSG00000071350 | 302.145861 | -0.170758828 | 0.184140948 | -0.92732676 | 0.353756904 | 0.79449673 | Setdb2 | SET domain, bifurcated 2 [Source:MGI Syr | 330.08 | 357.72 | 272.69 | 329.68 | 298.83 | 223.88 |
| 20035 | ENSMUSG00000071356 | 0 | NA | NA | NA | NA | NA | Reg3b | regenerating islet-derived 3 beta [Source: | 0 | 0 | 0 | 0 | 0 | 0 |
| 20036 | ENSMUSG00000071359 | 126.0390721 | -0.073642339 | 0.222723246 | -0.330645052 | 0.740912612 | 0.948792385 | Tbpl1 | TATA box binding protein-like 1 [Source:M | 118.06 | 128.49 | 141.88 | 148.45 | 107.97 | 111.38 |
| 20037 | ENSMUSG00000071361 | 0 | NA | NA | NA | NA | NA | Mcpt9 | mast cell protease 9 [Source:MGI Symbol; | 0 | 0 | 0 | 0 | 0 | 0 |
| 20038 | ENSMUSG00000071362 | 0 | NA | NA | NA | NA | NA | Gm10330 | predicted gene 10330 [Source:MGI Symbo | 0 | 0 | 0 | 0 | 0 | 0 |
| 20039 | ENSMUSG00000071369 | 888.9630173 | -0.078648656 | 0.159304189 | -0.493701119 | 0.621517275 | 0.918339551 | Map3k5 | mitogen-activated protein kinase kinase k | 705.12 | 1030 | 1006.2 | 973.62 | 765.61 | 853.2 |
| 20040 | ENSMUSG00000071379 | 232.1322075 | -0.452160576 | 0.174355353 | -2.593327752 | 0.009505214 | 0.104661272 | Hpcal1 | hippocalcin-like 1 [Source:MGI Symbol;Ac | 297.95 | 251.84 | 254.58 | 211.11 | 170.14 | 207.17 |
| 20041 | ENSMUSG00000071392 | 1.711104161 | -1.916517388 | 1.775392458 | -1.079489427 | 0.280369604 | NA | Ect2l | epithelial cell transforming sequence 2 on | 4.0155 | 1.0279 | 3.0187 | 0 | 1.0906 | 1.1138 |
| 20042 | ENSMUSG00000071398 | 0 | NA | NA | NA | NA | NA | 2410004P03 | RIKEN cDNA 2410004P03 gene [Source:MC | 0 | 0 | 0 | 0 | 0 | 0 |
| 20043 | ENSMUSG00000071414 | 0.688942901 | 0.052211695 | 2.79596103 | 0.018673971 | 0.985101193 | NA | Gm6736 | predicted gene 6736 [Source:MGI Symbol | 0 | 2.0558 | 0 | 0.964 | 0 | 1.1138 |
| 20044 | ENSMUSG00000071415 | 3429.618854 | 0.065213207 | 0.124281748 | 0.524720714 | 0.599777355 | 0.915217938 | Rpl23 | ribosomal protein L23 [Source:MGI Symbo | 3036.5 | 3639.9 | 3380 | 2986.4 | 3566.3 | 3968.6 |

| | | | | | | | | | | | | | | | |
|---|---|---|---|---|---|---|---|---|---|---|---|---|---|---|---|
| 20045 | ENSMUSG00000071419 | 0.710049642 | 0.116753315 | 2.770146132 | 0.042146988 | 0.966381523 | NA | Rps15-ps2 | ribosomal protein S15, pseudogene 2 [Sou | 0 | 2.0558 | 0 | 0 | 1.0906 | 1.1138 |
| 20046 | ENSMUSG00000071424 | 0.167707431 | -0.796629176 | 4.080472857 | -0.195229623 | 0.84521319 | NA | Grid2 | glutamate receptor, ionotropic, delta 2 [S | 0 | 0 | 1.0062 | 0 | 0 | 0 |
| 20047 | ENSMUSG00000071428 | 0 | NA | NA | NA | NA | NA | Vmn1r27 | vomeronasal 1 receptor 27 [Source:MGI S | 0 | 0 | 0 | 0 | 0 | 0 |
| 20048 | ENSMUSG00000071434 | 0 | NA | NA | NA | NA | NA | 9230019H11 | RIKEN cDNA 9230019H11 gene [Source:M | 0 | 0 | 0 | 0 | 0 | 0 |
| 20049 | ENSMUSG00000071451 | 131.1672093 | -0.250545205 | 0.209066127 | -1.198401718 | 0.230760664 | 0.677910147 | Psmg4 | proteasome (prosome, macropain) assem | 144.56 | 138.77 | 143.89 | 106.04 | 114.51 | 139.23 |
| 20050 | ENSMUSG00000071452 | 0 | NA | NA | NA | NA | NA | Serpinb9h | serine (or cysteine) peptidase inhibitor, cl | 0 | 0 | 0 | 0 | 0 | 0 |
| 20051 | ENSMUSG00000071454 | 407.6955207 | 0.200401613 | 0.147523219 | 1.358441159 | 0.174323742 | 0.604151393 | Dtnb | dystrobrevin, beta [Source:MGI Symbol;A | 346.94 | 384.44 | 407.53 | 429.93 | 488.6 | 388.73 |
| 20052 | ENSMUSG00000071456 | 38.9368286 | -0.07552244 | 0.402057781 | -0.187839767 | 0.851002261 | 0.973568625 | 1110002L01 | RIKEN cDNA 1110002L01 gene [Source:M | 49.792 | 29.81 | 40.25 | 53.019 | 23.994 | 36.757 |
| 20053 | ENSMUSG00000071470 | 0 | NA | NA | NA | NA | NA | Ccnb1ip1 | cyclin B1 interacting protein 1 [Source:M | 0 | 0 | 0 | 0 | 0 | 0 |
| 20054 | ENSMUSG00000071471 | 0 | NA | NA | NA | NA | NA | Krtap26-1 | keratin associated protein 26-1 [Source:M | 0 | 0 | 0 | 0 | 0 | 0 |
| 20055 | ENSMUSG00000071475 | 0 | NA | NA | NA | NA | NA | Rpl21-ps5 | ribosomal protein L21, pseudogene 5 [Sou | 0 | 0 | 0 | 0 | 0 | 0 |
| 20056 | ENSMUSG00000071477 | 246.6444301 | 0.055619648 | 0.182918088 | 0.304068606 | 0.761075624 | 0.95402891 | Zfp777 | zinc finger protein 777 [Source:MGI Symb | 273.86 | 215.86 | 235.46 | 257.38 | 289.01 | 208.29 |
| 20057 | ENSMUSG00000071478 | 0.462220778 | -0.685400455 | 3.191475091 | -0.21475977 | 0.82995462 | NA | H2ac7 | H2A clustered histone 7 [Source:MGI Sym | 0.8031 | 0 | 1.0062 | 0.964 | 0 | 0 |
| 20058 | ENSMUSG00000071481 | 0 | NA | NA | NA | NA | NA | Olfr437 | olfactory receptor 437 [Source:MGI Symb | 0 | 0 | 0 | 0 | 0 | 0 |
| 20059 | ENSMUSG00000071489 | 0 | NA | NA | NA | NA | NA | Ptgdr | prostaglandin D receptor [Source:MGI Syr | 0 | 0 | 0 | 0 | 0 | 0 |
| 20060 | ENSMUSG00000071490 | 0 | NA | NA | NA | NA | NA | Vmn1r212 | vomeronasal 1 receptor 212 [Source:MGI | 0 | 0 | 0 | 0 | 0 | 0 |
| 20061 | ENSMUSG00000071491 | 0 | NA | NA | NA | NA | NA | Vmn1r209 | vomeronasal 1 receptor 209 [Source:MGI | 0 | 0 | 0 | 0 | 0 | 0 |
| 20062 | ENSMUSG00000071493 | 0 | NA | NA | NA | NA | NA | Vmn1r208 | vomeronasal 1 receptor 208 [Source:MGI | 0 | 0 | 0 | 0 | 0 | 0 |
| 20063 | ENSMUSG00000071494 | 0 | NA | NA | NA | NA | NA | Olfr455 | olfactory receptor 455 [Source:MGI Symb | 0 | 0 | 0 | 0 | 0 | 0 |
| 20064 | ENSMUSG00000071497 | 1.673737746 | -1.038170307 | 1.728080672 | -0.600764955 | 0.547996549 | NA | Nutf2-ps1 | nuclear transport factor 2, pseudogene 1 | 1.6062 | 4.1117 | 1.0062 | 0 | 1.0906 | 2.2277 |
| 20065 | ENSMUSG00000071506 | 1.804240148 | -1.392121677 | 1.803836581 | -0.771755985 | 0.440258965 | NA | Tmem139 | transmembrane protein 139 [Source:MGI | 0.8031 | 4.1117 | 3.0187 | 2.8919 | 0 | 0 |
| 20066 | ENSMUSG00000071510 | 0 | NA | NA | NA | NA | NA | Olfr172 | olfactory receptor 172 [Source:MGI Symb | 0 | 0 | 0 | 0 | 0 | 0 |
| 20067 | ENSMUSG00000071516 | 0 | NA | NA | NA | NA | NA | H2ac13 | H2A clustered histone 13 [Source:MGI Syr | 0 | 0 | 0 | 0 | 0 | 0 |
| 20068 | ENSMUSG00000071517 | 0 | NA | NA | NA | NA | NA | Gm10334 | predicted gene 10334 [Source:MGI Symbo | 0 | 0 | 0 | 0 | 0 | 0 |
| 20069 | ENSMUSG00000071519 | 0 | NA | NA | NA | NA | NA | Prss3 | protease, serine 3 [Source:MGI Symbol;Ac | 0 | 0 | 0 | 0 | 0 | 0 |
| 20070 | ENSMUSG00000071521 | 0 | NA | NA | NA | NA | NA | Try10 | trypsin 10 [Source:MGI Symbol;Acc:MGI:3 | 0 | 0 | 0 | 0 | 0 | 0 |
| 20071 | ENSMUSG00000071522 | 0 | NA | NA | NA | NA | NA | Olfr263 | olfactory receptor 263 [Source:MGI Symb | 0 | 0 | 0 | 0 | 0 | 0 |
| 20072 | ENSMUSG00000071528 | 2532.773759 | -0.010830526 | 0.116525663 | -0.092945413 | 0.925946928 | 0.987546253 | Atp5md | ATP synthase membrane subunit DAPIT [S | 2585.2 | 2532.8 | 2507.6 | 2138.1 | 2630.6 | 2802.4 |
| 20073 | ENSMUSG00000071531 | 2.585609833 | -2.713224818 | 1.763540546 | -1.53851003 | 0.123923958 | NA | Gprin2 | G protein regulated inducer of neurite ou | 10.44 | 0 | 3.0187 | 0.964 | 1.0906 | 0 |
| 20074 | ENSMUSG00000071532 | 3.307635803 | 0.189385698 | 1.261331516 | 0.15014744 | 0.880648293 | NA | Gm10335 | predicted gene 10335 [Source:MGI Symbo | 3.2124 | 0 | 6.0375 | 2.8919 | 4.3625 | 3.3415 |
| 20075 | ENSMUSG00000071533 | 994.481181 | 0.02539618 | 0.112106778 | 0.226535633 | 0.820784848 | 0.969160361 | Pcnp | PEST proteolytic signal containing nuclear | 995.04 | 921.01 | 1041.5 | 1073.9 | 958.65 | 976.84 |
| 20076 | ENSMUSG00000071537 | 1.223363286 | -0.304298652 | 2.246052434 | -0.135481544 | 0.892231155 | NA | Klrg2 | killer cell lectin-like receptor subfamily G, | 0 | 2.0558 | 2.0125 | 0 | 3.2719 | 0 |
| 20077 | ENSMUSG00000071540 | 0 | NA | NA | NA | NA | NA | 3425401B19 | RIKEN cDNA 3425401B19 gene [Source:M | 0 | 0 | 0 | 0 | 0 | 0 |
| 20078 | ENSMUSG00000071547 | 121.3640039 | -0.617121854 | 0.227633573 | -2.711031792 | 0.006707419 | 0.08160729 | Nt5dc2 | 5'-nucleotidase domain containing 2 [Sou | 165.44 | 125.41 | 149.93 | 107.97 | 99.246 | 80.197 |
| 20079 | ENSMUSG00000071550 | 0.401551119 | -2.072982726 | 4.008251979 | -0.517178744 | 0.605031388 | NA | Cfap44 | cilia and flagella associated protein 44 [So | 2.4093 | 0 | 0 | 0 | 0 | 0 |
| 20080 | ENSMUSG00000071551 | 1524.29948 | 0.104596063 | 0.323121667 | 0.323704887 | 0.74616147 | 0.949997329 | Akr1c19 | aldo-keto reductase family 1, member C1 | 1163.7 | 1571.7 | 1672.4 | 1976.2 | 1747.2 | 1014.7 |
| 20081 | ENSMUSG00000071552 | 0.133850373 | -0.796629176 | 4.080472857 | -0.195229623 | 0.84521319 | NA | Tigit | T cell immunoreceptor with Ig and ITIM d | 0.8031 | 0 | 0 | 0 | 0 | 0 |
| 20082 | ENSMUSG00000071553 | 0 | NA | NA | NA | NA | NA | Cpa2 | carboxypeptidase A2, pancreatic [Source: | 0 | 0 | 0 | 0 | 0 | 0 |
| 20083 | ENSMUSG00000071561 | 0 | NA | NA | NA | NA | NA | Cstdc5 | cystatin domain containing 5 [Source:MGI | 0 | 0 | 0 | 0 | 0 | 0 |
| 20084 | ENSMUSG00000071562 | 0 | NA | NA | NA | NA | NA | Stfa1 | stefin A1 [Source:MGI Symbol;Acc:MGI:1 | 0 | 0 | 0 | 0 | 0 | 0 |
| 20085 | ENSMUSG00000071568 | 0 | NA | NA | NA | NA | NA | Gm5874 | predicted gene 5874 [Source:MGI Symbol | 0 | 0 | 0 | 0 | 0 | 0 |
| 20086 | ENSMUSG00000071573 | 29.39785373 | -0.098270706 | 0.429687363 | -0.228702806 | 0.819099913 | 0.969160361 | Rnls | renalase, FAD-dependent amine oxidase [ | 40.155 | 28.782 | 22.137 | 34.703 | 29.447 | 21.163 |
| 20087 | ENSMUSG00000071586 | 0 | NA | NA | NA | NA | NA | Gm10337 | predicted gene 10337 [Source:MGI Symbo | 0 | 0 | 0 | 0 | 0 | 0 |
| 20088 | ENSMUSG00000071591 | 0 | NA | NA | NA | NA | NA | Gm10339 | predicted gene 10339 [Source:MGI Symbo | 0 | 0 | 0 | 0 | 0 | 0 |
| 20089 | ENSMUSG00000071604 | 11.76831687 | -0.613314452 | 0.670420087 | -0.914821116 | 0.360285547 | 0.799598236 | Fam189a2 | family with sequence similarity 189, meml | 10.44 | 12.335 | 20.125 | 14.46 | 5.4531 | 7.7969 |
| 20090 | ENSMUSG00000071613 | 0 | NA | NA | NA | NA | NA | Gm10340 | predicted gene 10340 [Source:MGI Symbo | 0 | 0 | 0 | 0 | 0 | 0 |
| 20091 | ENSMUSG00000071629 | 0 | NA | NA | NA | NA | NA | Olfr1477 | olfactory receptor 1477 [Source:MGI Syml | 0 | 0 | 0 | 0 | 0 | 0 |
| 20092 | ENSMUSG00000071632 | 218.6117689 | 0.316211044 | 0.188383855 | 1.678546408 | 0.093240479 | 0.449843323 | 2510002D24 | RIKEN cDNA 2510002D24 gene [Source:M | 206.4 | 176.8 | 200.24 | 196.65 | 248.66 | 282.92 |
| 20093 | ENSMUSG00000071633 | 1879.483103 | -0.167916946 | 0.105367391 | -1.593632954 | 0.111018269 | 0.489438373 | Gm4952 | predicted gene 4952 [Source:MGI Symbol | 1953.1 | 2110.3 | 1903.8 | 1906.7 | 1774.4 | 1628.4 |
| 20094 | ENSMUSG00000071636 | 0 | NA | NA | NA | NA | NA | Rimbp3 | RIMS binding protein 3 [Source:MGI Symb | 0 | 0 | 0 | 0 | 0 | 0 |
| 20095 | ENSMUSG00000071637 | 345.6475298 | -1.222638868 | 0.251275151 | -4.865737266 | 1.14E-06 | 9.11E-05 | Cebpd | CCAAT/enhancer binding protein (C/EBP), | 479.45 | 584.89 | 387.4 | 183.16 | 307.55 | 131.43 |
| 20096 | ENSMUSG00000071640 | 0.449470461 | -0.545136533 | 3.562105917 | -0.15303771 | 0.878368537 | NA | Stxbp3-ps | syntaxin-binding protein 3, pseudogene [S | 1.6062 | 0 | 0 | 0 | 1.0906 | 0 |
| 20097 | ENSMUSG00000071644 | 2598.972384 | 0.075288329 | 0.105400237 | 0.714308917 | 0.47503618 | 0.869368495 | Eef1g | eukaryotic translation elongation factor 1 | 2410.9 | 2689 | 2493.5 | 2431.2 | 2665.5 | 2903.8 |
| 20098 | ENSMUSG00000071645 | 234.8398122 | -0.119612178 | 0.175420665 | -0.681859106 | 0.495328046 | 0.877937486 | Tut1 | terminal uridylyl transferase 1, U6 snRNA- | 215.23 | 243.62 | 275.71 | 247.74 | 202.86 | 223.88 |
| 20099 | ENSMUSG00000071646 | 960.6704668 | -0.057124755 | 0.111019238 | -0.514548251 | 0.606868735 | 0.916376001 | Mta2 | metastasis-associated gene family, memb | 938.02 | 943.63 | 1057.6 | 904.21 | 978.28 | 942.31 |
| 20100 | ENSMUSG00000071647 | 417.6043852 | -0.084577645 | 0.171061123 | -0.494429381 | 0.621002968 | 0.918069106 | Eml3 | echinoderm microtubule associated prote | 350.15 | 416.31 | 524.25 | 446.32 | 364.27 | 404.33 |

| | | | | | | | | | | | | | | | |
|---|---|---|---|---|---|---|---|---|---|---|---|---|---|---|---|
| 20101 | ENSMUSG00000071648 | 54.90144645 | -0.036118308 | 0.327123678 | -0.110411781 | 0.91208281 | 0.985976515 | Rom1 | rod outer segment membrane protein 1 [! | 52.202 | 54.48 | 60.375 | 68.442 | 35.99 | 57.92 |
| 20102 | ENSMUSG00000071649 | 674.8845949 | 0.187528238 | 0.151312359 | 1.239345145 | 0.215217707 | 0.659728602 | B3gat3 | beta-1,3-glucuronyltransferase 3 (glucuro | 515.59 | 671.23 | 707.39 | 676.71 | 665.28 | 813.11 |
| 20103 | ENSMUSG00000071650 | 3305.888977 | 0.101451559 | 0.10477011 | 0.968325408 | 0.332881879 | 0.77523112 | Ganab | alpha glucosidase 2 alpha neutral subunit | 2838.2 | 3295.5 | 3437.3 | 3564.8 | 3410.4 | 3289.2 |
| 20104 | ENSMUSG00000071652 | 190.2905245 | 0.191929537 | 0.181246071 | 1.058944539 | 0.289625039 | 0.739999393 | Ints5 | integrator complex subunit 5 [Source:MG | 157.41 | 190.16 | 186.16 | 218.82 | 187.59 | 201.61 |
| 20105 | ENSMUSG00000071653 | 2.023711223 | 0.50709869 | 1.644641832 | 0.308333815 | 0.757828338 | NA | 1810009A15 | RIKEN cDNA 1810009A15 gene [Source:M | 0 | 4.1117 | 1.0062 | 4.8199 | 1.0906 | 1.1138 |
| 20106 | ENSMUSG00000071654 | 373.4970637 | -0.086602634 | 0.143373724 | -0.604034212 | 0.545820897 | 0.89549457 | Uqcc3 | ubiquinol-cytochrome c reductase comple | 395.93 | 375.19 | 382.37 | 337.39 | 343.54 | 406.55 |
| 20107 | ENSMUSG00000071655 | 1580.363097 | 0.23367297 | 0.136038936 | 1.717691836 | 0.085852839 | 0.431795087 | Ubxn1 | UBX domain protein 1 [Source:MGI Symbo | 1509.8 | 1351.7 | 1495.3 | 1477.8 | 1581.4 | 2066.2 |
| 20108 | ENSMUSG00000071656 | 2.97289172 | 2.090545375 | 1.456975433 | 1.434852865 | 0.15132903 | NA | Lrrn4cl | LRRN4 C-terminal like [Source:MGI Symbo | 3.2124 | 0 | 0 | 5.7839 | 3.2719 | 5.5692 |
| 20109 | ENSMUSG00000071657 | 779.2615337 | -0.061597628 | 0.12925039 | -0.476575956 | 0.633664109 | 0.922736909 | Bscl2 | Berardinelli-Seip congenital lipodystrophy | 905.1 | 754.49 | 726.51 | 710.45 | 803.79 | 775.24 |
| 20110 | ENSMUSG00000071658 | 1.266067585 | 2.695958699 | 2.278896902 | 1.183010384 | 0.236805032 | NA | Gng3 | guanine nucleotide binding protein (G pro | 0 | 0 | 1.0062 | 0 | 4.3625 | 2.2277 |
| 20111 | ENSMUSG00000071659 | 1914.694002 | 0.012324086 | 0.104000306 | 0.118500476 | 0.905671118 | 0.983972587 | Hnrnpul2 | heterogeneous nuclear ribonucleoprotein | 1790.1 | 2029.1 | 1900.8 | 1848.9 | 1848.6 | 2070.6 |
| 20112 | ENSMUSG00000071660 | 334.9000771 | 0.190864504 | 0.151545255 | 1.259455489 | 0.207865858 | 0.653034701 | Ttc9c | tetratricopeptide repeat domain 9C [Sour | 344.53 | 287.82 | 304.89 | 323.9 | 377.35 | 370.91 |
| 20113 | ENSMUSG00000071661 | 3.628360026 | 0.781008143 | 1.2297141 | 0.635113595 | 0.525354331 | NA | Zbtb3 | zinc finger and BTB domain containing 3 [! | 4.8186 | 3.0838 | 0 | 2.8919 | 7.6343 | 3.3415 |
| 20114 | ENSMUSG00000071662 | 214.6042098 | 0.095911416 | 0.174396271 | 0.54996254 | 0.582345067 | 0.908645447 | Polr2g | polymerase (RNA) II (DNA directed) polyp | 212.82 | 223.06 | 186.16 | 199.54 | 239.94 | 226.11 |
| 20115 | ENSMUSG00000071665 | 0 | NA | NA | NA | NA | NA | Foxr2 | forkhead box R2 [Source:MGI Symbol;Acc | 0 | 0 | 0 | 0 | 0 | 0 |
| 20116 | ENSMUSG00000071669 | 90.00321496 | -0.070328892 | 0.277959362 | -0.253018613 | 0.80025383 | 0.962424862 | Snx29 | sorting nexin 29 [Source:MGI Symbol;Acc: | 81.916 | 69.898 | 124.77 | 77.118 | 90.521 | 95.791 |
| 20117 | ENSMUSG00000071671 | 0.185640647 | 1.126924175 | 4.080472857 | 0.276174898 | 0.782413736 | NA | Gm10343 | predicted gene 10343 [Source:MGI Symbo | 0 | 0 | 0 | 0 | 0 | 1.1138 |
| 20118 | ENSMUSG00000071679 | 0 | NA | NA | NA | NA | NA | Rtl4 | retrotransposon Gag like 4 [Source:MGI S | 0 | 0 | 0 | 0 | 0 | 0 |
| 20119 | ENSMUSG00000071680 | 0 | NA | NA | NA | NA | NA | Gm7123 | predicted pseudogene 7123 [Source:MGI | 0 | 0 | 0 | 0 | 0 | 0 |
| 20120 | ENSMUSG00000071686 | 0 | NA | NA | NA | NA | NA | Tex13a | testis expressed 13A [Source:MGI Symbol | 0 | 0 | 0 | 0 | 0 | 0 |
| 20121 | ENSMUSG00000071691 | 8.710520132 | 0.900399977 | 0.788151753 | 1.142419557 | 0.253279666 | 0.706545243 | Gm960 | predicted gene 960 [Source:MGI Symbol;A | 3.2124 | 6.1675 | 9.0562 | 17.352 | 10.906 | 5.5692 |
| 20122 | ENSMUSG00000071708 | 115.4017376 | -0.135232746 | 0.213340156 | -0.633883222 | 0.526157036 | 0.888334265 | Sms | spermine synthase [Source:MGI Symbol;A | 115.65 | 116.15 | 130.81 | 112.79 | 113.42 | 103.59 |
| 20123 | ENSMUSG00000071711 | 2621.633158 | 0.038486541 | 0.106065407 | 0.362856675 | 0.716711953 | 0.9447719 | Mpst | mercaptopyruvate sulfurtransferase [Sour | 2733 | 2454.7 | 2571 | 2433.1 | 2923.9 | 2614.2 |
| 20124 | ENSMUSG00000071713 | 111.9354016 | -0.214294271 | 0.242709368 | -0.882925422 | 0.377276567 | 0.81030298 | Csf2rb | colony stimulating factor 2 receptor, beta | 133.31 | 108.96 | 117.73 | 80.974 | 101.43 | 129.21 |
| 20125 | ENSMUSG00000071714 | 16.31870542 | -1.860981497 | 0.603326804 | -3.084533102 | 0.002038719 | 0.03544235 | Csf2rb2 | colony stimulating factor 2 receptor, beta | 29.715 | 31.865 | 15.094 | 5.7839 | 6.5437 | 8.9108 |
| 20126 | ENSMUSG00000071715 | 16.49567984 | -0.892317817 | 0.565543999 | -1.577804411 | 0.11461055 | 0.497367836 | Ncf4 | neutrophil cytosolic factor 4 [Source:MGI | 22.487 | 27.754 | 14.087 | 12.532 | 7.6343 | 14.48 |
| 20127 | ENSMUSG00000071716 | 2.44990507 | -1.930968836 | 1.535891772 | -1.257229755 | 0.208670452 | NA | Apol7e | apolipoprotein L 7e [Source:MGI Symbol;A | 5.6217 | 1.0279 | 5.0312 | 1.928 | 1.0906 | 0 |
| 20128 | ENSMUSG00000071719 | 146.3063138 | 0.082228596 | 0.305340012 | 0.269301739 | 0.787697494 | 0.960870081 | Tmem28 | transmembrane protein 28 [Source:MGI S | 183.11 | 89.429 | 152.95 | 88.686 | 183.22 | 180.44 |
| 20129 | ENSMUSG00000071721 | 0 | NA | NA | NA | NA | NA | 1700010D01 | RIKEN cDNA 1700010D01 gene [Source:M | 0 | 0 | 0 | 0 | 0 | 0 |
| 20130 | ENSMUSG00000071722 | 8.546852038 | 0.302033479 | 0.726138212 | 0.415944891 | 0.67745033 | 0.93325821 | Spin4 | spindlin family, member 4 [Source:MGI Sy | 9.6372 | 8.2233 | 5.0312 | 9.6398 | 8.7249 | 10.025 |
| 20131 | ENSMUSG00000071723 | 6.937986868 | -0.058773806 | 0.838068514 | -0.070130073 | 0.944090131 | 0.990077979 | Gspt2 | G1 to S phase transition 2 [Source:MGI Sy | 4.0155 | 7.1954 | 10.062 | 3.8559 | 9.8156 | 6.6831 |
| 20132 | ENSMUSG00000071724 | 1.5524311 | 0.404206833 | 1.731706262 | 0.233415356 | 0.81543889 | NA | Smpd5 | sphingomyelin phosphodiesterase 5 [Sour | 0 | 2.0558 | 2.0125 | 1.928 | 1.0906 | 2.2277 |
| 20133 | ENSMUSG00000071726 | 0 | NA | NA | NA | NA | NA | Gm5941 | predicted gene 5941 [Source:MGI Symbol | 0 | 0 | 0 | 0 | 0 | 0 |
| 20134 | ENSMUSG00000071738 | 0 | NA | NA | NA | NA | NA | Cldn34b3 | claudin 34B3 [Source:MGI Symbol;Acc:MG | 0 | 0 | 0 | 0 | 0 | 0 |
| 20135 | ENSMUSG00000071745 | 0.267700746 | -1.497481649 | 4.057001436 | -0.369110456 | 0.712045396 | NA | Pwwp4b | PWWP domain containing 4B [Source:MG | 1.6062 | 0 | 0 | 0 | 0 | 0 |
| 20136 | ENSMUSG00000071748 | 9.081325683 | -0.401078329 | 0.719407202 | -0.557512252 | 0.57717749 | 0.907838603 | Gm14698 | predicted gene 14698 [Source:MGI Symbo | 9.6372 | 8.2233 | 13.081 | 4.8199 | 9.8156 | 8.9108 |
| 20137 | ENSMUSG00000071749 | 0 | NA | NA | NA | NA | NA | 4933412E24 | RIKEN cDNA 4933412E24 gene [Source:M | 0 | 0 | 0 | 0 | 0 | 0 |
| 20138 | ENSMUSG00000071750 | 0 | NA | NA | NA | NA | NA | Gm7713 | predicted gene 7713 [Source:MGI Symbol | 0 | 0 | 0 | 0 | 0 | 0 |
| 20139 | ENSMUSG00000071753 | 0 | NA | NA | NA | NA | NA | Cdr1os | cerebellar degeneration related antigen 1 | 0 | 0 | 0 | 0 | 0 | 0 |
| 20140 | ENSMUSG00000071757 | 285.8625475 | 0.183613833 | 0.172159634 | 1.066532428 | 0.286183028 | 0.73856672 | Zhx2 | zinc fingers and homeoboxes 2 [Source:M | 261.81 | 272.4 | 269.67 | 369.2 | 278.11 | 263.98 |
| 20141 | ENSMUSG00000071763 | 0 | NA | NA | NA | NA | NA | Gm53064 | 53064 [Source:MGI Symbol;Acc:MGI:6436 | 0 | 0 | 0 | 0 | 0 | 0 |
| 20142 | ENSMUSG00000071764 | 0 | NA | NA | NA | NA | NA | Olfr1320 | olfactory receptor 1320 [Source:MGI Sym | 0 | 0 | 0 | 0 | 0 | 0 |
| 20143 | ENSMUSG00000071766 | 0 | NA | NA | NA | NA | NA | Rhox12 | reproductive homeobox 12 [Source:MGI S | 0 | 0 | 0 | 0 | 0 | 0 |
| 20144 | ENSMUSG00000071767 | 0 | NA | NA | NA | NA | NA | Rhox7a | reproductive homeobox 7A [Source:MGI S | 0 | 0 | 0 | 0 | 0 | 0 |
| 20145 | ENSMUSG00000071769 | 0 | NA | NA | NA | NA | NA | Rhox3h | reproductive homeobox 3H [Source:MGI S | 0 | 0 | 0 | 0 | 0 | 0 |
| 20146 | ENSMUSG00000071770 | 0 | NA | NA | NA | NA | NA | Rhox4e | reproductive homeobox 4E [Source:MGI S | 0 | 0 | 0 | 0 | 0 | 0 |
| 20147 | ENSMUSG00000071771 | 0 | NA | NA | NA | NA | NA | Rhox4b | reproductive homeobox 4B [Source:MGI S | 0 | 0 | 0 | 0 | 0 | 0 |
| 20148 | ENSMUSG00000071772 | 0 | NA | NA | NA | NA | NA | Rhox3a | reproductive homeobox 3A [Source:MGI S | 0 | 0 | 0 | 0 | 0 | 0 |
| 20149 | ENSMUSG00000071773 | 0 | NA | NA | NA | NA | NA | Rhox1 | reproductive homeobox 1 [Source:MGI Sy | 0 | 0 | 0 | 0 | 0 | 0 |
| 20150 | ENSMUSG00000071788 | 0 | NA | NA | NA | NA | NA | Gm14525 | predicted gene 14525 [Source:MGI Symbo | 0 | 0 | 0 | 0 | 0 | 0 |
| 20151 | ENSMUSG00000071793 | 70.59600237 | 0.305228745 | 0.358955195 | 0.850325471 | 0.395144159 | 0.822933206 | 2610005L07I | RIKEN cDNA 2610005L07 gene [Source:M | 49.792 | 70.926 | 69.431 | 123.39 | 63.256 | 46.781 |
| 20152 | ENSMUSG00000071796 | 66.99329066 | -0.108815672 | 0.294166996 | -0.36991122 | 0.711448641 | 0.944239215 | 6820431F20 | RIKEN cDNA 6820431F20 gene [Source:M | 75.492 | 66.815 | 66.412 | 82.902 | 49.078 | 61.261 |
| 20153 | ENSMUSG00000071815 | 0 | NA | NA | NA | NA | NA | Fthl17e | ferritin, heavy polypeptide-like 17, memb | 0 | 0 | 0 | 0 | 0 | 0 |
| 20154 | ENSMUSG00000071816 | 0.321325949 | 1.911575114 | 4.048267893 | 0.472195804 | 0.636787034 | NA | Ssxb5 | synovial sarcoma, X member B5 [Source:N | 0 | 0 | 0 | 1.928 | 0 | 0 |
| 20155 | ENSMUSG00000071818 | 0 | NA | NA | NA | NA | NA | 4930417O22 | RIKEN cDNA 4930417O22 gene [Source:M | 0 | 0 | 0 | 0 | 0 | 0 |
| 20156 | ENSMUSG00000071847 | 1.000898647 | -2.10615299 | 2.475207832 | -0.850899453 | 0.394825209 | NA | Apcdd1 | adenomatosis polyposis coli down-regulat | 0.8031 | 4.1117 | 0 | 0 | 1.0906 | 0 |

| | | | | | | | | | | | | | | | |
|---|---|---|---|---|---|---|---|---|---|---|---|---|---|---|---|
| 20157 | ENSMUSG00000071855 | 0.859823856 | -1.825223727 | 2.563005556 | -0.712141931 | 0.476376891 | NA | Ccdc112 | coiled-coil domain containing 112 [Source | 0 | 2.0558 | 2.0125 | 0 | 1.0906 | 0 |
| 20158 | ENSMUSG00000071856 | 669.5569237 | -0.626407185 | 0.215374694 | -2.908453047 | 0.003632217 | 0.053773172 | Mcc | mutated in colorectal cancers [Source:MG | 489.89 | 955.96 | 993.16 | 513.8 | 590.02 | 474.5 |
| 20159 | ENSMUSG00000071858 | 0 | NA | NA | NA | NA | NA | Gm94 | predicted gene 94 [Source:MGI Symbol;Ac | 0 | 0 | 0 | 0 | 0 | 0 |
| 20160 | ENSMUSG00000071860 | 0.371281295 | 2.09253362 | 4.031852165 | 0.519000582 | 0.603760335 | NA | NA | NA | 0 | 0 | 0 | 0 | 0 | 2.2277 |
| 20161 | ENSMUSG00000071862 | 0.167707431 | -0.796629176 | 4.080472857 | -0.195229623 | 0.84521319 | NA | Lrrtm2 | leucine rich repeat transmembrane neuro | 0 | 0 | 1.0062 | 0 | 0 | 0 |
| 20162 | ENSMUSG00000071866 | 8122.611665 | 0.136042076 | 0.08561427 | 1.589011696 | 0.112057752 | 0.491152524 | Ppia | peptidylprolyl isomerase A [Source:MGI S | 7937.9 | 7778.3 | 7502.6 | 8089.7 | 8782.7 | 8644.5 |
| 20163 | ENSMUSG00000071867 | 1.536558863 | -2.847469017 | 1.981342673 | -1.437141114 | 0.150677901 | NA | Mrpl27-ps | mitochondrial ribosomal protein L27, pseu | 4.0155 | 3.0838 | 1.0062 | 0 | 0 | 1.1138 |
| 20164 | ENSMUSG00000071890 | 0 | NA | NA | NA | NA | NA | Mroh9 | maestro heat-like repeat family member 9 | 0 | 0 | 0 | 0 | 0 | 0 |
| 20165 | ENSMUSG00000071893 | 8.440411604 | -0.578779102 | 0.81272761 | -0.712144013 | 0.476375602 | 0.86953366 | Vmn1r4 | vomeronasal 1 receptor 4 [Source:MGI Sy | 8.031 | 6.1675 | 16.1 | 4.8199 | 3.2719 | 12.252 |
| 20166 | ENSMUSG00000071909 | 0 | NA | NA | NA | NA | NA | Nutm2 | NUT family member 2 [Source:MGI Symbo | 0 | 0 | 0 | 0 | 0 | 0 |
| 20167 | ENSMUSG00000071937 | 0 | NA | NA | NA | NA | NA | Adam25 | a disintegrin and metallopeptidase domai | 0 | 0 | 0 | 0 | 0 | 0 |
| 20168 | ENSMUSG00000071952 | 0 | NA | NA | NA | NA | NA | Gm10351 | predicted gene 10351 [Source:MGI Symbo | 0 | 0 | 0 | 0 | 0 | 0 |
| 20169 | ENSMUSG00000071984 | 3.077919588 | -1.799144981 | 1.335777596 | -1.346889622 | 0.178015785 | NA | Fndc1 | fibronectin type III domain containing 1 [S | 7.2279 | 4.1117 | 3.0187 | 1.928 | 2.1812 | 0 |
| 20170 | ENSMUSG00000072049 | 0 | NA | NA | NA | NA | NA | Vmn2r121 | vomeronasal 2, receptor 121 [Source:MGI | 0 | 0 | 0 | 0 | 0 | 0 |
| 20171 | ENSMUSG00000072066 | 1.461780829 | -0.307486614 | 2.13761504 | -0.143845645 | 0.885622358 | NA | 6720489N17 | RIKEN cDNA 6720489N17 gene [Source:M | 0.8031 | 4.1117 | 0 | 3.8559 | 0 | 0 |
| 20172 | ENSMUSG00000072068 | 0 | NA | NA | NA | NA | NA | Gm16261 | predicted gene 16261 [Source:MGI Symbo | 0 | 0 | 0 | 0 | 0 | 0 |
| 20173 | ENSMUSG00000072082 | 471.1118657 | -0.528482137 | 0.195347886 | -2.705338397 | 0.006823484 | 0.082520796 | Ccnf | cyclin F [Source:MGI Symbol;Acc:MGI:102 | 527.64 | 489.29 | 652.05 | 306.54 | 514.77 | 336.38 |
| 20174 | ENSMUSG00000072100 | 0 | NA | NA | NA | NA | NA | Cldn34b1 | claudin 34B1 [Source:MGI Symbol;Acc:MG | 0 | 0 | 0 | 0 | 0 | 0 |
| 20175 | ENSMUSG00000072109 | 2.820850532 | -1.107610686 | 1.277956593 | -0.866704466 | 0.386103958 | NA | A530040E14 | RIKEN cDNA A530040E14 gene [Source:M | 2.4093 | 4.1117 | 5.0312 | 0.964 | 2.1812 | 2.2277 |
| 20176 | ENSMUSG00000072115 | 2786.956043 | 0.611095809 | 0.115389062 | 5.295959579 | 1.18E-07 | 1.28E-05 | Ang | angiogenin, ribonuclease, RNase A family, | 2001.3 | 2141.2 | 2474.4 | 3150.3 | 3253.3 | 3701.3 |
| 20177 | ENSMUSG00000072145 | 0 | NA | NA | NA | NA | NA | BC061237 | cDNA sequence BC061237 [Source:MGI Sy | 0 | 0 | 0 | 0 | 0 | 0 |
| 20178 | ENSMUSG00000072188 | 0 | NA | NA | NA | NA | NA | Gm10354 | predicted gene 10354 [Source:MGI Symbo | 0 | 0 | 0 | 0 | 0 | 0 |
| 20179 | ENSMUSG00000072214 | 1.144113709 | 0.823834277 | 2.553104362 | 0.322679437 | 0.746938027 | NA | Septin5 | septin 5 [Source:MGI Symbol;Acc:MGI:11 | 2.4093 | 0 | 0 | 0 | 0 | 4.4554 |
| 20180 | ENSMUSG00000072235 | 22.46664866 | -0.363070358 | 0.577393143 | -0.628809613 | 0.529473706 | 0.888334265 | Tuba1a | tubulin, alpha 1A [Source:MGI Symbol;Ac | 32.927 | 23.642 | 19.119 | 20.244 | 5.4531 | 33.415 |
| 20181 | ENSMUSG00000072244 | 2.316648367 | 0.121300107 | 1.453296563 | 0.083465488 | 0.933481418 | NA | Trim6 | tripartite motif-containing 6 [Source:MGI | 1.6062 | 3.0838 | 2.0125 | 3.8559 | 0 | 3.3415 |
| 20182 | ENSMUSG00000072249 | 0 | NA | NA | NA | NA | NA | Fthl17f | ferritin, heavy polypeptide-like 17, memb | 0 | 0 | 0 | 0 | 0 | 0 |
| 20183 | ENSMUSG00000072255 | 0 | NA | NA | NA | NA | NA | Rpl31-ps17 | ribosomal protein L31, pseudogene 17 [So | 0 | 0 | 0 | 0 | 0 | 0 |
| 20184 | ENSMUSG00000072258 | 104.2051274 | 0.285771193 | 0.23434448 | 1.219449222 | 0.222673738 | 0.668407528 | Taf1a | TATA-box binding protein associated facto | 96.372 | 100.74 | 84.525 | 107 | 135.24 | 101.36 |
| 20185 | ENSMUSG00000072259 | 0 | NA | NA | NA | NA | NA | Gm5592 | predicted gene 5592 [Source:MGI Symbol | 0 | 0 | 0 | 0 | 0 | 0 |
| 20186 | ENSMUSG00000072294 | 320.4730152 | 0.459877743 | 0.151329325 | 3.038920196 | 0.002374277 | 0.039642097 | Klf12 | Kruppel-like factor 12 [Source:MGI Symbo | 240.93 | 273.43 | 295.84 | 362.46 | 391.53 | 358.66 |
| 20187 | ENSMUSG00000072295 | 0.36353943 | 2.065989254 | 4.034137301 | 0.512126658 | 0.608562368 | NA | C2cd6 | C2 calcium dependent domain containing | 0 | 0 | 0 | 0 | 2.1812 | 0 |
| 20188 | ENSMUSG00000072324 | 0 | NA | NA | NA | NA | NA | Gm8420 | predicted gene 8420 [Source:MGI Symbol | 0 | 0 | 0 | 0 | 0 | 0 |
| 20189 | ENSMUSG00000072381 | 0 | NA | NA | NA | NA | NA | Gm10355 | predicted gene 10355 [Source:MGI Symbo | 0 | 0 | 0 | 0 | 0 | 0 |
| 20190 | ENSMUSG00000072387 | 0 | NA | NA | NA | NA | NA | Gm10356 | predicted gene 10356 [Source:MGI Symbo | 0 | 0 | 0 | 0 | 0 | 0 |
| 20191 | ENSMUSG00000072407 | 0 | NA | NA | NA | NA | NA | Gm6419 | predicted gene 6419 [Source:MGI Symbol | 0 | 0 | 0 | 0 | 0 | 0 |
| 20192 | ENSMUSG00000072419 | 0 | NA | NA | NA | NA | NA | Dppa2 | developmental pluripotency associated 2 | 0 | 0 | 0 | 0 | 0 | 0 |
| 20193 | ENSMUSG00000072421 | 3.259380628 | -0.028272055 | 1.218535034 | -0.023201676 | 0.981489402 | NA | Gm10357 | predicted gene 10357 [Source:MGI Symbo | 0.8031 | 3.0838 | 6.0375 | 1.928 | 4.3625 | 3.3415 |
| 20194 | ENSMUSG00000072423 | 0 | NA | NA | NA | NA | NA | Psmb11 | proteasome (prosome, macropain) subun | 0 | 0 | 0 | 0 | 0 | 0 |
| 20195 | ENSMUSG00000072437 | 0.824622689 | 0.984749497 | 2.871267556 | 0.342966818 | 0.731623423 | NA | Nanos1 | nanos C2HC-type zinc finger 1 [Source:MG | 1.6062 | 0 | 0 | 0 | 0 | 3.3415 |
| 20196 | ENSMUSG00000072460 | 1.474144596 | 0.297359805 | 2.354681888 | 0.126284491 | 0.899506732 | NA | Atp5l-ps1 | ATP synthase, H+ transporting, mitochonc | 0 | 0 | 4.025 | 4.8199 | 0 | 0 |
| 20197 | ENSMUSG00000072473 | 0 | NA | NA | NA | NA | NA | 1700024G13 | RIKEN cDNA 1700024G13 gene [Source:M | 0 | 0 | 0 | 0 | 0 | 0 |
| 20198 | ENSMUSG00000072476 | 0.17131964 | -0.796629176 | 4.080472857 | -0.195229623 | 0.84521319 | NA | Gm9008 | predicted pseudogene 9008 [Source:MGI | 0 | 1.0279 | 0 | 0 | 0 | 0 |
| 20199 | ENSMUSG00000072477 | 0 | NA | NA | NA | NA | NA | Gm10360 | predicted gene 10360 [Source:MGI Symbo | 0 | 0 | 0 | 0 | 0 | 0 |
| 20200 | ENSMUSG00000072479 | 0 | NA | NA | NA | NA | NA | Xlr5b | X-linked lymphocyte-regulated 5B [Source | 0 | 0 | 0 | 0 | 0 | 0 |
| 20201 | ENSMUSG00000072487 | 0.167707431 | -0.796629176 | 4.080472857 | -0.195229623 | 0.84521319 | NA | Mroh5 | maestro heat-like repeat family member 5 | 0 | 0 | 1.0062 | 0 | 0 | 0 |
| 20202 | ENSMUSG00000072492 | 1.149405861 | -1.242804601 | 2.372406571 | -0.523858185 | 0.60037718 | NA | Gm10364 | predicted pseudogene 10364 [Source:MG | 4.8186 | 0 | 0 | 0.964 | 0 | 1.1138 |
| 20203 | ENSMUSG00000072494 | 151.0534394 | 0.237566408 | 0.237635127 | 0.999710823 | 0.317450473 | 0.764248275 | Ppp1r3e | protein phosphatase 1, regulatory subunit | 126.89 | 141.85 | 147.92 | 209.18 | 166.86 | 113.61 |
| 20204 | ENSMUSG00000072501 | 697.9323884 | -0.021365805 | 0.138732167 | -0.154007578 | 0.877603761 | 0.98072613 | Phf20l1 | PHD finger protein 20-like 1 [Source:MGI S | 632.04 | 788.41 | 690.28 | 766.36 | 703.45 | 607.04 |
| 20205 | ENSMUSG00000072511 | 0 | NA | NA | NA | NA | NA | Hhla1 | HERV-H LTR-associating 1 [Source:MGI Sy | 0 | 0 | 0 | 0 | 0 | 0 |
| 20206 | ENSMUSG00000072541 | 0 | NA | NA | NA | NA | NA | Gm5046 | predicted gene 5046 [Source:MGI Symbol | 0 | 0 | 0 | 0 | 0 | 0 |
| 20207 | ENSMUSG00000072553 | 0 | NA | NA | NA | NA | NA | Gm525 | predicted gene 525 [Source:MGI Symbol;A | 0 | 0 | 0 | 0 | 0 | 0 |
| 20208 | ENSMUSG00000072568 | 121.957984 | -0.140681926 | 0.29179333 | -0.482128656 | 0.629714554 | 0.92057725 | Lratd2 | LRAT domain containing 1 [Source:MGI Sy | 143.76 | 135.69 | 104.65 | 174.48 | 79.615 | 93.563 |
| 20209 | ENSMUSG00000072571 | 24.32287338 | 0.645903956 | 0.465061442 | 1.388857252 | 0.164876162 | 0.590031077 | Tmem253 | transmembrane protein 253 [Source:MGI | 18.471 | 16.447 | 22.137 | 41.451 | 21.812 | 25.618 |
| 20210 | ENSMUSG00000072572 | 17.25409195 | 0.277771161 | 0.541022141 | 0.513419211 | 0.60765811 | 0.916376001 | Slc39a2 | solute carrier family 39 (zinc transporter), | 10.44 | 16.447 | 20.125 | 20.244 | 22.903 | 13.366 |
| 20211 | ENSMUSG00000072573 | 1.168941911 | -2.435929877 | 2.172449565 | -1.121282591 | 0.262167593 | NA | Gm10369 | predicted gene 10369 [Source:MGI Symbo | 4.0155 | 1.0279 | 1.0062 | 0.964 | 0 | 0 |
| 20212 | ENSMUSG00000072574 | 0 | NA | NA | NA | NA | NA | Gm17184 | predicted gene 17184 [Source:MGI Symbo | 0 | 0 | 0 | 0 | 0 | 0 |

| | | | | | | | | | | | | | | | |
|---|---|---|---|---|---|---|---|---|---|---|---|---|---|---|---|
| 20213 | ENSMUSG00000072575 | 0 | NA | NA | NA | NA | NA | Eddm3b | epididymal protein 3B [Source:MGI Symb | 0 | 0 | 0 | 0 | 0 | 0 |
| 20214 | ENSMUSG00000072576 | 0 | NA | NA | NA | NA | NA | Gm16020 | predicted gene 16020 [Source:MGI Symbc | 0 | 0 | 0 | 0 | 0 | 0 |
| 20215 | ENSMUSG00000072582 | 350.1069898 | -0.083366973 | 0.169878198 | -0.490745568 | 0.623606413 | 0.919503423 | Ptrh2 | peptidyl-tRNA hydrolase 2 [Source:MGI S | 407.17 | 361.83 | 310.93 | 321.97 | 401.35 | 297.4 |
| 20216 | ENSMUSG00000072584 | 0 | NA | NA | NA | NA | NA | Gm7489 | predicted gene 7489 [Source:MGI Symbol | 0 | 0 | 0 | 0 | 0 | 0 |
| 20217 | ENSMUSG00000072589 | 0.319491021 | 0.16514334 | 4.044653141 | 0.040830038 | 0.967431393 | NA | Gm10371 | predicted gene 10371 [Source:MGI Symbc | 0.8031 | 0 | 0 | 0 | 0 | 1.1138 |
| 20218 | ENSMUSG00000072591 | 0 | NA | NA | NA | NA | NA | Fzd10os | frizzled class receptor 10, opposite strand | 0 | 0 | 0 | 0 | 0 | 0 |
| 20219 | ENSMUSG00000072592 | 0 | NA | NA | NA | NA | NA | Gm10373 | predicted gene 10373 [Source:MGI Symbc | 0 | 0 | 0 | 0 | 0 | 0 |
| 20220 | ENSMUSG00000072594 | 0 | NA | NA | NA | NA | NA | Gm16439 | predicted pseudogene 16439 [Source:MG | 0 | 0 | 0 | 0 | 0 | 0 |
| 20221 | ENSMUSG00000072595 | 0 | NA | NA | NA | NA | NA | 4930503E14 | RIKEN cDNA 4930503E14 gene [Source:M | 0 | 0 | 0 | 0 | 0 | 0 |
| 20222 | ENSMUSG00000072596 | 17.99468298 | -1.343212376 | 0.657173785 | -2.043922638 | 0.04096119 | 0.277370158 | Ear2 | eosinophil-associated, ribonuclease A fam | 35.336 | 35.977 | 6.0375 | 9.6398 | 8.7249 | 12.252 |
| 20223 | ENSMUSG00000072598 | 0 | NA | NA | NA | NA | NA | Ang6 | angiogenin, ribonuclease A family, memb | 0 | 0 | 0 | 0 | 0 | 0 |
| 20224 | ENSMUSG00000072599 | 0.294513347 | 0.165143496 | 4.080472857 | 0.040471657 | 0.967717103 | NA | Ear-ps2 | eosinophil-associated, ribonuclease A fam | 0.8031 | 0 | 0 | 0.964 | 0 | 0 |
| 20225 | ENSMUSG00000072600 | 0 | NA | NA | NA | NA | NA | Ear-ps9 | eosinophil-associated, ribonuclease A fam | 0 | 0 | 0 | 0 | 0 | 0 |
| 20226 | ENSMUSG00000072601 | 0 | NA | NA | NA | NA | NA | Ear1 | eosinophil-associated, ribonuclease A fam | 0 | 0 | 0 | 0 | 0 | 0 |
| 20227 | ENSMUSG00000072605 | 0 | NA | NA | NA | NA | NA | Gm10376 | predicted gene 10376 [Source:MGI Symbc | 0 | 0 | 0 | 0 | 0 | 0 |
| 20228 | ENSMUSG00000072612 | 48.52314026 | -0.153962465 | 0.434253539 | -0.354545102 | 0.72293041 | 0.9447719 | Gm10382 | predicted gene 10382 [Source:MGI Symbc | 56.217 | 38.033 | 59.368 | 77.118 | 40.353 | 20.049 |
| 20229 | ENSMUSG00000072616 | 0 | NA | NA | NA | NA | NA | Gm7853 | predicted gene 7853 [Source:MGI Symbol | 0 | 0 | 0 | 0 | 0 | 0 |
| 20230 | ENSMUSG00000072618 | 0.455176322 | 0.955156269 | 3.546674842 | 0.269310357 | 0.787690862 | NA | Gm10384 | predicted gene 10384 [Source:MGI Symbc | 0.8031 | 0 | 0 | 1.928 | 0 | 0 |
| 20231 | ENSMUSG00000072620 | 40.36311217 | -1.128015024 | 0.404494763 | -2.788701185 | 0.005291987 | 0.069144802 | Slfn2 | schlafen 2 [Source:MGI Symbol;Acc:MGI: | 57.823 | 60.647 | 47.293 | 12.532 | 33.809 | 30.074 |
| 20232 | ENSMUSG00000072621 | 0 | NA | NA | NA | NA | NA | Slfn10-ps | schlafen 10, pseudogene [Source:MGI Sy | 0 | 0 | 0 | 0 | 0 | 0 |
| 20233 | ENSMUSG00000072623 | 18.39784325 | -0.455371094 | 0.497797525 | -0.914771712 | 0.360311488 | 0.799598236 | Zfp9 | zinc finger protein 9 [Source:MGI Symbol; | 20.078 | 21.586 | 22.137 | 13.496 | 15.269 | 17.822 |
| 20234 | ENSMUSG00000072624 | 0 | NA | NA | NA | NA | NA | Gm5460 | predicted gene 5460 [Source:MGI Symbol | 0 | 0 | 0 | 0 | 0 | 0 |
| 20235 | ENSMUSG00000072625 | 230.935854 | -1.251934914 | 0.198325454 | -6.312527669 | 2.75E-10 | 6.73E-08 | Gdf2 | growth differentiation factor 2 [Source:M | 363.81 | 313.51 | 297.85 | 124.35 | 112.33 | 173.76 |
| 20236 | ENSMUSG00000072640 | 128.4125345 | 0.015926934 | 0.231656137 | 0.068752482 | 0.945186643 | 0.990309554 | Lyrm9 | LYR motif containing 9 [Source:MGI Symb | 105.21 | 134.66 | 143.89 | 135.92 | 148.32 | 102.47 |
| 20237 | ENSMUSG00000072647 | 8.398961396 | -1.227515907 | 0.856648808 | -1.432927819 | 0.151878466 | 0.567340036 | Adam1a | a disintegrin and metallopeptidase domai | 12.85 | 9.2513 | 13.081 | 0.964 | 10.906 | 3.3415 |
| 20238 | ENSMUSG00000072653 | 6.61942921 | 0.568533024 | 0.904077539 | 0.628854273 | 0.529444465 | 0.888334265 | Zfp783 | zinc finger protein 783 [Source:MGI Symb | 4.8186 | 9.2513 | 2.0125 | 11.568 | 8.7249 | 3.3415 |
| 20239 | ENSMUSG00000072660 | 3.065588494 | -0.673740558 | 1.320388811 | -0.510259215 | 0.609869872 | NA | Gm6288 | predicted gene 6288 [Source:MGI Symbol | 3.2124 | 6.1675 | 2.0125 | 4.8199 | 2.1812 | 0 |
| 20240 | ENSMUSG00000072662 | 14.10006452 | 0.136437363 | 0.607580176 | 0.224558615 | 0.822322663 | 0.969197142 | Mansc4 | MANSC domain containing 4 [Source:MGI | 16.865 | 6.1675 | 17.106 | 12.532 | 17.45 | 14.48 |
| 20241 | ENSMUSG00000072663 | 1.318612387 | -0.657242253 | 1.881088587 | -0.349394631 | 0.726793063 | NA | Spef2 | sperm flagellar 2 [Source:MGI Symbol;Acc | 0.8031 | 3.0838 | 1.0062 | 1.928 | 1.0906 | 0 |
| 20242 | ENSMUSG00000072664 | 1429.017874 | -0.057701162 | 0.132142185 | -0.436659666 | 0.662358185 | 0.9288455 | Ugt3a1 | UDP glycosyltransferases 3 family, polype | 1338 | 1380.5 | 1655.3 | 1512.5 | 1194.2 | 1493.7 |
| 20243 | ENSMUSG00000072672 | 0 | NA | NA | NA | NA | NA | NA | NA | 0 | 0 | 0 | 0 | 0 | 0 |
| 20244 | ENSMUSG00000072673 | 17.55792538 | -0.365764137 | 0.547438723 | -0.668137129 | 0.504046063 | 0.881510796 | Gm10392 | predicted gene 10392 [Source:MGI Symbc | 16.865 | 14.391 | 28.175 | 17.352 | 18.541 | 10.025 |
| 20245 | ENSMUSG00000072674 | 0 | NA | NA | NA | NA | NA | NA | NA | 0 | 0 | 0 | 0 | 0 | 0 |
| 20246 | ENSMUSG00000072675 | 0 | NA | NA | NA | NA | NA | NA | NA | 0 | 0 | 0 | 0 | 0 | 0 |
| 20247 | ENSMUSG00000072676 | 0 | NA | NA | NA | NA | NA | Tmem254a | transmembrane protein 254a [Source:MG | 0 | 0 | 0 | 0 | 0 | 0 |
| 20248 | ENSMUSG00000072677 | 0 | NA | NA | NA | NA | NA | Sox5it | SRY (sex determining region Y)-box 5, intr | 0 | 0 | 0 | 0 | 0 | 0 |
| 20249 | ENSMUSG00000072679 | 0 | NA | NA | NA | NA | NA | D6Ertd474e | DNA segment, Chr 6, ERATO Doi 474, expr | 0 | 0 | 0 | 0 | 0 | 0 |
| 20250 | ENSMUSG00000072680 | 0 | NA | NA | NA | NA | NA | NA | NA | 0 | 0 | 0 | 0 | 0 | 0 |
| 20251 | ENSMUSG00000072683 | 0.167707431 | -0.796629176 | 4.080472857 | -0.195229623 | 0.84521319 | NA | Gm7457 | predicted gene 7457 [Source:MGI Symbol | 0 | 0 | 1.0062 | 0 | 0 | 0 |
| 20252 | ENSMUSG00000072686 | 0 | NA | NA | NA | NA | NA | Gm10398 | predicted gene 10398 [Source:MGI Symbc | 0 | 0 | 0 | 0 | 0 | 0 |
| 20253 | ENSMUSG00000072688 | 0.294513347 | 0.165143496 | 4.080472857 | 0.040471657 | 0.967717103 | NA | Gm10399 | predicted gene 10399 [Source:MGI Symbc | 0.8031 | 0 | 0 | 0.964 | 0 | 0 |
| 20254 | ENSMUSG00000072692 | 33.1975878 | 0.044850574 | 0.40839652 | 0.109821146 | 0.912551221 | 0.986248982 | Rpl37rt | ribosomal protein L37, retrotransposed [S | 23.29 | 33.921 | 41.256 | 43.379 | 27.265 | 30.074 |
| 20255 | ENSMUSG00000072693 | 0 | NA | NA | NA | NA | NA | Gm10401 | predicted gene 10401 [Source:MGI Symbc | 0 | 0 | 0 | 0 | 0 | 0 |
| 20256 | ENSMUSG00000072694 | 12.78891191 | -0.824339652 | 0.65701852 | -1.254667299 | 0.209599562 | 0.653550636 | 1500011B03 | RIKEN cDNA 1500011B03 gene [Source:M | 15.259 | 24.67 | 9.0562 | 6.7478 | 7.6343 | 13.366 |
| 20257 | ENSMUSG00000072704 | 1720.670382 | -0.145256424 | 0.108018034 | -1.344742342 | 0.178708456 | 0.610813333 | Smim10l1 | small integral membrane protein 10 like 1 | 1751.6 | 1919.1 | 1751.9 | 1701.4 | 1727.5 | 1472.5 |
| 20258 | ENSMUSG00000072707 | 0 | NA | NA | NA | NA | NA | Olfr31 | olfactory receptor 31 [Source:MGI Symbo | 0 | 0 | 0 | 0 | 0 | 0 |
| 20259 | ENSMUSG00000072708 | 0 | NA | NA | NA | NA | NA | Olfr381 | olfactory receptor 381 [Source:MGI Symb | 0 | 0 | 0 | 0 | 0 | 0 |
| 20260 | ENSMUSG00000072709 | 0 | NA | NA | NA | NA | NA | Olfr380 | olfactory receptor 380 [Source:MGI Symb | 0 | 0 | 0 | 0 | 0 | 0 |
| 20261 | ENSMUSG00000072714 | 0 | NA | NA | NA | NA | NA | Rpl21-ps4 | ribosomal protein L21, pseudogene 4 [Sou | 0 | 0 | 0 | 0 | 0 | 0 |
| 20262 | ENSMUSG00000072717 | 0 | NA | NA | NA | NA | NA | Klra11-ps | killer cell lectin-like receptor subfamily A, | 0 | 0 | 0 | 0 | 0 | 0 |
| 20263 | ENSMUSG00000072718 | 0 | NA | NA | NA | NA | NA | Klra10 | killer cell lectin-like receptor subfamily A, | 0 | 0 | 0 | 0 | 0 | 0 |
| 20264 | ENSMUSG00000072720 | 2.30487507 | -1.144765697 | 1.670580622 | -0.685250195 | 0.493186054 | NA | Myo18b | myosin XVIIIb [Source:MGI Symbol;Acc:M | 6.4248 | 0 | 3.0187 | 0 | 3.2719 | 1.1138 |
| 20265 | ENSMUSG00000072721 | 0.17131964 | -0.796629176 | 4.080472857 | -0.195229623 | 0.84521319 | NA | Klra14-ps | killer cell lectin-like receptor subfamily A, | 0 | 1.0279 | 0 | 0 | 0 | 0 |
| 20266 | ENSMUSG00000072722 | 0 | NA | NA | NA | NA | NA | Gm6588 | predicted gene 6588 [Source:MGI Symbol | 0 | 0 | 0 | 0 | 0 | 0 |
| 20267 | ENSMUSG00000072723 | 1.898502627 | -0.715159168 | 1.65150961 | -0.43303361 | 0.664990353 | NA | Gm10044 | predicted gene 10044 [Source:MGI Symbc | 0 | 3.0838 | 4.025 | 0.964 | 1.0906 | 2.2277 |
| 20268 | ENSMUSG00000072726 | 0 | NA | NA | NA | NA | NA | Gm5797 | predicted gene 5797 [Source:MGI Symbol | 0 | 0 | 0 | 0 | 0 | 0 |

| | | | | | | | | | | | | | | | |
|---|---|---|---|---|---|---|---|---|---|---|---|---|---|---|---|
| 20269 | ENSMUSG00000072731 | 0 | NA | NA | NA | NA | NA | Gm16525 | predicted gene, 16525 [Source:MGI Symb | 0 | 0 | 0 | 0 | 0 | 0 |
| 20270 | ENSMUSG00000072738 | 0 | NA | NA | NA | NA | NA | Gm16440 | predicted gene 16440 [Source:MGI Symbo | 0 | 0 | 0 | 0 | 0 | 0 |
| 20271 | ENSMUSG00000072739 | 0 | NA | NA | NA | NA | NA | Gm10408 | predicted gene 10408 [Source:MGI Symbo | 0 | 0 | 0 | 0 | 0 | 0 |
| 20272 | ENSMUSG00000072753 | 0 | NA | NA | NA | NA | NA | 9230020A06 | RIKEN cDNA 9230020A06 gene [Source:M | 0 | 0 | 0 | 0 | 0 | 0 |
| 20273 | ENSMUSG00000072754 | 0 | NA | NA | NA | NA | NA | Lrcol1 | leucine rich colipase-like 1 [Source:MGI Sy | 0 | 0 | 0 | 0 | 0 | 0 |
| 20274 | ENSMUSG00000072761 | 17.79211783 | 0.461806594 | 0.552426651 | 0.835960019 | 0.403177395 | 0.828953151 | Gm6712 | predicted gene 6712 [Source:MGI Symbol | 10.44 | 16.447 | 18.112 | 16.388 | 14.178 | 31.188 |
| 20275 | ENSMUSG00000072762 | 43.79500671 | 0.317560039 | 0.378948748 | 0.83800261 | 0.402029238 | 0.82893099 | 4930522L14I | RIKEN cDNA 4930522L14 gene [Source:M( | 34.533 | 38.033 | 44.275 | 27.955 | 56.712 | 61.261 |
| 20276 | ENSMUSG00000072763 | 39.69415179 | 0.19694885 | 0.364403231 | 0.540469549 | 0.588873256 | 0.911039598 | 5430403G16 | RIKEN cDNA 5430403G16 gene [Source:M | 33.73 | 34.949 | 42.262 | 36.631 | 34.9 | 55.692 |
| 20277 | ENSMUSG00000072768 | 0 | NA | NA | NA | NA | NA | Gm10418 | predicted gene 10418 [Source:MGI Symbo | 0 | 0 | 0 | 0 | 0 | 0 |
| 20278 | ENSMUSG00000072769 | 0 | NA | NA | NA | NA | NA | Gm10419 | predicted gene 10419 [Source:MGI Symbo | 0 | 0 | 0 | 0 | 0 | 0 |
| 20279 | ENSMUSG00000072770 | 36.92882483 | 0.125852097 | 0.379813844 | 0.331352053 | 0.740378577 | 0.948792385 | Acrbp | proacrosin binding protein [Source:MGI S | 46.58 | 29.81 | 29.181 | 35.667 | 45.806 | 34.529 |
| 20280 | ENSMUSG00000072772 | 103.7551723 | 0.101299847 | 0.232466484 | 0.435761085 | 0.663010081 | 0.92906822 | Grcc10 | gene rich cluster, C10 gene [Source:MGI S | 85.129 | 112.04 | 103.64 | 107 | 116.7 | 98.018 |
| 20281 | ENSMUSG00000072774 | 34.93683909 | -0.26643021 | 0.430997559 | -0.618171041 | 0.536462597 | 0.890560262 | Zfp951 | zinc finger protein 951 [Source:MGI Symb | 39.352 | 41.117 | 34.212 | 43.379 | 37.081 | 14.48 |
| 20282 | ENSMUSG00000072778 | 0 | NA | NA | NA | NA | NA | Vmn2r27 | vomeronasal 2, receptor27 [Source:MGI S | 0 | 0 | 0 | 0 | 0 | 0 |
| 20283 | ENSMUSG00000072780 | 0 | NA | NA | NA | NA | NA | Vmn2r24 | vomeronasal 2, receptor 24 [Source:MGI S | 0 | 0 | 0 | 0 | 0 | 0 |
| 20284 | ENSMUSG00000072789 | 0.606727817 | -2.651277273 | 2.811524265 | -0.943003518 | 0.345679103 | NA | Gm10420 | predicted gene 10420 [Source:MGI Symbo | 1.6062 | 1.0279 | 1.0062 | 0 | 0 | 0 |
| 20285 | ENSMUSG00000072791 | 0 | NA | NA | NA | NA | NA | Abcb5 | ATP-binding cassette, sub-family B (MDR/ | 0 | 0 | 0 | 0 | 0 | 0 |
| 20286 | ENSMUSG00000072809 | 3.711971053 | -0.343391321 | 1.223275869 | -0.280714539 | 0.778929357 | NA | 9330160F10 | RIKEN cDNA 9330160F10 gene [Source:M( | 3.2124 | 7.1954 | 2.0125 | 0.964 | 1.0906 | 7.7969 |
| 20287 | ENSMUSG00000072812 | 9.377446251 | 0.196200738 | 0.722707736 | 0.271480058 | 0.786021836 | 0.960870081 | NA | NA | 8.031 | 8.2233 | 10.062 | 13.496 | 11.997 | 4.4554 |
| 20288 | ENSMUSG00000072813 | 0 | NA | NA | NA | NA | NA | E330014E10 | RIKEN cDNA E330014E10 gene [Source:M( | 0 | 0 | 0 | 0 | 0 | 0 |
| 20289 | ENSMUSG00000072814 | 0 | NA | NA | NA | NA | NA | Gm7982 | predicted gene 7982 [Source:MGI Symbol | 0 | 0 | 0 | 0 | 0 | 0 |
| 20290 | ENSMUSG00000072821 | 0 | NA | NA | NA | NA | NA | Gm6351 | predicted gene 6351 [Source:MGI Symbol | 0 | 0 | 0 | 0 | 0 | 0 |
| 20291 | ENSMUSG00000072822 | 0 | NA | NA | NA | NA | NA | Pramel37 | PRAME like 37 [Source:MGI Symbol;Acc:M | 0 | 0 | 0 | 0 | 0 | 0 |
| 20292 | ENSMUSG00000072825 | 729.8057093 | 0.254434396 | 0.136611339 | 1.86246909 | 0.062536998 | 0.357100334 | Cep170b | centrosomal protein 170B [Source:MGI Sy | 619.19 | 705.15 | 673.18 | 818.42 | 671.82 | 891.08 |
| 20293 | ENSMUSG00000072834 | 0 | NA | NA | NA | NA | NA | Gm12298 | predicted gene 12298 [Source:MGI Symbo | 0 | 0 | 0 | 0 | 0 | 0 |
| 20294 | ENSMUSG00000072837 | 0 | NA | NA | NA | NA | NA | Mir546 | microRNA 546 [Source:MGI Symbol;Acc:M | 0 | 0 | 0 | 0 | 0 | 0 |
| 20295 | ENSMUSG00000072844 | 3.207583186 | 0.000345198 | 1.265312519 | 0.000272817 | 0.999782324 | NA | G530011O06 | RIKEN cDNA G530011O06 gene [Source:M | 6.4248 | 2.0558 | 1.0062 | 0.964 | 5.4531 | 3.3415 |
| 20296 | ENSMUSG00000072845 | 0.133850373 | -0.796629176 | 4.080472857 | -0.195229623 | 0.84521319 | NA | Tmprss11a | transmembrane protease, serine 11a [Sou | 0.8031 | 0 | 0 | 0 | 0 | 0 |
| 20297 | ENSMUSG00000072847 | 50.97042447 | 0.003244715 | 0.325339887 | 0.009973308 | 0.992042583 | 0.99859176 | A530017D24 | RIKEN cDNA A530017D24 gene [Source:M | 40.155 | 63.731 | 49.306 | 57.839 | 46.897 | 47.895 |
| 20298 | ENSMUSG00000072849 | 28308.64278 | 3.099374859 | 0.364389674 | 8.505660516 | 1.81E-17 | 1.74E-14 | Serpina1e | serine (or cysteine) peptidase inhibitor, cl | 5912.4 | 6516 | 5319 | 26401 | 58049 | 67654 |
| 20299 | ENSMUSG00000072852 | 0.353348078 | 0.165142701 | 3.895042568 | 0.042398176 | 0.966181282 | NA | 2310040G07 | RIKEN cDNA 2310040G07 gene [Source:M | 0 | 0 | 1.0062 | 0 | 0 | 1.1138 |
| 20300 | ENSMUSG00000072872 | 127.3956893 | 0.132890849 | 0.211061531 | 0.629630842 | 0.52893614 | 0.888334265 | Rybp | RING1 and YY1 binding protein [Source:M | 106.81 | 119.24 | 138.86 | 127.25 | 136.33 | 135.89 |
| 20301 | ENSMUSG00000072874 | 0 | NA | NA | NA | NA | NA | Gm6116 | predicted gene 6116 [Source:MGI Symbol | 0 | 0 | 0 | 0 | 0 | 0 |
| 20302 | ENSMUSG00000072875 | 0 | NA | NA | NA | NA | NA | Gpr27 | G protein-coupled receptor 27 [Source:M( | 0 | 0 | 0 | 0 | 0 | 0 |
| 20303 | ENSMUSG00000072878 | 0 | NA | NA | NA | NA | NA | 1700123L14I | RIKEN cDNA 1700123L14 gene [Source:M( | 0 | 0 | 0 | 0 | 0 | 0 |
| 20304 | ENSMUSG00000072884 | 19.40888306 | -0.063371686 | 0.493193346 | -0.128492581 | 0.897759169 | 0.982798755 | Gm10433 | predicted gene 10433 [Source:MGI Symbo | 20.881 | 14.391 | 24.15 | 17.352 | 19.631 | 20.049 |
| 20305 | ENSMUSG00000072888 | 0 | NA | NA | NA | NA | NA | Samt2 | spermatogenesis associated multipass tra | 0 | 0 | 0 | 0 | 0 | 0 |
| 20306 | ENSMUSG00000072889 | 165.9707406 | -0.269155686 | 0.20812386 | -1.29324762 | 0.19592543 | 0.636484845 | Nfxl1 | nuclear transcription factor, X-box binding | 141.35 | 204.56 | 199.24 | 161.95 | 145.05 | 143.69 |
| 20307 | ENSMUSG00000072893 | 64.02417363 | 0.163223072 | 0.286818414 | 0.569081566 | 0.569300788 | 0.903735951 | 4933439C10 | RIKEN cDNA 4933439C10 gene [Source:M( | 60.233 | 67.843 | 53.331 | 78.082 | 56.712 | 67.944 |
| 20308 | ENSMUSG00000072898 | 1.503782206 | 1.374619617 | 2.077060455 | 0.661810114 | 0.508092923 | NA | Gm10434 | predicted gene 10434 [Source:MGI Symbo | 2.4093 | 0 | 0 | 0 | 3.2719 | 3.3415 |
| 20309 | ENSMUSG00000072900 | 0 | NA | NA | NA | NA | NA | Mir540 | microRNA 540 [Source:MGI Symbol;Acc:M | 0 | 0 | 0 | 0 | 0 | 0 |
| 20310 | ENSMUSG00000072902 | 0 | NA | NA | NA | NA | NA | Gm10435 | predicted gene 10435 [Source:MGI Symbo | 0 | 0 | 0 | 0 | 0 | 0 |
| 20311 | ENSMUSG00000072905 | 0 | NA | NA | NA | NA | NA | Eif1ad3 | eukaryotic translation initiation factor 1A | 0 | 0 | 0 | 0 | 0 | 0 |
| 20312 | ENSMUSG00000072915 | 38.68946462 | 0.073634691 | 0.355968338 | 0.206857416 | 0.836121214 | 0.971073158 | Gm12258 | predicted gene 12258 [Source:MGI Symbo | 32.927 | 40.089 | 40.25 | 38.559 | 46.897 | 33.415 |
| 20313 | ENSMUSG00000072919 | 1.987373963 | 0.649495865 | 1.467759283 | 0.442508436 | 0.658121327 | NA | Noxred1 | NADP+ dependent oxidoreductase domai | 1.6062 | 1.0279 | 2.0125 | 2.8919 | 3.2719 | 1.1138 |
| 20314 | ENSMUSG00000072923 | 0 | NA | NA | NA | NA | NA | Gm10439 | predicted gene 10439 [Source:MGI Symbo | 0 | 0 | 0 | 0 | 0 | 0 |
| 20315 | ENSMUSG00000072929 | 0 | NA | NA | NA | NA | NA | Gm15109 | predicted gene 15109 [Source:MGI Symbo | 0 | 0 | 0 | 0 | 0 | 0 |
| 20316 | ENSMUSG00000072930 | 0 | NA | NA | NA | NA | NA | Gm15107 | predicted gene 15107 [Source:MGI Symbo | 0 | 0 | 0 | 0 | 0 | 0 |
| 20317 | ENSMUSG00000072931 | 0 | NA | NA | NA | NA | NA | Gm15080 | predicted gene 15080 [Source:MGI Symbo | 0 | 0 | 0 | 0 | 0 | 0 |
| 20318 | ENSMUSG00000072934 | 0 | NA | NA | NA | NA | NA | Trpc5os | transient receptor potential cation chann | 0 | 0 | 0 | 0 | 0 | 0 |
| 20319 | ENSMUSG00000072940 | 5.644815815 | -0.132742595 | 0.91747952 | -0.144681808 | 0.884962103 | 0.981795936 | Gm10443 | predicted pseudogene 10443 [Source:MG | 6.4248 | 6.1675 | 5.0312 | 1.928 | 7.6343 | 6.6831 |
| 20320 | ENSMUSG00000072941 | 389.7061797 | -1.331880221 | 0.172650435 | -7.714317201 | 1.22E-14 | 7.33E-12 | Sod3 | superoxide dismutase 3, extracellular [Sou | 628.83 | 532.46 | 511.17 | 197.62 | 193.04 | 275.12 |
| 20321 | ENSMUSG00000072944 | 10.19356713 | 0.069583011 | 0.766076013 | 0.090830426 | 0.927627334 | 0.987866423 | Nup62cl | nucleoporin 62 C-terminal like [Source:M( | 9.6372 | 12.335 | 8.05 | 21.208 | 4.3625 | 5.5692 |
| 20322 | ENSMUSG00000072945 | 10.37816186 | 0.177075976 | 0.711449969 | 0.248894488 | 0.803442399 | 0.963596811 | Ripply1 | ripply transcriptional repressor 1 [Source: | 12.85 | 12.335 | 4.025 | 15.424 | 8.7249 | 8.9108 |
| 20323 | ENSMUSG00000072946 | 894.810525 | 0.090166919 | 0.126808694 | 0.711046825 | 0.47705522 | 0.869671045 | Ptgr2 | prostaglandin reductase 2 [Source:MGI Sy | 802.3 | 886.07 | 912.66 | 866.62 | 1051.4 | 849.86 |
| 20324 | ENSMUSG00000072949 | 1908.671206 | -0.834059903 | 0.58806906 | -1.418302644 | 0.156102425 | 0.574894692 | Acot1 | acyl-CoA thioesterase 1 [Source:MGI Syml | 2140.3 | 2691.1 | 2505.5 | 2429.2 | 1120.1 | 565.83 |

| | | | | | | | | | | | | | | | |
|---|---|---|---|---|---|---|---|---|---|---|---|---|---|---|---|
| 20325 | ENSMUSG00000072950 | 0 | NA | NA | NA | NA | NA | Gm10444 | predicted pseudogene 10444 [Source:MG | 0 | 0 | 0 | 0 | 0 | 0 |
| 20326 | ENSMUSG00000072952 | 0 | NA | NA | NA | NA | NA | Gm5878 | predicted gene 5878 [Source:MGI Symbol | 0 | 0 | 0 | 0 | 0 | 0 |
| 20327 | ENSMUSG00000072955 | 0 | NA | NA | NA | NA | NA | Tmsb15l | thymosin beta 15b like [Source:MGI Symb | 0 | 0 | 0 | 0 | 0 | 0 |
| 20328 | ENSMUSG00000072962 | 0 | NA | NA | NA | NA | NA | Gm16401 | predicted gene 16401 [Source:MGI Symbo | 0 | 0 | 0 | 0 | 0 | 0 |
| 20329 | ENSMUSG00000072963 | 30.08819512 | -0.624043395 | 0.403279381 | -1.54742202 | 0.121761515 | 0.511642546 | Gm10447 | predicted gene 10447 [Source:MGI Symbo | 39.352 | 35.977 | 34.212 | 26.991 | 26.175 | 17.822 |
| 20330 | ENSMUSG00000072964 | 49.36178692 | -1.010657336 | 0.338955617 | -2.981680447 | 0.002866711 | 0.045266486 | Bhlhb9 | basic helix-loop-helix domain containing, | 70.673 | 76.066 | 51.318 | 37.595 | 34.9 | 25.618 |
| 20331 | ENSMUSG00000072966 | 2.150337179 | 0.296289133 | 1.468331986 | 0.201786201 | 0.84008387 | NA | Gprasp2 | G protein-coupled receptor associated so | 0.8031 | 2.0558 | 3.0187 | 4.8199 | 1.0906 | 1.1138 |
| 20332 | ENSMUSG00000072968 | 0 | NA | NA | NA | NA | NA | Gm17728 | predicted gene, 17728 [Source:MGI Symb | 0 | 0 | 0 | 0 | 0 | 0 |
| 20333 | ENSMUSG00000072969 | 46.197236 | -0.039647691 | 0.36103311 | -0.109817327 | 0.912554249 | 0.986248982 | Armcx5 | armadillo repeat containing, X-linked 5 [S | 36.14 | 40.089 | 64.4 | 40.487 | 39.262 | 56.806 |
| 20334 | ENSMUSG00000072972 | 4.844168036 | 0.021520949 | 0.957542205 | 0.022475196 | 0.982068898 | NA | Adam4 | a disintegrin and metallopeptidase domai | 7.2279 | 3.0838 | 4.025 | 4.8199 | 5.4531 | 4.4554 |
| 20335 | ENSMUSG00000072974 | 9.281524945 | 0.058161746 | 0.688438278 | 0.084483603 | 0.932671939 | 0.988420305 | Gm4787 | predicted gene 4787 [Source:MGI Symbol | 8.031 | 8.2233 | 11.069 | 9.6398 | 9.8156 | 8.9108 |
| 20336 | ENSMUSG00000072978 | 0.859823856 | -1.825223727 | 2.563005556 | -0.712141931 | 0.476376891 | NA | Gm5830 | predicted pseudogene 5830 [Source:MGI | 0 | 2.0558 | 2.0125 | 0 | 1.0906 | 0 |
| 20337 | ENSMUSG00000072980 | 4.470095909 | -0.313641513 | 1.083655885 | -0.289429068 | 0.772253053 | NA | Oip5 | Opa interacting protein 5 [Source:MGI Sy | 5.6217 | 4.1117 | 5.0312 | 0.964 | 2.1812 | 8.9108 |
| 20338 | ENSMUSG00000072982 | 0 | NA | NA | NA | NA | NA | 9230009I02F | RIKEN cDNA 9230009I02 gene [Source:MC | 0 | 0 | 0 | 0 | 0 | 0 |
| 20339 | ENSMUSG00000072983 | 0 | NA | NA | NA | NA | NA | 4933414I15F | RIKEN cDNA 4933414I15 gene [Source:MC | 0 | 0 | 0 | 0 | 0 | 0 |
| 20340 | ENSMUSG00000072995 | 0 | NA | NA | NA | NA | NA | Cpxcr1 | CPX chromosome region, candidate 1 [Sou | 0 | 0 | 0 | 0 | 0 | 0 |
| 20341 | ENSMUSG00000072999 | 3.647670612 | -1.292067312 | 1.299516507 | -0.994267718 | 0.320092548 | NA | Gm15401 | predicted gene 15401 [Source:MGI Symbo | 6.4248 | 1.0279 | 8.05 | 1.928 | 0 | 4.4554 |
| 20342 | ENSMUSG00000073000 | 6.687613615 | -0.133898807 | 1.001391922 | -0.133712689 | 0.893629771 | 0.982562372 | Gm10451 | predicted gene 10451 [Source:MGI Symbo | 8.8341 | 7.1954 | 5.0312 | 13.496 | 0 | 5.5692 |
| 20343 | ENSMUSG00000073001 | 0 | NA | NA | NA | NA | NA | Cylc1 | cylicin, basic protein of sperm head cytos | 0 | 0 | 0 | 0 | 0 | 0 |
| 20344 | ENSMUSG00000073002 | 123.1332029 | 0.04101842 | 0.246695138 | 0.1662717 | 0.867943136 | 0.978326885 | Vamp5 | vesicle-associated membrane protein 5 [S | 114.04 | 106.9 | 142.89 | 92.542 | 127.6 | 154.82 |
| 20345 | ENSMUSG00000073006 | 0 | NA | NA | NA | NA | NA | Wmp | WAVE homology in membrane protrusion | 0 | 0 | 0 | 0 | 0 | 0 |
| 20346 | ENSMUSG00000073007 | 0.160662974 | 1.126924175 | 4.080472857 | 0.276174898 | 0.782413736 | NA | Tent5d | terminal nucleotidyltransferase 5D [Sourc | 0 | 0 | 0 | 0.964 | 0 | 0 |
| 20347 | ENSMUSG00000073008 | 1.836210144 | -4.231555844 | 1.999440877 | -2.116369577 | 0.034313387 | NA | Gpr174 | G protein-coupled receptor 174 [Source:N | 0.8031 | 7.1954 | 3.0187 | 0 | 0 | 0 |
| 20348 | ENSMUSG00000073010 | 0.160662974 | 1.126924175 | 4.080472857 | 0.276174898 | 0.782413736 | NA | Gm5127 | predicted gene 5127 [Source:MGI Symbol | 0 | 0 | 0 | 0.964 | 0 | 0 |
| 20349 | ENSMUSG00000073012 | 0 | NA | NA | NA | NA | NA | Fnd3c2 | fibronectin type III domain containing 3C2 | 0 | 0 | 0 | 0 | 0 | 0 |
| 20350 | ENSMUSG00000073016 | 25.36562014 | -0.382573502 | 0.428448713 | -0.892927183 | 0.371896169 | 0.807685567 | Uprt | uracil phosphoribosyltransferase [Source: | 27.305 | 25.698 | 33.206 | 24.099 | 20.722 | 21.163 |
| 20351 | ENSMUSG00000073018 | 0.481988923 | 2.492401104 | 3.837077151 | 0.649557203 | 0.515978285 | NA | Gm1070 | predicted gene 1070 [Source:MGI Symbol | 0 | 0 | 0 | 2.8919 | 0 | 0 |
| 20352 | ENSMUSG00000073027 | 0 | NA | NA | NA | NA | NA | Dmrtc1b | DMRT-like family C1b [Source:MGI Symbo | 0 | 0 | 0 | 0 | 0 | 0 |
| 20353 | ENSMUSG00000073028 | 0 | NA | NA | NA | NA | NA | Igkv4-71 | immunoglobulin kappa chain variable 4-7 | 0 | 0 | 0 | 0 | 0 | 0 |
| 20354 | ENSMUSG00000073043 | 0 | NA | NA | NA | NA | NA | Atoh1 | atonal bHLH transcription factor 1 [Source | 0 | 0 | 0 | 0 | 0 | 0 |
| 20355 | ENSMUSG00000073045 | 0 | NA | NA | NA | NA | NA | Gm5570 | predicted gene 5570 [Source:MGI Symbol | 0 | 0 | 0 | 0 | 0 | 0 |
| 20356 | ENSMUSG00000073051 | 0.185640647 | 1.126924175 | 4.080472857 | 0.276174898 | 0.782413736 | NA | Gm14812 | predicted gene 14812 [Source:MGI Symbo | 0 | 0 | 0 | 0 | 0 | 1.1138 |
| 20357 | ENSMUSG00000073052 | 0 | NA | NA | NA | NA | NA | D130052B06 | RIKEN cDNA D130052B06 gene [Source:M | 0 | 0 | 0 | 0 | 0 | 0 |
| 20358 | ENSMUSG00000073060 | 0.371281295 | 2.09253362 | 4.031852165 | 0.519000582 | 0.603760335 | NA | Zxda | zinc finger, X-linked, duplicated A [Source | 0 | 0 | 0 | 0 | 0 | 2.2277 |
| 20359 | ENSMUSG00000073062 | 32.60115716 | 0.422192573 | 0.409621274 | 1.030690053 | 0.302686191 | 0.752484788 | Zxdb | zinc finger, X-linked, duplicated B [Source | 33.73 | 16.447 | 33.206 | 39.523 | 38.172 | 34.529 |
| 20360 | ENSMUSG00000073063 | 0 | NA | NA | NA | NA | NA | Hbq1b | hemoglobin, theta 1B [Source:MGI Symbo | 0 | 0 | 0 | 0 | 0 | 0 |
| 20361 | ENSMUSG00000073067 | 5.601192142 | 0.150553544 | 1.009977438 | 0.149066245 | 0.881501361 | 0.981676444 | 9130019P16 | RIKEN cDNA 9130019P16 gene [Source:M | 0.8031 | 10.279 | 5.0312 | 8.6758 | 4.3625 | 4.4554 |
| 20362 | ENSMUSG00000073069 | 0 | NA | NA | NA | NA | NA | Mageb2 | MAGE family member B2 [Source:MGI Syr | 0 | 0 | 0 | 0 | 0 | 0 |
| 20363 | ENSMUSG00000073073 | 0.17131964 | -0.796629176 | 4.080472857 | -0.195229623 | 0.84521319 | NA | Gm8098 | predicted gene 8098 [Source:MGI Symbol | 0 | 1.0279 | 0 | 0 | 0 | 0 |
| 20364 | ENSMUSG00000073077 | 0.356960287 | 0.16514264 | 3.880370363 | 0.042558474 | 0.966053499 | NA | Cfap47 | cilia and flagella associated protein 47 [So | 0 | 1.0279 | 0 | 0 | 0 | 1.1138 |
| 20365 | ENSMUSG00000073079 | 71.61924952 | 0.341849554 | 0.281405188 | 1.214794779 | 0.224444372 | 0.670547386 | Srp54a | signal recognition particle 54A [Source:M | 56.217 | 77.094 | 56.35 | 69.406 | 93.793 | 76.855 |
| 20366 | ENSMUSG00000073085 | 0 | NA | NA | NA | NA | NA | Cldn34b4 | claudin 34B4 [Source:MGI Symbol;Acc:M | 0 | 0 | 0 | 0 | 0 | 0 |
| 20367 | ENSMUSG00000073094 | 2.434394938 | 0.007418854 | 1.444198993 | 0.005137002 | 0.995901283 | NA | Smim9 | small integral membrane protein 9 [Sourc | 3.2124 | 1.0279 | 3.0187 | 2.8919 | 0 | 4.4554 |
| 20368 | ENSMUSG00000073096 | 70.81859777 | -0.081581139 | 0.277958177 | -0.293501489 | 0.769138862 | 0.956045284 | Lrrc61 | leucine rich repeat containing 61 [Source: | 82.72 | 65.787 | 69.431 | 55.911 | 71.981 | 79.083 |
| 20369 | ENSMUSG00000073102 | 14.26379893 | 0.272347575 | 0.615448662 | 0.442518754 | 0.658113862 | 0.927303282 | Drc1 | dynein regulatory complex subunit 1 [Sou | 10.44 | 7.1954 | 21.131 | 11.568 | 18.541 | 16.708 |
| 20370 | ENSMUSG00000073103 | 0 | NA | NA | NA | NA | NA | Gm10466 | predicted gene 10466 [Source:MGI Symbo | 0 | 0 | 0 | 0 | 0 | 0 |
| 20371 | ENSMUSG00000073110 | 0 | NA | NA | NA | NA | NA | Olfr444 | olfactory receptor 444 [Source:MGI Symb | 0 | 0 | 0 | 0 | 0 | 0 |
| 20372 | ENSMUSG00000073111 | 0 | NA | NA | NA | NA | NA | Olfr446 | olfactory receptor 446 [Source:MGI Symb | 0 | 0 | 0 | 0 | 0 | 0 |
| 20373 | ENSMUSG00000073116 | 0 | NA | NA | NA | NA | NA | Gm10471 | predicted gene 10471 [Source:MGI Symbo | 0 | 0 | 0 | 0 | 0 | 0 |
| 20374 | ENSMUSG00000073117 | 0 | NA | NA | NA | NA | NA | Gm7347 | predicted gene 7347 [Source:MGI Symbol | 0 | 0 | 0 | 0 | 0 | 0 |
| 20375 | ENSMUSG00000073119 | 0 | NA | NA | NA | NA | NA | Speer4a | spermatogenesis associated glutamate (E) | 0 | 0 | 0 | 0 | 0 | 0 |
| 20376 | ENSMUSG00000073125 | 5.325612053 | -1.746623925 | 1.278256507 | -1.366411135 | 0.171809956 | 0.599531773 | Xlr3b | X-linked lymphocyte-regulated 3B [Source | 11.243 | 13.363 | 0 | 2.8919 | 0 | 4.4554 |
| 20377 | ENSMUSG00000073130 | 0.663785267 | -1.400021131 | 3.106754789 | -0.450637796 | 0.652250621 | NA | Gm1141 | predicted gene 1141 [Source:MGI Symbol | 0 | 0 | 3.0187 | 0.964 | 0 | 0 |
| 20378 | ENSMUSG00000073131 | 708.7915544 | -0.163212209 | 0.120333763 | -1.356329303 | 0.174994416 | 0.604952016 | Vma21 | VMA21 vacuolar H+-ATPase homolog (S. c | 677.02 | 792.52 | 777.83 | 659.36 | 656.55 | 689.47 |
| 20379 | ENSMUSG00000073139 | 348.6205595 | -0.217117139 | 0.160586121 | -1.352029287 | 0.176365947 | 0.607400048 | Tmem185a | transmembrane protein 185A [Source:MG | 366.21 | 432.75 | 326.02 | 349.92 | 326.09 | 290.71 |
| 20380 | ENSMUSG00000073141 | 0 | NA | NA | NA | NA | NA | 4930567H17 | RIKEN cDNA 4930567H17 gene [Source:M | 0 | 0 | 0 | 0 | 0 | 0 |

| | | | | | | | | | | | | | | | |
|---|---|---|---|---|---|---|---|---|---|---|---|---|---|---|---|
| 20381 | ENSMUSG00000073144 | 8.397098561 | -0.310055249 | 0.84631532 | -0.366359017 | 0.714097196 | 0.94455746 | 4930599N23 | RIKEN cDNA 4930599N23 gene [Source:M | 4.8186 | 6.1675 | 17.106 | 13.496 | 5.4531 | 3.3415 |
| 20382 | ENSMUSG00000073145 | 0 | NA | NA | NA | NA | NA | Gm12000 | predicted gene 12000 [Source:MGI Symbo | 0 | 0 | 0 | 0 | 0 | 0 |
| 20383 | ENSMUSG00000073147 | 209.1297185 | 0.10741235 | 0.195659224 | 0.548976671 | 0.58302146 | 0.908645447 | 5031425E22 | RIKEN cDNA 5031425E22 gene [Source:M | 190.34 | 182.97 | 231.44 | 263.17 | 191.95 | 194.92 |
| 20384 | ENSMUSG00000073152 | 0 | NA | NA | NA | NA | NA | Gm10475 | predicted gene 10475 [Source:MGI Symbo | 0 | 0 | 0 | 0 | 0 | 0 |
| 20385 | ENSMUSG00000073154 | 0 | NA | NA | NA | NA | NA | 9330158H04 | RIKEN cDNA 9330158H04 gene [Source:M | 0 | 0 | 0 | 0 | 0 | 0 |
| 20386 | ENSMUSG00000073155 | 1327.800683 | 0.16286983 | 0.130255244 | 1.250389807 | 0.211157186 | 0.655691903 | 1810058I24F | RIKEN cDNA 1810058I24 gene [Source:M | 1461.6 | 1197.5 | 1097.8 | 1313.9 | 1520.3 | 1375.6 |
| 20387 | ENSMUSG00000073158 | 0.335414862 | -1.779047242 | 4.030804179 | -0.441362855 | 0.658950329 | NA | 9030624G23 | RIKEN cDNA 9030624G23 gene [Source:M | 0 | 0 | 2.0125 | 0 | 0 | 0 |
| 20388 | ENSMUSG00000073164 | 0 | NA | NA | NA | NA | NA | 2410018L13I | RIKEN cDNA 2410018L13 gene [Source:M | 0 | 0 | 0 | 0 | 0 | 0 |
| 20389 | ENSMUSG00000073173 | 0 | NA | NA | NA | NA | NA | Gm10477 | predicted gene 10477 [Source:MGI Symbo | 0 | 0 | 0 | 0 | 0 | 0 |
| 20390 | ENSMUSG00000073174 | 0 | NA | NA | NA | NA | NA | Gm29254 | predicted gene 29254 [Source:MGI Symbo | 0 | 0 | 0 | 0 | 0 | 0 |
| 20391 | ENSMUSG00000073176 | 5.174374496 | -1.427757899 | 0.967938528 | -1.475050179 | 0.140199053 | 0.547324995 | Zfp449 | zinc finger protein 449 [Source:MGI Symb | 6.4248 | 6.1675 | 10.062 | 2.8919 | 3.2719 | 2.2277 |
| 20392 | ENSMUSG00000073177 | 0 | NA | NA | NA | NA | NA | Gm773 | predicted gene 773 [Source:MGI Symbol;A | 0 | 0 | 0 | 0 | 0 | 0 |
| 20393 | ENSMUSG00000073179 | 8.699950675 | 0.178429918 | 0.758562568 | 0.235221095 | 0.814037134 | 0.967157956 | Gm10478 | predicted gene 10478 [Source:MGI Symbo | 7.2279 | 6.1675 | 11.069 | 5.7839 | 15.269 | 6.6831 |
| 20394 | ENSMUSG00000073184 | 0 | NA | NA | NA | NA | NA | Gm49957 | predicted gene, 49957 [Source:MGI Symb | 0 | 0 | 0 | 0 | 0 | 0 |
| 20395 | ENSMUSG00000073197 | 3.808740666 | 0.655316046 | 1.132869471 | 0.578456797 | 0.562955757 | NA | 5730507C01 | RIKEN cDNA 5730507C01 gene [Source:M | 0.8031 | 5.1396 | 3.0187 | 2.8919 | 6.5437 | 4.4554 |
| 20396 | ENSMUSG00000073198 | 6.77533451 | 0.334359764 | 0.915363174 | 0.365275526 | 0.714905746 | 0.944595913 | Bnip3l-ps | BCL2/adenovirus E1B interacting protein 3 | 4.8186 | 2.0558 | 11.069 | 2.8919 | 10.906 | 8.9108 |
| 20397 | ENSMUSG00000073207 | 4.343449963 | -1.594179029 | 1.148185892 | -1.388432866 | 0.165005274 | NA | Ccdc160 | coiled-coil domain containing 160 [Source | 3.2124 | 11.307 | 5.0312 | 0.964 | 1.0906 | 4.4554 |
| 20398 | ENSMUSG00000073208 | 0 | NA | NA | NA | NA | NA | Speer4c | spermatogenesis associated glutamate (E) | 0 | 0 | 0 | 0 | 0 | 0 |
| 20399 | ENSMUSG00000073209 | 0 | NA | NA | NA | NA | NA | Klf14 | Kruppel-like factor 14 [Source:MGI Symbo | 0 | 0 | 0 | 0 | 0 | 0 |
| 20400 | ENSMUSG00000073226 | 0 | NA | NA | NA | NA | NA | Gm10482 | predicted gene 10482 [Source:MGI Symbo | 0 | 0 | 0 | 0 | 0 | 0 |
| 20401 | ENSMUSG00000073233 | 1.506341172 | 0.559890583 | 1.913681709 | 0.292572469 | 0.76984896 | NA | Nfyc-ps | nuclear transcription factor-Y gamma, pse | 1.6062 | 0 | 2.0125 | 0.964 | 0 | 4.4554 |
| 20402 | ENSMUSG00000073234 | 0 | NA | NA | NA | NA | NA | Gm8773 | predicted gene 8773 [Source:MGI Symbol | 0 | 0 | 0 | 0 | 0 | 0 |
| 20403 | ENSMUSG00000073236 | 22.32023629 | -0.505011879 | 0.457554469 | -1.103719696 | 0.269714749 | 0.725011151 | 2500004C02 | RIKEN cDNA 2500004C02 gene [Source:M | 29.715 | 25.698 | 23.144 | 21.208 | 17.45 | 16.708 |
| 20404 | ENSMUSG00000073242 | 0.760346335 | -0.431432104 | 2.662343224 | -0.162049769 | 0.871266662 | NA | Dnmt3aos | DNA methyltransferase 3A, opposite strar | 1.6062 | 1.0279 | 0 | 1.928 | 0 | 0 |
| 20405 | ENSMUSG00000073243 | 0 | NA | NA | NA | NA | NA | Gm9 | predicted gene 9 [Source:MGI Symbol;Acc | 0 | 0 | 0 | 0 | 0 | 0 |
| 20406 | ENSMUSG00000073245 | 0 | NA | NA | NA | NA | NA | Gm14819 | predicted gene 14819 [Source:MGI Symbo | 0 | 0 | 0 | 0 | 0 | 0 |
| 20407 | ENSMUSG00000073247 | 0 | NA | NA | NA | NA | NA | Gm10486 | predicted gene 10486 [Source:MGI Symbo | 0 | 0 | 0 | 0 | 0 | 0 |
| 20408 | ENSMUSG00000073255 | 0 | NA | NA | NA | NA | NA | Gm14632 | predicted gene 14632 [Source:MGI Symbo | 0 | 0 | 0 | 0 | 0 | 0 |
| 20409 | ENSMUSG00000073257 | 0 | NA | NA | NA | NA | NA | Gm10488 | predicted gene 10488 [Source:MGI Symbo | 0 | 0 | 0 | 0 | 0 | 0 |
| 20410 | ENSMUSG00000073267 | 0 | NA | NA | NA | NA | NA | Gm1993 | predicted gene 1993 [Source:MGI Symbol | 0 | 0 | 0 | 0 | 0 | 0 |
| 20411 | ENSMUSG00000073274 | 0.342432689 | 1.990653086 | 3.900281263 | 0.51038706 | 0.609780321 | NA | Gm14636 | predicted gene 14636 [Source:MGI Symbo | 0 | 0 | 0 | 0.964 | 1.0906 | 0 |
| 20412 | ENSMUSG00000073290 | 0 | NA | NA | NA | NA | NA | Gm10490 | predicted gene 10490 [Source:MGI Symbo | 0 | 0 | 0 | 0 | 0 | 0 |
| 20413 | ENSMUSG00000073291 | 0 | NA | NA | NA | NA | NA | Gm10491 | predicted gene 10491 [Source:MGI Symbo | 0 | 0 | 0 | 0 | 0 | 0 |
| 20414 | ENSMUSG00000073293 | 0.514011053 | 1.044022102 | 3.082659575 | 0.338675769 | 0.734853997 | NA | Nudt10 | nudix (nucleoside diphosphate linked moi | 0 | 0 | 1.0062 | 0.964 | 0 | 1.1138 |
| 20415 | ENSMUSG00000073294 | 0 | NA | NA | NA | NA | NA | Ezhip | EZH inhibitory protein [Source:MGI Symbo | 0 | 0 | 0 | 0 | 0 | 0 |
| 20416 | ENSMUSG00000073295 | 2.565402645 | 1.198512313 | 1.329566531 | 0.901430868 | 0.367359275 | NA | Nudt11 | nudix (nucleoside diphosphate linked moi | 1.6062 | 1.0279 | 2.0125 | 1.928 | 4.3625 | 4.4554 |
| 20417 | ENSMUSG00000073367 | 0 | NA | NA | NA | NA | NA | Gm10493 | predicted gene 10493 [Source:MGI Symbo | 0 | 0 | 0 | 0 | 0 | 0 |
| 20418 | ENSMUSG00000073371 | 0 | NA | NA | NA | NA | NA | Gm6594 | predicted pseudogene 6594 [Source:MGI | 0 | 0 | 0 | 0 | 0 | 0 |
| 20419 | ENSMUSG00000073374 | 16.87212296 | 0.294894282 | 0.564117835 | 0.522752984 | 0.601146162 | 0.915461118 | C030034I22F | RIKEN cDNA C030034I22 gene [Source:M | 16.062 | 7.1954 | 22.137 | 18.316 | 16.359 | 21.163 |
| 20420 | ENSMUSG00000073375 | 0 | NA | NA | NA | NA | NA | Lrrc30 | leucine rich repeat containing 30 [Source: | 0 | 0 | 0 | 0 | 0 | 0 |
| 20421 | ENSMUSG00000073377 | 0 | NA | NA | NA | NA | NA | AU016765 | expressed sequence AU016765 [Source:M | 0 | 0 | 0 | 0 | 0 | 0 |
| 20422 | ENSMUSG00000073380 | 0.267700746 | -1.497481649 | 4.057001436 | -0.369110456 | 0.712045396 | NA | Arrdc5 | arrestin domain containing 5 [Source:MGI | 1.6062 | 0 | 0 | 0 | 0 | 0 |
| 20423 | ENSMUSG00000073386 | 0 | NA | NA | NA | NA | NA | 9830107B12 | RIKEN cDNA 9830107B12 gene [Source:M | 0 | 0 | 0 | 0 | 0 | 0 |
| 20424 | ENSMUSG00000073394 | 0 | NA | NA | NA | NA | NA | Runx2os1 | runt related transcription factor 2, opposi | 0 | 0 | 0 | 0 | 0 | 0 |
| 20425 | ENSMUSG00000073396 | 0 | NA | NA | NA | NA | NA | Esp1 | exocrine gland secreted peptide 1 [Source | 0 | 0 | 0 | 0 | 0 | 0 |
| 20426 | ENSMUSG00000073399 | 0 | NA | NA | NA | NA | NA | Trim40 | tripartite motif-containing 40 [Source:MG | 0 | 0 | 0 | 0 | 0 | 0 |
| 20427 | ENSMUSG00000073400 | 3.347089469 | 1.6097171 | 1.43100067 | 1.124889131 | 0.260636018 | NA | Trim10 | tripartite motif-containing 10 [Source:MG | 0 | 0 | 5.0312 | 2.8919 | 4.3625 | 7.7969 |
| 20428 | ENSMUSG00000073402 | 0.335414862 | -1.779047242 | 4.030804179 | -0.441362855 | 0.658950329 | NA | Gm8909 | predicted gene 8909 [Source:MGI Symbol | 0 | 0 | 2.0125 | 0 | 0 | 0 |
| 20429 | ENSMUSG00000073403 | 0.346303622 | 2.004339653 | 3.884003423 | 0.516049919 | 0.605819544 | NA | Gm10499 | predicted gene 10499 [Source:MGI Symbo | 0 | 0 | 0 | 0.964 | 0 | 1.1138 |
| 20430 | ENSMUSG00000073405 | 11.12276194 | -1.563380148 | 0.709449731 | -2.20365176 | 0.027548844 | 0.215503776 | H2-T-ps | histocompatibility 2, T region locus, pseuc | 23.29 | 13.363 | 13.081 | 4.8199 | 3.2719 | 8.9108 |
| 20431 | ENSMUSG00000073406 | 57.43505608 | 0.182225798 | 0.422768195 | 0.431030055 | 0.666446517 | 0.929596279 | H2-Bl | histocompatibility 2, blastocyst [Source:M | 72.279 | 27.754 | 61.381 | 97.362 | 49.078 | 36.757 |
| 20432 | ENSMUSG00000073407 | 2.937950174 | -0.495787496 | 1.514370648 | -0.327388475 | 0.743374083 | NA | Gm6034 | predicted gene 6034 [Source:MGI Symbol | 7.2279 | 0 | 3.0187 | 1.928 | 5.4531 | 0 |
| 20433 | ENSMUSG00000073408 | 0 | NA | NA | NA | NA | NA | Mucl3 | mucin like 3 [Source:MGI Symbol;Acc:MG | 0 | 0 | 0 | 0 | 0 | 0 |
| 20434 | ENSMUSG00000073409 | 620.1532796 | -0.062584263 | 0.141935701 | -0.440933905 | 0.659260847 | 0.927763076 | H2-Q6 | histocompatibility 2, Q region locus 6 [Sou | 609.55 | 687.68 | 604.75 | 709.49 | 572.57 | 536.87 |
| 20435 | ENSMUSG00000073411 | 11601.87899 | -0.316372174 | 0.097024212 | -3.260754912 | 0.00111116 | 0.022572786 | H2-D1 | histocompatibility 2, D region locus 1 [Sou | 12699 | 13902 | 12008 | 10930 | 10396 | 9678.2 |
| 20436 | ENSMUSG00000073412 | 14.11442041 | -1.886049104 | 0.639149805 | -2.950871751 | 0.003168785 | 0.048599236 | Lst1 | leukocyte specific transcript 1 [Source:M | 17.668 | 31.865 | 17.106 | 4.8199 | 6.5437 | 6.6831 |

| | | | | | | | | | | | | | | | |
|---|---|---|---|---|---|---|---|---|---|---|---|---|---|---|---|
| 20437 | ENSMUSG00000073413 | 0.806689472 | -0.274753491 | 2.612595279 | -0.105164965 | 0.91624491 | NA | Ly6g6d | lymphocyte antigen 6 complex, locus G6D | 1.6062 | 0 | 1.0062 | 0 | 0 | 2.2277 |
| 20438 | ENSMUSG00000073414 | 0.315620088 | 0.165143421 | 4.063300985 | 0.040642675 | 0.967580763 | NA | Mpig6b | megakaryocyte and platelet inhibitory rec | 0.8031 | 0 | 0 | 0 | 1.0906 | 0 |
| 20439 | ENSMUSG00000073415 | 2.976794796 | 0.76557384 | 1.262249554 | 0.606515437 | 0.544172535 | NA | Gm10501 | predicted gene 10501 [Source:MGI Symbo | 1.6062 | 2.0558 | 3.0187 | 6.7478 | 1.0906 | 3.3415 |
| 20440 | ENSMUSG00000073418 | 22745.4421 | -0.019051821 | 0.089405145 | -0.213095357 | 0.831252586 | 0.97052742 | C4b | complement component 4B (Chido blood | 23663 | 23846 | 21178 | 23072 | 22962 | 21752 |
| 20441 | ENSMUSG00000073420 | 0 | NA | NA | NA | NA | NA | Btnl5-ps | butyrophilin-like 5, pseudogene [Source:N | 0 | 0 | 0 | 0 | 0 | 0 |
| 20442 | ENSMUSG00000073421 | 289.9179888 | -1.131651691 | 0.219487441 | -5.155883573 | 2.52E-07 | 2.43E-05 | H2-Ab1 | histocompatibility 2, class II antigen A, bet | 368.62 | 517.04 | 308.92 | 187.01 | 139.6 | 218.31 |
| 20443 | ENSMUSG00000073422 | 7.390381492 | 0.437593776 | 0.844086204 | 0.518423088 | 0.60416311 | 0.916322307 | H2-Ke6 | H2-K region expressed gene 6 [Source:MG | 9.6372 | 4.1117 | 5.0312 | 13.496 | 8.7249 | 3.3415 |
| 20444 | ENSMUSG00000073423 | 44.6957404 | 0.501255814 | 0.349829299 | 1.4328583 | 0.151898336 | 0.567340036 | Zfp414 | zinc finger protein 414 [Source:MGI Symb | 41.761 | 32.893 | 36.225 | 57.839 | 37.081 | 62.375 |
| 20445 | ENSMUSG00000073424 | 3527.694926 | 0.001135661 | 0.109212977 | 0.010398588 | 0.991703276 | 0.99859176 | Cyp4f15 | cytochrome P450, family 4, subfamily f, po | 3220.4 | 3392.1 | 3967.6 | 3625.5 | 3659 | 3301.4 |
| 20446 | ENSMUSG00000073427 | 20.01167347 | -0.143693663 | 0.526331366 | -0.27300988 | 0.784845621 | 0.960670212 | Gm4924 | predicted gene 4924 [Source:MGI Symbol | 32.124 | 16.447 | 14.087 | 14.46 | 22.903 | 20.049 |
| 20447 | ENSMUSG00000073430 | 0.160662974 | 1.126924175 | 4.080472857 | 0.276174898 | 0.782413736 | NA | Gm10505 | predicted gene 10505 [Source:MGI Symbo | 0 | 0 | 0 | 0.964 | 0 | 0 |
| 20448 | ENSMUSG00000073433 | 0 | NA | NA | NA | NA | NA | Arhgdig | Rho GDP dissociation inhibitor (GDI) gamr | 0 | 0 | 0 | 0 | 0 | 0 |
| 20449 | ENSMUSG00000073434 | 115.2122659 | 0.015700876 | 0.236264157 | 0.066454751 | 0.947015782 | 0.990677354 | Wdr90 | WD repeat domain 90 [Source:MGI Symbo | 99.585 | 113.07 | 131.82 | 140.74 | 104.7 | 101.36 |
| 20450 | ENSMUSG00000073435 | 0 | NA | NA | NA | NA | NA | Nme3 | NME/NM23 nucleoside diphosphate kinas | 0 | 0 | 0 | 0 | 0 | 0 |
| 20451 | ENSMUSG00000073436 | 56.27938994 | -0.198176516 | 0.301728767 | -0.656803521 | 0.51130726 | 0.883679955 | Eme2 | essential meiotic structure-specific endon | 58.626 | 55.508 | 66.412 | 55.911 | 58.893 | 42.326 |
| 20452 | ENSMUSG00000073437 | 19.9128542 | 0.319210413 | 0.560943774 | 0.569059553 | 0.569315726 | 0.903735951 | D330041H03 | RIKEN cDNA D330041H03 gene [Source:M | 30.518 | 8.2233 | 14.087 | 26.991 | 20.722 | 18.935 |
| 20453 | ENSMUSG00000073448 | 10.89401794 | -0.332888221 | 0.684766145 | -0.48613417 | 0.626872042 | 0.920273791 | Gm10509 | predicted gene 10509 [Source:MGI Symbo | 12.85 | 15.419 | 8.05 | 4.8199 | 13.087 | 11.138 |
| 20454 | ENSMUSG00000073457 | 0 | NA | NA | NA | NA | NA | Smok2b | sperm motility kinase 2B [Source:MGI Sym | 0 | 0 | 0 | 0 | 0 | 0 |
| 20455 | ENSMUSG00000073458 | 0 | NA | NA | NA | NA | NA | Smok2a | sperm motility kinase 2A [Source:MGI Sym | 0 | 0 | 0 | 0 | 0 | 0 |
| 20456 | ENSMUSG00000073460 | 177.6326357 | -0.277464976 | 0.320300008 | -0.866265906 | 0.386344359 | 0.817115886 | Pnldc1 | poly(A)-specific ribonuclease (PARN)-like | 304.38 | 165.49 | 113.71 | 206.29 | 171.23 | 104.7 |
| 20457 | ENSMUSG00000073468 | 665.0130634 | 0.102821252 | 0.118971647 | 0.864250054 | 0.387450541 | 0.817618864 | Sft2d1 | SFT2 domain containing 1 [Source:MGI Sy | 671.39 | 611.61 | 639.97 | 642.97 | 702.36 | 721.77 |
| 20458 | ENSMUSG00000073471 | 116.7300919 | 0.014450043 | 0.232295807 | 0.062205353 | 0.9503993 | 0.990815588 | Rsph3a | radial spoke 3A homolog (Chlamydomona | 119.66 | 117.18 | 111.69 | 125.32 | 137.42 | 89.108 |
| 20459 | ENSMUSG00000073478 | 30.30453431 | -0.150401788 | 0.401058517 | -0.375012079 | 0.707651484 | 0.943361388 | D730003I15I | RIKEN cDNA D730003I15 gene [Source:MG | 38.549 | 24.67 | 32.2 | 27.955 | 27.265 | 31.188 |
| 20460 | ENSMUSG00000073481 | 8846.499569 | -0.017830932 | 0.099333914 | -0.179504979 | 0.857541208 | 0.975116041 | Mtarc2 | mitochondrial amidoxime reducing compo | 8105.7 | 8994.3 | 9604.6 | 8542.8 | 9357.5 | 8474.1 |
| 20461 | ENSMUSG00000073482 | 0.765090549 | -0.195763385 | 2.933488969 | -0.066733977 | 0.946793485 | NA | Gm10517 | predicted gene 10517 [Source:MGI Symbo | 2.4093 | 0 | 0 | 0 | 2.1812 | 0 |
| 20462 | ENSMUSG00000073485 | 9.85370344 | 1.527155607 | 0.743439193 | 2.054176887 | 0.039958575 | 0.273655693 | H3f3aos | H3.3 histone A, opposite strand [Source:N | 3.2124 | 4.1117 | 8.05 | 18.316 | 8.7249 | 16.708 |
| 20463 | ENSMUSG00000073486 | 0 | NA | NA | NA | NA | NA | Gm10518 | predicted gene 10518 [Source:MGI Symbo | 0 | 0 | 0 | 0 | 0 | 0 |
| 20464 | ENSMUSG00000073489 | 82.31442139 | -1.108296467 | 0.278505013 | -3.979448906 | 6.91E-05 | 0.002683563 | Ifi204 | interferon activated gene 204 [Source:MG | 116.45 | 123.35 | 97.606 | 55.911 | 37.081 | 63.489 |
| 20465 | ENSMUSG00000073490 | 70.55499011 | -1.1226238 | 0.309319246 | -3.629337053 | 0.00028415 | 0.008319729 | Ifi207 | interferon activated gene 207 [Source:MG | 105.21 | 118.21 | 66.412 | 37.595 | 46.897 | 49.009 |
| 20466 | ENSMUSG00000073491 | 23.83836126 | -0.649272576 | 0.492597259 | -1.318059661 | 0.187483677 | 0.623967764 | Ifi213 | interferon activated gene 213 [Source:MG | 26.502 | 37.005 | 24.15 | 28.919 | 13.087 | 13.366 |
| 20467 | ENSMUSG00000073492 | 0.774435248 | -2.999089927 | 2.565713408 | -1.168910728 | 0.242439601 | NA | Gm10521 | predicted gene 10521 [Source:MGI Symbo | 1.6062 | 1.0279 | 2.0125 | 0 | 0 | 0 |
| 20468 | ENSMUSG00000073494 | 0.185640647 | 1.126924175 | 4.080472857 | 0.276174898 | 0.782413736 | NA | Sh2d1b2 | SH2 domain containing 1B2 [Source:MGI S | 0 | 0 | 0 | 0 | 0 | 1.1138 |
| 20469 | ENSMUSG00000073497 | 0 | NA | NA | NA | NA | NA | AA792892 | expressed sequence AA792892 [Source:M | 0 | 0 | 0 | 0 | 0 | 0 |
| 20470 | ENSMUSG00000073510 | 0 | NA | NA | NA | NA | NA | Gm7647 | predicted gene 7647 [Source:MGI Symbol | 0 | 0 | 0 | 0 | 0 | 0 |
| 20471 | ENSMUSG00000073514 | 0 | NA | NA | NA | NA | NA | Dok6 | docking protein 6 [Source:MGI Symbol;Ac | 0 | 0 | 0 | 0 | 0 | 0 |
| 20472 | ENSMUSG00000073528 | 0 | NA | NA | NA | NA | NA | Gm10530 | predicted gene 10530 [Source:MGI Symbo | 0 | 0 | 0 | 0 | 0 | 0 |
| 20473 | ENSMUSG00000073529 | 0.476489653 | -2.293616034 | 3.421506946 | -0.670352587 | 0.502633051 | NA | F830208F22I | RIKEN cDNA F830208F22 gene [Source:MG | 0.8031 | 2.0558 | 0 | 0 | 0 | 0 |
| 20474 | ENSMUSG00000073530 | 0.267700746 | -1.497481649 | 4.057001436 | -0.369110456 | 0.712045396 | NA | Pappa2 | pappalysin 2 [Source:MGI Symbol;Acc:MG | 1.6062 | 0 | 0 | 0 | 0 | 0 |
| 20475 | ENSMUSG00000073532 | 0 | NA | NA | NA | NA | NA | Gm7276 | predicted gene 7276 [Source:MGI Symbol | 0 | 0 | 0 | 0 | 0 | 0 |
| 20476 | ENSMUSG00000073535 | 3.383446192 | 0.815240389 | 1.205418541 | 0.676313133 | 0.498841857 | NA | Gm5532 | predicted gene 5532 [Source:MGI Symbol | 3.2124 | 2.0558 | 2.0125 | 1.928 | 2.1812 | 8.9108 |
| 20477 | ENSMUSG00000073538 | 1.566958698 | -0.226238548 | 1.890802846 | -0.119652109 | 0.904758739 | NA | E330020D12 | Riken cDNA E330020D12 gene [Source:MG | 0 | 3.0838 | 2.0125 | 0.964 | 0 | 3.3415 |
| 20478 | ENSMUSG00000073542 | 126.9747033 | 0.075631331 | 0.218723384 | 0.3457853 | 0.729504073 | 0.946534338 | Cep76 | centrosomal protein 76 [Source:MGI Syml | 118.86 | 136.71 | 115.72 | 145.56 | 106.88 | 138.12 |
| 20479 | ENSMUSG00000073551 | 0 | NA | NA | NA | NA | NA | Spink13 | serine peptidase inhibitor, Kazal type 13 [S | 0 | 0 | 0 | 0 | 0 | 0 |
| 20480 | ENSMUSG00000073554 | 0.160662974 | 1.126924175 | 4.080472857 | 0.276174898 | 0.782413736 | NA | Gm6960 | predicted pseudogene 6960 [Source:MGI | 0 | 0 | 0 | 0.964 | 0 | 0 |
| 20481 | ENSMUSG00000073555 | 1635.62657 | -0.520266738 | 0.163739951 | -3.177396453 | 0.001486037 | 0.02817044 | Gm4951 | predicted gene 4951 [Source:MGI Symbol | 1785.3 | 1808.1 | 2188.6 | 1138.5 | 1747.2 | 1146.1 |
| 20482 | ENSMUSG00000073557 | 125.7483979 | -0.126922531 | 0.213202349 | -0.59531488 | 0.551633011 | 0.895935901 | Ppp1r12b | protein phosphatase 1, regulatory subunit | 139.74 | 124.38 | 129.81 | 135.92 | 121.06 | 103.59 |
| 20483 | ENSMUSG00000073559 | 0 | NA | NA | NA | NA | NA | Gm8618 | predicted gene 8618 [Source:MGI Symbol | 0 | 0 | 0 | 0 | 0 | 0 |
| 20484 | ENSMUSG00000073563 | 833.0817324 | -0.29461939 | 0.123784381 | -2.380101502 | 0.01730787 | 0.156658967 | Csnk1g3 | casein kinase 1, gamma 3 [Source:MGI Syı | 909.11 | 1017.6 | 827.13 | 757.69 | 787.43 | 699.49 |
| 20485 | ENSMUSG00000073565 | 726.0995171 | 0.226965065 | 0.125707554 | 1.805500602 | 0.070996381 | 0.385759452 | Prr16 | proline rich 16 [Source:MGI Symbol;Acc:N | 635.25 | 614.69 | 757.7 | 777.93 | 761.25 | 809.76 |
| 20486 | ENSMUSG00000073568 | 0 | NA | NA | NA | NA | NA | Arl14epl | ADP-ribosylation factor-like 14 effector pr | 0 | 0 | 0 | 0 | 0 | 0 |
| 20487 | ENSMUSG00000073569 | 0.160662974 | 1.126924175 | 4.080472857 | 0.276174898 | 0.782413736 | NA | Gm10540 | predicted gene 10540 [Source:MGI Symbo | 0 | 0 | 0 | 0.964 | 0 | 0 |
| 20488 | ENSMUSG00000073572 | 0 | NA | NA | NA | NA | NA | Gm10542 | predicted gene 10542 [Source:MGI Symbo | 0 | 0 | 0 | 0 | 0 | 0 |
| 20489 | ENSMUSG00000073573 | 0 | NA | NA | NA | NA | NA | Spink11 | serine peptidase inhibitor, Kazal type 11 [S | 0 | 0 | 0 | 0 | 0 | 0 |
| 20490 | ENSMUSG00000073574 | 0 | NA | NA | NA | NA | NA | Grxcr2 | glutaredoxin, cysteine rich 2 [Source:MGI | 0 | 0 | 0 | 0 | 0 | 0 |
| 20491 | ENSMUSG00000073590 | 5.27994419 | -0.877876495 | 1.03549407 | -0.84778515 | 0.396557635 | 0.824110579 | 3222401L13I | RIKEN cDNA 3222401L13 gene [Source:MG | 10.44 | 2.0558 | 8.05 | 6.7478 | 3.2719 | 1.1138 |
| 20492 | ENSMUSG00000073591 | 3.613684311 | -1.982039098 | 1.365841609 | -1.45114857 | 0.146738494 | NA | Pcdhb22 | protocadherin beta 22 [Source:MGI Symb | 4.0155 | 6.1675 | 7.0437 | 0 | 0 | 4.4554 |

| | | | | | | | | | | | | | | | |
|---|---|---|---|---|---|---|---|---|---|---|---|---|---|---|---|
| 20493 | ENSMUSG00000073594 | 0 | NA | NA | NA | NA | NA | Gm10545 | predicted gene 10545 [Source:MGI Symbo | 0 | 0 | 0 | 0 | 0 | 0 |
| 20494 | ENSMUSG00000073598 | 0.503302254 | -0.837877584 | 3.426698643 | -0.244514523 | 0.806832356 | NA | 1700066B19 | RIKEN cDNA 1700066B19 gene [Source:M | 0 | 2.0558 | 0 | 0.964 | 0 | 0 |
| 20495 | ENSMUSG00000073599 | 17.91084973 | -0.518567387 | 0.511259816 | -1.014293263 | 0.310442838 | 0.759154604 | Ecscr | endothelial cell surface expressed chemot | 22.487 | 24.67 | 16.1 | 14.46 | 15.269 | 14.48 |
| 20496 | ENSMUSG00000073600 | 472.1105808 | -0.680041138 | 0.194170892 | -3.502281578 | 0.000461292 | 0.011705483 | Prob1 | proline rich basic protein 1 [Source:MGI S | 625.62 | 627.03 | 491.05 | 322.93 | 484.23 | 281.8 |
| 20497 | ENSMUSG00000073601 | 0 | NA | NA | NA | NA | NA | Serpinb3c | serine (or cysteine) peptidase inhibitor, cl | 0 | 0 | 0 | 0 | 0 | 0 |
| 20498 | ENSMUSG00000073602 | 0 | NA | NA | NA | NA | NA | Serpinb3b | serine (or cysteine) peptidase inhibitor, cl | 0 | 0 | 0 | 0 | 0 | 0 |
| 20499 | ENSMUSG00000073607 | 0.961910823 | -0.938554428 | 2.269016656 | -0.413639285 | 0.679138298 | NA | Gm10548 | predicted gene 10548 [Source:MGI Symbo | 0.8031 | 1.0279 | 2.0125 | 1.928 | 0 | 0 |
| 20500 | ENSMUSG00000073608 | 0.335414862 | -1.779047242 | 4.030804179 | -0.441362855 | 0.658950329 | NA | Gal3st2c | galactose-3-O-sulfotransferase 2C [Source | 0 | 0 | 2.0125 | 0 | 0 | 0 |
| 20501 | ENSMUSG00000073609 | 534.414289 | -0.028188101 | 0.130877515 | -0.21537772 | 0.829472839 | 0.969955221 | D2hgdh | D-2-hydroxyglutarate dehydrogenase [Sou | 512.38 | 547.88 | 558.47 | 472.35 | 554.03 | 561.38 |
| 20502 | ENSMUSG00000073610 | 3.750630273 | 1.886454082 | 1.357420082 | 1.389734915 | 0.164609388 | NA | NA | NA | 0.8031 | 1.0279 | 3.0187 | 6.7478 | 10.906 | 0 |
| 20503 | ENSMUSG00000073616 | 523.4898018 | -0.003134192 | 0.12324428 | -0.025430728 | 0.979711402 | 0.99820894 | Cops9 | COP9 signalosome subunit 9 [Source:MGI | 522.02 | 542.74 | 507.15 | 495.48 | 532.22 | 541.33 |
| 20504 | ENSMUSG00000073627 | 5.804812626 | 0.836948436 | 0.911373532 | 0.91833744 | 0.358442232 | 0.798191951 | C130036L24 | RIKEN cDNA C130036L24 gene [Source:M( | 2.4093 | 5.1396 | 5.0312 | 6.7478 | 4.3625 | 11.138 |
| 20505 | ENSMUSG00000073628 | 0.371281295 | 2.09253362 | 4.031852165 | 0.519000582 | 0.603760335 | NA | Gm10552 | predicted gene 10552 [Source:MGI Symbo | 0 | 0 | 0 | 0 | 0 | 2.2277 |
| 20506 | ENSMUSG00000073631 | 0 | NA | NA | NA | NA | NA | Gm10553 | predicted gene 10553 [Source:MGI Symbo | 0 | 0 | 0 | 0 | 0 | 0 |
| 20507 | ENSMUSG00000073633 | 103.6018657 | 0.041343784 | 0.246750182 | 0.167553207 | 0.866934786 | 0.977930657 | Fbxo36 | F-box protein 36 [Source:MGI Symbol;Acc | 111.63 | 111.02 | 83.518 | 103.15 | 125.42 | 86.88 |
| 20508 | ENSMUSG00000073639 | 1933.012811 | -0.006570032 | 0.100567031 | -0.065329876 | 0.947911355 | 0.990677354 | Rab18 | RAB18, member RAS oncogene family [So | 1918.6 | 1950 | 1943.1 | 1767 | 2058 | 1961.5 |
| 20509 | ENSMUSG00000073640 | 1.889675376 | 0.43256143 | 1.678006472 | 0.257782933 | 0.796574437 | NA | Rpl27-ps3 | ribosomal protein L27, pseudogene 3 [Sou | 0.8031 | 0 | 4.025 | 0.964 | 1.0906 | 4.4554 |
| 20510 | ENSMUSG00000073643 | 193.4432318 | 0.178360389 | 0.227123886 | 0.785300005 | 0.432277693 | 0.845264432 | Wdfy1 | WD repeat and FYVE domain containing 1 | 205.59 | 190.16 | 148.92 | 267.02 | 194.13 | 154.82 |
| 20511 | ENSMUSG00000073647 | 18.9641963 | 0.69296212 | 0.514023688 | 1.3481132 | 0.177621976 | 0.608962907 | Gm10557 | predicted gene 10557 [Source:MGI Symbo | 16.062 | 14.391 | 13.081 | 32.775 | 18.541 | 18.935 |
| 20512 | ENSMUSG00000073650 | 0.562214094 | -1.122836648 | 3.276281221 | -0.34271681 | 0.731811515 | NA | Catip | ciliogenesis associated TTC17 interacting p | 2.4093 | 0 | 0 | 0.964 | 0 | 0 |
| 20513 | ENSMUSG00000073652 | 6.395797989 | 0.456071913 | 0.841464207 | 0.541997994 | 0.587819888 | 0.910798643 | Apol7d | apolipoprotein L 7d [Source:MGI Symbol;A | 4.0155 | 7.1954 | 5.0312 | 6.7478 | 9.8156 | 5.5692 |
| 20514 | ENSMUSG00000073656 | 0 | NA | NA | NA | NA | NA | Gm10558 | predicted gene 10558 [Source:MGI Symbo | 0 | 0 | 0 | 0 | 0 | 0 |
| 20515 | ENSMUSG00000073658 | 0 | NA | NA | NA | NA | NA | Crygb | crystallin, gamma B [Source:MGI Symbol;A | 0 | 0 | 0 | 0 | 0 | 0 |
| 20516 | ENSMUSG00000073664 | 554.624202 | 0.100682519 | 0.134811522 | 0.746839124 | 0.455160676 | 0.857337687 | Nbeal1 | neurobeachin like 1 [Source:MGI Symbol;A | 507.56 | 586.94 | 512.18 | 603.45 | 607.47 | 510.14 |
| 20517 | ENSMUSG00000073676 | 2380.825055 | 0.097087908 | 0.094898028 | 1.023076145 | 0.306271845 | 0.755442696 | Hspe1 | heat shock protein 1 (chaperonin 10) [Sou | 2226.2 | 2249.1 | 2427.1 | 2383 | 2424.4 | 2575.2 |
| 20518 | ENSMUSG00000073678 | 347.3424862 | 0.681414951 | 0.177067286 | 3.848339051 | 0.000118921 | 0.00416484 | Pgap1 | post-GPI attachment to proteins 1 [Source | 329.27 | 247.73 | 222.38 | 477.17 | 419.89 | 387.62 |
| 20519 | ENSMUSG00000073679 | 0 | NA | NA | NA | NA | NA | Mxra8os | matrix-remodelling associated 8, opposite | 0 | 0 | 0 | 0 | 0 | 0 |
| 20520 | ENSMUSG00000073680 | 51.47901096 | -0.118829168 | 0.378795524 | -0.313702672 | 0.753746871 | 0.952989051 | Tmem88b | transmembrane protein 88B [Source:MGI | 54.611 | 34.949 | 71.443 | 66.514 | 50.168 | 31.188 |
| 20521 | ENSMUSG00000073682 | 1.358014806 | -2.646870292 | 2.21080923 | -1.197240475 | 0.231212838 | NA | Gm10563 | predicted gene 10563 [Source:MGI Symbo | 4.0155 | 0 | 3.0187 | 0 | 0 | 1.1138 |
| 20522 | ENSMUSG00000073684 | 130.2934133 | 0.259852409 | 0.215847028 | 1.203873003 | 0.228638668 | 0.675041782 | Faap20 | Fanconi anemia core complex associated p | 133.31 | 113.07 | 108.67 | 123.39 | 140.69 | 162.62 |
| 20523 | ENSMUSG00000073686 | 0 | NA | NA | NA | NA | NA | Gm10564 | predicted gene 10564 [Source:MGI Symbo | 0 | 0 | 0 | 0 | 0 | 0 |
| 20524 | ENSMUSG00000073699 | 0 | NA | NA | NA | NA | NA | Gm833 | predicted gene 833 [Source:MGI Symbol;A | 0 | 0 | 0 | 0 | 0 | 0 |
| 20525 | ENSMUSG00000073700 | 1103.759504 | 0.170574215 | 0.139475358 | 1.222970258 | 0.22134093 | 0.667727967 | Klhl21 | kelch-like 21 [Source:MGI Symbol;Acc:MG | 1076.2 | 1091.6 | 946.88 | 979.4 | 1148.4 | 1380.1 |
| 20526 | ENSMUSG00000073702 | 2319.779389 | 0.082385107 | 0.111491016 | 0.738939424 | 0.459943782 | 0.859753653 | Rpl31 | ribosomal protein L31 [Source:MGI Symbo | 2155.5 | 2452.6 | 2152.4 | 2143.9 | 2416.8 | 2597.5 |
| 20527 | ENSMUSG00000073705 | 3.162254954 | -1.866679885 | 1.324304743 | -1.409554632 | 0.158671231 | NA | Cenps | centromere protein S [Source:MGI Symbo | 4.8186 | 4.1117 | 6.0375 | 2.8919 | 0 | 1.1138 |
| 20528 | ENSMUSG00000073716 | 2.402987366 | 0.435705866 | 1.393593995 | 0.31264907 | 0.754547293 | NA | Gm13241 | predicted gene 13241 [Source:MGI Symbo | 4.0155 | 1.0279 | 1.0062 | 2.8919 | 4.3625 | 1.1138 |
| 20529 | ENSMUSG00000073721 | 0 | NA | NA | NA | NA | NA | Pramel15 | PRAME like 15 [Source:MGI Symbol;Acc:M | 0 | 0 | 0 | 0 | 0 | 0 |
| 20530 | ENSMUSG00000073722 | 0 | NA | NA | NA | NA | NA | 4931408C20 | RIKEN cDNA 4931408C20 gene [Source:M( | 0 | 0 | 0 | 0 | 0 | 0 |
| 20531 | ENSMUSG00000073723 | 0 | NA | NA | NA | NA | NA | Gm13102 | predicted gene 13102 [Source:MGI Symbo | 0 | 0 | 0 | 0 | 0 | 0 |
| 20532 | ENSMUSG00000073724 | 0 | NA | NA | NA | NA | NA | Pramel4 | PRAME like 4 [Source:MGI Symbol;Acc:M( | 0 | 0 | 0 | 0 | 0 | 0 |
| 20533 | ENSMUSG00000073725 | 1472.68461 | -0.078000616 | 0.103961338 | -0.75028484 | 0.453083171 | 0.857073339 | Lmbrd1 | LMBR1 domain containing 1 [Source:MGI | 1490.6 | 1547 | 1499.3 | 1312 | 1468 | 1519.3 |
| 20534 | ENSMUSG00000073728 | 2.853421128 | 0.260747942 | 1.346569488 | 0.193638683 | 0.846458807 | NA | Tmem51os1 | Tmem51 opposite strand 1 [Source:MGI S | 0.8031 | 1.0279 | 6.0375 | 4.8199 | 1.0906 | 3.3415 |
| 20535 | ENSMUSG00000073730 | 0 | NA | NA | NA | NA | NA | Ppp1r14bl | protein phosphatase 1, regulatory inhibito | 0 | 0 | 0 | 0 | 0 | 0 |
| 20536 | ENSMUSG00000073731 | 0 | NA | NA | NA | NA | NA | AI507597 | expressed sequence AI507597 [Source:M( | 0 | 0 | 0 | 0 | 0 | 0 |
| 20537 | ENSMUSG00000073733 | 4.382827708 | -0.078609087 | 1.069649622 | -0.073490502 | 0.941415802 | NA | Cplane2 | ciliogenesis and planar polarity effector 2 | 2.4093 | 3.0838 | 8.05 | 2.8919 | 7.6343 | 2.2277 |
| 20538 | ENSMUSG00000073735 | 0 | NA | NA | NA | NA | NA | Defb18 | defensin beta 18 [Source:MGI Symbol;Acc | 0 | 0 | 0 | 0 | 0 | 0 |
| 20539 | ENSMUSG00000073737 | 0.167707431 | -0.796629176 | 4.080472857 | -0.195229623 | 0.84521319 | NA | Gm10566 | predicted gene 10566 [Source:MGI Symbo | 0 | 0 | 1.0062 | 0 | 0 | 0 |
| 20540 | ENSMUSG00000073739 | 0 | NA | NA | NA | NA | NA | Gm16287 | predicted gene 16287 [Source:MGI Symbo | 0 | 0 | 0 | 0 | 0 | 0 |
| 20541 | ENSMUSG00000073747 | 0 | NA | NA | NA | NA | NA | Zpld2 | zona pellucida like domain containing 2 [S | 0 | 0 | 0 | 0 | 0 | 0 |
| 20542 | ENSMUSG00000073752 | 0 | NA | NA | NA | NA | NA | Gm10570 | predicted gene 10570 [Source:MGI Symbo | 0 | 0 | 0 | 0 | 0 | 0 |
| 20543 | ENSMUSG00000073753 | 0 | NA | NA | NA | NA | NA | Gm10571 | predicted gene 10571 [Source:MGI Symbo | 0 | 0 | 0 | 0 | 0 | 0 |
| 20544 | ENSMUSG00000073755 | 157.6508985 | -0.277535309 | 0.191104961 | -1.452266372 | 0.146427554 | 0.556855766 | 5730409E04 | RIKEN cDNA 5730409E04Rik gene [Source | 187.12 | 175.77 | 154.96 | 135.92 | 142.87 | 149.26 |
| 20545 | ENSMUSG00000073758 | 31.14102409 | 0.019338228 | 0.427892847 | 0.04519409 | 0.963952605 | 0.993245949 | Sh3d21 | SH3 domain containing 21 [Source:MGI Sy | 42.564 | 31.865 | 18.112 | 33.739 | 32.719 | 27.846 |
| 20546 | ENSMUSG00000073759 | 0 | NA | NA | NA | NA | NA | 4933407E24 | RIKEN cDNA 4933407E24 gene [Source:M( | 0 | 0 | 0 | 0 | 0 | 0 |
| 20547 | ENSMUSG00000073761 | 0 | NA | NA | NA | NA | NA | 4933427I04F | Riken cDNA 4933427I04 gene [Source:MG | 0 | 0 | 0 | 0 | 0 | 0 |
| 20548 | ENSMUSG00000073764 | 0 | NA | NA | NA | NA | NA | Gm12888 | predicted gene 12888 [Source:MGI Symbo | 0 | 0 | 0 | 0 | 0 | 0 |

| | | | | | | | | | | | | | | | |
|---|---|---|---|---|---|---|---|---|---|---|---|---|---|---|---|
| 20549 | ENSMUSG00000073765 | 0 | NA | NA | NA | NA | NA | Gm12863 | predicted gene 12863 [Source:MGI Symbo | 0 | 0 | 0 | 0 | 0 | 0 |
| 20550 | ENSMUSG00000073768 | 0 | NA | NA | NA | NA | NA | Olfr1330 | olfactory receptor 1330 [Source:MGI Sym | 0 | 0 | 0 | 0 | 0 | 0 |
| 20551 | ENSMUSG00000073769 | 0 | NA | NA | NA | NA | NA | Olfr1331 | olfactory receptor 1331 [Source:MGI Sym | 0 | 0 | 0 | 0 | 0 | 0 |
| 20552 | ENSMUSG00000073771 | 28.65041053 | 0.33958636 | 0.456955637 | 0.743149514 | 0.457391165 | 0.858923498 | Btbd19 | BTB (POZ) domain containing 19 [Source:I | 19.274 | 31.865 | 25.156 | 48.199 | 22.903 | 24.505 |
| 20553 | ENSMUSG00000073774 | 0 | NA | NA | NA | NA | NA | Kncn | kinocilin [Source:MGI Symbol;Acc:MGI:36 | 0 | 0 | 0 | 0 | 0 | 0 |
| 20554 | ENSMUSG00000073775 | 164.83877 | 0.065652795 | 0.197384027 | 0.332614528 | 0.739425275 | 0.948310621 | Kti12 | KTI12 homolog, chromatin associated [So | 177.49 | 153.16 | 151.94 | 147.49 | 195.22 | 163.74 |
| 20555 | ENSMUSG00000073778 | 0 | NA | NA | NA | NA | NA | Faddos | Fas (TNFRSF6)-associated via death doma | 0 | 0 | 0 | 0 | 0 | 0 |
| 20556 | ENSMUSG00000073779 | 0 | NA | NA | NA | NA | NA | Lrp8os2 | low density lipoprotein receptor-related p | 0 | 0 | 0 | 0 | 0 | 0 |
| 20557 | ENSMUSG00000073781 | 0 | NA | NA | NA | NA | NA | Gm6471 | predicted gene 6471 [Source:MGI Symbol | 0 | 0 | 0 | 0 | 0 | 0 |
| 20558 | ENSMUSG00000073785 | 0 | NA | NA | NA | NA | NA | Krtap5-5 | keratin associated protein 5-5 [Source:MG | 0 | 0 | 0 | 0 | 0 | 0 |
| 20559 | ENSMUSG00000073786 | 0 | NA | NA | NA | NA | NA | Gm7579 | predicted gene 7579 [Source:MGI Symbol | 0 | 0 | 0 | 0 | 0 | 0 |
| 20560 | ENSMUSG00000073787 | 2.787070583 | -1.152035608 | 1.300831428 | -0.885614833 | 0.375825116 | NA | Gm10575 | predicted gene 10575 [Source:MGI Symbo | 6.4248 | 2.0558 | 3.0187 | 1.928 | 2.1812 | 1.1138 |
| 20561 | ENSMUSG00000073789 | 0 | NA | NA | NA | NA | NA | Gm10577 | predicted gene 10577 [Source:MGI Symbo | 0 | 0 | 0 | 0 | 0 | 0 |
| 20562 | ENSMUSG00000073791 | 5.858489734 | 1.276864943 | 0.941171984 | 1.356675469 | 0.17488435 | 0.604860651 | Efcab7 | EF-hand calcium binding domain 7 [Sourc | 3.2124 | 2.0558 | 5.0312 | 10.604 | 10.906 | 3.3415 |
| 20563 | ENSMUSG00000073792 | 211.0793042 | 0.258580823 | 0.17884746 | 1.4458177 | 0.14822834 | 0.560006817 | Alg6 | asparagine-linked glycosylation 6 (alpha-1 | 213.63 | 184 | 178.11 | 203.4 | 233.39 | 253.96 |
| 20564 | ENSMUSG00000073794 | 0 | NA | NA | NA | NA | NA | I0C0044D17I | RIKEN cDNA I0C0044D17 gene [Source:MG | 0 | 0 | 0 | 0 | 0 | 0 |
| 20565 | ENSMUSG00000073795 | 0.167707431 | -0.796629176 | 4.080472857 | -0.195229623 | 0.84521319 | NA | Spef1l | sperm flagellar 1 like [Source:MGI Symbol | 0 | 0 | 1.0062 | 0 | 0 | 0 |
| 20566 | ENSMUSG00000073797 | 0 | NA | NA | NA | NA | NA | Gm12650 | predicted gene 12650 [Source:MGI Symbo | 0 | 0 | 0 | 0 | 0 | 0 |
| 20567 | ENSMUSG00000073802 | 2.218308364 | 0.533160896 | 1.56771313 | 0.340088302 | 0.733790031 | NA | Cdkn2b | cyclin dependent kinase inhibitor 2B [Sou | 2.4093 | 2.0558 | 1.0062 | 6.7478 | 1.0906 | 0 |
| 20568 | ENSMUSG00000073803 | 0 | NA | NA | NA | NA | NA | 4930544L04I | RIKEN cDNA 4930544L04 gene [Source:MG | 0 | 0 | 0 | 0 | 0 | 0 |
| 20569 | ENSMUSG00000073804 | 0 | NA | NA | NA | NA | NA | Nps | neuropeptide S [Source:MGI Symbol;Acc:I | 0 | 0 | 0 | 0 | 0 | 0 |
| 20570 | ENSMUSG00000073805 | 0.524408994 | -0.837196744 | 3.377823722 | -0.247850928 | 0.804249749 | NA | Insyn2a | inhibitory synaptic factor 2A [Source:MGI | 0 | 2.0558 | 0 | 0 | 1.0906 | 0 |
| 20571 | ENSMUSG00000073807 | 0 | NA | NA | NA | NA | NA | Gm10581 | predicted gene 10581 [Source:MGI Symbo | 0 | 0 | 0 | 0 | 0 | 0 |
| 20572 | ENSMUSG00000073809 | 0 | NA | NA | NA | NA | NA | Gm10583 | predicted gene 10583 [Source:MGI Symbo | 0 | 0 | 0 | 0 | 0 | 0 |
| 20573 | ENSMUSG00000073811 | 0 | NA | NA | NA | NA | NA | Ifna12 | interferon alpha 12 [Source:MGI Symbol;A | 0 | 0 | 0 | 0 | 0 | 0 |
| 20574 | ENSMUSG00000073821 | 0 | NA | NA | NA | NA | NA | 8030451A03 | RIKEN cDNA 8030451A03 gene [Source:M | 0 | 0 | 0 | 0 | 0 | 0 |
| 20575 | ENSMUSG00000073830 | 2013.12189 | 1.186178397 | 0.714420974 | 1.66033535 | 0.096847008 | 0.458530879 | Mup14 | major urinary protein 14 [Source:MGI Sym | 746.89 | 505.74 | 2435.1 | 3036.5 | 1820.2 | 3534.2 |
| 20576 | ENSMUSG00000073834 | 3231.188701 | 2.214070207 | 0.576646628 | 3.839561529 | 0.000123254 | 0.004275178 | Mup11 | major urinary protein 11 [Source:MGI Sym | 1215.9 | 750.38 | 1471.1 | 2121.7 | 6868.7 | 6959.3 |
| 20577 | ENSMUSG00000073835 | 79.4342805 | 0.913940474 | 0.447110286 | 2.044105228 | 0.040943153 | 0.277370158 | Mup-ps12 | major urinary protein, pseudogene 12 [So | 88.341 | 30.838 | 45.281 | 40.487 | 162.5 | 109.16 |
| 20578 | ENSMUSG00000073838 | 1736.470861 | -0.101067998 | 0.096108871 | -1.051599059 | 0.29298354 | 0.743850082 | Tufm | Tu translation elongation factor, mitochor | 1770.8 | 1819.4 | 1801.2 | 1581.9 | 1707.9 | 1737.6 |
| 20579 | ENSMUSG00000073840 | 7.074699898 | 0.155969255 | 0.860222751 | 0.181312637 | 0.856122188 | 0.974319798 | Mup-ps2 | major urinary protein, pseudogene 2 [Sou | 8.8341 | 2.0558 | 9.0562 | 4.8199 | 6.5437 | 11.138 |
| 20580 | ENSMUSG00000073842 | 24928.0033 | 1.935208985 | 0.629298296 | 3.075185482 | 0.002103717 | 0.036180927 | Mup7 | major urinary protein 7 [Source:MGI Sym | 12175 | 3279.1 | 15548 | 22551 | 57392 | 38623 |
| 20581 | ENSMUSG00000073843 | 0 | NA | NA | NA | NA | NA | Gm12542 | predicted gene 12542 [Source:MGI Symbo | 0 | 0 | 0 | 0 | 0 | 0 |
| 20582 | ENSMUSG00000073844 | 0 | NA | NA | NA | NA | NA | Gm21957 | predicted gene, 21957 [Source:MGI Symb | 0 | 0 | 0 | 0 | 0 | 0 |
| 20583 | ENSMUSG00000073846 | 0 | NA | NA | NA | NA | NA | Gm12526 | predicted gene 12526 [Source:MGI Symbo | 0 | 0 | 0 | 0 | 0 | 0 |
| 20584 | ENSMUSG00000073856 | 1.68999742 | -1.958997631 | 1.783516978 | -1.098390234 | 0.272034125 | NA | Iqck | IQ motif containing K [Source:MGI Symbo | 4.0155 | 1.0279 | 3.0187 | 0.964 | 0 | 1.1138 |
| 20585 | ENSMUSG00000073860 | 0 | NA | NA | NA | NA | NA | Gm829 | predicted gene 829 [Source:MGI Symbol;A | 0 | 0 | 0 | 0 | 0 | 0 |
| 20586 | ENSMUSG00000073862 | 0 | NA | NA | NA | NA | NA | Gm5600 | predicted gene 5600 [Source:MGI Symbol | 0 | 0 | 0 | 0 | 0 | 0 |
| 20587 | ENSMUSG00000073867 | 17.8787603 | 0.319561828 | 0.543827337 | 0.587616338 | 0.556789843 | 0.898167794 | AA474408 | expressed sequence AA474408 [Source:M | 16.062 | 18.503 | 13.081 | 13.496 | 30.537 | 15.594 |
| 20588 | ENSMUSG00000073876 | 0.17131964 | -0.796629176 | 4.080472857 | -0.195229623 | 0.84521319 | NA | Gm13305 | predicted gene 13305 [Source:MGI Symbo | 0 | 1.0279 | 0 | 0 | 0 | 0 |
| 20589 | ENSMUSG00000073877 | 0.185640647 | 1.126924175 | 4.080472857 | 0.276174898 | 0.782413736 | NA | Gm13306 | predicted gene 13306 [Source:MGI Symbo | 0 | 0 | 0 | 0 | 0 | 1.1138 |
| 20590 | ENSMUSG00000073878 | 0 | NA | NA | NA | NA | NA | Gm13304 | predicted gene 13304 [Source:MGI Symbo | 0 | 0 | 0 | 0 | 0 | 0 |
| 20591 | ENSMUSG00000073879 | 0 | NA | NA | NA | NA | NA | Gm5859 | predicted pseudogene 5859 [Source:MGI | 0 | 0 | 0 | 0 | 0 | 0 |
| 20592 | ENSMUSG00000073888 | 174.7837999 | 0.032573122 | 0.205427741 | 0.158562429 | 0.874013631 | 0.979979315 | Ccl27a | chemokine (C-C motif) ligand 27A [Source | 165.44 | 160.36 | 193.2 | 207.26 | 183.22 | 139.23 |
| 20593 | ENSMUSG00000073889 | 521.1618785 | 0.130951989 | 0.148311577 | 0.882951902 | 0.377262259 | 0.81030298 | Il11ra1 | interleukin 11 receptor, alpha chain 1 [Sou | 432.87 | 501.62 | 558.47 | 484.88 | 539.86 | 609.27 |
| 20594 | ENSMUSG00000073893 | 0 | NA | NA | NA | NA | NA | Olfr472 | olfactory receptor 472 [Source:MGI Symb | 0 | 0 | 0 | 0 | 0 | 0 |
| 20595 | ENSMUSG00000073894 | 0 | NA | NA | NA | NA | NA | Rbmxl2 | RNA binding motif protein, X-linked-like 2 | 0 | 0 | 0 | 0 | 0 | 0 |
| 20596 | ENSMUSG00000073896 | 0 | NA | NA | NA | NA | NA | Olfr716 | olfactory receptor 716 [Source:MGI Symb | 0 | 0 | 0 | 0 | 0 | 0 |
| 20597 | ENSMUSG00000073897 | 0 | NA | NA | NA | NA | NA | Olfr17 | olfactory receptor 17 [Source:MGI Symbo | 0 | 0 | 0 | 0 | 0 | 0 |
| 20598 | ENSMUSG00000073898 | 0 | NA | NA | NA | NA | NA | Olfr713 | olfactory receptor 713 [Source:MGI Symb | 0 | 0 | 0 | 0 | 0 | 0 |
| 20599 | ENSMUSG00000073899 | 0 | NA | NA | NA | NA | NA | Olfr1532-ps1 | olfactory receptor 1532, pseudogene 1 [So | 0 | 0 | 0 | 0 | 0 | 0 |
| 20600 | ENSMUSG00000073900 | 0 | NA | NA | NA | NA | NA | Olfr704 | olfactory receptor 704 [Source:MGI Symb | 0 | 0 | 0 | 0 | 0 | 0 |
| 20601 | ENSMUSG00000073901 | 0.167707431 | -0.796629176 | 4.080472857 | -0.195229623 | 0.84521319 | NA | Olfr703 | olfactory receptor 703 [Source:MGI Symb | 0 | 0 | 1.0062 | 0 | 0 | 0 |
| 20602 | ENSMUSG00000073902 | 14.00751905 | -1.691062391 | 0.630585478 | -2.681733801 | 0.007324172 | 0.086502219 | Gvin3 | GTPase, very large interferon inducible, fa | 14.456 | 27.754 | 22.137 | 8.6758 | 5.4531 | 5.5692 |
| 20603 | ENSMUSG00000073906 | 0 | NA | NA | NA | NA | NA | Olfr692 | olfactory receptor 692 [Source:MGI Symb | 0 | 0 | 0 | 0 | 0 | 0 |
| 20604 | ENSMUSG00000073907 | 0 | NA | NA | NA | NA | NA | Olfr689 | olfactory receptor 689 [Source:MGI Symb | 0 | 0 | 0 | 0 | 0 | 0 |

| | | | | | | | | | | | | | | | |
|---|---|---|---|---|---|---|---|---|---|---|---|---|---|---|---|
| 20605 | ENSMUSG00000073908 | 0 | NA | NA | NA | NA | NA | Olfr687 | olfactory receptor 687 [Source:MGI Symb | 0 | 0 | 0 | 0 | 0 | 0 |
| 20606 | ENSMUSG00000073909 | 0 | NA | NA | NA | NA | NA | Olfr688 | olfactory receptor 688 [Source:MGI Symb | 0 | 0 | 0 | 0 | 0 | 0 |
| 20607 | ENSMUSG00000073910 | 610.6778026 | 0.156859479 | 0.125674465 | 1.248141213 | 0.211979348 | 0.656735971 | Mob3b | MOB kinase activator 3B [Source:MGI Syn | 589.48 | 602.36 | 540.35 | 631.41 | 598.75 | 701.72 |
| 20608 | ENSMUSG00000073913 | 0 | NA | NA | NA | NA | NA | Olfr678 | olfactory receptor 678 [Source:MGI Symb | 0 | 0 | 0 | 0 | 0 | 0 |
| 20609 | ENSMUSG00000073914 | 0 | NA | NA | NA | NA | NA | Olfr677 | olfactory receptor 677 [Source:MGI Symb | 0 | 0 | 0 | 0 | 0 | 0 |
| 20610 | ENSMUSG00000073915 | 0 | NA | NA | NA | NA | NA | Olfr676 | olfactory receptor 676 [Source:MGI Symb | 0 | 0 | 0 | 0 | 0 | 0 |
| 20611 | ENSMUSG00000073916 | 0 | NA | NA | NA | NA | NA | Olfr669 | olfactory receptor 669 [Source:MGI Symb | 0 | 0 | 0 | 0 | 0 | 0 |
| 20612 | ENSMUSG00000073917 | 0 | NA | NA | NA | NA | NA | Olfr665 | olfactory receptor 665 [Source:MGI Symb | 0 | 0 | 0 | 0 | 0 | 0 |
| 20613 | ENSMUSG00000073920 | 0 | NA | NA | NA | NA | NA | Olfr661 | olfactory receptor 661 [Source:MGI Symb | 0 | 0 | 0 | 0 | 0 | 0 |
| 20614 | ENSMUSG00000073922 | 0 | NA | NA | NA | NA | NA | Olfr659 | olfactory receptor 659 [Source:MGI Symb | 0 | 0 | 0 | 0 | 0 | 0 |
| 20615 | ENSMUSG00000073923 | 0 | NA | NA | NA | NA | NA | Olfr657 | olfactory receptor 657 [Source:MGI Symb | 0 | 0 | 0 | 0 | 0 | 0 |
| 20616 | ENSMUSG00000073924 | 0 | NA | NA | NA | NA | NA | Olfr656 | olfactory receptor 656 [Source:MGI Symb | 0 | 0 | 0 | 0 | 0 | 0 |
| 20617 | ENSMUSG00000073925 | 0 | NA | NA | NA | NA | NA | Olfr654 | olfactory receptor 654 [Source:MGI Symb | 0 | 0 | 0 | 0 | 0 | 0 |
| 20618 | ENSMUSG00000073926 | 0 | NA | NA | NA | NA | NA | Olfr653 | olfactory receptor 653 [Source:MGI Symb | 0 | 0 | 0 | 0 | 0 | 0 |
| 20619 | ENSMUSG00000073927 | 0 | NA | NA | NA | NA | NA | Olfr652 | olfactory receptor 652 [Source:MGI Symb | 0 | 0 | 0 | 0 | 0 | 0 |
| 20620 | ENSMUSG00000073928 | 0 | NA | NA | NA | NA | NA | Olfr651 | olfactory receptor 651 [Source:MGI Symb | 0 | 0 | 0 | 0 | 0 | 0 |
| 20621 | ENSMUSG00000073929 | 0 | NA | NA | NA | NA | NA | Trim30e-ps1 | tripartite motif-containing 30E, pseudoge | 0 | 0 | 0 | 0 | 0 | 0 |
| 20622 | ENSMUSG00000073931 | 0 | NA | NA | NA | NA | NA | Olfr646 | olfactory receptor 646 [Source:MGI Symb | 0 | 0 | 0 | 0 | 0 | 0 |
| 20623 | ENSMUSG00000073932 | 0 | NA | NA | NA | NA | NA | Olfr641 | olfactory receptor 641 [Source:MGI Symb | 0 | 0 | 0 | 0 | 0 | 0 |
| 20624 | ENSMUSG00000073937 | 0 | NA | NA | NA | NA | NA | Olfr633 | olfactory receptor 633 [Source:MGI Symb | 0 | 0 | 0 | 0 | 0 | 0 |
| 20625 | ENSMUSG00000073938 | 0 | NA | NA | NA | NA | NA | Olfr632 | olfactory receptor 632 [Source:MGI Symb | 0 | 0 | 0 | 0 | 0 | 0 |
| 20626 | ENSMUSG00000073940 | 304.8712742 | 0.388465461 | 0.209346197 | 1.855612694 | 0.063508783 | 0.36045804 | Hbb-bt | hemoglobin, beta adult t chain [Source:M | 212.82 | 243.62 | 336.09 | 288.23 | 315.19 | 433.29 |
| 20627 | ENSMUSG00000073943 | 0 | NA | NA | NA | NA | NA | Olfr625-ps1 | olfactory receptor 625, pseudogene 1 [So | 0 | 0 | 0 | 0 | 0 | 0 |
| 20628 | ENSMUSG00000073944 | 0 | NA | NA | NA | NA | NA | Olfr619 | olfactory receptor 619 [Source:MGI Symb | 0 | 0 | 0 | 0 | 0 | 0 |
| 20629 | ENSMUSG00000073945 | 0 | NA | NA | NA | NA | NA | Olfr618 | olfactory receptor 618 [Source:MGI Symb | 0 | 0 | 0 | 0 | 0 | 0 |
| 20630 | ENSMUSG00000073946 | 0 | NA | NA | NA | NA | NA | Olfr617 | olfactory receptor 617 [Source:MGI Symb | 0 | 0 | 0 | 0 | 0 | 0 |
| 20631 | ENSMUSG00000073947 | 0 | NA | NA | NA | NA | NA | Olfr615 | olfactory receptor 615 [Source:MGI Symb | 0 | 0 | 0 | 0 | 0 | 0 |
| 20632 | ENSMUSG00000073948 | 0 | NA | NA | NA | NA | NA | Olfr608 | olfactory receptor 608 [Source:MGI Symb | 0 | 0 | 0 | 0 | 0 | 0 |
| 20633 | ENSMUSG00000073949 | 0 | NA | NA | NA | NA | NA | Olfr606 | olfactory receptor 606 [Source:MGI Symb | 0 | 0 | 0 | 0 | 0 | 0 |
| 20634 | ENSMUSG00000073950 | 0 | NA | NA | NA | NA | NA | Olfr599 | olfactory receptor 599 [Source:MGI Symb | 0 | 0 | 0 | 0 | 0 | 0 |
| 20635 | ENSMUSG00000073951 | 0 | NA | NA | NA | NA | NA | Olfr598 | olfactory receptor 598 [Source:MGI Symb | 0 | 0 | 0 | 0 | 0 | 0 |
| 20636 | ENSMUSG00000073952 | 0 | NA | NA | NA | NA | NA | Olfr597 | olfactory receptor 597 [Source:MGI Symb | 0 | 0 | 0 | 0 | 0 | 0 |
| 20637 | ENSMUSG00000073953 | 0 | NA | NA | NA | NA | NA | Olfr596 | olfactory receptor 596 [Source:MGI Symb | 0 | 0 | 0 | 0 | 0 | 0 |
| 20638 | ENSMUSG00000073954 | 0 | NA | NA | NA | NA | NA | Olfr594 | olfactory receptor 594 [Source:MGI Symb | 0 | 0 | 0 | 0 | 0 | 0 |
| 20639 | ENSMUSG00000073955 | 0 | NA | NA | NA | NA | NA | Olfr593 | olfactory receptor 593 [Source:MGI Symb | 0 | 0 | 0 | 0 | 0 | 0 |
| 20640 | ENSMUSG00000073956 | 0 | NA | NA | NA | NA | NA | Olfr592 | olfactory receptor 592 [Source:MGI Symb | 0 | 0 | 0 | 0 | 0 | 0 |
| 20641 | ENSMUSG00000073959 | 0 | NA | NA | NA | NA | NA | Olfr584 | olfactory receptor 584 [Source:MGI Symb | 0 | 0 | 0 | 0 | 0 | 0 |
| 20642 | ENSMUSG00000073960 | 0 | NA | NA | NA | NA | NA | Olfr583 | olfactory receptor 583 [Source:MGI Symb | 0 | 0 | 0 | 0 | 0 | 0 |
| 20643 | ENSMUSG00000073961 | 0 | NA | NA | NA | NA | NA | Olfr582 | olfactory receptor 582 [Source:MGI Symb | 0 | 0 | 0 | 0 | 0 | 0 |
| 20644 | ENSMUSG00000073962 | 0 | NA | NA | NA | NA | NA | Olfr576 | olfactory receptor 576 [Source:MGI Symb | 0 | 0 | 0 | 0 | 0 | 0 |
| 20645 | ENSMUSG00000073963 | 0 | NA | NA | NA | NA | NA | Olfr572 | olfactory receptor 572 [Source:MGI Symb | 0 | 0 | 0 | 0 | 0 | 0 |
| 20646 | ENSMUSG00000073964 | 0 | NA | NA | NA | NA | NA | Olfr570 | olfactory receptor 570 [Source:MGI Symb | 0 | 0 | 0 | 0 | 0 | 0 |
| 20647 | ENSMUSG00000073965 | 0 | NA | NA | NA | NA | NA | Olfr568 | olfactory receptor 568 [Source:MGI Symb | 0 | 0 | 0 | 0 | 0 | 0 |
| 20648 | ENSMUSG00000073966 | 0.267700746 | -1.497481649 | 4.057001436 | -0.369110456 | 0.712045396 | NA | Olfr561 | olfactory receptor 561 [Source:MGI Symb | 1.6062 | 0 | 0 | 0 | 0 | 0 |
| 20649 | ENSMUSG00000073967 | 0 | NA | NA | NA | NA | NA | Olfr557 | olfactory receptor 557 [Source:MGI Symb | 0 | 0 | 0 | 0 | 0 | 0 |
| 20650 | ENSMUSG00000073968 | 23.37221747 | 0.78449122 | 0.501640853 | 1.563850344 | 0.117852718 | 0.502838264 | Trim68 | tripartite motif-containing 68 [Source:MG | 20.881 | 10.279 | 20.125 | 26.991 | 19.631 | 42.326 |
| 20651 | ENSMUSG00000073969 | 0 | NA | NA | NA | NA | NA | Olfr556 | olfactory receptor 556 [Source:MGI Symb | 0 | 0 | 0 | 0 | 0 | 0 |
| 20652 | ENSMUSG00000073970 | 0 | NA | NA | NA | NA | NA | Olfr555 | olfactory receptor 555 [Source:MGI Symb | 0 | 0 | 0 | 0 | 0 | 0 |
| 20653 | ENSMUSG00000073971 | 0 | NA | NA | NA | NA | NA | Olfr554 | olfactory receptor 554 [Source:MGI Symb | 0 | 0 | 0 | 0 | 0 | 0 |
| 20654 | ENSMUSG00000073972 | 0 | NA | NA | NA | NA | NA | Olfr553 | olfactory receptor 553 [Source:MGI Symb | 0 | 0 | 0 | 0 | 0 | 0 |
| 20655 | ENSMUSG00000073973 | 0 | NA | NA | NA | NA | NA | Olfr552 | olfactory receptor 552 [Source:MGI Symb | 0 | 0 | 0 | 0 | 0 | 0 |
| 20656 | ENSMUSG00000073974 | 0 | NA | NA | NA | NA | NA | Olfr551 | olfactory receptor 551 [Source:MGI Symb | 0 | 0 | 0 | 0 | 0 | 0 |
| 20657 | ENSMUSG00000073975 | 0 | NA | NA | NA | NA | NA | Olfr550 | olfactory receptor 550 [Source:MGI Symb | 0 | 0 | 0 | 0 | 0 | 0 |
| 20658 | ENSMUSG00000073976 | 0 | NA | NA | NA | NA | NA | 1700026J12I | RIKEN cDNA 1700026J12 gene [Source:MC | 0 | 0 | 0 | 0 | 0 | 0 |
| 20659 | ENSMUSG00000073977 | 0.321325949 | 1.911575114 | 4.048267893 | 0.472195804 | 0.636787034 | NA | Olfr549 | olfactory receptor 549 [Source:MGI Symb | 0 | 0 | 0 | 1.928 | 0 | 0 |
| 20660 | ENSMUSG00000073978 | 0 | NA | NA | NA | NA | NA | Olfr548-ps1 | olfactory receptor 548, pseudogene 1 [So | 0 | 0 | 0 | 0 | 0 | 0 |

| | | | | | | | | | | | | | | | |
|---|---|---|---|---|---|---|---|---|---|---|---|---|---|---|---|
| 20661 | ENSMUSG00000073979 | 0 | NA | NA | NA | NA | NA | Olfr547 | olfactory receptor 547 [Source:MGI Symb | 0 | 0 | 0 | 0 | 0 | 0 |
| 20662 | ENSMUSG00000073980 | 1.873704914 | -0.228153256 | 1.689548342 | -0.135038016 | 0.892581817 | NA | Gm9574 | predicted gene 9574 [Source:MGI Symbol | 0 | 5.1396 | 1.0062 | 2.8919 | 1.0906 | 1.1138 |
| 20663 | ENSMUSG00000073982 | 246.7197086 | -0.098745624 | 0.180439421 | -0.547250834 | 0.584206417 | 0.908955096 | Rhog | ras homolog family member G [Source:M( | 279.48 | 249.78 | 235.46 | 218.82 | 209.4 | 287.37 |
| 20664 | ENSMUSG00000073985 | 12.43282251 | 0.218646498 | 0.677992803 | 0.322490884 | 0.747080843 | 0.950561663 | Gm10602 | predicted gene 10602 [Source:MGI Symbc | 7.2279 | 16.447 | 11.069 | 22.171 | 6.5437 | 11.138 |
| 20665 | ENSMUSG00000073987 | 155.7521825 | -0.187584141 | 0.192185612 | -0.976057151 | 0.329036144 | 0.773866742 | Ggh | gamma-glutamyl hydrolase [Source:MGI S | 150.18 | 181.94 | 166.03 | 154.24 | 141.78 | 140.34 |
| 20666 | ENSMUSG00000073988 | 10874.69673 | -0.416895075 | 0.087251008 | -4.778111847 | 1.77E-06 | 0.000129183 | Ttpa | tocopherol (alpha) transfer protein [Sourc | 12461 | 12653 | 12193 | 9949.2 | 9063 | 8929.7 |
| 20667 | ENSMUSG00000073991 | 0 | NA | NA | NA | NA | NA | Cnbd1 | cyclic nucleotide binding domain containi | 0 | 0 | 0 | 0 | 0 | 0 |
| 20668 | ENSMUSG00000073995 | 0 | NA | NA | NA | NA | NA | NA | NA | 0 | 0 | 0 | 0 | 0 | 0 |
| 20669 | ENSMUSG00000073997 | 0 | NA | NA | NA | NA | NA | Olfr521 | olfactory receptor 521 [Source:MGI Symb | 0 | 0 | 0 | 0 | 0 | 0 |
| 20670 | ENSMUSG00000073998 | 0 | NA | NA | NA | NA | NA | Olfr520 | olfactory receptor 520 [Source:MGI Symb | 0 | 0 | 0 | 0 | 0 | 0 |
| 20671 | ENSMUSG00000074001 | 0 | NA | NA | NA | NA | NA | Klhl40 | kelch-like 40 [Source:MGI Symbol;Acc:MG | 0 | 0 | 0 | 0 | 0 | 0 |
| 20672 | ENSMUSG00000074003 | 0 | NA | NA | NA | NA | NA | Gucy2d | guanylate cyclase 2d [Source:MGI Symbol | 0 | 0 | 0 | 0 | 0 | 0 |
| 20673 | ENSMUSG00000074004 | 0 | NA | NA | NA | NA | NA | B3gnt6 | UDP-GlcNAc:betaGal beta-1,3-N-acetylglu | 0 | 0 | 0 | 0 | 0 | 0 |
| 20674 | ENSMUSG00000074006 | 1.261865794 | 0.252247416 | 2.009952904 | 0.125499168 | 0.900128384 | NA | Omp | olfactory marker protein [Source:MGI Syn | 2.4093 | 0 | 1.0062 | 1.928 | 0 | 2.2277 |
| 20675 | ENSMUSG00000074011 | 0 | NA | NA | NA | NA | NA | Pramel41 | PRAME like 41 [Source:MGI Symbol;Acc:N | 0 | 0 | 0 | 0 | 0 | 0 |
| 20676 | ENSMUSG00000074024 | 35.06202474 | 0.090413932 | 0.491809677 | 0.183839269 | 0.854139549 | 0.973825032 | 4632427E13 | RIKEN cDNA 4632427E13 gene [Source:M( | 40.155 | 26.726 | 35.219 | 67.478 | 19.631 | 21.163 |
| 20677 | ENSMUSG00000074026 | 0.160662974 | 1.126924175 | 4.080472857 | 0.276174898 | 0.782413736 | NA | Gm6091 | predicted pseudogene 6091 [Source:MGI | 0 | 0 | 0 | 0.964 | 0 | 0 |
| 20678 | ENSMUSG00000074028 | 0.319491021 | 0.16514334 | 4.044653141 | 0.040830038 | 0.967431393 | NA | Slc22a13 | solute carrier family 22 (organic cation tra | 0.8031 | 0 | 0 | 0 | 0 | 1.1138 |
| 20679 | ENSMUSG00000074029 | 0.36353943 | 2.065989254 | 4.034137301 | 0.512126658 | 0.608562368 | NA | Gm47289 | predicted gene, 47289 [Source:MGI Symb | 0 | 0 | 0 | 0 | 2.1812 | 0 |
| 20680 | ENSMUSG00000074030 | 230.8255421 | 0.009992094 | 0.182182168 | 0.054846719 | 0.95626058 | 0.991678478 | Exoc8 | exocyst complex component 8 [Source:M( | 227.28 | 212.78 | 249.55 | 196.65 | 223.58 | 275.12 |
| 20681 | ENSMUSG00000074034 | 0.331982614 | 0.16514309 | 3.986751476 | 0.041422971 | 0.966958701 | NA | Gm5921 | predicted gene 5921 [Source:MGI Symbol | 0 | 1.0279 | 0 | 0.964 | 0 | 0 |
| 20682 | ENSMUSG00000074037 | 0 | NA | NA | NA | NA | NA | Mc1r | melanocortin 1 receptor [Source:MGI Sym | 0 | 0 | 0 | 0 | 0 | 0 |
| 20683 | ENSMUSG00000074039 | 1.124144488 | 0.633232731 | 1.971100216 | 0.321258517 | 0.74801449 | NA | 4930520O04 | RIKEN cDNA 4930520O04 gene [Source:M | 1.6062 | 0 | 1.0062 | 1.928 | 1.0906 | 1.1138 |
| 20684 | ENSMUSG00000074052 | 76.06051476 | 0.183064464 | 0.279900085 | 0.654035044 | 0.513089229 | 0.884860213 | BC048644 | cDNA sequence BC048644 [Source:MGI Sy | 85.932 | 69.898 | 57.356 | 70.37 | 97.065 | 75.741 |
| 20685 | ENSMUSG00000074059 | 0 | NA | NA | NA | NA | NA | Fbxw18 | F-box and WD-40 domain protein 18 [Sou | 0 | 0 | 0 | 0 | 0 | 0 |
| 20686 | ENSMUSG00000074060 | 0.513752329 | 1.031892132 | 3.082998058 | 0.334704114 | 0.737848298 | NA | Fbxw15 | F-box and WD-40 domain protein 15 [Sou | 0 | 1.0279 | 0 | 0.964 | 1.0906 | 0 |
| 20687 | ENSMUSG00000074061 | 0 | NA | NA | NA | NA | NA | Fbxw19 | F-box and WD-40 domain protein 19 [Sou | 0 | 0 | 0 | 0 | 0 | 0 |
| 20688 | ENSMUSG00000074062 | 0 | NA | NA | NA | NA | NA | Fbxw16 | F-box and WD-40 domain protein 16 [Sou | 0 | 0 | 0 | 0 | 0 | 0 |
| 20689 | ENSMUSG00000074063 | 1875.247081 | 1.161502548 | 0.552669354 | 2.101622862 | 0.035586326 | 0.253307388 | Osgin1 | oxidative stress induced growth inhibitor | 1099.4 | 615.72 | 1760.9 | 3909.9 | 1861.7 | 2003.8 |
| 20690 | ENSMUSG00000074064 | 2536.621239 | 0.159542735 | 0.098571687 | 1.618545244 | 0.105545145 | 0.478709618 | Mlycd | malonyl-CoA decarboxylase [Source:MGI S | 2569.1 | 2269.6 | 2349.6 | 2663.5 | 2810.5 | 2557.4 |
| 20691 | ENSMUSG00000074065 | 0.496077836 | -0.812642266 | 3.444120572 | -0.235950586 | 0.813471014 | NA | Gm10617 | predicted gene 10617 [Source:MGI Symbc | 0 | 0 | 2.0125 | 0.964 | 0 | 0 |
| 20692 | ENSMUSG00000074067 | 0.133850373 | -0.796629176 | 4.080472857 | -0.195229623 | 0.84521319 | NA | Gm10619 | predicted gene 10619 [Source:MGI Symbc | 0.8031 | 0 | 0 | 0 | 0 | 0 |
| 20693 | ENSMUSG00000074071 | 508.3922162 | -0.00686582 | 0.136073943 | -0.050456537 | 0.959758584 | 0.992148164 | Fam169b | family with sequence similarity 169, meml | 561.37 | 446.12 | 520.23 | 487.77 | 518.04 | 516.82 |
| 20694 | ENSMUSG00000074072 | 0 | NA | NA | NA | NA | NA | Gm10620 | predicted gene 10620 [Source:MGI Symbc | 0 | 0 | 0 | 0 | 0 | 0 |
| 20695 | ENSMUSG00000074088 | 149.0802086 | -0.029056356 | 0.198841913 | -0.146127923 | 0.883820404 | 0.981769403 | Snrnp40 | small nuclear ribonucleoprotein 40 (U5) [S | 131.71 | 158.3 | 162.01 | 134.96 | 152.69 | 154.82 |
| 20696 | ENSMUSG00000074092 | 0 | NA | NA | NA | NA | NA | Gm6181 | predicted pseudogene 6181 [Source:MGI | 0 | 0 | 0 | 0 | 0 | 0 |
| 20697 | ENSMUSG00000074093 | 259.6256118 | 0.040255996 | 0.158498485 | 0.253983478 | 0.799508322 | 0.962424862 | Svip | small VCP/p97-interacting protein [Source | 240.13 | 272.4 | 255.59 | 243.89 | 262.84 | 282.92 |
| 20698 | ENSMUSG00000074102 | 309.1048791 | 0.04311312 | 0.148599123 | 0.29013038 | 0.771716495 | 0.957083104 | Rbm15b | RNA binding motif protein 15B [Source:M | 319.63 | 314.54 | 278.73 | 296.91 | 318.46 | 326.36 |
| 20699 | ENSMUSG00000074108 | 0 | NA | NA | NA | NA | NA | Rpl10-ps2 | ribosomal protein L10, pseudogene 2 [Sou | 0 | 0 | 0 | 0 | 0 | 0 |
| 20700 | ENSMUSG00000074109 | 0 | NA | NA | NA | NA | NA | Mrgprx2 | MAS-related GPR, member X2 [Source:M( | 0 | 0 | 0 | 0 | 0 | 0 |
| 20701 | ENSMUSG00000074111 | 0 | NA | NA | NA | NA | NA | Mrgpra9 | MAS-related GPR, member A9 [Source:M( | 0 | 0 | 0 | 0 | 0 | 0 |
| 20702 | ENSMUSG00000074113 | 0 | NA | NA | NA | NA | NA | Gm10629 | predicted gene 10629 [Source:MGI Symbc | 0 | 0 | 0 | 0 | 0 | 0 |
| 20703 | ENSMUSG00000074115 | 991.5016559 | -0.848945985 | 0.503707297 | -1.685395448 | 0.091912301 | 0.446623824 | Saa1 | serum amyloid A 1 [Source:MGI Symbol;A | 1423.1 | 1768 | 633.93 | 453.07 | 778.7 | 892.19 |
| 20704 | ENSMUSG00000074121 | 2.60004086 | -2.009643844 | 1.467462767 | -1.36946837 | 0.170852915 | NA | Ntf5 | neurotrophin 5 [Source:MGI Symbol;Acc:I | 2.4093 | 4.1117 | 6.0375 | 1.928 | 0 | 1.1138 |
| 20705 | ENSMUSG00000074123 | 0 | NA | NA | NA | NA | NA | 7420426K07 | RIKEN cDNA 7420426K07 gene [Source:M( | 0 | 0 | 0 | 0 | 0 | 0 |
| 20706 | ENSMUSG00000074127 | 0 | NA | NA | NA | NA | NA | Cmtm2a | CKLF-like MARVEL transmembrane domai | 0 | 0 | 0 | 0 | 0 | 0 |
| 20707 | ENSMUSG00000074128 | 0 | NA | NA | NA | NA | NA | Gm10631 | predicted gene 10631 [Source:MGI Symbc | 0 | 0 | 0 | 0 | 0 | 0 |
| 20708 | ENSMUSG00000074129 | 163.7594925 | 0.154148417 | 0.184299764 | 0.836400513 | 0.402929624 | 0.828953151 | Rpl13a | ribosomal protein L13A [Source:MGI Syml | 146.16 | 160.36 | 158.99 | 183.16 | 169.05 | 164.85 |
| 20709 | ENSMUSG00000074136 | 8.55285774 | 0.140396542 | 0.756898363 | 0.185489293 | 0.852845278 | 0.973781977 | 4930513N1C | RIKEN cDNA 4930513N10 gene [Source:M | 8.031 | 9.2513 | 7.0437 | 3.8559 | 11.997 | 11.138 |
| 20710 | ENSMUSG00000074139 | 4.503855101 | -2.920988303 | 1.212715691 | -2.408634048 | 0.016012345 | NA | 1700057G04 | RIKEN cDNA 1700057G04 gene [Source:M | 9.6372 | 6.1675 | 8.05 | 0.964 | 1.0906 | 1.1138 |
| 20711 | ENSMUSG00000074141 | 1.736468913 | 2.209297504 | 1.799024268 | 1.228053197 | 0.219426999 | NA | Il4i1 | interleukin 4 induced 1 [Source:MGI Symt | 0.8031 | 1.0279 | 0 | 1.928 | 1.0906 | 5.5692 |
| 20712 | ENSMUSG00000074146 | 0 | NA | NA | NA | NA | NA | 4930579C12 | RIKEN cDNA 4930579C12 gene [Source:M( | 0 | 0 | 0 | 0 | 0 | 0 |
| 20713 | ENSMUSG00000074149 | 0.803102239 | -3.069617558 | 3.178309912 | -0.965801839 | 0.334143348 | NA | Gm10634 | predicted gene 10634 [Source:MGI Symbc | 4.8186 | 0 | 0 | 0 | 0 | 0 |
| 20714 | ENSMUSG00000074151 | 422.3597682 | 0.23075183 | 0.171445755 | 1.345917429 | 0.178329147 | 0.610642232 | Nlrc5 | NLR family, CARD domain containing 5 [Sc | 358.18 | 427.61 | 381.37 | 568.75 | 383.9 | 414.35 |
| 20715 | ENSMUSG00000074155 | 0 | NA | NA | NA | NA | NA | Klk5 | kallikrein related-peptidase 5 [Source:MG | 0 | 0 | 0 | 0 | 0 | 0 |
| 20716 | ENSMUSG00000074156 | 0.267700746 | -1.497481649 | 4.057001436 | -0.369110456 | 0.712045396 | NA | Ces1h | carboxylesterase 1H [Source:MGI Symbol; | 1.6062 | 0 | 0 | 0 | 0 | 0 |

| | | | | | | | | | | | | | | | |
|---|---|---|---|---|---|---|---|---|---|---|---|---|---|---|---|
| 20717 | ENSMUSG00000074158 | 67.80508957 | -0.177923879 | 0.31019144 | -0.573593775 | 0.566242725 | 0.902076435 | Zfp976 | zinc finger protein 976 [Source:MGI Symb | 74.689 | 81.206 | 60.375 | 83.866 | 63.256 | 43.44 |
| 20718 | ENSMUSG00000074164 | 0 | NA | NA | NA | NA | NA | Vmn2r-ps54 | vomeronasal 2, receptor, pseudogene 54 | 0 | 0 | 0 | 0 | 0 | 0 |
| 20719 | ENSMUSG00000074165 | 74.69362552 | -0.366490592 | 0.263167408 | -1.392613907 | 0.163736575 | 0.587809835 | Zfp788 | zinc finger protein 788 [Source:MGI Symb | 96.372 | 80.178 | 75.468 | 61.695 | 67.618 | 66.831 |
| 20720 | ENSMUSG00000074166 | 87.02953271 | 0.200761328 | 0.250882818 | 0.80021952 | 0.423583622 | 0.84018348 | AW146154 | expressed sequence AW146154 [Source:N | 85.932 | 80.178 | 76.475 | 78.082 | 109.06 | 92.449 |
| 20721 | ENSMUSG00000074170 | 300.2991876 | -0.127999344 | 0.187733836 | -0.681812864 | 0.495357289 | 0.877937486 | Plekhf1 | pleckstrin homology domain containing, fa | 346.14 | 362.85 | 231.44 | 277.63 | 309.74 | 274.01 |
| 20722 | ENSMUSG00000074171 | 27.2156417 | -0.372076866 | 0.471156837 | -0.789709152 | 0.429697646 | 0.84307112 | Gm6658 | predicted gene 6658 [Source:MGI Symbol | 24.896 | 40.089 | 27.169 | 17.352 | 37.081 | 16.708 |
| 20723 | ENSMUSG00000074178 | 5.023732513 | -1.23083712 | 1.143337263 | -1.076530224 | 0.281690191 | NA | Gm10638 | predicted gene 10638 [Source:MGI Symbo | 4.0155 | 9.2513 | 8.05 | 7.7118 | 0 | 1.1138 |
| 20724 | ENSMUSG00000074179 | 0.476489653 | -2.293616034 | 3.421506946 | -0.670352587 | 0.502633051 | NA | Gsta5 | glutathione S-transferase alpha 5 [Source: | 0.8031 | 2.0558 | 0 | 0 | 0 | 0 |
| 20725 | ENSMUSG00000074182 | 140.1623294 | 0.094287001 | 0.224142558 | 0.420656398 | 0.674006005 | 0.932009833 | Znhit6 | zinc finger, HIT type 6 [Source:MGI Symbo | 117.25 | 129.52 | 159.99 | 115.68 | 158.14 | 160.39 |
| 20726 | ENSMUSG00000074183 | 16.44348851 | -2.166630498 | 0.681522812 | -3.179101945 | 0.001477321 | 0.028041961 | Gsta1 | glutathione S-transferase, alpha 1 (Ya) [So | 45.777 | 23.642 | 11.069 | 3.8559 | 7.6343 | 6.6831 |
| 20727 | ENSMUSG00000074184 | 0.160662974 | 1.126924175 | 4.080472857 | 0.276174898 | 0.782413736 | NA | Smt3h2-ps2 | SMT3 suppressor of mif two 3 homolog 2, | 0 | 0 | 0 | 0.964 | 0 | 0 |
| 20728 | ENSMUSG00000074194 | 19.89635365 | -0.513622277 | 0.547802108 | -0.937605513 | 0.348447177 | 0.789711685 | Zfp791 | zinc finger protein 791 [Source:MGI Symb | 31.321 | 14.391 | 24.15 | 8.6758 | 17.45 | 23.391 |
| 20729 | ENSMUSG00000074195 | 0 | NA | NA | NA | NA | NA | Clca4b | chloride channel accessory 4B [Source:MG | 0 | 0 | 0 | 0 | 0 | 0 |
| 20730 | ENSMUSG00000074196 | 0 | NA | NA | NA | NA | NA | Clca4c-ps | chloride channel accessory 4C, pseudoger | 0 | 0 | 0 | 0 | 0 | 0 |
| 20731 | ENSMUSG00000074199 | 0 | NA | NA | NA | NA | NA | Krtdap | keratinocyte differentiation associated pr | 0 | 0 | 0 | 0 | 0 | 0 |
| 20732 | ENSMUSG00000074203 | 3.295976085 | -3.067387045 | 1.499629053 | -2.045430528 | 0.040812432 | NA | G430095P16 | RIKEN cDNA G430095P16 gene [Source:M | 2.4093 | 9.2513 | 6.0375 | 0.964 | 0 | 1.1138 |
| 20733 | ENSMUSG00000074206 | 1.737813023 | 0.671249826 | 1.665698391 | 0.402984016 | 0.686959984 | NA | Adh6b | alcohol dehydrogenase 6B (class V) [Sourc | 0 | 3.0838 | 1.0062 | 1.928 | 2.1812 | 2.2277 |
| 20734 | ENSMUSG00000074207 | 32518.90724 | -0.140351628 | 0.112041992 | -1.252669883 | 0.210325871 | 0.654367261 | Adh1 | alcohol dehydrogenase 1 (class I) [Source: | 32280 | 34235 | 35785 | 33648 | 32910 | 26256 |
| 20735 | ENSMUSG00000074210 | 0 | NA | NA | NA | NA | NA | E130208F15 | RIKEN cDNA E130208F15 gene [Source:MG | 0 | 0 | 0 | 0 | 0 | 0 |
| 20736 | ENSMUSG00000074211 | 97.55114099 | -0.159310088 | 0.243843384 | -0.65332955 | 0.513543846 | 0.885004045 | Sdhaf1 | succinate dehydrogenase complex asseml | 104.4 | 91.485 | 112.7 | 81.938 | 111.24 | 83.538 |
| 20737 | ENSMUSG00000074212 | 460.1681353 | 0.040626066 | 0.133606019 | 0.304073621 | 0.761071804 | 0.95402891 | Dnajb14 | DnaJ heat shock protein family (Hsp40) m | 425.64 | 468.73 | 466.9 | 442.47 | 448.24 | 509.03 |
| 20738 | ENSMUSG00000074213 | 4.502995708 | 1.240651532 | 1.048387398 | 1.183390352 | 0.236654479 | NA | Gm10642 | predicted gene 10642 [Source:MGI Symbo | 4.8186 | 2.0558 | 1.0062 | 4.8199 | 7.6343 | 6.6831 |
| 20739 | ENSMUSG00000074215 | 7.417369666 | -0.320297167 | 0.774482338 | -0.413562906 | 0.679194244 | 0.934062537 | Gm10643 | predicted gene 10643 [Source:MGI Symbo | 10.44 | 7.1954 | 7.0437 | 7.7118 | 6.5437 | 5.5692 |
| 20740 | ENSMUSG00000074217 | 8.340556904 | -0.315288347 | 0.814633577 | -0.387030876 | 0.69873335 | 0.940837756 | Misp3 | MISP family member 3 [Source:MGI Symb | 9.6372 | 6.1675 | 12.075 | 14.46 | 4.3625 | 3.3415 |
| 20741 | ENSMUSG00000074218 | 26.14675246 | -0.748427582 | 0.44019211 | -1.700229436 | 0.089087777 | 0.439824707 | Cox7a1 | cytochrome c oxidase subunit 7A1 [Sourc | 32.124 | 29.81 | 36.225 | 13.496 | 20.722 | 24.505 |
| 20742 | ENSMUSG00000074219 | 27.36586809 | 0.389950946 | 0.431113301 | 0.904520796 | 0.365719312 | 0.804235439 | Gm10644 | predicted gene 10644 [Source:MGI Symbo | 22.487 | 17.475 | 31.194 | 34.703 | 32.719 | 25.618 |
| 20743 | ENSMUSG00000074220 | 0.979611946 | -3.327922556 | 2.381869301 | -1.397189407 | 0.162356621 | NA | Zfp382 | zinc finger protein 382 [Source:MGI Symb | 0.8031 | 2.0558 | 3.0187 | 0 | 0 | 0 |
| 20744 | ENSMUSG00000074221 | 227.4515676 | 0.209563577 | 0.165287401 | 1.267873873 | 0.204842996 | 0.650460832 | Zfp568 | zinc finger protein 568 [Source:MGI Symb | 191.14 | 222.03 | 220.37 | 251.6 | 232.3 | 247.27 |
| 20745 | ENSMUSG00000074224 | 0 | NA | NA | NA | NA | NA | 4932431P20 | RIKEN cDNA 4932431P20 gene [Source:M | 0 | 0 | 0 | 0 | 0 | 0 |
| 20746 | ENSMUSG00000074227 | 1504.748361 | 0.262215571 | 0.120149128 | 2.182417593 | 0.029078727 | 0.223720589 | Spint2 | serine protease inhibitor, Kunitz type 2 [So | 1407.8 | 1258.2 | 1437.9 | 1440.2 | 1661 | 1823.4 |
| 20747 | ENSMUSG00000074228 | 0 | NA | NA | NA | NA | NA | Gm10645 | predicted gene 10645 [Source:MGI Symbo | 0 | 0 | 0 | 0 | 0 | 0 |
| 20748 | ENSMUSG00000074232 | 2.619461018 | 2.998724662 | 1.677074832 | 1.78806849 | 0.073764959 | NA | Gm10647 | predicted gene 10647 [Source:MGI Symbo | 1.6062 | 0 | 0 | 1.928 | 3.2719 | 8.9108 |
| 20749 | ENSMUSG00000074235 | 7.456875476 | -0.146417526 | 0.867862673 | -0.168710478 | 0.866024377 | 0.977767277 | Gm10649 | predicted gene 10649 [Source:MGI Symbo | 3.2124 | 13.363 | 7.0437 | 5.7839 | 11.997 | 3.3415 |
| 20750 | ENSMUSG00000074236 | 0 | NA | NA | NA | NA | NA | Gm9387 | predicted pseudogene 9387 [Source:MGI | 0 | 0 | 0 | 0 | 0 | 0 |
| 20751 | ENSMUSG00000074237 | 0 | NA | NA | NA | NA | NA | Gm10650 | predicted gene 10650 [Source:MGI Symbo | 0 | 0 | 0 | 0 | 0 | 0 |
| 20752 | ENSMUSG00000074238 | 408.4375106 | 0.079086606 | 0.136482041 | 0.579465289 | 0.562275261 | 0.900565925 | Ap1ar | adaptor-related protein complex 1 associa | 359.79 | 413.22 | 419.6 | 427.04 | 416.62 | 414.35 |
| 20753 | ENSMUSG00000074240 | 58.83122635 | 1.426989246 | 0.451426404 | 3.161067308 | 0.001571922 | 0.029454792 | Cib3 | calcium and integrin binding family memb | 44.171 | 33.921 | 17.106 | 41.451 | 145.05 | 71.286 |
| 20754 | ENSMUSG00000074243 | 0.902269431 | 2.107869146 | 2.547671313 | 0.827370915 | 0.40802686 | NA | Gm10651 | predicted pseudogene 10651 [Source:MG | 0 | 1.0279 | 0 | 0 | 3.2719 | 1.1138 |
| 20755 | ENSMUSG00000074247 | 361.9437902 | -0.059544008 | 0.138882207 | -0.428737488 | 0.668114276 | 0.930136128 | Dda1 | DET1 and DDB1 associated 1 [Source:MGI | 391.91 | 362.85 | 353.19 | 369.2 | 339.18 | 355.32 |
| 20756 | ENSMUSG00000074248 | 0.331982614 | 0.16514309 | 3.986751476 | 0.041422971 | 0.966958701 | NA | 4930432M17 | RIKEN cDNA 4930432M17 gene [Source:M | 0 | 1.0279 | 0 | 0.964 | 0 | 0 |
| 20757 | ENSMUSG00000074250 | 0 | NA | NA | NA | NA | NA | Gm10653 | predicted gene 10653 [Source:MGI Symbo | 0 | 0 | 0 | 0 | 0 | 0 |
| 20758 | ENSMUSG00000074252 | 0 | NA | NA | NA | NA | NA | Gm10654 | predicted gene 10654 [Source:MGI Symbo | 0 | 0 | 0 | 0 | 0 | 0 |
| 20759 | ENSMUSG00000074254 | 34.81351102 | -3.207623555 | 0.548436328 | -5.848670828 | 4.96E-09 | 8.53E-07 | Cyp2a4 | cytochrome P450, family 2, subfamily a, p | 65.051 | 58.591 | 65.406 | 15.424 | 2.1812 | 2.2277 |
| 20760 | ENSMUSG00000074256 | 0.160662974 | 1.126924175 | 4.080472857 | 0.276174898 | 0.782413736 | NA | Gm10655 | predicted gene 10655 [Source:MGI Symbo | 0 | 0 | 0 | 0.964 | 0 | 0 |
| 20761 | ENSMUSG00000074259 | 0.644197084 | -2.725354879 | 2.762261801 | -0.986638876 | 0.323819705 | NA | Gramd2 | GRAM domain containing 2 [Source:MGI S | 0.8031 | 2.0558 | 1.0062 | 0 | 0 | 0 |
| 20762 | ENSMUSG00000074261 | 0.185640647 | 1.126924175 | 4.080472857 | 0.276174898 | 0.782413736 | NA | Erich4 | glutamate rich 4 [Source:MGI Symbol;Acc | 0 | 0 | 0 | 0 | 0 | 1.1138 |
| 20763 | ENSMUSG00000074264 | 5462.051183 | -0.066754211 | 0.106327678 | -0.627815936 | 0.530124525 | 0.888501493 | Amy1 | amylase 1, salivary [Source:MGI Symbol;A | 5000.1 | 5769.7 | 5997.2 | 5605.5 | 5502.2 | 4897.6 |
| 20764 | ENSMUSG00000074268 | 0 | NA | NA | NA | NA | NA | Amy2a5 | amylase 2a5 [Source:MGI Symbol;Acc:MG | 0 | 0 | 0 | 0 | 0 | 0 |
| 20765 | ENSMUSG00000074269 | 11.58036826 | -0.646919875 | 0.826526343 | -0.782697225 | 0.433804932 | 0.845264432 | Rec114 | REC114 meiotic recombination protein [So | 8.031 | 5.1396 | 29.181 | 3.8559 | 5.4531 | 17.822 |
| 20766 | ENSMUSG00000074272 | 2519.564885 | 0.013010306 | 0.095300099 | 0.136519336 | 0.891410739 | 0.982489666 | Ceacam1 | carcinoembryonic antigen-related cell adh | 2351.5 | 2539 | 2635.4 | 2584.4 | 2437.5 | 2569.6 |
| 20767 | ENSMUSG00000074274 | 0 | NA | NA | NA | NA | NA | D930028M14 | RIKEN cDNA D930028M14 gene [Source:M | 0 | 0 | 0 | 0 | 0 | 0 |
| 20768 | ENSMUSG00000074277 | 202.9962416 | 0.059587129 | 0.212927023 | 0.279847658 | 0.779594386 | 0.959870443 | Phldb3 | pleckstrin homology like domain, family B | 159.82 | 176.8 | 260.62 | 228.46 | 199.58 | 192.69 |
| 20769 | ENSMUSG00000074280 | 0 | NA | NA | NA | NA | NA | Gm6166 | predicted gene 6166 [Source:MGI Symbol | 0 | 0 | 0 | 0 | 0 | 0 |
| 20770 | ENSMUSG00000074281 | 0.339027071 | -1.79174149 | 3.919585585 | -0.457125237 | 0.647581032 | NA | Gm12522 | predicted gene 12522 [Source:MGI Symbo | 0 | 1.0279 | 1.0062 | 0 | 0 | 0 |
| 20771 | ENSMUSG00000074282 | 42.76470972 | -0.110057533 | 0.357962079 | -0.307455843 | 0.758496428 | 0.953824756 | Zfp94 | zinc finger protein 94 [Source:MGI Symbo | 40.958 | 41.117 | 51.318 | 46.271 | 49.078 | 27.846 |
| 20772 | ENSMUSG00000074283 | 48.55223499 | -0.247596439 | 0.317066829 | -0.780896696 | 0.434863255 | 0.845325456 | Zfp109 | zinc finger protein 109 [Source:MGI Symb | 50.595 | 47.284 | 60.375 | 48.199 | 42.534 | 42.326 |

| | | | | | | | | | | | | | | | |
|---|---|---|---|---|---|---|---|---|---|---|---|---|---|---|---|
| 20773 | ENSMUSG00000074284 | 124.8254014 | 0.225576612 | 0.231768024 | 0.973286166 | 0.330411096 | 0.774747658 | Gm10658 | predicted gene 10658 [Source:MGI Symbo | 103.6 | 107.93 | 133.83 | 125.32 | 113.42 | 164.85 |
| 20774 | ENSMUSG00000074291 | 0 | NA | NA | NA | NA | NA | Vmn1r168 | vomeronasal 1 receptor 168 [Source:MGI | 0 | 0 | 0 | 0 | 0 | 0 |
| 20775 | ENSMUSG00000074292 | 0 | NA | NA | NA | NA | NA | Gm10660 | predicted gene 10660 [Source:MGI Symbo | 0 | 0 | 0 | 0 | 0 | 0 |
| 20776 | ENSMUSG00000074300 | 0.353089354 | 0.165142706 | 3.896102244 | 0.042386646 | 0.966190474 | NA | Smim31 | small integral membrane protein 31 [Sour | 0 | 1.0279 | 0 | 0 | 1.0906 | 0 |
| 20777 | ENSMUSG00000074302 | 0 | NA | NA | NA | NA | NA | Gm10663 | predicted gene 10663 [Source:MGI Symbo | 0 | 0 | 0 | 0 | 0 | 0 |
| 20778 | ENSMUSG00000074305 | 192.5265542 | -0.241102159 | 0.191158118 | -1.261270834 | 0.207211286 | 0.652364421 | Peak1 | pseudopodium-enriched atypical kinase 1 | 208 | 232.31 | 185.15 | 160.02 | 162.5 | 207.17 |
| 20779 | ENSMUSG00000074311 | 0 | NA | NA | NA | NA | NA | Vmn1r139 | vomeronasal 1 receptor 139 [Source:MGI | 0 | 0 | 0 | 0 | 0 | 0 |
| 20780 | ENSMUSG00000074322 | 0 | NA | NA | NA | NA | NA | Vmn1r132 | vomeronasal 1 receptor 132 [Source:MGI | 0 | 0 | 0 | 0 | 0 | 0 |
| 20781 | ENSMUSG00000074336 | 10738.79174 | 0.039178699 | 0.100103927 | 0.391380243 | 0.695516194 | 0.939992732 | Apoc4 | apolipoprotein C-IV [Source:MGI Symbol;/ | 11784 | 9731.3 | 10262 | 10483 | 11226 | 10947 |
| 20782 | ENSMUSG00000074340 | 2.712292549 | -0.042726359 | 1.375758926 | -0.031056574 | 0.975224422 | NA | Ovgp1 | oviductal glycoprotein 1 [Source:MGI Sym | 3.2124 | 3.0838 | 2.0125 | 5.7839 | 2.1812 | 0 |
| 20783 | ENSMUSG00000074342 | 9.81384421 | -0.555452703 | 0.777839847 | -0.714096488 | 0.475167517 | 0.869368495 | I830077J02R | RIKEN cDNA I830077J02 gene [Source:MG | 14.456 | 12.335 | 8.05 | 1.928 | 7.6343 | 14.48 |
| 20784 | ENSMUSG00000074343 | 0 | NA | NA | NA | NA | NA | Gldnos | gliomedin, opposite strand [Source:MGI S | 0 | 0 | 0 | 0 | 0 | 0 |
| 20785 | ENSMUSG00000074344 | 0 | NA | NA | NA | NA | NA | Tmigd3 | transmembrane and immunoglobulin don | 0 | 0 | 0 | 0 | 0 | 0 |
| 20786 | ENSMUSG00000074345 | 15.18283255 | 0.687993075 | 0.599289921 | 1.14801376 | 0.250962883 | 0.702929347 | Tnfaip8l3 | tumor necrosis factor, alpha-induced prot | 9.6372 | 11.307 | 14.087 | 25.063 | 8.7249 | 22.277 |
| 20787 | ENSMUSG00000074346 | 0 | NA | NA | NA | NA | NA | Kcnd3os | potassium voltage-gated channel, Shal-rel | 0 | 0 | 0 | 0 | 0 | 0 |
| 20788 | ENSMUSG00000074354 | 0 | NA | NA | NA | NA | NA | Arhgap20os | Rho GTPase activating protein 20, opposit | 0 | 0 | 0 | 0 | 0 | 0 |
| 20789 | ENSMUSG00000074355 | 2.637343528 | 1.1769987 | 1.505156231 | 0.781977762 | 0.434227643 | NA | Gm10676 | predicted gene 10676 [Source:MGI Symbo | 0.8031 | 2.0558 | 2.0125 | 0 | 8.7249 | 2.2277 |
| 20790 | ENSMUSG00000074357 | 7.548726417 | 0.818132048 | 0.883560311 | 0.925949296 | 0.354472331 | 0.794633876 | AA386476 | expressed sequence AA386476 [Source:M | 2.4093 | 3.0838 | 11.069 | 14.46 | 9.8156 | 4.4554 |
| 20791 | ENSMUSG00000074358 | 99.10204198 | 0.273089861 | 0.24284971 | 1.12452208 | 0.260791609 | 0.714817099 | Ccdc61 | coiled-coil domain containing 61 [Source:I | 86.735 | 85.317 | 97.606 | 131.1 | 102.52 | 91.335 |
| 20792 | ENSMUSG00000074359 | 0 | NA | NA | NA | NA | NA | Psg23 | pregnancy-specific glycoprotein 23 [Sourc | 0 | 0 | 0 | 0 | 0 | 0 |
| 20793 | ENSMUSG00000074361 | 19.84963096 | -0.622434992 | 0.602034068 | -1.03388666 | 0.301189152 | 0.7521219 | C5ar2 | complement component 5a receptor 2 [S | 20.078 | 37.005 | 15.094 | 12.532 | 6.5437 | 27.846 |
| 20794 | ENSMUSG00000074364 | 46.98112466 | -0.705153109 | 0.350150072 | -2.013859669 | 0.044024269 | 0.289702924 | Ehd2 | EH-domain containing 2 [Source:MGI Sym | 78.704 | 48.312 | 47.293 | 33.739 | 37.081 | 36.757 |
| 20795 | ENSMUSG00000074365 | 0 | NA | NA | NA | NA | NA | Crxos | cone-rod homeobox, opposite strand [Sou | 0 | 0 | 0 | 0 | 0 | 0 |
| 20796 | ENSMUSG00000074366 | 0 | NA | NA | NA | NA | NA | Obox5 | oocyte specific homeobox 5 [Source:MGI | 0 | 0 | 0 | 0 | 0 | 0 |
| 20797 | ENSMUSG00000074369 | 0 | NA | NA | NA | NA | NA | Obox2 | oocyte specific homeobox 2 [Source:MGI | 0 | 0 | 0 | 0 | 0 | 0 |
| 20798 | ENSMUSG00000074373 | 0 | NA | NA | NA | NA | NA | Gm10680 | predicted gene 10680 [Source:MGI Symbo | 0 | 0 | 0 | 0 | 0 | 0 |
| 20799 | ENSMUSG00000074375 | 0 | NA | NA | NA | NA | NA | Sult2a3 | sulfotransferase family 2A, dehydroepian | 0 | 0 | 0 | 0 | 0 | 0 |
| 20800 | ENSMUSG00000074377 | 0 | NA | NA | NA | NA | NA | Sult2a4 | sulfotransferase family 2A, dehydroepian | 0 | 0 | 0 | 0 | 0 | 0 |
| 20801 | ENSMUSG00000074378 | 0 | NA | NA | NA | NA | NA | Bsph1 | binder of sperm protein homolog 1 [Sourc | 0 | 0 | 0 | 0 | 0 | 0 |
| 20802 | ENSMUSG00000074384 | 16.52425109 | 0.025731031 | 0.543642229 | 0.047330817 | 0.962249568 | 0.992956886 | AI429214 | expressed sequence AI429214 [Source:M | 13.653 | 21.586 | 14.087 | 21.208 | 16.359 | 12.252 |
| 20803 | ENSMUSG00000074385 | 0 | NA | NA | NA | NA | NA | Gm10684 | predicted gene 10684 [Source:MGI Symbo | 0 | 0 | 0 | 0 | 0 | 0 |
| 20804 | ENSMUSG00000074388 | 2.692183058 | -0.543549384 | 1.363468631 | -0.398651917 | 0.690149702 | NA | Gm5544 | predicted gene 5544 [Source:MGI Symbol | 2.4093 | 4.1117 | 3.0187 | 0 | 3.2719 | 3.3415 |
| 20805 | ENSMUSG00000074397 | 0 | NA | NA | NA | NA | NA | Foxr1 | forkhead box R1 [Source:MGI Symbol;Acc | 0 | 0 | 0 | 0 | 0 | 0 |
| 20806 | ENSMUSG00000074398 | 16.76777298 | -0.714653053 | 0.701996864 | -1.018028839 | 0.308664251 | 0.75728287 | Gm15441 | predicted gene 15441 [Source:MGI Symbo | 16.062 | 24.67 | 22.137 | 28.919 | 4.3625 | 4.4554 |
| 20807 | ENSMUSG00000074401 | 0 | NA | NA | NA | NA | NA | NA | NA | 0 | 0 | 0 | 0 | 0 | 0 |
| 20808 | ENSMUSG00000074402 | 0 | NA | NA | NA | NA | NA | NA | NA | 0 | 0 | 0 | 0 | 0 | 0 |
| 20809 | ENSMUSG00000074403 | 0 | NA | NA | NA | NA | NA | H3c13 | H3 clustered histone 13 [Source:MGI Sym | 0 | 0 | 0 | 0 | 0 | 0 |
| 20810 | ENSMUSG00000074404 | 0 | NA | NA | NA | NA | NA | Gm1698 | predicted gene 1698 [Source:MGI Symbol | 0 | 0 | 0 | 0 | 0 | 0 |
| 20811 | ENSMUSG00000074405 | 224.2656223 | 0.027034895 | 0.219987885 | 0.122892653 | 0.902192104 | 0.983821605 | Zfp865 | zinc finger protein 865 [Source:MGI Symb | 208.81 | 209.7 | 248.54 | 278.59 | 152.69 | 247.27 |
| 20812 | ENSMUSG00000074406 | 128.9789179 | 0.091155215 | 0.217613319 | 0.418886195 | 0.675299306 | 0.932446243 | Zfp628 | zinc finger protein 628 [Source:MGI Symb | 114.84 | 138.77 | 121.76 | 157.13 | 119.97 | 121.41 |
| 20813 | ENSMUSG00000074412 | 0.339027071 | -1.79174149 | 3.919585585 | -0.457125237 | 0.647581032 | NA | Gm10689 | predicted gene 10689 [Source:MGI Symbo | 0 | 1.0279 | 1.0062 | 0 | 0 | 0 |
| 20814 | ENSMUSG00000074415 | 8.460320018 | -0.219198853 | 0.82763993 | -0.264848088 | 0.791126491 | 0.961019028 | Mir100hg | Mir100 Mirlet7a-2 Mir125b-1 cluster host | 7.2279 | 5.1396 | 15.094 | 14.46 | 3.2719 | 5.5692 |
| 20815 | ENSMUSG00000074417 | 0.685278559 | -2.809870862 | 3.376578027 | -0.832165239 | 0.405315682 | NA | Gm14548 | predicted gene 14548 [Source:MGI Symbo | 0 | 4.1117 | 0 | 0 | 0 | 0 |
| 20816 | ENSMUSG00000074418 | 0 | NA | NA | NA | NA | NA | Gm15930 | predicted gene 15930 [Source:MGI Symbo | 0 | 0 | 0 | 0 | 0 | 0 |
| 20817 | ENSMUSG00000074419 | 0.465832987 | -0.698295221 | 3.18348759 | -0.219349126 | 0.826378096 | NA | Gm15448 | predicted gene 15448 [Source:MGI Symbo | 0.8031 | 1.0279 | 0 | 0.964 | 0 | 0 |
| 20818 | ENSMUSG00000074420 | 0 | NA | NA | NA | NA | NA | Gm10693 | predicted pseudogene 10693 [Source:MG | 0 | 0 | 0 | 0 | 0 | 0 |
| 20819 | ENSMUSG00000074424 | 0 | NA | NA | NA | NA | NA | Spopfm2 | speckle-type BTB/POZ protein family mem | 0 | 0 | 0 | 0 | 0 | 0 |
| 20820 | ENSMUSG00000074433 | 0 | NA | NA | NA | NA | NA | Lce3e | late cornified envelope 3E [Source:MGI Sy | 0 | 0 | 0 | 0 | 0 | 0 |
| 20821 | ENSMUSG00000074434 | 0 | NA | NA | NA | NA | NA | Defa28 | defensin, alpha, 28 [Source:MGI Symbol;A | 0 | 0 | 0 | 0 | 0 | 0 |
| 20822 | ENSMUSG00000074435 | 0 | NA | NA | NA | NA | NA | Smcp | sperm mitochondria-associated cysteine-r | 0 | 0 | 0 | 0 | 0 | 0 |
| 20823 | ENSMUSG00000074436 | 0 | NA | NA | NA | NA | NA | Defa-ps18 | defensin, alpha, pseudogene 18 [Source:M | 0 | 0 | 0 | 0 | 0 | 0 |
| 20824 | ENSMUSG00000074437 | 0 | NA | NA | NA | NA | NA | Defa29 | defensin, alpha, 29 [Source:MGI Symbol;A | 0 | 0 | 0 | 0 | 0 | 0 |
| 20825 | ENSMUSG00000074439 | 0 | NA | NA | NA | NA | NA | Defa5 | defensin, alpha, 5 [Source:MGI Symbol;Ac | 0 | 0 | 0 | 0 | 0 | 0 |
| 20826 | ENSMUSG00000074440 | 0 | NA | NA | NA | NA | NA | Defa3 | defensin, alpha, 3 [Source:MGI Symbol;Ac | 0 | 0 | 0 | 0 | 0 | 0 |
| 20827 | ENSMUSG00000074441 | 0 | NA | NA | NA | NA | NA | Defa40 | defensin, alpha, 40 [Source:MGI Symbol;A | 0 | 0 | 0 | 0 | 0 | 0 |
| 20828 | ENSMUSG00000074442 | 0 | NA | NA | NA | NA | NA | Defa31 | defensin, alpha, 31 [Source:MGI Symbol;A | 0 | 0 | 0 | 0 | 0 | 0 |

| | | | | | | | | | | | | | | | |
|---|---|---|---|---|---|---|---|---|---|---|---|---|---|---|---|
| 20829 | ENSMUSG00000074443 | 0 | NA | NA | NA | NA | NA | Defa22 | defensin, alpha, 22 [Source:MGI Symbol;A | 0 | 0 | 0 | 0 | 0 | 0 |
| 20830 | ENSMUSG00000074444 | 0 | NA | NA | NA | NA | NA | Defa30 | defensin, alpha, 30 [Source:MGI Symbol;A | 0 | 0 | 0 | 0 | 0 | 0 |
| 20831 | ENSMUSG00000074445 | 0 | NA | NA | NA | NA | NA | Sprr2a3 | small proline-rich protein 2A3 [Source:MG | 0 | 0 | 0 | 0 | 0 | 0 |
| 20832 | ENSMUSG00000074446 | 0 | NA | NA | NA | NA | NA | Defa23 | defensin, alpha, 23 [Source:MGI Symbol;A | 0 | 0 | 0 | 0 | 0 | 0 |
| 20833 | ENSMUSG00000074447 | 0 | NA | NA | NA | NA | NA | Defa21 | defensin, alpha, 21 [Source:MGI Symbol;A | 0 | 0 | 0 | 0 | 0 | 0 |
| 20834 | ENSMUSG00000074448 | 0 | NA | NA | NA | NA | NA | Pate9 | prostate and testis expressed 9 [Source:M | 0 | 0 | 0 | 0 | 0 | 0 |
| 20835 | ENSMUSG00000074449 | 0 | NA | NA | NA | NA | NA | Gm15319 | predicted gene 15319 [Source:MGI Symbo | 0 | 0 | 0 | 0 | 0 | 0 |
| 20836 | ENSMUSG00000074452 | 0 | NA | NA | NA | NA | NA | Pate2 | prostate and testis expressed 2 [Source:M | 0 | 0 | 0 | 0 | 0 | 0 |
| 20837 | ENSMUSG00000074454 | 0 | NA | NA | NA | NA | NA | Defb33 | defensin beta 33 [Source:MGI Symbol;Acc | 0 | 0 | 0 | 0 | 0 | 0 |
| 20838 | ENSMUSG00000074457 | 396.2250668 | 0.005235229 | 0.135600858 | 0.038607638 | 0.969203212 | 0.994602269 | S100a16 | S100 calcium binding protein A16 [Source | 423.23 | 376.22 | 386.4 | 389.45 | 414.43 | 387.62 |
| 20839 | ENSMUSG00000074461 | 7.653667815 | -0.600311602 | 0.895165413 | -0.670615278 | 0.502465647 | 0.881220881 | Gm10699 | predicted gene 10699 [Source:MGI Symbo | 18.471 | 4.1117 | 5.0312 | 10.604 | 4.3625 | 3.3415 |
| 20840 | ENSMUSG00000074465 | 1.618412372 | -1.905872387 | 1.858460436 | -1.025511413 | 0.305121943 | NA | Gm10701 | predicted gene 10701 [Source:MGI Symbo | 5.6217 | 1.0279 | 1.0062 | 0.964 | 1.0906 | 0 |
| 20841 | ENSMUSG00000074466 | 148.3971253 | 0.221660125 | 0.227688091 | 0.973525332 | 0.330292276 | 0.774747658 | Gm15417 | predicted gene 15417 [Source:MGI Symbo | 150.18 | 132.6 | 127.79 | 149.42 | 124.33 | 206.06 |
| 20842 | ENSMUSG00000074467 | 0 | NA | NA | NA | NA | NA | Gm10702 | predicted gene 10702 [Source:MGI Symbo | 0 | 0 | 0 | 0 | 0 | 0 |
| 20843 | ENSMUSG00000074469 | 0.349477146 | 0.165142768 | 3.911021834 | 0.042224967 | 0.966319359 | NA | Gm15348 | predicted gene 15348 [Source:MGI Symbo | 0 | 0 | 1.0062 | 0 | 1.0906 | 0 |
| 20844 | ENSMUSG00000074472 | 15.15721457 | 0.575504848 | 0.705795541 | 0.815398815 | 0.414844062 | 0.836001897 | Zfp872 | zinc finger protein 872 [Source:MGI Symb | 15.259 | 5.1396 | 16.1 | 28.919 | 4.3625 | 21.163 |
| 20845 | ENSMUSG00000074473 | 0 | NA | NA | NA | NA | NA | A230072I06I | RIKEN cDNA A230072I06 gene [Source:MG | 0 | 0 | 0 | 0 | 0 | 0 |
| 20846 | ENSMUSG00000074476 | 364.5262937 | 0.211297168 | 0.157101988 | 1.34496814 | 0.178635523 | 0.610813333 | Spc24 | SPC24, NDC80 kinetochore complex comp | 363.81 | 298.1 | 351.18 | 382.7 | 444.97 | 346.41 |
| 20847 | ENSMUSG00000074479 | 0.356960287 | 0.16514264 | 3.880370363 | 0.042558474 | 0.966053499 | NA | Rtraf-ps | RNA transcription, translation and transpo | 0 | 1.0279 | 0 | 0 | 0 | 1.1138 |
| 20848 | ENSMUSG00000074480 | 1.004562989 | 0.155751777 | 2.077846764 | 0.07495826 | 0.940247922 | NA | Mex3a | mex3 RNA binding family member A [Sour | 0.8031 | 2.0558 | 0 | 0.964 | 1.0906 | 1.1138 |
| 20849 | ENSMUSG00000074482 | 0.160662974 | 1.126924175 | 4.080472857 | 0.276174898 | 0.782413736 | NA | Gm16589 | predicted gene 16589 [Source:MGI Symbo | 0 | 0 | 0 | 0.964 | 0 | 0 |
| 20850 | ENSMUSG00000074483 | 0 | NA | NA | NA | NA | NA | Bglap | bone gamma carboxyglutamate protein [S | 0 | 0 | 0 | 0 | 0 | 0 |
| 20851 | ENSMUSG00000074485 | 0 | NA | NA | NA | NA | NA | Gm7381 | predicted gene 7381 [Source:MGI Symbol | 0 | 0 | 0 | 0 | 0 | 0 |
| 20852 | ENSMUSG00000074486 | 0 | NA | NA | NA | NA | NA | Bglap2 | bone gamma-carboxyglutamate protein 2 | 0 | 0 | 0 | 0 | 0 | 0 |
| 20853 | ENSMUSG00000074489 | 7.134492966 | 0.442635645 | 0.989833981 | 0.447181703 | 0.654743871 | 0.925797231 | Bglap3 | bone gamma-carboxyglutamate protein 3 | 12.85 | 5.1396 | 0 | 11.568 | 5.4531 | 7.7969 |
| 20854 | ENSMUSG00000074491 | 473.6845271 | -0.512649249 | 0.175575146 | -2.919828123 | 0.003502245 | 0.052331064 | Clec4g | C-type lectin domain family 4, member g [ | 555.75 | 495.46 | 619.85 | 423.19 | 291.2 | 456.68 |
| 20855 | ENSMUSG00000074497 | 1.955585053 | -1.48964766 | 1.607208819 | -0.926853836 | 0.354002429 | NA | NA | NA | 1.6062 | 3.0838 | 4.025 | 1.928 | 1.0906 | 0 |
| 20856 | ENSMUSG00000074500 | 8.997450418 | 0.086746737 | 0.828176708 | 0.104744237 | 0.916578759 | 0.986374671 | Zfp558 | zinc finger protein 558 [Source:MGI Symb | 12.85 | 3.0838 | 10.062 | 3.8559 | 17.45 | 6.6831 |
| 20857 | ENSMUSG00000074501 | 0 | NA | NA | NA | NA | NA | Gm5612 | predicted gene 5612 [Source:MGI Symbol | 0 | 0 | 0 | 0 | 0 | 0 |
| 20858 | ENSMUSG00000074502 | 0 | NA | NA | NA | NA | NA | Ubtfl1 | upstream binding transcription factor, RN | 0 | 0 | 0 | 0 | 0 | 0 |
| 20859 | ENSMUSG00000074505 | 0 | NA | NA | NA | NA | NA | Fat3 | FAT atypical cadherin 3 [Source:MGI Symb | 0 | 0 | 0 | 0 | 0 | 0 |
| 20860 | ENSMUSG00000074506 | 1.053024754 | -1.051058068 | 2.223622046 | -0.472678381 | 0.636442653 | NA | Gm10705 | predicted gene 10705 [Source:MGI Symbo | 3.2124 | 1.0279 | 0 | 0.964 | 0 | 1.1138 |
| 20861 | ENSMUSG00000074507 | 0 | NA | NA | NA | NA | NA | Gm14340 | predicted gene 14340 [Source:MGI Symbo | 0 | 0 | 0 | 0 | 0 | 0 |
| 20862 | ENSMUSG00000074508 | 0 | NA | NA | NA | NA | NA | Gm10706 | predicted gene 10706 [Source:MGI Symbo | 0 | 0 | 0 | 0 | 0 | 0 |
| 20863 | ENSMUSG00000074513 | 66.40452186 | -0.375166808 | 0.292606062 | -1.282156648 | 0.1997877 | 0.642000421 | Arfip1 | ADP-ribosylation factor interacting protei | 58.626 | 83.261 | 83.518 | 62.659 | 47.987 | 62.375 |
| 20864 | ENSMUSG00000074516 | 3.284506118 | 1.351699977 | 1.336834783 | 1.011119694 | 0.311959146 | NA | Gm10709 | predicted gene 10709 [Source:MGI Symbo | 2.4093 | 3.0838 | 0 | 0.964 | 5.4531 | 7.7969 |
| 20865 | ENSMUSG00000074517 | 0 | NA | NA | NA | NA | NA | Gm10710 | predicted gene 10710 [Source:MGI Symbo | 0 | 0 | 0 | 0 | 0 | 0 |
| 20866 | ENSMUSG00000074519 | 60.61682227 | 0.339735083 | 0.315080558 | 1.078248321 | 0.280922953 | 0.733710968 | Zfp971 | zinc finger protein 971 [Source:MGI Symb | 44.171 | 43.173 | 73.456 | 67.478 | 74.162 | 61.261 |
| 20867 | ENSMUSG00000074521 | 88.02716453 | -0.214460238 | 0.261332721 | -0.820640587 | 0.411851022 | 0.834198737 | Gm14327 | predicted gene 14327 [Source:MGI Symbo | 83.523 | 109.99 | 90.562 | 90.614 | 63.256 | 90.221 |
| 20868 | ENSMUSG00000074527 | 3.482825847 | -2.454856271 | 1.32673307 | -1.850301561 | 0.064270098 | NA | Gm14296 | predicted gene 14296 [Source:MGI Symbo | 6.4248 | 5.1396 | 6.0375 | 0 | 2.1812 | 1.1138 |
| 20869 | ENSMUSG00000074529 | 20.82240679 | 0.373224092 | 0.644686983 | 0.578922954 | 0.562641161 | 0.900565925 | Zfp972 | zinc finger protein 972 [Source:MGI Symb | 16.865 | 24.67 | 13.081 | 41.451 | 4.3625 | 24.505 |
| 20870 | ENSMUSG00000074547 | 0 | NA | NA | NA | NA | NA | Rps8-ps4 | ribosomal protein S8, pseudogene 4 [Sour | 0 | 0 | 0 | 0 | 0 | 0 |
| 20871 | ENSMUSG00000074553 | 0.881628005 | -0.457461187 | 2.549924572 | -0.17940185 | 0.857622178 | NA | Eif1-ps1 | eukaryotic translation initiation factor 1, p | 0 | 2.0558 | 1.0062 | 0 | 0 | 2.2277 |
| 20872 | ENSMUSG00000074555 | 0 | NA | NA | NA | NA | NA | Gm10714 | predicted gene 10714 [Source:MGI Symbo | 0 | 0 | 0 | 0 | 0 | 0 |
| 20873 | ENSMUSG00000074564 | 0 | NA | NA | NA | NA | NA | Gm10720 | predicted gene 10720 [Source:MGI Symbo | 0 | 0 | 0 | 0 | 0 | 0 |
| 20874 | ENSMUSG00000074569 | 6.302569877 | -0.839974411 | 0.847171997 | -0.991503985 | 0.321439548 | 0.76866499 | Gcnt7 | glucosaminyl (N-acetyl) transferase family | 8.031 | 8.2233 | 8.05 | 5.7839 | 3.2719 | 4.4554 |
| 20875 | ENSMUSG00000074570 | 0 | NA | NA | NA | NA | NA | Cass4 | Cas scaffolding protein family member 4 [ | 0 | 0 | 0 | 0 | 0 | 0 |
| 20876 | ENSMUSG00000074575 | 0 | NA | NA | NA | NA | NA | Kcng1 | potassium voltage-gated channel, subfam | 0 | 0 | 0 | 0 | 0 | 0 |
| 20877 | ENSMUSG00000074576 | 205.5547195 | 0.056394274 | 0.180795625 | 0.311922782 | 0.755099208 | 0.953085143 | Mocs3 | molybdenum cofactor synthesis 3 [Source | 211.22 | 201.47 | 192.19 | 242.92 | 202.86 | 182.67 |
| 20878 | ENSMUSG00000074577 | 31.09473842 | 0.303287137 | 0.480475331 | 0.631223119 | 0.527894648 | 0.888334265 | Ripor3 | RIPOR family member 3 [Source:MGI Sym | 43.368 | 17.475 | 22.137 | 25.063 | 26.175 | 52.351 |
| 20879 | ENSMUSG00000074578 | 195.954885 | 0.203862257 | 0.187230464 | 1.088830595 | 0.276228597 | 0.730038567 | Zfas1 | zinc finger, NFX1-type containing 1, antise | 193.55 | 184 | 169.05 | 245.81 | 201.76 | 181.56 |
| 20880 | ENSMUSG00000074579 | 3.370991727 | -3.157290127 | 1.520041963 | -2.077107214 | 0.037791665 | NA | Lekr1 | leucine, glutamate and lysine rich 1 [Sourc | 4.0155 | 9.2513 | 5.0312 | 1.928 | 0 | 0 |
| 20881 | ENSMUSG00000074580 | 6.941505826 | -0.019382721 | 0.826797893 | -0.023443119 | 0.981296811 | 0.99820894 | 4931440P22 | RIKEN cDNA 4931440P22 gene [Source:M | 4.8186 | 9.2513 | 7.0437 | 10.604 | 4.3625 | 5.5692 |
| 20882 | ENSMUSG00000074582 | 1478.83795 | -0.060407483 | 0.103909452 | -0.581347335 | 0.561006385 | 0.900199284 | Arfgef2 | ADP-ribosylation factor guanine nucleotid | 1513.8 | 1588.1 | 1426.9 | 1361.1 | 1456 | 1527.1 |
| 20883 | ENSMUSG00000074589 | 0 | NA | NA | NA | NA | NA | 4930449A18 | RIKEN cDNA 4930449A18 gene [Source:M | 0 | 0 | 0 | 0 | 0 | 0 |
| 20884 | ENSMUSG00000074591 | 6.175872087 | -0.22097727 | 0.948368108 | -0.233007909 | 0.815755264 | 0.967767553 | Ankub1 | ankrin repeat and ubiquitin domain conta | 8.8341 | 5.1396 | 6.0375 | 11.568 | 4.3625 | 1.1138 |

| | | | | | | | | | | | | | | | |
|---|---|---|---|---|---|---|---|---|---|---|---|---|---|---|---|
| 20885 | ENSMUSG00000074593 | 0 | NA | NA | NA | NA | NA | Spint5 | serine protease inhibitor, Kunitz type 5 [So | 0 | 0 | 0 | 0 | 0 | 0 |
| 20886 | ENSMUSG00000074594 | 0 | NA | NA | NA | NA | NA | Wfdc9 | WAP four-disulfide core domain 9 [Source | 0 | 0 | 0 | 0 | 0 | 0 |
| 20887 | ENSMUSG00000074595 | 1.178849562 | 0.581643986 | 1.973500221 | 0.294727095 | 0.768202364 | NA | Wfdc6a | WAP four-disulfide core domain 6A [Sourc | 0.8031 | 1.0279 | 1.0062 | 0.964 | 3.2719 | 0 |
| 20888 | ENSMUSG00000074596 | 0 | NA | NA | NA | NA | NA | Spint3 | serine peptidase inhibitor, Kunitz type, 3 [ | 0 | 0 | 0 | 0 | 0 | 0 |
| 20889 | ENSMUSG00000074603 | 0.181769715 | 1.126924175 | 4.080472857 | 0.276174898 | 0.782413736 | NA | Gm10729 | predicted gene 10729 [Source:MGI Symbo | 0 | 0 | 0 | 0 | 1.0906 | 0 |
| 20890 | ENSMUSG00000074604 | 1.855978288 | -2.185132062 | 2.037853264 | -1.072271542 | 0.283598085 | NA | Mgst2 | microsomal glutathione S-transferase 2 [S | 0 | 7.1954 | 2.0125 | 1.928 | 0 | 0 |
| 20891 | ENSMUSG00000074607 | 3.282165885 | -0.089446579 | 1.269186114 | -0.070475542 | 0.943815167 | NA | Tox2 | TOX high mobility group box family memb | 4.0155 | 3.0838 | 3.0187 | 2.8919 | 0 | 6.6831 |
| 20892 | ENSMUSG00000074619 | 1.379617879 | -0.182671712 | 2.126686134 | -0.085895003 | 0.931549884 | NA | 1700034I23R | RIKEN cDNA 1700034I23 gene [Source:MG | 2.4093 | 0 | 2.0125 | 3.8559 | 0 | 0 |
| 20893 | ENSMUSG00000074620 | 0 | NA | NA | NA | NA | NA | Gm10731 | predicted gene 10731 [Source:MGI Symbo | 0 | 0 | 0 | 0 | 0 | 0 |
| 20894 | ENSMUSG00000074622 | 1945.652816 | -0.846679388 | 0.18696885 | -4.528451596 | 5.94E-06 | 0.000358089 | Mafb | v-maf musculoaponeurotic fibrosarcoma o | 1880.9 | 3319.1 | 2302.3 | 1098 | 1543.2 | 1530.4 |
| 20895 | ENSMUSG00000074623 | 64.13485553 | -0.683388369 | 0.290131095 | -2.355446832 | 0.018500444 | 0.163849644 | Gm826 | predicted gene 826 [Source:MGI Symbol;A | 82.72 | 72.982 | 81.506 | 53.019 | 56.712 | 37.871 |
| 20896 | ENSMUSG00000074625 | 0 | NA | NA | NA | NA | NA | Arhgap40 | Rho GTPase activating protein 40 [Source: | 0 | 0 | 0 | 0 | 0 | 0 |
| 20897 | ENSMUSG00000074627 | 1.757426182 | -0.101786716 | 1.629413579 | -0.062468312 | 0.950189896 | NA | Mroh8 | maestro heat-like repeat family member 8 | 2.4093 | 2.0558 | 1.0062 | 2.8919 | 2.1812 | 0 |
| 20898 | ENSMUSG00000074628 | 0 | NA | NA | NA | NA | NA | Tldc2 | TBC/LysM associated domain containing 2 | 0 | 0 | 0 | 0 | 0 | 0 |
| 20899 | ENSMUSG00000074629 | 10.96677151 | 0.505091941 | 0.667667292 | 0.756502449 | 0.449348005 | 0.854879451 | 4930518I15R | RIKEN cDNA 4930518I15 gene [Source:MG | 12.85 | 5.1396 | 9.0562 | 14.46 | 9.8156 | 14.48 |
| 20900 | ENSMUSG00000074634 | 73.71740191 | -0.781611807 | 0.323241353 | -2.41804398 | 0.015604191 | 0.145989013 | Tmem267 | transmembrane protein 267 [Source:MGI | 122.87 | 86.345 | 70.437 | 73.262 | 39.262 | 50.123 |
| 20901 | ENSMUSG00000074635 | 2.996449072 | -0.445078588 | 1.364401678 | -0.326207887 | 0.744267075 | NA | 3110070M22 | RIKEN cDNA 3110070M22 gene [Source:M | 3.2124 | 3.0838 | 4.025 | 0 | 6.5437 | 1.1138 |
| 20902 | ENSMUSG00000074637 | 0 | NA | NA | NA | NA | NA | Sox2 | SRY (sex determining region Y)-box 2 [Sou | 0 | 0 | 0 | 0 | 0 | 0 |
| 20903 | ENSMUSG00000074639 | 1711.031458 | -0.106962351 | 0.172670404 | -0.619459669 | 0.535613583 | 0.890063748 | Rdh16f2 | RDH16 family member 2 [Source:MGI Sym | 1306.6 | 1672.4 | 2345.6 | 1735.2 | 1533.4 | 1673 |
| 20904 | ENSMUSG00000074643 | 1.638182171 | -1.058788624 | 1.880806833 | -0.562943842 | 0.573473125 | NA | Cpne1 | copine I [Source:MGI Symbol;Acc:MGI:23 | 1.6062 | 0 | 5.0312 | 0.964 | 0 | 2.2277 |
| 20905 | ENSMUSG00000074646 | 38.65856663 | 0.00074777 | 0.361915894 | 0.002066143 | 0.998351457 | 0.999764475 | 6430550D23 | RIKEN cDNA 6430550D23 gene [Source:M | 36.943 | 31.865 | 47.293 | 45.307 | 34.9 | 35.643 |
| 20906 | ENSMUSG00000074647 | 7.40003652 | -0.522621874 | 0.862222879 | -0.606133155 | 0.544426336 | 0.894940621 | Fam83c | family with sequence similarity 83, memb | 12.047 | 5.1396 | 9.0562 | 11.568 | 4.3625 | 2.2277 |
| 20907 | ENSMUSG00000074649 | 195.8210797 | 0.071219626 | 0.192917171 | 0.369172043 | 0.711999492 | 0.944239215 | BC029722 | cDNA sequence BC029722 [Source:MGI Sy | 226.47 | 158.3 | 187.16 | 184.12 | 206.13 | 212.74 |
| 20908 | ENSMUSG00000074650 | 0 | NA | NA | NA | NA | NA | Gm10735 | predicted gene 10735 [Source:MGI Symbo | 0 | 0 | 0 | 0 | 0 | 0 |
| 20909 | ENSMUSG00000074651 | 0 | NA | NA | NA | NA | NA | Mcidas | multiciliate differentiation and DNA synth | 0 | 0 | 0 | 0 | 0 | 0 |
| 20910 | ENSMUSG00000074652 | 2.741084036 | 2.348377895 | 1.469613832 | 1.597955765 | 0.110052822 | NA | Myh7b | myosin, heavy chain 7B, cardiac muscle, b | 1.6062 | 0 | 1.0062 | 3.8559 | 2.1812 | 7.7969 |
| 20911 | ENSMUSG00000074653 | 1.584944048 | 1.931604541 | 2.097974855 | 0.920699567 | 0.357207302 | NA | Lrrc31 | leucine rich repeat containing 31 [Source: | 0 | 0 | 2.0125 | 1.928 | 0 | 5.5692 |
| 20912 | ENSMUSG00000074655 | 0 | NA | NA | NA | NA | NA | Gm1527 | predicted gene 1527 [Source:MGI Symbol | 0 | 0 | 0 | 0 | 0 | 0 |
| 20913 | ENSMUSG00000074656 | 1373.624146 | 0.123107816 | 0.10310576 | 1.193995531 | 0.232479716 | 0.679847566 | Eif2s2 | eukaryotic translation initiation factor 2, s | 1391 | 1295.2 | 1257.8 | 1379.5 | 1409.1 | 1509.3 |
| 20914 | ENSMUSG00000074657 | 62.94196437 | 0.269936396 | 0.325189782 | 0.830088799 | 0.406488579 | 0.830399297 | Kif5a | kinesin family member 5A [Source:MGI Sy | 66.657 | 35.977 | 68.425 | 80.01 | 70.89 | 55.692 |
| 20915 | ENSMUSG00000074664 | 0 | NA | NA | NA | NA | NA | A830092H15 | RIKEN cDNA A830092H15 gene [Source:M | 0 | 0 | 0 | 0 | 0 | 0 |
| 20916 | ENSMUSG00000074665 | 0 | NA | NA | NA | NA | NA | Bpifb4 | BPI fold containing family B, member 4 [So | 0 | 0 | 0 | 0 | 0 | 0 |
| 20917 | ENSMUSG00000074671 | 4.861615948 | -0.43962122 | 1.073261458 | -0.409612417 | 0.682090287 | NA | Tspyl3 | TSPY-like 3 [Source:MGI Symbol;Acc:MGI: | 5.6217 | 7.1954 | 4.025 | 5.7839 | 6.5437 | 0 |
| 20918 | ENSMUSG00000074673 | 0 | NA | NA | NA | NA | NA | Ttll9 | tubulin tyrosine ligase-like family, membe | 0 | 0 | 0 | 0 | 0 | 0 |
| 20919 | ENSMUSG00000074676 | 2.320312709 | 1.001020219 | 1.500161011 | 0.667275187 | 0.504596373 | NA | Foxs1 | forkhead box S1 [Source:MGI Symbol;Acc | 1.6062 | 1.0279 | 2.0125 | 4.8199 | 0 | 4.4554 |
| 20920 | ENSMUSG00000074677 | 1.975379827 | -2.286161101 | 1.827472728 | -1.250996015 | 0.210935931 | NA | Sirpb1c | signal-regulatory protein beta 1C [Source: | 0.8031 | 3.0838 | 6.0375 | 1.928 | 0 | 0 |
| 20921 | ENSMUSG00000074678 | 0.167707431 | -0.796629176 | 4.080472857 | -0.195229623 | 0.84521319 | NA | Defb25 | defensin beta 25 [Source:MGI Symbol;Acc | 0 | 0 | 1.0062 | 0 | 0 | 0 |
| 20922 | ENSMUSG00000074679 | 0 | NA | NA | NA | NA | NA | Defb28 | defensin beta 28 [Source:MGI Symbol;Acc | 0 | 0 | 0 | 0 | 0 | 0 |
| 20923 | ENSMUSG00000074680 | 0 | NA | NA | NA | NA | NA | Defb26 | defensin beta 26 [Source:MGI Symbol;Acc | 0 | 0 | 0 | 0 | 0 | 0 |
| 20924 | ENSMUSG00000074681 | 0 | NA | NA | NA | NA | NA | Defb23 | defensin beta 23 [Source:MGI Symbol;Acc | 0 | 0 | 0 | 0 | 0 | 0 |
| 20925 | ENSMUSG00000074682 | 7.979326162 | -1.481144107 | 0.78378278 | -1.889738004 | 0.058793009 | 0.344306009 | Zcchc3 | zinc finger, CCHC domain containing 3 [So | 12.85 | 11.307 | 11.069 | 3.8559 | 5.4531 | 3.3415 |
| 20926 | ENSMUSG00000074695 | 0 | NA | NA | NA | NA | NA | Il22 | interleukin 22 [Source:MGI Symbol;Acc:M | 0 | 0 | 0 | 0 | 0 | 0 |
| 20927 | ENSMUSG00000074698 | 1653.83706 | 0.123827994 | 0.103699971 | 1.194098634 | 0.232439387 | 0.679847566 | Csnk2a1 | casein kinase 2, alpha 1 polypeptide [Sour | 1579.7 | 1611.8 | 1556.7 | 1585.7 | 1867.1 | 1722 |
| 20928 | ENSMUSG00000074704 | 0 | NA | NA | NA | NA | NA | Rad21l | RAD21-like (S. pombe) [Source:MGI Symb | 0 | 0 | 0 | 0 | 0 | 0 |
| 20929 | ENSMUSG00000074715 | 1.789448312 | -0.615349653 | 1.531017275 | -0.401922083 | 0.68774137 | NA | Ccl28 | chemokine (C-C motif) ligand 28 [Source:M | 2.4093 | 2.0558 | 2.0125 | 0.964 | 2.1812 | 1.1138 |
| 20930 | ENSMUSG00000074720 | 0 | NA | NA | NA | NA | NA | | | 0 | 0 | 0 | 0 | 0 | 0 |
| 20931 | ENSMUSG00000074731 | 0.167707431 | -0.796629176 | 4.080472857 | -0.195229623 | 0.84521319 | NA | Zfp345 | zinc finger protein 345 [Source:MGI Symb | 0 | 0 | 1.0062 | 0 | 0 | 0 |
| 20932 | ENSMUSG00000074733 | 235.550005 | -0.113970672 | 0.163001915 | -0.699198365 | 0.484428072 | 0.87256358 | Zfp950 | zinc finger protein 950 [Source:MGI Symb | 252.17 | 246.7 | 235.46 | 219.79 | 248.66 | 210.52 |
| 20933 | ENSMUSG00000074734 | 0 | NA | NA | NA | NA | NA | Taf7l2 | Taf7l2 [Source:MGI Symbol;Acc:MGI:6274 | 0 | 0 | 0 | 0 | 0 | 0 |
| 20934 | ENSMUSG00000074735 | 0 | NA | NA | NA | NA | NA | Gm21994 | predicted gene 21994 [Source:MGI Symbo | 0 | 0 | 0 | 0 | 0 | 0 |
| 20935 | ENSMUSG00000074736 | 0 | NA | NA | NA | NA | NA | Syndig1 | synapse differentiation inducing 1 [Source | 0 | 0 | 0 | 0 | 0 | 0 |
| 20936 | ENSMUSG00000074737 | 0 | NA | NA | NA | NA | NA | C530025M09 | RIKEN cDNA C530025M09 gene [Source:M | 0 | 0 | 0 | 0 | 0 | 0 |
| 20937 | ENSMUSG00000074738 | 10.97206865 | -0.376570563 | 0.697508032 | -0.539879895 | 0.589279864 | 0.911039598 | Fndc10 | fibronectin type III domain containing 10 [ | 16.865 | 11.307 | 9.0562 | 15.424 | 8.7249 | 4.4554 |
| 20938 | ENSMUSG00000074739 | 0 | NA | NA | NA | NA | NA | Gm10750 | predicted gene 10750 [Source:MGI Symbo | 0 | 0 | 0 | 0 | 0 | 0 |
| 20939 | ENSMUSG00000074743 | 26.9641448 | -1.469870408 | 0.446144823 | -3.29460375 | 0.000985606 | 0.020720637 | Thbd | thrombomodulin [Source:MGI Symbol;Acc | 34.533 | 48.312 | 36.225 | 17.352 | 11.997 | 13.366 |
| 20940 | ENSMUSG00000074746 | 747.3166421 | 0.135269553 | 0.110699376 | 1.221954067 | 0.221724998 | 0.667949912 | Pdzd8 | PDZ domain containing 8 [Source:MGI Syr | 718.78 | 710.29 | 707.39 | 748.05 | 788.52 | 810.88 |

| | | | | | | | | | | | | | | | |
|---|---|---|---|---|---|---|---|---|---|---|---|---|---|---|---|
| 20941 | ENSMUSG00000074748 | 1865.727118 | 0.023075301 | 0.091065588 | 0.253392101 | 0.799965231 | 0.962424862 | Atxn7l3b | ataxin 7-like 3B [Source:MGI Symbol;Acc:I | 1864 | 1829.7 | 1858.5 | 1878.8 | 1846.4 | 1916.9 |
| 20942 | ENSMUSG00000074749 | 304.6960879 | -0.009763567 | 0.145114469 | -0.067281831 | 0.946357341 | 0.990456772 | Kiz | kizuna centrosomal protein [Source:MGI S | 318.03 | 297.07 | 301.87 | 307.51 | 314.1 | 289.6 |
| 20943 | ENSMUSG00000074753 | 0 | NA | NA | NA | NA | NA | Gm14092 | predicted gene 14092 [Source:MGI Symbo | 0 | 0 | 0 | 0 | 0 | 0 |
| 20944 | ENSMUSG00000074754 | 171.1805366 | -0.043592421 | 0.206968412 | -0.210623549 | 0.833181035 | 0.970611245 | Smim26 | small integral membrane protein 26 [Sour | 203.99 | 184 | 132.82 | 173.52 | 170.14 | 162.62 |
| 20945 | ENSMUSG00000074758 | 0 | NA | NA | NA | NA | NA | Gm5535 | predicted gene 5535 [Source:MGI Symbol | 0 | 0 | 0 | 0 | 0 | 0 |
| 20946 | ENSMUSG00000074763 | 0 | NA | NA | NA | NA | NA | Macrod2os2 | mono-ADP ribosylhydrolase 2, opposite st | 0 | 0 | 0 | 0 | 0 | 0 |
| 20947 | ENSMUSG00000074764 | 0 | NA | NA | NA | NA | NA | Sel1l2 | sel-1 suppressor of lin-12-like 2 (C. elegan | 0 | 0 | 0 | 0 | 0 | 0 |
| 20948 | ENSMUSG00000074766 | 1.165329702 | -2.428778977 | 2.321199838 | -1.04634635 | 0.295401153 | NA | Ism1 | isthmin 1, angiogenesis inhibitor [Source:I | 4.0155 | 0 | 2.0125 | 0.964 | 0 | 0 |
| 20949 | ENSMUSG00000074768 | 65683.00091 | -0.554612826 | 0.171180679 | -3.239926544 | 0.001195605 | 0.023787111 | Bhmt | betaine-homocysteine methyltransferase | 79529 | 82728 | 72207 | 36627 | 68798 | 54209 |
| 20950 | ENSMUSG00000074771 | 0 | NA | NA | NA | NA | NA | Ankef1 | ankyrin repeat and EF-hand domain conta | 0 | 0 | 0 | 0 | 0 | 0 |
| 20951 | ENSMUSG00000074776 | 0 | NA | NA | NA | NA | NA | Gm10754 | predicted gene 10754 [Source:MGI Symbo | 0 | 0 | 0 | 0 | 0 | 0 |
| 20952 | ENSMUSG00000074780 | 0 | NA | NA | NA | NA | NA | Anapc15-ps | anaphase promoting complex C subunit 1 | 0 | 0 | 0 | 0 | 0 | 0 |
| 20953 | ENSMUSG00000074781 | 699.0941586 | -0.04758168 | 0.118022355 | -0.403158201 | 0.686831847 | 0.936582996 | Ube2n | ubiquitin-conjugating enzyme E2N [Sourc | 752.51 | 711.32 | 667.14 | 655.5 | 723.08 | 685.01 |
| 20954 | ENSMUSG00000074782 | 26.85779311 | -0.490405856 | 0.41537268 | -1.180640614 | 0.237745522 | 0.686377492 | 4833422C13 | RIKEN cDNA 4833422C13 gene [Source:M | 32.124 | 32.893 | 29.181 | 25.063 | 20.722 | 21.163 |
| 20955 | ENSMUSG00000074783 | 0.160662974 | 1.126924175 | 4.080472857 | 0.276174898 | 0.782413736 | NA | AU019990 | expressed sequence AU019990 [Source:M | 0 | 0 | 0 | 0.964 | 0 | 0 |
| 20956 | ENSMUSG00000074785 | 162.6600531 | -0.610389961 | 0.215853097 | -2.827802655 | 0.004686869 | 0.064014043 | Plxnc1 | plexin C1 [Source:MGI Symbol;Acc:MGI:1 | 205.59 | 158.3 | 225.4 | 119.53 | 113.42 | 153.71 |
| 20957 | ENSMUSG00000074793 | 13.14374193 | -0.675113383 | 0.602136998 | -1.12119565 | 0.262204591 | 0.716133499 | Hspa12b | heat shock protein 12B [Source:MGI Symb | 16.062 | 12.335 | 20.125 | 11.568 | 7.6343 | 11.138 |
| 20958 | ENSMUSG00000074794 | 2629.948047 | 1.275668049 | 0.501474568 | 2.543833985 | 0.010964318 | 0.115926826 | Arrdc3 | arrestin domain containing 3 [Source:MGI | 2288.8 | 706.18 | 1617 | 4378.4 | 3957.9 | 2831.4 |
| 20959 | ENSMUSG00000074796 | 2.988298113 | 1.57101473 | 1.332572082 | 1.178934147 | 0.238424399 | NA | Slc4a11 | solute carrier family 4, sodium bicarbonat | 2.4093 | 2.0558 | 0 | 5.7839 | 5.4531 | 2.2277 |
| 20960 | ENSMUSG00000074797 | 461.303003 | 0.06757762 | 0.130244166 | 0.518853337 | 0.60386302 | 0.916214699 | Itpa | inosine triphosphatase (nucleoside tripho | 458.57 | 425.56 | 466.9 | 453.07 | 459.15 | 504.57 |
| 20961 | ENSMUSG00000074800 | 10.02490726 | 0.289979297 | 0.71049964 | 0.408134334 | 0.683175055 | 0.935211432 | Gm4149 | predicted pseudogene 4149 [Source:MGI | 9.6372 | 11.307 | 6.0375 | 7.7118 | 7.6343 | 17.822 |
| 20962 | ENSMUSG00000074802 | 35.1933627 | 0.675752273 | 0.385023116 | 1.75509533 | 0.079243003 | 0.413332418 | Gas2l3 | growth arrest-specific 2 like 3 [Source:MG | 36.14 | 23.642 | 21.131 | 44.343 | 45.806 | 40.098 |
| 20963 | ENSMUSG00000074804 | 0 | NA | NA | NA | NA | NA | AU015228 | expressed sequence AU015228 [Source:M | 0 | 0 | 0 | 0 | 0 | 0 |
| 20964 | ENSMUSG00000074805 | 0.321325949 | 1.911575114 | 4.048267893 | 0.472195804 | 0.636787034 | NA | Il1bos | interleukin 1 beta, opposite strand [Sourc | 0 | 0 | 0 | 1.928 | 0 | 0 |
| 20965 | ENSMUSG00000074807 | 85.82784653 | -0.334728884 | 0.276315002 | -1.211403225 | 0.225740904 | 0.6725584 | Gm10762 | predicted gene 10762 [Source:MGI Symbo | 88.341 | 105.88 | 93.581 | 97.362 | 76.343 | 53.465 |
| 20966 | ENSMUSG00000074811 | 105.5993266 | -0.317205891 | 0.228423406 | -1.388675077 | 0.164931576 | 0.590049547 | Hps6 | HPS6, biogenesis of lysosomal organelles | 112.43 | 117.18 | 121.76 | 85.794 | 86.159 | 110.27 |
| 20967 | ENSMUSG00000074812 | 0.881705114 | -0.66605326 | 2.559736336 | -0.260203854 | 0.794706531 | NA | Spdye4c | speedy/RINGO cell cycle regulator family, | 3.2124 | 0 | 0 | 0.964 | 0 | 1.1138 |
| 20968 | ENSMUSG00000074813 | 5.244381686 | -1.458022486 | 0.958302148 | -1.521464279 | 0.128143369 | 0.524051388 | Morrbid | myeloid RNA regulator of BCL2L11 induce | 8.8341 | 6.1675 | 8.05 | 2.8919 | 2.1812 | 3.3415 |
| 20969 | ENSMUSG00000074817 | 0 | NA | NA | NA | NA | NA | Papolb | poly (A) polymerase beta (testis specific) [ | 0 | 0 | 0 | 0 | 0 | 0 |
| 20970 | ENSMUSG00000074818 | 2.446712084 | -0.576320464 | 1.65850064 | -0.34749487 | 0.728219572 | NA | Pdzd7 | PDZ domain containing 7 [Source:MGI Syr | 0.8031 | 2.0558 | 6.0375 | 5.7839 | 0 | 0 |
| 20971 | ENSMUSG00000074824 | 14.25947585 | 0.132720477 | 0.593362322 | 0.223675269 | 0.823009991 | 0.969382451 | Rslcan18 | regulator of sex-limitation candidate 18 [S | 10.44 | 18.503 | 12.075 | 20.244 | 9.8156 | 14.48 |
| 20972 | ENSMUSG00000074825 | 117.6214267 | 0.099228609 | 0.227251876 | 0.436645939 | 0.662368141 | 0.9288455 | Itpripl1 | inositol 1,4,5-triphosphate receptor intera | 110.83 | 108.96 | 120.75 | 107.97 | 107.97 | 149.26 |
| 20973 | ENSMUSG00000074826 | 22.06208435 | 0.007209587 | 0.494284592 | 0.014585903 | 0.988362546 | 0.99859176 | Gm10767 | predicted gene 10767 [Source:MGI Symbo | 14.456 | 22.614 | 29.181 | 22.171 | 28.356 | 15.594 |
| 20974 | ENSMUSG00000074829 | 454.4974075 | 0.194406055 | 0.144236858 | 1.347825084 | 0.177714648 | 0.609100512 | 2010315B03 | RIKEN cDNA 2010315B03 gene [Source:M | 416.81 | 397.8 | 456.84 | 434.75 | 551.85 | 468.93 |
| 20975 | ENSMUSG00000074830 | 0 | NA | NA | NA | NA | NA | Zfp640 | zinc finger protein 640 [Source:MGI Symb | 0 | 0 | 0 | 0 | 0 | 0 |
| 20976 | ENSMUSG00000074832 | 0 | NA | NA | NA | NA | NA | Zfp998 | zinc finger protein 998 [Source:MGI Symb | 0 | 0 | 0 | 0 | 0 | 0 |
| 20977 | ENSMUSG00000074837 | 0 | NA | NA | NA | NA | NA | Gm10770 | predicted gene 10770 [Source:MGI Symbo | 0 | 0 | 0 | 0 | 0 | 0 |
| 20978 | ENSMUSG00000074841 | 0 | NA | NA | NA | NA | NA | Cbx3-ps1 | chromobox 3, pseudogene 1 [Source:MGI | 0 | 0 | 0 | 0 | 0 | 0 |
| 20979 | ENSMUSG00000074846 | 0 | NA | NA | NA | NA | NA | Gm10774 | predicted pseudogene 10774 [Source:MG | 0 | 0 | 0 | 0 | 0 | 0 |
| 20980 | ENSMUSG00000074847 | 0 | NA | NA | NA | NA | NA | Gm10775 | predicted gene 10775 [Source:MGI Symbo | 0 | 0 | 0 | 0 | 0 | 0 |
| 20981 | ENSMUSG00000074849 | 0 | NA | NA | NA | NA | NA | Spata31d1c | spermatogenesis associated 31 subfamily | 0 | 0 | 0 | 0 | 0 | 0 |
| 20982 | ENSMUSG00000074852 | 0 | NA | NA | NA | NA | NA | Hpse2 | heparanase 2 [Source:MGI Symbol;Acc:M | 0 | 0 | 0 | 0 | 0 | 0 |
| 20983 | ENSMUSG00000074862 | 5.820864056 | 1.068244129 | 0.980693394 | 1.089274319 | 0.276032934 | 0.730038567 | BC025920 | cDNA sequence BC025920 [Source:MGI S | 8.031 | 1.0279 | 2.0125 | 10.604 | 5.4531 | 7.7969 |
| 20984 | ENSMUSG00000074863 | 12.17725636 | -0.352185099 | 0.627720867 | -0.561053674 | 0.574760947 | 0.906163061 | Platr25 | pluripotency associated transcript 25 [Sou | 10.44 | 15.419 | 15.094 | 6.7478 | 11.997 | 13.366 |
| 20985 | ENSMUSG00000074865 | 6.576616545 | -0.018218938 | 0.851022113 | -0.021408302 | 0.982919951 | 0.998580735 | Zfp934 | zinc finger protein 934 [Source:MGI Symb | 9.6372 | 5.1396 | 5.0312 | 8.6758 | 7.6343 | 3.3415 |
| 20986 | ENSMUSG00000074867 | 2.095315734 | -0.479403659 | 1.532438122 | -0.312837205 | 0.754404347 | NA | Zfp808 | zinc finger protein 808 [Source:MGI Symb | 3.2124 | 3.0838 | 1.0062 | 1.928 | 0 | 3.3415 |
| 20987 | ENSMUSG00000074868 | 0 | NA | NA | NA | NA | NA | Gm6478 | predicted pseudogene 6478 [Source:MGI | 0 | 0 | 0 | 0 | 0 | 0 |
| 20988 | ENSMUSG00000074870 | 0 | NA | NA | NA | NA | NA | Cts3 | cathepsin 3 [Source:MGI Symbol;Acc:MGI | 0 | 0 | 0 | 0 | 0 | 0 |
| 20989 | ENSMUSG00000074871 | 0 | NA | NA | NA | NA | NA | Ctsm | cathepsin M [Source:MGI Symbol;Acc:MG | 0 | 0 | 0 | 0 | 0 | 0 |
| 20990 | ENSMUSG00000074872 | 0 | NA | NA | NA | NA | NA | Ctxn2 | cortexin 2 [Source:MGI Symbol;Acc:MGI:2 | 0 | 0 | 0 | 0 | 0 | 0 |
| 20991 | ENSMUSG00000074873 | 41.84261798 | 0.162366879 | 0.341917126 | 0.474872028 | 0.63487819 | 0.923478764 | NA | NA | 39.352 | 37.005 | 42.262 | 52.055 | 43.625 | 36.757 |
| 20992 | ENSMUSG00000074874 | 0.435408177 | -2.179616019 | 3.531869654 | -0.617128103 | 0.537150234 | NA | Ctla2b | cytotoxic T lymphocyte-associated proteir | 1.6062 | 0 | 1.0062 | 0 | 0 | 0 |
| 20993 | ENSMUSG00000074876 | 103.1984355 | -0.183160024 | 0.232660082 | -0.787243015 | 0.431139621 | 0.84474037 | NA | NA | 106.01 | 101.76 | 121.76 | 109.89 | 83.978 | 95.791 |
| 20994 | ENSMUSG00000074877 | 0 | NA | NA | NA | NA | NA | Gm10780 | predicted gene 10780 [Source:MGI Symbo | 0 | 0 | 0 | 0 | 0 | 0 |
| 20995 | ENSMUSG00000074881 | 0.181769715 | 1.126924175 | 4.080472857 | 0.276174898 | 0.782413736 | NA | Mageb3 | MAGE family member B3 [Source:MGI Syr | 0 | 0 | 0 | 0 | 1.0906 | 0 |
| 20996 | ENSMUSG00000074882 | 2855.728266 | -0.1477425 | 0.16036154 | -0.921308809 | 0.356889223 | 0.796166767 | Cyp2c68 | cytochrome P450, family 2, subfamily c, p | 2404.5 | 2844.2 | 3758.3 | 3139.7 | 2308.8 | 2678.8 |

| | | | | | | | | | | | | | | | |
|---|---|---|---|---|---|---|---|---|---|---|---|---|---|---|---|
| 20997 | ENSMUSG00000074884 | 1154.488138 | -0.012688727 | 0.127037784 | -0.099881517 | 0.92043839 | 0.986685804 | Serf2 | small EDRK-rich factor 2 [Source:MGI Sym | 1306.6 | 1042.3 | 1129 | 1252.2 | 1033.9 | 1162.9 |
| 20998 | ENSMUSG00000074885 | 0 | NA | NA | NA | NA | NA | Gm10782 | predicted gene 10782 [Source:MGI Symbo | 0 | 0 | 0 | 0 | 0 | 0 |
| 20999 | ENSMUSG00000074886 | 351.3218285 | -0.020968906 | 0.153545249 | -0.136564992 | 0.89137465 | 0.982489666 | Grk6 | G protein-coupled receptor kinase 6 [Sour | 329.27 | 394.72 | 338.1 | 351.85 | 310.83 | 383.16 |
| 21000 | ENSMUSG00000074887 | 0 | NA | NA | NA | NA | NA | EU599041 | expressed sequence EU599041 [Source:M | 0 | 0 | 0 | 0 | 0 | 0 |
| 21001 | ENSMUSG00000074890 | 197.9111703 | -0.066094154 | 0.18162964 | -0.3638952 | 0.715936269 | 0.9447719 | Lcmt2 | leucine carboxyl methyltransferase 2 [Sou | 234.51 | 186.05 | 186.16 | 199.54 | 196.31 | 184.9 |
| 21002 | ENSMUSG00000074892 | 0 | NA | NA | NA | NA | NA | B3galt5 | UDP-Gal:betaGlcNAc beta 1,3-galactosyltr | 0 | 0 | 0 | 0 | 0 | 0 |
| 21003 | ENSMUSG00000074895 | 0 | NA | NA | NA | NA | NA | Eif4e1b | eukaryotic translation initiation factor 4E f | 0 | 0 | 0 | 0 | 0 | 0 |
| 21004 | ENSMUSG00000074896 | 284.2573669 | -0.082750617 | 0.251607129 | -0.328888203 | 0.742240191 | 0.949043964 | Ifit3 | interferon-induced protein with tetratrico | 403.96 | 251.84 | 220.37 | 230.39 | 378.44 | 220.54 |
| 21005 | ENSMUSG00000074899 | 1.145457819 | -1.111308189 | 1.995909703 | -0.556792818 | 0.577668991 | NA | Sptbn5 | spectrin beta, non-erythrocytic 5 [Source: | 1.6062 | 2.0558 | 1.0062 | 0 | 1.0906 | 1.1138 |
| 21006 | ENSMUSG00000074903 | 0 | NA | NA | NA | NA | NA | Gm2058 | predicted gene 2058 [Source:MGI Symbol | 0 | 0 | 0 | 0 | 0 | 0 |
| 21007 | ENSMUSG00000074909 | 379.0370271 | 0.190877666 | 0.144150169 | 1.324158463 | 0.185450423 | 0.621059253 | Ranbp6 | RAN binding protein 6 [Source:MGI Symb | 348.55 | 322.77 | 390.42 | 375.95 | 416.62 | 419.92 |
| 21008 | ENSMUSG00000074912 | 4.035798621 | 0.668071885 | 1.043482991 | 0.640232654 | 0.522021356 | NA | Gm14207 | predicted gene 14207 [Source:MGI Symbo | 3.2124 | 4.1117 | 2.0125 | 3.8559 | 5.4531 | 5.5692 |
| 21009 | ENSMUSG00000074913 | 0 | NA | NA | NA | NA | NA | Gm815 | predicted gene 815 [Source:MGI Symbol;A | 0 | 0 | 0 | 0 | 0 | 0 |
| 21010 | ENSMUSG00000074916 | 17.20756748 | -0.66634061 | 0.541526212 | -1.230486345 | 0.218515037 | 0.663735712 | Chst14 | carbohydrate sulfotransferase 14 [Source | 21.684 | 18.503 | 23.144 | 14.46 | 7.6343 | 17.822 |
| 21011 | ENSMUSG00000074918 | 250.4657517 | -0.172162041 | 0.158066753 | -1.08917301 | 0.276077599 | 0.730038567 | Inafm2 | InaF motif containing 2 [Source:MGI Symb | 268.24 | 252.87 | 274.7 | 220.75 | 233.39 | 252.84 |
| 21012 | ENSMUSG00000074919 | 2.172317201 | 0.387375938 | 1.539211634 | 0.251671654 | 0.801294868 | NA | Gm10787 | predicted gene 10787 [Source:MGI Symbo | 1.6062 | 1.0279 | 3.0187 | 1.928 | 5.4531 | 0 |
| 21013 | ENSMUSG00000074922 | 182.4607415 | -0.138705108 | 0.185352321 | -0.748332188 | 0.454259814 | 0.857337687 | Fam122a | family with sequence similarity 122, meml | 166.24 | 207.64 | 200.24 | 164.84 | 186.5 | 169.3 |
| 21014 | ENSMUSG00000074923 | 4.155722593 | -1.29544874 | 1.087191355 | -1.19155541 | 0.233435618 | NA | Pak6 | p21 (RAC1) activated kinase 6 [Source:MG | 5.6217 | 4.1117 | 8.05 | 3.8559 | 2.1812 | 1.1138 |
| 21015 | ENSMUSG00000074925 | 34.66248661 | 0.178722598 | 0.372397303 | 0.479924522 | 0.631281064 | 0.921377326 | Ptar1 | protein prenyltransferase alpha subunit re | 30.518 | 33.921 | 33.206 | 36.631 | 43.625 | 30.074 |
| 21016 | ENSMUSG00000074928 | 0 | NA | NA | NA | NA | NA | Krtap14 | keratin associated protein 14 [Source:MG | 0 | 0 | 0 | 0 | 0 | 0 |
| 21017 | ENSMUSG00000074930 | 0 | NA | NA | NA | NA | NA | Gm13981 | predicted gene 13981 [Source:MGI Symbo | 0 | 0 | 0 | 0 | 0 | 0 |
| 21018 | ENSMUSG00000074934 | 0 | NA | NA | NA | NA | NA | Grem1 | gremlin 1, DAN family BMP antagonist [So | 0 | 0 | 0 | 0 | 0 | 0 |
| 21019 | ENSMUSG00000074939 | 0 | NA | NA | NA | NA | NA | Chrm5 | cholinergic receptor, muscarinic 5 [Source | 0 | 0 | 0 | 0 | 0 | 0 |
| 21020 | ENSMUSG00000074945 | 0 | NA | NA | NA | NA | NA | Olfr1314 | olfactory receptor 1314 [Source:MGI Syml | 0 | 0 | 0 | 0 | 0 | 0 |
| 21021 | ENSMUSG00000074946 | 0 | NA | NA | NA | NA | NA | Olfr1313 | olfactory receptor 1313 [Source:MGI Syml | 0 | 0 | 0 | 0 | 0 | 0 |
| 21022 | ENSMUSG00000074947 | 0 | NA | NA | NA | NA | NA | Olfr1312 | olfactory receptor 1312 [Source:MGI Syml | 0 | 0 | 0 | 0 | 0 | 0 |
| 21023 | ENSMUSG00000074952 | 0 | NA | NA | NA | NA | NA | Olfr1308 | olfactory receptor 1308 [Source:MGI Syml | 0 | 0 | 0 | 0 | 0 | 0 |
| 21024 | ENSMUSG00000074955 | 0 | NA | NA | NA | NA | NA | Olfr1305 | olfactory receptor 1305 [Source:MGI Syml | 0 | 0 | 0 | 0 | 0 | 0 |
| 21025 | ENSMUSG00000074965 | 0 | NA | NA | NA | NA | NA | Olfr1277 | olfactory receptor 1277 [Source:MGI Syml | 0 | 0 | 0 | 0 | 0 | 0 |
| 21026 | ENSMUSG00000074966 | 0 | NA | NA | NA | NA | NA | Olfr1275 | olfactory receptor 1275 [Source:MGI Syml | 0 | 0 | 0 | 0 | 0 | 0 |
| 21027 | ENSMUSG00000074968 | 0 | NA | NA | NA | NA | NA | Ano3 | anoctamin 3 [Source:MGI Symbol;Acc:MG | 0 | 0 | 0 | 0 | 0 | 0 |
| 21028 | ENSMUSG00000074970 | 0 | NA | NA | NA | NA | NA | Gm10794 | predicted gene 10794 [Source:MGI Symbo | 0 | 0 | 0 | 0 | 0 | 0 |
| 21029 | ENSMUSG00000074971 | 22.16199588 | -0.617848721 | 0.467975635 | -1.320258309 | 0.18674879 | 0.623054278 | Fibin | fin bud initiation factor homolog (zebrafis | 30.518 | 29.81 | 20.125 | 18.316 | 15.269 | 18.935 |
| 21030 | ENSMUSG00000074973 | 0 | NA | NA | NA | NA | NA | Gm11382 | predicted pseudogene 11382 [Source:MG | 0 | 0 | 0 | 0 | 0 | 0 |
| 21031 | ENSMUSG00000074978 | 0 | NA | NA | NA | NA | NA | Actg-ps1 | actin, gamma, pseudogene 1 [Source:MGI | 0 | 0 | 0 | 0 | 0 | 0 |
| 21032 | ENSMUSG00000074981 | 0 | NA | NA | NA | NA | NA | Dcdc5 | doublecortin domain containing 5 [Source | 0 | 0 | 0 | 0 | 0 | 0 |
| 21033 | ENSMUSG00000074987 | 0.133850373 | -0.796629176 | 4.080472857 | -0.195229623 | 0.84521319 | NA | Wt1os | Wilms tumor 1 homolog, opposite strand | 0.8031 | 0 | 0 | 0 | 0 | 0 |
| 21034 | ENSMUSG00000074991 | 0 | NA | NA | NA | NA | NA | Gabrr3 | gamma-aminobutyric acid (GABA) recepto | 0 | 0 | 0 | 0 | 0 | 0 |
| 21035 | ENSMUSG00000074994 | 301.7916062 | -0.030748966 | 0.15391533 | -0.199778452 | 0.841653854 | 0.971854408 | Qser1 | glutamine and serine rich 1 [Source:MGI S | 300.36 | 306.32 | 307.91 | 262.2 | 305.37 | 328.58 |
| 21036 | ENSMUSG00000074995 | 0 | NA | NA | NA | NA | NA | Olfr201 | olfactory receptor 201 [Source:MGI Symb | 0 | 0 | 0 | 0 | 0 | 0 |
| 21037 | ENSMUSG00000074996 | 0 | NA | NA | NA | NA | NA | Olfr199 | olfactory receptor 199 [Source:MGI Symb | 0 | 0 | 0 | 0 | 0 | 0 |
| 21038 | ENSMUSG00000074997 | 0 | NA | NA | NA | NA | NA | Pin1rt1 | protein (peptidyl-prolyl cis/trans isomeras | 0 | 0 | 0 | 0 | 0 | 0 |
| 21039 | ENSMUSG00000074999 | 0 | NA | NA | NA | NA | NA | Gm10797 | predicted gene 10797 [Source:MGI Symbo | 0 | 0 | 0 | 0 | 0 | 0 |
| 21040 | ENSMUSG00000075000 | 468.9615986 | -0.374093566 | 0.130668423 | -2.862922489 | 0.004197532 | 0.059406172 | Nrbf2 | nuclear receptor binding factor 2 [Source: | 509.97 | 527.32 | 551.42 | 414.51 | 434.07 | 376.48 |
| 21041 | ENSMUSG00000075002 | 0.535401493 | -2.48596965 | 3.684117979 | -0.674780141 | 0.499815459 | NA | Ftdc1 | ferritin domain containing 1 [Source:MGI | 3.2124 | 0 | 0 | 0 | 0 | 0 |
| 21042 | ENSMUSG00000075006 | 0 | NA | NA | NA | NA | NA | Gm10799 | predicted gene 10799 [Source:MGI Symbo | 0 | 0 | 0 | 0 | 0 | 0 |
| 21043 | ENSMUSG00000075010 | 1048.974821 | -0.136207834 | 0.107598824 | -1.2658859 | 0.205553935 | 0.650907725 | AW112010 | expressed sequence AW112010 [Source:N | 1149.2 | 1038.2 | 1106.9 | 952.41 | 999.01 | 1048.1 |
| 21044 | ENSMUSG00000075012 | 2.211155781 | -1.052467226 | 1.569296879 | -0.670661645 | 0.502436102 | NA | Fjx1 | four jointed box 1 [Source:MGI Symbol;Ac | 0.8031 | 5.1396 | 3.0187 | 0.964 | 0 | 3.3415 |
| 21045 | ENSMUSG00000075014 | 0.315620088 | 0.165143421 | 4.063300985 | 0.040642675 | 0.967580763 | NA | Gm10800 | predicted gene 10800 [Source:MGI Symbo | 0.8031 | 0 | 0 | 0 | 1.0906 | 0 |
| 21046 | ENSMUSG00000075015 | 0 | NA | NA | NA | NA | NA | Gm10801 | predicted gene 10801 [Source:MGI Symbo | 0 | 0 | 0 | 0 | 0 | 0 |
| 21047 | ENSMUSG00000075020 | 17.25660714 | -1.542647214 | 0.6279749 | -2.456542792 | 0.014028109 | 0.136450958 | Mir670hg | MIR670 host gene (non-protein coding) [S | 30.518 | 25.698 | 21.131 | 17.352 | 3.2719 | 5.5692 |
| 21048 | ENSMUSG00000075023 | 0 | NA | NA | NA | NA | NA | Accsl | 1-aminocyclopropane-1-carboxylate synth | 0 | 0 | 0 | 0 | 0 | 0 |
| 21049 | ENSMUSG00000075024 | 0 | NA | NA | NA | NA | NA | NA | NA | 0 | 0 | 0 | 0 | 0 | 0 |
| 21050 | ENSMUSG00000075025 | 35.60713311 | -0.532936422 | 0.376253279 | -1.416429974 | 0.156649653 | 0.575786484 | Gm10804 | predicted gene 10804 [Source:MGI Symbo | 41.761 | 51.396 | 33.206 | 28.919 | 31.628 | 26.732 |
| 21051 | ENSMUSG00000075027 | 0.353089354 | 0.165142706 | 3.896102244 | 0.042386646 | 0.966190474 | NA | 4631405J19I | RIKEN cDNA 4631405J19 gene [Source:MC | 0 | 1.0279 | 0 | 0 | 1.0906 | 0 |
| 21052 | ENSMUSG00000075028 | 20.16670085 | -0.61669126 | 0.499384056 | -1.234903783 | 0.216866316 | 0.662741261 | Prdm11 | PR domain containing 11 [Source:MGI Syr | 19.274 | 31.865 | 22.137 | 13.496 | 14.178 | 20.049 |

| | | | | | | | | | | | | | | | |
|---|---|---|---|---|---|---|---|---|---|---|---|---|---|---|---|
| 21053 | ENSMUSG00000075029 | 0.160662974 | 1.126924175 | 4.080472857 | 0.276174898 | 0.782413736 | NA | 4930558J22I | RIKEN cDNA 4930558J22 gene [Source:M( | 0 | 0 | 0 | 0.964 | 0 | 0 |
| 21054 | ENSMUSG00000075031 | 0.89478629 | 2.103406 | 2.814836302 | 0.747256953 | 0.454908471 | NA | H2bc3 | H2B clustered histone 3 [Source:MGI Sym | 0 | 0 | 1.0062 | 0 | 4.3625 | 0 |
| 21055 | ENSMUSG00000075033 | 7.893636047 | -1.131674089 | 0.862789388 | -1.311645814 | 0.189639668 | 0.626663447 | Nxpe3 | neurexophilin and PC-esterase domain fai | 3.2124 | 10.279 | 19.119 | 4.8199 | 4.3625 | 5.5692 |
| 21056 | ENSMUSG00000075036 | 0 | NA | NA | NA | NA | NA | Gm10805 | predicted gene 10805 [Source:MGI Symbo | 0 | 0 | 0 | 0 | 0 | 0 |
| 21057 | ENSMUSG00000075040 | 84.31419847 | 0.302593226 | 0.274995584 | 1.100356673 | 0.271176749 | 0.72689038 | Zfp408 | zinc finger protein 408 [Source:MGI Symb | 75.492 | 67.843 | 83.518 | 120.5 | 87.249 | 71.286 |
| 21058 | ENSMUSG00000075042 | 0 | NA | NA | NA | NA | NA | 4930431P03 | RIKEN cDNA 4930431P03 gene [Source:M | 0 | 0 | 0 | 0 | 0 | 0 |
| 21059 | ENSMUSG00000075044 | 49.35472403 | -2.084582174 | 0.436192019 | -4.779047034 | 1.76E-06 | 0.000129183 | Slc22a29 | solute carrier family 22. member 29 [Sour | 104.4 | 84.289 | 51.318 | 30.847 | 16.359 | 8.9108 |
| 21060 | ENSMUSG00000075045 | 0.167707431 | -0.796629176 | 4.080472857 | -0.195229623 | 0.84521319 | NA | Gm4981 | predicted gene 4981 [Source:MGI Symbol | 0 | 0 | 1.0062 | 0 | 0 | 0 |
| 21061 | ENSMUSG00000075046 | 0 | NA | NA | NA | NA | NA | Duxf3 | double homeobox family member 3 [Sour | 0 | 0 | 0 | 0 | 0 | 0 |
| 21062 | ENSMUSG00000075053 | 4.414415241 | -0.102242752 | 1.036802429 | -0.098613535 | 0.921445123 | NA | Vdac3-ps1 | voltage-dependent anion channel 3, pseu | 1.6062 | 5.1396 | 7.0437 | 3.8559 | 3.2719 | 5.5692 |
| 21063 | ENSMUSG00000075054 | 288.654414 | -0.099962038 | 0.15851947 | -0.630597857 | 0.5283035 | 0.888334265 | Yae1d1 | Yae1 domain containing 1 [Source:MGI Sy | 267.43 | 301.18 | 328.04 | 282.45 | 296.65 | 256.18 |
| 21064 | ENSMUSG00000075061 | 0 | NA | NA | NA | NA | NA | Olfr1272 | olfactory receptor 1272 [Source:MGI Syml | 0 | 0 | 0 | 0 | 0 | 0 |
| 21065 | ENSMUSG00000075062 | 0 | NA | NA | NA | NA | NA | Olfr1271 | olfactory receptor 1271 [Source:MGI Syml | 0 | 0 | 0 | 0 | 0 | 0 |
| 21066 | ENSMUSG00000075063 | 0 | NA | NA | NA | NA | NA | Olfr142 | olfactory receptor 142 [Source:MGI Symb | 0 | 0 | 0 | 0 | 0 | 0 |
| 21067 | ENSMUSG00000075064 | 0 | NA | NA | NA | NA | NA | Olfr1506 | olfactory receptor 1506 [Source:MGI Syml | 0 | 0 | 0 | 0 | 0 | 0 |
| 21068 | ENSMUSG00000075065 | 0 | NA | NA | NA | NA | NA | Olfr1270 | olfactory receptor 1270 [Source:MGI Syml | 0 | 0 | 0 | 0 | 0 | 0 |
| 21069 | ENSMUSG00000075066 | 0 | NA | NA | NA | NA | NA | Olfr32 | olfactory receptor 32 [Source:MGI Symbo | 0 | 0 | 0 | 0 | 0 | 0 |
| 21070 | ENSMUSG00000075067 | 0 | NA | NA | NA | NA | NA | Olfr1267-ps1 | olfactory receptor 1267, pseudogene 1 [S( | 0 | 0 | 0 | 0 | 0 | 0 |
| 21071 | ENSMUSG00000075068 | 0 | NA | NA | NA | NA | NA | Olfr140 | olfactory receptor 140 [Source:MGI Symb | 0 | 0 | 0 | 0 | 0 | 0 |
| 21072 | ENSMUSG00000075069 | 0 | NA | NA | NA | NA | NA | Olfr1264 | olfactory receptor 1264 [Source:MGI Syml | 0 | 0 | 0 | 0 | 0 | 0 |
| 21073 | ENSMUSG00000075070 | 0 | NA | NA | NA | NA | NA | 4932412D23 | RIKEN cDNA 4932412D23 gene [Source:M | 0 | 0 | 0 | 0 | 0 | 0 |
| 21074 | ENSMUSG00000075072 | 0 | NA | NA | NA | NA | NA | Olfr48 | olfactory receptor 48 [Source:MGI Symbo | 0 | 0 | 0 | 0 | 0 | 0 |
| 21075 | ENSMUSG00000075073 | 0 | NA | NA | NA | NA | NA | Olfr1256 | olfactory receptor 1256 [Source:MGI Syml | 0 | 0 | 0 | 0 | 0 | 0 |
| 21076 | ENSMUSG00000075074 | 0 | NA | NA | NA | NA | NA | Olfr1254 | olfactory receptor 1254 [Source:MGI Syml | 0 | 0 | 0 | 0 | 0 | 0 |
| 21077 | ENSMUSG00000075075 | 0 | NA | NA | NA | NA | NA | Olfr1253 | olfactory receptor 1253 [Source:MGI Syml | 0 | 0 | 0 | 0 | 0 | 0 |
| 21078 | ENSMUSG00000075078 | 0 | NA | NA | NA | NA | NA | Olfr1250 | olfactory receptor 1250 [Source:MGI Syml | 0 | 0 | 0 | 0 | 0 | 0 |
| 21079 | ENSMUSG00000075079 | 0 | NA | NA | NA | NA | NA | Olfr1249 | olfactory receptor 1249 [Source:MGI Syml | 0 | 0 | 0 | 0 | 0 | 0 |
| 21080 | ENSMUSG00000075081 | 0 | NA | NA | NA | NA | NA | Olfr1247 | olfactory receptor 1247 [Source:MGI Syml | 0 | 0 | 0 | 0 | 0 | 0 |
| 21081 | ENSMUSG00000075084 | 0 | NA | NA | NA | NA | NA | Olfr1243 | olfactory receptor 1243 [Source:MGI Syml | 0 | 0 | 0 | 0 | 0 | 0 |
| 21082 | ENSMUSG00000075085 | 0 | NA | NA | NA | NA | NA | Olfr1242 | olfactory receptor 1242 [Source:MGI Syml | 0 | 0 | 0 | 0 | 0 | 0 |
| 21083 | ENSMUSG00000075086 | 0 | NA | NA | NA | NA | NA | Olfr1241 | olfactory receptor 1241 [Source:MGI Syml | 0 | 0 | 0 | 0 | 0 | 0 |
| 21084 | ENSMUSG00000075088 | 0 | NA | NA | NA | NA | NA | Olfr1239 | olfactory receptor 1239 [Source:MGI Syml | 0 | 0 | 0 | 0 | 0 | 0 |
| 21085 | ENSMUSG00000075090 | 0 | NA | NA | NA | NA | NA | Olfr1234 | olfactory receptor 1234 [Source:MGI Syml | 0 | 0 | 0 | 0 | 0 | 0 |
| 21086 | ENSMUSG00000075091 | 0 | NA | NA | NA | NA | NA | Olfr1233 | olfactory receptor 1233 [Source:MGI Syml | 0 | 0 | 0 | 0 | 0 | 0 |
| 21087 | ENSMUSG00000075092 | 0 | NA | NA | NA | NA | NA | Olfr1232 | olfactory receptor 1232 [Source:MGI Syml | 0 | 0 | 0 | 0 | 0 | 0 |
| 21088 | ENSMUSG00000075093 | 0 | NA | NA | NA | NA | NA | Olfr1231 | olfactory receptor 1231 [Source:MGI Syml | 0 | 0 | 0 | 0 | 0 | 0 |
| 21089 | ENSMUSG00000075094 | 0 | NA | NA | NA | NA | NA | Olfr1230 | olfactory receptor 1230 [Source:MGI Syml | 0 | 0 | 0 | 0 | 0 | 0 |
| 21090 | ENSMUSG00000075095 | 0 | NA | NA | NA | NA | NA | Olfr1229 | olfactory receptor 1229 [Source:MGI Syml | 0 | 0 | 0 | 0 | 0 | 0 |
| 21091 | ENSMUSG00000075097 | 0 | NA | NA | NA | NA | NA | Olfr1226 | olfactory receptor 1226 [Source:MGI Syml | 0 | 0 | 0 | 0 | 0 | 0 |
| 21092 | ENSMUSG00000075099 | 0 | NA | NA | NA | NA | NA | Olfr1224-ps1 | olfactory receptor 1224, pseudogene 1 [S( | 0 | 0 | 0 | 0 | 0 | 0 |
| 21093 | ENSMUSG00000075100 | 0 | NA | NA | NA | NA | NA | Olfr1223 | olfactory receptor 1223 [Source:MGI Syml | 0 | 0 | 0 | 0 | 0 | 0 |
| 21094 | ENSMUSG00000075101 | 0 | NA | NA | NA | NA | NA | Olfr1222 | olfactory receptor 1222 [Source:MGI Syml | 0 | 0 | 0 | 0 | 0 | 0 |
| 21095 | ENSMUSG00000075102 | 0 | NA | NA | NA | NA | NA | Olfr1221 | olfactory receptor 1221 [Source:MGI Syml | 0 | 0 | 0 | 0 | 0 | 0 |
| 21096 | ENSMUSG00000075104 | 0 | NA | NA | NA | NA | NA | Olfr1219 | olfactory receptor 1219 [Source:MGI Syml | 0 | 0 | 0 | 0 | 0 | 0 |
| 21097 | ENSMUSG00000075105 | 0 | NA | NA | NA | NA | NA | Olfr1218 | olfactory receptor 1218 [Source:MGI Syml | 0 | 0 | 0 | 0 | 0 | 0 |
| 21098 | ENSMUSG00000075107 | 0 | NA | NA | NA | NA | NA | Olfr1216 | olfactory receptor 1216 [Source:MGI Syml | 0 | 0 | 0 | 0 | 0 | 0 |
| 21099 | ENSMUSG00000075110 | 0 | NA | NA | NA | NA | NA | Olfr1214 | olfactory receptor 1214 [Source:MGI Syml | 0 | 0 | 0 | 0 | 0 | 0 |
| 21100 | ENSMUSG00000075111 | 0 | NA | NA | NA | NA | NA | Olfr1213 | olfactory receptor 1213 [Source:MGI Syml | 0 | 0 | 0 | 0 | 0 | 0 |
| 21101 | ENSMUSG00000075112 | 0 | NA | NA | NA | NA | NA | Olfr1211 | olfactory receptor 1211 [Source:MGI Syml | 0 | 0 | 0 | 0 | 0 | 0 |
| 21102 | ENSMUSG00000075113 | 0 | NA | NA | NA | NA | NA | Olfr1209 | olfactory receptor 1209 [Source:MGI Syml | 0 | 0 | 0 | 0 | 0 | 0 |
| 21103 | ENSMUSG00000075114 | 0 | NA | NA | NA | NA | NA | Olfr1208 | olfactory receptor 1208 [Source:MGI Syml | 0 | 0 | 0 | 0 | 0 | 0 |
| 21104 | ENSMUSG00000075115 | 0 | NA | NA | NA | NA | NA | Olfr1200 | olfactory receptor 1200 [Source:MGI Syml | 0 | 0 | 0 | 0 | 0 | 0 |
| 21105 | ENSMUSG00000075117 | 0 | NA | NA | NA | NA | NA | Olfr1198 | olfactory receptor 1198 [Source:MGI Syml | 0 | 0 | 0 | 0 | 0 | 0 |
| 21106 | ENSMUSG00000075119 | 0 | NA | NA | NA | NA | NA | Olfr1197 | olfactory receptor 1197 [Source:MGI Syml | 0 | 0 | 0 | 0 | 0 | 0 |
| 21107 | ENSMUSG00000075120 | 0 | NA | NA | NA | NA | NA | Olfr1196 | olfactory receptor 1196 [Source:MGI Syml | 0 | 0 | 0 | 0 | 0 | 0 |
| 21108 | ENSMUSG00000075121 | 0 | NA | NA | NA | NA | NA | Olfr1195 | olfactory receptor 1195 [Source:MGI Syml | 0 | 0 | 0 | 0 | 0 | 0 |

| | | | | | | | | | | | | | | | |
|---|---|---|---|---|---|---|---|---|---|---|---|---|---|---|---|
| 21109 | ENSMUSG00000075122 | 8.135989489 | -0.642941254 | 0.757847535 | -0.848378103 | 0.396227434 | 0.823764863 | Cd80 | CD80 antigen [Source:MGI Symbol;Acc:M | 6.4248 | 12.335 | 11.069 | 5.7839 | 7.6343 | 5.5692 |
| 21110 | ENSMUSG00000075125 | 0 | NA | NA | NA | NA | NA | Olfr1181 | olfactory receptor 1181 [Source:MGI Sym | 0 | 0 | 0 | 0 | 0 | 0 |
| 21111 | ENSMUSG00000075126 | 0 | NA | NA | NA | NA | NA | Olfr475-ps1 | olfactory receptor 475, pseudogene 1 [So | 0 | 0 | 0 | 0 | 0 | 0 |
| 21112 | ENSMUSG00000075127 | 0 | NA | NA | NA | NA | NA | Olfr1179 | olfactory receptor 1179 [Source:MGI Sym | 0 | 0 | 0 | 0 | 0 | 0 |
| 21113 | ENSMUSG00000075128 | 0 | NA | NA | NA | NA | NA | Olfr1177-ps | olfactory receptor 1177, pseudogene [Sou | 0 | 0 | 0 | 0 | 0 | 0 |
| 21114 | ENSMUSG00000075132 | 0 | NA | NA | NA | NA | NA | Olfr1173 | olfactory receptor 1173 [Source:MGI Sym | 0 | 0 | 0 | 0 | 0 | 0 |
| 21115 | ENSMUSG00000075133 | 0 | NA | NA | NA | NA | NA | Olfr1170 | olfactory receptor 1170 [Source:MGI Sym | 0 | 0 | 0 | 0 | 0 | 0 |
| 21116 | ENSMUSG00000075136 | 0 | NA | NA | NA | NA | NA | Olfr1164 | olfactory receptor 1164 [Source:MGI Sym | 0 | 0 | 0 | 0 | 0 | 0 |
| 21117 | ENSMUSG00000075137 | 0 | NA | NA | NA | NA | NA | Olfr1163 | olfactory receptor 1163 [Source:MGI Sym | 0 | 0 | 0 | 0 | 0 | 0 |
| 21118 | ENSMUSG00000075139 | 0 | NA | NA | NA | NA | NA | Olfr1162 | olfactory receptor 1162 [Source:MGI Sym | 0 | 0 | 0 | 0 | 0 | 0 |
| 21119 | ENSMUSG00000075140 | 0 | NA | NA | NA | NA | NA | Olfr73 | olfactory receptor 73 [Source:MGI Symbo | 0 | 0 | 0 | 0 | 0 | 0 |
| 21120 | ENSMUSG00000075141 | 0 | NA | NA | NA | NA | NA | Olfr1160 | olfactory receptor 1160 [Source:MGI Sym | 0 | 0 | 0 | 0 | 0 | 0 |
| 21121 | ENSMUSG00000075142 | 0 | NA | NA | NA | NA | NA | Olfr74 | olfactory receptor 74 [Source:MGI Symbo | 0 | 0 | 0 | 0 | 0 | 0 |
| 21122 | ENSMUSG00000075143 | 0 | NA | NA | NA | NA | NA | Olfr1157 | olfactory receptor 1157 [Source:MGI Sym | 0 | 0 | 0 | 0 | 0 | 0 |
| 21123 | ENSMUSG00000075144 | 0 | NA | NA | NA | NA | NA | Olfr1156 | olfactory receptor 1156 [Source:MGI Sym | 0 | 0 | 0 | 0 | 0 | 0 |
| 21124 | ENSMUSG00000075145 | 0 | NA | NA | NA | NA | NA | Olfr1155 | olfactory receptor 1155 [Source:MGI Sym | 0 | 0 | 0 | 0 | 0 | 0 |
| 21125 | ENSMUSG00000075146 | 0 | NA | NA | NA | NA | NA | Olfr1154 | olfactory receptor 1154 [Source:MGI Sym | 0 | 0 | 0 | 0 | 0 | 0 |
| 21126 | ENSMUSG00000075148 | 0 | NA | NA | NA | NA | NA | Olfr1141 | olfactory receptor 1141 [Source:MGI Sym | 0 | 0 | 0 | 0 | 0 | 0 |
| 21127 | ENSMUSG00000075149 | 0 | NA | NA | NA | NA | NA | Olfr1138 | olfactory receptor 1138 [Source:MGI Sym | 0 | 0 | 0 | 0 | 0 | 0 |
| 21128 | ENSMUSG00000075150 | 0 | NA | NA | NA | NA | NA | Olfr1137 | olfactory receptor 1137 [Source:MGI Sym | 0 | 0 | 0 | 0 | 0 | 0 |
| 21129 | ENSMUSG00000075151 | 0 | NA | NA | NA | NA | NA | Olfr1136 | olfactory receptor 1136 [Source:MGI Sym | 0 | 0 | 0 | 0 | 0 | 0 |
| 21130 | ENSMUSG00000075153 | 0 | NA | NA | NA | NA | NA | Olfr1135 | olfactory receptor 1135 [Source:MGI Sym | 0 | 0 | 0 | 0 | 0 | 0 |
| 21131 | ENSMUSG00000075155 | 0 | NA | NA | NA | NA | NA | Olfr1133 | olfactory receptor 1133 [Source:MGI Sym | 0 | 0 | 0 | 0 | 0 | 0 |
| 21132 | ENSMUSG00000075156 | 0 | NA | NA | NA | NA | NA | Olfr1128 | olfactory receptor 1128 [Source:MGI Sym | 0 | 0 | 0 | 0 | 0 | 0 |
| 21133 | ENSMUSG00000075158 | 0 | NA | NA | NA | NA | NA | Olfr1111 | olfactory receptor 1111 [Source:MGI Sym | 0 | 0 | 0 | 0 | 0 | 0 |
| 21134 | ENSMUSG00000075159 | 0 | NA | NA | NA | NA | NA | Olfr1110 | olfactory receptor 1110 [Source:MGI Sym | 0 | 0 | 0 | 0 | 0 | 0 |
| 21135 | ENSMUSG00000075160 | 0 | NA | NA | NA | NA | NA | Olfr259 | olfactory receptor 259 [Source:MGI Symb | 0 | 0 | 0 | 0 | 0 | 0 |
| 21136 | ENSMUSG00000075161 | 0 | NA | NA | NA | NA | NA | Olfr1109 | olfactory receptor 1109 [Source:MGI Sym | 0 | 0 | 0 | 0 | 0 | 0 |
| 21137 | ENSMUSG00000075162 | 0 | NA | NA | NA | NA | NA | Gm10812 | predicted gene 10812 [Source:MGI Symbo | 0 | 0 | 0 | 0 | 0 | 0 |
| 21138 | ENSMUSG00000075163 | 0 | NA | NA | NA | NA | NA | Olfr1107 | olfactory receptor 1107 [Source:MGI Sym | 0 | 0 | 0 | 0 | 0 | 0 |
| 21139 | ENSMUSG00000075164 | 0 | NA | NA | NA | NA | NA | Olfr1106 | olfactory receptor 1106 [Source:MGI Sym | 0 | 0 | 0 | 0 | 0 | 0 |
| 21140 | ENSMUSG00000075165 | 0 | NA | NA | NA | NA | NA | Olfr1105 | olfactory receptor 1105 [Source:MGI Sym | 0 | 0 | 0 | 0 | 0 | 0 |
| 21141 | ENSMUSG00000075166 | 0 | NA | NA | NA | NA | NA | Olfr1104 | olfactory receptor 1104 [Source:MGI Sym | 0 | 0 | 0 | 0 | 0 | 0 |
| 21142 | ENSMUSG00000075167 | 0 | NA | NA | NA | NA | NA | Olfr1101 | olfactory receptor 1101 [Source:MGI Sym | 0 | 0 | 0 | 0 | 0 | 0 |
| 21143 | ENSMUSG00000075168 | 0 | NA | NA | NA | NA | NA | Olfr1099 | olfactory receptor 1099 [Source:MGI Sym | 0 | 0 | 0 | 0 | 0 | 0 |
| 21144 | ENSMUSG00000075169 | 0 | NA | NA | NA | NA | NA | Olfr1098 | olfactory receptor 1098 [Source:MGI Sym | 0 | 0 | 0 | 0 | 0 | 0 |
| 21145 | ENSMUSG00000075170 | 0 | NA | NA | NA | NA | NA | Olfr1097 | olfactory receptor 1097 [Source:MGI Sym | 0 | 0 | 0 | 0 | 0 | 0 |
| 21146 | ENSMUSG00000075171 | 0 | NA | NA | NA | NA | NA | Olfr1095 | olfactory receptor 1095 [Source:MGI Sym | 0 | 0 | 0 | 0 | 0 | 0 |
| 21147 | ENSMUSG00000075172 | 0 | NA | NA | NA | NA | NA | Olfr1090 | olfactory receptor 1090 [Source:MGI Sym | 0 | 0 | 0 | 0 | 0 | 0 |
| 21148 | ENSMUSG00000075174 | 0 | NA | NA | NA | NA | NA | Olfr1087 | olfactory receptor 1087 [Source:MGI Sym | 0 | 0 | 0 | 0 | 0 | 0 |
| 21149 | ENSMUSG00000075175 | 0 | NA | NA | NA | NA | NA | Olfr1086 | olfactory receptor 1086 [Source:MGI Sym | 0 | 0 | 0 | 0 | 0 | 0 |
| 21150 | ENSMUSG00000075176 | 0 | NA | NA | NA | NA | NA | Olfr1085 | olfactory receptor 1085 [Source:MGI Sym | 0 | 0 | 0 | 0 | 0 | 0 |
| 21151 | ENSMUSG00000075179 | 0 | NA | NA | NA | NA | NA | Olfr1079 | olfactory receptor 1079 [Source:MGI Sym | 0 | 0 | 0 | 0 | 0 | 0 |
| 21152 | ENSMUSG00000075181 | 0 | NA | NA | NA | NA | NA | Olfr1066 | olfactory receptor 1066 [Source:MGI Sym | 0 | 0 | 0 | 0 | 0 | 0 |
| 21153 | ENSMUSG00000075184 | 1.117074527 | -3.52147603 | 2.24739728 | -1.566913007 | 0.117135027 | NA | F930017D23 | RIKEN cDNA F930017D23 gene [Source:M | 1.6062 | 3.0838 | 2.0125 | 0 | 0 | 0 |
| 21154 | ENSMUSG00000075185 | 0 | NA | NA | NA | NA | NA | Olfr1061 | olfactory receptor 1061 [Source:MGI Sym | 0 | 0 | 0 | 0 | 0 | 0 |
| 21155 | ENSMUSG00000075186 | 0 | NA | NA | NA | NA | NA | Olfr1058 | olfactory receptor 1058 [Source:MGI Sym | 0 | 0 | 0 | 0 | 0 | 0 |
| 21156 | ENSMUSG00000075187 | 0 | NA | NA | NA | NA | NA | Olfr1057 | olfactory receptor 1057 [Source:MGI Sym | 0 | 0 | 0 | 0 | 0 | 0 |
| 21157 | ENSMUSG00000075188 | 0 | NA | NA | NA | NA | NA | Olfr1056 | olfactory receptor 1056 [Source:MGI Sym | 0 | 0 | 0 | 0 | 0 | 0 |
| 21158 | ENSMUSG00000075189 | 0 | NA | NA | NA | NA | NA | Olfr1055 | olfactory receptor 1055 [Source:MGI Sym | 0 | 0 | 0 | 0 | 0 | 0 |
| 21159 | ENSMUSG00000075190 | 0 | NA | NA | NA | NA | NA | Olfr1054 | olfactory receptor 1054 [Source:MGI Sym | 0 | 0 | 0 | 0 | 0 | 0 |
| 21160 | ENSMUSG00000075192 | 0 | NA | NA | NA | NA | NA | Olfr1053 | olfactory receptor 1053 [Source:MGI Sym | 0 | 0 | 0 | 0 | 0 | 0 |
| 21161 | ENSMUSG00000075193 | 0 | NA | NA | NA | NA | NA | Olfr1051 | olfactory receptor 1051 [Source:MGI Sym | 0 | 0 | 0 | 0 | 0 | 0 |
| 21162 | ENSMUSG00000075194 | 0 | NA | NA | NA | NA | NA | Olfr1049 | olfactory receptor 1049 [Source:MGI Sym | 0 | 0 | 0 | 0 | 0 | 0 |
| 21163 | ENSMUSG00000075195 | 0 | NA | NA | NA | NA | NA | Olfr1048 | olfactory receptor 1048 [Source:MGI Sym | 0 | 0 | 0 | 0 | 0 | 0 |
| 21164 | ENSMUSG00000075196 | 0 | NA | NA | NA | NA | NA | Olfr1047 | olfactory receptor 1047 [Source:MGI Sym | 0 | 0 | 0 | 0 | 0 | 0 |

| | | | | | | | | | | | | | | | |
|---|---|---|---|---|---|---|---|---|---|---|---|---|---|---|---|
| 21165 | ENSMUSG00000075197 | 0 | NA | NA | NA | NA | NA | Olfr1046 | olfactory receptor 1046 [Source:MGI Symb | 0 | 0 | 0 | 0 | 0 | 0 |
| 21166 | ENSMUSG00000075198 | 0 | NA | NA | NA | NA | NA | Olfr1045 | olfactory receptor 1045 [Source:MGI Symb | 0 | 0 | 0 | 0 | 0 | 0 |
| 21167 | ENSMUSG00000075199 | 0 | NA | NA | NA | NA | NA | Olfr52 | olfactory receptor 52 [Source:MGI Symbo | 0 | 0 | 0 | 0 | 0 | 0 |
| 21168 | ENSMUSG00000075200 | 0 | NA | NA | NA | NA | NA | Olfr1044 | olfactory receptor 1044 [Source:MGI Symb | 0 | 0 | 0 | 0 | 0 | 0 |
| 21169 | ENSMUSG00000075201 | 0 | NA | NA | NA | NA | NA | Olfr1043 | olfactory receptor 1043 [Source:MGI Symb | 0 | 0 | 0 | 0 | 0 | 0 |
| 21170 | ENSMUSG00000075202 | 0 | NA | NA | NA | NA | NA | Olfr1042 | olfactory receptor 1042 [Source:MGI Symb | 0 | 0 | 0 | 0 | 0 | 0 |
| 21171 | ENSMUSG00000075203 | 0 | NA | NA | NA | NA | NA | Olfr1040 | olfactory receptor 1040 [Source:MGI Symb | 0 | 0 | 0 | 0 | 0 | 0 |
| 21172 | ENSMUSG00000075204 | 0 | NA | NA | NA | NA | NA | Olfr1039 | olfactory receptor 1039 [Source:MGI Symb | 0 | 0 | 0 | 0 | 0 | 0 |
| 21173 | ENSMUSG00000075205 | 0 | NA | NA | NA | NA | NA | Olfr1037 | olfactory receptor 1037 [Source:MGI Symb | 0 | 0 | 0 | 0 | 0 | 0 |
| 21174 | ENSMUSG00000075206 | 0 | NA | NA | NA | NA | NA | Olfr1024 | olfactory receptor 1024 [Source:MGI Symb | 0 | 0 | 0 | 0 | 0 | 0 |
| 21175 | ENSMUSG00000075208 | 0 | NA | NA | NA | NA | NA | Olfr1019 | olfactory receptor 1019 [Source:MGI Symb | 0 | 0 | 0 | 0 | 0 | 0 |
| 21176 | ENSMUSG00000075209 | 0 | NA | NA | NA | NA | NA | Olfr1016 | olfactory receptor 1016 [Source:MGI Symb | 0 | 0 | 0 | 0 | 0 | 0 |
| 21177 | ENSMUSG00000075210 | 0 | NA | NA | NA | NA | NA | Olfr1012 | olfactory receptor 1012 [Source:MGI Symb | 0 | 0 | 0 | 0 | 0 | 0 |
| 21178 | ENSMUSG00000075211 | 0 | NA | NA | NA | NA | NA | Olfr1006 | olfactory receptor 1006 [Source:MGI Symb | 0 | 0 | 0 | 0 | 0 | 0 |
| 21179 | ENSMUSG00000075212 | 0 | NA | NA | NA | NA | NA | Olfr154 | olfactory receptor 154 [Source:MGI Symb | 0 | 0 | 0 | 0 | 0 | 0 |
| 21180 | ENSMUSG00000075214 | 0 | NA | NA | NA | NA | NA | Olfr1002 | olfactory receptor 1002 [Source:MGI Symb | 0 | 0 | 0 | 0 | 0 | 0 |
| 21181 | ENSMUSG00000075215 | 0 | NA | NA | NA | NA | NA | Olfr1000 | olfactory receptor 1000 [Source:MGI Symb | 0 | 0 | 0 | 0 | 0 | 0 |
| 21182 | ENSMUSG00000075217 | 0 | NA | NA | NA | NA | NA | Fads2b | fatty acid desaturase 2B [Source:MGI Sym | 0 | 0 | 0 | 0 | 0 | 0 |
| 21183 | ENSMUSG00000075218 | 0 | NA | NA | NA | NA | NA | Olfr995 | olfactory receptor 995 [Source:MGI Symb | 0 | 0 | 0 | 0 | 0 | 0 |
| 21184 | ENSMUSG00000075219 | 0 | NA | NA | NA | NA | NA | Olfr994 | olfactory receptor 994 [Source:MGI Symb | 0 | 0 | 0 | 0 | 0 | 0 |
| 21185 | ENSMUSG00000075220 | 0 | NA | NA | NA | NA | NA | Olfr993 | olfactory receptor 993 [Source:MGI Symb | 0 | 0 | 0 | 0 | 0 | 0 |
| 21186 | ENSMUSG00000075221 | 0 | NA | NA | NA | NA | NA | Olfr992 | olfactory receptor 992 [Source:MGI Symb | 0 | 0 | 0 | 0 | 0 | 0 |
| 21187 | ENSMUSG00000075222 | 0 | NA | NA | NA | NA | NA | Olfr988 | olfactory receptor 988 [Source:MGI Symb | 0 | 0 | 0 | 0 | 0 | 0 |
| 21188 | ENSMUSG00000075223 | 0 | NA | NA | NA | NA | NA | Olfr987 | olfactory receptor 987 [Source:MGI Symb | 0 | 0 | 0 | 0 | 0 | 0 |
| 21189 | ENSMUSG00000075224 | 0 | NA | NA | NA | NA | NA | Lrrc55 | leucine rich repeat containing 55 [Source: | 0 | 0 | 0 | 0 | 0 | 0 |
| 21190 | ENSMUSG00000075225 | 154.4959595 | 0.204097674 | 0.235916536 | 0.865126615 | 0.386969298 | 0.8175758 | Ccdc162 | coiled-coil domain containing 162 [Source | 162.23 | 143.91 | 123.77 | 115.68 | 187.59 | 193.81 |
| 21191 | ENSMUSG00000075227 | 482.299519 | 0.05257127 | 0.149004387 | 0.352816924 | 0.724225695 | 0.944786345 | Znhit2 | zinc finger, HIT domain containing 2 [Sour | 509.97 | 398.83 | 511.17 | 504.16 | 441.7 | 527.96 |
| 21192 | ENSMUSG00000075228 | 0 | NA | NA | NA | NA | NA | Gm10813 | predicted gene 10813 [Source:MGI Symbo | 0 | 0 | 0 | 0 | 0 | 0 |
| 21193 | ENSMUSG00000075229 | 427.6468808 | 0.12686125 | 0.129427502 | 0.980172282 | 0.327001084 | 0.772772049 | Ccdc58 | coiled-coil domain containing 58 [Source:I | 411.19 | 408.08 | 407.53 | 467.53 | 446.06 | 425.49 |
| 21194 | ENSMUSG00000075232 | 589.1128645 | -0.036006165 | 0.127896882 | -0.281524962 | 0.778307785 | 0.959559604 | Amd1 | S-adenosylmethionine decarboxylase 1 [S | 554.94 | 613.67 | 620.85 | 526.33 | 611.84 | 607.04 |
| 21195 | ENSMUSG00000075245 | 0 | NA | NA | NA | NA | NA | Gm6043 | predicted gene 6043 [Source:MGI Symbol | 0 | 0 | 0 | 0 | 0 | 0 |
| 21196 | ENSMUSG00000075249 | 0 | NA | NA | NA | NA | NA | Fsip2 | fibrous sheath-interacting protein 2 [Sour | 0 | 0 | 0 | 0 | 0 | 0 |
| 21197 | ENSMUSG00000075254 | 150.6706085 | -0.575086823 | 0.202091159 | -2.845680262 | 0.004431667 | 0.061582136 | Heg1 | heart development protein with EGF-like | 195.96 | 186.05 | 158.99 | 135.92 | 110.15 | 116.95 |
| 21198 | ENSMUSG00000075256 | 54.47540149 | -0.57741104 | 0.308661757 | -1.870691874 | 0.061387802 | 0.353912206 | Cerkl | ceramide kinase-like [Source:MGI Symbol | 57.823 | 60.647 | 77.481 | 48.199 | 39.262 | 43.44 |
| 21199 | ENSMUSG00000075265 | 1.779314608 | -0.23717903 | 1.844262451 | -0.12860373 | 0.897671215 | NA | Tnk2os | tyrosine kinase, non-receptor 2, opposite | 0.8031 | 1.0279 | 4.025 | 4.8199 | 0 | 0 |
| 21200 | ENSMUSG00000075266 | 32.23403916 | -0.452475338 | 0.425550664 | -1.063270196 | 0.287659437 | 0.73951397 | Cenpw | centromere protein W [Source:MGI Symb | 37.746 | 27.754 | 46.287 | 28.919 | 35.99 | 16.708 |
| 21201 | ENSMUSG00000075267 | 0.339027071 | -1.79174149 | 3.919585585 | -0.457125237 | 0.647581032 | NA | Pjvk | pejvakin [Source:MGI Symbol;Acc:MGI:26 | 0 | 1.0279 | 1.0062 | 0 | 0 | 0 |
| 21202 | ENSMUSG00000075268 | 3.262863356 | 0.503545331 | 1.264096721 | 0.398343989 | 0.690376639 | NA | Gm10819 | predicted gene 10819 [Source:MGI Symbo | 4.0155 | 0 | 4.025 | 3.8559 | 5.4531 | 2.2277 |
| 21203 | ENSMUSG00000075269 | 0 | NA | NA | NA | NA | NA | Bex6 | brain expressed family member 6 [Source | 0 | 0 | 0 | 0 | 0 | 0 |
| 21204 | ENSMUSG00000075270 | 0 | NA | NA | NA | NA | NA | Pde11a | phosphodiesterase 11A [Source:MGI Sym | 0 | 0 | 0 | 0 | 0 | 0 |
| 21205 | ENSMUSG00000075271 | 38.78073411 | 0.016679211 | 0.366219155 | 0.045544344 | 0.96367343 | 0.993245949 | Ttc30a1 | tetratricopeptide repeat domain 30A1 [So | 43.368 | 29.81 | 42.262 | 36.631 | 32.719 | 47.895 |
| 21206 | ENSMUSG00000075272 | 4.382594187 | -0.276975176 | 1.054916253 | -0.262556554 | 0.79289239 | NA | Ttc30a2 | tetratricopeptide repeat domain 30A2 [So | 3.2124 | 5.1396 | 6.0375 | 1.928 | 2.1812 | 7.7969 |
| 21207 | ENSMUSG00000075273 | 152.8572739 | 0.057304649 | 0.198161313 | 0.289181821 | 0.772442242 | 0.957706548 | Ttc30b | tetratricopeptide repeat domain 30B [Sou | 149.38 | 151.1 | 148.92 | 147.49 | 182.13 | 138.12 |
| 21208 | ENSMUSG00000075276 | 0 | NA | NA | NA | NA | NA | Gm13653 | predicted gene 13653 [Source:MGI Symbo | 0 | 0 | 0 | 0 | 0 | 0 |
| 21209 | ENSMUSG00000075277 | 0 | NA | NA | NA | NA | NA | Haglr | Hoxd antisense growth associated long nc | 0 | 0 | 0 | 0 | 0 | 0 |
| 21210 | ENSMUSG00000075279 | 5.245280471 | -0.387645363 | 0.973829067 | -0.398063044 | 0.690583715 | 0.937606823 | Mrpl23-ps1 | mitichondrial ribosomal protein L23, pseu | 9.6372 | 2.0558 | 6.0375 | 3.8559 | 6.5437 | 3.3415 |
| 21211 | ENSMUSG00000075284 | 62.79280124 | -0.648018766 | 0.336985329 | -1.922988063 | 0.054481549 | 0.328615982 | Wipf1 | WAS/WASL interacting protein family, me | 73.885 | 82.233 | 73.456 | 27.955 | 50.168 | 69.058 |
| 21212 | ENSMUSG00000075286 | 0.506966596 | 2.558866737 | 3.339834942 | 0.766165628 | 0.443577757 | NA | Gm1968 | predicted gene 1968 [Source:MGI Symbol | 0 | 0 | 0 | 1.928 | 0 | 1.1138 |
| 21213 | ENSMUSG00000075288 | 0 | NA | NA | NA | NA | NA | Gm6913 | predicted gene 6913 [Source:MGI Symbol | 0 | 0 | 0 | 0 | 0 | 0 |
| 21214 | ENSMUSG00000075289 | 19.49538806 | 0.354262067 | 0.529861146 | 0.668594159 | 0.503754399 | 0.881510796 | Carns1 | carnosine synthase 1 [Source:MGI Symbo | 20.881 | 8.2233 | 22.137 | 26.027 | 18.541 | 21.163 |
| 21215 | ENSMUSG00000075296 | 0.845528352 | 3.296539104 | 2.749625613 | 1.198904712 | 0.230564999 | NA | Aldh3b2 | aldehyde dehydrogenase 3 family, membe | 0 | 0 | 0 | 2.8919 | 2.1812 | 0 |
| 21216 | ENSMUSG00000075297 | 0.267700746 | -1.497481649 | 4.057001436 | -0.369110456 | 0.712045396 | NA | H60b | histocompatibility 60b [Source:MGI Symb | 1.6062 | 0 | 0 | 0 | 0 | 0 |
| 21217 | ENSMUSG00000075302 | 0 | NA | NA | NA | NA | NA | Erich2 | glutamate rich 2 [Source:MGI Symbol;Acc | 0 | 0 | 0 | 0 | 0 | 0 |
| 21218 | ENSMUSG00000075304 | 58.08377462 | 0.667009491 | 0.406374081 | 1.641368194 | 0.100721008 | 0.46725495 | Sp5 | trans-acting transcription factor 5 [Source | 64.248 | 18.503 | 51.318 | 54.947 | 94.884 | 64.603 |
| 21219 | ENSMUSG00000075307 | 2.517695408 | -0.117497293 | 1.371694465 | -0.085658502 | 0.931737892 | NA | Klhl41 | kelch-like 41 [Source:MGI Symbol;Acc:MG | 0.8031 | 5.1396 | 2.0125 | 3.8559 | 2.1812 | 1.1138 |
| 21220 | ENSMUSG00000075312 | 0.342639279 | -1.805976596 | 4.028547714 | -0.448294702 | 0.653940524 | NA | Gm13597 | predicted gene 13597 [Source:MGI Symbo | 0 | 2.0558 | 0 | 0 | 0 | 0 |

| | | | | | | | | | | | | | | | |
|---|---|---|---|---|---|---|---|---|---|---|---|---|---|---|---|
| 21221 | ENSMUSG00000075314 | 0 | NA | NA | NA | NA | NA | NA | NA | 0 | 0 | 0 | 0 | 0 | 0 |
| 21222 | ENSMUSG00000075316 | 0 | NA | NA | NA | NA | NA | Scn9a | sodium channel, voltage-gated, type IX, al | 0 | 0 | 0 | 0 | 0 | 0 |
| 21223 | ENSMUSG00000075318 | 0.342432689 | 1.990653086 | 3.900281263 | 0.51038706 | 0.609780321 | NA | Scn2a | sodium channel, voltage-gated, type II, al | 0 | 0 | 0 | 0.964 | 1.0906 | 0 |
| 21224 | ENSMUSG00000075324 | 125.9976001 | 0.033440077 | 0.229512986 | 0.145700154 | 0.8841581 | 0.981769403 | Fign | fidgetin [Source:MGI Symbol;Acc:MGI:18 | 126.89 | 129.52 | 116.72 | 99.29 | 126.51 | 157.05 |
| 21225 | ENSMUSG00000075325 | 0.859617266 | 0.673351556 | 2.564846152 | 0.262530973 | 0.792912109 | NA | Gm13582 | predicted gene 13582 [Source:MGI Symbo | 0 | 0 | 2.0125 | 0.964 | 2.1812 | 0 |
| 21226 | ENSMUSG00000075327 | 113.5628395 | 0.366641744 | 0.248826266 | 1.473484895 | 0.140620334 | 0.548229787 | Zbtb2 | zinc finger and BTB domain containing 2 [ | 82.72 | 102.79 | 112.7 | 144.6 | 94.884 | 143.69 |
| 21227 | ENSMUSG00000075330 | 0 | NA | NA | NA | NA | NA | A930003A15 | RIKEN cDNA A930003A15 gene [Source:M | 0 | 0 | 0 | 0 | 0 | 0 |
| 21228 | ENSMUSG00000075334 | 0 | NA | NA | NA | NA | NA | Rprm | reprimo, TP53 dependent G2 arrest media | 0 | 0 | 0 | 0 | 0 | 0 |
| 21229 | ENSMUSG00000075335 | 0 | NA | NA | NA | NA | NA | 4930588K23 | RIKEN cDNA 4930588K23 gene [Source:M | 0 | 0 | 0 | 0 | 0 | 0 |
| 21230 | ENSMUSG00000075359 | 0.185640647 | 1.126924175 | 4.080472857 | 0.276174898 | 0.782413736 | NA | Gm20775 | predicted gene, 20775 [Source:MGI Symb | 0 | 0 | 0 | 0 | 0 | 1.1138 |
| 21231 | ENSMUSG00000075370 | 0 | NA | NA | NA | NA | NA | Igll1 | immunoglobulin lambda-like polypeptide | 0 | 0 | 0 | 0 | 0 | 0 |
| 21232 | ENSMUSG00000075376 | 292.249068 | -0.078939289 | 0.146798111 | -0.537740495 | 0.590756218 | 0.911823491 | Rc3h2 | ring finger and CCCH-type zinc finger dom | 304.38 | 301.18 | 294.83 | 268.95 | 290.1 | 294.05 |
| 21233 | ENSMUSG00000075377 | 0 | NA | NA | NA | NA | NA | Olfr362 | olfactory receptor 362 [Source:MGI Symb | 0 | 0 | 0 | 0 | 0 | 0 |
| 21234 | ENSMUSG00000075378 | 0 | NA | NA | NA | NA | NA | Olfr361 | olfactory receptor 361 [Source:MGI Symb | 0 | 0 | 0 | 0 | 0 | 0 |
| 21235 | ENSMUSG00000075379 | 0 | NA | NA | NA | NA | NA | Olfr358 | olfactory receptor 358 [Source:MGI Symb | 0 | 0 | 0 | 0 | 0 | 0 |
| 21236 | ENSMUSG00000075380 | 0 | NA | NA | NA | NA | NA | Olfr355 | olfactory receptor 355 [Source:MGI Symb | 0 | 0 | 0 | 0 | 0 | 0 |
| 21237 | ENSMUSG00000075382 | 0 | NA | NA | NA | NA | NA | Olfr353 | olfactory receptor 353 [Source:MGI Symb | 0 | 0 | 0 | 0 | 0 | 0 |
| 21238 | ENSMUSG00000075383 | 0 | NA | NA | NA | NA | NA | Olfr351 | olfactory receptor 351 [Source:MGI Symb | 0 | 0 | 0 | 0 | 0 | 0 |
| 21239 | ENSMUSG00000075384 | 0 | NA | NA | NA | NA | NA | Olfr3 | olfactory receptor 3 [Source:MGI Symbol; | 0 | 0 | 0 | 0 | 0 | 0 |
| 21240 | ENSMUSG00000075387 | 0 | NA | NA | NA | NA | NA | Olfr341 | olfactory receptor 341 [Source:MGI Symb | 0 | 0 | 0 | 0 | 0 | 0 |
| 21241 | ENSMUSG00000075389 | 109.8158061 | 0.114327972 | 0.261112906 | 0.437848799 | 0.661495894 | 0.928379255 | 2810410L24 | RIKEN cDNA 2810410L24 gene [Source:M | 111.63 | 78.122 | 126.79 | 139.78 | 111.24 | 91.335 |
| 21242 | ENSMUSG00000075390 | 0 | NA | NA | NA | NA | NA | Rpl36-ps8 | ribosomal protein L36, pseudogene 8 [Sou | 0 | 0 | 0 | 0 | 0 | 0 |
| 21243 | ENSMUSG00000075391 | 14.51438892 | 0.729053399 | 0.581103279 | 1.254602108 | 0.209623238 | 0.653550636 | Glo1-ps | glyoxalase 1, pseudogene [Source:MGI Sy | 15.259 | 10.279 | 7.0437 | 19.28 | 19.631 | 15.594 |
| 21244 | ENSMUSG00000075394 | 1.000898647 | -2.10615299 | 2.475207832 | -0.850899453 | 0.394825209 | NA | Hoxc4 | homeobox C4 [Source:MGI Symbol;Acc:M | 0.8031 | 4.1117 | 0 | 0 | 1.0906 | 0 |
| 21245 | ENSMUSG00000075395 | 0 | NA | NA | NA | NA | NA | A630010A05 | RIKEN cDNA A630010A05 gene [Source:M | 0 | 0 | 0 | 0 | 0 | 0 |
| 21246 | ENSMUSG00000075402 | 0 | NA | NA | NA | NA | NA | Krt76 | keratin 76 [Source:MGI Symbol;Acc:MGI:1 | 0 | 0 | 0 | 0 | 0 | 0 |
| 21247 | ENSMUSG00000075405 | 0 | NA | NA | NA | NA | NA | 9430097D07 | RIKEN cDNA 9430097D07 gene [Source:M | 0 | 0 | 0 | 0 | 0 | 0 |
| 21248 | ENSMUSG00000075408 | 0 | NA | NA | NA | NA | NA | Smim41 | small integral membrane protein 41 [Sour | 0 | 0 | 0 | 0 | 0 | 0 |
| 21249 | ENSMUSG00000075410 | 3.125089916 | 0.019168043 | 1.291047053 | 0.014846897 | 0.988154325 | NA | Prcd | photoreceptor disc component [Source:M | 3.2124 | 3.0838 | 3.0187 | 2.8919 | 6.5437 | 0 |
| 21250 | ENSMUSG00000075415 | 252.1035975 | -0.118613569 | 0.160192935 | -0.740441947 | 0.459031874 | 0.859322431 | Fnbp1 | formin binding protein 1 [Source:MGI Sym | 260.21 | 244.64 | 282.75 | 256.42 | 224.67 | 243.93 |
| 21251 | ENSMUSG00000075416 | 0.181769715 | 1.126924175 | 4.080472857 | 0.276174898 | 0.782413736 | NA | Gm14488 | predicted gene 14488 [Source:MGI Symbo | 0 | 0 | 0 | 0 | 1.0906 | 0 |
| 21252 | ENSMUSG00000075419 | 305.3042275 | 0.114121998 | 0.153549176 | 0.743227681 | 0.457343847 | 0.858923498 | Dolk | dolichol kinase [Source:MGI Symbol;Acc:M | 331.68 | 274.45 | 272.69 | 318.11 | 314.1 | 320.79 |
| 21253 | ENSMUSG00000075420 | 16.98282977 | -0.743867759 | 0.530268958 | -1.402812192 | 0.16067285 | 0.583512653 | Smim6 | small integral membrane protein 6 [Sourc | 16.062 | 24.67 | 23.144 | 11.568 | 13.087 | 13.366 |
| 21254 | ENSMUSG00000075425 | 0.167707431 | -0.796629176 | 4.080472857 | -0.195229623 | 0.84521319 | NA | Gm13547 | predicted gene 13547 [Source:MGI Symbo | 0 | 0 | 1.0062 | 0 | 0 | 0 |
| 21255 | ENSMUSG00000075427 | 0 | NA | NA | NA | NA | NA | Olfr288 | olfactory receptor 288 [Source:MGI Symb | 0 | 0 | 0 | 0 | 0 | 0 |
| 21256 | ENSMUSG00000075431 | 0 | NA | NA | NA | NA | NA | Gm11691 | predicted gene 11691 [Source:MGI Symbo | 0 | 0 | 0 | 0 | 0 | 0 |
| 21257 | ENSMUSG00000075437 | 0 | NA | NA | NA | NA | NA | Gm11681 | predicted gene 11681 [Source:MGI Symbo | 0 | 0 | 0 | 0 | 0 | 0 |
| 21258 | ENSMUSG00000075463 | 15.41374009 | 0.771591831 | 0.600085747 | 1.285802629 | 0.198511953 | 0.64076699 | 4930594M22 | RIKEN cDNA 4930594M22 gene [Source:M | 12.85 | 12.335 | 9.0562 | 31.811 | 14.178 | 12.252 |
| 21259 | ENSMUSG00000075465 | 0 | NA | NA | NA | NA | NA | Gm10837 | predicted gene 10837 [Source:MGI Symbo | 0 | 0 | 0 | 0 | 0 | 0 |
| 21260 | ENSMUSG00000075467 | 487.0560417 | -0.035385939 | 0.129676979 | -0.272877569 | 0.78494733 | 0.960670212 | Dnlz | DNL-type zinc finger [Source:MGI Symbol; | 475.44 | 488.26 | 515.2 | 441.5 | 495.14 | 506.8 |
| 21261 | ENSMUSG00000075469 | 0 | NA | NA | NA | NA | NA | Gm17205 | predicted gene 17205 [Source:MGI Symbo | 0 | 0 | 0 | 0 | 0 | 0 |
| 21262 | ENSMUSG00000075470 | 794.397993 | -0.068000189 | 0.124544054 | -0.545993063 | 0.585070706 | 0.909160151 | Alg10b | asparagine-linked glycosylation 10B (alph | 713.96 | 860.37 | 866.38 | 793.35 | 799.42 | 732.91 |
| 21263 | ENSMUSG00000075478 | 0 | NA | NA | NA | NA | NA | Slitrk1 | SLIT and NTRK-like family, member 1 [Sou | 0 | 0 | 0 | 0 | 0 | 0 |
| 21264 | ENSMUSG00000075480 | 0 | NA | NA | NA | NA | NA | Gm10840 | predicted gene 10840 [Source:MGI Symbo | 0 | 0 | 0 | 0 | 0 | 0 |
| 21265 | ENSMUSG00000075485 | 0 | NA | NA | NA | NA | NA | NA | NA | 0 | 0 | 0 | 0 | 0 | 0 |
| 21266 | ENSMUSG00000075486 | 542.9468386 | -0.069936736 | 0.126645602 | -0.552223962 | 0.580794919 | 0.908645447 | Commd6 | COMM domain containing 6 [Source:MGI | 595.1 | 553.02 | 519.22 | 499.34 | 549.67 | 541.33 |
| 21267 | ENSMUSG00000075502 | 2.1105943 | -0.509025831 | 1.549084959 | -0.328597749 | 0.74245975 | NA | Kbtbd6 | kelch repeat and BTB (POZ) domain conta | 2.4093 | 0 | 5.0312 | 1.928 | 2.1812 | 1.1138 |
| 21268 | ENSMUSG00000075510 | 0 | NA | NA | NA | NA | NA | Fam187a | family with sequence similarity 187, mem | 0 | 0 | 0 | 0 | 0 | 0 |
| 21269 | ENSMUSG00000075511 | 255.4527112 | 0.364799059 | 0.178002806 | 2.04940061 | 0.040422958 | 0.274883714 | 1700001L05 | RIKEN cDNA 1700001L05 gene [Source:M | 199.97 | 204.56 | 265.65 | 255.45 | 311.92 | 295.17 |
| 21270 | ENSMUSG00000075512 | 0 | NA | NA | NA | NA | NA | Gm1587 | predicted gene 1587 [Source:MGI Symbol | 0 | 0 | 0 | 0 | 0 | 0 |
| 21271 | ENSMUSG00000075514 | 42.96169639 | 0.502627295 | 0.351116764 | 1.431510387 | 0.152283992 | 0.568128879 | Gm13375 | predicted gene 13375 [Source:MGI Symbo | 40.155 | 40.089 | 26.162 | 44.343 | 47.987 | 59.034 |
| 21272 | ENSMUSG00000075517 | 2733.246475 | 0.163451167 | 0.143745664 | 1.137085898 | 0.255502379 | 0.708894389 | Cyp2d37-ps | cytochrome P450, family 2, subfamily d, p | 2986.7 | 2249.1 | 2499.5 | 3402.8 | 2847.6 | 2413.7 |
| 21273 | ENSMUSG00000075520 | 0.160662974 | 1.126924175 | 4.080472857 | 0.276174898 | 0.782413736 | NA | Malrd1 | MAM and LDL receptor class A domain co | 0 | 0 | 0 | 0.964 | 0 | 0 |
| 21274 | ENSMUSG00000075524 | 0 | NA | NA | NA | NA | NA | 4930407I10 | RIKEN cDNA 4930407I10 gene [Source:M | 0 | 0 | 0 | 0 | 0 | 0 |
| 21275 | ENSMUSG00000075525 | 0 | NA | NA | NA | NA | NA | Gm10849 | predicted gene 10849 [Source:MGI Symbo | 0 | 0 | 0 | 0 | 0 | 0 |
| 21276 | ENSMUSG00000075528 | 6.490319192 | 0.619997507 | 0.851935872 | 0.727751381 | 0.466765791 | 0.863779479 | Aarsd1 | alanyl-tRNA synthetase domain containin | 3.2124 | 5.1396 | 7.0437 | 4.8199 | 9.8156 | 8.9108 |

| | | | | | | | | | | | | | | | |
|---|---|---|---|---|---|---|---|---|---|---|---|---|---|---|---|
| 21277 | ENSMUSG00000075534 | 0 | NA | NA | NA | NA | NA | Gm13262 | predicted gene 13262 [Source:MGI Symbo | 0 | 0 | 0 | 0 | 0 | 0 |
| 21278 | ENSMUSG00000075538 | 0 | NA | NA | NA | NA | NA | Gm10855 | predicted gene 10855 [Source:MGI Symbo | 0 | 0 | 0 | 0 | 0 | 0 |
| 21279 | ENSMUSG00000075543 | 961.3705967 | 0.195879473 | 0.105874166 | 1.850115851 | 0.064296854 | 0.36329674 | Urad | ureidoimidazoline (2-oxo-4-hydroxy-4-car | 866.55 | 917.93 | 904.61 | 1018 | 1072.1 | 989.09 |
| 21280 | ENSMUSG00000075549 | 0 | NA | NA | NA | NA | NA | Gm6878 | predicted gene 6878 [Source:MGI Symbol | 0 | 0 | 0 | 0 | 0 | 0 |
| 21281 | ENSMUSG00000075551 | 0 | NA | NA | NA | NA | NA | Cyp3a41a | cytochrome P450, family 3, subfamily a, p | 0 | 0 | 0 | 0 | 0 | 0 |
| 21282 | ENSMUSG00000075552 | 0 | NA | NA | NA | NA | NA | Cyp3a41b | cytochrome P450, family 3, subfamily a, p | 0 | 0 | 0 | 0 | 0 | 0 |
| 21283 | ENSMUSG00000075553 | 0.17131964 | -0.796629176 | 4.080472857 | -0.195229623 | 0.84521319 | NA | Gm5464 | predicted gene 5464 [Source:MGI Symbol | 0 | 1.0279 | 0 | 0 | 0 | 0 |
| 21284 | ENSMUSG00000075555 | 1.345678198 | 0.272738895 | 2.169909302 | 0.12569138 | 0.899976226 | NA | Gm10863 | predicted gene 10863 [Source:MGI Symbo | 1.6062 | 0 | 2.0125 | 0 | 0 | 4.4554 |
| 21285 | ENSMUSG00000075558 | 0.48903338 | 0.95637552 | 3.456690196 | 0.276673773 | 0.782030612 | NA | Gm17151 | predicted gene 17151 [Source:MGI Symbo | 0 | 0 | 1.0062 | 1.928 | 0 | 0 |
| 21286 | ENSMUSG00000075566 | 0 | NA | NA | NA | NA | NA | Krtap4-6 | keratin associated protein 4-6 [Source:MG | 0 | 0 | 0 | 0 | 0 | 0 |
| 21287 | ENSMUSG00000075567 | 0 | NA | NA | NA | NA | NA | Krtap1-4 | keratin associated protein 1-4 [Source:MG | 0 | 0 | 0 | 0 | 0 | 0 |
| 21288 | ENSMUSG00000075569 | 1.393338816 | 0.950037025 | 1.855657742 | 0.511967807 | 0.60867354 | NA | Rsph10b | radial spoke head 10 homolog B (Chlamyd | 0.8031 | 1.0279 | 1.0062 | 0 | 2.1812 | 3.3415 |
| 21289 | ENSMUSG00000075570 | 0 | NA | NA | NA | NA | NA | Krt26 | keratin 26 [Source:MGI Symbol;Acc:MGI:2 | 0 | 0 | 0 | 0 | 0 | 0 |
| 21290 | ENSMUSG00000075571 | 0 | NA | NA | NA | NA | NA | Defb30 | defensin beta 30 [Source:MGI Symbol;Acc | 0 | 0 | 0 | 0 | 0 | 0 |
| 21291 | ENSMUSG00000075572 | 0 | NA | NA | NA | NA | NA | Defb43 | defensin beta 43 [Source:MGI Symbol;Acc | 0 | 0 | 0 | 0 | 0 | 0 |
| 21292 | ENSMUSG00000075573 | 0.481988923 | 2.492401104 | 3.837077151 | 0.649557203 | 0.515978285 | NA | Defb47 | defensin beta 47 [Source:MGI Symbol;Acc | 0 | 0 | 0 | 2.8919 | 0 | 0 |
| 21293 | ENSMUSG00000075574 | 0 | NA | NA | NA | NA | NA | Defb48 | defensin beta 48 [Source:MGI Symbol;Acc | 0 | 0 | 0 | 0 | 0 | 0 |
| 21294 | ENSMUSG00000075576 | 9.2850187 | 0.171160663 | 0.824476773 | 0.207599133 | 0.835541982 | 0.971073158 | Gm12359 | predicted gene 12359 [Source:MGI Symbo | 8.031 | 9.2513 | 9.0562 | 17.352 | 10.906 | 1.1138 |
| 21295 | ENSMUSG00000075581 | 1.68261713 | -1.111715545 | 1.648617335 | -0.674332073 | 0.500100218 | NA | Gm16409 | predicted gene 16409 [Source:MGI Symbo | 0.8031 | 4.1117 | 2.0125 | 0.964 | 1.0906 | 1.1138 |
| 21296 | ENSMUSG00000075585 | 26.69353566 | 0.268139823 | 0.442246939 | 0.606312447 | 0.544307295 | 0.894846638 | 6330403L08I | RIKEN cDNA 6330403L08 gene [Source:MG | 33.73 | 20.558 | 18.112 | 33.739 | 26.175 | 27.846 |
| 21297 | ENSMUSG00000075586 | 0 | NA | NA | NA | NA | NA | Gm11529 | predicted gene 11529 [Source:MGI Symbo | 0 | 0 | 0 | 0 | 0 | 0 |
| 21298 | ENSMUSG00000075587 | 0 | NA | NA | NA | NA | NA | Gm11531 | predicted gene 11531 [Source:MGI Symbo | 0 | 0 | 0 | 0 | 0 | 0 |
| 21299 | ENSMUSG00000075588 | 2.412125475 | 0.39126411 | 1.443210004 | 0.271106844 | 0.786308858 | NA | Hoxb2 | homeobox B2 [Source:MGI Symbol;Acc:M | 3.2124 | 0 | 3.0187 | 3.8559 | 3.2719 | 1.1138 |
| 21300 | ENSMUSG00000075589 | 0.760346335 | -0.431432104 | 2.662343224 | -0.162049769 | 0.871266662 | NA | Gm11536 | predicted gene 11536 [Source:MGI Symbo | 1.6062 | 1.0279 | 0 | 1.928 | 0 | 0 |
| 21301 | ENSMUSG00000075590 | 1025.716782 | -0.197899033 | 0.142057835 | -1.393087775 | 0.163593249 | 0.58776904 | Nrbp2 | nuclear receptor binding protein 2 [Sourc | 934.01 | 1148.2 | 1207.5 | 1098.9 | 930.3 | 835.38 |
| 21302 | ENSMUSG00000075591 | 4.753616758 | -0.033184086 | 1.016445695 | -0.032647181 | 0.973955945 | NA | Gm10874 | predicted gene 10874 [Source:MGI Symbo | 3.2124 | 6.1675 | 5.0312 | 1.928 | 3.2719 | 8.9108 |
| 21303 | ENSMUSG00000075592 | 7.147234221 | 0.539802344 | 0.865907865 | 0.623394665 | 0.533025203 | 0.889143097 | Nynrin | NYN domain and retroviral integrase cont | 7.2279 | 5.1396 | 5.0312 | 5.7839 | 16.359 | 3.3415 |
| 21304 | ENSMUSG00000075593 | 0 | NA | NA | NA | NA | NA | Gal3st4 | galactose-3-O-sulfotransferase 4 [Source:I | 0 | 0 | 0 | 0 | 0 | 0 |
| 21305 | ENSMUSG00000075595 | 704.0840227 | 0.011400553 | 0.135399874 | 0.084199139 | 0.932898103 | 0.988420305 | Zfp652 | zinc finger protein 652 [Source:MGI Symb | 610.36 | 737.02 | 757.7 | 731.66 | 631.47 | 756.3 |
| 21306 | ENSMUSG00000075596 | 1.847151037 | -0.712820295 | 1.635079079 | -0.43595463 | 0.662869647 | NA | NA | NA | 0.8031 | 4.1117 | 2.0125 | 1.928 | 0 | 2.2277 |
| 21307 | ENSMUSG00000075598 | 0 | NA | NA | NA | NA | NA | Smok3c | sperm motility kinase 3C [Source:MGI Sym | 0 | 0 | 0 | 0 | 0 | 0 |
| 21308 | ENSMUSG00000075599 | 0 | NA | NA | NA | NA | NA | Smok3a | sperm motility kinase 3A [Source:MGI Syn | 0 | 0 | 0 | 0 | 0 | 0 |
| 21309 | ENSMUSG00000075600 | 88.78465576 | -0.62983754 | 0.268763437 | -2.343464377 | 0.019105587 | 0.168091974 | Zc3h3 | zinc finger CCCH type containing 3 [Source | 116.45 | 101.76 | 105.66 | 86.758 | 73.071 | 49.009 |
| 21310 | ENSMUSG00000075602 | 68.22957347 | -0.625741398 | 0.285322631 | -2.193101177 | 0.02830009 | 0.219952985 | Ly6a | lymphocyte antigen 6 complex, locus A [S | 88.341 | 80.178 | 79.493 | 44.343 | 49.078 | 67.944 |
| 21311 | ENSMUSG00000075604 | 0 | NA | NA | NA | NA | NA | Cyp11b1 | cytochrome P450, family 11, subfamily b, | 0 | 0 | 0 | 0 | 0 | 0 |
| 21312 | ENSMUSG00000075605 | 0 | NA | NA | NA | NA | NA | Slurp2 | secreted Ly6/Plaur domain containing 2 [S | 0 | 0 | 0 | 0 | 0 | 0 |
| 21313 | ENSMUSG00000075609 | 1.077718728 | 1.149008664 | 2.221862456 | 0.517137621 | 0.605060093 | NA | Gm7935 | predicted pseudogene 7935 [Source:MGI | 0 | 1.0279 | 1.0062 | 0 | 1.0906 | 3.3415 |
| 21314 | ENSMUSG00000075610 | 0 | NA | NA | NA | NA | NA | Tmem92 | transmembrane protein 92 [Source:MGI S | 0 | 0 | 0 | 0 | 0 | 0 |
| 21315 | ENSMUSG00000075611 | 0.356960287 | 0.16514264 | 3.880370363 | 0.042558474 | 0.966053499 | NA | Gm11545 | predicted gene 11545 [Source:MGI Symbo | 0 | 1.0279 | 0 | 0 | 0 | 1.1138 |
| 21316 | ENSMUSG00000075700 | 3713.607357 | -0.026727349 | 0.093809621 | -0.284910531 | 0.775712687 | 0.95893551 | Selenot | selenoprotein T [Source:MGI Symbol;Acc: | 3754.5 | 3935.9 | 3553 | 3482.9 | 3707 | 3848.3 |
| 21317 | ENSMUSG00000075701 | 728.3836437 | 0.362389739 | 0.117521335 | 3.083608079 | 0.002045068 | 0.035510042 | Selenos | selenoprotein S [Source:MGI Symbol;Acc: | 615.18 | 695.9 | 601.73 | 839.62 | 810.33 | 807.54 |
| 21318 | ENSMUSG00000075702 | 25.91099895 | -0.710245002 | 0.528180446 | -1.344701433 | 0.178721672 | 0.610813333 | Selenom | selenoprotein M [Source:MGI Symbol;Acc | 47.383 | 35.977 | 13.081 | 26.027 | 19.631 | 13.366 |
| 21319 | ENSMUSG00000075703 | 886.2450307 | 0.031582035 | 0.112858014 | 0.279838654 | 0.779601294 | 0.959870443 | Selenoi | selenoprotein I [Source:MGI Symbol;Acc:I | 815.15 | 901.48 | 913.67 | 869.51 | 947.75 | 869.91 |
| 21320 | ENSMUSG00000075704 | 656.4873568 | -0.143950643 | 0.122616421 | -1.173991552 | 0.240398409 | 0.690453256 | Txnrd2 | thioredoxin reductase 2 [Source:MGI Sym | 649.71 | 721.6 | 696.32 | 569.71 | 651.1 | 650.48 |
| 21321 | ENSMUSG00000075705 | 6686.004915 | 0.205793619 | 0.089006845 | 2.312110016 | 0.020771621 | 0.177565444 | Msrb1 | methionine sulfoxide reductase B1 [Sourc | 5983.1 | 6008.2 | 6639.2 | 7145 | 7332.2 | 7008.3 |
| 21322 | ENSMUSG00000075706 | 5072.889525 | -0.180737241 | 0.109287906 | -1.653771642 | 0.09817392 | 0.462085121 | Gpx4 | glutathione peroxidase 4 [Source:MGI Syr | 5976.7 | 5487 | 4706.2 | 4965.4 | 4747.5 | 4554.5 |
| 21323 | ENSMUSG00000075707 | 3.194566826 | -0.494251181 | 1.350103444 | -0.366083935 | 0.714302445 | NA | Dio3 | deiodinase, iodothyronine type III [Source | 3.2124 | 2.0558 | 6.0375 | 6.7478 | 0 | 1.1138 |
| 21324 | ENSMUSG00000075719 | 0 | NA | NA | NA | NA | NA | Gm23741 | predicted gene, 23741 [Source:MGI Symb | 0 | 0 | 0 | 0 | 0 | 0 |
| 21325 | ENSMUSG00000075726 | 0 | NA | NA | NA | NA | NA | Gm23254 | predicted gene, 23254 [Source:MGI Symb | 0 | 0 | 0 | 0 | 0 | 0 |
| 21326 | ENSMUSG00000075760 | 0 | NA | NA | NA | NA | NA | NA | NA | 0 | 0 | 0 | 0 | 0 | 0 |
| 21327 | ENSMUSG00000075770 | 0 | NA | NA | NA | NA | NA | Gm22589 | predicted gene, 22589 [Source:MGI Symb | 0 | 0 | 0 | 0 | 0 | 0 |
| 21328 | ENSMUSG00000075826 | 0 | NA | NA | NA | NA | NA | Gm23691 | predicted gene, 23691 [Source:MGI Symb | 0 | 0 | 0 | 0 | 0 | 0 |
| 21329 | ENSMUSG00000075829 | 0 | NA | NA | NA | NA | NA | Gm23692 | predicted gene, 23692 [Source:MGI Symb | 0 | 0 | 0 | 0 | 0 | 0 |
| 21330 | ENSMUSG00000075832 | 0 | NA | NA | NA | NA | NA | Gm22004 | predicted gene, 22004 [Source:MGI Symb | 0 | 0 | 0 | 0 | 0 | 0 |
| 21331 | ENSMUSG00000075833 | 0 | NA | NA | NA | NA | NA | NA | NA | 0 | 0 | 0 | 0 | 0 | 0 |
| 21332 | ENSMUSG00000075837 | 0 | NA | NA | NA | NA | NA | Gm22005 | predicted gene, 22005 [Source:MGI Symb | 0 | 0 | 0 | 0 | 0 | 0 |

| | | | | | | | | | | | | | | | |
|---|---|---|---|---|---|---|---|---|---|---|---|---|---|---|---|
| 21333 | ENSMUSG00000075864 | 0 | NA | NA | NA | NA | NA | NA | NA | 0 | 0 | 0 | 0 | 0 | 0 |
| 21334 | ENSMUSG00000075874 | 0 | NA | NA | NA | NA | NA | n-R5s1 | nuclear encoded rRNA 5S 1 [Source:MGI S | 0 | 0 | 0 | 0 | 0 | 0 |
| 21335 | ENSMUSG00000075893 | 0 | NA | NA | NA | NA | NA | NA | NA | 0 | 0 | 0 | 0 | 0 | 0 |
| 21336 | ENSMUSG00000075903 | 0 | NA | NA | NA | NA | NA | Gm23941 | predicted gene, 23941 [Source:MGI Symb | 0 | 0 | 0 | 0 | 0 | 0 |
| 21337 | ENSMUSG00000075906 | 0 | NA | NA | NA | NA | NA | NA | NA | 0 | 0 | 0 | 0 | 0 | 0 |
| 21338 | ENSMUSG00000075908 | 0.36353943 | 2.065989254 | 4.034137301 | 0.512126658 | 0.608562368 | NA | Gm23940 | predicted gene, 23940 [Source:MGI Symb | 0 | 0 | 0 | 0 | 2.1812 | 0 |
| 21339 | ENSMUSG00000075918 | 0 | NA | NA | NA | NA | NA | NA | NA | 0 | 0 | 0 | 0 | 0 | 0 |
| 21340 | ENSMUSG00000075934 | 0 | NA | NA | NA | NA | NA | Gm24423 | predicted gene, 24423 [Source:MGI Symb | 0 | 0 | 0 | 0 | 0 | 0 |
| 21341 | ENSMUSG00000075952 | 0 | NA | NA | NA | NA | NA | Gm23422 | predicted gene, 23422 [Source:MGI Symb | 0 | 0 | 0 | 0 | 0 | 0 |
| 21342 | ENSMUSG00000075958 | 0 | NA | NA | NA | NA | NA | Gm23423 | predicted gene, 23423 [Source:MGI Symb | 0 | 0 | 0 | 0 | 0 | 0 |
| 21343 | ENSMUSG00000075968 | 0 | NA | NA | NA | NA | NA | Gm25089 | predicted gene, 25089 [Source:MGI Symb | 0 | 0 | 0 | 0 | 0 | 0 |
| 21344 | ENSMUSG00000075973 | 0 | NA | NA | NA | NA | NA | NA | NA | 0 | 0 | 0 | 0 | 0 | 0 |
| 21345 | ENSMUSG00000075983 | 0 | NA | NA | NA | NA | NA | Gm24121 | predicted gene, 24121 [Source:MGI Symb | 0 | 0 | 0 | 0 | 0 | 0 |
| 21346 | ENSMUSG00000076005 | 0 | NA | NA | NA | NA | NA | Mir547 | microRNA 547 [Source:MGI Symbol;Acc:M | 0 | 0 | 0 | 0 | 0 | 0 |
| 21347 | ENSMUSG00000076006 | 0 | NA | NA | NA | NA | NA | Mir376c | microRNA 376c [Source:MGI Symbol;Acc:I | 0 | 0 | 0 | 0 | 0 | 0 |
| 21348 | ENSMUSG00000076009 | 0 | NA | NA | NA | NA | NA | Mir688 | microRNA 688 [Source:MGI Symbol;Acc:M | 0 | 0 | 0 | 0 | 0 | 0 |
| 21349 | ENSMUSG00000076010 | 0 | NA | NA | NA | NA | NA | Mir615 | microRNA 615 [Source:MGI Symbol;Acc:M | 0 | 0 | 0 | 0 | 0 | 0 |
| 21350 | ENSMUSG00000076011 | 0 | NA | NA | NA | NA | NA | Mir652 | microRNA 652 [Source:MGI Symbol;Acc:M | 0 | 0 | 0 | 0 | 0 | 0 |
| 21351 | ENSMUSG00000076012 | 0 | NA | NA | NA | NA | NA | Mir705 | microRNA 705 [Source:MGI Symbol;Acc:M | 0 | 0 | 0 | 0 | 0 | 0 |
| 21352 | ENSMUSG00000076014 | 0 | NA | NA | NA | NA | NA | Mir669a-2 | microRNA 669a-2 [Source:MGI Symbol;Ac | 0 | 0 | 0 | 0 | 0 | 0 |
| 21353 | ENSMUSG00000076028 | 0 | NA | NA | NA | NA | NA | Mir669a-3 | microRNA 669a-3 [Source:MGI Symbol;Ac | 0 | 0 | 0 | 0 | 0 | 0 |
| 21354 | ENSMUSG00000076030 | 0 | NA | NA | NA | NA | NA | NA | NA | 0 | 0 | 0 | 0 | 0 | 0 |
| 21355 | ENSMUSG00000076031 | 0.133850373 | -0.796629176 | 4.080472857 | -0.195229623 | 0.84521319 | NA | NA | NA | 0.8031 | 0 | 0 | 0 | 0 | 0 |
| 21356 | ENSMUSG00000076033 | 0 | NA | NA | NA | NA | NA | Mir692-3 | microRNA 692-3 [Source:MGI Symbol;Acc | 0 | 0 | 0 | 0 | 0 | 0 |
| 21357 | ENSMUSG00000076036 | 0 | NA | NA | NA | NA | NA | NA | NA | 0 | 0 | 0 | 0 | 0 | 0 |
| 21358 | ENSMUSG00000076037 | 0 | NA | NA | NA | NA | NA | NA | NA | 0 | 0 | 0 | 0 | 0 | 0 |
| 21359 | ENSMUSG00000076038 | 7.506474989 | -0.148611751 | 0.839540067 | -0.177015674 | 0.859496078 | 0.975502162 | NA | NA | 2.4093 | 12.335 | 9.0562 | 5.7839 | 6.5437 | 8.9108 |
| 21360 | ENSMUSG00000076039 | 0.160662974 | 1.126924175 | 4.080472857 | 0.276174898 | 0.782413736 | NA | NA | NA | 0 | 0 | 0 | 0.964 | 0 | 0 |
| 21361 | ENSMUSG00000076041 | 0 | NA | NA | NA | NA | NA | Mir20b | microRNA 20b [Source:MGI Symbol;Acc:M | 0 | 0 | 0 | 0 | 0 | 0 |
| 21362 | ENSMUSG00000076042 | 0 | NA | NA | NA | NA | NA | Mir710 | microRNA 710 [Source:MGI Symbol;Acc:M | 0 | 0 | 0 | 0 | 0 | 0 |
| 21363 | ENSMUSG00000076049 | 0 | NA | NA | NA | NA | NA | Mir598 | microRNA 598 [Source:MGI Symbol;Acc:M | 0 | 0 | 0 | 0 | 0 | 0 |
| 21364 | ENSMUSG00000076050 | 0 | NA | NA | NA | NA | NA | Mir450b | microRNA 450b [Source:MGI Symbol;Acc: | 0 | 0 | 0 | 0 | 0 | 0 |
| 21365 | ENSMUSG00000076051 | 0 | NA | NA | NA | NA | NA | Mir707 | microRNA 707 [Source:MGI Symbol;Acc:M | 0 | 0 | 0 | 0 | 0 | 0 |
| 21366 | ENSMUSG00000076052 | 0 | NA | NA | NA | NA | NA | Mir541 | microRNA 541 [Source:MGI Symbol;Acc:M | 0 | 0 | 0 | 0 | 0 | 0 |
| 21367 | ENSMUSG00000076060 | 0 | NA | NA | NA | NA | NA | Mir704 | microRNA 704 [Source:MGI Symbol;Acc:M | 0 | 0 | 0 | 0 | 0 | 0 |
| 21368 | ENSMUSG00000076062 | 0 | NA | NA | NA | NA | NA | Mir92-1 | microRNA 92-1 [Source:MGI Symbol;Acc:I | 0 | 0 | 0 | 0 | 0 | 0 |
| 21369 | ENSMUSG00000076063 | 0 | NA | NA | NA | NA | NA | Mir539 | microRNA 539 [Source:MGI Symbol;Acc:M | 0 | 0 | 0 | 0 | 0 | 0 |
| 21370 | ENSMUSG00000076064 | 0 | NA | NA | NA | NA | NA | Mir690 | microRNA 690 [Source:MGI Symbol;Acc:M | 0 | 0 | 0 | 0 | 0 | 0 |
| 21371 | ENSMUSG00000076065 | 0 | NA | NA | NA | NA | NA | Mir681 | microRNA 681 [Source:MGI Symbol;Acc:M | 0 | 0 | 0 | 0 | 0 | 0 |
| 21372 | ENSMUSG00000076066 | 0 | NA | NA | NA | NA | NA | Mir223 | microRNA 223 [Source:MGI Symbol;Acc:M | 0 | 0 | 0 | 0 | 0 | 0 |
| 21373 | ENSMUSG00000076067 | 0 | NA | NA | NA | NA | NA | Mir592 | microRNA 592 [Source:MGI Symbol;Acc:M | 0 | 0 | 0 | 0 | 0 | 0 |
| 21374 | ENSMUSG00000076068 | 0 | NA | NA | NA | NA | NA | Mir678 | microRNA 678 [Source:MGI Symbol;Acc:M | 0 | 0 | 0 | 0 | 0 | 0 |
| 21375 | ENSMUSG00000076117 | 0 | NA | NA | NA | NA | NA | NA | NA | 0 | 0 | 0 | 0 | 0 | 0 |
| 21376 | ENSMUSG00000076118 | 0 | NA | NA | NA | NA | NA | Mir669c | microRNA 669c [Source:MGI Symbol;Acc:I | 0 | 0 | 0 | 0 | 0 | 0 |
| 21377 | ENSMUSG00000076119 | 0 | NA | NA | NA | NA | NA | Mir698 | microRNA 698 [Source:MGI Symbol;Acc:M | 0 | 0 | 0 | 0 | 0 | 0 |
| 21378 | ENSMUSG00000076120 | 0 | NA | NA | NA | NA | NA | Mir551b | microRNA 551b [Source:MGI Symbol;Acc: | 0 | 0 | 0 | 0 | 0 | 0 |
| 21379 | ENSMUSG00000076121 | 0 | NA | NA | NA | NA | NA | Mir687 | microRNA 687 [Source:MGI Symbol;Acc:M | 0 | 0 | 0 | 0 | 0 | 0 |
| 21380 | ENSMUSG00000076122 | 0 | NA | NA | NA | NA | NA | Mir503 | microRNA 503 [Source:MGI Symbol;Acc:M | 0 | 0 | 0 | 0 | 0 | 0 |
| 21381 | ENSMUSG00000076123 | 0 | NA | NA | NA | NA | NA | Mir700 | microRNA 700 [Source:MGI Symbol;Acc:M | 0 | 0 | 0 | 0 | 0 | 0 |
| 21382 | ENSMUSG00000076126 | 0 | NA | NA | NA | NA | NA | Mir669b | microRNA 669b [Source:MGI Symbol;Acc: | 0 | 0 | 0 | 0 | 0 | 0 |
| 21383 | ENSMUSG00000076127 | 0 | NA | NA | NA | NA | NA | Mir718 | microRNA 718 [Source:MGI Symbol;Acc:M | 0 | 0 | 0 | 0 | 0 | 0 |
| 21384 | ENSMUSG00000076128 | 0 | NA | NA | NA | NA | NA | Mir686 | microRNA 686 [Source:MGI Symbol;Acc:M | 0 | 0 | 0 | 0 | 0 | 0 |
| 21385 | ENSMUSG00000076129 | 0 | NA | NA | NA | NA | NA | NA | NA | 0 | 0 | 0 | 0 | 0 | 0 |
| 21386 | ENSMUSG00000076135 | 0 | NA | NA | NA | NA | NA | NA | NA | 0 | 0 | 0 | 0 | 0 | 0 |
| 21387 | ENSMUSG00000076136 | 0 | NA | NA | NA | NA | NA | NA | NA | 0 | 0 | 0 | 0 | 0 | 0 |
| 21388 | ENSMUSG00000076137 | 4.186482349 | -0.035063514 | 1.101957541 | -0.031819297 | 0.974616158 | NA | NA | NA | 1.6062 | 8.2233 | 3.0187 | 6.7478 | 2.1812 | 3.3415 |

| | | | | | | | | | | | | | | | |
|---|---|---|---|---|---|---|---|---|---|---|---|---|---|---|---|
| 21389 | ENSMUSG00000076138 | 0 | NA | NA | NA | NA | NA | Gm26377 | predicted gene, 26377 [Source:MGI Symb | 0 | 0 | 0 | 0 | 0 | 0 |
| 21390 | ENSMUSG00000076140 | 0 | NA | NA | NA | NA | NA | Mir542 | microRNA 542 [Source:MGI Symbol;Acc:M | 0 | 0 | 0 | 0 | 0 | 0 |
| 21391 | ENSMUSG00000076141 | 0 | NA | NA | NA | NA | NA | Mir693 | microRNA 693 [Source:MGI Symbol;Acc:M | 0 | 0 | 0 | 0 | 0 | 0 |
| 21392 | ENSMUSG00000076142 | 0 | NA | NA | NA | NA | NA | Mir711 | microRNA 711 [Source:MGI Symbol;Acc:M | 0 | 0 | 0 | 0 | 0 | 0 |
| 21393 | ENSMUSG00000076143 | 0 | NA | NA | NA | NA | NA | Mir708 | microRNA 708 [Source:MGI Symbol;Acc:M | 0 | 0 | 0 | 0 | 0 | 0 |
| 21394 | ENSMUSG00000076144 | 0.496077836 | -0.812642266 | 3.444120572 | -0.235950586 | 0.813471014 | NA | Mir709 | microRNA 709 [Source:MGI Symbol;Acc:M | 0 | 0 | 2.0125 | 0.964 | 0 | 0 |
| 21395 | ENSMUSG00000076145 | 0 | NA | NA | NA | NA | NA | Mir679 | microRNA 679 [Source:MGI Symbol;Acc:M | 0 | 0 | 0 | 0 | 0 | 0 |
| 21396 | ENSMUSG00000076146 | 0 | NA | NA | NA | NA | NA | Mir449c | microRNA 449c [Source:MGI Symbol;Acc:I | 0 | 0 | 0 | 0 | 0 | 0 |
| 21397 | ENSMUSG00000076147 | 0 | NA | NA | NA | NA | NA | Mir694 | microRNA 694 [Source:MGI Symbol;Acc:M | 0 | 0 | 0 | 0 | 0 | 0 |
| 21398 | ENSMUSG00000076163 | 0 | NA | NA | NA | NA | NA | Mir701 | microRNA 701 [Source:MGI Symbol;Acc:M | 0 | 0 | 0 | 0 | 0 | 0 |
| 21399 | ENSMUSG00000076180 | 0 | NA | NA | NA | NA | NA | NA | NA | 0 | 0 | 0 | 0 | 0 | 0 |
| 21400 | ENSMUSG00000076190 | 0 | NA | NA | NA | NA | NA | NA | NA | 0 | 0 | 0 | 0 | 0 | 0 |
| 21401 | ENSMUSG00000076199 | 0 | NA | NA | NA | NA | NA | NA | NA | 0 | 0 | 0 | 0 | 0 | 0 |
| 21402 | ENSMUSG00000076212 | 0 | NA | NA | NA | NA | NA | Mir680-1 | microRNA 680-1 [Source:MGI Symbol;Acc | 0 | 0 | 0 | 0 | 0 | 0 |
| 21403 | ENSMUSG00000076214 | 0 | NA | NA | NA | NA | NA | Mir717 | microRNA 717 [Source:MGI Symbol;Acc:M | 0 | 0 | 0 | 0 | 0 | 0 |
| 21404 | ENSMUSG00000076216 | 0 | NA | NA | NA | NA | NA | Mir695 | microRNA 695 [Source:MGI Symbol;Acc:M | 0 | 0 | 0 | 0 | 0 | 0 |
| 21405 | ENSMUSG00000076217 | 0 | NA | NA | NA | NA | NA | Mir706 | microRNA 706 [Source:MGI Symbol;Acc:M | 0 | 0 | 0 | 0 | 0 | 0 |
| 21406 | ENSMUSG00000076218 | 1.048817988 | -0.908682365 | 2.659333108 | -0.341695579 | 0.73258 | NA | Mir692-1 | microRNA 692-1 [Source:MGI Symbol;Acc | 0 | 4.1117 | 0 | 0 | 2.1812 | 0 |
| 21407 | ENSMUSG00000076219 | 0 | NA | NA | NA | NA | NA | Mir487b | microRNA 487b [Source:MGI Symbol;Acc: | 0 | 0 | 0 | 0 | 0 | 0 |
| 21408 | ENSMUSG00000076222 | 0 | NA | NA | NA | NA | NA | Mir297b | microRNA 297b [Source:MGI Symbol;Acc: | 0 | 0 | 0 | 0 | 0 | 0 |
| 21409 | ENSMUSG00000076227 | 0 | NA | NA | NA | NA | NA | NA | NA | 0 | 0 | 0 | 0 | 0 | 0 |
| 21410 | ENSMUSG00000076228 | 0 | NA | NA | NA | NA | NA | NA | NA | 0 | 0 | 0 | 0 | 0 | 0 |
| 21411 | ENSMUSG00000076229 | 0 | NA | NA | NA | NA | NA | NA | NA | 0 | 0 | 0 | 0 | 0 | 0 |
| 21412 | ENSMUSG00000076230 | 0 | NA | NA | NA | NA | NA | NA | NA | 0 | 0 | 0 | 0 | 0 | 0 |
| 21413 | ENSMUSG00000076231 | 0 | NA | NA | NA | NA | NA | NA | NA | 0 | 0 | 0 | 0 | 0 | 0 |
| 21414 | ENSMUSG00000076232 | 0 | NA | NA | NA | NA | NA | NA | NA | 0 | 0 | 0 | 0 | 0 | 0 |
| 21415 | ENSMUSG00000076233 | 0 | NA | NA | NA | NA | NA | NA | NA | 0 | 0 | 0 | 0 | 0 | 0 |
| 21416 | ENSMUSG00000076234 | 0 | NA | NA | NA | NA | NA | NA | NA | 0 | 0 | 0 | 0 | 0 | 0 |
| 21417 | ENSMUSG00000076236 | 0 | NA | NA | NA | NA | NA | Mir682 | microRNA 682 [Source:MGI Symbol;Acc:M | 0 | 0 | 0 | 0 | 0 | 0 |
| 21418 | ENSMUSG00000076237 | 0 | NA | NA | NA | NA | NA | NA | NA | 0 | 0 | 0 | 0 | 0 | 0 |
| 21419 | ENSMUSG00000076238 | 0 | NA | NA | NA | NA | NA | NA | NA | 0 | 0 | 0 | 0 | 0 | 0 |
| 21420 | ENSMUSG00000076240 | 0 | NA | NA | NA | NA | NA | Mir702 | microRNA 702 [Source:MGI Symbol;Acc:M | 0 | 0 | 0 | 0 | 0 | 0 |
| 21421 | ENSMUSG00000076241 | 0 | NA | NA | NA | NA | NA | Mir543 | microRNA 543 [Source:MGI Symbol;Acc:M | 0 | 0 | 0 | 0 | 0 | 0 |
| 21422 | ENSMUSG00000076246 | 0 | NA | NA | NA | NA | NA | NA | NA | 0 | 0 | 0 | 0 | 0 | 0 |
| 21423 | ENSMUSG00000076251 | 0 | NA | NA | NA | NA | NA | Mir501 | microRNA 501 [Source:MGI Symbol;Acc:M | 0 | 0 | 0 | 0 | 0 | 0 |
| 21424 | ENSMUSG00000076252 | 0 | NA | NA | NA | NA | NA | Mir302b | microRNA 302b [Source:MGI Symbol;Acc: | 0 | 0 | 0 | 0 | 0 | 0 |
| 21425 | ENSMUSG00000076253 | 0 | NA | NA | NA | NA | NA | Mir680-3 | microRNA 680-3 [Source:MGI Symbol;Acc | 0 | 0 | 0 | 0 | 0 | 0 |
| 21426 | ENSMUSG00000076255 | 0 | NA | NA | NA | NA | NA | Mir92b | microRNA 92b [Source:MGI Symbol;Acc:M | 0 | 0 | 0 | 0 | 0 | 0 |
| 21427 | ENSMUSG00000076256 | 0 | NA | NA | NA | NA | NA | Mir19b-1 | microRNA 19b-1 [Source:MGI Symbol;Acc | 0 | 0 | 0 | 0 | 0 | 0 |
| 21428 | ENSMUSG00000076258 | 20693.59827 | 0.803178936 | 1.023808046 | 0.784501488 | 0.432745909 | 0.845264432 | NA | NA | 12657 | 16973 | 15603 | 10357 | 63271 | 5300.8 |
| 21429 | ENSMUSG00000076269 | 0 | NA | NA | NA | NA | NA | Mir374b | microRNA 374b [Source:MGI Symbol;Acc: | 0 | 0 | 0 | 0 | 0 | 0 |
| 21430 | ENSMUSG00000076270 | 0 | NA | NA | NA | NA | NA | Mir719 | microRNA 719 [Source:MGI Symbol;Acc:M | 0 | 0 | 0 | 0 | 0 | 0 |
| 21431 | ENSMUSG00000076271 | 0.160662974 | 1.126924175 | 4.080472857 | 0.276174898 | 0.782413736 | NA | Mir697 | microRNA 697 [Source:MGI Symbol;Acc:M | 0 | 0 | 0 | 0.964 | 0 | 0 |
| 21432 | ENSMUSG00000076272 | 0 | NA | NA | NA | NA | NA | Mir666 | microRNA 666 [Source:MGI Symbol;Acc:M | 0 | 0 | 0 | 0 | 0 | 0 |
| 21433 | ENSMUSG00000076274 | 0 | NA | NA | NA | NA | NA | NA | NA | 0 | 0 | 0 | 0 | 0 | 0 |
| 21434 | ENSMUSG00000076275 | 0 | NA | NA | NA | NA | NA | Mir675 | microRNA 675 [Source:MGI Symbol;Acc:M | 0 | 0 | 0 | 0 | 0 | 0 |
| 21435 | ENSMUSG00000076281 | 175.4390295 | 1.142558117 | 1.173495513 | 0.973636545 | 0.330237034 | 0.774747658 | NA | NA | 93.16 | 121.29 | 113.71 | 92.542 | 609.66 | 22.277 |
| 21436 | ENSMUSG00000076282 | 0 | NA | NA | NA | NA | NA | NA | NA | 0 | 0 | 0 | 0 | 0 | 0 |
| 21437 | ENSMUSG00000076284 | 0 | NA | NA | NA | NA | NA | NA | NA | 0 | 0 | 0 | 0 | 0 | 0 |
| 21438 | ENSMUSG00000076286 | 0 | NA | NA | NA | NA | NA | Mir672 | microRNA 672 [Source:MGI Symbol;Acc:M | 0 | 0 | 0 | 0 | 0 | 0 |
| 21439 | ENSMUSG00000076287 | 0.267700746 | -1.497481649 | 4.057001436 | -0.369110456 | 0.712045396 | NA | NA | NA | 1.6062 | 0 | 0 | 0 | 0 | 0 |
| 21440 | ENSMUSG00000076288 | 0 | NA | NA | NA | NA | NA | Mir301b | microRNA 301b [Source:MGI Symbol;Acc: | 0 | 0 | 0 | 0 | 0 | 0 |
| 21441 | ENSMUSG00000076294 | 0 | NA | NA | NA | NA | NA | NA | NA | 0 | 0 | 0 | 0 | 0 | 0 |
| 21442 | ENSMUSG00000076300 | 0 | NA | NA | NA | NA | NA | NA | NA | 0 | 0 | 0 | 0 | 0 | 0 |
| 21443 | ENSMUSG00000076303 | 0 | NA | NA | NA | NA | NA | NA | NA | 0 | 0 | 0 | 0 | 0 | 0 |
| 21444 | ENSMUSG00000076304 | 0 | NA | NA | NA | NA | NA | NA | NA | 0 | 0 | 0 | 0 | 0 | 0 |

| | | | | | | | | | | | | | | | |
|---|---|---|---|---|---|---|---|---|---|---|---|---|---|---|---|
| 21445 | ENSMUSG00000076305 | 0 | NA | NA | NA | NA | NA | NA | NA | 0 | 0 | 0 | 0 | 0 | 0 |
| 21446 | ENSMUSG00000076309 | 0 | NA | NA | NA | NA | NA | Mir147 | microRNA 147 [Source:MGI Symbol;Acc:M | 0 | 0 | 0 | 0 | 0 | 0 |
| 21447 | ENSMUSG00000076312 | 0 | NA | NA | NA | NA | NA | Mir499 | microRNA 499 [Source:MGI Symbol;Acc:M | 0 | 0 | 0 | 0 | 0 | 0 |
| 21448 | ENSMUSG00000076313 | 0 | NA | NA | NA | NA | NA | Mir665 | microRNA 665 [Source:MGI Symbol;Acc:M | 0 | 0 | 0 | 0 | 0 | 0 |
| 21449 | ENSMUSG00000076315 | 1.388594601 | 0.820146163 | 1.971024992 | 0.416101351 | 0.677335841 | NA | Mir343 | microRNA 343 [Source:MGI Symbol;Acc:M | 0 | 2.0558 | 1.0062 | 1.928 | 0 | 3.3415 |
| 21450 | ENSMUSG00000076316 | 0 | NA | NA | NA | NA | NA | Mir673 | microRNA 673 [Source:MGI Symbol;Acc:M | 0 | 0 | 0 | 0 | 0 | 0 |
| 21451 | ENSMUSG00000076318 | 0 | NA | NA | NA | NA | NA | Mir216b | microRNA 216b [Source:MGI Symbol;Acc: | 0 | 0 | 0 | 0 | 0 | 0 |
| 21452 | ENSMUSG00000076320 | 0 | NA | NA | NA | NA | NA | NA | NA | 0 | 0 | 0 | 0 | 0 | 0 |
| 21453 | ENSMUSG00000076323 | 0 | NA | NA | NA | NA | NA | NA | NA | 0 | 0 | 0 | 0 | 0 | 0 |
| 21454 | ENSMUSG00000076327 | 0 | NA | NA | NA | NA | NA | NA | NA | 0 | 0 | 0 | 0 | 0 | 0 |
| 21455 | ENSMUSG00000076332 | 0 | NA | NA | NA | NA | NA | NA | NA | 0 | 0 | 0 | 0 | 0 | 0 |
| 21456 | ENSMUSG00000076333 | 0.181769715 | 1.126924175 | 4.080472857 | 0.276174898 | 0.782413736 | NA | Mir599 | microRNA 599 [Source:MGI Symbol;Acc:M | 0 | 0 | 0 | 0 | 1.0906 | 0 |
| 21457 | ENSMUSG00000076335 | 0 | NA | NA | NA | NA | NA | Mir670 | microRNA 670 [Source:MGI Symbol;Acc:M | 0 | 0 | 0 | 0 | 0 | 0 |
| 21458 | ENSMUSG00000076337 | 0 | NA | NA | NA | NA | NA | NA | NA | 0 | 0 | 0 | 0 | 0 | 0 |
| 21459 | ENSMUSG00000076338 | 0 | NA | NA | NA | NA | NA | Mir181d | microRNA 181d [Source:MGI Symbol;Acc: | 0 | 0 | 0 | 0 | 0 | 0 |
| 21460 | ENSMUSG00000076343 | 0 | NA | NA | NA | NA | NA | NA | NA | 0 | 0 | 0 | 0 | 0 | 0 |
| 21461 | ENSMUSG00000076355 | 0 | NA | NA | NA | NA | NA | NA | NA | 0 | 0 | 0 | 0 | 0 | 0 |
| 21462 | ENSMUSG00000076357 | 0 | NA | NA | NA | NA | NA | Mir653 | microRNA 653 [Source:MGI Symbol;Acc:M | 0 | 0 | 0 | 0 | 0 | 0 |
| 21463 | ENSMUSG00000076360 | 0 | NA | NA | NA | NA | NA | NA | NA | 0 | 0 | 0 | 0 | 0 | 0 |
| 21464 | ENSMUSG00000076361 | 0 | NA | NA | NA | NA | NA | Mir182 | microRNA 182 [Source:MGI Symbol;Acc:M | 0 | 0 | 0 | 0 | 0 | 0 |
| 21465 | ENSMUSG00000076364 | 0 | NA | NA | NA | NA | NA | NA | NA | 0 | 0 | 0 | 0 | 0 | 0 |
| 21466 | ENSMUSG00000076366 | 0 | NA | NA | NA | NA | NA | NA | NA | 0 | 0 | 0 | 0 | 0 | 0 |
| 21467 | ENSMUSG00000076370 | 0 | NA | NA | NA | NA | NA | NA | NA | 0 | 0 | 0 | 0 | 0 | 0 |
| 21468 | ENSMUSG00000076371 | 0 | NA | NA | NA | NA | NA | NA | NA | 0 | 0 | 0 | 0 | 0 | 0 |
| 21469 | ENSMUSG00000076372 | 0 | NA | NA | NA | NA | NA | Mir568 | microRNA 568 [Source:MGI Symbol;Acc:M | 0 | 0 | 0 | 0 | 0 | 0 |
| 21470 | ENSMUSG00000076376 | 0.167707431 | -0.796629176 | 4.080472857 | -0.195229623 | 0.84521319 | NA | Mir674 | microRNA 674 [Source:MGI Symbol;Acc:M | 0 | 0 | 1.0062 | 0 | 0 | 0 |
| 21471 | ENSMUSG00000076377 | 0.160662974 | 1.126924175 | 4.080472857 | 0.276174898 | 0.782413736 | NA | Gm22370 | predicted gene, 22370 [Source:MGI Symb | 0 | 0 | 0 | 0.964 | 0 | 0 |
| 21472 | ENSMUSG00000076379 | 0 | NA | NA | NA | NA | NA | Mir190a | microRNA 190a [Source:MGI Symbol;Acc: | 0 | 0 | 0 | 0 | 0 | 0 |
| 21473 | ENSMUSG00000076382 | 0 | NA | NA | NA | NA | NA | NA | NA | 0 | 0 | 0 | 0 | 0 | 0 |
| 21474 | ENSMUSG00000076385 | 0 | NA | NA | NA | NA | NA | NA | NA | 0 | 0 | 0 | 0 | 0 | 0 |
| 21475 | ENSMUSG00000076387 | 0 | NA | NA | NA | NA | NA | Mir671 | microRNA 671 [Source:MGI Symbol;Acc:M | 0 | 0 | 0 | 0 | 0 | 0 |
| 21476 | ENSMUSG00000076389 | 0 | NA | NA | NA | NA | NA | NA | NA | 0 | 0 | 0 | 0 | 0 | 0 |
| 21477 | ENSMUSG00000076392 | 0 | NA | NA | NA | NA | NA | NA | NA | 0 | 0 | 0 | 0 | 0 | 0 |
| 21478 | ENSMUSG00000076394 | 0 | NA | NA | NA | NA | NA | NA | NA | 0 | 0 | 0 | 0 | 0 | 0 |
| 21479 | ENSMUSG00000076395 | 0 | NA | NA | NA | NA | NA | NA | NA | 0 | 0 | 0 | 0 | 0 | 0 |
| 21480 | ENSMUSG00000076396 | 0 | NA | NA | NA | NA | NA | Mir667 | microRNA 667 [Source:MGI Symbol;Acc:M | 0 | 0 | 0 | 0 | 0 | 0 |
| 21481 | ENSMUSG00000076398 | 0 | NA | NA | NA | NA | NA | Mir676 | microRNA 676 [Source:MGI Symbol;Acc:M | 0 | 0 | 0 | 0 | 0 | 0 |
| 21482 | ENSMUSG00000076399 | 0 | NA | NA | NA | NA | NA | Mir450-2 | microRNA 450-2 [Source:MGI Symbol;Acc | 0 | 0 | 0 | 0 | 0 | 0 |
| 21483 | ENSMUSG00000076407 | 0 | NA | NA | NA | NA | NA | NA | NA | 0 | 0 | 0 | 0 | 0 | 0 |
| 21484 | ENSMUSG00000076424 | 0 | NA | NA | NA | NA | NA | Gm25986 | predicted gene, 25986 [Source:MGI Symb | 0 | 0 | 0 | 0 | 0 | 0 |
| 21485 | ENSMUSG00000076430 | 0.36353943 | 2.065989254 | 4.034137301 | 0.512126658 | 0.608562368 | NA | Hus1b | HUS1 checkpoint clamp component B [Sou | 0 | 0 | 0 | 0 | 2.1812 | 0 |
| 21486 | ENSMUSG00000076431 | 12.27454751 | 0.09918901 | 0.628219366 | 0.157889131 | 0.874544163 | 0.980188144 | Sox4 | SRY (sex determining region Y)-box 4 [Sou | 7.2279 | 14.391 | 14.087 | 12.532 | 9.8156 | 15.594 |
| 21487 | ENSMUSG00000076432 | 1013.549329 | 0.066073673 | 0.104121174 | 0.634584405 | 0.525699502 | 0.888334265 | Ywhaq | tyrosine 3-monooxygenase/tryptophan 5- | 1033.6 | 944.66 | 992.16 | 1042.1 | 1026.3 | 1042.6 |
| 21488 | ENSMUSG00000076433 | 0.342639279 | -1.805976596 | 4.028547714 | -0.448294702 | 0.653940524 | NA | Cep295nl | CEP295 N-terminal like [Source:MGI Symb | 0 | 2.0558 | 0 | 0 | 0 | 0 |
| 21489 | ENSMUSG00000076434 | 1.846996052 | -3.162536966 | 1.841939344 | -1.716960429 | 0.085986402 | NA | Wfdc3 | WAP four-disulfide core domain 3 [Source | 4.0155 | 3.0838 | 3.0187 | 0.964 | 0 | 0 |
| 21490 | ENSMUSG00000076435 | 4167.440107 | -0.083154497 | 0.086752815 | -0.958522177 | 0.337799514 | 0.780126485 | Acsf2 | acyl-CoA synthetase family member 2 [Sou | 4401 | 4123 | 4337.9 | 4009.2 | 4171.6 | 3961.9 |
| 21491 | ENSMUSG00000076436 | 0.160662974 | 1.126924175 | 4.080472857 | 0.276174898 | 0.782413736 | NA | Oxct2a | 3-oxoacid CoA transferase 2A [Source:MG | 0 | 0 | 0 | 0.964 | 0 | 0 |
| 21492 | ENSMUSG00000076437 | 155.2691607 | -0.017397698 | 0.212637772 | -0.081818473 | 0.934791066 | 0.988420305 | Selenoh | selenoprotein H [Source:MGI Symbol;Acc: | 159.01 | 148.02 | 161 | 120.5 | 155.96 | 187.13 |
| 21493 | ENSMUSG00000076438 | 0.160662974 | 1.126924175 | 4.080472857 | 0.276174898 | 0.782413736 | NA | Oxct2b | 3-oxoacid CoA transferase 2B [Source:MG | 0 | 0 | 0 | 0.964 | 0 | 0 |
| 21494 | ENSMUSG00000076439 | 0.185640647 | 1.126924175 | 4.080472857 | 0.276174898 | 0.782413736 | NA | Mog | myelin oligodendrocyte glycoprotein [Sou | 0 | 0 | 0 | 0 | 0 | 1.1138 |
| 21495 | ENSMUSG00000076441 | 53762.47686 | 0.109505198 | 0.124778253 | 0.877598427 | 0.380161683 | 0.812628154 | Ass1 | argininosuccinate synthetase 1 [Source:M | 51892 | 48941 | 54335 | 46444 | 55023 | 65940 |
| 21496 | ENSMUSG00000076444 | 0 | NA | NA | NA | NA | NA | Mir761 | microRNA 761 [Source:MGI Symbol;Acc:M | 0 | 0 | 0 | 0 | 0 | 0 |
| 21497 | ENSMUSG00000076445 | 0 | NA | NA | NA | NA | NA | NA | NA | 0 | 0 | 0 | 0 | 0 | 0 |
| 21498 | ENSMUSG00000076448 | 0 | NA | NA | NA | NA | NA | Mir763 | microRNA 763 [Source:MGI Symbol;Acc:M | 0 | 0 | 0 | 0 | 0 | 0 |
| 21499 | ENSMUSG00000076450 | 0 | NA | NA | NA | NA | NA | Mir804 | microRNA 804 [Source:MGI Symbol;Acc:M | 0 | 0 | 0 | 0 | 0 | 0 |
| 21500 | ENSMUSG00000076451 | 0 | NA | NA | NA | NA | NA | Mir770 | microRNA 770 [Source:MGI Symbol;Acc:M | 0 | 0 | 0 | 0 | 0 | 0 |

| | | | | | | | | | | | | | | | |
|---|---|---|---|---|---|---|---|---|---|---|---|---|---|---|---|
| 21501 | ENSMUSG00000076453 | 0 | NA | NA | NA | NA | NA | Mir759 | microRNA 759 [Source:MGI Symbol;Acc:M | 0 | 0 | 0 | 0 | 0 | 0 |
| 21502 | ENSMUSG00000076454 | 0 | NA | NA | NA | NA | NA | Mir762 | microRNA 762 [Source:MGI Symbol;Acc:M | 0 | 0 | 0 | 0 | 0 | 0 |
| 21503 | ENSMUSG00000076455 | 0 | NA | NA | NA | NA | NA | Mir767 | microRNA 767 [Source:MGI Symbol;Acc:M | 0 | 0 | 0 | 0 | 0 | 0 |
| 21504 | ENSMUSG00000076456 | 0 | NA | NA | NA | NA | NA | Mir760 | microRNA 760 [Source:MGI Symbol;Acc:M | 0 | 0 | 0 | 0 | 0 | 0 |
| 21505 | ENSMUSG00000076457 | 0.17131964 | -0.796629176 | 4.080472857 | -0.195229623 | 0.84521319 | NA | Mir802 | microRNA 802 [Source:MGI Symbol;Acc:M | 0 | 1.0279 | 0 | 0 | 0 | 0 |
| 21506 | ENSMUSG00000076458 | 0 | NA | NA | NA | NA | NA | Mir764 | microRNA 764 [Source:MGI Symbol;Acc:M | 0 | 0 | 0 | 0 | 0 | 0 |
| 21507 | ENSMUSG00000076459 | 0 | NA | NA | NA | NA | NA | Mir758 | microRNA 758 [Source:MGI Symbol;Acc:M | 0 | 0 | 0 | 0 | 0 | 0 |
| 21508 | ENSMUSG00000076460 | 0 | NA | NA | NA | NA | NA | Mir744 | microRNA 744 [Source:MGI Symbol;Acc:M | 0 | 0 | 0 | 0 | 0 | 0 |
| 21509 | ENSMUSG00000076461 | 0.335414862 | -1.779047242 | 4.030804179 | -0.441362855 | 0.658950329 | NA | Trbv1 | T cell receptor beta, variable 1 [Source:M | 0 | 0 | 2.0125 | 0 | 0 | 0 |
| 21510 | ENSMUSG00000076462 | 0.301557804 | -1.64335916 | 4.042814367 | -0.406488899 | 0.684383412 | NA | Trbv2 | T cell receptor beta, variable 2 [Source:M | 0.8031 | 0 | 1.0062 | 0 | 0 | 0 |
| 21511 | ENSMUSG00000076463 | 0 | NA | NA | NA | NA | NA | Trbv3 | T cell receptor beta, variable 3 [Source:M | 0 | 0 | 0 | 0 | 0 | 0 |
| 21512 | ENSMUSG00000076465 | 0 | NA | NA | NA | NA | NA | Trbv5 | T cell receptor beta, variable 5 [Source:M | 0 | 0 | 0 | 0 | 0 | 0 |
| 21513 | ENSMUSG00000076467 | 0.483327519 | -0.685704547 | 3.145069396 | -0.218025252 | 0.827409436 | NA | Trbv13-1 | T cell receptor beta, variable 13-1 [Source | 0.8031 | 0 | 1.0062 | 0 | 1.0906 | 0 |
| 21514 | ENSMUSG00000076469 | 1.409907932 | -0.615129367 | 2.179673303 | -0.282211727 | 0.777781167 | NA | Trbv13-2 | T cell receptor beta, variable 13-2 [Source | 0 | 4.1117 | 1.0062 | 0 | 0 | 3.3415 |
| 21515 | ENSMUSG00000076470 | 0.401551119 | -2.072982726 | 4.008251979 | -0.517178744 | 0.605031388 | NA | Trbv13-3 | T cell receptor beta, variable 13-3 [Source | 2.4093 | 0 | 0 | 0 | 0 | 0 |
| 21516 | ENSMUSG00000076471 | 0 | NA | NA | NA | NA | NA | Trbv14 | T cell receptor beta, variable 14 [Source:M | 0 | 0 | 0 | 0 | 0 | 0 |
| 21517 | ENSMUSG00000076472 | 0 | NA | NA | NA | NA | NA | Trbv15 | T cell receptor beta, variable 15 [Source:M | 0 | 0 | 0 | 0 | 0 | 0 |
| 21518 | ENSMUSG00000076473 | 0 | NA | NA | NA | NA | NA | Trbv16 | T cell receptor beta, variable 16 [Source:M | 0 | 0 | 0 | 0 | 0 | 0 |
| 21519 | ENSMUSG00000076474 | 0.321325949 | 1.911575114 | 4.048267893 | 0.472195804 | 0.636787034 | NA | Trbv17 | T cell receptor beta, variable 17 [Source:M | 0 | 0 | 0 | 1.928 | 0 | 0 |
| 21520 | ENSMUSG00000076475 | 0.133850373 | -0.796629176 | 4.080472857 | -0.195229623 | 0.84521319 | NA | Trbv19 | T cell receptor beta, variable 19 [Source:M | 0.8031 | 0 | 0 | 0 | 0 | 0 |
| 21521 | ENSMUSG00000076476 | 0 | NA | NA | NA | NA | NA | Trbv20 | T cell receptor beta, variable 20 [Source:M | 0 | 0 | 0 | 0 | 0 | 0 |
| 21522 | ENSMUSG00000076477 | 0 | NA | NA | NA | NA | NA | Trbv21 | T cell receptor beta, variable 21 [Source:M | 0 | 0 | 0 | 0 | 0 | 0 |
| 21523 | ENSMUSG00000076478 | 0 | NA | NA | NA | NA | NA | Trbv23 | T cell receptor beta, variable V23 [Source: | 0 | 0 | 0 | 0 | 0 | 0 |
| 21524 | ENSMUSG00000076479 | 0 | NA | NA | NA | NA | NA | Trbv26 | T cell receptor beta, variable 26 [Source:M | 0 | 0 | 0 | 0 | 0 | 0 |
| 21525 | ENSMUSG00000076480 | 0 | NA | NA | NA | NA | NA | Trbv29 | T cell receptor beta, variable 29 [Source:M | 0 | 0 | 0 | 0 | 0 | 0 |
| 21526 | ENSMUSG00000076481 | 0 | NA | NA | NA | NA | NA | Trbv30 | T cell receptor beta, variable 30 [Source:M | 0 | 0 | 0 | 0 | 0 | 0 |
| 21527 | ENSMUSG00000076483 | 0 | NA | NA | NA | NA | NA | Trbj1-1 | T cell receptor beta joining 1-1 [Source:M | 0 | 0 | 0 | 0 | 0 | 0 |
| 21528 | ENSMUSG00000076484 | 0 | NA | NA | NA | NA | NA | Trbj1-2 | T cell receptor beta joining 1-2 [Source:M | 0 | 0 | 0 | 0 | 0 | 0 |
| 21529 | ENSMUSG00000076485 | 0 | NA | NA | NA | NA | NA | Trbj1-3 | T cell receptor beta joining 1-3 [Source:M | 0 | 0 | 0 | 0 | 0 | 0 |
| 21530 | ENSMUSG00000076486 | 0 | NA | NA | NA | NA | NA | Trbj1-4 | T cell receptor beta joining 1-4 [Source:M | 0 | 0 | 0 | 0 | 0 | 0 |
| 21531 | ENSMUSG00000076487 | 0 | NA | NA | NA | NA | NA | Trbj1-5 | T cell receptor beta joining 1-5 [Source:M | 0 | 0 | 0 | 0 | 0 | 0 |
| 21532 | ENSMUSG00000076488 | 0 | NA | NA | NA | NA | NA | Trbj1-6 | T cell receptor beta joining 1-6 [Source:M | 0 | 0 | 0 | 0 | 0 | 0 |
| 21533 | ENSMUSG00000076489 | 0 | NA | NA | NA | NA | NA | Trbj1-7 | T cell receptor beta joining 1-7 [Source:M | 0 | 0 | 0 | 0 | 0 | 0 |
| 21534 | ENSMUSG00000076490 | 4.667478608 | -0.88118531 | 1.101117052 | -0.800264884 | 0.423557344 | NA | Trbc1 | T cell receptor beta, constant region 1 [So | 4.8186 | 8.2233 | 5.0312 | 0 | 4.3625 | 5.5692 |
| 21535 | ENSMUSG00000076492 | 0 | NA | NA | NA | NA | NA | Trbj2-1 | T cell receptor beta joining 2-1 [Source:M | 0 | 0 | 0 | 0 | 0 | 0 |
| 21536 | ENSMUSG00000076493 | 0 | NA | NA | NA | NA | NA | Trbj2-2 | T cell receptor beta joining 2-2 [Source:M | 0 | 0 | 0 | 0 | 0 | 0 |
| 21537 | ENSMUSG00000076494 | 0 | NA | NA | NA | NA | NA | Trbj2-3 | T cell receptor beta joining 2-3 [Source:M | 0 | 0 | 0 | 0 | 0 | 0 |
| 21538 | ENSMUSG00000076495 | 0 | NA | NA | NA | NA | NA | Trbj2-4 | T cell receptor beta joining 2-4 [Source:M | 0 | 0 | 0 | 0 | 0 | 0 |
| 21539 | ENSMUSG00000076496 | 0.167707431 | -0.796629176 | 4.080472857 | -0.195229623 | 0.84521319 | NA | Trbj2-5 | T cell receptor beta joining 2-5 [Source:M | 0 | 0 | 1.0062 | 0 | 0 | 0 |
| 21540 | ENSMUSG00000076497 | 0 | NA | NA | NA | NA | NA | Trbj2-7 | T cell receptor beta joining 2-7 [Source:M | 0 | 0 | 0 | 0 | 0 | 0 |
| 21541 | ENSMUSG00000076498 | 7.35729569 | -1.670509389 | 0.86633341 | -1.928252298 | 0.053823752 | 0.326281119 | Trbc2 | T cell receptor beta, constant 2 [Source:M | 15.259 | 6.1675 | 12.075 | 2.8919 | 2.1812 | 5.5692 |
| 21542 | ENSMUSG00000076499 | 0 | NA | NA | NA | NA | NA | Trbv31 | T cell receptor beta, variable 31 [Source:M | 0 | 0 | 0 | 0 | 0 | 0 |
| 21543 | ENSMUSG00000076500 | 0 | NA | NA | NA | NA | NA | Gm20730 | predicted gene, 20730 [Source:MGI Symb | 0 | 0 | 0 | 0 | 0 | 0 |
| 21544 | ENSMUSG00000076501 | 0 | NA | NA | NA | NA | NA | Igkv2-137 | immunoglobulin kappa chain variable 2-1 | 0 | 0 | 0 | 0 | 0 | 0 |
| 21545 | ENSMUSG00000076505 | 0 | NA | NA | NA | NA | NA | Igkv1-131 | immunoglobulin kappa variable 1-131 [So | 0 | 0 | 0 | 0 | 0 | 0 |
| 21546 | ENSMUSG00000076508 | 0 | NA | NA | NA | NA | NA | Igkv17-127 | immunoglobulin kappa variable 17-127 [S | 0 | 0 | 0 | 0 | 0 | 0 |
| 21547 | ENSMUSG00000076512 | 0 | NA | NA | NA | NA | NA | Igkv9-123 | immunoglobulin kappa variable 9-123 [So | 0 | 0 | 0 | 0 | 0 | 0 |
| 21548 | ENSMUSG00000076514 | 0 | NA | NA | NA | NA | NA | Igkv17-121 | immunoglobulin kappa variable 17-121 [S | 0 | 0 | 0 | 0 | 0 | 0 |
| 21549 | ENSMUSG00000076518 | 0.267700746 | -1.497481649 | 4.057001436 | -0.369110456 | 0.712045396 | NA | Igkv2-112 | immunoglobulin kappa variable 2-112 [So | 1.6062 | 0 | 0 | 0 | 0 | 0 |
| 21550 | ENSMUSG00000076522 | 0.946860264 | 0.386550567 | 2.714838108 | 0.142384389 | 0.886776392 | NA | Igkv16-104 | immunoglobulin kappa variable 16-104 [S | 2.4093 | 0 | 0 | 0 | 3.2719 | 0 |
| 21551 | ENSMUSG00000076523 | 0.267700746 | -1.497481649 | 4.057001436 | -0.369110456 | 0.712045396 | NA | Igkv15-103 | immunoglobulin kappa chain variable 15- | 1.6062 | 0 | 0 | 0 | 0 | 0 |
| 21552 | ENSMUSG00000076525 | 0 | NA | NA | NA | NA | NA | Igkv1-99 | immunoglobulin kappa variable 1-99 [Sou | 0 | 0 | 0 | 0 | 0 | 0 |
| 21553 | ENSMUSG00000076526 | 0.185640647 | 1.126924175 | 4.080472857 | 0.276174898 | 0.782413736 | NA | Igkv12-98 | immunoglobulin kappa variable 12-98 [So | 0 | 0 | 0 | 0 | 0 | 1.1138 |
| 21554 | ENSMUSG00000076530 | 0 | NA | NA | NA | NA | NA | Igkv11-106 | immunoglobulin kappa variable 11-106 [S | 0 | 0 | 0 | 0 | 0 | 0 |
| 21555 | ENSMUSG00000076531 | 0 | NA | NA | NA | NA | NA | Igkv4-92 | immunoglobulin kappa variable 4-92 [Sou | 0 | 0 | 0 | 0 | 0 | 0 |
| 21556 | ENSMUSG00000076532 | 0 | NA | NA | NA | NA | NA | Igkv4-91 | immunoglobulin kappa chain variable 4-9 | 0 | 0 | 0 | 0 | 0 | 0 |

| | | | | | | | | | | | | | | | |
|---|---|---|---|---|---|---|---|---|---|---|---|---|---|---|---|
| 21557 | ENSMUSG00000076533 | 0 | NA | NA | NA | NA | NA | Igkv4-90 | immunoglobulin kappa chain variable 4-9( | 0 | 0 | 0 | 0 | 0 | 0 |
| 21558 | ENSMUSG00000076534 | 0.305170013 | -1.657836477 | 4.041480335 | -0.410205257 | 0.681655385 | NA | Igkv12-89 | immunoglobulin kappa chain variable 12-{ | 0.8031 | 1.0279 | 0 | 0 | 0 | 0 |
| 21559 | ENSMUSG00000076535 | 0 | NA | NA | NA | NA | NA | Igkv1-88 | immunoglobulin kappa chain variable 1-8{ | 0 | 0 | 0 | 0 | 0 | 0 |
| 21560 | ENSMUSG00000076536 | 0 | NA | NA | NA | NA | NA | Igkv4-86 | immunoglobulin kappa variable 4-86 [Sou | 0 | 0 | 0 | 0 | 0 | 0 |
| 21561 | ENSMUSG00000076538 | 0 | NA | NA | NA | NA | NA | Igkv13-84 | immunoglobulin kappa chain variable 13-{ | 0 | 0 | 0 | 0 | 0 | 0 |
| 21562 | ENSMUSG00000076539 | 0 | NA | NA | NA | NA | NA | Igkv4-81 | immunoglobulin kappa variable 4-81 [Sou | 0 | 0 | 0 | 0 | 0 | 0 |
| 21563 | ENSMUSG00000076540 | 0 | NA | NA | NA | NA | NA | Igkv4-80 | immunoglobulin kappa variable 4-80 [Sou | 0 | 0 | 0 | 0 | 0 | 0 |
| 21564 | ENSMUSG00000076541 | 0 | NA | NA | NA | NA | NA | Igkv4-79 | immunoglobulin kappa variable 4-79 [Sou | 0 | 0 | 0 | 0 | 0 | 0 |
| 21565 | ENSMUSG00000076543 | 0 | NA | NA | NA | NA | NA | Igkv4-74 | immunoglobulin kappa variable 4-74 [Sou | 0 | 0 | 0 | 0 | 0 | 0 |
| 21566 | ENSMUSG00000076545 | 0.305170013 | -1.657836477 | 4.041480335 | -0.410205257 | 0.681655385 | NA | Igkv4-72 | immunoglobulin kappa chain variable 4-7: | 0.8031 | 1.0279 | 0 | 0 | 0 | 0 |
| 21567 | ENSMUSG00000076547 | 0 | NA | NA | NA | NA | NA | Igkv4-70 | immunoglobulin kappa chain variable 4-7( | 0 | 0 | 0 | 0 | 0 | 0 |
| 21568 | ENSMUSG00000076548 | 0 | NA | NA | NA | NA | NA | Igkv4-69 | immunoglobulin kappa variable 4-69 [Sou | 0 | 0 | 0 | 0 | 0 | 0 |
| 21569 | ENSMUSG00000076549 | 0 | NA | NA | NA | NA | NA | Igkv4-68 | immunoglobulin kappa variable 4-68 [Sou | 0 | 0 | 0 | 0 | 0 | 0 |
| 21570 | ENSMUSG00000076550 | 0.181769715 | 1.126924175 | 4.080472857 | 0.276174898 | 0.782413736 | NA | Igkv4-63 | immunoglobulin kappa variable 4-63 [Sou | 0 | 0 | 0 | 0 | 1.0906 | 0 |
| 21571 | ENSMUSG00000076552 | 0 | NA | NA | NA | NA | NA | Igkv4-61 | immunoglobulin kappa chain variable 4-6: | 0 | 0 | 0 | 0 | 0 | 0 |
| 21572 | ENSMUSG00000076555 | 0.401551119 | -2.072982726 | 4.008251979 | -0.517178744 | 0.605031388 | NA | Igkv4-57-1 | immunoglobulin kappa variable 4-57-1 [Sc | 2.4093 | 0 | 0 | 0 | 0 | 0 |
| 21573 | ENSMUSG00000076556 | 0.267700746 | -1.497481649 | 4.057001436 | -0.369110456 | 0.712045396 | NA | Igkv4-57 | immunoglobulin kappa variable 4-57 [Sou | 1.6062 | 0 | 0 | 0 | 0 | 0 |
| 21574 | ENSMUSG00000076557 | 0 | NA | NA | NA | NA | NA | Igkv4-83 | immunoglobulin kappa chain variable 4-8: | 0 | 0 | 0 | 0 | 0 | 0 |
| 21575 | ENSMUSG00000076562 | 0.181769715 | 1.126924175 | 4.080472857 | 0.276174898 | 0.782413736 | NA | Igkv4-50 | immunoglobulin kappa variable 4-50 [Sou | 0 | 0 | 0 | 0 | 1.0906 | 0 |
| 21576 | ENSMUSG00000076563 | 0.17131964 | -0.796629176 | 4.080472857 | -0.195229623 | 0.84521319 | NA | Igkv5-48 | immunoglobulin kappa variable 5-48 [Sou | 0 | 1.0279 | 0 | 0 | 0 | 0 |
| 21577 | ENSMUSG00000076564 | 0 | NA | NA | NA | NA | NA | Igkv12-46 | immunoglobulin kappa variable 12-46 [So | 0 | 0 | 0 | 0 | 0 | 0 |
| 21578 | ENSMUSG00000076569 | 0.804680097 | -3.048447205 | 2.843448763 | -1.072095001 | 0.283677364 | NA | Igkv5-39 | immunoglobulin kappa variable 5-39 [Sou | 0.8031 | 0 | 4.025 | 0 | 0 | 0 |
| 21579 | ENSMUSG00000076570 | 0 | NA | NA | NA | NA | NA | Igkv12-38 | immunoglobulin kappa chain variable 12-: | 0 | 0 | 0 | 0 | 0 | 0 |
| 21580 | ENSMUSG00000076571 | 0 | NA | NA | NA | NA | NA | Igkv5-37 | immunoglobulin kappa variable 5-37 [Sou | 0 | 0 | 0 | 0 | 0 | 0 |
| 21581 | ENSMUSG00000076572 | 0 | NA | NA | NA | NA | NA | Igkv18-36 | immunoglobulin kappa chain variable 18-: | 0 | 0 | 0 | 0 | 0 | 0 |
| 21582 | ENSMUSG00000076573 | 0 | NA | NA | NA | NA | NA | Igkv1-35 | immunoglobulin kappa variable 1-35 [Sou | 0 | 0 | 0 | 0 | 0 | 0 |
| 21583 | ENSMUSG00000076575 | 0.321325949 | 1.911575114 | 4.048267893 | 0.472195804 | 0.636787034 | NA | Igkv7-33 | immunoglobulin kappa chain variable 7-3: | 0 | 0 | 0 | 1.928 | 0 | 0 |
| 21584 | ENSMUSG00000076576 | 0.167707431 | -0.796629176 | 4.080472857 | -0.195229623 | 0.84521319 | NA | Igkv6-32 | immunoglobulin kappa variable 6-32 [Sou | 0 | 0 | 1.0062 | 0 | 0 | 0 |
| 21585 | ENSMUSG00000076577 | 0.319491021 | 0.16514334 | 4.044653141 | 0.040830038 | 0.967431393 | NA | Igkv8-30 | immunoglobulin kappa chain variable 8-3( | 0.8031 | 0 | 0 | 0 | 0 | 1.1138 |
| 21586 | ENSMUSG00000076578 | 0 | NA | NA | NA | NA | NA | Igkv6-29 | immunoglobulin kappa chain variable 6-2! | 0 | 0 | 0 | 0 | 0 | 0 |
| 21587 | ENSMUSG00000076580 | 0 | NA | NA | NA | NA | NA | Igkv8-27 | immunoglobulin kappa chain variable 8-2: | 0 | 0 | 0 | 0 | 0 | 0 |
| 21588 | ENSMUSG00000076581 | 0 | NA | NA | NA | NA | NA | Igkv8-26 | immunoglobulin kappa variable 8-26 [Sou | 0 | 0 | 0 | 0 | 0 | 0 |
| 21589 | ENSMUSG00000076583 | 0 | NA | NA | NA | NA | NA | Igkv8-24 | immunoglobulin kappa chain variable 8-2‹ | 0 | 0 | 0 | 0 | 0 | 0 |
| 21590 | ENSMUSG00000076586 | 0 | NA | NA | NA | NA | NA | Igkv8-21 | immunoglobulin kappa variable 8-21 [Sou | 0 | 0 | 0 | 0 | 0 | 0 |
| 21591 | ENSMUSG00000076587 | 0 | NA | NA | NA | NA | NA | Igkv6-20 | immunoglobulin kappa variable 6-20 [Sou | 0 | 0 | 0 | 0 | 0 | 0 |
| 21592 | ENSMUSG00000076589 | 0 | NA | NA | NA | NA | NA | Igkv8-18 | immunoglobulin kappa variable 8-18 [Sou | 0 | 0 | 0 | 0 | 0 | 0 |
| 21593 | ENSMUSG00000076591 | 0 | NA | NA | NA | NA | NA | Igkv8-16 | immunoglobulin kappa variable 8-16 [Sou | 0 | 0 | 0 | 0 | 0 | 0 |
| 21594 | ENSMUSG00000076594 | 0.267700746 | -1.497481649 | 4.057001436 | -0.369110456 | 0.712045396 | NA | Igkv6-13 | immunoglobulin kappa variable 6-13 [Sou | 1.6062 | 0 | 0 | 0 | 0 | 0 |
| 21595 | ENSMUSG00000076596 | 0.513958919 | -2.394274821 | 3.756096013 | -0.637437066 | 0.523840191 | NA | Igkv3-10 | immunoglobulin kappa variable 3-10 [Sou | 0 | 3.0838 | 0 | 0 | 0 | 0 |
| 21596 | ENSMUSG00000076598 | 0 | NA | NA | NA | NA | NA | Igkv3-7 | immunoglobulin kappa variable 3-7 [Sourc | 0 | 0 | 0 | 0 | 0 | 0 |
| 21597 | ENSMUSG00000076604 | 0 | NA | NA | NA | NA | NA | Igkj1 | immunoglobulin kappa joining 1 [Source:I | 0 | 0 | 0 | 0 | 0 | 0 |
| 21598 | ENSMUSG00000076605 | 0 | NA | NA | NA | NA | NA | Igkj2 | immunoglobulin kappa joining 2 [Source:I | 0 | 0 | 0 | 0 | 0 | 0 |
| 21599 | ENSMUSG00000076606 | 0.335414862 | -1.779047242 | 4.030804179 | -0.441362855 | 0.658950329 | NA | Igkj3 | immunoglobulin kappa joining 3 [Source:I | 0 | 0 | 2.0125 | 0 | 0 | 0 |
| 21600 | ENSMUSG00000076607 | 0.167707431 | -0.796629176 | 4.080472857 | -0.195229623 | 0.84521319 | NA | Igkj4 | immunoglobulin kappa joining 4 [Source:I | 0 | 0 | 1.0062 | 0 | 0 | 0 |
| 21601 | ENSMUSG00000076608 | 1.017133346 | -0.93167857 | 2.468499519 | -0.377427082 | 0.705856244 | NA | Igkj5 | immunoglobulin kappa joining 5 [Source:I | 0 | 0 | 4.025 | 0.964 | 0 | 1.1138 |
| 21602 | ENSMUSG00000076609 | 41.3990722 | -0.602407168 | 0.404147063 | -1.49056426 | 0.136075933 | 0.538744881 | Igkc | immunoglobulin kappa constant [Source:I | 58.626 | 61.675 | 29.181 | 24.099 | 43.625 | 31.188 |
| 21603 | ENSMUSG00000076612 | 10.09560589 | 0.128205415 | 0.854391702 | 0.150054612 | 0.880721529 | 0.981448736 | Ighg2c | immunoglobulin heavy constant gamma 2 | 18.471 | 9.2513 | 1.0062 | 7.7118 | 17.45 | 6.6831 |
| 21604 | ENSMUSG00000076613 | 13.6240476 | -0.26180353 | 0.613892127 | -0.426465038 | 0.669769018 | 0.930867597 | Ighg2b | immunoglobulin heavy constant gamma 2 | 20.078 | 15.419 | 9.0562 | 17.352 | 9.8156 | 10.025 |
| 21605 | ENSMUSG00000076614 | 3.591511059 | 1.197498742 | 1.223071984 | 0.979090976 | 0.32753503 | NA | Ighg1 | immunoglobulin heavy constant gamma 1 | 2.4093 | 4.1117 | 0 | 2.8919 | 5.4531 | 6.6831 |
| 21606 | ENSMUSG00000076615 | 0.339027071 | -1.79174149 | 3.919585585 | -0.457125237 | 0.647581032 | NA | Ighg3 | Immunoglobulin heavy constant gamma 3 | 0 | 1.0279 | 1.0062 | 0 | 0 | 0 |
| 21607 | ENSMUSG00000076617 | 67.76231301 | -0.473796076 | 0.338522892 | -1.399598334 | 0.161633633 | 0.584894158 | Ighm | immunoglobulin heavy constant mu [Sour | 89.144 | 83.261 | 63.393 | 32.775 | 57.803 | 80.197 |
| 21608 | ENSMUSG00000076618 | 0.267700746 | -1.497481649 | 4.057001436 | -0.369110456 | 0.712045396 | NA | Ighj4 | immunoglobulin heavy joining 4 [Source:I | 1.6062 | 0 | 0 | 0 | 0 | 0 |
| 21609 | ENSMUSG00000076619 | 0 | NA | NA | NA | NA | NA | Ighj3 | immunoglobulin heavy joining 3 [Source:I | 0 | 0 | 0 | 0 | 0 | 0 |
| 21610 | ENSMUSG00000076620 | 0 | NA | NA | NA | NA | NA | Ighj2 | immunoglobulin heavy joining 2 [Source:I | 0 | 0 | 0 | 0 | 0 | 0 |
| 21611 | ENSMUSG00000076621 | 0 | NA | NA | NA | NA | NA | Ighj1 | immunoglobulin heavy joining 1 [Source:I | 0 | 0 | 0 | 0 | 0 | 0 |
| 21612 | ENSMUSG00000076630 | 0 | NA | NA | NA | NA | NA | Ighd1-1 | immunoglobulin heavy diversity 1-1 [Sour | 0 | 0 | 0 | 0 | 0 | 0 |

| | | | | | | | | | | | | | | | |
|---|---|---|---|---|---|---|---|---|---|---|---|---|---|---|---|
| 21613 | ENSMUSG00000076632 | 0 | NA | NA | NA | NA | NA | Gm16968 | predicted gene, 16968 [Source:MGI Symb | 0 | 0 | 0 | 0 | 0 | 0 |
| 21614 | ENSMUSG00000076633 | 0 | NA | NA | NA | NA | NA | Ighv5-2 | immunoglobulin heavy variable 5-2 [Sourc | 0 | 0 | 0 | 0 | 0 | 0 |
| 21615 | ENSMUSG00000076646 | 0 | NA | NA | NA | NA | NA | Ighv2-6-8 | immunoglobulin heavy variable 2-6-8 [Sou | 0 | 0 | 0 | 0 | 0 | 0 |
| 21616 | ENSMUSG00000076652 | 0.167707431 | -0.796629176 | 4.080472857 | -0.195229623 | 0.84521319 | NA | Ighv7-3 | immunoglobulin heavy variable 7-3 [Sourc | 0 | 0 | 1.0062 | 0 | 0 | 0 |
| 21617 | ENSMUSG00000076653 | 0 | NA | NA | NA | NA | NA | Ighv7-2 | immunoglobulin heavy variable 7-2 [Sourc | 0 | 0 | 0 | 0 | 0 | 0 |
| 21618 | ENSMUSG00000076655 | 0 | NA | NA | NA | NA | NA | Ighv4-1 | immunoglobulin heavy variable 4-1 [Sourc | 0 | 0 | 0 | 0 | 0 | 0 |
| 21619 | ENSMUSG00000076661 | 0 | NA | NA | NA | NA | NA | Ighv16-1 | immunoglobulin heavy variable 16-1 [Sour | 0 | 0 | 0 | 0 | 0 | 0 |
| 21620 | ENSMUSG00000076665 | 0.703108924 | -2.875459482 | 2.96384054 | -0.970180225 | 0.331956666 | NA | Ighv7-1 | immunoglobulin heavy variable 7-1 [Sourc | 3.2124 | 0 | 1.0062 | 0 | 0 | 0 |
| 21621 | ENSMUSG00000076666 | 0 | NA | NA | NA | NA | NA | Ighv14-4 | immunoglobulin heavy variable 14-4 [Sour | 0 | 0 | 0 | 0 | 0 | 0 |
| 21622 | ENSMUSG00000076668 | 0 | NA | NA | NA | NA | NA | Ighv7-4 | immunoglobulin heavy variable 7-4 [Sourc | 0 | 0 | 0 | 0 | 0 | 0 |
| 21623 | ENSMUSG00000076670 | 0 | NA | NA | NA | NA | NA | Ighv3-5 | immunoglobulin heavy variable 3-5 [Sourc | 0 | 0 | 0 | 0 | 0 | 0 |
| 21624 | ENSMUSG00000076671 | 0 | NA | NA | NA | NA | NA | Ighv13-2 | immunoglobulin heavy variable 13-2 [Sour | 0 | 0 | 0 | 0 | 0 | 0 |
| 21625 | ENSMUSG00000076672 | 0.503095663 | 2.549494177 | 3.348715477 | 0.761334964 | 0.446457022 | NA | Ighv3-6 | immunoglobulin heavy variable 3-6 [Sourc | 0 | 0 | 0 | 1.928 | 1.0906 | 0 |
| 21626 | ENSMUSG00000076674 | 0 | NA | NA | NA | NA | NA | Ighv3-8 | immunoglobulin heavy variable V3-8 [Sou | 0 | 0 | 0 | 0 | 0 | 0 |
| 21627 | ENSMUSG00000076676 | 0 | NA | NA | NA | NA | NA | Ighv12-3 | immunoglobulin heavy variable V12-3 [So | 0 | 0 | 0 | 0 | 0 | 0 |
| 21628 | ENSMUSG00000076677 | 0.315620088 | 0.165143421 | 4.063300985 | 0.040642675 | 0.967580763 | NA | Ighv6-3 | immunoglobulin heavy variable 6-3 [Sourc | 0.8031 | 0 | 0 | 0 | 1.0906 | 0 |
| 21629 | ENSMUSG00000076680 | 0.811904515 | -3.057764991 | 2.533063159 | -1.207141236 | 0.227377757 | NA | Ighv6-6 | immunoglobulin heavy variable 6-6 [Sourc | 0.8031 | 2.0558 | 2.0125 | 0 | 0 | 0 |
| 21630 | ENSMUSG00000076688 | 0 | NA | NA | NA | NA | NA | Ighv15-2 | immunoglobulin heavy variable V15-2 [So | 0 | 0 | 0 | 0 | 0 | 0 |
| 21631 | ENSMUSG00000076695 | 0.695728634 | -1.425610641 | 3.05889997 | -0.46605337 | 0.641177299 | NA | Ighv1-18 | immunoglobulin heavy variable V1-18 [So | 0 | 3.0838 | 0 | 0 | 1.0906 | 0 |
| 21632 | ENSMUSG00000076709 | 0 | NA | NA | NA | NA | NA | Ighv1-47 | immunoglobulin heavy variable 1-47 [Sour | 0 | 0 | 0 | 0 | 0 | 0 |
| 21633 | ENSMUSG00000076710 | 0 | NA | NA | NA | NA | NA | Ighv1-49 | immunoglobulin heavy variable 1-49 [Sour | 0 | 0 | 0 | 0 | 0 | 0 |
| 21634 | ENSMUSG00000076731 | 0.481988923 | 2.492401104 | 3.837077151 | 0.649557203 | 0.515978285 | NA | Ighv8-12 | immunoglobulin heavy variable V8-12 [So | 0 | 0 | 0 | 2.8919 | 0 | 0 |
| 21635 | ENSMUSG00000076733 | 0 | NA | NA | NA | NA | NA | Ighv8-13 | immunoglobulin heavy variable 8-13 [Sour | 0 | 0 | 0 | 0 | 0 | 0 |
| 21636 | ENSMUSG00000076744 | 0 | NA | NA | NA | NA | NA | Trgv7 | T cell receptor gamma, variable 7 [Source: | 0 | 0 | 0 | 0 | 0 | 0 |
| 21637 | ENSMUSG00000076745 | 0 | NA | NA | NA | NA | NA | Trgv4 | T cell receptor gamma, variable 4 [Source: | 0 | 0 | 0 | 0 | 0 | 0 |
| 21638 | ENSMUSG00000076746 | 0 | NA | NA | NA | NA | NA | Trgv6 | T cell receptor gamma, variable 6 [Source: | 0 | 0 | 0 | 0 | 0 | 0 |
| 21639 | ENSMUSG00000076747 | 0 | NA | NA | NA | NA | NA | Trgv5 | T cell receptor gamma, variable 5 [Source: | 0 | 0 | 0 | 0 | 0 | 0 |
| 21640 | ENSMUSG00000076749 | 0.774435248 | -2.999089927 | 2.565713408 | -1.168910728 | 0.242439601 | NA | Trgc1 | T cell receptor gamma, constant 1 [Source | 1.6062 | 1.0279 | 2.0125 | 0 | 0 | 0 |
| 21641 | ENSMUSG00000076750 | 0 | NA | NA | NA | NA | NA | Trgv3 | T cell receptor gamma, variable 3 [Source: | 0 | 0 | 0 | 0 | 0 | 0 |
| 21642 | ENSMUSG00000076752 | 1.998779642 | -2.254462364 | 1.679887861 | -1.342031463 | 0.179585794 | NA | Trgc2 | T cell receptor gamma, constant 2 [Source | 4.8186 | 3.0838 | 2.0125 | 0.964 | 0 | 1.1138 |
| 21643 | ENSMUSG00000076754 | 0 | NA | NA | NA | NA | NA | Trgv2 | T cell receptor gamma variable 2 [Source:I | 0 | 0 | 0 | 0 | 0 | 0 |
| 21644 | ENSMUSG00000076755 | 0 | NA | NA | NA | NA | NA | Trgv1 | T cell receptor gamma, variable 1 [Source: | 0 | 0 | 0 | 0 | 0 | 0 |
| 21645 | ENSMUSG00000076757 | 0.356960287 | 0.16514264 | 3.880370363 | 0.042558474 | 0.966053499 | NA | Trgc4 | T cell receptor gamma, constant 4 [Source | 0 | 1.0279 | 0 | 0 | 0 | 1.1138 |
| 21646 | ENSMUSG00000076758 | 0 | NA | NA | NA | NA | NA | Trav1 | T cell receptor alpha variable 1 [Source:M | 0 | 0 | 0 | 0 | 0 | 0 |
| 21647 | ENSMUSG00000076759 | 0 | NA | NA | NA | NA | NA | Trav2 | T cell receptor alpha variable 2 [Source:M | 0 | 0 | 0 | 0 | 0 | 0 |
| 21648 | ENSMUSG00000076760 | 0 | NA | NA | NA | NA | NA | Trav3-1 | T cell receptor alpha variable 3-1 [Source: | 0 | 0 | 0 | 0 | 0 | 0 |
| 21649 | ENSMUSG00000076761 | 0 | NA | NA | NA | NA | NA | Trav5-1 | T cell receptor alpha variable 5-1 [Source: | 0 | 0 | 0 | 0 | 0 | 0 |
| 21650 | ENSMUSG00000076762 | 0 | NA | NA | NA | NA | NA | Trav6-1 | T cell receptor alpha variable 6-1 [Source: | 0 | 0 | 0 | 0 | 0 | 0 |
| 21651 | ENSMUSG00000076764 | 0 | NA | NA | NA | NA | NA | Trav6-2 | T cell receptor alpha variable 6-2 [Source: | 0 | 0 | 0 | 0 | 0 | 0 |
| 21652 | ENSMUSG00000076770 | 0 | NA | NA | NA | NA | NA | Trav8d-1 | T cell receptor alpha variable 8D-1 [Source | 0 | 0 | 0 | 0 | 0 | 0 |
| 21653 | ENSMUSG00000076778 | 0 | NA | NA | NA | NA | NA | Trav13d-1 | T cell receptor alpha variable 13D-1 [Sour | 0 | 0 | 0 | 0 | 0 | 0 |
| 21654 | ENSMUSG00000076795 | 0 | NA | NA | NA | NA | NA | Trav8d-2 | T cell receptor alpha variable 8D-2 [Sourc | 0 | 0 | 0 | 0 | 0 | 0 |
| 21655 | ENSMUSG00000076796 | 0 | NA | NA | NA | NA | NA | Trav16d-dv1 | T cell receptor alpha variable 16D-DV11 [S | 0 | 0 | 0 | 0 | 0 | 0 |
| 21656 | ENSMUSG00000076800 | 0 | NA | NA | NA | NA | NA | Trav6n-5 | T cell receptor alpha variable 6N-5 [Source | 0 | 0 | 0 | 0 | 0 | 0 |
| 21657 | ENSMUSG00000076802 | 0 | NA | NA | NA | NA | NA | Trav6n-6 | T cell receptor alpha variable 6N-6 [Source | 0 | 0 | 0 | 0 | 0 | 0 |
| 21658 | ENSMUSG00000076821 | 0 | NA | NA | NA | NA | NA | Trav8n-2 | T cell receptor alpha variable 8n-2 [Sourc | 0 | 0 | 0 | 0 | 0 | 0 |
| 21659 | ENSMUSG00000076822 | 0 | NA | NA | NA | NA | NA | Trav16n | T cell receptor alpha variable 16n [Source: | 0 | 0 | 0 | 0 | 0 | 0 |
| 21660 | ENSMUSG00000076823 | 0 | NA | NA | NA | NA | NA | Trav13n-4 | T cell receptor alpha variable 13N-4 [Sour | 0 | 0 | 0 | 0 | 0 | 0 |
| 21661 | ENSMUSG00000076824 | 0 | NA | NA | NA | NA | NA | Trav14n-3 | T cell receptor alpha variable 14N-3 [Sour | 0 | 0 | 0 | 0 | 0 | 0 |
| 21662 | ENSMUSG00000076826 | 0 | NA | NA | NA | NA | NA | Trav4-2 | T cell receptor alpha variable 4-2 [Source: | 0 | 0 | 0 | 0 | 0 | 0 |
| 21663 | ENSMUSG00000076831 | 0 | NA | NA | NA | NA | NA | Trav8-1 | T cell receptor alpha variable 8-1 [Source: | 0 | 0 | 0 | 0 | 0 | 0 |
| 21664 | ENSMUSG00000076839 | 0.535401493 | -2.48596965 | 3.684117979 | -0.674780141 | 0.499815459 | NA | Trav13-1 | T cell receptor alpha variable 13-1 [Source | 3.2124 | 0 | 0 | 0 | 0 | 0 |
| 21665 | ENSMUSG00000076840 | 0 | NA | NA | NA | NA | NA | Trav14-1 | T cell receptor alpha variable 14-1 [Source | 0 | 0 | 0 | 0 | 0 | 0 |
| 21666 | ENSMUSG00000076846 | 0 | NA | NA | NA | NA | NA | Trav13-2 | T cell receptor alpha variable 13-2 [Source | 0 | 0 | 0 | 0 | 0 | 0 |
| 21667 | ENSMUSG00000076848 | 0 | NA | NA | NA | NA | NA | Trav15-2-dv( | T cell receptor alpha variable 15-2-DV6-2 [ | 0 | 0 | 0 | 0 | 0 | 0 |
| 21668 | ENSMUSG00000076855 | 0 | NA | NA | NA | NA | NA | Trav16 | T cell receptor alpha variable 16 [Source:M | 0 | 0 | 0 | 0 | 0 | 0 |

| | | | | | | | | | | | | | | | |
|---|---|---|---|---|---|---|---|---|---|---|---|---|---|---|---|
| 21669 | ENSMUSG00000076858 | 0 | NA | NA | NA | NA | NA | Trav3-4 | T cell receptor alpha variable 3-4 [Source: | 0 | 0 | 0 | 0 | 0 | 0 |
| 21670 | ENSMUSG00000076861 | 0 | NA | NA | NA | NA | NA | Trav18 | T cell receptor alpha variable 18 [Source:N | 0 | 0 | 0 | 0 | 0 | 0 |
| 21671 | ENSMUSG00000076862 | 0 | NA | NA | NA | NA | NA | Trav19 | T cell receptor alpha variable 19 [Source:N | 0 | 0 | 0 | 0 | 0 | 0 |
| 21672 | ENSMUSG00000076863 | 0 | NA | NA | NA | NA | NA | Trav21-dv12 | T cell receptor alpha variable 21-DV12 [So | 0 | 0 | 0 | 0 | 0 | 0 |
| 21673 | ENSMUSG00000076864 | 0 | NA | NA | NA | NA | NA | Trdv1 | T cell receptor delta variable 1 [Source:M( | 0 | 0 | 0 | 0 | 0 | 0 |
| 21674 | ENSMUSG00000076865 | 0 | NA | NA | NA | NA | NA | Trdv2-1 | T cell receptor delta variable 2-1 [Source:I | 0 | 0 | 0 | 0 | 0 | 0 |
| 21675 | ENSMUSG00000076866 | 0.133850373 | -0.796629176 | 4.080472857 | -0.195229623 | 0.84521319 | NA | Trdv2-2 | T cell receptor delta variable 2-2 [Source:I | 0.8031 | 0 | 0 | 0 | 0 | 0 |
| 21676 | ENSMUSG00000076867 | 0 | NA | NA | NA | NA | NA | Trdv4 | T cell receptor delta variable 4 [Source:M( | 0 | 0 | 0 | 0 | 0 | 0 |
| 21677 | ENSMUSG00000076870 | 0 | NA | NA | NA | NA | NA | Trdj1 | T cell receptor delta joining 1 [Source:MG | 0 | 0 | 0 | 0 | 0 | 0 |
| 21678 | ENSMUSG00000076871 | 0 | NA | NA | NA | NA | NA | Trdj2 | T cell receptor delta joining 2 [Source:MG | 0 | 0 | 0 | 0 | 0 | 0 |
| 21679 | ENSMUSG00000076873 | 0 | NA | NA | NA | NA | NA | Trdv5 | T cell receptor delta variable 5 [Source:M( | 0 | 0 | 0 | 0 | 0 | 0 |
| 21680 | ENSMUSG00000076874 | 0 | NA | NA | NA | NA | NA | Traj59 | T cell receptor alpha joining 59 [Source:M | 0 | 0 | 0 | 0 | 0 | 0 |
| 21681 | ENSMUSG00000076875 | 0 | NA | NA | NA | NA | NA | Traj58 | T cell receptor alpha joining 58 [Source:M | 0 | 0 | 0 | 0 | 0 | 0 |
| 21682 | ENSMUSG00000076876 | 0 | NA | NA | NA | NA | NA | Traj57 | T cell receptor alpha joining 57 [Source:M | 0 | 0 | 0 | 0 | 0 | 0 |
| 21683 | ENSMUSG00000076877 | 0 | NA | NA | NA | NA | NA | Traj56 | T cell receptor alpha joining 56 [Source:M | 0 | 0 | 0 | 0 | 0 | 0 |
| 21684 | ENSMUSG00000076878 | 0 | NA | NA | NA | NA | NA | Traj53 | T cell receptor alpha joining 53 [Source:M | 0 | 0 | 0 | 0 | 0 | 0 |
| 21685 | ENSMUSG00000076879 | 0 | NA | NA | NA | NA | NA | Traj52 | T cell receptor alpha joining 52 [Source:M | 0 | 0 | 0 | 0 | 0 | 0 |
| 21686 | ENSMUSG00000076880 | 0 | NA | NA | NA | NA | NA | Traj51 | T cell receptor alpha joining 51 [Source:M | 0 | 0 | 0 | 0 | 0 | 0 |
| 21687 | ENSMUSG00000076881 | 0 | NA | NA | NA | NA | NA | Traj50 | T cell receptor alpha joining 50 [Source:M | 0 | 0 | 0 | 0 | 0 | 0 |
| 21688 | ENSMUSG00000076882 | 0 | NA | NA | NA | NA | NA | Traj49 | T cell receptor alpha joining 49 [Source:M | 0 | 0 | 0 | 0 | 0 | 0 |
| 21689 | ENSMUSG00000076883 | 0 | NA | NA | NA | NA | NA | Traj48 | T cell receptor alpha joining 48 [Source:M | 0 | 0 | 0 | 0 | 0 | 0 |
| 21690 | ENSMUSG00000076884 | 0 | NA | NA | NA | NA | NA | Traj46 | T cell receptor alpha joining 46 [Source:M | 0 | 0 | 0 | 0 | 0 | 0 |
| 21691 | ENSMUSG00000076885 | 0 | NA | NA | NA | NA | NA | Traj45 | T cell receptor alpha joining 45 [Source:M | 0 | 0 | 0 | 0 | 0 | 0 |
| 21692 | ENSMUSG00000076886 | 0 | NA | NA | NA | NA | NA | Traj44 | T cell receptor alpha joining 44 [Source:M | 0 | 0 | 0 | 0 | 0 | 0 |
| 21693 | ENSMUSG00000076887 | 0 | NA | NA | NA | NA | NA | Traj43 | T cell receptor alpha joining 43 [Source:M | 0 | 0 | 0 | 0 | 0 | 0 |
| 21694 | ENSMUSG00000076888 | 0 | NA | NA | NA | NA | NA | Traj42 | T cell receptor alpha joining 42 [Source:M | 0 | 0 | 0 | 0 | 0 | 0 |
| 21695 | ENSMUSG00000076889 | 0 | NA | NA | NA | NA | NA | Traj41 | T cell receptor alpha joining 41 [Source:M | 0 | 0 | 0 | 0 | 0 | 0 |
| 21696 | ENSMUSG00000076890 | 0 | NA | NA | NA | NA | NA | Traj40 | T cell receptor alpha joining 40 [Source:M | 0 | 0 | 0 | 0 | 0 | 0 |
| 21697 | ENSMUSG00000076891 | 0 | NA | NA | NA | NA | NA | Traj39 | T cell receptor alpha joining 39 [Source:M | 0 | 0 | 0 | 0 | 0 | 0 |
| 21698 | ENSMUSG00000076892 | 0 | NA | NA | NA | NA | NA | Traj38 | T cell receptor alpha joining 38 [Source:M | 0 | 0 | 0 | 0 | 0 | 0 |
| 21699 | ENSMUSG00000076893 | 0 | NA | NA | NA | NA | NA | Traj37 | T cell receptor alpha joining 37 [Source:M | 0 | 0 | 0 | 0 | 0 | 0 |
| 21700 | ENSMUSG00000076894 | 0 | NA | NA | NA | NA | NA | Traj35 | T cell receptor alpha joining 35 [Source:M | 0 | 0 | 0 | 0 | 0 | 0 |
| 21701 | ENSMUSG00000076895 | 0 | NA | NA | NA | NA | NA | Traj34 | T cell receptor alpha joining 34 [Source:M | 0 | 0 | 0 | 0 | 0 | 0 |
| 21702 | ENSMUSG00000076896 | 0 | NA | NA | NA | NA | NA | Traj33 | T cell receptor alpha joining 33 [Source:M | 0 | 0 | 0 | 0 | 0 | 0 |
| 21703 | ENSMUSG00000076897 | 0 | NA | NA | NA | NA | NA | Traj32 | T cell receptor alpha joining 32 [Source:M | 0 | 0 | 0 | 0 | 0 | 0 |
| 21704 | ENSMUSG00000076898 | 0 | NA | NA | NA | NA | NA | Traj31 | T cell receptor alpha joining 31 [Source:M | 0 | 0 | 0 | 0 | 0 | 0 |
| 21705 | ENSMUSG00000076899 | 0 | NA | NA | NA | NA | NA | Traj30 | T cell receptor alpha joining 30 [Source:M | 0 | 0 | 0 | 0 | 0 | 0 |
| 21706 | ENSMUSG00000076900 | 0 | NA | NA | NA | NA | NA | Traj29 | T cell receptor alpha joining 29 [Source:M | 0 | 0 | 0 | 0 | 0 | 0 |
| 21707 | ENSMUSG00000076901 | 0 | NA | NA | NA | NA | NA | Traj28 | T cell receptor alpha joining 28 [Source:M | 0 | 0 | 0 | 0 | 0 | 0 |
| 21708 | ENSMUSG00000076902 | 0 | NA | NA | NA | NA | NA | Traj27 | T cell receptor alpha joining 27 [Source:M | 0 | 0 | 0 | 0 | 0 | 0 |
| 21709 | ENSMUSG00000076903 | 0 | NA | NA | NA | NA | NA | Traj26 | T cell receptor alpha joining 26 [Source:M | 0 | 0 | 0 | 0 | 0 | 0 |
| 21710 | ENSMUSG00000076904 | 0 | NA | NA | NA | NA | NA | Traj25 | T cell receptor alpha joining 25 [Source:M | 0 | 0 | 0 | 0 | 0 | 0 |
| 21711 | ENSMUSG00000076905 | 0 | NA | NA | NA | NA | NA | Traj24 | T cell receptor alpha joining 24 [Source:M | 0 | 0 | 0 | 0 | 0 | 0 |
| 21712 | ENSMUSG00000076906 | 0 | NA | NA | NA | NA | NA | Traj23 | T cell receptor alpha joining 23 [Source:M | 0 | 0 | 0 | 0 | 0 | 0 |
| 21713 | ENSMUSG00000076907 | 0 | NA | NA | NA | NA | NA | Traj22 | T cell receptor alpha joining 22 [Source:M | 0 | 0 | 0 | 0 | 0 | 0 |
| 21714 | ENSMUSG00000076908 | 0 | NA | NA | NA | NA | NA | Traj21 | T cell receptor alpha joining 21 [Source:M | 0 | 0 | 0 | 0 | 0 | 0 |
| 21715 | ENSMUSG00000076909 | 0 | NA | NA | NA | NA | NA | Traj20 | T cell receptor alpha joining 20 [Source:M | 0 | 0 | 0 | 0 | 0 | 0 |
| 21716 | ENSMUSG00000076910 | 0.133850373 | -0.796629176 | 4.080472857 | -0.195229623 | 0.84521319 | NA | Traj19 | T cell receptor alpha joining 19 [Source:M | 0.8031 | 0 | 0 | 0 | 0 | 0 |
| 21717 | ENSMUSG00000076911 | 0 | NA | NA | NA | NA | NA | Traj18 | T cell receptor alpha joining 18 [Source:M | 0 | 0 | 0 | 0 | 0 | 0 |
| 21718 | ENSMUSG00000076912 | 0 | NA | NA | NA | NA | NA | Traj17 | T cell receptor alpha joining 17 [Source:M | 0 | 0 | 0 | 0 | 0 | 0 |
| 21719 | ENSMUSG00000076913 | 0 | NA | NA | NA | NA | NA | Traj16 | T cell receptor alpha joining 16 [Source:M | 0 | 0 | 0 | 0 | 0 | 0 |
| 21720 | ENSMUSG00000076914 | 0 | NA | NA | NA | NA | NA | Traj15 | T cell receptor alpha joining 15 [Source:M | 0 | 0 | 0 | 0 | 0 | 0 |
| 21721 | ENSMUSG00000076915 | 0 | NA | NA | NA | NA | NA | Traj14 | T cell receptor alpha joining 14 [Source:M | 0 | 0 | 0 | 0 | 0 | 0 |
| 21722 | ENSMUSG00000076916 | 0 | NA | NA | NA | NA | NA | Traj13 | T cell receptor alpha joining 13 [Source:M | 0 | 0 | 0 | 0 | 0 | 0 |
| 21723 | ENSMUSG00000076917 | 0.167707431 | -0.796629176 | 4.080472857 | -0.195229623 | 0.84521319 | NA | Traj12 | T cell receptor alpha joining 12 [Source:M | 0 | 0 | 1.0062 | 0 | 0 | 0 |
| 21724 | ENSMUSG00000076918 | 0 | NA | NA | NA | NA | NA | Traj11 | T cell receptor alpha joining 11 [Source:M | 0 | 0 | 0 | 0 | 0 | 0 |

| | | | | | | | | | | | | | | | |
|---|---|---|---|---|---|---|---|---|---|---|---|---|---|---|---|
| 21725 | ENSMUSG00000076919 | 0 | NA | NA | NA | NA | NA | Traj9 | T cell receptor alpha joining 9 [Source:MG | 0 | 0 | 0 | 0 | 0 | 0 |
| 21726 | ENSMUSG00000076920 | 0 | NA | NA | NA | NA | NA | Traj8 | T cell receptor alpha joining 8 [Source:MG | 0 | 0 | 0 | 0 | 0 | 0 |
| 21727 | ENSMUSG00000076921 | 0 | NA | NA | NA | NA | NA | Traj7 | T cell receptor alpha joining 7 [Source:MG | 0 | 0 | 0 | 0 | 0 | 0 |
| 21728 | ENSMUSG00000076922 | 0 | NA | NA | NA | NA | NA | Traj6 | T cell receptor alpha joining 6 [Source:MG | 0 | 0 | 0 | 0 | 0 | 0 |
| 21729 | ENSMUSG00000076923 | 0 | NA | NA | NA | NA | NA | Traj5 | T cell receptor alpha joining 5 [Source:MG | 0 | 0 | 0 | 0 | 0 | 0 |
| 21730 | ENSMUSG00000076924 | 0 | NA | NA | NA | NA | NA | Traj4 | T cell receptor alpha joining 4 [Source:MG | 0 | 0 | 0 | 0 | 0 | 0 |
| 21731 | ENSMUSG00000076925 | 0 | NA | NA | NA | NA | NA | Traj3 | T cell receptor alpha joining 3 [Source:MG | 0 | 0 | 0 | 0 | 0 | 0 |
| 21732 | ENSMUSG00000076926 | 0 | NA | NA | NA | NA | NA | Traj2 | T cell receptor alpha joining 2 [Source:MG | 0 | 0 | 0 | 0 | 0 | 0 |
| 21733 | ENSMUSG00000076927 | 0 | NA | NA | NA | NA | NA | Traj1 | T cell receptor alpha joining 1 [Source:MG | 0 | 0 | 0 | 0 | 0 | 0 |
| 21734 | ENSMUSG00000076928 | 6.631995179 | -0.740702981 | 0.836532215 | -0.885444658 | 0.375916856 | 0.809690069 | Trac | T cell receptor alpha constant [Source:MG | 5.6217 | 8.2233 | 11.069 | 3.8559 | 5.4531 | 5.5692 |
| 21735 | ENSMUSG00000076934 | 3.420947073 | -3.08355871 | 1.506621954 | -2.046670501 | 0.040690448 | NA | Iglv1 | immunoglobulin lambda variable 1 [Sourc | 4.0155 | 9.2513 | 5.0312 | 0 | 0 | 2.2277 |
| 21736 | ENSMUSG00000076937 | 4.636491146 | -0.964652346 | 1.104792615 | -0.87315242 | 0.382580005 | NA | Iglc2 | immunoglobulin lambda constant 2 [Sour | 7.2279 | 4.1117 | 7.0437 | 2.8919 | 6.5437 | 0 |
| 21737 | ENSMUSG00000076939 | 0.624661033 | -1.234026074 | 2.854658177 | -0.432285058 | 0.665534246 | NA | Iglv3 | immunoglobulin lambda variable 3 [Sourc | 1.6062 | 1.0279 | 0 | 0 | 0 | 1.1138 |
| 21738 | ENSMUSG00000076940 | 0.51034671 | -2.381743948 | 3.331081356 | -0.715006238 | 0.474605189 | NA | Iglv2 | immunoglobulin lambda variable 2 [Sourc | 0 | 2.0558 | 1.0062 | 0 | 0 | 0 |
| 21739 | ENSMUSG00000076941 | 0 | NA | NA | NA | NA | NA | NA | NA | 0 | 0 | 0 | 0 | 0 | 0 |
| 21740 | ENSMUSG00000076944 | 0 | NA | NA | NA | NA | NA | NA | NA | 0 | 0 | 0 | 0 | 0 | 0 |
| 21741 | ENSMUSG00000076946 | 0 | NA | NA | NA | NA | NA | Mir582 | microRNA 582 [Source:MGI Symbol;Acc:M | 0 | 0 | 0 | 0 | 0 | 0 |
| 21742 | ENSMUSG00000076948 | 0 | NA | NA | NA | NA | NA | Mir467e | microRNA 467e [Source:MGI Symbol;Acc: | 0 | 0 | 0 | 0 | 0 | 0 |
| 21743 | ENSMUSG00000076949 | 0 | NA | NA | NA | NA | NA | NA | NA | 0 | 0 | 0 | 0 | 0 | 0 |
| 21744 | ENSMUSG00000076950 | 0 | NA | NA | NA | NA | NA | NA | NA | 0 | 0 | 0 | 0 | 0 | 0 |
| 21745 | ENSMUSG00000076951 | 0 | NA | NA | NA | NA | NA | NA | NA | 0 | 0 | 0 | 0 | 0 | 0 |
| 21746 | ENSMUSG00000076952 | 0 | NA | NA | NA | NA | NA | NA | NA | 0 | 0 | 0 | 0 | 0 | 0 |
| 21747 | ENSMUSG00000076956 | 0 | NA | NA | NA | NA | NA | Mir297a-3 | microRNA 297a-3 [Source:MGI Symbol;Ac | 0 | 0 | 0 | 0 | 0 | 0 |
| 21748 | ENSMUSG00000076958 | 0 | NA | NA | NA | NA | NA | NA | NA | 0 | 0 | 0 | 0 | 0 | 0 |
| 21749 | ENSMUSG00000076959 | 0 | NA | NA | NA | NA | NA | Mir669l | microRNA 669l [Source:MGI Symbol;Acc:M | 0 | 0 | 0 | 0 | 0 | 0 |
| 21750 | ENSMUSG00000076961 | 0.356960287 | 0.16514264 | 3.880370363 | 0.042558474 | 0.966053499 | NA | NA | NA | 0 | 1.0279 | 0 | 0 | 0 | 1.1138 |
| 21751 | ENSMUSG00000076964 | 0 | NA | NA | NA | NA | NA | NA | NA | 0 | 0 | 0 | 0 | 0 | 0 |
| 21752 | ENSMUSG00000076966 | 0 | NA | NA | NA | NA | NA | NA | NA | 0 | 0 | 0 | 0 | 0 | 0 |
| 21753 | ENSMUSG00000076972 | 0 | NA | NA | NA | NA | NA | Mir193b | microRNA 193b [Source:MGI Symbol;Acc: | 0 | 0 | 0 | 0 | 0 | 0 |
| 21754 | ENSMUSG00000076976 | 0 | NA | NA | NA | NA | NA | Mir297c | microRNA 297c [Source:MGI Symbol;Acc:I | 0 | 0 | 0 | 0 | 0 | 0 |
| 21755 | ENSMUSG00000076977 | 0 | NA | NA | NA | NA | NA | NA | NA | 0 | 0 | 0 | 0 | 0 | 0 |
| 21756 | ENSMUSG00000076983 | 0 | NA | NA | NA | NA | NA | Mir297a-2 | microRNA 297a-2 [Source:MGI Symbol;Ac | 0 | 0 | 0 | 0 | 0 | 0 |
| 21757 | ENSMUSG00000076987 | 0 | NA | NA | NA | NA | NA | NA | NA | 0 | 0 | 0 | 0 | 0 | 0 |
| 21758 | ENSMUSG00000076994 | 0 | NA | NA | NA | NA | NA | Mir466b-3 | microRNA 466b-3 [Source:MGI Symbol;Ac | 0 | 0 | 0 | 0 | 0 | 0 |
| 21759 | ENSMUSG00000076999 | 0 | NA | NA | NA | NA | NA | NA | NA | 0 | 0 | 0 | 0 | 0 | 0 |
| 21760 | ENSMUSG00000077001 | 0 | NA | NA | NA | NA | NA | Mir544 | microRNA 544 [Source:MGI Symbol;Acc:M | 0 | 0 | 0 | 0 | 0 | 0 |
| 21761 | ENSMUSG00000077004 | 0 | NA | NA | NA | NA | NA | NA | NA | 0 | 0 | 0 | 0 | 0 | 0 |
| 21762 | ENSMUSG00000077010 | 0 | NA | NA | NA | NA | NA | NA | NA | 0 | 0 | 0 | 0 | 0 | 0 |
| 21763 | ENSMUSG00000077015 | 0 | NA | NA | NA | NA | NA | NA | NA | 0 | 0 | 0 | 0 | 0 | 0 |
| 21764 | ENSMUSG00000077019 | 0 | NA | NA | NA | NA | NA | NA | NA | 0 | 0 | 0 | 0 | 0 | 0 |
| 21765 | ENSMUSG00000077021 | 0 | NA | NA | NA | NA | NA | Mir467d | microRNA 467d [Source:MGI Symbol;Acc: | 0 | 0 | 0 | 0 | 0 | 0 |
| 21766 | ENSMUSG00000077022 | 0 | NA | NA | NA | NA | NA | NA | NA | 0 | 0 | 0 | 0 | 0 | 0 |
| 21767 | ENSMUSG00000077025 | 0 | NA | NA | NA | NA | NA | Mir493 | microRNA 493 [Source:MGI Symbol;Acc:M | 0 | 0 | 0 | 0 | 0 | 0 |
| 21768 | ENSMUSG00000077038 | 0 | NA | NA | NA | NA | NA | Mir669f | microRNA 669f [Source:MGI Symbol;Acc:I | 0 | 0 | 0 | 0 | 0 | 0 |
| 21769 | ENSMUSG00000077039 | 0 | NA | NA | NA | NA | NA | NA | NA | 0 | 0 | 0 | 0 | 0 | 0 |
| 21770 | ENSMUSG00000077040 | 0 | NA | NA | NA | NA | NA | NA | NA | 0 | 0 | 0 | 0 | 0 | 0 |
| 21771 | ENSMUSG00000077042 | 0.321325949 | 1.911575114 | 4.048267893 | 0.472195804 | 0.636787034 | NA | Mir574 | microRNA 574 [Source:MGI Symbol;Acc:M | 0 | 0 | 0 | 1.928 | 0 | 0 |
| 21772 | ENSMUSG00000077049 | 0 | NA | NA | NA | NA | NA | Mir467c | microRNA 467c [Source:MGI Symbol;Acc:I | 0 | 0 | 0 | 0 | 0 | 0 |
| 21773 | ENSMUSG00000077050 | 0 | NA | NA | NA | NA | NA | NA | NA | 0 | 0 | 0 | 0 | 0 | 0 |
| 21774 | ENSMUSG00000077055 | 0 | NA | NA | NA | NA | NA | NA | NA | 0 | 0 | 0 | 0 | 0 | 0 |
| 21775 | ENSMUSG00000077058 | 0 | NA | NA | NA | NA | NA | NA | NA | 0 | 0 | 0 | 0 | 0 | 0 |
| 21776 | ENSMUSG00000077059 | 0 | NA | NA | NA | NA | NA | NA | NA | 0 | 0 | 0 | 0 | 0 | 0 |
| 21777 | ENSMUSG00000077065 | 0 | NA | NA | NA | NA | NA | NA | NA | 0 | 0 | 0 | 0 | 0 | 0 |
| 21778 | ENSMUSG00000077081 | 0 | NA | NA | NA | NA | NA | NA | NA | 0 | 0 | 0 | 0 | 0 | 0 |
| 21779 | ENSMUSG00000077082 | 0 | NA | NA | NA | NA | NA | NA | NA | 0 | 0 | 0 | 0 | 0 | 0 |
| 21780 | ENSMUSG00000077085 | 0 | NA | NA | NA | NA | NA | NA | NA | 0 | 0 | 0 | 0 | 0 | 0 |

| | | | | | | | | | | | | | | | |
|---|---|---|---|---|---|---|---|---|---|---|---|---|---|---|---|
| 21781 | ENSMUSG00000077086 | 0 | NA | NA | NA | NA | NA | Mir669o | microRNA 669o [Source:MGI Symbol;Acc: | 0 | 0 | 0 | 0 | 0 | 0 |
| 21782 | ENSMUSG00000077090 | 0 | NA | NA | NA | NA | NA | NA | NA | 0 | 0 | 0 | 0 | 0 | 0 |
| 21783 | ENSMUSG00000077092 | 0 | NA | NA | NA | NA | NA | Mir742 | microRNA 742 [Source:MGI Symbol;Acc:M | 0 | 0 | 0 | 0 | 0 | 0 |
| 21784 | ENSMUSG00000077095 | 0 | NA | NA | NA | NA | NA | NA | NA | 0 | 0 | 0 | 0 | 0 | 0 |
| 21785 | ENSMUSG00000077096 | 0 | NA | NA | NA | NA | NA | NA | NA | 0 | 0 | 0 | 0 | 0 | 0 |
| 21786 | ENSMUSG00000077104 | 0 | NA | NA | NA | NA | NA | NA | NA | 0 | 0 | 0 | 0 | 0 | 0 |
| 21787 | ENSMUSG00000077107 | 0 | NA | NA | NA | NA | NA | Mir344c | microRNA 344c [Source:MGI Symbol;Acc:I | 0 | 0 | 0 | 0 | 0 | 0 |
| 21788 | ENSMUSG00000077110 | 0 | NA | NA | NA | NA | NA | NA | NA | 0 | 0 | 0 | 0 | 0 | 0 |
| 21789 | ENSMUSG00000077111 | 0 | NA | NA | NA | NA | NA | Mir18b | microRNA 18b [Source:MGI Symbol;Acc:M | 0 | 0 | 0 | 0 | 0 | 0 |
| 21790 | ENSMUSG00000077113 | 0 | NA | NA | NA | NA | NA | Mir466p | microRNA 466p [Source:MGI Symbol;Acc: | 0 | 0 | 0 | 0 | 0 | 0 |
| 21791 | ENSMUSG00000077119 | 0 | NA | NA | NA | NA | NA | Mir741 | microRNA 741 [Source:MGI Symbol;Acc:M | 0 | 0 | 0 | 0 | 0 | 0 |
| 21792 | ENSMUSG00000077125 | 0 | NA | NA | NA | NA | NA | NA | NA | 0 | 0 | 0 | 0 | 0 | 0 |
| 21793 | ENSMUSG00000077127 | 0 | NA | NA | NA | NA | NA | Mir743 | microRNA 743 [Source:MGI Symbol;Acc:M | 0 | 0 | 0 | 0 | 0 | 0 |
| 21794 | ENSMUSG00000077131 | 0 | NA | NA | NA | NA | NA | NA | NA | 0 | 0 | 0 | 0 | 0 | 0 |
| 21795 | ENSMUSG00000077138 | 0 | NA | NA | NA | NA | NA | NA | NA | 0 | 0 | 0 | 0 | 0 | 0 |
| 21796 | ENSMUSG00000077142 | 0 | NA | NA | NA | NA | NA | Gm22937 | predicted gene, 22937 [Source:MGI Symb | 0 | 0 | 0 | 0 | 0 | 0 |
| 21797 | ENSMUSG00000077143 | 0 | NA | NA | NA | NA | NA | Gm22936 | predicted gene, 22936 [Source:MGI Symb | 0 | 0 | 0 | 0 | 0 | 0 |
| 21798 | ENSMUSG00000077144 | 0 | NA | NA | NA | NA | NA | Gm22938 | predicted gene, 22938 [Source:MGI Symb | 0 | 0 | 0 | 0 | 0 | 0 |
| 21799 | ENSMUSG00000077146 | 0 | NA | NA | NA | NA | NA | Gm22939 | predicted gene, 22939 [Source:MGI Symb | 0 | 0 | 0 | 0 | 0 | 0 |
| 21800 | ENSMUSG00000077148 | 1.131136811 | -2.325642949 | 2.16414089 | -1.074626407 | 0.282542024 | NA | Gm22935 | predicted gene, 22935 [Source:MGI Symb | 1.6062 | 3.0838 | 1.0062 | 0 | 1.0906 | 0 |
| 21801 | ENSMUSG00000077149 | 0 | NA | NA | NA | NA | NA | Gm22934 | predicted gene, 22934 [Source:MGI Symb | 0 | 0 | 0 | 0 | 0 | 0 |
| 21802 | ENSMUSG00000077150 | 0 | NA | NA | NA | NA | NA | Gm24581 | predicted gene, 24581 [Source:MGI Symb | 0 | 0 | 0 | 0 | 0 | 0 |
| 21803 | ENSMUSG00000077152 | 0 | NA | NA | NA | NA | NA | Gm24579 | predicted gene, 24579 [Source:MGI Symb | 0 | 0 | 0 | 0 | 0 | 0 |
| 21804 | ENSMUSG00000077153 | 0 | NA | NA | NA | NA | NA | Gm24580 | predicted gene, 24580 [Source:MGI Symb | 0 | 0 | 0 | 0 | 0 | 0 |
| 21805 | ENSMUSG00000077155 | 0 | NA | NA | NA | NA | NA | Gm24582 | predicted gene, 24582 [Source:MGI Symb | 0 | 0 | 0 | 0 | 0 | 0 |
| 21806 | ENSMUSG00000077156 | 0 | NA | NA | NA | NA | NA | NA | NA | 0 | 0 | 0 | 0 | 0 | 0 |
| 21807 | ENSMUSG00000077158 | 0 | NA | NA | NA | NA | NA | Gm24578 | predicted gene, 24578 [Source:MGI Symb | 0 | 0 | 0 | 0 | 0 | 0 |
| 21808 | ENSMUSG00000077160 | 0 | NA | NA | NA | NA | NA | Gm24118 | predicted gene, 24118 [Source:MGI Symb | 0 | 0 | 0 | 0 | 0 | 0 |
| 21809 | ENSMUSG00000077161 | 0 | NA | NA | NA | NA | NA | Gm24117 | predicted gene, 24117 [Source:MGI Symb | 0 | 0 | 0 | 0 | 0 | 0 |
| 21810 | ENSMUSG00000077163 | 0 | NA | NA | NA | NA | NA | Gm24116 | predicted gene, 24116 [Source:MGI Symb | 0 | 0 | 0 | 0 | 0 | 0 |
| 21811 | ENSMUSG00000077166 | 0 | NA | NA | NA | NA | NA | Gm24120 | predicted gene, 24120 [Source:MGI Symb | 0 | 0 | 0 | 0 | 0 | 0 |
| 21812 | ENSMUSG00000077167 | 5.182419463 | 1.258269536 | 0.961923913 | 1.308075949 | 0.19084755 | 0.628409928 | Gm24119 | predicted gene, 24119 [Source:MGI Symb | 4.0155 | 3.0838 | 2.0125 | 7.7118 | 9.8156 | 4.4554 |
| 21813 | ENSMUSG00000077169 | 0 | NA | NA | NA | NA | NA | Gm25396 | predicted gene, 25396 [Source:MGI Symb | 0 | 0 | 0 | 0 | 0 | 0 |
| 21814 | ENSMUSG00000077171 | 0 | NA | NA | NA | NA | NA | Gm25761 | predicted gene, 25761 [Source:MGI Symb | 0 | 0 | 0 | 0 | 0 | 0 |
| 21815 | ENSMUSG00000077172 | 0 | NA | NA | NA | NA | NA | Gm25762 | predicted gene, 25762 [Source:MGI Symb | 0 | 0 | 0 | 0 | 0 | 0 |
| 21816 | ENSMUSG00000077174 | 0 | NA | NA | NA | NA | NA | Gm25757 | predicted gene, 25757 [Source:MGI Symb | 0 | 0 | 0 | 0 | 0 | 0 |
| 21817 | ENSMUSG00000077175 | 0 | NA | NA | NA | NA | NA | Gm25758 | predicted gene, 25758 [Source:MGI Symb | 0 | 0 | 0 | 0 | 0 | 0 |
| 21818 | ENSMUSG00000077176 | 0 | NA | NA | NA | NA | NA | Gm25759 | predicted gene, 25759 [Source:MGI Symb | 0 | 0 | 0 | 0 | 0 | 0 |
| 21819 | ENSMUSG00000077177 | 0 | NA | NA | NA | NA | NA | Gm25760 | predicted gene, 25760 [Source:MGI Symb | 0 | 0 | 0 | 0 | 0 | 0 |
| 21820 | ENSMUSG00000077178 | 0 | NA | NA | NA | NA | NA | Gm25756 | predicted gene, 25756 [Source:MGI Symb | 0 | 0 | 0 | 0 | 0 | 0 |
| 21821 | ENSMUSG00000077180 | 0 | NA | NA | NA | NA | NA | Gm25094 | predicted gene, 25094 [Source:MGI Symb | 0 | 0 | 0 | 0 | 0 | 0 |
| 21822 | ENSMUSG00000077181 | 0 | NA | NA | NA | NA | NA | Gm25093 | predicted gene, 25093 [Source:MGI Symb | 0 | 0 | 0 | 0 | 0 | 0 |
| 21823 | ENSMUSG00000077184 | 0 | NA | NA | NA | NA | NA | Gm25092 | predicted gene, 25092 [Source:MGI Symb | 0 | 0 | 0 | 0 | 0 | 0 |
| 21824 | ENSMUSG00000077185 | 0 | NA | NA | NA | NA | NA | Gm25091 | predicted gene, 25091 [Source:MGI Symb | 0 | 0 | 0 | 0 | 0 | 0 |
| 21825 | ENSMUSG00000077188 | 0 | NA | NA | NA | NA | NA | NA | NA | 0 | 0 | 0 | 0 | 0 | 0 |
| 21826 | ENSMUSG00000077190 | 0 | NA | NA | NA | NA | NA | NA | NA | 0 | 0 | 0 | 0 | 0 | 0 |
| 21827 | ENSMUSG00000077191 | 0 | NA | NA | NA | NA | NA | Snord64 | small nucleolar RNA, C/D box 64 [Source:I | 0 | 0 | 0 | 0 | 0 | 0 |
| 21828 | ENSMUSG00000077192 | 3.23886679 | 0.11683794 | 1.334013767 | 0.087583759 | 0.930207511 | NA | Snora17 | small nucleolar RNA, H/ACA box 17 [Sourc | 7.2279 | 0 | 2.0125 | 5.7839 | 2.1812 | 2.2277 |
| 21829 | ENSMUSG00000077194 | 0 | NA | NA | NA | NA | NA | Gm23431 | predicted gene, 23431 [Source:MGI Symb | 0 | 0 | 0 | 0 | 0 | 0 |
| 21830 | ENSMUSG00000077196 | 0 | NA | NA | NA | NA | NA | Gm22267 | predicted gene, 22267 [Source:MGI Symb | 0 | 0 | 0 | 0 | 0 | 0 |
| 21831 | ENSMUSG00000077197 | 0 | NA | NA | NA | NA | NA | Gm22268 | predicted gene, 22268 [Source:MGI Symb | 0 | 0 | 0 | 0 | 0 | 0 |
| 21832 | ENSMUSG00000077199 | 0 | NA | NA | NA | NA | NA | Gm22266 | predicted gene, 22266 [Source:MGI Symb | 0 | 0 | 0 | 0 | 0 | 0 |
| 21833 | ENSMUSG00000077200 | 0 | NA | NA | NA | NA | NA | Gm25613 | predicted gene, 25613 [Source:MGI Symb | 0 | 0 | 0 | 0 | 0 | 0 |
| 21834 | ENSMUSG00000077201 | 0 | NA | NA | NA | NA | NA | Gm25614 | predicted gene, 25614 [Source:MGI Symb | 0 | 0 | 0 | 0 | 0 | 0 |
| 21835 | ENSMUSG00000077202 | 1.325656843 | -1.489779841 | 1.921596063 | -0.775282522 | 0.438172735 | NA | Gm25612 | predicted gene, 25612 [Source:MGI Symb | 0.8031 | 3.0838 | 2.0125 | 0.964 | 1.0906 | 0 |
| 21836 | ENSMUSG00000077204 | 0 | NA | NA | NA | NA | NA | Gm25610 | predicted gene, 25610 [Source:MGI Symb | 0 | 0 | 0 | 0 | 0 | 0 |

| | | | | | | | | | | | | | | | |
|---|---|---|---|---|---|---|---|---|---|---|---|---|---|---|---|
| 21837 | ENSMUSG00000077205 | 0 | NA | NA | NA | NA | NA | Gm25611 | predicted gene, 25611 [Source:MGI Symb | 0 | 0 | 0 | 0 | 0 | 0 |
| 21838 | ENSMUSG00000077207 | 0 | NA | NA | NA | NA | NA | Gm25609 | predicted gene, 25609 [Source:MGI Symb | 0 | 0 | 0 | 0 | 0 | 0 |
| 21839 | ENSMUSG00000077208 | 0 | NA | NA | NA | NA | NA | Gm25607 | predicted gene, 25607 [Source:MGI Symb | 0 | 0 | 0 | 0 | 0 | 0 |
| 21840 | ENSMUSG00000077209 | 0 | NA | NA | NA | NA | NA | Gm25608 | predicted gene, 25608 [Source:MGI Symb | 0 | 0 | 0 | 0 | 0 | 0 |
| 21841 | ENSMUSG00000077212 | 0 | NA | NA | NA | NA | NA | Snord69 | small nucleolar RNA, C/D box 69 [Source:I | 0 | 0 | 0 | 0 | 0 | 0 |
| 21842 | ENSMUSG00000077214 | 0 | NA | NA | NA | NA | NA | Gm23950 | predicted gene, 23950 [Source:MGI Symb | 0 | 0 | 0 | 0 | 0 | 0 |
| 21843 | ENSMUSG00000077215 | 0 | NA | NA | NA | NA | NA | Gm23949 | predicted gene, 23949 [Source:MGI Symb | 0 | 0 | 0 | 0 | 0 | 0 |
| 21844 | ENSMUSG00000077217 | 0 | NA | NA | NA | NA | NA | Gm23951 | predicted gene, 23951 [Source:MGI Symb | 0 | 0 | 0 | 0 | 0 | 0 |
| 21845 | ENSMUSG00000077218 | 0 | NA | NA | NA | NA | NA | NA | NA | 0 | 0 | 0 | 0 | 0 | 0 |
| 21846 | ENSMUSG00000077220 | 0 | NA | NA | NA | NA | NA | Snord78 | small nucleolar RNA, C/D box 78 [Source:I | 0 | 0 | 0 | 0 | 0 | 0 |
| 21847 | ENSMUSG00000077221 | 0 | NA | NA | NA | NA | NA | Snord67 | small nucleolar RNA, C/D box 67 [Source:I | 0 | 0 | 0 | 0 | 0 | 0 |
| 21848 | ENSMUSG00000077222 | 6.307753304 | 0.482079559 | 0.838026266 | 0.575255906 | 0.565118238 | 0.901542552 | Gm22270 | predicted gene, 22270 [Source:MGI Symb | 5.6217 | 4.1117 | 6.0375 | 7.7118 | 5.4531 | 8.9108 |
| 21849 | ENSMUSG00000077223 | 0.975999737 | -3.32375764 | 2.421946067 | -1.372349981 | 0.169954513 | NA | Gm22271 | predicted gene, 22271 [Source:MGI Symb | 0.8031 | 1.0279 | 4.025 | 0 | 0 | 0 |
| 21850 | ENSMUSG00000077224 | 0 | NA | NA | NA | NA | NA | Gm23437 | predicted gene, 23437 [Source:MGI Symb | 0 | 0 | 0 | 0 | 0 | 0 |
| 21851 | ENSMUSG00000077227 | 0 | NA | NA | NA | NA | NA | Gm22272 | predicted gene, 22272 [Source:MGI Symb | 0 | 0 | 0 | 0 | 0 | 0 |
| 21852 | ENSMUSG00000077228 | 0 | NA | NA | NA | NA | NA | Gm22273 | predicted gene, 22273 [Source:MGI Symb | 0 | 0 | 0 | 0 | 0 | 0 |
| 21853 | ENSMUSG00000077230 | 0 | NA | NA | NA | NA | NA | Gm25097 | predicted gene, 25097 [Source:MGI Symb | 0 | 0 | 0 | 0 | 0 | 0 |
| 21854 | ENSMUSG00000077231 | 0 | NA | NA | NA | NA | NA | Gm25096 | predicted gene, 25096 [Source:MGI Symb | 0 | 0 | 0 | 0 | 0 | 0 |
| 21855 | ENSMUSG00000077235 | 0 | NA | NA | NA | NA | NA | Gm25095 | predicted gene, 25095 [Source:MGI Symb | 0 | 0 | 0 | 0 | 0 | 0 |
| 21856 | ENSMUSG00000077236 | 0 | NA | NA | NA | NA | NA | NA | NA | 0 | 0 | 0 | 0 | 0 | 0 |
| 21857 | ENSMUSG00000077237 | 0 | NA | NA | NA | NA | NA | Gm22438 | predicted gene, 22438 [Source:MGI Symb | 0 | 0 | 0 | 0 | 0 | 0 |
| 21858 | ENSMUSG00000077239 | 0 | NA | NA | NA | NA | NA | Snord66 | small nucleolar RNA, C/D box 66 [Source:I | 0 | 0 | 0 | 0 | 0 | 0 |
| 21859 | ENSMUSG00000077240 | 0 | NA | NA | NA | NA | NA | Gm23271 | predicted gene, 23271 [Source:MGI Symb | 0 | 0 | 0 | 0 | 0 | 0 |
| 21860 | ENSMUSG00000077241 | 0 | NA | NA | NA | NA | NA | Gm23272 | predicted gene, 23272 [Source:MGI Symb | 0 | 0 | 0 | 0 | 0 | 0 |
| 21861 | ENSMUSG00000077243 | 0 | NA | NA | NA | NA | NA | Gm23270 | predicted gene, 23270 [Source:MGI Symb | 0 | 0 | 0 | 0 | 0 | 0 |
| 21862 | ENSMUSG00000077244 | 0 | NA | NA | NA | NA | NA | Gm23274 | predicted gene, 23274 [Source:MGI Symb | 0 | 0 | 0 | 0 | 0 | 0 |
| 21863 | ENSMUSG00000077246 | 0 | NA | NA | NA | NA | NA | Gm23273 | predicted gene, 23273 [Source:MGI Symb | 0 | 0 | 0 | 0 | 0 | 0 |
| 21864 | ENSMUSG00000077248 | 0 | NA | NA | NA | NA | NA | Gm23275 | predicted gene, 23275 [Source:MGI Symb | 0 | 0 | 0 | 0 | 0 | 0 |
| 21865 | ENSMUSG00000077250 | 0 | NA | NA | NA | NA | NA | Gm26075 | predicted gene, 26075 [Source:MGI Symb | 0 | 0 | 0 | 0 | 0 | 0 |
| 21866 | ENSMUSG00000077251 | 0 | NA | NA | NA | NA | NA | Gm26074 | predicted gene, 26074 [Source:MGI Symb | 0 | 0 | 0 | 0 | 0 | 0 |
| 21867 | ENSMUSG00000077252 | 0 | NA | NA | NA | NA | NA | Gm26077 | predicted gene, 26077 [Source:MGI Symb | 0 | 0 | 0 | 0 | 0 | 0 |
| 21868 | ENSMUSG00000077253 | 0 | NA | NA | NA | NA | NA | NA | NA | 0 | 0 | 0 | 0 | 0 | 0 |
| 21869 | ENSMUSG00000077254 | 0 | NA | NA | NA | NA | NA | Gm26079 | predicted gene, 26079 [Source:MGI Symb | 0 | 0 | 0 | 0 | 0 | 0 |
| 21870 | ENSMUSG00000077255 | 0 | NA | NA | NA | NA | NA | Gm26078 | predicted gene, 26078 [Source:MGI Symb | 0 | 0 | 0 | 0 | 0 | 0 |
| 21871 | ENSMUSG00000077256 | 0 | NA | NA | NA | NA | NA | Gm26081 | predicted gene, 26081 [Source:MGI Symb | 0 | 0 | 0 | 0 | 0 | 0 |
| 21872 | ENSMUSG00000077257 | 0 | NA | NA | NA | NA | NA | Gm26080 | predicted gene, 26080 [Source:MGI Symb | 0 | 0 | 0 | 0 | 0 | 0 |
| 21873 | ENSMUSG00000077258 | 0 | NA | NA | NA | NA | NA | Gm26082 | predicted gene, 26082 [Source:MGI Symb | 0 | 0 | 0 | 0 | 0 | 0 |
| 21874 | ENSMUSG00000077259 | 0 | NA | NA | NA | NA | NA | Gm23979 | predicted gene, 23979 [Source:MGI Symb | 0 | 0 | 0 | 0 | 0 | 0 |
| 21875 | ENSMUSG00000077260 | 0 | NA | NA | NA | NA | NA | Gm24869 | predicted gene, 24869 [Source:MGI Symb | 0 | 0 | 0 | 0 | 0 | 0 |
| 21876 | ENSMUSG00000077261 | 0 | NA | NA | NA | NA | NA | NA | NA | 0 | 0 | 0 | 0 | 0 | 0 |
| 21877 | ENSMUSG00000077262 | 0 | NA | NA | NA | NA | NA | NA | NA | 0 | 0 | 0 | 0 | 0 | 0 |
| 21878 | ENSMUSG00000077263 | 0 | NA | NA | NA | NA | NA | Gm24432 | predicted gene, 24432 [Source:MGI Symb | 0 | 0 | 0 | 0 | 0 | 0 |
| 21879 | ENSMUSG00000077264 | 0 | NA | NA | NA | NA | NA | Gm24429 | predicted gene, 24429 [Source:MGI Symb | 0 | 0 | 0 | 0 | 0 | 0 |
| 21880 | ENSMUSG00000077265 | 1.080168213 | 2.425859534 | 2.660049518 | 0.911960292 | 0.361789621 | NA | Gm24430 | predicted gene, 24430 [Source:MGI Symb | 0 | 1.0279 | 0 | 0 | 5.4531 | 0 |
| 21881 | ENSMUSG00000077268 | 0 | NA | NA | NA | NA | NA | Gm24433 | predicted gene, 24433 [Source:MGI Symb | 0 | 0 | 0 | 0 | 0 | 0 |
| 21882 | ENSMUSG00000077269 | 0 | NA | NA | NA | NA | NA | Gm24434 | predicted gene, 24434 [Source:MGI Symb | 0 | 0 | 0 | 0 | 0 | 0 |
| 21883 | ENSMUSG00000077270 | 0 | NA | NA | NA | NA | NA | NA | NA | 0 | 0 | 0 | 0 | 0 | 0 |
| 21884 | ENSMUSG00000077271 | 0.688942901 | 0.052211695 | 2.79596103 | 0.018673971 | 0.985101193 | NA | Gm22781 | predicted gene, 22781 [Source:MGI Symb | 0 | 2.0558 | 0 | 0.964 | 0 | 1.1138 |
| 21885 | ENSMUSG00000077272 | 0 | NA | NA | NA | NA | NA | NA | NA | 0 | 0 | 0 | 0 | 0 | 0 |
| 21886 | ENSMUSG00000077273 | 0 | NA | NA | NA | NA | NA | Gm22779 | predicted gene, 22779 [Source:MGI Symb | 0 | 0 | 0 | 0 | 0 | 0 |
| 21887 | ENSMUSG00000077274 | 0.17131964 | -0.796629176 | 4.080472857 | -0.195229623 | 0.84521319 | NA | Gm22786 | predicted gene, 22786 [Source:MGI Symb | 0 | 1.0279 | 0 | 0 | 0 | 0 |
| 21888 | ENSMUSG00000077275 | 0 | NA | NA | NA | NA | NA | Gm22785 | predicted gene, 22785 [Source:MGI Symb | 0 | 0 | 0 | 0 | 0 | 0 |
| 21889 | ENSMUSG00000077276 | 0 | NA | NA | NA | NA | NA | Gm22784 | predicted gene, 22784 [Source:MGI Symb | 0 | 0 | 0 | 0 | 0 | 0 |
| 21890 | ENSMUSG00000077277 | 0 | NA | NA | NA | NA | NA | Gm22783 | predicted gene, 22783 [Source:MGI Symb | 0 | 0 | 0 | 0 | 0 | 0 |
| 21891 | ENSMUSG00000077278 | 0 | NA | NA | NA | NA | NA | Gm22778 | predicted gene, 22778 [Source:MGI Symb | 0 | 0 | 0 | 0 | 0 | 0 |
| 21892 | ENSMUSG00000077279 | 0 | NA | NA | NA | NA | NA | Gm22777 | predicted gene, 22777 [Source:MGI Symb | 0 | 0 | 0 | 0 | 0 | 0 |

| | | | | | | | | | | | | | | | |
|---|---|---|---|---|---|---|---|---|---|---|---|---|---|---|---|
| 21893 | ENSMUSG00000077280 | 0 | NA | NA | NA | NA | NA | Gm25764 | predicted gene, 25764 [Source:MGI Symb | 0 | 0 | 0 | 0 | 0 | 0 |
| 21894 | ENSMUSG00000077281 | 0 | NA | NA | NA | NA | NA | NA | NA | 0 | 0 | 0 | 0 | 0 | 0 |
| 21895 | ENSMUSG00000077282 | 0 | NA | NA | NA | NA | NA | Gm26395 | predicted gene, 26395 [Source:MGI Symb | 0 | 0 | 0 | 0 | 0 | 0 |
| 21896 | ENSMUSG00000077283 | 0.181769715 | 1.126924175 | 4.080472857 | 0.276174898 | 0.782413736 | NA | NA | NA | 0 | 0 | 0 | 0 | 1.0906 | 0 |
| 21897 | ENSMUSG00000077285 | 0 | NA | NA | NA | NA | NA | NA | NA | 0 | 0 | 0 | 0 | 0 | 0 |
| 21898 | ENSMUSG00000077287 | 0 | NA | NA | NA | NA | NA | Gm25763 | predicted gene, 25763 [Source:MGI Symb | 0 | 0 | 0 | 0 | 0 | 0 |
| 21899 | ENSMUSG00000077288 | 0 | NA | NA | NA | NA | NA | Gm25765 | predicted gene, 25765 [Source:MGI Symb | 0 | 0 | 0 | 0 | 0 | 0 |
| 21900 | ENSMUSG00000077289 | 0 | NA | NA | NA | NA | NA | Gm25766 | predicted gene, 25766 [Source:MGI Symb | 0 | 0 | 0 | 0 | 0 | 0 |
| 21901 | ENSMUSG00000077290 | 0 | NA | NA | NA | NA | NA | Gm25407 | predicted gene, 25407 [Source:MGI Symb | 0 | 0 | 0 | 0 | 0 | 0 |
| 21902 | ENSMUSG00000077291 | 0 | NA | NA | NA | NA | NA | Gm24124 | predicted gene, 24124 [Source:MGI Symb | 0 | 0 | 0 | 0 | 0 | 0 |
| 21903 | ENSMUSG00000077292 | 0 | NA | NA | NA | NA | NA | Gm25408 | predicted gene, 25408 [Source:MGI Symb | 0 | 0 | 0 | 0 | 0 | 0 |
| 21904 | ENSMUSG00000077293 | 0 | NA | NA | NA | NA | NA | Gm24125 | predicted gene, 24125 [Source:MGI Symb | 0 | 0 | 0 | 0 | 0 | 0 |
| 21905 | ENSMUSG00000077294 | 0 | NA | NA | NA | NA | NA | Gm24123 | predicted gene, 24123 [Source:MGI Symb | 0 | 0 | 0 | 0 | 0 | 0 |
| 21906 | ENSMUSG00000077295 | 0 | NA | NA | NA | NA | NA | Gm24122 | predicted gene, 24122 [Source:MGI Symb | 0 | 0 | 0 | 0 | 0 | 0 |
| 21907 | ENSMUSG00000077298 | 0 | NA | NA | NA | NA | NA | Gm24127 | predicted gene, 24127 [Source:MGI Symb | 0 | 0 | 0 | 0 | 0 | 0 |
| 21908 | ENSMUSG00000077299 | 0 | NA | NA | NA | NA | NA | Gm24126 | predicted gene, 24126 [Source:MGI Symb | 0 | 0 | 0 | 0 | 0 | 0 |
| 21909 | ENSMUSG00000077302 | 0 | NA | NA | NA | NA | NA | Gm22473 | predicted gene, 22473 [Source:MGI Symb | 0 | 0 | 0 | 0 | 0 | 0 |
| 21910 | ENSMUSG00000077303 | 0 | NA | NA | NA | NA | NA | Gm22472 | predicted gene, 22472 [Source:MGI Symb | 0 | 0 | 0 | 0 | 0 | 0 |
| 21911 | ENSMUSG00000077304 | 0 | NA | NA | NA | NA | NA | Gm22471 | predicted gene, 22471 [Source:MGI Symb | 0 | 0 | 0 | 0 | 0 | 0 |
| 21912 | ENSMUSG00000077305 | 0 | NA | NA | NA | NA | NA | Gm22470 | predicted gene, 22470 [Source:MGI Symb | 0 | 0 | 0 | 0 | 0 | 0 |
| 21913 | ENSMUSG00000077306 | 0 | NA | NA | NA | NA | NA | Gm22469 | predicted gene, 22469 [Source:MGI Symb | 0 | 0 | 0 | 0 | 0 | 0 |
| 21914 | ENSMUSG00000077309 | 0 | NA | NA | NA | NA | NA | NA | NA | 0 | 0 | 0 | 0 | 0 | 0 |
| 21915 | ENSMUSG00000077312 | 0 | NA | NA | NA | NA | NA | Gm24170 | predicted gene, 24170 [Source:MGI Symb | 0 | 0 | 0 | 0 | 0 | 0 |
| 21916 | ENSMUSG00000077314 | 0 | NA | NA | NA | NA | NA | Gm24171 | predicted gene, 24171 [Source:MGI Symb | 0 | 0 | 0 | 0 | 0 | 0 |
| 21917 | ENSMUSG00000077315 | 0 | NA | NA | NA | NA | NA | Gm24172 | predicted gene, 24172 [Source:MGI Symb | 0 | 0 | 0 | 0 | 0 | 0 |
| 21918 | ENSMUSG00000077316 | 0.535401493 | -2.48596965 | 3.684117979 | -0.674780141 | 0.499815459 | NA | NA | NA | 3.2124 | 0 | 0 | 0 | 0 | 0 |
| 21919 | ENSMUSG00000077317 | 0 | NA | NA | NA | NA | NA | Gm25785 | predicted gene, 25785 [Source:MGI Symb | 0 | 0 | 0 | 0 | 0 | 0 |
| 21920 | ENSMUSG00000077318 | 0 | NA | NA | NA | NA | NA | Gm24173 | predicted gene, 24173 [Source:MGI Symb | 0 | 0 | 0 | 0 | 0 | 0 |
| 21921 | ENSMUSG00000077320 | 0 | NA | NA | NA | NA | NA | Gm22284 | predicted gene, 22284 [Source:MGI Symb | 0 | 0 | 0 | 0 | 0 | 0 |
| 21922 | ENSMUSG00000077323 | 0 | NA | NA | NA | NA | NA | Rnu11 | U11 small nuclear RNA [Source:MGI Symb | 0 | 0 | 0 | 0 | 0 | 0 |
| 21923 | ENSMUSG00000077325 | 0 | NA | NA | NA | NA | NA | Gm25833 | predicted gene, 25833 [Source:MGI Symb | 0 | 0 | 0 | 0 | 0 | 0 |
| 21924 | ENSMUSG00000077327 | 0 | NA | NA | NA | NA | NA | Gm25938 | predicted gene, 25938 [Source:MGI Symb | 0 | 0 | 0 | 0 | 0 | 0 |
| 21925 | ENSMUSG00000077328 | 0 | NA | NA | NA | NA | NA | Gm25832 | predicted gene, 25832 [Source:MGI Symb | 0 | 0 | 0 | 0 | 0 | 0 |
| 21926 | ENSMUSG00000077332 | 0 | NA | NA | NA | NA | NA | Gm23014 | predicted gene, 23014 [Source:MGI Symb | 0 | 0 | 0 | 0 | 0 | 0 |
| 21927 | ENSMUSG00000077333 | 0 | NA | NA | NA | NA | NA | Gm23015 | predicted gene, 23015 [Source:MGI Symb | 0 | 0 | 0 | 0 | 0 | 0 |
| 21928 | ENSMUSG00000077334 | 0 | NA | NA | NA | NA | NA | Gm23013 | predicted gene, 23013 [Source:MGI Symb | 0 | 0 | 0 | 0 | 0 | 0 |
| 21929 | ENSMUSG00000077336 | 0 | NA | NA | NA | NA | NA | Gm23011 | predicted gene, 23011 [Source:MGI Symb | 0 | 0 | 0 | 0 | 0 | 0 |
| 21930 | ENSMUSG00000077337 | 0 | NA | NA | NA | NA | NA | Gm23012 | predicted gene, 23012 [Source:MGI Symb | 0 | 0 | 0 | 0 | 0 | 0 |
| 21931 | ENSMUSG00000077338 | 0 | NA | NA | NA | NA | NA | Gm23294 | predicted gene, 23294 [Source:MGI Symb | 0 | 0 | 0 | 0 | 0 | 0 |
| 21932 | ENSMUSG00000077339 | 0.167707431 | -0.796629176 | 4.080472857 | -0.195229623 | 0.84521319 | NA | Gm23010 | predicted gene, 23010 [Source:MGI Symb | 0 | 0 | 1.0062 | 0 | 0 | 0 |
| 21933 | ENSMUSG00000077342 | 0 | NA | NA | NA | NA | NA | Gm24670 | predicted gene, 24670 [Source:MGI Symb | 0 | 0 | 0 | 0 | 0 | 0 |
| 21934 | ENSMUSG00000077344 | 0 | NA | NA | NA | NA | NA | Gm24326 | predicted gene, 24326 [Source:MGI Symb | 0 | 0 | 0 | 0 | 0 | 0 |
| 21935 | ENSMUSG00000077345 | 0.342639279 | -1.805976596 | 4.028547714 | -0.448294702 | 0.653940524 | NA | Snord70 | small nucleolar RNA, C/D box 70 [Source:I | 0 | 2.0558 | 0 | 0 | 0 | 0 |
| 21936 | ENSMUSG00000077347 | 0 | NA | NA | NA | NA | NA | Gm24671 | predicted gene, 24671 [Source:MGI Symb | 0 | 0 | 0 | 0 | 0 | 0 |
| 21937 | ENSMUSG00000077348 | 0 | NA | NA | NA | NA | NA | Gm24665 | predicted gene, 24665 [Source:MGI Symb | 0 | 0 | 0 | 0 | 0 | 0 |
| 21938 | ENSMUSG00000077349 | 0 | NA | NA | NA | NA | NA | Gm24867 | predicted gene, 24867 [Source:MGI Symb | 0 | 0 | 0 | 0 | 0 | 0 |
| 21939 | ENSMUSG00000077350 | 0 | NA | NA | NA | NA | NA | NA | NA | 0 | 0 | 0 | 0 | 0 | 0 |
| 21940 | ENSMUSG00000077351 | 0 | NA | NA | NA | NA | NA | NA | NA | 0 | 0 | 0 | 0 | 0 | 0 |
| 21941 | ENSMUSG00000077354 | 0.133850373 | -0.796629176 | 4.080472857 | -0.195229623 | 0.84521319 | NA | Gm26286 | predicted gene, 26286 [Source:MGI Symb | 0.8031 | 0 | 0 | 0 | 0 | 0 |
| 21942 | ENSMUSG00000077356 | 0 | NA | NA | NA | NA | NA | NA | NA | 0 | 0 | 0 | 0 | 0 | 0 |
| 21943 | ENSMUSG00000077357 | 0 | NA | NA | NA | NA | NA | Gm26289 | predicted gene, 26289 [Source:MGI Symb | 0 | 0 | 0 | 0 | 0 | 0 |
| 21944 | ENSMUSG00000077358 | 0 | NA | NA | NA | NA | NA | Gm26293 | predicted gene, 26293 [Source:MGI Symb | 0 | 0 | 0 | 0 | 0 | 0 |
| 21945 | ENSMUSG00000077359 | 0 | NA | NA | NA | NA | NA | Gm25308 | predicted gene, 25308 [Source:MGI Symb | 0 | 0 | 0 | 0 | 0 | 0 |
| 21946 | ENSMUSG00000077360 | 0.17131964 | -0.796629176 | 4.080472857 | -0.195229623 | 0.84521319 | NA | Gm23482 | predicted gene, 23482 [Source:MGI Symb | 0 | 1.0279 | 0 | 0 | 0 | 0 |
| 21947 | ENSMUSG00000077361 | 0 | NA | NA | NA | NA | NA | Gm23481 | predicted gene, 23481 [Source:MGI Symb | 0 | 0 | 0 | 0 | 0 | 0 |
| 21948 | ENSMUSG00000077366 | 0 | NA | NA | NA | NA | NA | Gm23484 | predicted gene, 23484 [Source:MGI Symb | 0 | 0 | 0 | 0 | 0 | 0 |

| | | | | | | | | | | | | | | | |
|---|---|---|---|---|---|---|---|---|---|---|---|---|---|---|---|
| 21949 | ENSMUSG00000077367 | 0 | NA | NA | NA | NA | NA | Gm23483 | predicted gene, 23483 [Source:MGI Symb | 0 | 0 | 0 | 0 | 0 | 0 |
| 21950 | ENSMUSG00000077368 | 0 | NA | NA | NA | NA | NA | Gm26273 | predicted gene, 26273 [Source:MGI Symb | 0 | 0 | 0 | 0 | 0 | 0 |
| 21951 | ENSMUSG00000077369 | 0 | NA | NA | NA | NA | NA | Gm23485 | predicted gene, 23485 [Source:MGI Symb | 0 | 0 | 0 | 0 | 0 | 0 |
| 21952 | ENSMUSG00000077376 | 0 | NA | NA | NA | NA | NA | Gm25167 | predicted gene, 25167 [Source:MGI Symb | 0 | 0 | 0 | 0 | 0 | 0 |
| 21953 | ENSMUSG00000077377 | 0 | NA | NA | NA | NA | NA | Gm25168 | predicted gene, 25168 [Source:MGI Symb | 0 | 0 | 0 | 0 | 0 | 0 |
| 21954 | ENSMUSG00000077378 | 0 | NA | NA | NA | NA | NA | Gm25169 | predicted gene, 25169 [Source:MGI Symb | 0 | 0 | 0 | 0 | 0 | 0 |
| 21955 | ENSMUSG00000077379 | 0 | NA | NA | NA | NA | NA | Gm25170 | predicted gene, 25170 [Source:MGI Symb | 0 | 0 | 0 | 0 | 0 | 0 |
| 21956 | ENSMUSG00000077380 | 0.86008258 | -1.819272683 | 2.588418657 | -0.702851016 | 0.482148601 | NA | Gm22661 | predicted gene, 22661 [Source:MGI Symb | 0 | 1.0279 | 3.0187 | 0 | 0 | 1.1138 |
| 21957 | ENSMUSG00000077381 | 0 | NA | NA | NA | NA | NA | Gm22660 | predicted gene, 22660 [Source:MGI Symb | 0 | 0 | 0 | 0 | 0 | 0 |
| 21958 | ENSMUSG00000077382 | 0.160662974 | 1.126924175 | 4.080472857 | 0.276174898 | 0.782413736 | NA | NA | NA | 0 | 0 | 0 | 0.964 | 0 | 0 |
| 21959 | ENSMUSG00000077383 | 0 | NA | NA | NA | NA | NA | Gm22659 | predicted gene, 22659 [Source:MGI Symb | 0 | 0 | 0 | 0 | 0 | 0 |
| 21960 | ENSMUSG00000077384 | 0 | NA | NA | NA | NA | NA | Gm22658 | predicted gene, 22658 [Source:MGI Symb | 0 | 0 | 0 | 0 | 0 | 0 |
| 21961 | ENSMUSG00000077385 | 0 | NA | NA | NA | NA | NA | Gm22657 | predicted gene, 22657 [Source:MGI Symb | 0 | 0 | 0 | 0 | 0 | 0 |
| 21962 | ENSMUSG00000077387 | 0.167707431 | -0.796629176 | 4.080472857 | -0.195229623 | 0.84521319 | NA | Gm22656 | predicted gene, 22656 [Source:MGI Symb | 0 | 0 | 1.0062 | 0 | 0 | 0 |
| 21963 | ENSMUSG00000077391 | 1.429055777 | -0.12521544 | 1.898409786 | -0.065958067 | 0.94741121 | NA | Gm24336 | predicted gene, 24336 [Source:MGI Symb | 2.4093 | 2.0558 | 0 | 1.928 | 2.1812 | 0 |
| 21964 | ENSMUSG00000077392 | 0 | NA | NA | NA | NA | NA | Gm24337 | predicted gene, 24337 [Source:MGI Symb | 0 | 0 | 0 | 0 | 0 | 0 |
| 21965 | ENSMUSG00000077393 | 0 | NA | NA | NA | NA | NA | Gm24338 | predicted gene, 24338 [Source:MGI Symb | 0 | 0 | 0 | 0 | 0 | 0 |
| 21966 | ENSMUSG00000077394 | 0 | NA | NA | NA | NA | NA | Gm24339 | predicted gene, 24339 [Source:MGI Symb | 0 | 0 | 0 | 0 | 0 | 0 |
| 21967 | ENSMUSG00000077396 | 0 | NA | NA | NA | NA | NA | Gm24340 | predicted gene, 24340 [Source:MGI Symb | 0 | 0 | 0 | 0 | 0 | 0 |
| 21968 | ENSMUSG00000077397 | 0 | NA | NA | NA | NA | NA | Gm24341 | predicted gene, 24341 [Source:MGI Symb | 0 | 0 | 0 | 0 | 0 | 0 |
| 21969 | ENSMUSG00000077398 | 0 | NA | NA | NA | NA | NA | Gm24334 | predicted gene, 24334 [Source:MGI Symb | 0 | 0 | 0 | 0 | 0 | 0 |
| 21970 | ENSMUSG00000077399 | 0 | NA | NA | NA | NA | NA | Gm24335 | predicted gene, 24335 [Source:MGI Symb | 0 | 0 | 0 | 0 | 0 | 0 |
| 21971 | ENSMUSG00000077400 | 0 | NA | NA | NA | NA | NA | Gm25417 | predicted gene, 25417 [Source:MGI Symb | 0 | 0 | 0 | 0 | 0 | 0 |
| 21972 | ENSMUSG00000077401 | 0 | NA | NA | NA | NA | NA | NA | NA | 0 | 0 | 0 | 0 | 0 | 0 |
| 21973 | ENSMUSG00000077402 | 0 | NA | NA | NA | NA | NA | Gm25418 | predicted gene, 25418 [Source:MGI Symb | 0 | 0 | 0 | 0 | 0 | 0 |
| 21974 | ENSMUSG00000077403 | 0 | NA | NA | NA | NA | NA | Gm25419 | predicted gene, 25419 [Source:MGI Symb | 0 | 0 | 0 | 0 | 0 | 0 |
| 21975 | ENSMUSG00000077404 | 0 | NA | NA | NA | NA | NA | Gm25420 | predicted gene, 25420 [Source:MGI Symb | 0 | 0 | 0 | 0 | 0 | 0 |
| 21976 | ENSMUSG00000077406 | 0 | NA | NA | NA | NA | NA | Gm25421 | predicted gene, 25421 [Source:MGI Symb | 0 | 0 | 0 | 0 | 0 | 0 |
| 21977 | ENSMUSG00000077407 | 0 | NA | NA | NA | NA | NA | Gm25422 | predicted gene, 25422 [Source:MGI Symb | 0 | 0 | 0 | 0 | 0 | 0 |
| 21978 | ENSMUSG00000077408 | 0 | NA | NA | NA | NA | NA | Gm25415 | predicted gene, 25415 [Source:MGI Symb | 0 | 0 | 0 | 0 | 0 | 0 |
| 21979 | ENSMUSG00000077409 | 0 | NA | NA | NA | NA | NA | Gm25416 | predicted gene, 25416 [Source:MGI Symb | 0 | 0 | 0 | 0 | 0 | 0 |
| 21980 | ENSMUSG00000077410 | 0 | NA | NA | NA | NA | NA | Gm23750 | predicted gene, 23750 [Source:MGI Symb | 0 | 0 | 0 | 0 | 0 | 0 |
| 21981 | ENSMUSG00000077411 | 0 | NA | NA | NA | NA | NA | Gm23749 | predicted gene, 23749 [Source:MGI Symb | 0 | 0 | 0 | 0 | 0 | 0 |
| 21982 | ENSMUSG00000077413 | 0 | NA | NA | NA | NA | NA | Gm23389 | predicted gene, 23389 [Source:MGI Symb | 0 | 0 | 0 | 0 | 0 | 0 |
| 21983 | ENSMUSG00000077415 | 0 | NA | NA | NA | NA | NA | Gm23748 | predicted gene, 23748 [Source:MGI Symb | 0 | 0 | 0 | 0 | 0 | 0 |
| 21984 | ENSMUSG00000077416 | 0.503122293 | -2.366675966 | 3.785596405 | -0.625179156 | 0.531853481 | NA | Gm23747 | predicted gene, 23747 [Source:MGI Symb | 0 | 0 | 3.0187 | 0 | 0 | 0 |
| 21985 | ENSMUSG00000077417 | 0 | NA | NA | NA | NA | NA | Gm23746 | predicted gene, 23746 [Source:MGI Symb | 0 | 0 | 0 | 0 | 0 | 0 |
| 21986 | ENSMUSG00000077418 | 0 | NA | NA | NA | NA | NA | Gm23745 | predicted gene, 23745 [Source:MGI Symb | 0 | 0 | 0 | 0 | 0 | 0 |
| 21987 | ENSMUSG00000077419 | 0 | NA | NA | NA | NA | NA | Gm23744 | predicted gene, 23744 [Source:MGI Symb | 0 | 0 | 0 | 0 | 0 | 0 |
| 21988 | ENSMUSG00000077420 | 0.696064467 | -1.534444742 | 3.043667477 | -0.504143358 | 0.61416064 | NA | NA | NA | 3.2124 | 0 | 0 | 0.964 | 0 | 0 |
| 21989 | ENSMUSG00000077421 | 0 | NA | NA | NA | NA | NA | NA | NA | 0 | 0 | 0 | 0 | 0 | 0 |
| 21990 | ENSMUSG00000077422 | 0 | NA | NA | NA | NA | NA | Gm24281 | predicted gene, 24281 [Source:MGI Symb | 0 | 0 | 0 | 0 | 0 | 0 |
| 21991 | ENSMUSG00000077424 | 0 | NA | NA | NA | NA | NA | Gm24279 | predicted gene, 24279 [Source:MGI Symb | 0 | 0 | 0 | 0 | 0 | 0 |
| 21992 | ENSMUSG00000077425 | 0 | NA | NA | NA | NA | NA | Gm24280 | predicted gene, 24280 [Source:MGI Symb | 0 | 0 | 0 | 0 | 0 | 0 |
| 21993 | ENSMUSG00000077426 | 2.804957708 | 0.988773085 | 1.31896537 | 0.749658109 | 0.453460644 | NA | Gm26387 | predicted gene, 26387 [Source:MGI Symb | 1.6062 | 3.0838 | 1.0062 | 6.7478 | 3.2719 | 1.1138 |
| 21994 | ENSMUSG00000077427 | 0 | NA | NA | NA | NA | NA | Gm24278 | predicted gene, 24278 [Source:MGI Symb | 0 | 0 | 0 | 0 | 0 | 0 |
| 21995 | ENSMUSG00000077428 | 0 | NA | NA | NA | NA | NA | NA | NA | 0 | 0 | 0 | 0 | 0 | 0 |
| 21996 | ENSMUSG00000077430 | 0 | NA | NA | NA | NA | NA | Gm22592 | predicted gene, 22592 [Source:MGI Symb | 0 | 0 | 0 | 0 | 0 | 0 |
| 21997 | ENSMUSG00000077431 | 0 | NA | NA | NA | NA | NA | Gm22591 | predicted gene, 22591 [Source:MGI Symb | 0 | 0 | 0 | 0 | 0 | 0 |
| 21998 | ENSMUSG00000077433 | 0 | NA | NA | NA | NA | NA | Gm22593 | predicted gene, 22593 [Source:MGI Symb | 0 | 0 | 0 | 0 | 0 | 0 |
| 21999 | ENSMUSG00000077435 | 0 | NA | NA | NA | NA | NA | Gm22590 | predicted gene, 22590 [Source:MGI Symb | 0 | 0 | 0 | 0 | 0 | 0 |
| 22000 | ENSMUSG00000077437 | 0 | NA | NA | NA | NA | NA | NA | NA | 0 | 0 | 0 | 0 | 0 | 0 |
| 22001 | ENSMUSG00000077438 | 0 | NA | NA | NA | NA | NA | Gm22595 | predicted gene, 22595 [Source:MGI Symb | 0 | 0 | 0 | 0 | 0 | 0 |
| 22002 | ENSMUSG00000077439 | 0 | NA | NA | NA | NA | NA | Gm22594 | predicted gene, 22594 [Source:MGI Symb | 0 | 0 | 0 | 0 | 0 | 0 |
| 22003 | ENSMUSG00000077440 | 0 | NA | NA | NA | NA | NA | Gm23130 | predicted gene, 23130 [Source:MGI Symb | 0 | 0 | 0 | 0 | 0 | 0 |
| 22004 | ENSMUSG00000077441 | 0 | NA | NA | NA | NA | NA | Gm23131 | predicted gene, 23131 [Source:MGI Symb | 0 | 0 | 0 | 0 | 0 | 0 |

| | | | | | | | | | | | | | | | |
|---|---|---|---|---|---|---|---|---|---|---|---|---|---|---|---|
| 22005 | ENSMUSG00000077442 | 0 | NA | NA | NA | NA | NA | NA | NA | 0 | 0 | 0 | 0 | 0 | 0 |
| 22006 | ENSMUSG00000077443 | 0 | NA | NA | NA | NA | NA | NA | NA | 0 | 0 | 0 | 0 | 0 | 0 |
| 22007 | ENSMUSG00000077445 | 0 | NA | NA | NA | NA | NA | NA | NA | 0 | 0 | 0 | 0 | 0 | 0 |
| 22008 | ENSMUSG00000077447 | 0 | NA | NA | NA | NA | NA | Gm23129 | predicted gene, 23129 [Source:MGI Symb | 0 | 0 | 0 | 0 | 0 | 0 |
| 22009 | ENSMUSG00000077448 | 0 | NA | NA | NA | NA | NA | Gm23126 | predicted gene, 23126 [Source:MGI Symb | 0 | 0 | 0 | 0 | 0 | 0 |
| 22010 | ENSMUSG00000077449 | 0 | NA | NA | NA | NA | NA | Gm23127 | predicted gene, 23127 [Source:MGI Symb | 0 | 0 | 0 | 0 | 0 | 0 |
| 22011 | ENSMUSG00000077450 | 965.3078385 | -0.062682752 | 0.106095362 | -0.590815196 | 0.554644252 | 0.897558119 | Rab11b | RAB11B, member RAS oncogene family [S | 958.1 | 1009.4 | 992.16 | 993.86 | 919.39 | 918.92 |
| 22012 | ENSMUSG00000077451 | 0 | NA | NA | NA | NA | NA | Gm25916 | predicted gene, 25916 [Source:MGI Symb | 0 | 0 | 0 | 0 | 0 | 0 |
| 22013 | ENSMUSG00000077453 | 0 | NA | NA | NA | NA | NA | Gm25915 | predicted gene, 25915 [Source:MGI Symb | 0 | 0 | 0 | 0 | 0 | 0 |
| 22014 | ENSMUSG00000077454 | 0 | NA | NA | NA | NA | NA | Gm25918 | predicted gene, 25918 [Source:MGI Symb | 0 | 0 | 0 | 0 | 0 | 0 |
| 22015 | ENSMUSG00000077455 | 0 | NA | NA | NA | NA | NA | Gm25917 | predicted gene, 25917 [Source:MGI Symb | 0 | 0 | 0 | 0 | 0 | 0 |
| 22016 | ENSMUSG00000077456 | 0 | NA | NA | NA | NA | NA | Gm22927 | predicted gene, 22927 [Source:MGI Symb | 0 | 0 | 0 | 0 | 0 | 0 |
| 22017 | ENSMUSG00000077457 | 1.55449208 | -1.714554891 | 2.084436349 | -0.82255085 | 0.410763459 | NA | Snord65 | small nucleolar RNA, C/D box 65 [Source:I | 4.0155 | 3.0838 | 0 | 0 | 0 | 2.2277 |
| 22018 | ENSMUSG00000077459 | 0 | NA | NA | NA | NA | NA | Gm25919 | predicted gene, 25919 [Source:MGI Symb | 0 | 0 | 0 | 0 | 0 | 0 |
| 22019 | ENSMUSG00000077460 | 0.133850373 | -0.796629176 | 4.080472857 | -0.195229623 | 0.84521319 | NA | Gm26419 | predicted gene, 26419 [Source:MGI Symb | 0.8031 | 0 | 0 | 0 | 0 | 0 |
| 22020 | ENSMUSG00000077461 | 0 | NA | NA | NA | NA | NA | Gm26420 | predicted gene, 26420 [Source:MGI Symb | 0 | 0 | 0 | 0 | 0 | 0 |
| 22021 | ENSMUSG00000077462 | 0 | NA | NA | NA | NA | NA | Gm26417 | predicted gene, 26417 [Source:MGI Symb | 0 | 0 | 0 | 0 | 0 | 0 |
| 22022 | ENSMUSG00000077463 | 0 | NA | NA | NA | NA | NA | Gm26418 | predicted gene, 26418 [Source:MGI Symb | 0 | 0 | 0 | 0 | 0 | 0 |
| 22023 | ENSMUSG00000077464 | 0 | NA | NA | NA | NA | NA | NA | NA | 0 | 0 | 0 | 0 | 0 | 0 |
| 22024 | ENSMUSG00000077465 | 0 | NA | NA | NA | NA | NA | Gm26423 | predicted gene, 26423 [Source:MGI Symb | 0 | 0 | 0 | 0 | 0 | 0 |
| 22025 | ENSMUSG00000077467 | 0 | NA | NA | NA | NA | NA | NA | NA | 0 | 0 | 0 | 0 | 0 | 0 |
| 22026 | ENSMUSG00000077468 | 0 | NA | NA | NA | NA | NA | Gm26416 | predicted gene, 26416 [Source:MGI Symb | 0 | 0 | 0 | 0 | 0 | 0 |
| 22027 | ENSMUSG00000077469 | 0 | NA | NA | NA | NA | NA | Gm25922 | predicted gene, 25922 [Source:MGI Symb | 0 | 0 | 0 | 0 | 0 | 0 |
| 22028 | ENSMUSG00000077470 | 0 | NA | NA | NA | NA | NA | Gm24789 | predicted gene, 24789 [Source:MGI Symb | 0 | 0 | 0 | 0 | 0 | 0 |
| 22029 | ENSMUSG00000077471 | 0 | NA | NA | NA | NA | NA | Gm24788 | predicted gene, 24788 [Source:MGI Symb | 0 | 0 | 0 | 0 | 0 | 0 |
| 22030 | ENSMUSG00000077472 | 0 | NA | NA | NA | NA | NA | Gm24791 | predicted gene, 24791 [Source:MGI Symb | 0 | 0 | 0 | 0 | 0 | 0 |
| 22031 | ENSMUSG00000077473 | 0 | NA | NA | NA | NA | NA | Gm24790 | predicted gene, 24790 [Source:MGI Symb | 0 | 0 | 0 | 0 | 0 | 0 |
| 22032 | ENSMUSG00000077474 | 0 | NA | NA | NA | NA | NA | Gm24792 | predicted gene, 24792 [Source:MGI Symb | 0 | 0 | 0 | 0 | 0 | 0 |
| 22033 | ENSMUSG00000077476 | 0 | NA | NA | NA | NA | NA | Gm24794 | predicted gene, 24794 [Source:MGI Symb | 0 | 0 | 0 | 0 | 0 | 0 |
| 22034 | ENSMUSG00000077477 | 0 | NA | NA | NA | NA | NA | Gm24793 | predicted gene, 24793 [Source:MGI Symb | 0 | 0 | 0 | 0 | 0 | 0 |
| 22035 | ENSMUSG00000077480 | 0 | NA | NA | NA | NA | NA | NA | NA | 0 | 0 | 0 | 0 | 0 | 0 |
| 22036 | ENSMUSG00000077481 | 0 | NA | NA | NA | NA | NA | Gm25272 | predicted gene, 25272 [Source:MGI Symb | 0 | 0 | 0 | 0 | 0 | 0 |
| 22037 | ENSMUSG00000077483 | 0 | NA | NA | NA | NA | NA | NA | NA | 0 | 0 | 0 | 0 | 0 | 0 |
| 22038 | ENSMUSG00000077484 | 0 | NA | NA | NA | NA | NA | Gm25274 | predicted gene, 25274 [Source:MGI Symb | 0 | 0 | 0 | 0 | 0 | 0 |
| 22039 | ENSMUSG00000077485 | 0 | NA | NA | NA | NA | NA | Gm25275 | predicted gene, 25275 [Source:MGI Symb | 0 | 0 | 0 | 0 | 0 | 0 |
| 22040 | ENSMUSG00000077487 | 0 | NA | NA | NA | NA | NA | NA | NA | 0 | 0 | 0 | 0 | 0 | 0 |
| 22041 | ENSMUSG00000077488 | 0 | NA | NA | NA | NA | NA | Gm25276 | predicted gene, 25276 [Source:MGI Symb | 0 | 0 | 0 | 0 | 0 | 0 |
| 22042 | ENSMUSG00000077492 | 0 | NA | NA | NA | NA | NA | Gm23596 | predicted gene, 23596 [Source:MGI Symb | 0 | 0 | 0 | 0 | 0 | 0 |
| 22043 | ENSMUSG00000077493 | 9.112629288 | -0.319044206 | 0.735144578 | -0.433988382 | 0.664296878 | 0.929301301 | Snord91a | small nucleolar RNA, C/D box 91A [Source | 8.031 | 10.279 | 12.075 | 6.7478 | 13.087 | 4.4554 |
| 22044 | ENSMUSG00000077494 | 0 | NA | NA | NA | NA | NA | Gm23595 | predicted gene, 23595 [Source:MGI Symb | 0 | 0 | 0 | 0 | 0 | 0 |
| 22045 | ENSMUSG00000077495 | 0 | NA | NA | NA | NA | NA | Gm23594 | predicted gene, 23594 [Source:MGI Symb | 0 | 0 | 0 | 0 | 0 | 0 |
| 22046 | ENSMUSG00000077497 | 0.181769715 | 1.126924175 | 4.080472857 | 0.276174898 | 0.782413736 | NA | Gm23593 | predicted gene, 23593 [Source:MGI Symb | 0 | 0 | 0 | 0 | 1.0906 | 0 |
| 22047 | ENSMUSG00000077499 | 0 | NA | NA | NA | NA | NA | NA | NA | 0 | 0 | 0 | 0 | 0 | 0 |
| 22048 | ENSMUSG00000077501 | 0 | NA | NA | NA | NA | NA | Gm24230 | predicted gene, 24230 [Source:MGI Symb | 0 | 0 | 0 | 0 | 0 | 0 |
| 22049 | ENSMUSG00000077502 | 0 | NA | NA | NA | NA | NA | Gm24232 | predicted gene, 24232 [Source:MGI Symb | 0 | 0 | 0 | 0 | 0 | 0 |
| 22050 | ENSMUSG00000077503 | 0 | NA | NA | NA | NA | NA | Gm24231 | predicted gene, 24231 [Source:MGI Symb | 0 | 0 | 0 | 0 | 0 | 0 |
| 22051 | ENSMUSG00000077504 | 0 | NA | NA | NA | NA | NA | Gm24234 | predicted gene, 24234 [Source:MGI Symb | 0 | 0 | 0 | 0 | 0 | 0 |
| 22052 | ENSMUSG00000077505 | 5.504010776 | -0.174130017 | 0.991633165 | -0.175599227 | 0.86060881 | 0.975943719 | Gm24233 | predicted gene, 24233 [Source:MGI Symb | 2.4093 | 7.1954 | 8.05 | 7.7118 | 6.5437 | 1.1138 |
| 22053 | ENSMUSG00000077506 | 0 | NA | NA | NA | NA | NA | Scarna9 | small Cajal body-specific RNA 9 [Source:M | 0 | 0 | 0 | 0 | 0 | 0 |
| 22054 | ENSMUSG00000077507 | 0 | NA | NA | NA | NA | NA | Gm24235 | predicted gene, 24235 [Source:MGI Symb | 0 | 0 | 0 | 0 | 0 | 0 |
| 22055 | ENSMUSG00000077511 | 0 | NA | NA | NA | NA | NA | Gm25875 | predicted gene, 25875 [Source:MGI Symb | 0 | 0 | 0 | 0 | 0 | 0 |
| 22056 | ENSMUSG00000077514 | 0 | NA | NA | NA | NA | NA | Gm25877 | predicted gene, 25877 [Source:MGI Symb | 0 | 0 | 0 | 0 | 0 | 0 |
| 22057 | ENSMUSG00000077515 | 0 | NA | NA | NA | NA | NA | Gm25878 | predicted gene, 25878 [Source:MGI Symb | 0 | 0 | 0 | 0 | 0 | 0 |
| 22058 | ENSMUSG00000077516 | 0 | NA | NA | NA | NA | NA | NA | NA | 0 | 0 | 0 | 0 | 0 | 0 |
| 22059 | ENSMUSG00000077517 | 0 | NA | NA | NA | NA | NA | NA | NA | 0 | 0 | 0 | 0 | 0 | 0 |
| 22060 | ENSMUSG00000077518 | 0 | NA | NA | NA | NA | NA | Gm25879 | predicted gene, 25879 [Source:MGI Symb | 0 | 0 | 0 | 0 | 0 | 0 |

| | | | | | | | | | | | | | | | |
|---|---|---|---|---|---|---|---|---|---|---|---|---|---|---|---|
| 22061 | ENSMUSG00000077519 | 0 | NA | NA | NA | NA | NA | Gm25880 | predicted gene, 25880 [Source:MGI Symb | 0 | 0 | 0 | 0 | 0 | 0 |
| 22062 | ENSMUSG00000077521 | 0 | NA | NA | NA | NA | NA | Gm25378 | predicted gene, 25378 [Source:MGI Symb | 0 | 0 | 0 | 0 | 0 | 0 |
| 22063 | ENSMUSG00000077522 | 0 | NA | NA | NA | NA | NA | Gm25377 | predicted gene, 25377 [Source:MGI Symb | 0 | 0 | 0 | 0 | 0 | 0 |
| 22064 | ENSMUSG00000077524 | 0.506966596 | 2.558866737 | 3.339834942 | 0.766165628 | 0.443577757 | NA | NA | NA | 0 | 0 | 0 | 1.928 | 0 | 1.1138 |
| 22065 | ENSMUSG00000077526 | 0 | NA | NA | NA | NA | NA | Gm25379 | predicted gene, 25379 [Source:MGI Symb | 0 | 0 | 0 | 0 | 0 | 0 |
| 22066 | ENSMUSG00000077527 | 0 | NA | NA | NA | NA | NA | Gm25506 | predicted gene, 25506 [Source:MGI Symb | 0 | 0 | 0 | 0 | 0 | 0 |
| 22067 | ENSMUSG00000077528 | 0 | NA | NA | NA | NA | NA | Gm25376 | predicted gene, 25376 [Source:MGI Symb | 0 | 0 | 0 | 0 | 0 | 0 |
| 22068 | ENSMUSG00000077535 | 0 | NA | NA | NA | NA | NA | Gm22543 | predicted gene, 22543 [Source:MGI Symb | 0 | 0 | 0 | 0 | 0 | 0 |
| 22069 | ENSMUSG00000077537 | 0 | NA | NA | NA | NA | NA | Gm22544 | predicted gene, 22544 [Source:MGI Symb | 0 | 0 | 0 | 0 | 0 | 0 |
| 22070 | ENSMUSG00000077538 | 0 | NA | NA | NA | NA | NA | Gm22545 | predicted gene, 22545 [Source:MGI Symb | 0 | 0 | 0 | 0 | 0 | 0 |
| 22071 | ENSMUSG00000077539 | 0 | NA | NA | NA | NA | NA | Gm22546 | predicted gene, 22546 [Source:MGI Symb | 0 | 0 | 0 | 0 | 0 | 0 |
| 22072 | ENSMUSG00000077541 | 0 | NA | NA | NA | NA | NA | Gm22021 | predicted gene, 22021 [Source:MGI Symb | 0 | 0 | 0 | 0 | 0 | 0 |
| 22073 | ENSMUSG00000077542 | 0 | NA | NA | NA | NA | NA | Gm22023 | predicted gene, 22023 [Source:MGI Symb | 0 | 0 | 0 | 0 | 0 | 0 |
| 22074 | ENSMUSG00000077543 | 0 | NA | NA | NA | NA | NA | Gm22022 | predicted gene, 22022 [Source:MGI Symb | 0 | 0 | 0 | 0 | 0 | 0 |
| 22075 | ENSMUSG00000077545 | 0 | NA | NA | NA | NA | NA | Gm22018 | predicted gene, 22018 [Source:MGI Symb | 0 | 0 | 0 | 0 | 0 | 0 |
| 22076 | ENSMUSG00000077546 | 0 | NA | NA | NA | NA | NA | Gm22020 | predicted gene, 22020 [Source:MGI Symb | 0 | 0 | 0 | 0 | 0 | 0 |
| 22077 | ENSMUSG00000077547 | 0 | NA | NA | NA | NA | NA | Gm22019 | predicted gene, 22019 [Source:MGI Symb | 0 | 0 | 0 | 0 | 0 | 0 |
| 22078 | ENSMUSG00000077549 | 1.225999361 | -1.452919977 | 2.243334418 | -0.647660895 | 0.517204302 | NA | Snord71 | small nucleolar RNA, C/D box 71 [Source:I | 2.4093 | 0 | 3.0187 | 1.928 | 0 | 0 |
| 22079 | ENSMUSG00000077552 | 0 | NA | NA | NA | NA | NA | Gm23709 | predicted gene, 23709 [Source:MGI Symb | 0 | 0 | 0 | 0 | 0 | 0 |
| 22080 | ENSMUSG00000077553 | 0 | NA | NA | NA | NA | NA | Gm23176 | predicted gene, 23176 [Source:MGI Symb | 0 | 0 | 0 | 0 | 0 | 0 |
| 22081 | ENSMUSG00000077554 | 0 | NA | NA | NA | NA | NA | Gm23707 | predicted gene, 23707 [Source:MGI Symb | 0 | 0 | 0 | 0 | 0 | 0 |
| 22082 | ENSMUSG00000077555 | 0 | NA | NA | NA | NA | NA | Gm23708 | predicted gene, 23708 [Source:MGI Symb | 0 | 0 | 0 | 0 | 0 | 0 |
| 22083 | ENSMUSG00000077557 | 0.160662974 | 1.126924175 | 4.080472857 | 0.276174898 | 0.782413736 | NA | Gm23706 | predicted gene, 23706 [Source:MGI Symb | 0 | 0 | 0 | 0.964 | 0 | 0 |
| 22084 | ENSMUSG00000077558 | 0 | NA | NA | NA | NA | NA | Gm23705 | predicted gene, 23705 [Source:MGI Symb | 0 | 0 | 0 | 0 | 0 | 0 |
| 22085 | ENSMUSG00000077559 | 0 | NA | NA | NA | NA | NA | NA | NA | 0 | 0 | 0 | 0 | 0 | 0 |
| 22086 | ENSMUSG00000077561 | 0 | NA | NA | NA | NA | NA | Gm24607 | predicted gene, 24607 [Source:MGI Symb | 0 | 0 | 0 | 0 | 0 | 0 |
| 22087 | ENSMUSG00000077562 | 0 | NA | NA | NA | NA | NA | Gm23203 | predicted gene, 23203 [Source:MGI Symb | 0 | 0 | 0 | 0 | 0 | 0 |
| 22088 | ENSMUSG00000077563 | 0 | NA | NA | NA | NA | NA | Snora68 | small nucleolar RNA, H/ACA box 68 [Sourc | 0 | 0 | 0 | 0 | 0 | 0 |
| 22089 | ENSMUSG00000077564 | 0 | NA | NA | NA | NA | NA | Gm23202 | predicted gene, 23202 [Source:MGI Symb | 0 | 0 | 0 | 0 | 0 | 0 |
| 22090 | ENSMUSG00000077565 | 0 | NA | NA | NA | NA | NA | Gm23201 | predicted gene, 23201 [Source:MGI Symb | 0 | 0 | 0 | 0 | 0 | 0 |
| 22091 | ENSMUSG00000077567 | 0 | NA | NA | NA | NA | NA | Gm23200 | predicted gene, 23200 [Source:MGI Symb | 0 | 0 | 0 | 0 | 0 | 0 |
| 22092 | ENSMUSG00000077569 | 0.538988726 | 1.126003851 | 3.343747182 | 0.336749099 | 0.736306047 | NA | Gm24608 | predicted gene, 24608 [Source:MGI Symb | 0 | 0 | 1.0062 | 0 | 0 | 2.2277 |
| 22093 | ENSMUSG00000077571 | 0 | NA | NA | NA | NA | NA | NA | NA | 0 | 0 | 0 | 0 | 0 | 0 |
| 22094 | ENSMUSG00000077572 | 0 | NA | NA | NA | NA | NA | Gm24886 | predicted gene, 24886 [Source:MGI Symb | 0 | 0 | 0 | 0 | 0 | 0 |
| 22095 | ENSMUSG00000077574 | 0 | NA | NA | NA | NA | NA | Gm24887 | predicted gene, 24887 [Source:MGI Symb | 0 | 0 | 0 | 0 | 0 | 0 |
| 22096 | ENSMUSG00000077575 | 2.396254893 | -1.768497719 | 1.673381441 | -1.056840763 | 0.290584267 | NA | Gm24888 | predicted gene, 24888 [Source:MGI Symb | 4.0155 | 0 | 7.0437 | 0 | 1.0906 | 2.2277 |
| 22097 | ENSMUSG00000077576 | 0 | NA | NA | NA | NA | NA | Gm24889 | predicted gene, 24889 [Source:MGI Symb | 0 | 0 | 0 | 0 | 0 | 0 |
| 22098 | ENSMUSG00000077577 | 0 | NA | NA | NA | NA | NA | Gm24890 | predicted gene, 24890 [Source:MGI Symb | 0 | 0 | 0 | 0 | 0 | 0 |
| 22099 | ENSMUSG00000077578 | 0 | NA | NA | NA | NA | NA | Gm25631 | predicted gene, 25631 [Source:MGI Symb | 0 | 0 | 0 | 0 | 0 | 0 |
| 22100 | ENSMUSG00000077579 | 0 | NA | NA | NA | NA | NA | Gm24891 | predicted gene, 24891 [Source:MGI Symb | 0 | 0 | 0 | 0 | 0 | 0 |
| 22101 | ENSMUSG00000077581 | 0 | NA | NA | NA | NA | NA | Gm24373 | predicted gene, 24373 [Source:MGI Symb | 0 | 0 | 0 | 0 | 0 | 0 |
| 22102 | ENSMUSG00000077582 | 0 | NA | NA | NA | NA | NA | Gm24374 | predicted gene, 24374 [Source:MGI Symb | 0 | 0 | 0 | 0 | 0 | 0 |
| 22103 | ENSMUSG00000077585 | 0 | NA | NA | NA | NA | NA | Gm24375 | predicted gene, 24375 [Source:MGI Symb | 0 | 0 | 0 | 0 | 0 | 0 |
| 22104 | ENSMUSG00000077586 | 0 | NA | NA | NA | NA | NA | Gm24377 | predicted gene, 24377 [Source:MGI Symb | 0 | 0 | 0 | 0 | 0 | 0 |
| 22105 | ENSMUSG00000077587 | 0 | NA | NA | NA | NA | NA | Gm24376 | predicted gene, 24376 [Source:MGI Symb | 0 | 0 | 0 | 0 | 0 | 0 |
| 22106 | ENSMUSG00000077588 | 0 | NA | NA | NA | NA | NA | NA | NA | 0 | 0 | 0 | 0 | 0 | 0 |
| 22107 | ENSMUSG00000077590 | 0 | NA | NA | NA | NA | NA | Gm26015 | predicted gene, 26015 [Source:MGI Symb | 0 | 0 | 0 | 0 | 0 | 0 |
| 22108 | ENSMUSG00000077591 | 0 | NA | NA | NA | NA | NA | Gm26016 | predicted gene, 26016 [Source:MGI Symb | 0 | 0 | 0 | 0 | 0 | 0 |
| 22109 | ENSMUSG00000077592 | 0 | NA | NA | NA | NA | NA | Gm26013 | predicted gene, 26013 [Source:MGI Symb | 0 | 0 | 0 | 0 | 0 | 0 |
| 22110 | ENSMUSG00000077593 | 0 | NA | NA | NA | NA | NA | Gm26014 | predicted gene, 26014 [Source:MGI Symb | 0 | 0 | 0 | 0 | 0 | 0 |
| 22111 | ENSMUSG00000077594 | 0 | NA | NA | NA | NA | NA | Gm26018 | predicted gene, 26018 [Source:MGI Symb | 0 | 0 | 0 | 0 | 0 | 0 |
| 22112 | ENSMUSG00000077595 | 0 | NA | NA | NA | NA | NA | Gm26019 | predicted gene, 26019 [Source:MGI Symb | 0 | 0 | 0 | 0 | 0 | 0 |
| 22113 | ENSMUSG00000077597 | 0 | NA | NA | NA | NA | NA | Gm26017 | predicted gene, 26017 [Source:MGI Symb | 0 | 0 | 0 | 0 | 0 | 0 |
| 22114 | ENSMUSG00000077598 | 0 | NA | NA | NA | NA | NA | Gm26011 | predicted gene, 26011 [Source:MGI Symb | 0 | 0 | 0 | 0 | 0 | 0 |
| 22115 | ENSMUSG00000077599 | 0 | NA | NA | NA | NA | NA | Gm26012 | predicted gene, 26012 [Source:MGI Symb | 0 | 0 | 0 | 0 | 0 | 0 |
| 22116 | ENSMUSG00000077601 | 0 | NA | NA | NA | NA | NA | Gm22674 | predicted gene, 22674 [Source:MGI Symb | 0 | 0 | 0 | 0 | 0 | 0 |

| | | | | | | | | | | | | | | | |
|---|---|---|---|---|---|---|---|---|---|---|---|---|---|---|---|
| 22117 | ENSMUSG00000077602 | 0 | NA | NA | NA | NA | NA | Gm22672 | predicted gene, 22672 [Source:MGI Symb | 0 | 0 | 0 | 0 | 0 | 0 |
| 22118 | ENSMUSG00000077603 | 0 | NA | NA | NA | NA | NA | Gm22673 | predicted gene, 22673 [Source:MGI Symb | 0 | 0 | 0 | 0 | 0 | 0 |
| 22119 | ENSMUSG00000077604 | 0 | NA | NA | NA | NA | NA | Snord1b | small nucleolar RNA, C/D box 1B [Source:I | 0 | 0 | 0 | 0 | 0 | 0 |
| 22120 | ENSMUSG00000077607 | 0 | NA | NA | NA | NA | NA | Gm25803 | predicted gene, 25803 [Source:MGI Symb | 0 | 0 | 0 | 0 | 0 | 0 |
| 22121 | ENSMUSG00000077608 | 0 | NA | NA | NA | NA | NA | Gm22671 | predicted gene, 22671 [Source:MGI Symb | 0 | 0 | 0 | 0 | 0 | 0 |
| 22122 | ENSMUSG00000077609 | 0 | NA | NA | NA | NA | NA | Gm25795 | predicted gene, 25795 [Source:MGI Symb | 0 | 0 | 0 | 0 | 0 | 0 |
| 22123 | ENSMUSG00000077611 | 0 | NA | NA | NA | NA | NA | Gm23946 | predicted gene, 23946 [Source:MGI Symb | 0 | 0 | 0 | 0 | 0 | 0 |
| 22124 | ENSMUSG00000077612 | 0 | NA | NA | NA | NA | NA | Gm25498 | predicted gene, 25498 [Source:MGI Symb | 0 | 0 | 0 | 0 | 0 | 0 |
| 22125 | ENSMUSG00000077613 | 0 | NA | NA | NA | NA | NA | NA | NA | 0 | 0 | 0 | 0 | 0 | 0 |
| 22126 | ENSMUSG00000077614 | 0 | NA | NA | NA | NA | NA | NA | NA | 0 | 0 | 0 | 0 | 0 | 0 |
| 22127 | ENSMUSG00000077615 | 0 | NA | NA | NA | NA | NA | Gm25499 | predicted gene, 25499 [Source:MGI Symb | 0 | 0 | 0 | 0 | 0 | 0 |
| 22128 | ENSMUSG00000077616 | 0 | NA | NA | NA | NA | NA | Gm25502 | predicted gene, 25502 [Source:MGI Symb | 0 | 0 | 0 | 0 | 0 | 0 |
| 22129 | ENSMUSG00000077617 | 6.663759113 | 0.210232532 | 0.878955781 | 0.239184424 | 0.81096258 | 0.966452061 | Gm25501 | predicted gene, 25501 [Source:MGI Symb | 2.4093 | 5.1396 | 11.069 | 4.8199 | 7.6343 | 8.9108 |
| 22130 | ENSMUSG00000077618 | 0 | NA | NA | NA | NA | NA | Gm25496 | predicted gene, 25496 [Source:MGI Symb | 0 | 0 | 0 | 0 | 0 | 0 |
| 22131 | ENSMUSG00000077619 | 0 | NA | NA | NA | NA | NA | Gm25495 | predicted gene, 25495 [Source:MGI Symb | 0 | 0 | 0 | 0 | 0 | 0 |
| 22132 | ENSMUSG00000077620 | 0 | NA | NA | NA | NA | NA | NA | NA | 0 | 0 | 0 | 0 | 0 | 0 |
| 22133 | ENSMUSG00000077621 | 0.133850373 | -0.796629176 | 4.080472857 | -0.195229623 | 0.84521319 | NA | NA | NA | 0.8031 | 0 | 0 | 0 | 0 | 0 |
| 22134 | ENSMUSG00000077622 | 0 | NA | NA | NA | NA | NA | NA | NA | 0 | 0 | 0 | 0 | 0 | 0 |
| 22135 | ENSMUSG00000077624 | 0 | NA | NA | NA | NA | NA | Gm23834 | predicted gene, 23834 [Source:MGI Symb | 0 | 0 | 0 | 0 | 0 | 0 |
| 22136 | ENSMUSG00000077625 | 0 | NA | NA | NA | NA | NA | Snord4a | small nucleolar RNA, C/D box 4A [Source:I | 0 | 0 | 0 | 0 | 0 | 0 |
| 22137 | ENSMUSG00000077627 | 0 | NA | NA | NA | NA | NA | Gm23835 | predicted gene, 23835 [Source:MGI Symb | 0 | 0 | 0 | 0 | 0 | 0 |
| 22138 | ENSMUSG00000077628 | 0 | NA | NA | NA | NA | NA | Gm23768 | predicted gene, 23768 [Source:MGI Symb | 0 | 0 | 0 | 0 | 0 | 0 |
| 22139 | ENSMUSG00000077629 | 0 | NA | NA | NA | NA | NA | Gm23769 | predicted gene, 23769 [Source:MGI Symb | 0 | 0 | 0 | 0 | 0 | 0 |
| 22140 | ENSMUSG00000077630 | 0 | NA | NA | NA | NA | NA | Gm22149 | predicted gene, 22149 [Source:MGI Symb | 0 | 0 | 0 | 0 | 0 | 0 |
| 22141 | ENSMUSG00000077633 | 0 | NA | NA | NA | NA | NA | Gm22148 | predicted gene, 22148 [Source:MGI Symb | 0 | 0 | 0 | 0 | 0 | 0 |
| 22142 | ENSMUSG00000077635 | 0 | NA | NA | NA | NA | NA | Gm22150 | predicted gene, 22150 [Source:MGI Symb | 0 | 0 | 0 | 0 | 0 | 0 |
| 22143 | ENSMUSG00000077637 | 3.895079665 | 0.845867091 | 1.156114888 | 0.731646223 | 0.464384528 | NA | Gm22771 | predicted gene, 22771 [Source:MGI Symb | 3.2124 | 3.0838 | 2.0125 | 1.928 | 10.906 | 2.2277 |
| 22144 | ENSMUSG00000077639 | 1.37068636 | -1.664899446 | 2.166676757 | -0.76841155 | 0.44224272 | NA | Gm22772 | predicted gene, 22772 [Source:MGI Symb | 3.2124 | 3.0838 | 0 | 1.928 | 0 | 0 |
| 22145 | ENSMUSG00000077640 | 0 | NA | NA | NA | NA | NA | NA | NA | 0 | 0 | 0 | 0 | 0 | 0 |
| 22146 | ENSMUSG00000077641 | 0 | NA | NA | NA | NA | NA | Gm24987 | predicted gene, 24987 [Source:MGI Symb | 0 | 0 | 0 | 0 | 0 | 0 |
| 22147 | ENSMUSG00000077642 | 0 | NA | NA | NA | NA | NA | Gm24984 | predicted gene, 24984 [Source:MGI Symb | 0 | 0 | 0 | 0 | 0 | 0 |
| 22148 | ENSMUSG00000077643 | 0 | NA | NA | NA | NA | NA | Gm24985 | predicted gene, 24985 [Source:MGI Symb | 0 | 0 | 0 | 0 | 0 | 0 |
| 22149 | ENSMUSG00000077644 | 0 | NA | NA | NA | NA | NA | NA | NA | 0 | 0 | 0 | 0 | 0 | 0 |
| 22150 | ENSMUSG00000077645 | 0 | NA | NA | NA | NA | NA | Gm24983 | predicted gene, 24983 [Source:MGI Symb | 0 | 0 | 0 | 0 | 0 | 0 |
| 22151 | ENSMUSG00000077647 | 0 | NA | NA | NA | NA | NA | Gm24981 | predicted gene, 24981 [Source:MGI Symb | 0 | 0 | 0 | 0 | 0 | 0 |
| 22152 | ENSMUSG00000077648 | 0 | NA | NA | NA | NA | NA | Gm24988 | predicted gene, 24988 [Source:MGI Symb | 0 | 0 | 0 | 0 | 0 | 0 |
| 22153 | ENSMUSG00000077649 | 0 | NA | NA | NA | NA | NA | Gm24989 | predicted gene, 24989 [Source:MGI Symb | 0 | 0 | 0 | 0 | 0 | 0 |
| 22154 | ENSMUSG00000077650 | 0 | NA | NA | NA | NA | NA | Gm23320 | predicted gene, 23320 [Source:MGI Symb | 0 | 0 | 0 | 0 | 0 | 0 |
| 22155 | ENSMUSG00000077652 | 0.331982614 | 0.16514309 | 3.986751476 | 0.041422971 | 0.966958701 | NA | Gm23322 | predicted gene, 23322 [Source:MGI Symb | 0 | 1.0279 | 0 | 0.964 | 0 | 0 |
| 22156 | ENSMUSG00000077653 | 0 | NA | NA | NA | NA | NA | Gm23321 | predicted gene, 23321 [Source:MGI Symb | 0 | 0 | 0 | 0 | 0 | 0 |
| 22157 | ENSMUSG00000077655 | 0 | NA | NA | NA | NA | NA | Gm23319 | predicted gene, 23319 [Source:MGI Symb | 0 | 0 | 0 | 0 | 0 | 0 |
| 22158 | ENSMUSG00000077659 | 0 | NA | NA | NA | NA | NA | Gm23323 | predicted gene, 23323 [Source:MGI Symb | 0 | 0 | 0 | 0 | 0 | 0 |
| 22159 | ENSMUSG00000077660 | 0 | NA | NA | NA | NA | NA | Gm24257 | predicted gene, 24257 [Source:MGI Symb | 0 | 0 | 0 | 0 | 0 | 0 |
| 22160 | ENSMUSG00000077661 | 0 | NA | NA | NA | NA | NA | NA | NA | 0 | 0 | 0 | 0 | 0 | 0 |
| 22161 | ENSMUSG00000077663 | 0 | NA | NA | NA | NA | NA | Gm26144 | predicted gene, 26144 [Source:MGI Symb | 0 | 0 | 0 | 0 | 0 | 0 |
| 22162 | ENSMUSG00000077664 | 0 | NA | NA | NA | NA | NA | Gm26145 | predicted gene, 26145 [Source:MGI Symb | 0 | 0 | 0 | 0 | 0 | 0 |
| 22163 | ENSMUSG00000077666 | 0 | NA | NA | NA | NA | NA | Gm26146 | predicted gene, 26146 [Source:MGI Symb | 0 | 0 | 0 | 0 | 0 | 0 |
| 22164 | ENSMUSG00000077667 | 0.160662974 | 1.126924175 | 4.080472857 | 0.276174898 | 0.782413736 | NA | Gm26147 | predicted gene, 26147 [Source:MGI Symb | 0 | 0 | 0 | 0.964 | 0 | 0 |
| 22165 | ENSMUSG00000077668 | 0 | NA | NA | NA | NA | NA | Gm26143 | predicted gene, 26143 [Source:MGI Symb | 0 | 0 | 0 | 0 | 0 | 0 |
| 22166 | ENSMUSG00000077669 | 0 | NA | NA | NA | NA | NA | Gm24255 | predicted gene, 24255 [Source:MGI Symb | 0 | 0 | 0 | 0 | 0 | 0 |
| 22167 | ENSMUSG00000077671 | 0 | NA | NA | NA | NA | NA | Gm24473 | predicted gene, 24473 [Source:MGI Symb | 0 | 0 | 0 | 0 | 0 | 0 |
| 22168 | ENSMUSG00000077672 | 0 | NA | NA | NA | NA | NA | Gm24472 | predicted gene, 24472 [Source:MGI Symb | 0 | 0 | 0 | 0 | 0 | 0 |
| 22169 | ENSMUSG00000077673 | 0 | NA | NA | NA | NA | NA | Gm24471 | predicted gene, 24471 [Source:MGI Symb | 0 | 0 | 0 | 0 | 0 | 0 |
| 22170 | ENSMUSG00000077674 | 0 | NA | NA | NA | NA | NA | Gm24470 | predicted gene, 24470 [Source:MGI Symb | 0 | 0 | 0 | 0 | 0 | 0 |
| 22171 | ENSMUSG00000077675 | 0 | NA | NA | NA | NA | NA | Gm24469 | predicted gene, 24469 [Source:MGI Symb | 0 | 0 | 0 | 0 | 0 | 0 |
| 22172 | ENSMUSG00000077677 | 0.17131964 | -0.796629176 | 4.080472857 | -0.195229623 | 0.84521319 | NA | Gm24468 | predicted gene, 24468 [Source:MGI Symb | 0 | 1.0279 | 0 | 0 | 0 | 0 |

| | | | | | | | | | | | | | | | |
|---|---|---|---|---|---|---|---|---|---|---|---|---|---|---|---|
| 22173 | ENSMUSG00000077678 | 0 | NA | NA | NA | NA | NA | Gm24318 | predicted gene, 24318 [Source:MGI Symb | 0 | 0 | 0 | 0 | 0 | 0 |
| 22174 | ENSMUSG00000077680 | 0 | NA | NA | NA | NA | NA | Gm22495 | predicted gene, 22495 [Source:MGI Symb | 0 | 0 | 0 | 0 | 0 | 0 |
| 22175 | ENSMUSG00000077681 | 0 | NA | NA | NA | NA | NA | Gm22496 | predicted gene, 22496 [Source:MGI Symb | 0 | 0 | 0 | 0 | 0 | 0 |
| 22176 | ENSMUSG00000077682 | 0 | NA | NA | NA | NA | NA | Gm22493 | predicted gene, 22493 [Source:MGI Symb | 0 | 0 | 0 | 0 | 0 | 0 |
| 22177 | ENSMUSG00000077683 | 0 | NA | NA | NA | NA | NA | Gm22494 | predicted gene, 22494 [Source:MGI Symb | 0 | 0 | 0 | 0 | 0 | 0 |
| 22178 | ENSMUSG00000077685 | 0 | NA | NA | NA | NA | NA | Gm22498 | predicted gene, 22498 [Source:MGI Symb | 0 | 0 | 0 | 0 | 0 | 0 |
| 22179 | ENSMUSG00000077687 | 0 | NA | NA | NA | NA | NA | Gm22497 | predicted gene, 22497 [Source:MGI Symb | 0 | 0 | 0 | 0 | 0 | 0 |
| 22180 | ENSMUSG00000077689 | 0 | NA | NA | NA | NA | NA | Gm22499 | predicted gene, 22499 [Source:MGI Symb | 0 | 0 | 0 | 0 | 0 | 0 |
| 22181 | ENSMUSG00000077691 | 0 | NA | NA | NA | NA | NA | Gm25344 | predicted gene, 25344 [Source:MGI Symb | 0 | 0 | 0 | 0 | 0 | 0 |
| 22182 | ENSMUSG00000077693 | 0 | NA | NA | NA | NA | NA | Gm25345 | predicted gene, 25345 [Source:MGI Symb | 0 | 0 | 0 | 0 | 0 | 0 |
| 22183 | ENSMUSG00000077696 | 0 | NA | NA | NA | NA | NA | NA | NA | 0 | 0 | 0 | 0 | 0 | 0 |
| 22184 | ENSMUSG00000077697 | 0 | NA | NA | NA | NA | NA | Gm25346 | predicted gene, 25346 [Source:MGI Symb | 0 | 0 | 0 | 0 | 0 | 0 |
| 22185 | ENSMUSG00000077698 | 0 | NA | NA | NA | NA | NA | Snord12 | small nucleolar RNA, C/D box 12 [Source:I | 0 | 0 | 0 | 0 | 0 | 0 |
| 22186 | ENSMUSG00000077700 | 0 | NA | NA | NA | NA | NA | Gm24069 | predicted gene, 24069 [Source:MGI Symb | 0 | 0 | 0 | 0 | 0 | 0 |
| 22187 | ENSMUSG00000077701 | 0 | NA | NA | NA | NA | NA | Gm24068 | predicted gene, 24068 [Source:MGI Symb | 0 | 0 | 0 | 0 | 0 | 0 |
| 22188 | ENSMUSG00000077702 | 0 | NA | NA | NA | NA | NA | Gm24067 | predicted gene, 24067 [Source:MGI Symb | 0 | 0 | 0 | 0 | 0 | 0 |
| 22189 | ENSMUSG00000077703 | 0 | NA | NA | NA | NA | NA | Gm24066 | predicted gene, 24066 [Source:MGI Symb | 0 | 0 | 0 | 0 | 0 | 0 |
| 22190 | ENSMUSG00000077704 | 3.667746543 | -0.753640565 | 1.387059492 | -0.543336871 | 0.586897882 | NA | Snord89 | small nucleolar RNA, C/D box 89 [Source:I | 5.6217 | 8.2233 | 0 | 4.8199 | 0 | 3.3415 |
| 22191 | ENSMUSG00000077705 | 0 | NA | NA | NA | NA | NA | NA | NA | 0 | 0 | 0 | 0 | 0 | 0 |
| 22192 | ENSMUSG00000077706 | 0 | NA | NA | NA | NA | NA | NA | NA | 0 | 0 | 0 | 0 | 0 | 0 |
| 22193 | ENSMUSG00000077707 | 0 | NA | NA | NA | NA | NA | Gm24070 | predicted gene, 24070 [Source:MGI Symb | 0 | 0 | 0 | 0 | 0 | 0 |
| 22194 | ENSMUSG00000077708 | 0 | NA | NA | NA | NA | NA | Gm24071 | predicted gene, 24071 [Source:MGI Symb | 0 | 0 | 0 | 0 | 0 | 0 |
| 22195 | ENSMUSG00000077709 | 0 | NA | NA | NA | NA | NA | Snora64 | small nucleolar RNA, H/ACA box 64 [Sourc | 0 | 0 | 0 | 0 | 0 | 0 |
| 22196 | ENSMUSG00000077710 | 0 | NA | NA | NA | NA | NA | NA | NA | 0 | 0 | 0 | 0 | 0 | 0 |
| 22197 | ENSMUSG00000077711 | 5.260200134 | -0.83992825 | 0.995808458 | -0.843463663 | 0.39896918 | 0.826036389 | AF357399 | snoRNA AF357399 [Source:MGI Symbol;A | 8.031 | 9.2513 | 3.0187 | 5.7839 | 4.3625 | 1.1138 |
| 22198 | ENSMUSG00000077713 | 0 | NA | NA | NA | NA | NA | Gm25728 | predicted gene, 25728 [Source:MGI Symb | 0 | 0 | 0 | 0 | 0 | 0 |
| 22199 | ENSMUSG00000077714 | 0.371281295 | 2.09253362 | 4.031852165 | 0.519000582 | 0.603760335 | NA | Snord17 | small nucleolar RNA, C/D box 17 [Source:I | 0 | 0 | 0 | 0 | 0 | 2.2277 |
| 22200 | ENSMUSG00000077716 | 0 | NA | NA | NA | NA | NA | Gm25725 | predicted gene, 25725 [Source:MGI Symb | 0 | 0 | 0 | 0 | 0 | 0 |
| 22201 | ENSMUSG00000077717 | 0 | NA | NA | NA | NA | NA | Gm25726 | predicted gene, 25726 [Source:MGI Symb | 0 | 0 | 0 | 0 | 0 | 0 |
| 22202 | ENSMUSG00000077718 | 0 | NA | NA | NA | NA | NA | Gm25723 | predicted gene, 25723 [Source:MGI Symb | 0 | 0 | 0 | 0 | 0 | 0 |
| 22203 | ENSMUSG00000077719 | 0 | NA | NA | NA | NA | NA | Gm25724 | predicted gene, 25724 [Source:MGI Symb | 0 | 0 | 0 | 0 | 0 | 0 |
| 22204 | ENSMUSG00000077726 | 0 | NA | NA | NA | NA | NA | NA | NA | 0 | 0 | 0 | 0 | 0 | 0 |
| 22205 | ENSMUSG00000077728 | 0 | NA | NA | NA | NA | NA | NA | NA | 0 | 0 | 0 | 0 | 0 | 0 |
| 22206 | ENSMUSG00000077730 | 0 | NA | NA | NA | NA | NA | Gm23609 | predicted gene, 23609 [Source:MGI Symb | 0 | 0 | 0 | 0 | 0 | 0 |
| 22207 | ENSMUSG00000077733 | 0.535401493 | -2.48596965 | 3.684117979 | -0.674780141 | 0.499815459 | NA | Gm23608 | predicted gene, 23608 [Source:MGI Symb | 3.2124 | 0 | 0 | 0 | 0 | 0 |
| 22208 | ENSMUSG00000077734 | 1.337936333 | 0.255247689 | 1.957898427 | 0.130368197 | 0.896275127 | NA | Snord83b | small nucleolar RNA, C/D box 83B [Source | 1.6062 | 0 | 2.0125 | 0 | 2.1812 | 2.2277 |
| 22209 | ENSMUSG00000077735 | 0 | NA | NA | NA | NA | NA | Gm24556 | predicted gene, 24556 [Source:MGI Symb | 0 | 0 | 0 | 0 | 0 | 0 |
| 22210 | ENSMUSG00000077736 | 0 | NA | NA | NA | NA | NA | Gm24555 | predicted gene, 24555 [Source:MGI Symb | 0 | 0 | 0 | 0 | 0 | 0 |
| 22211 | ENSMUSG00000077737 | 3.958680354 | 0.300545125 | 1.239561174 | 0.242460906 | 0.808423046 | NA | Snord72 | small nucleolar RNA, C/D box 72 [Source:I | 1.6062 | 5.1396 | 4.025 | 9.6398 | 0 | 3.3415 |
| 22212 | ENSMUSG00000077738 | 0 | NA | NA | NA | NA | NA | Gm23607 | predicted gene, 23607 [Source:MGI Symb | 0 | 0 | 0 | 0 | 0 | 0 |
| 22213 | ENSMUSG00000077741 | 0 | NA | NA | NA | NA | NA | NA | NA | 0 | 0 | 0 | 0 | 0 | 0 |
| 22214 | ENSMUSG00000077743 | 0 | NA | NA | NA | NA | NA | NA | NA | 0 | 0 | 0 | 0 | 0 | 0 |
| 22215 | ENSMUSG00000077744 | 0 | NA | NA | NA | NA | NA | Gm26343 | predicted gene, 26343 [Source:MGI Symb | 0 | 0 | 0 | 0 | 0 | 0 |
| 22216 | ENSMUSG00000077745 | 0 | NA | NA | NA | NA | NA | Gm26342 | predicted gene, 26342 [Source:MGI Symb | 0 | 0 | 0 | 0 | 0 | 0 |
| 22217 | ENSMUSG00000077749 | 0 | NA | NA | NA | NA | NA | Gm26341 | predicted gene, 26341 [Source:MGI Symb | 0 | 0 | 0 | 0 | 0 | 0 |
| 22218 | ENSMUSG00000077750 | 0 | NA | NA | NA | NA | NA | NA | NA | 0 | 0 | 0 | 0 | 0 | 0 |
| 22219 | ENSMUSG00000077752 | 0 | NA | NA | NA | NA | NA | Gm23530 | predicted gene, 23530 [Source:MGI Symb | 0 | 0 | 0 | 0 | 0 | 0 |
| 22220 | ENSMUSG00000077755 | 0 | NA | NA | NA | NA | NA | NA | NA | 0 | 0 | 0 | 0 | 0 | 0 |
| 22221 | ENSMUSG00000077756 | 0 | NA | NA | NA | NA | NA | Snord90 | small nucleolar RNA, C/D box 90 [Source:I | 0 | 0 | 0 | 0 | 0 | 0 |
| 22222 | ENSMUSG00000077757 | 0 | NA | NA | NA | NA | NA | Gm23532 | predicted gene, 23532 [Source:MGI Symb | 0 | 0 | 0 | 0 | 0 | 0 |
| 22223 | ENSMUSG00000077758 | 0 | NA | NA | NA | NA | NA | Gm23527 | predicted gene, 23527 [Source:MGI Symb | 0 | 0 | 0 | 0 | 0 | 0 |
| 22224 | ENSMUSG00000077759 | 0 | NA | NA | NA | NA | NA | Gm23528 | predicted gene, 23528 [Source:MGI Symb | 0 | 0 | 0 | 0 | 0 | 0 |
| 22225 | ENSMUSG00000077761 | 0 | NA | NA | NA | NA | NA | Gm25214 | predicted gene, 25214 [Source:MGI Symb | 0 | 0 | 0 | 0 | 0 | 0 |
| 22226 | ENSMUSG00000077767 | 0 | NA | NA | NA | NA | NA | Snora35 | small nucleolar RNA, H/ACA box 35 [Sourc | 0 | 0 | 0 | 0 | 0 | 0 |
| 22227 | ENSMUSG00000077768 | 0 | NA | NA | NA | NA | NA | Gm25216 | predicted gene, 25216 [Source:MGI Symb | 0 | 0 | 0 | 0 | 0 | 0 |
| 22228 | ENSMUSG00000077769 | 0 | NA | NA | NA | NA | NA | Gm25215 | predicted gene, 25215 [Source:MGI Symb | 0 | 0 | 0 | 0 | 0 | 0 |

| | | | | | | | | | | | | | | | |
|---|---|---|---|---|---|---|---|---|---|---|---|---|---|---|---|
| 22229 | ENSMUSG00000077771 | 0 | NA | NA | NA | NA | NA | NA | NA | 0 | 0 | 0 | 0 | 0 | 0 |
| 22230 | ENSMUSG00000077772 | 0 | NA | NA | NA | NA | NA | Gm22373 | predicted gene, 22373 [Source:MGI Symb | 0 | 0 | 0 | 0 | 0 | 0 |
| 22231 | ENSMUSG00000077776 | 0.17131964 | -0.796629176 | 4.080472857 | -0.195229623 | 0.84521319 | NA | Gm22371 | predicted gene, 22371 [Source:MGI Symb | 0 | 1.0279 | 0 | 0 | 0 | 0 |
| 22232 | ENSMUSG00000077777 | 0 | NA | NA | NA | NA | NA | Gm22372 | predicted gene, 22372 [Source:MGI Symb | 0 | 0 | 0 | 0 | 0 | 0 |
| 22233 | ENSMUSG00000077780 | 0 | NA | NA | NA | NA | NA | Gm24161 | predicted gene, 24161 [Source:MGI Symb | 0 | 0 | 0 | 0 | 0 | 0 |
| 22234 | ENSMUSG00000077782 | 0 | NA | NA | NA | NA | NA | Gm24160 | predicted gene, 24160 [Source:MGI Symb | 0 | 0 | 0 | 0 | 0 | 0 |
| 22235 | ENSMUSG00000077783 | 0 | NA | NA | NA | NA | NA | Gm23907 | predicted gene, 23907 [Source:MGI Symb | 0 | 0 | 0 | 0 | 0 | 0 |
| 22236 | ENSMUSG00000077784 | 0 | NA | NA | NA | NA | NA | Gm23911 | predicted gene, 23911 [Source:MGI Symb | 0 | 0 | 0 | 0 | 0 | 0 |
| 22237 | ENSMUSG00000077785 | 0 | NA | NA | NA | NA | NA | NA | NA | 0 | 0 | 0 | 0 | 0 | 0 |
| 22238 | ENSMUSG00000077786 | 0 | NA | NA | NA | NA | NA | Gm23909 | predicted gene, 23909 [Source:MGI Symb | 0 | 0 | 0 | 0 | 0 | 0 |
| 22239 | ENSMUSG00000077787 | 0.353348078 | 0.165142701 | 3.895042568 | 0.042398176 | 0.966181282 | NA | Gm23908 | predicted gene, 23908 [Source:MGI Symb | 0 | 0 | 1.0062 | 0 | 0 | 1.1138 |
| 22240 | ENSMUSG00000077789 | 0 | NA | NA | NA | NA | NA | NA | NA | 0 | 0 | 0 | 0 | 0 | 0 |
| 22241 | ENSMUSG00000077791 | 0 | NA | NA | NA | NA | NA | Gm25563 | predicted gene, 25563 [Source:MGI Symb | 0 | 0 | 0 | 0 | 0 | 0 |
| 22242 | ENSMUSG00000077792 | 0 | NA | NA | NA | NA | NA | NA | NA | 0 | 0 | 0 | 0 | 0 | 0 |
| 22243 | ENSMUSG00000077793 | 0 | NA | NA | NA | NA | NA | Gm25565 | predicted gene, 25565 [Source:MGI Symb | 0 | 0 | 0 | 0 | 0 | 0 |
| 22244 | ENSMUSG00000077794 | 0 | NA | NA | NA | NA | NA | Gm25560 | predicted gene, 25560 [Source:MGI Symb | 0 | 0 | 0 | 0 | 0 | 0 |
| 22245 | ENSMUSG00000077795 | 0.133850373 | -0.796629176 | 4.080472857 | -0.195229623 | 0.84521319 | NA | Gm25561 | predicted gene, 25561 [Source:MGI Symb | 0.8031 | 0 | 0 | 0 | 0 | 0 |
| 22246 | ENSMUSG00000077796 | 0 | NA | NA | NA | NA | NA | Gm25562 | predicted gene, 25562 [Source:MGI Symb | 0 | 0 | 0 | 0 | 0 | 0 |
| 22247 | ENSMUSG00000077797 | 0 | NA | NA | NA | NA | NA | Snord19 | small nucleolar RNA, C/D box 19 [Source:I | 0 | 0 | 0 | 0 | 0 | 0 |
| 22248 | ENSMUSG00000077799 | 0 | NA | NA | NA | NA | NA | Gm25148 | predicted gene, 25148 [Source:MGI Symb | 0 | 0 | 0 | 0 | 0 | 0 |
| 22249 | ENSMUSG00000077802 | 0 | NA | NA | NA | NA | NA | NA | NA | 0 | 0 | 0 | 0 | 0 | 0 |
| 22250 | ENSMUSG00000077804 | 0 | NA | NA | NA | NA | NA | Gm24403 | predicted gene, 24403 [Source:MGI Symb | 0 | 0 | 0 | 0 | 0 | 0 |
| 22251 | ENSMUSG00000077815 | 1.353988001 | -1.451133495 | 2.240453263 | -0.647696392 | 0.517181338 | NA | Mir290a | microRNA 290a [Source:MGI Symbol;Acc: | 0.8031 | 5.1396 | 0 | 0 | 2.1812 | 0 |
| 22252 | ENSMUSG00000077821 | 0 | NA | NA | NA | NA | NA | NA | NA | 0 | 0 | 0 | 0 | 0 | 0 |
| 22253 | ENSMUSG00000077829 | 0 | NA | NA | NA | NA | NA | NA | NA | 0 | 0 | 0 | 0 | 0 | 0 |
| 22254 | ENSMUSG00000077830 | 0.160662974 | 1.126924175 | 4.080472857 | 0.276174898 | 0.782413736 | NA | NA | NA | 0 | 0 | 0 | 0.964 | 0 | 0 |
| 22255 | ENSMUSG00000077832 | 0 | NA | NA | NA | NA | NA | NA | NA | 0 | 0 | 0 | 0 | 0 | 0 |
| 22256 | ENSMUSG00000077834 | 0 | NA | NA | NA | NA | NA | Mir669d | microRNA 669d [Source:MGI Symbol;Acc: | 0 | 0 | 0 | 0 | 0 | 0 |
| 22257 | ENSMUSG00000077836 | 0 | NA | NA | NA | NA | NA | NA | NA | 0 | 0 | 0 | 0 | 0 | 0 |
| 22258 | ENSMUSG00000077844 | 0 | NA | NA | NA | NA | NA | NA | NA | 0 | 0 | 0 | 0 | 0 | 0 |
| 22259 | ENSMUSG00000077846 | 0 | NA | NA | NA | NA | NA | NA | NA | 0 | 0 | 0 | 0 | 0 | 0 |
| 22260 | ENSMUSG00000077847 | 0 | NA | NA | NA | NA | NA | Mir105 | microRNA 105 [Source:MGI Symbol;Acc:N | 0 | 0 | 0 | 0 | 0 | 0 |
| 22261 | ENSMUSG00000077851 | 0 | NA | NA | NA | NA | NA | Mir876 | microRNA 876 [Source:MGI Symbol;Acc:N | 0 | 0 | 0 | 0 | 0 | 0 |
| 22262 | ENSMUSG00000077857 | 0 | NA | NA | NA | NA | NA | NA | NA | 0 | 0 | 0 | 0 | 0 | 0 |
| 22263 | ENSMUSG00000077858 | 0 | NA | NA | NA | NA | NA | NA | NA | 0 | 0 | 0 | 0 | 0 | 0 |
| 22264 | ENSMUSG00000077869 | 0 | NA | NA | NA | NA | NA | Mir344e | microRNA 344e [Source:MGI Symbol;Acc: | 0 | 0 | 0 | 0 | 0 | 0 |
| 22265 | ENSMUSG00000077872 | 0 | NA | NA | NA | NA | NA | Mir878 | microRNA 878 [Source:MGI Symbol;Acc:N | 0 | 0 | 0 | 0 | 0 | 0 |
| 22266 | ENSMUSG00000077879 | 0 | NA | NA | NA | NA | NA | NA | NA | 0 | 0 | 0 | 0 | 0 | 0 |
| 22267 | ENSMUSG00000077881 | 0 | NA | NA | NA | NA | NA | Mir882 | microRNA 882 [Source:MGI Symbol;Acc:N | 0 | 0 | 0 | 0 | 0 | 0 |
| 22268 | ENSMUSG00000077884 | 0 | NA | NA | NA | NA | NA | NA | NA | 0 | 0 | 0 | 0 | 0 | 0 |
| 22269 | ENSMUSG00000077886 | 0 | NA | NA | NA | NA | NA | Mir295 | microRNA 295 [Source:MGI Symbol;Acc:N | 0 | 0 | 0 | 0 | 0 | 0 |
| 22270 | ENSMUSG00000077890 | 0 | NA | NA | NA | NA | NA | Mir875 | microRNA 875 [Source:MGI Symbol;Acc:N | 0 | 0 | 0 | 0 | 0 | 0 |
| 22271 | ENSMUSG00000077892 | 0 | NA | NA | NA | NA | NA | NA | NA | 0 | 0 | 0 | 0 | 0 | 0 |
| 22272 | ENSMUSG00000077896 | 0 | NA | NA | NA | NA | NA | Mir879 | microRNA 879 [Source:MGI Symbol;Acc:N | 0 | 0 | 0 | 0 | 0 | 0 |
| 22273 | ENSMUSG00000077897 | 0 | NA | NA | NA | NA | NA | Mir511 | microRNA 511 [Source:MGI Symbol;Acc:N | 0 | 0 | 0 | 0 | 0 | 0 |
| 22274 | ENSMUSG00000077902 | 0 | NA | NA | NA | NA | NA | NA | NA | 0 | 0 | 0 | 0 | 0 | 0 |
| 22275 | ENSMUSG00000077903 | 0 | NA | NA | NA | NA | NA | Mir294 | microRNA 294 [Source:MGI Symbol;Acc:N | 0 | 0 | 0 | 0 | 0 | 0 |
| 22276 | ENSMUSG00000077921 | 0.160662974 | 1.126924175 | 4.080472857 | 0.276174898 | 0.782413736 | NA | Mir872 | microRNA 872 [Source:MGI Symbol;Acc:N | 0 | 0 | 0 | 0.964 | 0 | 0 |
| 22277 | ENSMUSG00000077925 | 0 | NA | NA | NA | NA | NA | Mir453 | microRNA 453 [Source:MGI Symbol;Acc:N | 0 | 0 | 0 | 0 | 0 | 0 |
| 22278 | ENSMUSG00000077928 | 0 | NA | NA | NA | NA | NA | Mir208b | microRNA 208b [Source:MGI Symbol;Acc: | 0 | 0 | 0 | 0 | 0 | 0 |
| 22279 | ENSMUSG00000077931 | 0 | NA | NA | NA | NA | NA | Mir877 | microRNA 877 [Source:MGI Symbol;Acc:N | 0 | 0 | 0 | 0 | 0 | 0 |
| 22280 | ENSMUSG00000077938 | 0 | NA | NA | NA | NA | NA | NA | NA | 0 | 0 | 0 | 0 | 0 | 0 |
| 22281 | ENSMUSG00000077939 | 0 | NA | NA | NA | NA | NA | NA | NA | 0 | 0 | 0 | 0 | 0 | 0 |
| 22282 | ENSMUSG00000077941 | 0 | NA | NA | NA | NA | NA | Mir466h | microRNA 466h [Source:MGI Symbol;Acc: | 0 | 0 | 0 | 0 | 0 | 0 |
| 22283 | ENSMUSG00000077948 | 0 | NA | NA | NA | NA | NA | NA | NA | 0 | 0 | 0 | 0 | 0 | 0 |
| 22284 | ENSMUSG00000077954 | 0 | NA | NA | NA | NA | NA | Mir297a-4 | microRNA 297a-4 [Source:MGI Symbol;Ac | 0 | 0 | 0 | 0 | 0 | 0 |

| | | | | | | | | | | | | | | | |
|---|---|---|---|---|---|---|---|---|---|---|---|---|---|---|---|
| 22285 | ENSMUSG00000077962 | 0.133850373 | -0.796629176 | 4.080472857 | -0.195229623 | 0.84521319 | NA | Mir874 | microRNA 874 [Source:MGI Symbol;Acc:M | 0.8031 | 0 | 0 | 0 | 0 | 0 |
| 22286 | ENSMUSG00000077965 | 0 | NA | NA | NA | NA | NA | NA | NA | 0 | 0 | 0 | 0 | 0 | 0 |
| 22287 | ENSMUSG00000077971 | 0 | NA | NA | NA | NA | NA | NA | NA | 0 | 0 | 0 | 0 | 0 | 0 |
| 22288 | ENSMUSG00000077972 | 0 | NA | NA | NA | NA | NA | Mir466f-1 | microRNA 466f-1 [Source:MGI Symbol;Ac | 0 | 0 | 0 | 0 | 0 | 0 |
| 22289 | ENSMUSG00000077980 | 0 | NA | NA | NA | NA | NA | Mir654 | microRNA 654 [Source:MGI Symbol;Acc:M | 0 | 0 | 0 | 0 | 0 | 0 |
| 22290 | ENSMUSG00000077981 | 0 | NA | NA | NA | NA | NA | Mir466f-3 | microRNA 466f-3 [Source:MGI Symbol;Ac | 0 | 0 | 0 | 0 | 0 | 0 |
| 22291 | ENSMUSG00000077990 | 0 | NA | NA | NA | NA | NA | NA | NA | 0 | 0 | 0 | 0 | 0 | 0 |
| 22292 | ENSMUSG00000077991 | 0 | NA | NA | NA | NA | NA | Mir466n | microRNA 466n [Source:MGI Symbol;Acc: | 0 | 0 | 0 | 0 | 0 | 0 |
| 22293 | ENSMUSG00000077996 | 0 | NA | NA | NA | NA | NA | Mir190b | microRNA 190b [Source:MGI Symbol;Acc: | 0 | 0 | 0 | 0 | 0 | 0 |
| 22294 | ENSMUSG00000077998 | 0 | NA | NA | NA | NA | NA | Mir466f-2 | microRNA 466f-2 [Source:MGI Symbol;Ac | 0 | 0 | 0 | 0 | 0 | 0 |
| 22295 | ENSMUSG00000078002 | 0 | NA | NA | NA | NA | NA | Mir880 | microRNA 880 [Source:MGI Symbol;Acc:M | 0 | 0 | 0 | 0 | 0 | 0 |
| 22296 | ENSMUSG00000078006 | 0 | NA | NA | NA | NA | NA | NA | NA | 0 | 0 | 0 | 0 | 0 | 0 |
| 22297 | ENSMUSG00000078008 | 0 | NA | NA | NA | NA | NA | Mir291a | microRNA 291a [Source:MGI Symbol;Acc: | 0 | 0 | 0 | 0 | 0 | 0 |
| 22298 | ENSMUSG00000078017 | 0 | NA | NA | NA | NA | NA | Mir883a | microRNA 883a [Source:MGI Symbol;Acc: | 0 | 0 | 0 | 0 | 0 | 0 |
| 22299 | ENSMUSG00000078024 | 0 | NA | NA | NA | NA | NA | Mir883b | microRNA 883b [Source:MGI Symbol;Acc: | 0 | 0 | 0 | 0 | 0 | 0 |
| 22300 | ENSMUSG00000078025 | 0 | NA | NA | NA | NA | NA | Mir466g | microRNA 466g [Source:MGI Symbol;Acc: | 0 | 0 | 0 | 0 | 0 | 0 |
| 22301 | ENSMUSG00000078026 | 0 | NA | NA | NA | NA | NA | Mir466o | microRNA 466o [Source:MGI Symbol;Acc: | 0 | 0 | 0 | 0 | 0 | 0 |
| 22302 | ENSMUSG00000078027 | 0 | NA | NA | NA | NA | NA | Mir449b | microRNA 449b [Source:MGI Symbol;Acc: | 0 | 0 | 0 | 0 | 0 | 0 |
| 22303 | ENSMUSG00000078031 | 0 | NA | NA | NA | NA | NA | Mir466d | microRNA 466d [Source:MGI Symbol;Acc: | 0 | 0 | 0 | 0 | 0 | 0 |
| 22304 | ENSMUSG00000078032 | 0 | NA | NA | NA | NA | NA | Mir291b | microRNA 291b [Source:MGI Symbol;Acc: | 0 | 0 | 0 | 0 | 0 | 0 |
| 22305 | ENSMUSG00000078034 | 0 | NA | NA | NA | NA | NA | NA | NA | 0 | 0 | 0 | 0 | 0 | 0 |
| 22306 | ENSMUSG00000078035 | 0 | NA | NA | NA | NA | NA | Mir293 | microRNA 293 [Source:MGI Symbol;Acc:M | 0 | 0 | 0 | 0 | 0 | 0 |
| 22307 | ENSMUSG00000078038 | 0 | NA | NA | NA | NA | NA | Mir881 | microRNA 881 [Source:MGI Symbol;Acc:M | 0 | 0 | 0 | 0 | 0 | 0 |
| 22308 | ENSMUSG00000078041 | 0 | NA | NA | NA | NA | NA | Mir292 | microRNA 292 [Source:MGI Symbol;Acc:M | 0 | 0 | 0 | 0 | 0 | 0 |
| 22309 | ENSMUSG00000078042 | 1.563495732 | 1.202074448 | 1.991445035 | 0.603619195 | 0.546096848 | NA | NA | NA | 0.8031 | 1.0279 | 1.0062 | 0 | 6.5437 | 0 |
| 22310 | ENSMUSG00000078057 | 0 | NA | NA | NA | NA | NA | Mir873a | microRNA 873a [Source:MGI Symbol;Acc: | 0 | 0 | 0 | 0 | 0 | 0 |
| 22311 | ENSMUSG00000078060 | 0 | NA | NA | NA | NA | NA | Mir466l | microRNA 466l [Source:MGI Symbol;Acc:M | 0 | 0 | 0 | 0 | 0 | 0 |
| 22312 | ENSMUSG00000078061 | 0 | NA | NA | NA | NA | NA | NA | NA | 0 | 0 | 0 | 0 | 0 | 0 |
| 22313 | ENSMUSG00000078080 | 0 | NA | NA | NA | NA | NA | Olfr585 | olfactory receptor 585 [Source:MGI Symb | 0 | 0 | 0 | 0 | 0 | 0 |
| 22314 | ENSMUSG00000078087 | 0 | NA | NA | NA | NA | NA | Rps12l1 | ribosomal protein S12-like 1 [Source:MGI | 0 | 0 | 0 | 0 | 0 | 0 |
| 22315 | ENSMUSG00000078091 | 0.328370405 | 0.16514316 | 4.003149536 | 0.041253308 | 0.967093956 | NA | Gm10912 | predicted gene 10912 [Source:MGI Symbo | 0 | 0 | 1.0062 | 0.964 | 0 | 0 |
| 22316 | ENSMUSG00000078097 | 0 | NA | NA | NA | NA | NA | Gm10913 | predicted pseudogene 10913 [Source:MG | 0 | 0 | 0 | 0 | 0 | 0 |
| 22317 | ENSMUSG00000078116 | 0 | NA | NA | NA | NA | NA | Olfr828 | olfactory receptor 828 [Source:MGI Symb | 0 | 0 | 0 | 0 | 0 | 0 |
| 22318 | ENSMUSG00000078117 | 0.788497532 | -1.691662715 | 2.429158853 | -0.696398555 | 0.486179268 | NA | Gm16485 | predicted gene 16485 [Source:MGI Symbo | 1.6062 | 1.0279 | 1.0062 | 0 | 1.0906 | 0 |
| 22319 | ENSMUSG00000078118 | 0 | NA | NA | NA | NA | NA | Olfr483 | olfactory receptor 483 [Source:MGI Symb | 0 | 0 | 0 | 0 | 0 | 0 |
| 22320 | ENSMUSG00000078122 | 6.441757606 | -1.406093225 | 0.89225414 | -1.575888709 | 0.115051454 | 0.498085671 | F630028O10 | RIKEN cDNA F630028O10 gene [Source:M | 9.6372 | 13.363 | 5.0312 | 2.8919 | 3.2719 | 4.4554 |
| 22321 | ENSMUSG00000078123 | 0 | NA | NA | NA | NA | NA | Gm5071 | predicted pseudogene 5071 [Source:MGI | 0 | 0 | 0 | 0 | 0 | 0 |
| 22322 | ENSMUSG00000078125 | 1.33249471 | -0.610195187 | 1.87565474 | -0.325323832 | 0.744935996 | NA | Gm10916 | predicted gene 10916 [Source:MGI Symbo | 0.8031 | 1.0279 | 3.0187 | 0.964 | 2.1812 | 0 |
| 22323 | ENSMUSG00000078126 | 7.811722639 | 0.640016498 | 0.778920455 | 0.821671191 | 0.41126406 | 0.833705705 | Rpl23a-ps3 | ribosomal protein L23A, pseudogene 3 [S | 3.2124 | 7.1954 | 8.05 | 9.6398 | 7.6343 | 11.138 |
| 22324 | ENSMUSG00000078127 | 0.294513347 | 0.165143496 | 4.080472857 | 0.040471657 | 0.967717103 | NA | Fam170b | family with sequence similarity 170, mem | 0.8031 | 0 | 0 | 0.964 | 0 | 0 |
| 22325 | ENSMUSG00000078128 | 0.160662974 | 1.126924175 | 4.080472857 | 0.276174898 | 0.782413736 | NA | Gm2178 | predicted gene 2178 [Source:MGI Symbol | 0 | 0 | 0 | 0.964 | 0 | 0 |
| 22326 | ENSMUSG00000078129 | 0 | NA | NA | NA | NA | NA | Actl10 | actin-like 10 [Source:MGI Symbol;Acc:MG | 0 | 0 | 0 | 0 | 0 | 0 |
| 22327 | ENSMUSG00000078130 | 0 | NA | NA | NA | NA | NA | Gm11555 | predicted gene 11555 [Source:MGI Symbo | 0 | 0 | 0 | 0 | 0 | 0 |
| 22328 | ENSMUSG00000078131 | 0 | NA | NA | NA | NA | NA | Krtap1-3 | keratin associated protein 1-3 [Source:MG | 0 | 0 | 0 | 0 | 0 | 0 |
| 22329 | ENSMUSG00000078132 | 0 | NA | NA | NA | NA | NA | Gm11939 | predicted gene 11939 [Source:MGI Symbo | 0 | 0 | 0 | 0 | 0 | 0 |
| 22330 | ENSMUSG00000078134 | 0.660353019 | -0.01500701 | 2.83315355 | -0.005296928 | 0.995773683 | NA | Srsf3-ps | serine and arginine rich splicing factor 3, p | 0 | 1.0279 | 1.0062 | 1.928 | 0 | 0 |
| 22331 | ENSMUSG00000078137 | 0 | NA | NA | NA | NA | NA | Ankrd63 | ankyrin repeat domain 63 [Source:MGI Sy | 0 | 0 | 0 | 0 | 0 | 0 |
| 22332 | ENSMUSG00000078139 | 3.081144359 | 0.098715165 | 1.296665762 | 0.07613 | 0.939315672 | NA | AK157302 | cDNA sequence AK157302 [Source:MGI Sy | 0.8031 | 7.1954 | 1.0062 | 2.8919 | 4.3625 | 2.2277 |
| 22333 | ENSMUSG00000078141 | 0.503302254 | -0.837877584 | 3.426698643 | -0.244514523 | 0.806832356 | NA | Gm2399 | predicted gene 2399 [Source:MGI Symbol | 0 | 2.0558 | 0 | 0.964 | 0 | 0 |
| 22334 | ENSMUSG00000078143 | 0.435408177 | -2.179616019 | 3.531869654 | -0.617128103 | 0.537150234 | NA | Gm17344 | predicted gene, 17344 [Source:MGI Symb | 1.6062 | 0 | 1.0062 | 0 | 0 | 0 |
| 22335 | ENSMUSG00000078144 | 0 | NA | NA | NA | NA | NA | Capns2 | calpain, small subunit 2 [Source:MGI Sym | 0 | 0 | 0 | 0 | 0 | 0 |
| 22336 | ENSMUSG00000078151 | 0.785324008 | -0.358914107 | 2.435960446 | -0.147339875 | 0.88286376 | NA | Gm12226 | predicted pseudogene 12226 [Source:MG | 1.6062 | 1.0279 | 0 | 0.964 | 0 | 1.1138 |
| 22337 | ENSMUSG00000078153 | 60.57741601 | -0.155108152 | 0.285400953 | -0.54347454 | 0.586803116 | 0.910448513 | Psme2b | protease (prosome, macropain) activator | 55.414 | 67.843 | 68.425 | 54.947 | 57.803 | 59.034 |
| 22338 | ENSMUSG00000078154 | 6.397528639 | 0.726454309 | 0.870624579 | 0.834405926 | 0.404052279 | 0.829083864 | Gm12184 | predicted gene 12184 [Source:MGI Symbo | 7.2279 | 4.1117 | 3.0187 | 9.6398 | 4.3625 | 10.025 |
| 22339 | ENSMUSG00000078157 | 0 | NA | NA | NA | NA | NA | Fem1al | fem-1 homolog A like [Source:MGI Symbo | 0 | 0 | 0 | 0 | 0 | 0 |
| 22340 | ENSMUSG00000078160 | 0.321325949 | 1.911575114 | 4.048267893 | 0.472195804 | 0.636787034 | NA | NA | NA | 0 | 0 | 0 | 1.928 | 0 | 0 |

| | | | | | | | | | | | | | | | |
|---|---|---|---|---|---|---|---|---|---|---|---|---|---|---|---|
| 22341 | ENSMUSG00000078161 | 0.82806045 | -0.59635427 | 2.600266913 | -0.229343483 | 0.81860196 | NA | Erich3 | glutamate rich 3 [Source:MGI Symbol;Acc | 0 | 1.0279 | 2.0125 | 1.928 | 0 | 0 |
| 22342 | ENSMUSG00000078162 | 0.517184577 | -0.812115922 | 3.393570547 | -0.239310163 | 0.810865085 | NA | Gm2574 | predicted pseudogene 2574 [Source:MGI | 0 | 0 | 2.0125 | 0 | 1.0906 | 0 |
| 22343 | ENSMUSG00000078173 | 1.049360412 | -3.442031943 | 2.546201196 | -1.351830307 | 0.176429607 | NA | Lenep | lens epithelial protein [Source:MGI Symbo | 3.2124 | 3.0838 | 0 | 0 | 0 | 0 |
| 22344 | ENSMUSG00000078177 | 0 | NA | NA | NA | NA | NA | Gm4877 | predicted gene 4877 [Source:MGI Symbol | 0 | 0 | 0 | 0 | 0 | 0 |
| 22345 | ENSMUSG00000078179 | 0.669251866 | -2.807208097 | 3.391018549 | -0.827836255 | 0.407763239 | NA | Rnf148 | ring finger protein 148 [Source:MGI Symb | 4.0155 | 0 | 0 | 0 | 0 | 0 |
| 22346 | ENSMUSG00000078180 | 2.20795154 | -0.032580689 | 1.538393536 | -0.021178384 | 0.983103358 | NA | Gm20900 | predicted gene, 20900 [Source:MGI Symb | 1.6062 | 2.0558 | 3.0187 | 0 | 5.4531 | 1.1138 |
| 22347 | ENSMUSG00000078182 | 0 | NA | NA | NA | NA | NA | Gm6083 | predicted gene 6083 [Source:MGI Symbol | 0 | 0 | 0 | 0 | 0 | 0 |
| 22348 | ENSMUSG00000078183 | 7.972262772 | -0.124861353 | 0.779592458 | -0.16016234 | 0.872753194 | 0.979477204 | Gm15610 | predicted gene 15610 [Source:MGI Symbo | 4.8186 | 8.2233 | 12.075 | 10.604 | 6.5437 | 5.5692 |
| 22349 | ENSMUSG00000078184 | 0.133850373 | -0.796629176 | 4.080472857 | -0.195229623 | 0.84521319 | NA | Rbm8a2 | RNA binding motif protein 8A2 [Source:M | 0.8031 | 0 | 0 | 0 | 0 | 0 |
| 22350 | ENSMUSG00000078185 | 35.60471848 | -0.268837126 | 0.375377273 | -0.716178483 | 0.473881147 | 0.868952133 | Chml | choroideremia-like [Source:MGI Symbol;A | 30.518 | 44.2 | 42.262 | 33.739 | 27.265 | 35.643 |
| 22351 | ENSMUSG00000078190 | 7.116335949 | -1.453221344 | 0.847616037 | -1.714480709 | 0.086440477 | 0.433134324 | Dnm3os | dynamin 3, opposite strand [Source:MGI S | 8.031 | 10.279 | 13.081 | 5.7839 | 2.1812 | 3.3415 |
| 22352 | ENSMUSG00000078192 | 0 | NA | NA | NA | NA | NA | Gm7290 | predicted pseudogene 7290 [Source:MGI | 0 | 0 | 0 | 0 | 0 | 0 |
| 22353 | ENSMUSG00000078193 | 11.8897708 | -1.076174742 | 0.639168653 | -1.68371014 | 0.092237701 | 0.447243079 | Gm2000 | predicted gene 2000 [Source:MGI Symbol | 16.865 | 18.503 | 13.081 | 9.6398 | 5.4531 | 7.7969 |
| 22354 | ENSMUSG00000078197 | 0 | NA | NA | NA | NA | NA | Gm17374 | predicted gene, 17374 [Source:MGI Symb | 0 | 0 | 0 | 0 | 0 | 0 |
| 22355 | ENSMUSG00000078198 | 0 | NA | NA | NA | NA | NA | Olfr364-ps1 | olfactory receptor 364, pseudogene 1 [So | 0 | 0 | 0 | 0 | 0 | 0 |
| 22356 | ENSMUSG00000078201 | 141.7274675 | -0.113423857 | 0.205696013 | -0.55141495 | 0.581349255 | 0.908645447 | Tmem203 | transmembrane protein 203 [Source:MGI | 134.12 | 151.1 | 156.97 | 133.03 | 155.96 | 119.18 |
| 22357 | ENSMUSG00000078202 | 3.932966558 | -4.319152591 | 1.60005305 | -2.699380868 | 0.006946862 | NA | Nrarp | Notch-regulated ankyrin repeat protein [S | 11.243 | 9.2513 | 2.0125 | 0 | 1.0906 | 0 |
| 22358 | ENSMUSG00000078206 | 0 | NA | NA | NA | NA | NA | Fthl17d | ferritin, heavy polypeptide-like 17, memb | 0 | 0 | 0 | 0 | 0 | 0 |
| 22359 | ENSMUSG00000078208 | 0 | NA | NA | NA | NA | NA | Fthl17b | ferritin, heavy polypeptide-like 17, memb | 0 | 0 | 0 | 0 | 0 | 0 |
| 22360 | ENSMUSG00000078210 | 0 | NA | NA | NA | NA | NA | Gm5924 | predicted gene 5924 [Source:MGI Symbol | 0 | 0 | 0 | 0 | 0 | 0 |
| 22361 | ENSMUSG00000078212 | 0 | NA | NA | NA | NA | NA | Gm3757 | predicted pseudogene 3757 [Source:MGI | 0 | 0 | 0 | 0 | 0 | 0 |
| 22362 | ENSMUSG00000078213 | 0 | NA | NA | NA | NA | NA | Btbd35f4 | BTB domain containing 35, family membe | 0 | 0 | 0 | 0 | 0 | 0 |
| 22363 | ENSMUSG00000078214 | 0 | NA | NA | NA | NA | NA | Gm5925 | predicted gene 5925 [Source:MGI Symbol | 0 | 0 | 0 | 0 | 0 | 0 |
| 22364 | ENSMUSG00000078218 | 0 | NA | NA | NA | NA | NA | Btbd35f3 | BTB domain containing 35, family membe | 0 | 0 | 0 | 0 | 0 | 0 |
| 22365 | ENSMUSG00000078234 | 992.2687523 | 0.824467178 | 0.176806592 | 4.663102032 | 3.11E-06 | 0.000204816 | Klhdc7a | kelch domain containing 7A [Source:MGI S | 881.81 | 676.37 | 588.65 | 1052.7 | 1170.2 | 1583.9 |
| 22366 | ENSMUSG00000078235 | 0 | NA | NA | NA | NA | NA | Fam43b | family with sequence similarity 43, memb | 0 | 0 | 0 | 0 | 0 | 0 |
| 22367 | ENSMUSG00000078238 | 5.241750526 | 1.907401375 | 1.007312688 | 1.893554404 | 0.058284181 | 0.342880809 | Gm12854 | predicted gene 12854 [Source:MGI Symbo | 1.6062 | 3.0838 | 2.0125 | 11.568 | 8.7249 | 4.4554 |
| 22368 | ENSMUSG00000078240 | 0.342432689 | 1.990653086 | 3.900281263 | 0.51038706 | 0.609780321 | NA | Gm3550 | predicted gene 3550 [Source:MGI Symbol | 0 | 0 | 0 | 0.964 | 1.0906 | 0 |
| 22369 | ENSMUSG00000078247 | 6.306123851 | 0.505064996 | 1.015425064 | 0.497392682 | 0.618912165 | 0.917509899 | Airn | antisense Igf2r RNA [Source:MGI Symbol;A | 2.4093 | 12.335 | 1.0062 | 6.7478 | 11.997 | 3.3415 |
| 22370 | ENSMUSG00000078249 | 0.68145976 | 0.058694068 | 2.590797836 | 0.022654824 | 0.981925612 | NA | Hmga1b | high mobility group AT-hook 1B [Source:M | 0 | 1.0279 | 1.0062 | 0.964 | 1.0906 | 0 |
| 22371 | ENSMUSG00000078251 | 0.481988923 | 2.492401104 | 3.837077151 | 0.649557203 | 0.515978285 | NA | Gm16487 | predicted gene 16487 [Source:MGI Symbo | 0 | 0 | 0 | 2.8919 | 0 | 0 |
| 22372 | ENSMUSG00000078252 | 0 | NA | NA | NA | NA | NA | Krtap17-1 | keratin associated protein 17-1 [Source:M | 0 | 0 | 0 | 0 | 0 | 0 |
| 22373 | ENSMUSG00000078253 | 0 | NA | NA | NA | NA | NA | Krtap16-1 | keratin associated protein 16-1 [Source:M | 0 | 0 | 0 | 0 | 0 | 0 |
| 22374 | ENSMUSG00000078254 | 0 | NA | NA | NA | NA | NA | Krtap29-1 | keratin associated protein 29-1 [Source:M | 0 | 0 | 0 | 0 | 0 | 0 |
| 22375 | ENSMUSG00000078255 | 0 | NA | NA | NA | NA | NA | Krtap9-5 | keratin associated protein 9-5 [Source:MG | 0 | 0 | 0 | 0 | 0 | 0 |
| 22376 | ENSMUSG00000078256 | 0 | NA | NA | NA | NA | NA | Gm11565 | predicted gene 11565 [Source:MGI Symbo | 0 | 0 | 0 | 0 | 0 | 0 |
| 22377 | ENSMUSG00000078257 | 0 | NA | NA | NA | NA | NA | 2300003K06 | RIKEN cDNA 2300003K06 gene [Source:M | 0 | 0 | 0 | 0 | 0 | 0 |
| 22378 | ENSMUSG00000078258 | 0 | NA | NA | NA | NA | NA | Gm11564 | predicted gene 11564 [Source:MGI Symbo | 0 | 0 | 0 | 0 | 0 | 0 |
| 22379 | ENSMUSG00000078259 | 0 | NA | NA | NA | NA | NA | Gm11554 | predicted gene 11554 [Source:MGI Symbo | 0 | 0 | 0 | 0 | 0 | 0 |
| 22380 | ENSMUSG00000078260 | 0 | NA | NA | NA | NA | NA | Gm11569 | predicted gene 11569 [Source:MGI Symbo | 0 | 0 | 0 | 0 | 0 | 0 |
| 22381 | ENSMUSG00000078261 | 0 | NA | NA | NA | NA | NA | Gm11596 | predicted gene 11596 [Source:MGI Symbo | 0 | 0 | 0 | 0 | 0 | 0 |
| 22382 | ENSMUSG00000078262 | 0 | NA | NA | NA | NA | NA | Krtap4-9 | keratin associated protein 4-9 [Source:MG | 0 | 0 | 0 | 0 | 0 | 0 |
| 22383 | ENSMUSG00000078263 | 0 | NA | NA | NA | NA | NA | Gm14188 | predicted gene 14188 [Source:MGI Symbo | 0 | 0 | 0 | 0 | 0 | 0 |
| 22384 | ENSMUSG00000078269 | 0 | NA | NA | NA | NA | NA | Gm14180 | predicted gene 14180 [Source:MGI Symbo | 0 | 0 | 0 | 0 | 0 | 0 |
| 22385 | ENSMUSG00000078270 | 0 | NA | NA | NA | NA | NA | Gm14193 | predicted gene 14193 [Source:MGI Symbo | 0 | 0 | 0 | 0 | 0 | 0 |
| 22386 | ENSMUSG00000078276 | 0 | NA | NA | NA | NA | NA | Gm14190 | predicted gene 14190 [Source:MGI Symbo | 0 | 0 | 0 | 0 | 0 | 0 |
| 22387 | ENSMUSG00000078280 | 0 | NA | NA | NA | NA | NA | Tas2r122 | taste receptor, type 2, member 122 [Sour | 0 | 0 | 0 | 0 | 0 | 0 |
| 22388 | ENSMUSG00000078289 | 0 | NA | NA | NA | NA | NA | Gm10923 | predicted gene 10923 [Source:MGI Symbo | 0 | 0 | 0 | 0 | 0 | 0 |
| 22389 | ENSMUSG00000078291 | 0 | NA | NA | NA | NA | NA | Gm3940 | predicted gene 3940 [Source:MGI Symbol | 0 | 0 | 0 | 0 | 0 | 0 |
| 22390 | ENSMUSG00000078300 | 0.506734502 | -2.372847156 | 3.338060886 | -0.710845978 | 0.477179685 | NA | Gm2606 | predicted pseudogene 2606 [Source:MGI | 0 | 1.0279 | 2.0125 | 0 | 0 | 0 |
| 22391 | ENSMUSG00000078302 | 0 | NA | NA | NA | NA | NA | Foxd1 | forkhead box D1 [Source:MGI Symbol;Acc | 0 | 0 | 0 | 0 | 0 | 0 |
| 22392 | ENSMUSG00000078307 | 0 | NA | NA | NA | NA | NA | AI593442 | expressed sequence AI593442 [Source:MG | 0 | 0 | 0 | 0 | 0 | 0 |
| 22393 | ENSMUSG00000078308 | 0 | NA | NA | NA | NA | NA | Gm47854 | predicted gene, 47854 [Source:MGI Symb | 0 | 0 | 0 | 0 | 0 | 0 |
| 22394 | ENSMUSG00000078314 | 0.305170013 | -1.657836477 | 4.041480335 | -0.410205257 | 0.681655385 | NA | Gm14762 | predicted gene 14762 [Source:MGI Symbo | 0.8031 | 1.0279 | 0 | 0 | 0 | 0 |
| 22395 | ENSMUSG00000078315 | 0 | NA | NA | NA | NA | NA | Fam47c | family with sequence similarity 47, memb | 0 | 0 | 0 | 0 | 0 | 0 |
| 22396 | ENSMUSG00000078317 | 179.5085178 | 0.101649382 | 0.205273263 | 0.495190562 | 0.620465612 | 0.917977899 | F8a | factor 8-associated gene A [Source:MGI S | 203.18 | 173.72 | 141.88 | 176.41 | 165.77 | 216.09 |

| | | | | | | | | | | | | | | | |
|---|---|---|---|---|---|---|---|---|---|---|---|---|---|---|---|
| 22397 | ENSMUSG00000078320 | 0 | NA | NA | NA | NA | NA | Tex13c1 | TEX13 family member C1 [Source:MGI Syr | 0 | 0 | 0 | 0 | 0 | 0 |
| 22398 | ENSMUSG00000078324 | 0 | NA | NA | NA | NA | NA | Btbd35f14 | BTB domain containing 35, family membe | 0 | 0 | 0 | 0 | 0 | 0 |
| 22399 | ENSMUSG00000078346 | 0 | NA | NA | NA | NA | NA | H2al1n | H2A histone family member L1N [Source:I | 0 | 0 | 0 | 0 | 0 | 0 |
| 22400 | ENSMUSG00000078348 | 468.6850897 | 0.00489025 | 0.134058754 | 0.036478408 | 0.970900895 | 0.994966572 | Sf3b5 | splicing factor 3b, subunit 5 [Source:MGI S | 494.71 | 416.31 | 492.05 | 463.67 | 484.23 | 461.13 |
| 22401 | ENSMUSG00000078349 | 32.78596489 | 0.104480655 | 0.411551615 | 0.253870112 | 0.799595905 | 0.962424862 | AW011738 | expressed sequence AW011738 [Source:N | 37.746 | 23.642 | 33.206 | 31.811 | 45.806 | 24.505 |
| 22402 | ENSMUSG00000078350 | 42.25332286 | -0.419874564 | 0.38638316 | -1.086679252 | 0.277178586 | 0.730041765 | Smim1 | small integral membrane protein 1 [Sourc | 34.533 | 63.731 | 47.293 | 47.235 | 25.084 | 35.643 |
| 22403 | ENSMUSG00000078354 | 0 | NA | NA | NA | NA | NA | Ifna2 | interferon alpha 2 [Source:MGI Symbol;Ac | 0 | 0 | 0 | 0 | 0 | 0 |
| 22404 | ENSMUSG00000078355 | 0 | NA | NA | NA | NA | NA | Ifna16 | interferon alpha 16 [Source:MGI Symbol;A | 0 | 0 | 0 | 0 | 0 | 0 |
| 22405 | ENSMUSG00000078365 | 0 | NA | NA | NA | NA | NA | Mos | Moloney sarcoma oncogene [Source:MGI | 0 | 0 | 0 | 0 | 0 | 0 |
| 22406 | ENSMUSG00000078370 | 0.328370405 | 0.16514316 | 4.003149536 | 0.041253308 | 0.967093956 | NA | Gm6316 | predicted pseudogene 6316 [Source:MGI | 0 | 0 | 1.0062 | 0.964 | 0 | 0 |
| 22407 | ENSMUSG00000078377 | 6.814223569 | 0.277228457 | 0.827970858 | 0.334828761 | 0.737754263 | 0.948137969 | Gm4294 | predicted gene 4294 [Source:MGI Symbol | 7.2279 | 4.1117 | 7.0437 | 4.8199 | 6.5437 | 11.138 |
| 22408 | ENSMUSG00000078380 | 0 | NA | NA | NA | NA | NA | Gm6096 | predicted gene 6096 [Source:MGI Symbol | 0 | 0 | 0 | 0 | 0 | 0 |
| 22409 | ENSMUSG00000078420 | 0 | NA | NA | NA | NA | NA | Olfr141 | olfactory receptor 141 [Source:MGI Symb | 0 | 0 | 0 | 0 | 0 | 0 |
| 22410 | ENSMUSG00000078427 | 336.9833286 | 0.09875602 | 0.143434683 | 0.688508646 | 0.491132528 | 0.876368691 | Sarnp | SAP domain containing ribonucleoprotein | 301.97 | 323.79 | 351.18 | 353.78 | 338.09 | 353.09 |
| 22411 | ENSMUSG00000078429 | 979.9471234 | 0.016141118 | 0.11023836 | 0.146420161 | 0.883589713 | 0.981769403 | Ctdsp2 | CTD (carboxy-terminal domain, RNA polyr | 909.91 | 1050.5 | 963.98 | 1006.4 | 958.65 | 990.21 |
| 22412 | ENSMUSG00000078430 | 0.181769715 | 1.126924175 | 4.080472857 | 0.276174898 | 0.782413736 | NA | Gm10936 | predicted gene 10936 [Source:MGI Symbo | 0 | 0 | 0 | 0 | 1.0906 | 0 |
| 22413 | ENSMUSG00000078435 | 44.93302335 | -0.082496964 | 0.325456352 | -0.253480884 | 0.799896631 | 0.962424862 | AU041133 | expressed sequence AU041133 [Source:M | 48.186 | 44.2 | 46.287 | 48.199 | 38.172 | 44.554 |
| 22414 | ENSMUSG00000078439 | 11.1447776 | -0.659399993 | 0.699433415 | -0.942763068 | 0.345802106 | 0.786899236 | Smim24 | small integral membrane protein 24 [Sour | 14.456 | 19.53 | 7.0437 | 11.568 | 9.8156 | 4.4554 |
| 22415 | ENSMUSG00000078440 | 171.4345491 | 0.180680538 | 0.199830784 | 0.904167692 | 0.365906489 | 0.804235439 | Dohh | deoxyhypusine hydroxylase/monooxygen | 142.15 | 152.13 | 188.17 | 169.66 | 208.31 | 168.19 |
| 22416 | ENSMUSG00000078442 | 0.181769715 | 1.126924175 | 4.080472857 | 0.276174898 | 0.782413736 | NA | Ccdc105 | coiled-coil domain containing 105 [Source | 0 | 0 | 0 | 0 | 1.0906 | 0 |
| 22417 | ENSMUSG00000078444 | 3.529401079 | -1.165981494 | 1.245110558 | -0.936448163 | 0.34904249 | NA | Gm10941 | predicted gene 10941 [Source:MGI Symbo | 6.4248 | 3.0838 | 5.0312 | 0 | 2.1812 | 4.4554 |
| 22418 | ENSMUSG00000078451 | 8.485697662 | -1.449583997 | 0.820874042 | -1.765903077 | 0.077412111 | 0.40712138 | Ppil6 | peptidylprolyl isomerase (cyclophilin)-like | 16.865 | 10.279 | 10.062 | 4.8199 | 1.0906 | 7.7969 |
| 22419 | ENSMUSG00000078452 | 133.6340287 | -0.172727663 | 0.637585371 | -0.270909075 | 0.786460965 | 0.960870081 | Raet1d | retinoic acid early transcript delta [Source | 258.6 | 97.652 | 68.425 | 163.88 | 133.06 | 80.197 |
| 22420 | ENSMUSG00000078453 | 129.0784455 | -0.021751617 | 0.222154344 | -0.097912184 | 0.922002025 | 0.986685804 | Abracl | ABRA C-terminal like [Source:MGI Symbol | 146.16 | 138.77 | 104.65 | 114.71 | 127.6 | 142.57 |
| 22421 | ENSMUSG00000078480 | 0.487198452 | -0.685759572 | 3.136894395 | -0.218610985 | 0.826953095 | NA | Mrpl48-ps | mitochondrial ribosomal protein L48 pseu | 0.8031 | 0 | 1.0062 | 0 | 0 | 1.1138 |
| 22422 | ENSMUSG00000078481 | 0 | NA | NA | NA | NA | NA | Gm10944 | predicted gene 10944 [Source:MGI Symbo | 0 | 0 | 0 | 0 | 0 | 0 |
| 22423 | ENSMUSG00000078485 | 216.5863102 | 0.241170649 | 0.170817737 | 1.411859527 | 0.157991327 | 0.578096473 | Plekhn1 | pleckstrin homology domain containing, fa | 178.29 | 204.56 | 213.32 | 239.07 | 218.12 | 246.16 |
| 22424 | ENSMUSG00000078486 | 3.537117741 | -0.883052399 | 1.304578192 | -0.676887291 | 0.498477468 | NA | Perm1 | PPARGC1 and ESRR induced regulator, mu | 5.6217 | 2.0558 | 6.0375 | 0.964 | 6.5437 | 0 |
| 22425 | ENSMUSG00000078487 | 0 | NA | NA | NA | NA | NA | Ankrd65 | ankyrin repeat domain 65 [Source:MGI Sy | 0 | 0 | 0 | 0 | 0 | 0 |
| 22426 | ENSMUSG00000078488 | 0 | NA | NA | NA | NA | NA | Gm10945 | predicted gene 10945 [Source:MGI Symbo | 0 | 0 | 0 | 0 | 0 | 0 |
| 22427 | ENSMUSG00000078489 | 0.315620088 | 0.165143421 | 4.063300985 | 0.040642675 | 0.967580763 | NA | Gm17106 | predicted gene 17106 [Source:MGI Symbo | 0.8031 | 0 | 0 | 0 | 1.0906 | 0 |
| 22428 | ENSMUSG00000078490 | 3.397597737 | -1.595212696 | 1.291021056 | -1.235620974 | 0.216599485 | NA | Cfap74 | cilia and flagella associated protein 74 [So | 3.2124 | 7.1954 | 5.0312 | 3.8559 | 1.0906 | 0 |
| 22429 | ENSMUSG00000078491 | 0 | NA | NA | NA | NA | NA | Gm13090 | predicted gene 13090 [Source:MGI Symbo | 0 | 0 | 0 | 0 | 0 | 0 |
| 22430 | ENSMUSG00000078492 | 9.142691693 | -2.039378252 | 0.802311149 | -2.541879487 | 0.011025818 | 0.11632198 | 1700045H11 | RIKEN cDNA 1700045H11 gene [Source:M | 14.456 | 16.447 | 13.081 | 0.964 | 5.4531 | 4.4554 |
| 22431 | ENSMUSG00000078494 | 0 | NA | NA | NA | NA | NA | Gm13201 | predicted gene 13201 [Source:MGI Symbo | 0 | 0 | 0 | 0 | 0 | 0 |
| 22432 | ENSMUSG00000078495 | 38.79447032 | 0.028957564 | 0.38056631 | 0.076090718 | 0.939346925 | 0.988812319 | Zfp984 | zinc finger protein 984 [Source:MGI Symb | 42.564 | 47.284 | 25.156 | 31.811 | 43.625 | 42.326 |
| 22433 | ENSMUSG00000078496 | 91.65985017 | -0.380500201 | 0.248811963 | -1.529268112 | 0.126197991 | 0.521094669 | Zfp982 | zinc finger protein 982 [Source:MGI Symb | 110.03 | 92.513 | 108.67 | 95.434 | 69.8 | 73.514 |
| 22434 | ENSMUSG00000078497 | 1.633477947 | 3.112778638 | 2.118118316 | 1.469596205 | 0.141671149 | NA | Zfp978 | zinc finger protein 978 [Source:MGI Symb | 0 | 0 | 1.0062 | 0 | 5.4531 | 3.3415 |
| 22435 | ENSMUSG00000078498 | 0.906682787 | -0.583665192 | 2.768799895 | -0.210800785 | 0.833042726 | NA | Zfp988 | zinc finger protein 988 [Source:MGI Symb | 3.2124 | 0 | 0 | 0 | 0 | 2.2277 |
| 22436 | ENSMUSG00000078500 | 0 | NA | NA | NA | NA | NA | Zfp986 | zinc finger protein 986 [Source:MGI Symb | 0 | 0 | 0 | 0 | 0 | 0 |
| 22437 | ENSMUSG00000078502 | 4.883135869 | -1.318590008 | 1.1030942 | -1.195355762 | 0.231948062 | NA | Zfp268 | zinc finger protein 268 [Source:MGI Symb | 5.6217 | 11.307 | 4.025 | 2.8919 | 5.4531 | 0 |
| 22438 | ENSMUSG00000078503 | 0 | NA | NA | NA | NA | NA | Zfp990 | zinc finger protein 990 [Source:MGI Symb | 0 | 0 | 0 | 0 | 0 | 0 |
| 22439 | ENSMUSG00000078504 | 0 | NA | NA | NA | NA | NA | Aadacl4fm5 | AADACL4 family member 5 [Source:MGI S | 0 | 0 | 0 | 0 | 0 | 0 |
| 22440 | ENSMUSG00000078505 | 0 | NA | NA | NA | NA | NA | Aadacl4fm4 | AADACL4 family member 4 [Source:MGI S | 0 | 0 | 0 | 0 | 0 | 0 |
| 22441 | ENSMUSG00000078506 | 0 | NA | NA | NA | NA | NA | Aadacl4fm2 | AADACL4 family member 2 [Source:MGI S | 0 | 0 | 0 | 0 | 0 | 0 |
| 22442 | ENSMUSG00000078507 | 0 | NA | NA | NA | NA | NA | Aadacl3 | arylacetamide deacetylase like 3 [Source:I | 0 | 0 | 0 | 0 | 0 | 0 |
| 22443 | ENSMUSG00000078508 | 0 | NA | NA | NA | NA | NA | Pramel30 | PRAME like 30 [Source:MGI Symbol;Acc:M | 0 | 0 | 0 | 0 | 0 | 0 |
| 22444 | ENSMUSG00000078509 | 0 | NA | NA | NA | NA | NA | Pramel14 | PRAME like 14 [Source:MGI Symbol;Acc:M | 0 | 0 | 0 | 0 | 0 | 0 |
| 22445 | ENSMUSG00000078510 | 0 | NA | NA | NA | NA | NA | Pramel28 | PRAME like 28 [Source:MGI Symbol;Acc:M | 0 | 0 | 0 | 0 | 0 | 0 |
| 22446 | ENSMUSG00000078511 | 0 | NA | NA | NA | NA | NA | Pramel16 | PRAME like 16 [Source:MGI Symbol;Acc:M | 0 | 0 | 0 | 0 | 0 | 0 |
| 22447 | ENSMUSG00000078512 | 0 | NA | NA | NA | NA | NA | Pramel11 | PRAME like 11 [Source:MGI Symbol;Acc:M | 0 | 0 | 0 | 0 | 0 | 0 |
| 22448 | ENSMUSG00000078513 | 0 | NA | NA | NA | NA | NA | Pramel22 | PRAME like 22 [Source:MGI Symbol;Acc:M | 0 | 0 | 0 | 0 | 0 | 0 |
| 22449 | ENSMUSG00000078515 | 3802.05246 | -0.040252107 | 0.113773555 | -0.353791411 | 0.723495211 | 0.9447719 | Ddi2 | DNA-damage inducible protein 2 [Source: | 3401.9 | 3798.2 | 4366.1 | 3618.8 | 3620.9 | 4006.5 |
| 22450 | ENSMUSG00000078517 | 434.4442159 | -0.101933221 | 0.151069121 | -0.674745577 | 0.499837422 | 0.880716237 | Emc1 | ER membrane protein complex subunit 1 [ | 508.36 | 401.92 | 438.72 | 464.64 | 419.89 | 373.14 |
| 22451 | ENSMUSG00000078518 | 0 | NA | NA | NA | NA | NA | Gm13030 | predicted gene 13030 [Source:MGI Symbo | 0 | 0 | 0 | 0 | 0 | 0 |
| 22452 | ENSMUSG00000078519 | 7.980285266 | -0.673991352 | 0.914136921 | -0.73729803 | 0.46094113 | 0.859821403 | 2310026L22I | RIKEN cDNA 2310026L22 gene [Source:MG | 8.031 | 19.53 | 2.0125 | 10.604 | 4.3625 | 3.3415 |

| | | | | | | | | | | | | | | | |
|---|---|---|---|---|---|---|---|---|---|---|---|---|---|---|---|
| 22453 | ENSMUSG00000078520 | 0 | NA | NA | NA | NA | NA | Cela3a | chymotrypsin-like elastase family, membe | 0 | 0 | 0 | 0 | 0 | 0 |
| 22454 | ENSMUSG00000078521 | 1.801752777 | -0.597647822 | 2.021676579 | -0.295619897 | 0.767520377 | NA | Aunip | aurora kinase A and ninein interacting prc | 6.4248 | 0 | 0 | 0 | 3.2719 | 1.1138 |
| 22455 | ENSMUSG00000078528 | 0.160662974 | 1.126924175 | 4.080472857 | 0.276174898 | 0.782413736 | NA | Gm17068 | predicted gene 17068 [Source:MGI Symbo | 0 | 0 | 0 | 0.964 | 0 | 0 |
| 22456 | ENSMUSG00000078531 | 0 | NA | NA | NA | NA | NA | Gm831 | predicted gene 831 [Source:MGI Symbol;/ | 0 | 0 | 0 | 0 | 0 | 0 |
| 22457 | ENSMUSG00000078532 | 1.80790449 | -0.201353257 | 1.658277662 | -0.121423126 | 0.903355901 | NA | Nkain1 | Na+/K+ transporting ATPase interacting 1 | 0.8031 | 2.0558 | 3.0187 | 3.8559 | 0 | 1.1138 |
| 22458 | ENSMUSG00000078537 | 0 | NA | NA | NA | NA | NA | B020031M17 | RIKEN cDNA B020031M17 gene [Source:N | 0 | 0 | 0 | 0 | 0 | 0 |
| 22459 | ENSMUSG00000078546 | 38.95495007 | -0.794076037 | 0.380813626 | -2.085209098 | 0.037050324 | 0.260269982 | Zfp995 | zinc finger protein 995 [Source:MGI Symb | 62.642 | 47.284 | 38.237 | 32.775 | 31.628 | 21.163 |
| 22460 | ENSMUSG00000078552 | 32.61287892 | -0.007781482 | 0.376426698 | -0.020671971 | 0.983507328 | 0.998580735 | Dcdc2b | doublecortin domain containing 2b [Sourc | 32.124 | 29.81 | 36.225 | 34.703 | 31.628 | 31.188 |
| 22461 | ENSMUSG00000078554 | 0.17131964 | -0.796629176 | 4.080472857 | -0.195229623 | 0.84521319 | NA | Fam229a | family with sequence similarity 229, meml | 0 | 1.0279 | 0 | 0 | 0 | 0 |
| 22462 | ENSMUSG00000078566 | 4529.501668 | -0.211450209 | 0.113500114 | -1.86299557 | 0.06246289 | 0.356959002 | Bnip3 | BCL2/adenovirus E1B interacting protein 3 | 4774.4 | 4978.2 | 4831 | 4589.5 | 4455.2 | 3548.7 |
| 22463 | ENSMUSG00000078570 | 210.9168091 | 0.159428541 | 0.196451824 | 0.811540145 | 0.417055549 | 0.836878476 | 1110065P20 | RIKEN cDNA 1110065P20 gene [Source:M | 208.81 | 192.22 | 197.22 | 280.52 | 198.49 | 188.24 |
| 22464 | ENSMUSG00000078572 | 315.4029065 | 0.021410968 | 0.155470123 | 0.137717574 | 0.89046363 | 0.982489666 | Ndufaf8 | NADH:ubiquinone oxidoreductase comple | 295.54 | 308.38 | 335.08 | 277.63 | 328.28 | 347.52 |
| 22465 | ENSMUSG00000078575 | 0 | NA | NA | NA | NA | NA | Gm12887 | predicted gene 12887 [Source:MGI Symbo | 0 | 0 | 0 | 0 | 0 | 0 |
| 22466 | ENSMUSG00000078576 | 0 | NA | NA | NA | NA | NA | Gm12886 | predicted gene 12886 [Source:MGI Symbo | 0 | 0 | 0 | 0 | 0 | 0 |
| 22467 | ENSMUSG00000078577 | 0.167707431 | -0.796629176 | 4.080472857 | -0.195229623 | 0.84521319 | NA | Tmco2 | transmembrane and coiled-coil domains 2 | 0 | 0 | 1.0062 | 0 | 0 | 0 |
| 22468 | ENSMUSG00000078578 | 4532.717595 | 0.023431875 | 0.084563159 | 0.27709318 | 0.781708559 | 0.960079945 | Ube2d3 | ubiquitin-conjugating enzyme E2D 3 [Sour | 4366.5 | 4594.8 | 4527.1 | 4554.8 | 4674.4 | 4478.8 |
| 22469 | ENSMUSG00000078580 | 105.3834955 | -0.193083758 | 0.233572626 | -0.826654051 | 0.408433172 | 0.832151413 | E430018J23F | RIKEN cDNA E430018J23 gene [Source:MC | 107.62 | 126.43 | 103.64 | 113.75 | 85.068 | 95.791 |
| 22470 | ENSMUSG00000078584 | 1132.947038 | 0.046569826 | 0.152093028 | 0.306193036 | 0.759457674 | 0.95402891 | AU022252 | expressed sequence AU022252 [Source:M | 1027.2 | 1020.7 | 1296 | 1045.9 | 1007.7 | 1400.1 |
| 22471 | ENSMUSG00000078586 | 0 | NA | NA | NA | NA | NA | Gm11735 | predicted gene 11735 [Source:MGI Symbo | 0 | 0 | 0 | 0 | 0 | 0 |
| 22472 | ENSMUSG00000078588 | 3.623078363 | 0.618450839 | 1.305302977 | 0.473798689 | 0.63564347 | NA | Ccdc24 | coiled-coil domain containing 24 [Source:I | 6.4248 | 2.0558 | 0 | 7.7118 | 1.0906 | 4.4554 |
| 22473 | ENSMUSG00000078590 | 0.531246861 | 1.101376188 | 3.36099124 | 0.327693858 | 0.743143147 | NA | Gm10959 | predicted gene 10959 [Source:MGI Symbo | 0 | 0 | 1.0062 | 0 | 2.1812 | 0 |
| 22474 | ENSMUSG00000078591 | 0 | NA | NA | NA | NA | NA | Hs3st4 | heparan sulfate (glucosamine) 3-O-sulfotr | 0 | 0 | 0 | 0 | 0 | 0 |
| 22475 | ENSMUSG00000078592 | 0.514011053 | 1.044022102 | 3.082659575 | 0.338675769 | 0.734853997 | NA | Gm4609 | predicted gene 4609 [Source:MGI Symbol | 0 | 0 | 1.0062 | 0.964 | 0 | 1.1138 |
| 22476 | ENSMUSG00000078593 | 0 | NA | NA | NA | NA | NA | 1700042G07 | RIKEN cDNA 1700042G07 gene [Source:M | 0 | 0 | 0 | 0 | 0 | 0 |
| 22477 | ENSMUSG00000078597 | 549.2138792 | 2.241358258 | 0.460174129 | 4.8706742 | 1.11E-06 | 8.94E-05 | Cyp4a12b | cytochrome P450, family 4, subfamily a, p | 255.39 | 164.47 | 154.96 | 532.12 | 1376.4 | 811.99 |
| 22478 | ENSMUSG00000078598 | 0 | NA | NA | NA | NA | NA | Skint5 | selection and upkeep of intraepithelial T c | 0 | 0 | 0 | 0 | 0 | 0 |
| 22479 | ENSMUSG00000078599 | 0 | NA | NA | NA | NA | NA | Skint8 | selection and upkeep of intraepithelial T c | 0 | 0 | 0 | 0 | 0 | 0 |
| 22480 | ENSMUSG00000078600 | 0 | NA | NA | NA | NA | NA | 1700092K14 | RIKEN cDNA 1700092K14 gene [Source:M | 0 | 0 | 0 | 0 | 0 | 0 |
| 22481 | ENSMUSG00000078601 | 0 | NA | NA | NA | NA | NA | Gm12525 | predicted gene 12525 [Source:MGI Symbo | 0 | 0 | 0 | 0 | 0 | 0 |
| 22482 | ENSMUSG00000078604 | 0.736965982 | -2.935629304 | 2.883220453 | -1.018177192 | 0.308593757 | NA | Gm10961 | predicted gene 10961 [Source:MGI Symbo | 2.4093 | 0 | 2.0125 | 0 | 0 | 0 |
| 22483 | ENSMUSG00000078605 | 0 | NA | NA | NA | NA | NA | E030025P04 | RIKEN cDNA E030025P04 gene [Source:M | 0 | 0 | 0 | 0 | 0 | 0 |
| 22484 | ENSMUSG00000078606 | 15.36767406 | 0.058436868 | 0.593821238 | 0.098408181 | 0.921608179 | 0.986685804 | Gvin2 | GTPase, very large interferon inducible, fa | 12.85 | 10.279 | 22.137 | 19.28 | 8.7249 | 18.935 |
| 22485 | ENSMUSG00000078607 | 75.2166207 | -0.04895101 | 0.25973531 | -0.18846498 | 0.850512166 | 0.973432436 | 1810010H24 | RIKEN cDNA 1810010H24 gene [Source:M | 77.901 | 77.094 | 74.462 | 73.262 | 83.978 | 64.603 |
| 22486 | ENSMUSG00000078611 | 0 | NA | NA | NA | NA | NA | Gm5901 | predicted gene 5901 [Source:MGI Symbol | 0 | 0 | 0 | 0 | 0 | 0 |
| 22487 | ENSMUSG00000078612 | 86.23643346 | -0.385334838 | 0.247106736 | -1.559386218 | 0.118904996 | 0.504648435 | Fyb2 | FYN binding protein 2 [Source:MGI Symbo | 110.83 | 89.429 | 92.575 | 79.046 | 69.8 | 75.741 |
| 22488 | ENSMUSG00000078616 | 1.028636663 | 1.245965894 | 2.457526307 | 0.507000023 | 0.612154802 | NA | Trim30c | tripartite motif-containing 30C [Source:M | 0.8031 | 0 | 1.0062 | 0 | 4.3625 | 0 |
| 22489 | ENSMUSG00000078618 | 0 | NA | NA | NA | NA | NA | 4930456L15 | RIKEN cDNA 4930456L15 gene [Source:M | 0 | 0 | 0 | 0 | 0 | 0 |
| 22490 | ENSMUSG00000078619 | 1872.041527 | 0.034472644 | 0.115467214 | 0.298549196 | 0.765284034 | 0.954980327 | Smarcd2 | SWI/SNF related, matrix associated, actin | 1829.5 | 1747.5 | 1971.2 | 1660 | 1944.6 | 2079.5 |
| 22491 | ENSMUSG00000078620 | 0.589026695 | 0.241472922 | 3.228838524 | 0.074786311 | 0.940384733 | NA | Gm10964 | predicted gene 10964 [Source:MGI Symbo | 1.6062 | 0 | 0 | 1.928 | 0 | 0 |
| 22492 | ENSMUSG00000078621 | 0 | NA | NA | NA | NA | NA | Hbb-bh2 | hemoglobin beta, bh2 [Source:MGI Symbo | 0 | 0 | 0 | 0 | 0 | 0 |
| 22493 | ENSMUSG00000078622 | 1784.534955 | 0.053394746 | 0.101419684 | 0.526473204 | 0.598559465 | 0.915055389 | Ccdc47 | coiled-coil domain containing 47 [Source:I | 1614.2 | 1806.1 | 1835.4 | 1802.6 | 1885.7 | 1763.2 |
| 22494 | ENSMUSG00000078624 | 0 | NA | NA | NA | NA | NA | Olfr613 | olfactory receptor 613 [Source:MGI Symb | 0 | 0 | 0 | 0 | 0 | 0 |
| 22495 | ENSMUSG00000078625 | 0 | NA | NA | NA | NA | NA | Gm12789 | predicted gene 12789 [Source:MGI Symbo | 0 | 0 | 0 | 0 | 0 | 0 |
| 22496 | ENSMUSG00000078626 | 0 | NA | NA | NA | NA | NA | Gm12790 | predicted gene 12790 [Source:MGI Symbo | 0 | 0 | 0 | 0 | 0 | 0 |
| 22497 | ENSMUSG00000078627 | 0 | NA | NA | NA | NA | NA | Marchf10 | membrane associated ring-CH-type finger | 0 | 0 | 0 | 0 | 0 | 0 |
| 22498 | ENSMUSG00000078630 | 2.01101304 | 1.719077906 | 1.535676119 | 1.119427387 | 0.262957852 | NA | Tomt | transmembrane O-methyltransferase [Sou | 0.8031 | 1.0279 | 1.0062 | 4.8199 | 2.1812 | 2.2277 |
| 22499 | ENSMUSG00000078632 | 0 | NA | NA | NA | NA | NA | Lrrc37a | leucine rich repeat containing 37A [Source | 0 | 0 | 0 | 0 | 0 | 0 |
| 22500 | ENSMUSG00000078635 | 0 | NA | NA | NA | NA | NA | 1700028N14 | RIKEN cDNA 1700028N14 gene [Source:M | 0 | 0 | 0 | 0 | 0 | 0 |
| 22501 | ENSMUSG00000078636 | 4.710007334 | 1.257624376 | 0.986398504 | 1.274965818 | 0.202321341 | NA | Gm7336 | predicted gene 7336 [Source:MGI Symbol | 3.2124 | 3.0838 | 2.0125 | 6.7478 | 7.6343 | 5.5692 |
| 22502 | ENSMUSG00000078639 | 0 | NA | NA | NA | NA | NA | Gm12695 | predicted gene 12695 [Source:MGI Symbo | 0 | 0 | 0 | 0 | 0 | 0 |
| 22503 | ENSMUSG00000078640 | 0.371281295 | 2.09253362 | 4.031852165 | 0.519000582 | 0.603760335 | NA | Gm11627 | predicted gene 11627 [Source:MGI Symbo | 0 | 0 | 0 | 0 | 0 | 2.2277 |
| 22504 | ENSMUSG00000078648 | 0 | NA | NA | NA | NA | NA | Gm17546 | predicted gene, 17546 [Source:MGI Symb | 0 | 0 | 0 | 0 | 0 | 0 |
| 22505 | ENSMUSG00000078650 | 5650.266485 | -1.249683936 | 0.394672756 | -3.166380037 | 0.00154349 | 0.029101522 | G6pc | glucose-6-phosphatase, catalytic [Source: | 7855.1 | 8967.6 | 7042.7 | 3510.8 | 4788.9 | 1736.5 |
| 22506 | ENSMUSG00000078651 | 17.55113159 | -0.00664925 | 0.509549595 | -0.013049269 | 0.989588485 | 0.99859176 | Aoc2 | amine oxidase, copper containing 2 (retin | 15.259 | 18.503 | 19.119 | 18.316 | 19.631 | 14.48 |
| 22507 | ENSMUSG00000078652 | 984.3221589 | -0.039090841 | 0.112181651 | -0.348460205 | 0.727494595 | 0.945907265 | Psme3 | proteaseome (prosome, macropain) activa | 913.13 | 1008.4 | 1072.7 | 1000.6 | 951.02 | 960.13 |
| 22508 | ENSMUSG00000078653 | 2.229243216 | -0.709261846 | 1.705379027 | -0.415896897 | 0.67748545 | NA | Cntd1 | cyclin N-terminal domain containing 1 [So | 7.2279 | 1.0279 | 0 | 2.8919 | 0 | 2.2277 |

| | | | | | | | | | | | | | | | |
|---|---|---|---|---|---|---|---|---|---|---|---|---|---|---|---|
| 22509 | ENSMUSG00000078655 | 0 | NA | NA | NA | NA | NA | NA | NA | 0 | 0 | 0 | 0 | 0 | 0 |
| 22510 | ENSMUSG00000078656 | 4.54816022 | 0.663948455 | 1.138334278 | 0.583263166 | 0.559716155 | NA | Vps25 | vacuolar protein sorting 25 [Source:MGI S | 2.4093 | 8.2233 | 0 | 6.7478 | 5.4531 | 4.4554 |
| 22511 | ENSMUSG00000078657 | 0 | NA | NA | NA | NA | NA | Crnn | cornulin [Source:MGI Symbol;Acc:MGI:26 | 0 | 0 | 0 | 0 | 0 | 0 |
| 22512 | ENSMUSG00000078664 | 0 | NA | NA | NA | NA | NA | Sprr2a1 | small proline-rich protein 2A1 [Source:MG | 0 | 0 | 0 | 0 | 0 | 0 |
| 22513 | ENSMUSG00000078668 | 0 | NA | NA | NA | NA | NA | Gm11595 | predicted gene 11595 [Source:MGI Symbo | 0 | 0 | 0 | 0 | 0 | 0 |
| 22514 | ENSMUSG00000078670 | 50.50060039 | -0.591278046 | 0.344282236 | -1.717422465 | 0.08590201 | 0.431868847 | Fam174b | family with sequence similarity 174, meml | 61.839 | 62.703 | 57.356 | 33.739 | 29.447 | 57.92 |
| 22515 | ENSMUSG00000078671 | 565.3679369 | -0.035281169 | 0.138587378 | -0.254577075 | 0.799049767 | 0.962424862 | Chd2 | chromodomain helicase DNA binding prot | 499.53 | 585.91 | 631.92 | 512.84 | 562.76 | 599.25 |
| 22516 | ENSMUSG00000078672 | 56014.22056 | 0.787085438 | 0.174072677 | 4.521590933 | 6.14E-06 | 0.000368361 | Mup20 | major urinary protein 20 [Source:MGI Syn | 49652 | 30484 | 43170 | 54494 | 74590 | 83696 |
| 22517 | ENSMUSG00000078673 | 325.6163959 | 0.966492893 | 0.696834509 | 1.38697622 | 0.165449014 | 0.591023596 | Mup19 | major urinary protein 19 [Source:MGI Syn | 277.07 | 83.261 | 300.87 | 291.12 | 730.71 | 270.66 |
| 22518 | ENSMUSG00000078674 | 90.21382471 | 3.11567305 | 1.013586058 | 3.073910721 | 0.002112726 | 0.036292727 | Mup18 | major urinary protein 18 [Source:MGI Syn | 30.518 | 6.1675 | 19.119 | 34.703 | 329.37 | 121.41 |
| 22519 | ENSMUSG00000078675 | 650.6320161 | 2.316513669 | 0.776697915 | 2.982515626 | 0.002858901 | 0.0451925 | Mup16 | major urinary protein 16 [Source:MGI Syn | 237.72 | 108.96 | 305.9 | 310.4 | 1818.1 | 1122.8 |
| 22520 | ENSMUSG00000078676 | 369.2779458 | -0.100852489 | 0.163217957 | -0.617900695 | 0.536640801 | 0.890560262 | Casc3 | cancer susceptibility candidate 3 [Source:I | 345.33 | 403.97 | 398.47 | 425.11 | 309.74 | 333.04 |
| 22521 | ENSMUSG00000078677 | 0 | NA | NA | NA | NA | NA | Gm10974 | predicted gene 10974 [Source:MGI Symbo | 0 | 0 | 0 | 0 | 0 | 0 |
| 22522 | ENSMUSG00000078680 | 15.88167313 | 0.471534196 | 0.676960757 | 0.696545836 | 0.486087063 | 0.873111725 | Mup10 | major urinary protein 10 [Source:MGI Syn | 17.668 | 8.2233 | 14.087 | 36.631 | 11.997 | 6.6831 |
| 22523 | ENSMUSG00000078681 | 329.0001379 | 0.248581614 | 0.143171969 | 1.736244994 | 0.082520525 | 0.423554602 | Tm2d3 | TM2 domain containing 3 [Source:MGI Sy | 289.12 | 299.12 | 313.95 | 334.5 | 368.63 | 368.68 |
| 22524 | ENSMUSG00000078683 | 335.9457331 | 1.312809582 | 0.683188407 | 1.921592299 | 0.054657078 | 0.329262799 | Mup1 | major urinary protein 1 [Source:MGI Syml | 126.89 | 90.457 | 361.24 | 285.34 | 678.36 | 473.38 |
| 22525 | ENSMUSG00000078684 | 50.9938927 | 0.629016319 | 0.351682248 | 1.788592749 | 0.073680426 | 0.394708773 | 5830417I10F | RIKEN cDNA 5830417I10 gene [Source:MC | 28.912 | 46.256 | 45.281 | 49.163 | 82.887 | 53.465 |
| 22526 | ENSMUSG00000078685 | 9.499779183 | 0.986096541 | 1.136329205 | 0.86779125 | 0.385508621 | 0.816437825 | Mup-ps4 | major urinary protein, pseudogene 4 [Sou | 12.85 | 2.0558 | 4.025 | 0.964 | 35.99 | 1.1138 |
| 22527 | ENSMUSG00000078686 | 45.59073657 | 3.033545649 | 1.100911713 | 2.755484944 | 0.00586052 | 0.074161384 | Mup9 | major urinary protein 9 [Source:MGI Syml | 14.456 | 5.1396 | 10.062 | 16.388 | 196.31 | 31.188 |
| 22528 | ENSMUSG00000078687 | 10.37546653 | 1.211966724 | 0.761867538 | 1.590784046 | 0.111658185 | 0.490292648 | Mup8 | major urinary protein 8 [Source:MGI Syml | 5.6217 | 3.0838 | 10.062 | 10.604 | 25.084 | 7.7969 |
| 22529 | ENSMUSG00000078688 | 175.0438943 | 0.384376963 | 0.19552727 | 1.965848354 | 0.04931613 | 0.309919254 | Mup2 | major urinary protein 2 [Source:MGI Syml | 150.98 | 125.41 | 179.11 | 193.76 | 208.31 | 192.69 |
| 22530 | ENSMUSG00000078689 | 5.752452231 | -0.073249797 | 1.100025648 | -0.066589171 | 0.946908767 | 0.990677354 | Mup6 | major urinary protein 6 [Source:MGI Syml | 10.44 | 5.1396 | 2.0125 | 5.7839 | 0 | 11.138 |
| 22531 | ENSMUSG00000078691 | 0 | NA | NA | NA | NA | NA | Gm6821 | predicted gene 6821 [Source:MGI Symbol | 0 | 0 | 0 | 0 | 0 | 0 |
| 22532 | ENSMUSG00000078695 | 1043.840618 | -0.163676888 | 0.128034242 | -1.278383694 | 0.20111417 | 0.645135365 | Cisd3 | CDGSH iron sulfur domain 3 [Source:MGI | 1260.1 | 1082.4 | 964.99 | 927.35 | 1049.2 | 979.07 |
| 22533 | ENSMUSG00000078698 | 0 | NA | NA | NA | NA | NA | Mrgpra3 | MAS-related GPR, member A3 [Source:M( | 0 | 0 | 0 | 0 | 0 | 0 |
| 22534 | ENSMUSG00000078700 | 13.54825246 | 0.252314814 | 0.57342566 | 0.440013121 | 0.659927604 | 0.927787797 | D030028A08 | RIKEN cDNA D030028A08 gene [Source:M | 13.653 | 11.307 | 12.075 | 14.46 | 13.087 | 16.708 |
| 22535 | ENSMUSG00000078706 | 0 | NA | NA | NA | NA | NA | Gm53 | predicted gene 53 [Source:MGI Symbol;A( | 0 | 0 | 0 | 0 | 0 | 0 |
| 22536 | ENSMUSG00000078713 | 423.6192979 | -0.038650055 | 0.152442357 | -0.253538818 | 0.799851868 | 0.962424862 | Tomm5 | translocase of outer mitochondrial memb | 444.92 | 416.31 | 425.64 | 343.18 | 444.97 | 466.7 |
| 22537 | ENSMUSG00000078716 | 47.89775876 | 0.557493809 | 0.372084214 | 1.498300085 | 0.134055302 | 0.53726126 | Tmem8b | transmembrane protein 8B [Source:MGI S | 42.564 | 43.173 | 30.187 | 35.667 | 56.712 | 79.083 |
| 22538 | ENSMUSG00000078719 | 0 | NA | NA | NA | NA | NA | Msmp | microseminoprotein, prostate associated | 0 | 0 | 0 | 0 | 0 | 0 |
| 22539 | ENSMUSG00000078721 | 0.267700746 | -1.497481649 | 4.057001436 | -0.369110456 | 0.712045396 | NA | Fam205a1 | family with sequence similarity 205, meml | 1.6062 | 0 | 0 | 0 | 0 | 0 |
| 22540 | ENSMUSG00000078722 | 0 | NA | NA | NA | NA | NA | Gm12394 | predicted gene 12394 [Source:MGI Symbo | 0 | 0 | 0 | 0 | 0 | 0 |
| 22541 | ENSMUSG00000078735 | 0 | NA | NA | NA | NA | NA | Il11ra2 | interleukin 11 receptor, alpha chain 2 [Sou | 0 | 0 | 0 | 0 | 0 | 0 |
| 22542 | ENSMUSG00000078737 | 0 | NA | NA | NA | NA | NA | Gm2128 | predicted gene 2128 [Source:MGI Symbol | 0 | 0 | 0 | 0 | 0 | 0 |
| 22543 | ENSMUSG00000078742 | 0 | NA | NA | NA | NA | NA | Gm10985 | predicted gene 10985 [Source:MGI Symbo | 0 | 0 | 0 | 0 | 0 | 0 |
| 22544 | ENSMUSG00000078746 | 0 | NA | NA | NA | NA | NA | Fam205a4 | family with sequence similarity 205, meml | 0 | 0 | 0 | 0 | 0 | 0 |
| 22545 | ENSMUSG00000078747 | 0 | NA | NA | NA | NA | NA | Gm20878 | predicted gene, 20878 [Source:MGI Symb | 0 | 0 | 0 | 0 | 0 | 0 |
| 22546 | ENSMUSG00000078752 | 0 | NA | NA | NA | NA | NA | Scgb1b30 | secretoglobin, family 1B, member 30 [Sou | 0 | 0 | 0 | 0 | 0 | 0 |
| 22547 | ENSMUSG00000078753 | 0 | NA | NA | NA | NA | NA | Scgb1b24 | secretoglobin, family 1B, member 24 [Sou | 0 | 0 | 0 | 0 | 0 | 0 |
| 22548 | ENSMUSG00000078754 | 0 | NA | NA | NA | NA | NA | Scgb2b3 | secretoglobin, family 2B, member 3 [Sour | 0 | 0 | 0 | 0 | 0 | 0 |
| 22549 | ENSMUSG00000078757 | 0 | NA | NA | NA | NA | NA | Scgb1b29 | secretoglobin, family 1B, member 29 [Sou | 0 | 0 | 0 | 0 | 0 | 0 |
| 22550 | ENSMUSG00000078759 | 0 | NA | NA | NA | NA | NA | Scgb1b7 | secretoglobin, family 1B, member 7 [Sour | 0 | 0 | 0 | 0 | 0 | 0 |
| 22551 | ENSMUSG00000078762 | 20.51250309 | 0.778064409 | 0.503915961 | 1.544036047 | 0.1225796 | 0.513018327 | Haus5 | HAUS augmin-like complex, subunit 5 [Sou | 12.85 | 14.391 | 18.112 | 20.244 | 20.722 | 36.757 |
| 22552 | ENSMUSG00000078763 | 2.335868046 | -1.13265752 | 1.544396947 | -0.733397927 | 0.463315763 | NA | Slfn1 | schlafen 1 [Source:MGI Symbol;Acc:MGI:1 | 2.4093 | 6.1675 | 1.0062 | 0 | 1.0906 | 3.3415 |
| 22553 | ENSMUSG00000078765 | 0.965523032 | -0.944628791 | 2.267086424 | -0.416670834 | 0.67691919 | NA | U2af1l4 | U2 small nuclear RNA auxiliary factor 1-lik | 0.8031 | 2.0558 | 1.0062 | 1.928 | 0 | 0 |
| 22554 | ENSMUSG00000078768 | 6.275654065 | 0.677920421 | 0.86722683 | 0.781710618 | 0.43438466 | 0.845264432 | Zfp566 | zinc finger protein 566 [Source:MGI Symb | 2.4093 | 6.1675 | 6.0375 | 8.6758 | 5.4531 | 8.9108 |
| 22555 | ENSMUSG00000078771 | 8.949353787 | -0.24441156 | 0.754181225 | -0.324075371 | 0.745880972 | 0.949997329 | Evi2a | ecotropic viral integration site 2a [Source: | 9.6372 | 14.391 | 5.0312 | 4.8199 | 10.906 | 8.9108 |
| 22556 | ENSMUSG00000078772 | 1.521540975 | 1.167057869 | 1.77177728 | 0.658693326 | 0.51009272 | NA | Gm12353 | predicted gene 12353 [Source:MGI Symbo | 0.8031 | 0 | 2.0125 | 1.928 | 3.2719 | 1.1138 |
| 22557 | ENSMUSG00000078773 | 11.0463004 | -0.4897623 | 0.659657357 | -0.742449538 | 0.457815015 | 0.859309158 | Rad54b | RAD54 homolog B (S. cerevisiae) [Source:I | 15.259 | 8.2233 | 15.094 | 6.7478 | 9.8156 | 11.138 |
| 22558 | ENSMUSG00000078776 | 0.315620088 | 0.165143421 | 4.063300985 | 0.040642675 | 0.967580763 | NA | 9530053A07 | RIKEN cDNA 9530053A07 gene [Source:M | 0.8031 | 0 | 0 | 0 | 1.0906 | 0 |
| 22559 | ENSMUSG00000078779 | 42.48660245 | 0.367857348 | 0.346939892 | 1.060291297 | 0.289012097 | 0.739603195 | Zfp59 | zinc finger protein 59 [Source:MGI Symbo | 41.761 | 41.117 | 28.175 | 42.415 | 47.987 | 53.465 |
| 22560 | ENSMUSG00000078780 | 5.57684463 | -0.236687623 | 1.039385501 | -0.227718804 | 0.819864853 | 0.969160361 | Gm5150 | predicted gene 5150 [Source:MGI Symbol | 4.8186 | 6.1675 | 7.0437 | 0 | 7.6343 | 7.7969 |
| 22561 | ENSMUSG00000078783 | 0 | NA | NA | NA | NA | NA | Gm9733 | predicted gene 9733 [Source:MGI Symbol | 0 | 0 | 0 | 0 | 0 | 0 |
| 22562 | ENSMUSG00000078784 | 315.3350896 | 0.027144853 | 0.158136675 | 0.171654383 | 0.863709251 | 0.976954351 | Rbis | ribosomal biogenesis factor [Source:MGI ! | 342.12 | 299.12 | 294.83 | 285.34 | 310.83 | 359.77 |
| 22563 | ENSMUSG00000078786 | 136.7390229 | -0.201422745 | 0.217403737 | -0.92649164 | 0.35419054 | 0.79449673 | BC024978 | cDNA sequence BC024978 [Source:MGI Sy | 138.94 | 169.61 | 130.81 | 146.52 | 126.51 | 108.04 |
| 22564 | ENSMUSG00000078787 | 5.07391444 | -0.49748126 | 1.054032555 | -0.471979027 | 0.636941759 | NA | Cyp2t4 | cytochrome P450, family 2, subfamily t, pc | 5.6217 | 6.1675 | 6.0375 | 4.8199 | 0 | 7.7969 |

| | | | | | | | | | | | | | | | |
|---|---|---|---|---|---|---|---|---|---|---|---|---|---|---|---|
| 22565 | ENSMUSG00000078789 | 214.6997174 | 0.012576968 | 0.180858105 | 0.069540526 | 0.944559376 | 0.990079485 | Dph1 | diphthamide biosynthesis 1 [Source:MGI ! | 201.58 | 196.33 | 243.51 | 205.33 | 244.3 | 197.15 |
| 22566 | ENSMUSG00000078793 | 0 | NA | NA | NA | NA | NA | Gm5155 | predicted gene 5155 [Source:MGI Symbol | 0 | 0 | 0 | 0 | 0 | 0 |
| 22567 | ENSMUSG00000078794 | 8.866299509 | 0.283620855 | 0.785374459 | 0.361128188 | 0.718003622 | 0.9447719 | Dact3 | dishevelled-binding antagonist of beta-ca | 8.8341 | 2.0558 | 13.081 | 11.568 | 7.6343 | 10.025 |
| 22568 | ENSMUSG00000078795 | 0 | NA | NA | NA | NA | NA | Ceacam15 | carcinoembryonic antigen-related cell adh | 0 | 0 | 0 | 0 | 0 | 0 |
| 22569 | ENSMUSG00000078796 | 0.342639279 | -1.805976596 | 4.028547714 | -0.448294702 | 0.653940524 | NA | Zfp541 | zinc finger protein 541 [Source:MGI Symb | 0 | 2.0558 | 0 | 0 | 0 | 0 |
| 22570 | ENSMUSG00000078798 | 0 | NA | NA | NA | NA | NA | Sult2a1 | sulfotransferase family 2A, dehydroepian | 0 | 0 | 0 | 0 | 0 | 0 |
| 22571 | ENSMUSG00000078799 | 0 | NA | NA | NA | NA | NA | Sult2a5 | sulfotransferase family 2A, dehydroepian | 0 | 0 | 0 | 0 | 0 | 0 |
| 22572 | ENSMUSG00000078800 | 0 | NA | NA | NA | NA | NA | Bsph2 | binder of sperm protein homolog 2 [Sourc | 0 | 0 | 0 | 0 | 0 | 0 |
| 22573 | ENSMUSG00000078808 | 0 | NA | NA | NA | NA | NA | Vmn1r58 | vomeronasal 1 receptor 58 [Source:MGI S | 0 | 0 | 0 | 0 | 0 | 0 |
| 22574 | ENSMUSG00000078810 | 0.439020386 | -2.189747911 | 3.52148864 | -0.621824499 | 0.534057275 | NA | Gp6 | glycoprotein 6 (platelet) [Source:MGI Sym | 1.6062 | 1.0279 | 0 | 0 | 0 | 0 |
| 22575 | ENSMUSG00000078812 | 6622.17949 | 0.04109309 | 0.088088788 | 0.466496259 | 0.640860325 | 0.923932929 | Eif5a | eukaryotic translation initiation factor 5A | 6336.5 | 6526.3 | 6720.7 | 6325.6 | 6846.9 | 6977.1 |
| 22576 | ENSMUSG00000078813 | 160.0643071 | 0.344200268 | 0.194116591 | 1.773162543 | 0.076201777 | 0.403877794 | Leng1 | leukocyte receptor cluster (LRC) member | 129.3 | 152.13 | 141.88 | 158.09 | 196.31 | 182.67 |
| 22577 | ENSMUSG00000078815 | 0 | NA | NA | NA | NA | NA | Cacng6 | calcium channel, voltage-dependent, gam | 0 | 0 | 0 | 0 | 0 | 0 |
| 22578 | ENSMUSG00000078816 | 1.195418679 | -1.279722102 | 2.166649774 | -0.590645575 | 0.554757921 | NA | Prkcg | protein kinase C, gamma [Source:MGI Syn | 0 | 4.1117 | 1.0062 | 0.964 | 1.0906 | 0 |
| 22579 | ENSMUSG00000078817 | 1503.992684 | -0.505946327 | 0.136071234 | -3.718246045 | 0.000200611 | 0.006321645 | Nlrp12 | NLR family, pyrin domain containing 12 [S | 1810.2 | 1721.8 | 1763.9 | 1460.4 | 1001.2 | 1266.4 |
| 22580 | ENSMUSG00000078838 | 0 | NA | NA | NA | NA | NA | Gm17382 | predicted gene, 17382 [Source:MGI Symb | 0 | 0 | 0 | 0 | 0 | 0 |
| 22581 | ENSMUSG00000078840 | 0 | NA | NA | NA | NA | NA | Gm10999 | predicted gene 10999 [Source:MGI Symbo | 0 | 0 | 0 | 0 | 0 | 0 |
| 22582 | ENSMUSG00000078847 | 0 | NA | NA | NA | NA | NA | Gm6746 | predicted pseudogene 6746 [Source:MGI | 0 | 0 | 0 | 0 | 0 | 0 |
| 22583 | ENSMUSG00000078850 | 0 | NA | NA | NA | NA | NA | Gm2348 | predicted pseudogene 2348 [Source:MGI | 0 | 0 | 0 | 0 | 0 | 0 |
| 22584 | ENSMUSG00000078851 | 28.76207108 | 0.182216386 | 0.428150672 | 0.425589396 | 0.670407069 | 0.93116767 | H2aw | H2A.W histone [Source:MGI Symbol;Acc:I | 29.715 | 25.698 | 25.156 | 19.28 | 37.081 | 35.643 |
| 22585 | ENSMUSG00000078853 | 678.5461451 | -0.22005883 | 0.167902194 | -1.310637013 | 0.189980428 | 0.627369158 | Igtp | interferon gamma induced GTPase [Sourc | 824.79 | 746.27 | 619.85 | 783.71 | 545.31 | 551.35 |
| 22586 | ENSMUSG00000078861 | 104.6800162 | -0.24138429 | 0.25930555 | -0.930887482 | 0.35191177 | 0.792889192 | Zfp931 | zinc finger protein 931 [Source:MGI Symb | 119.66 | 86.345 | 133.83 | 82.902 | 86.159 | 119.18 |
| 22587 | ENSMUSG00000078862 | 72.47304782 | 0.009247434 | 0.280138895 | 0.033010176 | 0.973666473 | 0.996075612 | Gm14326 | predicted gene 14326 [Source:MGI Symbo | 55.414 | 85.317 | 76.475 | 74.226 | 65.437 | 77.969 |
| 22588 | ENSMUSG00000078864 | 14.94723444 | -1.324774987 | 0.646168961 | -2.050199046 | 0.040345011 | 0.274714427 | Gm14322 | predicted gene 14322 [Source:MGI Symbo | 22.487 | 27.754 | 14.087 | 15.424 | 4.3625 | 5.5692 |
| 22589 | ENSMUSG00000078865 | 0.160662974 | 1.126924175 | 4.080472857 | 0.276174898 | 0.782413736 | NA | Gm14406 | predicted gene 14406 [Source:MGI Symbo | 0 | 0 | 0 | 0.964 | 0 | 0 |
| 22590 | ENSMUSG00000078866 | 521.409691 | 0.234897218 | 0.216819068 | 1.083378969 | 0.278640247 | 0.731329747 | Zfp970 | zinc finger protein 970 [Source:MGI Symb | 449.74 | 539.66 | 448.79 | 802.99 | 387.17 | 500.12 |
| 22591 | ENSMUSG00000078867 | 25.57225165 | -0.095086346 | 0.432990135 | -0.219603956 | 0.82617961 | 0.96948417 | Gm14418 | predicted gene 14418 [Source:MGI Symbo | 23.29 | 29.81 | 26.162 | 20.244 | 30.537 | 23.391 |
| 22592 | ENSMUSG00000078868 | 1.700188771 | -1.181027616 | 1.709372862 | -0.690912815 | 0.489620332 | NA | Gm14412 | predicted gene 14412 [Source:MGI Symbo | 4.0155 | 1.0279 | 2.0125 | 0.964 | 2.1812 | 0 |
| 22593 | ENSMUSG00000078869 | 0 | NA | NA | NA | NA | NA | Gm14409 | predicted gene 14409 [Source:MGI Symbo | 0 | 0 | 0 | 0 | 0 | 0 |
| 22594 | ENSMUSG00000078870 | 44.39750248 | -0.161797812 | 0.328200357 | -0.492984875 | 0.622023274 | 0.918602439 | Gm14410 | predicted gene 14410 [Source:MGI Symbo | 48.989 | 43.173 | 48.3 | 36.631 | 43.625 | 45.668 |
| 22595 | ENSMUSG00000078872 | 0.82806045 | -0.59635427 | 2.600266913 | -0.229343483 | 0.81860196 | NA | Gm14401 | predicted gene 14401 [Source:MGI Symbo | 0 | 1.0279 | 2.0125 | 1.928 | 0 | 0 |
| 22596 | ENSMUSG00000078875 | 1.438400476 | -1.731166102 | 1.949644859 | -0.887939203 | 0.374573456 | NA | Gm14419 | predicted gene 14419 [Source:MGI Symbo | 1.6062 | 3.0838 | 2.0125 | 1.928 | 0 | 0 |
| 22597 | ENSMUSG00000078876 | 0 | NA | NA | NA | NA | NA | Gm14408 | predicted gene 14408 [Source:MGI Symbo | 0 | 0 | 0 | 0 | 0 | 0 |
| 22598 | ENSMUSG00000078877 | 46.62436933 | -0.49553458 | 0.337745175 | -1.467184781 | 0.142325796 | 0.55198239 | Gm14295 | predicted gene 14295 [Source:MGI Symbo | 40.958 | 57.563 | 65.406 | 37.595 | 40.353 | 37.871 |
| 22599 | ENSMUSG00000078878 | 47.05445761 | 0.196141944 | 0.339631653 | 0.577513732 | 0.563592465 | 0.900821918 | Gm14305 | predicted gene 14305 [Source:MGI Symbo | 40.958 | 47.284 | 43.269 | 41.451 | 42.534 | 66.831 |
| 22600 | ENSMUSG00000078879 | 0.68166635 | -2.79976525 | 3.012029182 | -0.92952793 | 0.352615558 | NA | Zfp973 | zinc finger protein 973 [Source:MGI Symb | 0 | 3.0838 | 1.0062 | 0 | 0 | 0 |
| 22601 | ENSMUSG00000078880 | 0 | NA | NA | NA | NA | NA | Gm14308 | predicted gene 14308 [Source:MGI Symbo | 0 | 0 | 0 | 0 | 0 | 0 |
| 22602 | ENSMUSG00000078881 | 0 | NA | NA | NA | NA | NA | Gm14434 | predicted gene 14434 [Source:MGI Symbo | 0 | 0 | 0 | 0 | 0 | 0 |
| 22603 | ENSMUSG00000078886 | 0.936952612 | -3.291460614 | 3.015666682 | -1.091453719 | 0.275073286 | NA | Gm2026 | predicted gene 2026 [Source:MGI Symbol | 5.6217 | 0 | 0 | 0 | 0 | 0 |
| 22604 | ENSMUSG00000078887 | 0.167707431 | -0.796629176 | 4.080472857 | -0.195229623 | 0.84521319 | NA | Gm6710 | predicted gene 6710 [Source:MGI Symbol | 0 | 0 | 1.0062 | 0 | 0 | 0 |
| 22605 | ENSMUSG00000078889 | 2.606878727 | -1.458229691 | 1.381630879 | -1.055440866 | 0.291223741 | NA | Gm14288 | predicted gene 14288 [Source:MGI Symbo | 2.4093 | 2.0558 | 7.0437 | 1.928 | 1.0906 | 1.1138 |
| 22606 | ENSMUSG00000078891 | 0.133850373 | -0.796629176 | 4.080472857 | -0.195229623 | 0.84521319 | NA | Gm11008 | predicted gene 11008 [Source:MGI Symbo | 0.8031 | 0 | 0 | 0 | 0 | 0 |
| 22607 | ENSMUSG00000078894 | 2.68364502 | -0.682160753 | 1.28719195 | -0.529960394 | 0.596139391 | NA | 2210418O1C | RIKEN cDNA 2210418O10 gene [Source:M | 4.8186 | 3.0838 | 2.0125 | 2.8919 | 2.1812 | 1.1138 |
| 22608 | ENSMUSG00000078895 | 0 | NA | NA | NA | NA | NA | Gm11009 | predicted gene 11009 [Source:MGI Symbo | 0 | 0 | 0 | 0 | 0 | 0 |
| 22609 | ENSMUSG00000078896 | 0.833502073 | 0.798571965 | 2.398861853 | 0.332896188 | 0.739212647 | NA | Zfp965 | zinc finger protein 965 [Source:MGI Symb | 0.8031 | 0 | 1.0062 | 0.964 | 0 | 2.2277 |
| 22610 | ENSMUSG00000078897 | 6.314570054 | 1.090664552 | 0.865145268 | 1.260672158 | 0.207426989 | 0.652364421 | Gm4724 | predicted gene 4724 [Source:MGI Symbol | 4.0155 | 4.1117 | 4.025 | 12.532 | 7.6343 | 5.5692 |
| 22611 | ENSMUSG00000078898 | 0 | NA | NA | NA | NA | NA | Zfp968 | zinc finger protein 968 [Source:MGI Symb | 0 | 0 | 0 | 0 | 0 | 0 |
| 22612 | ENSMUSG00000078899 | 0 | NA | NA | NA | NA | NA | Gm4631 | predicted gene 4631 [Source:MGI Symbol | 0 | 0 | 0 | 0 | 0 | 0 |
| 22613 | ENSMUSG00000078901 | 0.342432689 | 1.990653086 | 3.900281263 | 0.51038706 | 0.609780321 | NA | Gm14440 | predicted gene 14440 [Source:MGI Symbo | 0 | 0 | 0 | 0.964 | 1.0906 | 0 |
| 22614 | ENSMUSG00000078902 | 4.937279048 | 0.590741699 | 1.016565419 | 0.581115281 | 0.56116276 | NA | Gm14443 | predicted gene 14443 [Source:MGI Symbo | 0.8031 | 5.1396 | 6.0375 | 7.7118 | 4.3625 | 5.5692 |
| 22615 | ENSMUSG00000078903 | 0.497389803 | 1.101799453 | 3.439268845 | 0.320358629 | 0.748696483 | NA | Gm14391 | predicted gene 14391 [Source:MGI Symbo | 0.8031 | 0 | 0 | 0 | 2.1812 | 0 |
| 22616 | ENSMUSG00000078905 | 1.766564291 | -0.162897681 | 1.663770919 | -0.09790872 | 0.922004776 | NA | Gm14393 | predicted gene 14393 [Source:MGI Symbo | 1.6062 | 1.0279 | 3.0187 | 3.8559 | 1.0906 | 0 |
| 22617 | ENSMUSG00000078906 | 0 | NA | NA | NA | NA | NA | Gm14444 | predicted gene 14444 [Source:MGI Symbo | 0 | 0 | 0 | 0 | 0 | 0 |
| 22618 | ENSMUSG00000078907 | 3.528069651 | -0.931499208 | 1.323504728 | -0.703812528 | 0.481549531 | NA | Fam186b | family with sequence similarity 186, meml | 0.8031 | 4.1117 | 9.0562 | 3.8559 | 0 | 3.3415 |
| 22619 | ENSMUSG00000078908 | 220.2203702 | -0.04831501 | 0.186002488 | -0.259754645 | 0.795053039 | 0.962189806 | Mon1b | MON1 homolog B, secretory traffciking as | 197.56 | 259.04 | 215.34 | 193.76 | 206.13 | 249.5 |
| 22620 | ENSMUSG00000078912 | 0 | NA | NA | NA | NA | NA | Gm11011 | predicted gene 11011 [Source:MGI Symbo | 0 | 0 | 0 | 0 | 0 | 0 |

| | | | | | | | | | | | | | | | |
|---|---|---|---|---|---|---|---|---|---|---|---|---|---|---|---|
| 22621 | ENSMUSG00000078915 | 0.335414862 | -1.779047242 | 4.030804179 | -0.441362855 | 0.658950329 | NA | Hsp25-ps1 | heat shock protein 25, pseudogene 1 [Sou | 0 | 0 | 2.0125 | 0 | 0 | 0 |
| 22622 | ENSMUSG00000078917 | 0 | NA | NA | NA | NA | NA | Gm11281 | predicted gene 11281 [Source:MGI Symbo | 0 | 0 | 0 | 0 | 0 | 0 |
| 22623 | ENSMUSG00000078919 | 92.07824172 | 0.443777414 | 0.276542621 | 1.604734244 | 0.108552302 | 0.485347914 | Dpm1 | dolichol-phosphate (beta-D) mannosyltra | 95.569 | 70.926 | 67.418 | 136.88 | 99.246 | 82.424 |
| 22624 | ENSMUSG00000078920 | 342.396211 | 0.121304085 | 0.144067257 | 0.841996213 | 0.399790073 | 0.82676382 | Ifi47 | interferon gamma inducible protein 47 [S | 344.53 | 344.35 | 294.83 | 358.6 | 351.18 | 360.89 |
| 22625 | ENSMUSG00000078921 | 25.41962863 | -0.992850615 | 0.453502417 | -2.189295092 | 0.028575399 | 0.221616391 | Tgtp2 | T cell specific GTPase 2 [Source:MGI Symb | 27.305 | 33.921 | 40.25 | 13.496 | 15.269 | 22.277 |
| 22626 | ENSMUSG00000078922 | 46.92357245 | -0.398638299 | 0.340508003 | -1.170716389 | 0.241712795 | 0.691791181 | Tgtp1 | T cell specific GTPase 1 [Source:MGI Symb | 52.202 | 66.815 | 41.256 | 46.271 | 34.9 | 40.098 |
| 22627 | ENSMUSG00000078923 | 8.265579298 | -0.210077004 | 0.755794293 | -0.277955266 | 0.781046699 | 0.959993512 | Ube2v1 | ubiquitin-conjugating enzyme E2 variant 1 | 11.243 | 10.279 | 5.0312 | 8.6758 | 5.4531 | 8.9108 |
| 22628 | ENSMUSG00000078924 | 0 | NA | NA | NA | NA | NA | Gm12169 | predicted gene 12169 [Source:MGI Symbo | 0 | 0 | 0 | 0 | 0 | 0 |
| 22629 | ENSMUSG00000078925 | 0 | NA | NA | NA | NA | NA | D030018L15 | RIKEN cDNA D030018L15 gene [Source:M | 0 | 0 | 0 | 0 | 0 | 0 |
| 22630 | ENSMUSG00000078926 | 0 | NA | NA | NA | NA | NA | Cdc20b | cell division cycle 20B [Source:MGI Symbo | 0 | 0 | 0 | 0 | 0 | 0 |
| 22631 | ENSMUSG00000078931 | 6.986111146 | -0.106626187 | 0.832920946 | -0.128014775 | 0.89813728 | 0.982798755 | Pdf | peptide deformylase (mitochondrial) [Sou | 6.4248 | 10.279 | 5.0312 | 4.8199 | 10.906 | 4.4554 |
| 22632 | ENSMUSG00000078932 | 0 | NA | NA | NA | NA | NA | CN725425 | cDNA sequence CN725425 [Source:MGI S | 0 | 0 | 0 | 0 | 0 | 0 |
| 22633 | ENSMUSG00000078934 | 0 | NA | NA | NA | NA | NA | Pate13 | prostate and testis expressed 13 [Source: | 0 | 0 | 0 | 0 | 0 | 0 |
| 22634 | ENSMUSG00000078935 | 0 | NA | NA | NA | NA | NA | 1700025C18 | RIKEN cDNA 1700025C18 gene [Source:M | 0 | 0 | 0 | 0 | 0 | 0 |
| 22635 | ENSMUSG00000078937 | 5.096779299 | 0.874004159 | 1.045996362 | 0.835570936 | 0.403396325 | 0.828953151 | Cpt1b | carnitine palmitoyltransferase 1b, muscle | 0.8031 | 4.1117 | 6.0375 | 8.6758 | 8.7249 | 2.2277 |
| 22636 | ENSMUSG00000078938 | 1.038523786 | -3.429083216 | 2.553540547 | -1.342874003 | 0.179312776 | NA | Syce3 | synaptonemal complex central element p | 3.2124 | 0 | 3.0187 | 0 | 0 | 0 |
| 22637 | ENSMUSG00000078940 | 0 | NA | NA | NA | NA | NA | Wfdc11 | WAP four-disulfide core domain 11 [Sour | 0 | 0 | 0 | 0 | 0 | 0 |
| 22638 | ENSMUSG00000078941 | 127.1660775 | -0.021258789 | 0.211735444 | -0.100402599 | 0.920024706 | 0.986685804 | Ak6 | adenylate kinase 6 [Source:MGI Symbol;A | 134.12 | 124.38 | 125.78 | 131.1 | 140.69 | 106.93 |
| 22639 | ENSMUSG00000078942 | 14.28780962 | -2.030682077 | 0.620766685 | -3.271248482 | 0.001070738 | 0.022061471 | Naip6 | NLR family, apoptosis inhibitory protein 6 | 20.078 | 24.67 | 24.15 | 5.7839 | 4.3625 | 6.6831 |
| 22640 | ENSMUSG00000078945 | 89.69717053 | -0.552132875 | 0.273312848 | -2.020149727 | 0.043367859 | 0.287081334 | Naip2 | NLR family, apoptosis inhibitory protein 2 | 94.766 | 119.24 | 106.66 | 97.362 | 58.893 | 61.261 |
| 22641 | ENSMUSG00000078949 | 46.80253946 | 0.517307093 | 0.388229633 | 1.332477092 | 0.182703464 | 0.616714797 | R3hdml | R3H domain containing-like [Source:MGI | 38.549 | 32.893 | 44.275 | 76.154 | 59.984 | 28.96 |
| 22642 | ENSMUSG00000078952 | 0 | NA | NA | NA | NA | NA | Lncenc1 | long non-coding RNA, embryonic stem cel | 0 | 0 | 0 | 0 | 0 | 0 |
| 22643 | ENSMUSG00000078953 | 0 | NA | NA | NA | NA | NA | Gm11020 | predicted gene 11020 [Source:MGI Symbo | 0 | 0 | 0 | 0 | 0 | 0 |
| 22644 | ENSMUSG00000078954 | 1.214277311 | 1.568214921 | 2.15339429 | 0.728252568 | 0.466458992 | NA | Arhgap8 | Rho GTPase activating protein 8 [Source:N | 0.8031 | 0 | 1.0062 | 0 | 4.3625 | 1.1138 |
| 22645 | ENSMUSG00000078955 | 0 | NA | NA | NA | NA | NA | Gm14222 | predicted gene 14222 [Source:MGI Symbo | 0 | 0 | 0 | 0 | 0 | 0 |
| 22646 | ENSMUSG00000078956 | 8.081774704 | -1.204929877 | 0.770466713 | -1.563896085 | 0.117841974 | 0.502838264 | Gm14221 | predicted gene 14221 [Source:MGI Symbo | 10.44 | 13.363 | 10.062 | 5.7839 | 3.2719 | 5.5692 |
| 22647 | ENSMUSG00000078957 | 0 | NA | NA | NA | NA | NA | 1700060C20 | RIKEN cDNA 1700060C20 gene [Source:M | 0 | 0 | 0 | 0 | 0 | 0 |
| 22648 | ENSMUSG00000078958 | 0 | NA | NA | NA | NA | NA | Atp6ap1l | ATPase, H+ transporting, lysosomal acces | 0 | 0 | 0 | 0 | 0 | 0 |
| 22649 | ENSMUSG00000078962 | 0 | NA | NA | NA | NA | NA | 4932414J04I | RIKEN cDNA 4932414J04 gene [Source:M | 0 | 0 | 0 | 0 | 0 | 0 |
| 22650 | ENSMUSG00000078963 | 33.84068883 | 0.339870458 | 0.459970642 | 0.738895979 | 0.459970164 | 0.859753653 | Hsbp1l1 | heat shock factor binding protein 1-like 1 | 38.549 | 19.53 | 31.194 | 31.811 | 21.812 | 60.148 |
| 22651 | ENSMUSG00000078964 | 1661.502457 | -0.222331928 | 0.138866644 | -1.601046312 | 0.109366662 | 0.486233674 | Ces1b | carboxylesterase 1B [Source:MGI Symbol; | 1459.2 | 1924.3 | 1986.3 | 1727.4 | 1452.7 | 1419 |
| 22652 | ENSMUSG00000078965 | 1.099373405 | -1.223504443 | 2.147555505 | -0.56971959 | 0.568867901 | NA | Gm12033 | predicted gene 12033 [Source:MGI Symbo | 1.6062 | 2.0558 | 1.0062 | 1.928 | 0 | 0 |
| 22653 | ENSMUSG00000078967 | 0.36353943 | 2.065989254 | 4.034137301 | 0.512126658 | 0.608562368 | NA | Gapdh-ps16 | glyceraldehyde-3-phosphate dehydrogen | 0 | 0 | 0 | 0 | 2.1812 | 0 |
| 22654 | ENSMUSG00000078970 | 204.2427798 | 0.167275836 | 0.174733817 | 0.957318044 | 0.338406751 | 0.780780576 | Wdr92 | WD repeat domain 92 [Source:MGI Symbo | 184.71 | 193.25 | 199.24 | 201.47 | 202.86 | 243.93 |
| 22655 | ENSMUSG00000078972 | 0 | NA | NA | NA | NA | NA | Gm15557 | predicted gene 15557 [Source:MGI Symbo | 0 | 0 | 0 | 0 | 0 | 0 |
| 22656 | ENSMUSG00000078974 | 722.3036673 | 0.458571786 | 0.152135908 | 3.014224528 | 0.002576371 | 0.042202293 | Sec61g | SEC61, gamma subunit [Source:MGI Symb | 659.35 | 653.76 | 511.17 | 708.52 | 839.78 | 961.25 |
| 22657 | ENSMUSG00000078982 | 0.181769715 | 1.126924175 | 4.080472857 | 0.276174898 | 0.782413736 | NA | Gm11025 | predicted gene 11025 [Source:MGI Symbo | 0 | 0 | 0 | 0 | 1.0906 | 0 |
| 22658 | ENSMUSG00000078984 | 0 | NA | NA | NA | NA | NA | Gm11027 | predicted gene 11027 [Source:MGI Symbo | 0 | 0 | 0 | 0 | 0 | 0 |
| 22659 | ENSMUSG00000078994 | 21.45516754 | -0.452610856 | 0.541921551 | -0.835196265 | 0.403607212 | 0.828953151 | Zfp429 | zinc finger protein 429 [Source:MGI Symb | 21.684 | 32.893 | 20.125 | 30.847 | 9.8156 | 13.366 |
| 22660 | ENSMUSG00000078995 | 6.565852282 | 0.552244816 | 0.845508996 | 0.653150728 | 0.513659112 | 0.885004045 | Zfp456 | zinc finger protein 456 [Source:MGI Symb | 4.8186 | 3.0838 | 8.05 | 5.7839 | 7.6343 | 10.025 |
| 22661 | ENSMUSG00000078998 | 0 | NA | NA | NA | NA | NA | Bpifa6 | BPI fold containing family A, member 6 [S | 0 | 0 | 0 | 0 | 0 | 0 |
| 22662 | ENSMUSG00000079000 | 0 | NA | NA | NA | NA | NA | Dnmt3bos | DNA methyltransferase 3B, opposite stra | 0 | 0 | 0 | 0 | 0 | 0 |
| 22663 | ENSMUSG00000079001 | 0 | NA | NA | NA | NA | NA | 4930404H24 | RIKEN cDNA 4930404H24 gene [Source:M | 0 | 0 | 0 | 0 | 0 | 0 |
| 22664 | ENSMUSG00000079003 | 291.8164014 | 0.145837875 | 0.193748174 | 0.752718704 | 0.451618968 | 0.856234992 | Samd1 | sterile alpha motif domain containing 1 [S | 255.39 | 224.09 | 352.19 | 334.5 | 262.84 | 321.9 |
| 22665 | ENSMUSG00000079005 | 0 | NA | NA | NA | NA | NA | Gm14147 | predicted gene 14147 [Source:MGI Symbo | 0 | 0 | 0 | 0 | 0 | 0 |
| 22666 | ENSMUSG00000079006 | 0 | NA | NA | NA | NA | NA | Gm14151 | predicted gene 14151 [Source:MGI Symbo | 0 | 0 | 0 | 0 | 0 | 0 |
| 22667 | ENSMUSG00000079007 | 0 | NA | NA | NA | NA | NA | Tcl1b4 | T cell leukemia/lymphoma 1B, 4 [Source:N | 0 | 0 | 0 | 0 | 0 | 0 |
| 22668 | ENSMUSG00000079008 | 0.167707431 | -0.796629176 | 4.080472857 | -0.195229623 | 0.84521319 | NA | Gm14124 | predicted gene 14124 [Source:MGI Symbo | 0 | 0 | 1.0062 | 0 | 0 | 0 |
| 22669 | ENSMUSG00000079009 | 0 | NA | NA | NA | NA | NA | Gm14139 | predicted gene 14139 [Source:MGI Symbo | 0 | 0 | 0 | 0 | 0 | 0 |
| 22670 | ENSMUSG00000079010 | 0 | NA | NA | NA | NA | NA | Gm11032 | predicted gene 11032 [Source:MGI Symbo | 0 | 0 | 0 | 0 | 0 | 0 |
| 22671 | ENSMUSG00000079011 | 0 | NA | NA | NA | NA | NA | Gm11033 | predicted gene 11033 [Source:MGI Symbo | 0 | 0 | 0 | 0 | 0 | 0 |
| 22672 | ENSMUSG00000079012 | 6340.673319 | -0.538817995 | 0.11992762 | -4.492859899 | 7.03E-06 | 0.000404554 | Serpina3m | serine (or cysteine) peptidase inhibitor, cl | 6352.5 | 7895.4 | 8287.4 | 5623.8 | 4786.7 | 5098.1 |
| 22673 | ENSMUSG00000079013 | 0 | NA | NA | NA | NA | NA | Serpina3j | serine (or cysteine) peptidase inhibitor, cl | 0 | 0 | 0 | 0 | 0 | 0 |
| 22674 | ENSMUSG00000079014 | 0.658052777 | 1.6393978 | 2.811180186 | 0.583170658 | 0.559778421 | NA | Serpina3i | serine (or cysteine) peptidase inhibitor, cl | 0.8031 | 0 | 0 | 0.964 | 2.1812 | 0 |
| 22675 | ENSMUSG00000079015 | 25895.657 | 0.371484399 | 0.152130029 | 2.441887392 | 0.014610703 | 0.140429803 | Serpina1c | serine (or cysteine) peptidase inhibitor, cl | 22478 | 17189 | 28073 | 31947 | 26307 | 29380 |
| 22676 | ENSMUSG00000079016 | 0 | NA | NA | NA | NA | NA | Gm11034 | predicted gene 11034 [Source:MGI Symbo | 0 | 0 | 0 | 0 | 0 | 0 |

| | | | | | | | | | | | | | | | |
|---|---|---|---|---|---|---|---|---|---|---|---|---|---|---|---|
| 22677 | ENSMUSG00000079017 | 62.89122635 | -1.296123565 | 0.299328685 | -4.330101419 | 1.49E-05 | 0.000730754 | Ifi27l2a | interferon, alpha-inducible protein 27 like | 86.735 | 96.624 | 84.525 | 28.919 | 35.99 | 44.554 |
| 22678 | ENSMUSG00000079018 | 4.628774484 | -1.182892577 | 1.014217942 | -1.166310049 | 0.243489112 | NA | Ly6c1 | lymphocyte antigen 6 complex, locus C1 [ | 8.031 | 5.1396 | 6.0375 | 1.928 | 2.1812 | 4.4554 |
| 22679 | ENSMUSG00000079019 | 7.388606992 | -0.60252256 | 0.854254766 | -0.705319518 | 0.480611417 | 0.870278888 | Insl3 | insulin-like 3 [Source:MGI Symbol;Acc:MG | 5.6217 | 5.1396 | 16.1 | 8.6758 | 5.4531 | 3.3415 |
| 22680 | ENSMUSG00000079020 | 207.4284569 | 0.209233235 | 0.201133065 | 1.040272693 | 0.298213227 | 0.748976579 | Slc45a4 | solute carrier family 45, member 4 [Sourc | 216.03 | 191.19 | 169.05 | 177.37 | 222.49 | 268.44 |
| 22681 | ENSMUSG00000079022 | 0.556921942 | 2.679043159 | 3.638243063 | 0.736356289 | 0.461513899 | NA | Col22a1 | collagen, type XXII, alpha 1 [Source:MGI S | 0 | 0 | 0 | 0 | 0 | 3.3415 |
| 22682 | ENSMUSG00000079024 | 0 | NA | NA | NA | NA | NA | Gm21961 | predicted gene, 21961 [Source:MGI Symb | 0 | 0 | 0 | 0 | 0 | 0 |
| 22683 | ENSMUSG00000079025 | 0 | NA | NA | NA | NA | NA | Gsdmc | gasdermin C [Source:MGI Symbol;Acc:MG | 0 | 0 | 0 | 0 | 0 | 0 |
| 22684 | ENSMUSG00000079029 | 0 | NA | NA | NA | NA | NA | Eif1ad7 | eukaryotic translation initiation factor 1A | 0 | 0 | 0 | 0 | 0 | 0 |
| 22685 | ENSMUSG00000079031 | 0 | NA | NA | NA | NA | NA | Eif1ad2 | eukaryotic translation initiation factor 1A | 0 | 0 | 0 | 0 | 0 | 0 |
| 22686 | ENSMUSG00000079033 | 0.535401493 | -2.48596965 | 3.684117979 | -0.674780141 | 0.499815459 | NA | Mef2b | myocyte enhancer factor 2B [Source:MGI | 3.2124 | 0 | 0 | 0 | 0 | 0 |
| 22687 | ENSMUSG00000079034 | 0 | NA | NA | NA | NA | NA | Eif1ad8 | eukaryotic translation initiation factor 1A | 0 | 0 | 0 | 0 | 0 | 0 |
| 22688 | ENSMUSG00000079036 | 122.4611298 | 0.189339464 | 0.218018989 | 0.868454005 | 0.385145842 | 0.816437825 | Alkbh1 | alkB homolog 1, histone H2A dioxygenase | 110.03 | 103.82 | 129.81 | 147.49 | 118.88 | 124.75 |
| 22689 | ENSMUSG00000079037 | 895.7338359 | -0.207538932 | 0.155577993 | -1.333986438 | 0.182208301 | 0.615722956 | Prnp | prion protein [Source:MGI Symbol;Acc:M | 987.01 | 964.19 | 928.76 | 792.39 | 1041.5 | 660.51 |
| 22690 | ENSMUSG00000079038 | 1.142077704 | 1.613415074 | 2.128062572 | 0.758161482 | 0.448354317 | NA | D130040H23 | RIKEN cDNA D130040H23 gene [Source:M | 1.6062 | 0 | 0 | 1.928 | 1.0906 | 2.2277 |
| 22691 | ENSMUSG00000079039 | 0 | NA | NA | NA | NA | NA | Gm11037 | predicted gene 11037 [Source:MGI Symbo | 0 | 0 | 0 | 0 | 0 | 0 |
| 22692 | ENSMUSG00000079042 | 0 | NA | NA | NA | NA | NA | Apela | apelin receptor early endogenous ligand [ | 0 | 0 | 0 | 0 | 0 | 0 |
| 22693 | ENSMUSG00000079043 | 72.10130481 | -0.073588804 | 0.273232986 | -0.269326208 | 0.787678666 | 0.960870081 | Fastkd5 | FAST kinase domains 5 [Source:MGI Symb | 89.144 | 70.926 | 61.381 | 72.298 | 69.8 | 69.058 |
| 22694 | ENSMUSG00000079045 | 143.1679851 | 0.114553829 | 0.255665313 | 0.448061678 | 0.654108684 | 0.925797231 | Prox1os | prospero homeobox 1, opposite strand [S | 97.175 | 179.89 | 135.84 | 116.64 | 167.96 | 161.51 |
| 22695 | ENSMUSG00000079048 | 0 | NA | NA | NA | NA | NA | 4933413L06 | RIKEN cDNA 4933413L06 gene [Source:M | 0 | 0 | 0 | 0 | 0 | 0 |
| 22696 | ENSMUSG00000079049 | 0.321325949 | 1.911575114 | 4.048267893 | 0.472195804 | 0.636787034 | NA | Serpinb1c | serine (or cysteine) peptidase inhibitor, cl | 0 | 0 | 0 | 1.928 | 0 | 0 |
| 22697 | ENSMUSG00000079051 | 3.800662428 | 0.099446788 | 1.093873673 | 0.090912498 | 0.927562119 | NA | Vinac1 | vinculin/alpha-catenin family member 1 [ | 4.8186 | 3.0838 | 3.0187 | 1.928 | 3.2719 | 6.6831 |
| 22698 | ENSMUSG00000079053 | 0 | NA | NA | NA | NA | NA | Gm14010 | predicted gene 14010 [Source:MGI Symbo | 0 | 0 | 0 | 0 | 0 | 0 |
| 22699 | ENSMUSG00000079055 | 2.045897836 | -0.370160517 | 1.646460378 | -0.224822001 | 0.82211775 | NA | Slc8a3 | solute carrier family 8 (sodium/calcium ex | 0.8031 | 5.1396 | 1.0062 | 0.964 | 4.3625 | 0 |
| 22700 | ENSMUSG00000079056 | 3.779742288 | -2.137008899 | 1.269866691 | -1.682860819 | 0.092402038 | NA | Kcnip3 | Kv channel interacting protein 3, calsenilir | 3.2124 | 10.279 | 5.0312 | 1.928 | 0 | 2.2277 |
| 22701 | ENSMUSG00000079057 | 8756.966447 | 0.226450748 | 0.143376914 | 1.579408718 | 0.114242339 | 0.496868018 | Cyp4v3 | cytochrome P450, family 4, subfamily v, p | 7385.3 | 7412.3 | 9415.4 | 7556.6 | 10878 | 9894.3 |
| 22702 | ENSMUSG00000079058 | 0 | NA | NA | NA | NA | NA | Adam34 | a disintegrin and metallopeptidase domai | 0 | 0 | 0 | 0 | 0 | 0 |
| 22703 | ENSMUSG00000079061 | 0 | NA | NA | NA | NA | NA | Gm11042 | predicted gene 11042 [Source:MGI Symbo | 0 | 0 | 0 | 0 | 0 | 0 |
| 22704 | ENSMUSG00000079064 | 0 | NA | NA | NA | NA | NA | Vmn1r-ps12 | vomeronasal 1 receptor, pseudogene 123 | 0 | 0 | 0 | 0 | 0 | 0 |
| 22705 | ENSMUSG00000079065 | 151.2752056 | 0.029328721 | 0.201130975 | 0.145819018 | 0.884064262 | 0.981769403 | NA | NA | 152.59 | 149.05 | 147.92 | 180.26 | 134.15 | 143.69 |
| 22706 | ENSMUSG00000079067 | 2.197733559 | 0.862871338 | 1.407361389 | 0.61311284 | 0.539801731 | NA | Hmgn2-ps1 | high mobility group nucleosomal binding | 1.6062 | 2.0558 | 1.0062 | 1.928 | 4.3625 | 2.2277 |
| 22707 | ENSMUSG00000079069 | 0.335414862 | -1.779047242 | 4.030804179 | -0.441362855 | 0.658950329 | NA | 8430423G03 | RIKEN cDNA 8430423G03 gene [Source:M | 0 | 0 | 2.0125 | 0 | 0 | 0 |
| 22708 | ENSMUSG00000079070 | 0 | NA | NA | NA | NA | NA | Gm3985 | predicted gene 3985 [Source:MGI Symbol | 0 | 0 | 0 | 0 | 0 | 0 |
| 22709 | ENSMUSG00000079071 | 0.301557804 | -1.64335916 | 4.042814367 | -0.406488899 | 0.684383412 | NA | Slc28a2b | solute carrier family 28 member 2b [Sourc | 0.8031 | 0 | 1.0062 | 0 | 0 | 0 |
| 22710 | ENSMUSG00000079076 | 1.988277961 | 0.141323054 | 1.564345252 | 0.090340067 | 0.928016982 | NA | Gm3086 | predicted gene 3086 [Source:MGI Symbol | 1.6062 | 3.0838 | 1.0062 | 2.8919 | 0 | 3.3415 |
| 22711 | ENSMUSG00000079083 | 76.24455235 | 0.123535167 | 0.258658304 | 0.477599847 | 0.63293504 | 0.922107984 | Jrkl | Jrk-like [Source:MGI Symbol;Acc:MGI:192 | 76.295 | 72.982 | 69.431 | 73.262 | 74.162 | 91.335 |
| 22712 | ENSMUSG00000079084 | 107.0454931 | -0.129584982 | 0.22697607 | -0.570919135 | 0.568054459 | 0.903352921 | Ccdc82 | coiled-coil domain containing 82 [Source: | 106.01 | 121.29 | 108.67 | 116.64 | 90.521 | 99.132 |
| 22713 | ENSMUSG00000079091 | 0 | NA | NA | NA | NA | NA | Gm3404 | predicted gene 3404 [Source:MGI Symbol | 0 | 0 | 0 | 0 | 0 | 0 |
| 22714 | ENSMUSG00000079092 | 0 | NA | NA | NA | NA | NA | Prl2c2 | prolactin family 2, subfamily c, member 2 | 0 | 0 | 0 | 0 | 0 | 0 |
| 22715 | ENSMUSG00000079093 | 0 | NA | NA | NA | NA | NA | 4930449I24 | RIKEN cDNA 4930449I24 gene [Source:M | 0 | 0 | 0 | 0 | 0 | 0 |
| 22716 | ENSMUSG00000079101 | 0.6621303 | -1.329905799 | 2.807279375 | -0.473734752 | 0.635689068 | NA | Esd-ps | esterase D/formylglutathione hydrolase, | 0.8031 | 2.0558 | 0 | 0 | 0 | 1.1138 |
| 22717 | ENSMUSG00000079103 | 1.584685324 | 1.929790969 | 1.830011038 | 1.054524224 | 0.291642976 | NA | Tgm7 | transglutaminase 7 [Source:MGI Symbol; | 0 | 1.0279 | 1.0062 | 1.928 | 1.0906 | 4.4554 |
| 22718 | ENSMUSG00000079104 | 477.7584303 | -0.067438445 | 0.13122661 | -0.513908311 | 0.607316096 | 0.916376001 | Prps1l3 | phosphoribosyl pyrophosphate synthetas | 514.79 | 483.12 | 468.91 | 506.09 | 454.79 | 438.85 |
| 22719 | ENSMUSG00000079105 | 6.029008813 | -0.209020515 | 0.874684026 | -0.238966883 | 0.811131263 | 0.966452061 | C7 | complement component 7 [Source:MGI S | 7.2279 | 3.0838 | 9.0562 | 5.7839 | 5.4531 | 5.5692 |
| 22720 | ENSMUSG00000079108 | 129.0229009 | -0.168854941 | 0.225801412 | -0.747802857 | 0.454579078 | 0.857337687 | Srp54c | signal recognition particle 54C [Source:M | 146.97 | 132.6 | 129.81 | 119.53 | 95.974 | 149.26 |
| 22721 | ENSMUSG00000079109 | 38.45323241 | 0.063157001 | 0.354291781 | 0.178262675 | 0.85851669 | 0.975439379 | Pms2 | PMS1 homolog2, mismatch repair system | 37.746 | 37.005 | 38.237 | 47.235 | 37.081 | 33.415 |
| 22722 | ENSMUSG00000079110 | 25.17504634 | 0.567836498 | 0.478833296 | 1.185875131 | 0.235671615 | 0.683802256 | Capn3 | calpain 3 [Source:MGI Symbol;Acc:MGI:1 | 24.093 | 13.363 | 23.144 | 21.208 | 43.625 | 25.618 |
| 22723 | ENSMUSG00000079111 | 1435.961926 | 0.064601528 | 0.097752506 | 0.660868248 | 0.508696811 | 0.882793133 | Kdelr2 | KDEL (Lys-Asp-Glu-Leu) endoplasmic retic | 1411.9 | 1341.4 | 1458 | 1470.1 | 1493.1 | 1441.3 |
| 22724 | ENSMUSG00000079112 | 0 | NA | NA | NA | NA | NA | Fam90a1a | family with sequence similarity 90, memb | 0 | 0 | 0 | 0 | 0 | 0 |
| 22725 | ENSMUSG00000079113 | 0 | NA | NA | NA | NA | NA | Defa43 | defensin, alpha, 43 [Source:MGI Symbol; | 0 | 0 | 0 | 0 | 0 | 0 |
| 22726 | ENSMUSG00000079114 | 0 | NA | NA | NA | NA | NA | Defa42 | defensin, alpha, 42 [Source:MGI Symbol; | 0 | 0 | 0 | 0 | 0 | 0 |
| 22727 | ENSMUSG00000079116 | 0 | NA | NA | NA | NA | NA | Defa41 | defensin, alpha, 41 [Source:MGI Symbol; | 0 | 0 | 0 | 0 | 0 | 0 |
| 22728 | ENSMUSG00000079120 | 0 | NA | NA | NA | NA | NA | AY761185 | cDNA sequence AY761185 [Source:MGI S | 0 | 0 | 0 | 0 | 0 | 0 |
| 22729 | ENSMUSG00000079137 | 0.492645588 | 0.9565034 | 3.447648281 | 0.277436479 | 0.781444974 | NA | Rpl27-ps1 | ribosomal protein L27, pseudogene 1 [Sou | 0 | 1.0279 | 0 | 1.928 | 0 | 0 |
| 22730 | ENSMUSG00000079138 | 0 | NA | NA | NA | NA | NA | Gm8818 | predicted pseudogene 8818 [Source:MGI | 0 | 0 | 0 | 0 | 0 | 0 |
| 22731 | ENSMUSG00000079139 | 69.04213928 | -0.095720981 | 0.286485905 | -0.334121084 | 0.738288191 | 0.948137969 | Gm4204 | predicted gene 4204 [Source:MGI Symbol | 65.051 | 68.87 | 80.5 | 82.902 | 53.44 | 63.489 |
| 22732 | ENSMUSG00000079140 | 0 | NA | NA | NA | NA | NA | Gm6374 | predicted gene 6374 [Source:MGI Symbol | 0 | 0 | 0 | 0 | 0 | 0 |

| | | | | | | | | | | | | | | | |
|---|---|---|---|---|---|---|---|---|---|---|---|---|---|---|---|
| 22733 | ENSMUSG00000079143 | 0.17131964 | -0.796629176 | 4.080472857 | -0.195229623 | 0.84521319 | NA | Atxn7l1os1 | ataxin 7-like 1, opposite strand 1 [Source:I | 0 | 1.0279 | 0 | 0 | 0 | 0 |
| 22734 | ENSMUSG00000079144 | 103.1249818 | 0.095085262 | 0.246749062 | 0.385352071 | 0.699976587 | 0.941372511 | A130010J15I | RIKEN cDNA A130010J15 gene [Source:M( | 85.129 | 89.429 | 124.77 | 94.47 | 106.88 | 118.07 |
| 22735 | ENSMUSG00000079146 | 0 | NA | NA | NA | NA | NA | Gm15353 | predicted gene 15353 [Source:MGI Symbo | 0 | 0 | 0 | 0 | 0 | 0 |
| 22736 | ENSMUSG00000079156 | 0 | NA | NA | NA | NA | NA | Smok3b | sperm motility kinase 3B [Source:MGI Sym | 0 | 0 | 0 | 0 | 0 | 0 |
| 22737 | ENSMUSG00000079157 | 0 | NA | NA | NA | NA | NA | Fam155a | family with sequence similarity 155, meml | 0 | 0 | 0 | 0 | 0 | 0 |
| 22738 | ENSMUSG00000079162 | 0 | NA | NA | NA | NA | NA | Trim43b | tripartite motif-containing 43B [Source:M | 0 | 0 | 0 | 0 | 0 | 0 |
| 22739 | ENSMUSG00000079163 | 0 | NA | NA | NA | NA | NA | Gm15498 | predicted gene 15498 [Source:MGI Symbo | 0 | 0 | 0 | 0 | 0 | 0 |
| 22740 | ENSMUSG00000079164 | 263.4396146 | -0.058629343 | 0.219288777 | -0.267361347 | 0.789190957 | 0.960994657 | Tlr5 | toll-like receptor 5 [Source:MGI Symbol;A | 310 | 301.18 | 195.21 | 305.58 | 273.75 | 194.92 |
| 22741 | ENSMUSG00000079165 | 0 | NA | NA | NA | NA | NA | Sap25 | sin3 associated polypeptide [Source:MGI : | 0 | 0 | 0 | 0 | 0 | 0 |
| 22742 | ENSMUSG00000079168 | 0.133850373 | -0.796629176 | 4.080472857 | -0.195229623 | 0.84521319 | NA | Cd209g | CD209g antigen [Source:MGI Symbol;Acc: | 0.8031 | 0 | 0 | 0 | 0 | 0 |
| 22743 | ENSMUSG00000079169 | 0 | NA | NA | NA | NA | NA | Gm15130 | predicted gene 15130 [Source:MGI Symbo | 0 | 0 | 0 | 0 | 0 | 0 |
| 22744 | ENSMUSG00000079170 | 0 | NA | NA | NA | NA | NA | Gm13941 | predicted gene 13941 [Source:MGI Symbo | 0 | 0 | 0 | 0 | 0 | 0 |
| 22745 | ENSMUSG00000079171 | 0 | NA | NA | NA | NA | NA | Gm13942 | predicted gene 13942 [Source:MGI Symbo | 0 | 0 | 0 | 0 | 0 | 0 |
| 22746 | ENSMUSG00000079173 | 0 | NA | NA | NA | NA | NA | Zan | zonadhesin [Source:MGI Symbol;Acc:MGI | 0 | 0 | 0 | 0 | 0 | 0 |
| 22747 | ENSMUSG00000079174 | 0 | NA | NA | NA | NA | NA | NA | NA | 0 | 0 | 0 | 0 | 0 | 0 |
| 22748 | ENSMUSG00000079175 | 0 | NA | NA | NA | NA | NA | Gm11060 | predicted gene 11060 [Source:MGI Symbo | 0 | 0 | 0 | 0 | 0 | 0 |
| 22749 | ENSMUSG00000079177 | 5.140490808 | -0.76358383 | 0.937338761 | -0.814629525 | 0.415284405 | 0.836001897 | Fam228a | family with sequence similarity 228, meml | 7.2279 | 6.1675 | 6.0375 | 4.8199 | 4.3625 | 2.2277 |
| 22750 | ENSMUSG00000079179 | 153.6358889 | 0.365462542 | 0.221309363 | 1.651365029 | 0.098664061 | 0.463186297 | Rab10os | RAB10, member RAS oncogene family, op | 153.39 | 132.6 | 116.72 | 210.15 | 137.42 | 171.53 |
| 22751 | ENSMUSG00000079180 | 0 | NA | NA | NA | NA | NA | Mptx2 | mucosal pentraxin 2 [Source:MGI Symbol; | 0 | 0 | 0 | 0 | 0 | 0 |
| 22752 | ENSMUSG00000079183 | 0 | NA | NA | NA | NA | NA | C030005K15 | RIKEN cDNA C030005K15 gene [Source:M | 0 | 0 | 0 | 0 | 0 | 0 |
| 22753 | ENSMUSG00000079184 | 261.9543018 | -0.084885565 | 0.181198865 | -0.468466314 | 0.639451155 | 0.923932929 | Mphosph8 | M-phase phosphoprotein 8 [Source:MGI : | 265.83 | 300.15 | 242.5 | 205.33 | 267.2 | 290.71 |
| 22754 | ENSMUSG00000079186 | 0.167707431 | -0.796629176 | 4.080472857 | -0.195229623 | 0.84521319 | NA | Gzmc | granzyme C [Source:MGI Symbol;Acc:MGI | 0 | 0 | 1.0062 | 0 | 0 | 0 |
| 22755 | ENSMUSG00000079190 | 0 | NA | NA | NA | NA | NA | | | 0 | 0 | 0 | 0 | 0 | 0 |
| 22756 | ENSMUSG00000079192 | 0 | NA | NA | NA | NA | NA | | | 0 | 0 | 0 | 0 | 0 | 0 |
| 22757 | ENSMUSG00000079197 | 1293.490589 | 0.126390339 | 0.113702364 | 1.111589366 | 0.266314747 | 0.7224262 | Psme2 | proteasome (prosome, macropain) activa | 1286.6 | 1187.2 | 1235.7 | 1254.1 | 1285.8 | 1511.5 |
| 22758 | ENSMUSG00000079215 | 537.2527584 | -0.014020004 | 0.132878414 | -0.105510022 | 0.915971118 | 0.986374671 | Zfp664 | zinc finger protein 664 [Source:MGI Symb | 481.86 | 585.91 | 552.43 | 501.27 | 554.03 | 548.01 |
| 22759 | ENSMUSG00000079222 | 0 | NA | NA | NA | NA | NA | | | 0 | 0 | 0 | 0 | 0 | 0 |
| 22760 | ENSMUSG00000079223 | 0 | NA | NA | NA | NA | NA | Gm8778 | predicted gene 8778 [Source:MGI Symbol | 0 | 0 | 0 | 0 | 0 | 0 |
| 22761 | ENSMUSG00000079224 | 0.517184577 | -0.812115922 | 3.393570547 | -0.239310163 | 0.810865085 | NA | Gm6565 | predicted gene 6565 [Source:MGI Symbol | 0 | 0 | 2.0125 | 0 | 1.0906 | 0 |
| 22762 | ENSMUSG00000079225 | 6.352737856 | -0.042670578 | 0.876330677 | -0.048692325 | 0.961164493 | 0.992594832 | Gm9531 | predicted gene 9531 [Source:MGI Symbol | 3.2124 | 8.2233 | 8.05 | 8.6758 | 3.2719 | 6.6831 |
| 22763 | ENSMUSG00000079227 | 43.82942535 | -1.023617733 | 0.392461636 | -2.608198204 | 0.009102023 | 0.101114951 | Ccr5 | chemokine (C-C motif) receptor 5 [Source | 61.839 | 62.703 | 51.318 | 16.388 | 26.175 | 44.554 |
| 22764 | ENSMUSG00000079230 | 0 | NA | NA | NA | NA | NA | Gm11065 | predicted gene 11065 [Source:MGI Symbo | 0 | 0 | 0 | 0 | 0 | 0 |
| 22765 | ENSMUSG00000079235 | 0.17131964 | -0.796629176 | 4.080472857 | -0.195229623 | 0.84521319 | NA | Ccdc13 | coiled-coil domain containing 13 [Source:I | 0 | 1.0279 | 0 | 0 | 0 | 0 |
| 22766 | ENSMUSG00000079242 | 16.98467407 | -0.688788922 | 0.635223722 | -1.084324936 | 0.278220753 | 0.73110838 | C730034F03 | RIKEN cDNA C730034F03 gene [Source:M( | 15.259 | 29.81 | 18.112 | 21.208 | 14.178 | 3.3415 |
| 22767 | ENSMUSG00000079243 | 29.52308382 | 0.788379798 | 0.451522712 | 1.746046824 | 0.080802816 | 0.41758625 | Xirp1 | xin actin-binding repeat containing 1 [Sou | 24.093 | 21.586 | 19.119 | 28.919 | 57.803 | 25.618 |
| 22768 | ENSMUSG00000079244 | 0.677847551 | 0.063310748 | 2.809814301 | 0.022532004 | 0.982023583 | NA | Gm5622 | predicted gene 5622 [Source:MGI Symbol | 0 | 0 | 2.0125 | 0.964 | 1.0906 | 0 |
| 22769 | ENSMUSG00000079247 | 0 | NA | NA | NA | NA | NA | Gm13691 | predicted gene 13691 [Source:MGI Symbo | 0 | 0 | 0 | 0 | 0 | 0 |
| 22770 | ENSMUSG00000079258 | 0 | NA | NA | NA | NA | NA | Calhm1 | calcium homeostasis modulator 1 [Source | 0 | 0 | 0 | 0 | 0 | 0 |
| 22771 | ENSMUSG00000079259 | 0.17131964 | -0.796629176 | 4.080472857 | -0.195229623 | 0.84521319 | NA | Trim71 | tripartite motif-containing 71 [Source:MG | 0 | 1.0279 | 0 | 0 | 0 | 0 |
| 22772 | ENSMUSG00000079260 | 47.28317265 | 0.255138828 | 0.327596894 | 0.778819436 | 0.436086087 | 0.84624805 | Tmppe | transmembrane protein with metallophos | 40.155 | 44.2 | 45.281 | 62.659 | 49.078 | 42.326 |
| 22773 | ENSMUSG00000079261 | 0 | NA | NA | NA | NA | NA | Gm15217 | predicted gene 15217 [Source:MGI Symbo | 0 | 0 | 0 | 0 | 0 | 0 |
| 22774 | ENSMUSG00000079262 | 0.342639279 | -1.805976596 | 4.028547714 | -0.448294702 | 0.653940524 | NA | Slco1a6 | solute carrier organic anion transporter fa | 0 | 2.0558 | 0 | 0 | 0 | 0 |
| 22775 | ENSMUSG00000079263 | 11.71119864 | 0.45521163 | 0.83021732 | 0.548304184 | 0.583483055 | 0.908645447 | Gm6614 | predicted gene 6614 [Source:MGI Symbol | 3.2124 | 23.642 | 3.0187 | 17.352 | 16.359 | 6.6831 |
| 22776 | ENSMUSG00000079264 | 0 | NA | NA | NA | NA | NA | Gm11077 | predicted gene 11077 [Source:MGI Symbo | 0 | 0 | 0 | 0 | 0 | 0 |
| 22777 | ENSMUSG00000079265 | 0 | NA | NA | NA | NA | NA | Gm8257 | predicted pseudogene 8257 [Source:MGI | 0 | 0 | 0 | 0 | 0 | 0 |
| 22778 | ENSMUSG00000079266 | 0 | NA | NA | NA | NA | NA | Gm6490 | predicted gene 6490 [Source:MGI Symbol | 0 | 0 | 0 | 0 | 0 | 0 |
| 22779 | ENSMUSG00000079267 | 0 | NA | NA | NA | NA | NA | Gm5930 | predicted gene 5930 [Source:MGI Symbol | 0 | 0 | 0 | 0 | 0 | 0 |
| 22780 | ENSMUSG00000079269 | 0 | NA | NA | NA | NA | NA | Gm3676 | predicted gene 3676 [Source:MGI Symbol | 0 | 0 | 0 | 0 | 0 | 0 |
| 22781 | ENSMUSG00000079271 | 0 | NA | NA | NA | NA | NA | 1700049E17 | RIKEN cDNA 1700049E17 gene, gene 1 [Sc | 0 | 0 | 0 | 0 | 0 | 0 |
| 22782 | ENSMUSG00000079272 | 0 | NA | NA | NA | NA | NA | Gm7152 | predicted gene 7152 [Source:MGI Symbol | 0 | 0 | 0 | 0 | 0 | 0 |
| 22783 | ENSMUSG00000079273 | 0 | NA | NA | NA | NA | NA | Gm17165 | predicted gene 17165 [Source:MGI Symbo | 0 | 0 | 0 | 0 | 0 | 0 |
| 22784 | ENSMUSG00000079276 | 0 | NA | NA | NA | NA | NA | Gm4736 | predicted gene 4736 [Source:MGI Symbol | 0 | 0 | 0 | 0 | 0 | 0 |
| 22785 | ENSMUSG00000079277 | 0 | NA | NA | NA | NA | NA | Hoxd3 | homeobox D3 [Source:MGI Symbol;Acc:M | 0 | 0 | 0 | 0 | 0 | 0 |
| 22786 | ENSMUSG00000079278 | 0 | NA | NA | NA | NA | NA | Tmem233 | transmembrane protein 233 [Source:MGI | 0 | 0 | 0 | 0 | 0 | 0 |
| 22787 | ENSMUSG00000079283 | 84.30025419 | -0.190726108 | 0.271142458 | -0.703416609 | 0.481796159 | 0.871305282 | 2310009B15 | RIKEN cDNA 2310009B15 gene [Source:M | 71.476 | 100.74 | 97.606 | 70.37 | 68.709 | 96.904 |
| 22788 | ENSMUSG00000079286 | 0.874403588 | -0.441200908 | 2.817192576 | -0.156610134 | 0.875552115 | NA | Gm11084 | predicted gene 11084 [Source:MGI Symbo | 0 | 0 | 3.0187 | 0 | 0 | 2.2277 |

| | | | | | | | | | | | | | | | |
|---|---|---|---|---|---|---|---|---|---|---|---|---|---|---|---|
| 22789 | ENSMUSG00000079293 | 46.27629135 | -1.148315194 | 0.396338873 | -2.897306507 | 0.003763818 | 0.054823634 | Clec7a | C-type lectin domain family 7, member a [ | 69.87 | 64.759 | 56.35 | 20.244 | 18.541 | 47.895 |
| 22790 | ENSMUSG00000079295 | 0 | NA | NA | NA | NA | NA | Klrb1-ps1 | killer cell lectin-like receptor subfamily B r | 0 | 0 | 0 | 0 | 0 | 0 |
| 22791 | ENSMUSG00000079297 | 5.330566995 | 0.231132672 | 0.99058415 | 0.233329669 | 0.815505421 | 0.967767553 | Gm2223 | predicted pseudogene 2223 [Source:MGI | 1.6062 | 4.1117 | 9.0562 | 2.8919 | 7.6343 | 6.6831 |
| 22792 | ENSMUSG00000079298 | 0.644197084 | -2.725354879 | 2.762261801 | -0.986638876 | 0.323819705 | NA | Klrb1b | killer cell lectin-like receptor subfamily B r | 0.8031 | 2.0558 | 1.0062 | 0 | 0 | 0 |
| 22793 | ENSMUSG00000079299 | 0.319491021 | 0.16514334 | 4.044653141 | 0.040830038 | 0.967431393 | NA | Klrb1 | killer cell lectin-like receptor subfamily B r | 0.8031 | 0 | 0 | 0 | 0 | 1.1138 |
| 22794 | ENSMUSG00000079304 | 0 | NA | NA | NA | NA | NA | Tex52 | testis expressed 52 [Source:MGI Symbol;A | 0 | 0 | 0 | 0 | 0 | 0 |
| 22795 | ENSMUSG00000079311 | 1.199083021 | 0.480008596 | 1.941368705 | 0.24725267 | 0.804712685 | NA | Gm3222 | predicted pseudogene 3222 [Source:MGI | 0 | 2.0558 | 1.0062 | 1.928 | 1.0906 | 1.1138 |
| 22796 | ENSMUSG00000079316 | 874.6128087 | 0.035389861 | 0.11420703 | 0.309874628 | 0.756656298 | 0.953085143 | Rab9 | RAB9, member RAS oncogene family [Sou | 929.19 | 868.59 | 792.92 | 867.58 | 915.03 | 874.37 |
| 22797 | ENSMUSG00000079317 | 90.7575488 | -0.228899155 | 0.25361828 | -0.902534135 | 0.366773197 | 0.805239112 | Trappc2 | trafficking protein particle complex 2 [Sou | 110.83 | 76.066 | 106.66 | 85.794 | 88.34 | 76.855 |
| 22798 | ENSMUSG00000079323 | 0 | NA | NA | NA | NA | NA | Gm20661 | predicted gene 20661 [Source:MGI Symbo | 0 | 0 | 0 | 0 | 0 | 0 |
| 22799 | ENSMUSG00000079324 | 0 | NA | NA | NA | NA | NA | 4932414N04 | RIKEN cDNA 4932414N04 gene [Source:M | 0 | 0 | 0 | 0 | 0 | 0 |
| 22800 | ENSMUSG00000079330 | 0 | NA | NA | NA | NA | NA | Lemd1 | LEM domain containing 1 [Source:MGI Sy | 0 | 0 | 0 | 0 | 0 | 0 |
| 22801 | ENSMUSG00000079333 | 0 | NA | NA | NA | NA | NA | Gm11096 | predicted gene 11096 [Source:MGI Symbo | 0 | 0 | 0 | 0 | 0 | 0 |
| 22802 | ENSMUSG00000079334 | 691.8072199 | 0.236167691 | 0.115302028 | 2.048252699 | 0.040535245 | 0.275368634 | Naa80 | N(alpha)-acetyltransferase 80, NatH catal | 622.4 | 611.61 | 672.17 | 759.61 | 718.72 | 766.32 |
| 22803 | ENSMUSG00000079339 | 4.885405623 | -0.99853205 | 1.037387657 | -0.962544757 | 0.335776032 | NA | Ifit1bl1 | interferon induced protein with tetratricp | 3.2124 | 9.2513 | 7.0437 | 0.964 | 3.2719 | 5.5692 |
| 22804 | ENSMUSG00000079342 | 0 | NA | NA | NA | NA | NA | Lipo1 | lipase, member O1 [Source:MGI Symbol;A | 0 | 0 | 0 | 0 | 0 | 0 |
| 22805 | ENSMUSG00000079343 | 0.717584917 | 3.048786435 | 2.951298099 | 1.033032358 | 0.301588756 | NA | C1s2 | complement component 1, s subcompone | 0 | 0 | 0 | 0.964 | 0 | 3.3415 |
| 22806 | ENSMUSG00000079344 | 0 | NA | NA | NA | NA | NA | Lipo4 | lipase, member O4 [Source:MGI Symbol;A | 0 | 0 | 0 | 0 | 0 | 0 |
| 22807 | ENSMUSG00000079346 | 0 | NA | NA | NA | NA | NA | 1700013D24 | RIKEN cDNA 1700013D24 gene [Source:M | 0 | 0 | 0 | 0 | 0 | 0 |
| 22808 | ENSMUSG00000079349 | 0 | NA | NA | NA | NA | NA | Magea5 | MAGE family member A5 [Source:MGI Sy | 0 | 0 | 0 | 0 | 0 | 0 |
| 22809 | ENSMUSG00000079350 | 0 | NA | NA | NA | NA | NA | Magea8 | MAGE family member A8 [Source:MGI Sy | 0 | 0 | 0 | 0 | 0 | 0 |
| 22810 | ENSMUSG00000079353 | 0 | NA | NA | NA | NA | NA | Gm11099 | predicted gene 11099 [Source:MGI Symbo | 0 | 0 | 0 | 0 | 0 | 0 |
| 22811 | ENSMUSG00000079355 | 27.55201743 | -0.326869024 | 0.420441129 | -0.777443026 | 0.436897436 | 0.846852749 | Ackr4 | atypical chemokine receptor 4 [Source:M( | 36.943 | 26.726 | 28.175 | 25.063 | 28.356 | 20.049 |
| 22812 | ENSMUSG00000079356 | 0 | NA | NA | NA | NA | NA | Mettl4-ps1 | methyltransferase like 4, pseudogene 1 [S | 0 | 0 | 0 | 0 | 0 | 0 |
| 22813 | ENSMUSG00000079357 | 0.496077836 | -0.812642266 | 3.444120572 | -0.235950586 | 0.813471014 | NA | Gm11100 | predicted gene 11100 [Source:MGI Symbo | 0 | 0 | 2.0125 | 0.964 | 0 | 0 |
| 22814 | ENSMUSG00000079359 | 0 | NA | NA | NA | NA | NA | Gm9006 | predicted gene 9006 [Source:MGI Symbol | 0 | 0 | 0 | 0 | 0 | 0 |
| 22815 | ENSMUSG00000079362 | 0 | NA | NA | NA | NA | NA | Gm43302 | predicted gene 43302 [Source:MGI Symbo | 0 | 0 | 0 | 0 | 0 | 0 |
| 22816 | ENSMUSG00000079363 | 33.75560122 | -0.278357431 | 0.414335105 | -0.671817154 | 0.501700107 | 0.880974509 | Gbp4 | guanylate binding protein 4 [Source:MGI ! | 28.109 | 37.005 | 46.287 | 41.451 | 20.722 | 28.96 |
| 22817 | ENSMUSG00000079364 | 0.644197084 | -2.725354879 | 2.762261801 | -0.986638876 | 0.323819705 | NA | Gm3558 | predicted gene 3558 [Source:MGI Symbol | 0.8031 | 2.0558 | 1.0062 | 0 | 0 | 0 |
| 22818 | ENSMUSG00000079371 | 0 | NA | NA | NA | NA | NA | Gm3476 | predicted gene 3476 [Source:MGI Symbol | 0 | 0 | 0 | 0 | 0 | 0 |
| 22819 | ENSMUSG00000079374 | 0 | NA | NA | NA | NA | NA | Gm8334 | predicted gene 8334 [Source:MGI Symbol | 0 | 0 | 0 | 0 | 0 | 0 |
| 22820 | ENSMUSG00000079378 | 0 | NA | NA | NA | NA | NA | Gm8279 | predicted gene 8279 [Source:MGI Symbol | 0 | 0 | 0 | 0 | 0 | 0 |
| 22821 | ENSMUSG00000079379 | 0 | NA | NA | NA | NA | NA | Gm3242 | predicted gene 3242 [Source:MGI Symbol | 0 | 0 | 0 | 0 | 0 | 0 |
| 22822 | ENSMUSG00000079380 | 0 | NA | NA | NA | NA | NA | Gm8271 | predicted gene 8271 [Source:MGI Symbol | 0 | 0 | 0 | 0 | 0 | 0 |
| 22823 | ENSMUSG00000079382 | 0 | NA | NA | NA | NA | NA | Gm3443 | predicted gene 3443 [Source:MGI Symbol | 0 | 0 | 0 | 0 | 0 | 0 |
| 22824 | ENSMUSG00000079383 | 0 | NA | NA | NA | NA | NA | Gm3298 | predicted gene 3298 [Source:MGI Symbol | 0 | 0 | 0 | 0 | 0 | 0 |
| 22825 | ENSMUSG00000079386 | 0 | NA | NA | NA | NA | NA | Gm3173 | predicted gene 3173 [Source:MGI Symbol | 0 | 0 | 0 | 0 | 0 | 0 |
| 22826 | ENSMUSG00000079387 | 0 | NA | NA | NA | NA | NA | Luzp4 | leucine zipper protein 4 [Source:MGI Sym | 0 | 0 | 0 | 0 | 0 | 0 |
| 22827 | ENSMUSG00000079388 | 0 | NA | NA | NA | NA | NA | 2610042L04I | RIKEN cDNA 2610042L04 gene [Source:M( | 0 | 0 | 0 | 0 | 0 | 0 |
| 22828 | ENSMUSG00000079389 | 0 | NA | NA | NA | NA | NA | Gm3149 | predicted gene 3149 [Source:MGI Symbol | 0 | 0 | 0 | 0 | 0 | 0 |
| 22829 | ENSMUSG00000079391 | 0 | NA | NA | NA | NA | NA | Gm2974 | predicted gene 2974 [Source:MGI Symbol | 0 | 0 | 0 | 0 | 0 | 0 |
| 22830 | ENSMUSG00000079392 | 0 | NA | NA | NA | NA | NA | Gm17170 | predicted gene 17170 [Source:MGI Symbo | 0 | 0 | 0 | 0 | 0 | 0 |
| 22831 | ENSMUSG00000079395 | 0 | NA | NA | NA | NA | NA | Gm15114 | predicted gene 15114 [Source:MGI Symbo | 0 | 0 | 0 | 0 | 0 | 0 |
| 22832 | ENSMUSG00000079396 | 0 | NA | NA | NA | NA | NA | Gm3411 | predicted gene 3411 [Source:MGI Symbol | 0 | 0 | 0 | 0 | 0 | 0 |
| 22833 | ENSMUSG00000079402 | 0 | NA | NA | NA | NA | NA | Gm3020 | predicted gene 3020 [Source:MGI Symbol | 0 | 0 | 0 | 0 | 0 | 0 |
| 22834 | ENSMUSG00000079407 | 0 | NA | NA | NA | NA | NA | 1700110I01F | RIKEN cDNA 1700110I01 gene [Source:M( | 0 | 0 | 0 | 0 | 0 | 0 |
| 22835 | ENSMUSG00000079409 | 0 | NA | NA | NA | NA | NA | Gm5795 | predicted gene 5795 [Source:MGI Symbol | 0 | 0 | 0 | 0 | 0 | 0 |
| 22836 | ENSMUSG00000079410 | 0 | NA | NA | NA | NA | NA | Gm2897 | predicted gene 2897 [Source:MGI Symbol | 0 | 0 | 0 | 0 | 0 | 0 |
| 22837 | ENSMUSG00000079414 | 1.171831736 | -1.161928617 | 2.135676568 | -0.544056453 | 0.586402626 | NA | Gm11110 | predicted gene 11110 [Source:MGI Symbo | 0.8031 | 1.0279 | 3.0187 | 0 | 2.1812 | 0 |
| 22838 | ENSMUSG00000079415 | 3.840489585 | 0.221178834 | 1.134855528 | 0.194896027 | 0.845474344 | NA | Cntf | ciliary neurotrophic factor [Source:MGI Sy | 1.6062 | 4.1117 | 5.0312 | 6.7478 | 1.0906 | 4.4554 |
| 22839 | ENSMUSG00000079416 | 0 | NA | NA | NA | NA | NA | Gm11111 | predicted gene 11111 [Source:MGI Symbo | 0 | 0 | 0 | 0 | 0 | 0 |
| 22840 | ENSMUSG00000079418 | 257.8308831 | 0.011795597 | 0.194495378 | 0.060647184 | 0.951640195 | 0.990883058 | Atg4a | autophagy related 4A, cysteine peptidase | 316.42 | 246.7 | 206.28 | 262.2 | 224.67 | 290.71 |
| 22841 | ENSMUSG00000079419 | 11.2551118 | -2.09087438 | 0.793013551 | -2.636618729 | 0.008373689 | 0.095217798 | Ms4a6c | membrane-spanning 4-domains, subfamil | 24.896 | 20.558 | 9.0562 | 1.928 | 2.1812 | 8.9108 |
| 22842 | ENSMUSG00000079421 | 0 | NA | NA | NA | NA | NA | 4930402F06 | RIKEN cDNA 4930402F06 gene [Source:M( | 0 | 0 | 0 | 0 | 0 | 0 |
| 22843 | ENSMUSG00000079423 | 0 | NA | NA | NA | NA | NA | Gm3286 | predicted gene 3286 [Source:MGI Symbol | 0 | 0 | 0 | 0 | 0 | 0 |
| 22844 | ENSMUSG00000079424 | 0 | NA | NA | NA | NA | NA | Pramel47 | PRAME like 47 [Source:MGI Symbol;Acc:N | 0 | 0 | 0 | 0 | 0 | 0 |

| | | | | | | | | | | | | | | | |
|---|---|---|---|---|---|---|---|---|---|---|---|---|---|---|---|
| 22845 | ENSMUSG00000079426 | 576.0791968 | -0.144215048 | 0.133811527 | -1.077747572 | 0.281146422 | 0.733710968 | Arpc4 | actin related protein 2/3 complex, subunit | 618.39 | 584.89 | 610.79 | 531.15 | 487.51 | 623.75 |
| 22846 | ENSMUSG00000079427 | 328.9209654 | 0.058910834 | 0.147878165 | 0.398374117 | 0.690354434 | 0.937497562 | Mthfsl | 5, 10-methenyltetrahydrofolate synthetas | 350.96 | 289.87 | 325.02 | 329.68 | 329.37 | 348.63 |
| 22847 | ENSMUSG00000079428 | 0 | NA | NA | NA | NA | NA | Tceal7 | transcription elongation factor A (SII)-like | 0 | 0 | 0 | 0 | 0 | 0 |
| 22848 | ENSMUSG00000079429 | 175.0980469 | -0.143147547 | 0.206112215 | -0.69451268 | 0.487360751 | 0.873910634 | Mroh2a | maestro heat-like repeat family member 2 | 224.87 | 161.38 | 164.02 | 147.49 | 169.05 | 183.78 |
| 22849 | ENSMUSG00000079432 | 0 | NA | NA | NA | NA | NA | Gm15023 | predicted gene 15023 [Source:MGI Symbo | 0 | 0 | 0 | 0 | 0 | 0 |
| 22850 | ENSMUSG00000079433 | 0 | NA | NA | NA | NA | NA | Gm11114 | predicted gene 11114 [Source:MGI Symbo | 0 | 0 | 0 | 0 | 0 | 0 |
| 22851 | ENSMUSG00000079434 | 57.79374124 | -0.315204405 | 0.307042662 | -1.026581789 | 0.304617433 | 0.753546528 | Neu2 | neuraminidase 2 [Source:MGI Symbol;Acc | 52.202 | 72.982 | 67.418 | 55.911 | 41.443 | 56.806 |
| 22852 | ENSMUSG00000079435 | 1386.859486 | 0.000851782 | 0.115785546 | 0.00735655 | 0.994130375 | 0.998618081 | Rpl36a | ribosomal protein L36A [Source:MGI Symb | 1238.4 | 1503.8 | 1417.8 | 1283.1 | 1416.7 | 1461.4 |
| 22853 | ENSMUSG00000079436 | 0.301557804 | -1.64335916 | 4.042814367 | -0.406488899 | 0.684383412 | NA | Kcnj13 | potassium inwardly-rectifying channel, su | 0.8031 | 0 | 1.0062 | 0 | 0 | 0 |
| 22854 | ENSMUSG00000079438 | 0 | NA | NA | NA | NA | NA | Gm11115 | predicted gene 11115 [Source:MGI Symbo | 0 | 0 | 0 | 0 | 0 | 0 |
| 22855 | ENSMUSG00000079439 | 0 | NA | NA | NA | NA | NA | Gm11116 | predicted gene 11116 [Source:MGI Symbo | 0 | 0 | 0 | 0 | 0 | 0 |
| 22856 | ENSMUSG00000079440 | 0 | NA | NA | NA | NA | NA | Alpi | alkaline phosphatase, intestinal [Source:N | 0 | 0 | 0 | 0 | 0 | 0 |
| 22857 | ENSMUSG00000079442 | 17.82898466 | -0.642415261 | 0.620314377 | -1.035628522 | 0.30037548 | 0.751875664 | St6galnac4 | ST6 (alpha-N-acetyl-neuraminyl-2,3-beta-g | 31.321 | 17.475 | 16.1 | 7.7118 | 7.6343 | 26.732 |
| 22858 | ENSMUSG00000079444 | 0.808265676 | 0.728755068 | 2.41725015 | 0.301481031 | 0.763047712 | NA | Gm21981 | predicted gene 21981 [Source:MGI Symbo | 0.8031 | 1.0279 | 0 | 1.928 | 1.0906 | 0 |
| 22859 | ENSMUSG00000079445 | 1.186875127 | -1.301275081 | 2.153802803 | -0.604175591 | 0.545726908 | NA | B3gnt7 | UDP-GlcNAc:betaGal beta-1,3-N-acetylglu | 4.0155 | 1.0279 | 0 | 0.964 | 0 | 1.1138 |
| 22860 | ENSMUSG00000079450 | 5.704566988 | 0.07081927 | 0.932919985 | 0.075911408 | 0.939489581 | 0.988812319 | Cldn34c1 | claudin 34C1 [Source:MGI Symbol;Acc:MG | 5.6217 | 2.0558 | 9.0562 | 8.6758 | 4.3625 | 4.4554 |
| 22861 | ENSMUSG00000079451 | 0 | NA | NA | NA | NA | NA | Tmprss11g | transmembrane protease, serine 11g [Sou | 0 | 0 | 0 | 0 | 0 | 0 |
| 22862 | ENSMUSG00000079455 | 0.17131964 | -0.796629176 | 4.080472857 | -0.195229623 | 0.84521319 | NA | Gm16026 | predicted pseudogene 16026 [Source:MG | 0 | 1.0279 | 0 | 0 | 0 | 0 |
| 22863 | ENSMUSG00000079457 | 18.20506858 | -0.271707596 | 0.58908643 | -0.461235537 | 0.644629628 | 0.925643431 | Gm7609 | predicted pseudogene 7609 [Source:MGI | 20.881 | 20.558 | 18.112 | 8.6758 | 9.8156 | 31.188 |
| 22864 | ENSMUSG00000079460 | 0 | NA | NA | NA | NA | NA | 4933403O08 | RIKEN cDNA 4933403O08 gene [Source:M | 0 | 0 | 0 | 0 | 0 | 0 |
| 22865 | ENSMUSG00000079462 | 0 | NA | NA | NA | NA | NA | Gm15737 | predicted gene 15737 [Source:MGI Symbo | 0 | 0 | 0 | 0 | 0 | 0 |
| 22866 | ENSMUSG00000079465 | 10.89015251 | 0.084644123 | 0.66481255 | 0.127320285 | 0.898686906 | 0.982798755 | Col4a3 | collagen, type IV, alpha 3 [Source:MGI Syr | 15.259 | 8.2233 | 8.05 | 10.604 | 8.7249 | 14.48 |
| 22867 | ENSMUSG00000079466 | 0 | NA | NA | NA | NA | NA | Prdm12 | PR domain containing 12 [Source:MGI Syr | 0 | 0 | 0 | 0 | 0 | 0 |
| 22868 | ENSMUSG00000079467 | 5.583175536 | -0.571533035 | 0.898938781 | -0.635786381 | 0.524915665 | 0.888334265 | Gm14966 | predicted gene 14966 [Source:MGI Symbo | 8.8341 | 5.1396 | 6.0375 | 5.7839 | 4.3625 | 3.3415 |
| 22869 | ENSMUSG00000079469 | 133.4208623 | 0.033339595 | 0.244275553 | 0.136483552 | 0.891439026 | 0.982489666 | Pigb | phosphatidylinositol glycan anchor biosyn | 143.76 | 106.9 | 144.9 | 145.56 | 162.5 | 96.904 |
| 22870 | ENSMUSG00000079470 | 102.8043098 | 0.205435312 | 0.265557379 | 0.773600463 | 0.439167098 | 0.848147242 | Utp14b | UTP14B small subunit processome compo | 85.129 | 99.708 | 101.63 | 82.902 | 148.32 | 99.132 |
| 22871 | ENSMUSG00000079471 | 0.524202404 | 2.600864256 | 3.301637836 | 0.787749713 | 0.430843121 | NA | Mymx | myomixer, myoblast fusion factor [Source | 0 | 0 | 0 | 0.964 | 2.1812 | 0 |
| 22872 | ENSMUSG00000079476 | 0 | NA | NA | NA | NA | NA | 1700011M02 | RIKEN cDNA 1700011M02 gene [Source:M | 0 | 0 | 0 | 0 | 0 | 0 |
| 22873 | ENSMUSG00000079477 | 3234.307894 | -0.118565909 | 0.094243913 | -1.258074972 | 0.208364644 | 0.653455953 | Rab7 | RAB7, member RAS oncogene family [Sou | 3184.3 | 3401.4 | 3516.8 | 3144.5 | 3223.9 | 2935 |
| 22874 | ENSMUSG00000079478 | 156.46496 | -0.119258644 | 0.200856334 | -0.593750978 | 0.552678682 | 0.896081656 | Znrd2 | zinc ribbon domain containing 2 [Source:N | 184.71 | 138.77 | 165.02 | 164.84 | 142.87 | 142.57 |
| 22875 | ENSMUSG00000079479 | 0 | NA | NA | NA | NA | NA | Gm9112 | predicted gene 9112 [Source:MGI Symbol | 0 | 0 | 0 | 0 | 0 | 0 |
| 22876 | ENSMUSG00000079480 | 244.23108 | 0.201371134 | 0.159005635 | 1.266440237 | 0.205355513 | 0.650805014 | Pin4 | protein (peptidyl-prolyl cis/trans isomeras | 225.67 | 243.62 | 212.32 | 253.53 | 258.48 | 271.78 |
| 22877 | ENSMUSG00000079481 | 10.40822646 | -0.97531146 | 0.711434644 | -1.370908021 | 0.170403631 | 0.597556794 | Nhsl2 | NHS-like 2 [Source:MGI Symbol;Acc:MGI:3 | 8.8341 | 22.614 | 10.062 | 7.7118 | 6.5437 | 6.6831 |
| 22878 | ENSMUSG00000079484 | 99.73998022 | 0.295906673 | 0.243492799 | 1.215258415 | 0.224267547 | 0.67048487 | Phyhd1 | phytanoyl-CoA dioxygenase domain conta | 98.782 | 84.289 | 85.531 | 130.14 | 89.431 | 110.27 |
| 22879 | ENSMUSG00000079487 | 311.9508594 | 0.268392929 | 0.215902375 | 1.243121709 | 0.213822987 | 0.658589371 | Med12 | mediator complex subunit 12 [Source:MG | 223.26 | 347.44 | 279.74 | 448.25 | 290.1 | 282.92 |
| 22880 | ENSMUSG00000079491 | 70.04036423 | -0.070903447 | 0.306369171 | -0.231431402 | 0.816979672 | 0.967910123 | H2-T10 | histocompatibility 2, T region locus 10 [So | 97.978 | 50.368 | 66.412 | 73.262 | 67.618 | 64.603 |
| 22881 | ENSMUSG00000079492 | 0 | NA | NA | NA | NA | NA | Gm11127 | predicted gene 11127 [Source:MGI Symbo | 0 | 0 | 0 | 0 | 0 | 0 |
| 22882 | ENSMUSG00000079494 | 144.7278158 | -0.289133167 | 0.325750785 | -0.887590084 | 0.37476129 | 0.809332934 | Nat8f5 | N-acetyltransferase 8 (GCN5-related) fam | 139.74 | 140.82 | 197.22 | 133.03 | 197.4 | 60.148 |
| 22883 | ENSMUSG00000079495 | 11.33996077 | 0.05309091 | 0.718142843 | 0.073928064 | 0.941067625 | 0.989159676 | Nat8f6 | N-acetyltransferase 8 (GCN5-related) fam | 12.047 | 9.2513 | 12.075 | 10.604 | 20.722 | 3.3415 |
| 22884 | ENSMUSG00000079497 | 0.919174381 | -0.746991182 | 2.31347519 | -0.322887051 | 0.746780783 | NA | Gm13420 | predicted gene 13420 [Source:MGI Symbo | 2.4093 | 1.0279 | 0 | 0.964 | 0 | 1.1138 |
| 22885 | ENSMUSG00000079499 | 13.73530971 | -0.526249441 | 0.623554511 | -0.843950981 | 0.398696798 | 0.82588279 | 6530402F18 | RIKEN cDNA 6530402F18 gene [Source:M | 20.078 | 12.335 | 16.1 | 8.6758 | 18.541 | 6.6831 |
| 22886 | ENSMUSG00000079501 | 3.746551922 | -1.325430407 | 1.165363469 | -1.13735366 | 0.255390471 | NA | Gm5138 | predicted gene 5138 [Source:MGI Symbol | 4.8186 | 7.1954 | 4.025 | 0.964 | 4.3625 | 1.1138 |
| 22887 | ENSMUSG00000079502 | 2.74067584 | 0.689960775 | 1.309187542 | 0.527014467 | 0.598183542 | NA | Cfap77 | cilia and flagella associated protein 77 [So | 3.2124 | 2.0558 | 1.0062 | 5.7839 | 3.2719 | 1.1138 |
| 22888 | ENSMUSG00000079505 | 33.90726667 | 0.545579754 | 0.39268567 | 1.389354885 | 0.164724862 | 0.589893637 | Gm11131 | predicted gene 11131 [Source:MGI Symbo | 28.912 | 25.698 | 28.175 | 51.091 | 28.356 | 41.212 |
| 22889 | ENSMUSG00000079507 | 228.4409168 | -0.225935357 | 0.557682455 | -0.405132625 | 0.685380026 | 0.93583845 | H2-Q1 | histocompatibility 2, Q region locus 1 [Sou | 347.74 | 271.37 | 119.74 | 310.4 | 128.69 | 192.69 |
| 22890 | ENSMUSG00000079508 | 395.1927732 | -0.155378918 | 0.146558445 | -1.060183996 | 0.2890609 | 0.739603195 | Apoo | apolipoprotein O [Source:MGI Symbol;Acc | 419.22 | 409.11 | 420.61 | 338.36 | 431.88 | 351.97 |
| 22891 | ENSMUSG00000079509 | 284.6984714 | -0.186503146 | 0.168487514 | -1.106925619 | 0.268326089 | 0.724039098 | Zfx | zinc finger protein X-linked [Source:MGI S | 301.97 | 348.46 | 258.6 | 243.89 | 286.83 | 268.44 |
| 22892 | ENSMUSG00000079511 | 13.90903462 | -0.091325443 | 0.595075579 | -0.153468646 | 0.878028714 | 0.980817031 | Gm42688 | predicted gene 42688 [Source:MGI Symbo | 9.6372 | 13.363 | 20.125 | 10.604 | 16.359 | 13.366 |
| 22893 | ENSMUSG00000079513 | 0 | NA | NA | NA | NA | NA | 4932429P05 | RIKEN cDNA 4932429P05 gene [Source:M | 0 | 0 | 0 | 0 | 0 | 0 |
| 22894 | ENSMUSG00000079516 | 0 | NA | NA | NA | NA | NA | Reg3a | regenerating islet-derived 3 alpha [Source | 0 | 0 | 0 | 0 | 0 | 0 |
| 22895 | ENSMUSG00000079519 | 0 | NA | NA | NA | NA | NA | Gm14743 | predicted gene 14743 [Source:MGI Symbo | 0 | 0 | 0 | 0 | 0 | 0 |
| 22896 | ENSMUSG00000079521 | 0 | NA | NA | NA | NA | NA | Gm5938 | predicted gene 5938 [Source:MGI Symbol | 0 | 0 | 0 | 0 | 0 | 0 |
| 22897 | ENSMUSG00000079522 | 0 | NA | NA | NA | NA | NA | Gm14744 | predicted gene 14744 [Source:MGI Symbo | 0 | 0 | 0 | 0 | 0 | 0 |
| 22898 | ENSMUSG00000079523 | 117.4002198 | -0.769867533 | 0.290614953 | -2.649098148 | 0.008070689 | 0.092572914 | Tmsb10 | thymosin, beta 10 [Source:MGI Symbol;Ac | 199.97 | 138.77 | 104.65 | 81.938 | 64.346 | 114.73 |
| 22899 | ENSMUSG00000079524 | 0 | NA | NA | NA | NA | NA | Gm15402 | predicted gene 15402 [Source:MGI Symbo | 0 | 0 | 0 | 0 | 0 | 0 |
| 22900 | ENSMUSG00000079525 | 0 | NA | NA | NA | NA | NA | Cldn34d | claudin 34D [Source:MGI Symbol;Acc:MGI | 0 | 0 | 0 | 0 | 0 | 0 |

| | | | | | | | | | | | | | | | |
|---|---|---|---|---|---|---|---|---|---|---|---|---|---|---|---|
| 22901 | ENSMUSG00000079528 | 0 | NA | NA | NA | NA | NA | Clnkos | cytokine-dependent hematopoietic cell lin | 0 | 0 | 0 | 0 | 0 | 0 |
| 22902 | ENSMUSG00000079531 | 0.751028265 | -1.614680879 | 2.682726358 | -0.601880573 | 0.547253636 | NA | Pwwp4d | PWWP domain containing 4D [Source:MG | 2.4093 | 0 | 1.0062 | 0 | 1.0906 | 0 |
| 22903 | ENSMUSG00000079532 | 0 | NA | NA | NA | NA | NA | Ctag2l2 | CTAG2 like 2 [Source:MGI Symbol;Acc:MG | 0 | 0 | 0 | 0 | 0 | 0 |
| 22904 | ENSMUSG00000079534 | 3.458545582 | -1.992244959 | 1.26659584 | -1.572912918 | 0.115738984 | NA | Pwwp4c | PWWP domain containing 4C [Source:MG | 6.4248 | 5.1396 | 5.0312 | 1.928 | 0 | 2.2277 |
| 22905 | ENSMUSG00000079536 | 0 | NA | NA | NA | NA | NA | Ctag2l1 | CTAG2 like 1 [Source:MGI Symbol;Acc:MG | 0 | 0 | 0 | 0 | 0 | 0 |
| 22906 | ENSMUSG00000079539 | 0.160662974 | 1.126924175 | 4.080472857 | 0.276174898 | 0.782413736 | NA | Obp2b | odorant binding protein 2B [Source:MGI S | 0 | 0 | 0 | 0.964 | 0 | 0 |
| 22907 | ENSMUSG00000079542 | 0 | NA | NA | NA | NA | NA | Gm42606 | predicted gene 42606 [Source:MGI Symbo | 0 | 0 | 0 | 0 | 0 | 0 |
| 22908 | ENSMUSG00000079543 | 0 | NA | NA | NA | NA | NA | Igkv13-85 | immunoglobulin kappa chain variable 13-8 | 0 | 0 | 0 | 0 | 0 | 0 |
| 22909 | ENSMUSG00000079544 | 0 | NA | NA | NA | NA | NA | Igkv13-87 | immunoglobulin kappa variable 13-87 [So | 0 | 0 | 0 | 0 | 0 | 0 |
| 22910 | ENSMUSG00000079546 | 0 | NA | NA | NA | NA | NA | Gm11146 | predicted gene 11146 [Source:MGI Symbo | 0 | 0 | 0 | 0 | 0 | 0 |
| 22911 | ENSMUSG00000079547 | 38.03161649 | -1.006303817 | 0.423069891 | -2.378575829 | 0.017379662 | 0.15701401 | H2-DMb1 | histocompatibility 2, class II, locus Mb1 [S | 52.202 | 71.954 | 28.175 | 25.063 | 20.722 | 30.074 |
| 22912 | ENSMUSG00000079550 | 32.11525429 | 0.46807159 | 0.463102311 | 1.010730413 | 0.312145477 | 0.759830222 | Mpp4 | membrane protein, palmitoylated 4 (MAG | 24.093 | 35.977 | 21.131 | 59.767 | 29.447 | 22.277 |
| 22913 | ENSMUSG00000079553 | 5.90882505 | -0.464298943 | 1.053997925 | -0.440512198 | 0.65956618 | 0.927787797 | Kifc1 | kinesin family member C1 [Source:MGI Sy | 15.259 | 3.0838 | 2.0125 | 2.8919 | 2.1812 | 10.025 |
| 22914 | ENSMUSG00000079554 | 1.685764025 | 0.102391998 | 1.840331574 | 0.055637799 | 0.955630352 | NA | Aox2 | aldehyde oxidase 2 [Source:MGI Symbol;A | 0.8031 | 4.1117 | 0 | 1.928 | 3.2719 | 0 |
| 22915 | ENSMUSG00000079555 | 55.08898804 | -0.066713619 | 0.304221845 | -0.219292663 | 0.826422077 | 0.969648181 | Haus3 | HAUS augmin-like complex, subunit 3 [Sou | 65.051 | 57.563 | 46.287 | 55.911 | 56.712 | 49.009 |
| 22916 | ENSMUSG00000079557 | 1290.832414 | -0.044844644 | 0.107572495 | -0.416878344 | 0.676767394 | 0.932891189 | Marchf2 | membrane associated ring-CH-type finger | 1313.1 | 1270.5 | 1348.4 | 1180.9 | 1247.7 | 1384.5 |
| 22917 | ENSMUSG00000079559 | 1.536558863 | -2.847469017 | 1.981342673 | -1.437141114 | 0.150677901 | NA | Colca2 | COLCA2 homolog [Source:MGI Symbol;Ac | 4.0155 | 3.0838 | 1.0062 | 0 | 0 | 1.1138 |
| 22918 | ENSMUSG00000079560 | 3.005384149 | -1.775032966 | 1.381764293 | -1.284613429 | 0.198927404 | NA | Hoxa3 | homeobox A3 [Source:MGI Symbol;Acc:M | 4.8186 | 7.1954 | 2.0125 | 2.8919 | 0 | 1.1138 |
| 22919 | ENSMUSG00000079562 | 602.6315287 | 0.110585311 | 0.120747934 | 0.91583605 | 0.359752892 | 0.799452417 | Maea | macrophage erythroblast attacher [Source | 602.33 | 546.85 | 588.65 | 594.77 | 626.01 | 657.17 |
| 22920 | ENSMUSG00000079563 | 1816.347088 | 0.453632512 | 0.125278603 | 3.620989544 | 0.000293478 | 0.008540988 | Pglyrp2 | peptidoglycan recognition protein 2 [Sour | 1625.5 | 1700.2 | 1272.9 | 1992.5 | 2103.8 | 2203.2 |
| 22921 | ENSMUSG00000079564 | 0 | NA | NA | NA | NA | NA | Gm11149 | predicted gene 11149 [Source:MGI Symbo | 0 | 0 | 0 | 0 | 0 | 0 |
| 22922 | ENSMUSG00000079566 | 0 | NA | NA | NA | NA | NA | Spin2d | spindlin family, member 2D [Source:MGI S | 0 | 0 | 0 | 0 | 0 | 0 |
| 22923 | ENSMUSG00000079575 | 0 | NA | NA | NA | NA | NA | Rbpj-ps3 | recombination signal binding protein for i | 0 | 0 | 0 | 0 | 0 | 0 |
| 22924 | ENSMUSG00000079577 | 0 | NA | NA | NA | NA | NA | Gm14692 | predicted gene 14692 [Source:MGI Symbo | 0 | 0 | 0 | 0 | 0 | 0 |
| 22925 | ENSMUSG00000079578 | 0 | NA | NA | NA | NA | NA | Gm1140 | predicted gene 1140 [Source:MGI Symbol | 0 | 0 | 0 | 0 | 0 | 0 |
| 22926 | ENSMUSG00000079579 | 0 | NA | NA | NA | NA | NA | Gm6760 | predicted gene 6760 [Source:MGI Symbol | 0 | 0 | 0 | 0 | 0 | 0 |
| 22927 | ENSMUSG00000079580 | 0 | NA | NA | NA | NA | NA | Tmem217 | transmembrane protein 217 [Source:MGI | 0 | 0 | 0 | 0 | 0 | 0 |
| 22928 | ENSMUSG00000079583 | 0 | NA | NA | NA | NA | NA | Gm7073 | predicted gene 7073 [Source:MGI Symbol | 0 | 0 | 0 | 0 | 0 | 0 |
| 22929 | ENSMUSG00000079584 | 0 | NA | NA | NA | NA | NA | Tm9sf5 | transmembrane 9 superfamily member 5 | 0 | 0 | 0 | 0 | 0 | 0 |
| 22930 | ENSMUSG00000079588 | 0 | NA | NA | NA | NA | NA | Tmem182 | transmembrane protein 182 [Source:MGI | 0 | 0 | 0 | 0 | 0 | 0 |
| 22931 | ENSMUSG00000079592 | 0 | NA | NA | NA | NA | NA | C1qtnf5 | C1q and tumor necrosis factor related pro | 0 | 0 | 0 | 0 | 0 | 0 |
| 22932 | ENSMUSG00000079593 | 5.201825204 | -1.19974794 | 0.990903954 | -1.210761079 | 0.225986986 | 0.67256703 | 4933416I08R | RIKEN cDNA 4933416I08 gene [Source:MG | 10.44 | 8.2233 | 3.0187 | 2.8919 | 2.1812 | 4.4554 |
| 22933 | ENSMUSG00000079594 | 0 | NA | NA | NA | NA | NA | Cstdc6 | cystatin domain containing 6 [Source:MGI | 0 | 0 | 0 | 0 | 0 | 0 |
| 22934 | ENSMUSG00000079595 | 0 | NA | NA | NA | NA | NA | Gm7260 | predicted gene 7260 [Source:MGI Symbol | 0 | 0 | 0 | 0 | 0 | 0 |
| 22935 | ENSMUSG00000079597 | 0 | NA | NA | NA | NA | NA | Cstdc4 | cystatin domain containing 4 [Source:MGI | 0 | 0 | 0 | 0 | 0 | 0 |
| 22936 | ENSMUSG00000079598 | 0.181769715 | 1.126924175 | 4.080472857 | 0.276174898 | 0.782413736 | NA | Clec2l | C-type lectin domain family 2, member L [ | 0 | 0 | 0 | 0 | 1.0906 | 0 |
| 22937 | ENSMUSG00000079600 | 0 | NA | NA | NA | NA | NA | Gm17604 | predicted gene, 17604 [Source:MGI Symb | 0 | 0 | 0 | 0 | 0 | 0 |
| 22938 | ENSMUSG00000079602 | 0 | NA | NA | NA | NA | NA | 9230102O04 | RIKEN cDNA 9230102O04 gene [Source:M | 0 | 0 | 0 | 0 | 0 | 0 |
| 22939 | ENSMUSG00000079603 | 0 | NA | NA | NA | NA | NA | Dreg1 | distal regulatory enhancer RNA of Gata3 [ | 0 | 0 | 0 | 0 | 0 | 0 |
| 22940 | ENSMUSG00000079604 | 0 | NA | NA | NA | NA | NA | Gm13219 | predicted gene 13219 [Source:MGI Symbo | 0 | 0 | 0 | 0 | 0 | 0 |
| 22941 | ENSMUSG00000079605 | 0 | NA | NA | NA | NA | NA | Zbtb9 | zinc finger and BTB domain containing 9 [S | 0 | 0 | 0 | 0 | 0 | 0 |
| 22942 | ENSMUSG00000079606 | 0 | NA | NA | NA | NA | NA | Fsip2l | fibrous sheath-interacting protein 2-like [S | 0 | 0 | 0 | 0 | 0 | 0 |
| 22943 | ENSMUSG00000079608 | 1.352548208 | -1.519193148 | 2.288634739 | -0.663798868 | 0.506819051 | NA | Stard6 | StAR-related lipid transfer (START) domain | 0 | 0 | 6.0375 | 0.964 | 0 | 1.1138 |
| 22944 | ENSMUSG00000079610 | 51.36559621 | -0.176279081 | 0.321375585 | -0.548514228 | 0.583338862 | 0.908645447 | Ankrd39 | ankyrin repeat domain 39 [Source:MGI Sy | 45.777 | 54.48 | 63.393 | 46.271 | 40.353 | 57.92 |
| 22945 | ENSMUSG00000079614 | 829.8662427 | 0.12194377 | 0.12033088 | 1.013403791 | 0.31086733 | 0.759594595 | Seh1l | SEH1-like (S. cerevisiae [Source:MGI Symb | 802.3 | 875.79 | 706.38 | 866.62 | 863.77 | 864.34 |
| 22946 | ENSMUSG00000079619 | 0 | NA | NA | NA | NA | NA | Cypt15 | cysteine-rich perinuclear theca 15 [Source | 0 | 0 | 0 | 0 | 0 | 0 |
| 22947 | ENSMUSG00000079620 | 0 | NA | NA | NA | NA | NA | NA | NA | 0 | 0 | 0 | 0 | 0 | 0 |
| 22948 | ENSMUSG00000079625 | 0.17131964 | -0.796629176 | 4.080472857 | -0.195229623 | 0.84521319 | NA | Tm4sf19 | transmembrane 4 L six family member 19 | 0 | 1.0279 | 0 | 0 | 0 | 0 |
| 22949 | ENSMUSG00000079626 | 0 | NA | NA | NA | NA | NA | Rhox7b | reproductive homeobox 7B [Source:MGI S | 0 | 0 | 0 | 0 | 0 | 0 |
| 22950 | ENSMUSG00000079627 | 0 | NA | NA | NA | NA | NA | Rhox2h | reproductive homeobox 2H [Source:MGI S | 0 | 0 | 0 | 0 | 0 | 0 |
| 22951 | ENSMUSG00000079628 | 0 | NA | NA | NA | NA | NA | Rhox4g | reproductive homeobox 4G [Source:MGI S | 0 | 0 | 0 | 0 | 0 | 0 |
| 22952 | ENSMUSG00000079629 | 0 | NA | NA | NA | NA | NA | Rhox2g | reproductive homeobox 2G [Source:MGI S | 0 | 0 | 0 | 0 | 0 | 0 |
| 22953 | ENSMUSG00000079630 | 0 | NA | NA | NA | NA | NA | Rhox4f | reproductive homeobox 4F [Source:MGI S | 0 | 0 | 0 | 0 | 0 | 0 |
| 22954 | ENSMUSG00000079633 | 0 | NA | NA | NA | NA | NA | Rhox4d | reproductive homeobox 4D [Source:MGI S | 0 | 0 | 0 | 0 | 0 | 0 |
| 22955 | ENSMUSG00000079635 | 0 | NA | NA | NA | NA | NA | Rhox4c | reproductive homeobox 4C [Source:MGI S | 0 | 0 | 0 | 0 | 0 | 0 |
| 22956 | ENSMUSG00000079636 | 0 | NA | NA | NA | NA | NA | Rhox3c | reproductive homeobox 3C [Source:MGI S | 0 | 0 | 0 | 0 | 0 | 0 |

| | | | | | | | | | | | | | | | |
|---|---|---|---|---|---|---|---|---|---|---|---|---|---|---|---|
| 22957 | ENSMUSG00000079637 | 0 | NA | NA | NA | NA | NA | Rhox2c | reproductive homeobox 2C [Source:MGI S | 0 | 0 | 0 | 0 | 0 | 0 |
| 22958 | ENSMUSG00000079638 | 0 | NA | NA | NA | NA | NA | Rhox2b | reproductive homeobox 2B [Source:MGI S | 0 | 0 | 0 | 0 | 0 | 0 |
| 22959 | ENSMUSG00000079639 | 0 | NA | NA | NA | NA | NA | Rhox4a | reproductive homeobox 4A [Source:MGI S | 0 | 0 | 0 | 0 | 0 | 0 |
| 22960 | ENSMUSG00000079641 | 1107.096292 | 0.144717884 | 0.116169927 | 1.245743087 | 0.21285873 | 0.657720288 | Rpl39 | ribosomal protein L39 [Source:MGI Symbo | 1028 | 1110.2 | 1016.3 | 1046.9 | 1163.7 | 1277.6 |
| 22961 | ENSMUSG00000079642 | 0 | NA | NA | NA | NA | NA | Gm6268 | predicted gene 6268 [Source:MGI Symbol | 0 | 0 | 0 | 0 | 0 | 0 |
| 22962 | ENSMUSG00000079644 | 0 | NA | NA | NA | NA | NA | Gm1110 | predicted gene 1110 [Source:MGI Symbol | 0 | 0 | 0 | 0 | 0 | 0 |
| 22963 | ENSMUSG00000079645 | 11.81121999 | 0.844360667 | 0.671544312 | 1.2573417 | 0.20862993 | 0.653455953 | Gm17193 | predicted gene 17193 [Source:MGI Symbo | 12.047 | 4.1117 | 9.0562 | 20.244 | 9.8156 | 15.594 |
| 22964 | ENSMUSG00000079649 | 0 | NA | NA | NA | NA | NA | Gm2892 | predicted pseudogene 2892 [Source:MGI | 0 | 0 | 0 | 0 | 0 | 0 |
| 22965 | ENSMUSG00000079652 | 0.160662974 | 1.126924175 | 4.080472857 | 0.276174898 | 0.782413736 | NA | Fam71f2 | family with sequence similarity 71, memb | 0 | 0 | 0 | 0.964 | 0 | 0 |
| 22966 | ENSMUSG00000079654 | 0 | NA | NA | NA | NA | NA | Prrt4 | proline-rich transmembrane protein 4 [So | 0 | 0 | 0 | 0 | 0 | 0 |
| 22967 | ENSMUSG00000079655 | 0 | NA | NA | NA | NA | NA | Gm5168 | predicted gene 5168 [Source:MGI Symbol | 0 | 0 | 0 | 0 | 0 | 0 |
| 22968 | ENSMUSG00000079657 | 0.321325949 | 1.911575114 | 4.048267893 | 0.472195804 | 0.636787034 | NA | Rab26 | RAB26, member RAS oncogene family [So | 0 | 0 | 0 | 1.928 | 0 | 0 |
| 22969 | ENSMUSG00000079658 | 579.3389977 | 0.086997257 | 0.122451946 | 0.710460386 | 0.477418688 | 0.869706396 | Eloc | elongin C [Source:MGI Symbol;Acc:MGI:1 | 607.95 | 531.43 | 545.38 | 598.63 | 588.93 | 603.7 |
| 22970 | ENSMUSG00000079659 | 1633.02125 | 0.008065195 | 0.105117105 | 0.076725809 | 0.938841672 | 0.988721002 | Tmem243 | transmembrane protein 243, mitochondri | 1688.1 | 1558.3 | 1638.2 | 1616.6 | 1519.2 | 1777.7 |
| 22971 | ENSMUSG00000079666 | 0 | NA | NA | NA | NA | NA | Pttg1ip2 | PTTG1IP family member 2 [Source:MGI Sy | 0 | 0 | 0 | 0 | 0 | 0 |
| 22972 | ENSMUSG00000079669 | 2.403536957 | -0.903639602 | 1.512533915 | -0.597434274 | 0.550217476 | NA | Gm17396 | predicted gene, 17396 [Source:MGI Symb | 2.4093 | 1.0279 | 6.0375 | 3.8559 | 1.0906 | 0 |
| 22973 | ENSMUSG00000079671 | 9.904775777 | 0.644086303 | 0.780453851 | 0.825271478 | 0.409217481 | 0.833065722 | 2610203C22 | RIKEN cDNA 2610203C22 gene [Source:M | 8.031 | 4.1117 | 11.069 | 17.352 | 3.2719 | 15.594 |
| 22974 | ENSMUSG00000079677 | 0.85143529 | 2.085519423 | 2.590856022 | 0.804953809 | 0.420846336 | NA | Fdx2 | ferredoxin 2 [Source:MGI Symbol;Acc:MG | 0.8031 | 0 | 0 | 0.964 | 0 | 3.3415 |
| 22975 | ENSMUSG00000079679 | 0 | NA | NA | NA | NA | NA | Vwde | von Willebrand factor D and EGF domains | 0 | 0 | 0 | 0 | 0 | 0 |
| 22976 | ENSMUSG00000079681 | 0 | NA | NA | NA | NA | NA | Zglp1 | zinc finger, GATA-like protein 1 [Source:M | 0 | 0 | 0 | 0 | 0 | 0 |
| 22977 | ENSMUSG00000079685 | 5.282419647 | -0.171615872 | 1.042191182 | -0.164668321 | 0.869205053 | 0.978605211 | Ulbp1 | UL16 binding protein 1 [Source:MGI Symb | 1.6062 | 8.2233 | 7.0437 | 4.8199 | 1.0906 | 8.9108 |
| 22978 | ENSMUSG00000079693 | 0 | NA | NA | NA | NA | NA | Vmn1r-ps15 | vomeronasal 1 receptor, pseudogene 150 | 0 | 0 | 0 | 0 | 0 | 0 |
| 22979 | ENSMUSG00000079694 | 0 | NA | NA | NA | NA | NA | Gm14862 | predicted gene 14862 [Source:MGI Symbo | 0 | 0 | 0 | 0 | 0 | 0 |
| 22980 | ENSMUSG00000079697 | 3.614878352 | 0.890644803 | 1.286041678 | 0.692547386 | 0.488593638 | NA | Gm5751 | predicted gene 5751 [Source:MGI Symbol | 1.6062 | 2.0558 | 4.025 | 9.6398 | 4.3625 | 0 |
| 22981 | ENSMUSG00000079698 | 0.133850373 | -0.796629176 | 4.080472857 | -0.195229623 | 0.84521319 | NA | Vmn2r93 | vomeronasal 2, receptor 93 [Source:MGI S | 0.8031 | 0 | 0 | 0 | 0 | 0 |
| 22982 | ENSMUSG00000079699 | 0.781659665 | -3.010437285 | 2.830777252 | -1.063466679 | 0.287570368 | NA | Gm6592 | predicted gene 6592 [Source:MGI Symbol | 1.6062 | 3.0838 | 0 | 0 | 0 | 0 |
| 22983 | ENSMUSG00000079700 | 0 | NA | NA | NA | NA | NA | Fpr3 | formyl peptide receptor 3 [Source:MGI Sy | 0 | 0 | 0 | 0 | 0 | 0 |
| 22984 | ENSMUSG00000079701 | 0 | NA | NA | NA | NA | NA | Ssxb3 | synovial sarcoma, X member B3 [Source:N | 0 | 0 | 0 | 0 | 0 | 0 |
| 22985 | ENSMUSG00000079702 | 0 | NA | NA | NA | NA | NA | Ssxb6 | synovial sarcoma, X member B6 [Source:N | 0 | 0 | 0 | 0 | 0 | 0 |
| 22986 | ENSMUSG00000079703 | 0 | NA | NA | NA | NA | NA | Ssxb8 | synovial sarcoma, X member B8 [Source:N | 0 | 0 | 0 | 0 | 0 | 0 |
| 22987 | ENSMUSG00000079704 | 0 | NA | NA | NA | NA | NA | Gm14459 | predicted gene 14459 [Source:MGI Symbo | 0 | 0 | 0 | 0 | 0 | 0 |
| 22988 | ENSMUSG00000079705 | 0 | NA | NA | NA | NA | NA | Ssxb1 | synovial sarcoma, X member B1 [Source:N | 0 | 0 | 0 | 0 | 0 | 0 |
| 22989 | ENSMUSG00000079707 | 0 | NA | NA | NA | NA | NA | Dynlt2a1 | dynein light chain Tctex-type 2A1 [Source: | 0 | 0 | 0 | 0 | 0 | 0 |
| 22990 | ENSMUSG00000079710 | 0 | NA | NA | NA | NA | NA | Dynlt2a2 | dynein light chain Tctex-type 2A2 [Source: | 0 | 0 | 0 | 0 | 0 | 0 |
| 22991 | ENSMUSG00000079711 | 0 | NA | NA | NA | NA | NA | Smok4a | sperm motility kinase 4A [Source:MGI Syn | 0 | 0 | 0 | 0 | 0 | 0 |
| 22992 | ENSMUSG00000079722 | 0.503095663 | 2.549494177 | 3.348715477 | 0.761334964 | 0.446457022 | NA | Ttll2 | tubulin tyrosine ligase-like family, membe | 0 | 0 | 0 | 1.928 | 1.0906 | 0 |
| 22993 | ENSMUSG00000079737 | 37.41375905 | 0.625779467 | 0.380506671 | 1.64459526 | 0.100053305 | 0.4657776 | 3110001I22R | RIKEN cDNA 3110001I22 gene [Source:MG | 30.518 | 22.614 | 35.219 | 57.839 | 37.081 | 41.212 |
| 22994 | ENSMUSG00000079740 | 0 | NA | NA | NA | NA | NA | Gm11172 | predicted gene 11172 [Source:MGI Symbo | 0 | 0 | 0 | 0 | 0 | 0 |
| 22995 | ENSMUSG00000079741 | 0 | NA | NA | NA | NA | NA | Nlrp4g | NLR family, pyrin domain containing 4G [S | 0 | 0 | 0 | 0 | 0 | 0 |
| 22996 | ENSMUSG00000079764 | 0 | NA | NA | NA | NA | NA | NA | NA | 0 | 0 | 0 | 0 | 0 | 0 |
| 22997 | ENSMUSG00000079773 | 0 | NA | NA | NA | NA | NA | NA | NA | 0 | 0 | 0 | 0 | 0 | 0 |
| 22998 | ENSMUSG00000079774 | 0 | NA | NA | NA | NA | NA | NA | NA | 0 | 0 | 0 | 0 | 0 | 0 |
| 22999 | ENSMUSG00000079794 | 0 | NA | NA | NA | NA | NA | | | 0 | 0 | 0 | 0 | 0 | 0 |
| 23000 | ENSMUSG00000079800 | 0 | NA | NA | NA | NA | NA | | | 0 | 0 | 0 | 0 | 0 | 0 |
| 23001 | ENSMUSG00000079806 | 0 | NA | NA | NA | NA | NA | Gm21719 | predicted gene, 21719 [Source:MGI Symb | 0 | 0 | 0 | 0 | 0 | 0 |
| 23002 | ENSMUSG00000079808 | 0.185640647 | 1.126924175 | 4.080472857 | 0.276174898 | 0.782413736 | NA | | | 0 | 0 | 0 | 0 | 0 | 1.1138 |
| 23003 | ENSMUSG00000079834 | 272.2626133 | 0.024933785 | 0.184749281 | 0.134960118 | 0.892643407 | 0.982512308 | NA | NA | 254.58 | 306.32 | 248.54 | 215.93 | 311.92 | 296.28 |
| 23004 | ENSMUSG00000079842 | 0 | NA | NA | NA | NA | NA | Spag11a | sperm associated antigen 11A [Source:MG | 0 | 0 | 0 | 0 | 0 | 0 |
| 23005 | ENSMUSG00000079845 | 4.726331903 | 0.752917997 | 3.097419864 | 0.243079088 | 0.80794413 | NA | Xlr4a | X-linked lymphocyte-regulated 4A [Source | 6.4248 | 4.1117 | 0 | 0 | 0 | 17.822 |
| 23006 | ENSMUSG00000079852 | 0.167707431 | -0.796629176 | 4.080472857 | -0.195229623 | 0.84521319 | NA | Klra4 | killer cell lectin-like receptor, subfamily A, | 0 | 0 | 1.0062 | 0 | 0 | 0 |
| 23007 | ENSMUSG00000079853 | 0.321325949 | 1.911575114 | 4.048267893 | 0.472195804 | 0.636787034 | NA | Klra1 | killer cell lectin-like receptor, subfamily A, | 0 | 0 | 0 | 1.928 | 0 | 0 |
| 23008 | ENSMUSG00000079871 | 0 | NA | NA | NA | NA | NA | Gm4984 | predicted pseudogene 4984 [Source:MGI | 0 | 0 | 0 | 0 | 0 | 0 |
| 23009 | ENSMUSG00000079875 | 0 | NA | NA | NA | NA | NA | Olfr22-ps1 | olfactory receptor 22, pseudogene 1 [Sou | 0 | 0 | 0 | 0 | 0 | 0 |
| 23010 | ENSMUSG00000079884 | 5.887239576 | 1.38268545 | 0.969021512 | 1.426888292 | 0.153612096 | 0.570925073 | Gm10698 | predicted gene 10698 [Source:MGI Symbo | 5.6217 | 1.0279 | 3.0187 | 13.496 | 4.3625 | 7.7969 |
| 23011 | ENSMUSG00000079889 | 0.160662974 | 1.126924175 | 4.080472857 | 0.276174898 | 0.782413736 | NA | Gm11951 | predicted gene 11951 [Source:MGI Symbo | 0 | 0 | 0 | 0.964 | 0 | 0 |
| 23012 | ENSMUSG00000079890 | 0 | NA | NA | NA | NA | NA | Gm12106 | predicted gene 12106 [Source:MGI Symbo | 0 | 0 | 0 | 0 | 0 | 0 |

| | | | | | | | | | | | | | | | |
|---|---|---|---|---|---|---|---|---|---|---|---|---|---|---|---|
| 23013 | ENSMUSG00000079892 | 0 | NA | NA | NA | NA | NA | Scgb1b22-ps | secretoglobin, family 1B, member 22, pseu | 0 | 0 | 0 | 0 | 0 | 0 |
| 23014 | ENSMUSG00000079893 | 0 | NA | NA | NA | NA | NA | Scgb1b23-ps | secretoglobin, family 1B, member 23, pseu | 0 | 0 | 0 | 0 | 0 | 0 |
| 23015 | ENSMUSG00000079906 | 2.712551273 | -0.035737836 | 1.314582306 | -0.027185697 | 0.978311624 | NA | Gm15846 | predicted gene 15846 [Source:MGI Symbo | 3.2124 | 2.0558 | 3.0187 | 5.7839 | 1.0906 | 1.1138 |
| 23016 | ENSMUSG00000079915 | 1.195212088 | 0.46832086 | 2.069471296 | 0.226299761 | 0.820968284 | NA | Gm16073 | predicted gene 16073 [Source:MGI Symbo | 0 | 2.0558 | 1.0062 | 1.928 | 2.1812 | 0 |
| 23017 | ENSMUSG00000079941 | 10.38086621 | 0.235314175 | 0.652380742 | 0.360700677 | 0.718323218 | 0.9447719 | Cox5b-ps | cytochrome c oxidase subunit 5B, pseudo | 11.243 | 8.2233 | 9.0562 | 10.604 | 10.906 | 12.252 |
| 23018 | ENSMUSG00000079942 | 4.515287351 | -0.20843923 | 1.050037679 | -0.198506429 | 0.842648855 | NA | Rpl28-ps3 | ribosomal protein L28, pseudogene 3 [Sou | 2.4093 | 6.1675 | 6.0375 | 4.8199 | 6.5437 | 1.1138 |
| 23019 | ENSMUSG00000079962 | 0 | NA | NA | NA | NA | NA | Gm12643 | predicted gene 12643 [Source:MGI Symbo | 0 | 0 | 0 | 0 | 0 | 0 |
| 23020 | ENSMUSG00000079963 | 0 | NA | NA | NA | NA | NA | Gm7146 | predicted pseudogene 7146 [Source:MGI | 0 | 0 | 0 | 0 | 0 | 0 |
| 23021 | ENSMUSG00000079965 | 0.160662974 | 1.126924175 | 4.080472857 | 0.276174898 | 0.782413736 | NA | Gm14853 | predicted gene 14853 [Source:MGI Symbo | 0 | 0 | 0 | 0.964 | 0 | 0 |
| 23022 | ENSMUSG00000079966 | 0 | NA | NA | NA | NA | NA | Gm14058 | predicted gene 14058 [Source:MGI Symbo | 0 | 0 | 0 | 0 | 0 | 0 |
| 23023 | ENSMUSG00000079988 | 0 | NA | NA | NA | NA | NA | Gm11970 | predicted gene 11970 [Source:MGI Symbo | 0 | 0 | 0 | 0 | 0 | 0 |
| 23024 | ENSMUSG00000079993 | 3.244774267 | 0.561136974 | 1.258757508 | 0.445786397 | 0.655751549 | NA | Gm11799 | predicted gene 11799 [Source:MGI Symbo | 0.8031 | 5.1396 | 2.0125 | 3.8559 | 6.5437 | 1.1138 |
| 23025 | ENSMUSG00000079998 | 0 | NA | NA | NA | NA | NA | Gm12947 | predicted gene 12947 [Source:MGI Symbo | 0 | 0 | 0 | 0 | 0 | 0 |
| 23026 | ENSMUSG00000080002 | 1.971942065 | -1.374933004 | 1.711235585 | -0.803473827 | 0.421700919 | NA | Gm5566 | predicted pseudogene 5566 [Source:MGI | 2.4093 | 2.0558 | 4.025 | 0 | 0 | 3.3415 |
| 23027 | ENSMUSG00000080006 | 1.048870122 | 1.109222282 | 2.077678687 | 0.533875757 | 0.593427496 | NA | Rps19-ps7 | ribosomal protein S19, pseudogene 7 [Sou | 0 | 1.0279 | 1.0062 | 0.964 | 2.1812 | 1.1138 |
| 23028 | ENSMUSG00000080021 | 0.160662974 | 1.126924175 | 4.080472857 | 0.276174898 | 0.782413736 | NA | Gm5915 | predicted pseudogene 5915 [Source:MGI | 0 | 0 | 0 | 0.964 | 0 | 0 |
| 23029 | ENSMUSG00000080046 | 0 | NA | NA | NA | NA | NA | Gm15108 | predicted gene 15108 [Source:MGI Symbo | 0 | 0 | 0 | 0 | 0 | 0 |
| 23030 | ENSMUSG00000080050 | 0 | NA | NA | NA | NA | NA | Gm14864 | predicted gene 14864 [Source:MGI Symbo | 0 | 0 | 0 | 0 | 0 | 0 |
| 23031 | ENSMUSG00000080051 | 0 | NA | NA | NA | NA | NA | Gm14510 | predicted gene 14510 [Source:MGI Symbo | 0 | 0 | 0 | 0 | 0 | 0 |
| 23032 | ENSMUSG00000080054 | 0.160662974 | 1.126924175 | 4.080472857 | 0.276174898 | 0.782413736 | NA | Rpl12-ps1 | ribosomal protein L12, pseudogene 1 [Sou | 0 | 0 | 0 | 0.964 | 0 | 0 |
| 23033 | ENSMUSG00000080057 | 0 | NA | NA | NA | NA | NA | Gm12378 | predicted gene 12378 [Source:MGI Symbo | 0 | 0 | 0 | 0 | 0 | 0 |
| 23034 | ENSMUSG00000080058 | 0 | NA | NA | NA | NA | NA | Gm11175 | predicted gene 11175 [Source:MGI Symbo | 0 | 0 | 0 | 0 | 0 | 0 |
| 23035 | ENSMUSG00000080059 | 15.0950229 | 0.179747881 | 0.577227826 | 0.311398504 | 0.755497691 | 0.953085143 | Rps19-ps3 | ribosomal protein S19, pseudogene 3 [Sou | 12.85 | 18.503 | 11.069 | 10.604 | 15.269 | 22.277 |
| 23036 | ENSMUSG00000080065 | 0 | NA | NA | NA | NA | NA | Gm11864 | predicted gene 11864 [Source:MGI Symbo | 0 | 0 | 0 | 0 | 0 | 0 |
| 23037 | ENSMUSG00000080069 | 0 | NA | NA | NA | NA | NA | Gm41 | predicted pseudogene 41 [Source:MGI Sy | 0 | 0 | 0 | 0 | 0 | 0 |
| 23038 | ENSMUSG00000080071 | 0 | NA | NA | NA | NA | NA | Gm15125 | predicted gene 15125 [Source:MGI Symbo | 0 | 0 | 0 | 0 | 0 | 0 |
| 23039 | ENSMUSG00000080072 | 0 | NA | NA | NA | NA | NA | Spin2-ps2 | spindlin family, member 2, pseudogene 2 | 0 | 0 | 0 | 0 | 0 | 0 |
| 23040 | ENSMUSG00000080073 | 0 | NA | NA | NA | NA | NA | Gm9427 | predicted pseudogene 9427 [Source:MGI | 0 | 0 | 0 | 0 | 0 | 0 |
| 23041 | ENSMUSG00000080076 | 0 | NA | NA | NA | NA | NA | H2ac14 | H2A clustered histone 14 [Source:MGI Syr | 0 | 0 | 0 | 0 | 0 | 0 |
| 23042 | ENSMUSG00000080078 | 0 | NA | NA | NA | NA | NA | Gm15583 | predicted gene 15583 [Source:MGI Symbo | 0 | 0 | 0 | 0 | 0 | 0 |
| 23043 | ENSMUSG00000080115 | 0.371281295 | 2.09253362 | 4.031852165 | 0.519000582 | 0.603760335 | NA | Eef1akmt3 | EEF1A lysine methyltransferase 3 [Source: | 0 | 0 | 0 | 0 | 0 | 2.2277 |
| 23044 | ENSMUSG00000080142 | 0 | NA | NA | NA | NA | NA | Gm12838 | predicted gene 12838 [Source:MGI Symbo | 0 | 0 | 0 | 0 | 0 | 0 |
| 23045 | ENSMUSG00000080152 | 0.496077836 | -0.812642266 | 3.444120572 | -0.235950586 | 0.813471014 | NA | H3f4 | H3.4 histone [Source:MGI Symbol;Acc:MG | 0 | 0 | 2.0125 | 0.964 | 0 | 0 |
| 23046 | ENSMUSG00000080155 | 0 | NA | NA | NA | NA | NA | Gm14777 | predicted gene 14777 [Source:MGI Symbo | 0 | 0 | 0 | 0 | 0 | 0 |
| 23047 | ENSMUSG00000080158 | 0 | NA | NA | NA | NA | NA | Gm14385 | predicted gene 14385 [Source:MGI Symbo | 0 | 0 | 0 | 0 | 0 | 0 |
| 23048 | ENSMUSG00000080175 | 0 | NA | NA | NA | NA | NA | Tpm3-rs2 | tropomyosin 3, related sequence 2 [Sourc | 0 | 0 | 0 | 0 | 0 | 0 |
| 23049 | ENSMUSG00000080180 | 0.185640647 | 1.126924175 | 4.080472857 | 0.276174898 | 0.782413736 | NA | Gm17235 | predicted gene 17235 [Source:MGI Symbo | 0 | 0 | 0 | 0 | 0 | 1.1138 |
| 23050 | ENSMUSG00000080186 | 0 | NA | NA | NA | NA | NA | Gm14448 | predicted gene 14448 [Source:MGI Symbo | 0 | 0 | 0 | 0 | 0 | 0 |
| 23051 | ENSMUSG00000080193 | 0 | NA | NA | NA | NA | NA | Gm14756 | predicted gene 14756 [Source:MGI Symbo | 0 | 0 | 0 | 0 | 0 | 0 |
| 23052 | ENSMUSG00000080198 | 2.203232829 | 2.633637754 | 1.640464342 | 1.605422127 | 0.108400938 | NA | Rpl10a-ps4 | ribosomal protein L10A, pseudogene 4 [So | 0.8031 | 0 | 1.0062 | 4.8199 | 4.3625 | 2.2277 |
| 23053 | ENSMUSG00000080205 | 1.295411994 | -1.45671837 | 2.07740264 | -0.701221007 | 0.483165102 | NA | Gm15526 | predicted gene 15526 [Source:MGI Symbo | 1.6062 | 4.1117 | 0 | 0.964 | 1.0906 | 0 |
| 23054 | ENSMUSG00000080222 | 0 | NA | NA | NA | NA | NA | Gm11486 | predicted gene 11486 [Source:MGI Symbo | 0 | 0 | 0 | 0 | 0 | 0 |
| 23055 | ENSMUSG00000080223 | 0.181769715 | 1.126924175 | 4.080472857 | 0.276174898 | 0.782413736 | NA | Gm13768 | predicted gene 13768 [Source:MGI Symbo | 0 | 0 | 0 | 0 | 1.0906 | 0 |
| 23056 | ENSMUSG00000080227 | 0 | NA | NA | NA | NA | NA | Gm13343 | predicted gene 13343 [Source:MGI Symbo | 0 | 0 | 0 | 0 | 0 | 0 |
| 23057 | ENSMUSG00000080237 | 3.140625781 | 0.666062155 | 1.284005495 | 0.518737776 | 0.603943615 | NA | Gm14239 | predicted gene 14239 [Source:MGI Symbo | 3.2124 | 0 | 4.025 | 3.8559 | 2.1812 | 5.5692 |
| 23058 | ENSMUSG00000080241 | 0 | NA | NA | NA | NA | NA | Dppa3-ps | developmental pluripotency-associated 3, | 0 | 0 | 0 | 0 | 0 | 0 |
| 23059 | ENSMUSG00000080242 | 5.110117244 | 0.302553041 | 0.974386622 | 0.310506152 | 0.756176082 | 0.953085143 | Atp6v0c-ps2 | ATPase, H+ transporting, lysosomal V0 sul | 1.6062 | 8.2233 | 4.025 | 5.7839 | 5.4531 | 5.5692 |
| 23060 | ENSMUSG00000080250 | 0 | NA | NA | NA | NA | NA | Gm16400 | predicted gene 16400 [Source:MGI Symbo | 0 | 0 | 0 | 0 | 0 | 0 |
| 23061 | ENSMUSG00000080254 | 0.531944269 | 2.618916564 | 3.285123409 | 0.797204926 | 0.425332026 | NA | Fam220-ps | family with sequence similarity 220, pseu | 0 | 0 | 0 | 0.964 | 0 | 2.2277 |
| 23062 | ENSMUSG00000080266 | 0 | NA | NA | NA | NA | NA | Gm14772 | predicted gene 14772 [Source:MGI Symbo | 0 | 0 | 0 | 0 | 0 | 0 |
| 23063 | ENSMUSG00000080268 | 377.1174999 | -0.003635544 | 0.150232756 | -0.024199412 | 0.980693547 | 0.99820894 | Brms1 | breast cancer metastasis-suppressor 1 [Sc | 369.43 | 386.5 | 376.34 | 318.11 | 402.44 | 409.89 |
| 23064 | ENSMUSG00000080283 | 0 | NA | NA | NA | NA | NA | Gm14806 | predicted gene 14806 [Source:MGI Symbo | 0 | 0 | 0 | 0 | 0 | 0 |
| 23065 | ENSMUSG00000080298 | 0 | NA | NA | NA | NA | NA | Gm14807 | predicted gene 14807 [Source:MGI Symbo | 0 | 0 | 0 | 0 | 0 | 0 |
| 23066 | ENSMUSG00000080299 | 0 | NA | NA | NA | NA | NA | Gm14350 | predicted gene 14350 [Source:MGI Symbo | 0 | 0 | 0 | 0 | 0 | 0 |
| 23067 | ENSMUSG00000080300 | 0 | NA | NA | NA | NA | NA | Spin2-ps1 | spindlin family, member 2, pseudogene 1 | 0 | 0 | 0 | 0 | 0 | 0 |
| 23068 | ENSMUSG00000080303 | 0 | NA | NA | NA | NA | NA | Gm12419 | predicted gene 12419 [Source:MGI Symbo | 0 | 0 | 0 | 0 | 0 | 0 |

| | | | | | | | | | | | | | | | |
|---|---|---|---|---|---|---|---|---|---|---|---|---|---|---|---|
| 23069 | ENSMUSG00000080316 | 27.42700438 | 0.71023625 | 0.610932855 | 1.162543878 | 0.245014613 | 0.695017707 | Spaca6 | sperm acrosome associated 6 [Source:MG | 20.078 | 15.419 | 27.169 | 73.262 | 15.269 | 13.366 |
| 23070 | ENSMUSG00000080317 | 0 | NA | NA | NA | NA | NA | Vmn2r-ps94 | vomeronasal 2, receptor, pseudogene 94 | 0 | 0 | 0 | 0 | 0 | 0 |
| 23071 | ENSMUSG00000080321 | 0 | NA | NA | NA | NA | NA | Gm14644 | predicted gene 14644 [Source:MGI Symbo | 0 | 0 | 0 | 0 | 0 | 0 |
| 23072 | ENSMUSG00000080330 | 0 | NA | NA | NA | NA | NA | NA | NA | 0 | 0 | 0 | 0 | 0 | 0 |
| 23073 | ENSMUSG00000080331 | 0 | NA | NA | NA | NA | NA | Mir1298 | microRNA 1298 [Source:MGI Symbol;Acc: | 0 | 0 | 0 | 0 | 0 | 0 |
| 23074 | ENSMUSG00000080334 | 0 | NA | NA | NA | NA | NA | NA | NA | 0 | 0 | 0 | 0 | 0 | 0 |
| 23075 | ENSMUSG00000080336 | 0 | NA | NA | NA | NA | NA | NA | NA | 0 | 0 | 0 | 0 | 0 | 0 |
| 23076 | ENSMUSG00000080342 | 0 | NA | NA | NA | NA | NA | NA | NA | 0 | 0 | 0 | 0 | 0 | 0 |
| 23077 | ENSMUSG00000080345 | 0 | NA | NA | NA | NA | NA | NA | NA | 0 | 0 | 0 | 0 | 0 | 0 |
| 23078 | ENSMUSG00000080346 | 0 | NA | NA | NA | NA | NA | NA | NA | 0 | 0 | 0 | 0 | 0 | 0 |
| 23079 | ENSMUSG00000080347 | 0 | NA | NA | NA | NA | NA | NA | NA | 0 | 0 | 0 | 0 | 0 | 0 |
| 23080 | ENSMUSG00000080348 | 0 | NA | NA | NA | NA | NA | NA | NA | 0 | 0 | 0 | 0 | 0 | 0 |
| 23081 | ENSMUSG00000080349 | 0 | NA | NA | NA | NA | NA | NA | NA | 0 | 0 | 0 | 0 | 0 | 0 |
| 23082 | ENSMUSG00000080350 | 0 | NA | NA | NA | NA | NA | NA | NA | 0 | 0 | 0 | 0 | 0 | 0 |
| 23083 | ENSMUSG00000080352 | 0 | NA | NA | NA | NA | NA | Gm26247 | predicted gene, 26247 [Source:MGI Symb | 0 | 0 | 0 | 0 | 0 | 0 |
| 23084 | ENSMUSG00000080356 | 0 | NA | NA | NA | NA | NA | Mir1247 | microRNA 1247 [Source:MGI Symbol;Acc: | 0 | 0 | 0 | 0 | 0 | 0 |
| 23085 | ENSMUSG00000080358 | 0 | NA | NA | NA | NA | NA | Gm26249 | predicted gene, 26249 [Source:MGI Symb | 0 | 0 | 0 | 0 | 0 | 0 |
| 23086 | ENSMUSG00000080359 | 0 | NA | NA | NA | NA | NA | NA | NA | 0 | 0 | 0 | 0 | 0 | 0 |
| 23087 | ENSMUSG00000080360 | 0 | NA | NA | NA | NA | NA | NA | NA | 0 | 0 | 0 | 0 | 0 | 0 |
| 23088 | ENSMUSG00000080363 | 0 | NA | NA | NA | NA | NA | NA | NA | 0 | 0 | 0 | 0 | 0 | 0 |
| 23089 | ENSMUSG00000080364 | 0.963481514 | 0.243735673 | 2.098497405 | 0.116147712 | 0.907535476 | NA | Gm25777 | predicted gene, 25777 [Source:MGI Symb | 1.6062 | 0 | 1.0062 | 0.964 | 1.0906 | 1.1138 |
| 23090 | ENSMUSG00000080365 | 0.506734502 | -2.372847156 | 3.338060886 | -0.710845978 | 0.477179685 | NA | Gm25776 | predicted gene, 25776 [Source:MGI Symb | 0 | 1.0279 | 2.0125 | 0 | 0 | 0 |
| 23091 | ENSMUSG00000080366 | 0 | NA | NA | NA | NA | NA | NA | NA | 0 | 0 | 0 | 0 | 0 | 0 |
| 23092 | ENSMUSG00000080367 | 0 | NA | NA | NA | NA | NA | NA | NA | 0 | 0 | 0 | 0 | 0 | 0 |
| 23093 | ENSMUSG00000080368 | 0 | NA | NA | NA | NA | NA | NA | NA | 0 | 0 | 0 | 0 | 0 | 0 |
| 23094 | ENSMUSG00000080370 | 0.738691657 | 3.085611452 | 2.924644825 | 1.055038008 | 0.291407942 | NA | NA | NA | 0 | 0 | 0 | 0 | 1.0906 | 3.3415 |
| 23095 | ENSMUSG00000080374 | 0 | NA | NA | NA | NA | NA | Gm22953 | predicted gene, 22953 [Source:MGI Symb | 0 | 0 | 0 | 0 | 0 | 0 |
| 23096 | ENSMUSG00000080378 | 0 | NA | NA | NA | NA | NA | Gm25309 | predicted gene, 25309 [Source:MGI Symb | 0 | 0 | 0 | 0 | 0 | 0 |
| 23097 | ENSMUSG00000080380 | 0 | NA | NA | NA | NA | NA | NA | NA | 0 | 0 | 0 | 0 | 0 | 0 |
| 23098 | ENSMUSG00000080383 | 0 | NA | NA | NA | NA | NA | NA | NA | 0 | 0 | 0 | 0 | 0 | 0 |
| 23099 | ENSMUSG00000080384 | 0 | NA | NA | NA | NA | NA | Gm22105 | predicted gene, 22105 [Source:MGI Symb | 0 | 0 | 0 | 0 | 0 | 0 |
| 23100 | ENSMUSG00000080391 | 0 | NA | NA | NA | NA | NA | NA | NA | 0 | 0 | 0 | 0 | 0 | 0 |
| 23101 | ENSMUSG00000080394 | 0 | NA | NA | NA | NA | NA | NA | NA | 0 | 0 | 0 | 0 | 0 | 0 |
| 23102 | ENSMUSG00000080396 | 0 | NA | NA | NA | NA | NA | Snord111 | small nucleolar RNA, C/D box 111 [Source | 0 | 0 | 0 | 0 | 0 | 0 |
| 23103 | ENSMUSG00000080401 | 0 | NA | NA | NA | NA | NA | NA | NA | 0 | 0 | 0 | 0 | 0 | 0 |
| 23104 | ENSMUSG00000080402 | 0 | NA | NA | NA | NA | NA | NA | NA | 0 | 0 | 0 | 0 | 0 | 0 |
| 23105 | ENSMUSG00000080405 | 0 | NA | NA | NA | NA | NA | NA | NA | 0 | 0 | 0 | 0 | 0 | 0 |
| 23106 | ENSMUSG00000080406 | 0 | NA | NA | NA | NA | NA | Mir1188 | microRNA 1188 [Source:MGI Symbol;Acc: | 0 | 0 | 0 | 0 | 0 | 0 |
| 23107 | ENSMUSG00000080408 | 0.321325949 | 1.911575114 | 4.048267893 | 0.472195804 | 0.636787034 | NA | NA | NA | 0 | 0 | 0 | 1.928 | 0 | 0 |
| 23108 | ENSMUSG00000080409 | 0 | NA | NA | NA | NA | NA | Mir467h | microRNA 467h [Source:MGI Symbol;Acc: | 0 | 0 | 0 | 0 | 0 | 0 |
| 23109 | ENSMUSG00000080410 | 0 | NA | NA | NA | NA | NA | NA | NA | 0 | 0 | 0 | 0 | 0 | 0 |
| 23110 | ENSMUSG00000080411 | 0 | NA | NA | NA | NA | NA | Mir1193 | microRNA 1193 [Source:MGI Symbol;Acc: | 0 | 0 | 0 | 0 | 0 | 0 |
| 23111 | ENSMUSG00000080412 | 0 | NA | NA | NA | NA | NA | NA | NA | 0 | 0 | 0 | 0 | 0 | 0 |
| 23112 | ENSMUSG00000080413 | 0 | NA | NA | NA | NA | NA | Gm26473 | predicted gene, 26473 [Source:MGI Symb | 0 | 0 | 0 | 0 | 0 | 0 |
| 23113 | ENSMUSG00000080415 | 0 | NA | NA | NA | NA | NA | NA | NA | 0 | 0 | 0 | 0 | 0 | 0 |
| 23114 | ENSMUSG00000080419 | 0 | NA | NA | NA | NA | NA | NA | NA | 0 | 0 | 0 | 0 | 0 | 0 |
| 23115 | ENSMUSG00000080424 | 0 | NA | NA | NA | NA | NA | Mir466k | microRNA 466k [Source:MGI Symbol;Acc:I | 0 | 0 | 0 | 0 | 0 | 0 |
| 23116 | ENSMUSG00000080425 | 0 | NA | NA | NA | NA | NA | Gm24833 | predicted gene, 24833 [Source:MGI Symb | 0 | 0 | 0 | 0 | 0 | 0 |
| 23117 | ENSMUSG00000080427 | 0 | NA | NA | NA | NA | NA | NA | NA | 0 | 0 | 0 | 0 | 0 | 0 |
| 23118 | ENSMUSG00000080428 | 0 | NA | NA | NA | NA | NA | Gm24831 | predicted gene, 24831 [Source:MGI Symb | 0 | 0 | 0 | 0 | 0 | 0 |
| 23119 | ENSMUSG00000080430 | 0 | NA | NA | NA | NA | NA | NA | NA | 0 | 0 | 0 | 0 | 0 | 0 |
| 23120 | ENSMUSG00000080432 | 0 | NA | NA | NA | NA | NA | NA | NA | 0 | 0 | 0 | 0 | 0 | 0 |
| 23121 | ENSMUSG00000080434 | 0 | NA | NA | NA | NA | NA | NA | NA | 0 | 0 | 0 | 0 | 0 | 0 |
| 23122 | ENSMUSG00000080437 | 0 | NA | NA | NA | NA | NA | Gm24274 | predicted gene, 24274 [Source:MGI Symb | 0 | 0 | 0 | 0 | 0 | 0 |
| 23123 | ENSMUSG00000080440 | 0.17131964 | -0.796629176 | 4.080472857 | -0.195229623 | 0.84521319 | NA | Gm25848 | predicted gene, 25848 [Source:MGI Symb | 0 | 1.0279 | 0 | 0 | 0 | 0 |
| 23124 | ENSMUSG00000080441 | 0 | NA | NA | NA | NA | NA | Mir1249 | microRNA 1249 [Source:MGI Symbol;Acc: | 0 | 0 | 0 | 0 | 0 | 0 |

| | | | | | | | | | | | | | | | |
|---|---|---|---|---|---|---|---|---|---|---|---|---|---|---|---|
| 23125 | ENSMUSG00000080443 | 0 | NA | NA | NA | NA | NA | NA | NA | 0 | 0 | 0 | 0 | 0 | 0 |
| 23126 | ENSMUSG00000080451 | 0 | NA | NA | NA | NA | NA | NA | NA | 0 | 0 | 0 | 0 | 0 | 0 |
| 23127 | ENSMUSG00000080452 | 0 | NA | NA | NA | NA | NA | Mir1306 | microRNA 1306 [Source:MGI Symbol;Acc: | 0 | 0 | 0 | 0 | 0 | 0 |
| 23128 | ENSMUSG00000080454 | 0.596071152 | -1.224343926 | 2.897583865 | -0.4225396 | 0.672631203 | NA | Gm24195 | predicted gene, 24195 [Source:MGI Symb | 1.6062 | 0 | 1.0062 | 0.964 | 0 | 0 |
| 23129 | ENSMUSG00000080455 | 0 | NA | NA | NA | NA | NA | NA | NA | 0 | 0 | 0 | 0 | 0 | 0 |
| 23130 | ENSMUSG00000080459 | 0 | NA | NA | NA | NA | NA | Mir669g | microRNA 669g [Source:MGI Symbol;Acc: | 0 | 0 | 0 | 0 | 0 | 0 |
| 23131 | ENSMUSG00000080461 | 0 | NA | NA | NA | NA | NA | Gm22488 | predicted gene, 22488 [Source:MGI Symb | 0 | 0 | 0 | 0 | 0 | 0 |
| 23132 | ENSMUSG00000080463 | 0 | NA | NA | NA | NA | NA | Gm22489 | predicted gene, 22489 [Source:MGI Symb | 0 | 0 | 0 | 0 | 0 | 0 |
| 23133 | ENSMUSG00000080465 | 2.227951778 | 2.640129933 | 1.63788473 | 1.61191437 | 0.106980575 | NA | Snord94 | small nucleolar RNA, C/D box 94 [Source:I | 0.8031 | 1.0279 | 0 | 3.8559 | 5.4531 | 2.2277 |
| 23134 | ENSMUSG00000080468 | 0 | NA | NA | NA | NA | NA | Mir1195 | microRNA 1195 [Source:MGI Symbol;Acc: | 0 | 0 | 0 | 0 | 0 | 0 |
| 23135 | ENSMUSG00000080469 | 0.788497532 | -1.691662715 | 2.429158853 | -0.696398555 | 0.486179268 | NA | Snord98 | small nucleolar RNA, C/D box 98 [Source:I | 1.6062 | 1.0279 | 1.0062 | 0 | 1.0906 | 0 |
| 23136 | ENSMUSG00000080473 | 0 | NA | NA | NA | NA | NA | NA | NA | 0 | 0 | 0 | 0 | 0 | 0 |
| 23137 | ENSMUSG00000080474 | 0 | NA | NA | NA | NA | NA | NA | NA | 0 | 0 | 0 | 0 | 0 | 0 |
| 23138 | ENSMUSG00000080476 | 0 | NA | NA | NA | NA | NA | NA | NA | 0 | 0 | 0 | 0 | 0 | 0 |
| 23139 | ENSMUSG00000080478 | 1.047732603 | 1.259689577 | 2.445764001 | 0.515049521 | 0.606518416 | NA | Snord23 | small nucleolar RNA, C/D box 23 [Source:I | 0.8031 | 1.0279 | 0 | 0 | 0 | 4.4554 |
| 23140 | ENSMUSG00000080480 | 0 | NA | NA | NA | NA | NA | Gm23494 | predicted gene, 23494 [Source:MGI Symb | 0 | 0 | 0 | 0 | 0 | 0 |
| 23141 | ENSMUSG00000080485 | 0.133850373 | -0.796629176 | 4.080472857 | -0.195229623 | 0.84521319 | NA | NA | NA | 0.8031 | 0 | 0 | 0 | 0 | 0 |
| 23142 | ENSMUSG00000080486 | 0 | NA | NA | NA | NA | NA | NA | NA | 0 | 0 | 0 | 0 | 0 | 0 |
| 23143 | ENSMUSG00000080487 | 0 | NA | NA | NA | NA | NA | NA | NA | 0 | 0 | 0 | 0 | 0 | 0 |
| 23144 | ENSMUSG00000080495 | 0 | NA | NA | NA | NA | NA | NA | NA | 0 | 0 | 0 | 0 | 0 | 0 |
| 23145 | ENSMUSG00000080498 | 0 | NA | NA | NA | NA | NA | NA | NA | 0 | 0 | 0 | 0 | 0 | 0 |
| 23146 | ENSMUSG00000080499 | 0 | NA | NA | NA | NA | NA | Gm25338 | predicted gene, 25338 [Source:MGI Symb | 0 | 0 | 0 | 0 | 0 | 0 |
| 23147 | ENSMUSG00000080500 | 0 | NA | NA | NA | NA | NA | NA | NA | 0 | 0 | 0 | 0 | 0 | 0 |
| 23148 | ENSMUSG00000080504 | 0 | NA | NA | NA | NA | NA | Mir669d-2 | microRNA 669d-2 [Source:MGI Symbol;Ac | 0 | 0 | 0 | 0 | 0 | 0 |
| 23149 | ENSMUSG00000080506 | 0 | NA | NA | NA | NA | NA | NA | NA | 0 | 0 | 0 | 0 | 0 | 0 |
| 23150 | ENSMUSG00000080507 | 0 | NA | NA | NA | NA | NA | NA | NA | 0 | 0 | 0 | 0 | 0 | 0 |
| 23151 | ENSMUSG00000080508 | 0 | NA | NA | NA | NA | NA | NA | NA | 0 | 0 | 0 | 0 | 0 | 0 |
| 23152 | ENSMUSG00000080511 | 0 | NA | NA | NA | NA | NA | NA | NA | 0 | 0 | 0 | 0 | 0 | 0 |
| 23153 | ENSMUSG00000080515 | 0 | NA | NA | NA | NA | NA | NA | NA | 0 | 0 | 0 | 0 | 0 | 0 |
| 23154 | ENSMUSG00000080516 | 0 | NA | NA | NA | NA | NA | NA | NA | 0 | 0 | 0 | 0 | 0 | 0 |
| 23155 | ENSMUSG00000080517 | 0 | NA | NA | NA | NA | NA | Gm23109 | predicted gene, 23109 [Source:MGI Symb | 0 | 0 | 0 | 0 | 0 | 0 |
| 23156 | ENSMUSG00000080518 | 0 | NA | NA | NA | NA | NA | Gm22193 | predicted gene, 22193 [Source:MGI Symb | 0 | 0 | 0 | 0 | 0 | 0 |
| 23157 | ENSMUSG00000080519 | 0 | NA | NA | NA | NA | NA | NA | NA | 0 | 0 | 0 | 0 | 0 | 0 |
| 23158 | ENSMUSG00000080522 | 0 | NA | NA | NA | NA | NA | NA | NA | 0 | 0 | 0 | 0 | 0 | 0 |
| 23159 | ENSMUSG00000080524 | 0 | NA | NA | NA | NA | NA | Gm23879 | predicted gene, 23879 [Source:MGI Symb | 0 | 0 | 0 | 0 | 0 | 0 |
| 23160 | ENSMUSG00000080526 | 0 | NA | NA | NA | NA | NA | Mir466j | microRNA 466j [Source:MGI Symbol;Acc:I | 0 | 0 | 0 | 0 | 0 | 0 |
| 23161 | ENSMUSG00000080527 | 0 | NA | NA | NA | NA | NA | NA | NA | 0 | 0 | 0 | 0 | 0 | 0 |
| 23162 | ENSMUSG00000080529 | 0 | NA | NA | NA | NA | NA | NA | NA | 0 | 0 | 0 | 0 | 0 | 0 |
| 23163 | ENSMUSG00000080530 | 1.51511629 | -2.80177921 | 2.062042808 | -1.358739595 | 0.17422912 | NA | NA | NA | 0.8031 | 6.1675 | 1.0062 | 0 | 0 | 1.1138 |
| 23164 | ENSMUSG00000080533 | 0 | NA | NA | NA | NA | NA | Mir1191 | microRNA 1191 [Source:MGI Symbol;Acc: | 0 | 0 | 0 | 0 | 0 | 0 |
| 23165 | ENSMUSG00000080536 | 0 | NA | NA | NA | NA | NA | Gm25539 | predicted gene, 25539 [Source:MGI Symb | 0 | 0 | 0 | 0 | 0 | 0 |
| 23166 | ENSMUSG00000080538 | 0 | NA | NA | NA | NA | NA | Gm25541 | predicted gene, 25541 [Source:MGI Symb | 0 | 0 | 0 | 0 | 0 | 0 |
| 23167 | ENSMUSG00000080539 | 0 | NA | NA | NA | NA | NA | Gm25542 | predicted gene, 25542 [Source:MGI Symb | 0 | 0 | 0 | 0 | 0 | 0 |
| 23168 | ENSMUSG00000080540 | 0 | NA | NA | NA | NA | NA | Gm22711 | predicted gene, 22711 [Source:MGI Symb | 0 | 0 | 0 | 0 | 0 | 0 |
| 23169 | ENSMUSG00000080542 | 0 | NA | NA | NA | NA | NA | Rnu4atac | RNA, U4atac small nuclear (U12-depende | 0 | 0 | 0 | 0 | 0 | 0 |
| 23170 | ENSMUSG00000080543 | 0 | NA | NA | NA | NA | NA | NA | NA | 0 | 0 | 0 | 0 | 0 | 0 |
| 23171 | ENSMUSG00000080544 | 0 | NA | NA | NA | NA | NA | NA | NA | 0 | 0 | 0 | 0 | 0 | 0 |
| 23172 | ENSMUSG00000080545 | 0 | NA | NA | NA | NA | NA | Gm22713 | predicted gene, 22713 [Source:MGI Symb | 0 | 0 | 0 | 0 | 0 | 0 |
| 23173 | ENSMUSG00000080549 | 0 | NA | NA | NA | NA | NA | NA | NA | 0 | 0 | 0 | 0 | 0 | 0 |
| 23174 | ENSMUSG00000080555 | 0 | NA | NA | NA | NA | NA | NA | NA | 0 | 0 | 0 | 0 | 0 | 0 |
| 23175 | ENSMUSG00000080556 | 0 | NA | NA | NA | NA | NA | NA | NA | 0 | 0 | 0 | 0 | 0 | 0 |
| 23176 | ENSMUSG00000080558 | 0 | NA | NA | NA | NA | NA | NA | NA | 0 | 0 | 0 | 0 | 0 | 0 |
| 23177 | ENSMUSG00000080561 | 0 | NA | NA | NA | NA | NA | NA | NA | 0 | 0 | 0 | 0 | 0 | 0 |
| 23178 | ENSMUSG00000080562 | 0 | NA | NA | NA | NA | NA | NA | NA | 0 | 0 | 0 | 0 | 0 | 0 |
| 23179 | ENSMUSG00000080566 | 0 | NA | NA | NA | NA | NA | Gm26044 | predicted gene, 26044 [Source:MGI Symb | 0 | 0 | 0 | 0 | 0 | 0 |
| 23180 | ENSMUSG00000080573 | 0.49081066 | -0.698504554 | 3.1295234 | -0.22319838 | 0.823381113 | NA | Mir467f | microRNA 467f [Source:MGI Symbol;Acc:I | 0.8031 | 1.0279 | 0 | 0 | 0 | 1.1138 |

| | | | | | | | | | | | | | | | |
|---|---|---|---|---|---|---|---|---|---|---|---|---|---|---|---|
| 23181 | ENSMUSG00000080574 | 2.48437526 | -0.843339317 | 1.519153859 | -0.555137527 | 0.578800596 | NA | Mir466i | microRNA 466i [Source:MGI Symbol;Acc:N | 6.4248 | 2.0558 | 1.0062 | 0.964 | 0 | 4.4554 |
| 23182 | ENSMUSG00000080575 | 0 | NA | NA | NA | NA | NA | Mir1190 | microRNA 1190 [Source:MGI Symbol;Acc: | 0 | 0 | 0 | 0 | 0 | 0 |
| 23183 | ENSMUSG00000080577 | 0 | NA | NA | NA | NA | NA | Gm23226 | predicted gene, 23226 [Source:MGI Symb | 0 | 0 | 0 | 0 | 0 | 0 |
| 23184 | ENSMUSG00000080580 | 0 | NA | NA | NA | NA | NA | Mir466f-4 | microRNA 466f-4 [Source:MGI Symbol;Acc | 0 | 0 | 0 | 0 | 0 | 0 |
| 23185 | ENSMUSG00000080582 | 0 | NA | NA | NA | NA | NA | Gm24907 | predicted gene, 24907 [Source:MGI Symb | 0 | 0 | 0 | 0 | 0 | 0 |
| 23186 | ENSMUSG00000080586 | 0 | NA | NA | NA | NA | NA | Mir1187 | microRNA 1187 [Source:MGI Symbol;Acc: | 0 | 0 | 0 | 0 | 0 | 0 |
| 23187 | ENSMUSG00000080590 | 0 | NA | NA | NA | NA | NA | NA | NA | 0 | 0 | 0 | 0 | 0 | 0 |
| 23188 | ENSMUSG00000080592 | 0 | NA | NA | NA | NA | NA | NA | NA | 0 | 0 | 0 | 0 | 0 | 0 |
| 23189 | ENSMUSG00000080594 | 0 | NA | NA | NA | NA | NA | Mir669h | microRNA 669h [Source:MGI Symbol;Acc: | 0 | 0 | 0 | 0 | 0 | 0 |
| 23190 | ENSMUSG00000080595 | 0 | NA | NA | NA | NA | NA | Gm22031 | predicted gene, 22031 [Source:MGI Symb | 0 | 0 | 0 | 0 | 0 | 0 |
| 23191 | ENSMUSG00000080596 | 0 | NA | NA | NA | NA | NA | NA | NA | 0 | 0 | 0 | 0 | 0 | 0 |
| 23192 | ENSMUSG00000080597 | 0 | NA | NA | NA | NA | NA | NA | NA | 0 | 0 | 0 | 0 | 0 | 0 |
| 23193 | ENSMUSG00000080600 | 0 | NA | NA | NA | NA | NA | NA | NA | 0 | 0 | 0 | 0 | 0 | 0 |
| 23194 | ENSMUSG00000080601 | 0 | NA | NA | NA | NA | NA | NA | NA | 0 | 0 | 0 | 0 | 0 | 0 |
| 23195 | ENSMUSG00000080607 | 0 | NA | NA | NA | NA | NA | Gm23839 | predicted gene, 23839 [Source:MGI Symb | 0 | 0 | 0 | 0 | 0 | 0 |
| 23196 | ENSMUSG00000080609 | 0.133850373 | -0.796629176 | 4.080472857 | -0.195229623 | 0.84521319 | NA | NA | NA | 0.8031 | 0 | 0 | 0 | 0 | 0 |
| 23197 | ENSMUSG00000080610 | 0 | NA | NA | NA | NA | NA | Snord110 | small nucleolar RNA, C/D box 110 [Source | 0 | 0 | 0 | 0 | 0 | 0 |
| 23198 | ENSMUSG00000080613 | 0 | NA | NA | NA | NA | NA | NA | NA | 0 | 0 | 0 | 0 | 0 | 0 |
| 23199 | ENSMUSG00000080615 | 0 | NA | NA | NA | NA | NA | Snord99 | small nucleolar RNA, C/D box 99 [Source:I | 0 | 0 | 0 | 0 | 0 | 0 |
| 23200 | ENSMUSG00000080616 | 0 | NA | NA | NA | NA | NA | Gm22152 | predicted gene, 22152 [Source:MGI Symb | 0 | 0 | 0 | 0 | 0 | 0 |
| 23201 | ENSMUSG00000080622 | 0 | NA | NA | NA | NA | NA | Gm22676 | predicted gene, 22676 [Source:MGI Symb | 0 | 0 | 0 | 0 | 0 | 0 |
| 23202 | ENSMUSG00000080624 | 0 | NA | NA | NA | NA | NA | NA | NA | 0 | 0 | 0 | 0 | 0 | 0 |
| 23203 | ENSMUSG00000080625 | 0 | NA | NA | NA | NA | NA | NA | NA | 0 | 0 | 0 | 0 | 0 | 0 |
| 23204 | ENSMUSG00000080626 | 1.158285245 | -1.361361327 | 2.327755416 | -0.584838647 | 0.558656216 | NA | Mir1192 | microRNA 1192 [Source:MGI Symbol;Acc: | 4.0155 | 0 | 1.0062 | 1.928 | 0 | 0 |
| 23205 | ENSMUSG00000080627 | 0 | NA | NA | NA | NA | NA | NA | NA | 0 | 0 | 0 | 0 | 0 | 0 |
| 23206 | ENSMUSG00000080631 | 0 | NA | NA | NA | NA | NA | NA | NA | 0 | 0 | 0 | 0 | 0 | 0 |
| 23207 | ENSMUSG00000080635 | 0 | NA | NA | NA | NA | NA | NA | NA | 0 | 0 | 0 | 0 | 0 | 0 |
| 23208 | ENSMUSG00000080636 | 0 | NA | NA | NA | NA | NA | NA | NA | 0 | 0 | 0 | 0 | 0 | 0 |
| 23209 | ENSMUSG00000080645 | 1.125921769 | -0.10056922 | 2.125718962 | -0.047310685 | 0.962265612 | NA | Mir1198 | microRNA 1198 [Source:MGI Symbol;Acc: | 2.4093 | 1.0279 | 0 | 0 | 1.0906 | 2.2277 |
| 23210 | ENSMUSG00000080649 | 0 | NA | NA | NA | NA | NA | NA | NA | 0 | 0 | 0 | 0 | 0 | 0 |
| 23211 | ENSMUSG00000080653 | 0 | NA | NA | NA | NA | NA | Mir669e | microRNA 669e [Source:MGI Symbol;Acc: | 0 | 0 | 0 | 0 | 0 | 0 |
| 23212 | ENSMUSG00000080655 | 0 | NA | NA | NA | NA | NA | NA | NA | 0 | 0 | 0 | 0 | 0 | 0 |
| 23213 | ENSMUSG00000080657 | 0 | NA | NA | NA | NA | NA | Mir669i | microRNA 669i [Source:MGI Symbol;Acc:N | 0 | 0 | 0 | 0 | 0 | 0 |
| 23214 | ENSMUSG00000080659 | 0.17131964 | -0.796629176 | 4.080472857 | -0.195229623 | 0.84521319 | NA | NA | NA | 0 | 1.0279 | 0 | 0 | 0 | 0 |
| 23215 | ENSMUSG00000080662 | 0 | NA | NA | NA | NA | NA | Mir1b | microRNA 1b [Source:MGI Symbol;Acc:M( | 0 | 0 | 0 | 0 | 0 | 0 |
| 23216 | ENSMUSG00000080663 | 0 | NA | NA | NA | NA | NA | NA | NA | 0 | 0 | 0 | 0 | 0 | 0 |
| 23217 | ENSMUSG00000080665 | 0 | NA | NA | NA | NA | NA | Mir1199 | microRNA 1199 [Source:MGI Symbol;Acc: | 0 | 0 | 0 | 0 | 0 | 0 |
| 23218 | ENSMUSG00000080666 | 0 | NA | NA | NA | NA | NA | NA | NA | 0 | 0 | 0 | 0 | 0 | 0 |
| 23219 | ENSMUSG00000080668 | 0 | NA | NA | NA | NA | NA | NA | NA | 0 | 0 | 0 | 0 | 0 | 0 |
| 23220 | ENSMUSG00000080669 | 0 | NA | NA | NA | NA | NA | Mir1224 | microRNA 1224 [Source:MGI Symbol;Acc: | 0 | 0 | 0 | 0 | 0 | 0 |
| 23221 | ENSMUSG00000080671 | 0 | NA | NA | NA | NA | NA | NA | NA | 0 | 0 | 0 | 0 | 0 | 0 |
| 23222 | ENSMUSG00000080673 | 0 | NA | NA | NA | NA | NA | Gm23336 | predicted gene, 23336 [Source:MGI Symb | 0 | 0 | 0 | 0 | 0 | 0 |
| 23223 | ENSMUSG00000080676 | 0 | NA | NA | NA | NA | NA | NA | NA | 0 | 0 | 0 | 0 | 0 | 0 |
| 23224 | ENSMUSG00000080678 | 0 | NA | NA | NA | NA | NA | NA | NA | 0 | 0 | 0 | 0 | 0 | 0 |
| 23225 | ENSMUSG00000080680 | 0 | NA | NA | NA | NA | NA | NA | NA | 0 | 0 | 0 | 0 | 0 | 0 |
| 23226 | ENSMUSG00000080681 | 0 | NA | NA | NA | NA | NA | NA | NA | 0 | 0 | 0 | 0 | 0 | 0 |
| 23227 | ENSMUSG00000080683 | 0 | NA | NA | NA | NA | NA | Gm23991 | predicted gene, 23991 [Source:MGI Symb | 0 | 0 | 0 | 0 | 0 | 0 |
| 23228 | ENSMUSG00000080684 | 0 | NA | NA | NA | NA | NA | NA | NA | 0 | 0 | 0 | 0 | 0 | 0 |
| 23229 | ENSMUSG00000080686 | 0 | NA | NA | NA | NA | NA | NA | NA | 0 | 0 | 0 | 0 | 0 | 0 |
| 23230 | ENSMUSG00000080687 | 0 | NA | NA | NA | NA | NA | Mir1197 | microRNA 1197 [Source:MGI Symbol;Acc: | 0 | 0 | 0 | 0 | 0 | 0 |
| 23231 | ENSMUSG00000080689 | 0 | NA | NA | NA | NA | NA | Gm23987 | predicted gene, 23987 [Source:MGI Symb | 0 | 0 | 0 | 0 | 0 | 0 |
| 23232 | ENSMUSG00000080694 | 0 | NA | NA | NA | NA | NA | Gm11621 | predicted gene 11621 [Source:MGI Symbo | 0 | 0 | 0 | 0 | 0 | 0 |
| 23233 | ENSMUSG00000080695 | 0 | NA | NA | NA | NA | NA | Gm5394 | predicted gene 5394 [Source:MGI Symbol | 0 | 0 | 0 | 0 | 0 | 0 |
| 23234 | ENSMUSG00000080696 | 0 | NA | NA | NA | NA | NA | Gm12825 | predicted gene 12825 [Source:MGI Symbo | 0 | 0 | 0 | 0 | 0 | 0 |
| 23235 | ENSMUSG00000080697 | 0 | NA | NA | NA | NA | NA | Vmn1r-ps10 | vomeronasal 1 receptor, pseudogene 107 | 0 | 0 | 0 | 0 | 0 | 0 |
| 23236 | ENSMUSG00000080698 | 0 | NA | NA | NA | NA | NA | Gm11333 | predicted gene 11333 [Source:MGI Symbo | 0 | 0 | 0 | 0 | 0 | 0 |

| | | | | | | | | | | | | | | | |
|---|---|---|---|---|---|---|---|---|---|---|---|---|---|---|---|
| 23237 | ENSMUSG00000080700 | 2.272930816 | 0.779585779 | 1.47847752 | 0.527289572 | 0.597992515 | NA | Gm15981 | predicted gene 15981 [Source:MGI Symbo | 0 | 3.0838 | 2.0125 | 1.928 | 3.2719 | 3.3415 |
| 23238 | ENSMUSG00000080701 | 0.503122293 | -2.366675966 | 3.785596405 | -0.625179156 | 0.531853481 | NA | Gm12340 | predicted gene 12340 [Source:MGI Symbo | 0 | 0 | 3.0187 | 0 | 0 | 0 |
| 23239 | ENSMUSG00000080703 | 0 | NA | NA | NA | NA | NA | Gm13183 | predicted gene 13183 [Source:MGI Symbo | 0 | 0 | 0 | 0 | 0 | 0 |
| 23240 | ENSMUSG00000080704 | 0 | NA | NA | NA | NA | NA | Gm6031 | predicted gene 6031 [Source:MGI Symbol | 0 | 0 | 0 | 0 | 0 | 0 |
| 23241 | ENSMUSG00000080705 | 0 | NA | NA | NA | NA | NA | Sult2a-ps4 | sulfotransferase family 2A, dehydroepiand | 0 | 0 | 0 | 0 | 0 | 0 |
| 23242 | ENSMUSG00000080706 | 0 | NA | NA | NA | NA | NA | Gm12890 | predicted gene 12890 [Source:MGI Symbo | 0 | 0 | 0 | 0 | 0 | 0 |
| 23243 | ENSMUSG00000080707 | 0 | NA | NA | NA | NA | NA | Gm13823 | predicted gene 13823 [Source:MGI Symbo | 0 | 0 | 0 | 0 | 0 | 0 |
| 23244 | ENSMUSG00000080708 | 0.503122293 | -2.366675966 | 3.785596405 | -0.625179156 | 0.531853481 | NA | Ccnd3-ps | cyclin D3, pseudogene [Source:MGI Symb | 0 | 0 | 3.0187 | 0 | 0 | 0 |
| 23245 | ENSMUSG00000080709 | 0 | NA | NA | NA | NA | NA | Gm14324 | predicted gene 14324 [Source:MGI Symbo | 0 | 0 | 0 | 0 | 0 | 0 |
| 23246 | ENSMUSG00000080710 | 0 | NA | NA | NA | NA | NA | Gm8107 | predicted gene 8107 [Source:MGI Symbol | 0 | 0 | 0 | 0 | 0 | 0 |
| 23247 | ENSMUSG00000080711 | 0 | NA | NA | NA | NA | NA | Gm11686 | predicted gene 11686 [Source:MGI Symbo | 0 | 0 | 0 | 0 | 0 | 0 |
| 23248 | ENSMUSG00000080712 | 1.152993094 | 0.681708558 | 2.114908884 | 0.322334718 | 0.747199135 | NA | H2bu1-ps | H2B.U histone 1, pseudogene [Source:MG | 1.6062 | 0 | 1.0062 | 0.964 | 0 | 3.3415 |
| 23249 | ENSMUSG00000080713 | 0 | NA | NA | NA | NA | NA | Olfr1174 | olfactory receptor 1174 [Source:MGI Sym | 0 | 0 | 0 | 0 | 0 | 0 |
| 23250 | ENSMUSG00000080714 | 0 | NA | NA | NA | NA | NA | Gm14602 | predicted gene 14602 [Source:MGI Symbo | 0 | 0 | 0 | 0 | 0 | 0 |
| 23251 | ENSMUSG00000080715 | 0 | NA | NA | NA | NA | NA | Gm5406 | predicted gene 5406 [Source:MGI Symbol | 0 | 0 | 0 | 0 | 0 | 0 |
| 23252 | ENSMUSG00000080716 | 0 | NA | NA | NA | NA | NA | Gm13503 | predicted gene 13503 [Source:MGI Symbo | 0 | 0 | 0 | 0 | 0 | 0 |
| 23253 | ENSMUSG00000080717 | 39.32333986 | 0.044318457 | 0.405106679 | 0.109399473 | 0.912885652 | 0.98625062 | B230307C23 | RIKEN cDNA B230307C23 gene [Source:M | 53.005 | 23.642 | 39.244 | 45.307 | 46.897 | 27.846 |
| 23254 | ENSMUSG00000080718 | 0 | NA | NA | NA | NA | NA | Gm8019 | predicted gene 8019 [Source:MGI Symbol | 0 | 0 | 0 | 0 | 0 | 0 |
| 23255 | ENSMUSG00000080719 | 0 | NA | NA | NA | NA | NA | Gm15532 | predicted gene 15532 [Source:MGI Symbo | 0 | 0 | 0 | 0 | 0 | 0 |
| 23256 | ENSMUSG00000080720 | 0 | NA | NA | NA | NA | NA | Gm16077 | predicted gene 16077 [Source:MGI Symbo | 0 | 0 | 0 | 0 | 0 | 0 |
| 23257 | ENSMUSG00000080722 | 0 | NA | NA | NA | NA | NA | Defa-ps7 | defensin, alpha, pseudogene 7 [Source:M | 0 | 0 | 0 | 0 | 0 | 0 |
| 23258 | ENSMUSG00000080723 | 0 | NA | NA | NA | NA | NA | Gm12564 | predicted gene 12564 [Source:MGI Symbo | 0 | 0 | 0 | 0 | 0 | 0 |
| 23259 | ENSMUSG00000080725 | 0 | NA | NA | NA | NA | NA | Gm6121 | predicted gene 6121 [Source:MGI Symbol | 0 | 0 | 0 | 0 | 0 | 0 |
| 23260 | ENSMUSG00000080726 | 0 | NA | NA | NA | NA | NA | Gm15366 | predicted gene 15366 [Source:MGI Symbo | 0 | 0 | 0 | 0 | 0 | 0 |
| 23261 | ENSMUSG00000080727 | 13.57371341 | -0.116444685 | 0.720097669 | -0.161706794 | 0.871536754 | 0.979227205 | C920021L13 | RIKEN cDNA C920021L13 gene [Source:M | 8.031 | 17.475 | 17.106 | 20.244 | 16.359 | 2.2277 |
| 23262 | ENSMUSG00000080728 | 0 | NA | NA | NA | NA | NA | Gm13616 | predicted gene 13616 [Source:MGI Symbo | 0 | 0 | 0 | 0 | 0 | 0 |
| 23263 | ENSMUSG00000080729 | 0 | NA | NA | NA | NA | NA | Gm8084 | predicted gene 8084 [Source:MGI Symbol | 0 | 0 | 0 | 0 | 0 | 0 |
| 23264 | ENSMUSG00000080730 | 0 | NA | NA | NA | NA | NA | Gm11313 | predicted gene 11313 [Source:MGI Symbo | 0 | 0 | 0 | 0 | 0 | 0 |
| 23265 | ENSMUSG00000080731 | 0.804653467 | 0.739261853 | 2.420504647 | 0.305416416 | 0.760049021 | NA | Rps2-ps9 | ribosomal protein S2, pseudogene 9 [Sour | 0.8031 | 0 | 1.0062 | 1.928 | 1.0906 | 0 |
| 23266 | ENSMUSG00000080732 | 0 | NA | NA | NA | NA | NA | Gm13222 | predicted gene 13222 [Source:MGI Symbo | 0 | 0 | 0 | 0 | 0 | 0 |
| 23267 | ENSMUSG00000080733 | 0 | NA | NA | NA | NA | NA | Gm14867 | predicted gene 14867 [Source:MGI Symbo | 0 | 0 | 0 | 0 | 0 | 0 |
| 23268 | ENSMUSG00000080734 | 0 | NA | NA | NA | NA | NA | Gm12006 | predicted gene 12006 [Source:MGI Symbo | 0 | 0 | 0 | 0 | 0 | 0 |
| 23269 | ENSMUSG00000080735 | 0 | NA | NA | NA | NA | NA | Gm12075 | predicted gene 12075 [Source:MGI Symbo | 0 | 0 | 0 | 0 | 0 | 0 |
| 23270 | ENSMUSG00000080736 | 0 | NA | NA | NA | NA | NA | Gm15617 | predicted gene 15617 [Source:MGI Symbo | 0 | 0 | 0 | 0 | 0 | 0 |
| 23271 | ENSMUSG00000080737 | 0 | NA | NA | NA | NA | NA | Gm6982 | predicted gene 6982 [Source:MGI Symbol | 0 | 0 | 0 | 0 | 0 | 0 |
| 23272 | ENSMUSG00000080738 | 0 | NA | NA | NA | NA | NA | Mup-ps22 | major urinary protein, pseudogene 22 [So | 0 | 0 | 0 | 0 | 0 | 0 |
| 23273 | ENSMUSG00000080741 | 0 | NA | NA | NA | NA | NA | Gm11398 | predicted gene 11398 [Source:MGI Symbo | 0 | 0 | 0 | 0 | 0 | 0 |
| 23274 | ENSMUSG00000080742 | 0 | NA | NA | NA | NA | NA | Gm14035 | predicted gene 14035 [Source:MGI Symbo | 0 | 0 | 0 | 0 | 0 | 0 |
| 23275 | ENSMUSG00000080743 | 0 | NA | NA | NA | NA | NA | Gm7150 | predicted gene 7150 [Source:MGI Symbol | 0 | 0 | 0 | 0 | 0 | 0 |
| 23276 | ENSMUSG00000080744 | 0 | NA | NA | NA | NA | NA | Gm12710 | predicted gene 12710 [Source:MGI Symbo | 0 | 0 | 0 | 0 | 0 | 0 |
| 23277 | ENSMUSG00000080746 | 0.52105551 | -0.812028029 | 3.385364865 | -0.239864257 | 0.810435491 | NA | Rpsa-ps12 | ribosomal protein SA, pseudogene 12 [So | 0 | 0 | 2.0125 | 0 | 0 | 1.1138 |
| 23278 | ENSMUSG00000080747 | 0 | NA | NA | NA | NA | NA | Gm14016 | predicted gene 14016 [Source:MGI Symbo | 0 | 0 | 0 | 0 | 0 | 0 |
| 23279 | ENSMUSG00000080748 | 0 | NA | NA | NA | NA | NA | Gm15661 | predicted gene 15661 [Source:MGI Symbo | 0 | 0 | 0 | 0 | 0 | 0 |
| 23280 | ENSMUSG00000080749 | 0 | NA | NA | NA | NA | NA | Gm12134 | predicted gene 12134 [Source:MGI Symbo | 0 | 0 | 0 | 0 | 0 | 0 |
| 23281 | ENSMUSG00000080750 | 0 | NA | NA | NA | NA | NA | Gm15553 | predicted gene 15553 [Source:MGI Symbo | 0 | 0 | 0 | 0 | 0 | 0 |
| 23282 | ENSMUSG00000080751 | 0 | NA | NA | NA | NA | NA | Olfr634-ps1 | olfactory receptor 634, pseudogene 1 [So | 0 | 0 | 0 | 0 | 0 | 0 |
| 23283 | ENSMUSG00000080752 | 0 | NA | NA | NA | NA | NA | Olfr1069-ps1 | olfactory receptor 1069, pseudogene 1 [S | 0 | 0 | 0 | 0 | 0 | 0 |
| 23284 | ENSMUSG00000080753 | 0.17131964 | -0.796629176 | 4.080472857 | -0.195229623 | 0.84521319 | NA | Gm15613 | predicted gene 15613 [Source:MGI Symbo | 0 | 1.0279 | 0 | 0 | 0 | 0 |
| 23285 | ENSMUSG00000080754 | 0 | NA | NA | NA | NA | NA | Rpl36-ps7 | ribosomal protein L36, pseudogene 7 [Sou | 0 | 0 | 0 | 0 | 0 | 0 |
| 23286 | ENSMUSG00000080755 | 0 | NA | NA | NA | NA | NA | Gm15062 | predicted gene 15062 [Source:MGI Symbo | 0 | 0 | 0 | 0 | 0 | 0 |
| 23287 | ENSMUSG00000080756 | 0 | NA | NA | NA | NA | NA | Gm12633 | predicted gene 12633 [Source:MGI Symbo | 0 | 0 | 0 | 0 | 0 | 0 |
| 23288 | ENSMUSG00000080757 | 0 | NA | NA | NA | NA | NA | Gm15029 | predicted gene 15029 [Source:MGI Symbo | 0 | 0 | 0 | 0 | 0 | 0 |
| 23289 | ENSMUSG00000080758 | 0 | NA | NA | NA | NA | NA | Gm5392 | predicted gene 5392 [Source:MGI Symbol | 0 | 0 | 0 | 0 | 0 | 0 |
| 23290 | ENSMUSG00000080759 | 0 | NA | NA | NA | NA | NA | Gm15573 | predicted gene 15573 [Source:MGI Symbo | 0 | 0 | 0 | 0 | 0 | 0 |
| 23291 | ENSMUSG00000080760 | 0 | NA | NA | NA | NA | NA | Gm7571 | predicted gene 7571 [Source:MGI Symbol | 0 | 0 | 0 | 0 | 0 | 0 |
| 23292 | ENSMUSG00000080762 | 0 | NA | NA | NA | NA | NA | Gm11879 | predicted gene 11879 [Source:MGI Symbo | 0 | 0 | 0 | 0 | 0 | 0 |

| | | | | | | | | | | | | | | | |
|---|---|---|---|---|---|---|---|---|---|---|---|---|---|---|---|
| 23293 | ENSMUSG00000080763 | 0 | NA | NA | NA | NA | NA | Gm12341 | predicted gene 12341 [Source:MGI Symbo | 0 | 0 | 0 | 0 | 0 | 0 |
| 23294 | ENSMUSG00000080764 | 0 | NA | NA | NA | NA | NA | Gm14606 | predicted gene 14606 [Source:MGI Symbo | 0 | 0 | 0 | 0 | 0 | 0 |
| 23295 | ENSMUSG00000080765 | 0 | NA | NA | NA | NA | NA | Gm8908 | predicted gene 8908 [Source:MGI Symbol | 0 | 0 | 0 | 0 | 0 | 0 |
| 23296 | ENSMUSG00000080766 | 0 | NA | NA | NA | NA | NA | Gm13593 | predicted gene 13593 [Source:MGI Symbo | 0 | 0 | 0 | 0 | 0 | 0 |
| 23297 | ENSMUSG00000080767 | 0 | NA | NA | NA | NA | NA | Gm15595 | predicted gene 15595 [Source:MGI Symbo | 0 | 0 | 0 | 0 | 0 | 0 |
| 23298 | ENSMUSG00000080768 | 0.860314674 | 2.043280159 | 2.552981052 | 0.800350695 | 0.423507639 | NA | Gm12219 | predicted gene 12219 [Source:MGI Symbo | 0 | 0 | 1.0062 | 1.928 | 0 | 2.2277 |
| 23299 | ENSMUSG00000080770 | 0 | NA | NA | NA | NA | NA | Gm11278 | predicted gene 11278 [Source:MGI Symbo | 0 | 0 | 0 | 0 | 0 | 0 |
| 23300 | ENSMUSG00000080771 | 0 | NA | NA | NA | NA | NA | Gm11925 | predicted gene 11925 [Source:MGI Symbo | 0 | 0 | 0 | 0 | 0 | 0 |
| 23301 | ENSMUSG00000080772 | 0 | NA | NA | NA | NA | NA | Gm12543 | predicted gene 12543 [Source:MGI Symbo | 0 | 0 | 0 | 0 | 0 | 0 |
| 23302 | ENSMUSG00000080773 | 0.878015797 | -0.449419083 | 2.552723878 | -0.176054718 | 0.860250955 | NA | Gm12955 | predicted gene 12955 [Source:MGI Symbo | 0 | 1.0279 | 2.0125 | 0 | 0 | 2.2277 |
| 23303 | ENSMUSG00000080774 | 0 | NA | NA | NA | NA | NA | Gm9143 | predicted gene 9143 [Source:MGI Symbol | 0 | 0 | 0 | 0 | 0 | 0 |
| 23304 | ENSMUSG00000080775 | 0 | NA | NA | NA | NA | NA | Gm6368 | predicted gene 6368 [Source:MGI Symbol | 0 | 0 | 0 | 0 | 0 | 0 |
| 23305 | ENSMUSG00000080776 | 3.172135976 | 0.262173315 | 1.205378266 | 0.217502939 | 0.827816417 | NA | Gm12174 | predicted gene 12174 [Source:MGI Symbo | 1.6062 | 4.1117 | 3.0187 | 4.8199 | 4.3625 | 1.1138 |
| 23306 | ENSMUSG00000080777 | 0 | NA | NA | NA | NA | NA | Gm15365 | predicted gene 15365 [Source:MGI Symbo | 0 | 0 | 0 | 0 | 0 | 0 |
| 23307 | ENSMUSG00000080778 | 0 | NA | NA | NA | NA | NA | Gm7199 | predicted gene 7199 [Source:MGI Symbol | 0 | 0 | 0 | 0 | 0 | 0 |
| 23308 | ENSMUSG00000080779 | 0 | NA | NA | NA | NA | NA | Gm8731 | predicted gene 8731 [Source:MGI Symbol | 0 | 0 | 0 | 0 | 0 | 0 |
| 23309 | ENSMUSG00000080780 | 0 | NA | NA | NA | NA | NA | Gm11252 | predicted gene 11252 [Source:MGI Symbo | 0 | 0 | 0 | 0 | 0 | 0 |
| 23310 | ENSMUSG00000080781 | 0 | NA | NA | NA | NA | NA | Olfr388-ps1 | olfactory receptor 388, pseudogene 1 [Sou | 0 | 0 | 0 | 0 | 0 | 0 |
| 23311 | ENSMUSG00000080782 | 0 | NA | NA | NA | NA | NA | Gm13495 | predicted gene 13495 [Source:MGI Symbo | 0 | 0 | 0 | 0 | 0 | 0 |
| 23312 | ENSMUSG00000080783 | 3.092861424 | 1.518908553 | 1.320254285 | 1.150466672 | 0.249951713 | NA | Gm8250 | predicted gene 8250 [Source:MGI Symbol | 0.8031 | 1.0279 | 3.0187 | 4.8199 | 1.0906 | 7.7969 |
| 23313 | ENSMUSG00000080785 | 0 | NA | NA | NA | NA | NA | Gm11591 | predicted gene 11591 [Source:MGI Symbo | 0 | 0 | 0 | 0 | 0 | 0 |
| 23314 | ENSMUSG00000080788 | 0 | NA | NA | NA | NA | NA | Gm7841 | predicted gene 7841 [Source:MGI Symbol | 0 | 0 | 0 | 0 | 0 | 0 |
| 23315 | ENSMUSG00000080789 | 0 | NA | NA | NA | NA | NA | Gm14007 | predicted gene 14007 [Source:MGI Symbo | 0 | 0 | 0 | 0 | 0 | 0 |
| 23316 | ENSMUSG00000080790 | 1.024099039 | -0.954604393 | 2.258409355 | -0.422688823 | 0.672522311 | NA | Gm7638 | predicted gene 7638 [Source:MGI Symbol | 0 | 3.0838 | 1.0062 | 0.964 | 1.0906 | 0 |
| 23317 | ENSMUSG00000080791 | 0 | NA | NA | NA | NA | NA | Akirin1-ps | akirin 1, pseudogene [Source:MGI Symbol | 0 | 0 | 0 | 0 | 0 | 0 |
| 23318 | ENSMUSG00000080792 | 0 | NA | NA | NA | NA | NA | Gm14367 | predicted gene 14367 [Source:MGI Symbo | 0 | 0 | 0 | 0 | 0 | 0 |
| 23319 | ENSMUSG00000080793 | 0.669251866 | -2.807208097 | 3.391018549 | -0.827836255 | 0.407763239 | NA | Cbx3-ps3 | chromobox 3, pseudogene 3 [Source:MGI | 4.0155 | 0 | 0 | 0 | 0 | 0 |
| 23320 | ENSMUSG00000080794 | 0 | NA | NA | NA | NA | NA | Vmn1r-ps13 | vomeronasal 1 receptor, pseudogene 132 | 0 | 0 | 0 | 0 | 0 | 0 |
| 23321 | ENSMUSG00000080795 | 0 | NA | NA | NA | NA | NA | Gm6783 | predicted gene 6783 [Source:MGI Symbol | 0 | 0 | 0 | 0 | 0 | 0 |
| 23322 | ENSMUSG00000080796 | 0 | NA | NA | NA | NA | NA | Gm9159 | predicted gene 9159 [Source:MGI Symbol | 0 | 0 | 0 | 0 | 0 | 0 |
| 23323 | ENSMUSG00000080797 | 19.60557523 | -0.609702094 | 0.522081677 | -1.167828946 | 0.24287577 | 0.692615513 | Gm15760 | predicted gene 15760 [Source:MGI Symbo | 31.321 | 23.642 | 16.1 | 21.208 | 11.997 | 13.366 |
| 23324 | ENSMUSG00000080798 | 0 | NA | NA | NA | NA | NA | Gm5135 | predicted pseudogene 5135 [Source:MGI | 0 | 0 | 0 | 0 | 0 | 0 |
| 23325 | ENSMUSG00000080799 | 0 | NA | NA | NA | NA | NA | Gm15166 | predicted gene 15166 [Source:MGI Symbo | 0 | 0 | 0 | 0 | 0 | 0 |
| 23326 | ENSMUSG00000080800 | 0 | NA | NA | NA | NA | NA | Gm11582 | predicted gene 11582 [Source:MGI Symbo | 0 | 0 | 0 | 0 | 0 | 0 |
| 23327 | ENSMUSG00000080801 | 0.342639279 | -1.805976596 | 4.028547714 | -0.448294702 | 0.653940524 | NA | Gm13310 | predicted gene 13310 [Source:MGI Symbo | 0 | 2.0558 | 0 | 0 | 0 | 0 |
| 23328 | ENSMUSG00000080802 | 0 | NA | NA | NA | NA | NA | Gm13804 | predicted gene 13804 [Source:MGI Symbo | 0 | 0 | 0 | 0 | 0 | 0 |
| 23329 | ENSMUSG00000080803 | 0 | NA | NA | NA | NA | NA | Gm12683 | predicted gene 12683 [Source:MGI Symbo | 0 | 0 | 0 | 0 | 0 | 0 |
| 23330 | ENSMUSG00000080805 | 0 | NA | NA | NA | NA | NA | Gm11219 | predicted gene 11219 [Source:MGI Symbo | 0 | 0 | 0 | 0 | 0 | 0 |
| 23331 | ENSMUSG00000080806 | 0.181769715 | 1.126924175 | 4.080472857 | 0.276174898 | 0.782413736 | NA | Gm13005 | predicted gene 13005 [Source:MGI Symbo | 0 | 0 | 0 | 0 | 1.0906 | 0 |
| 23332 | ENSMUSG00000080807 | 0 | NA | NA | NA | NA | NA | Gm13117 | predicted gene 13117 [Source:MGI Symbo | 0 | 0 | 0 | 0 | 0 | 0 |
| 23333 | ENSMUSG00000080808 | 0 | NA | NA | NA | NA | NA | Gm12608 | predicted gene 12608 [Source:MGI Symbo | 0 | 0 | 0 | 0 | 0 | 0 |
| 23334 | ENSMUSG00000080809 | 0 | NA | NA | NA | NA | NA | Olfr1273-ps | olfactory receptor 1273, pseudogene [Sou | 0 | 0 | 0 | 0 | 0 | 0 |
| 23335 | ENSMUSG00000080810 | 0.305170013 | -1.657836477 | 4.041480335 | -0.410205257 | 0.681655385 | NA | Gm11737 | predicted gene 11737 [Source:MGI Symbo | 0.8031 | 1.0279 | 0 | 0 | 0 | 0 |
| 23336 | ENSMUSG00000080811 | 0 | NA | NA | NA | NA | NA | Gm14513 | predicted gene 14513 [Source:MGI Symbo | 0 | 0 | 0 | 0 | 0 | 0 |
| 23337 | ENSMUSG00000080812 | 0 | NA | NA | NA | NA | NA | Gm13526 | predicted gene 13526 [Source:MGI Symbo | 0 | 0 | 0 | 0 | 0 | 0 |
| 23338 | ENSMUSG00000080813 | 0 | NA | NA | NA | NA | NA | Gm14828 | predicted gene 14828 [Source:MGI Symbo | 0 | 0 | 0 | 0 | 0 | 0 |
| 23339 | ENSMUSG00000080814 | 0 | NA | NA | NA | NA | NA | Gm12314 | predicted gene 12314 [Source:MGI Symbo | 0 | 0 | 0 | 0 | 0 | 0 |
| 23340 | ENSMUSG00000080815 | 0 | NA | NA | NA | NA | NA | Hmgb1-rs18 | high mobility group box 1, related sequen | 0 | 0 | 0 | 0 | 0 | 0 |
| 23341 | ENSMUSG00000080816 | 0 | NA | NA | NA | NA | NA | Gm12447 | predicted gene 12447 [Source:MGI Symbo | 0 | 0 | 0 | 0 | 0 | 0 |
| 23342 | ENSMUSG00000080817 | 0 | NA | NA | NA | NA | NA | Gm13751 | predicted gene 13751 [Source:MGI Symbo | 0 | 0 | 0 | 0 | 0 | 0 |
| 23343 | ENSMUSG00000080818 | 0 | NA | NA | NA | NA | NA | Gm9434 | predicted gene 9434 [Source:MGI Symbol | 0 | 0 | 0 | 0 | 0 | 0 |
| 23344 | ENSMUSG00000080819 | 0 | NA | NA | NA | NA | NA | Gm2005 | predicted pseudogene 2005 [Source:MGI | 0 | 0 | 0 | 0 | 0 | 0 |
| 23345 | ENSMUSG00000080820 | 0 | NA | NA | NA | NA | NA | Gm13197 | predicted gene 13197 [Source:MGI Symbo | 0 | 0 | 0 | 0 | 0 | 0 |
| 23346 | ENSMUSG00000080822 | 0 | NA | NA | NA | NA | NA | Gm14694 | predicted gene 14694 [Source:MGI Symbo | 0 | 0 | 0 | 0 | 0 | 0 |
| 23347 | ENSMUSG00000080823 | 0 | NA | NA | NA | NA | NA | Atp5l-ps2 | ATP synthase, H+ transporting, mitochonc | 0 | 0 | 0 | 0 | 0 | 0 |
| 23348 | ENSMUSG00000080824 | 0 | NA | NA | NA | NA | NA | Gm9001 | predicted gene 9001 [Source:MGI Symbol | 0 | 0 | 0 | 0 | 0 | 0 |

| | | | | | | | | | | | | | | | |
|---|---|---|---|---|---|---|---|---|---|---|---|---|---|---|---|
| 23349 | ENSMUSG00000080825 | 0 | NA | NA | NA | NA | NA | Gm14892 | predicted gene 14892 [Source:MGI Symbo | 0 | 0 | 0 | 0 | 0 | 0 |
| 23350 | ENSMUSG00000080826 | 0 | NA | NA | NA | NA | NA | Gm12019 | predicted gene 12019 [Source:MGI Symbo | 0 | 0 | 0 | 0 | 0 | 0 |
| 23351 | ENSMUSG00000080827 | 0 | NA | NA | NA | NA | NA | Gm12086 | predicted gene 12086 [Source:MGI Symbo | 0 | 0 | 0 | 0 | 0 | 0 |
| 23352 | ENSMUSG00000080828 | 0 | NA | NA | NA | NA | NA | Gm14669 | predicted gene 14669 [Source:MGI Symbo | 0 | 0 | 0 | 0 | 0 | 0 |
| 23353 | ENSMUSG00000080829 | 0 | NA | NA | NA | NA | NA | Gm8545 | predicted gene 8545 [Source:MGI Symbol | 0 | 0 | 0 | 0 | 0 | 0 |
| 23354 | ENSMUSG00000080831 | 0.492645588 | 0.9565034 | 3.447648281 | 0.277436479 | 0.781444974 | NA | Gm12529 | predicted gene 12529 [Source:MGI Symbo | 0 | 1.0279 | 0 | 1.928 | 0 | 0 |
| 23355 | ENSMUSG00000080832 | 0.185640647 | 1.126924175 | 4.080472857 | 0.276174898 | 0.782413736 | NA | M6pr-ps | mannose-6-phosphate receptor, pseudog | 0 | 0 | 0 | 0 | 0 | 1.1138 |
| 23356 | ENSMUSG00000080833 | 0 | NA | NA | NA | NA | NA | Gm11976 | predicted gene 11976 [Source:MGI Symbo | 0 | 0 | 0 | 0 | 0 | 0 |
| 23357 | ENSMUSG00000080834 | 0 | NA | NA | NA | NA | NA | Gm15216 | predicted gene 15216 [Source:MGI Symbo | 0 | 0 | 0 | 0 | 0 | 0 |
| 23358 | ENSMUSG00000080835 | 0 | NA | NA | NA | NA | NA | Gm12137 | predicted gene 12137 [Source:MGI Symbo | 0 | 0 | 0 | 0 | 0 | 0 |
| 23359 | ENSMUSG00000080836 | 0.185640647 | 1.126924175 | 4.080472857 | 0.276174898 | 0.782413736 | NA | Gm6444 | predicted gene 6444 [Source:MGI Symbol | 0 | 0 | 0 | 0 | 0 | 1.1138 |
| 23360 | ENSMUSG00000080839 | 0 | NA | NA | NA | NA | NA | Gm11625 | predicted gene 11625 [Source:MGI Symbo | 0 | 0 | 0 | 0 | 0 | 0 |
| 23361 | ENSMUSG00000080840 | 0.133850373 | -0.796629176 | 4.080472857 | -0.195229623 | 0.84521319 | NA | Gm11349 | predicted gene 11349 [Source:MGI Symbo | 0.8031 | 0 | 0 | 0 | 0 | 0 |
| 23362 | ENSMUSG00000080841 | 0 | NA | NA | NA | NA | NA | Gm14880 | predicted gene 14880 [Source:MGI Symbo | 0 | 0 | 0 | 0 | 0 | 0 |
| 23363 | ENSMUSG00000080842 | 0 | NA | NA | NA | NA | NA | Gm15400 | predicted gene 15400 [Source:MGI Symbo | 0 | 0 | 0 | 0 | 0 | 0 |
| 23364 | ENSMUSG00000080843 | 0 | NA | NA | NA | NA | NA | Gm14726 | predicted gene 14726 [Source:MGI Symbo | 0 | 0 | 0 | 0 | 0 | 0 |
| 23365 | ENSMUSG00000080844 | 0 | NA | NA | NA | NA | NA | Gm14107 | predicted gene 14107 [Source:MGI Symbo | 0 | 0 | 0 | 0 | 0 | 0 |
| 23366 | ENSMUSG00000080845 | 0 | NA | NA | NA | NA | NA | Gm9115 | predicted gene 9115 [Source:MGI Symbol | 0 | 0 | 0 | 0 | 0 | 0 |
| 23367 | ENSMUSG00000080846 | 0 | NA | NA | NA | NA | NA | Gm13379 | predicted gene 13379 [Source:MGI Symbo | 0 | 0 | 0 | 0 | 0 | 0 |
| 23368 | ENSMUSG00000080847 | 0 | NA | NA | NA | NA | NA | Gm4773 | predicted gene 4773 [Source:MGI Symbol | 0 | 0 | 0 | 0 | 0 | 0 |
| 23369 | ENSMUSG00000080848 | 9.947538321 | -0.39055063 | 0.799785512 | -0.488319211 | 0.625323756 | 0.919638611 | Gm9385 | predicted pseudogene 9385 [Source:MGI | 9.6372 | 8.2233 | 16.1 | 13.496 | 1.0906 | 11.138 |
| 23370 | ENSMUSG00000080849 | 0 | NA | NA | NA | NA | NA | Gm15702 | predicted gene 15702 [Source:MGI Symbo | 0 | 0 | 0 | 0 | 0 | 0 |
| 23371 | ENSMUSG00000080850 | 0 | NA | NA | NA | NA | NA | Gm12439 | predicted gene 12439 [Source:MGI Symbo | 0 | 0 | 0 | 0 | 0 | 0 |
| 23372 | ENSMUSG00000080851 | 0 | NA | NA | NA | NA | NA | Gm12094 | predicted gene 12094 [Source:MGI Symbo | 0 | 0 | 0 | 0 | 0 | 0 |
| 23373 | ENSMUSG00000080852 | 0 | NA | NA | NA | NA | NA | Gm14918 | predicted gene 14918 [Source:MGI Symbo | 0 | 0 | 0 | 0 | 0 | 0 |
| 23374 | ENSMUSG00000080855 | 0 | NA | NA | NA | NA | NA | Rpl35-ps1 | ribosomal protein L35, pseudogene 1 [Sou | 0 | 0 | 0 | 0 | 0 | 0 |
| 23375 | ENSMUSG00000080857 | 0.160662974 | 1.126924175 | 4.080472857 | 0.276174898 | 0.782413736 | NA | Gm12859 | predicted gene 12859 [Source:MGI Symbo | 0 | 0 | 0 | 0.964 | 0 | 0 |
| 23376 | ENSMUSG00000080859 | 0 | NA | NA | NA | NA | NA | Rpl10-ps1 | ribosomal protein L10, pseudogene 1 [Sou | 0 | 0 | 0 | 0 | 0 | 0 |
| 23377 | ENSMUSG00000080860 | 0 | NA | NA | NA | NA | NA | Gm13815 | predicted gene 13815 [Source:MGI Symbo | 0 | 0 | 0 | 0 | 0 | 0 |
| 23378 | ENSMUSG00000080861 | 0 | NA | NA | NA | NA | NA | Gm11325 | predicted gene 11325 [Source:MGI Symbo | 0 | 0 | 0 | 0 | 0 | 0 |
| 23379 | ENSMUSG00000080862 | 0 | NA | NA | NA | NA | NA | Gm14523 | predicted gene 14523 [Source:MGI Symbo | 0 | 0 | 0 | 0 | 0 | 0 |
| 23380 | ENSMUSG00000080863 | 0 | NA | NA | NA | NA | NA | Gm14787 | predicted gene 14787 [Source:MGI Symbo | 0 | 0 | 0 | 0 | 0 | 0 |
| 23381 | ENSMUSG00000080865 | 0 | NA | NA | NA | NA | NA | Eif1-ps3 | eukaryotic translation initiation factor 1, p | 0 | 0 | 0 | 0 | 0 | 0 |
| 23382 | ENSMUSG00000080866 | 0 | NA | NA | NA | NA | NA | Gm5687 | predicted gene 5687 [Source:MGI Symbol | 0 | 0 | 0 | 0 | 0 | 0 |
| 23383 | ENSMUSG00000080868 | 0 | NA | NA | NA | NA | NA | Gm15745 | predicted gene 15745 [Source:MGI Symbo | 0 | 0 | 0 | 0 | 0 | 0 |
| 23384 | ENSMUSG00000080869 | 0 | NA | NA | NA | NA | NA | Gm11856 | predicted gene 11856 [Source:MGI Symbo | 0 | 0 | 0 | 0 | 0 | 0 |
| 23385 | ENSMUSG00000080870 | 0 | NA | NA | NA | NA | NA | Gm14111 | predicted gene 14111 [Source:MGI Symbo | 0 | 0 | 0 | 0 | 0 | 0 |
| 23386 | ENSMUSG00000080871 | 0 | NA | NA | NA | NA | NA | Gm12875 | predicted gene 12875 [Source:MGI Symbo | 0 | 0 | 0 | 0 | 0 | 0 |
| 23387 | ENSMUSG00000080872 | 0 | NA | NA | NA | NA | NA | Gm11506 | predicted gene 11506 [Source:MGI Symbo | 0 | 0 | 0 | 0 | 0 | 0 |
| 23388 | ENSMUSG00000080873 | 0 | NA | NA | NA | NA | NA | Rpl30-ps3 | ribosomal protein L30, pseudogene 3 [Sou | 0 | 0 | 0 | 0 | 0 | 0 |
| 23389 | ENSMUSG00000080874 | 0 | NA | NA | NA | NA | NA | Gm15010 | predicted gene 15010 [Source:MGI Symbo | 0 | 0 | 0 | 0 | 0 | 0 |
| 23390 | ENSMUSG00000080875 | 2.78518405 | 0.417894463 | 1.269837845 | 0.329092777 | 0.742085563 | NA | Gm7332 | predicted gene 7332 [Source:MGI Symbol | 4.0155 | 1.0279 | 2.0125 | 1.928 | 3.2719 | 4.4554 |
| 23391 | ENSMUSG00000080876 | 0 | NA | NA | NA | NA | NA | Gm14332 | predicted gene 14332 [Source:MGI Symbo | 0 | 0 | 0 | 0 | 0 | 0 |
| 23392 | ENSMUSG00000080877 | 0.367410362 | 2.078255025 | 3.800070157 | 0.54689912 | 0.58444804 | NA | Rpl22-ps1 | ribosomal protein L22, pseudogene 1 [Sou | 0 | 0 | 0 | 0 | 1.0906 | 1.1138 |
| 23393 | ENSMUSG00000080878 | 0 | NA | NA | NA | NA | NA | Btf3-ps14 | basic transcription factor 3, pseudogene 1 | 0 | 0 | 0 | 0 | 0 | 0 |
| 23394 | ENSMUSG00000080880 | 0 | NA | NA | NA | NA | NA | Gm14278 | predicted gene 14278 [Source:MGI Symbo | 0 | 0 | 0 | 0 | 0 | 0 |
| 23395 | ENSMUSG00000080884 | 0 | NA | NA | NA | NA | NA | Cyp2j7-ps3 | cytochrome P450, family 2, subfamily j, pc | 0 | 0 | 0 | 0 | 0 | 0 |
| 23396 | ENSMUSG00000080885 | 3.412125336 | -1.995411004 | 1.312726902 | -1.520050363 | 0.128498318 | NA | Rpl10-ps6 | ribosomal protein L10, pseudogene 6 [Sou | 3.2124 | 8.2233 | 5.0312 | 2.8919 | 0 | 1.1138 |
| 23397 | ENSMUSG00000080886 | 0 | NA | NA | NA | NA | NA | Gm14847 | predicted gene 14847 [Source:MGI Symbo | 0 | 0 | 0 | 0 | 0 | 0 |
| 23398 | ENSMUSG00000080887 | 0 | NA | NA | NA | NA | NA | Gm14831 | predicted gene 14831 [Source:MGI Symbo | 0 | 0 | 0 | 0 | 0 | 0 |
| 23399 | ENSMUSG00000080888 | 1.004769579 | -2.106122861 | 2.47240066 | -0.851853381 | 0.394295474 | NA | Gm14387 | predicted gene 14387 [Source:MGI Symbo | 0.8031 | 4.1117 | 0 | 0 | 0 | 1.1138 |
| 23400 | ENSMUSG00000080889 | 0 | NA | NA | NA | NA | NA | Gm7974 | predicted gene 7974 [Source:MGI Symbol | 0 | 0 | 0 | 0 | 0 | 0 |
| 23401 | ENSMUSG00000080890 | 0 | NA | NA | NA | NA | NA | Gm16458 | predicted pseudogene 16458 [Source:MG | 0 | 0 | 0 | 0 | 0 | 0 |
| 23402 | ENSMUSG00000080891 | 0 | NA | NA | NA | NA | NA | Olfr587-ps1 | olfactory receptor 587, pseudogene 1 [So | 0 | 0 | 0 | 0 | 0 | 0 |
| 23403 | ENSMUSG00000080892 | 0 | NA | NA | NA | NA | NA | Gm9266 | predicted gene 9266 [Source:MGI Symbol | 0 | 0 | 0 | 0 | 0 | 0 |
| 23404 | ENSMUSG00000080893 | 1.99462501 | -0.85187238 | 1.479696346 | -0.575707565 | 0.564812861 | NA | Ndufa12-ps | NADH:ubiquinone oxidoreductase subuni | 1.6062 | 3.0838 | 3.0187 | 0.964 | 2.1812 | 1.1138 |

| | | | | | | | | | | | | | | | |
|---|---|---|---|---|---|---|---|---|---|---|---|---|---|---|---|
| 23405 | ENSMUSG00000080894 | 0 | NA | NA | NA | NA | NA | Defa-ps15 | defensin, alpha, pseudogene 15 [Source:N | 0 | 0 | 0 | 0 | 0 | 0 |
| 23406 | ENSMUSG00000080895 | 0 | NA | NA | NA | NA | NA | Gm11645 | predicted gene 11645 [Source:MGI Symbo | 0 | 0 | 0 | 0 | 0 | 0 |
| 23407 | ENSMUSG00000080896 | 1.948973767 | -0.238208026 | 1.615908122 | -0.147414338 | 0.882804989 | NA | Gm14567 | predicted gene 14567 [Source:MGI Symbo | 3.2124 | 2.0558 | 1.0062 | 0.964 | 0 | 4.4554 |
| 23408 | ENSMUSG00000080897 | 0 | NA | NA | NA | NA | NA | Gm14500 | predicted gene 14500 [Source:MGI Symbo | 0 | 0 | 0 | 0 | 0 | 0 |
| 23409 | ENSMUSG00000080898 | 0 | NA | NA | NA | NA | NA | Gm14836 | predicted gene 14836 [Source:MGI Symbo | 0 | 0 | 0 | 0 | 0 | 0 |
| 23410 | ENSMUSG00000080899 | 0 | NA | NA | NA | NA | NA | Gm9109 | predicted gene 9109 [Source:MGI Symbol | 0 | 0 | 0 | 0 | 0 | 0 |
| 23411 | ENSMUSG00000080902 | 0.401551119 | -2.072982726 | 4.008251979 | -0.517178744 | 0.605031388 | NA | Ywhaq-ps3 | tyrosine 3-monooxygenase/tryptophan 5- | 2.4093 | 0 | 0 | 0 | 0 | 0 |
| 23412 | ENSMUSG00000080903 | 0 | NA | NA | NA | NA | NA | Gm16383 | predicted gene 16383 [Source:MGI Symbo | 0 | 0 | 0 | 0 | 0 | 0 |
| 23413 | ENSMUSG00000080904 | 0.353348078 | 0.165142701 | 3.895042568 | 0.042398176 | 0.966181282 | NA | Gm11966 | predicted gene 11966 [Source:MGI Symbo | 0 | 0 | 1.0062 | 0 | 0 | 1.1138 |
| 23414 | ENSMUSG00000080906 | 0 | NA | NA | NA | NA | NA | Gm12095 | predicted gene 12095 [Source:MGI Symbo | 0 | 0 | 0 | 0 | 0 | 0 |
| 23415 | ENSMUSG00000080907 | 0.342639279 | -1.805976596 | 4.028547714 | -0.448294702 | 0.653940524 | NA | 4930455H04 | RIKEN cDNA 4930455H04 gene [Source:M | 0 | 2.0558 | 0 | 0 | 0 | 0 |
| 23416 | ENSMUSG00000080910 | 0 | NA | NA | NA | NA | NA | Gm14821 | predicted gene 14821 [Source:MGI Symbo | 0 | 0 | 0 | 0 | 0 | 0 |
| 23417 | ENSMUSG00000080911 | 0 | NA | NA | NA | NA | NA | Gm11918 | predicted gene 11918 [Source:MGI Symbo | 0 | 0 | 0 | 0 | 0 | 0 |
| 23418 | ENSMUSG00000080912 | 0 | NA | NA | NA | NA | NA | Gm12269 | predicted gene 12269 [Source:MGI Symbo | 0 | 0 | 0 | 0 | 0 | 0 |
| 23419 | ENSMUSG00000080913 | 0 | NA | NA | NA | NA | NA | Gm12719 | predicted gene 12719 [Source:MGI Symbo | 0 | 0 | 0 | 0 | 0 | 0 |
| 23420 | ENSMUSG00000080914 | 0 | NA | NA | NA | NA | NA | Gm7129 | predicted gene 7129 [Source:MGI Symbol | 0 | 0 | 0 | 0 | 0 | 0 |
| 23421 | ENSMUSG00000080915 | 0 | NA | NA | NA | NA | NA | Gm6446 | predicted gene 6446 [Source:MGI Symbol | 0 | 0 | 0 | 0 | 0 | 0 |
| 23422 | ENSMUSG00000080916 | 1.049412545 | -1.045567553 | 2.225303659 | -0.469853878 | 0.638459418 | NA | Gm12463 | predicted gene 12463 [Source:MGI Symbo | 3.2124 | 0 | 1.0062 | 0.964 | 0 | 1.1138 |
| 23423 | ENSMUSG00000080917 | 0 | NA | NA | NA | NA | NA | Gm15925 | predicted gene 15925 [Source:MGI Symbo | 0 | 0 | 0 | 0 | 0 | 0 |
| 23424 | ENSMUSG00000080918 | 0 | NA | NA | NA | NA | NA | Gm12202 | predicted gene 12202 [Source:MGI Symbo | 0 | 0 | 0 | 0 | 0 | 0 |
| 23425 | ENSMUSG00000080919 | 0 | NA | NA | NA | NA | NA | Gm11854 | predicted gene 11854 [Source:MGI Symbo | 0 | 0 | 0 | 0 | 0 | 0 |
| 23426 | ENSMUSG00000080921 | 75.30766789 | 0.293948924 | 0.293644835 | 1.001035569 | 0.316809613 | 0.763467884 | Rpl38-ps2 | ribosomal protein L38, pseudogene 2 [Sou | 50.595 | 82.233 | 70.437 | 66.514 | 80.706 | 101.36 |
| 23427 | ENSMUSG00000080922 | 0.133850373 | -0.796629176 | 4.080472857 | -0.195229623 | 0.84521319 | NA | Gm2214 | predicted gene 2214 [Source:MGI Symbol | 0.8031 | 0 | 0 | 0 | 0 | 0 |
| 23428 | ENSMUSG00000080923 | 0 | NA | NA | NA | NA | NA | Gm5400 | predicted gene 5400 [Source:MGI Symbol | 0 | 0 | 0 | 0 | 0 | 0 |
| 23429 | ENSMUSG00000080924 | 0 | NA | NA | NA | NA | NA | Olfr1078-ps1 | olfactory receptor 1078, pseudogene 1 [S | 0 | 0 | 0 | 0 | 0 | 0 |
| 23430 | ENSMUSG00000080925 | 0 | NA | NA | NA | NA | NA | Gm13682 | predicted gene 13682 [Source:MGI Symbo | 0 | 0 | 0 | 0 | 0 | 0 |
| 23431 | ENSMUSG00000080927 | 0 | NA | NA | NA | NA | NA | Gm11430 | predicted gene 11430 [Source:MGI Symbo | 0 | 0 | 0 | 0 | 0 | 0 |
| 23432 | ENSMUSG00000080928 | 0.685071968 | 0.040605487 | 2.800847395 | 0.014497572 | 0.988433016 | NA | Hmgb1-ps6 | high mobility group box 1, pseudogene 6 [ | 0 | 2.0558 | 0 | 0.964 | 1.0906 | 0 |
| 23433 | ENSMUSG00000080929 | 0 | NA | NA | NA | NA | NA | Gm12891 | predicted gene 12891 [Source:MGI Symbo | 0 | 0 | 0 | 0 | 0 | 0 |
| 23434 | ENSMUSG00000080930 | 0 | NA | NA | NA | NA | NA | Gm13367 | predicted gene 13367 [Source:MGI Symbo | 0 | 0 | 0 | 0 | 0 | 0 |
| 23435 | ENSMUSG00000080932 | 1.573226982 | -0.909763205 | 2.333189705 | -0.389922518 | 0.696593842 | NA | Gm10224 | predicted pseudogene 10224 [Source:MG | 0 | 6.1675 | 0 | 0 | 3.2719 | 0 |
| 23436 | ENSMUSG00000080933 | 0 | NA | NA | NA | NA | NA | Rhox3g | reproductive homeobox 3G [Source:MGI S | 0 | 0 | 0 | 0 | 0 | 0 |
| 23437 | ENSMUSG00000080934 | 0 | NA | NA | NA | NA | NA | Gm12931 | predicted gene 12931 [Source:MGI Symbo | 0 | 0 | 0 | 0 | 0 | 0 |
| 23438 | ENSMUSG00000080935 | 4.308766544 | 0.401181928 | 0.994537373 | 0.403385472 | 0.686664672 | NA | Got2-ps1 | glutamatic-oxaloacetic transaminase 2, m | 4.0155 | 3.0838 | 4.025 | 4.8199 | 5.4531 | 4.4554 |
| 23439 | ENSMUSG00000080936 | 0 | NA | NA | NA | NA | NA | Olfr337-ps1 | olfactory receptor 337, pseudogene 1 [So | 0 | 0 | 0 | 0 | 0 | 0 |
| 23440 | ENSMUSG00000080937 | 0 | NA | NA | NA | NA | NA | Gm13510 | predicted gene 13510 [Source:MGI Symbo | 0 | 0 | 0 | 0 | 0 | 0 |
| 23441 | ENSMUSG00000080938 | 0 | NA | NA | NA | NA | NA | Gm6062 | predicted gene 6062 [Source:MGI Symbol | 0 | 0 | 0 | 0 | 0 | 0 |
| 23442 | ENSMUSG00000080939 | 0 | NA | NA | NA | NA | NA | Gm9208 | predicted gene 9208 [Source:MGI Symbol | 0 | 0 | 0 | 0 | 0 | 0 |
| 23443 | ENSMUSG00000080941 | 0 | NA | NA | NA | NA | NA | Gm16056 | predicted gene 16056 [Source:MGI Symbo | 0 | 0 | 0 | 0 | 0 | 0 |
| 23444 | ENSMUSG00000080942 | 12.2106681 | 0.23289456 | 0.629130379 | 0.370184889 | 0.711244735 | 0.944239215 | Zfp36l1-ps | zinc finger protein 36, C3H type-like 1, pse | 11.243 | 9.2513 | 13.081 | 7.7118 | 15.269 | 16.708 |
| 23445 | ENSMUSG00000080943 | 0 | NA | NA | NA | NA | NA | Gm13130 | predicted gene 13130 [Source:MGI Symbo | 0 | 0 | 0 | 0 | 0 | 0 |
| 23446 | ENSMUSG00000080944 | 0 | NA | NA | NA | NA | NA | Gm12967 | predicted gene 12967 [Source:MGI Symbo | 0 | 0 | 0 | 0 | 0 | 0 |
| 23447 | ENSMUSG00000080946 | 0 | NA | NA | NA | NA | NA | Gm12985 | predicted gene 12985 [Source:MGI Symbo | 0 | 0 | 0 | 0 | 0 | 0 |
| 23448 | ENSMUSG00000080947 | 0 | NA | NA | NA | NA | NA | Gm12779 | predicted gene 12779 [Source:MGI Symbo | 0 | 0 | 0 | 0 | 0 | 0 |
| 23449 | ENSMUSG00000080948 | 0 | NA | NA | NA | NA | NA | Gm14481 | predicted gene 14481 [Source:MGI Symbo | 0 | 0 | 0 | 0 | 0 | 0 |
| 23450 | ENSMUSG00000080949 | 0 | NA | NA | NA | NA | NA | Gm13683 | predicted gene 13683 [Source:MGI Symbo | 0 | 0 | 0 | 0 | 0 | 0 |
| 23451 | ENSMUSG00000080950 | 0 | NA | NA | NA | NA | NA | Gm7278 | predicted gene 7278 [Source:MGI Symbol | 0 | 0 | 0 | 0 | 0 | 0 |
| 23452 | ENSMUSG00000080951 | 1.492025678 | -0.350050528 | 2.090670314 | -0.167434591 | 0.86702811 | NA | Gm12208 | predicted gene 12208 [Source:MGI Symbo | 0 | 3.0838 | 2.0125 | 3.8559 | 0 | 0 |
| 23453 | ENSMUSG00000080952 | 0 | NA | NA | NA | NA | NA | Gm5379 | predicted gene 5379 [Source:MGI Symbol | 0 | 0 | 0 | 0 | 0 | 0 |
| 23454 | ENSMUSG00000080953 | 0 | NA | NA | NA | NA | NA | Gm15025 | predicted gene 15025 [Source:MGI Symbo | 0 | 0 | 0 | 0 | 0 | 0 |
| 23455 | ENSMUSG00000080954 | 0 | NA | NA | NA | NA | NA | Gm14919 | predicted gene 14919 [Source:MGI Symbo | 0 | 0 | 0 | 0 | 0 | 0 |
| 23456 | ENSMUSG00000080955 | 0 | NA | NA | NA | NA | NA | Gm14263 | predicted gene 14263 [Source:MGI Symbo | 0 | 0 | 0 | 0 | 0 | 0 |
| 23457 | ENSMUSG00000080956 | 0.503302254 | -0.837877584 | 3.426698643 | -0.244514523 | 0.806832356 | NA | Gm16179 | predicted gene 16179 [Source:MGI Symbo | 0 | 2.0558 | 0 | 0.964 | 0 | 0 |
| 23458 | ENSMUSG00000080957 | 0.133850373 | -0.796629176 | 4.080472857 | -0.195229623 | 0.84521319 | NA | Gm15739 | predicted gene 15739 [Source:MGI Symbo | 0.8031 | 0 | 0 | 0 | 0 | 0 |
| 23459 | ENSMUSG00000080958 | 0 | NA | NA | NA | NA | NA | Gm11902 | predicted gene 11902 [Source:MGI Symbo | 0 | 0 | 0 | 0 | 0 | 0 |
| 23460 | ENSMUSG00000080959 | 0 | NA | NA | NA | NA | NA | Gm2003 | predicted pseudogene 2003 [Source:MGI | 0 | 0 | 0 | 0 | 0 | 0 |

| | | | | | | | | | | | | | | | |
|---|---|---|---|---|---|---|---|---|---|---|---|---|---|---|---|
| 23461 | ENSMUSG00000080960 | 0 | NA | NA | NA | NA | NA | Olfr343-ps1 | olfactory receptor 343, pseudogene 1 [Sou | 0 | 0 | 0 | 0 | 0 | 0 |
| 23462 | ENSMUSG00000080961 | 4.88775137 | -0.116725277 | 1.207864544 | -0.096637721 | 0.923014099 | NA | Gm11798 | predicted gene 11798 [Source:MGI Symbo | 0 | 11.307 | 4.025 | 1.928 | 8.7249 | 3.3415 |
| 23463 | ENSMUSG00000080962 | 0.17131964 | -0.796629176 | 4.080472857 | -0.195229623 | 0.84521319 | NA | Gm11453 | predicted gene 11453 [Source:MGI Symbo | 0 | 1.0279 | 0 | 0 | 0 | 0 |
| 23464 | ENSMUSG00000080964 | 0 | NA | NA | NA | NA | NA | Ifna-ps1 | interferon alpha gene, pseudogene 1 [Sou | 0 | 0 | 0 | 0 | 0 | 0 |
| 23465 | ENSMUSG00000080965 | 0 | NA | NA | NA | NA | NA | Gm6461 | predicted gene 6461 [Source:MGI Symbol | 0 | 0 | 0 | 0 | 0 | 0 |
| 23466 | ENSMUSG00000080966 | 0.36353943 | 2.065989254 | 4.034137301 | 0.512126658 | 0.608562368 | NA | Gm6263 | predicted gene 6263 [Source:MGI Symbol | 0 | 0 | 0 | 0 | 2.1812 | 0 |
| 23467 | ENSMUSG00000080967 | 0 | NA | NA | NA | NA | NA | Gm12706 | predicted gene 12706 [Source:MGI Symbo | 0 | 0 | 0 | 0 | 0 | 0 |
| 23468 | ENSMUSG00000080968 | 0.353089354 | 0.165142706 | 3.896102244 | 0.042386646 | 0.966190474 | NA | Atf1-ps | activating transcription factor 1, pseudoge | 0 | 1.0279 | 0 | 0 | 1.0906 | 0 |
| 23469 | ENSMUSG00000080969 | 0.643990493 | 0.179747576 | 2.633236711 | 0.068261078 | 0.945577807 | NA | Taf9-ps | TATA-box binding protein associated facto | 0.8031 | 0 | 1.0062 | 0.964 | 1.0906 | 0 |
| 23470 | ENSMUSG00000080970 | 0 | NA | NA | NA | NA | NA | Gm12173 | predicted gene 12173 [Source:MGI Symbo | 0 | 0 | 0 | 0 | 0 | 0 |
| 23471 | ENSMUSG00000080971 | 0.842149364 | -3.107408315 | 2.805524882 | -1.107603192 | 0.268033224 | NA | Gm16128 | predicted gene 16128 [Source:MGI Symbo | 0 | 1.0279 | 4.025 | 0 | 0 | 0 |
| 23472 | ENSMUSG00000080972 | 3.885234874 | -1.176563773 | 1.196549779 | -0.983296971 | 0.3254613 | NA | Gm16061 | predicted gene 16061 [Source:MGI Symbo | 4.0155 | 8.2233 | 4.025 | 4.8199 | 0 | 2.2277 |
| 23473 | ENSMUSG00000080974 | 0.17131964 | -0.796629176 | 4.080472857 | -0.195229623 | 0.84521319 | NA | Gm9009 | predicted pseudogene 9009 [Source:MGI | 0 | 1.0279 | 0 | 0 | 0 | 0 |
| 23474 | ENSMUSG00000080976 | 0 | NA | NA | NA | NA | NA | Gm11490 | predicted gene 11490 [Source:MGI Symbo | 0 | 0 | 0 | 0 | 0 | 0 |
| 23475 | ENSMUSG00000080977 | 1.542388992 | 1.178831019 | 1.840581277 | 0.640466701 | 0.521869231 | NA | Gm13772 | predicted gene 13772 [Source:MGI Symbo | 0.8031 | 1.0279 | 1.0062 | 0.964 | 5.4531 | 0 |
| 23476 | ENSMUSG00000080978 | 0 | NA | NA | NA | NA | NA | Gm12297 | predicted gene 12297 [Source:MGI Symbo | 0 | 0 | 0 | 0 | 0 | 0 |
| 23477 | ENSMUSG00000080979 | 0.17131964 | -0.796629176 | 4.080472857 | -0.195229623 | 0.84521319 | NA | Gm13675 | predicted gene 13675 [Source:MGI Symbo | 0 | 1.0279 | 0 | 0 | 0 | 0 |
| 23478 | ENSMUSG00000080980 | 0 | NA | NA | NA | NA | NA | 1700071K01 | RIKEN cDNA 1700071K01 gene [Source:M | 0 | 0 | 0 | 0 | 0 | 0 |
| 23479 | ENSMUSG00000080981 | 0.315620088 | 0.165143421 | 4.063300985 | 0.040642675 | 0.967580763 | NA | Gm12161 | predicted gene 12161 [Source:MGI Symbo | 0.8031 | 0 | 0 | 0 | 1.0906 | 0 |
| 23480 | ENSMUSG00000080982 | 0 | NA | NA | NA | NA | NA | Gm13549 | predicted gene 13549 [Source:MGI Symbo | 0 | 0 | 0 | 0 | 0 | 0 |
| 23481 | ENSMUSG00000080983 | 0 | NA | NA | NA | NA | NA | Gm5419 | predicted gene 5419 [Source:MGI Symbol | 0 | 0 | 0 | 0 | 0 | 0 |
| 23482 | ENSMUSG00000080984 | 0 | NA | NA | NA | NA | NA | Gm14469 | predicted gene 14469 [Source:MGI Symbo | 0 | 0 | 0 | 0 | 0 | 0 |
| 23483 | ENSMUSG00000080985 | 1.392000219 | 2.859136945 | 2.037875014 | 1.402999166 | 0.160617088 | NA | Gm13559 | predicted gene 13559 [Source:MGI Symbo | 0 | 1.0279 | 0 | 2.8919 | 1.0906 | 3.3415 |
| 23484 | ENSMUSG00000080986 | 0 | NA | NA | NA | NA | NA | Olfr1081-ps1 | olfactory receptor 1081, pseudogene 1 [So | 0 | 0 | 0 | 0 | 0 | 0 |
| 23485 | ENSMUSG00000080987 | 0 | NA | NA | NA | NA | NA | Gm12073 | predicted gene 12073 [Source:MGI Symbo | 0 | 0 | 0 | 0 | 0 | 0 |
| 23486 | ENSMUSG00000080989 | 0 | NA | NA | NA | NA | NA | Gm14048 | predicted gene 14048 [Source:MGI Symbo | 0 | 0 | 0 | 0 | 0 | 0 |
| 23487 | ENSMUSG00000080990 | 0 | NA | NA | NA | NA | NA | Olfr118 | olfactory receptor 118 [Source:MGI Symb | 0 | 0 | 0 | 0 | 0 | 0 |
| 23488 | ENSMUSG00000080992 | 0 | NA | NA | NA | NA | NA | Psmb5-ps | proteasome (prosome, macropain) subun | 0 | 0 | 0 | 0 | 0 | 0 |
| 23489 | ENSMUSG00000080993 | 0 | NA | NA | NA | NA | NA | Gm12903 | predicted gene 12903 [Source:MGI Symbo | 0 | 0 | 0 | 0 | 0 | 0 |
| 23490 | ENSMUSG00000080994 | 0.936410189 | -0.694018354 | 2.480552701 | -0.279783757 | 0.779643414 | NA | Gm13464 | predicted gene 13464 [Source:MGI Symbo | 2.4093 | 1.0279 | 0 | 0 | 2.1812 | 0 |
| 23491 | ENSMUSG00000080995 | 0 | NA | NA | NA | NA | NA | Gm15165 | predicted gene 15165 [Source:MGI Symbo | 0 | 0 | 0 | 0 | 0 | 0 |
| 23492 | ENSMUSG00000080996 | 0 | NA | NA | NA | NA | NA | Gm4986 | predicted pseudogene 4986 [Source:MGI | 0 | 0 | 0 | 0 | 0 | 0 |
| 23493 | ENSMUSG00000080997 | 0 | NA | NA | NA | NA | NA | Gm11885 | predicted gene 11885 [Source:MGI Symbo | 0 | 0 | 0 | 0 | 0 | 0 |
| 23494 | ENSMUSG00000080998 | 0 | NA | NA | NA | NA | NA | Gm5384 | predicted gene 5384 [Source:MGI Symbol | 0 | 0 | 0 | 0 | 0 | 0 |
| 23495 | ENSMUSG00000080999 | 0 | NA | NA | NA | NA | NA | H2ac5-ps | H2A clustered histone 5, pseudogene [Sou | 0 | 0 | 0 | 0 | 0 | 0 |
| 23496 | ENSMUSG00000081000 | 0 | NA | NA | NA | NA | NA | Gm11200 | predicted gene 11200 [Source:MGI Symbo | 0 | 0 | 0 | 0 | 0 | 0 |
| 23497 | ENSMUSG00000081001 | 0 | NA | NA | NA | NA | NA | Gm14678 | predicted gene 14678 [Source:MGI Symbo | 0 | 0 | 0 | 0 | 0 | 0 |
| 23498 | ENSMUSG00000081002 | 0 | NA | NA | NA | NA | NA | Gm14240 | predicted gene 14240 [Source:MGI Symbo | 0 | 0 | 0 | 0 | 0 | 0 |
| 23499 | ENSMUSG00000081003 | 0 | NA | NA | NA | NA | NA | Gm14301 | predicted gene 14301 [Source:MGI Symbo | 0 | 0 | 0 | 0 | 0 | 0 |
| 23500 | ENSMUSG00000081004 | 3.876014442 | 0.527858022 | 1.147856108 | 0.45986428 | 0.645613641 | NA | Gm9157 | predicted gene 9157 [Source:MGI Symbol | 2.4093 | 5.1396 | 2.0125 | 3.8559 | 8.7249 | 1.1138 |
| 23501 | ENSMUSG00000081005 | 0 | NA | NA | NA | NA | NA | Gm13198 | predicted gene 13198 [Source:MGI Symbo | 0 | 0 | 0 | 0 | 0 | 0 |
| 23502 | ENSMUSG00000081006 | 0 | NA | NA | NA | NA | NA | Cyp4b1-ps1 | cytochrome P450, family 4, subfamily b, p | 0 | 0 | 0 | 0 | 0 | 0 |
| 23503 | ENSMUSG00000081007 | 0 | NA | NA | NA | NA | NA | Gm11543 | predicted gene 11543 [Source:MGI Symbo | 0 | 0 | 0 | 0 | 0 | 0 |
| 23504 | ENSMUSG00000081008 | 0 | NA | NA | NA | NA | NA | Gm13314 | predicted gene 13314 [Source:MGI Symbo | 0 | 0 | 0 | 0 | 0 | 0 |
| 23505 | ENSMUSG00000081009 | 0 | NA | NA | NA | NA | NA | Gm14065 | predicted gene 14065 [Source:MGI Symbo | 0 | 0 | 0 | 0 | 0 | 0 |
| 23506 | ENSMUSG00000081010 | 7.653903467 | -1.714292517 | 0.992859196 | -1.726621986 | 0.084235575 | 0.427953408 | Gm13880 | predicted gene 13880 [Source:MGI Symbo | 1.6062 | 19.53 | 14.087 | 1.928 | 6.5437 | 2.2277 |
| 23507 | ENSMUSG00000081011 | 0 | NA | NA | NA | NA | NA | Gm11913 | predicted gene 11913 [Source:MGI Symbo | 0 | 0 | 0 | 0 | 0 | 0 |
| 23508 | ENSMUSG00000081012 | 0 | NA | NA | NA | NA | NA | Gm15224 | predicted gene 15224 [Source:MGI Symbo | 0 | 0 | 0 | 0 | 0 | 0 |
| 23509 | ENSMUSG00000081013 | 0 | NA | NA | NA | NA | NA | Gm14497 | predicted gene 14497 [Source:MGI Symbo | 0 | 0 | 0 | 0 | 0 | 0 |
| 23510 | ENSMUSG00000081014 | 0 | NA | NA | NA | NA | NA | Gm13714 | predicted gene 13714 [Source:MGI Symbo | 0 | 0 | 0 | 0 | 0 | 0 |
| 23511 | ENSMUSG00000081016 | 0 | NA | NA | NA | NA | NA | Olfr1397-ps1 | olfactory receptor 1397, pseudogene 1 [So | 0 | 0 | 0 | 0 | 0 | 0 |
| 23512 | ENSMUSG00000081017 | 0 | NA | NA | NA | NA | NA | Gm15000 | predicted gene 15000 [Source:MGI Symbo | 0 | 0 | 0 | 0 | 0 | 0 |
| 23513 | ENSMUSG00000081018 | 0 | NA | NA | NA | NA | NA | Gm15012 | predicted gene 15012 [Source:MGI Symbo | 0 | 0 | 0 | 0 | 0 | 0 |
| 23514 | ENSMUSG00000081020 | 0 | NA | NA | NA | NA | NA | Gm14509 | predicted gene 14509 [Source:MGI Symbo | 0 | 0 | 0 | 0 | 0 | 0 |
| 23515 | ENSMUSG00000081021 | 0.610340026 | -2.657561537 | 3.094276798 | -0.858863544 | 0.390415805 | NA | Gm11964 | predicted gene 11964 [Source:MGI Symbo | 1.6062 | 2.0558 | 0 | 0 | 0 | 0 |
| 23516 | ENSMUSG00000081022 | 0 | NA | NA | NA | NA | NA | Gm15936 | predicted gene 15936 [Source:MGI Symbo | 0 | 0 | 0 | 0 | 0 | 0 |

| | | | | | | | | | | | | | | | |
|---|---|---|---|---|---|---|---|---|---|---|---|---|---|---|---|
| 23517 | ENSMUSG00000081023 | 0 | NA | NA | NA | NA | NA | Gm15766 | predicted gene 15766 [Source:MGI Symbo | 0 | 0 | 0 | 0 | 0 | 0 |
| 23518 | ENSMUSG00000081024 | 0 | NA | NA | NA | NA | NA | Gm14627 | predicted gene 14627 [Source:MGI Symbo | 0 | 0 | 0 | 0 | 0 | 0 |
| 23519 | ENSMUSG00000081025 | 0 | NA | NA | NA | NA | NA | Gm11943 | predicted gene 11943 [Source:MGI Symbo | 0 | 0 | 0 | 0 | 0 | 0 |
| 23520 | ENSMUSG00000081026 | 0 | NA | NA | NA | NA | NA | Gm11855 | predicted gene 11855 [Source:MGI Symbo | 0 | 0 | 0 | 0 | 0 | 0 |
| 23521 | ENSMUSG00000081029 | 0 | NA | NA | NA | NA | NA | Gm15077 | predicted gene 15077 [Source:MGI Symbo | 0 | 0 | 0 | 0 | 0 | 0 |
| 23522 | ENSMUSG00000081030 | 0 | NA | NA | NA | NA | NA | Gm12704 | predicted gene 12704 [Source:MGI Symbo | 0 | 0 | 0 | 0 | 0 | 0 |
| 23523 | ENSMUSG00000081033 | 0 | NA | NA | NA | NA | NA | Gm14797 | predicted gene 14797 [Source:MGI Symbo | 0 | 0 | 0 | 0 | 0 | 0 |
| 23524 | ENSMUSG00000081034 | 0 | NA | NA | NA | NA | NA | Gm5944 | predicted gene 5944 [Source:MGI Symbol | 0 | 0 | 0 | 0 | 0 | 0 |
| 23525 | ENSMUSG00000081036 | 0 | NA | NA | NA | NA | NA | Gm14628 | predicted gene 14628 [Source:MGI Symbo | 0 | 0 | 0 | 0 | 0 | 0 |
| 23526 | ENSMUSG00000081037 | 0 | NA | NA | NA | NA | NA | Gm13929 | predicted gene 13929 [Source:MGI Symbo | 0 | 0 | 0 | 0 | 0 | 0 |
| 23527 | ENSMUSG00000081038 | 0 | NA | NA | NA | NA | NA | Olfr626-ps1 | olfactory receptor 626, pseudogene 1 [Sou | 0 | 0 | 0 | 0 | 0 | 0 |
| 23528 | ENSMUSG00000081041 | 0 | NA | NA | NA | NA | NA | Nlk-ps1 | nemo like kinase, pseudogene 1 [Source:N | 0 | 0 | 0 | 0 | 0 | 0 |
| 23529 | ENSMUSG00000081042 | 0.17131964 | -0.796629176 | 4.080472857 | -0.195229623 | 0.84521319 | NA | Psmb7-ps2 | proteasome (prosome, macropain) subun | 0 | 1.0279 | 0 | 0 | 0 | 0 |
| 23530 | ENSMUSG00000081043 | 0 | NA | NA | NA | NA | NA | Gm11512 | predicted gene 11512 [Source:MGI Symbo | 0 | 0 | 0 | 0 | 0 | 0 |
| 23531 | ENSMUSG00000081044 | 0 | NA | NA | NA | NA | NA | AU015836 | expressed sequence AU015836 [Source:M | 0 | 0 | 0 | 0 | 0 | 0 |
| 23532 | ENSMUSG00000081045 | 0 | NA | NA | NA | NA | NA | Gm12797 | predicted gene 12797 [Source:MGI Symbo | 0 | 0 | 0 | 0 | 0 | 0 |
| 23533 | ENSMUSG00000081046 | 0 | NA | NA | NA | NA | NA | Gm12846 | predicted gene 12846 [Source:MGI Symbo | 0 | 0 | 0 | 0 | 0 | 0 |
| 23534 | ENSMUSG00000081047 | 0 | NA | NA | NA | NA | NA | Olfr997-ps1 | olfactory receptor 997, pseudogene 1 [Sou | 0 | 0 | 0 | 0 | 0 | 0 |
| 23535 | ENSMUSG00000081048 | 0 | NA | NA | NA | NA | NA | Gm11760 | predicted gene 11760 [Source:MGI Symbo | 0 | 0 | 0 | 0 | 0 | 0 |
| 23536 | ENSMUSG00000081049 | 2.914316839 | -0.807413832 | 1.337852631 | -0.603514777 | 0.546166288 | NA | Rps24-ps3 | ribosomal protein S24, pseudogene 3 [Sou | 4.0155 | 2.0558 | 5.0312 | 1.928 | 0 | 4.4554 |
| 23537 | ENSMUSG00000081050 | 0 | NA | NA | NA | NA | NA | Gm12461 | predicted gene 12461 [Source:MGI Symbo | 0 | 0 | 0 | 0 | 0 | 0 |
| 23538 | ENSMUSG00000081051 | 15.21746351 | 0.375283373 | 0.587987147 | 0.63825098 | 0.523310316 | 0.888334265 | Gm15427 | predicted pseudogene 15427 [Source:MG | 11.243 | 13.363 | 15.094 | 8.6758 | 23.994 | 18.935 |
| 23539 | ENSMUSG00000081052 | 0.692116425 | -1.412606127 | 2.773087916 | -0.509398248 | 0.610473104 | NA | Gm6305 | predicted gene 6305 [Source:MGI Symbol | 0 | 2.0558 | 1.0062 | 0 | 1.0906 | 0 |
| 23540 | ENSMUSG00000081053 | 1.027324697 | 0.082525906 | 2.406406141 | 0.034294255 | 0.972642506 | NA | Mup-ps17 | major urinary protein, pseudogene 17 [So | 0 | 0 | 3.0187 | 0.964 | 2.1812 | 0 |
| 23541 | ENSMUSG00000081054 | 0 | NA | NA | NA | NA | NA | Gm12025 | predicted gene 12025 [Source:MGI Symbo | 0 | 0 | 0 | 0 | 0 | 0 |
| 23542 | ENSMUSG00000081055 | 0 | NA | NA | NA | NA | NA | Gm6987 | predicted pseudogene 6987 [Source:MGI | 0 | 0 | 0 | 0 | 0 | 0 |
| 23543 | ENSMUSG00000081056 | 0 | NA | NA | NA | NA | NA | Gm13998 | predicted gene 13998 [Source:MGI Symbo | 0 | 0 | 0 | 0 | 0 | 0 |
| 23544 | ENSMUSG00000081057 | 0 | NA | NA | NA | NA | NA | Olfr1017-ps1 | olfactory receptor 1017, pseudogene 1 [S | 0 | 0 | 0 | 0 | 0 | 0 |
| 23545 | ENSMUSG00000081058 | 11.0096409 | 0.455177668 | 0.780406009 | 0.583257513 | 0.55971996 | 0.899475387 | H3c15 | H3 clustered histone 15 [Source:MGI Sym | 10.44 | 7.1954 | 10.062 | 1.928 | 15.269 | 21.163 |
| 23546 | ENSMUSG00000081059 | 0.185640647 | 1.126924175 | 4.080472857 | 0.276174898 | 0.782413736 | NA | Gm11945 | predicted gene 11945 [Source:MGI Symbo | 0 | 0 | 0 | 0 | 0 | 1.1138 |
| 23547 | ENSMUSG00000081060 | 0 | NA | NA | NA | NA | NA | Gm11318 | predicted gene 11318 [Source:MGI Symbo | 0 | 0 | 0 | 0 | 0 | 0 |
| 23548 | ENSMUSG00000081061 | 0 | NA | NA | NA | NA | NA | Gm14712 | predicted gene 14712 [Source:MGI Symbo | 0 | 0 | 0 | 0 | 0 | 0 |
| 23549 | ENSMUSG00000081062 | 0 | NA | NA | NA | NA | NA | Gm13640 | predicted gene 13640 [Source:MGI Symbo | 0 | 0 | 0 | 0 | 0 | 0 |
| 23550 | ENSMUSG00000081063 | 0 | NA | NA | NA | NA | NA | Gm11825 | predicted gene 11825 [Source:MGI Symbo | 0 | 0 | 0 | 0 | 0 | 0 |
| 23551 | ENSMUSG00000081064 | 0 | NA | NA | NA | NA | NA | Gm6071 | predicted gene 6071 [Source:MGI Symbol | 0 | 0 | 0 | 0 | 0 | 0 |
| 23552 | ENSMUSG00000081065 | 0 | NA | NA | NA | NA | NA | Gm13576 | predicted gene 13576 [Source:MGI Symbo | 0 | 0 | 0 | 0 | 0 | 0 |
| 23553 | ENSMUSG00000081066 | 0 | NA | NA | NA | NA | NA | Gm12817 | predicted gene 12817 [Source:MGI Symbo | 0 | 0 | 0 | 0 | 0 | 0 |
| 23554 | ENSMUSG00000081067 | 0 | NA | NA | NA | NA | NA | Gm12912 | predicted gene 12912 [Source:MGI Symbo | 0 | 0 | 0 | 0 | 0 | 0 |
| 23555 | ENSMUSG00000081068 | 0 | NA | NA | NA | NA | NA | Gm12804 | predicted gene 12804 [Source:MGI Symbo | 0 | 0 | 0 | 0 | 0 | 0 |
| 23556 | ENSMUSG00000081069 | 0 | NA | NA | NA | NA | NA | Gm13928 | predicted gene 13928 [Source:MGI Symbo | 0 | 0 | 0 | 0 | 0 | 0 |
| 23557 | ENSMUSG00000081070 | 0 | NA | NA | NA | NA | NA | Gm13181 | predicted gene 13181 [Source:MGI Symbo | 0 | 0 | 0 | 0 | 0 | 0 |
| 23558 | ENSMUSG00000081071 | 1.024357763 | -0.938111495 | 2.232331503 | -0.420238434 | 0.674311281 | NA | Gm9836 | predicted pseudogene 9836 [Source:MGI | 0 | 2.0558 | 2.0125 | 0.964 | 0 | 1.1138 |
| 23559 | ENSMUSG00000081072 | 0 | NA | NA | NA | NA | NA | Gm14977 | predicted gene 14977 [Source:MGI Symbo | 0 | 0 | 0 | 0 | 0 | 0 |
| 23560 | ENSMUSG00000081073 | 0 | NA | NA | NA | NA | NA | Gm13758 | predicted gene 13758 [Source:MGI Symbo | 0 | 0 | 0 | 0 | 0 | 0 |
| 23561 | ENSMUSG00000081076 | 1.8046162 | 3.302415052 | 2.330988609 | 1.416744397 | 0.156557672 | NA | Rpsa-ps4 | ribosomal protein S4, pseudogene 4 [Sour | 0.8031 | 0 | 0 | 0 | 0 | 10.025 |
| 23562 | ENSMUSG00000081077 | 0 | NA | NA | NA | NA | NA | Gm13899 | predicted gene 13899 [Source:MGI Symbo | 0 | 0 | 0 | 0 | 0 | 0 |
| 23563 | ENSMUSG00000081078 | 0 | NA | NA | NA | NA | NA | Gm12568 | predicted gene 12568 [Source:MGI Symbo | 0 | 0 | 0 | 0 | 0 | 0 |
| 23564 | ENSMUSG00000081079 | 0 | NA | NA | NA | NA | NA | Gm13059 | predicted gene 13059 [Source:MGI Symbo | 0 | 0 | 0 | 0 | 0 | 0 |
| 23565 | ENSMUSG00000081080 | 0 | NA | NA | NA | NA | NA | Vmn1r-ps10 | vomeronasal 1 receptor, pseudogene 102 | 0 | 0 | 0 | 0 | 0 | 0 |
| 23566 | ENSMUSG00000081081 | 0 | NA | NA | NA | NA | NA | Gm14856 | predicted gene 14856 [Source:MGI Symbo | 0 | 0 | 0 | 0 | 0 | 0 |
| 23567 | ENSMUSG00000081082 | 0 | NA | NA | NA | NA | NA | Gm14937 | predicted gene 14937 [Source:MGI Symbo | 0 | 0 | 0 | 0 | 0 | 0 |
| 23568 | ENSMUSG00000081083 | 0.185640647 | 1.126924175 | 4.080472857 | 0.276174898 | 0.782413736 | NA | Gm6079 | predicted gene 6079 [Source:MGI Symbol | 0 | 0 | 0 | 0 | 0 | 1.1138 |
| 23569 | ENSMUSG00000081084 | 0.167707431 | -0.796629176 | 4.080472857 | -0.195229623 | 0.84521319 | NA | Gm5638 | predicted gene 5638 [Source:MGI Symbol | 0 | 0 | 1.0062 | 0 | 0 | 0 |
| 23570 | ENSMUSG00000081085 | 0 | NA | NA | NA | NA | NA | Gm13463 | predicted gene 13463 [Source:MGI Symbo | 0 | 0 | 0 | 0 | 0 | 0 |
| 23571 | ENSMUSG00000081086 | 0 | NA | NA | NA | NA | NA | Gm9506 | predicted gene 9506 [Source:MGI Symbol | 0 | 0 | 0 | 0 | 0 | 0 |
| 23572 | ENSMUSG00000081087 | 2.19151688 | -1.062793289 | 1.679100697 | -0.632953873 | 0.526763766 | NA | Rps15a-ps7 | ribosomal protein S15A, pseudogene 7 [So | 4.8186 | 0 | 4.025 | 0.964 | 0 | 3.3415 |

| | | | | | | | | | | | | | | | |
|---|---|---|---|---|---|---|---|---|---|---|---|---|---|---|---|
| 23573 | ENSMUSG00000081088 | 0 | NA | NA | NA | NA | NA | Gm11679 | predicted gene 11679 [Source:MGI Symbo | 0 | 0 | 0 | 0 | 0 | 0 |
| 23574 | ENSMUSG00000081089 | 0 | NA | NA | NA | NA | NA | Gm14624 | predicted gene 14624 [Source:MGI Symbo | 0 | 0 | 0 | 0 | 0 | 0 |
| 23575 | ENSMUSG00000081092 | 0 | NA | NA | NA | NA | NA | Gm14128 | predicted gene 14128 [Source:MGI Symbo | 0 | 0 | 0 | 0 | 0 | 0 |
| 23576 | ENSMUSG00000081093 | 0.353089354 | 0.165142706 | 3.896102244 | 0.042386646 | 0.966190474 | NA | Gm12565 | predicted gene 12565 [Source:MGI Symbo | 0 | 1.0279 | 0 | 0 | 1.0906 | 0 |
| 23577 | ENSMUSG00000081094 | 27.97345797 | -0.269913644 | 0.425982809 | -0.633625674 | 0.526325142 | 0.888334265 | Rpl19-ps11 | ribosomal protein L19, pseudogene 11 [Sc | 24.896 | 35.977 | 31.194 | 32.775 | 20.722 | 22.277 |
| 23578 | ENSMUSG00000081095 | 0 | NA | NA | NA | NA | NA | Gm14946 | predicted gene 14946 [Source:MGI Symbo | 0 | 0 | 0 | 0 | 0 | 0 |
| 23579 | ENSMUSG00000081096 | 0.133850373 | -0.796629176 | 4.080472857 | -0.195229623 | 0.84521319 | NA | Gm12691 | predicted gene 12691 [Source:MGI Symbo | 0.8031 | 0 | 0 | 0 | 0 | 0 |
| 23580 | ENSMUSG00000081097 | 0 | NA | NA | NA | NA | NA | Gm12135 | predicted gene 12135 [Source:MGI Symbo | 0 | 0 | 0 | 0 | 0 | 0 |
| 23581 | ENSMUSG00000081098 | 0 | NA | NA | NA | NA | NA | Gm12220 | predicted gene 12220 [Source:MGI Symbo | 0 | 0 | 0 | 0 | 0 | 0 |
| 23582 | ENSMUSG00000081099 | 0 | NA | NA | NA | NA | NA | Gm5270 | predicted gene 5270 [Source:MGI Symbol | 0 | 0 | 0 | 0 | 0 | 0 |
| 23583 | ENSMUSG00000081100 | 0 | NA | NA | NA | NA | NA | Gm14630 | predicted gene 14630 [Source:MGI Symbo | 0 | 0 | 0 | 0 | 0 | 0 |
| 23584 | ENSMUSG00000081101 | 0 | NA | NA | NA | NA | NA | Gm12024 | predicted gene 12024 [Source:MGI Symbo | 0 | 0 | 0 | 0 | 0 | 0 |
| 23585 | ENSMUSG00000081103 | 0.339027071 | -1.79174149 | 3.919585585 | -0.457125237 | 0.647581032 | NA | Rps12-ps26 | ribosomal protein S12, pseudogene 26 [Sc | 0 | 1.0279 | 1.0062 | 0 | 0 | 0 |
| 23586 | ENSMUSG00000081104 | 0 | NA | NA | NA | NA | NA | Gm15989 | predicted gene 15989 [Source:MGI Symbo | 0 | 0 | 0 | 0 | 0 | 0 |
| 23587 | ENSMUSG00000081106 | 0 | NA | NA | NA | NA | NA | Gm7155 | predicted gene 7155 [Source:MGI Symbol | 0 | 0 | 0 | 0 | 0 | 0 |
| 23588 | ENSMUSG00000081107 | 0 | NA | NA | NA | NA | NA | Gm12235 | predicted gene 12235 [Source:MGI Symbo | 0 | 0 | 0 | 0 | 0 | 0 |
| 23589 | ENSMUSG00000081108 | 0 | NA | NA | NA | NA | NA | Gm4234 | predicted pseudogene 4234 [Source:MGI | 0 | 0 | 0 | 0 | 0 | 0 |
| 23590 | ENSMUSG00000081110 | 0.556921942 | 2.679043159 | 3.638243063 | 0.736356289 | 0.461513899 | NA | Gm12745 | predicted gene 12745 [Source:MGI Symbo | 0 | 0 | 0 | 0 | 0 | 3.3415 |
| 23591 | ENSMUSG00000081111 | 0.635111108 | 0.379830552 | 2.868137728 | 0.132431071 | 0.89464334 | NA | Gm5913 | predicted gene 5913 [Source:MGI Symbol | 1.6062 | 0 | 0 | 0 | 1.0906 | 1.1138 |
| 23592 | ENSMUSG00000081112 | 0 | NA | NA | NA | NA | NA | Gm12140 | predicted gene 12140 [Source:MGI Symbo | 0 | 0 | 0 | 0 | 0 | 0 |
| 23593 | ENSMUSG00000081113 | 0 | NA | NA | NA | NA | NA | Gm7308 | predicted pseudogene 7308 [Source:MGI | 0 | 0 | 0 | 0 | 0 | 0 |
| 23594 | ENSMUSG00000081114 | 0 | NA | NA | NA | NA | NA | Gm15390 | predicted gene 15390 [Source:MGI Symbo | 0 | 0 | 0 | 0 | 0 | 0 |
| 23595 | ENSMUSG00000081115 | 0 | NA | NA | NA | NA | NA | Gm15913 | predicted gene 15913 [Source:MGI Symbo | 0 | 0 | 0 | 0 | 0 | 0 |
| 23596 | ENSMUSG00000081116 | 0 | NA | NA | NA | NA | NA | Gm13329 | predicted gene 13329 [Source:MGI Symbo | 0 | 0 | 0 | 0 | 0 | 0 |
| 23597 | ENSMUSG00000081117 | 0 | NA | NA | NA | NA | NA | Gm11233 | predicted gene 11233 [Source:MGI Symbo | 0 | 0 | 0 | 0 | 0 | 0 |
| 23598 | ENSMUSG00000081118 | 0.349477146 | 0.165142768 | 3.911021834 | 0.042224967 | 0.966319359 | NA | Gm6758 | predicted gene 6758 [Source:MGI Symbol | 0 | 0 | 1.0062 | 0 | 1.0906 | 0 |
| 23599 | ENSMUSG00000081119 | 0 | NA | NA | NA | NA | NA | Gm14601 | predicted gene 14601 [Source:MGI Symbo | 0 | 0 | 0 | 0 | 0 | 0 |
| 23600 | ENSMUSG00000081120 | 0 | NA | NA | NA | NA | NA | Gm12458 | predicted gene 12458 [Source:MGI Symbo | 0 | 0 | 0 | 0 | 0 | 0 |
| 23601 | ENSMUSG00000081121 | 0.328370405 | 0.16514316 | 4.003149536 | 0.041253308 | 0.967093956 | NA | Gm12791 | predicted gene 12791 [Source:MGI Symbo | 0 | 0 | 1.0062 | 0.964 | 0 | 0 |
| 23602 | ENSMUSG00000081122 | 0 | NA | NA | NA | NA | NA | Gm13678 | predicted gene 13678 [Source:MGI Symbo | 0 | 0 | 0 | 0 | 0 | 0 |
| 23603 | ENSMUSG00000081124 | 0 | NA | NA | NA | NA | NA | Gm14906 | predicted gene 14906 [Source:MGI Symbo | 0 | 0 | 0 | 0 | 0 | 0 |
| 23604 | ENSMUSG00000081125 | 0 | NA | NA | NA | NA | NA | Rpl26-ps5 | ribosomal protein L26, pseudogene 5 [Sou | 0 | 0 | 0 | 0 | 0 | 0 |
| 23605 | ENSMUSG00000081126 | 0 | NA | NA | NA | NA | NA | Gm15784 | predicted gene 15784 [Source:MGI Symbo | 0 | 0 | 0 | 0 | 0 | 0 |
| 23606 | ENSMUSG00000081127 | 0 | NA | NA | NA | NA | NA | Gm12677 | predicted gene 12677 [Source:MGI Symbo | 0 | 0 | 0 | 0 | 0 | 0 |
| 23607 | ENSMUSG00000081128 | 1.833114257 | 1.540649022 | 1.703903167 | 0.904188132 | 0.365895652 | NA | Gm13328 | predicted gene 13328 [Source:MGI Symbo | 0.8031 | 1.0279 | 1.0062 | 4.8199 | 0 | 3.3415 |
| 23608 | ENSMUSG00000081129 | 0 | NA | NA | NA | NA | NA | Gm14605 | predicted gene 14605 [Source:MGI Symbo | 0 | 0 | 0 | 0 | 0 | 0 |
| 23609 | ENSMUSG00000081130 | 0 | NA | NA | NA | NA | NA | Gm8217 | predicted gene 8217 [Source:MGI Symbol | 0 | 0 | 0 | 0 | 0 | 0 |
| 23610 | ENSMUSG00000081131 | 0 | NA | NA | NA | NA | NA | Gm14833 | predicted gene 14833 [Source:MGI Symbo | 0 | 0 | 0 | 0 | 0 | 0 |
| 23611 | ENSMUSG00000081132 | 0 | NA | NA | NA | NA | NA | Gm13744 | predicted gene 13744 [Source:MGI Symbo | 0 | 0 | 0 | 0 | 0 | 0 |
| 23612 | ENSMUSG00000081133 | 0 | NA | NA | NA | NA | NA | Rhox11-ps2 | reproductive homeobox 11, pseudogene : | 0 | 0 | 0 | 0 | 0 | 0 |
| 23613 | ENSMUSG00000081134 | 0 | NA | NA | NA | NA | NA | Gm14913 | predicted gene 14913 [Source:MGI Symbo | 0 | 0 | 0 | 0 | 0 | 0 |
| 23614 | ENSMUSG00000081135 | 0 | NA | NA | NA | NA | NA | Gm14800 | predicted gene 14800 [Source:MGI Symbo | 0 | 0 | 0 | 0 | 0 | 0 |
| 23615 | ENSMUSG00000081136 | 0 | NA | NA | NA | NA | NA | Gm12990 | predicted gene 12990 [Source:MGI Symbo | 0 | 0 | 0 | 0 | 0 | 0 |
| 23616 | ENSMUSG00000081137 | 1.748521822 | 1.334906653 | 1.674037031 | 0.797417637 | 0.42520852 | NA | BC022960 | cDNA sequence BC022960 [Source:MGI Sy | 0 | 1.0279 | 2.0125 | 1.928 | 2.1812 | 3.3415 |
| 23617 | ENSMUSG00000081138 | 0 | NA | NA | NA | NA | NA | Gm12834 | predicted gene 12834 [Source:MGI Symbo | 0 | 0 | 0 | 0 | 0 | 0 |
| 23618 | ENSMUSG00000081139 | 0 | NA | NA | NA | NA | NA | Gm13333 | predicted gene 13333 [Source:MGI Symbo | 0 | 0 | 0 | 0 | 0 | 0 |
| 23619 | ENSMUSG00000081141 | 0 | NA | NA | NA | NA | NA | Gm11501 | predicted gene 11501 [Source:MGI Symbo | 0 | 0 | 0 | 0 | 0 | 0 |
| 23620 | ENSMUSG00000081142 | 0.401551119 | -2.072982726 | 4.008251979 | -0.517178744 | 0.605031388 | NA | Gm15497 | predicted gene 15497 [Source:MGI Symbo | 2.4093 | 0 | 0 | 0 | 0 | 0 |
| 23621 | ENSMUSG00000081143 | 2.708622693 | -0.661293395 | 1.26715814 | -0.521871244 | 0.601759981 | NA | Gm15823 | predicted gene 15823 [Source:MGI Symbo | 4.8186 | 3.0838 | 2.0125 | 1.928 | 2.1812 | 2.2277 |
| 23622 | ENSMUSG00000081144 | 0 | NA | NA | NA | NA | NA | Vmn2r-ps2 | vomeronasal 2, receptor, pseudogene 2 [S | 0 | 0 | 0 | 0 | 0 | 0 |
| 23623 | ENSMUSG00000081145 | 0 | NA | NA | NA | NA | NA | Gm14585 | predicted gene 14585 [Source:MGI Symbo | 0 | 0 | 0 | 0 | 0 | 0 |
| 23624 | ENSMUSG00000081146 | 0 | NA | NA | NA | NA | NA | Gm6207 | predicted gene 6207 [Source:MGI Symbol | 0 | 0 | 0 | 0 | 0 | 0 |
| 23625 | ENSMUSG00000081147 | 0 | NA | NA | NA | NA | NA | Gm6024 | predicted gene 6024 [Source:MGI Symbol | 0 | 0 | 0 | 0 | 0 | 0 |
| 23626 | ENSMUSG00000081148 | 0 | NA | NA | NA | NA | NA | Gm11891 | predicted gene 11891 [Source:MGI Symbo | 0 | 0 | 0 | 0 | 0 | 0 |
| 23627 | ENSMUSG00000081150 | 0 | NA | NA | NA | NA | NA | Gm12684 | predicted gene 12684 [Source:MGI Symbo | 0 | 0 | 0 | 0 | 0 | 0 |
| 23628 | ENSMUSG00000081151 | 0.185640647 | 1.126924175 | 4.080472857 | 0.276174898 | 0.782413736 | NA | Gm11448 | predicted gene 11448 [Source:MGI Symbo | 0 | 0 | 0 | 0 | 0 | 1.1138 |

| | | | | | | | | | | | | | | | |
|---|---|---|---|---|---|---|---|---|---|---|---|---|---|---|---|
| 23629 | ENSMUSG00000081152 | 0 | NA | NA | NA | NA | NA | Gm12430 | predicted gene 12430 [Source:MGI Symbo | 0 | 0 | 0 | 0 | 0 | 0 |
| 23630 | ENSMUSG00000081153 | 0 | NA | NA | NA | NA | NA | Gm11401 | predicted gene 11401 [Source:MGI Symbo | 0 | 0 | 0 | 0 | 0 | 0 |
| 23631 | ENSMUSG00000081154 | 0 | NA | NA | NA | NA | NA | Defa-ps13 | defensin, alpha, pseudogene 13 [Source:N | 0 | 0 | 0 | 0 | 0 | 0 |
| 23632 | ENSMUSG00000081155 | 0 | NA | NA | NA | NA | NA | Gm12771 | predicted gene 12771 [Source:MGI Symbo | 0 | 0 | 0 | 0 | 0 | 0 |
| 23633 | ENSMUSG00000081156 | 1.955558423 | 0.108512568 | 1.579109327 | 0.068717578 | 0.945214426 | NA | Gm14425 | predicted gene 14425 [Source:MGI Symbo | 1.6062 | 3.0838 | 1.0062 | 3.8559 | 2.1812 | 0 |
| 23634 | ENSMUSG00000081157 | 1.172218286 | -2.379491135 | 2.373447105 | -1.00254652 | 0.316079711 | NA | Gm13552 | predicted gene 13552 [Source:MGI Symbo | 0.8031 | 5.1396 | 0 | 0 | 1.0906 | 0 |
| 23635 | ENSMUSG00000081158 | 0.68166635 | -2.79976525 | 3.012029182 | -0.92952793 | 0.352615558 | NA | Gm13521 | predicted gene 13521 [Source:MGI Symbo | 0 | 3.0838 | 1.0062 | 0 | 0 | 0 |
| 23636 | ENSMUSG00000081159 | 0 | NA | NA | NA | NA | NA | Gm6023 | predicted gene 6023 [Source:MGI Symbol | 0 | 0 | 0 | 0 | 0 | 0 |
| 23637 | ENSMUSG00000081160 | 0 | NA | NA | NA | NA | NA | Gm14897 | predicted gene 14897 [Source:MGI Symbo | 0 | 0 | 0 | 0 | 0 | 0 |
| 23638 | ENSMUSG00000081161 | 0.706696157 | 0.149852971 | 3.037999338 | 0.049326203 | 0.960659338 | NA | Gm12783 | predicted gene 12783 [Source:MGI Symbo | 0 | 0 | 2.0125 | 0 | 0 | 2.2277 |
| 23639 | ENSMUSG00000081162 | 0 | NA | NA | NA | NA | NA | Olfr400-ps1 | olfactory receptor 400, pseudogene 1 [Sou | 0 | 0 | 0 | 0 | 0 | 0 |
| 23640 | ENSMUSG00000081163 | 0 | NA | NA | NA | NA | NA | Defa-ps12 | defensin, alpha, pseudogene 12 [Source:N | 0 | 0 | 0 | 0 | 0 | 0 |
| 23641 | ENSMUSG00000081164 | 0 | NA | NA | NA | NA | NA | Gm8722 | predicted gene 8722 [Source:MGI Symbol | 0 | 0 | 0 | 0 | 0 | 0 |
| 23642 | ENSMUSG00000081165 | 0.160662974 | 1.126924175 | 4.080472857 | 0.276174898 | 0.782413736 | NA | Gm11783 | predicted gene 11783 [Source:MGI Symbo | 0 | 0 | 0 | 0.964 | 0 | 0 |
| 23643 | ENSMUSG00000081166 | 0 | NA | NA | NA | NA | NA | Gm12819 | predicted gene 12819 [Source:MGI Symbo | 0 | 0 | 0 | 0 | 0 | 0 |
| 23644 | ENSMUSG00000081167 | 0 | NA | NA | NA | NA | NA | Gm8880 | predicted gene 8880 [Source:MGI Symbol | 0 | 0 | 0 | 0 | 0 | 0 |
| 23645 | ENSMUSG00000081168 | 0 | NA | NA | NA | NA | NA | Gm13730 | predicted gene 13730 [Source:MGI Symbo | 0 | 0 | 0 | 0 | 0 | 0 |
| 23646 | ENSMUSG00000081169 | 0.305170013 | -1.657836477 | 4.041480335 | -0.410205257 | 0.681655385 | NA | Gm12551 | predicted gene 12551 [Source:MGI Symbo | 0.8031 | 1.0279 | 0 | 0 | 0 | 0 |
| 23647 | ENSMUSG00000081170 | 0.481988923 | 2.492401104 | 3.837077151 | 0.649557203 | 0.515978285 | NA | Gm12922 | predicted gene 12922 [Source:MGI Symbo | 0 | 0 | 0 | 2.8919 | 0 | 0 |
| 23648 | ENSMUSG00000081171 | 0 | NA | NA | NA | NA | NA | Gm7485 | predicted gene 7485 [Source:MGI Symbol | 0 | 0 | 0 | 0 | 0 | 0 |
| 23649 | ENSMUSG00000081172 | 0 | NA | NA | NA | NA | NA | Gm12849 | predicted gene 12849 [Source:MGI Symbo | 0 | 0 | 0 | 0 | 0 | 0 |
| 23650 | ENSMUSG00000081173 | 0 | NA | NA | NA | NA | NA | Gm13500 | predicted gene 13500 [Source:MGI Symbo | 0 | 0 | 0 | 0 | 0 | 0 |
| 23651 | ENSMUSG00000081174 | 0 | NA | NA | NA | NA | NA | Gm13439 | predicted gene 13439 [Source:MGI Symbo | 0 | 0 | 0 | 0 | 0 | 0 |
| 23652 | ENSMUSG00000081175 | 0 | NA | NA | NA | NA | NA | Gm5793 | predicted pseudogene 5793 [Source:MGI | 0 | 0 | 0 | 0 | 0 | 0 |
| 23653 | ENSMUSG00000081176 | 1.591291096 | 0.465286826 | 2.057426799 | 0.226149881 | 0.82108485 | NA | H3f3a-ps2 | H3.3 histone A, pseudogene 2 [Source:MG | 0 | 0 | 4.025 | 0 | 2.1812 | 3.3415 |
| 23654 | ENSMUSG00000081177 | 0 | NA | NA | NA | NA | NA | Gm12509 | predicted gene 12509 [Source:MGI Symbo | 0 | 0 | 0 | 0 | 0 | 0 |
| 23655 | ENSMUSG00000081179 | 0.556921942 | 2.679043159 | 3.638243063 | 0.736356289 | 0.461513899 | NA | Gm13136 | predicted gene 13136 [Source:MGI Symbo | 0 | 0 | 0 | 0 | 0 | 3.3415 |
| 23656 | ENSMUSG00000081180 | 0 | NA | NA | NA | NA | NA | Gm7212 | predicted gene 7212 [Source:MGI Symbol | 0 | 0 | 0 | 0 | 0 | 0 |
| 23657 | ENSMUSG00000081181 | 0 | NA | NA | NA | NA | NA | Gm13353 | predicted gene 13353 [Source:MGI Symbo | 0 | 0 | 0 | 0 | 0 | 0 |
| 23658 | ENSMUSG00000081182 | 0 | NA | NA | NA | NA | NA | Gm14935 | predicted gene 14935 [Source:MGI Symbo | 0 | 0 | 0 | 0 | 0 | 0 |
| 23659 | ENSMUSG00000081183 | 0 | NA | NA | NA | NA | NA | Gm12449 | predicted gene 12449 [Source:MGI Symbo | 0 | 0 | 0 | 0 | 0 | 0 |
| 23660 | ENSMUSG00000081184 | 0 | NA | NA | NA | NA | NA | Gm12685 | predicted gene 12685 [Source:MGI Symbo | 0 | 0 | 0 | 0 | 0 | 0 |
| 23661 | ENSMUSG00000081185 | 0.133850373 | -0.796629176 | 4.080472857 | -0.195229623 | 0.84521319 | NA | Gm4852 | predicted pseudogene 4852 [Source:MGI | 0.8031 | 0 | 0 | 0 | 0 | 0 |
| 23662 | ENSMUSG00000081186 | 0 | NA | NA | NA | NA | NA | Gm12578 | predicted gene 12578 [Source:MGI Symbo | 0 | 0 | 0 | 0 | 0 | 0 |
| 23663 | ENSMUSG00000081187 | 0.83344994 | -1.76917134 | 2.609239342 | -0.678041034 | 0.497745676 | NA | 3110067C02 | RIKEN cDNA 3110067C02 gene [Source:M | 0.8031 | 3.0838 | 0 | 0 | 0 | 1.1138 |
| 23664 | ENSMUSG00000081188 | 0 | NA | NA | NA | NA | NA | Gm14621 | predicted gene 14621 [Source:MGI Symbo | 0 | 0 | 0 | 0 | 0 | 0 |
| 23665 | ENSMUSG00000081189 | 3.963211936 | -0.880116053 | 1.067997027 | -0.824080995 | 0.409893538 | NA | Hspd1-ps4 | heat shock protein 1 (chaperonin), pseud | 7.2279 | 4.1117 | 4.025 | 2.8919 | 2.1812 | 3.3415 |
| 23666 | ENSMUSG00000081191 | 0.167707431 | -0.796629176 | 4.080472857 | -0.195229623 | 0.84521319 | NA | Gm11271 | predicted gene 11271 [Source:MGI Symbo | 0 | 0 | 1.0062 | 0 | 0 | 0 |
| 23667 | ENSMUSG00000081194 | 0 | NA | NA | NA | NA | NA | Gm8424 | predicted gene 8424 [Source:MGI Symbol | 0 | 0 | 0 | 0 | 0 | 0 |
| 23668 | ENSMUSG00000081195 | 0 | NA | NA | NA | NA | NA | Rhox2-ps | reproductive homeobox 2, pseudogene [S | 0 | 0 | 0 | 0 | 0 | 0 |
| 23669 | ENSMUSG00000081196 | 0 | NA | NA | NA | NA | NA | Gm13858 | predicted gene 13858 [Source:MGI Symbo | 0 | 0 | 0 | 0 | 0 | 0 |
| 23670 | ENSMUSG00000081197 | 0 | NA | NA | NA | NA | NA | Gm15076 | predicted gene 15076 [Source:MGI Symbo | 0 | 0 | 0 | 0 | 0 | 0 |
| 23671 | ENSMUSG00000081199 | 0 | NA | NA | NA | NA | NA | Gm16407 | predicted pseudogene 16407 [Source:MG | 0 | 0 | 0 | 0 | 0 | 0 |
| 23672 | ENSMUSG00000081200 | 0 | NA | NA | NA | NA | NA | Gm15615 | predicted gene 15615 [Source:MGI Symbo | 0 | 0 | 0 | 0 | 0 | 0 |
| 23673 | ENSMUSG00000081201 | 0 | NA | NA | NA | NA | NA | Smt3h2-ps4 | SMT3 suppressor of mif two 3 homolog 2, | 0 | 0 | 0 | 0 | 0 | 0 |
| 23674 | ENSMUSG00000081202 | 0 | NA | NA | NA | NA | NA | Gm13220 | predicted gene 13220 [Source:MGI Symbo | 0 | 0 | 0 | 0 | 0 | 0 |
| 23675 | ENSMUSG00000081205 | 0.160662974 | 1.126924175 | 4.080472857 | 0.276174898 | 0.782413736 | NA | Gm5940 | predicted gene 5940 [Source:MGI Symbol | 0 | 0 | 0 | 0.964 | 0 | 0 |
| 23676 | ENSMUSG00000081206 | 0 | NA | NA | NA | NA | NA | Gm11935 | predicted gene 11935 [Source:MGI Symbo | 0 | 0 | 0 | 0 | 0 | 0 |
| 23677 | ENSMUSG00000081207 | 201.194486 | 0.164476122 | 0.260071793 | 0.63242584 | 0.527108654 | 0.888334265 | Gm13775 | predicted gene 13775 [Source:MGI Symbo | 238.52 | 128.49 | 201.25 | 157.13 | 285.74 | 196.04 |
| 23678 | ENSMUSG00000081208 | 12.85461207 | -0.475208362 | 0.75955859 | -0.625637532 | 0.531552715 | 0.888947136 | Gm11400 | predicted gene 11400 [Source:MGI Symbo | 10.44 | 23.642 | 11.069 | 23.135 | 3.2719 | 5.5692 |
| 23679 | ENSMUSG00000081209 | 0 | NA | NA | NA | NA | NA | Gm14890 | predicted gene 14890 [Source:MGI Symbo | 0 | 0 | 0 | 0 | 0 | 0 |
| 23680 | ENSMUSG00000081210 | 0.353089354 | 0.165142706 | 3.896102244 | 0.042386646 | 0.966190474 | NA | Gm12933 | predicted gene 12933 [Source:MGI Symbo | 0 | 1.0279 | 0 | 0 | 1.0906 | 0 |
| 23681 | ENSMUSG00000081211 | 0 | NA | NA | NA | NA | NA | Gm14886 | predicted gene 14886 [Source:MGI Symbo | 0 | 0 | 0 | 0 | 0 | 0 |
| 23682 | ENSMUSG00000081212 | 0.331982614 | 0.16514309 | 3.986751476 | 0.041422971 | 0.966958701 | NA | Gm6964 | predicted gene 6964 [Source:MGI Symbol | 0 | 1.0279 | 0 | 0.964 | 0 | 0 |
| 23683 | ENSMUSG00000081214 | 6.738714915 | -0.306567435 | 1.004514657 | -0.305189608 | 0.760221747 | 0.95402891 | Rpl35a-ps2 | ribosomal protein L35A, pseudogene 2 [So | 16.062 | 2.0558 | 4.025 | 3.8559 | 2.1812 | 12.252 |
| 23684 | ENSMUSG00000081215 | 0 | NA | NA | NA | NA | NA | Gm11446 | predicted gene 11446 [Source:MGI Symbo | 0 | 0 | 0 | 0 | 0 | 0 |

| | | | | | | | | | | | | | | | |
|---|---|---|---|---|---|---|---|---|---|---|---|---|---|---|---|
| 23685 | ENSMUSG00000081216 | 0 | NA | NA | NA | NA | NA | Gm6744 | predicted gene 6744 [Source:MGI Symbol | 0 | 0 | 0 | 0 | 0 | 0 |
| 23686 | ENSMUSG00000081217 | 0.185640647 | 1.126924175 | 4.080472857 | 0.276174898 | 0.782413736 | NA | Gm15268 | predicted gene 15268 [Source:MGI Symbo | 0 | 0 | 0 | 0 | 0 | 1.1138 |
| 23687 | ENSMUSG00000081218 | 0 | NA | NA | NA | NA | NA | Gm4297 | predicted gene 4297 [Source:MGI Symbol | 0 | 0 | 0 | 0 | 0 | 0 |
| 23688 | ENSMUSG00000081219 | 0.167707431 | -0.796629176 | 4.080472857 | -0.195229623 | 0.84521319 | NA | Bambi-ps1 | BMP and activin membrane-bound inhibit | 0 | 0 | 1.0062 | 0 | 0 | 0 |
| 23689 | ENSMUSG00000081220 | 0 | NA | NA | NA | NA | NA | Gm14542 | predicted gene 14542 [Source:MGI Symbo | 0 | 0 | 0 | 0 | 0 | 0 |
| 23690 | ENSMUSG00000081221 | 0.938710431 | -2.042881906 | 2.283279595 | -0.894713863 | 0.370940066 | NA | Gm14760 | predicted gene 14760 [Source:MGI Symbo | 1.6062 | 2.0558 | 1.0062 | 0.964 | 0 | 0 |
| 23691 | ENSMUSG00000081222 | 0 | NA | NA | NA | NA | NA | Gm15173 | predicted gene 15173 [Source:MGI Symbo | 0 | 0 | 0 | 0 | 0 | 0 |
| 23692 | ENSMUSG00000081223 | 37.45155718 | 0.267346838 | 0.391213841 | 0.683377757 | 0.494368171 | 0.877473722 | Gm12247 | predicted gene 12247 [Source:MGI Symbo | 41.761 | 21.586 | 38.237 | 33.739 | 39.262 | 50.123 |
| 23693 | ENSMUSG00000081224 | 0 | NA | NA | NA | NA | NA | Gm2066 | predicted pseudogene 2066 [Source:MGI | 0 | 0 | 0 | 0 | 0 | 0 |
| 23694 | ENSMUSG00000081225 | 0 | NA | NA | NA | NA | NA | Cyp2j12 | cytochrome P450, family 2, subfamily j, po | 0 | 0 | 0 | 0 | 0 | 0 |
| 23695 | ENSMUSG00000081226 | 0 | NA | NA | NA | NA | NA | Gm8802 | predicted gene 8802 [Source:MGI Symbol | 0 | 0 | 0 | 0 | 0 | 0 |
| 23696 | ENSMUSG00000081227 | 0 | NA | NA | NA | NA | NA | Gm13082 | predicted gene 13082 [Source:MGI Symbo | 0 | 0 | 0 | 0 | 0 | 0 |
| 23697 | ENSMUSG00000081228 | 3.606253303 | -1.230842573 | 1.226344916 | -1.00366753 | 0.315538893 | NA | Gm16089 | predicted gene 16089 [Source:MGI Symbo | 4.0155 | 2.0558 | 9.0562 | 0.964 | 1.0906 | 4.4554 |
| 23698 | ENSMUSG00000081229 | 0 | NA | NA | NA | NA | NA | Lamr1-ps1 | laminin receptor 1 (ribosomal protein SA), | 0 | 0 | 0 | 0 | 0 | 0 |
| 23699 | ENSMUSG00000081230 | 0 | NA | NA | NA | NA | NA | Gm9097 | predicted gene 9097 [Source:MGI Symbol | 0 | 0 | 0 | 0 | 0 | 0 |
| 23700 | ENSMUSG00000081231 | 0 | NA | NA | NA | NA | NA | Tcp1-ps1 | t-complex protein 1, pseudogene 1 [Sourc | 0 | 0 | 0 | 0 | 0 | 0 |
| 23701 | ENSMUSG00000081232 | 0.346303622 | 2.004339653 | 3.884003423 | 0.516049919 | 0.605819544 | NA | Gm14373 | predicted gene 14373 [Source:MGI Symbo | 0 | 0 | 0 | 0.964 | 0 | 1.1138 |
| 23702 | ENSMUSG00000081233 | 0 | NA | NA | NA | NA | NA | Gm12857 | predicted gene 12857 [Source:MGI Symbo | 0 | 0 | 0 | 0 | 0 | 0 |
| 23703 | ENSMUSG00000081234 | 0 | NA | NA | NA | NA | NA | Olfr1120 | olfactory receptor 1120 [Source:MGI Sym | 0 | 0 | 0 | 0 | 0 | 0 |
| 23704 | ENSMUSG00000081235 | 0 | NA | NA | NA | NA | NA | Gm14788 | predicted gene 14788 [Source:MGI Symbo | 0 | 0 | 0 | 0 | 0 | 0 |
| 23705 | ENSMUSG00000081236 | 0 | NA | NA | NA | NA | NA | Gm13574 | predicted gene 13574 [Source:MGI Symbo | 0 | 0 | 0 | 0 | 0 | 0 |
| 23706 | ENSMUSG00000081237 | 0 | NA | NA | NA | NA | NA | Gm11385 | predicted gene 11385 [Source:MGI Symbo | 0 | 0 | 0 | 0 | 0 | 0 |
| 23707 | ENSMUSG00000081238 | 0.506734502 | -2.372847156 | 3.338060886 | -0.710845978 | 0.477179685 | NA | Gm15836 | predicted gene 15836 [Source:MGI Symbo | 0 | 1.0279 | 2.0125 | 0 | 0 | 0 |
| 23708 | ENSMUSG00000081239 | 0 | NA | NA | NA | NA | NA | Gm11836 | predicted gene 11836 [Source:MGI Symbo | 0 | 0 | 0 | 0 | 0 | 0 |
| 23709 | ENSMUSG00000081240 | 0 | NA | NA | NA | NA | NA | Gm16511 | predicted gene 16511 [Source:MGI Symbo | 0 | 0 | 0 | 0 | 0 | 0 |
| 23710 | ENSMUSG00000081242 | 0.342639279 | -1.805976596 | 4.028547714 | -0.448294702 | 0.653940524 | NA | Gm15719 | predicted gene 15719 [Source:MGI Symbo | 0 | 2.0558 | 0 | 0 | 0 | 0 |
| 23711 | ENSMUSG00000081243 | 0 | NA | NA | NA | NA | NA | Gm14037 | predicted gene 14037 [Source:MGI Symbo | 0 | 0 | 0 | 0 | 0 | 0 |
| 23712 | ENSMUSG00000081244 | 0 | NA | NA | NA | NA | NA | Gm13529 | predicted gene 13529 [Source:MGI Symbo | 0 | 0 | 0 | 0 | 0 | 0 |
| 23713 | ENSMUSG00000081245 | 0 | NA | NA | NA | NA | NA | Gm6587 | predicted gene 6587 [Source:MGI Symbol | 0 | 0 | 0 | 0 | 0 | 0 |
| 23714 | ENSMUSG00000081248 | 0 | NA | NA | NA | NA | NA | Gm13787 | predicted gene 13787 [Source:MGI Symbo | 0 | 0 | 0 | 0 | 0 | 0 |
| 23715 | ENSMUSG00000081249 | 2.56450386 | -0.876429423 | 1.492002445 | -0.587418221 | 0.556922859 | NA | Gm11517 | predicted gene 11517 [Source:MGI Symbo | 0.8031 | 5.1396 | 4.025 | 0.964 | 0 | 4.4554 |
| 23716 | ENSMUSG00000081250 | 0 | NA | NA | NA | NA | NA | Gm15554 | predicted gene 15554 [Source:MGI Symbo | 0 | 0 | 0 | 0 | 0 | 0 |
| 23717 | ENSMUSG00000081251 | 0.167707431 | -0.796629176 | 4.080472857 | -0.195229623 | 0.84521319 | NA | Gm12164 | predicted gene 12164 [Source:MGI Symbo | 0 | 0 | 1.0062 | 0 | 0 | 0 |
| 23718 | ENSMUSG00000081252 | 0 | NA | NA | NA | NA | NA | Gm16032 | predicted gene 16032 [Source:MGI Symbo | 0 | 0 | 0 | 0 | 0 | 0 |
| 23719 | ENSMUSG00000081253 | 0 | NA | NA | NA | NA | NA | Gm8501 | predicted gene 8501 [Source:MGI Symbol | 0 | 0 | 0 | 0 | 0 | 0 |
| 23720 | ENSMUSG00000081254 | 2.085796587 | -0.085485834 | 1.50781082 | -0.056695331 | 0.954787893 | NA | Gm12112 | predicted gene 12112 [Source:MGI Symbo | 2.4093 | 2.0558 | 2.0125 | 3.8559 | 2.1812 | 0 |
| 23721 | ENSMUSG00000081255 | 0.167707431 | -0.796629176 | 4.080472857 | -0.195229623 | 0.84521319 | NA | Gm5380 | predicted gene 5380 [Source:MGI Symbol | 0 | 0 | 1.0062 | 0 | 0 | 0 |
| 23722 | ENSMUSG00000081256 | 0 | NA | NA | NA | NA | NA | Gm15791 | predicted gene 15791 [Source:MGI Symbo | 0 | 0 | 0 | 0 | 0 | 0 |
| 23723 | ENSMUSG00000081257 | 0.346303622 | 2.004339653 | 3.884003423 | 0.516049919 | 0.605819544 | NA | Gm13607 | predicted gene 13607 [Source:MGI Symbo | 0 | 0 | 0 | 0.964 | 0 | 1.1138 |
| 23724 | ENSMUSG00000081258 | 0 | NA | NA | NA | NA | NA | Gm14013 | predicted gene 14013 [Source:MGI Symbo | 0 | 0 | 0 | 0 | 0 | 0 |
| 23725 | ENSMUSG00000081259 | 0 | NA | NA | NA | NA | NA | Gm6893 | predicted gene 6893 [Source:MGI Symbol | 0 | 0 | 0 | 0 | 0 | 0 |
| 23726 | ENSMUSG00000081260 | 0 | NA | NA | NA | NA | NA | Gm13149 | predicted gene 13149 [Source:MGI Symbo | 0 | 0 | 0 | 0 | 0 | 0 |
| 23727 | ENSMUSG00000081261 | 0.743983808 | -0.269067535 | 2.702716736 | -0.099554471 | 0.920698041 | NA | Gm15370 | predicted gene 15370 [Source:MGI Symbo | 2.4093 | 0 | 0 | 0.964 | 1.0906 | 0 |
| 23728 | ENSMUSG00000081262 | 0 | NA | NA | NA | NA | NA | Gm12261 | predicted gene 12261 [Source:MGI Symbo | 0 | 0 | 0 | 0 | 0 | 0 |
| 23729 | ENSMUSG00000081263 | 0 | NA | NA | NA | NA | NA | Gm14859 | predicted gene 14859 [Source:MGI Symbo | 0 | 0 | 0 | 0 | 0 | 0 |
| 23730 | ENSMUSG00000081264 | 0 | NA | NA | NA | NA | NA | Gm14021 | predicted gene 14021 [Source:MGI Symbo | 0 | 0 | 0 | 0 | 0 | 0 |
| 23731 | ENSMUSG00000081265 | 0 | NA | NA | NA | NA | NA | Gm11282 | predicted gene 11282 [Source:MGI Symbo | 0 | 0 | 0 | 0 | 0 | 0 |
| 23732 | ENSMUSG00000081268 | 0 | NA | NA | NA | NA | NA | Gm14909 | predicted gene 14909 [Source:MGI Symbo | 0 | 0 | 0 | 0 | 0 | 0 |
| 23733 | ENSMUSG00000081269 | 0 | NA | NA | NA | NA | NA | Gm14659 | predicted gene 14659 [Source:MGI Symbo | 0 | 0 | 0 | 0 | 0 | 0 |
| 23734 | ENSMUSG00000081270 | 0 | NA | NA | NA | NA | NA | Gm11653 | predicted gene 11653 [Source:MGI Symbo | 0 | 0 | 0 | 0 | 0 | 0 |
| 23735 | ENSMUSG00000081271 | 0 | NA | NA | NA | NA | NA | Gm14883 | predicted gene 14883 [Source:MGI Symbo | 0 | 0 | 0 | 0 | 0 | 0 |
| 23736 | ENSMUSG00000081272 | 18.43822286 | 0.279292576 | 0.518541676 | 0.538611628 | 0.590154863 | 0.911690353 | Ap2m1-ps | adaptor-related protein complex 2, mu 1 s | 14.456 | 13.363 | 22.137 | 15.424 | 19.631 | 25.618 |
| 23737 | ENSMUSG00000081273 | 0 | NA | NA | NA | NA | NA | Gm15896 | predicted gene 15896 [Source:MGI Symbo | 0 | 0 | 0 | 0 | 0 | 0 |
| 23738 | ENSMUSG00000081274 | 0 | NA | NA | NA | NA | NA | Gm15727 | predicted gene 15727 [Source:MGI Symbo | 0 | 0 | 0 | 0 | 0 | 0 |
| 23739 | ENSMUSG00000081275 | 0 | NA | NA | NA | NA | NA | Gm13875 | predicted gene 13875 [Source:MGI Symbo | 0 | 0 | 0 | 0 | 0 | 0 |
| 23740 | ENSMUSG00000081276 | 0 | NA | NA | NA | NA | NA | Gm13132 | predicted gene 13132 [Source:MGI Symbo | 0 | 0 | 0 | 0 | 0 | 0 |

| | | | | | | | | | | | | | | | |
|---|---|---|---|---|---|---|---|---|---|---|---|---|---|---|---|
| 23741 | ENSMUSG00000081277 | 0.185640647 | 1.126924175 | 4.080472857 | 0.276174898 | 0.782413736 | NA | Gm15285 | predicted gene 15285 [Source:MGI Symbo | 0 | 0 | 0 | 0 | 0 | 1.1138 |
| 23742 | ENSMUSG00000081278 | 0 | NA | NA | NA | NA | NA | Gm13186 | predicted gene 13186 [Source:MGI Symbo | 0 | 0 | 0 | 0 | 0 | 0 |
| 23743 | ENSMUSG00000081279 | 0 | NA | NA | NA | NA | NA | Vmn1r-ps2 | vomeronasal 1 receptor, pseudogene 2 [S | 0 | 0 | 0 | 0 | 0 | 0 |
| 23744 | ENSMUSG00000081280 | 0 | NA | NA | NA | NA | NA | Gm9516 | predicted gene 9516 [Source:MGI Symbol | 0 | 0 | 0 | 0 | 0 | 0 |
| 23745 | ENSMUSG00000081281 | 0.967300313 | -2.048316886 | 2.458008888 | -0.833323629 | 0.404662234 | NA | Rpl7-ps9 | ribosomal protein L7, pseudogene 9 [Sour | 1.6062 | 3.0838 | 0 | 0 | 0 | 1.1138 |
| 23746 | ENSMUSG00000081282 | 0 | NA | NA | NA | NA | NA | Gm13836 | predicted gene 13836 [Source:MGI Symbo | 0 | 0 | 0 | 0 | 0 | 0 |
| 23747 | ENSMUSG00000081283 | 0 | NA | NA | NA | NA | NA | Vmn1r-ps10 | vomeronasal 1 receptor, pseudogene 104 | 0 | 0 | 0 | 0 | 0 | 0 |
| 23748 | ENSMUSG00000081284 | 0.160662974 | 1.126924175 | 4.080472857 | 0.276174898 | 0.782413736 | NA | Gm13673 | predicted gene 13673 [Source:MGI Symbo | 0 | 0 | 0 | 0.964 | 0 | 0 |
| 23749 | ENSMUSG00000081285 | 0 | NA | NA | NA | NA | NA | Gm18859 | predicted gene, 18859 [Source:MGI Symb | 0 | 0 | 0 | 0 | 0 | 0 |
| 23750 | ENSMUSG00000081286 | 0 | NA | NA | NA | NA | NA | Gm14995 | predicted gene 14995 [Source:MGI Symbo | 0 | 0 | 0 | 0 | 0 | 0 |
| 23751 | ENSMUSG00000081287 | 0 | NA | NA | NA | NA | NA | Olfr333-ps1 | olfactory receptor 333, pseudogene 1 [Sou | 0 | 0 | 0 | 0 | 0 | 0 |
| 23752 | ENSMUSG00000081288 | 0 | NA | NA | NA | NA | NA | Gm11386 | predicted gene 11386 [Source:MGI Symbo | 0 | 0 | 0 | 0 | 0 | 0 |
| 23753 | ENSMUSG00000081289 | 0 | NA | NA | NA | NA | NA | Gm14857 | predicted gene 14857 [Source:MGI Symbo | 0 | 0 | 0 | 0 | 0 | 0 |
| 23754 | ENSMUSG00000081290 | 0 | NA | NA | NA | NA | NA | Gm12520 | predicted gene 12520 [Source:MGI Symbo | 0 | 0 | 0 | 0 | 0 | 0 |
| 23755 | ENSMUSG00000081292 | 0 | NA | NA | NA | NA | NA | Gm15018 | predicted gene 15018 [Source:MGI Symbo | 0 | 0 | 0 | 0 | 0 | 0 |
| 23756 | ENSMUSG00000081293 | 0 | NA | NA | NA | NA | NA | Gm11673 | predicted gene 11673 [Source:MGI Symbo | 0 | 0 | 0 | 0 | 0 | 0 |
| 23757 | ENSMUSG00000081295 | 0 | NA | NA | NA | NA | NA | Gm6275 | predicted gene 6275 [Source:MGI Symbol | 0 | 0 | 0 | 0 | 0 | 0 |
| 23758 | ENSMUSG00000081296 | 0 | NA | NA | NA | NA | NA | Gm12203 | predicted gene 12203 [Source:MGI Symbo | 0 | 0 | 0 | 0 | 0 | 0 |
| 23759 | ENSMUSG00000081297 | 0 | NA | NA | NA | NA | NA | Gm12209 | predicted gene 12209 [Source:MGI Symbo | 0 | 0 | 0 | 0 | 0 | 0 |
| 23760 | ENSMUSG00000081298 | 0 | NA | NA | NA | NA | NA | Gm14622 | predicted gene 14622 [Source:MGI Symbo | 0 | 0 | 0 | 0 | 0 | 0 |
| 23761 | ENSMUSG00000081299 | 0 | NA | NA | NA | NA | NA | Gm13603 | predicted gene 13603 [Source:MGI Symbo | 0 | 0 | 0 | 0 | 0 | 0 |
| 23762 | ENSMUSG00000081300 | 0 | NA | NA | NA | NA | NA | Gm14587 | predicted gene 14587 [Source:MGI Symbo | 0 | 0 | 0 | 0 | 0 | 0 |
| 23763 | ENSMUSG00000081301 | 0 | NA | NA | NA | NA | NA | Gm15951 | predicted gene 15951 [Source:MGI Symbo | 0 | 0 | 0 | 0 | 0 | 0 |
| 23764 | ENSMUSG00000081302 | 0 | NA | NA | NA | NA | NA | Gm12020 | predicted gene 12020 [Source:MGI Symbo | 0 | 0 | 0 | 0 | 0 | 0 |
| 23765 | ENSMUSG00000081303 | 0.167707431 | -0.796629176 | 4.080472857 | -0.195229623 | 0.84521319 | NA | Gm16011 | predicted gene 16011 [Source:MGI Symbo | 0 | 0 | 1.0062 | 0 | 0 | 0 |
| 23766 | ENSMUSG00000081304 | 0 | NA | NA | NA | NA | NA | Gm11509 | predicted gene 11509 [Source:MGI Symbo | 0 | 0 | 0 | 0 | 0 | 0 |
| 23767 | ENSMUSG00000081305 | 0 | NA | NA | NA | NA | NA | Gm12879 | predicted gene 12879 [Source:MGI Symbo | 0 | 0 | 0 | 0 | 0 | 0 |
| 23768 | ENSMUSG00000081306 | 0 | NA | NA | NA | NA | NA | Gm15266 | predicted gene 15266 [Source:MGI Symbo | 0 | 0 | 0 | 0 | 0 | 0 |
| 23769 | ENSMUSG00000081307 | 0 | NA | NA | NA | NA | NA | Gm13627 | predicted gene 13627 [Source:MGI Symbo | 0 | 0 | 0 | 0 | 0 | 0 |
| 23770 | ENSMUSG00000081308 | 0 | NA | NA | NA | NA | NA | Gm14480 | predicted gene 14480 [Source:MGI Symbo | 0 | 0 | 0 | 0 | 0 | 0 |
| 23771 | ENSMUSG00000081309 | 0 | NA | NA | NA | NA | NA | Gm12397 | predicted gene 12397 [Source:MGI Symbo | 0 | 0 | 0 | 0 | 0 | 0 |
| 23772 | ENSMUSG00000081310 | 0 | NA | NA | NA | NA | NA | Gm14054 | predicted gene 14054 [Source:MGI Symbo | 0 | 0 | 0 | 0 | 0 | 0 |
| 23773 | ENSMUSG00000081312 | 0 | NA | NA | NA | NA | NA | Gm14520 | predicted gene 14520 [Source:MGI Symbo | 0 | 0 | 0 | 0 | 0 | 0 |
| 23774 | ENSMUSG00000081313 | 0 | NA | NA | NA | NA | NA | Gm14213 | predicted gene 14213 [Source:MGI Symbo | 0 | 0 | 0 | 0 | 0 | 0 |
| 23775 | ENSMUSG00000081314 | 0.167707431 | -0.796629176 | 4.080472857 | -0.195229623 | 0.84521319 | NA | Gm14132 | predicted gene 14132 [Source:MGI Symbo | 0 | 0 | 1.0062 | 0 | 0 | 0 |
| 23776 | ENSMUSG00000081315 | 0 | NA | NA | NA | NA | NA | Olfr621-ps1 | olfactory receptor 621, pseudogene 1 [Sou | 0 | 0 | 0 | 0 | 0 | 0 |
| 23777 | ENSMUSG00000081316 | 0 | NA | NA | NA | NA | NA | Gm16205 | predicted gene 16205 [Source:MGI Symbo | 0 | 0 | 0 | 0 | 0 | 0 |
| 23778 | ENSMUSG00000081317 | 0 | NA | NA | NA | NA | NA | Gm12757 | predicted gene 12757 [Source:MGI Symbo | 0 | 0 | 0 | 0 | 0 | 0 |
| 23779 | ENSMUSG00000081318 | 0 | NA | NA | NA | NA | NA | Rpl36-ps1 | ribosomal protein L36, pseudogene 1 [Sou | 0 | 0 | 0 | 0 | 0 | 0 |
| 23780 | ENSMUSG00000081319 | 0 | NA | NA | NA | NA | NA | Gm12936 | predicted gene 12936 [Source:MGI Symbo | 0 | 0 | 0 | 0 | 0 | 0 |
| 23781 | ENSMUSG00000081320 | 0 | NA | NA | NA | NA | NA | Gm15034 | predicted gene 15034 [Source:MGI Symbo | 0 | 0 | 0 | 0 | 0 | 0 |
| 23782 | ENSMUSG00000081321 | 0 | NA | NA | NA | NA | NA | Gm15304 | predicted gene 15304 [Source:MGI Symbo | 0 | 0 | 0 | 0 | 0 | 0 |
| 23783 | ENSMUSG00000081323 | 0 | NA | NA | NA | NA | NA | Rpl7l1-ps1 | ribosomal protein L7-like 1, pseudogene 1 | 0 | 0 | 0 | 0 | 0 | 0 |
| 23784 | ENSMUSG00000081324 | 0 | NA | NA | NA | NA | NA | Gm14550 | predicted gene 14550 [Source:MGI Symbo | 0 | 0 | 0 | 0 | 0 | 0 |
| 23785 | ENSMUSG00000081325 | 0 | NA | NA | NA | NA | NA | Gm12599 | predicted gene 12599 [Source:MGI Symbo | 0 | 0 | 0 | 0 | 0 | 0 |
| 23786 | ENSMUSG00000081326 | 0 | NA | NA | NA | NA | NA | Gm14417 | predicted gene 14417 [Source:MGI Symbo | 0 | 0 | 0 | 0 | 0 | 0 |
| 23787 | ENSMUSG00000081327 | 0 | NA | NA | NA | NA | NA | Gm11819 | predicted gene 11819 [Source:MGI Symbo | 0 | 0 | 0 | 0 | 0 | 0 |
| 23788 | ENSMUSG00000081329 | 0 | NA | NA | NA | NA | NA | Gm12492 | predicted gene 12492 [Source:MGI Symbo | 0 | 0 | 0 | 0 | 0 | 0 |
| 23789 | ENSMUSG00000081330 | 0 | NA | NA | NA | NA | NA | Gm13013 | predicted gene 13013 [Source:MGI Symbo | 0 | 0 | 0 | 0 | 0 | 0 |
| 23790 | ENSMUSG00000081331 | 0 | NA | NA | NA | NA | NA | Gm15151 | predicted gene 15151 [Source:MGI Symbo | 0 | 0 | 0 | 0 | 0 | 0 |
| 23791 | ENSMUSG00000081332 | 0.428363721 | -0.543730382 | 3.624677344 | -0.15000794 | 0.880758351 | NA | Gm12664 | predicted gene 12664 [Source:MGI Symbo | 1.6062 | 0 | 0 | 0.964 | 0 | 0 |
| 23792 | ENSMUSG00000081333 | 0 | NA | NA | NA | NA | NA | Gm12307 | predicted gene 12307 [Source:MGI Symbo | 0 | 0 | 0 | 0 | 0 | 0 |
| 23793 | ENSMUSG00000081334 | 0 | NA | NA | NA | NA | NA | Vmn1r-ps94 | vomeronasal 1 receptor, pseudogene 94 [ | 0 | 0 | 0 | 0 | 0 | 0 |
| 23794 | ENSMUSG00000081335 | 0 | NA | NA | NA | NA | NA | Gm15786 | predicted gene 15786 [Source:MGI Symbo | 0 | 0 | 0 | 0 | 0 | 0 |
| 23795 | ENSMUSG00000081336 | 0 | NA | NA | NA | NA | NA | Gm15073 | predicted gene 15073 [Source:MGI Symbo | 0 | 0 | 0 | 0 | 0 | 0 |
| 23796 | ENSMUSG00000081339 | 0.695522043 | 1.639439662 | 2.762392557 | 0.593485404 | 0.552856349 | NA | Gm16044 | predicted gene 16044 [Source:MGI Symbo | 0 | 1.0279 | 0 | 0.964 | 2.1812 | 0 |

| | | | | | | | | | | | | | | | |
|---|---|---|---|---|---|---|---|---|---|---|---|---|---|---|---|
| 23797 | ENSMUSG00000081340 | 1.250977034 | -1.387274498 | 2.079707712 | -0.66705263 | 0.504738517 | NA | Gm12275 | predicted gene 12275 [Source:MGI Symbo | 2.4093 | 0 | 3.0187 | 0.964 | 0 | 1.1138 |
| 23798 | ENSMUSG00000081341 | 0 | NA | NA | NA | NA | NA | Gm14891 | predicted gene 14891 [Source:MGI Symbo | 0 | 0 | 0 | 0 | 0 | 0 |
| 23799 | ENSMUSG00000081342 | 0.181769715 | 1.126924175 | 4.080472857 | 0.276174898 | 0.782413736 | NA | Gm11434 | predicted gene 11434 [Source:MGI Symbo | 0 | 0 | 0 | 0 | 1.0906 | 0 |
| 23800 | ENSMUSG00000081343 | 0 | NA | NA | NA | NA | NA | Gm16071 | predicted gene 16071 [Source:MGI Symbo | 0 | 0 | 0 | 0 | 0 | 0 |
| 23801 | ENSMUSG00000081344 | 9.859153508 | 0.926162201 | 0.732187393 | 1.264925086 | 0.205898184 | 0.651085207 | Gm14303 | predicted gene 14303 [Source:MGI Symbo | 8.031 | 7.1954 | 5.0312 | 5.7839 | 14.178 | 18.935 |
| 23802 | ENSMUSG00000081345 | 0 | NA | NA | NA | NA | NA | Olfr1159-ps1 | olfactory receptor 1159, pseudogene 1 [S | 0 | 0 | 0 | 0 | 0 | 0 |
| 23803 | ENSMUSG00000081346 | 0 | NA | NA | NA | NA | NA | Gm12030 | predicted gene 12030 [Source:MGI Symbo | 0 | 0 | 0 | 0 | 0 | 0 |
| 23804 | ENSMUSG00000081348 | 0 | NA | NA | NA | NA | NA | Gm13878 | predicted gene 13878 [Source:MGI Symbo | 0 | 0 | 0 | 0 | 0 | 0 |
| 23805 | ENSMUSG00000081349 | 0 | NA | NA | NA | NA | NA | Gm15041 | predicted gene 15041 [Source:MGI Symbo | 0 | 0 | 0 | 0 | 0 | 0 |
| 23806 | ENSMUSG00000081350 | 0 | NA | NA | NA | NA | NA | Gm5388 | predicted gene 5388 [Source:MGI Symbol | 0 | 0 | 0 | 0 | 0 | 0 |
| 23807 | ENSMUSG00000081352 | 0 | NA | NA | NA | NA | NA | Gm12389 | predicted gene 12389 [Source:MGI Symbo | 0 | 0 | 0 | 0 | 0 | 0 |
| 23808 | ENSMUSG00000081353 | 0 | NA | NA | NA | NA | NA | Gm13818 | predicted gene 13818 [Source:MGI Symbo | 0 | 0 | 0 | 0 | 0 | 0 |
| 23809 | ENSMUSG00000081354 | 0 | NA | NA | NA | NA | NA | Gm14894 | predicted gene 14894 [Source:MGI Symbo | 0 | 0 | 0 | 0 | 0 | 0 |
| 23810 | ENSMUSG00000081355 | 0.133850373 | -0.796629176 | 4.080472857 | -0.195229623 | 0.84521319 | NA | Gm15264 | predicted gene 15264 [Source:MGI Symbo | 0.8031 | 0 | 0 | 0 | 0 | 0 |
| 23811 | ENSMUSG00000081357 | 0 | NA | NA | NA | NA | NA | Xlr4e-ps | X-linked lymphocyte-regulated 4E, pseudo | 0 | 0 | 0 | 0 | 0 | 0 |
| 23812 | ENSMUSG00000081358 | 0 | NA | NA | NA | NA | NA | Gm14746 | predicted gene 14746 [Source:MGI Symbo | 0 | 0 | 0 | 0 | 0 | 0 |
| 23813 | ENSMUSG00000081359 | 0 | NA | NA | NA | NA | NA | Cox20b | cytochrome c oxidase assembly protein 2( | 0 | 0 | 0 | 0 | 0 | 0 |
| 23814 | ENSMUSG00000081360 | 0 | NA | NA | NA | NA | NA | Gm11718 | predicted gene 11718 [Source:MGI Symbo | 0 | 0 | 0 | 0 | 0 | 0 |
| 23815 | ENSMUSG00000081361 | 0 | NA | NA | NA | NA | NA | Gm11813 | predicted gene 11813 [Source:MGI Symbo | 0 | 0 | 0 | 0 | 0 | 0 |
| 23816 | ENSMUSG00000081362 | 0 | NA | NA | NA | NA | NA | Cyp2j7 | cytochrome P450, family 2, subfamily j, pc | 0 | 0 | 0 | 0 | 0 | 0 |
| 23817 | ENSMUSG00000081363 | 0 | NA | NA | NA | NA | NA | Gm11500 | predicted gene 11500 [Source:MGI Symbo | 0 | 0 | 0 | 0 | 0 | 0 |
| 23818 | ENSMUSG00000081364 | 0 | NA | NA | NA | NA | NA | Gm8595 | predicted gene 8595 [Source:MGI Symbol | 0 | 0 | 0 | 0 | 0 | 0 |
| 23819 | ENSMUSG00000081365 | 0 | NA | NA | NA | NA | NA | Gm15031 | predicted gene 15031 [Source:MGI Symbo | 0 | 0 | 0 | 0 | 0 | 0 |
| 23820 | ENSMUSG00000081367 | 0 | NA | NA | NA | NA | NA | Gm13265 | predicted gene 13265 [Source:MGI Symbo | 0 | 0 | 0 | 0 | 0 | 0 |
| 23821 | ENSMUSG00000081369 | 0 | NA | NA | NA | NA | NA | Gm11896 | predicted gene 11896 [Source:MGI Symbo | 0 | 0 | 0 | 0 | 0 | 0 |
| 23822 | ENSMUSG00000081370 | 0.133850373 | -0.796629176 | 4.080472857 | -0.195229623 | 0.84521319 | NA | Gm9105 | predicted gene 9105 [Source:MGI Symbol | 0.8031 | 0 | 0 | 0 | 0 | 0 |
| 23823 | ENSMUSG00000081372 | 0.621048825 | -1.224891493 | 2.859518283 | -0.428355888 | 0.668392035 | NA | Gm16171 | predicted gene 16171 [Source:MGI Symbo | 1.6062 | 0 | 1.0062 | 0 | 0 | 1.1138 |
| 23824 | ENSMUSG00000081373 | 0 | NA | NA | NA | NA | NA | Gm16436 | predicted gene 16436 [Source:MGI Symbo | 0 | 0 | 0 | 0 | 0 | 0 |
| 23825 | ENSMUSG00000081374 | 0.185640647 | 1.126924175 | 4.080472857 | 0.276174898 | 0.782413736 | NA | Gm15011 | predicted gene 15011 [Source:MGI Symbo | 0 | 0 | 0 | 0 | 0 | 1.1138 |
| 23826 | ENSMUSG00000081375 | 0 | NA | NA | NA | NA | NA | Gm14686 | predicted gene 14686 [Source:MGI Symbo | 0 | 0 | 0 | 0 | 0 | 0 |
| 23827 | ENSMUSG00000081376 | 0.181769715 | 1.126924175 | 4.080472857 | 0.276174898 | 0.782413736 | NA | Gm11694 | predicted gene 11694 [Source:MGI Symbo | 0 | 0 | 0 | 0 | 1.0906 | 0 |
| 23828 | ENSMUSG00000081377 | 0 | NA | NA | NA | NA | NA | Gm12877 | predicted gene 12877 [Source:MGI Symbo | 0 | 0 | 0 | 0 | 0 | 0 |
| 23829 | ENSMUSG00000081378 | 1.184840777 | -1.255103961 | 2.367539016 | -0.530130212 | 0.596021653 | NA | Rps13-ps4 | ribosomal protein S13, pseudogene 4 [Sou | 0 | 0 | 5.0312 | 0.964 | 0 | 1.1138 |
| 23830 | ENSMUSG00000081379 | 0 | NA | NA | NA | NA | NA | Gm14771 | predicted gene 14771 [Source:MGI Symbo | 0 | 0 | 0 | 0 | 0 | 0 |
| 23831 | ENSMUSG00000081381 | 0 | NA | NA | NA | NA | NA | Gm5752 | predicted gene 5752 [Source:MGI Symbol | 0 | 0 | 0 | 0 | 0 | 0 |
| 23832 | ENSMUSG00000081382 | 1.634491198 | -0.442467688 | 1.583430466 | -0.279436134 | 0.779910144 | NA | Rpl18-ps1 | ribosomal protein L18, pseudogene 1 [Sou | 1.6062 | 2.0558 | 2.0125 | 1.928 | 1.0906 | 1.1138 |
| 23833 | ENSMUSG00000081383 | 0 | NA | NA | NA | NA | NA | Gm5742 | predicted gene 5742 [Source:MGI Symbol | 0 | 0 | 0 | 0 | 0 | 0 |
| 23834 | ENSMUSG00000081384 | 0 | NA | NA | NA | NA | NA | Gm12752 | predicted gene 12752 [Source:MGI Symbo | 0 | 0 | 0 | 0 | 0 | 0 |
| 23835 | ENSMUSG00000081386 | 0 | NA | NA | NA | NA | NA | Gm15160 | predicted gene 15160 [Source:MGI Symbo | 0 | 0 | 0 | 0 | 0 | 0 |
| 23836 | ENSMUSG00000081388 | 0 | NA | NA | NA | NA | NA | Gm11801 | predicted gene 11801 [Source:MGI Symbo | 0 | 0 | 0 | 0 | 0 | 0 |
| 23837 | ENSMUSG00000081389 | 0 | NA | NA | NA | NA | NA | Gm13368 | predicted gene 13368 [Source:MGI Symbo | 0 | 0 | 0 | 0 | 0 | 0 |
| 23838 | ENSMUSG00000081390 | 0.321325949 | 1.911575114 | 4.048267893 | 0.472195804 | 0.636787034 | NA | Gm14538 | predicted gene 14538 [Source:MGI Symbo | 0 | 0 | 0 | 1.928 | 0 | 0 |
| 23839 | ENSMUSG00000081391 | 0 | NA | NA | NA | NA | NA | Gm11765 | predicted gene 11765 [Source:MGI Symbo | 0 | 0 | 0 | 0 | 0 | 0 |
| 23840 | ENSMUSG00000081392 | 0 | NA | NA | NA | NA | NA | Gm11668 | predicted gene 11668 [Source:MGI Symbo | 0 | 0 | 0 | 0 | 0 | 0 |
| 23841 | ENSMUSG00000081394 | 0.321325949 | 1.911575114 | 4.048267893 | 0.472195804 | 0.636787034 | NA | Gm13215 | predicted gene 13215 [Source:MGI Symbo | 0 | 0 | 0 | 1.928 | 0 | 0 |
| 23842 | ENSMUSG00000081395 | 0 | NA | NA | NA | NA | NA | Gm14879 | predicted gene 14879 [Source:MGI Symbo | 0 | 0 | 0 | 0 | 0 | 0 |
| 23843 | ENSMUSG00000081396 | 0.562214094 | -1.122836648 | 3.276281221 | -0.34271681 | 0.731811515 | NA | Gm15132 | predicted gene 15132 [Source:MGI Symbo | 2.4093 | 0 | 0 | 0.964 | 0 | 0 |
| 23844 | ENSMUSG00000081397 | 0 | NA | NA | NA | NA | NA | Gm14815 | predicted gene 14815 [Source:MGI Symbo | 0 | 0 | 0 | 0 | 0 | 0 |
| 23845 | ENSMUSG00000081398 | 0 | NA | NA | NA | NA | NA | Gm7657 | predicted gene 7657 [Source:MGI Symbol | 0 | 0 | 0 | 0 | 0 | 0 |
| 23846 | ENSMUSG00000081399 | 0 | NA | NA | NA | NA | NA | Gm14921 | predicted gene 14921 [Source:MGI Symbo | 0 | 0 | 0 | 0 | 0 | 0 |
| 23847 | ENSMUSG00000081400 | 1.276134668 | -0.503793086 | 2.011628704 | -0.250440394 | 0.802246795 | NA | Gm13680 | predicted gene 13680 [Source:MGI Symbo | 2.4093 | 2.0558 | 0 | 0.964 | 0 | 2.2277 |
| 23848 | ENSMUSG00000081402 | 0.716887508 | 1.693828953 | 2.739549946 | 0.618287305 | 0.536385969 | NA | Gm15455 | predicted gene 15455 [Source:MGI Symbo | 0 | 0 | 1.0062 | 0 | 2.1812 | 1.1138 |
| 23849 | ENSMUSG00000081404 | 1.158001546 | 0.562584638 | 1.95700862 | 0.287471722 | 0.773751151 | NA | Gm15694 | predicted gene 15694 [Source:MGI Symbo | 0.8031 | 0 | 2.0125 | 1.928 | 1.0906 | 1.1138 |
| 23850 | ENSMUSG00000081405 | 0 | NA | NA | NA | NA | NA | Gm13142 | predicted gene 13142 [Source:MGI Symbo | 0 | 0 | 0 | 0 | 0 | 0 |
| 23851 | ENSMUSG00000081406 | 19.29165139 | 0.210540039 | 0.486142284 | 0.433083166 | 0.664954352 | 0.929426323 | Rps6-ps4 | ribosomal protein S6, pseudogene 4 [Sour | 16.062 | 21.586 | 16.1 | 21.208 | 19.631 | 21.163 |
| 23852 | ENSMUSG00000081407 | 0 | NA | NA | NA | NA | NA | Gm14568 | predicted gene 14568 [Source:MGI Symbo | 0 | 0 | 0 | 0 | 0 | 0 |

| | | | | | | | | | | | | | | | |
|---|---|---|---|---|---|---|---|---|---|---|---|---|---|---|---|
| 23853 | ENSMUSG00000081408 | 0 | NA | NA | NA | NA | NA | Gm15047 | predicted gene 15047 [Source:MGI Symbo | 0 | 0 | 0 | 0 | 0 | 0 |
| 23854 | ENSMUSG00000081409 | 0 | NA | NA | NA | NA | NA | Gm8215 | predicted gene 8215 [Source:MGI Symbol | 0 | 0 | 0 | 0 | 0 | 0 |
| 23855 | ENSMUSG00000081410 | 0 | NA | NA | NA | NA | NA | Gm6797 | predicted pseudogene 6797 [Source:MGI | 0 | 0 | 0 | 0 | 0 | 0 |
| 23856 | ENSMUSG00000081412 | 0 | NA | NA | NA | NA | NA | Gm15403 | predicted gene 15403 [Source:MGI Symbo | 0 | 0 | 0 | 0 | 0 | 0 |
| 23857 | ENSMUSG00000081413 | 0 | NA | NA | NA | NA | NA | Uqcrh-ps2 | ubiquinol-cytochrome c reductase hinge p | 0 | 0 | 0 | 0 | 0 | 0 |
| 23858 | ENSMUSG00000081414 | 0 | NA | NA | NA | NA | NA | Gm3943 | predicted gene 3943 [Source:MGI Symbol | 0 | 0 | 0 | 0 | 0 | 0 |
| 23859 | ENSMUSG00000081415 | 0 | NA | NA | NA | NA | NA | Gm15176 | predicted gene 15176 [Source:MGI Symbo | 0 | 0 | 0 | 0 | 0 | 0 |
| 23860 | ENSMUSG00000081416 | 0 | NA | NA | NA | NA | NA | Gm14383 | predicted gene 14383 [Source:MGI Symbo | 0 | 0 | 0 | 0 | 0 | 0 |
| 23861 | ENSMUSG00000081417 | 0 | NA | NA | NA | NA | NA | Gm15603 | predicted gene 15603 [Source:MGI Symbo | 0 | 0 | 0 | 0 | 0 | 0 |
| 23862 | ENSMUSG00000081418 | 0.503095663 | 2.549494177 | 3.348715477 | 0.761334964 | 0.446457022 | NA | Gm14830 | predicted gene 14830 [Source:MGI Symbo | 0 | 0 | 0 | 1.928 | 1.0906 | 0 |
| 23863 | ENSMUSG00000081421 | 0 | NA | NA | NA | NA | NA | Gm13445 | predicted gene 13445 [Source:MGI Symbo | 0 | 0 | 0 | 0 | 0 | 0 |
| 23864 | ENSMUSG00000081422 | 0 | NA | NA | NA | NA | NA | Gm14282 | predicted gene 14282 [Source:MGI Symbo | 0 | 0 | 0 | 0 | 0 | 0 |
| 23865 | ENSMUSG00000081423 | 0 | NA | NA | NA | NA | NA | Gm12882 | predicted gene 12882 [Source:MGI Symbo | 0 | 0 | 0 | 0 | 0 | 0 |
| 23866 | ENSMUSG00000081424 | 0 | NA | NA | NA | NA | NA | Gm14774 | predicted gene 14774 [Source:MGI Symbo | 0 | 0 | 0 | 0 | 0 | 0 |
| 23867 | ENSMUSG00000081427 | 0 | NA | NA | NA | NA | NA | Gm12382 | predicted gene 12382 [Source:MGI Symbo | 0 | 0 | 0 | 0 | 0 | 0 |
| 23868 | ENSMUSG00000081428 | 0 | NA | NA | NA | NA | NA | Tpt1-ps2 | tumor protein, translationally-controlled, | 0 | 0 | 0 | 0 | 0 | 0 |
| 23869 | ENSMUSG00000081429 | 0.678054141 | -2.792799308 | 2.990673428 | -0.933836266 | 0.350388359 | NA | Rps27-ps1 | ribosomal protein S27, pseudogene 1 [Sou | 0 | 2.0558 | 2.0125 | 0 | 0 | 0 |
| 23870 | ENSMUSG00000081431 | 0.681718484 | 0.074918168 | 2.804856827 | 0.026710158 | 0.978690911 | NA | Gm15483 | predicted gene 15483 [Source:MGI Symbo | 0 | 0 | 2.0125 | 0.964 | 0 | 1.1138 |
| 23871 | ENSMUSG00000081432 | 0 | NA | NA | NA | NA | NA | Gm12593 | predicted gene 12593 [Source:MGI Symbo | 0 | 0 | 0 | 0 | 0 | 0 |
| 23872 | ENSMUSG00000081433 | 0 | NA | NA | NA | NA | NA | Serpinb1-ps: | serine (or cysteine) peptidase inhibitor, cl | 0 | 0 | 0 | 0 | 0 | 0 |
| 23873 | ENSMUSG00000081434 | 0.160662974 | 1.126924175 | 4.080472857 | 0.276174898 | 0.782413736 | NA | Gm14165 | predicted gene 14165 [Source:MGI Symbo | 0 | 0 | 0 | 0.964 | 0 | 0 |
| 23874 | ENSMUSG00000081435 | 0 | NA | NA | NA | NA | NA | Gm11403 | predicted gene 11403 [Source:MGI Symbo | 0 | 0 | 0 | 0 | 0 | 0 |
| 23875 | ENSMUSG00000081436 | 0 | NA | NA | NA | NA | NA | Gm5912 | predicted gene 5912 [Source:MGI Symbol | 0 | 0 | 0 | 0 | 0 | 0 |
| 23876 | ENSMUSG00000081437 | 0 | NA | NA | NA | NA | NA | Gm12402 | predicted gene 12402 [Source:MGI Symbo | 0 | 0 | 0 | 0 | 0 | 0 |
| 23877 | ENSMUSG00000081438 | 0 | NA | NA | NA | NA | NA | Gm11254 | predicted gene 11254 [Source:MGI Symbo | 0 | 0 | 0 | 0 | 0 | 0 |
| 23878 | ENSMUSG00000081439 | 0 | NA | NA | NA | NA | NA | Gm7840 | predicted gene 7840 [Source:MGI Symbol | 0 | 0 | 0 | 0 | 0 | 0 |
| 23879 | ENSMUSG00000081440 | 0 | NA | NA | NA | NA | NA | Gm12464 | predicted gene 12464 [Source:MGI Symbo | 0 | 0 | 0 | 0 | 0 | 0 |
| 23880 | ENSMUSG00000081441 | 4.266997261 | 0.532367601 | 1.122356853 | 0.474330067 | 0.635264554 | NA | Gm6161 | predicted gene 6161 [Source:MGI Symbol | 2.4093 | 6.1675 | 2.0125 | 10.604 | 2.1812 | 2.2277 |
| 23881 | ENSMUSG00000081442 | 0 | NA | NA | NA | NA | NA | Gm14682 | predicted gene 14682 [Source:MGI Symbo | 0 | 0 | 0 | 0 | 0 | 0 |
| 23882 | ENSMUSG00000081443 | 0 | NA | NA | NA | NA | NA | Gm3880 | predicted pseudogene 3880 [Source:MGI | 0 | 0 | 0 | 0 | 0 | 0 |
| 23883 | ENSMUSG00000081444 | 0 | NA | NA | NA | NA | NA | Gm15803 | predicted gene 15803 [Source:MGI Symbo | 0 | 0 | 0 | 0 | 0 | 0 |
| 23884 | ENSMUSG00000081445 | 0.449470461 | -0.545136533 | 3.562105917 | -0.15303771 | 0.878368537 | NA | Rpl23a-ps4 | ribosomal protein L23A, pseudogene 4 [Sc | 1.6062 | 0 | 0 | 0 | 1.0906 | 0 |
| 23885 | ENSMUSG00000081447 | 0 | NA | NA | NA | NA | NA | Gm13870 | predicted gene 13870 [Source:MGI Symbo | 0 | 0 | 0 | 0 | 0 | 0 |
| 23886 | ENSMUSG00000081449 | 0 | NA | NA | NA | NA | NA | Gm11432 | predicted gene 11432 [Source:MGI Symbo | 0 | 0 | 0 | 0 | 0 | 0 |
| 23887 | ENSMUSG00000081450 | 0 | NA | NA | NA | NA | NA | Gm11700 | predicted gene 11700 [Source:MGI Symbo | 0 | 0 | 0 | 0 | 0 | 0 |
| 23888 | ENSMUSG00000081451 | 1.271825051 | -1.34972375 | 2.043905257 | -0.660365125 | 0.509019547 | NA | Gm12734 | predicted gene 12734 [Source:MGI Symbo | 2.4093 | 1.0279 | 2.0125 | 0 | 2.1812 | 0 |
| 23889 | ENSMUSG00000081452 | 0 | NA | NA | NA | NA | NA | Gm14761 | predicted gene 14761 [Source:MGI Symbo | 0 | 0 | 0 | 0 | 0 | 0 |
| 23890 | ENSMUSG00000081453 | 4.770180661 | 0.819943457 | 0.988748495 | 0.829274038 | 0.406949358 | NA | Gm6767 | predicted gene 6767 [Source:MGI Symbol | 3.2124 | 3.0838 | 4.025 | 2.8919 | 8.7249 | 6.6831 |
| 23891 | ENSMUSG00000081455 | 0.342639279 | -1.805976596 | 4.028547714 | -0.448294702 | 0.653940524 | NA | Hmgb1-ps3 | high mobility group box 1, pseudogene 3 [ | 0 | 2.0558 | 0 | 0 | 0 | 0 |
| 23892 | ENSMUSG00000081456 | 0 | NA | NA | NA | NA | NA | Gm7781 | predicted gene 7781 [Source:MGI Symbol | 0 | 0 | 0 | 0 | 0 | 0 |
| 23893 | ENSMUSG00000081457 | 0.133850373 | -0.796629176 | 4.080472857 | -0.195229623 | 0.84521319 | NA | Gm13488 | predicted gene 13488 [Source:MGI Symbo | 0.8031 | 0 | 0 | 0 | 0 | 0 |
| 23894 | ENSMUSG00000081458 | 0 | NA | NA | NA | NA | NA | Gm14796 | predicted gene 14796 [Source:MGI Symbo | 0 | 0 | 0 | 0 | 0 | 0 |
| 23895 | ENSMUSG00000081459 | 0.455176322 | 0.955156269 | 3.546674842 | 0.269310357 | 0.787690862 | NA | Llph-ps1 | LLP homolog, pseudogene 1 [Source:MGI | 0.8031 | 0 | 0 | 1.928 | 0 | 0 |
| 23896 | ENSMUSG00000081460 | 0.167707431 | -0.796629176 | 4.080472857 | -0.195229623 | 0.84521319 | NA | Gm13168 | predicted gene 13168 [Source:MGI Symbo | 0 | 0 | 1.0062 | 0 | 0 | 0 |
| 23897 | ENSMUSG00000081461 | 0 | NA | NA | NA | NA | NA | Rpl30-ps11 | ribosomal protein L30, pseudogene 11 [Sc | 0 | 0 | 0 | 0 | 0 | 0 |
| 23898 | ENSMUSG00000081463 | 0 | NA | NA | NA | NA | NA | Pou5f1-rs13 | POU domain, class 5, transcription factor : | 0 | 0 | 0 | 0 | 0 | 0 |
| 23899 | ENSMUSG00000081464 | 0 | NA | NA | NA | NA | NA | Gm14732 | predicted gene 14732 [Source:MGI Symbo | 0 | 0 | 0 | 0 | 0 | 0 |
| 23900 | ENSMUSG00000081465 | 0 | NA | NA | NA | NA | NA | Gm12884 | predicted gene 12884 [Source:MGI Symbo | 0 | 0 | 0 | 0 | 0 | 0 |
| 23901 | ENSMUSG00000081466 | 0 | NA | NA | NA | NA | NA | Gm14912 | predicted gene 14912 [Source:MGI Symbo | 0 | 0 | 0 | 0 | 0 | 0 |
| 23902 | ENSMUSG00000081467 | 0 | NA | NA | NA | NA | NA | Gm13337 | predicted gene 13337 [Source:MGI Symbo | 0 | 0 | 0 | 0 | 0 | 0 |
| 23903 | ENSMUSG00000081469 | 0 | NA | NA | NA | NA | NA | Vmn1r-ps95 | vomeronasal 1 receptor, pseudogene 95 [ | 0 | 0 | 0 | 0 | 0 | 0 |
| 23904 | ENSMUSG00000081470 | 0 | NA | NA | NA | NA | NA | Gm14131 | predicted gene 14131 [Source:MGI Symbo | 0 | 0 | 0 | 0 | 0 | 0 |
| 23905 | ENSMUSG00000081471 | 0 | NA | NA | NA | NA | NA | Gm14735 | predicted gene 14735 [Source:MGI Symbo | 0 | 0 | 0 | 0 | 0 | 0 |
| 23906 | ENSMUSG00000081473 | 1.66700966 | 0.594725401 | 1.860118244 | 0.319724514 | 0.749177175 | NA | Gm4853 | predicted pseudogene 4853 [Source:MGI | 0 | 2.0558 | 2.0125 | 4.8199 | 0 | 1.1138 |
| 23907 | ENSMUSG00000081474 | 0 | NA | NA | NA | NA | NA | Gm12855 | predicted gene 12855 [Source:MGI Symbo | 0 | 0 | 0 | 0 | 0 | 0 |
| 23908 | ENSMUSG00000081475 | 0 | NA | NA | NA | NA | NA | Gm14558 | predicted gene 14558 [Source:MGI Symbo | 0 | 0 | 0 | 0 | 0 | 0 |

| | | | | | | | | | | | | | | |
|---|---|---|---|---|---|---|---|---|---|---|---|---|---|---|
| 23909 | ENSMUSG00000081476 | 23.08671117 | -0.286838146 | 0.451365343 | -0.635489965 | 0.525108909 | 0.888334265 | Itpa-ps1 | inosine triphosphatase (nucleoside tripho | 28.109 | 26.726 | 21.131 | 17.352 | 21.812 | 23.391 |
| 23910 | ENSMUSG00000081477 | 0 | NA | NA | NA | NA | NA | Gm7916 | predicted gene 7916 [Source:MGI Symbol | 0 | 0 | 0 | 0 | 0 | 0 |
| 23911 | ENSMUSG00000081478 | 0 | NA | NA | NA | NA | NA | AA623943 | expressed sequence AA623943 [Source:M | 0 | 0 | 0 | 0 | 0 | 0 |
| 23912 | ENSMUSG00000081480 | 0.167707431 | -0.796629176 | 4.080472857 | -0.195229623 | 0.84521319 | NA | Gm13608 | predicted gene 13608 [Source:MGI Symbo | 0 | 0 | 1.0062 | 0 | 0 | 0 |
| 23913 | ENSMUSG00000081481 | 0 | NA | NA | NA | NA | NA | Gm14293 | predicted gene 14293 [Source:MGI Symbo | 0 | 0 | 0 | 0 | 0 | 0 |
| 23914 | ENSMUSG00000081483 | 0 | NA | NA | NA | NA | NA | Gm12445 | predicted gene 12445 [Source:MGI Symbo | 0 | 0 | 0 | 0 | 0 | 0 |
| 23915 | ENSMUSG00000081484 | 0 | NA | NA | NA | NA | NA | Gm13492 | predicted gene 13492 [Source:MGI Symbo | 0 | 0 | 0 | 0 | 0 | 0 |
| 23916 | ENSMUSG00000081485 | 30.81080568 | 0.075185638 | 0.413392504 | 0.181874701 | 0.855681059 | 0.974158771 | Gm12338 | predicted gene 12338 [Source:MGI Symbo | 32.124 | 19.53 | 38.237 | 29.883 | 29.447 | 35.643 |
| 23917 | ENSMUSG00000081486 | 0 | NA | NA | NA | NA | NA | Gm14452 | predicted gene 14452 [Source:MGI Symbo | 0 | 0 | 0 | 0 | 0 | 0 |
| 23918 | ENSMUSG00000081487 | 0.542600935 | 1.125931187 | 3.335923981 | 0.33751704 | 0.735727169 | NA | Gm13689 | predicted gene 13689 [Source:MGI Symbo | 0 | 1.0279 | 0 | 0 | 0 | 2.2277 |
| 23919 | ENSMUSG00000081488 | 3.483115588 | -0.676854661 | 1.238770946 | -0.546392102 | 0.584796438 | NA | Gm15368 | predicted gene 15368 [Source:MGI Symbo | 4.8186 | 3.0838 | 5.0312 | 5.7839 | 2.1812 | 0 |
| 23920 | ENSMUSG00000081490 | 0 | NA | NA | NA | NA | NA | Gm11830 | predicted gene 11830 [Source:MGI Symbo | 0 | 0 | 0 | 0 | 0 | 0 |
| 23921 | ENSMUSG00000081491 | 0 | NA | NA | NA | NA | NA | Gm12778 | predicted gene 12778 [Source:MGI Symbo | 0 | 0 | 0 | 0 | 0 | 0 |
| 23922 | ENSMUSG00000081492 | 0.513752329 | 1.031892132 | 3.082998058 | 0.334704114 | 0.737848298 | NA | Gm9311 | predicted gene 9311 [Source:MGI Symbol | 0 | 1.0279 | 0 | 0.964 | 1.0906 | 0 |
| 23923 | ENSMUSG00000081494 | 0 | NA | NA | NA | NA | NA | Gm14130 | predicted gene 14130 [Source:MGI Symbo | 0 | 0 | 0 | 0 | 0 | 0 |
| 23924 | ENSMUSG00000081495 | 0 | NA | NA | NA | NA | NA | Vmn1r-ps10 | vomeronasal 1 receptor, pseudogene 108 | 0 | 0 | 0 | 0 | 0 | 0 |
| 23925 | ENSMUSG00000081496 | 0 | NA | NA | NA | NA | NA | Gm13428 | predicted gene 13428 [Source:MGI Symbo | 0 | 0 | 0 | 0 | 0 | 0 |
| 23926 | ENSMUSG00000081497 | 1.042084389 | 2.377088107 | 2.219862216 | 1.070826869 | 0.284247283 | NA | Gm15560 | predicted gene 15560 [Source:MGI Symbo | 0 | 0 | 1.0062 | 1.928 | 1.0906 | 2.2277 |
| 23927 | ENSMUSG00000081498 | 0 | NA | NA | NA | NA | NA | Olfr407-ps1 | olfactory receptor 407, pseudogene 1 [So | 0 | 0 | 0 | 0 | 0 | 0 |
| 23928 | ENSMUSG00000081499 | 0.401551119 | -2.072982726 | 4.008251979 | -0.517178744 | 0.605031388 | NA | Gm12517 | predicted gene 12517 [Source:MGI Symbo | 2.4093 | 0 | 0 | 0 | 0 | 0 |
| 23929 | ENSMUSG00000081500 | 0 | NA | NA | NA | NA | NA | Gm15233 | predicted gene 15233 [Source:MGI Symbo | 0 | 0 | 0 | 0 | 0 | 0 |
| 23930 | ENSMUSG00000081501 | 0 | NA | NA | NA | NA | NA | Gm13286 | predicted gene 13286 [Source:MGI Symbo | 0 | 0 | 0 | 0 | 0 | 0 |
| 23931 | ENSMUSG00000081502 | 0 | NA | NA | NA | NA | NA | Gm5646 | predicted pseudogene 5646 [Source:MGI | 0 | 0 | 0 | 0 | 0 | 0 |
| 23932 | ENSMUSG00000081504 | 0 | NA | NA | NA | NA | NA | Gm12508 | predicted gene 12508 [Source:MGI Symbo | 0 | 0 | 0 | 0 | 0 | 0 |
| 23933 | ENSMUSG00000081505 | 0 | NA | NA | NA | NA | NA | Gm8644 | predicted gene 8644 [Source:MGI Symbol | 0 | 0 | 0 | 0 | 0 | 0 |
| 23934 | ENSMUSG00000081506 | 0 | NA | NA | NA | NA | NA | Amy2-ps1 | amylase 2, pseudogene 1 [Source:MGI Sy | 0 | 0 | 0 | 0 | 0 | 0 |
| 23935 | ENSMUSG00000081507 | 0 | NA | NA | NA | NA | NA | Gm14993 | predicted gene 14993 [Source:MGI Symbo | 0 | 0 | 0 | 0 | 0 | 0 |
| 23936 | ENSMUSG00000081508 | 0 | NA | NA | NA | NA | NA | Gm12654 | predicted gene 12654 [Source:MGI Symbo | 0 | 0 | 0 | 0 | 0 | 0 |
| 23937 | ENSMUSG00000081510 | 0 | NA | NA | NA | NA | NA | Gm12768 | predicted gene 12768 [Source:MGI Symbo | 0 | 0 | 0 | 0 | 0 | 0 |
| 23938 | ENSMUSG00000081511 | 0 | NA | NA | NA | NA | NA | Gm11475 | predicted gene 11475 [Source:MGI Symbo | 0 | 0 | 0 | 0 | 0 | 0 |
| 23939 | ENSMUSG00000081512 | 2.445102321 | -0.764605199 | 1.88883569 | -0.404802389 | 0.685622773 | NA | Gm15821 | predicted gene 15821 [Source:MGI Symbo | 0 | 9.2513 | 0 | 0.964 | 0 | 4.4554 |
| 23940 | ENSMUSG00000081513 | 0 | NA | NA | NA | NA | NA | Gm11623 | predicted gene 11623 [Source:MGI Symbo | 0 | 0 | 0 | 0 | 0 | 0 |
| 23941 | ENSMUSG00000081514 | 0 | NA | NA | NA | NA | NA | Gm14673 | predicted gene 14673 [Source:MGI Symbo | 0 | 0 | 0 | 0 | 0 | 0 |
| 23942 | ENSMUSG00000081515 | 0 | NA | NA | NA | NA | NA | Gm12311 | predicted gene 12311 [Source:MGI Symbo | 0 | 0 | 0 | 0 | 0 | 0 |
| 23943 | ENSMUSG00000081516 | 0 | NA | NA | NA | NA | NA | Gm12470 | predicted gene 12470 [Source:MGI Symbo | 0 | 0 | 0 | 0 | 0 | 0 |
| 23944 | ENSMUSG00000081517 | 0 | NA | NA | NA | NA | NA | Gm16204 | predicted gene 16204 [Source:MGI Symbo | 0 | 0 | 0 | 0 | 0 | 0 |
| 23945 | ENSMUSG00000081519 | 0 | NA | NA | NA | NA | NA | Gm13331 | predicted gene 13331 [Source:MGI Symbo | 0 | 0 | 0 | 0 | 0 | 0 |
| 23946 | ENSMUSG00000081520 | 1.361316686 | 1.776258021 | 1.88191201 | 0.943858167 | 0.345242132 | NA | Gm16200 | predicted gene 16200 [Source:MGI Symbo | 0.8031 | 1.0279 | 0 | 1.928 | 2.1812 | 2.2277 |
| 23947 | ENSMUSG00000081522 | 0 | NA | NA | NA | NA | NA | Gm13170 | predicted gene 13170 [Source:MGI Symbo | 0 | 0 | 0 | 0 | 0 | 0 |
| 23948 | ENSMUSG00000081523 | 0 | NA | NA | NA | NA | NA | Gm13058 | predicted gene 13058 [Source:MGI Symbo | 0 | 0 | 0 | 0 | 0 | 0 |
| 23949 | ENSMUSG00000081524 | 0 | NA | NA | NA | NA | NA | Gm11658 | predicted gene 11658 [Source:MGI Symbo | 0 | 0 | 0 | 0 | 0 | 0 |
| 23950 | ENSMUSG00000081525 | 0 | NA | NA | NA | NA | NA | Gm13325 | predicted gene 13325 [Source:MGI Symbo | 0 | 0 | 0 | 0 | 0 | 0 |
| 23951 | ENSMUSG00000081526 | 0 | NA | NA | NA | NA | NA | Gm7219 | predicted pseudogene 7219 [Source:MGI | 0 | 0 | 0 | 0 | 0 | 0 |
| 23952 | ENSMUSG00000081528 | 0 | NA | NA | NA | NA | NA | Gm12924 | predicted gene 12924 [Source:MGI Symbo | 0 | 0 | 0 | 0 | 0 | 0 |
| 23953 | ENSMUSG00000081529 | 0 | NA | NA | NA | NA | NA | Gm14884 | predicted gene 14884 [Source:MGI Symbo | 0 | 0 | 0 | 0 | 0 | 0 |
| 23954 | ENSMUSG00000081530 | 0 | NA | NA | NA | NA | NA | Gm12646 | predicted gene 12646 [Source:MGI Symbo | 0 | 0 | 0 | 0 | 0 | 0 |
| 23955 | ENSMUSG00000081531 | 0 | NA | NA | NA | NA | NA | Vmn1r-ps11 | vomeronasal 1 receptor, pseudogene 113 | 0 | 0 | 0 | 0 | 0 | 0 |
| 23956 | ENSMUSG00000081532 | 0 | NA | NA | NA | NA | NA | Gm15781 | predicted gene 15781 [Source:MGI Symbo | 0 | 0 | 0 | 0 | 0 | 0 |
| 23957 | ENSMUSG00000081533 | 0 | NA | NA | NA | NA | NA | Gm15111 | predicted gene 15111 [Source:MGI Symbo | 0 | 0 | 0 | 0 | 0 | 0 |
| 23958 | ENSMUSG00000081534 | 713.3941058 | -0.080617611 | 0.111295091 | -0.724359088 | 0.46884531 | 0.865855282 | Slc48a1 | solute carrier family 48 (heme transporter | 749.29 | 725.71 | 724.5 | 678.64 | 684.91 | 717.32 |
| 23959 | ENSMUSG00000081536 | 0 | NA | NA | NA | NA | NA | Gm15193 | predicted gene 15193 [Source:MGI Symbo | 0 | 0 | 0 | 0 | 0 | 0 |
| 23960 | ENSMUSG00000081537 | 0 | NA | NA | NA | NA | NA | Gm37832 | predicted gene, 37832 [Source:MGI Symb | 0 | 0 | 0 | 0 | 0 | 0 |
| 23961 | ENSMUSG00000081538 | 0 | NA | NA | NA | NA | NA | Rps13-ps8 | ribosmal protein S13, pseudogene 8 [Sour | 0 | 0 | 0 | 0 | 0 | 0 |
| 23962 | ENSMUSG00000081539 | 0 | NA | NA | NA | NA | NA | Gm11990 | predicted gene 11990 [Source:MGI Symbo | 0 | 0 | 0 | 0 | 0 | 0 |
| 23963 | ENSMUSG00000081540 | 0 | NA | NA | NA | NA | NA | Gm12538 | predicted gene 12538 [Source:MGI Symbo | 0 | 0 | 0 | 0 | 0 | 0 |
| 23964 | ENSMUSG00000081541 | 0 | NA | NA | NA | NA | NA | Gm13940 | predicted gene 13940 [Source:MGI Symbo | 0 | 0 | 0 | 0 | 0 | 0 |

| | | | | | | | | | | | | | | | |
|---|---|---|---|---|---|---|---|---|---|---|---|---|---|---|---|
| 23965 | ENSMUSG00000081542 | 0 | NA | NA | NA | NA | NA | Gm12512 | predicted gene 12512 [Source:MGI Symbo | 0 | 0 | 0 | 0 | 0 | 0 |
| 23966 | ENSMUSG00000081543 | 0 | NA | NA | NA | NA | NA | Gm12733 | predicted gene 12733 [Source:MGI Symbo | 0 | 0 | 0 | 0 | 0 | 0 |
| 23967 | ENSMUSG00000081544 | 0 | NA | NA | NA | NA | NA | Gm13077 | predicted gene 13077 [Source:MGI Symbo | 0 | 0 | 0 | 0 | 0 | 0 |
| 23968 | ENSMUSG00000081546 | 0 | NA | NA | NA | NA | NA | Gm14927 | predicted gene 14927 [Source:MGI Symbo | 0 | 0 | 0 | 0 | 0 | 0 |
| 23969 | ENSMUSG00000081547 | 0 | NA | NA | NA | NA | NA | Gm11553 | predicted gene 11553 [Source:MGI Symbo | 0 | 0 | 0 | 0 | 0 | 0 |
| 23970 | ENSMUSG00000081548 | 0.167707431 | -0.796629176 | 4.080472857 | -0.195229623 | 0.84521319 | NA | Gm11692 | predicted gene 11692 [Source:MGI Symbo | 0 | 0 | 1.0062 | 0 | 0 | 0 |
| 23971 | ENSMUSG00000081550 | 0 | NA | NA | NA | NA | NA | Gm15877 | predicted gene 15877 [Source:MGI Symbo | 0 | 0 | 0 | 0 | 0 | 0 |
| 23972 | ENSMUSG00000081551 | 0 | NA | NA | NA | NA | NA | Gm14654 | predicted gene 14654 [Source:MGI Symbo | 0 | 0 | 0 | 0 | 0 | 0 |
| 23973 | ENSMUSG00000081552 | 0 | NA | NA | NA | NA | NA | Gm12328 | predicted gene 12328 [Source:MGI Symbo | 0 | 0 | 0 | 0 | 0 | 0 |
| 23974 | ENSMUSG00000081553 | 0 | NA | NA | NA | NA | NA | Ccdc50-ps | Ccdc50 retrotransposed pseudogene [Sou | 0 | 0 | 0 | 0 | 0 | 0 |
| 23975 | ENSMUSG00000081555 | 0 | NA | NA | NA | NA | NA | Gm6427 | predicted gene 6427 [Source:MGI Symbol | 0 | 0 | 0 | 0 | 0 | 0 |
| 23976 | ENSMUSG00000081556 | 0 | NA | NA | NA | NA | NA | Gm15778 | predicted gene 15778 [Source:MGI Symbo | 0 | 0 | 0 | 0 | 0 | 0 |
| 23977 | ENSMUSG00000081557 | 0 | NA | NA | NA | NA | NA | Gm5697 | predicted gene 5697 [Source:MGI Symbol | 0 | 0 | 0 | 0 | 0 | 0 |
| 23978 | ENSMUSG00000081559 | 0.840287806 | -0.383646848 | 2.239474228 | -0.171311124 | 0.863979134 | NA | Gm12411 | predicted gene 12411 [Source:MGI Symbo | 0.8031 | 1.0279 | 1.0062 | 0 | 1.0906 | 1.1138 |
| 23979 | ENSMUSG00000081560 | 0 | NA | NA | NA | NA | NA | Gm9218 | predicted gene 9218 [Source:MGI Symbol | 0 | 0 | 0 | 0 | 0 | 0 |
| 23980 | ENSMUSG00000081561 | 0 | NA | NA | NA | NA | NA | Gm12425 | predicted gene 12425 [Source:MGI Symbo | 0 | 0 | 0 | 0 | 0 | 0 |
| 23981 | ENSMUSG00000081562 | 0.342639279 | -1.805976596 | 4.028547714 | -0.448294702 | 0.653940524 | NA | Gm11575 | predicted gene 11575 [Source:MGI Symbo | 0 | 2.0558 | 0 | 0 | 0 | 0 |
| 23982 | ENSMUSG00000081563 | 0 | NA | NA | NA | NA | NA | Gm12635 | predicted gene 12635 [Source:MGI Symbo | 0 | 0 | 0 | 0 | 0 | 0 |
| 23983 | ENSMUSG00000081564 | 0.342432689 | 1.990653086 | 3.900281263 | 0.51038706 | 0.609780321 | NA | Gm13717 | predicted gene 13717 [Source:MGI Symbo | 0 | 0 | 0 | 0.964 | 1.0906 | 0 |
| 23984 | ENSMUSG00000081565 | 0 | NA | NA | NA | NA | NA | Rps12-ps5 | ribosomal protein S12, pseudogene 5 [Sou | 0 | 0 | 0 | 0 | 0 | 0 |
| 23985 | ENSMUSG00000081566 | 0 | NA | NA | NA | NA | NA | Gm13756 | predicted gene 13756 [Source:MGI Symbo | 0 | 0 | 0 | 0 | 0 | 0 |
| 23986 | ENSMUSG00000081567 | 0 | NA | NA | NA | NA | NA | Gm14838 | predicted gene 14838 [Source:MGI Symbo | 0 | 0 | 0 | 0 | 0 | 0 |
| 23987 | ENSMUSG00000081568 | 0 | NA | NA | NA | NA | NA | Gm15032 | predicted gene 15032 [Source:MGI Symbo | 0 | 0 | 0 | 0 | 0 | 0 |
| 23988 | ENSMUSG00000081569 | 0 | NA | NA | NA | NA | NA | Gm12490 | predicted gene 12490 [Source:MGI Symbo | 0 | 0 | 0 | 0 | 0 | 0 |
| 23989 | ENSMUSG00000081570 | 0 | NA | NA | NA | NA | NA | Gm12192 | predicted gene 12192 [Source:MGI Symbo | 0 | 0 | 0 | 0 | 0 | 0 |
| 23990 | ENSMUSG00000081571 | 0 | NA | NA | NA | NA | NA | Gm12125 | predicted gene 12125 [Source:MGI Symbo | 0 | 0 | 0 | 0 | 0 | 0 |
| 23991 | ENSMUSG00000081572 | 0.167707431 | -0.796629176 | 4.080472857 | -0.195229623 | 0.84521319 | NA | Gm14138 | predicted gene 14138 [Source:MGI Symbo | 0 | 0 | 1.0062 | 0 | 0 | 0 |
| 23992 | ENSMUSG00000081573 | 0 | NA | NA | NA | NA | NA | Gm15164 | predicted gene 15164 [Source:MGI Symbo | 0 | 0 | 0 | 0 | 0 | 0 |
| 23993 | ENSMUSG00000081574 | 2.127453158 | 0.650304564 | 1.934650377 | 0.336135444 | 0.736768731 | NA | Gm14429 | predicted gene 14429 [Source:MGI Symbo | 0 | 1.0279 | 4.025 | 7.7118 | 0 | 0 |
| 23994 | ENSMUSG00000081575 | 0 | NA | NA | NA | NA | NA | Gm12090 | predicted gene 12090 [Source:MGI Symbo | 0 | 0 | 0 | 0 | 0 | 0 |
| 23995 | ENSMUSG00000081576 | 0 | NA | NA | NA | NA | NA | Gm11279 | predicted gene 11279 [Source:MGI Symbo | 0 | 0 | 0 | 0 | 0 | 0 |
| 23996 | ENSMUSG00000081577 | 0 | NA | NA | NA | NA | NA | Gm13455 | predicted gene 13455 [Source:MGI Symbo | 0 | 0 | 0 | 0 | 0 | 0 |
| 23997 | ENSMUSG00000081578 | 4.851827051 | -0.139931142 | 1.048059992 | -0.133514439 | 0.893786546 | NA | Gm12611 | predicted gene 12611 [Source:MGI Symbo | 8.031 | 3.0838 | 4.025 | 1.928 | 9.8156 | 2.2277 |
| 23998 | ENSMUSG00000081579 | 0 | NA | NA | NA | NA | NA | Gm11909 | predicted gene 11909 [Source:MGI Symbo | 0 | 0 | 0 | 0 | 0 | 0 |
| 23999 | ENSMUSG00000081580 | 0 | NA | NA | NA | NA | NA | Gm14129 | predicted gene 14129 [Source:MGI Symbo | 0 | 0 | 0 | 0 | 0 | 0 |
| 24000 | ENSMUSG00000081583 | 0 | NA | NA | NA | NA | NA | Gm14769 | predicted gene 14769 [Source:MGI Symbo | 0 | 0 | 0 | 0 | 0 | 0 |
| 24001 | ENSMUSG00000081584 | 0 | NA | NA | NA | NA | NA | Gm12716 | predicted gene 12716 [Source:MGI Symbo | 0 | 0 | 0 | 0 | 0 | 0 |
| 24002 | ENSMUSG00000081585 | 0.55305101 | 2.667977666 | 3.241982584 | 0.822946329 | 0.410538515 | NA | Gm12926 | predicted gene 12926 [Source:MGI Symbo | 0 | 0 | 0 | 0 | 1.0906 | 2.2277 |
| 24003 | ENSMUSG00000081586 | 0 | NA | NA | NA | NA | NA | Gm11544 | predicted gene 11544 [Source:MGI Symbo | 0 | 0 | 0 | 0 | 0 | 0 |
| 24004 | ENSMUSG00000081587 | 0 | NA | NA | NA | NA | NA | Gm14926 | predicted gene 14926 [Source:MGI Symbo | 0 | 0 | 0 | 0 | 0 | 0 |
| 24005 | ENSMUSG00000081588 | 0 | NA | NA | NA | NA | NA | Gm11661 | predicted gene 11661 [Source:MGI Symbo | 0 | 0 | 0 | 0 | 0 | 0 |
| 24006 | ENSMUSG00000081589 | 0 | NA | NA | NA | NA | NA | Gm14353 | predicted gene 14353 [Source:MGI Symbo | 0 | 0 | 0 | 0 | 0 | 0 |
| 24007 | ENSMUSG00000081590 | 0.167707431 | -0.796629176 | 4.080472857 | -0.195229623 | 0.84521319 | NA | Gm14413 | predicted gene 14413 [Source:MGI Symbo | 0 | 0 | 1.0062 | 0 | 0 | 0 |
| 24008 | ENSMUSG00000081591 | 0 | NA | NA | NA | NA | NA | Btf3-ps9 | basic transcription factor 3, pseudogene 9 | 0 | 0 | 0 | 0 | 0 | 0 |
| 24009 | ENSMUSG00000081592 | 0 | NA | NA | NA | NA | NA | Gm11936 | predicted gene 11936 [Source:MGI Symbo | 0 | 0 | 0 | 0 | 0 | 0 |
| 24010 | ENSMUSG00000081593 | 0.849167191 | -0.527085716 | 2.389288686 | -0.220603613 | 0.825401088 | NA | Gm11841 | predicted gene 11841 [Source:MGI Symbo | 0 | 1.0279 | 2.0125 | 0.964 | 1.0906 | 0 |
| 24011 | ENSMUSG00000081594 | 0 | NA | NA | NA | NA | NA | Gm15467 | predicted gene 15467 [Source:MGI Symbo | 0 | 0 | 0 | 0 | 0 | 0 |
| 24012 | ENSMUSG00000081595 | 0 | NA | NA | NA | NA | NA | Gm13036 | predicted gene 13036 [Source:MGI Symbo | 0 | 0 | 0 | 0 | 0 | 0 |
| 24013 | ENSMUSG00000081596 | 0 | NA | NA | NA | NA | NA | Gm15968 | predicted gene 15968 [Source:MGI Symbo | 0 | 0 | 0 | 0 | 0 | 0 |
| 24014 | ENSMUSG00000081597 | 0.267700746 | -1.497481649 | 4.057001436 | -0.369110456 | 0.712045396 | NA | Gm14506 | predicted gene 14506 [Source:MGI Symbo | 1.6062 | 0 | 0 | 0 | 0 | 0 |
| 24015 | ENSMUSG00000081598 | 0 | NA | NA | NA | NA | NA | Gm8312 | predicted gene 8312 [Source:MGI Symbol | 0 | 0 | 0 | 0 | 0 | 0 |
| 24016 | ENSMUSG00000081599 | 0 | NA | NA | NA | NA | NA | Gm14390 | predicted gene 14390 [Source:MGI Symbo | 0 | 0 | 0 | 0 | 0 | 0 |
| 24017 | ENSMUSG00000081600 | 2.47160548 | -0.030941667 | 1.314839158 | -0.023532664 | 0.981225384 | NA | Gm12286 | predicted gene 12286 [Source:MGI Symbo | 2.4093 | 3.0838 | 2.0125 | 2.8919 | 1.0906 | 3.3415 |
| 24018 | ENSMUSG00000081601 | 0.133850373 | -0.796629176 | 4.080472857 | -0.195229623 | 0.84521319 | NA | Gm13065 | predicted gene 13065 [Source:MGI Symbo | 0.8031 | 0 | 0 | 0 | 0 | 0 |
| 24019 | ENSMUSG00000081602 | 0.610133435 | 0.304049584 | 2.904025954 | 0.104699334 | 0.91661439 | NA | Gm15369 | predicted gene 15369 [Source:MGI Symbo | 1.6062 | 0 | 0 | 0.964 | 1.0906 | 0 |
| 24020 | ENSMUSG00000081603 | 4.766521532 | -0.932892463 | 1.139204257 | -0.818898329 | 0.412844423 | NA | Gm14681 | predicted gene 14681 [Source:MGI Symbo | 2.4093 | 11.307 | 5.0312 | 0.964 | 1.0906 | 7.7969 |

| | | | | | | | | | | | | | | | |
|---|---|---|---|---|---|---|---|---|---|---|---|---|---|---|---|
| 24021 | ENSMUSG00000081604 | 7.187661148 | -0.268973914 | 0.812205108 | -0.331165012 | 0.740519847 | 0.948792385 | Gm11518 | predicted gene 11518 [Source:MGI Symbo | 6.4248 | 8.2233 | 9.0562 | 10.604 | 4.3625 | 4.4554 |
| 24022 | ENSMUSG00000081605 | 2.347584883 | 0.063599757 | 1.599990943 | 0.039750073 | 0.968292381 | NA | Gm15953 | predicted gene 15953 [Source:MGI Symbo | 4.8186 | 0 | 2.0125 | 2.8919 | 4.3625 | 0 |
| 24023 | ENSMUSG00000081606 | 0 | NA | NA | NA | NA | NA | Gm9059 | predicted gene 9059 [Source:MGI Symbol | 0 | 0 | 0 | 0 | 0 | 0 |
| 24024 | ENSMUSG00000081607 | 0 | NA | NA | NA | NA | NA | Gm15294 | predicted gene 15294 [Source:MGI Symbo | 0 | 0 | 0 | 0 | 0 | 0 |
| 24025 | ENSMUSG00000081608 | 0 | NA | NA | NA | NA | NA | Gm16210 | predicted gene 16210 [Source:MGI Symbo | 0 | 0 | 0 | 0 | 0 | 0 |
| 24026 | ENSMUSG00000081609 | 0 | NA | NA | NA | NA | NA | Gm12939 | predicted gene 12939 [Source:MGI Symbo | 0 | 0 | 0 | 0 | 0 | 0 |
| 24027 | ENSMUSG00000081610 | 0 | NA | NA | NA | NA | NA | Gm13733 | predicted gene 13733 [Source:MGI Symbo | 0 | 0 | 0 | 0 | 0 | 0 |
| 24028 | ENSMUSG00000081611 | 0.160662974 | 1.126924175 | 4.080472857 | 0.276174898 | 0.782413736 | NA | Gm13651 | predicted gene 13651 [Source:MGI Symbo | 0 | 0 | 0 | 0.964 | 0 | 0 |
| 24029 | ENSMUSG00000081612 | 0 | NA | NA | NA | NA | NA | Xlr5e-ps | X-linked lymphocyte-regulated 5E, pseudo | 0 | 0 | 0 | 0 | 0 | 0 |
| 24030 | ENSMUSG00000081613 | 0 | NA | NA | NA | NA | NA | Gm12109 | predicted gene 12109 [Source:MGI Symbo | 0 | 0 | 0 | 0 | 0 | 0 |
| 24031 | ENSMUSG00000081614 | 0 | NA | NA | NA | NA | NA | Gm9436 | predicted gene 9436 [Source:MGI Symbol | 0 | 0 | 0 | 0 | 0 | 0 |
| 24032 | ENSMUSG00000081615 | 0 | NA | NA | NA | NA | NA | Gm12539 | predicted gene 12539 [Source:MGI Symbo | 0 | 0 | 0 | 0 | 0 | 0 |
| 24033 | ENSMUSG00000081616 | 0 | NA | NA | NA | NA | NA | Gm14247 | predicted gene 14247 [Source:MGI Symbo | 0 | 0 | 0 | 0 | 0 | 0 |
| 24034 | ENSMUSG00000081617 | 0 | NA | NA | NA | NA | NA | Rps13-ps7 | ribosomal protein S13, pseudogene 7 [Sou | 0 | 0 | 0 | 0 | 0 | 0 |
| 24035 | ENSMUSG00000081618 | 0 | NA | NA | NA | NA | NA | 5033423K11 | RIKEN cDNA 5033423K11 gene [Source:M | 0 | 0 | 0 | 0 | 0 | 0 |
| 24036 | ENSMUSG00000081619 | 1.566597122 | -0.993370648 | 1.770883718 | -0.560946288 | 0.574834154 | NA | Gm13351 | predicted gene 13351 [Source:MGI Symbo | 3.2124 | 0 | 3.0187 | 0.964 | 1.0906 | 1.1138 |
| 24037 | ENSMUSG00000081620 | 0 | NA | NA | NA | NA | NA | Gm14633 | predicted gene 14633 [Source:MGI Symbo | 0 | 0 | 0 | 0 | 0 | 0 |
| 24038 | ENSMUSG00000081621 | 0 | NA | NA | NA | NA | NA | Gm12793 | predicted gene 12793 [Source:MGI Symbo | 0 | 0 | 0 | 0 | 0 | 0 |
| 24039 | ENSMUSG00000081622 | 0 | NA | NA | NA | NA | NA | Gm5937 | predicted gene 5937 [Source:MGI Symbol | 0 | 0 | 0 | 0 | 0 | 0 |
| 24040 | ENSMUSG00000081623 | 0 | NA | NA | NA | NA | NA | Gm8809 | predicted gene 8809 [Source:MGI Symbol | 0 | 0 | 0 | 0 | 0 | 0 |
| 24041 | ENSMUSG00000081624 | 0 | NA | NA | NA | NA | NA | Gm6912 | predicted gene 6912 [Source:MGI Symbol | 0 | 0 | 0 | 0 | 0 | 0 |
| 24042 | ENSMUSG00000081625 | 0 | NA | NA | NA | NA | NA | Rps12-ps7 | ribosomal protein S12, pseudogene 7 [Sou | 0 | 0 | 0 | 0 | 0 | 0 |
| 24043 | ENSMUSG00000081626 | 0 | NA | NA | NA | NA | NA | Gm14369 | predicted gene 14369 [Source:MGI Symbo | 0 | 0 | 0 | 0 | 0 | 0 |
| 24044 | ENSMUSG00000081627 | 0 | NA | NA | NA | NA | NA | Gm13158 | predicted gene 13158 [Source:MGI Symbo | 0 | 0 | 0 | 0 | 0 | 0 |
| 24045 | ENSMUSG00000081628 | 0 | NA | NA | NA | NA | NA | Gm14804 | predicted gene 14804 [Source:MGI Symbo | 0 | 0 | 0 | 0 | 0 | 0 |
| 24046 | ENSMUSG00000081629 | 0 | NA | NA | NA | NA | NA | Eif3s6-ps1 | eukaryotic translation initiation factor 3, s | 0 | 0 | 0 | 0 | 0 | 0 |
| 24047 | ENSMUSG00000081630 | 0 | NA | NA | NA | NA | NA | Gm11498 | predicted gene 11498 [Source:MGI Symbo | 0 | 0 | 0 | 0 | 0 | 0 |
| 24048 | ENSMUSG00000081631 | 0 | NA | NA | NA | NA | NA | Gm12005 | predicted gene 12005 [Source:MGI Symbo | 0 | 0 | 0 | 0 | 0 | 0 |
| 24049 | ENSMUSG00000081632 | 0 | NA | NA | NA | NA | NA | Gm11443 | predicted gene 11443 [Source:MGI Symbo | 0 | 0 | 0 | 0 | 0 | 0 |
| 24050 | ENSMUSG00000081633 | 1.385007368 | -0.792709412 | 1.958378789 | -0.404778389 | 0.685640415 | NA | Gm8522 | predicted gene 8522 [Source:MGI Symbol | 3.2124 | 2.0558 | 0 | 1.928 | 0 | 1.1138 |
| 24051 | ENSMUSG00000081635 | 0 | NA | NA | NA | NA | NA | Gm15071 | predicted gene 15071 [Source:MGI Symbo | 0 | 0 | 0 | 0 | 0 | 0 |
| 24052 | ENSMUSG00000081636 | 0 | NA | NA | NA | NA | NA | Gm15901 | predicted gene 15901 [Source:MGI Symbo | 0 | 0 | 0 | 0 | 0 | 0 |
| 24053 | ENSMUSG00000081638 | 0 | NA | NA | NA | NA | NA | Gm14594 | predicted gene 14594 [Source:MGI Symbo | 0 | 0 | 0 | 0 | 0 | 0 |
| 24054 | ENSMUSG00000081639 | 0 | NA | NA | NA | NA | NA | Gm14973 | predicted gene 14973 [Source:MGI Symbo | 0 | 0 | 0 | 0 | 0 | 0 |
| 24055 | ENSMUSG00000081640 | 0 | NA | NA | NA | NA | NA | Rplp0-ps1 | ribosomal protein, large, P0, pseudogene | 0 | 0 | 0 | 0 | 0 | 0 |
| 24056 | ENSMUSG00000081641 | 0 | NA | NA | NA | NA | NA | Gm9439 | predicted gene 9439 [Source:MGI Symbol | 0 | 0 | 0 | 0 | 0 | 0 |
| 24057 | ENSMUSG00000081642 | 1.241296502 | 0.539413051 | 2.077601579 | 0.259632576 | 0.795147205 | NA | Gm13532 | predicted gene 13532 [Source:MGI Symbo | 0 | 2.0558 | 1.0062 | 0 | 3.2719 | 1.1138 |
| 24058 | ENSMUSG00000081643 | 3.335119183 | 1.786091146 | 1.270592601 | 1.405715054 | 0.159808755 | NA | Gm11605 | predicted gene 11605 [Source:MGI Symbo | 2.4093 | 0 | 2.0125 | 6.7478 | 3.2719 | 5.5692 |
| 24059 | ENSMUSG00000081646 | 0 | NA | NA | NA | NA | NA | Gm15060 | predicted gene 15060 [Source:MGI Symbo | 0 | 0 | 0 | 0 | 0 | 0 |
| 24060 | ENSMUSG00000081647 | 0 | NA | NA | NA | NA | NA | Gm13022 | predicted gene 13022 [Source:MGI Symbo | 0 | 0 | 0 | 0 | 0 | 0 |
| 24061 | ENSMUSG00000081648 | 3.059651067 | 0.428799175 | 1.224493795 | 0.350184849 | 0.726199977 | NA | Gm13423 | predicted gene 13423 [Source:MGI Symbo | 0.8031 | 3.0838 | 4.025 | 3.8559 | 4.3625 | 2.2277 |
| 24062 | ENSMUSG00000081649 | 0 | NA | NA | NA | NA | NA | Olfr213 | olfactory receptor 213 [Source:MGI Symb | 0 | 0 | 0 | 0 | 0 | 0 |
| 24063 | ENSMUSG00000081650 | 0.817403785 | 0.568896211 | 2.872137803 | 0.198074135 | 0.842987061 | NA | Gm16181 | predicted gene 16181 [Source:MGI Symbo | 0 | 0 | 2.0125 | 2.8919 | 0 | 0 |
| 24064 | ENSMUSG00000081651 | 0.160662974 | 1.126924175 | 4.080472857 | 0.276174898 | 0.782413736 | NA | Gm15530 | predicted gene 15530 [Source:MGI Symbo | 0 | 0 | 0 | 0.964 | 0 | 0 |
| 24065 | ENSMUSG00000081652 | 0 | NA | NA | NA | NA | NA | Gm14283 | predicted gene 14283 [Source:MGI Symbo | 0 | 0 | 0 | 0 | 0 | 0 |
| 24066 | ENSMUSG00000081653 | 0 | NA | NA | NA | NA | NA | Gm11354 | predicted gene 11354 [Source:MGI Symbo | 0 | 0 | 0 | 0 | 0 | 0 |
| 24067 | ENSMUSG00000081654 | 0 | NA | NA | NA | NA | NA | Gm5383 | predicted gene 5383 [Source:MGI Symbol | 0 | 0 | 0 | 0 | 0 | 0 |
| 24068 | ENSMUSG00000081656 | 0 | NA | NA | NA | NA | NA | Gm11246 | predicted gene 11246 [Source:MGI Symbo | 0 | 0 | 0 | 0 | 0 | 0 |
| 24069 | ENSMUSG00000081657 | 0 | NA | NA | NA | NA | NA | Gm15466 | predicted gene 15466 [Source:MGI Symbo | 0 | 0 | 0 | 0 | 0 | 0 |
| 24070 | ENSMUSG00000081658 | 0 | NA | NA | NA | NA | NA | Gm12077 | predicted gene 12077 [Source:MGI Symbo | 0 | 0 | 0 | 0 | 0 | 0 |
| 24071 | ENSMUSG00000081659 | 0 | NA | NA | NA | NA | NA | Gm14917 | predicted gene 14917 [Source:MGI Symbo | 0 | 0 | 0 | 0 | 0 | 0 |
| 24072 | ENSMUSG00000081660 | 0 | NA | NA | NA | NA | NA | Gm11787 | predicted gene 11787 [Source:MGI Symbo | 0 | 0 | 0 | 0 | 0 | 0 |
| 24073 | ENSMUSG00000081661 | 0 | NA | NA | NA | NA | NA | Gm14450 | predicted gene 14450 [Source:MGI Symbo | 0 | 0 | 0 | 0 | 0 | 0 |
| 24074 | ENSMUSG00000081662 | 0 | NA | NA | NA | NA | NA | Gm14098 | predicted gene 14098 [Source:MGI Symbo | 0 | 0 | 0 | 0 | 0 | 0 |
| 24075 | ENSMUSG00000081663 | 0 | NA | NA | NA | NA | NA | Gm11207 | predicted gene 11207 [Source:MGI Symbo | 0 | 0 | 0 | 0 | 0 | 0 |
| 24076 | ENSMUSG00000081664 | 0 | NA | NA | NA | NA | NA | Gm15544 | predicted gene 15544 [Source:MGI Symbo | 0 | 0 | 0 | 0 | 0 | 0 |

| | | | | | | | | | | | | | | | |
|---|---|---|---|---|---|---|---|---|---|---|---|---|---|---|---|
| 24077 | ENSMUSG00000081665 | 2.544998298 | -1.514950187 | 1.611559356 | -0.940052367 | 0.3471907 | NA | Gm15922 | predicted gene 15922 [Source:MGI Symbo | 3.2124 | 7.1954 | 1.0062 | 3.8559 | 0 | 0 |
| 24078 | ENSMUSG00000081666 | 0.160662974 | 1.126924175 | 4.080472857 | 0.276174898 | 0.782413736 | NA | Gm11677 | predicted gene 11677 [Source:MGI Symbo | 0 | 0 | 0 | 0.964 | 0 | 0 |
| 24079 | ENSMUSG00000081667 | 0 | NA | NA | NA | NA | NA | Gm12883 | predicted gene 12883 [Source:MGI Symbo | 0 | 0 | 0 | 0 | 0 | 0 |
| 24080 | ENSMUSG00000081668 | 0 | NA | NA | NA | NA | NA | Gm7869 | predicted gene 7869 [Source:MGI Symbol | 0 | 0 | 0 | 0 | 0 | 0 |
| 24081 | ENSMUSG00000081669 | 0.620790101 | -1.233953725 | 2.860353486 | -0.431399032 | 0.666178254 | NA | Npm3-ps1 | nucleoplasmin 3, pseudogene 1 [Source:N | 1.6062 | 1.0279 | 0 | 0 | 1.0906 | 0 |
| 24082 | ENSMUSG00000081670 | 0.65103495 | -1.312578592 | 2.821398903 | -0.465222621 | 0.641772041 | NA | Gm15697 | predicted gene 15697 [Source:MGI Symbo | 0.8031 | 0 | 2.0125 | 0 | 1.0906 | 0 |
| 24083 | ENSMUSG00000081671 | 0 | NA | NA | NA | NA | NA | Gm13167 | predicted gene 13167 [Source:MGI Symbo | 0 | 0 | 0 | 0 | 0 | 0 |
| 24084 | ENSMUSG00000081672 | 0 | NA | NA | NA | NA | NA | Gm13000 | predicted gene 13000 [Source:MGI Symbo | 0 | 0 | 0 | 0 | 0 | 0 |
| 24085 | ENSMUSG00000081673 | 0 | NA | NA | NA | NA | NA | Gm14794 | predicted gene 14794 [Source:MGI Symbo | 0 | 0 | 0 | 0 | 0 | 0 |
| 24086 | ENSMUSG00000081674 | 0 | NA | NA | NA | NA | NA | Gm6184 | predicted gene 6184 [Source:MGI Symbol | 0 | 0 | 0 | 0 | 0 | 0 |
| 24087 | ENSMUSG00000081675 | 0.181769715 | 1.126924175 | 4.080472857 | 0.276174898 | 0.782413736 | NA | Gm12712 | predicted gene 12712 [Source:MGI Symbo | 0 | 0 | 0 | 0 | 1.0906 | 0 |
| 24088 | ENSMUSG00000081676 | 0 | NA | NA | NA | NA | NA | Gm11643 | predicted gene 11643 [Source:MGI Symbo | 0 | 0 | 0 | 0 | 0 | 0 |
| 24089 | ENSMUSG00000081677 | 0.321325949 | 1.911575114 | 4.048267893 | 0.472195804 | 0.636787034 | NA | Mdk-ps1 | midkine pseudogene 1 [Source:MGI Symb | 0 | 0 | 0 | 1.928 | 0 | 0 |
| 24090 | ENSMUSG00000081678 | 0 | NA | NA | NA | NA | NA | Gm12959 | predicted gene 12959 [Source:MGI Symbo | 0 | 0 | 0 | 0 | 0 | 0 |
| 24091 | ENSMUSG00000081679 | 1.27380945 | 2.704413443 | 2.275930377 | 1.188267212 | 0.234728151 | NA | Gm11874 | predicted gene 11874 [Source:MGI Symbo | 0 | 0 | 1.0062 | 0 | 2.1812 | 4.4554 |
| 24092 | ENSMUSG00000081680 | 0 | NA | NA | NA | NA | NA | Gm14473 | predicted gene 14473 [Source:MGI Symbo | 0 | 0 | 0 | 0 | 0 | 0 |
| 24093 | ENSMUSG00000081681 | 0 | NA | NA | NA | NA | NA | Gm14868 | predicted gene 14868 [Source:MGI Symbo | 0 | 0 | 0 | 0 | 0 | 0 |
| 24094 | ENSMUSG00000081683 | 0.133850373 | -0.796629176 | 4.080472857 | -0.195229623 | 0.84521319 | NA | Fzd10 | frizzled class receptor 10 [Source:MGI Syn | 0.8031 | 0 | 0 | 0 | 0 | 0 |
| 24095 | ENSMUSG00000081684 | 1.752134031 | 1.329022762 | 1.673282937 | 0.794260631 | 0.427043718 | NA | Rps2-ps13 | ribosomal protein S2, pseudogene 13 [Sou | 0 | 2.0558 | 1.0062 | 1.928 | 2.1812 | 3.3415 |
| 24096 | ENSMUSG00000081685 | 0 | NA | NA | NA | NA | NA | Gm12647 | predicted gene 12647 [Source:MGI Symbo | 0 | 0 | 0 | 0 | 0 | 0 |
| 24097 | ENSMUSG00000081686 | 0 | NA | NA | NA | NA | NA | Gm11774 | predicted gene 11774 [Source:MGI Symbo | 0 | 0 | 0 | 0 | 0 | 0 |
| 24098 | ENSMUSG00000081687 | 0 | NA | NA | NA | NA | NA | Gm13996 | predicted gene 13996 [Source:MGI Symbo | 0 | 0 | 0 | 0 | 0 | 0 |
| 24099 | ENSMUSG00000081688 | 0 | NA | NA | NA | NA | NA | Gm12499 | predicted gene 12499 [Source:MGI Symbo | 0 | 0 | 0 | 0 | 0 | 0 |
| 24100 | ENSMUSG00000081689 | 0 | NA | NA | NA | NA | NA | Gm9166 | predicted gene 9166 [Source:MGI Symbol | 0 | 0 | 0 | 0 | 0 | 0 |
| 24101 | ENSMUSG00000081690 | 0.321325949 | 1.911575114 | 4.048267893 | 0.472195804 | 0.636787034 | NA | Gm12317 | predicted gene 12317 [Source:MGI Symbo | 0 | 0 | 0 | 1.928 | 0 | 0 |
| 24102 | ENSMUSG00000081691 | 0 | NA | NA | NA | NA | NA | Gm11965 | predicted gene 11965 [Source:MGI Symbo | 0 | 0 | 0 | 0 | 0 | 0 |
| 24103 | ENSMUSG00000081692 | 0.678054141 | -2.792799308 | 2.990673428 | -0.933836266 | 0.350388359 | NA | Gm15971 | predicted gene 15971 [Source:MGI Symbo | 0 | 2.0558 | 2.0125 | 0 | 0 | 0 |
| 24104 | ENSMUSG00000081693 | 0 | NA | NA | NA | NA | NA | Gm13297 | predicted gene 13297 [Source:MGI Symbo | 0 | 0 | 0 | 0 | 0 | 0 |
| 24105 | ENSMUSG00000081694 | 0 | NA | NA | NA | NA | NA | Olfr660-ps1 | olfactory receptor 660, pseudogene 1 [Sou | 0 | 0 | 0 | 0 | 0 | 0 |
| 24106 | ENSMUSG00000081695 | 0 | NA | NA | NA | NA | NA | Rpsa-ps5 | ribosomal protein SA, pseudogene 5 [Sou | 0 | 0 | 0 | 0 | 0 | 0 |
| 24107 | ENSMUSG00000081696 | 0 | NA | NA | NA | NA | NA | Gm12717 | predicted gene 12717 [Source:MGI Symbo | 0 | 0 | 0 | 0 | 0 | 0 |
| 24108 | ENSMUSG00000081697 | 0 | NA | NA | NA | NA | NA | Gm14714 | predicted gene 14714 [Source:MGI Symbo | 0 | 0 | 0 | 0 | 0 | 0 |
| 24109 | ENSMUSG00000081698 | 0 | NA | NA | NA | NA | NA | Gm12329 | predicted gene 12329 [Source:MGI Symbo | 0 | 0 | 0 | 0 | 0 | 0 |
| 24110 | ENSMUSG00000081699 | 0 | NA | NA | NA | NA | NA | Gm15213 | predicted gene 15213 [Source:MGI Symbo | 0 | 0 | 0 | 0 | 0 | 0 |
| 24111 | ENSMUSG00000081700 | 0 | NA | NA | NA | NA | NA | Atp5k-ps2 | ATP synthase, H+ transporting, mitochonc | 0 | 0 | 0 | 0 | 0 | 0 |
| 24112 | ENSMUSG00000081701 | 0 | NA | NA | NA | NA | NA | Gm13890 | predicted gene 13890 [Source:MGI Symbo | 0 | 0 | 0 | 0 | 0 | 0 |
| 24113 | ENSMUSG00000081703 | 0.514011053 | 1.044022102 | 3.082659575 | 0.338675769 | 0.734853997 | NA | Gm6285 | predicted gene 6285 [Source:MGI Symbol | 0 | 0 | 1.0062 | 0.964 | 0 | 1.1138 |
| 24114 | ENSMUSG00000081704 | 0 | NA | NA | NA | NA | NA | Gm14603 | predicted gene 14603 [Source:MGI Symbo | 0 | 0 | 0 | 0 | 0 | 0 |
| 24115 | ENSMUSG00000081705 | 0 | NA | NA | NA | NA | NA | Gm12582 | predicted gene 12582 [Source:MGI Symbo | 0 | 0 | 0 | 0 | 0 | 0 |
| 24116 | ENSMUSG00000081706 | 0 | NA | NA | NA | NA | NA | Gm7927 | predicted gene 7927 [Source:MGI Symbol | 0 | 0 | 0 | 0 | 0 | 0 |
| 24117 | ENSMUSG00000081707 | 0 | NA | NA | NA | NA | NA | Gm11343 | predicted gene 11343 [Source:MGI Symbo | 0 | 0 | 0 | 0 | 0 | 0 |
| 24118 | ENSMUSG00000081709 | 0 | NA | NA | NA | NA | NA | Gm15022 | predicted gene 15022 [Source:MGI Symbo | 0 | 0 | 0 | 0 | 0 | 0 |
| 24119 | ENSMUSG00000081711 | 0 | NA | NA | NA | NA | NA | Gm11785 | predicted gene 11785 [Source:MGI Symbo | 0 | 0 | 0 | 0 | 0 | 0 |
| 24120 | ENSMUSG00000081712 | 0 | NA | NA | NA | NA | NA | Gm12096 | predicted gene 12096 [Source:MGI Symbo | 0 | 0 | 0 | 0 | 0 | 0 |
| 24121 | ENSMUSG00000081713 | 0 | NA | NA | NA | NA | NA | Gm12170 | predicted gene 12170 [Source:MGI Symbo | 0 | 0 | 0 | 0 | 0 | 0 |
| 24122 | ENSMUSG00000081714 | 0 | NA | NA | NA | NA | NA | Gm12782 | predicted gene 12782 [Source:MGI Symbo | 0 | 0 | 0 | 0 | 0 | 0 |
| 24123 | ENSMUSG00000081715 | 0.167707431 | -0.796629176 | 4.080472857 | -0.195229623 | 0.84521319 | NA | Rps11-ps3 | ribosomal protein S11, pseudogene 3 [Sou | 0 | 0 | 1.0062 | 0 | 0 | 0 |
| 24124 | ENSMUSG00000081716 | 0 | NA | NA | NA | NA | NA | Gm14839 | predicted gene 14839 [Source:MGI Symbo | 0 | 0 | 0 | 0 | 0 | 0 |
| 24125 | ENSMUSG00000081717 | 0 | NA | NA | NA | NA | NA | Vmn1r-ps10 | vomeronasal 1 receptor, pseudogene 106 | 0 | 0 | 0 | 0 | 0 | 0 |
| 24126 | ENSMUSG00000081718 | 0 | NA | NA | NA | NA | NA | Gm8261 | predicted gene 8261 [Source:MGI Symbol | 0 | 0 | 0 | 0 | 0 | 0 |
| 24127 | ENSMUSG00000081719 | 0 | NA | NA | NA | NA | NA | Gm13726 | predicted gene 13726 [Source:MGI Symbo | 0 | 0 | 0 | 0 | 0 | 0 |
| 24128 | ENSMUSG00000081720 | 0 | NA | NA | NA | NA | NA | Gm12391 | predicted gene 12391 [Source:MGI Symbo | 0 | 0 | 0 | 0 | 0 | 0 |
| 24129 | ENSMUSG00000081721 | 11.09796528 | -0.083681307 | 0.689550144 | -0.121356377 | 0.903408768 | 0.983821605 | Gm13882 | predicted gene 13882 [Source:MGI Symbo | 16.865 | 7.1954 | 10.062 | 12.532 | 5.4531 | 14.48 |
| 24130 | ENSMUSG00000081722 | 0 | NA | NA | NA | NA | NA | Gm11763 | predicted gene 11763 [Source:MGI Symbo | 0 | 0 | 0 | 0 | 0 | 0 |
| 24131 | ENSMUSG00000081723 | 0.133850373 | -0.796629176 | 4.080472857 | -0.195229623 | 0.84521319 | NA | Gm15931 | predicted gene 15931 [Source:MGI Symbo | 0.8031 | 0 | 0 | 0 | 0 | 0 |
| 24132 | ENSMUSG00000081724 | 0 | NA | NA | NA | NA | NA | Olfr129 | olfactory receptor 129 [Source:MGI Symb | 0 | 0 | 0 | 0 | 0 | 0 |

| | | | | | | | | | | | | | | | | |
|---|---|---|---|---|---|---|---|---|---|---|---|---|---|---|---|---|
| 24133 | ENSMUSG00000081728 | 0 | NA | NA | NA | NA | NA | Gm14930 | predicted gene 14930 [Source:MGI Symbo | 0 | 0 | 0 | 0 | 0 | 0 |
| 24134 | ENSMUSG00000081729 | 0 | NA | NA | NA | NA | NA | Hspa9-ps1 | heat shock protein 9, pseudogene 1 [Sour | 0 | 0 | 0 | 0 | 0 | 0 |
| 24135 | ENSMUSG00000081730 | 0.167707431 | -0.796629176 | 4.080472857 | -0.195229623 | 0.84521319 | NA | Gm13785 | predicted gene 13785 [Source:MGI Symbo | 0 | 0 | 1.0062 | 0 | 0 | 0 |
| 24136 | ENSMUSG00000081731 | 0 | NA | NA | NA | NA | NA | Calr-ps | calreticulin, pseudogene [Source:MGI Sym | 0 | 0 | 0 | 0 | 0 | 0 |
| 24137 | ENSMUSG00000081732 | 1.549025481 | -0.918540389 | 1.905431756 | -0.482064176 | 0.629760358 | NA | Gm15495 | predicted gene 15495 [Source:MGI Symbo | 0 | 3.0838 | 3.0187 | 0.964 | 0 | 2.2277 |
| 24138 | ENSMUSG00000081733 | 0 | NA | NA | NA | NA | NA | Gm12852 | predicted gene 12852 [Source:MGI Symbo | 0 | 0 | 0 | 0 | 0 | 0 |
| 24139 | ENSMUSG00000081734 | 0 | NA | NA | NA | NA | NA | Gm12740 | predicted gene 12740 [Source:MGI Symbo | 0 | 0 | 0 | 0 | 0 | 0 |
| 24140 | ENSMUSG00000081735 | 0 | NA | NA | NA | NA | NA | Gm14530 | predicted gene 14530 [Source:MGI Symbo | 0 | 0 | 0 | 0 | 0 | 0 |
| 24141 | ENSMUSG00000081736 | 0 | NA | NA | NA | NA | NA | Gm14971 | predicted gene 14971 [Source:MGI Symbo | 0 | 0 | 0 | 0 | 0 | 0 |
| 24142 | ENSMUSG00000081737 | 0 | NA | NA | NA | NA | NA | Olfr359-ps1 | olfactory receptor 359, pseudogene 1 [So | 0 | 0 | 0 | 0 | 0 | 0 |
| 24143 | ENSMUSG00000081738 | 0.68145976 | 0.058694068 | 2.590797836 | 0.022654824 | 0.981925612 | NA | Hmgb1-ps2 | high mobility group box 1, pseudogene 2 | | 0 | 1.0279 | 1.0062 | 0.964 | 1.0906 | 0 |
| 24144 | ENSMUSG00000081739 | 0.17131964 | -0.796629176 | 4.080472857 | -0.195229623 | 0.84521319 | NA | Mdm4-ps | transformed mouse 3T3 cell double minut | 0 | 1.0279 | 0 | 0 | 0 | 0 |
| 24145 | ENSMUSG00000081740 | 0.499690045 | -0.82080011 | 3.113553401 | -0.263621658 | 0.792071468 | NA | Gm14279 | predicted gene 14279 [Source:MGI Symbo | 0 | 1.0279 | 1.0062 | 0.964 | 0 | 0 |
| 24146 | ENSMUSG00000081741 | 0 | NA | NA | NA | NA | NA | Gm14106 | predicted gene 14106 [Source:MGI Symbo | 0 | 0 | 0 | 0 | 0 | 0 |
| 24147 | ENSMUSG00000081742 | 0 | NA | NA | NA | NA | NA | Gm9051 | predicted gene 9051 [Source:MGI Symbol | 0 | 0 | 0 | 0 | 0 | 0 |
| 24148 | ENSMUSG00000081743 | 0 | NA | NA | NA | NA | NA | Gm4911 | predicted pseudogene 4911 [Source:MGI | 0 | 0 | 0 | 0 | 0 | 0 |
| 24149 | ENSMUSG00000081744 | 0 | NA | NA | NA | NA | NA | Gm5721 | predicted gene 5721 [Source:MGI Symbol | 0 | 0 | 0 | 0 | 0 | 0 |
| 24150 | ENSMUSG00000081745 | 0 | NA | NA | NA | NA | NA | Gm15549 | predicted gene 15549 [Source:MGI Symbo | 0 | 0 | 0 | 0 | 0 | 0 |
| 24151 | ENSMUSG00000081746 | 0 | NA | NA | NA | NA | NA | Gm13925 | predicted gene 13925 [Source:MGI Symbo | 0 | 0 | 0 | 0 | 0 | 0 |
| 24152 | ENSMUSG00000081747 | 0.160662974 | 1.126924175 | 4.080472857 | 0.276174898 | 0.782413736 | NA | Rpl7a-ps9 | ribosomal protein L7A, pseudogene 9 [Sou | 0 | 0 | 0 | 0.964 | 0 | 0 |
| 24153 | ENSMUSG00000081748 | 0 | NA | NA | NA | NA | NA | Gm14870 | predicted gene 14870 [Source:MGI Symbo | 0 | 0 | 0 | 0 | 0 | 0 |
| 24154 | ENSMUSG00000081749 | 0 | NA | NA | NA | NA | NA | Gm12241 | predicted gene 12241 [Source:MGI Symbo | 0 | 0 | 0 | 0 | 0 | 0 |
| 24155 | ENSMUSG00000081750 | 0 | NA | NA | NA | NA | NA | Gm4995 | predicted pseudogene 4995 [Source:MGI | 0 | 0 | 0 | 0 | 0 | 0 |
| 24156 | ENSMUSG00000081751 | 0.696064467 | -1.534444742 | 3.043667477 | -0.504143358 | 0.61416064 | NA | Gm14251 | predicted gene 14251 [Source:MGI Symbo | 3.2124 | 0 | 0 | 0.964 | 0 | 0 |
| 24157 | ENSMUSG00000081752 | 1.2118528 | -1.230711982 | 2.294392741 | -0.536399876 | 0.591682217 | NA | Sms-ps | spermine synthase, pseudogene [Source:I | 4.0155 | 1.0279 | 0 | 0 | 0 | 2.2277 |
| 24158 | ENSMUSG00000081755 | 1.725166445 | -1.140617036 | 1.703421579 | -0.669603491 | 0.503110588 | NA | Mup-ps23 | major urinary protein, pseudogene 23 [So | 4.0155 | 1.0279 | 2.0125 | 0 | 2.1812 | 1.1138 |
| 24159 | ENSMUSG00000081756 | 0.674621893 | -1.426050892 | 3.091195043 | -0.461326727 | 0.644564212 | NA | Gm12291 | predicted gene 12291 [Source:MGI Symbo | 0 | 3.0838 | 0 | 0.964 | 0 | 0 |
| 24160 | ENSMUSG00000081757 | 0 | NA | NA | NA | NA | NA | Gm13798 | predicted gene 13798 [Source:MGI Symbo | 0 | 0 | 0 | 0 | 0 | 0 |
| 24161 | ENSMUSG00000081758 | 0 | NA | NA | NA | NA | NA | Gm11460 | predicted gene 11460 [Source:MGI Symbo | 0 | 0 | 0 | 0 | 0 | 0 |
| 24162 | ENSMUSG00000081761 | 0 | NA | NA | NA | NA | NA | Gm6298 | predicted gene 6298 [Source:MGI Symbol | 0 | 0 | 0 | 0 | 0 | 0 |
| 24163 | ENSMUSG00000081762 | 0 | NA | NA | NA | NA | NA | Gm13738 | predicted gene 13738 [Source:MGI Symbo | 0 | 0 | 0 | 0 | 0 | 0 |
| 24164 | ENSMUSG00000081763 | 0.727078859 | 3.067238961 | 3.29680021 | 0.930368468 | 0.352180339 | NA | Gm9435 | predicted gene 9435 [Source:MGI Symbol | 0 | 0 | 0 | 0 | 4.3625 | 0 |
| 24165 | ENSMUSG00000081765 | 0 | NA | NA | NA | NA | NA | Gm14677 | predicted gene 14677 [Source:MGI Symbo | 0 | 0 | 0 | 0 | 0 | 0 |
| 24166 | ENSMUSG00000081766 | 0 | NA | NA | NA | NA | NA | 2210409E12 | RIKEN cDNA 2210409E12 gene [Source:M | 0 | 0 | 0 | 0 | 0 | 0 |
| 24167 | ENSMUSG00000081767 | 0 | NA | NA | NA | NA | NA | Gm12067 | predicted gene 12067 [Source:MGI Symbo | 0 | 0 | 0 | 0 | 0 | 0 |
| 24168 | ENSMUSG00000081768 | 0 | NA | NA | NA | NA | NA | Gm12755 | predicted gene 12755 [Source:MGI Symbo | 0 | 0 | 0 | 0 | 0 | 0 |
| 24169 | ENSMUSG00000081769 | 22.46383445 | 0.444692666 | 0.550698044 | 0.807507254 | 0.419374297 | 0.837591673 | Gm12216 | predicted gene 12216 [Source:MGI Symbo | 14.456 | 20.558 | 22.137 | 22.171 | 10.906 | 44.554 |
| 24170 | ENSMUSG00000081770 | 0 | NA | NA | NA | NA | NA | Rbpj-ps1 | recombination signal binding protein for i | 0 | 0 | 0 | 0 | 0 | 0 |
| 24171 | ENSMUSG00000081771 | 0 | NA | NA | NA | NA | NA | Gm15381 | predicted gene 15381 [Source:MGI Symbo | 0 | 0 | 0 | 0 | 0 | 0 |
| 24172 | ENSMUSG00000081772 | 0 | NA | NA | NA | NA | NA | Gm13131 | predicted gene 13131 [Source:MGI Symbo | 0 | 0 | 0 | 0 | 0 | 0 |
| 24173 | ENSMUSG00000081773 | 0.722877068 | -0.331317305 | 2.989821669 | -0.110815073 | 0.911762993 | NA | Gm8302 | predicted gene 8302 [Source:MGI Symbol | 2.4093 | 0 | 0 | 1.928 | 0 | 0 |
| 24174 | ENSMUSG00000081774 | 0 | NA | NA | NA | NA | NA | Gm13866 | predicted gene 13866 [Source:MGI Symbo | 0 | 0 | 0 | 0 | 0 | 0 |
| 24175 | ENSMUSG00000081775 | 0 | NA | NA | NA | NA | NA | Gm12420 | predicted gene 12420 [Source:MGI Symbo | 0 | 0 | 0 | 0 | 0 | 0 |
| 24176 | ENSMUSG00000081777 | 0 | NA | NA | NA | NA | NA | Gm14159 | predicted gene 14159 [Source:MGI Symbo | 0 | 0 | 0 | 0 | 0 | 0 |
| 24177 | ENSMUSG00000081778 | 0 | NA | NA | NA | NA | NA | Gm6325 | predicted gene 6325 [Source:MGI Symbol | 0 | 0 | 0 | 0 | 0 | 0 |
| 24178 | ENSMUSG00000081779 | 0 | NA | NA | NA | NA | NA | Gm21998 | predicted gene 21998 [Source:MGI Symbo | 0 | 0 | 0 | 0 | 0 | 0 |
| 24179 | ENSMUSG00000081781 | 0 | NA | NA | NA | NA | NA | Gm14753 | predicted gene 14753 [Source:MGI Symbo | 0 | 0 | 0 | 0 | 0 | 0 |
| 24180 | ENSMUSG00000081782 | 0.305170013 | -1.657836477 | 4.041480335 | -0.410205257 | 0.681655385 | NA | Gm15152 | predicted gene 15152 [Source:MGI Symbo | 0.8031 | 1.0279 | 0 | 0 | 0 | 0 |
| 24181 | ENSMUSG00000081783 | 0.342432689 | 1.990653086 | 3.900281263 | 0.51038706 | 0.609780321 | NA | Gm15599 | predicted gene 15599 [Source:MGI Symbo | 0 | 0 | 0 | 0.964 | 1.0906 | 0 |
| 24182 | ENSMUSG00000081784 | 0 | NA | NA | NA | NA | NA | Gm13281 | predicted gene 13281 [Source:MGI Symbo | 0 | 0 | 0 | 0 | 0 | 0 |
| 24183 | ENSMUSG00000081786 | 0 | NA | NA | NA | NA | NA | Gm11897 | predicted gene 11897 [Source:MGI Symbo | 0 | 0 | 0 | 0 | 0 | 0 |
| 24184 | ENSMUSG00000081787 | 1.017313306 | 0.03731332 | 2.211598506 | 0.016871652 | 0.986539008 | NA | Gm13991 | predicted gene 13991 [Source:MGI Symbo | 0 | 2.0558 | 1.0062 | 1.928 | 0 | 1.1138 |
| 24185 | ENSMUSG00000081788 | 1.625669461 | 0.656220888 | 1.759409577 | 0.372977899 | 0.709164893 | NA | Gm5898 | predicted gene 5898 [Source:MGI Symbol | 0.8031 | 1.0279 | 2.0125 | 4.8199 | 1.0906 | 0 |
| 24186 | ENSMUSG00000081789 | 0.160662974 | 1.126924175 | 4.080472857 | 0.276174898 | 0.782413736 | NA | Gm11893 | predicted gene 11893 [Source:MGI Symbo | 0 | 0 | 0 | 0.964 | 0 | 0 |
| 24187 | ENSMUSG00000081791 | 0 | NA | NA | NA | NA | NA | Gm12190 | predicted gene 12190 [Source:MGI Symbo | 0 | 0 | 0 | 0 | 0 | 0 |
| 24188 | ENSMUSG00000081792 | 0 | NA | NA | NA | NA | NA | Anp32b-ps1 | Bacidic (leucine-rich) nuclear phosphopro | 0 | 0 | 0 | 0 | 0 | 0 |

| | | | | | | | | | | | | | | | |
|---|---|---|---|---|---|---|---|---|---|---|---|---|---|---|---|
| 24189 | ENSMUSG00000081793 | 0.506966596 | 2.558866737 | 3.339834942 | 0.766165628 | 0.443577757 | NA | 5730416F02 | RIKEN cDNA 5730416F02 gene [Source:M | 0 | 0 | 0 | 1.928 | 0 | 1.1138 |
| 24190 | ENSMUSG00000081796 | 0 | NA | NA | NA | NA | NA | Gm9005 | predicted gene 9005 [Source:MGI Symbol | 0 | 0 | 0 | 0 | 0 | 0 |
| 24191 | ENSMUSG00000081797 | 0 | NA | NA | NA | NA | NA | Gm8662 | predicted gene 8662 [Source:MGI Symbol | 0 | 0 | 0 | 0 | 0 | 0 |
| 24192 | ENSMUSG00000081798 | 0.636946036 | 1.591416711 | 2.83781177 | 0.560790087 | 0.574940645 | NA | Gm8152 | predicted gene 8152 [Source:MGI Symbol | 0.8031 | 0 | 0 | 1.928 | 1.0906 | 0 |
| 24193 | ENSMUSG00000081799 | 0 | NA | NA | NA | NA | NA | Gm14071 | predicted gene 14071 [Source:MGI Symbo | 0 | 0 | 0 | 0 | 0 | 0 |
| 24194 | ENSMUSG00000081800 | 0 | NA | NA | NA | NA | NA | Gm14200 | predicted gene 14200 [Source:MGI Symbo | 0 | 0 | 0 | 0 | 0 | 0 |
| 24195 | ENSMUSG00000081801 | 0 | NA | NA | NA | NA | NA | Dnmt3l-ps1 | DNA methyltransferase 3-like, pseudogen | 0 | 0 | 0 | 0 | 0 | 0 |
| 24196 | ENSMUSG00000081803 | 0 | NA | NA | NA | NA | NA | Gm15282 | predicted gene 15282 [Source:MGI Symbo | 0 | 0 | 0 | 0 | 0 | 0 |
| 24197 | ENSMUSG00000081804 | 0 | NA | NA | NA | NA | NA | Xlr5d-ps | X-linked lymphocyte-regulated 5D, pseud | 0 | 0 | 0 | 0 | 0 | 0 |
| 24198 | ENSMUSG00000081805 | 2.103988528 | 0.394457894 | 1.43375981 | 0.275121322 | 0.783223031 | NA | Gm14335 | predicted gene 14335 [Source:MGI Symbo | 2.4093 | 1.0279 | 2.0125 | 3.8559 | 1.0906 | 2.2277 |
| 24199 | ENSMUSG00000081806 | 0 | NA | NA | NA | NA | NA | Gm14901 | predicted gene 14901 [Source:MGI Symbo | 0 | 0 | 0 | 0 | 0 | 0 |
| 24200 | ENSMUSG00000081807 | 0 | NA | NA | NA | NA | NA | Gm14849 | predicted gene 14849 [Source:MGI Symbo | 0 | 0 | 0 | 0 | 0 | 0 |
| 24201 | ENSMUSG00000081809 | 0.465832987 | -0.698295221 | 3.18348759 | -0.219349126 | 0.826378096 | NA | Gm15539 | predicted gene 15539 [Source:MGI Symbo | 0.8031 | 1.0279 | 0 | 0.964 | 0 | 0 |
| 24202 | ENSMUSG00000081810 | 0.469265235 | -2.274672822 | 3.439794491 | -0.661281605 | 0.508431736 | NA | Gm11878 | predicted gene 11878 [Source:MGI Symbo | 0.8031 | 0 | 2.0125 | 0 | 0 | 0 |
| 24203 | ENSMUSG00000081811 | 0 | NA | NA | NA | NA | NA | Gm12713 | predicted gene 12713 [Source:MGI Symbo | 0 | 0 | 0 | 0 | 0 | 0 |
| 24204 | ENSMUSG00000081812 | 0 | NA | NA | NA | NA | NA | Gm4789 | predicted gene 4789 [Source:MGI Symbol | 0 | 0 | 0 | 0 | 0 | 0 |
| 24205 | ENSMUSG00000081813 | 0 | NA | NA | NA | NA | NA | Gm12806 | predicted gene 12806 [Source:MGI Symbo | 0 | 0 | 0 | 0 | 0 | 0 |
| 24206 | ENSMUSG00000081814 | 0 | NA | NA | NA | NA | NA | Gm14695 | predicted gene 14695 [Source:MGI Symbo | 0 | 0 | 0 | 0 | 0 | 0 |
| 24207 | ENSMUSG00000081816 | 0 | NA | NA | NA | NA | NA | Gm15528 | predicted gene 15528 [Source:MGI Symbo | 0 | 0 | 0 | 0 | 0 | 0 |
| 24208 | ENSMUSG00000081817 | 0.531944269 | 2.618916564 | 3.285123409 | 0.797204926 | 0.425332026 | NA | Gm13590 | predicted gene 13590 [Source:MGI Symbo | 0 | 0 | 0 | 0.964 | 0 | 2.2277 |
| 24209 | ENSMUSG00000081819 | 0 | NA | NA | NA | NA | NA | Gm12722 | predicted gene 12722 [Source:MGI Symbo | 0 | 0 | 0 | 0 | 0 | 0 |
| 24210 | ENSMUSG00000081820 | 0 | NA | NA | NA | NA | NA | Gm12029 | predicted gene 12029 [Source:MGI Symbo | 0 | 0 | 0 | 0 | 0 | 0 |
| 24211 | ENSMUSG00000081821 | 0 | NA | NA | NA | NA | NA | Gm15158 | predicted gene 15158 [Source:MGI Symbo | 0 | 0 | 0 | 0 | 0 | 0 |
| 24212 | ENSMUSG00000081822 | 0 | NA | NA | NA | NA | NA | Gm15626 | predicted gene 15626 [Source:MGI Symbo | 0 | 0 | 0 | 0 | 0 | 0 |
| 24213 | ENSMUSG00000081823 | 0.685278559 | -2.809870862 | 3.376578027 | -0.832165239 | 0.405315682 | NA | Cyp4a28-ps | cytochrome P450, family 4, subfamily a, p | 0 | 4.1117 | 0 | 0 | 0 | 0 |
| 24214 | ENSMUSG00000081824 | 4.281055157 | -1.025158543 | 1.060185563 | -0.966961426 | 0.333563318 | NA | Ndufs5-ps | NADH:ubiquinone oxidoreductase core su | 4.0155 | 7.1954 | 6.0375 | 2.8919 | 1.0906 | 4.4554 |
| 24215 | ENSMUSG00000081827 | 0 | NA | NA | NA | NA | NA | Gm15103 | predicted gene 15103 [Source:MGI Symbo | 0 | 0 | 0 | 0 | 0 | 0 |
| 24216 | ENSMUSG00000081828 | 0.167707431 | -0.796629176 | 4.080472857 | -0.195229623 | 0.84521319 | NA | Gm11420 | predicted gene 11420 [Source:MGI Symbo | 0 | 0 | 1.0062 | 0 | 0 | 0 |
| 24217 | ENSMUSG00000081829 | 0 | NA | NA | NA | NA | NA | Gm15460 | predicted gene 15460 [Source:MGI Symbo | 0 | 0 | 0 | 0 | 0 | 0 |
| 24218 | ENSMUSG00000081830 | 0 | NA | NA | NA | NA | NA | Gm13475 | predicted gene 13475 [Source:MGI Symbo | 0 | 0 | 0 | 0 | 0 | 0 |
| 24219 | ENSMUSG00000081831 | 0 | NA | NA | NA | NA | NA | Gm7405 | predicted gene 7405 [Source:MGI Symbol | 0 | 0 | 0 | 0 | 0 | 0 |
| 24220 | ENSMUSG00000081832 | 0 | NA | NA | NA | NA | NA | Gm6897 | predicted pseudogene 6897 [Source:MGI | 0 | 0 | 0 | 0 | 0 | 0 |
| 24221 | ENSMUSG00000081833 | 0 | NA | NA | NA | NA | NA | Gm13669 | predicted gene 13669 [Source:MGI Symbo | 0 | 0 | 0 | 0 | 0 | 0 |
| 24222 | ENSMUSG00000081834 | 0 | NA | NA | NA | NA | NA | Gm13770 | predicted gene 13770 [Source:MGI Symbo | 0 | 0 | 0 | 0 | 0 | 0 |
| 24223 | ENSMUSG00000081835 | 0 | NA | NA | NA | NA | NA | Gm11790 | predicted gene 11790 [Source:MGI Symbo | 0 | 0 | 0 | 0 | 0 | 0 |
| 24224 | ENSMUSG00000081836 | 0 | NA | NA | NA | NA | NA | Olfr1180 | olfactory receptor 1180 [Source:MGI Sym | 0 | 0 | 0 | 0 | 0 | 0 |
| 24225 | ENSMUSG00000081837 | 0 | NA | NA | NA | NA | NA | Gm14649 | predicted gene 14649 [Source:MGI Symbo | 0 | 0 | 0 | 0 | 0 | 0 |
| 24226 | ENSMUSG00000081838 | 0.185640647 | 1.126924175 | 4.080472857 | 0.276174898 | 0.782413736 | NA | Gm13038 | predicted gene 13038 [Source:MGI Symbo | 0 | 0 | 0 | 0 | 0 | 1.1138 |
| 24227 | ENSMUSG00000081839 | 0 | NA | NA | NA | NA | NA | Gm13454 | predicted gene 13454 [Source:MGI Symbo | 0 | 0 | 0 | 0 | 0 | 0 |
| 24228 | ENSMUSG00000081840 | 0 | NA | NA | NA | NA | NA | Gm15220 | predicted gene 15220 [Source:MGI Symbo | 0 | 0 | 0 | 0 | 0 | 0 |
| 24229 | ENSMUSG00000081841 | 0 | NA | NA | NA | NA | NA | Gm13444 | predicted gene 13444 [Source:MGI Symbo | 0 | 0 | 0 | 0 | 0 | 0 |
| 24230 | ENSMUSG00000081842 | 0 | NA | NA | NA | NA | NA | Rap1a-ps2 | RAS related protein 1a, pseudogene 2 [So | 0 | 0 | 0 | 0 | 0 | 0 |
| 24231 | ENSMUSG00000081843 | 0 | NA | NA | NA | NA | NA | Gm7851 | predicted gene 7851 [Source:MGI Symbol | 0 | 0 | 0 | 0 | 0 | 0 |
| 24232 | ENSMUSG00000081844 | 0 | NA | NA | NA | NA | NA | Gm11241 | predicted gene 11241 [Source:MGI Symbo | 0 | 0 | 0 | 0 | 0 | 0 |
| 24233 | ENSMUSG00000081845 | 0 | NA | NA | NA | NA | NA | Prkaca-ps1 | protein kinase, cAMP dependent, catalyti | 0 | 0 | 0 | 0 | 0 | 0 |
| 24234 | ENSMUSG00000081847 | 0 | NA | NA | NA | NA | NA | Btf3-ps7 | basic transcription factor 3, pseudogene 7 | 0 | 0 | 0 | 0 | 0 | 0 |
| 24235 | ENSMUSG00000081848 | 0 | NA | NA | NA | NA | NA | Gm14925 | predicted gene 14925 [Source:MGI Symbo | 0 | 0 | 0 | 0 | 0 | 0 |
| 24236 | ENSMUSG00000081849 | 0 | NA | NA | NA | NA | NA | Zar1-ps | zygote arrest 1, pseudogene [Source:MGI | 0 | 0 | 0 | 0 | 0 | 0 |
| 24237 | ENSMUSG00000081851 | 0 | NA | NA | NA | NA | NA | Gm13323 | predicted gene 13323 [Source:MGI Symbo | 0 | 0 | 0 | 0 | 0 | 0 |
| 24238 | ENSMUSG00000081852 | 0 | NA | NA | NA | NA | NA | Gm12281 | predicted gene 12281 [Source:MGI Symbo | 0 | 0 | 0 | 0 | 0 | 0 |
| 24239 | ENSMUSG00000081853 | 0 | NA | NA | NA | NA | NA | Gm13735 | predicted gene 13735 [Source:MGI Symbo | 0 | 0 | 0 | 0 | 0 | 0 |
| 24240 | ENSMUSG00000081854 | 0.294513347 | 0.165143496 | 4.080472857 | 0.040471657 | 0.967717103 | NA | Gm11447 | predicted gene 11447 [Source:MGI Symbo | 0.8031 | 0 | 0 | 0.964 | 0 | 0 |
| 24241 | ENSMUSG00000081855 | 6.067529896 | 0.299343787 | 0.926416636 | 0.323120047 | 0.746604329 | 0.950187858 | Rpl17-ps5 | ribosomal protein L17, pseudogene 5 [Sou | 8.031 | 6.1675 | 2.0125 | 5.7839 | 3.2719 | 11.138 |
| 24242 | ENSMUSG00000081856 | 0 | NA | NA | NA | NA | NA | Gm14580 | predicted gene 14580 [Source:MGI Symbo | 0 | 0 | 0 | 0 | 0 | 0 |
| 24243 | ENSMUSG00000081857 | 0 | NA | NA | NA | NA | NA | Gm13624 | predicted gene 13624 [Source:MGI Symbo | 0 | 0 | 0 | 0 | 0 | 0 |
| 24244 | ENSMUSG00000081858 | 0 | NA | NA | NA | NA | NA | Gm14514 | predicted gene 14514 [Source:MGI Symbo | 0 | 0 | 0 | 0 | 0 | 0 |

| | | | | | | | | | | | | | | | |
|---|---|---|---|---|---|---|---|---|---|---|---|---|---|---|---|
| 24245 | ENSMUSG00000081859 | 0 | NA | NA | NA | NA | NA | Olfr650-ps1 | olfactory receptor 650, pseudogene 1 [Sou | 0 | 0 | 0 | 0 | 0 | 0 |
| 24246 | ENSMUSG00000081860 | 0.670829724 | -2.782124174 | 3.401449285 | -0.817923168 | 0.41340106 | NA | Gm11533 | predicted gene 11533 [Source:MGI Symbo | 0 | 0 | 4.025 | 0 | 0 | 0 |
| 24247 | ENSMUSG00000081861 | 0.267700746 | -1.497481649 | 4.057001436 | -0.369110456 | 0.712045396 | NA | Gm14441 | predicted gene 14441 [Source:MGI Symbo | 1.6062 | 0 | 0 | 0 | 0 | 0 |
| 24248 | ENSMUSG00000081862 | 0 | NA | NA | NA | NA | NA | Gm12413 | predicted gene 12413 [Source:MGI Symbo | 0 | 0 | 0 | 0 | 0 | 0 |
| 24249 | ENSMUSG00000081863 | 0 | NA | NA | NA | NA | NA | Gm12780 | predicted gene 12780 [Source:MGI Symbo | 0 | 0 | 0 | 0 | 0 | 0 |
| 24250 | ENSMUSG00000081864 | 0 | NA | NA | NA | NA | NA | Olfr405-ps1 | olfactory receptor 405, pseudogene 1 [Sou | 0 | 0 | 0 | 0 | 0 | 0 |
| 24251 | ENSMUSG00000081865 | 0.685071968 | 0.040605487 | 2.800847395 | 0.014497572 | 0.988433016 | NA | Gm15484 | predicted gene 15484 [Source:MGI Symbo | 0 | 2.0558 | 0 | 0.964 | 1.0906 | 0 |
| 24252 | ENSMUSG00000081866 | 0 | NA | NA | NA | NA | NA | Gm12401 | predicted gene 12401 [Source:MGI Symbo | 0 | 0 | 0 | 0 | 0 | 0 |
| 24253 | ENSMUSG00000081868 | 0.328370405 | 0.16514316 | 4.003149536 | 0.041253308 | 0.967093956 | NA | Gm12039 | predicted gene 12039 [Source:MGI Symbo | 0 | 0 | 1.0062 | 0.964 | 0 | 0 |
| 24254 | ENSMUSG00000081869 | 0 | NA | NA | NA | NA | NA | Gm15002 | predicted gene 15002 [Source:MGI Symbo | 0 | 0 | 0 | 0 | 0 | 0 |
| 24255 | ENSMUSG00000081870 | 0 | NA | NA | NA | NA | NA | Gm15141 | predicted gene 15141 [Source:MGI Symbo | 0 | 0 | 0 | 0 | 0 | 0 |
| 24256 | ENSMUSG00000081871 | 0.866661723 | -0.464185162 | 2.825297661 | -0.16429602 | 0.869498114 | NA | Gm11488 | predicted gene 11488 [Source:MGI Symbo | 0 | 0 | 3.0187 | 0 | 2.1812 | 0 |
| 24257 | ENSMUSG00000081872 | 0 | NA | NA | NA | NA | NA | Gm15572 | predicted gene 15572 [Source:MGI Symbo | 0 | 0 | 0 | 0 | 0 | 0 |
| 24258 | ENSMUSG00000081873 | 0 | NA | NA | NA | NA | NA | Gm15197 | predicted gene 15197 [Source:MGI Symbo | 0 | 0 | 0 | 0 | 0 | 0 |
| 24259 | ENSMUSG00000081874 | 0 | NA | NA | NA | NA | NA | Gm14983 | predicted gene 14983 [Source:MGI Symbo | 0 | 0 | 0 | 0 | 0 | 0 |
| 24260 | ENSMUSG00000081875 | 0.160662974 | 1.126924175 | 4.080472857 | 0.276174898 | 0.782413736 | NA | Gm15223 | predicted gene 15223 [Source:MGI Symbo | 0 | 0 | 0 | 0.964 | 0 | 0 |
| 24261 | ENSMUSG00000081876 | 0 | NA | NA | NA | NA | NA | Gm12644 | predicted gene 12644 [Source:MGI Symbo | 0 | 0 | 0 | 0 | 0 | 0 |
| 24262 | ENSMUSG00000081877 | 0 | NA | NA | NA | NA | NA | Gm8858 | predicted gene 8858 [Source:MGI Symbol | 0 | 0 | 0 | 0 | 0 | 0 |
| 24263 | ENSMUSG00000081878 | 0 | NA | NA | NA | NA | NA | Gm13739 | predicted gene 13739 [Source:MGI Symbo | 0 | 0 | 0 | 0 | 0 | 0 |
| 24264 | ENSMUSG00000081879 | 0 | NA | NA | NA | NA | NA | Gm11933 | predicted gene 11933 [Source:MGI Symbo | 0 | 0 | 0 | 0 | 0 | 0 |
| 24265 | ENSMUSG00000081880 | 0 | NA | NA | NA | NA | NA | Gm5385 | predicted gene 5385 [Source:MGI Symbol | 0 | 0 | 0 | 0 | 0 | 0 |
| 24266 | ENSMUSG00000081881 | 0 | NA | NA | NA | NA | NA | Gm14752 | predicted gene 14752 [Source:MGI Symbo | 0 | 0 | 0 | 0 | 0 | 0 |
| 24267 | ENSMUSG00000081882 | 0 | NA | NA | NA | NA | NA | Gm14294 | predicted gene 14294 [Source:MGI Symbo | 0 | 0 | 0 | 0 | 0 | 0 |
| 24268 | ENSMUSG00000081883 | 0 | NA | NA | NA | NA | NA | Gm15574 | predicted gene 15574 [Source:MGI Symbo | 0 | 0 | 0 | 0 | 0 | 0 |
| 24269 | ENSMUSG00000081884 | 0 | NA | NA | NA | NA | NA | Gm11821 | predicted gene 11821 [Source:MGI Symbo | 0 | 0 | 0 | 0 | 0 | 0 |
| 24270 | ENSMUSG00000081885 | 0.167707431 | -0.796629176 | 4.080472857 | -0.195229623 | 0.84521319 | NA | Gm13231 | predicted gene 13231 [Source:MGI Symbo | 0 | 0 | 1.0062 | 0 | 0 | 0 |
| 24271 | ENSMUSG00000081886 | 0 | NA | NA | NA | NA | NA | Gm6175 | predicted gene 6175 [Source:MGI Symbol | 0 | 0 | 0 | 0 | 0 | 0 |
| 24272 | ENSMUSG00000081887 | 0.133850373 | -0.796629176 | 4.080472857 | -0.195229623 | 0.84521319 | NA | Gm12097 | predicted gene 12097 [Source:MGI Symbo | 0.8031 | 0 | 0 | 0 | 0 | 0 |
| 24273 | ENSMUSG00000081888 | 0 | NA | NA | NA | NA | NA | Spcs2-ps | signal peptidase complex subunit 2, pseud | 0 | 0 | 0 | 0 | 0 | 0 |
| 24274 | ENSMUSG00000081889 | 0 | NA | NA | NA | NA | NA | Gm13188 | predicted gene 13188 [Source:MGI Symbo | 0 | 0 | 0 | 0 | 0 | 0 |
| 24275 | ENSMUSG00000081890 | 0 | NA | NA | NA | NA | NA | Gm11243 | predicted gene 11243 [Source:MGI Symbo | 0 | 0 | 0 | 0 | 0 | 0 |
| 24276 | ENSMUSG00000081891 | 0 | NA | NA | NA | NA | NA | Gm14660 | predicted gene 14660 [Source:MGI Symbo | 0 | 0 | 0 | 0 | 0 | 0 |
| 24277 | ENSMUSG00000081892 | 0 | NA | NA | NA | NA | NA | Gm8864 | predicted gene 8864 [Source:MGI Symbol | 0 | 0 | 0 | 0 | 0 | 0 |
| 24278 | ENSMUSG00000081893 | 0 | NA | NA | NA | NA | NA | Gm13273 | predicted gene 13273 [Source:MGI Symbo | 0 | 0 | 0 | 0 | 0 | 0 |
| 24279 | ENSMUSG00000081894 | 0.167707431 | -0.796629176 | 4.080472857 | -0.195229623 | 0.84521319 | NA | Gm14212 | predicted gene 14212 [Source:MGI Symbo | 0 | 0 | 1.0062 | 0 | 0 | 0 |
| 24280 | ENSMUSG00000081895 | 21.75677927 | -0.057355796 | 0.51082536 | -0.112280635 | 0.910600896 | 0.985662791 | Rpl17-ps10 | ribosomal protein L17, pseudogene 10 [Sc | 16.865 | 19.53 | 30.187 | 16.388 | 15.269 | 32.301 |
| 24281 | ENSMUSG00000081896 | 0 | NA | NA | NA | NA | NA | Gm5389 | predicted gene 5389 [Source:MGI Symbol | 0 | 0 | 0 | 0 | 0 | 0 |
| 24282 | ENSMUSG00000081897 | 0 | NA | NA | NA | NA | NA | Gm15067 | predicted gene 15067 [Source:MGI Symbo | 0 | 0 | 0 | 0 | 0 | 0 |
| 24283 | ENSMUSG00000081898 | 0 | NA | NA | NA | NA | NA | Gm5754 | predicted gene 5754 [Source:MGI Symbol | 0 | 0 | 0 | 0 | 0 | 0 |
| 24284 | ENSMUSG00000081900 | 0 | NA | NA | NA | NA | NA | Gm14985 | predicted gene 14985 [Source:MGI Symbo | 0 | 0 | 0 | 0 | 0 | 0 |
| 24285 | ENSMUSG00000081901 | 0 | NA | NA | NA | NA | NA | Gm15357 | predicted gene 15357 [Source:MGI Symbo | 0 | 0 | 0 | 0 | 0 | 0 |
| 24286 | ENSMUSG00000081903 | 0 | NA | NA | NA | NA | NA | Gm11387 | predicted gene 11387 [Source:MGI Symbo | 0 | 0 | 0 | 0 | 0 | 0 |
| 24287 | ENSMUSG00000081904 | 0 | NA | NA | NA | NA | NA | Gm13419 | predicted gene 13419 [Source:MGI Symbo | 0 | 0 | 0 | 0 | 0 | 0 |
| 24288 | ENSMUSG00000081905 | 0 | NA | NA | NA | NA | NA | Gm11370 | predicted gene 11370 [Source:MGI Symbo | 0 | 0 | 0 | 0 | 0 | 0 |
| 24289 | ENSMUSG00000081906 | 0 | NA | NA | NA | NA | NA | Rpl9-ps1 | ribosomal protein L9, pseudogene 1 [Sour | 0 | 0 | 0 | 0 | 0 | 0 |
| 24290 | ENSMUSG00000081907 | 0 | NA | NA | NA | NA | NA | Gm13731 | predicted gene 13731 [Source:MGI Symbo | 0 | 0 | 0 | 0 | 0 | 0 |
| 24291 | ENSMUSG00000081908 | 0 | NA | NA | NA | NA | NA | Olfr1147-ps1 | olfactory receptor 1147, pseudogene 1 [So | 0 | 0 | 0 | 0 | 0 | 0 |
| 24292 | ENSMUSG00000081909 | 14.46328926 | 1.147552338 | 0.636744745 | 1.80221721 | 0.071511238 | 0.38710107 | Gm11341 | predicted gene 11341 [Source:MGI Symbo | 8.8341 | 5.1396 | 13.081 | 29.883 | 10.906 | 18.935 |
| 24293 | ENSMUSG00000081910 | 0 | NA | NA | NA | NA | NA | Gm15986 | predicted gene 15986 [Source:MGI Symbo | 0 | 0 | 0 | 0 | 0 | 0 |
| 24294 | ENSMUSG00000081911 | 0.181769715 | 1.126924175 | 4.080472857 | 0.276174898 | 0.782413736 | NA | Mrps36-ps2 | mitochondrial ribosomal protein S36, pse | 0 | 0 | 0 | 0 | 1.0906 | 0 |
| 24295 | ENSMUSG00000081912 | 0 | NA | NA | NA | NA | NA | Vmn1r-ps12 | vomeronasal 1 receptor, pseudogene 127 | 0 | 0 | 0 | 0 | 0 | 0 |
| 24296 | ENSMUSG00000081913 | 0 | NA | NA | NA | NA | NA | Gm11803 | predicted gene 11803 [Source:MGI Symbo | 0 | 0 | 0 | 0 | 0 | 0 |
| 24297 | ENSMUSG00000081914 | 0 | NA | NA | NA | NA | NA | Vmn1r-ps10 | vomeronasal 1 receptor, pseudogene 109 | 0 | 0 | 0 | 0 | 0 | 0 |
| 24298 | ENSMUSG00000081916 | 0 | NA | NA | NA | NA | NA | Gm12899 | predicted gene 12899 [Source:MGI Symbo | 0 | 0 | 0 | 0 | 0 | 0 |
| 24299 | ENSMUSG00000081917 | 0 | NA | NA | NA | NA | NA | Gm15209 | predicted gene 15209 [Source:MGI Symbo | 0 | 0 | 0 | 0 | 0 | 0 |
| 24300 | ENSMUSG00000081918 | 0 | NA | NA | NA | NA | NA | Gm14613 | predicted gene 14613 [Source:MGI Symbo | 0 | 0 | 0 | 0 | 0 | 0 |

| | | | | | | | | | | | | | | | |
|---|---|---|---|---|---|---|---|---|---|---|---|---|---|---|---|
| 24301 | ENSMUSG00000081919 | 0.160662974 | 1.126924175 | 4.080472857 | 0.276174898 | 0.782413736 | NA | Gm12399 | predicted gene 12399 [Source:MGI Symbo | 0 | 0 | 0 | 0.964 | 0 | 0 |
| 24302 | ENSMUSG00000081920 | 0 | NA | NA | NA | NA | NA | Selenot-ps | selenoprotein T, pseudogene [Source:MG | 0 | 0 | 0 | 0 | 0 | 0 |
| 24303 | ENSMUSG00000081921 | 0 | NA | NA | NA | NA | NA | Gm12858 | predicted gene 12858 [Source:MGI Symbo | 0 | 0 | 0 | 0 | 0 | 0 |
| 24304 | ENSMUSG00000081923 | 0 | NA | NA | NA | NA | NA | Gm9062 | predicted gene 9062 [Source:MGI Symbol | 0 | 0 | 0 | 0 | 0 | 0 |
| 24305 | ENSMUSG00000081924 | 0 | NA | NA | NA | NA | NA | Gm11191 | predicted gene 11191 [Source:MGI Symbo | 0 | 0 | 0 | 0 | 0 | 0 |
| 24306 | ENSMUSG00000081925 | 0 | NA | NA | NA | NA | NA | Rpl31-ps22 | ribosomal protein L31, pseudogene 22 [Sc | 0 | 0 | 0 | 0 | 0 | 0 |
| 24307 | ENSMUSG00000081926 | 0.545309144 | 2.651974629 | 3.666219369 | 0.72335405 | 0.469462394 | NA | Gm15536 | predicted gene 15536 [Source:MGI Symbo | 0 | 0 | 0 | 0 | 3.2719 | 0 |
| 24308 | ENSMUSG00000081928 | 0 | NA | NA | NA | NA | NA | Gm15105 | predicted gene 15105 [Source:MGI Symbo | 0 | 0 | 0 | 0 | 0 | 0 |
| 24309 | ENSMUSG00000081929 | 0.185640647 | 1.126924175 | 4.080472857 | 0.276174898 | 0.782413736 | NA | Rps11-ps2 | ribosomal protein S11, pseudogene 2 [Sou | 0 | 0 | 0 | 0 | 0 | 1.1138 |
| 24310 | ENSMUSG00000081931 | 0 | NA | NA | NA | NA | NA | Gm15211 | predicted gene 15211 [Source:MGI Symbo | 0 | 0 | 0 | 0 | 0 | 0 |
| 24311 | ENSMUSG00000081932 | 0 | NA | NA | NA | NA | NA | Gm12331 | predicted gene 12331 [Source:MGI Symbo | 0 | 0 | 0 | 0 | 0 | 0 |
| 24312 | ENSMUSG00000081933 | 1.123885764 | 0.622981963 | 2.101105212 | 0.296502031 | 0.766846716 | NA | Gm14767 | predicted gene 14767 [Source:MGI Symbo | 1.6062 | 1.0279 | 0 | 1.928 | 2.1812 | 0 |
| 24313 | ENSMUSG00000081934 | 0 | NA | NA | NA | NA | NA | Gm12193 | predicted gene 12193 [Source:MGI Symbo | 0 | 0 | 0 | 0 | 0 | 0 |
| 24314 | ENSMUSG00000081935 | 0 | NA | NA | NA | NA | NA | Gm16186 | predicted gene 16186 [Source:MGI Symbo | 0 | 0 | 0 | 0 | 0 | 0 |
| 24315 | ENSMUSG00000081936 | 0 | NA | NA | NA | NA | NA | Gm11244 | predicted gene 11244 [Source:MGI Symbo | 0 | 0 | 0 | 0 | 0 | 0 |
| 24316 | ENSMUSG00000081937 | 0 | NA | NA | NA | NA | NA | Gm14599 | predicted gene 14599 [Source:MGI Symbo | 0 | 0 | 0 | 0 | 0 | 0 |
| 24317 | ENSMUSG00000081938 | 0 | NA | NA | NA | NA | NA | Gm15039 | predicted gene 15039 [Source:MGI Symbo | 0 | 0 | 0 | 0 | 0 | 0 |
| 24318 | ENSMUSG00000081939 | 0 | NA | NA | NA | NA | NA | Gm15399 | predicted gene 15399 [Source:MGI Symbo | 0 | 0 | 0 | 0 | 0 | 0 |
| 24319 | ENSMUSG00000081940 | 0 | NA | NA | NA | NA | NA | Xlr3e-ps | X-linked lymphocyte-regulated 3E, pseudo | 0 | 0 | 0 | 0 | 0 | 0 |
| 24320 | ENSMUSG00000081941 | 0 | NA | NA | NA | NA | NA | Gm11218 | predicted gene 11218 [Source:MGI Symbo | 0 | 0 | 0 | 0 | 0 | 0 |
| 24321 | ENSMUSG00000081942 | 0 | NA | NA | NA | NA | NA | Gm15149 | predicted gene 15149 [Source:MGI Symbo | 0 | 0 | 0 | 0 | 0 | 0 |
| 24322 | ENSMUSG00000081943 | 0 | NA | NA | NA | NA | NA | Gm9078 | predicted gene 9078 [Source:MGI Symbol | 0 | 0 | 0 | 0 | 0 | 0 |
| 24323 | ENSMUSG00000081944 | 13.45700928 | 0.14062017 | 0.595153296 | 0.236275547 | 0.813218861 | 0.966802988 | Gm12276 | predicted gene 12276 [Source:MGI Symbo | 8.031 | 16.447 | 14.087 | 13.496 | 13.087 | 15.594 |
| 24324 | ENSMUSG00000081945 | 0 | NA | NA | NA | NA | NA | Olfr607 | olfactory receptor 607 [Source:MGI Symb | 0 | 0 | 0 | 0 | 0 | 0 |
| 24325 | ENSMUSG00000081946 | 0 | NA | NA | NA | NA | NA | Gm11472 | predicted gene 11472 [Source:MGI Symbo | 0 | 0 | 0 | 0 | 0 | 0 |
| 24326 | ENSMUSG00000081947 | 0 | NA | NA | NA | NA | NA | Gm14512 | predicted gene 14512 [Source:MGI Symbo | 0 | 0 | 0 | 0 | 0 | 0 |
| 24327 | ENSMUSG00000081948 | 0 | NA | NA | NA | NA | NA | Olfr1191-ps1 | olfactory receptor 1191, pseudogene 1 [So | 0 | 0 | 0 | 0 | 0 | 0 |
| 24328 | ENSMUSG00000081949 | 0 | NA | NA | NA | NA | NA | Gm14392 | predicted gene 14392 [Source:MGI Symbo | 0 | 0 | 0 | 0 | 0 | 0 |
| 24329 | ENSMUSG00000081950 | 0 | NA | NA | NA | NA | NA | Gm11600 | predicted gene 11600 [Source:MGI Symbo | 0 | 0 | 0 | 0 | 0 | 0 |
| 24330 | ENSMUSG00000081951 | 0 | NA | NA | NA | NA | NA | Gm15352 | predicted gene 15352 [Source:MGI Symbo | 0 | 0 | 0 | 0 | 0 | 0 |
| 24331 | ENSMUSG00000081952 | 0.481988923 | 2.492401104 | 3.837077151 | 0.649557203 | 0.515978285 | NA | Gm12178 | predicted gene 12178 [Source:MGI Symbo | 0 | 0 | 0 | 2.8919 | 0 | 0 |
| 24332 | ENSMUSG00000081953 | 4.947516491 | -0.19182133 | 0.944012673 | -0.203197834 | 0.838980406 | NA | Gm12435 | predicted gene 12435 [Source:MGI Symbo | 5.6217 | 4.1117 | 6.0375 | 2.8919 | 5.4531 | 5.5692 |
| 24333 | ENSMUSG00000081954 | 0.160662974 | 1.126924175 | 4.080472857 | 0.276174898 | 0.782413736 | NA | Gm13626 | predicted gene 13626 [Source:MGI Symbo | 0 | 0 | 0 | 0.964 | 0 | 0 |
| 24334 | ENSMUSG00000081955 | 0 | NA | NA | NA | NA | NA | Olfr999-ps1 | olfactory receptor 999, pseudogene 1 [Sou | 0 | 0 | 0 | 0 | 0 | 0 |
| 24335 | ENSMUSG00000081957 | 1.262894061 | 3.862452193 | 2.195331677 | 1.75939346 | 0.078510702 | NA | Ak3l2-ps | adenylate kinase 3-like 2, pseudogene [So | 0 | 0 | 0 | 0.964 | 3.2719 | 3.3415 |
| 24336 | ENSMUSG00000081958 | 0 | NA | NA | NA | NA | NA | Gm11594 | predicted gene 11594 [Source:MGI Symbo | 0 | 0 | 0 | 0 | 0 | 0 |
| 24337 | ENSMUSG00000081959 | 0 | NA | NA | NA | NA | NA | Gm16480 | predicted gene 16480 [Source:MGI Symbo | 0 | 0 | 0 | 0 | 0 | 0 |
| 24338 | ENSMUSG00000081960 | 0 | NA | NA | NA | NA | NA | Mageb17-ps | MAGE family member B17, pseudogene [S | 0 | 0 | 0 | 0 | 0 | 0 |
| 24339 | ENSMUSG00000081961 | 0 | NA | NA | NA | NA | NA | Olfr383-ps1 | olfactory receptor 383, pseudogene 1 [Sou | 0 | 0 | 0 | 0 | 0 | 0 |
| 24340 | ENSMUSG00000081962 | 0 | NA | NA | NA | NA | NA | Gm12773 | predicted gene 12773 [Source:MGI Symbo | 0 | 0 | 0 | 0 | 0 | 0 |
| 24341 | ENSMUSG00000081964 | 0 | NA | NA | NA | NA | NA | Gm11873 | predicted gene 11873 [Source:MGI Symbo | 0 | 0 | 0 | 0 | 0 | 0 |
| 24342 | ENSMUSG00000081965 | 0.66762957 | 2.958119316 | 3.019579549 | 0.979646096 | 0.327260843 | NA | Gm11620 | predicted gene 11620 [Source:MGI Symbo | 0 | 0 | 0 | 2.8919 | 0 | 1.1138 |
| 24343 | ENSMUSG00000081966 | 0 | NA | NA | NA | NA | NA | Gm13900 | predicted gene 13900 [Source:MGI Symbo | 0 | 0 | 0 | 0 | 0 | 0 |
| 24344 | ENSMUSG00000081967 | 0.876412963 | 2.129655479 | 2.836103289 | 0.750908998 | 0.452707425 | NA | Gm14017 | predicted gene 14017 [Source:MGI Symbo | 0.8031 | 0 | 0 | 0 | 0 | 4.4554 |
| 24345 | ENSMUSG00000081968 | 0.185640647 | 1.126924175 | 4.080472857 | 0.276174898 | 0.782413736 | NA | Rpl23a-ps2 | ribosomal protein L23A, pseudogene 2 [Sc | 0 | 0 | 0 | 0 | 0 | 1.1138 |
| 24346 | ENSMUSG00000081969 | 0 | NA | NA | NA | NA | NA | Gm16471 | predicted gene 16471 [Source:MGI Symbo | 0 | 0 | 0 | 0 | 0 | 0 |
| 24347 | ENSMUSG00000081970 | 0 | NA | NA | NA | NA | NA | Gm16057 | predicted gene 16057 [Source:MGI Symbo | 0 | 0 | 0 | 0 | 0 | 0 |
| 24348 | ENSMUSG00000081971 | 0 | NA | NA | NA | NA | NA | Gm14115 | predicted gene 14115 [Source:MGI Symbo | 0 | 0 | 0 | 0 | 0 | 0 |
| 24349 | ENSMUSG00000081972 | 0 | NA | NA | NA | NA | NA | Gm16397 | predicted pseudogene 16397 [Source:MG | 0 | 0 | 0 | 0 | 0 | 0 |
| 24350 | ENSMUSG00000081973 | 0 | NA | NA | NA | NA | NA | Gm13600 | predicted gene 13600 [Source:MGI Symbo | 0 | 0 | 0 | 0 | 0 | 0 |
| 24351 | ENSMUSG00000081974 | 1.05829193 | -1.166662428 | 2.355445425 | -0.495304377 | 0.620385282 | NA | Gm11960 | predicted gene 11960 [Source:MGI Symbo | 2.4093 | 0 | 2.0125 | 1.928 | 0 | 0 |
| 24352 | ENSMUSG00000081975 | 0 | NA | NA | NA | NA | NA | Gm12482 | predicted gene 12482 [Source:MGI Symbo | 0 | 0 | 0 | 0 | 0 | 0 |
| 24353 | ENSMUSG00000081976 | 0 | NA | NA | NA | NA | NA | Gm14823 | predicted gene 14823 [Source:MGI Symbo | 0 | 0 | 0 | 0 | 0 | 0 |
| 24354 | ENSMUSG00000081978 | 0 | NA | NA | NA | NA | NA | Gm14895 | predicted gene 14895 [Source:MGI Symbo | 0 | 0 | 0 | 0 | 0 | 0 |
| 24355 | ENSMUSG00000081979 | 0 | NA | NA | NA | NA | NA | Rps8-ps3 | ribosomal protein S8, pseudogene 3 [Sour | 0 | 0 | 0 | 0 | 0 | 0 |
| 24356 | ENSMUSG00000081980 | 0 | NA | NA | NA | NA | NA | Gm13702 | predicted gene 13702 [Source:MGI Symbo | 0 | 0 | 0 | 0 | 0 | 0 |

| | | | | | | | | | | | | | | | |
|---|---|---|---|---|---|---|---|---|---|---|---|---|---|---|---|
| 24357 | ENSMUSG00000081982 | 0 | NA | NA | NA | NA | NA | Sult2a-ps2 | sulfotransferase family 2A, dehydroepian | 0 | 0 | 0 | 0 | 0 | 0 |
| 24358 | ENSMUSG00000081984 | 5.476164394 | -0.900160835 | 0.915265137 | -0.983497347 | 0.32536272 | 0.771501263 | Dnajb3 | DnaJ heat shock protein family (Hsp40) m | 7.2279 | 5.1396 | 9.0562 | 4.8199 | 3.2719 | 3.3415 |
| 24359 | ENSMUSG00000081985 | 0 | NA | NA | NA | NA | NA | Gng2-ps1 | guanine nucleotide binding protein (G pro | 0 | 0 | 0 | 0 | 0 | 0 |
| 24360 | ENSMUSG00000081986 | 0 | NA | NA | NA | NA | NA | Gm14841 | predicted gene 14841 [Source:MGI Symbo | 0 | 0 | 0 | 0 | 0 | 0 |
| 24361 | ENSMUSG00000081987 | 0 | NA | NA | NA | NA | NA | Gm12639 | predicted gene 12639 [Source:MGI Symbo | 0 | 0 | 0 | 0 | 0 | 0 |
| 24362 | ENSMUSG00000081988 | 60.41118976 | 0.59863482 | 0.393467081 | 1.521435587 | 0.128150564 | 0.524051388 | Gm15746 | predicted gene 15746 [Source:MGI Symbo | 57.823 | 44.2 | 42.262 | 123.39 | 46.897 | 47.895 |
| 24363 | ENSMUSG00000081991 | 0 | NA | NA | NA | NA | NA | Gm13859 | predicted gene 13859 [Source:MGI Symbo | 0 | 0 | 0 | 0 | 0 | 0 |
| 24364 | ENSMUSG00000081992 | 1.634982017 | 1.430840227 | 1.857759571 | 0.770196666 | 0.441183241 | NA | Gm13408 | predicted gene 13408 [Source:MGI Symbo | 1.6062 | 0 | 1.0062 | 3.8559 | 0 | 3.3415 |
| 24365 | ENSMUSG00000081993 | 0 | NA | NA | NA | NA | NA | Gm8475 | predicted gene 8475 [Source:MGI Symbol | 0 | 0 | 0 | 0 | 0 | 0 |
| 24366 | ENSMUSG00000081994 | 0 | NA | NA | NA | NA | NA | Gm14517 | predicted gene 14517 [Source:MGI Symbo | 0 | 0 | 0 | 0 | 0 | 0 |
| 24367 | ENSMUSG00000081995 | 0.36353943 | 2.065989254 | 4.034137301 | 0.512126658 | 0.608562368 | NA | Gm11719 | predicted gene 11719 [Source:MGI Symbo | 0 | 0 | 0 | 0 | 2.1812 | 0 |
| 24368 | ENSMUSG00000081996 | 2.47431369 | 0.367682816 | 1.323063487 | 0.277902625 | 0.781087109 | NA | Gm7153 | predicted gene 7153 [Source:MGI Symbol | 2.4093 | 2.0558 | 2.0125 | 2.8919 | 4.3625 | 1.1138 |
| 24369 | ENSMUSG00000081998 | 1.744212205 | 0.69889292 | 1.863633191 | 0.375016351 | 0.707648306 | NA | Gm13245 | predicted gene 13245 [Source:MGI Symbo | 0 | 0 | 4.025 | 0.964 | 4.3625 | 1.1138 |
| 24370 | ENSMUSG00000081999 | 3.753621882 | -0.185206767 | 1.082931868 | -0.171023472 | 0.86420531 | NA | Gm13461 | predicted gene 13461 [Source:MGI Symbo | 4.8186 | 4.1117 | 3.0187 | 2.8919 | 5.4531 | 2.2277 |
| 24371 | ENSMUSG00000082000 | 0 | NA | NA | NA | NA | NA | Gm12488 | predicted gene 12488 [Source:MGI Symbo | 0 | 0 | 0 | 0 | 0 | 0 |
| 24372 | ENSMUSG00000082001 | 0 | NA | NA | NA | NA | NA | Gm15607 | predicted gene 15607 [Source:MGI Symbo | 0 | 0 | 0 | 0 | 0 | 0 |
| 24373 | ENSMUSG00000082002 | 0.17131964 | -0.796629176 | 4.080472857 | -0.195229623 | 0.84521319 | NA | Gm6568 | predicted pseudogene 6568 [Source:MGI | 0 | 1.0279 | 0 | 0 | 0 | 0 |
| 24374 | ENSMUSG00000082005 | 0 | NA | NA | NA | NA | NA | Gm7189 | predicted pseudogene 7189 [Source:MGI | 0 | 0 | 0 | 0 | 0 | 0 |
| 24375 | ENSMUSG00000082006 | 0 | NA | NA | NA | NA | NA | Gm13649 | predicted gene 13649 [Source:MGI Symbo | 0 | 0 | 0 | 0 | 0 | 0 |
| 24376 | ENSMUSG00000082007 | 0 | NA | NA | NA | NA | NA | Gm15206 | predicted gene 15206 [Source:MGI Symbo | 0 | 0 | 0 | 0 | 0 | 0 |
| 24377 | ENSMUSG00000082008 | 0.339027071 | -1.79174149 | 3.919585585 | -0.457125237 | 0.647581032 | NA | Gm12390 | predicted gene 12390 [Source:MGI Symbo | 0 | 1.0279 | 1.0062 | 0 | 0 | 0 |
| 24378 | ENSMUSG00000082009 | 0 | NA | NA | NA | NA | NA | Gm12954 | predicted gene 12954 [Source:MGI Symbo | 0 | 0 | 0 | 0 | 0 | 0 |
| 24379 | ENSMUSG00000082010 | 0 | NA | NA | NA | NA | NA | Olfr1171-ps1 | olfactory receptor 1171, pseudogene 1 [S | 0 | 0 | 0 | 0 | 0 | 0 |
| 24380 | ENSMUSG00000082011 | 0 | NA | NA | NA | NA | NA | Gm15043 | predicted gene 15043 [Source:MGI Symbo | 0 | 0 | 0 | 0 | 0 | 0 |
| 24381 | ENSMUSG00000082012 | 0 | NA | NA | NA | NA | NA | Gm5257 | predicted gene 5257 [Source:MGI Symbol | 0 | 0 | 0 | 0 | 0 | 0 |
| 24382 | ENSMUSG00000082013 | 0.133850373 | -0.796629176 | 4.080472857 | -0.195229623 | 0.84521319 | NA | Gm13216 | predicted gene 13216 [Source:MGI Symbo | 0.8031 | 0 | 0 | 0 | 0 | 0 |
| 24383 | ENSMUSG00000082014 | 0 | NA | NA | NA | NA | NA | Gm11371 | predicted gene 11371 [Source:MGI Symbo | 0 | 0 | 0 | 0 | 0 | 0 |
| 24384 | ENSMUSG00000082015 | 0 | NA | NA | NA | NA | NA | Gm13452 | predicted gene 13452 [Source:MGI Symbo | 0 | 0 | 0 | 0 | 0 | 0 |
| 24385 | ENSMUSG00000082016 | 5.697644316 | -1.184103752 | 0.927124091 | -1.277179359 | 0.201538932 | 0.645999739 | Pgam1-ps2 | phosphoglycerate mutase 1, pseudogene | 10.44 | 9.2513 | 4.025 | 3.8559 | 3.2719 | 3.3415 |
| 24386 | ENSMUSG00000082017 | 0 | NA | NA | NA | NA | NA | Gm11229 | predicted gene 11229 [Source:MGI Symbo | 0 | 0 | 0 | 0 | 0 | 0 |
| 24387 | ENSMUSG00000082018 | 0 | NA | NA | NA | NA | NA | Gm6322 | predicted gene 6322 [Source:MGI Symbol | 0 | 0 | 0 | 0 | 0 | 0 |
| 24388 | ENSMUSG00000082019 | 0 | NA | NA | NA | NA | NA | Gm15231 | predicted gene 15231 [Source:MGI Symbo | 0 | 0 | 0 | 0 | 0 | 0 |
| 24389 | ENSMUSG00000082020 | 0 | NA | NA | NA | NA | NA | Gm14182 | predicted gene 14182 [Source:MGI Symbo | 0 | 0 | 0 | 0 | 0 | 0 |
| 24390 | ENSMUSG00000082021 | 0 | NA | NA | NA | NA | NA | Gm14991 | predicted gene 14991 [Source:MGI Symbo | 0 | 0 | 0 | 0 | 0 | 0 |
| 24391 | ENSMUSG00000082022 | 0 | NA | NA | NA | NA | NA | Gm12386 | predicted gene 12386 [Source:MGI Symbo | 0 | 0 | 0 | 0 | 0 | 0 |
| 24392 | ENSMUSG00000082023 | 0 | NA | NA | NA | NA | NA | Gm14407 | predicted gene 14407 [Source:MGI Symbo | 0 | 0 | 0 | 0 | 0 | 0 |
| 24393 | ENSMUSG00000082024 | 0 | NA | NA | NA | NA | NA | Gm11794 | predicted gene 11794 [Source:MGI Symbo | 0 | 0 | 0 | 0 | 0 | 0 |
| 24394 | ENSMUSG00000082025 | 0 | NA | NA | NA | NA | NA | Gm14359 | predicted gene 14359 [Source:MGI Symbo | 0 | 0 | 0 | 0 | 0 | 0 |
| 24395 | ENSMUSG00000082026 | 0 | NA | NA | NA | NA | NA | Gm5764 | predicted gene 5764 [Source:MGI Symbol | 0 | 0 | 0 | 0 | 0 | 0 |
| 24396 | ENSMUSG00000082027 | 0 | NA | NA | NA | NA | NA | Gm11416 | predicted gene 11416 [Source:MGI Symbo | 0 | 0 | 0 | 0 | 0 | 0 |
| 24397 | ENSMUSG00000082028 | 0.353348078 | 0.165142701 | 3.895042568 | 0.042398176 | 0.966181282 | NA | Gm1826 | predicted gene 1826 [Source:MGI Symbol | 0 | 0 | 1.0062 | 0 | 0 | 1.1138 |
| 24398 | ENSMUSG00000082029 | 2.061201606 | -0.839438557 | 1.727497589 | -0.485927484 | 0.627018581 | NA | H3f3c | H3 histone, family 3C [Source:MGI Symbo | 0.8031 | 3.0838 | 4.025 | 0 | 0 | 4.4554 |
| 24399 | ENSMUSG00000082030 | 0 | NA | NA | NA | NA | NA | Gm13081 | predicted gene 13081 [Source:MGI Symbo | 0 | 0 | 0 | 0 | 0 | 0 |
| 24400 | ENSMUSG00000082031 | 0 | NA | NA | NA | NA | NA | Gm8097 | predicted gene 8097 [Source:MGI Symbol | 0 | 0 | 0 | 0 | 0 | 0 |
| 24401 | ENSMUSG00000082032 | 0 | NA | NA | NA | NA | NA | Gm12271 | predicted gene 12271 [Source:MGI Symbo | 0 | 0 | 0 | 0 | 0 | 0 |
| 24402 | ENSMUSG00000082033 | 0 | NA | NA | NA | NA | NA | Mageb10-ps | MAGE family member B10, pseudogene [S | 0 | 0 | 0 | 0 | 0 | 0 |
| 24403 | ENSMUSG00000082035 | 0.321325949 | 1.911575114 | 4.048267893 | 0.472195804 | 0.636787034 | NA | Rpl17-ps8 | ribosomal protein L17, pseudogene 8 [Sou | 0 | 0 | 0 | 1.928 | 0 | 0 |
| 24404 | ENSMUSG00000082036 | 0 | NA | NA | NA | NA | NA | Gm13724 | predicted gene 13724 [Source:MGI Symbo | 0 | 0 | 0 | 0 | 0 | 0 |
| 24405 | ENSMUSG00000082037 | 0 | NA | NA | NA | NA | NA | Vmn1r-ps11 | vomeronasal 1 receptor, pseudogene 116 | 0 | 0 | 0 | 0 | 0 | 0 |
| 24406 | ENSMUSG00000082038 | 0 | NA | NA | NA | NA | NA | Gm12071 | predicted gene 12071 [Source:MGI Symbo | 0 | 0 | 0 | 0 | 0 | 0 |
| 24407 | ENSMUSG00000082039 | 6.927242367 | 1.089424225 | 1.083870989 | 1.005123521 | 0.314837375 | 0.761124486 | Gm15653 | predicted gene 15653 [Source:MGI Symbo | 3.2124 | 0 | 10.062 | 1.928 | 17.45 | 8.9108 |
| 24408 | ENSMUSG00000082040 | 0 | NA | NA | NA | NA | NA | Gm12046 | predicted gene 12046 [Source:MGI Symbo | 0 | 0 | 0 | 0 | 0 | 0 |
| 24409 | ENSMUSG00000082042 | 0 | NA | NA | NA | NA | NA | Gm15923 | predicted gene 15923 [Source:MGI Symbo | 0 | 0 | 0 | 0 | 0 | 0 |
| 24410 | ENSMUSG00000082043 | 0 | NA | NA | NA | NA | NA | Gm12848 | predicted gene 12848 [Source:MGI Symbo | 0 | 0 | 0 | 0 | 0 | 0 |
| 24411 | ENSMUSG00000082044 | 16.52845891 | 0.894784976 | 0.545153799 | 1.641344109 | 0.100726005 | 0.46725495 | Snrpert | small nuclear ribonucleoprotein E, pseudo | 10.44 | 10.279 | 14.087 | 27.955 | 16.359 | 20.049 |
| 24412 | ENSMUSG00000082045 | 17.34838128 | 0.246524756 | 0.583695773 | 0.422351451 | 0.672768509 | 0.931454408 | Gm11585 | predicted gene 11585 [Source:MGI Symbo | 7.2279 | 15.419 | 25.156 | 15.424 | 16.359 | 24.505 |

| | | | | | | | | | | | | | | | |
|---|---|---|---|---|---|---|---|---|---|---|---|---|---|---|---|
| 24413 | ENSMUSG00000082046 | 0 | NA | NA | NA | NA | NA | Gm14464 | predicted gene 14464 [Source:MGI Symbo | 0 | 0 | 0 | 0 | 0 | 0 |
| 24414 | ENSMUSG00000082047 | 0 | NA | NA | NA | NA | NA | Gm12948 | predicted gene 12948 [Source:MGI Symbo | 0 | 0 | 0 | 0 | 0 | 0 |
| 24415 | ENSMUSG00000082048 | 0 | NA | NA | NA | NA | NA | Gm11481 | predicted gene 11481 [Source:MGI Symbo | 0 | 0 | 0 | 0 | 0 | 0 |
| 24416 | ENSMUSG00000082049 | 0.160662974 | 1.126924175 | 4.080472857 | 0.276174898 | 0.782413736 | NA | Gm12105 | predicted gene 12105 [Source:MGI Symbo | 0 | 0 | 0 | 0.964 | 0 | 0 |
| 24417 | ENSMUSG00000082051 | 0 | NA | NA | NA | NA | NA | Gm16072 | predicted gene 16072 [Source:MGI Symbo | 0 | 0 | 0 | 0 | 0 | 0 |
| 24418 | ENSMUSG00000082052 | 0.185640647 | 1.126924175 | 4.080472857 | 0.276174898 | 0.782413736 | NA | Ywhaq-ps2 | tyrosine 3-monooxygenase/tryptophan 5- | 0 | 0 | 0 | 0 | 0 | 1.1138 |
| 24419 | ENSMUSG00000082053 | 0 | NA | NA | NA | NA | NA | Gm11220 | predicted gene 11220 [Source:MGI Symbo | 0 | 0 | 0 | 0 | 0 | 0 |
| 24420 | ENSMUSG00000082054 | 0 | NA | NA | NA | NA | NA | Psmb6-ps | proteasome (prosome, macropain) subun | 0 | 0 | 0 | 0 | 0 | 0 |
| 24421 | ENSMUSG00000082056 | 0 | NA | NA | NA | NA | NA | Gm13745 | predicted gene 13745 [Source:MGI Symbo | 0 | 0 | 0 | 0 | 0 | 0 |
| 24422 | ENSMUSG00000082057 | 0.517623261 | 1.044040381 | 3.075707576 | 0.339447218 | 0.734272853 | NA | Gm15789 | predicted gene 15789 [Source:MGI Symbo | 0 | 1.0279 | 0 | 0.964 | 0 | 1.1138 |
| 24423 | ENSMUSG00000082058 | 0 | NA | NA | NA | NA | NA | Gm11923 | predicted gene 11923 [Source:MGI Symbo | 0 | 0 | 0 | 0 | 0 | 0 |
| 24424 | ENSMUSG00000082059 | 0 | NA | NA | NA | NA | NA | Gm15183 | predicted gene 15183 [Source:MGI Symbo | 0 | 0 | 0 | 0 | 0 | 0 |
| 24425 | ENSMUSG00000082060 | 0 | NA | NA | NA | NA | NA | Gm15048 | predicted gene 15048 [Source:MGI Symbo | 0 | 0 | 0 | 0 | 0 | 0 |
| 24426 | ENSMUSG00000082061 | 0 | NA | NA | NA | NA | NA | Gm12726 | predicted gene 12726 [Source:MGI Symbo | 0 | 0 | 0 | 0 | 0 | 0 |
| 24427 | ENSMUSG00000082062 | 1.364773909 | -0.710082016 | 2.005588113 | -0.354051767 | 0.723300088 | NA | Ftl2-ps | ferritin light polypeptide 2, pseudogene [S | 4.0155 | 1.0279 | 0 | 0.964 | 2.1812 | 0 |
| 24428 | ENSMUSG00000082063 | 3.898937236 | -1.451651702 | 1.135607868 | -1.278303666 | 0.201142376 | NA | Gm12993 | predicted gene 12993 [Source:MGI Symbo | 4.0155 | 4.1117 | 9.0562 | 2.8919 | 1.0906 | 2.2277 |
| 24429 | ENSMUSG00000082064 | 0.854608814 | 0.841289729 | 2.383906808 | 0.352903782 | 0.724160575 | NA | Rpl5-ps2 | ribisomal protein L5, pseudogene 2 [Sourc | 0.8031 | 0 | 1.0062 | 0 | 1.0906 | 2.2277 |
| 24430 | ENSMUSG00000082065 | 51.74663376 | 0.757744653 | 0.496918913 | 1.524885921 | 0.127287554 | 0.522739121 | Mup-ps14 | major urinary protein, pseudogene 14 [So | 73.082 | 13.363 | 28.175 | 45.307 | 95.974 | 54.578 |
| 24431 | ENSMUSG00000082066 | 0 | NA | NA | NA | NA | NA | Gm13904 | predicted gene 13904 [Source:MGI Symbo | 0 | 0 | 0 | 0 | 0 | 0 |
| 24432 | ENSMUSG00000082068 | 0.185640647 | 1.126924175 | 4.080472857 | 0.276174898 | 0.782413736 | NA | Gm2673 | predicted gene 2673 [Source:MGI Symbol | 0 | 0 | 0 | 0 | 0 | 1.1138 |
| 24433 | ENSMUSG00000082069 | 0 | NA | NA | NA | NA | NA | Gm16242 | predicted gene 16242 [Source:MGI Symbo | 0 | 0 | 0 | 0 | 0 | 0 |
| 24434 | ENSMUSG00000082070 | 0.167707431 | -0.796629176 | 4.080472857 | -0.195229623 | 0.84521319 | NA | Gm1866 | predicted gene 1866 [Source:MGI Symbol | 0 | 0 | 1.0062 | 0 | 0 | 0 |
| 24435 | ENSMUSG00000082071 | 0 | NA | NA | NA | NA | NA | Pramex2 | PRAME like, X-linked 2 [Source:MGI Symb | 0 | 0 | 0 | 0 | 0 | 0 |
| 24436 | ENSMUSG00000082072 | 1.511245357 | -2.801803125 | 2.063304422 | -1.35792038 | 0.17448895 | NA | Gm15785 | predicted gene 15785 [Source:MGI Symbo | 0.8031 | 6.1675 | 1.0062 | 0 | 1.0906 | 0 |
| 24437 | ENSMUSG00000082073 | 0 | NA | NA | NA | NA | NA | Gm13114 | predicted gene 13114 [Source:MGI Symbo | 0 | 0 | 0 | 0 | 0 | 0 |
| 24438 | ENSMUSG00000082074 | 0 | NA | NA | NA | NA | NA | Gm12661 | predicted gene 12661 [Source:MGI Symbo | 0 | 0 | 0 | 0 | 0 | 0 |
| 24439 | ENSMUSG00000082076 | 0 | NA | NA | NA | NA | NA | Gm8659 | predicted gene 8659 [Source:MGI Symbol | 0 | 0 | 0 | 0 | 0 | 0 |
| 24440 | ENSMUSG00000082077 | 0 | NA | NA | NA | NA | NA | Olfr260-ps1 | olfactory receptor 260, pseudogene 1 [Sou | 0 | 0 | 0 | 0 | 0 | 0 |
| 24441 | ENSMUSG00000082078 | 0.133850373 | -0.796629176 | 4.080472857 | -0.195229623 | 0.84521319 | NA | Gm12098 | predicted gene 12098 [Source:MGI Symbo | 0.8031 | 0 | 0 | 0 | 0 | 0 |
| 24442 | ENSMUSG00000082079 | 0 | NA | NA | NA | NA | NA | Dnmt3c | DNA methyltransferase 3C [Source:MGI Sy | 0 | 0 | 0 | 0 | 0 | 0 |
| 24443 | ENSMUSG00000082080 | 0 | NA | NA | NA | NA | NA | Gm9246 | predicted gene 9246 [Source:MGI Symbol | 0 | 0 | 0 | 0 | 0 | 0 |
| 24444 | ENSMUSG00000082081 | 0 | NA | NA | NA | NA | NA | Gm14358 | predicted gene 14358 [Source:MGI Symbo | 0 | 0 | 0 | 0 | 0 | 0 |
| 24445 | ENSMUSG00000082082 | 0 | NA | NA | NA | NA | NA | Gm13230 | predicted gene 13230 [Source:MGI Symbo | 0 | 0 | 0 | 0 | 0 | 0 |
| 24446 | ENSMUSG00000082083 | 0 | NA | NA | NA | NA | NA | Gm14728 | predicted gene 14728 [Source:MGI Symbo | 0 | 0 | 0 | 0 | 0 | 0 |
| 24447 | ENSMUSG00000082084 | 0 | NA | NA | NA | NA | NA | Usp17le-ps | ubiquitin specific peptidase 17-like E, pseu | 0 | 0 | 0 | 0 | 0 | 0 |
| 24448 | ENSMUSG00000082085 | 0 | NA | NA | NA | NA | NA | Gm6441 | predicted gene 6441 [Source:MGI Symbol | 0 | 0 | 0 | 0 | 0 | 0 |
| 24449 | ENSMUSG00000082087 | 0 | NA | NA | NA | NA | NA | Gm12138 | predicted gene 12138 [Source:MGI Symbo | 0 | 0 | 0 | 0 | 0 | 0 |
| 24450 | ENSMUSG00000082088 | 0.160662974 | 1.126924175 | 4.080472857 | 0.276174898 | 0.782413736 | NA | Gm15753 | predicted gene 15753 [Source:MGI Symbo | 0 | 0 | 0 | 0.964 | 0 | 0 |
| 24451 | ENSMUSG00000082089 | 0 | NA | NA | NA | NA | NA | Gm12431 | predicted gene 12431 [Source:MGI Symbo | 0 | 0 | 0 | 0 | 0 | 0 |
| 24452 | ENSMUSG00000082090 | 0.133850373 | -0.796629176 | 4.080472857 | -0.195229623 | 0.84521319 | NA | Gm16481 | predicted gene 16481 [Source:MGI Symbo | 0.8031 | 0 | 0 | 0 | 0 | 0 |
| 24453 | ENSMUSG00000082091 | 0 | NA | NA | NA | NA | NA | Gm13922 | predicted gene 13922 [Source:MGI Symbo | 0 | 0 | 0 | 0 | 0 | 0 |
| 24454 | ENSMUSG00000082092 | 0 | NA | NA | NA | NA | NA | Gm14665 | predicted gene 14665 [Source:MGI Symbo | 0 | 0 | 0 | 0 | 0 | 0 |
| 24455 | ENSMUSG00000082093 | 0 | NA | NA | NA | NA | NA | Gm15928 | predicted gene 15928 [Source:MGI Symbo | 0 | 0 | 0 | 0 | 0 | 0 |
| 24456 | ENSMUSG00000082095 | 0 | NA | NA | NA | NA | NA | Gm11991 | predicted gene 11991 [Source:MGI Symbo | 0 | 0 | 0 | 0 | 0 | 0 |
| 24457 | ENSMUSG00000082096 | 0 | NA | NA | NA | NA | NA | Gm13505 | predicted gene 13505 [Source:MGI Symbo | 0 | 0 | 0 | 0 | 0 | 0 |
| 24458 | ENSMUSG00000082097 | 0 | NA | NA | NA | NA | NA | Gm14039 | predicted gene 14039 [Source:MGI Symbo | 0 | 0 | 0 | 0 | 0 | 0 |
| 24459 | ENSMUSG00000082098 | 0 | NA | NA | NA | NA | NA | Gm15377 | predicted gene 15377 [Source:MGI Symbo | 0 | 0 | 0 | 0 | 0 | 0 |
| 24460 | ENSMUSG00000082099 | 0 | NA | NA | NA | NA | NA | Gm12013 | predicted gene 12013 [Source:MGI Symbo | 0 | 0 | 0 | 0 | 0 | 0 |
| 24461 | ENSMUSG00000082100 | 1.696886004 | -1.870516383 | 1.850956004 | -1.010567717 | 0.312223373 | NA | Glns-ps1 | glutamine synthetase pseudogene 1 [Sour | 0.8031 | 6.1675 | 1.0062 | 0 | 1.0906 | 1.1138 |
| 24462 | ENSMUSG00000082101 | 0 | NA | NA | NA | NA | NA | Slfn14 | schlafen 14 [Source:MGI Symbol;Acc:MGI | 0 | 0 | 0 | 0 | 0 | 0 |
| 24463 | ENSMUSG00000082103 | 0 | NA | NA | NA | NA | NA | Gm15320 | predicted gene 15320 [Source:MGI Symbo | 0 | 0 | 0 | 0 | 0 | 0 |
| 24464 | ENSMUSG00000082104 | 0 | NA | NA | NA | NA | NA | Gm15933 | predicted gene 15933 [Source:MGI Symbo | 0 | 0 | 0 | 0 | 0 | 0 |
| 24465 | ENSMUSG00000082106 | 0 | NA | NA | NA | NA | NA | Mageb16-ps | MAGE family member B16, pseudogene 1 | 0 | 0 | 0 | 0 | 0 | 0 |
| 24466 | ENSMUSG00000082107 | 0 | NA | NA | NA | NA | NA | Gm16216 | predicted gene 16216 [Source:MGI Symbo | 0 | 0 | 0 | 0 | 0 | 0 |
| 24467 | ENSMUSG00000082108 | 0.185640647 | 1.126924175 | 4.080472857 | 0.276174898 | 0.782413736 | NA | Gm13777 | predicted gene 13777 [Source:MGI Symbo | 0 | 0 | 0 | 0 | 0 | 1.1138 |
| 24468 | ENSMUSG00000082109 | 0.367410362 | 2.078255025 | 3.800070157 | 0.54689912 | 0.58444804 | NA | Rpl19-ps12 | ribosomal protein L19, pseudogene 12 [Sc | 0 | 0 | 0 | 0 | 1.0906 | 1.1138 |

| | | | | | | | | | | | | | | | |
|---|---|---|---|---|---|---|---|---|---|---|---|---|---|---|---|
| 24469 | ENSMUSG00000082110 | 0.328370405 | 0.16514316 | 4.003149536 | 0.041253308 | 0.967093956 | NA | Gm11196 | predicted gene 11196 [Source:MGI Symbo | 0 | 0 | 1.0062 | 0.964 | 0 | 0 |
| 24470 | ENSMUSG00000082111 | 0 | NA | NA | NA | NA | NA | Gm12479 | predicted gene 12479 [Source:MGI Symbo | 0 | 0 | 0 | 0 | 0 | 0 |
| 24471 | ENSMUSG00000082112 | 0 | NA | NA | NA | NA | NA | Gm14445 | predicted gene 14445 [Source:MGI Symbo | 0 | 0 | 0 | 0 | 0 | 0 |
| 24472 | ENSMUSG00000082113 | 0 | NA | NA | NA | NA | NA | Gm13915 | predicted gene 13915 [Source:MGI Symbo | 0 | 0 | 0 | 0 | 0 | 0 |
| 24473 | ENSMUSG00000082114 | 0 | NA | NA | NA | NA | NA | Gm13489 | predicted gene 13489 [Source:MGI Symbo | 0 | 0 | 0 | 0 | 0 | 0 |
| 24474 | ENSMUSG00000082115 | 0 | NA | NA | NA | NA | NA | Gm15490 | predicted gene 15490 [Source:MGI Symbo | 0 | 0 | 0 | 0 | 0 | 0 |
| 24475 | ENSMUSG00000082116 | 0.913882229 | 2.129306995 | 2.79764157 | 0.761107862 | 0.446592645 | NA | Gm12188 | predicted gene 12188 [Source:MGI Symbo | 0 | 1.0279 | 0 | 0 | 0 | 4.4554 |
| 24476 | ENSMUSG00000082117 | 0 | NA | NA | NA | NA | NA | Gm4994 | predicted gene 4994 [Source:MGI Symbol | 0 | 0 | 0 | 0 | 0 | 0 |
| 24477 | ENSMUSG00000082118 | 0.160662974 | 1.126924175 | 4.080472857 | 0.276174898 | 0.782413736 | NA | Gm13862 | predicted gene 13862 [Source:MGI Symbo | 0 | 0 | 0 | 0.964 | 0 | 0 |
| 24478 | ENSMUSG00000082119 | 0 | NA | NA | NA | NA | NA | Cyp2j14-ps | cytochrome P450, family 2, subfamily j, m | 0 | 0 | 0 | 0 | 0 | 0 |
| 24479 | ENSMUSG00000082120 | 0 | NA | NA | NA | NA | NA | Gm15720 | predicted gene 15720 [Source:MGI Symbo | 0 | 0 | 0 | 0 | 0 | 0 |
| 24480 | ENSMUSG00000082121 | 0 | NA | NA | NA | NA | NA | Gm8199 | predicted gene 8199 [Source:MGI Symbol | 0 | 0 | 0 | 0 | 0 | 0 |
| 24481 | ENSMUSG00000082122 | 0 | NA | NA | NA | NA | NA | Gm11809 | predicted gene 11809 [Source:MGI Symbo | 0 | 0 | 0 | 0 | 0 | 0 |
| 24482 | ENSMUSG00000082123 | 0 | NA | NA | NA | NA | NA | Gm16149 | predicted gene 16149 [Source:MGI Symbo | 0 | 0 | 0 | 0 | 0 | 0 |
| 24483 | ENSMUSG00000082125 | 0 | NA | NA | NA | NA | NA | Gm14679 | predicted gene 14679 [Source:MGI Symbo | 0 | 0 | 0 | 0 | 0 | 0 |
| 24484 | ENSMUSG00000082126 | 0 | NA | NA | NA | NA | NA | Gm14975 | predicted gene 14975 [Source:MGI Symbo | 0 | 0 | 0 | 0 | 0 | 0 |
| 24485 | ENSMUSG00000082127 | 51.75371569 | 0.106650305 | 0.302120214 | 0.353006188 | 0.724083801 | 0.944786345 | Gm13577 | predicted gene 13577 [Source:MGI Symbo | 49.792 | 53.452 | 46.287 | 53.019 | 55.622 | 52.351 |
| 24486 | ENSMUSG00000082128 | 0.17131964 | -0.796629176 | 4.080472857 | -0.195229623 | 0.84521319 | NA | Gm11824 | predicted gene 11824 [Source:MGI Symbo | 0 | 1.0279 | 0 | 0 | 0 | 0 |
| 24487 | ENSMUSG00000082129 | 0.497389803 | 1.101799453 | 3.439268845 | 0.320358629 | 0.748696483 | NA | Gm14502 | predicted gene 14502 [Source:MGI Symbo | 0.8031 | 0 | 0 | 0 | 2.1812 | 0 |
| 24488 | ENSMUSG00000082130 | 0.513958919 | -2.394274821 | 3.756096013 | -0.637437066 | 0.523840191 | NA | Gm15799 | predicted gene 15799 [Source:MGI Symbo | 0 | 3.0838 | 0 | 0 | 0 | 0 |
| 24489 | ENSMUSG00000082132 | 0 | NA | NA | NA | NA | NA | Gm13541 | predicted gene 13541 [Source:MGI Symbo | 0 | 0 | 0 | 0 | 0 | 0 |
| 24490 | ENSMUSG00000082134 | 0 | NA | NA | NA | NA | NA | Gm15236 | predicted gene 15236 [Source:MGI Symbo | 0 | 0 | 0 | 0 | 0 | 0 |
| 24491 | ENSMUSG00000082135 | 0 | NA | NA | NA | NA | NA | Gm14988 | predicted gene 14988 [Source:MGI Symbo | 0 | 0 | 0 | 0 | 0 | 0 |
| 24492 | ENSMUSG00000082136 | 0 | NA | NA | NA | NA | NA | Gm15660 | predicted gene 15660 [Source:MGI Symbo | 0 | 0 | 0 | 0 | 0 | 0 |
| 24493 | ENSMUSG00000082138 | 0 | NA | NA | NA | NA | NA | Gm11905 | predicted gene 11905 [Source:MGI Symbo | 0 | 0 | 0 | 0 | 0 | 0 |
| 24494 | ENSMUSG00000082139 | 0 | NA | NA | NA | NA | NA | Gm16463 | predicted gene 16463 [Source:MGI Symbo | 0 | 0 | 0 | 0 | 0 | 0 |
| 24495 | ENSMUSG00000082140 | 0 | NA | NA | NA | NA | NA | Gm13094 | predicted gene 13094 [Source:MGI Symbo | 0 | 0 | 0 | 0 | 0 | 0 |
| 24496 | ENSMUSG00000082141 | 1.04956753 | 2.382147202 | 2.398712306 | 0.993094168 | 0.320664066 | NA | Gm11212 | predicted gene 11212 [Source:MGI Symbo | 0 | 1.0279 | 0 | 1.928 | 0 | 3.3415 |
| 24497 | ENSMUSG00000082142 | 0 | NA | NA | NA | NA | NA | Gm13518 | predicted gene 13518 [Source:MGI Symbo | 0 | 0 | 0 | 0 | 0 | 0 |
| 24498 | ENSMUSG00000082143 | 0 | NA | NA | NA | NA | NA | Gm12864 | predicted gene 12864 [Source:MGI Symbo | 0 | 0 | 0 | 0 | 0 | 0 |
| 24499 | ENSMUSG00000082144 | 2.610541415 | -0.310188794 | 1.444827487 | -0.214689156 | 0.830009678 | NA | Gm12788 | predicted gene 12788 [Source:MGI Symbo | 5.6217 | 1.0279 | 2.0125 | 4.8199 | 2.1812 | 0 |
| 24500 | ENSMUSG00000082145 | 0 | NA | NA | NA | NA | NA | Gm12312 | predicted gene 12312 [Source:MGI Symbo | 0 | 0 | 0 | 0 | 0 | 0 |
| 24501 | ENSMUSG00000082146 | 0 | NA | NA | NA | NA | NA | Rpl23a-ps13 | ribosomal protein L23A, pseudogene 13 [S | 0 | 0 | 0 | 0 | 0 | 0 |
| 24502 | ENSMUSG00000082147 | 0 | NA | NA | NA | NA | NA | Gm14570 | predicted gene 14570 [Source:MGI Symbo | 0 | 0 | 0 | 0 | 0 | 0 |
| 24503 | ENSMUSG00000082148 | 0.181769715 | 1.126924175 | 4.080472857 | 0.276174898 | 0.782413736 | NA | Gm12266 | predicted gene 12266 [Source:MGI Symbo | 0 | 0 | 0 | 0 | 1.0906 | 0 |
| 24504 | ENSMUSG00000082149 | 0 | NA | NA | NA | NA | NA | Gm13002 | predicted gene 13002 [Source:MGI Symbo | 0 | 0 | 0 | 0 | 0 | 0 |
| 24505 | ENSMUSG00000082151 | 0 | NA | NA | NA | NA | NA | Gm11810 | predicted gene 11810 [Source:MGI Symbo | 0 | 0 | 0 | 0 | 0 | 0 |
| 24506 | ENSMUSG00000082152 | 0.321325949 | 1.911575114 | 4.048267893 | 0.472195804 | 0.636787034 | NA | Gm13655 | predicted gene 13655 [Source:MGI Symbo | 0 | 0 | 0 | 1.928 | 0 | 0 |
| 24507 | ENSMUSG00000082154 | 1.543345125 | 2.014016556 | 1.814397526 | 1.110019457 | 0.266990642 | NA | Gm16464 | predicted gene 16464 [Source:MGI Symbo | 0.8031 | 0 | 1.0062 | 1.928 | 2.1812 | 3.3415 |
| 24508 | ENSMUSG00000082155 | 0 | NA | NA | NA | NA | NA | Gm7164 | predicted gene 7164 [Source:MGI Symbol | 0 | 0 | 0 | 0 | 0 | 0 |
| 24509 | ENSMUSG00000082156 | 0 | NA | NA | NA | NA | NA | Gm12008 | predicted gene 12008 [Source:MGI Symbo | 0 | 0 | 0 | 0 | 0 | 0 |
| 24510 | ENSMUSG00000082157 | 0 | NA | NA | NA | NA | NA | Gm12038 | predicted gene 12038 [Source:MGI Symbo | 0 | 0 | 0 | 0 | 0 | 0 |
| 24511 | ENSMUSG00000082158 | 0 | NA | NA | NA | NA | NA | Gm14116 | predicted gene 14116 [Source:MGI Symbo | 0 | 0 | 0 | 0 | 0 | 0 |
| 24512 | ENSMUSG00000082159 | 0 | NA | NA | NA | NA | NA | Gm7910 | predicted gene 7910 [Source:MGI Symbol | 0 | 0 | 0 | 0 | 0 | 0 |
| 24513 | ENSMUSG00000082160 | 0 | NA | NA | NA | NA | NA | Gm11578 | predicted gene 11578 [Source:MGI Symbo | 0 | 0 | 0 | 0 | 0 | 0 |
| 24514 | ENSMUSG00000082161 | 0 | NA | NA | NA | NA | NA | Gm11350 | predicted gene 11350 [Source:MGI Symbo | 0 | 0 | 0 | 0 | 0 | 0 |
| 24515 | ENSMUSG00000082162 | 0 | NA | NA | NA | NA | NA | Gm14701 | predicted gene 14701 [Source:MGI Symbo | 0 | 0 | 0 | 0 | 0 | 0 |
| 24516 | ENSMUSG00000082163 | 3.681111418 | -0.74216544 | 1.280893901 | -0.579412112 | 0.562311134 | NA | Gm14276 | predicted gene 14276 [Source:MGI Symbo | 5.6217 | 8.2233 | 0 | 3.8559 | 3.2719 | 1.1138 |
| 24517 | ENSMUSG00000082164 | 0 | NA | NA | NA | NA | NA | Gm13360 | predicted gene 13360 [Source:MGI Symbo | 0 | 0 | 0 | 0 | 0 | 0 |
| 24518 | ENSMUSG00000082165 | 0 | NA | NA | NA | NA | NA | Gm7091 | predicted gene 7091 [Source:MGI Symbol | 0 | 0 | 0 | 0 | 0 | 0 |
| 24519 | ENSMUSG00000082167 | 0 | NA | NA | NA | NA | NA | Gm14360 | predicted gene 14360 [Source:MGI Symbo | 0 | 0 | 0 | 0 | 0 | 0 |
| 24520 | ENSMUSG00000082168 | 0 | NA | NA | NA | NA | NA | Gm14656 | predicted gene 14656 [Source:MGI Symbo | 0 | 0 | 0 | 0 | 0 | 0 |
| 24521 | ENSMUSG00000082169 | 0 | NA | NA | NA | NA | NA | Gm12369 | predicted gene 12369 [Source:MGI Symbo | 0 | 0 | 0 | 0 | 0 | 0 |
| 24522 | ENSMUSG00000082170 | 0 | NA | NA | NA | NA | NA | Gm8617 | predicted gene 8617 [Source:MGI Symbol | 0 | 0 | 0 | 0 | 0 | 0 |
| 24523 | ENSMUSG00000082171 | 0 | NA | NA | NA | NA | NA | Rpl27-ps2 | ribosomal protein L27, pseudogene 2 [Sou | 0 | 0 | 0 | 0 | 0 | 0 |
| 24524 | ENSMUSG00000082172 | 0 | NA | NA | NA | NA | NA | Gm14835 | predicted gene 14835 [Source:MGI Symbo | 0 | 0 | 0 | 0 | 0 | 0 |

| | | | | | | | | | | | | | | | |
|---|---|---|---|---|---|---|---|---|---|---|---|---|---|---|---|
| 24525 | ENSMUSG00000082173 | 0.167707431 | -0.796629176 | 4.080472857 | -0.195229623 | 0.84521319 | NA | Mup-ps10 | major urinary protein, pseudogene 10 [So | 0 | 0 | 1.0062 | 0 | 0 | 0 |
| 24526 | ENSMUSG00000082174 | 0 | NA | NA | NA | NA | NA | Gm14877 | predicted gene 14877 [Source:MGI Symbo | 0 | 0 | 0 | 0 | 0 | 0 |
| 24527 | ENSMUSG00000082175 | 0 | NA | NA | NA | NA | NA | Gm14563 | predicted gene 14563 [Source:MGI Symbo | 0 | 0 | 0 | 0 | 0 | 0 |
| 24528 | ENSMUSG00000082176 | 0 | NA | NA | NA | NA | NA | Gm13163 | predicted gene 13163 [Source:MGI Symbo | 0 | 0 | 0 | 0 | 0 | 0 |
| 24529 | ENSMUSG00000082177 | 0 | NA | NA | NA | NA | NA | Gm16005 | predicted gene 16005 [Source:MGI Symbo | 0 | 0 | 0 | 0 | 0 | 0 |
| 24530 | ENSMUSG00000082178 | 0 | NA | NA | NA | NA | NA | Gm11307 | predicted gene 11307 [Source:MGI Symbo | 0 | 0 | 0 | 0 | 0 | 0 |
| 24531 | ENSMUSG00000082179 | 0 | NA | NA | NA | NA | NA | Gm11407 | predicted gene 11407 [Source:MGI Symbo | 0 | 0 | 0 | 0 | 0 | 0 |
| 24532 | ENSMUSG00000082180 | 0.52105551 | -0.812028029 | 3.385364865 | -0.239864257 | 0.810435491 | NA | Gm12230 | predicted gene 12230 [Source:MGI Symbo | 0 | 0 | 2.0125 | 0 | 0 | 1.1138 |
| 24533 | ENSMUSG00000082181 | 0 | NA | NA | NA | NA | NA | Gm12252 | predicted gene 12252 [Source:MGI Symbo | 0 | 0 | 0 | 0 | 0 | 0 |
| 24534 | ENSMUSG00000082183 | 0.17131964 | -0.796629176 | 4.080472857 | -0.195229623 | 0.84521319 | NA | Gm15994 | predicted gene 15994 [Source:MGI Symbo | 0 | 1.0279 | 0 | 0 | 0 | 0 |
| 24535 | ENSMUSG00000082185 | 0 | NA | NA | NA | NA | NA | Gm12428 | predicted gene 12428 [Source:MGI Symbo | 0 | 0 | 0 | 0 | 0 | 0 |
| 24536 | ENSMUSG00000082186 | 0.713920574 | 0.125792037 | 3.028037396 | 0.041542432 | 0.966863466 | NA | Gm12403 | predicted gene 12403 [Source:MGI Symbo | 0 | 2.0558 | 0 | 0 | 0 | 2.2277 |
| 24537 | ENSMUSG00000082187 | 0.160662974 | 1.126924175 | 4.080472857 | 0.276174898 | 0.782413736 | NA | Gm13670 | predicted gene 13670 [Source:MGI Symbo | 0 | 0 | 0 | 0.964 | 0 | 0 |
| 24538 | ENSMUSG00000082188 | 0 | NA | NA | NA | NA | NA | Gm2768 | predicted gene 2768 [Source:MGI Symbol | 0 | 0 | 0 | 0 | 0 | 0 |
| 24539 | ENSMUSG00000082189 | 0 | NA | NA | NA | NA | NA | Gm4784 | predicted gene 4784 [Source:MGI Symbol | 0 | 0 | 0 | 0 | 0 | 0 |
| 24540 | ENSMUSG00000082190 | 0.181769715 | 1.126924175 | 4.080472857 | 0.276174898 | 0.782413736 | NA | Gm13641 | predicted gene 13641 [Source:MGI Symbo | 0 | 0 | 0 | 0 | 1.0906 | 0 |
| 24541 | ENSMUSG00000082191 | 0 | NA | NA | NA | NA | NA | Gm5166 | predicted pseudogene 5166 [Source:MGI | 0 | 0 | 0 | 0 | 0 | 0 |
| 24542 | ENSMUSG00000082192 | 0.160662974 | 1.126924175 | 4.080472857 | 0.276174898 | 0.782413736 | NA | Gm14719 | predicted gene 14719 [Source:MGI Symbo | 0 | 0 | 0 | 0.964 | 0 | 0 |
| 24543 | ENSMUSG00000082193 | 0.356960287 | 0.16514264 | 3.880370363 | 0.042558474 | 0.966053499 | NA | Rpl5-ps1 | ribosomal protein L5, pseudogene 1 [Sour | 0 | 1.0279 | 0 | 0 | 0 | 1.1138 |
| 24544 | ENSMUSG00000082194 | 0 | NA | NA | NA | NA | NA | Gm12444 | predicted gene 12444 [Source:MGI Symbo | 0 | 0 | 0 | 0 | 0 | 0 |
| 24545 | ENSMUSG00000082195 | 0.301557804 | -1.64335916 | 4.042814367 | -0.406488899 | 0.684383412 | NA | Gm13034 | predicted gene 13034 [Source:MGI Symbo | 0.8031 | 0 | 1.0062 | 0 | 0 | 0 |
| 24546 | ENSMUSG00000082196 | 0 | NA | NA | NA | NA | NA | Gm14231 | predicted gene 14231 [Source:MGI Symbo | 0 | 0 | 0 | 0 | 0 | 0 |
| 24547 | ENSMUSG00000082197 | 0 | NA | NA | NA | NA | NA | Gm8577 | predicted gene 8577 [Source:MGI Symbol | 0 | 0 | 0 | 0 | 0 | 0 |
| 24548 | ENSMUSG00000082198 | 0 | NA | NA | NA | NA | NA | Gm14504 | predicted gene 14504 [Source:MGI Symbo | 0 | 0 | 0 | 0 | 0 | 0 |
| 24549 | ENSMUSG00000082199 | 0 | NA | NA | NA | NA | NA | Gm15763 | predicted gene 15763 [Source:MGI Symbo | 0 | 0 | 0 | 0 | 0 | 0 |
| 24550 | ENSMUSG00000082200 | 0 | NA | NA | NA | NA | NA | Gm12871 | predicted gene 12871 [Source:MGI Symbo | 0 | 0 | 0 | 0 | 0 | 0 |
| 24551 | ENSMUSG00000082202 | 0 | NA | NA | NA | NA | NA | Gm4910 | predicted pseudogene 4910 [Source:MGI | 0 | 0 | 0 | 0 | 0 | 0 |
| 24552 | ENSMUSG00000082203 | 0 | NA | NA | NA | NA | NA | Gm15636 | predicted gene 15636 [Source:MGI Symbo | 0 | 0 | 0 | 0 | 0 | 0 |
| 24553 | ENSMUSG00000082204 | 0 | NA | NA | NA | NA | NA | Gm7722 | predicted gene 7722 [Source:MGI Symbol | 0 | 0 | 0 | 0 | 0 | 0 |
| 24554 | ENSMUSG00000082205 | 0 | NA | NA | NA | NA | NA | Gm7058 | predicted gene 7058 [Source:MGI Symbol | 0 | 0 | 0 | 0 | 0 | 0 |
| 24555 | ENSMUSG00000082207 | 0.17131964 | -0.796629176 | 4.080472857 | -0.195229623 | 0.84521319 | NA | Gm12998 | predicted gene 12998 [Source:MGI Symbo | 0 | 1.0279 | 0 | 0 | 0 | 0 |
| 24556 | ENSMUSG00000082208 | 0.133850373 | -0.796629176 | 4.080472857 | -0.195229623 | 0.84521319 | NA | Gm15379 | predicted gene 15379 [Source:MGI Symbo | 0.8031 | 0 | 0 | 0 | 0 | 0 |
| 24557 | ENSMUSG00000082210 | 0 | NA | NA | NA | NA | NA | Gm14382 | predicted gene 14382 [Source:MGI Symbo | 0 | 0 | 0 | 0 | 0 | 0 |
| 24558 | ENSMUSG00000082211 | 0 | NA | NA | NA | NA | NA | Defa27 | defensin, alpha, 27 [Source:MGI Symbol;A | 0 | 0 | 0 | 0 | 0 | 0 |
| 24559 | ENSMUSG00000082212 | 0.17131964 | -0.796629176 | 4.080472857 | -0.195229623 | 0.84521319 | NA | Gm15826 | predicted gene 15826 [Source:MGI Symbo | 0 | 1.0279 | 0 | 0 | 0 | 0 |
| 24560 | ENSMUSG00000082213 | 0.331982614 | 0.16514309 | 3.986751476 | 0.041422971 | 0.966958701 | NA | Gm12102 | predicted gene 12102 [Source:MGI Symbo | 0 | 1.0279 | 0 | 0.964 | 0 | 0 |
| 24561 | ENSMUSG00000082214 | 0 | NA | NA | NA | NA | NA | Gm15009 | predicted gene 15009 [Source:MGI Symbo | 0 | 0 | 0 | 0 | 0 | 0 |
| 24562 | ENSMUSG00000082215 | 0 | NA | NA | NA | NA | NA | Gm14314 | predicted gene 14314 [Source:MGI Symbo | 0 | 0 | 0 | 0 | 0 | 0 |
| 24563 | ENSMUSG00000082216 | 0 | NA | NA | NA | NA | NA | Olfr271-ps1 | olfactory receptor 271, pseudogene 1 [So | 0 | 0 | 0 | 0 | 0 | 0 |
| 24564 | ENSMUSG00000082217 | 0 | NA | NA | NA | NA | NA | Gm1401 | predicted gene 1401 [Source:MGI Symbol | 0 | 0 | 0 | 0 | 0 | 0 |
| 24565 | ENSMUSG00000082218 | 0 | NA | NA | NA | NA | NA | Gm13602 | predicted gene 13602 [Source:MGI Symbo | 0 | 0 | 0 | 0 | 0 | 0 |
| 24566 | ENSMUSG00000082220 | 0 | NA | NA | NA | NA | NA | Gm12943 | predicted gene 12943 [Source:MGI Symbo | 0 | 0 | 0 | 0 | 0 | 0 |
| 24567 | ENSMUSG00000082222 | 0 | NA | NA | NA | NA | NA | Gm14865 | predicted gene 14865 [Source:MGI Symbo | 0 | 0 | 0 | 0 | 0 | 0 |
| 24568 | ENSMUSG00000082223 | 0 | NA | NA | NA | NA | NA | Gm15079 | predicted gene 15079 [Source:MGI Symbo | 0 | 0 | 0 | 0 | 0 | 0 |
| 24569 | ENSMUSG00000082225 | 0 | NA | NA | NA | NA | NA | Gm13728 | predicted gene 13728 [Source:MGI Symbo | 0 | 0 | 0 | 0 | 0 | 0 |
| 24570 | ENSMUSG00000082226 | 2.194818759 | 1.369342142 | 1.434392773 | 0.954649359 | 0.339755047 | NA | Gm715 | predicted gene 715 [Source:MGI Symbol;A | 1.6062 | 1.0279 | 1.0062 | 2.8919 | 2.1812 | 4.4554 |
| 24571 | ENSMUSG00000082227 | 0 | NA | NA | NA | NA | NA | Olfr1144-ps1 | olfactory receptor 1144, pseudogene 1 [So | 0 | 0 | 0 | 0 | 0 | 0 |
| 24572 | ENSMUSG00000082228 | 0 | NA | NA | NA | NA | NA | Gm8344 | predicted gene 8344 [Source:MGI Symbol | 0 | 0 | 0 | 0 | 0 | 0 |
| 24573 | ENSMUSG00000082229 | 0 | NA | NA | NA | NA | NA | Nap1l2 | nucleosome assembly protein 1-like 2 [Sou | 0 | 0 | 0 | 0 | 0 | 0 |
| 24574 | ENSMUSG00000082233 | 0 | NA | NA | NA | NA | NA | Gm6506 | predicted gene 6506 [Source:MGI Symbol | 0 | 0 | 0 | 0 | 0 | 0 |
| 24575 | ENSMUSG00000082234 | 0 | NA | NA | NA | NA | NA | Gm16221 | predicted gene 16221 [Source:MGI Symbo | 0 | 0 | 0 | 0 | 0 | 0 |
| 24576 | ENSMUSG00000082235 | 0 | NA | NA | NA | NA | NA | Gm13326 | predicted gene 13326 [Source:MGI Symbo | 0 | 0 | 0 | 0 | 0 | 0 |
| 24577 | ENSMUSG00000082236 | 0 | NA | NA | NA | NA | NA | Gm11355 | predicted gene 11355 [Source:MGI Symbo | 0 | 0 | 0 | 0 | 0 | 0 |
| 24578 | ENSMUSG00000082237 | 0 | NA | NA | NA | NA | NA | Olfr1172-ps1 | olfactory receptor 1172, pseudogene 1 [So | 0 | 0 | 0 | 0 | 0 | 0 |
| 24579 | ENSMUSG00000082238 | 0 | NA | NA | NA | NA | NA | Gm12383 | predicted gene 12383 [Source:MGI Symbo | 0 | 0 | 0 | 0 | 0 | 0 |
| 24580 | ENSMUSG00000082239 | 0 | NA | NA | NA | NA | NA | Gm14234 | predicted gene 14234 [Source:MGI Symbo | 0 | 0 | 0 | 0 | 0 | 0 |

| | | | | | | | | | | | | | | | |
|---|---|---|---|---|---|---|---|---|---|---|---|---|---|---|---|
| 24581 | ENSMUSG00000082240 | 0 | NA | NA | NA | NA | NA | Gm14380 | predicted gene 14380 [Source:MGI Symbc | 0 | 0 | 0 | 0 | 0 | 0 |
| 24582 | ENSMUSG00000082241 | 0.321325949 | 1.911575114 | 4.048267893 | 0.472195804 | 0.636787034 | NA | Gng5-ps | G protein subunit gamma 5, pseudogene [ | 0 | 0 | 0 | 1.928 | 0 | 0 |
| 24583 | ENSMUSG00000082242 | 0 | NA | NA | NA | NA | NA | Gm12443 | predicted gene 12443 [Source:MGI Symbc | 0 | 0 | 0 | 0 | 0 | 0 |
| 24584 | ENSMUSG00000082243 | 0 | NA | NA | NA | NA | NA | Gm3810 | predicted gene 3810 [Source:MGI Symbol | 0 | 0 | 0 | 0 | 0 | 0 |
| 24585 | ENSMUSG00000082244 | 0 | NA | NA | NA | NA | NA | Gm8503 | predicted gene 8503 [Source:MGI Symbol | 0 | 0 | 0 | 0 | 0 | 0 |
| 24586 | ENSMUSG00000082245 | 0 | NA | NA | NA | NA | NA | Gm6292 | predicted gene 6292 [Source:MGI Symbol | 0 | 0 | 0 | 0 | 0 | 0 |
| 24587 | ENSMUSG00000082246 | 0.17131964 | -0.796629176 | 4.080472857 | -0.195229623 | 0.84521319 | NA | Gm13047 | predicted gene 13047 [Source:MGI Symbc | 0 | 1.0279 | 0 | 0 | 0 | 0 |
| 24588 | ENSMUSG00000082247 | 0 | NA | NA | NA | NA | NA | Gm16074 | predicted gene 16074 [Source:MGI Symbc | 0 | 0 | 0 | 0 | 0 | 0 |
| 24589 | ENSMUSG00000082248 | 9.727692999 | -1.300854498 | 0.711279216 | -1.828894292 | 0.067415446 | 0.373457298 | Gm13161 | predicted gene 13161 [Source:MGI Symbc | 16.062 | 11.307 | 14.087 | 4.8199 | 7.6343 | 4.4554 |
| 24590 | ENSMUSG00000082249 | 0 | NA | NA | NA | NA | NA | Gm12997 | predicted gene 12997 [Source:MGI Symbc | 0 | 0 | 0 | 0 | 0 | 0 |
| 24591 | ENSMUSG00000082250 | 0 | NA | NA | NA | NA | NA | Gm12181 | predicted gene 12181 [Source:MGI Symbc | 0 | 0 | 0 | 0 | 0 | 0 |
| 24592 | ENSMUSG00000082251 | 0 | NA | NA | NA | NA | NA | Gm13438 | predicted gene 13438 [Source:MGI Symbc | 0 | 0 | 0 | 0 | 0 | 0 |
| 24593 | ENSMUSG00000082252 | 0 | NA | NA | NA | NA | NA | Gm11204 | predicted gene 11204 [Source:MGI Symbc | 0 | 0 | 0 | 0 | 0 | 0 |
| 24594 | ENSMUSG00000082253 | 0 | NA | NA | NA | NA | NA | Gm14639 | predicted gene 14639 [Source:MGI Symbc | 0 | 0 | 0 | 0 | 0 | 0 |
| 24595 | ENSMUSG00000082254 | 0 | NA | NA | NA | NA | NA | Gm11922 | predicted gene 11922 [Source:MGI Symbc | 0 | 0 | 0 | 0 | 0 | 0 |
| 24596 | ENSMUSG00000082255 | 0 | NA | NA | NA | NA | NA | Gm11822 | predicted gene 11822 [Source:MGI Symbc | 0 | 0 | 0 | 0 | 0 | 0 |
| 24597 | ENSMUSG00000082256 | 0 | NA | NA | NA | NA | NA | Gm12287 | predicted gene 12287 [Source:MGI Symbc | 0 | 0 | 0 | 0 | 0 | 0 |
| 24598 | ENSMUSG00000082258 | 0 | NA | NA | NA | NA | NA | Gm15289 | predicted gene 15289 [Source:MGI Symbc | 0 | 0 | 0 | 0 | 0 | 0 |
| 24599 | ENSMUSG00000082259 | 0 | NA | NA | NA | NA | NA | Gm13808 | predicted gene 13808 [Source:MGI Symbc | 0 | 0 | 0 | 0 | 0 | 0 |
| 24600 | ENSMUSG00000082260 | 1.913003595 | 0.345665838 | 1.64151462 | 0.210577374 | 0.833217069 | NA | Gm7429 | predicted pseudogene 7429 [Source:MGI | 0 | 4.1117 | 1.0062 | 1.928 | 1.0906 | 3.3415 |
| 24601 | ENSMUSG00000082261 | 0 | NA | NA | NA | NA | NA | Gm14999 | predicted gene 14999 [Source:MGI Symbc | 0 | 0 | 0 | 0 | 0 | 0 |
| 24602 | ENSMUSG00000082262 | 0.167707431 | -0.796629176 | 4.080472857 | -0.195229623 | 0.84521319 | NA | Gm14730 | predicted gene 14730 [Source:MGI Symbc | 0 | 0 | 1.0062 | 0 | 0 | 0 |
| 24603 | ENSMUSG00000082263 | 0 | NA | NA | NA | NA | NA | Gm11188 | predicted gene 11188 [Source:MGI Symbc | 0 | 0 | 0 | 0 | 0 | 0 |
| 24604 | ENSMUSG00000082264 | 0.935330317 | 1.257565026 | 2.487133718 | 0.505628232 | 0.613117658 | NA | Gm12799 | predicted gene 12799 [Source:MGI Symbc | 1.6062 | 0 | 0 | 2.8919 | 0 | 1.1138 |
| 24605 | ENSMUSG00000082265 | 1.665199708 | -1.264794106 | 2.000078561 | -0.632372213 | 0.527143687 | NA | Gm15547 | predicted gene 15547 [Source:MGI Symbc | 4.0155 | 3.0838 | 0 | 2.8919 | 0 | 0 |
| 24606 | ENSMUSG00000082266 | 0.267700746 | -1.497481649 | 4.057001436 | -0.369110456 | 0.712045396 | NA | Gm12303 | predicted gene 12303 [Source:MGI Symbc | 1.6062 | 0 | 0 | 0 | 0 | 0 |
| 24607 | ENSMUSG00000082267 | 0 | NA | NA | NA | NA | NA | Gm12467 | predicted gene 12467 [Source:MGI Symbc | 0 | 0 | 0 | 0 | 0 | 0 |
| 24608 | ENSMUSG00000082268 | 0 | NA | NA | NA | NA | NA | Vmn1r-ps12 | vomeronasal 1 receptor, pseudogene 120 | 0 | 0 | 0 | 0 | 0 | 0 |
| 24609 | ENSMUSG00000082269 | 0 | NA | NA | NA | NA | NA | Gm14676 | predicted gene 14676 [Source:MGI Symbc | 0 | 0 | 0 | 0 | 0 | 0 |
| 24610 | ENSMUSG00000082270 | 0 | NA | NA | NA | NA | NA | Gm14375 | predicted gene 14375 [Source:MGI Symbc | 0 | 0 | 0 | 0 | 0 | 0 |
| 24611 | ENSMUSG00000082271 | 23.09620794 | -0.203962816 | 0.51280862 | -0.397736715 | 0.69082427 | 0.937606823 | Gm12217 | predicted gene 12217 [Source:MGI Symbc | 14.456 | 35.977 | 24.15 | 29.883 | 19.631 | 14.48 |
| 24612 | ENSMUSG00000082272 | 0.17131964 | -0.796629176 | 4.080472857 | -0.195229623 | 0.84521319 | NA | Gm11675 | predicted gene 11675 [Source:MGI Symbc | 0 | 1.0279 | 0 | 0 | 0 | 0 |
| 24613 | ENSMUSG00000082274 | 0.967093722 | 0.236383879 | 2.096580567 | 0.112747338 | 0.91023087 | NA | Gm14026 | predicted gene 14026 [Source:MGI Symbc | 1.6062 | 1.0279 | 0 | 0.964 | 1.0906 | 1.1138 |
| 24614 | ENSMUSG00000082275 | 0 | NA | NA | NA | NA | NA | Gm14854 | predicted gene 14854 [Source:MGI Symbc | 0 | 0 | 0 | 0 | 0 | 0 |
| 24615 | ENSMUSG00000082276 | 0.335414862 | -1.779047242 | 4.030804179 | -0.441362855 | 0.658950329 | NA | Gm13885 | predicted gene 13885 [Source:MGI Symbc | 0 | 0 | 2.0125 | 0 | 0 | 0 |
| 24616 | ENSMUSG00000082277 | 0 | NA | NA | NA | NA | NA | Gm15807 | predicted gene 15807 [Source:MGI Symbc | 0 | 0 | 0 | 0 | 0 | 0 |
| 24617 | ENSMUSG00000082278 | 0 | NA | NA | NA | NA | NA | Gm13471 | predicted gene 13471 [Source:MGI Symbc | 0 | 0 | 0 | 0 | 0 | 0 |
| 24618 | ENSMUSG00000082279 | 0.819128932 | -3.070895205 | 2.828495588 | -1.085699132 | 0.277612125 | NA | Gm12870 | predicted gene 12870 [Source:MGI Symbc | 0.8031 | 4.1117 | 0 | 0 | 0 | 0 |
| 24619 | ENSMUSG00000082280 | 0 | NA | NA | NA | NA | NA | Gm5847 | predicted gene 5847 [Source:MGI Symbol | 0 | 0 | 0 | 0 | 0 | 0 |
| 24620 | ENSMUSG00000082281 | 0 | NA | NA | NA | NA | NA | Gm4886 | predicted gene 4886 [Source:MGI Symbol | 0 | 0 | 0 | 0 | 0 | 0 |
| 24621 | ENSMUSG00000082283 | 0.185640647 | 1.126924175 | 4.080472857 | 0.276174898 | 0.782413736 | NA | Gm14472 | predicted gene 14472 [Source:MGI Symbc | 0 | 0 | 0 | 0 | 0 | 1.1138 |
| 24622 | ENSMUSG00000082284 | 20.67812518 | 0.005798401 | 0.49480699 | 0.011718511 | 0.990650195 | 0.99859176 | H3f3a-ps1 | H3.3 histone A, pseudogene 1 [Source:MC | 20.078 | 28.782 | 13.081 | 20.244 | 20.722 | 21.163 |
| 24623 | ENSMUSG00000082285 | 0 | NA | NA | NA | NA | NA | Gm14069 | predicted gene 14069 [Source:MGI Symbc | 0 | 0 | 0 | 0 | 0 | 0 |
| 24624 | ENSMUSG00000082286 | 60.72008111 | -0.130407568 | 0.295307517 | -0.441599218 | 0.65877925 | 0.927599242 | Pisd-ps1 | phosphatidylserine decarboxylase, pseudo | 67.461 | 63.731 | 59.368 | 71.334 | 54.531 | 47.895 |
| 24625 | ENSMUSG00000082287 | 1.257762767 | -2.513989027 | 2.089651093 | -1.203066404 | 0.228950625 | NA | Gm15333 | predicted gene 15333 [Source:MGI Symbc | 2.4093 | 1.0279 | 3.0187 | 0 | 1.0906 | 0 |
| 24626 | ENSMUSG00000082288 | 0 | NA | NA | NA | NA | NA | Gm14755 | predicted gene 14755 [Source:MGI Symbc | 0 | 0 | 0 | 0 | 0 | 0 |
| 24627 | ENSMUSG00000082289 | 0 | NA | NA | NA | NA | NA | Gm15596 | predicted gene 15596 [Source:MGI Symbc | 0 | 0 | 0 | 0 | 0 | 0 |
| 24628 | ENSMUSG00000082291 | 0 | NA | NA | NA | NA | NA | Gm5762 | predicted gene 5762 [Source:MGI Symbol | 0 | 0 | 0 | 0 | 0 | 0 |
| 24629 | ENSMUSG00000082292 | 28.4394777 | -0.264655305 | 0.425548836 | -0.621915238 | 0.533997605 | 0.889524514 | Gm12250 | predicted gene 12250 [Source:MGI Symbc | 36.943 | 34.949 | 21.131 | 27.955 | 21.812 | 27.846 |
| 24630 | ENSMUSG00000082294 | 0 | NA | NA | NA | NA | NA | Gm12072 | predicted gene 12072 [Source:MGI Symbc | 0 | 0 | 0 | 0 | 0 | 0 |
| 24631 | ENSMUSG00000082295 | 0 | NA | NA | NA | NA | NA | Gm12133 | predicted gene 12133 [Source:MGI Symbc | 0 | 0 | 0 | 0 | 0 | 0 |
| 24632 | ENSMUSG00000082296 | 0 | NA | NA | NA | NA | NA | Gm11450 | predicted gene 11450 [Source:MGI Symbc | 0 | 0 | 0 | 0 | 0 | 0 |
| 24633 | ENSMUSG00000082297 | 0 | NA | NA | NA | NA | NA | Rhox3d-ps | reproductive homeobox 3D, pseudogene | 0 | 0 | 0 | 0 | 0 | 0 |
| 24634 | ENSMUSG00000082301 | 0 | NA | NA | NA | NA | NA | Gm14032 | predicted gene 14032 [Source:MGI Symbc | 0 | 0 | 0 | 0 | 0 | 0 |
| 24635 | ENSMUSG00000082302 | 0 | NA | NA | NA | NA | NA | Gm12385 | predicted gene 12385 [Source:MGI Symbc | 0 | 0 | 0 | 0 | 0 | 0 |
| 24636 | ENSMUSG00000082303 | 0 | NA | NA | NA | NA | NA | Gm13633 | predicted gene 13633 [Source:MGI Symbc | 0 | 0 | 0 | 0 | 0 | 0 |

| | | | | | | | | | | | | | | | |
|---|---|---|---|---|---|---|---|---|---|---|---|---|---|---|---|
| 24637 | ENSMUSG00000082304 | 0.342639279 | -1.805976596 | 4.028547714 | -0.448294702 | 0.653940524 | NA | Gm11198 | predicted gene 11198 [Source:MGI Symbo | 0 | 2.0558 | 0 | 0 | 0 | 0 |
| 24638 | ENSMUSG00000082305 | 1.508071833 | -1.724607796 | 2.048149871 | -0.842032031 | 0.399770024 | NA | Gm2420 | predicted gene 2420 [Source:MGI Symbol | 0.8031 | 6.1675 | 0 | 0.964 | 0 | 1.1138 |
| 24639 | ENSMUSG00000082306 | 0 | NA | NA | NA | NA | NA | Gm11358 | predicted gene 11358 [Source:MGI Symbo | 0 | 0 | 0 | 0 | 0 | 0 |
| 24640 | ENSMUSG00000082307 | 0.449470461 | -0.545136533 | 3.562105917 | -0.15303771 | 0.878368537 | NA | Gm13418 | predicted gene 13418 [Source:MGI Symbo | 1.6062 | 0 | 0 | 0 | 1.0906 | 0 |
| 24641 | ENSMUSG00000082308 | 0.321325949 | 1.911575114 | 4.048267893 | 0.472195804 | 0.636787034 | NA | Gm15770 | predicted gene 15770 [Source:MGI Symbo | 0 | 0 | 0 | 1.928 | 0 | 0 |
| 24642 | ENSMUSG00000082309 | 0 | NA | NA | NA | NA | NA | Gm12123 | predicted gene 12123 [Source:MGI Symbo | 0 | 0 | 0 | 0 | 0 | 0 |
| 24643 | ENSMUSG00000082310 | 0 | NA | NA | NA | NA | NA | Gm12904 | predicted gene 12904 [Source:MGI Symbo | 0 | 0 | 0 | 0 | 0 | 0 |
| 24644 | ENSMUSG00000082311 | 0 | NA | NA | NA | NA | NA | Gm14791 | predicted gene 14791 [Source:MGI Symbo | 0 | 0 | 0 | 0 | 0 | 0 |
| 24645 | ENSMUSG00000082312 | 0 | NA | NA | NA | NA | NA | Gm11996 | predicted gene 11996 [Source:MGI Symbo | 0 | 0 | 0 | 0 | 0 | 0 |
| 24646 | ENSMUSG00000082313 | 0 | NA | NA | NA | NA | NA | Gm11701 | predicted gene 11701 [Source:MGI Symbo | 0 | 0 | 0 | 0 | 0 | 0 |
| 24647 | ENSMUSG00000082314 | 0 | NA | NA | NA | NA | NA | Gm13296 | predicted gene 13296 [Source:MGI Symbo | 0 | 0 | 0 | 0 | 0 | 0 |
| 24648 | ENSMUSG00000082315 | 0.17131964 | -0.796629176 | 4.080472857 | -0.195229623 | 0.84521319 | NA | Gm16523 | predicted gene, 16523 [Source:MGI Symb | 0 | 1.0279 | 0 | 0 | 0 | 0 |
| 24649 | ENSMUSG00000082316 | 0 | NA | NA | NA | NA | NA | Vmn1r221 | vomeronasal 1 receptor 221 [Source:MGI | 0 | 0 | 0 | 0 | 0 | 0 |
| 24650 | ENSMUSG00000082317 | 0 | NA | NA | NA | NA | NA | Gm14933 | predicted gene 14933 [Source:MGI Symbo | 0 | 0 | 0 | 0 | 0 | 0 |
| 24651 | ENSMUSG00000082318 | 0 | NA | NA | NA | NA | NA | Gm14162 | predicted gene 14162 [Source:MGI Symbo | 0 | 0 | 0 | 0 | 0 | 0 |
| 24652 | ENSMUSG00000082319 | 1.3207255 | -1.513936874 | 2.0972413 | -0.721870618 | 0.470374027 | NA | Actr3-ps | actin related protein 3, pseudogene [Sour | 4.8186 | 1.0279 | 0 | 0.964 | 0 | 1.1138 |
| 24653 | ENSMUSG00000082320 | 0 | NA | NA | NA | NA | NA | Gm11877 | predicted gene 11877 [Source:MGI Symbo | 0 | 0 | 0 | 0 | 0 | 0 |
| 24654 | ENSMUSG00000082321 | 0.614004368 | 0.315622926 | 2.898299649 | 0.108899342 | 0.913282328 | NA | Gm14253 | predicted gene 14253 [Source:MGI Symbo | 1.6062 | 0 | 0 | 0.964 | 0 | 1.1138 |
| 24655 | ENSMUSG00000082322 | 0.181769715 | 1.126924175 | 4.080472857 | 0.276174898 | 0.782413736 | NA | Gm9164 | predicted gene 9164 [Source:MGI Symbol | 0 | 0 | 0 | 0 | 1.0906 | 0 |
| 24656 | ENSMUSG00000082323 | 0 | NA | NA | NA | NA | NA | Gm15314 | predicted gene 15314 [Source:MGI Symbo | 0 | 0 | 0 | 0 | 0 | 0 |
| 24657 | ENSMUSG00000082324 | 0 | NA | NA | NA | NA | NA | Gm14615 | predicted gene 14615 [Source:MGI Symbo | 0 | 0 | 0 | 0 | 0 | 0 |
| 24658 | ENSMUSG00000082325 | 0 | NA | NA | NA | NA | NA | Gm12642 | predicted gene 12642 [Source:MGI Symbo | 0 | 0 | 0 | 0 | 0 | 0 |
| 24659 | ENSMUSG00000082326 | 0 | NA | NA | NA | NA | NA | Gm14366 | predicted gene 14366 [Source:MGI Symbo | 0 | 0 | 0 | 0 | 0 | 0 |
| 24660 | ENSMUSG00000082327 | 0 | NA | NA | NA | NA | NA | Gm13801 | predicted gene 13801 [Source:MGI Symbo | 0 | 0 | 0 | 0 | 0 | 0 |
| 24661 | ENSMUSG00000082328 | 0.181769715 | 1.126924175 | 4.080472857 | 0.276174898 | 0.782413736 | NA | Gm13921 | predicted gene 13921 [Source:MGI Symbo | 0 | 0 | 0 | 0 | 1.0906 | 0 |
| 24662 | ENSMUSG00000082329 | 0 | NA | NA | NA | NA | NA | Gm14287 | predicted gene 14287 [Source:MGI Symbo | 0 | 0 | 0 | 0 | 0 | 0 |
| 24663 | ENSMUSG00000082330 | 0 | NA | NA | NA | NA | NA | Ldha-ps | lactate dehydrogenase A, pseudogene [Sc | 0 | 0 | 0 | 0 | 0 | 0 |
| 24664 | ENSMUSG00000082332 | 0 | NA | NA | NA | NA | NA | Gm13384 | predicted gene 13384 [Source:MGI Symbo | 0 | 0 | 0 | 0 | 0 | 0 |
| 24665 | ENSMUSG00000082333 | 0 | NA | NA | NA | NA | NA | Gm11984 | predicted gene 11984 [Source:MGI Symbo | 0 | 0 | 0 | 0 | 0 | 0 |
| 24666 | ENSMUSG00000082334 | 18.57118313 | -0.288101171 | 0.626356901 | -0.459963274 | 0.645542582 | 0.925643431 | Gm13244 | predicted gene 13244 [Source:MGI Symbo | 31.321 | 21.586 | 8.05 | 10.604 | 10.906 | 28.96 |
| 24667 | ENSMUSG00000082336 | 0 | NA | NA | NA | NA | NA | Gm8401 | predicted gene 8401 [Source:MGI Symbol | 0 | 0 | 0 | 0 | 0 | 0 |
| 24668 | ENSMUSG00000082337 | 0 | NA | NA | NA | NA | NA | Gm14723 | predicted gene 14723 [Source:MGI Symbo | 0 | 0 | 0 | 0 | 0 | 0 |
| 24669 | ENSMUSG00000082338 | 0 | NA | NA | NA | NA | NA | Gm12327 | predicted gene 12327 [Source:MGI Symbo | 0 | 0 | 0 | 0 | 0 | 0 |
| 24670 | ENSMUSG00000082339 | 0 | NA | NA | NA | NA | NA | Gm14028 | predicted gene 14028 [Source:MGI Symbo | 0 | 0 | 0 | 0 | 0 | 0 |
| 24671 | ENSMUSG00000082340 | 0 | NA | NA | NA | NA | NA | Gm11321 | predicted gene 11321 [Source:MGI Symbo | 0 | 0 | 0 | 0 | 0 | 0 |
| 24672 | ENSMUSG00000082341 | 0 | NA | NA | NA | NA | NA | Gm14143 | predicted gene 14143 [Source:MGI Symbo | 0 | 0 | 0 | 0 | 0 | 0 |
| 24673 | ENSMUSG00000082342 | 0 | NA | NA | NA | NA | NA | Gm5395 | predicted gene 5395 [Source:MGI Symbol | 0 | 0 | 0 | 0 | 0 | 0 |
| 24674 | ENSMUSG00000082343 | 0 | NA | NA | NA | NA | NA | Gm14404 | predicted gene 14404 [Source:MGI Symbo | 0 | 0 | 0 | 0 | 0 | 0 |
| 24675 | ENSMUSG00000082344 | 0 | NA | NA | NA | NA | NA | 4930415H17 | RIKEN cDNA 4930415H17 gene [Source:M | 0 | 0 | 0 | 0 | 0 | 0 |
| 24676 | ENSMUSG00000082345 | 0.339027071 | -1.79174149 | 3.919585585 | -0.457125237 | 0.647581032 | NA | Gm13622 | predicted gene 13622 [Source:MGI Symbo | 0 | 1.0279 | 1.0062 | 0 | 0 | 0 |
| 24677 | ENSMUSG00000082346 | 0 | NA | NA | NA | NA | NA | Gm6952 | predicted gene 6952 [Source:MGI Symbol | 0 | 0 | 0 | 0 | 0 | 0 |
| 24678 | ENSMUSG00000082347 | 0 | NA | NA | NA | NA | NA | Gm15036 | predicted gene 15036 [Source:MGI Symbo | 0 | 0 | 0 | 0 | 0 | 0 |
| 24679 | ENSMUSG00000082348 | 0.181769715 | 1.126924175 | 4.080472857 | 0.276174898 | 0.782413736 | NA | Gm12807 | predicted gene 12807 [Source:MGI Symbo | 0 | 0 | 0 | 0 | 1.0906 | 0 |
| 24680 | ENSMUSG00000082350 | 0 | NA | NA | NA | NA | NA | Gm14008 | predicted gene 14008 [Source:MGI Symbo | 0 | 0 | 0 | 0 | 0 | 0 |
| 24681 | ENSMUSG00000082351 | 0 | NA | NA | NA | NA | NA | Gm11260 | predicted gene 11260 [Source:MGI Symbo | 0 | 0 | 0 | 0 | 0 | 0 |
| 24682 | ENSMUSG00000082352 | 0 | NA | NA | NA | NA | NA | Gm13228 | predicted gene 13228 [Source:MGI Symbo | 0 | 0 | 0 | 0 | 0 | 0 |
| 24683 | ENSMUSG00000082353 | 0 | NA | NA | NA | NA | NA | Vmn1r-ps1 | vomeronasal 1 receptor, pseudogene 1 [S | 0 | 0 | 0 | 0 | 0 | 0 |
| 24684 | ENSMUSG00000082354 | 0 | NA | NA | NA | NA | NA | Gm378 | predicted pseudogene 378 [Source:MGI S | 0 | 0 | 0 | 0 | 0 | 0 |
| 24685 | ENSMUSG00000082356 | 0 | NA | NA | NA | NA | NA | Gm15565 | predicted gene 15565 [Source:MGI Symbo | 0 | 0 | 0 | 0 | 0 | 0 |
| 24686 | ENSMUSG00000082357 | 0 | NA | NA | NA | NA | NA | Gm13542 | predicted gene 13542 [Source:MGI Symbo | 0 | 0 | 0 | 0 | 0 | 0 |
| 24687 | ENSMUSG00000082358 | 0 | NA | NA | NA | NA | NA | Gm14990 | predicted gene 14990 [Source:MGI Symbo | 0 | 0 | 0 | 0 | 0 | 0 |
| 24688 | ENSMUSG00000082359 | 0 | NA | NA | NA | NA | NA | Mif-ps11 | macrophage migration inhibitory factor, p | 0 | 0 | 0 | 0 | 0 | 0 |
| 24689 | ENSMUSG00000082360 | 0 | NA | NA | NA | NA | NA | Gm8814 | predicted gene 8814 [Source:MGI Symbol | 0 | 0 | 0 | 0 | 0 | 0 |
| 24690 | ENSMUSG00000082361 | 37.38164945 | -0.317763681 | 0.355230173 | -0.894528969 | 0.371038937 | 0.807214774 | Btc | betacellulin, epidermal growth factor fam | 40.155 | 39.061 | 45.281 | 34.703 | 29.447 | 35.643 |
| 24691 | ENSMUSG00000082362 | 0 | NA | NA | NA | NA | NA | Gm5187 | predicted gene 5187 [Source:MGI Symbol | 0 | 0 | 0 | 0 | 0 | 0 |
| 24692 | ENSMUSG00000082363 | 0 | NA | NA | NA | NA | NA | Mup-ps18 | major urinary protein, pseudogene 18 [So | 0 | 0 | 0 | 0 | 0 | 0 |

| | | | | | | | | | | | | | | | |
|---|---|---|---|---|---|---|---|---|---|---|---|---|---|---|---|
| 24693 | ENSMUSG00000082364 | 30.15326146 | -0.010357771 | 0.483718744 | -0.021412796 | 0.982916366 | 0.998580735 | Gm11955 | predicted gene 11955 [Source:MGI Symbo | 28.109 | 22.614 | 40.25 | 39.523 | 38.172 | 12.252 |
| 24694 | ENSMUSG00000082365 | 0 | NA | NA | NA | NA | NA | Btf3-ps11 | basic transcription factor 3, pseudogene 1 | 0 | 0 | 0 | 0 | 0 | 0 |
| 24695 | ENSMUSG00000082367 | 0 | NA | NA | NA | NA | NA | Gm12363 | predicted gene 12363 [Source:MGI Symbo | 0 | 0 | 0 | 0 | 0 | 0 |
| 24696 | ENSMUSG00000082368 | 0 | NA | NA | NA | NA | NA | Gm11225 | predicted gene 11225 [Source:MGI Symbo | 0 | 0 | 0 | 0 | 0 | 0 |
| 24697 | ENSMUSG00000082369 | 0 | NA | NA | NA | NA | NA | Gm14208 | predicted gene 14208 [Source:MGI Symbo | 0 | 0 | 0 | 0 | 0 | 0 |
| 24698 | ENSMUSG00000082370 | 0 | NA | NA | NA | NA | NA | Gm12313 | predicted gene 12313 [Source:MGI Symbo | 0 | 0 | 0 | 0 | 0 | 0 |
| 24699 | ENSMUSG00000082371 | 0 | NA | NA | NA | NA | NA | Gm3712 | predicted gene 3712 [Source:MGI Symbol | 0 | 0 | 0 | 0 | 0 | 0 |
| 24700 | ENSMUSG00000082372 | 0 | NA | NA | NA | NA | NA | Gm4772 | predicted gene 4772 [Source:MGI Symbol | 0 | 0 | 0 | 0 | 0 | 0 |
| 24701 | ENSMUSG00000082373 | 0 | NA | NA | NA | NA | NA | Gm13754 | predicted gene 13754 [Source:MGI Symbo | 0 | 0 | 0 | 0 | 0 | 0 |
| 24702 | ENSMUSG00000082374 | 0 | NA | NA | NA | NA | NA | Gm12741 | predicted gene 12741 [Source:MGI Symbo | 0 | 0 | 0 | 0 | 0 | 0 |
| 24703 | ENSMUSG00000082375 | 0 | NA | NA | NA | NA | NA | Gm14141 | predicted gene 14141 [Source:MGI Symbo | 0 | 0 | 0 | 0 | 0 | 0 |
| 24704 | ENSMUSG00000082376 | 0 | NA | NA | NA | NA | NA | Gm14442 | predicted gene 14442 [Source:MGI Symbo | 0 | 0 | 0 | 0 | 0 | 0 |
| 24705 | ENSMUSG00000082377 | 0.483327519 | -0.685704547 | 3.145069396 | -0.218025252 | 0.827409436 | NA | Gm14415 | predicted gene 14415 [Source:MGI Symbo | 0.8031 | 0 | 1.0062 | 0 | 1.0906 | 0 |
| 24706 | ENSMUSG00000082378 | 0 | NA | NA | NA | NA | NA | Olfr408-ps1 | olfactory receptor 408, pseudogene 1 [Sou | 0 | 0 | 0 | 0 | 0 | 0 |
| 24707 | ENSMUSG00000082379 | 0 | NA | NA | NA | NA | NA | Gm13884 | predicted gene 13884 [Source:MGI Symbo | 0 | 0 | 0 | 0 | 0 | 0 |
| 24708 | ENSMUSG00000082380 | 2.939629758 | 4.042324229 | 1.675814376 | 2.412155121 | 0.015858533 | NA | Gm12218 | predicted gene 12218 [Source:MGI Symbo | 0 | 1.0279 | 0 | 6.7478 | 7.6343 | 2.2277 |
| 24709 | ENSMUSG00000082381 | 0.167707431 | -0.796629176 | 4.080472857 | -0.195229623 | 0.84521319 | NA | Gm12928 | predicted gene 12928 [Source:MGI Symbo | 0 | 0 | 1.0062 | 0 | 0 | 0 |
| 24710 | ENSMUSG00000082382 | 0 | NA | NA | NA | NA | NA | Gm14970 | predicted gene 14970 [Source:MGI Symbo | 0 | 0 | 0 | 0 | 0 | 0 |
| 24711 | ENSMUSG00000082383 | 7.148914392 | 0.402396155 | 0.797331535 | 0.50467859 | 0.613784601 | 0.917353781 | Gm9670 | predicted gene 9670 [Source:MGI Symbol | 7.2279 | 4.1117 | 7.0437 | 5.7839 | 9.8156 | 8.9108 |
| 24712 | ENSMUSG00000082388 | 0 | NA | NA | NA | NA | NA | Gm12692 | predicted gene 12692 [Source:MGI Symbo | 0 | 0 | 0 | 0 | 0 | 0 |
| 24713 | ENSMUSG00000082389 | 0 | NA | NA | NA | NA | NA | Mkrn1-ps1 | makorin, ring finger protein 1, pseudogen | 0 | 0 | 0 | 0 | 0 | 0 |
| 24714 | ENSMUSG00000082390 | 0 | NA | NA | NA | NA | NA | Gm12136 | predicted gene 12136 [Source:MGI Symbo | 0 | 0 | 0 | 0 | 0 | 0 |
| 24715 | ENSMUSG00000082392 | 0 | NA | NA | NA | NA | NA | Gm14938 | predicted gene 14938 [Source:MGI Symbo | 0 | 0 | 0 | 0 | 0 | 0 |
| 24716 | ENSMUSG00000082393 | 0 | NA | NA | NA | NA | NA | Gm15028 | predicted gene 15028 [Source:MGI Symbo | 0 | 0 | 0 | 0 | 0 | 0 |
| 24717 | ENSMUSG00000082394 | 0.633540418 | -1.320719359 | 2.623870149 | -0.503347835 | 0.61471974 | NA | Gm4596 | predicted gene 4596 [Source:MGI Symbol | 0.8031 | 1.0279 | 1.0062 | 0.964 | 0 | 0 |
| 24718 | ENSMUSG00000082395 | 0.958478575 | 0.054139297 | 2.45225727 | 0.022077331 | 0.982386269 | NA | Gm15480 | predicted gene 15480 [Source:MGI Symbo | 0.8031 | 2.0558 | 0 | 2.8919 | 0 | 0 |
| 24719 | ENSMUSG00000082396 | 1.273945333 | -3.713031505 | 2.423959792 | -1.531804083 | 0.125570788 | NA | Zfp389 | zinc finger protein 389 [Source:MGI Symb | 1.6062 | 0 | 6.0375 | 0 | 0 | 0 |
| 24720 | ENSMUSG00000082397 | 0 | NA | NA | NA | NA | NA | Gm14803 | predicted gene 14803 [Source:MGI Symbo | 0 | 0 | 0 | 0 | 0 | 0 |
| 24721 | ENSMUSG00000082398 | 0 | NA | NA | NA | NA | NA | Gm11927 | predicted gene 11927 [Source:MGI Symbo | 0 | 0 | 0 | 0 | 0 | 0 |
| 24722 | ENSMUSG00000082399 | 0.181769715 | 1.126924175 | 4.080472857 | 0.276174898 | 0.782413736 | NA | Gm14036 | predicted gene 14036 [Source:MGI Symbo | 0 | 0 | 0 | 0 | 1.0906 | 0 |
| 24723 | ENSMUSG00000082400 | 0 | NA | NA | NA | NA | NA | Gm16017 | predicted gene 16017 [Source:MGI Symbo | 0 | 0 | 0 | 0 | 0 | 0 |
| 24724 | ENSMUSG00000082402 | 0 | NA | NA | NA | NA | NA | Gm15621 | predicted gene 15621 [Source:MGI Symbo | 0 | 0 | 0 | 0 | 0 | 0 |
| 24725 | ENSMUSG00000082403 | 0 | NA | NA | NA | NA | NA | Gm14916 | predicted gene 14916 [Source:MGI Symbo | 0 | 0 | 0 | 0 | 0 | 0 |
| 24726 | ENSMUSG00000082404 | 0 | NA | NA | NA | NA | NA | Gm14799 | predicted gene 14799 [Source:MGI Symbo | 0 | 0 | 0 | 0 | 0 | 0 |
| 24727 | ENSMUSG00000082405 | 0 | NA | NA | NA | NA | NA | Gm16021 | predicted gene 16021 [Source:MGI Symbo | 0 | 0 | 0 | 0 | 0 | 0 |
| 24728 | ENSMUSG00000082406 | 0 | NA | NA | NA | NA | NA | Gm5401 | predicted pseudogene 5401 [Source:MGI | 0 | 0 | 0 | 0 | 0 | 0 |
| 24729 | ENSMUSG00000082407 | 0 | NA | NA | NA | NA | NA | Gm13045 | predicted gene 13045 [Source:MGI Symbo | 0 | 0 | 0 | 0 | 0 | 0 |
| 24730 | ENSMUSG00000082408 | 0 | NA | NA | NA | NA | NA | Gm13545 | predicted gene 13545 [Source:MGI Symbo | 0 | 0 | 0 | 0 | 0 | 0 |
| 24731 | ENSMUSG00000082410 | 0 | NA | NA | NA | NA | NA | Gm12011 | predicted gene 12011 [Source:MGI Symbo | 0 | 0 | 0 | 0 | 0 | 0 |
| 24732 | ENSMUSG00000082411 | 0 | NA | NA | NA | NA | NA | Gm12016 | predicted gene 12016 [Source:MGI Symbo | 0 | 0 | 0 | 0 | 0 | 0 |
| 24733 | ENSMUSG00000082412 | 0 | NA | NA | NA | NA | NA | Fau-ps1 | Finkel-Biskis-Reilly murine sarcoma virus ( | 0 | 0 | 0 | 0 | 0 | 0 |
| 24734 | ENSMUSG00000082413 | 0 | NA | NA | NA | NA | NA | Gm13401 | predicted gene 13401 [Source:MGI Symbo | 0 | 0 | 0 | 0 | 0 | 0 |
| 24735 | ENSMUSG00000082414 | 0 | NA | NA | NA | NA | NA | Gm13303 | predicted gene, 21093 [Source:MGI Symb | 0 | 0 | 0 | 0 | 0 | 0 |
| 24736 | ENSMUSG00000082414 | 0 | NA | NA | NA | NA | NA | Gm21093 | predicted gene, 21093 [Source:MGI Symb | 0 | 0 | 0 | 0 | 0 | 0 |
| 24737 | ENSMUSG00000082415 | 0 | NA | NA | NA | NA | NA | Gm12486 | predicted gene 12486 [Source:MGI Symbo | 0 | 0 | 0 | 0 | 0 | 0 |
| 24738 | ENSMUSG00000082416 | 0 | NA | NA | NA | NA | NA | Gm11414 | predicted gene 11414 [Source:MGI Symbo | 0 | 0 | 0 | 0 | 0 | 0 |
| 24739 | ENSMUSG00000082417 | 0 | NA | NA | NA | NA | NA | Gm13740 | predicted gene 13740 [Source:MGI Symbo | 0 | 0 | 0 | 0 | 0 | 0 |
| 24740 | ENSMUSG00000082418 | 0 | NA | NA | NA | NA | NA | Gm13126 | predicted gene 13126 [Source:MGI Symbo | 0 | 0 | 0 | 0 | 0 | 0 |
| 24741 | ENSMUSG00000082419 | 0.506966596 | 2.558866737 | 3.339834942 | 0.766165628 | 0.443577757 | NA | Gm11425 | predicted gene 11425 [Source:MGI Symbo | 0 | 0 | 0 | 1.928 | 0 | 1.1138 |
| 24742 | ENSMUSG00000082420 | 0 | NA | NA | NA | NA | NA | Gm6517 | predicted gene 6517 [Source:MGI Symbol | 0 | 0 | 0 | 0 | 0 | 0 |
| 24743 | ENSMUSG00000082421 | 0 | NA | NA | NA | NA | NA | Gm15004 | predicted gene 15004 [Source:MGI Symbo | 0 | 0 | 0 | 0 | 0 | 0 |
| 24744 | ENSMUSG00000082422 | 0 | NA | NA | NA | NA | NA | Gm6973 | predicted gene 6973 [Source:MGI Symbol | 0 | 0 | 0 | 0 | 0 | 0 |
| 24745 | ENSMUSG00000082423 | 0.517623261 | 1.044040381 | 3.075707576 | 0.339447218 | 0.734272853 | NA | Gm15148 | predicted gene 15148 [Source:MGI Symbo | 0 | 1.0279 | 0 | 0.964 | 0 | 1.1138 |
| 24746 | ENSMUSG00000082424 | 1.111859485 | -1.041840268 | 2.15740327 | -0.482914012 | 0.629156792 | NA | Gm13292 | predicted gene 13292 [Source:MGI Symbo | 2.4093 | 1.0279 | 1.0062 | 0 | 0 | 2.2277 |
| 24747 | ENSMUSG00000082426 | 0 | NA | NA | NA | NA | NA | Gm14610 | predicted gene 14610 [Source:MGI Symbo | 0 | 0 | 0 | 0 | 0 | 0 |
| 24748 | ENSMUSG00000082427 | 0 | NA | NA | NA | NA | NA | Olfr1327-ps1 | olfactory receptor 1327, pseudogene 1 [So | 0 | 0 | 0 | 0 | 0 | 0 |

| | | | | | | | | | | | | | | | |
|---|---|---|---|---|---|---|---|---|---|---|---|---|---|---|---|
| 24749 | ENSMUSG00000082428 | 0 | NA | NA | NA | NA | NA | Gm13742 | predicted gene 13742 [Source:MGI Symbo | 0 | 0 | 0 | 0 | 0 | 0 |
| 24750 | ENSMUSG00000082429 | 0 | NA | NA | NA | NA | NA | Gm13171 | predicted gene 13171 [Source:MGI Symbo | 0 | 0 | 0 | 0 | 0 | 0 |
| 24751 | ENSMUSG00000082431 | 0.517623261 | 1.044040381 | 3.075707576 | 0.339447218 | 0.734272853 | NA | Prdx2-ps1 | peroxiredoxin 2, pseudogene 1 [Source:M | 0 | 1.0279 | 0 | 0.964 | 0 | 1.1138 |
| 24752 | ENSMUSG00000082432 | 0 | NA | NA | NA | NA | NA | Gm11231 | predicted gene 11231 [Source:MGI Symbo | 0 | 0 | 0 | 0 | 0 | 0 |
| 24753 | ENSMUSG00000082433 | 0 | NA | NA | NA | NA | NA | Gm9025 | predicted gene 9025 [Source:MGI Symbol | 0 | 0 | 0 | 0 | 0 | 0 |
| 24754 | ENSMUSG00000082435 | 0 | NA | NA | NA | NA | NA | Gm13813 | predicted gene 13813 [Source:MGI Symbo | 0 | 0 | 0 | 0 | 0 | 0 |
| 24755 | ENSMUSG00000082436 | 0.569258551 | -2.569337407 | 3.18860282 | -0.805787849 | 0.420365185 | NA | Gm11688 | predicted gene 11688 [Source:MGI Symbo | 2.4093 | 0 | 1.0062 | 0 | 0 | 0 |
| 24756 | ENSMUSG00000082437 | 0 | NA | NA | NA | NA | NA | Gm15136 | predicted gene 15136 [Source:MGI Symbo | 0 | 0 | 0 | 0 | 0 | 0 |
| 24757 | ENSMUSG00000082440 | 0 | NA | NA | NA | NA | NA | Gm4914 | predicted gene 4914 [Source:MGI Symbol | 0 | 0 | 0 | 0 | 0 | 0 |
| 24758 | ENSMUSG00000082442 | 0 | NA | NA | NA | NA | NA | Gm12686 | predicted gene 12686 [Source:MGI Symbo | 0 | 0 | 0 | 0 | 0 | 0 |
| 24759 | ENSMUSG00000082443 | 0 | NA | NA | NA | NA | NA | Gm12604 | predicted gene 12604 [Source:MGI Symbo | 0 | 0 | 0 | 0 | 0 | 0 |
| 24760 | ENSMUSG00000082444 | 0 | NA | NA | NA | NA | NA | Gm14709 | predicted gene 14709 [Source:MGI Symbo | 0 | 0 | 0 | 0 | 0 | 0 |
| 24761 | ENSMUSG00000082446 | 0 | NA | NA | NA | NA | NA | Gm13863 | predicted gene 13863 [Source:MGI Symbo | 0 | 0 | 0 | 0 | 0 | 0 |
| 24762 | ENSMUSG00000082447 | 0 | NA | NA | NA | NA | NA | Gm15364 | predicted gene 15364 [Source:MGI Symbo | 0 | 0 | 0 | 0 | 0 | 0 |
| 24763 | ENSMUSG00000082448 | 0 | NA | NA | NA | NA | NA | Gm11676 | predicted gene 11676 [Source:MGI Symbo | 0 | 0 | 0 | 0 | 0 | 0 |
| 24764 | ENSMUSG00000082449 | 0 | NA | NA | NA | NA | NA | Gm15244 | predicted gene 15244 [Source:MGI Symbo | 0 | 0 | 0 | 0 | 0 | 0 |
| 24765 | ENSMUSG00000082450 | 0 | NA | NA | NA | NA | NA | Mageb7-ps | MAGE family member B7, pseudogene [Sc | 0 | 0 | 0 | 0 | 0 | 0 |
| 24766 | ENSMUSG00000082451 | 0 | NA | NA | NA | NA | NA | Gm11894 | predicted gene 11894 [Source:MGI Symbo | 0 | 0 | 0 | 0 | 0 | 0 |
| 24767 | ENSMUSG00000082453 | 0 | NA | NA | NA | NA | NA | Gm11502 | predicted gene 11502 [Source:MGI Symbo | 0 | 0 | 0 | 0 | 0 | 0 |
| 24768 | ENSMUSG00000082454 | 3.358935259 | -0.416426543 | 1.222149354 | -0.340732941 | 0.733304638 | NA | Gm12183 | predicted gene 12183 [Source:MGI Symbo | 2.4093 | 2.0558 | 7.0437 | 0.964 | 5.4531 | 2.2277 |
| 24769 | ENSMUSG00000082455 | 0.17131964 | -0.796629176 | 4.080472857 | -0.195229623 | 0.84521319 | NA | Gm13665 | predicted gene 13665 [Source:MGI Symbo | 0 | 1.0279 | 0 | 0 | 0 | 0 |
| 24770 | ENSMUSG00000082456 | 0 | NA | NA | NA | NA | NA | Gm11598 | predicted gene 11598 [Source:MGI Symbo | 0 | 0 | 0 | 0 | 0 | 0 |
| 24771 | ENSMUSG00000082457 | 2.84918169 | -1.090407172 | 1.361334697 | -0.800983898 | 0.423140968 | NA | Gm13776 | predicted gene 13776 [Source:MGI Symbo | 2.4093 | 6.1675 | 3.0187 | 0 | 3.2719 | 2.2277 |
| 24772 | ENSMUSG00000082458 | 0 | NA | NA | NA | NA | NA | Gm9430 | predicted gene 9430 [Source:MGI Symbol | 0 | 0 | 0 | 0 | 0 | 0 |
| 24773 | ENSMUSG00000082459 | 0 | NA | NA | NA | NA | NA | Gm11988 | predicted gene 11988 [Source:MGI Symbo | 0 | 0 | 0 | 0 | 0 | 0 |
| 24774 | ENSMUSG00000082460 | 0 | NA | NA | NA | NA | NA | Gm13685 | predicted gene 13685 [Source:MGI Symbo | 0 | 0 | 0 | 0 | 0 | 0 |
| 24775 | ENSMUSG00000082461 | 0 | NA | NA | NA | NA | NA | Gm8844 | predicted gene 8844 [Source:MGI Symbol | 0 | 0 | 0 | 0 | 0 | 0 |
| 24776 | ENSMUSG00000082462 | 0 | NA | NA | NA | NA | NA | Gm15147 | predicted gene 15147 [Source:MGI Symbo | 0 | 0 | 0 | 0 | 0 | 0 |
| 24777 | ENSMUSG00000082463 | 0 | NA | NA | NA | NA | NA | Gm15765 | predicted gene 15765 [Source:MGI Symbo | 0 | 0 | 0 | 0 | 0 | 0 |
| 24778 | ENSMUSG00000082464 | 0 | NA | NA | NA | NA | NA | Rab9b-ps1 | RAB9B, member RAS oncogene family, ps | 0 | 0 | 0 | 0 | 0 | 0 |
| 24779 | ENSMUSG00000082465 | 5.060073762 | 0.460839215 | 1.071183723 | 0.430214916 | 0.667039313 | NA | Rps6-ps3 | ribosomal protein S6, pseudogene 3 [Sour | 1.6062 | 6.1675 | 5.0312 | 0.964 | 5.4531 | 11.138 |
| 24780 | ENSMUSG00000082467 | 0 | NA | NA | NA | NA | NA | Gm6743 | predicted pseudogene 6743 [Source:MGI | 0 | 0 | 0 | 0 | 0 | 0 |
| 24781 | ENSMUSG00000082468 | 0 | NA | NA | NA | NA | NA | Gm12980 | predicted gene 12980 [Source:MGI Symbo | 0 | 0 | 0 | 0 | 0 | 0 |
| 24782 | ENSMUSG00000082472 | 0 | NA | NA | NA | NA | NA | Gm13501 | predicted gene 13501 [Source:MGI Symbo | 0 | 0 | 0 | 0 | 0 | 0 |
| 24783 | ENSMUSG00000082473 | 0 | NA | NA | NA | NA | NA | Kif22-ps | kinesin family member 22, pseudogene [S | 0 | 0 | 0 | 0 | 0 | 0 |
| 24784 | ENSMUSG00000082474 | 3.2778491 | -0.055208211 | 1.41459297 | -0.03902763 | 0.96886836 | NA | Gm8213 | predicted pseudogene 8213 [Source:MGI | 8.8341 | 1.0279 | 0 | 0.964 | 3.2719 | 5.5692 |
| 24785 | ENSMUSG00000082475 | 1.94837921 | -0.151060143 | 1.951628925 | -0.077402083 | 0.938303684 | NA | Gm7206 | predicted pseudogene 7206 [Source:MGI | 0 | 6.1675 | 0 | 0 | 2.1812 | 3.3415 |
| 24786 | ENSMUSG00000082476 | 0 | NA | NA | NA | NA | NA | Gm12242 | predicted gene 12242 [Source:MGI Symbo | 0 | 0 | 0 | 0 | 0 | 0 |
| 24787 | ENSMUSG00000082478 | 0.342639279 | -1.805976596 | 4.028547714 | -0.448294702 | 0.653940524 | NA | Gm11619 | predicted gene 11619 [Source:MGI Symbo | 0 | 2.0558 | 0 | 0 | 0 | 0 |
| 24788 | ENSMUSG00000082479 | 0 | NA | NA | NA | NA | NA | Gm4746 | predicted gene 4746 [Source:MGI Symbol | 0 | 0 | 0 | 0 | 0 | 0 |
| 24789 | ENSMUSG00000082480 | 0.342432689 | 1.990653086 | 3.900281263 | 0.51038706 | 0.609780321 | NA | Gm11687 | predicted gene 11687 [Source:MGI Symbo | 0 | 0 | 0 | 0.964 | 1.0906 | 0 |
| 24790 | ENSMUSG00000082481 | 0.371281295 | 2.09253362 | 4.031852165 | 0.519000582 | 0.603760335 | NA | Rps12-ps19 | ribosomal protein S12, pseudogene 19 [Sc | 0 | 0 | 0 | 0 | 0 | 2.2277 |
| 24791 | ENSMUSG00000082482 | 0 | NA | NA | NA | NA | NA | H2ab2 | H2A.B variant histone 2 [Source:MGI Syml | 0 | 0 | 0 | 0 | 0 | 0 |
| 24792 | ENSMUSG00000082483 | 0.305170013 | -1.657836477 | 4.041480335 | -0.410205257 | 0.681655385 | NA | Gm13486 | predicted gene 13486 [Source:MGI Symbo | 0.8031 | 1.0279 | 0 | 0 | 0 | 0 |
| 24793 | ENSMUSG00000082484 | 1.247492652 | -2.525342597 | 2.270614284 | -1.112184758 | 0.26605872 | NA | Gm16177 | predicted gene 16177 [Source:MGI Symbo | 2.4093 | 4.1117 | 0 | 0.964 | 0 | 0 |
| 24794 | ENSMUSG00000082485 | 0 | NA | NA | NA | NA | NA | Gm15693 | predicted gene 15693 [Source:MGI Symbo | 0 | 0 | 0 | 0 | 0 | 0 |
| 24795 | ENSMUSG00000082486 | 0 | NA | NA | NA | NA | NA | Gm11919 | predicted gene 11919 [Source:MGI Symbo | 0 | 0 | 0 | 0 | 0 | 0 |
| 24796 | ENSMUSG00000082487 | 0.674441933 | -2.786326546 | 3.022203109 | -0.921952114 | 0.356553554 | NA | Gm11628 | predicted gene 11628 [Source:MGI Symbo | 0 | 1.0279 | 3.0187 | 0 | 0 | 0 |
| 24797 | ENSMUSG00000082488 | 0 | NA | NA | NA | NA | NA | 1700119I11F | RIKEN cDNA 1700119I11 gene [Source:MG | 0 | 0 | 0 | 0 | 0 | 0 |
| 24798 | ENSMUSG00000082489 | 0 | NA | NA | NA | NA | NA | Gm14356 | predicted gene 14356 [Source:MGI Symbo | 0 | 0 | 0 | 0 | 0 | 0 |
| 24799 | ENSMUSG00000082490 | 0 | NA | NA | NA | NA | NA | Gm11880 | predicted gene 11880 [Source:MGI Symbo | 0 | 0 | 0 | 0 | 0 | 0 |
| 24800 | ENSMUSG00000082491 | 0.167707431 | -0.796629176 | 4.080472857 | -0.195229623 | 0.84521319 | NA | Gm5909 | predicted gene 5909 [Source:MGI Symbol | 0 | 0 | 1.0062 | 0 | 0 | 0 |
| 24801 | ENSMUSG00000082492 | 0 | NA | NA | NA | NA | NA | Gm14871 | predicted gene 14871 [Source:MGI Symbo | 0 | 0 | 0 | 0 | 0 | 0 |
| 24802 | ENSMUSG00000082494 | 0.513958919 | -2.394274821 | 3.756096013 | -0.637437066 | 0.523840191 | NA | Gm13687 | predicted gene 13687 [Source:MGI Symbo | 0 | 3.0838 | 0 | 0 | 0 | 0 |
| 24803 | ENSMUSG00000082495 | 0 | NA | NA | NA | NA | NA | Gm12570 | predicted gene 12570 [Source:MGI Symbo | 0 | 0 | 0 | 0 | 0 | 0 |
| 24804 | ENSMUSG00000082496 | 0 | NA | NA | NA | NA | NA | Gm6798 | predicted gene 6798 [Source:MGI Symbol | 0 | 0 | 0 | 0 | 0 | 0 |

| | | | | | | | | | | | | | | | |
|---|---|---|---|---|---|---|---|---|---|---|---|---|---|---|---|
| 24805 | ENSMUSG00000082497 | 0 | NA | NA | NA | NA | NA | Gm40 | predicted pseudogene 40 [Source:MGI Sy | 0 | 0 | 0 | 0 | 0 | 0 |
| 24806 | ENSMUSG00000082498 | 0.342432689 | 1.990653086 | 3.900281263 | 0.51038706 | 0.609780321 | NA | Gm12937 | predicted gene 12937 [Source:MGI Symbo | 0 | 0 | 0 | 0.964 | 1.0906 | 0 |
| 24807 | ENSMUSG00000082499 | 0 | NA | NA | NA | NA | NA | Gm13615 | predicted gene 13615 [Source:MGI Symbo | 0 | 0 | 0 | 0 | 0 | 0 |
| 24808 | ENSMUSG00000082500 | 0 | NA | NA | NA | NA | NA | Gm15195 | predicted gene 15195 [Source:MGI Symbo | 0 | 0 | 0 | 0 | 0 | 0 |
| 24809 | ENSMUSG00000082501 | 0 | NA | NA | NA | NA | NA | Gm11835 | predicted gene 11835 [Source:MGI Symbo | 0 | 0 | 0 | 0 | 0 | 0 |
| 24810 | ENSMUSG00000082502 | 0 | NA | NA | NA | NA | NA | Gm12636 | predicted gene 12636 [Source:MGI Symbo | 0 | 0 | 0 | 0 | 0 | 0 |
| 24811 | ENSMUSG00000082503 | 0 | NA | NA | NA | NA | NA | Gm12229 | predicted gene 12229 [Source:MGI Symbo | 0 | 0 | 0 | 0 | 0 | 0 |
| 24812 | ENSMUSG00000082504 | 0 | NA | NA | NA | NA | NA | Gm15003 | predicted gene 15003 [Source:MGI Symbo | 0 | 0 | 0 | 0 | 0 | 0 |
| 24813 | ENSMUSG00000082505 | 0 | NA | NA | NA | NA | NA | Gm14816 | predicted gene 14816 [Source:MGI Symbo | 0 | 0 | 0 | 0 | 0 | 0 |
| 24814 | ENSMUSG00000082507 | 1.486113227 | 0.427102758 | 1.769508984 | 0.241367951 | 0.80926995 | NA | Gm9378 | predicted gene 9378 [Source:MGI Symbol | 0.8031 | 1.0279 | 2.0125 | 2.8919 | 2.1812 | 0 |
| 24815 | ENSMUSG00000082508 | 0.401551119 | -2.072982726 | 4.008251979 | -0.517178744 | 0.605031388 | NA | Rpl15-ps4 | ribosomal protein L15, pseudogene 4 [Sou | 2.4093 | 0 | 0 | 0 | 0 | 0 |
| 24816 | ENSMUSG00000082510 | 0.626495961 | 0.101295816 | 2.879413718 | 0.03517932 | 0.971936753 | NA | Gm11793 | predicted gene 11793 [Source:MGI Symbo | 0.8031 | 1.0279 | 0 | 1.928 | 0 | 0 |
| 24817 | ENSMUSG00000082511 | 0 | NA | NA | NA | NA | NA | Gm4915 | predicted gene 4915 [Source:MGI Symbol | 0 | 0 | 0 | 0 | 0 | 0 |
| 24818 | ENSMUSG00000082512 | 0 | NA | NA | NA | NA | NA | Gm13937 | predicted gene 13937 [Source:MGI Symbo | 0 | 0 | 0 | 0 | 0 | 0 |
| 24819 | ENSMUSG00000082513 | 1.366790452 | 0.064014805 | 1.832463861 | 0.034933734 | 0.972132581 | NA | Gm15374 | predicted gene 15374 [Source:MGI Symbo | 0 | 2.0558 | 2.0125 | 1.928 | 1.0906 | 1.1138 |
| 24820 | ENSMUSG00000082514 | 0 | NA | NA | NA | NA | NA | Gm11452 | predicted gene 11452 [Source:MGI Symbo | 0 | 0 | 0 | 0 | 0 | 0 |
| 24821 | ENSMUSG00000082515 | 1.37618563 | 0.485221616 | 2.169618527 | 0.223643747 | 0.823034521 | NA | Gm12902 | predicted gene 12902 [Source:MGI Symbo | 2.4093 | 1.0279 | 0 | 4.8199 | 0 | 0 |
| 24822 | ENSMUSG00000082516 | 0 | NA | NA | NA | NA | NA | Gm13342 | predicted gene 13342 [Source:MGI Symbo | 0 | 0 | 0 | 0 | 0 | 0 |
| 24823 | ENSMUSG00000082517 | 0 | NA | NA | NA | NA | NA | Gm5757 | predicted gene 5757 [Source:MGI Symbol | 0 | 0 | 0 | 0 | 0 | 0 |
| 24824 | ENSMUSG00000082519 | 0 | NA | NA | NA | NA | NA | Vamp7-ps | vesicle-associated membrane protein 7, p | 0 | 0 | 0 | 0 | 0 | 0 |
| 24825 | ENSMUSG00000082520 | 0.524202404 | 2.600864256 | 3.301637836 | 0.787749713 | 0.430843121 | NA | Gm13349 | predicted gene 13349 [Source:MGI Symbo | 0 | 0 | 0 | 0.964 | 2.1812 | 0 |
| 24826 | ENSMUSG00000082523 | 0 | NA | NA | NA | NA | NA | Gm12667 | predicted gene 12667 [Source:MGI Symbo | 0 | 0 | 0 | 0 | 0 | 0 |
| 24827 | ENSMUSG00000082524 | 0 | NA | NA | NA | NA | NA | Gm6272 | predicted pseudogene 6272 [Source:MGI | 0 | 0 | 0 | 0 | 0 | 0 |
| 24828 | ENSMUSG00000082526 | 0 | NA | NA | NA | NA | NA | Gm14067 | predicted gene 14067 [Source:MGI Symbo | 0 | 0 | 0 | 0 | 0 | 0 |
| 24829 | ENSMUSG00000082527 | 0.133850373 | -0.796629176 | 4.080472857 | -0.195229623 | 0.84521319 | NA | Gm8200 | predicted gene 8200 [Source:MGI Symbol | 0.8031 | 0 | 0 | 0 | 0 | 0 |
| 24830 | ENSMUSG00000082528 | 0 | NA | NA | NA | NA | NA | Gm15182 | predicted gene 15182 [Source:MGI Symbo | 0 | 0 | 0 | 0 | 0 | 0 |
| 24831 | ENSMUSG00000082529 | 0 | NA | NA | NA | NA | NA | Gm11199 | predicted gene 11199 [Source:MGI Symbo | 0 | 0 | 0 | 0 | 0 | 0 |
| 24832 | ENSMUSG00000082530 | 0.17131964 | -0.796629176 | 4.080472857 | -0.195229623 | 0.84521319 | NA | Gm12168 | predicted gene 12168 [Source:MGI Symbo | 0 | 1.0279 | 0 | 0 | 0 | 0 |
| 24833 | ENSMUSG00000082531 | 0 | NA | NA | NA | NA | NA | Gm11556 | predicted gene 11556 [Source:MGI Symbo | 0 | 0 | 0 | 0 | 0 | 0 |
| 24834 | ENSMUSG00000082532 | 0.133850373 | -0.796629176 | 4.080472857 | -0.195229623 | 0.84521319 | NA | Rpl31-ps6 | ribosomal protein L31, pseudogene 6 [Sou | 0.8031 | 0 | 0 | 0 | 0 | 0 |
| 24835 | ENSMUSG00000082533 | 0 | NA | NA | NA | NA | NA | Gm11253 | predicted gene 11253 [Source:MGI Symbo | 0 | 0 | 0 | 0 | 0 | 0 |
| 24836 | ENSMUSG00000082534 | 0 | NA | NA | NA | NA | NA | Gm13712 | predicted gene 13712 [Source:MGI Symbo | 0 | 0 | 0 | 0 | 0 | 0 |
| 24837 | ENSMUSG00000082535 | 0 | NA | NA | NA | NA | NA | Gm7860 | predicted gene 7860 [Source:MGI Symbol | 0 | 0 | 0 | 0 | 0 | 0 |
| 24838 | ENSMUSG00000082536 | 58.48802654 | 0.027757198 | 0.314027651 | 0.088390935 | 0.929565966 | 0.988405116 | Gm13456 | predicted gene 13456 [Source:MGI Symbo | 45.777 | 65.787 | 62.387 | 51.091 | 51.259 | 74.628 |
| 24839 | ENSMUSG00000082537 | 0 | NA | NA | NA | NA | NA | Gm14981 | predicted gene 14981 [Source:MGI Symbo | 0 | 0 | 0 | 0 | 0 | 0 |
| 24840 | ENSMUSG00000082538 | 0 | NA | NA | NA | NA | NA | Gm14704 | predicted gene 14704 [Source:MGI Symbo | 0 | 0 | 0 | 0 | 0 | 0 |
| 24841 | ENSMUSG00000082539 | 0 | NA | NA | NA | NA | NA | Gm15207 | predicted gene 15207 [Source:MGI Symbo | 0 | 0 | 0 | 0 | 0 | 0 |
| 24842 | ENSMUSG00000082540 | 0 | NA | NA | NA | NA | NA | Vmn1r-ps11 | vomeronasal 1 receptor, pseudogene 112 | 0 | 0 | 0 | 0 | 0 | 0 |
| 24843 | ENSMUSG00000082541 | 0 | NA | NA | NA | NA | NA | Gm9095 | predicted gene 9095 [Source:MGI Symbol | 0 | 0 | 0 | 0 | 0 | 0 |
| 24844 | ENSMUSG00000082542 | 0 | NA | NA | NA | NA | NA | Gm13311 | predicted gene 13311 [Source:MGI Symbo | 0 | 0 | 0 | 0 | 0 | 0 |
| 24845 | ENSMUSG00000082543 | 0 | NA | NA | NA | NA | NA | Gm14336 | predicted gene 14336 [Source:MGI Symbo | 0 | 0 | 0 | 0 | 0 | 0 |
| 24846 | ENSMUSG00000082544 | 0 | NA | NA | NA | NA | NA | Gm11667 | predicted gene 11667 [Source:MGI Symbo | 0 | 0 | 0 | 0 | 0 | 0 |
| 24847 | ENSMUSG00000082545 | 0 | NA | NA | NA | NA | NA | Gm12196 | predicted gene 12196 [Source:MGI Symbo | 0 | 0 | 0 | 0 | 0 | 0 |
| 24848 | ENSMUSG00000082546 | 0 | NA | NA | NA | NA | NA | Gm14898 | predicted gene 14898 [Source:MGI Symbo | 0 | 0 | 0 | 0 | 0 | 0 |
| 24849 | ENSMUSG00000082547 | 0 | NA | NA | NA | NA | NA | Olfr1073-ps1 | olfactory receptor 1073, pseudogene 1 [S | 0 | 0 | 0 | 0 | 0 | 0 |
| 24850 | ENSMUSG00000082548 | 0 | NA | NA | NA | NA | NA | Gm12377 | predicted gene 12377 [Source:MGI Symbo | 0 | 0 | 0 | 0 | 0 | 0 |
| 24851 | ENSMUSG00000082549 | 0 | NA | NA | NA | NA | NA | Gm14813 | predicted gene 14813 [Source:MGI Symbo | 0 | 0 | 0 | 0 | 0 | 0 |
| 24852 | ENSMUSG00000082550 | 0 | NA | NA | NA | NA | NA | Gm12344 | predicted gene 12344 [Source:MGI Symbo | 0 | 0 | 0 | 0 | 0 | 0 |
| 24853 | ENSMUSG00000082551 | 0 | NA | NA | NA | NA | NA | Gm15065 | predicted gene 15065 [Source:MGI Symbo | 0 | 0 | 0 | 0 | 0 | 0 |
| 24854 | ENSMUSG00000082552 | 0 | NA | NA | NA | NA | NA | Gm11850 | predicted gene 11850 [Source:MGI Symbo | 0 | 0 | 0 | 0 | 0 | 0 |
| 24855 | ENSMUSG00000082553 | 0 | NA | NA | NA | NA | NA | Gm11404 | predicted gene 11404 [Source:MGI Symbo | 0 | 0 | 0 | 0 | 0 | 0 |
| 24856 | ENSMUSG00000082554 | 0 | NA | NA | NA | NA | NA | Gm15806 | predicted gene 15806 [Source:MGI Symbo | 0 | 0 | 0 | 0 | 0 | 0 |
| 24857 | ENSMUSG00000082555 | 0 | NA | NA | NA | NA | NA | Gm11406 | predicted gene 11406 [Source:MGI Symbo | 0 | 0 | 0 | 0 | 0 | 0 |
| 24858 | ENSMUSG00000082556 | 0 | NA | NA | NA | NA | NA | Gm5387 | predicted gene 5387 [Source:MGI Symbol | 0 | 0 | 0 | 0 | 0 | 0 |
| 24859 | ENSMUSG00000082558 | 0 | NA | NA | NA | NA | NA | Snrp1c-ps2 | U1 small nuclear ribonucleoprotein 1C, ps | 0 | 0 | 0 | 0 | 0 | 0 |
| 24860 | ENSMUSG00000082560 | 0 | NA | NA | NA | NA | NA | Gm15157 | predicted gene 15157 [Source:MGI Symbo | 0 | 0 | 0 | 0 | 0 | 0 |

| | | | | | | | | | | | | | | | |
|---|---|---|---|---|---|---|---|---|---|---|---|---|---|---|---|
| 24861 | ENSMUSG00000082561 | 0.160662974 | 1.126924175 | 4.080472857 | 0.276174898 | 0.782413736 | NA | Gm15580 | predicted gene 15580 [Source:MGI Symbo | 0 | 0 | 0 | 0.964 | 0 | 0 |
| 24862 | ENSMUSG00000082562 | 0 | NA | NA | NA | NA | NA | Gm11726 | predicted gene 11726 [Source:MGI Symbo | 0 | 0 | 0 | 0 | 0 | 0 |
| 24863 | ENSMUSG00000082563 | 0 | NA | NA | NA | NA | NA | Gm15026 | predicted gene 15026 [Source:MGI Symbo | 0 | 0 | 0 | 0 | 0 | 0 |
| 24864 | ENSMUSG00000082564 | 0 | NA | NA | NA | NA | NA | Gm15586 | predicted gene 15586 [Source:MGI Symbo | 0 | 0 | 0 | 0 | 0 | 0 |
| 24865 | ENSMUSG00000082565 | 0 | NA | NA | NA | NA | NA | Gm5255 | predicted gene 5255 [Source:MGI Symbol | 0 | 0 | 0 | 0 | 0 | 0 |
| 24866 | ENSMUSG00000082566 | 0 | NA | NA | NA | NA | NA | Gm11485 | predicted gene 11485 [Source:MGI Symbo | 0 | 0 | 0 | 0 | 0 | 0 |
| 24867 | ENSMUSG00000082567 | 0 | NA | NA | NA | NA | NA | Gm14896 | predicted gene 14896 [Source:MGI Symbo | 0 | 0 | 0 | 0 | 0 | 0 |
| 24868 | ENSMUSG00000082569 | 0 | NA | NA | NA | NA | NA | Gm12634 | predicted gene 12634 [Source:MGI Symbo | 0 | 0 | 0 | 0 | 0 | 0 |
| 24869 | ENSMUSG00000082570 | 0.160662974 | 1.126924175 | 4.080472857 | 0.276174898 | 0.782413736 | NA | Gm15711 | predicted gene 15711 [Source:MGI Symbo | 0 | 0 | 0 | 0.964 | 0 | 0 |
| 24870 | ENSMUSG00000082571 | 0 | NA | NA | NA | NA | NA | Gm8737 | predicted gene 8737 [Source:MGI Symbol | 0 | 0 | 0 | 0 | 0 | 0 |
| 24871 | ENSMUSG00000082573 | 0 | NA | NA | NA | NA | NA | Gm11903 | predicted gene 11903 [Source:MGI Symbo | 0 | 0 | 0 | 0 | 0 | 0 |
| 24872 | ENSMUSG00000082574 | 0 | NA | NA | NA | NA | NA | Olfr1268-ps1 | olfactory receptor 1268, pseudogene 1 [S | 0 | 0 | 0 | 0 | 0 | 0 |
| 24873 | ENSMUSG00000082575 | 2.467004996 | 0.896117717 | 1.495525077 | 0.599199392 | 0.549039928 | NA | Eef2-ps2 | eukaryotic translation elongation factor 2, | 4.0155 | 1.0279 | 0 | 0.964 | 5.4531 | 3.3415 |
| 24874 | ENSMUSG00000082576 | 0 | NA | NA | NA | NA | NA | Gm12058 | predicted gene 12058 [Source:MGI Symbo | 0 | 0 | 0 | 0 | 0 | 0 |
| 24875 | ENSMUSG00000082577 | 0 | NA | NA | NA | NA | NA | Olfr1003-ps1 | olfactory receptor 1003, pseudogene 1 [S | 0 | 0 | 0 | 0 | 0 | 0 |
| 24876 | ENSMUSG00000082578 | 0 | NA | NA | NA | NA | NA | Gm13527 | predicted gene 13527 [Source:MGI Symbo | 0 | 0 | 0 | 0 | 0 | 0 |
| 24877 | ENSMUSG00000082579 | 0.17131964 | -0.796629176 | 4.080472857 | -0.195229623 | 0.84521319 | NA | Gm11979 | predicted gene 11979 [Source:MGI Symbo | 0 | 1.0279 | 0 | 0 | 0 | 0 |
| 24878 | ENSMUSG00000082580 | 0.181769715 | 1.126924175 | 4.080472857 | 0.276174898 | 0.782413736 | NA | Gm13182 | predicted gene 13182 [Source:MGI Symbo | 0 | 0 | 0 | 0 | 1.0906 | 0 |
| 24879 | ENSMUSG00000082581 | 0 | NA | NA | NA | NA | NA | Gm13761 | predicted gene 13761 [Source:MGI Symbo | 0 | 0 | 0 | 0 | 0 | 0 |
| 24880 | ENSMUSG00000082583 | 0 | NA | NA | NA | NA | NA | Olfr336-ps1 | olfactory receptor 336, pseudogene 1 [Sou | 0 | 0 | 0 | 0 | 0 | 0 |
| 24881 | ENSMUSG00000082584 | 0 | NA | NA | NA | NA | NA | Gm8569 | predicted gene 8569 [Source:MGI Symbol | 0 | 0 | 0 | 0 | 0 | 0 |
| 24882 | ENSMUSG00000082585 | 5.314823114 | 0.624264518 | 0.939111129 | 0.664739772 | 0.50621695 | 0.881592135 | Gm15387 | predicted gene 15387 [Source:MGI Symbo | 2.4093 | 6.1675 | 4.025 | 3.8559 | 7.6343 | 7.7969 |
| 24883 | ENSMUSG00000082586 | 0 | NA | NA | NA | NA | NA | Sult2a-ps1 | sulfotransferase family 2A, dehydroepian | 0 | 0 | 0 | 0 | 0 | 0 |
| 24884 | ENSMUSG00000082587 | 0 | NA | NA | NA | NA | NA | Gm12129 | predicted gene 12129 [Source:MGI Symbo | 0 | 0 | 0 | 0 | 0 | 0 |
| 24885 | ENSMUSG00000082588 | 0.346303622 | 2.004339653 | 3.884003423 | 0.516049919 | 0.605819544 | NA | Gm15443 | predicted gene 15443 [Source:MGI Symbo | 0 | 0 | 0 | 0.964 | 0 | 1.1138 |
| 24886 | ENSMUSG00000082589 | 0 | NA | NA | NA | NA | NA | Gm12433 | predicted gene 12433 [Source:MGI Symbo | 0 | 0 | 0 | 0 | 0 | 0 |
| 24887 | ENSMUSG00000082590 | 0 | NA | NA | NA | NA | NA | Gm14943 | predicted gene 14943 [Source:MGI Symbo | 0 | 0 | 0 | 0 | 0 | 0 |
| 24888 | ENSMUSG00000082591 | 0 | NA | NA | NA | NA | NA | Gm11558 | predicted gene 11558 [Source:MGI Symbo | 0 | 0 | 0 | 0 | 0 | 0 |
| 24889 | ENSMUSG00000082592 | 0 | NA | NA | NA | NA | NA | Gm12950 | predicted gene 12950 [Source:MGI Symbo | 0 | 0 | 0 | 0 | 0 | 0 |
| 24890 | ENSMUSG00000082593 | 0 | NA | NA | NA | NA | NA | Gm11331 | predicted gene 11331 [Source:MGI Symbo | 0 | 0 | 0 | 0 | 0 | 0 |
| 24891 | ENSMUSG00000082594 | 0 | NA | NA | NA | NA | NA | Gm15074 | predicted gene 15074 [Source:MGI Symbo | 0 | 0 | 0 | 0 | 0 | 0 |
| 24892 | ENSMUSG00000082595 | 0 | NA | NA | NA | NA | NA | Gm14793 | predicted gene 14793 [Source:MGI Symbo | 0 | 0 | 0 | 0 | 0 | 0 |
| 24893 | ENSMUSG00000082596 | 0.633540418 | -1.320719359 | 2.623870149 | -0.503347835 | 0.61471974 | NA | Gm14227 | predicted gene 14227 [Source:MGI Symbo | 0.8031 | 1.0279 | 1.0062 | 0.964 | 0 | 0 |
| 24894 | ENSMUSG00000082597 | 0 | NA | NA | NA | NA | NA | Gm12645 | predicted gene 12645 [Source:MGI Symbo | 0 | 0 | 0 | 0 | 0 | 0 |
| 24895 | ENSMUSG00000082598 | 0 | NA | NA | NA | NA | NA | Gm15212 | predicted gene 15212 [Source:MGI Symbo | 0 | 0 | 0 | 0 | 0 | 0 |
| 24896 | ENSMUSG00000082599 | 0.503122293 | -2.366675966 | 3.785596405 | -0.625179156 | 0.531853481 | NA | Gm13141 | predicted gene 13141 [Source:MGI Symbo | 0 | 0 | 3.0187 | 0 | 0 | 0 |
| 24897 | ENSMUSG00000082600 | 0 | NA | NA | NA | NA | NA | Gm9673 | predicted gene 9673 [Source:MGI Symbol | 0 | 0 | 0 | 0 | 0 | 0 |
| 24898 | ENSMUSG00000082601 | 0 | NA | NA | NA | NA | NA | Gm12700 | predicted gene 12700 [Source:MGI Symbo | 0 | 0 | 0 | 0 | 0 | 0 |
| 24899 | ENSMUSG00000082602 | 0 | NA | NA | NA | NA | NA | Gm15021 | predicted gene 15021 [Source:MGI Symbo | 0 | 0 | 0 | 0 | 0 | 0 |
| 24900 | ENSMUSG00000082604 | 0 | NA | NA | NA | NA | NA | Gm8482 | predicted pseudogene 8482 [Source:MGI | 0 | 0 | 0 | 0 | 0 | 0 |
| 24901 | ENSMUSG00000082605 | 0.596071152 | -1.224343926 | 2.897583865 | -0.4225396 | 0.672631203 | NA | Gm16148 | predicted gene 16148 [Source:MGI Symbo | 1.6062 | 0 | 1.0062 | 0.964 | 0 | 0 |
| 24902 | ENSMUSG00000082606 | 0 | NA | NA | NA | NA | NA | Gm14638 | predicted gene 14638 [Source:MGI Symbo | 0 | 0 | 0 | 0 | 0 | 0 |
| 24903 | ENSMUSG00000082607 | 0 | NA | NA | NA | NA | NA | Gm12045 | predicted gene 12045 [Source:MGI Symbo | 0 | 0 | 0 | 0 | 0 | 0 |
| 24904 | ENSMUSG00000082608 | 0 | NA | NA | NA | NA | NA | Gm7670 | predicted gene 7670 [Source:MGI Symbol | 0 | 0 | 0 | 0 | 0 | 0 |
| 24905 | ENSMUSG00000082609 | 0 | NA | NA | NA | NA | NA | Gm15464 | predicted gene 15464 [Source:MGI Symbo | 0 | 0 | 0 | 0 | 0 | 0 |
| 24906 | ENSMUSG00000082610 | 0.321325949 | 1.911575114 | 4.048267893 | 0.472195804 | 0.636787034 | NA | Gm13829 | predicted gene 13829 [Source:MGI Symbo | 0 | 0 | 0 | 1.928 | 0 | 0 |
| 24907 | ENSMUSG00000082611 | 0 | NA | NA | NA | NA | NA | Gm7746 | predicted gene 7746 [Source:MGI Symbol | 0 | 0 | 0 | 0 | 0 | 0 |
| 24908 | ENSMUSG00000082612 | 0 | NA | NA | NA | NA | NA | 4930503H13 | RIKEN cDNA 4930503H13 gene [Source:M | 0 | 0 | 0 | 0 | 0 | 0 |
| 24909 | ENSMUSG00000082613 | 0 | NA | NA | NA | NA | NA | Gm12518 | predicted gene 12518 [Source:MGI Symbo | 0 | 0 | 0 | 0 | 0 | 0 |
| 24910 | ENSMUSG00000082614 | 0 | NA | NA | NA | NA | NA | Gm12149 | predicted gene 12149 [Source:MGI Symbo | 0 | 0 | 0 | 0 | 0 | 0 |
| 24911 | ENSMUSG00000082615 | 0 | NA | NA | NA | NA | NA | Gm13239 | predicted gene 13239 [Source:MGI Symbo | 0 | 0 | 0 | 0 | 0 | 0 |
| 24912 | ENSMUSG00000082616 | 0 | NA | NA | NA | NA | NA | Gm6025 | predicted pseudogene 6025 [Source:MGI | 0 | 0 | 0 | 0 | 0 | 0 |
| 24913 | ENSMUSG00000082617 | 0 | NA | NA | NA | NA | NA | Smt3h2-ps | SMT3 suppressor of mif two 3 homolog 2, | 0 | 0 | 0 | 0 | 0 | 0 |
| 24914 | ENSMUSG00000082618 | 0 | NA | NA | NA | NA | NA | Gm14852 | predicted gene 14852 [Source:MGI Symbo | 0 | 0 | 0 | 0 | 0 | 0 |
| 24915 | ENSMUSG00000082619 | 0 | NA | NA | NA | NA | NA | Gm16219 | predicted gene 16219 [Source:MGI Symbo | 0 | 0 | 0 | 0 | 0 | 0 |
| 24916 | ENSMUSG00000082620 | 0 | NA | NA | NA | NA | NA | Gm8244 | predicted gene 8244 [Source:MGI Symbol | 0 | 0 | 0 | 0 | 0 | 0 |

| | | | | | | | | | | | | | | | |
|---|---|---|---|---|---|---|---|---|---|---|---|---|---|---|---|
| 24917 | ENSMUSG00000082621 | 0 | NA | NA | NA | NA | NA | Rpl7a-ps13 | ribosomal protein L7A, pseudogene 13 [So | 0 | 0 | 0 | 0 | 0 | 0 |
| 24918 | ENSMUSG00000082622 | 0 | NA | NA | NA | NA | NA | Gm12632 | predicted gene 12632 [Source:MGI Symbo | 0 | 0 | 0 | 0 | 0 | 0 |
| 24919 | ENSMUSG00000082623 | 0 | NA | NA | NA | NA | NA | Gm15493 | predicted gene 15493 [Source:MGI Symbo | 0 | 0 | 0 | 0 | 0 | 0 |
| 24920 | ENSMUSG00000082624 | 0 | NA | NA | NA | NA | NA | Gm12535 | predicted gene 12535 [Source:MGI Symbo | 0 | 0 | 0 | 0 | 0 | 0 |
| 24921 | ENSMUSG00000082625 | 0.979378725 | 2.336871961 | 2.444416672 | 0.956003936 | 0.339070246 | NA | Gm15204 | predicted gene 15204 [Source:MGI Symbo | 0.8031 | 0 | 0 | 2.8919 | 2.1812 | 0 |
| 24922 | ENSMUSG00000082626 | 0 | NA | NA | NA | NA | NA | Rps13-ps5 | ribosomal protein S13, pseudogene 5 [Sou | 0 | 0 | 0 | 0 | 0 | 0 |
| 24923 | ENSMUSG00000082628 | 0 | NA | NA | NA | NA | NA | Gm12566 | predicted gene 12566 [Source:MGI Symbo | 0 | 0 | 0 | 0 | 0 | 0 |
| 24924 | ENSMUSG00000082629 | 0 | NA | NA | NA | NA | NA | Pwwp4-ps | PWWP domain containing 4, pseudogene | 0 | 0 | 0 | 0 | 0 | 0 |
| 24925 | ENSMUSG00000082630 | 0 | NA | NA | NA | NA | NA | Gm14834 | predicted gene 14834 [Source:MGI Symbo | 0 | 0 | 0 | 0 | 0 | 0 |
| 24926 | ENSMUSG00000082632 | 0 | NA | NA | NA | NA | NA | Gm13365 | predicted gene 13365 [Source:MGI Symbo | 0 | 0 | 0 | 0 | 0 | 0 |
| 24927 | ENSMUSG00000082633 | 2.877116197 | 1.90734598 | 1.570207123 | 1.214709799 | 0.224476794 | NA | Gm15779 | predicted gene 15779 [Source:MGI Symbo | 1.6062 | 1.0279 | 1.0062 | 12.532 | 1.0906 | 0 |
| 24928 | ENSMUSG00000082634 | 0 | NA | NA | NA | NA | NA | Gm12900 | predicted gene 12900 [Source:MGI Symbo | 0 | 0 | 0 | 0 | 0 | 0 |
| 24929 | ENSMUSG00000082635 | 0 | NA | NA | NA | NA | NA | Gm14750 | predicted gene 14750 [Source:MGI Symbo | 0 | 0 | 0 | 0 | 0 | 0 |
| 24930 | ENSMUSG00000082636 | 0 | NA | NA | NA | NA | NA | Gm14923 | predicted gene 14923 [Source:MGI Symbo | 0 | 0 | 0 | 0 | 0 | 0 |
| 24931 | ENSMUSG00000082637 | 0 | NA | NA | NA | NA | NA | Gm11248 | predicted gene 11248 [Source:MGI Symbo | 0 | 0 | 0 | 0 | 0 | 0 |
| 24932 | ENSMUSG00000082638 | 0 | NA | NA | NA | NA | NA | Gm12815 | predicted gene 12815 [Source:MGI Symbo | 0 | 0 | 0 | 0 | 0 | 0 |
| 24933 | ENSMUSG00000082639 | 0 | NA | NA | NA | NA | NA | Gm2012 | predicted gene 2012 [Source:MGI Symbol | 0 | 0 | 0 | 0 | 0 | 0 |
| 24934 | ENSMUSG00000082641 | 0 | NA | NA | NA | NA | NA | Rpl30-ps5 | ribosomal protein L30, pseudogene 5 [Sou | 0 | 0 | 0 | 0 | 0 | 0 |
| 24935 | ENSMUSG00000082644 | 32.23208332 | 1.181078164 | 0.419737627 | 2.813848672 | 0.004895227 | 0.065864706 | Mup-ps19 | major urinary protein, pseudogene 19 [So | 23.29 | 16.447 | 19.119 | 29.883 | 54.531 | 50.123 |
| 24936 | ENSMUSG00000082645 | 0 | NA | NA | NA | NA | NA | Gm15481 | predicted gene 15481 [Source:MGI Symbo | 0 | 0 | 0 | 0 | 0 | 0 |
| 24937 | ENSMUSG00000082646 | 0 | NA | NA | NA | NA | NA | Gm12732 | predicted gene 12732 [Source:MGI Symbo | 0 | 0 | 0 | 0 | 0 | 0 |
| 24938 | ENSMUSG00000082647 | 0.167707431 | -0.796629176 | 4.080472857 | -0.195229623 | 0.84521319 | NA | Gm14056 | predicted gene 14056 [Source:MGI Symbo | 0 | 0 | 1.0062 | 0 | 0 | 0 |
| 24939 | ENSMUSG00000082648 | 0 | NA | NA | NA | NA | NA | Gm11366 | predicted gene 11366 [Source:MGI Symbo | 0 | 0 | 0 | 0 | 0 | 0 |
| 24940 | ENSMUSG00000082649 | 0 | NA | NA | NA | NA | NA | Gm12754 | predicted gene 12754 [Source:MGI Symbo | 0 | 0 | 0 | 0 | 0 | 0 |
| 24941 | ENSMUSG00000082650 | 0 | NA | NA | NA | NA | NA | Gm15358 | predicted gene 15358 [Source:MGI Symbo | 0 | 0 | 0 | 0 | 0 | 0 |
| 24942 | ENSMUSG00000082651 | 1.80975888 | -0.776848742 | 1.69052801 | -0.45953024 | 0.645853442 | NA | Gm14001 | predicted gene 14001 [Source:MGI Symbo | 4.8186 | 1.0279 | 1.0062 | 2.8919 | 0 | 1.1138 |
| 24943 | ENSMUSG00000082652 | 0 | NA | NA | NA | NA | NA | Gm14725 | predicted gene 14725 [Source:MGI Symbo | 0 | 0 | 0 | 0 | 0 | 0 |
| 24944 | ENSMUSG00000082653 | 0.992155673 | -1.012754205 | 2.706122858 | -0.374245464 | 0.708221702 | NA | Gm15897 | predicted gene 15897 [Source:MGI Symbo | 0 | 0 | 4.025 | 1.928 | 0 | 0 |
| 24945 | ENSMUSG00000082655 | 0 | NA | NA | NA | NA | NA | Gm7857 | predicted gene 7857 [Source:MGI Symbol | 0 | 0 | 0 | 0 | 0 | 0 |
| 24946 | ENSMUSG00000082656 | 0 | NA | NA | NA | NA | NA | Vmn1r-ps12 | vomeronasal 1 receptor, pseudogene 125 | 0 | 0 | 0 | 0 | 0 | 0 |
| 24947 | ENSMUSG00000082657 | 0 | NA | NA | NA | NA | NA | Gm14736 | predicted gene 14736 [Source:MGI Symbo | 0 | 0 | 0 | 0 | 0 | 0 |
| 24948 | ENSMUSG00000082658 | 1.15116368 | -0.31437627 | 1.962373926 | -0.160202022 | 0.872721937 | NA | Fau-ps2 | Finkel-Biskis-Reilly murine sarcoma virus ( | 0.8031 | 2.0558 | 1.0062 | 1.928 | 0 | 1.1138 |
| 24949 | ENSMUSG00000082660 | 0 | NA | NA | NA | NA | NA | Gm14612 | predicted gene 14612 [Source:MGI Symbo | 0 | 0 | 0 | 0 | 0 | 0 |
| 24950 | ENSMUSG00000082661 | 0 | NA | NA | NA | NA | NA | Gm11590 | predicted gene 11590 [Source:MGI Symbo | 0 | 0 | 0 | 0 | 0 | 0 |
| 24951 | ENSMUSG00000082662 | 0 | NA | NA | NA | NA | NA | Gm15516 | predicted gene 15516 [Source:MGI Symbo | 0 | 0 | 0 | 0 | 0 | 0 |
| 24952 | ENSMUSG00000082663 | 0 | NA | NA | NA | NA | NA | Trav13-3 | T cell receptor alpha variable 13-3 [Source | 0 | 0 | 0 | 0 | 0 | 0 |
| 24953 | ENSMUSG00000082664 | 0 | NA | NA | NA | NA | NA | Gm8761 | predicted gene 8761 [Source:MGI Symbol | 0 | 0 | 0 | 0 | 0 | 0 |
| 24954 | ENSMUSG00000082665 | 7.117249751 | 0.456361039 | 0.855080779 | 0.533705178 | 0.593545526 | 0.913011006 | Gm11470 | predicted gene 11470 [Source:MGI Symbo | 4.8186 | 4.1117 | 9.0562 | 3.8559 | 14.178 | 6.6831 |
| 24955 | ENSMUSG00000082666 | 0 | NA | NA | NA | NA | NA | Gm13677 | predicted gene 13677 [Source:MGI Symbo | 0 | 0 | 0 | 0 | 0 | 0 |
| 24956 | ENSMUSG00000082667 | 0 | NA | NA | NA | NA | NA | Gm11348 | predicted gene 11348 [Source:MGI Symbo | 0 | 0 | 0 | 0 | 0 | 0 |
| 24957 | ENSMUSG00000082669 | 0 | NA | NA | NA | NA | NA | Gm5393 | predicted pseudogene 5393 [Source:MGI | 0 | 0 | 0 | 0 | 0 | 0 |
| 24958 | ENSMUSG00000082670 | 0 | NA | NA | NA | NA | NA | Gm14050 | predicted gene 14050 [Source:MGI Symbo | 0 | 0 | 0 | 0 | 0 | 0 |
| 24959 | ENSMUSG00000082671 | 0 | NA | NA | NA | NA | NA | Gm14006 | predicted gene 14006 [Source:MGI Symbo | 0 | 0 | 0 | 0 | 0 | 0 |
| 24960 | ENSMUSG00000082672 | 0 | NA | NA | NA | NA | NA | Gm11876 | predicted gene 11876 [Source:MGI Symbo | 0 | 0 | 0 | 0 | 0 | 0 |
| 24961 | ENSMUSG00000082673 | 0 | NA | NA | NA | NA | NA | Gm11857 | predicted gene 11857 [Source:MGI Symbo | 0 | 0 | 0 | 0 | 0 | 0 |
| 24962 | ENSMUSG00000082674 | 68.80955928 | 0.208822605 | 0.291796491 | 0.715644677 | 0.474210779 | 0.86899591 | Gm11914 | predicted gene 11914 [Source:MGI Symbo | 50.595 | 67.843 | 73.456 | 75.19 | 58.893 | 86.88 |
| 24963 | ENSMUSG00000082675 | 0 | NA | NA | NA | NA | NA | Gm6382 | predicted gene 6382 [Source:MGI Symbol | 0 | 0 | 0 | 0 | 0 | 0 |
| 24964 | ENSMUSG00000082676 | 14.77657622 | -1.406432635 | 0.603305112 | -2.33121286 | 0.019742138 | 0.172018247 | Gm11843 | predicted gene 11843 [Source:MGI Symbo | 18.471 | 30.838 | 15.094 | 7.7118 | 7.6343 | 8.9108 |
| 24965 | ENSMUSG00000082677 | 0.428363721 | -0.543730382 | 3.624677344 | -0.15000794 | 0.880758351 | NA | Gm15371 | predicted gene 15371 [Source:MGI Symbo | 1.6062 | 0 | 0 | 0.964 | 0 | 0 |
| 24966 | ENSMUSG00000082678 | 0 | NA | NA | NA | NA | NA | Gm12818 | predicted gene 12818 [Source:MGI Symbo | 0 | 0 | 0 | 0 | 0 | 0 |
| 24967 | ENSMUSG00000082679 | 0 | NA | NA | NA | NA | NA | Gm13469 | predicted gene 13469 [Source:MGI Symbo | 0 | 0 | 0 | 0 | 0 | 0 |
| 24968 | ENSMUSG00000082680 | 0 | NA | NA | NA | NA | NA | Gm14885 | predicted gene 14885 [Source:MGI Symbo | 0 | 0 | 0 | 0 | 0 | 0 |
| 24969 | ENSMUSG00000082681 | 0 | NA | NA | NA | NA | NA | Cyp2j7-ps1 | cytochrome P450, family 2, subfamily j, pc | 0 | 0 | 0 | 0 | 0 | 0 |
| 24970 | ENSMUSG00000082682 | 0 | NA | NA | NA | NA | NA | Gm15349 | predicted gene 15349 [Source:MGI Symbo | 0 | 0 | 0 | 0 | 0 | 0 |
| 24971 | ENSMUSG00000082683 | 0 | NA | NA | NA | NA | NA | Gm12697 | predicted gene 12697 [Source:MGI Symbo | 0 | 0 | 0 | 0 | 0 | 0 |
| 24972 | ENSMUSG00000082684 | 0.722877068 | -0.331317305 | 2.989821669 | -0.110815073 | 0.911762993 | NA | Gm12774 | predicted gene 12774 [Source:MGI Symbo | 2.4093 | 0 | 0 | 1.928 | 0 | 0 |

| | | | | | | | | | | | | | | | |
|---|---|---|---|---|---|---|---|---|---|---|---|---|---|---|---|
| 24973 | ENSMUSG00000082686 | 0.133850373 | -0.796629176 | 4.080472857 | -0.195229623 | 0.84521319 | NA | Gm12961 | predicted gene 12961 [Source:MGI Symbo | 0.8031 | 0 | 0 | 0 | 0 | 0 |
| 24974 | ENSMUSG00000082687 | 0.160662974 | 1.126924175 | 4.080472857 | 0.276174898 | 0.782413736 | NA | Gm11474 | predicted gene 11474 [Source:MGI Symbo | 0 | 0 | 0 | 0.964 | 0 | 0 |
| 24975 | ENSMUSG00000082688 | 0 | NA | NA | NA | NA | NA | Gm12124 | predicted gene 12124 [Source:MGI Symbo | 0 | 0 | 0 | 0 | 0 | 0 |
| 24976 | ENSMUSG00000082689 | 0 | NA | NA | NA | NA | NA | Gm14156 | predicted gene 14156 [Source:MGI Symbo | 0 | 0 | 0 | 0 | 0 | 0 |
| 24977 | ENSMUSG00000082690 | 0 | NA | NA | NA | NA | NA | Gm14972 | predicted gene 14972 [Source:MGI Symbo | 0 | 0 | 0 | 0 | 0 | 0 |
| 24978 | ENSMUSG00000082691 | 0.535117793 | 1.111049431 | 3.044213722 | 0.364970903 | 0.715133126 | NA | Dynlt1-ps1 | dynein light chain Tctex-type 1, pseudog | 0 | 0 | 1.0062 | 0 | 1.0906 | 1.1138 |
| 24979 | ENSMUSG00000082692 | 0 | NA | NA | NA | NA | NA | H2af-ps1 | H2A histone family, pseudogene 1 [Source | 0 | 0 | 0 | 0 | 0 | 0 |
| 24980 | ENSMUSG00000082693 | 0.528279927 | -0.837071592 | 3.369085264 | -0.248456636 | 0.803781118 | NA | Gm15190 | predicted gene 15190 [Source:MGI Symbo | 0 | 2.0558 | 0 | 0 | 0 | 1.1138 |
| 24981 | ENSMUSG00000082696 | 0 | NA | NA | NA | NA | NA | Gm15040 | predicted gene 15040 [Source:MGI Symbo | 0 | 0 | 0 | 0 | 0 | 0 |
| 24982 | ENSMUSG00000082698 | 0 | NA | NA | NA | NA | NA | Gm15316 | predicted gene 15316 [Source:MGI Symbo | 0 | 0 | 0 | 0 | 0 | 0 |
| 24983 | ENSMUSG00000082699 | 0 | NA | NA | NA | NA | NA | Gm12736 | predicted gene 12736 [Source:MGI Symbo | 0 | 0 | 0 | 0 | 0 | 0 |
| 24984 | ENSMUSG00000082700 | 0 | NA | NA | NA | NA | NA | Gm13792 | predicted gene 13792 [Source:MGI Symbo | 0 | 0 | 0 | 0 | 0 | 0 |
| 24985 | ENSMUSG00000082701 | 0 | NA | NA | NA | NA | NA | Gm14460 | predicted gene 14460 [Source:MGI Symbo | 0 | 0 | 0 | 0 | 0 | 0 |
| 24986 | ENSMUSG00000082703 | 0 | NA | NA | NA | NA | NA | Gm11907 | predicted gene 11907 [Source:MGI Symbo | 0 | 0 | 0 | 0 | 0 | 0 |
| 24987 | ENSMUSG00000082704 | 0 | NA | NA | NA | NA | NA | Gm12515 | predicted gene 12515 [Source:MGI Symbo | 0 | 0 | 0 | 0 | 0 | 0 |
| 24988 | ENSMUSG00000082705 | 0.36353943 | 2.065989254 | 4.034137301 | 0.512126658 | 0.608562368 | NA | Gm15616 | predicted gene 15616 [Source:MGI Symbo | 0 | 0 | 0 | 0 | 2.1812 | 0 |
| 24989 | ENSMUSG00000082706 | 0 | NA | NA | NA | NA | NA | Gm11663 | predicted gene 11663 [Source:MGI Symbo | 0 | 0 | 0 | 0 | 0 | 0 |
| 24990 | ENSMUSG00000082707 | 0 | NA | NA | NA | NA | NA | Gm6881 | predicted gene 6881 [Source:MGI Symbol | 0 | 0 | 0 | 0 | 0 | 0 |
| 24991 | ENSMUSG00000082708 | 0 | NA | NA | NA | NA | NA | Gm15172 | predicted gene 15172 [Source:MGI Symbo | 0 | 0 | 0 | 0 | 0 | 0 |
| 24992 | ENSMUSG00000082709 | 0 | NA | NA | NA | NA | NA | Gm12362 | predicted gene 12362 [Source:MGI Symbo | 0 | 0 | 0 | 0 | 0 | 0 |
| 24993 | ENSMUSG00000082710 | 0 | NA | NA | NA | NA | NA | Defa-ps14 | defensin, alpha, pseudogene 14 [Source:N | 0 | 0 | 0 | 0 | 0 | 0 |
| 24994 | ENSMUSG00000082711 | 0 | NA | NA | NA | NA | NA | Gm12987 | predicted gene 12987 [Source:MGI Symbo | 0 | 0 | 0 | 0 | 0 | 0 |
| 24995 | ENSMUSG00000082712 | 0 | NA | NA | NA | NA | NA | Olfr990-ps1 | olfactory receptor 990, pseudogene 1 [Sou | 0 | 0 | 0 | 0 | 0 | 0 |
| 24996 | ENSMUSG00000082713 | 0 | NA | NA | NA | NA | NA | Gm14721 | predicted gene 14721 [Source:MGI Symbo | 0 | 0 | 0 | 0 | 0 | 0 |
| 24997 | ENSMUSG00000082714 | 0 | NA | NA | NA | NA | NA | Gm15120 | predicted gene 15120 [Source:MGI Symbo | 0 | 0 | 0 | 0 | 0 | 0 |
| 24998 | ENSMUSG00000082715 | 0 | NA | NA | NA | NA | NA | Gm11633 | predicted gene 11633 [Source:MGI Symbo | 0 | 0 | 0 | 0 | 0 | 0 |
| 24999 | ENSMUSG00000082716 | 0 | NA | NA | NA | NA | NA | Rpl31-ps5 | ribosomal protein L31, pseudogene 5 [Sou | 0 | 0 | 0 | 0 | 0 | 0 |
| 25000 | ENSMUSG00000082717 | 0 | NA | NA | NA | NA | NA | Gm13087 | predicted gene 13087 [Source:MGI Symbo | 0 | 0 | 0 | 0 | 0 | 0 |
| 25001 | ENSMUSG00000082718 | 0.160662974 | 1.126924175 | 4.080472857 | 0.276174898 | 0.782413736 | NA | Gm14928 | predicted gene 14928 [Source:MGI Symbo | 0 | 0 | 0 | 0.964 | 0 | 0 |
| 25002 | ENSMUSG00000082720 | 0 | NA | NA | NA | NA | NA | Gm12489 | predicted gene 12489 [Source:MGI Symbo | 0 | 0 | 0 | 0 | 0 | 0 |
| 25003 | ENSMUSG00000082721 | 0 | NA | NA | NA | NA | NA | Gm12087 | predicted gene 12087 [Source:MGI Symbo | 0 | 0 | 0 | 0 | 0 | 0 |
| 25004 | ENSMUSG00000082722 | 0 | NA | NA | NA | NA | NA | Gm11393 | predicted gene 11393 [Source:MGI Symbo | 0 | 0 | 0 | 0 | 0 | 0 |
| 25005 | ENSMUSG00000082723 | 0 | NA | NA | NA | NA | NA | Gm14564 | predicted gene 14564 [Source:MGI Symbo | 0 | 0 | 0 | 0 | 0 | 0 |
| 25006 | ENSMUSG00000082724 | 0.346303622 | 2.004339653 | 3.884003423 | 0.516049919 | 0.605819544 | NA | Gm14416 | predicted gene 14416 [Source:MGI Symbo | 0 | 0 | 0 | 0.964 | 0 | 1.1138 |
| 25007 | ENSMUSG00000082725 | 1.363616928 | 0.787807956 | 1.978397547 | 0.398205081 | 0.690479021 | NA | Gm12451 | predicted gene 12451 [Source:MGI Symbo | 0 | 2.0558 | 1.0062 | 2.8919 | 0 | 2.2277 |
| 25008 | ENSMUSG00000082726 | 0 | NA | NA | NA | NA | NA | Gm11424 | predicted gene 11424 [Source:MGI Symbo | 0 | 0 | 0 | 0 | 0 | 0 |
| 25009 | ENSMUSG00000082727 | 0.181769715 | 1.126924175 | 4.080472857 | 0.276174898 | 0.782413736 | NA | Gm13260 | predicted gene 13260 [Source:MGI Symbo | 0 | 0 | 0 | 0 | 1.0906 | 0 |
| 25010 | ENSMUSG00000082728 | 0 | NA | NA | NA | NA | NA | Supt20-ps | SPT20 SAGA complex component, pseudo | 0 | 0 | 0 | 0 | 0 | 0 |
| 25011 | ENSMUSG00000082729 | 0.185640647 | 1.126924175 | 4.080472857 | 0.276174898 | 0.782413736 | NA | Gm14845 | predicted gene 14845 [Source:MGI Symbo | 0 | 0 | 0 | 0 | 0 | 1.1138 |
| 25012 | ENSMUSG00000082730 | 0 | NA | NA | NA | NA | NA | Gm11363 | predicted gene 11363 [Source:MGI Symbo | 0 | 0 | 0 | 0 | 0 | 0 |
| 25013 | ENSMUSG00000082732 | 0 | NA | NA | NA | NA | NA | Rpl23a-ps6 | ribosomal protein 23A, pseudogene 6 [Sou | 0 | 0 | 0 | 0 | 0 | 0 |
| 25014 | ENSMUSG00000082733 | 0 | NA | NA | NA | NA | NA | Gm15153 | predicted gene 15153 [Source:MGI Symbo | 0 | 0 | 0 | 0 | 0 | 0 |
| 25015 | ENSMUSG00000082734 | 0 | NA | NA | NA | NA | NA | Rhox11-ps1 | reproductive homeobox 11, pseudogene 1 | 0 | 0 | 0 | 0 | 0 | 0 |
| 25016 | ENSMUSG00000082735 | 0 | NA | NA | NA | NA | NA | Gm16394 | predicted gene 16394 [Source:MGI Symbo | 0 | 0 | 0 | 0 | 0 | 0 |
| 25017 | ENSMUSG00000082736 | 0 | NA | NA | NA | NA | NA | Gm5381 | predicted gene 5381 [Source:MGI Symbol | 0 | 0 | 0 | 0 | 0 | 0 |
| 25018 | ENSMUSG00000082737 | 0 | NA | NA | NA | NA | NA | Gm12026 | predicted gene 12026 [Source:MGI Symbo | 0 | 0 | 0 | 0 | 0 | 0 |
| 25019 | ENSMUSG00000082738 | 0.267700746 | -1.497481649 | 4.057001436 | -0.369110456 | 0.712045396 | NA | Gm15422 | predicted gene 15422 [Source:MGI Symbo | 1.6062 | 0 | 0 | 0 | 0 | 0 |
| 25020 | ENSMUSG00000082739 | 0 | NA | NA | NA | NA | NA | Olfr602-ps1 | olfactory receptor 602, pseudogene 1 [Sou | 0 | 0 | 0 | 0 | 0 | 0 |
| 25021 | ENSMUSG00000082740 | 0.185640647 | 1.126924175 | 4.080472857 | 0.276174898 | 0.782413736 | NA | Gm14127 | predicted gene 14127 [Source:MGI Symbo | 0 | 0 | 0 | 0 | 0 | 1.1138 |
| 25022 | ENSMUSG00000082741 | 0.305170013 | -1.657836477 | 4.041480335 | -0.410205257 | 0.681655385 | NA | Gm9703 | predicted gene 9703 [Source:MGI Symbol | 0.8031 | 1.0279 | 0 | 0 | 0 | 0 |
| 25023 | ENSMUSG00000082743 | 0 | NA | NA | NA | NA | NA | Gm12988 | predicted gene 12988 [Source:MGI Symbo | 0 | 0 | 0 | 0 | 0 | 0 |
| 25024 | ENSMUSG00000082744 | 0 | NA | NA | NA | NA | NA | Gm15625 | predicted gene 15625 [Source:MGI Symbo | 0 | 0 | 0 | 0 | 0 | 0 |
| 25025 | ENSMUSG00000082745 | 0 | NA | NA | NA | NA | NA | Gm15181 | predicted gene 15181 [Source:MGI Symbo | 0 | 0 | 0 | 0 | 0 | 0 |
| 25026 | ENSMUSG00000082746 | 1.405623819 | 2.875628331 | 2.010832145 | 1.430068809 | 0.152697271 | NA | Rps12-ps1 | ribosomal protein S12, pseudogene 1 [Sou | 0 | 0 | 1.0062 | 1.928 | 3.2719 | 2.2277 |
| 25027 | ENSMUSG00000082749 | 0 | NA | NA | NA | NA | NA | Rpl9-ps2 | ribosomal protein L9, pseudogene 2 [Sour | 0 | 0 | 0 | 0 | 0 | 0 |
| 25028 | ENSMUSG00000082750 | 0 | NA | NA | NA | NA | NA | Gm12156 | predicted gene 12156 [Source:MGI Symbo | 0 | 0 | 0 | 0 | 0 | 0 |

| | | | | | | | | | | | | | | | |
|---|---|---|---|---|---|---|---|---|---|---|---|---|---|---|---|
| 25029 | ENSMUSG00000082751 | 0 | NA | NA | NA | NA | NA | Gm6620 | predicted gene 6620 [Source:MGI Symbol | 0 | 0 | 0 | 0 | 0 | 0 |
| 25030 | ENSMUSG00000082752 | 0 | NA | NA | NA | NA | NA | Gm14899 | predicted gene 14899 [Source:MGI Symbo | 0 | 0 | 0 | 0 | 0 | 0 |
| 25031 | ENSMUSG00000082754 | 0 | NA | NA | NA | NA | NA | Olfr1021-ps1 | olfactory receptor 1021, pseudogene 1 [S | 0 | 0 | 0 | 0 | 0 | 0 |
| 25032 | ENSMUSG00000082755 | 0 | NA | NA | NA | NA | NA | Gm8692 | predicted gene 8692 [Source:MGI Symbol | 0 | 0 | 0 | 0 | 0 | 0 |
| 25033 | ENSMUSG00000082757 | 0 | NA | NA | NA | NA | NA | Gm6923 | predicted pseudogene 6923 [Source:MGI | 0 | 0 | 0 | 0 | 0 | 0 |
| 25034 | ENSMUSG00000082758 | 0 | NA | NA | NA | NA | NA | Gm13024 | predicted gene 13024 [Source:MGI Symbo | 0 | 0 | 0 | 0 | 0 | 0 |
| 25035 | ENSMUSG00000082759 | 0 | NA | NA | NA | NA | NA | Gm14826 | predicted gene 14826 [Source:MGI Symbo | 0 | 0 | 0 | 0 | 0 | 0 |
| 25036 | ENSMUSG00000082760 | 0 | NA | NA | NA | NA | NA | Gm15546 | predicted gene 15546 [Source:MGI Symbo | 0 | 0 | 0 | 0 | 0 | 0 |
| 25037 | ENSMUSG00000082762 | 3.740232032 | 0.401779865 | 1.117692262 | 0.359472708 | 0.719241491 | NA | Gm12366 | predicted gene 12366 [Source:MGI Symbo | 1.6062 | 3.0838 | 5.0312 | 3.8559 | 2.1812 | 6.6831 |
| 25038 | ENSMUSG00000082763 | 0 | NA | NA | NA | NA | NA | Tra2b-ps | transformer 2 beta, pseudogene [Source:I | 0 | 0 | 0 | 0 | 0 | 0 |
| 25039 | ENSMUSG00000082765 | 1.195212088 | 0.46832086 | 2.069471296 | 0.226299761 | 0.820968284 | NA | Gm14411 | predicted gene 14411 [Source:MGI Symbo | 0 | 2.0558 | 1.0062 | 1.928 | 2.1812 | 0 |
| 25040 | ENSMUSG00000082766 | 0 | NA | NA | NA | NA | NA | 1700064H15 | RIKEN cDNA 1700064H15 gene [Source:M | 0 | 0 | 0 | 0 | 0 | 0 |
| 25041 | ENSMUSG00000082767 | 0 | NA | NA | NA | NA | NA | Gm11622 | predicted gene 11622 [Source:MGI Symbo | 0 | 0 | 0 | 0 | 0 | 0 |
| 25042 | ENSMUSG00000082768 | 0.640816969 | 1.600078776 | 2.832778663 | 0.564844263 | 0.572179697 | NA | Gm12727 | predicted gene 12727 [Source:MGI Symbo | 0.8031 | 0 | 0 | 1.928 | 0 | 1.1138 |
| 25043 | ENSMUSG00000082769 | 1.540280853 | 0.134906922 | 2.056817962 | 0.065590113 | 0.947704161 | NA | Gm15793 | predicted gene 15793 [Source:MGI Symbo | 2.4093 | 0 | 2.0125 | 4.8199 | 0 | 0 |
| 25044 | ENSMUSG00000082770 | 0 | NA | NA | NA | NA | NA | Gm14722 | predicted gene 14722 [Source:MGI Symbo | 0 | 0 | 0 | 0 | 0 | 0 |
| 25045 | ENSMUSG00000082771 | 1.028636663 | 1.245965894 | 2.457526307 | 0.507000023 | 0.612154802 | NA | Gm12803 | predicted gene 12803 [Source:MGI Symbo | 0.8031 | 0 | 1.0062 | 0 | 4.3625 | 0 |
| 25046 | ENSMUSG00000082772 | 0 | NA | NA | NA | NA | NA | Gm381 | predicted pseudogene 381 [Source:MGI S | 0 | 0 | 0 | 0 | 0 | 0 |
| 25047 | ENSMUSG00000082773 | 0 | NA | NA | NA | NA | NA | Gm15058 | predicted gene 15058 [Source:MGI Symbo | 0 | 0 | 0 | 0 | 0 | 0 |
| 25048 | ENSMUSG00000082774 | 0 | NA | NA | NA | NA | NA | Gm8805 | predicted gene 8805 [Source:MGI Symbol | 0 | 0 | 0 | 0 | 0 | 0 |
| 25049 | ENSMUSG00000082775 | 0 | NA | NA | NA | NA | NA | Itpa-ps3 | inosine triphosphatase (nucleoside tripho | 0 | 0 | 0 | 0 | 0 | 0 |
| 25050 | ENSMUSG00000082776 | 3.465661047 | 0.92647028 | 1.303532743 | 0.71073802 | 0.477246595 | NA | Gm7061 | predicted gene 7061 [Source:MGI Symbol | 4.0155 | 3.0838 | 0 | 3.8559 | 8.7249 | 1.1138 |
| 25051 | ENSMUSG00000082777 | 0 | NA | NA | NA | NA | NA | Gm15861 | predicted gene 15861 [Source:MGI Symbo | 0 | 0 | 0 | 0 | 0 | 0 |
| 25052 | ENSMUSG00000082778 | 0.36353943 | 2.065989254 | 4.034137301 | 0.512126658 | 0.608562368 | NA | Gm15191 | predicted gene 15191 [Source:MGI Symbo | 0 | 0 | 0 | 0 | 2.1812 | 0 |
| 25053 | ENSMUSG00000082779 | 0 | NA | NA | NA | NA | NA | Gm13162 | predicted pseudogene 13162 [Source:MG | 0 | 0 | 0 | 0 | 0 | 0 |
| 25054 | ENSMUSG00000082780 | 0 | NA | NA | NA | NA | NA | Rps11-ps5 | ribosomal protein S11, pseudogene 5 [Sou | 0 | 0 | 0 | 0 | 0 | 0 |
| 25055 | ENSMUSG00000082781 | 0 | NA | NA | NA | NA | NA | Rps19-ps14 | ribosomal protein S19, pseudogene 14 [Sc | 0 | 0 | 0 | 0 | 0 | 0 |
| 25056 | ENSMUSG00000082782 | 0 | NA | NA | NA | NA | NA | Gm5319 | predicted gene 5319 [Source:MGI Symbol | 0 | 0 | 0 | 0 | 0 | 0 |
| 25057 | ENSMUSG00000082783 | 0 | NA | NA | NA | NA | NA | Gm9126 | predicted pseudogene 9126 [Source:MGI | 0 | 0 | 0 | 0 | 0 | 0 |
| 25058 | ENSMUSG00000082784 | 0 | NA | NA | NA | NA | NA | Gm15957 | predicted gene 15957 [Source:MGI Symbo | 0 | 0 | 0 | 0 | 0 | 0 |
| 25059 | ENSMUSG00000082785 | 0 | NA | NA | NA | NA | NA | Vmn1r-ps10 | vomeronasal 1 receptor, pseudogene 100 | 0 | 0 | 0 | 0 | 0 | 0 |
| 25060 | ENSMUSG00000082786 | 0 | NA | NA | NA | NA | NA | Gm14489 | predicted gene 14489 [Source:MGI Symbo | 0 | 0 | 0 | 0 | 0 | 0 |
| 25061 | ENSMUSG00000082788 | 0 | NA | NA | NA | NA | NA | Gm13396 | predicted gene 13396 [Source:MGI Symbo | 0 | 0 | 0 | 0 | 0 | 0 |
| 25062 | ENSMUSG00000082789 | 0 | NA | NA | NA | NA | NA | Gm13284 | predicted gene 13284 [Source:MGI Symbo | 0 | 0 | 0 | 0 | 0 | 0 |
| 25063 | ENSMUSG00000082790 | 0 | NA | NA | NA | NA | NA | Gm12155 | predicted gene 12155 [Source:MGI Symbo | 0 | 0 | 0 | 0 | 0 | 0 |
| 25064 | ENSMUSG00000082791 | 0 | NA | NA | NA | NA | NA | Gm4875 | predicted gene 4875 [Source:MGI Symbol | 0 | 0 | 0 | 0 | 0 | 0 |
| 25065 | ENSMUSG00000082792 | 0 | NA | NA | NA | NA | NA | Gm12176 | predicted gene 12176 [Source:MGI Symbo | 0 | 0 | 0 | 0 | 0 | 0 |
| 25066 | ENSMUSG00000082793 | 0 | NA | NA | NA | NA | NA | Btf3-ps4 | basic transcription factor 3, pseudogene 4 | 0 | 0 | 0 | 0 | 0 | 0 |
| 25067 | ENSMUSG00000082794 | 0 | NA | NA | NA | NA | NA | Gm6806 | predicted gene 6806 [Source:MGI Symbol | 0 | 0 | 0 | 0 | 0 | 0 |
| 25068 | ENSMUSG00000082795 | 0.17131964 | -0.796629176 | 4.080472857 | -0.195229623 | 0.84521319 | NA | Gm13193 | predicted gene 13193 [Source:MGI Symbo | 0 | 1.0279 | 0 | 0 | 0 | 0 |
| 25069 | ENSMUSG00000082796 | 0 | NA | NA | NA | NA | NA | Gm13385 | predicted gene 13385 [Source:MGI Symbo | 0 | 0 | 0 | 0 | 0 | 0 |
| 25070 | ENSMUSG00000082797 | 0 | NA | NA | NA | NA | NA | Gm12015 | predicted gene 12015 [Source:MGI Symbo | 0 | 0 | 0 | 0 | 0 | 0 |
| 25071 | ENSMUSG00000082798 | 0 | NA | NA | NA | NA | NA | Gm11561 | predicted gene 11561 [Source:MGI Symbo | 0 | 0 | 0 | 0 | 0 | 0 |
| 25072 | ENSMUSG00000082799 | 0 | NA | NA | NA | NA | NA | Gm14554 | predicted gene 14554 [Source:MGI Symbo | 0 | 0 | 0 | 0 | 0 | 0 |
| 25073 | ENSMUSG00000082800 | 0 | NA | NA | NA | NA | NA | Gm13560 | predicted gene 13560 [Source:MGI Symbo | 0 | 0 | 0 | 0 | 0 | 0 |
| 25074 | ENSMUSG00000082802 | 0 | NA | NA | NA | NA | NA | Gm15465 | predicted gene 15465 [Source:MGI Symbo | 0 | 0 | 0 | 0 | 0 | 0 |
| 25075 | ENSMUSG00000082804 | 0 | NA | NA | NA | NA | NA | Rpl17-ps1 | ribosomal protein L17, pseudogene 1 [Sou | 0 | 0 | 0 | 0 | 0 | 0 |
| 25076 | ENSMUSG00000082806 | 0.492645588 | 0.9565034 | 3.447648281 | 0.277436479 | 0.781444974 | NA | Rpl13-ps1 | ribosomal protein L13, pseudogene 1 [Sou | 0 | 1.0279 | 0 | 1.928 | 0 | 0 |
| 25077 | ENSMUSG00000082807 | 0 | NA | NA | NA | NA | NA | Gm14782 | predicted gene 14782 [Source:MGI Symbo | 0 | 0 | 0 | 0 | 0 | 0 |
| 25078 | ENSMUSG00000082809 | 0.181769715 | 1.126924175 | 4.080472857 | 0.276174898 | 0.782413736 | NA | Gm14150 | predicted gene 14150 [Source:MGI Symbo | 0 | 0 | 0 | 0 | 1.0906 | 0 |
| 25079 | ENSMUSG00000082810 | 0 | NA | NA | NA | NA | NA | Gm8682 | predicted gene 8682 [Source:MGI Symbol | 0 | 0 | 0 | 0 | 0 | 0 |
| 25080 | ENSMUSG00000082812 | 0.321325949 | 1.911575114 | 4.048267893 | 0.472195804 | 0.636787034 | NA | Gm14860 | predicted gene 14860 [Source:MGI Symbo | 0 | 0 | 0 | 1.928 | 0 | 0 |
| 25081 | ENSMUSG00000082814 | 1.441516881 | 1.746670698 | 2.156450264 | 0.809974951 | 0.417954572 | NA | Gm12956 | predicted gene 12956 [Source:MGI Symbo | 0 | 0 | 2.0125 | 0 | 2.1812 | 4.4554 |
| 25082 | ENSMUSG00000082815 | 0 | NA | NA | NA | NA | NA | Gm5678 | predicted gene 5678 [Source:MGI Symbol | 0 | 0 | 0 | 0 | 0 | 0 |
| 25083 | ENSMUSG00000082816 | 0.267700746 | -1.497481649 | 4.057001436 | -0.369110456 | 0.712045396 | NA | Gm11953 | predicted gene 11953 [Source:MGI Symbo | 1.6062 | 0 | 0 | 0 | 0 | 0 |
| 25084 | ENSMUSG00000082817 | 0 | NA | NA | NA | NA | NA | Gm16428 | predicted gene 16428 [Source:MGI Symbo | 0 | 0 | 0 | 0 | 0 | 0 |

| | | | | | | | | | | | | | | | |
|---|---|---|---|---|---|---|---|---|---|---|---|---|---|---|---|
| 25085 | ENSMUSG00000082818 | 0 | NA | NA | NA | NA | NA | Gm12175 | predicted gene 12175 [Source:MGI Symbo | 0 | 0 | 0 | 0 | 0 | 0 |
| 25086 | ENSMUSG00000082819 | 0 | NA | NA | NA | NA | NA | Trav4d-2 | T cell receptor alpha variable 4D-2 [Source | 0 | 0 | 0 | 0 | 0 | 0 |
| 25087 | ENSMUSG00000082820 | 0.465832987 | -0.698295221 | 3.18348759 | -0.219349126 | 0.826378096 | NA | Gm13803 | predicted gene 13803 [Source:MGI Symbo | 0.8031 | 1.0279 | 0 | 0.964 | 0 | 0 |
| 25088 | ENSMUSG00000082821 | 0 | NA | NA | NA | NA | NA | Gm12880 | predicted gene 12880 [Source:MGI Symbo | 0 | 0 | 0 | 0 | 0 | 0 |
| 25089 | ENSMUSG00000082822 | 0 | NA | NA | NA | NA | NA | Gm14731 | predicted gene 14731 [Source:MGI Symbo | 0 | 0 | 0 | 0 | 0 | 0 |
| 25090 | ENSMUSG00000082823 | 0 | NA | NA | NA | NA | NA | Gm15828 | predicted gene 15828 [Source:MGI Symbo | 0 | 0 | 0 | 0 | 0 | 0 |
| 25091 | ENSMUSG00000082824 | 0 | NA | NA | NA | NA | NA | Gm14716 | predicted gene 14716 [Source:MGI Symbo | 0 | 0 | 0 | 0 | 0 | 0 |
| 25092 | ENSMUSG00000082825 | 0 | NA | NA | NA | NA | NA | Gm15741 | predicted gene 15741 [Source:MGI Symbo | 0 | 0 | 0 | 0 | 0 | 0 |
| 25093 | ENSMUSG00000082826 | 0 | NA | NA | NA | NA | NA | Mif-ps3 | macrophage migration inhibitory factor, p | 0 | 0 | 0 | 0 | 0 | 0 |
| 25094 | ENSMUSG00000082827 | 0 | NA | NA | NA | NA | NA | Gm5397 | predicted gene 5397 [Source:MGI Symbol | 0 | 0 | 0 | 0 | 0 | 0 |
| 25095 | ENSMUSG00000082828 | 0 | NA | NA | NA | NA | NA | Gm16106 | predicted gene 16106 [Source:MGI Symbo | 0 | 0 | 0 | 0 | 0 | 0 |
| 25096 | ENSMUSG00000082829 | 0.545309144 | 2.651974629 | 3.666219369 | 0.72335405 | 0.469462394 | NA | Gm15780 | predicted gene 15780 [Source:MGI Symbo | 0 | 0 | 0 | 0 | 3.2719 | 0 |
| 25097 | ENSMUSG00000082831 | 0 | NA | NA | NA | NA | NA | Gm12591 | predicted gene 12591 [Source:MGI Symbo | 0 | 0 | 0 | 0 | 0 | 0 |
| 25098 | ENSMUSG00000082832 | 0 | NA | NA | NA | NA | NA | Gm14786 | predicted gene 14786 [Source:MGI Symbo | 0 | 0 | 0 | 0 | 0 | 0 |
| 25099 | ENSMUSG00000082833 | 0 | NA | NA | NA | NA | NA | Gm12982 | predicted gene 12982 [Source:MGI Symbo | 0 | 0 | 0 | 0 | 0 | 0 |
| 25100 | ENSMUSG00000082835 | 0 | NA | NA | NA | NA | NA | Gm13118 | predicted gene 13118 [Source:MGI Symbo | 0 | 0 | 0 | 0 | 0 | 0 |
| 25101 | ENSMUSG00000082836 | 1.875354368 | -0.113329325 | 1.567157098 | -0.07231523 | 0.942351045 | NA | Gm13612 | predicted gene 13612 [Source:MGI Symbo | 0.8031 | 1.0279 | 4.025 | 0.964 | 1.0906 | 3.3415 |
| 25102 | ENSMUSG00000082837 | 0.185640647 | 1.126924175 | 4.080472857 | 0.276174898 | 0.782413736 | NA | Gm15373 | predicted gene 15373 [Source:MGI Symbo | 0 | 0 | 0 | 0 | 0 | 1.1138 |
| 25103 | ENSMUSG00000082838 | 0.185640647 | 1.126924175 | 4.080472857 | 0.276174898 | 0.782413736 | NA | Gm14519 | predicted gene 14519 [Source:MGI Symbo | 0 | 0 | 0 | 0 | 0 | 1.1138 |
| 25104 | ENSMUSG00000082839 | 0 | NA | NA | NA | NA | NA | Gm14588 | predicted gene 14588 [Source:MGI Symbo | 0 | 0 | 0 | 0 | 0 | 0 |
| 25105 | ENSMUSG00000082840 | 0 | NA | NA | NA | NA | NA | Gm16421 | predicted gene 16421 [Source:MGI Symbo | 0 | 0 | 0 | 0 | 0 | 0 |
| 25106 | ENSMUSG00000082841 | 0 | NA | NA | NA | NA | NA | Gm12994 | predicted gene 12994 [Source:MGI Symbo | 0 | 0 | 0 | 0 | 0 | 0 |
| 25107 | ENSMUSG00000082842 | 0 | NA | NA | NA | NA | NA | Gm14748 | predicted gene 14748 [Source:MGI Symbo | 0 | 0 | 0 | 0 | 0 | 0 |
| 25108 | ENSMUSG00000082843 | 0 | NA | NA | NA | NA | NA | 4930448K20 | RIKEN cDNA 4930448K20 gene [Source:M | 0 | 0 | 0 | 0 | 0 | 0 |
| 25109 | ENSMUSG00000082844 | 0 | NA | NA | NA | NA | NA | Gm13537 | predicted gene 13537 [Source:MGI Symbo | 0 | 0 | 0 | 0 | 0 | 0 |
| 25110 | ENSMUSG00000082845 | 0 | NA | NA | NA | NA | NA | Gm13879 | predicted gene 13879 [Source:MGI Symbo | 0 | 0 | 0 | 0 | 0 | 0 |
| 25111 | ENSMUSG00000082846 | 0 | NA | NA | NA | NA | NA | Gm13484 | predicted gene 13484 [Source:MGI Symbo | 0 | 0 | 0 | 0 | 0 | 0 |
| 25112 | ENSMUSG00000082847 | 0 | NA | NA | NA | NA | NA | Gm5759 | predicted gene 5759 [Source:MGI Symbol | 0 | 0 | 0 | 0 | 0 | 0 |
| 25113 | ENSMUSG00000082848 | 0.321325949 | 1.911575114 | 4.048267893 | 0.472195804 | 0.636787034 | NA | Gm16066 | predicted gene 16066 [Source:MGI Symbo | 0 | 0 | 0 | 1.928 | 0 | 0 |
| 25114 | ENSMUSG00000082849 | 0 | NA | NA | NA | NA | NA | Gm14053 | predicted gene 14053 [Source:MGI Symbo | 0 | 0 | 0 | 0 | 0 | 0 |
| 25115 | ENSMUSG00000082850 | 0 | NA | NA | NA | NA | NA | Gm14299 | predicted gene 14299 [Source:MGI Symbo | 0 | 0 | 0 | 0 | 0 | 0 |
| 25116 | ENSMUSG00000082851 | 0.797609011 | 1.992900111 | 2.636035165 | 0.756021823 | 0.449636112 | NA | Btf3-ps8 | basic transcription factor 3, pseudogene 8 | 0.8031 | 0 | 0 | 2.8919 | 1.0906 | 0 |
| 25117 | ENSMUSG00000082852 | 0 | NA | NA | NA | NA | NA | Gm12021 | predicted gene 12021 [Source:MGI Symbo | 0 | 0 | 0 | 0 | 0 | 0 |
| 25118 | ENSMUSG00000082853 | 0 | NA | NA | NA | NA | NA | Gm7134 | predicted gene 7134 [Source:MGI Symbol | 0 | 0 | 0 | 0 | 0 | 0 |
| 25119 | ENSMUSG00000082855 | 0.181769715 | 1.126924175 | 4.080472857 | 0.276174898 | 0.782413736 | NA | Gm14537 | predicted gene 14537 [Source:MGI Symbo | 0 | 0 | 0 | 0 | 1.0906 | 0 |
| 25120 | ENSMUSG00000082856 | 0 | NA | NA | NA | NA | NA | Gm15106 | predicted gene 15106 [Source:MGI Symbo | 0 | 0 | 0 | 0 | 0 | 0 |
| 25121 | ENSMUSG00000082857 | 0 | NA | NA | NA | NA | NA | Gm12293 | predicted gene 12293 [Source:MGI Symbo | 0 | 0 | 0 | 0 | 0 | 0 |
| 25122 | ENSMUSG00000082858 | 0 | NA | NA | NA | NA | NA | Gm11746 | predicted gene 11746 [Source:MGI Symbo | 0 | 0 | 0 | 0 | 0 | 0 |
| 25123 | ENSMUSG00000082859 | 0 | NA | NA | NA | NA | NA | Defa-ps9 | defensin, alpha, pseudogene 9 [Source:M | 0 | 0 | 0 | 0 | 0 | 0 |
| 25124 | ENSMUSG00000082860 | 5.449118573 | -0.148456904 | 0.921180543 | -0.1611594 | 0.871967857 | 0.979369708 | Gm13204 | predicted gene 13204 [Source:MGI Symbo | 8.031 | 3.0838 | 6.0375 | 6.7478 | 5.4531 | 3.3415 |
| 25125 | ENSMUSG00000082861 | 0 | NA | NA | NA | NA | NA | Gm11362 | predicted gene 11362 [Source:MGI Symbo | 0 | 0 | 0 | 0 | 0 | 0 |
| 25126 | ENSMUSG00000082862 | 0 | NA | NA | NA | NA | NA | Gm13806 | predicted gene 13806 [Source:MGI Symbo | 0 | 0 | 0 | 0 | 0 | 0 |
| 25127 | ENSMUSG00000082863 | 0 | NA | NA | NA | NA | NA | Gm14674 | predicted gene 14674 [Source:MGI Symbo | 0 | 0 | 0 | 0 | 0 | 0 |
| 25128 | ENSMUSG00000082864 | 0 | NA | NA | NA | NA | NA | Gm13358 | predicted gene 13358 [Source:MGI Symbo | 0 | 0 | 0 | 0 | 0 | 0 |
| 25129 | ENSMUSG00000082865 | 0 | NA | NA | NA | NA | NA | Gm14745 | predicted gene 14745 [Source:MGI Symbo | 0 | 0 | 0 | 0 | 0 | 0 |
| 25130 | ENSMUSG00000082867 | 0 | NA | NA | NA | NA | NA | Gm15730 | predicted gene 15730 [Source:MGI Symbo | 0 | 0 | 0 | 0 | 0 | 0 |
| 25131 | ENSMUSG00000082868 | 0 | NA | NA | NA | NA | NA | Mup-ps5 | major urinary protein, pseudogene 5 [Sou | 0 | 0 | 0 | 0 | 0 | 0 |
| 25132 | ENSMUSG00000082869 | 0 | NA | NA | NA | NA | NA | Vmn1r-ps93 | vomeronasal 1 receptor, pseudogene 93 [ | 0 | 0 | 0 | 0 | 0 | 0 |
| 25133 | ENSMUSG00000082870 | 0 | NA | NA | NA | NA | NA | Gm12165 | predicted gene 12165 [Source:MGI Symbo | 0 | 0 | 0 | 0 | 0 | 0 |
| 25134 | ENSMUSG00000082871 | 0 | NA | NA | NA | NA | NA | Gm13263 | predicted gene 13263 [Source:MGI Symbo | 0 | 0 | 0 | 0 | 0 | 0 |
| 25135 | ENSMUSG00000082872 | 0 | NA | NA | NA | NA | NA | Gm15773 | predicted gene 15773 [Source:MGI Symbo | 0 | 0 | 0 | 0 | 0 | 0 |
| 25136 | ENSMUSG00000082873 | 0 | NA | NA | NA | NA | NA | Gm15367 | predicted gene 15367 [Source:MGI Symbo | 0 | 0 | 0 | 0 | 0 | 0 |
| 25137 | ENSMUSG00000082874 | 0 | NA | NA | NA | NA | NA | Rpl15-ps1 | ribosomal protein L15, pseudogene 1 [Sou | 0 | 0 | 0 | 0 | 0 | 0 |
| 25138 | ENSMUSG00000082875 | 0 | NA | NA | NA | NA | NA | Gm12598 | predicted gene 12598 [Source:MGI Symbo | 0 | 0 | 0 | 0 | 0 | 0 |
| 25139 | ENSMUSG00000082876 | 1.055449264 | 2.395599456 | 2.43289538 | 0.984670149 | 0.32478612 | NA | Gm11889 | predicted gene 11889 [Source:MGI Symbo | 0 | 0 | 1.0062 | 0.964 | 4.3625 | 0 |
| 25140 | ENSMUSG00000082877 | 0 | NA | NA | NA | NA | NA | Gm13009 | predicted gene 13009 [Source:MGI Symbo | 0 | 0 | 0 | 0 | 0 | 0 |

| | | | | | | | | | | | | | | | |
|---|---|---|---|---|---|---|---|---|---|---|---|---|---|---|---|
| 25141 | ENSMUSG00000082878 | 0 | NA | NA | NA | NA | NA | Gm12583 | predicted gene 12583 [Source:MGI Symbo | 0 | 0 | 0 | 0 | 0 | 0 |
| 25142 | ENSMUSG00000082879 | 0 | NA | NA | NA | NA | NA | Gm13525 | predicted gene 13525 [Source:MGI Symbo | 0 | 0 | 0 | 0 | 0 | 0 |
| 25143 | ENSMUSG00000082882 | 0 | NA | NA | NA | NA | NA | Olfr1186 | olfactory receptor 1186 [Source:MGI Sym | 0 | 0 | 0 | 0 | 0 | 0 |
| 25144 | ENSMUSG00000082884 | 12.99313188 | 0.238227834 | 0.643549544 | 0.370177923 | 0.711249925 | 0.944239215 | Gm13339 | predicted gene 13339 [Source:MGI Symbo | 11.243 | 19.53 | 5.0312 | 13.496 | 14.178 | 14.48 |
| 25145 | ENSMUSG00000082886 | 0 | NA | NA | NA | NA | NA | Gm5386 | predicted pseudogene 5386 [Source:MGI | 0 | 0 | 0 | 0 | 0 | 0 |
| 25146 | ENSMUSG00000082887 | 0 | NA | NA | NA | NA | NA | Gm14607 | predicted gene 14607 [Source:MGI Symbo | 0 | 0 | 0 | 0 | 0 | 0 |
| 25147 | ENSMUSG00000082888 | 0.513958919 | -2.394274821 | 3.756096013 | -0.637437066 | 0.523840191 | NA | Gm12934 | predicted gene 12934 [Source:MGI Symbo | 0 | 3.0838 | 0 | 0 | 0 | 0 |
| 25148 | ENSMUSG00000082889 | 0 | NA | NA | NA | NA | NA | Cyp4a29-ps1 | cytochrome P450, family 4, subfamily a, p | 0 | 0 | 0 | 0 | 0 | 0 |
| 25149 | ENSMUSG00000082890 | 0 | NA | NA | NA | NA | NA | Gm15014 | predicted gene 15014 [Source:MGI Symbo | 0 | 0 | 0 | 0 | 0 | 0 |
| 25150 | ENSMUSG00000082891 | 0 | NA | NA | NA | NA | NA | Gm12881 | predicted gene 12881 [Source:MGI Symbo | 0 | 0 | 0 | 0 | 0 | 0 |
| 25151 | ENSMUSG00000082892 | 0 | NA | NA | NA | NA | NA | Olfr1027-ps1 | olfactory receptor 1027, pseudogene 1 [S | 0 | 0 | 0 | 0 | 0 | 0 |
| 25152 | ENSMUSG00000082893 | 0 | NA | NA | NA | NA | NA | Gm11383 | predicted gene 11383 [Source:MGI Symbo | 0 | 0 | 0 | 0 | 0 | 0 |
| 25153 | ENSMUSG00000082894 | 0 | NA | NA | NA | NA | NA | Gm6480 | predicted pseudogene 6480 [Source:MGI | 0 | 0 | 0 | 0 | 0 | 0 |
| 25154 | ENSMUSG00000082895 | 1.219724447 | 2.639236362 | 2.296847442 | 1.149069074 | 0.2505275 | NA | Rpsa-ps9 | ribosomal protein SA, pseudogene 9 [Sou | 0 | 1.0279 | 0 | 1.928 | 4.3625 | 0 |
| 25155 | ENSMUSG00000082896 | 2.301856531 | 0.66713975 | 1.396269054 | 0.477801716 | 0.63279134 | NA | Gm5844 | predicted gene 5844 [Source:MGI Symbol | 3.2124 | 1.0279 | 1.0062 | 1.928 | 2.1812 | 4.4554 |
| 25156 | ENSMUSG00000082897 | 0 | NA | NA | NA | NA | NA | Gm11982 | predicted gene 11982 [Source:MGI Symbo | 0 | 0 | 0 | 0 | 0 | 0 |
| 25157 | ENSMUSG00000082898 | 0 | NA | NA | NA | NA | NA | Gm12041 | predicted gene 12041 [Source:MGI Symbo | 0 | 0 | 0 | 0 | 0 | 0 |
| 25158 | ENSMUSG00000082899 | 0 | NA | NA | NA | NA | NA | Gm12379 | predicted gene 12379 [Source:MGI Symbo | 0 | 0 | 0 | 0 | 0 | 0 |
| 25159 | ENSMUSG00000082900 | 0 | NA | NA | NA | NA | NA | Gm11557 | predicted gene 11557 [Source:MGI Symbo | 0 | 0 | 0 | 0 | 0 | 0 |
| 25160 | ENSMUSG00000082901 | 0 | NA | NA | NA | NA | NA | Rps8-ps5 | ribosomal protein S8, pseudogene 5 [Sour | 0 | 0 | 0 | 0 | 0 | 0 |
| 25161 | ENSMUSG00000082902 | 1.288394167 | -3.725625981 | 2.154147746 | -1.729512745 | 0.083717367 | NA | Ccl19-ps1 | chemokine (C-C motif) ligand 19, pseudog | 1.6062 | 4.1117 | 2.0125 | 0 | 0 | 0 |
| 25162 | ENSMUSG00000082903 | 0 | NA | NA | NA | NA | NA | Gm14083 | predicted gene 14083 [Source:MGI Symbo | 0 | 0 | 0 | 0 | 0 | 0 |
| 25163 | ENSMUSG00000082905 | 0 | NA | NA | NA | NA | NA | Gm7823 | predicted gene 7823 [Source:MGI Symbol | 0 | 0 | 0 | 0 | 0 | 0 |
| 25164 | ENSMUSG00000082906 | 0 | NA | NA | NA | NA | NA | Smt3h1-ps | SMT3 suppressor of mif two 3 homolog 1, | 0 | 0 | 0 | 0 | 0 | 0 |
| 25165 | ENSMUSG00000082907 | 0 | NA | NA | NA | NA | NA | Gm14832 | predicted gene 14832 [Source:MGI Symbo | 0 | 0 | 0 | 0 | 0 | 0 |
| 25166 | ENSMUSG00000082908 | 1.381136435 | -0.799612225 | 1.959780848 | -0.408011041 | 0.683265569 | NA | Gm13736 | predicted gene 13736 [Source:MGI Symbo | 3.2124 | 2.0558 | 0 | 1.928 | 1.0906 | 0 |
| 25167 | ENSMUSG00000082910 | 0 | NA | NA | NA | NA | NA | Gm14758 | predicted gene 14758 [Source:MGI Symbo | 0 | 0 | 0 | 0 | 0 | 0 |
| 25168 | ENSMUSG00000082911 | 0 | NA | NA | NA | NA | NA | Gm14119 | predicted gene 14119 [Source:MGI Symbo | 0 | 0 | 0 | 0 | 0 | 0 |
| 25169 | ENSMUSG00000082912 | 0 | NA | NA | NA | NA | NA | Gm8770 | predicted gene 8770 [Source:MGI Symbol | 0 | 0 | 0 | 0 | 0 | 0 |
| 25170 | ENSMUSG00000082913 | 0 | NA | NA | NA | NA | NA | Gm14739 | predicted gene 14739 [Source:MGI Symbo | 0 | 0 | 0 | 0 | 0 | 0 |
| 25171 | ENSMUSG00000082915 | 0 | NA | NA | NA | NA | NA | Gm15240 | predicted gene 15240 [Source:MGI Symbo | 0 | 0 | 0 | 0 | 0 | 0 |
| 25172 | ENSMUSG00000082917 | 0 | NA | NA | NA | NA | NA | Gm11440 | predicted gene 11440 [Source:MGI Symbo | 0 | 0 | 0 | 0 | 0 | 0 |
| 25173 | ENSMUSG00000082918 | 0 | NA | NA | NA | NA | NA | Defa-ps16 | defensin, alpha, pseudogene 16 [Source:N | 0 | 0 | 0 | 0 | 0 | 0 |
| 25174 | ENSMUSG00000082919 | 0 | NA | NA | NA | NA | NA | Gm12641 | predicted gene 12641 [Source:MGI Symbo | 0 | 0 | 0 | 0 | 0 | 0 |
| 25175 | ENSMUSG00000082920 | 1.42224454 | -3.871861966 | 2.074950443 | -1.86600214 | 0.062041066 | NA | Gm13864 | predicted gene 13864 [Source:MGI Symbo | 2.4093 | 4.1117 | 2.0125 | 0 | 0 | 0 |
| 25176 | ENSMUSG00000082922 | 4.992112837 | -0.503203235 | 0.978844627 | -0.514078763 | 0.607196925 | NA | Gm6939 | predicted gene 6939 [Source:MGI Symbol | 6.4248 | 5.1396 | 6.0375 | 5.7839 | 5.4531 | 1.1138 |
| 25177 | ENSMUSG00000082924 | 0 | NA | NA | NA | NA | NA | Gm11462 | predicted gene 11462 [Source:MGI Symbo | 0 | 0 | 0 | 0 | 0 | 0 |
| 25178 | ENSMUSG00000082925 | 0.904673413 | -3.227537609 | 2.678727401 | -1.204877214 | 0.228250706 | NA | Gm13135 | predicted gene 13135 [Source:MGI Symbo | 2.4093 | 0 | 3.0187 | 0 | 0 | 0 |
| 25179 | ENSMUSG00000082926 | 0.795774083 | 0.899657507 | 2.433208733 | 0.369741196 | 0.711575334 | NA | Gm8172 | predicted pseudogene 8172 [Source:MGI | 1.6062 | 0 | 0 | 0.964 | 1.0906 | 1.1138 |
| 25180 | ENSMUSG00000082927 | 0 | NA | NA | NA | NA | NA | Gm5863 | predicted gene 5863 [Source:MGI Symbol | 0 | 0 | 0 | 0 | 0 | 0 |
| 25181 | ENSMUSG00000082928 | 0.181769715 | 1.126924175 | 4.080472857 | 0.276174898 | 0.782413736 | NA | Gm12785 | predicted gene 12785 [Source:MGI Symbo | 0 | 0 | 0 | 0 | 1.0906 | 0 |
| 25182 | ENSMUSG00000082929 | 0 | NA | NA | NA | NA | NA | Gm6067 | predicted gene 6067 [Source:MGI Symbol | 0 | 0 | 0 | 0 | 0 | 0 |
| 25183 | ENSMUSG00000082930 | 0 | NA | NA | NA | NA | NA | Gm13462 | predicted gene 13462 [Source:MGI Symbo | 0 | 0 | 0 | 0 | 0 | 0 |
| 25184 | ENSMUSG00000082931 | 0 | NA | NA | NA | NA | NA | Gm13586 | predicted gene 13586 [Source:MGI Symbo | 0 | 0 | 0 | 0 | 0 | 0 |
| 25185 | ENSMUSG00000082932 | 1.661587499 | -1.257330817 | 1.845442275 | -0.681316796 | 0.495671058 | NA | Cyp2j8 | cytochrome P450, family 2, subfamily j, pc | 4.0155 | 2.0558 | 1.0062 | 2.8919 | 0 | 0 |
| 25186 | ENSMUSG00000082933 | 0 | NA | NA | NA | NA | NA | Gm14811 | predicted gene 14811 [Source:MGI Symbo | 0 | 0 | 0 | 0 | 0 | 0 |
| 25187 | ENSMUSG00000082934 | 0 | NA | NA | NA | NA | NA | Gm13887 | predicted gene 13887 [Source:MGI Symbo | 0 | 0 | 0 | 0 | 0 | 0 |
| 25188 | ENSMUSG00000082935 | 0.995767881 | -1.008676959 | 2.44979061 | -0.411740071 | 0.680529951 | NA | Gm7658 | predicted gene 7658 [Source:MGI Symbol | 0 | 1.0279 | 3.0187 | 1.928 | 0 | 0 |
| 25189 | ENSMUSG00000082936 | 0.925960114 | -1.988843075 | 2.485090455 | -0.800310134 | 0.423531133 | NA | Gm16166 | predicted gene 16166 [Source:MGI Symbo | 2.4093 | 2.0558 | 0 | 0 | 1.0906 | 0 |
| 25190 | ENSMUSG00000082937 | 0 | NA | NA | NA | NA | NA | Gm15095 | predicted gene 15095 [Source:MGI Symbo | 0 | 0 | 0 | 0 | 0 | 0 |
| 25191 | ENSMUSG00000082938 | 3.237005471 | -0.333478036 | 1.371882978 | -0.243080526 | 0.807943017 | NA | Gm2810 | predicted pseudogene 2810 [Source:MGI | 1.6062 | 9.2513 | 0 | 1.928 | 2.1812 | 4.4554 |
| 25192 | ENSMUSG00000082939 | 0 | NA | NA | NA | NA | NA | Gm14881 | predicted gene 14881 [Source:MGI Symbo | 0 | 0 | 0 | 0 | 0 | 0 |
| 25193 | ENSMUSG00000082940 | 0 | NA | NA | NA | NA | NA | Gm14103 | predicted gene 14103 [Source:MGI Symbo | 0 | 0 | 0 | 0 | 0 | 0 |
| 25194 | ENSMUSG00000082941 | 0.17131964 | -0.796629176 | 4.080472857 | -0.195229623 | 0.84521319 | NA | Gm12850 | predicted gene 12850 [Source:MGI Symbo | 0 | 1.0279 | 0 | 0 | 0 | 0 |
| 25195 | ENSMUSG00000082943 | 0 | NA | NA | NA | NA | NA | Gm13355 | predicted gene 13355 [Source:MGI Symbo | 0 | 0 | 0 | 0 | 0 | 0 |
| 25196 | ENSMUSG00000082945 | 0.654905883 | -1.312574656 | 2.816093566 | -0.466097672 | 0.641145589 | NA | Gm11540 | predicted gene 11540 [Source:MGI Symbo | 0.8031 | 0 | 2.0125 | 0 | 0 | 1.1138 |

| | | | | | | | | | | | | | | | |
|---|---|---|---|---|---|---|---|---|---|---|---|---|---|---|---|
| 25197 | ENSMUSG00000082947 | 0 | NA | NA | NA | NA | NA | Gm12587 | predicted gene 12587 [Source:MGI Symbo | 0 | 0 | 0 | 0 | 0 | 0 |
| 25198 | ENSMUSG00000082949 | 0 | NA | NA | NA | NA | NA | Gm11920 | predicted gene 11920 [Source:MGI Symbo | 0 | 0 | 0 | 0 | 0 | 0 |
| 25199 | ENSMUSG00000082950 | 0 | NA | NA | NA | NA | NA | Btf3-ps10 | basic transcription factor 3, pseudogene 1 | 0 | 0 | 0 | 0 | 0 | 0 |
| 25200 | ENSMUSG00000082951 | 0 | NA | NA | NA | NA | NA | Gm13359 | predicted gene 13359 [Source:MGI Symbo | 0 | 0 | 0 | 0 | 0 | 0 |
| 25201 | ENSMUSG00000082952 | 0 | NA | NA | NA | NA | NA | Gm12507 | predicted gene 12507 [Source:MGI Symbo | 0 | 0 | 0 | 0 | 0 | 0 |
| 25202 | ENSMUSG00000082953 | 0.342432689 | 1.990653086 | 3.900281263 | 0.51038706 | 0.609780321 | NA | Gm13217 | predicted gene 13217 [Source:MGI Symbo | 0 | 0 | 0 | 0.964 | 1.0906 | 0 |
| 25203 | ENSMUSG00000082954 | 0 | NA | NA | NA | NA | NA | Gm15504 | predicted gene 15504 [Source:MGI Symbo | 0 | 0 | 0 | 0 | 0 | 0 |
| 25204 | ENSMUSG00000082955 | 0 | NA | NA | NA | NA | NA | Gm8036 | predicted gene 8036 [Source:MGI Symbol | 0 | 0 | 0 | 0 | 0 | 0 |
| 25205 | ENSMUSG00000082956 | 0 | NA | NA | NA | NA | NA | Naip3 | NLR family, apoptosis inhibitory protein 3 | 0 | 0 | 0 | 0 | 0 | 0 |
| 25206 | ENSMUSG00000082957 | 0 | NA | NA | NA | NA | NA | Gm15303 | predicted gene 15303 [Source:MGI Symbo | 0 | 0 | 0 | 0 | 0 | 0 |
| 25207 | ENSMUSG00000082958 | 0 | NA | NA | NA | NA | NA | Gm12731 | predicted gene 12731 [Source:MGI Symbo | 0 | 0 | 0 | 0 | 0 | 0 |
| 25208 | ENSMUSG00000082960 | 0 | NA | NA | NA | NA | NA | Gm6884 | predicted gene 6884 [Source:MGI Symbol | 0 | 0 | 0 | 0 | 0 | 0 |
| 25209 | ENSMUSG00000082961 | 0.160662974 | 1.126924175 | 4.080472857 | 0.276174898 | 0.782413736 | NA | Gm15131 | predicted gene 15131 [Source:MGI Symbo | 0 | 0 | 0 | 0.964 | 0 | 0 |
| 25210 | ENSMUSG00000082962 | 0 | NA | NA | NA | NA | NA | Gm10601 | predicted pseudogene 10601 [Source:MG | 0 | 0 | 0 | 0 | 0 | 0 |
| 25211 | ENSMUSG00000082963 | 0 | NA | NA | NA | NA | NA | Gm12465 | predicted gene 12465 [Source:MGI Symbo | 0 | 0 | 0 | 0 | 0 | 0 |
| 25212 | ENSMUSG00000082964 | 1.385338226 | 0.930866123 | 2.200087305 | 0.423104175 | 0.672219257 | NA | Rpl13-ps5 | ribosomal protein L13, pseudogene 5 [Sou | 0.8031 | 2.0558 | 0 | 0 | 5.4531 | 0 |
| 25213 | ENSMUSG00000082965 | 0 | NA | NA | NA | NA | NA | Gm6754 | predicted pseudogene 6754 [Source:MGI | 0 | 0 | 0 | 0 | 0 | 0 |
| 25214 | ENSMUSG00000082966 | 0 | NA | NA | NA | NA | NA | Gm14614 | predicted gene 14614 [Source:MGI Symbo | 0 | 0 | 0 | 0 | 0 | 0 |
| 25215 | ENSMUSG00000082967 | 0 | NA | NA | NA | NA | NA | Gm16082 | predicted gene 16082 [Source:MGI Symbo | 0 | 0 | 0 | 0 | 0 | 0 |
| 25216 | ENSMUSG00000082968 | 0 | NA | NA | NA | NA | NA | Gm14581 | predicted gene 14581 [Source:MGI Symbo | 0 | 0 | 0 | 0 | 0 | 0 |
| 25217 | ENSMUSG00000082969 | 0.845941533 | -1.843011426 | 2.87307209 | -0.641477613 | 0.521212422 | NA | Gm12630 | predicted gene 12630 [Source:MGI Symbo | 0 | 4.1117 | 0 | 0.964 | 0 | 0 |
| 25218 | ENSMUSG00000082970 | 0 | NA | NA | NA | NA | NA | Gm15214 | predicted gene 15214 [Source:MGI Symbo | 0 | 0 | 0 | 0 | 0 | 0 |
| 25219 | ENSMUSG00000082971 | 0 | NA | NA | NA | NA | NA | Gm13517 | predicted gene 13517 [Source:MGI Symbo | 0 | 0 | 0 | 0 | 0 | 0 |
| 25220 | ENSMUSG00000082973 | 0 | NA | NA | NA | NA | NA | Gm13672 | predicted gene 13672 [Source:MGI Symbo | 0 | 0 | 0 | 0 | 0 | 0 |
| 25221 | ENSMUSG00000082974 | 0.342639279 | -1.805976596 | 4.028547714 | -0.448294702 | 0.653940524 | NA | Gm15129 | predicted gene 15129 [Source:MGI Symbo | 0 | 2.0558 | 0 | 0 | 0 | 0 |
| 25222 | ENSMUSG00000082975 | 0 | NA | NA | NA | NA | NA | Gm14363 | predicted gene 14363 [Source:MGI Symbo | 0 | 0 | 0 | 0 | 0 | 0 |
| 25223 | ENSMUSG00000082976 | 0 | NA | NA | NA | NA | NA | Gm15056 | predicted gene 15056 [Source:MGI Symbo | 0 | 0 | 0 | 0 | 0 | 0 |
| 25224 | ENSMUSG00000082977 | 0 | NA | NA | NA | NA | NA | Gm11911 | predicted gene 11911 [Source:MGI Symbo | 0 | 0 | 0 | 0 | 0 | 0 |
| 25225 | ENSMUSG00000082978 | 0 | NA | NA | NA | NA | NA | Rpsa-ps11 | ribosomal protein SA, pseudogene 11 [Sou | 0 | 0 | 0 | 0 | 0 | 0 |
| 25226 | ENSMUSG00000082980 | 0 | NA | NA | NA | NA | NA | Olfr1274 | olfactory receptor 1274 [Source:MGI Syml | 0 | 0 | 0 | 0 | 0 | 0 |
| 25227 | ENSMUSG00000082981 | 0.133850373 | -0.796629176 | 4.080472857 | -0.195229623 | 0.84521319 | NA | Polr2k-ps | polymerase (RNA) II (DNA directed) polyp | 0.8031 | 0 | 0 | 0 | 0 | 0 |
| 25228 | ENSMUSG00000082982 | 0 | NA | NA | NA | NA | NA | Olfr1149-ps1 | olfactory receptor 1149, pseudogene 1 [So | 0 | 0 | 0 | 0 | 0 | 0 |
| 25229 | ENSMUSG00000082983 | 0.160662974 | 1.126924175 | 4.080472857 | 0.276174898 | 0.782413736 | NA | Gm12318 | predicted gene 12318 [Source:MGI Symbo | 0 | 0 | 0 | 0.964 | 0 | 0 |
| 25230 | ENSMUSG00000082985 | 0 | NA | NA | NA | NA | NA | Gm14042 | predicted gene 14042 [Source:MGI Symbo | 0 | 0 | 0 | 0 | 0 | 0 |
| 25231 | ENSMUSG00000082986 | 0 | NA | NA | NA | NA | NA | Gm11389 | predicted gene 11389 [Source:MGI Symbo | 0 | 0 | 0 | 0 | 0 | 0 |
| 25232 | ENSMUSG00000082987 | 0 | NA | NA | NA | NA | NA | Rps13-ps6 | ribosomal protein S13, pseudogene 6 [Sou | 0 | 0 | 0 | 0 | 0 | 0 |
| 25233 | ENSMUSG00000082988 | 0.367410362 | 2.078255025 | 3.800070157 | 0.54689912 | 0.58444804 | NA | Gm5903 | predicted gene 5903 [Source:MGI Symbol | 0 | 0 | 0 | 0 | 1.0906 | 1.1138 |
| 25234 | ENSMUSG00000082989 | 0 | NA | NA | NA | NA | NA | Gm13557 | predicted gene 13557 [Source:MGI Symbo | 0 | 0 | 0 | 0 | 0 | 0 |
| 25235 | ENSMUSG00000082990 | 0.455176322 | 0.955156269 | 3.546674842 | 0.269310357 | 0.787690862 | NA | Gm14052 | predicted gene 14052 [Source:MGI Symbo | 0.8031 | 0 | 0 | 1.928 | 0 | 0 |
| 25236 | ENSMUSG00000082991 | 0 | NA | NA | NA | NA | NA | Rps10-ps3 | ribosomal protein S10, pseudogene 3 [Sou | 0 | 0 | 0 | 0 | 0 | 0 |
| 25237 | ENSMUSG00000082993 | 0 | NA | NA | NA | NA | NA | Gm11566 | predicted gene 11566 [Source:MGI Symbo | 0 | 0 | 0 | 0 | 0 | 0 |
| 25238 | ENSMUSG00000082994 | 0 | NA | NA | NA | NA | NA | Gm11848 | predicted gene 11848 [Source:MGI Symbo | 0 | 0 | 0 | 0 | 0 | 0 |
| 25239 | ENSMUSG00000082995 | 0 | NA | NA | NA | NA | NA | Gm15126 | predicted gene 15126 [Source:MGI Symbo | 0 | 0 | 0 | 0 | 0 | 0 |
| 25240 | ENSMUSG00000082996 | 0 | NA | NA | NA | NA | NA | Gm6238 | predicted pseudogene 6238 [Source:MGI | 0 | 0 | 0 | 0 | 0 | 0 |
| 25241 | ENSMUSG00000082997 | 0 | NA | NA | NA | NA | NA | Gm13975 | predicted gene 13975 [Source:MGI Symbo | 0 | 0 | 0 | 0 | 0 | 0 |
| 25242 | ENSMUSG00000082998 | 0.465832987 | -0.698295221 | 3.18348759 | -0.219349126 | 0.826378096 | NA | Gm11221 | predicted gene 11221 [Source:MGI Symbo | 0.8031 | 1.0279 | 0 | 0.964 | 0 | 0 |
| 25243 | ENSMUSG00000083000 | 0 | NA | NA | NA | NA | NA | Gm15678 | predicted gene 15678 [Source:MGI Symbo | 0 | 0 | 0 | 0 | 0 | 0 |
| 25244 | ENSMUSG00000083001 | 0 | NA | NA | NA | NA | NA | Gm13250 | predicted gene 13250 [Source:MGI Symbo | 0 | 0 | 0 | 0 | 0 | 0 |
| 25245 | ENSMUSG00000083004 | 2.738025278 | 0.947735624 | 1.48613548 | 0.637718187 | 0.523657144 | NA | Gm13413 | predicted gene 13413 [Source:MGI Symbo | 1.6062 | 2.0558 | 2.0125 | 9.6398 | 0 | 1.1138 |
| 25246 | ENSMUSG00000083005 | 0 | NA | NA | NA | NA | NA | Gm12392 | predicted gene 12392 [Source:MGI Symbo | 0 | 0 | 0 | 0 | 0 | 0 |
| 25247 | ENSMUSG00000083006 | 0 | NA | NA | NA | NA | NA | Gm15715 | predicted gene 15715 [Source:MGI Symbo | 0 | 0 | 0 | 0 | 0 | 0 |
| 25248 | ENSMUSG00000083007 | 0 | NA | NA | NA | NA | NA | Gm13623 | predicted gene 13623 [Source:MGI Symbo | 0 | 0 | 0 | 0 | 0 | 0 |
| 25249 | ENSMUSG00000083008 | 0 | NA | NA | NA | NA | NA | Gm12053 | predicted gene 12053 [Source:MGI Symbo | 0 | 0 | 0 | 0 | 0 | 0 |
| 25250 | ENSMUSG00000083009 | 0 | NA | NA | NA | NA | NA | Gm1335 | predicted gene 1335 [Source:MGI Symbol | 0 | 0 | 0 | 0 | 0 | 0 |
| 25251 | ENSMUSG00000083010 | 0 | NA | NA | NA | NA | NA | Gm14687 | predicted gene 14687 [Source:MGI Symbo | 0 | 0 | 0 | 0 | 0 | 0 |
| 25252 | ENSMUSG00000083011 | 1.063656444 | 0.142299728 | 2.383976614 | 0.059690069 | 0.952402482 | NA | Gm12816 | predicted gene 12816 [Source:MGI Symbo | 0 | 1.0279 | 2.0125 | 0 | 0 | 3.3415 |

| | | | | | | | | | | | | | | | |
|---|---|---|---|---|---|---|---|---|---|---|---|---|---|---|---|
| 25253 | ENSMUSG00000083012 | 179.3474922 | 0.058428652 | 0.191716489 | 0.304765921 | 0.760544438 | 0.95402891 | Fam220a | family with sequence similarity 220, meml | 145.36 | 191.19 | 191.19 | 173.52 | 182.13 | 192.69 |
| 25254 | ENSMUSG00000083013 | 0 | NA | NA | NA | NA | NA | Gm12978 | predicted gene 12978 [Source:MGI Symbc | 0 | 0 | 0 | 0 | 0 | 0 |
| 25255 | ENSMUSG00000083014 | 0.133850373 | -0.796629176 | 4.080472857 | -0.195229623 | 0.84521319 | NA | Gm11764 | predicted gene 11764 [Source:MGI Symbc | 0.8031 | 0 | 0 | 0 | 0 | 0 |
| 25256 | ENSMUSG00000083015 | 0 | NA | NA | NA | NA | NA | Gm16518 | predicted gene, 16518 [Source:MGI Symb | 0 | 0 | 0 | 0 | 0 | 0 |
| 25257 | ENSMUSG00000083016 | 0.966655038 | -0.754805966 | 2.43372943 | -0.310143747 | 0.756451646 | NA | Gm12618 | predicted gene 12618 [Source:MGI Symbc | 1.6062 | 0 | 2.0125 | 0 | 2.1812 | 0 |
| 25258 | ENSMUSG00000083017 | 0 | NA | NA | NA | NA | NA | Gm12163 | predicted gene 12163 [Source:MGI Symbc | 0 | 0 | 0 | 0 | 0 | 0 |
| 25259 | ENSMUSG00000083018 | 0 | NA | NA | NA | NA | NA | Gm13555 | predicted gene 13555 [Source:MGI Symbc | 0 | 0 | 0 | 0 | 0 | 0 |
| 25260 | ENSMUSG00000083019 | 0.371281295 | 2.09253362 | 4.031852165 | 0.519000582 | 0.603760335 | NA | Gm6222 | predicted pseudogene 6222 [Source:MGI | 0 | 0 | 0 | 0 | 0 | 2.2277 |
| 25261 | ENSMUSG00000083020 | 0 | NA | NA | NA | NA | NA | Gm15331 | predicted gene 15331 [Source:MGI Symbc | 0 | 0 | 0 | 0 | 0 | 0 |
| 25262 | ENSMUSG00000083022 | 0 | NA | NA | NA | NA | NA | Rps15a-ps6 | ribosomal protein S15A, pseudogene 6 [So | 0 | 0 | 0 | 0 | 0 | 0 |
| 25263 | ENSMUSG00000083023 | 0 | NA | NA | NA | NA | NA | Gm11804 | predicted gene 11804 [Source:MGI Symbc | 0 | 0 | 0 | 0 | 0 | 0 |
| 25264 | ENSMUSG00000083024 | 0 | NA | NA | NA | NA | NA | Gm14982 | predicted gene 14982 [Source:MGI Symbc | 0 | 0 | 0 | 0 | 0 | 0 |
| 25265 | ENSMUSG00000083025 | 0 | NA | NA | NA | NA | NA | Gm11829 | predicted gene 11829 [Source:MGI Symbc | 0 | 0 | 0 | 0 | 0 | 0 |
| 25266 | ENSMUSG00000083026 | 0 | NA | NA | NA | NA | NA | Gm6373 | predicted gene 6373 [Source:MGI Symbol | 0 | 0 | 0 | 0 | 0 | 0 |
| 25267 | ENSMUSG00000083027 | 0.160662974 | 1.126924175 | 4.080472857 | 0.276174898 | 0.782413736 | NA | Gm13140 | predicted gene 13140 [Source:MGI Symbc | 0 | 0 | 0 | 0.964 | 0 | 0 |
| 25268 | ENSMUSG00000083029 | 0 | NA | NA | NA | NA | NA | Gm13720 | predicted gene 13720 [Source:MGI Symbc | 0 | 0 | 0 | 0 | 0 | 0 |
| 25269 | ENSMUSG00000083030 | 0 | NA | NA | NA | NA | NA | Gm14561 | predicted gene 14561 [Source:MGI Symbc | 0 | 0 | 0 | 0 | 0 | 0 |
| 25270 | ENSMUSG00000083031 | 0 | NA | NA | NA | NA | NA | Gm14910 | predicted gene 14910 [Source:MGI Symbc | 0 | 0 | 0 | 0 | 0 | 0 |
| 25271 | ENSMUSG00000083032 | 0 | NA | NA | NA | NA | NA | Gm8087 | predicted gene 8087 [Source:MGI Symbol | 0 | 0 | 0 | 0 | 0 | 0 |
| 25272 | ENSMUSG00000083033 | 1.029540662 | 0.20390886 | 2.204283627 | 0.092505727 | 0.926296241 | NA | Gm9083 | predicted gene 9083 [Source:MGI Symbol | 0.8031 | 2.0558 | 0 | 0 | 1.0906 | 2.2277 |
| 25273 | ENSMUSG00000083034 | 0.342639279 | -1.805976596 | 4.028547714 | -0.448294702 | 0.653940524 | NA | Olfr334-ps1 | olfactory receptor 334, pseudogene 1 [So | 0 | 2.0558 | 0 | 0 | 0 | 0 |
| 25274 | ENSMUSG00000083036 | 0 | NA | NA | NA | NA | NA | Gm15959 | predicted gene 15959 [Source:MGI Symbc | 0 | 0 | 0 | 0 | 0 | 0 |
| 25275 | ENSMUSG00000083037 | 0 | NA | NA | NA | NA | NA | Gm14900 | predicted gene 14900 [Source:MGI Symbc | 0 | 0 | 0 | 0 | 0 | 0 |
| 25276 | ENSMUSG00000083038 | 0.904853373 | -1.988674067 | 2.502700767 | -0.794611203 | 0.4268397 | NA | Gm13233 | predicted gene 13233 [Source:MGI Symbc | 2.4093 | 2.0558 | 0 | 0.964 | 0 | 0 |
| 25277 | ENSMUSG00000083039 | 0 | NA | NA | NA | NA | NA | Gm11606 | predicted gene 11606 [Source:MGI Symbc | 0 | 0 | 0 | 0 | 0 | 0 |
| 25278 | ENSMUSG00000083041 | 0.885033624 | 2.078055061 | 2.356585208 | 0.881807733 | 0.377880789 | NA | Gmfg-ps | glia maturation factor, gamma, pseudoger | 0 | 1.0279 | 0 | 0.964 | 1.0906 | 2.2277 |
| 25279 | ENSMUSG00000083042 | 0 | NA | NA | NA | NA | NA | Gm11388 | predicted gene 11388 [Source:MGI Symbc | 0 | 0 | 0 | 0 | 0 | 0 |
| 25280 | ENSMUSG00000083043 | 0.36353943 | 2.065989254 | 4.034137301 | 0.512126658 | 0.608562368 | NA | Gm13433 | predicted gene 13433 [Source:MGI Symbc | 0 | 0 | 0 | 0 | 2.1812 | 0 |
| 25281 | ENSMUSG00000083044 | 0.583320834 | -1.123770786 | 3.233988906 | -0.34748752 | 0.728225092 | NA | Gm12416 | predicted gene 12416 [Source:MGI Symbc | 2.4093 | 0 | 0 | 0 | 1.0906 | 0 |
| 25282 | ENSMUSG00000083045 | 0.133850373 | -0.796629176 | 4.080472857 | -0.195229623 | 0.84521319 | NA | Gm16100 | predicted gene 16100 [Source:MGI Symbc | 0.8031 | 0 | 0 | 0 | 0 | 0 |
| 25283 | ENSMUSG00000083046 | 0 | NA | NA | NA | NA | NA | Gm14699 | predicted gene 14699 [Source:MGI Symbc | 0 | 0 | 0 | 0 | 0 | 0 |
| 25284 | ENSMUSG00000083047 | 0 | NA | NA | NA | NA | NA | Gm14082 | predicted gene 14082 [Source:MGI Symbc | 0 | 0 | 0 | 0 | 0 | 0 |
| 25285 | ENSMUSG00000083049 | 0.167707431 | -0.796629176 | 4.080472857 | -0.195229623 | 0.84521319 | NA | Gm14822 | predicted gene 14822 [Source:MGI Symbc | 0 | 0 | 1.0062 | 0 | 0 | 0 |
| 25286 | ENSMUSG00000083050 | 0 | NA | NA | NA | NA | NA | Gm11242 | predicted gene 11242 [Source:MGI Symbc | 0 | 0 | 0 | 0 | 0 | 0 |
| 25287 | ENSMUSG00000083052 | 0 | NA | NA | NA | NA | NA | Gm12285 | predicted gene 12285 [Source:MGI Symbc | 0 | 0 | 0 | 0 | 0 | 0 |
| 25288 | ENSMUSG00000083053 | 0 | NA | NA | NA | NA | NA | Gm15001 | predicted gene 15001 [Source:MGI Symbc | 0 | 0 | 0 | 0 | 0 | 0 |
| 25289 | ENSMUSG00000083054 | 0 | NA | NA | NA | NA | NA | Gm12986 | predicted gene 12986 [Source:MGI Symbc | 0 | 0 | 0 | 0 | 0 | 0 |
| 25290 | ENSMUSG00000083055 | 0 | NA | NA | NA | NA | NA | Gm14140 | predicted gene 14140 [Source:MGI Symbc | 0 | 0 | 0 | 0 | 0 | 0 |
| 25291 | ENSMUSG00000083057 | 0 | NA | NA | NA | NA | NA | Gm13356 | predicted gene 13356 [Source:MGI Symbc | 0 | 0 | 0 | 0 | 0 | 0 |
| 25292 | ENSMUSG00000083058 | 0 | NA | NA | NA | NA | NA | Gm14795 | predicted gene 14795 [Source:MGI Symbc | 0 | 0 | 0 | 0 | 0 | 0 |
| 25293 | ENSMUSG00000083059 | 0 | NA | NA | NA | NA | NA | Olfr268-ps1 | olfactory receptor 268, pseudogene 1 [So | 0 | 0 | 0 | 0 | 0 | 0 |
| 25294 | ENSMUSG00000083060 | 0 | NA | NA | NA | NA | NA | Gm14533 | predicted gene 14533 [Source:MGI Symbc | 0 | 0 | 0 | 0 | 0 | 0 |
| 25295 | ENSMUSG00000083061 | 12.67131819 | 0.102773324 | 0.604656912 | 0.169969651 | 0.865034005 | 0.977227145 | Gm12191 | predicted gene 12191 [Source:MGI Symbc | 11.243 | 10.279 | 15.094 | 9.6398 | 14.178 | 15.594 |
| 25296 | ENSMUSG00000083062 | 0 | NA | NA | NA | NA | NA | Gm14980 | predicted gene 14980 [Source:MGI Symbc | 0 | 0 | 0 | 0 | 0 | 0 |
| 25297 | ENSMUSG00000083063 | 0 | NA | NA | NA | NA | NA | Gm14882 | predicted gene 14882 [Source:MGI Symbc | 0 | 0 | 0 | 0 | 0 | 0 |
| 25298 | ENSMUSG00000083064 | 1.992821099 | 1.095504097 | 1.614970396 | 0.678343145 | 0.497554149 | NA | Gm7785 | predicted gene 7785 [Source:MGI Symbol | 0.8031 | 2.0558 | 1.0062 | 4.8199 | 3.2719 | 0 |
| 25299 | ENSMUSG00000083065 | 0 | NA | NA | NA | NA | NA | Gm13195 | predicted gene 13195 [Source:MGI Symbc | 0 | 0 | 0 | 0 | 0 | 0 |
| 25300 | ENSMUSG00000083066 | 0 | NA | NA | NA | NA | NA | Gm12748 | predicted gene 12748 [Source:MGI Symbc | 0 | 0 | 0 | 0 | 0 | 0 |
| 25301 | ENSMUSG00000083067 | 0 | NA | NA | NA | NA | NA | Gm14751 | predicted gene 14751 [Source:MGI Symbc | 0 | 0 | 0 | 0 | 0 | 0 |
| 25302 | ENSMUSG00000083068 | 0 | NA | NA | NA | NA | NA | Gm11800 | predicted gene 11800 [Source:MGI Symbc | 0 | 0 | 0 | 0 | 0 | 0 |
| 25303 | ENSMUSG00000083069 | 0 | NA | NA | NA | NA | NA | Gm16431 | predicted gene 16431 [Source:MGI Symbc | 0 | 0 | 0 | 0 | 0 | 0 |
| 25304 | ENSMUSG00000083070 | 0 | NA | NA | NA | NA | NA | Gm13350 | predicted gene 13350 [Source:MGI Symbc | 0 | 0 | 0 | 0 | 0 | 0 |
| 25305 | ENSMUSG00000083072 | 0.185640647 | 1.126924175 | 4.080472857 | 0.276174898 | 0.782413736 | NA | Gm13668 | predicted gene 13668 [Source:MGI Symbc | 0 | 0 | 0 | 0 | 0 | 1.1138 |
| 25306 | ENSMUSG00000083073 | 0 | NA | NA | NA | NA | NA | Gm11845 | predicted gene 11845 [Source:MGI Symbc | 0 | 0 | 0 | 0 | 0 | 0 |
| 25307 | ENSMUSG00000083074 | 0 | NA | NA | NA | NA | NA | Gm14670 | predicted gene 14670 [Source:MGI Symbc | 0 | 0 | 0 | 0 | 0 | 0 |
| 25308 | ENSMUSG00000083075 | 0 | NA | NA | NA | NA | NA | Gm11980 | predicted gene 11980 [Source:MGI Symbc | 0 | 0 | 0 | 0 | 0 | 0 |

| | | | | | | | | | | | | | | | |
|---|---|---|---|---|---|---|---|---|---|---|---|---|---|---|---|
| 25309 | ENSMUSG00000083076 | 1.663287671 | -1.827066991 | 1.910213642 | -0.956472591 | 0.338833526 | NA | Gm11408 | predicted gene 11408 [Source:MGI Symbo | 1.6062 | 5.1396 | 1.0062 | 0 | 0 | 2.2277 |
| 25310 | ENSMUSG00000083077 | 0 | NA | NA | NA | NA | NA | Gm14126 | predicted gene 14126 [Source:MGI Symbo | 0 | 0 | 0 | 0 | 0 | 0 |
| 25311 | ENSMUSG00000083078 | 0 | NA | NA | NA | NA | NA | Gm13172 | predicted gene 13172 [Source:MGI Symbo | 0 | 0 | 0 | 0 | 0 | 0 |
| 25312 | ENSMUSG00000083079 | 0 | NA | NA | NA | NA | NA | Amy2b | amylase 2b [Source:MGI Symbol;Acc:MGI | 0 | 0 | 0 | 0 | 0 | 0 |
| 25313 | ENSMUSG00000083080 | 0 | NA | NA | NA | NA | NA | Gm11484 | predicted gene 11484 [Source:MGI Symbo | 0 | 0 | 0 | 0 | 0 | 0 |
| 25314 | ENSMUSG00000083081 | 0 | NA | NA | NA | NA | NA | Gm13223 | predicted gene 13223 [Source:MGI Symbo | 0 | 0 | 0 | 0 | 0 | 0 |
| 25315 | ENSMUSG00000083083 | 1.875695715 | -0.391512521 | 1.921930159 | -0.203707986 | 0.838581701 | NA | Gm15382 | predicted gene 15382 [Source:MGI Symbo | 2.4093 | 0 | 4.025 | 4.8199 | 0 | 0 |
| 25316 | ENSMUSG00000083084 | 0 | NA | NA | NA | NA | NA | Gm14667 | predicted gene 14667 [Source:MGI Symbo | 0 | 0 | 0 | 0 | 0 | 0 |
| 25317 | ENSMUSG00000083085 | 0 | NA | NA | NA | NA | NA | Gm12765 | predicted gene 12765 [Source:MGI Symbo | 0 | 0 | 0 | 0 | 0 | 0 |
| 25318 | ENSMUSG00000083086 | 0 | NA | NA | NA | NA | NA | Gm15531 | predicted gene 15531 [Source:MGI Symbo | 0 | 0 | 0 | 0 | 0 | 0 |
| 25319 | ENSMUSG00000083087 | 2.927998085 | -0.203338083 | 1.397939362 | -0.145455581 | 0.884351184 | NA | Gm11249 | predicted gene 11249 [Source:MGI Symbo | 2.4093 | 0 | 7.0437 | 4.8199 | 2.1812 | 1.1138 |
| 25320 | ENSMUSG00000083088 | 0 | NA | NA | NA | NA | NA | Gm14153 | predicted gene 14153 [Source:MGI Symbo | 0 | 0 | 0 | 0 | 0 | 0 |
| 25321 | ENSMUSG00000083089 | 0 | NA | NA | NA | NA | NA | Gm4760 | predicted gene 4760 [Source:MGI Symbol | 0 | 0 | 0 | 0 | 0 | 0 |
| 25322 | ENSMUSG00000083090 | 0 | NA | NA | NA | NA | NA | Gm11736 | predicted gene 11736 [Source:MGI Symbo | 0 | 0 | 0 | 0 | 0 | 0 |
| 25323 | ENSMUSG00000083091 | 0 | NA | NA | NA | NA | NA | Gm12438 | predicted gene 12438 [Source:MGI Symbo | 0 | 0 | 0 | 0 | 0 | 0 |
| 25324 | ENSMUSG00000083093 | 0.506966596 | 2.558866737 | 3.339834942 | 0.766165628 | 0.443577757 | NA | Gm15385 | predicted gene 15385 [Source:MGI Symbo | 0 | 0 | 0 | 1.928 | 0 | 1.1138 |
| 25325 | ENSMUSG00000083094 | 0 | NA | NA | NA | NA | NA | Gm4908 | predicted gene 4908 [Source:MGI Symbol | 0 | 0 | 0 | 0 | 0 | 0 |
| 25326 | ENSMUSG00000083096 | 0 | NA | NA | NA | NA | NA | Gm11356 | predicted gene 11356 [Source:MGI Symbo | 0 | 0 | 0 | 0 | 0 | 0 |
| 25327 | ENSMUSG00000083097 | 1.755591254 | -0.58112224 | 1.566000719 | -0.371086828 | 0.710572861 | NA | Gm14494 | predicted gene 14494 [Source:MGI Symbo | 3.2124 | 2.0558 | 1.0062 | 0.964 | 2.1812 | 1.1138 |
| 25328 | ENSMUSG00000083098 | 0 | NA | NA | NA | NA | NA | Gm14578 | predicted gene 14578 [Source:MGI Symbo | 0 | 0 | 0 | 0 | 0 | 0 |
| 25329 | ENSMUSG00000083099 | 0 | NA | NA | NA | NA | NA | Gm4732 | predicted gene 4732 [Source:MGI Symbol | 0 | 0 | 0 | 0 | 0 | 0 |
| 25330 | ENSMUSG00000083100 | 0.524408994 | -0.837196744 | 3.377823722 | -0.247850928 | 0.804249749 | NA | Gm6044 | predicted gene 6044 [Source:MGI Symbol | 0 | 2.0558 | 0 | 0 | 1.0906 | 0 |
| 25331 | ENSMUSG00000083101 | 0 | NA | NA | NA | NA | NA | Gm13430 | predicted gene 13430 [Source:MGI Symbo | 0 | 0 | 0 | 0 | 0 | 0 |
| 25332 | ENSMUSG00000083103 | 0 | NA | NA | NA | NA | NA | Gm6429 | predicted gene 6429 [Source:MGI Symbol | 0 | 0 | 0 | 0 | 0 | 0 |
| 25333 | ENSMUSG00000083104 | 2.383715553 | 0.773388847 | 1.505131547 | 0.51383472 | 0.607367551 | NA | Gm12917 | predicted gene 12917 [Source:MGI Symbo | 3.2124 | 1.0279 | 1.0062 | 5.7839 | 3.2719 | 0 |
| 25334 | ENSMUSG00000083105 | 0 | NA | NA | NA | NA | NA | Gm12393 | predicted gene 12393 [Source:MGI Symbo | 0 | 0 | 0 | 0 | 0 | 0 |
| 25335 | ENSMUSG00000083106 | 0 | NA | NA | NA | NA | NA | Gm15309 | predicted gene 15309 [Source:MGI Symbo | 0 | 0 | 0 | 0 | 0 | 0 |
| 25336 | ENSMUSG00000083107 | 0 | NA | NA | NA | NA | NA | Gm5303 | predicted gene 5303 [Source:MGI Symbol | 0 | 0 | 0 | 0 | 0 | 0 |
| 25337 | ENSMUSG00000083109 | 0 | NA | NA | NA | NA | NA | Gm12876 | predicted gene 12876 [Source:MGI Symbo | 0 | 0 | 0 | 0 | 0 | 0 |
| 25338 | ENSMUSG00000083110 | 0 | NA | NA | NA | NA | NA | Gm14073 | predicted gene 14073 [Source:MGI Symbo | 0 | 0 | 0 | 0 | 0 | 0 |
| 25339 | ENSMUSG00000083111 | 2.729522843 | 0.083129628 | 1.29339268 | 0.064272536 | 0.948753221 | NA | Gm14421 | predicted gene 14421 [Source:MGI Symbo | 4.8186 | 1.0279 | 2.0125 | 1.928 | 4.3625 | 2.2277 |
| 25340 | ENSMUSG00000083112 | 0 | NA | NA | NA | NA | NA | Gm14620 | predicted gene 14620 [Source:MGI Symbo | 0 | 0 | 0 | 0 | 0 | 0 |
| 25341 | ENSMUSG00000083113 | 0 | NA | NA | NA | NA | NA | Rpl10-ps4 | ribosomal protein L10, pseudogene 4 [Sou | 0 | 0 | 0 | 0 | 0 | 0 |
| 25342 | ENSMUSG00000083114 | 0 | NA | NA | NA | NA | NA | Gm14790 | predicted gene 14790 [Source:MGI Symbo | 0 | 0 | 0 | 0 | 0 | 0 |
| 25343 | ENSMUSG00000083115 | 0 | NA | NA | NA | NA | NA | Gm13386 | predicted gene 13386 [Source:MGI Symbo | 0 | 0 | 0 | 0 | 0 | 0 |
| 25344 | ENSMUSG00000083116 | 0.167707431 | -0.796629176 | 4.080472857 | -0.195229623 | 0.84521319 | NA | Gm13410 | predicted gene 13410 [Source:MGI Symbo | 0 | 0 | 1.0062 | 0 | 0 | 0 |
| 25345 | ENSMUSG00000083117 | 0 | NA | NA | NA | NA | NA | Gm15171 | predicted gene 15171 [Source:MGI Symbo | 0 | 0 | 0 | 0 | 0 | 0 |
| 25346 | ENSMUSG00000083118 | 0 | NA | NA | NA | NA | NA | Gm11609 | predicted gene 11609 [Source:MGI Symbo | 0 | 0 | 0 | 0 | 0 | 0 |
| 25347 | ENSMUSG00000083119 | 0 | NA | NA | NA | NA | NA | Gm14724 | predicted gene 14724 [Source:MGI Symbo | 0 | 0 | 0 | 0 | 0 | 0 |
| 25348 | ENSMUSG00000083120 | 0 | NA | NA | NA | NA | NA | Gm15854 | predicted gene 15854 [Source:MGI Symbo | 0 | 0 | 0 | 0 | 0 | 0 |
| 25349 | ENSMUSG00000083121 | 0.435408177 | -2.179616019 | 3.531869654 | -0.617128103 | 0.537150234 | NA | Gm6341 | predicted pseudogene 6341 [Source:MGI | 1.6062 | 0 | 1.0062 | 0 | 0 | 0 |
| 25350 | ENSMUSG00000083122 | 0 | NA | NA | NA | NA | NA | Gm15578 | predicted gene 15578 [Source:MGI Symbo | 0 | 0 | 0 | 0 | 0 | 0 |
| 25351 | ENSMUSG00000083123 | 0 | NA | NA | NA | NA | NA | Fthl17-ps2 | ferritin, heavy polypeptide-like 17, pseudo | 0 | 0 | 0 | 0 | 0 | 0 |
| 25352 | ENSMUSG00000083124 | 0 | NA | NA | NA | NA | NA | Vmn1r-ps12 | vomeronasal 1 receptor, pseudogene 126 | 0 | 0 | 0 | 0 | 0 | 0 |
| 25353 | ENSMUSG00000083126 | 0 | NA | NA | NA | NA | NA | Gm14109 | predicted gene 14109 [Source:MGI Symbo | 0 | 0 | 0 | 0 | 0 | 0 |
| 25354 | ENSMUSG00000083127 | 0 | NA | NA | NA | NA | NA | Gm15189 | predicted gene 15189 [Source:MGI Symbo | 0 | 0 | 0 | 0 | 0 | 0 |
| 25355 | ENSMUSG00000083128 | 0 | NA | NA | NA | NA | NA | Gm12723 | predicted gene 12723 [Source:MGI Symbo | 0 | 0 | 0 | 0 | 0 | 0 |
| 25356 | ENSMUSG00000083129 | 0 | NA | NA | NA | NA | NA | Gm12210 | predicted gene 12210 [Source:MGI Symbo | 0 | 0 | 0 | 0 | 0 | 0 |
| 25357 | ENSMUSG00000083130 | 0 | NA | NA | NA | NA | NA | Gm14560 | predicted gene 14560 [Source:MGI Symbo | 0 | 0 | 0 | 0 | 0 | 0 |
| 25358 | ENSMUSG00000083135 | 0 | NA | NA | NA | NA | NA | Gm11504 | predicted gene 11504 [Source:MGI Symbo | 0 | 0 | 0 | 0 | 0 | 0 |
| 25359 | ENSMUSG00000083136 | 0 | NA | NA | NA | NA | NA | Gm15170 | predicted gene 15170 [Source:MGI Symbo | 0 | 0 | 0 | 0 | 0 | 0 |
| 25360 | ENSMUSG00000083137 | 0 | NA | NA | NA | NA | NA | Gm11792 | predicted gene 11792 [Source:MGI Symbo | 0 | 0 | 0 | 0 | 0 | 0 |
| 25361 | ENSMUSG00000083138 | 0 | NA | NA | NA | NA | NA | Cyp4a29 | cytochrome P450, family 4, subfamily a, p | 0 | 0 | 0 | 0 | 0 | 0 |
| 25362 | ENSMUSG00000083139 | 0.506734502 | -2.372847156 | 3.338060886 | -0.710845978 | 0.477179685 | NA | Gm12418 | predicted gene 12418 [Source:MGI Symbo | 0 | 1.0279 | 2.0125 | 0 | 0 | 0 |
| 25363 | ENSMUSG00000083140 | 0 | NA | NA | NA | NA | NA | Gm5073 | predicted pseudogene 5073 [Source:MGI | 0 | 0 | 0 | 0 | 0 | 0 |
| 25364 | ENSMUSG00000083141 | 0 | NA | NA | NA | NA | NA | Gm15752 | predicted gene 15752 [Source:MGI Symbo | 0 | 0 | 0 | 0 | 0 | 0 |

| | | | | | | | | | | | | | | | |
|---|---|---|---|---|---|---|---|---|---|---|---|---|---|---|---|
| 25365 | ENSMUSG00000083142 | 0.335414862 | -1.779047242 | 4.030804179 | -0.441362855 | 0.658950329 | NA | Gm12288 | predicted gene 12288 [Source:MGI Symbo | 0 | 0 | 2.0125 | 0 | 0 | 0 |
| 25366 | ENSMUSG00000083143 | 0 | NA | NA | NA | NA | NA | Gm13424 | predicted gene 13424 [Source:MGI Symbo | 0 | 0 | 0 | 0 | 0 | 0 |
| 25367 | ENSMUSG00000083144 | 0 | NA | NA | NA | NA | NA | Gm14334 | predicted gene 14334 [Source:MGI Symbo | 0 | 0 | 0 | 0 | 0 | 0 |
| 25368 | ENSMUSG00000083145 | 0 | NA | NA | NA | NA | NA | Gm12579 | predicted gene 12579 [Source:MGI Symbo | 0 | 0 | 0 | 0 | 0 | 0 |
| 25369 | ENSMUSG00000083147 | 0 | NA | NA | NA | NA | NA | Olfr1103-ps1 | olfactory receptor 1103, pseudogene 1 [S | 0 | 0 | 0 | 0 | 0 | 0 |
| 25370 | ENSMUSG00000083149 | 0 | NA | NA | NA | NA | NA | Gm11380 | predicted gene 11380 [Source:MGI Symbo | 0 | 0 | 0 | 0 | 0 | 0 |
| 25371 | ENSMUSG00000083150 | 0 | NA | NA | NA | NA | NA | Gm11852 | predicted gene 11852 [Source:MGI Symbo | 0 | 0 | 0 | 0 | 0 | 0 |
| 25372 | ENSMUSG00000083151 | 0 | NA | NA | NA | NA | NA | Gm4913 | predicted gene 4913 [Source:MGI Symbol | 0 | 0 | 0 | 0 | 0 | 0 |
| 25373 | ENSMUSG00000083152 | 3.033742465 | -0.195414267 | 1.234750213 | -0.158262185 | 0.874250204 | NA | Apc-ps1 | adenomatosis polyposis coli, pseudogene | 1.6062 | 5.1396 | 3.0187 | 2.8919 | 1.0906 | 4.4554 |
| 25374 | ENSMUSG00000083153 | 0 | NA | NA | NA | NA | NA | Gm12459 | predicted gene 12459 [Source:MGI Symbo | 0 | 0 | 0 | 0 | 0 | 0 |
| 25375 | ENSMUSG00000083154 | 0.335414862 | -1.779047242 | 4.030804179 | -0.441362855 | 0.658950329 | NA | Gm11892 | predicted gene 11892 [Source:MGI Symbo | 0 | 0 | 2.0125 | 0 | 0 | 0 |
| 25376 | ENSMUSG00000083155 | 0 | NA | NA | NA | NA | NA | Gm11866 | predicted gene 11866 [Source:MGI Symbo | 0 | 0 | 0 | 0 | 0 | 0 |
| 25377 | ENSMUSG00000083156 | 0 | NA | NA | NA | NA | NA | Gm12201 | predicted gene 12201 [Source:MGI Symbo | 0 | 0 | 0 | 0 | 0 | 0 |
| 25378 | ENSMUSG00000083157 | 0.492645588 | 0.9565034 | 3.447648281 | 0.277436479 | 0.781444974 | NA | Gm13511 | predicted gene 13511 [Source:MGI Symbo | 0 | 1.0279 | 0 | 1.928 | 0 | 0 |
| 25379 | ENSMUSG00000083158 | 0 | NA | NA | NA | NA | NA | Gm14655 | predicted gene 14655 [Source:MGI Symbo | 0 | 0 | 0 | 0 | 0 | 0 |
| 25380 | ENSMUSG00000083159 | 0 | NA | NA | NA | NA | NA | Gm12180 | predicted gene 12180 [Source:MGI Symbo | 0 | 0 | 0 | 0 | 0 | 0 |
| 25381 | ENSMUSG00000083160 | 0 | NA | NA | NA | NA | NA | Gm15967 | predicted gene 15967 [Source:MGI Symbo | 0 | 0 | 0 | 0 | 0 | 0 |
| 25382 | ENSMUSG00000083161 | 2.376923778 | -1.835046552 | 1.526098691 | -1.202442911 | 0.229191972 | NA | Gm11427 | predicted gene 11427 [Source:MGI Symbo | 8.031 | 2.0558 | 1.0062 | 0.964 | 1.0906 | 1.1138 |
| 25383 | ENSMUSG00000083162 | 0 | NA | NA | NA | NA | NA | Gm4917 | predicted gene 4917 [Source:MGI Symbol | 0 | 0 | 0 | 0 | 0 | 0 |
| 25384 | ENSMUSG00000083163 | 0 | NA | NA | NA | NA | NA | Gm11429 | predicted gene 11429 [Source:MGI Symbo | 0 | 0 | 0 | 0 | 0 | 0 |
| 25385 | ENSMUSG00000083165 | 1.017261173 | -2.166287499 | 2.717865413 | -0.797054736 | 0.425419243 | NA | Gm11344 | predicted gene 11344 [Source:MGI Symbo | 0 | 5.1396 | 0 | 0.964 | 0 | 0 |
| 25386 | ENSMUSG00000083166 | 0 | NA | NA | NA | NA | NA | Gm8648 | predicted gene 8648 [Source:MGI Symbol | 0 | 0 | 0 | 0 | 0 | 0 |
| 25387 | ENSMUSG00000083167 | 0 | NA | NA | NA | NA | NA | Vmn1r-ps97 | vomeronasal 1 receptor, pseudogene 97 [ | 0 | 0 | 0 | 0 | 0 | 0 |
| 25388 | ENSMUSG00000083168 | 1.322276729 | 0.865032458 | 1.778635474 | 0.486346118 | 0.626721786 | NA | Gm14201 | predicted gene 14201 [Source:MGI Symbo | 0.8031 | 1.0279 | 1.0062 | 2.8919 | 1.0906 | 1.1138 |
| 25389 | ENSMUSG00000083169 | 1.072560804 | -2.278047819 | 2.189625656 | -1.04038232 | 0.298162313 | NA | Gm6580 | predicted gene 6580 [Source:MGI Symbol | 2.4093 | 2.0558 | 1.0062 | 0.964 | 0 | 0 |
| 25390 | ENSMUSG00000083170 | 0 | NA | NA | NA | NA | NA | Gm14405 | predicted gene 14405 [Source:MGI Symbo | 0 | 0 | 0 | 0 | 0 | 0 |
| 25391 | ENSMUSG00000083171 | 0 | NA | NA | NA | NA | NA | Gm12321 | predicted gene 12321 [Source:MGI Symbo | 0 | 0 | 0 | 0 | 0 | 0 |
| 25392 | ENSMUSG00000083172 | 0 | NA | NA | NA | NA | NA | Gm13409 | predicted gene 13409 [Source:MGI Symbo | 0 | 0 | 0 | 0 | 0 | 0 |
| 25393 | ENSMUSG00000083173 | 0 | NA | NA | NA | NA | NA | Gm13064 | predicted gene 13064 [Source:MGI Symbo | 0 | 0 | 0 | 0 | 0 | 0 |
| 25394 | ENSMUSG00000083174 | 0 | NA | NA | NA | NA | NA | Gm7190 | predicted gene 7190 [Source:MGI Symbol | 0 | 0 | 0 | 0 | 0 | 0 |
| 25395 | ENSMUSG00000083175 | 0 | NA | NA | NA | NA | NA | Gm12206 | predicted gene 12206 [Source:MGI Symbo | 0 | 0 | 0 | 0 | 0 | 0 |
| 25396 | ENSMUSG00000083176 | 0 | NA | NA | NA | NA | NA | D11Bhm181 | DNA segment, Chr 11, Boehm 181, expres | 0 | 0 | 0 | 0 | 0 | 0 |
| 25397 | ENSMUSG00000083177 | 0 | NA | NA | NA | NA | NA | Gm7820 | predicted gene 7820 [Source:MGI Symbol | 0 | 0 | 0 | 0 | 0 | 0 |
| 25398 | ENSMUSG00000083178 | 0 | NA | NA | NA | NA | NA | Gm12187 | predicted gene 12187 [Source:MGI Symbo | 0 | 0 | 0 | 0 | 0 | 0 |
| 25399 | ENSMUSG00000083179 | 0.797609011 | 1.992900111 | 2.636035165 | 0.756021823 | 0.449636112 | NA | Gm12693 | predicted gene 12693 [Source:MGI Symbo | 0.8031 | 0 | 0 | 2.8919 | 1.0906 | 0 |
| 25400 | ENSMUSG00000083180 | 0 | NA | NA | NA | NA | NA | Gm15007 | predicted gene 15007 [Source:MGI Symbo | 0 | 0 | 0 | 0 | 0 | 0 |
| 25401 | ENSMUSG00000083181 | 0 | NA | NA | NA | NA | NA | Gm13903 | predicted gene 13903 [Source:MGI Symbo | 0 | 0 | 0 | 0 | 0 | 0 |
| 25402 | ENSMUSG00000083182 | 0 | NA | NA | NA | NA | NA | Gm13828 | predicted gene 13828 [Source:MGI Symbo | 0 | 0 | 0 | 0 | 0 | 0 |
| 25403 | ENSMUSG00000083183 | 0 | NA | NA | NA | NA | NA | Gm15575 | predicted gene 15575 [Source:MGI Symbo | 0 | 0 | 0 | 0 | 0 | 0 |
| 25404 | ENSMUSG00000083185 | 0 | NA | NA | NA | NA | NA | Olfr1146-ps1 | olfactory receptor 1146, pseudogene 1 [S | 0 | 0 | 0 | 0 | 0 | 0 |
| 25405 | ENSMUSG00000083186 | 0.133850373 | -0.796629176 | 4.080472857 | -0.195229623 | 0.84521319 | NA | Gm11805 | predicted gene 11805 [Source:MGI Symbo | 0.8031 | 0 | 0 | 0 | 0 | 0 |
| 25406 | ENSMUSG00000083188 | 0 | NA | NA | NA | NA | NA | Olfr404-ps1 | olfactory receptor 404, pseudogene 1 [So | 0 | 0 | 0 | 0 | 0 | 0 |
| 25407 | ENSMUSG00000083190 | 0 | NA | NA | NA | NA | NA | Cyp4a30-ps | cytochrome P450, family 4, subfamily a, m | 0 | 0 | 0 | 0 | 0 | 0 |
| 25408 | ENSMUSG00000083191 | 0 | NA | NA | NA | NA | NA | Gm15571 | predicted gene 15571 [Source:MGI Symbo | 0 | 0 | 0 | 0 | 0 | 0 |
| 25409 | ENSMUSG00000083193 | 0 | NA | NA | NA | NA | NA | 4930595D18 | RIKEN cDNA 4930595D18 gene [Source:M | 0 | 0 | 0 | 0 | 0 | 0 |
| 25410 | ENSMUSG00000083195 | 0 | NA | NA | NA | NA | NA | Gm13567 | predicted gene 13567 [Source:MGI Symbo | 0 | 0 | 0 | 0 | 0 | 0 |
| 25411 | ENSMUSG00000083196 | 0 | NA | NA | NA | NA | NA | Gm15272 | predicted gene 15272 [Source:MGI Symbo | 0 | 0 | 0 | 0 | 0 | 0 |
| 25412 | ENSMUSG00000083197 | 0 | NA | NA | NA | NA | NA | Gm13581 | predicted gene 13581 [Source:MGI Symbo | 0 | 0 | 0 | 0 | 0 | 0 |
| 25413 | ENSMUSG00000083199 | 0.17131964 | -0.796629176 | 4.080472857 | -0.195229623 | 0.84521319 | NA | Gm9432 | predicted gene 9432 [Source:MGI Symbol | 0 | 1.0279 | 0 | 0 | 0 | 0 |
| 25414 | ENSMUSG00000083200 | 0 | NA | NA | NA | NA | NA | Gm15066 | predicted gene 15066 [Source:MGI Symbo | 0 | 0 | 0 | 0 | 0 | 0 |
| 25415 | ENSMUSG00000083201 | 0 | NA | NA | NA | NA | NA | Gm12080 | predicted gene 12080 [Source:MGI Symbo | 0 | 0 | 0 | 0 | 0 | 0 |
| 25416 | ENSMUSG00000083202 | 0 | NA | NA | NA | NA | NA | Gm12388 | predicted gene 12388 [Source:MGI Symbo | 0 | 0 | 0 | 0 | 0 | 0 |
| 25417 | ENSMUSG00000083203 | 0.160662974 | 1.126924175 | 4.080472857 | 0.276174898 | 0.782413736 | NA | Gm11421 | predicted gene 11421 [Source:MGI Symbo | 0 | 0 | 0 | 0.964 | 0 | 0 |
| 25418 | ENSMUSG00000083204 | 0 | NA | NA | NA | NA | NA | Gm13788 | predicted gene 13788 [Source:MGI Symbo | 0 | 0 | 0 | 0 | 0 | 0 |
| 25419 | ENSMUSG00000083205 | 0 | NA | NA | NA | NA | NA | Gm7821 | predicted gene 7821 [Source:MGI Symbol | 0 | 0 | 0 | 0 | 0 | 0 |
| 25420 | ENSMUSG00000083206 | 1.081124346 | 3.638382224 | 2.309068382 | 1.575692713 | 0.115096638 | NA | Gm11338 | predicted gene 11338 [Source:MGI Symbo | 0 | 0 | 0 | 0.964 | 2.1812 | 3.3415 |

| | | | | | | | | | | | | | | | |
|---|---|---|---|---|---|---|---|---|---|---|---|---|---|---|---|
| 25421 | ENSMUSG00000083207 | 0 | NA | NA | NA | NA | NA | Gm14780 | predicted gene 14780 [Source:MGI Symbo | 0 | 0 | 0 | 0 | 0 | 0 |
| 25422 | ENSMUSG00000083208 | 0 | NA | NA | NA | NA | NA | Gm11522 | predicted gene 11522 [Source:MGI Symbo | 0 | 0 | 0 | 0 | 0 | 0 |
| 25423 | ENSMUSG00000083209 | 0 | NA | NA | NA | NA | NA | Gm11969 | predicted gene 11969 [Source:MGI Symbo | 0 | 0 | 0 | 0 | 0 | 0 |
| 25424 | ENSMUSG00000083210 | 0.17131964 | -0.796629176 | 4.080472857 | -0.195229623 | 0.84521319 | NA | Gm14869 | predicted gene 14869 [Source:MGI Symbo | 0 | 1.0279 | 0 | 0 | 0 | 0 |
| 25425 | ENSMUSG00000083211 | 0 | NA | NA | NA | NA | NA | Gm13737 | predicted gene 13737 [Source:MGI Symbo | 0 | 0 | 0 | 0 | 0 | 0 |
| 25426 | ENSMUSG00000083212 | 0 | NA | NA | NA | NA | NA | Gm12274 | predicted gene 12274 [Source:MGI Symbo | 0 | 0 | 0 | 0 | 0 | 0 |
| 25427 | ENSMUSG00000083213 | 0 | NA | NA | NA | NA | NA | Gm14875 | predicted gene 14875 [Source:MGI Symbo | 0 | 0 | 0 | 0 | 0 | 0 |
| 25428 | ENSMUSG00000083214 | 0 | NA | NA | NA | NA | NA | Gm11912 | predicted gene 11912 [Source:MGI Symbo | 0 | 0 | 0 | 0 | 0 | 0 |
| 25429 | ENSMUSG00000083215 | 0 | NA | NA | NA | NA | NA | Rbmx2-ps | RNA binding motif protein, X-linked 2, pse | 0 | 0 | 0 | 0 | 0 | 0 |
| 25430 | ENSMUSG00000083216 | 0 | NA | NA | NA | NA | NA | Hbb-bh3 | hemoglobin beta, pseudogene bh3 [Sourc | 0 | 0 | 0 | 0 | 0 | 0 |
| 25431 | ENSMUSG00000083218 | 0 | NA | NA | NA | NA | NA | Gm16425 | predicted gene 16425 [Source:MGI Symbo | 0 | 0 | 0 | 0 | 0 | 0 |
| 25432 | ENSMUSG00000083219 | 0 | NA | NA | NA | NA | NA | Gm11410 | predicted gene 11410 [Source:MGI Symbo | 0 | 0 | 0 | 0 | 0 | 0 |
| 25433 | ENSMUSG00000083220 | 0 | NA | NA | NA | NA | NA | Gm12919 | predicted gene 12919 [Source:MGI Symbo | 0 | 0 | 0 | 0 | 0 | 0 |
| 25434 | ENSMUSG00000083221 | 0 | NA | NA | NA | NA | NA | Gm14528 | predicted gene 14528 [Source:MGI Symbo | 0 | 0 | 0 | 0 | 0 | 0 |
| 25435 | ENSMUSG00000083222 | 0 | NA | NA | NA | NA | NA | Gm12427 | predicted gene 12427 [Source:MGI Symbo | 0 | 0 | 0 | 0 | 0 | 0 |
| 25436 | ENSMUSG00000083223 | 0 | NA | NA | NA | NA | NA | Gm15016 | predicted gene 15016 [Source:MGI Symbo | 0 | 0 | 0 | 0 | 0 | 0 |
| 25437 | ENSMUSG00000083224 | 0 | NA | NA | NA | NA | NA | Csnk2a1-ps4 | casein kinase 2, alpha 1 polypeptide, pseu | 0 | 0 | 0 | 0 | 0 | 0 |
| 25438 | ENSMUSG00000083225 | 0 | NA | NA | NA | NA | NA | Gm15215 | predicted gene 15215 [Source:MGI Symbo | 0 | 0 | 0 | 0 | 0 | 0 |
| 25439 | ENSMUSG00000083226 | 0.181769715 | 1.126924175 | 4.080472857 | 0.276174898 | 0.782413736 | NA | Gm7831 | predicted gene 7831 [Source:MGI Symbol | 0 | 0 | 0 | 0 | 1.0906 | 0 |
| 25440 | ENSMUSG00000083227 | 0 | NA | NA | NA | NA | NA | Gm11546 | predicted gene 11546 [Source:MGI Symbo | 0 | 0 | 0 | 0 | 0 | 0 |
| 25441 | ENSMUSG00000083228 | 0 | NA | NA | NA | NA | NA | Gm11779 | predicted gene 11779 [Source:MGI Symbo | 0 | 0 | 0 | 0 | 0 | 0 |
| 25442 | ENSMUSG00000083229 | 0 | NA | NA | NA | NA | NA | Gm11862 | predicted gene 11862 [Source:MGI Symbo | 0 | 0 | 0 | 0 | 0 | 0 |
| 25443 | ENSMUSG00000083230 | 0 | NA | NA | NA | NA | NA | Gm14158 | predicted gene 14158 [Source:MGI Symbo | 0 | 0 | 0 | 0 | 0 | 0 |
| 25444 | ENSMUSG00000083231 | 0 | NA | NA | NA | NA | NA | Gm15096 | predicted gene 15096 [Source:MGI Symbo | 0 | 0 | 0 | 0 | 0 | 0 |
| 25445 | ENSMUSG00000083232 | 0 | NA | NA | NA | NA | NA | Gm11796 | predicted gene 11796 [Source:MGI Symbo | 0 | 0 | 0 | 0 | 0 | 0 |
| 25446 | ENSMUSG00000083233 | 0 | NA | NA | NA | NA | NA | Gm13406 | predicted gene 13406 [Source:MGI Symbo | 0 | 0 | 0 | 0 | 0 | 0 |
| 25447 | ENSMUSG00000083234 | 0 | NA | NA | NA | NA | NA | Gm12262 | predicted gene 12262 [Source:MGI Symbo | 0 | 0 | 0 | 0 | 0 | 0 |
| 25448 | ENSMUSG00000083235 | 0 | NA | NA | NA | NA | NA | Gm13246 | predicted gene 13246 [Source:MGI Symbo | 0 | 0 | 0 | 0 | 0 | 0 |
| 25449 | ENSMUSG00000083236 | 0 | NA | NA | NA | NA | NA | Gm7222 | predicted gene 7222 [Source:MGI Symbol | 0 | 0 | 0 | 0 | 0 | 0 |
| 25450 | ENSMUSG00000083237 | 0.167707431 | -0.796629176 | 4.080472857 | -0.195229623 | 0.84521319 | NA | Btf3-ps13 | basic transcription factor 3, pseudogene 1 | 0 | 0 | 1.0062 | 0 | 0 | 0 |
| 25451 | ENSMUSG00000083238 | 0 | NA | NA | NA | NA | NA | Hspe1-ps4 | heat shock protein 1 (chaperonin 10), pse | 0 | 0 | 0 | 0 | 0 | 0 |
| 25452 | ENSMUSG00000083239 | 2.714602293 | 3.910856509 | 1.753689532 | 2.230073476 | 0.025742565 | NA | Mup-ps1 | major urinary protein, pseudogene 1 [Sou | 0 | 0 | 1.0062 | 0.964 | 7.6343 | 6.6831 |
| 25453 | ENSMUSG00000083240 | 1.238640426 | 1.454958615 | 2.28972858 | 0.635428421 | 0.525149037 | NA | Gm13453 | predicted gene 13453 [Source:MGI Symbo | 0 | 0 | 2.0125 | 0.964 | 0 | 4.4554 |
| 25454 | ENSMUSG00000083241 | 0.181769715 | 1.126924175 | 4.080472857 | 0.276174898 | 0.782413736 | NA | Gm13259 | predicted gene 13259 [Source:MGI Symbo | 0 | 0 | 0 | 0 | 1.0906 | 0 |
| 25455 | ENSMUSG00000083242 | 0 | NA | NA | NA | NA | NA | Gm15659 | predicted gene 15659 [Source:MGI Symbo | 0 | 0 | 0 | 0 | 0 | 0 |
| 25456 | ENSMUSG00000083243 | 0 | NA | NA | NA | NA | NA | Gm14668 | predicted gene 14668 [Source:MGI Symbo | 0 | 0 | 0 | 0 | 0 | 0 |
| 25457 | ENSMUSG00000083244 | 0 | NA | NA | NA | NA | NA | Gm15180 | predicted gene 15180 [Source:MGI Symbo | 0 | 0 | 0 | 0 | 0 | 0 |
| 25458 | ENSMUSG00000083245 | 0 | NA | NA | NA | NA | NA | Gm11185 | predicted gene 11185 [Source:MGI Symbo | 0 | 0 | 0 | 0 | 0 | 0 |
| 25459 | ENSMUSG00000083246 | 4.996422454 | -0.307688647 | 0.949429382 | -0.324077444 | 0.745879404 | NA | Gm11839 | predicted gene 11839 [Source:MGI Symbo | 6.4248 | 6.1675 | 4.025 | 6.7478 | 3.2719 | 3.3415 |
| 25460 | ENSMUSG00000083248 | 0 | NA | NA | NA | NA | NA | Gm6574 | predicted gene 6574 [Source:MGI Symbol | 0 | 0 | 0 | 0 | 0 | 0 |
| 25461 | ENSMUSG00000083249 | 0 | NA | NA | NA | NA | NA | Gm14623 | predicted gene 14623 [Source:MGI Symbo | 0 | 0 | 0 | 0 | 0 | 0 |
| 25462 | ENSMUSG00000083250 | 0 | NA | NA | NA | NA | NA | Gm5403 | predicted gene 5403 [Source:MGI Symbol | 0 | 0 | 0 | 0 | 0 | 0 |
| 25463 | ENSMUSG00000083251 | 0 | NA | NA | NA | NA | NA | Gm15300 | predicted gene 15300 [Source:MGI Symbo | 0 | 0 | 0 | 0 | 0 | 0 |
| 25464 | ENSMUSG00000083252 | 0 | NA | NA | NA | NA | NA | Gm13210 | predicted gene 13210 [Source:MGI Symbo | 0 | 0 | 0 | 0 | 0 | 0 |
| 25465 | ENSMUSG00000083253 | 0 | NA | NA | NA | NA | NA | Gm7108 | predicted gene 7108 [Source:MGI Symbol | 0 | 0 | 0 | 0 | 0 | 0 |
| 25466 | ENSMUSG00000083254 | 0 | NA | NA | NA | NA | NA | Gm15116 | predicted gene 15116 [Source:MGI Symbo | 0 | 0 | 0 | 0 | 0 | 0 |
| 25467 | ENSMUSG00000083255 | 0 | NA | NA | NA | NA | NA | Gm12738 | predicted gene 12738 [Source:MGI Symbo | 0 | 0 | 0 | 0 | 0 | 0 |
| 25468 | ENSMUSG00000083256 | 0 | NA | NA | NA | NA | NA | Kars-ps1 | lysyl-tRNA synthetase, pseudogene 1 [Sou | 0 | 0 | 0 | 0 | 0 | 0 |
| 25469 | ENSMUSG00000083257 | 0.17131964 | -0.796629176 | 4.080472857 | -0.195229623 | 0.84521319 | NA | Gm15568 | predicted gene 15568 [Source:MGI Symbo | 0 | 1.0279 | 0 | 0 | 0 | 0 |
| 25470 | ENSMUSG00000083258 | 0 | NA | NA | NA | NA | NA | Gm5939 | predicted pseudogene 5939 [Source:MGI | 0 | 0 | 0 | 0 | 0 | 0 |
| 25471 | ENSMUSG00000083260 | 0 | NA | NA | NA | NA | NA | Gm13460 | predicted gene 13460 [Source:MGI Symbo | 0 | 0 | 0 | 0 | 0 | 0 |
| 25472 | ENSMUSG00000083261 | 4.586633137 | -0.661078756 | 1.014989828 | -0.651315646 | 0.51484275 | NA | Gm7816 | predicted pseudogene 7816 [Source:MGI | 5.6217 | 6.1675 | 5.0312 | 1.928 | 6.5437 | 2.2277 |
| 25473 | ENSMUSG00000083262 | 0 | NA | NA | NA | NA | NA | Gm14470 | predicted gene 14470 [Source:MGI Symbo | 0 | 0 | 0 | 0 | 0 | 0 |
| 25474 | ENSMUSG00000083263 | 0 | NA | NA | NA | NA | NA | Gm14874 | predicted gene 14874 [Source:MGI Symbo | 0 | 0 | 0 | 0 | 0 | 0 |
| 25475 | ENSMUSG00000083264 | 0.877757073 | -0.466619568 | 2.366442592 | -0.197181867 | 0.84368522 | NA | Gm11461 | predicted gene 11461 [Source:MGI Symbo | 0 | 2.0558 | 1.0062 | 0 | 1.0906 | 1.1138 |
| 25476 | ENSMUSG00000083267 | 0 | NA | NA | NA | NA | NA | Olfr1127-ps1 | olfactory receptor 1127, pseudogene 1 [S | 0 | 0 | 0 | 0 | 0 | 0 |

| | | | | | | | | | | | | | | | |
|---|---|---|---|---|---|---|---|---|---|---|---|---|---|---|---|
| 25477 | ENSMUSG00000083268 | 0 | NA | NA | NA | NA | NA | Gm11511 | predicted gene 11511 [Source:MGI Symbo | 0 | 0 | 0 | 0 | 0 | 0 |
| 25478 | ENSMUSG00000083269 | 0 | NA | NA | NA | NA | NA | Gm13254 | predicted gene 13254 [Source:MGI Symbo | 0 | 0 | 0 | 0 | 0 | 0 |
| 25479 | ENSMUSG00000083270 | 0 | NA | NA | NA | NA | NA | Gm13498 | predicted gene 13498 [Source:MGI Symbo | 0 | 0 | 0 | 0 | 0 | 0 |
| 25480 | ENSMUSG00000083271 | 0 | NA | NA | NA | NA | NA | Gm12384 | predicted gene 12384 [Source:MGI Symbo | 0 | 0 | 0 | 0 | 0 | 0 |
| 25481 | ENSMUSG00000083272 | 0 | NA | NA | NA | NA | NA | Gm12550 | predicted gene 12550 [Source:MGI Symbo | 0 | 0 | 0 | 0 | 0 | 0 |
| 25482 | ENSMUSG00000083274 | 0 | NA | NA | NA | NA | NA | Zcchc9-ps | zinc finger, CCHC domain containing 9, ps | 0 | 0 | 0 | 0 | 0 | 0 |
| 25483 | ENSMUSG00000083275 | 0 | NA | NA | NA | NA | NA | Gm4993 | predicted gene 4993 [Source:MGI Symbol | 0 | 0 | 0 | 0 | 0 | 0 |
| 25484 | ENSMUSG00000083276 | 0 | NA | NA | NA | NA | NA | Gm13551 | predicted gene 13551 [Source:MGI Symbo | 0 | 0 | 0 | 0 | 0 | 0 |
| 25485 | ENSMUSG00000083277 | 1.49183855 | -0.261732931 | 1.949908924 | -0.134228285 | 0.89322206 | NA | Gm15376 | predicted gene 15376 [Source:MGI Symbo | 4.8186 | 0 | 0 | 1.928 | 1.0906 | 1.1138 |
| 25486 | ENSMUSG00000083278 | 0 | NA | NA | NA | NA | NA | Gm14589 | predicted gene 14589 [Source:MGI Symbo | 0 | 0 | 0 | 0 | 0 | 0 |
| 25487 | ENSMUSG00000083279 | 0 | NA | NA | NA | NA | NA | Pex13-ps | peroxisomal biogenesis factor 13, pseudo | 0 | 0 | 0 | 0 | 0 | 0 |
| 25488 | ENSMUSG00000083280 | 0 | NA | NA | NA | NA | NA | Gm11838 | predicted gene 11838 [Source:MGI Symbo | 0 | 0 | 0 | 0 | 0 | 0 |
| 25489 | ENSMUSG00000083281 | 0 | NA | NA | NA | NA | NA | Gm13332 | predicted gene 13332 [Source:MGI Symbo | 0 | 0 | 0 | 0 | 0 | 0 |
| 25490 | ENSMUSG00000083282 | 488.6583481 | -0.175259315 | 0.157146808 | -1.115258508 | 0.26473966 | 0.720450507 | Ctsf | cathepsin F [Source:MGI Symbol;Acc:MGI | 420.02 | 580.77 | 555.45 | 477.17 | 486.42 | 412.12 |
| 25491 | ENSMUSG00000083283 | 0.665794642 | 1.656189793 | 2.801711575 | 0.591135007 | 0.554429965 | NA | Gm15361 | predicted pseudogene 15361 [Source:MG | 0.8031 | 0 | 0 | 0.964 | 0 | 2.2277 |
| 25492 | ENSMUSG00000083284 | 0.305170013 | -1.657836477 | 4.041480335 | -0.410205257 | 0.681655385 | NA | Gm13688 | predicted gene 13688 [Source:MGI Symbo | 0.8031 | 1.0279 | 0 | 0 | 0 | 0 |
| 25493 | ENSMUSG00000083285 | 0 | NA | NA | NA | NA | NA | Gm15815 | predicted gene 15815 [Source:MGI Symbo | 0 | 0 | 0 | 0 | 0 | 0 |
| 25494 | ENSMUSG00000083287 | 1.907168253 | 0.819806318 | 1.722131562 | 0.476041631 | 0.634044719 | NA | Idi1-ps1 | isopentenyl-diphosphate delta isomerase, | 4.0155 | 0 | 0 | 1.928 | 3.2719 | 2.2277 |
| 25495 | ENSMUSG00000083288 | 0 | NA | NA | NA | NA | NA | Csnk2a1-ps2 | casein kinase 2, alpha 1 polypeptide, pseu | 0 | 0 | 0 | 0 | 0 | 0 |
| 25496 | ENSMUSG00000083289 | 0 | NA | NA | NA | NA | NA | Gm8812 | predicted gene 8812 [Source:MGI Symbol | 0 | 0 | 0 | 0 | 0 | 0 |
| 25497 | ENSMUSG00000083290 | 0 | NA | NA | NA | NA | NA | Gm15030 | predicted gene 15030 [Source:MGI Symbo | 0 | 0 | 0 | 0 | 0 | 0 |
| 25498 | ENSMUSG00000083291 | 0 | NA | NA | NA | NA | NA | Gm13679 | predicted gene 13679 [Source:MGI Symbo | 0 | 0 | 0 | 0 | 0 | 0 |
| 25499 | ENSMUSG00000083292 | 0 | NA | NA | NA | NA | NA | Gm12010 | predicted gene 12010 [Source:MGI Symbo | 0 | 0 | 0 | 0 | 0 | 0 |
| 25500 | ENSMUSG00000083293 | 0 | NA | NA | NA | NA | NA | Mup-ps28 | major urinary protein, pseudogene 28 [So | 0 | 0 | 0 | 0 | 0 | 0 |
| 25501 | ENSMUSG00000083294 | 0 | NA | NA | NA | NA | NA | Gm14957 | predicted gene 14957 [Source:MGI Symbo | 0 | 0 | 0 | 0 | 0 | 0 |
| 25502 | ENSMUSG00000083296 | 0 | NA | NA | NA | NA | NA | Gm12638 | predicted gene 12638 [Source:MGI Symbo | 0 | 0 | 0 | 0 | 0 | 0 |
| 25503 | ENSMUSG00000083297 | 0 | NA | NA | NA | NA | NA | Vmn1r-ps11 | vomeronasal 1 receptor, pseudogene 114 | 0 | 0 | 0 | 0 | 0 | 0 |
| 25504 | ENSMUSG00000083298 | 0 | NA | NA | NA | NA | NA | Gm12085 | predicted gene 12085 [Source:MGI Symbo | 0 | 0 | 0 | 0 | 0 | 0 |
| 25505 | ENSMUSG00000083299 | 0 | NA | NA | NA | NA | NA | Gm12332 | predicted gene 12332 [Source:MGI Symbo | 0 | 0 | 0 | 0 | 0 | 0 |
| 25506 | ENSMUSG00000083300 | 0 | NA | NA | NA | NA | NA | Gm12309 | predicted gene 12309 [Source:MGI Symbo | 0 | 0 | 0 | 0 | 0 | 0 |
| 25507 | ENSMUSG00000083303 | 0 | NA | NA | NA | NA | NA | Gm12110 | predicted gene 12110 [Source:MGI Symbo | 0 | 0 | 0 | 0 | 0 | 0 |
| 25508 | ENSMUSG00000083304 | 15.42988401 | 1.839059898 | 0.605808268 | 3.035712772 | 0.002399678 | 0.039987268 | Mup-ps15 | major urinary protein, pseudogene 15 [So | 8.8341 | 4.1117 | 7.0437 | 16.388 | 28.356 | 27.846 |
| 25509 | ENSMUSG00000083305 | 0 | NA | NA | NA | NA | NA | Gm13315 | predicted gene 13315 [Source:MGI Symbo | 0 | 0 | 0 | 0 | 0 | 0 |
| 25510 | ENSMUSG00000083306 | 0.17131964 | -0.796629176 | 4.080472857 | -0.195229623 | 0.84521319 | NA | Gm13868 | predicted gene 13868 [Source:MGI Symbo | 0 | 1.0279 | 0 | 0 | 0 | 0 |
| 25511 | ENSMUSG00000083307 | 1.284755329 | -0.613014061 | 2.01101397 | -0.304828346 | 0.760496892 | NA | AA414768 | expressed sequence AA414768 [Source:M | 1.6062 | 3.0838 | 0 | 1.928 | 1.0906 | 0 |
| 25512 | ENSMUSG00000083309 | 0 | NA | NA | NA | NA | NA | Gm15926 | predicted gene 15926 [Source:MGI Symbo | 0 | 0 | 0 | 0 | 0 | 0 |
| 25513 | ENSMUSG00000083310 | 0 | NA | NA | NA | NA | NA | Gm5760 | predicted gene 5760 [Source:MGI Symbol | 0 | 0 | 0 | 0 | 0 | 0 |
| 25514 | ENSMUSG00000083311 | 0.181769715 | 1.126924175 | 4.080472857 | 0.276174898 | 0.782413736 | NA | Gm5643 | predicted gene 5643 [Source:MGI Symbol | 0 | 0 | 0 | 0 | 1.0906 | 0 |
| 25515 | ENSMUSG00000083312 | 0 | NA | NA | NA | NA | NA | Olfr1374-ps1 | olfactory receptor 1374, pseudogene 1 [S | 0 | 0 | 0 | 0 | 0 | 0 |
| 25516 | ENSMUSG00000083314 | 0 | NA | NA | NA | NA | NA | Gm11742 | predicted gene 11742 [Source:MGI Symbo | 0 | 0 | 0 | 0 | 0 | 0 |
| 25517 | ENSMUSG00000083315 | 0 | NA | NA | NA | NA | NA | Gm11871 | predicted gene 11871 [Source:MGI Symbo | 0 | 0 | 0 | 0 | 0 | 0 |
| 25518 | ENSMUSG00000083316 | 0 | NA | NA | NA | NA | NA | Gm15185 | predicted gene 15185 [Source:MGI Symbo | 0 | 0 | 0 | 0 | 0 | 0 |
| 25519 | ENSMUSG00000083317 | 0 | NA | NA | NA | NA | NA | Gm14402 | predicted gene 14402 [Source:MGI Symbo | 0 | 0 | 0 | 0 | 0 | 0 |
| 25520 | ENSMUSG00000083318 | 0 | NA | NA | NA | NA | NA | Gm15192 | predicted gene 15192 [Source:MGI Symbo | 0 | 0 | 0 | 0 | 0 | 0 |
| 25521 | ENSMUSG00000083319 | 0 | NA | NA | NA | NA | NA | Gm11284 | predicted gene 11284 [Source:MGI Symbo | 0 | 0 | 0 | 0 | 0 | 0 |
| 25522 | ENSMUSG00000083320 | 0 | NA | NA | NA | NA | NA | Gm13935 | predicted gene 13935 [Source:MGI Symbo | 0 | 0 | 0 | 0 | 0 | 0 |
| 25523 | ENSMUSG00000083321 | 0 | NA | NA | NA | NA | NA | Gm14931 | predicted gene 14931 [Source:MGI Symbo | 0 | 0 | 0 | 0 | 0 | 0 |
| 25524 | ENSMUSG00000083324 | 0 | NA | NA | NA | NA | NA | Gm14997 | predicted gene 14997 [Source:MGI Symbo | 0 | 0 | 0 | 0 | 0 | 0 |
| 25525 | ENSMUSG00000083325 | 0.85435009 | 0.823372753 | 2.383559862 | 0.345438252 | 0.729764924 | NA | Gm14121 | predicted gene 14121 [Source:MGI Symbo | 0.8031 | 1.0279 | 0 | 0 | 2.1812 | 1.1138 |
| 25526 | ENSMUSG00000083326 | 0.503302254 | -0.837877584 | 3.426698643 | -0.244514523 | 0.806832356 | NA | Rpl38-ps1 | ribosomal protein L38, pseudogene 1 [Sou | 0 | 2.0558 | 0 | 0.964 | 0 | 0 |
| 25527 | ENSMUSG00000083327 | 17.5179494 | 0.317254981 | 0.548044612 | 0.578885321 | 0.562666556 | 0.900565925 | Vcp-rs | valosin containing protein, related sequer | 15.259 | 21.586 | 10.062 | 25.063 | 13.087 | 20.049 |
| 25528 | ENSMUSG00000083328 | 0.867306998 | -1.831940351 | 2.582928097 | -0.709249457 | 0.478169687 | NA | Gm11826 | predicted gene 11826 [Source:MGI Symbo | 0 | 3.0838 | 1.0062 | 0 | 0 | 1.1138 |
| 25529 | ENSMUSG00000083329 | 0 | NA | NA | NA | NA | NA | Gm14220 | predicted gene 14220 [Source:MGI Symbo | 0 | 0 | 0 | 0 | 0 | 0 |
| 25530 | ENSMUSG00000083330 | 0 | NA | NA | NA | NA | NA | Gm11859 | predicted gene 11859 [Source:MGI Symbo | 0 | 0 | 0 | 0 | 0 | 0 |
| 25531 | ENSMUSG00000083331 | 0.663758638 | 2.950973782 | 3.025092005 | 0.975498853 | 0.32931287 | NA | Gm12400 | predicted gene 12400 [Source:MGI Symbo | 0 | 0 | 0 | 2.8919 | 1.0906 | 0 |
| 25532 | ENSMUSG00000083332 | 5.495557933 | 0.687508557 | 1.145066723 | 0.600409167 | 0.54823358 | 0.895600915 | Gm7599 | predicted gene 7599 [Source:MGI Symbol | 2.4093 | 10.279 | 0 | 3.8559 | 13.087 | 3.3415 |

| | | | | | | | | | | | | | | | |
|---|---|---|---|---|---|---|---|---|---|---|---|---|---|---|---|
| 25533 | ENSMUSG00000083333 | 0 | NA | NA | NA | NA | NA | Magea7-ps | MAGE family member A7, pseudogene [Sc | 0 | 0 | 0 | 0 | 0 | 0 |
| 25534 | ENSMUSG00000083334 | 0 | NA | NA | NA | NA | NA | Gm13579 | predicted gene 13579 [Source:MGI Symbc | 0 | 0 | 0 | 0 | 0 | 0 |
| 25535 | ENSMUSG00000083335 | 0 | NA | NA | NA | NA | NA | Gm13397 | predicted gene 13397 [Source:MGI Symbc | 0 | 0 | 0 | 0 | 0 | 0 |
| 25536 | ENSMUSG00000083336 | 0 | NA | NA | NA | NA | NA | Olfr1332-ps1 | olfactory receptor 1332, pseudogene 1 [Sc | 0 | 0 | 0 | 0 | 0 | 0 |
| 25537 | ENSMUSG00000083337 | 0.439020386 | -2.189747911 | 3.52148864 | -0.621824499 | 0.534057275 | NA | Gm11539 | predicted gene 11539 [Source:MGI Symbc | 1.6062 | 1.0279 | 0 | 0 | 0 | 0 |
| 25538 | ENSMUSG00000083338 | 0 | NA | NA | NA | NA | NA | Gm12475 | predicted gene 12475 [Source:MGI Symbc | 0 | 0 | 0 | 0 | 0 | 0 |
| 25539 | ENSMUSG00000083339 | 0.181769715 | 1.126924175 | 4.080472857 | 0.276174898 | 0.782413736 | NA | Gm11693 | predicted gene 11693 [Source:MGI Symbc | 0 | 0 | 0 | 0 | 1.0906 | 0 |
| 25540 | ENSMUSG00000083340 | 2.132837134 | 0.419872099 | 1.52472695 | 0.275375272 | 0.78302794 | NA | Gm13156 | predicted gene 13156 [Source:MGI Symbc | 2.4093 | 1.0279 | 2.0125 | 2.8919 | 0 | 4.4554 |
| 25541 | ENSMUSG00000083341 | 0 | NA | NA | NA | NA | NA | Gm14238 | predicted gene 14238 [Source:MGI Symbc | 0 | 0 | 0 | 0 | 0 | 0 |
| 25542 | ENSMUSG00000083342 | 0 | NA | NA | NA | NA | NA | Gm14237 | predicted gene 14237 [Source:MGI Symbc | 0 | 0 | 0 | 0 | 0 | 0 |
| 25543 | ENSMUSG00000083343 | 0 | NA | NA | NA | NA | NA | Gm14666 | predicted gene 14666 [Source:MGI Symbc | 0 | 0 | 0 | 0 | 0 | 0 |
| 25544 | ENSMUSG00000083344 | 0 | NA | NA | NA | NA | NA | Gm7363 | predicted gene 7363 [Source:MGI Symbol | 0 | 0 | 0 | 0 | 0 | 0 |
| 25545 | ENSMUSG00000083345 | 0 | NA | NA | NA | NA | NA | Gm6491 | predicted gene 6491 [Source:MGI Symbol | 0 | 0 | 0 | 0 | 0 | 0 |
| 25546 | ENSMUSG00000083346 | 0 | NA | NA | NA | NA | NA | Gm8260 | predicted gene 8260 [Source:MGI Symbol | 0 | 0 | 0 | 0 | 0 | 0 |
| 25547 | ENSMUSG00000083347 | 0 | NA | NA | NA | NA | NA | Gm13913 | predicted gene 13913 [Source:MGI Symbc | 0 | 0 | 0 | 0 | 0 | 0 |
| 25548 | ENSMUSG00000083348 | 0 | NA | NA | NA | NA | NA | Vmn1r-ps99 | vomeronasal 1 receptor, pseudogene 97 [ | 0 | 0 | 0 | 0 | 0 | 0 |
| 25549 | ENSMUSG00000083350 | 0 | NA | NA | NA | NA | NA | Gm9060 | predicted gene 9060 [Source:MGI Symbol | 0 | 0 | 0 | 0 | 0 | 0 |
| 25550 | ENSMUSG00000083351 | 0 | NA | NA | NA | NA | NA | Gm15783 | predicted gene 15783 [Source:MGI Symbc | 0 | 0 | 0 | 0 | 0 | 0 |
| 25551 | ENSMUSG00000083352 | 0 | NA | NA | NA | NA | NA | Gm14907 | predicted gene 14907 [Source:MGI Symbc | 0 | 0 | 0 | 0 | 0 | 0 |
| 25552 | ENSMUSG00000083353 | 0 | NA | NA | NA | NA | NA | Gm12540 | predicted gene 12540 [Source:MGI Symbc | 0 | 0 | 0 | 0 | 0 | 0 |
| 25553 | ENSMUSG00000083355 | 0.59968336 | -1.233555632 | 2.892386901 | -0.426483618 | 0.669755482 | NA | Gm11581 | predicted gene 11581 [Source:MGI Symbc | 1.6062 | 1.0279 | 0 | 0.964 | 0 | 0 |
| 25554 | ENSMUSG00000083356 | 0 | NA | NA | NA | NA | NA | Gm11708 | predicted gene 11708 [Source:MGI Symbc | 0 | 0 | 0 | 0 | 0 | 0 |
| 25555 | ENSMUSG00000083358 | 0 | NA | NA | NA | NA | NA | Olfr1004-ps1 | olfactory receptor 1004, pseudogene 1 [Sc | 0 | 0 | 0 | 0 | 0 | 0 |
| 25556 | ENSMUSG00000083359 | 0.897628956 | -1.977329804 | 2.507804229 | -0.78847056 | 0.430421512 | NA | Gm12656 | predicted gene 12656 [Source:MGI Symbc | 2.4093 | 0 | 2.0125 | 0.964 | 0 | 0 |
| 25557 | ENSMUSG00000083361 | 0 | NA | NA | NA | NA | NA | Olfr360 | olfactory receptor 360 [Source:MGI Symb | 0 | 0 | 0 | 0 | 0 | 0 |
| 25558 | ENSMUSG00000083362 | 0 | NA | NA | NA | NA | NA | Gm8606 | predicted gene 8606 [Source:MGI Symbol | 0 | 0 | 0 | 0 | 0 | 0 |
| 25559 | ENSMUSG00000083363 | 0 | NA | NA | NA | NA | NA | Gm12237 | predicted gene 12237 [Source:MGI Symbc | 0 | 0 | 0 | 0 | 0 | 0 |
| 25560 | ENSMUSG00000083364 | 5.04288986 | 0.573781644 | 0.959097529 | 0.598251613 | 0.549672058 | NA | Llph-ps2 | LLP homolog, pseudogene 2 [Source:MGI | 4.0155 | 3.0838 | 5.0312 | 3.8559 | 9.8156 | 4.4554 |
| 25561 | ENSMUSG00000083365 | 0 | NA | NA | NA | NA | NA | Gm12895 | predicted gene 12895 [Source:MGI Symbc | 0 | 0 | 0 | 0 | 0 | 0 |
| 25562 | ENSMUSG00000083366 | 0 | NA | NA | NA | NA | NA | Gm14887 | predicted gene 14887 [Source:MGI Symbc | 0 | 0 | 0 | 0 | 0 | 0 |
| 25563 | ENSMUSG00000083367 | 0.51034671 | -2.381743948 | 3.331081356 | -0.715006238 | 0.474605189 | NA | Gm8806 | predicted gene 8806 [Source:MGI Symbol | 0 | 2.0558 | 1.0062 | 0 | 0 | 0 |
| 25564 | ENSMUSG00000083368 | 0.481988923 | 2.492401104 | 3.837077151 | 0.649557203 | 0.515978285 | NA | Gm13214 | predicted gene 13214 [Source:MGI Symbc | 0 | 0 | 0 | 2.8919 | 0 | 0 |
| 25565 | ENSMUSG00000083369 | 0 | NA | NA | NA | NA | NA | Gm15332 | predicted gene 15332 [Source:MGI Symbc | 0 | 0 | 0 | 0 | 0 | 0 |
| 25566 | ENSMUSG00000083371 | 0 | NA | NA | NA | NA | NA | Gm11422 | predicted gene 11422 [Source:MGI Symbc | 0 | 0 | 0 | 0 | 0 | 0 |
| 25567 | ENSMUSG00000083372 | 0 | NA | NA | NA | NA | NA | Gm11235 | predicted gene 11235 [Source:MGI Symbc | 0 | 0 | 0 | 0 | 0 | 0 |
| 25568 | ENSMUSG00000083373 | 0.808265676 | 0.728755068 | 2.41725015 | 0.301481031 | 0.763047712 | NA | Gm13392 | predicted gene 13392 [Source:MGI Symbc | 0.8031 | 1.0279 | 0 | 1.928 | 1.0906 | 0 |
| 25569 | ENSMUSG00000083374 | 0 | NA | NA | NA | NA | NA | Gm11449 | predicted gene 11449 [Source:MGI Symbc | 0 | 0 | 0 | 0 | 0 | 0 |
| 25570 | ENSMUSG00000083375 | 0 | NA | NA | NA | NA | NA | Gm15057 | predicted gene 15057 [Source:MGI Symbc | 0 | 0 | 0 | 0 | 0 | 0 |
| 25571 | ENSMUSG00000083376 | 0 | NA | NA | NA | NA | NA | Gm11882 | predicted gene 11882 [Source:MGI Symbc | 0 | 0 | 0 | 0 | 0 | 0 |
| 25572 | ENSMUSG00000083377 | 0 | NA | NA | NA | NA | NA | Gm12653 | predicted gene 12653 [Source:MGI Symbc | 0 | 0 | 0 | 0 | 0 | 0 |
| 25573 | ENSMUSG00000083378 | 0 | NA | NA | NA | NA | NA | Gm13196 | predicted gene 13196 [Source:MGI Symbc | 0 | 0 | 0 | 0 | 0 | 0 |
| 25574 | ENSMUSG00000083379 | 0 | NA | NA | NA | NA | NA | Gm4919 | predicted gene 4919 [Source:MGI Symbol | 0 | 0 | 0 | 0 | 0 | 0 |
| 25575 | ENSMUSG00000083380 | 1.696292863 | 0.15334619 | 1.730260434 | 0.088626075 | 0.929379085 | NA | Ndufb4c | NADH:ubiquinone oxidoreductase subuni | 0.8031 | 0 | 4.025 | 0.964 | 3.2719 | 1.1138 |
| 25576 | ENSMUSG00000083381 | 0 | NA | NA | NA | NA | NA | Scgb2b22-ps | secretoglobin, family 2B, member 22, pseu | 0 | 0 | 0 | 0 | 0 | 0 |
| 25577 | ENSMUSG00000083382 | 0 | NA | NA | NA | NA | NA | Gm6433 | predicted gene 6433 [Source:MGI Symbol | 0 | 0 | 0 | 0 | 0 | 0 |
| 25578 | ENSMUSG00000083383 | 0 | NA | NA | NA | NA | NA | Olfr595-ps1 | olfactory receptor 595, pseudogene 1 [Sou | 0 | 0 | 0 | 0 | 0 | 0 |
| 25579 | ENSMUSG00000083384 | 0 | NA | NA | NA | NA | NA | Gm12813 | predicted gene 12813 [Source:MGI Symbc | 0 | 0 | 0 | 0 | 0 | 0 |
| 25580 | ENSMUSG00000083385 | 0 | NA | NA | NA | NA | NA | Gm11411 | predicted gene 11411 [Source:MGI Symbc | 0 | 0 | 0 | 0 | 0 | 0 |
| 25581 | ENSMUSG00000083386 | 0 | NA | NA | NA | NA | NA | Gm15426 | predicted gene 15426 [Source:MGI Symbc | 0 | 0 | 0 | 0 | 0 | 0 |
| 25582 | ENSMUSG00000083387 | 0 | NA | NA | NA | NA | NA | Gm11860 | predicted gene 11860 [Source:MGI Symbc | 0 | 0 | 0 | 0 | 0 | 0 |
| 25583 | ENSMUSG00000083388 | 0 | NA | NA | NA | NA | NA | Gm11550 | predicted gene 11550 [Source:MGI Symbc | 0 | 0 | 0 | 0 | 0 | 0 |
| 25584 | ENSMUSG00000083389 | 0 | NA | NA | NA | NA | NA | Gm13681 | predicted gene 13681 [Source:MGI Symbc | 0 | 0 | 0 | 0 | 0 | 0 |
| 25585 | ENSMUSG00000083390 | 0 | NA | NA | NA | NA | NA | Gm13121 | predicted gene 13121 [Source:MGI Symbc | 0 | 0 | 0 | 0 | 0 | 0 |
| 25586 | ENSMUSG00000083391 | 0 | NA | NA | NA | NA | NA | Gm14148 | predicted gene 14148 [Source:MGI Symbc | 0 | 0 | 0 | 0 | 0 | 0 |
| 25587 | ENSMUSG00000083392 | 0.677847551 | 0.063310748 | 2.809814301 | 0.022532004 | 0.982023583 | NA | Gm6335 | predicted gene 6335 [Source:MGI Symbol | 0 | 0 | 2.0125 | 0.964 | 1.0906 | 0 |
| 25588 | ENSMUSG00000083393 | 0 | NA | NA | NA | NA | NA | Gm15194 | predicted gene 15194 [Source:MGI Symbc | 0 | 0 | 0 | 0 | 0 | 0 |

| | | | | | | | | | | | | | | | | |
|---|---|---|---|---|---|---|---|---|---|---|---|---|---|---|---|---|
| 25589 | ENSMUSG00000083394 | 0.59968336 | -1.233555632 | 2.892386901 | -0.426483618 | 0.669755482 | NA | | Gm11703 | predicted gene 11703 [Source:MGI Symbo | 1.6062 | 1.0279 | 0 | 0.964 | 0 | 0 |
| 25590 | ENSMUSG00000083395 | 0 | NA | NA | NA | NA | NA | | Gm11203 | predicted gene 11203 [Source:MGI Symbo | 0 | 0 | 0 | 0 | 0 | 0 |
| 25591 | ENSMUSG00000083396 | 16.87514304 | 0.452013109 | 0.532747743 | 0.848456168 | 0.396183973 | 0.823764863 | | Gm15542 | predicted gene 15542 [Source:MGI Symbo | 15.259 | 16.447 | 11.069 | 23.135 | 14.178 | 21.163 |
| 25592 | ENSMUSG00000083397 | 0.181769715 | 1.126924175 | 4.080472857 | 0.276174898 | 0.782413736 | NA | | Gm5466 | predicted gene 5466 [Source:MGI Symbol | 0 | 0 | 0 | 0 | 1.0906 | 0 |
| 25593 | ENSMUSG00000083398 | 0 | NA | NA | NA | NA | NA | | Gm13458 | predicted gene 13458 [Source:MGI Symbo | 0 | 0 | 0 | 0 | 0 | 0 |
| 25594 | ENSMUSG00000083399 | 0 | NA | NA | NA | NA | NA | | Gm15033 | predicted gene 15033 [Source:MGI Symbo | 0 | 0 | 0 | 0 | 0 | 0 |
| 25595 | ENSMUSG00000083401 | 0 | NA | NA | NA | NA | NA | | Gm13506 | predicted gene 13506 [Source:MGI Symbo | 0 | 0 | 0 | 0 | 0 | 0 |
| 25596 | ENSMUSG00000083402 | 0 | NA | NA | NA | NA | NA | | Gm14394 | predicted gene 14394 [Source:MGI Symbo | 0 | 0 | 0 | 0 | 0 | 0 |
| 25597 | ENSMUSG00000083403 | 0 | NA | NA | NA | NA | NA | | Gm13041 | predicted gene 13041 [Source:MGI Symbo | 0 | 0 | 0 | 0 | 0 | 0 |
| 25598 | ENSMUSG00000083404 | 0.499690045 | -0.82080011 | 3.113553401 | -0.263621658 | 0.792071468 | NA | | Gm13019 | predicted gene 13019 [Source:MGI Symbo | 0 | 1.0279 | 1.0062 | 0.964 | 0 | 0 |
| 25599 | ENSMUSG00000083405 | 0 | NA | NA | NA | NA | NA | | Gm15725 | predicted gene 15725 [Source:MGI Symbo | 0 | 0 | 0 | 0 | 0 | 0 |
| 25600 | ENSMUSG00000083406 | 0 | NA | NA | NA | NA | NA | | Gm8410 | predicted gene 8410 [Source:MGI Symbol | 0 | 0 | 0 | 0 | 0 | 0 |
| 25601 | ENSMUSG00000083407 | 0 | NA | NA | NA | NA | NA | | Gm13176 | predicted gene 13176 [Source:MGI Symbo | 0 | 0 | 0 | 0 | 0 | 0 |
| 25602 | ENSMUSG00000083409 | 3.417205912 | 0.331646725 | 1.244589028 | 0.266470873 | 0.789876589 | NA | | Gm11340 | predicted gene 11340 [Source:MGI Symbo | 4.0155 | 0 | 5.0312 | 4.8199 | 2.1812 | 4.4554 |
| 25603 | ENSMUSG00000083410 | 0 | NA | NA | NA | NA | NA | | Gm14300 | predicted gene 14300 [Source:MGI Symbo | 0 | 0 | 0 | 0 | 0 | 0 |
| 25604 | ENSMUSG00000083411 | 0.885033624 | 2.078055061 | 2.356585208 | 0.881807733 | 0.377880789 | NA | | Rpl30-ps10 | ribosomal protein L30, pseudogene 10 [Sc | 0 | 1.0279 | 0 | 0.964 | 1.0906 | 2.2277 |
| 25605 | ENSMUSG00000083412 | 0 | NA | NA | NA | NA | NA | | Gm12268 | predicted gene 12268 [Source:MGI Symbo | 0 | 0 | 0 | 0 | 0 | 0 |
| 25606 | ENSMUSG00000083413 | 0 | NA | NA | NA | NA | NA | | Vmn1r-ps11 | vomeronasal 1 receptor, pseudogene 110 | 0 | 0 | 0 | 0 | 0 | 0 |
| 25607 | ENSMUSG00000083414 | 0 | NA | NA | NA | NA | NA | | Gm14840 | predicted gene 14840 [Source:MGI Symbo | 0 | 0 | 0 | 0 | 0 | 0 |
| 25608 | ENSMUSG00000083416 | 0.185640647 | 1.126924175 | 4.080472857 | 0.276174898 | 0.782413736 | NA | | Gm15378 | predicted gene 15378 [Source:MGI Symbo | 0 | 0 | 0 | 0 | 0 | 1.1138 |
| 25609 | ENSMUSG00000083417 | 0 | NA | NA | NA | NA | NA | | Olfr269-ps1 | olfactory receptor 269, pseudogene 1 [So | 0 | 0 | 0 | 0 | 0 | 0 |
| 25610 | ENSMUSG00000083418 | 0 | NA | NA | NA | NA | NA | | Olfr1035-ps1 | olfactory receptor 1035, pseudogene 1 [S | 0 | 0 | 0 | 0 | 0 | 0 |
| 25611 | ENSMUSG00000083419 | 0 | NA | NA | NA | NA | NA | | Gm14485 | predicted gene 14485 [Source:MGI Symbo | 0 | 0 | 0 | 0 | 0 | 0 |
| 25612 | ENSMUSG00000083420 | 0 | NA | NA | NA | NA | NA | | Gm15665 | predicted gene 15665 [Source:MGI Symbo | 0 | 0 | 0 | 0 | 0 | 0 |
| 25613 | ENSMUSG00000083421 | 0 | NA | NA | NA | NA | NA | | Gm14776 | predicted gene 14776 [Source:MGI Symbo | 0 | 0 | 0 | 0 | 0 | 0 |
| 25614 | ENSMUSG00000083422 | 0 | NA | NA | NA | NA | NA | | Gm15604 | predicted gene 15604 [Source:MGI Symbo | 0 | 0 | 0 | 0 | 0 | 0 |
| 25615 | ENSMUSG00000083423 | 0 | NA | NA | NA | NA | NA | | Gm13221 | predicted gene 13221 [Source:MGI Symbo | 0 | 0 | 0 | 0 | 0 | 0 |
| 25616 | ENSMUSG00000083424 | 0.59968336 | -1.233555632 | 2.892386901 | -0.426483618 | 0.669755482 | NA | | Rpl35a-ps4 | ribosomal protein 35A, pseudogene 4 [So | 1.6062 | 1.0279 | 0 | 0.964 | 0 | 0 |
| 25617 | ENSMUSG00000083425 | 0 | NA | NA | NA | NA | NA | | Gm13992 | predicted gene 13992 [Source:MGI Symbo | 0 | 0 | 0 | 0 | 0 | 0 |
| 25618 | ENSMUSG00000083427 | 0 | NA | NA | NA | NA | NA | | Gm12007 | predicted gene 12007 [Source:MGI Symbo | 0 | 0 | 0 | 0 | 0 | 0 |
| 25619 | ENSMUSG00000083428 | 0 | NA | NA | NA | NA | NA | | Gm14591 | predicted gene 14591 [Source:MGI Symbo | 0 | 0 | 0 | 0 | 0 | 0 |
| 25620 | ENSMUSG00000083429 | 0 | NA | NA | NA | NA | NA | | Gm15198 | predicted gene 15198 [Source:MGI Symbo | 0 | 0 | 0 | 0 | 0 | 0 |
| 25621 | ENSMUSG00000083430 | 0 | NA | NA | NA | NA | NA | | Gm13006 | predicted gene 13006 [Source:MGI Symbo | 0 | 0 | 0 | 0 | 0 | 0 |
| 25622 | ENSMUSG00000083431 | 0 | NA | NA | NA | NA | NA | | Gm13437 | predicted gene 13437 [Source:MGI Symbo | 0 | 0 | 0 | 0 | 0 | 0 |
| 25623 | ENSMUSG00000083432 | 0 | NA | NA | NA | NA | NA | | Gm14878 | predicted gene 14878 [Source:MGI Symbo | 0 | 0 | 0 | 0 | 0 | 0 |
| 25624 | ENSMUSG00000083434 | 0 | NA | NA | NA | NA | NA | | Gm12760 | predicted gene 12760 [Source:MGI Symbo | 0 | 0 | 0 | 0 | 0 | 0 |
| 25625 | ENSMUSG00000083435 | 0 | NA | NA | NA | NA | NA | | Gm15824 | predicted gene 15824 [Source:MGI Symbo | 0 | 0 | 0 | 0 | 0 | 0 |
| 25626 | ENSMUSG00000083436 | 0 | NA | NA | NA | NA | NA | | Gm11471 | predicted gene 11471 [Source:MGI Symbo | 0 | 0 | 0 | 0 | 0 | 0 |
| 25627 | ENSMUSG00000083437 | 0 | NA | NA | NA | NA | NA | | Gm15187 | predicted gene 15187 [Source:MGI Symbo | 0 | 0 | 0 | 0 | 0 | 0 |
| 25628 | ENSMUSG00000083438 | 0 | NA | NA | NA | NA | NA | | Gm5402 | predicted gene 5402 [Source:MGI Symbol | 0 | 0 | 0 | 0 | 0 | 0 |
| 25629 | ENSMUSG00000083439 | 0.181769715 | 1.126924175 | 4.080472857 | 0.276174898 | 0.782413736 | NA | | Gm14491 | predicted gene 14491 [Source:MGI Symbo | 0 | 0 | 0 | 0 | 1.0906 | 0 |
| 25630 | ENSMUSG00000083440 | 0 | NA | NA | NA | NA | NA | | Rpl7a-ps12 | ribosomal protein L7A, pseudogene 12 [Sc | 0 | 0 | 0 | 0 | 0 | 0 |
| 25631 | ENSMUSG00000083441 | 0 | NA | NA | NA | NA | NA | | Gm15329 | predicted gene 15329 [Source:MGI Symbo | 0 | 0 | 0 | 0 | 0 | 0 |
| 25632 | ENSMUSG00000083443 | 0.797635641 | -1.749522449 | 2.643029821 | -0.661938218 | 0.508010818 | NA | | Gm15519 | predicted gene 15519 [Source:MGI Symbo | 0.8031 | 0 | 3.0187 | 0.964 | 0 | 0 |
| 25633 | ENSMUSG00000083444 | 0 | NA | NA | NA | NA | NA | | Gm14768 | predicted gene 14768 [Source:MGI Symbo | 0 | 0 | 0 | 0 | 0 | 0 |
| 25634 | ENSMUSG00000083445 | 0 | NA | NA | NA | NA | NA | | Gm15038 | predicted gene 15038 [Source:MGI Symbo | 0 | 0 | 0 | 0 | 0 | 0 |
| 25635 | ENSMUSG00000083446 | 0 | NA | NA | NA | NA | NA | | Gm12826 | predicted gene 12826 [Source:MGI Symbo | 0 | 0 | 0 | 0 | 0 | 0 |
| 25636 | ENSMUSG00000083447 | 0 | NA | NA | NA | NA | NA | | Gm11904 | predicted gene 11904 [Source:MGI Symbo | 0 | 0 | 0 | 0 | 0 | 0 |
| 25637 | ENSMUSG00000083448 | 0 | NA | NA | NA | NA | NA | | Gm8749 | predicted gene 8749 [Source:MGI Symbol | 0 | 0 | 0 | 0 | 0 | 0 |
| 25638 | ENSMUSG00000083449 | 0 | NA | NA | NA | NA | NA | | Gm13487 | predicted gene 13487 [Source:MGI Symbo | 0 | 0 | 0 | 0 | 0 | 0 |
| 25639 | ENSMUSG00000083450 | 0 | NA | NA | NA | NA | NA | | Gm14619 | predicted gene 14619 [Source:MGI Symbo | 0 | 0 | 0 | 0 | 0 | 0 |
| 25640 | ENSMUSG00000083451 | 0 | NA | NA | NA | NA | NA | | Gm11445 | predicted gene 11445 [Source:MGI Symbo | 0 | 0 | 0 | 0 | 0 | 0 |
| 25641 | ENSMUSG00000083452 | 0 | NA | NA | NA | NA | NA | | Gm14905 | predicted gene 14905 [Source:MGI Symbo | 0 | 0 | 0 | 0 | 0 | 0 |
| 25642 | ENSMUSG00000083453 | 0 | NA | NA | NA | NA | NA | | Gm13338 | predicted gene 13338 [Source:MGI Symbo | 0 | 0 | 0 | 0 | 0 | 0 |
| 25643 | ENSMUSG00000083454 | 1.248418068 | -0.406672647 | 2.240634769 | -0.181498856 | 0.855976031 | NA | | Gm13226 | predicted gene 13226 [Source:MGI Symbo | 3.2124 | 0 | 1.0062 | 0 | 3.2719 | 0 |
| 25644 | ENSMUSG00000083455 | 0 | NA | NA | NA | NA | NA | | Gm15037 | predicted gene 15037 [Source:MGI Symbo | 0 | 0 | 0 | 0 | 0 | 0 |

| | | | | | | | | | | | | | | | |
|---|---|---|---|---|---|---|---|---|---|---|---|---|---|---|---|
| 25645 | ENSMUSG00000083456 | 0 | NA | NA | NA | NA | NA | Gm13108 | predicted gene 13108 [Source:MGI Symbo | 0 | 0 | 0 | 0 | 0 | 0 |
| 25646 | ENSMUSG00000083457 | 0.877550482 | 2.072039088 | 2.361583201 | 0.877394066 | 0.380272635 | NA | Cyp4b1-ps2 | cytochrome P450, family 4, subfamily b, p | 0 | 0 | 1.0062 | 0.964 | 2.1812 | 1.1138 |
| 25647 | ENSMUSG00000083458 | 0 | NA | NA | NA | NA | NA | Gm5510 | predicted pseudogene 5510 [Source:MGI | 0 | 0 | 0 | 0 | 0 | 0 |
| 25648 | ENSMUSG00000083460 | 0 | NA | NA | NA | NA | NA | Gm12182 | predicted gene 12182 [Source:MGI Symbo | 0 | 0 | 0 | 0 | 0 | 0 |
| 25649 | ENSMUSG00000083461 | 0 | NA | NA | NA | NA | NA | Gm12935 | predicted gene 12935 [Source:MGI Symbo | 0 | 0 | 0 | 0 | 0 | 0 |
| 25650 | ENSMUSG00000083462 | 0 | NA | NA | NA | NA | NA | Gm12501 | predicted gene 12501 [Source:MGI Symbo | 0 | 0 | 0 | 0 | 0 | 0 |
| 25651 | ENSMUSG00000083463 | 0 | NA | NA | NA | NA | NA | Gm4909 | predicted gene 4909 [Source:MGI Symbol | 0 | 0 | 0 | 0 | 0 | 0 |
| 25652 | ENSMUSG00000083465 | 0.133850373 | -0.796629176 | 4.080472857 | -0.195229623 | 0.84521319 | NA | Gm11652 | predicted gene 11652 [Source:MGI Symbo | 0.8031 | 0 | 0 | 0 | 0 | 0 |
| 25653 | ENSMUSG00000083466 | 0 | NA | NA | NA | NA | NA | Gm15225 | predicted gene 15225 [Source:MGI Symbo | 0 | 0 | 0 | 0 | 0 | 0 |
| 25654 | ENSMUSG00000083467 | 0 | NA | NA | NA | NA | NA | Gm15840 | predicted gene 15840 [Source:MGI Symbo | 0 | 0 | 0 | 0 | 0 | 0 |
| 25655 | ENSMUSG00000083468 | 0 | NA | NA | NA | NA | NA | Gm9431 | predicted gene 9431 [Source:MGI Symbol | 0 | 0 | 0 | 0 | 0 | 0 |
| 25656 | ENSMUSG00000083469 | 0 | NA | NA | NA | NA | NA | Gm14281 | predicted gene 14281 [Source:MGI Symbo | 0 | 0 | 0 | 0 | 0 | 0 |
| 25657 | ENSMUSG00000083470 | 0 | NA | NA | NA | NA | NA | Nsa2-ps2 | NSA2 ribosome biogenesis homolog, pseu | 0 | 0 | 0 | 0 | 0 | 0 |
| 25658 | ENSMUSG00000083471 | 0 | NA | NA | NA | NA | NA | Gm12118 | predicted gene 12118 [Source:MGI Symbo | 0 | 0 | 0 | 0 | 0 | 0 |
| 25659 | ENSMUSG00000083472 | 0 | NA | NA | NA | NA | NA | Gm13491 | predicted gene 13491 [Source:MGI Symbo | 0 | 0 | 0 | 0 | 0 | 0 |
| 25660 | ENSMUSG00000083473 | 0 | NA | NA | NA | NA | NA | Gm13207 | predicted gene 13207 [Source:MGI Symbo | 0 | 0 | 0 | 0 | 0 | 0 |
| 25661 | ENSMUSG00000083474 | 0.670829724 | -2.782124174 | 3.401449285 | -0.817923168 | 0.41340106 | NA | Gm15267 | predicted gene 15267 [Source:MGI Symbo | 0 | 0 | 4.025 | 0 | 0 | 0 |
| 25662 | ENSMUSG00000083475 | 0 | NA | NA | NA | NA | NA | Gm9440 | predicted gene 9440 [Source:MGI Symbol | 0 | 0 | 0 | 0 | 0 | 0 |
| 25663 | ENSMUSG00000083477 | 0.55305101 | 2.667977666 | 3.241982584 | 0.822946329 | 0.410538515 | NA | Gm5555 | predicted pseudogene 5555 [Source:MGI | 0 | 0 | 0 | 0 | 1.0906 | 2.2277 |
| 25664 | ENSMUSG00000083478 | 3.693268595 | 0.047544687 | 1.129175526 | 0.042105665 | 0.966414464 | NA | Gm15375 | predicted gene 15375 [Source:MGI Symbo | 4.8186 | 2.0558 | 4.025 | 5.7839 | 4.3625 | 1.1138 |
| 25665 | ENSMUSG00000083481 | 0 | NA | NA | NA | NA | NA | Rps8-ps2 | ribosomal protein S8, pseudogene 2 [Sour | 0 | 0 | 0 | 0 | 0 | 0 |
| 25666 | ENSMUSG00000083482 | 0 | NA | NA | NA | NA | NA | Gm14706 | predicted gene 14706 [Source:MGI Symbo | 0 | 0 | 0 | 0 | 0 | 0 |
| 25667 | ENSMUSG00000083483 | 0.535117793 | 1.111049431 | 3.044213722 | 0.364970903 | 0.715133126 | NA | Gm14044 | predicted gene 14044 [Source:MGI Symbo | 0 | 0 | 1.0062 | 0 | 1.0906 | 1.1138 |
| 25668 | ENSMUSG00000083484 | 0 | NA | NA | NA | NA | NA | Gm14574 | predicted gene 14574 [Source:MGI Symbo | 0 | 0 | 0 | 0 | 0 | 0 |
| 25669 | ENSMUSG00000083485 | 0.160662974 | 1.126924175 | 4.080472857 | 0.276174898 | 0.782413736 | NA | Gm12308 | predicted gene 12308 [Source:MGI Symbo | 0 | 0 | 0 | 0.964 | 0 | 0 |
| 25670 | ENSMUSG00000083486 | 0 | NA | NA | NA | NA | NA | Gm14521 | predicted gene 14521 [Source:MGI Symbo | 0 | 0 | 0 | 0 | 0 | 0 |
| 25671 | ENSMUSG00000083487 | 0 | NA | NA | NA | NA | NA | Gm13888 | predicted gene 13888 [Source:MGI Symbo | 0 | 0 | 0 | 0 | 0 | 0 |
| 25672 | ENSMUSG00000083488 | 0 | NA | NA | NA | NA | NA | Gm8534 | predicted gene 8534 [Source:MGI Symbol | 0 | 0 | 0 | 0 | 0 | 0 |
| 25673 | ENSMUSG00000083489 | 0 | NA | NA | NA | NA | NA | Gm11784 | predicted gene 11784 [Source:MGI Symbo | 0 | 0 | 0 | 0 | 0 | 0 |
| 25674 | ENSMUSG00000083490 | 0 | NA | NA | NA | NA | NA | Gm14940 | predicted gene 14940 [Source:MGI Symbo | 0 | 0 | 0 | 0 | 0 | 0 |
| 25675 | ENSMUSG00000083491 | 0 | NA | NA | NA | NA | NA | Gm12423 | predicted gene 12423 [Source:MGI Symbo | 0 | 0 | 0 | 0 | 0 | 0 |
| 25676 | ENSMUSG00000083492 | 0 | NA | NA | NA | NA | NA | Gm14515 | predicted gene 14515 [Source:MGI Symbo | 0 | 0 | 0 | 0 | 0 | 0 |
| 25677 | ENSMUSG00000083493 | 0 | NA | NA | NA | NA | NA | Gm15044 | predicted gene 15044 [Source:MGI Symbo | 0 | 0 | 0 | 0 | 0 | 0 |
| 25678 | ENSMUSG00000083494 | 0 | NA | NA | NA | NA | NA | Gm12873 | predicted gene 12873 [Source:MGI Symbo | 0 | 0 | 0 | 0 | 0 | 0 |
| 25679 | ENSMUSG00000083495 | 0 | NA | NA | NA | NA | NA | Gm13877 | predicted gene 13877 [Source:MGI Symbo | 0 | 0 | 0 | 0 | 0 | 0 |
| 25680 | ENSMUSG00000083496 | 2.270125269 | -0.337888932 | 1.510945737 | -0.223627443 | 0.823047208 | NA | Gm11263 | predicted gene 11263 [Source:MGI Symbo | 1.6062 | 2.0558 | 4.025 | 4.8199 | 0 | 1.1138 |
| 25681 | ENSMUSG00000083498 | 0 | NA | NA | NA | NA | NA | Gm14172 | predicted gene 14172 [Source:MGI Symbo | 0 | 0 | 0 | 0 | 0 | 0 |
| 25682 | ENSMUSG00000083499 | 0 | NA | NA | NA | NA | NA | Gm14609 | predicted gene 14609 [Source:MGI Symbo | 0 | 0 | 0 | 0 | 0 | 0 |
| 25683 | ENSMUSG00000083500 | 0.342639279 | -1.805976596 | 4.028547714 | -0.448294702 | 0.653940524 | NA | Gm15470 | predicted gene 15470 [Source:MGI Symbo | 0 | 2.0558 | 0 | 0 | 0 | 0 |
| 25684 | ENSMUSG00000083501 | 0 | NA | NA | NA | NA | NA | Gm12255 | predicted gene 12255 [Source:MGI Symbo | 0 | 0 | 0 | 0 | 0 | 0 |
| 25685 | ENSMUSG00000083502 | 0 | NA | NA | NA | NA | NA | Gm14861 | predicted gene 14861 [Source:MGI Symbo | 0 | 0 | 0 | 0 | 0 | 0 |
| 25686 | ENSMUSG00000083503 | 0.167707431 | -0.796629176 | 4.080472857 | -0.195229623 | 0.84521319 | NA | Gm13407 | predicted gene 13407 [Source:MGI Symbo | 0 | 0 | 1.0062 | 0 | 0 | 0 |
| 25687 | ENSMUSG00000083505 | 0.17131964 | -0.796629176 | 4.080472857 | -0.195229623 | 0.84521319 | NA | Gm7541 | predicted gene 7541 [Source:MGI Symbol | 0 | 1.0279 | 0 | 0 | 0 | 0 |
| 25688 | ENSMUSG00000083507 | 0 | NA | NA | NA | NA | NA | Gm12842 | predicted gene 12842 [Source:MGI Symbo | 0 | 0 | 0 | 0 | 0 | 0 |
| 25689 | ENSMUSG00000083508 | 0.335414862 | -1.779047242 | 4.030804179 | -0.441362855 | 0.658950329 | NA | Gm6027 | predicted gene 6027 [Source:MGI Symbol | 0 | 0 | 2.0125 | 0 | 0 | 0 |
| 25690 | ENSMUSG00000083509 | 0 | NA | NA | NA | NA | NA | Defa-ps4 | defensin, alpha, pseudogene 4 [Source:M | 0 | 0 | 0 | 0 | 0 | 0 |
| 25691 | ENSMUSG00000083510 | 0 | NA | NA | NA | NA | NA | Gm13478 | predicted gene 13478 [Source:MGI Symbo | 0 | 0 | 0 | 0 | 0 | 0 |
| 25692 | ENSMUSG00000083512 | 0.9173971 | 0.153222854 | 2.48201849 | 0.061733164 | 0.950775329 | NA | Gm12749 | predicted gene 12749 [Source:MGI Symbo | 1.6062 | 0 | 1.0062 | 2.8919 | 0 | 0 |
| 25693 | ENSMUSG00000083513 | 0 | NA | NA | NA | NA | NA | Olfr647-ps1 | olfactory receptor 647, pseudogene 1 [So | 0 | 0 | 0 | 0 | 0 | 0 |
| 25694 | ENSMUSG00000083514 | 0 | NA | NA | NA | NA | NA | Gm11881 | predicted gene 11881 [Source:MGI Symbo | 0 | 0 | 0 | 0 | 0 | 0 |
| 25695 | ENSMUSG00000083515 | 0 | NA | NA | NA | NA | NA | Gm14577 | predicted gene 14577 [Source:MGI Symbo | 0 | 0 | 0 | 0 | 0 | 0 |
| 25696 | ENSMUSG00000083516 | 0 | NA | NA | NA | NA | NA | Gm15196 | predicted gene 15196 [Source:MGI Symbo | 0 | 0 | 0 | 0 | 0 | 0 |
| 25697 | ENSMUSG00000083517 | 0 | NA | NA | NA | NA | NA | Gm14998 | predicted gene 14998 [Source:MGI Symbo | 0 | 0 | 0 | 0 | 0 | 0 |
| 25698 | ENSMUSG00000083518 | 0.181769715 | 1.126924175 | 4.080472857 | 0.276174898 | 0.782413736 | NA | Gm9429 | predicted gene 9429 [Source:MGI Symbol | 0 | 0 | 0 | 0 | 1.0906 | 0 |
| 25699 | ENSMUSG00000083519 | 0 | NA | NA | NA | NA | NA | Gm13674 | predicted gene 13674 [Source:MGI Symbo | 0 | 0 | 0 | 0 | 0 | 0 |
| 25700 | ENSMUSG00000083520 | 0 | NA | NA | NA | NA | NA | Gm6418 | predicted gene 6418 [Source:MGI Symbol | 0 | 0 | 0 | 0 | 0 | 0 |

| | | | | | | | | | | | | | | | |
|---|---|---|---|---|---|---|---|---|---|---|---|---|---|---|---|
| 25701 | ENSMUSG00000083522 | 0 | NA | NA | NA | NA | NA | Gm15321 | predicted gene 15321 [Source:MGI Symbo | 0 | 0 | 0 | 0 | 0 | 0 |
| 25702 | ENSMUSG00000083524 | 0 | NA | NA | NA | NA | NA | Mup-ps21 | major urinary protein, pseudogene 21 [So | 0 | 0 | 0 | 0 | 0 | 0 |
| 25703 | ENSMUSG00000083527 | 0 | NA | NA | NA | NA | NA | Gm12001 | predicted gene 12001 [Source:MGI Symbo | 0 | 0 | 0 | 0 | 0 | 0 |
| 25704 | ENSMUSG00000083528 | 0 | NA | NA | NA | NA | NA | Gm13432 | predicted gene 13432 [Source:MGI Symbo | 0 | 0 | 0 | 0 | 0 | 0 |
| 25705 | ENSMUSG00000083530 | 0 | NA | NA | NA | NA | NA | Gm13867 | predicted gene 13867 [Source:MGI Symbo | 0 | 0 | 0 | 0 | 0 | 0 |
| 25706 | ENSMUSG00000083533 | 0 | NA | NA | NA | NA | NA | Gm13450 | predicted gene 13450 [Source:MGI Symbo | 0 | 0 | 0 | 0 | 0 | 0 |
| 25707 | ENSMUSG00000083534 | 3.428863686 | -0.204472562 | 1.187545902 | -0.172180765 | 0.863295419 | NA | H2-M6-ps | histocompatibility 2, M region locus 6, pse | 4.8186 | 5.1396 | 1.0062 | 1.928 | 5.4531 | 2.2277 |
| 25708 | ENSMUSG00000083535 | 0 | NA | NA | NA | NA | NA | Gm14381 | predicted gene 14381 [Source:MGI Symbo | 0 | 0 | 0 | 0 | 0 | 0 |
| 25709 | ENSMUSG00000083536 | 0 | NA | NA | NA | NA | NA | Gm15808 | predicted gene 15808 [Source:MGI Symbo | 0 | 0 | 0 | 0 | 0 | 0 |
| 25710 | ENSMUSG00000083537 | 0 | NA | NA | NA | NA | NA | Gm14942 | predicted gene 14942 [Source:MGI Symbo | 0 | 0 | 0 | 0 | 0 | 0 |
| 25711 | ENSMUSG00000083538 | 0.301557804 | -1.64335916 | 4.042814367 | -0.406488899 | 0.684383412 | NA | Gm16016 | predicted gene 16016 [Source:MGI Symbo | 0.8031 | 0 | 1.0062 | 0 | 0 | 0 |
| 25712 | ENSMUSG00000083539 | 0 | NA | NA | NA | NA | NA | Gm12466 | predicted gene 12466 [Source:MGI Symbo | 0 | 0 | 0 | 0 | 0 | 0 |
| 25713 | ENSMUSG00000083541 | 0 | NA | NA | NA | NA | NA | Gm12585 | predicted gene 12585 [Source:MGI Symbo | 0 | 0 | 0 | 0 | 0 | 0 |
| 25714 | ENSMUSG00000083542 | 0 | NA | NA | NA | NA | NA | Olfr1379-ps1 | olfactory receptor 1379, pseudogene 1 [S | 0 | 0 | 0 | 0 | 0 | 0 |
| 25715 | ENSMUSG00000083543 | 0 | NA | NA | NA | NA | NA | Gm14663 | predicted gene 14663 [Source:MGI Symbo | 0 | 0 | 0 | 0 | 0 | 0 |
| 25716 | ENSMUSG00000083544 | 0 | NA | NA | NA | NA | NA | Oat-rs1 | ornithine aminotransferase related seque | 0 | 0 | 0 | 0 | 0 | 0 |
| 25717 | ENSMUSG00000083545 | 0 | NA | NA | NA | NA | NA | Gm13320 | predicted gene 13320 [Source:MGI Symbo | 0 | 0 | 0 | 0 | 0 | 0 |
| 25718 | ENSMUSG00000083546 | 0 | NA | NA | NA | NA | NA | Tpt1-ps1 | tumor protein, translationally-controlled, | 0 | 0 | 0 | 0 | 0 | 0 |
| 25719 | ENSMUSG00000083547 | 0 | NA | NA | NA | NA | NA | Gm13321 | predicted gene 13321 [Source:MGI Symbo | 0 | 0 | 0 | 0 | 0 | 0 |
| 25720 | ENSMUSG00000083548 | 0 | NA | NA | NA | NA | NA | Gm11514 | predicted gene 11514 [Source:MGI Symbo | 0 | 0 | 0 | 0 | 0 | 0 |
| 25721 | ENSMUSG00000083549 | 0 | NA | NA | NA | NA | NA | Gm11227 | predicted gene 11227 [Source:MGI Symbo | 0 | 0 | 0 | 0 | 0 | 0 |
| 25722 | ENSMUSG00000083550 | 0 | NA | NA | NA | NA | NA | Gm13729 | predicted gene 13729 [Source:MGI Symbo | 0 | 0 | 0 | 0 | 0 | 0 |
| 25723 | ENSMUSG00000083551 | 0 | NA | NA | NA | NA | NA | Gm14516 | predicted gene 14516 [Source:MGI Symbo | 0 | 0 | 0 | 0 | 0 | 0 |
| 25724 | ENSMUSG00000083553 | 0 | NA | NA | NA | NA | NA | Gm11989 | predicted gene 11989 [Source:MGI Symbo | 0 | 0 | 0 | 0 | 0 | 0 |
| 25725 | ENSMUSG00000083554 | 0 | NA | NA | NA | NA | NA | Gm6645 | predicted gene 6645 [Source:MGI Symbol | 0 | 0 | 0 | 0 | 0 | 0 |
| 25726 | ENSMUSG00000083555 | 0.160662974 | 1.126924175 | 4.080472857 | 0.276174898 | 0.782413736 | NA | Gm12035 | predicted gene 12035 [Source:MGI Symbo | 0 | 0 | 0 | 0.964 | 0 | 0 |
| 25727 | ENSMUSG00000083556 | 0.181769715 | 1.126924175 | 4.080472857 | 0.276174898 | 0.782413736 | NA | Gm9043 | predicted gene 9043 [Source:MGI Symbol | 0 | 0 | 0 | 0 | 1.0906 | 0 |
| 25728 | ENSMUSG00000083557 | 0.453341394 | -0.545391191 | 3.551113277 | -0.153583158 | 0.877938417 | NA | Gm6877 | predicted pseudogene 6877 [Source:MGI | 1.6062 | 0 | 0 | 0 | 0 | 1.1138 |
| 25729 | ENSMUSG00000083558 | 0 | NA | NA | NA | NA | NA | Gm12896 | predicted gene 12896 [Source:MGI Symbo | 0 | 0 | 0 | 0 | 0 | 0 |
| 25730 | ENSMUSG00000083560 | 0 | NA | NA | NA | NA | NA | Gm14939 | predicted gene 14939 [Source:MGI Symbo | 0 | 0 | 0 | 0 | 0 | 0 |
| 25731 | ENSMUSG00000083561 | 0 | NA | NA | NA | NA | NA | Gm5942 | predicted gene 5942 [Source:MGI Symbol | 0 | 0 | 0 | 0 | 0 | 0 |
| 25732 | ENSMUSG00000083562 | 0 | NA | NA | NA | NA | NA | Gm13234 | predicted gene 13234 [Source:MGI Symbo | 0 | 0 | 0 | 0 | 0 | 0 |
| 25733 | ENSMUSG00000083563 | 213.1947672 | 0.037038373 | 0.211022762 | 0.175518379 | 0.860672331 | 0.975943719 | Gm13340 | predicted gene 13340 [Source:MGI Symbo | 172.67 | 237.45 | 221.37 | 167.73 | 214.85 | 265.09 |
| 25734 | ENSMUSG00000083564 | 0 | NA | NA | NA | NA | NA | Gm11929 | predicted gene 11929 [Source:MGI Symbo | 0 | 0 | 0 | 0 | 0 | 0 |
| 25735 | ENSMUSG00000083565 | 0 | NA | NA | NA | NA | NA | Gm11941 | predicted gene 11941 [Source:MGI Symbo | 0 | 0 | 0 | 0 | 0 | 0 |
| 25736 | ENSMUSG00000083566 | 0 | NA | NA | NA | NA | NA | Gm14711 | predicted gene 14711 [Source:MGI Symbo | 0 | 0 | 0 | 0 | 0 | 0 |
| 25737 | ENSMUSG00000083567 | 20.09502186 | 0.487082663 | 0.497031555 | 0.979983381 | 0.327094322 | 0.772772049 | Gm11451 | predicted gene 11451 [Source:MGI Symbo | 18.471 | 16.447 | 15.094 | 15.424 | 26.175 | 28.96 |
| 25738 | ENSMUSG00000083568 | 0 | NA | NA | NA | NA | NA | Gm12770 | predicted gene 12770 [Source:MGI Symbo | 0 | 0 | 0 | 0 | 0 | 0 |
| 25739 | ENSMUSG00000083569 | 0 | NA | NA | NA | NA | NA | Gm13666 | predicted gene 13666 [Source:MGI Symbo | 0 | 0 | 0 | 0 | 0 | 0 |
| 25740 | ENSMUSG00000083570 | 0 | NA | NA | NA | NA | NA | Gm14671 | predicted gene 14671 [Source:MGI Symbo | 0 | 0 | 0 | 0 | 0 | 0 |
| 25741 | ENSMUSG00000083572 | 0 | NA | NA | NA | NA | NA | Gm11849 | predicted gene 11849 [Source:MGI Symbo | 0 | 0 | 0 | 0 | 0 | 0 |
| 25742 | ENSMUSG00000083573 | 0 | NA | NA | NA | NA | NA | Gm7312 | predicted gene 7312 [Source:MGI Symbol | 0 | 0 | 0 | 0 | 0 | 0 |
| 25743 | ENSMUSG00000083574 | 0 | NA | NA | NA | NA | NA | Gm15709 | predicted gene 15709 [Source:MGI Symbo | 0 | 0 | 0 | 0 | 0 | 0 |
| 25744 | ENSMUSG00000083576 | 0 | NA | NA | NA | NA | NA | Gm16411 | predicted gene 16411 [Source:MGI Symbo | 0 | 0 | 0 | 0 | 0 | 0 |
| 25745 | ENSMUSG00000083577 | 0 | NA | NA | NA | NA | NA | Gm12432 | predicted gene 12432 [Source:MGI Symbo | 0 | 0 | 0 | 0 | 0 | 0 |
| 25746 | ENSMUSG00000083579 | 0.910270021 | 2.129340375 | 2.801245283 | 0.760140637 | 0.447170524 | NA | Gm15538 | predicted gene 15538 [Source:MGI Symbo | 0 | 0 | 1.0062 | 0 | 0 | 4.4554 |
| 25747 | ENSMUSG00000083580 | 0 | NA | NA | NA | NA | NA | Rpsa-ps3 | ribosomal protein SA, pseudogene 3 [Sou | 0 | 0 | 0 | 0 | 0 | 0 |
| 25748 | ENSMUSG00000083581 | 0 | NA | NA | NA | NA | NA | Gm14186 | predicted gene 14186 [Source:MGI Symbo | 0 | 0 | 0 | 0 | 0 | 0 |
| 25749 | ENSMUSG00000083582 | 0 | NA | NA | NA | NA | NA | Gm15393 | predicted gene 15393 [Source:MGI Symbo | 0 | 0 | 0 | 0 | 0 | 0 |
| 25750 | ENSMUSG00000083584 | 0 | NA | NA | NA | NA | NA | Gm15394 | predicted gene 15394 [Source:MGI Symbo | 0 | 0 | 0 | 0 | 0 | 0 |
| 25751 | ENSMUSG00000083585 | 0 | NA | NA | NA | NA | NA | Gm15885 | predicted gene 15885 [Source:MGI Symbo | 0 | 0 | 0 | 0 | 0 | 0 |
| 25752 | ENSMUSG00000083586 | 0 | NA | NA | NA | NA | NA | Gm8688 | predicted gene 8688 [Source:MGI Symbol | 0 | 0 | 0 | 0 | 0 | 0 |
| 25753 | ENSMUSG00000083587 | 0 | NA | NA | NA | NA | NA | Gm14741 | predicted gene 14741 [Source:MGI Symbo | 0 | 0 | 0 | 0 | 0 | 0 |
| 25754 | ENSMUSG00000083588 | 0 | NA | NA | NA | NA | NA | Gm8051 | predicted gene 8051 [Source:MGI Symbol | 0 | 0 | 0 | 0 | 0 | 0 |
| 25755 | ENSMUSG00000083590 | 0 | NA | NA | NA | NA | NA | Gm13916 | predicted gene 13916 [Source:MGI Symbo | 0 | 0 | 0 | 0 | 0 | 0 |
| 25756 | ENSMUSG00000083592 | 1.474266909 | -0.167907102 | 1.8867113 | -0.088994592 | 0.929086208 | NA | Gm13614 | predicted gene 13614 [Source:MGI Symbo | 1.6062 | 3.0838 | 0 | 1.928 | 0 | 2.2277 |

| | | | | | | | | | | | | | | | |
|---|---|---|---|---|---|---|---|---|---|---|---|---|---|---|---|
| 25757 | ENSMUSG00000083594 | 0 | NA | NA | NA | NA | NA | Gm13722 | predicted gene 13722 [Source:MGI Symbo | 0 | 0 | 0 | 0 | 0 | 0 |
| 25758 | ENSMUSG00000083595 | 0 | NA | NA | NA | NA | NA | Gm9200 | predicted gene 9200 [Source:MGI Symbol | 0 | 0 | 0 | 0 | 0 | 0 |
| 25759 | ENSMUSG00000083596 | 1.073382481 | 3.629918471 | 2.351148199 | 1.543891819 | 0.122614542 | NA | Rpl21-ps15 | ribosomal protein L21, pseudogene 15 [So | 0 | 0 | 0 | 0.964 | 4.3625 | 1.1138 |
| 25760 | ENSMUSG00000083597 | 0 | NA | NA | NA | NA | NA | Gm14805 | predicted gene 14805 [Source:MGI Symbo | 0 | 0 | 0 | 0 | 0 | 0 |
| 25761 | ENSMUSG00000083598 | 0 | NA | NA | NA | NA | NA | Gm14579 | predicted gene 14579 [Source:MGI Symbo | 0 | 0 | 0 | 0 | 0 | 0 |
| 25762 | ENSMUSG00000083599 | 0 | NA | NA | NA | NA | NA | Gm12821 | predicted gene 12821 [Source:MGI Symbo | 0 | 0 | 0 | 0 | 0 | 0 |
| 25763 | ENSMUSG00000083601 | 0 | NA | NA | NA | NA | NA | Gm12028 | predicted gene 12028 [Source:MGI Symbo | 0 | 0 | 0 | 0 | 0 | 0 |
| 25764 | ENSMUSG00000083602 | 0 | NA | NA | NA | NA | NA | Gm14911 | predicted gene 14911 [Source:MGI Symbo | 0 | 0 | 0 | 0 | 0 | 0 |
| 25765 | ENSMUSG00000083603 | 0.401551119 | -2.072982726 | 4.008251979 | -0.517178744 | 0.605031388 | NA | Gm14810 | predicted gene 14810 [Source:MGI Symbo | 2.4093 | 0 | 0 | 0 | 0 | 0 |
| 25766 | ENSMUSG00000083605 | 0.294513347 | 0.165143496 | 4.080472857 | 0.040471657 | 0.967717103 | NA | Gm14893 | predicted gene 14893 [Source:MGI Symbo | 0.8031 | 0 | 0 | 0.964 | 0 | 0 |
| 25767 | ENSMUSG00000083606 | 0 | NA | NA | NA | NA | NA | Gm15916 | predicted gene 15916 [Source:MGI Symbo | 0 | 0 | 0 | 0 | 0 | 0 |
| 25768 | ENSMUSG00000083607 | 0 | NA | NA | NA | NA | NA | Gm12640 | predicted gene 12640 [Source:MGI Symbo | 0 | 0 | 0 | 0 | 0 | 0 |
| 25769 | ENSMUSG00000083608 | 0.503095663 | 2.549494177 | 3.348715477 | 0.761334964 | 0.446457022 | NA | Gm14863 | predicted gene 14863 [Source:MGI Symbo | 0 | 0 | 0 | 1.928 | 1.0906 | 0 |
| 25770 | ENSMUSG00000083609 | 0 | NA | NA | NA | NA | NA | Gm16007 | predicted gene 16007 [Source:MGI Symbo | 0 | 0 | 0 | 0 | 0 | 0 |
| 25771 | ENSMUSG00000083610 | 0.492645588 | 0.9565034 | 3.447648281 | 0.277436479 | 0.781444974 | NA | Oaz2-ps | ornithine decarboxylase antizyme 2, pseu | 0 | 1.0279 | 0 | 1.928 | 0 | 0 |
| 25772 | ENSMUSG00000083611 | 0 | NA | NA | NA | NA | NA | Gm8832 | predicted gene 8832 [Source:MGI Symbol | 0 | 0 | 0 | 0 | 0 | 0 |
| 25773 | ENSMUSG00000083613 | 0 | NA | NA | NA | NA | NA | Pou2f3-rs1 | POU domain, class 2, transcription factor : | 0 | 0 | 0 | 0 | 0 | 0 |
| 25774 | ENSMUSG00000083614 | 8.709271536 | -0.75141255 | 0.823416816 | -0.912554292 | 0.361477003 | 0.800421803 | Gm12795 | predicted gene 12795 [Source:MGI Symbo | 18.471 | 9.2513 | 5.0312 | 9.6398 | 7.6343 | 2.2277 |
| 25775 | ENSMUSG00000083615 | 0 | NA | NA | NA | NA | NA | Gm12878 | predicted gene 12878 [Source:MGI Symbo | 0 | 0 | 0 | 0 | 0 | 0 |
| 25776 | ENSMUSG00000083616 | 0 | NA | NA | NA | NA | NA | H2ab3 | H2A.B variant histone 3 [Source:MGI Sym | 0 | 0 | 0 | 0 | 0 | 0 |
| 25777 | ENSMUSG00000083617 | 0 | NA | NA | NA | NA | NA | Gm5390 | predicted gene 5390 [Source:MGI Symbol | 0 | 0 | 0 | 0 | 0 | 0 |
| 25778 | ENSMUSG00000083618 | 0 | NA | NA | NA | NA | NA | Gm11771 | predicted gene 11771 [Source:MGI Symbo | 0 | 0 | 0 | 0 | 0 | 0 |
| 25779 | ENSMUSG00000083619 | 0 | NA | NA | NA | NA | NA | Gm14414 | predicted gene 14414 [Source:MGI Symbo | 0 | 0 | 0 | 0 | 0 | 0 |
| 25780 | ENSMUSG00000083620 | 0.167707431 | -0.796629176 | 4.080472857 | -0.195229623 | 0.84521319 | NA | Gm15529 | predicted gene 15529 [Source:MGI Symbo | 0 | 0 | 1.0062 | 0 | 0 | 0 |
| 25781 | ENSMUSG00000083621 | 9.249703892 | -0.023399702 | 0.691047867 | -0.033861188 | 0.972987843 | 0.995820834 | Gm14586 | predicted gene 14586 [Source:MGI Symbo | 9.6372 | 8.2233 | 10.062 | 7.7118 | 8.7249 | 11.138 |
| 25782 | ENSMUSG00000083622 | 0 | NA | NA | NA | NA | NA | Gm15570 | predicted gene 15570 [Source:MGI Symbo | 0 | 0 | 0 | 0 | 0 | 0 |
| 25783 | ENSMUSG00000083623 | 0.181769715 | 1.126924175 | 4.080472857 | 0.276174898 | 0.782413736 | NA | Gm7224 | predicted gene 7224 [Source:MGI Symbol | 0 | 0 | 0 | 0 | 1.0906 | 0 |
| 25784 | ENSMUSG00000083624 | 0 | NA | NA | NA | NA | NA | Gm12469 | predicted gene 12469 [Source:MGI Symbo | 0 | 0 | 0 | 0 | 0 | 0 |
| 25785 | ENSMUSG00000083625 | 0 | NA | NA | NA | NA | NA | Gm15188 | predicted gene 15188 [Source:MGI Symbo | 0 | 0 | 0 | 0 | 0 | 0 |
| 25786 | ENSMUSG00000083626 | 2.960406766 | -0.875911435 | 1.237040491 | -0.708070141 | 0.478901701 | NA | Gm4459 | predicted gene 4459 [Source:MGI Symbol | 2.4093 | 4.1117 | 5.0312 | 2.8919 | 1.0906 | 2.2277 |
| 25787 | ENSMUSG00000083627 | 0 | NA | NA | NA | NA | NA | Gm12502 | predicted gene 12502 [Source:MGI Symbo | 0 | 0 | 0 | 0 | 0 | 0 |
| 25788 | ENSMUSG00000083628 | 0 | NA | NA | NA | NA | NA | Gm2030 | predicted gene 2030 [Source:MGI Symbol | 0 | 0 | 0 | 0 | 0 | 0 |
| 25789 | ENSMUSG00000083629 | 0 | NA | NA | NA | NA | NA | Gm11262 | predicted gene 11262 [Source:MGI Symbo | 0 | 0 | 0 | 0 | 0 | 0 |
| 25790 | ENSMUSG00000083630 | 0 | NA | NA | NA | NA | NA | Gm15602 | predicted gene 15602 [Source:MGI Symbo | 0 | 0 | 0 | 0 | 0 | 0 |
| 25791 | ENSMUSG00000083631 | 0 | NA | NA | NA | NA | NA | Gm13366 | predicted gene 13366 [Source:MGI Symbo | 0 | 0 | 0 | 0 | 0 | 0 |
| 25792 | ENSMUSG00000083632 | 0 | NA | NA | NA | NA | NA | Gm11786 | predicted gene 11786 [Source:MGI Symbo | 0 | 0 | 0 | 0 | 0 | 0 |
| 25793 | ENSMUSG00000083633 | 0 | NA | NA | NA | NA | NA | Gm13312 | predicted gene 13312 [Source:MGI Symbo | 0 | 0 | 0 | 0 | 0 | 0 |
| 25794 | ENSMUSG00000083635 | 0 | NA | NA | NA | NA | NA | Gm11480 | predicted gene 11480 [Source:MGI Symbo | 0 | 0 | 0 | 0 | 0 | 0 |
| 25795 | ENSMUSG00000083636 | 0.160662974 | 1.126924175 | 4.080472857 | 0.276174898 | 0.782413736 | NA | Gm15175 | predicted gene 15175 [Source:MGI Symbo | 0 | 0 | 0 | 0.964 | 0 | 0 |
| 25796 | ENSMUSG00000083637 | 0 | NA | NA | NA | NA | NA | Gm14754 | predicted gene 14754 [Source:MGI Symbo | 0 | 0 | 0 | 0 | 0 | 0 |
| 25797 | ENSMUSG00000083640 | 0 | NA | NA | NA | NA | NA | Gm8876 | predicted gene 8876 [Source:MGI Symbol | 0 | 0 | 0 | 0 | 0 | 0 |
| 25798 | ENSMUSG00000083641 | 0 | NA | NA | NA | NA | NA | Gm9225 | predicted gene 9225 [Source:MGI Symbol | 0 | 0 | 0 | 0 | 0 | 0 |
| 25799 | ENSMUSG00000083642 | 0 | NA | NA | NA | NA | NA | Gm13148 | predicted gene 13148 [Source:MGI Symbo | 0 | 0 | 0 | 0 | 0 | 0 |
| 25800 | ENSMUSG00000083643 | 0 | NA | NA | NA | NA | NA | Gm13876 | predicted gene 13876 [Source:MGI Symbo | 0 | 0 | 0 | 0 | 0 | 0 |
| 25801 | ENSMUSG00000083644 | 0 | NA | NA | NA | NA | NA | Gm12787 | predicted gene 12787 [Source:MGI Symbo | 0 | 0 | 0 | 0 | 0 | 0 |
| 25802 | ENSMUSG00000083645 | 0 | NA | NA | NA | NA | NA | Gm14086 | predicted gene 14086 [Source:MGI Symbo | 0 | 0 | 0 | 0 | 0 | 0 |
| 25803 | ENSMUSG00000083646 | 0 | NA | NA | NA | NA | NA | Gm6088 | predicted gene 6088 [Source:MGI Symbol | 0 | 0 | 0 | 0 | 0 | 0 |
| 25804 | ENSMUSG00000083648 | 0 | NA | NA | NA | NA | NA | Gm14229 | predicted gene 14229 [Source:MGI Symbo | 0 | 0 | 0 | 0 | 0 | 0 |
| 25805 | ENSMUSG00000083649 | 19.51619412 | -0.167434811 | 0.60397761 | -0.277220229 | 0.781611008 | 0.960079945 | Rasl2-9 | RAS-like, family 2, locus 9 [Source:MGI Sy | 10.44 | 15.419 | 36.225 | 17.352 | 9.8156 | 27.846 |
| 25806 | ENSMUSG00000083650 | 0.339027071 | -1.79174149 | 3.919585585 | -0.457125237 | 0.647581032 | NA | Gm13357 | predicted gene 13357 [Source:MGI Symbo | 0 | 1.0279 | 1.0062 | 0 | 0 | 0 |
| 25807 | ENSMUSG00000083651 | 0 | NA | NA | NA | NA | NA | Gm11524 | predicted gene 11524 [Source:MGI Symbo | 0 | 0 | 0 | 0 | 0 | 0 |
| 25808 | ENSMUSG00000083653 | 0 | NA | NA | NA | NA | NA | Gm11384 | predicted gene 11384 [Source:MGI Symbo | 0 | 0 | 0 | 0 | 0 | 0 |
| 25809 | ENSMUSG00000083654 | 0 | NA | NA | NA | NA | NA | Gm11875 | predicted gene 11875 [Source:MGI Symbo | 0 | 0 | 0 | 0 | 0 | 0 |
| 25810 | ENSMUSG00000083655 | 0 | NA | NA | NA | NA | NA | Gm12099 | predicted gene 12099 [Source:MGI Symbo | 0 | 0 | 0 | 0 | 0 | 0 |
| 25811 | ENSMUSG00000083657 | 0 | NA | NA | NA | NA | NA | Gm12245 | predicted gene 12245 [Source:MGI Symbo | 0 | 0 | 0 | 0 | 0 | 0 |
| 25812 | ENSMUSG00000083658 | 0 | NA | NA | NA | NA | NA | Gm15798 | predicted gene 15798 [Source:MGI Symbo | 0 | 0 | 0 | 0 | 0 | 0 |

| | | | | | | | | | | | | | | | |
|---|---|---|---|---|---|---|---|---|---|---|---|---|---|---|---|
| 25813 | ENSMUSG00000083659 | 0 | NA | NA | NA | NA | NA | Gm11258 | predicted gene 11258 [Source:MGI Symbo | 0 | 0 | 0 | 0 | 0 | 0 |
| 25814 | ENSMUSG00000083660 | 0 | NA | NA | NA | NA | NA | Gm11853 | predicted gene 11853 [Source:MGI Symbo | 0 | 0 | 0 | 0 | 0 | 0 |
| 25815 | ENSMUSG00000083661 | 0 | NA | NA | NA | NA | NA | Gm11473 | predicted gene 11473 [Source:MGI Symbo | 0 | 0 | 0 | 0 | 0 | 0 |
| 25816 | ENSMUSG00000083662 | 0 | NA | NA | NA | NA | NA | Gm12152 | predicted gene 12152 [Source:MGI Symbo | 0 | 0 | 0 | 0 | 0 | 0 |
| 25817 | ENSMUSG00000083663 | 0 | NA | NA | NA | NA | NA | Gm14903 | predicted gene 14903 [Source:MGI Symbo | 0 | 0 | 0 | 0 | 0 | 0 |
| 25818 | ENSMUSG00000083664 | 0 | NA | NA | NA | NA | NA | Gm11433 | predicted gene 11433 [Source:MGI Symbo | 0 | 0 | 0 | 0 | 0 | 0 |
| 25819 | ENSMUSG00000083665 | 0 | NA | NA | NA | NA | NA | Gm14575 | predicted gene 14575 [Source:MGI Symbo | 0 | 0 | 0 | 0 | 0 | 0 |
| 25820 | ENSMUSG00000083666 | 0 | NA | NA | NA | NA | NA | Gm12358 | predicted gene 12358 [Source:MGI Symbo | 0 | 0 | 0 | 0 | 0 | 0 |
| 25821 | ENSMUSG00000083667 | 0 | NA | NA | NA | NA | NA | Gm11334 | predicted gene 11334 [Source:MGI Symbo | 0 | 0 | 0 | 0 | 0 | 0 |
| 25822 | ENSMUSG00000083668 | 0 | NA | NA | NA | NA | NA | Gm5648 | predicted pseudogene 5648 [Source:MGI | 0 | 0 | 0 | 0 | 0 | 0 |
| 25823 | ENSMUSG00000083669 | 0 | NA | NA | NA | NA | NA | Mif-ps4 | macrophage migration inhibitory factor, p | 0 | 0 | 0 | 0 | 0 | 0 |
| 25824 | ENSMUSG00000083670 | 4.434287124 | -0.393715633 | 1.022903593 | -0.384900039 | 0.700311477 | NA | Gm6829 | predicted pseudogene 6829 [Source:MGI | 4.0155 | 3.0838 | 8.05 | 4.8199 | 2.1812 | 4.4554 |
| 25825 | ENSMUSG00000083671 | 0 | NA | NA | NA | NA | NA | Gm14304 | predicted gene 14304 [Source:MGI Symbo | 0 | 0 | 0 | 0 | 0 | 0 |
| 25826 | ENSMUSG00000083672 | 0 | NA | NA | NA | NA | NA | Kpna2-ps | Kpna2 retrotransposed pseudogene [Sour | 0 | 0 | 0 | 0 | 0 | 0 |
| 25827 | ENSMUSG00000083673 | 0 | NA | NA | NA | NA | NA | Vmn1r-ps11 | vomeronasal 1 receptor, pseudogene 111 | 0 | 0 | 0 | 0 | 0 | 0 |
| 25828 | ENSMUSG00000083674 | 4.111687545 | 0.593654811 | 1.10988377 | 0.534880162 | 0.592732727 | NA | Zfp133-ps | zinc finger protein 133, pseudogene [Sour | 5.6217 | 3.0838 | 1.0062 | 2.8919 | 8.7249 | 3.3415 |
| 25829 | ENSMUSG00000083676 | 0 | NA | NA | NA | NA | NA | Gm11367 | predicted gene 11367 [Source:MGI Symbo | 0 | 0 | 0 | 0 | 0 | 0 |
| 25830 | ENSMUSG00000083677 | 0 | NA | NA | NA | NA | NA | Gm11699 | predicted gene 11699 [Source:MGI Symbo | 0 | 0 | 0 | 0 | 0 | 0 |
| 25831 | ENSMUSG00000083678 | 0.699392976 | 1.647556484 | 2.55058628 | 0.645952069 | 0.518310397 | NA | Gm12989 | predicted gene 12989 [Source:MGI Symbo | 0 | 1.0279 | 0 | 0.964 | 1.0906 | 1.1138 |
| 25832 | ENSMUSG00000083679 | 0.133850373 | -0.796629176 | 4.080472857 | -0.195229623 | 0.84521319 | NA | Gm12892 | predicted gene 12892 [Source:MGI Symbo | 0.8031 | 0 | 0 | 0 | 0 | 0 |
| 25833 | ENSMUSG00000083680 | 0 | NA | NA | NA | NA | NA | Gm9119 | predicted pseudogene 9119 [Source:MGI | 0 | 0 | 0 | 0 | 0 | 0 |
| 25834 | ENSMUSG00000083681 | 0 | NA | NA | NA | NA | NA | Gm6640 | predicted gene 6640 [Source:MGI Symbol | 0 | 0 | 0 | 0 | 0 | 0 |
| 25835 | ENSMUSG00000083682 | 0.160662974 | 1.126924175 | 4.080472857 | 0.276174898 | 0.782413736 | NA | Pcna-ps1 | proliferating cell nuclear antigen pseudog | 0 | 0 | 0 | 0.964 | 0 | 0 |
| 25836 | ENSMUSG00000083683 | 0 | NA | NA | NA | NA | NA | Pou5f1-rs8 | POU domain, class 5, transcription factor : | 0 | 0 | 0 | 0 | 0 | 0 |
| 25837 | ENSMUSG00000083684 | 0 | NA | NA | NA | NA | NA | Gm14034 | predicted gene 14034 [Source:MGI Symbo | 0 | 0 | 0 | 0 | 0 | 0 |
| 25838 | ENSMUSG00000083685 | 0 | NA | NA | NA | NA | NA | Gm12412 | predicted gene 12412 [Source:MGI Symbo | 0 | 0 | 0 | 0 | 0 | 0 |
| 25839 | ENSMUSG00000083686 | 0 | NA | NA | NA | NA | NA | Gm13504 | predicted gene 13504 [Source:MGI Symbo | 0 | 0 | 0 | 0 | 0 | 0 |
| 25840 | ENSMUSG00000083687 | 0 | NA | NA | NA | NA | NA | Gm15154 | predicted gene 15154 [Source:MGI Symbo | 0 | 0 | 0 | 0 | 0 | 0 |
| 25841 | ENSMUSG00000083688 | 0 | NA | NA | NA | NA | NA | Gm13743 | predicted gene 13743 [Source:MGI Symbo | 0 | 0 | 0 | 0 | 0 | 0 |
| 25842 | ENSMUSG00000083689 | 0 | NA | NA | NA | NA | NA | Gm14094 | predicted gene 14094 [Source:MGI Symbo | 0 | 0 | 0 | 0 | 0 | 0 |
| 25843 | ENSMUSG00000083691 | 0 | NA | NA | NA | NA | NA | Gm14465 | predicted gene 14465 [Source:MGI Symbo | 0 | 0 | 0 | 0 | 0 | 0 |
| 25844 | ENSMUSG00000083692 | 0 | NA | NA | NA | NA | NA | Gm9575 | predicted gene 9575 [Source:MGI Symbol | 0 | 0 | 0 | 0 | 0 | 0 |
| 25845 | ENSMUSG00000083693 | 0 | NA | NA | NA | NA | NA | Gm11971 | predicted gene 11971 [Source:MGI Symbo | 0 | 0 | 0 | 0 | 0 | 0 |
| 25846 | ENSMUSG00000083695 | 0 | NA | NA | NA | NA | NA | Rnf138rt1 | ring finger protein 138, retrogene 1 [Sourc | 0 | 0 | 0 | 0 | 0 | 0 |
| 25847 | ENSMUSG00000083697 | 0 | NA | NA | NA | NA | NA | Gm13914 | predicted gene 13914 [Source:MGI Symbo | 0 | 0 | 0 | 0 | 0 | 0 |
| 25848 | ENSMUSG00000083698 | 0 | NA | NA | NA | NA | NA | Gm12687 | predicted gene 12687 [Source:MGI Symbo | 0 | 0 | 0 | 0 | 0 | 0 |
| 25849 | ENSMUSG00000083699 | 0 | NA | NA | NA | NA | NA | Gm14932 | predicted gene 14932 [Source:MGI Symbo | 0 | 0 | 0 | 0 | 0 | 0 |
| 25850 | ENSMUSG00000083700 | 0 | NA | NA | NA | NA | NA | Gm2986 | predicted gene 2986 [Source:MGI Symbol | 0 | 0 | 0 | 0 | 0 | 0 |
| 25851 | ENSMUSG00000083701 | 0 | NA | NA | NA | NA | NA | Gm4912 | predicted pseudogene 4912 [Source:MGI | 0 | 0 | 0 | 0 | 0 | 0 |
| 25852 | ENSMUSG00000083702 | 0 | NA | NA | NA | NA | NA | Gm14428 | predicted gene 14428 [Source:MGI Symbo | 0 | 0 | 0 | 0 | 0 | 0 |
| 25853 | ENSMUSG00000083704 | 0.160662974 | 1.126924175 | 4.080472857 | 0.276174898 | 0.782413736 | NA | Gm13153 | predicted gene 13153 [Source:MGI Symbo | 0 | 0 | 0 | 0.964 | 0 | 0 |
| 25854 | ENSMUSG00000083705 | 0 | NA | NA | NA | NA | NA | Gm8624 | predicted gene 8624 [Source:MGI Symbol | 0 | 0 | 0 | 0 | 0 | 0 |
| 25855 | ENSMUSG00000083706 | 0 | NA | NA | NA | NA | NA | Olfr1118 | olfactory receptor 1118 [Source:MGI Sym | 0 | 0 | 0 | 0 | 0 | 0 |
| 25856 | ENSMUSG00000083707 | 0 | NA | NA | NA | NA | NA | Gm13592 | predicted gene 13592 [Source:MGI Symbo | 0 | 0 | 0 | 0 | 0 | 0 |
| 25857 | ENSMUSG00000083708 | 0.17131964 | -0.796629176 | 4.080472857 | -0.195229623 | 0.84521319 | NA | Gm13123 | predicted gene 13123 [Source:MGI Symbo | 0 | 1.0279 | 0 | 0 | 0 | 0 |
| 25858 | ENSMUSG00000083709 | 0 | NA | NA | NA | NA | NA | Gm14688 | predicted gene 14688 [Source:MGI Symbo | 0 | 0 | 0 | 0 | 0 | 0 |
| 25859 | ENSMUSG00000083710 | 0 | NA | NA | NA | NA | NA | Gm15078 | predicted gene 15078 [Source:MGI Symbo | 0 | 0 | 0 | 0 | 0 | 0 |
| 25860 | ENSMUSG00000083711 | 0 | NA | NA | NA | NA | NA | Gm3931 | predicted gene 3931 [Source:MGI Symbol | 0 | 0 | 0 | 0 | 0 | 0 |
| 25861 | ENSMUSG00000083713 | 0 | NA | NA | NA | NA | NA | Gm5883 | predicted gene 5883 [Source:MGI Symbol | 0 | 0 | 0 | 0 | 0 | 0 |
| 25862 | ENSMUSG00000083714 | 0 | NA | NA | NA | NA | NA | Fthl17-ps1 | ferritin, heavy polypeptide-like 17, pseudo | 0 | 0 | 0 | 0 | 0 | 0 |
| 25863 | ENSMUSG00000083715 | 1.03119563 | 0.094051261 | 2.229235409 | 0.042189919 | 0.966347299 | NA | Gm14202 | predicted gene 14202 [Source:MGI Symbo | 0 | 0 | 3.0187 | 0.964 | 1.0906 | 1.1138 |
| 25864 | ENSMUSG00000083716 | 6.973315853 | 0.177447315 | 0.80272105 | 0.221057258 | 0.82504785 | 0.96943211 | Gm13436 | predicted gene 13436 [Source:MGI Symbo | 6.4248 | 5.1396 | 8.05 | 6.7478 | 5.4531 | 10.025 |
| 25865 | ENSMUSG00000083717 | 0 | NA | NA | NA | NA | NA | Gm13865 | predicted gene 13865 [Source:MGI Symbo | 0 | 0 | 0 | 0 | 0 | 0 |
| 25866 | ENSMUSG00000083718 | 16.62622032 | 0.002329455 | 0.568274872 | 0.00409917 | 0.996729345 | 0.999154389 | Ccnb2-ps | cyclin B2, pseudogene [Source:MGI Symb | 19.274 | 9.2513 | 21.131 | 12.532 | 14.178 | 23.391 |
| 25867 | ENSMUSG00000083719 | 0 | NA | NA | NA | NA | NA | Gm15161 | predicted gene 15161 [Source:MGI Symbo | 0 | 0 | 0 | 0 | 0 | 0 |
| 25868 | ENSMUSG00000083720 | 0 | NA | NA | NA | NA | NA | Gm12901 | predicted gene 12901 [Source:MGI Symbo | 0 | 0 | 0 | 0 | 0 | 0 |

| | | | | | | | | | | | | | | | |
|---|---|---|---|---|---|---|---|---|---|---|---|---|---|---|---|
| 25869 | ENSMUSG00000083721 | 0 | NA | NA | NA | NA | NA | Gm7700 | predicted gene 7700 [Source:MGI Symbol | 0 | 0 | 0 | 0 | 0 | 0 |
| 25870 | ENSMUSG00000083722 | 0 | NA | NA | NA | NA | NA | Adgrd2-ps | adhesion G protein-coupled receptor D2, | 0 | 0 | 0 | 0 | 0 | 0 |
| 25871 | ENSMUSG00000083723 | 0 | NA | NA | NA | NA | NA | Rpl23a-ps14 | ribosomal protein L23A, pseudogene 14 [S | 0 | 0 | 0 | 0 | 0 | 0 |
| 25872 | ENSMUSG00000083725 | 0.185640647 | 1.126924175 | 4.080472857 | 0.276174898 | 0.782413736 | NA | Gm11865 | predicted gene 11865 [Source:MGI Symbo | 0 | 0 | 0 | 0 | 0 | 1.1138 |
| 25873 | ENSMUSG00000083728 | 0 | NA | NA | NA | NA | NA | Gm13854 | predicted gene 13854 [Source:MGI Symbo | 0 | 0 | 0 | 0 | 0 | 0 |
| 25874 | ENSMUSG00000083729 | 0 | NA | NA | NA | NA | NA | Sh3d2c-ps1 | SH3 domain protein 2C, pseudogene 1 [Sc | 0 | 0 | 0 | 0 | 0 | 0 |
| 25875 | ENSMUSG00000083731 | 0.331982614 | 0.16514309 | 3.986751476 | 0.041422971 | 0.966958701 | NA | Gm16202 | predicted gene 16202 [Source:MGI Symbo | 0 | 1.0279 | 0 | 0.964 | 0 | 0 |
| 25876 | ENSMUSG00000083732 | 0.665097234 | 0.245106807 | 2.82617131 | 0.086727512 | 0.930888107 | NA | Gm14197 | predicted gene 14197 [Source:MGI Symbo | 0.8031 | 0 | 1.0062 | 0 | 2.1812 | 0 |
| 25877 | ENSMUSG00000083733 | 0 | NA | NA | NA | NA | NA | Gm13035 | predicted gene 13035 [Source:MGI Symbo | 0 | 0 | 0 | 0 | 0 | 0 |
| 25878 | ENSMUSG00000083734 | 4.631952396 | 0.277772874 | 1.0629314 | 0.261327188 | 0.793840202 | NA | Gm16209 | predicted gene 16209 [Source:MGI Symbo | 6.4248 | 4.1117 | 2.0125 | 8.6758 | 5.4531 | 1.1138 |
| 25879 | ENSMUSG00000083735 | 0.167707431 | -0.796629176 | 4.080472857 | -0.195229623 | 0.84521319 | NA | Gm12495 | predicted gene 12495 [Source:MGI Symbo | 0 | 0 | 1.0062 | 0 | 0 | 0 |
| 25880 | ENSMUSG00000083736 | 0.133850373 | -0.796629176 | 4.080472857 | -0.195229623 | 0.84521319 | NA | Gm6039 | predicted gene 6039 [Source:MGI Symbol | 0.8031 | 0 | 0 | 0 | 0 | 0 |
| 25881 | ENSMUSG00000083737 | 0 | NA | NA | NA | NA | NA | Prdx6-ps2 | peroxiredoxin 6 pseudogene 2 [Source:M | 0 | 0 | 0 | 0 | 0 | 0 |
| 25882 | ENSMUSG00000083738 | 0 | NA | NA | NA | NA | NA | Gm14592 | predicted gene 14592 [Source:MGI Symbo | 0 | 0 | 0 | 0 | 0 | 0 |
| 25883 | ENSMUSG00000083740 | 0 | NA | NA | NA | NA | NA | Gm11456 | predicted gene 11456 [Source:MGI Symbo | 0 | 0 | 0 | 0 | 0 | 0 |
| 25884 | ENSMUSG00000083741 | 0 | NA | NA | NA | NA | NA | Gm15297 | predicted gene 15297 [Source:MGI Symbo | 0 | 0 | 0 | 0 | 0 | 0 |
| 25885 | ENSMUSG00000083742 | 0 | NA | NA | NA | NA | NA | Vmn1r-ps3 | vomeronasal 1 receptor, pseudogene 3 [S | 0 | 0 | 0 | 0 | 0 | 0 |
| 25886 | ENSMUSG00000083743 | 0 | NA | NA | NA | NA | NA | Gm12421 | predicted gene 12421 [Source:MGI Symbo | 0 | 0 | 0 | 0 | 0 | 0 |
| 25887 | ENSMUSG00000083744 | 0 | NA | NA | NA | NA | NA | Gm14824 | predicted gene 14824 [Source:MGI Symbo | 0 | 0 | 0 | 0 | 0 | 0 |
| 25888 | ENSMUSG00000083745 | 0.17131964 | -0.796629176 | 4.080472857 | -0.195229623 | 0.84521319 | NA | Gm13543 | predicted gene 13543 [Source:MGI Symbo | 0 | 1.0279 | 0 | 0 | 0 | 0 |
| 25889 | ENSMUSG00000083747 | 0 | NA | NA | NA | NA | NA | Gm14727 | predicted gene 14727 [Source:MGI Symbo | 0 | 0 | 0 | 0 | 0 | 0 |
| 25890 | ENSMUSG00000083748 | 0 | NA | NA | NA | NA | NA | Gm11662 | predicted gene 11662 [Source:MGI Symbo | 0 | 0 | 0 | 0 | 0 | 0 |
| 25891 | ENSMUSG00000083749 | 0 | NA | NA | NA | NA | NA | Gm13100 | predicted gene 13100 [Source:MGI Symbo | 0 | 0 | 0 | 0 | 0 | 0 |
| 25892 | ENSMUSG00000083750 | 0 | NA | NA | NA | NA | NA | Gm14108 | predicted gene 14108 [Source:MGI Symbo | 0 | 0 | 0 | 0 | 0 | 0 |
| 25893 | ENSMUSG00000083751 | 0 | NA | NA | NA | NA | NA | Gm14534 | predicted gene 14534 [Source:MGI Symbo | 0 | 0 | 0 | 0 | 0 | 0 |
| 25894 | ENSMUSG00000083752 | 0 | NA | NA | NA | NA | NA | Gm12014 | predicted gene 12014 [Source:MGI Symbo | 0 | 0 | 0 | 0 | 0 | 0 |
| 25895 | ENSMUSG00000083753 | 0 | NA | NA | NA | NA | NA | Gm12814 | predicted gene 12814 [Source:MGI Symbo | 0 | 0 | 0 | 0 | 0 | 0 |
| 25896 | ENSMUSG00000083754 | 0.874144864 | -0.458978284 | 2.368814316 | -0.193758659 | 0.846364859 | NA | Gm7803 | predicted gene 7803 [Source:MGI Symbol | 0 | 1.0279 | 2.0125 | 0 | 1.0906 | 1.1138 |
| 25897 | ENSMUSG00000083755 | 0 | NA | NA | NA | NA | NA | Gm14559 | predicted gene 14559 [Source:MGI Symbo | 0 | 0 | 0 | 0 | 0 | 0 |
| 25898 | ENSMUSG00000083757 | 0.439020386 | -2.189747911 | 3.52148864 | -0.621824499 | 0.534057275 | NA | Rps15a-ps4 | ribosomal protein S15A, pseudogene 4 [S | 1.6062 | 1.0279 | 0 | 0 | 0 | 0 |
| 25899 | ENSMUSG00000083760 | 0 | NA | NA | NA | NA | NA | Gm14524 | predicted gene 14524 [Source:MGI Symbo | 0 | 0 | 0 | 0 | 0 | 0 |
| 25900 | ENSMUSG00000083761 | 0 | NA | NA | NA | NA | NA | Pgam1-ps1 | phosphoglycerate mutase 1, pseudogene | 0 | 0 | 0 | 0 | 0 | 0 |
| 25901 | ENSMUSG00000083763 | 0 | NA | NA | NA | NA | NA | Gm9050 | predicted gene 9050 [Source:MGI Symbol | 0 | 0 | 0 | 0 | 0 | 0 |
| 25902 | ENSMUSG00000083764 | 0 | NA | NA | NA | NA | NA | Gm11516 | predicted gene 11516 [Source:MGI Symbo | 0 | 0 | 0 | 0 | 0 | 0 |
| 25903 | ENSMUSG00000083766 | 0 | NA | NA | NA | NA | NA | Gm15335 | predicted gene 15335 [Source:MGI Symbo | 0 | 0 | 0 | 0 | 0 | 0 |
| 25904 | ENSMUSG00000083767 | 0 | NA | NA | NA | NA | NA | Gm11405 | predicted gene 11405 [Source:MGI Symbo | 0 | 0 | 0 | 0 | 0 | 0 |
| 25905 | ENSMUSG00000083768 | 0 | NA | NA | NA | NA | NA | Gm15362 | predicted gene 15362 [Source:MGI Symbo | 0 | 0 | 0 | 0 | 0 | 0 |
| 25906 | ENSMUSG00000083769 | 0 | NA | NA | NA | NA | NA | Gm11689 | predicted gene 11689 [Source:MGI Symbo | 0 | 0 | 0 | 0 | 0 | 0 |
| 25907 | ENSMUSG00000083771 | 0 | NA | NA | NA | NA | NA | Gm15988 | predicted gene 15988 [Source:MGI Symbo | 0 | 0 | 0 | 0 | 0 | 0 |
| 25908 | ENSMUSG00000083772 | 0 | NA | NA | NA | NA | NA | Gm15566 | predicted gene 15566 [Source:MGI Symbo | 0 | 0 | 0 | 0 | 0 | 0 |
| 25909 | ENSMUSG00000083773 | 1.707181623 | 1.398765519 | 1.682955976 | 0.831136131 | 0.405896731 | NA | Gm13394 | predicted gene 13394 [Source:MGI Symbo | 0.8031 | 0 | 2.0125 | 1.928 | 3.2719 | 2.2277 |
| 25910 | ENSMUSG00000083774 | 0.52105551 | -0.812028029 | 3.385364865 | -0.239864257 | 0.810435491 | NA | Gm7180 | predicted pseudogene 7180 [Source:MGI | 0 | 0 | 2.0125 | 0 | 0 | 1.1138 |
| 25911 | ENSMUSG00000083775 | 0 | NA | NA | NA | NA | NA | Gm14837 | predicted gene 14837 [Source:MGI Symbo | 0 | 0 | 0 | 0 | 0 | 0 |
| 25912 | ENSMUSG00000083776 | 0 | NA | NA | NA | NA | NA | Gm13473 | predicted gene 13473 [Source:MGI Symbo | 0 | 0 | 0 | 0 | 0 | 0 |
| 25913 | ENSMUSG00000083777 | 0 | NA | NA | NA | NA | NA | Gm13361 | predicted gene 13361 [Source:MGI Symbo | 0 | 0 | 0 | 0 | 0 | 0 |
| 25914 | ENSMUSG00000083778 | 0 | NA | NA | NA | NA | NA | Gm15094 | predicted gene 15094 [Source:MGI Symbo | 0 | 0 | 0 | 0 | 0 | 0 |
| 25915 | ENSMUSG00000083779 | 0 | NA | NA | NA | NA | NA | Gm14775 | predicted gene 14775 [Source:MGI Symbo | 0 | 0 | 0 | 0 | 0 | 0 |
| 25916 | ENSMUSG00000083780 | 0 | NA | NA | NA | NA | NA | Gm12728 | predicted gene 12728 [Source:MGI Symbo | 0 | 0 | 0 | 0 | 0 | 0 |
| 25917 | ENSMUSG00000083781 | 0 | NA | NA | NA | NA | NA | Gm14842 | predicted gene 14842 [Source:MGI Symbo | 0 | 0 | 0 | 0 | 0 | 0 |
| 25918 | ENSMUSG00000083782 | 0 | NA | NA | NA | NA | NA | Gm11660 | predicted gene 11660 [Source:MGI Symbo | 0 | 0 | 0 | 0 | 0 | 0 |
| 25919 | ENSMUSG00000083783 | 0.321325949 | 1.911575114 | 4.048267893 | 0.472195804 | 0.636787034 | NA | Gm12739 | predicted gene 12739 [Source:MGI Symbo | 0 | 0 | 0 | 1.928 | 0 | 0 |
| 25920 | ENSMUSG00000083784 | 0 | NA | NA | NA | NA | NA | Gm14729 | predicted gene 14729 [Source:MGI Symbo | 0 | 0 | 0 | 0 | 0 | 0 |
| 25921 | ENSMUSG00000083785 | 0 | NA | NA | NA | NA | NA | Gm11888 | predicted gene 11888 [Source:MGI Symbo | 0 | 0 | 0 | 0 | 0 | 0 |
| 25922 | ENSMUSG00000083786 | 0 | NA | NA | NA | NA | NA | Rps18-ps2 | ribosomal protein S18, pseudogene 2 [Sou | 0 | 0 | 0 | 0 | 0 | 0 |
| 25923 | ENSMUSG00000083787 | 0 | NA | NA | NA | NA | NA | Gm11644 | predicted gene 11644 [Source:MGI Symbo | 0 | 0 | 0 | 0 | 0 | 0 |
| 25924 | ENSMUSG00000083789 | 0 | NA | NA | NA | NA | NA | Gm6092 | predicted gene 6092 [Source:MGI Symbol | 0 | 0 | 0 | 0 | 0 | 0 |

| | | | | | | | | | | | | | | | |
|---|---|---|---|---|---|---|---|---|---|---|---|---|---|---|---|
| 25925 | ENSMUSG00000083791 | 0.520796785 | -0.820103173 | 3.072141206 | -0.266948398 | 0.789508892 | NA | Gm12721 | predicted gene 12721 [Source:MGI Symbo | 0 | 1.0279 | 1.0062 | 0 | 1.0906 | 0 |
| 25926 | ENSMUSG00000083792 | 0 | NA | NA | NA | NA | NA | Rpl12-ps2 | ribosomal proteimn L12-ps2 [Source:MGI | 0 | 0 | 0 | 0 | 0 | 0 |
| 25927 | ENSMUSG00000083793 | 0 | NA | NA | NA | NA | NA | Gm14274 | predicted gene 14274 [Source:MGI Symbo | 0 | 0 | 0 | 0 | 0 | 0 |
| 25928 | ENSMUSG00000083794 | 0 | NA | NA | NA | NA | NA | Gm16123 | predicted gene 16123 [Source:MGI Symbo | 0 | 0 | 0 | 0 | 0 | 0 |
| 25929 | ENSMUSG00000083795 | 0 | NA | NA | NA | NA | NA | Gm15670 | predicted gene 15670 [Source:MGI Symbo | 0 | 0 | 0 | 0 | 0 | 0 |
| 25930 | ENSMUSG00000083796 | 0.371281295 | 2.09253362 | 4.031852165 | 0.519000582 | 0.603760335 | NA | Gm13369 | predicted gene 13369 [Source:MGI Symbo | 0 | 0 | 0 | 0 | 0 | 2.2277 |
| 25931 | ENSMUSG00000083798 | 1.088168803 | 2.43902648 | 2.378968265 | 1.025245488 | 0.30524737 | NA | Gm14584 | predicted gene 14584 [Source:MGI Symbo | 0 | 0 | 1.0062 | 0 | 2.1812 | 3.3415 |
| 25932 | ENSMUSG00000083801 | 0 | NA | NA | NA | NA | NA | Cyp2j7-ps2 | cytochrome P450, family 2, subfamily j, pc | 0 | 0 | 0 | 0 | 0 | 0 |
| 25933 | ENSMUSG00000083803 | 0 | NA | NA | NA | NA | NA | Gm12455 | predicted gene 12455 [Source:MGI Symbo | 0 | 0 | 0 | 0 | 0 | 0 |
| 25934 | ENSMUSG00000083804 | 0 | NA | NA | NA | NA | NA | Gm13676 | predicted gene 13676 [Source:MGI Symbo | 0 | 0 | 0 | 0 | 0 | 0 |
| 25935 | ENSMUSG00000083805 | 0 | NA | NA | NA | NA | NA | Gm14848 | predicted gene 14848 [Source:MGI Symbo | 0 | 0 | 0 | 0 | 0 | 0 |
| 25936 | ENSMUSG00000083806 | 0 | NA | NA | NA | NA | NA | Gm12131 | predicted gene 12131 [Source:MGI Symbo | 0 | 0 | 0 | 0 | 0 | 0 |
| 25937 | ENSMUSG00000083807 | 0 | NA | NA | NA | NA | NA | Gm13466 | predicted gene 13466 [Source:MGI Symbo | 0 | 0 | 0 | 0 | 0 | 0 |
| 25938 | ENSMUSG00000083808 | 0 | NA | NA | NA | NA | NA | Gm3828 | predicted gene 3828 [Source:MGI Symbol | 0 | 0 | 0 | 0 | 0 | 0 |
| 25939 | ENSMUSG00000083809 | 0 | NA | NA | NA | NA | NA | Gm15110 | predicted gene 15110 [Source:MGI Symbo | 0 | 0 | 0 | 0 | 0 | 0 |
| 25940 | ENSMUSG00000083810 | 0 | NA | NA | NA | NA | NA | Gm14713 | predicted gene 14713 [Source:MGI Symbo | 0 | 0 | 0 | 0 | 0 | 0 |
| 25941 | ENSMUSG00000083811 | 5.532233569 | -0.31749696 | 0.927819151 | -0.342197032 | 0.732202618 | 0.947515796 | Gm13071 | predicted gene 13071 [Source:MGI Symbo | 7.2279 | 3.0838 | 8.05 | 3.8559 | 7.6343 | 3.3415 |
| 25942 | ENSMUSG00000083812 | 0 | NA | NA | NA | NA | NA | Gm5054 | predicted gene 5054 [Source:MGI Symbol | 0 | 0 | 0 | 0 | 0 | 0 |
| 25943 | ENSMUSG00000083813 | 29.76194909 | -0.641892025 | 0.40900048 | -1.569416311 | 0.116550966 | 0.501814922 | Gm15502 | predicted gene 15502 [Source:MGI Symbo | 37.746 | 38.033 | 33.206 | 29.883 | 18.541 | 21.163 |
| 25944 | ENSMUSG00000083814 | 0 | NA | NA | NA | NA | NA | Gm5133 | predicted pseudogene 5133 [Source:MGI | 0 | 0 | 0 | 0 | 0 | 0 |
| 25945 | ENSMUSG00000083815 | 0 | NA | NA | NA | NA | NA | Gm13435 | predicted gene 13435 [Source:MGI Symbo | 0 | 0 | 0 | 0 | 0 | 0 |
| 25946 | ENSMUSG00000083816 | 5.054321043 | 0.520479291 | 0.969147747 | 0.537048445 | 0.591234151 | NA | Gm13033 | predicted gene 13033 [Source:MGI Symbo | 7.2279 | 2.0558 | 3.0187 | 4.8199 | 7.6343 | 5.5692 |
| 25947 | ENSMUSG00000083817 | 0 | NA | NA | NA | NA | NA | Gm14400 | predicted gene 14400 [Source:MGI Symbo | 0 | 0 | 0 | 0 | 0 | 0 |
| 25948 | ENSMUSG00000083818 | 0 | NA | NA | NA | NA | NA | Gm14908 | predicted gene 14908 [Source:MGI Symbo | 0 | 0 | 0 | 0 | 0 | 0 |
| 25949 | ENSMUSG00000083819 | 0 | NA | NA | NA | NA | NA | Gm11977 | predicted gene 11977 [Source:MGI Symbo | 0 | 0 | 0 | 0 | 0 | 0 |
| 25950 | ENSMUSG00000083820 | 6.650421168 | 0.638167494 | 0.997552075 | 0.639733514 | 0.522345863 | 0.888334265 | Ndufs6b | NADH:ubiquinone oxidoreductase subuni | 1.6062 | 1.0279 | 13.081 | 7.7118 | 10.906 | 5.5692 |
| 25951 | ENSMUSG00000083821 | 0 | NA | NA | NA | NA | NA | Gm14631 | predicted gene 14631 [Source:MGI Symbo | 0 | 0 | 0 | 0 | 0 | 0 |
| 25952 | ENSMUSG00000083822 | 0 | NA | NA | NA | NA | NA | Hmgb1-ps5 | high mobility group box 1, pseudogene 5 [ | 0 | 0 | 0 | 0 | 0 | 0 |
| 25953 | ENSMUSG00000083823 | 0 | NA | NA | NA | NA | NA | Amd-ps1 | S-adenosylmethionine decarboxylase, pse | 0 | 0 | 0 | 0 | 0 | 0 |
| 25954 | ENSMUSG00000083825 | 0 | NA | NA | NA | NA | NA | Olfr1376-ps1 | olfactory receptor 1376, pseudogene 1 [So | 0 | 0 | 0 | 0 | 0 | 0 |
| 25955 | ENSMUSG00000083826 | 0.328370405 | 0.16514316 | 4.003149536 | 0.041253308 | 0.967093956 | NA | Gm13039 | predicted gene 13039 [Source:MGI Symbo | 0 | 0 | 1.0062 | 0.964 | 0 | 0 |
| 25956 | ENSMUSG00000083827 | 0 | NA | NA | NA | NA | NA | Gm15712 | predicted gene 15712 [Source:MGI Symbo | 0 | 0 | 0 | 0 | 0 | 0 |
| 25957 | ENSMUSG00000083828 | 0 | NA | NA | NA | NA | NA | Gm15006 | predicted gene 15006 [Source:MGI Symbo | 0 | 0 | 0 | 0 | 0 | 0 |
| 25958 | ENSMUSG00000083829 | 0 | NA | NA | NA | NA | NA | Gm2199 | predicted gene 2199 [Source:MGI Symbol | 0 | 0 | 0 | 0 | 0 | 0 |
| 25959 | ENSMUSG00000083830 | 0 | NA | NA | NA | NA | NA | Gm8839 | predicted gene 8839 [Source:MGI Symbol | 0 | 0 | 0 | 0 | 0 | 0 |
| 25960 | ENSMUSG00000083831 | 0 | NA | NA | NA | NA | NA | Gm14248 | predicted gene 14248 [Source:MGI Symbo | 0 | 0 | 0 | 0 | 0 | 0 |
| 25961 | ENSMUSG00000083832 | 0 | NA | NA | NA | NA | NA | Gm13771 | predicted gene 13771 [Source:MGI Symbo | 0 | 0 | 0 | 0 | 0 | 0 |
| 25962 | ENSMUSG00000083833 | 5.677370507 | -0.097308709 | 1.098848557 | -0.088555159 | 0.929435446 | 0.988405116 | Gm13841 | predicted gene 13841 [Source:MGI Symbo | 2.4093 | 11.307 | 4.025 | 9.6398 | 0 | 6.6831 |
| 25963 | ENSMUSG00000083834 | 0 | NA | NA | NA | NA | NA | Gm12577 | predicted gene 12577 [Source:MGI Symbo | 0 | 0 | 0 | 0 | 0 | 0 |
| 25964 | ENSMUSG00000083835 | 0 | NA | NA | NA | NA | NA | Gm11898 | predicted gene 11898 [Source:MGI Symbo | 0 | 0 | 0 | 0 | 0 | 0 |
| 25965 | ENSMUSG00000083836 | 2.286171423 | -1.187997355 | 1.483341895 | -0.80089247 | 0.4231939 | NA | Ldha-ps2 | lactate dehydrogenase A, pseudogene 2 [ | 2.4093 | 5.1396 | 2.0125 | 1.928 | 0 | 2.2277 |
| 25966 | ENSMUSG00000083837 | 0.160662974 | 1.126924175 | 4.080472857 | 0.276174898 | 0.782413736 | NA | Gm14427 | predicted gene 14427 [Source:MGI Symbo | 0 | 0 | 0 | 0.964 | 0 | 0 |
| 25967 | ENSMUSG00000083838 | 1.209843426 | -3.647354608 | 2.210792771 | -1.649794886 | 0.098984895 | NA | Gm11957 | predicted gene 11957 [Source:MGI Symbo | 3.2124 | 1.0279 | 3.0187 | 0 | 0 | 0 |
| 25968 | ENSMUSG00000083839 | 0 | NA | NA | NA | NA | NA | Gm14765 | predicted gene 14765 [Source:MGI Symbo | 0 | 0 | 0 | 0 | 0 | 0 |
| 25969 | ENSMUSG00000083841 | 0 | NA | NA | NA | NA | NA | Gm1947 | predicted pseudogene 1947 [Source:MGI | 0 | 0 | 0 | 0 | 0 | 0 |
| 25970 | ENSMUSG00000083842 | 0 | NA | NA | NA | NA | NA | Gm12485 | predicted gene 12485 [Source:MGI Symbo | 0 | 0 | 0 | 0 | 0 | 0 |
| 25971 | ENSMUSG00000083843 | 0 | NA | NA | NA | NA | NA | Gm7071 | predicted gene 7071 [Source:MGI Symbol | 0 | 0 | 0 | 0 | 0 | 0 |
| 25972 | ENSMUSG00000083844 | 208.3017094 | 0.106541816 | 0.20076645 | 0.530675401 | 0.595643735 | 0.913808971 | Ube2d-ps | ubiquitin-conjugating enzyme E2D, pseud | 218.44 | 205.58 | 177.1 | 186.05 | 190.86 | 271.78 |
| 25973 | ENSMUSG00000083845 | 0 | NA | NA | NA | NA | NA | Gm14571 | predicted gene 14571 [Source:MGI Symbo | 0 | 0 | 0 | 0 | 0 | 0 |
| 25974 | ENSMUSG00000083846 | 0 | NA | NA | NA | NA | NA | Gm13797 | predicted gene 13797 [Source:MGI Symbo | 0 | 0 | 0 | 0 | 0 | 0 |
| 25975 | ENSMUSG00000083847 | 0 | NA | NA | NA | NA | NA | Xlr3d-ps | X-linked lymphocyte-regulated 3D, pseud | 0 | 0 | 0 | 0 | 0 | 0 |
| 25976 | ENSMUSG00000083848 | 0 | NA | NA | NA | NA | NA | Gm7309 | predicted gene 7309 [Source:MGI Symbol | 0 | 0 | 0 | 0 | 0 | 0 |
| 25977 | ENSMUSG00000083849 | 0.797635641 | -1.749522449 | 2.643029821 | -0.661938218 | 0.508010818 | NA | Gm13477 | predicted gene 13477 [Source:MGI Symbo | 0.8031 | 0 | 3.0187 | 0.964 | 0 | 0 |
| 25978 | ENSMUSG00000083851 | 0 | NA | NA | NA | NA | NA | Gm12365 | predicted gene 12365 [Source:MGI Symbo | 0 | 0 | 0 | 0 | 0 | 0 |
| 25979 | ENSMUSG00000083852 | 0 | NA | NA | NA | NA | NA | Gm13007 | predicted gene 13007 [Source:MGI Symbo | 0 | 0 | 0 | 0 | 0 | 0 |
| 25980 | ENSMUSG00000083853 | 1.544947958 | 0.403903114 | 1.757006713 | 0.229881372 | 0.818183952 | NA | Gm15696 | predicted gene 15696 [Source:MGI Symbo | 0 | 1.0279 | 3.0187 | 1.928 | 2.1812 | 1.1138 |

| | | | | | | | | | | | | | | | |
|---|---|---|---|---|---|---|---|---|---|---|---|---|---|---|---|
| 25981 | ENSMUSG00000083854 | 1.027711248 | -0.96742891 | 2.462926793 | -0.392796454 | 0.694469824 | NA | Dnajc19-ps | DnaJ heat shock protein family (Hsp40) m | 0 | 4.1117 | 0 | 0.964 | 1.0906 | 0 |
| 25982 | ENSMUSG00000083855 | 0 | NA | NA | NA | NA | NA | Olfr1175 | olfactory receptor 1175 [Source:MGI Syml | 0 | 0 | 0 | 0 | 0 | 0 |
| 25983 | ENSMUSG00000083856 | 2.004253937 | 0.593935302 | 1.72781418 | 0.343749524 | 0.731034663 | NA | Gm13886 | predicted gene 13886 [Source:MGI Symbo | 0.8031 | 0 | 4.025 | 3.8559 | 0 | 3.3415 |
| 25984 | ENSMUSG00000083857 | 0 | NA | NA | NA | NA | NA | Gm12584 | predicted gene 12584 [Source:MGI Symbo | 0 | 0 | 0 | 0 | 0 | 0 |
| 25985 | ENSMUSG00000083858 | 0 | NA | NA | NA | NA | NA | Gm14979 | predicted gene 14979 [Source:MGI Symbo | 0 | 0 | 0 | 0 | 0 | 0 |
| 25986 | ENSMUSG00000083859 | 0 | NA | NA | NA | NA | NA | Gm12003 | predicted gene 12003 [Source:MGI Symbo | 0 | 0 | 0 | 0 | 0 | 0 |
| 25987 | ENSMUSG00000083860 | 1.87463033 | 0.959021347 | 1.547614372 | 0.619677204 | 0.535470327 | NA | Gm13313 | predicted gene 13313 [Source:MGI Symbo | 0.8031 | 1.0279 | 2.0125 | 1.928 | 4.3625 | 1.1138 |
| 25988 | ENSMUSG00000083861 | 0 | NA | NA | NA | NA | NA | Gm9599 | predicted gene 9599 [Source:MGI Symbol | 0 | 0 | 0 | 0 | 0 | 0 |
| 25989 | ENSMUSG00000083863 | 64.66340767 | 0.573945223 | 0.327012506 | 1.755117046 | 0.079239289 | 0.413332418 | Gm13341 | predicted gene 13341 [Source:MGI Symbo | 40.958 | 63.731 | 51.318 | 51.091 | 83.978 | 96.904 |
| 25990 | ENSMUSG00000083864 | 0 | NA | NA | NA | NA | NA | Gm13719 | predicted gene 13719 [Source:MGI Symbo | 0 | 0 | 0 | 0 | 0 | 0 |
| 25991 | ENSMUSG00000083866 | 0.513958919 | -2.394274821 | 3.756096013 | -0.637437066 | 0.523840191 | NA | Gm13472 | predicted gene 13472 [Source:MGI Symbo | 0 | 3.0838 | 0 | 0 | 0 | 0 |
| 25992 | ENSMUSG00000083867 | 0 | NA | NA | NA | NA | NA | Gm12893 | predicted gene 12893 [Source:MGI Symbo | 0 | 0 | 0 | 0 | 0 | 0 |
| 25993 | ENSMUSG00000083868 | 0 | NA | NA | NA | NA | NA | Gm12477 | predicted gene 12477 [Source:MGI Symbo | 0 | 0 | 0 | 0 | 0 | 0 |
| 25994 | ENSMUSG00000083869 | 0 | NA | NA | NA | NA | NA | Gm12442 | predicted gene 12442 [Source:MGI Symbo | 0 | 0 | 0 | 0 | 0 | 0 |
| 25995 | ENSMUSG00000083870 | 0.160662974 | 1.126924175 | 4.080472857 | 0.276174898 | 0.782413736 | NA | Gm11761 | predicted gene 11761 [Source:MGI Symbo | 0 | 0 | 0 | 0.964 | 0 | 0 |
| 25996 | ENSMUSG00000083871 | 0 | NA | NA | NA | NA | NA | Gm16479 | predicted gene 16479 [Source:MGI Symbo | 0 | 0 | 0 | 0 | 0 | 0 |
| 25997 | ENSMUSG00000083872 | 0 | NA | NA | NA | NA | NA | Gm11928 | predicted gene 11928 [Source:MGI Symbo | 0 | 0 | 0 | 0 | 0 | 0 |
| 25998 | ENSMUSG00000083873 | 0 | NA | NA | NA | NA | NA | Vmn1r-ps92 | vomeronasal 1 receptor, pseudogene 92 [ | 0 | 0 | 0 | 0 | 0 | 0 |
| 25999 | ENSMUSG00000083874 | 0 | NA | NA | NA | NA | NA | Gm7494 | predicted gene 7494 [Source:MGI Symbol | 0 | 0 | 0 | 0 | 0 | 0 |
| 26000 | ENSMUSG00000083875 | 0.706178709 | 0.10198885 | 3.03884937 | 0.033561667 | 0.97322669 | NA | Gm13831 | predicted gene 13831 [Source:MGI Symbo | 0 | 2.0558 | 0 | 0 | 2.1812 | 0 |
| 26001 | ENSMUSG00000083876 | 0 | NA | NA | NA | NA | NA | Gm11439 | predicted gene 11439 [Source:MGI Symbo | 0 | 0 | 0 | 0 | 0 | 0 |
| 26002 | ENSMUSG00000083877 | 0 | NA | NA | NA | NA | NA | Gm14740 | predicted gene 14740 [Source:MGI Symbo | 0 | 0 | 0 | 0 | 0 | 0 |
| 26003 | ENSMUSG00000083878 | 0 | NA | NA | NA | NA | NA | Gm12034 | predicted gene 12034 [Source:MGI Symbo | 0 | 0 | 0 | 0 | 0 | 0 |
| 26004 | ENSMUSG00000083879 | 0.713016575 | 1.687601183 | 3.028998059 | 0.557148321 | 0.577426095 | NA | Gm14038 | predicted gene 14038 [Source:MGI Symbo | 0 | 0 | 1.0062 | 0 | 3.2719 | 0 |
| 26005 | ENSMUSG00000083880 | 0 | NA | NA | NA | NA | NA | Hspe1-ps6 | heat shock protein 1 (chaperonin 10), pse | 0 | 0 | 0 | 0 | 0 | 0 |
| 26006 | ENSMUSG00000083882 | 0 | NA | NA | NA | NA | NA | Gm11369 | predicted gene 11369 [Source:MGI Symbo | 0 | 0 | 0 | 0 | 0 | 0 |
| 26007 | ENSMUSG00000083884 | 0 | NA | NA | NA | NA | NA | Gm15388 | predicted gene 15388 [Source:MGI Symbo | 0 | 0 | 0 | 0 | 0 | 0 |
| 26008 | ENSMUSG00000083885 | 0.185640647 | 1.126924175 | 4.080472857 | 0.276174898 | 0.782413736 | NA | Gm13213 | predicted gene 13213 [Source:MGI Symbo | 0 | 0 | 0 | 0 | 0 | 1.1138 |
| 26009 | ENSMUSG00000083886 | 0 | NA | NA | NA | NA | NA | Olfr409-ps1 | olfactory receptor 409, pseudogene 1 [So | 0 | 0 | 0 | 0 | 0 | 0 |
| 26010 | ENSMUSG00000083887 | 0 | NA | NA | NA | NA | NA | Gm11930 | predicted gene 11930 [Source:MGI Symbo | 0 | 0 | 0 | 0 | 0 | 0 |
| 26011 | ENSMUSG00000083888 | 0 | NA | NA | NA | NA | NA | Gm12441 | predicted gene 12441 [Source:MGI Symbo | 0 | 0 | 0 | 0 | 0 | 0 |
| 26012 | ENSMUSG00000083889 | 0 | NA | NA | NA | NA | NA | E530001F21 | RIKEN cDNA E530001F21 gene [Source:M( | 0 | 0 | 0 | 0 | 0 | 0 |
| 26013 | ENSMUSG00000083890 | 0 | NA | NA | NA | NA | NA | Gm15703 | predicted gene 15703 [Source:MGI Symbo | 0 | 0 | 0 | 0 | 0 | 0 |
| 26014 | ENSMUSG00000083892 | 0.133850373 | -0.796629176 | 4.080472857 | -0.195229623 | 0.84521319 | NA | Gm1848 | predicted gene 1848 [Source:MGI Symbol | 0.8031 | 0 | 0 | 0 | 0 | 0 |
| 26015 | ENSMUSG00000083893 | 0 | NA | NA | NA | NA | NA | Gm13347 | predicted gene 13347 [Source:MGI Symbo | 0 | 0 | 0 | 0 | 0 | 0 |
| 26016 | ENSMUSG00000083894 | 0 | NA | NA | NA | NA | NA | Gm11489 | predicted gene 11489 [Source:MGI Symbo | 0 | 0 | 0 | 0 | 0 | 0 |
| 26017 | ENSMUSG00000083895 | 0 | NA | NA | NA | NA | NA | Gm12074 | predicted gene 12074 [Source:MGI Symbo | 0 | 0 | 0 | 0 | 0 | 0 |
| 26018 | ENSMUSG00000083896 | 0 | NA | NA | NA | NA | NA | Scgb2b23-ps | secretoglobin, family 2B, member 23, pse | 0 | 0 | 0 | 0 | 0 | 0 |
| 26019 | ENSMUSG00000083897 | 0.835104907 | -1.8196403 | 2.6116807 | -0.696731534 | 0.48597082 | NA | Gm14843 | predicted gene 14843 [Source:MGI Symbo | 0 | 1.0279 | 3.0187 | 0.964 | 0 | 0 |
| 26020 | ENSMUSG00000083898 | 0 | NA | NA | NA | NA | NA | Gm7340 | predicted gene 7340 [Source:MGI Symbol | 0 | 0 | 0 | 0 | 0 | 0 |
| 26021 | ENSMUSG00000083899 | 8.404367944 | 0.049699002 | 0.779563514 | 0.063752345 | 0.949167425 | 0.990731019 | Gm12346 | predicted gene 12346 [Source:MGI Symbo | 9.6372 | 4.1117 | 11.069 | 13.496 | 6.5437 | 5.5692 |
| 26022 | ENSMUSG00000083900 | 0 | NA | NA | NA | NA | NA | Gm13294 | predicted gene 13294 [Source:MGI Symbo | 0 | 0 | 0 | 0 | 0 | 0 |
| 26023 | ENSMUSG00000083901 | 0 | NA | NA | NA | NA | NA | Gm12885 | predicted gene 12885 [Source:MGI Symbo | 0 | 0 | 0 | 0 | 0 | 0 |
| 26024 | ENSMUSG00000083902 | 0.17131964 | -0.796629176 | 4.080472857 | -0.195229623 | 0.84521319 | NA | Tent2-ps1 | terminal nucleotidyltransferase 2, pseudo | 0 | 1.0279 | 0 | 0 | 0 | 0 |
| 26025 | ENSMUSG00000083903 | 2.889519126 | -0.488305172 | 1.321388591 | -0.369539419 | 0.711725697 | NA | Gm5405 | predicted gene 5405 [Source:MGI Symbol | 4.0155 | 4.1117 | 2.0125 | 3.8559 | 0 | 3.3415 |
| 26026 | ENSMUSG00000083904 | 0 | NA | NA | NA | NA | NA | Gm12805 | predicted gene 12805 [Source:MGI Symbo | 0 | 0 | 0 | 0 | 0 | 0 |
| 26027 | ENSMUSG00000083905 | 0 | NA | NA | NA | NA | NA | Gm15355 | predicted gene 15355 [Source:MGI Symbo | 0 | 0 | 0 | 0 | 0 | 0 |
| 26028 | ENSMUSG00000083906 | 0.826457617 | 2.043192078 | 2.584773689 | 0.790472329 | 0.429251976 | NA | Gm15186 | predicted gene 15186 [Source:MGI Symbo | 0.8031 | 0 | 0 | 1.928 | 0 | 2.2277 |
| 26029 | ENSMUSG00000083907 | 3.760873458 | 0.963759763 | 1.098862121 | 0.877052493 | 0.380458127 | NA | Plk-ps1 | polo like kinase, pseudogene 1 [Source:M | 1.6062 | 2.0558 | 4.025 | 3.8559 | 5.4531 | 5.5692 |
| 26030 | ENSMUSG00000083908 | 0 | NA | NA | NA | NA | NA | Gm6447 | predicted gene 6447 [Source:MGI Symbol | 0 | 0 | 0 | 0 | 0 | 0 |
| 26031 | ENSMUSG00000083909 | 1.909622952 | -2.164997245 | 1.863079921 | -1.162052803 | 0.245214017 | NA | Gm15842 | predicted gene 15842 [Source:MGI Symbo | 3.2124 | 6.1675 | 0 | 0.964 | 0 | 1.1138 |
| 26032 | ENSMUSG00000083910 | 0 | NA | NA | NA | NA | NA | Gm13604 | predicted gene 13604 [Source:MGI Symbo | 0 | 0 | 0 | 0 | 0 | 0 |
| 26033 | ENSMUSG00000083911 | 0.160662974 | 1.126924175 | 4.080472857 | 0.276174898 | 0.782413736 | NA | Gm4342 | predicted gene 4342 [Source:MGI Symbol | 0 | 0 | 0 | 0.964 | 0 | 0 |
| 26034 | ENSMUSG00000083912 | 0 | NA | NA | NA | NA | NA | Gm5391 | predicted gene 5391 [Source:MGI Symbol | 0 | 0 | 0 | 0 | 0 | 0 |
| 26035 | ENSMUSG00000083913 | 0 | NA | NA | NA | NA | NA | Gm14219 | predicted gene 14219 [Source:MGI Symbo | 0 | 0 | 0 | 0 | 0 | 0 |
| 26036 | ENSMUSG00000083914 | 0.342639279 | -1.805976596 | 4.028547714 | -0.448294702 | 0.653940524 | NA | Rps18-ps1 | ribosomal protein S18, pseudogene 1 [Sou | 0 | 2.0558 | 0 | 0 | 0 | 0 |

| | | | | | | | | | | | | | | | |
|---|---|---|---|---|---|---|---|---|---|---|---|---|---|---|---|
| 26037 | ENSMUSG00000083915 | 0 | NA | NA | NA | NA | NA | Gm13932 | predicted gene 13932 [Source:MGI Symbc | 0 | 0 | 0 | 0 | 0 | 0 |
| 26038 | ENSMUSG00000083916 | 0 | NA | NA | NA | NA | NA | Gm15476 | predicted gene 15476 [Source:MGI Symbc | 0 | 0 | 0 | 0 | 0 | 0 |
| 26039 | ENSMUSG00000083917 | 0 | NA | NA | NA | NA | NA | Gm12093 | predicted gene 12093 [Source:MGI Symbc | 0 | 0 | 0 | 0 | 0 | 0 |
| 26040 | ENSMUSG00000083918 | 0 | NA | NA | NA | NA | NA | Gm14352 | predicted gene 14352 [Source:MGI Symbc | 0 | 0 | 0 | 0 | 0 | 0 |
| 26041 | ENSMUSG00000083919 | 0 | NA | NA | NA | NA | NA | Tpi-rs8 | triosephosphate isomerase related seque | 0 | 0 | 0 | 0 | 0 | 0 |
| 26042 | ENSMUSG00000083920 | 0 | NA | NA | NA | NA | NA | Gm14987 | predicted gene 14987 [Source:MGI Symbc | 0 | 0 | 0 | 0 | 0 | 0 |
| 26043 | ENSMUSG00000083921 | 0.167707431 | -0.796629176 | 4.080472857 | -0.195229623 | 0.84521319 | NA | Gm15750 | predicted gene 15750 [Source:MGI Symbc | 0 | 0 | 1.0062 | 0 | 0 | 0 |
| 26044 | ENSMUSG00000083922 | 0 | NA | NA | NA | NA | NA | Gm3835 | predicted pseudogene 3835 [Source:MGI | 0 | 0 | 0 | 0 | 0 | 0 |
| 26045 | ENSMUSG00000083923 | 0 | NA | NA | NA | NA | NA | Gm649 | predicted gene 649 [Source:MGI Symbol;/ | 0 | 0 | 0 | 0 | 0 | 0 |
| 26046 | ENSMUSG00000083924 | 0 | NA | NA | NA | NA | NA | Gm12304 | predicted gene 12304 [Source:MGI Symbc | 0 | 0 | 0 | 0 | 0 | 0 |
| 26047 | ENSMUSG00000083925 | 0 | NA | NA | NA | NA | NA | Gm12012 | predicted gene 12012 [Source:MGI Symbc | 0 | 0 | 0 | 0 | 0 | 0 |
| 26048 | ENSMUSG00000083926 | 0 | NA | NA | NA | NA | NA | Gm14783 | predicted gene 14783 [Source:MGI Symbc | 0 | 0 | 0 | 0 | 0 | 0 |
| 26049 | ENSMUSG00000083927 | 0 | NA | NA | NA | NA | NA | Gm14749 | predicted gene 14749 [Source:MGI Symbc | 0 | 0 | 0 | 0 | 0 | 0 |
| 26050 | ENSMUSG00000083928 | 0 | NA | NA | NA | NA | NA | Gm4991 | predicted gene 4991 [Source:MGI Symbol | 0 | 0 | 0 | 0 | 0 | 0 |
| 26051 | ENSMUSG00000083929 | 0 | NA | NA | NA | NA | NA | Gm10600 | predicted gene 10600 [Source:MGI Symbc | 0 | 0 | 0 | 0 | 0 | 0 |
| 26052 | ENSMUSG00000083930 | 0 | NA | NA | NA | NA | NA | Gm13042 | predicted gene 13042 [Source:MGI Symbc | 0 | 0 | 0 | 0 | 0 | 0 |
| 26053 | ENSMUSG00000083931 | 0 | NA | NA | NA | NA | NA | Gm14518 | predicted gene 14518 [Source:MGI Symbc | 0 | 0 | 0 | 0 | 0 | 0 |
| 26054 | ENSMUSG00000083932 | 0 | NA | NA | NA | NA | NA | Atp5k-ps3 | ATP synthase, H+ transporting, mitochonc | 0 | 0 | 0 | 0 | 0 | 0 |
| 26055 | ENSMUSG00000083933 | 0 | NA | NA | NA | NA | NA | Gm15515 | predicted gene 15515 [Source:MGI Symbc | 0 | 0 | 0 | 0 | 0 | 0 |
| 26056 | ENSMUSG00000083934 | 0 | NA | NA | NA | NA | NA | Gm12368 | predicted gene 12368 [Source:MGI Symbc | 0 | 0 | 0 | 0 | 0 | 0 |
| 26057 | ENSMUSG00000083935 | 0 | NA | NA | NA | NA | NA | Gm12294 | predicted gene 12294 [Source:MGI Symbc | 0 | 0 | 0 | 0 | 0 | 0 |
| 26058 | ENSMUSG00000083936 | 0 | NA | NA | NA | NA | NA | Gm14829 | predicted gene 14829 [Source:MGI Symbc | 0 | 0 | 0 | 0 | 0 | 0 |
| 26059 | ENSMUSG00000083937 | 0 | NA | NA | NA | NA | NA | Cct3-ps1 | chaperonin containing Tcp1, subunit 3 (ga | 0 | 0 | 0 | 0 | 0 | 0 |
| 26060 | ENSMUSG00000083938 | 0 | NA | NA | NA | NA | NA | Gm12448 | predicted gene 12448 [Source:MGI Symbc | 0 | 0 | 0 | 0 | 0 | 0 |
| 26061 | ENSMUSG00000083939 | 0 | NA | NA | NA | NA | NA | Gm14451 | predicted gene 14451 [Source:MGI Symbc | 0 | 0 | 0 | 0 | 0 | 0 |
| 26062 | ENSMUSG00000083940 | 0 | NA | NA | NA | NA | NA | Gm11298 | predicted gene 11298 [Source:MGI Symbc | 0 | 0 | 0 | 0 | 0 | 0 |
| 26063 | ENSMUSG00000083941 | 0 | NA | NA | NA | NA | NA | Gm12889 | predicted gene 12889 [Source:MGI Symbc | 0 | 0 | 0 | 0 | 0 | 0 |
| 26064 | ENSMUSG00000083942 | 0 | NA | NA | NA | NA | NA | Scgb1b26-ps | secretoglobin, family 1B, member 26, pseu | 0 | 0 | 0 | 0 | 0 | 0 |
| 26065 | ENSMUSG00000083943 | 0 | NA | NA | NA | NA | NA | Gm11483 | predicted gene 11483 [Source:MGI Symbc | 0 | 0 | 0 | 0 | 0 | 0 |
| 26066 | ENSMUSG00000083944 | 0 | NA | NA | NA | NA | NA | Gm14733 | predicted gene 14733 [Source:MGI Symbc | 0 | 0 | 0 | 0 | 0 | 0 |
| 26067 | ENSMUSG00000083945 | 0 | NA | NA | NA | NA | NA | Gm15242 | predicted gene 15242 [Source:MGI Symbc | 0 | 0 | 0 | 0 | 0 | 0 |
| 26068 | ENSMUSG00000083946 | 0 | NA | NA | NA | NA | NA | Gm11359 | predicted gene 11359 [Source:MGI Symbc | 0 | 0 | 0 | 0 | 0 | 0 |
| 26069 | ENSMUSG00000083947 | 0 | NA | NA | NA | NA | NA | Olfr122 | olfactory receptor 122 [Source:MGI Symb | 0 | 0 | 0 | 0 | 0 | 0 |
| 26070 | ENSMUSG00000083948 | 0 | NA | NA | NA | NA | NA | Gm7158 | predicted gene 7158 [Source:MGI Symbol | 0 | 0 | 0 | 0 | 0 | 0 |
| 26071 | ENSMUSG00000083949 | 0 | NA | NA | NA | NA | NA | Gm12897 | predicted gene 12897 [Source:MGI Symbc | 0 | 0 | 0 | 0 | 0 | 0 |
| 26072 | ENSMUSG00000083950 | 0 | NA | NA | NA | NA | NA | Gm14466 | predicted gene 14466 [Source:MGI Symbc | 0 | 0 | 0 | 0 | 0 | 0 |
| 26073 | ENSMUSG00000083951 | 0.133850373 | -0.796629176 | 4.080472857 | -0.195229623 | 0.84521319 | NA | Gm13786 | predicted gene 13786 [Source:MGI Symbc | 0.8031 | 0 | 0 | 0 | 0 | 0 |
| 26074 | ENSMUSG00000083953 | 1.099193445 | -2.312770509 | 2.368100974 | -0.976635091 | 0.32874984 | NA | Gm14049 | predicted gene 14049 [Source:MGI Symbc | 1.6062 | 0 | 4.025 | 0.964 | 0 | 0 |
| 26075 | ENSMUSG00000083954 | 0 | NA | NA | NA | NA | NA | Gm12179 | predicted gene 12179 [Source:MGI Symbc | 0 | 0 | 0 | 0 | 0 | 0 |
| 26076 | ENSMUSG00000083955 | 0 | NA | NA | NA | NA | NA | Gm14785 | predicted gene 14785 [Source:MGI Symbc | 0 | 0 | 0 | 0 | 0 | 0 |
| 26077 | ENSMUSG00000083956 | 0 | NA | NA | NA | NA | NA | Gm12361 | predicted gene 12361 [Source:MGI Symbc | 0 | 0 | 0 | 0 | 0 | 0 |
| 26078 | ENSMUSG00000083957 | 0 | NA | NA | NA | NA | NA | Gm12742 | predicted gene 12742 [Source:MGI Symbc | 0 | 0 | 0 | 0 | 0 | 0 |
| 26079 | ENSMUSG00000083958 | 0 | NA | NA | NA | NA | NA | Gm12601 | predicted gene 12601 [Source:MGI Symbc | 0 | 0 | 0 | 0 | 0 | 0 |
| 26080 | ENSMUSG00000083960 | 0 | NA | NA | NA | NA | NA | Gm4918 | predicted gene 4918 [Source:MGI Symbol | 0 | 0 | 0 | 0 | 0 | 0 |
| 26081 | ENSMUSG00000083961 | 0 | NA | NA | NA | NA | NA | Gm13725 | predicted gene 13725 [Source:MGI Symbc | 0 | 0 | 0 | 0 | 0 | 0 |
| 26082 | ENSMUSG00000083962 | 0.339027071 | -1.79174149 | 3.919585585 | -0.457125237 | 0.647581032 | NA | Gm15707 | predicted gene 15707 [Source:MGI Symbc | 0 | 1.0279 | 1.0062 | 0 | 0 | 0 |
| 26083 | ENSMUSG00000083963 | 0 | NA | NA | NA | NA | NA | Gm15203 | predicted gene 15203 [Source:MGI Symbc | 0 | 0 | 0 | 0 | 0 | 0 |
| 26084 | ENSMUSG00000083964 | 3.299608283 | -0.69189814 | 1.22122363 | -0.566561376 | 0.571012223 | NA | Gm14122 | predicted gene 14122 [Source:MGI Symbc | 4.0155 | 5.1396 | 3.0187 | 0.964 | 1.0906 | 5.5692 |
| 26085 | ENSMUSG00000083966 | 0 | NA | NA | NA | NA | NA | Gm13482 | predicted gene 13482 [Source:MGI Symbc | 0 | 0 | 0 | 0 | 0 | 0 |
| 26086 | ENSMUSG00000083967 | 0 | NA | NA | NA | NA | NA | Gm15115 | predicted gene 15115 [Source:MGI Symbc | 0 | 0 | 0 | 0 | 0 | 0 |
| 26087 | ENSMUSG00000083969 | 0 | NA | NA | NA | NA | NA | Olfr1050-ps1 | olfactory receptor 1050, pseudogene 1 [S | 0 | 0 | 0 | 0 | 0 | 0 |
| 26088 | ENSMUSG00000083970 | 0 | NA | NA | NA | NA | NA | Gm15081 | predicted gene 15081 [Source:MGI Symbc | 0 | 0 | 0 | 0 | 0 | 0 |
| 26089 | ENSMUSG00000083971 | 0 | NA | NA | NA | NA | NA | Gm5943 | predicted pseudogene 5943 [Source:MGI | 0 | 0 | 0 | 0 | 0 | 0 |
| 26090 | ENSMUSG00000083972 | 0 | NA | NA | NA | NA | NA | Gm14802 | predicted gene 14802 [Source:MGI Symbc | 0 | 0 | 0 | 0 | 0 | 0 |
| 26091 | ENSMUSG00000083973 | 0 | NA | NA | NA | NA | NA | Gm13459 | predicted gene 13459 [Source:MGI Symbc | 0 | 0 | 0 | 0 | 0 | 0 |
| 26092 | ENSMUSG00000083974 | 1.503581129 | 3.029179708 | 2.149011099 | 1.409569131 | 0.158666947 | NA | Gm16053 | predicted gene 16053 [Source:MGI Symbc | 0.8031 | 0 | 0 | 3.8559 | 4.3625 | 0 |

| | | | | | | | | | | | | | | | |
|---|---|---|---|---|---|---|---|---|---|---|---|---|---|---|---|
| 26093 | ENSMUSG00000083975 | 0 | NA | NA | NA | NA | NA | Gm14215 | predicted gene 14215 [Source:MGI Symbo | 0 | 0 | 0 | 0 | 0 | 0 |
| 26094 | ENSMUSG00000083976 | 0 | NA | NA | NA | NA | NA | Gm13664 | predicted gene 13664 [Source:MGI Symbo | 0 | 0 | 0 | 0 | 0 | 0 |
| 26095 | ENSMUSG00000083977 | 0 | NA | NA | NA | NA | NA | Mageb8-ps | MAGE family member B8, pseudogene [So | 0 | 0 | 0 | 0 | 0 | 0 |
| 26096 | ENSMUSG00000083978 | 0 | NA | NA | NA | NA | NA | Gm5060 | predicted gene 5060 [Source:MGI Symbol | 0 | 0 | 0 | 0 | 0 | 0 |
| 26097 | ENSMUSG00000083979 | 0 | NA | NA | NA | NA | NA | Gm12527 | predicted gene 12527 [Source:MGI Symbo | 0 | 0 | 0 | 0 | 0 | 0 |
| 26098 | ENSMUSG00000083981 | 0 | NA | NA | NA | NA | NA | Gm15363 | predicted gene 15363 [Source:MGI Symbo | 0 | 0 | 0 | 0 | 0 | 0 |
| 26099 | ENSMUSG00000083982 | 0 | NA | NA | NA | NA | NA | Gm14650 | predicted gene 14650 [Source:MGI Symbo | 0 | 0 | 0 | 0 | 0 | 0 |
| 26100 | ENSMUSG00000083983 | 0 | NA | NA | NA | NA | NA | Gm12375 | predicted gene 12375 [Source:MGI Symbo | 0 | 0 | 0 | 0 | 0 | 0 |
| 26101 | ENSMUSG00000083985 | 0 | NA | NA | NA | NA | NA | Gm12468 | predicted gene 12468 [Source:MGI Symbo | 0 | 0 | 0 | 0 | 0 | 0 |
| 26102 | ENSMUSG00000083986 | 0 | NA | NA | NA | NA | NA | Gm12213 | predicted gene 12213 [Source:MGI Symbo | 0 | 0 | 0 | 0 | 0 | 0 |
| 26103 | ENSMUSG00000083987 | 0 | NA | NA | NA | NA | NA | Gm15311 | predicted gene 15311 [Source:MGI Symbo | 0 | 0 | 0 | 0 | 0 | 0 |
| 26104 | ENSMUSG00000083988 | 0.676451308 | 0.255551166 | 2.811516859 | 0.09089441 | 0.927576492 | NA | Rps10-ps4 | ribosomal protein S10, pseudogene 4 [Sou | 0.8031 | 1.0279 | 0 | 0 | 0 | 2.2277 |
| 26105 | ENSMUSG00000083989 | 0 | NA | NA | NA | NA | NA | Gm12607 | predicted gene 12607 [Source:MGI Symbo | 0 | 0 | 0 | 0 | 0 | 0 |
| 26106 | ENSMUSG00000083990 | 0 | NA | NA | NA | NA | NA | Gm12101 | predicted gene 12101 [Source:MGI Symbo | 0 | 0 | 0 | 0 | 0 | 0 |
| 26107 | ENSMUSG00000083991 | 0.514011053 | 1.044022102 | 3.082659575 | 0.338675769 | 0.734853997 | NA | Gm11618 | predicted gene 11618 [Source:MGI Symbo | 0 | 0 | 1.0062 | 0.964 | 0 | 1.1138 |
| 26108 | ENSMUSG00000083992 | 7.032361561 | 0.990705502 | 0.817489134 | 1.211888282 | 0.225555148 | 0.672301295 | Gm11478 | predicted gene 11478 [Source:MGI Symbo | 4.0155 | 6.1675 | 4.025 | 12.532 | 6.5437 | 8.9108 |
| 26109 | ENSMUSG00000083993 | 0 | NA | NA | NA | NA | NA | Gm14398 | predicted gene 14398 [Source:MGI Symbo | 0 | 0 | 0 | 0 | 0 | 0 |
| 26110 | ENSMUSG00000083994 | 0 | NA | NA | NA | NA | NA | Gm16465 | predicted gene 16465 [Source:MGI Symbo | 0 | 0 | 0 | 0 | 0 | 0 |
| 26111 | ENSMUSG00000083995 | 0 | NA | NA | NA | NA | NA | Gm12808 | predicted gene 12808 [Source:MGI Symbo | 0 | 0 | 0 | 0 | 0 | 0 |
| 26112 | ENSMUSG00000083996 | 0 | NA | NA | NA | NA | NA | Gm11649 | predicted gene 11649 [Source:MGI Symbo | 0 | 0 | 0 | 0 | 0 | 0 |
| 26113 | ENSMUSG00000083997 | 0 | NA | NA | NA | NA | NA | Gm12920 | predicted gene 12920 [Source:MGI Symbo | 0 | 0 | 0 | 0 | 0 | 0 |
| 26114 | ENSMUSG00000083998 | 0 | NA | NA | NA | NA | NA | Prdx2-ps2 | peroxiredoxin 2, pseudogene 2 [Source:M | 0 | 0 | 0 | 0 | 0 | 0 |
| 26115 | ENSMUSG00000083999 | 0 | NA | NA | NA | NA | NA | Gm7790 | predicted gene 7790 [Source:MGI Symbol | 0 | 0 | 0 | 0 | 0 | 0 |
| 26116 | ENSMUSG00000084000 | 0 | NA | NA | NA | NA | NA | Gm14653 | predicted gene 14653 [Source:MGI Symbo | 0 | 0 | 0 | 0 | 0 | 0 |
| 26117 | ENSMUSG00000084001 | 0 | NA | NA | NA | NA | NA | Gm15113 | predicted gene 15113 [Source:MGI Symbo | 0 | 0 | 0 | 0 | 0 | 0 |
| 26118 | ENSMUSG00000084004 | 0 | NA | NA | NA | NA | NA | Gm12047 | predicted gene 12047 [Source:MGI Symbo | 0 | 0 | 0 | 0 | 0 | 0 |
| 26119 | ENSMUSG00000084006 | 0 | NA | NA | NA | NA | NA | Gm14245 | predicted gene 14245 [Source:MGI Symbo | 0 | 0 | 0 | 0 | 0 | 0 |
| 26120 | ENSMUSG00000084007 | 0.346303622 | 2.004339653 | 3.884003423 | 0.516049919 | 0.605819544 | NA | Gm9003 | predicted pseudogene 9003 [Source:MGI | 0 | 0 | 0 | 0.964 | 0 | 1.1138 |
| 26121 | ENSMUSG00000084009 | 0 | NA | NA | NA | NA | NA | Rhox7-ps1 | reproductive homeobox 7, pseudogene 1 | 0 | 0 | 0 | 0 | 0 | 0 |
| 26122 | ENSMUSG00000084010 | 0 | NA | NA | NA | NA | NA | Gm13302 | predicted gene 13302 [Source:MGI Symbo | 0 | 0 | 0 | 0 | 0 | 0 |
| 26123 | ENSMUSG00000084011 | 0 | NA | NA | NA | NA | NA | Gm12969 | predicted gene 12969 [Source:MGI Symbo | 0 | 0 | 0 | 0 | 0 | 0 |
| 26124 | ENSMUSG00000084012 | 0 | NA | NA | NA | NA | NA | Gm11956 | predicted gene 11956 [Source:MGI Symbo | 0 | 0 | 0 | 0 | 0 | 0 |
| 26125 | ENSMUSG00000084013 | 3.926311364 | 0.777322424 | 1.144401582 | 0.679239208 | 0.496986308 | NA | Gm14270 | predicted gene 14270 [Source:MGI Symbo | 1.6062 | 4.1117 | 3.0187 | 4.8199 | 1.0906 | 8.9108 |
| 26126 | ENSMUSG00000084014 | 0 | NA | NA | NA | NA | NA | Gm7209 | predicted pseudogene 7209 [Source:MGI | 0 | 0 | 0 | 0 | 0 | 0 |
| 26127 | ENSMUSG00000084015 | 0 | NA | NA | NA | NA | NA | Gm14175 | predicted gene 14175 [Source:MGI Symbo | 0 | 0 | 0 | 0 | 0 | 0 |
| 26128 | ENSMUSG00000084017 | 0 | NA | NA | NA | NA | NA | Rpl21-ps9 | ribosomal protein L21, pseudogene 9 [Sou | 0 | 0 | 0 | 0 | 0 | 0 |
| 26129 | ENSMUSG00000084018 | 0 | NA | NA | NA | NA | NA | Gm11895 | predicted gene 11895 [Source:MGI Symbo | 0 | 0 | 0 | 0 | 0 | 0 |
| 26130 | ENSMUSG00000084019 | 0 | NA | NA | NA | NA | NA | Gm8754 | predicted gene 8754 [Source:MGI Symbol | 0 | 0 | 0 | 0 | 0 | 0 |
| 26131 | ENSMUSG00000084020 | 0 | NA | NA | NA | NA | NA | Wsb2-ps | WD repeat and SOCS box-containing 2, ps | 0 | 0 | 0 | 0 | 0 | 0 |
| 26132 | ENSMUSG00000084021 | 0.267700746 | -1.497481649 | 4.057001436 | -0.369110456 | 0.712045396 | NA | Gm5832 | predicted gene 5832 [Source:MGI Symbol | 1.6062 | 0 | 0 | 0 | 0 | 0 |
| 26133 | ENSMUSG00000084022 | 0 | NA | NA | NA | NA | NA | Gm16459 | predicted gene 16459 [Source:MGI Symbo | 0 | 0 | 0 | 0 | 0 | 0 |
| 26134 | ENSMUSG00000084024 | 0.476489653 | -2.293616034 | 3.421506946 | -0.670352587 | 0.502633051 | NA | Gm15937 | predicted gene 15937 [Source:MGI Symbo | 0.8031 | 2.0558 | 0 | 0 | 0 | 0 |
| 26135 | ENSMUSG00000084025 | 0 | NA | NA | NA | NA | NA | Gm6276 | predicted pseudogene 6276 [Source:MGI | 0 | 0 | 0 | 0 | 0 | 0 |
| 26136 | ENSMUSG00000084026 | 0 | NA | NA | NA | NA | NA | Gm11259 | predicted gene 11259 [Source:MGI Symbo | 0 | 0 | 0 | 0 | 0 | 0 |
| 26137 | ENSMUSG00000084027 | 0 | NA | NA | NA | NA | NA | Gm12376 | predicted gene 12376 [Source:MGI Symbo | 0 | 0 | 0 | 0 | 0 | 0 |
| 26138 | ENSMUSG00000084028 | 0 | NA | NA | NA | NA | NA | Gm13983 | predicted gene 13983 [Source:MGI Symbo | 0 | 0 | 0 | 0 | 0 | 0 |
| 26139 | ENSMUSG00000084029 | 0 | NA | NA | NA | NA | NA | Gm8855 | predicted gene 8855 [Source:MGI Symbol | 0 | 0 | 0 | 0 | 0 | 0 |
| 26140 | ENSMUSG00000084030 | 0 | NA | NA | NA | NA | NA | Gm16423 | predicted gene 16423 [Source:MGI Symbo | 0 | 0 | 0 | 0 | 0 | 0 |
| 26141 | ENSMUSG00000084031 | 0 | NA | NA | NA | NA | NA | Gm7416 | predicted gene 7416 [Source:MGI Symbol | 0 | 0 | 0 | 0 | 0 | 0 |
| 26142 | ENSMUSG00000084032 | 1.593436353 | -2.89780663 | 2.131204121 | -1.359703935 | 0.173923632 | NA | Gm12991 | predicted gene 12991 [Source:MGI Symbo | 2.4093 | 0 | 6.0375 | 0 | 0 | 1.1138 |
| 26143 | ENSMUSG00000084033 | 2.905500615 | -0.295368626 | 1.527208624 | -0.193404242 | 0.846642394 | NA | Gm7645 | predicted gene 7645 [Source:MGI Symbol | 1.6062 | 3.0838 | 5.0312 | 7.7118 | 0 | 0 |
| 26144 | ENSMUSG00000084034 | 0 | NA | NA | NA | NA | NA | Gm12617 | predicted gene 12617 [Source:MGI Symbo | 0 | 0 | 0 | 0 | 0 | 0 |
| 26145 | ENSMUSG00000084038 | 0 | NA | NA | NA | NA | NA | Gm11295 | predicted gene 11295 [Source:MGI Symbo | 0 | 0 | 0 | 0 | 0 | 0 |
| 26146 | ENSMUSG00000084039 | 0 | NA | NA | NA | NA | NA | Gm14551 | predicted gene 14551 [Source:MGI Symbo | 0 | 0 | 0 | 0 | 0 | 0 |
| 26147 | ENSMUSG00000084041 | 0 | NA | NA | NA | NA | NA | Gm15312 | predicted gene 15312 [Source:MGI Symbo | 0 | 0 | 0 | 0 | 0 | 0 |
| 26148 | ENSMUSG00000084043 | 0 | NA | NA | NA | NA | NA | Gm13732 | predicted gene 13732 [Source:MGI Symbo | 0 | 0 | 0 | 0 | 0 | 0 |

| | | | | | | | | | | | | | | | |
|---|---|---|---|---|---|---|---|---|---|---|---|---|---|---|---|
| 26149 | ENSMUSG00000084044 | 0 | NA | NA | NA | NA | NA | Gm14647 | predicted gene 14647 [Source:MGI Symbo | 0 | 0 | 0 | 0 | 0 | 0 |
| 26150 | ENSMUSG00000084045 | 0 | NA | NA | NA | NA | NA | Gm11997 | predicted gene 11997 [Source:MGI Symbo | 0 | 0 | 0 | 0 | 0 | 0 |
| 26151 | ENSMUSG00000084046 | 0 | NA | NA | NA | NA | NA | Gm11833 | predicted gene 11833 [Source:MGI Symbo | 0 | 0 | 0 | 0 | 0 | 0 |
| 26152 | ENSMUSG00000084047 | 0 | NA | NA | NA | NA | NA | Gm14984 | predicted gene 14984 [Source:MGI Symbo | 0 | 0 | 0 | 0 | 0 | 0 |
| 26153 | ENSMUSG00000084048 | 0 | NA | NA | NA | NA | NA | Gm12042 | predicted gene 12042 [Source:MGI Symbo | 0 | 0 | 0 | 0 | 0 | 0 |
| 26154 | ENSMUSG00000084050 | 2.797733291 | 0.995418334 | 1.319175087 | 0.754576358 | 0.450503213 | NA | Gm14439 | predicted gene 14439 [Source:MGI Symbo | 1.6062 | 1.0279 | 3.0187 | 6.7478 | 3.2719 | 1.1138 |
| 26155 | ENSMUSG00000084052 | 0 | NA | NA | NA | NA | NA | Gm11745 | predicted gene 11745 [Source:MGI Symbo | 0 | 0 | 0 | 0 | 0 | 0 |
| 26156 | ENSMUSG00000084053 | 0 | NA | NA | NA | NA | NA | Gm14904 | predicted gene 14904 [Source:MGI Symbo | 0 | 0 | 0 | 0 | 0 | 0 |
| 26157 | ENSMUSG00000084054 | 0 | NA | NA | NA | NA | NA | Gm14945 | predicted gene 14945 [Source:MGI Symbo | 0 | 0 | 0 | 0 | 0 | 0 |
| 26158 | ENSMUSG00000084055 | 0 | NA | NA | NA | NA | NA | Gm12004 | predicted gene 12004 [Source:MGI Symbo | 0 | 0 | 0 | 0 | 0 | 0 |
| 26159 | ENSMUSG00000084056 | 0 | NA | NA | NA | NA | NA | Rhox3b-ps | reproductive homeobox 3B, pseudogene | | 0 | 0 | 0 | 0 | 0 | 0 |
| 26160 | ENSMUSG00000084057 | 0 | NA | NA | NA | NA | NA | Gm8916 | predicted gene 8916 [Source:MGI Symbol | 0 | 0 | 0 | 0 | 0 | 0 |
| 26161 | ENSMUSG00000084058 | 0 | NA | NA | NA | NA | NA | Gm15263 | predicted gene 15263 [Source:MGI Symbo | 0 | 0 | 0 | 0 | 0 | 0 |
| 26162 | ENSMUSG00000084059 | 0 | NA | NA | NA | NA | NA | Gm13046 | predicted gene 13046 [Source:MGI Symbo | 0 | 0 | 0 | 0 | 0 | 0 |
| 26163 | ENSMUSG00000084060 | 0 | NA | NA | NA | NA | NA | Gm16412 | predicted pseudogene 16412 [Source:MG | 0 | 0 | 0 | 0 | 0 | 0 |
| 26164 | ENSMUSG00000084061 | 0 | NA | NA | NA | NA | NA | Gm4996 | predicted gene 4996 [Source:MGI Symbol | 0 | 0 | 0 | 0 | 0 | 0 |
| 26165 | ENSMUSG00000084063 | 0 | NA | NA | NA | NA | NA | Gm5934 | predicted gene 5934 [Source:MGI Symbol | 0 | 0 | 0 | 0 | 0 | 0 |
| 26166 | ENSMUSG00000084064 | 0 | NA | NA | NA | NA | NA | Gm12079 | predicted gene 12079 [Source:MGI Symbo | 0 | 0 | 0 | 0 | 0 | 0 |
| 26167 | ENSMUSG00000084065 | 0 | NA | NA | NA | NA | NA | Gm15159 | predicted gene 15159 [Source:MGI Symbo | 0 | 0 | 0 | 0 | 0 | 0 |
| 26168 | ENSMUSG00000084066 | 0 | NA | NA | NA | NA | NA | Xlr4d-ps | X-linked lymphocyte-regulated 4D, pseud | 0 | 0 | 0 | 0 | 0 | 0 |
| 26169 | ENSMUSG00000084067 | 1.176141353 | -0.268928376 | 1.952237713 | -0.137753909 | 0.890434912 | NA | Gm14269 | predicted gene 14269 [Source:MGI Symbo | 0.8031 | 2.0558 | 1.0062 | 0.964 | 0 | 2.2277 |
| 26170 | ENSMUSG00000084068 | 0 | NA | NA | NA | NA | NA | Gm11573 | predicted gene 11573 [Source:MGI Symbo | 0 | 0 | 0 | 0 | 0 | 0 |
| 26171 | ENSMUSG00000084069 | 0.17131964 | -0.796629176 | 4.080472857 | -0.195229623 | 0.84521319 | NA | Gm12306 | predicted gene 12306 [Source:MGI Symbo | 0 | 1.0279 | 0 | 0 | 0 | 0 |
| 26172 | ENSMUSG00000084070 | 0 | NA | NA | NA | NA | NA | Gm6591 | predicted gene 6591 [Source:MGI Symbol | 0 | 0 | 0 | 0 | 0 | 0 |
| 26173 | ENSMUSG00000084071 | 0.353089354 | 0.165142706 | 3.896102244 | 0.042386646 | 0.966190474 | NA | Gm13378 | predicted gene 13378 [Source:MGI Symbo | 0 | 1.0279 | 0 | 0 | 1.0906 | 0 |
| 26174 | ENSMUSG00000084072 | 0 | NA | NA | NA | NA | NA | Gm11285 | predicted gene 11285 [Source:MGI Symbo | 0 | 0 | 0 | 0 | 0 | 0 |
| 26175 | ENSMUSG00000084073 | 0 | NA | NA | NA | NA | NA | Gm11910 | predicted gene 11910 [Source:MGI Symbo | 0 | 0 | 0 | 0 | 0 | 0 |
| 26176 | ENSMUSG00000084074 | 0 | NA | NA | NA | NA | NA | Gm8310 | predicted gene 8310 [Source:MGI Symbol | 0 | 0 | 0 | 0 | 0 | 0 |
| 26177 | ENSMUSG00000084076 | 0 | NA | NA | NA | NA | NA | Gm14600 | predicted gene 14600 [Source:MGI Symbo | 0 | 0 | 0 | 0 | 0 | 0 |
| 26178 | ENSMUSG00000084077 | 0 | NA | NA | NA | NA | NA | Gm11952 | predicted gene 11952 [Source:MGI Symbo | 0 | 0 | 0 | 0 | 0 | 0 |
| 26179 | ENSMUSG00000084078 | 0 | NA | NA | NA | NA | NA | Olfr349-ps1 | olfactory receptor 349, pseudogene 1 [So | 0 | 0 | 0 | 0 | 0 | 0 |
| 26180 | ENSMUSG00000084079 | 0 | NA | NA | NA | NA | NA | Gm11631 | predicted gene 11631 [Source:MGI Symbo | 0 | 0 | 0 | 0 | 0 | 0 |
| 26181 | ENSMUSG00000084080 | 0 | NA | NA | NA | NA | NA | Gm15227 | predicted gene 15227 [Source:MGI Symbo | 0 | 0 | 0 | 0 | 0 | 0 |
| 26182 | ENSMUSG00000084081 | 0 | NA | NA | NA | NA | NA | Gm12057 | predicted gene 12057 [Source:MGI Symbo | 0 | 0 | 0 | 0 | 0 | 0 |
| 26183 | ENSMUSG00000084082 | 0 | NA | NA | NA | NA | NA | Gm12872 | predicted gene 12872 [Source:MGI Symbo | 0 | 0 | 0 | 0 | 0 | 0 |
| 26184 | ENSMUSG00000084083 | 0 | NA | NA | NA | NA | NA | Gm15782 | predicted gene 15782 [Source:MGI Symbo | 0 | 0 | 0 | 0 | 0 | 0 |
| 26185 | ENSMUSG00000084085 | 0 | NA | NA | NA | NA | NA | Gm16140 | predicted gene 16140 [Source:MGI Symbo | 0 | 0 | 0 | 0 | 0 | 0 |
| 26186 | ENSMUSG00000084086 | 0 | NA | NA | NA | NA | NA | Gm5761 | predicted gene 5761 [Source:MGI Symbol | 0 | 0 | 0 | 0 | 0 | 0 |
| 26187 | ENSMUSG00000084087 | 0 | NA | NA | NA | NA | NA | Gm13650 | predicted gene 13650 [Source:MGI Symbo | 0 | 0 | 0 | 0 | 0 | 0 |
| 26188 | ENSMUSG00000084088 | 2.311433324 | 1.06206186 | 1.523799643 | 0.696982615 | 0.485813674 | NA | Gm12941 | predicted gene 12941 [Source:MGI Symbo | 2.4093 | 1.0279 | 1.0062 | 3.8559 | 0 | 5.5692 |
| 26189 | ENSMUSG00000084089 | 0 | NA | NA | NA | NA | NA | Gm14099 | predicted gene 14099 [Source:MGI Symbo | 0 | 0 | 0 | 0 | 0 | 0 |
| 26190 | ENSMUSG00000084090 | 0 | NA | NA | NA | NA | NA | Gm11245 | predicted gene 11245 [Source:MGI Symbo | 0 | 0 | 0 | 0 | 0 | 0 |
| 26191 | ENSMUSG00000084092 | 0 | NA | NA | NA | NA | NA | Gm5843 | predicted gene 5843 [Source:MGI Symbol | 0 | 0 | 0 | 0 | 0 | 0 |
| 26192 | ENSMUSG00000084093 | 3.241265797 | -0.411179952 | 1.275421018 | -0.322387624 | 0.747159059 | NA | Gm16418 | predicted pseudogene 16418 [Source:MG | 4.0155 | 2.0558 | 5.0312 | 2.8919 | 5.4531 | 0 |
| 26193 | ENSMUSG00000084094 | 0 | NA | NA | NA | NA | NA | Gm13961 | predicted gene 13961 [Source:MGI Symbo | 0 | 0 | 0 | 0 | 0 | 0 |
| 26194 | ENSMUSG00000084095 | 0 | NA | NA | NA | NA | NA | Gm11834 | predicted gene 11834 [Source:MGI Symbo | 0 | 0 | 0 | 0 | 0 | 0 |
| 26195 | ENSMUSG00000084096 | 0 | NA | NA | NA | NA | NA | Vmn1r-ps13 | vomeronasal 1 receptor, pseudogene 135 | 0 | 0 | 0 | 0 | 0 | 0 |
| 26196 | ENSMUSG00000084097 | 0 | NA | NA | NA | NA | NA | Gm14922 | predicted gene 14922 [Source:MGI Symbo | 0 | 0 | 0 | 0 | 0 | 0 |
| 26197 | ENSMUSG00000084098 | 0.401551119 | -2.072982726 | 4.008251979 | -0.517178744 | 0.605031388 | NA | Gm13422 | predicted gene 13422 [Source:MGI Symbo | 2.4093 | 0 | 0 | 0 | 0 | 0 |
| 26198 | ENSMUSG00000084099 | 0 | NA | NA | NA | NA | NA | Gm12929 | predicted gene 12929 [Source:MGI Symbo | 0 | 0 | 0 | 0 | 0 | 0 |
| 26199 | ENSMUSG00000084101 | 0 | NA | NA | NA | NA | NA | Gm17014 | predicted gene 17014 [Source:MGI Symbo | 0 | 0 | 0 | 0 | 0 | 0 |
| 26200 | ENSMUSG00000084102 | 0 | NA | NA | NA | NA | NA | Sult2a-ps3 | sulfotransferase family 2A, dehydroepian | 0 | 0 | 0 | 0 | 0 | 0 |
| 26201 | ENSMUSG00000084103 | 0 | NA | NA | NA | NA | NA | Gm14629 | predicted gene 14629 [Source:MGI Symbo | 0 | 0 | 0 | 0 | 0 | 0 |
| 26202 | ENSMUSG00000084104 | 0.181769715 | 1.126924175 | 4.080472857 | 0.276174898 | 0.782413736 | NA | Gm13578 | predicted gene 13578 [Source:MGI Symbo | 0 | 0 | 0 | 0 | 1.0906 | 0 |
| 26203 | ENSMUSG00000084105 | 0.335414862 | -1.779047242 | 4.030804179 | -0.441362855 | 0.658950329 | NA | Gm13252 | predicted gene 13252 [Source:MGI Symbo | 0 | 0 | 2.0125 | 0 | 0 | 0 |
| 26204 | ENSMUSG00000084106 | 16.25727438 | -0.043124578 | 0.547709464 | -0.078736229 | 0.937242428 | 0.988420305 | Gm6136 | predicted pseudogene 6136 [Source:MGI | 12.047 | 21.586 | 16.1 | 20.244 | 13.087 | 14.48 |

| | | | | | | | | | | | | | | | |
|---|---|---|---|---|---|---|---|---|---|---|---|---|---|---|---|
| 26205 | ENSMUSG00000084107 | 0 | NA | NA | NA | NA | NA | Gm13700 | predicted gene 13700 [Source:MGI Symbo | 0 | 0 | 0 | 0 | 0 | 0 |
| 26206 | ENSMUSG00000084108 | 0 | NA | NA | NA | NA | NA | Gm15794 | predicted gene 15794 [Source:MGI Symbo | 0 | 0 | 0 | 0 | 0 | 0 |
| 26207 | ENSMUSG00000084110 | 0 | NA | NA | NA | NA | NA | Gm4992 | predicted pseudogene 4992 [Source:MGI | 0 | 0 | 0 | 0 | 0 | 0 |
| 26208 | ENSMUSG00000084111 | 9.887147387 | -0.438572251 | 0.680739857 | -0.644258224 | 0.519408001 | 0.887731952 | Gm15710 | predicted gene 15710 [Source:MGI Symbo | 8.8341 | 11.307 | 14.087 | 9.6398 | 6.5437 | 8.9108 |
| 26209 | ENSMUSG00000084112 | 0 | NA | NA | NA | NA | NA | Gm15112 | predicted gene 15112 [Source:MGI Symbo | 0 | 0 | 0 | 0 | 0 | 0 |
| 26210 | ENSMUSG00000084113 | 0 | NA | NA | NA | NA | NA | Gm14277 | predicted gene 14277 [Source:MGI Symbo | 0 | 0 | 0 | 0 | 0 | 0 |
| 26211 | ENSMUSG00000084114 | 0 | NA | NA | NA | NA | NA | Gm8421 | predicted gene 8421 [Source:MGI Symbol | 0 | 0 | 0 | 0 | 0 | 0 |
| 26212 | ENSMUSG00000084115 | 0 | NA | NA | NA | NA | NA | Gm13566 | predicted gene 13566 [Source:MGI Symbo | 0 | 0 | 0 | 0 | 0 | 0 |
| 26213 | ENSMUSG00000084116 | 0 | NA | NA | NA | NA | NA | Gm12460 | predicted gene 12460 [Source:MGI Symbo | 0 | 0 | 0 | 0 | 0 | 0 |
| 26214 | ENSMUSG00000084117 | 0 | NA | NA | NA | NA | NA | Olfr1063-ps1 | olfactory receptor 1063, pseudogene 1 [So | 0 | 0 | 0 | 0 | 0 | 0 |
| 26215 | ENSMUSG00000084118 | 0 | NA | NA | NA | NA | NA | Gm14989 | predicted gene 14989 [Source:MGI Symbo | 0 | 0 | 0 | 0 | 0 | 0 |
| 26216 | ENSMUSG00000084119 | 0 | NA | NA | NA | NA | NA | Gm12351 | predicted gene 12351 [Source:MGI Symbo | 0 | 0 | 0 | 0 | 0 | 0 |
| 26217 | ENSMUSG00000084120 | 0 | NA | NA | NA | NA | NA | Gm11624 | predicted gene 11624 [Source:MGI Symbo | 0 | 0 | 0 | 0 | 0 | 0 |
| 26218 | ENSMUSG00000084123 | 0 | NA | NA | NA | NA | NA | Gm13106 | predicted gene 13106 [Source:MGI Symbo | 0 | 0 | 0 | 0 | 0 | 0 |
| 26219 | ENSMUSG00000084124 | 0 | NA | NA | NA | NA | NA | Gm13588 | predicted gene 13588 [Source:MGI Symbo | 0 | 0 | 0 | 0 | 0 | 0 |
| 26220 | ENSMUSG00000084125 | 0 | NA | NA | NA | NA | NA | Gm8757 | predicted gene 8757 [Source:MGI Symbol | 0 | 0 | 0 | 0 | 0 | 0 |
| 26221 | ENSMUSG00000084126 | 0 | NA | NA | NA | NA | NA | Gm15844 | predicted gene 15844 [Source:MGI Symbo | 0 | 0 | 0 | 0 | 0 | 0 |
| 26222 | ENSMUSG00000084127 | 7.607690241 | 0.64143094 | 0.982950858 | 0.652556468 | 0.514042258 | 0.885004045 | Gm13169 | predicted gene 13169 [Source:MGI Symbo | 14.456 | 2.0558 | 1.0062 | 3.8559 | 10.906 | 13.366 |
| 26223 | ENSMUSG00000084128 | 1752.025706 | 0.222577062 | 0.121606356 | 1.830307808 | 0.067203924 | 0.373143019 | Esrp2 | epithelial splicing regulatory protein 2 [So | 1487.3 | 1497.7 | 1867.6 | 2010.9 | 1895.5 | 1753.2 |
| 26224 | ENSMUSG00000084129 | 0.371281295 | 2.09253362 | 4.031852165 | 0.519000582 | 0.603760335 | NA | Hmgb1-ps1 | high mobility group box 1, pseudogene 1 [ | 0 | 0 | 0 | 0 | 0 | 2.2277 |
| 26225 | ENSMUSG00000084130 | 0 | NA | NA | NA | NA | NA | Gm11704 | predicted gene 11704 [Source:MGI Symbo | 0 | 0 | 0 | 0 | 0 | 0 |
| 26226 | ENSMUSG00000084131 | 0.356960287 | 0.16514264 | 3.880370363 | 0.042558474 | 0.966053499 | NA | Rpl3-ps2 | ribosomal protein L3, pseudogene 2 [Sour | 0 | 1.0279 | 0 | 0 | 0 | 1.1138 |
| 26227 | ENSMUSG00000084132 | 8.249102845 | 1.310623573 | 0.862621505 | 1.519349524 | 0.12867454 | 0.524858587 | Gm15982 | predicted gene 15982 [Source:MGI Symbo | 4.0155 | 8.2233 | 2.0125 | 6.7478 | 21.812 | 6.6831 |
| 26228 | ENSMUSG00000084133 | 0 | NA | NA | NA | NA | NA | Gm12856 | predicted gene 12856 [Source:MGI Symbo | 0 | 0 | 0 | 0 | 0 | 0 |
| 26229 | ENSMUSG00000084134 | 0 | NA | NA | NA | NA | NA | Gm13930 | predicted gene 13930 [Source:MGI Symbo | 0 | 0 | 0 | 0 | 0 | 0 |
| 26230 | ENSMUSG00000084135 | 0 | NA | NA | NA | NA | NA | Pom121l12 | POM121 membrane glycoprotein-like 12 [ | 0 | 0 | 0 | 0 | 0 | 0 |
| 26231 | ENSMUSG00000084136 | 0 | NA | NA | NA | NA | NA | Vmn1r207 | vomeronasal 1 receptor 207 [Source:MGI | 0 | 0 | 0 | 0 | 0 | 0 |
| 26232 | ENSMUSG00000084137 | 0.133850373 | -0.796629176 | 4.080472857 | -0.195229623 | 0.84521319 | NA | Gm9085 | predicted gene 9085 [Source:MGI Symbol | 0.8031 | 0 | 0 | 0 | 0 | 0 |
| 26233 | ENSMUSG00000084138 | 0 | NA | NA | NA | NA | NA | Gm14915 | predicted gene 14915 [Source:MGI Symbo | 0 | 0 | 0 | 0 | 0 | 0 |
| 26234 | ENSMUSG00000084139 | 0.880724006 | 0.715724162 | 2.808295084 | 0.25486074 | 0.79883066 | NA | Gm12322 | predicted gene 12322 [Source:MGI Symbo | 0 | 0 | 2.0125 | 0 | 3.2719 | 0 |
| 26235 | ENSMUSG00000084140 | 2.166383334 | -0.548756928 | 1.452133391 | -0.377897052 | 0.705507073 | NA | Gm14000 | predicted gene 14000 [Source:MGI Symbo | 1.6062 | 5.1396 | 1.0062 | 1.928 | 1.0906 | 2.2277 |
| 26236 | ENSMUSG00000084142 | 0.36353943 | 2.065989254 | 4.034137301 | 0.512126658 | 0.608562368 | NA | Gm9103 | predicted gene 9103 [Source:MGI Symbol | 0 | 0 | 0 | 0 | 2.1812 | 0 |
| 26237 | ENSMUSG00000084143 | 0.335414862 | -1.779047242 | 4.030804179 | -0.441362855 | 0.658950329 | NA | Gm13400 | predicted gene 13400 [Source:MGI Symbo | 0 | 0 | 2.0125 | 0 | 0 | 0 |
| 26238 | ENSMUSG00000084144 | 0 | NA | NA | NA | NA | NA | Gm12516 | predicted gene 12516 [Source:MGI Symbo | 0 | 0 | 0 | 0 | 0 | 0 |
| 26239 | ENSMUSG00000084145 | 1.751979046 | -0.57772173 | 1.566421279 | -0.368816319 | 0.71226464 | NA | Gm12263 | predicted gene 12263 [Source:MGI Symbo | 3.2124 | 1.0279 | 2.0125 | 0.964 | 2.1812 | 1.1138 |
| 26240 | ENSMUSG00000084146 | 0.614004368 | 0.315622926 | 2.898299649 | 0.108899342 | 0.913282328 | NA | Gm12104 | predicted gene 12104 [Source:MGI Symbo | 1.6062 | 0 | 0 | 0.964 | 0 | 1.1138 |
| 26241 | ENSMUSG00000084147 | 0 | NA | NA | NA | NA | NA | Gm15075 | predicted gene 15075 [Source:MGI Symbo | 0 | 0 | 0 | 0 | 0 | 0 |
| 26242 | ENSMUSG00000084148 | 0 | NA | NA | NA | NA | NA | Gm12380 | predicted gene 12380 [Source:MGI Symbo | 0 | 0 | 0 | 0 | 0 | 0 |
| 26243 | ENSMUSG00000084149 | 0.346303622 | 2.004339653 | 3.884003423 | 0.516049919 | 0.605819544 | NA | Gm12944 | predicted gene 12944 [Source:MGI Symbo | 0 | 0 | 0 | 0.964 | 0 | 1.1138 |
| 26244 | ENSMUSG00000084150 | 0 | NA | NA | NA | NA | NA | Gm14395 | predicted gene 14395 [Source:MGI Symbo | 0 | 0 | 0 | 0 | 0 | 0 |
| 26245 | ENSMUSG00000084151 | 0 | NA | NA | NA | NA | NA | Gm11577 | predicted gene 11577 [Source:MGI Symbo | 0 | 0 | 0 | 0 | 0 | 0 |
| 26246 | ENSMUSG00000084152 | 0 | NA | NA | NA | NA | NA | Gm11617 | predicted gene 11617 [Source:MGI Symbo | 0 | 0 | 0 | 0 | 0 | 0 |
| 26247 | ENSMUSG00000084153 | 0.695522043 | 1.639439662 | 2.762392557 | 0.593485404 | 0.552856349 | NA | Gm12186 | predicted gene 12186 [Source:MGI Symbo | 0 | 1.0279 | 0 | 0.964 | 2.1812 | 0 |
| 26248 | ENSMUSG00000084154 | 0 | NA | NA | NA | NA | NA | Gm15644 | predicted gene 15644 [Source:MGI Symbo | 0 | 0 | 0 | 0 | 0 | 0 |
| 26249 | ENSMUSG00000084155 | 0 | NA | NA | NA | NA | NA | Gm7113 | predicted gene 7113 [Source:MGI Symbol | 0 | 0 | 0 | 0 | 0 | 0 |
| 26250 | ENSMUSG00000084157 | 0 | NA | NA | NA | NA | NA | Gm15576 | predicted gene 15576 [Source:MGI Symbo | 0 | 0 | 0 | 0 | 0 | 0 |
| 26251 | ENSMUSG00000084158 | 0 | NA | NA | NA | NA | NA | Gm14652 | predicted gene 14652 [Source:MGI Symbo | 0 | 0 | 0 | 0 | 0 | 0 |
| 26252 | ENSMUSG00000084159 | 0.997286438 | -2.098554371 | 2.276243307 | -0.921937635 | 0.356561107 | NA | Gm12696 | predicted gene 12696 [Source:MGI Symbo | 0.8031 | 3.0838 | 1.0062 | 0 | 1.0906 | 0 |
| 26253 | ENSMUSG00000084160 | 0 | NA | NA | NA | NA | NA | Gm13345 | predicted gene 13345 [Source:MGI Symbo | 0 | 0 | 0 | 0 | 0 | 0 |
| 26254 | ENSMUSG00000084161 | 0 | NA | NA | NA | NA | NA | Gm15061 | predicted gene 15061 [Source:MGI Symbo | 0 | 0 | 0 | 0 | 0 | 0 |
| 26255 | ENSMUSG00000084162 | 0 | NA | NA | NA | NA | NA | Gm11251 | predicted gene 11251 [Source:MGI Symbo | 0 | 0 | 0 | 0 | 0 | 0 |
| 26256 | ENSMUSG00000084163 | 0 | NA | NA | NA | NA | NA | Gm11901 | predicted gene 11901 [Source:MGI Symbo | 0 | 0 | 0 | 0 | 0 | 0 |
| 26257 | ENSMUSG00000084164 | 0 | NA | NA | NA | NA | NA | Gm13902 | predicted gene 13902 [Source:MGI Symbo | 0 | 0 | 0 | 0 | 0 | 0 |
| 26258 | ENSMUSG00000084166 | 8.76960589 | 0.175325832 | 0.733826411 | 0.238920035 | 0.81116759 | 0.966452061 | Gm6451 | predicted gene 6451 [Source:MGI Symbol | 10.44 | 8.2233 | 6.0375 | 12.532 | 9.8156 | 5.5692 |
| 26259 | ENSMUSG00000084167 | 0 | NA | NA | NA | NA | NA | Gm13924 | predicted gene 13924 [Source:MGI Symbo | 0 | 0 | 0 | 0 | 0 | 0 |
| 26260 | ENSMUSG00000084168 | 1.371069352 | 1.803135641 | 2.08749324 | 0.86378035 | 0.387708565 | NA | Atp5l2-ps | ATP synthase, H+ transporting, mitochonc | 0.8031 | 0 | 1.0062 | 0.964 | 5.4531 | 0 |

| | | | | | | | | | | | | | | | |
|---|---|---|---|---|---|---|---|---|---|---|---|---|---|---|---|
| 26261 | ENSMUSG00000084169 | 0.335414862 | -1.779047242 | 4.030804179 | -0.441362855 | 0.658950329 | NA | Gm12240 | predicted gene 12240 [Source:MGI Symbo | 0 | 0 | 2.0125 | 0 | 0 | 0 |
| 26262 | ENSMUSG00000084170 | 0 | NA | NA | NA | NA | NA | Gm11954 | predicted gene 11954 [Source:MGI Symbo | 0 | 0 | 0 | 0 | 0 | 0 |
| 26263 | ENSMUSG00000084171 | 0 | NA | NA | NA | NA | NA | Gm13457 | predicted gene 13457 [Source:MGI Symbo | 0 | 0 | 0 | 0 | 0 | 0 |
| 26264 | ENSMUSG00000084172 | 0 | NA | NA | NA | NA | NA | Gm16420 | predicted gene 16420 [Source:MGI Symbo | 0 | 0 | 0 | 0 | 0 | 0 |
| 26265 | ENSMUSG00000084173 | 0 | NA | NA | NA | NA | NA | Gm5287 | predicted gene 5287 [Source:MGI Symbol | 0 | 0 | 0 | 0 | 0 | 0 |
| 26266 | ENSMUSG00000084174 | 0 | NA | NA | NA | NA | NA | Sycn | syncollin [Source:MGI Symbol;Acc:MGI:19 | 0 | 0 | 0 | 0 | 0 | 0 |
| 26267 | ENSMUSG00000084175 | 0 | NA | NA | NA | NA | NA | Gm11372 | predicted gene 11372 [Source:MGI Symbo | 0 | 0 | 0 | 0 | 0 | 0 |
| 26268 | ENSMUSG00000084177 | 0 | NA | NA | NA | NA | NA | Gm13508 | predicted gene 13508 [Source:MGI Symbo | 0 | 0 | 0 | 0 | 0 | 0 |
| 26269 | ENSMUSG00000084178 | 0 | NA | NA | NA | NA | NA | Gm11759 | predicted gene 11759 [Source:MGI Symbo | 0 | 0 | 0 | 0 | 0 | 0 |
| 26270 | ENSMUSG00000084179 | 0 | NA | NA | NA | NA | NA | Gm11890 | predicted gene 11890 [Source:MGI Symbo | 0 | 0 | 0 | 0 | 0 | 0 |
| 26271 | ENSMUSG00000084180 | 0 | NA | NA | NA | NA | NA | Gm15237 | predicted gene 15237 [Source:MGI Symbo | 0 | 0 | 0 | 0 | 0 | 0 |
| 26272 | ENSMUSG00000084181 | 0 | NA | NA | NA | NA | NA | Gm11858 | predicted gene 11858 [Source:MGI Symbo | 0 | 0 | 0 | 0 | 0 | 0 |
| 26273 | ENSMUSG00000084182 | 0 | NA | NA | NA | NA | NA | Taar7c-ps | trace amine-associated receptor 7C, pseu | 0 | 0 | 0 | 0 | 0 | 0 |
| 26274 | ENSMUSG00000084183 | 0 | NA | NA | NA | NA | NA | Gm12009 | predicted gene 12009 [Source:MGI Symbo | 0 | 0 | 0 | 0 | 0 | 0 |
| 26275 | ENSMUSG00000084184 | 0 | NA | NA | NA | NA | NA | Gm13474 | predicted gene 13474 [Source:MGI Symbo | 0 | 0 | 0 | 0 | 0 | 0 |
| 26276 | ENSMUSG00000084185 | 0 | NA | NA | NA | NA | NA | Rplp1-ps1 | ribosomal protein, large, P1, pseudogene | 0 | 0 | 0 | 0 | 0 | 0 |
| 26277 | ENSMUSG00000084186 | 0 | NA | NA | NA | NA | NA | Rpl36-ps10 | ribosomal protein L36, pseudogene 10 [So | 0 | 0 | 0 | 0 | 0 | 0 |
| 26278 | ENSMUSG00000084188 | 0 | NA | NA | NA | NA | NA | Gm12851 | predicted gene 12851 [Source:MGI Symbo | 0 | 0 | 0 | 0 | 0 | 0 |
| 26279 | ENSMUSG00000084189 | 0 | NA | NA | NA | NA | NA | Gm8089 | predicted gene 8089 [Source:MGI Symbol | 0 | 0 | 0 | 0 | 0 | 0 |
| 26280 | ENSMUSG00000084190 | 0.481988923 | 2.492401104 | 3.837077151 | 0.649557203 | 0.515978285 | NA | Gm11869 | predicted gene 11869 [Source:MGI Symbo | 0 | 0 | 0 | 2.8919 | 0 | 0 |
| 26281 | ENSMUSG00000084191 | 0 | NA | NA | NA | NA | NA | Gm15167 | predicted gene 15167 [Source:MGI Symbo | 0 | 0 | 0 | 0 | 0 | 0 |
| 26282 | ENSMUSG00000084192 | 0 | NA | NA | NA | NA | NA | Gm13282 | predicted gene 13282 [Source:MGI Symbo | 0 | 0 | 0 | 0 | 0 | 0 |
| 26283 | ENSMUSG00000084193 | 0 | NA | NA | NA | NA | NA | Gm6823 | predicted gene 6823 [Source:MGI Symbol | 0 | 0 | 0 | 0 | 0 | 0 |
| 26284 | ENSMUSG00000084195 | 2.011298393 | 0.110962363 | 1.735784681 | 0.063926341 | 0.949028878 | NA | Gm15372 | predicted gene 15372 [Source:MGI Symbo | 0.8031 | 0 | 5.0312 | 2.8919 | 0 | 3.3415 |
| 26285 | ENSMUSG00000084196 | 0 | NA | NA | NA | NA | NA | Gm12874 | predicted gene 12874 [Source:MGI Symbo | 0 | 0 | 0 | 0 | 0 | 0 |
| 26286 | ENSMUSG00000084197 | 0 | NA | NA | NA | NA | NA | Gm9089 | predicted gene 9089 [Source:MGI Symbol | 0 | 0 | 0 | 0 | 0 | 0 |
| 26287 | ENSMUSG00000084198 | 0 | NA | NA | NA | NA | NA | Gm13711 | predicted gene 13711 [Source:MGI Symbo | 0 | 0 | 0 | 0 | 0 | 0 |
| 26288 | ENSMUSG00000084199 | 0 | NA | NA | NA | NA | NA | Gm12422 | predicted gene 12422 [Source:MGI Symbo | 0 | 0 | 0 | 0 | 0 | 0 |
| 26289 | ENSMUSG00000084200 | 0 | NA | NA | NA | NA | NA | Gm11584 | predicted gene 11584 [Source:MGI Symbo | 0 | 0 | 0 | 0 | 0 | 0 |
| 26290 | ENSMUSG00000084201 | 0 | NA | NA | NA | NA | NA | Gm11926 | predicted gene 11926 [Source:MGI Symbo | 0 | 0 | 0 | 0 | 0 | 0 |
| 26291 | ENSMUSG00000084202 | 0 | NA | NA | NA | NA | NA | Gm5398 | predicted gene 5398 [Source:MGI Symbol | 0 | 0 | 0 | 0 | 0 | 0 |
| 26292 | ENSMUSG00000084203 | 0 | NA | NA | NA | NA | NA | Gm15593 | predicted gene 15593 [Source:MGI Symbo | 0 | 0 | 0 | 0 | 0 | 0 |
| 26293 | ENSMUSG00000084204 | 0 | NA | NA | NA | NA | NA | Gm6584 | predicted gene 6584 [Source:MGI Symbol | 0 | 0 | 0 | 0 | 0 | 0 |
| 26294 | ENSMUSG00000084205 | 0 | NA | NA | NA | NA | NA | Gm8061 | predicted gene 8061 [Source:MGI Symbol | 0 | 0 | 0 | 0 | 0 | 0 |
| 26295 | ENSMUSG00000084206 | 0 | NA | NA | NA | NA | NA | Gm12062 | predicted gene 12062 [Source:MGI Symbo | 0 | 0 | 0 | 0 | 0 | 0 |
| 26296 | ENSMUSG00000084207 | 0 | NA | NA | NA | NA | NA | Gm14070 | predicted gene 14070 [Source:MGI Symbo | 0 | 0 | 0 | 0 | 0 | 0 |
| 26297 | ENSMUSG00000084208 | 0 | NA | NA | NA | NA | NA | Gm12792 | predicted gene 12792 [Source:MGI Symbo | 0 | 0 | 0 | 0 | 0 | 0 |
| 26298 | ENSMUSG00000084209 | 0 | NA | NA | NA | NA | NA | Gm13811 | predicted gene 13811 [Source:MGI Symbo | 0 | 0 | 0 | 0 | 0 | 0 |
| 26299 | ENSMUSG00000084210 | 0 | NA | NA | NA | NA | NA | Gm15168 | predicted gene 15168 [Source:MGI Symbo | 0 | 0 | 0 | 0 | 0 | 0 |
| 26300 | ENSMUSG00000084211 | 0 | NA | NA | NA | NA | NA | Gm11949 | predicted gene 11949 [Source:MGI Symbo | 0 | 0 | 0 | 0 | 0 | 0 |
| 26301 | ENSMUSG00000084212 | 0 | NA | NA | NA | NA | NA | Gm12225 | predicted gene 12225 [Source:MGI Symbo | 0 | 0 | 0 | 0 | 0 | 0 |
| 26302 | ENSMUSG00000084213 | 0.17131964 | -0.796629176 | 4.080472857 | -0.195229623 | 0.84521319 | NA | Hspe1-ps5 | heat shock protein 1 (chaperonin 10), pse | 0 | 1.0279 | 0 | 0 | 0 | 0 |
| 26303 | ENSMUSG00000084214 | 0 | NA | NA | NA | NA | NA | Gm14825 | predicted gene 14825 [Source:MGI Symbo | 0 | 0 | 0 | 0 | 0 | 0 |
| 26304 | ENSMUSG00000084215 | 0.185640647 | 1.126924175 | 4.080472857 | 0.276174898 | 0.782413736 | NA | Gm11814 | predicted gene 11814 [Source:MGI Symbo | 0 | 0 | 0 | 0 | 0 | 1.1138 |
| 26305 | ENSMUSG00000084216 | 0 | NA | NA | NA | NA | NA | Btf3-ps3 | basic transcription factor 3, pseudogene 3 | 0 | 0 | 0 | 0 | 0 | 0 |
| 26306 | ENSMUSG00000084217 | 0.692607243 | 3.004183657 | 2.957557172 | 1.015765202 | 0.309741211 | NA | Setd9-ps | SET domain containing 9, pseudogene [So | 0 | 0 | 0 | 1.928 | 0 | 2.2277 |
| 26307 | ENSMUSG00000084218 | 0 | NA | NA | NA | NA | NA | Gm8145 | predicted gene 8145 [Source:MGI Symbol | 0 | 0 | 0 | 0 | 0 | 0 |
| 26308 | ENSMUSG00000084219 | 0 | NA | NA | NA | NA | NA | Gm14876 | predicted gene 14876 [Source:MGI Symbo | 0 | 0 | 0 | 0 | 0 | 0 |
| 26309 | ENSMUSG00000084220 | 0 | NA | NA | NA | NA | NA | Gm12387 | predicted gene 12387 [Source:MGI Symbo | 0 | 0 | 0 | 0 | 0 | 0 |
| 26310 | ENSMUSG00000084221 | 0 | NA | NA | NA | NA | NA | Hmgb1-ps4 | high mobility group box 1, pseudogene 4 [ | 0 | 0 | 0 | 0 | 0 | 0 |
| 26311 | ENSMUSG00000084222 | 0 | NA | NA | NA | NA | NA | Gm13973 | predicted gene 13973 [Source:MGI Symbo | 0 | 0 | 0 | 0 | 0 | 0 |
| 26312 | ENSMUSG00000084223 | 0 | NA | NA | NA | NA | NA | Gm12244 | predicted gene 12244 [Source:MGI Symbo | 0 | 0 | 0 | 0 | 0 | 0 |
| 26313 | ENSMUSG00000084224 | 10.6829137 | 1.19794722 | 0.726072429 | 1.64990044 | 0.098963301 | 0.463570426 | Gm12337 | predicted gene 12337 [Source:MGI Symbo | 12.047 | 3.0838 | 4.025 | 17.352 | 11.997 | 15.594 |
| 26314 | ENSMUSG00000084226 | 0 | NA | NA | NA | NA | NA | Gm13426 | predicted gene 13426 [Source:MGI Symbo | 0 | 0 | 0 | 0 | 0 | 0 |
| 26315 | ENSMUSG00000084228 | 0 | NA | NA | NA | NA | NA | Gm8080 | predicted gene 8080 [Source:MGI Symbol | 0 | 0 | 0 | 0 | 0 | 0 |
| 26316 | ENSMUSG00000084230 | 1.117100031 | 1.582896982 | 2.163697584 | 0.731570342 | 0.464430856 | NA | Gm14388 | predicted gene 14388 [Source:MGI Symbo | 1.6062 | 0 | 0 | 2.8919 | 1.0906 | 1.1138 |

| | | | | | | | | | | | | | | | |
|---|---|---|---|---|---|---|---|---|---|---|---|---|---|---|---|
| 26317 | ENSMUSG00000084231 | 0.160662974 | 1.126924175 | 4.080472857 | 0.276174898 | 0.782413736 | NA | Gm11975 | predicted gene 11975 [Source:MGI Symbo | 0 | 0 | 0 | 0.964 | 0 | 0 |
| 26318 | ENSMUSG00000084234 | 0 | NA | NA | NA | NA | NA | 4933405O20 | RIKEN cDNA 4933405O20 gene [Source:M | 0 | 0 | 0 | 0 | 0 | 0 |
| 26319 | ENSMUSG00000084235 | 1.277737502 | -2.558404199 | 2.061944144 | -1.240772795 | 0.214689694 | NA | Gm15421 | predicted gene 15421 [Source:MGI Symbo | 1.6062 | 3.0838 | 2.0125 | 0.964 | 0 | 0 |
| 26320 | ENSMUSG00000084236 | 0 | NA | NA | NA | NA | NA | Olfr637-ps1 | olfactory receptor 637, pseudogene 1 [Sou | 0 | 0 | 0 | 0 | 0 | 0 |
| 26321 | ENSMUSG00000084237 | 0 | NA | NA | NA | NA | NA | Gm11781 | predicted gene 11781 [Source:MGI Symbo | 0 | 0 | 0 | 0 | 0 | 0 |
| 26322 | ENSMUSG00000084238 | 0 | NA | NA | NA | NA | NA | Gm13076 | predicted gene 13076 [Source:MGI Symbo | 0 | 0 | 0 | 0 | 0 | 0 |
| 26323 | ENSMUSG00000084239 | 0 | NA | NA | NA | NA | NA | Gm14463 | predicted gene 14463 [Source:MGI Symbo | 0 | 0 | 0 | 0 | 0 | 0 |
| 26324 | ENSMUSG00000084240 | 0 | NA | NA | NA | NA | NA | Gm15383 | predicted gene 15383 [Source:MGI Symbo | 0 | 0 | 0 | 0 | 0 | 0 |
| 26325 | ENSMUSG00000084241 | 0 | NA | NA | NA | NA | NA | Gm13416 | predicted gene 13416 [Source:MGI Symbo | 0 | 0 | 0 | 0 | 0 | 0 |
| 26326 | ENSMUSG00000084243 | 0.331982614 | 0.16514309 | 3.986751476 | 0.041422971 | 0.966958701 | NA | Gm13784 | predicted gene 13784 [Source:MGI Symbo | 0 | 1.0279 | 0 | 0.964 | 0 | 0 |
| 26327 | ENSMUSG00000084244 | 0 | NA | NA | NA | NA | NA | Gm14020 | predicted gene 14020 [Source:MGI Symbo | 0 | 0 | 0 | 0 | 0 | 0 |
| 26328 | ENSMUSG00000084245 | 0 | NA | NA | NA | NA | NA | Gm14994 | predicted gene 14994 [Source:MGI Symbo | 0 | 0 | 0 | 0 | 0 | 0 |
| 26329 | ENSMUSG00000084246 | 0 | NA | NA | NA | NA | NA | Gm15162 | predicted gene 15162 [Source:MGI Symbo | 0 | 0 | 0 | 0 | 0 | 0 |
| 26330 | ENSMUSG00000084247 | 0 | NA | NA | NA | NA | NA | Gm14566 | predicted gene 14566 [Source:MGI Symbo | 0 | 0 | 0 | 0 | 0 | 0 |
| 26331 | ENSMUSG00000084248 | 0 | NA | NA | NA | NA | NA | Gm11409 | predicted gene 11409 [Source:MGI Symbo | 0 | 0 | 0 | 0 | 0 | 0 |
| 26332 | ENSMUSG00000084249 | 0.481988923 | 2.492401104 | 3.837077151 | 0.649557203 | 0.515978285 | NA | Gm12267 | predicted gene 12267 [Source:MGI Symbo | 0 | 0 | 0 | 2.8919 | 0 | 0 |
| 26333 | ENSMUSG00000084250 | 0 | NA | NA | NA | NA | NA | Gm13550 | predicted gene 13550 [Source:MGI Symbo | 0 | 0 | 0 | 0 | 0 | 0 |
| 26334 | ENSMUSG00000084251 | 0 | NA | NA | NA | NA | NA | Gm12310 | predicted gene 12310 [Source:MGI Symbo | 0 | 0 | 0 | 0 | 0 | 0 |
| 26335 | ENSMUSG00000084252 | 0.499690045 | -0.82080011 | 3.113553401 | -0.263621658 | 0.792071468 | NA | Gm12036 | predicted gene 12036 [Source:MGI Symbo | 0 | 1.0279 | 1.0062 | 0.964 | 0 | 0 |
| 26336 | ENSMUSG00000084253 | 0 | NA | NA | NA | NA | NA | Gm11256 | predicted gene 11256 [Source:MGI Symbo | 0 | 0 | 0 | 0 | 0 | 0 |
| 26337 | ENSMUSG00000084254 | 0 | NA | NA | NA | NA | NA | Defa-ps11 | defensin, alpha, pseudogene 11 [Source:N | 0 | 0 | 0 | 0 | 0 | 0 |
| 26338 | ENSMUSG00000084256 | 0 | NA | NA | NA | NA | NA | Gm14976 | predicted gene 14976 [Source:MGI Symbo | 0 | 0 | 0 | 0 | 0 | 0 |
| 26339 | ENSMUSG00000084257 | 0 | NA | NA | NA | NA | NA | Gm11597 | predicted gene 11597 [Source:MGI Symbo | 0 | 0 | 0 | 0 | 0 | 0 |
| 26340 | ENSMUSG00000084259 | 0 | NA | NA | NA | NA | NA | Gm12040 | predicted gene 12040 [Source:MGI Symbo | 0 | 0 | 0 | 0 | 0 | 0 |
| 26341 | ENSMUSG00000084260 | 0.505131668 | 1.126696199 | 3.421159819 | 0.329331647 | 0.741905026 | NA | Gm12844 | predicted gene 12844 [Source:MGI Symbo | 0.8031 | 0 | 0 | 0 | 0 | 2.2277 |
| 26342 | ENSMUSG00000084261 | 0 | NA | NA | NA | NA | NA | Rpl36a-ps4 | ribosomal protein L36A, pseudogene 4 [So | 0 | 0 | 0 | 0 | 0 | 0 |
| 26343 | ENSMUSG00000084262 | 0 | NA | NA | NA | NA | NA | Olfr588-ps1 | olfactory receptor 588, pseudogene 1 [Sou | 0 | 0 | 0 | 0 | 0 | 0 |
| 26344 | ENSMUSG00000084264 | 0 | NA | NA | NA | NA | NA | Gm14142 | predicted gene 14142 [Source:MGI Symbo | 0 | 0 | 0 | 0 | 0 | 0 |
| 26345 | ENSMUSG00000084265 | 0 | NA | NA | NA | NA | NA | Gm14541 | predicted gene 14541 [Source:MGI Symbo | 0 | 0 | 0 | 0 | 0 | 0 |
| 26346 | ENSMUSG00000084267 | 0 | NA | NA | NA | NA | NA | Gm12417 | predicted gene 12417 [Source:MGI Symbo | 0 | 0 | 0 | 0 | 0 | 0 |
| 26347 | ENSMUSG00000084269 | 0 | NA | NA | NA | NA | NA | Gm14784 | predicted gene 14784 [Source:MGI Symbo | 0 | 0 | 0 | 0 | 0 | 0 |
| 26348 | ENSMUSG00000084270 | 0 | NA | NA | NA | NA | NA | Gm15177 | predicted gene 15177 [Source:MGI Symbo | 0 | 0 | 0 | 0 | 0 | 0 |
| 26349 | ENSMUSG00000084271 | 0 | NA | NA | NA | NA | NA | Gm14846 | predicted gene 14846 [Source:MGI Symbo | 0 | 0 | 0 | 0 | 0 | 0 |
| 26350 | ENSMUSG00000084272 | 0 | NA | NA | NA | NA | NA | Gm13224 | predicted gene 13224 [Source:MGI Symbo | 0 | 0 | 0 | 0 | 0 | 0 |
| 26351 | ENSMUSG00000084273 | 0 | NA | NA | NA | NA | NA | Gm12628 | predicted gene 12628 [Source:MGI Symbo | 0 | 0 | 0 | 0 | 0 | 0 |
| 26352 | ENSMUSG00000084274 | 0.349477146 | 0.165142768 | 3.911021834 | 0.042224967 | 0.966319359 | NA | Ptma-ps2 | prothymosin alpha, pseudogene 2 [Source | 0 | 0 | 1.0062 | 0 | 1.0906 | 0 |
| 26353 | ENSMUSG00000084275 | 0 | NA | NA | NA | NA | NA | Gm5396 | predicted pseudogene 5396 [Source:MGI | 0 | 0 | 0 | 0 | 0 | 0 |
| 26354 | ENSMUSG00000084276 | 0 | NA | NA | NA | NA | NA | Gm8317 | predicted gene 8317 [Source:MGI Symbol | 0 | 0 | 0 | 0 | 0 | 0 |
| 26355 | ENSMUSG00000084277 | 0 | NA | NA | NA | NA | NA | Gm11702 | predicted gene 11702 [Source:MGI Symbo | 0 | 0 | 0 | 0 | 0 | 0 |
| 26356 | ENSMUSG00000084278 | 0 | NA | NA | NA | NA | NA | Gm12189 | predicted gene 12189 [Source:MGI Symbo | 0 | 0 | 0 | 0 | 0 | 0 |
| 26357 | ENSMUSG00000084279 | 0 | NA | NA | NA | NA | NA | Gm13232 | predicted gene 13232 [Source:MGI Symbo | 0 | 0 | 0 | 0 | 0 | 0 |
| 26358 | ENSMUSG00000084280 | 0 | NA | NA | NA | NA | NA | Gm11972 | predicted gene 11972 [Source:MGI Symbo | 0 | 0 | 0 | 0 | 0 | 0 |
| 26359 | ENSMUSG00000084281 | 0 | NA | NA | NA | NA | NA | Gm14770 | predicted gene 14770 [Source:MGI Symbo | 0 | 0 | 0 | 0 | 0 | 0 |
| 26360 | ENSMUSG00000084282 | 0 | NA | NA | NA | NA | NA | Gm15054 | predicted gene 15054 [Source:MGI Symbo | 0 | 0 | 0 | 0 | 0 | 0 |
| 26361 | ENSMUSG00000084283 | 0 | NA | NA | NA | NA | NA | Gm14914 | predicted gene 14914 [Source:MGI Symbo | 0 | 0 | 0 | 0 | 0 | 0 |
| 26362 | ENSMUSG00000084284 | 1.681014297 | 0.194163177 | 1.69802288 | 0.11434662 | 0.908963027 | NA | Tpt1-ps6 | tumor protein, translationally-controlled, | 1.6062 | 3.0838 | 0 | 0.964 | 1.0906 | 3.3415 |
| 26363 | ENSMUSG00000084285 | 0 | NA | NA | NA | NA | NA | Gm13451 | predicted gene 13451 [Source:MGI Symbo | 0 | 0 | 0 | 0 | 0 | 0 |
| 26364 | ENSMUSG00000084286 | 0 | NA | NA | NA | NA | NA | Cyp2j15-ps | cytochrome P450, family 2, subfamily j, m | 0 | 0 | 0 | 0 | 0 | 0 |
| 26365 | ENSMUSG00000084287 | 0 | NA | NA | NA | NA | NA | Gm14576 | predicted gene 14576 [Source:MGI Symbo | 0 | 0 | 0 | 0 | 0 | 0 |
| 26366 | ENSMUSG00000084288 | 0.371281295 | 2.09253362 | 4.031852165 | 0.519000582 | 0.603760335 | NA | Mif-ps8 | macrophage migration inhibitory factor, p | 0 | 0 | 0 | 0 | 0 | 2.2277 |
| 26367 | ENSMUSG00000084289 | 0 | NA | NA | NA | NA | NA | Fth1-ps | ferritin heavy polypeptide 1, pseudogene | 0 | 0 | 0 | 0 | 0 | 0 |
| 26368 | ENSMUSG00000084291 | 0 | NA | NA | NA | NA | NA | Gm6654 | predicted pseudogene 6654 [Source:MGI | 0 | 0 | 0 | 0 | 0 | 0 |
| 26369 | ENSMUSG00000084292 | 1.168192897 | 1.507867275 | 2.134947591 | 0.706278356 | 0.48001505 | NA | Gm13535 | predicted gene 13535 [Source:MGI Symbo | 0.8031 | 0 | 1.0062 | 1.928 | 3.2719 | 0 |
| 26370 | ENSMUSG00000084293 | 0 | NA | NA | NA | NA | NA | Gm12143 | predicted gene 12143 [Source:MGI Symbo | 0 | 0 | 0 | 0 | 0 | 0 |
| 26371 | ENSMUSG00000084294 | 0 | NA | NA | NA | NA | NA | Gm13533 | predicted gene 13533 [Source:MGI Symbo | 0 | 0 | 0 | 0 | 0 | 0 |
| 26372 | ENSMUSG00000084295 | 0 | NA | NA | NA | NA | NA | Olfr1235-ps1 | olfactory receptor 1235, pseudogene 1 [So | 0 | 0 | 0 | 0 | 0 | 0 |

| | | | | | | | | | | | | | | | |
|---|---|---|---|---|---|---|---|---|---|---|---|---|---|---|---|
| 26373 | ENSMUSG00000084296 | 0 | NA | NA | NA | NA | NA | Scgb2b25-ps | secretoglobin, family 2B, member 25, pseu | 0 | 0 | 0 | 0 | 0 | 0 |
| 26374 | ENSMUSG00000084297 | 0 | NA | NA | NA | NA | NA | Gm13755 | predicted gene 13755 [Source:MGI Symbo | 0 | 0 | 0 | 0 | 0 | 0 |
| 26375 | ENSMUSG00000084298 | 0 | NA | NA | NA | NA | NA | Gm14331 | predicted gene 14331 [Source:MGI Symbo | 0 | 0 | 0 | 0 | 0 | 0 |
| 26376 | ENSMUSG00000084299 | 0 | NA | NA | NA | NA | NA | Gm6630 | predicted pseudogene 6630 [Source:MGI | 0 | 0 | 0 | 0 | 0 | 0 |
| 26377 | ENSMUSG00000084300 | 0 | NA | NA | NA | NA | NA | Gm6954 | predicted pseudogene 6954 [Source:MGI | 0 | 0 | 0 | 0 | 0 | 0 |
| 26378 | ENSMUSG00000084301 | 0 | NA | NA | NA | NA | NA | Gm7856 | predicted gene 7856 [Source:MGI Symbol | 0 | 0 | 0 | 0 | 0 | 0 |
| 26379 | ENSMUSG00000084302 | 0.36353943 | 2.065989254 | 4.034137301 | 0.512126658 | 0.608562368 | NA | Gm12194 | predicted gene 12194 [Source:MGI Symbo | 0 | 0 | 0 | 0 | 2.1812 | 0 |
| 26380 | ENSMUSG00000084303 | 0 | NA | NA | NA | NA | NA | Gm8216 | predicted gene 8216 [Source:MGI Symbol | 0 | 0 | 0 | 0 | 0 | 0 |
| 26381 | ENSMUSG00000084304 | 1.671753875 | 0.709685837 | 1.595041209 | 0.444932603 | 0.65636846 | NA | Gm6142 | predicted pseudogene 6142 [Source:MGI | 0.8031 | 1.0279 | 2.0125 | 2.8919 | 2.1812 | 1.1138 |
| 26382 | ENSMUSG00000084305 | 0 | NA | NA | NA | NA | NA | Gm14645 | predicted gene 14645 [Source:MGI Symbo | 0 | 0 | 0 | 0 | 0 | 0 |
| 26383 | ENSMUSG00000084306 | 0.496077836 | -0.812642266 | 3.444120572 | -0.235950586 | 0.813471014 | NA | Gm11743 | predicted gene 11743 [Source:MGI Symbo | 0 | 0 | 2.0125 | 0.964 | 0 | 0 |
| 26384 | ENSMUSG00000084307 | 0 | NA | NA | NA | NA | NA | Gm12151 | predicted gene 12151 [Source:MGI Symbo | 0 | 0 | 0 | 0 | 0 | 0 |
| 26385 | ENSMUSG00000084308 | 0 | NA | NA | NA | NA | NA | Gm14593 | predicted gene 14593 [Source:MGI Symbo | 0 | 0 | 0 | 0 | 0 | 0 |
| 26386 | ENSMUSG00000084309 | 38.42360438 | 3.969008464 | 1.126990782 | 3.521775447 | 0.000428667 | 0.01119177 | Mup-ps20 | major urinary protein, pseudogene 20 [So | 0.8031 | 5.1396 | 8.05 | 12.532 | 95.974 | 108.04 |
| 26387 | ENSMUSG00000084310 | 0 | NA | NA | NA | NA | NA | Gm11950 | predicted gene 11950 [Source:MGI Symbo | 0 | 0 | 0 | 0 | 0 | 0 |
| 26388 | ENSMUSG00000084311 | 0 | NA | NA | NA | NA | NA | Gm11257 | predicted gene 11257 [Source:MGI Symbo | 0 | 0 | 0 | 0 | 0 | 0 |
| 26389 | ENSMUSG00000084312 | 0 | NA | NA | NA | NA | NA | Gm13122 | predicted gene 13122 [Source:MGI Symbo | 0 | 0 | 0 | 0 | 0 | 0 |
| 26390 | ENSMUSG00000084313 | 0 | NA | NA | NA | NA | NA | Gm13809 | predicted gene 13809 [Source:MGI Symbo | 0 | 0 | 0 | 0 | 0 | 0 |
| 26391 | ENSMUSG00000084314 | 0 | NA | NA | NA | NA | NA | Rps15a-ps3 | ribosomal protein S15A, pseudogene 3 [So | 0 | 0 | 0 | 0 | 0 | 0 |
| 26392 | ENSMUSG00000084315 | 0 | NA | NA | NA | NA | NA | Vmn1r-ps12 | vomeronasal 1 receptor, pseudogene 128 | 0 | 0 | 0 | 0 | 0 | 0 |
| 26393 | ENSMUSG00000084316 | 0 | NA | NA | NA | NA | NA | Gm8668 | predicted gene 8668 [Source:MGI Symbol | 0 | 0 | 0 | 0 | 0 | 0 |
| 26394 | ENSMUSG00000084318 | 0 | NA | NA | NA | NA | NA | Gm11987 | predicted gene 11987 [Source:MGI Symbo | 0 | 0 | 0 | 0 | 0 | 0 |
| 26395 | ENSMUSG00000084319 | 115.5582092 | -0.122490588 | 0.219150542 | -0.558933536 | 0.576207081 | 0.90747596 | Tpt1-ps3 | tumor protein, translationally-controlled, | 113.24 | 130.55 | 117.73 | 108.93 | 98.156 | 124.75 |
| 26396 | ENSMUSG00000084320 | 0 | NA | NA | NA | NA | NA | Gm14125 | predicted gene 14125 [Source:MGI Symbo | 0 | 0 | 0 | 0 | 0 | 0 |
| 26397 | ENSMUSG00000084323 | 0.17131964 | -0.796629176 | 4.080472857 | -0.195229623 | 0.84521319 | NA | Gm14438 | predicted gene 14438 [Source:MGI Symbo | 0 | 1.0279 | 0 | 0 | 0 | 0 |
| 26398 | ENSMUSG00000084324 | 0 | NA | NA | NA | NA | NA | Gm6500 | predicted gene 6500 [Source:MGI Symbol | 0 | 0 | 0 | 0 | 0 | 0 |
| 26399 | ENSMUSG00000084325 | 0.538988726 | 1.126003851 | 3.343747182 | 0.336749099 | 0.736306047 | NA | Gm15550 | predicted gene 15550 [Source:MGI Symbo | 0 | 0 | 1.0062 | 0 | 0 | 2.2277 |
| 26400 | ENSMUSG00000084326 | 0 | NA | NA | NA | NA | NA | Gm12820 | predicted gene 12820 [Source:MGI Symbo | 0 | 0 | 0 | 0 | 0 | 0 |
| 26401 | ENSMUSG00000084327 | 0 | NA | NA | NA | NA | NA | Gm14531 | predicted gene 14531 [Source:MGI Symbo | 0 | 0 | 0 | 0 | 0 | 0 |
| 26402 | ENSMUSG00000084328 | 0.167707431 | -0.796629176 | 4.080472857 | -0.195229623 | 0.84521319 | NA | Gm14046 | predicted gene 14046 [Source:MGI Symbo | 0 | 0 | 1.0062 | 0 | 0 | 0 |
| 26403 | ENSMUSG00000084329 | 1.353781411 | 0.135944945 | 2.00516922 | 0.067797243 | 0.945947038 | NA | Gm6733 | predicted gene 6733 [Source:MGI Symbol | 0.8031 | 3.0838 | 0 | 0.964 | 3.2719 | 0 |
| 26404 | ENSMUSG00000084330 | 0 | NA | NA | NA | NA | NA | Gm15019 | predicted gene 15019 [Source:MGI Symbo | 0 | 0 | 0 | 0 | 0 | 0 |
| 26405 | ENSMUSG00000084331 | 0 | NA | NA | NA | NA | NA | Gm8163 | predicted gene 8163 [Source:MGI Symbol | 0 | 0 | 0 | 0 | 0 | 0 |
| 26406 | ENSMUSG00000084332 | 0 | NA | NA | NA | NA | NA | Gm8840 | predicted gene 8840 [Source:MGI Symbol | 0 | 0 | 0 | 0 | 0 | 0 |
| 26407 | ENSMUSG00000084333 | 0 | NA | NA | NA | NA | NA | Gm11599 | predicted gene 11599 [Source:MGI Symbo | 0 | 0 | 0 | 0 | 0 | 0 |
| 26408 | ENSMUSG00000084334 | 0 | NA | NA | NA | NA | NA | Gm13494 | predicted gene 13494 [Source:MGI Symbo | 0 | 0 | 0 | 0 | 0 | 0 |
| 26409 | ENSMUSG00000084335 | 0 | NA | NA | NA | NA | NA | Gm12619 | predicted gene 12619 [Source:MGI Symbo | 0 | 0 | 0 | 0 | 0 | 0 |
| 26410 | ENSMUSG00000084336 | 0 | NA | NA | NA | NA | NA | Olfr1269 | olfactory receptor 1269 [Source:MGI Sym | 0 | 0 | 0 | 0 | 0 | 0 |
| 26411 | ENSMUSG00000084337 | 0 | NA | NA | NA | NA | NA | Gm14986 | predicted gene 14986 [Source:MGI Symbo | 0 | 0 | 0 | 0 | 0 | 0 |
| 26412 | ENSMUSG00000084338 | 0 | NA | NA | NA | NA | NA | Gm13531 | predicted gene 13531 [Source:MGI Symbo | 0 | 0 | 0 | 0 | 0 | 0 |
| 26413 | ENSMUSG00000084339 | 0 | NA | NA | NA | NA | NA | Gm6141 | predicted pseudogene 6141 [Source:MGI | 0 | 0 | 0 | 0 | 0 | 0 |
| 26414 | ENSMUSG00000084340 | 0 | NA | NA | NA | NA | NA | Gm14766 | predicted gene 14766 [Source:MGI Symbo | 0 | 0 | 0 | 0 | 0 | 0 |
| 26415 | ENSMUSG00000084341 | 0 | NA | NA | NA | NA | NA | Gm3928 | predicted gene 3928 [Source:MGI Symbol | 0 | 0 | 0 | 0 | 0 | 0 |
| 26416 | ENSMUSG00000084342 | 0 | NA | NA | NA | NA | NA | Gm11224 | predicted gene 11224 [Source:MGI Symbo | 0 | 0 | 0 | 0 | 0 | 0 |
| 26417 | ENSMUSG00000084343 | 0 | NA | NA | NA | NA | NA | Gm16198 | predicted gene 16198 [Source:MGI Symbo | 0 | 0 | 0 | 0 | 0 | 0 |
| 26418 | ENSMUSG00000084344 | 0 | NA | NA | NA | NA | NA | Gm15305 | predicted gene 15305 [Source:MGI Symbo | 0 | 0 | 0 | 0 | 0 | 0 |
| 26419 | ENSMUSG00000084345 | 5.622165542 | 0.408977316 | 0.942421696 | 0.433964241 | 0.664314409 | 0.929301301 | Rpl17-ps4 | ribosomal protein L17, pseudogene 4 [Sou | 2.4093 | 3.0838 | 9.0562 | 4.8199 | 5.4531 | 8.9108 |
| 26420 | ENSMUSG00000084346 | 0 | NA | NA | NA | NA | NA | Cyp4a30b | cytochrome P450, family 4, subfamily a, p | 0 | 0 | 0 | 0 | 0 | 0 |
| 26421 | ENSMUSG00000084347 | 0.328370405 | 0.16514316 | 4.003149536 | 0.041253308 | 0.967093956 | NA | Akt2-ps | thymoma viral proto-oncogene 2, pseudo | 0 | 0 | 1.0062 | 0.964 | 0 | 0 |
| 26422 | ENSMUSG00000084348 | 0.181769715 | 1.126924175 | 4.080472857 | 0.276174898 | 0.782413736 | NA | Gm14608 | predicted gene 14608 [Source:MGI Symbo | 0 | 0 | 0 | 0 | 1.0906 | 0 |
| 26423 | ENSMUSG00000084349 | 19.98220941 | 0.077685553 | 0.574046193 | 0.135329794 | 0.89235113 | 0.982489666 | Rpl3-ps1 | ribosomal protein L3, pseudogene 1 [Sour | 9.6372 | 22.614 | 26.162 | 9.6398 | 23.994 | 27.846 |
| 26424 | ENSMUSG00000084350 | 20.98327497 | -0.116483909 | 0.493270957 | -0.236145891 | 0.813319465 | 0.966802988 | Znf41-ps | ZNF41, pseudogene [Source:MGI Symbol; | 25.699 | 25.698 | 14.087 | 24.099 | 20.722 | 15.594 |
| 26425 | ENSMUSG00000084351 | 0 | NA | NA | NA | NA | NA | Gm13085 | predicted gene 13085 [Source:MGI Symbo | 0 | 0 | 0 | 0 | 0 | 0 |
| 26426 | ENSMUSG00000084352 | 0 | NA | NA | NA | NA | NA | Gm12370 | predicted gene 12370 [Source:MGI Symbo | 0 | 0 | 0 | 0 | 0 | 0 |
| 26427 | ENSMUSG00000084353 | 1.004356399 | 2.372094719 | 2.241352889 | 1.058331658 | 0.289904265 | NA | Gm15452 | predicted gene 15452 [Source:MGI Symbo | 0.8031 | 0 | 0 | 1.928 | 2.1812 | 1.1138 |
| 26428 | ENSMUSG00000084355 | 0 | NA | NA | NA | NA | NA | Gm14792 | predicted gene 14792 [Source:MGI Symbo | 0 | 0 | 0 | 0 | 0 | 0 |

| | | | | | | | | | | | | | | | |
|---|---|---|---|---|---|---|---|---|---|---|---|---|---|---|---|
| 26429 | ENSMUSG00000084356 | 0 | NA | NA | NA | NA | NA | Olfr590-ps1 | olfactory receptor 590, pseudogene 1 [Sou | 0 | 0 | 0 | 0 | 0 | 0 |
| 26430 | ENSMUSG00000084357 | 0 | NA | NA | NA | NA | NA | Gm11268 | predicted gene 11268 [Source:MGI Symbo | 0 | 0 | 0 | 0 | 0 | 0 |
| 26431 | ENSMUSG00000084358 | 0 | NA | NA | NA | NA | NA | Gm11352 | predicted gene 11352 [Source:MGI Symbo | 0 | 0 | 0 | 0 | 0 | 0 |
| 26432 | ENSMUSG00000084359 | 0 | NA | NA | NA | NA | NA | Gm13692 | predicted gene 13692 [Source:MGI Symbo | 0 | 0 | 0 | 0 | 0 | 0 |
| 26433 | ENSMUSG00000084360 | 0 | NA | NA | NA | NA | NA | Gm11797 | predicted gene 11797 [Source:MGI Symbo | 0 | 0 | 0 | 0 | 0 | 0 |
| 26434 | ENSMUSG00000084361 | 0 | NA | NA | NA | NA | NA | Gm8202 | predicted gene 8202 [Source:MGI Symbol | 0 | 0 | 0 | 0 | 0 | 0 |
| 26435 | ENSMUSG00000084362 | 0 | NA | NA | NA | NA | NA | 2610024D14 | RIKEN cDNA 2610024D14 gene [Source:M | 0 | 0 | 0 | 0 | 0 | 0 |
| 26436 | ENSMUSG00000084363 | 0 | NA | NA | NA | NA | NA | Gm2288 | predicted gene 2288 [Source:MGI Symbol | 0 | 0 | 0 | 0 | 0 | 0 |
| 26437 | ENSMUSG00000084366 | 0 | NA | NA | NA | NA | NA | Gm15313 | predicted gene 15313 [Source:MGI Symbo | 0 | 0 | 0 | 0 | 0 | 0 |
| 26438 | ENSMUSG00000084367 | 0 | NA | NA | NA | NA | NA | Gm15142 | predicted gene 15142 [Source:MGI Symbo | 0 | 0 | 0 | 0 | 0 | 0 |
| 26439 | ENSMUSG00000084368 | 0 | NA | NA | NA | NA | NA | Gm11572 | predicted gene 11572 [Source:MGI Symbo | 0 | 0 | 0 | 0 | 0 | 0 |
| 26440 | ENSMUSG00000084369 | 0 | NA | NA | NA | NA | NA | Gm14626 | predicted gene 14626 [Source:MGI Symbo | 0 | 0 | 0 | 0 | 0 | 0 |
| 26441 | ENSMUSG00000084370 | 0 | NA | NA | NA | NA | NA | Gm15442 | predicted gene 15442 [Source:MGI Symbo | 0 | 0 | 0 | 0 | 0 | 0 |
| 26442 | ENSMUSG00000084371 | 0 | NA | NA | NA | NA | NA | Gm14113 | predicted gene 14113 [Source:MGI Symbo | 0 | 0 | 0 | 0 | 0 | 0 |
| 26443 | ENSMUSG00000084372 | 0 | NA | NA | NA | NA | NA | Gm13988 | predicted gene 13988 [Source:MGI Symbo | 0 | 0 | 0 | 0 | 0 | 0 |
| 26444 | ENSMUSG00000084373 | 0 | NA | NA | NA | NA | NA | Rps2-ps4 | ribosomal protein S2, pseudogene 4 [Sour | 0 | 0 | 0 | 0 | 0 | 0 |
| 26445 | ENSMUSG00000084374 | 0 | NA | NA | NA | NA | NA | Gm13255 | predicted gene 13255 [Source:MGI Symbo | 0 | 0 | 0 | 0 | 0 | 0 |
| 26446 | ENSMUSG00000084375 | 0 | NA | NA | NA | NA | NA | Gm11637 | predicted gene 11637 [Source:MGI Symbo | 0 | 0 | 0 | 0 | 0 | 0 |
| 26447 | ENSMUSG00000084376 | 0 | NA | NA | NA | NA | NA | Gm14855 | predicted gene 14855 [Source:MGI Symbo | 0 | 0 | 0 | 0 | 0 | 0 |
| 26448 | ENSMUSG00000084377 | 0 | NA | NA | NA | NA | NA | Gm15302 | predicted gene 15302 [Source:MGI Symbo | 0 | 0 | 0 | 0 | 0 | 0 |
| 26449 | ENSMUSG00000084378 | 0 | NA | NA | NA | NA | NA | Gm15238 | predicted gene 15238 [Source:MGI Symbo | 0 | 0 | 0 | 0 | 0 | 0 |
| 26450 | ENSMUSG00000084379 | 0.884878638 | -1.920491624 | 2.550705096 | -0.75292578 | 0.451494515 | NA | Gm12957 | predicted gene 12957 [Source:MGI Symbo | 3.2124 | 0 | 1.0062 | 0 | 1.0906 | 0 |
| 26451 | ENSMUSG00000084380 | 0 | NA | NA | NA | NA | NA | Tmc4-ps | Tmc4 retrotransposed pseudogene [Sourc | 0 | 0 | 0 | 0 | 0 | 0 |
| 26452 | ENSMUSG00000084381 | 2.290740065 | 2.049373752 | 1.691919377 | 1.211271518 | 0.225791361 | NA | AA413626 | expressed sequence AA413626 [Source:M | 1.6062 | 1.0279 | 0 | 6.7478 | 4.3625 | 0 |
| 26453 | ENSMUSG00000084382 | 0 | NA | NA | NA | NA | NA | Gm3267 | predicted gene 3267 [Source:MGI Symbol | 0 | 0 | 0 | 0 | 0 | 0 |
| 26454 | ENSMUSG00000084383 | 0.778047457 | -3.004661392 | 2.562427823 | -1.172583815 | 0.240962743 | NA | Gm13370 | predicted gene 13370 [Source:MGI Symbo | 1.6062 | 2.0558 | 1.0062 | 0 | 0 | 0 |
| 26455 | ENSMUSG00000084384 | 3.584550351 | -0.955303135 | 1.147002608 | -0.832869192 | 0.404918507 | NA | Gm12251 | predicted gene 12251 [Source:MGI Symbo | 4.0155 | 7.1954 | 3.0187 | 2.8919 | 3.2719 | 1.1138 |
| 26456 | ENSMUSG00000084385 | 0 | NA | NA | NA | NA | NA | Gm13873 | predicted gene 13873 [Source:MGI Symbo | 0 | 0 | 0 | 0 | 0 | 0 |
| 26457 | ENSMUSG00000084386 | 0 | NA | NA | NA | NA | NA | Gm13852 | predicted gene 13852 [Source:MGI Symbo | 0 | 0 | 0 | 0 | 0 | 0 |
| 26458 | ENSMUSG00000084387 | 0 | NA | NA | NA | NA | NA | Olfr379-ps1 | olfactory receptor 379, pseudogene 1 [Sou | 0 | 0 | 0 | 0 | 0 | 0 |
| 26459 | ENSMUSG00000084390 | 0 | NA | NA | NA | NA | NA | Gm15425 | predicted gene 15425 [Source:MGI Symbo | 0 | 0 | 0 | 0 | 0 | 0 |
| 26460 | ENSMUSG00000084391 | 0 | NA | NA | NA | NA | NA | Gm13967 | predicted gene 13967 [Source:MGI Symbo | 0 | 0 | 0 | 0 | 0 | 0 |
| 26461 | ENSMUSG00000084392 | 0.133850373 | -0.796629176 | 4.080472857 | -0.195229623 | 0.84521319 | NA | Rps29-ps | ribosomal protein S29, pseudogene [Sour | 0.8031 | 0 | 0 | 0 | 0 | 0 |
| 26462 | ENSMUSG00000084393 | 0 | NA | NA | NA | NA | NA | Gm14101 | predicted gene 14101 [Source:MGI Symbo | 0 | 0 | 0 | 0 | 0 | 0 |
| 26463 | ENSMUSG00000084394 | 0 | NA | NA | NA | NA | NA | Gm8111 | predicted gene 8111 [Source:MGI Symbol | 0 | 0 | 0 | 0 | 0 | 0 |
| 26464 | ENSMUSG00000084395 | 0 | NA | NA | NA | NA | NA | Gm15087 | predicted gene 15087 [Source:MGI Symbo | 0 | 0 | 0 | 0 | 0 | 0 |
| 26465 | ENSMUSG00000084396 | 0 | NA | NA | NA | NA | NA | Gm14657 | predicted gene 14657 [Source:MGI Symbo | 0 | 0 | 0 | 0 | 0 | 0 |
| 26466 | ENSMUSG00000084397 | 0 | NA | NA | NA | NA | NA | Gm9133 | predicted pseudogene 9133 [Source:MGI | 0 | 0 | 0 | 0 | 0 | 0 |
| 26467 | ENSMUSG00000084399 | 0.487198452 | -0.685759572 | 3.136894395 | -0.218610985 | 0.826953095 | NA | Gm15666 | predicted gene 15666 [Source:MGI Symbo | 0.8031 | 0 | 1.0062 | 0 | 0 | 1.1138 |
| 26468 | ENSMUSG00000084400 | 0.133850373 | -0.796629176 | 4.080472857 | -0.195229623 | 0.84521319 | NA | Gm11576 | predicted gene 11576 [Source:MGI Symbo | 0.8031 | 0 | 0 | 0 | 0 | 0 |
| 26469 | ENSMUSG00000084401 | 0 | NA | NA | NA | NA | NA | Gm12586 | predicted gene 12586 [Source:MGI Symbo | 0 | 0 | 0 | 0 | 0 | 0 |
| 26470 | ENSMUSG00000084403 | 0 | NA | NA | NA | NA | NA | Rps15a-ps8 | ribosomal protein S15A, pseudogene 8 [S | 0 | 0 | 0 | 0 | 0 | 0 |
| 26471 | ENSMUSG00000084404 | 0 | NA | NA | NA | NA | NA | Gm12513 | predicted gene 12513 [Source:MGI Symbo | 0 | 0 | 0 | 0 | 0 | 0 |
| 26472 | ENSMUSG00000084405 | 0 | NA | NA | NA | NA | NA | Olfr1169-ps1 | olfactory receptor 1169, pseudogene 1 [S | 0 | 0 | 0 | 0 | 0 | 0 |
| 26473 | ENSMUSG00000084407 | 11.77933857 | -0.486041561 | 0.736413773 | -0.660011503 | 0.509246448 | 0.882793133 | Gm14018 | predicted gene 14018 [Source:MGI Symbo | 9.6372 | 14.391 | 17.106 | 1.928 | 10.906 | 16.708 |
| 26474 | ENSMUSG00000084408 | 0.538988726 | 1.126003851 | 3.343747182 | 0.336749099 | 0.736306047 | NA | Gm11870 | predicted gene 11870 [Source:MGI Symbo | 0 | 0 | 1.0062 | 0 | 0 | 2.2277 |
| 26475 | ENSMUSG00000084411 | 0.401551119 | -2.072982726 | 4.008251979 | -0.517178744 | 0.605031388 | NA | Gm11510 | predicted gene 11510 [Source:MGI Symbo | 2.4093 | 0 | 0 | 0 | 0 | 0 |
| 26476 | ENSMUSG00000084412 | 0 | NA | NA | NA | NA | NA | Gm12044 | predicted gene 12044 [Source:MGI Symbo | 0 | 0 | 0 | 0 | 0 | 0 |
| 26477 | ENSMUSG00000084413 | 0.535117793 | 1.111049431 | 3.044213722 | 0.364970903 | 0.715133126 | NA | Gm12017 | predicted gene 12017 [Source:MGI Symbo | 0 | 0 | 1.0062 | 0 | 1.0906 | 1.1138 |
| 26478 | ENSMUSG00000084414 | 0.371281295 | 2.09253362 | 4.031852165 | 0.519000582 | 0.603760335 | NA | Gm11270 | predicted gene 11270 [Source:MGI Symbo | 0 | 0 | 0 | 0 | 0 | 2.2277 |
| 26479 | ENSMUSG00000084415 | 0.531944269 | 2.618916564 | 3.285123409 | 0.797204926 | 0.425332026 | NA | Gm12837 | predicted gene 12837 [Source:MGI Symbo | 0 | 0 | 0 | 0.964 | 0 | 2.2277 |
| 26480 | ENSMUSG00000084416 | 16.80286367 | 0.23922566 | 0.532493508 | 0.449255544 | 0.653247324 | 0.925756399 | Rpl10a-ps1 | ribosomal protein L10A, pseudogene 1 [S | 12.85 | 14.391 | 19.119 | 19.28 | 21.812 | 13.366 |
| 26481 | ENSMUSG00000084417 | 0 | NA | NA | NA | NA | NA | Gm23463 | predicted gene, 23463 [Source:MGI Symb | 0 | 0 | 0 | 0 | 0 | 0 |
| 26482 | ENSMUSG00000084418 | 0 | NA | NA | NA | NA | NA | NA | NA | 0 | 0 | 0 | 0 | 0 | 0 |
| 26483 | ENSMUSG00000084419 | 0.160662974 | 1.126924175 | 4.080472857 | 0.276174898 | 0.782413736 | NA | NA | NA | 0 | 0 | 0 | 0.964 | 0 | 0 |
| 26484 | ENSMUSG00000084421 | 0 | NA | NA | NA | NA | NA | Gm25107 | predicted gene, 25107 [Source:MGI Symb | 0 | 0 | 0 | 0 | 0 | 0 |

| | | | | | | | | | | | | | | |
|---|---|---|---|---|---|---|---|---|---|---|---|---|---|---|
| 26485 | ENSMUSG00000084422 | 0.492645588 | 0.9565034 | 3.447648281 | 0.277436479 | 0.781444974 | NA | NA | NA | 0 | 1.0279 | 0 | 1.928 | 0 | 0 |
| 26486 | ENSMUSG00000084423 | 0 | NA | NA | NA | NA | NA | NA | NA | 0 | 0 | 0 | 0 | 0 | 0 |
| 26487 | ENSMUSG00000084425 | 0 | NA | NA | NA | NA | NA | Gm25104 | predicted gene, 25104 [Source:MGI Symb | 0 | 0 | 0 | 0 | 0 | 0 |
| 26488 | ENSMUSG00000084426 | 0 | NA | NA | NA | NA | NA | NA | NA | 0 | 0 | 0 | 0 | 0 | 0 |
| 26489 | ENSMUSG00000084428 | 0 | NA | NA | NA | NA | NA | Gm25102 | predicted gene, 25102 [Source:MGI Symb | 0 | 0 | 0 | 0 | 0 | 0 |
| 26490 | ENSMUSG00000084431 | 0 | NA | NA | NA | NA | NA | NA | NA | 0 | 0 | 0 | 0 | 0 | 0 |
| 26491 | ENSMUSG00000084432 | 0 | NA | NA | NA | NA | NA | NA | NA | 0 | 0 | 0 | 0 | 0 | 0 |
| 26492 | ENSMUSG00000084433 | 3.912105122 | -0.482679719 | 1.411606339 | -0.341936492 | 0.732398687 | NA | NA | NA | 1.6062 | 7.1954 | 5.0312 | 9.6398 | 0 | 0 |
| 26493 | ENSMUSG00000084435 | 0 | NA | NA | NA | NA | NA | Gm23429 | predicted gene, 23429 [Source:MGI Symb | 0 | 0 | 0 | 0 | 0 | 0 |
| 26494 | ENSMUSG00000084437 | 0 | NA | NA | NA | NA | NA | Gm23428 | predicted gene, 23428 [Source:MGI Symb | 0 | 0 | 0 | 0 | 0 | 0 |
| 26495 | ENSMUSG00000084439 | 0 | NA | NA | NA | NA | NA | NA | NA | 0 | 0 | 0 | 0 | 0 | 0 |
| 26496 | ENSMUSG00000084440 | 0.878040772 | -3.188971767 | 2.713192499 | -1.175357727 | 0.239851629 | NA | Gm26254 | predicted gene, 26254 [Source:MGI Symb | 3.2124 | 2.0558 | 0 | 0 | 0 | 0 |
| 26497 | ENSMUSG00000084441 | 0 | NA | NA | NA | NA | NA | Gm26255 | predicted gene, 26255 [Source:MGI Symb | 0 | 0 | 0 | 0 | 0 | 0 |
| 26498 | ENSMUSG00000084442 | 0 | NA | NA | NA | NA | NA | Gm26253 | predicted gene, 26253 [Source:MGI Symb | 0 | 0 | 0 | 0 | 0 | 0 |
| 26499 | ENSMUSG00000084443 | 0 | NA | NA | NA | NA | NA | NA | NA | 0 | 0 | 0 | 0 | 0 | 0 |
| 26500 | ENSMUSG00000084444 | 0 | NA | NA | NA | NA | NA | NA | NA | 0 | 0 | 0 | 0 | 0 | 0 |
| 26501 | ENSMUSG00000084445 | 0 | NA | NA | NA | NA | NA | NA | NA | 0 | 0 | 0 | 0 | 0 | 0 |
| 26502 | ENSMUSG00000084446 | 0 | NA | NA | NA | NA | NA | NA | NA | 0 | 0 | 0 | 0 | 0 | 0 |
| 26503 | ENSMUSG00000084447 | 0 | NA | NA | NA | NA | NA | Gm26251 | predicted gene, 26251 [Source:MGI Symb | 0 | 0 | 0 | 0 | 0 | 0 |
| 26504 | ENSMUSG00000084448 | 0 | NA | NA | NA | NA | NA | Gm26256 | predicted gene, 26256 [Source:MGI Symb | 0 | 0 | 0 | 0 | 0 | 0 |
| 26505 | ENSMUSG00000084449 | 0 | NA | NA | NA | NA | NA | Gm26257 | predicted gene, 26257 [Source:MGI Symb | 0 | 0 | 0 | 0 | 0 | 0 |
| 26506 | ENSMUSG00000084450 | 0 | NA | NA | NA | NA | NA | Gm24595 | predicted gene, 24595 [Source:MGI Symb | 0 | 0 | 0 | 0 | 0 | 0 |
| 26507 | ENSMUSG00000084451 | 0 | NA | NA | NA | NA | NA | NA | NA | 0 | 0 | 0 | 0 | 0 | 0 |
| 26508 | ENSMUSG00000084452 | 17.5221384 | -0.251338411 | 0.633695881 | -0.396623079 | 0.691645431 | 0.938105731 | NA | NA | 18.471 | 16.447 | 22.137 | 8.6758 | 32.719 | 6.6831 |
| 26509 | ENSMUSG00000084453 | 0.342432689 | 1.990653086 | 3.900281263 | 0.51038706 | 0.609780321 | NA | Gm24596 | predicted gene, 24596 [Source:MGI Symb | 0 | 0 | 0 | 0.964 | 1.0906 | 0 |
| 26510 | ENSMUSG00000084454 | 0.36353943 | 2.065989254 | 4.034137301 | 0.512126658 | 0.608562368 | NA | Gm24592 | predicted gene, 24592 [Source:MGI Symb | 0 | 0 | 0 | 0 | 2.1812 | 0 |
| 26511 | ENSMUSG00000084456 | 0 | NA | NA | NA | NA | NA | Gm24593 | predicted gene, 24593 [Source:MGI Symb | 0 | 0 | 0 | 0 | 0 | 0 |
| 26512 | ENSMUSG00000084457 | 0.160662974 | 1.126924175 | 4.080472857 | 0.276174898 | 0.782413736 | NA | NA | NA | 0 | 0 | 0 | 0.964 | 0 | 0 |
| 26513 | ENSMUSG00000084458 | 0 | NA | NA | NA | NA | NA | Mir1907 | microRNA 1907 [Source:MGI Symbol;Acc: | 0 | 0 | 0 | 0 | 0 | 0 |
| 26514 | ENSMUSG00000084459 | 0 | NA | NA | NA | NA | NA | Gm24598 | predicted gene, 24598 [Source:MGI Symb | 0 | 0 | 0 | 0 | 0 | 0 |
| 26515 | ENSMUSG00000084460 | 0 | NA | NA | NA | NA | NA | NA | NA | 0 | 0 | 0 | 0 | 0 | 0 |
| 26516 | ENSMUSG00000084461 | 0 | NA | NA | NA | NA | NA | Gm22958 | predicted gene, 22958 [Source:MGI Symb | 0 | 0 | 0 | 0 | 0 | 0 |
| 26517 | ENSMUSG00000084462 | 0 | NA | NA | NA | NA | NA | NA | NA | 0 | 0 | 0 | 0 | 0 | 0 |
| 26518 | ENSMUSG00000084463 | 0 | NA | NA | NA | NA | NA | NA | NA | 0 | 0 | 0 | 0 | 0 | 0 |
| 26519 | ENSMUSG00000084465 | 0 | NA | NA | NA | NA | NA | NA | NA | 0 | 0 | 0 | 0 | 0 | 0 |
| 26520 | ENSMUSG00000084466 | 0 | NA | NA | NA | NA | NA | Gm22960 | predicted gene, 22960 [Source:MGI Symb | 0 | 0 | 0 | 0 | 0 | 0 |
| 26521 | ENSMUSG00000084467 | 0 | NA | NA | NA | NA | NA | NA | NA | 0 | 0 | 0 | 0 | 0 | 0 |
| 26522 | ENSMUSG00000084468 | 0 | NA | NA | NA | NA | NA | NA | NA | 0 | 0 | 0 | 0 | 0 | 0 |
| 26523 | ENSMUSG00000084469 | 0 | NA | NA | NA | NA | NA | Gm22956 | predicted gene, 22956 [Source:MGI Symb | 0 | 0 | 0 | 0 | 0 | 0 |
| 26524 | ENSMUSG00000084472 | 0.615839296 | 1.537131441 | 3.17677869 | 0.483864817 | 0.628481811 | NA | NA | NA | 0.8031 | 0 | 0 | 2.8919 | 0 | 0 |
| 26525 | ENSMUSG00000084473 | 0 | NA | NA | NA | NA | NA | Gm25784 | predicted gene, 25784 [Source:MGI Symb | 0 | 0 | 0 | 0 | 0 | 0 |
| 26526 | ENSMUSG00000084474 | 0 | NA | NA | NA | NA | NA | Gm25783 | predicted gene, 25783 [Source:MGI Symb | 0 | 0 | 0 | 0 | 0 | 0 |
| 26527 | ENSMUSG00000084475 | 0 | NA | NA | NA | NA | NA | NA | NA | 0 | 0 | 0 | 0 | 0 | 0 |
| 26528 | ENSMUSG00000084476 | 0 | NA | NA | NA | NA | NA | NA | NA | 0 | 0 | 0 | 0 | 0 | 0 |
| 26529 | ENSMUSG00000084477 | 0.167707431 | -0.796629176 | 4.080472857 | -0.195229623 | 0.84521319 | NA | NA | NA | 0 | 0 | 1.0062 | 0 | 0 | 0 |
| 26530 | ENSMUSG00000084478 | 0 | NA | NA | NA | NA | NA | Gm26425 | predicted gene, 26425 [Source:MGI Symb | 0 | 0 | 0 | 0 | 0 | 0 |
| 26531 | ENSMUSG00000084479 | 9.246280826 | -1.44976214 | 0.780870839 | -1.856596592 | 0.063368569 | 0.360000344 | Gm26424 | predicted gene, 26424 [Source:MGI Symb | 11.243 | 7.1954 | 22.137 | 3.8559 | 4.3625 | 6.6831 |
| 26532 | ENSMUSG00000084480 | 0 | NA | NA | NA | NA | NA | NA | NA | 0 | 0 | 0 | 0 | 0 | 0 |
| 26533 | ENSMUSG00000084484 | 0 | NA | NA | NA | NA | NA | Gm23788 | predicted gene, 23788 [Source:MGI Symb | 0 | 0 | 0 | 0 | 0 | 0 |
| 26534 | ENSMUSG00000084485 | 0 | NA | NA | NA | NA | NA | NA | NA | 0 | 0 | 0 | 0 | 0 | 0 |
| 26535 | ENSMUSG00000084487 | 0 | NA | NA | NA | NA | NA | Gm23787 | predicted gene, 23787 [Source:MGI Symb | 0 | 0 | 0 | 0 | 0 | 0 |
| 26536 | ENSMUSG00000084492 | 0 | NA | NA | NA | NA | NA | NA | NA | 0 | 0 | 0 | 0 | 0 | 0 |
| 26537 | ENSMUSG00000084493 | 0.185640647 | 1.126924175 | 4.080472857 | 0.276174898 | 0.782413736 | NA | Gm22106 | predicted gene, 22106 [Source:MGI Symb | 0 | 0 | 0 | 0 | 0 | 1.1138 |
| 26538 | ENSMUSG00000084496 | 0 | NA | NA | NA | NA | NA | NA | NA | 0 | 0 | 0 | 0 | 0 | 0 |
| 26539 | ENSMUSG00000084497 | 1.593920542 | 1.913431964 | 2.140701436 | 0.89383411 | 0.371410656 | NA | Gm22107 | predicted gene, 22107 [Source:MGI Symb | 0 | 2.0558 | 0 | 0.964 | 6.5437 | 0 |
| 26540 | ENSMUSG00000084498 | 0 | NA | NA | NA | NA | NA | NA | NA | 0 | 0 | 0 | 0 | 0 | 0 |

| | | | | | | | | | | | | | | | |
|---|---|---|---|---|---|---|---|---|---|---|---|---|---|---|---|
| 26541 | ENSMUSG00000084500 | 0 | NA | NA | NA | NA | NA | NA | NA | 0 | 0 | 0 | 0 | 0 | 0 |
| 26542 | ENSMUSG00000084501 | 0 | NA | NA | NA | NA | NA | NA | NA | 0 | 0 | 0 | 0 | 0 | 0 |
| 26543 | ENSMUSG00000084502 | 0 | NA | NA | NA | NA | NA | NA | NA | 0 | 0 | 0 | 0 | 0 | 0 |
| 26544 | ENSMUSG00000084503 | 0 | NA | NA | NA | NA | NA | NA | NA | 0 | 0 | 0 | 0 | 0 | 0 |
| 26545 | ENSMUSG00000084504 | 0.160662974 | 1.126924175 | 4.080472857 | 0.276174898 | 0.782413736 | NA | Gm25323 | predicted gene, 25323 [Source:MGI Symb | 0 | 0 | 0 | 0.964 | 0 | 0 |
| 26546 | ENSMUSG00000084505 | 0 | NA | NA | NA | NA | NA | Gm25322 | predicted gene, 25322 [Source:MGI Symb | 0 | 0 | 0 | 0 | 0 | 0 |
| 26547 | ENSMUSG00000084506 | 0 | NA | NA | NA | NA | NA | Gm23792 | predicted gene, 23792 [Source:MGI Symb | 0 | 0 | 0 | 0 | 0 | 0 |
| 26548 | ENSMUSG00000084507 | 1.391250677 | 0.06749928 | 2.180657966 | 0.03095363 | 0.97530652 | NA | Gm25321 | predicted gene, 25321 [Source:MGI Symb | 0 | 4.1117 | 0 | 0.964 | 3.2719 | 0 |
| 26549 | ENSMUSG00000084508 | 0 | NA | NA | NA | NA | NA | NA | NA | 0 | 0 | 0 | 0 | 0 | 0 |
| 26550 | ENSMUSG00000084511 | 0 | NA | NA | NA | NA | NA | NA | NA | 0 | 0 | 0 | 0 | 0 | 0 |
| 26551 | ENSMUSG00000084512 | 0.321325949 | 1.911575114 | 4.048267893 | 0.472195804 | 0.636787034 | NA | NA | NA | 0 | 0 | 0 | 1.928 | 0 | 0 |
| 26552 | ENSMUSG00000084513 | 0 | NA | NA | NA | NA | NA | NA | NA | 0 | 0 | 0 | 0 | 0 | 0 |
| 26553 | ENSMUSG00000084514 | 0 | NA | NA | NA | NA | NA | Mir1893 | microRNA 1893 [Source:MGI Symbol;Acc: | 0 | 0 | 0 | 0 | 0 | 0 |
| 26554 | ENSMUSG00000084515 | 0 | NA | NA | NA | NA | NA | Gm22479 | predicted gene, 22479 [Source:MGI Symb | 0 | 0 | 0 | 0 | 0 | 0 |
| 26555 | ENSMUSG00000084516 | 0 | NA | NA | NA | NA | NA | Mir1902 | microRNA 1902 [Source:MGI Symbol;Acc: | 0 | 0 | 0 | 0 | 0 | 0 |
| 26556 | ENSMUSG00000084517 | 0 | NA | NA | NA | NA | NA | Gm22480 | predicted gene, 22480 [Source:MGI Symb | 0 | 0 | 0 | 0 | 0 | 0 |
| 26557 | ENSMUSG00000084518 | 0 | NA | NA | NA | NA | NA | NA | NA | 0 | 0 | 0 | 0 | 0 | 0 |
| 26558 | ENSMUSG00000084519 | 3.398987341 | 0.898523523 | 1.185033032 | 0.758226563 | 0.448315362 | NA | NA | NA | 4.0155 | 1.0279 | 2.0125 | 6.7478 | 4.3625 | 2.2277 |
| 26559 | ENSMUSG00000084520 | 0.513958919 | -2.394274821 | 3.756096013 | -0.637437066 | 0.523840191 | NA | NA | NA | 0 | 3.0838 | 0 | 0 | 0 | 0 |
| 26560 | ENSMUSG00000084521 | 0 | NA | NA | NA | NA | NA | Gm24183 | predicted gene, 24183 [Source:MGI Symb | 0 | 0 | 0 | 0 | 0 | 0 |
| 26561 | ENSMUSG00000084522 | 0.160662974 | 1.126924175 | 4.080472857 | 0.276174898 | 0.782413736 | NA | NA | NA | 0 | 0 | 0 | 0.964 | 0 | 0 |
| 26562 | ENSMUSG00000084523 | 0 | NA | NA | NA | NA | NA | Mir1903 | microRNA 1903 [Source:MGI Symbol;Acc: | 0 | 0 | 0 | 0 | 0 | 0 |
| 26563 | ENSMUSG00000084524 | 0 | NA | NA | NA | NA | NA | NA | NA | 0 | 0 | 0 | 0 | 0 | 0 |
| 26564 | ENSMUSG00000084526 | 0 | NA | NA | NA | NA | NA | NA | NA | 0 | 0 | 0 | 0 | 0 | 0 |
| 26565 | ENSMUSG00000084527 | 0 | NA | NA | NA | NA | NA | Gm24185 | predicted gene, 24185 [Source:MGI Symb | 0 | 0 | 0 | 0 | 0 | 0 |
| 26566 | ENSMUSG00000084530 | 0 | NA | NA | NA | NA | NA | NA | NA | 0 | 0 | 0 | 0 | 0 | 0 |
| 26567 | ENSMUSG00000084531 | 0 | NA | NA | NA | NA | NA | NA | NA | 0 | 0 | 0 | 0 | 0 | 0 |
| 26568 | ENSMUSG00000084532 | 0 | NA | NA | NA | NA | NA | NA | NA | 0 | 0 | 0 | 0 | 0 | 0 |
| 26569 | ENSMUSG00000084535 | 0 | NA | NA | NA | NA | NA | Mir1906-1 | microRNA 1906-1 [Source:MGI Symbol;Ac | 0 | 0 | 0 | 0 | 0 | 0 |
| 26570 | ENSMUSG00000084536 | 0 | NA | NA | NA | NA | NA | NA | NA | 0 | 0 | 0 | 0 | 0 | 0 |
| 26571 | ENSMUSG00000084537 | 0 | NA | NA | NA | NA | NA | NA | NA | 0 | 0 | 0 | 0 | 0 | 0 |
| 26572 | ENSMUSG00000084539 | 0.335414862 | -1.779047242 | 4.030804179 | -0.441362855 | 0.658950329 | NA | NA | NA | 0 | 0 | 2.0125 | 0 | 0 | 0 |
| 26573 | ENSMUSG00000084540 | 0.167707431 | -0.796629176 | 4.080472857 | -0.195229623 | 0.84521319 | NA | NA | NA | 0 | 0 | 1.0062 | 0 | 0 | 0 |
| 26574 | ENSMUSG00000084541 | 0 | NA | NA | NA | NA | NA | NA | NA | 0 | 0 | 0 | 0 | 0 | 0 |
| 26575 | ENSMUSG00000084543 | 0.528073336 | 2.60973587 | 2.979250867 | 0.875970499 | 0.381046072 | NA | NA | NA | 0 | 0 | 0 | 0.964 | 1.0906 | 1.1138 |
| 26576 | ENSMUSG00000084544 | 0 | NA | NA | NA | NA | NA | NA | NA | 0 | 0 | 0 | 0 | 0 | 0 |
| 26577 | ENSMUSG00000084545 | 0 | NA | NA | NA | NA | NA | NA | NA | 0 | 0 | 0 | 0 | 0 | 0 |
| 26578 | ENSMUSG00000084546 | 0 | NA | NA | NA | NA | NA | NA | NA | 0 | 0 | 0 | 0 | 0 | 0 |
| 26579 | ENSMUSG00000084547 | 0 | NA | NA | NA | NA | NA | NA | NA | 0 | 0 | 0 | 0 | 0 | 0 |
| 26580 | ENSMUSG00000084548 | 0 | NA | NA | NA | NA | NA | NA | NA | 0 | 0 | 0 | 0 | 0 | 0 |
| 26581 | ENSMUSG00000084549 | 0 | NA | NA | NA | NA | NA | DQ267100 | snoRNA DQ267100 [Source:MGI Symbol;A | 0 | 0 | 0 | 0 | 0 | 0 |
| 26582 | ENSMUSG00000084550 | 0 | NA | NA | NA | NA | NA | Gm25269 | predicted gene, 25269 [Source:MGI Symb | 0 | 0 | 0 | 0 | 0 | 0 |
| 26583 | ENSMUSG00000084551 | 0 | NA | NA | NA | NA | NA | Gm24839 | predicted gene, 24839 [Source:MGI Symb | 0 | 0 | 0 | 0 | 0 | 0 |
| 26584 | ENSMUSG00000084552 | 0.596071152 | -1.224343926 | 2.897583865 | -0.4225396 | 0.672631203 | NA | Gm24840 | predicted gene, 24840 [Source:MGI Symb | 1.6062 | 0 | 1.0062 | 0.964 | 0 | 0 |
| 26585 | ENSMUSG00000084555 | 0 | NA | NA | NA | NA | NA | Gm25270 | predicted gene, 25270 [Source:MGI Symb | 0 | 0 | 0 | 0 | 0 | 0 |
| 26586 | ENSMUSG00000084557 | 0 | NA | NA | NA | NA | NA | Gm25279 | predicted gene, 25279 [Source:MGI Symb | 0 | 0 | 0 | 0 | 0 | 0 |
| 26587 | ENSMUSG00000084558 | 0 | NA | NA | NA | NA | NA | Gm24836 | predicted gene, 24836 [Source:MGI Symb | 0 | 0 | 0 | 0 | 0 | 0 |
| 26588 | ENSMUSG00000084559 | 0 | NA | NA | NA | NA | NA | Mir1906-2 | microRNA 1906-2 [Source:MGI Symbol;Ac | 0 | 0 | 0 | 0 | 0 | 0 |
| 26589 | ENSMUSG00000084560 | 0 | NA | NA | NA | NA | NA | Gm26478 | predicted gene, 26478 [Source:MGI Symb | 0 | 0 | 0 | 0 | 0 | 0 |
| 26590 | ENSMUSG00000084562 | 0.696064467 | -1.534444742 | 3.043667477 | -0.504143358 | 0.61416064 | NA | NA | NA | 3.2124 | 0 | 0 | 0.964 | 0 | 0 |
| 26591 | ENSMUSG00000084564 | 0 | NA | NA | NA | NA | NA | Mir1895 | microRNA 1895 [Source:MGI Symbol;Acc: | 0 | 0 | 0 | 0 | 0 | 0 |
| 26592 | ENSMUSG00000084565 | 0 | NA | NA | NA | NA | NA | Mir1901 | microRNA 1901 [Source:MGI Symbol;Acc: | 0 | 0 | 0 | 0 | 0 | 0 |
| 26593 | ENSMUSG00000084566 | 0 | NA | NA | NA | NA | NA | NA | NA | 0 | 0 | 0 | 0 | 0 | 0 |
| 26594 | ENSMUSG00000084567 | 0 | NA | NA | NA | NA | NA | NA | NA | 0 | 0 | 0 | 0 | 0 | 0 |
| 26595 | ENSMUSG00000084568 | 0 | NA | NA | NA | NA | NA | NA | NA | 0 | 0 | 0 | 0 | 0 | 0 |
| 26596 | ENSMUSG00000084569 | 0 | NA | NA | NA | NA | NA | Gm26479 | predicted gene, 26479 [Source:MGI Symb | 0 | 0 | 0 | 0 | 0 | 0 |

| | | | | | | | | | | | | | | | |
|---|---|---|---|---|---|---|---|---|---|---|---|---|---|---|---|
| 26597 | ENSMUSG00000084571 | 0.160662974 | 1.126924175 | 4.080472857 | 0.276174898 | 0.782413736 | NA | NA | NA | 0 | 0 | 0 | 0.964 | 0 | 0 |
| 26598 | ENSMUSG00000084572 | 0.160662974 | 1.126924175 | 4.080472857 | 0.276174898 | 0.782413736 | NA | NA | NA | 0 | 0 | 0 | 0.964 | 0 | 0 |
| 26599 | ENSMUSG00000084573 | 0 | NA | NA | NA | NA | NA | NA | NA | 0 | 0 | 0 | 0 | 0 | 0 |
| 26600 | ENSMUSG00000084574 | 0 | NA | NA | NA | NA | NA | NA | NA | 0 | 0 | 0 | 0 | 0 | 0 |
| 26601 | ENSMUSG00000084575 | 0.160662974 | 1.126924175 | 4.080472857 | 0.276174898 | 0.782413736 | NA | Gm23656 | predicted gene, 23656 [Source:MGI Symb | 0 | 0 | 0 | 0.964 | 0 | 0 |
| 26602 | ENSMUSG00000084576 | 0 | NA | NA | NA | NA | NA | Gm23655 | predicted gene, 23655 [Source:MGI Symb | 0 | 0 | 0 | 0 | 0 | 0 |
| 26603 | ENSMUSG00000084579 | 0 | NA | NA | NA | NA | NA | Gm23661 | predicted gene, 23661 [Source:MGI Symb | 0 | 0 | 0 | 0 | 0 | 0 |
| 26604 | ENSMUSG00000084581 | 0 | NA | NA | NA | NA | NA | NA | NA | 0 | 0 | 0 | 0 | 0 | 0 |
| 26605 | ENSMUSG00000084582 | 0 | NA | NA | NA | NA | NA | NA | NA | 0 | 0 | 0 | 0 | 0 | 0 |
| 26606 | ENSMUSG00000084583 | 0 | NA | NA | NA | NA | NA | Gm25177 | predicted gene, 25177 [Source:MGI Symb | 0 | 0 | 0 | 0 | 0 | 0 |
| 26607 | ENSMUSG00000084584 | 0 | NA | NA | NA | NA | NA | NA | NA | 0 | 0 | 0 | 0 | 0 | 0 |
| 26608 | ENSMUSG00000084587 | 0.160662974 | 1.126924175 | 4.080472857 | 0.276174898 | 0.782413736 | NA | NA | NA | 0 | 0 | 0 | 0.964 | 0 | 0 |
| 26609 | ENSMUSG00000084588 | 0 | NA | NA | NA | NA | NA | NA | NA | 0 | 0 | 0 | 0 | 0 | 0 |
| 26610 | ENSMUSG00000084589 | 0 | NA | NA | NA | NA | NA | Gm25176 | predicted gene, 25176 [Source:MGI Symb | 0 | 0 | 0 | 0 | 0 | 0 |
| 26611 | ENSMUSG00000084590 | 0 | NA | NA | NA | NA | NA | Mir1896 | microRNA 1896 [Source:MGI Symbol;Acc: | 0 | 0 | 0 | 0 | 0 | 0 |
| 26612 | ENSMUSG00000084591 | 0 | NA | NA | NA | NA | NA | NA | NA | 0 | 0 | 0 | 0 | 0 | 0 |
| 26613 | ENSMUSG00000084592 | 2.749648476 | 0.558513098 | 1.501760382 | 0.371905601 | 0.709963135 | NA | Mir1898 | microRNA 1898 [Source:MGI Symbol;Acc: | 1.6062 | 1.0279 | 4.025 | 0 | 8.7249 | 1.1138 |
| 26614 | ENSMUSG00000084593 | 0.685071968 | 0.040605487 | 2.800847395 | 0.014497572 | 0.988433016 | NA | Gm22345 | predicted gene, 22345 [Source:MGI Symb | 0 | 2.0558 | 0 | 0.964 | 1.0906 | 0 |
| 26615 | ENSMUSG00000084595 | 0 | NA | NA | NA | NA | NA | NA | NA | 0 | 0 | 0 | 0 | 0 | 0 |
| 26616 | ENSMUSG00000084596 | 0 | NA | NA | NA | NA | NA | NA | NA | 0 | 0 | 0 | 0 | 0 | 0 |
| 26617 | ENSMUSG00000084597 | 0 | NA | NA | NA | NA | NA | NA | NA | 0 | 0 | 0 | 0 | 0 | 0 |
| 26618 | ENSMUSG00000084598 | 0 | NA | NA | NA | NA | NA | NA | NA | 0 | 0 | 0 | 0 | 0 | 0 |
| 26619 | ENSMUSG00000084599 | 0 | NA | NA | NA | NA | NA | NA | NA | 0 | 0 | 0 | 0 | 0 | 0 |
| 26620 | ENSMUSG00000084600 | 0 | NA | NA | NA | NA | NA | NA | NA | 0 | 0 | 0 | 0 | 0 | 0 |
| 26621 | ENSMUSG00000084601 | 0 | NA | NA | NA | NA | NA | Gm23111 | predicted gene, 23111 [Source:MGI Symb | 0 | 0 | 0 | 0 | 0 | 0 |
| 26622 | ENSMUSG00000084602 | 0 | NA | NA | NA | NA | NA | NA | NA | 0 | 0 | 0 | 0 | 0 | 0 |
| 26623 | ENSMUSG00000084603 | 0.656740811 | -0.013525372 | 3.112905782 | -0.004344934 | 0.996533255 | NA | NA | NA | 0 | 0 | 2.0125 | 1.928 | 0 | 0 |
| 26624 | ENSMUSG00000084604 | 0 | NA | NA | NA | NA | NA | Gm22205 | predicted gene, 22205 [Source:MGI Symb | 0 | 0 | 0 | 0 | 0 | 0 |
| 26625 | ENSMUSG00000084605 | 0 | NA | NA | NA | NA | NA | NA | NA | 0 | 0 | 0 | 0 | 0 | 0 |
| 26626 | ENSMUSG00000084606 | 0 | NA | NA | NA | NA | NA | Gm23118 | predicted gene, 23118 [Source:MGI Symb | 0 | 0 | 0 | 0 | 0 | 0 |
| 26627 | ENSMUSG00000084607 | 1.028022105 | 1.080690112 | 2.389621529 | 0.45224321 | 0.651093782 | NA | Gm22208 | predicted gene, 22208 [Source:MGI Symb | 0 | 0 | 2.0125 | 1.928 | 0 | 2.2277 |
| 26628 | ENSMUSG00000084608 | 0 | NA | NA | NA | NA | NA | NA | NA | 0 | 0 | 0 | 0 | 0 | 0 |
| 26629 | ENSMUSG00000084609 | 0 | NA | NA | NA | NA | NA | Gm22199 | predicted gene, 22199 [Source:MGI Symb | 0 | 0 | 0 | 0 | 0 | 0 |
| 26630 | ENSMUSG00000084610 | 0 | NA | NA | NA | NA | NA | Gm25052 | predicted gene, 25052 [Source:MGI Symb | 0 | 0 | 0 | 0 | 0 | 0 |
| 26631 | ENSMUSG00000084611 | 0 | NA | NA | NA | NA | NA | NA | NA | 0 | 0 | 0 | 0 | 0 | 0 |
| 26632 | ENSMUSG00000084612 | 0 | NA | NA | NA | NA | NA | NA | NA | 0 | 0 | 0 | 0 | 0 | 0 |
| 26633 | ENSMUSG00000084613 | 0 | NA | NA | NA | NA | NA | NA | NA | 0 | 0 | 0 | 0 | 0 | 0 |
| 26634 | ENSMUSG00000084616 | 8.16254888 | -0.575799482 | 0.876791718 | -0.656711817 | 0.511366235 | 0.883679955 | NA | NA | 4.0155 | 15.419 | 10.062 | 9.6398 | 8.7249 | 1.1138 |
| 26635 | ENSMUSG00000084617 | 0 | NA | NA | NA | NA | NA | Gm22073 | predicted gene, 22073 [Source:MGI Symb | 0 | 0 | 0 | 0 | 0 | 0 |
| 26636 | ENSMUSG00000084619 | 0 | NA | NA | NA | NA | NA | Gm25045 | predicted gene, 25045 [Source:MGI Symb | 0 | 0 | 0 | 0 | 0 | 0 |
| 26637 | ENSMUSG00000084620 | 0 | NA | NA | NA | NA | NA | NA | NA | 0 | 0 | 0 | 0 | 0 | 0 |
| 26638 | ENSMUSG00000084621 | 0 | NA | NA | NA | NA | NA | Gm25106 | predicted gene, 25106 [Source:MGI Symb | 0 | 0 | 0 | 0 | 0 | 0 |
| 26639 | ENSMUSG00000084622 | 3.474010151 | 0.459262721 | 1.350748846 | 0.340006006 | 0.733852005 | NA | NA | NA | 0.8031 | 2.0558 | 6.0375 | 8.6758 | 3.2719 | 0 |
| 26640 | ENSMUSG00000084624 | 0 | NA | NA | NA | NA | NA | NA | NA | 0 | 0 | 0 | 0 | 0 | 0 |
| 26641 | ENSMUSG00000084625 | 0 | NA | NA | NA | NA | NA | NA | NA | 0 | 0 | 0 | 0 | 0 | 0 |
| 26642 | ENSMUSG00000084627 | 0 | NA | NA | NA | NA | NA | NA | NA | 0 | 0 | 0 | 0 | 0 | 0 |
| 26643 | ENSMUSG00000084628 | 0 | NA | NA | NA | NA | NA | NA | NA | 0 | 0 | 0 | 0 | 0 | 0 |
| 26644 | ENSMUSG00000084629 | 0 | NA | NA | NA | NA | NA | Gm25112 | predicted gene, 25112 [Source:MGI Symb | 0 | 0 | 0 | 0 | 0 | 0 |
| 26645 | ENSMUSG00000084630 | 0 | NA | NA | NA | NA | NA | NA | NA | 0 | 0 | 0 | 0 | 0 | 0 |
| 26646 | ENSMUSG00000084631 | 0 | NA | NA | NA | NA | NA | NA | NA | 0 | 0 | 0 | 0 | 0 | 0 |
| 26647 | ENSMUSG00000084632 | 0.856391608 | -0.549592269 | 2.599657283 | -0.211409509 | 0.832567737 | NA | Gm23887 | predicted gene, 23887 [Source:MGI Symb | 0 | 3.0838 | 0 | 0.964 | 1.0906 | 0 |
| 26648 | ENSMUSG00000084633 | 0 | NA | NA | NA | NA | NA | NA | NA | 0 | 0 | 0 | 0 | 0 | 0 |
| 26649 | ENSMUSG00000084635 | 0 | NA | NA | NA | NA | NA | NA | NA | 0 | 0 | 0 | 0 | 0 | 0 |
| 26650 | ENSMUSG00000084636 | 0 | NA | NA | NA | NA | NA | NA | NA | 0 | 0 | 0 | 0 | 0 | 0 |
| 26651 | ENSMUSG00000084637 | 0 | NA | NA | NA | NA | NA | NA | NA | 0 | 0 | 0 | 0 | 0 | 0 |
| 26652 | ENSMUSG00000084638 | 0.339027071 | -1.79174149 | 3.919585585 | -0.457125237 | 0.647581032 | NA | Gm23889 | predicted gene, 23889 [Source:MGI Symb | 0 | 1.0279 | 1.0062 | 0 | 0 | 0 |

| | | | | | | | | | | | | | | | |
|---|---|---|---|---|---|---|---|---|---|---|---|---|---|---|---|
| 26653 | ENSMUSG00000084639 | 0 | NA | NA | NA | NA | NA | NA | NA | 0 | 0 | 0 | 0 | 0 | 0 |
| 26654 | ENSMUSG00000084640 | 0 | NA | NA | NA | NA | NA | NA | NA | 0 | 0 | 0 | 0 | 0 | 0 |
| 26655 | ENSMUSG00000084641 | 0.328370405 | 0.16514316 | 4.003149536 | 0.041253308 | 0.967093956 | NA | NA | NA | 0 | 0 | 1.0062 | 0.964 | 0 | 0 |
| 26656 | ENSMUSG00000084642 | 0.36353943 | 2.065989254 | 4.034137301 | 0.512126658 | 0.608562368 | NA | NA | NA | 0 | 0 | 0 | 0 | 2.1812 | 0 |
| 26657 | ENSMUSG00000084643 | 0 | NA | NA | NA | NA | NA | NA | NA | 0 | 0 | 0 | 0 | 0 | 0 |
| 26658 | ENSMUSG00000084644 | 0.531246861 | 1.101376188 | 3.36099124 | 0.327693858 | 0.743143147 | NA | Gm24379 | predicted gene, 24379 [Source:MGI Symb | 0 | 0 | 1.0062 | 0 | 2.1812 | 0 |
| 26659 | ENSMUSG00000084645 | 0 | NA | NA | NA | NA | NA | NA | NA | 0 | 0 | 0 | 0 | 0 | 0 |
| 26660 | ENSMUSG00000084646 | 0 | NA | NA | NA | NA | NA | Gm24381 | predicted gene, 24381 [Source:MGI Symb | 0 | 0 | 0 | 0 | 0 | 0 |
| 26661 | ENSMUSG00000084647 | 0 | NA | NA | NA | NA | NA | Gm24382 | predicted gene, 24382 [Source:MGI Symb | 0 | 0 | 0 | 0 | 0 | 0 |
| 26662 | ENSMUSG00000084648 | 0 | NA | NA | NA | NA | NA | Gm24378 | predicted gene, 24378 [Source:MGI Symb | 0 | 0 | 0 | 0 | 0 | 0 |
| 26663 | ENSMUSG00000084649 | 0 | NA | NA | NA | NA | NA | NA | NA | 0 | 0 | 0 | 0 | 0 | 0 |
| 26664 | ENSMUSG00000084650 | 0 | NA | NA | NA | NA | NA | NA | NA | 0 | 0 | 0 | 0 | 0 | 0 |
| 26665 | ENSMUSG00000084651 | 0 | NA | NA | NA | NA | NA | Mir1904 | microRNA 1904 [Source:MGI Symbol;Acc: | 0 | 0 | 0 | 0 | 0 | 0 |
| 26666 | ENSMUSG00000084652 | 0 | NA | NA | NA | NA | NA | Gm22717 | predicted gene, 22717 [Source:MGI Symb | 0 | 0 | 0 | 0 | 0 | 0 |
| 26667 | ENSMUSG00000084653 | 0 | NA | NA | NA | NA | NA | NA | NA | 0 | 0 | 0 | 0 | 0 | 0 |
| 26668 | ENSMUSG00000084654 | 0 | NA | NA | NA | NA | NA | NA | NA | 0 | 0 | 0 | 0 | 0 | 0 |
| 26669 | ENSMUSG00000084658 | 0 | NA | NA | NA | NA | NA | NA | NA | 0 | 0 | 0 | 0 | 0 | 0 |
| 26670 | ENSMUSG00000084659 | 0.17131964 | -0.796629176 | 4.080472857 | -0.195229623 | 0.84521319 | NA | Gm22720 | predicted gene, 22720 [Source:MGI Symb | 0 | 1.0279 | 0 | 0 | 0 | 0 |
| 26671 | ENSMUSG00000084660 | 0 | NA | NA | NA | NA | NA | NA | NA | 0 | 0 | 0 | 0 | 0 | 0 |
| 26672 | ENSMUSG00000084661 | 0 | NA | NA | NA | NA | NA | Gm24635 | predicted gene, 24635 [Source:MGI Symb | 0 | 0 | 0 | 0 | 0 | 0 |
| 26673 | ENSMUSG00000084663 | 0 | NA | NA | NA | NA | NA | Mir1900 | microRNA 1900 [Source:MGI Symbol;Acc: | 0 | 0 | 0 | 0 | 0 | 0 |
| 26674 | ENSMUSG00000084664 | 0 | NA | NA | NA | NA | NA | NA | NA | 0 | 0 | 0 | 0 | 0 | 0 |
| 26675 | ENSMUSG00000084665 | 0 | NA | NA | NA | NA | NA | Gm23214 | predicted gene, 23214 [Source:MGI Symb | 0 | 0 | 0 | 0 | 0 | 0 |
| 26676 | ENSMUSG00000084666 | 0.167707431 | -0.796629176 | 4.080472857 | -0.195229623 | 0.84521319 | NA | NA | NA | 0 | 0 | 1.0062 | 0 | 0 | 0 |
| 26677 | ENSMUSG00000084667 | 0 | NA | NA | NA | NA | NA | Gm23212 | predicted gene, 23212 [Source:MGI Symb | 0 | 0 | 0 | 0 | 0 | 0 |
| 26678 | ENSMUSG00000084668 | 0 | NA | NA | NA | NA | NA | NA | NA | 0 | 0 | 0 | 0 | 0 | 0 |
| 26679 | ENSMUSG00000084669 | 0 | NA | NA | NA | NA | NA | Gm23206 | predicted gene, 23206 [Source:MGI Symb | 0 | 0 | 0 | 0 | 0 | 0 |
| 26680 | ENSMUSG00000084671 | 0 | NA | NA | NA | NA | NA | NA | NA | 0 | 0 | 0 | 0 | 0 | 0 |
| 26681 | ENSMUSG00000084673 | 0 | NA | NA | NA | NA | NA | NA | NA | 0 | 0 | 0 | 0 | 0 | 0 |
| 26682 | ENSMUSG00000084674 | 0.167707431 | -0.796629176 | 4.080472857 | -0.195229623 | 0.84521319 | NA | NA | NA | 0 | 0 | 1.0062 | 0 | 0 | 0 |
| 26683 | ENSMUSG00000084675 | 0 | NA | NA | NA | NA | NA | NA | NA | 0 | 0 | 0 | 0 | 0 | 0 |
| 26684 | ENSMUSG00000084678 | 3.555954956 | -0.882891579 | 1.139373655 | -0.774892043 | 0.438403455 | NA | Gm26020 | predicted gene, 26020 [Source:MGI Symb | 5.6217 | 4.1117 | 4.025 | 0.964 | 3.2719 | 3.3415 |
| 26685 | ENSMUSG00000084682 | 0 | NA | NA | NA | NA | NA | NA | NA | 0 | 0 | 0 | 0 | 0 | 0 |
| 26686 | ENSMUSG00000084683 | 0 | NA | NA | NA | NA | NA | NA | NA | 0 | 0 | 0 | 0 | 0 | 0 |
| 26687 | ENSMUSG00000084684 | 0 | NA | NA | NA | NA | NA | NA | NA | 0 | 0 | 0 | 0 | 0 | 0 |
| 26688 | ENSMUSG00000084686 | 1.896775826 | 1.195032785 | 1.771216077 | 0.674696216 | 0.499868789 | NA | Gm22027 | predicted gene, 22027 [Source:MGI Symb | 2.4093 | 0 | 1.0062 | 5.7839 | 2.1812 | 0 |
| 26689 | ENSMUSG00000084688 | 0 | NA | NA | NA | NA | NA | NA | NA | 0 | 0 | 0 | 0 | 0 | 0 |
| 26690 | ENSMUSG00000084689 | 0 | NA | NA | NA | NA | NA | NA | NA | 0 | 0 | 0 | 0 | 0 | 0 |
| 26691 | ENSMUSG00000084690 | 0 | NA | NA | NA | NA | NA | Gm24896 | predicted gene, 24896 [Source:MGI Symb | 0 | 0 | 0 | 0 | 0 | 0 |
| 26692 | ENSMUSG00000084691 | 0 | NA | NA | NA | NA | NA | Gm24895 | predicted gene, 24895 [Source:MGI Symb | 0 | 0 | 0 | 0 | 0 | 0 |
| 26693 | ENSMUSG00000084693 | 0 | NA | NA | NA | NA | NA | NA | NA | 0 | 0 | 0 | 0 | 0 | 0 |
| 26694 | ENSMUSG00000084694 | 0 | NA | NA | NA | NA | NA | NA | NA | 0 | 0 | 0 | 0 | 0 | 0 |
| 26695 | ENSMUSG00000084696 | 0 | NA | NA | NA | NA | NA | Gm24892 | predicted gene, 24892 [Source:MGI Symb | 0 | 0 | 0 | 0 | 0 | 0 |
| 26696 | ENSMUSG00000084700 | 0 | NA | NA | NA | NA | NA | Gm22984 | predicted gene, 22984 [Source:MGI Symb | 0 | 0 | 0 | 0 | 0 | 0 |
| 26697 | ENSMUSG00000084701 | 0 | NA | NA | NA | NA | NA | NA | NA | 0 | 0 | 0 | 0 | 0 | 0 |
| 26698 | ENSMUSG00000084702 | 0 | NA | NA | NA | NA | NA | NA | NA | 0 | 0 | 0 | 0 | 0 | 0 |
| 26699 | ENSMUSG00000084703 | 0 | NA | NA | NA | NA | NA | Gm22985 | predicted gene, 22985 [Source:MGI Symb | 0 | 0 | 0 | 0 | 0 | 0 |
| 26700 | ENSMUSG00000084704 | 0 | NA | NA | NA | NA | NA | NA | NA | 0 | 0 | 0 | 0 | 0 | 0 |
| 26701 | ENSMUSG00000084705 | 0 | NA | NA | NA | NA | NA | NA | NA | 0 | 0 | 0 | 0 | 0 | 0 |
| 26702 | ENSMUSG00000084706 | 0 | NA | NA | NA | NA | NA | NA | NA | 0 | 0 | 0 | 0 | 0 | 0 |
| 26703 | ENSMUSG00000084708 | 0 | NA | NA | NA | NA | NA | NA | NA | 0 | 0 | 0 | 0 | 0 | 0 |
| 26704 | ENSMUSG00000084710 | 0 | NA | NA | NA | NA | NA | NA | NA | 0 | 0 | 0 | 0 | 0 | 0 |
| 26705 | ENSMUSG00000084711 | 0 | NA | NA | NA | NA | NA | Gm24629 | predicted gene, 24629 [Source:MGI Symb | 0 | 0 | 0 | 0 | 0 | 0 |
| 26706 | ENSMUSG00000084713 | 0 | NA | NA | NA | NA | NA | Gm24627 | predicted gene, 24627 [Source:MGI Symb | 0 | 0 | 0 | 0 | 0 | 0 |
| 26707 | ENSMUSG00000084714 | 0 | NA | NA | NA | NA | NA | Gm24631 | predicted gene, 24631 [Source:MGI Symb | 0 | 0 | 0 | 0 | 0 | 0 |
| 26708 | ENSMUSG00000084717 | 0 | NA | NA | NA | NA | NA | Gm24630 | predicted gene, 24630 [Source:MGI Symb | 0 | 0 | 0 | 0 | 0 | 0 |

| | | | | | | | | | | | | | | | |
|---|---|---|---|---|---|---|---|---|---|---|---|---|---|---|---|
| 26709 | ENSMUSG00000084718 | 0 | NA | NA | NA | NA | NA | NA | NA | 0 | 0 | 0 | 0 | 0 | 0 |
| 26710 | ENSMUSG00000084719 | 0 | NA | NA | NA | NA | NA | NA | NA | 0 | 0 | 0 | 0 | 0 | 0 |
| 26711 | ENSMUSG00000084722 | 0.160662974 | 1.126924175 | 4.080472857 | 0.276174898 | 0.782413736 | NA | NA | NA | 0 | 0 | 0 | 0.964 | 0 | 0 |
| 26712 | ENSMUSG00000084725 | 0 | NA | NA | NA | NA | NA | Gm24141 | predicted gene, 24141 [Source:MGI Symb | 0 | 0 | 0 | 0 | 0 | 0 |
| 26713 | ENSMUSG00000084728 | 0 | NA | NA | NA | NA | NA | Mir1905 | microRNA 1905 [Source:MGI Symbol;Acc: | 0 | 0 | 0 | 0 | 0 | 0 |
| 26714 | ENSMUSG00000084729 | 0 | NA | NA | NA | NA | NA | NA | NA | 0 | 0 | 0 | 0 | 0 | 0 |
| 26715 | ENSMUSG00000084731 | 0 | NA | NA | NA | NA | NA | Gm25806 | predicted gene, 25806 [Source:MGI Symb | 0 | 0 | 0 | 0 | 0 | 0 |
| 26716 | ENSMUSG00000084732 | 0 | NA | NA | NA | NA | NA | NA | NA | 0 | 0 | 0 | 0 | 0 | 0 |
| 26717 | ENSMUSG00000084733 | 0 | NA | NA | NA | NA | NA | NA | NA | 0 | 0 | 0 | 0 | 0 | 0 |
| 26718 | ENSMUSG00000084734 | 0 | NA | NA | NA | NA | NA | NA | NA | 0 | 0 | 0 | 0 | 0 | 0 |
| 26719 | ENSMUSG00000084737 | 0 | NA | NA | NA | NA | NA | Gm25805 | predicted gene, 25805 [Source:MGI Symb | 0 | 0 | 0 | 0 | 0 | 0 |
| 26720 | ENSMUSG00000084738 | 0.321325949 | 1.911575114 | 4.048267893 | 0.472195804 | 0.636787034 | NA | NA | NA | 0 | 0 | 0 | 1.928 | 0 | 0 |
| 26721 | ENSMUSG00000084739 | 0 | NA | NA | NA | NA | NA | Gm22067 | predicted gene, 22067 [Source:MGI Symb | 0 | 0 | 0 | 0 | 0 | 0 |
| 26722 | ENSMUSG00000084740 | 0 | NA | NA | NA | NA | NA | Gm25294 | predicted gene, 25294 [Source:MGI Symb | 0 | 0 | 0 | 0 | 0 | 0 |
| 26723 | ENSMUSG00000084741 | 0 | NA | NA | NA | NA | NA | NA | NA | 0 | 0 | 0 | 0 | 0 | 0 |
| 26724 | ENSMUSG00000084742 | 0 | NA | NA | NA | NA | NA | NA | NA | 0 | 0 | 0 | 0 | 0 | 0 |
| 26725 | ENSMUSG00000084743 | 0 | NA | NA | NA | NA | NA | NA | NA | 0 | 0 | 0 | 0 | 0 | 0 |
| 26726 | ENSMUSG00000084744 | 2.440741986 | -0.801607753 | 1.378532165 | -0.581493688 | 0.560907771 | NA | NA | NA | 3.2124 | 1.0279 | 5.0312 | 0.964 | 2.1812 | 2.2277 |
| 26727 | ENSMUSG00000084745 | 0.353348078 | 0.165142701 | 3.895042568 | 0.042398176 | 0.966181282 | NA | Gm25290 | predicted gene, 25290 [Source:MGI Symb | 0 | 0 | 1.0062 | 0 | 0 | 1.1138 |
| 26728 | ENSMUSG00000084746 | 0 | NA | NA | NA | NA | NA | NA | NA | 0 | 0 | 0 | 0 | 0 | 0 |
| 26729 | ENSMUSG00000084747 | 0 | NA | NA | NA | NA | NA | Mir1899 | microRNA 1899 [Source:MGI Symbol;Acc: | 0 | 0 | 0 | 0 | 0 | 0 |
| 26730 | ENSMUSG00000084748 | 0 | NA | NA | NA | NA | NA | Gm25289 | predicted gene, 25289 [Source:MGI Symb | 0 | 0 | 0 | 0 | 0 | 0 |
| 26731 | ENSMUSG00000084749 | 0 | NA | NA | NA | NA | NA | NA | NA | 0 | 0 | 0 | 0 | 0 | 0 |
| 26732 | ENSMUSG00000084750 | 0 | NA | NA | NA | NA | NA | NA | NA | 0 | 0 | 0 | 0 | 0 | 0 |
| 26733 | ENSMUSG00000084751 | 0 | NA | NA | NA | NA | NA | Mir1894 | microRNA 1894 [Source:MGI Symbol;Acc: | 0 | 0 | 0 | 0 | 0 | 0 |
| 26734 | ENSMUSG00000084752 | 0 | NA | NA | NA | NA | NA | Gm22451 | predicted gene, 22451 [Source:MGI Symb | 0 | 0 | 0 | 0 | 0 | 0 |
| 26735 | ENSMUSG00000084753 | 0 | NA | NA | NA | NA | NA | Gm22452 | predicted gene, 22452 [Source:MGI Symb | 0 | 0 | 0 | 0 | 0 | 0 |
| 26736 | ENSMUSG00000084754 | 0 | NA | NA | NA | NA | NA | Gm12532 | predicted gene 12532 [Source:MGI Symbo | 0 | 0 | 0 | 0 | 0 | 0 |
| 26737 | ENSMUSG00000084755 | 3.428657095 | 0.365753977 | 1.17905985 | 0.310208152 | 0.756402672 | NA | Gm16276 | predicted gene 16276 [Source:MGI Symbo | 4.8186 | 3.0838 | 1.0062 | 2.8919 | 6.5437 | 2.2277 |
| 26738 | ENSMUSG00000084756 | 0 | NA | NA | NA | NA | NA | Gm13881 | predicted gene 13881 [Source:MGI Symbo | 0 | 0 | 0 | 0 | 0 | 0 |
| 26739 | ENSMUSG00000084757 | 0 | NA | NA | NA | NA | NA | 1700057H15 | RIKEN cDNA 1700057H15 gene [Source:M | 0 | 0 | 0 | 0 | 0 | 0 |
| 26740 | ENSMUSG00000084758 | 6.204946984 | -1.534877031 | 0.963717373 | -1.592663029 | 0.111235806 | 0.489450944 | Gm12798 | predicted gene 12798 [Source:MGI Symbo | 7.2279 | 15.419 | 5.0312 | 2.8919 | 1.0906 | 5.5692 |
| 26741 | ENSMUSG00000084760 | 0.167707431 | -0.796629176 | 4.080472857 | -0.195229623 | 0.84521319 | NA | Gm15472 | predicted gene 15472 [Source:MGI Symbo | 0 | 0 | 1.0062 | 0 | 0 | 0 |
| 26742 | ENSMUSG00000084761 | 2.54705572 | -0.488197876 | 1.482367601 | -0.32933658 | 0.741901298 | NA | Gm12406 | predicted gene 12406 [Source:MGI Symbo | 4.8186 | 1.0279 | 3.0187 | 0.964 | 5.4531 | 0 |
| 26743 | ENSMUSG00000084762 | 0 | NA | NA | NA | NA | NA | Platr3 | pluripotency associated transcript 3 [Sour | 0 | 0 | 0 | 0 | 0 | 0 |
| 26744 | ENSMUSG00000084764 | 0 | NA | NA | NA | NA | NA | Grip1os2 | glutamate receptor interacting protein 1, | 0 | 0 | 0 | 0 | 0 | 0 |
| 26745 | ENSMUSG00000084766 | 0 | NA | NA | NA | NA | NA | Gm12714 | predicted gene 12714 [Source:MGI Symbo | 0 | 0 | 0 | 0 | 0 | 0 |
| 26746 | ENSMUSG00000084768 | 0 | NA | NA | NA | NA | NA | Gm13605 | predicted gene 13605 [Source:MGI Symbo | 0 | 0 | 0 | 0 | 0 | 0 |
| 26747 | ENSMUSG00000084769 | 0.133850373 | -0.796629176 | 4.080472857 | -0.195229623 | 0.84521319 | NA | Gm12061 | predicted gene 12061 [Source:MGI Symbo | 0.8031 | 0 | 0 | 0 | 0 | 0 |
| 26748 | ENSMUSG00000084771 | 3.825181427 | -1.684771723 | 1.222625545 | -1.377994865 | 0.168204873 | NA | A230072E10 | RIKEN cDNA A230072E10 gene [Source:M | 3.2124 | 7.1954 | 7.0437 | 0 | 3.2719 | 2.2277 |
| 26749 | ENSMUSG00000084772 | 0 | NA | NA | NA | NA | NA | Gm15473 | predicted gene 15473 [Source:MGI Symbo | 0 | 0 | 0 | 0 | 0 | 0 |
| 26750 | ENSMUSG00000084773 | 0.449470461 | -0.545136533 | 3.562105917 | -0.15303771 | 0.878368537 | NA | Gm12159 | predicted gene 12159 [Source:MGI Symbo | 1.6062 | 0 | 0 | 0 | 1.0906 | 0 |
| 26751 | ENSMUSG00000084774 | 0 | NA | NA | NA | NA | NA | Gm14110 | predicted gene 14110 [Source:MGI Symbo | 0 | 0 | 0 | 0 | 0 | 0 |
| 26752 | ENSMUSG00000084775 | 0.572870759 | -2.577107685 | 3.182795569 | -0.809699407 | 0.418112959 | NA | Gm16741 | predicted gene, 16741 [Source:MGI Symb | 2.4093 | 1.0279 | 0 | 0 | 0 | 0 |
| 26753 | ENSMUSG00000084776 | 0 | NA | NA | NA | NA | NA | Gm14951 | predicted gene 14951 [Source:MGI Symbo | 0 | 0 | 0 | 0 | 0 | 0 |
| 26754 | ENSMUSG00000084780 | 0 | NA | NA | NA | NA | NA | Gm15350 | predicted gene 15350 [Source:MGI Symbo | 0 | 0 | 0 | 0 | 0 | 0 |
| 26755 | ENSMUSG00000084781 | 12.37181933 | -0.341639558 | 0.713585149 | -0.47876495 | 0.632105854 | 0.921791304 | D930015M0 | RIKEN cDNA D930015M05 gene [Source:N | 11.243 | 12.335 | 18.112 | 18.316 | 11.997 | 2.2277 |
| 26756 | ENSMUSG00000084782 | 0 | NA | NA | NA | NA | NA | Gm11213 | predicted gene 11213 [Source:MGI Symbo | 0 | 0 | 0 | 0 | 0 | 0 |
| 26757 | ENSMUSG00000084783 | 0 | NA | NA | NA | NA | NA | Gm15419 | predicted gene 15419 [Source:MGI Symbo | 0 | 0 | 0 | 0 | 0 | 0 |
| 26758 | ENSMUSG00000084785 | 0.133850373 | -0.796629176 | 4.080472857 | -0.195229623 | 0.84521319 | NA | Gm13972 | predicted gene 13972 [Source:MGI Symbo | 0.8031 | 0 | 0 | 0 | 0 | 0 |
| 26759 | ENSMUSG00000084786 | 950.727584 | 0.02896476 | 0.109529716 | 0.264446593 | 0.791435813 | 0.961152611 | Ubl5 | ubiquitin-like 5 [Source:MGI Symbol;Acc:N | 970.15 | 903.54 | 948.89 | 894.57 | 988.1 | 999.12 |
| 26760 | ENSMUSG00000084787 | 0 | NA | NA | NA | NA | NA | Gm12436 | predicted gene 12436 [Source:MGI Symbo | 0 | 0 | 0 | 0 | 0 | 0 |
| 26761 | ENSMUSG00000084788 | 15.52350079 | 0.891860822 | 0.555524513 | 1.605439185 | 0.108397187 | 0.48522218 | Gm11342 | predicted gene 11342 [Source:MGI Symbo | 11.243 | 13.363 | 8.05 | 24.099 | 17.45 | 18.935 |
| 26762 | ENSMUSG00000084789 | 0.36353943 | 2.065989254 | 4.034137301 | 0.512126658 | 0.608562368 | NA | Gm12974 | predicted gene 12974 [Source:MGI Symbo | 0 | 0 | 0 | 0 | 2.1812 | 0 |
| 26763 | ENSMUSG00000084790 | 2.532244422 | 0.762756572 | 1.605657407 | 0.475043162 | 0.634756209 | NA | Gm15879 | predicted gene 15879 [Source:MGI Symbo | 1.6062 | 0 | 4.025 | 1.928 | 7.6343 | 0 |
| 26764 | ENSMUSG00000084791 | 0 | NA | NA | NA | NA | NA | Gm13667 | predicted gene 13667 [Source:MGI Symbo | 0 | 0 | 0 | 0 | 0 | 0 |

| | | | | | | | | | | | | | | | |
|---|---|---|---|---|---|---|---|---|---|---|---|---|---|---|---|
| 26765 | ENSMUSG00000084792 | 4.259399354 | 1.546308706 | 1.078138351 | 1.43423959 | 0.151503903 | NA | 1700056N10 | RIKEN cDNA 1700056N10 gene [Source:M | 2.4093 | 3.0838 | 1.0062 | 5.7839 | 4.3625 | 8.9108 |
| 26766 | ENSMUSG00000084793 | 13.24483399 | -0.208097089 | 0.610089876 | -0.341092513 | 0.733033938 | 0.947915087 | Gm2568 | predicted gene 2568 [Source:MGI Symbol | 16.062 | 14.391 | 12.075 | 10.604 | 18.541 | 7.7969 |
| 26767 | ENSMUSG00000084794 | 0 | NA | NA | NA | NA | NA | Gm13662 | predicted gene 13662 [Source:MGI Symbo | 0 | 0 | 0 | 0 | 0 | 0 |
| 26768 | ENSMUSG00000084795 | 0 | NA | NA | NA | NA | NA | Gm14617 | predicted gene 14617 [Source:MGI Symbo | 0 | 0 | 0 | 0 | 0 | 0 |
| 26769 | ENSMUSG00000084796 | 3.986826038 | 0.157682046 | 1.092383229 | 0.144346821 | 0.885226608 | NA | Mir142hg | Mir142 host gene (non-protein coding) [S | 3.2124 | 3.0838 | 5.0312 | 4.8199 | 1.0906 | 6.6831 |
| 26770 | ENSMUSG00000084797 | 5.593837715 | -0.811112923 | 1.015105139 | -0.799043263 | 0.424265326 | 0.840450984 | Gm14321 | predicted gene 14321 [Source:MGI Symbo | 7.2279 | 8.2233 | 6.0375 | 7.7118 | 4.3625 | 0 |
| 26771 | ENSMUSG00000084798 | 0.181769715 | 1.126924175 | 4.080472857 | 0.276174898 | 0.782413736 | NA | 4930533B01 | RIKEN cDNA 4930533B01 gene [Source:M | 0 | 0 | 0 | 0 | 1.0906 | 0 |
| 26772 | ENSMUSG00000084799 | 18.38768576 | -0.216132769 | 0.567141125 | -0.381091688 | 0.703135215 | 0.942506832 | Ino80dos | INO80 complex subunit D, opposite stranc | 20.881 | 9.2513 | 29.181 | 20.244 | 19.631 | 11.138 |
| 26773 | ENSMUSG00000084800 | 0.524408994 | -0.837196744 | 3.377823722 | -0.247850928 | 0.804249749 | NA | Gm12205 | predicted gene 12205 [Source:MGI Symbo | 0 | 2.0558 | 0 | 0 | 1.0906 | 0 |
| 26774 | ENSMUSG00000084802 | 0 | NA | NA | NA | NA | NA | Gm14089 | predicted gene 14089 [Source:MGI Symbo | 0 | 0 | 0 | 0 | 0 | 0 |
| 26775 | ENSMUSG00000084803 | 8.436623822 | 0.204505703 | 0.83305385 | 0.245489176 | 0.806077688 | 0.964428432 | 5830444B04 | RIKEN cDNA 5830444B04 gene [Source:M | 6.4248 | 2.0558 | 15.094 | 11.568 | 5.4531 | 10.025 |
| 26776 | ENSMUSG00000084804 | 0 | NA | NA | NA | NA | NA | 1700095J12I | RIKEN cDNA 1700095J12 gene [Source:M( | 0 | 0 | 0 | 0 | 0 | 0 |
| 26777 | ENSMUSG00000084805 | 0 | NA | NA | NA | NA | NA | Gm12273 | predicted gene 12273 [Source:MGI Symbo | 0 | 0 | 0 | 0 | 0 | 0 |
| 26778 | ENSMUSG00000084806 | 0 | NA | NA | NA | NA | NA | Gm15232 | predicted gene 15232 [Source:MGI Symbo | 0 | 0 | 0 | 0 | 0 | 0 |
| 26779 | ENSMUSG00000084807 | 0.181769715 | 1.126924175 | 4.080472857 | 0.276174898 | 0.782413736 | NA | Gm13073 | predicted gene 13073 [Source:MGI Symbo | 0 | 0 | 0 | 0 | 1.0906 | 0 |
| 26780 | ENSMUSG00000084808 | 51.03215615 | 0.415800569 | 0.329383795 | 1.262358912 | 0.206819667 | 0.652194904 | 9430091E24 | RIKEN cDNA 9430091E24 gene [Source:M | 40.958 | 32.893 | 57.356 | 54.947 | 64.346 | 55.692 |
| 26781 | ENSMUSG00000084809 | 0.160662974 | 1.126924175 | 4.080472857 | 0.276174898 | 0.782413736 | NA | Gm12631 | predicted gene 12631 [Source:MGI Symbo | 0 | 0 | 0 | 0.964 | 0 | 0 |
| 26782 | ENSMUSG00000084811 | 0 | NA | NA | NA | NA | NA | 1700092E19 | RIKEN cDNA 1700092E19 gene [Source:M | 0 | 0 | 0 | 0 | 0 | 0 |
| 26783 | ENSMUSG00000084816 | 3.015447673 | -0.939498517 | 1.235445443 | -0.760453262 | 0.446983697 | NA | Platr29 | pluripotency associated transcript 29 [Sou | 4.8186 | 2.0558 | 5.0312 | 2.8919 | 2.1812 | 1.1138 |
| 26784 | ENSMUSG00000084817 | 5.664608935 | -0.194655467 | 0.936980938 | -0.207747521 | 0.835426111 | 0.971073158 | Gm5526 | predicted pseudogene 5526 [Source:MGI | 8.8341 | 7.1954 | 2.0125 | 3.8559 | 7.6343 | 4.4554 |
| 26785 | ENSMUSG00000084818 | 0 | NA | NA | NA | NA | NA | Gm14640 | predicted gene 14640 [Source:MGI Symbo | 0 | 0 | 0 | 0 | 0 | 0 |
| 26786 | ENSMUSG00000084819 | 38.5276391 | -0.128251789 | 0.374651027 | -0.342323334 | 0.732107577 | 0.947494989 | Gm11967 | predicted gene 11967 [Source:MGI Symbo | 40.155 | 52.424 | 28.175 | 37.595 | 32.719 | 40.098 |
| 26787 | ENSMUSG00000084821 | 1.021592206 | 2.3956617 | 2.456185367 | 0.975358673 | 0.329382376 | NA | Gm15880 | predicted gene 15880 [Source:MGI Symbo | 0.8031 | 0 | 0 | 0.964 | 4.3625 | 0 |
| 26788 | ENSMUSG00000084822 | 16.55570617 | -0.156365052 | 0.55823618 | -0.280105551 | 0.779396526 | 0.959870443 | Myadml2os | myeloid-associated differentiation marker | 20.881 | 10.279 | 21.131 | 18.316 | 10.906 | 17.822 |
| 26789 | ENSMUSG00000084823 | 0 | NA | NA | NA | NA | NA | Gm17216 | predicted gene 17216 [Source:MGI Symbo | 0 | 0 | 0 | 0 | 0 | 0 |
| 26790 | ENSMUSG00000084824 | 0 | NA | NA | NA | NA | NA | Gm16344 | predicted gene 16344 [Source:MGI Symbo | 0 | 0 | 0 | 0 | 0 | 0 |
| 26791 | ENSMUSG00000084825 | 0 | NA | NA | NA | NA | NA | 2410152P15 | RIKEN cDNA 2410152P15 gene [Source:M | 0 | 0 | 0 | 0 | 0 | 0 |
| 26792 | ENSMUSG00000084826 | 0.185640647 | 1.126924175 | 4.080472857 | 0.276174898 | 0.782413736 | NA | AI847159 | expressed sequence AI847159 [Source:M( | 0 | 0 | 0 | 0 | 0 | 1.1138 |
| 26793 | ENSMUSG00000084828 | 0 | NA | NA | NA | NA | NA | Gm12367 | predicted gene 12367 [Source:MGI Symbo | 0 | 0 | 0 | 0 | 0 | 0 |
| 26794 | ENSMUSG00000084829 | 0.133850373 | -0.796629176 | 4.080472857 | -0.195229623 | 0.84521319 | NA | Gm14266 | predicted gene 14266 [Source:MGI Symbo | 0.8031 | 0 | 0 | 0 | 0 | 0 |
| 26795 | ENSMUSG00000084830 | 21.14028423 | -0.174765384 | 0.491748828 | -0.355395628 | 0.722293224 | 0.9447719 | Gm14539 | predicted gene 14539 [Source:MGI Symbo | 23.29 | 26.726 | 17.106 | 13.496 | 26.175 | 20.049 |
| 26796 | ENSMUSG00000084831 | 0 | NA | NA | NA | NA | NA | Gm14710 | predicted gene 14710 [Source:MGI Symbo | 0 | 0 | 0 | 0 | 0 | 0 |
| 26797 | ENSMUSG00000084833 | 0 | NA | NA | NA | NA | NA | Gm12649 | predicted gene 12649 [Source:MGI Symbo | 0 | 0 | 0 | 0 | 0 | 0 |
| 26798 | ENSMUSG00000084834 | 22.4984114 | 0.109467675 | 0.499934096 | 0.218964211 | 0.826677927 | 0.969648181 | 4930565N06 | RIKEN cDNA 4930565N06 gene [Source:M | 18.471 | 22.614 | 24.15 | 35.667 | 20.722 | 13.366 |
| 26799 | ENSMUSG00000084835 | 2.051941184 | -0.406175939 | 1.633486021 | -0.248655901 | 0.803626963 | NA | Gm12352 | predicted gene 12352 [Source:MGI Symbo | 0 | 1.0279 | 6.0375 | 1.928 | 1.0906 | 2.2277 |
| 26800 | ENSMUSG00000084836 | 0 | NA | NA | NA | NA | NA | Gm16274 | predicted gene 16274 [Source:MGI Symbo | 0 | 0 | 0 | 0 | 0 | 0 |
| 26801 | ENSMUSG00000084837 | 0 | NA | NA | NA | NA | NA | 1700108N11 | RIKEN cDNA 1700108N11 gene [Source:M | 0 | 0 | 0 | 0 | 0 | 0 |
| 26802 | ENSMUSG00000084838 | 1.402476924 | 0.860466798 | 1.992844265 | 0.431778244 | 0.665902593 | NA | Gm10241 | predicted pseudogene 10241 [Source:MG | 0 | 0 | 3.0187 | 0.964 | 1.0906 | 3.3415 |
| 26803 | ENSMUSG00000084839 | 336.1472876 | 0.005375826 | 0.22258105 | 0.02415222 | 0.98073119 | 0.99820894 | Gm14097 | predicted gene 14097 [Source:MGI Symbo | 448.93 | 279.59 | 276.72 | 275.7 | 432.98 | 302.97 |
| 26804 | ENSMUSG00000084840 | 0 | NA | NA | NA | NA | NA | Gm11468 | predicted gene 11468 [Source:MGI Symbo | 0 | 0 | 0 | 0 | 0 | 0 |
| 26805 | ENSMUSG00000084842 | 0 | NA | NA | NA | NA | NA | Pabpc1l2b | poly(A) binding protein, cytoplasmic 1-like | 0 | 0 | 0 | 0 | 0 | 0 |
| 26806 | ENSMUSG00000084843 | 0 | NA | NA | NA | NA | NA | B230312C02 | RIKEN cDNA B230312C02 gene [Source:M | 0 | 0 | 0 | 0 | 0 | 0 |
| 26807 | ENSMUSG00000084844 | 2.676910892 | -2.810879032 | 1.631629628 | -1.722743314 | 0.084934954 | NA | Hoxb3os | homeobox B3 and homeobox B2, opposite | 8.031 | 3.0838 | 3.0187 | 1.928 | 0 | 0 |
| 26808 | ENSMUSG00000084845 | 4.794214345 | -0.144245108 | 0.995853028 | -0.144845779 | 0.884832637 | NA | Tmem240 | transmembrane protein 240 [Source:MGI | 4.0155 | 2.0558 | 9.0562 | 4.8199 | 4.3625 | 4.4554 |
| 26809 | ENSMUSG00000084846 | 4.530763914 | 0.025363941 | 1.051068669 | 0.024131574 | 0.980747658 | NA | A730011C13 | RIKEN cDNA A730011C13 gene [Source:M | 7.2279 | 5.1396 | 1.0062 | 3.8559 | 3.2719 | 6.6831 |
| 26810 | ENSMUSG00000084847 | 0 | NA | NA | NA | NA | NA | Gm15507 | predicted gene 15507 [Source:MGI Symbo | 0 | 0 | 0 | 0 | 0 | 0 |
| 26811 | ENSMUSG00000084848 | 0 | NA | NA | NA | NA | NA | 4933408N05 | RIKEN cDNA 4933408N05 gene [Source:M | 0 | 0 | 0 | 0 | 0 | 0 |
| 26812 | ENSMUSG00000084849 | 0 | NA | NA | NA | NA | NA | Gm16105 | predicted gene 16105 [Source:MGI Symbo | 0 | 0 | 0 | 0 | 0 | 0 |
| 26813 | ENSMUSG00000084850 | 0 | NA | NA | NA | NA | NA | Gm12092 | predicted gene 12092 [Source:MGI Symbo | 0 | 0 | 0 | 0 | 0 | 0 |
| 26814 | ENSMUSG00000084851 | 0 | NA | NA | NA | NA | NA | Gm11267 | predicted gene 11267 [Source:MGI Symbo | 0 | 0 | 0 | 0 | 0 | 0 |
| 26815 | ENSMUSG00000084852 | 0 | NA | NA | NA | NA | NA | Gm16359 | predicted gene 16359 [Source:MGI Symbo | 0 | 0 | 0 | 0 | 0 | 0 |
| 26816 | ENSMUSG00000084853 | 0 | NA | NA | NA | NA | NA | Gm11791 | predicted gene 11791 [Source:MGI Symbo | 0 | 0 | 0 | 0 | 0 | 0 |
| 26817 | ENSMUSG00000084854 | 0.17131964 | -0.796629176 | 4.080472857 | -0.195229623 | 0.84521319 | NA | Gm12678 | predicted gene 12678 [Source:MGI Symbo | 0 | 1.0279 | 0 | 0 | 0 | 0 |
| 26818 | ENSMUSG00000084855 | 0.335414862 | -1.779047242 | 4.030804179 | -0.441362855 | 0.658950329 | NA | Gm9850 | predicted gene 9850 [Source:MGI Symbol | 0 | 0 | 2.0125 | 0 | 0 | 0 |
| 26819 | ENSMUSG00000084856 | 0 | NA | NA | NA | NA | NA | Gm11528 | predicted gene 11528 [Source:MGI Symbo | 0 | 0 | 0 | 0 | 0 | 0 |
| 26820 | ENSMUSG00000084858 | 0 | NA | NA | NA | NA | NA | Gm1980 | predicted gene 1980 [Source:MGI Symbol | 0 | 0 | 0 | 0 | 0 | 0 |

| | | | | | | | | | | | | | | | |
|---|---|---|---|---|---|---|---|---|---|---|---|---|---|---|---|
| 26821 | ENSMUSG00000084859 | 0 | NA | NA | NA | NA | NA | 1700080N15 | RIKEN cDNA 1700080N15 gene [Source:M | 0 | 0 | 0 | 0 | 0 | 0 |
| 26822 | ENSMUSG00000084860 | 0 | NA | NA | NA | NA | NA | 2610204G07 | RIKEN cDNA 2610204G07 gene [Source:M | 0 | 0 | 0 | 0 | 0 | 0 |
| 26823 | ENSMUSG00000084861 | 0 | NA | NA | NA | NA | NA | 1700095J07I | RIKEN cDNA 1700095J07 gene [Source:MC | 0 | 0 | 0 | 0 | 0 | 0 |
| 26824 | ENSMUSG00000084862 | 4.292951955 | -0.145435934 | 1.062863872 | -0.136834018 | 0.891161994 | NA | Gm16278 | predicted gene 16278 [Source:MGI Symbo | 6.4248 | 3.0838 | 4.025 | 6.7478 | 4.3625 | 1.1138 |
| 26825 | ENSMUSG00000084863 | 0 | NA | NA | NA | NA | NA | Gm12523 | predicted gene 12523 [Source:MGI Symbo | 0 | 0 | 0 | 0 | 0 | 0 |
| 26826 | ENSMUSG00000084864 | 0.17131964 | -0.796629176 | 4.080472857 | -0.195229623 | 0.84521319 | NA | 1700027A07 | RIKEN cDNA 1700027A07 gene [Source:M | 0 | 1.0279 | 0 | 0 | 0 | 0 |
| 26827 | ENSMUSG00000084865 | 0.505131668 | 1.126696199 | 3.421159819 | 0.329331647 | 0.741905026 | NA | Gm15723 | predicted gene 15723 [Source:MGI Symbo | 0.8031 | 0 | 0 | 0 | 0 | 2.2277 |
| 26828 | ENSMUSG00000084866 | 3.060890899 | 0.027130025 | 1.297674282 | 0.020906652 | 0.98332012 | NA | A930006K02 | RIKEN cDNA A930006K02 gene [Source:M | 4.0155 | 2.0558 | 3.0187 | 4.8199 | 0 | 4.4554 |
| 26829 | ENSMUSG00000084869 | 0 | NA | NA | NA | NA | NA | Gm13063 | predicted gene 13063 [Source:MGI Symbo | 0 | 0 | 0 | 0 | 0 | 0 |
| 26830 | ENSMUSG00000084870 | 0.321325949 | 1.911575114 | 4.048267893 | 0.472195804 | 0.636787034 | NA | Gm13954 | predicted gene 13954 [Source:MGI Symbo | 0 | 0 | 0 | 1.928 | 0 | 0 |
| 26831 | ENSMUSG00000084871 | 0 | NA | NA | NA | NA | NA | Gm2800 | predicted gene 2800 [Source:MGI Symbol | 0 | 0 | 0 | 0 | 0 | 0 |
| 26832 | ENSMUSG00000084873 | 0 | NA | NA | NA | NA | NA | C030047K22 | RIKEN cDNA C030047K22 gene [Source:M | 0 | 0 | 0 | 0 | 0 | 0 |
| 26833 | ENSMUSG00000084874 | 0.36353943 | 2.065989254 | 4.034137301 | 0.512126658 | 0.608562368 | NA | Gm12906 | predicted gene 12906 [Source:MGI Symbo | 0 | 0 | 0 | 0 | 2.1812 | 0 |
| 26834 | ENSMUSG00000084875 | 0 | NA | NA | NA | NA | NA | 4930570D08 | RIKEN cDNA 4930570D08 gene [Source:M | 0 | 0 | 0 | 0 | 0 | 0 |
| 26835 | ENSMUSG00000084876 | 0 | NA | NA | NA | NA | NA | Gm14965 | predicted gene 14965 [Source:MGI Symbo | 0 | 0 | 0 | 0 | 0 | 0 |
| 26836 | ENSMUSG00000084878 | 2.854267779 | 1.431558654 | 1.353961313 | 1.057311343 | 0.290369518 | NA | Lrrc8dos | leucine rich repeat containing 8D, opposit | 1.6062 | 0 | 3.0187 | 4.8199 | 5.4531 | 2.2277 |
| 26837 | ENSMUSG00000084880 | 9.327354494 | 0.106267722 | 0.765206588 | 0.138874552 | 0.88954928 | 0.982382091 | Tomm6os | translocase of outer mitochondrial memb | 9.6372 | 12.335 | 5.0312 | 12.532 | 13.087 | 3.3415 |
| 26838 | ENSMUSG00000084883 | 370.7623783 | 0.275628062 | 0.149321475 | 1.845870211 | 0.064911053 | 0.36546262 | Ccdc85c | coiled-coil domain containing 85C [Source | 324.45 | 315.57 | 366.27 | 387.52 | 374.08 | 456.68 |
| 26839 | ENSMUSG00000084884 | 0 | NA | NA | NA | NA | NA | Gm12289 | predicted gene 12289 [Source:MGI Symbo | 0 | 0 | 0 | 0 | 0 | 0 |
| 26840 | ENSMUSG00000084885 | 0.160662974 | 1.126924175 | 4.080472857 | 0.276174898 | 0.782413736 | NA | 3010001F23 | RIKEN cDNA 3010001F23 gene [Source:M | 0 | 0 | 0 | 0.964 | 0 | 0 |
| 26841 | ENSMUSG00000084887 | 0 | NA | NA | NA | NA | NA | Gm11749 | predicted gene 11749 [Source:MGI Symbo | 0 | 0 | 0 | 0 | 0 | 0 |
| 26842 | ENSMUSG00000084889 | 0 | NA | NA | NA | NA | NA | Gm12148 | predicted gene 12148 [Source:MGI Symbo | 0 | 0 | 0 | 0 | 0 | 0 |
| 26843 | ENSMUSG00000084890 | 0 | NA | NA | NA | NA | NA | A830036E02 | RIKEN cDNA A830036E02 gene [Source:M | 0 | 0 | 0 | 0 | 0 | 0 |
| 26844 | ENSMUSG00000084891 | 0 | NA | NA | NA | NA | NA | Gm12958 | predicted gene 12958 [Source:MGI Symbo | 0 | 0 | 0 | 0 | 0 | 0 |
| 26845 | ENSMUSG00000084892 | 0.52105551 | -0.812028029 | 3.385364865 | -0.239864257 | 0.810435491 | NA | Gm14471 | predicted gene 14471 [Source:MGI Symbo | 0 | 0 | 2.0125 | 0 | 0 | 1.1138 |
| 26846 | ENSMUSG00000084893 | 0.167707431 | -0.796629176 | 4.080472857 | -0.195229623 | 0.84521319 | NA | Hba-ps4 | hemoglobin alpha, pseudogene 4 [Source | 0 | 0 | 1.0062 | 0 | 0 | 0 |
| 26847 | ENSMUSG00000084895 | 0 | NA | NA | NA | NA | NA | AA672651 | expressed sequence AA672651 [Source:M | 0 | 0 | 0 | 0 | 0 | 0 |
| 26848 | ENSMUSG00000084896 | 0 | NA | NA | NA | NA | NA | Gm11632 | predicted gene 11632 [Source:MGI Symbo | 0 | 0 | 0 | 0 | 0 | 0 |
| 26849 | ENSMUSG00000084897 | 1.504331798 | -0.865885113 | 1.968269715 | -0.439921981 | 0.659993615 | NA | Gm14226 | predicted gene 14226 [Source:MGI Symbo | 0.8031 | 0 | 5.0312 | 0.964 | 0 | 2.2277 |
| 26850 | ENSMUSG00000084898 | 0 | NA | NA | NA | NA | NA | Gm12371 | predicted gene 12371 [Source:MGI Symbo | 0 | 0 | 0 | 0 | 0 | 0 |
| 26851 | ENSMUSG00000084899 | 16.42558458 | -0.24041493 | 0.552997907 | -0.434748355 | 0.663745096 | 0.929301301 | Gm15344 | predicted gene 15344 [Source:MGI Symbo | 25.699 | 12.335 | 15.094 | 13.496 | 17.45 | 14.48 |
| 26852 | ENSMUSG00000084901 | 0 | NA | NA | NA | NA | NA | Gm13266 | predicted gene 13266 [Source:MGI Symbo | 0 | 0 | 0 | 0 | 0 | 0 |
| 26853 | ENSMUSG00000084902 | 0 | NA | NA | NA | NA | NA | Gm281 | predicted gene 281 [Source:MGI Symbol;/ | 0 | 0 | 0 | 0 | 0 | 0 |
| 26854 | ENSMUSG00000084904 | 0.503302254 | -0.837877584 | 3.426698643 | -0.244514523 | 0.806832356 | NA | Gm14827 | predicted gene 14827 [Source:MGI Symbo | 0 | 2.0558 | 0 | 0.964 | 0 | 0 |
| 26855 | ENSMUSG00000084905 | 0 | NA | NA | NA | NA | NA | Gm12081 | predicted gene 12081 [Source:MGI Symbo | 0 | 0 | 0 | 0 | 0 | 0 |
| 26856 | ENSMUSG00000084906 | 0 | NA | NA | NA | NA | NA | Gm13794 | predicted gene 13794 [Source:MGI Symbo | 0 | 0 | 0 | 0 | 0 | 0 |
| 26857 | ENSMUSG00000084907 | 0 | NA | NA | NA | NA | NA | Gm14505 | predicted gene 14505 [Source:MGI Symbo | 0 | 0 | 0 | 0 | 0 | 0 |
| 26858 | ENSMUSG00000084908 | 0 | NA | NA | NA | NA | NA | C79798 | expressed sequence C79798 [Source:MGI | 0 | 0 | 0 | 0 | 0 | 0 |
| 26859 | ENSMUSG00000084909 | 0 | NA | NA | NA | NA | NA | 4930528G23 | RIKEN cDNA 4930528G23 gene [Source:M | 0 | 0 | 0 | 0 | 0 | 0 |
| 26860 | ENSMUSG00000084910 | 62.90325597 | 0.609009161 | 0.312786649 | 1.947043338 | 0.051529539 | 0.318923087 | C630043F03 | RIKEN cDNA C630043F03 gene [Source:M | 53.808 | 38.033 | 57.356 | 71.334 | 59.984 | 96.904 |
| 26861 | ENSMUSG00000084911 | 3.318524034 | -1.463485511 | 1.221326302 | -1.198275603 | 0.230809741 | NA | Gm16185 | predicted gene 16185 [Source:MGI Symbo | 6.4248 | 4.1117 | 4.025 | 0.964 | 3.2719 | 1.1138 |
| 26862 | ENSMUSG00000084913 | 0 | NA | NA | NA | NA | NA | Gm7616 | predicted gene 7616 [Source:MGI Symbol | 0 | 0 | 0 | 0 | 0 | 0 |
| 26863 | ENSMUSG00000084915 | 7.410583933 | -0.189785127 | 0.780341824 | -0.243207683 | 0.807844515 | 0.965116302 | C230037L18 | RIKEN cDNA C230037L18 gene [Source:M | 10.44 | 6.1675 | 7.0437 | 8.6758 | 5.4531 | 6.6831 |
| 26864 | ENSMUSG00000084916 | 0 | NA | NA | NA | NA | NA | Gm11656 | predicted gene 11656 [Source:MGI Symbo | 0 | 0 | 0 | 0 | 0 | 0 |
| 26865 | ENSMUSG00000084917 | 1.366531728 | 0.046504607 | 1.976116938 | 0.023533328 | 0.981224854 | NA | Gm17477 | predicted gene, 17477 [Source:MGI Symb | 0 | 3.0838 | 1.0062 | 1.928 | 2.1812 | 0 |
| 26866 | ENSMUSG00000084918 | 0.133850373 | -0.796629176 | 4.080472857 | -0.195229623 | 0.84521319 | NA | Gm12708 | predicted gene 12708 [Source:MGI Symbo | 0.8031 | 0 | 0 | 0 | 0 | 0 |
| 26867 | ENSMUSG00000084919 | 0 | NA | NA | NA | NA | NA | Ptprtos | protein tyrosine phosphatase, receptor ty | 0 | 0 | 0 | 0 | 0 | 0 |
| 26868 | ENSMUSG00000084920 | 0 | NA | NA | NA | NA | NA | Gm15230 | predicted gene 15230 [Source:MGI Symbo | 0 | 0 | 0 | 0 | 0 | 0 |
| 26869 | ENSMUSG00000084921 | 0 | NA | NA | NA | NA | NA | Gm13838 | predicted gene 13838 [Source:MGI Symbo | 0 | 0 | 0 | 0 | 0 | 0 |
| 26870 | ENSMUSG00000084923 | 9.882384618 | 0.895563937 | 0.741412709 | 1.207915546 | 0.227079749 | 0.6734635 | Gm15611 | predicted gene 15611 [Source:MGI Symbo | 6.4248 | 9.2513 | 5.0312 | 8.6758 | 7.6343 | 22.277 |
| 26871 | ENSMUSG00000084924 | 0 | NA | NA | NA | NA | NA | Vmn1r-ps10 | vomeronasal 1 receptor, pseudogene 103 | 0 | 0 | 0 | 0 | 0 | 0 |
| 26872 | ENSMUSG00000084925 | 4.946185063 | -0.043411679 | 1.140283197 | -0.038070962 | 0.969631103 | NA | 1810062O18 | RIKEN cDNA 1810062O18 gene [Source:M | 0 | 5.1396 | 10.062 | 6.7478 | 3.2719 | 4.4554 |
| 26873 | ENSMUSG00000084927 | 0 | NA | NA | NA | NA | NA | Gm5724 | predicted gene 5724 [Source:MGI Symbol | 0 | 0 | 0 | 0 | 0 | 0 |
| 26874 | ENSMUSG00000084929 | 0 | NA | NA | NA | NA | NA | Foxo6os | forkhead box O6, opposite strand [Source | 0 | 0 | 0 | 0 | 0 | 0 |
| 26875 | ENSMUSG00000084930 | 1.805138933 | 1.620780737 | 1.801828275 | 0.899520093 | 0.368375698 | NA | Gm11706 | predicted gene 11706 [Source:MGI Symbo | 1.6062 | 0 | 1.0062 | 3.8559 | 4.3625 | 0 |
| 26876 | ENSMUSG00000084932 | 0 | NA | NA | NA | NA | NA | Gm15156 | predicted gene 15156 [Source:MGI Symbo | 0 | 0 | 0 | 0 | 0 | 0 |

| | | | | | | | | | | | | | | | |
|---|---|---|---|---|---|---|---|---|---|---|---|---|---|---|---|
| 26877 | ENSMUSG00000084933 | 0 | NA | NA | NA | NA | NA | Gm13962 | predicted gene 13962 [Source:MGI Symbo | 0 | 0 | 0 | 0 | 0 | 0 |
| 26878 | ENSMUSG00000084934 | 13.1115564 | 0.437102162 | 0.630954153 | 0.692763746 | 0.488457827 | 0.874431125 | Gm16035 | predicted gene 16035 [Source:MGI Symbo | 8.031 | 18.503 | 7.0437 | 16.388 | 11.997 | 16.708 |
| 26879 | ENSMUSG00000084935 | 0 | NA | NA | NA | NA | NA | Gm14161 | predicted gene 14161 [Source:MGI Symbo | 0 | 0 | 0 | 0 | 0 | 0 |
| 26880 | ENSMUSG00000084936 | 0 | NA | NA | NA | NA | NA | Gm11583 | predicted gene 11583 [Source:MGI Symbo | 0 | 0 | 0 | 0 | 0 | 0 |
| 26881 | ENSMUSG00000084937 | 0 | NA | NA | NA | NA | NA | Gm11228 | predicted gene 11228 [Source:MGI Symbo | 0 | 0 | 0 | 0 | 0 | 0 |
| 26882 | ENSMUSG00000084938 | 0 | NA | NA | NA | NA | NA | BB557941 | expressed sequence BB557941 [Source:M | 0 | 0 | 0 | 0 | 0 | 0 |
| 26883 | ENSMUSG00000084939 | 3.48548601 | -2.043986684 | 1.384321403 | -1.476526101 | 0.139802714 | NA | Gm830 | predicted gene 830 [Source:MGI Symbol;A | 5.6217 | 10.279 | 1.0062 | 2.8919 | 0 | 1.1138 |
| 26884 | ENSMUSG00000084940 | 0 | NA | NA | NA | NA | NA | Gm15935 | predicted gene 15935 [Source:MGI Symbo | 0 | 0 | 0 | 0 | 0 | 0 |
| 26885 | ENSMUSG00000084941 | 4.187459898 | 0.982977583 | 1.068225752 | 0.920196486 | 0.35747009 | NA | Gm11944 | predicted gene 11944 [Source:MGI Symbo | 2.4093 | 1.0279 | 5.0312 | 6.7478 | 5.4531 | 4.4554 |
| 26886 | ENSMUSG00000084942 | 0 | NA | NA | NA | NA | NA | 7330404K18 | RIKEN cDNA 7330404K18 gene [Source:M | 0 | 0 | 0 | 0 | 0 | 0 |
| 26887 | ENSMUSG00000084944 | 0 | NA | NA | NA | NA | NA | Gm14684 | predicted gene 14684 [Source:MGI Symbo | 0 | 0 | 0 | 0 | 0 | 0 |
| 26888 | ENSMUSG00000084945 | 0.133850373 | -0.796629176 | 4.080472857 | -0.195229623 | 0.84521319 | NA | C030037F17 | RIKEN cDNA C030037F17 gene [Source:M | 0.8031 | 0 | 0 | 0 | 0 | 0 |
| 26889 | ENSMUSG00000084946 | 7.785427258 | 0.343975905 | 0.784884812 | 0.438250173 | 0.661204941 | 0.928341045 | Dlx1as | distal-less homeobox 1, antisense [Source | 6.4248 | 5.1396 | 9.0562 | 9.6398 | 11.997 | 4.4554 |
| 26890 | ENSMUSG00000084947 | 0 | NA | NA | NA | NA | NA | Gm15594 | predicted gene 15594 [Source:MGI Symbo | 0 | 0 | 0 | 0 | 0 | 0 |
| 26891 | ENSMUSG00000084948 | 0.937165244 | 2.270987635 | 2.769260373 | 0.820070101 | 0.412176145 | NA | 1700061H18 | RIKEN cDNA 1700061H18 gene [Source:M | 0.8031 | 0 | 0 | 4.8199 | 0 | 0 |
| 26892 | ENSMUSG00000084949 | 0 | NA | NA | NA | NA | NA | Gm16249 | predicted gene 16249 [Source:MGI Symbo | 0 | 0 | 0 | 0 | 0 | 0 |
| 26893 | ENSMUSG00000084950 | 1.277557541 | -3.715649068 | 2.202711627 | -1.686852252 | 0.091631765 | NA | Gm5577 | predicted gene 5577 [Source:MGI Symbol | 1.6062 | 1.0279 | 5.0312 | 0 | 0 | 0 |
| 26894 | ENSMUSG00000084951 | 0.321325949 | 1.911575114 | 4.048267893 | 0.472195804 | 0.636787034 | NA | Mipepos | mitochondrial intermediate peptidase, op | 0 | 0 | 0 | 1.928 | 0 | 0 |
| 26895 | ENSMUSG00000084954 | 0.36353943 | 2.065989254 | 4.034137301 | 0.512126658 | 0.608562368 | NA | Gm16351 | predicted gene 16351 [Source:MGI Symbo | 0 | 0 | 0 | 0 | 2.1812 | 0 |
| 26896 | ENSMUSG00000084955 | 0 | NA | NA | NA | NA | NA | Gm14349 | predicted gene 14349 [Source:MGI Symbo | 0 | 0 | 0 | 0 | 0 | 0 |
| 26897 | ENSMUSG00000084957 | 200.1532606 | 0.306604349 | 0.200699876 | 1.527675829 | 0.126593044 | 0.521187766 | Bbip1 | BBSome interacting protein 1 [Source:MG | 172.67 | 175.77 | 188.17 | 174.48 | 273.75 | 216.09 |
| 26898 | ENSMUSG00000084959 | 0 | NA | NA | NA | NA | NA | 4933407I08F | RIKEN cDNA 4933407I08 gene [Source:MG | 0 | 0 | 0 | 0 | 0 | 0 |
| 26899 | ENSMUSG00000084960 | 2.911014372 | 0.932773952 | 1.500033104 | 0.621835578 | 0.53404999 | NA | B430010I23F | RIKEN cDNA B430010I23 gene [Source:MG | 0 | 3.0838 | 3.0187 | 4.8199 | 6.5437 | 0 |
| 26900 | ENSMUSG00000084962 | 0 | NA | NA | NA | NA | NA | Gm11947 | predicted gene 11947 [Source:MGI Symbo | 0 | 0 | 0 | 0 | 0 | 0 |
| 26901 | ENSMUSG00000084963 | 0 | NA | NA | NA | NA | NA | Gm14936 | predicted gene 14936 [Source:MGI Symbo | 0 | 0 | 0 | 0 | 0 | 0 |
| 26902 | ENSMUSG00000084964 | 0 | NA | NA | NA | NA | NA | Gm15503 | predicted gene 15503 [Source:MGI Symbo | 0 | 0 | 0 | 0 | 0 | 0 |
| 26903 | ENSMUSG00000084965 | 0 | NA | NA | NA | NA | NA | Gm13857 | predicted gene 13857 [Source:MGI Symbo | 0 | 0 | 0 | 0 | 0 | 0 |
| 26904 | ENSMUSG00000084966 | 0 | NA | NA | NA | NA | NA | 2810471M0 | RIKEN cDNA 2810471M01 gene [Source:M | 0 | 0 | 0 | 0 | 0 | 0 |
| 26905 | ENSMUSG00000084967 | 0 | NA | NA | NA | NA | NA | Gm12296 | predicted gene 12296 [Source:MGI Symbo | 0 | 0 | 0 | 0 | 0 | 0 |
| 26906 | ENSMUSG00000084968 | 0.305170013 | -1.657836477 | 4.041480335 | -0.410205257 | 0.681655385 | NA | Gm12743 | predicted gene 12743 [Source:MGI Symbo | 0.8031 | 1.0279 | 0 | 0 | 0 | 0 |
| 26907 | ENSMUSG00000084969 | 2.269726575 | 1.935671343 | 1.657416972 | 1.16788435 | 0.242853418 | NA | Gm11205 | predicted gene 11205 [Source:MGI Symbo | 0.8031 | 0 | 2.0125 | 0.964 | 8.7249 | 1.1138 |
| 26908 | ENSMUSG00000084970 | 1.40643048 | 0.428278072 | 2.131914534 | 0.20088895 | 0.840785408 | NA | 1700060J05F | RIKEN cDNA 1700060J05 gene [Source:MG | 1.6062 | 0 | 2.0125 | 4.8199 | 0 | 0 |
| 26909 | ENSMUSG00000084971 | 0 | NA | NA | NA | NA | NA | 4930579D09 | RIKEN cDNA 4930579D09 gene [Source:M | 0 | 0 | 0 | 0 | 0 | 0 |
| 26910 | ENSMUSG00000084973 | 0 | NA | NA | NA | NA | NA | Gm13848 | predicted gene 13848 [Source:MGI Symbo | 0 | 0 | 0 | 0 | 0 | 0 |
| 26911 | ENSMUSG00000084974 | 2.253394766 | 1.334380375 | 1.429177617 | 0.933670077 | 0.350474105 | NA | Gm15567 | predicted gene 15567 [Source:MGI Symbo | 0.8031 | 2.0558 | 1.0062 | 1.928 | 3.2719 | 4.4554 |
| 26912 | ENSMUSG00000084978 | 0 | NA | NA | NA | NA | NA | Gm11655 | predicted gene 11655 [Source:MGI Symbo | 0 | 0 | 0 | 0 | 0 | 0 |
| 26913 | ENSMUSG00000084979 | 0 | NA | NA | NA | NA | NA | Gm16267 | predicted gene 16267 [Source:MGI Symbo | 0 | 0 | 0 | 0 | 0 | 0 |
| 26914 | ENSMUSG00000084980 | 0 | NA | NA | NA | NA | NA | Slc36a1os | solute carrier family 36 (proton/amino aci | 0 | 0 | 0 | 0 | 0 | 0 |
| 26915 | ENSMUSG00000084981 | 40.81011668 | 0.844166157 | 0.461029338 | 1.831046504 | 0.067093602 | 0.372673526 | Gm15962 | predicted gene 15962 [Source:MGI Symbo | 28.912 | 23.642 | 35.219 | 82.902 | 20.722 | 53.465 |
| 26916 | ENSMUSG00000084982 | 0 | NA | NA | NA | NA | NA | Gm15275 | predicted gene 15275 [Source:MGI Symbo | 0 | 0 | 0 | 0 | 0 | 0 |
| 26917 | ENSMUSG00000084983 | 689.2303197 | -0.045180983 | 0.135815038 | -0.332665541 | 0.739386764 | 0.948310621 | Gm11789 | predicted gene 11789 [Source:MGI Symbo | 802.3 | 595.16 | 701.35 | 678.64 | 672.91 | 685.01 |
| 26918 | ENSMUSG00000084984 | 0.335414862 | -1.779047242 | 4.030804179 | -0.441362855 | 0.658950329 | NA | Far1os | fatty acyl CoA reductase 1, opposite stran | 0 | 0 | 2.0125 | 0 | 0 | 0 |
| 26919 | ENSMUSG00000084985 | 0.481988923 | 2.492401104 | 3.837077151 | 0.649557203 | 0.515978285 | NA | Gm16135 | predicted gene 16135 [Source:MGI Symbo | 0 | 0 | 0 | 2.8919 | 0 | 0 |
| 26920 | ENSMUSG00000084986 | 0 | NA | NA | NA | NA | NA | Gm16224 | predicted gene 16224 [Source:MGI Symbo | 0 | 0 | 0 | 0 | 0 | 0 |
| 26921 | ENSMUSG00000084987 | 0 | NA | NA | NA | NA | NA | Gm13134 | predicted gene 13134 [Source:MGI Symbo | 0 | 0 | 0 | 0 | 0 | 0 |
| 26922 | ENSMUSG00000084989 | 0 | NA | NA | NA | NA | NA | Crocc2 | ciliary rootlet coiled-coil, rootletin family r | 0 | 0 | 0 | 0 | 0 | 0 |
| 26923 | ENSMUSG00000084990 | 0 | NA | NA | NA | NA | NA | Gm14549 | predicted gene 14549 [Source:MGI Symbo | 0 | 0 | 0 | 0 | 0 | 0 |
| 26924 | ENSMUSG00000084991 | 0.160662974 | 1.126924175 | 4.080472857 | 0.276174898 | 0.782413736 | NA | Gm16260 | predicted gene 16260 [Source:MGI Symbo | 0 | 0 | 0 | 0.964 | 0 | 0 |
| 26925 | ENSMUSG00000084992 | 0 | NA | NA | NA | NA | NA | Gm13842 | predicted gene 13842 [Source:MGI Symbo | 0 | 0 | 0 | 0 | 0 | 0 |
| 26926 | ENSMUSG00000084993 | 0 | NA | NA | NA | NA | NA | Gm12305 | predicted gene 12305 [Source:MGI Symbo | 0 | 0 | 0 | 0 | 0 | 0 |
| 26927 | ENSMUSG00000084995 | 0 | NA | NA | NA | NA | NA | Lyzl4os | lysozyme-like 4, opposite strand [Source:N | 0 | 0 | 0 | 0 | 0 | 0 |
| 26928 | ENSMUSG00000084996 | 16.65860683 | -0.829913635 | 0.540803538 | -1.534593576 | 0.124883692 | 0.518460883 | Gm11419 | predicted gene 11419 [Source:MGI Symbo | 24.093 | 23.642 | 16.1 | 9.6398 | 11.997 | 14.48 |
| 26929 | ENSMUSG00000084997 | 0 | NA | NA | NA | NA | NA | Gm15471 | predicted gene 15471 [Source:MGI Symbo | 0 | 0 | 0 | 0 | 0 | 0 |
| 26930 | ENSMUSG00000084998 | 13.71345857 | 0.417780769 | 0.57226394 | 0.730049092 | 0.465360178 | 0.863256867 | Gm16279 | predicted gene 16279 [Source:MGI Symbo | 12.85 | 10.279 | 12.075 | 17.352 | 16.359 | 13.366 |
| 26931 | ENSMUSG00000084999 | 0 | NA | NA | NA | NA | NA | Gm15839 | predicted gene 15839 [Source:MGI Symbo | 0 | 0 | 0 | 0 | 0 | 0 |
| 26932 | ENSMUSG00000085000 | 0 | NA | NA | NA | NA | NA | Gm14553 | predicted gene 14553 [Source:MGI Symbo | 0 | 0 | 0 | 0 | 0 | 0 |

| | | | | | | | | | | | | | | | |
|---|---|---|---|---|---|---|---|---|---|---|---|---|---|---|---|
| 26933 | ENSMUSG00000085001 | 10.6310157 | 0.569235796 | 0.806815425 | 0.705534101 | 0.480477918 | 0.870278888 | Rapgef4os2 | Rap guanine nucleotide exchange factor ( | 10.44 | 11.307 | 4.025 | 26.991 | 5.4531 | 5.5692 |
| 26934 | ENSMUSG00000085002 | 0 | NA | NA | NA | NA | NA | Gm12984 | predicted gene 12984 [Source:MGI Symbo | 0 | 0 | 0 | 0 | 0 | 0 |
| 26935 | ENSMUSG00000085003 | 0 | NA | NA | NA | NA | NA | Pip5k1bos | phosphatidylinositol-4-phosphate 5-kinas | 0 | 0 | 0 | 0 | 0 | 0 |
| 26936 | ENSMUSG00000085004 | 0 | NA | NA | NA | NA | NA | 5430427M07 | RIKEN cDNA 5430427M07 gene [Source:M | 0 | 0 | 0 | 0 | 0 | 0 |
| 26937 | ENSMUSG00000085006 | 39.65604752 | 1.257822949 | 0.403424057 | 3.117867978 | 0.001821644 | 0.032561965 | BC021767 | cDNA sequence BC021767 [Source:MGI Sy | 17.668 | 16.447 | 36.225 | 45.307 | 63.256 | 59.034 |
| 26938 | ENSMUSG00000085007 | 0 | NA | NA | NA | NA | NA | Smim43 | small integral membrane protein 43 [Sour | 0 | 0 | 0 | 0 | 0 | 0 |
| 26939 | ENSMUSG00000085008 | 0 | NA | NA | NA | NA | NA | Dbhos | dopamine beta hydroxylase, opposite stra | 0 | 0 | 0 | 0 | 0 | 0 |
| 26940 | ENSMUSG00000085009 | 0.658259367 | -1.329928433 | 2.812497761 | -0.472863819 | 0.636310339 | NA | Gm12977 | predicted gene 12977 [Source:MGI Symbo | 0.8031 | 2.0558 | 0 | 0 | 1.0906 | 0 |
| 26941 | ENSMUSG00000085010 | 0 | NA | NA | NA | NA | NA | Gssos1 | glutathione synthase, opposite strand 1 [S | 0 | 0 | 0 | 0 | 0 | 0 |
| 26942 | ENSMUSG00000085012 | 0.838717116 | -1.825569927 | 2.582619186 | -0.706867639 | 0.479648736 | NA | Gm15492 | predicted gene 15492 [Source:MGI Symbo | 0 | 2.0558 | 2.0125 | 0.964 | 0 | 0 |
| 26943 | ENSMUSG00000085014 | 0 | NA | NA | NA | NA | NA | Gm13490 | predicted gene 13490 [Source:MGI Symbo | 0 | 0 | 0 | 0 | 0 | 0 |
| 26944 | ENSMUSG00000085015 | 0 | NA | NA | NA | NA | NA | Gm14424 | predicted gene 14424 [Source:MGI Symbo | 0 | 0 | 0 | 0 | 0 | 0 |
| 26945 | ENSMUSG00000085016 | 1.69624073 | -1.081326345 | 1.840571232 | -0.587494972 | 0.556871326 | NA | Gm11335 | predicted gene 11335 [Source:MGI Symbo | 0.8031 | 3.0838 | 3.0187 | 0 | 3.2719 | 0 |
| 26946 | ENSMUSG00000085017 | 2.197785164 | -1.62890336 | 1.65128467 | -0.986446123 | 0.323914241 | NA | Gm13412 | predicted gene 13412 [Source:MGI Symbo | 4.8186 | 3.0838 | 2.0125 | 0 | 3.2719 | 0 |
| 26947 | ENSMUSG00000085019 | 0.160662974 | 1.126924175 | 4.080472857 | 0.276174898 | 0.782413736 | NA | Pard3bos2 | par-3 family cell polarity regulator beta, o | 0 | 0 | 0 | 0.964 | 0 | 0 |
| 26948 | ENSMUSG00000085020 | 0 | NA | NA | NA | NA | NA | 2310081O03 | RIKEN cDNA 2310081O03 gene [Source:M | 0 | 0 | 0 | 0 | 0 | 0 |
| 26949 | ENSMUSG00000085021 | 0.167707431 | -0.796629176 | 4.080472857 | -0.195229623 | 0.84521319 | NA | Gm12051 | predicted gene 12051 [Source:MGI Symbo | 0 | 0 | 1.0062 | 0 | 0 | 0 |
| 26950 | ENSMUSG00000085022 | 0 | NA | NA | NA | NA | NA | Gm5860 | predicted gene 5860 [Source:MGI Symbol | 0 | 0 | 0 | 0 | 0 | 0 |
| 26951 | ENSMUSG00000085023 | 0 | NA | NA | NA | NA | NA | Gm12744 | predicted gene 12744 [Source:MGI Symbo | 0 | 0 | 0 | 0 | 0 | 0 |
| 26952 | ENSMUSG00000085024 | 4.532709749 | -0.715158365 | 1.039432152 | -0.688027943 | 0.491435185 | NA | C230035I16I | RIKEN cDNA C230035I16 gene [Source:MG | 4.8186 | 4.1117 | 8.05 | 5.7839 | 1.0906 | 3.3415 |
| 26953 | ENSMUSG00000085025 | 0 | NA | NA | NA | NA | NA | Gm13715 | predicted gene 13715 [Source:MGI Symbo | 0 | 0 | 0 | 0 | 0 | 0 |
| 26954 | ENSMUSG00000085026 | 0 | NA | NA | NA | NA | NA | Srrm3os | serine/arginine repetitive matrix 3, opposi | 0 | 0 | 0 | 0 | 0 | 0 |
| 26955 | ENSMUSG00000085027 | 0 | NA | NA | NA | NA | NA | Gm11840 | predicted gene 11840 [Source:MGI Symbo | 0 | 0 | 0 | 0 | 0 | 0 |
| 26956 | ENSMUSG00000085028 | 126.7487131 | 0.241321741 | 0.211419189 | 1.141437268 | 0.253688003 | 0.706909089 | Slc2a4rg-ps | Slc2a4 regulator, pseudogene [Source:MG | 114.84 | 108.96 | 124.77 | 142.67 | 118.88 | 150.37 |
| 26957 | ENSMUSG00000085029 | 0 | NA | NA | NA | NA | NA | Gm13569 | predicted gene 13569 [Source:MGI Symbo | 0 | 0 | 0 | 0 | 0 | 0 |
| 26958 | ENSMUSG00000085030 | 1.506341172 | 0.559890583 | 1.913681709 | 0.292572469 | 0.76984896 | NA | 2810455O05 | RIKEN cDNA 2810455O05 gene [Source:M | 1.6062 | 0 | 2.0125 | 0.964 | 0 | 4.4554 |
| 26959 | ENSMUSG00000085032 | 0 | NA | NA | NA | NA | NA | Gm11217 | predicted gene 11217 [Source:MGI Symbo | 0 | 0 | 0 | 0 | 0 | 0 |
| 26960 | ENSMUSG00000085033 | 0.294513347 | 0.165143496 | 4.080472857 | 0.040471657 | 0.967717103 | NA | Gm11646 | predicted gene 11646 [Source:MGI Symbo | 0.8031 | 0 | 0 | 0.964 | 0 | 0 |
| 26961 | ENSMUSG00000085034 | 0 | NA | NA | NA | NA | NA | Gm11240 | predicted gene 11240 [Source:MGI Symbo | 0 | 0 | 0 | 0 | 0 | 0 |
| 26962 | ENSMUSG00000085035 | 0 | NA | NA | NA | NA | NA | Gm12031 | predicted gene 12031 [Source:MGI Symbo | 0 | 0 | 0 | 0 | 0 | 0 |
| 26963 | ENSMUSG00000085036 | 0 | NA | NA | NA | NA | NA | Gm14011 | predicted gene 14011 [Source:MGI Symbo | 0 | 0 | 0 | 0 | 0 | 0 |
| 26964 | ENSMUSG00000085037 | 78.24245579 | 0.137940795 | 0.288450682 | 0.478212755 | 0.632498785 | 0.922076476 | 4933421O10 | RIKEN cDNA 4933421O10 gene [Source:M | 89.947 | 75.038 | 58.362 | 99.29 | 62.165 | 84.652 |
| 26965 | ENSMUSG00000085039 | 2.658822332 | 0.480094663 | 1.308119002 | 0.367011459 | 0.713610471 | NA | Gm15927 | predicted gene 15927 [Source:MGI Symbo | 1.6062 | 4.1117 | 1.0062 | 4.8199 | 2.1812 | 2.2277 |
| 26966 | ENSMUSG00000085041 | 0 | NA | NA | NA | NA | NA | BB031773 | expressed sequence BB031773 [Source:M | 0 | 0 | 0 | 0 | 0 | 0 |
| 26967 | ENSMUSG00000085042 | 16.71950013 | -0.217599867 | 0.578064723 | -0.376428206 | 0.70659858 | 0.94295036 | Abhd11os | abhydrolase domain containing 11, oppos | 18.471 | 18.503 | 17.106 | 24.099 | 6.5437 | 15.594 |
| 26968 | ENSMUSG00000085043 | 0.167707431 | -0.796629176 | 4.080472857 | -0.195229623 | 0.84521319 | NA | Gm13184 | predicted gene 13184 [Source:MGI Symbo | 0 | 0 | 1.0062 | 0 | 0 | 0 |
| 26969 | ENSMUSG00000085044 | 0 | NA | NA | NA | NA | NA | 1700109G15 | RIKEN cDNA 1700109G15 gene [Source:M | 0 | 0 | 0 | 0 | 0 | 0 |
| 26970 | ENSMUSG00000085046 | 0 | NA | NA | NA | NA | NA | Gm11467 | predicted gene 11467 [Source:MGI Symbo | 0 | 0 | 0 | 0 | 0 | 0 |
| 26971 | ENSMUSG00000085047 | 0 | NA | NA | NA | NA | NA | Gm13026 | predicted gene 13026 [Source:MGI Symbo | 0 | 0 | 0 | 0 | 0 | 0 |
| 26972 | ENSMUSG00000085048 | 0.301557804 | -1.64335916 | 4.042814367 | -0.406488899 | 0.684383412 | NA | Gm15537 | predicted gene 15537 [Source:MGI Symbo | 0.8031 | 0 | 1.0062 | 0 | 0 | 0 |
| 26973 | ENSMUSG00000085049 | 0.328370405 | 0.16514316 | 4.003149536 | 0.041253308 | 0.967093956 | NA | Parvaos | parvin, alpha, opposite strand [Source:MG | 0 | 0 | 1.0062 | 0.964 | 0 | 0 |
| 26974 | ENSMUSG00000085051 | 0 | NA | NA | NA | NA | NA | Gm11542 | predicted gene 11542 [Source:MGI Symbo | 0 | 0 | 0 | 0 | 0 | 0 |
| 26975 | ENSMUSG00000085053 | 0 | NA | NA | NA | NA | NA | Gm14809 | predicted gene 14809 [Source:MGI Symbo | 0 | 0 | 0 | 0 | 0 | 0 |
| 26976 | ENSMUSG00000085054 | 0.562214094 | -1.122836648 | 3.276281221 | -0.34271681 | 0.731811515 | NA | Gm15834 | predicted gene 15834 [Source:MGI Symbo | 2.4093 | 0 | 0 | 0.964 | 0 | 0 |
| 26977 | ENSMUSG00000085055 | 1.233813361 | 0.537562785 | 2.27529665 | 0.236260527 | 0.813230515 | NA | Gm15958 | predicted gene 15958 [Source:MGI Symbo | 0 | 1.0279 | 2.0125 | 0 | 4.3625 | 0 |
| 26978 | ENSMUSG00000085057 | 0.990500705 | -0.87623328 | 2.111888074 | -0.41490517 | 0.678211325 | NA | Gm13415 | predicted gene 13415 [Source:MGI Symbo | 0.8031 | 2.0558 | 1.0062 | 0.964 | 0 | 1.1138 |
| 26979 | ENSMUSG00000085058 | 0.335414862 | -1.779047242 | 4.030804179 | -0.441362855 | 0.658950329 | NA | 8030453O22 | RIKEN cDNA 8030453O22 gene [Source:M | 0 | 0 | 2.0125 | 0 | 0 | 0 |
| 26980 | ENSMUSG00000085059 | 0 | NA | NA | NA | NA | NA | Gm11750 | predicted gene 11750 [Source:MGI Symbo | 0 | 0 | 0 | 0 | 0 | 0 |
| 26981 | ENSMUSG00000085060 | 0 | NA | NA | NA | NA | NA | E330016L19I | RIKEN cDNA E330016L19 gene [Source:MG | 0 | 0 | 0 | 0 | 0 | 0 |
| 26982 | ENSMUSG00000085063 | 2.756825734 | -1.132925838 | 1.347386225 | -0.840832285 | 0.400441898 | NA | Gm16024 | predicted gene 16024 [Source:MGI Symbo | 7.2279 | 3.0838 | 1.0062 | 1.928 | 2.1812 | 1.1138 |
| 26983 | ENSMUSG00000085066 | 0 | NA | NA | NA | NA | NA | Gm4989 | predicted pseudogene 4989 [Source:MGI | 0 | 0 | 0 | 0 | 0 | 0 |
| 26984 | ENSMUSG00000085067 | 0 | NA | NA | NA | NA | NA | Gm15631 | predicted gene 15631 [Source:MGI Symbo | 0 | 0 | 0 | 0 | 0 | 0 |
| 26985 | ENSMUSG00000085068 | 0.267700746 | -1.497481649 | 4.057001436 | -0.369110456 | 0.712045396 | NA | Gm15895 | predicted gene 15895 [Source:MGI Symbo | 1.6062 | 0 | 0 | 0 | 0 | 0 |
| 26986 | ENSMUSG00000085069 | 0.480153995 | 1.043847617 | 3.151673025 | 0.331204287 | 0.740490182 | NA | Prdm16os | Prdm16 opposite strand transcript [Sourc | 0.8031 | 0 | 0 | 0.964 | 0 | 1.1138 |
| 26987 | ENSMUSG00000085070 | 0 | NA | NA | NA | NA | NA | Gm13211 | predicted gene 13211 [Source:MGI Symbo | 0 | 0 | 0 | 0 | 0 | 0 |
| 26988 | ENSMUSG00000085071 | 0 | NA | NA | NA | NA | NA | Gm14066 | predicted gene 14066 [Source:MGI Symbo | 0 | 0 | 0 | 0 | 0 | 0 |

| | | | | | | | | | | | | | | | |
|---|---|---|---|---|---|---|---|---|---|---|---|---|---|---|---|
| 26989 | ENSMUSG00000085072 | 3.656032548 | -1.337813502 | 1.151483731 | -1.161817111 | 0.245309762 | NA | Ict1os | immature colon carcinoma transcript 1, o | 5.6217 | 3.0838 | 7.0437 | 2.8919 | 2.1812 | 1.1138 |
| 26990 | ENSMUSG00000085073 | 0 | NA | NA | NA | NA | NA | Gm14562 | predicted gene 14562 [Source:MGI Symbo | 0 | 0 | 0 | 0 | 0 | 0 |
| 26991 | ENSMUSG00000085074 | 0 | NA | NA | NA | NA | NA | Gm11494 | predicted gene 11494 [Source:MGI Symbo | 0 | 0 | 0 | 0 | 0 | 0 |
| 26992 | ENSMUSG00000085075 | 0 | NA | NA | NA | NA | NA | Gm15669 | predicted gene 15669 [Source:MGI Symbo | 0 | 0 | 0 | 0 | 0 | 0 |
| 26993 | ENSMUSG00000085077 | 0 | NA | NA | NA | NA | NA | 1700049E15 | RIKEN cDNA 1700049E15 gene [Source:M | 0 | 0 | 0 | 0 | 0 | 0 |
| 26994 | ENSMUSG00000085078 | 0 | NA | NA | NA | NA | NA | C030013C21 | RIKEN cDNA C030013C21 gene [Source:M | 0 | 0 | 0 | 0 | 0 | 0 |
| 26995 | ENSMUSG00000085079 | 0 | NA | NA | NA | NA | NA | Khdc1b | KH domain containing 1B [Source:MGI Syr | 0 | 0 | 0 | 0 | 0 | 0 |
| 26996 | ENSMUSG00000085080 | 0.17131964 | -0.796629176 | 4.080472857 | -0.195229623 | 0.84521319 | NA | 2310044K18 | RIKEN cDNA 2310044K18 gene [Source:M | 0 | 1.0279 | 0 | 0 | 0 | 0 |
| 26997 | ENSMUSG00000085081 | 0.670803094 | 1.59186814 | 2.791527899 | 0.570249769 | 0.568508305 | NA | Rptoros | regulatory associated protein of MTOR, co | 0 | 0 | 1.0062 | 1.928 | 1.0906 | 0 |
| 26998 | ENSMUSG00000085082 | 0 | NA | NA | NA | NA | NA | Gm16330 | predicted gene 16330 [Source:MGI Symbo | 0 | 0 | 0 | 0 | 0 | 0 |
| 26999 | ENSMUSG00000085083 | 0.167707431 | -0.796629176 | 4.080472857 | -0.195229623 | 0.84521319 | NA | Gm11615 | predicted gene 11615 [Source:MGI Symbo | 0 | 0 | 1.0062 | 0 | 0 | 0 |
| 27000 | ENSMUSG00000085084 | 0.678286236 | 1.600482029 | 2.782216593 | 0.575254289 | 0.565119331 | NA | 4930570G19 | RIKEN cDNA 4930570G19 gene [Source:M | 0 | 1.0279 | 0 | 1.928 | 0 | 1.1138 |
| 27001 | ENSMUSG00000085085 | 1.049153821 | -1.060647667 | 2.226169081 | -0.476445242 | 0.63375721 | NA | 1700086P04 | RIKEN cDNA 1700086P04 gene [Source:M | 3.2124 | 1.0279 | 0 | 0.964 | 1.0906 | 0 |
| 27002 | ENSMUSG00000085087 | 2.773446984 | -1.170252993 | 1.376971473 | -0.849874537 | 0.395394844 | NA | Gm13528 | predicted gene 13528 [Source:MGI Symbo | 6.4248 | 3.0838 | 2.0125 | 2.8919 | 0 | 2.2277 |
| 27003 | ENSMUSG00000085088 | 2.919866827 | -0.121389422 | 1.530369932 | -0.079320313 | 0.93677785 | NA | 4931413K12 | RIKEN cDNA 4931413K12 gene [Source:M | 0 | 7.1954 | 2.0125 | 3.8559 | 0 | 4.4554 |
| 27004 | ENSMUSG00000085089 | 1.72689212 | 1.507003766 | 1.839821313 | 0.819103331 | 0.412727462 | NA | Arhgap15os | Rho GTPase activating protein 15, opposit | 1.6062 | 1.0279 | 0 | 0 | 3.2719 | 4.4554 |
| 27005 | ENSMUSG00000085090 | 0 | NA | NA | NA | NA | NA | Gm12711 | predicted gene 12711 [Source:MGI Symbo | 0 | 0 | 0 | 0 | 0 | 0 |
| 27006 | ENSMUSG00000085091 | 3.104711602 | 2.03189016 | 1.526068948 | 1.331453708 | 0.183039767 | NA | Egfros | epidermal growth factor receptor, opposi | 1.6062 | 2.0558 | 0 | 10.604 | 4.3625 | 0 |
| 27007 | ENSMUSG00000085093 | 0.181769715 | 1.126924175 | 4.080472857 | 0.276174898 | 0.782413736 | NA | Gm12737 | predicted gene 12737 [Source:MGI Symbo | 0 | 0 | 0 | 0 | 1.0906 | 0 |
| 27008 | ENSMUSG00000085094 | 0 | NA | NA | NA | NA | NA | Gm15692 | predicted gene 15692 [Source:MGI Symbo | 0 | 0 | 0 | 0 | 0 | 0 |
| 27009 | ENSMUSG00000085095 | 33.60124155 | 0.050595629 | 0.403423512 | 0.125415669 | 0.900194484 | 0.982805511 | Gm15635 | predicted gene 15635 [Source:MGI Symbo | 40.958 | 32.893 | 25.156 | 45.307 | 29.447 | 27.846 |
| 27010 | ENSMUSG00000085097 | 0.36353943 | 2.065989254 | 4.034137301 | 0.512126658 | 0.608562368 | NA | Gm13049 | predicted gene 13049 [Source:MGI Symbo | 0 | 0 | 0 | 0 | 2.1812 | 0 |
| 27011 | ENSMUSG00000085099 | 0.695728634 | -1.425610641 | 3.05889997 | -0.46605337 | 0.641177299 | NA | Gm16288 | predicted gene 16288 [Source:MGI Symbo | 0 | 3.0838 | 0 | 0 | 1.0906 | 0 |
| 27012 | ENSMUSG00000085100 | 0 | NA | NA | NA | NA | NA | Gm13324 | predicted gene 13324 [Source:MGI Symbo | 0 | 0 | 0 | 0 | 0 | 0 |
| 27013 | ENSMUSG00000085101 | 0 | NA | NA | NA | NA | NA | Platr16 | pluripotency associated transcript 16 [Sou | 0 | 0 | 0 | 0 | 0 | 0 |
| 27014 | ENSMUSG00000085102 | 0.160662974 | 1.126924175 | 4.080472857 | 0.276174898 | 0.782413736 | NA | 1700010K24 | RIKEN cDNA 1700010K24 gene [Source:M | 0 | 0 | 0 | 0.964 | 0 | 0 |
| 27015 | ENSMUSG00000085103 | 0 | NA | NA | NA | NA | NA | Gm16357 | predicted gene 16357 [Source:MGI Symbo | 0 | 0 | 0 | 0 | 0 | 0 |
| 27016 | ENSMUSG00000085104 | 0 | NA | NA | NA | NA | NA | Gm11731 | predicted gene 11731 [Source:MGI Symbo | 0 | 0 | 0 | 0 | 0 | 0 |
| 27017 | ENSMUSG00000085105 | 5.216541346 | -0.812697938 | 1.096857079 | -0.740933302 | 0.458733882 | 0.859322431 | Gm12758 | predicted gene 12758 [Source:MGI Symbo | 8.8341 | 2.0558 | 9.0562 | 5.7839 | 0 | 5.5692 |
| 27018 | ENSMUSG00000085106 | 0.185640647 | 1.126924175 | 4.080472857 | 0.276174898 | 0.782413736 | NA | Gm16000 | predicted gene 16000 [Source:MGI Symbo | 0 | 0 | 0 | 0 | 0 | 1.1138 |
| 27019 | ENSMUSG00000085107 | 0 | NA | NA | NA | NA | NA | Gm4764 | predicted pseudogene 4764 [Source:MGI | 0 | 0 | 0 | 0 | 0 | 0 |
| 27020 | ENSMUSG00000085108 | 0 | NA | NA | NA | NA | NA | Saxo1os | stabilizer of axonemal microtubules 1, op | 0 | 0 | 0 | 0 | 0 | 0 |
| 27021 | ENSMUSG00000085109 | 0.167707431 | -0.796629176 | 4.080472857 | -0.195229623 | 0.84521319 | NA | Gm15356 | predicted gene 15356 [Source:MGI Symbo | 0 | 0 | 1.0062 | 0 | 0 | 0 |
| 27022 | ENSMUSG00000085110 | 0 | NA | NA | NA | NA | NA | Gm13920 | predicted gene 13920 [Source:MGI Symbo | 0 | 0 | 0 | 0 | 0 | 0 |
| 27023 | ENSMUSG00000085111 | 0.167707431 | -0.796629176 | 4.080472857 | -0.195229623 | 0.84521319 | NA | Ascl4 | achaete-scute family bHLH transcription fa | 0 | 0 | 1.0062 | 0 | 0 | 0 |
| 27024 | ENSMUSG00000085112 | 12.9147561 | -0.295206315 | 0.664414514 | -0.444310455 | 0.656818143 | 0.9268505 | A530072M1 | RIKEN cDNA gene A530072M11 [Source:M | 14.456 | 10.279 | 18.112 | 19.28 | 10.906 | 4.4554 |
| 27025 | ENSMUSG00000085113 | 0 | NA | NA | NA | NA | NA | B130011K05 | RIKEN cDNA B130011K05 gene [Source:M | 0 | 0 | 0 | 0 | 0 | 0 |
| 27026 | ENSMUSG00000085114 | 0 | NA | NA | NA | NA | NA | 1600025M17 | RIKEN cDNA 1600025M17 gene [Source:M | 0 | 0 | 0 | 0 | 0 | 0 |
| 27027 | ENSMUSG00000085115 | 0 | NA | NA | NA | NA | NA | Gm11197 | predicted gene 11197 [Source:MGI Symbo | 0 | 0 | 0 | 0 | 0 | 0 |
| 27028 | ENSMUSG00000085116 | 0 | NA | NA | NA | NA | NA | Gm2735 | predicted gene 2735 [Source:MGI Symbol | 0 | 0 | 0 | 0 | 0 | 0 |
| 27029 | ENSMUSG00000085118 | 1.417851174 | 2.92126731 | 2.197497013 | 1.32936122 | 0.183728826 | NA | Gm15774 | predicted gene 15774 [Source:MGI Symbo | 0.8031 | 0 | 0 | 0 | 4.3625 | 3.3415 |
| 27030 | ENSMUSG00000085120 | 0 | NA | NA | NA | NA | NA | Gm16086 | predicted gene 16086 [Source:MGI Symbo | 0 | 0 | 0 | 0 | 0 | 0 |
| 27031 | ENSMUSG00000085121 | 0 | NA | NA | NA | NA | NA | 5730437C11 | RIKEN cDNA 5730437C11 gene [Source:M | 0 | 0 | 0 | 0 | 0 | 0 |
| 27032 | ENSMUSG00000085122 | 0 | NA | NA | NA | NA | NA | 4933404K08 | RIKEN cDNA 4933404K08 gene [Source:M | 0 | 0 | 0 | 0 | 0 | 0 |
| 27033 | ENSMUSG00000085123 | 0 | NA | NA | NA | NA | NA | Rubie | RNA upstream of Bmp4 expressed in inne | 0 | 0 | 0 | 0 | 0 | 0 |
| 27034 | ENSMUSG00000085124 | 0 | NA | NA | NA | NA | NA | Gm12766 | predicted gene 12766 [Source:MGI Symbo | 0 | 0 | 0 | 0 | 0 | 0 |
| 27035 | ENSMUSG00000085125 | 0 | NA | NA | NA | NA | NA | Gm16070 | predicted gene 16070 [Source:MGI Symbo | 0 | 0 | 0 | 0 | 0 | 0 |
| 27036 | ENSMUSG00000085126 | 0 | NA | NA | NA | NA | NA | Gm12589 | predicted gene 12589 [Source:MGI Symbo | 0 | 0 | 0 | 0 | 0 | 0 |
| 27037 | ENSMUSG00000085127 | 0 | NA | NA | NA | NA | NA | 4930444E06 | RIKEN cDNA 4930444E06 gene [Source:M | 0 | 0 | 0 | 0 | 0 | 0 |
| 27038 | ENSMUSG00000085129 | 2.303802366 | -0.800962607 | 1.741500716 | -0.45992666 | 0.645568864 | NA | 5031425F14 | RIKEN cDNA 5031425F14 gene [Source:M | 0.8031 | 0 | 8.05 | 3.8559 | 0 | 1.1138 |
| 27039 | ENSMUSG00000085130 | 0 | NA | NA | NA | NA | NA | Gm11417 | predicted gene 11417 [Source:MGI Symbo | 0 | 0 | 0 | 0 | 0 | 0 |
| 27040 | ENSMUSG00000085132 | 14.33388768 | -0.429883034 | 0.637328125 | -0.674508181 | 0.499988286 | 0.880716237 | Gm12265 | predicted gene 12265 [Source:MGI Symbo | 24.896 | 6.1675 | 18.112 | 12.532 | 9.8156 | 14.48 |
| 27041 | ENSMUSG00000085133 | 41.88661877 | 0.510528804 | 0.340558769 | 1.499091641 | 0.13384986 | 0.53673534 | B930095G15 | RIKEN cDNA B930095G15 gene [Source:M | 34.533 | 29.81 | 39.244 | 45.307 | 54.531 | 47.895 |
| 27042 | ENSMUSG00000085134 | 0.342639279 | -1.805976596 | 4.028547714 | -0.448294702 | 0.653940524 | NA | Gm11659 | predicted gene 11659 [Source:MGI Symbo | 0 | 2.0558 | 0 | 0 | 0 | 0 |
| 27043 | ENSMUSG00000085135 | 0 | NA | NA | NA | NA | NA | Lrrc55os | leucine rich repeat containing 55, opposit | 0 | 0 | 0 | 0 | 0 | 0 |
| 27044 | ENSMUSG00000085139 | 5.747037777 | -0.405143395 | 0.932257667 | -0.434583066 | 0.663865089 | 0.929301301 | A730046J19 | RIKEN cDNA A730046J19 gene [Source:M | 2.4093 | 11.307 | 6.0375 | 4.8199 | 5.4531 | 4.4554 |

| | | | | | | | | | | | | | | | |
|---|---|---|---|---|---|---|---|---|---|---|---|---|---|---|---|
| 27045 | ENSMUSG00000085141 | 0.17131964 | -0.796629176 | 4.080472857 | -0.195229623 | 0.84521319 | NA | Gm13429 | predicted gene 13429 [Source:MGI Symbo | 0 | 1.0279 | 0 | 0 | 0 | 0 |
| 27046 | ENSMUSG00000085142 | 0 | NA | NA | NA | NA | NA | Gm12702 | predicted gene 12702 [Source:MGI Symbo | 0 | 0 | 0 | 0 | 0 | 0 |
| 27047 | ENSMUSG00000085143 | 1.48314078 | -0.033064259 | 1.946149588 | -0.016989578 | 0.98644493 | NA | Gm11520 | predicted gene 11520 [Source:MGI Symbo | 2.4093 | 1.0279 | 1.0062 | 0 | 0 | 4.4554 |
| 27048 | ENSMUSG00000085145 | 2.340747016 | -0.365199847 | 1.479437217 | -0.246850521 | 0.80502391 | NA | Gm11614 | predicted gene 11614 [Source:MGI Symbo | 4.8186 | 2.0558 | 1.0062 | 2.8919 | 3.2719 | 0 |
| 27049 | ENSMUSG00000085146 | 0 | NA | NA | NA | NA | NA | Eif2c5 | eukaryotic translation initiation factor 2C, | 0 | 0 | 0 | 0 | 0 | 0 |
| 27050 | ENSMUSG00000085147 | 0 | NA | NA | NA | NA | NA | Gm12609 | predicted gene 12609 [Source:MGI Symbo | 0 | 0 | 0 | 0 | 0 | 0 |
| 27051 | ENSMUSG00000085148 | 665.4043783 | 0.273323489 | 0.175214962 | 1.559932362 | 0.118775866 | 0.504308497 | Mir22hg | Mir22 host gene (non-protein coding) [So | 565.38 | 554.05 | 689.28 | 907.1 | 725.26 | 551.35 |
| 27052 | ENSMUSG00000085150 | 0 | NA | NA | NA | NA | NA | Gm16346 | predicted gene 16346 [Source:MGI Symbo | 0 | 0 | 0 | 0 | 0 | 0 |
| 27053 | ENSMUSG00000085151 | 9.561645151 | -0.283072031 | 0.730961232 | -0.387259979 | 0.698563751 | 0.940696964 | 1110018N2C | RIKEN cDNA 1110018N20 gene [Source:M | 11.243 | 8.2233 | 12.075 | 12.532 | 3.2719 | 10.025 |
| 27054 | ENSMUSG00000085152 | 0.535401493 | -2.48596965 | 3.684117979 | -0.674780141 | 0.499815459 | NA | Gm11496 | predicted gene 11496 [Source:MGI Symbo | 3.2124 | 0 | 0 | 0 | 0 | 0 |
| 27055 | ENSMUSG00000085154 | 8.148293294 | 0.0527246 | 0.947693501 | 0.055634654 | 0.955632857 | 0.991678478 | C130046K22 | RIKEN cDNA C130046K22 gene [Source:M | 1.6062 | 16.447 | 6.0375 | 4.8199 | 3.2719 | 16.708 |
| 27056 | ENSMUSG00000085155 | 0 | NA | NA | NA | NA | NA | Gm11780 | predicted gene 11780 [Source:MGI Symbo | 0 | 0 | 0 | 0 | 0 | 0 |
| 27057 | ENSMUSG00000085156 | 24.74489387 | 0.285341402 | 0.47478108 | 0.60099573 | 0.54784283 | 0.895600915 | Snhg15 | small nucleolar RNA host gene 15 [Source | 32.124 | 12.335 | 22.137 | 28.919 | 23.994 | 28.96 |
| 27058 | ENSMUSG00000085157 | 0 | NA | NA | NA | NA | NA | 4930515L19I | RIKEN cDNA 4930515L19 gene [Source:M( | 0 | 0 | 0 | 0 | 0 | 0 |
| 27059 | ENSMUSG00000085158 | 0 | NA | NA | NA | NA | NA | Gm14264 | predicted gene 14264 [Source:MGI Symbo | 0 | 0 | 0 | 0 | 0 | 0 |
| 27060 | ENSMUSG00000085160 | 0 | NA | NA | NA | NA | NA | Gm13617 | predicted gene 13617 [Source:MGI Symbo | 0 | 0 | 0 | 0 | 0 | 0 |
| 27061 | ENSMUSG00000085161 | 0 | NA | NA | NA | NA | NA | Gm7580 | predicted gene 7580 [Source:MGI Symbol | 0 | 0 | 0 | 0 | 0 | 0 |
| 27062 | ENSMUSG00000085162 | 0 | NA | NA | NA | NA | NA | Gm12295 | predicted gene 12295 [Source:MGI Symbo | 0 | 0 | 0 | 0 | 0 | 0 |
| 27063 | ENSMUSG00000085165 | 0.401551119 | -2.072982726 | 4.008251979 | -0.517178744 | 0.605031388 | NA | Gm12089 | predicted gene 12089 [Source:MGI Symbo | 2.4093 | 0 | 0 | 0 | 0 | 0 |
| 27064 | ENSMUSG00000085166 | 0 | NA | NA | NA | NA | NA | Gm15522 | predicted gene 15522 [Source:MGI Symbo | 0 | 0 | 0 | 0 | 0 | 0 |
| 27065 | ENSMUSG00000085167 | 0 | NA | NA | NA | NA | NA | Gm12063 | predicted gene 12063 [Source:MGI Symbo | 0 | 0 | 0 | 0 | 0 | 0 |
| 27066 | ENSMUSG00000085168 | 0 | NA | NA | NA | NA | NA | Gm16252 | predicted gene 16252 [Source:MGI Symbo | 0 | 0 | 0 | 0 | 0 | 0 |
| 27067 | ENSMUSG00000085169 | 5.939517179 | -0.867771119 | 0.916906895 | -0.94641138 | 0.343938803 | 0.785649218 | Gm10785 | predicted gene 10785 [Source:MGI Symbo | 8.8341 | 3.0838 | 11.069 | 3.8559 | 5.4531 | 3.3415 |
| 27068 | ENSMUSG00000085170 | 0 | NA | NA | NA | NA | NA | Lrrc75aos1 | leucine rich repeat containing 75A, oppos | 0 | 0 | 0 | 0 | 0 | 0 |
| 27069 | ENSMUSG00000085171 | 1.14024829 | 0.506576368 | 2.118787831 | 0.239087822 | 0.811037485 | NA | D830026I12I | RIKEN cDNA D830026I12 gene [Source:M( | 0.8031 | 2.0558 | 0 | 2.8919 | 1.0906 | 0 |
| 27070 | ENSMUSG00000085172 | 1.736030229 | 1.424630206 | 1.712542178 | 0.831880361 | 0.405476478 | NA | Gm6542 | predicted gene 6542 [Source:MGI Symbol | 0.8031 | 0 | 2.0125 | 0.964 | 2.1812 | 4.4554 |
| 27071 | ENSMUSG00000085173 | 0 | NA | NA | NA | NA | NA | Gm2869 | predicted gene 2869 [Source:MGI Symbol | 0 | 0 | 0 | 0 | 0 | 0 |
| 27072 | ENSMUSG00000085174 | 9.224854933 | -0.608024762 | 0.719291779 | -0.845310316 | 0.397937603 | 0.825555003 | Gm16206 | predicted gene 16206 [Source:MGI Symbo | 16.062 | 7.1954 | 10.062 | 7.7118 | 7.6343 | 6.6831 |
| 27073 | ENSMUSG00000085175 | 40.18259861 | -0.013917525 | 0.401228784 | -0.034687255 | 0.972329124 | 0.995715135 | Gm11423 | predicted gene 11423 [Source:MGI Symbo | 36.14 | 39.061 | 46.287 | 55.911 | 42.534 | 21.163 |
| 27074 | ENSMUSG00000085176 | 0 | NA | NA | NA | NA | NA | Gm15397 | predicted gene 15397 [Source:MGI Symbo | 0 | 0 | 0 | 0 | 0 | 0 |
| 27075 | ENSMUSG00000085177 | 0.486939728 | -0.698472448 | 3.137608162 | -0.222613026 | 0.823836697 | NA | Gm11209 | predicted gene 11209 [Source:MGI Symbo | 0.8031 | 1.0279 | 0 | 0 | 1.0906 | 0 |
| 27076 | ENSMUSG00000085178 | 3.29532417 | 0.513688371 | 1.165998333 | 0.440556694 | 0.65953396 | NA | Kdm6bos | KDM1 lysine (K)-specific demethylase 6B, | 4.0155 | 1.0279 | 3.0187 | 2.8919 | 4.3625 | 4.4554 |
| 27077 | ENSMUSG00000085179 | 0 | NA | NA | NA | NA | NA | Gm13645 | predicted gene 13645 [Source:MGI Symbo | 0 | 0 | 0 | 0 | 0 | 0 |
| 27078 | ENSMUSG00000085180 | 12.92481322 | 0.217468692 | 0.615160146 | 0.353515574 | 0.723701955 | 0.944786345 | AI838599 | expressed sequence AI838599 [Source:M( | 9.6372 | 9.2513 | 17.106 | 17.352 | 14.178 | 10.025 |
| 27079 | ENSMUSG00000085181 | 0.133850373 | -0.796629176 | 4.080472857 | -0.195229623 | 0.84521319 | NA | Gm12709 | predicted gene 12709 [Source:MGI Symbo | 0.8031 | 0 | 0 | 0 | 0 | 0 |
| 27080 | ENSMUSG00000085182 | 0.167707431 | -0.796629176 | 4.080472857 | -0.195229623 | 0.84521319 | NA | Gm13596 | predicted gene 13596 [Source:MGI Symbo | 0 | 0 | 1.0062 | 0 | 0 | 0 |
| 27081 | ENSMUSG00000085183 | 0 | NA | NA | NA | NA | NA | Wincr1 | WNT induced non-coding RNA 1 [Source:I | 0 | 0 | 0 | 0 | 0 | 0 |
| 27082 | ENSMUSG00000085184 | 0.990320745 | -2.084832509 | 2.480930004 | -0.84034314 | 0.40071602 | NA | 4933439K11 | RIKEN cDNA 4933439K11 gene [Source:M( | 0.8031 | 0 | 4.025 | 0 | 0 | 1.1138 |
| 27083 | ENSMUSG00000085185 | 7.721533067 | -0.932813033 | 0.798402133 | -1.168349876 | 0.242665664 | 0.692154638 | BC028777 | cDNA sequence BC028777 [Source:MGI Sy | 8.031 | 11.307 | 11.069 | 3.8559 | 8.7249 | 3.3415 |
| 27084 | ENSMUSG00000085187 | 0 | NA | NA | NA | NA | NA | Gm13860 | predicted gene 13860 [Source:MGI Symbo | 0 | 0 | 0 | 0 | 0 | 0 |
| 27085 | ENSMUSG00000085188 | 0 | NA | NA | NA | NA | NA | Gm11368 | predicted gene 11368 [Source:MGI Symbo | 0 | 0 | 0 | 0 | 0 | 0 |
| 27086 | ENSMUSG00000085189 | 64.8301768 | 0.856993732 | 0.295411797 | 2.901013905 | 0.003719574 | 0.054619202 | Gm11963 | predicted gene 11963 [Source:MGI Symbo | 46.58 | 53.452 | 38.237 | 75.19 | 74.162 | 101.36 |
| 27087 | ENSMUSG00000085192 | 16.90965132 | 1.25254027 | 0.550810323 | 2.273995633 | 0.022966249 | 0.19080059 | Gm12195 | predicted gene 12195 [Source:MGI Symbo | 9.6372 | 8.2233 | 12.075 | 16.388 | 26.175 | 28.96 |
| 27088 | ENSMUSG00000085193 | 0 | NA | NA | NA | NA | NA | Gm13322 | predicted gene 13322 [Source:MGI Symbo | 0 | 0 | 0 | 0 | 0 | 0 |
| 27089 | ENSMUSG00000085194 | 0 | NA | NA | NA | NA | NA | Platr32 | pluripotency associated transcript 32 [Sou | 0 | 0 | 0 | 0 | 0 | 0 |
| 27090 | ENSMUSG00000085196 | 2.072115341 | -0.935412708 | 1.707532924 | -0.547815328 | 0.583818713 | NA | Gm14963 | predicted gene 14963 [Source:MGI Symbo | 4.0155 | 4.1117 | 0 | 0.964 | 0 | 3.3415 |
| 27091 | ENSMUSG00000085197 | 0 | NA | NA | NA | NA | NA | 1700125H03 | RIKEN cDNA 1700125H03 gene [Source:M | 0 | 0 | 0 | 0 | 0 | 0 |
| 27092 | ENSMUSG00000085198 | 0 | NA | NA | NA | NA | NA | 4933416E03 | RIKEN cDNA 4933416E03 gene [Source:M( | 0 | 0 | 0 | 0 | 0 | 0 |
| 27093 | ENSMUSG00000085199 | 0.167707431 | -0.796629176 | 4.080472857 | -0.195229623 | 0.84521319 | NA | Gm15346 | predicted gene 15346 [Source:MGI Symbo | 0 | 0 | 1.0062 | 0 | 0 | 0 |
| 27094 | ENSMUSG00000085201 | 0.476489653 | -2.293616034 | 3.421506946 | -0.670352587 | 0.502633051 | NA | Nr6a1os | nuclear receptor subfamily 6, group A, me | 0.8031 | 2.0558 | 0 | 0 | 0 | 0 |
| 27095 | ENSMUSG00000085202 | 0.696064467 | -1.534444742 | 3.043667477 | -0.504143358 | 0.61416064 | NA | Gm15620 | predicted gene 15620 [Source:MGI Symbo | 3.2124 | 0 | 0 | 0.964 | 0 | 0 |
| 27096 | ENSMUSG00000085203 | 0 | NA | NA | NA | NA | NA | Gm12927 | predicted gene 12927 [Source:MGI Symbo | 0 | 0 | 0 | 0 | 0 | 0 |
| 27097 | ENSMUSG00000085204 | 3.367347903 | 0.762807872 | 1.156444784 | 0.659614608 | 0.509501178 | NA | Gm15327 | predicted gene 15327 [Source:MGI Symbo | 2.4093 | 2.0558 | 3.0187 | 3.8559 | 2.1812 | 6.6831 |
| 27098 | ENSMUSG00000085205 | 0.524408994 | -0.837196744 | 3.377823722 | -0.247850928 | 0.804249749 | NA | Gm13185 | predicted gene 13185 [Source:MGI Symbo | 0 | 2.0558 | 0 | 0 | 1.0906 | 0 |
| 27099 | ENSMUSG00000085206 | 0 | NA | NA | NA | NA | NA | Gm13380 | predicted gene 13380 [Source:MGI Symbo | 0 | 0 | 0 | 0 | 0 | 0 |
| 27100 | ENSMUSG00000085207 | 7.900647234 | 0.211292325 | 0.779399408 | 0.271096338 | 0.786316938 | 0.960870081 | Gm11767 | predicted gene 11767 [Source:MGI Symbo | 4.8186 | 5.1396 | 12.075 | 7.7118 | 7.6343 | 10.025 |

| | | | | | | | | | | | | | | | |
|---|---|---|---|---|---|---|---|---|---|---|---|---|---|---|---|
| 27101 | ENSMUSG00000085208 | 486.9060541 | 0.067752067 | 0.141395022 | 0.479168687 | 0.631818628 | 0.921696888 | Brip1os | BRCA1 interacting protein C-terminal helic | 485.07 | 422.47 | 518.22 | 450.18 | 542.04 | 503.46 |
| 27102 | ENSMUSG00000085209 | 0 | NA | NA | NA | NA | NA | Gm12637 | predicted gene 12637 [Source:MGI Symbo | 0 | 0 | 0 | 0 | 0 | 0 |
| 27103 | ENSMUSG00000085211 | 59.18881117 | -0.578281426 | 0.915400558 | -0.631725009 | 0.527566583 | 0.888334265 | B430219N15 | RIKEN cDNA B430219N15 gene [Source:M | 155.8 | 16.447 | 40.25 | 40.487 | 67.618 | 34.529 |
| 27104 | ENSMUSG00000085213 | 2.524895737 | -0.970683403 | 1.48487782 | -0.653712642 | 0.513296957 | NA | Gm13091 | predicted gene 13091 [Source:MGI Symbo | 4.0155 | 0 | 6.0375 | 2.8919 | 1.0906 | 1.1138 |
| 27105 | ENSMUSG00000085217 | 0 | NA | NA | NA | NA | NA | 6030471H07 | RIKEN cDNA 6030471H07 gene [Source:M | 0 | 0 | 0 | 0 | 0 | 0 |
| 27106 | ENSMUSG00000085218 | 2.47807537 | 2.074411004 | 1.548387283 | 1.339723612 | 0.180335218 | NA | BB218582 | expressed sequence BB218582 [Source:M | 0.8031 | 2.0558 | 0 | 0.964 | 4.3625 | 6.6831 |
| 27107 | ENSMUSG00000085221 | 0 | NA | NA | NA | NA | NA | Gm11309 | predicted gene 11309 [Source:MGI Symbo | 0 | 0 | 0 | 0 | 0 | 0 |
| 27108 | ENSMUSG00000085222 | 0 | NA | NA | NA | NA | NA | Gm13974 | predicted gene 13974 [Source:MGI Symbo | 0 | 0 | 0 | 0 | 0 | 0 |
| 27109 | ENSMUSG00000085223 | 2.091496935 | 0.458711786 | 1.528192385 | 0.300166255 | 0.764050343 | NA | Gm11210 | predicted gene 11210 [Source:MGI Symbo | 3.2124 | 0 | 2.0125 | 2.8919 | 1.0906 | 3.3415 |
| 27110 | ENSMUSG00000085224 | 0 | NA | NA | NA | NA | NA | Gm13425 | predicted gene 13425 [Source:MGI Symbo | 0 | 0 | 0 | 0 | 0 | 0 |
| 27111 | ENSMUSG00000085225 | 0 | NA | NA | NA | NA | NA | Gm14637 | predicted gene 14637 [Source:MGI Symbo | 0 | 0 | 0 | 0 | 0 | 0 |
| 27112 | ENSMUSG00000085226 | 0.181769715 | 1.126924175 | 4.080472857 | 0.276174898 | 0.782413736 | NA | Gm7855 | predicted gene 7855 [Source:MGI Symbol | 0 | 0 | 0 | 0 | 1.0906 | 0 |
| 27113 | ENSMUSG00000085227 | 3.586483505 | -1.613952071 | 1.190016123 | -1.356243869 | 0.175021588 | NA | 6330418K02 | RIKEN cDNA 6330418K02 gene [Source:M | 8.031 | 3.0838 | 5.0312 | 0.964 | 2.1812 | 2.2277 |
| 27114 | ENSMUSG00000085228 | 0.167707431 | -0.796629176 | 4.080472857 | -0.195229623 | 0.84521319 | NA | Gm14376 | predicted gene 14376 [Source:MGI Symbo | 0 | 0 | 1.0062 | 0 | 0 | 0 |
| 27115 | ENSMUSG00000085229 | 0.36353943 | 2.065989254 | 4.034137301 | 0.512126658 | 0.608562368 | NA | Gm11672 | predicted gene 11672 [Source:MGI Symbo | 0 | 0 | 0 | 0 | 2.1812 | 0 |
| 27116 | ENSMUSG00000085230 | 0 | NA | NA | NA | NA | NA | Gm16013 | predicted gene 16013 [Source:MGI Symbo | 0 | 0 | 0 | 0 | 0 | 0 |
| 27117 | ENSMUSG00000085232 | 14.75775599 | 0.952392354 | 0.625973987 | 1.52145676 | 0.128145255 | 0.524051388 | Gm11962 | predicted gene 11962 [Source:MGI Symbo | 8.031 | 10.279 | 12.075 | 32.775 | 10.906 | 14.48 |
| 27118 | ENSMUSG00000085234 | 0.160662974 | 1.126924175 | 4.080472857 | 0.276174898 | 0.782413736 | NA | Gm15614 | predicted gene 15614 [Source:MGI Symbo | 0 | 0 | 0 | 0.964 | 0 | 0 |
| 27119 | ENSMUSG00000085235 | 19.22071416 | -0.094048927 | 0.486124238 | -0.193466854 | 0.846593362 | 0.972803041 | Gm12576 | predicted gene 12576 [Source:MGI Symbo | 20.881 | 17.475 | 21.131 | 18.316 | 16.359 | 21.163 |
| 27120 | ENSMUSG00000085236 | 0.894196708 | -0.816209142 | 2.51745277 | -0.32422024 | 0.7457713 | NA | 2610206C17 | RIKEN cDNA 2610206C17 gene [Source:M | 2.4093 | 1.0279 | 0 | 1.928 | 0 | 0 |
| 27121 | ENSMUSG00000085237 | 0.267700746 | -1.497481649 | 4.057001436 | -0.369110456 | 0.712045396 | NA | Gm15406 | predicted gene 15406 [Source:MGI Symbo | 1.6062 | 0 | 0 | 0 | 0 | 0 |
| 27122 | ENSMUSG00000085238 | 0.68145976 | 0.058694068 | 2.590797836 | 0.022654824 | 0.981925612 | NA | 4930479D17 | RIKEN cDNA 4930479D17 gene [Source:M | 0 | 1.0279 | 1.0062 | 0.964 | 1.0906 | 0 |
| 27123 | ENSMUSG00000085239 | 0 | NA | NA | NA | NA | NA | Gm12150 | predicted gene 12150 [Source:MGI Symbo | 0 | 0 | 0 | 0 | 0 | 0 |
| 27124 | ENSMUSG00000085240 | 0 | NA | NA | NA | NA | NA | Gm12119 | predicted gene 12119 [Source:MGI Symbo | 0 | 0 | 0 | 0 | 0 | 0 |
| 27125 | ENSMUSG00000085241 | 1.076173013 | -2.281611141 | 2.368826644 | -0.963181982 | 0.335456206 | NA | Snhg3 | small nucleolar RNA host gene 3 [Source:I | 2.4093 | 3.0838 | 0 | 0.964 | 0 | 0 |
| 27126 | ENSMUSG00000085242 | 1.22362201 | -0.289417703 | 2.087230194 | -0.138661133 | 0.889717933 | NA | Gm11537 | predicted gene 11537 [Source:MGI Symbo | 0 | 1.0279 | 3.0187 | 0 | 2.1812 | 1.1138 |
| 27127 | ENSMUSG00000085244 | 0 | NA | NA | NA | NA | NA | Spag17os | sperm associated antigen 17, opposite str | 0 | 0 | 0 | 0 | 0 | 0 |
| 27128 | ENSMUSG00000085245 | 0 | NA | NA | NA | NA | NA | Gm11713 | predicted gene 11713 [Source:MGI Symbo | 0 | 0 | 0 | 0 | 0 | 0 |
| 27129 | ENSMUSG00000085246 | 1.08321747 | -3.483110813 | 2.291372099 | -1.520098291 | 0.128486274 | NA | Gm15893 | predicted gene 15893 [Source:MGI Symbo | 2.4093 | 3.0838 | 1.0062 | 0 | 0 | 0 |
| 27130 | ENSMUSG00000085247 | 1.006191327 | 3.548856365 | 2.592453122 | 1.36891824 | 0.171024834 | NA | 4930545L23I | RIKEN cDNA 4930545L23 gene [Source:M( | 0 | 0 | 0 | 3.8559 | 2.1812 | 0 |
| 27131 | ENSMUSG00000085248 | 0 | NA | NA | NA | NA | NA | Gm11587 | predicted gene 11587 [Source:MGI Symbo | 0 | 0 | 0 | 0 | 0 | 0 |
| 27132 | ENSMUSG00000085249 | 1.762170397 | -0.006795748 | 1.697425725 | -0.004003561 | 0.996805629 | NA | NA | NA | 3.2124 | 1.0279 | 1.0062 | 0.964 | 4.3625 | 0 |
| 27133 | ENSMUSG00000085250 | 1.238122978 | 1.432230464 | 2.076151604 | 0.68984869 | 0.490289346 | NA | Gm15234 | predicted gene 15234 [Source:MGI Symbo | 0 | 2.0558 | 0 | 0.964 | 2.1812 | 2.2277 |
| 27134 | ENSMUSG00000085251 | 0 | NA | NA | NA | NA | NA | Gm12326 | predicted gene 12326 [Source:MGI Symbo | 0 | 0 | 0 | 0 | 0 | 0 |
| 27135 | ENSMUSG00000085252 | 0 | NA | NA | NA | NA | NA | 8430437L04I | RIKEN cDNA 8430437L04 gene [Source:M( | 0 | 0 | 0 | 0 | 0 | 0 |
| 27136 | ENSMUSG00000085253 | 0.133850373 | -0.796629176 | 4.080472857 | -0.195229623 | 0.84521319 | NA | Gm16833 | predicted gene, 16833 [Source:MGI Symb | 0.8031 | 0 | 0 | 0 | 0 | 0 |
| 27137 | ENSMUSG00000085254 | 0 | NA | NA | NA | NA | NA | Gm15881 | predicted gene 15881 [Source:MGI Symbo | 0 | 0 | 0 | 0 | 0 | 0 |
| 27138 | ENSMUSG00000085255 | 6.547323382 | 1.831188238 | 0.887404202 | 2.06353343 | 0.039061975 | 0.270232919 | Taco1os | translational activator of mitochondrially ( | 1.6062 | 4.1117 | 3.0187 | 9.6398 | 11.997 | 8.9108 |
| 27139 | ENSMUSG00000085256 | 0 | NA | NA | NA | NA | NA | Dmrta2os | doublesex and mab-3 related transcriptio | 0 | 0 | 0 | 0 | 0 | 0 |
| 27140 | ENSMUSG00000085257 | 0 | NA | NA | NA | NA | NA | Gm13264 | predicted gene 13264 [Source:MGI Symbo | 0 | 0 | 0 | 0 | 0 | 0 |
| 27141 | ENSMUSG00000085260 | 0.181769715 | 1.126924175 | 4.080472857 | 0.276174898 | 0.782413736 | NA | Med9os | mediator complex subunit 9, opposite stra | 0 | 0 | 0 | 0 | 1.0906 | 0 |
| 27142 | ENSMUSG00000085261 | 0.160662974 | 1.126924175 | 4.080472857 | 0.276174898 | 0.782413736 | NA | Gm13814 | predicted gene 13814 [Source:MGI Symbo | 0 | 0 | 0 | 0.964 | 0 | 0 |
| 27143 | ENSMUSG00000085262 | 0.321325949 | 1.911575114 | 4.048267893 | 0.472195804 | 0.636787034 | NA | Gm11574 | predicted gene 11574 [Source:MGI Symbo | 0 | 0 | 0 | 1.928 | 0 | 0 |
| 27144 | ENSMUSG00000085263 | 0 | NA | NA | NA | NA | NA | 1700125G02 | RIKEN cDNA 1700125G02 gene [Source:M | 0 | 0 | 0 | 0 | 0 | 0 |
| 27145 | ENSMUSG00000085264 | 0 | NA | NA | NA | NA | NA | Gm15581 | predicted gene 15581 [Source:MGI Symbo | 0 | 0 | 0 | 0 | 0 | 0 |
| 27146 | ENSMUSG00000085265 | 0 | NA | NA | NA | NA | NA | Nell1os | NEL-like 1, opposite strand [Source:MGI S | 0 | 0 | 0 | 0 | 0 | 0 |
| 27147 | ENSMUSG00000085267 | 0 | NA | NA | NA | NA | NA | Gm14235 | predicted gene 14235 [Source:MGI Symbo | 0 | 0 | 0 | 0 | 0 | 0 |
| 27148 | ENSMUSG00000085269 | 7.889889622 | -0.397373405 | 0.840255544 | -0.472919707 | 0.636270465 | 0.923478764 | Gm15777 | predicted gene 15777 [Source:MGI Symbo | 13.653 | 8.2233 | 5.0312 | 11.568 | 2.1812 | 6.6831 |
| 27149 | ENSMUSG00000085271 | 0 | NA | NA | NA | NA | NA | E130006D01 | RIKEN cDNA E130006D01 gene [Source:M | 0 | 0 | 0 | 0 | 0 | 0 |
| 27150 | ENSMUSG00000085272 | 0 | NA | NA | NA | NA | NA | Sbk3 | SH3 domain binding kinase family, membe | 0 | 0 | 0 | 0 | 0 | 0 |
| 27151 | ENSMUSG00000085274 | 2.836798223 | 0.543717218 | 1.505019237 | 0.361269281 | 0.717898155 | NA | BC046401 | cDNA sequence BC046401 [Source:MGI Sy | 4.8186 | 2.0558 | 0 | 5.7839 | 4.3625 | 0 |
| 27152 | ENSMUSG00000085275 | 0 | NA | NA | NA | NA | NA | Gm14487 | predicted gene 14487 [Source:MGI Symbo | 0 | 0 | 0 | 0 | 0 | 0 |
| 27153 | ENSMUSG00000085276 | 0 | NA | NA | NA | NA | NA | Gm15812 | predicted gene 15812 [Source:MGI Symbo | 0 | 0 | 0 | 0 | 0 | 0 |
| 27154 | ENSMUSG00000085277 | 0 | NA | NA | NA | NA | NA | Gm13943 | predicted gene 13943 [Source:MGI Symbo | 0 | 0 | 0 | 0 | 0 | 0 |
| 27155 | ENSMUSG00000085278 | 0 | NA | NA | NA | NA | NA | Gm12841 | predicted gene 12841 [Source:MGI Symbo | 0 | 0 | 0 | 0 | 0 | 0 |
| 27156 | ENSMUSG00000085279 | 0 | NA | NA | NA | NA | NA | Gm15965 | predicted gene 15965 [Source:MGI Symbo | 0 | 0 | 0 | 0 | 0 | 0 |

| | | | | | | | | | | | | | | | |
|---|---|---|---|---|---|---|---|---|---|---|---|---|---|---|---|
| 27157 | ENSMUSG00000085280 | 0.692607243 | 3.004183657 | 2.957557172 | 1.015765202 | 0.309741211 | NA | Gm16151 | predicted gene 16151 [Source:MGI Symbo | 0 | 0 | 0 | 1.928 | 0 | 2.2277 |
| 27158 | ENSMUSG00000085281 | 0 | NA | NA | NA | NA | NA | Gm13316 | predicted gene 13316 [Source:MGI Symbo | 0 | 0 | 0 | 0 | 0 | 0 |
| 27159 | ENSMUSG00000085282 | 8.904336413 | -0.62007891 | 0.72798014 | -0.851779981 | 0.394336219 | 0.822827574 | Gm15663 | predicted gene 15663 [Source:MGI Symbo | 8.031 | 12.335 | 12.075 | 7.7118 | 4.3625 | 8.9108 |
| 27160 | ENSMUSG00000085284 | 0 | NA | NA | NA | NA | NA | Gm16189 | predicted gene 16189 [Source:MGI Symbo | 0 | 0 | 0 | 0 | 0 | 0 |
| 27161 | ENSMUSG00000085285 | 0 | NA | NA | NA | NA | NA | Gm14968 | predicted gene 14968 [Source:MGI Symbo | 0 | 0 | 0 | 0 | 0 | 0 |
| 27162 | ENSMUSG00000085286 | 8.318166632 | -0.569489556 | 0.771299053 | -0.738351166 | 0.460301081 | 0.859817816 | Ube4bos3 | ubiquitination factor E4B, opposite strand | 14.456 | 5.1396 | 10.062 | 4.8199 | 7.6343 | 7.7969 |
| 27163 | ENSMUSG00000085287 | 108.8050788 | 0.111058491 | 0.235609448 | 0.47136688 | 0.637378763 | 0.923478764 | 4833418N02 | RIKEN cDNA 4833418N02 gene [Source:M | 111.63 | 100.74 | 101.63 | 133.03 | 116.7 | 89.108 |
| 27164 | ENSMUSG00000085288 | 0 | NA | NA | NA | NA | NA | Gm16356 | predicted gene 16356 [Source:MGI Symbo | 0 | 0 | 0 | 0 | 0 | 0 |
| 27165 | ENSMUSG00000085289 | 3.713444934 | 1.614750796 | 1.210991759 | 1.333411878 | 0.182396676 | NA | Gm15337 | predicted gene 15337 [Source:MGI Symbo | 2.4093 | 0 | 3.0187 | 5.7839 | 3.2719 | 7.7969 |
| 27166 | ENSMUSG00000085290 | 0 | NA | NA | NA | NA | NA | Gm16129 | predicted gene 16129 [Source:MGI Symbo | 0 | 0 | 0 | 0 | 0 | 0 |
| 27167 | ENSMUSG00000085292 | 0 | NA | NA | NA | NA | NA | Gm12853 | predicted gene 12853 [Source:MGI Symbo | 0 | 0 | 0 | 0 | 0 | 0 |
| 27168 | ENSMUSG00000085293 | 0 | NA | NA | NA | NA | NA | Gm13817 | predicted gene 13817 [Source:MGI Symbo | 0 | 0 | 0 | 0 | 0 | 0 |
| 27169 | ENSMUSG00000085295 | 0.342639279 | -1.805976596 | 4.028547714 | -0.448294702 | 0.653940524 | NA | 4930430E12 | RIKEN cDNA 4930430E12 gene [Source:M | 0 | 2.0558 | 0 | 0 | 0 | 0 |
| 27170 | ENSMUSG00000085297 | 0 | NA | NA | NA | NA | NA | Gm11651 | predicted gene 11651 [Source:MGI Symbo | 0 | 0 | 0 | 0 | 0 | 0 |
| 27171 | ENSMUSG00000085298 | 3.005281298 | -2.333935301 | 1.492344586 | -1.563938599 | 0.117831989 | NA | F730035M0 | RIKEN cDNA F730035M05 gene [Source:M | 8.031 | 3.0838 | 4.025 | 2.8919 | 0 | 0 |
| 27172 | ENSMUSG00000085300 | 0 | NA | NA | NA | NA | NA | Gm16345 | predicted gene 16345 [Source:MGI Symbo | 0 | 0 | 0 | 0 | 0 | 0 |
| 27173 | ENSMUSG00000085301 | 0 | NA | NA | NA | NA | NA | Gm12132 | predicted gene 12132 [Source:MGI Symbo | 0 | 0 | 0 | 0 | 0 | 0 |
| 27174 | ENSMUSG00000085302 | 0 | NA | NA | NA | NA | NA | Trav8-2 | T-cell receptor alpha variable 8-2 [Source: | 0 | 0 | 0 | 0 | 0 | 0 |
| 27175 | ENSMUSG00000085303 | 1.542854306 | -0.154089419 | 1.75946098 | -0.087577628 | 0.930212384 | NA | Gm16751 | predicted gene, 16751 [Source:MGI Symb | 0.8031 | 2.0558 | 2.0125 | 0 | 3.2719 | 1.1138 |
| 27176 | ENSMUSG00000085304 | 0 | NA | NA | NA | NA | NA | Gm12802 | predicted gene 12802 [Source:MGI Symbo | 0 | 0 | 0 | 0 | 0 | 0 |
| 27177 | ENSMUSG00000085305 | 0 | NA | NA | NA | NA | NA | Gm12410 | predicted gene 12410 [Source:MGI Symbo | 0 | 0 | 0 | 0 | 0 | 0 |
| 27178 | ENSMUSG00000085307 | 0.321325949 | 1.911575114 | 4.048267893 | 0.472195804 | 0.636787034 | NA | Gm11525 | predicted gene 11525 [Source:MGI Symbo | 0 | 0 | 0 | 1.928 | 0 | 0 |
| 27179 | ENSMUSG00000085310 | 0.160662974 | 1.126924175 | 4.080472857 | 0.276174898 | 0.782413736 | NA | Gm11491 | predicted gene 11491 [Source:MGI Symbo | 0 | 0 | 0 | 0.964 | 0 | 0 |
| 27180 | ENSMUSG00000085311 | 2.640910232 | -0.919029312 | 1.427990541 | -0.643582212 | 0.519846388 | NA | Gm15445 | predicted gene 15445 [Source:MGI Symbo | 3.2124 | 2.0558 | 5.0312 | 0 | 1.0906 | 4.4554 |
| 27181 | ENSMUSG00000085312 | 0 | NA | NA | NA | NA | NA | Gm12763 | predicted gene 12763 [Source:MGI Symbo | 0 | 0 | 0 | 0 | 0 | 0 |
| 27182 | ENSMUSG00000085314 | 0 | NA | NA | NA | NA | NA | Gm14866 | predicted gene 14866 [Source:MGI Symbo | 0 | 0 | 0 | 0 | 0 | 0 |
| 27183 | ENSMUSG00000085315 | 0.353348078 | 0.165142701 | 3.895042568 | 0.042398176 | 0.966181282 | NA | A430018G1 | RIKEN cDNA A430018G15 gene [Source:M | 0 | 0 | 1.0062 | 0 | 0 | 1.1138 |
| 27184 | ENSMUSG00000085316 | 0.167707431 | -0.796629176 | 4.080472857 | -0.195229623 | 0.84521319 | NA | D330050G2 | RIKEN cDNA D330050G23 gene [Source:M | 0 | 0 | 1.0062 | 0 | 0 | 0 |
| 27185 | ENSMUSG00000085317 | 1.010062259 | 3.555195121 | 2.385557701 | 1.490299362 | 0.13614554 | NA | Gssos2 | glutathione synthase, opposite strand 2 [S | 0 | 0 | 0 | 3.8559 | 1.0906 | 1.1138 |
| 27186 | ENSMUSG00000085318 | 0 | NA | NA | NA | NA | NA | Gm12524 | predicted gene 12524 [Source:MGI Symbo | 0 | 0 | 0 | 0 | 0 | 0 |
| 27187 | ENSMUSG00000085319 | 0 | NA | NA | NA | NA | NA | Gm13330 | predicted gene 13330 [Source:MGI Symbo | 0 | 0 | 0 | 0 | 0 | 0 |
| 27188 | ENSMUSG00000085320 | 0 | NA | NA | NA | NA | NA | Gm11548 | predicted gene 11548 [Source:MGI Symbo | 0 | 0 | 0 | 0 | 0 | 0 |
| 27189 | ENSMUSG00000085321 | 0 | NA | NA | NA | NA | NA | 4930412F09 | RIKEN cDNA 4930412F09 gene [Source:M | 0 | 0 | 0 | 0 | 0 | 0 |
| 27190 | ENSMUSG00000085322 | 0 | NA | NA | NA | NA | NA | Gm14261 | predicted gene 14261 [Source:MGI Symbo | 0 | 0 | 0 | 0 | 0 | 0 |
| 27191 | ENSMUSG00000085323 | 0 | NA | NA | NA | NA | NA | 9130410C08 | RIKEN cDNA 9130410C08 gene [Source:M | 0 | 0 | 0 | 0 | 0 | 0 |
| 27192 | ENSMUSG00000085324 | 0 | NA | NA | NA | NA | NA | Gm11680 | predicted gene 11680 [Source:MGI Symbo | 0 | 0 | 0 | 0 | 0 | 0 |
| 27193 | ENSMUSG00000085325 | 0 | NA | NA | NA | NA | NA | Gm14974 | predicted gene 14974 [Source:MGI Symbo | 0 | 0 | 0 | 0 | 0 | 0 |
| 27194 | ENSMUSG00000085326 | 0 | NA | NA | NA | NA | NA | Gm15642 | predicted gene 15642 [Source:MGI Symbo | 0 | 0 | 0 | 0 | 0 | 0 |
| 27195 | ENSMUSG00000085327 | 1.402656885 | 1.683059303 | 1.993626824 | 0.844219833 | 0.398546574 | NA | Gm16104 | predicted gene 16104 [Source:MGI Symbo | 0 | 2.0558 | 0 | 1.928 | 1.0906 | 3.3415 |
| 27196 | ENSMUSG00000085328 | 0 | NA | NA | NA | NA | NA | Gm17131 | predicted gene 17131 [Source:MGI Symbo | 0 | 0 | 0 | 0 | 0 | 0 |
| 27197 | ENSMUSG00000085329 | 0 | NA | NA | NA | NA | NA | 2810404F17 | RIKEN cDNA 2810404F17 gene [Source:M | 0 | 0 | 0 | 0 | 0 | 0 |
| 27198 | ENSMUSG00000085330 | 0 | NA | NA | NA | NA | NA | Gm13417 | predicted gene 13417 [Source:MGI Symbo | 0 | 0 | 0 | 0 | 0 | 0 |
| 27199 | ENSMUSG00000085331 | 0.772832414 | -0.17263844 | 2.923794788 | -0.059046018 | 0.952915455 | NA | Gm11274 | predicted gene 11274 [Source:MGI Symbo | 2.4093 | 0 | 0 | 0 | 0 | 2.2277 |
| 27200 | ENSMUSG00000085333 | 0.167707431 | -0.796629176 | 4.080472857 | -0.195229623 | 0.84521319 | NA | 1700030A11 | RIKEN cDNA 1700030A11 gene [Source:M | 0 | 0 | 1.0062 | 0 | 0 | 0 |
| 27201 | ENSMUSG00000085334 | 82.58566977 | -0.225529624 | 0.259797517 | -0.868097689 | 0.385340856 | 0.816437825 | Gm12940 | predicted gene 12940 [Source:MGI Symbo | 75.492 | 91.485 | 100.62 | 86.758 | 66.528 | 74.628 |
| 27202 | ENSMUSG00000085335 | 0.663758638 | 2.950973782 | 3.025092005 | 0.975498853 | 0.32931287 | NA | Gm13684 | predicted gene 13684 [Source:MGI Symbo | 0 | 0 | 0 | 2.8919 | 1.0906 | 0 |
| 27203 | ENSMUSG00000085336 | 0 | NA | NA | NA | NA | NA | Gm11732 | predicted gene 11732 [Source:MGI Symbo | 0 | 0 | 0 | 0 | 0 | 0 |
| 27204 | ENSMUSG00000085337 | 7.163938322 | 0.641619093 | 0.782542673 | 0.819915788 | 0.412264116 | 0.834424863 | Gm15964 | predicted gene 15964 [Source:MGI Symbo | 5.6217 | 5.1396 | 6.0375 | 9.6398 | 7.6343 | 8.9108 |
| 27205 | ENSMUSG00000085338 | 0.133850373 | -0.796629176 | 4.080472857 | -0.195229623 | 0.84521319 | NA | 2410004I01 | RIKEN cDNA 2410004I01 gene [Source:MG | 0.8031 | 0 | 0 | 0 | 0 | 0 |
| 27206 | ENSMUSG00000085340 | 0 | NA | NA | NA | NA | NA | Gm16335 | predicted gene 16335 [Source:MGI Symbo | 0 | 0 | 0 | 0 | 0 | 0 |
| 27207 | ENSMUSG00000085342 | 6.121853933 | 0.40636964 | 0.838776388 | 0.484479112 | 0.628045885 | 0.920328819 | Gm12254 | predicted gene 12254 [Source:MGI Symbo | 5.6217 | 5.1396 | 5.0312 | 7.7118 | 6.5437 | 6.6831 |
| 27208 | ENSMUSG00000085345 | 0 | NA | NA | NA | NA | NA | Gm16064 | predicted gene 16064 [Source:MGI Symbo | 0 | 0 | 0 | 0 | 0 | 0 |
| 27209 | ENSMUSG00000085347 | 0.339027071 | -1.79174149 | 3.919585585 | -0.457125237 | 0.647581032 | NA | Hk1os | hexokinase 1, opposite strand [Source:MG | 0 | 1.0279 | 1.0062 | 0 | 0 | 0 |
| 27210 | ENSMUSG00000085348 | 0 | NA | NA | NA | NA | NA | Myhas | myosin heavy chain gene antisense RNA [ | 0 | 0 | 0 | 0 | 0 | 0 |
| 27211 | ENSMUSG00000085350 | 0 | NA | NA | NA | NA | NA | 4930529I22 | RIKEN cDNA 4930529I22 gene [Source:MG | 0 | 0 | 0 | 0 | 0 | 0 |
| 27212 | ENSMUSG00000085351 | 0.481988923 | 2.492401104 | 3.837077151 | 0.649557203 | 0.515978285 | NA | Gm12472 | predicted gene 12472 [Source:MGI Symbo | 0 | 0 | 0 | 2.8919 | 0 | 0 |

| | | | | | | | | | | | | | | | |
|---|---|---|---|---|---|---|---|---|---|---|---|---|---|---|---|
| 27213 | ENSMUSG00000085352 | 0 | NA | NA | NA | NA | NA | Gm4128 | predicted gene 4128 [Source:MGI Symbol | 0 | 0 | 0 | 0 | 0 | 0 |
| 27214 | ENSMUSG00000085353 | 0 | NA | NA | NA | NA | NA | 1700092C17 | RIKEN cDNA 1700092C17 gene [Source:M | 0 | 0 | 0 | 0 | 0 | 0 |
| 27215 | ENSMUSG00000085354 | 0 | NA | NA | NA | NA | NA | Gm2044 | predicted gene 2044 [Source:MGI Symbol | 0 | 0 | 0 | 0 | 0 | 0 |
| 27216 | ENSMUSG00000085355 | 8.419271594 | 0.744680099 | 0.854749175 | 0.871226461 | 0.383630515 | 0.816002185 | 3010003L21 | RIKEN cDNA 3010003L21 gene [Source:M | 9.6372 | 1.0279 | 8.05 | 9.6398 | 5.4531 | 16.708 |
| 27217 | ENSMUSG00000085357 | 0 | NA | NA | NA | NA | NA | Gm13480 | predicted gene 13480 [Source:MGI Symbo | 0 | 0 | 0 | 0 | 0 | 0 |
| 27218 | ENSMUSG00000085358 | 0 | NA | NA | NA | NA | NA | Gm14437 | predicted gene 14437 [Source:MGI Symbo | 0 | 0 | 0 | 0 | 0 | 0 |
| 27219 | ENSMUSG00000085359 | 1.908022073 | 2.272219232 | 1.977790709 | 1.148867381 | 0.25061067 | NA | Gm12336 | predicted gene 12336 [Source:MGI Symbo | 0 | 0 | 2.0125 | 2.8919 | 6.5437 | 0 |
| 27220 | ENSMUSG00000085360 | 0 | NA | NA | NA | NA | NA | Arhgap27os2 | Rho GTPase activating protein 27, opposit | 0 | 0 | 0 | 0 | 0 | 0 |
| 27221 | ENSMUSG00000085361 | 0.294513347 | 0.165143496 | 4.080472857 | 0.040471657 | 0.967717103 | NA | Gm16305 | predicted gene 16305 [Source:MGI Symbo | 0.8031 | 0 | 0 | 0.964 | 0 | 0 |
| 27222 | ENSMUSG00000085363 | 0 | NA | NA | NA | NA | NA | Gm15478 | predicted gene 15478 [Source:MGI Symbo | 0 | 0 | 0 | 0 | 0 | 0 |
| 27223 | ENSMUSG00000085366 | 0 | NA | NA | NA | NA | NA | Gm16358 | predicted gene 16358 [Source:MGI Symbo | 0 | 0 | 0 | 0 | 0 | 0 |
| 27224 | ENSMUSG00000085368 | 1.202076585 | -1.206969044 | 2.552319891 | -0.472890976 | 0.636290964 | NA | Gm16188 | predicted gene 16188 [Source:MGI Symbo | 0 | 0 | 5.0312 | 0 | 2.1812 | 0 |
| 27225 | ENSMUSG00000085369 | 0.870712616 | 0.665767941 | 2.367602424 | 0.281199214 | 0.778557607 | NA | 9430093N23 | RIKEN cDNA 9430093N23 gene [Source:M | 0 | 2.0558 | 0 | 0.964 | 1.0906 | 1.1138 |
| 27226 | ENSMUSG00000085371 | 0 | NA | NA | NA | NA | NA | Gm3160 | predicted pseudogene 3160 [Source:MGI | 0 | 0 | 0 | 0 | 0 | 0 |
| 27227 | ENSMUSG00000085372 | 0 | NA | NA | NA | NA | NA | Gm16145 | predicted gene 16145 [Source:MGI Symbo | 0 | 0 | 0 | 0 | 0 | 0 |
| 27228 | ENSMUSG00000085375 | 0 | NA | NA | NA | NA | NA | Gm12506 | predicted gene 12506 [Source:MGI Symbo | 0 | 0 | 0 | 0 | 0 | 0 |
| 27229 | ENSMUSG00000085376 | 6.237700382 | -0.529873214 | 0.950927061 | -0.557217516 | 0.577378824 | 0.907838603 | Gm14508 | predicted gene 14508 [Source:MGI Symbo | 8.031 | 3.0838 | 11.069 | 8.6758 | 5.4531 | 1.1138 |
| 27230 | ENSMUSG00000085377 | 0 | NA | NA | NA | NA | NA | 4930551L18 | RIKEN cDNA 4930551L18 gene [Source:M | 0 | 0 | 0 | 0 | 0 | 0 |
| 27231 | ENSMUSG00000085378 | 0 | NA | NA | NA | NA | NA | Gm11415 | predicted gene 11415 [Source:MGI Symbo | 0 | 0 | 0 | 0 | 0 | 0 |
| 27232 | ENSMUSG00000085379 | 17.93570065 | 0.066210024 | 0.522843961 | 0.126634386 | 0.89922978 | 0.982798755 | 2310058D17 | RIKEN cDNA 2310058D17 gene [Source:M | 12.047 | 20.558 | 20.125 | 16.388 | 22.903 | 15.594 |
| 27233 | ENSMUSG00000085380 | 0 | NA | NA | NA | NA | NA | E130120K24 | RIKEN cDNA E130120K24 gene [Source:M | 0 | 0 | 0 | 0 | 0 | 0 |
| 27234 | ENSMUSG00000085382 | 0 | NA | NA | NA | NA | NA | Gm13861 | predicted gene 13861 [Source:MGI Symbo | 0 | 0 | 0 | 0 | 0 | 0 |
| 27235 | ENSMUSG00000085383 | 0 | NA | NA | NA | NA | NA | Gm14218 | predicted gene 14218 [Source:MGI Symbo | 0 | 0 | 0 | 0 | 0 | 0 |
| 27236 | ENSMUSG00000085385 | 207.3860668 | 0.143730226 | 0.207014429 | 0.69430052 | 0.487493765 | 0.873910634 | Snhg17 | small nucleolar RNA host gene 17 [Source | 231.29 | 168.58 | 191.19 | 269.91 | 200.67 | 182.67 |
| 27237 | ENSMUSG00000085386 | 0 | NA | NA | NA | NA | NA | Gm13630 | predicted gene 13630 [Source:MGI Symbo | 0 | 0 | 0 | 0 | 0 | 0 |
| 27238 | ENSMUSG00000085387 | 0 | NA | NA | NA | NA | NA | A630050E04 | RIKEN cDNA A630050E04 gene [Source:M | 0 | 0 | 0 | 0 | 0 | 0 |
| 27239 | ENSMUSG00000085388 | 0 | NA | NA | NA | NA | NA | Ptf1aos | pancreas specific transcription factor 1a, c | 0 | 0 | 0 | 0 | 0 | 0 |
| 27240 | ENSMUSG00000085389 | 0 | NA | NA | NA | NA | NA | 1700003M07 | RIKEN cDNA 1700003M07 gene [Source:M | 0 | 0 | 0 | 0 | 0 | 0 |
| 27241 | ENSMUSG00000085390 | 0 | NA | NA | NA | NA | NA | Gm13115 | predicted gene 13115 [Source:MGI Symbo | 0 | 0 | 0 | 0 | 0 | 0 |
| 27242 | ENSMUSG00000085391 | 0 | NA | NA | NA | NA | NA | Gm16150 | predicted gene 16150 [Source:MGI Symbo | 0 | 0 | 0 | 0 | 0 | 0 |
| 27243 | ENSMUSG00000085393 | 0 | NA | NA | NA | NA | NA | Gm11280 | predicted gene 11280 [Source:MGI Symbo | 0 | 0 | 0 | 0 | 0 | 0 |
| 27244 | ENSMUSG00000085394 | 0 | NA | NA | NA | NA | NA | 2210414B05 | RIKEN cDNA 2210414B05 gene [Source:M | 0 | 0 | 0 | 0 | 0 | 0 |
| 27245 | ENSMUSG00000085395 | 4.99228177 | -0.182729639 | 1.054014361 | -0.173365417 | 0.862364207 | NA | Gm13056 | predicted gene 13056 [Source:MGI Symbo | 9.6372 | 3.0838 | 3.0187 | 0.964 | 5.4531 | 7.7969 |
| 27246 | ENSMUSG00000085396 | 832.3588757 | 0.03788559 | 0.12462722 | 0.303991293 | 0.761134525 | 0.95402891 | Firre | functional intergenic repeating RNA elem | 796.68 | 874.76 | 793.93 | 935.06 | 845.23 | 748.5 |
| 27247 | ENSMUSG00000085397 | 0 | NA | NA | NA | NA | NA | Gm13524 | predicted gene 13524 [Source:MGI Symbo | 0 | 0 | 0 | 0 | 0 | 0 |
| 27248 | ENSMUSG00000085398 | 0.315620088 | 0.165143421 | 4.063300985 | 0.040642675 | 0.967580763 | NA | 4931406G06 | RIKEN cDNA 4931406G06 gene [Source:M | 0.8031 | 0 | 0 | 0 | 1.0906 | 0 |
| 27249 | ENSMUSG00000085399 | 1.277737502 | -2.558404199 | 2.061944144 | -1.240772795 | 0.214689694 | NA | Foxd2os | forkhead box D2, opposite strand [Source | 1.6062 | 3.0838 | 2.0125 | 0.964 | 0 | 0 |
| 27250 | ENSMUSG00000085400 | 4.482380912 | 0.096585286 | 1.009289474 | 0.095696317 | 0.923761766 | NA | Pard3bos3 | par-3 family cell polarity regulator beta, o | 4.8186 | 3.0838 | 5.0312 | 2.8919 | 3.2719 | 7.7969 |
| 27251 | ENSMUSG00000085401 | 0.167707431 | -0.796629176 | 4.080472857 | -0.195229623 | 0.84521319 | NA | Slc39a1-ps | solute carrier family 39 (zinc transporter), | 0 | 0 | 1.0062 | 0 | 0 | 0 |
| 27252 | ENSMUSG00000085402 | 0 | NA | NA | NA | NA | NA | Gm12111 | predicted gene 12111 [Source:MGI Symbo | 0 | 0 | 0 | 0 | 0 | 0 |
| 27253 | ENSMUSG00000085403 | 2.9500338 | -1.331175243 | 1.349120141 | -0.986698814 | 0.323790312 | NA | Gm13068 | predicted gene 13068 [Source:MGI Symbo | 5.6217 | 3.0838 | 4.025 | 3.8559 | 0 | 1.1138 |
| 27254 | ENSMUSG00000085404 | 128.9432508 | 0.184347342 | 0.241340706 | 0.763846866 | 0.444958505 | 0.852095831 | Gm12909 | predicted gene 12909 [Source:MGI Symbo | 122.87 | 92.513 | 146.91 | 157.13 | 143.96 | 110.27 |
| 27255 | ENSMUSG00000085405 | 0 | NA | NA | NA | NA | NA | 4930441J16 | RIKEN cDNA 4930441J16 gene [Source:M | 0 | 0 | 0 | 0 | 0 | 0 |
| 27256 | ENSMUSG00000085407 | 0 | NA | NA | NA | NA | NA | 1700095J03 | RIKEN cDNA 1700095J03 gene [Source:M | 0 | 0 | 0 | 0 | 0 | 0 |
| 27257 | ENSMUSG00000085408 | 8.729843249 | -0.460592292 | 0.747565822 | -0.616122726 | 0.537813524 | 0.891159905 | C530005A16 | RIKEN cDNA C530005A16 gene [Source:M | 8.031 | 11.307 | 11.069 | 8.6758 | 3.2719 | 10.025 |
| 27258 | ENSMUSG00000085409 | 0.133850373 | -0.796629176 | 4.080472857 | -0.195229623 | 0.84521319 | NA | Gm16155 | predicted gene 16155 [Source:MGI Symbo | 0.8031 | 0 | 0 | 0 | 0 | 0 |
| 27259 | ENSMUSG00000085411 | 5.407228482 | -0.605391128 | 1.038954504 | -0.582692626 | 0.560100238 | 0.899475387 | Gm14319 | predicted gene 14319 [Source:MGI Symbo | 11.243 | 6.1675 | 2.0125 | 1.928 | 2.1812 | 8.9108 |
| 27260 | ENSMUSG00000085412 | 0 | NA | NA | NA | NA | NA | Halr1 | Hoxa adjacent long noncoding RNA 1 [Sou | 0 | 0 | 0 | 0 | 0 | 0 |
| 27261 | ENSMUSG00000085413 | 0 | NA | NA | NA | NA | NA | Kbtbd8os | kelch repeat and BTB (POZ) domain conta | 0 | 0 | 0 | 0 | 0 | 0 |
| 27262 | ENSMUSG00000085414 | 0 | NA | NA | NA | NA | NA | Sspnos | sarcospan, opposite strand [Source:MGI S | 0 | 0 | 0 | 0 | 0 | 0 |
| 27263 | ENSMUSG00000085415 | 0.562214094 | -1.122836648 | 3.276281221 | -0.34271681 | 0.731811515 | NA | Selenok-ps1 | selenoprotein K, pseudogene 1 [Source:M | 2.4093 | 0 | 0 | 0.964 | 0 | 0 |
| 27264 | ENSMUSG00000085416 | 0 | NA | NA | NA | NA | NA | 1700019G24 | RIKEN cDNA 1700019G24 gene [Source:M | 0 | 0 | 0 | 0 | 0 | 0 |
| 27265 | ENSMUSG00000085417 | 0 | NA | NA | NA | NA | NA | Gm13919 | predicted gene 13919 [Source:MGI Symbo | 0 | 0 | 0 | 0 | 0 | 0 |
| 27266 | ENSMUSG00000085418 | 0 | NA | NA | NA | NA | NA | Grip1os1 | glutamate receptor interacting protein 1, | 0 | 0 | 0 | 0 | 0 | 0 |
| 27267 | ENSMUSG00000085419 | 0 | NA | NA | NA | NA | NA | Gm11734 | predicted gene 11734 [Source:MGI Symbo | 0 | 0 | 0 | 0 | 0 | 0 |
| 27268 | ENSMUSG00000085420 | 0 | NA | NA | NA | NA | NA | 4930453O03 | RIKEN cDNA 4930453O03 gene [Source:M | 0 | 0 | 0 | 0 | 0 | 0 |

| | | | | | | | | | | | | | | | |
|---|---|---|---|---|---|---|---|---|---|---|---|---|---|---|---|
| 27269 | ENSMUSG00000085421 | 0 | NA | NA | NA | NA | NA | 4732490B19 | RIKEN cDNA 4732490B19 gene [Source:M | 0 | 0 | 0 | 0 | 0 | 0 |
| 27270 | ENSMUSG00000085423 | 0.133850373 | -0.796629176 | 4.080472857 | -0.195229623 | 0.84521319 | NA | 4930594M2: | RIKEN cDNA 4930594M22 gene [Source:M | 0.8031 | 0 | 0 | 0 | 0 | 0 |
| 27271 | ENSMUSG00000085424 | 0 | NA | NA | NA | NA | NA | Fhad1os1 | forkhead-associated (FHA) phosphopeptic | 0 | 0 | 0 | 0 | 0 | 0 |
| 27272 | ENSMUSG00000085426 | 0 | NA | NA | NA | NA | NA | Gm14272 | predicted gene 14272 [Source:MGI Symbo | 0 | 0 | 0 | 0 | 0 | 0 |
| 27273 | ENSMUSG00000085427 | 5.639766774 | 1.696755027 | 1.092825757 | 1.552630889 | 0.120511342 | 0.508333638 | 6430710C18 | RIKEN cDNA 6430710C18 gene [Source:M | 4.8186 | 0 | 3.0187 | 11.568 | 2.1812 | 12.252 |
| 27274 | ENSMUSG00000085428 | 0 | NA | NA | NA | NA | NA | Gm14344 | predicted gene 14344 [Source:MGI Symbo | 0 | 0 | 0 | 0 | 0 | 0 |
| 27275 | ENSMUSG00000085429 | 0 | NA | NA | NA | NA | NA | Gm13485 | predicted gene 13485 [Source:MGI Symbo | 0 | 0 | 0 | 0 | 0 | 0 |
| 27276 | ENSMUSG00000085430 | 0 | NA | NA | NA | NA | NA | Gm15619 | predicted gene 15619 [Source:MGI Symbo | 0 | 0 | 0 | 0 | 0 | 0 |
| 27277 | ENSMUSG00000085431 | 0 | NA | NA | NA | NA | NA | 4930440I19F | RIKEN cDNA 4930440I19 gene [Source:MC | 0 | 0 | 0 | 0 | 0 | 0 |
| 27278 | ENSMUSG00000085433 | 13.87494343 | -0.191405312 | 0.708762173 | -0.270055767 | 0.787117351 | 0.960870081 | Gm16001 | predicted gene 16001 [Source:MGI Symbo | 11.243 | 6.1675 | 27.169 | 22.171 | 9.8156 | 6.6831 |
| 27279 | ENSMUSG00000085434 | 1.465265211 | 0.405379506 | 1.949922552 | 0.207895183 | 0.83531081 | NA | Gm11725 | predicted gene 11725 [Source:MGI Symbo | 0.8031 | 0 | 3.0187 | 3.8559 | 0 | 1.1138 |
| 27280 | ENSMUSG00000085436 | 13.36902006 | -0.200434825 | 0.602326019 | -0.332768001 | 0.739309414 | 0.948310621 | Zfp335os | zinc finger protein 335, opposite strand [S | 14.456 | 17.475 | 11.069 | 17.352 | 8.7249 | 11.138 |
| 27281 | ENSMUSG00000085437 | 0 | NA | NA | NA | NA | NA | 1700047A11 | RIKEN cDNA 1700047A11 gene [Source:M | 0 | 0 | 0 | 0 | 0 | 0 |
| 27282 | ENSMUSG00000085438 | 1442.462687 | 0.167660574 | 0.097707862 | 1.715937389 | 0.086173502 | 0.432482144 | Oip5os1 | Opa interacting protein 5, opposite strand | 1333.1 | 1381.5 | 1361.4 | 1455.6 | 1577 | 1546 |
| 27283 | ENSMUSG00000085439 | 7.026934415 | 1.745113798 | 0.948341627 | 1.840174203 | 0.065742666 | 0.368337165 | Rapgef4os1 | Rap guanine nucleotide exchange factor ( | 6.4248 | 0 | 3.0187 | 10.604 | 7.6343 | 14.48 |
| 27284 | ENSMUSG00000085440 | 3.111827892 | 0.385333191 | 1.374248804 | 0.280395508 | 0.779174083 | NA | Sorbs2os | sorbin and SH3 domain containing 2, oppo | 0 | 7.1954 | 1.0062 | 3.8559 | 3.2719 | 3.3415 |
| 27285 | ENSMUSG00000085441 | 0 | NA | NA | NA | NA | NA | Gm8289 | predicted gene 8289 [Source:MGI Symbol | 0 | 0 | 0 | 0 | 0 | 0 |
| 27286 | ENSMUSG00000085442 | 2.371993202 | -0.188915262 | 1.815639313 | -0.104048894 | 0.917130548 | NA | Gm3362 | predicted pseudogene 3362 [Source:MGI | 2.4093 | 5.1396 | 0 | 0 | 0 | 6.6831 |
| 27287 | ENSMUSG00000085443 | 0 | NA | NA | NA | NA | NA | Gm15416 | predicted gene 15416 [Source:MGI Symbo | 0 | 0 | 0 | 0 | 0 | 0 |
| 27288 | ENSMUSG00000085444 | 0 | NA | NA | NA | NA | NA | Gm13936 | predicted gene 13936 [Source:MGI Symbo | 0 | 0 | 0 | 0 | 0 | 0 |
| 27289 | ENSMUSG00000085445 | 72.67788475 | 0.809316653 | 0.392783802 | 2.060463416 | 0.03935426 | 0.271444928 | Gm16348 | predicted gene 16348 [Source:MGI Symbo | 57.823 | 57.563 | 43.269 | 158.09 | 45.806 | 73.514 |
| 27290 | ENSMUSG00000085447 | 0 | NA | NA | NA | NA | NA | 9530026F06 | RIKEN cDNA 9530026F06 gene [Source:M | 0 | 0 | 0 | 0 | 0 | 0 |
| 27291 | ENSMUSG00000085448 | 0 | NA | NA | NA | NA | NA | Gm13963 | predicted gene 13963 [Source:MGI Symbo | 0 | 0 | 0 | 0 | 0 | 0 |
| 27292 | ENSMUSG00000085449 | 0 | NA | NA | NA | NA | NA | Gm15520 | predicted gene 15520 [Source:MGI Symbo | 0 | 0 | 0 | 0 | 0 | 0 |
| 27293 | ENSMUSG00000085450 | 0 | NA | NA | NA | NA | NA | Gm13686 | predicted gene 13686 [Source:MGI Symbo | 0 | 0 | 0 | 0 | 0 | 0 |
| 27294 | ENSMUSG00000085451 | 0 | NA | NA | NA | NA | NA | Gm40614 | predicted gene, 40614 [Source:MGI Symb | 0 | 0 | 0 | 0 | 0 | 0 |
| 27295 | ENSMUSG00000085452 | 0 | NA | NA | NA | NA | NA | 4930554G24 | RIKEN cDNA 4930554G24 gene [Source:M | 0 | 0 | 0 | 0 | 0 | 0 |
| 27296 | ENSMUSG00000085453 | 0 | NA | NA | NA | NA | NA | Gm12018 | predicted gene 12018 [Source:MGI Symbo | 0 | 0 | 0 | 0 | 0 | 0 |
| 27297 | ENSMUSG00000085454 | 0.342639279 | -1.805976596 | 4.028547714 | -0.448294702 | 0.653940524 | NA | Grip1os3 | glutamate receptor interacting protein 1, | 0 | 2.0558 | 0 | 0 | 0 | 0 |
| 27298 | ENSMUSG00000085456 | 0 | NA | NA | NA | NA | NA | Gm15398 | predicted gene 15398 [Source:MGI Symbo | 0 | 0 | 0 | 0 | 0 | 0 |
| 27299 | ENSMUSG00000085457 | 3.856039936 | 0.959594251 | 1.179157354 | 0.813796604 | 0.415761481 | NA | 1110046J04F | RIKEN cDNA 1110046J04 gene [Source:MC | 0.8031 | 3.0838 | 4.025 | 1.928 | 3.2719 | 10.025 |
| 27300 | ENSMUSG00000085459 | 0 | NA | NA | NA | NA | NA | Gm12272 | predicted gene 12272 [Source:MGI Symbo | 0 | 0 | 0 | 0 | 0 | 0 |
| 27301 | ENSMUSG00000085463 | 0 | NA | NA | NA | NA | NA | 9430024E24 | RIKEN cDNA 9430024E24 gene [Source:M | 0 | 0 | 0 | 0 | 0 | 0 |
| 27302 | ENSMUSG00000085464 | 0.335414862 | -1.779047242 | 4.030804179 | -0.441362855 | 0.658950329 | NA | Gm16208 | predicted gene 16208 [Source:MGI Symbo | 0 | 0 | 2.0125 | 0 | 0 | 0 |
| 27303 | ENSMUSG00000085465 | 1.507944006 | -0.86875514 | 1.814643171 | -0.478747092 | 0.63211856 | NA | Gm15347 | predicted gene 15347 [Source:MGI Symbo | 0.8031 | 1.0279 | 4.025 | 0.964 | 0 | 2.2277 |
| 27304 | ENSMUSG00000085466 | 0 | NA | NA | NA | NA | NA | Hmgn2l6 | high-mobility group nucleosomal binding | 0 | 0 | 0 | 0 | 0 | 0 |
| 27305 | ENSMUSG00000085467 | 0.535401493 | -2.48596965 | 3.684117979 | -0.674780141 | 0.499815459 | NA | Gm11842 | predicted gene 11842 [Source:MGI Symbo | 3.2124 | 0 | 0 | 0 | 0 | 0 |
| 27306 | ENSMUSG00000085468 | 55.45598522 | -0.057532162 | 0.31818886 | -0.180811365 | 0.856515642 | 0.974562795 | Gm15343 | predicted gene 15343 [Source:MGI Symbo | 45.777 | 62.703 | 61.381 | 46.271 | 68.709 | 47.895 |
| 27307 | ENSMUSG00000085469 | 0 | NA | NA | NA | NA | NA | Gm14242 | predicted gene 14242 [Source:MGI Symbo | 0 | 0 | 0 | 0 | 0 | 0 |
| 27308 | ENSMUSG00000085470 | 0 | NA | NA | NA | NA | NA | 1700012C14 | RIKEN cDNA 1700012C14 gene [Source:M | 0 | 0 | 0 | 0 | 0 | 0 |
| 27309 | ENSMUSG00000085471 | 1.681175683 | -0.670357496 | 1.982721922 | -0.338099604 | 0.735288129 | NA | 4933423P22 | RIKEN cDNA 4933423P22 gene [Source:M | 3.2124 | 0 | 3.0187 | 3.8559 | 0 | 0 |
| 27310 | ENSMUSG00000085472 | 0 | NA | NA | NA | NA | NA | Gm15691 | predicted gene 15691 [Source:MGI Symbo | 0 | 0 | 0 | 0 | 0 | 0 |
| 27311 | ENSMUSG00000085476 | 0 | NA | NA | NA | NA | NA | Gm12162 | predicted gene 12162 [Source:MGI Symbo | 0 | 0 | 0 | 0 | 0 | 0 |
| 27312 | ENSMUSG00000085477 | 0 | NA | NA | NA | NA | NA | Gm13335 | predicted gene 13335 [Source:MGI Symbo | 0 | 0 | 0 | 0 | 0 | 0 |
| 27313 | ENSMUSG00000085478 | 0 | NA | NA | NA | NA | NA | Gm11851 | predicted gene 11851 [Source:MGI Symbo | 0 | 0 | 0 | 0 | 0 | 0 |
| 27314 | ENSMUSG00000085479 | 0 | NA | NA | NA | NA | NA | 9430073C21 | Riken cDNA 9430073C21 gene [Source:MC | 0 | 0 | 0 | 0 | 0 | 0 |
| 27315 | ENSMUSG00000085481 | 0 | NA | NA | NA | NA | NA | Gm16304 | predicted gene 16304 [Source:MGI Symbo | 0 | 0 | 0 | 0 | 0 | 0 |
| 27316 | ENSMUSG00000085482 | 4.00522434 | -0.416710544 | 1.149397043 | -0.362547082 | 0.716943246 | NA | A230083N12 | RIKEN cDNA A230083N12 gene [Source:M | 5.6217 | 4.1117 | 4.025 | 4.8199 | 5.4531 | 0 |
| 27317 | ENSMUSG00000085483 | 0.133850373 | -0.796629176 | 4.080472857 | -0.195229623 | 0.84521319 | NA | Gm14198 | predicted gene 14198 [Source:MGI Symbo | 0.8031 | 0 | 0 | 0 | 0 | 0 |
| 27318 | ENSMUSG00000085484 | 0 | NA | NA | NA | NA | NA | Gm13539 | predicted gene 13539 [Source:MGI Symbo | 0 | 0 | 0 | 0 | 0 | 0 |
| 27319 | ENSMUSG00000085485 | 0 | NA | NA | NA | NA | NA | Gm14024 | predicted gene 14024 [Source:MGI Symbo | 0 | 0 | 0 | 0 | 0 | 0 |
| 27320 | ENSMUSG00000085486 | 1.075966422 | -0.197496223 | 2.150415569 | -0.091840957 | 0.926824403 | NA | Gm11634 | predicted gene 11634 [Source:MGI Symbo | 2.4093 | 1.0279 | 0 | 1.928 | 1.0906 | 0 |
| 27321 | ENSMUSG00000085487 | 0 | NA | NA | NA | NA | NA | Gm15654 | predicted gene 15654 [Source:MGI Symbo | 0 | 0 | 0 | 0 | 0 | 0 |
| 27322 | ENSMUSG00000085488 | 0 | NA | NA | NA | NA | NA | 4930557F10 | RIKEN cDNA 4930557F10 gene [Source:M | 0 | 0 | 0 | 0 | 0 | 0 |
| 27323 | ENSMUSG00000085489 | 0 | NA | NA | NA | NA | NA | Gm3669 | predicted gene 3669 [Source:MGI Symbol | 0 | 0 | 0 | 0 | 0 | 0 |
| 27324 | ENSMUSG00000085491 | 0 | NA | NA | NA | NA | NA | 4930527E20 | RIKEN cDNA 4930527E20 gene [Source:M | 0 | 0 | 0 | 0 | 0 | 0 |

| | | | | | | | | | | | | | | | |
|---|---|---|---|---|---|---|---|---|---|---|---|---|---|---|---|
| 27325 | ENSMUSG00000085492 | 57.3136789 | -0.157440823 | 0.290260343 | -0.542412448 | 0.587534406 | 0.910765033 | Trmt61b | tRNA methyltransferase 61B [Source:MGI | 59.43 | 59.619 | 62.387 | 57.839 | 56.712 | 47.895 |
| 27326 | ENSMUSG00000085495 | 0.267700746 | -1.497481649 | 4.057001436 | -0.369110456 | 0.712045396 | NA | Gm16796 | predicted gene, 16796 [Source:MGI Symb | 1.6062 | 0 | 0 | 0 | 0 | 0 |
| 27327 | ENSMUSG00000085496 | 0 | NA | NA | NA | NA | NA | 4930517E11 | RIKEN cDNA 4930517E11 gene [Source:M | 0 | 0 | 0 | 0 | 0 | 0 |
| 27328 | ENSMUSG00000085497 | 0 | NA | NA | NA | NA | NA | Gm15985 | predicted gene 15985 [Source:MGI Symbc | 0 | 0 | 0 | 0 | 0 | 0 |
| 27329 | ENSMUSG00000085498 | 15.85666073 | 0.334348815 | 0.629998651 | 0.530713542 | 0.595617301 | 0.913808971 | Gm14023 | predicted gene 14023 [Source:MGI Symbc | 26.502 | 6.1675 | 9.0562 | 19.28 | 18.541 | 15.594 |
| 27330 | ENSMUSG00000085499 | 0.692607243 | 3.004183657 | 2.957557172 | 1.015765202 | 0.309741211 | NA | Gm15713 | predicted gene 15713 [Source:MGI Symbc | 0 | 0 | 0 | 1.928 | 0 | 2.2277 |
| 27331 | ENSMUSG00000085500 | 0 | NA | NA | NA | NA | NA | Gm16976 | predicted gene, 16976 [Source:MGI Symb | 0 | 0 | 0 | 0 | 0 | 0 |
| 27332 | ENSMUSG00000085501 | 14.56034906 | 0.429335726 | 0.586965379 | 0.731449829 | 0.464504439 | 0.862553639 | Gm11772 | predicted gene 11772 [Source:MGI Symbc | 17.668 | 12.335 | 7.0437 | 18.316 | 14.178 | 17.822 |
| 27333 | ENSMUSG00000085502 | 0.160662974 | 1.126924175 | 4.080472857 | 0.276174898 | 0.782413736 | NA | Gm12320 | predicted gene 12320 [Source:MGI Symbc | 0 | 0 | 0 | 0.964 | 0 | 0 |
| 27334 | ENSMUSG00000085504 | 0 | NA | NA | NA | NA | NA | Gm14055 | predicted gene 14055 [Source:MGI Symbc | 0 | 0 | 0 | 0 | 0 | 0 |
| 27335 | ENSMUSG00000085505 | 0.160662974 | 1.126924175 | 4.080472857 | 0.276174898 | 0.782413736 | NA | Gm12868 | predicted gene 12868 [Source:MGI Symbc | 0 | 0 | 0 | 0.964 | 0 | 0 |
| 27336 | ENSMUSG00000085507 | 0 | NA | NA | NA | NA | NA | Gm2709 | predicted gene 2709 [Source:MGI Symbol | 0 | 0 | 0 | 0 | 0 | 0 |
| 27337 | ENSMUSG00000085508 | 0 | NA | NA | NA | NA | NA | NA | NA | 0 | 0 | 0 | 0 | 0 | 0 |
| 27338 | ENSMUSG00000085509 | 0 | NA | NA | NA | NA | NA | Gm11521 | predicted gene 11521 [Source:MGI Symbc | 0 | 0 | 0 | 0 | 0 | 0 |
| 27339 | ENSMUSG00000085510 | 0 | NA | NA | NA | NA | NA | Mir217hg | microRNA 217 host gene [Source:MGI Syn | 0 | 0 | 0 | 0 | 0 | 0 |
| 27340 | ENSMUSG00000085511 | 0 | NA | NA | NA | NA | NA | Gm11738 | predicted gene 11738 [Source:MGI Symbc | 0 | 0 | 0 | 0 | 0 | 0 |
| 27341 | ENSMUSG00000085512 | 0 | NA | NA | NA | NA | NA | Gm11958 | predicted gene 11958 [Source:MGI Symbc | 0 | 0 | 0 | 0 | 0 | 0 |
| 27342 | ENSMUSG00000085513 | 0 | NA | NA | NA | NA | NA | Gm16363 | predicted gene 16363 [Source:MGI Symbc | 0 | 0 | 0 | 0 | 0 | 0 |
| 27343 | ENSMUSG00000085514 | 1.165432554 | -1.22162459 | 2.176801979 | -0.561201525 | 0.574660164 | NA | Bcas3os2 | breast carcinoma amplified sequence 3 op | 0.8031 | 4.1117 | 0 | 0.964 | 0 | 1.1138 |
| 27344 | ENSMUSG00000085515 | 0 | NA | NA | NA | NA | NA | C630028M0 | RIKEN cDNA C630028M04 gene [Source:M | 0 | 0 | 0 | 0 | 0 | 0 |
| 27345 | ENSMUSG00000085516 | 0 | NA | NA | NA | NA | NA | 4930538E20 | RIKEN cDNA 4930538E20 gene [Source:M | 0 | 0 | 0 | 0 | 0 | 0 |
| 27346 | ENSMUSG00000085517 | 1.211910448 | 0.174485504 | 2.253456452 | 0.077430165 | 0.938281345 | NA | Gm12963 | predicted gene 12963 [Source:MGI Symbc | 2.4093 | 0 | 1.0062 | 3.8559 | 0 | 0 |
| 27347 | ENSMUSG00000085519 | 0.767390791 | -1.69152783 | 2.44773922 | -0.691057207 | 0.489529591 | NA | Gm13703 | predicted gene 13703 [Source:MGI Symbc | 1.6062 | 1.0279 | 1.0062 | 0.964 | 0 | 0 |
| 27348 | ENSMUSG00000085521 | 0.528073336 | 2.60973587 | 2.979250867 | 0.875970499 | 0.381046072 | NA | 4930526A20 | RIKEN cDNA 4930526A20 gene [Source:M | 0 | 0 | 0 | 0.964 | 1.0906 | 1.1138 |
| 27349 | ENSMUSG00000085522 | 0 | NA | NA | NA | NA | NA | 4930556G01 | RIKEN cDNA 4930556G01 gene [Source:M | 0 | 0 | 0 | 0 | 0 | 0 |
| 27350 | ENSMUSG00000085523 | 0.96895528 | -2.08853317 | 2.292908757 | -0.910866237 | 0.362365857 | NA | Gm15945 | predicted gene 15945 [Source:MGI Symbc | 0.8031 | 1.0279 | 3.0187 | 0.964 | 0 | 0 |
| 27351 | ENSMUSG00000085524 | 0 | NA | NA | NA | NA | NA | Gm14224 | predicted gene 14224 [Source:MGI Symbc | 0 | 0 | 0 | 0 | 0 | 0 |
| 27352 | ENSMUSG00000085525 | 5.086802386 | 2.459545068 | 1.145615357 | 2.146920476 | 0.031799613 | 0.237577275 | Gm13166 | predicted gene 13166 [Source:MGI Symbc | 1.6062 | 3.0838 | 0 | 3.8559 | 14.178 | 7.7969 |
| 27353 | ENSMUSG00000085526 | 0 | NA | NA | NA | NA | NA | Gm16083 | predicted gene 16083 [Source:MGI Symbc | 0 | 0 | 0 | 0 | 0 | 0 |
| 27354 | ENSMUSG00000085527 | 18.94993904 | 0.172447431 | 0.568513124 | 0.303330607 | 0.761637923 | 0.954216198 | Gm15535 | predicted gene 15535 [Source:MGI Symbc | 20.078 | 14.391 | 19.119 | 34.703 | 9.8156 | 15.594 |
| 27355 | ENSMUSG00000085528 | 0 | NA | NA | NA | NA | NA | 4930401O1C | RIKEN cDNA 4930401O10 gene [Source:M | 0 | 0 | 0 | 0 | 0 | 0 |
| 27356 | ENSMUSG00000085530 | 0 | NA | NA | NA | NA | NA | Gm13901 | predicted gene 13901 [Source:MGI Symbc | 0 | 0 | 0 | 0 | 0 | 0 |
| 27357 | ENSMUSG00000085531 | 0.17131964 | -0.796629176 | 4.080472857 | -0.195229623 | 0.84521319 | NA | Slc36a3os | solute carrier family 36 (proton/amino aci | 0 | 1.0279 | 0 | 0 | 0 | 0 |
| 27358 | ENSMUSG00000085532 | 0 | NA | NA | NA | NA | NA | B430319H21 | RIKEN cDNA B430319H21 gene [Source:M | 0 | 0 | 0 | 0 | 0 | 0 |
| 27359 | ENSMUSG00000085533 | 0 | NA | NA | NA | NA | NA | Gm12146 | predicted gene 12146 [Source:MGI Symbc | 0 | 0 | 0 | 0 | 0 | 0 |
| 27360 | ENSMUSG00000085534 | 0 | NA | NA | NA | NA | NA | Gm2117 | predicted gene 2117 [Source:MGI Symbol | 0 | 0 | 0 | 0 | 0 | 0 |
| 27361 | ENSMUSG00000085536 | 0 | NA | NA | NA | NA | NA | Gm14950 | predicted gene 14950 [Source:MGI Symbc | 0 | 0 | 0 | 0 | 0 | 0 |
| 27362 | ENSMUSG00000085538 | 0.730949792 | 3.072456834 | 2.934263219 | 1.04709653 | 0.29505506 | NA | Gm11455 | predicted gene 11455 [Source:MGI Symbc | 0 | 0 | 0 | 0 | 3.2719 | 1.1138 |
| 27363 | ENSMUSG00000085539 | 0 | NA | NA | NA | NA | NA | Gm15561 | predicted gene 15561 [Source:MGI Symbc | 0 | 0 | 0 | 0 | 0 | 0 |
| 27364 | ENSMUSG00000085540 | 0 | NA | NA | NA | NA | NA | Gm16036 | predicted gene 16036 [Source:MGI Symbc | 0 | 0 | 0 | 0 | 0 | 0 |
| 27365 | ENSMUSG00000085541 | 0 | NA | NA | NA | NA | NA | Gm16010 | predicted gene 16010 [Source:MGI Symbc | 0 | 0 | 0 | 0 | 0 | 0 |
| 27366 | ENSMUSG00000085542 | 0 | NA | NA | NA | NA | NA | 1700037C06 | RIKEN cDNA 1700037C06 gene [Source:M | 0 | 0 | 0 | 0 | 0 | 0 |
| 27367 | ENSMUSG00000085543 | 0.294513347 | 0.165143496 | 4.080472857 | 0.040471657 | 0.967717103 | NA | Gm13568 | predicted gene 13568 [Source:MGI Symbc | 0.8031 | 0 | 0 | 0.964 | 0 | 0 |
| 27368 | ENSMUSG00000085545 | 0 | NA | NA | NA | NA | NA | Gm13553 | predicted gene 13553 [Source:MGI Symbc | 0 | 0 | 0 | 0 | 0 | 0 |
| 27369 | ENSMUSG00000085546 | 0.401551119 | -2.072982726 | 4.008251979 | -0.517178744 | 0.605031388 | NA | Gm14252 | predicted gene 14252 [Source:MGI Symbc | 2.4093 | 0 | 0 | 0 | 0 | 0 |
| 27370 | ENSMUSG00000085547 | 0 | NA | NA | NA | NA | NA | Gm13767 | predicted gene 13767 [Source:MGI Symbc | 0 | 0 | 0 | 0 | 0 | 0 |
| 27371 | ENSMUSG00000085548 | 0.133850373 | -0.796629176 | 4.080472857 | -0.195229623 | 0.84521319 | NA | Gm11753 | predicted gene 11753 [Source:MGI Symbc | 0.8031 | 0 | 0 | 0 | 0 | 0 |
| 27372 | ENSMUSG00000085549 | 0 | NA | NA | NA | NA | NA | 1700047F07 | RIKEN cDNA 1700047F07 gene [Source:M | 0 | 0 | 0 | 0 | 0 | 0 |
| 27373 | ENSMUSG00000085551 | 0 | NA | NA | NA | NA | NA | 1810063I02F | RIKEN cDNA 1810063I02 gene [Source:MC | 0 | 0 | 0 | 0 | 0 | 0 |
| 27374 | ENSMUSG00000085552 | 0 | NA | NA | NA | NA | NA | Gm21990 | predicted gene 21990 [Source:MGI Symbc | 0 | 0 | 0 | 0 | 0 | 0 |
| 27375 | ENSMUSG00000085553 | 0 | NA | NA | NA | NA | NA | Gm14808 | predicted gene 14808 [Source:MGI Symbc | 0 | 0 | 0 | 0 | 0 | 0 |
| 27376 | ENSMUSG00000085555 | 0 | NA | NA | NA | NA | NA | 2610035F20 | RIKEN cDNA 2610035F20 gene [Source:M | 0 | 0 | 0 | 0 | 0 | 0 |
| 27377 | ENSMUSG00000085556 | 0 | NA | NA | NA | NA | NA | Gm15020 | predicted gene 15020 [Source:MGI Symbc | 0 | 0 | 0 | 0 | 0 | 0 |
| 27378 | ENSMUSG00000085558 | 113.4996988 | 0.076313704 | 0.212141252 | 0.359730619 | 0.719048592 | 0.9447719 | 4930412C18 | RIKEN cDNA 4930412C18 gene [Source:M | 115.65 | 108.96 | 106.66 | 112.79 | 118.88 | 118.07 |
| 27379 | ENSMUSG00000085559 | 0 | NA | NA | NA | NA | NA | Gm11959 | predicted gene 11959 [Source:MGI Symbc | 0 | 0 | 0 | 0 | 0 | 0 |
| 27380 | ENSMUSG00000085560 | 0.728241582 | 1.712995752 | 3.00895953 | 0.569298369 | 0.569153674 | NA | C230038L03 | RIKEN cDNA C230038L03 gene [Source:M | 0 | 1.0279 | 0 | 0 | 0 | 3.3415 |

| | | | | | | | | | | | | | | | |
|---|---|---|---|---|---|---|---|---|---|---|---|---|---|---|---|
| 27381 | ENSMUSG00000085561 | 0 | NA | NA | NA | NA | NA | 1700046C09 | RIKEN cDNA 1700046C09 gene [Source:M | 0 | 0 | 0 | 0 | 0 | 0 |
| 27382 | ENSMUSG00000085562 | 0.181769715 | 1.126924175 | 4.080472857 | 0.276174898 | 0.782413736 | NA | 2610028E06 | RIKEN cDNA 2610028E06 gene [Source:M | 0 | 0 | 0 | 0 | 1.0906 | 0 |
| 27383 | ENSMUSG00000085563 | 0 | NA | NA | NA | NA | NA | 2210411M0! | RIKEN cDNA 2210411M09 gene [Source:M | 0 | 0 | 0 | 0 | 0 | 0 |
| 27384 | ENSMUSG00000085564 | 0 | NA | NA | NA | NA | NA | Gm12198 | predicted gene 12198 [Source:MGI Symbo | 0 | 0 | 0 | 0 | 0 | 0 |
| 27385 | ENSMUSG00000085565 | 0 | NA | NA | NA | NA | NA | Gm15721 | predicted gene 15721 [Source:MGI Symbo | 0 | 0 | 0 | 0 | 0 | 0 |
| 27386 | ENSMUSG00000085566 | 141.1897864 | -0.053390461 | 0.207241069 | -0.257624905 | 0.796696407 | 0.962236316 | A730017L22 | RIKEN cDNA A730017L22 gene [Source:M | 133.31 | 154.19 | 143.89 | 119.53 | 160.32 | 135.89 |
| 27387 | ENSMUSG00000085568 | 0.267700746 | -1.497481649 | 4.057001436 | -0.369110456 | 0.712045396 | NA | Gm14858 | predicted gene 14858 [Source:MGI Symbo | 1.6062 | 0 | 0 | 0 | 0 | 0 |
| 27388 | ENSMUSG00000085569 | 3.966333855 | 0.182786786 | 1.031958922 | 0.177126029 | 0.859409397 | NA | Gm12602 | predicted gene 12602 [Source:MGI Symbo | 4.0155 | 3.0838 | 4.025 | 3.8559 | 4.3625 | 4.4554 |
| 27389 | ENSMUSG00000085572 | 0.339027071 | -1.79174149 | 3.919585585 | -0.457125237 | 0.647581032 | NA | Gm16234 | predicted gene 16234 [Source:MGI Symbo | 0 | 1.0279 | 1.0062 | 0 | 0 | 0 |
| 27390 | ENSMUSG00000085573 | 0.589026695 | 0.241472922 | 3.228838524 | 0.074786311 | 0.940384733 | NA | Gm15418 | predicted gene 15418 [Source:MGI Symbo | 1.6062 | 0 | 0 | 1.928 | 0 | 0 |
| 27391 | ENSMUSG00000085574 | 0 | NA | NA | NA | NA | NA | Gm13053 | predicted gene 13053 [Source:MGI Symbo | 0 | 0 | 0 | 0 | 0 | 0 |
| 27392 | ENSMUSG00000085575 | 0.17131964 | -0.796629176 | 4.080472857 | -0.195229623 | 0.84521319 | NA | Gm15485 | predicted gene 15485 [Source:MGI Symbo | 0 | 1.0279 | 0 | 0 | 0 | 0 |
| 27393 | ENSMUSG00000085576 | 0.167707431 | -0.796629176 | 4.080472857 | -0.195229623 | 0.84521319 | NA | Dpy19l2 | dpy-19-like 2 (C. elegans) [Source:MGI Syr | 0 | 0 | 1.0062 | 0 | 0 | 0 |
| 27394 | ENSMUSG00000085577 | 0 | NA | NA | NA | NA | NA | Mageb6-ps | MAGE family member B6, pseudogene [So | 0 | 0 | 0 | 0 | 0 | 0 |
| 27395 | ENSMUSG00000085578 | 0 | NA | NA | NA | NA | NA | Gm11527 | predicted gene 11527 [Source:MGI Symbo | 0 | 0 | 0 | 0 | 0 | 0 |
| 27396 | ENSMUSG00000085579 | 0 | NA | NA | NA | NA | NA | Gm12066 | predicted gene 12066 [Source:MGI Symbo | 0 | 0 | 0 | 0 | 0 | 0 |
| 27397 | ENSMUSG00000085580 | 0 | NA | NA | NA | NA | NA | 1700061F12 | RIKEN cDNA 1700061F12 gene [Source:M | 0 | 0 | 0 | 0 | 0 | 0 |
| 27398 | ENSMUSG00000085581 | 2.656573695 | -1.051970545 | 1.42529735 | -0.738070933 | 0.460471346 | NA | Gm12786 | predicted gene 12786 [Source:MGI Symbo | 5.6217 | 4.1117 | 1.0062 | 1.928 | 3.2719 | 0 |
| 27399 | ENSMUSG00000085582 | 0 | NA | NA | NA | NA | NA | 3110099E03 | RIKEN cDNA 3110099E03 gene [Source:M | 0 | 0 | 0 | 0 | 0 | 0 |
| 27400 | ENSMUSG00000085584 | 0 | NA | NA | NA | NA | NA | Rtl9 | retrotransposon Gag like 9 [Source:MGI S | 0 | 0 | 0 | 0 | 0 | 0 |
| 27401 | ENSMUSG00000085585 | 0 | NA | NA | NA | NA | NA | Gm12223 | predicted gene 12223 [Source:MGI Symbo | 0 | 0 | 0 | 0 | 0 | 0 |
| 27402 | ENSMUSG00000085586 | 3.832251388 | -0.47316529 | 1.078547131 | -0.438706178 | 0.66087445 | NA | Gm11613 | predicted gene 11613 [Source:MGI Symbo | 3.2124 | 4.1117 | 6.0375 | 1.928 | 4.3625 | 3.3415 |
| 27403 | ENSMUSG00000085587 | 0 | NA | NA | NA | NA | NA | Gm14493 | predicted gene 14493 [Source:MGI Symbo | 0 | 0 | 0 | 0 | 0 | 0 |
| 27404 | ENSMUSG00000085588 | 0 | NA | NA | NA | NA | NA | 3110004A20 | RIKEN cDNA 3110004A20 gene [Source:M | 0 | 0 | 0 | 0 | 0 | 0 |
| 27405 | ENSMUSG00000085589 | 0.499690045 | -0.82080011 | 3.113553401 | -0.263621658 | 0.792071468 | NA | A430078I02I | RIKEN cDNA A430078I02 gene [Source:M | 0 | 1.0279 | 1.0062 | 0.964 | 0 | 0 |
| 27406 | ENSMUSG00000085591 | 0 | NA | NA | NA | NA | NA | Gm13479 | predicted gene 13479 [Source:MGI Symbo | 0 | 0 | 0 | 0 | 0 | 0 |
| 27407 | ENSMUSG00000085592 | 0.342639279 | -1.805976596 | 4.028547714 | -0.448294702 | 0.653940524 | NA | Gm16321 | predicted gene 16321 [Source:MGI Symbo | 0 | 2.0558 | 0 | 0 | 0 | 0 |
| 27408 | ENSMUSG00000085593 | 0 | NA | NA | NA | NA | NA | 4930589P08 | RIKEN cDNA 4930589P08 gene [Source:M | 0 | 0 | 0 | 0 | 0 | 0 |
| 27409 | ENSMUSG00000085594 | 0 | NA | NA | NA | NA | NA | Gm11551 | predicted gene 11551 [Source:MGI Symbo | 0 | 0 | 0 | 0 | 0 | 0 |
| 27410 | ENSMUSG00000085595 | 4.333847956 | -0.308567109 | 1.067533374 | -0.289046804 | 0.772545561 | NA | Gm16090 | predicted gene 16090 [Source:MGI Symbo | 7.2279 | 1.0279 | 6.0375 | 2.8919 | 4.3625 | 4.4554 |
| 27411 | ENSMUSG00000085596 | 0.517623261 | 1.044040381 | 3.075707576 | 0.339447218 | 0.734272853 | NA | Gm11476 | predicted gene 11476 [Source:MGI Symbo | 0 | 1.0279 | 0 | 0.964 | 0 | 1.1138 |
| 27412 | ENSMUSG00000085598 | 0 | NA | NA | NA | NA | NA | Gm14715 | predicted gene 14715 [Source:MGI Symbo | 0 | 0 | 0 | 0 | 0 | 0 |
| 27413 | ENSMUSG00000085599 | 13.06995739 | 0.95821228 | 0.752426332 | 1.273496474 | 0.202841923 | 0.647682949 | Gm13449 | predicted gene 13449 [Source:MGI Symbo | 5.6217 | 5.1396 | 16.1 | 31.811 | 14.178 | 5.5692 |
| 27414 | ENSMUSG00000085600 | 0 | NA | NA | NA | NA | NA | E330012B07 | RIKEN cDNA E330012B07 gene [Source:M | 0 | 0 | 0 | 0 | 0 | 0 |
| 27415 | ENSMUSG00000085601 | 1.199289611 | -1.270320185 | 2.16443215 | -0.58690691 | 0.557266229 | NA | Meiosin | meiosis initiator [Source:MGI Symbol;Acc: | 0 | 4.1117 | 1.0062 | 0.964 | 0 | 1.1138 |
| 27416 | ENSMUSG00000085603 | 2.58047818 | -0.203716528 | 1.297048808 | -0.157061574 | 0.875196321 | NA | Gm11346 | predicted gene 11346 [Source:MGI Symbo | 3.2124 | 3.0838 | 2.0125 | 3.8559 | 1.0906 | 2.2277 |
| 27417 | ENSMUSG00000085604 | 1.004356399 | 2.372094719 | 2.241352889 | 1.058331658 | 0.289904265 | NA | Dhx58os | DEAH (Asp-Glu-Ala-His) box polypeptide 5 | 0.8031 | 0 | 0 | 1.928 | 2.1812 | 1.1138 |
| 27418 | ENSMUSG00000085606 | 0 | NA | NA | NA | NA | NA | Gm15792 | predicted gene 15792 [Source:MGI Symbo | 0 | 0 | 0 | 0 | 0 | 0 |
| 27419 | ENSMUSG00000085608 | 0 | NA | NA | NA | NA | NA | Gm16354 | predicted gene 16354 [Source:MGI Symbo | 0 | 0 | 0 | 0 | 0 | 0 |
| 27420 | ENSMUSG00000085609 | 0 | NA | NA | NA | NA | NA | 1700016P03 | RIKEN cDNA 1700016P03 gene [Source:M | 0 | 0 | 0 | 0 | 0 | 0 |
| 27421 | ENSMUSG00000085610 | 0 | NA | NA | NA | NA | NA | Gm14747 | predicted gene 14747 [Source:MGI Symbo | 0 | 0 | 0 | 0 | 0 | 0 |
| 27422 | ENSMUSG00000085611 | 0.294513347 | 0.165143496 | 4.080472857 | 0.040471657 | 0.967717103 | NA | Ap3s1-ps1 | adaptor-related protein complex 3, sigma | 0.8031 | 0 | 0 | 0.964 | 0 | 0 |
| 27423 | ENSMUSG00000085612 | 1.489958656 | -3.930900617 | 2.094800059 | -1.876503965 | 0.060586111 | NA | Gm15868 | predicted gene 15868 [Source:MGI Symbo | 0.8031 | 4.1117 | 4.025 | 0 | 0 | 0 |
| 27424 | ENSMUSG00000085614 | 0.185640647 | 1.126924175 | 4.080472857 | 0.276174898 | 0.782413736 | NA | 1700123M0 | RIKEN cDNA 1700123M08 gene [Source:M | 0 | 0 | 0 | 0 | 0 | 1.1138 |
| 27425 | ENSMUSG00000085615 | 52.11554947 | 0.352850516 | 0.346029807 | 1.019711335 | 0.307865384 | 0.756343859 | A330035P11 | RIKEN cDNA A330035P11 gene [Source:M | 28.109 | 55.508 | 54.337 | 64.587 | 57.803 | 52.351 |
| 27426 | ENSMUSG00000085616 | 0 | NA | NA | NA | NA | NA | Gm13850 | predicted gene 13850 [Source:MGI Symbo | 0 | 0 | 0 | 0 | 0 | 0 |
| 27427 | ENSMUSG00000085618 | 0.353089354 | 0.165142706 | 3.896102244 | 0.042386646 | 0.966190474 | NA | Gm11466 | predicted gene 11466 [Source:MGI Symbo | 0 | 1.0279 | 0 | 0 | 1.0906 | 0 |
| 27428 | ENSMUSG00000085619 | 0 | NA | NA | NA | NA | NA | 4930555B12 | RIKEN cDNA 4930555B12 gene [Source:M | 0 | 0 | 0 | 0 | 0 | 0 |
| 27429 | ENSMUSG00000085620 | 0 | NA | NA | NA | NA | NA | G630018N1 | RIKEN cDNA G630018N14 gene [Source:M | 0 | 0 | 0 | 0 | 0 | 0 |
| 27430 | ENSMUSG00000085622 | 53.30577968 | -0.189648902 | 0.315642494 | -0.600834506 | 0.547950219 | 0.895600915 | 3110056K07 | RIKEN cDNA 3110056K07 gene [Source:M | 57.823 | 48.312 | 64.4 | 56.875 | 53.44 | 38.985 |
| 27431 | ENSMUSG00000085623 | 0 | NA | NA | NA | NA | NA | Gm16041 | predicted gene 16041 [Source:MGI Symbo | 0 | 0 | 0 | 0 | 0 | 0 |
| 27432 | ENSMUSG00000085624 | 0.55305101 | 2.667977666 | 3.241982584 | 0.822946329 | 0.410538515 | NA | Gm14573 | predicted gene 14573 [Source:MGI Symbo | 0 | 0 | 0 | 0 | 1.0906 | 2.2277 |
| 27433 | ENSMUSG00000085625 | 0 | NA | NA | NA | NA | NA | Gm8813 | predicted gene 8813 [Source:MGI Symbol | 0 | 0 | 0 | 0 | 0 | 0 |
| 27434 | ENSMUSG00000085626 | 0 | NA | NA | NA | NA | NA | Gm12898 | predicted gene 12898 [Source:MGI Symbo | 0 | 0 | 0 | 0 | 0 | 0 |
| 27435 | ENSMUSG00000085627 | 0 | NA | NA | NA | NA | NA | Gm11222 | predicted gene 11222 [Source:MGI Symbo | 0 | 0 | 0 | 0 | 0 | 0 |
| 27436 | ENSMUSG00000085628 | 15.69921136 | 0.204146916 | 0.549575509 | 0.371462907 | 0.710292779 | 0.944239215 | Appbp2os | amyloid beta precursor protein (cytoplasr | 10.44 | 14.391 | 19.119 | 18.316 | 17.45 | 14.48 |

| | | | | | | | | | | | | | | | |
|---|---|---|---|---|---|---|---|---|---|---|---|---|---|---|---|
| 27437 | ENSMUSG00000085629 | 0.692375149 | -1.404276945 | 2.772265464 | -0.506544905 | 0.612474175 | NA | Gm11697 | predicted gene 11697 [Source:MGI Symbo | 0 | 1.0279 | 2.0125 | 0 | 0 | 1.1138 |
| 27438 | ENSMUSG00000085630 | 0 | NA | NA | NA | NA | NA | Gm11795 | predicted gene 11795 [Source:MGI Symbo | 0 | 0 | 0 | 0 | 0 | 0 |
| 27439 | ENSMUSG00000085631 | 0 | NA | NA | NA | NA | NA | 9630028H03 | RIKEN cDNA 9630028H03 gene [Source:M | 0 | 0 | 0 | 0 | 0 | 0 |
| 27440 | ENSMUSG00000085632 | 0 | NA | NA | NA | NA | NA | Gm12381 | predicted gene 12381 [Source:MGI Symbo | 0 | 0 | 0 | 0 | 0 | 0 |
| 27441 | ENSMUSG00000085633 | 1.371534667 | 0.184953453 | 1.854502525 | 0.099732112 | 0.920557006 | NA | Bloc1s6os | biogenesis of lysosomal organelles comple | 0.8031 | 1.0279 | 2.0125 | 0 | 3.2719 | 1.1138 |
| 27442 | ENSMUSG00000085634 | 0 | NA | NA | NA | NA | NA | Gm15290 | predicted gene 15290 [Source:MGI Symbo | 0 | 0 | 0 | 0 | 0 | 0 |
| 27443 | ENSMUSG00000085635 | 0 | NA | NA | NA | NA | NA | Gm14565 | predicted gene 14565 [Source:MGI Symbo | 0 | 0 | 0 | 0 | 0 | 0 |
| 27444 | ENSMUSG00000085636 | 2.405907908 | 0.47336767 | 1.601148072 | 0.295642656 | 0.767502994 | NA | Gm11769 | predicted gene 11769 [Source:MGI Symbo | 0 | 4.1117 | 2.0125 | 3.8559 | 0 | 4.4554 |
| 27445 | ENSMUSG00000085637 | 6.134655856 | -0.445860617 | 0.848614409 | -0.525398358 | 0.599306295 | 0.915117178 | 1700048M1 | RIKEN cDNA 1700048M11 gene [Source:M | 8.031 | 6.1675 | 7.0437 | 6.7478 | 4.3625 | 4.4554 |
| 27446 | ENSMUSG00000085638 | 0 | NA | NA | NA | NA | NA | Gm15521 | predicted gene 15521 [Source:MGI Symbo | 0 | 0 | 0 | 0 | 0 | 0 |
| 27447 | ENSMUSG00000085639 | 0.583320834 | -1.123770786 | 3.233988906 | -0.34748752 | 0.728225092 | NA | Gm15410 | predicted gene 15410 [Source:MGI Symbo | 2.4093 | 0 | 0 | 0 | 1.0906 | 0 |
| 27448 | ENSMUSG00000085640 | 0 | NA | NA | NA | NA | NA | Gm14064 | predicted gene 14064 [Source:MGI Symbo | 0 | 0 | 0 | 0 | 0 | 0 |
| 27449 | ENSMUSG00000085641 | 0 | NA | NA | NA | NA | NA | 4930465M2 | RIKEN cDNA 4930465M20 gene [Source:M | 0 | 0 | 0 | 0 | 0 | 0 |
| 27450 | ENSMUSG00000085642 | 13.47505151 | 0.216367074 | 0.610929175 | 0.354160651 | 0.723218491 | 0.9447719 | 3110053B16 | RIKEN cDNA 3110053B16 gene [Source:M | 16.062 | 7.1954 | 14.087 | 19.28 | 13.087 | 11.138 |
| 27451 | ENSMUSG00000085643 | 0 | NA | NA | NA | NA | NA | Gm12519 | predicted gene 12519 [Source:MGI Symbo | 0 | 0 | 0 | 0 | 0 | 0 |
| 27452 | ENSMUSG00000085645 | 0.167707431 | -0.796629176 | 4.080472857 | -0.195229623 | 0.84521319 | NA | Hoxb5os | homeobox B5 and homeobox B6, opposit | 0 | 0 | 1.0062 | 0 | 0 | 0 |
| 27453 | ENSMUSG00000085646 | 0 | NA | NA | NA | NA | NA | Gm11493 | predicted gene 11493 [Source:MGI Symbo | 0 | 0 | 0 | 0 | 0 | 0 |
| 27454 | ENSMUSG00000085648 | 0 | NA | NA | NA | NA | NA | Gm13173 | predicted gene 13173 [Source:MGI Symbo | 0 | 0 | 0 | 0 | 0 | 0 |
| 27455 | ENSMUSG00000085649 | 0 | NA | NA | NA | NA | NA | A730032A03 | RIKEN cDNA A730032A03 gene [Source:M | 0 | 0 | 0 | 0 | 0 | 0 |
| 27456 | ENSMUSG00000085651 | 15.55253914 | -0.905866107 | 0.681203626 | -1.329802238 | 0.183583438 | 0.617851749 | Gm11695 | predicted gene 11695 [Source:MGI Symbo | 31.321 | 20.558 | 9.0562 | 20.244 | 5.4531 | 6.6831 |
| 27457 | ENSMUSG00000085652 | 0 | NA | NA | NA | NA | NA | Gm11999 | predicted gene 11999 [Source:MGI Symbo | 0 | 0 | 0 | 0 | 0 | 0 |
| 27458 | ENSMUSG00000085653 | 0 | NA | NA | NA | NA | NA | Gm15179 | predicted gene 15179 [Source:MGI Symbo | 0 | 0 | 0 | 0 | 0 | 0 |
| 27459 | ENSMUSG00000085654 | 0.524408994 | -0.837196744 | 3.377823722 | -0.247850928 | 0.804249749 | NA | Gm12536 | predicted gene 12536 [Source:MGI Symbo | 0 | 2.0558 | 0 | 0 | 1.0906 | 0 |
| 27460 | ENSMUSG00000085655 | 13.83729346 | 0.25367214 | 0.586668546 | 0.432394309 | 0.665454854 | 0.929426323 | Gm15952 | predicted gene 15952 [Source:MGI Symbo | 14.456 | 9.2513 | 14.087 | 14.46 | 19.631 | 11.138 |
| 27461 | ENSMUSG00000085656 | 0 | NA | NA | NA | NA | NA | Gm13180 | predicted gene 13180 [Source:MGI Symbo | 0 | 0 | 0 | 0 | 0 | 0 |
| 27462 | ENSMUSG00000085657 | 0 | NA | NA | NA | NA | NA | Gm15984 | predicted gene 15984 [Source:MGI Symbo | 0 | 0 | 0 | 0 | 0 | 0 |
| 27463 | ENSMUSG00000085658 | 0.556921942 | 2.679043159 | 3.638243063 | 0.736356289 | 0.461513899 | NA | Gm15704 | predicted gene 15704 [Source:MGI Symbo | 0 | 0 | 0 | 0 | 0 | 3.3415 |
| 27464 | ENSMUSG00000085659 | 0 | NA | NA | NA | NA | NA | Gm12325 | predicted gene 12325 [Source:MGI Symbo | 0 | 0 | 0 | 0 | 0 | 0 |
| 27465 | ENSMUSG00000085660 | 0 | NA | NA | NA | NA | NA | Gm12434 | predicted gene 12434 [Source:MGI Symbo | 0 | 0 | 0 | 0 | 0 | 0 |
| 27466 | ENSMUSG00000085662 | 0 | NA | NA | NA | NA | NA | Gm16220 | predicted gene 16220 [Source:MGI Symbo | 0 | 0 | 0 | 0 | 0 | 0 |
| 27467 | ENSMUSG00000085663 | 5.037829274 | 0.201202974 | 0.982559505 | 0.20477434 | 0.837748436 | NA | Gm15718 | predicted gene 15718 [Source:MGI Symbo | 4.8186 | 6.1675 | 3.0187 | 1.928 | 8.7249 | 5.5692 |
| 27468 | ENSMUSG00000085664 | 3.773323934 | 1.603101306 | 1.33556662 | 1.200315493 | 0.230016835 | NA | Atxn7l1os2 | ataxin 7-like 1, opposite strand 2 [Source:I | 1.6062 | 1.0279 | 3.0187 | 12.532 | 0 | 4.4554 |
| 27469 | ENSMUSG00000085665 | 4.441639896 | 1.010682269 | 1.091909319 | 0.925610077 | 0.354648655 | NA | Gm12059 | predicted gene 12059 [Source:MGI Symbo | 0.8031 | 6.1675 | 2.0125 | 7.7118 | 3.2719 | 6.6831 |
| 27470 | ENSMUSG00000085666 | 0.133850373 | -0.796629176 | 4.080472857 | -0.195229623 | 0.84521319 | NA | Tdg-ps2 | thymine DNA glycosylase, pseudogene 2 [ | 0.8031 | 0 | 0 | 0 | 0 | 0 |
| 27471 | ENSMUSG00000085667 | 7.185180945 | -0.24395242 | 0.806662759 | -0.30242182 | 0.762330523 | 0.954414324 | Gm12992 | predicted gene 12992 [Source:MGI Symbo | 7.2279 | 5.1396 | 11.069 | 8.6758 | 6.5437 | 4.4554 |
| 27472 | ENSMUSG00000085668 | 0 | NA | NA | NA | NA | NA | Gm35202 | predicted gene, 35202 [Source:MGI Symb | 0 | 0 | 0 | 0 | 0 | 0 |
| 27473 | ENSMUSG00000085669 | 0.319491021 | 0.16514334 | 4.044653141 | 0.040830038 | 0.967431393 | NA | Gm14817 | predicted gene 14817 [Source:MGI Symbo | 0.8031 | 0 | 0 | 0 | 0 | 1.1138 |
| 27474 | ENSMUSG00000085672 | 0.492645588 | 0.9565034 | 3.447648281 | 0.277436479 | 0.781444974 | NA | Gm16193 | predicted gene 16193 [Source:MGI Symbo | 0 | 1.0279 | 0 | 1.928 | 0 | 0 |
| 27475 | ENSMUSG00000085673 | 0.331982614 | 0.16514309 | 3.986751476 | 0.041422971 | 0.966958701 | NA | Gm16122 | predicted gene 16122 [Source:MGI Symbo | 0 | 1.0279 | 0 | 0.964 | 0 | 0 |
| 27476 | ENSMUSG00000085675 | 0 | NA | NA | NA | NA | NA | Gm12022 | predicted gene 12022 [Source:MGI Symbo | 0 | 0 | 0 | 0 | 0 | 0 |
| 27477 | ENSMUSG00000085676 | 0 | NA | NA | NA | NA | NA | 4930555O08 | RIKEN cDNA 4930555O08 gene [Source:M | 0 | 0 | 0 | 0 | 0 | 0 |
| 27478 | ENSMUSG00000085677 | 0 | NA | NA | NA | NA | NA | 4930594O21 | RIKEN cDNA 4930594O21 gene [Source:M | 0 | 0 | 0 | 0 | 0 | 0 |
| 27479 | ENSMUSG00000085678 | 0 | NA | NA | NA | NA | NA | Gm16349 | predicted gene 16349 [Source:MGI Symbo | 0 | 0 | 0 | 0 | 0 | 0 |
| 27480 | ENSMUSG00000085679 | 0 | NA | NA | NA | NA | NA | Gm16328 | predicted gene 16328 [Source:MGI Symbo | 0 | 0 | 0 | 0 | 0 | 0 |
| 27481 | ENSMUSG00000085680 | 0 | NA | NA | NA | NA | NA | Gm13327 | predicted gene 13327 [Source:MGI Symbo | 0 | 0 | 0 | 0 | 0 | 0 |
| 27482 | ENSMUSG00000085681 | 0 | NA | NA | NA | NA | NA | Gm13982 | predicted gene 13982 [Source:MGI Symbo | 0 | 0 | 0 | 0 | 0 | 0 |
| 27483 | ENSMUSG00000085682 | 0 | NA | NA | NA | NA | NA | Gm14267 | predicted gene 14267 [Source:MGI Symbo | 0 | 0 | 0 | 0 | 0 | 0 |
| 27484 | ENSMUSG00000085683 | 0 | NA | NA | NA | NA | NA | Tmem238l | transmembrane protein 238 like [Source:I | 0 | 0 | 0 | 0 | 0 | 0 |
| 27485 | ENSMUSG00000085684 | 0.465832987 | -0.698295221 | 3.18348759 | -0.219349126 | 0.826378096 | NA | 4930469K13 | RIKEN cDNA 4930469K13 gene [Source:M | 0.8031 | 1.0279 | 0 | 0.964 | 0 | 0 |
| 27486 | ENSMUSG00000085686 | 0 | NA | NA | NA | NA | NA | Gm9498 | predicted gene 9498 [Source:MGI Symbol | 0 | 0 | 0 | 0 | 0 | 0 |
| 27487 | ENSMUSG00000085687 | 28.37992746 | 0.468645392 | 0.417029771 | 1.123769631 | 0.261110768 | 0.715149811 | Gm16153 | predicted gene 16153 [Source:MGI Symbo | 25.699 | 23.642 | 22.137 | 40.487 | 33.809 | 24.505 |
| 27488 | ENSMUSG00000085688 | 0 | NA | NA | NA | NA | NA | Gm15395 | predicted gene 15395 [Source:MGI Symbo | 0 | 0 | 0 | 0 | 0 | 0 |
| 27489 | ENSMUSG00000085689 | 0 | NA | NA | NA | NA | NA | Gm16838 | predicted gene, 16838 [Source:MGI Symb | 0 | 0 | 0 | 0 | 0 | 0 |
| 27490 | ENSMUSG00000085690 | 0 | NA | NA | NA | NA | NA | Gm12629 | predicted gene 12629 [Source:MGI Symbo | 0 | 0 | 0 | 0 | 0 | 0 |
| 27491 | ENSMUSG00000085691 | 0 | NA | NA | NA | NA | NA | Gm14216 | predicted gene 14216 [Source:MGI Symbo | 0 | 0 | 0 | 0 | 0 | 0 |
| 27492 | ENSMUSG00000085693 | 0 | NA | NA | NA | NA | NA | Gm13371 | predicted gene 13371 [Source:MGI Symbo | 0 | 0 | 0 | 0 | 0 | 0 |

| | | | | | | | | | | | | | | | |
|---|---|---|---|---|---|---|---|---|---|---|---|---|---|---|---|
| 27493 | ENSMUSG00000085694 | 0 | NA | NA | NA | NA | NA | NA | NA | 0 | 0 | 0 | 0 | 0 | 0 |
| 27494 | ENSMUSG00000085695 | 0 | NA | NA | NA | NA | NA | Gm15243 | predicted gene 15243 [Source:MGI Symbc | 0 | 0 | 0 | 0 | 0 | 0 |
| 27495 | ENSMUSG00000085696 | 0.785324008 | -0.358914107 | 2.435960446 | -0.147339875 | 0.88286376 | NA | Hoxaas3 | Hoxa cluster antisense RNA 3 [Source:MG | 1.6062 | 1.0279 | 0 | 0.964 | 0 | 1.1138 |
| 27496 | ENSMUSG00000085699 | 0 | NA | NA | NA | NA | NA | Gm14298 | predicted gene 14298 [Source:MGI Symbc | 0 | 0 | 0 | 0 | 0 | 0 |
| 27497 | ENSMUSG00000085700 | 0 | NA | NA | NA | NA | NA | Hbb-bh0 | hemoglobin, beta, pseudogene bh0 [Sour | 0 | 0 | 0 | 0 | 0 | 0 |
| 27498 | ENSMUSG00000085701 | 0 | NA | NA | NA | NA | NA | Gm12968 | predicted gene 12968 [Source:MGI Symbc | 0 | 0 | 0 | 0 | 0 | 0 |
| 27499 | ENSMUSG00000085702 | 0 | NA | NA | NA | NA | NA | Mecomos | MDS1 and EVI1 complex locus, opposite s | 0 | 0 | 0 | 0 | 0 | 0 |
| 27500 | ENSMUSG00000085704 | 31.65538169 | -0.005423255 | 0.405855134 | -0.01336254 | 0.989338553 | 0.99859176 | 4921531C22 | RIKEN cDNA 4921531C22 gene [Source:M | 32.124 | 27.754 | 35.219 | 31.811 | 21.812 | 41.212 |
| 27501 | ENSMUSG00000085705 | 0 | NA | NA | NA | NA | NA | Gm16046 | predicted gene 16046 [Source:MGI Symbc | 0 | 0 | 0 | 0 | 0 | 0 |
| 27502 | ENSMUSG00000085706 | 0 | NA | NA | NA | NA | NA | Gm9428 | predicted gene 9428 [Source:MGI Symbol | 0 | 0 | 0 | 0 | 0 | 0 |
| 27503 | ENSMUSG00000085707 | 1.94309346 | -1.421698022 | 1.536588294 | -0.925230283 | 0.354846134 | NA | Gm12212 | predicted gene 12212 [Source:MGI Symbc | 2.4093 | 2.0558 | 4.025 | 0.964 | 1.0906 | 1.1138 |
| 27504 | ENSMUSG00000085708 | 7.559111915 | 0.39058029 | 0.833874575 | 0.468392132 | 0.639504193 | 0.923932929 | Gm16063 | predicted gene 16063 [Source:MGI Symbc | 2.4093 | 11.307 | 6.0375 | 5.7839 | 10.906 | 8.9108 |
| 27505 | ENSMUSG00000085709 | 0 | NA | NA | NA | NA | NA | Gm14211 | predicted gene 14211 [Source:MGI Symbc | 0 | 0 | 0 | 0 | 0 | 0 |
| 27506 | ENSMUSG00000085710 | 0.315620088 | 0.165143421 | 4.063300985 | 0.040642675 | 0.967580763 | NA | Gm12676 | predicted gene 12676 [Source:MGI Symbc | 0.8031 | 0 | 0 | 0 | 1.0906 | 0 |
| 27507 | ENSMUSG00000085711 | 12.35239889 | -0.406844717 | 0.68163542 | -0.596865575 | 0.550597133 | 0.895703112 | Gm15163 | predicted gene 15163 [Source:MGI Symbc | 9.6372 | 22.614 | 10.062 | 9.6398 | 5.4531 | 16.708 |
| 27508 | ENSMUSG00000085712 | 0 | NA | NA | NA | NA | NA | Gm15124 | predicted gene 15124 [Source:MGI Symbc | 0 | 0 | 0 | 0 | 0 | 0 |
| 27509 | ENSMUSG00000085713 | 0 | NA | NA | NA | NA | NA | Gm15736 | predicted gene 15736 [Source:MGI Symbc | 0 | 0 | 0 | 0 | 0 | 0 |
| 27510 | ENSMUSG00000085714 | 0 | NA | NA | NA | NA | NA | Gm13008 | predicted gene 13008 [Source:MGI Symbc | 0 | 0 | 0 | 0 | 0 | 0 |
| 27511 | ENSMUSG00000085715 | 0.671009685 | -1.413087452 | 2.800223337 | -0.504633839 | 0.613816038 | NA | Tsix | X (inactive)-specific transcript, opposite st | 0 | 2.0558 | 1.0062 | 0.964 | 0 | 0 |
| 27512 | ENSMUSG00000085716 | 0 | NA | NA | NA | NA | NA | Gm13388 | predicted gene 13388 [Source:MGI Symbc | 0 | 0 | 0 | 0 | 0 | 0 |
| 27513 | ENSMUSG00000085717 | 0 | NA | NA | NA | NA | NA | Hmgb4os | high-mobility group box 4, opposite stran | 0 | 0 | 0 | 0 | 0 | 0 |
| 27514 | ENSMUSG00000085718 | 0 | NA | NA | NA | NA | NA | Rpl9-ps3 | ribosomal protein L9, pseudogene 3 [Sour | 0 | 0 | 0 | 0 | 0 | 0 |
| 27515 | ENSMUSG00000085719 | 0 | NA | NA | NA | NA | NA | 4933424L21I | RIKEN cDNA 4933424L21 gene [Source:M | 0 | 0 | 0 | 0 | 0 | 0 |
| 27516 | ENSMUSG00000085720 | 11.84377851 | -0.413176543 | 0.623609362 | -0.662556671 | 0.507614528 | 0.882258527 | Gm7854 | predicted gene 7854 [Source:MGI Symbol | 16.062 | 14.391 | 10.062 | 9.6398 | 11.997 | 8.9108 |
| 27517 | ENSMUSG00000085721 | 5.479150431 | 1.092678953 | 0.916481937 | 1.19225367 | 0.233161795 | 0.680673483 | Gm12796 | predicted gene 12796 [Source:MGI Symbc | 2.4093 | 4.1117 | 4.025 | 5.7839 | 7.6343 | 8.9108 |
| 27518 | ENSMUSG00000085722 | 0 | NA | NA | NA | NA | NA | Gm4916 | predicted gene 4916 [Source:MGI Symbol | 0 | 0 | 0 | 0 | 0 | 0 |
| 27519 | ENSMUSG00000085723 | 0 | NA | NA | NA | NA | NA | Gm15915 | predicted gene 15915 [Source:MGI Symbc | 0 | 0 | 0 | 0 | 0 | 0 |
| 27520 | ENSMUSG00000085724 | 0 | NA | NA | NA | NA | NA | Gm13096 | predicted gene 13096 [Source:MGI Symbc | 0 | 0 | 0 | 0 | 0 | 0 |
| 27521 | ENSMUSG00000085725 | 0 | NA | NA | NA | NA | NA | Gm15873 | predicted gene 15873 [Source:MGI Symbc | 0 | 0 | 0 | 0 | 0 | 0 |
| 27522 | ENSMUSG00000085728 | 0 | NA | NA | NA | NA | NA | Gm14339 | predicted gene 14339 [Source:MGI Symbc | 0 | 0 | 0 | 0 | 0 | 0 |
| 27523 | ENSMUSG00000085730 | 0 | NA | NA | NA | NA | NA | Gm13307 | predicted gene 13307 [Source:MGI Symbc | 0 | 0 | 0 | 0 | 0 | 0 |
| 27524 | ENSMUSG00000085731 | 0 | NA | NA | NA | NA | NA | Gm16229 | predicted gene 16229 [Source:MGI Symbc | 0 | 0 | 0 | 0 | 0 | 0 |
| 27525 | ENSMUSG00000085732 | 0.319491021 | 0.16514334 | 4.044653141 | 0.040830038 | 0.967431393 | NA | Gm4279 | predicted gene 4279 [Source:MGI Symbol | 0.8031 | 0 | 0 | 0 | 0 | 1.1138 |
| 27526 | ENSMUSG00000085733 | 0 | NA | NA | NA | NA | NA | Arhgap27os: | Rho GTPase activating protein 27, opposit | 0 | 0 | 0 | 0 | 0 | 0 |
| 27527 | ENSMUSG00000085734 | 0.371281295 | 2.09253362 | 4.031852165 | 0.519000582 | 0.603760335 | NA | Gm11684 | predicted gene 11684 [Source:MGI Symbc | 0 | 0 | 0 | 0 | 0 | 2.2277 |
| 27528 | ENSMUSG00000085735 | 3.356618875 | -0.055620706 | 1.463960821 | -0.037993303 | 0.969693022 | NA | Mettl5os | methyltransferase like 5, opposite strand | 7.2279 | 0 | 3.0187 | 7.7118 | 2.1812 | 0 |
| 27529 | ENSMUSG00000085737 | 0 | NA | NA | NA | NA | NA | Gm13442 | predicted gene 13442 [Source:MGI Symbc | 0 | 0 | 0 | 0 | 0 | 0 |
| 27530 | ENSMUSG00000085738 | 0.321325949 | 1.911575114 | 4.048267893 | 0.472195804 | 0.636787034 | NA | Gm12335 | predicted gene 12335 [Source:MGI Symbc | 0 | 0 | 0 | 1.928 | 0 | 0 |
| 27531 | ENSMUSG00000085739 | 0 | NA | NA | NA | NA | NA | Gm15862 | predicted gene 15862 [Source:MGI Symbc | 0 | 0 | 0 | 0 | 0 | 0 |
| 27532 | ENSMUSG00000085740 | 0 | NA | NA | NA | NA | NA | 4930505K14 | RIKEN cDNA 4930505K14 gene [Source:M | 0 | 0 | 0 | 0 | 0 | 0 |
| 27533 | ENSMUSG00000085741 | 14.67811637 | -0.04371208 | 0.613230247 | -0.071281676 | 0.943173579 | 0.990077979 | 5430405H02 | RIKEN cDNA 5430405H02 gene [Source:M | 14.456 | 8.2233 | 22.137 | 21.208 | 10.906 | 11.138 |
| 27534 | ENSMUSG00000085742 | 0.795774083 | 0.899657507 | 2.433208733 | 0.369741196 | 0.711575334 | NA | Gm13536 | predicted gene 13536 [Source:MGI Symbc | 1.6062 | 0 | 0 | 0.964 | 1.0906 | 1.1138 |
| 27535 | ENSMUSG00000085743 | 0 | NA | NA | NA | NA | NA | 8430419K02 | RIKEN cDNA 8430419K02 gene [Source:M | 0 | 0 | 0 | 0 | 0 | 0 |
| 27536 | ENSMUSG00000085744 | 0 | NA | NA | NA | NA | NA | Gm12259 | predicted gene 12259 [Source:MGI Symbc | 0 | 0 | 0 | 0 | 0 | 0 |
| 27537 | ENSMUSG00000085745 | 0.48903338 | 0.95637552 | 3.456690196 | 0.276673773 | 0.782030612 | NA | Gm14372 | predicted gene 14372 [Source:MGI Symbc | 0 | 0 | 1.0062 | 1.928 | 0 | 0 |
| 27538 | ENSMUSG00000085747 | 0.321325949 | 1.911575114 | 4.048267893 | 0.472195804 | 0.636787034 | NA | Slc13a2os | solute carrier family 13 (sodium-depender | 0 | 0 | 0 | 1.928 | 0 | 0 |
| 27539 | ENSMUSG00000085748 | 0 | NA | NA | NA | NA | NA | Gm12280 | predicted gene 12280 [Source:MGI Symbc | 0 | 0 | 0 | 0 | 0 | 0 |
| 27540 | ENSMUSG00000085749 | 3.37947152 | 2.023065152 | 1.434736166 | 1.410060749 | 0.158521746 | NA | Gm12843 | predicted gene 12843 [Source:MGI Symbc | 0 | 4.1117 | 0 | 2.8919 | 4.3625 | 8.9108 |
| 27541 | ENSMUSG00000085750 | 0 | NA | NA | NA | NA | NA | Gm14009 | predicted gene 14009 [Source:MGI Symbc | 0 | 0 | 0 | 0 | 0 | 0 |
| 27542 | ENSMUSG00000085751 | 0 | NA | NA | NA | NA | NA | Gm15412 | predicted gene 15412 [Source:MGI Symbc | 0 | 0 | 0 | 0 | 0 | 0 |
| 27543 | ENSMUSG00000085753 | 0 | NA | NA | NA | NA | NA | Gm16131 | predicted gene 16131 [Source:MGI Symbc | 0 | 0 | 0 | 0 | 0 | 0 |
| 27544 | ENSMUSG00000085754 | 0.481988923 | 2.492401104 | 3.837077151 | 0.649557203 | 0.515978285 | NA | Gm15886 | predicted gene 15886 [Source:MGI Symbc | 0 | 0 | 0 | 2.8919 | 0 | 0 |
| 27545 | ENSMUSG00000085755 | 1.416487074 | 0.12239615 | 1.960786118 | 0.062421979 | 0.950226792 | NA | Gm13496 | predicted gene 13496 [Source:MGI Symbc | 0 | 3.0838 | 1.0062 | 0 | 2.1812 | 2.2277 |
| 27546 | ENSMUSG00000085756 | 0 | NA | NA | NA | NA | NA | Gm15279 | predicted gene 15279 [Source:MGI Symbc | 0 | 0 | 0 | 0 | 0 | 0 |
| 27547 | ENSMUSG00000085757 | 0 | NA | NA | NA | NA | NA | Tslrn1 | testis specific long noncoding RNA 1 [Sour | 0 | 0 | 0 | 0 | 0 | 0 |
| 27548 | ENSMUSG00000085759 | 4.186300734 | -0.090887355 | 1.144032717 | -0.079444716 | 0.936678902 | NA | 1700061E18 | RIKEN cDNA 1700061E18 gene [Source:M | 4.8186 | 7.1954 | 1.0062 | 7.7118 | 3.2719 | 1.1138 |

| | | | | | | | | | | | | | | | |
|---|---|---|---|---|---|---|---|---|---|---|---|---|---|---|---|
| 27549 | ENSMUSG00000085760 | 1.167319615 | 1.349049689 | 2.163487966 | 0.623553128 | 0.532921101 | NA | Gm14302 | predicted gene 14302 [Source:MGI Symbo | 0 | 1.0279 | 1.0062 | 3.8559 | 0 | 1.1138 |
| 27550 | ENSMUSG00000085761 | 1.858758094 | -1.302346802 | 1.792288887 | -0.726638887 | 0.4674472 | NA | 4930455G09 | RIKEN cDNA 4930455G09 gene [Source:M | 4.8186 | 2.0558 | 1.0062 | 0 | 3.2719 | 0 |
| 27551 | ENSMUSG00000085762 | 0 | NA | NA | NA | NA | NA | Lrrc75aos2 | leucine rich repeat containing 75A, oppos | 0 | 0 | 0 | 0 | 0 | 0 |
| 27552 | ENSMUSG00000085763 | 1.431176059 | -1.725849062 | 1.98770212 | -0.868263431 | 0.385250137 | NA | Smc2os | structural maintenance of chromosomes 2 | 1.6062 | 1.0279 | 4.025 | 1.928 | 0 | 0 |
| 27553 | ENSMUSG00000085765 | 0.469265235 | -2.274672822 | 3.439794491 | -0.661281605 | 0.508431736 | NA | Gm12333 | predicted gene 12333 [Source:MGI Symbo | 0.8031 | 0 | 2.0125 | 0 | 0 | 0 |
| 27554 | ENSMUSG00000085766 | 0.973672865 | 1.321976834 | 2.461749649 | 0.537007016 | 0.591262767 | NA | 2810430I11F | RIKEN cDNA 2810430I11 gene [Source:MG | 1.6062 | 0 | 0 | 0.964 | 3.2719 | 0 |
| 27555 | ENSMUSG00000085767 | 8.061025383 | 1.67150282 | 0.84426824 | 1.97982435 | 0.047723269 | 0.303548534 | Gm13563 | predicted gene 13563 [Source:MGI Symbo | 1.6062 | 2.0558 | 8.05 | 13.496 | 10.906 | 12.252 |
| 27556 | ENSMUSG00000085768 | 0.804680097 | -3.048447205 | 2.843448763 | -1.072095001 | 0.283677364 | NA | Gm13707 | predicted gene 13707 [Source:MGI Symbo | 0.8031 | 0 | 4.025 | 0 | 0 | 0 |
| 27557 | ENSMUSG00000085770 | 0 | NA | NA | NA | NA | NA | Gm11381 | predicted gene 11381 [Source:MGI Symbo | 0 | 0 | 0 | 0 | 0 | 0 |
| 27558 | ENSMUSG00000085771 | 0 | NA | NA | NA | NA | NA | C230066G23 | RIKEN cDNA C230066G23 gene [Source:M | 0 | 0 | 0 | 0 | 0 | 0 |
| 27559 | ENSMUSG00000085772 | 15.36852246 | 1.656868032 | 0.587414928 | 2.82060934 | 0.004793254 | 0.064976213 | D630024D03 | RIKEN cDNA D630024D03 gene [Source:M | 8.8341 | 6.1675 | 7.0437 | 17.352 | 30.537 | 22.277 |
| 27560 | ENSMUSG00000085773 | 0 | NA | NA | NA | NA | NA | Gm16233 | predicted gene 16233 [Source:MGI Symbo | 0 | 0 | 0 | 0 | 0 | 0 |
| 27561 | ENSMUSG00000085774 | 0.167707431 | -0.796629176 | 4.080472857 | -0.195229623 | 0.84521319 | NA | Gm13055 | predicted gene 13055 [Source:MGI Symbo | 0 | 0 | 1.0062 | 0 | 0 | 0 |
| 27562 | ENSMUSG00000085775 | 0 | NA | NA | NA | NA | NA | Gm14364 | predicted gene 14364 [Source:MGI Symbo | 0 | 0 | 0 | 0 | 0 | 0 |
| 27563 | ENSMUSG00000085776 | 21.50920648 | -0.272871159 | 0.508749745 | -0.536356355 | 0.591712289 | 0.912678231 | 5430402O13 | RIKEN cDNA 5430402O13 gene [Source:M | 32.927 | 12.335 | 25.156 | 21.208 | 20.722 | 16.708 |
| 27564 | ENSMUSG00000085779 | 0.17131964 | -0.796629176 | 4.080472857 | -0.195229623 | 0.84521319 | NA | Atcayos | ataxia, cerebellar, Cayman type, opposite | 0 | 1.0279 | 0 | 0 | 0 | 0 |
| 27565 | ENSMUSG00000085781 | 2.407272008 | 2.040403008 | 1.453037327 | 1.40423303 | 0.160249469 | NA | Gm15640 | predicted gene 15640 [Source:MGI Symbo | 0.8031 | 1.0279 | 1.0062 | 3.8559 | 2.1812 | 5.5692 |
| 27566 | ENSMUSG00000085782 | 1.806916214 | 1.06222317 | 1.65810419 | 0.640625104 | 0.521766285 | NA | Gm15624 | predicted gene 15624 [Source:MGI Symbo | 2.4093 | 1.0279 | 0 | 1.928 | 4.3625 | 1.1138 |
| 27567 | ENSMUSG00000085783 | 4.709708619 | 0.010640017 | 1.044127652 | 0.010190341 | 0.991869425 | NA | Gm9816 | predicted pseudogene 9816 [Source:MGI | 4.0155 | 5.1396 | 5.0312 | 9.6398 | 1.0906 | 3.3415 |
| 27568 | ENSMUSG00000085784 | 0 | NA | NA | NA | NA | NA | Gm12158 | predicted gene 12158 [Source:MGI Symbo | 0 | 0 | 0 | 0 | 0 | 0 |
| 27569 | ENSMUSG00000085785 | 0 | NA | NA | NA | NA | NA | Sox5os3 | SRY (sex determining region Y)-box 5, opp | 0 | 0 | 0 | 0 | 0 | 0 |
| 27570 | ENSMUSG00000085786 | 0.68166635 | -2.79976525 | 3.012029182 | -0.92952793 | 0.352615558 | NA | Gm15987 | predicted gene 15987 [Source:MGI Symbo | 0 | 3.0838 | 1.0062 | 0 | 0 | 0 |
| 27571 | ENSMUSG00000085787 | 0.356960287 | 0.16514264 | 3.880370363 | 0.042558474 | 0.966053499 | NA | Gm13092 | predicted gene 13092 [Source:MGI Symbo | 0 | 1.0279 | 0 | 0 | 0 | 1.1138 |
| 27572 | ENSMUSG00000085788 | 0.160662974 | 1.126924175 | 4.080472857 | 0.276174898 | 0.782413736 | NA | 4930470G03 | RIKEN cDNA 4930470G03 gene [Source:M | 0 | 0 | 0 | 0.964 | 0 | 0 |
| 27573 | ENSMUSG00000085790 | 0 | NA | NA | NA | NA | NA | Gm12729 | predicted gene 12729 [Source:MGI Symbo | 0 | 0 | 0 | 0 | 0 | 0 |
| 27574 | ENSMUSG00000085791 | 0 | NA | NA | NA | NA | NA | Rpl30-ps9 | ribosomal protein L30, pseudogene 9 [Sou | 0 | 0 | 0 | 0 | 0 | 0 |
| 27575 | ENSMUSG00000085792 | 0 | NA | NA | NA | NA | NA | Gm15414 | predicted gene 15414 [Source:MGI Symbo | 0 | 0 | 0 | 0 | 0 | 0 |
| 27576 | ENSMUSG00000085793 | 147.644099 | 0.396224801 | 0.253471885 | 1.563190331 | 0.118007834 | 0.503351609 | Lin52 | lin-52 homolog (C. elegans) [Source:MGI S | 131.71 | 129.52 | 120.75 | 120.5 | 146.14 | 237.25 |
| 27577 | ENSMUSG00000085794 | 0 | NA | NA | NA | NA | NA | Vax2os | ventral anterior homeobox 2, opposite str | 0 | 0 | 0 | 0 | 0 | 0 |
| 27578 | ENSMUSG00000085795 | 175.1882627 | 0.125681805 | 0.208276197 | 0.603438156 | 0.546217245 | 0.89549457 | Zfp703 | zinc finger protein 703 [Source:MGI Symb | 167.85 | 156.24 | 179.11 | 230.39 | 152.69 | 164.85 |
| 27579 | ENSMUSG00000085796 | 0 | NA | NA | NA | NA | NA | 4933406D12 | RIKEN cDNA 4933406D12 gene [Source:M | 0 | 0 | 0 | 0 | 0 | 0 |
| 27580 | ENSMUSG00000085797 | 0 | NA | NA | NA | NA | NA | A530010F05 | RIKEN cDNA A530010F05 gene [Source:M | 0 | 0 | 0 | 0 | 0 | 0 |
| 27581 | ENSMUSG00000085798 | 0 | NA | NA | NA | NA | NA | Gm2018 | predicted gene 2018 [Source:MGI Symbol | 0 | 0 | 0 | 0 | 0 | 0 |
| 27582 | ENSMUSG00000085799 | 0 | NA | NA | NA | NA | NA | Gm6963 | predicted gene 6963 [Source:MGI Symbol | 0 | 0 | 0 | 0 | 0 | 0 |
| 27583 | ENSMUSG00000085800 | 0 | NA | NA | NA | NA | NA | 1700066O22 | RIKEN cDNA 1700066O22 gene [Source:M | 0 | 0 | 0 | 0 | 0 | 0 |
| 27584 | ENSMUSG00000085801 | 0 | NA | NA | NA | NA | NA | 1700013N06 | RIKEN cDNA 1700013N06 gene [Source:M | 0 | 0 | 0 | 0 | 0 | 0 |
| 27585 | ENSMUSG00000085802 | 0 | NA | NA | NA | NA | NA | Gm16059 | predicted gene 16059 [Source:MGI Symbo | 0 | 0 | 0 | 0 | 0 | 0 |
| 27586 | ENSMUSG00000085803 | 0 | NA | NA | NA | NA | NA | Gm12056 | predicted gene 12056 [Source:MGI Symbo | 0 | 0 | 0 | 0 | 0 | 0 |
| 27587 | ENSMUSG00000085804 | 0 | NA | NA | NA | NA | NA | Gm12973 | predicted gene 12973 [Source:MGI Symbo | 0 | 0 | 0 | 0 | 0 | 0 |
| 27588 | ENSMUSG00000085805 | 0 | NA | NA | NA | NA | NA | Gm14145 | predicted gene 14145 [Source:MGI Symbo | 0 | 0 | 0 | 0 | 0 | 0 |
| 27589 | ENSMUSG00000085806 | 0 | NA | NA | NA | NA | NA | Gm12023 | predicted gene 12023 [Source:MGI Symbo | 0 | 0 | 0 | 0 | 0 | 0 |
| 27590 | ENSMUSG00000085807 | 1.413365684 | 2.88291135 | 2.043675042 | 1.410650564 | 0.158347674 | NA | Ube4bos2 | ubiquitination factor E4B, opposite strand | 0 | 0 | 1.0062 | 1.928 | 1.0906 | 4.4554 |
| 27591 | ENSMUSG00000085808 | 0 | NA | NA | NA | NA | NA | Gm14718 | predicted gene 14718 [Source:MGI Symbo | 0 | 0 | 0 | 0 | 0 | 0 |
| 27592 | ENSMUSG00000085810 | 0.133850373 | -0.796629176 | 4.080472857 | -0.195229623 | 0.84521319 | NA | Gm16325 | predicted gene 16325 [Source:MGI Symbo | 0.8031 | 0 | 0 | 0 | 0 | 0 |
| 27593 | ENSMUSG00000085811 | 0.503302254 | -0.837877584 | 3.426698643 | -0.244514523 | 0.806832356 | NA | Cep112it | centrosomal protein 112, intronic transcri | 0 | 2.0558 | 0 | 0.964 | 0 | 0 |
| 27594 | ENSMUSG00000085812 | 0 | NA | NA | NA | NA | NA | Gm15843 | predicted gene 15843 [Source:MGI Symbo | 0 | 0 | 0 | 0 | 0 | 0 |
| 27595 | ENSMUSG00000085813 | 0 | NA | NA | NA | NA | NA | Gm15870 | predicted gene 15870 [Source:MGI Symbo | 0 | 0 | 0 | 0 | 0 | 0 |
| 27596 | ENSMUSG00000085815 | 0 | NA | NA | NA | NA | NA | Gm13548 | predicted gene 13548 [Source:MGI Symbo | 0 | 0 | 0 | 0 | 0 | 0 |
| 27597 | ENSMUSG00000085816 | 0 | NA | NA | NA | NA | NA | Gm13052 | predicted gene 13052 [Source:MGI Symbo | 0 | 0 | 0 | 0 | 0 | 0 |
| 27598 | ENSMUSG00000085817 | 0 | NA | NA | NA | NA | NA | Gm12668 | predicted gene 12668 [Source:MGI Symbo | 0 | 0 | 0 | 0 | 0 | 0 |
| 27599 | ENSMUSG00000085818 | 4.858986502 | -0.851222231 | 0.972012755 | -0.875731544 | 0.381175993 | NA | Gm13267 | predicted gene 13267 [Source:MGI Symbo | 5.6217 | 5.1396 | 8.05 | 4.8199 | 2.1812 | 3.3415 |
| 27600 | ENSMUSG00000085819 | 1.463455258 | -1.794018729 | 2.008227707 | -0.893334318 | 0.371678165 | NA | Ube4bos1 | ubiquitination factor E4B, opposite strand | 4.8186 | 1.0279 | 1.0062 | 1.928 | 0 | 0 |
| 27601 | ENSMUSG00000085820 | 0 | NA | NA | NA | NA | NA | 4933406K04 | RIKEN cDNA 4933406K04 gene [Source:M | 0 | 0 | 0 | 0 | 0 | 0 |
| 27602 | ENSMUSG00000085822 | 0 | NA | NA | NA | NA | NA | Gm15584 | predicted gene 15584 [Source:MGI Symbo | 0 | 0 | 0 | 0 | 0 | 0 |
| 27603 | ENSMUSG00000085823 | 3.211459632 | -0.096475478 | 1.219602444 | -0.079104038 | 0.936949872 | NA | Gm13012 | predicted gene 13012 [Source:MGI Symbo | 4.8186 | 4.1117 | 1.0062 | 3.8559 | 4.3625 | 1.1138 |
| 27604 | ENSMUSG00000085824 | 0 | NA | NA | NA | NA | NA | Platr9 | pluripotency associated transcript 9 [Sour | 0 | 0 | 0 | 0 | 0 | 0 |

| | | | | | | | | | | | | | | | |
|---|---|---|---|---|---|---|---|---|---|---|---|---|---|---|---|
| 27605 | ENSMUSG00000085826 | 130.609068 | 0.669044033 | 0.216057319 | 3.096604353 | 0.001957509 | 0.034277745 | Gm15638 | predicted gene 15638 [Source:MGI Symbo | 82.72 | 108.96 | 111.69 | 171.59 | 149.41 | 159.28 |
| 27606 | ENSMUSG00000085827 | 0 | NA | NA | NA | NA | NA | Gm11290 | predicted gene 11290 [Source:MGI Symbo | 0 | 0 | 0 | 0 | 0 | 0 |
| 27607 | ENSMUSG00000085828 | 0.663965228 | -0.037580459 | 3.10181903 | -0.012115619 | 0.990333371 | NA | Gm15612 | predicted gene 15612 [Source:MGI Symbo | 0 | 2.0558 | 0 | 1.928 | 0 | 0 |
| 27608 | ENSMUSG00000085829 | 53.01742063 | 0.424605914 | 0.33794599 | 1.256431284 | 0.208959645 | 0.65350014 | Gm4285 | predicted gene 4285 [Source:MGI Symbol | 42.564 | 39.061 | 54.337 | 64.587 | 76.343 | 41.212 |
| 27609 | ENSMUSG00000085830 | 0 | NA | NA | NA | NA | NA | Grin1os | glutamate receptor, ionotropic, NMDA1 (a | 0 | 0 | 0 | 0 | 0 | 0 |
| 27610 | ENSMUSG00000085831 | 0 | NA | NA | NA | NA | NA | Lncbate3 | brown adipose tissue enriched long nonco | 0 | 0 | 0 | 0 | 0 | 0 |
| 27611 | ENSMUSG00000085833 | 0 | NA | NA | NA | NA | NA | Gm13003 | predicted gene 13003 [Source:MGI Symbo | 0 | 0 | 0 | 0 | 0 | 0 |
| 27612 | ENSMUSG00000085834 | 184.3623008 | -1.344524127 | 0.974653477 | -1.379489386 | 0.167743916 | 0.593804353 | Gm15622 | predicted gene 15622 [Source:MGI Symbo | 230.49 | 222.03 | 341.12 | 72.298 | 225.76 | 14.48 |
| 27613 | ENSMUSG00000085835 | 0 | NA | NA | NA | NA | NA | Gm12707 | predicted gene 12707 [Source:MGI Symbo | 0 | 0 | 0 | 0 | 0 | 0 |
| 27614 | ENSMUSG00000085836 | 3.761389251 | 1.152158118 | 1.149157644 | 1.00261102 | 0.316048577 | NA | Gm13074 | predicted gene 13074 [Source:MGI Symbo | 4.8186 | 1.0279 | 1.0062 | 5.7839 | 4.3625 | 5.5692 |
| 27615 | ENSMUSG00000085837 | 0 | NA | NA | NA | NA | NA | Kcnmb4os2 | potassium large conductance calcium-acti | 0 | 0 | 0 | 0 | 0 | 0 |
| 27616 | ENSMUSG00000085838 | 0 | NA | NA | NA | NA | NA | Chn1os1 | chimerin 1, opposite strand 1 [Source:MG | 0 | 0 | 0 | 0 | 0 | 0 |
| 27617 | ENSMUSG00000085839 | 0 | NA | NA | NA | NA | NA | Gm15949 | predicted gene 15949 [Source:MGI Symbo | 0 | 0 | 0 | 0 | 0 | 0 |
| 27618 | ENSMUSG00000085840 | 0 | NA | NA | NA | NA | NA | Gm11261 | predicted gene 11261 [Source:MGI Symbo | 0 | 0 | 0 | 0 | 0 | 0 |
| 27619 | ENSMUSG00000085841 | 0 | NA | NA | NA | NA | NA | Gm4954 | predicted gene 4954 [Source:MGI Symbol | 0 | 0 | 0 | 0 | 0 | 0 |
| 27620 | ENSMUSG00000085842 | 0 | NA | NA | NA | NA | NA | Pard3bos1 | par-3 family cell polarity regulator beta, o | 0 | 0 | 0 | 0 | 0 | 0 |
| 27621 | ENSMUSG00000085843 | 0 | NA | NA | NA | NA | NA | Kank4os | KN motif and ankyrin repeat domains 4, o | 0 | 0 | 0 | 0 | 0 | 0 |
| 27622 | ENSMUSG00000085844 | 0 | NA | NA | NA | NA | NA | Gm11690 | predicted gene 11690 [Source:MGI Symbo | 0 | 0 | 0 | 0 | 0 | 0 |
| 27623 | ENSMUSG00000085845 | 0.353089354 | 0.165142706 | 3.896102244 | 0.042386646 | 0.966190474 | NA | Gm13944 | predicted gene 13944 [Source:MGI Symbo | 0 | 1.0279 | 0 | 0 | 1.0906 | 0 |
| 27624 | ENSMUSG00000085846 | 0.267700746 | -1.497481649 | 4.057001436 | -0.369110456 | 0.712045396 | NA | Gm15597 | predicted gene 15597 [Source:MGI Symbo | 1.6062 | 0 | 0 | 0 | 0 | 0 |
| 27625 | ENSMUSG00000085847 | 0.328370405 | 0.16514316 | 4.003149536 | 0.041253308 | 0.967093956 | NA | Gm14540 | predicted gene 14540 [Source:MGI Symbo | 0 | 0 | 1.0062 | 0.964 | 0 | 0 |
| 27626 | ENSMUSG00000085851 | 0.339027071 | -1.79174149 | 3.919585585 | -0.457125237 | 0.647581032 | NA | 4921518K17 | RIKEN cDNA 4921518K17 gene [Source:M | 0 | 1.0279 | 1.0062 | 0 | 0 | 0 |
| 27627 | ENSMUSG00000085852 | 8.090622774 | 0.83907661 | 0.788077426 | 1.064713418 | 0.287005636 | 0.73951397 | Gm13807 | predicted gene 13807 [Source:MGI Symbo | 7.2279 | 3.0838 | 7.0437 | 12.532 | 13.087 | 5.5692 |
| 27628 | ENSMUSG00000085853 | 0 | NA | NA | NA | NA | NA | Gm12167 | predicted gene 12167 [Source:MGI Symbo | 0 | 0 | 0 | 0 | 0 | 0 |
| 27629 | ENSMUSG00000085856 | 0.321325949 | 1.911575114 | 4.048267893 | 0.472195804 | 0.636787034 | NA | Gm12050 | predicted gene 12050 [Source:MGI Symbo | 0 | 0 | 0 | 1.928 | 0 | 0 |
| 27630 | ENSMUSG00000085857 | 0.133850373 | -0.796629176 | 4.080472857 | -0.195229623 | 0.84521319 | NA | Gm15392 | predicted gene 15392 [Source:MGI Symbo | 0.8031 | 0 | 0 | 0 | 0 | 0 |
| 27631 | ENSMUSG00000085858 | 0 | NA | NA | NA | NA | NA | Gm15831 | predicted gene 15831 [Source:MGI Symbo | 0 | 0 | 0 | 0 | 0 | 0 |
| 27632 | ENSMUSG00000085860 | 1.797314445 | 2.155800046 | 2.034641238 | 1.059547996 | 0.289350283 | NA | 2410003L11 | RIKEN cDNA 2410003L11 gene [Source:M | 0 | 0 | 2.0125 | 0 | 6.5437 | 2.2277 |
| 27633 | ENSMUSG00000085861 | 0.439020386 | -2.189747911 | 3.52148864 | -0.621824499 | 0.534057275 | NA | Armh2 | armadillo-like helical domain containing 2 | 1.6062 | 1.0279 | 0 | 0 | 0 | 0 |
| 27634 | ENSMUSG00000085862 | 4.589397568 | 0.378252248 | 0.989553877 | 0.382245229 | 0.702279476 | NA | Gm13483 | predicted gene 13483 [Source:MGI Symbo | 4.8186 | 5.1396 | 2.0125 | 6.7478 | 4.3625 | 4.4554 |
| 27635 | ENSMUSG00000085863 | 0 | NA | NA | NA | NA | NA | Gm12801 | predicted gene 12801 [Source:MGI Symbo | 0 | 0 | 0 | 0 | 0 | 0 |
| 27636 | ENSMUSG00000085864 | 0 | NA | NA | NA | NA | NA | C330019F10 | RIKEN cDNA C330019F10 gene [Source:M | 0 | 0 | 0 | 0 | 0 | 0 |
| 27637 | ENSMUSG00000085865 | 0 | NA | NA | NA | NA | NA | Gm15966 | predicted gene 15966 [Source:MGI Symbo | 0 | 0 | 0 | 0 | 0 | 0 |
| 27638 | ENSMUSG00000085866 | 0 | NA | NA | NA | NA | NA | Gm15218 | predicted gene 15218 [Source:MGI Symbo | 0 | 0 | 0 | 0 | 0 | 0 |
| 27639 | ENSMUSG00000085867 | 101.0609109 | 0.292162856 | 0.230766676 | 1.266053059 | 0.205494087 | 0.650907725 | Gm5834 | predicted pseudogene 5834 [Source:MGI | 82.72 | 93.541 | 96.599 | 108.93 | 124.33 | 100.25 |
| 27640 | ENSMUSG00000085868 | 0 | NA | NA | NA | NA | NA | Gm13632 | predicted gene 13632 [Source:MGI Symbo | 0 | 0 | 0 | 0 | 0 | 0 |
| 27641 | ENSMUSG00000085869 | 0 | NA | NA | NA | NA | NA | Gm16079 | predicted gene 16079 [Source:MGI Symbo | 0 | 0 | 0 | 0 | 0 | 0 |
| 27642 | ENSMUSG00000085870 | 0 | NA | NA | NA | NA | NA | Gm12829 | predicted gene 12829 [Source:MGI Symbo | 0 | 0 | 0 | 0 | 0 | 0 |
| 27643 | ENSMUSG00000085871 | 0 | NA | NA | NA | NA | NA | Ube2uos | ubiquitin-conjugating enzyme E2U (putati | 0 | 0 | 0 | 0 | 0 | 0 |
| 27644 | ENSMUSG00000085872 | 0 | NA | NA | NA | NA | NA | Gm11505 | predicted gene 11505 [Source:MGI Symbo | 0 | 0 | 0 | 0 | 0 | 0 |
| 27645 | ENSMUSG00000085873 | 10.98991907 | -2.258210687 | 0.778718125 | -2.899907701 | 0.003732726 | 0.054645736 | Ttc39aos1 | Ttc39a opposite strand RNA 1 [Source:MG | 28.109 | 13.363 | 13.081 | 4.8199 | 5.4531 | 1.1138 |
| 27646 | ENSMUSG00000085875 | 2.792963572 | -0.874956142 | 1.568646692 | -0.557777699 | 0.576996193 | NA | Gm12905 | predicted gene 12905 [Source:MGI Symbo | 0.8031 | 5.1396 | 5.0312 | 5.7839 | 0 | 0 |
| 27647 | ENSMUSG00000085876 | 0 | NA | NA | NA | NA | NA | Gm12409 | predicted gene 12409 [Source:MGI Symbo | 0 | 0 | 0 | 0 | 0 | 0 |
| 27648 | ENSMUSG00000085877 | 0 | NA | NA | NA | NA | NA | Spin2-ps4 | spindlin family, member 2, pseudogene 4 | 0 | 0 | 0 | 0 | 0 | 0 |
| 27649 | ENSMUSG00000085878 | 0 | NA | NA | NA | NA | NA | Gm12823 | predicted gene 12823 [Source:MGI Symbo | 0 | 0 | 0 | 0 | 0 | 0 |
| 27650 | ENSMUSG00000085880 | 0 | NA | NA | NA | NA | NA | Gm15229 | predicted gene 15229 [Source:MGI Symbo | 0 | 0 | 0 | 0 | 0 | 0 |
| 27651 | ENSMUSG00000085881 | 4.994970518 | 0.387585796 | 0.95272386 | 0.406818609 | 0.684141218 | NA | Gm15912 | predicted gene 15912 [Source:MGI Symbo | 4.8186 | 3.0838 | 5.0312 | 3.8559 | 8.7249 | 4.4554 |
| 27652 | ENSMUSG00000085882 | 143.4722742 | 0.31435594 | 0.203315772 | 1.546146357 | 0.122069225 | 0.511805228 | 2610507I01 | RIKEN cDNA 2610507I01 gene [Source:MG | 120.47 | 146.99 | 116.72 | 172.55 | 155.96 | 148.14 |
| 27653 | ENSMUSG00000085883 | 0 | NA | NA | NA | NA | NA | NA | NA | 0 | 0 | 0 | 0 | 0 | 0 |
| 27654 | ENSMUSG00000085884 | 0.831492699 | -1.816502146 | 2.887260166 | -0.629143909 | 0.529254847 | NA | Gm15342 | predicted gene 15342 [Source:MGI Symbo | 0 | 0 | 4.025 | 0.964 | 0 | 0 |
| 27655 | ENSMUSG00000085885 | 0.66762957 | 2.958119316 | 3.019579549 | 0.979646096 | 0.327260843 | NA | Gm11906 | predicted gene 11906 [Source:MGI Symbo | 0 | 0 | 0 | 2.8919 | 0 | 1.1138 |
| 27656 | ENSMUSG00000085886 | 0 | NA | NA | NA | NA | NA | D030047H15 | RIKEN cDNA D030047H15 gene [Source:M | 0 | 0 | 0 | 0 | 0 | 0 |
| 27657 | ENSMUSG00000085887 | 2.287954218 | -1.84248748 | 1.724773914 | -1.068248693 | 0.285408345 | NA | Arhgap27os | Rho GTPase activating protein 27, opposit | 1.6062 | 8.2233 | 1.0062 | 2.8919 | 0 | 0 |
| 27658 | ENSMUSG00000085888 | 0 | NA | NA | NA | NA | NA | Gm12224 | predicted gene 12224 [Source:MGI Symbo | 0 | 0 | 0 | 0 | 0 | 0 |
| 27659 | ENSMUSG00000085889 | 0.528073336 | 2.60973587 | 2.979250867 | 0.875970499 | 0.381046072 | NA | 1700054M1 | RIKEN cDNA 1700054M17 gene [Source:M | 0 | 0 | 0 | 0.964 | 1.0906 | 1.1138 |
| 27660 | ENSMUSG00000085890 | 8.455739534 | 0.086188159 | 0.807438119 | 0.106742742 | 0.914993073 | 0.98625062 | Tnfsf13os | tumor necrosis factor (ligand) superfamily | 6.4248 | 7.1954 | 11.069 | 9.6398 | 14.178 | 2.2277 |

| | | | | | | | | | | | | | | | |
|---|---|---|---|---|---|---|---|---|---|---|---|---|---|---|---|
| 27661 | ENSMUSG00000085891 | 3.759327743 | 0.038023121 | 1.072047449 | 0.03546776 | 0.971706754 | NA | Gm14634 | predicted gene 14634 [Source:MGI Symbo | 4.0155 | 4.1117 | 3.0187 | 4.8199 | 4.3625 | 2.2277 |
| 27662 | ENSMUSG00000085892 | 0 | NA | NA | NA | NA | NA | Gm15339 | predicted gene 15339 [Source:MGI Symbo | 0 | 0 | 0 | 0 | 0 | 0 |
| 27663 | ENSMUSG00000085893 | 0.160662974 | 1.126924175 | 4.080472857 | 0.276174898 | 0.782413736 | NA | Gm12091 | predicted gene 12091 [Source:MGI Symbo | 0 | 0 | 0 | 0.964 | 0 | 0 |
| 27664 | ENSMUSG00000085894 | 28.48321629 | -0.346943066 | 0.449543359 | -0.771767749 | 0.440251997 | 0.84892746 | Gm15832 | predicted gene 15832 [Source:MGI Symbo | 32.927 | 28.782 | 34.212 | 37.595 | 22.903 | 14.48 |
| 27665 | ENSMUSG00000085895 | 0 | NA | NA | NA | NA | NA | Gm12861 | predicted gene 12861 [Source:MGI Symbo | 0 | 0 | 0 | 0 | 0 | 0 |
| 27666 | ENSMUSG00000085896 | 0 | NA | NA | NA | NA | NA | 5330429C05 | RIKEN cDNA 5330429C05 gene [Source:M | 0 | 0 | 0 | 0 | 0 | 0 |
| 27667 | ENSMUSG00000085897 | 0.36353943 | 2.065989254 | 4.034137301 | 0.512126658 | 0.608562368 | NA | Gm11604 | predicted gene 11604 [Source:MGI Symbo | 0 | 0 | 0 | 0 | 2.1812 | 0 |
| 27668 | ENSMUSG00000085898 | 0 | NA | NA | NA | NA | NA | Gm6689 | predicted gene 6689 [Source:MGI Symbol | 0 | 0 | 0 | 0 | 0 | 0 |
| 27669 | ENSMUSG00000085899 | 0 | NA | NA | NA | NA | NA | Gm15338 | predicted gene 15338 [Source:MGI Symbo | 0 | 0 | 0 | 0 | 0 | 0 |
| 27670 | ENSMUSG00000085900 | 0.636972666 | -2.712703855 | 3.077964703 | -0.881330397 | 0.378139018 | NA | A930041C12 | RIKEN cDNA A930041C12 gene [Source:M | 0.8031 | 0 | 3.0187 | 0 | 0 | 0 |
| 27671 | ENSMUSG00000085902 | 0 | NA | NA | NA | NA | NA | Gm12200 | predicted gene 12200 [Source:MGI Symbo | 0 | 0 | 0 | 0 | 0 | 0 |
| 27672 | ENSMUSG00000085903 | 0 | NA | NA | NA | NA | NA | Gm15340 | predicted gene 15340 [Source:MGI Symbo | 0 | 0 | 0 | 0 | 0 | 0 |
| 27673 | ENSMUSG00000085905 | 0 | NA | NA | NA | NA | NA | Gm14273 | predicted gene 14273 [Source:MGI Symbo | 0 | 0 | 0 | 0 | 0 | 0 |
| 27674 | ENSMUSG00000085908 | 0 | NA | NA | NA | NA | NA | Gm14209 | predicted gene 14209 [Source:MGI Symbo | 0 | 0 | 0 | 0 | 0 | 0 |
| 27675 | ENSMUSG00000085909 | 0 | NA | NA | NA | NA | NA | Gm11846 | predicted gene 11846 [Source:MGI Symbo | 0 | 0 | 0 | 0 | 0 | 0 |
| 27676 | ENSMUSG00000085910 | 0 | NA | NA | NA | NA | NA | Gm12395 | predicted gene 12395 [Source:MGI Symbo | 0 | 0 | 0 | 0 | 0 | 0 |
| 27677 | ENSMUSG00000085911 | 0 | NA | NA | NA | NA | NA | Gm14061 | predicted gene 14061 [Source:MGI Symbo | 0 | 0 | 0 | 0 | 0 | 0 |
| 27678 | ENSMUSG00000085912 | 1.234355784 | -1.29456175 | 2.242850757 | -0.57719478 | 0.563807883 | NA | Trp53cor1 | tumor protein p53 pathway corepressor 1 | 3.2124 | 0 | 2.0125 | 0 | 2.1812 | 0 |
| 27679 | ENSMUSG00000085913 | 8.345314179 | 0.027854408 | 0.732783707 | 0.038011773 | 0.969678295 | 0.994674947 | Gm15601 | predicted gene 15601 [Source:MGI Symbo | 10.44 | 8.2233 | 6.0375 | 7.7118 | 7.6343 | 10.025 |
| 27680 | ENSMUSG00000085914 | 1.902088206 | 0.870192199 | 1.74023998 | 0.500041494 | 0.617045861 | NA | Gm14320 | predicted gene 14320 [Source:MGI Symbo | 0 | 4.1117 | 0 | 2.8919 | 2.1812 | 2.2277 |
| 27681 | ENSMUSG00000085915 | 0 | NA | NA | NA | NA | NA | Gm8091 | predicted gene 8091 [Source:MGI Symbol | 0 | 0 | 0 | 0 | 0 | 0 |
| 27682 | ENSMUSG00000085917 | 0.503302254 | -0.837877584 | 3.426698643 | -0.244514523 | 0.806832356 | NA | Gm15899 | predicted gene 15899 [Source:MGI Symbo | 0 | 2.0558 | 0 | 0.964 | 0 | 0 |
| 27683 | ENSMUSG00000085918 | 0 | NA | NA | NA | NA | NA | Gm13032 | predicted gene 13032 [Source:MGI Symbo | 0 | 0 | 0 | 0 | 0 | 0 |
| 27684 | ENSMUSG00000085919 | 0 | NA | NA | NA | NA | NA | Gm16019 | predicted gene 16019 [Source:MGI Symbo | 0 | 0 | 0 | 0 | 0 | 0 |
| 27685 | ENSMUSG00000085920 | 0 | NA | NA | NA | NA | NA | Gm12951 | predicted gene 12951 [Source:MGI Symbo | 0 | 0 | 0 | 0 | 0 | 0 |
| 27686 | ENSMUSG00000085921 | 0 | NA | NA | NA | NA | NA | Gm13752 | predicted gene 13752 [Source:MGI Symbo | 0 | 0 | 0 | 0 | 0 | 0 |
| 27687 | ENSMUSG00000085922 | 0 | NA | NA | NA | NA | NA | Fhad1os2 | forkhead-associated (FHA) phosphopeptic | 0 | 0 | 0 | 0 | 0 | 0 |
| 27688 | ENSMUSG00000085923 | 106.2774909 | -0.207746364 | 0.229055046 | -0.90697135 | 0.364421948 | 0.802897466 | Gm12781 | predicted gene 12781 [Source:MGI Symbo | 102.8 | 131.57 | 107.67 | 99.29 | 90.521 | 105.82 |
| 27689 | ENSMUSG00000085924 | 0 | NA | NA | NA | NA | NA | Gm8980 | predicted gene 8980 [Source:MGI Symbol | 0 | 0 | 0 | 0 | 0 | 0 |
| 27690 | ENSMUSG00000085925 | 0 | NA | NA | NA | NA | NA | Rtl1 | retrotransposon Gaglike 1 [Source:MGI Sy | 0 | 0 | 0 | 0 | 0 | 0 |
| 27691 | ENSMUSG00000085926 | 0 | NA | NA | NA | NA | NA | Gm12299 | predicted gene 12299 [Source:MGI Symbo | 0 | 0 | 0 | 0 | 0 | 0 |
| 27692 | ENSMUSG00000085927 | 0 | NA | NA | NA | NA | NA | Gm15888 | predicted gene 15888 [Source:MGI Symbo | 0 | 0 | 0 | 0 | 0 | 0 |
| 27693 | ENSMUSG00000085928 | 0 | NA | NA | NA | NA | NA | 4933427I22R | RIKEN cDNA 4933427I22 gene [Source:MG | 0 | 0 | 0 | 0 | 0 | 0 |
| 27694 | ENSMUSG00000085929 | 29.66443409 | 0.960851931 | 0.403672203 | 2.380277672 | 0.017299597 | 0.156658967 | Gm13421 | predicted gene 13421 [Source:MGI Symbo | 21.684 | 19.53 | 19.119 | 39.523 | 44.715 | 33.415 |
| 27695 | ENSMUSG00000085931 | 0 | NA | NA | NA | NA | NA | Gm12648 | predicted gene 12648 [Source:MGI Symbo | 0 | 0 | 0 | 0 | 0 | 0 |
| 27696 | ENSMUSG00000085932 | 0.160662974 | 1.126924175 | 4.080472857 | 0.276174898 | 0.782413736 | NA | Gm15556 | predicted gene 15556 [Source:MGI Symbo | 0 | 0 | 0 | 0.964 | 0 | 0 |
| 27697 | ENSMUSG00000085933 | 0 | NA | NA | NA | NA | NA | Tmem61 | transmembrane protein 61 [Source:MGI S | 0 | 0 | 0 | 0 | 0 | 0 |
| 27698 | ENSMUSG00000085934 | 0 | NA | NA | NA | NA | NA | Gm8988 | predicted gene 8988 [Source:MGI Symbol | 0 | 0 | 0 | 0 | 0 | 0 |
| 27699 | ENSMUSG00000085936 | 0.356960287 | 0.16514264 | 3.880370363 | 0.042558474 | 0.966053499 | NA | 2610307P16 | RIKEN cDNA 2610307P16 gene [Source:M | 0 | 1.0279 | 0 | 0 | 0 | 1.1138 |
| 27700 | ENSMUSG00000085937 | 0 | NA | NA | NA | NA | NA | Ccdc42os | coiled-coil domain containing 42, opposite | 0 | 0 | 0 | 0 | 0 | 0 |
| 27701 | ENSMUSG00000085938 | 0 | NA | NA | NA | NA | NA | Gm14371 | predicted gene 14371 [Source:MGI Symbo | 0 | 0 | 0 | 0 | 0 | 0 |
| 27702 | ENSMUSG00000085939 | 0.52105551 | -0.812028029 | 3.385364865 | -0.239864257 | 0.810435491 | NA | Cd63-ps | CD63 antigen, pseudogene [Source:MGI S | 0 | 0 | 2.0125 | 0 | 0 | 1.1138 |
| 27703 | ENSMUSG00000085940 | 0 | NA | NA | NA | NA | NA | 4930405D11 | RIKEN cDNA 4930405D11 gene [Source:M | 0 | 0 | 0 | 0 | 0 | 0 |
| 27704 | ENSMUSG00000085941 | 0 | NA | NA | NA | NA | NA | Gm11201 | predicted gene 11201 [Source:MGI Symbo | 0 | 0 | 0 | 0 | 0 | 0 |
| 27705 | ENSMUSG00000085944 | 0.321325949 | 1.911575114 | 4.048267893 | 0.472195804 | 0.636787034 | NA | 1700003D09 | RIKEN cDNA 1700003D09 gene [Source:M | 0 | 0 | 0 | 1.928 | 0 | 0 |
| 27706 | ENSMUSG00000085945 | 1.464742249 | 0.602336868 | 1.775288764 | 0.339289517 | 0.734391639 | NA | 2310014F06 | RIKEN cDNA 2310014F06 gene [Source:M | 2.4093 | 0 | 1.0062 | 0.964 | 2.1812 | 2.2277 |
| 27707 | ENSMUSG00000085946 | 0 | NA | NA | NA | NA | NA | AA387200 | expressed sequence AA387200 [Source:M | 0 | 0 | 0 | 0 | 0 | 0 |
| 27708 | ENSMUSG00000085949 | 0 | NA | NA | NA | NA | NA | Gm14275 | predicted gene 14275 [Source:MGI Symbo | 0 | 0 | 0 | 0 | 0 | 0 |
| 27709 | ENSMUSG00000085950 | 0 | NA | NA | NA | NA | NA | Gm13589 | predicted gene 13589 [Source:MGI Symbo | 0 | 0 | 0 | 0 | 0 | 0 |
| 27710 | ENSMUSG00000085951 | 0 | NA | NA | NA | NA | NA | Gm12145 | predicted gene 12145 [Source:MGI Symbo | 0 | 0 | 0 | 0 | 0 | 0 |
| 27711 | ENSMUSG00000085952 | 5.271384278 | 2.242547751 | 1.151433785 | 1.947613299 | 0.051461251 | 0.318636789 | Gm12315 | predicted gene 12315 [Source:MGI Symbo | 2.4093 | 0 | 3.0187 | 1.928 | 10.906 | 13.366 |
| 27712 | ENSMUSG00000085953 | 0 | NA | NA | NA | NA | NA | Gm15496 | predicted gene 15496 [Source:MGI Symbo | 0 | 0 | 0 | 0 | 0 | 0 |
| 27713 | ENSMUSG00000085955 | 1.675599304 | -2.966598134 | 1.950974718 | -1.52057231 | 0.1283672 | NA | Gm12356 | predicted gene 12356 [Source:MGI Symbo | 0.8031 | 4.1117 | 4.025 | 0 | 0 | 1.1138 |
| 27714 | ENSMUSG00000085956 | 15.40197712 | 0.224044811 | 0.548469108 | 0.408491213 | 0.68291308 | 0.935181411 | 4930481B07 | RIKEN cDNA 4930481B07 gene [Source:M | 16.062 | 12.335 | 14.087 | 13.496 | 15.269 | 21.163 |
| 27715 | ENSMUSG00000085957 | 0 | NA | NA | NA | NA | NA | Syna | syncytin a [Source:MGI Symbol;Acc:MGI:2 | 0 | 0 | 0 | 0 | 0 | 0 |
| 27716 | ENSMUSG00000085958 | 1.584969023 | 0.277469063 | 1.860840428 | 0.149109541 | 0.881467197 | NA | A930019D19 | RIKEN cDNA A930019D19 gene [Source:M | 3.2124 | 1.0279 | 0 | 1.928 | 0 | 3.3415 |

| | | | | | | | | | | | | | | | |
|---|---|---|---|---|---|---|---|---|---|---|---|---|---|---|---|
| 27717 | ENSMUSG00000085961 | 0 | NA | NA | NA | NA | NA | Gm15904 | predicted gene 15904 [Source:MGI Symbo | 0 | 0 | 0 | 0 | 0 | 0 |
| 27718 | ENSMUSG00000085962 | 38.66590598 | -0.035046964 | 0.488795846 | -0.071700617 | 0.942840165 | 0.990074885 | Gm16984 | predicted gene, 16984 [Source:MGI Symb | 42.564 | 31.865 | 43.269 | 70.37 | 29.447 | 14.48 |
| 27719 | ENSMUSG00000085965 | 0 | NA | NA | NA | NA | NA | 2310016D23 | RIKEN cDNA 2310016D23 gene [Source:M | 0 | 0 | 0 | 0 | 0 | 0 |
| 27720 | ENSMUSG00000085966 | 0 | NA | NA | NA | NA | NA | Gm14368 | predicted gene 14368 [Source:MGI Symbo | 0 | 0 | 0 | 0 | 0 | 0 |
| 27721 | ENSMUSG00000085967 | 0 | NA | NA | NA | NA | NA | Gm12530 | predicted gene 12530 [Source:MGI Symbo | 0 | 0 | 0 | 0 | 0 | 0 |
| 27722 | ENSMUSG00000085968 | 0 | NA | NA | NA | NA | NA | Gm13027 | predicted gene 13027 [Source:MGI Symbo | 0 | 0 | 0 | 0 | 0 | 0 |
| 27723 | ENSMUSG00000085969 | 0 | NA | NA | NA | NA | NA | Gm15600 | predicted gene 15600 [Source:MGI Symbo | 0 | 0 | 0 | 0 | 0 | 0 |
| 27724 | ENSMUSG00000085970 | 0 | NA | NA | NA | NA | NA | Gm15643 | predicted gene 15643 [Source:MGI Symbo | 0 | 0 | 0 | 0 | 0 | 0 |
| 27725 | ENSMUSG00000085971 | 20.48273087 | -0.020633808 | 0.564154909 | -0.036574721 | 0.9708241 | 0.994966572 | Gm15411 | predicted gene 15411 [Source:MGI Symbo | 36.14 | 15.419 | 10.062 | 26.027 | 18.541 | 16.708 |
| 27726 | ENSMUSG00000085972 | 25.14980633 | 0.389270801 | 0.485561436 | 0.801692169 | 0.422731047 | 0.84005258 | 1110028F11 | RIKEN cDNA 1110028F11 gene [Source:M | 31.321 | 20.558 | 13.081 | 22.171 | 39.262 | 24.505 |
| 27727 | ENSMUSG00000085973 | 0 | NA | NA | NA | NA | NA | Gm14742 | predicted gene 14742 [Source:MGI Symbo | 0 | 0 | 0 | 0 | 0 | 0 |
| 27728 | ENSMUSG00000085975 | 0.665097234 | 0.245106807 | 2.82617131 | 0.086727512 | 0.930888107 | NA | Gm13572 | predicted gene 13572 [Source:MGI Symbo | 0.8031 | 0 | 1.0062 | 0 | 2.1812 | 0 |
| 27729 | ENSMUSG00000085976 | 0.506966596 | 2.558866737 | 3.339834942 | 0.766165628 | 0.443577757 | NA | Gm13816 | predicted gene 13816 [Source:MGI Symbo | 0 | 0 | 0 | 1.928 | 0 | 1.1138 |
| 27730 | ENSMUSG00000085977 | 125.807211 | 2.806301593 | 0.282591991 | 9.930577239 | 3.06E-23 | 7.39E-20 | Gm5970 | predicted gene 5970 [Source:MGI Symbol | 19.274 | 38.033 | 38.237 | 241.96 | 173.41 | 243.93 |
| 27731 | ENSMUSG00000085978 | 0 | NA | NA | NA | NA | NA | Gm15258 | predicted gene 15258 [Source:MGI Symbo | 0 | 0 | 0 | 0 | 0 | 0 |
| 27732 | ENSMUSG00000085979 | 0 | NA | NA | NA | NA | NA | Gm12055 | predicted gene 12055 [Source:MGI Symbo | 0 | 0 | 0 | 0 | 0 | 0 |
| 27733 | ENSMUSG00000085980 | 0 | NA | NA | NA | NA | NA | Gm12408 | predicted gene 12408 [Source:MGI Symbo | 0 | 0 | 0 | 0 | 0 | 0 |
| 27734 | ENSMUSG00000085982 | 1.006424547 | -2.146212304 | 2.438584396 | -0.880105814 | 0.37880199 | NA | 9530051G07 | RIKEN cDNA 9530051G07 gene [Source:M | 0 | 2.0558 | 3.0187 | 0.964 | 0 | 0 |
| 27735 | ENSMUSG00000085984 | 0.160662974 | 1.126924175 | 4.080472857 | 0.276174898 | 0.782413736 | NA | 1700001G11 | RIKEN cDNA 1700001G11 gene [Source:M | 0 | 0 | 0 | 0.964 | 0 | 0 |
| 27736 | ENSMUSG00000085985 | 0 | NA | NA | NA | NA | NA | Gm11682 | predicted gene 11682 [Source:MGI Symbo | 0 | 0 | 0 | 0 | 0 | 0 |
| 27737 | ENSMUSG00000085987 | 0 | NA | NA | NA | NA | NA | Gm13403 | predicted gene 13403 [Source:MGI Symbo | 0 | 0 | 0 | 0 | 0 | 0 |
| 27738 | ENSMUSG00000085989 | 0 | NA | NA | NA | NA | NA | Gm7437 | predicted gene 7437 [Source:MGI Symbol | 0 | 0 | 0 | 0 | 0 | 0 |
| 27739 | ENSMUSG00000085990 | 11.43411884 | 1.458426991 | 0.702517547 | 2.076000802 | 0.037893877 | 0.26432283 | Gm16731 | predicted gene, 16731 [Source:MGI Symb | 3.2124 | 7.1954 | 8.05 | 12.532 | 11.997 | 25.618 |
| 27740 | ENSMUSG00000085991 | 0 | NA | NA | NA | NA | NA | Gm11479 | predicted gene 11479 [Source:MGI Symbo | 0 | 0 | 0 | 0 | 0 | 0 |
| 27741 | ENSMUSG00000085992 | 0 | NA | NA | NA | NA | NA | Gm11515 | predicted gene 11515 [Source:MGI Symbo | 0 | 0 | 0 | 0 | 0 | 0 |
| 27742 | ENSMUSG00000085993 | 0 | NA | NA | NA | NA | NA | Gm15905 | predicted gene 15905 [Source:MGI Symbo | 0 | 0 | 0 | 0 | 0 | 0 |
| 27743 | ENSMUSG00000085995 | 758.5898747 | 0.107859216 | 0.183563906 | 0.587584009 | 0.556811547 | 0.898167794 | Gm2788 | predicted gene 2788 [Source:MGI Symbol | 766.16 | 831.59 | 593.68 | 1021.8 | 705.63 | 632.66 |
| 27744 | ENSMUSG00000085996 | 0 | NA | NA | NA | NA | NA | A830012C17 | RIKEN cDNA A830012C17 gene [Source:M | 0 | 0 | 0 | 0 | 0 | 0 |
| 27745 | ENSMUSG00000085998 | 0 | NA | NA | NA | NA | NA | AW822252 | expressed sequence AW822252 [Source:N | 0 | 0 | 0 | 0 | 0 | 0 |
| 27746 | ENSMUSG00000085999 | 0 | NA | NA | NA | NA | NA | Gm13411 | predicted gene 13411 [Source:MGI Symbo | 0 | 0 | 0 | 0 | 0 | 0 |
| 27747 | ENSMUSG00000086000 | 1.16117507 | -0.27256011 | 2.112400613 | -0.129028608 | 0.897335012 | NA | Gm12493 | predicted gene 12493 [Source:MGI Symbo | 0.8031 | 0 | 3.0187 | 0.964 | 2.1812 | 0 |
| 27748 | ENSMUSG00000086001 | 0 | NA | NA | NA | NA | NA | Gm13448 | predicted gene 13448 [Source:MGI Symbo | 0 | 0 | 0 | 0 | 0 | 0 |
| 27749 | ENSMUSG00000086003 | 40.43271927 | 0.117447626 | 0.43140541 | 0.272244213 | 0.785434248 | 0.960720589 | B230206L02 | RIKEN cDNA B230206L02 gene [Source:M | 30.518 | 38.033 | 48.3 | 67.478 | 35.99 | 22.277 |
| 27750 | ENSMUSG00000086004 | 0 | NA | NA | NA | NA | NA | Gm7805 | predicted gene 7805 [Source:MGI Symbol | 0 | 0 | 0 | 0 | 0 | 0 |
| 27751 | ENSMUSG00000086006 | 0.181769715 | 1.126924175 | 4.080472857 | 0.276174898 | 0.782413736 | NA | Gm13293 | predicted gene 13293 [Source:MGI Symbo | 0 | 0 | 0 | 0 | 1.0906 | 0 |
| 27752 | ENSMUSG00000086007 | 0 | NA | NA | NA | NA | NA | 1700057H21 | RIKEN cDNA 1700057H21 gene [Source:M | 0 | 0 | 0 | 0 | 0 | 0 |
| 27753 | ENSMUSG00000086008 | 0 | NA | NA | NA | NA | NA | Gm8817 | predicted gene 8817 [Source:MGI Symbol | 0 | 0 | 0 | 0 | 0 | 0 |
| 27754 | ENSMUSG00000086009 | 0 | NA | NA | NA | NA | NA | 4930587A21 | RIKEN cDNA 4930587A21 gene [Source:M | 0 | 0 | 0 | 0 | 0 | 0 |
| 27755 | ENSMUSG00000086010 | 99.77448034 | 0.558975089 | 0.256115603 | 2.182510876 | 0.029071849 | 0.223720589 | Gm15318 | predicted gene 15318 [Source:MGI Symbo | 85.129 | 77.094 | 79.493 | 97.362 | 154.87 | 104.7 |
| 27756 | ENSMUSG00000086012 | 0.51014012 | 1.031855138 | 3.09002894 | 0.333930574 | 0.738431948 | NA | Gm15902 | predicted gene 15902 [Source:MGI Symbo | 0 | 0 | 1.0062 | 0.964 | 1.0906 | 0 |
| 27757 | ENSMUSG00000086013 | 11.50111782 | -0.619464434 | 0.668447533 | -0.926721102 | 0.354071358 | 0.79449673 | Gm15706 | predicted gene 15706 [Source:MGI Symbo | 15.259 | 18.503 | 8.05 | 10.604 | 5.4531 | 11.138 |
| 27758 | ENSMUSG00000086014 | 0 | NA | NA | NA | NA | NA | Gm16095 | predicted gene 16095 [Source:MGI Symbo | 0 | 0 | 0 | 0 | 0 | 0 |
| 27759 | ENSMUSG00000086015 | 5.371968701 | 0.589750501 | 1.166391745 | 0.505619577 | 0.613123736 | 0.916813074 | 4833417C18 | RIKEN cDNA 4833417C18 gene [Source:M | 0.8031 | 7.1954 | 5.0312 | 11.568 | 7.6343 | 0 |
| 27760 | ENSMUSG00000086016 | 0 | NA | NA | NA | NA | NA | 1700084C06 | RIKEN cDNA 1700084C06 gene [Source:M | 0 | 0 | 0 | 0 | 0 | 0 |
| 27761 | ENSMUSG00000086017 | 0 | NA | NA | NA | NA | NA | Gm11872 | predicted gene 11872 [Source:MGI Symbo | 0 | 0 | 0 | 0 | 0 | 0 |
| 27762 | ENSMUSG00000086018 | 0 | NA | NA | NA | NA | NA | Gm13179 | predicted gene 13179 [Source:MGI Symbo | 0 | 0 | 0 | 0 | 0 | 0 |
| 27763 | ENSMUSG00000086019 | 0 | NA | NA | NA | NA | NA | Gm15762 | predicted gene 15762 [Source:MGI Symbo | 0 | 0 | 0 | 0 | 0 | 0 |
| 27764 | ENSMUSG00000086020 | 0 | NA | NA | NA | NA | NA | Gm12239 | predicted gene 12239 [Source:MGI Symbo | 0 | 0 | 0 | 0 | 0 | 0 |
| 27765 | ENSMUSG00000086021 | 0 | NA | NA | NA | NA | NA | Gm15767 | predicted gene 15767 [Source:MGI Symbo | 0 | 0 | 0 | 0 | 0 | 0 |
| 27766 | ENSMUSG00000086022 | 0.17131964 | -0.796629176 | 4.080472857 | -0.195229623 | 0.84521319 | NA | Rad51ap2 | RAD51 associated protein 2 [Source:MGI S | 0 | 1.0279 | 0 | 0 | 0 | 0 |
| 27767 | ENSMUSG00000086025 | 0 | NA | NA | NA | NA | NA | 4933406G16 | RIKEN cDNA 4933406G16 gene [Source:M | 0 | 0 | 0 | 0 | 0 | 0 |
| 27768 | ENSMUSG00000086026 | 0 | NA | NA | NA | NA | NA | Gm12374 | predicted gene 12374 [Source:MGI Symbo | 0 | 0 | 0 | 0 | 0 | 0 |
| 27769 | ENSMUSG00000086027 | 0 | NA | NA | NA | NA | NA | Gm11250 | predicted gene 11250 [Source:MGI Symbo | 0 | 0 | 0 | 0 | 0 | 0 |
| 27770 | ENSMUSG00000086028 | 0 | NA | NA | NA | NA | NA | Gm12243 | predicted gene 12243 [Source:MGI Symbo | 0 | 0 | 0 | 0 | 0 | 0 |
| 27771 | ENSMUSG00000086029 | 0 | NA | NA | NA | NA | NA | Pax6os1 | paired box 6 opposite strand 1 [Source:M | 0 | 0 | 0 | 0 | 0 | 0 |
| 27772 | ENSMUSG00000086030 | 0 | NA | NA | NA | NA | NA | Gm14255 | predicted gene 14255 [Source:MGI Symbo | 0 | 0 | 0 | 0 | 0 | 0 |

| | | | | | | | | | | | | | | | |
|---|---|---|---|---|---|---|---|---|---|---|---|---|---|---|---|
| 27773 | ENSMUSG00000086031 | 0 | NA | NA | NA | NA | NA | Gm12348 | predicted gene 12348 [Source:MGI Symbo | 0 | 0 | 0 | 0 | 0 | 0 |
| 27774 | ENSMUSG00000086032 | 0.331982614 | 0.16514309 | 3.986751476 | 0.041422971 | 0.966958701 | NA | Gm15929 | predicted gene 15929 [Source:MGI Symbo | 0 | 1.0279 | 0 | 0.964 | 0 | 0 |
| 27775 | ENSMUSG00000086034 | 0 | NA | NA | NA | NA | NA | Gm15201 | predicted gene 15201 [Source:MGI Symbo | 0 | 0 | 0 | 0 | 0 | 0 |
| 27776 | ENSMUSG00000086035 | 0 | NA | NA | NA | NA | NA | Gm12610 | predicted gene 12610 [Source:MGI Symbo | 0 | 0 | 0 | 0 | 0 | 0 |
| 27777 | ENSMUSG00000086036 | 0 | NA | NA | NA | NA | NA | Gm14254 | predicted gene 14254 [Source:MGI Symbo | 0 | 0 | 0 | 0 | 0 | 0 |
| 27778 | ENSMUSG00000086037 | 0 | NA | NA | NA | NA | NA | Gm16085 | predicted gene 16085 [Source:MGI Symbo | 0 | 0 | 0 | 0 | 0 | 0 |
| 27779 | ENSMUSG00000086039 | 0.17131964 | -0.796629176 | 4.080472857 | -0.195229623 | 0.84521319 | NA | Gm12227 | predicted gene 12227 [Source:MGI Symbo | 0 | 1.0279 | 0 | 0 | 0 | 0 |
| 27780 | ENSMUSG00000086040 | 323.3411368 | -0.00497856 | 0.234027035 | -0.02127344 | 0.983027531 | 0.998580735 | Wipf3 | WAS/WASL interacting protein family, me | 452.95 | 292.96 | 224.39 | 267.99 | 311.92 | 389.85 |
| 27781 | ENSMUSG00000086041 | 0.435408177 | -2.179616019 | 3.531869654 | -0.617128103 | 0.537150234 | NA | Gm12869 | predicted gene 12869 [Source:MGI Symbo | 1.6062 | 0 | 1.0062 | 0 | 0 | 0 |
| 27782 | ENSMUSG00000086042 | 0 | NA | NA | NA | NA | NA | Gm12670 | predicted gene 12670 [Source:MGI Symbo | 0 | 0 | 0 | 0 | 0 | 0 |
| 27783 | ENSMUSG00000086043 | 56.77607211 | 0.127514025 | 0.358824942 | 0.355365556 | 0.722315749 | 0.9447719 | Gm12473 | predicted gene 12473 [Source:MGI Symbo | 47.383 | 33.921 | 81.506 | 50.127 | 69.8 | 57.92 |
| 27784 | ENSMUSG00000086044 | 0 | NA | NA | NA | NA | NA | A330043C09 | RIKEN cDNA A330043C09 gene [Source:M | 0 | 0 | 0 | 0 | 0 | 0 |
| 27785 | ENSMUSG00000086046 | 1.214974719 | 2.675163135 | 2.126070915 | 1.258266183 | 0.208295507 | NA | 1700095A21 | RIKEN cDNA 1700095A21 gene [Source:M | 0.8031 | 0 | 0 | 0.964 | 2.1812 | 3.3415 |
| 27786 | ENSMUSG00000086047 | 9.148656509 | 1.470001591 | 0.861252912 | 1.706817557 | 0.087855968 | 0.43690328 | 9530046B11 | RIKEN cDNA 9530046B11 gene [Source:M | 6.4248 | 0 | 8.05 | 17.352 | 15.269 | 7.7969 |
| 27787 | ENSMUSG00000086048 | 0 | NA | NA | NA | NA | NA | Drr1 | developmentally regulated repeat elemen | 0 | 0 | 0 | 0 | 0 | 0 |
| 27788 | ENSMUSG00000086049 | 3.358789875 | -0.947706697 | 1.40859097 | -0.672804751 | 0.501071515 | NA | Gm12735 | predicted gene 12735 [Source:MGI Symbo | 3.2124 | 6.1675 | 4.025 | 6.7478 | 0 | 0 |
| 27789 | ENSMUSG00000086050 | 0.481988923 | 2.492401104 | 3.837077151 | 0.649557203 | 0.515978285 | NA | Gm16045 | predicted gene 16045 [Source:MGI Symbo | 0 | 0 | 0 | 2.8919 | 0 | 0 |
| 27790 | ENSMUSG00000086052 | 0.545309144 | 2.651974629 | 3.666219369 | 0.72335405 | 0.469462394 | NA | Gm11802 | predicted gene 11802 [Source:MGI Symbo | 0 | 0 | 0 | 0 | 3.2719 | 0 |
| 27791 | ENSMUSG00000086053 | 0 | NA | NA | NA | NA | NA | Gm15178 | predicted gene 15178 [Source:MGI Symbo | 0 | 0 | 0 | 0 | 0 | 0 |
| 27792 | ENSMUSG00000086054 | 173.2664221 | 0.288217305 | 0.21108496 | 1.365409005 | 0.172124535 | 0.60005044 | Hnf1aos1 | HNF1 homeobox A, opposite strand 1 [Sou | 189.53 | 152.13 | 125.78 | 209.18 | 163.59 | 199.38 |
| 27793 | ENSMUSG00000086055 | 0 | NA | NA | NA | NA | NA | Gm13837 | predicted gene 13837 [Source:MGI Symbo | 0 | 0 | 0 | 0 | 0 | 0 |
| 27794 | ENSMUSG00000086056 | 0 | NA | NA | NA | NA | NA | Gm4846 | predicted gene 4846 [Source:MGI Symbol | 0 | 0 | 0 | 0 | 0 | 0 |
| 27795 | ENSMUSG00000086057 | 0 | NA | NA | NA | NA | NA | Gm15324 | predicted gene 15324 [Source:MGI Symbo | 0 | 0 | 0 | 0 | 0 | 0 |
| 27796 | ENSMUSG00000086058 | 1.870196445 | -2.203275093 | 1.788360242 | -1.232008541 | 0.217945893 | NA | Unc45bos | unc-45 myosin chaperone B, opposite stra | 3.2124 | 2.0558 | 4.025 | 1.928 | 0 | 0 |
| 27797 | ENSMUSG00000086060 | 0 | NA | NA | NA | NA | NA | Gm16084 | predicted gene 16084 [Source:MGI Symbo | 0 | 0 | 0 | 0 | 0 | 0 |
| 27798 | ENSMUSG00000086061 | 0 | NA | NA | NA | NA | NA | Gm13075 | predicted gene 13075 [Source:MGI Symbo | 0 | 0 | 0 | 0 | 0 | 0 |
| 27799 | ENSMUSG00000086063 | 0.321325949 | 1.911575114 | 4.048267893 | 0.472195804 | 0.636787034 | NA | Gm12257 | predicted gene 12257 [Source:MGI Symbo | 0 | 0 | 0 | 1.928 | 0 | 0 |
| 27800 | ENSMUSG00000086064 | 0.821015994 | 0.567017314 | 2.624106235 | 0.216080167 | 0.828925259 | NA | 4930439A04 | RIKEN cDNA 4930439A04 gene [Source:M | 0 | 1.0279 | 1.0062 | 2.8919 | 0 | 0 |
| 27801 | ENSMUSG00000086065 | 0 | NA | NA | NA | NA | NA | Gm16180 | predicted gene 16180 [Source:MGI Symbo | 0 | 0 | 0 | 0 | 0 | 0 |
| 27802 | ENSMUSG00000086066 | 0 | NA | NA | NA | NA | NA | 4930423M02 | RIKEN cDNA 4930423M02 gene [Source:M | 0 | 0 | 0 | 0 | 0 | 0 |
| 27803 | ENSMUSG00000086067 | 0 | NA | NA | NA | NA | NA | Gm16183 | predicted gene 16183 [Source:MGI Symbo | 0 | 0 | 0 | 0 | 0 | 0 |
| 27804 | ENSMUSG00000086068 | 0 | NA | NA | NA | NA | NA | 1700120G11 | RIKEN cDNA 1700120G11 gene [Source:M | 0 | 0 | 0 | 0 | 0 | 0 |
| 27805 | ENSMUSG00000086069 | 0.321325949 | 1.911575114 | 4.048267893 | 0.472195804 | 0.636787034 | NA | Gm13708 | predicted gene 13708 [Source:MGI Symbo | 0 | 0 | 0 | 1.928 | 0 | 0 |
| 27806 | ENSMUSG00000086070 | 0 | NA | NA | NA | NA | NA | 8430436N08 | RIKEN cDNA 8430436N08 gene [Source:M | 0 | 0 | 0 | 0 | 0 | 0 |
| 27807 | ENSMUSG00000086071 | 69.12246161 | 0.010538921 | 0.278469413 | 0.037845882 | 0.969810562 | 0.994705713 | Gm12354 | predicted gene 12354 [Source:MGI Symbo | 77.098 | 61.675 | 67.418 | 58.803 | 81.796 | 67.944 |
| 27808 | ENSMUSG00000086072 | 0 | NA | NA | NA | NA | NA | Gm11861 | predicted gene 11861 [Source:MGI Symbo | 0 | 0 | 0 | 0 | 0 | 0 |
| 27809 | ENSMUSG00000086074 | 0 | NA | NA | NA | NA | NA | Gm13274 | predicted gene 13274 [Source:MGI Symbo | 0 | 0 | 0 | 0 | 0 | 0 |
| 27810 | ENSMUSG00000086075 | 0 | NA | NA | NA | NA | NA | Gm15728 | predicted gene 15728 [Source:MGI Symbo | 0 | 0 | 0 | 0 | 0 | 0 |
| 27811 | ENSMUSG00000086076 | 0.133850373 | -0.796629176 | 4.080472857 | -0.195229623 | 0.84521319 | NA | Ncr3-ps | natural cytotoxicity triggering receptor 3, | 0.8031 | 0 | 0 | 0 | 0 | 0 |
| 27812 | ENSMUSG00000086077 | 0 | NA | NA | NA | NA | NA | Gm14396 | predicted gene 14396 [Source:MGI Symbo | 0 | 0 | 0 | 0 | 0 | 0 |
| 27813 | ENSMUSG00000086078 | 0 | NA | NA | NA | NA | NA | Gm11657 | predicted gene 11657 [Source:MGI Symbo | 0 | 0 | 0 | 0 | 0 | 0 |
| 27814 | ENSMUSG00000086080 | 0 | NA | NA | NA | NA | NA | 1700123O12 | RIKEN cDNA 1700123O12 gene [Source:M | 0 | 0 | 0 | 0 | 0 | 0 |
| 27815 | ENSMUSG00000086081 | 0 | NA | NA | NA | NA | NA | Spin2-ps3 | spindlin family, member 2, pseudogene 3 | 0 | 0 | 0 | 0 | 0 | 0 |
| 27816 | ENSMUSG00000086083 | 1.351125335 | 0.898520505 | 1.866150402 | 0.481483434 | 0.63017295 | NA | Gm15829 | predicted gene 15829 [Source:MGI Symbo | 0.8031 | 1.0279 | 1.0062 | 1.928 | 0 | 3.3415 |
| 27817 | ENSMUSG00000086087 | 0 | NA | NA | NA | NA | NA | Gm12916 | predicted gene 12916 [Source:MGI Symbo | 0 | 0 | 0 | 0 | 0 | 0 |
| 27818 | ENSMUSG00000086089 | 0 | NA | NA | NA | NA | NA | Gm11640 | predicted gene 11640 [Source:MGI Symbo | 0 | 0 | 0 | 0 | 0 | 0 |
| 27819 | ENSMUSG00000086090 | 0.167707431 | -0.796629176 | 4.080472857 | -0.195229623 | 0.84521319 | NA | Gm12531 | predicted gene 12531 [Source:MGI Symbo | 0 | 0 | 1.0062 | 0 | 0 | 0 |
| 27820 | ENSMUSG00000086092 | 0 | NA | NA | NA | NA | NA | Gm15323 | predicted gene 15323 [Source:MGI Symbo | 0 | 0 | 0 | 0 | 0 | 0 |
| 27821 | ENSMUSG00000086095 | 2.78030538 | 2.292614126 | 1.426169076 | 1.607533191 | 0.107937455 | NA | Gm15328 | predicted gene 15328 [Source:MGI Symbo | 0.8031 | 1.0279 | 1.0062 | 2.8919 | 8.7249 | 2.2277 |
| 27822 | ENSMUSG00000086096 | 0 | NA | NA | NA | NA | NA | Gm12688 | predicted gene 12688 [Source:MGI Symbo | 0 | 0 | 0 | 0 | 0 | 0 |
| 27823 | ENSMUSG00000086097 | 0 | NA | NA | NA | NA | NA | Gm16250 | predicted gene 16250 [Source:MGI Symbo | 0 | 0 | 0 | 0 | 0 | 0 |
| 27824 | ENSMUSG00000086098 | 0 | NA | NA | NA | NA | NA | Gm14291 | predicted gene 14291 [Source:MGI Symbo | 0 | 0 | 0 | 0 | 0 | 0 |
| 27825 | ENSMUSG00000086103 | 5.7706717 | -2.441909536 | 1.06335761 | -2.296414219 | 0.021652209 | 0.183082289 | Gm11832 | predicted gene 11832 [Source:MGI Symbo | 12.85 | 7.1954 | 9.0562 | 0 | 2.1812 | 3.3415 |
| 27826 | ENSMUSG00000086104 | 0 | NA | NA | NA | NA | NA | Gm14707 | predicted gene 14707 [Source:MGI Symbo | 0 | 0 | 0 | 0 | 0 | 0 |
| 27827 | ENSMUSG00000086105 | 0 | NA | NA | NA | NA | NA | Gm11636 | predicted gene 11636 [Source:MGI Symbo | 0 | 0 | 0 | 0 | 0 | 0 |
| 27828 | ENSMUSG00000086107 | 0 | NA | NA | NA | NA | NA | Gm16312 | predicted gene 16312 [Source:MGI Symbo | 0 | 0 | 0 | 0 | 0 | 0 |

| | | | | | | | | | | | | | | | |
|---|---|---|---|---|---|---|---|---|---|---|---|---|---|---|---|
| 27829 | ENSMUSG00000086108 | 3.038569302 | -0.312596534 | 1.298521712 | -0.24073262 | 0.809762355 | NA | Gm5602 | predicted gene 5602 [Source:MGI Symbol | 4.0155 | 4.1117 | 2.0125 | 4.8199 | 3.2719 | 0 |
| 27830 | ENSMUSG00000086109 | 0.852599439 | -1.816262765 | 2.863514483 | -0.634277485 | 0.525899747 | NA | Gm13391 | predicted gene 13391 [Source:MGI Symbo | 0 | 0 | 4.025 | 0 | 1.0906 | 0 |
| 27831 | ENSMUSG00000086111 | 0 | NA | NA | NA | NA | NA | Gm15326 | predicted gene 15326 [Source:MGI Symbo | 0 | 0 | 0 | 0 | 0 | 0 |
| 27832 | ENSMUSG00000086112 | 0 | NA | NA | NA | NA | NA | 4930563I02R | RIKEN cDNA 4930563I02 gene [Source:MG | 0 | 0 | 0 | 0 | 0 | 0 |
| 27833 | ENSMUSG00000086114 | 0 | NA | NA | NA | NA | NA | Gm14642 | predicted gene 14642 [Source:MGI Symbo | 0 | 0 | 0 | 0 | 0 | 0 |
| 27834 | ENSMUSG00000086115 | 0 | NA | NA | NA | NA | NA | Gm15733 | predicted gene 15733 [Source:MGI Symbo | 0 | 0 | 0 | 0 | 0 | 0 |
| 27835 | ENSMUSG00000086117 | 0 | NA | NA | NA | NA | NA | Gm12065 | predicted gene 12065 [Source:MGI Symbo | 0 | 0 | 0 | 0 | 0 | 0 |
| 27836 | ENSMUSG00000086118 | 0 | NA | NA | NA | NA | NA | Gm14169 | predicted gene 14169 [Source:MGI Symbo | 0 | 0 | 0 | 0 | 0 | 0 |
| 27837 | ENSMUSG00000086119 | 13.9167359 | 0.587204603 | 0.64430122 | 0.911382105 | 0.362094079 | 0.800872989 | Gm2415 | predicted gene 2415 [Source:MGI Symbol | 7.2279 | 7.1954 | 19.119 | 21.208 | 9.8156 | 18.935 |
| 27838 | ENSMUSG00000086121 | 1.010036755 | -2.151347842 | 2.436507427 | -0.882963794 | 0.377255834 | NA | Gm16068 | predicted gene 16068 [Source:MGI Symbo | 0 | 3.0838 | 2.0125 | 0.964 | 0 | 0 |
| 27839 | ENSMUSG00000086122 | 0 | NA | NA | NA | NA | NA | Gm14507 | predicted gene 14507 [Source:MGI Symbo | 0 | 0 | 0 | 0 | 0 | 0 |
| 27840 | ENSMUSG00000086123 | 0 | NA | NA | NA | NA | NA | Gm16060 | predicted pseudogene 16060 [Source:MG | 0 | 0 | 0 | 0 | 0 | 0 |
| 27841 | ENSMUSG00000086124 | 0.17131964 | -0.796629176 | 4.080472857 | -0.195229623 | 0.84521319 | NA | A530076I17R | RIKEN cDNA A530076I17 gene [Source:MG | 0 | 1.0279 | 0 | 0 | 0 | 0 |
| 27842 | ENSMUSG00000086125 | 0 | NA | NA | NA | NA | NA | Gm15942 | predicted gene 15942 [Source:MGI Symbo | 0 | 0 | 0 | 0 | 0 | 0 |
| 27843 | ENSMUSG00000086126 | 0 | NA | NA | NA | NA | NA | Evx1os | even skipped homeotic gene 1, opposite s | 0 | 0 | 0 | 0 | 0 | 0 |
| 27844 | ENSMUSG00000086127 | 0.160662974 | 1.126924175 | 4.080472857 | 0.276174898 | 0.782413736 | NA | Gm11934 | predicted gene 11934 [Source:MGI Symbo | 0 | 0 | 0 | 0.964 | 0 | 0 |
| 27845 | ENSMUSG00000086128 | 5.766128503 | 0.67012006 | 0.884327589 | 0.757773554 | 0.448586557 | 0.854238022 | St8sia3os | ST8 alpha-N-acetyl-neuraminide alpha-2,8 | 3.2124 | 6.1675 | 4.025 | 5.7839 | 8.7249 | 6.6831 |
| 27846 | ENSMUSG00000086129 | 0.321325949 | 1.911575114 | 4.048267893 | 0.472195804 | 0.636787034 | NA | Gm13558 | predicted gene 13558 [Source:MGI Symbo | 0 | 0 | 0 | 1.928 | 0 | 0 |
| 27847 | ENSMUSG00000086130 | 0.185640647 | 1.126924175 | 4.080472857 | 0.276174898 | 0.782413736 | NA | Gm16211 | predicted gene 16211 [Source:MGI Symbo | 0 | 0 | 0 | 0 | 0 | 1.1138 |
| 27848 | ENSMUSG00000086132 | 0 | NA | NA | NA | NA | NA | Gm13631 | predicted gene 13631 [Source:MGI Symbo | 0 | 0 | 0 | 0 | 0 | 0 |
| 27849 | ENSMUSG00000086133 | 0 | NA | NA | NA | NA | NA | Gm16331 | predicted gene 16331 [Source:MGI Symbo | 0 | 0 | 0 | 0 | 0 | 0 |
| 27850 | ENSMUSG00000086134 | 0.167707431 | -0.796629176 | 4.080472857 | -0.195229623 | 0.84521319 | NA | Gm16159 | predicted gene 16159 [Source:MGI Symbo | 0 | 0 | 1.0062 | 0 | 0 | 0 |
| 27851 | ENSMUSG00000086135 | 0 | NA | NA | NA | NA | NA | 1700001J04R | RIKEN cDNA 1700001J04 gene [Source:MG | 0 | 0 | 0 | 0 | 0 | 0 |
| 27852 | ENSMUSG00000086136 | 40.29216665 | 1.452448625 | 0.528220408 | 2.749701835 | 0.005964951 | 0.075023529 | Gm12718 | predicted gene 12718 [Source:MGI Symbo | 28.912 | 7.1954 | 28.175 | 23.135 | 75.253 | 79.083 |
| 27853 | ENSMUSG00000086137 | 0 | NA | NA | NA | NA | NA | Gm16248 | predicted gene 16248 [Source:MGI Symbo | 0 | 0 | 0 | 0 | 0 | 0 |
| 27854 | ENSMUSG00000086138 | 0.503302254 | -0.837877584 | 3.426698643 | -0.244514523 | 0.806832356 | NA | NA | NA | 0 | 2.0558 | 0 | 0.964 | 0 | 0 |
| 27855 | ENSMUSG00000086139 | 0.321325949 | 1.911575114 | 4.048267893 | 0.472195804 | 0.636787034 | NA | Gm11211 | predicted gene 11211 [Source:MGI Symbo | 0 | 0 | 0 | 1.928 | 0 | 0 |
| 27856 | ENSMUSG00000086140 | 11.86356597 | -0.652334363 | 0.700153345 | -0.931702131 | 0.351490485 | 0.792511048 | Hnf1aos2 | HNF1 homeobox A, opposite strand 2 [Sou | 11.243 | 13.363 | 19.119 | 16.388 | 3.2719 | 7.7969 |
| 27857 | ENSMUSG00000086141 | 260.3081787 | 0.782696031 | 0.260216786 | 3.007861422 | 0.002630931 | 0.042853359 | 9030622O22 | RIKEN cDNA 9030622O22 gene [Source:M | 185.52 | 181.94 | 207.29 | 486.81 | 305.37 | 194.92 |
| 27858 | ENSMUSG00000086142 | 0 | NA | NA | NA | NA | NA | Ppp4r3c-ps | protein phosphatase 4 regulatory subunit | 0 | 0 | 0 | 0 | 0 | 0 |
| 27859 | ENSMUSG00000086144 | 0 | NA | NA | NA | NA | NA | Gm11379 | predicted gene 11379 [Source:MGI Symbo | 0 | 0 | 0 | 0 | 0 | 0 |
| 27860 | ENSMUSG00000086145 | 0.842329324 | -1.832359871 | 2.605893773 | -0.703159848 | 0.481956138 | NA | Gm15479 | predicted gene 15479 [Source:MGI Symbo | 0 | 3.0838 | 1.0062 | 0.964 | 0 | 0 |
| 27861 | ENSMUSG00000086146 | 0 | NA | NA | NA | NA | NA | Gm15729 | predicted gene 15729 [Source:MGI Symbo | 0 | 0 | 0 | 0 | 0 | 0 |
| 27862 | ENSMUSG00000086147 | 0 | NA | NA | NA | NA | NA | Zfp989 | zinc finger protein 989 [Source:MGI Symb | 0 | 0 | 0 | 0 | 0 | 0 |
| 27863 | ENSMUSG00000086148 | 0.321325949 | 1.911575114 | 4.048267893 | 0.472195804 | 0.636787034 | NA | Gm15271 | predicted gene 15271 [Source:MGI Symbo | 0 | 0 | 0 | 1.928 | 0 | 0 |
| 27864 | ENSMUSG00000086150 | 8.863816295 | -2.188532834 | 0.854132136 | -2.562288365 | 0.010398494 | 0.111804391 | Bach2os | BTB and CNC homology 2, opposite stranc | 20.078 | 12.335 | 11.069 | 1.928 | 1.0906 | 6.6831 |
| 27865 | ENSMUSG00000086152 | 1.106102019 | 3.668747043 | 2.525738057 | 1.452544547 | 0.146350252 | NA | Gm11454 | predicted gene 11454 [Source:MGI Symbo | 0 | 0 | 0 | 0 | 2.1812 | 4.4554 |
| 27866 | ENSMUSG00000086153 | 0.167707431 | -0.796629176 | 4.080472857 | -0.195229623 | 0.84521319 | NA | Gm12923 | predicted gene 12923 [Source:MGI Symbo | 0 | 0 | 1.0062 | 0 | 0 | 0 |
| 27867 | ENSMUSG00000086154 | 0.181769715 | 1.126924175 | 4.080472857 | 0.276174898 | 0.782413736 | NA | Gm16196 | predicted gene 16196 [Source:MGI Symbo | 0 | 0 | 0 | 0 | 1.0906 | 0 |
| 27868 | ENSMUSG00000086155 | 0.335414862 | -1.779047242 | 4.030804179 | -0.441362855 | 0.658950329 | NA | 9430041J12R | RIKEN cDNA 9430041J12 gene [Source:MG | 0 | 0 | 2.0125 | 0 | 0 | 0 |
| 27869 | ENSMUSG00000086156 | 0.267700746 | -1.497481649 | 4.057001436 | -0.369110456 | 0.712045396 | NA | Gm15354 | predicted gene 15354 [Source:MGI Symbo | 1.6062 | 0 | 0 | 0 | 0 | 0 |
| 27870 | ENSMUSG00000086157 | 0 | NA | NA | NA | NA | NA | Gm14643 | predicted gene 14643 [Source:MGI Symbo | 0 | 0 | 0 | 0 | 0 | 0 |
| 27871 | ENSMUSG00000086158 | 104.5069968 | -0.033944218 | 0.243385975 | -0.139466615 | 0.889081436 | 0.982382091 | Ccpg1os | cell cycle progression 1, opposite strand [S | 115.65 | 85.317 | 115.72 | 86.758 | 117.79 | 105.82 |
| 27872 | ENSMUSG00000086159 | 0 | NA | NA | NA | NA | NA | Gm13025 | predicted gene 13025 [Source:MGI Symbo | 0 | 0 | 0 | 0 | 0 | 0 |
| 27873 | ENSMUSG00000086160 | 0 | NA | NA | NA | NA | NA | 4933438A12 | RIKEN cDNA 4933438A12 gene [Source:M | 0 | 0 | 0 | 0 | 0 | 0 |
| 27874 | ENSMUSG00000086161 | 0 | NA | NA | NA | NA | NA | Gm15389 | predicted gene 15389 [Source:MGI Symbo | 0 | 0 | 0 | 0 | 0 | 0 |
| 27875 | ENSMUSG00000086162 | 0.181769715 | 1.126924175 | 4.080472857 | 0.276174898 | 0.782413736 | NA | Gm16342 | predicted gene 16342 [Source:MGI Symbo | 0 | 0 | 0 | 0 | 1.0906 | 0 |
| 27876 | ENSMUSG00000086163 | 0 | NA | NA | NA | NA | NA | Gm14206 | predicted gene 14206 [Source:MGI Symbo | 0 | 0 | 0 | 0 | 0 | 0 |
| 27877 | ENSMUSG00000086164 | 0 | NA | NA | NA | NA | NA | Gm13029 | predicted gene 13029 [Source:MGI Symbo | 0 | 0 | 0 | 0 | 0 | 0 |
| 27878 | ENSMUSG00000086165 | 79.81423059 | -1.633671779 | 0.340682576 | -4.79529008 | 1.62E-06 | 0.000121737 | Gm15690 | predicted gene 15690 [Source:MGI Symbo | 98.782 | 161.38 | 101.63 | 21.208 | 47.987 | 47.895 |
| 27879 | ENSMUSG00000086166 | 0 | NA | NA | NA | NA | NA | Gm14342 | predicted gene 14342 [Source:MGI Symbo | 0 | 0 | 0 | 0 | 0 | 0 |
| 27880 | ENSMUSG00000086167 | 0.401551119 | -2.072982726 | 4.008251979 | -0.517178744 | 0.605031388 | NA | Gm13827 | predicted gene 13827 [Source:MGI Symbo | 2.4093 | 0 | 0 | 0 | 0 | 0 |
| 27881 | ENSMUSG00000086168 | 0 | NA | NA | NA | NA | NA | Gm15221 | predicted gene 15221 [Source:MGI Symbo | 0 | 0 | 0 | 0 | 0 | 0 |
| 27882 | ENSMUSG00000086170 | 0 | NA | NA | NA | NA | NA | Gm12144 | predicted gene 12144 [Source:MGI Symbo | 0 | 0 | 0 | 0 | 0 | 0 |
| 27883 | ENSMUSG00000086171 | 0 | NA | NA | NA | NA | NA | Pcsk2os1 | proprotein convertase subtilisin/kexin typ | 0 | 0 | 0 | 0 | 0 | 0 |
| 27884 | ENSMUSG00000086172 | 0.663758638 | 2.950973782 | 3.025092005 | 0.975498853 | 0.32931287 | NA | 2700068H02 | RIKEN cDNA 2700068H02 gene [Source:M | 0 | 0 | 0 | 2.8919 | 1.0906 | 0 |

| | | | | | | | | | | | | | | | |
|---|---|---|---|---|---|---|---|---|---|---|---|---|---|---|---|
| 27885 | ENSMUSG00000086173 | 0 | NA | NA | NA | NA | NA | Kis2 | Kaplan integration site 2 [Source:MGI Syn | 0 | 0 | 0 | 0 | 0 | 0 |
| 27886 | ENSMUSG00000086174 | 0 | NA | NA | NA | NA | NA | Gm6425 | predicted pseudogene 6425 [Source:MGI | 0 | 0 | 0 | 0 | 0 | 0 |
| 27887 | ENSMUSG00000086175 | 0 | NA | NA | NA | NA | NA | Gm15802 | predicted gene 15802 [Source:MGI Symbo | 0 | 0 | 0 | 0 | 0 | 0 |
| 27888 | ENSMUSG00000086176 | 0 | NA | NA | NA | NA | NA | Gm15534 | predicted gene 15534 [Source:MGI Symbo | 0 | 0 | 0 | 0 | 0 | 0 |
| 27889 | ENSMUSG00000086177 | 0 | NA | NA | NA | NA | NA | Gm14616 | predicted gene 14616 [Source:MGI Symbo | 0 | 0 | 0 | 0 | 0 | 0 |
| 27890 | ENSMUSG00000086179 | 0 | NA | NA | NA | NA | NA | Gm14317 | predicted gene 14317 [Source:MGI Symbo | 0 | 0 | 0 | 0 | 0 | 0 |
| 27891 | ENSMUSG00000086181 | 0.346303622 | 2.004339653 | 3.884003423 | 0.516049919 | 0.605819544 | NA | C230034O21 | RIKEN cDNA C230034O21 gene [Source:M | 0 | 0 | 0 | 0.964 | 0 | 1.1138 |
| 27892 | ENSMUSG00000086184 | 0 | NA | NA | NA | NA | NA | Gm12764 | predicted gene 12764 [Source:MGI Symbo | 0 | 0 | 0 | 0 | 0 | 0 |
| 27893 | ENSMUSG00000086185 | 0 | NA | NA | NA | NA | NA | 1700036O09 | RIKEN cDNA 1700036O09 gene [Source:M | 0 | 0 | 0 | 0 | 0 | 0 |
| 27894 | ENSMUSG00000086186 | 0 | NA | NA | NA | NA | NA | Ccdc34os | coiled-coil domain containing 34, opposite | 0 | 0 | 0 | 0 | 0 | 0 |
| 27895 | ENSMUSG00000086187 | 0 | NA | NA | NA | NA | NA | Gm12860 | predicted gene 12860 [Source:MGI Symbo | 0 | 0 | 0 | 0 | 0 | 0 |
| 27896 | ENSMUSG00000086188 | 0 | NA | NA | NA | NA | NA | Gm15169 | predicted gene 15169 [Source:MGI Symbo | 0 | 0 | 0 | 0 | 0 | 0 |
| 27897 | ENSMUSG00000086189 | 0 | NA | NA | NA | NA | NA | Gm15462 | predicted gene 15462 [Source:MGI Symbo | 0 | 0 | 0 | 0 | 0 | 0 |
| 27898 | ENSMUSG00000086190 | 0 | NA | NA | NA | NA | NA | 4930579M01 | RIKEN cDNA 4930579M01 gene [Source:M | 0 | 0 | 0 | 0 | 0 | 0 |
| 27899 | ENSMUSG00000086191 | 1.534059199 | -0.925135918 | 1.961985055 | -0.471530563 | 0.637261899 | NA | Zfp652os | zinc finger protein 652, opposite strand [S | 0 | 1.0279 | 5.0312 | 0.964 | 2.1812 | 0 |
| 27900 | ENSMUSG00000086192 | 0.321325949 | 1.911575114 | 4.048267893 | 0.472195804 | 0.636787034 | NA | Gm13609 | predicted gene 13609 [Source:MGI Symbo | 0 | 0 | 0 | 1.928 | 0 | 0 |
| 27901 | ENSMUSG00000086193 | 0 | NA | NA | NA | NA | NA | Gm11508 | predicted gene 11508 [Source:MGI Symbo | 0 | 0 | 0 | 0 | 0 | 0 |
| 27902 | ENSMUSG00000086194 | 0 | NA | NA | NA | NA | NA | Gm16329 | predicted gene 16329 [Source:MGI Symbo | 0 | 0 | 0 | 0 | 0 | 0 |
| 27903 | ENSMUSG00000086195 | 0 | NA | NA | NA | NA | NA | Gm6152 | predicted gene 6152 [Source:MGI Symbol | 0 | 0 | 0 | 0 | 0 | 0 |
| 27904 | ENSMUSG00000086196 | 0 | NA | NA | NA | NA | NA | Gm13571 | predicted gene 13571 [Source:MGI Symbo | 0 | 0 | 0 | 0 | 0 | 0 |
| 27905 | ENSMUSG00000086197 | 0 | NA | NA | NA | NA | NA | Gm13976 | predicted gene 13976 [Source:MGI Symbo | 0 | 0 | 0 | 0 | 0 | 0 |
| 27906 | ENSMUSG00000086199 | 0.503095663 | 2.549494177 | 3.348715477 | 0.761334964 | 0.446457022 | NA | Bcas3os1 | breast carcinoma amplified sequence 3, o | 0 | 0 | 0 | 1.928 | 1.0906 | 0 |
| 27907 | ENSMUSG00000086201 | 0 | NA | NA | NA | NA | NA | Gm6270 | predicted gene 6270 [Source:MGI Symbol | 0 | 0 | 0 | 0 | 0 | 0 |
| 27908 | ENSMUSG00000086202 | 0 | NA | NA | NA | NA | NA | 1700019E08 | RIKEN cDNA 1700019E08 gene [Source:M | 0 | 0 | 0 | 0 | 0 | 0 |
| 27909 | ENSMUSG00000086203 | 0 | NA | NA | NA | NA | NA | Gm13985 | predicted gene 13985 [Source:MGI Symbo | 0 | 0 | 0 | 0 | 0 | 0 |
| 27910 | ENSMUSG00000086204 | 0 | NA | NA | NA | NA | NA | Gm15668 | predicted gene 15668 [Source:MGI Symbo | 0 | 0 | 0 | 0 | 0 | 0 |
| 27911 | ENSMUSG00000086205 | 0.729921525 | -1.61423457 | 2.706831497 | -0.596355766 | 0.550937584 | NA | Gm12679 | predicted gene 12679 [Source:MGI Symbo | 2.4093 | 0 | 1.0062 | 0.964 | 0 | 0 |
| 27912 | ENSMUSG00000086206 | 2.699568861 | -1.113930955 | 1.354665986 | -0.822291965 | 0.410910749 | NA | Gm13783 | predicted gene 13783 [Source:MGI Symbo | 4.0155 | 3.0838 | 4.025 | 2.8919 | 2.1812 | 0 |
| 27913 | ENSMUSG00000086207 | 0 | NA | NA | NA | NA | NA | Gm14717 | predicted gene 14717 [Source:MGI Symbo | 0 | 0 | 0 | 0 | 0 | 0 |
| 27914 | ENSMUSG00000086209 | 0 | NA | NA | NA | NA | NA | 4933405E24 | RIKEN cDNA 4933405E24 gene [Source:M | 0 | 0 | 0 | 0 | 0 | 0 |
| 27915 | ENSMUSG00000086211 | 0 | NA | NA | NA | NA | NA | Gm12462 | predicted gene 12462 [Source:MGI Symbo | 0 | 0 | 0 | 0 | 0 | 0 |
| 27916 | ENSMUSG00000086212 | 10.82851277 | 0.065741746 | 0.658852922 | 0.099782126 | 0.920517298 | 0.986685804 | Mkln1os | muskelin 1, intracellular mediator contain | 10.44 | 12.335 | 9.0562 | 14.46 | 11.997 | 6.6831 |
| 27917 | ENSMUSG00000086213 | 46.64326771 | 0.807192052 | 0.40201154 | 2.007882787 | 0.044655749 | 0.291866583 | A330040F15 | RIKEN cDNA A330040F15 gene [Source:M | 24.093 | 37.005 | 41.256 | 91.578 | 44.715 | 41.212 |
| 27918 | ENSMUSG00000086214 | 0 | NA | NA | NA | NA | NA | Gm14062 | predicted gene 14062 [Source:MGI Symbo | 0 | 0 | 0 | 0 | 0 | 0 |
| 27919 | ENSMUSG00000086216 | 1.895684937 | -0.114973682 | 1.82309441 | -0.063065128 | 0.949714643 | NA | Gm13610 | predicted gene 13610 [Source:MGI Symbo | 0.8031 | 4.1117 | 1.0062 | 0 | 5.4531 | 0 |
| 27920 | ENSMUSG00000086219 | 20.97445556 | 0.402969336 | 0.544568173 | 0.739979594 | 0.459312376 | 0.859442977 | Srrm4os | serine/arginine repetitive matrix 4, opposi | 27.305 | 19.53 | 7.0437 | 20.244 | 29.447 | 22.277 |
| 27921 | ENSMUSG00000086220 | 0 | NA | NA | NA | NA | NA | Gm16599 | predicted gene, 16599 [Source:MGI Symb | 0 | 0 | 0 | 0 | 0 | 0 |
| 27922 | ENSMUSG00000086221 | 0 | NA | NA | NA | NA | NA | 4930412B13 | RIKEN cDNA 4930412B13 gene [Source:M | 0 | 0 | 0 | 0 | 0 | 0 |
| 27923 | ENSMUSG00000086222 | 0 | NA | NA | NA | NA | NA | Gm11665 | predicted gene 11665 [Source:MGI Symbo | 0 | 0 | 0 | 0 | 0 | 0 |
| 27924 | ENSMUSG00000086224 | 3.690627006 | 0.498307839 | 1.156782284 | 0.430770635 | 0.666635153 | NA | 2700069I18R | RIKEN cDNA 2700069I18 gene [Source:MG | 4.0155 | 2.0558 | 3.0187 | 0.964 | 7.6343 | 4.4554 |
| 27925 | ENSMUSG00000086225 | 0 | NA | NA | NA | NA | NA | Gm8661 | predicted gene 8661 [Source:MGI Symbol | 0 | 0 | 0 | 0 | 0 | 0 |
| 27926 | ENSMUSG00000086226 | 0 | NA | NA | NA | NA | NA | Gm12660 | predicted gene 12660 [Source:MGI Symbo | 0 | 0 | 0 | 0 | 0 | 0 |
| 27927 | ENSMUSG00000086227 | 0 | NA | NA | NA | NA | NA | Gm11820 | predicted gene 11820 [Source:MGI Symbo | 0 | 0 | 0 | 0 | 0 | 0 |
| 27928 | ENSMUSG00000086228 | 1.766222944 | 0.132158881 | 1.788353031 | 0.073899772 | 0.941090138 | NA | Ubap1l | ubiquitin-associated protein 1-like [Source | 0 | 2.0558 | 3.0187 | 0 | 2.1812 | 3.3415 |
| 27929 | ENSMUSG00000086229 | 0.353348078 | 0.165142701 | 3.895042568 | 0.042398176 | 0.966181282 | NA | Gm4887 | predicted gene 4887 [Source:MGI Symbol | 0 | 0 | 1.0062 | 0 | 0 | 1.1138 |
| 27930 | ENSMUSG00000086231 | 48.95658623 | 0.839795427 | 0.572229447 | 1.46758513 | 0.14221695 | 0.551925399 | Rapgef4os3 | Rap guanine nucleotide exchange factor ( | 43.368 | 31.865 | 30.187 | 137.85 | 35.99 | 14.48 |
| 27931 | ENSMUSG00000086232 | 0 | NA | NA | NA | NA | NA | Gm7546 | predicted gene 7546 [Source:MGI Symbol | 0 | 0 | 0 | 0 | 0 | 0 |
| 27932 | ENSMUSG00000086233 | 0 | NA | NA | NA | NA | NA | Gm11816 | predicted gene 11816 [Source:MGI Symbo | 0 | 0 | 0 | 0 | 0 | 0 |
| 27933 | ENSMUSG00000086234 | 0 | NA | NA | NA | NA | NA | Usp46os2 | ubiquitin specific peptidase 46, opposite s | 0 | 0 | 0 | 0 | 0 | 0 |
| 27934 | ENSMUSG00000086235 | 0 | NA | NA | NA | NA | NA | Gm16284 | predicted gene 16284 [Source:MGI Symbo | 0 | 0 | 0 | 0 | 0 | 0 |
| 27935 | ENSMUSG00000086236 | 0 | NA | NA | NA | NA | NA | 5830418P13 | RIKEN cDNA 5830418P13 gene [Source:M | 0 | 0 | 0 | 0 | 0 | 0 |
| 27936 | ENSMUSG00000086237 | 1.316570868 | 0.208290484 | 1.749968478 | 0.119025277 | 0.90525533 | NA | Gm15591 | predicted gene 15591 [Source:MGI Symbo | 1.6062 | 1.0279 | 1.0062 | 0.964 | 2.1812 | 1.1138 |
| 27937 | ENSMUSG00000086238 | 0 | NA | NA | NA | NA | NA | Gm14258 | predicted gene 14258 [Source:MGI Symbo | 0 | 0 | 0 | 0 | 0 | 0 |
| 27938 | ENSMUSG00000086239 | 0 | NA | NA | NA | NA | NA | Gm14329 | predicted gene 14329 [Source:MGI Symbo | 0 | 0 | 0 | 0 | 0 | 0 |
| 27939 | ENSMUSG00000086240 | 8.103724538 | -0.096052048 | 0.825256059 | -0.1163906 | 0.907342985 | 0.984526042 | Gm7846 | predicted gene 7846 [Source:MGI Symbol | 8.8341 | 5.1396 | 11.069 | 3.8559 | 15.269 | 4.4554 |
| 27940 | ENSMUSG00000086241 | 0.51014012 | 1.031855138 | 3.09002894 | 0.333930574 | 0.738431948 | NA | 4930483K19 | RIKEN cDNA 4930483K19 gene [Source:M | 0 | 0 | 1.0062 | 0.964 | 1.0906 | 0 |

| | | | | | | | | | | | | | | | |
|---|---|---|---|---|---|---|---|---|---|---|---|---|---|---|---|
| 27941 | ENSMUSG00000086243 | 0.181769715 | 1.126924175 | 4.080472857 | 0.276174898 | 0.782413736 | NA | Gm12121 | predicted gene 12121 [Source:MGI Symbo | 0 | 0 | 0 | 0 | 1.0906 | 0 |
| 27942 | ENSMUSG00000086244 | 0 | NA | NA | NA | NA | NA | 4930558G05 | RIKEN cDNA 4930558G05 gene [Source:M | 0 | 0 | 0 | 0 | 0 | 0 |
| 27943 | ENSMUSG00000086245 | 40.92988015 | -0.051990316 | 0.389381703 | -0.133520182 | 0.893782004 | 0.982562372 | Gm16170 | predicted gene 16170 [Source:MGI Symbo | 49.792 | 41.117 | 34.212 | 58.803 | 33.809 | 27.846 |
| 27944 | ENSMUSG00000086246 | 0 | NA | NA | NA | NA | NA | Gm15274 | predicted gene 15274 [Source:MGI Symbo | 0 | 0 | 0 | 0 | 0 | 0 |
| 27945 | ENSMUSG00000086247 | 14.59932495 | -0.352813533 | 0.561974115 | -0.627811003 | 0.530127758 | 0.888501493 | Gm15787 | predicted gene 15787 [Source:MGI Symbo | 17.668 | 15.419 | 16.1 | 16.388 | 11.997 | 10.025 |
| 27946 | ENSMUSG00000086248 | 0 | NA | NA | NA | NA | NA | Gm16127 | predicted gene 16127 [Source:MGI Symbo | 0 | 0 | 0 | 0 | 0 | 0 |
| 27947 | ENSMUSG00000086249 | 0.160662974 | 1.126924175 | 4.080472857 | 0.276174898 | 0.782413736 | NA | Gm12724 | predicted gene 12724 [Source:MGI Symbo | 0 | 0 | 0 | 0.964 | 0 | 0 |
| 27948 | ENSMUSG00000086250 | 0 | NA | NA | NA | NA | NA | Gm14095 | predicted gene 14095 [Source:MGI Symbo | 0 | 0 | 0 | 0 | 0 | 0 |
| 27949 | ENSMUSG00000086251 | 0.685278559 | -2.809870862 | 3.376578027 | -0.832165239 | 0.405315682 | NA | Gm12694 | predicted gene 12694 [Source:MGI Symbo | 0 | 4.1117 | 0 | 0 | 0 | 0 |
| 27950 | ENSMUSG00000086253 | 6.094876387 | 1.514994473 | 0.998327264 | 1.517532905 | 0.129132192 | 0.525983675 | Gm13773 | predicted gene 13773 [Source:MGI Symbo | 2.4093 | 1.0279 | 6.0375 | 2.8919 | 14.178 | 10.025 |
| 27951 | ENSMUSG00000086254 | 0 | NA | NA | NA | NA | NA | 3100003L05I | RIKEN cDNA 3100003L05 gene [Source:M( | 0 | 0 | 0 | 0 | 0 | 0 |
| 27952 | ENSMUSG00000086255 | 0 | NA | NA | NA | NA | NA | Gm11534 | predicted gene 11534 [Source:MGI Symbo | 0 | 0 | 0 | 0 | 0 | 0 |
| 27953 | ENSMUSG00000086256 | 1.411587875 | -2.726783388 | 1.993845459 | -1.36760017 | 0.171437263 | NA | Gm12052 | predicted gene 12052 [Source:MGI Symbo | 2.4093 | 3.0838 | 2.0125 | 0.964 | 0 | 0 |
| 27954 | ENSMUSG00000086257 | 0 | NA | NA | NA | NA | NA | 4930548K13 | RIKEN cDNA 4930548K13 gene [Source:M | 0 | 0 | 0 | 0 | 0 | 0 |
| 27955 | ENSMUSG00000086258 | 0 | NA | NA | NA | NA | NA | Gm15239 | predicted gene 15239 [Source:MGI Symbo | 0 | 0 | 0 | 0 | 0 | 0 |
| 27956 | ENSMUSG00000086259 | 0 | NA | NA | NA | NA | NA | Tmem150co | transmembrane protein 150C, opposite st | 0 | 0 | 0 | 0 | 0 | 0 |
| 27957 | ENSMUSG00000086260 | 0 | NA | NA | NA | NA | NA | Gm16259 | predicted gene 16259 [Source:MGI Symbo | 0 | 0 | 0 | 0 | 0 | 0 |
| 27958 | ENSMUSG00000086261 | 0 | NA | NA | NA | NA | NA | Gm3765 | predicted gene 3765 [Source:MGI Symbol | 0 | 0 | 0 | 0 | 0 | 0 |
| 27959 | ENSMUSG00000086262 | 0 | NA | NA | NA | NA | NA | A930031H19 | RIKEN cDNA A930031H19 gene [Source:M | 0 | 0 | 0 | 0 | 0 | 0 |
| 27960 | ENSMUSG00000086263 | 0 | NA | NA | NA | NA | NA | 4930519D14 | RIKEN cDNA 4930519D14 gene [Source:M | 0 | 0 | 0 | 0 | 0 | 0 |
| 27961 | ENSMUSG00000086264 | 0 | NA | NA | NA | NA | NA | Gm15850 | predicted gene 15850 [Source:MGI Symbo | 0 | 0 | 0 | 0 | 0 | 0 |
| 27962 | ENSMUSG00000086265 | 0 | NA | NA | NA | NA | NA | Marcksl1-ps | MARCKS-like 1, pseudogene 4 [Source:MC | 0 | 0 | 0 | 0 | 0 | 0 |
| 27963 | ENSMUSG00000086266 | 0 | NA | NA | NA | NA | NA | Igf2os | insulin-like growth factor 2, opposite strar | 0 | 0 | 0 | 0 | 0 | 0 |
| 27964 | ENSMUSG00000086267 | 0 | NA | NA | NA | NA | NA | Gm12496 | predicted gene 12496 [Source:MGI Symbo | 0 | 0 | 0 | 0 | 0 | 0 |
| 27965 | ENSMUSG00000086268 | 0.810359328 | 1.950931649 | 2.89643804 | 0.673562363 | 0.500589588 | NA | Gm11670 | predicted gene 11670 [Source:MGI Symbo | 0 | 0 | 1.0062 | 3.8559 | 0 | 0 |
| 27966 | ENSMUSG00000086269 | 0.856727441 | -0.738450537 | 2.821005776 | -0.26176853 | 0.793499902 | NA | Tmcc3os | transmembrane and coiled coil domains 3 | 3.2124 | 0 | 0 | 1.928 | 0 | 0 |
| 27967 | ENSMUSG00000086272 | 0 | NA | NA | NA | NA | NA | BC039966 | cDNA sequence BC039966 [Source:MGI Sy | 0 | 0 | 0 | 0 | 0 | 0 |
| 27968 | ENSMUSG00000086273 | 0 | NA | NA | NA | NA | NA | Rbm46os | RNA binding motif protein 46, opposite st | 0 | 0 | 0 | 0 | 0 | 0 |
| 27969 | ENSMUSG00000086275 | 0 | NA | NA | NA | NA | NA | 1700121C08 | RIKEN cDNA 1700121C08 gene [Source:M | 0 | 0 | 0 | 0 | 0 | 0 |
| 27970 | ENSMUSG00000086277 | 0.695987358 | -1.412518496 | 2.768247659 | -0.510257271 | 0.609871234 | NA | 4930558K02 | RIKEN cDNA 4930558K02 gene [Source:M | 0 | 2.0558 | 1.0062 | 0 | 0 | 1.1138 |
| 27971 | ENSMUSG00000086278 | 0 | NA | NA | NA | NA | NA | Olfr75-ps1 | olfactory receptor 75, pseudogene 1 [Sou | 0 | 0 | 0 | 0 | 0 | 0 |
| 27972 | ENSMUSG00000086279 | 5.492696154 | 0.433398715 | 1.06055831 | 0.408651472 | 0.682795451 | 0.935166147 | Gm15634 | predicted gene 15634 [Source:MGI Symbo | 8.8341 | 0 | 5.0312 | 4.8199 | 9.8156 | 4.4554 |
| 27973 | ENSMUSG00000086282 | 3.347191025 | -1.491929709 | 1.279378773 | -1.16613605 | 0.243559443 | NA | Sox5os4 | SRY (sex determining region Y)-box 5, opp | 9.6372 | 3.0838 | 2.0125 | 0.964 | 3.2719 | 1.1138 |
| 27974 | ENSMUSG00000086283 | 0.133850373 | -0.796629176 | 4.080472857 | -0.195229623 | 0.84521319 | NA | 2810433D01 | RIKEN cDNA 2810433D01 gene [Source:M | 0.8031 | 0 | 0 | 0 | 0 | 0 |
| 27975 | ENSMUSG00000086284 | 0 | NA | NA | NA | NA | NA | Frmpd1os | FERM and PDZ domain containing 1, oppc | 0 | 0 | 0 | 0 | 0 | 0 |
| 27976 | ENSMUSG00000086286 | 0 | NA | NA | NA | NA | NA | Gm15138 | predicted gene 15138 [Source:MGI Symbo | 0 | 0 | 0 | 0 | 0 | 0 |
| 27977 | ENSMUSG00000086287 | 0.838537155 | -3.104405377 | 3.144633814 | -0.987207275 | 0.323541036 | NA | Gm15972 | predicted gene 15972 [Source:MGI Symbo | 0 | 0 | 5.0312 | 0 | 0 | 0 |
| 27978 | ENSMUSG00000086288 | 12.36315988 | 0.588967558 | 0.658221688 | 0.89478601 | 0.37090149 | 0.807214774 | Gm15265 | predicted gene 15265 [Source:MGI Symbo | 15.259 | 4.1117 | 10.062 | 18.316 | 14.178 | 12.252 |
| 27979 | ENSMUSG00000086289 | 0 | NA | NA | NA | NA | NA | Gm16933 | predicted gene, 16933 [Source:MGI Symb | 0 | 0 | 0 | 0 | 0 | 0 |
| 27980 | ENSMUSG00000086290 | 146.8267015 | 0.201855243 | 0.208069149 | 0.970135382 | 0.331979015 | 0.774849842 | Snhg12 | small nucleolar RNA host gene 12 [Source | 144.56 | 129.52 | 135.84 | 188.94 | 142.87 | 139.23 |
| 27981 | ENSMUSG00000086291 | 0.17131964 | -0.796629176 | 4.080472857 | -0.195229623 | 0.84521319 | NA | Gm15513 | predicted gene 15513 [Source:MGI Symbo | 0 | 1.0279 | 0 | 0 | 0 | 0 |
| 27982 | ENSMUSG00000086293 | 0 | NA | NA | NA | NA | NA | Gm11269 | predicted gene 11269 [Source:MGI Symbo | 0 | 0 | 0 | 0 | 0 | 0 |
| 27983 | ENSMUSG00000086296 | 37.69649269 | -0.453476078 | 0.442797997 | -1.024115018 | 0.305780952 | 0.755004385 | D030055H07 | RIKEN cDNA D030055H07 gene [Source:M | 54.611 | 37.005 | 39.244 | 51.091 | 15.269 | 28.96 |
| 27984 | ENSMUSG00000086297 | 0 | NA | NA | NA | NA | NA | Gm15632 | predicted gene 15632 [Source:MGI Symbo | 0 | 0 | 0 | 0 | 0 | 0 |
| 27985 | ENSMUSG00000086298 | 0 | NA | NA | NA | NA | NA | Gm11716 | predicted gene 11716 [Source:MGI Symbo | 0 | 0 | 0 | 0 | 0 | 0 |
| 27986 | ENSMUSG00000086302 | 0.160662974 | 1.126924175 | 4.080472857 | 0.276174898 | 0.782413736 | NA | Gm13790 | predicted gene 13790 [Source:MGI Symbo | 0 | 0 | 0 | 0.964 | 0 | 0 |
| 27987 | ENSMUSG00000086306 | 0.167707431 | -0.796629176 | 4.080472857 | -0.195229623 | 0.84521319 | NA | Gm11754 | predicted gene 11754 [Source:MGI Symbo | 0 | 0 | 1.0062 | 0 | 0 | 0 |
| 27988 | ENSMUSG00000086308 | 1.309294317 | -1.39726663 | 2.007141872 | -0.696147417 | 0.486336514 | NA | G630016G05 | RIKEN cDNA G630016G05 gene [Source:M | 1.6062 | 2.0558 | 2.0125 | 0 | 2.1812 | 0 |
| 27989 | ENSMUSG00000086310 | 0 | NA | NA | NA | NA | NA | Gm12278 | predicted gene 12278 [Source:MGI Symbo | 0 | 0 | 0 | 0 | 0 | 0 |
| 27990 | ENSMUSG00000086311 | 0 | NA | NA | NA | NA | NA | Gm12122 | predicted gene 12122 [Source:MGI Symbo | 0 | 0 | 0 | 0 | 0 | 0 |
| 27991 | ENSMUSG00000086312 | 0.17131964 | -0.796629176 | 4.080472857 | -0.195229623 | 0.84521319 | NA | Gm15336 | predicted gene 15336 [Source:MGI Symbo | 0 | 1.0279 | 0 | 0 | 0 | 0 |
| 27992 | ENSMUSG00000086313 | 0 | NA | NA | NA | NA | NA | Gm15940 | predicted gene 15940 [Source:MGI Symbo | 0 | 0 | 0 | 0 | 0 | 0 |
| 27993 | ENSMUSG00000086315 | 0.181769715 | 1.126924175 | 4.080472857 | 0.276174898 | 0.782413736 | NA | Arl4aos | ADP-ribosylation factor-like 4A, opposite s | 0 | 0 | 0 | 0 | 1.0906 | 0 |
| 27994 | ENSMUSG00000086316 | 14.13452 | -0.198476132 | 0.686911841 | -0.288939745 | 0.772627488 | 0.957772025 | Nbdy | negative regulator of P-body association [ | 16.865 | 14.391 | 14.087 | 15.424 | 21.812 | 2.2277 |
| 27995 | ENSMUSG00000086319 | 0 | NA | NA | NA | NA | NA | Gm16087 | predicted gene 16087 [Source:MGI Symbo | 0 | 0 | 0 | 0 | 0 | 0 |
| 27996 | ENSMUSG00000086320 | 1.448741299 | 1.727639027 | 2.153019475 | 0.802426103 | 0.422306519 | NA | Gm12840 | predicted gene 12840 [Source:MGI Symbo | 0 | 2.0558 | 0 | 0 | 2.1812 | 4.4554 |

| | | | | | | | | | | | | | | | |
|---|---|---|---|---|---|---|---|---|---|---|---|---|---|---|---|
| 27997 | ENSMUSG00000086321 | 0 | NA | NA | NA | NA | NA | Gm11413 | predicted gene 11413 [Source:MGI Symbo | 0 | 0 | 0 | 0 | 0 | 0 |
| 27998 | ENSMUSG00000086322 | 0 | NA | NA | NA | NA | NA | E130218I03F | RIKEN cDNA E130218I03 gene [Source:MG | 0 | 0 | 0 | 0 | 0 | 0 |
| 27999 | ENSMUSG00000086324 | 18.16500989 | 0.873452716 | 0.621240099 | 1.405982513 | 0.159729318 | 0.581652782 | Gm15564 | predicted gene 15564 [Source:MGI Symbo | 8.031 | 19.53 | 11.069 | 14.46 | 42.534 | 13.366 |
| 28000 | ENSMUSG00000086326 | 0 | NA | NA | NA | NA | NA | Gm13200 | predicted gene 13200 [Source:MGI Symbo | 0 | 0 | 0 | 0 | 0 | 0 |
| 28001 | ENSMUSG00000086327 | 0 | NA | NA | NA | NA | NA | Slfn5os | schlafen 5, opposite strand [Source:MGI S | 0 | 0 | 0 | 0 | 0 | 0 |
| 28002 | ENSMUSG00000086328 | 4.987692313 | 0.13571358 | 1.087275186 | 0.1248199 | 0.900666133 | NA | 2700033N17 | RIKEN cDNA 2700033N17 gene [Source:M | 8.031 | 5.1396 | 1.0062 | 4.8199 | 9.8156 | 1.1138 |
| 28003 | ENSMUSG00000086329 | 0 | NA | NA | NA | NA | NA | Gm14233 | predicted gene 14233 [Source:MGI Symbo | 0 | 0 | 0 | 0 | 0 | 0 |
| 28004 | ENSMUSG00000086330 | 3.115854169 | -2.990841933 | 1.48453779 | -2.01466204 | 0.043940072 | NA | 1700007J10I | RIKEN cDNA 1700007J10 gene [Source:MG | 6.4248 | 6.1675 | 4.025 | 0.964 | 0 | 1.1138 |
| 28005 | ENSMUSG00000086331 | 1.381395159 | -0.788978462 | 1.843451663 | -0.427989775 | 0.668658565 | NA | Gm16310 | predicted gene 16310 [Source:MGI Symbo | 3.2124 | 1.0279 | 1.0062 | 1.928 | 0 | 1.1138 |
| 28006 | ENSMUSG00000086332 | 66.55361269 | -1.122667474 | 0.316427914 | -3.547940704 | 0.000388256 | 0.010477107 | 4930480G23 | RIKEN cDNA 4930480G23 gene [Source:M | 85.932 | 98.68 | 89.556 | 55.911 | 43.625 | 25.618 |
| 28007 | ENSMUSG00000086333 | 0 | NA | NA | NA | NA | NA | 1700120E14 | RIKEN cDNA 1700120E14 gene [Source:M | 0 | 0 | 0 | 0 | 0 | 0 |
| 28008 | ENSMUSG00000086335 | 0.524202404 | 2.600864256 | 3.301637836 | 0.787749713 | 0.430843121 | NA | Gm12107 | predicted gene 12107 [Source:MGI Symbo | 0 | 0 | 0 | 0.964 | 2.1812 | 0 |
| 28009 | ENSMUSG00000086336 | 0 | NA | NA | NA | NA | NA | Gm12979 | predicted gene 12979 [Source:MGI Symbo | 0 | 0 | 0 | 0 | 0 | 0 |
| 28010 | ENSMUSG00000086337 | 0 | NA | NA | NA | NA | NA | Gm11535 | predicted gene 11535 [Source:MGI Symbo | 0 | 0 | 0 | 0 | 0 | 0 |
| 28011 | ENSMUSG00000086338 | 0 | NA | NA | NA | NA | NA | Olfr1096 | olfactory receptor 1096 [Source:MGI Sym | 0 | 0 | 0 | 0 | 0 | 0 |
| 28012 | ENSMUSG00000086339 | 0 | NA | NA | NA | NA | NA | Gm11916 | predicted gene 11916 [Source:MGI Symbo | 0 | 0 | 0 | 0 | 0 | 0 |
| 28013 | ENSMUSG00000086340 | 2.913824894 | 0.882505493 | 1.263652835 | 0.698376538 | 0.484941745 | NA | 1810059C17 | RIKEN cDNA 1810059C17 gene [Source:M | 4.0155 | 1.0279 | 1.0062 | 4.8199 | 3.2719 | 3.3415 |
| 28014 | ENSMUSG00000086341 | 9.293163258 | 0.06131328 | 0.758924018 | 0.080789748 | 0.935609161 | 0.988420305 | Gm15932 | predicted gene 15932 [Source:MGI Symbo | 4.0155 | 14.391 | 9.0562 | 9.6398 | 13.087 | 5.5692 |
| 28015 | ENSMUSG00000086342 | 1.328855871 | 1.755398778 | 2.048295649 | 0.857004593 | 0.391442346 | NA | Gm12932 | predicted gene 12932 [Source:MGI Symbo | 0.8031 | 0 | 1.0062 | 2.8919 | 3.2719 | 0 |
| 28016 | ENSMUSG00000086346 | 0 | NA | NA | NA | NA | NA | Gm12510 | predicted gene 12510 [Source:MGI Symbo | 0 | 0 | 0 | 0 | 0 | 0 |
| 28017 | ENSMUSG00000086347 | 0.133850373 | -0.796629176 | 4.080472857 | -0.195229623 | 0.84521319 | NA | Gm11626 | predicted gene 11626 [Source:MGI Symbo | 0.8031 | 0 | 0 | 0 | 0 | 0 |
| 28018 | ENSMUSG00000086348 | 0 | NA | NA | NA | NA | NA | 1700017J07I | RIKEN cDNA 1700017J07 gene [Source:MG | 0 | 0 | 0 | 0 | 0 | 0 |
| 28019 | ENSMUSG00000086349 | 0 | NA | NA | NA | NA | NA | Gm13497 | predicted gene 13497 [Source:MGI Symbo | 0 | 0 | 0 | 0 | 0 | 0 |
| 28020 | ENSMUSG00000086350 | 22.24245504 | 0.420244139 | 0.47453828 | 0.885585328 | 0.37584102 | 0.809690069 | B230369F24 | RIKEN cDNA B230369F24 gene [Source:M | 18.471 | 21.586 | 17.106 | 27.955 | 31.628 | 16.708 |
| 28021 | ENSMUSG00000086352 | 0 | NA | NA | NA | NA | NA | 4930401G09 | RIKEN cDNA 4930401G09 gene [Source:M | 0 | 0 | 0 | 0 | 0 | 0 |
| 28022 | ENSMUSG00000086353 | 0.74256259 | 3.09446134 | 3.273353887 | 0.945348852 | 0.344480805 | NA | Gm13481 | predicted gene 13481 [Source:MGI Symbo | 0 | 0 | 0 | 0 | 0 | 4.4554 |
| 28023 | ENSMUSG00000086354 | 0 | NA | NA | NA | NA | NA | Gm13938 | predicted gene 13938 [Source:MGI Symbo | 0 | 0 | 0 | 0 | 0 | 0 |
| 28024 | ENSMUSG00000086355 | 0 | NA | NA | NA | NA | NA | 4933428C19 | RIKEN cDNA 4933428C19 gene [Source:M | 0 | 0 | 0 | 0 | 0 | 0 |
| 28025 | ENSMUSG00000086356 | 0 | NA | NA | NA | NA | NA | Gm13441 | predicted gene 13441 [Source:MGI Symbo | 0 | 0 | 0 | 0 | 0 | 0 |
| 28026 | ENSMUSG00000086358 | 4.239947052 | -0.28341145 | 1.015757202 | -0.279014955 | 0.780233348 | NA | Gm13270 | predicted gene 13270 [Source:MGI Symbo | 4.8186 | 5.1396 | 4.025 | 4.8199 | 2.1812 | 4.4554 |
| 28027 | ENSMUSG00000086359 | 4.181938972 | 0.371129069 | 1.37912194 | 0.269105333 | 0.787848627 | NA | 9630013K17 | RIKEN cDNA 9630013K17 gene [Source:M | 8.8341 | 0 | 2.0125 | 8.6758 | 0 | 5.5692 |
| 28028 | ENSMUSG00000086360 | 0.160662974 | 1.126924175 | 4.080472857 | 0.276174898 | 0.782413736 | NA | Gm16214 | predicted gene 16214 [Source:MGI Symbo | 0 | 0 | 0 | 0.964 | 0 | 0 |
| 28029 | ENSMUSG00000086361 | 0 | NA | NA | NA | NA | NA | Gm6569 | predicted gene 6569 [Source:MGI Symbol | 0 | 0 | 0 | 0 | 0 | 0 |
| 28030 | ENSMUSG00000086363 | 0 | NA | NA | NA | NA | NA | A330102I10I | RIKEN cDNA A330102I10 gene [Source:MG | 0 | 0 | 0 | 0 | 0 | 0 |
| 28031 | ENSMUSG00000086364 | 0 | NA | NA | NA | NA | NA | Gm11751 | predicted gene 11751 [Source:MGI Symbo | 0 | 0 | 0 | 0 | 0 | 0 |
| 28032 | ENSMUSG00000086365 | 0 | NA | NA | NA | NA | NA | 4930593C16 | RIKEN cDNA 4930593C16 gene [Source:M | 0 | 0 | 0 | 0 | 0 | 0 |
| 28033 | ENSMUSG00000086366 | 0.133850373 | -0.796629176 | 4.080472857 | -0.195229623 | 0.84521319 | NA | 1700061J23I | RIKEN cDNA 1700061J23 gene [Source:MG | 0.8031 | 0 | 0 | 0 | 0 | 0 |
| 28034 | ENSMUSG00000086367 | 0 | NA | NA | NA | NA | NA | Gm6182 | predicted gene 6182 [Source:MGI Symbol | 0 | 0 | 0 | 0 | 0 | 0 |
| 28035 | ENSMUSG00000086368 | 0 | NA | NA | NA | NA | NA | Gm13830 | predicted gene 13830 [Source:MGI Symbo | 0 | 0 | 0 | 0 | 0 | 0 |
| 28036 | ENSMUSG00000086369 | 0.663758638 | 2.950973782 | 3.025092005 | 0.975498853 | 0.32931287 | NA | E330017L17I | RIKEN cDNA E330017L17 gene [Source:MG | 0 | 0 | 0 | 2.8919 | 1.0906 | 0 |
| 28037 | ENSMUSG00000086370 | 50.78533233 | 0.370225359 | 0.345426845 | 1.071790928 | 0.283813949 | 0.736651636 | Ftx | Ftx transcript, Xist regulator (non-protein | 32.927 | 38.033 | 62.387 | 65.55 | 52.35 | 53.465 |
| 28038 | ENSMUSG00000086371 | 0 | NA | NA | NA | NA | NA | Gm12478 | predicted gene 12478 [Source:MGI Symbo | 0 | 0 | 0 | 0 | 0 | 0 |
| 28039 | ENSMUSG00000086372 | 0.822586684 | 2.037574431 | 2.401368532 | 0.84850551 | 0.396156505 | NA | Gm16246 | predicted gene 16246 [Source:MGI Symbo | 0.8031 | 0 | 0 | 1.928 | 1.0906 | 1.1138 |
| 28040 | ENSMUSG00000086373 | 0 | NA | NA | NA | NA | NA | Gm11978 | predicted gene 11978 [Source:MGI Symbo | 0 | 0 | 0 | 0 | 0 | 0 |
| 28041 | ENSMUSG00000086375 | 0 | NA | NA | NA | NA | NA | Gm12127 | predicted gene 12127 [Source:MGI Symbo | 0 | 0 | 0 | 0 | 0 | 0 |
| 28042 | ENSMUSG00000086376 | 0 | NA | NA | NA | NA | NA | 1700071G01 | RIKEN cDNA 1700071G01 gene [Source:M | 0 | 0 | 0 | 0 | 0 | 0 |
| 28043 | ENSMUSG00000086377 | 0 | NA | NA | NA | NA | NA | 4930529C04 | RIKEN cDNA 4930529C04 gene [Source:M | 0 | 0 | 0 | 0 | 0 | 0 |
| 28044 | ENSMUSG00000086379 | 0 | NA | NA | NA | NA | NA | 1700026D11 | RIKEN cDNA 1700026D11 gene [Source:M | 0 | 0 | 0 | 0 | 0 | 0 |
| 28045 | ENSMUSG00000086380 | 0 | NA | NA | NA | NA | NA | Gm11195 | predicted gene 11195 [Source:MGI Symbo | 0 | 0 | 0 | 0 | 0 | 0 |
| 28046 | ENSMUSG00000086381 | 0 | NA | NA | NA | NA | NA | AV064505 | expressed sequence AV064505 [Source:M | 0 | 0 | 0 | 0 | 0 | 0 |
| 28047 | ENSMUSG00000086382 | 49.41736114 | 0.165544374 | 0.317821229 | 0.520872615 | 0.602455514 | 0.915541976 | Chrna1os | cholinergic receptor, nicotinic, alpha poly | 44.171 | 52.424 | 43.269 | 53.983 | 43.625 | 59.034 |
| 28048 | ENSMUSG00000086384 | 0 | NA | NA | NA | NA | NA | Banf2os | barrier to autointegration factor 2, opposi | 0 | 0 | 0 | 0 | 0 | 0 |
| 28049 | ENSMUSG00000086385 | 0.669251866 | -2.807208097 | 3.391018549 | -0.827836255 | 0.407763239 | NA | Gm12701 | predicted gene 12701 [Source:MGI Symbo | 4.0155 | 0 | 0 | 0 | 0 | 0 |
| 28050 | ENSMUSG00000086386 | 0 | NA | NA | NA | NA | NA | Gm12108 | predicted gene 12108 [Source:MGI Symbo | 0 | 0 | 0 | 0 | 0 | 0 |
| 28051 | ENSMUSG00000086387 | 2.153742797 | 1.270474454 | 1.491784348 | 0.851647529 | 0.394409751 | NA | Gm15563 | predicted gene 15563 [Source:MGI Symbo | 0.8031 | 1.0279 | 2.0125 | 5.7839 | 2.1812 | 1.1138 |
| 28052 | ENSMUSG00000086389 | 6.40983387 | 1.37711896 | 0.941879975 | 1.462096018 | 0.1437149 | 0.55387485 | Gm15998 | predicted gene 15998 [Source:MGI Symbo | 1.6062 | 7.1954 | 2.0125 | 7.7118 | 5.4531 | 14.48 |

| | | | | | | | | | | | | | | | |
|---|---|---|---|---|---|---|---|---|---|---|---|---|---|---|---|
| 28053 | ENSMUSG00000086390 | 603.0385811 | 0.178843799 | 0.201083583 | 0.889400304 | 0.373787982 | 0.808708253 | 1810019D21 | RIKEN cDNA 1810019D21 gene [Source:M | 541.29 | 443.03 | 713.43 | 780.82 | 670.73 | 468.93 |
| 28054 | ENSMUSG00000086391 | 0.747854741 | -0.257859511 | 2.698313412 | -0.095563217 | 0.92386748 | NA | 1700042O10 | RIKEN cDNA 1700042O10 gene [Source:M | 2.4093 | 0 | 0 | 0.964 | 0 | 1.1138 |
| 28055 | ENSMUSG00000086392 | 3.365859835 | 0.246011575 | 1.239355702 | 0.198499571 | 0.842654221 | NA | Mccc1os | methylcrotonoyl-Coenzyme A carboxylase | 3.2124 | 3.0838 | 3.0187 | 8.6758 | 1.0906 | 1.1138 |
| 28056 | ENSMUSG00000086395 | 0.958298615 | -0.940364986 | 2.473304036 | -0.380205981 | 0.70379252 | NA | A630014C17 | RIKEN cDNA A630014C17 gene [Source:M | 0.8031 | 0 | 3.0187 | 1.928 | 0 | 0 |
| 28057 | ENSMUSG00000086396 | 0 | NA | NA | NA | NA | NA | 4930555B11 | RIKEN cDNA 4930555B11 gene [Source:M | 0 | 0 | 0 | 0 | 0 | 0 |
| 28058 | ENSMUSG00000086399 | 0 | NA | NA | NA | NA | NA | Gm16144 | predicted gene 16144 [Source:MGI Symbo | 0 | 0 | 0 | 0 | 0 | 0 |
| 28059 | ENSMUSG00000086401 | 3.060011876 | -0.354040415 | 1.37460473 | -0.257557978 | 0.796748064 | NA | Gm15559 | predicted gene 15559 [Source:MGI Symbo | 7.2279 | 1.0279 | 2.0125 | 4.8199 | 3.2719 | 0 |
| 28060 | ENSMUSG00000086402 | 0 | NA | NA | NA | NA | NA | Gm15322 | predicted gene 15322 [Source:MGI Symbo | 0 | 0 | 0 | 0 | 0 | 0 |
| 28061 | ENSMUSG00000086404 | 3.043002887 | -0.478374584 | 1.437299837 | -0.332828664 | 0.73926362 | NA | Gm6787 | predicted gene 6787 [Source:MGI Symbol | 2.4093 | 7.1954 | 1.0062 | 0.964 | 0 | 6.6831 |
| 28062 | ENSMUSG00000086405 | 0.342639279 | -1.805976596 | 4.028547714 | -0.448294702 | 0.653940524 | NA | 9330198N18 | RIKEN cDNA 9330198N18 gene [Source:M | 0 | 2.0558 | 0 | 0 | 0 | 0 |
| 28063 | ENSMUSG00000086407 | 0.181769715 | 1.126924175 | 4.080472857 | 0.276174898 | 0.782413736 | NA | Gm14123 | predicted gene 14123 [Source:MGI Symbo | 0 | 0 | 0 | 0 | 1.0906 | 0 |
| 28064 | ENSMUSG00000086408 | 0 | NA | NA | NA | NA | NA | Gm11588 | predicted gene 11588 [Source:MGI Symbo | 0 | 0 | 0 | 0 | 0 | 0 |
| 28065 | ENSMUSG00000086409 | 0 | NA | NA | NA | NA | NA | Gm13898 | predicted gene 13898 [Source:MGI Symbo | 0 | 0 | 0 | 0 | 0 | 0 |
| 28066 | ENSMUSG00000086410 | 8.687924396 | -0.031148487 | 0.76597989 | -0.040664888 | 0.967563053 | 0.994231936 | Gm16158 | predicted gene 16158 [Source:MGI Symbo | 8.031 | 6.1675 | 12.075 | 3.8559 | 13.087 | 8.9108 |
| 28067 | ENSMUSG00000086411 | 0.185640647 | 1.126924175 | 4.080472857 | 0.276174898 | 0.782413736 | NA | Gm14236 | predicted gene 14236 [Source:MGI Symbo | 0 | 0 | 0 | 0 | 0 | 1.1138 |
| 28068 | ENSMUSG00000086413 | 0 | NA | NA | NA | NA | NA | Gm12415 | predicted gene 12415 [Source:MGI Symbo | 0 | 0 | 0 | 0 | 0 | 0 |
| 28069 | ENSMUSG00000086414 | 0 | NA | NA | NA | NA | NA | Gm15645 | predicted gene 15645 [Source:MGI Symbo | 0 | 0 | 0 | 0 | 0 | 0 |
| 28070 | ENSMUSG00000086415 | 0 | NA | NA | NA | NA | NA | Gm4487 | predicted gene 4487 [Source:MGI Symbol | 0 | 0 | 0 | 0 | 0 | 0 |
| 28071 | ENSMUSG00000086416 | 0 | NA | NA | NA | NA | NA | Gm14002 | predicted gene 14002 [Source:MGI Symbo | 0 | 0 | 0 | 0 | 0 | 0 |
| 28072 | ENSMUSG00000086417 | 4.421510176 | 0.287328642 | 0.985175001 | 0.291652388 | 0.770552416 | NA | Gm12996 | predicted gene 12996 [Source:MGI Symbo | 4.8186 | 3.0838 | 4.025 | 5.7839 | 4.3625 | 4.4554 |
| 28073 | ENSMUSG00000086418 | 0 | NA | NA | NA | NA | NA | Gm13362 | predicted gene 13362 [Source:MGI Symbo | 0 | 0 | 0 | 0 | 0 | 0 |
| 28074 | ENSMUSG00000086420 | 0 | NA | NA | NA | NA | NA | Gm8865 | predicted gene 8865 [Source:MGI Symbol | 0 | 0 | 0 | 0 | 0 | 0 |
| 28075 | ENSMUSG00000086421 | 0.133850373 | -0.796629176 | 4.080472857 | -0.195229623 | 0.84521319 | NA | Gm14091 | predicted gene 14091 [Source:MGI Symbo | 0.8031 | 0 | 0 | 0 | 0 | 0 |
| 28076 | ENSMUSG00000086422 | 0 | NA | NA | NA | NA | NA | 4930421P07 | RIKEN cDNA 4930421P07 gene [Source:M | 0 | 0 | 0 | 0 | 0 | 0 |
| 28077 | ENSMUSG00000086423 | 0 | NA | NA | NA | NA | NA | Gm11364 | predicted gene 11364 [Source:MGI Symbo | 0 | 0 | 0 | 0 | 0 | 0 |
| 28078 | ENSMUSG00000086424 | 0 | NA | NA | NA | NA | NA | Gm15569 | predicted gene 15569 [Source:MGI Symbo | 0 | 0 | 0 | 0 | 0 | 0 |
| 28079 | ENSMUSG00000086425 | 0 | NA | NA | NA | NA | NA | F730016J06I | RIKEN cDNA F730016J06 gene [Source:M | 0 | 0 | 0 | 0 | 0 | 0 |
| 28080 | ENSMUSG00000086426 | 0 | NA | NA | NA | NA | NA | Gm15295 | predicted gene 15295 [Source:MGI Symbo | 0 | 0 | 0 | 0 | 0 | 0 |
| 28081 | ENSMUSG00000086427 | 0 | NA | NA | NA | NA | NA | Hoxa11os | homeobox A11, opposite strand [Source:I | 0 | 0 | 0 | 0 | 0 | 0 |
| 28082 | ENSMUSG00000086428 | 1.562907805 | -0.87216067 | 1.953251246 | -0.446517401 | 0.655223546 | NA | Gm15199 | predicted gene 15199 [Source:MGI Symbo | 0 | 1.0279 | 5.0312 | 0 | 1.0906 | 2.2277 |
| 28083 | ENSMUSG00000086429 | 124.8028337 | -0.152545986 | 0.235210008 | -0.648552273 | 0.516627813 | 0.886716042 | Gt(ROSA)26 | gene trap ROSA 26, Philippe Soriano [Sour | 130.1 | 161.38 | 102.64 | 110.86 | 109.06 | 134.78 |
| 28084 | ENSMUSG00000086430 | 7.992669551 | 1.02620248 | 0.776397545 | 1.321748745 | 0.186251831 | 0.62230226 | 4930551O13 | RIKEN cDNA 4930551O13 gene [Source:M | 5.6217 | 6.1675 | 4.025 | 14.46 | 6.5437 | 11.138 |
| 28085 | ENSMUSG00000086431 | 0 | NA | NA | NA | NA | NA | Gm14929 | predicted gene 14929 [Source:MGI Symbo | 0 | 0 | 0 | 0 | 0 | 0 |
| 28086 | ENSMUSG00000086432 | 412.3501917 | 0.33343405 | 0.411317143 | 0.810649533 | 0.417566963 | 0.837107483 | B430119L08 | RIKEN cDNA B430119L08 gene [Source:M | 236.92 | 376.22 | 481.99 | 540.79 | 286.83 | 551.35 |
| 28087 | ENSMUSG00000086433 | 0 | NA | NA | NA | NA | NA | Rsf1os1 | remodeling and spacing factor 1, opposite | 0 | 0 | 0 | 0 | 0 | 0 |
| 28088 | ENSMUSG00000086434 | 0 | NA | NA | NA | NA | NA | Gm15200 | predicted gene 15200 [Source:MGI Symbo | 0 | 0 | 0 | 0 | 0 | 0 |
| 28089 | ENSMUSG00000086436 | 0.538730002 | 1.11098254 | 3.037664833 | 0.365735722 | 0.714562288 | NA | Gm13690 | predicted gene 13690 [Source:MGI Symbo | 0 | 1.0279 | 0 | 0 | 1.0906 | 1.1138 |
| 28090 | ENSMUSG00000086437 | 33.34370531 | -0.801180447 | 0.430681409 | -1.860262439 | 0.062848405 | 0.358595396 | Gm13021 | predicted gene 13021 [Source:MGI Symbo | 42.564 | 57.563 | 27.169 | 29.883 | 26.175 | 16.708 |
| 28091 | ENSMUSG00000086438 | 0 | NA | NA | NA | NA | NA | Asb17os | ankyrin repeat and SOCS box-containing 1 | 0 | 0 | 0 | 0 | 0 | 0 |
| 28092 | ENSMUSG00000086439 | 0 | NA | NA | NA | NA | NA | Gm12069 | predicted gene 12069 [Source:MGI Symbo | 0 | 0 | 0 | 0 | 0 | 0 |
| 28093 | ENSMUSG00000086441 | 0 | NA | NA | NA | NA | NA | Gm15046 | predicted gene 15046 [Source:MGI Symbo | 0 | 0 | 0 | 0 | 0 | 0 |
| 28094 | ENSMUSG00000086442 | 0.328370405 | 0.16514316 | 4.003149536 | 0.041253308 | 0.967093956 | NA | Gm11722 | predicted gene 11722 [Source:MGI Symbo | 0 | 0 | 1.0062 | 0.964 | 0 | 0 |
| 28095 | ENSMUSG00000086443 | 19.46640364 | -0.450932178 | 0.485318252 | -0.929147371 | 0.352812718 | 0.793259753 | 4933421A08 | RIKEN cDNA 4933421A08 gene [Source:M | 25.699 | 19.53 | 22.137 | 16.388 | 17.45 | 15.594 |
| 28096 | ENSMUSG00000086444 | 0 | NA | NA | NA | NA | NA | Gm15961 | predicted gene 15961 [Source:MGI Symbo | 0 | 0 | 0 | 0 | 0 | 0 |
| 28097 | ENSMUSG00000086445 | 0 | NA | NA | NA | NA | NA | Gm13191 | predicted gene 13191 [Source:MGI Symbo | 0 | 0 | 0 | 0 | 0 | 0 |
| 28098 | ENSMUSG00000086446 | 4.923577573 | -0.108870788 | 0.998763982 | -0.109005521 | 0.913198112 | NA | Prkag2os1 | protein kinase, AMP-activated, gamma 2 r | 7.2279 | 3.0838 | 5.0312 | 8.6758 | 2.1812 | 3.3415 |
| 28099 | ENSMUSG00000086447 | 51.22230647 | 1.477541898 | 0.367498946 | 4.020533698 | 5.81E-05 | 0.002370229 | Gm13522 | predicted gene 13522 [Source:MGI Symbo | 32.124 | 22.614 | 26.162 | 69.406 | 105.79 | 51.237 |
| 28100 | ENSMUSG00000086448 | 95.94448208 | 0.13556727 | 0.240625814 | 0.563394541 | 0.573166253 | 0.905249693 | 9330162012 | cDNA RIKEN 9330162012 gene [Source:M | 87.538 | 78.122 | 108.67 | 97.362 | 98.156 | 105.82 |
| 28101 | ENSMUSG00000086449 | 0 | NA | NA | NA | NA | NA | 9030204H09 | RIKEN cDNA 9030204H09 gene [Source:M | 0 | 0 | 0 | 0 | 0 | 0 |
| 28102 | ENSMUSG00000086450 | 0.160662974 | 1.126924175 | 4.080472857 | 0.276174898 | 0.782413736 | NA | Macrod2os1 | mono-ADP ribosylhydrolase 2, opposite st | 0 | 0 | 0 | 0.964 | 0 | 0 |
| 28103 | ENSMUSG00000086451 | 0.160662974 | 1.126924175 | 4.080472857 | 0.276174898 | 0.782413736 | NA | 4933431K23 | RIKEN cDNA 4933431K23 gene [Source:M | 0 | 0 | 0 | 0.964 | 0 | 0 |
| 28104 | ENSMUSG00000086452 | 0 | NA | NA | NA | NA | NA | Gm14947 | predicted gene 14947 [Source:MGI Symbo | 0 | 0 | 0 | 0 | 0 | 0 |
| 28105 | ENSMUSG00000086453 | 1.664993117 | -0.039351357 | 1.865004704 | -0.02109987 | 0.983165989 | NA | Gm11457 | predicted gene 11457 [Source:MGI Symbo | 4.0155 | 1.0279 | 0 | 3.8559 | 1.0906 | 0 |
| 28106 | ENSMUSG00000086454 | 0 | NA | NA | NA | NA | NA | Platr14 | pluripotency associated transcript 14 [Sou | 0 | 0 | 0 | 0 | 0 | 0 |
| 28107 | ENSMUSG00000086455 | 0.167707431 | -0.796629176 | 4.080472857 | -0.195229623 | 0.84521319 | NA | Gm11815 | predicted gene 11815 [Source:MGI Symbo | 0 | 0 | 1.0062 | 0 | 0 | 0 |
| 28108 | ENSMUSG00000086458 | 0 | NA | NA | NA | NA | NA | Gm2639 | predicted gene 2639 [Source:MGI Symbol | 0 | 0 | 0 | 0 | 0 | 0 |

| | | | | | | | | | | | | | | | |
|---|---|---|---|---|---|---|---|---|---|---|---|---|---|---|---|
| 28109 | ENSMUSG00000086459 | 0.371281295 | 2.09253362 | 4.031852165 | 0.519000582 | 0.603760335 | NA | 1700030C12 | RIKEN cDNA 1700030C12 gene [Source:M | 0 | 0 | 0 | 0 | 0 | 2.2277 |
| 28110 | ENSMUSG00000086460 | 0 | NA | NA | NA | NA | NA | Gm12236 | predicted gene 12236 [Source:MGI Symbo | 0 | 0 | 0 | 0 | 0 | 0 |
| 28111 | ENSMUSG00000086462 | 0 | NA | NA | NA | NA | NA | Gm13990 | predicted gene 13990 [Source:MGI Symbo | 0 | 0 | 0 | 0 | 0 | 0 |
| 28112 | ENSMUSG00000086464 | 8.073671264 | -1.374132309 | 0.816088189 | -1.683803697 | 0.092219613 | 0.447243079 | Gm15835 | predicted gene 15835 [Source:MGI Symbo | 13.653 | 10.279 | 11.069 | 5.7839 | 6.5437 | 1.1138 |
| 28113 | ENSMUSG00000086467 | 0 | NA | NA | NA | NA | NA | 4930571N24 | RIKEN cDNA 4930571N24 gene [Source:M | 0 | 0 | 0 | 0 | 0 | 0 |
| 28114 | ENSMUSG00000086468 | 14.50143389 | 0.94150828 | 0.59114022 | 1.592698733 | 0.111227792 | 0.489450944 | Etaa1os | Ewing tumor-associated antigen 1, opposi | 10.44 | 7.1954 | 12.075 | 16.388 | 14.178 | 26.732 |
| 28115 | ENSMUSG00000086470 | 0 | NA | NA | NA | NA | NA | Gm12580 | predicted gene 12580 [Source:MGI Symbo | 0 | 0 | 0 | 0 | 0 | 0 |
| 28116 | ENSMUSG00000086472 | 7.843936644 | 0.000107474 | 0.805450142 | 0.000133434 | 0.999893535 | 0.999897338 | Gm16172 | predicted gene 16172 [Source:MGI Symbo | 6.4248 | 7.1954 | 10.062 | 13.496 | 6.5437 | 3.3415 |
| 28117 | ENSMUSG00000086473 | 0 | NA | NA | NA | NA | NA | Gm16265 | predicted gene 16265 [Source:MGI Symbo | 0 | 0 | 0 | 0 | 0 | 0 |
| 28118 | ENSMUSG00000086474 | 6.698771726 | 0.202542505 | 0.864826962 | 0.234200035 | 0.814829688 | 0.967547542 | 9130204K15 | RIKEN cDNA 9130204K15 gene [Source:M | 6.4248 | 5.1396 | 7.0437 | 3.8559 | 4.3625 | 13.366 |
| 28119 | ENSMUSG00000086475 | 0 | NA | NA | NA | NA | NA | Platr13 | pluripotency associated transcript 13 [Sou | 0 | 0 | 0 | 0 | 0 | 0 |
| 28120 | ENSMUSG00000086477 | 11.339826 | 0.356385011 | 0.705178084 | 0.505382993 | 0.613289862 | 0.916813074 | Gm15506 | predicted gene 15506 [Source:MGI Symbo | 14.456 | 3.0838 | 12.075 | 8.6758 | 15.269 | 14.48 |
| 28121 | ENSMUSG00000086478 | 0 | NA | NA | NA | NA | NA | Gm14102 | predicted gene 14102 [Source:MGI Symbo | 0 | 0 | 0 | 0 | 0 | 0 |
| 28122 | ENSMUSG00000086479 | 0.449470461 | -0.545136533 | 3.562105917 | -0.15303771 | 0.878368537 | NA | Gm16014 | predicted gene 16014 [Source:MGI Symbo | 1.6062 | 0 | 0 | 0 | 1.0906 | 0 |
| 28123 | ENSMUSG00000086480 | 0.528279927 | -0.837071592 | 3.369085264 | -0.248456636 | 0.803781118 | NA | Gm15287 | predicted gene 15287 [Source:MGI Symbo | 0 | 2.0558 | 0 | 0 | 0 | 1.1138 |
| 28124 | ENSMUSG00000086481 | 4.784562875 | 0.235423269 | 1.056885881 | 0.222751835 | 0.823728656 | NA | Gm11707 | predicted gene 11707 [Source:MGI Symbo | 4.0155 | 8.2233 | 1.0062 | 7.7118 | 2.1812 | 5.5692 |
| 28125 | ENSMUSG00000086482 | 0 | NA | NA | NA | NA | NA | 1700012C08 | RIKEN cDNA 1700012C08 gene [Source:M | 0 | 0 | 0 | 0 | 0 | 0 |
| 28126 | ENSMUSG00000086483 | 0 | NA | NA | NA | NA | NA | 8030443G20 | RIKEN cDNA 8030443G20 gene [Source:M | 0 | 0 | 0 | 0 | 0 | 0 |
| 28127 | ENSMUSG00000086484 | 0 | NA | NA | NA | NA | NA | Nron | non-protein coding RNA, repressor of NFA | 0 | 0 | 0 | 0 | 0 | 0 |
| 28128 | ENSMUSG00000086485 | 0 | NA | NA | NA | NA | NA | Gm15458 | predicted gene 15458 [Source:MGI Symbo | 0 | 0 | 0 | 0 | 0 | 0 |
| 28129 | ENSMUSG00000086486 | 1.275954708 | -1.321633042 | 2.213943522 | -0.596958788 | 0.550534896 | NA | Ift88os | intraflagellar transport 88, opposite strand | 2.4093 | 0 | 3.0187 | 0 | 0 | 2.2277 |
| 28130 | ENSMUSG00000086487 | 0 | NA | NA | NA | NA | NA | Gm11638 | predicted gene 11638 [Source:MGI Symbo | 0 | 0 | 0 | 0 | 0 | 0 |
| 28131 | ENSMUSG00000086489 | 0 | NA | NA | NA | NA | NA | Gm16362 | predicted gene 16362 [Source:MGI Symbo | 0 | 0 | 0 | 0 | 0 | 0 |
| 28132 | ENSMUSG00000086490 | 0 | NA | NA | NA | NA | NA | Gm12292 | predicted gene 12292 [Source:MGI Symbo | 0 | 0 | 0 | 0 | 0 | 0 |
| 28133 | ENSMUSG00000086491 | 0 | NA | NA | NA | NA | NA | Gm13291 | predicted gene 13291 [Source:MGI Symbo | 0 | 0 | 0 | 0 | 0 | 0 |
| 28134 | ENSMUSG00000086493 | 0.267700746 | -1.497481649 | 4.057001436 | -0.369110456 | 0.712045396 | NA | Gm12204 | predicted gene 12204 [Source:MGI Symbo | 1.6062 | 0 | 0 | 0 | 0 | 0 |
| 28135 | ENSMUSG00000086494 | 25.82098027 | -0.112561967 | 0.461101656 | -0.244115296 | 0.807141526 | 0.964718338 | 2210417A02 | RIKEN cDNA 2210417A02 gene [Source:M | 33.73 | 21.586 | 25.156 | 34.703 | 16.359 | 23.391 |
| 28136 | ENSMUSG00000086495 | 4.475923855 | -0.586641768 | 1.2472313 | -0.470355232 | 0.638101243 | NA | Gm13778 | predicted gene 13778 [Source:MGI Symbo | 8.8341 | 7.1954 | 0 | 0.964 | 7.6343 | 2.2277 |
| 28137 | ENSMUSG00000086496 | 0 | NA | NA | NA | NA | NA | Gm14204 | predicted gene 14204 [Source:MGI Symbo | 0 | 0 | 0 | 0 | 0 | 0 |
| 28138 | ENSMUSG00000086497 | 0 | NA | NA | NA | NA | NA | Gm15259 | predicted gene 15259 [Source:MGI Symbo | 0 | 0 | 0 | 0 | 0 | 0 |
| 28139 | ENSMUSG00000086498 | 0 | NA | NA | NA | NA | NA | Gm12681 | predicted gene 12681 [Source:MGI Symbo | 0 | 0 | 0 | 0 | 0 | 0 |
| 28140 | ENSMUSG00000086499 | 1.272580107 | 0.800386714 | 2.244168912 | 0.356651725 | 0.72135255 | NA | Gm16217 | predicted gene 16217 [Source:MGI Symbo | 0.8031 | 0 | 2.0125 | 4.8199 | 0 | 0 |
| 28141 | ENSMUSG00000086500 | 0 | NA | NA | NA | NA | NA | Gm15875 | predicted gene 15875 [Source:MGI Symbo | 0 | 0 | 0 | 0 | 0 | 0 |
| 28142 | ENSMUSG00000086501 | 0 | NA | NA | NA | NA | NA | 4930597A21 | RIKEN cDNA 4930597A21 gene [Source:M | 0 | 0 | 0 | 0 | 0 | 0 |
| 28143 | ENSMUSG00000086502 | 26.0068233 | 0.066841681 | 0.439557519 | 0.152065835 | 0.87913501 | 0.981233798 | B130055M24 | RIKEN cDNA B130055M24 gene [Source:M | 18.471 | 31.865 | 26.162 | 29.883 | 21.812 | 27.846 |
| 28144 | ENSMUSG00000086503 | 2.871478083 | -0.36872344 | 1.411919446 | -0.261150479 | 0.793976465 | NA | Xist | inactive X specific transcripts [Source:MGI | 1.6062 | 3.0838 | 5.0312 | 0.964 | 6.5437 | 0 |
| 28145 | ENSMUSG00000086504 | 0 | NA | NA | NA | NA | NA | Kif16bos | kinesin family member 16B, opposite strar | 0 | 0 | 0 | 0 | 0 | 0 |
| 28146 | ENSMUSG00000086505 | 0 | NA | NA | NA | NA | NA | Gm16058 | predicted gene 16058 [Source:MGI Symbo | 0 | 0 | 0 | 0 | 0 | 0 |
| 28147 | ENSMUSG00000086506 | 4.204391477 | -0.289002035 | 1.130206427 | -0.255707301 | 0.798176857 | NA | Gm15911 | predicted gene 15911 [Source:MGI Symbo | 4.8186 | 1.0279 | 8.05 | 5.7839 | 1.0906 | 4.4554 |
| 28148 | ENSMUSG00000086507 | 78.85675987 | -0.316135736 | 0.290489003 | -1.088288136 | 0.276467926 | 0.730038567 | Adap2os | ArfGAP with dual PH domains 2, opposite | 73.885 | 81.206 | 107.67 | 75.19 | 85.068 | 50.123 |
| 28149 | ENSMUSG00000086509 | 0 | NA | NA | NA | NA | NA | Nkx2-2os | NK2 homeobox 2, opposite strand [Source | 0 | 0 | 0 | 0 | 0 | 0 |
| 28150 | ENSMUSG00000086512 | 0 | NA | NA | NA | NA | NA | Gm12600 | predicted gene 12600 [Source:MGI Symbo | 0 | 0 | 0 | 0 | 0 | 0 |
| 28151 | ENSMUSG00000086513 | 0.331982614 | 0.16514309 | 3.986751476 | 0.041422971 | 0.966958701 | NA | Gvin-ps1 | GTPase, very large interferon inducible, ps | 0 | 1.0279 | 0 | 0.964 | 0 | 0 |
| 28152 | ENSMUSG00000086514 | 0 | NA | NA | NA | NA | NA | Gm11747 | predicted gene 11747 [Source:MGI Symbo | 0 | 0 | 0 | 0 | 0 | 0 |
| 28153 | ENSMUSG00000086515 | 0 | NA | NA | NA | NA | NA | Erich2os | glutamate rich 2, opposite strand [Source: | 0 | 0 | 0 | 0 | 0 | 0 |
| 28154 | ENSMUSG00000086517 | 0 | NA | NA | NA | NA | NA | Gm12534 | predicted gene 12534 [Source:MGI Symbo | 0 | 0 | 0 | 0 | 0 | 0 |
| 28155 | ENSMUSG00000086518 | 0 | NA | NA | NA | NA | NA | Gm14503 | predicted gene 14503 [Source:MGI Symbo | 0 | 0 | 0 | 0 | 0 | 0 |
| 28156 | ENSMUSG00000086519 | 0 | NA | NA | NA | NA | NA | Gm13393 | predicted gene 13393 [Source:MGI Symbo | 0 | 0 | 0 | 0 | 0 | 0 |
| 28157 | ENSMUSG00000086521 | 0 | NA | NA | NA | NA | NA | Olfr1343-ps1 | olfactory receptor 1343, pseudogene 1 [So | 0 | 0 | 0 | 0 | 0 | 0 |
| 28158 | ENSMUSG00000086522 | 0 | NA | NA | NA | NA | NA | Gm4473 | predicted gene 4473 [Source:MGI Symbol | 0 | 0 | 0 | 0 | 0 | 0 |
| 28159 | ENSMUSG00000086523 | 0 | NA | NA | NA | NA | NA | Gm13499 | predicted gene 13499 [Source:MGI Symbo | 0 | 0 | 0 | 0 | 0 | 0 |
| 28160 | ENSMUSG00000086524 | 0 | NA | NA | NA | NA | NA | Pabpc1l2a | poly(A) binding protein, cytoplasmic 1-like | 0 | 0 | 0 | 0 | 0 | 0 |
| 28161 | ENSMUSG00000086525 | 0 | NA | NA | NA | NA | NA | Gm15281 | predicted gene 15281 [Source:MGI Symbo | 0 | 0 | 0 | 0 | 0 | 0 |
| 28162 | ENSMUSG00000086526 | 2.039681157 | -3.297486011 | 1.793114232 | -1.838971524 | 0.065919375 | NA | Gm12762 | predicted gene 12762 [Source:MGI Symbo | 4.0155 | 3.0838 | 4.025 | 0 | 0 | 1.1138 |
| 28163 | ENSMUSG00000086527 | 0 | NA | NA | NA | NA | NA | Gm15856 | predicted gene 15856 [Source:MGI Symbo | 0 | 0 | 0 | 0 | 0 | 0 |
| 28164 | ENSMUSG00000086528 | 0 | NA | NA | NA | NA | NA | Gm15731 | predicted gene 15731 [Source:MGI Symbo | 0 | 0 | 0 | 0 | 0 | 0 |

| | | | | | | | | | | | | | | | |
|---|---|---|---|---|---|---|---|---|---|---|---|---|---|---|---|
| 28165 | ENSMUSG00000086529 | 2.737553921 | -0.818002118 | 1.451153474 | -0.563690976 | 0.57296446 | NA | Acss2os | acyl-CoA synthetase short-chain family me | 6.4248 | 3.0838 | 1.0062 | 4.8199 | 1.0906 | 0 |
| 28166 | ENSMUSG00000086530 | 1.006191327 | 3.548856365 | 2.592453122 | 1.36891824 | 0.171024834 | NA | Gm12910 | predicted gene 12910 [Source:MGI Symbo | 0 | 0 | 0 | 3.8559 | 2.1812 | 0 |
| 28167 | ENSMUSG00000086531 | 0 | NA | NA | NA | NA | NA | Gm11351 | predicted gene 11351 [Source:MGI Symbo | 0 | 0 | 0 | 0 | 0 | 0 |
| 28168 | ENSMUSG00000086533 | 12.00403903 | -0.573063568 | 0.646177507 | -0.886851619 | 0.375158793 | 0.809332934 | Mypopos | Myb-related transcription factor, partner | 13.653 | 18.503 | 11.069 | 14.46 | 6.5437 | 7.7969 |
| 28169 | ENSMUSG00000086534 | 0 | NA | NA | NA | NA | NA | Gm16758 | predicted gene, 16758 [Source:MGI Symb | 0 | 0 | 0 | 0 | 0 | 0 |
| 28170 | ENSMUSG00000086536 | 3.057097076 | -0.041063766 | 1.396181314 | -0.029411485 | 0.976536413 | NA | Gm12264 | predicted gene 12264 [Source:MGI Symbo | 7.2279 | 0 | 2.0125 | 5.7839 | 1.0906 | 2.2277 |
| 28171 | ENSMUSG00000086537 | 0 | NA | NA | NA | NA | NA | Gnasas1 | GNAS antisense RNA 1 [Source:MGI Symb | 0 | 0 | 0 | 0 | 0 | 0 |
| 28172 | ENSMUSG00000086539 | 0 | NA | NA | NA | NA | NA | Gm16759 | predicted gene, 16759 [Source:MGI Symb | 0 | 0 | 0 | 0 | 0 | 0 |
| 28173 | ENSMUSG00000086540 | 0 | NA | NA | NA | NA | NA | Scml1 | Scm polycomb group protein like 1 [Sourc | 0 | 0 | 0 | 0 | 0 | 0 |
| 28174 | ENSMUSG00000086541 | 0 | NA | NA | NA | NA | NA | Has2os | hyaluronan synthase 2, opposite strand [S | 0 | 0 | 0 | 0 | 0 | 0 |
| 28175 | ENSMUSG00000086542 | 0 | NA | NA | NA | NA | NA | Gm13782 | predicted gene 13782 [Source:MGI Symbo | 0 | 0 | 0 | 0 | 0 | 0 |
| 28176 | ENSMUSG00000086544 | 45.97001094 | -0.157211631 | 0.364948481 | -0.430777601 | 0.666630088 | 0.929596279 | Chn1os3 | chimerin 1, opposite strand 3 [Source:MG | 62.642 | 51.396 | 31.194 | 49.163 | 46.897 | 34.529 |
| 28177 | ENSMUSG00000086545 | 0 | NA | NA | NA | NA | NA | Lrrc3c | leucine rich repeat containing 3C [Source: | 0 | 0 | 0 | 0 | 0 | 0 |
| 28178 | ENSMUSG00000086546 | 0 | NA | NA | NA | NA | NA | Gm13709 | predicted gene 13709 [Source:MGI Symbo | 0 | 0 | 0 | 0 | 0 | 0 |
| 28179 | ENSMUSG00000086547 | 0.346303622 | 2.004339653 | 3.884003423 | 0.516049919 | 0.605819544 | NA | Gm11755 | predicted gene 11755 [Source:MGI Symbo | 0 | 0 | 0 | 0.964 | 0 | 1.1138 |
| 28180 | ENSMUSG00000086548 | 0 | NA | NA | NA | NA | NA | Gm12972 | predicted gene 12972 [Source:MGI Symbo | 0 | 0 | 0 | 0 | 0 | 0 |
| 28181 | ENSMUSG00000086549 | 0.321325949 | 1.911575114 | 4.048267893 | 0.472195804 | 0.636787034 | NA | Gm13648 | predicted gene 13648 [Source:MGI Symbo | 0 | 0 | 0 | 1.928 | 0 | 0 |
| 28182 | ENSMUSG00000086550 | 0 | NA | NA | NA | NA | NA | Pla2g10os | phospholipase A2, group X, opposite strar | 0 | 0 | 0 | 0 | 0 | 0 |
| 28183 | ENSMUSG00000086552 | 1.042698947 | 2.426023578 | 2.689838701 | 0.901921583 | 0.367098525 | NA | Dlx4os | distal-less homeobox 4, opposite strand [S | 0.8031 | 0 | 0 | 0 | 5.4531 | 0 |
| 28184 | ENSMUSG00000086554 | 1.898993445 | 0.892245359 | 1.766229822 | 0.505169456 | 0.613439821 | NA | 9530034E10 | RIKEN cDNA 9530034E10 gene [Source:M( | 0 | 1.0279 | 3.0187 | 2.8919 | 0 | 4.4554 |
| 28185 | ENSMUSG00000086555 | 0 | NA | NA | NA | NA | NA | Gm13446 | predicted gene 13446 [Source:MGI Symbo | 0 | 0 | 0 | 0 | 0 | 0 |
| 28186 | ENSMUSG00000086556 | 0 | NA | NA | NA | NA | NA | Gm15444 | predicted gene 15444 [Source:MGI Symbo | 0 | 0 | 0 | 0 | 0 | 0 |
| 28187 | ENSMUSG00000086557 | 0 | NA | NA | NA | NA | NA | Gm15999 | predicted gene 15999 [Source:MGI Symbo | 0 | 0 | 0 | 0 | 0 | 0 |
| 28188 | ENSMUSG00000086558 | 0 | NA | NA | NA | NA | NA | Gm14223 | predicted gene 14223 [Source:MGI Symbo | 0 | 0 | 0 | 0 | 0 | 0 |
| 28189 | ENSMUSG00000086559 | 0 | NA | NA | NA | NA | NA | Gm14426 | predicted gene 14426 [Source:MGI Symbo | 0 | 0 | 0 | 0 | 0 | 0 |
| 28190 | ENSMUSG00000086560 | 0.663758638 | 2.950973782 | 3.025092005 | 0.975498853 | 0.32931287 | NA | Gm13372 | predicted gene 13372 [Source:MGI Symbo | 0 | 0 | 0 | 2.8919 | 1.0906 | 0 |
| 28191 | ENSMUSG00000086561 | 6.920310193 | -1.18297423 | 0.900070274 | -1.314313187 | 0.188740831 | 0.625984911 | Gm15540 | predicted gene 15540 [Source:MGI Symbo | 10.44 | 11.307 | 7.0437 | 2.8919 | 8.7249 | 1.1138 |
| 28192 | ENSMUSG00000086562 | 0 | NA | NA | NA | NA | NA | Gm14651 | predicted gene 14651 [Source:MGI Symbo | 0 | 0 | 0 | 0 | 0 | 0 |
| 28193 | ENSMUSG00000086563 | 0 | NA | NA | NA | NA | NA | Platr15 | pluripotency associated transcript 15 [Sou | 0 | 0 | 0 | 0 | 0 | 0 |
| 28194 | ENSMUSG00000086564 | 0.305170013 | -1.657836477 | 4.041480335 | -0.410205257 | 0.681655385 | NA | Cd101 | CD101 antigen [Source:MGI Symbol;Acc:M | 0.8031 | 1.0279 | 0 | 0 | 0 | 0 |
| 28195 | ENSMUSG00000086565 | 0 | NA | NA | NA | NA | NA | Gm13133 | predicted gene 13133 [Source:MGI Symbo | 0 | 0 | 0 | 0 | 0 | 0 |
| 28196 | ENSMUSG00000086567 | 3.620386295 | 0.343660064 | 1.265076423 | 0.271651624 | 0.785889902 | NA | Gm2830 | predicted gene 2830 [Source:MGI Symbol | 2.4093 | 4.1117 | 3.0187 | 0 | 3.2719 | 8.9108 |
| 28197 | ENSMUSG00000086569 | 0 | NA | NA | NA | NA | NA | Rab11fip4os | RAB11 family interacting protein 4 (class I | 0 | 0 | 0 | 0 | 0 | 0 |
| 28198 | ENSMUSG00000086573 | 0.492645588 | 0.9565034 | 3.447648281 | 0.277436479 | 0.781444974 | NA | Gm15384 | predicted gene 15384 [Source:MGI Symbo | 0 | 1.0279 | 0 | 1.928 | 0 | 0 |
| 28199 | ENSMUSG00000086575 | 0.771003 | -1.69715066 | 2.640552792 | -0.642725518 | 0.52040222 | NA | Gm14104 | predicted gene 14104 [Source:MGI Symbo | 1.6062 | 2.0558 | 0 | 0.964 | 0 | 0 |
| 28200 | ENSMUSG00000086576 | 0 | NA | NA | NA | NA | NA | Gm13660 | predicted gene 13660 [Source:MGI Symbo | 0 | 0 | 0 | 0 | 0 | 0 |
| 28201 | ENSMUSG00000086577 | 0 | NA | NA | NA | NA | NA | Gm7076 | predicted pseudogene 7076 [Source:MGI | 0 | 0 | 0 | 0 | 0 | 0 |
| 28202 | ENSMUSG00000086578 | 0 | NA | NA | NA | NA | NA | Gm13583 | predicted gene 13583 [Source:MGI Symbo | 0 | 0 | 0 | 0 | 0 | 0 |
| 28203 | ENSMUSG00000086580 | 0 | NA | NA | NA | NA | NA | Gm15280 | predicted gene 15280 [Source:MGI Symbo | 0 | 0 | 0 | 0 | 0 | 0 |
| 28204 | ENSMUSG00000086582 | 0.356960287 | 0.16514264 | 3.880370363 | 0.042558474 | 0.966053499 | NA | Gm16272 | predicted gene 16272 [Source:MGI Symbo | 0 | 1.0279 | 0 | 0 | 0 | 1.1138 |
| 28205 | ENSMUSG00000086583 | 255.1865887 | 0.007316427 | 0.178397227 | 0.041012001 | 0.967286328 | 0.994231936 | Gm15500 | predicted pseudogene 15500 [Source:MG | 255.39 | 273.43 | 234.45 | 231.35 | 226.85 | 309.65 |
| 28206 | ENSMUSG00000086584 | 0 | NA | NA | NA | NA | NA | Gm12002 | predicted gene 12002 [Source:MGI Symbo | 0 | 0 | 0 | 0 | 0 | 0 |
| 28207 | ENSMUSG00000086585 | 0 | NA | NA | NA | NA | NA | Gm16126 | predicted gene 16126 [Source:MGI Symbo | 0 | 0 | 0 | 0 | 0 | 0 |
| 28208 | ENSMUSG00000086586 | 1.662435805 | 0.199041593 | 1.701221328 | 0.116999235 | 0.90686066 | NA | Gm11192 | predicted gene 11192 [Source:MGI Symbo | 1.6062 | 0 | 3.0187 | 0.964 | 3.2719 | 1.1138 |
| 28209 | ENSMUSG00000086587 | 21.23147232 | 0.329786866 | 0.529899843 | 0.622356981 | 0.533707161 | 0.889143097 | Gm11837 | predicted gene 11837 [Source:MGI Symbo | 12.047 | 13.363 | 31.194 | 22.171 | 18.541 | 30.074 |
| 28210 | ENSMUSG00000086588 | 0 | NA | NA | NA | NA | NA | Gm11729 | predicted gene 11729 [Source:MGI Symbo | 0 | 0 | 0 | 0 | 0 | 0 |
| 28211 | ENSMUSG00000086589 | 10.40845997 | -0.385573344 | 0.660234286 | -0.583994731 | 0.559223857 | 0.899423311 | Gm12915 | predicted gene 12915 [Source:MGI Symbo | 12.047 | 11.307 | 12.075 | 11.568 | 6.5437 | 8.9108 |
| 28212 | ENSMUSG00000086591 | 1.787877622 | -1.31455666 | 1.649753357 | -0.796820115 | 0.425555511 | NA | Dnah2os | dynein, axonemal, heavy chain 2, opposite | 1.6062 | 3.0838 | 3.0187 | 1.928 | 1.0906 | 0 |
| 28213 | ENSMUSG00000086592 | 0 | NA | NA | NA | NA | NA | Gm12682 | predicted gene 12682 [Source:MGI Symbo | 0 | 0 | 0 | 0 | 0 | 0 |
| 28214 | ENSMUSG00000086593 | 78.28498575 | 0.616220093 | 0.289402632 | 2.129282961 | 0.033230857 | 0.243995238 | Gm16548 | predicted gene 16548 [Source:MGI Symbo | 69.067 | 52.424 | 63.393 | 66.514 | 104.7 | 113.61 |
| 28215 | ENSMUSG00000086594 | 2.699955941 | 1.242631806 | 1.334160932 | 0.931395738 | 0.351648895 | NA | Nudt12os | nudix (nucleoside diphosphate linked moi | 0.8031 | 3.0838 | 1.0062 | 5.7839 | 2.1812 | 3.3415 |
| 28216 | ENSMUSG00000086595 | 0 | NA | NA | NA | NA | NA | Gm12214 | predicted gene 12214 [Source:MGI Symbo | 0 | 0 | 0 | 0 | 0 | 0 |
| 28217 | ENSMUSG00000086596 | 0 | NA | NA | NA | NA | NA | Susd5 | sushi domain containing 5 [Source:MGI Sy | 0 | 0 | 0 | 0 | 0 | 0 |
| 28218 | ENSMUSG00000086598 | 1.327304643 | -0.651539922 | 2.215152027 | -0.294128761 | 0.768659514 | NA | Btbd18 | BTB (POZ) domain containing 18 [Source:I | 4.8186 | 0 | 0 | 0.964 | 2.1812 | 0 |
| 28219 | ENSMUSG00000086600 | 0.301557804 | -1.64335916 | 4.042814367 | -0.406488899 | 0.684383412 | NA | C030005K06 | RIKEN cDNA C030005K06 gene [Source:M | 0.8031 | 0 | 1.0062 | 0 | 0 | 0 |
| 28220 | ENSMUSG00000086601 | 0 | NA | NA | NA | NA | NA | Rapgef3os1 | Rap guanine nucleotide exchange factor ( | 0 | 0 | 0 | 0 | 0 | 0 |

| | | | | | | | | | | | | | | | |
|---|---|---|---|---|---|---|---|---|---|---|---|---|---|---|---|
| 28221 | ENSMUSG00000086602 | 0.267700746 | -1.497481649 | 4.057001436 | -0.369110456 | 0.712045396 | NA | Gm15609 | predicted gene 15609 [Source:MGI Symbo | 1.6062 | 0 | 0 | 0 | 0 | 0 |
| 28222 | ENSMUSG00000086603 | 1.413365684 | 2.88291135 | 2.043675042 | 1.410650564 | 0.158347674 | NA | Gm13620 | predicted gene 13620 [Source:MGI Symbo | 0 | 0 | 1.0062 | 1.928 | 1.0906 | 4.4554 |
| 28223 | ENSMUSG00000086604 | 5.070615532 | -0.594931912 | 1.326994842 | -0.448330237 | 0.653914882 | NA | Gm15510 | predicted gene 15510 [Source:MGI Symbo | 4.0155 | 14.391 | 0 | 8.6758 | 0 | 3.3415 |
| 28224 | ENSMUSG00000086606 | 0.294513347 | 0.165143496 | 4.080472857 | 0.040471657 | 0.967717103 | NA | Gm13205 | predicted gene 13205 [Source:MGI Symbo | 0.8031 | 0 | 0 | 0.964 | 0 | 0 |
| 28225 | ENSMUSG00000086607 | 0.603115608 | -2.642822926 | 3.106511347 | -0.850736608 | 0.394915683 | NA | 4930511M06 | RIKEN cDNA 4930511M06 gene [Source:M | 1.6062 | 0 | 2.0125 | 0 | 0 | 0 |
| 28226 | ENSMUSG00000086608 | 0 | NA | NA | NA | NA | NA | 4930562D21 | RIKEN cDNA 4930562D21 gene [Source:M | 0 | 0 | 0 | 0 | 0 | 0 |
| 28227 | ENSMUSG00000086610 | 97.20170354 | 0.418720574 | 0.300833713 | 1.391867188 | 0.16396262 | 0.588267729 | Gm15408 | predicted gene 15408 [Source:MGI Symbo | 123.68 | 53.452 | 71.443 | 103.15 | 113.42 | 118.07 |
| 28228 | ENSMUSG00000086612 | 0 | NA | NA | NA | NA | NA | Gm12064 | predicted gene 12064 [Source:MGI Symbo | 0 | 0 | 0 | 0 | 0 | 0 |
| 28229 | ENSMUSG00000086613 | 0 | NA | NA | NA | NA | NA | 1700007M16 | RIKEN cDNA 1700007M16 gene [Source:M | 0 | 0 | 0 | 0 | 0 | 0 |
| 28230 | ENSMUSG00000086614 | 0 | NA | NA | NA | NA | NA | Gm14330 | predicted gene 14330 [Source:MGI Symbo | 0 | 0 | 0 | 0 | 0 | 0 |
| 28231 | ENSMUSG00000086617 | 0 | NA | NA | NA | NA | NA | Gm11635 | predicted gene 11635 [Source:MGI Symbo | 0 | 0 | 0 | 0 | 0 | 0 |
| 28232 | ENSMUSG00000086618 | 0 | NA | NA | NA | NA | NA | Gm13256 | predicted gene 13256 [Source:MGI Symbo | 0 | 0 | 0 | 0 | 0 | 0 |
| 28233 | ENSMUSG00000086620 | 0 | NA | NA | NA | NA | NA | Rspo4os | R-spondin 4, opposite strand [Source:MGI | 0 | 0 | 0 | 0 | 0 | 0 |
| 28234 | ENSMUSG00000086621 | 0 | NA | NA | NA | NA | NA | Gm13348 | predicted gene 13348 [Source:MGI Symbo | 0 | 0 | 0 | 0 | 0 | 0 |
| 28235 | ENSMUSG00000086624 | 0 | NA | NA | NA | NA | NA | Gm16130 | predicted gene 16130 [Source:MGI Symbo | 0 | 0 | 0 | 0 | 0 | 0 |
| 28236 | ENSMUSG00000086625 | 0 | NA | NA | NA | NA | NA | Gm11831 | predicted gene 11831 [Source:MGI Symbo | 0 | 0 | 0 | 0 | 0 | 0 |
| 28237 | ENSMUSG00000086628 | 269.8664971 | -0.201018449 | 0.158683299 | -1.266790202 | 0.205230317 | 0.650728757 | Gm16157 | predicted gene 16157 [Source:MGI Symbo | 290.72 | 277.54 | 297.85 | 263.17 | 269.38 | 220.54 |
| 28238 | ENSMUSG00000086629 | 18.44224891 | -0.240799839 | 0.532608622 | -0.452114045 | 0.651186826 | 0.925643431 | 2810403D21 | RIKEN cDNA 2810403D21 gene [Source:M | 23.29 | 12.335 | 24.15 | 13.496 | 22.903 | 14.48 |
| 28239 | ENSMUSG00000086630 | 0 | NA | NA | NA | NA | NA | Gm14397 | predicted gene 14397 [Source:MGI Symbo | 0 | 0 | 0 | 0 | 0 | 0 |
| 28240 | ENSMUSG00000086631 | 0 | NA | NA | NA | NA | NA | Gm12784 | predicted gene 12784 [Source:MGI Symbo | 0 | 0 | 0 | 0 | 0 | 0 |
| 28241 | ENSMUSG00000086632 | 0 | NA | NA | NA | NA | NA | 1700112J05I | RIKEN cDNA 1700112J05 gene [Source:M( | 0 | 0 | 0 | 0 | 0 | 0 |
| 28242 | ENSMUSG00000086633 | 0 | NA | NA | NA | NA | NA | Gm16081 | predicted gene 16081 [Source:MGI Symbo | 0 | 0 | 0 | 0 | 0 | 0 |
| 28243 | ENSMUSG00000086636 | 0.160662974 | 1.126924175 | 4.080472857 | 0.276174898 | 0.782413736 | NA | Gm12949 | predicted gene 12949 [Source:MGI Symbo | 0 | 0 | 0 | 0.964 | 0 | 0 |
| 28244 | ENSMUSG00000086637 | 0 | NA | NA | NA | NA | NA | Gm14040 | predicted gene 14040 [Source:MGI Symbo | 0 | 0 | 0 | 0 | 0 | 0 |
| 28245 | ENSMUSG00000086638 | 3.063652369 | -1.729526459 | 1.357416532 | -1.274130982 | 0.202617 | NA | 4930405A21 | RIKEN cDNA 4930405A21 gene [Source:M | 4.0155 | 1.0279 | 9.0562 | 0.964 | 1.0906 | 2.2277 |
| 28246 | ENSMUSG00000086641 | 0 | NA | NA | NA | NA | NA | Gm13663 | predicted gene 13663 [Source:MGI Symbo | 0 | 0 | 0 | 0 | 0 | 0 |
| 28247 | ENSMUSG00000086643 | 0 | NA | NA | NA | NA | NA | Gm15687 | predicted gene 15687 [Source:MGI Symbo | 0 | 0 | 0 | 0 | 0 | 0 |
| 28248 | ENSMUSG00000086644 | 0.167707431 | -0.796629176 | 4.080472857 | -0.195229623 | 0.84521319 | NA | Gm13470 | predicted gene 13470 [Source:MGI Symbo | 0 | 0 | 1.0062 | 0 | 0 | 0 |
| 28249 | ENSMUSG00000086645 | 5.715403085 | -1.227983029 | 0.944898271 | -1.299592841 | 0.193740555 | 0.631851946 | Gm15743 | predicted gene 15743 [Source:MGI Symbo | 8.8341 | 9.2513 | 6.0375 | 5.7839 | 3.2719 | 1.1138 |
| 28250 | ENSMUSG00000086646 | 0 | NA | NA | NA | NA | NA | Platr20 | pluripotency associated transcript 20 [Sou | 0 | 0 | 0 | 0 | 0 | 0 |
| 28251 | ENSMUSG00000086648 | 0.133850373 | -0.796629176 | 4.080472857 | -0.195229623 | 0.84521319 | NA | Gm15511 | predicted gene 15511 [Source:MGI Symbo | 0.8031 | 0 | 0 | 0 | 0 | 0 |
| 28252 | ENSMUSG00000086649 | 0 | NA | NA | NA | NA | NA | Gm15286 | predicted gene 15286 [Source:MGI Symbo | 0 | 0 | 0 | 0 | 0 | 0 |
| 28253 | ENSMUSG00000086650 | 0 | NA | NA | NA | NA | NA | A430048G15 | RIKEN cDNA A430048G15 gene [Source:M | 0 | 0 | 0 | 0 | 0 | 0 |
| 28254 | ENSMUSG00000086652 | 0.572870759 | -2.577107685 | 3.182795569 | -0.809699407 | 0.418112959 | NA | Gm14029 | predicted gene 14029 [Source:MGI Symbo | 2.4093 | 1.0279 | 0 | 0 | 0 | 0 |
| 28255 | ENSMUSG00000086653 | 0 | NA | NA | NA | NA | NA | 9130015L21I | RIKEN cDNA 9130015L21 gene [Source:M( | 0 | 0 | 0 | 0 | 0 | 0 |
| 28256 | ENSMUSG00000086654 | 1.20408596 | 0.63264855 | 2.067078626 | 0.306059258 | 0.759559528 | NA | Gm13165 | predicted gene 13165 [Source:MGI Symbo | 0.8031 | 0 | 2.0125 | 0 | 2.1812 | 2.2277 |
| 28257 | ENSMUSG00000086655 | 0 | NA | NA | NA | NA | NA | C330018A13 | RIKEN cDNA C330018A13 gene [Source:M | 0 | 0 | 0 | 0 | 0 | 0 |
| 28258 | ENSMUSG00000086656 | 0 | NA | NA | NA | NA | NA | Gm15701 | predicted gene 15701 [Source:MGI Symbo | 0 | 0 | 0 | 0 | 0 | 0 |
| 28259 | ENSMUSG00000086657 | 9.202703337 | 0.465513287 | 0.749919023 | 0.620751405 | 0.534763201 | 0.889858671 | Stamos | signal transducing adaptor molecule (SH3 | 8.8341 | 8.2233 | 6.0375 | 5.7839 | 18.541 | 7.7969 |
| 28260 | ENSMUSG00000086658 | 0 | NA | NA | NA | NA | NA | Gm14260 | predicted gene 14260 [Source:MGI Symbo | 0 | 0 | 0 | 0 | 0 | 0 |
| 28261 | ENSMUSG00000086659 | 0 | NA | NA | NA | NA | NA | Gm14146 | predicted gene 14146 [Source:MGI Symbo | 0 | 0 | 0 | 0 | 0 | 0 |
| 28262 | ENSMUSG00000086660 | 0 | NA | NA | NA | NA | NA | Gm12594 | predicted gene 12594 [Source:MGI Symbo | 0 | 0 | 0 | 0 | 0 | 0 |
| 28263 | ENSMUSG00000086662 | 0 | NA | NA | NA | NA | NA | BC046251 | cDNA sequence BC046251 [Source:MGI Sy | 0 | 0 | 0 | 0 | 0 | 0 |
| 28264 | ENSMUSG00000086665 | 0.321325949 | 1.911575114 | 4.048267893 | 0.472195804 | 0.636787034 | NA | Gm13067 | predicted gene 13067 [Source:MGI Symbo | 0 | 0 | 0 | 1.928 | 0 | 0 |
| 28265 | ENSMUSG00000086666 | 0 | NA | NA | NA | NA | NA | Gm12249 | predicted gene 12249 [Source:MGI Symbo | 0 | 0 | 0 | 0 | 0 | 0 |
| 28266 | ENSMUSG00000086668 | 0 | NA | NA | NA | NA | NA | Gm15273 | predicted gene 15273 [Source:MGI Symbo | 0 | 0 | 0 | 0 | 0 | 0 |
| 28267 | ENSMUSG00000086669 | 1.38268215 | 1.810549411 | 1.900767638 | 0.952535899 | 0.340825268 | NA | AA645442 | expressed sequence AA645442 [Source:M | 0.8031 | 0 | 1.0062 | 0.964 | 2.1812 | 3.3415 |
| 28268 | ENSMUSG00000086670 | 36.03459223 | -0.090165523 | 0.372445006 | -0.242090837 | 0.808709778 | 0.965682062 | Gm13194 | predicted gene 13194 [Source:MGI Symbo | 40.155 | 42.145 | 29.181 | 37.595 | 38.172 | 28.96 |
| 28269 | ENSMUSG00000086671 | 0.17131964 | -0.796629176 | 4.080472857 | -0.195229623 | 0.84521319 | NA | Gm13618 | predicted gene 13618 [Source:MGI Symbo | 0 | 1.0279 | 0 | 0 | 0 | 0 |
| 28270 | ENSMUSG00000086674 | 0 | NA | NA | NA | NA | NA | Zfp286os | zinc finger protein 286, opposite strand [S | 0 | 0 | 0 | 0 | 0 | 0 |
| 28271 | ENSMUSG00000086675 | 0 | NA | NA | NA | NA | NA | Plxna4os2 | plexin A4, opposite strand 2 [Source:MGI | 0 | 0 | 0 | 0 | 0 | 0 |
| 28272 | ENSMUSG00000086676 | 0 | NA | NA | NA | NA | NA | Gm14705 | predicted gene 14705 [Source:MGI Symbo | 0 | 0 | 0 | 0 | 0 | 0 |
| 28273 | ENSMUSG00000086677 | 0 | NA | NA | NA | NA | NA | Tvp23bos | trans-golgi network vesicle protein 23B, o | 0 | 0 | 0 | 0 | 0 | 0 |
| 28274 | ENSMUSG00000086679 | 2.786605269 | -0.40239735 | 1.410237566 | -0.285340115 | 0.775383582 | NA | Gm15551 | predicted gene 15551 [Source:MGI Symbo | 6.4248 | 1.0279 | 2.0125 | 2.8919 | 4.3625 | 0 |
| 28275 | ENSMUSG00000086680 | 0.692607243 | 3.004183657 | 2.957557172 | 1.015765202 | 0.309741211 | NA | Gm11592 | predicted gene 11592 [Source:MGI Symbo | 0 | 0 | 0 | 1.928 | 0 | 2.2277 |
| 28276 | ENSMUSG00000086681 | 0.185640647 | 1.126924175 | 4.080472857 | 0.276174898 | 0.782413736 | NA | Gm16178 | predicted gene 16178 [Source:MGI Symbo | 0 | 0 | 0 | 0 | 0 | 1.1138 |

| | | | | | | | | | | | | | | | |
|---|---|---|---|---|---|---|---|---|---|---|---|---|---|---|---|
| 28277 | ENSMUSG00000086682 | 14.76712616 | 0.079211693 | 0.569477823 | 0.139095307 | 0.889374837 | 0.982382091 | Gm16023 | predicted gene 16023 [Source:MGI Symbo | 13.653 | 10.279 | 19.119 | 12.532 | 18.541 | 14.48 |
| 28278 | ENSMUSG00000086683 | 2.086618264 | 3.511705996 | 1.84646263 | 1.901855981 | 0.057189985 | NA | Gm12867 | predicted gene 12867 [Source:MGI Symbo | 0 | 0 | 1.0062 | 3.8559 | 6.5437 | 1.1138 |
| 28279 | ENSMUSG00000086684 | 0 | NA | NA | NA | NA | NA | Gm12590 | predicted gene 12590 [Source:MGI Symbo | 0 | 0 | 0 | 0 | 0 | 0 |
| 28280 | ENSMUSG00000086685 | 0 | NA | NA | NA | NA | NA | Gm15015 | predicted gene 15015 [Source:MGI Symbo | 0 | 0 | 0 | 0 | 0 | 0 |
| 28281 | ENSMUSG00000086686 | 0.133850373 | -0.796629176 | 4.080472857 | -0.195229623 | 0.84521319 | NA | F630206G17 | RIKEN cDNA F630206G17 gene [Source:M | 0.8031 | 0 | 0 | 0 | 0 | 0 |
| 28282 | ENSMUSG00000086687 | 0 | NA | NA | NA | NA | NA | 4930547E08 | RIKEN cDNA 4930547E08 gene [Source:M | 0 | 0 | 0 | 0 | 0 | 0 |
| 28283 | ENSMUSG00000086688 | 1.364024895 | 2.868628407 | 2.231715633 | 1.285391545 | 0.198655494 | NA | Gm11560 | predicted gene 11560 [Source:MGI Symbo | 0.8031 | 0 | 0 | 1.928 | 5.4531 | 0 |
| 28284 | ENSMUSG00000086689 | 0 | NA | NA | NA | NA | NA | Gm16876 | predicted gene, 16876 [Source:MGI Symb | 0 | 0 | 0 | 0 | 0 | 0 |
| 28285 | ENSMUSG00000086690 | 0 | NA | NA | NA | NA | NA | 4933406J10I | RIKEN cDNA 4933406J10 gene [Source:M( | 0 | 0 | 0 | 0 | 0 | 0 |
| 28286 | ENSMUSG00000086691 | 0.160662974 | 1.126924175 | 4.080472857 | 0.276174898 | 0.782413736 | NA | Gm15432 | predicted gene 15432 [Source:MGI Symbo | 0 | 0 | 0 | 0.964 | 0 | 0 |
| 28287 | ENSMUSG00000086693 | 2.496588667 | -0.137373311 | 1.409214877 | -0.097482161 | 0.9223435 | NA | A730081D07 | RIKEN cDNA A730081D07 gene [Source:M | 0.8031 | 5.1396 | 2.0125 | 4.8199 | 1.0906 | 1.1138 |
| 28288 | ENSMUSG00000086694 | 0 | NA | NA | NA | NA | NA | Gm16237 | predicted gene 16237 [Source:MGI Symbo | 0 | 0 | 0 | 0 | 0 | 0 |
| 28289 | ENSMUSG00000086695 | 1.971534398 | 0.562300619 | 1.74989673 | 0.321333602 | 0.747957595 | NA | Gm15247 | predicted gene 15247 [Source:MGI Symbo | 0.8031 | 0 | 4.025 | 4.8199 | 2.1812 | 0 |
| 28290 | ENSMUSG00000086696 | 0 | NA | NA | NA | NA | NA | Gm11234 | predicted gene 11234 [Source:MGI Symbo | 0 | 0 | 0 | 0 | 0 | 0 |
| 28291 | ENSMUSG00000086697 | 1.646982792 | -0.522551125 | 1.710761906 | -0.305449357 | 0.760023935 | NA | 4933427G23 | RIKEN cDNA 4933427G23 gene [Source:M | 0.8031 | 3.0838 | 2.0125 | 2.8919 | 1.0906 | 0 |
| 28292 | ENSMUSG00000086698 | 0 | NA | NA | NA | NA | NA | 4930511O05 | RIKEN cDNA 4930511O05 gene [Source:M | 0 | 0 | 0 | 0 | 0 | 0 |
| 28293 | ENSMUSG00000086699 | 0 | NA | NA | NA | NA | NA | Gm16510 | predicted pseudogene 16510 [Source:MG | 0 | 0 | 0 | 0 | 0 | 0 |
| 28294 | ENSMUSG00000086700 | 0.17131964 | -0.796629176 | 4.080472857 | -0.195229623 | 0.84521319 | NA | Gm15747 | predicted gene 15747 [Source:MGI Symbo | 0 | 1.0279 | 0 | 0 | 0 | 0 |
| 28295 | ENSMUSG00000086701 | 0 | NA | NA | NA | NA | NA | Gm13595 | predicted gene 13595 [Source:MGI Symbo | 0 | 0 | 0 | 0 | 0 | 0 |
| 28296 | ENSMUSG00000086702 | 0 | NA | NA | NA | NA | NA | Gm12347 | predicted gene 12347 [Source:MGI Symbo | 0 | 0 | 0 | 0 | 0 | 0 |
| 28297 | ENSMUSG00000086703 | 0 | NA | NA | NA | NA | NA | 4930507D10 | RIKEN cDNA 4930507D10 gene [Source:M | 0 | 0 | 0 | 0 | 0 | 0 |
| 28298 | ENSMUSG00000086704 | 0.133850373 | -0.796629176 | 4.080472857 | -0.195229623 | 0.84521319 | NA | Gm14582 | predicted gene 14582 [Source:MGI Symbo | 0.8031 | 0 | 0 | 0 | 0 | 0 |
| 28299 | ENSMUSG00000086706 | 4.65981438 | 0.627483333 | 0.987996837 | 0.635106621 | 0.525358879 | NA | Gm15848 | predicted gene 15848 [Source:MGI Symbo | 4.8186 | 2.0558 | 4.025 | 3.8559 | 7.6343 | 5.5692 |
| 28300 | ENSMUSG00000086707 | 0 | NA | NA | NA | NA | NA | Gm12088 | predicted gene 12088 [Source:MGI Symbo | 0 | 0 | 0 | 0 | 0 | 0 |
| 28301 | ENSMUSG00000086708 | 0 | NA | NA | NA | NA | NA | Gm15577 | predicted gene 15577 [Source:MGI Symbo | 0 | 0 | 0 | 0 | 0 | 0 |
| 28302 | ENSMUSG00000086709 | 0 | NA | NA | NA | NA | NA | Gm16263 | predicted gene 16263 [Source:MGI Symbo | 0 | 0 | 0 | 0 | 0 | 0 |
| 28303 | ENSMUSG00000086710 | 0 | NA | NA | NA | NA | NA | Gm13465 | predicted gene 13465 [Source:MGI Symbo | 0 | 0 | 0 | 0 | 0 | 0 |
| 28304 | ENSMUSG00000086711 | 0 | NA | NA | NA | NA | NA | Gm15482 | predicted gene 15482 [Source:MGI Symbo | 0 | 0 | 0 | 0 | 0 | 0 |
| 28305 | ENSMUSG00000086712 | 2.352070601 | 0.727568739 | 1.680971427 | 0.432826357 | 0.665140924 | NA | Mexis | macrophage expressed LXRa(NR1H3)-dep | 3.2124 | 0 | 2.0125 | 0 | 1.0906 | 7.7969 |
| 28306 | ENSMUSG00000086713 | 0 | NA | NA | NA | NA | NA | 4930407G08 | RIKEN cDNA 4930407G08 gene [Source:M | 0 | 0 | 0 | 0 | 0 | 0 |
| 28307 | ENSMUSG00000086714 | 7.138671436 | 1.021479291 | 0.818022529 | 1.248717798 | 0.211768309 | 0.656595245 | 0610009E02 | RIKEN cDNA 0610009E02 gene [Source:M( | 4.0155 | 3.0838 | 7.0437 | 7.7118 | 8.7249 | 12.252 |
| 28308 | ENSMUSG00000086715 | 0 | NA | NA | NA | NA | NA | B230112J18I | RIKEN cDNA B230112J18 gene [Source:M( | 0 | 0 | 0 | 0 | 0 | 0 |
| 28309 | ENSMUSG00000086716 | 0 | NA | NA | NA | NA | NA | Gm11629 | predicted gene 11629 [Source:MGI Symbo | 0 | 0 | 0 | 0 | 0 | 0 |
| 28310 | ENSMUSG00000086717 | 0 | NA | NA | NA | NA | NA | Gm15655 | predicted gene 15655 [Source:MGI Symbo | 0 | 0 | 0 | 0 | 0 | 0 |
| 28311 | ENSMUSG00000086718 | 1.263468628 | -1.494299982 | 2.056380425 | -0.726665146 | 0.46743111 | NA | Gm16069 | predicted gene 16069 [Source:MGI Symbo | 1.6062 | 1.0279 | 3.0187 | 1.928 | 0 | 0 |
| 28312 | ENSMUSG00000086719 | 0 | NA | NA | NA | NA | NA | Gm16275 | predicted gene 16275 [Source:MGI Symbo | 0 | 0 | 0 | 0 | 0 | 0 |
| 28313 | ENSMUSG00000086720 | 0 | NA | NA | NA | NA | NA | Plxna4os3 | plexin A4, opposite strand 3 [Source:MGI | 0 | 0 | 0 | 0 | 0 | 0 |
| 28314 | ENSMUSG00000086721 | 0 | NA | NA | NA | NA | NA | Gm13750 | predicted gene 13750 [Source:MGI Symbo | 0 | 0 | 0 | 0 | 0 | 0 |
| 28315 | ENSMUSG00000086722 | 0 | NA | NA | NA | NA | NA | Gm15499 | predicted gene 15499 [Source:MGI Symbo | 0 | 0 | 0 | 0 | 0 | 0 |
| 28316 | ENSMUSG00000086724 | 0 | NA | NA | NA | NA | NA | Gm12833 | predicted gene 12833 [Source:MGI Symbo | 0 | 0 | 0 | 0 | 0 | 0 |
| 28317 | ENSMUSG00000086725 | 44.62960725 | 0.323566603 | 0.343751326 | 0.941281032 | 0.34656086 | 0.787534373 | A630052C17 | RIKEN cDNA A630052C17 gene [Source:M | 46.58 | 40.089 | 32.2 | 60.731 | 43.625 | 44.554 |
| 28318 | ENSMUSG00000086726 | 0 | NA | NA | NA | NA | NA | Gm9458 | predicted gene 9458 [Source:MGI Symbol | 0 | 0 | 0 | 0 | 0 | 0 |
| 28319 | ENSMUSG00000086727 | 19.88178881 | -0.58001499 | 0.519580848 | -1.11631326 | 0.264288068 | 0.719492304 | 4931428L18I | RIKEN cDNA 4931428L18 gene [Source:M( | 29.715 | 23.642 | 18.112 | 19.28 | 19.631 | 8.9108 |
| 28320 | ENSMUSG00000086728 | 9.562537007 | -0.016875918 | 0.700057058 | -0.024106489 | 0.980767667 | 0.99820894 | Man2c1os | mannosidase, alpha, class 2C, member 1, | 10.44 | 12.335 | 6.0375 | 8.6758 | 7.6343 | 12.252 |
| 28321 | ENSMUSG00000086729 | 0 | NA | NA | NA | NA | NA | Gm15589 | predicted gene 15589 [Source:MGI Symbo | 0 | 0 | 0 | 0 | 0 | 0 |
| 28322 | ENSMUSG00000086730 | 0 | NA | NA | NA | NA | NA | Platr12 | pluripotency associated transcript 12 [Sou | 0 | 0 | 0 | 0 | 0 | 0 |
| 28323 | ENSMUSG00000086731 | 0 | NA | NA | NA | NA | NA | Sox5os5 | SRY (sex determining region Y)-box 5, opp | 0 | 0 | 0 | 0 | 0 | 0 |
| 28324 | ENSMUSG00000086732 | 0 | NA | NA | NA | NA | NA | Gm6117 | predicted gene 6117 [Source:MGI Symbol | 0 | 0 | 0 | 0 | 0 | 0 |
| 28325 | ENSMUSG00000086733 | 0 | NA | NA | NA | NA | NA | Gm11685 | predicted gene 11685 [Source:MGI Symbo | 0 | 0 | 0 | 0 | 0 | 0 |
| 28326 | ENSMUSG00000086735 | 0 | NA | NA | NA | NA | NA | Gm13977 | predicted gene 13977 [Source:MGI Symbo | 0 | 0 | 0 | 0 | 0 | 0 |
| 28327 | ENSMUSG00000086736 | 0 | NA | NA | NA | NA | NA | Gm14902 | predicted gene 14902 [Source:MGI Symbo | 0 | 0 | 0 | 0 | 0 | 0 |
| 28328 | ENSMUSG00000086738 | 0 | NA | NA | NA | NA | NA | Gm12154 | predicted gene 12154 [Source:MGI Symbo | 0 | 0 | 0 | 0 | 0 | 0 |
| 28329 | ENSMUSG00000086739 | 0 | NA | NA | NA | NA | NA | Gm4409 | predicted gene 4409 [Source:MGI Symbol | 0 | 0 | 0 | 0 | 0 | 0 |
| 28330 | ENSMUSG00000086740 | 14.39722643 | 0.602555043 | 0.656511116 | 0.917813923 | 0.358716292 | 0.79836408 | Gm17029 | predicted gene 17029 [Source:MGI Symbo | 8.8341 | 18.503 | 7.0437 | 12.532 | 29.447 | 10.025 |
| 28331 | ENSMUSG00000086741 | 0 | NA | NA | NA | NA | NA | Gm15816 | predicted gene 15816 [Source:MGI Symbo | 0 | 0 | 0 | 0 | 0 | 0 |
| 28332 | ENSMUSG00000086742 | 0.353348078 | 0.165142701 | 3.895042568 | 0.042398176 | 0.966181282 | NA | Gm16201 | predicted gene 16201 [Source:MGI Symbo | 0 | 0 | 1.0062 | 0 | 0 | 1.1138 |

| | | | | | | | | | | | | | | | |
|---|---|---|---|---|---|---|---|---|---|---|---|---|---|---|---|
| 28333 | ENSMUSG00000086743 | 0 | NA | NA | NA | NA | NA | Hnrnpa1l2-p | heterogeneous nuclear ribonucleoprotein | 0 | 0 | 0 | 0 | 0 | 0 |
| 28334 | ENSMUSG00000086745 | 0 | NA | NA | NA | NA | NA | Tspan2os | tetraspanin 2, opposite strand [Source:M( | 0 | 0 | 0 | 0 | 0 | 0 |
| 28335 | ENSMUSG00000086746 | 4.595182192 | 0.359057753 | 1.022132942 | 0.35128283 | 0.725376175 | NA | Gm15222 | predicted gene 15222 [Source:MGI Symbo | 4.0155 | 2.0558 | 6.0375 | 7.7118 | 2.1812 | 5.5692 |
| 28336 | ENSMUSG00000086748 | 5.191241429 | 1.181794338 | 1.019775038 | 1.158877492 | 0.246506125 | 0.696725814 | Gm13261 | predicted gene 13261 [Source:MGI Symbo | 2.4093 | 4.1117 | 3.0187 | 3.8559 | 3.2719 | 14.48 |
| 28337 | ENSMUSG00000086749 | 0 | NA | NA | NA | NA | NA | Gm12037 | predicted gene 12037 [Source:MGI Symbo | 0 | 0 | 0 | 0 | 0 | 0 |
| 28338 | ENSMUSG00000086750 | 0 | NA | NA | NA | NA | NA | Gm11770 | predicted gene 11770 [Source:MGI Symbo | 0 | 0 | 0 | 0 | 0 | 0 |
| 28339 | ENSMUSG00000086752 | 0 | NA | NA | NA | NA | NA | Gm11674 | predicted gene 11674 [Source:MGI Symbo | 0 | 0 | 0 | 0 | 0 | 0 |
| 28340 | ENSMUSG00000086753 | 15.24425212 | -0.142506115 | 0.571894081 | -0.249182707 | 0.803219455 | 0.963596811 | Gm15751 | predicted gene 15751 [Source:MGI Symbo | 13.653 | 15.419 | 19.119 | 21.208 | 9.8156 | 12.252 |
| 28341 | ENSMUSG00000086754 | 1.174105348 | 0.433231377 | 2.10389636 | 0.205918592 | 0.836854502 | NA | Gm16098 | predicted gene 16098 [Source:MGI Symbo | 0 | 2.0558 | 1.0062 | 2.8919 | 1.0906 | 0 |
| 28342 | ENSMUSG00000086755 | 0 | NA | NA | NA | NA | NA | Gm11216 | predicted gene 11216 [Source:MGI Symbo | 0 | 0 | 0 | 0 | 0 | 0 |
| 28343 | ENSMUSG00000086756 | 0 | NA | NA | NA | NA | NA | Gm14051 | predicted gene 14051 [Source:MGI Symbo | 0 | 0 | 0 | 0 | 0 | 0 |
| 28344 | ENSMUSG00000086757 | 0 | NA | NA | NA | NA | NA | Gm2309 | predicted gene 2309 [Source:MGI Symbol | 0 | 0 | 0 | 0 | 0 | 0 |
| 28345 | ENSMUSG00000086758 | 0 | NA | NA | NA | NA | NA | Gm13912 | predicted gene 13912 [Source:MGI Symbo | 0 | 0 | 0 | 0 | 0 | 0 |
| 28346 | ENSMUSG00000086760 | 0.160662974 | 1.126924175 | 4.080472857 | 0.276174898 | 0.782413736 | NA | 1700025D23 | RIKEN cDNA 1700025D23 gene [Source:M | 0 | 0 | 0 | 0.964 | 0 | 0 |
| 28347 | ENSMUSG00000086761 | 0 | NA | NA | NA | NA | NA | Cep112os1 | centrosomal protein 112, opposite strand | 0 | 0 | 0 | 0 | 0 | 0 |
| 28348 | ENSMUSG00000086762 | 0 | NA | NA | NA | NA | NA | Gm14546 | predicted gene 14546 [Source:MGI Symbo | 0 | 0 | 0 | 0 | 0 | 0 |
| 28349 | ENSMUSG00000086763 | 2.883665448 | -2.911047528 | 1.587667808 | -1.833536911 | 0.066722765 | NA | Plxna4os1 | plexin A4, opposite strand 1 [Source:MGI | 3.2124 | 4.1117 | 8.05 | 1.928 | 0 | 0 |
| 28350 | ENSMUSG00000086764 | 0 | NA | NA | NA | NA | NA | 4930527A07 | RIKEN cDNA 4930527A07 gene [Source:M | 0 | 0 | 0 | 0 | 0 | 0 |
| 28351 | ENSMUSG00000086765 | 22.95654732 | -2.333307665 | 0.540498624 | -4.316953938 | 1.58E-05 | 0.000773028 | Gm11827 | predicted gene 11827 [Source:MGI Symbo | 37.746 | 46.256 | 31.194 | 11.568 | 7.6343 | 3.3415 |
| 28352 | ENSMUSG00000086766 | 0 | NA | NA | NA | NA | NA | Gm15969 | predicted gene 15969 [Source:MGI Symbo | 0 | 0 | 0 | 0 | 0 | 0 |
| 28353 | ENSMUSG00000086767 | 0 | NA | NA | NA | NA | NA | Gm13070 | predicted gene 13070 [Source:MGI Symbo | 0 | 0 | 0 | 0 | 0 | 0 |
| 28354 | ENSMUSG00000086769 | 0.824421612 | 3.264491806 | 2.808832384 | 1.162223786 | 0.245144575 | NA | Gm15587 | predicted gene 15587 [Source:MGI Symbo | 0 | 0 | 0 | 3.8559 | 1.0906 | 0 |
| 28355 | ENSMUSG00000086770 | 0.181769715 | 1.126924175 | 4.080472857 | 0.276174898 | 0.782413736 | NA | Gm15351 | predicted gene 15351 [Source:MGI Symbo | 0 | 0 | 0 | 0 | 1.0906 | 0 |
| 28356 | ENSMUSG00000086771 | 84.01654235 | 0.227395598 | 0.286761377 | 0.792978468 | 0.427790368 | 0.842753542 | NA | NA | 104.4 | 63.731 | 63.393 | 87.722 | 106.88 | 77.969 |
| 28357 | ENSMUSG00000086773 | 0.267700746 | -1.497481649 | 4.057001436 | -0.369110456 | 0.712045396 | NA | Gm16192 | predicted gene 16192 [Source:MGI Symbo | 1.6062 | 0 | 0 | 0 | 0 | 0 |
| 28358 | ENSMUSG00000086774 | 0 | NA | NA | NA | NA | NA | Gm11915 | predicted gene 11915 [Source:MGI Symbo | 0 | 0 | 0 | 0 | 0 | 0 |
| 28359 | ENSMUSG00000086775 | 0.294513347 | 0.165143496 | 4.080472857 | 0.040471657 | 0.967717103 | NA | Snhg7os | small nucleolar RNA host gene 7, opposite | 0.8031 | 0 | 0 | 0.964 | 0 | 0 |
| 28360 | ENSMUSG00000086777 | 0 | NA | NA | NA | NA | NA | Far2os2 | fatty acyl CoA reductase 2, opposite stran | 0 | 0 | 0 | 0 | 0 | 0 |
| 28361 | ENSMUSG00000086779 | 3.68914468 | 0.319581897 | 1.286487333 | 0.248414336 | 0.803813842 | NA | Gm13562 | predicted gene 13562 [Source:MGI Symbo | 0.8031 | 5.1396 | 4.025 | 7.7118 | 0 | 4.4554 |
| 28362 | ENSMUSG00000086781 | 0 | NA | NA | NA | NA | NA | 1700092C02 | RIKEN cDNA 1700092C02 gene [Source:M | 0 | 0 | 0 | 0 | 0 | 0 |
| 28363 | ENSMUSG00000086782 | 93.44501083 | -0.103160003 | 0.323348936 | -0.31903616 | 0.749699092 | 0.951447545 | E130102H24 | RIKEN cDNA E130102H24 gene [Source:M | 96.372 | 106.9 | 87.543 | 128.21 | 95.974 | 45.668 |
| 28364 | ENSMUSG00000086783 | 0 | NA | NA | NA | NA | NA | Gm14262 | predicted gene 14262 [Source:MGI Symbo | 0 | 0 | 0 | 0 | 0 | 0 |
| 28365 | ENSMUSG00000086784 | 3410.196538 | 0.039192726 | 0.104212332 | 0.376085295 | 0.706853486 | 0.943006403 | Isoc2a | isochorismatase domain containing 2a [So | 3504.7 | 3470.3 | 3115.3 | 3130 | 3674.3 | 3566.5 |
| 28366 | ENSMUSG00000086785 | 0 | NA | NA | NA | NA | NA | Gm6081 | predicted gene 6081 [Source:MGI Symbol | 0 | 0 | 0 | 0 | 0 | 0 |
| 28367 | ENSMUSG00000086786 | 0 | NA | NA | NA | NA | NA | Gm15908 | predicted gene 15908 [Source:MGI Symbo | 0 | 0 | 0 | 0 | 0 | 0 |
| 28368 | ENSMUSG00000086788 | 0 | NA | NA | NA | NA | NA | 1700029M2( | RIKEN cDNA 1700029M20 gene [Source:M | 0 | 0 | 0 | 0 | 0 | 0 |
| 28369 | ENSMUSG00000086789 | 0 | NA | NA | NA | NA | NA | Lyrm7os | LYR motif containing 7, opposite strand [S | 0 | 0 | 0 | 0 | 0 | 0 |
| 28370 | ENSMUSG00000086790 | 0 | NA | NA | NA | NA | NA | Gm12911 | predicted gene 12911 [Source:MGI Symbo | 0 | 0 | 0 | 0 | 0 | 0 |
| 28371 | ENSMUSG00000086791 | 0 | NA | NA | NA | NA | NA | Gm12147 | predicted gene 12147 [Source:MGI Symbo | 0 | 0 | 0 | 0 | 0 | 0 |
| 28372 | ENSMUSG00000086792 | 0.301557804 | -1.64335916 | 4.042814367 | -0.406488899 | 0.684383412 | NA | Gm12364 | predicted gene 12364 [Source:MGI Symbo | 0.8031 | 0 | 1.0062 | 0 | 0 | 0 |
| 28373 | ENSMUSG00000086794 | 1.566958698 | -0.226238548 | 1.890802846 | -0.119652109 | 0.904758739 | NA | Gm11642 | predicted gene 11642 [Source:MGI Symbo | 0 | 3.0838 | 2.0125 | 0.964 | 0 | 3.3415 |
| 28374 | ENSMUSG00000086795 | 0 | NA | NA | NA | NA | NA | Gm11579 | predicted gene 11579 [Source:MGI Symbo | 0 | 0 | 0 | 0 | 0 | 0 |
| 28375 | ENSMUSG00000086796 | 8.983604854 | 0.710810467 | 0.703185215 | 1.010843874 | 0.312091161 | 0.759830222 | 4932702P03 | RIKEN cDNA 4932702P03 gene [Source:M | 7.2279 | 6.1675 | 7.0437 | 12.532 | 10.906 | 10.025 |
| 28376 | ENSMUSG00000086798 | 0 | NA | NA | NA | NA | NA | Gm16222 | predicted gene 16222 [Source:MGI Symbo | 0 | 0 | 0 | 0 | 0 | 0 |
| 28377 | ENSMUSG00000086799 | 0.688762941 | -1.399623551 | 3.067774249 | -0.456234207 | 0.64822157 | NA | 4930502E09 | RIKEN cDNA 4930502E09 gene [Source:M | 0 | 0 | 3.0187 | 0 | 0 | 1.1138 |
| 28378 | ENSMUSG00000086800 | 0 | NA | NA | NA | NA | NA | 4930556L07I | RIKEN cDNA 4930556L07 gene [Source:M | 0 | 0 | 0 | 0 | 0 | 0 |
| 28379 | ENSMUSG00000086801 | 0 | NA | NA | NA | NA | NA | Gm15943 | predicted gene 15943 [Source:MGI Symbo | 0 | 0 | 0 | 0 | 0 | 0 |
| 28380 | ENSMUSG00000086804 | 0.267700746 | -1.497481649 | 4.057001436 | -0.369110456 | 0.712045396 | NA | Gm43154 | predicted gene 43154 [Source:MGI Symbo | 1.6062 | 0 | 0 | 0 | 0 | 0 |
| 28381 | ENSMUSG00000086805 | 0.915510039 | -3.242382683 | 2.669421139 | -1.214638873 | 0.224503856 | NA | 4932443L11I | RIKEN cDNA 4932443L11 gene [Source:M | 2.4093 | 3.0838 | 0 | 0 | 0 | 0 |
| 28382 | ENSMUSG00000086806 | 0 | NA | NA | NA | NA | NA | Gm13054 | predicted gene 13054 [Source:MGI Symbo | 0 | 0 | 0 | 0 | 0 | 0 |
| 28383 | ENSMUSG00000086807 | 0 | NA | NA | NA | NA | NA | Platr21 | pluripotency associated transcript 21 [Sou | 0 | 0 | 0 | 0 | 0 | 0 |
| 28384 | ENSMUSG00000086808 | 0 | NA | NA | NA | NA | NA | Gm11981 | predicted gene 11981 [Source:MGI Symbo | 0 | 0 | 0 | 0 | 0 | 0 |
| 28385 | ENSMUSG00000086810 | 0 | NA | NA | NA | NA | NA | Gm13110 | predicted gene 13110 [Source:MGI Symbo | 0 | 0 | 0 | 0 | 0 | 0 |
| 28386 | ENSMUSG00000086812 | 0 | NA | NA | NA | NA | NA | Gm8978 | predicted gene 8978 [Source:MGI Symbol | 0 | 0 | 0 | 0 | 0 | 0 |
| 28387 | ENSMUSG00000086813 | 13.97613123 | -0.837450974 | 0.608914519 | -1.375317796 | 0.169032942 | 0.596320535 | Gm13657 | predicted gene 13657 [Source:MGI Symbo | 23.29 | 13.363 | 17.106 | 12.532 | 11.997 | 5.5692 |
| 28388 | ENSMUSG00000086814 | 0 | NA | NA | NA | NA | NA | Gm15677 | predicted gene 15677 [Source:MGI Symbo | 0 | 0 | 0 | 0 | 0 | 0 |

| | | | | | | | | | | | | | | | |
|---|---|---|---|---|---|---|---|---|---|---|---|---|---|---|---|
| 28389 | ENSMUSG00000086815 | 0 | NA | NA | NA | NA | NA | 3110082J24I | RIKEN cDNA 3110082J24 gene [Source:MG | 0 | 0 | 0 | 0 | 0 | 0 |
| 28390 | ENSMUSG00000086816 | 0.513958919 | -2.394274821 | 3.756096013 | -0.637437066 | 0.523840191 | NA | Gm16726 | predicted gene, 16726 [Source:MGI Symb | 0 | 3.0838 | 0 | 0 | 0 | 0 |
| 28391 | ENSMUSG00000086817 | 0 | NA | NA | NA | NA | NA | Gm11482 | predicted gene 11482 [Source:MGI Symbo | 0 | 0 | 0 | 0 | 0 | 0 |
| 28392 | ENSMUSG00000086818 | 0 | NA | NA | NA | NA | NA | Ctrcos | chymotrypsin C (caldecrin), opposite strar | 0 | 0 | 0 | 0 | 0 | 0 |
| 28393 | ENSMUSG00000086819 | 2.086261902 | -1.05519918 | 1.529757687 | -0.68978191 | 0.490331346 | NA | Gm13613 | predicted gene 13613 [Source:MGI Symbo | 2.4093 | 3.0838 | 3.0187 | 2.8919 | 0 | 1.1138 |
| 28394 | ENSMUSG00000086820 | 0.17131964 | -0.796629176 | 4.080472857 | -0.195229623 | 0.84521319 | NA | Gm11465 | predicted gene 11465 [Source:MGI Symbo | 0 | 1.0279 | 0 | 0 | 0 | 0 |
| 28395 | ENSMUSG00000086821 | 0 | NA | NA | NA | NA | NA | Gm13377 | predicted gene 13377 [Source:MGI Symbo | 0 | 0 | 0 | 0 | 0 | 0 |
| 28396 | ENSMUSG00000086822 | 0 | NA | NA | NA | NA | NA | 5330413P13 | RIKEN cDNA 5330413P13 gene [Source:M | 0 | 0 | 0 | 0 | 0 | 0 |
| 28397 | ENSMUSG00000086823 | 0 | NA | NA | NA | NA | NA | 1700028P15 | RIKEN cDNA 1700028P15 gene [Source:M | 0 | 0 | 0 | 0 | 0 | 0 |
| 28398 | ENSMUSG00000086824 | 0.610133435 | 0.304049584 | 2.904025954 | 0.104699334 | 0.91661439 | NA | 4930448D08 | RIKEN cDNA 4930448D08 gene [Source:M | 1.6062 | 0 | 0 | 0.964 | 1.0906 | 0 |
| 28399 | ENSMUSG00000086825 | 54.37918632 | -0.498859393 | 0.316073517 | -1.578301775 | 0.114496298 | 0.497223548 | Gm15675 | predicted gene 15675 [Source:MGI Symbo | 61.036 | 66.815 | 63.393 | 48.199 | 54.531 | 32.301 |
| 28400 | ENSMUSG00000086826 | 20.7095455 | 0.335089795 | 0.501465769 | 0.668220677 | 0.503992739 | 0.881510796 | Gm11739 | predicted gene 11739 [Source:MGI Symbo | 18.471 | 18.503 | 18.112 | 31.811 | 23.994 | 13.366 |
| 28401 | ENSMUSG00000086827 | 0 | NA | NA | NA | NA | NA | 4930461C15 | RIKEN cDNA 4930461C15 gene [Source:M | 0 | 0 | 0 | 0 | 0 | 0 |
| 28402 | ENSMUSG00000086828 | 0 | NA | NA | NA | NA | NA | Gm15579 | predicted gene 15579 [Source:MGI Symbo | 0 | 0 | 0 | 0 | 0 | 0 |
| 28403 | ENSMUSG00000086830 | 0 | NA | NA | NA | NA | NA | Gm13467 | predicted gene 13467 [Source:MGI Symbo | 0 | 0 | 0 | 0 | 0 | 0 |
| 28404 | ENSMUSG00000086831 | 0 | NA | NA | NA | NA | NA | Gm14280 | predicted gene 14280 [Source:MGI Symbo | 0 | 0 | 0 | 0 | 0 | 0 |
| 28405 | ENSMUSG00000086833 | 0 | NA | NA | NA | NA | NA | Gm12440 | predicted gene 12440 [Source:MGI Symbo | 0 | 0 | 0 | 0 | 0 | 0 |
| 28406 | ENSMUSG00000086834 | 0 | NA | NA | NA | NA | NA | Gm14547 | predicted gene 14547 [Source:MGI Symbo | 0 | 0 | 0 | 0 | 0 | 0 |
| 28407 | ENSMUSG00000086835 | 0 | NA | NA | NA | NA | NA | Gm11775 | predicted gene 11775 [Source:MGI Symbo | 0 | 0 | 0 | 0 | 0 | 0 |
| 28408 | ENSMUSG00000086836 | 0 | NA | NA | NA | NA | NA | Gm13748 | predicted gene 13748 [Source:MGI Symbo | 0 | 0 | 0 | 0 | 0 | 0 |
| 28409 | ENSMUSG00000086837 | 3.439398038 | -0.272499228 | 1.22960169 | -0.221615854 | 0.82461294 | NA | Gm16618 | predicted gene, 16618 [Source:MGI Symb | 3.2124 | 3.0838 | 5.0312 | 3.8559 | 5.4531 | 0 |
| 28410 | ENSMUSG00000086838 | 0 | NA | NA | NA | NA | NA | Gm16212 | predicted gene 16212 [Source:MGI Symbo | 0 | 0 | 0 | 0 | 0 | 0 |
| 28411 | ENSMUSG00000086839 | 1.902399064 | 2.274882612 | 1.816128472 | 1.252600048 | 0.210351297 | NA | Gm11973 | predicted gene 11973 [Source:MGI Symbo | 0 | 0 | 2.0125 | 3.8559 | 1.0906 | 4.4554 |
| 28412 | ENSMUSG00000086840 | 0 | NA | NA | NA | NA | NA | Gm14250 | predicted gene 14250 [Source:MGI Symbo | 0 | 0 | 0 | 0 | 0 | 0 |
| 28413 | ENSMUSG00000086841 | 180.6771068 | 0.243284972 | 0.176035079 | 1.382025526 | 0.166963863 | 0.593319485 | 2410006H16 | RIKEN cDNA 2410006H16 gene [Source:M | 155 | 172.69 | 169.05 | 192.8 | 198.49 | 196.04 |
| 28414 | ENSMUSG00000086842 | 0 | NA | NA | NA | NA | NA | Gm15681 | predicted gene 15681 [Source:MGI Symbo | 0 | 0 | 0 | 0 | 0 | 0 |
| 28415 | ENSMUSG00000086843 | 1.270222217 | 0.391045348 | 2.200220979 | 0.177730033 | 0.858934997 | NA | E030013I19F | RIKEN cDNA E030013I19 gene [Source:MG | 3.2124 | 0 | 0 | 0 | 2.1812 | 2.2277 |
| 28416 | ENSMUSG00000086844 | 7.005343496 | 0.004169297 | 0.800569454 | 0.005207914 | 0.995844704 | 0.999003684 | B230206H07 | RIKEN cDNA B230206H07 gene [Source:M | 4.8186 | 7.1954 | 9.0562 | 7.7118 | 5.4531 | 7.7969 |
| 28417 | ENSMUSG00000086845 | 0 | NA | NA | NA | NA | NA | Gm13010 | predicted gene 13010 [Source:MGI Symbo | 0 | 0 | 0 | 0 | 0 | 0 |
| 28418 | ENSMUSG00000086846 | 0.428363721 | -0.543730382 | 3.624677344 | -0.15000794 | 0.880758351 | NA | Gm13642 | predicted gene 13642 [Source:MGI Symbo | 1.6062 | 0 | 0 | 0.964 | 0 | 0 |
| 28419 | ENSMUSG00000086847 | 0.342639279 | -1.805976596 | 4.028547714 | -0.448294702 | 0.653940524 | NA | Tbx3os2 | T-box 3, opposite strand 2 [Source:MGI Sy | 0 | 2.0558 | 0 | 0 | 0 | 0 |
| 28420 | ENSMUSG00000086848 | 0 | NA | NA | NA | NA | NA | Lce6a | late cornified envelope 6A [Source:MGI Sy | 0 | 0 | 0 | 0 | 0 | 0 |
| 28421 | ENSMUSG00000086849 | 2.725625509 | 1.861467464 | 1.615680759 | 1.152125786 | 0.249269389 | NA | Gm13840 | predicted gene 13840 [Source:MGI Symbo | 2.4093 | 1.0279 | 0 | 2.8919 | 0 | 10.025 |
| 28422 | ENSMUSG00000086850 | 0 | NA | NA | NA | NA | NA | Gm13257 | predicted gene 13257 [Source:MGI Symbo | 0 | 0 | 0 | 0 | 0 | 0 |
| 28423 | ENSMUSG00000086851 | 0 | NA | NA | NA | NA | NA | A430108G06 | RIKEN cDNA A430108G06 gene [Source:M | 0 | 0 | 0 | 0 | 0 | 0 |
| 28424 | ENSMUSG00000086854 | 0 | NA | NA | NA | NA | NA | Gm13781 | predicted gene 13781 [Source:MGI Symbo | 0 | 0 | 0 | 0 | 0 | 0 |
| 28425 | ENSMUSG00000086855 | 1.534239159 | -0.273786383 | 1.873238264 | -0.146156732 | 0.883797661 | NA | Gm13844 | predicted gene 13844 [Source:MGI Symbo | 0 | 3.0838 | 2.0125 | 1.928 | 2.1812 | 0 |
| 28426 | ENSMUSG00000086856 | 0 | NA | NA | NA | NA | NA | Gm16080 | predicted gene 16080 [Source:MGI Symbo | 0 | 0 | 0 | 0 | 0 | 0 |
| 28427 | ENSMUSG00000086858 | 0 | NA | NA | NA | NA | NA | Trav20 | T cell receptor alpha variable 20 [Source:N | 0 | 0 | 0 | 0 | 0 | 0 |
| 28428 | ENSMUSG00000086859 | 188.8112805 | 0.244115104 | 0.175902011 | 1.387790294 | 0.165200911 | 0.590574884 | Snhg20 | small nucleolar RNA host gene 20 [Source | 170.26 | 161.38 | 187.16 | 213.04 | 207.22 | 193.81 |
| 28429 | ENSMUSG00000086860 | 0 | NA | NA | NA | NA | NA | Gm1720 | predicted gene 1720 [Source:MGI Symbol | 0 | 0 | 0 | 0 | 0 | 0 |
| 28430 | ENSMUSG00000086861 | 0 | NA | NA | NA | NA | NA | Gm15325 | predicted gene 15325 [Source:MGI Symbo | 0 | 0 | 0 | 0 | 0 | 0 |
| 28431 | ENSMUSG00000086862 | 0 | NA | NA | NA | NA | NA | Gm13546 | predicted gene 13546 [Source:MGI Symbo | 0 | 0 | 0 | 0 | 0 | 0 |
| 28432 | ENSMUSG00000086863 | 0 | NA | NA | NA | NA | NA | Gm15543 | predicted gene 15543 [Source:MGI Symbo | 0 | 0 | 0 | 0 | 0 | 0 |
| 28433 | ENSMUSG00000086866 | 1.275230669 | 0.274142193 | 2.031337397 | 0.134956504 | 0.892646265 | NA | 4930512H18 | RIKEN cDNA 4930512H18 gene [Source:M | 2.4093 | 0 | 1.0062 | 0.964 | 3.2719 | 0 |
| 28434 | ENSMUSG00000086867 | 0 | NA | NA | NA | NA | NA | Gm4577 | predicted gene 4577 [Source:MGI Symbol | 0 | 0 | 0 | 0 | 0 | 0 |
| 28435 | ENSMUSG00000086868 | 145.1591114 | 0.311780669 | 0.277435479 | 1.123795232 | 0.261099905 | 0.715149811 | Gm15883 | predicted gene 15883 [Source:MGI Symbo | 138.13 | 102.79 | 147.92 | 214 | 172.32 | 95.791 |
| 28436 | ENSMUSG00000086870 | 0 | NA | NA | NA | NA | NA | Gm15288 | predicted gene 15288 [Source:MGI Symbo | 0 | 0 | 0 | 0 | 0 | 0 |
| 28437 | ENSMUSG00000086872 | 0 | NA | NA | NA | NA | NA | Gm14225 | predicted gene 14225 [Source:MGI Symbo | 0 | 0 | 0 | 0 | 0 | 0 |
| 28438 | ENSMUSG00000086873 | 0.342639279 | -1.805976596 | 4.028547714 | -0.448294702 | 0.653940524 | NA | Gm15672 | predicted gene 15672 [Source:MGI Symbo | 0 | 2.0558 | 0 | 0 | 0 | 0 |
| 28439 | ENSMUSG00000086874 | 0 | NA | NA | NA | NA | NA | Gm13175 | predicted gene 13175 [Source:MGI Symbo | 0 | 0 | 0 | 0 | 0 | 0 |
| 28440 | ENSMUSG00000086875 | 0 | NA | NA | NA | NA | NA | Lipo5 | lipase, member O5 [Source:MGI Symbol;A | 0 | 0 | 0 | 0 | 0 | 0 |
| 28441 | ENSMUSG00000086876 | 0 | NA | NA | NA | NA | NA | Gm12828 | predicted gene 12828 [Source:MGI Symbo | 0 | 0 | 0 | 0 | 0 | 0 |
| 28442 | ENSMUSG00000086877 | 48.70855675 | 0.271625654 | 0.319400981 | 0.850422104 | 0.395090451 | 0.822933206 | A230072C01 | RIKEN cDNA A230072C01 gene [Source:M | 45.777 | 42.145 | 44.275 | 45.307 | 51.259 | 63.489 |
| 28443 | ENSMUSG00000086879 | 0 | NA | NA | NA | NA | NA | NA | NA | 0 | 0 | 0 | 0 | 0 | 0 |
| 28444 | ENSMUSG00000086881 | 12.35053242 | 0.732340703 | 0.652144624 | 1.122972844 | 0.261449028 | 0.71553796 | Gm13594 | predicted gene 13594 [Source:MGI Symbo | 10.44 | 12.335 | 5.0312 | 14.46 | 21.812 | 10.025 |

| | | | | | | | | | | | | | | | |
|---|---|---|---|---|---|---|---|---|---|---|---|---|---|---|---|
| 28445 | ENSMUSG00000086882 | 0 | NA | NA | NA | NA | NA | Gm13352 | predicted gene 13352 [Source:MGI Symbo | 0 | 0 | 0 | 0 | 0 | 0 |
| 28446 | ENSMUSG00000086884 | 0 | NA | NA | NA | NA | NA | Gm16225 | predicted gene 16225 [Source:MGI Symbo | 0 | 0 | 0 | 0 | 0 | 0 |
| 28447 | ENSMUSG00000086887 | 0.453341394 | -0.545391191 | 3.551113277 | -0.153583158 | 0.877938417 | NA | 1700060C16 | RIKEN cDNA 1700060C16 gene [Source:M | 1.6062 | 0 | 0 | 0 | 0 | 1.1138 |
| 28448 | ENSMUSG00000086889 | 7.628524537 | 1.054804454 | 0.818538063 | 1.288644355 | 0.197521756 | 0.639425845 | Phf2os1 | PHD finger protein 2, opposite strand 1 [S | 5.6217 | 6.1675 | 3.0187 | 7.7118 | 6.5437 | 16.708 |
| 28449 | ENSMUSG00000086890 | 2.974494553 | 1.176002001 | 1.235875552 | 0.951553737 | 0.341323353 | NA | 1700021J08I | RIKEN cDNA 1700021J08 gene [Source:M( | 2.4093 | 1.0279 | 2.0125 | 5.7839 | 3.2719 | 3.3415 |
| 28450 | ENSMUSG00000086891 | 0 | NA | NA | NA | NA | NA | Gm15963 | predicted gene 15963 [Source:MGI Symbo | 0 | 0 | 0 | 0 | 0 | 0 |
| 28451 | ENSMUSG00000086892 | 0 | NA | NA | NA | NA | NA | 4933430M04 | RIKEN cDNA 4933430M04 gene [Source:M | 0 | 0 | 0 | 0 | 0 | 0 |
| 28452 | ENSMUSG00000086893 | 0.185640647 | 1.126924175 | 4.080472857 | 0.276174898 | 0.782413736 | NA | Gm11365 | predicted gene 11365 [Source:MGI Symbo | 0 | 0 | 0 | 0 | 0 | 1.1138 |
| 28453 | ENSMUSG00000086894 | 11.34893665 | -0.530705844 | 0.697899262 | -0.760433307 | 0.44699562 | 0.853398186 | Gm15708 | predicted gene 15708 [Source:MGI Symbo | 17.668 | 10.279 | 12.075 | 4.8199 | 6.5437 | 16.708 |
| 28454 | ENSMUSG00000086896 | 0 | NA | NA | NA | NA | NA | Cyp4x1os | cytochrome P450, family 4, subfamily x, p | 0 | 0 | 0 | 0 | 0 | 0 |
| 28455 | ENSMUSG00000086898 | 0 | NA | NA | NA | NA | NA | Itpr3os | inositol 1,4,5-triphosphate receptor 3, op | 0 | 0 | 0 | 0 | 0 | 0 |
| 28456 | ENSMUSG00000086900 | 0 | NA | NA | NA | NA | NA | Kcnab3os | potassium voltage-gated channel, shaker- | 0 | 0 | 0 | 0 | 0 | 0 |
| 28457 | ENSMUSG00000086902 | 0 | NA | NA | NA | NA | NA | Gm11728 | predicted gene 11728 [Source:MGI Symbo | 0 | 0 | 0 | 0 | 0 | 0 |
| 28458 | ENSMUSG00000086903 | 0 | NA | NA | NA | NA | NA | Hotair | HOX transcript antisense RNA (non-protei | 0 | 0 | 0 | 0 | 0 | 0 |
| 28459 | ENSMUSG00000086904 | 0 | NA | NA | NA | NA | NA | Gm13404 | predicted gene 13404 [Source:MGI Symbo | 0 | 0 | 0 | 0 | 0 | 0 |
| 28460 | ENSMUSG00000086905 | 0 | NA | NA | NA | NA | NA | Gm13716 | predicted gene 13716 [Source:MGI Symbo | 0 | 0 | 0 | 0 | 0 | 0 |
| 28461 | ENSMUSG00000086906 | 0 | NA | NA | NA | NA | NA | Gm15276 | predicted gene 15276 [Source:MGI Symbo | 0 | 0 | 0 | 0 | 0 | 0 |
| 28462 | ENSMUSG00000086907 | 0 | NA | NA | NA | NA | NA | Gm15298 | predicted gene 15298 [Source:MGI Symbo | 0 | 0 | 0 | 0 | 0 | 0 |
| 28463 | ENSMUSG00000086908 | 0 | NA | NA | NA | NA | NA | Gm13598 | predicted gene 13598 [Source:MGI Symbo | 0 | 0 | 0 | 0 | 0 | 0 |
| 28464 | ENSMUSG00000086909 | 0.17131964 | -0.796629176 | 4.080472857 | -0.195229623 | 0.84521319 | NA | Gm4926 | predicted gene 4926 [Source:MGI Symbol | 0 | 1.0279 | 0 | 0 | 0 | 0 |
| 28465 | ENSMUSG00000086910 | 0 | NA | NA | NA | NA | NA | Gm15755 | predicted gene 15755 [Source:MGI Symbo | 0 | 0 | 0 | 0 | 0 | 0 |
| 28466 | ENSMUSG00000086911 | 0 | NA | NA | NA | NA | NA | Gm12027 | predicted gene 12027 [Source:MGI Symbo | 0 | 0 | 0 | 0 | 0 | 0 |
| 28467 | ENSMUSG00000086912 | 0 | NA | NA | NA | NA | NA | 4930505O20 | RIKEN cDNA 4930505O20 gene [Source:M | 0 | 0 | 0 | 0 | 0 | 0 |
| 28468 | ENSMUSG00000086913 | 0 | NA | NA | NA | NA | NA | Gm14019 | predicted gene 14019 [Source:MGI Symbo | 0 | 0 | 0 | 0 | 0 | 0 |
| 28469 | ENSMUSG00000086914 | 62.85949437 | 0.023361479 | 0.29480229 | 0.079244563 | 0.9368381 | 0.988420305 | Gm16124 | predicted gene 16124 [Source:MGI Symbo | 60.233 | 61.675 | 65.406 | 73.262 | 69.8 | 46.781 |
| 28470 | ENSMUSG00000086915 | 0.185640647 | 1.126924175 | 4.080472857 | 0.276174898 | 0.782413736 | NA | Gm16364 | predicted gene 16364 [Source:MGI Symbo | 0 | 0 | 0 | 0 | 0 | 1.1138 |
| 28471 | ENSMUSG00000086916 | 2.975011473 | -1.305981644 | 1.315459918 | -0.992794707 | 0.32081001 | NA | Gm15903 | predicted gene 15903 [Source:MGI Symbo | 5.6217 | 3.0838 | 4.025 | 2.8919 | 0 | 2.2277 |
| 28472 | ENSMUSG00000086917 | 0 | NA | NA | NA | NA | NA | Gm11630 | predicted gene 11630 [Source:MGI Symbo | 0 | 0 | 0 | 0 | 0 | 0 |
| 28473 | ENSMUSG00000086918 | 2.96282985 | 0.957896561 | 1.516308634 | 0.63172928 | 0.527563792 | NA | 4930429F24 | RIKEN cDNA 4930429F24 gene [Source:M( | 0 | 4.1117 | 2.0125 | 3.8559 | 0 | 7.7969 |
| 28474 | ENSMUSG00000086919 | 0 | NA | NA | NA | NA | NA | Gm11497 | predicted gene 11497 [Source:MGI Symbo | 0 | 0 | 0 | 0 | 0 | 0 |
| 28475 | ENSMUSG00000086920 | 4.126563658 | -0.018564735 | 1.166861993 | -0.015909966 | 0.987306219 | NA | Gm12207 | predicted gene 12207 [Source:MGI Symbo | 2.4093 | 4.1117 | 6.0375 | 6.7478 | 5.4531 | 0 |
| 28476 | ENSMUSG00000086921 | 0 | NA | NA | NA | NA | NA | Gm13189 | predicted gene 13189 [Source:MGI Symbo | 0 | 0 | 0 | 0 | 0 | 0 |
| 28477 | ENSMUSG00000086922 | 13.01704325 | -0.051061778 | 0.609432429 | -0.08378579 | 0.933226746 | 0.988420305 | Gm13835 | predicted gene 13835 [Source:MGI Symbo | 10.44 | 15.419 | 14.087 | 18.316 | 9.8156 | 10.025 |
| 28478 | ENSMUSG00000086923 | 0 | NA | NA | NA | NA | NA | 4930406D18 | RIKEN cDNA 4930406D18 gene [Source:M | 0 | 0 | 0 | 0 | 0 | 0 |
| 28479 | ENSMUSG00000086924 | 9.106288869 | -0.463856966 | 0.758680546 | -0.611399579 | 0.540935081 | 0.893466372 | Gm11766 | predicted gene 11766 [Source:MGI Symbo | 10.44 | 6.1675 | 15.094 | 9.6398 | 3.2719 | 10.025 |
| 28480 | ENSMUSG00000086925 | 0.797609011 | 1.992900111 | 2.636035165 | 0.756021823 | 0.449636112 | NA | Gm6286 | predicted gene 6286 [Source:MGI Symbol | 0.8031 | 0 | 0 | 2.8919 | 1.0906 | 0 |
| 28481 | ENSMUSG00000086926 | 0 | NA | NA | NA | NA | NA | Gm6226 | predicted gene 6226 [Source:MGI Symbol | 0 | 0 | 0 | 0 | 0 | 0 |
| 28482 | ENSMUSG00000086927 | 0 | NA | NA | NA | NA | NA | Gm11823 | predicted gene 11823 [Source:MGI Symbo | 0 | 0 | 0 | 0 | 0 | 0 |
| 28483 | ENSMUSG00000086928 | 0.371281295 | 2.09253362 | 4.031852165 | 0.519000582 | 0.603760335 | NA | Gm14210 | predicted gene 14210 [Source:MGI Symbo | 0 | 0 | 0 | 0 | 0 | 2.2277 |
| 28484 | ENSMUSG00000086929 | 25.42294906 | -0.040481743 | 0.54125352 | -0.074792572 | 0.940379752 | 0.988839835 | Gm11788 | predicted gene 11788 [Source:MGI Symbo | 24.896 | 13.363 | 39.244 | 36.631 | 27.265 | 11.138 |
| 28485 | ENSMUSG00000086930 | 4.291117027 | -0.338150601 | 1.029548317 | -0.328445587 | 0.742574779 | NA | Frs3os | fibroblast growth factor receptor substrat | 7.2279 | 3.0838 | 4.025 | 4.8199 | 4.3625 | 2.2277 |
| 28486 | ENSMUSG00000086931 | 0.17131964 | -0.796629176 | 4.080472857 | -0.195229623 | 0.84521319 | NA | Gm14532 | predicted gene 14532 [Source:MGI Symbo | 0 | 1.0279 | 0 | 0 | 0 | 0 |
| 28487 | ENSMUSG00000086932 | 0 | NA | NA | NA | NA | NA | Gm22000 | predicted gene 22000 [Source:MGI Symbo | 0 | 0 | 0 | 0 | 0 | 0 |
| 28488 | ENSMUSG00000086933 | 0.294513347 | 0.165143496 | 4.080472857 | 0.040471657 | 0.967717103 | NA | 4933426K07 | RIKEN cDNA 4933426K07 gene [Source:M | 0.8031 | 0 | 0 | 0.964 | 0 | 0 |
| 28489 | ENSMUSG00000086935 | 5.725492352 | -0.587424908 | 0.936797464 | -0.627056467 | 0.530622222 | 0.888794797 | Gm15956 | predicted gene 15956 [Source:MGI Symbo | 2.4093 | 10.279 | 8.05 | 4.8199 | 5.4531 | 3.3415 |
| 28490 | ENSMUSG00000086936 | 0 | NA | NA | NA | NA | NA | Gm9697 | predicted gene 9697 [Source:MGI Symbol | 0 | 0 | 0 | 0 | 0 | 0 |
| 28491 | ENSMUSG00000086937 | 0 | NA | NA | NA | NA | NA | Gm15063 | predicted gene 15063 [Source:MGI Symbo | 0 | 0 | 0 | 0 | 0 | 0 |
| 28492 | ENSMUSG00000086938 | 107.0813093 | 0.16064482 | 0.241712203 | 0.664611958 | 0.506298719 | 0.881592135 | 4930481A15 | RIKEN cDNA 4930481A15 gene [Source:M | 100.39 | 78.122 | 124.77 | 109.89 | 111.24 | 118.07 |
| 28493 | ENSMUSG00000086939 | 0 | NA | NA | NA | NA | NA | Gm13530 | predicted gene 13530 [Source:MGI Symbo | 0 | 0 | 0 | 0 | 0 | 0 |
| 28494 | ENSMUSG00000086940 | 0.185640647 | 1.126924175 | 4.080472857 | 0.276174898 | 0.782413736 | NA | Gm12746 | predicted gene 12746 [Source:MGI Symbo | 0 | 0 | 0 | 0 | 0 | 1.1138 |
| 28495 | ENSMUSG00000086941 | 0 | NA | NA | NA | NA | NA | Gm2020 | predicted gene 2020 [Source:MGI Symbol | 0 | 0 | 0 | 0 | 0 | 0 |
| 28496 | ENSMUSG00000086942 | 1.216836277 | 0.526797018 | 2.086953666 | 0.252423917 | 0.800713416 | NA | Gm15489 | predicted gene 15489 [Source:MGI Symbo | 0 | 0 | 3.0187 | 0.964 | 1.0906 | 2.2277 |
| 28497 | ENSMUSG00000086943 | 0 | NA | NA | NA | NA | NA | 4732414G09 | RIKEN cDNA 4732414G09 gene [Source:M | 0 | 0 | 0 | 0 | 0 | 0 |
| 28498 | ENSMUSG00000086944 | 0.305170013 | -1.657836477 | 4.041480335 | -0.410205257 | 0.681655385 | NA | Gm15859 | predicted gene 15859 [Source:MGI Symbo | 0.8031 | 1.0279 | 0 | 0 | 0 | 0 |
| 28499 | ENSMUSG00000086945 | 0 | NA | NA | NA | NA | NA | 4930562A09 | RIKEN cDNA 4930562A09 gene [Source:M | 0 | 0 | 0 | 0 | 0 | 0 |
| 28500 | ENSMUSG00000086946 | 1.794922079 | -2.055900096 | 1.734060458 | -1.185598856 | 0.235780754 | NA | Gm15527 | predicted gene 15527 [Source:MGI Symbo | 1.6062 | 3.0838 | 4.025 | 0.964 | 1.0906 | 0 |

| | | | | | | | | | | | | | | | |
|---|---|---|---|---|---|---|---|---|---|---|---|---|---|---|---|
| 28501 | ENSMUSG00000086947 | 0 | NA | NA | NA | NA | NA | 4930522O17 | RIKEN cDNA 4930522O17 gene [Source:M | 0 | 0 | 0 | 0 | 0 | 0 |
| 28502 | ENSMUSG00000086948 | 0 | NA | NA | NA | NA | NA | Gm14697 | predicted gene 14697 [Source:MGI Symbo | 0 | 0 | 0 | 0 | 0 | 0 |
| 28503 | ENSMUSG00000086949 | 0 | NA | NA | NA | NA | NA | Gm13066 | predicted gene 13066 [Source:MGI Symbo | 0 | 0 | 0 | 0 | 0 | 0 |
| 28504 | ENSMUSG00000086951 | 0.167707431 | -0.796629176 | 4.080472857 | -0.195229623 | 0.84521319 | NA | 1700051A21 | RIKEN cDNA 1700051A21 gene [Source:M | 0 | 0 | 1.0062 | 0 | 0 | 0 |
| 28505 | ENSMUSG00000086952 | 0.531944269 | 2.618916564 | 3.285123409 | 0.797204926 | 0.425332026 | NA | Gm12596 | predicted gene 12596 [Source:MGI Symbo | 0 | 0 | 0 | 0.964 | 0 | 2.2277 |
| 28506 | ENSMUSG00000086953 | 0 | NA | NA | NA | NA | NA | Aknaos | AT-hook transcription factor, opposite str | 0 | 0 | 0 | 0 | 0 | 0 |
| 28507 | ENSMUSG00000086954 | 0 | NA | NA | NA | NA | NA | 4930583P06 | RIKEN cDNA 4930583P06 gene [Source:M | 0 | 0 | 0 | 0 | 0 | 0 |
| 28508 | ENSMUSG00000086955 | 0 | NA | NA | NA | NA | NA | 1700021L23I | RIKEN cDNA 1700021L23 gene [Source:M | 0 | 0 | 0 | 0 | 0 | 0 |
| 28509 | ENSMUSG00000086957 | 0.670829724 | -2.782124174 | 3.401449285 | -0.817923168 | 0.41340106 | NA | Gm13869 | predicted gene 13869 [Source:MGI Symbo | 0 | 0 | 4.025 | 0 | 0 | 0 |
| 28510 | ENSMUSG00000086959 | 0 | NA | NA | NA | NA | NA | E330010L02I | RIKEN cDNA E330010L02 gene [Source:M | 0 | 0 | 0 | 0 | 0 | 0 |
| 28511 | ENSMUSG00000086960 | 1.562829041 | -0.239127084 | 1.790649738 | -0.133542076 | 0.89376469 | NA | Gm13017 | predicted gene 13017 [Source:MGI Symbo | 0 | 4.1117 | 1.0062 | 0.964 | 2.1812 | 1.1138 |
| 28512 | ENSMUSG00000086961 | 3.110154351 | -0.350508272 | 1.208137343 | -0.290122868 | 0.771722241 | NA | Gm12946 | predicted gene 12946 [Source:MGI Symbo | 2.4093 | 4.1117 | 4.025 | 4.8199 | 2.1812 | 1.1138 |
| 28513 | ENSMUSG00000086962 | 0.695728634 | -1.425610641 | 3.05889997 | -0.46605337 | 0.641177299 | NA | Gm12248 | predicted gene 12248 [Source:MGI Symbo | 0 | 3.0838 | 0 | 0 | 1.0906 | 0 |
| 28514 | ENSMUSG00000086963 | 0 | NA | NA | NA | NA | NA | Gm14135 | predicted gene 14135 [Source:MGI Symbo | 0 | 0 | 0 | 0 | 0 | 0 |
| 28515 | ENSMUSG00000086964 | 0 | NA | NA | NA | NA | NA | Gm15948 | predicted gene 15948 [Source:MGI Symbo | 0 | 0 | 0 | 0 | 0 | 0 |
| 28516 | ENSMUSG00000086965 | 7.28921964 | -0.11384741 | 0.792599256 | -0.143638048 | 0.885786294 | 0.982139744 | Rtl10 | retrotransposon Gag like 10 [Source:MGI | 10.44 | 5.1396 | 7.0437 | 6.7478 | 5.4531 | 8.9108 |
| 28517 | ENSMUSG00000086968 | 29.30279273 | -0.448256766 | 0.453928149 | -0.98750599 | 0.323394648 | 0.770475661 | 4933431E20 | RIKEN cDNA 4933431E20 gene [Source:M | 25.699 | 33.921 | 42.262 | 37.595 | 19.631 | 16.708 |
| 28518 | ENSMUSG00000086969 | 0.476489653 | -2.293616034 | 3.421506946 | -0.670352587 | 0.502633051 | NA | 4930443O20 | RIKEN cDNA 4930443O20 gene [Source:M | 0.8031 | 2.0558 | 0 | 0 | 0 | 0 |
| 28519 | ENSMUSG00000086970 | 0 | NA | NA | NA | NA | NA | Bcas1os1 | breast carcinoma amplified sequence 1, o | 0 | 0 | 0 | 0 | 0 | 0 |
| 28520 | ENSMUSG00000086971 | 0 | NA | NA | NA | NA | NA | 4930527B05 | RIKEN cDNA 4930527B05 gene [Source:M | 0 | 0 | 0 | 0 | 0 | 0 |
| 28521 | ENSMUSG00000086974 | 0 | NA | NA | NA | NA | NA | Gm15092 | predicted gene 15092 [Source:MGI Symbo | 0 | 0 | 0 | 0 | 0 | 0 |
| 28522 | ENSMUSG00000086975 | 0 | NA | NA | NA | NA | NA | Gm15208 | predicted gene 15208 [Source:MGI Symbo | 0 | 0 | 0 | 0 | 0 | 0 |
| 28523 | ENSMUSG00000086976 | 1.136842672 | -1.279029039 | 2.158329997 | -0.592601243 | 0.553448047 | NA | Gm12316 | predicted gene 12316 [Source:MGI Symbo | 0.8031 | 3.0838 | 1.0062 | 1.928 | 0 | 0 |
| 28524 | ENSMUSG00000086978 | 0 | NA | NA | NA | NA | NA | 4933435F18 | RIKEN cDNA 4933435F18 gene [Source:M | 0 | 0 | 0 | 0 | 0 | 0 |
| 28525 | ENSMUSG00000086980 | 0 | NA | NA | NA | NA | NA | Gm13791 | predicted gene 13791 [Source:MGI Symbo | 0 | 0 | 0 | 0 | 0 | 0 |
| 28526 | ENSMUSG00000086981 | 0 | NA | NA | NA | NA | NA | Gm12171 | predicted gene 12171 [Source:MGI Symbo | 0 | 0 | 0 | 0 | 0 | 0 |
| 28527 | ENSMUSG00000086984 | 0 | NA | NA | NA | NA | NA | Gm6762 | predicted pseudogene 6762 [Source:MGI | 0 | 0 | 0 | 0 | 0 | 0 |
| 28528 | ENSMUSG00000086986 | 0.185640647 | 1.126924175 | 4.080472857 | 0.276174898 | 0.782413736 | NA | Gm12103 | predicted gene 12103 [Source:MGI Symbo | 0 | 0 | 0 | 0 | 0 | 1.1138 |
| 28529 | ENSMUSG00000086987 | 0 | NA | NA | NA | NA | NA | Gm12484 | predicted gene 12484 [Source:MGI Symbo | 0 | 0 | 0 | 0 | 0 | 0 |
| 28530 | ENSMUSG00000086989 | 0 | NA | NA | NA | NA | NA | Gm16441 | predicted pseudogene 16441 [Source:MG | 0 | 0 | 0 | 0 | 0 | 0 |
| 28531 | ENSMUSG00000086991 | 0.167707431 | -0.796629176 | 4.080472857 | -0.195229623 | 0.84521319 | NA | Gm15334 | predicted gene 15334 [Source:MGI Symbo | 0 | 0 | 1.0062 | 0 | 0 | 0 |
| 28532 | ENSMUSG00000086992 | 0.160662974 | 1.126924175 | 4.080472857 | 0.276174898 | 0.782413736 | NA | Gm15941 | predicted gene 15941 [Source:MGI Symbo | 0 | 0 | 0 | 0.964 | 0 | 0 |
| 28533 | ENSMUSG00000086993 | 2.78312307 | 1.691893526 | 1.498177699 | 1.129300969 | 0.258770892 | NA | Rsf1os2 | remodeling and spacing factor 1, opposite | 0 | 0 | 4.025 | 3.8559 | 4.3625 | 4.4554 |
| 28534 | ENSMUSG00000086994 | 0 | NA | NA | NA | NA | NA | Gm13402 | predicted gene 13402 [Source:MGI Symbo | 0 | 0 | 0 | 0 | 0 | 0 |
| 28535 | ENSMUSG00000086995 | 0 | NA | NA | NA | NA | NA | Gm13544 | predicted gene 13544 [Source:MGI Symbo | 0 | 0 | 0 | 0 | 0 | 0 |
| 28536 | ENSMUSG00000086996 | 0 | NA | NA | NA | NA | NA | 4933416E14 | RIKEN cDNA 4933416E14 gene [Source:M | 0 | 0 | 0 | 0 | 0 | 0 |
| 28537 | ENSMUSG00000086997 | 0.401551119 | -2.072982726 | 4.008251979 | -0.517178744 | 0.605031388 | NA | Gm12981 | predicted gene 12981 [Source:MGI Symbo | 2.4093 | 0 | 0 | 0 | 0 | 0 |
| 28538 | ENSMUSG00000086998 | 0 | NA | NA | NA | NA | NA | Gm16033 | predicted gene 16033 [Source:MGI Symbo | 0 | 0 | 0 | 0 | 0 | 0 |
| 28539 | ENSMUSG00000086999 | 0 | NA | NA | NA | NA | NA | Bcas1os2 | breast carcinoma amplified sequence 1, o | 0 | 0 | 0 | 0 | 0 | 0 |
| 28540 | ENSMUSG00000087001 | 0.17131964 | -0.796629176 | 4.080472857 | -0.195229623 | 0.84521319 | NA | Gm15475 | predicted gene 15475 [Source:MGI Symbo | 0 | 1.0279 | 0 | 0 | 0 | 0 |
| 28541 | ENSMUSG00000087002 | 0 | NA | NA | NA | NA | NA | Gm16277 | predicted gene 16277 [Source:MGI Symbo | 0 | 0 | 0 | 0 | 0 | 0 |
| 28542 | ENSMUSG00000087003 | 0.185640647 | 1.126924175 | 4.080472857 | 0.276174898 | 0.782413736 | NA | Cntrobos | centrobin, centrosomal BRCA2 interacting | 0 | 0 | 0 | 0 | 0 | 1.1138 |
| 28543 | ENSMUSG00000087004 | 0 | NA | NA | NA | NA | NA | Gm14154 | predicted gene 14154 [Source:MGI Symbo | 0 | 0 | 0 | 0 | 0 | 0 |
| 28544 | ENSMUSG00000087006 | 23.3823237 | -0.558076693 | 0.449279638 | -1.242158883 | 0.214177947 | 0.658874792 | Gm13889 | predicted gene 13889 [Source:MGI Symbo | 25.699 | 26.726 | 31.194 | 20.244 | 15.269 | 21.163 |
| 28545 | ENSMUSG00000087007 | 0 | NA | NA | NA | NA | NA | Gm2231 | predicted gene 2231 [Source:MGI Symbol | 0 | 0 | 0 | 0 | 0 | 0 |
| 28546 | ENSMUSG00000087008 | 0.305170013 | -1.657836477 | 4.041480335 | -0.410205257 | 0.681655385 | NA | Gm5530 | predicted gene 5530 [Source:MGI Symbol | 0.8031 | 1.0279 | 0 | 0 | 0 | 0 |
| 28547 | ENSMUSG00000087009 | 0 | NA | NA | NA | NA | NA | Nphs1os | nephrosis 1 homolog, nephrin, opposite s | 0 | 0 | 0 | 0 | 0 | 0 |
| 28548 | ENSMUSG00000087010 | 0 | NA | NA | NA | NA | NA | 1700041M05 | RIKEN cDNA 1700041M05 gene [Source:M | 0 | 0 | 0 | 0 | 0 | 0 |
| 28549 | ENSMUSG00000087011 | 0 | NA | NA | NA | NA | NA | Gm12930 | predicted gene 12930 [Source:MGI Symbo | 0 | 0 | 0 | 0 | 0 | 0 |
| 28550 | ENSMUSG00000087013 | 3.934905754 | -1.525608476 | 1.179600703 | -1.293326184 | 0.195898268 | NA | 2610027K06 | RIKEN cDNA 2610027K06 gene [Source:M | 10.44 | 3.0838 | 4.025 | 3.8559 | 1.0906 | 1.1138 |
| 28551 | ENSMUSG00000087014 | 0.87367955 | 2.066316271 | 2.569175551 | 0.804272121 | 0.421239835 | NA | Gm16364 | predicted gene 16364 [Source:MGI Symbo | 0 | 0 | 1.0062 | 0.964 | 3.2719 | 0 |
| 28552 | ENSMUSG00000087015 | 0 | NA | NA | NA | NA | NA | Gm14041 | predicted gene 14041 [Source:MGI Symbo | 0 | 0 | 0 | 0 | 0 | 0 |
| 28553 | ENSMUSG00000087016 | 0 | NA | NA | NA | NA | NA | Gm15979 | predicted gene 15979 [Source:MGI Symbo | 0 | 0 | 0 | 0 | 0 | 0 |
| 28554 | ENSMUSG00000087017 | 5.973146758 | 0.133001011 | 0.979260062 | 0.135817865 | 0.891965267 | 0.982489666 | 4930417H01 | RIKEN cDNA 4930417H01 gene [Source:M | 4.0155 | 3.0838 | 10.062 | 8.6758 | 1.0906 | 8.9108 |
| 28555 | ENSMUSG00000087018 | 0 | NA | NA | NA | NA | NA | 2900072N19 | RIKEN cDNA 2900072N19 gene [Source:M | 0 | 0 | 0 | 0 | 0 | 0 |
| 28556 | ENSMUSG00000087020 | 10.19969657 | -0.339253177 | 0.746351065 | -0.454549063 | 0.649433687 | 0.925643431 | Gm16283 | predicted gene 16283 [Source:MGI Symbo | 8.8341 | 15.419 | 10.062 | 11.568 | 13.087 | 2.2277 |

| | | | | | | | | | | | | | | | |
|---|---|---|---|---|---|---|---|---|---|---|---|---|---|---|---|
| 28557 | ENSMUSG00000087021 | 0 | NA | NA | NA | NA | NA | Fthl17-ps3 | ferritin, heavy polypeptide-like 17, pseudo | 0 | 0 | 0 | 0 | 0 | 0 |
| 28558 | ENSMUSG00000087022 | 3.749426133 | 1.021309568 | 1.170099694 | 0.872839787 | 0.38275041 | NA | 9130024F11 | RIKEN cDNA 9130024F11 gene [Source:M( | 2.4093 | 2.0558 | 3.0187 | 10.604 | 2.1812 | 2.2277 |
| 28559 | ENSMUSG00000087024 | 0 | NA | NA | NA | NA | NA | B230119M0! | RIKEN cDNA B230119M05 gene [Source:M | 0 | 0 | 0 | 0 | 0 | 0 |
| 28560 | ENSMUSG00000087026 | 4.380867386 | 0.526342815 | 1.023508232 | 0.514253621 | 0.607074683 | NA | A230103J11I | RIKEN cDNA A230103J11 gene [Source:M( | 5.6217 | 2.0558 | 3.0187 | 6.7478 | 3.2719 | 5.5692 |
| 28561 | ENSMUSG00000087027 | 0 | NA | NA | NA | NA | NA | Gm13206 | predicted gene 13206 [Source:MGI Symbo | 0 | 0 | 0 | 0 | 0 | 0 |
| 28562 | ENSMUSG00000087028 | 0.888207148 | 0.719020891 | 2.356182394 | 0.305163511 | 0.760241622 | NA | Gm13387 | predicted gene 13387 [Source:MGI Symbo | 0 | 1.0279 | 1.0062 | 0 | 2.1812 | 1.1138 |
| 28563 | ENSMUSG00000087029 | 0 | NA | NA | NA | NA | NA | Gm14133 | predicted gene 14133 [Source:MGI Symbo | 0 | 0 | 0 | 0 | 0 | 0 |
| 28564 | ENSMUSG00000087030 | 14.04040918 | 0.429337723 | 0.597578407 | 0.718462578 | 0.472472114 | 0.868340109 | Gm16143 | predicted gene 16143 [Source:MGI Symbo | 9.6372 | 9.2513 | 17.106 | 16.388 | 20.722 | 11.138 |
| 28565 | ENSMUSG00000087031 | 0 | NA | NA | NA | NA | NA | Gm14241 | predicted gene 14241 [Source:MGI Symbo | 0 | 0 | 0 | 0 | 0 | 0 |
| 28566 | ENSMUSG00000087032 | 0 | NA | NA | NA | NA | NA | Gm13874 | predicted gene 13874 [Source:MGI Symbo | 0 | 0 | 0 | 0 | 0 | 0 |
| 28567 | ENSMUSG00000087033 | 0 | NA | NA | NA | NA | NA | Gm14155 | predicted gene 14155 [Source:MGI Symbo | 0 | 0 | 0 | 0 | 0 | 0 |
| 28568 | ENSMUSG00000087034 | 0 | NA | NA | NA | NA | NA | Cbfa2t2-ps1 | CBFA2/RUNX1 translocation partner 2, ps | 0 | 0 | 0 | 0 | 0 | 0 |
| 28569 | ENSMUSG00000087035 | 0 | NA | NA | NA | NA | NA | Tmem74bos | transmembrane 74B, opposite strand [Sou | 0 | 0 | 0 | 0 | 0 | 0 |
| 28570 | ENSMUSG00000087036 | 2.675082006 | -0.007350798 | 1.35205861 | -0.005436745 | 0.995662126 | NA | Gm11748 | predicted gene 11748 [Source:MGI Symbo | 4.0155 | 1.0279 | 3.0187 | 5.7839 | 1.0906 | 1.1138 |
| 28571 | ENSMUSG00000087037 | 0 | NA | NA | NA | NA | NA | 4930432L08I | RIKEN cDNA 4930432L08 gene [Source:M( | 0 | 0 | 0 | 0 | 0 | 0 |
| 28572 | ENSMUSG00000087038 | 0 | NA | NA | NA | NA | NA | 2900079G21 | RIKEN cDNA 2900079G21 gene [Source:M | 0 | 0 | 0 | 0 | 0 | 0 |
| 28573 | ENSMUSG00000087039 | 0.185640647 | 1.126924175 | 4.080472857 | 0.276174898 | 0.782413736 | NA | Cdrt4os2 | CMT1A duplicated region transcript 4, op | 0 | 0 | 0 | 0 | 0 | 1.1138 |
| 28574 | ENSMUSG00000087040 | 0 | NA | NA | NA | NA | NA | Gm14033 | predicted gene 14033 [Source:MGI Symbo | 0 | 0 | 0 | 0 | 0 | 0 |
| 28575 | ENSMUSG00000087041 | 0 | NA | NA | NA | NA | NA | Gm16295 | predicted gene 16295 [Source:MGI Symbo | 0 | 0 | 0 | 0 | 0 | 0 |
| 28576 | ENSMUSG00000087042 | 0.845761573 | -3.112447952 | 2.7647732 | -1.125751636 | 0.260270661 | NA | Gm11611 | predicted gene 11611 [Source:MGI Symbo | 0 | 2.0558 | 3.0187 | 0 | 0 | 0 |
| 28577 | ENSMUSG00000087043 | 0 | NA | NA | NA | NA | NA | Gm13599 | predicted gene 13599 [Source:MGI Symbo | 0 | 0 | 0 | 0 | 0 | 0 |
| 28578 | ENSMUSG00000087044 | 0 | NA | NA | NA | NA | NA | 1700042G15 | RIKEN cDNA 1700042G15 gene [Source:M | 0 | 0 | 0 | 0 | 0 | 0 |
| 28579 | ENSMUSG00000087045 | 0 | NA | NA | NA | NA | NA | 4930515B02 | RIKEN cDNA 4930515B02 gene [Source:M | 0 | 0 | 0 | 0 | 0 | 0 |
| 28580 | ENSMUSG00000087047 | 0.481988923 | 2.492401104 | 3.837077151 | 0.649557203 | 0.515978285 | NA | 1700110K17 | RIKEN cDNA 1700110K17 gene [Source:M( | 0 | 0 | 0 | 2.8919 | 0 | 0 |
| 28581 | ENSMUSG00000087049 | 2.330789414 | 0.151221756 | 1.615989399 | 0.093578433 | 0.925444042 | NA | Bach2it1 | BTB and CNC homology 2, intronic transcr | 1.6062 | 0 | 5.0312 | 2.8919 | 0 | 4.4554 |
| 28582 | ENSMUSG00000087050 | 0 | NA | NA | NA | NA | NA | Dhrs13os | dehydrogenase/reductase (SDR family) m | 0 | 0 | 0 | 0 | 0 | 0 |
| 28583 | ENSMUSG00000087051 | 0.642651897 | 2.906894826 | 3.440535777 | 0.844895974 | 0.398168923 | NA | Gm12730 | predicted gene 12730 [Source:MGI Symbo | 0 | 0 | 0 | 3.8559 | 0 | 0 |
| 28584 | ENSMUSG00000087052 | 0.181769715 | 1.126924175 | 4.080472857 | 0.276174898 | 0.782413736 | NA | Gm8093 | predicted gene 8093 [Source:MGI Symbol | 0 | 0 | 0 | 0 | 1.0906 | 0 |
| 28585 | ENSMUSG00000087053 | 0 | NA | NA | NA | NA | NA | Gm15082 | predicted gene 15082 [Source:MGI Symbo | 0 | 0 | 0 | 0 | 0 | 0 |
| 28586 | ENSMUSG00000087054 | 1.997714785 | 4.523214684 | 1.933691375 | 2.339160603 | 0.019327123 | NA | Gm12405 | predicted gene 12405 [Source:MGI Symbo | 0 | 0 | 0 | 0.964 | 5.4531 | 5.5692 |
| 28587 | ENSMUSG00000087055 | 1.031479329 | -2.220845979 | 2.403244684 | -0.924103148 | 0.355432614 | NA | Gm11948 | predicted gene 11948 [Source:MGI Symbo | 3.2124 | 0 | 2.0125 | 0.964 | 0 | 0 |
| 28588 | ENSMUSG00000087056 | 0 | NA | NA | NA | NA | NA | Gm14004 | predicted gene 14004 [Source:MGI Symbo | 0 | 0 | 0 | 0 | 0 | 0 |
| 28589 | ENSMUSG00000087057 | 0 | NA | NA | NA | NA | NA | Gm11730 | predicted gene 11730 [Source:MGI Symbo | 0 | 0 | 0 | 0 | 0 | 0 |
| 28590 | ENSMUSG00000087058 | 0 | NA | NA | NA | NA | NA | Gm12301 | predicted gene 12301 [Source:MGI Symbo | 0 | 0 | 0 | 0 | 0 | 0 |
| 28591 | ENSMUSG00000087059 | 0 | NA | NA | NA | NA | NA | Gm12339 | predicted gene 12339 [Source:MGI Symbo | 0 | 0 | 0 | 0 | 0 | 0 |
| 28592 | ENSMUSG00000087060 | 1.802275739 | -0.778147066 | 1.7939284 | -0.43376707 | 0.664457597 | NA | Eldr | Egfr long non-coding downstream RNA [S | 4.8186 | 0 | 2.0125 | 2.8919 | 1.0906 | 0 |
| 28593 | ENSMUSG00000087061 | 0 | NA | NA | NA | NA | NA | A130030D18 | RIKEN cDNA A130030D18 gene [Source:M | 0 | 0 | 0 | 0 | 0 | 0 |
| 28594 | ENSMUSG00000087062 | 0 | NA | NA | NA | NA | NA | NA | NA | 0 | 0 | 0 | 0 | 0 | 0 |
| 28595 | ENSMUSG00000087063 | 0 | NA | NA | NA | NA | NA | Gm15857 | predicted gene 15857 [Source:MGI Symbo | 0 | 0 | 0 | 0 | 0 | 0 |
| 28596 | ENSMUSG00000087064 | 1.808957732 | 0.404348583 | 1.744387565 | 0.23179974 | 0.816693559 | NA | Sap30bpos | SAP30 binding protein, opposite strand [S | 1.6062 | 3.0838 | 0 | 2.8919 | 3.2719 | 0 |
| 28597 | ENSMUSG00000087066 | 0 | NA | NA | NA | NA | NA | Gm15518 | predicted gene 15518 [Source:MGI Symbo | 0 | 0 | 0 | 0 | 0 | 0 |
| 28598 | ENSMUSG00000087067 | 0 | NA | NA | NA | NA | NA | Gm11532 | predicted gene 11532 [Source:MGI Symbo | 0 | 0 | 0 | 0 | 0 | 0 |
| 28599 | ENSMUSG00000087069 | 0 | NA | NA | NA | NA | NA | Gm13793 | predicted gene 13793 [Source:MGI Symbo | 0 | 0 | 0 | 0 | 0 | 0 |
| 28600 | ENSMUSG00000087070 | 0 | NA | NA | NA | NA | NA | Gm12505 | predicted gene 12505 [Source:MGI Symbo | 0 | 0 | 0 | 0 | 0 | 0 |
| 28601 | ENSMUSG00000087071 | 0 | NA | NA | NA | NA | NA | Trp73os | transformation related protein 73, opposi | 0 | 0 | 0 | 0 | 0 | 0 |
| 28602 | ENSMUSG00000087072 | 0.294513347 | 0.165143496 | 4.080472857 | 0.040471657 | 0.967717103 | NA | Recql5os1 | RecQ-like 5, opposite strand 1 [Source:M( | 0.8031 | 0 | 0 | 0.964 | 0 | 0 |
| 28603 | ENSMUSG00000087074 | 0 | NA | NA | NA | NA | NA | Gm14249 | predicted gene 14249 [Source:MGI Symbo | 0 | 0 | 0 | 0 | 0 | 0 |
| 28604 | ENSMUSG00000087075 | 0 | NA | NA | NA | NA | NA | Lbhd2 | LBH domain containing 2 [Source:MGI Syr | 0 | 0 | 0 | 0 | 0 | 0 |
| 28605 | ENSMUSG00000087076 | 0 | NA | NA | NA | NA | NA | Gm13229 | predicted gene 13229 [Source:MGI Symbo | 0 | 0 | 0 | 0 | 0 | 0 |
| 28606 | ENSMUSG00000087077 | 0 | NA | NA | NA | NA | NA | Gm12480 | predicted gene 12480 [Source:MGI Symbo | 0 | 0 | 0 | 0 | 0 | 0 |
| 28607 | ENSMUSG00000087079 | 0 | NA | NA | NA | NA | NA | Gm13389 | predicted gene 13389 [Source:MGI Symbo | 0 | 0 | 0 | 0 | 0 | 0 |
| 28608 | ENSMUSG00000087080 | 0 | NA | NA | NA | NA | NA | Gm12199 | predicted gene 12199 [Source:MGI Symbo | 0 | 0 | 0 | 0 | 0 | 0 |
| 28609 | ENSMUSG00000087082 | 0.845528352 | 3.296539104 | 2.749625613 | 1.198904712 | 0.230564999 | NA | Gm15423 | predicted gene 15423 [Source:MGI Symbo | 0 | 0 | 0 | 2.8919 | 2.1812 | 0 |
| 28610 | ENSMUSG00000087083 | 0 | NA | NA | NA | NA | NA | Gm15083 | predicted gene 15083 [Source:MGI Symbo | 0 | 0 | 0 | 0 | 0 | 0 |
| 28611 | ENSMUSG00000087084 | 0 | NA | NA | NA | NA | NA | Gm13016 | predicted gene 13016 [Source:MGI Symbo | 0 | 0 | 0 | 0 | 0 | 0 |
| 28612 | ENSMUSG00000087085 | 0 | NA | NA | NA | NA | NA | Gm12068 | predicted gene 12068 [Source:MGI Symbo | 0 | 0 | 0 | 0 | 0 | 0 |

| | | | | | | | | | | | | | | | |
|---|---|---|---|---|---|---|---|---|---|---|---|---|---|---|---|
| 28613 | ENSMUSG00000087086 | 0 | NA | NA | NA | NA | NA | Gm13269 | predicted gene 13269 [Source:MGI Symbo | 0 | 0 | 0 | 0 | 0 | 0 |
| 28614 | ENSMUSG00000087088 | 6.023249165 | -0.559438913 | 0.929499233 | -0.601871302 | 0.547259808 | 0.895518681 | Gm16638 | predicted gene, 16638 [Source:MGI Symb | 11.243 | 7.1954 | 3.0187 | 4.8199 | 7.6343 | 2.2277 |
| 28615 | ENSMUSG00000087089 | 0 | NA | NA | NA | NA | NA | Gm12160 | predicted gene 12160 [Source:MGI Symbo | 0 | 0 | 0 | 0 | 0 | 0 |
| 28616 | ENSMUSG00000087090 | 0.513958919 | -2.394274821 | 3.756096013 | -0.637437066 | 0.523840191 | NA | Nctc1 | non-coding transcript 1 [Source:MGI Syml | 0 | 3.0838 | 0 | 0 | 0 | 0 |
| 28617 | ENSMUSG00000087093 | 0 | NA | NA | NA | NA | NA | Gm15486 | predicted gene 15486 [Source:MGI Symbo | 0 | 0 | 0 | 0 | 0 | 0 |
| 28618 | ENSMUSG00000087094 | 1.756211554 | 0.100173744 | 1.862579119 | 0.053782276 | 0.957108631 | NA | Gm13093 | predicted gene 13093 [Source:MGI Symbo | 0 | 4.1117 | 1.0062 | 0.964 | 0 | 4.4554 |
| 28619 | ENSMUSG00000087095 | 0 | NA | NA | NA | NA | NA | Emx2os | Emx2 opposite strand/antisense transcrip | 0 | 0 | 0 | 0 | 0 | 0 |
| 28620 | ENSMUSG00000087097 | 0 | NA | NA | NA | NA | NA | 4930586N03 | RIKEN cDNA 4930586N03 gene [Source:M | 0 | 0 | 0 | 0 | 0 | 0 |
| 28621 | ENSMUSG00000087098 | 0 | NA | NA | NA | NA | NA | Gm11884 | predicted gene 11884 [Source:MGI Symbo | 0 | 0 | 0 | 0 | 0 | 0 |
| 28622 | ENSMUSG00000087099 | 0 | NA | NA | NA | NA | NA | Gm11782 | predicted gene 11782 [Source:MGI Symbo | 0 | 0 | 0 | 0 | 0 | 0 |
| 28623 | ENSMUSG00000087100 | 0 | NA | NA | NA | NA | NA | Gm14641 | predicted gene 14641 [Source:MGI Symbo | 0 | 0 | 0 | 0 | 0 | 0 |
| 28624 | ENSMUSG00000087102 | 0 | NA | NA | NA | NA | NA | Gm13661 | predicted gene 13661 [Source:MGI Symbo | 0 | 0 | 0 | 0 | 0 | 0 |
| 28625 | ENSMUSG00000087103 | 0 | NA | NA | NA | NA | NA | Gm13344 | predicted gene 13344 [Source:MGI Symbo | 0 | 0 | 0 | 0 | 0 | 0 |
| 28626 | ENSMUSG00000087104 | 0 | NA | NA | NA | NA | NA | Tmem132co | transmembrane protein 132C, opposite st | 0 | 0 | 0 | 0 | 0 | 0 |
| 28627 | ENSMUSG00000087106 | 0.695522043 | 1.639439662 | 2.762392557 | 0.593485404 | 0.552856349 | NA | Gm13964 | predicted gene 13964 [Source:MGI Symbo | 0 | 1.0279 | 0 | 0.964 | 2.1812 | 0 |
| 28628 | ENSMUSG00000087107 | 8.599527447 | -1.307731101 | 0.806642133 | -1.621203564 | 0.104973989 | 0.477616789 | AI662270 | expressed sequence AI662270 [Source:M( | 7.2279 | 18.503 | 11.069 | 4.8199 | 2.1812 | 7.7969 |
| 28629 | ENSMUSG00000087108 | 0.486939728 | -0.698472448 | 3.137608162 | -0.222613026 | 0.823836697 | NA | Gm16093 | predicted gene 16093 [Source:MGI Symbo | 0.8031 | 1.0279 | 0 | 0 | 1.0906 | 0 |
| 28630 | ENSMUSG00000087109 | 0.181769715 | 1.126924175 | 4.080472857 | 0.276174898 | 0.782413736 | NA | 4930474H06 | RIKEN cDNA 4930474H06 gene [Source:M | 0 | 0 | 0 | 0 | 1.0906 | 0 |
| 28631 | ENSMUSG00000087110 | 0 | NA | NA | NA | NA | NA | Gm11264 | predicted gene 11264 [Source:MGI Symbo | 0 | 0 | 0 | 0 | 0 | 0 |
| 28632 | ENSMUSG00000087111 | 0.301557804 | -1.64335916 | 4.042814367 | -0.406488899 | 0.684383412 | NA | Gm11399 | predicted gene 11399 [Source:MGI Symbo | 0.8031 | 0 | 1.0062 | 0 | 0 | 0 |
| 28633 | ENSMUSG00000087112 | 0 | NA | NA | NA | NA | NA | Gm7237 | predicted gene 7237 [Source:MGI Symbol | 0 | 0 | 0 | 0 | 0 | 0 |
| 28634 | ENSMUSG00000087113 | 0 | NA | NA | NA | NA | NA | Gm11714 | predicted gene 11714 [Source:MGI Symbo | 0 | 0 | 0 | 0 | 0 | 0 |
| 28635 | ENSMUSG00000087114 | 0 | NA | NA | NA | NA | NA | Gm16099 | predicted gene 16099 [Source:MGI Symbo | 0 | 0 | 0 | 0 | 0 | 0 |
| 28636 | ENSMUSG00000087115 | 0 | NA | NA | NA | NA | NA | Pcsk2os2 | proprotein convertase subtilisin/kexin typ | 0 | 0 | 0 | 0 | 0 | 0 |
| 28637 | ENSMUSG00000087116 | 0 | NA | NA | NA | NA | NA | Gm12324 | predicted gene 12324 [Source:MGI Symbo | 0 | 0 | 0 | 0 | 0 | 0 |
| 28638 | ENSMUSG00000087117 | 0 | NA | NA | NA | NA | NA | Gm11523 | predicted gene 11523 [Source:MGI Symbo | 0 | 0 | 0 | 0 | 0 | 0 |
| 28639 | ENSMUSG00000087119 | 0.644197084 | -2.725354879 | 2.762261801 | -0.986638876 | 0.323819705 | NA | Atg4a-ps | autophagy related 4A, pseudogene [Sourc | 0.8031 | 2.0558 | 1.0062 | 0 | 0 | 0 |
| 28640 | ENSMUSG00000087120 | 5.100370979 | -0.611249509 | 1.017490522 | -0.600742214 | 0.548011698 | 0.895600915 | Gm12279 | predicted gene 12279 [Source:MGI Symbo | 6.4248 | 5.1396 | 7.0437 | 8.6758 | 1.0906 | 2.2277 |
| 28641 | ENSMUSG00000087121 | 0 | NA | NA | NA | NA | NA | Gm7847 | predicted gene 7847 [Source:MGI Symbol | 0 | 0 | 0 | 0 | 0 | 0 |
| 28642 | ENSMUSG00000087122 | 0 | NA | NA | NA | NA | NA | 4930403D09 | RIKEN cDNA 4930403D09 gene [Source:M | 0 | 0 | 0 | 0 | 0 | 0 |
| 28643 | ENSMUSG00000087123 | 0 | NA | NA | NA | NA | NA | Gm6706 | predicted gene 6706 [Source:MGI Symbol | 0 | 0 | 0 | 0 | 0 | 0 |
| 28644 | ENSMUSG00000087124 | 0 | NA | NA | NA | NA | NA | Gm13839 | predicted gene 13839 [Source:MGI Symbo | 0 | 0 | 0 | 0 | 0 | 0 |
| 28645 | ENSMUSG00000087125 | 1.400751249 | -2.716509929 | 2.200550022 | -1.234468611 | 0.217028336 | NA | A230108P19 | RIKEN cDNA A230108P19 gene [Source:M | 2.4093 | 0 | 5.0312 | 0.964 | 0 | 0 |
| 28646 | ENSMUSG00000087127 | 0 | NA | NA | NA | NA | NA | Gm12756 | predicted gene 12756 [Source:MGI Symbo | 0 | 0 | 0 | 0 | 0 | 0 |
| 28647 | ENSMUSG00000087128 | 6.115053723 | -0.351312341 | 1.077395391 | -0.326075593 | 0.744367163 | 0.949997329 | Gm12655 | predicted gene 12655 [Source:MGI Symbo | 2.4093 | 11.307 | 7.0437 | 11.568 | 4.3625 | 0 |
| 28648 | ENSMUSG00000087129 | 0.997622271 | -2.171875158 | 2.465204635 | -0.881012119 | 0.378311261 | NA | Gm16316 | predicted gene 16316 [Source:MGI Symbo | 4.0155 | 0 | 1.0062 | 0.964 | 0 | 0 |
| 28649 | ENSMUSG00000087131 | 0 | NA | NA | NA | NA | NA | Gm16152 | predicted gene 16152 [Source:MGI Symbo | 0 | 0 | 0 | 0 | 0 | 0 |
| 28650 | ENSMUSG00000087132 | 18.88059815 | 0.352289024 | 0.564544486 | 0.624023497 | 0.532612154 | 0.889143097 | A930001C03 | RIKEN cDNA A930001C03 gene [Source:M | 19.274 | 6.1675 | 24.15 | 17.352 | 20.722 | 25.618 |
| 28651 | ENSMUSG00000087133 | 0 | NA | NA | NA | NA | NA | Gm27192 | predicted gene 27192 [Source:MGI Symbo | 0 | 0 | 0 | 0 | 0 | 0 |
| 28652 | ENSMUSG00000087135 | 0 | NA | NA | NA | NA | NA | Gm16096 | predicted gene 16096 [Source:MGI Symbo | 0 | 0 | 0 | 0 | 0 | 0 |
| 28653 | ENSMUSG00000087136 | 0 | NA | NA | NA | NA | NA | Gm15864 | predicted gene 15864 [Source:MGI Symbo | 0 | 0 | 0 | 0 | 0 | 0 |
| 28654 | ENSMUSG00000087137 | 0 | NA | NA | NA | NA | NA | 4930509K18 | RIKEN cDNA 4930509K18 gene [Source:M( | 0 | 0 | 0 | 0 | 0 | 0 |
| 28655 | ENSMUSG00000087138 | 45.2215933 | -0.059579948 | 0.345790208 | -0.172300854 | 0.863201013 | 0.976790757 | Gm15545 | predicted gene 15545 [Source:MGI Symbo | 53.005 | 39.061 | 46.287 | 47.235 | 53.44 | 32.301 |
| 28656 | ENSMUSG00000087139 | 0 | NA | NA | NA | NA | NA | Gm11683 | predicted gene 11683 [Source:MGI Symbo | 0 | 0 | 0 | 0 | 0 | 0 |
| 28657 | ENSMUSG00000087140 | 0 | NA | NA | NA | NA | NA | Gm11376 | predicted gene 11376 [Source:MGI Symbo | 0 | 0 | 0 | 0 | 0 | 0 |
| 28658 | ENSMUSG00000087141 | 2100.034185 | 0.908620853 | 0.182472987 | 4.97948145 | 6.38E-07 | 5.46E-05 | Plcxd2 | phosphatidylinositol-specific phospholipa | 1523.5 | 1643.6 | 1212.5 | 3568.6 | 1964.2 | 2687.7 |
| 28659 | ENSMUSG00000087142 | 3.993328071 | 0.470518563 | 1.19758253 | 0.392890303 | 0.694400504 | NA | Gm12454 | predicted gene 12454 [Source:MGI Symbo | 0 | 4.1117 | 6.0375 | 3.8559 | 3.2719 | 6.6831 |
| 28660 | ENSMUSG00000087143 | 3.892092202 | -1.435529613 | 1.14398862 | -1.254846061 | 0.209534648 | NA | A830082K12 | RIKEN cDNA A830082K12 gene [Source:M | 8.8341 | 5.1396 | 3.0187 | 1.928 | 1.0906 | 3.3415 |
| 28661 | ENSMUSG00000087144 | 0 | NA | NA | NA | NA | NA | Gm5958 | predicted pseudogene 5958 [Source:MGI | 0 | 0 | 0 | 0 | 0 | 0 |
| 28662 | ENSMUSG00000087145 | 0 | NA | NA | NA | NA | NA | Gm12665 | predicted gene 12665 [Source:MGI Symbo | 0 | 0 | 0 | 0 | 0 | 0 |
| 28663 | ENSMUSG00000087146 | 0 | NA | NA | NA | NA | NA | NA | NA | 0 | 0 | 0 | 0 | 0 | 0 |
| 28664 | ENSMUSG00000087147 | 0 | NA | NA | NA | NA | NA | Gm14228 | predicted gene 14228 [Source:MGI Symbo | 0 | 0 | 0 | 0 | 0 | 0 |
| 28665 | ENSMUSG00000087148 | 0.68303045 | 1.69421863 | 2.781340055 | 0.609137537 | 0.542433279 | NA | 1700030C14 | RIKEN cDNA 1700030C14 gene [Source:M | 0.8031 | 0 | 0 | 0 | 2.1812 | 1.1138 |
| 28666 | ENSMUSG00000087149 | 0 | NA | NA | NA | NA | NA | Itih5l-ps | inter-alpha (globulin) inhibitor H5-like, pse | 0 | 0 | 0 | 0 | 0 | 0 |
| 28667 | ENSMUSG00000087150 | 1.011607446 | -0.82475717 | 2.098656838 | -0.392992868 | 0.694324749 | NA | BC064078 | cDNA sequence BC064078 [Source:MGI Sy | 0.8031 | 2.0558 | 1.0062 | 0 | 1.0906 | 1.1138 |
| 28668 | ENSMUSG00000087151 | 4.318420446 | 0.538449133 | 1.074189753 | 0.501260724 | 0.616187643 | NA | Gm14022 | predicted gene 14022 [Source:MGI Symbo | 6.4248 | 1.0279 | 3.0187 | 7.7118 | 3.2719 | 4.4554 |

| | | | | | | | | | | | | | | | |
|---|---|---|---|---|---|---|---|---|---|---|---|---|---|---|---|
| 28669 | ENSMUSG00000087152 | 0 | NA | NA | NA | NA | NA | Cldn34-ps | claudin 34, pseudogene [Source:MGI Sym | 0 | 0 | 0 | 0 | 0 | 0 |
| 28670 | ENSMUSG00000087153 | 13.44495472 | 0.272799398 | 0.60361541 | 0.451942402 | 0.651310477 | 0.925643431 | Gm6483 | predicted gene 6483 [Source:MGI Symbol | 12.047 | 17.475 | 7.0437 | 15.424 | 13.087 | 15.594 |
| 28671 | ENSMUSG00000087155 | 0 | NA | NA | NA | NA | NA | Gm15008 | predicted gene 15008 [Source:MGI Symbo | 0 | 0 | 0 | 0 | 0 | 0 |
| 28672 | ENSMUSG00000087157 | 0 | NA | NA | NA | NA | NA | Sox5os2 | SRY (sex determining region Y)-box 5, opp | 0 | 0 | 0 | 0 | 0 | 0 |
| 28673 | ENSMUSG00000087158 | 0 | NA | NA | NA | NA | NA | Gm14093 | predicted gene 14093 [Source:MGI Symbo | 0 | 0 | 0 | 0 | 0 | 0 |
| 28674 | ENSMUSG00000087159 | 0.342432689 | 1.990653086 | 3.900281263 | 0.51038706 | 0.609780321 | NA | Gm15246 | predicted gene 15246 [Source:MGI Symbo | 0 | 0 | 0 | 0.964 | 1.0906 | 0 |
| 28675 | ENSMUSG00000087160 | 0 | NA | NA | NA | NA | NA | Gm16336 | predicted gene 16336 [Source:MGI Symbo | 0 | 0 | 0 | 0 | 0 | 0 |
| 28676 | ENSMUSG00000087162 | 0 | NA | NA | NA | NA | NA | Gm14244 | predicted gene 14244 [Source:MGI Symbo | 0 | 0 | 0 | 0 | 0 | 0 |
| 28677 | ENSMUSG00000087163 | 6.702693677 | 0.978216071 | 0.868311267 | 1.126573049 | 0.259923039 | 0.713382702 | Gm16230 | predicted gene 16230 [Source:MGI Symbo | 2.4093 | 8.2233 | 3.0187 | 6.7478 | 10.906 | 8.9108 |
| 28678 | ENSMUSG00000087164 | 0 | NA | NA | NA | NA | NA | Nr5a1os | nuclear receptor subfamily 5, group A, me | 0 | 0 | 0 | 0 | 0 | 0 |
| 28679 | ENSMUSG00000087165 | 79.53551401 | -0.003167668 | 0.25301024 | -0.01251992 | 0.99001081 | 0.99859176 | 2010001A14 | RIKEN cDNA 2010001A14 gene [Source:M | 85.932 | 80.178 | 72.45 | 72.298 | 86.159 | 80.197 |
| 28680 | ENSMUSG00000087166 | 0 | NA | NA | NA | NA | NA | L1td1 | LINE-1 type transposase domain containir | 0 | 0 | 0 | 0 | 0 | 0 |
| 28681 | ENSMUSG00000087167 | 0 | NA | NA | NA | NA | NA | Gm15891 | predicted gene 15891 [Source:MGI Symbo | 0 | 0 | 0 | 0 | 0 | 0 |
| 28682 | ENSMUSG00000087168 | 0.181769715 | 1.126924175 | 4.080472857 | 0.276174898 | 0.782413736 | NA | Gm15983 | predicted gene 15983 [Source:MGI Symbo | 0 | 0 | 0 | 0 | 1.0906 | 0 |
| 28683 | ENSMUSG00000087170 | 0 | NA | NA | NA | NA | NA | 1700001G01 | RIKEN cDNA 1700001G01 gene [Source:M | 0 | 0 | 0 | 0 | 0 | 0 |
| 28684 | ENSMUSG00000087171 | 0 | NA | NA | NA | NA | NA | Gm12962 | predicted gene 12962 [Source:MGI Symbo | 0 | 0 | 0 | 0 | 0 | 0 |
| 28685 | ENSMUSG00000087172 | 0 | NA | NA | NA | NA | NA | Gm12153 | predicted gene 12153 [Source:MGI Symbo | 0 | 0 | 0 | 0 | 0 | 0 |
| 28686 | ENSMUSG00000087173 | 14.20853672 | 0.284322071 | 0.658233986 | 0.431946811 | 0.665780071 | 0.929596279 | Gm11337 | predicted gene 11337 [Source:MGI Symbo | 11.243 | 5.1396 | 22.137 | 19.28 | 18.541 | 8.9108 |
| 28687 | ENSMUSG00000087174 | 78.31712995 | 0.033032163 | 0.259780127 | 0.127154311 | 0.898818266 | 0.982798755 | 5530601H04 | RIKEN cDNA 5530601H04 gene [Source:M | 77.901 | 64.759 | 89.556 | 80.01 | 75.253 | 82.424 |
| 28688 | ENSMUSG00000087175 | 1.879405261 | 0.39787096 | 1.645441944 | 0.241801883 | 0.80893368 | NA | Gm15133 | predicted gene 15133 [Source:MGI Symbo | 0.8031 | 3.0838 | 1.0062 | 1.928 | 0 | 4.4554 |
| 28689 | ENSMUSG00000087176 | 3.566378401 | 0.239682059 | 1.147077401 | 0.208950206 | 0.83448711 | NA | D230022J07 | RIKEN cDNA D230022J07 gene [Source:M( | 5.6217 | 3.0838 | 1.0062 | 2.8919 | 5.4531 | 3.3415 |
| 28690 | ENSMUSG00000087177 | 14.37294517 | -0.086952188 | 0.66454419 | -0.130844855 | 0.895898039 | 0.982798755 | E130307A14 | RIKEN cDNA E130307A14 gene [Source:M | 8.031 | 17.475 | 19.119 | 18.316 | 4.3625 | 18.935 |
| 28691 | ENSMUSG00000087178 | 10.50522264 | 0.192505899 | 0.675942261 | 0.284796365 | 0.775800157 | 0.95893551 | A230056P14 | RIKEN cDNA A230056P14 gene [Source:M | 12.047 | 8.2233 | 9.0562 | 11.568 | 6.5437 | 15.594 |
| 28692 | ENSMUSG00000087179 | 1.224164433 | -2.469837792 | 2.276305006 | -1.085020586 | 0.277912538 | NA | 5730471H19 | RIKEN cDNA 5730471H19 gene [Source:M | 3.2124 | 0 | 3.0187 | 0 | 0 | 1.1138 |
| 28693 | ENSMUSG00000087180 | 0 | NA | NA | NA | NA | NA | Gm11924 | predicted gene 11924 [Source:MGI Symbo | 0 | 0 | 0 | 0 | 0 | 0 |
| 28694 | ENSMUSG00000087181 | 0 | NA | NA | NA | NA | NA | Gm13570 | predicted gene 13570 [Source:MGI Symbo | 0 | 0 | 0 | 0 | 0 | 0 |
| 28695 | ENSMUSG00000087184 | 0 | NA | NA | NA | NA | NA | Gm11650 | predicted gene 11650 [Source:MGI Symbo | 0 | 0 | 0 | 0 | 0 | 0 |
| 28696 | ENSMUSG00000087185 | 0.853038123 | -0.516856556 | 2.386032976 | -0.216617524 | 0.828506428 | NA | Gm13872 | predicted gene 13872 [Source:MGI Symbo | 0 | 1.0279 | 2.0125 | 0.964 | 0 | 1.1138 |
| 28697 | ENSMUSG00000087187 | 2.548605522 | 1.654135311 | 1.47255559 | 1.123309246 | 0.261306178 | NA | Gm13431 | predicted gene 13431 [Source:MGI Symbo | 1.6062 | 2.0558 | 0 | 3.8559 | 1.0906 | 6.6831 |
| 28698 | ENSMUSG00000087190 | 0 | NA | NA | NA | NA | NA | D430001F17 | RIKEN cDNA D430001F17 gene [Source:M | 0 | 0 | 0 | 0 | 0 | 0 |
| 28699 | ENSMUSG00000087192 | 0 | NA | NA | NA | NA | NA | Lexis1 | lipid responsive LXR-induced inhibitor of c | 0 | 0 | 0 | 0 | 0 | 0 |
| 28700 | ENSMUSG00000087193 | 0 | NA | NA | NA | NA | NA | Gm14820 | predicted gene 14820 [Source:MGI Symbo | 0 | 0 | 0 | 0 | 0 | 0 |
| 28701 | ENSMUSG00000087194 | 0 | NA | NA | NA | NA | NA | Skint6 | selection and upkeep of intraepithelial T c | 0 | 0 | 0 | 0 | 0 | 0 |
| 28702 | ENSMUSG00000087195 | 0 | NA | NA | NA | NA | NA | Gm12907 | predicted gene 12907 [Source:MGI Symbo | 0 | 0 | 0 | 0 | 0 | 0 |
| 28703 | ENSMUSG00000087196 | 0 | NA | NA | NA | NA | NA | Gm13373 | predicted gene 13373 [Source:MGI Symbo | 0 | 0 | 0 | 0 | 0 | 0 |
| 28704 | ENSMUSG00000087197 | 0 | NA | NA | NA | NA | NA | Gm13780 | predicted gene 13780 [Source:MGI Symbo | 0 | 0 | 0 | 0 | 0 | 0 |
| 28705 | ENSMUSG00000087198 | 1.353781411 | 0.135944945 | 2.00516922 | 0.067797243 | 0.945947038 | NA | Tmem274 | transmembrane protein 274 [Source:MGI | 0.8031 | 3.0838 | 0 | 0.964 | 3.2719 | 0 |
| 28706 | ENSMUSG00000087199 | 0 | NA | NA | NA | NA | NA | Gm15818 | predicted gene 15818 [Source:MGI Symbo | 0 | 0 | 0 | 0 | 0 | 0 |
| 28707 | ENSMUSG00000087200 | 0 | NA | NA | NA | NA | NA | Lrp8os3 | low density lipoprotein receptor-related p | 0 | 0 | 0 | 0 | 0 | 0 |
| 28708 | ENSMUSG00000087201 | 11.58065855 | -0.079504307 | 0.670502909 | -0.118574141 | 0.905612753 | 0.983972587 | Gm15261 | predicted gene 15261 [Source:MGI Symbo | 14.456 | 8.2233 | 13.081 | 18.316 | 8.7249 | 6.6831 |
| 28709 | ENSMUSG00000087202 | 0 | NA | NA | NA | NA | NA | Gm15813 | predicted gene 15813 [Source:MGI Symbo | 0 | 0 | 0 | 0 | 0 | 0 |
| 28710 | ENSMUSG00000087203 | 0 | NA | NA | NA | NA | NA | Gm13986 | predicted gene 13986 [Source:MGI Symbo | 0 | 0 | 0 | 0 | 0 | 0 |
| 28711 | ENSMUSG00000087204 | 0 | NA | NA | NA | NA | NA | 1700073E17 | RIKEN cDNA 1700073E17 gene [Source:M( | 0 | 0 | 0 | 0 | 0 | 0 |
| 28712 | ENSMUSG00000087205 | 0 | NA | NA | NA | NA | NA | 4930445N08 | RIKEN cDNA 4930445N08 gene [Source:M | 0 | 0 | 0 | 0 | 0 | 0 |
| 28713 | ENSMUSG00000087206 | 1.661380909 | -0.034870331 | 1.86576186 | -0.018689593 | 0.98508873 | NA | Gm16175 | predicted gene 16175 [Source:MGI Symbo | 4.0155 | 0 | 1.0062 | 3.8559 | 1.0906 | 0 |
| 28714 | ENSMUSG00000087207 | 0 | NA | NA | NA | NA | NA | Gm13147 | predicted gene 13147 [Source:MGI Symbo | 0 | 0 | 0 | 0 | 0 | 0 |
| 28715 | ENSMUSG00000087208 | 0.401551119 | -2.072982726 | 4.008251979 | -0.517178744 | 0.605031388 | NA | Gm5125 | predicted gene 5125 [Source:MGI Symbol | 2.4093 | 0 | 0 | 0 | 0 | 0 |
| 28716 | ENSMUSG00000087210 | 0 | NA | NA | NA | NA | NA | Gm15391 | predicted gene 15391 [Source:MGI Symbo | 0 | 0 | 0 | 0 | 0 | 0 |
| 28717 | ENSMUSG00000087211 | 0.133850373 | -0.796629176 | 4.080472857 | -0.195229623 | 0.84521319 | NA | Lhx1os | LIM homeobox 1, opposite strand [Source | 0.8031 | 0 | 0 | 0 | 0 | 0 |
| 28718 | ENSMUSG00000087213 | 60.3700704 | -0.148724379 | 0.337979703 | -0.440039382 | 0.659908585 | 0.927787797 | 2810408I11F | RIKEN cDNA 2810408I11 gene [Source:MC | 87.538 | 44.2 | 58.362 | 66.514 | 62.165 | 43.44 |
| 28719 | ENSMUSG00000087214 | 0 | NA | NA | NA | NA | NA | Gm15555 | predicted gene 15555 [Source:MGI Symbo | 0 | 0 | 0 | 0 | 0 | 0 |
| 28720 | ENSMUSG00000087215 | 0 | NA | NA | NA | NA | NA | Gm14618 | predicted gene 14618 [Source:MGI Symbo | 0 | 0 | 0 | 0 | 0 | 0 |
| 28721 | ENSMUSG00000087216 | 0 | NA | NA | NA | NA | NA | E130111B04 | RIKEN cDNA E130111B04 gene [Source:M | 0 | 0 | 0 | 0 | 0 | 0 |
| 28722 | ENSMUSG00000087217 | 0 | NA | NA | NA | NA | NA | Gm13209 | predicted gene 13209 [Source:MGI Symbo | 0 | 0 | 0 | 0 | 0 | 0 |
| 28723 | ENSMUSG00000087219 | 0 | NA | NA | NA | NA | NA | Gm13376 | predicted gene 13376 [Source:MGI Symbo | 0 | 0 | 0 | 0 | 0 | 0 |
| 28724 | ENSMUSG00000087220 | 0 | NA | NA | NA | NA | NA | Gm11377 | predicted gene 11377 [Source:MGI Symbo | 0 | 0 | 0 | 0 | 0 | 0 |

| | | | | | | | | | | | | | | | |
|---|---|---|---|---|---|---|---|---|---|---|---|---|---|---|---|
| 28725 | ENSMUSG00000087221 | 0.321325949 | 1.911575114 | 4.048267893 | 0.472195804 | 0.636787034 | NA | BC037032 | cDNA Sequence BC037032 [Source:MGI S\ | 0 | 0 | 0 | 1.928 | 0 | 0 |
| 28726 | ENSMUSG00000087222 | 32.57271029 | -0.250503728 | 0.401035317 | -0.624642562 | 0.532205679 | 0.889143097 | E030042O20 | RIKEN cDNA E030042O20 gene [Source:M | 36.14 | 39.061 | 31.194 | 39.523 | 28.356 | 21.163 |
| 28727 | ENSMUSG00000087223 | 1.929980679 | 0.349641944 | 1.815950336 | 0.192539375 | 0.847319728 | NA | 4930442L01I | RIKEN cDNA 4930442L01 gene [Source:MC | 0 | 5.1396 | 0 | 0.964 | 4.3625 | 1.1138 |
| 28728 | ENSMUSG00000087224 | 0 | NA | NA | NA | NA | NA | Gm2182 | predicted gene 2182 [Source:MGI Symbol | 0 | 0 | 0 | 0 | 0 | 0 |
| 28729 | ENSMUSG00000087225 | 0 | NA | NA | NA | NA | NA | Gm14014 | predicted gene 14014 [Source:MGI Symbo | 0 | 0 | 0 | 0 | 0 | 0 |
| 28730 | ENSMUSG00000087226 | 0 | NA | NA | NA | NA | NA | Gm14015 | predicted gene 14015 [Source:MGI Symbo | 0 | 0 | 0 | 0 | 0 | 0 |
| 28731 | ENSMUSG00000087228 | 0 | NA | NA | NA | NA | NA | Gm12827 | predicted gene 12827 [Source:MGI Symbo | 0 | 0 | 0 | 0 | 0 | 0 |
| 28732 | ENSMUSG00000087230 | 0 | NA | NA | NA | NA | NA | Mroh3 | maestro heat-like repeat family member 3 | 0 | 0 | 0 | 0 | 0 | 0 |
| 28733 | ENSMUSG00000087231 | 41.32106412 | 1.214435714 | 0.398241293 | 3.049497218 | 0.002292248 | 0.038552406 | E230016M11 | RIKEN cDNA E230016M11 gene [Source:M | 33.73 | 15.419 | 25.156 | 71.334 | 61.075 | 41.212 |
| 28734 | ENSMUSG00000087232 | 0.349477146 | 0.165142768 | 3.911021834 | 0.042224967 | 0.966319359 | NA | Gm14764 | predicted gene 14764 [Source:MGI Symbo | 0 | 0 | 1.0062 | 0 | 1.0906 | 0 |
| 28735 | ENSMUSG00000087233 | 0 | NA | NA | NA | NA | NA | Gm43213 | predicted gene 43213 [Source:MGI Symbo | 0 | 0 | 0 | 0 | 0 | 0 |
| 28736 | ENSMUSG00000087235 | 0 | NA | NA | NA | NA | NA | Gm4750 | predicted gene 4750 [Source:MGI Symbol | 0 | 0 | 0 | 0 | 0 | 0 |
| 28737 | ENSMUSG00000087236 | 0 | NA | NA | NA | NA | NA | Kif28 | kinesin family member 28 [Source:MGI Sy | 0 | 0 | 0 | 0 | 0 | 0 |
| 28738 | ENSMUSG00000087238 | 0 | NA | NA | NA | NA | NA | Gm12865 | predicted gene 12865 [Source:MGI Symbo | 0 | 0 | 0 | 0 | 0 | 0 |
| 28739 | ENSMUSG00000087239 | 0 | NA | NA | NA | NA | NA | Btf3-ps2 | basic transcription factor 3, pseudogene 2 | 0 | 0 | 0 | 0 | 0 | 0 |
| 28740 | ENSMUSG00000087240 | 0 | NA | NA | NA | NA | NA | Gm15976 | predicted gene 15976 [Source:MGI Symbo | 0 | 0 | 0 | 0 | 0 | 0 |
| 28741 | ENSMUSG00000087241 | 209.4165362 | -0.192066712 | 0.197695529 | -0.971527846 | 0.331285494 | 0.774747658 | Gm11844 | predicted gene 11844 [Source:MGI Symbo | 221.66 | 244.64 | 203.26 | 154.24 | 236.66 | 196.04 |
| 28742 | ENSMUSG00000087242 | 0 | NA | NA | NA | NA | NA | C78197 | expressed sequence C78197 [Source:MGI | 0 | 0 | 0 | 0 | 0 | 0 |
| 28743 | ENSMUSG00000087244 | 0 | NA | NA | NA | NA | NA | Gm7284 | predicted gene 7284 [Source:MGI Symbol | 0 | 0 | 0 | 0 | 0 | 0 |
| 28744 | ENSMUSG00000087245 | 0 | NA | NA | NA | NA | NA | Gm12126 | predicted gene 12126 [Source:MGI Symbo | 0 | 0 | 0 | 0 | 0 | 0 |
| 28745 | ENSMUSG00000087246 | 0 | NA | NA | NA | NA | NA | Gm13346 | predicted gene 13346 [Source:MGI Symbo | 0 | 0 | 0 | 0 | 0 | 0 |
| 28746 | ENSMUSG00000087247 | 0 | NA | NA | NA | NA | NA | Alkal1 | ALK and LTK ligand 1 [Source:MGI Symbol | 0 | 0 | 0 | 0 | 0 | 0 |
| 28747 | ENSMUSG00000087248 | 0.167707431 | -0.796629176 | 4.080472857 | -0.195229623 | 0.84521319 | NA | Gm16120 | predicted gene 16120 [Source:MGI Symbo | 0 | 0 | 1.0062 | 0 | 0 | 0 |
| 28748 | ENSMUSG00000087249 | 2.780387702 | -0.350208581 | 1.293257597 | -0.270795688 | 0.786548176 | NA | Gm16062 | predicted gene 16062 [Source:MGI Symbo | 3.2124 | 5.1396 | 1.0062 | 2.8919 | 1.0906 | 3.3415 |
| 28749 | ENSMUSG00000087250 | 0 | NA | NA | NA | NA | NA | Usp46os1 | ubiquitin specific peptidase 46, opposite s | 0 | 0 | 0 | 0 | 0 | 0 |
| 28750 | ENSMUSG00000087252 | 3.153013106 | -0.653166476 | 1.214253782 | -0.537915949 | 0.590635077 | NA | Gm14379 | predicted gene 14379 [Source:MGI Symbo | 2.4093 | 7.1954 | 2.0125 | 2.8919 | 2.1812 | 2.2277 |
| 28751 | ENSMUSG00000087253 | 5.057840529 | 2.321750621 | 1.254266511 | 1.851082366 | 0.064157705 | NA | Gm12043 | predicted gene 12043 [Source:MGI Symbo | 4.8186 | 0 | 0 | 14.46 | 3.2719 | 7.7969 |
| 28752 | ENSMUSG00000087254 | 4.913795315 | -1.604277101 | 1.031175931 | -1.555774387 | 0.119761751 | NA | 2310005A03 | RIKEN cDNA 2310005A03 gene [Source:M | 8.031 | 4.1117 | 10.062 | 2.8919 | 3.2719 | 1.1138 |
| 28753 | ENSMUSG00000087255 | 0 | NA | NA | NA | NA | NA | Gm6985 | predicted pseudogene 6985 [Source:MGI | 0 | 0 | 0 | 0 | 0 | 0 |
| 28754 | ENSMUSG00000087256 | 0.815310133 | -0.44746426 | 2.258708535 | -0.198106242 | 0.842961941 | NA | Gm15990 | predicted gene 15990 [Source:MGI Symbo | 0.8031 | 1.0279 | 1.0062 | 0.964 | 1.0906 | 0 |
| 28755 | ENSMUSG00000087258 | 0.181769715 | 1.126924175 | 4.080472857 | 0.276174898 | 0.782413736 | NA | Gm15461 | predicted gene 15461 [Source:MGI Symbo | 0 | 0 | 0 | 0 | 1.0906 | 0 |
| 28756 | ENSMUSG00000087259 | 11.16392762 | 0.41261809 | 0.729111619 | 0.565918961 | 0.571448871 | 0.904651508 | 2610035D17 | RIKEN cDNA 2610035D17 gene [Source:M | 14.456 | 3.0838 | 11.069 | 17.352 | 6.5437 | 14.48 |
| 28757 | ENSMUSG00000087260 | 455.1560584 | -0.050975661 | 0.138078945 | -0.369177655 | 0.71199531 | 0.944239215 | Lamtor5 | late endosomal/lysosomal adaptor, MAPK | 506.76 | 409.11 | 472.93 | 445.36 | 464.6 | 432.17 |
| 28758 | ENSMUSG00000087261 | 0 | NA | NA | NA | NA | NA | Gm15477 | predicted gene 15477 [Source:MGI Symbo | 0 | 0 | 0 | 0 | 0 | 0 |
| 28759 | ENSMUSG00000087263 | 0.531246861 | 1.101376188 | 3.36099124 | 0.327693858 | 0.743143147 | NA | Gm15726 | predicted gene 15726 [Source:MGI Symbo | 0 | 0 | 1.0062 | 0 | 2.1812 | 0 |
| 28760 | ENSMUSG00000087264 | 0 | NA | NA | NA | NA | NA | Gad1os | glutamate decarboxylase 1, opposite strar | 0 | 0 | 0 | 0 | 0 | 0 |
| 28761 | ENSMUSG00000087265 | 0.663785267 | -1.400021131 | 3.106754789 | -0.450637796 | 0.652250621 | NA | Gm12349 | predicted gene 12349 [Source:MGI Symbo | 0 | 0 | 3.0187 | 0.964 | 0 | 0 |
| 28762 | ENSMUSG00000087266 | 0.492645588 | 0.9565034 | 3.447648281 | 0.277436479 | 0.781444974 | NA | Gm15991 | predicted gene 15991 [Source:MGI Symbo | 0 | 1.0279 | 0 | 1.928 | 0 | 0 |
| 28763 | ENSMUSG00000087267 | 0 | NA | NA | NA | NA | NA | 4933427J07I | RIKEN cDNA 4933427J07 gene [Source:MC | 0 | 0 | 0 | 0 | 0 | 0 |
| 28764 | ENSMUSG00000087268 | 0.185640647 | 1.126924175 | 4.080472857 | 0.276174898 | 0.782413736 | NA | Gm14486 | predicted gene 14486 [Source:MGI Symbo | 0 | 0 | 0 | 0 | 0 | 1.1138 |
| 28765 | ENSMUSG00000087269 | 33.17171842 | 0.048652875 | 0.38001844 | 0.128027668 | 0.898127077 | 0.982798755 | D330023K18 | RIKEN cDNA D330023K18 gene [Source:M | 32.927 | 29.81 | 35.219 | 40.487 | 31.628 | 28.96 |
| 28766 | ENSMUSG00000087270 | 0 | NA | NA | NA | NA | NA | Gm12983 | predicted gene 12983 [Source:MGI Symbo | 0 | 0 | 0 | 0 | 0 | 0 |
| 28767 | ENSMUSG00000087272 | 4.943139127 | 0.65356233 | 1.053757486 | 0.620220818 | 0.535112417 | NA | Gm15409 | predicted gene 15409 [Source:MGI Symbo | 6.4248 | 1.0279 | 4.025 | 11.568 | 3.2719 | 3.3415 |
| 28768 | ENSMUSG00000087273 | 9.242203323 | 0.563107822 | 0.947107257 | 0.594555493 | 0.552140638 | 0.895935901 | Gm13203 | predicted gene 13203 [Source:MGI Symbo | 8.031 | 7.1954 | 7.0437 | 0 | 10.906 | 22.277 |
| 28769 | ENSMUSG00000087276 | 0 | NA | NA | NA | NA | NA | Gm11378 | predicted gene 11378 [Source:MGI Symbo | 0 | 0 | 0 | 0 | 0 | 0 |
| 28770 | ENSMUSG00000087277 | 0 | NA | NA | NA | NA | NA | 2010013B24 | RIKEN cDNA 2010013B24 gene [Source:M | 0 | 0 | 0 | 0 | 0 | 0 |
| 28771 | ENSMUSG00000087278 | 0 | NA | NA | NA | NA | NA | A930006I01I | RIKEN cDNA A930006I01 gene [Source:MC | 0 | 0 | 0 | 0 | 0 | 0 |
| 28772 | ENSMUSG00000087279 | 0 | NA | NA | NA | NA | NA | 4930544D05 | RIKEN cDNA 4930544D05 gene [Source:M | 0 | 0 | 0 | 0 | 0 | 0 |
| 28773 | ENSMUSG00000087281 | 3.022988462 | -0.364786909 | 1.369376444 | -0.266389064 | 0.789939587 | NA | Gm16015 | predicted gene 16015 [Source:MGI Symbo | 3.2124 | 2.0558 | 5.0312 | 6.7478 | 1.0906 | 0 |
| 28774 | ENSMUSG00000087283 | 0 | NA | NA | NA | NA | NA | Gm13580 | predicted gene 13580 [Source:MGI Symbo | 0 | 0 | 0 | 0 | 0 | 0 |
| 28775 | ENSMUSG00000087286 | 0 | NA | NA | NA | NA | NA | Gm9013 | predicted gene 9013 [Source:MGI Symbol | 0 | 0 | 0 | 0 | 0 | 0 |
| 28776 | ENSMUSG00000087288 | 0 | NA | NA | NA | NA | NA | Gm13061 | predicted gene 13061 [Source:MGI Symbo | 0 | 0 | 0 | 0 | 0 | 0 |
| 28777 | ENSMUSG00000087289 | 0 | NA | NA | NA | NA | NA | 4933424M12 | RIKEN cDNA 4933424M12 gene [Source:M | 0 | 0 | 0 | 0 | 0 | 0 |
| 28778 | ENSMUSG00000087290 | 0 | NA | NA | NA | NA | NA | Gm15866 | predicted gene 15866 [Source:MGI Symbo | 0 | 0 | 0 | 0 | 0 | 0 |
| 28779 | ENSMUSG00000087291 | 0.17131964 | -0.796629176 | 4.080472857 | -0.195229623 | 0.84521319 | NA | Gm11946 | predicted gene 11946 [Source:MGI Symbo | 0 | 1.0279 | 0 | 0 | 0 | 0 |
| 28780 | ENSMUSG00000087292 | 0 | NA | NA | NA | NA | NA | Gm13832 | predicted gene 13832 [Source:MGI Symbo | 0 | 0 | 0 | 0 | 0 | 0 |

| | | | | | | | | | | | | | | | |
|---|---|---|---|---|---|---|---|---|---|---|---|---|---|---|---|
| 28781 | ENSMUSG00000087293 | 0 | NA | NA | NA | NA | NA | Gm14341 | predicted gene 14341 [Source:MGI Symbo | 0 | 0 | 0 | 0 | 0 | 0 |
| 28782 | ENSMUSG00000087294 | 0.181769715 | 1.126924175 | 4.080472857 | 0.276174898 | 0.782413736 | NA | Gm13556 | predicted gene 13556 [Source:MGI Symbo | 0 | 0 | 0 | 0 | 1.0906 | 0 |
| 28783 | ENSMUSG00000087295 | 0 | NA | NA | NA | NA | NA | Gm3948 | predicted gene 3948 [Source:MGI Symbol | 0 | 0 | 0 | 0 | 0 | 0 |
| 28784 | ENSMUSG00000087296 | 0 | NA | NA | NA | NA | NA | Gm12130 | predicted gene 12130 [Source:MGI Symbo | 0 | 0 | 0 | 0 | 0 | 0 |
| 28785 | ENSMUSG00000087298 | 8.263958429 | 0.020224927 | 0.77144875 | 0.026216812 | 0.979084406 | 0.998055255 | Gm9392 | predicted pseudogene 9392 [Source:MGI | 6.4248 | 9.2513 | 9.0562 | 10.604 | 10.906 | 3.3415 |
| 28786 | ENSMUSG00000087299 | 0.815516723 | -3.06336425 | 2.557025255 | -1.19801877 | 0.23090971 | NA | Gm12953 | predicted gene 12953 [Source:MGI Symbo | 0.8031 | 3.0838 | 1.0062 | 0 | 0 | 0 |
| 28787 | ENSMUSG00000087300 | 0 | NA | NA | NA | NA | NA | Gm16725 | predicted gene, 16725 [Source:MGI Symb | 0 | 0 | 0 | 0 | 0 | 0 |
| 28788 | ENSMUSG00000087301 | 0 | NA | NA | NA | NA | NA | Gm13629 | predicted gene 13629 [Source:MGI Symbo | 0 | 0 | 0 | 0 | 0 | 0 |
| 28789 | ENSMUSG00000087302 | 0 | NA | NA | NA | NA | NA | Sox5os1 | SRY (sex determining region Y)-box 5, opp | 0 | 0 | 0 | 0 | 0 | 0 |
| 28790 | ENSMUSG00000087303 | 0 | NA | NA | NA | NA | NA | Lipo2 | lipase, member O2 [Source:MGI Symbol;A | 0 | 0 | 0 | 0 | 0 | 0 |
| 28791 | ENSMUSG00000087305 | 35.32575556 | -0.08369327 | 0.418550134 | -0.199959966 | 0.841511892 | 0.971854408 | A430035B10 | RIKEN cDNA A430035B10 gene [Source:M | 45.777 | 39.061 | 24.15 | 41.451 | 40.353 | 21.163 |
| 28792 | ENSMUSG00000087306 | 0 | NA | NA | NA | NA | NA | A230004M1 | RIKEN cDNA A230004M16 gene [Source:M | 0 | 0 | 0 | 0 | 0 | 0 |
| 28793 | ENSMUSG00000087307 | 0.679159517 | 1.687875387 | 3.079592583 | 0.548083989 | 0.583634234 | NA | Gm12925 | predicted gene 12925 [Source:MGI Symbo | 0.8031 | 0 | 0 | 0 | 3.2719 | 0 |
| 28794 | ENSMUSG00000087308 | 0 | NA | NA | NA | NA | NA | Gm16195 | predicted gene 16195 [Source:MGI Symbo | 0 | 0 | 0 | 0 | 0 | 0 |
| 28795 | ENSMUSG00000087309 | 0 | NA | NA | NA | NA | NA | 4930528P14 | RIKEN cDNA 4930528P14 gene [Source:M | 0 | 0 | 0 | 0 | 0 | 0 |
| 28796 | ENSMUSG00000087312 | 0.931459384 | 1.251134958 | 2.489841943 | 0.502495735 | 0.615318852 | NA | Gm13833 | predicted gene 13833 [Source:MGI Symbo | 1.6062 | 0 | 0 | 2.8919 | 1.0906 | 0 |
| 28797 | ENSMUSG00000087314 | 15.83098679 | -0.531079847 | 0.649128713 | -0.81814259 | 0.413275772 | 0.8353299 | Prkag2os2 | protein kinase, AMP-activated, gamma 2 r | 22.487 | 18.503 | 15.094 | 11.568 | 23.994 | 3.3415 |
| 28798 | ENSMUSG00000087316 | 0 | NA | NA | NA | NA | NA | Gm11917 | predicted gene 11917 [Source:MGI Symbo | 0 | 0 | 0 | 0 | 0 | 0 |
| 28799 | ENSMUSG00000087317 | 0 | NA | NA | NA | NA | NA | 4930466I24R | RIKEN cDNA 4930466I24 gene [Source:MG | 0 | 0 | 0 | 0 | 0 | 0 |
| 28800 | ENSMUSG00000087318 | 0 | NA | NA | NA | NA | NA | Gm12114 | predicted gene 12114 [Source:MGI Symbo | 0 | 0 | 0 | 0 | 0 | 0 |
| 28801 | ENSMUSG00000087319 | 0 | NA | NA | NA | NA | NA | Gm15907 | predicted gene 15907 [Source:MGI Symbo | 0 | 0 | 0 | 0 | 0 | 0 |
| 28802 | ENSMUSG00000087321 | 0.49081066 | -0.698504554 | 3.1295234 | -0.22319838 | 0.823381113 | NA | Gm5353 | predicted gene 5353 [Source:MGI Symbol | 0.8031 | 1.0279 | 0 | 0 | 0 | 1.1138 |
| 28803 | ENSMUSG00000087322 | 0 | NA | NA | NA | NA | NA | Gm16075 | predicted gene 16075 [Source:MGI Symbo | 0 | 0 | 0 | 0 | 0 | 0 |
| 28804 | ENSMUSG00000087323 | 0.160662974 | 1.126924175 | 4.080472857 | 0.276174898 | 0.782413736 | NA | Gm14170 | predicted gene 14170 [Source:MGI Symbo | 0 | 0 | 0 | 0.964 | 0 | 0 |
| 28805 | ENSMUSG00000087325 | 0 | NA | NA | NA | NA | NA | Gm6038 | predicted pseudogene 6038 [Source:MGI | 0 | 0 | 0 | 0 | 0 | 0 |
| 28806 | ENSMUSG00000087326 | 0.688736311 | 2.998073904 | 2.697618692 | 1.11137794 | 0.266405704 | NA | Gm12503 | predicted gene 12503 [Source:MGI Symbo | 0 | 0 | 0 | 1.928 | 1.0906 | 1.1138 |
| 28807 | ENSMUSG00000087327 | 2.244748902 | 1.919112783 | 1.745950475 | 1.099179393 | 0.271689825 | NA | Gm15884 | predicted gene 15884 [Source:MGI Symbo | 0.8031 | 0 | 2.0125 | 1.928 | 8.7249 | 0 |
| 28808 | ENSMUSG00000087328 | 0 | NA | NA | NA | NA | NA | Gm12120 | predicted gene 12120 [Source:MGI Symbo | 0 | 0 | 0 | 0 | 0 | 0 |
| 28809 | ENSMUSG00000087329 | 0 | NA | NA | NA | NA | NA | Gm15671 | predicted gene 15671 [Source:MGI Symbo | 0 | 0 | 0 | 0 | 0 | 0 |
| 28810 | ENSMUSG00000087330 | 1.481311366 | -0.836652103 | 1.78562457 | -0.468548718 | 0.63939224 | NA | Bloodlinc | Bloodlinc, erythroid developmental long i | 1.6062 | 3.0838 | 1.0062 | 0.964 | 0 | 2.2277 |
| 28811 | ENSMUSG00000087331 | 50.35094444 | 0.124369876 | 0.320272866 | 0.388324737 | 0.697775733 | 0.940502785 | 1810021B22 | RIKEN cDNA 1810021B22 gene [Source:M | 48.989 | 54.48 | 41.256 | 62.659 | 50.168 | 44.554 |
| 28812 | ENSMUSG00000087332 | 0 | NA | NA | NA | NA | NA | Gm12690 | predicted gene 12690 [Source:MGI Symbo | 0 | 0 | 0 | 0 | 0 | 0 |
| 28813 | ENSMUSG00000087333 | 0 | NA | NA | NA | NA | NA | Gm13652 | predicted gene 13652 [Source:MGI Symbo | 0 | 0 | 0 | 0 | 0 | 0 |
| 28814 | ENSMUSG00000087334 | 5.035024255 | 1.172491669 | 0.981812356 | 1.194211564 | 0.232395221 | NA | AW495222 | expressed sequence AW495222 [Source:M | 3.2124 | 1.0279 | 5.0312 | 7.7118 | 6.5437 | 6.6831 |
| 28815 | ENSMUSG00000087335 | 0 | NA | NA | NA | NA | NA | 4930526F13 | RIKEN cDNA 4930526F13 gene [Source:M | 0 | 0 | 0 | 0 | 0 | 0 |
| 28816 | ENSMUSG00000087336 | 7.00949096 | -0.158856149 | 0.859545271 | -0.184814173 | 0.853374791 | 0.973825032 | Gm15860 | predicted gene 15860 [Source:MGI Symbo | 12.85 | 6.1675 | 3.0187 | 6.7478 | 4.3625 | 8.9108 |
| 28817 | ENSMUSG00000087337 | 0 | NA | NA | NA | NA | NA | Gm14144 | predicted gene 14144 [Source:MGI Symbo | 0 | 0 | 0 | 0 | 0 | 0 |
| 28818 | ENSMUSG00000087338 | 1.34359006 | -0.611265528 | 1.777582845 | -0.343874565 | 0.730940621 | NA | Scpep1os | serine carboxypeptidase 1, opposite stran | 0.8031 | 3.0838 | 1.0062 | 0.964 | 1.0906 | 1.1138 |
| 28819 | ENSMUSG00000087339 | 5.914564481 | -1.291780844 | 1.01625517 | -1.271118595 | 0.203686461 | 0.64863958 | Gm11586 | predicted gene 11586 [Source:MGI Symbo | 12.047 | 4.1117 | 9.0562 | 4.8199 | 5.4531 | 0 |
| 28820 | ENSMUSG00000087340 | 0.305170013 | -1.657836477 | 4.041480335 | -0.410205257 | 0.681655385 | NA | Gm15228 | predicted gene 15228 [Source:MGI Symbo | 0.8031 | 1.0279 | 0 | 0 | 0 | 0 |
| 28821 | ENSMUSG00000087341 | 2.02370571 | 0.672774429 | 1.47497065 | 0.456127333 | 0.648298416 | NA | 0610040F04 | RIKEN cDNA 0610040F04 gene [Source:M | 1.6062 | 2.0558 | 1.0062 | 1.928 | 1.0906 | 4.4554 |
| 28822 | ENSMUSG00000087342 | 0 | NA | NA | NA | NA | NA | Gm12238 | predicted gene 12238 [Source:MGI Symbo | 0 | 0 | 0 | 0 | 0 | 0 |
| 28823 | ENSMUSG00000087343 | 0.401551119 | -2.072982726 | 4.008251979 | -0.517178744 | 0.605031388 | NA | 1700021N21 | RIKEN cDNA 1700021N21 gene [Source:M | 2.4093 | 0 | 0 | 0 | 0 | 0 |
| 28824 | ENSMUSG00000087344 | 0.294513347 | 0.165143496 | 4.080472857 | 0.040471657 | 0.967717103 | NA | Gm15627 | predicted gene 15627 [Source:MGI Symbo | 0.8031 | 0 | 0 | 0.964 | 0 | 0 |
| 28825 | ENSMUSG00000087345 | 0 | NA | NA | NA | NA | NA | Gm13746 | predicted gene 13746 [Source:MGI Symbo | 0 | 0 | 0 | 0 | 0 | 0 |
| 28826 | ENSMUSG00000087347 | 1.857413984 | -0.078773297 | 1.95784029 | -0.040234792 | 0.96790594 | NA | 1700047K16 | RIKEN cDNA 1700047K16 gene [Source:M | 5.6217 | 0 | 0 | 0 | 2.1812 | 3.3415 |
| 28827 | ENSMUSG00000087348 | 1.041825665 | 2.372145788 | 2.219541612 | 1.068754817 | 0.285180164 | NA | Gm15582 | predicted gene 15582 [Source:MGI Symbo | 0 | 1.0279 | 0 | 1.928 | 2.1812 | 1.1138 |
| 28828 | ENSMUSG00000087350 | 0 | NA | NA | NA | NA | NA | Gm16254 | predicted gene 16254 [Source:MGI Symbo | 0 | 0 | 0 | 0 | 0 | 0 |
| 28829 | ENSMUSG00000087351 | 0.503122293 | -2.366675966 | 3.785596405 | -0.625179156 | 0.531853481 | NA | Gm11464 | predicted gene 11464 [Source:MGI Symbo | 0 | 0 | 3.0187 | 0 | 0 | 0 |
| 28830 | ENSMUSG00000087352 | 0 | NA | NA | NA | NA | NA | Gm12999 | predicted gene 12999 [Source:MGI Symbo | 0 | 0 | 0 | 0 | 0 | 0 |
| 28831 | ENSMUSG00000087353 | 0 | NA | NA | NA | NA | NA | Gm13727 | predicted gene 13727 [Source:MGI Symbo | 0 | 0 | 0 | 0 | 0 | 0 |
| 28832 | ENSMUSG00000087354 | 5.435216788 | 0.184262824 | 0.961801422 | 0.191580944 | 0.848070471 | 0.972803041 | 4930404I05R | RIKEN cDNA 4930404I05 gene [Source:MG | 3.2124 | 6.1675 | 6.0375 | 10.604 | 4.3625 | 2.2277 |
| 28833 | ENSMUSG00000087355 | 0 | NA | NA | NA | NA | NA | Gm13187 | predicted gene 13187 [Source:MGI Symbo | 0 | 0 | 0 | 0 | 0 | 0 |
| 28834 | ENSMUSG00000087356 | 0.160662974 | 1.126924175 | 4.080472857 | 0.276174898 | 0.782413736 | NA | Gm13856 | predicted gene 13856 [Source:MGI Symbo | 0 | 0 | 0 | 0.964 | 0 | 0 |
| 28835 | ENSMUSG00000087357 | 0.267700746 | -1.497481649 | 4.057001436 | -0.369110456 | 0.712045396 | NA | Gm12498 | predicted gene 12498 [Source:MGI Symbo | 1.6062 | 0 | 0 | 0 | 0 | 0 |
| 28836 | ENSMUSG00000087358 | 0 | NA | NA | NA | NA | NA | 4930453H23 | RIKEN cDNA 4930453H23 gene [Source:M | 0 | 0 | 0 | 0 | 0 | 0 |

| | | | | | | | | | | | | | | | |
|---|---|---|---|---|---|---|---|---|---|---|---|---|---|---|---|
| 28837 | ENSMUSG00000087360 | 0.17131964 | -0.796629176 | 4.080472857 | -0.195229623 | 0.84521319 | NA | Gm15104 | predicted gene 15104 [Source:MGI Symbo | 0 | 1.0279 | 0 | 0 | 0 | 0 |
| 28838 | ENSMUSG00000087361 | 38.11766613 | -0.261744043 | 0.388372367 | -0.673951253 | 0.500342306 | 0.880716237 | 0610043K17 | RIKEN cDNA 0610043K17 gene [Source:M | 39.352 | 52.424 | 33.206 | 44.343 | 35.99 | 23.391 |
| 28839 | ENSMUSG00000087362 | 4.296812399 | 0.529763466 | 1.10370251 | 0.479987552 | 0.631236245 | NA | Gm13710 | predicted gene 13710 [Source:MGI Symbo | 2.4093 | 4.1117 | 4.025 | 0.964 | 9.8156 | 4.4554 |
| 28840 | ENSMUSG00000087363 | 0 | NA | NA | NA | NA | NA | Gm16347 | predicted gene 16347 [Source:MGI Symbo | 0 | 0 | 0 | 0 | 0 | 0 |
| 28841 | ENSMUSG00000087364 | 0 | NA | NA | NA | NA | NA | Gm13398 | predicted gene 13398 [Source:MGI Symbo | 0 | 0 | 0 | 0 | 0 | 0 |
| 28842 | ENSMUSG00000087365 | 2.048631249 | -0.292504249 | 1.743656719 | -0.167753346 | 0.866777327 | NA | C430049B03 | RIKEN cDNA C430049B03 gene [Source:M | 1.6062 | 5.1396 | 0 | 0 | 1.0906 | 4.4554 |
| 28843 | ENSMUSG00000087366 | 7.407921228 | 0.379178465 | 0.798549856 | 0.474833803 | 0.634905437 | 0.923478764 | Junos | jun proto-oncogene, opposite strand [Sou | 8.031 | 8.2233 | 3.0187 | 8.6758 | 9.8156 | 6.6831 |
| 28844 | ENSMUSG00000087367 | 0 | NA | NA | NA | NA | NA | Gm15491 | predicted gene 15491 [Source:MGI Symbo | 0 | 0 | 0 | 0 | 0 | 0 |
| 28845 | ENSMUSG00000087368 | 17.97208756 | -0.518300374 | 0.527718783 | -0.982152599 | 0.326024676 | 0.772018857 | BC065397 | cDNA sequence BC065397 [Source:MGI Sy | 17.668 | 29.81 | 16.1 | 14.46 | 13.087 | 16.708 |
| 28846 | ENSMUSG00000087369 | 0 | NA | NA | NA | NA | NA | Gm14696 | predicted gene 14696 [Source:MGI Symbo | 0 | 0 | 0 | 0 | 0 | 0 |
| 28847 | ENSMUSG00000087370 | 352.2731746 | 0.028580248 | 0.159593497 | 0.179081534 | 0.857873681 | 0.975277277 | Tmem170b | transmembrane protein 170B [Source:MG | 348.55 | 407.06 | 290.8 | 349.92 | 366.45 | 350.86 |
| 28848 | ENSMUSG00000087371 | 0.133850373 | -0.796629176 | 4.080472857 | -0.195229623 | 0.84521319 | NA | Gm15541 | predicted gene 15541 [Source:MGI Symbo | 0.8031 | 0 | 0 | 0 | 0 | 0 |
| 28849 | ENSMUSG00000087373 | 0.321325949 | 1.911575114 | 4.048267893 | 0.472195804 | 0.636787034 | NA | Gm15892 | predicted gene 15892 [Source:MGI Symbo | 0 | 0 | 0 | 1.928 | 0 | 0 |
| 28850 | ENSMUSG00000087374 | 0 | NA | NA | NA | NA | NA | Gm15457 | predicted gene 15457 [Source:MGI Symbo | 0 | 0 | 0 | 0 | 0 | 0 |
| 28851 | ENSMUSG00000087376 | 0 | NA | NA | NA | NA | NA | Gm15517 | predicted gene 15517 [Source:MGI Symbo | 0 | 0 | 0 | 0 | 0 | 0 |
| 28852 | ENSMUSG00000087377 | 6.110783558 | 0.046399479 | 0.87737179 | 0.052884626 | 0.957823834 | 0.991861398 | AV099323 | expressed sequence AV099323 [Source:M | 8.8341 | 4.1117 | 5.0312 | 7.7118 | 7.6343 | 3.3415 |
| 28853 | ENSMUSG00000087378 | 0 | NA | NA | NA | NA | NA | Gm12414 | predicted gene 12414 [Source:MGI Symbo | 0 | 0 | 0 | 0 | 0 | 0 |
| 28854 | ENSMUSG00000087379 | 0 | NA | NA | NA | NA | NA | Gm13849 | predicted gene 13849 [Source:MGI Symbo | 0 | 0 | 0 | 0 | 0 | 0 |
| 28855 | ENSMUSG00000087380 | 168.5334299 | 0.203523598 | 0.199382463 | 1.020769802 | 0.307363515 | 0.756237058 | 2210408F21 | RIKEN cDNA 2210408F21 gene [Source:M | 154.2 | 162.41 | 153.96 | 217.86 | 167.96 | 154.82 |
| 28856 | ENSMUSG00000087381 | 18.59350249 | 0.346136184 | 0.626515097 | 0.552478601 | 0.580620491 | 0.908645447 | Gm16008 | predicted gene 16008 [Source:MGI Symbo | 8.8341 | 18.503 | 22.137 | 36.631 | 7.6343 | 17.822 |
| 28857 | ENSMUSG00000087382 | 347.2702726 | -0.599640795 | 0.216053777 | -2.775423806 | 0.005512982 | 0.071069309 | Ctcflos | CCCTC-binding factor (zinc finger protein)- | 462.59 | 406.03 | 387.4 | 391.38 | 227.94 | 208.29 |
| 28858 | ENSMUSG00000087383 | 0 | NA | NA | NA | NA | NA | Gm12446 | predicted gene 12446 [Source:MGI Symbo | 0 | 0 | 0 | 0 | 0 | 0 |
| 28859 | ENSMUSG00000087384 | 0.653308562 | 1.538013299 | 3.112409277 | 0.494155223 | 0.62119656 | NA | Gm15558 | predicted gene 15558 [Source:MGI Symbo | 0 | 1.0279 | 0 | 2.8919 | 0 | 0 |
| 28860 | ENSMUSG00000087385 | 116.2325175 | 0.03802102 | 0.233905013 | 0.162548977 | 0.870873563 | 0.97888679 | Frg2f1 | FSHD region gene 2 family member 1 [Sou | 142.15 | 92.513 | 108.67 | 113.75 | 117.79 | 122.52 |
| 28861 | ENSMUSG00000087386 | 0 | NA | NA | NA | NA | NA | Gm15637 | predicted gene 15637 [Source:MGI Symbo | 0 | 0 | 0 | 0 | 0 | 0 |
| 28862 | ENSMUSG00000087387 | 1.027504657 | 1.047266995 | 2.389110177 | 0.43835023 | 0.661132419 | NA | Gm15420 | predicted gene 15420 [Source:MGI Symbo | 0 | 2.0558 | 0 | 1.928 | 2.1812 | 0 |
| 28863 | ENSMUSG00000087388 | 0 | NA | NA | NA | NA | NA | Gm12128 | predicted gene 12128 [Source:MGI Symbo | 0 | 0 | 0 | 0 | 0 | 0 |
| 28864 | ENSMUSG00000087389 | 0 | NA | NA | NA | NA | NA | Gm15592 | predicted gene 15592 [Source:MGI Symbo | 0 | 0 | 0 | 0 | 0 | 0 |
| 28865 | ENSMUSG00000087390 | 2.265731374 | -0.216318405 | 1.370698966 | -0.157816129 | 0.874601689 | NA | Gm7598 | predicted gene 7598 [Source:MGI Symbol | 3.2124 | 2.0558 | 2.0125 | 1.928 | 3.2719 | 1.1138 |
| 28866 | ENSMUSG00000087393 | 0 | NA | NA | NA | NA | NA | 1700023C21 | RIKEN cDNA 1700023C21 gene [Source:M | 0 | 0 | 0 | 0 | 0 | 0 |
| 28867 | ENSMUSG00000087394 | 0 | NA | NA | NA | NA | NA | Gm12945 | predicted gene 12945 [Source:MGI Symbo | 0 | 0 | 0 | 0 | 0 | 0 |
| 28868 | ENSMUSG00000087395 | 0 | NA | NA | NA | NA | NA | Gm12705 | predicted gene 12705 [Source:MGI Symbo | 0 | 0 | 0 | 0 | 0 | 0 |
| 28869 | ENSMUSG00000087396 | 17.45390383 | -0.164261629 | 0.522989795 | -0.314081901 | 0.753458835 | 0.952989051 | 4933407K13 | RIKEN cDNA 4933407K13 gene [Source:M | 12.85 | 20.558 | 22.137 | 18.316 | 15.269 | 15.594 |
| 28870 | ENSMUSG00000087397 | 0.506734502 | -2.372847156 | 3.338060886 | -0.710845978 | 0.477179685 | NA | Rapgef3os2 | Rap guanine nucleotide exchange factor ( | 0 | 1.0279 | 2.0125 | 0 | 0 | 0 |
| 28871 | ENSMUSG00000087398 | 0.342639279 | -1.805976596 | 4.028547714 | -0.448294702 | 0.653940524 | NA | Gm12343 | predicted gene 12343 [Source:MGI Symbo | 0 | 2.0558 | 0 | 0 | 0 | 0 |
| 28872 | ENSMUSG00000087399 | 0 | NA | NA | NA | NA | NA | Gm11899 | predicted gene 11899 [Source:MGI Symbo | 0 | 0 | 0 | 0 | 0 | 0 |
| 28873 | ENSMUSG00000087400 | 0.840571505 | -3.133009828 | 2.792076153 | -1.122107585 | 0.261816699 | NA | Gm15270 | predicted gene 15270 [Source:MGI Symbo | 4.0155 | 1.0279 | 0 | 0 | 0 | 0 |
| 28874 | ENSMUSG00000087401 | 0 | NA | NA | NA | NA | NA | Gm14664 | predicted gene 14664 [Source:MGI Symbo | 0 | 0 | 0 | 0 | 0 | 0 |
| 28875 | ENSMUSG00000087402 | 0 | NA | NA | NA | NA | NA | Gm11867 | predicted gene 11867 [Source:MGI Symbo | 0 | 0 | 0 | 0 | 0 | 0 |
| 28876 | ENSMUSG00000087403 | 28.45655092 | 0.877155007 | 0.432091237 | 2.030022673 | 0.042354235 | 0.283485262 | Kantr | Kdm5c adjacent non-coding transcript [So | 13.653 | 23.642 | 23.144 | 30.847 | 34.9 | 44.554 |
| 28877 | ENSMUSG00000087404 | 84.59471701 | -0.402792593 | 0.35612788 | -1.131033585 | 0.258040956 | 0.711580339 | Gm11752 | predicted gene 11752 [Source:MGI Symbo | 148.57 | 84.289 | 55.343 | 52.055 | 93.793 | 73.514 |
| 28878 | ENSMUSG00000087405 | 0.17131964 | -0.796629176 | 4.080472857 | -0.195229623 | 0.84521319 | NA | Gm14232 | predicted gene 14232 [Source:MGI Symbo | 0 | 1.0279 | 0 | 0 | 0 | 0 |
| 28879 | ENSMUSG00000087406 | 0 | NA | NA | NA | NA | NA | E130215H24 | RIKEN cDNA E130215H24 gene [Source:M | 0 | 0 | 0 | 0 | 0 | 0 |
| 28880 | ENSMUSG00000087407 | 0 | NA | NA | NA | NA | NA | Gm11431 | predicted gene 11431 [Source:MGI Symbo | 0 | 0 | 0 | 0 | 0 | 0 |
| 28881 | ENSMUSG00000087408 | 0.17131964 | -0.796629176 | 4.080472857 | -0.195229623 | 0.84521319 | NA | Cers1 | ceramide synthase 1 [Source:MGI Symbol | 0 | 1.0279 | 0 | 0 | 0 | 0 |
| 28882 | ENSMUSG00000087409 | 0 | NA | NA | NA | NA | NA | Gm14271 | predicted gene 14271 [Source:MGI Symbo | 0 | 0 | 0 | 0 | 0 | 0 |
| 28883 | ENSMUSG00000087410 | 0 | NA | NA | NA | NA | NA | 2310065F04 | RIKEN cDNA 2310065F04 gene [Source:M | 0 | 0 | 0 | 0 | 0 | 0 |
| 28884 | ENSMUSG00000087411 | 0 | NA | NA | NA | NA | NA | Gm16048 | predicted gene 16048 [Source:MGI Symbo | 0 | 0 | 0 | 0 | 0 | 0 |
| 28885 | ENSMUSG00000087412 | 6.894075648 | 0.367789351 | 0.950628854 | 0.386890582 | 0.698837214 | 0.940854732 | Gm15501 | predicted pseudogene 15501 [Source:MG | 8.8341 | 8.2233 | 1.0062 | 14.46 | 3.2719 | 5.5692 |
| 28886 | ENSMUSG00000087413 | 67.40170889 | 0.16281015 | 0.325137435 | 0.500742555 | 0.616552319 | 0.917509899 | Gm11266 | predicted gene 11266 [Source:MGI Symbo | 61.839 | 51.396 | 77.481 | 58.803 | 101.43 | 53.465 |
| 28887 | ENSMUSG00000087415 | 0 | NA | NA | NA | NA | NA | Gm14529 | predicted gene 14529 [Source:MGI Symbo | 0 | 0 | 0 | 0 | 0 | 0 |
| 28888 | ENSMUSG00000087416 | 0 | NA | NA | NA | NA | NA | Gm15906 | predicted gene 15906 [Source:MGI Symbo | 0 | 0 | 0 | 0 | 0 | 0 |
| 28889 | ENSMUSG00000087417 | 0 | NA | NA | NA | NA | NA | Gm13001 | predicted gene 13001 [Source:MGI Symbo | 0 | 0 | 0 | 0 | 0 | 0 |
| 28890 | ENSMUSG00000087422 | 0 | NA | NA | NA | NA | NA | Gm13564 | predicted gene 13564 [Source:MGI Symbo | 0 | 0 | 0 | 0 | 0 | 0 |
| 28891 | ENSMUSG00000087424 | 14.81505546 | -0.530664391 | 0.56946246 | -0.931868961 | 0.35140425 | 0.792511048 | 5730405O15 | RIKEN cDNA 5730405O15 gene [Source:M | 20.078 | 13.363 | 19.119 | 15.424 | 11.997 | 8.9108 |
| 28892 | ENSMUSG00000087425 | 0 | NA | NA | NA | NA | NA | Gm15404 | predicted gene 15404 [Source:MGI Symbo | 0 | 0 | 0 | 0 | 0 | 0 |

| | | | | | | | | | | | | | | | |
|---|---|---|---|---|---|---|---|---|---|---|---|---|---|---|---|
| 28893 | ENSMUSG00000087428 | 0 | NA | NA | NA | NA | NA | Gm15017 | predicted gene 15017 [Source:MGI Symbo | 0 | 0 | 0 | 0 | 0 | 0 |
| 28894 | ENSMUSG00000087429 | 0.301557804 | -1.64335916 | 4.042814367 | -0.406488899 | 0.684383412 | NA | Gm16235 | predicted gene 16235 [Source:MGI Symbo | 0.8031 | 0 | 1.0062 | 0 | 0 | 0 |
| 28895 | ENSMUSG00000087430 | 0 | NA | NA | NA | NA | NA | Gm13405 | predicted gene 13405 [Source:MGI Symbo | 0 | 0 | 0 | 0 | 0 | 0 |
| 28896 | ENSMUSG00000087431 | 0 | NA | NA | NA | NA | NA | Gm16054 | predicted gene 16054 [Source:MGI Symbo | 0 | 0 | 0 | 0 | 0 | 0 |
| 28897 | ENSMUSG00000087432 | 0 | NA | NA | NA | NA | NA | Gm15123 | predicted gene 15123 [Source:MGI Symbo | 0 | 0 | 0 | 0 | 0 | 0 |
| 28898 | ENSMUSG00000087433 | 2.330536204 | -0.003744611 | 1.664054768 | -0.002250293 | 0.998204527 | NA | Gm14167 | predicted gene 14167 [Source:MGI Symbo | 0 | 3.0838 | 4.025 | 5.7839 | 1.0906 | 0 |
| 28899 | ENSMUSG00000087434 | 0 | NA | NA | NA | NA | NA | Rab11fip4os | RAB11 family interacting protein 4 (class I | 0 | 0 | 0 | 0 | 0 | 0 |
| 28900 | ENSMUSG00000087435 | 0 | NA | NA | NA | NA | NA | Gm16323 | predicted gene 16323 [Source:MGI Symbo | 0 | 0 | 0 | 0 | 0 | 0 |
| 28901 | ENSMUSG00000087436 | 0.160662974 | 1.126924175 | 4.080472857 | 0.276174898 | 0.782413736 | NA | Gm16156 | predicted gene 16156 [Source:MGI Symbo | 0 | 0 | 0 | 0.964 | 0 | 0 |
| 28902 | ENSMUSG00000087437 | 0 | NA | NA | NA | NA | NA | Gm13112 | predicted gene 13112 [Source:MGI Symbo | 0 | 0 | 0 | 0 | 0 | 0 |
| 28903 | ENSMUSG00000087439 | 33.95981515 | -0.081595581 | 0.393133955 | -0.207551599 | 0.8355791 | 0.971073158 | Gm15788 | predicted gene 15788 [Source:MGI Symbo | 28.912 | 33.921 | 42.262 | 42.415 | 26.175 | 30.074 |
| 28904 | ENSMUSG00000087440 | 9.115554037 | -0.997191866 | 0.833686874 | -1.196122786 | 0.231648647 | 0.67889946 | 4930577N17 | RIKEN cDNA 4930577N17 gene [Source:M | 19.274 | 5.1396 | 12.075 | 11.568 | 2.1812 | 4.4554 |
| 28905 | ENSMUSG00000087441 | 0 | NA | NA | NA | NA | NA | Gm13853 | predicted gene 13853 [Source:MGI Symbo | 0 | 0 | 0 | 0 | 0 | 0 |
| 28906 | ENSMUSG00000087442 | 0.622883753 | 0.113321311 | 2.884614835 | 0.039284729 | 0.968663382 | NA | Gm14453 | predicted gene 14453 [Source:MGI Symbo | 0.8031 | 0 | 1.0062 | 1.928 | 0 | 0 |
| 28907 | ENSMUSG00000087443 | 0 | NA | NA | NA | NA | NA | Ppp1r18os | protein phosphatase 1, regulatory subunit | 0 | 0 | 0 | 0 | 0 | 0 |
| 28908 | ENSMUSG00000087444 | 0.481988923 | 2.492401104 | 3.837077151 | 0.649557203 | 0.515978285 | NA | Gm5475 | predicted gene 5475 [Source:MGI Symbol | 0 | 0 | 0 | 2.8919 | 0 | 0 |
| 28909 | ENSMUSG00000087445 | 9.491744433 | 0.265914843 | 0.821600786 | 0.323654562 | 0.746199575 | 0.949997329 | Gm14286 | predicted gene 14286 [Source:MGI Symbo | 13.653 | 6.1675 | 6.0375 | 21.208 | 6.5437 | 3.3415 |
| 28910 | ENSMUSG00000087449 | 1.061904139 | -1.168441417 | 2.189134959 | -0.533745721 | 0.593517472 | NA | Gm12971 | predicted gene 12971 [Source:MGI Symbo | 2.4093 | 1.0279 | 1.0062 | 1.928 | 0 | 0 |
| 28911 | ENSMUSG00000087450 | 0.83344994 | -1.76917134 | 2.609239342 | -0.678041034 | 0.497745676 | NA | Gm13994 | predicted gene 13994 [Source:MGI Symbo | 0.8031 | 3.0838 | 0 | 0 | 0 | 1.1138 |
| 28912 | ENSMUSG00000087451 | 0.819128932 | -3.070895205 | 2.828495588 | -1.085699132 | 0.277612125 | NA | Gm12970 | predicted gene 12970 [Source:MGI Symbo | 0.8031 | 4.1117 | 0 | 0 | 0 | 0 |
| 28913 | ENSMUSG00000087452 | 0 | NA | NA | NA | NA | NA | Gm11998 | predicted gene 11998 [Source:MGI Symbo | 0 | 0 | 0 | 0 | 0 | 0 |
| 28914 | ENSMUSG00000087454 | 0 | NA | NA | NA | NA | NA | Gm13174 | predicted gene 13174 [Source:MGI Symbo | 0 | 0 | 0 | 0 | 0 | 0 |
| 28915 | ENSMUSG00000087455 | 0.305170013 | -1.657836477 | 4.041480335 | -0.410205257 | 0.681655385 | NA | 5330411J11R | RIKEN cDNA 5330411J11 gene [Source:MG | 0.8031 | 1.0279 | 0 | 0 | 0 | 0 |
| 28916 | ENSMUSG00000087456 | 0.480153995 | 1.043847617 | 3.151673025 | 0.331204287 | 0.740490182 | NA | Gm16322 | predicted gene 16322 [Source:MGI Symbo | 0.8031 | 0 | 0 | 0.964 | 0 | 1.1138 |
| 28917 | ENSMUSG00000087457 | 0 | NA | NA | NA | NA | NA | Gm7799 | predicted gene 7799 [Source:MGI Symbol | 0 | 0 | 0 | 0 | 0 | 0 |
| 28918 | ENSMUSG00000087458 | 0 | NA | NA | NA | NA | NA | Gm13999 | predicted gene 13999 [Source:MGI Symbo | 0 | 0 | 0 | 0 | 0 | 0 |
| 28919 | ENSMUSG00000087459 | 0 | NA | NA | NA | NA | NA | Csmd2os | CUB and Sushi multiple domains 2, oppos | 0 | 0 | 0 | 0 | 0 | 0 |
| 28920 | ENSMUSG00000087460 | 0 | NA | NA | NA | NA | NA | 4930401O12 | RIKEN cDNA 4930401O12 gene [Source:M | 0 | 0 | 0 | 0 | 0 | 0 |
| 28921 | ENSMUSG00000087461 | 0 | NA | NA | NA | NA | NA | C230014O12 | RIKEN cDNA C230014O12 gene [Source:M | 0 | 0 | 0 | 0 | 0 | 0 |
| 28922 | ENSMUSG00000087462 | 0 | NA | NA | NA | NA | NA | Gm2200 | predicted gene 2200 [Source:MGI Symbol | 0 | 0 | 0 | 0 | 0 | 0 |
| 28923 | ENSMUSG00000087464 | 0 | NA | NA | NA | NA | NA | Gm12862 | predicted gene 12862 [Source:MGI Symbo | 0 | 0 | 0 | 0 | 0 | 0 |
| 28924 | ENSMUSG00000087465 | 0 | NA | NA | NA | NA | NA | Gm13905 | predicted gene 13905 [Source:MGI Symbo | 0 | 0 | 0 | 0 | 0 | 0 |
| 28925 | ENSMUSG00000087466 | 4.831915177 | -1.061988024 | 0.970942658 | -1.093770074 | 0.274055832 | NA | A330041J22I | RIKEN cDNA A330041J22 gene [Source:MG | 6.4248 | 6.1675 | 7.0437 | 3.8559 | 3.2719 | 2.2277 |
| 28926 | ENSMUSG00000087467 | 0 | NA | NA | NA | NA | NA | Gm13601 | predicted gene 13601 [Source:MGI Symbo | 0 | 0 | 0 | 0 | 0 | 0 |
| 28927 | ENSMUSG00000087468 | 0 | NA | NA | NA | NA | NA | Gm7023 | predicted gene 7023 [Source:MGI Symbol | 0 | 0 | 0 | 0 | 0 | 0 |
| 28928 | ENSMUSG00000087469 | 0 | NA | NA | NA | NA | NA | Gm16142 | predicted gene 16142 [Source:MGI Symbo | 0 | 0 | 0 | 0 | 0 | 0 |
| 28929 | ENSMUSG00000087470 | 0 | NA | NA | NA | NA | NA | A630031M0 | RIKEN cDNA A630031M04 gene [Source:N | 0 | 0 | 0 | 0 | 0 | 0 |
| 28930 | ENSMUSG00000087473 | 0 | NA | NA | NA | NA | NA | Gm13883 | predicted gene 13883 [Source:MGI Symbo | 0 | 0 | 0 | 0 | 0 | 0 |
| 28931 | ENSMUSG00000087474 | 0 | NA | NA | NA | NA | NA | Gm12100 | predicted gene 12100 [Source:MGI Symbo | 0 | 0 | 0 | 0 | 0 | 0 |
| 28932 | ENSMUSG00000087475 | 8.855978687 | -0.179494432 | 0.785277611 | -0.228574493 | 0.819199651 | 0.969160361 | 4933406I18R | RIKEN cDNA 4933406I18 gene [Source:MG | 8.031 | 7.1954 | 13.081 | 10.604 | 11.997 | 2.2277 |
| 28933 | ENSMUSG00000087476 | 1.588117572 | 1.132025104 | 1.923403777 | 0.588553021 | 0.556161156 | NA | Rap1gapos | RAP1 GTPase activating protein, opposite | 0 | 0 | 3.0187 | 0.964 | 1.0906 | 4.4554 |
| 28934 | ENSMUSG00000087477 | 1.714948464 | 0.017561858 | 1.686947635 | 0.010410435 | 0.991693825 | NA | Gm13822 | predicted gene 13822 [Source:MGI Symbo | 4.0155 | 1.0279 | 0 | 1.928 | 1.0906 | 2.2277 |
| 28935 | ENSMUSG00000087478 | 25.47610359 | 0.227696372 | 0.5264069 | 0.432548228 | 0.665343008 | 0.929426323 | 4930506C21 | RIKEN cDNA 4930506C21 gene [Source:M | 33.73 | 10.279 | 26.162 | 28.919 | 38.172 | 15.594 |
| 28936 | ENSMUSG00000087479 | 0 | NA | NA | NA | NA | NA | Gm16835 | predicted gene, 16835 [Source:MGI Symb | 0 | 0 | 0 | 0 | 0 | 0 |
| 28937 | ENSMUSG00000087480 | 0.339027071 | -1.79174149 | 3.919585585 | -0.457125237 | 0.647581032 | NA | Gm15910 | predicted gene 15910 [Source:MGI Symbo | 0 | 1.0279 | 1.0062 | 0 | 0 | 0 |
| 28938 | ENSMUSG00000087482 | 0 | NA | NA | NA | NA | NA | Gm12720 | predicted gene 12720 [Source:MGI Symbo | 0 | 0 | 0 | 0 | 0 | 0 |
| 28939 | ENSMUSG00000087484 | 21.53444109 | 0.350300233 | 0.49110666 | 0.713287481 | 0.475667883 | 0.869462941 | 2900089D17 | RIKEN cDNA 2900089D17 gene [Source:M | 24.093 | 14.391 | 18.112 | 25.063 | 16.359 | 31.188 |
| 28940 | ENSMUSG00000087485 | 0 | NA | NA | NA | NA | NA | Gm13383 | predicted gene 13383 [Source:MGI Symbo | 0 | 0 | 0 | 0 | 0 | 0 |
| 28941 | ENSMUSG00000087486 | 0 | NA | NA | NA | NA | NA | Gm11723 | predicted gene 11723 [Source:MGI Symbo | 0 | 0 | 0 | 0 | 0 | 0 |
| 28942 | ENSMUSG00000087487 | 0 | NA | NA | NA | NA | NA | Ppp2r2cos | protein phosphatase 2, regulatory subunit | 0 | 0 | 0 | 0 | 0 | 0 |
| 28943 | ENSMUSG00000087489 | 0 | NA | NA | NA | NA | NA | Gm15656 | predicted gene 15656 [Source:MGI Symbo | 0 | 0 | 0 | 0 | 0 | 0 |
| 28944 | ENSMUSG00000087491 | 0 | NA | NA | NA | NA | NA | Gm14552 | predicted gene 14552 [Source:MGI Symbo | 0 | 0 | 0 | 0 | 0 | 0 |
| 28945 | ENSMUSG00000087492 | 0 | NA | NA | NA | NA | NA | Gm15700 | predicted gene 15700 [Source:MGI Symbo | 0 | 0 | 0 | 0 | 0 | 0 |
| 28946 | ENSMUSG00000087495 | 0 | NA | NA | NA | NA | NA | Gm14152 | predicted gene 14152 [Source:MGI Symbo | 0 | 0 | 0 | 0 | 0 | 0 |
| 28947 | ENSMUSG00000087496 | 0.669251866 | -2.807208097 | 3.391018549 | -0.827836255 | 0.407763239 | NA | Gm13523 | predicted gene 13523 [Source:MGI Symbo | 4.0155 | 0 | 0 | 0 | 0 | 0 |
| 28948 | ENSMUSG00000087497 | 45.8495243 | 0.133977774 | 0.324277214 | 0.413158152 | 0.679490746 | 0.93418523 | 2810001G20 | RIKEN cDNA 2810001G20 gene [Source:M | 50.595 | 39.061 | 41.256 | 47.235 | 50.168 | 46.781 |

| | | | | | | | | | | | | | | | |
|---|---|---|---|---|---|---|---|---|---|---|---|---|---|---|---|
| 28949 | ENSMUSG00000087500 | 0 | NA | NA | NA | NA | NA | Gm12426 | predicted gene 12426 [Source:MGI Symbo | 0 | 0 | 0 | 0 | 0 | 0 |
| 28950 | ENSMUSG00000087501 | 0.160662974 | 1.126924175 | 4.080472857 | 0.276174898 | 0.782413736 | NA | Gm12116 | predicted gene 12116 [Source:MGI Symbo | 0 | 0 | 0 | 0.964 | 0 | 0 |
| 28951 | ENSMUSG00000087502 | 7.850231787 | -0.328055379 | 0.840215453 | -0.390441973 | 0.696209756 | 0.939992732 | Gm16091 | predicted gene 16091 [Source:MGI Symbo | 4.0155 | 10.279 | 12.075 | 9.6398 | 2.1812 | 8.9108 |
| 28952 | ENSMUSG00000087504 | 0 | NA | NA | NA | NA | NA | Gm13658 | predicted gene 13658 [Source:MGI Symbo | 0 | 0 | 0 | 0 | 0 | 0 |
| 28953 | ENSMUSG00000087505 | 0 | NA | NA | NA | NA | NA | Gm15241 | predicted gene 15241 [Source:MGI Symbo | 0 | 0 | 0 | 0 | 0 | 0 |
| 28954 | ENSMUSG00000087507 | 0.167707431 | -0.796629176 | 4.080472857 | -0.195229623 | 0.84521319 | NA | Gm16141 | predicted gene 16141 [Source:MGI Symbo | 0 | 0 | 1.0062 | 0 | 0 | 0 |
| 28955 | ENSMUSG00000087508 | 0 | NA | NA | NA | NA | NA | Gm14174 | predicted gene 14174 [Source:MGI Symbo | 0 | 0 | 0 | 0 | 0 | 0 |
| 28956 | ENSMUSG00000087510 | 0 | NA | NA | NA | NA | NA | 1700112K13 | RIKEN cDNA 1700112K13 gene [Source:M | 0 | 0 | 0 | 0 | 0 | 0 |
| 28957 | ENSMUSG00000087512 | 0 | NA | NA | NA | NA | NA | Ccdc201 | coiled coil domain 201 [Source:MGI Symb | 0 | 0 | 0 | 0 | 0 | 0 |
| 28958 | ENSMUSG00000087513 | 0 | NA | NA | NA | NA | NA | Gm13236 | predicted gene 13236 [Source:MGI Symbo | 0 | 0 | 0 | 0 | 0 | 0 |
| 28959 | ENSMUSG00000087514 | 0.654905883 | -1.312574656 | 2.816093566 | -0.466097672 | 0.641145589 | NA | Gm16076 | predicted gene 16076 [Source:MGI Symbo | 0.8031 | 0 | 2.0125 | 0 | 0 | 1.1138 |
| 28960 | ENSMUSG00000087515 | 0 | NA | NA | NA | NA | NA | Gm13554 | predicted gene 13554 [Source:MGI Symbo | 0 | 0 | 0 | 0 | 0 | 0 |
| 28961 | ENSMUSG00000087516 | 114.3374593 | 0.452568544 | 0.242106288 | 1.869296942 | 0.061581513 | 0.354584001 | Tbx3os1 | T-box 3, opposite strand 1 [Source:MGI Sy | 81.916 | 105.88 | 102.64 | 160.02 | 130.87 | 104.7 |
| 28962 | ENSMUSG00000087517 | 0 | NA | NA | NA | NA | NA | 4930577H14 | RIKEN cDNA 4930577H14 gene [Source:M | 0 | 0 | 0 | 0 | 0 | 0 |
| 28963 | ENSMUSG00000087518 | 3.230039778 | -0.322130455 | 1.235191796 | -0.260793875 | 0.794251468 | NA | Gm13561 | predicted gene 13561 [Source:MGI Symbo | 1.6062 | 6.1675 | 3.0187 | 1.928 | 1.0906 | 5.5692 |
| 28964 | ENSMUSG00000087519 | 1.575915191 | -0.430254688 | 1.72216694 | -0.249833322 | 0.802716249 | NA | AV039307 | expressed sequence AV039307 [Source:M | 2.4093 | 1.0279 | 2.0125 | 2.8919 | 0 | 1.1138 |
| 28965 | ENSMUSG00000087520 | 0 | NA | NA | NA | NA | NA | Gm14389 | predicted gene 14389 [Source:MGI Symbo | 0 | 0 | 0 | 0 | 0 | 0 |
| 28966 | ENSMUSG00000087521 | 0 | NA | NA | NA | NA | NA | AI646519 | expressed sequence AI646519 [Source:M( | 0 | 0 | 0 | 0 | 0 | 0 |
| 28967 | ENSMUSG00000087522 | 0 | NA | NA | NA | NA | NA | Gm371 | predicted pseudogene 371 [Source:MGI S | 0 | 0 | 0 | 0 | 0 | 0 |
| 28968 | ENSMUSG00000087523 | 27.6574699 | 0.022595689 | 0.437747183 | 0.051618126 | 0.958832976 | 0.992027194 | Gm12319 | predicted gene 12319 [Source:MGI Symbo | 26.502 | 27.754 | 28.175 | 33.739 | 16.359 | 33.415 |
| 28969 | ENSMUSG00000087524 | 0 | NA | NA | NA | NA | NA | Gm14285 | predicted gene 14285 [Source:MGI Symbo | 0 | 0 | 0 | 0 | 0 | 0 |
| 28970 | ENSMUSG00000087525 | 0 | NA | NA | NA | NA | NA | Gm13062 | predicted gene 13062 [Source:MGI Symbo | 0 | 0 | 0 | 0 | 0 | 0 |
| 28971 | ENSMUSG00000087526 | 1.020745555 | -0.932908796 | 2.259280719 | -0.412922922 | 0.679663086 | NA | Gm15738 | predicted gene 15738 [Source:MGI Symbo | 0 | 1.0279 | 3.0187 | 0.964 | 0 | 1.1138 |
| 28972 | ENSMUSG00000087527 | 0 | NA | NA | NA | NA | NA | Gm13656 | predicted gene 13656 [Source:MGI Symbo | 0 | 0 | 0 | 0 | 0 | 0 |
| 28973 | ENSMUSG00000087528 | 17.67105337 | -0.529921568 | 0.547360585 | -0.968139802 | 0.332974554 | 0.77523112 | 9830144P21 | RIKEN cDNA 9830144P21 gene [Source:M | 28.109 | 14.391 | 20.125 | 20.244 | 10.906 | 12.252 |
| 28974 | ENSMUSG00000087530 | 0.167707431 | -0.796629176 | 4.080472857 | -0.195229623 | 0.84521319 | NA | Gm15533 | predicted gene 15533 [Source:MGI Symbo | 0 | 0 | 1.0062 | 0 | 0 | 0 |
| 28975 | ENSMUSG00000087531 | 0 | NA | NA | NA | NA | NA | Gm15606 | predicted gene 15606 [Source:MGI Symbo | 0 | 0 | 0 | 0 | 0 | 0 |
| 28976 | ENSMUSG00000087532 | 0.435408177 | -2.179616019 | 3.531869654 | -0.617128103 | 0.537150234 | NA | Gm15938 | predicted gene 15938 [Source:MGI Symbo | 1.6062 | 0 | 1.0062 | 0 | 0 | 0 |
| 28977 | ENSMUSG00000087534 | 0 | NA | NA | NA | NA | NA | Gm11418 | predicted gene 11418 [Source:MGI Symbo | 0 | 0 | 0 | 0 | 0 | 0 |
| 28978 | ENSMUSG00000087535 | 7.159271217 | 0.578638415 | 0.816632414 | 0.708566554 | 0.478593498 | 0.86997315 | Zmiz1os1 | Zmiz1 opposite strand 1 [Source:MGI Sym | 8.031 | 4.1117 | 5.0312 | 12.532 | 5.4531 | 7.7969 |
| 28979 | ENSMUSG00000087536 | 0 | NA | NA | NA | NA | NA | Gm14246 | predicted gene 14246 [Source:MGI Symbo | 0 | 0 | 0 | 0 | 0 | 0 |
| 28980 | ENSMUSG00000087537 | 0 | NA | NA | NA | NA | NA | Gm13799 | predicted gene 13799 [Source:MGI Symbo | 0 | 0 | 0 | 0 | 0 | 0 |
| 28981 | ENSMUSG00000087538 | 0 | NA | NA | NA | NA | NA | Gm16341 | predicted gene 16341 [Source:MGI Symbo | 0 | 0 | 0 | 0 | 0 | 0 |
| 28982 | ENSMUSG00000087539 | 0 | NA | NA | NA | NA | NA | Gm13584 | predicted gene 13584 [Source:MGI Symbo | 0 | 0 | 0 | 0 | 0 | 0 |
| 28983 | ENSMUSG00000087540 | 0 | NA | NA | NA | NA | NA | Gm11412 | predicted gene 11412 [Source:MGI Symbo | 0 | 0 | 0 | 0 | 0 | 0 |
| 28984 | ENSMUSG00000087541 | 5.19179488 | 0.575399119 | 1.035807081 | 0.555507999 | 0.57854724 | 0.908137944 | Hopxos | HOP homeobox, opposite strand [Source: | 2.4093 | 7.1954 | 3.0187 | 9.6398 | 1.0906 | 7.7969 |
| 28985 | ENSMUSG00000087542 | 0 | NA | NA | NA | NA | NA | Gm15878 | predicted gene 15878 [Source:MGI Symbo | 0 | 0 | 0 | 0 | 0 | 0 |
| 28986 | ENSMUSG00000087543 | 21.58723147 | 0.143652238 | 0.497032699 | 0.289019692 | 0.772566308 | 0.957772025 | Gm16576 | predicted gene 16576 [Source:MGI Symbo | 20.881 | 13.363 | 27.169 | 16.388 | 29.447 | 22.277 |
| 28987 | ENSMUSG00000087544 | 0 | NA | NA | NA | NA | NA | Gm13923 | predicted gene 13923 [Source:MGI Symbo | 0 | 0 | 0 | 0 | 0 | 0 |
| 28988 | ENSMUSG00000087545 | 0 | NA | NA | NA | NA | NA | Gm13966 | predicted gene 13966 [Source:MGI Symbo | 0 | 0 | 0 | 0 | 0 | 0 |
| 28989 | ENSMUSG00000087547 | 0 | NA | NA | NA | NA | NA | Platr27 | pluripotency associated transcript 27 [Sou | 0 | 0 | 0 | 0 | 0 | 0 |
| 28990 | ENSMUSG00000087549 | 2.48252087 | -0.416501075 | 1.40924192 | -0.295549734 | 0.767573966 | NA | Gm13427 | predicted gene 13427 [Source:MGI Symbo | 2.4093 | 3.0838 | 3.0187 | 1.928 | 0 | 4.4554 |
| 28991 | ENSMUSG00000087550 | 0 | NA | NA | NA | NA | NA | Gm14343 | predicted gene 14343 [Source:MGI Symbo | 0 | 0 | 0 | 0 | 0 | 0 |
| 28992 | ENSMUSG00000087552 | 0.688762941 | -1.399623551 | 3.067774249 | -0.456234207 | 0.64822157 | NA | Gm15819 | predicted gene 15819 [Source:MGI Symbo | 0 | 0 | 3.0187 | 0 | 0 | 1.1138 |
| 28993 | ENSMUSG00000087553 | 0 | NA | NA | NA | NA | NA | 5730412P04 | RIKEN cDNA 5730412P04 gene [Source:M | 0 | 0 | 0 | 0 | 0 | 0 |
| 28994 | ENSMUSG00000087555 | 0 | NA | NA | NA | NA | NA | Gm13997 | predicted gene 13997 [Source:MGI Symbo | 0 | 0 | 0 | 0 | 0 | 0 |
| 28995 | ENSMUSG00000087556 | 30.27271607 | 0.048711945 | 0.458812329 | 0.106169652 | 0.915447749 | 0.98625062 | Gm15764 | predicted gene 15764 [Source:MGI Symbo | 30.518 | 24.67 | 34.212 | 42.415 | 14.178 | 35.643 |
| 28996 | ENSMUSG00000087559 | 0 | NA | NA | NA | NA | NA | Gm13802 | predicted gene 13802 [Source:MGI Symbo | 0 | 0 | 0 | 0 | 0 | 0 |
| 28997 | ENSMUSG00000087561 | 0 | NA | NA | NA | NA | NA | Gm11608 | predicted gene 11608 [Source:MGI Symbo | 0 | 0 | 0 | 0 | 0 | 0 |
| 28998 | ENSMUSG00000087563 | 0 | NA | NA | NA | NA | NA | Gm14205 | predicted gene 14205 [Source:MGI Symbo | 0 | 0 | 0 | 0 | 0 | 0 |
| 28999 | ENSMUSG00000087564 | 0 | NA | NA | NA | NA | NA | Gm11817 | predicted gene 11817 [Source:MGI Symbo | 0 | 0 | 0 | 0 | 0 | 0 |
| 29000 | ENSMUSG00000087565 | 0 | NA | NA | NA | NA | NA | Gm14662 | predicted gene 14662 [Source:MGI Symbo | 0 | 0 | 0 | 0 | 0 | 0 |
| 29001 | ENSMUSG00000087568 | 0 | NA | NA | NA | NA | NA | Maats1os | MYCBP-associated, testis expressed 1, op | 0 | 0 | 0 | 0 | 0 | 0 |
| 29002 | ENSMUSG00000087569 | 0 | NA | NA | NA | NA | NA | Actb-ps1 | actin, beta, pseudogene 1 [Source:MGI Sy | 0 | 0 | 0 | 0 | 0 | 0 |
| 29003 | ENSMUSG00000087570 | 0 | NA | NA | NA | NA | NA | Gm13647 | predicted gene 13647 [Source:MGI Symbo | 0 | 0 | 0 | 0 | 0 | 0 |
| 29004 | ENSMUSG00000087571 | 0 | NA | NA | NA | NA | NA | Gm15202 | predicted gene 15202 [Source:MGI Symbo | 0 | 0 | 0 | 0 | 0 | 0 |

| | | | | | | | | | | | | | | |
|---|---|---|---|---|---|---|---|---|---|---|---|---|---|---|
| 29005 ENSMUSG00000087574 | 11.65492156 | -0.09369176 | 0.679027711 | -0.137979287 | 0.890256787 | 0.982489666 | C030037D0 | RIKEN cDNA C030037D09 gene [Source:M | 4.8186 | 13.363 | 18.112 | 11.568 | 9.8156 | 12.252 |
| 29006 ENSMUSG00000087575 | 0.160662974 | 1.126924175 | 4.080472857 | 0.276174898 | 0.782413736 | NA | Gm12976 | predicted gene 12976 [Source:MGI Symbo | 0 | 0 | 0 | 0.964 | 0 | 0 |
| 29007 ENSMUSG00000087577 | 0 | NA | NA | NA | NA | NA | Gm13299 | predicted gene 13299 [Source:MGI Symbo | 0 | 0 | 0 | 0 | 0 | 0 |
| 29008 ENSMUSG00000087578 | 6.524253359 | 0.562780523 | 0.840366162 | 0.669684892 | 0.503058684 | 0.881220881 | Ccdc142os | coiled-coil domain containing 142, opposi | 5.6217 | 3.0838 | 7.0437 | 5.7839 | 9.8156 | 7.7969 |
| 29009 ENSMUSG00000087579 | 450.8697353 | 0.542275492 | 0.216739765 | 2.501965858 | 0.012350583 | 0.125184891 | Hectd2os | Hectd2, opposite strand [Source:MGI Sym | 460.98 | 328.93 | 309.92 | 364.38 | 567.12 | 673.88 |
| 29010 ENSMUSG00000087581 | 0 | NA | NA | NA | NA | NA | Gm11571 | predicted gene 11571 [Source:MGI Symbo | 0 | 0 | 0 | 0 | 0 | 0 |
| 29011 ENSMUSG00000087582 | 0 | NA | NA | NA | NA | NA | Ighv6-7 | immunoglobulin heavy variable V6-7 [Sou | 0 | 0 | 0 | 0 | 0 | 0 |
| 29012 ENSMUSG00000087584 | 0 | NA | NA | NA | NA | NA | Gm14457 | predicted gene 14457 [Source:MGI Symbo | 0 | 0 | 0 | 0 | 0 | 0 |
| 29013 ENSMUSG00000087585 | 0.688762941 | -1.399623551 | 3.067774249 | -0.456234207 | 0.64822157 | NA | Gm14100 | predicted gene 14100 [Source:MGI Symbo | 0 | 0 | 3.0187 | 0 | 0 | 1.1138 |
| 29014 ENSMUSG00000087586 | 0 | NA | NA | NA | NA | NA | Gm6938 | predicted gene 6938 [Source:MGI Symbol | 0 | 0 | 0 | 0 | 0 | 0 |
| 29015 ENSMUSG00000087587 | 0 | NA | NA | NA | NA | NA | Gm15245 | predicted gene 15245 [Source:MGI Symbo | 0 | 0 | 0 | 0 | 0 | 0 |
| 29016 ENSMUSG00000087588 | 0 | NA | NA | NA | NA | NA | Gm16355 | predicted gene 16355 [Source:MGI Symbo | 0 | 0 | 0 | 0 | 0 | 0 |
| 29017 ENSMUSG00000087589 | 3.769369851 | -0.256502256 | 1.186074049 | -0.216261587 | 0.82878385 | NA | D430040D2 | RIKEN cDNA D430040D24 gene [Source:M | 3.2124 | 5.1396 | 4.025 | 5.7839 | 0 | 4.4554 |
| 29018 ENSMUSG00000087590 | 142.1868284 | 0.178361707 | 0.200771354 | 0.888382249 | 0.374335171 | 0.809332934 | Epb41l4aos | erythrocyte membrane protein band 4.1 l | 126.89 | 146.99 | 126.79 | 163.88 | 152.69 | 135.89 |
| 29019 ENSMUSG00000087591 | 0 | NA | NA | NA | NA | NA | Gm14635 | predicted gene 14635 [Source:MGI Symbo | 0 | 0 | 0 | 0 | 0 | 0 |
| 29020 ENSMUSG00000087592 | 0 | NA | NA | NA | NA | NA | Gm11184 | predicted gene 11184 [Source:MGI Symbo | 0 | 0 | 0 | 0 | 0 | 0 |
| 29021 ENSMUSG00000087593 | 26.03033137 | 0.545053165 | 0.442379849 | 1.232093112 | 0.217914304 | 0.663697152 | Gm16174 | predicted gene 16174 [Source:MGI Symbo | 21.684 | 19.53 | 22.137 | 22.171 | 29.447 | 41.212 |
| 29022 ENSMUSG00000087595 | 17.27195913 | -0.398358152 | 0.558128904 | -0.713738617 | 0.475388823 | 0.86938592 | 1810012K08 | RIKEN cDNA 1810012K08 gene [Source:M | 18.471 | 15.419 | 25.156 | 20.244 | 7.6343 | 16.708 |
| 29023 ENSMUSG00000087596 | 0.301557804 | -1.64335916 | 4.042814367 | -0.406488899 | 0.684383412 | NA | Gm11715 | predicted gene 11715 [Source:MGI Symbo | 0.8031 | 0 | 1.0062 | 0 | 0 | 0 |
| 29024 ENSMUSG00000087597 | 0.342432689 | 1.990653086 | 3.900281263 | 0.51038706 | 0.609780321 | NA | Gm15345 | predicted gene 15345 [Source:MGI Symbo | 0 | 0 | 0 | 0.964 | 1.0906 | 0 |
| 29025 ENSMUSG00000087598 | 78.72393211 | -0.442149665 | 0.286676807 | -1.542327993 | 0.12299391 | 0.513267141 | Zfp111 | zinc finger protein 111 [Source:MGI Symb | 65.051 | 96.624 | 110.69 | 58.803 | 65.437 | 75.741 |
| 29026 ENSMUSG00000087600 | 0 | NA | NA | NA | NA | NA | Pmepa1os | prostate transmembrane protein, androg | 0 | 0 | 0 | 0 | 0 | 0 |
| 29027 ENSMUSG00000087601 | 0 | NA | NA | NA | NA | NA | Uchl1os | ubiquitin carboxy-terminal hydrolase L1, c | 0 | 0 | 0 | 0 | 0 | 0 |
| 29028 ENSMUSG00000087602 | 0 | NA | NA | NA | NA | NA | Gm12753 | predicted gene 12753 [Source:MGI Symbo | 0 | 0 | 0 | 0 | 0 | 0 |
| 29029 ENSMUSG00000087603 | 0 | NA | NA | NA | NA | NA | Gm11339 | predicted gene 11339 [Source:MGI Symbo | 0 | 0 | 0 | 0 | 0 | 0 |
| 29030 ENSMUSG00000087604 | 0.133850373 | -0.796629176 | 4.080472857 | -0.195229623 | 0.84521319 | NA | Cdrt4os1 | CMT1A duplicated region transcript 4, op | 0.8031 | 0 | 0 | 0 | 0 | 0 |
| 29031 ENSMUSG00000087605 | 3.825051946 | -1.079898412 | 1.127212697 | -0.958025415 | 0.338049943 | NA | Gm15742 | predicted gene 15742 [Source:MGI Symbo | 6.4248 | 3.0838 | 6.0375 | 1.928 | 4.3625 | 1.1138 |
| 29032 ENSMUSG00000087606 | 7.98350221 | -1.316074988 | 0.901106048 | -1.460510659 | 0.14414978 | 0.554380949 | Gm16121 | predicted gene 16121 [Source:MGI Symbo | 16.865 | 5.1396 | 12.075 | 2.8919 | 9.8156 | 1.1138 |
| 29033 ENSMUSG00000087607 | 0 | NA | NA | NA | NA | NA | Gm16034 | predicted gene 16034 [Source:MGI Symbo | 0 | 0 | 0 | 0 | 0 | 0 |
| 29034 ENSMUSG00000087608 | 0 | NA | NA | NA | NA | NA | Gm12914 | predicted gene 12914 [Source:MGI Symbo | 0 | 0 | 0 | 0 | 0 | 0 |
| 29035 ENSMUSG00000087609 | 0 | NA | NA | NA | NA | NA | 4930442J19 | RIKEN cDNA 4930442J19 gene [Source:M | 0 | 0 | 0 | 0 | 0 | 0 |
| 29036 ENSMUSG00000087610 | 1.484716984 | 0.510652645 | 1.746072406 | 0.292457886 | 0.769936555 | NA | Gm16253 | predicted gene 16253 [Source:MGI Symbo | 1.6062 | 2.0558 | 0 | 1.928 | 1.0906 | 2.2277 |
| 29037 ENSMUSG00000087611 | 13.99265336 | 0.02285443 | 0.613363787 | 0.037260807 | 0.970277055 | 0.994966572 | 4930458D05 | RIKEN cDNA 4930458D05 gene [Source:M | 15.259 | 14.391 | 12.075 | 21.208 | 6.5437 | 14.48 |
| 29038 ENSMUSG00000087612 | 11.34525836 | -0.571255127 | 0.639583329 | -0.893167631 | 0.371767409 | 0.807685567 | A230005M1 | RIKEN cDNA A230005M16 gene [Source:N | 11.243 | 15.419 | 14.087 | 9.6398 | 6.5437 | 11.138 |
| 29039 ENSMUSG00000087613 | 5.43766046 | -0.642270279 | 0.964872352 | -0.665653107 | 0.505632853 | 0.881592135 | Gm13855 | predicted gene 13855 [Source:MGI Symbo | 5.6217 | 8.2233 | 6.0375 | 3.8559 | 1.0906 | 7.7969 |
| 29040 ENSMUSG00000087614 | 0.349477146 | 0.165142768 | 3.911021834 | 0.042224967 | 0.966319359 | NA | Gm12514 | predicted gene 12514 [Source:MGI Symbo | 0 | 0 | 1.0062 | 0 | 1.0906 | 0 |
| 29041 ENSMUSG00000087615 | 0 | NA | NA | NA | NA | NA | Pnpla1os | patatin-like phospholipase domain contai | 0 | 0 | 0 | 0 | 0 | 0 |
| 29042 ENSMUSG00000087616 | 0.858479746 | 0.854607444 | 2.595754194 | 0.329232809 | 0.741979726 | NA | Gm14257 | predicted gene 14257 [Source:MGI Symbo | 0.8031 | 0 | 1.0062 | 0 | 0 | 3.3415 |
| 29043 ENSMUSG00000087617 | 0 | NA | NA | NA | NA | NA | Gm14268 | predicted gene 14268 [Source:MGI Symbo | 0 | 0 | 0 | 0 | 0 | 0 |
| 29044 ENSMUSG00000087618 | 0 | NA | NA | NA | NA | NA | Gm14114 | predicted gene 14114 [Source:MGI Symbo | 0 | 0 | 0 | 0 | 0 | 0 |
| 29045 ENSMUSG00000087620 | 0 | NA | NA | NA | NA | NA | 5330434G04 | RIKEN cDNA 5330434G04 gene [Source:M | 0 | 0 | 0 | 0 | 0 | 0 |
| 29046 ENSMUSG00000087621 | 0.606727817 | -2.651277273 | 2.811524265 | -0.943003518 | 0.345679103 | NA | 1700003G18 | RIKEN cDNA 1700003G18 gene [Source:M | 1.6062 | 1.0279 | 1.0062 | 0 | 0 | 0 |
| 29047 ENSMUSG00000087622 | 0 | NA | NA | NA | NA | NA | Gm12290 | predicted gene 12290 [Source:MGI Symbo | 0 | 0 | 0 | 0 | 0 | 0 |
| 29048 ENSMUSG00000087623 | 3.29083868 | 0.469209127 | 1.211845397 | 0.387185633 | 0.698618786 | NA | Gm12404 | predicted gene 12404 [Source:MGI Symbo | 3.2124 | 1.0279 | 4.025 | 4.8199 | 1.0906 | 5.5692 |
| 29049 ENSMUSG00000087624 | 3.623229261 | 1.034636629 | 1.203916208 | 0.859392557 | 0.390123974 | NA | 9230111E07 | RIKEN cDNA 9230111E07 gene [Source:M | 4.0155 | 0 | 3.0187 | 4.8199 | 6.5437 | 3.3415 |
| 29050 ENSMUSG00000087625 | 0 | NA | NA | NA | NA | NA | 4930419G24 | RIKEN cDNA 4930419G24 gene [Source:M | 0 | 0 | 0 | 0 | 0 | 0 |
| 29051 ENSMUSG00000087626 | 0.160662974 | 1.126924175 | 4.080472857 | 0.276174898 | 0.782413736 | NA | Gm15050 | predicted gene 15050 [Source:MGI Symbo | 0 | 0 | 0 | 0.964 | 0 | 0 |
| 29052 ENSMUSG00000087627 | 0.185640647 | 1.126924175 | 4.080472857 | 0.276174898 | 0.782413736 | NA | A230059L01 | RIKEN cDNA A230059L01 gene [Source:M | 0 | 0 | 0 | 0 | 0 | 1.1138 |
| 29053 ENSMUSG00000087628 | 0 | NA | NA | NA | NA | NA | Gm13028 | predicted gene 13028 [Source:MGI Symbo | 0 | 0 | 0 | 0 | 0 | 0 |
| 29054 ENSMUSG00000087629 | 0 | NA | NA | NA | NA | NA | Gm12246 | predicted gene 12246 [Source:MGI Symbo | 0 | 0 | 0 | 0 | 0 | 0 |
| 29055 ENSMUSG00000087630 | 1.694902133 | 0.035219933 | 1.801970698 | 0.019545231 | 0.984406155 | NA | Gm11513 | predicted gene 11513 [Source:MGI Symbo | 0 | 3.0838 | 2.0125 | 2.8919 | 2.1812 | 0 |
| 29056 ENSMUSG00000087632 | 0.17131964 | -0.796629176 | 4.080472857 | -0.195229623 | 0.84521319 | NA | Gm6058 | predicted gene 6058 [Source:MGI Symbol | 0 | 1.0279 | 0 | 0 | 0 | 0 |
| 29057 ENSMUSG00000087633 | 0 | NA | NA | NA | NA | NA | Gm14455 | predicted gene 14455 [Source:MGI Symbo | 0 | 0 | 0 | 0 | 0 | 0 |
| 29058 ENSMUSG00000087635 | 0.993467639 | 0.162634227 | 2.224152761 | 0.073121878 | 0.941709132 | NA | Gm13414 | predicted gene 13414 [Source:MGI Symbo | 0.8031 | 0 | 2.0125 | 0.964 | 2.1812 | 0 |
| 29059 ENSMUSG00000087636 | 0 | NA | NA | NA | NA | NA | Gm4660 | predicted gene 4660 [Source:MGI Symbol | 0 | 0 | 0 | 0 | 0 | 0 |
| 29060 ENSMUSG00000087637 | 0 | NA | NA | NA | NA | NA | Gm11986 | predicted gene 11986 [Source:MGI Symbo | 0 | 0 | 0 | 0 | 0 | 0 |

| | | | | | | | | | | | | | | | |
|---|---|---|---|---|---|---|---|---|---|---|---|---|---|---|---|
| 29061 | ENSMUSG00000087638 | 24.00811101 | -0.619477768 | 0.535095853 | -1.157694953 | 0.246988544 | 0.696925928 | Gm11773 | predicted gene 11773 [Source:MGI Symbo | 47.383 | 23.642 | 16.1 | 25.063 | 20.722 | 11.138 |
| 29062 | ENSMUSG00000087639 | 0 | NA | NA | NA | NA | NA | Gm15512 | predicted gene 15512 [Source:MGI Symbo | 0 | 0 | 0 | 0 | 0 | 0 |
| 29063 | ENSMUSG00000087640 | 0 | NA | NA | NA | NA | NA | Gm14798 | predicted gene 14798 [Source:MGI Symbo | 0 | 0 | 0 | 0 | 0 | 0 |
| 29064 | ENSMUSG00000087641 | 0 | NA | NA | NA | NA | NA | Gm12811 | predicted gene 12811 [Source:MGI Symbo | 0 | 0 | 0 | 0 | 0 | 0 |
| 29065 | ENSMUSG00000087642 | 0 | NA | NA | NA | NA | NA | Ighe | Immunoglobulin heavy constant epsilon [ | 0 | 0 | 0 | 0 | 0 | 0 |
| 29066 | ENSMUSG00000087643 | 0 | NA | NA | NA | NA | NA | Gm11314 | predicted gene 11314 [Source:MGI Symbo | 0 | 0 | 0 | 0 | 0 | 0 |
| 29067 | ENSMUSG00000087644 | 7.003889377 | -0.412055488 | 0.852018873 | -0.483622488 | 0.628653812 | 0.920328819 | Gm14703 | predicted gene 14703 [Source:MGI Symbo | 9.6372 | 9.2513 | 5.0312 | 3.8559 | 10.906 | 3.3415 |
| 29068 | ENSMUSG00000087646 | 0 | NA | NA | NA | NA | NA | Gm1667 | predicted gene 1667 [Source:MGI Symbol | 0 | 0 | 0 | 0 | 0 | 0 |
| 29069 | ENSMUSG00000087647 | 0 | NA | NA | NA | NA | NA | Gm15407 | predicted gene 15407 [Source:MGI Symbo | 0 | 0 | 0 | 0 | 0 | 0 |
| 29070 | ENSMUSG00000087648 | 2.296880222 | -0.688986042 | 1.447112656 | -0.476110854 | 0.633995405 | NA | E130018N17 | RIKEN cDNA E130018N17 gene [Source:M | 2.4093 | 3.0838 | 3.0187 | 1.928 | 0 | 3.3415 |
| 29071 | ENSMUSG00000087649 | 0 | NA | NA | NA | NA | NA | Gm21979 | predicted gene 21979 [Source:MGI Symbo | 0 | 0 | 0 | 0 | 0 | 0 |
| 29072 | ENSMUSG00000087651 | 0 | NA | NA | NA | NA | NA | 1500009L16 | RIKEN cDNA 1500009L16 gene [Source:M | 0 | 0 | 0 | 0 | 0 | 0 |
| 29073 | ENSMUSG00000087652 | 0.342639279 | -1.805976596 | 4.028547714 | -0.448294702 | 0.653940524 | NA | Gm15918 | predicted gene 15918 [Source:MGI Symbo | 0 | 2.0558 | 0 | 0 | 0 | 0 |
| 29074 | ENSMUSG00000087654 | 0 | NA | NA | NA | NA | NA | Gm12256 | predicted gene 12256 [Source:MGI Symbo | 0 | 0 | 0 | 0 | 0 | 0 |
| 29075 | ENSMUSG00000087657 | 0 | NA | NA | NA | NA | NA | Gm16764 | predicted gene, 16764 [Source:MGI Symb | 0 | 0 | 0 | 0 | 0 | 0 |
| 29076 | ENSMUSG00000087658 | 4.974838844 | -1.28712643 | 1.137625307 | -1.131415082 | 0.257880427 | NA | Hotairm1 | Hoxa transcript antisense RNA, myeloid-s | 12.85 | 5.1396 | 3.0187 | 0 | 3.2719 | 5.5692 |
| 29077 | ENSMUSG00000087659 | 0 | NA | NA | NA | NA | NA | Gm12606 | predicted gene 12606 [Source:MGI Symbo | 0 | 0 | 0 | 0 | 0 | 0 |
| 29078 | ENSMUSG00000087660 | 0.647809292 | -2.733819264 | 3.061311498 | -0.893022244 | 0.371845261 | NA | B230398E01 | RIKEN cDNA B230398E01 gene [Source:M | 0.8031 | 3.0838 | 0 | 0 | 0 | 0 |
| 29079 | ENSMUSG00000087662 | 0 | NA | NA | NA | NA | NA | Gm11373 | predicted gene 11373 [Source:MGI Symbo | 0 | 0 | 0 | 0 | 0 | 0 |
| 29080 | ENSMUSG00000087663 | 0 | NA | NA | NA | NA | NA | Defa-ps3 | defensin, alpha, pseudogene 3 [Source:M | 0 | 0 | 0 | 0 | 0 | 0 |
| 29081 | ENSMUSG00000087666 | 0 | NA | NA | NA | NA | NA | Trav13-5 | T cell receptor alpha variable 13-5 [Source | 0 | 0 | 0 | 0 | 0 | 0 |
| 29082 | ENSMUSG00000087667 | 0.481988923 | 2.492401104 | 3.837077151 | 0.649557203 | 0.515978285 | NA | Gm13381 | predicted gene 13381 [Source:MGI Symbo | 0 | 0 | 0 | 2.8919 | 0 | 0 |
| 29083 | ENSMUSG00000087668 | 0 | NA | NA | NA | NA | NA | Gm11186 | predicted gene 11186 [Source:MGI Symbo | 0 | 0 | 0 | 0 | 0 | 0 |
| 29084 | ENSMUSG00000087669 | 2.313934415 | -0.755967453 | 1.507273738 | -0.501546225 | 0.615986754 | NA | Gm11724 | predicted gene 11724 [Source:MGI Symbo | 5.6217 | 2.0558 | 1.0062 | 1.928 | 3.2719 | 0 |
| 29085 | ENSMUSG00000087670 | 0.339027071 | -1.79174149 | 3.919585585 | -0.457125237 | 0.647581032 | NA | 9530036M11 | RIKEN cDNA 9530036M11 gene [Source:M | 0 | 1.0279 | 1.0062 | 0 | 0 | 0 |
| 29086 | ENSMUSG00000087672 | 11.92478039 | 0.006832072 | 0.637579238 | 0.010715643 | 0.991450317 | 0.99859176 | Gm15122 | predicted gene 15122 [Source:MGI Symbo | 15.259 | 7.1954 | 13.081 | 10.604 | 9.8156 | 15.594 |
| 29087 | ENSMUSG00000087674 | 0 | NA | NA | NA | NA | NA | 4930447M23 | RIKEN cDNA 4930447M23 gene [Source:M | 0 | 0 | 0 | 0 | 0 | 0 |
| 29088 | ENSMUSG00000087675 | 0 | NA | NA | NA | NA | NA | Gm11762 | predicted gene 11762 [Source:MGI Symbo | 0 | 0 | 0 | 0 | 0 | 0 |
| 29089 | ENSMUSG00000087677 | 0 | NA | NA | NA | NA | NA | Gm15226 | predicted gene 15226 [Source:MGI Symbo | 0 | 0 | 0 | 0 | 0 | 0 |
| 29090 | ENSMUSG00000087678 | 0 | NA | NA | NA | NA | NA | Gm14120 | predicted gene 14120 [Source:MGI Symbo | 0 | 0 | 0 | 0 | 0 | 0 |
| 29091 | ENSMUSG00000087679 | 1183.782968 | 0.022468805 | 0.118751802 | 0.189208115 | 0.849929708 | 0.973277966 | Tmem250-ps | transmembrane protein 250, pseudogene | 1156.5 | 1095.8 | 1270.9 | 1088.3 | 1170.2 | 1321 |
| 29092 | ENSMUSG00000087681 | 0.453341394 | -0.545391191 | 3.551113277 | -0.153583158 | 0.877938417 | NA | Gm13565 | predicted gene 13565 [Source:MGI Symbo | 1.6062 | 0 | 0 | 0 | 0 | 1.1138 |
| 29093 | ENSMUSG00000087682 | 0 | NA | NA | NA | NA | NA | Gm14003 | predicted gene 14003 [Source:MGI Symbo | 0 | 0 | 0 | 0 | 0 | 0 |
| 29094 | ENSMUSG00000087683 | 0.185640647 | 1.126924175 | 4.080472857 | 0.276174898 | 0.782413736 | NA | Gm11712 | predicted gene 11712 [Source:MGI Symbo | 0 | 0 | 0 | 0 | 0 | 1.1138 |
| 29095 | ENSMUSG00000087684 | 3.411447917 | -0.49656254 | 1.126400941 | -0.440839955 | 0.659328866 | NA | 1200007C13 | RIKEN cDNA 1200007C13 gene [Source:M | 4.8186 | 3.0838 | 4.025 | 1.928 | 3.2719 | 3.3415 |
| 29096 | ENSMUSG00000087685 | 3.044600508 | 0.85269312 | 1.242702661 | 0.686160211 | 0.492612085 | NA | 1700122E12 | RIKEN cDNA 1700122E12 gene [Source:M | 2.4093 | 2.0558 | 2.0125 | 1.928 | 7.6343 | 2.2277 |
| 29097 | ENSMUSG00000087687 | 181.8118737 | -0.049170207 | 0.204646112 | -0.240269443 | 0.810121382 | 0.966157155 | Pet100 | PET100 homolog [Source:MGI Symbol;Acc | 167.85 | 207.64 | 179.11 | 144.6 | 175.59 | 216.09 |
| 29098 | ENSMUSG00000087688 | 0 | NA | NA | NA | NA | NA | Gm11300 | predicted gene 11300 [Source:MGI Symbo | 0 | 0 | 0 | 0 | 0 | 0 |
| 29099 | ENSMUSG00000087689 | 0 | NA | NA | NA | NA | NA | Gm15845 | predicted gene 15845 [Source:MGI Symbo | 0 | 0 | 0 | 0 | 0 | 0 |
| 29100 | ENSMUSG00000087690 | 0.349477146 | 0.165142768 | 3.911021834 | 0.042224967 | 0.966319359 | NA | Gm16031 | predicted gene 16031 [Source:MGI Symbo | 0 | 0 | 1.0062 | 0 | 1.0906 | 0 |
| 29101 | ENSMUSG00000087691 | 10.74615395 | -0.326952872 | 0.710681934 | -0.460055133 | 0.645476648 | 0.925643431 | Cd55os | CD55 molecule, opposite strand sequence | 9.6372 | 14.391 | 12.075 | 17.352 | 5.4531 | 5.5692 |
| 29102 | ENSMUSG00000087692 | 0 | NA | NA | NA | NA | NA | Gm11940 | predicted gene 11940 [Source:MGI Symbo | 0 | 0 | 0 | 0 | 0 | 0 |
| 29103 | ENSMUSG00000087693 | 0 | NA | NA | NA | NA | NA | Gm16191 | predicted gene 16191 [Source:MGI Symbo | 0 | 0 | 0 | 0 | 0 | 0 |
| 29104 | ENSMUSG00000087694 | 0 | NA | NA | NA | NA | NA | A530058N18 | RIKEN cDNA A530058N18 gene [Source:M | 0 | 0 | 0 | 0 | 0 | 0 |
| 29105 | ENSMUSG00000087695 | 8.975461132 | -1.310669536 | 0.788543278 | -1.66214027 | 0.096484653 | 0.458530879 | Gm16291 | predicted gene 16291 [Source:MGI Symbo | 8.031 | 17.475 | 13.081 | 8.6758 | 4.3625 | 2.2277 |
| 29106 | ENSMUSG00000087698 | 0.160662974 | 1.126924175 | 4.080472857 | 0.276174898 | 0.782413736 | NA | Gm13031 | predicted gene 13031 [Source:MGI Symbo | 0 | 0 | 0 | 0.964 | 0 | 0 |
| 29107 | ENSMUSG00000087700 | 6.112192334 | -0.692023769 | 1.048817549 | -0.659813301 | 0.509373647 | 0.882793133 | Gm15283 | predicted gene 15283 [Source:MGI Symbo | 11.243 | 10.279 | 1.0062 | 0.964 | 8.7249 | 4.4554 |
| 29108 | ENSMUSG00000087701 | 6.936854334 | -1.173043496 | 0.853227974 | -1.374830094 | 0.169184126 | 0.596414836 | Gm13493 | predicted gene 13493 [Source:MGI Symbo | 6.4248 | 13.363 | 9.0562 | 2.8919 | 6.5437 | 3.3415 |
| 29109 | ENSMUSG00000087702 | 0 | NA | NA | NA | NA | NA | Gm13575 | predicted gene 13575 [Source:MGI Symbo | 0 | 0 | 0 | 0 | 0 | 0 |
| 29110 | ENSMUSG00000087703 | 0.669251866 | -2.807208097 | 3.391018549 | -0.827836255 | 0.407763239 | NA | Gm15650 | predicted gene 15650 [Source:MGI Symbo | 4.0155 | 0 | 0 | 0 | 0 | 0 |
| 29111 | ENSMUSG00000087704 | 0 | NA | NA | NA | NA | NA | Gm14978 | predicted gene 14978 [Source:MGI Symbo | 0 | 0 | 0 | 0 | 0 | 0 |
| 29112 | ENSMUSG00000087707 | 0 | NA | NA | NA | NA | NA | Gm22999 | predicted gene, 22999 [Source:MGI Symb | 0 | 0 | 0 | 0 | 0 | 0 |
| 29113 | ENSMUSG00000087708 | 0 | NA | NA | NA | NA | NA | Gm23000 | predicted gene, 23000 [Source:MGI Symb | 0 | 0 | 0 | 0 | 0 | 0 |
| 29114 | ENSMUSG00000087710 | 11.72997975 | -0.105088819 | 0.741805811 | -0.141666212 | 0.887343665 | 0.982382091 | NA | NA | 24.093 | 5.1396 | 7.0437 | 15.424 | 11.997 | 6.6831 |
| 29115 | ENSMUSG00000087712 | 0 | NA | NA | NA | NA | NA | Gm25828 | predicted gene, 25828 [Source:MGI Symb | 0 | 0 | 0 | 0 | 0 | 0 |
| 29116 | ENSMUSG00000087714 | 0.367410362 | 2.078255025 | 3.800070157 | 0.54689912 | 0.58444804 | NA | Gm25827 | predicted gene, 25827 [Source:MGI Symb | 0 | 0 | 0 | 0 | 1.0906 | 1.1138 |

| | | | | | | | | | | | | | | | |
|---|---|---|---|---|---|---|---|---|---|---|---|---|---|---|---|
| 29117 | ENSMUSG00000087715 | 0 | NA | NA | NA | NA | NA | Gm25826 | predicted gene, 25826 [Source:MGI Symb | 0 | 0 | 0 | 0 | 0 | 0 |
| 29118 | ENSMUSG00000087716 | 0 | NA | NA | NA | NA | NA | NA | NA | 0 | 0 | 0 | 0 | 0 | 0 |
| 29119 | ENSMUSG00000087717 | 0 | NA | NA | NA | NA | NA | Gm25824 | predicted gene, 25824 [Source:MGI Symb | 0 | 0 | 0 | 0 | 0 | 0 |
| 29120 | ENSMUSG00000087718 | 0 | NA | NA | NA | NA | NA | Gm22236 | predicted gene, 22236 [Source:MGI Symb | 0 | 0 | 0 | 0 | 0 | 0 |
| 29121 | ENSMUSG00000087719 | 1.117100031 | 1.582896982 | 2.163697584 | 0.731570342 | 0.464430856 | NA | Gm26464 | predicted gene, 26464 [Source:MGI Symb | 1.6062 | 0 | 0 | 2.8919 | 1.0906 | 1.1138 |
| 29122 | ENSMUSG00000087720 | 0 | NA | NA | NA | NA | NA | Gm26279 | predicted gene, 26279 [Source:MGI Symb | 0 | 0 | 0 | 0 | 0 | 0 |
| 29123 | ENSMUSG00000087721 | 0.353089354 | 0.165142706 | 3.896102244 | 0.042386646 | 0.966190474 | NA | NA | NA | 0 | 1.0279 | 0 | 0 | 1.0906 | 0 |
| 29124 | ENSMUSG00000087722 | 0 | NA | NA | NA | NA | NA | Gm26277 | predicted gene, 26277 [Source:MGI Symb | 0 | 0 | 0 | 0 | 0 | 0 |
| 29125 | ENSMUSG00000087723 | 0 | NA | NA | NA | NA | NA | Gm26278 | predicted gene, 26278 [Source:MGI Symb | 0 | 0 | 0 | 0 | 0 | 0 |
| 29126 | ENSMUSG00000087724 | 0 | NA | NA | NA | NA | NA | Gm26276 | predicted gene, 26276 [Source:MGI Symb | 0 | 0 | 0 | 0 | 0 | 0 |
| 29127 | ENSMUSG00000087726 | 0 | NA | NA | NA | NA | NA | NA | NA | 0 | 0 | 0 | 0 | 0 | 0 |
| 29128 | ENSMUSG00000087728 | 0 | NA | NA | NA | NA | NA | NA | NA | 0 | 0 | 0 | 0 | 0 | 0 |
| 29129 | ENSMUSG00000087730 | 0.133850373 | -0.796629176 | 4.080472857 | -0.195229623 | 0.84521319 | NA | NA | NA | 0.8031 | 0 | 0 | 0 | 0 | 0 |
| 29130 | ENSMUSG00000087731 | 0 | NA | NA | NA | NA | NA | NA | NA | 0 | 0 | 0 | 0 | 0 | 0 |
| 29131 | ENSMUSG00000087732 | 0 | NA | NA | NA | NA | NA | NA | NA | 0 | 0 | 0 | 0 | 0 | 0 |
| 29132 | ENSMUSG00000087734 | 0.915510039 | -3.242382683 | 2.669421139 | -1.214638873 | 0.224503856 | NA | Gm24646 | predicted gene, 24646 [Source:MGI Symb | 2.4093 | 3.0838 | 0 | 0 | 0 | 0 |
| 29133 | ENSMUSG00000087736 | 0 | NA | NA | NA | NA | NA | Gm24648 | predicted gene, 24648 [Source:MGI Symb | 0 | 0 | 0 | 0 | 0 | 0 |
| 29134 | ENSMUSG00000087737 | 0 | NA | NA | NA | NA | NA | NA | NA | 0 | 0 | 0 | 0 | 0 | 0 |
| 29135 | ENSMUSG00000087739 | 6.317343667 | -0.284362532 | 1.091154828 | -0.26060695 | 0.794395629 | 0.962046762 | Gm24644 | predicted gene, 24644 [Source:MGI Symb | 4.8186 | 1.0279 | 15.094 | 12.532 | 1.0906 | 3.3415 |
| 29136 | ENSMUSG00000087741 | 0 | NA | NA | NA | NA | NA | NA | NA | 0 | 0 | 0 | 0 | 0 | 0 |
| 29137 | ENSMUSG00000087742 | 0 | NA | NA | NA | NA | NA | Gm25152 | predicted gene, 25152 [Source:MGI Symb | 0 | 0 | 0 | 0 | 0 | 0 |
| 29138 | ENSMUSG00000087743 | 0 | NA | NA | NA | NA | NA | NA | NA | 0 | 0 | 0 | 0 | 0 | 0 |
| 29139 | ENSMUSG00000087744 | 2.327252898 | -1.219369313 | 1.548770205 | -0.787314547 | 0.431097756 | NA | NA | NA | 1.6062 | 7.1954 | 1.0062 | 1.928 | 0 | 2.2277 |
| 29140 | ENSMUSG00000087746 | 0 | NA | NA | NA | NA | NA | Gm25150 | predicted gene, 25150 [Source:MGI Symb | 0 | 0 | 0 | 0 | 0 | 0 |
| 29141 | ENSMUSG00000087747 | 0 | NA | NA | NA | NA | NA | Gm25151 | predicted gene, 25151 [Source:MGI Symb | 0 | 0 | 0 | 0 | 0 | 0 |
| 29142 | ENSMUSG00000087748 | 0.133850373 | -0.796629176 | 4.080472857 | -0.195229623 | 0.84521319 | NA | NA | NA | 0.8031 | 0 | 0 | 0 | 0 | 0 |
| 29143 | ENSMUSG00000087750 | 0.133850373 | -0.796629176 | 4.080472857 | -0.195229623 | 0.84521319 | NA | NA | NA | 0.8031 | 0 | 0 | 0 | 0 | 0 |
| 29144 | ENSMUSG00000087752 | 0 | NA | NA | NA | NA | NA | Gm23467 | predicted gene, 23467 [Source:MGI Symb | 0 | 0 | 0 | 0 | 0 | 0 |
| 29145 | ENSMUSG00000087754 | 0 | NA | NA | NA | NA | NA | Gm23470 | predicted gene, 23470 [Source:MGI Symb | 0 | 0 | 0 | 0 | 0 | 0 |
| 29146 | ENSMUSG00000087756 | 0 | NA | NA | NA | NA | NA | NA | NA | 0 | 0 | 0 | 0 | 0 | 0 |
| 29147 | ENSMUSG00000087758 | 0 | NA | NA | NA | NA | NA | NA | NA | 0 | 0 | 0 | 0 | 0 | 0 |
| 29148 | ENSMUSG00000087759 | 0 | NA | NA | NA | NA | NA | NA | NA | 0 | 0 | 0 | 0 | 0 | 0 |
| 29149 | ENSMUSG00000087760 | 0 | NA | NA | NA | NA | NA | Gm23980 | predicted gene, 23980 [Source:MGI Symb | 0 | 0 | 0 | 0 | 0 | 0 |
| 29150 | ENSMUSG00000087761 | 0 | NA | NA | NA | NA | NA | Gm23981 | predicted gene, 23981 [Source:MGI Symb | 0 | 0 | 0 | 0 | 0 | 0 |
| 29151 | ENSMUSG00000087764 | 0 | NA | NA | NA | NA | NA | Gm23984 | predicted gene, 23984 [Source:MGI Symb | 0 | 0 | 0 | 0 | 0 | 0 |
| 29152 | ENSMUSG00000087765 | 0 | NA | NA | NA | NA | NA | Gm23985 | predicted gene, 23985 [Source:MGI Symb | 0 | 0 | 0 | 0 | 0 | 0 |
| 29153 | ENSMUSG00000087766 | 0 | NA | NA | NA | NA | NA | NA | NA | 0 | 0 | 0 | 0 | 0 | 0 |
| 29154 | ENSMUSG00000087767 | 0 | NA | NA | NA | NA | NA | NA | NA | 0 | 0 | 0 | 0 | 0 | 0 |
| 29155 | ENSMUSG00000087768 | 0 | NA | NA | NA | NA | NA | NA | NA | 0 | 0 | 0 | 0 | 0 | 0 |
| 29156 | ENSMUSG00000087769 | 0 | NA | NA | NA | NA | NA | Gm23986 | predicted gene, 23986 [Source:MGI Symb | 0 | 0 | 0 | 0 | 0 | 0 |
| 29157 | ENSMUSG00000087772 | 0 | NA | NA | NA | NA | NA | Gm22319 | predicted gene, 22319 [Source:MGI Symb | 0 | 0 | 0 | 0 | 0 | 0 |
| 29158 | ENSMUSG00000087774 | 0 | NA | NA | NA | NA | NA | NA | NA | 0 | 0 | 0 | 0 | 0 | 0 |
| 29159 | ENSMUSG00000087775 | 0 | NA | NA | NA | NA | NA | Rprl2 | ribonuclease P RNA-like 2 [Source:MGI Sy | 0 | 0 | 0 | 0 | 0 | 0 |
| 29160 | ENSMUSG00000087776 | 0 | NA | NA | NA | NA | NA | Gm22322 | predicted gene, 22322 [Source:MGI Symb | 0 | 0 | 0 | 0 | 0 | 0 |
| 29161 | ENSMUSG00000087777 | 0 | NA | NA | NA | NA | NA | Gm22321 | predicted gene, 22321 [Source:MGI Symb | 0 | 0 | 0 | 0 | 0 | 0 |
| 29162 | ENSMUSG00000087779 | 0 | NA | NA | NA | NA | NA | NA | NA | 0 | 0 | 0 | 0 | 0 | 0 |
| 29163 | ENSMUSG00000087780 | 0 | NA | NA | NA | NA | NA | Gm23167 | predicted gene, 23167 [Source:MGI Symb | 0 | 0 | 0 | 0 | 0 | 0 |
| 29164 | ENSMUSG00000087781 | 0 | NA | NA | NA | NA | NA | Gm23168 | predicted gene, 23168 [Source:MGI Symb | 0 | 0 | 0 | 0 | 0 | 0 |
| 29165 | ENSMUSG00000087782 | 0 | NA | NA | NA | NA | NA | Gm23169 | predicted gene, 23169 [Source:MGI Symb | 0 | 0 | 0 | 0 | 0 | 0 |
| 29166 | ENSMUSG00000087783 | 0.160662974 | 1.126924175 | 4.080472857 | 0.276174898 | 0.782413736 | NA | NA | NA | 0 | 0 | 0 | 0.964 | 0 | 0 |
| 29167 | ENSMUSG00000087784 | 0.335414862 | -1.779047242 | 4.030804179 | -0.441362855 | 0.658950329 | NA | NA | NA | 0 | 0 | 2.0125 | 0 | 0 | 0 |
| 29168 | ENSMUSG00000087785 | 1.496563302 | 1.139410575 | 1.910177139 | 0.596494719 | 0.550844781 | NA | NA | NA | 0.8031 | 0 | 2.0125 | 2.8919 | 3.2719 | 0 |
| 29169 | ENSMUSG00000087786 | 0 | NA | NA | NA | NA | NA | NA | NA | 0 | 0 | 0 | 0 | 0 | 0 |
| 29170 | ENSMUSG00000087789 | 0 | NA | NA | NA | NA | NA | NA | NA | 0 | 0 | 0 | 0 | 0 | 0 |
| 29171 | ENSMUSG00000087790 | 0.98432953 | 0.280713436 | 2.433774132 | 0.115340792 | 0.908175007 | NA | Gm25970 | predicted gene, 25970 [Source:MGI Symb | 1.6062 | 1.0279 | 0 | 0 | 3.2719 | 0 |
| 29172 | ENSMUSG00000087791 | 0 | NA | NA | NA | NA | NA | NA | NA | 0 | 0 | 0 | 0 | 0 | 0 |

| | | | | | | | | | | | | | | | |
|---|---|---|---|---|---|---|---|---|---|---|---|---|---|---|---|
| 29173 | ENSMUSG00000087792 | 0.185640647 | 1.126924175 | 4.080472857 | 0.276174898 | 0.782413736 | NA | Gm25968 | predicted gene, 25968 [Source:MGI Symb | 0 | 0 | 0 | 0 | 0 | 1.1138 |
| 29174 | ENSMUSG00000087793 | 0.160662974 | 1.126924175 | 4.080472857 | 0.276174898 | 0.782413736 | NA | Gm25967 | predicted gene, 25967 [Source:MGI Symb | 0 | 0 | 0 | 0.964 | 0 | 0 |
| 29175 | ENSMUSG00000087794 | 0.881628005 | -0.457461187 | 2.549924572 | -0.17940185 | 0.857622178 | NA | NA | NA | 0 | 2.0558 | 1.0062 | 0 | 0 | 2.2277 |
| 29176 | ENSMUSG00000087795 | 0.951254158 | 0.070446961 | 2.457323444 | 0.028668168 | 0.977129244 | NA | Gm25965 | predicted gene, 25965 [Source:MGI Symb | 0.8031 | 0 | 2.0125 | 2.8919 | 0 | 0 |
| 29177 | ENSMUSG00000087796 | 0 | NA | NA | NA | NA | NA | Gm25964 | predicted gene, 25964 [Source:MGI Symb | 0 | 0 | 0 | 0 | 0 | 0 |
| 29178 | ENSMUSG00000087798 | 0 | NA | NA | NA | NA | NA | NA | NA | 0 | 0 | 0 | 0 | 0 | 0 |
| 29179 | ENSMUSG00000087799 | 0 | NA | NA | NA | NA | NA | NA | NA | 0 | 0 | 0 | 0 | 0 | 0 |
| 29180 | ENSMUSG00000087800 | 0 | NA | NA | NA | NA | NA | Gm25569 | predicted gene, 25569 [Source:MGI Symb | 0 | 0 | 0 | 0 | 0 | 0 |
| 29181 | ENSMUSG00000087801 | 0 | NA | NA | NA | NA | NA | NA | NA | 0 | 0 | 0 | 0 | 0 | 0 |
| 29182 | ENSMUSG00000087802 | 0 | NA | NA | NA | NA | NA | Gm25979 | predicted gene, 25979 [Source:MGI Symb | 0 | 0 | 0 | 0 | 0 | 0 |
| 29183 | ENSMUSG00000087804 | 0 | NA | NA | NA | NA | NA | Gm23434 | predicted gene, 23434 [Source:MGI Symb | 0 | 0 | 0 | 0 | 0 | 0 |
| 29184 | ENSMUSG00000087805 | 0 | NA | NA | NA | NA | NA | Gm27926 | predicted gene, 27926 [Source:MGI Symb | 0 | 0 | 0 | 0 | 0 | 0 |
| 29185 | ENSMUSG00000087806 | 0 | NA | NA | NA | NA | NA | Gm23432 | predicted gene, 23432 [Source:MGI Symb | 0 | 0 | 0 | 0 | 0 | 0 |
| 29186 | ENSMUSG00000087807 | 0 | NA | NA | NA | NA | NA | NA | NA | 0 | 0 | 0 | 0 | 0 | 0 |
| 29187 | ENSMUSG00000087808 | 0 | NA | NA | NA | NA | NA | Gm23435 | predicted gene, 23435 [Source:MGI Symb | 0 | 0 | 0 | 0 | 0 | 0 |
| 29188 | ENSMUSG00000087810 | 0 | NA | NA | NA | NA | NA | Gm25113 | predicted gene, 25113 [Source:MGI Symb | 0 | 0 | 0 | 0 | 0 | 0 |
| 29189 | ENSMUSG00000087812 | 0 | NA | NA | NA | NA | NA | Gm25115 | predicted gene, 25115 [Source:MGI Symb | 0 | 0 | 0 | 0 | 0 | 0 |
| 29190 | ENSMUSG00000087813 | 0 | NA | NA | NA | NA | NA | Gm22490 | predicted gene, 22490 [Source:MGI Symb | 0 | 0 | 0 | 0 | 0 | 0 |
| 29191 | ENSMUSG00000087815 | 0 | NA | NA | NA | NA | NA | Gm22491 | predicted gene, 22491 [Source:MGI Symb | 0 | 0 | 0 | 0 | 0 | 0 |
| 29192 | ENSMUSG00000087816 | 0 | NA | NA | NA | NA | NA | Gm25116 | predicted gene, 25116 [Source:MGI Symb | 0 | 0 | 0 | 0 | 0 | 0 |
| 29193 | ENSMUSG00000087817 | 0 | NA | NA | NA | NA | NA | NA | NA | 0 | 0 | 0 | 0 | 0 | 0 |
| 29194 | ENSMUSG00000087819 | 0.335414862 | -1.779047242 | 4.030804179 | -0.441362855 | 0.658950329 | NA | Gm25117 | predicted gene, 25117 [Source:MGI Symb | 0 | 0 | 2.0125 | 0 | 0 | 0 |
| 29195 | ENSMUSG00000087820 | 0 | NA | NA | NA | NA | NA | Mir1928 | microRNA 1928 [Source:MGI Symbol;Acc: | 0 | 0 | 0 | 0 | 0 | 0 |
| 29196 | ENSMUSG00000087823 | 0.631240176 | 0.380643921 | 3.15447101 | 0.120668068 | 0.903953952 | NA | Gm22281 | predicted gene, 22281 [Source:MGI Symb | 1.6062 | 0 | 0 | 0 | 2.1812 | 0 |
| 29197 | ENSMUSG00000087825 | 0 | NA | NA | NA | NA | NA | Gm22279 | predicted gene, 22279 [Source:MGI Symb | 0 | 0 | 0 | 0 | 0 | 0 |
| 29198 | ENSMUSG00000087829 | 0 | NA | NA | NA | NA | NA | NA | NA | 0 | 0 | 0 | 0 | 0 | 0 |
| 29199 | ENSMUSG00000087830 | 0 | NA | NA | NA | NA | NA | NA | NA | 0 | 0 | 0 | 0 | 0 | 0 |
| 29200 | ENSMUSG00000087831 | 0 | NA | NA | NA | NA | NA | Gm23960 | predicted gene, 23960 [Source:MGI Symb | 0 | 0 | 0 | 0 | 0 | 0 |
| 29201 | ENSMUSG00000087832 | 0 | NA | NA | NA | NA | NA | Gm23958 | predicted gene, 23958 [Source:MGI Symb | 0 | 0 | 0 | 0 | 0 | 0 |
| 29202 | ENSMUSG00000087833 | 0 | NA | NA | NA | NA | NA | NA | NA | 0 | 0 | 0 | 0 | 0 | 0 |
| 29203 | ENSMUSG00000087834 | 0 | NA | NA | NA | NA | NA | NA | NA | 0 | 0 | 0 | 0 | 0 | 0 |
| 29204 | ENSMUSG00000087835 | 0 | NA | NA | NA | NA | NA | Gm24481 | predicted gene, 24481 [Source:MGI Symb | 0 | 0 | 0 | 0 | 0 | 0 |
| 29205 | ENSMUSG00000087836 | 0 | NA | NA | NA | NA | NA | NA | NA | 0 | 0 | 0 | 0 | 0 | 0 |
| 29206 | ENSMUSG00000087838 | 0.133850373 | -0.796629176 | 4.080472857 | -0.195229623 | 0.84521319 | NA | NA | NA | 0.8031 | 0 | 0 | 0 | 0 | 0 |
| 29207 | ENSMUSG00000087839 | 0 | NA | NA | NA | NA | NA | Gm23955 | predicted gene, 23955 [Source:MGI Symb | 0 | 0 | 0 | 0 | 0 | 0 |
| 29208 | ENSMUSG00000087841 | 0 | NA | NA | NA | NA | NA | Gm25623 | predicted gene, 25623 [Source:MGI Symb | 0 | 0 | 0 | 0 | 0 | 0 |
| 29209 | ENSMUSG00000087842 | 0 | NA | NA | NA | NA | NA | Gm25622 | predicted gene, 25622 [Source:MGI Symb | 0 | 0 | 0 | 0 | 0 | 0 |
| 29210 | ENSMUSG00000087843 | 0 | NA | NA | NA | NA | NA | NA | NA | 0 | 0 | 0 | 0 | 0 | 0 |
| 29211 | ENSMUSG00000087844 | 0 | NA | NA | NA | NA | NA | Mir1958 | microRNA 1958 [Source:MGI Symbol;Acc: | 0 | 0 | 0 | 0 | 0 | 0 |
| 29212 | ENSMUSG00000087845 | 0 | NA | NA | NA | NA | NA | NA | NA | 0 | 0 | 0 | 0 | 0 | 0 |
| 29213 | ENSMUSG00000087846 | 0 | NA | NA | NA | NA | NA | NA | NA | 0 | 0 | 0 | 0 | 0 | 0 |
| 29214 | ENSMUSG00000087847 | 0 | NA | NA | NA | NA | NA | Gm25624 | predicted gene, 25624 [Source:MGI Symb | 0 | 0 | 0 | 0 | 0 | 0 |
| 29215 | ENSMUSG00000087848 | 0 | NA | NA | NA | NA | NA | NA | NA | 0 | 0 | 0 | 0 | 0 | 0 |
| 29216 | ENSMUSG00000087850 | 0 | NA | NA | NA | NA | NA | Gm22792 | predicted gene, 22792 [Source:MGI Symb | 0 | 0 | 0 | 0 | 0 | 0 |
| 29217 | ENSMUSG00000087852 | 0 | NA | NA | NA | NA | NA | NA | NA | 0 | 0 | 0 | 0 | 0 | 0 |
| 29218 | ENSMUSG00000087853 | 0 | NA | NA | NA | NA | NA | NA | NA | 0 | 0 | 0 | 0 | 0 | 0 |
| 29219 | ENSMUSG00000087854 | 0 | NA | NA | NA | NA | NA | NA | NA | 0 | 0 | 0 | 0 | 0 | 0 |
| 29220 | ENSMUSG00000087855 | 0 | NA | NA | NA | NA | NA | Mir432 | microRNA 432 [Source:MGI Symbol;Acc:M | 0 | 0 | 0 | 0 | 0 | 0 |
| 29221 | ENSMUSG00000087857 | 0 | NA | NA | NA | NA | NA | Gm22791 | predicted gene, 22791 [Source:MGI Symb | 0 | 0 | 0 | 0 | 0 | 0 |
| 29222 | ENSMUSG00000087858 | 0 | NA | NA | NA | NA | NA | Gm22795 | predicted gene, 22795 [Source:MGI Symb | 0 | 0 | 0 | 0 | 0 | 0 |
| 29223 | ENSMUSG00000087859 | 0 | NA | NA | NA | NA | NA | NA | NA | 0 | 0 | 0 | 0 | 0 | 0 |
| 29224 | ENSMUSG00000087860 | 0 | NA | NA | NA | NA | NA | NA | NA | 0 | 0 | 0 | 0 | 0 | 0 |
| 29225 | ENSMUSG00000087861 | 0.181769715 | 1.126924175 | 4.080472857 | 0.276174898 | 0.782413736 | NA | Gm24440 | predicted gene, 24440 [Source:MGI Symb | 0 | 0 | 0 | 0 | 1.0906 | 0 |
| 29226 | ENSMUSG00000087862 | 0 | NA | NA | NA | NA | NA | Gm23048 | predicted gene, 23048 [Source:MGI Symb | 0 | 0 | 0 | 0 | 0 | 0 |
| 29227 | ENSMUSG00000087863 | 0 | NA | NA | NA | NA | NA | NA | NA | 0 | 0 | 0 | 0 | 0 | 0 |
| 29228 | ENSMUSG00000087867 | 0 | NA | NA | NA | NA | NA | Gm24443 | predicted gene, 24443 [Source:MGI Symb | 0 | 0 | 0 | 0 | 0 | 0 |

| | | | | | | | | | | | | | | | |
|---|---|---|---|---|---|---|---|---|---|---|---|---|---|---|---|
| 29229 | ENSMUSG00000087869 | 0 | NA | NA | NA | NA | NA | NA | NA | 0 | 0 | 0 | 0 | 0 | 0 |
| 29230 | ENSMUSG00000087870 | 0 | NA | NA | NA | NA | NA | Gm26100 | predicted gene, 26100 [Source:MGI Symb | 0 | 0 | 0 | 0 | 0 | 0 |
| 29231 | ENSMUSG00000087871 | 0 | NA | NA | NA | NA | NA | Gm26101 | predicted gene, 26101 [Source:MGI Symb | 0 | 0 | 0 | 0 | 0 | 0 |
| 29232 | ENSMUSG00000087872 | 0 | NA | NA | NA | NA | NA | NA | NA | 0 | 0 | 0 | 0 | 0 | 0 |
| 29233 | ENSMUSG00000087873 | 0 | NA | NA | NA | NA | NA | Gm26099 | predicted gene, 26099 [Source:MGI Symb | 0 | 0 | 0 | 0 | 0 | 0 |
| 29234 | ENSMUSG00000087875 | 0 | NA | NA | NA | NA | NA | Gm26104 | predicted gene, 26104 [Source:MGI Symb | 0 | 0 | 0 | 0 | 0 | 0 |
| 29235 | ENSMUSG00000087876 | 0 | NA | NA | NA | NA | NA | NA | NA | 0 | 0 | 0 | 0 | 0 | 0 |
| 29236 | ENSMUSG00000087877 | 0 | NA | NA | NA | NA | NA | NA | NA | 0 | 0 | 0 | 0 | 0 | 0 |
| 29237 | ENSMUSG00000087878 | 0 | NA | NA | NA | NA | NA | Gm26105 | predicted gene, 26105 [Source:MGI Symb | 0 | 0 | 0 | 0 | 0 | 0 |
| 29238 | ENSMUSG00000087879 | 0 | NA | NA | NA | NA | NA | NA | NA | 0 | 0 | 0 | 0 | 0 | 0 |
| 29239 | ENSMUSG00000087880 | 0 | NA | NA | NA | NA | NA | Gm23605 | predicted gene, 23605 [Source:MGI Symb | 0 | 0 | 0 | 0 | 0 | 0 |
| 29240 | ENSMUSG00000087881 | 5.194344017 | -0.17814959 | 1.109667208 | -0.160543259 | 0.872453148 | 0.979419285 | Gm22442 | predicted gene, 22442 [Source:MGI Symb | 6.4248 | 2.0558 | 8.05 | 4.8199 | 9.8156 | 0 |
| 29241 | ENSMUSG00000087883 | 0 | NA | NA | NA | NA | NA | Gm23602 | predicted gene, 23602 [Source:MGI Symb | 0 | 0 | 0 | 0 | 0 | 0 |
| 29242 | ENSMUSG00000087884 | 0 | NA | NA | NA | NA | NA | Gm23601 | predicted gene, 23601 [Source:MGI Symb | 0 | 0 | 0 | 0 | 0 | 0 |
| 29243 | ENSMUSG00000087885 | 0 | NA | NA | NA | NA | NA | NA | NA | 0 | 0 | 0 | 0 | 0 | 0 |
| 29244 | ENSMUSG00000087886 | 0 | NA | NA | NA | NA | NA | NA | NA | 0 | 0 | 0 | 0 | 0 | 0 |
| 29245 | ENSMUSG00000087887 | 0 | NA | NA | NA | NA | NA | Gm23598 | predicted gene, 23598 [Source:MGI Symb | 0 | 0 | 0 | 0 | 0 | 0 |
| 29246 | ENSMUSG00000087890 | 0 | NA | NA | NA | NA | NA | NA | NA | 0 | 0 | 0 | 0 | 0 | 0 |
| 29247 | ENSMUSG00000087891 | 0 | NA | NA | NA | NA | NA | NA | NA | 0 | 0 | 0 | 0 | 0 | 0 |
| 29248 | ENSMUSG00000087892 | 0 | NA | NA | NA | NA | NA | Gm25278 | predicted gene, 25278 [Source:MGI Symb | 0 | 0 | 0 | 0 | 0 | 0 |
| 29249 | ENSMUSG00000087893 | 0 | NA | NA | NA | NA | NA | Gm23441 | predicted gene, 23441 [Source:MGI Symb | 0 | 0 | 0 | 0 | 0 | 0 |
| 29250 | ENSMUSG00000087894 | 0 | NA | NA | NA | NA | NA | Gm25280 | predicted gene, 25280 [Source:MGI Symb | 0 | 0 | 0 | 0 | 0 | 0 |
| 29251 | ENSMUSG00000087895 | 0 | NA | NA | NA | NA | NA | NA | NA | 0 | 0 | 0 | 0 | 0 | 0 |
| 29252 | ENSMUSG00000087896 | 0.698954292 | 0.126053087 | 3.048972385 | 0.04134281 | 0.967022605 | NA | NA | NA | 0 | 0 | 2.0125 | 0 | 2.1812 | 0 |
| 29253 | ENSMUSG00000087897 | 0 | NA | NA | NA | NA | NA | Gm25283 | predicted gene, 25283 [Source:MGI Symb | 0 | 0 | 0 | 0 | 0 | 0 |
| 29254 | ENSMUSG00000087898 | 0 | NA | NA | NA | NA | NA | NA | NA | 0 | 0 | 0 | 0 | 0 | 0 |
| 29255 | ENSMUSG00000087899 | 0 | NA | NA | NA | NA | NA | Mir1942 | microRNA 1942 [Source:MGI Symbol;Acc: | 0 | 0 | 0 | 0 | 0 | 0 |
| 29256 | ENSMUSG00000087902 | 0 | NA | NA | NA | NA | NA | NA | NA | 0 | 0 | 0 | 0 | 0 | 0 |
| 29257 | ENSMUSG00000087903 | 0.160662974 | 1.126924175 | 4.080472857 | 0.276174898 | 0.782413736 | NA | Gm22039 | predicted gene, 22039 [Source:MGI Symb | 0 | 0 | 0 | 0.964 | 0 | 0 |
| 29258 | ENSMUSG00000087904 | 3.763464039 | -2.687765245 | 1.414831028 | -1.899707592 | 0.057471504 | NA | NA | NA | 2.4093 | 8.2233 | 9.0562 | 2.8919 | 0 | 0 |
| 29259 | ENSMUSG00000087905 | 0.17131964 | -0.796629176 | 4.080472857 | -0.195229623 | 0.84521319 | NA | NA | NA | 0 | 1.0279 | 0 | 0 | 0 | 0 |
| 29260 | ENSMUSG00000087908 | 0 | NA | NA | NA | NA | NA | Gm22034 | predicted gene, 22034 [Source:MGI Symb | 0 | 0 | 0 | 0 | 0 | 0 |
| 29261 | ENSMUSG00000087909 | 0 | NA | NA | NA | NA | NA | NA | NA | 0 | 0 | 0 | 0 | 0 | 0 |
| 29262 | ENSMUSG00000087910 | 0 | NA | NA | NA | NA | NA | NA | NA | 0 | 0 | 0 | 0 | 0 | 0 |
| 29263 | ENSMUSG00000087911 | 0 | NA | NA | NA | NA | NA | NA | NA | 0 | 0 | 0 | 0 | 0 | 0 |
| 29264 | ENSMUSG00000087912 | 0 | NA | NA | NA | NA | NA | Gm24916 | predicted gene, 24916 [Source:MGI Symb | 0 | 0 | 0 | 0 | 0 | 0 |
| 29265 | ENSMUSG00000087913 | 0 | NA | NA | NA | NA | NA | NA | NA | 0 | 0 | 0 | 0 | 0 | 0 |
| 29266 | ENSMUSG00000087914 | 0 | NA | NA | NA | NA | NA | NA | NA | 0 | 0 | 0 | 0 | 0 | 0 |
| 29267 | ENSMUSG00000087915 | 0 | NA | NA | NA | NA | NA | NA | NA | 0 | 0 | 0 | 0 | 0 | 0 |
| 29268 | ENSMUSG00000087916 | 0 | NA | NA | NA | NA | NA | Gm24912 | predicted gene, 24912 [Source:MGI Symb | 0 | 0 | 0 | 0 | 0 | 0 |
| 29269 | ENSMUSG00000087917 | 0 | NA | NA | NA | NA | NA | Gm24911 | predicted gene, 24911 [Source:MGI Symb | 0 | 0 | 0 | 0 | 0 | 0 |
| 29270 | ENSMUSG00000087919 | 0 | NA | NA | NA | NA | NA | Gm24908 | predicted gene, 24908 [Source:MGI Symb | 0 | 0 | 0 | 0 | 0 | 0 |
| 29271 | ENSMUSG00000087921 | 0 | NA | NA | NA | NA | NA | NA | NA | 0 | 0 | 0 | 0 | 0 | 0 |
| 29272 | ENSMUSG00000087922 | 0 | NA | NA | NA | NA | NA | Gm23229 | predicted gene, 23229 [Source:MGI Symb | 0 | 0 | 0 | 0 | 0 | 0 |
| 29273 | ENSMUSG00000087923 | 0 | NA | NA | NA | NA | NA | Gm23230 | predicted gene, 23230 [Source:MGI Symb | 0 | 0 | 0 | 0 | 0 | 0 |
| 29274 | ENSMUSG00000087924 | 0 | NA | NA | NA | NA | NA | Gm23231 | predicted gene, 23231 [Source:MGI Symb | 0 | 0 | 0 | 0 | 0 | 0 |
| 29275 | ENSMUSG00000087925 | 0 | NA | NA | NA | NA | NA | NA | NA | 0 | 0 | 0 | 0 | 0 | 0 |
| 29276 | ENSMUSG00000087926 | 0 | NA | NA | NA | NA | NA | Gm23233 | predicted gene, 23233 [Source:MGI Symb | 0 | 0 | 0 | 0 | 0 | 0 |
| 29277 | ENSMUSG00000087929 | 0 | NA | NA | NA | NA | NA | Gm23234 | predicted gene, 23234 [Source:MGI Symb | 0 | 0 | 0 | 0 | 0 | 0 |
| 29278 | ENSMUSG00000087930 | 0 | NA | NA | NA | NA | NA | Gm26048 | predicted gene, 26048 [Source:MGI Symb | 0 | 0 | 0 | 0 | 0 | 0 |
| 29279 | ENSMUSG00000087932 | 0 | NA | NA | NA | NA | NA | Mir1971 | microRNA 1971 [Source:MGI Symbol;Acc: | 0 | 0 | 0 | 0 | 0 | 0 |
| 29280 | ENSMUSG00000087933 | 0 | NA | NA | NA | NA | NA | Gm26047 | predicted gene, 26047 [Source:MGI Symb | 0 | 0 | 0 | 0 | 0 | 0 |
| 29281 | ENSMUSG00000087935 | 0 | NA | NA | NA | NA | NA | Snora81 | small nucleolar RNA, H/ACA box 81 [Sourc | 0 | 0 | 0 | 0 | 0 | 0 |
| 29282 | ENSMUSG00000087936 | 0 | NA | NA | NA | NA | NA | Gm23675 | predicted gene, 23675 [Source:MGI Symb | 0 | 0 | 0 | 0 | 0 | 0 |
| 29283 | ENSMUSG00000087937 | 0 | NA | NA | NA | NA | NA | Gm26046 | predicted gene, 26046 [Source:MGI Symb | 0 | 0 | 0 | 0 | 0 | 0 |
| 29284 | ENSMUSG00000087938 | 1.108272252 | -3.532687305 | 2.562527587 | -1.378594839 | 0.168019709 | NA | Gm26049 | predicted gene, 26049 [Source:MGI Symb | 5.6217 | 1.0279 | 0 | 0 | 0 | 0 |

| | | | | | | | | | | | | | | | |
|---|---|---|---|---|---|---|---|---|---|---|---|---|---|---|---|
| 29285 | ENSMUSG00000087939 | 0.160662974 | 1.126924175 | 4.080472857 | 0.276174898 | 0.782413736 | NA | Gm23676 | predicted gene, 23676 [Source:MGI Symb | 0 | 0 | 0 | 0.964 | 0 | 0 |
| 29286 | ENSMUSG00000087940 | 0 | NA | NA | NA | NA | NA | Gm24246 | predicted gene, 24246 [Source:MGI Symb | 0 | 0 | 0 | 0 | 0 | 0 |
| 29287 | ENSMUSG00000087941 | 0 | NA | NA | NA | NA | NA | NA | NA | 0 | 0 | 0 | 0 | 0 | 0 |
| 29288 | ENSMUSG00000087942 | 0 | NA | NA | NA | NA | NA | Gm24244 | predicted gene, 24244 [Source:MGI Symb | 0 | 0 | 0 | 0 | 0 | 0 |
| 29289 | ENSMUSG00000087943 | 1.273918703 | -0.597986764 | 2.014559695 | -0.296832487 | 0.766594402 | NA | NA | NA | 1.6062 | 0 | 3.0187 | 1.928 | 1.0906 | 0 |
| 29290 | ENSMUSG00000087945 | 0 | NA | NA | NA | NA | NA | NA | NA | 0 | 0 | 0 | 0 | 0 | 0 |
| 29291 | ENSMUSG00000087946 | 0 | NA | NA | NA | NA | NA | NA | NA | 0 | 0 | 0 | 0 | 0 | 0 |
| 29292 | ENSMUSG00000087947 | 0 | NA | NA | NA | NA | NA | Gm24249 | predicted gene, 24249 [Source:MGI Symb | 0 | 0 | 0 | 0 | 0 | 0 |
| 29293 | ENSMUSG00000087949 | 0 | NA | NA | NA | NA | NA | Gm24251 | predicted gene, 24251 [Source:MGI Symb | 0 | 0 | 0 | 0 | 0 | 0 |
| 29294 | ENSMUSG00000087950 | 5.099976373 | 0.634761529 | 0.989176723 | 0.6417069 | 0.521063508 | 0.887887809 | Mir1948 | microRNA 1948 [Source:MGI Symbol;Acc: | 4.8186 | 6.1675 | 1.0062 | 8.6758 | 4.3625 | 5.5692 |
| 29295 | ENSMUSG00000087951 | 0.185640647 | 1.126924175 | 4.080472857 | 0.276174898 | 0.782413736 | NA | NA | NA | 0 | 0 | 0 | 0 | 0 | 1.1138 |
| 29296 | ENSMUSG00000087954 | 0 | NA | NA | NA | NA | NA | Gm22559 | predicted gene, 22559 [Source:MGI Symb | 0 | 0 | 0 | 0 | 0 | 0 |
| 29297 | ENSMUSG00000087956 | 0 | NA | NA | NA | NA | NA | NA | NA | 0 | 0 | 0 | 0 | 0 | 0 |
| 29298 | ENSMUSG00000087957 | 0 | NA | NA | NA | NA | NA | NA | NA | 0 | 0 | 0 | 0 | 0 | 0 |
| 29299 | ENSMUSG00000087958 | 0 | NA | NA | NA | NA | NA | NA | NA | 0 | 0 | 0 | 0 | 0 | 0 |
| 29300 | ENSMUSG00000087959 | 0 | NA | NA | NA | NA | NA | NA | NA | 0 | 0 | 0 | 0 | 0 | 0 |
| 29301 | ENSMUSG00000087960 | 0 | NA | NA | NA | NA | NA | NA | NA | 0 | 0 | 0 | 0 | 0 | 0 |
| 29302 | ENSMUSG00000087961 | 0 | NA | NA | NA | NA | NA | NA | NA | 0 | 0 | 0 | 0 | 0 | 0 |
| 29303 | ENSMUSG00000087963 | 0 | NA | NA | NA | NA | NA | Gm25394 | predicted gene, 25394 [Source:MGI Symb | 0 | 0 | 0 | 0 | 0 | 0 |
| 29304 | ENSMUSG00000087964 | 0 | NA | NA | NA | NA | NA | Gm25388 | predicted gene, 25388 [Source:MGI Symb | 0 | 0 | 0 | 0 | 0 | 0 |
| 29305 | ENSMUSG00000087965 | 0 | NA | NA | NA | NA | NA | NA | NA | 0 | 0 | 0 | 0 | 0 | 0 |
| 29306 | ENSMUSG00000087966 | 0 | NA | NA | NA | NA | NA | Gm25390 | predicted gene, 25390 [Source:MGI Symb | 0 | 0 | 0 | 0 | 0 | 0 |
| 29307 | ENSMUSG00000087967 | 0 | NA | NA | NA | NA | NA | Gm25391 | predicted gene, 25391 [Source:MGI Symb | 0 | 0 | 0 | 0 | 0 | 0 |
| 29308 | ENSMUSG00000087968 | 0.904646783 | 0.28560953 | 2.491718272 | 0.114623524 | 0.908743533 | NA | Gm25395 | predicted gene, 25395 [Source:MGI Symb | 2.4093 | 0 | 0 | 1.928 | 1.0906 | 0 |
| 29309 | ENSMUSG00000087973 | 0 | NA | NA | NA | NA | NA | Gm23716 | predicted gene, 23716 [Source:MGI Symb | 0 | 0 | 0 | 0 | 0 | 0 |
| 29310 | ENSMUSG00000087975 | 0 | NA | NA | NA | NA | NA | NA | NA | 0 | 0 | 0 | 0 | 0 | 0 |
| 29311 | ENSMUSG00000087979 | 0 | NA | NA | NA | NA | NA | NA | NA | 0 | 0 | 0 | 0 | 0 | 0 |
| 29312 | ENSMUSG00000087980 | 0 | NA | NA | NA | NA | NA | Gm22226 | predicted gene, 22226 [Source:MGI Symb | 0 | 0 | 0 | 0 | 0 | 0 |
| 29313 | ENSMUSG00000087981 | 0 | NA | NA | NA | NA | NA | NA | NA | 0 | 0 | 0 | 0 | 0 | 0 |
| 29314 | ENSMUSG00000087982 | 0 | NA | NA | NA | NA | NA | NA | NA | 0 | 0 | 0 | 0 | 0 | 0 |
| 29315 | ENSMUSG00000087983 | 0 | NA | NA | NA | NA | NA | NA | NA | 0 | 0 | 0 | 0 | 0 | 0 |
| 29316 | ENSMUSG00000087985 | 0 | NA | NA | NA | NA | NA | Gm22223 | predicted gene, 22223 [Source:MGI Symb | 0 | 0 | 0 | 0 | 0 | 0 |
| 29317 | ENSMUSG00000087987 | 0 | NA | NA | NA | NA | NA | Gm22221 | predicted gene, 22221 [Source:MGI Symb | 0 | 0 | 0 | 0 | 0 | 0 |
| 29318 | ENSMUSG00000087989 | 0 | NA | NA | NA | NA | NA | NA | NA | 0 | 0 | 0 | 0 | 0 | 0 |
| 29319 | ENSMUSG00000087990 | 0 | NA | NA | NA | NA | NA | Gm25058 | predicted gene, 25058 [Source:MGI Symb | 0 | 0 | 0 | 0 | 0 | 0 |
| 29320 | ENSMUSG00000087991 | 0 | NA | NA | NA | NA | NA | NA | NA | 0 | 0 | 0 | 0 | 0 | 0 |
| 29321 | ENSMUSG00000087992 | 0 | NA | NA | NA | NA | NA | Gm25059 | predicted gene, 25059 [Source:MGI Symb | 0 | 0 | 0 | 0 | 0 | 0 |
| 29322 | ENSMUSG00000087993 | 0 | NA | NA | NA | NA | NA | Mir1982 | microRNA 1982 [Source:MGI Symbol;Acc: | 0 | 0 | 0 | 0 | 0 | 0 |
| 29323 | ENSMUSG00000087994 | 0 | NA | NA | NA | NA | NA | NA | NA | 0 | 0 | 0 | 0 | 0 | 0 |
| 29324 | ENSMUSG00000087995 | 0 | NA | NA | NA | NA | NA | Gm25054 | predicted gene, 25054 [Source:MGI Symb | 0 | 0 | 0 | 0 | 0 | 0 |
| 29325 | ENSMUSG00000087997 | 0 | NA | NA | NA | NA | NA | Gm25056 | predicted gene, 25056 [Source:MGI Symb | 0 | 0 | 0 | 0 | 0 | 0 |
| 29326 | ENSMUSG00000087999 | 0 | NA | NA | NA | NA | NA | NA | NA | 0 | 0 | 0 | 0 | 0 | 0 |
| 29327 | ENSMUSG00000088000 | 0 | NA | NA | NA | NA | NA | NA | NA | 0 | 0 | 0 | 0 | 0 | 0 |
| 29328 | ENSMUSG00000088001 | 0 | NA | NA | NA | NA | NA | Gm22883 | predicted gene, 22883 [Source:MGI Symb | 0 | 0 | 0 | 0 | 0 | 0 |
| 29329 | ENSMUSG00000088002 | 0 | NA | NA | NA | NA | NA | Gm24775 | predicted gene, 24775 [Source:MGI Symb | 0 | 0 | 0 | 0 | 0 | 0 |
| 29330 | ENSMUSG00000088004 | 0 | NA | NA | NA | NA | NA | NA | NA | 0 | 0 | 0 | 0 | 0 | 0 |
| 29331 | ENSMUSG00000088005 | 0 | NA | NA | NA | NA | NA | Gm24787 | predicted gene, 24787 [Source:MGI Symb | 0 | 0 | 0 | 0 | 0 | 0 |
| 29332 | ENSMUSG00000088008 | 1.496563302 | 1.139410575 | 1.910177139 | 0.596494719 | 0.550844781 | NA | Gm25492 | predicted gene, 25492 [Source:MGI Symb | 0.8031 | 0 | 2.0125 | 2.8919 | 3.2719 | 0 |
| 29333 | ENSMUSG00000088009 | 0 | NA | NA | NA | NA | NA | Gm24774 | predicted gene, 24774 [Source:MGI Symb | 0 | 0 | 0 | 0 | 0 | 0 |
| 29334 | ENSMUSG00000088010 | 0 | NA | NA | NA | NA | NA | NA | NA | 0 | 0 | 0 | 0 | 0 | 0 |
| 29335 | ENSMUSG00000088014 | 0 | NA | NA | NA | NA | NA | NA | NA | 0 | 0 | 0 | 0 | 0 | 0 |
| 29336 | ENSMUSG00000088015 | 1.131627629 | 0.634716303 | 2.097733039 | 0.302572487 | 0.762215684 | NA | Mir1932 | microRNA 1932 [Source:MGI Symbol;Acc: | 1.6062 | 1.0279 | 0 | 1.928 | 0 | 2.2277 |
| 29337 | ENSMUSG00000088016 | 0 | NA | NA | NA | NA | NA | NA | NA | 0 | 0 | 0 | 0 | 0 | 0 |
| 29338 | ENSMUSG00000088017 | 0 | NA | NA | NA | NA | NA | NA | NA | 0 | 0 | 0 | 0 | 0 | 0 |
| 29339 | ENSMUSG00000088018 | 0.133850373 | -0.796629176 | 4.080472857 | -0.195229623 | 0.84521319 | NA | Gm23390 | predicted gene, 23390 [Source:MGI Symb | 0.8031 | 0 | 0 | 0 | 0 | 0 |
| 29340 | ENSMUSG00000088019 | 0 | NA | NA | NA | NA | NA | Gm23826 | predicted gene, 23826 [Source:MGI Symb | 0 | 0 | 0 | 0 | 0 | 0 |

| | | | | | | | | | | | | | | | |
|---|---|---|---|---|---|---|---|---|---|---|---|---|---|---|---|
| 29341 | ENSMUSG00000088020 | 0 | NA | NA | NA | NA | NA | Gm22762 | predicted gene, 22762 [Source:MGI Symb | 0 | 0 | 0 | 0 | 0 | 0 |
| 29342 | ENSMUSG00000088021 | 0 | NA | NA | NA | NA | NA | Gm22145 | predicted gene, 22145 [Source:MGI Symb | 0 | 0 | 0 | 0 | 0 | 0 |
| 29343 | ENSMUSG00000088022 | 2.625507697 | -0.308209516 | 1.549940449 | -0.198852489 | 0.842378136 | NA | NA | NA | 5.6217 | 3.0838 | 0 | 4.8199 | 0 | 2.2277 |
| 29344 | ENSMUSG00000088025 | 0 | NA | NA | NA | NA | NA | Rprl3 | ribonuclease P RNA-like 3 [Source:MGI Sy | 0 | 0 | 0 | 0 | 0 | 0 |
| 29345 | ENSMUSG00000088026 | 0 | NA | NA | NA | NA | NA | NA | NA | 0 | 0 | 0 | 0 | 0 | 0 |
| 29346 | ENSMUSG00000088027 | 0 | NA | NA | NA | NA | NA | Gm22144 | predicted gene, 22144 [Source:MGI Symb | 0 | 0 | 0 | 0 | 0 | 0 |
| 29347 | ENSMUSG00000088028 | 0 | NA | NA | NA | NA | NA | NA | NA | 0 | 0 | 0 | 0 | 0 | 0 |
| 29348 | ENSMUSG00000088030 | 0 | NA | NA | NA | NA | NA | NA | NA | 0 | 0 | 0 | 0 | 0 | 0 |
| 29349 | ENSMUSG00000088031 | 0 | NA | NA | NA | NA | NA | NA | NA | 0 | 0 | 0 | 0 | 0 | 0 |
| 29350 | ENSMUSG00000088036 | 0 | NA | NA | NA | NA | NA | NA | NA | 0 | 0 | 0 | 0 | 0 | 0 |
| 29351 | ENSMUSG00000088037 | 0 | NA | NA | NA | NA | NA | Gm24992 | predicted gene, 24992 [Source:MGI Symb | 0 | 0 | 0 | 0 | 0 | 0 |
| 29352 | ENSMUSG00000088039 | 0 | NA | NA | NA | NA | NA | Gm24994 | predicted gene, 24994 [Source:MGI Symb | 0 | 0 | 0 | 0 | 0 | 0 |
| 29353 | ENSMUSG00000088040 | 0 | NA | NA | NA | NA | NA | Gm23329 | predicted gene, 23329 [Source:MGI Symb | 0 | 0 | 0 | 0 | 0 | 0 |
| 29354 | ENSMUSG00000088041 | 0 | NA | NA | NA | NA | NA | Gm25261 | predicted gene, 25261 [Source:MGI Symb | 0 | 0 | 0 | 0 | 0 | 0 |
| 29355 | ENSMUSG00000088043 | 0 | NA | NA | NA | NA | NA | Gm23328 | predicted gene, 23328 [Source:MGI Symb | 0 | 0 | 0 | 0 | 0 | 0 |
| 29356 | ENSMUSG00000088044 | 0 | NA | NA | NA | NA | NA | Gm23326 | predicted gene, 23326 [Source:MGI Symb | 0 | 0 | 0 | 0 | 0 | 0 |
| 29357 | ENSMUSG00000088045 | 0 | NA | NA | NA | NA | NA | Gm23327 | predicted gene, 23327 [Source:MGI Symb | 0 | 0 | 0 | 0 | 0 | 0 |
| 29358 | ENSMUSG00000088046 | 0 | NA | NA | NA | NA | NA | Gm23324 | predicted gene, 23324 [Source:MGI Symb | 0 | 0 | 0 | 0 | 0 | 0 |
| 29359 | ENSMUSG00000088047 | 0 | NA | NA | NA | NA | NA | Gm23325 | predicted gene, 23325 [Source:MGI Symb | 0 | 0 | 0 | 0 | 0 | 0 |
| 29360 | ENSMUSG00000088048 | 0 | NA | NA | NA | NA | NA | Gm23331 | predicted gene, 23331 [Source:MGI Symb | 0 | 0 | 0 | 0 | 0 | 0 |
| 29361 | ENSMUSG00000088049 | 0 | NA | NA | NA | NA | NA | NA | NA | 0 | 0 | 0 | 0 | 0 | 0 |
| 29362 | ENSMUSG00000088050 | 0 | NA | NA | NA | NA | NA | Gm26151 | predicted gene, 26151 [Source:MGI Symb | 0 | 0 | 0 | 0 | 0 | 0 |
| 29363 | ENSMUSG00000088051 | 0 | NA | NA | NA | NA | NA | Gm26150 | predicted gene, 26150 [Source:MGI Symb | 0 | 0 | 0 | 0 | 0 | 0 |
| 29364 | ENSMUSG00000088052 | 0 | NA | NA | NA | NA | NA | NA | NA | 0 | 0 | 0 | 0 | 0 | 0 |
| 29365 | ENSMUSG00000088054 | 7.631342528 | 1.813193515 | 0.841039543 | 2.155895677 | 0.031091812 | 0.234103056 | Mir1968 | microRNA 1968 [Source:MGI Symbol;Acc: | 4.0155 | 1.0279 | 5.0312 | 12.532 | 9.8156 | 13.366 |
| 29366 | ENSMUSG00000088055 | 0 | NA | NA | NA | NA | NA | Gm26148 | predicted gene, 26148 [Source:MGI Symb | 0 | 0 | 0 | 0 | 0 | 0 |
| 29367 | ENSMUSG00000088056 | 0.535401493 | -2.48596965 | 3.684117979 | -0.674780141 | 0.499815459 | NA | NA | NA | 3.2124 | 0 | 0 | 0 | 0 | 0 |
| 29368 | ENSMUSG00000088057 | 0 | NA | NA | NA | NA | NA | NA | NA | 0 | 0 | 0 | 0 | 0 | 0 |
| 29369 | ENSMUSG00000088058 | 4.430802742 | -0.634072718 | 1.010706778 | -0.627355759 | 0.530426061 | NA | Gm26153 | predicted gene, 26153 [Source:MGI Symb | 4.0155 | 7.1954 | 5.0312 | 4.8199 | 2.1812 | 3.3415 |
| 29370 | ENSMUSG00000088059 | 0 | NA | NA | NA | NA | NA | NA | NA | 0 | 0 | 0 | 0 | 0 | 0 |
| 29371 | ENSMUSG00000088060 | 0 | NA | NA | NA | NA | NA | Gm24475 | predicted gene, 24475 [Source:MGI Symb | 0 | 0 | 0 | 0 | 0 | 0 |
| 29372 | ENSMUSG00000088062 | 0 | NA | NA | NA | NA | NA | NA | NA | 0 | 0 | 0 | 0 | 0 | 0 |
| 29373 | ENSMUSG00000088063 | 0 | NA | NA | NA | NA | NA | NA | NA | 0 | 0 | 0 | 0 | 0 | 0 |
| 29374 | ENSMUSG00000088064 | 0 | NA | NA | NA | NA | NA | NA | NA | 0 | 0 | 0 | 0 | 0 | 0 |
| 29375 | ENSMUSG00000088066 | 0.181769715 | 1.126924175 | 4.080472857 | 0.276174898 | 0.782413736 | NA | NA | NA | 0 | 0 | 0 | 0 | 1.0906 | 0 |
| 29376 | ENSMUSG00000088067 | 0 | NA | NA | NA | NA | NA | NA | NA | 0 | 0 | 0 | 0 | 0 | 0 |
| 29377 | ENSMUSG00000088068 | 0 | NA | NA | NA | NA | NA | Gm47290 | predicted gene, 47290 [Source:MGI Symb | 0 | 0 | 0 | 0 | 0 | 0 |
| 29378 | ENSMUSG00000088069 | 0.428363721 | -0.543730382 | 3.624677344 | -0.15000794 | 0.880758351 | NA | NA | NA | 1.6062 | 0 | 0 | 0.964 | 0 | 0 |
| 29379 | ENSMUSG00000088070 | 0 | NA | NA | NA | NA | NA | NA | NA | 0 | 0 | 0 | 0 | 0 | 0 |
| 29380 | ENSMUSG00000088071 | 0 | NA | NA | NA | NA | NA | Gm22818 | predicted gene, 22818 [Source:MGI Symb | 0 | 0 | 0 | 0 | 0 | 0 |
| 29381 | ENSMUSG00000088073 | 0 | NA | NA | NA | NA | NA | Gm22817 | predicted gene, 22817 [Source:MGI Symb | 0 | 0 | 0 | 0 | 0 | 0 |
| 29382 | ENSMUSG00000088075 | 0 | NA | NA | NA | NA | NA | NA | NA | 0 | 0 | 0 | 0 | 0 | 0 |
| 29383 | ENSMUSG00000088076 | 0 | NA | NA | NA | NA | NA | Gm22815 | predicted gene, 22815 [Source:MGI Symb | 0 | 0 | 0 | 0 | 0 | 0 |
| 29384 | ENSMUSG00000088077 | 0 | NA | NA | NA | NA | NA | NA | NA | 0 | 0 | 0 | 0 | 0 | 0 |
| 29385 | ENSMUSG00000088083 | 0 | NA | NA | NA | NA | NA | NA | NA | 0 | 0 | 0 | 0 | 0 | 0 |
| 29386 | ENSMUSG00000088084 | 0 | NA | NA | NA | NA | NA | Gm25342 | predicted gene, 25342 [Source:MGI Symb | 0 | 0 | 0 | 0 | 0 | 0 |
| 29387 | ENSMUSG00000088085 | 0 | NA | NA | NA | NA | NA | NA | NA | 0 | 0 | 0 | 0 | 0 | 0 |
| 29388 | ENSMUSG00000088086 | 0 | NA | NA | NA | NA | NA | NA | NA | 0 | 0 | 0 | 0 | 0 | 0 |
| 29389 | ENSMUSG00000088088 | 0 | NA | NA | NA | NA | NA | Rmrp | RNA component of mitochondrial RNAase | 0 | 0 | 0 | 0 | 0 | 0 |
| 29390 | ENSMUSG00000088090 | 0 | NA | NA | NA | NA | NA | NA | NA | 0 | 0 | 0 | 0 | 0 | 0 |
| 29391 | ENSMUSG00000088091 | 0 | NA | NA | NA | NA | NA | NA | NA | 0 | 0 | 0 | 0 | 0 | 0 |
| 29392 | ENSMUSG00000088093 | 0 | NA | NA | NA | NA | NA | Gm22811 | predicted gene, 22811 [Source:MGI Symb | 0 | 0 | 0 | 0 | 0 | 0 |
| 29393 | ENSMUSG00000088094 | 0 | NA | NA | NA | NA | NA | Gm23670 | predicted gene, 23670 [Source:MGI Symb | 0 | 0 | 0 | 0 | 0 | 0 |
| 29394 | ENSMUSG00000088095 | 0 | NA | NA | NA | NA | NA | Gm23669 | predicted gene, 23669 [Source:MGI Symb | 0 | 0 | 0 | 0 | 0 | 0 |
| 29395 | ENSMUSG00000088096 | 0 | NA | NA | NA | NA | NA | NA | NA | 0 | 0 | 0 | 0 | 0 | 0 |
| 29396 | ENSMUSG00000088097 | 0 | NA | NA | NA | NA | NA | NA | NA | 0 | 0 | 0 | 0 | 0 | 0 |

| | | | | | | | | | | | | | | | |
|---|---|---|---|---|---|---|---|---|---|---|---|---|---|---|---|
| 29397 | ENSMUSG00000088098 | 0 | NA | NA | NA | NA | NA | Gm23674 | predicted gene, 23674 [Source:MGI Symb | 0 | 0 | 0 | 0 | 0 | 0 |
| 29398 | ENSMUSG00000088099 | 0 | NA | NA | NA | NA | NA | NA | NA | 0 | 0 | 0 | 0 | 0 | 0 |
| 29399 | ENSMUSG00000088102 | 0 | NA | NA | NA | NA | NA | NA | NA | 0 | 0 | 0 | 0 | 0 | 0 |
| 29400 | ENSMUSG00000088103 | 0 | NA | NA | NA | NA | NA | Gm23993 | predicted gene, 23993 [Source:MGI Symb | 0 | 0 | 0 | 0 | 0 | 0 |
| 29401 | ENSMUSG00000088104 | 0 | NA | NA | NA | NA | NA | Gm22380 | predicted gene, 22380 [Source:MGI Symb | 0 | 0 | 0 | 0 | 0 | 0 |
| 29402 | ENSMUSG00000088105 | 0 | NA | NA | NA | NA | NA | Gm22379 | predicted gene, 22379 [Source:MGI Symb | 0 | 0 | 0 | 0 | 0 | 0 |
| 29403 | ENSMUSG00000088106 | 3.062533724 | 1.217243862 | 1.258669562 | 0.967087708 | 0.33350019 | NA | Gm22378 | predicted gene, 22378 [Source:MGI Symb | 2.4093 | 2.0558 | 1.0062 | 1.928 | 7.6343 | 3.3415 |
| 29404 | ENSMUSG00000088108 | 1.091781011 | 2.43899895 | 2.376659209 | 1.026229987 | 0.30478319 | NA | Snora47 | small nucleolar RNA, H/ACA box 47 [Sourc | 0 | 1.0279 | 0 | 0 | 2.1812 | 3.3415 |
| 29405 | ENSMUSG00000088111 | 0 | NA | NA | NA | NA | NA | Gm24064 | predicted gene, 24064 [Source:MGI Symb | 0 | 0 | 0 | 0 | 0 | 0 |
| 29406 | ENSMUSG00000088112 | 0 | NA | NA | NA | NA | NA | NA | NA | 0 | 0 | 0 | 0 | 0 | 0 |
| 29407 | ENSMUSG00000088113 | 0.167707431 | -0.796629176 | 4.080472857 | -0.195229623 | 0.84521319 | NA | Mir1960 | microRNA 1960 [Source:MGI Symbol;Acc: | 0 | 0 | 1.0062 | 0 | 0 | 0 |
| 29408 | ENSMUSG00000088114 | 0 | NA | NA | NA | NA | NA | Gm24061 | predicted gene, 24061 [Source:MGI Symb | 0 | 0 | 0 | 0 | 0 | 0 |
| 29409 | ENSMUSG00000088115 | 0.503302254 | -0.837877584 | 3.426698643 | -0.244514523 | 0.806832356 | NA | NA | NA | 0 | 2.0558 | 0 | 0.964 | 0 | 0 |
| 29410 | ENSMUSG00000088116 | 0 | NA | NA | NA | NA | NA | NA | NA | 0 | 0 | 0 | 0 | 0 | 0 |
| 29411 | ENSMUSG00000088118 | 0 | NA | NA | NA | NA | NA | NA | NA | 0 | 0 | 0 | 0 | 0 | 0 |
| 29412 | ENSMUSG00000088119 | 7.38197459 | 0.057056447 | 0.816931446 | 0.069842393 | 0.944319105 | 0.990077979 | NA | NA | 5.6217 | 6.1675 | 10.062 | 12.532 | 5.4531 | 4.4554 |
| 29413 | ENSMUSG00000088121 | 0 | NA | NA | NA | NA | NA | Gm25721 | predicted gene, 25721 [Source:MGI Symb | 0 | 0 | 0 | 0 | 0 | 0 |
| 29414 | ENSMUSG00000088122 | 0 | NA | NA | NA | NA | NA | NA | NA | 0 | 0 | 0 | 0 | 0 | 0 |
| 29415 | ENSMUSG00000088123 | 0 | NA | NA | NA | NA | NA | NA | NA | 0 | 0 | 0 | 0 | 0 | 0 |
| 29416 | ENSMUSG00000088124 | 0 | NA | NA | NA | NA | NA | NA | NA | 0 | 0 | 0 | 0 | 0 | 0 |
| 29417 | ENSMUSG00000088125 | 0 | NA | NA | NA | NA | NA | Gm26070 | predicted gene, 26070 [Source:MGI Symb | 0 | 0 | 0 | 0 | 0 | 0 |
| 29418 | ENSMUSG00000088126 | 0 | NA | NA | NA | NA | NA | Gm22870 | predicted gene, 22870 [Source:MGI Symb | 0 | 0 | 0 | 0 | 0 | 0 |
| 29419 | ENSMUSG00000088128 | 0 | NA | NA | NA | NA | NA | NA | NA | 0 | 0 | 0 | 0 | 0 | 0 |
| 29420 | ENSMUSG00000088132 | 0 | NA | NA | NA | NA | NA | NA | NA | 0 | 0 | 0 | 0 | 0 | 0 |
| 29421 | ENSMUSG00000088133 | 0 | NA | NA | NA | NA | NA | NA | NA | 0 | 0 | 0 | 0 | 0 | 0 |
| 29422 | ENSMUSG00000088134 | 0 | NA | NA | NA | NA | NA | Gm22310 | predicted gene, 22310 [Source:MGI Symb | 0 | 0 | 0 | 0 | 0 | 0 |
| 29423 | ENSMUSG00000088135 | 0 | NA | NA | NA | NA | NA | Gm22588 | predicted gene, 22588 [Source:MGI Symb | 0 | 0 | 0 | 0 | 0 | 0 |
| 29424 | ENSMUSG00000088136 | 0 | NA | NA | NA | NA | NA | NA | NA | 0 | 0 | 0 | 0 | 0 | 0 |
| 29425 | ENSMUSG00000088138 | 0 | NA | NA | NA | NA | NA | NA | NA | 0 | 0 | 0 | 0 | 0 | 0 |
| 29426 | ENSMUSG00000088139 | 0.401551119 | -2.072982726 | 4.008251979 | -0.517178744 | 0.605031388 | NA | Gm27343 | predicted gene, 27343 [Source:MGI Symb | 2.4093 | 0 | 0 | 0 | 0 | 0 |
| 29427 | ENSMUSG00000088141 | 0 | NA | NA | NA | NA | NA | Mir1962 | microRNA 1962 [Source:MGI Symbol;Acc: | 0 | 0 | 0 | 0 | 0 | 0 |
| 29428 | ENSMUSG00000088143 | 0 | NA | NA | NA | NA | NA | NA | NA | 0 | 0 | 0 | 0 | 0 | 0 |
| 29429 | ENSMUSG00000088144 | 0 | NA | NA | NA | NA | NA | NA | NA | 0 | 0 | 0 | 0 | 0 | 0 |
| 29430 | ENSMUSG00000088145 | 0 | NA | NA | NA | NA | NA | Gm24719 | predicted gene, 24719 [Source:MGI Symb | 0 | 0 | 0 | 0 | 0 | 0 |
| 29431 | ENSMUSG00000088146 | 0 | NA | NA | NA | NA | NA | Gm24718 | predicted gene, 24718 [Source:MGI Symb | 0 | 0 | 0 | 0 | 0 | 0 |
| 29432 | ENSMUSG00000088147 | 0 | NA | NA | NA | NA | NA | Gm24717 | predicted gene, 24717 [Source:MGI Symb | 0 | 0 | 0 | 0 | 0 | 0 |
| 29433 | ENSMUSG00000088148 | 0 | NA | NA | NA | NA | NA | Mir1983 | microRNA 1983 [Source:MGI Symbol;Acc: | 0 | 0 | 0 | 0 | 0 | 0 |
| 29434 | ENSMUSG00000088149 | 0 | NA | NA | NA | NA | NA | Gm24716 | predicted gene, 24716 [Source:MGI Symb | 0 | 0 | 0 | 0 | 0 | 0 |
| 29435 | ENSMUSG00000088150 | 0 | NA | NA | NA | NA | NA | NA | NA | 0 | 0 | 0 | 0 | 0 | 0 |
| 29436 | ENSMUSG00000088151 | 1.216525419 | -1.227051403 | 2.308900077 | -0.531444135 | 0.595111044 | NA | Gm26347 | predicted gene, 26347 [Source:MGI Symb | 0 | 4.1117 | 1.0062 | 0 | 2.1812 | 0 |
| 29437 | ENSMUSG00000088152 | 0 | NA | NA | NA | NA | NA | Gm25572 | predicted gene, 25572 [Source:MGI Symb | 0 | 0 | 0 | 0 | 0 | 0 |
| 29438 | ENSMUSG00000088153 | 0 | NA | NA | NA | NA | NA | NA | NA | 0 | 0 | 0 | 0 | 0 | 0 |
| 29439 | ENSMUSG00000088154 | 0 | NA | NA | NA | NA | NA | Gm25637 | predicted gene, 25637 [Source:MGI Symb | 0 | 0 | 0 | 0 | 0 | 0 |
| 29440 | ENSMUSG00000088155 | 0 | NA | NA | NA | NA | NA | Gm26349 | predicted gene, 26349 [Source:MGI Symb | 0 | 0 | 0 | 0 | 0 | 0 |
| 29441 | ENSMUSG00000088156 | 0 | NA | NA | NA | NA | NA | NA | NA | 0 | 0 | 0 | 0 | 0 | 0 |
| 29442 | ENSMUSG00000088157 | 0 | NA | NA | NA | NA | NA | Gm26351 | predicted gene, 26351 [Source:MGI Symb | 0 | 0 | 0 | 0 | 0 | 0 |
| 29443 | ENSMUSG00000088158 | 0 | NA | NA | NA | NA | NA | NA | NA | 0 | 0 | 0 | 0 | 0 | 0 |
| 29444 | ENSMUSG00000088159 | 0 | NA | NA | NA | NA | NA | Gm26345 | predicted gene, 26345 [Source:MGI Symb | 0 | 0 | 0 | 0 | 0 | 0 |
| 29445 | ENSMUSG00000088160 | 0 | NA | NA | NA | NA | NA | Gm23538 | predicted gene, 23538 [Source:MGI Symb | 0 | 0 | 0 | 0 | 0 | 0 |
| 29446 | ENSMUSG00000088161 | 0 | NA | NA | NA | NA | NA | NA | NA | 0 | 0 | 0 | 0 | 0 | 0 |
| 29447 | ENSMUSG00000088162 | 0 | NA | NA | NA | NA | NA | NA | NA | 0 | 0 | 0 | 0 | 0 | 0 |
| 29448 | ENSMUSG00000088164 | 0 | NA | NA | NA | NA | NA | Gm23534 | predicted gene, 23534 [Source:MGI Symb | 0 | 0 | 0 | 0 | 0 | 0 |
| 29449 | ENSMUSG00000088165 | 0 | NA | NA | NA | NA | NA | NA | NA | 0 | 0 | 0 | 0 | 0 | 0 |
| 29450 | ENSMUSG00000088166 | 0 | NA | NA | NA | NA | NA | NA | NA | 0 | 0 | 0 | 0 | 0 | 0 |
| 29451 | ENSMUSG00000088167 | 0 | NA | NA | NA | NA | NA | Gm23535 | predicted gene, 23535 [Source:MGI Symb | 0 | 0 | 0 | 0 | 0 | 0 |
| 29452 | ENSMUSG00000088168 | 0 | NA | NA | NA | NA | NA | Gm23540 | predicted gene, 23540 [Source:MGI Symb | 0 | 0 | 0 | 0 | 0 | 0 |

| | | | | | | | | | | | | | | | |
|---|---|---|---|---|---|---|---|---|---|---|---|---|---|---|---|
| 29453 | ENSMUSG00000088169 | 0 | NA | NA | NA | NA | NA | Gm22059 | predicted gene, 22059 [Source:MGI Symb | 0 | 0 | 0 | 0 | 0 | 0 |
| 29454 | ENSMUSG00000088170 | 1.339512537 | 0.884662434 | 1.869203337 | 0.473283145 | 0.636011186 | NA | NA | NA | 0.8031 | 1.0279 | 1.0062 | 1.928 | 3.2719 | 0 |
| 29455 | ENSMUSG00000088171 | 1.042135994 | -3.435243139 | 2.323779668 | -1.478299852 | 0.139327537 | NA | NA | NA | 3.2124 | 1.0279 | 2.0125 | 0 | 0 | 0 |
| 29456 | ENSMUSG00000088172 | 0.267700746 | -1.497481649 | 4.057001436 | -0.369110456 | 0.712045396 | NA | Gm25217 | predicted gene, 25217 [Source:MGI Symb | 1.6062 | 0 | 0 | 0 | 0 | 0 |
| 29457 | ENSMUSG00000088175 | 0 | NA | NA | NA | NA | NA | Gm23101 | predicted gene, 23101 [Source:MGI Symb | 0 | 0 | 0 | 0 | 0 | 0 |
| 29458 | ENSMUSG00000088176 | 0 | NA | NA | NA | NA | NA | NA | NA | 0 | 0 | 0 | 0 | 0 | 0 |
| 29459 | ENSMUSG00000088177 | 0 | NA | NA | NA | NA | NA | NA | NA | 0 | 0 | 0 | 0 | 0 | 0 |
| 29460 | ENSMUSG00000088179 | 0 | NA | NA | NA | NA | NA | NA | NA | 0 | 0 | 0 | 0 | 0 | 0 |
| 29461 | ENSMUSG00000088180 | 0 | NA | NA | NA | NA | NA | Gm22218 | predicted gene, 22218 [Source:MGI Symb | 0 | 0 | 0 | 0 | 0 | 0 |
| 29462 | ENSMUSG00000088181 | 0 | NA | NA | NA | NA | NA | NA | NA | 0 | 0 | 0 | 0 | 0 | 0 |
| 29463 | ENSMUSG00000088182 | 0 | NA | NA | NA | NA | NA | NA | NA | 0 | 0 | 0 | 0 | 0 | 0 |
| 29464 | ENSMUSG00000088183 | 1.306068659 | -2.567602762 | 2.262929625 | -1.134636594 | 0.256527614 | NA | Gm22215 | predicted gene, 22215 [Source:MGI Symb | 1.6062 | 5.1396 | 0 | 0 | 1.0906 | 0 |
| 29465 | ENSMUSG00000088184 | 0 | NA | NA | NA | NA | NA | NA | NA | 0 | 0 | 0 | 0 | 0 | 0 |
| 29466 | ENSMUSG00000088185 | 6.026288461 | 0.217487839 | 0.913289745 | 0.238136736 | 0.811775043 | 0.96653525 | Scarna2 | small Cajal body-specific RNA 2 [Source:M | 6.4248 | 6.1675 | 4.025 | 1.928 | 9.8156 | 7.7969 |
| 29467 | ENSMUSG00000088187 | 0 | NA | NA | NA | NA | NA | NA | NA | 0 | 0 | 0 | 0 | 0 | 0 |
| 29468 | ENSMUSG00000088188 | 0 | NA | NA | NA | NA | NA | NA | NA | 0 | 0 | 0 | 0 | 0 | 0 |
| 29469 | ENSMUSG00000088189 | 0.167707431 | -0.796629176 | 4.080472857 | -0.195229623 | 0.84521319 | NA | NA | NA | 0 | 0 | 1.0062 | 0 | 0 | 0 |
| 29470 | ENSMUSG00000088191 | 0 | NA | NA | NA | NA | NA | Gm23903 | predicted gene, 23903 [Source:MGI Symb | 0 | 0 | 0 | 0 | 0 | 0 |
| 29471 | ENSMUSG00000088192 | 0 | NA | NA | NA | NA | NA | Gm23904 | predicted gene, 23904 [Source:MGI Symb | 0 | 0 | 0 | 0 | 0 | 0 |
| 29472 | ENSMUSG00000088193 | 0 | NA | NA | NA | NA | NA | NA | NA | 0 | 0 | 0 | 0 | 0 | 0 |
| 29473 | ENSMUSG00000088194 | 0 | NA | NA | NA | NA | NA | NA | NA | 0 | 0 | 0 | 0 | 0 | 0 |
| 29474 | ENSMUSG00000088195 | 0 | NA | NA | NA | NA | NA | Gm23901 | predicted gene, 23901 [Source:MGI Symb | 0 | 0 | 0 | 0 | 0 | 0 |
| 29475 | ENSMUSG00000088196 | 0 | NA | NA | NA | NA | NA | Gm23902 | predicted gene, 23902 [Source:MGI Symb | 0 | 0 | 0 | 0 | 0 | 0 |
| 29476 | ENSMUSG00000088197 | 0 | NA | NA | NA | NA | NA | Gm24138 | predicted gene, 24138 [Source:MGI Symb | 0 | 0 | 0 | 0 | 0 | 0 |
| 29477 | ENSMUSG00000088198 | 0 | NA | NA | NA | NA | NA | Gm23906 | predicted gene, 23906 [Source:MGI Symb | 0 | 0 | 0 | 0 | 0 | 0 |
| 29478 | ENSMUSG00000088200 | 0.133850373 | -0.796629176 | 4.080472857 | -0.195229623 | 0.84521319 | NA | NA | NA | 0.8031 | 0 | 0 | 0 | 0 | 0 |
| 29479 | ENSMUSG00000088201 | 0 | NA | NA | NA | NA | NA | Gm23754 | predicted gene, 23754 [Source:MGI Symb | 0 | 0 | 0 | 0 | 0 | 0 |
| 29480 | ENSMUSG00000088203 | 0 | NA | NA | NA | NA | NA | NA | NA | 0 | 0 | 0 | 0 | 0 | 0 |
| 29481 | ENSMUSG00000088204 | 0 | NA | NA | NA | NA | NA | NA | NA | 0 | 0 | 0 | 0 | 0 | 0 |
| 29482 | ENSMUSG00000088205 | 0 | NA | NA | NA | NA | NA | NA | NA | 0 | 0 | 0 | 0 | 0 | 0 |
| 29483 | ENSMUSG00000088207 | 0 | NA | NA | NA | NA | NA | NA | NA | 0 | 0 | 0 | 0 | 0 | 0 |
| 29484 | ENSMUSG00000088208 | 0.462220778 | -0.685400455 | 3.191475091 | -0.21475977 | 0.82995462 | NA | Gm23751 | predicted gene, 23751 [Source:MGI Symb | 0.8031 | 0 | 1.0062 | 0.964 | 0 | 0 |
| 29485 | ENSMUSG00000088209 | 0 | NA | NA | NA | NA | NA | NA | NA | 0 | 0 | 0 | 0 | 0 | 0 |
| 29486 | ENSMUSG00000088210 | 0 | NA | NA | NA | NA | NA | NA | NA | 0 | 0 | 0 | 0 | 0 | 0 |
| 29487 | ENSMUSG00000088211 | 0 | NA | NA | NA | NA | NA | NA | NA | 0 | 0 | 0 | 0 | 0 | 0 |
| 29488 | ENSMUSG00000088212 | 0 | NA | NA | NA | NA | NA | Gm22082 | predicted gene, 22082 [Source:MGI Symb | 0 | 0 | 0 | 0 | 0 | 0 |
| 29489 | ENSMUSG00000088213 | 0 | NA | NA | NA | NA | NA | NA | NA | 0 | 0 | 0 | 0 | 0 | 0 |
| 29490 | ENSMUSG00000088216 | 0 | NA | NA | NA | NA | NA | NA | NA | 0 | 0 | 0 | 0 | 0 | 0 |
| 29491 | ENSMUSG00000088217 | 0.315620088 | 0.165143421 | 4.063300985 | 0.040642675 | 0.967580763 | NA | Gm22079 | predicted gene, 22079 [Source:MGI Symb | 0.8031 | 0 | 0 | 0 | 1.0906 | 0 |
| 29492 | ENSMUSG00000088218 | 0 | NA | NA | NA | NA | NA | Gm22078 | predicted gene, 22078 [Source:MGI Symb | 0 | 0 | 0 | 0 | 0 | 0 |
| 29493 | ENSMUSG00000088219 | 0 | NA | NA | NA | NA | NA | NA | NA | 0 | 0 | 0 | 0 | 0 | 0 |
| 29494 | ENSMUSG00000088220 | 0 | NA | NA | NA | NA | NA | NA | NA | 0 | 0 | 0 | 0 | 0 | 0 |
| 29495 | ENSMUSG00000088221 | 0 | NA | NA | NA | NA | NA | NA | NA | 0 | 0 | 0 | 0 | 0 | 0 |
| 29496 | ENSMUSG00000088222 | 0 | NA | NA | NA | NA | NA | NA | NA | 0 | 0 | 0 | 0 | 0 | 0 |
| 29497 | ENSMUSG00000088223 | 0 | NA | NA | NA | NA | NA | NA | NA | 0 | 0 | 0 | 0 | 0 | 0 |
| 29498 | ENSMUSG00000088224 | 0.367410362 | 2.078255025 | 3.800070157 | 0.54689912 | 0.58444804 | NA | Gm25406 | predicted gene, 25406 [Source:MGI Symb | 0 | 0 | 0 | 0 | 1.0906 | 1.1138 |
| 29499 | ENSMUSG00000088225 | 0 | NA | NA | NA | NA | NA | Gm22598 | predicted gene, 22598 [Source:MGI Symb | 0 | 0 | 0 | 0 | 0 | 0 |
| 29500 | ENSMUSG00000088226 | 2.230123666 | -1.16262684 | 1.520663567 | -0.764552308 | 0.444538178 | NA | NA | NA | 3.2124 | 1.0279 | 5.0312 | 1.928 | 2.1812 | 0 |
| 29501 | ENSMUSG00000088227 | 0 | NA | NA | NA | NA | NA | Gm22597 | predicted gene, 22597 [Source:MGI Symb | 0 | 0 | 0 | 0 | 0 | 0 |
| 29502 | ENSMUSG00000088228 | 0 | NA | NA | NA | NA | NA | Gm22603 | predicted gene, 22603 [Source:MGI Symb | 0 | 0 | 0 | 0 | 0 | 0 |
| 29503 | ENSMUSG00000088229 | 0 | NA | NA | NA | NA | NA | NA | NA | 0 | 0 | 0 | 0 | 0 | 0 |
| 29504 | ENSMUSG00000088230 | 0 | NA | NA | NA | NA | NA | NA | NA | 0 | 0 | 0 | 0 | 0 | 0 |
| 29505 | ENSMUSG00000088231 | 0 | NA | NA | NA | NA | NA | NA | NA | 0 | 0 | 0 | 0 | 0 | 0 |
| 29506 | ENSMUSG00000088232 | 0 | NA | NA | NA | NA | NA | Gm25429 | predicted gene, 25429 [Source:MGI Symb | 0 | 0 | 0 | 0 | 0 | 0 |
| 29507 | ENSMUSG00000088233 | 0.185640647 | 1.126924175 | 4.080472857 | 0.276174898 | 0.782413736 | NA | NA | NA | 0 | 0 | 0 | 0 | 0 | 1.1138 |
| 29508 | ENSMUSG00000088234 | 0 | NA | NA | NA | NA | NA | NA | NA | 0 | 0 | 0 | 0 | 0 | 0 |

| | | | | | | | | | | | | | | | |
|---|---|---|---|---|---|---|---|---|---|---|---|---|---|---|---|
| 29509 | ENSMUSG00000088236 | 0 | NA | NA | NA | NA | NA | Gm25425 | predicted gene, 25425 [Source:MGI Symb | 0 | 0 | 0 | 0 | 0 | 0 |
| 29510 | ENSMUSG00000088237 | 0 | NA | NA | NA | NA | NA | NA | NA | 0 | 0 | 0 | 0 | 0 | 0 |
| 29511 | ENSMUSG00000088239 | 0 | NA | NA | NA | NA | NA | NA | NA | 0 | 0 | 0 | 0 | 0 | 0 |
| 29512 | ENSMUSG00000088240 | 0 | NA | NA | NA | NA | NA | Gm22915 | predicted gene, 22915 [Source:MGI Symb | 0 | 0 | 0 | 0 | 0 | 0 |
| 29513 | ENSMUSG00000088241 | 0 | NA | NA | NA | NA | NA | Gm22916 | predicted gene, 22916 [Source:MGI Symb | 0 | 0 | 0 | 0 | 0 | 0 |
| 29514 | ENSMUSG00000088242 | 0 | NA | NA | NA | NA | NA | Gm25913 | predicted gene, 25913 [Source:MGI Symb | 0 | 0 | 0 | 0 | 0 | 0 |
| 29515 | ENSMUSG00000088243 | 0 | NA | NA | NA | NA | NA | NA | NA | 0 | 0 | 0 | 0 | 0 | 0 |
| 29516 | ENSMUSG00000088244 | 0 | NA | NA | NA | NA | NA | Gm25909 | predicted gene, 25909 [Source:MGI Symb | 0 | 0 | 0 | 0 | 0 | 0 |
| 29517 | ENSMUSG00000088245 | 0 | NA | NA | NA | NA | NA | Gm25910 | predicted gene, 25910 [Source:MGI Symb | 0 | 0 | 0 | 0 | 0 | 0 |
| 29518 | ENSMUSG00000088246 | 0.133850373 | -0.796629176 | 4.080472857 | -0.195229623 | 0.84521319 | NA | NA | NA | 0.8031 | 0 | 0 | 0 | 0 | 0 |
| 29519 | ENSMUSG00000088248 | 0 | NA | NA | NA | NA | NA | Mir1969 | microRNA 1969 [Source:MGI Symbol;Acc: | 0 | 0 | 0 | 0 | 0 | 0 |
| 29520 | ENSMUSG00000088249 | 0.667397476 | -1.404797968 | 2.804497115 | -0.500909044 | 0.616435137 | NA | NA | NA | 0 | 1.0279 | 2.0125 | 0.964 | 0 | 0 |
| 29521 | ENSMUSG00000088250 | 0.167707431 | -0.796629176 | 4.080472857 | -0.195229623 | 0.84521319 | NA | NA | NA | 0 | 0 | 1.0062 | 0 | 0 | 0 |
| 29522 | ENSMUSG00000088252 | 0 | NA | NA | NA | NA | NA | Snord13 | small nucleolar RNA, C/D box 13 [Source:I | 0 | 0 | 0 | 0 | 0 | 0 |
| 29523 | ENSMUSG00000088253 | 0 | NA | NA | NA | NA | NA | Gm24285 | predicted gene, 24285 [Source:MGI Symb | 0 | 0 | 0 | 0 | 0 | 0 |
| 29524 | ENSMUSG00000088254 | 0 | NA | NA | NA | NA | NA | Gm24289 | predicted gene, 24289 [Source:MGI Symb | 0 | 0 | 0 | 0 | 0 | 0 |
| 29525 | ENSMUSG00000088255 | 0 | NA | NA | NA | NA | NA | Gm24288 | predicted gene, 24288 [Source:MGI Symb | 0 | 0 | 0 | 0 | 0 | 0 |
| 29526 | ENSMUSG00000088257 | 0 | NA | NA | NA | NA | NA | Gm24286 | predicted gene, 24286 [Source:MGI Symb | 0 | 0 | 0 | 0 | 0 | 0 |
| 29527 | ENSMUSG00000088258 | 0 | NA | NA | NA | NA | NA | Gm24291 | predicted gene, 24291 [Source:MGI Symb | 0 | 0 | 0 | 0 | 0 | 0 |
| 29528 | ENSMUSG00000088259 | 0 | NA | NA | NA | NA | NA | NA | NA | 0 | 0 | 0 | 0 | 0 | 0 |
| 29529 | ENSMUSG00000088260 | 0 | NA | NA | NA | NA | NA | Gm24780 | predicted gene, 24780 [Source:MGI Symb | 0 | 0 | 0 | 0 | 0 | 0 |
| 29530 | ENSMUSG00000088261 | 0 | NA | NA | NA | NA | NA | Gm24781 | predicted gene, 24781 [Source:MGI Symb | 0 | 0 | 0 | 0 | 0 | 0 |
| 29531 | ENSMUSG00000088264 | 0 | NA | NA | NA | NA | NA | NA | NA | 0 | 0 | 0 | 0 | 0 | 0 |
| 29532 | ENSMUSG00000088265 | 0 | NA | NA | NA | NA | NA | Gm24784 | predicted gene, 24784 [Source:MGI Symb | 0 | 0 | 0 | 0 | 0 | 0 |
| 29533 | ENSMUSG00000088266 | 0 | NA | NA | NA | NA | NA | Gm24782 | predicted gene, 24782 [Source:MGI Symb | 0 | 0 | 0 | 0 | 0 | 0 |
| 29534 | ENSMUSG00000088268 | 1.586339763 | -4.028767891 | 1.995558398 | -2.018867449 | 0.043500998 | NA | NA | NA | 2.4093 | 3.0838 | 4.025 | 0 | 0 | 0 |
| 29535 | ENSMUSG00000088269 | 0 | NA | NA | NA | NA | NA | NA | NA | 0 | 0 | 0 | 0 | 0 | 0 |
| 29536 | ENSMUSG00000088270 | 0 | NA | NA | NA | NA | NA | NA | NA | 0 | 0 | 0 | 0 | 0 | 0 |
| 29537 | ENSMUSG00000088273 | 0.462220778 | -0.685400455 | 3.191475091 | -0.21475977 | 0.82995462 | NA | Gm23123 | predicted gene, 23123 [Source:MGI Symb | 0.8031 | 0 | 1.0062 | 0.964 | 0 | 0 |
| 29538 | ENSMUSG00000088275 | 0 | NA | NA | NA | NA | NA | Gm23124 | predicted gene, 23124 [Source:MGI Symb | 0 | 0 | 0 | 0 | 0 | 0 |
| 29539 | ENSMUSG00000088276 | 0 | NA | NA | NA | NA | NA | Gm23125 | predicted gene, 23125 [Source:MGI Symb | 0 | 0 | 0 | 0 | 0 | 0 |
| 29540 | ENSMUSG00000088278 | 0 | NA | NA | NA | NA | NA | NA | NA | 0 | 0 | 0 | 0 | 0 | 0 |
| 29541 | ENSMUSG00000088279 | 0 | NA | NA | NA | NA | NA | NA | NA | 0 | 0 | 0 | 0 | 0 | 0 |
| 29542 | ENSMUSG00000088280 | 0 | NA | NA | NA | NA | NA | NA | NA | 0 | 0 | 0 | 0 | 0 | 0 |
| 29543 | ENSMUSG00000088281 | 0 | NA | NA | NA | NA | NA | Gm23585 | predicted gene, 23585 [Source:MGI Symb | 0 | 0 | 0 | 0 | 0 | 0 |
| 29544 | ENSMUSG00000088282 | 0 | NA | NA | NA | NA | NA | Gm23586 | predicted gene, 23586 [Source:MGI Symb | 0 | 0 | 0 | 0 | 0 | 0 |
| 29545 | ENSMUSG00000088283 | 0 | NA | NA | NA | NA | NA | NA | NA | 0 | 0 | 0 | 0 | 0 | 0 |
| 29546 | ENSMUSG00000088284 | 0 | NA | NA | NA | NA | NA | NA | NA | 0 | 0 | 0 | 0 | 0 | 0 |
| 29547 | ENSMUSG00000088285 | 0 | NA | NA | NA | NA | NA | NA | NA | 0 | 0 | 0 | 0 | 0 | 0 |
| 29548 | ENSMUSG00000088286 | 0 | NA | NA | NA | NA | NA | NA | NA | 0 | 0 | 0 | 0 | 0 | 0 |
| 29549 | ENSMUSG00000088288 | 0 | NA | NA | NA | NA | NA | NA | NA | 0 | 0 | 0 | 0 | 0 | 0 |
| 29550 | ENSMUSG00000088291 | 0 | NA | NA | NA | NA | NA | NA | NA | 0 | 0 | 0 | 0 | 0 | 0 |
| 29551 | ENSMUSG00000088293 | 0 | NA | NA | NA | NA | NA | Gm26411 | predicted gene, 26411 [Source:MGI Symb | 0 | 0 | 0 | 0 | 0 | 0 |
| 29552 | ENSMUSG00000088294 | 0 | NA | NA | NA | NA | NA | NA | NA | 0 | 0 | 0 | 0 | 0 | 0 |
| 29553 | ENSMUSG00000088295 | 0 | NA | NA | NA | NA | NA | NA | NA | 0 | 0 | 0 | 0 | 0 | 0 |
| 29554 | ENSMUSG00000088296 | 0 | NA | NA | NA | NA | NA | Gm26409 | predicted gene, 26409 [Source:MGI Symb | 0 | 0 | 0 | 0 | 0 | 0 |
| 29555 | ENSMUSG00000088297 | 0 | NA | NA | NA | NA | NA | Gm26408 | predicted gene, 26408 [Source:MGI Symb | 0 | 0 | 0 | 0 | 0 | 0 |
| 29556 | ENSMUSG00000088298 | 0.167707431 | -0.796629176 | 4.080472857 | -0.195229623 | 0.84521319 | NA | NA | NA | 0 | 0 | 1.0062 | 0 | 0 | 0 |
| 29557 | ENSMUSG00000088299 | 0 | NA | NA | NA | NA | NA | NA | NA | 0 | 0 | 0 | 0 | 0 | 0 |
| 29558 | ENSMUSG00000088301 | 0 | NA | NA | NA | NA | NA | NA | NA | 0 | 0 | 0 | 0 | 0 | 0 |
| 29559 | ENSMUSG00000088302 | 0 | NA | NA | NA | NA | NA | NA | NA | 0 | 0 | 0 | 0 | 0 | 0 |
| 29560 | ENSMUSG00000088303 | 0.17131964 | -0.796629176 | 4.080472857 | -0.195229623 | 0.84521319 | NA | Gm23382 | predicted gene, 23382 [Source:MGI Symb | 0 | 1.0279 | 0 | 0 | 0 | 0 |
| 29561 | ENSMUSG00000088304 | 0 | NA | NA | NA | NA | NA | Gm24504 | predicted gene, 24504 [Source:MGI Symb | 0 | 0 | 0 | 0 | 0 | 0 |
| 29562 | ENSMUSG00000088307 | 0 | NA | NA | NA | NA | NA | Gm24505 | predicted gene, 24505 [Source:MGI Symb | 0 | 0 | 0 | 0 | 0 | 0 |
| 29563 | ENSMUSG00000088308 | 0.439020386 | -2.189747911 | 3.52148864 | -0.621824499 | 0.534057275 | NA | Gm24507 | predicted gene, 24507 [Source:MGI Symb | 1.6062 | 1.0279 | 0 | 0 | 0 | 0 |
| 29564 | ENSMUSG00000088310 | 0 | NA | NA | NA | NA | NA | Gm26183 | predicted gene, 26183 [Source:MGI Symb | 0 | 0 | 0 | 0 | 0 | 0 |

| | | | | | | | | | | | | | | | |
|---|---|---|---|---|---|---|---|---|---|---|---|---|---|---|---|
| 29565 | ENSMUSG00000088311 | 0 | NA | NA | NA | NA | NA | Gm26184 | predicted gene, 26184 [Source:MGI Symb | 0 | 0 | 0 | 0 | 0 | 0 |
| 29566 | ENSMUSG00000088312 | 0 | NA | NA | NA | NA | NA | Gm26181 | predicted gene, 26181 [Source:MGI Symb | 0 | 0 | 0 | 0 | 0 | 0 |
| 29567 | ENSMUSG00000088313 | 0 | NA | NA | NA | NA | NA | Gm26182 | predicted gene, 26182 [Source:MGI Symb | 0 | 0 | 0 | 0 | 0 | 0 |
| 29568 | ENSMUSG00000088314 | 0 | NA | NA | NA | NA | NA | NA | NA | 0 | 0 | 0 | 0 | 0 | 0 |
| 29569 | ENSMUSG00000088317 | 0 | NA | NA | NA | NA | NA | NA | NA | 0 | 0 | 0 | 0 | 0 | 0 |
| 29570 | ENSMUSG00000088318 | 0 | NA | NA | NA | NA | NA | Gm26187 | predicted gene, 26187 [Source:MGI Symb | 0 | 0 | 0 | 0 | 0 | 0 |
| 29571 | ENSMUSG00000088320 | 0 | NA | NA | NA | NA | NA | Gm25665 | predicted gene, 25665 [Source:MGI Symb | 0 | 0 | 0 | 0 | 0 | 0 |
| 29572 | ENSMUSG00000088322 | 0 | NA | NA | NA | NA | NA | NA | NA | 0 | 0 | 0 | 0 | 0 | 0 |
| 29573 | ENSMUSG00000088323 | 0 | NA | NA | NA | NA | NA | Gm25663 | predicted gene, 25663 [Source:MGI Symb | 0 | 0 | 0 | 0 | 0 | 0 |
| 29574 | ENSMUSG00000088324 | 0 | NA | NA | NA | NA | NA | NA | NA | 0 | 0 | 0 | 0 | 0 | 0 |
| 29575 | ENSMUSG00000088325 | 0.321325949 | 1.911575114 | 4.048267893 | 0.472195804 | 0.636787034 | NA | Gm25668 | predicted gene, 25668 [Source:MGI Symb | 0 | 0 | 0 | 1.928 | 0 | 0 |
| 29576 | ENSMUSG00000088326 | 1.09031406 | -2.267649464 | 2.358096776 | -0.961643935 | 0.336228494 | NA | NA | NA | 2.4093 | 0 | 3.0187 | 0 | 0 | 1.1138 |
| 29577 | ENSMUSG00000088327 | 0 | NA | NA | NA | NA | NA | NA | NA | 0 | 0 | 0 | 0 | 0 | 0 |
| 29578 | ENSMUSG00000088329 | 0.636972666 | -2.712703855 | 3.077964703 | -0.881330397 | 0.378139018 | NA | Gm25661 | predicted gene, 25661 [Source:MGI Symb | 0.8031 | 0 | 3.0187 | 0 | 0 | 0 |
| 29579 | ENSMUSG00000088331 | 0 | NA | NA | NA | NA | NA | NA | NA | 0 | 0 | 0 | 0 | 0 | 0 |
| 29580 | ENSMUSG00000088332 | 0.85435009 | 0.823372753 | 2.383559862 | 0.345438252 | 0.729764924 | NA | Gm22847 | predicted gene, 22847 [Source:MGI Symb | 0.8031 | 1.0279 | 0 | 0 | 2.1812 | 1.1138 |
| 29581 | ENSMUSG00000088333 | 0 | NA | NA | NA | NA | NA | Gm27396 | predicted gene, 27396 [Source:MGI Symb | 0 | 0 | 0 | 0 | 0 | 0 |
| 29582 | ENSMUSG00000088334 | 0 | NA | NA | NA | NA | NA | NA | NA | 0 | 0 | 0 | 0 | 0 | 0 |
| 29583 | ENSMUSG00000088335 | 0 | NA | NA | NA | NA | NA | NA | NA | 0 | 0 | 0 | 0 | 0 | 0 |
| 29584 | ENSMUSG00000088337 | 0 | NA | NA | NA | NA | NA | Gm22845 | predicted gene, 22845 [Source:MGI Symb | 0 | 0 | 0 | 0 | 0 | 0 |
| 29585 | ENSMUSG00000088338 | 0.749689669 | 0.81510874 | 2.956253303 | 0.275723579 | 0.782760384 | NA | NA | NA | 1.6062 | 0 | 0 | 2.8919 | 0 | 0 |
| 29586 | ENSMUSG00000088339 | 0 | NA | NA | NA | NA | NA | Gm22850 | predicted gene, 22850 [Source:MGI Symb | 0 | 0 | 0 | 0 | 0 | 0 |
| 29587 | ENSMUSG00000088340 | 0 | NA | NA | NA | NA | NA | NA | NA | 0 | 0 | 0 | 0 | 0 | 0 |
| 29588 | ENSMUSG00000088342 | 0.36353943 | 2.065989254 | 4.034137301 | 0.512126658 | 0.608562368 | NA | NA | NA | 0 | 0 | 0 | 0 | 2.1812 | 0 |
| 29589 | ENSMUSG00000088343 | 0 | NA | NA | NA | NA | NA | Gm22339 | predicted gene, 22339 [Source:MGI Symb | 0 | 0 | 0 | 0 | 0 | 0 |
| 29590 | ENSMUSG00000088344 | 0 | NA | NA | NA | NA | NA | Gm22336 | predicted gene, 22336 [Source:MGI Symb | 0 | 0 | 0 | 0 | 0 | 0 |
| 29591 | ENSMUSG00000088346 | 0 | NA | NA | NA | NA | NA | Gm22337 | predicted gene, 22337 [Source:MGI Symb | 0 | 0 | 0 | 0 | 0 | 0 |
| 29592 | ENSMUSG00000088348 | 0 | NA | NA | NA | NA | NA | NA | NA | 0 | 0 | 0 | 0 | 0 | 0 |
| 29593 | ENSMUSG00000088349 | 0.267700746 | -1.497481649 | 4.057001436 | -0.369110456 | 0.712045396 | NA | Gm22335 | predicted gene, 22335 [Source:MGI Symb | 1.6062 | 0 | 0 | 0 | 0 | 0 |
| 29594 | ENSMUSG00000088350 | 0 | NA | NA | NA | NA | NA | NA | NA | 0 | 0 | 0 | 0 | 0 | 0 |
| 29595 | ENSMUSG00000088351 | 0 | NA | NA | NA | NA | NA | Gm24016 | predicted gene, 24016 [Source:MGI Symb | 0 | 0 | 0 | 0 | 0 | 0 |
| 29596 | ENSMUSG00000088352 | 0 | NA | NA | NA | NA | NA | Gm24013 | predicted gene, 24013 [Source:MGI Symb | 0 | 0 | 0 | 0 | 0 | 0 |
| 29597 | ENSMUSG00000088353 | 0 | NA | NA | NA | NA | NA | NA | NA | 0 | 0 | 0 | 0 | 0 | 0 |
| 29598 | ENSMUSG00000088354 | 0 | NA | NA | NA | NA | NA | NA | NA | 0 | 0 | 0 | 0 | 0 | 0 |
| 29599 | ENSMUSG00000088356 | 0 | NA | NA | NA | NA | NA | NA | NA | 0 | 0 | 0 | 0 | 0 | 0 |
| 29600 | ENSMUSG00000088357 | 0.342639279 | -1.805976596 | 4.028547714 | -0.448294702 | 0.653940524 | NA | NA | NA | 0 | 2.0558 | 0 | 0 | 0 | 0 |
| 29601 | ENSMUSG00000088358 | 0 | NA | NA | NA | NA | NA | Gm24008 | predicted gene, 24008 [Source:MGI Symb | 0 | 0 | 0 | 0 | 0 | 0 |
| 29602 | ENSMUSG00000088359 | 0 | NA | NA | NA | NA | NA | NA | NA | 0 | 0 | 0 | 0 | 0 | 0 |
| 29603 | ENSMUSG00000088360 | 0 | NA | NA | NA | NA | NA | NA | NA | 0 | 0 | 0 | 0 | 0 | 0 |
| 29604 | ENSMUSG00000088361 | 0 | NA | NA | NA | NA | NA | NA | NA | 0 | 0 | 0 | 0 | 0 | 0 |
| 29605 | ENSMUSG00000088362 | 0 | NA | NA | NA | NA | NA | Gm23498 | predicted gene, 23498 [Source:MGI Symb | 0 | 0 | 0 | 0 | 0 | 0 |
| 29606 | ENSMUSG00000088363 | 0 | NA | NA | NA | NA | NA | Gm23497 | predicted gene, 23497 [Source:MGI Symb | 0 | 0 | 0 | 0 | 0 | 0 |
| 29607 | ENSMUSG00000088364 | 0 | NA | NA | NA | NA | NA | Gm28020 | predicted gene, 28020 [Source:MGI Symb | 0 | 0 | 0 | 0 | 0 | 0 |
| 29608 | ENSMUSG00000088365 | 0 | NA | NA | NA | NA | NA | NA | NA | 0 | 0 | 0 | 0 | 0 | 0 |
| 29609 | ENSMUSG00000088366 | 0 | NA | NA | NA | NA | NA | NA | NA | 0 | 0 | 0 | 0 | 0 | 0 |
| 29610 | ENSMUSG00000088367 | 0 | NA | NA | NA | NA | NA | NA | NA | 0 | 0 | 0 | 0 | 0 | 0 |
| 29611 | ENSMUSG00000088368 | 0 | NA | NA | NA | NA | NA | NA | NA | 0 | 0 | 0 | 0 | 0 | 0 |
| 29612 | ENSMUSG00000088369 | 0 | NA | NA | NA | NA | NA | NA | NA | 0 | 0 | 0 | 0 | 0 | 0 |
| 29613 | ENSMUSG00000088370 | 0 | NA | NA | NA | NA | NA | Gm22880 | predicted gene, 22880 [Source:MGI Symb | 0 | 0 | 0 | 0 | 0 | 0 |
| 29614 | ENSMUSG00000088371 | 0 | NA | NA | NA | NA | NA | NA | NA | 0 | 0 | 0 | 0 | 0 | 0 |
| 29615 | ENSMUSG00000088372 | 3.272148753 | -0.414271279 | 1.324635989 | -0.312743488 | 0.754475553 | NA | Gm22881 | predicted gene, 22881 [Source:MGI Symb | 8.031 | 3.0838 | 0 | 1.928 | 4.3625 | 2.2277 |
| 29616 | ENSMUSG00000088373 | 0 | NA | NA | NA | NA | NA | Mir1956 | microRNA 1956 [Source:MGI Symbol;Acc: | 0 | 0 | 0 | 0 | 0 | 0 |
| 29617 | ENSMUSG00000088374 | 0 | NA | NA | NA | NA | NA | Gm22882 | predicted gene, 22882 [Source:MGI Symb | 0 | 0 | 0 | 0 | 0 | 0 |
| 29618 | ENSMUSG00000088377 | 0 | NA | NA | NA | NA | NA | Gm25183 | predicted gene, 25183 [Source:MGI Symb | 0 | 0 | 0 | 0 | 0 | 0 |
| 29619 | ENSMUSG00000088378 | 0.160662974 | 1.126924175 | 4.080472857 | 0.276174898 | 0.782413736 | NA | Gm25184 | predicted gene, 25184 [Source:MGI Symb | 0 | 0 | 0 | 0.964 | 0 | 0 |
| 29620 | ENSMUSG00000088379 | 0 | NA | NA | NA | NA | NA | Gm25185 | predicted gene, 25185 [Source:MGI Symb | 0 | 0 | 0 | 0 | 0 | 0 |

| | | | | | | | | | | | | | | | |
|---|---|---|---|---|---|---|---|---|---|---|---|---|---|---|---|
| 29621 | ENSMUSG00000088380 | 0 | NA | NA | NA | NA | NA | Gm24686 | predicted gene, 24686 [Source:MGI Symb | 0 | 0 | 0 | 0 | 0 | 0 |
| 29622 | ENSMUSG00000088383 | 0 | NA | NA | NA | NA | NA | Gm24687 | predicted gene, 24687 [Source:MGI Symb | 0 | 0 | 0 | 0 | 0 | 0 |
| 29623 | ENSMUSG00000088384 | 0 | NA | NA | NA | NA | NA | NA | NA | 0 | 0 | 0 | 0 | 0 | 0 |
| 29624 | ENSMUSG00000088385 | 0 | NA | NA | NA | NA | NA | NA | NA | 0 | 0 | 0 | 0 | 0 | 0 |
| 29625 | ENSMUSG00000088386 | 0 | NA | NA | NA | NA | NA | NA | NA | 0 | 0 | 0 | 0 | 0 | 0 |
| 29626 | ENSMUSG00000088387 | 0 | NA | NA | NA | NA | NA | Gm24691 | predicted gene, 24691 [Source:MGI Symb | 0 | 0 | 0 | 0 | 0 | 0 |
| 29627 | ENSMUSG00000088389 | 0 | NA | NA | NA | NA | NA | NA | NA | 0 | 0 | 0 | 0 | 0 | 0 |
| 29628 | ENSMUSG00000088391 | 0 | NA | NA | NA | NA | NA | Gm26310 | predicted gene, 26310 [Source:MGI Symb | 0 | 0 | 0 | 0 | 0 | 0 |
| 29629 | ENSMUSG00000088392 | 0 | NA | NA | NA | NA | NA | Gm26307 | predicted gene, 26307 [Source:MGI Symb | 0 | 0 | 0 | 0 | 0 | 0 |
| 29630 | ENSMUSG00000088393 | 0 | NA | NA | NA | NA | NA | NA | NA | 0 | 0 | 0 | 0 | 0 | 0 |
| 29631 | ENSMUSG00000088394 | 0 | NA | NA | NA | NA | NA | Gm26312 | predicted gene, 26312 [Source:MGI Symb | 0 | 0 | 0 | 0 | 0 | 0 |
| 29632 | ENSMUSG00000088395 | 0 | NA | NA | NA | NA | NA | NA | NA | 0 | 0 | 0 | 0 | 0 | 0 |
| 29633 | ENSMUSG00000088397 | 0 | NA | NA | NA | NA | NA | NA | NA | 0 | 0 | 0 | 0 | 0 | 0 |
| 29634 | ENSMUSG00000088398 | 0 | NA | NA | NA | NA | NA | Gm26306 | predicted gene, 26306 [Source:MGI Symb | 0 | 0 | 0 | 0 | 0 | 0 |
| 29635 | ENSMUSG00000088400 | 0.133850373 | -0.796629176 | 4.080472857 | -0.195229623 | 0.84521319 | NA | NA | NA | 0.8031 | 0 | 0 | 0 | 0 | 0 |
| 29636 | ENSMUSG00000088402 | 0 | NA | NA | NA | NA | NA | Gm25887 | predicted gene, 25887 [Source:MGI Symb | 0 | 0 | 0 | 0 | 0 | 0 |
| 29637 | ENSMUSG00000088403 | 0 | NA | NA | NA | NA | NA | NA | NA | 0 | 0 | 0 | 0 | 0 | 0 |
| 29638 | ENSMUSG00000088404 | 0 | NA | NA | NA | NA | NA | NA | NA | 0 | 0 | 0 | 0 | 0 | 0 |
| 29639 | ENSMUSG00000088405 | 0 | NA | NA | NA | NA | NA | Gm25886 | predicted gene, 25886 [Source:MGI Symb | 0 | 0 | 0 | 0 | 0 | 0 |
| 29640 | ENSMUSG00000088406 | 0 | NA | NA | NA | NA | NA | NA | NA | 0 | 0 | 0 | 0 | 0 | 0 |
| 29641 | ENSMUSG00000088407 | 0 | NA | NA | NA | NA | NA | Gm25884 | predicted gene, 25884 [Source:MGI Symb | 0 | 0 | 0 | 0 | 0 | 0 |
| 29642 | ENSMUSG00000088408 | 3.868516814 | 0.345036718 | 1.237563739 | 0.27880319 | 0.780395866 | NA | Gm25881 | predicted gene, 25881 [Source:MGI Symb | 3.2124 | 2.0558 | 5.0312 | 9.6398 | 3.2719 | 0 |
| 29643 | ENSMUSG00000088409 | 0 | NA | NA | NA | NA | NA | NA | NA | 0 | 0 | 0 | 0 | 0 | 0 |
| 29644 | ENSMUSG00000088410 | 0.185640647 | 1.126924175 | 4.080472857 | 0.276174898 | 0.782413736 | NA | Gm24240 | predicted gene, 24240 [Source:MGI Symb | 0 | 0 | 0 | 0 | 0 | 1.1138 |
| 29645 | ENSMUSG00000088412 | 0.160662974 | 1.126924175 | 4.080472857 | 0.276174898 | 0.782413736 | NA | Gm24241 | predicted gene, 24241 [Source:MGI Symb | 0 | 0 | 0 | 0.964 | 0 | 0 |
| 29646 | ENSMUSG00000088414 | 0 | NA | NA | NA | NA | NA | Mir1967 | microRNA 1967 [Source:MGI Symbol;Acc: | 0 | 0 | 0 | 0 | 0 | 0 |
| 29647 | ENSMUSG00000088415 | 0 | NA | NA | NA | NA | NA | NA | NA | 0 | 0 | 0 | 0 | 0 | 0 |
| 29648 | ENSMUSG00000088416 | 0 | NA | NA | NA | NA | NA | NA | NA | 0 | 0 | 0 | 0 | 0 | 0 |
| 29649 | ENSMUSG00000088417 | 0 | NA | NA | NA | NA | NA | NA | NA | 0 | 0 | 0 | 0 | 0 | 0 |
| 29650 | ENSMUSG00000088418 | 0 | NA | NA | NA | NA | NA | Gm26162 | predicted gene, 26162 [Source:MGI Symb | 0 | 0 | 0 | 0 | 0 | 0 |
| 29651 | ENSMUSG00000088419 | 0 | NA | NA | NA | NA | NA | Gm24236 | predicted gene, 24236 [Source:MGI Symb | 0 | 0 | 0 | 0 | 0 | 0 |
| 29652 | ENSMUSG00000088420 | 0 | NA | NA | NA | NA | NA | Gm22551 | predicted gene, 22551 [Source:MGI Symb | 0 | 0 | 0 | 0 | 0 | 0 |
| 29653 | ENSMUSG00000088421 | 0 | NA | NA | NA | NA | NA | NA | NA | 0 | 0 | 0 | 0 | 0 | 0 |
| 29654 | ENSMUSG00000088422 | 0 | NA | NA | NA | NA | NA | NA | NA | 0 | 0 | 0 | 0 | 0 | 0 |
| 29655 | ENSMUSG00000088423 | 0 | NA | NA | NA | NA | NA | Gm22554 | predicted gene, 22554 [Source:MGI Symb | 0 | 0 | 0 | 0 | 0 | 0 |
| 29656 | ENSMUSG00000088425 | 0 | NA | NA | NA | NA | NA | Gm24422 | predicted gene, 24422 [Source:MGI Symb | 0 | 0 | 0 | 0 | 0 | 0 |
| 29657 | ENSMUSG00000088426 | 0 | NA | NA | NA | NA | NA | NA | NA | 0 | 0 | 0 | 0 | 0 | 0 |
| 29658 | ENSMUSG00000088428 | 0 | NA | NA | NA | NA | NA | NA | NA | 0 | 0 | 0 | 0 | 0 | 0 |
| 29659 | ENSMUSG00000088429 | 0 | NA | NA | NA | NA | NA | Gm22557 | predicted gene, 22557 [Source:MGI Symb | 0 | 0 | 0 | 0 | 0 | 0 |
| 29660 | ENSMUSG00000088432 | 0 | NA | NA | NA | NA | NA | Gm25384 | predicted gene, 25384 [Source:MGI Symb | 0 | 0 | 0 | 0 | 0 | 0 |
| 29661 | ENSMUSG00000088433 | 0 | NA | NA | NA | NA | NA | NA | NA | 0 | 0 | 0 | 0 | 0 | 0 |
| 29662 | ENSMUSG00000088435 | 0 | NA | NA | NA | NA | NA | Gm25383 | predicted gene, 25383 [Source:MGI Symb | 0 | 0 | 0 | 0 | 0 | 0 |
| 29663 | ENSMUSG00000088436 | 0 | NA | NA | NA | NA | NA | NA | NA | 0 | 0 | 0 | 0 | 0 | 0 |
| 29664 | ENSMUSG00000088437 | 0 | NA | NA | NA | NA | NA | Gm24198 | predicted gene, 24198 [Source:MGI Symb | 0 | 0 | 0 | 0 | 0 | 0 |
| 29665 | ENSMUSG00000088438 | 0 | NA | NA | NA | NA | NA | NA | NA | 0 | 0 | 0 | 0 | 0 | 0 |
| 29666 | ENSMUSG00000088439 | 0 | NA | NA | NA | NA | NA | NA | NA | 0 | 0 | 0 | 0 | 0 | 0 |
| 29667 | ENSMUSG00000088440 | 0 | NA | NA | NA | NA | NA | Gm23555 | predicted gene, 23555 [Source:MGI Symb | 0 | 0 | 0 | 0 | 0 | 0 |
| 29668 | ENSMUSG00000088441 | 0 | NA | NA | NA | NA | NA | Gm25599 | predicted gene, 25599 [Source:MGI Symb | 0 | 0 | 0 | 0 | 0 | 0 |
| 29669 | ENSMUSG00000088442 | 0.658518091 | -1.320623475 | 2.595005939 | -0.508909616 | 0.61081558 | NA | NA | NA | 0.8031 | 1.0279 | 1.0062 | 0 | 0 | 1.1138 |
| 29670 | ENSMUSG00000088443 | 0 | NA | NA | NA | NA | NA | Gm23554 | predicted gene, 23554 [Source:MGI Symb | 0 | 0 | 0 | 0 | 0 | 0 |
| 29671 | ENSMUSG00000088444 | 0 | NA | NA | NA | NA | NA | Gm22187 | predicted gene, 22187 [Source:MGI Symb | 0 | 0 | 0 | 0 | 0 | 0 |
| 29672 | ENSMUSG00000088445 | 0 | NA | NA | NA | NA | NA | Gm23557 | predicted gene, 23557 [Source:MGI Symb | 0 | 0 | 0 | 0 | 0 | 0 |
| 29673 | ENSMUSG00000088446 | 0 | NA | NA | NA | NA | NA | NA | NA | 0 | 0 | 0 | 0 | 0 | 0 |
| 29674 | ENSMUSG00000088447 | 0 | NA | NA | NA | NA | NA | Gm22186 | predicted gene, 22186 [Source:MGI Symb | 0 | 0 | 0 | 0 | 0 | 0 |
| 29675 | ENSMUSG00000088448 | 0 | NA | NA | NA | NA | NA | NA | NA | 0 | 0 | 0 | 0 | 0 | 0 |
| 29676 | ENSMUSG00000088449 | 0 | NA | NA | NA | NA | NA | NA | NA | 0 | 0 | 0 | 0 | 0 | 0 |

| | | | | | | | | | | | | | | | |
|---|---|---|---|---|---|---|---|---|---|---|---|---|---|---|---|
| 29677 | ENSMUSG00000088450 | 0 | NA | NA | NA | NA | NA | Gm26369 | predicted gene, 26369 [Source:MGI Symb | 0 | 0 | 0 | 0 | 0 | 0 |
| 29678 | ENSMUSG00000088451 | 0 | NA | NA | NA | NA | NA | Gm26368 | predicted gene, 26368 [Source:MGI Symb | 0 | 0 | 0 | 0 | 0 | 0 |
| 29679 | ENSMUSG00000088453 | 0 | NA | NA | NA | NA | NA | NA | NA | 0 | 0 | 0 | 0 | 0 | 0 |
| 29680 | ENSMUSG00000088454 | 0 | NA | NA | NA | NA | NA | Gm26371 | predicted gene, 26371 [Source:MGI Symb | 0 | 0 | 0 | 0 | 0 | 0 |
| 29681 | ENSMUSG00000088456 | 0.339027071 | -1.79174149 | 3.919585585 | -0.457125237 | 0.647581032 | NA | NA | NA | 0 | 1.0279 | 1.0062 | 0 | 0 | 0 |
| 29682 | ENSMUSG00000088458 | 0 | NA | NA | NA | NA | NA | Gm26373 | predicted gene, 26373 [Source:MGI Symb | 0 | 0 | 0 | 0 | 0 | 0 |
| 29683 | ENSMUSG00000088459 | 0 | NA | NA | NA | NA | NA | Mir1941 | microRNA 1941 [Source:MGI Symbol;Acc: | 0 | 0 | 0 | 0 | 0 | 0 |
| 29684 | ENSMUSG00000088460 | 0 | NA | NA | NA | NA | NA | NA | NA | 0 | 0 | 0 | 0 | 0 | 0 |
| 29685 | ENSMUSG00000088461 | 0 | NA | NA | NA | NA | NA | NA | NA | 0 | 0 | 0 | 0 | 0 | 0 |
| 29686 | ENSMUSG00000088463 | 0 | NA | NA | NA | NA | NA | NA | NA | 0 | 0 | 0 | 0 | 0 | 0 |
| 29687 | ENSMUSG00000088464 | 0.549180077 | 2.659271622 | 3.249668912 | 0.818320787 | 0.41317404 | NA | Mir1970 | microRNA 1970 [Source:MGI Symbol;Acc: | 0 | 0 | 0 | 0 | 2.1812 | 1.1138 |
| 29688 | ENSMUSG00000088465 | 0 | NA | NA | NA | NA | NA | Gm24745 | predicted gene, 24745 [Source:MGI Symb | 0 | 0 | 0 | 0 | 0 | 0 |
| 29689 | ENSMUSG00000088468 | 0 | NA | NA | NA | NA | NA | Gm24750 | predicted gene, 24750 [Source:MGI Symb | 0 | 0 | 0 | 0 | 0 | 0 |
| 29690 | ENSMUSG00000088469 | 0 | NA | NA | NA | NA | NA | Gm24751 | predicted gene, 24751 [Source:MGI Symb | 0 | 0 | 0 | 0 | 0 | 0 |
| 29691 | ENSMUSG00000088470 | 0 | NA | NA | NA | NA | NA | NA | NA | 0 | 0 | 0 | 0 | 0 | 0 |
| 29692 | ENSMUSG00000088471 | 0 | NA | NA | NA | NA | NA | Gm23080 | predicted gene, 23080 [Source:MGI Symb | 0 | 0 | 0 | 0 | 0 | 0 |
| 29693 | ENSMUSG00000088472 | 0 | NA | NA | NA | NA | NA | Gm23079 | predicted gene, 23079 [Source:MGI Symb | 0 | 0 | 0 | 0 | 0 | 0 |
| 29694 | ENSMUSG00000088473 | 0 | NA | NA | NA | NA | NA | NA | NA | 0 | 0 | 0 | 0 | 0 | 0 |
| 29695 | ENSMUSG00000088474 | 0 | NA | NA | NA | NA | NA | NA | NA | 0 | 0 | 0 | 0 | 0 | 0 |
| 29696 | ENSMUSG00000088475 | 1.26003638 | -0.648637764 | 2.020901123 | -0.320964621 | 0.748237202 | NA | NA | NA | 1.6062 | 2.0558 | 1.0062 | 2.8919 | 0 | 0 |
| 29697 | ENSMUSG00000088476 | 0 | NA | NA | NA | NA | NA | Gm23083 | predicted gene, 23083 [Source:MGI Symb | 0 | 0 | 0 | 0 | 0 | 0 |
| 29698 | ENSMUSG00000088477 | 0 | NA | NA | NA | NA | NA | Gm23082 | predicted gene, 23082 [Source:MGI Symb | 0 | 0 | 0 | 0 | 0 | 0 |
| 29699 | ENSMUSG00000088478 | 0.615839296 | 1.537131441 | 3.17677869 | 0.483864817 | 0.628481811 | NA | Mir1946b | microRNA 1946b [Source:MGI Symbol;Acc | 0.8031 | 0 | 0 | 2.8919 | 0 | 0 |
| 29700 | ENSMUSG00000088479 | 0.133850373 | -0.796629176 | 4.080472857 | -0.195229623 | 0.84521319 | NA | Gm23077 | predicted gene, 23077 [Source:MGI Symb | 0.8031 | 0 | 0 | 0 | 0 | 0 |
| 29701 | ENSMUSG00000088480 | 0 | NA | NA | NA | NA | NA | NA | NA | 0 | 0 | 0 | 0 | 0 | 0 |
| 29702 | ENSMUSG00000088481 | 0 | NA | NA | NA | NA | NA | Gm23644 | predicted gene, 23644 [Source:MGI Symb | 0 | 0 | 0 | 0 | 0 | 0 |
| 29703 | ENSMUSG00000088482 | 0 | NA | NA | NA | NA | NA | Gm23641 | predicted gene, 23641 [Source:MGI Symb | 0 | 0 | 0 | 0 | 0 | 0 |
| 29704 | ENSMUSG00000088483 | 0 | NA | NA | NA | NA | NA | NA | NA | 0 | 0 | 0 | 0 | 0 | 0 |
| 29705 | ENSMUSG00000088484 | 0 | NA | NA | NA | NA | NA | Gm23592 | predicted gene, 23592 [Source:MGI Symb | 0 | 0 | 0 | 0 | 0 | 0 |
| 29706 | ENSMUSG00000088485 | 0.17131964 | -0.796629176 | 4.080472857 | -0.195229623 | 0.84521319 | NA | NA | NA | 0 | 1.0279 | 0 | 0 | 0 | 0 |
| 29707 | ENSMUSG00000088487 | 0 | NA | NA | NA | NA | NA | NA | NA | 0 | 0 | 0 | 0 | 0 | 0 |
| 29708 | ENSMUSG00000088490 | 1.829217821 | -1.343574369 | 1.678238561 | -0.800586043 | 0.423371333 | NA | Gm24400 | predicted gene, 24400 [Source:MGI Symb | 0.8031 | 4.1117 | 3.0187 | 1.928 | 0 | 1.1138 |
| 29709 | ENSMUSG00000088491 | 0 | NA | NA | NA | NA | NA | Mir1953 | microRNA 1953 [Source:MGI Symbol;Acc: | 0 | 0 | 0 | 0 | 0 | 0 |
| 29710 | ENSMUSG00000088493 | 0 | NA | NA | NA | NA | NA | Gm24401 | predicted gene, 24401 [Source:MGI Symb | 0 | 0 | 0 | 0 | 0 | 0 |
| 29711 | ENSMUSG00000088494 | 0 | NA | NA | NA | NA | NA | Gm24399 | predicted gene, 24399 [Source:MGI Symb | 0 | 0 | 0 | 0 | 0 | 0 |
| 29712 | ENSMUSG00000088495 | 0 | NA | NA | NA | NA | NA | Gm24398 | predicted gene, 24398 [Source:MGI Symb | 0 | 0 | 0 | 0 | 0 | 0 |
| 29713 | ENSMUSG00000088497 | 0 | NA | NA | NA | NA | NA | NA | NA | 0 | 0 | 0 | 0 | 0 | 0 |
| 29714 | ENSMUSG00000088498 | 0.994779605 | 1.370136611 | 2.701221166 | 0.507228593 | 0.611994435 | NA | NA | NA | 1.6062 | 0 | 0 | 0 | 4.3625 | 0 |
| 29715 | ENSMUSG00000088499 | 0 | NA | NA | NA | NA | NA | Gm22654 | predicted gene, 22654 [Source:MGI Symb | 0 | 0 | 0 | 0 | 0 | 0 |
| 29716 | ENSMUSG00000088500 | 0 | NA | NA | NA | NA | NA | NA | NA | 0 | 0 | 0 | 0 | 0 | 0 |
| 29717 | ENSMUSG00000088501 | 0 | NA | NA | NA | NA | NA | Gm22788 | predicted gene, 22788 [Source:MGI Symb | 0 | 0 | 0 | 0 | 0 | 0 |
| 29718 | ENSMUSG00000088502 | 0 | NA | NA | NA | NA | NA | NA | NA | 0 | 0 | 0 | 0 | 0 | 0 |
| 29719 | ENSMUSG00000088504 | 1.299484532 | 3.902703431 | 2.722060246 | 1.433731468 | 0.151648909 | NA | Gm26490 | predicted gene, 26490 [Source:MGI Symb | 0 | 0 | 0 | 0 | 0 | 7.7969 |
| 29720 | ENSMUSG00000088505 | 0 | NA | NA | NA | NA | NA | Gm26489 | predicted gene, 26489 [Source:MGI Symb | 0 | 0 | 0 | 0 | 0 | 0 |
| 29721 | ENSMUSG00000088506 | 0.480153995 | 1.043847617 | 3.151673025 | 0.331204287 | 0.740490182 | NA | Gm24409 | predicted gene, 24409 [Source:MGI Symb | 0.8031 | 0 | 0 | 0.964 | 0 | 1.1138 |
| 29722 | ENSMUSG00000088509 | 0 | NA | NA | NA | NA | NA | Gm22789 | predicted gene, 22789 [Source:MGI Symb | 0 | 0 | 0 | 0 | 0 | 0 |
| 29723 | ENSMUSG00000088510 | 0 | NA | NA | NA | NA | NA | Gm23036 | predicted gene, 23036 [Source:MGI Symb | 0 | 0 | 0 | 0 | 0 | 0 |
| 29724 | ENSMUSG00000088511 | 0 | NA | NA | NA | NA | NA | Gm24435 | predicted gene, 24435 [Source:MGI Symb | 0 | 0 | 0 | 0 | 0 | 0 |
| 29725 | ENSMUSG00000088512 | 0 | NA | NA | NA | NA | NA | Gm24467 | predicted gene, 24467 [Source:MGI Symb | 0 | 0 | 0 | 0 | 0 | 0 |
| 29726 | ENSMUSG00000088514 | 0 | NA | NA | NA | NA | NA | NA | NA | 0 | 0 | 0 | 0 | 0 | 0 |
| 29727 | ENSMUSG00000088515 | 0 | NA | NA | NA | NA | NA | Gm23043 | predicted gene, 23043 [Source:MGI Symb | 0 | 0 | 0 | 0 | 0 | 0 |
| 29728 | ENSMUSG00000088516 | 0.185640647 | 1.126924175 | 4.080472857 | 0.276174898 | 0.782413736 | NA | Gm23044 | predicted gene, 23044 [Source:MGI Symb | 0 | 0 | 0 | 0 | 0 | 1.1138 |
| 29729 | ENSMUSG00000088517 | 0 | NA | NA | NA | NA | NA | Gm24437 | predicted gene, 24437 [Source:MGI Symb | 0 | 0 | 0 | 0 | 0 | 0 |
| 29730 | ENSMUSG00000088518 | 0 | NA | NA | NA | NA | NA | NA | NA | 0 | 0 | 0 | 0 | 0 | 0 |
| 29731 | ENSMUSG00000088519 | 0 | NA | NA | NA | NA | NA | NA | NA | 0 | 0 | 0 | 0 | 0 | 0 |
| 29732 | ENSMUSG00000088521 | 0 | NA | NA | NA | NA | NA | Gm26088 | predicted gene, 26088 [Source:MGI Symb | 0 | 0 | 0 | 0 | 0 | 0 |

| | | | | | | | | | | | | | | | | |
|---|---|---|---|---|---|---|---|---|---|---|---|---|---|---|---|---|
| 29733 | ENSMUSG00000088522 | 0 | NA | NA | NA | NA | NA | NA | NA | NA | 0 | 0 | 0 | 0 | 0 | 0 |
| 29734 | ENSMUSG00000088523 | 0 | NA | NA | NA | NA | NA | NA | NA | NA | 0 | 0 | 0 | 0 | 0 | 0 |
| 29735 | ENSMUSG00000088524 | 0 | NA | NA | NA | NA | NA | NA | Snord2 | small nucleolar RNA, C/D box 2 [Source:M | 0 | 0 | 0 | 0 | 0 | 0 |
| 29736 | ENSMUSG00000088525 | 0 | NA | NA | NA | NA | NA | NA | Gm26085 | predicted gene, 26085 [Source:MGI Symb | 0 | 0 | 0 | 0 | 0 | 0 |
| 29737 | ENSMUSG00000088526 | 0 | NA | NA | NA | NA | NA | NA | Gm26087 | predicted gene, 26087 [Source:MGI Symb | 0 | 0 | 0 | 0 | 0 | 0 |
| 29738 | ENSMUSG00000088527 | 0 | NA | NA | NA | NA | NA | NA | NA | NA | 0 | 0 | 0 | 0 | 0 | 0 |
| 29739 | ENSMUSG00000088528 | 0.36353943 | 2.065989254 | 4.034137301 | 0.512126658 | 0.608562368 | NA | Gm26084 | predicted gene, 26084 [Source:MGI Symb | 0 | 0 | 0 | 0 | 2.1812 | 0 |
| 29740 | ENSMUSG00000088529 | 0 | NA | NA | NA | NA | NA | NA | NA | NA | 0 | 0 | 0 | 0 | 0 | 0 |
| 29741 | ENSMUSG00000088530 | 0 | NA | NA | NA | NA | NA | NA | Gm23280 | predicted gene, 23280 [Source:MGI Symb | 0 | 0 | 0 | 0 | 0 | 0 |
| 29742 | ENSMUSG00000088532 | 0 | NA | NA | NA | NA | NA | NA | NA | NA | 0 | 0 | 0 | 0 | 0 | 0 |
| 29743 | ENSMUSG00000088533 | 0 | NA | NA | NA | NA | NA | NA | Gm24998 | predicted gene, 24998 [Source:MGI Symb | 0 | 0 | 0 | 0 | 0 | 0 |
| 29744 | ENSMUSG00000088535 | 0 | NA | NA | NA | NA | NA | NA | NA | NA | 0 | 0 | 0 | 0 | 0 | 0 |
| 29745 | ENSMUSG00000088536 | 0 | NA | NA | NA | NA | NA | NA | Gm23278 | predicted gene, 23278 [Source:MGI Symb | 0 | 0 | 0 | 0 | 0 | 0 |
| 29746 | ENSMUSG00000088537 | 0 | NA | NA | NA | NA | NA | NA | NA | NA | 0 | 0 | 0 | 0 | 0 | 0 |
| 29747 | ENSMUSG00000088538 | 0 | NA | NA | NA | NA | NA | NA | Gm23276 | predicted gene, 23276 [Source:MGI Symb | 0 | 0 | 0 | 0 | 0 | 0 |
| 29748 | ENSMUSG00000088539 | 0 | NA | NA | NA | NA | NA | NA | Gm23277 | predicted gene, 23277 [Source:MGI Symb | 0 | 0 | 0 | 0 | 0 | 0 |
| 29749 | ENSMUSG00000088541 | 0 | NA | NA | NA | NA | NA | NA | Gm24963 | predicted gene, 24963 [Source:MGI Symb | 0 | 0 | 0 | 0 | 0 | 0 |
| 29750 | ENSMUSG00000088542 | 0 | NA | NA | NA | NA | NA | NA | NA | NA | 0 | 0 | 0 | 0 | 0 | 0 |
| 29751 | ENSMUSG00000088543 | 0 | NA | NA | NA | NA | NA | NA | NA | NA | 0 | 0 | 0 | 0 | 0 | 0 |
| 29752 | ENSMUSG00000088544 | 0.698954292 | 0.126053087 | 3.048972385 | 0.04134281 | 0.967022605 | NA | Mir1945 | microRNA 1945 [Source:MGI Symbol;Acc: | 0 | 0 | 2.0125 | 0 | 2.1812 | 0 |
| 29753 | ENSMUSG00000088545 | 0.787158936 | 0.679629554 | 2.65851484 | 0.255642566 | 0.798226848 | NA | Gm25982 | predicted gene, 25982 [Source:MGI Symb | 0.8031 | 1.0279 | 0 | 2.8919 | 0 | 0 |
| 29754 | ENSMUSG00000088546 | 0 | NA | NA | NA | NA | NA | NA | NA | NA | 0 | 0 | 0 | 0 | 0 | 0 |
| 29755 | ENSMUSG00000088547 | 0 | NA | NA | NA | NA | NA | NA | Gm24965 | predicted gene, 24965 [Source:MGI Symb | 0 | 0 | 0 | 0 | 0 | 0 |
| 29756 | ENSMUSG00000088548 | 0 | NA | NA | NA | NA | NA | NA | NA | NA | 0 | 0 | 0 | 0 | 0 | 0 |
| 29757 | ENSMUSG00000088549 | 0 | NA | NA | NA | NA | NA | NA | NA | NA | 0 | 0 | 0 | 0 | 0 | 0 |
| 29758 | ENSMUSG00000088550 | 0 | NA | NA | NA | NA | NA | NA | Gm22099 | predicted gene, 22099 [Source:MGI Symb | 0 | 0 | 0 | 0 | 0 | 0 |
| 29759 | ENSMUSG00000088551 | 0 | NA | NA | NA | NA | NA | NA | Gm22100 | predicted gene, 22100 [Source:MGI Symb | 0 | 0 | 0 | 0 | 0 | 0 |
| 29760 | ENSMUSG00000088552 | 0.301557804 | -1.64335916 | 4.042814367 | -0.406488899 | 0.684383412 | NA | Mir1957a | microRNA 1957a [Source:MGI Symbol;Acc | 0.8031 | 0 | 1.0062 | 0 | 0 | 0 |
| 29761 | ENSMUSG00000088553 | 0 | NA | NA | NA | NA | NA | NA | NA | NA | 0 | 0 | 0 | 0 | 0 | 0 |
| 29762 | ENSMUSG00000088554 | 0 | NA | NA | NA | NA | NA | NA | NA | NA | 0 | 0 | 0 | 0 | 0 | 0 |
| 29763 | ENSMUSG00000088555 | 0.48903338 | 0.95637552 | 3.456690196 | 0.276673773 | 0.782030612 | NA | NA | NA | 0 | 0 | 1.0062 | 1.928 | 0 | 0 |
| 29764 | ENSMUSG00000088558 | 0 | NA | NA | NA | NA | NA | NA | Gm22506 | predicted gene, 22506 [Source:MGI Symb | 0 | 0 | 0 | 0 | 0 | 0 |
| 29765 | ENSMUSG00000088559 | 0 | NA | NA | NA | NA | NA | NA | Mir1981 | microRNA 1981 [Source:MGI Symbol;Acc: | 0 | 0 | 0 | 0 | 0 | 0 |
| 29766 | ENSMUSG00000088562 | 0 | NA | NA | NA | NA | NA | NA | NA | NA | 0 | 0 | 0 | 0 | 0 | 0 |
| 29767 | ENSMUSG00000088563 | 0.133850373 | -0.796629176 | 4.080472857 | -0.195229623 | 0.84521319 | NA | Gm23779 | predicted gene, 23779 [Source:MGI Symb | 0.8031 | 0 | 0 | 0 | 0 | 0 |
| 29768 | ENSMUSG00000088564 | 0 | NA | NA | NA | NA | NA | NA | NA | NA | 0 | 0 | 0 | 0 | 0 | 0 |
| 29769 | ENSMUSG00000088565 | 0 | NA | NA | NA | NA | NA | NA | NA | NA | 0 | 0 | 0 | 0 | 0 | 0 |
| 29770 | ENSMUSG00000088566 | 0 | NA | NA | NA | NA | NA | NA | NA | NA | 0 | 0 | 0 | 0 | 0 | 0 |
| 29771 | ENSMUSG00000088570 | 0 | NA | NA | NA | NA | NA | NA | NA | NA | 0 | 0 | 0 | 0 | 0 | 0 |
| 29772 | ENSMUSG00000088571 | 0 | NA | NA | NA | NA | NA | NA | NA | NA | 0 | 0 | 0 | 0 | 0 | 0 |
| 29773 | ENSMUSG00000088572 | 0 | NA | NA | NA | NA | NA | NA | NA | NA | 0 | 0 | 0 | 0 | 0 | 0 |
| 29774 | ENSMUSG00000088573 | 0.160662974 | 1.126924175 | 4.080472857 | 0.276174898 | 0.782413736 | NA | NA | NA | 0 | 0 | 0 | 0.964 | 0 | 0 |
| 29775 | ENSMUSG00000088574 | 0 | NA | NA | NA | NA | NA | NA | NA | NA | 0 | 0 | 0 | 0 | 0 | 0 |
| 29776 | ENSMUSG00000088575 | 0 | NA | NA | NA | NA | NA | NA | Gm24537 | predicted gene, 24537 [Source:MGI Symb | 0 | 0 | 0 | 0 | 0 | 0 |
| 29777 | ENSMUSG00000088576 | 0 | NA | NA | NA | NA | NA | NA | NA | NA | 0 | 0 | 0 | 0 | 0 | 0 |
| 29778 | ENSMUSG00000088577 | 0 | NA | NA | NA | NA | NA | NA | Gm25456 | predicted gene, 25456 [Source:MGI Symb | 0 | 0 | 0 | 0 | 0 | 0 |
| 29779 | ENSMUSG00000088578 | 0 | NA | NA | NA | NA | NA | NA | Gm25459 | predicted gene, 25459 [Source:MGI Symb | 0 | 0 | 0 | 0 | 0 | 0 |
| 29780 | ENSMUSG00000088579 | 0 | NA | NA | NA | NA | NA | NA | Gm25460 | predicted gene, 25460 [Source:MGI Symb | 0 | 0 | 0 | 0 | 0 | 0 |
| 29781 | ENSMUSG00000088580 | 0 | NA | NA | NA | NA | NA | NA | NA | NA | 0 | 0 | 0 | 0 | 0 | 0 |
| 29782 | ENSMUSG00000088581 | 0 | NA | NA | NA | NA | NA | NA | NA | NA | 0 | 0 | 0 | 0 | 0 | 0 |
| 29783 | ENSMUSG00000088582 | 0 | NA | NA | NA | NA | NA | NA | Gm22966 | predicted gene, 22966 [Source:MGI Symb | 0 | 0 | 0 | 0 | 0 | 0 |
| 29784 | ENSMUSG00000088583 | 0.556921942 | 2.679043159 | 3.638243063 | 0.736356289 | 0.461513899 | NA | Gm22965 | predicted gene, 22965 [Source:MGI Symb | 0 | 0 | 0 | 0 | 0 | 3.3415 |
| 29785 | ENSMUSG00000088585 | 0 | NA | NA | NA | NA | NA | NA | NA | NA | 0 | 0 | 0 | 0 | 0 | 0 |
| 29786 | ENSMUSG00000088587 | 0 | NA | NA | NA | NA | NA | NA | NA | NA | 0 | 0 | 0 | 0 | 0 | 0 |
| 29787 | ENSMUSG00000088589 | 0 | NA | NA | NA | NA | NA | NA | NA | NA | 0 | 0 | 0 | 0 | 0 | 0 |
| 29788 | ENSMUSG00000088591 | 0.167707431 | -0.796629176 | 4.080472857 | -0.195229623 | 0.84521319 | NA | NA | NA | 0 | 0 | 1.0062 | 0 | 0 | 0 |

| | | | | | | | | | | | | | | | |
|---|---|---|---|---|---|---|---|---|---|---|---|---|---|---|---|
| 29789 | ENSMUSG00000088594 | 0 | NA | NA | NA | NA | NA | Gm23965 | predicted gene, 23965 [Source:MGI Symb | 0 | 0 | 0 | 0 | 0 | 0 |
| 29790 | ENSMUSG00000088595 | 0 | NA | NA | NA | NA | NA | NA | NA | 0 | 0 | 0 | 0 | 0 | 0 |
| 29791 | ENSMUSG00000088596 | 0 | NA | NA | NA | NA | NA | NA | NA | 0 | 0 | 0 | 0 | 0 | 0 |
| 29792 | ENSMUSG00000088597 | 0 | NA | NA | NA | NA | NA | NA | NA | 0 | 0 | 0 | 0 | 0 | 0 |
| 29793 | ENSMUSG00000088598 | 0 | NA | NA | NA | NA | NA | NA | NA | 0 | 0 | 0 | 0 | 0 | 0 |
| 29794 | ENSMUSG00000088600 | 0 | NA | NA | NA | NA | NA | NA | NA | 0 | 0 | 0 | 0 | 0 | 0 |
| 29795 | ENSMUSG00000088601 | 0 | NA | NA | NA | NA | NA | NA | NA | 0 | 0 | 0 | 0 | 0 | 0 |
| 29796 | ENSMUSG00000088602 | 0.181769715 | 1.126924175 | 4.080472857 | 0.276174898 | 0.782413736 | NA | NA | NA | 0 | 0 | 0 | 0 | 1.0906 | 0 |
| 29797 | ENSMUSG00000088603 | 0 | NA | NA | NA | NA | NA | Gm24192 | predicted gene, 24192 [Source:MGI Symb | 0 | 0 | 0 | 0 | 0 | 0 |
| 29798 | ENSMUSG00000088605 | 0 | NA | NA | NA | NA | NA | NA | NA | 0 | 0 | 0 | 0 | 0 | 0 |
| 29799 | ENSMUSG00000088606 | 0 | NA | NA | NA | NA | NA | Mir1951 | microRNA 1951 [Source:MGI Symbol;Acc: | 0 | 0 | 0 | 0 | 0 | 0 |
| 29800 | ENSMUSG00000088608 | 0 | NA | NA | NA | NA | NA | NA | NA | 0 | 0 | 0 | 0 | 0 | 0 |
| 29801 | ENSMUSG00000088609 | 21.60780019 | 0.132700352 | 0.700450606 | 0.189449978 | 0.849740157 | 0.973207825 | NA | NA | 8.031 | 31.865 | 22.137 | 17.352 | 45.806 | 4.4554 |
| 29802 | ENSMUSG00000088611 | 0 | NA | NA | NA | NA | NA | NA | NA | 0 | 0 | 0 | 0 | 0 | 0 |
| 29803 | ENSMUSG00000088614 | 0 | NA | NA | NA | NA | NA | NA | NA | 0 | 0 | 0 | 0 | 0 | 0 |
| 29804 | ENSMUSG00000088615 | 0.813009891 | 0.960077285 | 2.883567268 | 0.332947768 | 0.739173711 | NA | NA | NA | 1.6062 | 0 | 0 | 0 | 3.2719 | 0 |
| 29805 | ENSMUSG00000088616 | 0 | NA | NA | NA | NA | NA | Gm26000 | predicted gene, 26000 [Source:MGI Symb | 0 | 0 | 0 | 0 | 0 | 0 |
| 29806 | ENSMUSG00000088617 | 0 | NA | NA | NA | NA | NA | NA | NA | 0 | 0 | 0 | 0 | 0 | 0 |
| 29807 | ENSMUSG00000088618 | 0 | NA | NA | NA | NA | NA | NA | NA | 0 | 0 | 0 | 0 | 0 | 0 |
| 29808 | ENSMUSG00000088620 | 0 | NA | NA | NA | NA | NA | Gm23032 | predicted gene, 23032 [Source:MGI Symb | 0 | 0 | 0 | 0 | 0 | 0 |
| 29809 | ENSMUSG00000088621 | 0 | NA | NA | NA | NA | NA | Gm23033 | predicted gene, 23033 [Source:MGI Symb | 0 | 0 | 0 | 0 | 0 | 0 |
| 29810 | ENSMUSG00000088622 | 0 | NA | NA | NA | NA | NA | Gm23316 | predicted gene, 23316 [Source:MGI Symb | 0 | 0 | 0 | 0 | 0 | 0 |
| 29811 | ENSMUSG00000088623 | 0.181769715 | 1.126924175 | 4.080472857 | 0.276174898 | 0.782413736 | NA | Gm23031 | predicted gene, 23031 [Source:MGI Symb | 0 | 0 | 0 | 0 | 1.0906 | 0 |
| 29812 | ENSMUSG00000088624 | 0 | NA | NA | NA | NA | NA | NA | NA | 0 | 0 | 0 | 0 | 0 | 0 |
| 29813 | ENSMUSG00000088625 | 2.023782819 | 0.549356353 | 1.718506742 | 0.319670758 | 0.749217929 | NA | Gm23318 | predicted gene, 23318 [Source:MGI Symb | 4.8186 | 0 | 0 | 2.8919 | 1.0906 | 3.3415 |
| 29814 | ENSMUSG00000088626 | 1.636268479 | -0.999810553 | 1.723127554 | -0.580230147 | 0.561759426 | NA | Gm23034 | predicted gene, 23034 [Source:MGI Symb | 2.4093 | 3.0838 | 1.0062 | 0 | 1.0906 | 2.2277 |
| 29815 | ENSMUSG00000088627 | 0 | NA | NA | NA | NA | NA | Gm23035 | predicted gene, 23035 [Source:MGI Symb | 0 | 0 | 0 | 0 | 0 | 0 |
| 29816 | ENSMUSG00000088628 | 0 | NA | NA | NA | NA | NA | Gm23315 | predicted gene, 23315 [Source:MGI Symb | 0 | 0 | 0 | 0 | 0 | 0 |
| 29817 | ENSMUSG00000088629 | 0 | NA | NA | NA | NA | NA | NA | NA | 0 | 0 | 0 | 0 | 0 | 0 |
| 29818 | ENSMUSG00000088630 | 0 | NA | NA | NA | NA | NA | Gm22312 | predicted gene, 22312 [Source:MGI Symb | 0 | 0 | 0 | 0 | 0 | 0 |
| 29819 | ENSMUSG00000088631 | 0.524408994 | -0.837196744 | 3.377823722 | -0.247850928 | 0.804249749 | NA | NA | NA | 0 | 2.0558 | 0 | 0 | 1.0906 | 0 |
| 29820 | ENSMUSG00000088632 | 0 | NA | NA | NA | NA | NA | NA | NA | 0 | 0 | 0 | 0 | 0 | 0 |
| 29821 | ENSMUSG00000088633 | 0 | NA | NA | NA | NA | NA | Gm22313 | predicted gene, 22313 [Source:MGI Symb | 0 | 0 | 0 | 0 | 0 | 0 |
| 29822 | ENSMUSG00000088634 | 0 | NA | NA | NA | NA | NA | Gm24958 | predicted gene, 24958 [Source:MGI Symb | 0 | 0 | 0 | 0 | 0 | 0 |
| 29823 | ENSMUSG00000088635 | 0 | NA | NA | NA | NA | NA | Gm22315 | predicted gene, 22315 [Source:MGI Symb | 0 | 0 | 0 | 0 | 0 | 0 |
| 29824 | ENSMUSG00000088637 | 0 | NA | NA | NA | NA | NA | NA | NA | 0 | 0 | 0 | 0 | 0 | 0 |
| 29825 | ENSMUSG00000088638 | 0 | NA | NA | NA | NA | NA | Gm25846 | predicted gene, 25846 [Source:MGI Symb | 0 | 0 | 0 | 0 | 0 | 0 |
| 29826 | ENSMUSG00000088639 | 0.167707431 | -0.796629176 | 4.080472857 | -0.195229623 | 0.84521319 | NA | Mir1955 | microRNA 1955 [Source:MGI Symbol;Acc: | 0 | 0 | 1.0062 | 0 | 0 | 0 |
| 29827 | ENSMUSG00000088640 | 0 | NA | NA | NA | NA | NA | Gm26482 | predicted gene, 26482 [Source:MGI Symb | 0 | 0 | 0 | 0 | 0 | 0 |
| 29828 | ENSMUSG00000088641 | 0 | NA | NA | NA | NA | NA | NA | NA | 0 | 0 | 0 | 0 | 0 | 0 |
| 29829 | ENSMUSG00000088643 | 0.531944269 | 2.618916564 | 3.285123409 | 0.797204926 | 0.425332026 | NA | NA | NA | 0 | 0 | 0 | 0.964 | 0 | 2.2277 |
| 29830 | ENSMUSG00000088644 | 0 | NA | NA | NA | NA | NA | Gm26485 | predicted gene, 26485 [Source:MGI Symb | 0 | 0 | 0 | 0 | 0 | 0 |
| 29831 | ENSMUSG00000088646 | 0 | NA | NA | NA | NA | NA | NA | NA | 0 | 0 | 0 | 0 | 0 | 0 |
| 29832 | ENSMUSG00000088647 | 0 | NA | NA | NA | NA | NA | Gm26487 | predicted gene, 26487 [Source:MGI Symb | 0 | 0 | 0 | 0 | 0 | 0 |
| 29833 | ENSMUSG00000088649 | 0 | NA | NA | NA | NA | NA | Gm26262 | predicted gene, 26262 [Source:MGI Symb | 0 | 0 | 0 | 0 | 0 | 0 |
| 29834 | ENSMUSG00000088650 | 0.890584499 | -0.80930233 | 2.520154863 | -0.321131983 | 0.748110374 | NA | NA | NA | 2.4093 | 0 | 1.0062 | 1.928 | 0 | 0 |
| 29835 | ENSMUSG00000088651 | 0 | NA | NA | NA | NA | NA | Gm24853 | predicted gene, 24853 [Source:MGI Symb | 0 | 0 | 0 | 0 | 0 | 0 |
| 29836 | ENSMUSG00000088653 | 0 | NA | NA | NA | NA | NA | Gm24851 | predicted gene, 24851 [Source:MGI Symb | 0 | 0 | 0 | 0 | 0 | 0 |
| 29837 | ENSMUSG00000088654 | 0 | NA | NA | NA | NA | NA | Gm24850 | predicted gene, 24850 [Source:MGI Symb | 0 | 0 | 0 | 0 | 0 | 0 |
| 29838 | ENSMUSG00000088656 | 0 | NA | NA | NA | NA | NA | Gm24848 | predicted gene, 24848 [Source:MGI Symb | 0 | 0 | 0 | 0 | 0 | 0 |
| 29839 | ENSMUSG00000088657 | 0 | NA | NA | NA | NA | NA | NA | NA | 0 | 0 | 0 | 0 | 0 | 0 |
| 29840 | ENSMUSG00000088658 | 0 | NA | NA | NA | NA | NA | Gm24846 | predicted gene, 24846 [Source:MGI Symb | 0 | 0 | 0 | 0 | 0 | 0 |
| 29841 | ENSMUSG00000088659 | 0 | NA | NA | NA | NA | NA | NA | NA | 0 | 0 | 0 | 0 | 0 | 0 |
| 29842 | ENSMUSG00000088660 | 0 | NA | NA | NA | NA | NA | NA | NA | 0 | 0 | 0 | 0 | 0 | 0 |
| 29843 | ENSMUSG00000088661 | 0 | NA | NA | NA | NA | NA | NA | NA | 0 | 0 | 0 | 0 | 0 | 0 |
| 29844 | ENSMUSG00000088663 | 0 | NA | NA | NA | NA | NA | NA | NA | 0 | 0 | 0 | 0 | 0 | 0 |

| | | | | | | | | | | | | | | | |
|---|---|---|---|---|---|---|---|---|---|---|---|---|---|---|---|
| 29845 | ENSMUSG00000088666 | 0 | NA | NA | NA | NA | NA | NA | NA | 0 | 0 | 0 | 0 | 0 | 0 |
| 29846 | ENSMUSG00000088668 | 0 | NA | NA | NA | NA | NA | Gm25335 | predicted gene, 25335 [Source:MGI Symb | 0 | 0 | 0 | 0 | 0 | 0 |
| 29847 | ENSMUSG00000088669 | 0.160662974 | 1.126924175 | 4.080472857 | 0.276174898 | 0.782413736 | NA | Gm25336 | predicted gene, 25336 [Source:MGI Symb | 0 | 0 | 0 | 0.964 | 0 | 0 |
| 29848 | ENSMUSG00000088670 | 0.36353943 | 2.065989254 | 4.034137301 | 0.512126658 | 0.608562368 | NA | NA | NA | 0 | 0 | 0 | 0 | 2.1812 | 0 |
| 29849 | ENSMUSG00000088671 | 0 | NA | NA | NA | NA | NA | NA | NA | 0 | 0 | 0 | 0 | 0 | 0 |
| 29850 | ENSMUSG00000088672 | 0 | NA | NA | NA | NA | NA | Gm23664 | predicted gene, 23664 [Source:MGI Symb | 0 | 0 | 0 | 0 | 0 | 0 |
| 29851 | ENSMUSG00000088674 | 0 | NA | NA | NA | NA | NA | NA | NA | 0 | 0 | 0 | 0 | 0 | 0 |
| 29852 | ENSMUSG00000088675 | 0 | NA | NA | NA | NA | NA | Rprl1 | ribonuclease P RNA-like 1 [Source:MGI Sy | 0 | 0 | 0 | 0 | 0 | 0 |
| 29853 | ENSMUSG00000088677 | 0 | NA | NA | NA | NA | NA | NA | NA | 0 | 0 | 0 | 0 | 0 | 0 |
| 29854 | ENSMUSG00000088678 | 0 | NA | NA | NA | NA | NA | Gm23666 | predicted gene, 23666 [Source:MGI Symb | 0 | 0 | 0 | 0 | 0 | 0 |
| 29855 | ENSMUSG00000088679 | 0 | NA | NA | NA | NA | NA | Gm23665 | predicted gene, 23665 [Source:MGI Symb | 0 | 0 | 0 | 0 | 0 | 0 |
| 29856 | ENSMUSG00000088680 | 0 | NA | NA | NA | NA | NA | NA | NA | 0 | 0 | 0 | 0 | 0 | 0 |
| 29857 | ENSMUSG00000088682 | 0 | NA | NA | NA | NA | NA | NA | NA | 0 | 0 | 0 | 0 | 0 | 0 |
| 29858 | ENSMUSG00000088683 | 0 | NA | NA | NA | NA | NA | NA | NA | 0 | 0 | 0 | 0 | 0 | 0 |
| 29859 | ENSMUSG00000088684 | 0 | NA | NA | NA | NA | NA | Gm23994 | predicted gene, 23994 [Source:MGI Symb | 0 | 0 | 0 | 0 | 0 | 0 |
| 29860 | ENSMUSG00000088685 | 0 | NA | NA | NA | NA | NA | Gm23995 | predicted gene, 23995 [Source:MGI Symb | 0 | 0 | 0 | 0 | 0 | 0 |
| 29861 | ENSMUSG00000088686 | 0 | NA | NA | NA | NA | NA | NA | NA | 0 | 0 | 0 | 0 | 0 | 0 |
| 29862 | ENSMUSG00000088687 | 0 | NA | NA | NA | NA | NA | Gm23996 | predicted gene, 23996 [Source:MGI Symb | 0 | 0 | 0 | 0 | 0 | 0 |
| 29863 | ENSMUSG00000088688 | 0 | NA | NA | NA | NA | NA | Gm24000 | predicted gene, 24000 [Source:MGI Symb | 0 | 0 | 0 | 0 | 0 | 0 |
| 29864 | ENSMUSG00000088689 | 2.326839718 | 0.552728311 | 1.364238704 | 0.405155131 | 0.685363484 | NA | Scarna17 | small Cajal body-specific RNA 17 [Source:I | 1.6062 | 3.0838 | 1.0062 | 3.8559 | 2.1812 | 2.2277 |
| 29865 | ENSMUSG00000088691 | 0 | NA | NA | NA | NA | NA | Gm22331 | predicted gene, 22331 [Source:MGI Symb | 0 | 0 | 0 | 0 | 0 | 0 |
| 29866 | ENSMUSG00000088692 | 0 | NA | NA | NA | NA | NA | Gm23838 | predicted gene, 23838 [Source:MGI Symb | 0 | 0 | 0 | 0 | 0 | 0 |
| 29867 | ENSMUSG00000088693 | 0 | NA | NA | NA | NA | NA | Gm22330 | predicted gene, 22330 [Source:MGI Symb | 0 | 0 | 0 | 0 | 0 | 0 |
| 29868 | ENSMUSG00000088695 | 0.335414862 | -1.779047242 | 4.030804179 | -0.441362855 | 0.658950329 | NA | Gm22334 | predicted gene, 22334 [Source:MGI Symb | 0 | 0 | 2.0125 | 0 | 0 | 0 |
| 29869 | ENSMUSG00000088696 | 0 | NA | NA | NA | NA | NA | Gm22333 | predicted gene, 22333 [Source:MGI Symb | 0 | 0 | 0 | 0 | 0 | 0 |
| 29870 | ENSMUSG00000088697 | 0 | NA | NA | NA | NA | NA | Gm22332 | predicted gene, 22332 [Source:MGI Symb | 0 | 0 | 0 | 0 | 0 | 0 |
| 29871 | ENSMUSG00000088699 | 0 | NA | NA | NA | NA | NA | NA | NA | 0 | 0 | 0 | 0 | 0 | 0 |
| 29872 | ENSMUSG00000088700 | 0 | NA | NA | NA | NA | NA | NA | NA | 0 | 0 | 0 | 0 | 0 | 0 |
| 29873 | ENSMUSG00000088701 | 0 | NA | NA | NA | NA | NA | NA | NA | 0 | 0 | 0 | 0 | 0 | 0 |
| 29874 | ENSMUSG00000088702 | 0 | NA | NA | NA | NA | NA | NA | NA | 0 | 0 | 0 | 0 | 0 | 0 |
| 29875 | ENSMUSG00000088703 | 0.133850373 | -0.796629176 | 4.080472857 | -0.195229623 | 0.84521319 | NA | NA | NA | 0.8031 | 0 | 0 | 0 | 0 | 0 |
| 29876 | ENSMUSG00000088704 | 0 | NA | NA | NA | NA | NA | Gm25550 | predicted gene, 25550 [Source:MGI Symb | 0 | 0 | 0 | 0 | 0 | 0 |
| 29877 | ENSMUSG00000088705 | 0.167707431 | -0.796629176 | 4.080472857 | -0.195229623 | 0.84521319 | NA | Gm25549 | predicted gene, 25549 [Source:MGI Symb | 0 | 0 | 1.0062 | 0 | 0 | 0 |
| 29878 | ENSMUSG00000088706 | 0 | NA | NA | NA | NA | NA | Gm25552 | predicted gene, 25552 [Source:MGI Symb | 0 | 0 | 0 | 0 | 0 | 0 |
| 29879 | ENSMUSG00000088707 | 0 | NA | NA | NA | NA | NA | NA | NA | 0 | 0 | 0 | 0 | 0 | 0 |
| 29880 | ENSMUSG00000088708 | 0.671009685 | -1.413087452 | 2.800223337 | -0.504633839 | 0.613816038 | NA | Gm25558 | predicted gene, 25558 [Source:MGI Symb | 0 | 2.0558 | 1.0062 | 0.964 | 0 | 0 |
| 29881 | ENSMUSG00000088709 | 0 | NA | NA | NA | NA | NA | NA | NA | 0 | 0 | 0 | 0 | 0 | 0 |
| 29882 | ENSMUSG00000088710 | 0 | NA | NA | NA | NA | NA | NA | NA | 0 | 0 | 0 | 0 | 0 | 0 |
| 29883 | ENSMUSG00000088711 | 0 | NA | NA | NA | NA | NA | NA | NA | 0 | 0 | 0 | 0 | 0 | 0 |
| 29884 | ENSMUSG00000088712 | 0.342432689 | 1.990653086 | 3.900281263 | 0.51038706 | 0.609780321 | NA | Gm22728 | predicted gene, 22728 [Source:MGI Symb | 0 | 0 | 0 | 0.964 | 1.0906 | 0 |
| 29885 | ENSMUSG00000088713 | 0 | NA | NA | NA | NA | NA | NA | NA | 0 | 0 | 0 | 0 | 0 | 0 |
| 29886 | ENSMUSG00000088714 | 0 | NA | NA | NA | NA | NA | NA | NA | 0 | 0 | 0 | 0 | 0 | 0 |
| 29887 | ENSMUSG00000088715 | 0 | NA | NA | NA | NA | NA | Gm22727 | predicted gene, 22727 [Source:MGI Symb | 0 | 0 | 0 | 0 | 0 | 0 |
| 29888 | ENSMUSG00000088716 | 0 | NA | NA | NA | NA | NA | Gm22724 | predicted gene, 22724 [Source:MGI Symb | 0 | 0 | 0 | 0 | 0 | 0 |
| 29889 | ENSMUSG00000088717 | 0 | NA | NA | NA | NA | NA | Gm22725 | predicted gene, 22725 [Source:MGI Symb | 0 | 0 | 0 | 0 | 0 | 0 |
| 29890 | ENSMUSG00000088718 | 0 | NA | NA | NA | NA | NA | Gm22732 | predicted gene, 22732 [Source:MGI Symb | 0 | 0 | 0 | 0 | 0 | 0 |
| 29891 | ENSMUSG00000088719 | 0.17131964 | -0.796629176 | 4.080472857 | -0.195229623 | 0.84521319 | NA | Gm22733 | predicted gene, 22733 [Source:MGI Symb | 0 | 1.0279 | 0 | 0 | 0 | 0 |
| 29892 | ENSMUSG00000088721 | 0 | NA | NA | NA | NA | NA | Gm22212 | predicted gene, 22212 [Source:MGI Symb | 0 | 0 | 0 | 0 | 0 | 0 |
| 29893 | ENSMUSG00000088722 | 0 | NA | NA | NA | NA | NA | NA | NA | 0 | 0 | 0 | 0 | 0 | 0 |
| 29894 | ENSMUSG00000088726 | 0 | NA | NA | NA | NA | NA | Gm22210 | predicted gene, 22210 [Source:MGI Symb | 0 | 0 | 0 | 0 | 0 | 0 |
| 29895 | ENSMUSG00000088730 | 0 | NA | NA | NA | NA | NA | Gm23892 | predicted gene, 23892 [Source:MGI Symb | 0 | 0 | 0 | 0 | 0 | 0 |
| 29896 | ENSMUSG00000088731 | 0 | NA | NA | NA | NA | NA | NA | NA | 0 | 0 | 0 | 0 | 0 | 0 |
| 29897 | ENSMUSG00000088732 | 0 | NA | NA | NA | NA | NA | NA | NA | 0 | 0 | 0 | 0 | 0 | 0 |
| 29898 | ENSMUSG00000088733 | 0 | NA | NA | NA | NA | NA | Gm23895 | predicted gene, 23895 [Source:MGI Symb | 0 | 0 | 0 | 0 | 0 | 0 |
| 29899 | ENSMUSG00000088734 | 0 | NA | NA | NA | NA | NA | Gm23896 | predicted gene, 23896 [Source:MGI Symb | 0 | 0 | 0 | 0 | 0 | 0 |
| 29900 | ENSMUSG00000088735 | 0 | NA | NA | NA | NA | NA | Gm23897 | predicted gene, 23897 [Source:MGI Symb | 0 | 0 | 0 | 0 | 0 | 0 |

| | | | | | | | | | | | | | | | |
|---|---|---|---|---|---|---|---|---|---|---|---|---|---|---|---|
| 29901 | ENSMUSG00000088736 | 0 | NA | NA | NA | NA | NA | NA | NA | 0 | 0 | 0 | 0 | 0 | 0 |
| 29902 | ENSMUSG00000088737 | 0 | NA | NA | NA | NA | NA | NA | NA | 0 | 0 | 0 | 0 | 0 | 0 |
| 29903 | ENSMUSG00000088738 | 0 | NA | NA | NA | NA | NA | Gm27579 | predicted gene, 27579 [Source:MGI Symb | 0 | 0 | 0 | 0 | 0 | 0 |
| 29904 | ENSMUSG00000088739 | 0 | NA | NA | NA | NA | NA | Gm23891 | predicted gene, 23891 [Source:MGI Symb | 0 | 0 | 0 | 0 | 0 | 0 |
| 29905 | ENSMUSG00000088740 | 0 | NA | NA | NA | NA | NA | Gm23220 | predicted gene, 23220 [Source:MGI Symb | 0 | 0 | 0 | 0 | 0 | 0 |
| 29906 | ENSMUSG00000088742 | 0 | NA | NA | NA | NA | NA | NA | NA | 0 | 0 | 0 | 0 | 0 | 0 |
| 29907 | ENSMUSG00000088743 | 0 | NA | NA | NA | NA | NA | NA | NA | 0 | 0 | 0 | 0 | 0 | 0 |
| 29908 | ENSMUSG00000088744 | 0 | NA | NA | NA | NA | NA | Gm23223 | predicted gene, 23223 [Source:MGI Symb | 0 | 0 | 0 | 0 | 0 | 0 |
| 29909 | ENSMUSG00000088746 | 0 | NA | NA | NA | NA | NA | Gm23225 | predicted gene, 23225 [Source:MGI Symb | 0 | 0 | 0 | 0 | 0 | 0 |
| 29910 | ENSMUSG00000088747 | 0 | NA | NA | NA | NA | NA | NA | NA | 0 | 0 | 0 | 0 | 0 | 0 |
| 29911 | ENSMUSG00000088748 | 0 | NA | NA | NA | NA | NA | Gm23219 | predicted gene, 23219 [Source:MGI Symb | 0 | 0 | 0 | 0 | 0 | 0 |
| 29912 | ENSMUSG00000088749 | 2.824334914 | -0.696711379 | 1.435492546 | -0.485346567 | 0.627430529 | NA | NA | NA | 2.4093 | 0 | 8.05 | 0.964 | 2.1812 | 3.3415 |
| 29913 | ENSMUSG00000088750 | 0 | NA | NA | NA | NA | NA | Gm24901 | predicted gene, 24901 [Source:MGI Symb | 0 | 0 | 0 | 0 | 0 | 0 |
| 29914 | ENSMUSG00000088751 | 0 | NA | NA | NA | NA | NA | Gm24902 | predicted gene, 24902 [Source:MGI Symb | 0 | 0 | 0 | 0 | 0 | 0 |
| 29915 | ENSMUSG00000088752 | 0.904646783 | 0.28560953 | 2.491718272 | 0.114623524 | 0.908743533 | NA | Mir1934 | microRNA 1934 [Source:MGI Symbol;Acc: | 2.4093 | 0 | 0 | 1.928 | 1.0906 | 0 |
| 29916 | ENSMUSG00000088753 | 0 | NA | NA | NA | NA | NA | NA | NA | 0 | 0 | 0 | 0 | 0 | 0 |
| 29917 | ENSMUSG00000088754 | 0 | NA | NA | NA | NA | NA | Gm24905 | predicted gene, 24905 [Source:MGI Symb | 0 | 0 | 0 | 0 | 0 | 0 |
| 29918 | ENSMUSG00000088756 | 0 | NA | NA | NA | NA | NA | Gm24903 | predicted gene, 24903 [Source:MGI Symb | 0 | 0 | 0 | 0 | 0 | 0 |
| 29919 | ENSMUSG00000088758 | 1.19159988 | 0.475438562 | 2.070964431 | 0.229573505 | 0.818423197 | NA | NA | NA | 0 | 1.0279 | 2.0125 | 1.928 | 2.1812 | 0 |
| 29920 | ENSMUSG00000088760 | 0 | NA | NA | NA | NA | NA | Gm24393 | predicted gene, 24393 [Source:MGI Symb | 0 | 0 | 0 | 0 | 0 | 0 |
| 29921 | ENSMUSG00000088761 | 0 | NA | NA | NA | NA | NA | Gm24392 | predicted gene, 24392 [Source:MGI Symb | 0 | 0 | 0 | 0 | 0 | 0 |
| 29922 | ENSMUSG00000088763 | 0 | NA | NA | NA | NA | NA | NA | NA | 0 | 0 | 0 | 0 | 0 | 0 |
| 29923 | ENSMUSG00000088764 | 0 | NA | NA | NA | NA | NA | Gm24395 | predicted gene, 24395 [Source:MGI Symb | 0 | 0 | 0 | 0 | 0 | 0 |
| 29924 | ENSMUSG00000088766 | 0 | NA | NA | NA | NA | NA | Gm24394 | predicted gene, 24394 [Source:MGI Symb | 0 | 0 | 0 | 0 | 0 | 0 |
| 29925 | ENSMUSG00000088767 | 0 | NA | NA | NA | NA | NA | NA | NA | 0 | 0 | 0 | 0 | 0 | 0 |
| 29926 | ENSMUSG00000088768 | 0 | NA | NA | NA | NA | NA | Gm25494 | predicted gene, 25494 [Source:MGI Symb | 0 | 0 | 0 | 0 | 0 | 0 |
| 29927 | ENSMUSG00000088769 | 3.364718229 | -0.222697856 | 1.295854013 | -0.171854124 | 0.863552214 | NA | NA | NA | 4.8186 | 0 | 6.0375 | 3.8559 | 4.3625 | 1.1138 |
| 29928 | ENSMUSG00000088772 | 0 | NA | NA | NA | NA | NA | Gm26040 | predicted gene, 26040 [Source:MGI Symb | 0 | 0 | 0 | 0 | 0 | 0 |
| 29929 | ENSMUSG00000088773 | 0 | NA | NA | NA | NA | NA | NA | NA | 0 | 0 | 0 | 0 | 0 | 0 |
| 29930 | ENSMUSG00000088774 | 0 | NA | NA | NA | NA | NA | Gm26036 | predicted gene, 26036 [Source:MGI Symb | 0 | 0 | 0 | 0 | 0 | 0 |
| 29931 | ENSMUSG00000088775 | 0 | NA | NA | NA | NA | NA | NA | NA | 0 | 0 | 0 | 0 | 0 | 0 |
| 29932 | ENSMUSG00000088776 | 0 | NA | NA | NA | NA | NA | Gm26038 | predicted gene, 26038 [Source:MGI Symb | 0 | 0 | 0 | 0 | 0 | 0 |
| 29933 | ENSMUSG00000088777 | 0.663785267 | -1.400021131 | 3.106754789 | -0.450637796 | 0.652250621 | NA | NA | NA | 0 | 0 | 3.0187 | 0.964 | 0 | 0 |
| 29934 | ENSMUSG00000088779 | 0 | NA | NA | NA | NA | NA | NA | NA | 0 | 0 | 0 | 0 | 0 | 0 |
| 29935 | ENSMUSG00000088781 | 0 | NA | NA | NA | NA | NA | Gm25373 | predicted gene, 25373 [Source:MGI Symb | 0 | 0 | 0 | 0 | 0 | 0 |
| 29936 | ENSMUSG00000088782 | 0 | NA | NA | NA | NA | NA | Gm25375 | predicted gene, 25375 [Source:MGI Symb | 0 | 0 | 0 | 0 | 0 | 0 |
| 29937 | ENSMUSG00000088783 | 0 | NA | NA | NA | NA | NA | NA | NA | 0 | 0 | 0 | 0 | 0 | 0 |
| 29938 | ENSMUSG00000088784 | 0 | NA | NA | NA | NA | NA | Gm25370 | predicted gene, 25370 [Source:MGI Symb | 0 | 0 | 0 | 0 | 0 | 0 |
| 29939 | ENSMUSG00000088786 | 0 | NA | NA | NA | NA | NA | NA | NA | 0 | 0 | 0 | 0 | 0 | 0 |
| 29940 | ENSMUSG00000088787 | 0 | NA | NA | NA | NA | NA | NA | NA | 0 | 0 | 0 | 0 | 0 | 0 |
| 29941 | ENSMUSG00000088789 | 1.523789084 | -0.975344162 | 1.924518288 | -0.506799114 | 0.612295778 | NA | NA | NA | 0 | 4.1117 | 2.0125 | 1.928 | 1.0906 | 0 |
| 29942 | ENSMUSG00000088790 | 0 | NA | NA | NA | NA | NA | NA | NA | 0 | 0 | 0 | 0 | 0 | 0 |
| 29943 | ENSMUSG00000088791 | 0 | NA | NA | NA | NA | NA | Gm22542 | predicted gene, 22542 [Source:MGI Symb | 0 | 0 | 0 | 0 | 0 | 0 |
| 29944 | ENSMUSG00000088792 | 0 | NA | NA | NA | NA | NA | Gm22539 | predicted gene, 22539 [Source:MGI Symb | 0 | 0 | 0 | 0 | 0 | 0 |
| 29945 | ENSMUSG00000088793 | 0 | NA | NA | NA | NA | NA | Gm22540 | predicted gene, 22540 [Source:MGI Symb | 0 | 0 | 0 | 0 | 0 | 0 |
| 29946 | ENSMUSG00000088794 | 0 | NA | NA | NA | NA | NA | NA | NA | 0 | 0 | 0 | 0 | 0 | 0 |
| 29947 | ENSMUSG00000088796 | 0 | NA | NA | NA | NA | NA | Gm22536 | predicted gene, 22536 [Source:MGI Symb | 0 | 0 | 0 | 0 | 0 | 0 |
| 29948 | ENSMUSG00000088797 | 0 | NA | NA | NA | NA | NA | NA | NA | 0 | 0 | 0 | 0 | 0 | 0 |
| 29949 | ENSMUSG00000088798 | 0 | NA | NA | NA | NA | NA | Gm22534 | predicted gene, 22534 [Source:MGI Symb | 0 | 0 | 0 | 0 | 0 | 0 |
| 29950 | ENSMUSG00000088799 | 0 | NA | NA | NA | NA | NA | NA | NA | 0 | 0 | 0 | 0 | 0 | 0 |
| 29951 | ENSMUSG00000088800 | 0 | NA | NA | NA | NA | NA | Gm24075 | predicted gene, 24075 [Source:MGI Symb | 0 | 0 | 0 | 0 | 0 | 0 |
| 29952 | ENSMUSG00000088802 | 0 | NA | NA | NA | NA | NA | NA | NA | 0 | 0 | 0 | 0 | 0 | 0 |
| 29953 | ENSMUSG00000088803 | 0 | NA | NA | NA | NA | NA | NA | NA | 0 | 0 | 0 | 0 | 0 | 0 |
| 29954 | ENSMUSG00000088804 | 0 | NA | NA | NA | NA | NA | NA | NA | 0 | 0 | 0 | 0 | 0 | 0 |
| 29955 | ENSMUSG00000088808 | 0 | NA | NA | NA | NA | NA | NA | NA | 0 | 0 | 0 | 0 | 0 | 0 |
| 29956 | ENSMUSG00000088809 | 0 | NA | NA | NA | NA | NA | NA | NA | 0 | 0 | 0 | 0 | 0 | 0 |

| | | | | | | | | | | | | | | | |
|---|---|---|---|---|---|---|---|---|---|---|---|---|---|---|---|
| 29957 | ENSMUSG00000088810 | 0 | NA | NA | NA | NA | NA | Gm22382 | predicted gene, 22382 [Source:MGI Symb | 0 | 0 | 0 | 0 | 0 | 0 |
| 29958 | ENSMUSG00000088811 | 0 | NA | NA | NA | NA | NA | Gm22381 | predicted gene, 22381 [Source:MGI Symb | 0 | 0 | 0 | 0 | 0 | 0 |
| 29959 | ENSMUSG00000088812 | 0 | NA | NA | NA | NA | NA | NA | NA | 0 | 0 | 0 | 0 | 0 | 0 |
| 29960 | ENSMUSG00000088813 | 0.185640647 | 1.126924175 | 4.080472857 | 0.276174898 | 0.782413736 | NA | NA | NA | 0 | 0 | 0 | 0 | 0 | 1.1138 |
| 29961 | ENSMUSG00000088814 | 0 | NA | NA | NA | NA | NA | NA | NA | 0 | 0 | 0 | 0 | 0 | 0 |
| 29962 | ENSMUSG00000088816 | 0 | NA | NA | NA | NA | NA | Gm22387 | predicted gene, 22387 [Source:MGI Symb | 0 | 0 | 0 | 0 | 0 | 0 |
| 29963 | ENSMUSG00000088817 | 0 | NA | NA | NA | NA | NA | Gm22386 | predicted gene, 22386 [Source:MGI Symb | 0 | 0 | 0 | 0 | 0 | 0 |
| 29964 | ENSMUSG00000088818 | 0 | NA | NA | NA | NA | NA | NA | NA | 0 | 0 | 0 | 0 | 0 | 0 |
| 29965 | ENSMUSG00000088819 | 0 | NA | NA | NA | NA | NA | NA | NA | 0 | 0 | 0 | 0 | 0 | 0 |
| 29966 | ENSMUSG00000088820 | 0 | NA | NA | NA | NA | NA | Gm25224 | predicted gene, 25224 [Source:MGI Symb | 0 | 0 | 0 | 0 | 0 | 0 |
| 29967 | ENSMUSG00000088822 | 0 | NA | NA | NA | NA | NA | NA | NA | 0 | 0 | 0 | 0 | 0 | 0 |
| 29968 | ENSMUSG00000088823 | 0 | NA | NA | NA | NA | NA | Gm25226 | predicted gene, 25226 [Source:MGI Symb | 0 | 0 | 0 | 0 | 0 | 0 |
| 29969 | ENSMUSG00000088824 | 0 | NA | NA | NA | NA | NA | Gm25222 | predicted gene, 25222 [Source:MGI Symb | 0 | 0 | 0 | 0 | 0 | 0 |
| 29970 | ENSMUSG00000088825 | 0 | NA | NA | NA | NA | NA | NA | NA | 0 | 0 | 0 | 0 | 0 | 0 |
| 29971 | ENSMUSG00000088829 | 0 | NA | NA | NA | NA | NA | Gm25227 | predicted gene, 25227 [Source:MGI Symb | 0 | 0 | 0 | 0 | 0 | 0 |
| 29972 | ENSMUSG00000088830 | 0 | NA | NA | NA | NA | NA | Gm23545 | predicted gene, 23545 [Source:MGI Symb | 0 | 0 | 0 | 0 | 0 | 0 |
| 29973 | ENSMUSG00000088832 | 0 | NA | NA | NA | NA | NA | NA | NA | 0 | 0 | 0 | 0 | 0 | 0 |
| 29974 | ENSMUSG00000088833 | 0 | NA | NA | NA | NA | NA | NA | NA | 0 | 0 | 0 | 0 | 0 | 0 |
| 29975 | ENSMUSG00000088835 | 0 | NA | NA | NA | NA | NA | Gm23547 | predicted gene, 23547 [Source:MGI Symb | 0 | 0 | 0 | 0 | 0 | 0 |
| 29976 | ENSMUSG00000088836 | 0.185640647 | 1.126924175 | 4.080472857 | 0.276174898 | 0.782413736 | NA | Gm23546 | predicted gene, 23546 [Source:MGI Symb | 0 | 0 | 0 | 0 | 0 | 1.1138 |
| 29977 | ENSMUSG00000088839 | 0 | NA | NA | NA | NA | NA | Gm27510 | predicted gene, 27510 [Source:MGI Symb | 0 | 0 | 0 | 0 | 0 | 0 |
| 29978 | ENSMUSG00000088840 | 0 | NA | NA | NA | NA | NA | Gm26358 | predicted gene, 26358 [Source:MGI Symb | 0 | 0 | 0 | 0 | 0 | 0 |
| 29979 | ENSMUSG00000088842 | 0 | NA | NA | NA | NA | NA | NA | NA | 0 | 0 | 0 | 0 | 0 | 0 |
| 29980 | ENSMUSG00000088843 | 0 | NA | NA | NA | NA | NA | Gm26357 | predicted gene, 26357 [Source:MGI Symb | 0 | 0 | 0 | 0 | 0 | 0 |
| 29981 | ENSMUSG00000088845 | 0 | NA | NA | NA | NA | NA | NA | NA | 0 | 0 | 0 | 0 | 0 | 0 |
| 29982 | ENSMUSG00000088846 | 0 | NA | NA | NA | NA | NA | Gm26354 | predicted gene, 26354 [Source:MGI Symb | 0 | 0 | 0 | 0 | 0 | 0 |
| 29983 | ENSMUSG00000088848 | 0 | NA | NA | NA | NA | NA | Gm26352 | predicted gene, 26352 [Source:MGI Symb | 0 | 0 | 0 | 0 | 0 | 0 |
| 29984 | ENSMUSG00000088849 | 0 | NA | NA | NA | NA | NA | NA | NA | 0 | 0 | 0 | 0 | 0 | 0 |
| 29985 | ENSMUSG00000088850 | 0 | NA | NA | NA | NA | NA | Gm24729 | predicted gene, 24729 [Source:MGI Symb | 0 | 0 | 0 | 0 | 0 | 0 |
| 29986 | ENSMUSG00000088851 | 0 | NA | NA | NA | NA | NA | Gm24728 | predicted gene, 24728 [Source:MGI Symb | 0 | 0 | 0 | 0 | 0 | 0 |
| 29987 | ENSMUSG00000088852 | 0 | NA | NA | NA | NA | NA | Gm24731 | predicted gene, 24731 [Source:MGI Symb | 0 | 0 | 0 | 0 | 0 | 0 |
| 29988 | ENSMUSG00000088853 | 0 | NA | NA | NA | NA | NA | Gm24730 | predicted gene, 24730 [Source:MGI Symb | 0 | 0 | 0 | 0 | 0 | 0 |
| 29989 | ENSMUSG00000088854 | 0 | NA | NA | NA | NA | NA | Gm24725 | predicted gene, 24725 [Source:MGI Symb | 0 | 0 | 0 | 0 | 0 | 0 |
| 29990 | ENSMUSG00000088855 | 0 | NA | NA | NA | NA | NA | NA | NA | 0 | 0 | 0 | 0 | 0 | 0 |
| 29991 | ENSMUSG00000088856 | 1.615039425 | -0.558084326 | 2.077429868 | -0.268641716 | 0.788205407 | NA | NA | NA | 0.8031 | 0 | 5.0312 | 3.8559 | 0 | 0 |
| 29992 | ENSMUSG00000088857 | 0.870273931 | -0.470854203 | 2.55968665 | -0.183949939 | 0.854052728 | NA | Gm24726 | predicted gene, 24726 [Source:MGI Symb | 0 | 1.0279 | 2.0125 | 0 | 2.1812 | 0 |
| 29993 | ENSMUSG00000088858 | 0 | NA | NA | NA | NA | NA | Gm24723 | predicted gene, 24723 [Source:MGI Symb | 0 | 0 | 0 | 0 | 0 | 0 |
| 29994 | ENSMUSG00000088860 | 0.160662974 | 1.126924175 | 4.080472857 | 0.276174898 | 0.782413736 | NA | Mir1947 | microRNA 1947 [Source:MGI Symbol;Acc: | 0 | 0 | 0 | 0.964 | 0 | 0 |
| 29995 | ENSMUSG00000088861 | 0 | NA | NA | NA | NA | NA | NA | NA | 0 | 0 | 0 | 0 | 0 | 0 |
| 29996 | ENSMUSG00000088862 | 0 | NA | NA | NA | NA | NA | NA | NA | 0 | 0 | 0 | 0 | 0 | 0 |
| 29997 | ENSMUSG00000088863 | 0 | NA | NA | NA | NA | NA | Gm23067 | predicted gene, 23067 [Source:MGI Symb | 0 | 0 | 0 | 0 | 0 | 0 |
| 29998 | ENSMUSG00000088864 | 0 | NA | NA | NA | NA | NA | NA | NA | 0 | 0 | 0 | 0 | 0 | 0 |
| 29999 | ENSMUSG00000088865 | 0 | NA | NA | NA | NA | NA | NA | NA | 0 | 0 | 0 | 0 | 0 | 0 |
| 30000 | ENSMUSG00000088867 | 0 | NA | NA | NA | NA | NA | NA | NA | 0 | 0 | 0 | 0 | 0 | 0 |
| 30001 | ENSMUSG00000088868 | 0 | NA | NA | NA | NA | NA | Gm23072 | predicted gene, 23072 [Source:MGI Symb | 0 | 0 | 0 | 0 | 0 | 0 |
| 30002 | ENSMUSG00000088870 | 0 | NA | NA | NA | NA | NA | Gm25866 | predicted gene, 25866 [Source:MGI Symb | 0 | 0 | 0 | 0 | 0 | 0 |
| 30003 | ENSMUSG00000088872 | 0 | NA | NA | NA | NA | NA | NA | NA | 0 | 0 | 0 | 0 | 0 | 0 |
| 30004 | ENSMUSG00000088873 | 0.483327519 | -0.685704547 | 3.145069396 | -0.218025252 | 0.827409436 | NA | NA | NA | 0.8031 | 0 | 1.0062 | 0 | 1.0906 | 0 |
| 30005 | ENSMUSG00000088874 | 0 | NA | NA | NA | NA | NA | Gm25863 | predicted gene, 25863 [Source:MGI Symb | 0 | 0 | 0 | 0 | 0 | 0 |
| 30006 | ENSMUSG00000088875 | 0.328370405 | 0.16514316 | 4.003149536 | 0.041253308 | 0.967093956 | NA | Gm25862 | predicted gene, 25862 [Source:MGI Symb | 0 | 0 | 1.0062 | 0.964 | 0 | 0 |
| 30007 | ENSMUSG00000088876 | 0 | NA | NA | NA | NA | NA | Gm25861 | predicted gene, 25861 [Source:MGI Symb | 0 | 0 | 0 | 0 | 0 | 0 |
| 30008 | ENSMUSG00000088877 | 0 | NA | NA | NA | NA | NA | Gm25860 | predicted gene, 25860 [Source:MGI Symb | 0 | 0 | 0 | 0 | 0 | 0 |
| 30009 | ENSMUSG00000088878 | 0 | NA | NA | NA | NA | NA | NA | NA | 0 | 0 | 0 | 0 | 0 | 0 |
| 30010 | ENSMUSG00000088879 | 0 | NA | NA | NA | NA | NA | Gm25867 | predicted gene, 25867 [Source:MGI Symb | 0 | 0 | 0 | 0 | 0 | 0 |
| 30011 | ENSMUSG00000088880 | 0 | NA | NA | NA | NA | NA | NA | NA | 0 | 0 | 0 | 0 | 0 | 0 |
| 30012 | ENSMUSG00000088883 | 0 | NA | NA | NA | NA | NA | Gm24212 | predicted gene, 24212 [Source:MGI Symb | 0 | 0 | 0 | 0 | 0 | 0 |

| | | | | | | | | | | | | | | | |
|---|---|---|---|---|---|---|---|---|---|---|---|---|---|---|---|
| 30013 | ENSMUSG00000088884 | 0 | NA | NA | NA | NA | NA | Gm24216 | predicted gene, 24216 [Source:MGI Symb | 0 | 0 | 0 | 0 | 0 | 0 |
| 30014 | ENSMUSG00000088885 | 0 | NA | NA | NA | NA | NA | NA | NA | 0 | 0 | 0 | 0 | 0 | 0 |
| 30015 | ENSMUSG00000088888 | 0 | NA | NA | NA | NA | NA | Gm24210 | predicted gene, 24210 [Source:MGI Symb | 0 | 0 | 0 | 0 | 0 | 0 |
| 30016 | ENSMUSG00000088889 | 0 | NA | NA | NA | NA | NA | Gm24211 | predicted gene, 24211 [Source:MGI Symb | 0 | 0 | 0 | 0 | 0 | 0 |
| 30017 | ENSMUSG00000088890 | 0 | NA | NA | NA | NA | NA | NA | NA | 0 | 0 | 0 | 0 | 0 | 0 |
| 30018 | ENSMUSG00000088891 | 0.520796785 | -0.820103173 | 3.072141206 | -0.266948398 | 0.789508892 | NA | Gm22516 | predicted gene, 22516 [Source:MGI Symb | 0 | 1.0279 | 1.0062 | 0 | 1.0906 | 0 |
| 30019 | ENSMUSG00000088892 | 0 | NA | NA | NA | NA | NA | NA | NA | 0 | 0 | 0 | 0 | 0 | 0 |
| 30020 | ENSMUSG00000088894 | 0 | NA | NA | NA | NA | NA | Gm22520 | predicted gene, 22520 [Source:MGI Symb | 0 | 0 | 0 | 0 | 0 | 0 |
| 30021 | ENSMUSG00000088895 | 0 | NA | NA | NA | NA | NA | Gm22519 | predicted gene, 22519 [Source:MGI Symb | 0 | 0 | 0 | 0 | 0 | 0 |
| 30022 | ENSMUSG00000088896 | 0 | NA | NA | NA | NA | NA | Gm22522 | predicted gene, 22522 [Source:MGI Symb | 0 | 0 | 0 | 0 | 0 | 0 |
| 30023 | ENSMUSG00000088897 | 0 | NA | NA | NA | NA | NA | Gm22521 | predicted gene, 22521 [Source:MGI Symb | 0 | 0 | 0 | 0 | 0 | 0 |
| 30024 | ENSMUSG00000088898 | 0 | NA | NA | NA | NA | NA | NA | NA | 0 | 0 | 0 | 0 | 0 | 0 |
| 30025 | ENSMUSG00000088900 | 0 | NA | NA | NA | NA | NA | Gm25436 | predicted gene, 25436 [Source:MGI Symb | 0 | 0 | 0 | 0 | 0 | 0 |
| 30026 | ENSMUSG00000088901 | 0.663758638 | 2.950973782 | 3.025092005 | 0.975498853 | 0.32931287 | NA | Mir1943 | microRNA 1943 [Source:MGI Symbol;Acc: | 0 | 0 | 0 | 2.8919 | 1.0906 | 0 |
| 30027 | ENSMUSG00000088902 | 0 | NA | NA | NA | NA | NA | NA | NA | 0 | 0 | 0 | 0 | 0 | 0 |
| 30028 | ENSMUSG00000088903 | 0 | NA | NA | NA | NA | NA | Gm25434 | predicted gene, 25434 [Source:MGI Symb | 0 | 0 | 0 | 0 | 0 | 0 |
| 30029 | ENSMUSG00000088904 | 0 | NA | NA | NA | NA | NA | NA | NA | 0 | 0 | 0 | 0 | 0 | 0 |
| 30030 | ENSMUSG00000088905 | 0 | NA | NA | NA | NA | NA | Gm25439 | predicted gene, 25439 [Source:MGI Symb | 0 | 0 | 0 | 0 | 0 | 0 |
| 30031 | ENSMUSG00000088906 | 0 | NA | NA | NA | NA | NA | Gm25438 | predicted gene, 25438 [Source:MGI Symb | 0 | 0 | 0 | 0 | 0 | 0 |
| 30032 | ENSMUSG00000088907 | 0 | NA | NA | NA | NA | NA | Gm25437 | predicted gene, 25437 [Source:MGI Symb | 0 | 0 | 0 | 0 | 0 | 0 |
| 30033 | ENSMUSG00000088908 | 0 | NA | NA | NA | NA | NA | NA | NA | 0 | 0 | 0 | 0 | 0 | 0 |
| 30034 | ENSMUSG00000088909 | 0 | NA | NA | NA | NA | NA | NA | NA | 0 | 0 | 0 | 0 | 0 | 0 |
| 30035 | ENSMUSG00000088910 | 0 | NA | NA | NA | NA | NA | NA | NA | 0 | 0 | 0 | 0 | 0 | 0 |
| 30036 | ENSMUSG00000088911 | 0 | NA | NA | NA | NA | NA | Gm22609 | predicted gene, 22609 [Source:MGI Symb | 0 | 0 | 0 | 0 | 0 | 0 |
| 30037 | ENSMUSG00000088913 | 0 | NA | NA | NA | NA | NA | NA | NA | 0 | 0 | 0 | 0 | 0 | 0 |
| 30038 | ENSMUSG00000088914 | 0 | NA | NA | NA | NA | NA | Gm22605 | predicted gene, 22605 [Source:MGI Symb | 0 | 0 | 0 | 0 | 0 | 0 |
| 30039 | ENSMUSG00000088915 | 0.670829724 | -2.782124174 | 3.401449285 | -0.817923168 | 0.41340106 | NA | Gm22606 | predicted gene, 22606 [Source:MGI Symb | 0 | 0 | 4.025 | 0 | 0 | 0 |
| 30040 | ENSMUSG00000088916 | 0.167707431 | -0.796629176 | 4.080472857 | -0.195229623 | 0.84521319 | NA | NA | NA | 0 | 0 | 1.0062 | 0 | 0 | 0 |
| 30041 | ENSMUSG00000088918 | 0 | NA | NA | NA | NA | NA | NA | NA | 0 | 0 | 0 | 0 | 0 | 0 |
| 30042 | ENSMUSG00000088920 | 0 | NA | NA | NA | NA | NA | NA | NA | 0 | 0 | 0 | 0 | 0 | 0 |
| 30043 | ENSMUSG00000088922 | 0 | NA | NA | NA | NA | NA | Gm24296 | predicted gene, 24296 [Source:MGI Symb | 0 | 0 | 0 | 0 | 0 | 0 |
| 30044 | ENSMUSG00000088923 | 0 | NA | NA | NA | NA | NA | Gm24295 | predicted gene, 24295 [Source:MGI Symb | 0 | 0 | 0 | 0 | 0 | 0 |
| 30045 | ENSMUSG00000088924 | 0 | NA | NA | NA | NA | NA | Gm24298 | predicted gene, 24298 [Source:MGI Symb | 0 | 0 | 0 | 0 | 0 | 0 |
| 30046 | ENSMUSG00000088925 | 0 | NA | NA | NA | NA | NA | NA | NA | 0 | 0 | 0 | 0 | 0 | 0 |
| 30047 | ENSMUSG00000088926 | 0 | NA | NA | NA | NA | NA | NA | NA | 0 | 0 | 0 | 0 | 0 | 0 |
| 30048 | ENSMUSG00000088928 | 0 | NA | NA | NA | NA | NA | Gm25898 | predicted gene, 25898 [Source:MGI Symb | 0 | 0 | 0 | 0 | 0 | 0 |
| 30049 | ENSMUSG00000088929 | 0 | NA | NA | NA | NA | NA | Gm24299 | predicted gene, 24299 [Source:MGI Symb | 0 | 0 | 0 | 0 | 0 | 0 |
| 30050 | ENSMUSG00000088930 | 0 | NA | NA | NA | NA | NA | Gm25921 | predicted gene, 25921 [Source:MGI Symb | 0 | 0 | 0 | 0 | 0 | 0 |
| 30051 | ENSMUSG00000088932 | 0 | NA | NA | NA | NA | NA | NA | NA | 0 | 0 | 0 | 0 | 0 | 0 |
| 30052 | ENSMUSG00000088933 | 0 | NA | NA | NA | NA | NA | NA | NA | 0 | 0 | 0 | 0 | 0 | 0 |
| 30053 | ENSMUSG00000088935 | 0 | NA | NA | NA | NA | NA | Gm25924 | predicted gene, 25924 [Source:MGI Symb | 0 | 0 | 0 | 0 | 0 | 0 |
| 30054 | ENSMUSG00000088937 | 0 | NA | NA | NA | NA | NA | NA | NA | 0 | 0 | 0 | 0 | 0 | 0 |
| 30055 | ENSMUSG00000088938 | 0 | NA | NA | NA | NA | NA | NA | NA | 0 | 0 | 0 | 0 | 0 | 0 |
| 30056 | ENSMUSG00000088939 | 0 | NA | NA | NA | NA | NA | NA | NA | 0 | 0 | 0 | 0 | 0 | 0 |
| 30057 | ENSMUSG00000088940 | 0 | NA | NA | NA | NA | NA | NA | NA | 0 | 0 | 0 | 0 | 0 | 0 |
| 30058 | ENSMUSG00000088941 | 0 | NA | NA | NA | NA | NA | NA | NA | 0 | 0 | 0 | 0 | 0 | 0 |
| 30059 | ENSMUSG00000088942 | 0 | NA | NA | NA | NA | NA | Gm23260 | predicted gene, 23260 [Source:MGI Symb | 0 | 0 | 0 | 0 | 0 | 0 |
| 30060 | ENSMUSG00000088943 | 0 | NA | NA | NA | NA | NA | NA | NA | 0 | 0 | 0 | 0 | 0 | 0 |
| 30061 | ENSMUSG00000088944 | 0 | NA | NA | NA | NA | NA | Gm23258 | predicted gene, 23258 [Source:MGI Symb | 0 | 0 | 0 | 0 | 0 | 0 |
| 30062 | ENSMUSG00000088947 | 0.503302254 | -0.837877584 | 3.426698643 | -0.244514523 | 0.806832356 | NA | NA | NA | 0 | 2.0558 | 0 | 0.964 | 0 | 0 |
| 30063 | ENSMUSG00000088948 | 0 | NA | NA | NA | NA | NA | Gm23262 | predicted gene, 23262 [Source:MGI Symb | 0 | 0 | 0 | 0 | 0 | 0 |
| 30064 | ENSMUSG00000088949 | 0.185640647 | 1.126924175 | 4.080472857 | 0.276174898 | 0.782413736 | NA | NA | NA | 0 | 0 | 0 | 0 | 0 | 1.1138 |
| 30065 | ENSMUSG00000088950 | 0 | NA | NA | NA | NA | NA | NA | NA | 0 | 0 | 0 | 0 | 0 | 0 |
| 30066 | ENSMUSG00000088951 | 0 | NA | NA | NA | NA | NA | Gm24940 | predicted gene, 24940 [Source:MGI Symb | 0 | 0 | 0 | 0 | 0 | 0 |
| 30067 | ENSMUSG00000088953 | 0 | NA | NA | NA | NA | NA | Gm24942 | predicted gene, 24942 [Source:MGI Symb | 0 | 0 | 0 | 0 | 0 | 0 |
| 30068 | ENSMUSG00000088954 | 0 | NA | NA | NA | NA | NA | Gm24943 | predicted gene, 24943 [Source:MGI Symb | 0 | 0 | 0 | 0 | 0 | 0 |

| | | | | | | | | | | | | | | | |
|---|---|---|---|---|---|---|---|---|---|---|---|---|---|---|---|
| 30069 | ENSMUSG00000088955 | 0.885240214 | -0.466577448 | 2.807084628 | -0.166214244 | 0.867988351 | NA | NA | NA | 0 | 3.0838 | 0 | 0 | 0 | 2.2277 |
| 30070 | ENSMUSG00000088956 | 0 | NA | NA | NA | NA | NA | NA | NA | 0 | 0 | 0 | 0 | 0 | 0 |
| 30071 | ENSMUSG00000088957 | 0 | NA | NA | NA | NA | NA | NA | NA | 0 | 0 | 0 | 0 | 0 | 0 |
| 30072 | ENSMUSG00000088958 | 0 | NA | NA | NA | NA | NA | Scarna8 | small Cajal body-specific RNA 8 [Source:M | 0 | 0 | 0 | 0 | 0 | 0 |
| 30073 | ENSMUSG00000088959 | 0 | NA | NA | NA | NA | NA | Gm24947 | predicted gene, 24947 [Source:MGI Symb | 0 | 0 | 0 | 0 | 0 | 0 |
| 30074 | ENSMUSG00000088960 | 0 | NA | NA | NA | NA | NA | Gm22090 | predicted gene, 22090 [Source:MGI Symb | 0 | 0 | 0 | 0 | 0 | 0 |
| 30075 | ENSMUSG00000088961 | 0.167707431 | -0.796629176 | 4.080472857 | -0.195229623 | 0.84521319 | NA | Gm22089 | predicted gene, 22089 [Source:MGI Symb | 0 | 0 | 1.0062 | 0 | 0 | 0 |
| 30076 | ENSMUSG00000088964 | 0 | NA | NA | NA | NA | NA | Gm22087 | predicted gene, 22087 [Source:MGI Symb | 0 | 0 | 0 | 0 | 0 | 0 |
| 30077 | ENSMUSG00000088966 | 0 | NA | NA | NA | NA | NA | NA | NA | 0 | 0 | 0 | 0 | 0 | 0 |
| 30078 | ENSMUSG00000088969 | 0 | NA | NA | NA | NA | NA | Gm22085 | predicted gene, 22085 [Source:MGI Symb | 0 | 0 | 0 | 0 | 0 | 0 |
| 30079 | ENSMUSG00000088970 | 0 | NA | NA | NA | NA | NA | Gm23763 | predicted gene, 23763 [Source:MGI Symb | 0 | 0 | 0 | 0 | 0 | 0 |
| 30080 | ENSMUSG00000088971 | 0 | NA | NA | NA | NA | NA | NA | NA | 0 | 0 | 0 | 0 | 0 | 0 |
| 30081 | ENSMUSG00000088972 | 0 | NA | NA | NA | NA | NA | NA | NA | 0 | 0 | 0 | 0 | 0 | 0 |
| 30082 | ENSMUSG00000088973 | 0 | NA | NA | NA | NA | NA | NA | NA | 0 | 0 | 0 | 0 | 0 | 0 |
| 30083 | ENSMUSG00000088975 | 0 | NA | NA | NA | NA | NA | Gm23760 | predicted gene, 23760 [Source:MGI Symb | 0 | 0 | 0 | 0 | 0 | 0 |
| 30084 | ENSMUSG00000088976 | 0.305170013 | -1.657836477 | 4.041480335 | -0.410205257 | 0.681655385 | NA | NA | NA | 0.8031 | 1.0279 | 0 | 0 | 0 | 0 |
| 30085 | ENSMUSG00000088978 | 0.133850373 | -0.796629176 | 4.080472857 | -0.195229623 | 0.84521319 | NA | NA | NA | 0.8031 | 0 | 0 | 0 | 0 | 0 |
| 30086 | ENSMUSG00000088979 | 0.133850373 | -0.796629176 | 4.080472857 | -0.195229623 | 0.84521319 | NA | Gm23758 | predicted gene, 23758 [Source:MGI Symb | 0.8031 | 0 | 0 | 0 | 0 | 0 |
| 30087 | ENSMUSG00000088980 | 0 | NA | NA | NA | NA | NA | Mir669m-2 | microRNA 669m-2 [Source:MGI Symbol;A | 0 | 0 | 0 | 0 | 0 | 0 |
| 30088 | ENSMUSG00000088981 | 0 | NA | NA | NA | NA | NA | NA | NA | 0 | 0 | 0 | 0 | 0 | 0 |
| 30089 | ENSMUSG00000088982 | 0.321325949 | 1.911575114 | 4.048267893 | 0.472195804 | 0.636787034 | NA | Gm25600 | predicted gene, 25600 [Source:MGI Symb | 0 | 0 | 0 | 1.928 | 0 | 0 |
| 30090 | ENSMUSG00000088983 | 0 | NA | NA | NA | NA | NA | Mir1952 | microRNA 1952 [Source:MGI Symbol;Acc: | 0 | 0 | 0 | 0 | 0 | 0 |
| 30091 | ENSMUSG00000088984 | 1.618412372 | -1.905872387 | 1.858460436 | -1.025511413 | 0.305121943 | NA | Gm25604 | predicted gene, 25604 [Source:MGI Symb | 5.6217 | 1.0279 | 1.0062 | 0.964 | 1.0906 | 0 |
| 30092 | ENSMUSG00000088985 | 0 | NA | NA | NA | NA | NA | Mir1946a | microRNA 1946a [Source:MGI Symbol;Acc | 0 | 0 | 0 | 0 | 0 | 0 |
| 30093 | ENSMUSG00000088986 | 0 | NA | NA | NA | NA | NA | NA | NA | 0 | 0 | 0 | 0 | 0 | 0 |
| 30094 | ENSMUSG00000088987 | 0 | NA | NA | NA | NA | NA | Gm25602 | predicted gene, 25602 [Source:MGI Symb | 0 | 0 | 0 | 0 | 0 | 0 |
| 30095 | ENSMUSG00000088988 | 0 | NA | NA | NA | NA | NA | NA | NA | 0 | 0 | 0 | 0 | 0 | 0 |
| 30096 | ENSMUSG00000088989 | 0 | NA | NA | NA | NA | NA | NA | NA | 0 | 0 | 0 | 0 | 0 | 0 |
| 30097 | ENSMUSG00000088990 | 0 | NA | NA | NA | NA | NA | Gm22767 | predicted gene, 22767 [Source:MGI Symb | 0 | 0 | 0 | 0 | 0 | 0 |
| 30098 | ENSMUSG00000088991 | 0 | NA | NA | NA | NA | NA | NA | NA | 0 | 0 | 0 | 0 | 0 | 0 |
| 30099 | ENSMUSG00000088992 | 0 | NA | NA | NA | NA | NA | Gm22769 | predicted gene, 22769 [Source:MGI Symb | 0 | 0 | 0 | 0 | 0 | 0 |
| 30100 | ENSMUSG00000088993 | 0 | NA | NA | NA | NA | NA | NA | NA | 0 | 0 | 0 | 0 | 0 | 0 |
| 30101 | ENSMUSG00000088994 | 0 | NA | NA | NA | NA | NA | NA | NA | 0 | 0 | 0 | 0 | 0 | 0 |
| 30102 | ENSMUSG00000088995 | 0 | NA | NA | NA | NA | NA | Gm22764 | predicted gene, 22764 [Source:MGI Symb | 0 | 0 | 0 | 0 | 0 | 0 |
| 30103 | ENSMUSG00000088996 | 0 | NA | NA | NA | NA | NA | NA | NA | 0 | 0 | 0 | 0 | 0 | 0 |
| 30104 | ENSMUSG00000088997 | 0.133850373 | -0.796629176 | 4.080472857 | -0.195229623 | 0.84521319 | NA | Gm22766 | predicted gene, 22766 [Source:MGI Symb | 0.8031 | 0 | 0 | 0 | 0 | 0 |
| 30105 | ENSMUSG00000088999 | 0 | NA | NA | NA | NA | NA | Gm22763 | predicted gene, 22763 [Source:MGI Symb | 0 | 0 | 0 | 0 | 0 | 0 |
| 30106 | ENSMUSG00000089000 | 0 | NA | NA | NA | NA | NA | NA | NA | 0 | 0 | 0 | 0 | 0 | 0 |
| 30107 | ENSMUSG00000089001 | 0 | NA | NA | NA | NA | NA | Gm23852 | predicted gene, 23852 [Source:MGI Symb | 0 | 0 | 0 | 0 | 0 | 0 |
| 30108 | ENSMUSG00000089002 | 0 | NA | NA | NA | NA | NA | Gm23855 | predicted gene, 23855 [Source:MGI Symb | 0 | 0 | 0 | 0 | 0 | 0 |
| 30109 | ENSMUSG00000089003 | 0 | NA | NA | NA | NA | NA | NA | NA | 0 | 0 | 0 | 0 | 0 | 0 |
| 30110 | ENSMUSG00000089004 | 0 | NA | NA | NA | NA | NA | NA | NA | 0 | 0 | 0 | 0 | 0 | 0 |
| 30111 | ENSMUSG00000089005 | 0 | NA | NA | NA | NA | NA | Gm23856 | predicted gene, 23856 [Source:MGI Symb | 0 | 0 | 0 | 0 | 0 | 0 |
| 30112 | ENSMUSG00000089007 | 0 | NA | NA | NA | NA | NA | Gm23877 | predicted gene, 23877 [Source:MGI Symb | 0 | 0 | 0 | 0 | 0 | 0 |
| 30113 | ENSMUSG00000089008 | 0 | NA | NA | NA | NA | NA | Gm23859 | predicted gene, 23859 [Source:MGI Symb | 0 | 0 | 0 | 0 | 0 | 0 |
| 30114 | ENSMUSG00000089009 | 0.181769715 | 1.126924175 | 4.080472857 | 0.276174898 | 0.782413736 | NA | Gm23858 | predicted gene, 23858 [Source:MGI Symb | 0 | 0 | 0 | 0 | 1.0906 | 0 |
| 30115 | ENSMUSG00000089011 | 0 | NA | NA | NA | NA | NA | Gm24879 | predicted gene, 24879 [Source:MGI Symb | 0 | 0 | 0 | 0 | 0 | 0 |
| 30116 | ENSMUSG00000089014 | 0 | NA | NA | NA | NA | NA | Gm25520 | predicted gene, 25520 [Source:MGI Symb | 0 | 0 | 0 | 0 | 0 | 0 |
| 30117 | ENSMUSG00000089015 | 1.011684555 | -0.99386545 | 2.458038153 | -0.404332801 | 0.685968008 | NA | NA | NA | 4.0155 | 0 | 0 | 0.964 | 1.0906 | 0 |
| 30118 | ENSMUSG00000089016 | 0 | NA | NA | NA | NA | NA | NA | NA | 0 | 0 | 0 | 0 | 0 | 0 |
| 30119 | ENSMUSG00000089017 | 0 | NA | NA | NA | NA | NA | Gm25519 | predicted gene, 25519 [Source:MGI Symb | 0 | 0 | 0 | 0 | 0 | 0 |
| 30120 | ENSMUSG00000089018 | 0 | NA | NA | NA | NA | NA | Gm25521 | predicted gene, 25521 [Source:MGI Symb | 0 | 0 | 0 | 0 | 0 | 0 |
| 30121 | ENSMUSG00000089019 | 0 | NA | NA | NA | NA | NA | Gm25522 | predicted gene, 25522 [Source:MGI Symb | 0 | 0 | 0 | 0 | 0 | 0 |
| 30122 | ENSMUSG00000089021 | 0 | NA | NA | NA | NA | NA | NA | NA | 0 | 0 | 0 | 0 | 0 | 0 |
| 30123 | ENSMUSG00000089023 | 0 | NA | NA | NA | NA | NA | NA | NA | 0 | 0 | 0 | 0 | 0 | 0 |
| 30124 | ENSMUSG00000089024 | 0 | NA | NA | NA | NA | NA | Gm25016 | predicted gene, 25016 [Source:MGI Symb | 0 | 0 | 0 | 0 | 0 | 0 |

| | | | | | | | | | | | | | | | |
|---|---|---|---|---|---|---|---|---|---|---|---|---|---|---|---|
| 30125 | ENSMUSG00000089025 | 0 | NA | NA | NA | NA | NA | Gm25015 | predicted gene, 25015 [Source:MGI Symb | 0 | 0 | 0 | 0 | 0 | 0 |
| 30126 | ENSMUSG00000089026 | 0.36353943 | 2.065989254 | 4.034137301 | 0.512126658 | 0.608562368 | NA | Gm25014 | predicted gene, 25014 [Source:MGI Symb | 0 | 0 | 0 | 0 | 2.1812 | 0 |
| 30127 | ENSMUSG00000089027 | 0 | NA | NA | NA | NA | NA | Gm25013 | predicted gene, 25013 [Source:MGI Symb | 0 | 0 | 0 | 0 | 0 | 0 |
| 30128 | ENSMUSG00000089028 | 0 | NA | NA | NA | NA | NA | Gm25010 | predicted gene, 25010 [Source:MGI Symb | 0 | 0 | 0 | 0 | 0 | 0 |
| 30129 | ENSMUSG00000089029 | 0 | NA | NA | NA | NA | NA | Gm25009 | predicted gene, 25009 [Source:MGI Symb | 0 | 0 | 0 | 0 | 0 | 0 |
| 30130 | ENSMUSG00000089030 | 0 | NA | NA | NA | NA | NA | NA | NA | 0 | 0 | 0 | 0 | 0 | 0 |
| 30131 | ENSMUSG00000089033 | 0 | NA | NA | NA | NA | NA | Gm22163 | predicted gene, 22163 [Source:MGI Symb | 0 | 0 | 0 | 0 | 0 | 0 |
| 30132 | ENSMUSG00000089034 | 0 | NA | NA | NA | NA | NA | Gm22156 | predicted gene, 22156 [Source:MGI Symb | 0 | 0 | 0 | 0 | 0 | 0 |
| 30133 | ENSMUSG00000089035 | 0 | NA | NA | NA | NA | NA | Gm22157 | predicted gene, 22157 [Source:MGI Symb | 0 | 0 | 0 | 0 | 0 | 0 |
| 30134 | ENSMUSG00000089036 | 0 | NA | NA | NA | NA | NA | Gm22158 | predicted gene, 22158 [Source:MGI Symb | 0 | 0 | 0 | 0 | 0 | 0 |
| 30135 | ENSMUSG00000089037 | 0 | NA | NA | NA | NA | NA | Gm22159 | predicted gene, 22159 [Source:MGI Symb | 0 | 0 | 0 | 0 | 0 | 0 |
| 30136 | ENSMUSG00000089038 | 0 | NA | NA | NA | NA | NA | NA | NA | 0 | 0 | 0 | 0 | 0 | 0 |
| 30137 | ENSMUSG00000089039 | 0.181769715 | 1.126924175 | 4.080472857 | 0.276174898 | 0.782413736 | NA | Gm22165 | predicted gene, 22165 [Source:MGI Symb | 0 | 0 | 0 | 0 | 1.0906 | 0 |
| 30138 | ENSMUSG00000089040 | 0 | NA | NA | NA | NA | NA | Gm26174 | predicted gene, 26174 [Source:MGI Symb | 0 | 0 | 0 | 0 | 0 | 0 |
| 30139 | ENSMUSG00000089041 | 0 | NA | NA | NA | NA | NA | NA | NA | 0 | 0 | 0 | 0 | 0 | 0 |
| 30140 | ENSMUSG00000089043 | 0 | NA | NA | NA | NA | NA | NA | NA | 0 | 0 | 0 | 0 | 0 | 0 |
| 30141 | ENSMUSG00000089044 | 0 | NA | NA | NA | NA | NA | NA | NA | 0 | 0 | 0 | 0 | 0 | 0 |
| 30142 | ENSMUSG00000089045 | 0 | NA | NA | NA | NA | NA | Gm26169 | predicted gene, 26169 [Source:MGI Symb | 0 | 0 | 0 | 0 | 0 | 0 |
| 30143 | ENSMUSG00000089046 | 0 | NA | NA | NA | NA | NA | Gm26172 | predicted gene, 26172 [Source:MGI Symb | 0 | 0 | 0 | 0 | 0 | 0 |
| 30144 | ENSMUSG00000089047 | 0 | NA | NA | NA | NA | NA | NA | NA | 0 | 0 | 0 | 0 | 0 | 0 |
| 30145 | ENSMUSG00000089048 | 0.979611946 | -3.327922556 | 2.381869301 | -1.397189407 | 0.162356621 | NA | NA | NA | 0.8031 | 2.0558 | 3.0187 | 0 | 0 | 0 |
| 30146 | ENSMUSG00000089049 | 0 | NA | NA | NA | NA | NA | NA | NA | 0 | 0 | 0 | 0 | 0 | 0 |
| 30147 | ENSMUSG00000089050 | 0 | NA | NA | NA | NA | NA | NA | NA | 0 | 0 | 0 | 0 | 0 | 0 |
| 30148 | ENSMUSG00000089051 | 1.305940833 | -1.378756532 | 2.068830352 | -0.666442529 | 0.505128286 | NA | Gm23354 | predicted gene, 23354 [Source:MGI Symb | 1.6062 | 0 | 4.025 | 0 | 1.0906 | 1.1138 |
| 30149 | ENSMUSG00000089052 | 0 | NA | NA | NA | NA | NA | Gm23351 | predicted gene, 23351 [Source:MGI Symb | 0 | 0 | 0 | 0 | 0 | 0 |
| 30150 | ENSMUSG00000089053 | 0 | NA | NA | NA | NA | NA | Gm23352 | predicted gene, 23352 [Source:MGI Symb | 0 | 0 | 0 | 0 | 0 | 0 |
| 30151 | ENSMUSG00000089054 | 0.342639279 | -1.805976596 | 4.028547714 | -0.448294702 | 0.653940524 | NA | Gm23350 | predicted gene, 23350 [Source:MGI Symb | 0 | 2.0558 | 0 | 0 | 0 | 0 |
| 30152 | ENSMUSG00000089055 | 0 | NA | NA | NA | NA | NA | Gm22675 | predicted gene, 22675 [Source:MGI Symb | 0 | 0 | 0 | 0 | 0 | 0 |
| 30153 | ENSMUSG00000089057 | 0 | NA | NA | NA | NA | NA | Gm23349 | predicted gene, 23349 [Source:MGI Symb | 0 | 0 | 0 | 0 | 0 | 0 |
| 30154 | ENSMUSG00000089060 | 0 | NA | NA | NA | NA | NA | Gm22361 | predicted gene, 22361 [Source:MGI Symb | 0 | 0 | 0 | 0 | 0 | 0 |
| 30155 | ENSMUSG00000089061 | 0 | NA | NA | NA | NA | NA | NA | NA | 0 | 0 | 0 | 0 | 0 | 0 |
| 30156 | ENSMUSG00000089062 | 0 | NA | NA | NA | NA | NA | NA | NA | 0 | 0 | 0 | 0 | 0 | 0 |
| 30157 | ENSMUSG00000089065 | 0 | NA | NA | NA | NA | NA | NA | NA | 0 | 0 | 0 | 0 | 0 | 0 |
| 30158 | ENSMUSG00000089066 | 0 | NA | NA | NA | NA | NA | Gm22836 | predicted gene, 22836 [Source:MGI Symb | 0 | 0 | 0 | 0 | 0 | 0 |
| 30159 | ENSMUSG00000089068 | 0 | NA | NA | NA | NA | NA | Gm22841 | predicted gene, 22841 [Source:MGI Symb | 0 | 0 | 0 | 0 | 0 | 0 |
| 30160 | ENSMUSG00000089069 | 0 | NA | NA | NA | NA | NA | Gm22840 | predicted gene, 22840 [Source:MGI Symb | 0 | 0 | 0 | 0 | 0 | 0 |
| 30161 | ENSMUSG00000089070 | 0 | NA | NA | NA | NA | NA | NA | NA | 0 | 0 | 0 | 0 | 0 | 0 |
| 30162 | ENSMUSG00000089071 | 0 | NA | NA | NA | NA | NA | Gm24499 | predicted gene, 24499 [Source:MGI Symb | 0 | 0 | 0 | 0 | 0 | 0 |
| 30163 | ENSMUSG00000089072 | 1.432720119 | 1.285700338 | 1.931877464 | 0.665518576 | 0.505718867 | NA | NA | NA | 2.4093 | 0 | 0 | 2.8919 | 2.1812 | 1.1138 |
| 30164 | ENSMUSG00000089073 | 0 | NA | NA | NA | NA | NA | NA | NA | 0 | 0 | 0 | 0 | 0 | 0 |
| 30165 | ENSMUSG00000089074 | 0 | NA | NA | NA | NA | NA | Gm23817 | predicted gene, 23817 [Source:MGI Symb | 0 | 0 | 0 | 0 | 0 | 0 |
| 30166 | ENSMUSG00000089075 | 0 | NA | NA | NA | NA | NA | NA | NA | 0 | 0 | 0 | 0 | 0 | 0 |
| 30167 | ENSMUSG00000089077 | 0.321325949 | 1.911575114 | 4.048267893 | 0.472195804 | 0.636787034 | NA | NA | NA | 0 | 0 | 0 | 1.928 | 0 | 0 |
| 30168 | ENSMUSG00000089078 | 0 | NA | NA | NA | NA | NA | Gm24502 | predicted gene, 24502 [Source:MGI Symb | 0 | 0 | 0 | 0 | 0 | 0 |
| 30169 | ENSMUSG00000089079 | 0 | NA | NA | NA | NA | NA | Gm24503 | predicted gene, 24503 [Source:MGI Symb | 0 | 0 | 0 | 0 | 0 | 0 |
| 30170 | ENSMUSG00000089080 | 0 | NA | NA | NA | NA | NA | Gm24003 | predicted gene, 24003 [Source:MGI Symb | 0 | 0 | 0 | 0 | 0 | 0 |
| 30171 | ENSMUSG00000089081 | 0 | NA | NA | NA | NA | NA | Gm24002 | predicted gene, 24002 [Source:MGI Symb | 0 | 0 | 0 | 0 | 0 | 0 |
| 30172 | ENSMUSG00000089082 | 0 | NA | NA | NA | NA | NA | NA | NA | 0 | 0 | 0 | 0 | 0 | 0 |
| 30173 | ENSMUSG00000089083 | 0 | NA | NA | NA | NA | NA | Rnu7 | U7 small nuclear RNA [Source:MGI Symbc | 0 | 0 | 0 | 0 | 0 | 0 |
| 30174 | ENSMUSG00000089084 | 0 | NA | NA | NA | NA | NA | NA | NA | 0 | 0 | 0 | 0 | 0 | 0 |
| 30175 | ENSMUSG00000089085 | 0 | NA | NA | NA | NA | NA | Gm24004 | predicted gene, 24004 [Source:MGI Symb | 0 | 0 | 0 | 0 | 0 | 0 |
| 30176 | ENSMUSG00000089087 | 0 | NA | NA | NA | NA | NA | Gm24006 | predicted gene, 24006 [Source:MGI Symb | 0 | 0 | 0 | 0 | 0 | 0 |
| 30177 | ENSMUSG00000089088 | 0 | NA | NA | NA | NA | NA | NA | NA | 0 | 0 | 0 | 0 | 0 | 0 |
| 30178 | ENSMUSG00000089090 | 0 | NA | NA | NA | NA | NA | NA | NA | 0 | 0 | 0 | 0 | 0 | 0 |
| 30179 | ENSMUSG00000089091 | 0 | NA | NA | NA | NA | NA | Gm25656 | predicted gene, 25656 [Source:MGI Symb | 0 | 0 | 0 | 0 | 0 | 0 |
| 30180 | ENSMUSG00000089092 | 3.122744169 | -1.372257798 | 1.286915442 | -1.066315434 | 0.286281075 | NA | NA | NA | 6.4248 | 1.0279 | 6.0375 | 1.928 | 1.0906 | 2.2277 |

| | | | | | | | | | | | | | | | |
|---|---|---|---|---|---|---|---|---|---|---|---|---|---|---|---|
| 30181 | ENSMUSG00000089093 | 0 | NA | NA | NA | NA | NA | Snord11 | small nucleolar RNA, C/D box 11 [Source:I | 0 | 0 | 0 | 0 | 0 | 0 |
| 30182 | ENSMUSG00000089094 | 0 | NA | NA | NA | NA | NA | NA | NA | 0 | 0 | 0 | 0 | 0 | 0 |
| 30183 | ENSMUSG00000089095 | 0 | NA | NA | NA | NA | NA | NA | NA | 0 | 0 | 0 | 0 | 0 | 0 |
| 30184 | ENSMUSG00000089096 | 0 | NA | NA | NA | NA | NA | Gm25657 | predicted gene, 25657 [Source:MGI Symb | 0 | 0 | 0 | 0 | 0 | 0 |
| 30185 | ENSMUSG00000089097 | 0 | NA | NA | NA | NA | NA | Gm25658 | predicted gene, 25658 [Source:MGI Symb | 0 | 0 | 0 | 0 | 0 | 0 |
| 30186 | ENSMUSG00000089098 | 0.160662974 | 1.126924175 | 4.080472857 | 0.276174898 | 0.782413736 | NA | Gm25653 | predicted gene, 25653 [Source:MGI Symb | 0 | 0 | 0 | 0.964 | 0 | 0 |
| 30187 | ENSMUSG00000089102 | 0 | NA | NA | NA | NA | NA | Gm23113 | predicted gene, 23113 [Source:MGI Symb | 0 | 0 | 0 | 0 | 0 | 0 |
| 30188 | ENSMUSG00000089103 | 0 | NA | NA | NA | NA | NA | NA | NA | 0 | 0 | 0 | 0 | 0 | 0 |
| 30189 | ENSMUSG00000089105 | 0 | NA | NA | NA | NA | NA | NA | NA | 0 | 0 | 0 | 0 | 0 | 0 |
| 30190 | ENSMUSG00000089106 | 0 | NA | NA | NA | NA | NA | Gm23116 | predicted gene, 23116 [Source:MGI Symb | 0 | 0 | 0 | 0 | 0 | 0 |
| 30191 | ENSMUSG00000089107 | 0 | NA | NA | NA | NA | NA | Gm23117 | predicted gene, 23117 [Source:MGI Symb | 0 | 0 | 0 | 0 | 0 | 0 |
| 30192 | ENSMUSG00000089109 | 0 | NA | NA | NA | NA | NA | Gm23916 | predicted gene, 23916 [Source:MGI Symb | 0 | 0 | 0 | 0 | 0 | 0 |
| 30193 | ENSMUSG00000089111 | 0 | NA | NA | NA | NA | NA | NA | NA | 0 | 0 | 0 | 0 | 0 | 0 |
| 30194 | ENSMUSG00000089113 | 0 | NA | NA | NA | NA | NA | Gm22902 | predicted gene, 22902 [Source:MGI Symb | 0 | 0 | 0 | 0 | 0 | 0 |
| 30195 | ENSMUSG00000089114 | 0 | NA | NA | NA | NA | NA | Gm25906 | predicted gene, 25906 [Source:MGI Symb | 0 | 0 | 0 | 0 | 0 | 0 |
| 30196 | ENSMUSG00000089115 | 0 | NA | NA | NA | NA | NA | NA | NA | 0 | 0 | 0 | 0 | 0 | 0 |
| 30197 | ENSMUSG00000089116 | 0 | NA | NA | NA | NA | NA | Gm25904 | predicted gene, 25904 [Source:MGI Symb | 0 | 0 | 0 | 0 | 0 | 0 |
| 30198 | ENSMUSG00000089117 | 0 | NA | NA | NA | NA | NA | Gm25903 | predicted gene, 25903 [Source:MGI Symb | 0 | 0 | 0 | 0 | 0 | 0 |
| 30199 | ENSMUSG00000089118 | 0 | NA | NA | NA | NA | NA | NA | NA | 0 | 0 | 0 | 0 | 0 | 0 |
| 30200 | ENSMUSG00000089119 | 0 | NA | NA | NA | NA | NA | Gm25901 | predicted gene, 25901 [Source:MGI Symb | 0 | 0 | 0 | 0 | 0 | 0 |
| 30201 | ENSMUSG00000089120 | 0 | NA | NA | NA | NA | NA | NA | NA | 0 | 0 | 0 | 0 | 0 | 0 |
| 30202 | ENSMUSG00000089123 | 0 | NA | NA | NA | NA | NA | Gm26404 | predicted gene, 26404 [Source:MGI Symb | 0 | 0 | 0 | 0 | 0 | 0 |
| 30203 | ENSMUSG00000089124 | 0 | NA | NA | NA | NA | NA | Gm26402 | predicted gene, 26402 [Source:MGI Symb | 0 | 0 | 0 | 0 | 0 | 0 |
| 30204 | ENSMUSG00000089125 | 0.342639279 | -1.805976596 | 4.028547714 | -0.448294702 | 0.653940524 | NA | Gm26403 | predicted gene, 26403 [Source:MGI Symb | 0 | 2.0558 | 0 | 0 | 0 | 0 |
| 30205 | ENSMUSG00000089127 | 0 | NA | NA | NA | NA | NA | Gm26401 | predicted gene, 26401 [Source:MGI Symb | 0 | 0 | 0 | 0 | 0 | 0 |
| 30206 | ENSMUSG00000089128 | 0 | NA | NA | NA | NA | NA | Gm26406 | predicted gene, 26406 [Source:MGI Symb | 0 | 0 | 0 | 0 | 0 | 0 |
| 30207 | ENSMUSG00000089129 | 0.6996517 | 1.656076801 | 2.758183935 | 0.600422901 | 0.548224429 | NA | NA | NA | 0 | 0 | 1.0062 | 0.964 | 0 | 2.2277 |
| 30208 | ENSMUSG00000089130 | 0 | NA | NA | NA | NA | NA | Gm24770 | predicted gene, 24770 [Source:MGI Symb | 0 | 0 | 0 | 0 | 0 | 0 |
| 30209 | ENSMUSG00000089131 | 0 | NA | NA | NA | NA | NA | NA | NA | 0 | 0 | 0 | 0 | 0 | 0 |
| 30210 | ENSMUSG00000089132 | 0 | NA | NA | NA | NA | NA | NA | NA | 0 | 0 | 0 | 0 | 0 | 0 |
| 30211 | ENSMUSG00000089133 | 0 | NA | NA | NA | NA | NA | Gm24771 | predicted gene, 24771 [Source:MGI Symb | 0 | 0 | 0 | 0 | 0 | 0 |
| 30212 | ENSMUSG00000089134 | 0 | NA | NA | NA | NA | NA | NA | NA | 0 | 0 | 0 | 0 | 0 | 0 |
| 30213 | ENSMUSG00000089136 | 0 | NA | NA | NA | NA | NA | Gm24922 | predicted gene, 24922 [Source:MGI Symb | 0 | 0 | 0 | 0 | 0 | 0 |
| 30214 | ENSMUSG00000089137 | 0 | NA | NA | NA | NA | NA | NA | NA | 0 | 0 | 0 | 0 | 0 | 0 |
| 30215 | ENSMUSG00000089138 | 0 | NA | NA | NA | NA | NA | NA | NA | 0 | 0 | 0 | 0 | 0 | 0 |
| 30216 | ENSMUSG00000089139 | 0.185640647 | 1.126924175 | 4.080472857 | 0.276174898 | 0.782413736 | NA | Mir1964 | microRNA 1964 [Source:MGI Symbol;Acc: | 0 | 0 | 0 | 0 | 0 | 1.1138 |
| 30217 | ENSMUSG00000089141 | 0 | NA | NA | NA | NA | NA | NA | NA | 0 | 0 | 0 | 0 | 0 | 0 |
| 30218 | ENSMUSG00000089143 | 0 | NA | NA | NA | NA | NA | Gm25268 | predicted gene, 25268 [Source:MGI Symb | 0 | 0 | 0 | 0 | 0 | 0 |
| 30219 | ENSMUSG00000089144 | 0 | NA | NA | NA | NA | NA | Gm25264 | predicted gene, 25264 [Source:MGI Symb | 0 | 0 | 0 | 0 | 0 | 0 |
| 30220 | ENSMUSG00000089145 | 0 | NA | NA | NA | NA | NA | NA | NA | 0 | 0 | 0 | 0 | 0 | 0 |
| 30221 | ENSMUSG00000089146 | 0 | NA | NA | NA | NA | NA | NA | NA | 0 | 0 | 0 | 0 | 0 | 0 |
| 30222 | ENSMUSG00000089147 | 0 | NA | NA | NA | NA | NA | Gm25267 | predicted gene, 25267 [Source:MGI Symb | 0 | 0 | 0 | 0 | 0 | 0 |
| 30223 | ENSMUSG00000089148 | 0 | NA | NA | NA | NA | NA | NA | NA | 0 | 0 | 0 | 0 | 0 | 0 |
| 30224 | ENSMUSG00000089149 | 0 | NA | NA | NA | NA | NA | NA | NA | 0 | 0 | 0 | 0 | 0 | 0 |
| 30225 | ENSMUSG00000089151 | 0 | NA | NA | NA | NA | NA | Mir1950 | microRNA 1950 [Source:MGI Symbol;Acc: | 0 | 0 | 0 | 0 | 0 | 0 |
| 30226 | ENSMUSG00000089152 | 0 | NA | NA | NA | NA | NA | Gm22424 | predicted gene, 22424 [Source:MGI Symb | 0 | 0 | 0 | 0 | 0 | 0 |
| 30227 | ENSMUSG00000089153 | 0 | NA | NA | NA | NA | NA | NA | NA | 0 | 0 | 0 | 0 | 0 | 0 |
| 30228 | ENSMUSG00000089155 | 0 | NA | NA | NA | NA | NA | Gm23582 | predicted gene, 23582 [Source:MGI Symb | 0 | 0 | 0 | 0 | 0 | 0 |
| 30229 | ENSMUSG00000089156 | 0 | NA | NA | NA | NA | NA | Gm23581 | predicted gene, 23581 [Source:MGI Symb | 0 | 0 | 0 | 0 | 0 | 0 |
| 30230 | ENSMUSG00000089157 | 0 | NA | NA | NA | NA | NA | Gm22425 | predicted gene, 22425 [Source:MGI Symb | 0 | 0 | 0 | 0 | 0 | 0 |
| 30231 | ENSMUSG00000089158 | 0 | NA | NA | NA | NA | NA | NA | NA | 0 | 0 | 0 | 0 | 0 | 0 |
| 30232 | ENSMUSG00000089161 | 0 | NA | NA | NA | NA | NA | Gm24130 | predicted gene, 24130 [Source:MGI Symb | 0 | 0 | 0 | 0 | 0 | 0 |
| 30233 | ENSMUSG00000089162 | 0 | NA | NA | NA | NA | NA | Gm24129 | predicted gene, 24129 [Source:MGI Symb | 0 | 0 | 0 | 0 | 0 | 0 |
| 30234 | ENSMUSG00000089163 | 0 | NA | NA | NA | NA | NA | NA | NA | 0 | 0 | 0 | 0 | 0 | 0 |
| 30235 | ENSMUSG00000089164 | 0 | NA | NA | NA | NA | NA | Gm24131 | predicted gene, 24131 [Source:MGI Symb | 0 | 0 | 0 | 0 | 0 | 0 |
| 30236 | ENSMUSG00000089168 | 0 | NA | NA | NA | NA | NA | Gm24128 | predicted gene, 24128 [Source:MGI Symb | 0 | 0 | 0 | 0 | 0 | 0 |

| | | | | | | | | | | | | | | | |
|---|---|---|---|---|---|---|---|---|---|---|---|---|---|---|---|
| 30237 | ENSMUSG00000089169 | 0 | NA | NA | NA | NA | NA | NA | NA | 0 | 0 | 0 | 0 | 0 | 0 |
| 30238 | ENSMUSG00000089170 | 0 | NA | NA | NA | NA | NA | NA | NA | 0 | 0 | 0 | 0 | 0 | 0 |
| 30239 | ENSMUSG00000089171 | 0 | NA | NA | NA | NA | NA | Gm22430 | predicted gene, 22430 [Source:MGI Symb | 0 | 0 | 0 | 0 | 0 | 0 |
| 30240 | ENSMUSG00000089174 | 0 | NA | NA | NA | NA | NA | Gm22434 | predicted gene, 22434 [Source:MGI Symb | 0 | 0 | 0 | 0 | 0 | 0 |
| 30241 | ENSMUSG00000089175 | 0 | NA | NA | NA | NA | NA | Gm22433 | predicted gene, 22433 [Source:MGI Symb | 0 | 0 | 0 | 0 | 0 | 0 |
| 30242 | ENSMUSG00000089177 | 0 | NA | NA | NA | NA | NA | NA | NA | 0 | 0 | 0 | 0 | 0 | 0 |
| 30243 | ENSMUSG00000089178 | 0 | NA | NA | NA | NA | NA | Gm22429 | predicted gene, 22429 [Source:MGI Symb | 0 | 0 | 0 | 0 | 0 | 0 |
| 30244 | ENSMUSG00000089179 | 0 | NA | NA | NA | NA | NA | Gm22428 | predicted gene, 22428 [Source:MGI Symb | 0 | 0 | 0 | 0 | 0 | 0 |
| 30245 | ENSMUSG00000089182 | 0 | NA | NA | NA | NA | NA | Gm22945 | predicted gene, 22945 [Source:MGI Symb | 0 | 0 | 0 | 0 | 0 | 0 |
| 30246 | ENSMUSG00000089183 | 0 | NA | NA | NA | NA | NA | Gm22946 | predicted gene, 22946 [Source:MGI Symb | 0 | 0 | 0 | 0 | 0 | 0 |
| 30247 | ENSMUSG00000089184 | 0 | NA | NA | NA | NA | NA | Gm22947 | predicted gene, 22947 [Source:MGI Symb | 0 | 0 | 0 | 0 | 0 | 0 |
| 30248 | ENSMUSG00000089185 | 0 | NA | NA | NA | NA | NA | NA | NA | 0 | 0 | 0 | 0 | 0 | 0 |
| 30249 | ENSMUSG00000089186 | 0.321325949 | 1.911575114 | 4.048267893 | 0.472195804 | 0.636787034 | NA | NA | NA | 0 | 0 | 0 | 1.928 | 0 | 0 |
| 30250 | ENSMUSG00000089187 | 0 | NA | NA | NA | NA | NA | NA | NA | 0 | 0 | 0 | 0 | 0 | 0 |
| 30251 | ENSMUSG00000089188 | 0.401551119 | -2.072982726 | 4.008251979 | -0.517178744 | 0.605031388 | NA | Gm22942 | predicted gene, 22942 [Source:MGI Symb | 2.4093 | 0 | 0 | 0 | 0 | 0 |
| 30252 | ENSMUSG00000089189 | 0 | NA | NA | NA | NA | NA | NA | NA | 0 | 0 | 0 | 0 | 0 | 0 |
| 30253 | ENSMUSG00000089190 | 0 | NA | NA | NA | NA | NA | Gm25771 | predicted gene, 25771 [Source:MGI Symb | 0 | 0 | 0 | 0 | 0 | 0 |
| 30254 | ENSMUSG00000089191 | 0 | NA | NA | NA | NA | NA | Gm25770 | predicted gene, 25770 [Source:MGI Symb | 0 | 0 | 0 | 0 | 0 | 0 |
| 30255 | ENSMUSG00000089194 | 0 | NA | NA | NA | NA | NA | Gm25769 | predicted gene, 25769 [Source:MGI Symb | 0 | 0 | 0 | 0 | 0 | 0 |
| 30256 | ENSMUSG00000089195 | 0 | NA | NA | NA | NA | NA | Gm25768 | predicted gene, 25768 [Source:MGI Symb | 0 | 0 | 0 | 0 | 0 | 0 |
| 30257 | ENSMUSG00000089197 | 0 | NA | NA | NA | NA | NA | Gm25767 | predicted gene, 25767 [Source:MGI Symb | 0 | 0 | 0 | 0 | 0 | 0 |
| 30258 | ENSMUSG00000089199 | 0.133850373 | -0.796629176 | 4.080472857 | -0.195229623 | 0.84521319 | NA | NA | NA | 0.8031 | 0 | 0 | 0 | 0 | 0 |
| 30259 | ENSMUSG00000089203 | 0 | NA | NA | NA | NA | NA | Gm26211 | predicted gene, 26211 [Source:MGI Symb | 0 | 0 | 0 | 0 | 0 | 0 |
| 30260 | ENSMUSG00000089204 | 0 | NA | NA | NA | NA | NA | Mir1927 | microRNA 1927 [Source:MGI Symbol;Acc: | 0 | 0 | 0 | 0 | 0 | 0 |
| 30261 | ENSMUSG00000089205 | 0 | NA | NA | NA | NA | NA | Gm24577 | predicted gene, 24577 [Source:MGI Symb | 0 | 0 | 0 | 0 | 0 | 0 |
| 30262 | ENSMUSG00000089206 | 0 | NA | NA | NA | NA | NA | NA | NA | 0 | 0 | 0 | 0 | 0 | 0 |
| 30263 | ENSMUSG00000089207 | 0 | NA | NA | NA | NA | NA | NA | NA | 0 | 0 | 0 | 0 | 0 | 0 |
| 30264 | ENSMUSG00000089208 | 0 | NA | NA | NA | NA | NA | NA | NA | 0 | 0 | 0 | 0 | 0 | 0 |
| 30265 | ENSMUSG00000089209 | 0 | NA | NA | NA | NA | NA | NA | NA | 0 | 0 | 0 | 0 | 0 | 0 |
| 30266 | ENSMUSG00000089210 | 0 | NA | NA | NA | NA | NA | Gm23386 | predicted gene, 23386 [Source:MGI Symb | 0 | 0 | 0 | 0 | 0 | 0 |
| 30267 | ENSMUSG00000089211 | 0 | NA | NA | NA | NA | NA | NA | NA | 0 | 0 | 0 | 0 | 0 | 0 |
| 30268 | ENSMUSG00000089213 | 0 | NA | NA | NA | NA | NA | NA | NA | 0 | 0 | 0 | 0 | 0 | 0 |
| 30269 | ENSMUSG00000089214 | 0.321325949 | 1.911575114 | 4.048267893 | 0.472195804 | 0.636787034 | NA | Mir1931 | microRNA 1931 [Source:MGI Symbol;Acc: | 0 | 0 | 0 | 1.928 | 0 | 0 |
| 30270 | ENSMUSG00000089217 | 0 | NA | NA | NA | NA | NA | NA | NA | 0 | 0 | 0 | 0 | 0 | 0 |
| 30271 | ENSMUSG00000089218 | 0 | NA | NA | NA | NA | NA | NA | NA | 0 | 0 | 0 | 0 | 0 | 0 |
| 30272 | ENSMUSG00000089219 | 0 | NA | NA | NA | NA | NA | Mir1963 | microRNA 1963 [Source:MGI Symbol;Acc: | 0 | 0 | 0 | 0 | 0 | 0 |
| 30273 | ENSMUSG00000089220 | 0 | NA | NA | NA | NA | NA | Gm22074 | predicted gene, 22074 [Source:MGI Symb | 0 | 0 | 0 | 0 | 0 | 0 |
| 30274 | ENSMUSG00000089222 | 0 | NA | NA | NA | NA | NA | Gm25049 | predicted gene, 25049 [Source:MGI Symb | 0 | 0 | 0 | 0 | 0 | 0 |
| 30275 | ENSMUSG00000089223 | 0 | NA | NA | NA | NA | NA | NA | NA | 0 | 0 | 0 | 0 | 0 | 0 |
| 30276 | ENSMUSG00000089224 | 0 | NA | NA | NA | NA | NA | NA | NA | 0 | 0 | 0 | 0 | 0 | 0 |
| 30277 | ENSMUSG00000089226 | 0 | NA | NA | NA | NA | NA | NA | NA | 0 | 0 | 0 | 0 | 0 | 0 |
| 30278 | ENSMUSG00000089227 | 0 | NA | NA | NA | NA | NA | NA | NA | 0 | 0 | 0 | 0 | 0 | 0 |
| 30279 | ENSMUSG00000089228 | 0 | NA | NA | NA | NA | NA | NA | NA | 0 | 0 | 0 | 0 | 0 | 0 |
| 30280 | ENSMUSG00000089229 | 0 | NA | NA | NA | NA | NA | Gm25043 | predicted gene, 25043 [Source:MGI Symb | 0 | 0 | 0 | 0 | 0 | 0 |
| 30281 | ENSMUSG00000089230 | 0 | NA | NA | NA | NA | NA | Gm22203 | predicted gene, 22203 [Source:MGI Symb | 0 | 0 | 0 | 0 | 0 | 0 |
| 30282 | ENSMUSG00000089231 | 0 | NA | NA | NA | NA | NA | Gm23112 | predicted gene, 23112 [Source:MGI Symb | 0 | 0 | 0 | 0 | 0 | 0 |
| 30283 | ENSMUSG00000089233 | 0 | NA | NA | NA | NA | NA | NA | NA | 0 | 0 | 0 | 0 | 0 | 0 |
| 30284 | ENSMUSG00000089234 | 0 | NA | NA | NA | NA | NA | NA | NA | 0 | 0 | 0 | 0 | 0 | 0 |
| 30285 | ENSMUSG00000089235 | 8.13384808 | 0.537712431 | 0.781435162 | 0.688108824 | 0.491384254 | 0.876368691 | Gm23119 | predicted gene, 23119 [Source:MGI Symb | 9.6372 | 6.1675 | 4.025 | 13.496 | 5.4531 | 10.025 |
| 30286 | ENSMUSG00000089237 | 0 | NA | NA | NA | NA | NA | Gm22206 | predicted gene, 22206 [Source:MGI Symb | 0 | 0 | 0 | 0 | 0 | 0 |
| 30287 | ENSMUSG00000089239 | 3.342918167 | -0.00971577 | 1.283592648 | -0.0075692 | 0.99396071 | NA | Gm22200 | predicted gene, 22200 [Source:MGI Symb | 4.0155 | 3.0838 | 3.0187 | 7.7118 | 0 | 2.2277 |
| 30288 | ENSMUSG00000089241 | 0 | NA | NA | NA | NA | NA | NA | NA | 0 | 0 | 0 | 0 | 0 | 0 |
| 30289 | ENSMUSG00000089242 | 0 | NA | NA | NA | NA | NA | Gm24056 | predicted gene, 24056 [Source:MGI Symb | 0 | 0 | 0 | 0 | 0 | 0 |
| 30290 | ENSMUSG00000089244 | 0 | NA | NA | NA | NA | NA | NA | NA | 0 | 0 | 0 | 0 | 0 | 0 |
| 30291 | ENSMUSG00000089245 | 0 | NA | NA | NA | NA | NA | NA | NA | 0 | 0 | 0 | 0 | 0 | 0 |
| 30292 | ENSMUSG00000089247 | 0 | NA | NA | NA | NA | NA | NA | NA | 0 | 0 | 0 | 0 | 0 | 0 |

| | | | | | | | | | | | | | | | |
|---|---|---|---|---|---|---|---|---|---|---|---|---|---|---|---|
| 30293 | ENSMUSG00000089248 | 0 | NA | NA | NA | NA | NA | NA | NA | 0 | 0 | 0 | 0 | 0 | 0 |
| 30294 | ENSMUSG00000089250 | 0 | NA | NA | NA | NA | NA | NA | NA | 0 | 0 | 0 | 0 | 0 | 0 |
| 30295 | ENSMUSG00000089251 | 0 | NA | NA | NA | NA | NA | NA | NA | 0 | 0 | 0 | 0 | 0 | 0 |
| 30296 | ENSMUSG00000089252 | 0 | NA | NA | NA | NA | NA | Gm25715 | predicted gene, 25715 [Source:MGI Symb | 0 | 0 | 0 | 0 | 0 | 0 |
| 30297 | ENSMUSG00000089253 | 1.117100031 | 1.582896982 | 2.163697584 | 0.731570342 | 0.464430856 | NA | NA | NA | 1.6062 | 0 | 0 | 2.8919 | 1.0906 | 1.1138 |
| 30298 | ENSMUSG00000089255 | 1.930576891 | -0.121026649 | 1.990913088 | -0.060789519 | 0.951526838 | NA | Snora78 | small nucleolar RNA, H/ACA box 7 [Source | 0 | 0 | 6.0375 | 0 | 1.0906 | 4.4554 |
| 30299 | ENSMUSG00000089256 | 0 | NA | NA | NA | NA | NA | NA | NA | 0 | 0 | 0 | 0 | 0 | 0 |
| 30300 | ENSMUSG00000089257 | 0 | NA | NA | NA | NA | NA | Gm25718 | predicted gene, 25718 [Source:MGI Symb | 0 | 0 | 0 | 0 | 0 | 0 |
| 30301 | ENSMUSG00000089258 | 0 | NA | NA | NA | NA | NA | Gm25711 | predicted gene, 25711 [Source:MGI Symb | 0 | 0 | 0 | 0 | 0 | 0 |
| 30302 | ENSMUSG00000089259 | 0 | NA | NA | NA | NA | NA | NA | NA | 0 | 0 | 0 | 0 | 0 | 0 |
| 30303 | ENSMUSG00000089260 | 0.17131964 | -0.796629176 | 4.080472857 | -0.195229623 | 0.84521319 | NA | Mir1929 | microRNA 1929 [Source:MGI Symbol;Acc: | 0 | 1.0279 | 0 | 0 | 0 | 0 |
| 30304 | ENSMUSG00000089261 | 0.160662974 | 1.126924175 | 4.080472857 | 0.276174898 | 0.782413736 | NA | Mir669n | microRNA 669n [Source:MGI Symbol;Acc: | 0 | 0 | 0 | 0.964 | 0 | 0 |
| 30305 | ENSMUSG00000089262 | 0 | NA | NA | NA | NA | NA | Gm22893 | predicted gene, 22893 [Source:MGI Symb | 0 | 0 | 0 | 0 | 0 | 0 |
| 30306 | ENSMUSG00000089263 | 0 | NA | NA | NA | NA | NA | NA | NA | 0 | 0 | 0 | 0 | 0 | 0 |
| 30307 | ENSMUSG00000089264 | 0 | NA | NA | NA | NA | NA | Gm22889 | predicted gene, 22889 [Source:MGI Symb | 0 | 0 | 0 | 0 | 0 | 0 |
| 30308 | ENSMUSG00000089265 | 0.959817171 | -2.042882424 | 2.2694873 | -0.900151512 | 0.368039626 | NA | Gm22888 | predicted gene, 22888 [Source:MGI Symb | 1.6062 | 2.0558 | 1.0062 | 0 | 1.0906 | 0 |
| 30309 | ENSMUSG00000089266 | 0 | NA | NA | NA | NA | NA | NA | NA | 0 | 0 | 0 | 0 | 0 | 0 |
| 30310 | ENSMUSG00000089267 | 0 | NA | NA | NA | NA | NA | Gm22890 | predicted gene, 22890 [Source:MGI Symb | 0 | 0 | 0 | 0 | 0 | 0 |
| 30311 | ENSMUSG00000089268 | 0 | NA | NA | NA | NA | NA | Mir1961 | microRNA 1961 [Source:MGI Symbol;Acc: | 0 | 0 | 0 | 0 | 0 | 0 |
| 30312 | ENSMUSG00000089269 | 0 | NA | NA | NA | NA | NA | Gm26468 | predicted gene, 26468 [Source:MGI Symb | 0 | 0 | 0 | 0 | 0 | 0 |
| 30313 | ENSMUSG00000089270 | 0 | NA | NA | NA | NA | NA | NA | NA | 0 | 0 | 0 | 0 | 0 | 0 |
| 30314 | ENSMUSG00000089271 | 0 | NA | NA | NA | NA | NA | Gm24552 | predicted gene, 24552 [Source:MGI Symb | 0 | 0 | 0 | 0 | 0 | 0 |
| 30315 | ENSMUSG00000089272 | 0 | NA | NA | NA | NA | NA | Gm24549 | predicted gene, 24549 [Source:MGI Symb | 0 | 0 | 0 | 0 | 0 | 0 |
| 30316 | ENSMUSG00000089273 | 0 | NA | NA | NA | NA | NA | NA | NA | 0 | 0 | 0 | 0 | 0 | 0 |
| 30317 | ENSMUSG00000089274 | 0 | NA | NA | NA | NA | NA | Gm24547 | predicted gene, 24547 [Source:MGI Symb | 0 | 0 | 0 | 0 | 0 | 0 |
| 30318 | ENSMUSG00000089275 | 0 | NA | NA | NA | NA | NA | Gm24548 | predicted gene, 24548 [Source:MGI Symb | 0 | 0 | 0 | 0 | 0 | 0 |
| 30319 | ENSMUSG00000089276 | 0 | NA | NA | NA | NA | NA | NA | NA | 0 | 0 | 0 | 0 | 0 | 0 |
| 30320 | ENSMUSG00000089277 | 0 | NA | NA | NA | NA | NA | Gm24546 | predicted gene, 24546 [Source:MGI Symb | 0 | 0 | 0 | 0 | 0 | 0 |
| 30321 | ENSMUSG00000089278 | 0 | NA | NA | NA | NA | NA | Gm24553 | predicted gene, 24553 [Source:MGI Symb | 0 | 0 | 0 | 0 | 0 | 0 |
| 30322 | ENSMUSG00000089279 | 0 | NA | NA | NA | NA | NA | Gm24554 | predicted gene, 24554 [Source:MGI Symb | 0 | 0 | 0 | 0 | 0 | 0 |
| 30323 | ENSMUSG00000089280 | 0 | NA | NA | NA | NA | NA | NA | NA | 0 | 0 | 0 | 0 | 0 | 0 |
| 30324 | ENSMUSG00000089281 | 1.88036282 | -0.186378774 | 1.665199407 | -0.111925799 | 0.910882241 | NA | Scarna6 | small Cajal body-specific RNA 6 [Source:M | 0 | 1.0279 | 5.0312 | 1.928 | 2.1812 | 1.1138 |
| 30325 | ENSMUSG00000089282 | 0 | NA | NA | NA | NA | NA | NA | NA | 0 | 0 | 0 | 0 | 0 | 0 |
| 30326 | ENSMUSG00000089284 | 0 | NA | NA | NA | NA | NA | Gm26029 | predicted gene, 26029 [Source:MGI Symb | 0 | 0 | 0 | 0 | 0 | 0 |
| 30327 | ENSMUSG00000089285 | 0 | NA | NA | NA | NA | NA | NA | NA | 0 | 0 | 0 | 0 | 0 | 0 |
| 30328 | ENSMUSG00000089286 | 0 | NA | NA | NA | NA | NA | NA | NA | 0 | 0 | 0 | 0 | 0 | 0 |
| 30329 | ENSMUSG00000089287 | 0 | NA | NA | NA | NA | NA | Gm22944 | predicted gene, 22944 [Source:MGI Symb | 0 | 0 | 0 | 0 | 0 | 0 |
| 30330 | ENSMUSG00000089288 | 0 | NA | NA | NA | NA | NA | Gm26022 | predicted gene, 26022 [Source:MGI Symb | 0 | 0 | 0 | 0 | 0 | 0 |
| 30331 | ENSMUSG00000089289 | 0 | NA | NA | NA | NA | NA | Gm26021 | predicted gene, 26021 [Source:MGI Symb | 0 | 0 | 0 | 0 | 0 | 0 |
| 30332 | ENSMUSG00000089290 | 0 | NA | NA | NA | NA | NA | Gm23207 | predicted gene, 23207 [Source:MGI Symb | 0 | 0 | 0 | 0 | 0 | 0 |
| 30333 | ENSMUSG00000089291 | 0 | NA | NA | NA | NA | NA | NA | NA | 0 | 0 | 0 | 0 | 0 | 0 |
| 30334 | ENSMUSG00000089292 | 0 | NA | NA | NA | NA | NA | NA | NA | 0 | 0 | 0 | 0 | 0 | 0 |
| 30335 | ENSMUSG00000089293 | 0 | NA | NA | NA | NA | NA | Gm23210 | predicted gene, 23210 [Source:MGI Symb | 0 | 0 | 0 | 0 | 0 | 0 |
| 30336 | ENSMUSG00000089294 | 0 | NA | NA | NA | NA | NA | Mir2139 | microRNA 2139 [Source:MGI Symbol;Acc: | 0 | 0 | 0 | 0 | 0 | 0 |
| 30337 | ENSMUSG00000089296 | 1.820156821 | -1.378936396 | 1.801638616 | -0.765379019 | 0.444045883 | NA | Gm23205 | predicted gene, 23205 [Source:MGI Symb | 4.8186 | 3.0838 | 0 | 1.928 | 1.0906 | 0 |
| 30338 | ENSMUSG00000089297 | 0 | NA | NA | NA | NA | NA | NA | NA | 0 | 0 | 0 | 0 | 0 | 0 |
| 30339 | ENSMUSG00000089298 | 0.160662974 | 1.126924175 | 4.080472857 | 0.276174898 | 0.782413736 | NA | NA | NA | 0 | 0 | 0 | 0.964 | 0 | 0 |
| 30340 | ENSMUSG00000089299 | 0.267700746 | -1.497481649 | 4.057001436 | -0.369110456 | 0.712045396 | NA | Gm24637 | predicted gene, 24637 [Source:MGI Symb | 1.6062 | 0 | 0 | 0 | 0 | 0 |
| 30341 | ENSMUSG00000089301 | 0 | NA | NA | NA | NA | NA | NA | NA | 0 | 0 | 0 | 0 | 0 | 0 |
| 30342 | ENSMUSG00000089302 | 0 | NA | NA | NA | NA | NA | Gm26208 | predicted gene, 26208 [Source:MGI Symb | 0 | 0 | 0 | 0 | 0 | 0 |
| 30343 | ENSMUSG00000089304 | 0 | NA | NA | NA | NA | NA | Gm24844 | predicted gene, 24844 [Source:MGI Symb | 0 | 0 | 0 | 0 | 0 | 0 |
| 30344 | ENSMUSG00000089305 | 0 | NA | NA | NA | NA | NA | NA | NA | 0 | 0 | 0 | 0 | 0 | 0 |
| 30345 | ENSMUSG00000089306 | 0.321325949 | 1.911575114 | 4.048267893 | 0.472195804 | 0.636787034 | NA | NA | NA | 0 | 0 | 0 | 1.928 | 0 | 0 |
| 30346 | ENSMUSG00000089307 | 0 | NA | NA | NA | NA | NA | Gm24843 | predicted gene, 24843 [Source:MGI Symb | 0 | 0 | 0 | 0 | 0 | 0 |
| 30347 | ENSMUSG00000089308 | 0.658052777 | 1.6393978 | 2.811180186 | 0.583170658 | 0.559778421 | NA | Gm24837 | predicted gene, 24837 [Source:MGI Symb | 0.8031 | 0 | 0 | 0.964 | 2.1812 | 0 |
| 30348 | ENSMUSG00000089309 | 0 | NA | NA | NA | NA | NA | Gm24838 | predicted gene, 24838 [Source:MGI Symb | 0 | 0 | 0 | 0 | 0 | 0 |

| | | | | | | | | | | | | | | | |
|---|---|---|---|---|---|---|---|---|---|---|---|---|---|---|---|
| 30349 | ENSMUSG00000089310 | 0 | NA | NA | NA | NA | NA | Gm23181 | predicted gene, 23181 [Source:MGI Symb | 0 | 0 | 0 | 0 | 0 | 0 |
| 30350 | ENSMUSG00000089311 | 0 | NA | NA | NA | NA | NA | Gm23180 | predicted gene, 23180 [Source:MGI Symb | 0 | 0 | 0 | 0 | 0 | 0 |
| 30351 | ENSMUSG00000089312 | 0 | NA | NA | NA | NA | NA | NA | NA | 0 | 0 | 0 | 0 | 0 | 0 |
| 30352 | ENSMUSG00000089313 | 0 | NA | NA | NA | NA | NA | Gm23182 | predicted gene, 23182 [Source:MGI Symb | 0 | 0 | 0 | 0 | 0 | 0 |
| 30353 | ENSMUSG00000089314 | 0 | NA | NA | NA | NA | NA | Gm23185 | predicted gene, 23185 [Source:MGI Symb | 0 | 0 | 0 | 0 | 0 | 0 |
| 30354 | ENSMUSG00000089316 | 0 | NA | NA | NA | NA | NA | NA | NA | 0 | 0 | 0 | 0 | 0 | 0 |
| 30355 | ENSMUSG00000089317 | 0.181769715 | 1.126924175 | 4.080472857 | 0.276174898 | 0.782413736 | NA | Snord123 | small nucleolar RNA, C/D box 123 [Source | 0 | 0 | 0 | 0 | 1.0906 | 0 |
| 30356 | ENSMUSG00000089318 | 0 | NA | NA | NA | NA | NA | NA | NA | 0 | 0 | 0 | 0 | 0 | 0 |
| 30357 | ENSMUSG00000089320 | 0 | NA | NA | NA | NA | NA | Gm25976 | predicted gene, 25976 [Source:MGI Symb | 0 | 0 | 0 | 0 | 0 | 0 |
| 30358 | ENSMUSG00000089321 | 0.17131964 | -0.796629176 | 4.080472857 | -0.195229623 | 0.84521319 | NA | Gm22523 | predicted gene, 22523 [Source:MGI Symb | 0 | 1.0279 | 0 | 0 | 0 | 0 |
| 30359 | ENSMUSG00000089322 | 0 | NA | NA | NA | NA | NA | Gm25977 | predicted gene, 25977 [Source:MGI Symb | 0 | 0 | 0 | 0 | 0 | 0 |
| 30360 | ENSMUSG00000089323 | 0 | NA | NA | NA | NA | NA | NA | NA | 0 | 0 | 0 | 0 | 0 | 0 |
| 30361 | ENSMUSG00000089324 | 0 | NA | NA | NA | NA | NA | Mir1936 | microRNA 1936 [Source:MGI Symbol;Acc: | 0 | 0 | 0 | 0 | 0 | 0 |
| 30362 | ENSMUSG00000089326 | 0 | NA | NA | NA | NA | NA | NA | NA | 0 | 0 | 0 | 0 | 0 | 0 |
| 30363 | ENSMUSG00000089327 | 0 | NA | NA | NA | NA | NA | Gm25975 | predicted gene, 25975 [Source:MGI Symb | 0 | 0 | 0 | 0 | 0 | 0 |
| 30364 | ENSMUSG00000089331 | 0 | NA | NA | NA | NA | NA | Gm24345 | predicted gene, 24345 [Source:MGI Symb | 0 | 0 | 0 | 0 | 0 | 0 |
| 30365 | ENSMUSG00000089333 | 0 | NA | NA | NA | NA | NA | Gm24343 | predicted gene, 24343 [Source:MGI Symb | 0 | 0 | 0 | 0 | 0 | 0 |
| 30366 | ENSMUSG00000089334 | 0 | NA | NA | NA | NA | NA | Gm24349 | predicted gene, 24349 [Source:MGI Symb | 0 | 0 | 0 | 0 | 0 | 0 |
| 30367 | ENSMUSG00000089335 | 0 | NA | NA | NA | NA | NA | Gm24348 | predicted gene, 24348 [Source:MGI Symb | 0 | 0 | 0 | 0 | 0 | 0 |
| 30368 | ENSMUSG00000089336 | 0.160662974 | 1.126924175 | 4.080472857 | 0.276174898 | 0.782413736 | NA | Gm24347 | predicted gene, 24347 [Source:MGI Symb | 0 | 0 | 0 | 0.964 | 0 | 0 |
| 30369 | ENSMUSG00000089337 | 0.698954292 | 0.126053087 | 3.048972385 | 0.04134281 | 0.967022605 | NA | NA | NA | 0 | 0 | 2.0125 | 0 | 2.1812 | 0 |
| 30370 | ENSMUSG00000089338 | 0 | NA | NA | NA | NA | NA | NA | NA | 0 | 0 | 0 | 0 | 0 | 0 |
| 30371 | ENSMUSG00000089341 | 0 | NA | NA | NA | NA | NA | NA | NA | 0 | 0 | 0 | 0 | 0 | 0 |
| 30372 | ENSMUSG00000089342 | 0 | NA | NA | NA | NA | NA | NA | NA | 0 | 0 | 0 | 0 | 0 | 0 |
| 30373 | ENSMUSG00000089343 | 0 | NA | NA | NA | NA | NA | NA | NA | 0 | 0 | 0 | 0 | 0 | 0 |
| 30374 | ENSMUSG00000089344 | 0 | NA | NA | NA | NA | NA | NA | NA | 0 | 0 | 0 | 0 | 0 | 0 |
| 30375 | ENSMUSG00000089345 | 0 | NA | NA | NA | NA | NA | NA | NA | 0 | 0 | 0 | 0 | 0 | 0 |
| 30376 | ENSMUSG00000089346 | 0.133850373 | -0.796629176 | 4.080472857 | -0.195229623 | 0.84521319 | NA | Gm22663 | predicted gene, 22663 [Source:MGI Symb | 0.8031 | 0 | 0 | 0 | 0 | 0 |
| 30377 | ENSMUSG00000089348 | 0 | NA | NA | NA | NA | NA | NA | NA | 0 | 0 | 0 | 0 | 0 | 0 |
| 30378 | ENSMUSG00000089349 | 0 | NA | NA | NA | NA | NA | Gm22670 | predicted gene, 22670 [Source:MGI Symb | 0 | 0 | 0 | 0 | 0 | 0 |
| 30379 | ENSMUSG00000089350 | 0.160662974 | 1.126924175 | 4.080472857 | 0.276174898 | 0.782413736 | NA | Gm25487 | predicted gene, 25487 [Source:MGI Symb | 0 | 0 | 0 | 0.964 | 0 | 0 |
| 30380 | ENSMUSG00000089351 | 0 | NA | NA | NA | NA | NA | Gm25486 | predicted gene, 25486 [Source:MGI Symb | 0 | 0 | 0 | 0 | 0 | 0 |
| 30381 | ENSMUSG00000089352 | 0 | NA | NA | NA | NA | NA | Gm25489 | predicted gene, 25489 [Source:MGI Symb | 0 | 0 | 0 | 0 | 0 | 0 |
| 30382 | ENSMUSG00000089353 | 0 | NA | NA | NA | NA | NA | NA | NA | 0 | 0 | 0 | 0 | 0 | 0 |
| 30383 | ENSMUSG00000089357 | 0 | NA | NA | NA | NA | NA | Mir2137 | microRNA 2137 [Source:MGI Symbol;Acc: | 0 | 0 | 0 | 0 | 0 | 0 |
| 30384 | ENSMUSG00000089358 | 0 | NA | NA | NA | NA | NA | NA | NA | 0 | 0 | 0 | 0 | 0 | 0 |
| 30385 | ENSMUSG00000089359 | 0 | NA | NA | NA | NA | NA | Gm25490 | predicted gene, 25490 [Source:MGI Symb | 0 | 0 | 0 | 0 | 0 | 0 |
| 30386 | ENSMUSG00000089363 | 0 | NA | NA | NA | NA | NA | NA | NA | 0 | 0 | 0 | 0 | 0 | 0 |
| 30387 | ENSMUSG00000089364 | 0 | NA | NA | NA | NA | NA | Gm23822 | predicted gene, 23822 [Source:MGI Symb | 0 | 0 | 0 | 0 | 0 | 0 |
| 30388 | ENSMUSG00000089365 | 0 | NA | NA | NA | NA | NA | NA | NA | 0 | 0 | 0 | 0 | 0 | 0 |
| 30389 | ENSMUSG00000089366 | 0 | NA | NA | NA | NA | NA | NA | NA | 0 | 0 | 0 | 0 | 0 | 0 |
| 30390 | ENSMUSG00000089369 | 0 | NA | NA | NA | NA | NA | Gm23819 | predicted gene, 23819 [Source:MGI Symb | 0 | 0 | 0 | 0 | 0 | 0 |
| 30391 | ENSMUSG00000089370 | 0 | NA | NA | NA | NA | NA | Gm22140 | predicted gene, 22140 [Source:MGI Symb | 0 | 0 | 0 | 0 | 0 | 0 |
| 30392 | ENSMUSG00000089371 | 0 | NA | NA | NA | NA | NA | Mir1938 | microRNA 1938 [Source:MGI Symbol;Acc: | 0 | 0 | 0 | 0 | 0 | 0 |
| 30393 | ENSMUSG00000089372 | 0 | NA | NA | NA | NA | NA | Gm22139 | predicted gene, 22139 [Source:MGI Symb | 0 | 0 | 0 | 0 | 0 | 0 |
| 30394 | ENSMUSG00000089373 | 0 | NA | NA | NA | NA | NA | Gm22138 | predicted gene, 22138 [Source:MGI Symb | 0 | 0 | 0 | 0 | 0 | 0 |
| 30395 | ENSMUSG00000089375 | 0 | NA | NA | NA | NA | NA | Snord107 | small nucleolar RNA, C/D box 107 [Source | 0 | 0 | 0 | 0 | 0 | 0 |
| 30396 | ENSMUSG00000089376 | 0 | NA | NA | NA | NA | NA | NA | NA | 0 | 0 | 0 | 0 | 0 | 0 |
| 30397 | ENSMUSG00000089377 | 0 | NA | NA | NA | NA | NA | NA | NA | 0 | 0 | 0 | 0 | 0 | 0 |
| 30398 | ENSMUSG00000089380 | 0 | NA | NA | NA | NA | NA | Gm24677 | predicted gene, 24677 [Source:MGI Symb | 0 | 0 | 0 | 0 | 0 | 0 |
| 30399 | ENSMUSG00000089381 | 0 | NA | NA | NA | NA | NA | Gm24678 | predicted gene, 24678 [Source:MGI Symb | 0 | 0 | 0 | 0 | 0 | 0 |
| 30400 | ENSMUSG00000089382 | 0 | NA | NA | NA | NA | NA | NA | NA | 0 | 0 | 0 | 0 | 0 | 0 |
| 30401 | ENSMUSG00000089383 | 0 | NA | NA | NA | NA | NA | NA | NA | 0 | 0 | 0 | 0 | 0 | 0 |
| 30402 | ENSMUSG00000089384 | 0 | NA | NA | NA | NA | NA | Gm24680 | predicted gene, 24680 [Source:MGI Symb | 0 | 0 | 0 | 0 | 0 | 0 |
| 30403 | ENSMUSG00000089385 | 0 | NA | NA | NA | NA | NA | Gm24681 | predicted gene, 24681 [Source:MGI Symb | 0 | 0 | 0 | 0 | 0 | 0 |
| 30404 | ENSMUSG00000089387 | 0 | NA | NA | NA | NA | NA | Gm24679 | predicted gene, 24679 [Source:MGI Symb | 0 | 0 | 0 | 0 | 0 | 0 |

| | | | | | | | | | | | | | | | |
|---|---|---|---|---|---|---|---|---|---|---|---|---|---|---|---|
| 30405 | ENSMUSG00000089389 | 0 | NA | NA | NA | NA | NA | NA | NA | 0 | 0 | 0 | 0 | 0 | 0 |
| 30406 | ENSMUSG00000089390 | 0 | NA | NA | NA | NA | NA | Gm23021 | predicted gene, 23021 [Source:MGI Symb | 0 | 0 | 0 | 0 | 0 | 0 |
| 30407 | ENSMUSG00000089391 | 0 | NA | NA | NA | NA | NA | NA | NA | 0 | 0 | 0 | 0 | 0 | 0 |
| 30408 | ENSMUSG00000089392 | 0 | NA | NA | NA | NA | NA | NA | NA | 0 | 0 | 0 | 0 | 0 | 0 |
| 30409 | ENSMUSG00000089393 | 0 | NA | NA | NA | NA | NA | Gm23022 | predicted gene, 23022 [Source:MGI Symb | 0 | 0 | 0 | 0 | 0 | 0 |
| 30410 | ENSMUSG00000089394 | 0 | NA | NA | NA | NA | NA | Gm23025 | predicted gene, 23025 [Source:MGI Symb | 0 | 0 | 0 | 0 | 0 | 0 |
| 30411 | ENSMUSG00000089395 | 0 | NA | NA | NA | NA | NA | NA | NA | 0 | 0 | 0 | 0 | 0 | 0 |
| 30412 | ENSMUSG00000089396 | 0.979378725 | 2.336871961 | 2.444416672 | 0.956003936 | 0.339070246 | NA | NA | NA | 0.8031 | 0 | 0 | 2.8919 | 2.1812 | 0 |
| 30413 | ENSMUSG00000089397 | 0 | NA | NA | NA | NA | NA | NA | NA | 0 | 0 | 0 | 0 | 0 | 0 |
| 30414 | ENSMUSG00000089398 | 0 | NA | NA | NA | NA | NA | NA | NA | 0 | 0 | 0 | 0 | 0 | 0 |
| 30415 | ENSMUSG00000089399 | 0 | NA | NA | NA | NA | NA | Gm23312 | predicted gene, 23312 [Source:MGI Symb | 0 | 0 | 0 | 0 | 0 | 0 |
| 30416 | ENSMUSG00000089401 | 0 | NA | NA | NA | NA | NA | NA | NA | 0 | 0 | 0 | 0 | 0 | 0 |
| 30417 | ENSMUSG00000089404 | 0.160662974 | 1.126924175 | 4.080472857 | 0.276174898 | 0.782413736 | NA | NA | NA | 0 | 0 | 0 | 0.964 | 0 | 0 |
| 30418 | ENSMUSG00000089405 | 0 | NA | NA | NA | NA | NA | NA | NA | 0 | 0 | 0 | 0 | 0 | 0 |
| 30419 | ENSMUSG00000089406 | 0 | NA | NA | NA | NA | NA | Mir2136 | microRNA 2136 [Source:MGI Symbol;Acc: | 0 | 0 | 0 | 0 | 0 | 0 |
| 30420 | ENSMUSG00000089407 | 0 | NA | NA | NA | NA | NA | Gm24878 | predicted gene, 24878 [Source:MGI Symb | 0 | 0 | 0 | 0 | 0 | 0 |
| 30421 | ENSMUSG00000089408 | 0 | NA | NA | NA | NA | NA | Gm24885 | predicted gene, 24885 [Source:MGI Symb | 0 | 0 | 0 | 0 | 0 | 0 |
| 30422 | ENSMUSG00000089409 | 0 | NA | NA | NA | NA | NA | Gm24884 | predicted gene, 24884 [Source:MGI Symb | 0 | 0 | 0 | 0 | 0 | 0 |
| 30423 | ENSMUSG00000089410 | 0 | NA | NA | NA | NA | NA | NA | NA | 0 | 0 | 0 | 0 | 0 | 0 |
| 30424 | ENSMUSG00000089413 | 0 | NA | NA | NA | NA | NA | Gm22013 | predicted gene, 22013 [Source:MGI Symb | 0 | 0 | 0 | 0 | 0 | 0 |
| 30425 | ENSMUSG00000089414 | 0 | NA | NA | NA | NA | NA | NA | NA | 0 | 0 | 0 | 0 | 0 | 0 |
| 30426 | ENSMUSG00000089415 | 0 | NA | NA | NA | NA | NA | Gm22011 | predicted gene, 22011 [Source:MGI Symb | 0 | 0 | 0 | 0 | 0 | 0 |
| 30427 | ENSMUSG00000089416 | 0 | NA | NA | NA | NA | NA | Gm22116 | predicted gene, 22116 [Source:MGI Symb | 0 | 0 | 0 | 0 | 0 | 0 |
| 30428 | ENSMUSG00000089417 | 2.911059037 | -0.020175926 | 1.56889793 | -0.012859935 | 0.989739539 | NA | Gm22009 | predicted gene, 22009 [Source:MGI Symb | 5.6217 | 3.0838 | 0 | 0.964 | 0 | 7.7969 |
| 30429 | ENSMUSG00000089418 | 0 | NA | NA | NA | NA | NA | NA | NA | 0 | 0 | 0 | 0 | 0 | 0 |
| 30430 | ENSMUSG00000089420 | 0 | NA | NA | NA | NA | NA | Gm27934 | predicted gene, 27934 [Source:MGI Symb | 0 | 0 | 0 | 0 | 0 | 0 |
| 30431 | ENSMUSG00000089422 | 0 | NA | NA | NA | NA | NA | Gm26008 | predicted gene, 26008 [Source:MGI Symb | 0 | 0 | 0 | 0 | 0 | 0 |
| 30432 | ENSMUSG00000089423 | 0 | NA | NA | NA | NA | NA | Gm26007 | predicted gene, 26007 [Source:MGI Symb | 0 | 0 | 0 | 0 | 0 | 0 |
| 30433 | ENSMUSG00000089424 | 0 | NA | NA | NA | NA | NA | Mir1930 | microRNA 1930 [Source:MGI Symbol;Acc: | 0 | 0 | 0 | 0 | 0 | 0 |
| 30434 | ENSMUSG00000089425 | 0 | NA | NA | NA | NA | NA | Gm26006 | predicted gene, 26006 [Source:MGI Symb | 0 | 0 | 0 | 0 | 0 | 0 |
| 30435 | ENSMUSG00000089427 | 0 | NA | NA | NA | NA | NA | Gm26005 | predicted gene, 26005 [Source:MGI Symb | 0 | 0 | 0 | 0 | 0 | 0 |
| 30436 | ENSMUSG00000089429 | 0 | NA | NA | NA | NA | NA | NA | NA | 0 | 0 | 0 | 0 | 0 | 0 |
| 30437 | ENSMUSG00000089431 | 0 | NA | NA | NA | NA | NA | Gm23196 | predicted gene, 23196 [Source:MGI Symb | 0 | 0 | 0 | 0 | 0 | 0 |
| 30438 | ENSMUSG00000089432 | 0.167707431 | -0.796629176 | 4.080472857 | -0.195229623 | 0.84521319 | NA | Mir1966 | microRNA 1966 [Source:MGI Symbol;Acc: | 0 | 0 | 1.0062 | 0 | 0 | 0 |
| 30439 | ENSMUSG00000089434 | 0 | NA | NA | NA | NA | NA | NA | NA | 0 | 0 | 0 | 0 | 0 | 0 |
| 30440 | ENSMUSG00000089436 | 0 | NA | NA | NA | NA | NA | NA | NA | 0 | 0 | 0 | 0 | 0 | 0 |
| 30441 | ENSMUSG00000089437 | 0 | NA | NA | NA | NA | NA | Gm23199 | predicted gene, 23199 [Source:MGI Symb | 0 | 0 | 0 | 0 | 0 | 0 |
| 30442 | ENSMUSG00000089438 | 0 | NA | NA | NA | NA | NA | Gm23194 | predicted gene, 23194 [Source:MGI Symb | 0 | 0 | 0 | 0 | 0 | 0 |
| 30443 | ENSMUSG00000089439 | 0 | NA | NA | NA | NA | NA | Gm23195 | predicted gene, 23195 [Source:MGI Symb | 0 | 0 | 0 | 0 | 0 | 0 |
| 30444 | ENSMUSG00000089441 | 0 | NA | NA | NA | NA | NA | NA | NA | 0 | 0 | 0 | 0 | 0 | 0 |
| 30445 | ENSMUSG00000089442 | 0 | NA | NA | NA | NA | NA | Gm22529 | predicted gene, 22529 [Source:MGI Symb | 0 | 0 | 0 | 0 | 0 | 0 |
| 30446 | ENSMUSG00000089443 | 0 | NA | NA | NA | NA | NA | NA | NA | 0 | 0 | 0 | 0 | 0 | 0 |
| 30447 | ENSMUSG00000089444 | 0 | NA | NA | NA | NA | NA | Gm22531 | predicted gene, 22531 [Source:MGI Symb | 0 | 0 | 0 | 0 | 0 | 0 |
| 30448 | ENSMUSG00000089445 | 0 | NA | NA | NA | NA | NA | Gm22530 | predicted gene, 22530 [Source:MGI Symb | 0 | 0 | 0 | 0 | 0 | 0 |
| 30449 | ENSMUSG00000089446 | 0 | NA | NA | NA | NA | NA | NA | NA | 0 | 0 | 0 | 0 | 0 | 0 |
| 30450 | ENSMUSG00000089447 | 0 | NA | NA | NA | NA | NA | NA | NA | 0 | 0 | 0 | 0 | 0 | 0 |
| 30451 | ENSMUSG00000089450 | 0 | NA | NA | NA | NA | NA | NA | NA | 0 | 0 | 0 | 0 | 0 | 0 |
| 30452 | ENSMUSG00000089451 | 0 | NA | NA | NA | NA | NA | NA | NA | 0 | 0 | 0 | 0 | 0 | 0 |
| 30453 | ENSMUSG00000089452 | 0 | NA | NA | NA | NA | NA | Gm24223 | predicted gene, 24223 [Source:MGI Symb | 0 | 0 | 0 | 0 | 0 | 0 |
| 30454 | ENSMUSG00000089453 | 0 | NA | NA | NA | NA | NA | NA | NA | 0 | 0 | 0 | 0 | 0 | 0 |
| 30455 | ENSMUSG00000089454 | 0 | NA | NA | NA | NA | NA | Gm24228 | predicted gene, 24228 [Source:MGI Symb | 0 | 0 | 0 | 0 | 0 | 0 |
| 30456 | ENSMUSG00000089455 | 0 | NA | NA | NA | NA | NA | Gm24229 | predicted gene, 24229 [Source:MGI Symb | 0 | 0 | 0 | 0 | 0 | 0 |
| 30457 | ENSMUSG00000089458 | 0 | NA | NA | NA | NA | NA | Gm24221 | predicted gene, 24221 [Source:MGI Symb | 0 | 0 | 0 | 0 | 0 | 0 |
| 30458 | ENSMUSG00000089459 | 0 | NA | NA | NA | NA | NA | NA | NA | 0 | 0 | 0 | 0 | 0 | 0 |
| 30459 | ENSMUSG00000089460 | 0 | NA | NA | NA | NA | NA | Gm23699 | predicted gene, 23699 [Source:MGI Symb | 0 | 0 | 0 | 0 | 0 | 0 |
| 30460 | ENSMUSG00000089461 | 0 | NA | NA | NA | NA | NA | Gm23698 | predicted gene, 23698 [Source:MGI Symb | 0 | 0 | 0 | 0 | 0 | 0 |

| | | | | | | | | | | | | | | | |
|---|---|---|---|---|---|---|---|---|---|---|---|---|---|---|---|
| 30461 | ENSMUSG00000089462 | 0 | NA | NA | NA | NA | NA | NA | NA | 0 | 0 | 0 | 0 | 0 | 0 |
| 30462 | ENSMUSG00000089464 | 0 | NA | NA | NA | NA | NA | Gm23703 | predicted gene, 23703 [Source:MGI Symb | 0 | 0 | 0 | 0 | 0 | 0 |
| 30463 | ENSMUSG00000089465 | 0 | NA | NA | NA | NA | NA | NA | NA | 0 | 0 | 0 | 0 | 0 | 0 |
| 30464 | ENSMUSG00000089467 | 0 | NA | NA | NA | NA | NA | NA | NA | 0 | 0 | 0 | 0 | 0 | 0 |
| 30465 | ENSMUSG00000089468 | 0 | NA | NA | NA | NA | NA | Gm27649 | predicted gene, 27649 [Source:MGI Symb | 0 | 0 | 0 | 0 | 0 | 0 |
| 30466 | ENSMUSG00000089469 | 0 | NA | NA | NA | NA | NA | NA | NA | 0 | 0 | 0 | 0 | 0 | 0 |
| 30467 | ENSMUSG00000089470 | 0 | NA | NA | NA | NA | NA | NA | NA | 0 | 0 | 0 | 0 | 0 | 0 |
| 30468 | ENSMUSG00000089471 | 0 | NA | NA | NA | NA | NA | NA | NA | 0 | 0 | 0 | 0 | 0 | 0 |
| 30469 | ENSMUSG00000089472 | 0 | NA | NA | NA | NA | NA | NA | NA | 0 | 0 | 0 | 0 | 0 | 0 |
| 30470 | ENSMUSG00000089473 | 0 | NA | NA | NA | NA | NA | Gm25369 | predicted gene, 25369 [Source:MGI Symb | 0 | 0 | 0 | 0 | 0 | 0 |
| 30471 | ENSMUSG00000089475 | 0.656740811 | -0.013525372 | 3.112905782 | -0.004344934 | 0.996533255 | NA | NA | NA | 0 | 0 | 2.0125 | 1.928 | 0 | 0 |
| 30472 | ENSMUSG00000089476 | 0.614004368 | 0.315622926 | 2.898299649 | 0.108899342 | 0.913282328 | NA | Gm25364 | predicted gene, 25364 [Source:MGI Symb | 1.6062 | 0 | 0 | 0.964 | 0 | 1.1138 |
| 30473 | ENSMUSG00000089477 | 0 | NA | NA | NA | NA | NA | NA | NA | 0 | 0 | 0 | 0 | 0 | 0 |
| 30474 | ENSMUSG00000089479 | 0 | NA | NA | NA | NA | NA | Gm25362 | predicted gene, 25362 [Source:MGI Symb | 0 | 0 | 0 | 0 | 0 | 0 |
| 30475 | ENSMUSG00000089480 | 0 | NA | NA | NA | NA | NA | NA | NA | 0 | 0 | 0 | 0 | 0 | 0 |
| 30476 | ENSMUSG00000089481 | 0 | NA | NA | NA | NA | NA | NA | NA | 0 | 0 | 0 | 0 | 0 | 0 |
| 30477 | ENSMUSG00000089482 | 0 | NA | NA | NA | NA | NA | NA | NA | 0 | 0 | 0 | 0 | 0 | 0 |
| 30478 | ENSMUSG00000089484 | 0 | NA | NA | NA | NA | NA | Gm24735 | predicted gene, 24735 [Source:MGI Symb | 0 | 0 | 0 | 0 | 0 | 0 |
| 30479 | ENSMUSG00000089485 | 0 | NA | NA | NA | NA | NA | NA | NA | 0 | 0 | 0 | 0 | 0 | 0 |
| 30480 | ENSMUSG00000089486 | 0 | NA | NA | NA | NA | NA | Gm24737 | predicted gene, 24737 [Source:MGI Symb | 0 | 0 | 0 | 0 | 0 | 0 |
| 30481 | ENSMUSG00000089487 | 0 | NA | NA | NA | NA | NA | NA | NA | 0 | 0 | 0 | 0 | 0 | 0 |
| 30482 | ENSMUSG00000089488 | 0 | NA | NA | NA | NA | NA | NA | NA | 0 | 0 | 0 | 0 | 0 | 0 |
| 30483 | ENSMUSG00000089489 | 0 | NA | NA | NA | NA | NA | Gm24732 | predicted gene, 24732 [Source:MGI Symb | 0 | 0 | 0 | 0 | 0 | 0 |
| 30484 | ENSMUSG00000089490 | 0 | NA | NA | NA | NA | NA | NA | NA | 0 | 0 | 0 | 0 | 0 | 0 |
| 30485 | ENSMUSG00000089491 | 0.335414862 | -1.779047242 | 4.030804179 | -0.441362855 | 0.658950329 | NA | NA | NA | 0 | 0 | 2.0125 | 0 | 0 | 0 |
| 30486 | ENSMUSG00000089493 | 0 | NA | NA | NA | NA | NA | Gm26361 | predicted gene, 26361 [Source:MGI Symb | 0 | 0 | 0 | 0 | 0 | 0 |
| 30487 | ENSMUSG00000089497 | 0 | NA | NA | NA | NA | NA | NA | NA | 0 | 0 | 0 | 0 | 0 | 0 |
| 30488 | ENSMUSG00000089498 | 0.185640647 | 1.126924175 | 4.080472857 | 0.276174898 | 0.782413736 | NA | Gm26359 | predicted gene, 26359 [Source:MGI Symb | 0 | 0 | 0 | 0 | 0 | 1.1138 |
| 30489 | ENSMUSG00000089499 | 0 | NA | NA | NA | NA | NA | NA | NA | 0 | 0 | 0 | 0 | 0 | 0 |
| 30490 | ENSMUSG00000089501 | 0 | NA | NA | NA | NA | NA | Gm23479 | predicted gene, 23479 [Source:MGI Symb | 0 | 0 | 0 | 0 | 0 | 0 |
| 30491 | ENSMUSG00000089504 | 0 | NA | NA | NA | NA | NA | Gm23475 | predicted gene, 23475 [Source:MGI Symb | 0 | 0 | 0 | 0 | 0 | 0 |
| 30492 | ENSMUSG00000089506 | 0 | NA | NA | NA | NA | NA | NA | NA | 0 | 0 | 0 | 0 | 0 | 0 |
| 30493 | ENSMUSG00000089507 | 0 | NA | NA | NA | NA | NA | Gm23477 | predicted gene, 23477 [Source:MGI Symb | 0 | 0 | 0 | 0 | 0 | 0 |
| 30494 | ENSMUSG00000089508 | 0 | NA | NA | NA | NA | NA | NA | NA | 0 | 0 | 0 | 0 | 0 | 0 |
| 30495 | ENSMUSG00000089509 | 0 | NA | NA | NA | NA | NA | NA | NA | 0 | 0 | 0 | 0 | 0 | 0 |
| 30496 | ENSMUSG00000089510 | 0 | NA | NA | NA | NA | NA | NA | NA | 0 | 0 | 0 | 0 | 0 | 0 |
| 30497 | ENSMUSG00000089512 | 0.17131964 | -0.796629176 | 4.080472857 | -0.195229623 | 0.84521319 | NA | Gm26287 | predicted gene, 26287 [Source:MGI Symb | 0 | 1.0279 | 0 | 0 | 0 | 0 |
| 30498 | ENSMUSG00000089513 | 0.17131964 | -0.796629176 | 4.080472857 | -0.195229623 | 0.84521319 | NA | Gm24875 | predicted gene, 24875 [Source:MGI Symb | 0 | 1.0279 | 0 | 0 | 0 | 0 |
| 30499 | ENSMUSG00000089514 | 0 | NA | NA | NA | NA | NA | Gm26291 | predicted gene, 26291 [Source:MGI Symb | 0 | 0 | 0 | 0 | 0 | 0 |
| 30500 | ENSMUSG00000089516 | 0 | NA | NA | NA | NA | NA | NA | NA | 0 | 0 | 0 | 0 | 0 | 0 |
| 30501 | ENSMUSG00000089517 | 0 | NA | NA | NA | NA | NA | NA | NA | 0 | 0 | 0 | 0 | 0 | 0 |
| 30502 | ENSMUSG00000089520 | 0 | NA | NA | NA | NA | NA | NA | NA | 0 | 0 | 0 | 0 | 0 | 0 |
| 30503 | ENSMUSG00000089522 | 0 | NA | NA | NA | NA | NA | NA | NA | 0 | 0 | 0 | 0 | 0 | 0 |
| 30504 | ENSMUSG00000089524 | 0 | NA | NA | NA | NA | NA | Gm22327 | predicted gene, 22327 [Source:MGI Symb | 0 | 0 | 0 | 0 | 0 | 0 |
| 30505 | ENSMUSG00000089525 | 0 | NA | NA | NA | NA | NA | NA | NA | 0 | 0 | 0 | 0 | 0 | 0 |
| 30506 | ENSMUSG00000089528 | 0 | NA | NA | NA | NA | NA | Gm22324 | predicted gene, 22324 [Source:MGI Symb | 0 | 0 | 0 | 0 | 0 | 0 |
| 30507 | ENSMUSG00000089529 | 0 | NA | NA | NA | NA | NA | NA | NA | 0 | 0 | 0 | 0 | 0 | 0 |
| 30508 | ENSMUSG00000089530 | 0 | NA | NA | NA | NA | NA | Gm25163 | predicted gene, 25163 [Source:MGI Symb | 0 | 0 | 0 | 0 | 0 | 0 |
| 30509 | ENSMUSG00000089531 | 0 | NA | NA | NA | NA | NA | Gm25162 | predicted gene, 25162 [Source:MGI Symb | 0 | 0 | 0 | 0 | 0 | 0 |
| 30510 | ENSMUSG00000089533 | 0 | NA | NA | NA | NA | NA | NA | NA | 0 | 0 | 0 | 0 | 0 | 0 |
| 30511 | ENSMUSG00000089534 | 0 | NA | NA | NA | NA | NA | Gm25166 | predicted gene, 25166 [Source:MGI Symb | 0 | 0 | 0 | 0 | 0 | 0 |
| 30512 | ENSMUSG00000089536 | 0.17131964 | -0.796629176 | 4.080472857 | -0.195229623 | 0.84521319 | NA | NA | NA | 0 | 1.0279 | 0 | 0 | 0 | 0 |
| 30513 | ENSMUSG00000089538 | 0 | NA | NA | NA | NA | NA | Gm25161 | predicted gene, 25161 [Source:MGI Symb | 0 | 0 | 0 | 0 | 0 | 0 |
| 30514 | ENSMUSG00000089539 | 0 | NA | NA | NA | NA | NA | Gm25160 | predicted gene, 25160 [Source:MGI Symb | 0 | 0 | 0 | 0 | 0 | 0 |
| 30515 | ENSMUSG00000089541 | 0 | NA | NA | NA | NA | NA | Gm25834 | predicted gene, 25834 [Source:MGI Symb | 0 | 0 | 0 | 0 | 0 | 0 |
| 30516 | ENSMUSG00000089542 | 3.455345957 | 1.529473908 | 1.260510909 | 1.213376178 | 0.22498603 | NA | Gm25835 | predicted gene, 25835 [Source:MGI Symb | 3.2124 | 1.0279 | 1.0062 | 7.7118 | 1.0906 | 6.6831 |

| | | | | | | | | | | | | | | | |
|---|---|---|---|---|---|---|---|---|---|---|---|---|---|---|---|
| 30517 | ENSMUSG00000089543 | 0 | NA | NA | NA | NA | NA | Gm25836 | predicted gene, 25836 [Source:MGI Symb | 0 | 0 | 0 | 0 | 0 | 0 |
| 30518 | ENSMUSG00000089544 | 0 | NA | NA | NA | NA | NA | NA | NA | 0 | 0 | 0 | 0 | 0 | 0 |
| 30519 | ENSMUSG00000089545 | 0 | NA | NA | NA | NA | NA | NA | NA | 0 | 0 | 0 | 0 | 0 | 0 |
| 30520 | ENSMUSG00000089546 | 0 | NA | NA | NA | NA | NA | NA | NA | 0 | 0 | 0 | 0 | 0 | 0 |
| 30521 | ENSMUSG00000089548 | 0 | NA | NA | NA | NA | NA | Gm25830 | predicted gene, 25830 [Source:MGI Symb | 0 | 0 | 0 | 0 | 0 | 0 |
| 30522 | ENSMUSG00000089549 | 0 | NA | NA | NA | NA | NA | Gm25831 | predicted gene, 25831 [Source:MGI Symb | 0 | 0 | 0 | 0 | 0 | 0 |
| 30523 | ENSMUSG00000089550 | 0 | NA | NA | NA | NA | NA | Gm24179 | predicted gene, 24179 [Source:MGI Symb | 0 | 0 | 0 | 0 | 0 | 0 |
| 30524 | ENSMUSG00000089551 | 0 | NA | NA | NA | NA | NA | Gm24178 | predicted gene, 24178 [Source:MGI Symb | 0 | 0 | 0 | 0 | 0 | 0 |
| 30525 | ENSMUSG00000089552 | 0 | NA | NA | NA | NA | NA | Gm24177 | predicted gene, 24177 [Source:MGI Symb | 0 | 0 | 0 | 0 | 0 | 0 |
| 30526 | ENSMUSG00000089553 | 0 | NA | NA | NA | NA | NA | Gm24176 | predicted gene, 24176 [Source:MGI Symb | 0 | 0 | 0 | 0 | 0 | 0 |
| 30527 | ENSMUSG00000089555 | 0 | NA | NA | NA | NA | NA | Mir1933 | microRNA 1933 [Source:MGI Symbol;Acc: | 0 | 0 | 0 | 0 | 0 | 0 |
| 30528 | ENSMUSG00000089556 | 0 | NA | NA | NA | NA | NA | Gm24175 | predicted gene, 24175 [Source:MGI Symb | 0 | 0 | 0 | 0 | 0 | 0 |
| 30529 | ENSMUSG00000089558 | 0 | NA | NA | NA | NA | NA | NA | NA | 0 | 0 | 0 | 0 | 0 | 0 |
| 30530 | ENSMUSG00000089560 | 0 | NA | NA | NA | NA | NA | Gm24668 | predicted gene, 24668 [Source:MGI Symb | 0 | 0 | 0 | 0 | 0 | 0 |
| 30531 | ENSMUSG00000089561 | 0 | NA | NA | NA | NA | NA | Gm24669 | predicted gene, 24669 [Source:MGI Symb | 0 | 0 | 0 | 0 | 0 | 0 |
| 30532 | ENSMUSG00000089562 | 0 | NA | NA | NA | NA | NA | Gm24666 | predicted gene, 24666 [Source:MGI Symb | 0 | 0 | 0 | 0 | 0 | 0 |
| 30533 | ENSMUSG00000089563 | 0 | NA | NA | NA | NA | NA | Gm24667 | predicted gene, 24667 [Source:MGI Symb | 0 | 0 | 0 | 0 | 0 | 0 |
| 30534 | ENSMUSG00000089564 | 0 | NA | NA | NA | NA | NA | NA | NA | 0 | 0 | 0 | 0 | 0 | 0 |
| 30535 | ENSMUSG00000089570 | 0 | NA | NA | NA | NA | NA | Mir669m-1 | microRNA 669m-1 [Source:MGI Symbol;A | 0 | 0 | 0 | 0 | 0 | 0 |
| 30536 | ENSMUSG00000089571 | 0 | NA | NA | NA | NA | NA | Gm23008 | predicted gene, 23008 [Source:MGI Symb | 0 | 0 | 0 | 0 | 0 | 0 |
| 30537 | ENSMUSG00000089575 | 0 | NA | NA | NA | NA | NA | Gm23005 | predicted gene, 23005 [Source:MGI Symb | 0 | 0 | 0 | 0 | 0 | 0 |
| 30538 | ENSMUSG00000089576 | 0 | NA | NA | NA | NA | NA | NA | NA | 0 | 0 | 0 | 0 | 0 | 0 |
| 30539 | ENSMUSG00000089578 | 0 | NA | NA | NA | NA | NA | NA | NA | 0 | 0 | 0 | 0 | 0 | 0 |
| 30540 | ENSMUSG00000089580 | 2.039970898 | -0.131469121 | 1.719744176 | -0.076446906 | 0.939063553 | NA | Gm23340 | predicted gene, 23340 [Source:MGI Symb | 2.4093 | 1.0279 | 3.0187 | 5.7839 | 0 | 0 |
| 30541 | ENSMUSG00000089581 | 1.186152743 | -0.224486212 | 2.101231109 | -0.106835565 | 0.914919432 | NA | Gm23341 | predicted gene, 23341 [Source:MGI Symb | 0.8031 | 0 | 3.0187 | 0 | 2.1812 | 1.1138 |
| 30542 | ENSMUSG00000089583 | 0 | NA | NA | NA | NA | NA | NA | NA | 0 | 0 | 0 | 0 | 0 | 0 |
| 30543 | ENSMUSG00000089584 | 0 | NA | NA | NA | NA | NA | Gm23337 | predicted gene, 23337 [Source:MGI Symb | 0 | 0 | 0 | 0 | 0 | 0 |
| 30544 | ENSMUSG00000089585 | 0 | NA | NA | NA | NA | NA | Gm23338 | predicted gene, 23338 [Source:MGI Symb | 0 | 0 | 0 | 0 | 0 | 0 |
| 30545 | ENSMUSG00000089586 | 0 | NA | NA | NA | NA | NA | NA | NA | 0 | 0 | 0 | 0 | 0 | 0 |
| 30546 | ENSMUSG00000089587 | 0 | NA | NA | NA | NA | NA | Gm23339 | predicted gene, 23339 [Source:MGI Symb | 0 | 0 | 0 | 0 | 0 | 0 |
| 30547 | ENSMUSG00000089588 | 0 | NA | NA | NA | NA | NA | NA | NA | 0 | 0 | 0 | 0 | 0 | 0 |
| 30548 | ENSMUSG00000089589 | 0 | NA | NA | NA | NA | NA | Gm23343 | predicted gene, 23343 [Source:MGI Symb | 0 | 0 | 0 | 0 | 0 | 0 |
| 30549 | ENSMUSG00000089590 | 0 | NA | NA | NA | NA | NA | Gm26160 | predicted gene, 26160 [Source:MGI Symb | 0 | 0 | 0 | 0 | 0 | 0 |
| 30550 | ENSMUSG00000089591 | 0 | NA | NA | NA | NA | NA | Gm24352 | predicted gene, 24352 [Source:MGI Symb | 0 | 0 | 0 | 0 | 0 | 0 |
| 30551 | ENSMUSG00000089593 | 0 | NA | NA | NA | NA | NA | Gm26159 | predicted gene, 26159 [Source:MGI Symb | 0 | 0 | 0 | 0 | 0 | 0 |
| 30552 | ENSMUSG00000089596 | 0 | NA | NA | NA | NA | NA | NA | NA | 0 | 0 | 0 | 0 | 0 | 0 |
| 30553 | ENSMUSG00000089597 | 0 | NA | NA | NA | NA | NA | NA | NA | 0 | 0 | 0 | 0 | 0 | 0 |
| 30554 | ENSMUSG00000089598 | 0 | NA | NA | NA | NA | NA | NA | NA | 0 | 0 | 0 | 0 | 0 | 0 |
| 30555 | ENSMUSG00000089599 | 0 | NA | NA | NA | NA | NA | NA | NA | 0 | 0 | 0 | 0 | 0 | 0 |
| 30556 | ENSMUSG00000089601 | 0 | NA | NA | NA | NA | NA | NA | NA | 0 | 0 | 0 | 0 | 0 | 0 |
| 30557 | ENSMUSG00000089603 | 0 | NA | NA | NA | NA | NA | NA | NA | 0 | 0 | 0 | 0 | 0 | 0 |
| 30558 | ENSMUSG00000089605 | 0 | NA | NA | NA | NA | NA | NA | NA | 0 | 0 | 0 | 0 | 0 | 0 |
| 30559 | ENSMUSG00000089606 | 0 | NA | NA | NA | NA | NA | Gm22501 | predicted gene, 22501 [Source:MGI Symb | 0 | 0 | 0 | 0 | 0 | 0 |
| 30560 | ENSMUSG00000089607 | 0 | NA | NA | NA | NA | NA | Gm22500 | predicted gene, 22500 [Source:MGI Symb | 0 | 0 | 0 | 0 | 0 | 0 |
| 30561 | ENSMUSG00000089608 | 0 | NA | NA | NA | NA | NA | Gm22503 | predicted gene, 22503 [Source:MGI Symb | 0 | 0 | 0 | 0 | 0 | 0 |
| 30562 | ENSMUSG00000089609 | 0 | NA | NA | NA | NA | NA | NA | NA | 0 | 0 | 0 | 0 | 0 | 0 |
| 30563 | ENSMUSG00000089610 | 0 | NA | NA | NA | NA | NA | NA | NA | 0 | 0 | 0 | 0 | 0 | 0 |
| 30564 | ENSMUSG00000089612 | 0 | NA | NA | NA | NA | NA | Gm22708 | predicted gene, 22708 [Source:MGI Symb | 0 | 0 | 0 | 0 | 0 | 0 |
| 30565 | ENSMUSG00000089613 | 0.506734502 | -2.372847156 | 3.338060886 | -0.710845978 | 0.477179685 | NA | NA | NA | 0 | 1.0279 | 2.0125 | 0 | 0 | 0 |
| 30566 | ENSMUSG00000089614 | 0 | NA | NA | NA | NA | NA | Gm23505 | predicted gene, 23505 [Source:MGI Symb | 0 | 0 | 0 | 0 | 0 | 0 |
| 30567 | ENSMUSG00000089615 | 0 | NA | NA | NA | NA | NA | Gm23774 | predicted gene, 23774 [Source:MGI Symb | 0 | 0 | 0 | 0 | 0 | 0 |
| 30568 | ENSMUSG00000089616 | 0 | NA | NA | NA | NA | NA | NA | NA | 0 | 0 | 0 | 0 | 0 | 0 |
| 30569 | ENSMUSG00000089617 | 0 | NA | NA | NA | NA | NA | Scarna10 | small Cajal body-specific RNA 10 [Source:I | 0 | 0 | 0 | 0 | 0 | 0 |
| 30570 | ENSMUSG00000089618 | 0 | NA | NA | NA | NA | NA | NA | NA | 0 | 0 | 0 | 0 | 0 | 0 |
| 30571 | ENSMUSG00000089619 | 0 | NA | NA | NA | NA | NA | NA | NA | 0 | 0 | 0 | 0 | 0 | 0 |
| 30572 | ENSMUSG00000089621 | 0.401551119 | -2.072982726 | 4.008251979 | -0.517178744 | 0.605031388 | NA | NA | NA | 2.4093 | 0 | 0 | 0 | 0 | 0 |

| | | | | | | | | | | | | | | | |
|---|---|---|---|---|---|---|---|---|---|---|---|---|---|---|---|
| 30573 | ENSMUSG00000089622 | 0 | NA | NA | NA | NA | NA | NA | NA | 0 | 0 | 0 | 0 | 0 | 0 |
| 30574 | ENSMUSG00000089624 | 0 | NA | NA | NA | NA | NA | NA | NA | 0 | 0 | 0 | 0 | 0 | 0 |
| 30575 | ENSMUSG00000089625 | 0 | NA | NA | NA | NA | NA | Gm25443 | predicted gene, 25443 [Source:MGI Symb | 0 | 0 | 0 | 0 | 0 | 0 |
| 30576 | ENSMUSG00000089627 | 0 | NA | NA | NA | NA | NA | Gm16228 | predicted gene 16228 [Source:MGI Symbo | 0 | 0 | 0 | 0 | 0 | 0 |
| 30577 | ENSMUSG00000089628 | 0.160662974 | 1.126924175 | 4.080472857 | 0.276174898 | 0.782413736 | NA | Gm9727 | predicted gene 9727 [Source:MGI Symbol | 0 | 0 | 0 | 0.964 | 0 | 0 |
| 30578 | ENSMUSG00000089630 | 0 | NA | NA | NA | NA | NA | Gm6903 | predicted gene 6903 [Source:MGI Symbol | 0 | 0 | 0 | 0 | 0 | 0 |
| 30579 | ENSMUSG00000089631 | 0 | NA | NA | NA | NA | NA | Gm16326 | predicted gene 16326 [Source:MGI Symbo | 0 | 0 | 0 | 0 | 0 | 0 |
| 30580 | ENSMUSG00000089633 | 0 | NA | NA | NA | NA | NA | A230009B12 | RIKEN cDNA A230009B12 gene [Source:M | 0 | 0 | 0 | 0 | 0 | 0 |
| 30581 | ENSMUSG00000089634 | 1.650382897 | 0.870245044 | 1.705690166 | 0.510201126 | 0.609910563 | NA | Nat8b-ps | N-acetyltransferase 8B, pseudogene [Sou | 2.4093 | 0 | 1.0062 | 0.964 | 2.1812 | 3.3415 |
| 30582 | ENSMUSG00000089635 | 65.72058677 | 0.570749843 | 0.389508093 | 1.465309329 | 0.142836542 | 0.552550882 | Gm16559 | predicted gene 16559 [Source:MGI Symbo | 55.414 | 43.173 | 60.375 | 133.03 | 58.893 | 43.44 |
| 30583 | ENSMUSG00000089636 | 3.333645302 | -0.413435378 | 1.250554446 | -0.330601662 | 0.740945391 | NA | 1700058P15 | RIKEN cDNA 1700058P15 gene [Source:M | 3.2124 | 7.1954 | 1.0062 | 1.928 | 1.0906 | 5.5692 |
| 30584 | ENSMUSG00000089639 | 0.503302254 | -0.837877584 | 3.426698643 | -0.244514523 | 0.806832356 | NA | Gm2862 | predicted gene 2862 [Source:MGI Symbol | 0 | 2.0558 | 0 | 0.964 | 0 | 0 |
| 30585 | ENSMUSG00000089640 | 0 | NA | NA | NA | NA | NA | Gm6397 | predicted gene 6397 [Source:MGI Symbol | 0 | 0 | 0 | 0 | 0 | 0 |
| 30586 | ENSMUSG00000089642 | 0 | NA | NA | NA | NA | NA | NA | NA | 0 | 0 | 0 | 0 | 0 | 0 |
| 30587 | ENSMUSG00000089643 | 0.342639279 | -1.805976596 | 4.028547714 | -0.448294702 | 0.653940524 | NA | Gm16301 | predicted gene 16301 [Source:MGI Symbo | 0 | 2.0558 | 0 | 0 | 0 | 0 |
| 30588 | ENSMUSG00000089645 | 0 | NA | NA | NA | NA | NA | Gm5766 | predicted gene 5766 [Source:MGI Symbol | 0 | 0 | 0 | 0 | 0 | 0 |
| 30589 | ENSMUSG00000089646 | 0.167707431 | -0.796629176 | 4.080472857 | -0.195229623 | 0.84521319 | NA | Gm15796 | predicted gene 15796 [Source:MGI Symbo | 0 | 0 | 1.0062 | 0 | 0 | 0 |
| 30590 | ENSMUSG00000089647 | 0 | NA | NA | NA | NA | NA | Gm2245 | predicted gene 2245 [Source:MGI Symbol | 0 | 0 | 0 | 0 | 0 | 0 |
| 30591 | ENSMUSG00000089648 | 0.17131964 | -0.796629176 | 4.080472857 | -0.195229623 | 0.84521319 | NA | Gm15790 | predicted pseudogene 15790 [Source:MG | 0 | 1.0279 | 0 | 0 | 0 | 0 |
| 30592 | ENSMUSG00000089651 | 0 | NA | NA | NA | NA | NA | Gm16353 | predicted gene 16353 [Source:MGI Symbo | 0 | 0 | 0 | 0 | 0 | 0 |
| 30593 | ENSMUSG00000089652 | 0.82991484 | -1.853997749 | 2.875131703 | -0.644839242 | 0.519031369 | NA | Gm16025 | predicted gene 16025 [Source:MGI Symbo | 4.0155 | 0 | 0 | 0.964 | 0 | 0 |
| 30594 | ENSMUSG00000089653 | 0 | NA | NA | NA | NA | NA | Gm15841 | predicted gene 15841 [Source:MGI Symbo | 0 | 0 | 0 | 0 | 0 | 0 |
| 30595 | ENSMUSG00000089654 | 0 | NA | NA | NA | NA | NA | Ear-ps12 | eosinophil-associated, ribonuclease A fam | 0 | 0 | 0 | 0 | 0 | 0 |
| 30596 | ENSMUSG00000089655 | 0 | NA | NA | NA | NA | NA | Gm16004 | predicted gene 16004 [Source:MGI Symbo | 0 | 0 | 0 | 0 | 0 | 0 |
| 30597 | ENSMUSG00000089656 | 0.346303622 | 2.004339653 | 3.884003423 | 0.516049919 | 0.605819544 | NA | Gm16271 | predicted gene 16271 [Source:MGI Symbo | 0 | 0 | 0 | 0.964 | 0 | 1.1138 |
| 30598 | ENSMUSG00000089657 | 0.696064467 | -1.534444742 | 3.043667477 | -0.504143358 | 0.61416064 | NA | Gm16585 | predicted gene 16585 [Source:MGI Symbo | 3.2124 | 0 | 0 | 0.964 | 0 | 0 |
| 30599 | ENSMUSG00000089659 | 1.485590266 | 0.626680717 | 1.944843862 | 0.32222675 | 0.747280921 | NA | Gm3470 | predicted gene 3470 [Source:MGI Symbol | 2.4093 | 1.0279 | 0 | 0 | 4.3625 | 1.1138 |
| 30600 | ENSMUSG00000089661 | 0.349477146 | 0.165142768 | 3.911021834 | 0.042224967 | 0.966319359 | NA | Mia | melanoma inhibitory activity [Source:MGI | 0 | 0 | 1.0062 | 0 | 1.0906 | 0 |
| 30601 | ENSMUSG00000089662 | 9.895435534 | -0.616590874 | 0.745794264 | -0.826757329 | 0.40837462 | 0.832151413 | Gm14057 | predicted gene 14057 [Source:MGI Symbo | 13.653 | 13.363 | 9.0562 | 14.46 | 3.2719 | 5.5692 |
| 30602 | ENSMUSG00000089663 | 0 | NA | NA | NA | NA | NA | Gm15651 | predicted gene 15651 [Source:MGI Symbo | 0 | 0 | 0 | 0 | 0 | 0 |
| 30603 | ENSMUSG00000089664 | 1.266971584 | 1.463191283 | 2.276902105 | 0.642623712 | 0.520468293 | NA | Gm12494 | predicted gene 12494 [Source:MGI Symbo | 0 | 2.0558 | 0 | 0 | 1.0906 | 4.4554 |
| 30604 | ENSMUSG00000089665 | 8.771958266 | 0.272503722 | 0.735211751 | 0.370646581 | 0.710900783 | 0.944239215 | Fcor | Foxo1 corepressor [Source:MGI Symbol;A | 9.6372 | 6.1675 | 8.05 | 14.46 | 7.6343 | 6.6831 |
| 30605 | ENSMUSG00000089666 | 0 | NA | NA | NA | NA | NA | Gm16269 | predicted gene 16269 [Source:MGI Symbo | 0 | 0 | 0 | 0 | 0 | 0 |
| 30606 | ENSMUSG00000089667 | 0 | NA | NA | NA | NA | NA | Gm15914 | predicted gene 15914 [Source:MGI Symbo | 0 | 0 | 0 | 0 | 0 | 0 |
| 30607 | ENSMUSG00000089669 | 7.749513905 | -0.987735588 | 0.816032629 | -1.210411879 | 0.226120886 | 0.672682908 | Tnfsf13 | tumor necrosis factor (ligand) superfamily | 5.6217 | 14.391 | 11.069 | 7.7118 | 4.3625 | 3.3415 |
| 30608 | ENSMUSG00000089670 | 0.342639279 | -1.805976596 | 4.028547714 | -0.448294702 | 0.653940524 | NA | Gm16581 | predicted gene 16581 [Source:MGI Symbo | 0 | 2.0558 | 0 | 0 | 0 | 0 |
| 30609 | ENSMUSG00000089671 | 0.305170013 | -1.657836477 | 4.041480335 | -0.410205257 | 0.681655385 | NA | Gm16537 | predicted gene 16537 [Source:MGI Symbo | 0.8031 | 1.0279 | 0 | 0 | 0 | 0 |
| 30610 | ENSMUSG00000089673 | 0 | NA | NA | NA | NA | NA | Gm16546 | predicted gene 16546 [Source:MGI Symbo | 0 | 0 | 0 | 0 | 0 | 0 |
| 30611 | ENSMUSG00000089675 | 0.294513347 | 0.165143496 | 4.080472857 | 0.040471657 | 0.967717103 | NA | Ugt1a8 | UDP glucuronosyltransferase 1 family, pol | 0.8031 | 0 | 0 | 0.964 | 0 | 0 |
| 30612 | ENSMUSG00000089678 | 3467.890366 | -0.00017632 | 0.133568263 | -0.001320074 | 0.998946734 | 0.999764475 | Agxt2 | alanine-glyoxylate aminotransferase 2 [So | 3853.3 | 3464.1 | 3085.1 | 2856.3 | 3920.8 | 3627.8 |
| 30613 | ENSMUSG00000089679 | 1.777370428 | 1.357998065 | 1.742694439 | 0.779251965 | 0.435831305 | NA | Gm16299 | predicted gene 16299 [Source:MGI Symbo | 0 | 1.0279 | 2.0125 | 0.964 | 1.0906 | 5.5692 |
| 30614 | ENSMUSG00000089680 | 5.092288295 | 0.912099295 | 0.991525057 | 0.919895356 | 0.357627446 | 0.797029797 | Gm15722 | predicted gene 15722 [Source:MGI Symbo | 1.6062 | 2.0558 | 7.0437 | 7.7118 | 5.4531 | 6.6831 |
| 30615 | ENSMUSG00000089682 | 430.4225667 | 0.113077881 | 0.153623144 | 0.736073213 | 0.461686144 | 0.860323162 | Bcl2l2 | BCL2-like 2 [Source:MGI Symbol;Acc:MGI: | 380.67 | 450.23 | 410.55 | 479.1 | 371.9 | 490.09 |
| 30616 | ENSMUSG00000089683 | 1.828545916 | 0.915595374 | 1.653126031 | 0.55385697 | 0.579676736 | NA | 4930570N18 | RIKEN cDNA 4930570N18 gene [Source:M | 0.8031 | 1.0279 | 2.0125 | 3.8559 | 3.2719 | 0 |
| 30617 | ENSMUSG00000089684 | 0 | NA | NA | NA | NA | NA | Gm14163 | predicted gene 14163 [Source:MGI Symbo | 0 | 0 | 0 | 0 | 0 | 0 |
| 30618 | ENSMUSG00000089685 | 0 | NA | NA | NA | NA | NA | Gm9017 | predicted gene 9017 [Source:MGI Symbol | 0 | 0 | 0 | 0 | 0 | 0 |
| 30619 | ENSMUSG00000089686 | 0.356960287 | 0.16514264 | 3.880370363 | 0.042558474 | 0.966053499 | NA | Mnd1-ps | Mnd1 retrotransposed pseudogene [Sour | 0 | 1.0279 | 0 | 0 | 0 | 1.1138 |
| 30620 | ENSMUSG00000089687 | 1.131136811 | -2.325642949 | 2.16414089 | -1.074626407 | 0.282542024 | NA | Rab42 | RAB42, member RAS oncogene family [So | 1.6062 | 3.0838 | 1.0062 | 0 | 1.0906 | 0 |
| 30621 | ENSMUSG00000089689 | 0 | NA | NA | NA | NA | NA | Gm4982 | predicted gene 4982 [Source:MGI Symbol | 0 | 0 | 0 | 0 | 0 | 0 |
| 30622 | ENSMUSG00000089690 | 0 | NA | NA | NA | NA | NA | Gm16569 | predicted gene 16569 [Source:MGI Symbo | 0 | 0 | 0 | 0 | 0 | 0 |
| 30623 | ENSMUSG00000089694 | 4.868176745 | -0.448836348 | 1.103000171 | -0.40692319 | 0.684064403 | NA | Nat8f7 | N-acetyltransferase 8 (GCN5-related) fam | 4.8186 | 1.0279 | 11.069 | 6.7478 | 1.0906 | 4.4554 |
| 30624 | ENSMUSG00000089695 | 0.685071968 | 0.040605487 | 2.800847395 | 0.014497572 | 0.988433016 | NA | Gm15946 | predicted gene 15946 [Source:MGI Symbo | 0 | 2.0558 | 0 | 0.964 | 1.0906 | 0 |
| 30625 | ENSMUSG00000089696 | 0 | NA | NA | NA | NA | NA | Spopfm1 | speckle-type BTB/POZ protein family mem | 0 | 0 | 0 | 0 | 0 | 0 |
| 30626 | ENSMUSG00000089697 | 0 | NA | NA | NA | NA | NA | Gm15947 | predicted gene 15947 [Source:MGI Symbo | 0 | 0 | 0 | 0 | 0 | 0 |
| 30627 | ENSMUSG00000089698 | 0.945986982 | 0.197573854 | 2.252218024 | 0.087724124 | 0.930095945 | NA | Gm2541 | predicted pseudogene 2541 [Source:MGI | 1.6062 | 1.0279 | 0 | 1.928 | 0 | 1.1138 |
| 30628 | ENSMUSG00000089699 | 0 | NA | NA | NA | NA | NA | Gm1992 | predicted gene 1992 [Source:MGI Symbol | 0 | 0 | 0 | 0 | 0 | 0 |

| | | | | | | | | | | | | | | | |
|---|---|---|---|---|---|---|---|---|---|---|---|---|---|---|---|
| 30629 | ENSMUSG00000089700 | 0 | NA | NA | NA | NA | NA | Gm2343 | predicted gene 2343 [Source:MGI Symbol | 0 | 0 | 0 | 0 | 0 | 0 |
| 30630 | ENSMUSG00000089701 | 0.681718484 | 0.074918168 | 2.804856827 | 0.026710158 | 0.978690911 | NA | Gm15978 | predicted gene 15978 [Source:MGI Symbo | 0 | 0 | 2.0125 | 0.964 | 0 | 1.1138 |
| 30631 | ENSMUSG00000089702 | 0 | NA | NA | NA | NA | NA | Gm16568 | predicted gene 16568 [Source:MGI Symbo | 0 | 0 | 0 | 0 | 0 | 0 |
| 30632 | ENSMUSG00000089703 | 0 | NA | NA | NA | NA | NA | Gm15833 | predicted gene 15833 [Source:MGI Symbo | 0 | 0 | 0 | 0 | 0 | 0 |
| 30633 | ENSMUSG00000089704 | 15.44608952 | -0.51202829 | 0.583827073 | -0.877020463 | 0.380475524 | 0.812628154 | Galnt2 | polypeptide N-acetylgalactosaminyltransf | 12.85 | 20.558 | 21.131 | 11.568 | 7.6343 | 18.935 |
| 30634 | ENSMUSG00000089705 | 0 | NA | NA | NA | NA | NA | Gm15970 | predicted gene 15970 [Source:MGI Symbo | 0 | 0 | 0 | 0 | 0 | 0 |
| 30635 | ENSMUSG00000089706 | 1.656120901 | -0.606856099 | 2.023972734 | -0.299834128 | 0.764303682 | NA | B230216N24 | RIKEN cDNA B230216N24 gene [Source:M | 0 | 2.0558 | 4.025 | 3.8559 | 0 | 0 |
| 30636 | ENSMUSG00000089707 | 0 | NA | NA | NA | NA | NA | Slain1os | SLAIN motif family, member 1, opposite st | 0 | 0 | 0 | 0 | 0 | 0 |
| 30637 | ENSMUSG00000089708 | 0.17131964 | -0.796629176 | 4.080472857 | -0.195229623 | 0.84521319 | NA | 5330439A09 | RIKEN cDNA 5330439A09 gene [Source:M | 0 | 1.0279 | 0 | 0 | 0 | 0 |
| 30638 | ENSMUSG00000089709 | 0 | NA | NA | NA | NA | NA | Gm16262 | predicted gene 16262 [Source:MGI Symbo | 0 | 0 | 0 | 0 | 0 | 0 |
| 30639 | ENSMUSG00000089710 | 0 | NA | NA | NA | NA | NA | Gm15810 | predicted gene 15810 [Source:MGI Symbo | 0 | 0 | 0 | 0 | 0 | 0 |
| 30640 | ENSMUSG00000089711 | 0 | NA | NA | NA | NA | NA | Gm16240 | predicted gene 16240 [Source:MGI Symbo | 0 | 0 | 0 | 0 | 0 | 0 |
| 30641 | ENSMUSG00000089712 | 248.5636557 | -1.565060135 | 0.22824127 | -6.857042714 | 7.03E-12 | 2.54E-09 | Gm15889 | predicted gene 15889 [Source:MGI Symbo | 337.3 | 433.78 | 344.14 | 137.85 | 159.23 | 79.083 |
| 30642 | ENSMUSG00000089713 | 0 | NA | NA | NA | NA | NA | Gm16564 | predicted gene 16564 [Source:MGI Symbo | 0 | 0 | 0 | 0 | 0 | 0 |
| 30643 | ENSMUSG00000089714 | 0.160662974 | 1.126924175 | 4.080472857 | 0.276174898 | 0.782413736 | NA | Gm16092 | predicted gene 16092 [Source:MGI Symbo | 0 | 0 | 0 | 0.964 | 0 | 0 |
| 30644 | ENSMUSG00000089715 | 97.80260836 | -0.674249883 | 0.252890282 | -2.666175536 | 0.007671962 | 0.089130331 | Cbx6 | chromobox 6 [Source:MGI Symbol;Acc:MG | 130.1 | 136.71 | 93.581 | 66.514 | 75.253 | 84.652 |
| 30645 | ENSMUSG00000089716 | 0.17131964 | -0.796629176 | 4.080472857 | -0.195229623 | 0.84521319 | NA | Gm6264 | predicted gene 6264 [Source:MGI Symbol | 0 | 1.0279 | 0 | 0 | 0 | 0 |
| 30646 | ENSMUSG00000089717 | 0 | NA | NA | NA | NA | NA | Olfr275 | olfactory receptor 275 [Source:MGI Symb | 0 | 0 | 0 | 0 | 0 | 0 |
| 30647 | ENSMUSG00000089718 | 0 | NA | NA | NA | NA | NA | 2310075C17 | RIKEN cDNA 2310075C17 gene [Source:M | 0 | 0 | 0 | 0 | 0 | 0 |
| 30648 | ENSMUSG00000089719 | 0 | NA | NA | NA | NA | NA | Gm15758 | predicted gene 15758 [Source:MGI Symbo | 0 | 0 | 0 | 0 | 0 | 0 |
| 30649 | ENSMUSG00000089722 | 0 | NA | NA | NA | NA | NA | Cd300ld5 | CD300 molecule like family member D5 [S | 0 | 0 | 0 | 0 | 0 | 0 |
| 30650 | ENSMUSG00000089723 | 0 | NA | NA | NA | NA | NA | Gm16117 | predicted gene 16117 [Source:MGI Symbo | 0 | 0 | 0 | 0 | 0 | 0 |
| 30651 | ENSMUSG00000089724 | 0 | NA | NA | NA | NA | NA | Krtap4-8 | keratin associated protein 4-8 [Source:MG | 0 | 0 | 0 | 0 | 0 | 0 |
| 30652 | ENSMUSG00000089726 | 36.81456931 | -0.358032922 | 0.411784465 | -0.869466801 | 0.384591859 | 0.816437825 | Mir17hg | Mir17 host gene (non-protein coding) [So | 41.761 | 46.256 | 36.225 | 42.415 | 16.359 | 37.871 |
| 30653 | ENSMUSG00000089727 | 0.728241582 | 1.712995752 | 3.00895953 | 0.569298369 | 0.569153674 | NA | Klra8 | killer cell lectin-like receptor, subfamily A, | 0 | 1.0279 | 0 | 0 | 0 | 3.3415 |
| 30654 | ENSMUSG00000089728 | 0.185640647 | 1.126924175 | 4.080472857 | 0.276174898 | 0.782413736 | NA | Clec2f | C-type lectin domain family 2, member f [S | 0 | 0 | 0 | 0 | 0 | 1.1138 |
| 30655 | ENSMUSG00000089729 | 0.321325949 | 1.911575114 | 4.048267893 | 0.472195804 | 0.636787034 | NA | Gm16552 | predicted gene 16552 [Source:MGI Symbo | 0 | 0 | 0 | 1.928 | 0 | 0 |
| 30656 | ENSMUSG00000089730 | 0 | NA | NA | NA | NA | NA | 1700007P06 | RIKEN cDNA 1700007P06 gene [Source:M | 0 | 0 | 0 | 0 | 0 | 0 |
| 30657 | ENSMUSG00000089731 | 0 | NA | NA | NA | NA | NA | Gm10013 | predicted gene 10013 [Source:MGI Symbo | 0 | 0 | 0 | 0 | 0 | 0 |
| 30658 | ENSMUSG00000089732 | 0 | NA | NA | NA | NA | NA | Olfr1117-ps1 | olfactory receptor 1117, pseudogene 1 [So | 0 | 0 | 0 | 0 | 0 | 0 |
| 30659 | ENSMUSG00000089733 | 0 | NA | NA | NA | NA | NA | Gm15628 | predicted gene 15628 [Source:MGI Symbo | 0 | 0 | 0 | 0 | 0 | 0 |
| 30660 | ENSMUSG00000089735 | 0.167707431 | -0.796629176 | 4.080472857 | -0.195229623 | 0.84521319 | NA | Gm8428 | predicted gene 8428 [Source:MGI Symbol | 0 | 0 | 1.0062 | 0 | 0 | 0 |
| 30661 | ENSMUSG00000089736 | 42.23253874 | 0.075560709 | 0.368560933 | 0.205015514 | 0.837560004 | 0.971073158 | Tgfbr3l | transforming growth factor, beta receptor | 27.305 | 46.256 | 50.312 | 49.163 | 44.715 | 35.643 |
| 30662 | ENSMUSG00000089737 | 0 | NA | NA | NA | NA | NA | Gm15688 | predicted gene 15688 [Source:MGI Symbo | 0 | 0 | 0 | 0 | 0 | 0 |
| 30663 | ENSMUSG00000089739 | 0 | NA | NA | NA | NA | NA | Gm20431 | predicted gene 20431 [Source:MGI Symbo | 0 | 0 | 0 | 0 | 0 | 0 |
| 30664 | ENSMUSG00000089742 | 1.495373649 | -0.13711118 | 1.769211587 | -0.077498464 | 0.938227013 | NA | Gm15898 | predicted gene 15898 [Source:MGI Symbo | 1.6062 | 3.0838 | 0 | 0.964 | 1.0906 | 2.2277 |
| 30665 | ENSMUSG00000089743 | 0 | NA | NA | NA | NA | NA | Gm16226 | predicted gene 16226 [Source:MGI Symbo | 0 | 0 | 0 | 0 | 0 | 0 |
| 30666 | ENSMUSG00000089744 | 2.409593138 | -0.342767977 | 1.628136778 | -0.210527753 | 0.833255794 | NA | Gm16146 | predicted gene 16146 [Source:MGI Symbo | 4.0155 | 0 | 4.025 | 0.964 | 5.4531 | 0 |
| 30667 | ENSMUSG00000089745 | 0 | NA | NA | NA | NA | NA | 1810008B01 | RIKEN cDNA 1810008B01 gene [Source:M | 0 | 0 | 0 | 0 | 0 | 0 |
| 30668 | ENSMUSG00000089746 | 0 | NA | NA | NA | NA | NA | Gm3513 | predicted gene 3513 [Source:MGI Symbol | 0 | 0 | 0 | 0 | 0 | 0 |
| 30669 | ENSMUSG00000089748 | 0 | NA | NA | NA | NA | NA | Matr3-ps2 | matrin 3, pseudogene 2 [Source:MGI Sym | 0 | 0 | 0 | 0 | 0 | 0 |
| 30670 | ENSMUSG00000089749 | 0 | NA | NA | NA | NA | NA | Gm11357 | predicted gene 11357 [Source:MGI Symbo | 0 | 0 | 0 | 0 | 0 | 0 |
| 30671 | ENSMUSG00000089753 | 0.160662974 | 1.126924175 | 4.080472857 | 0.276174898 | 0.782413736 | NA | Cd300ld2 | CD300 molecule like family member D2 [S | 0 | 0 | 0 | 0.964 | 0 | 0 |
| 30672 | ENSMUSG00000089754 | 0 | NA | NA | NA | NA | NA | Ear-ps7 | eosinophil-associated, ribonuclease A fam | 0 | 0 | 0 | 0 | 0 | 0 |
| 30673 | ENSMUSG00000089755 | 1.147551471 | -0.308736011 | 1.963633334 | -0.157226915 | 0.875066017 | NA | 0610012D04 | RIKEN cDNA 0610012D04 gene [Source:M | 0.8031 | 1.0279 | 2.0125 | 1.928 | 0 | 1.1138 |
| 30674 | ENSMUSG00000089756 | 0.453341394 | -0.545391191 | 3.551113277 | -0.153583158 | 0.877938417 | NA | Zfp966 | zinc finger protein 966 [Source:MGI Symb | 1.6062 | 0 | 0 | 0 | 0 | 1.1138 |
| 30675 | ENSMUSG00000089757 | 0 | NA | NA | NA | NA | NA | NA | NA | 0 | 0 | 0 | 0 | 0 | 0 |
| 30676 | ENSMUSG00000089759 | 1.220266871 | 0.311146302 | 2.222383583 | 0.14000567 | 0.888655511 | NA | 3632454L22I | RIKEN cDNA 3632454L22 gene [Source:M | 3.2124 | 0 | 0 | 1.928 | 2.1812 | 0 |
| 30677 | ENSMUSG00000089760 | 0 | NA | NA | NA | NA | NA | D030046N08 | RIKEN cDNA D030046N08 gene [Source:M | 0 | 0 | 0 | 0 | 0 | 0 |
| 30678 | ENSMUSG00000089761 | 0 | NA | NA | NA | NA | NA | Vmn2r-ps11 | vomeronasal 2, receptor, pseudogene 111 | 0 | 0 | 0 | 0 | 0 | 0 |
| 30679 | ENSMUSG00000089762 | 22.12619446 | -0.115214734 | 0.522139226 | -0.220659027 | 0.825357937 | 0.96943211 | Ier5l | immediate early response 5-like [Source:N | 23.29 | 16.447 | 29.181 | 14.46 | 34.9 | 14.48 |
| 30680 | ENSMUSG00000089764 | 3.433852137 | -0.333407102 | 1.297990714 | -0.256864012 | 0.797283753 | NA | Gm16580 | predicted gene 16580 [Source:MGI Symbo | 6.4248 | 1.0279 | 4.025 | 5.7839 | 0 | 3.3415 |
| 30681 | ENSMUSG00000089766 | 0 | NA | NA | NA | NA | NA | Gm16538 | predicted gene 16538 [Source:MGI Symbo | 0 | 0 | 0 | 0 | 0 | 0 |
| 30682 | ENSMUSG00000089767 | 0 | NA | NA | NA | NA | NA | 4930556I23F | RIKEN cDNA 4930556I23 gene [Source:MG | 0 | 0 | 0 | 0 | 0 | 0 |
| 30683 | ENSMUSG00000089768 | 1.707646937 | 0.158341606 | 1.609530908 | 0.098377487 | 0.921632551 | NA | Tmsb15b1 | thymosin beta 15b1 [Source:MGI Symbol; | 0.8031 | 1.0279 | 3.0187 | 0.964 | 1.0906 | 3.3415 |
| 30684 | ENSMUSG00000089769 | 0.897628956 | -1.977329804 | 2.507804229 | -0.78847056 | 0.430421512 | NA | Gm16574 | predicted gene 16574 [Source:MGI Symbo | 2.4093 | 0 | 2.0125 | 0.964 | 0 | 0 |

| | | | | | | | | | | | | | | | |
|---|---|---|---|---|---|---|---|---|---|---|---|---|---|---|---|
| 30685 | ENSMUSG00000089771 | 0.160662974 | 1.126924175 | 4.080472857 | 0.276174898 | 0.782413736 | NA | Gm16311 | predicted gene 16311 [Source:MGI Symbo | 0 | 0 | 0 | 0.964 | 0 | 0 |
| 30686 | ENSMUSG00000089773 | 0 | NA | NA | NA | NA | NA | Skint1 | selection and upkeep of intraepithelial T c | 0 | 0 | 0 | 0 | 0 | 0 |
| 30687 | ENSMUSG00000089774 | 80.38464219 | 0.118722204 | 0.262606488 | 0.452091662 | 0.65120295 | 0.925643431 | Slc5a3 | solute carrier family 5 (inositol transporte | 73.082 | 69.898 | 88.55 | 94.47 | 87.249 | 69.058 |
| 30688 | ENSMUSG00000089775 | 0 | NA | NA | NA | NA | NA | Gm16270 | predicted gene 16270 [Source:MGI Symbo | 0 | 0 | 0 | 0 | 0 | 0 |
| 30689 | ENSMUSG00000089776 | 0 | NA | NA | NA | NA | NA | Gm15684 | predicted gene 15684 [Source:MGI Symbo | 0 | 0 | 0 | 0 | 0 | 0 |
| 30690 | ENSMUSG00000089779 | 0 | NA | NA | NA | NA | NA | Gm15641 | predicted gene 15641 [Source:MGI Symbo | 0 | 0 | 0 | 0 | 0 | 0 |
| 30691 | ENSMUSG00000089780 | 0.517623261 | 1.044040381 | 3.075707576 | 0.339447218 | 0.734272853 | NA | Gm16306 | predicted gene 16306 [Source:MGI Symbo | 0 | 1.0279 | 0 | 0.964 | 0 | 1.1138 |
| 30692 | ENSMUSG00000089781 | 10.09107158 | -0.737287737 | 0.849027562 | -0.868390816 | 0.385180421 | 0.816437825 | Gm15756 | predicted gene 15756 [Source:MGI Symbo | 7.2279 | 19.53 | 11.069 | 1.928 | 17.45 | 3.3415 |
| 30693 | ENSMUSG00000089782 | 6.246081421 | 1.004794156 | 0.889437768 | 1.129695851 | 0.258604406 | 0.712165697 | Btf3-ps1 | basic transcription factor 3, pseudogene 1 | 2.4093 | 6.1675 | 4.025 | 10.604 | 9.8156 | 4.4554 |
| 30694 | ENSMUSG00000089783 | 1.632397547 | -1.005803904 | 1.724074477 | -0.583387735 | 0.559632313 | NA | Gm454 | predicted gene 454 [Source:MGI Symbol;/ | 2.4093 | 3.0838 | 1.0062 | 0 | 2.1812 | 1.1138 |
| 30695 | ENSMUSG00000089785 | 3.966101761 | -0.561585395 | 1.058653296 | -0.530471494 | 0.595785068 | NA | Slc25a5-ps | Slc25a5 retrotransposed pseudogene [Sou | 4.0155 | 4.1117 | 6.0375 | 1.928 | 4.3625 | 3.3415 |
| 30696 | ENSMUSG00000089787 | 0 | NA | NA | NA | NA | NA | Gm15825 | predicted gene 15825 [Source:MGI Symbo | 0 | 0 | 0 | 0 | 0 | 0 |
| 30697 | ENSMUSG00000089789 | 4.593138729 | -1.73825497 | 1.13651671 | -1.529458348 | 0.126150857 | NA | Rdh1 | retinol dehydrogenase 1 (all trans) [Sourc | 4.8186 | 14.391 | 2.0125 | 1.928 | 2.1812 | 2.2277 |
| 30698 | ENSMUSG00000089790 | 1.305914203 | 0.963472305 | 1.857113917 | 0.518800864 | 0.603899615 | NA | Gm16588 | predicted gene 16588 [Source:MGI Symbo | 1.6062 | 0 | 1.0062 | 1.928 | 2.1812 | 1.1138 |
| 30699 | ENSMUSG00000089791 | 0.925753523 | 0.330822674 | 2.475764778 | 0.133624437 | 0.89369956 | NA | Gm3555 | predicted pseudogene 3555 [Source:MGI | 2.4093 | 0 | 0 | 0.964 | 2.1812 | 0 |
| 30700 | ENSMUSG00000089792 | 0 | NA | NA | NA | NA | NA | Gm16571 | predicted gene 16571 [Source:MGI Symbo | 0 | 0 | 0 | 0 | 0 | 0 |
| 30701 | ENSMUSG00000089793 | 0 | NA | NA | NA | NA | NA | Gm16112 | predicted gene 16112 [Source:MGI Symbo | 0 | 0 | 0 | 0 | 0 | 0 |
| 30702 | ENSMUSG00000089794 | 0 | NA | NA | NA | NA | NA | Gm16337 | predicted gene 16337 [Source:MGI Symbo | 0 | 0 | 0 | 0 | 0 | 0 |
| 30703 | ENSMUSG00000089795 | 0 | NA | NA | NA | NA | NA | Hmgb3-ps | Hmgb3 retrotransposed pseudogene [Sou | 0 | 0 | 0 | 0 | 0 | 0 |
| 30704 | ENSMUSG00000089796 | 1.857419497 | -0.267534294 | 1.733515604 | -0.154330479 | 0.877349166 | NA | Gm16575 | predicted gene 16575 [Source:MGI Symbo | 4.0155 | 2.0558 | 0 | 2.8919 | 2.1812 | 0 |
| 30705 | ENSMUSG00000089797 | 1.088866211 | 3.648835749 | 2.580470319 | 1.414019655 | 0.157356132 | NA | Gm16118 | predicted gene 16118 [Source:MGI Symbo | 0 | 0 | 0 | 0.964 | 0 | 5.5692 |
| 30706 | ENSMUSG00000089798 | 0 | NA | NA | NA | NA | NA | 1700028K03 | RIKEN cDNA 1700028K03 gene [Source:M | 0 | 0 | 0 | 0 | 0 | 0 |
| 30707 | ENSMUSG00000089800 | 1.268599393 | -2.525346601 | 2.259681915 | -1.117567293 | 0.263751844 | NA | Rps19-ps10 | ribosomal protein S19, pseudogene 10 [So | 2.4093 | 4.1117 | 0 | 0 | 1.0906 | 0 |
| 30708 | ENSMUSG00000089802 | 0.167707431 | -0.796629176 | 4.080472857 | -0.195229623 | 0.84521319 | NA | Gm16107 | predicted gene 16107 [Source:MGI Symbo | 0 | 0 | 1.0062 | 0 | 0 | 0 |
| 30709 | ENSMUSG00000089803 | 1.968044503 | 0.198905622 | 1.448504476 | 0.13731792 | 0.890779506 | NA | Gm10171 | predicted gene 10171 [Source:MGI Symbo | 2.4093 | 2.0558 | 1.0062 | 1.928 | 2.1812 | 2.2277 |
| 30710 | ENSMUSG00000089804 | 0.185640647 | 1.126924175 | 4.080472857 | 0.276174898 | 0.782413736 | NA | Gm16136 | predicted gene 16136 [Source:MGI Symbo | 0 | 0 | 0 | 0 | 0 | 1.1138 |
| 30711 | ENSMUSG00000089806 | 0 | NA | NA | NA | NA | NA | Gm16570 | predicted gene 16570 [Source:MGI Symbo | 0 | 0 | 0 | 0 | 0 | 0 |
| 30712 | ENSMUSG00000089807 | 0.267700746 | -1.497481649 | 4.057001436 | -0.369110456 | 0.712045396 | NA | Gm16298 | predicted gene 16298 [Source:MGI Symbo | 1.6062 | 0 | 0 | 0 | 0 | 0 |
| 30713 | ENSMUSG00000089808 | 0 | NA | NA | NA | NA | NA | Gm7241 | predicted pseudogene 7241 [Source:MGI | 0 | 0 | 0 | 0 | 0 | 0 |
| 30714 | ENSMUSG00000089809 | 158.4370313 | -0.926332395 | 0.303027037 | -3.056929855 | 0.002236166 | 0.037784935 | Rasgef1b | RasGEF domain family, member 1B [Sourc | 149.38 | 309.4 | 164.02 | 77.118 | 100.34 | 150.37 |
| 30715 | ENSMUSG00000089810 | 146.1484203 | 0.326574306 | 0.220765236 | 1.479283208 | 0.139064639 | 0.545066664 | Gm16536 | predicted gene 16536 [Source:MGI Symbo | 138.94 | 152.13 | 97.606 | 146.52 | 169.05 | 172.65 |
| 30716 | ENSMUSG00000089811 | 0 | NA | NA | NA | NA | NA | Gm16138 | predicted gene 16138 [Source:MGI Symbo | 0 | 0 | 0 | 0 | 0 | 0 |
| 30717 | ENSMUSG00000089812 | 0.642651897 | 2.906894826 | 3.440535777 | 0.844895974 | 0.398168923 | NA | Gm15867 | predicted gene 15867 [Source:MGI Symbo | 0 | 0 | 0 | 3.8559 | 0 | 0 |
| 30718 | ENSMUSG00000089814 | 0 | NA | NA | NA | NA | NA | Gm15310 | predicted gene 15310 [Source:MGI Symbo | 0 | 0 | 0 | 0 | 0 | 0 |
| 30719 | ENSMUSG00000089815 | 0 | NA | NA | NA | NA | NA | Gm5083 | predicted gene 5083 [Source:MGI Symbol | 0 | 0 | 0 | 0 | 0 | 0 |
| 30720 | ENSMUSG00000089816 | 0 | NA | NA | NA | NA | NA | Ear-ps8 | eosinophil-associated, ribonuclease A fan | 0 | 0 | 0 | 0 | 0 | 0 |
| 30721 | ENSMUSG00000089817 | 2.191773949 | -0.672186635 | 1.879472823 | -0.357646371 | 0.720607971 | NA | Gm7162 | predicted gene 7162 [Source:MGI Symbol | 8.031 | 0 | 0 | 2.8919 | 0 | 2.2277 |
| 30722 | ENSMUSG00000089818 | 2.337755107 | 0.14121728 | 1.355030123 | 0.104217078 | 0.916997082 | NA | Gm15950 | predicted gene 15950 [Source:MGI Symbo | 1.6062 | 3.0838 | 2.0125 | 2.8919 | 1.0906 | 3.3415 |
| 30723 | ENSMUSG00000089819 | 0 | NA | NA | NA | NA | NA | Gm15679 | predicted gene 15679 [Source:MGI Symbo | 0 | 0 | 0 | 0 | 0 | 0 |
| 30724 | ENSMUSG00000089820 | 1.670150512 | -2.98697967 | 2.089622154 | -1.4294353 | 0.152879159 | NA | Gm15775 | predicted gene 15775 [Source:MGI Symbo | 4.8186 | 4.1117 | 0 | 0 | 1.0906 | 0 |
| 30725 | ENSMUSG00000089821 | 0 | NA | NA | NA | NA | NA | Efhd1os | EF hand domain containing 1, opposite str | 0 | 0 | 0 | 0 | 0 | 0 |
| 30726 | ENSMUSG00000089822 | 0 | NA | NA | NA | NA | NA | Gm15759 | predicted gene 15759 [Source:MGI Symbo | 0 | 0 | 0 | 0 | 0 | 0 |
| 30727 | ENSMUSG00000089824 | 82.1433558 | -0.31231487 | 0.289517898 | -1.07874115 | 0.280703136 | 0.733530291 | Rbm12 | RNA binding motif protein 12 [Source:MG | 103.6 | 93.541 | 75.468 | 59.767 | 100.34 | 60.148 |
| 30728 | ENSMUSG00000089826 | 0 | NA | NA | NA | NA | NA | Gm16560 | predicted gene 16560 [Source:MGI Symbo | 0 | 0 | 0 | 0 | 0 | 0 |
| 30729 | ENSMUSG00000089827 | 20.0879725 | 1.149562136 | 0.752108754 | 1.52845201 | 0.12640035 | 0.521167236 | 1700023H06 | RIKEN cDNA 1700023H06 gene [Source:M | 15.259 | 2.0558 | 20.125 | 49.163 | 28.356 | 5.5692 |
| 30730 | ENSMUSG00000089828 | 0 | NA | NA | NA | NA | NA | Gm16300 | predicted gene 16300 [Source:MGI Symbo | 0 | 0 | 0 | 0 | 0 | 0 |
| 30731 | ENSMUSG00000089829 | 0.305170013 | -1.657836477 | 4.041480335 | -0.410205257 | 0.681655385 | NA | Gm16565 | predicted gene 16565 [Source:MGI Symbo | 0.8031 | 1.0279 | 0 | 0 | 0 | 0 |
| 30732 | ENSMUSG00000089830 | 0 | NA | NA | NA | NA | NA | Gm15749 | predicted gene 15749 [Source:MGI Symbo | 0 | 0 | 0 | 0 | 0 | 0 |
| 30733 | ENSMUSG00000089831 | 0 | NA | NA | NA | NA | NA | Gm15508 | predicted gene 15508 [Source:MGI Symbo | 0 | 0 | 0 | 0 | 0 | 0 |
| 30734 | ENSMUSG00000089832 | 123.1844262 | -0.113151047 | 0.233765383 | -0.484036795 | 0.628359756 | 0.920328819 | Shkbp1 | Sh3kbp1 binding protein 1 [Source:MGI Sy | 125.28 | 98.68 | 159.99 | 113.75 | 118.88 | 122.52 |
| 30735 | ENSMUSG00000089833 | 0 | NA | NA | NA | NA | NA | Fhitos | fragile histidine triad gene, opposite stran | 0 | 0 | 0 | 0 | 0 | 0 |
| 30736 | ENSMUSG00000089834 | 0 | NA | NA | NA | NA | NA | Gm16303 | predicted gene 16303 [Source:MGI Symbo | 0 | 0 | 0 | 0 | 0 | 0 |
| 30737 | ENSMUSG00000089835 | 0 | NA | NA | NA | NA | NA | Gm7097 | predicted gene 7097 [Source:MGI Symbol | 0 | 0 | 0 | 0 | 0 | 0 |
| 30738 | ENSMUSG00000089836 | 0 | NA | NA | NA | NA | NA | Gm16290 | predicted gene 16290 [Source:MGI Symbo | 0 | 0 | 0 | 0 | 0 | 0 |
| 30739 | ENSMUSG00000089837 | 0 | NA | NA | NA | NA | NA | Npcd | neuronal pentraxin chromo domain [Sour | 0 | 0 | 0 | 0 | 0 | 0 |
| 30740 | ENSMUSG00000089838 | 1.511038767 | -0.893514062 | 1.814760323 | -0.492359267 | 0.62246539 | NA | Gm2962 | predicted pseudogene 2962 [Source:MGI | 0.8031 | 4.1117 | 1.0062 | 0.964 | 2.1812 | 0 |

| | | | | | | | | | | | | | | | |
|---|---|---|---|---|---|---|---|---|---|---|---|---|---|---|---|
| 30741 | ENSMUSG00000089839 | 0 | NA | NA | NA | NA | NA | Rps19-ps11 | ribosomal protein S19, pseudogene 11 [Sc | 0 | 0 | 0 | 0 | 0 | 0 |
| 30742 | ENSMUSG00000089840 | 0.822586684 | 2.037574431 | 2.401368532 | 0.84850551 | 0.396156505 | NA | Gm4491 | predicted gene 4491 [Source:MGI Symbol | 0.8031 | 0 | 0 | 1.928 | 1.0906 | 1.1138 |
| 30743 | ENSMUSG00000089842 | 3.124397721 | -2.988551615 | 1.519575795 | -1.966701249 | 0.049217663 | NA | Pitpnm2os2 | phosphatidylinositol transfer protein, mer | 2.4093 | 9.2513 | 5.0312 | 0.964 | 1.0906 | 0 |
| 30744 | ENSMUSG00000089844 | 7.864086952 | -0.871164926 | 0.784726881 | -1.110150483 | 0.266934186 | 0.722778036 | A530032D15 | RIKEN cDNA A530032D15Rik gene [Source | 7.2279 | 12.335 | 11.069 | 7.7118 | 3.2719 | 5.5692 |
| 30745 | ENSMUSG00000089847 | 74.76151701 | -0.363174146 | 0.282498219 | -1.285580303 | 0.198589575 | 0.640874522 | Timm10b | translocase of inner mitochondrial membr | 102.8 | 68.87 | 80.5 | 76.154 | 54.531 | 65.717 |
| 30746 | ENSMUSG00000089849 | 0 | NA | NA | NA | NA | NA | Runx2os2 | runt related transcription factor 2, opposi | 0 | 0 | 0 | 0 | 0 | 0 |
| 30747 | ENSMUSG00000089851 | 0 | NA | NA | NA | NA | NA | Gm16579 | predicted gene 16579 [Source:MGI Symbo | 0 | 0 | 0 | 0 | 0 | 0 |
| 30748 | ENSMUSG00000089854 | 0 | NA | NA | NA | NA | NA | Gm16133 | predicted gene 16133 [Source:MGI Symbo | 0 | 0 | 0 | 0 | 0 | 0 |
| 30749 | ENSMUSG00000089857 | 38.13036918 | -0.080771291 | 0.358692574 | -0.225182502 | 0.821837302 | 0.969160361 | Zfp882 | zinc finger protein 882 [Source:MGI Symb | 48.186 | 33.921 | 35.219 | 37.595 | 35.99 | 37.871 |
| 30750 | ENSMUSG00000089859 | 0 | NA | NA | NA | NA | NA | Olfr1565 | olfactory receptor 1565 [Source:MGI Sym | 0 | 0 | 0 | 0 | 0 | 0 |
| 30751 | ENSMUSG00000089861 | 0.342432689 | 1.990653086 | 3.900281263 | 0.51038706 | 0.609780321 | NA | Gm16139 | predicted gene 16139 [Source:MGI Symbo | 0 | 0 | 0 | 0.964 | 1.0906 | 0 |
| 30752 | ENSMUSG00000089862 | 267.887394 | 0.127817875 | 0.164066854 | 0.779059706 | 0.435944545 | 0.84624805 | Umad1 | UMAP1-MVP12 associated (UMA) domair | 269.84 | 222.03 | 275.71 | 267.02 | 304.28 | 268.44 |
| 30753 | ENSMUSG00000089865 | 0 | NA | NA | NA | NA | NA | Gm44503 | predicted readthrough transcript (NMD ca | 0 | 0 | 0 | 0 | 0 | 0 |
| 30754 | ENSMUSG00000089866 | 0 | NA | NA | NA | NA | NA | Gm15699 | predicted gene 15699 [Source:MGI Symbo | 0 | 0 | 0 | 0 | 0 | 0 |
| 30755 | ENSMUSG00000089868 | 0 | NA | NA | NA | NA | NA | Gm16134 | predicted gene 16134 [Source:MGI Symbo | 0 | 0 | 0 | 0 | 0 | 0 |
| 30756 | ENSMUSG00000089870 | 0 | NA | NA | NA | NA | NA | Gm15562 | predicted gene 15562 [Source:MGI Symbo | 0 | 0 | 0 | 0 | 0 | 0 |
| 30757 | ENSMUSG00000089871 | 0 | NA | NA | NA | NA | NA | Speer4cos | spermatogenesis associated glutamate (E) | 0 | 0 | 0 | 0 | 0 | 0 |
| 30758 | ENSMUSG00000089872 | 167.7483366 | 0.048437327 | 0.194087191 | 0.249564779 | 0.802923939 | 0.963537682 | Rps6kc1 | ribosomal protein S6 kinase polypeptide 1 | 139.74 | 165.49 | 190.18 | 171.59 | 167.96 | 171.53 |
| 30759 | ENSMUSG00000089873 | 6.214812065 | 1.756276717 | 1.155931271 | 1.519360849 | 0.128671691 | 0.524858587 | Mup13 | major urinary protein 13 [Source:MGI Sym | 7.2279 | 1.0279 | 0 | 3.8559 | 20.722 | 4.4554 |
| 30760 | ENSMUSG00000089874 | 0 | NA | NA | NA | NA | NA | 9230117E06 | RIKEN cDNA 9230117E06 gene [Source:M | 0 | 0 | 0 | 0 | 0 | 0 |
| 30761 | ENSMUSG00000089875 | 47.78433411 | 0.027508184 | 0.34376656 | 0.080019954 | 0.936221386 | 0.988420305 | Etohd2 | ethanol decreased 2 [Source:MGI Symbol | 42.564 | 56.535 | 43.269 | 60.731 | 49.078 | 34.529 |
| 30762 | ENSMUSG00000089876 | 213.3285203 | 0.436396139 | 0.193231159 | 2.258414957 | 0.023919801 | 0.196800912 | Tmem102 | transmembrane protein 102 [Source:MGI | 191.14 | 144.94 | 207.29 | 223.64 | 233.39 | 279.57 |
| 30763 | ENSMUSG00000089877 | 0 | NA | NA | NA | NA | NA | Gm16102 | predicted gene 16102 [Source:MGI Symbo | 0 | 0 | 0 | 0 | 0 | 0 |
| 30764 | ENSMUSG00000089879 | 0.319491021 | 0.16514334 | 4.044653141 | 0.040830038 | 0.967431393 | NA | 4930448H16 | RIKEN cDNA 4930448H16 gene [Source:M | 0.8031 | 0 | 0 | 0 | 0 | 1.1138 |
| 30765 | ENSMUSG00000089881 | 0.17131964 | -0.796629176 | 4.080472857 | -0.195229623 | 0.84521319 | NA | Gm16280 | predicted gene 16280 [Source:MGI Symbo | 0 | 1.0279 | 0 | 0 | 0 | 0 |
| 30766 | ENSMUSG00000089883 | 0.167707431 | -0.796629176 | 4.080472857 | -0.195229623 | 0.84521319 | NA | Gm3355 | predicted gene 3355 [Source:MGI Symbol | 0 | 0 | 1.0062 | 0 | 0 | 0 |
| 30767 | ENSMUSG00000089885 | 0 | NA | NA | NA | NA | NA | Gm16296 | predicted gene 16296 [Source:MGI Symbo | 0 | 0 | 0 | 0 | 0 | 0 |
| 30768 | ENSMUSG00000089886 | 1.37019607 | 1.666070979 | 1.894315248 | 0.879510937 | 0.379124306 | NA | Gm16184 | predicted gene 16184 [Source:MGI Symbo | 0 | 1.0279 | 1.0062 | 2.8919 | 2.1812 | 1.1138 |
| 30769 | ENSMUSG00000089887 | 0 | NA | NA | NA | NA | NA | 4930428N03 | RIKEN cDNA 4930428N03 gene [Source:M | 0 | 0 | 0 | 0 | 0 | 0 |
| 30770 | ENSMUSG00000089888 | 0 | NA | NA | NA | NA | NA | Gm16547 | predicted gene 16547 [Source:MGI Symbo | 0 | 0 | 0 | 0 | 0 | 0 |
| 30771 | ENSMUSG00000089889 | 43.60272352 | -0.142719848 | 0.362164571 | -0.394074572 | 0.693525984 | 0.938983415 | 0610040B10 | RIKEN cDNA 0610040B10 gene [Source:M | 58.626 | 35.977 | 42.262 | 35.667 | 53.44 | 35.643 |
| 30772 | ENSMUSG00000089890 | 0 | NA | NA | NA | NA | NA | Gm16055 | predicted gene 16055 [Source:MGI Symbo | 0 | 0 | 0 | 0 | 0 | 0 |
| 30773 | ENSMUSG00000089891 | 0 | NA | NA | NA | NA | NA | Gm16561 | predicted gene 16561 [Source:MGI Symbo | 0 | 0 | 0 | 0 | 0 | 0 |
| 30774 | ENSMUSG00000089892 | 0 | NA | NA | NA | NA | NA | Olfr1199 | olfactory receptor 1199 [Source:MGI Sym | 0 | 0 | 0 | 0 | 0 | 0 |
| 30775 | ENSMUSG00000089893 | 0 | NA | NA | NA | NA | NA | Gm9141 | predicted gene 9141 [Source:MGI Symbol | 0 | 0 | 0 | 0 | 0 | 0 |
| 30776 | ENSMUSG00000089894 | 0 | NA | NA | NA | NA | NA | Gm15524 | predicted gene 15524 [Source:MGI Symbo | 0 | 0 | 0 | 0 | 0 | 0 |
| 30777 | ENSMUSG00000089898 | 0 | NA | NA | NA | NA | NA | Gm14199 | predicted gene 14199 [Source:MGI Symbo | 0 | 0 | 0 | 0 | 0 | 0 |
| 30778 | ENSMUSG00000089900 | 1.853259352 | 1.119533466 | 1.77475455 | 0.630810309 | 0.528164562 | NA | Tbc1d22bos | TBC1 domain family, member 22B, opposi | 2.4093 | 0 | 1.0062 | 0 | 4.3625 | 3.3415 |
| 30779 | ENSMUSG00000089901 | 5.529995658 | -2.822931071 | 1.116244283 | -2.52895456 | 0.011440284 | 0.119388358 | Gm8113 | predicted gene 8113 [Source:MGI Symbol | 8.8341 | 11.307 | 9.0562 | 2.8919 | 1.0906 | 0 |
| 30780 | ENSMUSG00000089902 | 1.354117244 | -0.004184291 | 2.152127421 | -0.001944258 | 0.998448708 | NA | Gm13625 | predicted gene 13625 [Source:MGI Symbo | 4.0155 | 0 | 0 | 1.928 | 2.1812 | 0 |
| 30781 | ENSMUSG00000089904 | 0.435408177 | -2.179616019 | 3.531869654 | -0.617128103 | 0.537150234 | NA | Gm15664 | predicted gene 15664 [Source:MGI Symbo | 1.6062 | 0 | 1.0062 | 0 | 0 | 0 |
| 30782 | ENSMUSG00000089905 | 0 | NA | NA | NA | NA | NA | Gm16563 | predicted gene 16563 [Source:MGI Symbo | 0 | 0 | 0 | 0 | 0 | 0 |
| 30783 | ENSMUSG00000089906 | 0 | NA | NA | NA | NA | NA | Gm6428 | predicted pseudogene 6428 [Source:MGI | 0 | 0 | 0 | 0 | 0 | 0 |
| 30784 | ENSMUSG00000089907 | 0 | NA | NA | NA | NA | NA | Gm16169 | predicted gene 16169 [Source:MGI Symbo | 0 | 0 | 0 | 0 | 0 | 0 |
| 30785 | ENSMUSG00000089908 | 0.181769715 | 1.126924175 | 4.080472857 | 0.276174898 | 0.782413736 | NA | Gm16558 | predicted gene 16558 [Source:MGI Symbo | 0 | 0 | 0 | 0 | 1.0906 | 0 |
| 30786 | ENSMUSG00000089909 | 0 | NA | NA | NA | NA | NA | Gm16164 | predicted gene 16164 [Source:MGI Symbo | 0 | 0 | 0 | 0 | 0 | 0 |
| 30787 | ENSMUSG00000089910 | 0 | NA | NA | NA | NA | NA | Gm16113 | predicted gene 16113 [Source:MGI Symbo | 0 | 0 | 0 | 0 | 0 | 0 |
| 30788 | ENSMUSG00000089911 | 357.8723136 | 0.101243692 | 0.141420169 | 0.715907024 | 0.474048761 | 0.86899591 | Mfsd14a | major facilitator superfamily domain cont | 322.85 | 336.13 | 377.34 | 361.49 | 375.17 | 374.25 |
| 30789 | ENSMUSG00000089914 | 0 | NA | NA | NA | NA | NA | 4930486I03F | RIKEN cDNA 4930486I03 gene [Source:M | 0 | 0 | 0 | 0 | 0 | 0 |
| 30790 | ENSMUSG00000089916 | 0 | NA | NA | NA | NA | NA | Gm16160 | predicted gene 16160 [Source:MGI Symbo | 0 | 0 | 0 | 0 | 0 | 0 |
| 30791 | ENSMUSG00000089917 | 352.6496686 | 0.22830113 | 0.166109728 | 1.374399522 | 0.169317685 | 0.5966686 | Uckl1 | uridine-cytidine kinase 1-like 1 [Source:M | 335.7 | 295.01 | 344.14 | 458.85 | 341.36 | 340.84 |
| 30792 | ENSMUSG00000089922 | 0 | NA | NA | NA | NA | NA | Gm43517 | predicted gene 43517 [Source:MGI Symbo | 0 | 0 | 0 | 0 | 0 | 0 |
| 30793 | ENSMUSG00000089924 | 0 | NA | NA | NA | NA | NA | Gm15689 | predicted gene 15689 [Source:MGI Symbo | 0 | 0 | 0 | 0 | 0 | 0 |
| 30794 | ENSMUSG00000089926 | 0 | NA | NA | NA | NA | NA | Gm15734 | predicted gene 15734 [Source:MGI Symbo | 0 | 0 | 0 | 0 | 0 | 0 |
| 30795 | ENSMUSG00000089927 | 0 | NA | NA | NA | NA | NA | Gm16539 | predicted gene 16539 [Source:MGI Symbo | 0 | 0 | 0 | 0 | 0 | 0 |
| 30796 | ENSMUSG00000089929 | 1.333192118 | 0.129315543 | 1.841092404 | 0.070238486 | 0.944003843 | NA | Bcl2a1b | B cell leukemia/lymphoma 2 related prote | 0.8031 | 1.0279 | 2.0125 | 1.928 | 0 | 2.2277 |

| | | | | | | | | | | | | | | | |
|---|---|---|---|---|---|---|---|---|---|---|---|---|---|---|---|
| 30797 | ENSMUSG00000089931 | 0 | NA | NA | NA | NA | NA | Gm8459 | predicted gene 8459 [Source:MGI Symbol | 0 | 0 | 0 | 0 | 0 | 0 |
| 30798 | ENSMUSG00000089932 | 0 | NA | NA | NA | NA | NA | Gm16114 | predicted gene 16114 [Source:MGI Symbo | 0 | 0 | 0 | 0 | 0 | 0 |
| 30799 | ENSMUSG00000089933 | 0 | NA | NA | NA | NA | NA | Gm16101 | predicted gene 16101 [Source:MGI Symbo | 0 | 0 | 0 | 0 | 0 | 0 |
| 30800 | ENSMUSG00000089934 | 0 | NA | NA | NA | NA | NA | 4930473D10 | RIKEN cDNA 4930473D10 gene [Source:M | 0 | 0 | 0 | 0 | 0 | 0 |
| 30801 | ENSMUSG00000089935 | 0.167707431 | -0.796629176 | 4.080472857 | -0.195229623 | 0.84521319 | NA | Gm16343 | predicted gene 16343 [Source:MGI Symbo | 0 | 0 | 1.0062 | 0 | 0 | 0 |
| 30802 | ENSMUSG00000089936 | 0.167707431 | -0.796629176 | 4.080472857 | -0.195229623 | 0.84521319 | NA | Gm16199 | predicted gene 16199 [Source:MGI Symbo | 0 | 0 | 1.0062 | 0 | 0 | 0 |
| 30803 | ENSMUSG00000089937 | 0 | NA | NA | NA | NA | NA | Gm7281 | predicted gene 7281 [Source:MGI Symbol | 0 | 0 | 0 | 0 | 0 | 0 |
| 30804 | ENSMUSG00000089938 | 0 | NA | NA | NA | NA | NA | Gm15744 | predicted gene 15744 [Source:MGI Symbo | 0 | 0 | 0 | 0 | 0 | 0 |
| 30805 | ENSMUSG00000089940 | 0.656740811 | -0.013525372 | 3.112905782 | -0.004344934 | 0.996533255 | NA | Gm4117 | predicted gene 4117 [Source:MGI Symbol | 0 | 0 | 2.0125 | 1.928 | 0 | 0 |
| 30806 | ENSMUSG00000089941 | 0.160662974 | 1.126924175 | 4.080472857 | 0.276174898 | 0.782413736 | NA | Gm16168 | predicted gene 16168 [Source:MGI Symbo | 0 | 0 | 0 | 0.964 | 0 | 0 |
| 30807 | ENSMUSG00000089942 | 4.785927573 | -2.979692288 | 1.239092585 | -2.404737405 | 0.01618409 | NA | Pira2 | paired-Ig-like receptor A2 [Source:MGI Sy | 8.031 | 12.335 | 5.0312 | 0 | 1.0906 | 2.2277 |
| 30808 | ENSMUSG00000089943 | 239.2639748 | 0.483928706 | 0.255329441 | 1.895311032 | 0.058051208 | 0.342435834 | Ugt1a5 | UDP glucuronosyltransferase 1 family, pol | 259.4 | 218.95 | 119.74 | 348.96 | 230.12 | 258.41 |
| 30809 | ENSMUSG00000089945 | 0 | NA | NA | NA | NA | NA | Pakap | paralemmin A kinase anchor protein [Sou | 0 | 0 | 0 | 0 | 0 | 0 |
| 30810 | ENSMUSG00000089947 | 0 | NA | NA | NA | NA | NA | Gm15509 | predicted gene 15509 [Source:MGI Symbo | 0 | 0 | 0 | 0 | 0 | 0 |
| 30811 | ENSMUSG00000089948 | 0 | NA | NA | NA | NA | NA | Far2os1 | fatty acyl CoA reductase 2, opposite stran | 0 | 0 | 0 | 0 | 0 | 0 |
| 30812 | ENSMUSG00000089949 | 0 | NA | NA | NA | NA | NA | Gm16137 | predicted gene 16137 [Source:MGI Symbo | 0 | 0 | 0 | 0 | 0 | 0 |
| 30813 | ENSMUSG00000089950 | 0 | NA | NA | NA | NA | NA | Gm16215 | predicted gene 16215 [Source:MGI Symbo | 0 | 0 | 0 | 0 | 0 | 0 |
| 30814 | ENSMUSG00000089951 | 0.36353943 | 2.065989254 | 4.034137301 | 0.512126658 | 0.608562368 | NA | Zfp968-ps | zinc finger protein 968, pseudogene [Sour | 0 | 0 | 0 | 0 | 2.1812 | 0 |
| 30815 | ENSMUSG00000089952 | 0 | NA | NA | NA | NA | NA | 4933413C19 | RIKEN cDNA 4933413C19 gene [Source:M | 0 | 0 | 0 | 0 | 0 | 0 |
| 30816 | ENSMUSG00000089953 | 0 | NA | NA | NA | NA | NA | Rnf224 | ring finger protein 224 [Source:MGI Symb | 0 | 0 | 0 | 0 | 0 | 0 |
| 30817 | ENSMUSG00000089956 | 2.161090415 | -1.121079927 | 1.794874826 | -0.624600619 | 0.532233213 | NA | Gm16553 | predicted gene 16553 [Source:MGI Symbo | 8.8341 | 0 | 0 | 1.928 | 1.0906 | 1.1138 |
| 30818 | ENSMUSG00000089957 | 0 | NA | NA | NA | NA | NA | A830011K09 | RIKEN cDNA A830011K09 gene [Source:M | 0 | 0 | 0 | 0 | 0 | 0 |
| 30819 | ENSMUSG00000089958 | 0 | NA | NA | NA | NA | NA | Gm16165 | predicted gene 16165 [Source:MGI Symbo | 0 | 0 | 0 | 0 | 0 | 0 |
| 30820 | ENSMUSG00000089959 | 3.770474698 | -1.403394675 | 1.16633469 | -1.203252108 | 0.228878776 | NA | Gm16268 | predicted gene 16268 [Source:MGI Symbo | 7.2279 | 7.1954 | 2.0125 | 2.8919 | 2.1812 | 1.1138 |
| 30821 | ENSMUSG00000089960 | 1533.435896 | -0.355261815 | 0.157903966 | -2.24985998 | 0.024457835 | 0.199836083 | Ugt1a1 | UDP glucuronosyltransferase 1 family, pol | 1444.8 | 1844.1 | 1876.6 | 1560.7 | 1437.4 | 1037 |
| 30822 | ENSMUSG00000089961 | 0 | NA | NA | NA | NA | NA | Gm16567 | predicted gene 16567 [Source:MGI Symbo | 0 | 0 | 0 | 0 | 0 | 0 |
| 30823 | ENSMUSG00000089963 | 0 | NA | NA | NA | NA | NA | Rhox11-ps3 | reproductive homeobox 11, pseudogene 3 | 0 | 0 | 0 | 0 | 0 | 0 |
| 30824 | ENSMUSG00000089964 | 0 | NA | NA | NA | NA | NA | Gm16582 | predicted gene 16582 [Source:MGI Symbo | 0 | 0 | 0 | 0 | 0 | 0 |
| 30825 | ENSMUSG00000089965 | 0 | NA | NA | NA | NA | NA | Gm16236 | predicted gene 16236 [Source:MGI Symbo | 0 | 0 | 0 | 0 | 0 | 0 |
| 30826 | ENSMUSG00000089966 | 0 | NA | NA | NA | NA | NA | Gm16030 | predicted gene 16030 [Source:MGI Symbo | 0 | 0 | 0 | 0 | 0 | 0 |
| 30827 | ENSMUSG00000089967 | 0 | NA | NA | NA | NA | NA | Gm16550 | predicted gene 16550 [Source:MGI Symbo | 0 | 0 | 0 | 0 | 0 | 0 |
| 30828 | ENSMUSG00000089968 | 0.305170013 | -1.657836477 | 4.041480335 | -0.410205257 | 0.681655385 | NA | Nckap5los | NCK-associated protein 5-like, opposite st | 0.8031 | 1.0279 | 0 | 0 | 0 | 0 |
| 30829 | ENSMUSG00000089972 | 0 | NA | NA | NA | NA | NA | Gm16318 | predicted gene 16318 [Source:MGI Symbo | 0 | 0 | 0 | 0 | 0 | 0 |
| 30830 | ENSMUSG00000089973 | 0 | NA | NA | NA | NA | NA | Vmn1r190-p | vomeronasal 1 receptor 190, pseudogene | 0 | 0 | 0 | 0 | 0 | 0 |
| 30831 | ENSMUSG00000089975 | 0.497389803 | 1.101799453 | 3.439268845 | 0.320358629 | 0.748696483 | NA | Gm7420 | predicted gene 7420 [Source:MGI Symbol | 0.8031 | 0 | 0 | 0 | 2.1812 | 0 |
| 30832 | ENSMUSG00000089976 | 0 | NA | NA | NA | NA | NA | Gm2389 | predicted pseudogene 2389 [Source:MGI | 0 | 0 | 0 | 0 | 0 | 0 |
| 30833 | ENSMUSG00000089978 | 0 | NA | NA | NA | NA | NA | Crb1-ps | crumbs family member 1, photoreceptor | 0 | 0 | 0 | 0 | 0 | 0 |
| 30834 | ENSMUSG00000089982 | 0.133850373 | -0.796629176 | 4.080472857 | -0.195229623 | 0.84521319 | NA | Gm7568 | predicted gene 7568 [Source:MGI Symbol | 0.8031 | 0 | 0 | 0 | 0 | 0 |
| 30835 | ENSMUSG00000089983 | 0.133850373 | -0.796629176 | 4.080472857 | -0.195229623 | 0.84521319 | NA | 2010320O07 | RIKEN cDNA 2010320O07 gene [Source:M | 0.8031 | 0 | 0 | 0 | 0 | 0 |
| 30836 | ENSMUSG00000089984 | 1.9932809 | 0.230787439 | 1.498539658 | 0.154008229 | 0.877603247 | NA | Fbxo24 | F-box protein 24 [Source:MGI Symbol;Acc | 2.4093 | 1.0279 | 2.0125 | 0.964 | 1.0906 | 4.4554 |
| 30837 | ENSMUSG00000089987 | 0 | NA | NA | NA | NA | NA | Gpx2-ps1 | glutathione peroxidase 2, pseudogene 1 [ | 0 | 0 | 0 | 0 | 0 | 0 |
| 30838 | ENSMUSG00000089988 | 0.695728634 | -1.425610641 | 3.05889997 | -0.46605337 | 0.641177299 | NA | Gm16238 | predicted gene 16238 [Source:MGI Symbo | 0 | 3.0838 | 0 | 0 | 1.0906 | 0 |
| 30839 | ENSMUSG00000089989 | 0 | NA | NA | NA | NA | NA | Gm45713 | predicted gene 45713 [Source:MGI Symbo | 0 | 0 | 0 | 0 | 0 | 0 |
| 30840 | ENSMUSG00000089990 | 0.160662974 | 1.126924175 | 4.080472857 | 0.276174898 | 0.782413736 | NA | Gm15852 | predicted gene 15852 [Source:MGI Symbo | 0 | 0 | 0 | 0.964 | 0 | 0 |
| 30841 | ENSMUSG00000089991 | 200.5822612 | 0.372743735 | 0.205718029 | 1.811915742 | 0.069999217 | 0.382496664 | Gm16332 | predicted gene 16332 [Source:MGI Symbo | 144.56 | 168.58 | 212.32 | 267.99 | 201.76 | 208.29 |
| 30842 | ENSMUSG00000089992 | 0 | NA | NA | NA | NA | NA | G6pd2 | glucose-6-phosphate dehydrogenase 2 [S | 0 | 0 | 0 | 0 | 0 | 0 |
| 30843 | ENSMUSG00000089993 | 0 | NA | NA | NA | NA | NA | Gm5822 | predicted gene 5822 [Source:MGI Symbol | 0 | 0 | 0 | 0 | 0 | 0 |
| 30844 | ENSMUSG00000089994 | 0.59968336 | -1.233555632 | 2.892386901 | -0.426483618 | 0.669755482 | NA | Gm16239 | predicted gene 16239 [Source:MGI Symbo | 1.6062 | 1.0279 | 0 | 0.964 | 0 | 0 |
| 30845 | ENSMUSG00000089995 | 0.160662974 | 1.126924175 | 4.080472857 | 0.276174898 | 0.782413736 | NA | Gm15716 | predicted gene 15716 [Source:MGI Symbo | 0 | 0 | 0 | 0.964 | 0 | 0 |
| 30846 | ENSMUSG00000089996 | 0 | NA | NA | NA | NA | NA | Tmsb15b2 | thymosin beta 15b2 [Source:MGI Symbol; | 0 | 0 | 0 | 0 | 0 | 0 |
| 30847 | ENSMUSG00000089997 | 0 | NA | NA | NA | NA | NA | 1810020O05 | Riken cDNA 1810020O05 gene [Source:M | 0 | 0 | 0 | 0 | 0 | 0 |
| 30848 | ENSMUSG00000089998 | 3.974309241 | 0.403355537 | 1.152053977 | 0.35011861 | 0.726249685 | NA | Phtf1os | putative homeodomain transcription fact | 3.2124 | 1.0279 | 6.0375 | 4.8199 | 7.6343 | 1.1138 |
| 30849 | ENSMUSG00000089999 | 1.150904955 | -0.332137203 | 2.117813663 | -0.156830229 | 0.875378648 | NA | Gm6485 | predicted gene 6485 [Source:MGI Symbol | 0.8031 | 3.0838 | 0 | 1.928 | 1.0906 | 0 |
| 30850 | ENSMUSG00000090000 | 159.6724922 | -0.088899417 | 0.201615754 | -0.440934874 | 0.659260146 | 0.927763076 | Ier3ip1 | immediate early response 3 interacting pr | 179.89 | 153.16 | 159.99 | 133.99 | 148.32 | 182.67 |
| 30851 | ENSMUSG00000090001 | 0 | NA | NA | NA | NA | NA | Gm9719 | predicted gene 9719 [Source:MGI Symbol | 0 | 0 | 0 | 0 | 0 | 0 |
| 30852 | ENSMUSG00000090002 | 10.9036661 | -0.363672033 | 0.745521169 | -0.48780913 | 0.625685044 | 0.919638611 | Gm16006 | predicted gene 16006 [Source:MGI Symbo | 19.274 | 11.307 | 6.0375 | 5.7839 | 17.45 | 5.5692 |

| | | | | | | | | | | | | | | | |
|---|---|---|---|---|---|---|---|---|---|---|---|---|---|---|---|
| 30853 | ENSMUSG00000090003 | 0 | NA | NA | NA | NA | NA | Gm15882 | predicted gene 15882 [Source:MGI Symbo | 0 | 0 | 0 | 0 | 0 | 0 |
| 30854 | ENSMUSG00000090004 | 0.48903338 | 0.95637552 | 3.456690196 | 0.276673773 | 0.782030612 | NA | Gm16125 | predicted gene 16125 [Source:MGI Symbo | 0 | 0 | 1.0062 | 1.928 | 0 | 0 |
| 30855 | ENSMUSG00000090005 | 0.952772715 | -0.827065859 | 2.269489396 | -0.364428166 | 0.715538306 | NA | Gm16540 | predicted gene 16540 [Source:MGI Symbo | 1.6062 | 2.0558 | 0 | 0.964 | 1.0906 | 0 |
| 30856 | ENSMUSG00000090006 | 0 | NA | NA | NA | NA | NA | Gm16227 | predicted gene 16227 [Source:MGI Symbo | 0 | 0 | 0 | 0 | 0 | 0 |
| 30857 | ENSMUSG00000090007 | 0.267700746 | -1.497481649 | 4.057001436 | -0.369110456 | 0.712045396 | NA | Rpl30-ps2 | ribosomal protein L30, pseudogene 2 [Sou | 1.6062 | 0 | 0 | 0 | 0 | 0 |
| 30858 | ENSMUSG00000090008 | 0.48903338 | 0.95637552 | 3.456690196 | 0.276673773 | 0.782030612 | NA | Gm16111 | predicted gene 16111 [Source:MGI Symbo | 0 | 0 | 1.0062 | 1.928 | 0 | 0 |
| 30859 | ENSMUSG00000090009 | 0 | NA | NA | NA | NA | NA | Gm16282 | predicted gene 16282 [Source:MGI Symbo | 0 | 0 | 0 | 0 | 0 | 0 |
| 30860 | ENSMUSG00000090012 | 0 | NA | NA | NA | NA | NA | Gm16555 | predicted gene 16555 [Source:MGI Symbo | 0 | 0 | 0 | 0 | 0 | 0 |
| 30861 | ENSMUSG00000090013 | 0 | NA | NA | NA | NA | NA | Gm6507 | predicted gene 6507 [Source:MGI Symbol | 0 | 0 | 0 | 0 | 0 | 0 |
| 30862 | ENSMUSG00000090015 | 68.61417416 | -0.061404921 | 0.28139117 | -0.218219076 | 0.827258422 | 0.969794868 | Gm15446 | predicted gene 15446 [Source:MGI Symbo | 64.248 | 61.675 | 84.525 | 73.262 | 57.803 | 70.172 |
| 30863 | ENSMUSG00000090016 | 0 | NA | NA | NA | NA | NA | Gm16578 | predicted gene 16578 [Source:MGI Symbo | 0 | 0 | 0 | 0 | 0 | 0 |
| 30864 | ENSMUSG00000090017 | 0 | NA | NA | NA | NA | NA | Gm16213 | predicted gene 16213 [Source:MGI Symbo | 0 | 0 | 0 | 0 | 0 | 0 |
| 30865 | ENSMUSG00000090019 | 45.1988876 | -0.241767016 | 0.355500989 | -0.680074102 | 0.496457542 | 0.878165817 | Gimap1 | GTPase, IMAP family member 1 [Source:M | 38.549 | 54.48 | 54.337 | 52.055 | 29.447 | 42.326 |
| 30866 | ENSMUSG00000090021 | 7.62971419 | 1.335472097 | 0.795887865 | 1.677965146 | 0.093353903 | 0.450090286 | Gm6493 | predicted gene 6493 [Source:MGI Symbol | 4.8186 | 3.0838 | 5.0312 | 9.6398 | 8.7249 | 14.48 |
| 30867 | ENSMUSG00000090023 | 0.185640647 | 1.126924175 | 4.080472857 | 0.276174898 | 0.782413736 | NA | Gm16302 | predicted gene 16302 [Source:MGI Symbo | 0 | 0 | 0 | 0 | 0 | 1.1138 |
| 30868 | ENSMUSG00000090024 | 0 | NA | NA | NA | NA | NA | Gm16350 | predicted gene 16350 [Source:MGI Symbo | 0 | 0 | 0 | 0 | 0 | 0 |
| 30869 | ENSMUSG00000090026 | 0.971022302 | 2.27143097 | 2.737623092 | 0.829709165 | 0.406703238 | NA | Gm15996 | predicted gene 15996 [Source:MGI Symbo | 0 | 0 | 1.0062 | 4.8199 | 0 | 0 |
| 30870 | ENSMUSG00000090027 | 0 | NA | NA | NA | NA | NA | Gm15740 | predicted gene 15740 [Source:MGI Symbo | 0 | 0 | 0 | 0 | 0 | 0 |
| 30871 | ENSMUSG00000090030 | 0 | NA | NA | NA | NA | NA | A430072P03 | RIKEN cDNA A430072P03 gene [Source:M | 0 | 0 | 0 | 0 | 0 | 0 |
| 30872 | ENSMUSG00000090031 | 34.39848734 | 0.684224705 | 0.458964075 | 1.490802312 | 0.136013405 | 0.538690549 | 4732440D04 | RIKEN cDNA 4732440D04 gene [Source:M | 30.518 | 19.53 | 29.181 | 67.478 | 21.812 | 37.871 |
| 30873 | ENSMUSG00000090033 | 0 | NA | NA | NA | NA | NA | Gm16243 | predicted gene 16243 [Source:MGI Symbo | 0 | 0 | 0 | 0 | 0 | 0 |
| 30874 | ENSMUSG00000090034 | 5.465848188 | -0.018932 | 1.001468009 | -0.018904248 | 0.984917491 | 0.99859176 | Gm15917 | predicted gene 15917 [Source:MGI Symbo | 2.4093 | 9.2513 | 5.0312 | 10.604 | 3.2719 | 2.2277 |
| 30875 | ENSMUSG00000090035 | 378.2536218 | 0.255922542 | 0.151875245 | 1.685083975 | 0.091972371 | 0.446623824 | Galnt4 | polypeptide N-acetylgalactosaminyltransf | 350.15 | 304.26 | 379.35 | 366.31 | 440.61 | 428.83 |
| 30876 | ENSMUSG00000090036 | 0 | NA | NA | NA | NA | NA | Gm15900 | predicted gene 15900 [Source:MGI Symbo | 0 | 0 | 0 | 0 | 0 | 0 |
| 30877 | ENSMUSG00000090038 | 365.2169725 | -0.866617185 | 0.176098604 | -4.921204171 | 8.60E-07 | 7.28E-05 | Gm16573 | predicted gene 16573 [Source:MGI Symbo | 389.5 | 501.62 | 524.25 | 225.57 | 308.64 | 241.7 |
| 30878 | ENSMUSG00000090039 | 0.133850373 | -0.796629176 | 4.080472857 | -0.195229623 | 0.84521319 | NA | Gm15894 | predicted gene 15894 [Source:MGI Symbo | 0.8031 | 0 | 0 | 0 | 0 | 0 |
| 30879 | ENSMUSG00000090040 | 0 | NA | NA | NA | NA | NA | Gm16232 | predicted gene 16232 [Source:MGI Symbo | 0 | 0 | 0 | 0 | 0 | 0 |
| 30880 | ENSMUSG00000090041 | 0 | NA | NA | NA | NA | NA | Gm16027 | predicted gene 16027 [Source:MGI Symbo | 0 | 0 | 0 | 0 | 0 | 0 |
| 30881 | ENSMUSG00000090042 | 0 | NA | NA | NA | NA | NA | Gm8550 | predicted gene 8550 [Source:MGI Symbol | 0 | 0 | 0 | 0 | 0 | 0 |
| 30882 | ENSMUSG00000090043 | 0.503122293 | -2.366675966 | 3.785596405 | -0.625179156 | 0.531853481 | NA | Gm16167 | predicted gene 16167 [Source:MGI Symbo | 0 | 0 | 3.0187 | 0 | 0 | 0 |
| 30883 | ENSMUSG00000090044 | 0 | NA | NA | NA | NA | NA | Gm16557 | predicted gene 16557 [Source:MGI Symbo | 0 | 0 | 0 | 0 | 0 | 0 |
| 30884 | ENSMUSG00000090047 | 0 | NA | NA | NA | NA | NA | Gm16297 | predicted gene 16297 [Source:MGI Symbo | 0 | 0 | 0 | 0 | 0 | 0 |
| 30885 | ENSMUSG00000090048 | 0 | NA | NA | NA | NA | NA | Runx2os3 | runt related transcription factor 2, opposi | 0 | 0 | 0 | 0 | 0 | 0 |
| 30886 | ENSMUSG00000090050 | 0 | NA | NA | NA | NA | NA | Gm16339 | predicted gene 16339 [Source:MGI Symbo | 0 | 0 | 0 | 0 | 0 | 0 |
| 30887 | ENSMUSG00000090051 | 0 | NA | NA | NA | NA | NA | Gm8194 | predicted gene 8194 [Source:MGI Symbol | 0 | 0 | 0 | 0 | 0 | 0 |
| 30888 | ENSMUSG00000090053 | 0 | NA | NA | NA | NA | NA | Pakap | paralemmin A kinase anchor protein [Sou | 0 | 0 | 0 | 0 | 0 | 0 |
| 30889 | ENSMUSG00000090054 | 0 | NA | NA | NA | NA | NA | Gm15629 | predicted gene 15629 [Source:MGI Symbo | 0 | 0 | 0 | 0 | 0 | 0 |
| 30890 | ENSMUSG00000090057 | 0 | NA | NA | NA | NA | NA | Gm16333 | predicted gene 16333 [Source:MGI Symbo | 0 | 0 | 0 | 0 | 0 | 0 |
| 30891 | ENSMUSG00000090059 | 0 | NA | NA | NA | NA | NA | Olfr1062 | olfactory receptor 1062 [Source:MGI Syml | 0 | 0 | 0 | 0 | 0 | 0 |
| 30892 | ENSMUSG00000090061 | 8.305371878 | -0.06188149 | 0.755318508 | -0.081927676 | 0.934704226 | 0.988420305 | Nwd2 | NACHT and WD repeat domain containing | 7.2279 | 5.1396 | 13.081 | 6.7478 | 8.7249 | 8.9108 |
| 30893 | ENSMUSG00000090062 | 0 | NA | NA | NA | NA | NA | Galnt6os | polypeptide N-acetylgalactosaminyltransf | 0 | 0 | 0 | 0 | 0 | 0 |
| 30894 | ENSMUSG00000090063 | 0 | NA | NA | NA | NA | NA | Dlx6os1 | distal-less homeobox 6, opposite strand 1 | 0 | 0 | 0 | 0 | 0 | 0 |
| 30895 | ENSMUSG00000090066 | 3.630539082 | -1.655750725 | 1.277608159 | -1.29597695 | 0.194983426 | NA | 1110002E22 | RIKEN cDNA 1110002E22 gene [Source:M | 2.4093 | 4.1117 | 10.062 | 1.928 | 3.2719 | 0 |
| 30896 | ENSMUSG00000090067 | 0 | NA | NA | NA | NA | NA | Gm2976 | predicted gene 2976 [Source:MGI Symbol | 0 | 0 | 0 | 0 | 0 | 0 |
| 30897 | ENSMUSG00000090069 | 2.242530982 | -0.620790015 | 1.432788207 | -0.433274096 | 0.664815655 | NA | E430024P14 | RIKEN cDNA E430024P14 gene [Source:M | 4.0155 | 3.0838 | 1.0062 | 0.964 | 3.2719 | 1.1138 |
| 30898 | ENSMUSG00000090070 | 2.112810265 | -0.455605729 | 1.437934158 | -0.316847421 | 0.75135938 | NA | Gm16577 | predicted gene 16577 [Source:MGI Symbo | 3.2124 | 2.0558 | 2.0125 | 0.964 | 1.0906 | 3.3415 |
| 30899 | ENSMUSG00000090071 | 0 | NA | NA | NA | NA | NA | Cdk5r2 | cyclin-dependent kinase 5, regulatory sub | 0 | 0 | 0 | 0 | 0 | 0 |
| 30900 | ENSMUSG00000090073 | 0 | NA | NA | NA | NA | NA | Gm13538 | predicted gene 13538 [Source:MGI Symbo | 0 | 0 | 0 | 0 | 0 | 0 |
| 30901 | ENSMUSG00000090074 | 0 | NA | NA | NA | NA | NA | Gm6921 | predicted pseudogene 6921 [Source:MGI | 0 | 0 | 0 | 0 | 0 | 0 |
| 30902 | ENSMUSG00000090075 | 0.160662974 | 1.126924175 | 4.080472857 | 0.276174898 | 0.782413736 | NA | Gm16251 | predicted gene 16251 [Source:MGI Symbo | 0 | 0 | 0 | 0.964 | 0 | 0 |
| 30903 | ENSMUSG00000090077 | 0.339027071 | -1.79174149 | 3.919585585 | -0.457125237 | 0.647581032 | NA | Lime1 | Lck interacting transmembrane adaptor 1 | 0 | 1.0279 | 1.0062 | 0 | 0 | 0 |
| 30904 | ENSMUSG00000090078 | 0 | NA | NA | NA | NA | NA | Gm16147 | predicted gene 16147 [Source:MGI Symbo | 0 | 0 | 0 | 0 | 0 | 0 |
| 30905 | ENSMUSG00000090079 | 0 | NA | NA | NA | NA | NA | Gm15849 | predicted gene 15849 [Source:MGI Symbo | 0 | 0 | 0 | 0 | 0 | 0 |
| 30906 | ENSMUSG00000090080 | 0 | NA | NA | NA | NA | NA | Gm15872 | predicted gene 15872 [Source:MGI Symbo | 0 | 0 | 0 | 0 | 0 | 0 |
| 30907 | ENSMUSG00000090081 | 6.469032491 | 0.462900621 | 0.884696646 | 0.523230899 | 0.600813581 | 0.915429015 | Gm16587 | predicted gene 16587 [Source:MGI Symbo | 3.2124 | 3.0838 | 10.062 | 4.8199 | 8.7249 | 8.9108 |
| 30908 | ENSMUSG00000090083 | 61.21550657 | 0.305049236 | 0.284526875 | 1.07212802 | 0.283662535 | 0.736605908 | Rnf8 | ring finger protein 8 [Source:MGI Symbol; | 59.43 | 48.312 | 56.35 | 65.55 | 74.162 | 63.489 |

| | | | | | | | | | | | | | | | |
|---|---|---|---|---|---|---|---|---|---|---|---|---|---|---|---|
| 30909 | ENSMUSG00000090084 | 3.470933437 | -0.900765361 | 1.194765138 | -0.753926719 | 0.450893226 | NA | Srpx | sushi-repeat-containing protein [Source:N | 2.4093 | 8.2233 | 3.0187 | 3.8559 | 1.0906 | 2.2277 |
| 30910 | ENSMUSG00000090086 | 158.6012201 | 0.405316474 | 0.251276257 | 1.613031325 | 0.106737702 | 0.482153069 | AI480526 | expressed sequence AI480526 [Source:M | 164.64 | 98.68 | 145.91 | 236.17 | 161.41 | 144.8 |
| 30911 | ENSMUSG00000090087 | 0 | NA | NA | NA | NA | NA | Gm15939 | predicted gene 15939 [Source:MGI Symbo | 0 | 0 | 0 | 0 | 0 | 0 |
| 30912 | ENSMUSG00000090088 | 0 | NA | NA | NA | NA | NA | Gm15934 | predicted gene 15934 [Source:MGI Symbo | 0 | 0 | 0 | 0 | 0 | 0 |
| 30913 | ENSMUSG00000090089 | 0 | NA | NA | NA | NA | NA | Magea9-ps | MAGE family member A9, pseudogene [S | 0 | 0 | 0 | 0 | 0 | 0 |
| 30914 | ENSMUSG00000090090 | 116.46079 | 0.282542552 | 0.306195312 | 0.922752706 | 0.356136092 | 0.795708016 | Mup-ps16 | major urinary protein, pseudogene 16 [So | 156.6 | 78.122 | 79.493 | 97.362 | 165.77 | 121.41 |
| 30915 | ENSMUSG00000090093 | 27.49880218 | 0.42195637 | 0.484775096 | 0.870416763 | 0.384072689 | 0.816413186 | Gm14399 | predicted gene 14399 [Source:MGI Symbo | 28.912 | 17.475 | 24.15 | 47.235 | 31.628 | 15.594 |
| 30916 | ENSMUSG00000090095 | 0.167707431 | -0.796629176 | 4.080472857 | -0.195229623 | 0.84521319 | NA | Rpl29-ps1 | ribosomal protein L29, pseudogene 1 [Sou | 0 | 0 | 1.0062 | 0 | 0 | 0 |
| 30917 | ENSMUSG00000090097 | 0 | NA | NA | NA | NA | NA | Olfr1182 | olfactory receptor 1182 [Source:MGI Sym | 0 | 0 | 0 | 0 | 0 | 0 |
| 30918 | ENSMUSG00000090098 | 0.524408994 | -0.837196744 | 3.377823722 | -0.247850928 | 0.804249749 | NA | Alms1-ps2 | ALMS1, centrosome and basal body assoc | 0 | 2.0558 | 0 | 0 | 1.0906 | 0 |
| 30919 | ENSMUSG00000090099 | 0 | NA | NA | NA | NA | NA | Gm8837 | predicted gene 8837 [Source:MGI Symbol | 0 | 0 | 0 | 0 | 0 | 0 |
| 30920 | ENSMUSG00000090100 | 69.79664871 | -0.169081409 | 0.272653316 | -0.620133331 | 0.535170009 | 0.889858671 | Ttbk2 | tau tubulin kinase 2 [Source:MGI Symbol; | 77.098 | 80.178 | 64.4 | 71.334 | 56.712 | 69.058 |
| 30921 | ENSMUSG00000090101 | 0 | NA | NA | NA | NA | NA | Snhg9 | small nucleolar RNA host gene 9 [Source:I | 0 | 0 | 0 | 0 | 0 | 0 |
| 30922 | ENSMUSG00000090102 | 0 | NA | NA | NA | NA | NA | Tesl2 | testin LIM domain protein like 2 [Source:N | 0 | 0 | 0 | 0 | 0 | 0 |
| 30923 | ENSMUSG00000090103 | 0.181769715 | 1.126924175 | 4.080472857 | 0.276174898 | 0.782413736 | NA | Gm16094 | predicted gene 16094 [Source:MGI Symbo | 0 | 0 | 0 | 0 | 1.0906 | 0 |
| 30924 | ENSMUSG00000090104 | 0.321325949 | 1.911575114 | 4.048267893 | 0.472195804 | 0.636787034 | NA | Slmapos2 | sarcolemma associated protein, opposite | 0 | 0 | 0 | 1.928 | 0 | 0 |
| 30925 | ENSMUSG00000090105 | 21.30032404 | -0.999896559 | 0.528681597 | -1.891301997 | 0.058584042 | 0.343617027 | Gm15890 | predicted gene 15890 [Source:MGI Symbo | 27.305 | 33.921 | 24.15 | 18.316 | 18.541 | 5.5692 |
| 30926 | ENSMUSG00000090106 | 0 | NA | NA | NA | NA | NA | Gm15838 | predicted gene 15838 [Source:MGI Symbo | 0 | 0 | 0 | 0 | 0 | 0 |
| 30927 | ENSMUSG00000090107 | 0 | NA | NA | NA | NA | NA | NA | NA | 0 | 0 | 0 | 0 | 0 | 0 |
| 30928 | ENSMUSG00000090109 | 0 | NA | NA | NA | NA | NA | Ear-ps10 | eosinophil-associated, ribonuclease A fam | 0 | 0 | 0 | 0 | 0 | 0 |
| 30929 | ENSMUSG00000090110 | 57.53627606 | -0.025115685 | 0.310315398 | -0.080935994 | 0.935492855 | 0.988420305 | Cmc4 | C-x(9)-C motif containing 4 [Source:MGI S | 61.839 | 65.787 | 46.287 | 52.055 | 49.078 | 70.172 |
| 30930 | ENSMUSG00000090111 | 0 | NA | NA | NA | NA | NA | Gm16584 | predicted gene 16584 [Source:MGI Symbo | 0 | 0 | 0 | 0 | 0 | 0 |
| 30931 | ENSMUSG00000090112 | 421.3810591 | -0.042665763 | 0.161059533 | -0.264906785 | 0.791081271 | 0.961019028 | Shprh | SNF2 histone linker PHD RING helicase [Sc | 450.54 | 447.14 | 385.39 | 461.75 | 450.43 | 333.04 |
| 30932 | ENSMUSG00000090113 | 0.133850373 | -0.796629176 | 4.080472857 | -0.195229623 | 0.84521319 | NA | Nhlrc4 | NHL repeat containing 4 [Source:MGI Sym | 0.8031 | 0 | 0 | 0 | 0 | 0 |
| 30933 | ENSMUSG00000090114 | 0.321325949 | 1.911575114 | 4.048267893 | 0.472195804 | 0.636787034 | NA | Gm3076 | predicted gene 3076 [Source:MGI Symbol | 0 | 0 | 0 | 1.928 | 0 | 0 |
| 30934 | ENSMUSG00000090115 | 43.66243641 | 0.442949316 | 0.35814869 | 1.236774915 | 0.216170657 | 0.66165144 | Usp49 | ubiquitin specific peptidase 49 [Source:M | 45.777 | 31.865 | 33.206 | 61.695 | 37.081 | 52.351 |
| 30935 | ENSMUSG00000090116 | 0 | NA | NA | NA | NA | NA | Gm15680 | predicted gene 15680 [Source:MGI Symbo | 0 | 0 | 0 | 0 | 0 | 0 |
| 30936 | ENSMUSG00000090117 | 0.160662974 | 1.126924175 | 4.080472857 | 0.276174898 | 0.782413736 | NA | Gm16541 | predicted gene 16541 [Source:MGI Symbo | 0 | 0 | 0 | 0.964 | 0 | 0 |
| 30937 | ENSMUSG00000090118 | 0.321325949 | 1.911575114 | 4.048267893 | 0.472195804 | 0.636787034 | NA | Gm16163 | predicted gene 16163 [Source:MGI Symbo | 0 | 0 | 0 | 1.928 | 0 | 0 |
| 30938 | ENSMUSG00000090119 | 0 | NA | NA | NA | NA | NA | Gm8185 | predicted gene 8185 [Source:MGI Symbol | 0 | 0 | 0 | 0 | 0 | 0 |
| 30939 | ENSMUSG00000090120 | 0.501260735 | 1.111687294 | 3.109072805 | 0.357562323 | 0.720670879 | NA | A730090N16 | RIKEN cDNA A730090N16 gene [Source:M | 0.8031 | 0 | 0 | 0 | 1.0906 | 1.1138 |
| 30940 | ENSMUSG00000090121 | 0 | NA | NA | NA | NA | NA | Abhd12b | abhydrolase domain containing 12B [Sour | 0 | 0 | 0 | 0 | 0 | 0 |
| 30941 | ENSMUSG00000090122 | 0 | NA | NA | NA | NA | NA | Kcne1l | potassium voltage-gated channel, Isk-rela | 0 | 0 | 0 | 0 | 0 | 0 |
| 30942 | ENSMUSG00000090124 | 7.170536926 | 0.507600123 | 0.833091843 | 0.609296715 | 0.542327784 | 0.894095573 | Ugt1a7c | UDP glucuronosyltransferase 1 family, pol | 10.44 | 3.0838 | 4.025 | 6.7478 | 9.8156 | 8.9108 |
| 30943 | ENSMUSG00000090125 | 0.342639279 | -1.805976596 | 4.028547714 | -0.448294702 | 0.653940524 | NA | Pou3f1 | POU domain, class 3, transcription factor | 0 | 2.0558 | 0 | 0 | 0 | 0 |
| 30944 | ENSMUSG00000090126 | 0 | NA | NA | NA | NA | NA | 4930519F09 | RIKEN cDNA 4930519F09 gene [Source:M | 0 | 0 | 0 | 0 | 0 | 0 |
| 30945 | ENSMUSG00000090128 | 0.185640647 | 1.126924175 | 4.080472857 | 0.276174898 | 0.782413736 | NA | Gm16038 | predicted gene 16038 [Source:MGI Symbo | 0 | 0 | 0 | 0 | 0 | 1.1138 |
| 30946 | ENSMUSG00000090129 | 0 | NA | NA | NA | NA | NA | Olfr287 | olfactory receptor 287 [Source:MGI Symb | 0 | 0 | 0 | 0 | 0 | 0 |
| 30947 | ENSMUSG00000090131 | 0 | NA | NA | NA | NA | NA | Gm16162 | predicted gene 16162 [Source:MGI Symbo | 0 | 0 | 0 | 0 | 0 | 0 |
| 30948 | ENSMUSG00000090132 | 0 | NA | NA | NA | NA | NA | Cypt2 | cysteine-rich perinuclear theca 2 [Source: | 0 | 0 | 0 | 0 | 0 | 0 |
| 30949 | ENSMUSG00000090135 | 0 | NA | NA | NA | NA | NA | Gm15809 | predicted gene 15809 [Source:MGI Symbo | 0 | 0 | 0 | 0 | 0 | 0 |
| 30950 | ENSMUSG00000090136 | 4.388273418 | 0.589904909 | 1.038431083 | 0.568073238 | 0.569985239 | NA | Gm10177 | predicted gene 10177 [Source:MGI Symbo | 2.4093 | 5.1396 | 3.0187 | 5.7839 | 2.1812 | 7.7969 |
| 30951 | ENSMUSG00000090137 | 213.0237161 | 0.052568542 | 0.169552757 | 0.310042391 | 0.75652872 | 0.953085143 | Uba52 | ubiquitin A-52 residue ribosomal protein f | 212.82 | 187.08 | 227.41 | 218.82 | 217.03 | 214.97 |
| 30952 | ENSMUSG00000090138 | 0 | NA | NA | NA | NA | NA | Gm3687 | predicted gene 3687 [Source:MGI Symbol | 0 | 0 | 0 | 0 | 0 | 0 |
| 30953 | ENSMUSG00000090141 | 0.17131964 | -0.796629176 | 4.080472857 | -0.195229623 | 0.84521319 | NA | Gm614 | predicted gene 614 [Source:MGI Symbol; | 0 | 1.0279 | 0 | 0 | 0 | 0 |
| 30954 | ENSMUSG00000090142 | 0.185640647 | 1.126924175 | 4.080472857 | 0.276174898 | 0.782413736 | NA | Gm15795 | predicted gene 15795 [Source:MGI Symbo | 0 | 0 | 0 | 0 | 0 | 1.1138 |
| 30955 | ENSMUSG00000090145 | 809.2892627 | -0.263750019 | 0.137601924 | -1.91676112 | 0.055268286 | 0.331563872 | Ugt1a6b | UDP glucuronosyltransferase 1 family, pol | 883.41 | 970.35 | 795.94 | 781.79 | 801.6 | 622.64 |
| 30956 | ENSMUSG00000090146 | 0 | NA | NA | NA | NA | NA | 4930405H06 | RIKEN cDNA 4930405H06 gene [Source:M | 0 | 0 | 0 | 0 | 0 | 0 |
| 30957 | ENSMUSG00000090147 | 0 | NA | NA | NA | NA | NA | Gm3329 | predicted gene 3329 [Source:MGI Symbol | 0 | 0 | 0 | 0 | 0 | 0 |
| 30958 | ENSMUSG00000090150 | 7222.197222 | -0.077467827 | 0.109083288 | -0.710171362 | 0.477597878 | 0.869794311 | Acad11 | acyl-Coenzyme A dehydrogenase family, r | 7098.6 | 7488.4 | 7662.6 | 7670.4 | 7358.4 | 6054.9 |
| 30959 | ENSMUSG00000090151 | 0.36353943 | 2.065989254 | 4.034137301 | 0.512126658 | 0.608562368 | NA | Gm16043 | predicted gene 16043 [Source:MGI Symbo | 0 | 0 | 0 | 0 | 2.1812 | 0 |
| 30960 | ENSMUSG00000090152 | 0 | NA | NA | NA | NA | NA | Gm16285 | predicted gene 16285 [Source:MGI Symbo | 0 | 0 | 0 | 0 | 0 | 0 |
| 30961 | ENSMUSG00000090157 | 0 | NA | NA | NA | NA | NA | Gm16534 | predicted gene 16534 [Source:MGI Symbo | 0 | 0 | 0 | 0 | 0 | 0 |
| 30962 | ENSMUSG00000090159 | 0 | NA | NA | NA | NA | NA | Gm8580 | predicted gene 8580 [Source:MGI Symbol | 0 | 0 | 0 | 0 | 0 | 0 |
| 30963 | ENSMUSG00000090160 | 0 | NA | NA | NA | NA | NA | 4930480K15 | RIKEN cDNA 4930480K15 gene [Source:M | 0 | 0 | 0 | 0 | 0 | 0 |
| 30964 | ENSMUSG00000090162 | 0 | NA | NA | NA | NA | NA | Gm6808 | predicted gene 6808 [Source:MGI Symbol | 0 | 0 | 0 | 0 | 0 | 0 |

| | | | | | | | | | | | | | | | |
|---|---|---|---|---|---|---|---|---|---|---|---|---|---|---|---|
| 30965 | ENSMUSG00000090163 | 0 | NA | NA | NA | NA | NA | Gm16231 | predicted gene 16231 [Source:MGI Symbo | 0 | 0 | 0 | 0 | 0 | 0 |
| 30966 | ENSMUSG00000090164 | 2.217966489 | -3.429344455 | 1.823384712 | -1.88075749 | 0.060004915 | NA | BC035044 | cDNA sequence BC035044 [Source:MGI Sy | 4.0155 | 7.1954 | 1.0062 | 0 | 1.0906 | 0 |
| 30967 | ENSMUSG00000090165 | 3.111931631 | -0.628998504 | 1.179669807 | -0.533198782 | 0.593895985 | NA | Ugt1a10 | UDP glycosyltransferase 1 family, polypep | 3.2124 | 5.1396 | 3.0187 | 2.8919 | 2.1812 | 2.2277 |
| 30968 | ENSMUSG00000090166 | 0 | NA | NA | NA | NA | NA | Ear10 | eosinophil-associated, ribonuclease A fam | 0 | 0 | 0 | 0 | 0 | 0 |
| 30969 | ENSMUSG00000090168 | 0 | NA | NA | NA | NA | NA | D630014O11 | RIKEN cDNA D630014O11 gene [Source:M | 0 | 0 | 0 | 0 | 0 | 0 |
| 30970 | ENSMUSG00000090171 | 2.928641705 | -0.302269895 | 1.542648237 | -0.195942204 | 0.844655404 | NA | Ugt1a2 | UDP glucuronosyltransferase 1 family, pol | 5.6217 | 4.1117 | 0 | 6.7478 | 1.0906 | 0 |
| 30971 | ENSMUSG00000090173 | 0.371281295 | 2.09253362 | 4.031852165 | 0.519000582 | 0.603760335 | NA | Fbxw10 | F-box and WD-40 domain protein 10 [Sou | 0 | 0 | 0 | 0 | 0 | 2.2277 |
| 30972 | ENSMUSG00000090174 | 0.656740811 | -0.013525372 | 3.112905782 | -0.004344934 | 0.996533255 | NA | Gm10612 | predicted gene 10612 [Source:MGI Symbo | 0 | 0 | 2.0125 | 1.928 | 0 | 0 |
| 30973 | ENSMUSG00000090175 | 147.7653881 | -0.833979996 | 0.208465585 | -4.000564396 | 6.32E-05 | 0.002510998 | Ugt1a9 | UDP glucuronosyltransferase 1 family, pol | 163.03 | 185.03 | 220.37 | 110.86 | 98.156 | 109.16 |
| 30974 | ENSMUSG00000090176 | 0.17131964 | -0.796629176 | 4.080472857 | -0.195229623 | 0.84521319 | NA | Cd200r2 | Cd200 receptor 2 [Source:MGI Symbol;Ac | 0 | 1.0279 | 0 | 0 | 0 | 0 |
| 30975 | ENSMUSG00000090179 | 0 | NA | NA | NA | NA | NA | Gm16307 | predicted gene 16307 [Source:MGI Symbo | 0 | 0 | 0 | 0 | 0 | 0 |
| 30976 | ENSMUSG00000090180 | 0 | NA | NA | NA | NA | NA | Ang-ps1 | angiogenin, ribonuclease A family, pseudo | 0 | 0 | 0 | 0 | 0 | 0 |
| 30977 | ENSMUSG00000090185 | 2.237141493 | -0.235731746 | 1.483127655 | -0.158942317 | 0.87371432 | NA | Gm15523 | predicted gene 15523 [Source:MGI Symbo | 3.2124 | 1.0279 | 3.0187 | 2.8919 | 3.2719 | 0 |
| 30978 | ENSMUSG00000090186 | 2.667799942 | -0.623556757 | 1.424309296 | -0.437795891 | 0.66153425 | NA | Gm52951 | predicted gene, 52951 [Source:MGI Symb | 5.6217 | 0 | 4.025 | 1.928 | 1.0906 | 3.3415 |
| 30979 | ENSMUSG00000090188 | 0 | NA | NA | NA | NA | NA | Gm7418 | predicted pseudogene 7418 [Source:MGI | 0 | 0 | 0 | 0 | 0 | 0 |
| 30980 | ENSMUSG00000090191 | 0 | NA | NA | NA | NA | NA | 9230105E05 | RIKEN cDNA 9230105E05 gene [Source:M | 0 | 0 | 0 | 0 | 0 | 0 |
| 30981 | ENSMUSG00000090192 | 0 | NA | NA | NA | NA | NA | Gm16556 | predicted gene 16556 [Source:MGI Symbo | 0 | 0 | 0 | 0 | 0 | 0 |
| 30982 | ENSMUSG00000090193 | 0 | NA | NA | NA | NA | NA | Gm15995 | predicted gene 15995 [Source:MGI Symbo | 0 | 0 | 0 | 0 | 0 | 0 |
| 30983 | ENSMUSG00000090194 | 3.274248447 | 2.196286303 | 1.365665718 | 1.608216619 | 0.107787745 | NA | Gm16161 | predicted gene 16161 [Source:MGI Symbo | 2.4093 | 0 | 1.0062 | 9.6398 | 4.3625 | 2.2277 |
| 30984 | ENSMUSG00000090197 | 0 | NA | NA | NA | NA | NA | Dnaja1-ps | DnaJ heat shock protein family (Hsp40) m | 0 | 0 | 0 | 0 | 0 | 0 |
| 30985 | ENSMUSG00000090199 | 0 | NA | NA | NA | NA | NA | Gm16029 | predicted gene 16029 [Source:MGI Symbo | 0 | 0 | 0 | 0 | 0 | 0 |
| 30986 | ENSMUSG00000090200 | 0 | NA | NA | NA | NA | NA | 1700025N21 | RIKEN cDNA 1700025N21 gene [Source:M | 0 | 0 | 0 | 0 | 0 | 0 |
| 30987 | ENSMUSG00000090202 | 0 | NA | NA | NA | NA | NA | 4930503B20 | RIKEN cDNA 4930503B20 gene [Source:M | 0 | 0 | 0 | 0 | 0 | 0 |
| 30988 | ENSMUSG00000090203 | 0 | NA | NA | NA | NA | NA | AU015336 | expressed sequence AU015336 [Source:M | 0 | 0 | 0 | 0 | 0 | 0 |
| 30989 | ENSMUSG00000090206 | 0.133850373 | -0.796629176 | 4.080472857 | -0.195229623 | 0.84521319 | NA | Tepp | testis, prostate and placenta expressed [S | 0.8031 | 0 | 0 | 0 | 0 | 0 |
| 30990 | ENSMUSG00000090207 | 12.64292607 | 0.080793104 | 0.644995883 | 0.125261425 | 0.90031659 | 0.982805974 | 4930524O07 | RIKEN cDNA 4930524O07 gene [Source:M | 19.274 | 11.307 | 6.0375 | 10.604 | 15.269 | 13.366 |
| 30991 | ENSMUSG00000090208 | 0 | NA | NA | NA | NA | NA | Gm15851 | predicted gene 15851 [Source:MGI Symbo | 0 | 0 | 0 | 0 | 0 | 0 |
| 30992 | ENSMUSG00000090210 | 25.40189441 | 0.283818529 | 0.453204548 | 0.626248192 | 0.531152162 | 0.888877582 | Itga10 | integrin, alpha 10 [Source:MGI Symbol;Ac | 16.062 | 25.698 | 27.169 | 26.027 | 21.812 | 35.643 |
| 30993 | ENSMUSG00000090211 | 0 | NA | NA | NA | NA | NA | Gm16050 | predicted gene 16050 [Source:MGI Symbo | 0 | 0 | 0 | 0 | 0 | 0 |
| 30994 | ENSMUSG00000090212 | 0 | NA | NA | NA | NA | NA | Gm15647 | predicted gene 15647 [Source:MGI Symbo | 0 | 0 | 0 | 0 | 0 | 0 |
| 30995 | ENSMUSG00000090213 | 458.1372833 | -0.45631045 | 0.184415031 | -2.474366908 | 0.013347254 | 0.131958089 | Tmem189 | transmembrane protein 189 [Source:MGI | 693.88 | 454.34 | 440.74 | 408.73 | 346.82 | 404.33 |
| 30996 | ENSMUSG00000090214 | 0 | NA | NA | NA | NA | NA | Gm15657 | predicted gene 15657 [Source:MGI Symbo | 0 | 0 | 0 | 0 | 0 | 0 |
| 30997 | ENSMUSG00000090215 | 1.031479329 | -2.220845979 | 2.403244684 | -0.924103148 | 0.355432614 | NA | Trim34b | tripartite motif-containing 34B [Source:M | 3.2124 | 0 | 2.0125 | 0.964 | 0 | 0 |
| 30998 | ENSMUSG00000090216 | 0 | NA | NA | NA | NA | NA | Gm16549 | predicted gene 16549 [Source:MGI Symbo | 0 | 0 | 0 | 0 | 0 | 0 |
| 30999 | ENSMUSG00000090217 | 9.62645374 | 0.502146794 | 0.73373758 | 0.684368373 | 0.493742582 | 0.877333256 | Gm16116 | predicted gene 16116 [Source:MGI Symbo | 9.6372 | 3.0838 | 11.069 | 8.6758 | 15.269 | 10.025 |
| 31000 | ENSMUSG00000090219 | 0 | NA | NA | NA | NA | NA | 4930516K23 | RIKEN cDNA 4930516K23 gene [Source:M | 0 | 0 | 0 | 0 | 0 | 0 |
| 31001 | ENSMUSG00000090220 | 1.607968339 | 2.112280823 | 2.006397327 | 1.052772945 | 0.292445068 | NA | Pitpnm2os1 | phosphatidylinositol transfer protein, mer | 0.8031 | 0 | 1.0062 | 6.7478 | 1.0906 | 0 |
| 31002 | ENSMUSG00000090222 | 25.47111576 | -0.784664944 | 0.449404553 | -1.746010223 | 0.080809176 | 0.41758625 | Ifi203-ps | interferon activated gene 203, pseudogen | 33.73 | 34.949 | 28.175 | 25.063 | 13.087 | 17.822 |
| 31003 | ENSMUSG00000090223 | 0 | NA | NA | NA | NA | NA | Pcp4 | Purkinje cell protein 4 [Source:MGI Symbo | 0 | 0 | 0 | 0 | 0 | 0 |
| 31004 | ENSMUSG00000090224 | 0 | NA | NA | NA | NA | NA | Gm15974 | predicted gene 15974 [Source:MGI Symbo | 0 | 0 | 0 | 0 | 0 | 0 |
| 31005 | ENSMUSG00000090225 | 0 | NA | NA | NA | NA | NA | Gm11559 | predicted gene 11559 [Source:MGI Symbo | 0 | 0 | 0 | 0 | 0 | 0 |
| 31006 | ENSMUSG00000090226 | 0 | NA | NA | NA | NA | NA | Gm16110 | predicted gene 16110 [Source:MGI Symbo | 0 | 0 | 0 | 0 | 0 | 0 |
| 31007 | ENSMUSG00000090227 | 0 | NA | NA | NA | NA | NA | Gm16554 | predicted gene 16554 [Source:MGI Symbo | 0 | 0 | 0 | 0 | 0 | 0 |
| 31008 | ENSMUSG00000090228 | 0 | NA | NA | NA | NA | NA | Gm16289 | predicted gene 16289 [Source:MGI Symbo | 0 | 0 | 0 | 0 | 0 | 0 |
| 31009 | ENSMUSG00000090230 | 0.883540042 | 0.243447205 | 2.772289218 | 0.087814505 | 0.930024109 | NA | Gm16315 | predicted gene 16315 [Source:MGI Symbo | 2.4093 | 0 | 0 | 2.8919 | 0 | 0 |
| 31010 | ENSMUSG00000090231 | 132.0505458 | 0.153095286 | 0.246806692 | 0.620304437 | 0.535057374 | 0.889858671 | Cfb | complement factor B [Source:MGI Symbo | 98.782 | 120.27 | 156.97 | 169.66 | 135.24 | 111.38 |
| 31011 | ENSMUSG00000090232 | 0 | NA | NA | NA | NA | NA | Gm16132 | predicted gene 16132 [Source:MGI Symbo | 0 | 0 | 0 | 0 | 0 | 0 |
| 31012 | ENSMUSG00000090235 | 0.910011297 | 2.120272794 | 2.542165232 | 0.834042086 | 0.404257268 | NA | Gm16244 | predicted gene 16244 [Source:MGI Symbo | 0 | 1.0279 | 0 | 0 | 1.0906 | 3.3415 |
| 31013 | ENSMUSG00000090236 | 1.827176303 | -0.643663442 | 1.527238673 | -0.421455699 | 0.673422354 | NA | Car15 | carbonic anhydrase 15 [Source:MGI Symb | 1.6062 | 2.0558 | 3.0187 | 0.964 | 1.0906 | 2.2277 |
| 31014 | ENSMUSG00000090237 | 0 | NA | NA | NA | NA | NA | Gm16308 | predicted gene 16308 [Source:MGI Symbo | 0 | 0 | 0 | 0 | 0 | 0 |
| 31015 | ENSMUSG00000090242 | 0 | NA | NA | NA | NA | NA | Rpl17-ps11 | ribosomal protein L17, pseudogene 11 [So | 0 | 0 | 0 | 0 | 0 | 0 |
| 31016 | ENSMUSG00000090243 | 0.133850373 | -0.796629176 | 4.080472857 | -0.195229623 | 0.84521319 | NA | Gm16103 | predicted gene 16103 [Source:MGI Symbo | 0.8031 | 0 | 0 | 0 | 0 | 0 |
| 31017 | ENSMUSG00000090244 | 0.435408177 | -2.179616019 | 3.531869654 | -0.617128103 | 0.537150234 | NA | Gm16572 | predicted gene 16572 [Source:MGI Symbo | 1.6062 | 0 | 1.0062 | 0 | 0 | 0 |
| 31018 | ENSMUSG00000090246 | 1.094158363 | 0.728962073 | 2.309869645 | 0.315585806 | 0.752316913 | NA | Gm17017 | predicted gene 17017 [Source:MGI Symbo | 2.4093 | 0 | 0 | 1.928 | 0 | 2.2277 |
| 31019 | ENSMUSG00000090247 | 10.650282 | 0.52803299 | 0.738039578 | 0.715453488 | 0.474328872 | 0.869056076 | Bloc1s1 | biogenesis of lysosomal organelles comple | 12.85 | 9.2513 | 4.025 | 21.208 | 6.5437 | 10.025 |
| 31020 | ENSMUSG00000090248 | 3.16553133 | -1.818041301 | 1.314589439 | -1.382972696 | 0.166673238 | NA | Gm14027 | predicted gene 14027 [Source:MGI Symbo | 1.6062 | 8.2233 | 5.0312 | 1.928 | 1.0906 | 1.1138 |

| | | | | | | | | | | | | | | | |
|---|---|---|---|---|---|---|---|---|---|---|---|---|---|---|---|
| 31021 | ENSMUSG00000090249 | 0 | NA | NA | NA | NA | NA | Gm3287 | predicted gene 3287 [Source:MGI Symbol | 0 | 0 | 0 | 0 | 0 | 0 |
| 31022 | ENSMUSG00000090252 | 0 | NA | NA | NA | NA | NA | Gm16028 | predicted gene 16028 [Source:MGI Symbo | 0 | 0 | 0 | 0 | 0 | 0 |
| 31023 | ENSMUSG00000090254 | 0 | NA | NA | NA | NA | NA | Gm1965 | predicted gene 1965 [Source:MGI Symbol | 0 | 0 | 0 | 0 | 0 | 0 |
| 31024 | ENSMUSG00000090255 | 0 | NA | NA | NA | NA | NA | 4921534H16 | RIKEN cDNA 4921534H16 gene [Source:M | 0 | 0 | 0 | 0 | 0 | 0 |
| 31025 | ENSMUSG00000090256 | 0.342639279 | -1.805976596 | 4.028547714 | -0.448294702 | 0.653940524 | NA | Alms1-ps1 | ALMS1, centrosome and basal body assoc | 0 | 2.0558 | 0 | 0 | 0 | 0 |
| 31026 | ENSMUSG00000090257 | 1.638362131 | -0.438886503 | 1.662862977 | -0.263934256 | 0.791830579 | NA | Gm4524 | predicted gene 4524 [Source:MGI Symbol | 1.6062 | 2.0558 | 2.0125 | 1.928 | 0 | 2.2277 |
| 31027 | ENSMUSG00000090258 | 209.5793876 | -0.02845704 | 0.196752439 | -0.144633732 | 0.885000063 | 0.981795936 | Churc1 | churchill domain containing 1 [Source:MG | 249.76 | 203.53 | 181.12 | 215.93 | 234.48 | 172.65 |
| 31028 | ENSMUSG00000090259 | 0 | NA | NA | NA | NA | NA | Gm16545 | predicted gene 16545 [Source:MGI Symbo | 0 | 0 | 0 | 0 | 0 | 0 |
| 31029 | ENSMUSG00000090260 | 0.935098222 | -2.037664976 | 2.285197225 | -0.891680138 | 0.372564403 | NA | 2810442N19 | RIKEN cDNA 2810442N19 gene [Source:M | 1.6062 | 1.0279 | 2.0125 | 0.964 | 0 | 0 |
| 31030 | ENSMUSG00000090263 | 0 | NA | NA | NA | NA | NA | D730045A05 | RIKEN cDNA D730045A05 gene [Source:M | 0 | 0 | 0 | 0 | 0 | 0 |
| 31031 | ENSMUSG00000090264 | 147.1904683 | -0.108396447 | 0.27152627 | -0.399211639 | 0.689737268 | 0.937261276 | Eif4ebp3 | eukaryotic translation initiation factor 4E I | 95.569 | 142.88 | 220.37 | 138.81 | 139.6 | 145.91 |
| 31032 | ENSMUSG00000090265 | 0 | NA | NA | NA | NA | NA | Gm9550 | predicted gene 9550 [Source:MGI Symbol | 0 | 0 | 0 | 0 | 0 | 0 |
| 31033 | ENSMUSG00000090266 | 294.2366775 | 0.099706838 | 0.153028652 | 0.651556667 | 0.51468721 | 0.885488593 | Mettl23 | methyltransferase like 23 [Source:MGI Syı | 267.43 | 268.29 | 316.97 | 314.26 | 297.74 | 300.74 |
| 31034 | ENSMUSG00000090268 | 0 | NA | NA | NA | NA | NA | Spopfm3 | speckle-type BTB/POZ protein family mem | 0 | 0 | 0 | 0 | 0 | 0 |
| 31035 | ENSMUSG00000090270 | 0 | NA | NA | NA | NA | NA | Gm17226 | predicted gene 17226 [Source:MGI Symbo | 0 | 0 | 0 | 0 | 0 | 0 |
| 31036 | ENSMUSG00000090272 | 102.1617286 | -0.516562649 | 0.233852922 | -2.208921074 | 0.027180131 | 0.21354341 | Mndal | myeloid nuclear differentiation antigen lik | 126.89 | 109.99 | 123.77 | 86.758 | 70.89 | 94.677 |
| 31037 | ENSMUSG00000090273 | 3.029717076 | 1.159576103 | 1.305970102 | 0.88790402 | 0.374592382 | NA | Prr22 | proline rich 22 [Source:MGI Symbol;Acc:M | 1.6062 | 0 | 4.025 | 4.8199 | 3.2719 | 4.4554 |
| 31038 | ENSMUSG00000090277 | 0.160662974 | 1.126924175 | 4.080472857 | 0.276174898 | 0.782413736 | NA | Gm8338 | predicted gene 8338 [Source:MGI Symbol | 0 | 0 | 0 | 0.964 | 0 | 0 |
| 31039 | ENSMUSG00000090284 | 0 | NA | NA | NA | NA | NA | Gm17613 | predicted gene, 17613 [Source:MGI Symb | 0 | 0 | 0 | 0 | 0 | 0 |
| 31040 | ENSMUSG00000090290 | 78.47440023 | 0.227684718 | 0.2872889 | 0.792528767 | 0.428052425 | 0.842753542 | Tarbp1 | TAR RNA binding protein 1 [Source:MGI S | 59.43 | 76.066 | 81.506 | 64.587 | 109.06 | 80.197 |
| 31041 | ENSMUSG00000090291 | 0.185640647 | 1.126924175 | 4.080472857 | 0.276174898 | 0.782413736 | NA | Lrrc10b | leucine rich repeat containing 10B [Source | 0 | 0 | 0 | 0 | 0 | 1.1138 |
| 31042 | ENSMUSG00000090292 | 0 | NA | NA | NA | NA | NA | Gm17152 | predicted gene 17152 [Source:MGI Symbo | 0 | 0 | 0 | 0 | 0 | 0 |
| 31043 | ENSMUSG00000090293 | 0 | NA | NA | NA | NA | NA | Gm17034 | predicted gene 17034 [Source:MGI Symbo | 0 | 0 | 0 | 0 | 0 | 0 |
| 31044 | ENSMUSG00000090298 | 0 | NA | NA | NA | NA | NA | Sult3a2 | sulfotransferase family 3A, member 2 [Sou | 0 | 0 | 0 | 0 | 0 | 0 |
| 31045 | ENSMUSG00000090300 | 0 | NA | NA | NA | NA | NA | Gm17119 | predicted gene 17119 [Source:MGI Symbo | 0 | 0 | 0 | 0 | 0 | 0 |
| 31046 | ENSMUSG00000090302 | 0 | NA | NA | NA | NA | NA | Smr2l | submaxillary gland androgen regulated pr | 0 | 0 | 0 | 0 | 0 | 0 |
| 31047 | ENSMUSG00000090303 | 0 | NA | NA | NA | NA | NA | Vmn2r-ps3 | vomeronasal 2, receptor, pseudogene 3 [S | 0 | 0 | 0 | 0 | 0 | 0 |
| 31048 | ENSMUSG00000090304 | 0 | NA | NA | NA | NA | NA | Vmn2r99 | vomeronasal 2, receptor 99 [Source:MGI S | 0 | 0 | 0 | 0 | 0 | 0 |
| 31049 | ENSMUSG00000090305 | 0 | NA | NA | NA | NA | NA | Gm5459 | predicted gene 5459 [Source:MGI Symbol | 0 | 0 | 0 | 0 | 0 | 0 |
| 31050 | ENSMUSG00000090306 | 770.9015981 | 0.85011575 | 0.118689025 | 7.162547251 | 7.92E-13 | 3.47E-10 | Adh6-ps1 | alcohol dehydrogenase 6 (class V), pseudo | 519.61 | 590.02 | 541.36 | 947.59 | 1063.4 | 963.47 |
| 31051 | ENSMUSG00000090307 | 0.167707431 | -0.796629176 | 4.080472857 | -0.195229623 | 0.84521319 | NA | 1700071M16 | RIKEN cDNA 1700071M16 gene [Source:M | 0 | 0 | 1.0062 | 0 | 0 | 0 |
| 31052 | ENSMUSG00000090308 | 0 | NA | NA | NA | NA | NA | Gm5174 | predicted gene 5174 [Source:MGI Symbol | 0 | 0 | 0 | 0 | 0 | 0 |
| 31053 | ENSMUSG00000090314 | 0 | NA | NA | NA | NA | NA | 2310050C09 | RIKEN cDNA 2310050C09 gene [Source:M | 0 | 0 | 0 | 0 | 0 | 0 |
| 31054 | ENSMUSG00000090315 | 0 | NA | NA | NA | NA | NA | Vmn2r104 | vomeronasal 2, receptor 104 [Source:MGI | 0 | 0 | 0 | 0 | 0 | 0 |
| 31055 | ENSMUSG00000090317 | 0.938503841 | 0.197878443 | 2.25690035 | 0.087677085 | 0.930133333 | NA | Gm17324 | predicted gene, 17324 [Source:MGI Symb | 1.6062 | 0 | 1.0062 | 1.928 | 1.0906 | 0 |
| 31056 | ENSMUSG00000090318 | 0 | NA | NA | NA | NA | NA | Gm17223 | predicted gene 17223 [Source:MGI Symbo | 0 | 0 | 0 | 0 | 0 | 0 |
| 31057 | ENSMUSG00000090319 | 0 | NA | NA | NA | NA | NA | Gm4462 | predicted gene 4462 [Source:MGI Symbol | 0 | 0 | 0 | 0 | 0 | 0 |
| 31058 | ENSMUSG00000090322 | 1.34359006 | -0.611265528 | 1.777582845 | -0.343874565 | 0.730940621 | NA | Gm17090 | predicted gene 17090 [Source:MGI Symbo | 0.8031 | 3.0838 | 1.0062 | 0.964 | 1.0906 | 1.1138 |
| 31059 | ENSMUSG00000090323 | 0 | NA | NA | NA | NA | NA | Gm5263 | predicted gene 5263 [Source:MGI Symbol | 0 | 0 | 0 | 0 | 0 | 0 |
| 31060 | ENSMUSG00000090326 | 0 | NA | NA | NA | NA | NA | Dthd1 | death domain containing 1 [Source:MGI S | 0 | 0 | 0 | 0 | 0 | 0 |
| 31061 | ENSMUSG00000090327 | 3.276601949 | -0.223089563 | 1.389920267 | -0.160505295 | 0.872483051 | NA | Gm17111 | predicted gene 17111 [Source:MGI Symbo | 1.6062 | 1.0279 | 8.05 | 6.7478 | 0 | 2.2277 |
| 31062 | ENSMUSG00000090328 | 0 | NA | NA | NA | NA | NA | Gm17073 | predicted gene 17073 [Source:MGI Symbo | 0 | 0 | 0 | 0 | 0 | 0 |
| 31063 | ENSMUSG00000090329 | 0 | NA | NA | NA | NA | NA | Gm17160 | predicted gene 17160 [Source:MGI Symbo | 0 | 0 | 0 | 0 | 0 | 0 |
| 31064 | ENSMUSG00000090330 | 21.97267179 | 0.77877305 | 0.500248685 | 1.556771807 | 0.119524673 | 0.505914517 | 9130221H12 | RIKEN cDNA 9130221H12 gene [Source:M | 12.047 | 17.475 | 19.119 | 18.316 | 39.262 | 25.618 |
| 31065 | ENSMUSG00000090331 | 0 | NA | NA | NA | NA | NA | Gm17199 | predicted gene 17199 [Source:MGI Symbo | 0 | 0 | 0 | 0 | 0 | 0 |
| 31066 | ENSMUSG00000090332 | 0 | NA | NA | NA | NA | NA | Gm3216 | predicted gene 3216 [Source:MGI Symbol | 0 | 0 | 0 | 0 | 0 | 0 |
| 31067 | ENSMUSG00000090333 | 0 | NA | NA | NA | NA | NA | Gm3025 | predicted gene 3025 [Source:MGI Symbol | 0 | 0 | 0 | 0 | 0 | 0 |
| 31068 | ENSMUSG00000090334 | 0.874428563 | -3.186291795 | 2.489702555 | -1.279788137 | 0.200619657 | NA | Gm17149 | predicted gene 17149 [Source:MGI Symbo | 3.2124 | 1.0279 | 1.0062 | 0 | 0 | 0 |
| 31069 | ENSMUSG00000090336 | 0 | NA | NA | NA | NA | NA | Cfap97d2 | CFAP97 domain containing 2 [Source:MGI | 0 | 0 | 0 | 0 | 0 | 0 |
| 31070 | ENSMUSG00000090338 | 0 | NA | NA | NA | NA | NA | Gm17081 | predicted gene 17081 [Source:MGI Symbo | 0 | 0 | 0 | 0 | 0 | 0 |
| 31071 | ENSMUSG00000090341 | 0 | NA | NA | NA | NA | NA | Gm17217 | predicted gene 17217 [Source:MGI Symbo | 0 | 0 | 0 | 0 | 0 | 0 |
| 31072 | ENSMUSG00000090342 | 0 | NA | NA | NA | NA | NA | NA | NA | 0 | 0 | 0 | 0 | 0 | 0 |
| 31073 | ENSMUSG00000090346 | 0.305170013 | -1.657836477 | 4.041480335 | -0.410205257 | 0.681655385 | NA | Vmn1r5 | vomeronasal 1 receptor 5 [Source:MGI Sy | 0.8031 | 1.0279 | 0 | 0 | 0 | 0 |
| 31074 | ENSMUSG00000090350 | 0 | NA | NA | NA | NA | NA | Gm17148 | predicted gene 17148 [Source:MGI Symbo | 0 | 0 | 0 | 0 | 0 | 0 |
| 31075 | ENSMUSG00000090351 | 0 | NA | NA | NA | NA | NA | Gm17204 | predicted gene 17204 [Source:MGI Symbo | 0 | 0 | 0 | 0 | 0 | 0 |
| 31076 | ENSMUSG00000090352 | 0 | NA | NA | NA | NA | NA | Gm3972 | predicted gene 3972 [Source:MGI Symbol | 0 | 0 | 0 | 0 | 0 | 0 |

| | | | | | | | | | | | | | | | |
|---|---|---|---|---|---|---|---|---|---|---|---|---|---|---|---|
| 31077 | ENSMUSG00000090353 | 0 | NA | NA | NA | NA | NA | Gm17555 | predicted gene, 17555 [Source:MGI Symb | 0 | 0 | 0 | 0 | 0 | 0 |
| 31078 | ENSMUSG00000090355 | 0 | NA | NA | NA | NA | NA | Vmn2r-ps12 | vomeronasal 2, receptor, pseudogene 12 | 0 | 0 | 0 | 0 | 0 | 0 |
| 31079 | ENSMUSG00000090356 | 0 | NA | NA | NA | NA | NA | Teddm3 | transmembrane epididymal family memb | 0 | 0 | 0 | 0 | 0 | 0 |
| 31080 | ENSMUSG00000090357 | 0 | NA | NA | NA | NA | NA | Gm17077 | predicted gene 17077 [Source:MGI Symbo | 0 | 0 | 0 | 0 | 0 | 0 |
| 31081 | ENSMUSG00000090358 | 0 | NA | NA | NA | NA | NA | Gm2822 | predicted gene 2822 [Source:MGI Symbol | 0 | 0 | 0 | 0 | 0 | 0 |
| 31082 | ENSMUSG00000090362 | 0 | NA | NA | NA | NA | NA | Vmn2r79 | vomeronasal 2, receptor 79 [Source:MGI S | 0 | 0 | 0 | 0 | 0 | 0 |
| 31083 | ENSMUSG00000090363 | 0.160662974 | 1.126924175 | 4.080472857 | 0.276174898 | 0.782413736 | NA | Gm3512 | predicted gene 3512 [Source:MGI Symbol | 0 | 0 | 0 | 0.964 | 0 | 0 |
| 31084 | ENSMUSG00000090364 | 0 | NA | NA | NA | NA | NA | Gm3043 | predicted gene 3043 [Source:MGI Symbol | 0 | 0 | 0 | 0 | 0 | 0 |
| 31085 | ENSMUSG00000090365 | 0 | NA | NA | NA | NA | NA | Gm17070 | predicted gene 17070 [Source:MGI Symbo | 0 | 0 | 0 | 0 | 0 | 0 |
| 31086 | ENSMUSG00000090369 | 67.17271287 | 0.345528724 | 0.300113545 | 1.151326657 | 0.249597874 | 0.701824194 | 4933411K16 | RIKEN cDNA 4933411K16 gene [Source:M | 61.839 | 41.117 | 74.462 | 78.082 | 80.706 | 66.831 |
| 31087 | ENSMUSG00000090370 | 0 | NA | NA | NA | NA | NA | Gm21989 | predicted gene 21989 [Source:MGI Symbo | 0 | 0 | 0 | 0 | 0 | 0 |
| 31088 | ENSMUSG00000090374 | 0 | NA | NA | NA | NA | NA | Gm17232 | predicted gene 17232 [Source:MGI Symbo | 0 | 0 | 0 | 0 | 0 | 0 |
| 31089 | ENSMUSG00000090379 | 0 | NA | NA | NA | NA | NA | Gm8229 | predicted gene 8229 [Source:MGI Symbol | 0 | 0 | 0 | 0 | 0 | 0 |
| 31090 | ENSMUSG00000090381 | 1.816260913 | -0.110580105 | 1.499801138 | -0.073729845 | 0.941225351 | NA | Gm6158 | predicted gene 6158 [Source:MGI Symbol | 1.6062 | 2.0558 | 2.0125 | 1.928 | 2.1812 | 1.1138 |
| 31091 | ENSMUSG00000090383 | 0 | NA | NA | NA | NA | NA | Vmn2r58 | vomeronasal 2, receptor 58 [Source:MGI S | 0 | 0 | 0 | 0 | 0 | 0 |
| 31092 | ENSMUSG00000090385 | 0 | NA | NA | NA | NA | NA | Gm7316 | predicted gene 7316 [Source:MGI Symbol | 0 | 0 | 0 | 0 | 0 | 0 |
| 31093 | ENSMUSG00000090386 | 26.93488604 | 0.07062452 | 0.42715207 | 0.16533812 | 0.868677856 | 0.97847181 | Mir99ahg | Mir99a and Mirlet7c-1 host gene (non-pro | 20.881 | 32.893 | 25.156 | 23.135 | 28.356 | 31.188 |
| 31094 | ENSMUSG00000090387 | 0.185640647 | 1.126924175 | 4.080472857 | 0.276174898 | 0.782413736 | NA | Gm17056 | predicted gene 17056 [Source:MGI Symbo | 0 | 0 | 0 | 0 | 0 | 1.1138 |
| 31095 | ENSMUSG00000090389 | 1.361368291 | -2.656489506 | 2.054860461 | -1.292783406 | 0.196085981 | NA | Cdv3-ps | Cdv3 retrotransposed pseudogene [Sourc | 4.0155 | 2.0558 | 1.0062 | 0 | 1.0906 | 0 |
| 31096 | ENSMUSG00000090394 | 198.912711 | 0.441744408 | 0.193491391 | 2.283018412 | 0.022429284 | 0.187573567 | 4930523C07 | RIKEN cDNA 4930523C07 gene [Source:M | 179.09 | 132.6 | 194.21 | 246.78 | 222.49 | 218.31 |
| 31097 | ENSMUSG00000090395 | 0 | NA | NA | NA | NA | NA | | | 0 | 0 | 0 | 0 | 0 | 0 |
| 31098 | ENSMUSG00000090396 | 0 | NA | NA | NA | NA | NA | Gm6536 | predicted gene 6536 [Source:MGI Symbol | 0 | 0 | 0 | 0 | 0 | 0 |
| 31099 | ENSMUSG00000090397 | 4.97704447 | -0.027020142 | 1.172107448 | -0.023052615 | 0.981608303 | NA | Gm17096 | predicted gene 17096 [Source:MGI Symbo | 0 | 4.1117 | 11.069 | 4.8199 | 7.6343 | 2.2277 |
| 31100 | ENSMUSG00000090399 | 1.218954904 | -0.591544171 | 2.219035622 | -0.266577141 | 0.789794758 | NA | NA | NA | 2.4093 | 0 | 2.0125 | 2.8919 | 0 | 0 |
| 31101 | ENSMUSG00000090404 | 0 | NA | NA | NA | NA | NA | Gm8362 | predicted gene 8362 [Source:MGI Symbol | 0 | 0 | 0 | 0 | 0 | 0 |
| 31102 | ENSMUSG00000090406 | 0.133850373 | -0.796629176 | 4.080472857 | -0.195229623 | 0.84521319 | NA | Gm17058 | predicted gene 17058 [Source:MGI Symbo | 0.8031 | 0 | 0 | 0 | 0 | 0 |
| 31103 | ENSMUSG00000090408 | 0 | NA | NA | NA | NA | NA | Gm17402 | predicted gene, 17402 [Source:MGI Symb | 0 | 0 | 0 | 0 | 0 | 0 |
| 31104 | ENSMUSG00000090411 | 0 | NA | NA | NA | NA | NA | Vmn1r174 | vomeronasal 1 receptor 174 [Source:MGI | 0 | 0 | 0 | 0 | 0 | 0 |
| 31105 | ENSMUSG00000090417 | 0 | NA | NA | NA | NA | NA | Vmn2r94 | vomeronasal 2, receptor 94 [Source:MGI S | 0 | 0 | 0 | 0 | 0 | 0 |
| 31106 | ENSMUSG00000090427 | 0 | NA | NA | NA | NA | NA | Gm17225 | predicted gene 17225 [Source:MGI Symbo | 0 | 0 | 0 | 0 | 0 | 0 |
| 31107 | ENSMUSG00000090429 | 0 | NA | NA | NA | NA | NA | Gm17271 | predicted gene, 17271 [Source:MGI Symb | 0 | 0 | 0 | 0 | 0 | 0 |
| 31108 | ENSMUSG00000090436 | 0 | NA | NA | NA | NA | NA | Vmn2r75 | vomeronasal 2, receptor 75 [Source:MGI S | 0 | 0 | 0 | 0 | 0 | 0 |
| 31109 | ENSMUSG00000090439 | 0.321325949 | 1.911575114 | 4.048267893 | 0.472195804 | 0.636787034 | NA | Gm17455 | predicted gene, 17455 [Source:MGI Symb | 0 | 0 | 0 | 1.928 | 0 | 0 |
| 31110 | ENSMUSG00000090440 | 0 | NA | NA | NA | NA | NA | 1700049E17 | tandem duplication of RIKEN cDNA 17000 | 0 | 0 | 0 | 0 | 0 | 0 |
| 31111 | ENSMUSG00000090441 | 0 | NA | NA | NA | NA | NA | Gm17651 | predicted gene, 17651 [Source:MGI Symb | 0 | 0 | 0 | 0 | 0 | 0 |
| 31112 | ENSMUSG00000090448 | 0 | NA | NA | NA | NA | NA | Gm17192 | predicted gene 17192 [Source:MGI Symbo | 0 | 0 | 0 | 0 | 0 | 0 |
| 31113 | ENSMUSG00000090449 | 0 | NA | NA | NA | NA | NA | Gm17076 | predicted gene 17076 [Source:MGI Symbo | 0 | 0 | 0 | 0 | 0 | 0 |
| 31114 | ENSMUSG00000090451 | 0 | NA | NA | NA | NA | NA | Gm6133 | predicted gene 6133 [Source:MGI Symbol | 0 | 0 | 0 | 0 | 0 | 0 |
| 31115 | ENSMUSG00000090455 | 0 | NA | NA | NA | NA | NA | Gm2431 | predicted gene 2431 [Source:MGI Symbol | 0 | 0 | 0 | 0 | 0 | 0 |
| 31116 | ENSMUSG00000090457 | 0 | NA | NA | NA | NA | NA | 4930571K23 | RIKEN cDNA 4930571K23 gene [Source:M | 0 | 0 | 0 | 0 | 0 | 0 |
| 31117 | ENSMUSG00000090458 | 0 | NA | NA | NA | NA | NA | Gm17122 | predicted gene 17122 [Source:MGI Symbo | 0 | 0 | 0 | 0 | 0 | 0 |
| 31118 | ENSMUSG00000090459 | 1.307955721 | 0.081029206 | 1.872596083 | 0.043271054 | 0.965485465 | NA | Gm17210 | predicted gene 17210 [Source:MGI Symbo | 0.8031 | 2.0558 | 1.0062 | 2.8919 | 1.0906 | 0 |
| 31119 | ENSMUSG00000090460 | 0.346303622 | 2.004339653 | 3.884003423 | 0.516049919 | 0.605819544 | NA | Gm17511 | predicted gene, 17511 [Source:MGI Symb | 0 | 0 | 0 | 0.964 | 0 | 1.1138 |
| 31120 | ENSMUSG00000090461 | 0 | NA | NA | NA | NA | NA | Iltifb | interleukin 10-related T cell-derived induc | 0 | 0 | 0 | 0 | 0 | 0 |
| 31121 | ENSMUSG00000090462 | 0 | NA | NA | NA | NA | NA | 1700091H14 | RIKEN cDNA 1700091H14 gene [Source:M | 0 | 0 | 0 | 0 | 0 | 0 |
| 31122 | ENSMUSG00000090464 | 0 | NA | NA | NA | NA | NA | Gm17140 | predicted gene 17140 [Source:MGI Symbo | 0 | 0 | 0 | 0 | 0 | 0 |
| 31123 | ENSMUSG00000090465 | 0 | NA | NA | NA | NA | NA | Gm3091 | predicted gene 3091 [Source:MGI Symbol | 0 | 0 | 0 | 0 | 0 | 0 |
| 31124 | ENSMUSG00000090466 | 0.160662974 | 1.126924175 | 4.080472857 | 0.276174898 | 0.782413736 | NA | Gm17043 | predicted gene 17043 [Source:MGI Symbo | 0 | 0 | 0 | 0.964 | 0 | 0 |
| 31125 | ENSMUSG00000090467 | 0.549180077 | 2.659271622 | 3.249668912 | 0.818320787 | 0.41317404 | NA | Gm4613 | predicted gene 4613 [Source:MGI Symbol | 0 | 0 | 0 | 0 | 2.1812 | 1.1138 |
| 31126 | ENSMUSG00000090468 | 0 | NA | NA | NA | NA | NA | Gm2840 | predicted gene 2840 [Source:MGI Symbol | 0 | 0 | 0 | 0 | 0 | 0 |
| 31127 | ENSMUSG00000090470 | 0 | NA | NA | NA | NA | NA | Prr23a3 | proline rich 23A, member 3 [Source:MGI S | 0 | 0 | 0 | 0 | 0 | 0 |
| 31128 | ENSMUSG00000090471 | 0 | NA | NA | NA | NA | NA | Gm4553 | predicted gene 4553 [Source:MGI Symbol | 0 | 0 | 0 | 0 | 0 | 0 |
| 31129 | ENSMUSG00000090472 | 0 | NA | NA | NA | NA | NA | Gm3047 | predicted gene 3047 [Source:MGI Symbol | 0 | 0 | 0 | 0 | 0 | 0 |
| 31130 | ENSMUSG00000090474 | 0 | NA | NA | NA | NA | NA | Gm9372 | predicted gene 9372 [Source:MGI Symbol | 0 | 0 | 0 | 0 | 0 | 0 |
| 31131 | ENSMUSG00000090475 | 0 | NA | NA | NA | NA | NA | Gm6245 | predicted gene 6245 [Source:MGI Symbol | 0 | 0 | 0 | 0 | 0 | 0 |
| 31132 | ENSMUSG00000090476 | 0 | NA | NA | NA | NA | NA | Gm17166 | predicted gene 17166 [Source:MGI Symbo | 0 | 0 | 0 | 0 | 0 | 0 |

| | | | | | | | | | | | | | | | |
|---|---|---|---|---|---|---|---|---|---|---|---|---|---|---|---|
| 31133 | ENSMUSG00000090478 | 0 | NA | NA | NA | NA | NA | Gm5656 | predicted gene 5656 [Source:MGI Symbol | 0 | 0 | 0 | 0 | 0 | 0 |
| 31134 | ENSMUSG00000090481 | 0 | NA | NA | NA | NA | NA | Gm2966 | predicted gene 2966 [Source:MGI Symbol | 0 | 0 | 0 | 0 | 0 | 0 |
| 31135 | ENSMUSG00000090482 | 0 | NA | NA | NA | NA | NA | Gm3675 | predicted gene 3675 [Source:MGI Symbol | 0 | 0 | 0 | 0 | 0 | 0 |
| 31136 | ENSMUSG00000090485 | 0 | NA | NA | NA | NA | NA | Gm17416 | predicted gene, 17416 [Source:MGI Symb | 0 | 0 | 0 | 0 | 0 | 0 |
| 31137 | ENSMUSG00000090486 | 0.669251866 | -2.807208097 | 3.391018549 | -0.827836255 | 0.407763239 | NA | BC035947 | cDNA sequence BC035947 [Source:MGI Sy | 4.0155 | 0 | 0 | 0 | 0 | 0 |
| 31138 | ENSMUSG00000090487 | 0 | NA | NA | NA | NA | NA | Gm2888 | predicted gene 2888 [Source:MGI Symbol | 0 | 0 | 0 | 0 | 0 | 0 |
| 31139 | ENSMUSG00000090489 | 2.377265892 | -0.391217793 | 1.63663095 | -0.239038492 | 0.811075736 | NA | Gm17415 | predicted gene, 17415 [Source:MGI Symb | 0 | 6.1675 | 2.0125 | 3.8559 | 0 | 2.2277 |
| 31140 | ENSMUSG00000090494 | 0 | NA | NA | NA | NA | NA | Traj36 | T cell receptor alpha joining 36 [Source:M | 0 | 0 | 0 | 0 | 0 | 0 |
| 31141 | ENSMUSG00000090497 | 0 | NA | NA | NA | NA | NA | Gm17611 | predicted gene, 17611 [Source:MGI Symb | 0 | 0 | 0 | 0 | 0 | 0 |
| 31142 | ENSMUSG00000090500 | 1.989880795 | -0.942410032 | 1.609255703 | -0.585618575 | 0.558131863 | NA | Gm17057 | predicted gene 17057 [Source:MGI Symbo | 0.8031 | 4.1117 | 3.0187 | 2.8919 | 0 | 1.1138 |
| 31143 | ENSMUSG00000090505 | 0 | NA | NA | NA | NA | NA | Gm3486 | predicted gene 3486 [Source:MGI Symbol | 0 | 0 | 0 | 0 | 0 | 0 |
| 31144 | ENSMUSG00000090509 | 0 | NA | NA | NA | NA | NA | Sfta2 | surfactant associated 2 [Source:MGI Symb | 0 | 0 | 0 | 0 | 0 | 0 |
| 31145 | ENSMUSG00000090512 | 0 | NA | NA | NA | NA | NA | Gm3424 | predicted gene 3424 [Source:MGI Symbol | 0 | 0 | 0 | 0 | 0 | 0 |
| 31146 | ENSMUSG00000090515 | 0 | NA | NA | NA | NA | NA | Krtap27-1 | keratin associated protein 27-1 [Source:M | 0 | 0 | 0 | 0 | 0 | 0 |
| 31147 | ENSMUSG00000090516 | 0.686901383 | 1.702364833 | 2.776876575 | 0.613050233 | 0.539843125 | NA | Rps11-ps1 | ribosomal protein S11, pseudogene 1 [Sou | 0.8031 | 0 | 0 | 0 | 1.0906 | 2.2277 |
| 31148 | ENSMUSG00000090519 | 0 | NA | NA | NA | NA | NA | Gm17061 | predicted gene 17061 [Source:MGI Symbo | 0 | 0 | 0 | 0 | 0 | 0 |
| 31149 | ENSMUSG00000090523 | 185.5051841 | -0.808315388 | 0.225219989 | -3.589003755 | 0.000331944 | 0.009479365 | Gypc | glycophorin C [Source:MGI Symbol;Acc:M | 203.18 | 281.65 | 224.39 | 154.24 | 153.78 | 95.791 |
| 31150 | ENSMUSG00000090527 | 0 | NA | NA | NA | NA | NA | Aadacl2fm2 | AADACL2 family member 2 [Source:MGI S | 0 | 0 | 0 | 0 | 0 | 0 |
| 31151 | ENSMUSG00000090533 | 0 | NA | NA | NA | NA | NA | Gm8214 | predicted gene 8214 [Source:MGI Symbol | 0 | 0 | 0 | 0 | 0 | 0 |
| 31152 | ENSMUSG00000090534 | 0 | NA | NA | NA | NA | NA | Gm4675 | predicted gene 4675 [Source:MGI Symbol | 0 | 0 | 0 | 0 | 0 | 0 |
| 31153 | ENSMUSG00000090535 | 0 | NA | NA | NA | NA | NA | Gm8540 | predicted gene 8540 [Source:MGI Symbol | 0 | 0 | 0 | 0 | 0 | 0 |
| 31154 | ENSMUSG00000090538 | 0 | NA | NA | NA | NA | NA | Gm4775 | predicted gene 4775 [Source:MGI Symbol | 0 | 0 | 0 | 0 | 0 | 0 |
| 31155 | ENSMUSG00000090539 | 0 | NA | NA | NA | NA | NA | Gm9602 | predicted gene 9602 [Source:MGI Symbol | 0 | 0 | 0 | 0 | 0 | 0 |
| 31156 | ENSMUSG00000090544 | 0 | NA | NA | NA | NA | NA | Gm6576 | predicted gene 6576 [Source:MGI Symbol | 0 | 0 | 0 | 0 | 0 | 0 |
| 31157 | ENSMUSG00000090546 | 0 | NA | NA | NA | NA | NA | Cdr1 | cerebellar degeneration related antigen 1 | 0 | 0 | 0 | 0 | 0 | 0 |
| 31158 | ENSMUSG00000090547 | 0 | NA | NA | NA | NA | NA | Gm8265 | predicted gene 8265 [Source:MGI Symbol | 0 | 0 | 0 | 0 | 0 | 0 |
| 31159 | ENSMUSG00000090549 | 0 | NA | NA | NA | NA | NA | Rpl28-ps4 | ribosomal protein L28, pseudogene 4 [Sou | 0 | 0 | 0 | 0 | 0 | 0 |
| 31160 | ENSMUSG00000090550 | 0 | NA | NA | NA | NA | NA | Prlh | prolactin releasing hormone [Source:MGI | 0 | 0 | 0 | 0 | 0 | 0 |
| 31161 | ENSMUSG00000090551 | 1.699206237 | 0.817222165 | 1.853809319 | 0.440833993 | 0.659333182 | NA | NA | NA | 1.6062 | 2.0558 | 0 | 0.964 | 0 | 5.5692 |
| 31162 | ENSMUSG00000090552 | 0 | NA | NA | NA | NA | NA | Gm17158 | predicted gene 17158 [Source:MGI Symbo | 0 | 0 | 0 | 0 | 0 | 0 |
| 31163 | ENSMUSG00000090553 | 316.8735894 | 0.08372111 | 0.150176023 | 0.557486528 | 0.577195061 | 0.907838603 | Snrpe | small nuclear ribonucleoprotein E [Source | 319.63 | 299.12 | 303.89 | 311.36 | 364.27 | 302.97 |
| 31164 | ENSMUSG00000090555 | 201.1529018 | 1.013735614 | 0.209417301 | 4.840744341 | 1.29E-06 | 0.00010059 | Gm8893 | predicted gene 8893 [Source:MGI Symbol | 122.87 | 107.93 | 169.05 | 248.71 | 246.48 | 311.88 |
| 31165 | ENSMUSG00000090556 | 0 | NA | NA | NA | NA | NA | Olfr753-ps1 | olfactory receptor 753, pseudogene 1 [So | 0 | 0 | 0 | 0 | 0 | 0 |
| 31166 | ENSMUSG00000090558 | 0 | NA | NA | NA | NA | NA | Gm17202 | predicted gene 17202 [Source:MGI Symbo | 0 | 0 | 0 | 0 | 0 | 0 |
| 31167 | ENSMUSG00000090559 | 1.256159933 | -0.400276649 | 2.054019058 | -0.194874847 | 0.845490926 | NA | Gm17137 | predicted gene 17137 [Source:MGI Symbo | 3.2124 | 0 | 1.0062 | 0 | 1.0906 | 2.2277 |
| 31168 | ENSMUSG00000090561 | 0 | NA | NA | NA | NA | NA | Gm17207 | predicted gene 17207 [Source:MGI Symbo | 0 | 0 | 0 | 0 | 0 | 0 |
| 31169 | ENSMUSG00000090564 | 1.463610243 | 0.484975761 | 1.887111255 | 0.25699373 | 0.797183613 | NA | A430057M0 | RIKEN cDNA A430057M04 gene [Source:M | 1.6062 | 2.0558 | 0 | 2.8919 | 0 | 2.2277 |
| 31170 | ENSMUSG00000090565 | 0.315620088 | 0.165143421 | 4.063300985 | 0.040642675 | 0.967580763 | NA | 4930405O22 | RIKEN cDNA 4930405O22 gene [Source:M | 0.8031 | 0 | 0 | 0 | 1.0906 | 0 |
| 31171 | ENSMUSG00000090570 | 0 | NA | NA | NA | NA | NA | Gm17041 | predicted gene 17041 [Source:MGI Symbo | 0 | 0 | 0 | 0 | 0 | 0 |
| 31172 | ENSMUSG00000090572 | 0 | NA | NA | NA | NA | NA | Vmn2r109 | vomeronasal 2, receptor 109 [Source:MGI | 0 | 0 | 0 | 0 | 0 | 0 |
| 31173 | ENSMUSG00000090574 | 0 | NA | NA | NA | NA | NA | Traj61 | T cell receptor alpha joining 61 [Source:M | 0 | 0 | 0 | 0 | 0 | 0 |
| 31174 | ENSMUSG00000090576 | 15.77905994 | -0.500153579 | 0.572361146 | -0.873842648 | 0.382203951 | 0.813997203 | Gm17055 | predicted gene 17055 [Source:MGI Symbo | 20.078 | 13.363 | 22.137 | 19.28 | 10.906 | 8.9108 |
| 31175 | ENSMUSG00000090577 | 0 | NA | NA | NA | NA | NA | Gm3027 | predicted gene 3027 [Source:MGI Symbol | 0 | 0 | 0 | 0 | 0 | 0 |
| 31176 | ENSMUSG00000090578 | 0 | NA | NA | NA | NA | NA | Gm8323 | predicted gene 8323 [Source:MGI Symbol | 0 | 0 | 0 | 0 | 0 | 0 |
| 31177 | ENSMUSG00000090581 | 0 | NA | NA | NA | NA | NA | Vmn2r6 | vomeronasal 2, receptor 6 [Source:MGI Sy | 0 | 0 | 0 | 0 | 0 | 0 |
| 31178 | ENSMUSG00000090582 | 14.80356244 | -0.242641485 | 0.642684361 | -0.377543784 | 0.705769533 | 0.94295036 | Gm17024 | predicted gene 17024 [Source:MGI Symbo | 18.471 | 23.642 | 6.0375 | 15.424 | 17.45 | 7.7969 |
| 31179 | ENSMUSG00000090584 | 0 | NA | NA | NA | NA | NA | Gm17169 | predicted gene 17169 [Source:MGI Symbo | 0 | 0 | 0 | 0 | 0 | 0 |
| 31180 | ENSMUSG00000090585 | 0 | NA | NA | NA | NA | NA | 4933406F09 | RIKEN cDNA 4933406F09 gene [Source:M | 0 | 0 | 0 | 0 | 0 | 0 |
| 31181 | ENSMUSG00000090588 | 0 | NA | NA | NA | NA | NA | Muc21 | mucin 21 [Source:MGI Symbol;Acc:MGI:3 | 0 | 0 | 0 | 0 | 0 | 0 |
| 31182 | ENSMUSG00000090589 | 0 | NA | NA | NA | NA | NA | Gm17180 | predicted gene 17180 [Source:MGI Symbo | 0 | 0 | 0 | 0 | 0 | 0 |
| 31183 | ENSMUSG00000090592 | 0 | NA | NA | NA | NA | NA | Gm17571 | predicted gene, 17571 [Source:MGI Symb | 0 | 0 | 0 | 0 | 0 | 0 |
| 31184 | ENSMUSG00000090594 | 0 | NA | NA | NA | NA | NA | Gm3327 | predicted gene 3327 [Source:MGI Symbol | 0 | 0 | 0 | 0 | 0 | 0 |
| 31185 | ENSMUSG00000090598 | 0 | NA | NA | NA | NA | NA | Gm8164 | predicted gene 8164 [Source:MGI Symbol | 0 | 0 | 0 | 0 | 0 | 0 |
| 31186 | ENSMUSG00000090602 | 0 | NA | NA | NA | NA | NA | Gm5611 | predicted gene 5611 [Source:MGI Symbol | 0 | 0 | 0 | 0 | 0 | 0 |
| 31187 | ENSMUSG00000090603 | 0 | NA | NA | NA | NA | NA | Gm17186 | predicted gene 17186 [Source:MGI Symbo | 0 | 0 | 0 | 0 | 0 | 0 |
| 31188 | ENSMUSG00000090607 | 0.133850373 | -0.796629176 | 4.080472857 | -0.195229623 | 0.84521319 | NA | Gm17052 | predicted gene 17052 [Source:MGI Symbo | 0.8031 | 0 | 0 | 0 | 0 | 0 |

| | | | | | | | | | | | | | | | |
|---|---|---|---|---|---|---|---|---|---|---|---|---|---|---|---|
| 31189 | ENSMUSG00000090608 | 0.658259367 | -1.329928433 | 2.812497761 | -0.472863819 | 0.636310339 | NA | Gm17200 | predicted gene 17200 [Source:MGI Symbo | 0.8031 | 2.0558 | 0 | 0 | 1.0906 | 0 |
| 31190 | ENSMUSG00000090610 | 2.603104006 | 2.892715456 | 1.557579396 | 1.857186519 | 0.063284622 | NA | Gm3571 | predicted gene 3571 [Source:MGI Symbol | 0.8031 | 1.0279 | 0 | 3.8559 | 4.3625 | 5.5692 |
| 31191 | ENSMUSG00000090618 | 0 | NA | NA | NA | NA | NA | Gm8379 | predicted gene 8379 [Source:MGI Symbol | 0 | 0 | 0 | 0 | 0 | 0 |
| 31192 | ENSMUSG00000090619 | 0 | NA | NA | NA | NA | NA | Vmn2r60 | vomeronasal 2, receptor 60 [Source:MGI S | 0 | 0 | 0 | 0 | 0 | 0 |
| 31193 | ENSMUSG00000090622 | 121.2109611 | -0.232670167 | 0.351171296 | -0.662554629 | 0.507615837 | 0.882258527 | A930033H14 | RIKEN cDNA A930033H14 gene [Source:M | 142.15 | 149.05 | 101.63 | 102.18 | 182.13 | 50.123 |
| 31194 | ENSMUSG00000090623 | 256.2518486 | 0.128925653 | 0.161439626 | 0.798599795 | 0.424522506 | 0.840450984 | Cfhr3 | complement factor H-related 3 [Source:M | 228.88 | 261.09 | 244.52 | 242.92 | 269.38 | 290.71 |
| 31195 | ENSMUSG00000090624 | 0 | NA | NA | NA | NA | NA | Gm17060 | predicted gene 17060 [Source:MGI Symbo | 0 | 0 | 0 | 0 | 0 | 0 |
| 31196 | ENSMUSG00000090625 | 0.160662974 | 1.126924175 | 4.080472857 | 0.276174898 | 0.782413736 | NA | Gm20721 | predicted gene, 20721 [Source:MGI Symb | 0 | 0 | 0 | 0.964 | 0 | 0 |
| 31197 | ENSMUSG00000090626 | 8.784142462 | 0.181908481 | 0.81104946 | 0.224287778 | 0.822533386 | 0.969219124 | Tex9 | testis expressed gene 9 [Source:MGI Sym | 11.243 | 9.2513 | 4.025 | 2.8919 | 15.269 | 10.025 |
| 31198 | ENSMUSG00000090627 | 0 | NA | NA | NA | NA | NA | Gm8356 | predicted gene 8356 [Source:MGI Symbol | 0 | 0 | 0 | 0 | 0 | 0 |
| 31199 | ENSMUSG00000090628 | 3.20351639 | -1.482530786 | 1.350132983 | -1.098062786 | 0.272177074 | NA | Gm17083 | predicted gene 17083 [Source:MGI Symbo | 4.0155 | 8.2233 | 2.0125 | 3.8559 | 0 | 1.1138 |
| 31200 | ENSMUSG00000090629 | 0 | NA | NA | NA | NA | NA | Olfr180 | olfactory receptor 180 [Source:MGI Symb | 0 | 0 | 0 | 0 | 0 | 0 |
| 31201 | ENSMUSG00000090631 | 0.17131964 | -0.796629176 | 4.080472857 | -0.195229623 | 0.84521319 | NA | Olfr456 | olfactory receptor 456 [Source:MGI Symb | 0 | 1.0279 | 0 | 0 | 0 | 0 |
| 31202 | ENSMUSG00000090634 | 0 | NA | NA | NA | NA | NA | Gm8126 | predicted gene 8126 [Source:MGI Symbol | 0 | 0 | 0 | 0 | 0 | 0 |
| 31203 | ENSMUSG00000090635 | 0 | NA | NA | NA | NA | NA | Gm2951 | predicted gene 2951 [Source:MGI Symbol | 0 | 0 | 0 | 0 | 0 | 0 |
| 31204 | ENSMUSG00000090636 | 0 | NA | NA | NA | NA | NA | Gm17069 | predicted gene 17069 [Source:MGI Symbo | 0 | 0 | 0 | 0 | 0 | 0 |
| 31205 | ENSMUSG00000090637 | 0 | NA | NA | NA | NA | NA | Gm6189 | predicted gene 6189 [Source:MGI Symbol | 0 | 0 | 0 | 0 | 0 | 0 |
| 31206 | ENSMUSG00000090638 | 0 | NA | NA | NA | NA | NA | Gm17028 | predicted gene 17028 [Source:MGI Symbo | 0 | 0 | 0 | 0 | 0 | 0 |
| 31207 | ENSMUSG00000090639 | 0 | NA | NA | NA | NA | NA | NA | NA | 0 | 0 | 0 | 0 | 0 | 0 |
| 31208 | ENSMUSG00000090641 | 16.30125246 | -0.007058268 | 0.553646232 | -0.012748696 | 0.989828288 | 0.99859176 | Zfp712 | zinc finger protein 712 [Source:MGI Symb | 19.274 | 19.53 | 10.062 | 12.532 | 16.359 | 20.049 |
| 31209 | ENSMUSG00000090642 | 0.481988923 | 2.492401104 | 3.837077151 | 0.649557203 | 0.515978285 | NA | Gm17182 | predicted gene 17182 [Source:MGI Symbo | 0 | 0 | 0 | 2.8919 | 0 | 0 |
| 31210 | ENSMUSG00000090643 | 0 | NA | NA | NA | NA | NA | Gm3453 | predicted gene 3453 [Source:MGI Symbol | 0 | 0 | 0 | 0 | 0 | 0 |
| 31211 | ENSMUSG00000090655 | 0 | NA | NA | NA | NA | NA | Vmn2r120 | vomeronasal 2, receptor 120 [Source:MGI | 0 | 0 | 0 | 0 | 0 | 0 |
| 31212 | ENSMUSG00000090657 | 0 | NA | NA | NA | NA | NA | Gm8114 | predicted gene 8114 [Source:MGI Symbol | 0 | 0 | 0 | 0 | 0 | 0 |
| 31213 | ENSMUSG00000090658 | 0 | NA | NA | NA | NA | NA | Prss47 | protease, serine 47 [Source:MGI Symbol;A | 0 | 0 | 0 | 0 | 0 | 0 |
| 31214 | ENSMUSG00000090659 | 34.41664426 | -0.252701186 | 0.396832467 | -0.636795643 | 0.524257964 | 0.888334265 | Zfp493 | zinc finger protein 493 [Source:MGI Symb | 35.336 | 37.005 | 40.25 | 44.343 | 26.175 | 23.391 |
| 31215 | ENSMUSG00000090660 | 0 | NA | NA | NA | NA | NA | Gm17030 | predicted gene 17030 [Source:MGI Symbo | 0 | 0 | 0 | 0 | 0 | 0 |
| 31216 | ENSMUSG00000090662 | 0 | NA | NA | NA | NA | NA | Vmn2r45 | vomeronasal 2, receptor 45 [Source:MGI S | 0 | 0 | 0 | 0 | 0 | 0 |
| 31217 | ENSMUSG00000090665 | 0 | NA | NA | NA | NA | NA | Gad1-ps | glutamate decarboxylase 1, pseudogene [ | 0 | 0 | 0 | 0 | 0 | 0 |
| 31218 | ENSMUSG00000090667 | 8.929435045 | 0.036905778 | 0.819073398 | 0.045057962 | 0.964061109 | 0.993245949 | Mdfic2 | MyoD family inhibitor domain containing | 12.047 | 12.335 | 2.0125 | 9.6398 | 13.087 | 4.4554 |
| 31219 | ENSMUSG00000090671 | 0 | NA | NA | NA | NA | NA | Gm5067 | predicted gene 5067 [Source:MGI Symbol | 0 | 0 | 0 | 0 | 0 | 0 |
| 31220 | ENSMUSG00000090673 | 53.83793963 | 0.567094441 | 0.319419947 | 1.775388317 | 0.075833792 | 0.403405651 | NA | NA | 54.611 | 37.005 | 38.237 | 74.226 | 63.256 | 55.692 |
| 31221 | ENSMUSG00000090674 | 1.574055288 | 0.442139132 | 2.0993284 | 0.210609799 | 0.833191765 | NA | Gm17082 | predicted gene 17082 [Source:MGI Symbo | 0 | 0 | 4.025 | 0.964 | 0 | 4.4554 |
| 31222 | ENSMUSG00000090675 | 0.160662974 | 1.126924175 | 4.080472857 | 0.276174898 | 0.782413736 | NA | Olfr111 | olfactory receptor 111 [Source:MGI Symb | 0 | 0 | 0 | 0.964 | 0 | 0 |
| 31223 | ENSMUSG00000090681 | 0 | NA | NA | NA | NA | NA | Gm3221 | predicted gene 3221 [Source:MGI Symbol | 0 | 0 | 0 | 0 | 0 | 0 |
| 31224 | ENSMUSG00000090682 | 0 | NA | NA | NA | NA | NA | Gm3852 | predicted gene 3852 [Source:MGI Symbol | 0 | 0 | 0 | 0 | 0 | 0 |
| 31225 | ENSMUSG00000090684 | 0 | NA | NA | NA | NA | NA | Gm17221 | predicted gene 17221 [Source:MGI Symbo | 0 | 0 | 0 | 0 | 0 | 0 |
| 31226 | ENSMUSG00000090685 | 0 | NA | NA | NA | NA | NA | Atp6v1fnb | Atp6v1f neighbor [Source:MGI Symbol;Ac | 0 | 0 | 0 | 0 | 0 | 0 |
| 31227 | ENSMUSG00000090688 | 0 | NA | NA | NA | NA | NA | Vmn2r12 | vomeronasal 2, receptor 12 [Source:MGI S | 0 | 0 | 0 | 0 | 0 | 0 |
| 31228 | ENSMUSG00000090690 | 0 | NA | NA | NA | NA | NA | Gm8020 | predicted gene 8020 [Source:MGI Symbol | 0 | 0 | 0 | 0 | 0 | 0 |
| 31229 | ENSMUSG00000090691 | 0 | NA | NA | NA | NA | NA | Gm3667 | predicted gene 3667 [Source:MGI Symbol | 0 | 0 | 0 | 0 | 0 | 0 |
| 31230 | ENSMUSG00000090692 | 0 | NA | NA | NA | NA | NA | Gm17172 | predicted gene 17172 [Source:MGI Symbo | 0 | 0 | 0 | 0 | 0 | 0 |
| 31231 | ENSMUSG00000090693 | 4.70363567 | -0.33891466 | 1.128550551 | -0.300309684 | 0.763940947 | NA | Trim43a | tripartite motif-containing 43A [Source:M | 9.6372 | 0 | 6.0375 | 4.8199 | 3.2719 | 4.4554 |
| 31232 | ENSMUSG00000090698 | 11.58424691 | -0.568167065 | 0.657342698 | -0.864339205 | 0.387401579 | 0.817618864 | Apold1 | apolipoprotein L domain containing 1 [Sou | 11.243 | 11.307 | 19.119 | 13.496 | 6.5437 | 7.7969 |
| 31233 | ENSMUSG00000090699 | 0 | NA | NA | NA | NA | NA | Gm9071 | predicted gene 9071 [Source:MGI Symbol | 0 | 0 | 0 | 0 | 0 | 0 |
| 31234 | ENSMUSG00000090700 | 0 | NA | NA | NA | NA | NA | Cyp4f40 | cytochrome P450, family 4, subfamily f, po | 0 | 0 | 0 | 0 | 0 | 0 |
| 31235 | ENSMUSG00000090701 | 0 | NA | NA | NA | NA | NA | Gm6486 | predicted gene 6486 [Source:MGI Symbol | 0 | 0 | 0 | 0 | 0 | 0 |
| 31236 | ENSMUSG00000090704 | 0 | NA | NA | NA | NA | NA | Trp53-ps | transformation related protein 53, pseudo | 0 | 0 | 0 | 0 | 0 | 0 |
| 31237 | ENSMUSG00000090706 | 0 | NA | NA | NA | NA | NA | Gm17233 | predicted gene 17233 [Source:MGI Symbo | 0 | 0 | 0 | 0 | 0 | 0 |
| 31238 | ENSMUSG00000090707 | 0 | NA | NA | NA | NA | NA | Gm8237 | predicted gene 8237 [Source:MGI Symbol | 0 | 0 | 0 | 0 | 0 | 0 |
| 31239 | ENSMUSG00000090708 | 6.768035416 | 0.745650495 | 0.819397684 | 0.909998295 | 0.362823409 | 0.801768464 | Gm17196 | predicted gene 17196 [Source:MGI Symbo | 4.0155 | 6.1675 | 5.0312 | 7.7118 | 6.5437 | 11.138 |
| 31240 | ENSMUSG00000090709 | 0 | NA | NA | NA | NA | NA | Gm17173 | predicted gene 17173 [Source:MGI Symbo | 0 | 0 | 0 | 0 | 0 | 0 |
| 31241 | ENSMUSG00000090710 | 0 | NA | NA | NA | NA | NA | Pate11 | prostate and testis expressed 11 [Source:I | 0 | 0 | 0 | 0 | 0 | 0 |
| 31242 | ENSMUSG00000090713 | 0 | NA | NA | NA | NA | NA | Gm8127 | predicted gene 8127 [Source:MGI Symbol | 0 | 0 | 0 | 0 | 0 | 0 |
| 31243 | ENSMUSG00000090714 | 0 | NA | NA | NA | NA | NA | Zscan4d | zinc finger and SCAN domain containing 4 | 0 | 0 | 0 | 0 | 0 | 0 |
| 31244 | ENSMUSG00000090715 | 0 | NA | NA | NA | NA | NA | Vmn1r167 | vomeronasal 1 receptor 167 [Source:MGI | 0 | 0 | 0 | 0 | 0 | 0 |

| | | | | | | | | | | | | | | | |
|---|---|---|---|---|---|---|---|---|---|---|---|---|---|---|---|
| 31245 | ENSMUSG00000090716 | 0 | NA | NA | NA | NA | NA | Gm3371 | predicted gene 3371 [Source:MGI Symbol | 0 | 0 | 0 | 0 | 0 | 0 |
| 31246 | ENSMUSG00000090718 | 0 | NA | NA | NA | NA | NA | Gm17376 | predicted gene, 17376 [Source:MGI Symb | 0 | 0 | 0 | 0 | 0 | 0 |
| 31247 | ENSMUSG00000090722 | 0.36353943 | 2.065989254 | 4.034137301 | 0.512126658 | 0.608562368 | NA | Gm8378 | predicted gene 8378 [Source:MGI Symbol | 0 | 0 | 0 | 0 | 2.1812 | 0 |
| 31248 | ENSMUSG00000090724 | 0 | NA | NA | NA | NA | NA | 4930542C12 | RIKEN cDNA 4930542C12 gene [Source:M | 0 | 0 | 0 | 0 | 0 | 0 |
| 31249 | ENSMUSG00000090726 | 0 | NA | NA | NA | NA | NA | Gm17421 | predicted gene, 17421 [Source:MGI Symb | 0 | 0 | 0 | 0 | 0 | 0 |
| 31250 | ENSMUSG00000090727 | 0 | NA | NA | NA | NA | NA | Gm3956 | predicted gene 3956 [Source:MGI Symbol | 0 | 0 | 0 | 0 | 0 | 0 |
| 31251 | ENSMUSG00000090728 | 0 | NA | NA | NA | NA | NA | Gm17420 | predicted gene, 17420 [Source:MGI Symb | 0 | 0 | 0 | 0 | 0 | 0 |
| 31252 | ENSMUSG00000090731 | 0 | NA | NA | NA | NA | NA | Gm2959 | predicted gene 2959 [Source:MGI Symbol | 0 | 0 | 0 | 0 | 0 | 0 |
| 31253 | ENSMUSG00000090732 | 0 | NA | NA | NA | NA | NA | 4930579O11 | RIKEN cDNA 4930579O11 gene [Source:M | 0 | 0 | 0 | 0 | 0 | 0 |
| 31254 | ENSMUSG00000090733 | 2037.312821 | 0.184977697 | 0.10221111 | 1.809761157 | 0.070332834 | 0.383450475 | Rps27 | ribosomal protein S27 [Source:MGI Symb | 1790.9 | 2047.6 | 1882.7 | 2051.3 | 2282.7 | 2168.7 |
| 31255 | ENSMUSG00000090736 | 0 | NA | NA | NA | NA | NA | Gm9593 | predicted gene 9593 [Source:MGI Symbol | 0 | 0 | 0 | 0 | 0 | 0 |
| 31256 | ENSMUSG00000090737 | 0 | NA | NA | NA | NA | NA | Gm6788 | predicted gene 6788 [Source:MGI Symbol | 0 | 0 | 0 | 0 | 0 | 0 |
| 31257 | ENSMUSG00000090738 | 0 | NA | NA | NA | NA | NA | Pate7 | prostate and testis expressed 7 [Source:M | 0 | 0 | 0 | 0 | 0 | 0 |
| 31258 | ENSMUSG00000090740 | 0 | NA | NA | NA | NA | NA | Gm17079 | predicted gene 17079 [Source:MGI Symb | 0 | 0 | 0 | 0 | 0 | 0 |
| 31259 | ENSMUSG00000090742 | 0 | NA | NA | NA | NA | NA | Gm3008 | predicted gene 3008 [Source:MGI Symbol | 0 | 0 | 0 | 0 | 0 | 0 |
| 31260 | ENSMUSG00000090743 | 0 | NA | NA | NA | NA | NA | Gm17213 | predicted gene 17213 [Source:MGI Symb | 0 | 0 | 0 | 0 | 0 | 0 |
| 31261 | ENSMUSG00000090744 | 0 | NA | NA | NA | NA | NA | Gm6871 | predicted gene 6871 [Source:MGI Symbol | 0 | 0 | 0 | 0 | 0 | 0 |
| 31262 | ENSMUSG00000090747 | 0 | NA | NA | NA | NA | NA | Esp8 | exocrine gland secreted peptide 8 [Source | 0 | 0 | 0 | 0 | 0 | 0 |
| 31263 | ENSMUSG00000090750 | 0 | NA | NA | NA | NA | NA | Gm21999 | predicted gene 21999 [Source:MGI Symb | 0 | 0 | 0 | 0 | 0 | 0 |
| 31264 | ENSMUSG00000090751 | 0 | NA | NA | NA | NA | NA | Vmn2r63 | vomeronasal 2, receptor 63 [Source:MGI S | 0 | 0 | 0 | 0 | 0 | 0 |
| 31265 | ENSMUSG00000090753 | 0 | NA | NA | NA | NA | NA | Rpl31-ps4 | ribosomal protein L31, pseudogene 4 [Sou | 0 | 0 | 0 | 0 | 0 | 0 |
| 31266 | ENSMUSG00000090758 | 0 | NA | NA | NA | NA | NA | Gm17178 | predicted gene 17178 [Source:MGI Symb | 0 | 0 | 0 | 0 | 0 | 0 |
| 31267 | ENSMUSG00000090761 | 39.21621099 | 0.279929287 | 0.358470511 | 0.780899064 | 0.434861863 | 0.845325456 | Gm17201 | predicted gene 17201 [Source:MGI Symb | 32.927 | 44.2 | 29.181 | 38.559 | 42.534 | 47.895 |
| 31268 | ENSMUSG00000090762 | 0 | NA | NA | NA | NA | NA | Vmn2r56 | vomeronasal 2, receptor 56 [Source:MGI S | 0 | 0 | 0 | 0 | 0 | 0 |
| 31269 | ENSMUSG00000090764 | 0 | NA | NA | NA | NA | NA | Gm3127 | predicted gene 3127 [Source:MGI Symbol | 0 | 0 | 0 | 0 | 0 | 0 |
| 31270 | ENSMUSG00000090765 | 0 | NA | NA | NA | NA | NA | | | 0 | 0 | 0 | 0 | 0 | 0 |
| 31271 | ENSMUSG00000090772 | 0 | NA | NA | NA | NA | NA | Gm2930 | predicted gene 2930 [Source:MGI Symbol | 0 | 0 | 0 | 0 | 0 | 0 |
| 31272 | ENSMUSG00000090773 | 0 | NA | NA | NA | NA | NA | Vmn2r-ps69 | vomeronasal 2, receptor, pseudogene 69 | 0 | 0 | 0 | 0 | 0 | 0 |
| 31273 | ENSMUSG00000090774 | 0 | NA | NA | NA | NA | NA | Vmn2r74 | vomeronasal 2, receptor 74 [Source:MGI S | 0 | 0 | 0 | 0 | 0 | 0 |
| 31274 | ENSMUSG00000090777 | 0 | NA | NA | NA | NA | NA | Ccdc188 | coiled-coil domain containing 188 [Source | 0 | 0 | 0 | 0 | 0 | 0 |
| 31275 | ENSMUSG00000090778 | 1.092328948 | -0.312353558 | 2.312443774 | -0.135075093 | 0.892552503 | NA | Gm3235 | predicted gene 3235 [Source:MGI Symbol | 1.6062 | 2.0558 | 0 | 2.8919 | 0 | 0 |
| 31276 | ENSMUSG00000090779 | 5.966298175 | 0.216755512 | 0.974503769 | 0.222426551 | 0.823981845 | 0.96943211 | Gm17110 | predicted gene 17110 [Source:MGI Symb | 2.4093 | 6.1675 | 8.05 | 3.8559 | 13.087 | 2.2277 |
| 31277 | ENSMUSG00000090781 | 0 | NA | NA | NA | NA | NA | Igkv2-95-2 | immunoglobulin kappa variable 2-95-2 [So | 0 | 0 | 0 | 0 | 0 | 0 |
| 31278 | ENSMUSG00000090782 | 0 | NA | NA | NA | NA | NA | Traj60 | T cell receptor alpha joining 60 [Source:M | 0 | 0 | 0 | 0 | 0 | 0 |
| 31279 | ENSMUSG00000090785 | 2.074875684 | 0.513799611 | 1.547425274 | 0.332035168 | 0.739862704 | NA | Gm17116 | predicted gene 17116 [Source:MGI Symb | 4.0155 | 0 | 1.0062 | 1.928 | 3.2719 | 2.2277 |
| 31280 | ENSMUSG00000090788 | 0 | NA | NA | NA | NA | NA | Gm9597 | predicted gene 9597 [Source:MGI Symbol | 0 | 0 | 0 | 0 | 0 | 0 |
| 31281 | ENSMUSG00000090790 | 0.160662974 | 1.126924175 | 4.080472857 | 0.276174898 | 0.782413736 | NA | Gm17377 | predicted gene, 17377 [Source:MGI Symb | 0 | 0 | 0 | 0.964 | 0 | 0 |
| 31282 | ENSMUSG00000090793 | 0 | NA | NA | NA | NA | NA | Gm6650 | predicted gene 6650 [Source:MGI Symbol | 0 | 0 | 0 | 0 | 0 | 0 |
| 31283 | ENSMUSG00000090794 | 0 | NA | NA | NA | NA | NA | NA | NA | 0 | 0 | 0 | 0 | 0 | 0 |
| 31284 | ENSMUSG00000090795 | 0 | NA | NA | NA | NA | NA | Gm7346 | predicted gene 7346 [Source:MGI Symbol | 0 | 0 | 0 | 0 | 0 | 0 |
| 31285 | ENSMUSG00000090796 | 0 | NA | NA | NA | NA | NA | Gm3141 | predicted gene 3141 [Source:MGI Symbol | 0 | 0 | 0 | 0 | 0 | 0 |
| 31286 | ENSMUSG00000090798 | 0 | NA | NA | NA | NA | NA | Gm8046 | predicted gene 8046 [Source:MGI Symbol | 0 | 0 | 0 | 0 | 0 | 0 |
| 31287 | ENSMUSG00000090799 | 2.028791271 | 0.474535976 | 1.58658496 | 0.299092698 | 0.764869318 | NA | Klhl33 | kelch-like 33 [Source:MGI Symbol;Acc:MG | 4.0155 | 0 | 1.0062 | 3.8559 | 2.1812 | 1.1138 |
| 31288 | ENSMUSG00000090801 | 0 | NA | NA | NA | NA | NA | Gm4862 | predicted gene 4862 [Source:MGI Symbol | 0 | 0 | 0 | 0 | 0 | 0 |
| 31289 | ENSMUSG00000090802 | 0 | NA | NA | NA | NA | NA | Gm8074 | predicted gene 8074 [Source:MGI Symbol | 0 | 0 | 0 | 0 | 0 | 0 |
| 31290 | ENSMUSG00000090805 | 0 | NA | NA | NA | NA | NA | NA | NA | 0 | 0 | 0 | 0 | 0 | 0 |
| 31291 | ENSMUSG00000090806 | 0 | NA | NA | NA | NA | NA | Vmn2r70 | vomeronasal 2, receptor 70 [Source:MGI S | 0 | 0 | 0 | 0 | 0 | 0 |
| 31292 | ENSMUSG00000090808 | 0 | NA | NA | NA | NA | NA | Gm1758 | predicted gene 1758 [Source:MGI Symbol | 0 | 0 | 0 | 0 | 0 | 0 |
| 31293 | ENSMUSG00000090812 | 0.319491021 | 0.16514334 | 4.044653141 | 0.040830038 | 0.967431393 | NA | Samd15 | sterile alpha motif domain containing 15 [ | 0.8031 | 0 | 0 | 0 | 0 | 1.1138 |
| 31294 | ENSMUSG00000090816 | 0 | NA | NA | NA | NA | NA | Gm6729 | predicted gene 6729 [Source:MGI Symbol | 0 | 0 | 0 | 0 | 0 | 0 |
| 31295 | ENSMUSG00000090817 | 0 | NA | NA | NA | NA | NA | Gm4450 | predicted gene 4450 [Source:MGI Symbol | 0 | 0 | 0 | 0 | 0 | 0 |
| 31296 | ENSMUSG00000090818 | 0 | NA | NA | NA | NA | NA | Trbj2-6 | T cell receptor beta joining 2-6 [Source:M | 0 | 0 | 0 | 0 | 0 | 0 |
| 31297 | ENSMUSG00000090821 | 0 | NA | NA | NA | NA | NA | Gm8161 | predicted gene 8161 [Source:MGI Symbol | 0 | 0 | 0 | 0 | 0 | 0 |
| 31298 | ENSMUSG00000090824 | 0 | NA | NA | NA | NA | NA | Olfr1344 | olfactory receptor 1344 [Source:MGI Sym | 0 | 0 | 0 | 0 | 0 | 0 |
| 31299 | ENSMUSG00000090827 | 0 | NA | NA | NA | NA | NA | Gm8297 | predicted gene 8297 [Source:MGI Symbol | 0 | 0 | 0 | 0 | 0 | 0 |
| 31300 | ENSMUSG00000090839 | 0 | NA | NA | NA | NA | NA | Gm17094 | predicted gene 17094 [Source:MGI Symb | 0 | 0 | 0 | 0 | 0 | 0 |

| | | | | | | | | | | | | | | | |
|---|---|---|---|---|---|---|---|---|---|---|---|---|---|---|---|
| 31301 | ENSMUSG00000090840 | 0.804860058 | -1.761048096 | 2.420315956 | -0.727610828 | 0.46685185 | NA | 1700092M07 | RIKEN cDNA 1700092M07 gene [Source:M | 0.8031 | 2.0558 | 1.0062 | 0.964 | 0 | 0 |
| 31302 | ENSMUSG00000090841 | 2912.181812 | 0.0397639 | 0.113086446 | 0.35162393 | 0.725120316 | 0.9448775 | Myl6 | myosin, light polypeptide 6, alkali, smooth | 3142.5 | 2706.5 | 2765.2 | 2653.8 | 2937 | 3268 |
| 31303 | ENSMUSG00000090843 | 0.185640647 | 1.126924175 | 4.080472857 | 0.276174898 | 0.782413736 | NA | Heatr4 | HEAT repeat containing 4 [Source:MGI Sy | 0 | 0 | 0 | 0 | 0 | 1.1138 |
| 31304 | ENSMUSG00000090849 | 0 | NA | NA | NA | NA | NA | Vmn2r-ps36 | vomeronasal 2, receptor, pseudogene 36 | 0 | 0 | 0 | 0 | 0 | 0 |
| 31305 | ENSMUSG00000090853 | 0 | NA | NA | NA | NA | NA | Gm17364 | predicted gene, 17364 [Source:MGI Symb | 0 | 0 | 0 | 0 | 0 | 0 |
| 31306 | ENSMUSG00000090854 | 0 | NA | NA | NA | NA | NA | Gm4340 | predicted gene 4340 [Source:MGI Symbol | 0 | 0 | 0 | 0 | 0 | 0 |
| 31307 | ENSMUSG00000090855 | 0 | NA | NA | NA | NA | NA | Gm17219 | predicted gene 17219 [Source:MGI Symbo | 0 | 0 | 0 | 0 | 0 | 0 |
| 31308 | ENSMUSG00000090860 | 0 | NA | NA | NA | NA | NA | Gm17134 | predicted gene 17134 [Source:MGI Symbo | 0 | 0 | 0 | 0 | 0 | 0 |
| 31309 | ENSMUSG00000090861 | 0.569258551 | -2.569337407 | 3.18860282 | -0.805787849 | 0.420365185 | NA | Arf4os | ADP-ribosylation factor 4, opposite strand | 2.4093 | 0 | 1.0062 | 0 | 0 | 0 |
| 31310 | ENSMUSG00000090862 | 1743.596762 | -0.089744543 | 0.09380205 | -0.956743938 | 0.338696515 | 0.780858357 | Rps13 | ribosomal protein S13 [Source:MGI Symbo | 1750.8 | 1822.5 | 1820.3 | 1638.8 | 1738.4 | 1690.8 |
| 31311 | ENSMUSG00000090863 | 0 | NA | NA | NA | NA | NA | A530084C06 | RIKEN cDNA A530084C06 gene [Source:M | 0 | 0 | 0 | 0 | 0 | 0 |
| 31312 | ENSMUSG00000090864 | 0 | NA | NA | NA | NA | NA | Vmn2r40 | vomeronasal 2, receptor 40 [Source:MGI S | 0 | 0 | 0 | 0 | 0 | 0 |
| 31313 | ENSMUSG00000090865 | 0 | NA | NA | NA | NA | NA | Gm17074 | predicted gene 17074 [Source:MGI Symbo | 0 | 0 | 0 | 0 | 0 | 0 |
| 31314 | ENSMUSG00000090868 | 0 | NA | NA | NA | NA | NA | Trav9d-4 | T cell receptor alpha variable 9D-4 [Source | 0 | 0 | 0 | 0 | 0 | 0 |
| 31315 | ENSMUSG00000090872 | 0 | NA | NA | NA | NA | NA | Gm3239 | predicted gene 3239 [Source:MGI Symbol | 0 | 0 | 0 | 0 | 0 | 0 |
| 31316 | ENSMUSG00000090873 | 12.5725491 | 0.215257162 | 0.610204265 | 0.352762468 | 0.724266524 | 0.944786345 | Gm17112 | predicted gene 17112 [Source:MGI Symbo | 9.6372 | 13.363 | 12.075 | 18.316 | 10.906 | 11.138 |
| 31317 | ENSMUSG00000090874 | 0 | NA | NA | NA | NA | NA | Olfr733 | olfactory receptor 733 [Source:MGI Symb | 0 | 0 | 0 | 0 | 0 | 0 |
| 31318 | ENSMUSG00000090877 | 140.3117268 | -0.990077095 | 0.243878442 | -4.059715524 | 4.91E-05 | 0.002077933 | Hspa1b | heat shock protein 1B [Source:MGI Symbo | 142.95 | 216.89 | 200.24 | 77.118 | 117.79 | 86.88 |
| 31319 | ENSMUSG00000090881 | 2.54496781 | -0.422454189 | 1.458244715 | -0.289700476 | 0.772045392 | NA | Phf11 | PHD finger protein 11 [Source:MGI Symbo | 1.6062 | 4.1117 | 3.0187 | 0.964 | 0 | 5.5692 |
| 31320 | ENSMUSG00000090882 | 0 | NA | NA | NA | NA | NA | Gm17065 | predicted gene 17065 [Source:MGI Symbo | 0 | 0 | 0 | 0 | 0 | 0 |
| 31321 | ENSMUSG00000090889 | 0 | NA | NA | NA | NA | NA | Gm17428 | predicted gene, 17428 [Source:MGI Symb | 0 | 0 | 0 | 0 | 0 | 0 |
| 31322 | ENSMUSG00000090891 | 0.160662974 | 1.126924175 | 4.080472857 | 0.276174898 | 0.782413736 | NA | D6Ertd527e | DNA segment, Chr 6, ERATO Doi 527, expr | 0 | 0 | 0 | 0.964 | 0 | 0 |
| 31323 | ENSMUSG00000090892 | 0 | NA | NA | NA | NA | NA | Vmn2r41 | vomeronasal 2, receptor 41 [Source:MGI S | 0 | 0 | 0 | 0 | 0 | 0 |
| 31324 | ENSMUSG00000090894 | 0 | NA | NA | NA | NA | NA | Olfr110 | olfactory receptor 110 [Source:MGI Symb | 0 | 0 | 0 | 0 | 0 | 0 |
| 31325 | ENSMUSG00000090897 | 0 | NA | NA | NA | NA | NA | Gm44501 | predicted readthrough transcript, 44501 [ | 0 | 0 | 0 | 0 | 0 | 0 |
| 31326 | ENSMUSG00000090900 | 0 | NA | NA | NA | NA | NA | Gm3326 | predicted gene 3326 [Source:MGI Symbol | 0 | 0 | 0 | 0 | 0 | 0 |
| 31327 | ENSMUSG00000090907 | 0 | NA | NA | NA | NA | NA | C030017D09 | RIKEN cDNA C030017D09 gene [Source:M | 0 | 0 | 0 | 0 | 0 | 0 |
| 31328 | ENSMUSG00000090919 | 4.465876083 | -0.90759836 | 1.096871137 | -0.827443014 | 0.407986009 | NA | Pabpc4l | poly(A) binding protein, cytoplasmic 4-like | 2.4093 | 4.1117 | 11.069 | 4.8199 | 3.2719 | 1.1138 |
| 31329 | ENSMUSG00000090925 | 21.59524169 | 0.943523478 | 0.555215003 | 1.699383973 | 0.08924686 | 0.440268274 | 1810064F22 | RIKEN cDNA 1810064F22 gene [Source:M | 12.047 | 15.419 | 17.106 | 46.271 | 11.997 | 26.732 |
| 31330 | ENSMUSG00000090927 | 0 | NA | NA | NA | NA | NA | Gm17047 | predicted gene 17047 [Source:MGI Symbo | 0 | 0 | 0 | 0 | 0 | 0 |
| 31331 | ENSMUSG00000090935 | 1341.478875 | -0.025661122 | 0.110561344 | -0.232098496 | 0.816461513 | 0.967767553 | Synj2bp | synaptojanin 2 binding protein [Source:M | 1267.3 | 1394.9 | 1398.7 | 1287.9 | 1456 | 1244.2 |
| 31332 | ENSMUSG00000090936 | 3.251358622 | -1.071039811 | 1.408003301 | -0.760679901 | 0.446848283 | NA | Gm17705 | predicted gene, 17705 [Source:MGI Symb | 0 | 10.279 | 3.0187 | 2.8919 | 1.0906 | 2.2277 |
| 31333 | ENSMUSG00000090939 | 0.305170013 | -1.657836477 | 4.041480335 | -0.410205257 | 0.681655385 | NA | Gm17147 | predicted gene 17147 [Source:MGI Symbo | 0.8031 | 1.0279 | 0 | 0 | 0 | 0 |
| 31334 | ENSMUSG00000090941 | 0 | NA | NA | NA | NA | NA | Gm17212 | predicted gene 17212 [Source:MGI Symbo | 0 | 0 | 0 | 0 | 0 | 0 |
| 31335 | ENSMUSG00000090942 | 318.4174117 | -1.929357738 | 0.893305546 | -2.159795993 | 0.030788466 | 0.232721383 | F830016B08 | RIKEN cDNA F830016B08 gene [Source:M | 505.95 | 426.59 | 580.6 | 60.731 | 302.1 | 34.529 |
| 31336 | ENSMUSG00000090946 | 142.0308532 | 0.058639262 | 0.196884303 | 0.297836145 | 0.765828226 | 0.954996635 | Ccdc71l | coiled-coil domain containing 71 like [Sou | 147.77 | 133.63 | 135.84 | 150.38 | 131.96 | 152.6 |
| 31337 | ENSMUSG00000090947 | 0 | NA | NA | NA | NA | NA | Gm3102 | predicted gene 3102 [Source:MGI Symbol | 0 | 0 | 0 | 0 | 0 | 0 |
| 31338 | ENSMUSG00000090949 | 0 | NA | NA | NA | NA | NA | Vmn2r77 | vomeronasal 2, receptor 77 [Source:MGI S | 0 | 0 | 0 | 0 | 0 | 0 |
| 31339 | ENSMUSG00000090951 | 0 | NA | NA | NA | NA | NA | Olfr181 | olfactory receptor 181 [Source:MGI Symb | 0 | 0 | 0 | 0 | 0 | 0 |
| 31340 | ENSMUSG00000090952 | 61.82726875 | 0.238274323 | 0.307341081 | 0.775276518 | 0.438176282 | 0.847566655 | Gm17251 | predicted gene, 17251 [Source:MGI Symb | 65.051 | 59.619 | 45.281 | 68.442 | 51.259 | 81.311 |
| 31341 | ENSMUSG00000090955 | 0 | NA | NA | NA | NA | NA | Gm17097 | predicted gene 17097 [Source:MGI Symbo | 0 | 0 | 0 | 0 | 0 | 0 |
| 31342 | ENSMUSG00000090957 | 0 | NA | NA | NA | NA | NA | Gm7535 | predicted gene 7535 [Source:MGI Symbol | 0 | 0 | 0 | 0 | 0 | 0 |
| 31343 | ENSMUSG00000090958 | 125.3499346 | -0.539852586 | 0.319489264 | -1.689736235 | 0.091078428 | 0.445203913 | Lrrc32 | leucine rich repeat containing 32 [Source: | 138.13 | 127.46 | 180.12 | 114.71 | 47.987 | 143.69 |
| 31344 | ENSMUSG00000090961 | 0.133850373 | -0.796629176 | 4.080472857 | -0.195229623 | 0.84521319 | NA | Vmn2r8 | vomeronasal 2, receptor 8 [Source:MGI Sy | 0.8031 | 0 | 0 | 0 | 0 | 0 |
| 31345 | ENSMUSG00000090963 | 11.35270633 | 0.838868503 | 0.703777328 | 1.191951587 | 0.233280229 | 0.680673483 | Gm17655 | predicted gene, 17655 [Source:MGI Symb | 7.2279 | 3.0838 | 14.087 | 10.604 | 14.178 | 18.935 |
| 31346 | ENSMUSG00000090964 | 0.428363721 | -0.543730382 | 3.624677344 | -0.15000794 | 0.880758351 | NA | Gm17206 | predicted gene 17206 [Source:MGI Symbo | 1.6062 | 0 | 0 | 0.964 | 0 | 0 |
| 31347 | ENSMUSG00000090965 | 0.620790101 | -1.233953725 | 2.860353486 | -0.431399032 | 0.666178254 | NA | Gm17203 | predicted gene 17203 [Source:MGI Symbo | 1.6062 | 1.0279 | 0 | 0 | 1.0906 | 0 |
| 31348 | ENSMUSG00000090966 | 0 | NA | NA | NA | NA | NA | Vmn2r116 | vomeronasal 2, receptor 116 [Source:MGI | 0 | 0 | 0 | 0 | 0 | 0 |
| 31349 | ENSMUSG00000090967 | 0 | NA | NA | NA | NA | NA | Vmn2r61 | vomeronasal 2, receptor 61 [Source:MGI S | 0 | 0 | 0 | 0 | 0 | 0 |
| 31350 | ENSMUSG00000090968 | 0 | NA | NA | NA | NA | NA | Gm3978 | predicted gene 3978 [Source:MGI Symbol | 0 | 0 | 0 | 0 | 0 | 0 |
| 31351 | ENSMUSG00000090975 | 0 | NA | NA | NA | NA | NA | Gm17162 | predicted gene 17162 [Source:MGI Symbo | 0 | 0 | 0 | 0 | 0 | 0 |
| 31352 | ENSMUSG00000090981 | 0 | NA | NA | NA | NA | NA | Olfr718-ps1 | olfactory receptor 718, pseudogene 1 [Sou | 0 | 0 | 0 | 0 | 0 | 0 |
| 31353 | ENSMUSG00000090989 | 0.401551119 | -2.072982726 | 4.008251979 | -0.517178744 | 0.605031388 | NA | Gm17042 | predicted gene 17042 [Source:MGI Symbo | 2.4093 | 0 | 0 | 0 | 0 | 0 |
| 31354 | ENSMUSG00000090990 | 0 | NA | NA | NA | NA | NA | Gm17197 | predicted gene 17197 [Source:MGI Symbo | 0 | 0 | 0 | 0 | 0 | 0 |
| 31355 | ENSMUSG00000090991 | 0 | NA | NA | NA | NA | NA | Gm6647 | predicted gene 6647 [Source:MGI Symbol | 0 | 0 | 0 | 0 | 0 | 0 |
| 31356 | ENSMUSG00000090994 | 0.167707431 | -0.796629176 | 4.080472857 | -0.195229623 | 0.84521319 | NA | Gm6153 | predicted gene 6153 [Source:MGI Symbol | 0 | 0 | 1.0062 | 0 | 0 | 0 |

| | | | | | | | | | | | | | | | |
|---|---|---|---|---|---|---|---|---|---|---|---|---|---|---|---|
| 31357 | ENSMUSG00000090996 | 0 | NA | NA | NA | NA | NA | Gm20458 | predicted gene 20458 [Source:MGI Symbc | 0 | 0 | 0 | 0 | 0 | 0 |
| 31358 | ENSMUSG00000091002 | 0 | NA | NA | NA | NA | NA | Tcerg1l | transcription elongation regulator 1-like [S | 0 | 0 | 0 | 0 | 0 | 0 |
| 31359 | ENSMUSG00000091003 | 0 | NA | NA | NA | NA | NA | Gm7244 | predicted gene 7244 [Source:MGI Symbol | 0 | 0 | 0 | 0 | 0 | 0 |
| 31360 | ENSMUSG00000091004 | 0 | NA | NA | NA | NA | NA | Gm6872 | predicted gene 6872 [Source:MGI Symbol | 0 | 0 | 0 | 0 | 0 | 0 |
| 31361 | ENSMUSG00000091005 | 0 | NA | NA | NA | NA | NA | Traj55 | T cell receptor alpha joining 55 [Source:M | 0 | 0 | 0 | 0 | 0 | 0 |
| 31362 | ENSMUSG00000091006 | 0 | NA | NA | NA | NA | NA | Vmn2r69 | vomeronasal 2, receptor 69 [Source:MGI S | 0 | 0 | 0 | 0 | 0 | 0 |
| 31363 | ENSMUSG00000091007 | 1.171831736 | -1.161928617 | 2.135676568 | -0.544056453 | 0.586402626 | NA | D630036H23 | RIKEN cDNA D630036H23 gene [Source:M | 0.8031 | 1.0279 | 3.0187 | 0 | 2.1812 | 0 |
| 31364 | ENSMUSG00000091008 | 0 | NA | NA | NA | NA | NA | Gm17472 | predicted gene, 17472 [Source:MGI Symb | 0 | 0 | 0 | 0 | 0 | 0 |
| 31365 | ENSMUSG00000091010 | 0 | NA | NA | NA | NA | NA | Gm17107 | predicted gene 17107 [Source:MGI Symbc | 0 | 0 | 0 | 0 | 0 | 0 |
| 31366 | ENSMUSG00000091011 | 0 | NA | NA | NA | NA | NA | Gm17187 | predicted gene 17187 [Source:MGI Symbc | 0 | 0 | 0 | 0 | 0 | 0 |
| 31367 | ENSMUSG00000091012 | 0 | NA | NA | NA | NA | NA | Vmn1r34 | vomeronasal 1 receptor 34 [Source:MGI S | 0 | 0 | 0 | 0 | 0 | 0 |
| 31368 | ENSMUSG00000091013 | 0 | NA | NA | NA | NA | NA | Vmn1r1 | vomeronasal 1 receptor 1 [Source:MGI Sy | 0 | 0 | 0 | 0 | 0 | 0 |
| 31369 | ENSMUSG00000091014 | 0 | NA | NA | NA | NA | NA | Gm5244 | predicted pseudogene 5244 [Source:MGI | 0 | 0 | 0 | 0 | 0 | 0 |
| 31370 | ENSMUSG00000091016 | 0 | NA | NA | NA | NA | NA | Gm2878 | predicted gene 2878 [Source:MGI Symbol | 0 | 0 | 0 | 0 | 0 | 0 |
| 31371 | ENSMUSG00000091017 | 0 | NA | NA | NA | NA | NA | Fam71a | family with sequence similarity 71, memb | 0 | 0 | 0 | 0 | 0 | 0 |
| 31372 | ENSMUSG00000091019 | 0 | NA | NA | NA | NA | NA | Gm7502 | predicted gene 7502 [Source:MGI Symbol | 0 | 0 | 0 | 0 | 0 | 0 |
| 31373 | ENSMUSG00000091020 | 0.685278559 | -2.809870862 | 3.376578027 | -0.832165239 | 0.405315682 | NA | Gm5828 | predicted gene 5828 [Source:MGI Symbol | 0 | 4.1117 | 0 | 0 | 0 | 0 |
| 31374 | ENSMUSG00000091021 | 10.2194321 | -2.871860536 | 0.828636017 | -3.465768416 | 0.000528718 | 0.012909222 | Gm17300 | predicted gene, 17300 [Source:MGI Symb | 23.29 | 19.53 | 11.069 | 1.928 | 3.2719 | 2.2277 |
| 31375 | ENSMUSG00000091022 | 0 | NA | NA | NA | NA | NA | Gm3115 | predicted gene 3115 [Source:MGI Symbol | 0 | 0 | 0 | 0 | 0 | 0 |
| 31376 | ENSMUSG00000091027 | 0 | NA | NA | NA | NA | NA | Gm11107 | predicted gene 11107 [Source:MGI Symbc | 0 | 0 | 0 | 0 | 0 | 0 |
| 31377 | ENSMUSG00000091028 | 0 | NA | NA | NA | NA | NA | Gm10722 | predicted gene 10722 [Source:MGI Symbc | 0 | 0 | 0 | 0 | 0 | 0 |
| 31378 | ENSMUSG00000091031 | 0 | NA | NA | NA | NA | NA | Gm17063 | predicted gene 17063 [Source:MGI Symbc | 0 | 0 | 0 | 0 | 0 | 0 |
| 31379 | ENSMUSG00000091034 | 0 | NA | NA | NA | NA | NA | Gm17660 | predicted gene, 17660 [Source:MGI Symb | 0 | 0 | 0 | 0 | 0 | 0 |
| 31380 | ENSMUSG00000091037 | 0 | NA | NA | NA | NA | NA | Gm17661 | predicted gene, 17661 [Source:MGI Symb | 0 | 0 | 0 | 0 | 0 | 0 |
| 31381 | ENSMUSG00000091039 | 0 | NA | NA | NA | NA | NA | Krtap20-2 | keratin associated protein 20-2 [Source:M | 0 | 0 | 0 | 0 | 0 | 0 |
| 31382 | ENSMUSG00000091040 | 0 | NA | NA | NA | NA | NA | Gm9359 | predicted gene 9359 [Source:MGI Symbol | 0 | 0 | 0 | 0 | 0 | 0 |
| 31383 | ENSMUSG00000091041 | 1.634491198 | -0.442467688 | 1.583430466 | -0.279436134 | 0.779910144 | NA | Gm17720 | predicted gene, 17720 [Source:MGI Symb | 1.6062 | 2.0558 | 2.0125 | 1.928 | 1.0906 | 1.1138 |
| 31384 | ENSMUSG00000091043 | 0 | NA | NA | NA | NA | NA | Glyatl3 | glycine-N-acyltransferase-like 3 [Source:M | 0 | 0 | 0 | 0 | 0 | 0 |
| 31385 | ENSMUSG00000091045 | 0 | NA | NA | NA | NA | NA | Vmn2r55 | vomeronasal 2, receptor 55 [Source:MGI S | 0 | 0 | 0 | 0 | 0 | 0 |
| 31386 | ENSMUSG00000091048 | 0 | NA | NA | NA | NA | NA | Gm8207 | predicted gene 8207 [Source:MGI Symbol | 0 | 0 | 0 | 0 | 0 | 0 |
| 31387 | ENSMUSG00000091049 | 0 | NA | NA | NA | NA | NA | Gm1979 | predicted gene 1979 [Source:MGI Symbol | 0 | 0 | 0 | 0 | 0 | 0 |
| 31388 | ENSMUSG00000091050 | 12.20587561 | 0.087696109 | 0.630073839 | 0.13918386 | 0.889304862 | 0.982382091 | 9330020H09 | RIKEN cDNA 9330020H09 gene [Source:M | 12.047 | 16.447 | 7.0437 | 13.496 | 14.178 | 10.025 |
| 31389 | ENSMUSG00000091055 | 0 | NA | NA | NA | NA | NA | Siglec15 | sialic acid binding Ig-like lectin 15 [Source: | 0 | 0 | 0 | 0 | 0 | 0 |
| 31390 | ENSMUSG00000091058 | 0.465832987 | -0.698295221 | 3.18348759 | -0.219349126 | 0.826378096 | NA | Gm17538 | predicted gene, 17538 [Source:MGI Symb | 0.8031 | 1.0279 | 0 | 0.964 | 0 | 0 |
| 31391 | ENSMUSG00000091059 | 0 | NA | NA | NA | NA | NA | Vmn2r14 | vomeronasal 2, receptor 14 [Source:MGI S | 0 | 0 | 0 | 0 | 0 | 0 |
| 31392 | ENSMUSG00000091060 | 0 | NA | NA | NA | NA | NA | Myocos | myocilin opposite strand [Source:MGI Syn | 0 | 0 | 0 | 0 | 0 | 0 |
| 31393 | ENSMUSG00000091061 | 0.133850373 | -0.796629176 | 4.080472857 | -0.195229623 | 0.84521319 | NA | 4930445B16 | RIKEN cDNA 4930445B16 gene [Source:M | 0.8031 | 0 | 0 | 0 | 0 | 0 |
| 31394 | ENSMUSG00000091071 | 2.947888543 | 0.173818617 | 1.197433348 | 0.145159325 | 0.884585079 | NA | 1700030C10 | RIKEN cDNA 1700030C10 gene [Source:M | 3.2124 | 3.0838 | 2.0125 | 3.8559 | 2.1812 | 3.3415 |
| 31395 | ENSMUSG00000091076 | 0 | NA | NA | NA | NA | NA | Vmn2r115 | vomeronasal 2, receptor 115 [Source:MGI | 0 | 0 | 0 | 0 | 0 | 0 |
| 31396 | ENSMUSG00000091077 | 0 | NA | NA | NA | NA | NA | Gm21292 | predicted gene, 21292 [Source:MGI Symb | 0 | 0 | 0 | 0 | 0 | 0 |
| 31397 | ENSMUSG00000091078 | 0.160662974 | 1.126924175 | 4.080472857 | 0.276174898 | 0.782413736 | NA | Gm17218 | predicted gene 17218 [Source:MGI Symbc | 0 | 0 | 0 | 0.964 | 0 | 0 |
| 31398 | ENSMUSG00000091079 | 0 | NA | NA | NA | NA | NA | Gm17105 | predicted gene 17105 [Source:MGI Symbc | 0 | 0 | 0 | 0 | 0 | 0 |
| 31399 | ENSMUSG00000091080 | 0 | NA | NA | NA | NA | NA | Prr23a1 | proline rich 23A, member 1 [Source:MGI S | 0 | 0 | 0 | 0 | 0 | 0 |
| 31400 | ENSMUSG00000091083 | 0 | NA | NA | NA | NA | NA | Gm4535 | predicted gene 4535 [Source:MGI Symbol | 0 | 0 | 0 | 0 | 0 | 0 |
| 31401 | ENSMUSG00000091086 | 4.465068296 | 1.565701035 | 1.106689234 | 1.414761242 | 0.157138511 | NA | Rpl6l | ribosomal protein L6-like [Source:MGI Syr | 0.8031 | 1.0279 | 5.0312 | 6.7478 | 8.7249 | 4.4554 |
| 31402 | ENSMUSG00000091087 | 0 | NA | NA | NA | NA | NA | Ighv8-14 | immunoglobulin heavy variable 8-14 [Sou | 0 | 0 | 0 | 0 | 0 | 0 |
| 31403 | ENSMUSG00000091091 | 0.167707431 | -0.796629176 | 4.080472857 | -0.195229623 | 0.84521319 | NA | Kcnmb3 | potassium large conductance calcium-acti | 0 | 0 | 1.0062 | 0 | 0 | 0 |
| 31404 | ENSMUSG00000091096 | 0 | NA | NA | NA | NA | NA | Gm17304 | predicted gene, 17304 [Source:MGI Symb | 0 | 0 | 0 | 0 | 0 | 0 |
| 31405 | ENSMUSG00000091099 | 0 | NA | NA | NA | NA | NA | Gm17164 | predicted gene 17164 [Source:MGI Symbc | 0 | 0 | 0 | 0 | 0 | 0 |
| 31406 | ENSMUSG00000091101 | 0 | NA | NA | NA | NA | NA | Gm4302 | predicted gene 4302 [Source:MGI Symbol | 0 | 0 | 0 | 0 | 0 | 0 |
| 31407 | ENSMUSG00000091102 | 4.085294167 | 1.133074586 | 1.111734663 | 1.019195159 | 0.308110325 | NA | 5830462I19F | RIKEN cDNA 5830462I19 gene [Source:MC | 1.6062 | 5.1396 | 1.0062 | 5.7839 | 7.6343 | 3.3415 |
| 31408 | ENSMUSG00000091105 | 0.181769715 | 1.126924175 | 4.080472857 | 0.276174898 | 0.782413736 | NA | Mrto4-ps2 | mRNA turnover 4, pseudogene 2 [Source: | 0 | 0 | 0 | 0 | 1.0906 | 0 |
| 31409 | ENSMUSG00000091109 | 0.17131964 | -0.796629176 | 4.080472857 | -0.195229623 | 0.84521319 | NA | Gm17622 | predicted gene, 17622 [Source:MGI Symb | 0 | 1.0279 | 0 | 0 | 0 | 0 |
| 31410 | ENSMUSG00000091110 | 0 | NA | NA | NA | NA | NA | Gm2832 | predicted gene 2832 [Source:MGI Symbol | 0 | 0 | 0 | 0 | 0 | 0 |
| 31411 | ENSMUSG00000091114 | 0 | NA | NA | NA | NA | NA | Gm3138 | predicted gene 3138 [Source:MGI Symbol | 0 | 0 | 0 | 0 | 0 | 0 |
| 31412 | ENSMUSG00000091116 | 0 | NA | NA | NA | NA | NA | Gm7991 | predicted gene 7991 [Source:MGI Symbol | 0 | 0 | 0 | 0 | 0 | 0 |

| | | | | | | | | | | | | | | | |
|---|---|---|---|---|---|---|---|---|---|---|---|---|---|---|---|
| 31413 | ENSMUSG00000091117 | 0.342639279 | -1.805976596 | 4.028547714 | -0.448294702 | 0.653940524 | NA | Gm9064 | predicted gene 9064 [Source:MGI Symbol | 0 | 2.0558 | 0 | 0 | 0 | 0 |
| 31414 | ENSMUSG00000091119 | 169.1761321 | -0.637496602 | 0.215344051 | -2.960363192 | 0.003072766 | 0.047432745 | Ccdc152 | coiled-coil domain containing 152 [Source | 211.22 | 161.38 | 245.52 | 152.31 | 124.33 | 120.3 |
| 31415 | ENSMUSG00000091122 | 0 | NA | NA | NA | NA | NA | Gm8247 | predicted gene 8247 [Source:MGI Symbol | 0 | 0 | 0 | 0 | 0 | 0 |
| 31416 | ENSMUSG00000091123 | 1.848779375 | 0.841800707 | 1.666703624 | 0.505069225 | 0.613510216 | NA | Gm17181 | predicted gene 17181 [Source:MGI Symbo | 0 | 2.0558 | 2.0125 | 4.8199 | 1.0906 | 1.1138 |
| 31417 | ENSMUSG00000091129 | 0 | NA | NA | NA | NA | NA | Iqcf6 | IQ motif containing F6 [Source:MGI Symb | 0 | 0 | 0 | 0 | 0 | 0 |
| 31418 | ENSMUSG00000091131 | 0 | NA | NA | NA | NA | NA | Gm7945 | predicted gene 7945 [Source:MGI Symbol | 0 | 0 | 0 | 0 | 0 | 0 |
| 31419 | ENSMUSG00000091132 | 0 | NA | NA | NA | NA | NA | Gm17109 | predicted gene 17109 [Source:MGI Symbo | 0 | 0 | 0 | 0 | 0 | 0 |
| 31420 | ENSMUSG00000091133 | 2.692783129 | -0.720493289 | 1.306852729 | -0.55131942 | 0.581414728 | NA | Gm17108 | predicted gene 17108 [Source:MGI Symbo | 4.0155 | 2.0558 | 4.025 | 3.8559 | 1.0906 | 1.1138 |
| 31421 | ENSMUSG00000091135 | 0 | NA | NA | NA | NA | NA | Lgals1-ps2 | lectin, galactose binding, soluble 1, pseud | 0 | 0 | 0 | 0 | 0 | 0 |
| 31422 | ENSMUSG00000091140 | 0 | NA | NA | NA | NA | NA | Gm17124 | predicted gene 17124 [Source:MGI Symbo | 0 | 0 | 0 | 0 | 0 | 0 |
| 31423 | ENSMUSG00000091142 | 0 | NA | NA | NA | NA | NA | Gm17175 | predicted gene 17175 [Source:MGI Symbo | 0 | 0 | 0 | 0 | 0 | 0 |
| 31424 | ENSMUSG00000091144 | 147.2133216 | -0.370508718 | 0.254753553 | -1.454380962 | 0.145840715 | 0.556690797 | Phf11c | PHD finger protein 11C [Source:MGI Symb | 148.57 | 229.23 | 120.75 | 143.63 | 133.06 | 108.04 |
| 31425 | ENSMUSG00000091146 | 0 | NA | NA | NA | NA | NA | Gm3077 | predicted gene 3077 [Source:MGI Symbol | 0 | 0 | 0 | 0 | 0 | 0 |
| 31426 | ENSMUSG00000091147 | 0 | NA | NA | NA | NA | NA | 4921509A06 | RIKEN cDNA 4921509A06 gene [Source:M | 0 | 0 | 0 | 0 | 0 | 0 |
| 31427 | ENSMUSG00000091148 | 0 | NA | NA | NA | NA | NA | Gm3182 | predicted gene 3182 [Source:MGI Symbol | 0 | 0 | 0 | 0 | 0 | 0 |
| 31428 | ENSMUSG00000091149 | 0.17131964 | -0.796629176 | 4.080472857 | -0.195229623 | 0.84521319 | NA | Gm17020 | predicted gene 17020 [Source:MGI Symbo | 0 | 1.0279 | 0 | 0 | 0 | 0 |
| 31429 | ENSMUSG00000091151 | 0 | NA | NA | NA | NA | NA | Vmn1r224 | vomeronasal 1 receptor 224 [Source:MGI | 0 | 0 | 0 | 0 | 0 | 0 |
| 31430 | ENSMUSG00000091154 | 0.954866367 | 0.067917755 | 2.273052348 | 0.029879539 | 0.976163124 | NA | Proscos | proline synthetase co-transcribed, opposi | 0.8031 | 1.0279 | 1.0062 | 2.8919 | 0 | 0 |
| 31431 | ENSMUSG00000091155 | 0.654647159 | -1.320638211 | 2.599368548 | -0.508061164 | 0.61141045 | NA | Serpine3 | serpin peptidase inhibitor, clade E (nexin, | 0.8031 | 1.0279 | 1.0062 | 0 | 1.0906 | 0 |
| 31432 | ENSMUSG00000091157 | 0 | NA | NA | NA | NA | NA | Serpina3l-ps | serine (or cysteine) peptidase inhibitor, cl | 0 | 0 | 0 | 0 | 0 | 0 |
| 31433 | ENSMUSG00000091159 | 0.66192371 | 1.647602865 | 2.590567345 | 0.636000785 | 0.52477591 | NA | Gm17545 | predicted gene, 17545 [Source:MGI Symb | 0.8031 | 0 | 0 | 0.964 | 1.0906 | 1.1138 |
| 31434 | ENSMUSG00000091160 | 6.33831951 | -0.439626671 | 0.956806882 | -0.45947273 | 0.64589473 | 0.925643431 | Gm17133 | predicted gene 17133 [Source:MGI Symbo | 4.8186 | 7.1954 | 10.062 | 11.568 | 3.2719 | 1.1138 |
| 31435 | ENSMUSG00000091163 | 0 | NA | NA | NA | NA | NA | Gm3962 | predicted gene 3962 [Source:MGI Symbol | 0 | 0 | 0 | 0 | 0 | 0 |
| 31436 | ENSMUSG00000091165 | 0 | NA | NA | NA | NA | NA | Gm17036 | predicted gene 17036 [Source:MGI Symbo | 0 | 0 | 0 | 0 | 0 | 0 |
| 31437 | ENSMUSG00000091172 | 0 | NA | NA | NA | NA | NA | Gm17120 | predicted gene 17120 [Source:MGI Symbo | 0 | 0 | 0 | 0 | 0 | 0 |
| 31438 | ENSMUSG00000091174 | 0 | NA | NA | NA | NA | NA | Pate10 | prostate and testis expressed 10 [Source:I | 0 | 0 | 0 | 0 | 0 | 0 |
| 31439 | ENSMUSG00000091175 | 0 | NA | NA | NA | NA | NA | S100a7l2 | S100 calcium binding protein A7 like 2 [So | 0 | 0 | 0 | 0 | 0 | 0 |
| 31440 | ENSMUSG00000091177 | 0 | NA | NA | NA | NA | NA | Gm15494 | predicted gene 15494 [Source:MGI Symbo | 0 | 0 | 0 | 0 | 0 | 0 |
| 31441 | ENSMUSG00000091183 | 2.598334288 | 0.268856381 | 1.45342342 | 0.184981456 | 0.85324358 | NA | Gm5141 | predicted gene 5141 [Source:MGI Symbol | 0 | 5.1396 | 2.0125 | 2.8919 | 1.0906 | 4.4554 |
| 31442 | ENSMUSG00000091184 | 0 | NA | NA | NA | NA | NA | Gm17059 | predicted gene 17059 [Source:MGI Symbo | 0 | 0 | 0 | 0 | 0 | 0 |
| 31443 | ENSMUSG00000091185 | 0 | NA | NA | NA | NA | NA | Gm3278 | predicted gene 3278 [Source:MGI Symbol | 0 | 0 | 0 | 0 | 0 | 0 |
| 31444 | ENSMUSG00000091189 | 0 | NA | NA | NA | NA | NA | Ear-ps3 | eosinophil-associated, ribonuclease A fam | 0 | 0 | 0 | 0 | 0 | 0 |
| 31445 | ENSMUSG00000091191 | 2.170383509 | 0.273933098 | 1.677871109 | 0.163262301 | 0.870311915 | NA | Gm17334 | predicted gene, 17334 [Source:MGI Symb | 4.8186 | 0 | 1.0062 | 3.8559 | 0 | 3.3415 |
| 31446 | ENSMUSG00000091192 | 1.803045281 | 1.064964002 | 1.800613135 | 0.591445203 | 0.55422216 | NA | Sardhos | sarcosine dehydrogenase, opposite stranc | 2.4093 | 1.0279 | 0 | 1.928 | 5.4531 | 0 |
| 31447 | ENSMUSG00000091193 | 0 | NA | NA | NA | NA | NA | Gm17333 | predicted gene, 17333 [Source:MGI Symb | 0 | 0 | 0 | 0 | 0 | 0 |
| 31448 | ENSMUSG00000091194 | 0.17131964 | -0.796629176 | 4.080472857 | -0.195229623 | 0.84521319 | NA | Gm19840 | predicted gene, 19840 [Source:MGI Symb | 0 | 1.0279 | 0 | 0 | 0 | 0 |
| 31449 | ENSMUSG00000091195 | 0 | NA | NA | NA | NA | NA | Gm17332 | predicted gene, 17332 [Source:MGI Symb | 0 | 0 | 0 | 0 | 0 | 0 |
| 31450 | ENSMUSG00000091198 | 0.160662974 | 1.126924175 | 4.080472857 | 0.276174898 | 0.782413736 | NA | Gm5218 | predicted gene 5218 [Source:MGI Symbol | 0 | 0 | 0 | 0.964 | 0 | 0 |
| 31451 | ENSMUSG00000091199 | 1.631137714 | -0.432376648 | 1.829183262 | -0.236376889 | 0.813140228 | NA | Gm2619 | predicted gene 2619 [Source:MGI Symbol | 1.6062 | 0 | 4.025 | 1.928 | 0 | 2.2277 |
| 31452 | ENSMUSG00000091204 | 0 | NA | NA | NA | NA | NA | Gm7271 | predicted gene 7271 [Source:MGI Symbol | 0 | 0 | 0 | 0 | 0 | 0 |
| 31453 | ENSMUSG00000091205 | 0 | NA | NA | NA | NA | NA | Vmn2r71 | vomeronasal 2, receptor 71 [Source:MGI ! | 0 | 0 | 0 | 0 | 0 | 0 |
| 31454 | ENSMUSG00000091206 | 0 | NA | NA | NA | NA | NA | Vmn2r91 | vomeronasal 2, receptor 91 [Source:MGI ! | 0 | 0 | 0 | 0 | 0 | 0 |
| 31455 | ENSMUSG00000091212 | 0 | NA | NA | NA | NA | NA | Krtap11-1 | keratin associated protein 11-1 [Source:M | 0 | 0 | 0 | 0 | 0 | 0 |
| 31456 | ENSMUSG00000091215 | 0 | NA | NA | NA | NA | NA | Pate1 | prostate and testis expressed 1 [Source:M | 0 | 0 | 0 | 0 | 0 | 0 |
| 31457 | ENSMUSG00000091216 | 0 | NA | NA | NA | NA | NA | Gm17208 | predicted gene 17208 [Source:MGI Symbo | 0 | 0 | 0 | 0 | 0 | 0 |
| 31458 | ENSMUSG00000091221 | 0 | NA | NA | NA | NA | NA | Gm3084 | predicted gene 3084 [Source:MGI Symbol | 0 | 0 | 0 | 0 | 0 | 0 |
| 31459 | ENSMUSG00000091223 | 0.874403588 | -0.441200908 | 2.817192576 | -0.156610134 | 0.875552115 | NA | Gm8775 | predicted gene 8775 [Source:MGI Symbol | 0 | 0 | 3.0187 | 0 | 0 | 2.2277 |
| 31460 | ENSMUSG00000091227 | 0 | NA | NA | NA | NA | NA | Gm3755 | predicted gene 3755 [Source:MGI Symbol | 0 | 0 | 0 | 0 | 0 | 0 |
| 31461 | ENSMUSG00000091228 | 0 | NA | NA | NA | NA | NA | Gm20390 | predicted gene 20390 [Source:MGI Symbo | 0 | 0 | 0 | 0 | 0 | 0 |
| 31462 | ENSMUSG00000091230 | 0 | NA | NA | NA | NA | NA | H1f11-ps | H1.11 linker histone, pseudogene [Source | 0 | 0 | 0 | 0 | 0 | 0 |
| 31463 | ENSMUSG00000091233 | 0 | NA | NA | NA | NA | NA | Gm6434 | predicted gene 6434 [Source:MGI Symbol | 0 | 0 | 0 | 0 | 0 | 0 |
| 31464 | ENSMUSG00000091237 | 0 | NA | NA | NA | NA | NA | Gm17114 | predicted gene 17114 [Source:MGI Symbo | 0 | 0 | 0 | 0 | 0 | 0 |
| 31465 | ENSMUSG00000091238 | 0.167707431 | -0.796629176 | 4.080472857 | -0.195229623 | 0.84521319 | NA | Gm17103 | predicted gene 17103 [Source:MGI Symbo | 0 | 0 | 1.0062 | 0 | 0 | 0 |
| 31466 | ENSMUSG00000091239 | 0 | NA | NA | NA | NA | NA | Vmn2r76 | vomeronasal 2, receptor 76 [Source:MGI ! | 0 | 0 | 0 | 0 | 0 | 0 |
| 31467 | ENSMUSG00000091243 | 4.330422876 | -0.329346249 | 1.05243903 | -0.312936179 | 0.75432915 | NA | Vgll3 | vestigial like family member 3 [Source:MG | 2.4093 | 3.0838 | 9.0562 | 4.8199 | 3.2719 | 3.3415 |
| 31468 | ENSMUSG00000091248 | 0 | NA | NA | NA | NA | NA | Pate8 | prostate and testis expressed 8 [Source:M | 0 | 0 | 0 | 0 | 0 | 0 |

| | | | | | | | | | | | | | | | |
|---|---|---|---|---|---|---|---|---|---|---|---|---|---|---|---|
| 31469 | ENSMUSG00000091255 | 0 | NA | NA | NA | NA | NA | Speer4e | spermatogenesis associated glutamate (E) | 0 | 0 | 0 | 0 | 0 | 0 |
| 31470 | ENSMUSG00000091259 | 0 | NA | NA | NA | NA | NA | Vmn2r110 | vomeronasal 2, receptor 110 [Source:MGI | 0 | 0 | 0 | 0 | 0 | 0 |
| 31471 | ENSMUSG00000091260 | 0 | NA | NA | NA | NA | NA | Vmn2r19 | vomeronasal 2, receptor 19 [Source:MGI S | 0 | 0 | 0 | 0 | 0 | 0 |
| 31472 | ENSMUSG00000091264 | 493.2681892 | 0.042779796 | 0.143244845 | 0.29864807 | 0.765208584 | 0.954980327 | Smim13 | small integral membrane protein 13 [Sour | 453.75 | 437.89 | 566.52 | 492.59 | 484.23 | 524.62 |
| 31473 | ENSMUSG00000091268 | 0 | NA | NA | NA | NA | NA | Igkv17-134 | immunoglobulin kappa chain variable 17-1 | 0 | 0 | 0 | 0 | 0 | 0 |
| 31474 | ENSMUSG00000091269 | 0 | NA | NA | NA | NA | NA | Gm6682 | predicted gene 6682 [Source:MGI Symbol | 0 | 0 | 0 | 0 | 0 | 0 |
| 31475 | ENSMUSG00000091272 | 0 | NA | NA | NA | NA | NA | Gm17641 | predicted gene, 17641 [Source:MGI Symb | 0 | 0 | 0 | 0 | 0 | 0 |
| 31476 | ENSMUSG00000091273 | 0 | NA | NA | NA | NA | NA | Gm6673 | predicted gene 6673 [Source:MGI Symbol | 0 | 0 | 0 | 0 | 0 | 0 |
| 31477 | ENSMUSG00000091275 | 0 | NA | NA | NA | NA | NA | Gm3248 | predicted gene 3248 [Source:MGI Symbol | 0 | 0 | 0 | 0 | 0 | 0 |
| 31478 | ENSMUSG00000091277 | 1.636320613 | 0.253984298 | 1.705583507 | 0.148913435 | 0.88162194 | NA | Gm1818 | predicted gene 1818 [Source:MGI Symbol | 2.4093 | 0 | 2.0125 | 0.964 | 1.0906 | 3.3415 |
| 31479 | ENSMUSG00000091280 | 0.185640647 | 1.126924175 | 4.080472857 | 0.276174898 | 0.782413736 | NA | Gm17032 | predicted gene 17032 [Source:MGI Symbo | 0 | 0 | 0 | 0 | 0 | 1.1138 |
| 31480 | ENSMUSG00000091281 | 0 | NA | NA | NA | NA | NA | Gm8065 | predicted gene 8065 [Source:MGI Symbol | 0 | 0 | 0 | 0 | 0 | 0 |
| 31481 | ENSMUSG00000091283 | 0 | NA | NA | NA | NA | NA | Gm17234 | predicted gene 17234 [Source:MGI Symbo | 0 | 0 | 0 | 0 | 0 | 0 |
| 31482 | ENSMUSG00000091285 | 2.799458438 | -0.037581159 | 1.292058275 | -0.029086272 | 0.976795784 | NA | Gm17430 | predicted gene, 17430 [Source:MGI Symb | 2.4093 | 5.1396 | 1.0062 | 3.8559 | 3.2719 | 1.1138 |
| 31483 | ENSMUSG00000091286 | 0 | NA | NA | NA | NA | NA | Gm3111 | predicted gene 3111 [Source:MGI Symbol | 0 | 0 | 0 | 0 | 0 | 0 |
| 31484 | ENSMUSG00000091288 | 0 | NA | NA | NA | NA | NA | Gm17075 | predicted gene 17075 [Source:MGI Symbo | 0 | 0 | 0 | 0 | 0 | 0 |
| 31485 | ENSMUSG00000091289 | 0 | NA | NA | NA | NA | NA | Gm17051 | predicted gene 17051 [Source:MGI Symbo | 0 | 0 | 0 | 0 | 0 | 0 |
| 31486 | ENSMUSG00000091290 | 0 | NA | NA | NA | NA | NA | Gm3275 | predicted gene 3275 [Source:MGI Symbol | 0 | 0 | 0 | 0 | 0 | 0 |
| 31487 | ENSMUSG00000091292 | 0 | NA | NA | NA | NA | NA | Gm3104 | predicted gene 3104 [Source:MGI Symbol | 0 | 0 | 0 | 0 | 0 | 0 |
| 31488 | ENSMUSG00000091293 | 0 | NA | NA | NA | NA | NA | Gm6047 | predicted gene 6047 [Source:MGI Symbol | 0 | 0 | 0 | 0 | 0 | 0 |
| 31489 | ENSMUSG00000091296 | 0 | NA | NA | NA | NA | NA | Gm4181 | predicted gene 4181 [Source:MGI Symbol | 0 | 0 | 0 | 0 | 0 | 0 |
| 31490 | ENSMUSG00000091297 | 0 | NA | NA | NA | NA | NA | Gm8439 | predicted gene 8439 [Source:MGI Symbol | 0 | 0 | 0 | 0 | 0 | 0 |
| 31491 | ENSMUSG00000091304 | 0 | NA | NA | NA | NA | NA | Speer6-ps1 | spermatogenesis associated glutamate (E) | 0 | 0 | 0 | 0 | 0 | 0 |
| 31492 | ENSMUSG00000091305 | 0 | NA | NA | NA | NA | NA | Gm17100 | predicted gene 17100 [Source:MGI Symbo | 0 | 0 | 0 | 0 | 0 | 0 |
| 31493 | ENSMUSG00000091306 | 0 | NA | NA | NA | NA | NA | Gm17606 | predicted gene, 17606 [Source:MGI Symb | 0 | 0 | 0 | 0 | 0 | 0 |
| 31494 | ENSMUSG00000091307 | 1.095993291 | 1.564303563 | 2.37013612 | 0.66000579 | 0.509250114 | NA | Gm8895 | predicted gene 8895 [Source:MGI Symbol | 1.6062 | 0 | 0 | 3.8559 | 0 | 1.1138 |
| 31495 | ENSMUSG00000091311 | 0 | NA | NA | NA | NA | NA | Spata31d1b | spermatogenesis associated 31 subfamily | 0 | 0 | 0 | 0 | 0 | 0 |
| 31496 | ENSMUSG00000091312 | 0 | NA | NA | NA | NA | NA | Gm17490 | predicted gene, 17490 [Source:MGI Symb | 0 | 0 | 0 | 0 | 0 | 0 |
| 31497 | ENSMUSG00000091313 | 0.649696354 | 1.537929172 | 3.118369374 | 0.493183773 | 0.621882743 | NA | Fth-ps2 | ferritin heavy chain, pseudogene 2 [Sourc | 0 | 0 | 1.0062 | 2.8919 | 0 | 0 |
| 31498 | ENSMUSG00000091315 | 0 | NA | NA | NA | NA | NA | Gm17214 | predicted gene 17214 [Source:MGI Symbo | 0 | 0 | 0 | 0 | 0 | 0 |
| 31499 | ENSMUSG00000091318 | 0 | NA | NA | NA | NA | NA | Gm5415 | predicted gene 5415 [Source:MGI Symbol | 0 | 0 | 0 | 0 | 0 | 0 |
| 31500 | ENSMUSG00000091325 | 0 | NA | NA | NA | NA | NA | Gm3760 | predicted gene 3760 [Source:MGI Symbol | 0 | 0 | 0 | 0 | 0 | 0 |
| 31501 | ENSMUSG00000091326 | 0 | NA | NA | NA | NA | NA | Traj54 | T cell receptor alpha joining 54 [Source:M | 0 | 0 | 0 | 0 | 0 | 0 |
| 31502 | ENSMUSG00000091329 | 5.042353537 | -1.104138682 | 0.984155166 | -1.121915243 | 0.261898479 | NA | 1700112D23 | RIKEN cDNA 1700112D23 gene [Source:M | 6.4248 | 4.1117 | 10.062 | 1.928 | 3.2719 | 4.4554 |
| 31503 | ENSMUSG00000091336 | 0 | NA | NA | NA | NA | NA | Gm17157 | predicted gene 17157 [Source:MGI Symbo | 0 | 0 | 0 | 0 | 0 | 0 |
| 31504 | ENSMUSG00000091337 | 66.64191957 | -0.042825127 | 0.294880121 | -0.14522894 | 0.884530118 | 0.981795936 | Eid1 | EP300 interacting inhibitor of differentiati | 68.264 | 68.87 | 65.406 | 46.271 | 73.071 | 77.969 |
| 31505 | ENSMUSG00000091345 | 1.067268653 | 0.134458805 | 2.381868772 | 0.056450971 | 0.954982553 | NA | Col6a5 | collagen, type VI, alpha 5 [Source:MGI Syr | 0 | 2.0558 | 1.0062 | 0 | 0 | 3.3415 |
| 31506 | ENSMUSG00000091347 | 0 | NA | NA | NA | NA | NA | Gm10772 | predicted gene 10772 [Source:MGI Symbo | 0 | 0 | 0 | 0 | 0 | 0 |
| 31507 | ENSMUSG00000091350 | 0 | NA | NA | NA | NA | NA | Vmn2r92 | vomeronasal 2, receptor 92 [Source:MGI S | 0 | 0 | 0 | 0 | 0 | 0 |
| 31508 | ENSMUSG00000091351 | 0 | NA | NA | NA | NA | NA | Gm17135 | predicted gene 17135 [Source:MGI Symbo | 0 | 0 | 0 | 0 | 0 | 0 |
| 31509 | ENSMUSG00000091355 | 0 | NA | NA | NA | NA | NA | Gm17142 | predicted gene 17142 [Source:MGI Symbo | 0 | 0 | 0 | 0 | 0 | 0 |
| 31510 | ENSMUSG00000091358 | 0 | NA | NA | NA | NA | NA | Speer5-ps1 | spermatogenesis associated glutamate (E) | 0 | 0 | 0 | 0 | 0 | 0 |
| 31511 | ENSMUSG00000091361 | 0 | NA | NA | NA | NA | NA | Gm5277 | predicted gene 5277 [Source:MGI Symbol | 0 | 0 | 0 | 0 | 0 | 0 |
| 31512 | ENSMUSG00000091363 | 0.513958919 | -2.394274821 | 3.756096013 | -0.637437066 | 0.523840191 | NA | Gm7701 | predicted gene 7701 [Source:MGI Symbol | 0 | 3.0838 | 0 | 0 | 0 | 0 |
| 31513 | ENSMUSG00000091366 | 10.67812066 | 0.843689723 | 0.721119413 | 1.169972279 | 0.242012125 | 0.691791181 | Gm17040 | predicted gene 17040 [Source:MGI Symbo | 4.8186 | 5.1396 | 13.081 | 14.46 | 7.6343 | 18.935 |
| 31514 | ENSMUSG00000091367 | 0 | NA | NA | NA | NA | NA | Gm17711 | predicted gene, 17711 [Source:MGI Symb | 0 | 0 | 0 | 0 | 0 | 0 |
| 31515 | ENSMUSG00000091368 | 0 | NA | NA | NA | NA | NA | Gm17085 | predicted gene 17085 [Source:MGI Symbo | 0 | 0 | 0 | 0 | 0 | 0 |
| 31516 | ENSMUSG00000091370 | 9.307318782 | 0.676849498 | 0.745222096 | 0.908252053 | 0.363745068 | 0.802534854 | 5730435O14 | RIKEN cDNA 5730435O14 gene [Source:M | 7.2279 | 4.1117 | 10.062 | 5.7839 | 14.178 | 14.48 |
| 31517 | ENSMUSG00000091373 | 0 | NA | NA | NA | NA | NA | Gm8810 | predicted gene 8810 [Source:MGI Symbol | 0 | 0 | 0 | 0 | 0 | 0 |
| 31518 | ENSMUSG00000091375 | 0 | NA | NA | NA | NA | NA | Vmn2r15 | vomeronasal 2, receptor 15 [Source:MGI S | 0 | 0 | 0 | 0 | 0 | 0 |
| 31519 | ENSMUSG00000091376 | 0 | NA | NA | NA | NA | NA | Aadacl2 | arylacetamide deacetylase like 2 [Source:I | 0 | 0 | 0 | 0 | 0 | 0 |
| 31520 | ENSMUSG00000091378 | 0 | NA | NA | NA | NA | NA | Gm4219 | predicted gene 4219 [Source:MGI Symbol | 0 | 0 | 0 | 0 | 0 | 0 |
| 31521 | ENSMUSG00000091381 | 0 | NA | NA | NA | NA | NA | Vmn2r83 | vomeronasal 2, receptor 83 [Source:MGI S | 0 | 0 | 0 | 0 | 0 | 0 |
| 31522 | ENSMUSG00000091382 | 0 | NA | NA | NA | NA | NA | Vmn1r18 | vomeronasal 1 receptor 18 [Source:MGI S | 0 | 0 | 0 | 0 | 0 | 0 |
| 31523 | ENSMUSG00000091383 | 0.160662974 | 1.126924175 | 4.080472857 | 0.276174898 | 0.782413736 | NA | H2af-ps2 | H2A histone family, pseudogene 2 [Source | 0 | 0 | 0 | 0.964 | 0 | 0 |
| 31524 | ENSMUSG00000091386 | 0 | NA | NA | NA | NA | NA | Olfr443-ps1 | olfactory receptor 443, pseudogene 1 [Sou | 0 | 0 | 0 | 0 | 0 | 0 |

| | | | | | | | | | | | | | | | |
|---|---|---|---|---|---|---|---|---|---|---|---|---|---|---|---|
| 31525 | ENSMUSG00000091387 | 106.5854002 | 0.576765168 | 0.25722525 | 2.242257193 | 0.024944755 | 0.202469658 | Gcnt4 | glucosaminyl (N-acetyl) transferase 4, cor | 64.248 | 88.401 | 104.65 | 121.46 | 151.6 | 109.16 |
| 31526 | ENSMUSG00000091390 | 5.206949479 | 0.836723256 | 1.07655123 | 0.777225674 | 0.437025637 | 0.846852749 | Gm17168 | predicted gene 17168 [Source:MGI Symbo | 3.2124 | 2.0558 | 6.0375 | 14.46 | 4.3625 | 1.1138 |
| 31527 | ENSMUSG00000091393 | 1.143912632 | 2.595556111 | 2.186080874 | 1.187310196 | 0.235105288 | NA | 5330438I03F | RIKEN cDNA 5330438I03 gene [Source:MC | 0.8031 | 0 | 0 | 3.8559 | 1.0906 | 1.1138 |
| 31528 | ENSMUSG00000091396 | 0 | NA | NA | NA | NA | NA | Spanxn4 | SPANX family, member N4 [Source:MGI Sy | 0 | 0 | 0 | 0 | 0 | 0 |
| 31529 | ENSMUSG00000091400 | 0 | NA | NA | NA | NA | NA | Gm6356 | predicted gene 6356 [Source:MGI Symbol | 0 | 0 | 0 | 0 | 0 | 0 |
| 31530 | ENSMUSG00000091401 | 0 | NA | NA | NA | NA | NA | Gm9252 | predicted gene 9252 [Source:MGI Symbol | 0 | 0 | 0 | 0 | 0 | 0 |
| 31531 | ENSMUSG00000091402 | 0 | NA | NA | NA | NA | NA | Rd3l | retinal degeneration 3-like [Source:MGI Sy | 0 | 0 | 0 | 0 | 0 | 0 |
| 31532 | ENSMUSG00000091404 | 0 | NA | NA | NA | NA | NA | Gm3140 | predicted gene 3140 [Source:MGI Symbol | 0 | 0 | 0 | 0 | 0 | 0 |
| 31533 | ENSMUSG00000091405 | 3.320018144 | 1.277990018 | 1.250200182 | 1.022228309 | 0.306672855 | NA | H4c14 | H4 clustered histone 14 [Source:MGI Sym | 0.8031 | 1.0279 | 4.025 | 1.928 | 5.4531 | 6.6831 |
| 31534 | ENSMUSG00000091407 | 0 | NA | NA | NA | NA | NA | Vmn2r117 | vomeronasal 2, receptor 117 [Source:MGI | 0 | 0 | 0 | 0 | 0 | 0 |
| 31535 | ENSMUSG00000091408 | 0.342639279 | -1.805976596 | 4.028547714 | -0.448294702 | 0.653940524 | NA | Gm6728 | predicted gene 6728 [Source:MGI Symbol | 0 | 2.0558 | 0 | 0 | 0 | 0 |
| 31536 | ENSMUSG00000091410 | 0 | NA | NA | NA | NA | NA | Gm17150 | predicted gene 17150 [Source:MGI Symbo | 0 | 0 | 0 | 0 | 0 | 0 |
| 31537 | ENSMUSG00000091411 | 0 | NA | NA | NA | NA | NA | Gm5916 | predicted gene 5916 [Source:MGI Symbol | 0 | 0 | 0 | 0 | 0 | 0 |
| 31538 | ENSMUSG00000091412 | 0 | NA | NA | NA | NA | NA | Gm2895 | predicted gene 2895 [Source:MGI Symbol | 0 | 0 | 0 | 0 | 0 | 0 |
| 31539 | ENSMUSG00000091415 | 0 | NA | NA | NA | NA | NA | Ak9 | adenylate kinase 9 [Source:MGI Symbol;A | 0 | 0 | 0 | 0 | 0 | 0 |
| 31540 | ENSMUSG00000091416 | 0 | NA | NA | NA | NA | NA | Gm6327 | predicted gene 6327 [Source:MGI Symbol | 0 | 0 | 0 | 0 | 0 | 0 |
| 31541 | ENSMUSG00000091418 | 0 | NA | NA | NA | NA | NA | Gm3164 | predicted gene 3164 [Source:MGI Symbol | 0 | 0 | 0 | 0 | 0 | 0 |
| 31542 | ENSMUSG00000091419 | 0 | NA | NA | NA | NA | NA | Gm17450 | predicted gene, 17450 [Source:MGI Symb | 0 | 0 | 0 | 0 | 0 | 0 |
| 31543 | ENSMUSG00000091421 | 3.256749528 | -0.401098122 | 1.186672456 | -0.338002386 | 0.735361389 | NA | Gm4202 | predicted gene 4202 [Source:MGI Symbol | 4.0155 | 2.0558 | 5.0312 | 2.8919 | 1.0906 | 4.4554 |
| 31544 | ENSMUSG00000091422 | 0 | NA | NA | NA | NA | NA | Gm6455 | predicted gene 6455 [Source:MGI Symbol | 0 | 0 | 0 | 0 | 0 | 0 |
| 31545 | ENSMUSG00000091423 | 0 | NA | NA | NA | NA | NA | Gm17509 | predicted gene, 17509 [Source:MGI Symb | 0 | 0 | 0 | 0 | 0 | 0 |
| 31546 | ENSMUSG00000091425 | 0 | NA | NA | NA | NA | NA | Gm17118 | predicted gene 17118 [Source:MGI Symbo | 0 | 0 | 0 | 0 | 0 | 0 |
| 31547 | ENSMUSG00000091426 | 0 | NA | NA | NA | NA | NA | Gm17228 | predicted gene 17228 [Source:MGI Symbo | 0 | 0 | 0 | 0 | 0 | 0 |
| 31548 | ENSMUSG00000091429 | 0 | NA | NA | NA | NA | NA | Gm17093 | predicted gene 17093 [Source:MGI Symbo | 0 | 0 | 0 | 0 | 0 | 0 |
| 31549 | ENSMUSG00000091430 | 1.170493139 | 0.44043163 | 2.105443608 | 0.209187094 | 0.834302187 | NA | Platr19 | pluripotency associated transcript 19 [Sou | 0 | 1.0279 | 2.0125 | 2.8919 | 1.0906 | 0 |
| 31550 | ENSMUSG00000091435 | 0 | NA | NA | NA | NA | NA | Vmn1r115 | vomeronasal 1 receptor 115 [Source:MGI | 0 | 0 | 0 | 0 | 0 | 0 |
| 31551 | ENSMUSG00000091438 | 8.928305581 | -0.542124267 | 0.735729199 | -0.736853 | 0.461211748 | 0.860071824 | Gm17088 | predicted gene 17088 [Source:MGI Symbo | 14.456 | 7.1954 | 10.062 | 8.6758 | 8.7249 | 4.4554 |
| 31552 | ENSMUSG00000091441 | 0 | NA | NA | NA | NA | NA | Gm17387 | predicted gene, 17387 [Source:MGI Symb | 0 | 0 | 0 | 0 | 0 | 0 |
| 31553 | ENSMUSG00000091442 | 0 | NA | NA | NA | NA | NA | Gm17031 | predicted gene 17031 [Source:MGI Symbo | 0 | 0 | 0 | 0 | 0 | 0 |
| 31554 | ENSMUSG00000091443 | 6.331264565 | 0.16257958 | 0.968574609 | 0.167854472 | 0.866697768 | 0.977930657 | Gm17023 | predicted gene 17023 [Source:MGI Symbo | 0.8031 | 8.2233 | 9.0562 | 6.7478 | 9.8156 | 3.3415 |
| 31555 | ENSMUSG00000091449 | 3.621296108 | 1.751882552 | 1.304897794 | 1.342543883 | 0.179419712 | NA | Gm10269 | predicted gene 10269 [Source:MGI Symbo | 0 | 4.1117 | 1.0062 | 6.7478 | 7.6343 | 2.2277 |
| 31556 | ENSMUSG00000091450 | 0 | NA | NA | NA | NA | NA | Vmn2r11 | vomeronasal 2, receptor 11 [Source:MGI S | 0 | 0 | 0 | 0 | 0 | 0 |
| 31557 | ENSMUSG00000091453 | 0 | NA | NA | NA | NA | NA | Gm6541 | predicted gene 6541 [Source:MGI Symbol | 0 | 0 | 0 | 0 | 0 | 0 |
| 31558 | ENSMUSG00000091455 | 0 | NA | NA | NA | NA | NA | Otogl | otogelin-like [Source:MGI Symbol;Acc:MG | 0 | 0 | 0 | 0 | 0 | 0 |
| 31559 | ENSMUSG00000091457 | 0 | NA | NA | NA | NA | NA | Gm17171 | predicted gene 17171 [Source:MGI Symbo | 0 | 0 | 0 | 0 | 0 | 0 |
| 31560 | ENSMUSG00000091459 | 0 | NA | NA | NA | NA | NA | Gm17054 | predicted gene 17054 [Source:MGI Symbo | 0 | 0 | 0 | 0 | 0 | 0 |
| 31561 | ENSMUSG00000091460 | 0 | NA | NA | NA | NA | NA | Gm7808 | predicted pseudogene 7808 [Source:MGI | 0 | 0 | 0 | 0 | 0 | 0 |
| 31562 | ENSMUSG00000091462 | 0 | NA | NA | NA | NA | NA | Gm17084 | predicted gene 17084 [Source:MGI Symbo | 0 | 0 | 0 | 0 | 0 | 0 |
| 31563 | ENSMUSG00000091466 | 0 | NA | NA | NA | NA | NA | Gm17183 | predicted gene 17183 [Source:MGI Symbo | 0 | 0 | 0 | 0 | 0 | 0 |
| 31564 | ENSMUSG00000091468 | 0 | NA | NA | NA | NA | NA | Vmn2r82 | vomeronasal 2, receptor 82 [Source:MGI S | 0 | 0 | 0 | 0 | 0 | 0 |
| 31565 | ENSMUSG00000091470 | 0 | NA | NA | NA | NA | NA | Gm17159 | predicted gene 17159 [Source:MGI Symbo | 0 | 0 | 0 | 0 | 0 | 0 |
| 31566 | ENSMUSG00000091471 | 0 | NA | NA | NA | NA | NA | Gm20538 | predicted gene 20538 [Source:MGI Symbo | 0 | 0 | 0 | 0 | 0 | 0 |
| 31567 | ENSMUSG00000091472 | 0.17131964 | -0.796629176 | 4.080472857 | -0.195229623 | 0.84521319 | NA | Gm3739 | predicted gene 3739 [Source:MGI Symbol | 0 | 1.0279 | 0 | 0 | 0 | 0 |
| 31568 | ENSMUSG00000091474 | 237.2383238 | 0.285338414 | 0.18196612 | 1.568085392 | 0.116861205 | 0.50225288 | 2610021A01 | RIKEN cDNA 2610021A01 gene [Source:M | 218.44 | 198.39 | 224.39 | 234.25 | 316.28 | 231.68 |
| 31569 | ENSMUSG00000091475 | 0.167707431 | -0.796629176 | 4.080472857 | -0.195229623 | 0.84521319 | NA | Cerox1 | cytoplasmic endogenous regulator of oxid | 0 | 0 | 1.0062 | 0 | 0 | 0 |
| 31570 | ENSMUSG00000091476 | 3.392123971 | -0.864602952 | 1.170019993 | -0.738964254 | 0.459928703 | NA | Catspere2 | cation channel sperm associated auxiliary | 4.0155 | 6.1675 | 3.0187 | 3.8559 | 2.1812 | 1.1138 |
| 31571 | ENSMUSG00000091477 | 0 | NA | NA | NA | NA | NA | Gm5799 | predicted gene 5799 [Source:MGI Symbol | 0 | 0 | 0 | 0 | 0 | 0 |
| 31572 | ENSMUSG00000091478 | 12.81823306 | -0.304826632 | 0.595081359 | -0.512243625 | 0.608480515 | 0.916376001 | Gm10039 | predicted pseudogene 10039 [Source:MG | 16.062 | 13.363 | 13.081 | 13.496 | 11.997 | 8.9108 |
| 31573 | ENSMUSG00000091479 | 0 | NA | NA | NA | NA | NA | Gm17035 | predicted gene 17035 [Source:MGI Symbo | 0 | 0 | 0 | 0 | 0 | 0 |
| 31574 | ENSMUSG00000091481 | 0 | NA | NA | NA | NA | NA | Gm17154 | predicted gene 17154 [Source:MGI Symbo | 0 | 0 | 0 | 0 | 0 | 0 |
| 31575 | ENSMUSG00000091483 | 0 | NA | NA | NA | NA | NA | Gm8715 | predicted gene 8715 [Source:MGI Symbol | 0 | 0 | 0 | 0 | 0 | 0 |
| 31576 | ENSMUSG00000091486 | 0 | NA | NA | NA | NA | NA | Gm7953 | predicted gene 7953 [Source:MGI Symbol | 0 | 0 | 0 | 0 | 0 | 0 |
| 31577 | ENSMUSG00000091487 | 0 | NA | NA | NA | NA | NA | Gm6586 | predicted gene 6586 [Source:MGI Symbol | 0 | 0 | 0 | 0 | 0 | 0 |
| 31578 | ENSMUSG00000091490 | 0 | NA | NA | NA | NA | NA | Triml2 | tripartite motif family-like 2 [Source:MGI S | 0 | 0 | 0 | 0 | 0 | 0 |
| 31579 | ENSMUSG00000091491 | 0.524408994 | -0.837196744 | 3.377823722 | -0.247850928 | 0.804249749 | NA | Vmn2r97 | vomeronasal 2, receptor 97 [Source:MGI S | 0 | 2.0558 | 0 | 0 | 1.0906 | 0 |
| 31580 | ENSMUSG00000091492 | 8.81545821 | -0.143490874 | 0.717379097 | -0.200020985 | 0.841464169 | 0.971854408 | Gm17025 | predicted gene 17025 [Source:MGI Symbo | 10.44 | 9.2513 | 8.05 | 8.6758 | 10.906 | 5.5692 |

| | | | | | | | | | | | | | | | |
|---|---|---|---|---|---|---|---|---|---|---|---|---|---|---|---|
| 31581 | ENSMUSG00000091494 | 0 | NA | NA | NA | NA | NA | Gm3269 | predicted gene 3269 [Source:MGI Symbol | 0 | 0 | 0 | 0 | 0 | 0 |
| 31582 | ENSMUSG00000091496 | 0 | NA | NA | NA | NA | NA | Gm8398 | predicted gene 8398 [Source:MGI Symbol | 0 | 0 | 0 | 0 | 0 | 0 |
| 31583 | ENSMUSG00000091497 | 0 | NA | NA | NA | NA | NA | Mageb11 | MAGE family member B11 [Source:MGI Sy | 0 | 0 | 0 | 0 | 0 | 0 |
| 31584 | ENSMUSG00000091498 | 343.0826821 | -0.319812085 | 0.148716613 | -2.150479882 | 0.031517276 | 0.236302595 | Mpc1-ps | mitochondrial pyruvate carrier 1, pseudog | 367.02 | 379.3 | 396.46 | 281.48 | 344.64 | 289.6 |
| 31585 | ENSMUSG00000091504 | 0 | NA | NA | NA | NA | NA | Vmn2r118 | vomeronasal 2, receptor 118 [Source:MGI | 0 | 0 | 0 | 0 | 0 | 0 |
| 31586 | ENSMUSG00000091507 | 1.186126113 | 2.643610322 | 2.135807954 | 1.237756568 | 0.215806337 | NA | Gm17194 | predicted gene 17194 [Source:MGI Symbo | 0.8031 | 0 | 0 | 1.928 | 3.2719 | 1.1138 |
| 31587 | ENSMUSG00000091509 | 63.79148834 | 0.186578558 | 0.34201966 | 0.545519981 | 0.585395942 | 0.909202452 | Gm17066 | predicted gene 17066 [Source:MGI Symbo | 83.523 | 60.647 | 34.212 | 58.803 | 68.709 | 76.855 |
| 31588 | ENSMUSG00000091510 | 0 | NA | NA | NA | NA | NA | Mtag2 | metastasis associated gene 2 [Source:MG | 0 | 0 | 0 | 0 | 0 | 0 |
| 31589 | ENSMUSG00000091511 | 0 | NA | NA | NA | NA | NA | Vmn2r87 | vomeronasal 2, receptor 87 [Source:MGI S | 0 | 0 | 0 | 0 | 0 | 0 |
| 31590 | ENSMUSG00000091512 | 254.0703838 | 0.111648963 | 0.166507874 | 0.670532632 | 0.502518311 | 0.881220881 | Lamtor3 | late endosomal/lysosomal adaptor, MAPK | 265.02 | 255.95 | 211.31 | 266.06 | 245.39 | 280.69 |
| 31591 | ENSMUSG00000091514 | 5.462530835 | 0.980823729 | 1.111235944 | 0.882642192 | 0.377429625 | 0.81030298 | Gm17484 | predicted gene, 17484 [Source:MGI Symb | 0 | 3.0838 | 8.05 | 2.8919 | 8.7249 | 10.025 |
| 31592 | ENSMUSG00000091515 | 18.53014731 | 0.15177801 | 0.495232311 | 0.306478408 | 0.759240417 | 0.95402891 | Zfp422-ps | zinc finger protein 422, pseudogene [Sour | 20.078 | 16.447 | 16.1 | 22.171 | 17.45 | 18.935 |
| 31593 | ENSMUSG00000091519 | 0 | NA | NA | NA | NA | NA | Skor2 | SKI family transcriptional corepressor 2 [S | 0 | 0 | 0 | 0 | 0 | 0 |
| 31594 | ENSMUSG00000091523 | 0 | NA | NA | NA | NA | NA | AU040972 | expressed sequence AU040972 [Source:M | 0 | 0 | 0 | 0 | 0 | 0 |
| 31595 | ENSMUSG00000091526 | 0 | NA | NA | NA | NA | NA | BB019430 | expressed sequence BB019430 [Source:M | 0 | 0 | 0 | 0 | 0 | 0 |
| 31596 | ENSMUSG00000091528 | 0 | NA | NA | NA | NA | NA | Vmn2r-ps15 | vomeronasal 2, receptor, pseudogene 158 | 0 | 0 | 0 | 0 | 0 | 0 |
| 31597 | ENSMUSG00000091530 | 5.167089879 | 0.269247272 | 1.159596156 | 0.232190552 | 0.816390016 | 0.967767553 | Cldn20 | claudin 20 [Source:MGI Symbol;Acc:MGI:3 | 8.8341 | 3.0838 | 2.0125 | 4.8199 | 0 | 12.252 |
| 31598 | ENSMUSG00000091531 | 0 | NA | NA | NA | NA | NA | Olfr102 | olfactory receptor 102 [Source:MGI Symb | 0 | 0 | 0 | 0 | 0 | 0 |
| 31599 | ENSMUSG00000091534 | 3.404842145 | 0.066228496 | 1.16366159 | 0.05691388 | 0.954613797 | NA | Gm17195 | predicted gene 17195 [Source:MGI Symbo | 4.8186 | 4.1117 | 1.0062 | 3.8559 | 2.1812 | 4.4554 |
| 31600 | ENSMUSG00000091537 | 797.3719423 | -0.024880966 | 0.110251976 | -0.225673652 | 0.821455252 | 0.969160361 | Tma7 | translational machinery associated 7 [Sou | 806.31 | 810 | 795.94 | 745.15 | 808.15 | 818.68 |
| 31601 | ENSMUSG00000091539 | 0 | NA | NA | NA | NA | NA | Vmn1r238 | vomeronasal 1 receptor, 238 [Source:MGI | 0 | 0 | 0 | 0 | 0 | 0 |
| 31602 | ENSMUSG00000091541 | 0 | NA | NA | NA | NA | NA | Vmn1r9 | vomeronasal 1 receptor 9 [Source:MGI Sy | 0 | 0 | 0 | 0 | 0 | 0 |
| 31603 | ENSMUSG00000091542 | 0.160662974 | 1.126924175 | 4.080472857 | 0.276174898 | 0.782413736 | NA | Gm17167 | predicted gene 17167 [Source:MGI Symbo | 0 | 0 | 0 | 0.964 | 0 | 0 |
| 31604 | ENSMUSG00000091548 | 0 | NA | NA | NA | NA | NA | Gm17572 | predicted gene, 17572 [Source:MGI Symb | 0 | 0 | 0 | 0 | 0 | 0 |
| 31605 | ENSMUSG00000091549 | 142.2333652 | 0.013994813 | 0.217701182 | 0.064284508 | 0.948743689 | 0.990731019 | Gm6548 | predicted gene 6548 [Source:MGI Symbol | 147.77 | 125.41 | 150.94 | 122.43 | 130.87 | 175.99 |
| 31606 | ENSMUSG00000091550 | 0 | NA | NA | NA | NA | NA | Gm6811 | predicted gene 6811 [Source:MGI Symbol | 0 | 0 | 0 | 0 | 0 | 0 |
| 31607 | ENSMUSG00000091552 | 0 | NA | NA | NA | NA | NA | Gm8711 | predicted gene 8711 [Source:MGI Symbol | 0 | 0 | 0 | 0 | 0 | 0 |
| 31608 | ENSMUSG00000091553 | 0.781659665 | -3.010437285 | 2.830777252 | -1.063466679 | 0.287570368 | NA | Serpina3e-ps | serine (or cysteine) peptidase inhibitor, cla | 1.6062 | 3.0838 | 0 | 0 | 0 | 0 |
| 31609 | ENSMUSG00000091556 | 0 | NA | NA | NA | NA | NA | Gm14569 | predicted gene 14569 [Source:MGI Symbo | 0 | 0 | 0 | 0 | 0 | 0 |
| 31610 | ENSMUSG00000091557 | 0 | NA | NA | NA | NA | NA | Gm8584 | predicted gene 8584 [Source:MGI Symbol | 0 | 0 | 0 | 0 | 0 | 0 |
| 31611 | ENSMUSG00000091558 | 0.835104907 | -1.8196403 | 2.6116807 | -0.696731534 | 0.48597082 | NA | Gm17227 | predicted gene 17227 [Source:MGI Symbo | 0 | 1.0279 | 3.0187 | 0.964 | 0 | 0 |
| 31612 | ENSMUSG00000091561 | 1.279049468 | -1.354713058 | 2.2140152 | -0.611880649 | 0.540616725 | NA | Gm6665 | predicted gene 6665 [Source:MGI Symbol | 2.4093 | 3.0838 | 0 | 0 | 2.1812 | 0 |
| 31613 | ENSMUSG00000091563 | 0 | NA | NA | NA | NA | NA | Gm8108 | predicted gene 8108 [Source:MGI Symbol | 0 | 0 | 0 | 0 | 0 | 0 |
| 31614 | ENSMUSG00000091568 | 0 | NA | NA | NA | NA | NA | Gm8206 | predicted gene 8206 [Source:MGI Symbol | 0 | 0 | 0 | 0 | 0 | 0 |
| 31615 | ENSMUSG00000091569 | 0 | NA | NA | NA | NA | NA | Gm8232 | predicted gene 8232 [Source:MGI Symbol | 0 | 0 | 0 | 0 | 0 | 0 |
| 31616 | ENSMUSG00000091570 | 0 | NA | NA | NA | NA | NA | Gm7707 | predicted gene 7707 [Source:MGI Symbol | 0 | 0 | 0 | 0 | 0 | 0 |
| 31617 | ENSMUSG00000091572 | 0 | NA | NA | NA | NA | NA | Vmn2r3 | vomeronasal 2, receptor 3 [Source:MGI Sy | 0 | 0 | 0 | 0 | 0 | 0 |
| 31618 | ENSMUSG00000091573 | 0 | NA | NA | NA | NA | NA | Serpina3d-ps | serine (or cysteine) peptidase inhibitor, cla | 0 | 0 | 0 | 0 | 0 | 0 |
| 31619 | ENSMUSG00000091575 | 9.71046732 | -0.355701269 | 0.798864638 | -0.445258499 | 0.656132956 | 0.92651636 | 2010016I18R | RIKEN cDNA 2010016I18 gene [Source:MG | 10.44 | 12.335 | 10.062 | 15.424 | 1.0906 | 8.9108 |
| 31620 | ENSMUSG00000091577 | 0 | NA | NA | NA | NA | NA | Gm6211 | predicted gene 6211 [Source:MGI Symbol | 0 | 0 | 0 | 0 | 0 | 0 |
| 31621 | ENSMUSG00000091583 | 0 | NA | NA | NA | NA | NA | Gm17033 | predicted gene 17033 [Source:MGI Symbo | 0 | 0 | 0 | 0 | 0 | 0 |
| 31622 | ENSMUSG00000091584 | 0 | NA | NA | NA | NA | NA | Gm8180 | predicted gene 8180 [Source:MGI Symbol | 0 | 0 | 0 | 0 | 0 | 0 |
| 31623 | ENSMUSG00000091585 | 0 | NA | NA | NA | NA | NA | | | 0 | 0 | 0 | 0 | 0 | 0 |
| 31624 | ENSMUSG00000091586 | 223.1990882 | -0.032290155 | 0.240354387 | -0.134343938 | 0.89313061 | 0.982562372 | Cyp4f17 | cytochrome P450, family 4, subfamily f, po | 218.44 | 222.03 | 237.47 | 321.97 | 177.77 | 161.51 |
| 31625 | ENSMUSG00000091587 | 0 | NA | NA | NA | NA | NA | Gm17191 | predicted gene 17191 [Source:MGI Symbo | 0 | 0 | 0 | 0 | 0 | 0 |
| 31626 | ENSMUSG00000091594 | 0.328370405 | 0.16514316 | 4.003149536 | 0.041253308 | 0.967093956 | NA | Gm17067 | predicted gene 17067 [Source:MGI Symbo | 0 | 0 | 1.0062 | 0.964 | 0 | 0 |
| 31627 | ENSMUSG00000091595 | 0 | NA | NA | NA | NA | NA | Gm3344 | predicted gene 3344 [Source:MGI Symbol | 0 | 0 | 0 | 0 | 0 | 0 |
| 31628 | ENSMUSG00000091601 | 0 | NA | NA | NA | NA | NA | Olfr93 | olfactory receptor 93 [Source:MGI Symbo | 0 | 0 | 0 | 0 | 0 | 0 |
| 31629 | ENSMUSG00000091604 | 0 | NA | NA | NA | NA | NA | Gm17349 | predicted gene, 17349 [Source:MGI Symb | 0 | 0 | 0 | 0 | 0 | 0 |
| 31630 | ENSMUSG00000091605 | 0 | NA | NA | NA | NA | NA | Gm8256 | predicted gene 8256 [Source:MGI Symbol | 0 | 0 | 0 | 0 | 0 | 0 |
| 31631 | ENSMUSG00000091606 | 0 | NA | NA | NA | NA | NA | Ighv2-1 | immunoglobulin heavy variable 2-1 [Sourc | 0 | 0 | 0 | 0 | 0 | 0 |
| 31632 | ENSMUSG00000091613 | 0.185640647 | 1.126924175 | 4.080472857 | 0.276174898 | 0.782413736 | NA | Gm17046 | predicted gene 17046 [Source:MGI Symbo | 0 | 0 | 0 | 0 | 0 | 1.1138 |
| 31633 | ENSMUSG00000091614 | 0 | NA | NA | NA | NA | NA | Gm17657 | predicted gene, 17657 [Source:MGI Symb | 0 | 0 | 0 | 0 | 0 | 0 |
| 31634 | ENSMUSG00000091615 | 0 | NA | NA | NA | NA | NA | Gm17044 | predicted gene 17044 [Source:MGI Symbo | 0 | 0 | 0 | 0 | 0 | 0 |
| 31635 | ENSMUSG00000091617 | 0 | NA | NA | NA | NA | NA | Gm3752 | predicted gene 3752 [Source:MGI Symbol | 0 | 0 | 0 | 0 | 0 | 0 |
| 31636 | ENSMUSG00000091618 | 0 | NA | NA | NA | NA | NA | H60c | histocompatibility 60c [Source:MGI Symbo | 0 | 0 | 0 | 0 | 0 | 0 |

| | | | | | | | | | | | | | | | |
|---|---|---|---|---|---|---|---|---|---|---|---|---|---|---|---|
| 31637 | ENSMUSG00000091619 | 0 | NA | NA | NA | NA | NA | NA | NA | 0 | 0 | 0 | 0 | 0 | 0 |
| 31638 | ENSMUSG00000091620 | 0 | NA | NA | NA | NA | NA | Vmn2r23 | vomeronasal 2, receptor 23 [Source:MGI S | 0 | 0 | 0 | 0 | 0 | 0 |
| 31639 | ENSMUSG00000091623 | 5.477358134 | -0.224224715 | 1.054470041 | -0.212642092 | 0.831606138 | 0.970533397 | Gm17092 | predicted gene 17092 [Source:MGI Symbo | 5.6217 | 5.1396 | 7.0437 | 10.604 | 0 | 4.4554 |
| 31640 | ENSMUSG00000091624 | 0 | NA | NA | NA | NA | NA | Vmn2r9 | vomeronasal 2, receptor 9 [Source:MGI Sy | 0 | 0 | 0 | 0 | 0 | 0 |
| 31641 | ENSMUSG00000091625 | 88.41731967 | 0.112454884 | 0.247700746 | 0.453994935 | 0.649832472 | 0.925643431 | Lsm5 | LSM5 homolog, U6 small nuclear RNA and | 98.782 | 81.206 | 74.462 | 89.65 | 86.159 | 100.25 |
| 31642 | ENSMUSG00000091627 | 0 | NA | NA | NA | NA | NA | Gm5929 | predicted gene 5929 [Source:MGI Symbol | 0 | 0 | 0 | 0 | 0 | 0 |
| 31643 | ENSMUSG00000091631 | 0 | NA | NA | NA | NA | NA | Vmn2r95 | vomeronasal 2, receptor 95 [Source:MGI S | 0 | 0 | 0 | 0 | 0 | 0 |
| 31644 | ENSMUSG00000091635 | 0 | NA | NA | NA | NA | NA | Vmn2r13 | vomeronasal 2, receptor 13 [Source:MGI S | 0 | 0 | 0 | 0 | 0 | 0 |
| 31645 | ENSMUSG00000091636 | 0 | NA | NA | NA | NA | NA | Akain1 | A kinase (PRKA) anchor inhibitor 1 [Source | 0 | 0 | 0 | 0 | 0 | 0 |
| 31646 | ENSMUSG00000091638 | 0 | NA | NA | NA | NA | NA | Vmn1r113 | vomeronasal 1 receptor 113 [Source:MGI | 0 | 0 | 0 | 0 | 0 | 0 |
| 31647 | ENSMUSG00000091639 | 2.242996296 | -1.648695868 | 1.533342985 | -1.075229667 | 0.282271916 | NA | Gm3756 | predicted gene 3756 [Source:MGI Symbol | 4.0155 | 4.1117 | 2.0125 | 0 | 1.0906 | 2.2277 |
| 31648 | ENSMUSG00000091648 | 52.60223948 | 0.33050541 | 0.340478475 | 0.970708676 | 0.33169337 | 0.774747658 | C2cd4d | C2 calcium-dependent domain containing | 61.036 | 37.005 | 41.256 | 52.055 | 75.253 | 49.009 |
| 31649 | ENSMUSG00000091649 | 10.10566126 | -1.159304242 | 0.711801506 | -1.628690346 | 0.103378588 | 0.473591415 | Phf11b | PHD finger protein 11B [Source:MGI Symb | 18.471 | 11.307 | 12.075 | 7.7118 | 3.2719 | 7.7969 |
| 31650 | ENSMUSG00000091650 | 0 | NA | NA | NA | NA | NA | Apol11a | apolipoprotein L 11a [Source:MGI Symbol | 0 | 0 | 0 | 0 | 0 | 0 |
| 31651 | ENSMUSG00000091651 | 0 | NA | NA | NA | NA | NA | Vmn2r36 | vomeronasal 2, receptor 36 [Source:MGI S | 0 | 0 | 0 | 0 | 0 | 0 |
| 31652 | ENSMUSG00000091652 | 0 | NA | NA | NA | NA | NA | Vmn1r57 | vomeronasal 1 receptor 57 [Source:MGI S | 0 | 0 | 0 | 0 | 0 | 0 |
| 31653 | ENSMUSG00000091655 | 0.733533733 | -1.621434111 | 2.70314781 | -0.599831835 | 0.548618315 | NA | Gm8741 | predicted gene 8741 [Source:MGI Symbol | 2.4093 | 1.0279 | 0 | 0.964 | 0 | 0 |
| 31654 | ENSMUSG00000091656 | 0 | NA | NA | NA | NA | NA | Vmn2r106 | vomeronasal 2, receptor 106 [Source:MGI | 0 | 0 | 0 | 0 | 0 | 0 |
| 31655 | ENSMUSG00000091657 | 0 | NA | NA | NA | NA | NA | Gm3072 | predicted gene 3072 [Source:MGI Symbol | 0 | 0 | 0 | 0 | 0 | 0 |
| 31656 | ENSMUSG00000091659 | 0 | NA | NA | NA | NA | NA | Gm17099 | predicted gene 17099 [Source:MGI Symbo | 0 | 0 | 0 | 0 | 0 | 0 |
| 31657 | ENSMUSG00000091661 | 0.160662974 | 1.126924175 | 4.080472857 | 0.276174898 | 0.782413736 | NA | Gm17021 | predicted gene 17021 [Source:MGI Symbo | 0 | 0 | 0 | 0.964 | 0 | 0 |
| 31658 | ENSMUSG00000091662 | 0 | NA | NA | NA | NA | NA | Vmn1r69 | vomeronasal 1 receptor 69 [Source:MGI S | 0 | 0 | 0 | 0 | 0 | 0 |
| 31659 | ENSMUSG00000091665 | 0 | NA | NA | NA | NA | NA | Gm17101 | predicted gene 17101 [Source:MGI Symbo | 0 | 0 | 0 | 0 | 0 | 0 |
| 31660 | ENSMUSG00000091668 | 0 | NA | NA | NA | NA | NA | Gm8358 | predicted gene 8358 [Source:MGI Symbol | 0 | 0 | 0 | 0 | 0 | 0 |
| 31661 | ENSMUSG00000091669 | 0 | NA | NA | NA | NA | NA | Gm17117 | predicted gene 17117 [Source:MGI Symbo | 0 | 0 | 0 | 0 | 0 | 0 |
| 31662 | ENSMUSG00000091670 | 0 | NA | NA | NA | NA | NA | Vmn2r105 | vomeronasal 2, receptor 105 [Source:MGI | 0 | 0 | 0 | 0 | 0 | 0 |
| 31663 | ENSMUSG00000091671 | 0 | NA | NA | NA | NA | NA | Trbv10 | T cell receptor beta variable 10 [Source:M | 0 | 0 | 0 | 0 | 0 | 0 |
| 31664 | ENSMUSG00000091672 | 0 | NA | NA | NA | NA | NA | Gm17129 | predicted gene 17129 [Source:MGI Symbo | 0 | 0 | 0 | 0 | 0 | 0 |
| 31665 | ENSMUSG00000091676 | 0 | NA | NA | NA | NA | NA | Gm8374 | predicted gene 8374 [Source:MGI Symbol | 0 | 0 | 0 | 0 | 0 | 0 |
| 31666 | ENSMUSG00000091679 | 0 | NA | NA | NA | NA | NA | Vmn2r96 | vomeronasal 2, receptor 96 [Source:MGI S | 0 | 0 | 0 | 0 | 0 | 0 |
| 31667 | ENSMUSG00000091680 | 11.0881861 | -0.972141584 | 0.739583606 | -1.314444475 | 0.188696672 | 0.625984911 | Klhdc7b | kelch domain containing 7B [Source:MGI S | 13.653 | 6.1675 | 24.15 | 3.8559 | 10.906 | 7.7969 |
| 31668 | ENSMUSG00000091682 | 0 | NA | NA | NA | NA | NA | Trgc3 | T cell receptor gamma, constant 3 [Source | 0 | 0 | 0 | 0 | 0 | 0 |
| 31669 | ENSMUSG00000091685 | 0.17131964 | -0.796629176 | 4.080472857 | -0.195229623 | 0.84521319 | NA | Gm17359 | predicted gene, 17359 [Source:MGI Symb | 0 | 1.0279 | 0 | 0 | 0 | 0 |
| 31670 | ENSMUSG00000091686 | 0 | NA | NA | NA | NA | NA | 4930471C04 | RIKEN cDNA 4930471C04 gene [Source:M | 0 | 0 | 0 | 0 | 0 | 0 |
| 31671 | ENSMUSG00000091692 | 0 | NA | NA | NA | NA | NA | 4930433I11F | RIKEN cDNA 4930433I11 gene [Source:MC | 0 | 0 | 0 | 0 | 0 | 0 |
| 31672 | ENSMUSG00000091694 | 1.461568725 | 1.316727391 | 1.924519225 | 0.684185107 | 0.493858286 | NA | Apol11b | apolipoprotein L 11b [Source:MGI Symbol | 2.4093 | 0 | 0 | 1.928 | 1.0906 | 3.3415 |
| 31673 | ENSMUSG00000091695 | 0.499690045 | -0.82080011 | 3.113553401 | -0.263621658 | 0.792071468 | NA | Gm17669 | predicted gene, 17669 [Source:MGI Symb | 0 | 1.0279 | 1.0062 | 0.964 | 0 | 0 |
| 31674 | ENSMUSG00000091697 | 0.185640647 | 1.126924175 | 4.080472857 | 0.276174898 | 0.782413736 | NA | Eif3s6-ps2 | eukaryotic translation initiation factor 3, s | 0 | 0 | 0 | 0 | 0 | 1.1138 |
| 31675 | ENSMUSG00000091698 | 0 | NA | NA | NA | NA | NA | Gm6526 | predicted gene 6526 [Source:MGI Symbol | 0 | 0 | 0 | 0 | 0 | 0 |
| 31676 | ENSMUSG00000091700 | 0 | NA | NA | NA | NA | NA | Gm7876 | predicted gene 7876 [Source:MGI Symbol | 0 | 0 | 0 | 0 | 0 | 0 |
| 31677 | ENSMUSG00000091703 | 0 | NA | NA | NA | NA | NA | Gm4222 | predicted gene 4222 [Source:MGI Symbol | 0 | 0 | 0 | 0 | 0 | 0 |
| 31678 | ENSMUSG00000091705 | 10.83117459 | 0.069995209 | 0.687120198 | 0.101867489 | 0.918861856 | 0.986374671 | H2-Q2 | histocompatibility 2, Q region locus 2 [Sou | 6.4248 | 16.447 | 9.0562 | 15.424 | 8.7249 | 8.9108 |
| 31679 | ENSMUSG00000091707 | 0 | NA | NA | NA | NA | NA | Gm8224 | predicted gene 8224 [Source:MGI Symbol | 0 | 0 | 0 | 0 | 0 | 0 |
| 31680 | ENSMUSG00000091709 | 0.167707431 | -0.796629176 | 4.080472857 | -0.195229623 | 0.84521319 | NA | Gm17189 | predicted gene 17189 [Source:MGI Symbo | 0 | 0 | 1.0062 | 0 | 0 | 0 |
| 31681 | ENSMUSG00000091712 | 1.676239365 | 0.800824098 | 1.751055378 | 0.457337962 | 0.647428148 | NA | Sec14l5 | SEC14-like lipid binding 5 [Source:MGI Syr | 1.6062 | 1.0279 | 1.0062 | 0.964 | 5.4531 | 0 |
| 31682 | ENSMUSG00000091713 | 0 | NA | NA | NA | NA | NA | Gm17126 | predicted gene 17126 [Source:MGI Symbo | 0 | 0 | 0 | 0 | 0 | 0 |
| 31683 | ENSMUSG00000091718 | 0 | NA | NA | NA | NA | NA | Gm16506 | predicted gene 16506 [Source:MGI Symbo | 0 | 0 | 0 | 0 | 0 | 0 |
| 31684 | ENSMUSG00000091721 | 39.40071726 | -0.209846699 | 0.364823475 | -0.575200649 | 0.565155604 | 0.901542552 | Gimd1 | GIMAP family P-loop NTPase domain cont | 40.155 | 44.2 | 42.262 | 26.991 | 34.9 | 47.895 |
| 31685 | ENSMUSG00000091722 | 0 | NA | NA | NA | NA | NA | Siah3 | siah E3 ubiquitin protein ligase family mer | 0 | 0 | 0 | 0 | 0 | 0 |
| 31686 | ENSMUSG00000091723 | 0 | NA | NA | NA | NA | NA | Gm17366 | predicted gene, 17366 [Source:MGI Symb | 0 | 0 | 0 | 0 | 0 | 0 |
| 31687 | ENSMUSG00000091725 | 0 | NA | NA | NA | NA | NA | Gm8220 | predicted gene 8220 [Source:MGI Symbol | 0 | 0 | 0 | 0 | 0 | 0 |
| 31688 | ENSMUSG00000091730 | 0.465832987 | -0.698295221 | 3.18348759 | -0.219349126 | 0.826378096 | NA | Gm17230 | predicted gene 17230 [Source:MGI Symbo | 0.8031 | 1.0279 | 0 | 0.964 | 0 | 0 |
| 31689 | ENSMUSG00000091731 | 0 | NA | NA | NA | NA | NA | Gm17542 | predicted gene, 17542 [Source:MGI Symb | 0 | 0 | 0 | 0 | 0 | 0 |
| 31690 | ENSMUSG00000091732 | 0 | NA | NA | NA | NA | NA | Gm17541 | predicted gene, 17541 [Source:MGI Symb | 0 | 0 | 0 | 0 | 0 | 0 |
| 31691 | ENSMUSG00000091733 | 0 | NA | NA | NA | NA | NA | Gm8094 | predicted gene 8094 [Source:MGI Symbol | 0 | 0 | 0 | 0 | 0 | 0 |
| 31692 | ENSMUSG00000091734 | 0 | NA | NA | NA | NA | NA | Vmn1r29 | vomeronasal 1 receptor 29 [Source:MGI S | 0 | 0 | 0 | 0 | 0 | 0 |

| | | | | | | | | | | | | | | | |
|---|---|---|---|---|---|---|---|---|---|---|---|---|---|---|---|
| 31693 | ENSMUSG00000091735 | 0.17131964 | -0.796629176 | 4.080472857 | -0.195229623 | 0.84521319 | NA | Gpr62 | G protein-coupled receptor 62 [Source:M | 0 | 1.0279 | 0 | 0 | 0 | 0 |
| 31694 | ENSMUSG00000091736 | 0 | NA | NA | NA | NA | NA | Yy2 | Yy2 transcription factor [Source:MGI Syml | 0 | 0 | 0 | 0 | 0 | 0 |
| 31695 | ENSMUSG00000091740 | 0 | NA | NA | NA | NA | NA | Gm7929 | predicted gene 7929 [Source:MGI Symbol | 0 | 0 | 0 | 0 | 0 | 0 |
| 31696 | ENSMUSG00000091741 | 0 | NA | NA | NA | NA | NA | Gm17161 | predicted gene 17161 [Source:MGI Symbo | 0 | 0 | 0 | 0 | 0 | 0 |
| 31697 | ENSMUSG00000091742 | 23.11467336 | 0.565457451 | 0.598270988 | 0.94515272 | 0.344580913 | 0.785874854 | Gm5093 | predicted gene 5093 [Source:MGI Symbol | 5.6217 | 13.363 | 37.231 | 25.063 | 23.994 | 33.415 |
| 31698 | ENSMUSG00000091743 | 0 | NA | NA | NA | NA | NA | Gm5705 | predicted gene 5705 [Source:MGI Symbol | 0 | 0 | 0 | 0 | 0 | 0 |
| 31699 | ENSMUSG00000091744 | 0 | NA | NA | NA | NA | NA | NA | NA | 0 | 0 | 0 | 0 | 0 | 0 |
| 31700 | ENSMUSG00000091745 | 0.956463687 | -2.031943053 | 2.464797407 | -0.824385423 | 0.409720595 | NA | Gm17098 | predicted gene 17098 [Source:MGI Symbo | 1.6062 | 0 | 3.0187 | 0 | 0 | 1.1138 |
| 31701 | ENSMUSG00000091749 | 0 | NA | NA | NA | NA | NA | Tspy-ps | testis specific protein-Y encoded, pseudog | 0 | 0 | 0 | 0 | 0 | 0 |
| 31702 | ENSMUSG00000091752 | 0 | NA | NA | NA | NA | NA | Gm3208 | predicted gene 3208 [Source:MGI Symbol | 0 | 0 | 0 | 0 | 0 | 0 |
| 31703 | ENSMUSG00000091754 | 0 | NA | NA | NA | NA | NA | Gm3636 | predicted gene 3636 [Source:MGI Symbol | 0 | 0 | 0 | 0 | 0 | 0 |
| 31704 | ENSMUSG00000091756 | 0 | NA | NA | NA | NA | NA | Gm3095 | predicted gene 3095 [Source:MGI Symbol | 0 | 0 | 0 | 0 | 0 | 0 |
| 31705 | ENSMUSG00000091757 | 0 | NA | NA | NA | NA | NA | Gm17064 | predicted gene 17064 [Source:MGI Symbo | 0 | 0 | 0 | 0 | 0 | 0 |
| 31706 | ENSMUSG00000091762 | 0 | NA | NA | NA | NA | NA | Gm3191 | predicted gene 3191 [Source:MGI Symbol | 0 | 0 | 0 | 0 | 0 | 0 |
| 31707 | ENSMUSG00000091764 | 27.00584489 | 0.062069112 | 0.44950767 | 0.138082432 | 0.89017527 | 0.982489666 | Zfp964 | zinc finger protein 964 [Source:MGI Symb | 21.684 | 19.53 | 38.237 | 31.811 | 21.812 | 28.96 |
| 31708 | ENSMUSG00000091768 | 0 | NA | NA | NA | NA | NA | Gm17617 | predicted gene, 17617 [Source:MGI Symb | 0 | 0 | 0 | 0 | 0 | 0 |
| 31709 | ENSMUSG00000091771 | 0 | NA | NA | NA | NA | NA | Vmn2r103 | vomeronasal 2, receptor 103 [Source:MGI | 0 | 0 | 0 | 0 | 0 | 0 |
| 31710 | ENSMUSG00000091775 | 0 | NA | NA | NA | NA | NA | Gm16494 | predicted gene 16494 [Source:MGI Symbo | 0 | 0 | 0 | 0 | 0 | 0 |
| 31711 | ENSMUSG00000091776 | 0 | NA | NA | NA | NA | NA | Gm8340 | predicted gene 8340 [Source:MGI Symbol | 0 | 0 | 0 | 0 | 0 | 0 |
| 31712 | ENSMUSG00000091779 | 0 | NA | NA | NA | NA | NA | Gm21293 | predicted gene, 21293 [Source:MGI Symb | 0 | 0 | 0 | 0 | 0 | 0 |
| 31713 | ENSMUSG00000091780 | 0 | NA | NA | NA | NA | NA | Sco2 | SCO2 cytochrome c oxidase assembly prot | 0 | 0 | 0 | 0 | 0 | 0 |
| 31714 | ENSMUSG00000091784 | 9.95742373 | 1.004117655 | 0.743060159 | 1.351327538 | 0.176590534 | 0.607421044 | Gm17022 | predicted gene 17022 [Source:MGI Symbo | 4.8186 | 7.1954 | 8.05 | 23.135 | 7.6343 | 8.9108 |
| 31715 | ENSMUSG00000091785 | 0 | NA | NA | NA | NA | NA | Rpl29-ps5 | ribosomal protein L29, pseudogene 5 [Sou | 0 | 0 | 0 | 0 | 0 | 0 |
| 31716 | ENSMUSG00000091786 | 0.181769715 | 1.126924175 | 4.080472857 | 0.276174898 | 0.782413736 | NA | Gm8425 | predicted gene 8425 [Source:MGI Symbol | 0 | 0 | 0 | 0 | 1.0906 | 0 |
| 31717 | ENSMUSG00000091788 | 0 | NA | NA | NA | NA | NA | Gm8032 | predicted gene 8032 [Source:MGI Symbol | 0 | 0 | 0 | 0 | 0 | 0 |
| 31718 | ENSMUSG00000091792 | 0 | NA | NA | NA | NA | NA | Gm3573 | predicted gene 3573 [Source:MGI Symbol | 0 | 0 | 0 | 0 | 0 | 0 |
| 31719 | ENSMUSG00000091794 | 0 | NA | NA | NA | NA | NA | Vmn2r18 | vomeronasal 2, receptor 18 [Source:MGI S | 0 | 0 | 0 | 0 | 0 | 0 |
| 31720 | ENSMUSG00000091800 | 0 | NA | NA | NA | NA | NA | Igkv1-115 | immunoglobulin kappa variable 1-115 [So | 0 | 0 | 0 | 0 | 0 | 0 |
| 31721 | ENSMUSG00000091803 | 215.0772497 | 0.071685432 | 0.175229856 | 0.409093711 | 0.682470891 | 0.93502957 | Cox16 | cytochrome c oxidase assembly protein 1( | 221.66 | 209.7 | 197.22 | 198.58 | 211.58 | 251.73 |
| 31722 | ENSMUSG00000091805 | 0 | NA | NA | NA | NA | NA | Vmn2r108 | vomeronasal 2, receptor 108 [Source:MGI | 0 | 0 | 0 | 0 | 0 | 0 |
| 31723 | ENSMUSG00000091806 | 0 | NA | NA | NA | NA | NA | Gm5258 | predicted gene 5258 [Source:MGI Symbol | 0 | 0 | 0 | 0 | 0 | 0 |
| 31724 | ENSMUSG00000091808 | 0 | NA | NA | NA | NA | NA | Gm9671 | predicted gene 9671 [Source:MGI Symbol | 0 | 0 | 0 | 0 | 0 | 0 |
| 31725 | ENSMUSG00000091809 | 0 | NA | NA | NA | NA | NA | Olfr721-ps1 | olfactory receptor 721, pseudogene 1 [So | 0 | 0 | 0 | 0 | 0 | 0 |
| 31726 | ENSMUSG00000091811 | 122.0285622 | 0.287941883 | 0.211275369 | 1.362874832 | 0.172921962 | 0.601945568 | Inafm1 | InaF motif containing 1 [Source:MGI Symb | 100.39 | 106.9 | 122.76 | 138.81 | 136.33 | 126.98 |
| 31727 | ENSMUSG00000091812 | 0 | NA | NA | NA | NA | NA | Ear-ps5 | eosinophil-associated, ribonuclease A fam | 0 | 0 | 0 | 0 | 0 | 0 |
| 31728 | ENSMUSG00000091813 | 11.28587457 | 0.345798513 | 0.630017566 | 0.548871224 | 0.583093827 | 0.908645447 | Ces2h | carboxylesterase 2H [Source:MGI Symbol; | 10.44 | 11.307 | 8.05 | 12.532 | 10.906 | 14.48 |
| 31729 | ENSMUSG00000091814 | 0 | NA | NA | NA | NA | NA | Gm8050 | predicted gene 8050 [Source:MGI Symbol | 0 | 0 | 0 | 0 | 0 | 0 |
| 31730 | ENSMUSG00000091815 | 0 | NA | NA | NA | NA | NA | Gm17153 | predicted gene 17153 [Source:MGI Symbo | 0 | 0 | 0 | 0 | 0 | 0 |
| 31731 | ENSMUSG00000091816 | 0 | NA | NA | NA | NA | NA | Gm17141 | predicted gene 17141 [Source:MGI Symbo | 0 | 0 | 0 | 0 | 0 | 0 |
| 31732 | ENSMUSG00000091817 | 0 | NA | NA | NA | NA | NA | Gm17091 | predicted gene 17091 [Source:MGI Symbo | 0 | 0 | 0 | 0 | 0 | 0 |
| 31733 | ENSMUSG00000091818 | 0 | NA | NA | NA | NA | NA | Rpl19-ps4 | ribosomal protein L19, pseudogene 4 [Sou | 0 | 0 | 0 | 0 | 0 | 0 |
| 31734 | ENSMUSG00000091821 | 0 | NA | NA | NA | NA | NA | Srsx | serine-rich, secreted, X-linked [Source:MG | 0 | 0 | 0 | 0 | 0 | 0 |
| 31735 | ENSMUSG00000091825 | 0 | NA | NA | NA | NA | NA | Gm5778 | predicted gene 5778 [Source:MGI Symbol | 0 | 0 | 0 | 0 | 0 | 0 |
| 31736 | ENSMUSG00000091827 | 0 | NA | NA | NA | NA | NA | Speer4f2 | spermatogenesis associated glutamate (E) | 0 | 0 | 0 | 0 | 0 | 0 |
| 31737 | ENSMUSG00000091831 | 0.653308562 | 1.538013299 | 3.112409277 | 0.494155223 | 0.62119656 | NA | Gm4707 | predicted gene 4707 [Source:MGI Symbol | 0 | 1.0279 | 0 | 2.8919 | 0 | 0 |
| 31738 | ENSMUSG00000091832 | 0 | NA | NA | NA | NA | NA | Gm17155 | predicted gene 17155 [Source:MGI Symbo | 0 | 0 | 0 | 0 | 0 | 0 |
| 31739 | ENSMUSG00000091834 | 0 | NA | NA | NA | NA | NA | Gm17177 | predicted gene 17177 [Source:MGI Symbo | 0 | 0 | 0 | 0 | 0 | 0 |
| 31740 | ENSMUSG00000091842 | 0 | NA | NA | NA | NA | NA | Gm16490 | predicted gene 16490 [Source:MGI Symbo | 0 | 0 | 0 | 0 | 0 | 0 |
| 31741 | ENSMUSG00000091844 | 2.099858871 | 0.384254109 | 1.527716582 | 0.251521855 | 0.801410667 | NA | Gm8251 | predicted gene 8251 [Source:MGI Symbol | 2.4093 | 2.0558 | 1.0062 | 3.8559 | 3.2719 | 0 |
| 31742 | ENSMUSG00000091845 | 7.924341775 | -0.152339411 | 0.790197885 | -0.192786407 | 0.847126249 | 0.972803041 | Rpl36-ps12 | ribosomal protein L36, pseudogene 12 [Sc | 8.8341 | 9.2513 | 7.0437 | 12.532 | 6.5437 | 3.3415 |
| 31743 | ENSMUSG00000091847 | 0 | NA | NA | NA | NA | NA | Gm2855 | predicted gene 2855 [Source:MGI Symbol | 0 | 0 | 0 | 0 | 0 | 0 |
| 31744 | ENSMUSG00000091849 | 0 | NA | NA | NA | NA | NA | Gm17188 | predicted gene 17188 [Source:MGI Symbo | 0 | 0 | 0 | 0 | 0 | 0 |
| 31745 | ENSMUSG00000091859 | 0 | NA | NA | NA | NA | NA | Vmn2r100 | vomeronasal 2, receptor 100 [Source:MGI | 0 | 0 | 0 | 0 | 0 | 0 |
| 31746 | ENSMUSG00000091863 | 0 | NA | NA | NA | NA | NA | Cldn34a | claudin 34A [Source:MGI Symbol;Acc:MGI | 0 | 0 | 0 | 0 | 0 | 0 |
| 31747 | ENSMUSG00000091864 | 1.591961346 | -1.753226202 | 2.068591819 | -0.847545749 | 0.396690999 | NA | Gm17102 | predicted gene 17102 [Source:MGI Symbo | 3.2124 | 4.1117 | 0 | 0 | 0 | 2.2277 |
| 31748 | ENSMUSG00000091866 | 0 | NA | NA | NA | NA | NA | Gm6257 | predicted gene 6257 [Source:MGI Symbol | 0 | 0 | 0 | 0 | 0 | 0 |

| | | | | | | | | | | | | | | | |
|---|---|---|---|---|---|---|---|---|---|---|---|---|---|---|---|
| 31749 | ENSMUSG00000091867 | 167.0271531 | -2.99536894 | 0.310979509 | -9.632046008 | 5.86E-22 | 1.21E-18 | Cyp2a22 | cytochrome P450, family 2, subfamily a, p | 261.01 | 405 | 225.4 | 56.875 | 32.719 | 21.163 |
| 31750 | ENSMUSG00000091873 | 0 | NA | NA | NA | NA | NA | Olfr732 | olfactory receptor 732 [Source:MGI Symb | 0 | 0 | 0 | 0 | 0 | 0 |
| 31751 | ENSMUSG00000091874 | 0 | NA | NA | NA | NA | NA | Vmn1r56 | vomeronasal 1 receptor 56 [Source:MGI S | 0 | 0 | 0 | 0 | 0 | 0 |
| 31752 | ENSMUSG00000091876 | 0.185640647 | 1.126924175 | 4.080472857 | 0.276174898 | 0.782413736 | NA | Gm3851 | predicted gene 3851 [Source:MGI Symbol | 0 | 0 | 0 | 0 | 0 | 1.1138 |
| 31753 | ENSMUSG00000091879 | 0 | NA | NA | NA | NA | NA | Vmn2r17 | vomeronasal 2, receptor 17 [Source:MGI ! | 0 | 0 | 0 | 0 | 0 | 0 |
| 31754 | ENSMUSG00000091881 | 0.696064467 | -1.534444742 | 3.043667477 | -0.504143358 | 0.61416064 | NA | Gm17146 | predicted gene 17146 [Source:MGI Symbo | 3.2124 | 0 | 0 | 0.964 | 0 | 0 |
| 31755 | ENSMUSG00000091882 | 0 | NA | NA | NA | NA | NA | Gm6337 | predicted gene 6337 [Source:MGI Symbol | 0 | 0 | 0 | 0 | 0 | 0 |
| 31756 | ENSMUSG00000091887 | 0 | NA | NA | NA | NA | NA | Gm17053 | predicted gene 17053 [Source:MGI Symbo | 0 | 0 | 0 | 0 | 0 | 0 |
| 31757 | ENSMUSG00000091888 | 0 | NA | NA | NA | NA | NA | Vmn2r80 | vomeronasal 2, receptor 80 [Source:MGI ! | 0 | 0 | 0 | 0 | 0 | 0 |
| 31758 | ENSMUSG00000091890 | 0 | NA | NA | NA | NA | NA | A830073O21 | RIKEN cDNA A830073O21 gene [Source:M | 0 | 0 | 0 | 0 | 0 | 0 |
| 31759 | ENSMUSG00000091894 | 0 | NA | NA | NA | NA | NA | Gm3030 | predicted gene 3030 [Source:MGI Symbol | 0 | 0 | 0 | 0 | 0 | 0 |
| 31760 | ENSMUSG00000091896 | 1016.137193 | 0.084946456 | 0.101873832 | 0.833839801 | 0.404371264 | 0.829271989 | Ube2d2a | ubiquitin-conjugating enzyme E2D 2A [Sou | 1005.5 | 995.02 | 957.94 | 1047.8 | 1053.5 | 1037 |
| 31761 | ENSMUSG00000091897 | 0 | NA | NA | NA | NA | NA | Gm17019 | predicted gene 17019 [Source:MGI Symbo | 0 | 0 | 0 | 0 | 0 | 0 |
| 31762 | ENSMUSG00000091898 | 27.99799821 | -0.087176059 | 0.62003157 | -0.140599387 | 0.888186433 | 0.982382091 | Tnnc1 | troponin C, cardiac/slow skeletal [Source: | 16.865 | 20.558 | 49.306 | 29.883 | 45.806 | 5.5692 |
| 31763 | ENSMUSG00000091900 | 0.349477146 | 0.165142768 | 3.911021834 | 0.042224967 | 0.966319359 | NA | Gm4353 | predicted gene 4353 [Source:MGI Symbol | 0 | 0 | 1.0062 | 0 | 1.0906 | 0 |
| 31764 | ENSMUSG00000091902 | 0 | NA | NA | NA | NA | NA | Gm6918 | predicted gene 6918 [Source:MGI Symbol | 0 | 0 | 0 | 0 | 0 | 0 |
| 31765 | ENSMUSG00000091903 | 0 | NA | NA | NA | NA | NA | Gm6460 | predicted gene 6460 [Source:MGI Symbol | 0 | 0 | 0 | 0 | 0 | 0 |
| 31766 | ENSMUSG00000091905 | 2.377136411 | 0.351640879 | 1.41583131 | 0.248363542 | 0.803853139 | NA | Dnajb6-ps | DnaJ heat shock protein family (Hsp40) m | 3.2124 | 2.0558 | 1.0062 | 5.7839 | 1.0906 | 1.1138 |
| 31767 | ENSMUSG00000091906 | 0.185640647 | 1.126924175 | 4.080472857 | 0.276174898 | 0.782413736 | NA | 1700099I09F | RIKEN cDNA 1700099I09 gene [Source:MG | 0 | 0 | 0 | 0 | 0 | 1.1138 |
| 31768 | ENSMUSG00000091908 | 0 | NA | NA | NA | NA | NA | Gm17231 | predicted gene 17231 [Source:MGI Symbo | 0 | 0 | 0 | 0 | 0 | 0 |
| 31769 | ENSMUSG00000091911 | 0 | NA | NA | NA | NA | NA | Gm9587 | predicted gene 9587 [Source:MGI Symbol | 0 | 0 | 0 | 0 | 0 | 0 |
| 31770 | ENSMUSG00000091922 | 0.353089354 | 0.165142706 | 3.896102244 | 0.042386646 | 0.966190474 | NA | Gm17409 | predicted gene, 17409 [Source:MGI Symb | 0 | 1.0279 | 0 | 0 | 1.0906 | 0 |
| 31771 | ENSMUSG00000091923 | 0 | NA | NA | NA | NA | NA | Gm8267 | predicted gene 8267 [Source:MGI Symbol | 0 | 0 | 0 | 0 | 0 | 0 |
| 31772 | ENSMUSG00000091924 | 0 | NA | NA | NA | NA | NA | Vmn1r185 | vomeronasal 1 receptor 185 [Source:MGI | 0 | 0 | 0 | 0 | 0 | 0 |
| 31773 | ENSMUSG00000091926 | 0 | NA | NA | NA | NA | NA | Vmn2r62 | vomeronasal 2, receptor 62 [Source:MGI ! | 0 | 0 | 0 | 0 | 0 | 0 |
| 31774 | ENSMUSG00000091929 | 0 | NA | NA | NA | NA | NA | Gm8582 | predicted gene 8582 [Source:MGI Symbol | 0 | 0 | 0 | 0 | 0 | 0 |
| 31775 | ENSMUSG00000091930 | 0.185640647 | 1.126924175 | 4.080472857 | 0.276174898 | 0.782413736 | NA | Vmn2r52 | vomeronasal 2, receptor 52 [Source:MGI ! | 0 | 0 | 0 | 0 | 0 | 1.1138 |
| 31776 | ENSMUSG00000091931 | 61.59194481 | 0.122563749 | 0.288504924 | 0.424823766 | 0.670965151 | 0.93116767 | Gon7 | GON7 subunit of KEOPS complex [Source: | 52.202 | 55.508 | 69.431 | 65.55 | 57.803 | 69.058 |
| 31777 | ENSMUSG00000091933 | 0 | NA | NA | NA | NA | NA | Gm8857 | predicted gene 8857 [Source:MGI Symbol | 0 | 0 | 0 | 0 | 0 | 0 |
| 31778 | ENSMUSG00000091936 | 0 | NA | NA | NA | NA | NA | Gm3053 | predicted gene 3053 [Source:MGI Symbol | 0 | 0 | 0 | 0 | 0 | 0 |
| 31779 | ENSMUSG00000091937 | 0 | NA | NA | NA | NA | NA | Gm3646 | predicted gene 3646 [Source:MGI Symbol | 0 | 0 | 0 | 0 | 0 | 0 |
| 31780 | ENSMUSG00000091938 | 0 | NA | NA | NA | NA | NA | Gm2564 | predicted gene 2564 [Source:MGI Symbol | 0 | 0 | 0 | 0 | 0 | 0 |
| 31781 | ENSMUSG00000091941 | 0.335414862 | -1.779047242 | 4.030804179 | -0.441362855 | 0.658950329 | NA | Gm7399 | predicted gene 7399 [Source:MGI Symbol | 0 | 0 | 2.0125 | 0 | 0 | 0 |
| 31782 | ENSMUSG00000091942 | 0 | NA | NA | NA | NA | NA | Gm17138 | predicted gene 17138 [Source:MGI Symbo | 0 | 0 | 0 | 0 | 0 | 0 |
| 31783 | ENSMUSG00000091945 | 0 | NA | NA | NA | NA | NA | Vmn2r114 | vomeronasal 2, receptor 114 [Source:MGI | 0 | 0 | 0 | 0 | 0 | 0 |
| 31784 | ENSMUSG00000091947 | 0 | NA | NA | NA | NA | NA | 6530401F13 | RIKEN cDNA 6530401F13 gene [Source:MG | 0 | 0 | 0 | 0 | 0 | 0 |
| 31785 | ENSMUSG00000091950 | 0 | NA | NA | NA | NA | NA | Olfr421-ps1 | olfactory receptor 421, pseudogene 1 [So | 0 | 0 | 0 | 0 | 0 | 0 |
| 31786 | ENSMUSG00000091952 | 5.779400128 | 0.713177229 | 0.944786808 | 0.754855194 | 0.450335873 | 0.855596751 | Gm17709 | predicted gene, 17709 [Source:MGI Symb | 4.0155 | 7.1954 | 2.0125 | 11.568 | 6.5437 | 3.3415 |
| 31787 | ENSMUSG00000091955 | 5.180021583 | 0.020552221 | 0.921139016 | 0.022311747 | 0.982199278 | 0.998580735 | Gm9844 | predicted pseudogene 9844 [Source:MGI | 7.2279 | 4.1117 | 4.025 | 5.7839 | 4.3625 | 5.5692 |
| 31788 | ENSMUSG00000091956 | 0 | NA | NA | NA | NA | NA | C2cd4b | C2 calcium-dependent domain containing | 0 | 0 | 0 | 0 | 0 | 0 |
| 31789 | ENSMUSG00000091957 | 8.343278174 | 0.126065996 | 0.728183801 | 0.173123868 | 0.862554064 | 0.976334255 | Rps2-ps10 | ribosomal protein S2, pseudogene 10 [Sou | 9.6372 | 8.2233 | 6.0375 | 9.6398 | 8.7249 | 7.7969 |
| 31790 | ENSMUSG00000091960 | 0 | NA | NA | NA | NA | NA | Gm17038 | predicted gene 17038 [Source:MGI Symbo | 0 | 0 | 0 | 0 | 0 | 0 |
| 31791 | ENSMUSG00000091961 | 0 | NA | NA | NA | NA | NA | Gm3123 | predicted gene 3123 [Source:MGI Symbol | 0 | 0 | 0 | 0 | 0 | 0 |
| 31792 | ENSMUSG00000091962 | 0 | NA | NA | NA | NA | NA | Vmn2r78 | vomeronasal 2, receptor 78 [Source:MGI ! | 0 | 0 | 0 | 0 | 0 | 0 |
| 31793 | ENSMUSG00000091967 | 0 | NA | NA | NA | NA | NA | Vmn2r128 | vomeronasal 2 receptor 128 [Source:MGI | 0 | 0 | 0 | 0 | 0 | 0 |
| 31794 | ENSMUSG00000091968 | 15.57470456 | 0.606125622 | 0.558560608 | 1.085156406 | 0.277852389 | 0.730834142 | Gm17115 | predicted gene 17115 [Source:MGI Symbo | 8.8341 | 12.335 | 16.1 | 23.135 | 17.45 | 15.594 |
| 31795 | ENSMUSG00000091970 | 0 | NA | NA | NA | NA | NA | Gm4868 | predicted gene 4868 [Source:MGI Symbol | 0 | 0 | 0 | 0 | 0 | 0 |
| 31796 | ENSMUSG00000091971 | 9.617914645 | -1.045537097 | 0.803417672 | -1.301361836 | 0.19313463 | 0.63128389 | Hspa1a | heat shock protein 1A [Source:MGI Symbo | 18.471 | 10.279 | 10.062 | 5.7839 | 11.997 | 1.1138 |
| 31797 | ENSMUSG00000091972 | 0 | NA | NA | NA | NA | NA | Tmem207 | transmembrane protein 207 [Source:MGI | 0 | 0 | 0 | 0 | 0 | 0 |
| 31798 | ENSMUSG00000091983 | 0 | NA | NA | NA | NA | NA | Olfr457 | olfactory receptor 457 [Source:MGI Symb | 0 | 0 | 0 | 0 | 0 | 0 |
| 31799 | ENSMUSG00000091985 | 44.43202715 | 0.023517165 | 0.33763632 | 0.069652355 | 0.944470365 | 0.990077979 | Gm17354 | predicted gene, 17354 [Source:MGI Symb | 44.974 | 41.117 | 46.287 | 54.947 | 43.625 | 35.643 |
| 31800 | ENSMUSG00000091987 | 0 | NA | NA | NA | NA | NA | Gm10352 | predicted gene 10352 [Source:MGI Symbo | 0 | 0 | 0 | 0 | 0 | 0 |
| 31801 | ENSMUSG00000091989 | 2.007838987 | -1.620778136 | 1.887808052 | -0.858550282 | 0.390588678 | NA | Ndufab1-ps | NADH:ubiquinone oxidoreductase subuni | 4.0155 | 5.1396 | 0 | 2.8919 | 0 | 0 |
| 31802 | ENSMUSG00000091990 | 0 | NA | NA | NA | NA | NA | Gm9672 | predicted gene 9672 [Source:MGI Symbol | 0 | 0 | 0 | 0 | 0 | 0 |
| 31803 | ENSMUSG00000091993 | 2.546472179 | 1.287523671 | 1.606046038 | 0.801672953 | 0.422742165 | NA | B930036N10 | RIKEN cDNA B930036N10 gene [Source:M | 2.4093 | 0 | 2.0125 | 8.6758 | 2.1812 | 0 |
| 31804 | ENSMUSG00000091994 | 6.075278701 | -0.088398313 | 0.859378817 | -0.102863034 | 0.918071676 | 0.986374671 | E130317F20 | RIKEN cDNA E130317F20 gene [Source:MG | 5.6217 | 7.1954 | 6.0375 | 7.7118 | 6.5437 | 3.3415 |

| | | | | | | | | | | | | | | | |
|---|---|---|---|---|---|---|---|---|---|---|---|---|---|---|---|
| 31805 | ENSMUSG00000091995 | 0 | NA | NA | NA | NA | NA | Gm3339 | predicted gene 3339 [Source:MGI Symbol | 0 | 0 | 0 | 0 | 0 | 0 |
| 31806 | ENSMUSG00000091996 | 5.227760964 | 0.018260801 | 0.939745704 | 0.019431641 | 0.98449677 | 0.998580735 | BC049352 | cDNA sequence BC049352 [Source:MGI Sy | 6.4248 | 2.0558 | 7.0437 | 4.8199 | 5.4531 | 5.5692 |
| 31807 | ENSMUSG00000091997 | 0.51014012 | 1.031855138 | 3.09002894 | 0.333930574 | 0.738431948 | NA | Gm6611 | predicted gene 6611 [Source:MGI Symbol | 0 | 0 | 1.0062 | 0.964 | 1.0906 | 0 |
| 31808 | ENSMUSG00000091998 | 0 | NA | NA | NA | NA | NA | Gm17072 | predicted gene 17072 [Source:MGI Symbo | 0 | 0 | 0 | 0 | 0 | 0 |
| 31809 | ENSMUSG00000092004 | 0 | NA | NA | NA | NA | NA | Gm17482 | predicted gene, 17482 [Source:MGI Symb | 0 | 0 | 0 | 0 | 0 | 0 |
| 31810 | ENSMUSG00000092006 | 0 | NA | NA | NA | NA | NA | Gm17139 | predicted gene 17139 [Source:MGI Symbo | 0 | 0 | 0 | 0 | 0 | 0 |
| 31811 | ENSMUSG00000092008 | 2.717218378 | 0.109734552 | 1.282683072 | 0.085550792 | 0.931823518 | NA | Cyp2c69 | cytochrome P450, family 2, subfamily c, p | 0.8031 | 3.0838 | 4.025 | 2.8919 | 3.2719 | 2.2277 |
| 31812 | ENSMUSG00000092009 | 0 | NA | NA | NA | NA | NA | Myh15 | myosin, heavy chain 15 [Source:MGI Symb | 0 | 0 | 0 | 0 | 0 | 0 |
| 31813 | ENSMUSG00000092014 | 0.656740811 | -0.013525372 | 3.112905782 | -0.004344934 | 0.996533255 | NA | Gm4468 | predicted gene 4468 [Source:MGI Symbol | 0 | 0 | 2.0125 | 1.928 | 0 | 0 |
| 31814 | ENSMUSG00000092019 | 0 | NA | NA | NA | NA | NA | Eif1ad6 | eukaryotic translation initiation factor 1A | 0 | 0 | 0 | 0 | 0 | 0 |
| 31815 | ENSMUSG00000092021 | 338.3252277 | 2.529759866 | 0.807664848 | 3.132190132 | 0.001735075 | 0.031567448 | Gbp11 | guanylate binding protein 11 [Source:MGI | 134.92 | 76.066 | 88.55 | 1136.5 | 147.23 | 446.65 |
| 31816 | ENSMUSG00000092027 | 0 | NA | NA | NA | NA | NA | Gm17062 | predicted gene 17062 [Source:MGI Symbo | 0 | 0 | 0 | 0 | 0 | 0 |
| 31817 | ENSMUSG00000092029 | 0.167707431 | -0.796629176 | 4.080472857 | -0.195229623 | 0.84521319 | NA | Gm9027 | predicted gene 9027 [Source:MGI Symbol | 0 | 0 | 1.0062 | 0 | 0 | 0 |
| 31818 | ENSMUSG00000092031 | 0 | NA | NA | NA | NA | NA | Gm2108 | predicted gene 2108 [Source:MGI Symbol | 0 | 0 | 0 | 0 | 0 | 0 |
| 31819 | ENSMUSG00000092032 | 0 | NA | NA | NA | NA | NA | Vmn2r59 | vomeronasal 2, receptor 59 [Source:MGI S | 0 | 0 | 0 | 0 | 0 | 0 |
| 31820 | ENSMUSG00000092035 | 7.218208799 | 0.749026797 | 0.818840618 | 0.914740647 | 0.3603278 | 0.799598236 | Peg10 | paternally expressed 10 [Source:MGI Sym | 4.0155 | 8.2233 | 4.025 | 11.568 | 5.4531 | 10.025 |
| 31821 | ENSMUSG00000092036 | 0 | NA | NA | NA | NA | NA | Gm2244 | predicted gene 2244 [Source:MGI Symbol | 0 | 0 | 0 | 0 | 0 | 0 |
| 31822 | ENSMUSG00000092039 | 0 | NA | NA | NA | NA | NA | Gm5501 | predicted gene 5501 [Source:MGI Symbol | 0 | 0 | 0 | 0 | 0 | 0 |
| 31823 | ENSMUSG00000092040 | 0 | NA | NA | NA | NA | NA | Gm17198 | predicted gene 17198 [Source:MGI Symbo | 0 | 0 | 0 | 0 | 0 | 0 |
| 31824 | ENSMUSG00000092042 | 0 | NA | NA | NA | NA | NA | Gm17163 | predicted gene 17163 [Source:MGI Symbo | 0 | 0 | 0 | 0 | 0 | 0 |
| 31825 | ENSMUSG00000092043 | 0 | NA | NA | NA | NA | NA | Esp6 | exocrine gland secreted peptide 6 [Source | 0 | 0 | 0 | 0 | 0 | 0 |
| 31826 | ENSMUSG00000092045 | 0 | NA | NA | NA | NA | NA | Gm17086 | predicted gene 17086 [Source:MGI Symbo | 0 | 0 | 0 | 0 | 0 | 0 |
| 31827 | ENSMUSG00000092047 | 0 | NA | NA | NA | NA | NA | Gm17631 | predicted gene, 17631 [Source:MGI Symb | 0 | 0 | 0 | 0 | 0 | 0 |
| 31828 | ENSMUSG00000092048 | 0 | NA | NA | NA | NA | NA | Vmn2r85 | vomeronasal 2, receptor 85 [Source:MGI S | 0 | 0 | 0 | 0 | 0 | 0 |
| 31829 | ENSMUSG00000092049 | 0 | NA | NA | NA | NA | NA | Vmn2r4 | vomeronasal 2, receptor 4 [Source:MGI Sy | 0 | 0 | 0 | 0 | 0 | 0 |
| 31830 | ENSMUSG00000092051 | 8.479041113 | 1.00689557 | 0.795528372 | 1.265694103 | 0.20562262 | 0.650907725 | Gm17229 | predicted gene 17229 [Source:MGI Symbo | 9.6372 | 2.0558 | 5.0312 | 15.424 | 9.8156 | 8.9108 |
| 31831 | ENSMUSG00000092054 | 0 | NA | NA | NA | NA | NA | Kif4-ps | kinesin family member 4, pseudogene [So | 0 | 0 | 0 | 0 | 0 | 0 |
| 31832 | ENSMUSG00000092055 | 0 | NA | NA | NA | NA | NA | Gm7925 | predicted gene 7925 [Source:MGI Symbol | 0 | 0 | 0 | 0 | 0 | 0 |
| 31833 | ENSMUSG00000092056 | 0 | NA | NA | NA | NA | NA | Gm7289 | predicted gene 7289 [Source:MGI Symbol | 0 | 0 | 0 | 0 | 0 | 0 |
| 31834 | ENSMUSG00000092060 | 6.335936117 | -1.390109267 | 0.877547581 | -1.584084211 | 0.113174557 | 0.493951959 | Bend4 | BEN domain containing 4 [Source:MGI Syr | 7.2279 | 10.279 | 10.062 | 3.8559 | 4.3625 | 2.2277 |
| 31835 | ENSMUSG00000092066 | 0 | NA | NA | NA | NA | NA | 1700121N20 | RIKEN cDNA 1700121N20 gene [Source:M | 0 | 0 | 0 | 0 | 0 | 0 |
| 31836 | ENSMUSG00000092071 | 0 | NA | NA | NA | NA | NA | A230065N10 | RIKEN cDNA A230065N10 gene [Source:M | 0 | 0 | 0 | 0 | 0 | 0 |
| 31837 | ENSMUSG00000092072 | 2.702710242 | -0.235934304 | 1.338469882 | -0.176271657 | 0.860080528 | NA | Gm4540 | predicted gene 4540 [Source:MGI Symbol | 5.6217 | 1.0279 | 2.0125 | 0.964 | 4.3625 | 2.2277 |
| 31838 | ENSMUSG00000092073 | 0 | NA | NA | NA | NA | NA | Gm6205 | predicted gene 6205 [Source:MGI Symbol | 0 | 0 | 0 | 0 | 0 | 0 |
| 31839 | ENSMUSG00000092074 | 7.357113188 | -1.000846141 | 0.805239222 | -1.242917773 | 0.213898136 | 0.658680569 | Dynlt1a | dynein light chain Tctex-type 1A [Source:N | 12.047 | 11.307 | 6.0375 | 4.8199 | 4.3625 | 5.5692 |
| 31840 | ENSMUSG00000092075 | 2344.99654 | 2.204300782 | 0.630252309 | 3.497489417 | 0.000469659 | 0.011855415 | Serpina4-ps1 | serine (or cysteine) peptidase inhibitor, cl | 811.94 | 778.13 | 918.7 | 7363.8 | 2491 | 1706.4 |
| 31841 | ENSMUSG00000092077 | 0 | NA | NA | NA | NA | NA | Olfr101 | olfactory receptor 101 [Source:MGI Symb | 0 | 0 | 0 | 0 | 0 | 0 |
| 31842 | ENSMUSG00000092080 | 0 | NA | NA | NA | NA | NA | Vmn2r16 | vomeronasal 2, receptor 16 [Source:MGI S | 0 | 0 | 0 | 0 | 0 | 0 |
| 31843 | ENSMUSG00000092083 | 0 | NA | NA | NA | NA | NA | Kcnb2 | potassium voltage gated channel, Shab-re | 0 | 0 | 0 | 0 | 0 | 0 |
| 31844 | ENSMUSG00000092085 | 0 | NA | NA | NA | NA | NA | Gm17224 | predicted gene 17224 [Source:MGI Symbo | 0 | 0 | 0 | 0 | 0 | 0 |
| 31845 | ENSMUSG00000092086 | 2.886728065 | -1.176334214 | 1.294809955 | -0.908499513 | 0.363614371 | NA | Gm6793 | predicted gene 6793 [Source:MGI Symbol | 4.8186 | 5.1396 | 2.0125 | 0.964 | 3.2719 | 1.1138 |
| 31846 | ENSMUSG00000092090 | 0.185640647 | 1.126924175 | 4.080472857 | 0.276174898 | 0.782413736 | NA | Gm3294 | predicted gene 3294 [Source:MGI Symbol | 0 | 0 | 0 | 0 | 0 | 1.1138 |
| 31847 | ENSMUSG00000092092 | 0 | NA | NA | NA | NA | NA | Gm17215 | predicted gene 17215 [Source:MGI Symbo | 0 | 0 | 0 | 0 | 0 | 0 |
| 31848 | ENSMUSG00000092093 | 0 | NA | NA | NA | NA | NA | Gm17528 | predicted gene, 17528 [Source:MGI Symb | 0 | 0 | 0 | 0 | 0 | 0 |
| 31849 | ENSMUSG00000092094 | 0.17131964 | -0.796629176 | 4.080472857 | -0.195229623 | 0.84521319 | NA | Zfp804b | zinc finger protein 804B [Source:MGI Sym | 0 | 1.0279 | 0 | 0 | 0 | 0 |
| 31850 | ENSMUSG00000092097 | 2.292285251 | 0.162535729 | 1.477665102 | 0.10999497 | 0.912413364 | NA | Gm5819 | predicted gene 5819 [Source:MGI Symbol | 2.4093 | 3.0838 | 1.0062 | 2.8919 | 4.3625 | 0 |
| 31851 | ENSMUSG00000092103 | 0.958473062 | 0.410959158 | 2.705027395 | 0.151924213 | 0.879246711 | NA | Gm17039 | predicted gene 17039 [Source:MGI Symbo | 2.4093 | 0 | 0 | 0 | 0 | 3.3415 |
| 31852 | ENSMUSG00000092107 | 0 | NA | NA | NA | NA | NA | Ear-ps4 | eosinophil-associated, ribonuclease A fam | 0 | 0 | 0 | 0 | 0 | 0 |
| 31853 | ENSMUSG00000092109 | 0 | NA | NA | NA | NA | NA | Gm9376 | predicted gene 9376 [Source:MGI Symbol | 0 | 0 | 0 | 0 | 0 | 0 |
| 31854 | ENSMUSG00000092111 | 0 | NA | NA | NA | NA | NA | Vmn2r113 | vomeronasal 2, receptor 113 [Source:MGI | 0 | 0 | 0 | 0 | 0 | 0 |
| 31855 | ENSMUSG00000092116 | 2.063009904 | 1.067414561 | 1.589887065 | 0.671377599 | 0.501980011 | NA | Gm10320 | predicted pseudogene 10320 [Source:MG | 0 | 3.0838 | 1.0062 | 3.8559 | 1.0906 | 3.3415 |
| 31856 | ENSMUSG00000092118 | 79.94912352 | 0.357512357 | 0.308547756 | 1.15869375 | 0.246581039 | 0.696725814 | Fancf | Fanconi anemia, complementation group | 60.233 | 72.982 | 77.481 | 110.86 | 53.44 | 104.7 |
| 31857 | ENSMUSG00000092120 | 0 | NA | NA | NA | NA | NA | Vmn2r90 | vomeronasal 2, receptor 90 [Source:MGI S | 0 | 0 | 0 | 0 | 0 | 0 |
| 31858 | ENSMUSG00000092123 | 0 | NA | NA | NA | NA | NA | Gm17581 | predicted gene, 17581 [Source:MGI Symb | 0 | 0 | 0 | 0 | 0 | 0 |
| 31859 | ENSMUSG00000092124 | 8.696007608 | -0.089379795 | 0.73184402 | -0.12212957 | 0.902796404 | 0.983821605 | B930094E09 | RIKEN cDNA B930094E09 gene [Source:M | 9.6372 | 5.1396 | 12.075 | 7.7118 | 9.8156 | 7.7969 |
| 31860 | ENSMUSG00000092125 | 0 | NA | NA | NA | NA | NA | AA543401 | cDNA sequence, AA543401 [Source:MGI S | 0 | 0 | 0 | 0 | 0 | 0 |

| | | | | | | | | | | | | | | | |
|---|---|---|---|---|---|---|---|---|---|---|---|---|---|---|---|
| 31861 | ENSMUSG00000092130 | 0 | NA | NA | NA | NA | NA | D030025P21 | RIKEN cDNA D030025P21 gene [Source:M | 0 | 0 | 0 | 0 | 0 | 0 |
| 31862 | ENSMUSG00000092131 | 0.160662974 | 1.126924175 | 4.080472857 | 0.276174898 | 0.782413736 | NA | BC050972 | cDNA sequence BC050972 [Source:MGI Sy | 0 | 0 | 0 | 0.964 | 0 | 0 |
| 31863 | ENSMUSG00000092134 | 1.048404808 | 3.600807465 | 2.562882816 | 1.404983264 | 0.160026255 | NA | Gm17089 | predicted gene 17089 [Source:MGI Symbo | 0 | 0 | 0 | 1.928 | 4.3625 | 0 |
| 31864 | ENSMUSG00000092142 | 0 | NA | NA | NA | NA | NA | Gm6482 | predicted gene 6482 [Source:MGI Symbol | 0 | 0 | 0 | 0 | 0 | 0 |
| 31865 | ENSMUSG00000092144 | 0 | NA | NA | NA | NA | NA | Gm3370 | predicted gene 3370 [Source:MGI Symbol | 0 | 0 | 0 | 0 | 0 | 0 |
| 31866 | ENSMUSG00000092147 | 0 | NA | NA | NA | NA | NA | Gm7052 | predicted gene 7052 [Source:MGI Symbol | 0 | 0 | 0 | 0 | 0 | 0 |
| 31867 | ENSMUSG00000092148 | 0 | NA | NA | NA | NA | NA | Gm8005 | predicted gene 8005 [Source:MGI Symbol | 0 | 0 | 0 | 0 | 0 | 0 |
| 31868 | ENSMUSG00000092152 | 0 | NA | NA | NA | NA | NA | Gm17026 | predicted gene 17026 [Source:MGI Symbo | 0 | 0 | 0 | 0 | 0 | 0 |
| 31869 | ENSMUSG00000092157 | 0 | NA | NA | NA | NA | NA | Gm17268 | predicted gene, 17268 [Source:MGI Symb | 0 | 0 | 0 | 0 | 0 | 0 |
| 31870 | ENSMUSG00000092162 | 0 | NA | NA | NA | NA | NA | Vmn2r86 | vomeronasal 2, receptor 86 [Source:MGI S | 0 | 0 | 0 | 0 | 0 | 0 |
| 31871 | ENSMUSG00000092164 | 0 | NA | NA | NA | NA | NA | Rergl | RERG/RAS-like [Source:MGI Symbol;Acc:N | 0 | 0 | 0 | 0 | 0 | 0 |
| 31872 | ENSMUSG00000092165 | 0 | NA | NA | NA | NA | NA | Gm5624 | predicted gene 5624 [Source:MGI Symbol | 0 | 0 | 0 | 0 | 0 | 0 |
| 31873 | ENSMUSG00000092166 | 0 | NA | NA | NA | NA | NA | Pramel45 | PRAME like 45 [Source:MGI Symbol;Acc:M | 0 | 0 | 0 | 0 | 0 | 0 |
| 31874 | ENSMUSG00000092167 | 0 | NA | NA | NA | NA | NA | Gm3696 | predicted gene 3696 [Source:MGI Symbol | 0 | 0 | 0 | 0 | 0 | 0 |
| 31875 | ENSMUSG00000092168 | 0 | NA | NA | NA | NA | NA | Gm9075 | predicted gene 9075 [Source:MGI Symbol | 0 | 0 | 0 | 0 | 0 | 0 |
| 31876 | ENSMUSG00000092171 | 0 | NA | NA | NA | NA | NA | 4833427F10 | RIKEN cDNA 4833427F10 gene [Source:M | 0 | 0 | 0 | 0 | 0 | 0 |
| 31877 | ENSMUSG00000092172 | 0 | NA | NA | NA | NA | NA | Gm20523 | predicted gene 20523 [Source:MGI Symbo | 0 | 0 | 0 | 0 | 0 | 0 |
| 31878 | ENSMUSG00000092173 | 0 | NA | NA | NA | NA | NA | Vmn2r-ps38 | vomeronasal 2, receptor, pseudogene 38 | 0 | 0 | 0 | 0 | 0 | 0 |
| 31879 | ENSMUSG00000092174 | 0 | NA | NA | NA | NA | NA | NA | NA | 0 | 0 | 0 | 0 | 0 | 0 |
| 31880 | ENSMUSG00000092175 | 0 | NA | NA | NA | NA | NA | Gm3584 | predicted gene 3584 [Source:MGI Symbol | 0 | 0 | 0 | 0 | 0 | 0 |
| 31881 | ENSMUSG00000092176 | 0 | NA | NA | NA | NA | NA | Gm20460 | predicted gene 20460 [Source:MGI Symbo | 0 | 0 | 0 | 0 | 0 | 0 |
| 31882 | ENSMUSG00000092178 | 1.413442265 | -3.880278124 | 2.342991152 | -1.656121544 | 0.097697207 | NA | Gm45351 | predicted gene 45351 [Source:MGI Symbo | 6.4248 | 2.0558 | 0 | 0 | 0 | 0 |
| 31883 | ENSMUSG00000092179 | 0 | NA | NA | NA | NA | NA | Vmn2r-ps75 | vomeronasal 2, receptor, pseudogene 75 | 0 | 0 | 0 | 0 | 0 | 0 |
| 31884 | ENSMUSG00000092181 | 0.160662974 | 1.126924175 | 4.080472857 | 0.276174898 | 0.782413736 | NA | Gm20432 | predicted gene 20432 [Source:MGI Symbo | 0 | 0 | 0 | 0.964 | 0 | 0 |
| 31885 | ENSMUSG00000092182 | 0.301557804 | -1.64335916 | 4.042814367 | -0.406488899 | 0.684383412 | NA | Gm20407 | predicted gene 20407 [Source:MGI Symbo | 0.8031 | 0 | 1.0062 | 0 | 0 | 0 |
| 31886 | ENSMUSG00000092183 | 3.697166157 | -0.744703498 | 1.123955505 | -0.662573825 | 0.507603539 | NA | 4930515G01 | RIKEN cDNA 4930515G01 gene [Source:M | 4.8186 | 2.0558 | 7.0437 | 3.8559 | 2.1812 | 2.2277 |
| 31887 | ENSMUSG00000092184 | 0 | NA | NA | NA | NA | NA | Gm18489 | predicted gene, 18489 [Source:MGI Symb | 0 | 0 | 0 | 0 | 0 | 0 |
| 31888 | ENSMUSG00000092185 | 0 | NA | NA | NA | NA | NA | Gm20416 | predicted gene 20416 [Source:MGI Symbo | 0 | 0 | 0 | 0 | 0 | 0 |
| 31889 | ENSMUSG00000092186 | 0 | NA | NA | NA | NA | NA | Gm5967 | predicted gene 5967 [Source:MGI Symbol | 0 | 0 | 0 | 0 | 0 | 0 |
| 31890 | ENSMUSG00000092187 | 20.8398361 | -0.583184196 | 0.478531072 | -1.218696613 | 0.222959364 | 0.668407528 | Gm20457 | predicted gene 20457 [Source:MGI Symbo | 23.29 | 25.698 | 26.162 | 21.208 | 13.087 | 15.594 |
| 31891 | ENSMUSG00000092188 | 0 | NA | NA | NA | NA | NA | Gm20471 | predicted gene 20471 [Source:MGI Symbo | 0 | 0 | 0 | 0 | 0 | 0 |
| 31892 | ENSMUSG00000092190 | 1.669660223 | 0.18897775 | 1.593280112 | 0.118609245 | 0.905584941 | NA | Gm20470 | predicted gene 20470 [Source:MGI Symbo | 1.6062 | 2.0558 | 1.0062 | 0.964 | 3.2719 | 1.1138 |
| 31893 | ENSMUSG00000092191 | 0 | NA | NA | NA | NA | NA | Gm7211 | predicted gene 7211 [Source:MGI Symbol | 0 | 0 | 0 | 0 | 0 | 0 |
| 31894 | ENSMUSG00000092192 | 0 | NA | NA | NA | NA | NA | Dnaaf4 | dynein axonemal assembly factor 4 [Sourc | 0 | 0 | 0 | 0 | 0 | 0 |
| 31895 | ENSMUSG00000092193 | 0 | NA | NA | NA | NA | NA | Cd9-ps | Cd9 antigen, pseudogene [Source:MGI Sy | 0 | 0 | 0 | 0 | 0 | 0 |
| 31896 | ENSMUSG00000092195 | 0 | NA | NA | NA | NA | NA | Vmn2r-ps87 | vomeronasal 2, receptor, pseudogene 87 | 0 | 0 | 0 | 0 | 0 | 0 |
| 31897 | ENSMUSG00000092196 | 0 | NA | NA | NA | NA | NA | Gm38505 | predicted gene, 38505 [Source:MGI Symb | 0 | 0 | 0 | 0 | 0 | 0 |
| 31898 | ENSMUSG00000092197 | 0 | NA | NA | NA | NA | NA | Gm20480 | predicted gene 20480 [Source:MGI Symbo | 0 | 0 | 0 | 0 | 0 | 0 |
| 31899 | ENSMUSG00000092199 | 0 | NA | NA | NA | NA | NA | Gm18358 | predicted gene, 18358 [Source:MGI Symb | 0 | 0 | 0 | 0 | 0 | 0 |
| 31900 | ENSMUSG00000092200 | 0.688736311 | 2.998073904 | 2.697618692 | 1.11137794 | 0.266405704 | NA | Tnxa | tenascin XA (pseudogene) [Source:MGI Sy | 0 | 0 | 0 | 1.928 | 1.0906 | 1.1138 |
| 31901 | ENSMUSG00000092201 | 0 | NA | NA | NA | NA | NA | A530058N18 | RIKEN cDNA A530058N18 gene [Source:M | 0 | 0 | 0 | 0 | 0 | 0 |
| 31902 | ENSMUSG00000092202 | 0.669251866 | -2.807208097 | 3.391018549 | -0.827836255 | 0.407763239 | NA | Stk19-ps1 | serine/threonine kinase 19, pseudogene 1 | 4.0155 | 0 | 0 | 0 | 0 | 0 |
| 31903 | ENSMUSG00000092203 | 146.4924297 | 0.189500937 | 0.216497323 | 0.875303837 | 0.381408607 | 0.8130053 | 1110038B12 | RIKEN cDNA 1110038B12 gene [Source:M | 168.65 | 119.24 | 121.76 | 139.78 | 164.68 | 164.85 |
| 31904 | ENSMUSG00000092204 | 0 | NA | NA | NA | NA | NA | Vmn2r-ps49 | vomeronasal 2, receptor, pseudogene 49 | 0 | 0 | 0 | 0 | 0 | 0 |
| 31905 | ENSMUSG00000092208 | 0 | NA | NA | NA | NA | NA | Vmn1r-ps58 | vomeronasal 1 receptor, pseudogene 58 [ | 0 | 0 | 0 | 0 | 0 | 0 |
| 31906 | ENSMUSG00000092210 | 0.672839099 | 0.267535205 | 2.816195809 | 0.094998794 | 0.924315785 | NA | A930009A15 | RIKEN cDNA A930009A15 gene [Source:M | 0.8031 | 0 | 1.0062 | 0 | 0 | 2.2277 |
| 31907 | ENSMUSG00000092211 | 0 | NA | NA | NA | NA | NA | Vmn2r-ps67 | vomeronasal 2, receptor, pseudogene 67 | 0 | 0 | 0 | 0 | 0 | 0 |
| 31908 | ENSMUSG00000092212 | 2.529101615 | 0.680583024 | 1.452802053 | 0.468462322 | 0.639454009 | NA | Slc22a13b | solute carrier family 22 (organic cation tra | 0.8031 | 3.0838 | 2.0125 | 4.8199 | 0 | 4.4554 |
| 31909 | ENSMUSG00000092213 | 0 | NA | NA | NA | NA | NA | Gm18522 | predicted gene, 18522 [Source:MGI Symb | 0 | 0 | 0 | 0 | 0 | 0 |
| 31910 | ENSMUSG00000092214 | 0 | NA | NA | NA | NA | NA | Gm18560 | predicted gene, 18560 [Source:MGI Symb | 0 | 0 | 0 | 0 | 0 | 0 |
| 31911 | ENSMUSG00000092216 | 0 | NA | NA | NA | NA | NA | Gm19345 | predicted gene, 19345 [Source:MGI Symb | 0 | 0 | 0 | 0 | 0 | 0 |
| 31912 | ENSMUSG00000092217 | 0 | NA | NA | NA | NA | NA | Vmn2r-ps62 | vomeronasal 2, receptor, pseudogene 62 | 0 | 0 | 0 | 0 | 0 | 0 |
| 31913 | ENSMUSG00000092218 | 0 | NA | NA | NA | NA | NA | Gm20529 | predicted gene 20529 [Source:MGI Symbo | 0 | 0 | 0 | 0 | 0 | 0 |
| 31914 | ENSMUSG00000092219 | 0 | NA | NA | NA | NA | NA | Gm20443 | predicted gene 20443 [Source:MGI Symbo | 0 | 0 | 0 | 0 | 0 | 0 |
| 31915 | ENSMUSG00000092220 | 0 | NA | NA | NA | NA | NA | Gm20528 | predicted gene 20528 [Source:MGI Symbo | 0 | 0 | 0 | 0 | 0 | 0 |
| 31916 | ENSMUSG00000092221 | 0 | NA | NA | NA | NA | NA | Gm7230 | predicted gene 7230 [Source:MGI Symbol | 0 | 0 | 0 | 0 | 0 | 0 |

| | | | | | | | | | | | | | | | |
|---|---|---|---|---|---|---|---|---|---|---|---|---|---|---|---|
| 31917 | ENSMUSG00000092222 | 0.185640647 | 1.126924175 | 4.080472857 | 0.276174898 | 0.782413736 | NA | NA | NA | 0 | 0 | 0 | 0 | 0 | 1.1138 |
| 31918 | ENSMUSG00000092223 | 0 | NA | NA | NA | NA | NA | Gm19807 | predicted gene, 19807 [Source:MGI Symb | 0 | 0 | 0 | 0 | 0 | 0 |
| 31919 | ENSMUSG00000092224 | 0 | NA | NA | NA | NA | NA | Gm8815 | predicted gene 8815 [Source:MGI Symbol | 0 | 0 | 0 | 0 | 0 | 0 |
| 31920 | ENSMUSG00000092225 | 0 | NA | NA | NA | NA | NA | Gm2381 | predicted gene 2381 [Source:MGI Symbol | 0 | 0 | 0 | 0 | 0 | 0 |
| 31921 | ENSMUSG00000092227 | 0 | NA | NA | NA | NA | NA | Gm7651 | predicted gene 7651 [Source:MGI Symbol | 0 | 0 | 0 | 0 | 0 | 0 |
| 31922 | ENSMUSG00000092228 | 0 | NA | NA | NA | NA | NA | Vmn1r-ps28 | vomeronasal 1 receptor, pseudogene 28 [ | 0 | 0 | 0 | 0 | 0 | 0 |
| 31923 | ENSMUSG00000092229 | 0.353348078 | 0.165142701 | 3.895042568 | 0.042398176 | 0.966181282 | NA | Gm5977 | predicted gene 5977 [Source:MGI Symbol | 0 | 0 | 1.0062 | 0 | 0 | 1.1138 |
| 31924 | ENSMUSG00000092230 | 0 | NA | NA | NA | NA | NA | Gm18889 | predicted gene, 18889 [Source:MGI Symb | 0 | 0 | 0 | 0 | 0 | 0 |
| 31925 | ENSMUSG00000092231 | 0 | NA | NA | NA | NA | NA | Lgals1-ps1 | lectin, galactose binding, soluble 1, pseud | 0 | 0 | 0 | 0 | 0 | 0 |
| 31926 | ENSMUSG00000092232 | 0 | NA | NA | NA | NA | NA | Gm20521 | predicted gene 20521 [Source:MGI Symbo | 0 | 0 | 0 | 0 | 0 | 0 |
| 31927 | ENSMUSG00000092233 | 1.277737502 | -2.558404199 | 2.061944144 | -1.240772795 | 0.214689694 | NA | Gm20403 | predicted gene 20403 [Source:MGI Symbo | 1.6062 | 3.0838 | 2.0125 | 0.964 | 0 | 0 |
| 31928 | ENSMUSG00000092237 | 0 | NA | NA | NA | NA | NA | Vmn1r-ps27 | vomeronasal 1 receptor, pseudogene 27 [ | 0 | 0 | 0 | 0 | 0 | 0 |
| 31929 | ENSMUSG00000092238 | 0 | NA | NA | NA | NA | NA | Gm6004 | predicted gene 6004 [Source:MGI Symbol | 0 | 0 | 0 | 0 | 0 | 0 |
| 31930 | ENSMUSG00000092239 | 0 | NA | NA | NA | NA | NA | 1700031A10 | RIKEN cDNA 1700031A10 gene [Source:M | 0 | 0 | 0 | 0 | 0 | 0 |
| 31931 | ENSMUSG00000092240 | 0 | NA | NA | NA | NA | NA | Gm4246 | predicted gene 4246 [Source:MGI Symbol | 0 | 0 | 0 | 0 | 0 | 0 |
| 31932 | ENSMUSG00000092241 | 14.78366148 | 0.20164564 | 0.561590319 | 0.359061816 | 0.719548846 | 0.9447719 | Gm20522 | predicted gene 20522 [Source:MGI Symbo | 17.668 | 12.335 | 11.069 | 13.496 | 18.541 | 15.594 |
| 31933 | ENSMUSG00000092242 | 0 | NA | NA | NA | NA | NA | Gm20515 | predicted gene 20515 [Source:MGI Symbo | 0 | 0 | 0 | 0 | 0 | 0 |
| 31934 | ENSMUSG00000092243 | 0.486939728 | -0.698472448 | 3.137608162 | -0.222613026 | 0.823836697 | NA | Gm7030 | predicted gene 7030 [Source:MGI Symbol | 0.8031 | 1.0279 | 0 | 0 | 1.0906 | 0 |
| 31935 | ENSMUSG00000092244 | 0 | NA | NA | NA | NA | NA | Esp34 | exocrine gland secreted peptide 34 [Sourc | 0 | 0 | 0 | 0 | 0 | 0 |
| 31936 | ENSMUSG00000092245 | 0 | NA | NA | NA | NA | NA | Gm17973 | predicted gene, 17973 [Source:MGI Symb | 0 | 0 | 0 | 0 | 0 | 0 |
| 31937 | ENSMUSG00000092246 | 0 | NA | NA | NA | NA | NA | Gm20493 | predicted gene 20493 [Source:MGI Symbo | 0 | 0 | 0 | 0 | 0 | 0 |
| 31938 | ENSMUSG00000092247 | 6.012009197 | -0.257007004 | 1.004013336 | -0.255979671 | 0.797966534 | 0.962377049 | Gm20426 | predicted gene 20426 [Source:MGI Symbo | 2.4093 | 14.391 | 3.0187 | 9.6398 | 3.2719 | 3.3415 |
| 31939 | ENSMUSG00000092248 | 0 | NA | NA | NA | NA | NA | Vmn1r-ps63 | vomeronasal 1 receptor, pseudogene 63 [ | 0 | 0 | 0 | 0 | 0 | 0 |
| 31940 | ENSMUSG00000092250 | 0 | NA | NA | NA | NA | NA | Gm20467 | predicted gene 20467 [Source:MGI Symbo | 0 | 0 | 0 | 0 | 0 | 0 |
| 31941 | ENSMUSG00000092251 | 0 | NA | NA | NA | NA | NA | Trav9n-1 | T cell receptor alpha variable 9N-1 [Sourc | 0 | 0 | 0 | 0 | 0 | 0 |
| 31942 | ENSMUSG00000092252 | 0 | NA | NA | NA | NA | NA | NA | NA | 0 | 0 | 0 | 0 | 0 | 0 |
| 31943 | ENSMUSG00000092253 | 0 | NA | NA | NA | NA | NA | H2-Q3 | histocompatibility 2, Q region locus 3 [Sou | 0 | 0 | 0 | 0 | 0 | 0 |
| 31944 | ENSMUSG00000092255 | 0 | NA | NA | NA | NA | NA | Gm20455 | predicted gene 20455 [Source:MGI Symbo | 0 | 0 | 0 | 0 | 0 | 0 |
| 31945 | ENSMUSG00000092256 | 0 | NA | NA | NA | NA | NA | Gm18359 | predicted gene, 18359 [Source:MGI Symb | 0 | 0 | 0 | 0 | 0 | 0 |
| 31946 | ENSMUSG00000092257 | 0.294513347 | 0.165143496 | 4.080472857 | 0.040471657 | 0.967717103 | NA | Gm2308 | predicted gene 2308 [Source:MGI Symbol | 0.8031 | 0 | 0 | 0.964 | 0 | 0 |
| 31947 | ENSMUSG00000092258 | 0 | NA | NA | NA | NA | NA | Gm20444 | predicted gene 20444 [Source:MGI Symbo | 0 | 0 | 0 | 0 | 0 | 0 |
| 31948 | ENSMUSG00000092259 | 0 | NA | NA | NA | NA | NA | Vmn2r-ps68 | vomeronasal 2, receptor, pseudogene 68 | 0 | 0 | 0 | 0 | 0 | 0 |
| 31949 | ENSMUSG00000092260 | 34.50326221 | -0.179368551 | 0.375764822 | -0.477342585 | 0.633118192 | 0.922107984 | Zfp963 | zinc finger protein 963 [Source:MGI Symb | 32.927 | 38.033 | 39.244 | 39.523 | 29.447 | 27.846 |
| 31950 | ENSMUSG00000092261 | 0.331982614 | 0.16514309 | 3.986751476 | 0.041422971 | 0.966958701 | NA | Gm20519 | predicted gene 20519 [Source:MGI Symbo | 0 | 1.0279 | 0 | 0.964 | 0 | 0 |
| 31951 | ENSMUSG00000092263 | 0 | NA | NA | NA | NA | NA | Vmn2r-ps79 | vomeronasal 2, receptor, pseudogene 79 | 0 | 0 | 0 | 0 | 0 | 0 |
| 31952 | ENSMUSG00000092264 | 0 | NA | NA | NA | NA | NA | Gm3585 | predicted gene 3585 [Source:MGI Symbol | 0 | 0 | 0 | 0 | 0 | 0 |
| 31953 | ENSMUSG00000092265 | 0 | NA | NA | NA | NA | NA | Gm5682 | predicted gene 5682 [Source:MGI Symbol | 0 | 0 | 0 | 0 | 0 | 0 |
| 31954 | ENSMUSG00000092266 | 0 | NA | NA | NA | NA | NA | Gm20500 | predicted gene 20500 [Source:MGI Symbo | 0 | 0 | 0 | 0 | 0 | 0 |
| 31955 | ENSMUSG00000092267 | 0 | NA | NA | NA | NA | NA | NA | NA | 0 | 0 | 0 | 0 | 0 | 0 |
| 31956 | ENSMUSG00000092268 | 5.120391218 | 0.379913178 | 1.131699663 | 0.335701415 | 0.737096039 | 0.948137969 | Gm20540 | predicted gene 20540 [Source:MGI Symbo | 3.2124 | 8.2233 | 2.0125 | 9.6398 | 7.6343 | 0 |
| 31957 | ENSMUSG00000092269 | 0 | NA | NA | NA | NA | NA | Gm9577 | predicted gene 9577 [Source:MGI Symbol | 0 | 0 | 0 | 0 | 0 | 0 |
| 31958 | ENSMUSG00000092270 | 0 | NA | NA | NA | NA | NA | Gm7031 | predicted gene 7031 [Source:MGI Symbol | 0 | 0 | 0 | 0 | 0 | 0 |
| 31959 | ENSMUSG00000092271 | 0 | NA | NA | NA | NA | NA | Vmn1r-ps46 | vomeronasal 1 receptor, pseudogene 46 [ | 0 | 0 | 0 | 0 | 0 | 0 |
| 31960 | ENSMUSG00000092272 | 0 | NA | NA | NA | NA | NA | Gm19301 | predicted gene, 19301 [Source:MGI Symb | 0 | 0 | 0 | 0 | 0 | 0 |
| 31961 | ENSMUSG00000092274 | 2080.364762 | 0.37479688 | 0.170046125 | 2.204089505 | 0.02751805 | 0.215379371 | Neat1 | nuclear paraspeckle assembly transcript 1 | 2346.7 | 1428.8 | 1657.3 | 1978.1 | 2666.6 | 2404.8 |
| 31962 | ENSMUSG00000092275 | 0 | NA | NA | NA | NA | NA | Gm20465 | predicted gene 20465 [Source:MGI Symbo | 0 | 0 | 0 | 0 | 0 | 0 |
| 31963 | ENSMUSG00000092276 | 0 | NA | NA | NA | NA | NA | 1700116B05 | RIKEN cDNA 1700116B05 gene [Source:M | 0 | 0 | 0 | 0 | 0 | 0 |
| 31964 | ENSMUSG00000092277 | 3.231021186 | 1.106668374 | 1.330600301 | 0.83170609 | 0.405574862 | NA | Gm19684 | predicted gene, 19684 [Source:MGI Symb | 4.0155 | 2.0558 | 0 | 6.7478 | 5.4531 | 1.1138 |
| 31965 | ENSMUSG00000092278 | 0 | NA | NA | NA | NA | NA | Gm8752 | predicted pseudogene 8752 [Source:MGI | 0 | 0 | 0 | 0 | 0 | 0 |
| 31966 | ENSMUSG00000092281 | 0.677847551 | 0.063310748 | 2.809814301 | 0.022532004 | 0.982023583 | NA | Hmgb1-ps7 | high-mobility group high mobility group b | 0 | 0 | 2.0125 | 0.964 | 1.0906 | 0 |
| 31967 | ENSMUSG00000092283 | 0 | NA | NA | NA | NA | NA | Gm20412 | predicted gene 20412 [Source:MGI Symbo | 0 | 0 | 0 | 0 | 0 | 0 |
| 31968 | ENSMUSG00000092284 | 0 | NA | NA | NA | NA | NA | Gm8801 | predicted gene 8801 [Source:MGI Symbol | 0 | 0 | 0 | 0 | 0 | 0 |
| 31969 | ENSMUSG00000092285 | 0 | NA | NA | NA | NA | NA | Vmn1r-ps56 | vomeronasal 1 receptor, pseudogene 56 [ | 0 | 0 | 0 | 0 | 0 | 0 |
| 31970 | ENSMUSG00000092286 | 0.339027071 | -1.79174149 | 3.919585585 | -0.457125237 | 0.647581032 | NA | Dtnbos | dystrobrevin, beta, opposite strand [Sourc | 0 | 1.0279 | 1.0062 | 0 | 0 | 0 |
| 31971 | ENSMUSG00000092287 | 0 | NA | NA | NA | NA | NA | Vmn2r-ps77 | vomeronasal 2, receptor, pseudogene 77 | 0 | 0 | 0 | 0 | 0 | 0 |
| 31972 | ENSMUSG00000092289 | 0 | NA | NA | NA | NA | NA | Gm6659 | predicted gene 6659 [Source:MGI Symbol | 0 | 0 | 0 | 0 | 0 | 0 |

| | | | | | | | | | | | | | | | |
|---|---|---|---|---|---|---|---|---|---|---|---|---|---|---|---|
| 31973 | ENSMUSG00000092290 | 0 | NA | NA | NA | NA | NA | Gm20452 | predicted gene 20452 [Source:MGI Symbo | 0 | 0 | 0 | 0 | 0 | 0 |
| 31974 | ENSMUSG00000092292 | 0 | NA | NA | NA | NA | NA | Olfr113 | olfactory receptor 113 [Source:MGI Symb | 0 | 0 | 0 | 0 | 0 | 0 |
| 31975 | ENSMUSG00000092293 | 0 | NA | NA | NA | NA | NA | Gm17919 | predicted gene, 17919 [Source:MGI Symb | 0 | 0 | 0 | 0 | 0 | 0 |
| 31976 | ENSMUSG00000092294 | 0 | NA | NA | NA | NA | NA | Gm18200 | predicted gene, 18200 [Source:MGI Symb | 0 | 0 | 0 | 0 | 0 | 0 |
| 31977 | ENSMUSG00000092295 | 0 | NA | NA | NA | NA | NA | Gm20543 | predicted gene 20543 [Source:MGI Symbo | 0 | 0 | 0 | 0 | 0 | 0 |
| 31978 | ENSMUSG00000092296 | 0 | NA | NA | NA | NA | NA | Gm20454 | predicted gene 20454 [Source:MGI Symbo | 0 | 0 | 0 | 0 | 0 | 0 |
| 31979 | ENSMUSG00000092297 | 0 | NA | NA | NA | NA | NA | Vmn1r258 | vomeronasal 1 receptor 258 [Source:MGI | 0 | 0 | 0 | 0 | 0 | 0 |
| 31980 | ENSMUSG00000092298 | 0 | NA | NA | NA | NA | NA | Gm8877 | predicted gene 8877 [Source:MGI Symbol | 0 | 0 | 0 | 0 | 0 | 0 |
| 31981 | ENSMUSG00000092300 | 4.063134184 | 0.82447414 | 1.101620883 | 0.748419127 | 0.454207389 | NA | Cdk3 | cyclin-dependent kinase 3 [Source:MGI Sy | 0.8031 | 4.1117 | 4.025 | 7.7118 | 3.2719 | 4.4554 |
| 31982 | ENSMUSG00000092301 | 0 | NA | NA | NA | NA | NA | Gm6887 | predicted gene 6887 [Source:MGI Symbol | 0 | 0 | 0 | 0 | 0 | 0 |
| 31983 | ENSMUSG00000092303 | 0 | NA | NA | NA | NA | NA | Vmn1r-ps29 | vomeronasal 1 receptor, pseudogene 29 [ | 0 | 0 | 0 | 0 | 0 | 0 |
| 31984 | ENSMUSG00000092304 | 0 | NA | NA | NA | NA | NA | Gm20530 | predicted gene 20530 [Source:MGI Symbo | 0 | 0 | 0 | 0 | 0 | 0 |
| 31985 | ENSMUSG00000092305 | 0.642651897 | 2.906894826 | 3.440535777 | 0.844895974 | 0.398168923 | NA | Prps1l1 | phosphoribosyl pyrophosphate synthetas | 0 | 0 | 0 | 3.8559 | 0 | 0 |
| 31986 | ENSMUSG00000092306 | 0 | NA | NA | NA | NA | NA | Gm20447 | predicted gene 20447 [Source:MGI Symbo | 0 | 0 | 0 | 0 | 0 | 0 |
| 31987 | ENSMUSG00000092307 | 0 | NA | NA | NA | NA | NA | Vmn2r-ps76 | vomeronasal 2, receptor, pseudogene 76 | 0 | 0 | 0 | 0 | 0 | 0 |
| 31988 | ENSMUSG00000092308 | 0 | NA | NA | NA | NA | NA | Gm3623 | predicted gene 3623 [Source:MGI Symbol | 0 | 0 | 0 | 0 | 0 | 0 |
| 31989 | ENSMUSG00000092309 | 0 | NA | NA | NA | NA | NA | 4930518C09 | RIKEN cDNA 4930518C09 gene [Source:M | 0 | 0 | 0 | 0 | 0 | 0 |
| 31990 | ENSMUSG00000092310 | 0 | NA | NA | NA | NA | NA | Gm20509 | predicted gene 20509 [Source:MGI Symbo | 0 | 0 | 0 | 0 | 0 | 0 |
| 31991 | ENSMUSG00000092311 | 0 | NA | NA | NA | NA | NA | Vmn1r-ps26 | vomeronasal 1 receptor, pseudogene 26 [ | 0 | 0 | 0 | 0 | 0 | 0 |
| 31992 | ENSMUSG00000092312 | 0 | NA | NA | NA | NA | NA | Zfp419 | zinc finger protein 419 [Source:MGI Symb | 0 | 0 | 0 | 0 | 0 | 0 |
| 31993 | ENSMUSG00000092313 | 0 | NA | NA | NA | NA | NA | Gm20415 | predicted gene 20415 [Source:MGI Symbo | 0 | 0 | 0 | 0 | 0 | 0 |
| 31994 | ENSMUSG00000092314 | 0 | NA | NA | NA | NA | NA | Gm3563 | predicted gene 3563 [Source:MGI Symbol | 0 | 0 | 0 | 0 | 0 | 0 |
| 31995 | ENSMUSG00000092316 | 0 | NA | NA | NA | NA | NA | Gm6726 | predicted gene 6726 [Source:MGI Symbol | 0 | 0 | 0 | 0 | 0 | 0 |
| 31996 | ENSMUSG00000092317 | 0 | NA | NA | NA | NA | NA | Gm6633 | predicted gene 6633 [Source:MGI Symbol | 0 | 0 | 0 | 0 | 0 | 0 |
| 31997 | ENSMUSG00000092318 | 0 | NA | NA | NA | NA | NA | Vmn1r-ps60 | vomeronasal 1 receptor, pseudogene 60 [ | 0 | 0 | 0 | 0 | 0 | 0 |
| 31998 | ENSMUSG00000092322 | 0 | NA | NA | NA | NA | NA | Esp36 | exocrine gland secreted peptide 36 [Sourc | 0 | 0 | 0 | 0 | 0 | 0 |
| 31999 | ENSMUSG00000092323 | 1.482707609 | -0.920791188 | 1.788393318 | -0.514870626 | 0.606643428 | NA | BB365896 | expressed sequence BB365896 [Source:M | 0.8031 | 2.0558 | 3.0187 | 1.928 | 1.0906 | 0 |
| 32000 | ENSMUSG00000092325 | 2.020325595 | 2.592831818 | 1.800420895 | 1.440125375 | 0.149831931 | NA | Gm18284 | predicted gene, 18284 [Source:MGI Symb | 1.6062 | 0 | 0 | 3.8559 | 1.0906 | 5.5692 |
| 32001 | ENSMUSG00000092326 | 0 | NA | NA | NA | NA | NA | Vmn1r-ps55 | vomeronasal 1 receptor, pseudogene 55 [ | 0 | 0 | 0 | 0 | 0 | 0 |
| 32002 | ENSMUSG00000092328 | 0 | NA | NA | NA | NA | NA | Vmn2r-ps57 | vomeronasal 2, receptor, pseudogene 57 | 0 | 0 | 0 | 0 | 0 | 0 |
| 32003 | ENSMUSG00000092329 | 0 | NA | NA | NA | NA | NA | Galnt2l | polypeptide N-acetylgalactosaminyltransf | 0 | 0 | 0 | 0 | 0 | 0 |
| 32004 | ENSMUSG00000092330 | 0 | NA | NA | NA | NA | NA | Vmn2r-ps82 | vomeronasal 2, receptor, pseudogene 82 | 0 | 0 | 0 | 0 | 0 | 0 |
| 32005 | ENSMUSG00000092331 | 0 | NA | NA | NA | NA | NA | Vmn1r-ps66 | vomeronasal 1 receptor, pseudogene 66 [ | 0 | 0 | 0 | 0 | 0 | 0 |
| 32006 | ENSMUSG00000092332 | 0 | NA | NA | NA | NA | NA | Gm8868 | predicted gene 8868 [Source:MGI Symbol | 0 | 0 | 0 | 0 | 0 | 0 |
| 32007 | ENSMUSG00000092335 | 11.64124559 | 0.035504283 | 0.639194525 | 0.055545349 | 0.955704002 | 0.991678478 | Zfp977 | zinc finger protein 977 [Source:MGI Symb | 11.243 | 8.2233 | 15.094 | 15.424 | 8.7249 | 11.138 |
| 32008 | ENSMUSG00000092336 | 0 | NA | NA | NA | NA | NA | Vmn1r-ps40 | vomeronasal 1 receptor, pseudogene 40 [ | 0 | 0 | 0 | 0 | 0 | 0 |
| 32009 | ENSMUSG00000092338 | 0 | NA | NA | NA | NA | NA | Gm26940 | predicted gene, 26940 [Source:MGI Symb | 0 | 0 | 0 | 0 | 0 | 0 |
| 32010 | ENSMUSG00000092339 | 0 | NA | NA | NA | NA | NA | Gm4831 | predicted gene 4831 [Source:MGI Symbol | 0 | 0 | 0 | 0 | 0 | 0 |
| 32011 | ENSMUSG00000092340 | 0 | NA | NA | NA | NA | NA | Olfr754-ps1 | olfactory receptor 754, pseudogene 1 [So | 0 | 0 | 0 | 0 | 0 | 0 |
| 32012 | ENSMUSG00000092341 | 6295.155898 | 0.183032267 | 0.337512474 | 0.542297786 | 0.587613381 | 0.910765033 | Malat1 | metastasis associated lung adenocarcinor | 5608.9 | 6189.1 | 5891.6 | 9629.2 | 6279.8 | 4172.5 |
| 32013 | ENSMUSG00000092342 | 0 | NA | NA | NA | NA | NA | Esp31 | exocrine gland secreted peptide 31 [Sourc | 0 | 0 | 0 | 0 | 0 | 0 |
| 32014 | ENSMUSG00000092343 | 0 | NA | NA | NA | NA | NA | Gm20525 | predicted gene 20525 [Source:MGI Symbo | 0 | 0 | 0 | 0 | 0 | 0 |
| 32015 | ENSMUSG00000092344 | 0.678054141 | -2.792799308 | 2.990673428 | -0.933836266 | 0.350388359 | NA | Hspe1-ps3 | heat shock protein 1 (chaperonin 10), pse | 0 | 2.0558 | 2.0125 | 0 | 0 | 0 |
| 32016 | ENSMUSG00000092345 | 0 | NA | NA | NA | NA | NA | Gm20503 | predicted gene 20503 [Source:MGI Symbo | 0 | 0 | 0 | 0 | 0 | 0 |
| 32017 | ENSMUSG00000092346 | 0 | NA | NA | NA | NA | NA | Tlx1os | T cell leukemia, homeobox 1, opposite str | 0 | 0 | 0 | 0 | 0 | 0 |
| 32018 | ENSMUSG00000092349 | 0.481988923 | 2.492401104 | 3.837077151 | 0.649557203 | 0.515978285 | NA | Smim40 | small integral membrane protein 40 [Sour | 0 | 0 | 0 | 2.8919 | 0 | 0 |
| 32019 | ENSMUSG00000092350 | 0 | NA | NA | NA | NA | NA | Vmn1r-ps62 | vomeronasal 1 receptor, pseudogene 62 [ | 0 | 0 | 0 | 0 | 0 | 0 |
| 32020 | ENSMUSG00000092351 | 0 | NA | NA | NA | NA | NA | Vmn1r-ps64 | vomeronasal 1 receptor, pseudogene 64 [ | 0 | 0 | 0 | 0 | 0 | 0 |
| 32021 | ENSMUSG00000092352 | 0 | NA | NA | NA | NA | NA | NA | NA | 0 | 0 | 0 | 0 | 0 | 0 |
| 32022 | ENSMUSG00000092353 | 0 | NA | NA | NA | NA | NA | Gm20539 | predicted gene 20539 [Source:MGI Symbo | 0 | 0 | 0 | 0 | 0 | 0 |
| 32023 | ENSMUSG00000092354 | 5.405954701 | -0.159905671 | 1.05418365 | -0.151686731 | 0.879434024 | 0.98128671 | Gm20548 | predicted gene 20548 [Source:MGI Symbo | 4.0155 | 6.1675 | 7.0437 | 9.6398 | 0 | 5.5692 |
| 32024 | ENSMUSG00000092356 | 0 | NA | NA | NA | NA | NA | Gm20532 | predicted gene 20532 [Source:MGI Symbo | 0 | 0 | 0 | 0 | 0 | 0 |
| 32025 | ENSMUSG00000092357 | 0 | NA | NA | NA | NA | NA | Vmn2r-ps83 | vomeronasal 2, receptor, pseudogene 83 | 0 | 0 | 0 | 0 | 0 | 0 |
| 32026 | ENSMUSG00000092358 | 0.80281854 | -0.292476728 | 2.419692615 | -0.120873505 | 0.903791228 | NA | Gm20430 | predicted gene 20430 [Source:MGI Symbo | 1.6062 | 0 | 1.0062 | 0 | 1.0906 | 1.1138 |
| 32027 | ENSMUSG00000092359 | 0 | NA | NA | NA | NA | NA | Vmn1r-ps68 | vomeronasal 1 receptor, pseudogene 68 [ | 0 | 0 | 0 | 0 | 0 | 0 |
| 32028 | ENSMUSG00000092360 | 0 | NA | NA | NA | NA | NA | Gm20441 | predicted gene 20441 [Source:MGI Symbo | 0 | 0 | 0 | 0 | 0 | 0 |

| | | | | | | | | | | | | | | | |
|---|---|---|---|---|---|---|---|---|---|---|---|---|---|---|---|
| 32029 | ENSMUSG00000092362 | 0 | NA | NA | NA | NA | NA | Gm20392 | predicted gene 20392 [Source:MGI Symbo | 0 | 0 | 0 | 0 | 0 | 0 |
| 32030 | ENSMUSG00000092363 | 0 | NA | NA | NA | NA | NA | Gm20391 | predicted gene 20391 [Source:MGI Symbo | 0 | 0 | 0 | 0 | 0 | 0 |
| 32031 | ENSMUSG00000092364 | 14.0888293 | -0.006777777 | 0.60835531 | -0.011141148 | 0.991110834 | 0.99859176 | Gm20476 | predicted gene 20476 [Source:MGI Symbo | 16.062 | 10.279 | 16.1 | 21.208 | 13.087 | 7.7969 |
| 32032 | ENSMUSG00000092365 | 0 | NA | NA | NA | NA | NA | BC023719 | cDNA sequence BC023719 [Source:MGI Sy | 0 | 0 | 0 | 0 | 0 | 0 |
| 32033 | ENSMUSG00000092366 | 0 | NA | NA | NA | NA | NA | Gm17782 | predicted gene, 17782 [Source:MGI Symb | 0 | 0 | 0 | 0 | 0 | 0 |
| 32034 | ENSMUSG00000092367 | 0 | NA | NA | NA | NA | NA | Gm20479 | predicted gene 20479 [Source:MGI Symbo | 0 | 0 | 0 | 0 | 0 | 0 |
| 32035 | ENSMUSG00000092368 | 0.674441933 | -2.786326546 | 3.022203109 | -0.921952114 | 0.356553554 | NA | A930015D03 | RIKEN cDNA A930015D03 gene [Source:M | 0 | 1.0279 | 3.0187 | 0 | 0 | 0 |
| 32036 | ENSMUSG00000092369 | 0 | NA | NA | NA | NA | NA | 1700039E22 | RIKEN cDNA 1700039E22 gene [Source:M | 0 | 0 | 0 | 0 | 0 | 0 |
| 32037 | ENSMUSG00000092370 | 0 | NA | NA | NA | NA | NA | Vmn2r-ps58 | vomeronasal 2, receptor, pseudogene 58 | 0 | 0 | 0 | 0 | 0 | 0 |
| 32038 | ENSMUSG00000092371 | 0.267700746 | -1.497481649 | 4.057001436 | -0.369110456 | 0.712045396 | NA | Gm20511 | predicted gene 20511 [Source:MGI Symbo | 1.6062 | 0 | 0 | 0 | 0 | 0 |
| 32039 | ENSMUSG00000092372 | 0 | NA | NA | NA | NA | NA | Olfr296-ps1 | olfactory receptor 296, pseudogene 1 [Sou | 0 | 0 | 0 | 0 | 0 | 0 |
| 32040 | ENSMUSG00000092373 | 1.177717557 | 0.416898694 | 2.290762114 | 0.181991264 | 0.855589581 | NA | Gm20535 | predicted gene 20535 [Source:MGI Symbo | 0 | 3.0838 | 0 | 2.8919 | 1.0906 | 0 |
| 32041 | ENSMUSG00000092374 | 0 | NA | NA | NA | NA | NA | Gm6623 | predicted gene 6623 [Source:MGI Symbol | 0 | 0 | 0 | 0 | 0 | 0 |
| 32042 | ENSMUSG00000092375 | 0 | NA | NA | NA | NA | NA | A730060N03 | RIKEN cDNA A730060N03 gene [Source:M | 0 | 0 | 0 | 0 | 0 | 0 |
| 32043 | ENSMUSG00000092376 | 0 | NA | NA | NA | NA | NA | Gm19203 | predicted gene, 19203 [Source:MGI Symb | 0 | 0 | 0 | 0 | 0 | 0 |
| 32044 | ENSMUSG00000092377 | 0 | NA | NA | NA | NA | NA | Gm20549 | predicted gene 20549 [Source:MGI Symbo | 0 | 0 | 0 | 0 | 0 | 0 |
| 32045 | ENSMUSG00000092381 | 1.167055378 | 1.64701604 | 2.143963744 | 0.768210771 | 0.442361976 | NA | Gm20512 | predicted gene 20512 [Source:MGI Symbo | 1.6062 | 0 | 0 | 0.964 | 1.0906 | 3.3415 |
| 32046 | ENSMUSG00000092382 | 0 | NA | NA | NA | NA | NA | Vmn2r-ps59 | vomeronasal 2, receptor, pseudogene 59 | 0 | 0 | 0 | 0 | 0 | 0 |
| 32047 | ENSMUSG00000092383 | 0 | NA | NA | NA | NA | NA | H2-Pb | histocompatibility 2, P region beta locus [S | 0 | 0 | 0 | 0 | 0 | 0 |
| 32048 | ENSMUSG00000092384 | 4.847814043 | -0.813663285 | 1.017531735 | -0.799644136 | 0.423917008 | NA | Gm4189 | predicted gene 4189 [Source:MGI Symbol | 2.4093 | 5.1396 | 11.069 | 3.8559 | 3.2719 | 3.3415 |
| 32049 | ENSMUSG00000092385 | 0 | NA | NA | NA | NA | NA | Gm20401 | predicted gene 20401 [Source:MGI Symbo | 0 | 0 | 0 | 0 | 0 | 0 |
| 32050 | ENSMUSG00000092386 | 0 | NA | NA | NA | NA | NA | Gm20536 | predicted gene 20536 [Source:MGI Symbo | 0 | 0 | 0 | 0 | 0 | 0 |
| 32051 | ENSMUSG00000092387 | 0 | NA | NA | NA | NA | NA | Gm7648 | predicted gene 7648 [Source:MGI Symbol | 0 | 0 | 0 | 0 | 0 | 0 |
| 32052 | ENSMUSG00000092388 | 0 | NA | NA | NA | NA | NA | Olfr306-ps1 | olfactory receptor 306, pseudogene 1 [Sou | 0 | 0 | 0 | 0 | 0 | 0 |
| 32053 | ENSMUSG00000092389 | 0 | NA | NA | NA | NA | NA | Gm20483 | predicted gene 20483 [Source:MGI Symbo | 0 | 0 | 0 | 0 | 0 | 0 |
| 32054 | ENSMUSG00000092390 | 0.839822492 | 2.066357436 | 2.600192809 | 0.794693928 | 0.426791566 | NA | Gm20541 | predicted gene 20541 [Source:MGI Symbo | 0.8031 | 0 | 0 | 0.964 | 3.2719 | 0 |
| 32055 | ENSMUSG00000092392 | 0 | NA | NA | NA | NA | NA | Gm20546 | predicted gene 20546 [Source:MGI Symbo | 0 | 0 | 0 | 0 | 0 | 0 |
| 32056 | ENSMUSG00000092393 | 0 | NA | NA | NA | NA | NA | Gm20399 | predicted gene 20399 [Source:MGI Symbo | 0 | 0 | 0 | 0 | 0 | 0 |
| 32057 | ENSMUSG00000092394 | 0.160662974 | 1.126924175 | 4.080472857 | 0.276174898 | 0.782413736 | NA | Gm20445 | predicted gene 20445 [Source:MGI Symbo | 0 | 0 | 0 | 0.964 | 0 | 0 |
| 32058 | ENSMUSG00000092395 | 0 | NA | NA | NA | NA | NA | Gm20463 | predicted gene 20463 [Source:MGI Symbo | 0 | 0 | 0 | 0 | 0 | 0 |
| 32059 | ENSMUSG00000092396 | 0 | NA | NA | NA | NA | NA | Olfr758-ps1 | olfactory receptor 758, pseudogene 1 [Sou | 0 | 0 | 0 | 0 | 0 | 0 |
| 32060 | ENSMUSG00000092397 | 0.636972666 | -2.712703855 | 3.077964703 | -0.881330397 | 0.378139018 | NA | C130080G10 | RIKEN cDNA C130080G10 gene [Source:M | 0.8031 | 0 | 3.0187 | 0 | 0 | 0 |
| 32061 | ENSMUSG00000092398 | 0 | NA | NA | NA | NA | NA | Vmn1r-ps59 | vomeronasal 1 receptor, pseudogene 59 [ | 0 | 0 | 0 | 0 | 0 | 0 |
| 32062 | ENSMUSG00000092399 | 13.53207102 | -0.244548557 | 0.580211069 | -0.421482061 | 0.673403108 | 0.931915649 | 4930556N13 | RIKEN cDNA 4930556N13 gene [Source:M | 14.456 | 15.419 | 14.087 | 9.6398 | 11.997 | 15.594 |
| 32063 | ENSMUSG00000092401 | 0 | NA | NA | NA | NA | NA | Olfr300-ps1 | olfactory receptor 300, pseudogene 1 [Sou | 0 | 0 | 0 | 0 | 0 | 0 |
| 32064 | ENSMUSG00000092402 | 0 | NA | NA | NA | NA | NA | Gm20485 | predicted gene 20485 [Source:MGI Symbo | 0 | 0 | 0 | 0 | 0 | 0 |
| 32065 | ENSMUSG00000092404 | 0 | NA | NA | NA | NA | NA | Gm9921 | predicted gene 9921 [Source:MGI Symbol | 0 | 0 | 0 | 0 | 0 | 0 |
| 32066 | ENSMUSG00000092405 | 0 | NA | NA | NA | NA | NA | Gm20402 | predicted gene 20402 [Source:MGI Symbo | 0 | 0 | 0 | 0 | 0 | 0 |
| 32067 | ENSMUSG00000092406 | 0 | NA | NA | NA | NA | NA | Gm20462 | predicted gene 20462 [Source:MGI Symbo | 0 | 0 | 0 | 0 | 0 | 0 |
| 32068 | ENSMUSG00000092408 | 0 | NA | NA | NA | NA | NA | Vhl-ps1 | von Hippel-Lindau tumor suppressor-like, | 0 | 0 | 0 | 0 | 0 | 0 |
| 32069 | ENSMUSG00000092409 | 0 | NA | NA | NA | NA | NA | Gm9613 | predicted gene 9613 [Source:MGI Symbol | 0 | 0 | 0 | 0 | 0 | 0 |
| 32070 | ENSMUSG00000092410 | 0 | NA | NA | NA | NA | NA | Gm18354 | predicted gene, 18354 [Source:MGI Symb | 0 | 0 | 0 | 0 | 0 | 0 |
| 32071 | ENSMUSG00000092411 | 0 | NA | NA | NA | NA | NA | Gm5594 | predicted gene 5594 [Source:MGI Symbol | 0 | 0 | 0 | 0 | 0 | 0 |
| 32072 | ENSMUSG00000092412 | 0 | NA | NA | NA | NA | NA | Gm20507 | predicted gene 20507 [Source:MGI Symbo | 0 | 0 | 0 | 0 | 0 | 0 |
| 32073 | ENSMUSG00000092413 | 0 | NA | NA | NA | NA | NA | Olfr115 | olfactory receptor 115 [Source:MGI Symb | 0 | 0 | 0 | 0 | 0 | 0 |
| 32074 | ENSMUSG00000092414 | 0 | NA | NA | NA | NA | NA | Gm20453 | predicted gene 20453 [Source:MGI Symbo | 0 | 0 | 0 | 0 | 0 | 0 |
| 32075 | ENSMUSG00000092415 | 0 | NA | NA | NA | NA | NA | Gm20513 | predicted gene 20513 [Source:MGI Symbo | 0 | 0 | 0 | 0 | 0 | 0 |
| 32076 | ENSMUSG00000092416 | 161.2844228 | 0.233004679 | 0.183937988 | 1.266756705 | 0.205242297 | 0.650728757 | Zfp141 | zinc finger protein 141 [Source:MGI Symb | 142.95 | 146.99 | 154.96 | 166.77 | 175.59 | 180.44 |
| 32077 | ENSMUSG00000092417 | 101.3635297 | 0.393258697 | 0.246873504 | 1.592956271 | 0.111170001 | 0.489450944 | Gpank1 | G patch domain and ankyrin repeats 1 [So | 86.735 | 106.9 | 69.431 | 123.39 | 101.43 | 120.3 |
| 32078 | ENSMUSG00000092418 | 0 | NA | NA | NA | NA | NA | Gm20406 | predicted gene 20406 [Source:MGI Symbo | 0 | 0 | 0 | 0 | 0 | 0 |
| 32079 | ENSMUSG00000092419 | 0 | NA | NA | NA | NA | NA | Vmn2r-ps86 | vomeronasal 2, receptor, pseudogene 86 | 0 | 0 | 0 | 0 | 0 | 0 |
| 32080 | ENSMUSG00000092422 | 0 | NA | NA | NA | NA | NA | Vmn2r-ps65 | vomeronasal 2, receptor, pseudogene 65 | 0 | 0 | 0 | 0 | 0 | 0 |
| 32081 | ENSMUSG00000092425 | 0 | NA | NA | NA | NA | NA | Tprgl-ps2 | transformation related protein 63 regulat | 0 | 0 | 0 | 0 | 0 | 0 |
| 32082 | ENSMUSG00000092426 | 0 | NA | NA | NA | NA | NA | Gm18779 | predicted gene, 18779 [Source:MGI Symb | 0 | 0 | 0 | 0 | 0 | 0 |
| 32083 | ENSMUSG00000092427 | 0 | NA | NA | NA | NA | NA | Vmn1r-ps69 | vomeronasal 1 receptor, pseudogene 69 [ | 0 | 0 | 0 | 0 | 0 | 0 |
| 32084 | ENSMUSG00000092428 | 0 | NA | NA | NA | NA | NA | Gm20545 | predicted gene 20545 [Source:MGI Symbo | 0 | 0 | 0 | 0 | 0 | 0 |

| | | | | | | | | | | | | | | | |
|---|---|---|---|---|---|---|---|---|---|---|---|---|---|---|---|
| 32085 | ENSMUSG00000092429 | 0 | NA | NA | NA | NA | NA | Tprgl-ps1 | transformation related protein 63 regulat | 0 | 0 | 0 | 0 | 0 | 0 |
| 32086 | ENSMUSG00000092431 | 0 | NA | NA | NA | NA | NA | Vmn2r-ps70 | vomeronasal 2, receptor, pseudogene 70 | 0 | 0 | 0 | 0 | 0 | 0 |
| 32087 | ENSMUSG00000092433 | 0 | NA | NA | NA | NA | NA | Vmn2r-ps40 | vomeronasal 2, receptor, pseudogene 40 | 0 | 0 | 0 | 0 | 0 | 0 |
| 32088 | ENSMUSG00000092435 | 0 | NA | NA | NA | NA | NA | Gm20526 | predicted gene 20526 [Source:MGI Symbo | 0 | 0 | 0 | 0 | 0 | 0 |
| 32089 | ENSMUSG00000092437 | 0 | NA | NA | NA | NA | NA | Rpl9-ps9 | ribosomal protein L9,pseudogene 9 [Sour | 0 | 0 | 0 | 0 | 0 | 0 |
| 32090 | ENSMUSG00000092438 | 0.428363721 | -0.543730382 | 3.624677344 | -0.15000794 | 0.880758351 | NA | Gm18734 | predicted gene, 18734 [Source:MGI Symb | 1.6062 | 0 | 0 | 0.964 | 0 | 0 |
| 32091 | ENSMUSG00000092439 | 0 | NA | NA | NA | NA | NA | Vmn2r-ps37 | vomeronasal 2, receptor, pseudogene 37 | 0 | 0 | 0 | 0 | 0 | 0 |
| 32092 | ENSMUSG00000092440 | 0 | NA | NA | NA | NA | NA | Vmn2r-ps78 | vomeronasal 2, receptor, pseudogene 78 | 0 | 0 | 0 | 0 | 0 | 0 |
| 32093 | ENSMUSG00000092441 | 0 | NA | NA | NA | NA | NA | Vmn2r-ps46 | vomeronasal 2, receptor, pseudogene 46 | 0 | 0 | 0 | 0 | 0 | 0 |
| 32094 | ENSMUSG00000092444 | 0 | NA | NA | NA | NA | NA | Gm18733 | predicted gene, 18733 [Source:MGI Symb | 0 | 0 | 0 | 0 | 0 | 0 |
| 32095 | ENSMUSG00000092447 | 0.556921942 | 2.679043159 | 3.638243063 | 0.736356289 | 0.461513899 | NA | Gm7696 | predicted gene 7696 [Source:MGI Symbol | 0 | 0 | 0 | 0 | 0 | 3.3415 |
| 32096 | ENSMUSG00000092448 | 0 | NA | NA | NA | NA | NA | Gm20387 | predicted gene 20387 [Source:MGI Symbo | 0 | 0 | 0 | 0 | 0 | 0 |
| 32097 | ENSMUSG00000092450 | 0 | NA | NA | NA | NA | NA | Vmn1r-ps31 | vomeronasal 1 receptor, pseudogene 31 [ | 0 | 0 | 0 | 0 | 0 | 0 |
| 32098 | ENSMUSG00000092451 | 0 | NA | NA | NA | NA | NA | Vmn2r-ps56 | vomeronasal 2, receptor, pseudogene 56 | 0 | 0 | 0 | 0 | 0 | 0 |
| 32099 | ENSMUSG00000092452 | 0 | NA | NA | NA | NA | NA | Vmn2r-ps45 | vomeronasal 2, receptor, pseudogene 45 | 0 | 0 | 0 | 0 | 0 | 0 |
| 32100 | ENSMUSG00000092454 | 1.634491198 | -0.442467688 | 1.583430466 | -0.279436134 | 0.779910144 | NA | Gm2991 | predicted pseudogene 2991 [Source:MGI | 1.6062 | 2.0558 | 2.0125 | 1.928 | 1.0906 | 1.1138 |
| 32101 | ENSMUSG00000092455 | 0.692607243 | 3.004183657 | 2.957557172 | 1.015765202 | 0.309741211 | NA | Gm20456 | predicted gene 20456 [Source:MGI Symbo | 0 | 0 | 0 | 1.928 | 0 | 2.2277 |
| 32102 | ENSMUSG00000092456 | 0 | NA | NA | NA | NA | NA | V1rd19 | vomeronasal 1 receptor, D19 [Source:MG | 0 | 0 | 0 | 0 | 0 | 0 |
| 32103 | ENSMUSG00000092457 | 0.462220778 | -0.685400455 | 3.191475091 | -0.21475977 | 0.82995462 | NA | Gm8835 | predicted gene 8835 [Source:MGI Symbol | 0.8031 | 0 | 1.0062 | 0.964 | 0 | 0 |
| 32104 | ENSMUSG00000092458 | 0 | NA | NA | NA | NA | NA | Gm18737 | predicted gene, 18737 [Source:MGI Symb | 0 | 0 | 0 | 0 | 0 | 0 |
| 32105 | ENSMUSG00000092460 | 0 | NA | NA | NA | NA | NA | Gm8878 | predicted gene 8878 [Source:MGI Symbol | 0 | 0 | 0 | 0 | 0 | 0 |
| 32106 | ENSMUSG00000092461 | 0 | NA | NA | NA | NA | NA | Gm18833 | predicted gene, 18833 [Source:MGI Symb | 0 | 0 | 0 | 0 | 0 | 0 |
| 32107 | ENSMUSG00000092462 | 0 | NA | NA | NA | NA | NA | Vmn1r-ps83 | vomeronasal 1 receptor, pseudogene 83 [ | 0 | 0 | 0 | 0 | 0 | 0 |
| 32108 | ENSMUSG00000092463 | 0 | NA | NA | NA | NA | NA | Gm20489 | predicted gene 20489 [Source:MGI Symbo | 0 | 0 | 0 | 0 | 0 | 0 |
| 32109 | ENSMUSG00000092464 | 0 | NA | NA | NA | NA | NA | Gm8902 | predicted gene 8902 [Source:MGI Symbol | 0 | 0 | 0 | 0 | 0 | 0 |
| 32110 | ENSMUSG00000092465 | 0 | NA | NA | NA | NA | NA | Gm20472 | predicted gene 20472 [Source:MGI Symbo | 0 | 0 | 0 | 0 | 0 | 0 |
| 32111 | ENSMUSG00000092466 | 0 | NA | NA | NA | NA | NA | Vmn1r-ps57 | vomeronasal 1 receptor, pseudogene 57 [ | 0 | 0 | 0 | 0 | 0 | 0 |
| 32112 | ENSMUSG00000092467 | 0 | NA | NA | NA | NA | NA | Gm18360 | predicted gene, 18360 [Source:MGI Symb | 0 | 0 | 0 | 0 | 0 | 0 |
| 32113 | ENSMUSG00000092468 | 0 | NA | NA | NA | NA | NA | Gm18520 | predicted gene, 18520 [Source:MGI Symb | 0 | 0 | 0 | 0 | 0 | 0 |
| 32114 | ENSMUSG00000092469 | 0 | NA | NA | NA | NA | NA | Gm20395 | predicted gene 20395 [Source:MGI Symbo | 0 | 0 | 0 | 0 | 0 | 0 |
| 32115 | ENSMUSG00000092470 | 0.849373781 | -3.117833363 | 2.761285217 | -1.129123983 | 0.258845535 | NA | Gm20518 | predicted gene 20518 [Source:MGI Symbo | 0 | 3.0838 | 2.0125 | 0 | 0 | 0 |
| 32116 | ENSMUSG00000092471 | 95.33599134 | 0.132485469 | 0.274931583 | 0.481885229 | 0.629887479 | 0.920736989 | Cyp21a2-ps | cytochrome P450, family 21, subfamily a, | 124.48 | 82.233 | 65.406 | 99.29 | 99.246 | 101.36 |
| 32117 | ENSMUSG00000092472 | 0 | NA | NA | NA | NA | NA | Gm20473 | predicted gene 20473 [Source:MGI Symbo | 0 | 0 | 0 | 0 | 0 | 0 |
| 32118 | ENSMUSG00000092473 | 0 | NA | NA | NA | NA | NA | Vmn1r180 | vomeronasal 1 receptor 180 [Source:MGI | 0 | 0 | 0 | 0 | 0 | 0 |
| 32119 | ENSMUSG00000092474 | 0.85143529 | 2.085519423 | 2.590856022 | 0.804953809 | 0.420846336 | NA | Gm20478 | predicted gene 20478 [Source:MGI Symbo | 0.8031 | 0 | 0 | 0.964 | 0 | 3.3415 |
| 32120 | ENSMUSG00000092476 | 0 | NA | NA | NA | NA | NA | Vmn1r-ps65 | vomeronasal 1 receptor, pseudogene 65 [ | 0 | 0 | 0 | 0 | 0 | 0 |
| 32121 | ENSMUSG00000092477 | 0 | NA | NA | NA | NA | NA | Gm18808 | predicted gene, 18808 [Source:MGI Symb | 0 | 0 | 0 | 0 | 0 | 0 |
| 32122 | ENSMUSG00000092480 | 0 | NA | NA | NA | NA | NA | Gm20495 | predicted gene 20495 [Source:MGI Symbo | 0 | 0 | 0 | 0 | 0 | 0 |
| 32123 | ENSMUSG00000092482 | 1.066803338 | 2.404617844 | 2.208738042 | 1.088684035 | 0.276293244 | NA | Gm20531 | predicted gene 20531 [Source:MGI Symbo | 0 | 1.0279 | 0 | 0.964 | 2.1812 | 2.2277 |
| 32124 | ENSMUSG00000092483 | 0 | NA | NA | NA | NA | NA | Gm20421 | predicted gene 20421 [Source:MGI Symbo | 0 | 0 | 0 | 0 | 0 | 0 |
| 32125 | ENSMUSG00000092485 | 0 | NA | NA | NA | NA | NA | Rpl9-ps5 | ribosomal protein L9, pseudogene 5 [Sour | 0 | 0 | 0 | 0 | 0 | 0 |
| 32126 | ENSMUSG00000092486 | 2.019421068 | -0.917860353 | 1.606609175 | -0.571302821 | 0.56779439 | NA | NA | NA | 4.8186 | 2.0558 | 1.0062 | 0.964 | 3.2719 | 0 |
| 32127 | ENSMUSG00000092487 | 0 | NA | NA | NA | NA | NA | Vmn1r-ps71 | vomeronasal 1 receptor, pseudogene 71 [ | 0 | 0 | 0 | 0 | 0 | 0 |
| 32128 | ENSMUSG00000092488 | 0 | NA | NA | NA | NA | NA | Gm18159 | predicted gene, 18159 [Source:MGI Symb | 0 | 0 | 0 | 0 | 0 | 0 |
| 32129 | ENSMUSG00000092489 | 0 | NA | NA | NA | NA | NA | Vmn1r-ps53 | vomeronasal 1 receptor, pseudogene 53 [ | 0 | 0 | 0 | 0 | 0 | 0 |
| 32130 | ENSMUSG00000092490 | 0.267700746 | -1.497481649 | 4.057001436 | -0.369110456 | 0.712045396 | NA | Gm20482 | predicted gene 20482 [Source:MGI Symbo | 1.6062 | 0 | 0 | 0 | 0 | 0 |
| 32131 | ENSMUSG00000092492 | 0 | NA | NA | NA | NA | NA | B230208B08 | RIKEN cDNA B230208B08 gene [Source:M | 0 | 0 | 0 | 0 | 0 | 0 |
| 32132 | ENSMUSG00000092493 | 0 | NA | NA | NA | NA | NA | Gm20451 | predicted gene 20451 [Source:MGI Symbo | 0 | 0 | 0 | 0 | 0 | 0 |
| 32133 | ENSMUSG00000092494 | 0.48903338 | 0.95637552 | 3.456690196 | 0.276673773 | 0.782030612 | NA | Gm20420 | predicted gene 20420 [Source:MGI Symbo | 0 | 0 | 1.0062 | 1.928 | 0 | 0 |
| 32134 | ENSMUSG00000092495 | 0 | NA | NA | NA | NA | NA | Gm20405 | predicted gene 20405 [Source:MGI Symbo | 0 | 0 | 0 | 0 | 0 | 0 |
| 32135 | ENSMUSG00000092496 | 0 | NA | NA | NA | NA | NA | Vmn1r-ps84 | vomeronasal 1 receptor, pseudogene 84 [ | 0 | 0 | 0 | 0 | 0 | 0 |
| 32136 | ENSMUSG00000092499 | 1.63658485 | 0.071686525 | 1.741232971 | 0.041169979 | 0.967160387 | NA | 1700092C10 | RIKEN cDNA 1700092C10 gene [Source:M | 0.8031 | 1.0279 | 3.0187 | 3.8559 | 0 | 1.1138 |
| 32137 | ENSMUSG00000092500 | 0.679159517 | 1.687875387 | 3.079592583 | 0.548083989 | 0.583634234 | NA | Gm20400 | predicted gene 20400 [Source:MGI Symbo | 0.8031 | 0 | 0 | 0 | 3.2719 | 0 |
| 32138 | ENSMUSG00000092502 | 0 | NA | NA | NA | NA | NA | Gm4252 | predicted gene 4252 [Source:MGI Symbol | 0 | 0 | 0 | 0 | 0 | 0 |
| 32139 | ENSMUSG00000092503 | 0 | NA | NA | NA | NA | NA | Gm20484 | predicted gene 20484 [Source:MGI Symbo | 0 | 0 | 0 | 0 | 0 | 0 |
| 32140 | ENSMUSG00000092504 | 0 | NA | NA | NA | NA | NA | Gm20396 | predicted gene 20396 [Source:MGI Symbo | 0 | 0 | 0 | 0 | 0 | 0 |

| | | | | | | | | | | | | | | | |
|---|---|---|---|---|---|---|---|---|---|---|---|---|---|---|---|
| 32141 | ENSMUSG00000092505 | 0 | NA | NA | NA | NA | NA | Gm19246 | predicted gene, 19246 [Source:MGI Symb | 0 | 0 | 0 | 0 | 0 | 0 |
| 32142 | ENSMUSG00000092506 | 0 | NA | NA | NA | NA | NA | Vmn1r-ps77 | vomeronasal 1 receptor, pseudogene 77 [ | 0 | 0 | 0 | 0 | 0 | 0 |
| 32143 | ENSMUSG00000092508 | 0 | NA | NA | NA | NA | NA | H2-M10.5-ps | histocompatibility 2, M region locus 10.5, | 0 | 0 | 0 | 0 | 0 | 0 |
| 32144 | ENSMUSG00000092509 | 0.160662974 | 1.126924175 | 4.080472857 | 0.276174898 | 0.782413736 | NA | Gm20394 | predicted gene 20394 [Source:MGI Symbo | 0 | 0 | 0 | 0.964 | 0 | 0 |
| 32145 | ENSMUSG00000092510 | 0.160662974 | 1.126924175 | 4.080472857 | 0.276174898 | 0.782413736 | NA | Gm18343 | predicted gene, 18343 [Source:MGI Symb | 0 | 0 | 0 | 0.964 | 0 | 0 |
| 32146 | ENSMUSG00000092511 | 0 | NA | NA | NA | NA | NA | Gm20547 | predicted gene 20547 [Source:MGI Symbo | 0 | 0 | 0 | 0 | 0 | 0 |
| 32147 | ENSMUSG00000092512 | 6.77198933 | -1.05068173 | 0.911181985 | -1.153097566 | 0.24887034 | 0.700732062 | Gm4152 | predicted gene 4152 [Source:MGI Symbol | 15.259 | 4.1117 | 8.05 | 7.7118 | 3.2719 | 2.2277 |
| 32148 | ENSMUSG00000092513 | 0 | NA | NA | NA | NA | NA | Gm20542 | predicted gene 20542 [Source:MGI Symbo | 0 | 0 | 0 | 0 | 0 | 0 |
| 32149 | ENSMUSG00000092514 | 0 | NA | NA | NA | NA | NA | Vmn1r-ps86 | vomeronasal 1 receptor, pseudogene 86 [ | 0 | 0 | 0 | 0 | 0 | 0 |
| 32150 | ENSMUSG00000092515 | 0 | NA | NA | NA | NA | NA | C87198 | expressed sequence C87198 [Source:MGI | 0 | 0 | 0 | 0 | 0 | 0 |
| 32151 | ENSMUSG00000092516 | 0 | NA | NA | NA | NA | NA | Gm20397 | predicted gene 20397 [Source:MGI Symbo | 0 | 0 | 0 | 0 | 0 | 0 |
| 32152 | ENSMUSG00000092517 | 2.183722881 | -2.357596848 | 1.697094616 | -1.38919588 | 0.164773194 | NA | Art2a | ADP-ribosyltransferase 2a [Source:MGI Sy | 4.8186 | 3.0838 | 3.0187 | 0 | 2.1812 | 0 |
| 32153 | ENSMUSG00000092518 | 0 | NA | NA | NA | NA | NA | Fam71e2 | family with sequence similarity 71, memb | 0 | 0 | 0 | 0 | 0 | 0 |
| 32154 | ENSMUSG00000092519 | 0.774667342 | 0.864398069 | 2.649064859 | 0.326303097 | 0.744195045 | NA | Actl9 | actin-like 9 [Source:MGI Symbol;Acc:MGI: | 1.6062 | 0 | 0 | 1.928 | 0 | 1.1138 |
| 32155 | ENSMUSG00000092520 | 0 | NA | NA | NA | NA | NA | Gm3611 | predicted gene 3611 [Source:MGI Symbol | 0 | 0 | 0 | 0 | 0 | 0 |
| 32156 | ENSMUSG00000092521 | 0 | NA | NA | NA | NA | NA | Gm4271 | predicted gene 4271 [Source:MGI Symbol | 0 | 0 | 0 | 0 | 0 | 0 |
| 32157 | ENSMUSG00000092522 | 0 | NA | NA | NA | NA | NA | Gm20389 | predicted gene 20389 [Source:MGI Symbo | 0 | 0 | 0 | 0 | 0 | 0 |
| 32158 | ENSMUSG00000092523 | 0 | NA | NA | NA | NA | NA | Gm18191 | predicted gene, 18191 [Source:MGI Symb | 0 | 0 | 0 | 0 | 0 | 0 |
| 32159 | ENSMUSG00000092524 | 0 | NA | NA | NA | NA | NA | Olfr756-ps1 | olfactory receptor 756, pseudogene 1 [So | 0 | 0 | 0 | 0 | 0 | 0 |
| 32160 | ENSMUSG00000092525 | 0 | NA | NA | NA | NA | NA | Gm20461 | predicted gene 20461 [Source:MGI Symbo | 0 | 0 | 0 | 0 | 0 | 0 |
| 32161 | ENSMUSG00000092526 | 1.849911381 | 0.943659187 | 1.639526827 | 0.575568006 | 0.564907211 | NA | Gm17907 | predicted gene, 17907 [Source:MGI Symb | 0.8031 | 0 | 3.0187 | 2.8919 | 3.2719 | 1.1138 |
| 32162 | ENSMUSG00000092528 | 1.850583286 | -1.300764674 | 1.673014246 | -0.777497668 | 0.43686521 | NA | Nlrp1c-ps | NLR family, pyrin domain containing 1C, p | 0.8031 | 3.0838 | 4.025 | 0.964 | 0 | 2.2277 |
| 32163 | ENSMUSG00000092529 | 0 | NA | NA | NA | NA | NA | Vmn2r-ps80 | vomeronasal 2, receptor, pseudogene 80 | 0 | 0 | 0 | 0 | 0 | 0 |
| 32164 | ENSMUSG00000092530 | 0 | NA | NA | NA | NA | NA | Vmn2r-ps64 | vomeronasal 2, receptor, pseudogene 64 | 0 | 0 | 0 | 0 | 0 | 0 |
| 32165 | ENSMUSG00000092532 | 0 | NA | NA | NA | NA | NA | Vmn2r-ps84 | vomeronasal 2, receptor, pseudogene 84 | 0 | 0 | 0 | 0 | 0 | 0 |
| 32166 | ENSMUSG00000092533 | 0 | NA | NA | NA | NA | NA | Vmn2r-ps72 | vomeronasal 2, receptor, pseudogene 72 | 0 | 0 | 0 | 0 | 0 | 0 |
| 32167 | ENSMUSG00000092534 | 0 | NA | NA | NA | NA | NA | Pagr1b | PAXIP1 associated glutamate rich protein | 0 | 0 | 0 | 0 | 0 | 0 |
| 32168 | ENSMUSG00000092535 | 0 | NA | NA | NA | NA | NA | Vmn1r-ps61 | vomeronasal 1 receptor, pseudogene 61 [ | 0 | 0 | 0 | 0 | 0 | 0 |
| 32169 | ENSMUSG00000092536 | 0 | NA | NA | NA | NA | NA | Gm20501 | predicted gene 20501 [Source:MGI Symbo | 0 | 0 | 0 | 0 | 0 | 0 |
| 32170 | ENSMUSG00000092538 | 0 | NA | NA | NA | NA | NA | Vmn2r-ps55 | vomeronasal 2, receptor, pseudogene 55 | 0 | 0 | 0 | 0 | 0 | 0 |
| 32171 | ENSMUSG00000092539 | 0 | NA | NA | NA | NA | NA | Gm20468 | predicted gene 20468 [Source:MGI Symbo | 0 | 0 | 0 | 0 | 0 | 0 |
| 32172 | ENSMUSG00000092540 | 0.513958919 | -2.394274821 | 3.756096013 | -0.637437066 | 0.523840191 | NA | Gm8738 | predicted gene 8738 [Source:MGI Symbol | 0 | 3.0838 | 0 | 0 | 0 | 0 |
| 32173 | ENSMUSG00000092541 | 0 | NA | NA | NA | NA | NA | Gm20537 | predicted gene 20537 [Source:MGI Symbo | 0 | 0 | 0 | 0 | 0 | 0 |
| 32174 | ENSMUSG00000092542 | 0 | NA | NA | NA | NA | NA | Olfr757-ps1 | olfactory receptor 757, pseudogene 1 [So | 0 | 0 | 0 | 0 | 0 | 0 |
| 32175 | ENSMUSG00000092543 | 0 | NA | NA | NA | NA | NA | Skiv2l-ps1 | superkiller viralicidic activity 2-like (S. cere | 0 | 0 | 0 | 0 | 0 | 0 |
| 32176 | ENSMUSG00000092544 | 0 | NA | NA | NA | NA | NA | Gm20422 | predicted gene 20422 [Source:MGI Symbo | 0 | 0 | 0 | 0 | 0 | 0 |
| 32177 | ENSMUSG00000092545 | 924.9143009 | 0.083323051 | 0.118230812 | 0.704749041 | 0.480966427 | 0.870348855 | Gm20319 | predicted gene, 20319 [Source:MGI Symb | 872.97 | 870.65 | 950.9 | 885.9 | 1055.7 | 913.35 |
| 32178 | ENSMUSG00000092547 | 0 | NA | NA | NA | NA | NA | Vmn1r-ps80 | vomeronasal 1 receptor, pseudogene 80 [ | 0 | 0 | 0 | 0 | 0 | 0 |
| 32179 | ENSMUSG00000092548 | 0 | NA | NA | NA | NA | NA | Vmn2r-ps41 | vomeronasal 2, receptor, pseudogene 41 | 0 | 0 | 0 | 0 | 0 | 0 |
| 32180 | ENSMUSG00000092549 | 0 | NA | NA | NA | NA | NA | Gm20491 | predicted gene 20491 [Source:MGI Symbo | 0 | 0 | 0 | 0 | 0 | 0 |
| 32181 | ENSMUSG00000092550 | 0.342432689 | 1.990653086 | 3.900281263 | 0.51038706 | 0.609780321 | NA | Gm20496 | predicted gene 20496 [Source:MGI Symbo | 0 | 0 | 0 | 0.964 | 1.0906 | 0 |
| 32182 | ENSMUSG00000092551 | 0 | NA | NA | NA | NA | NA | Gm20287 | predicted gene, 20287 [Source:MGI Symb | 0 | 0 | 0 | 0 | 0 | 0 |
| 32183 | ENSMUSG00000092552 | 0.993932953 | -2.088369984 | 2.277335805 | -0.917023295 | 0.359130433 | NA | Gm4287 | predicted gene 4287 [Source:MGI Symbol | 0.8031 | 1.0279 | 3.0187 | 0 | 0 | 1.1138 |
| 32184 | ENSMUSG00000092553 | 0 | NA | NA | NA | NA | NA | Gm20428 | predicted gene 20428 [Source:MGI Symbo | 0 | 0 | 0 | 0 | 0 | 0 |
| 32185 | ENSMUSG00000092554 | 0 | NA | NA | NA | NA | NA | Gm18128 | predicted gene, 18128 [Source:MGI Symb | 0 | 0 | 0 | 0 | 0 | 0 |
| 32186 | ENSMUSG00000092555 | 0 | NA | NA | NA | NA | NA | Stk-ps1 | serine/threonine kinase, pseudogene 1 [S | 0 | 0 | 0 | 0 | 0 | 0 |
| 32187 | ENSMUSG00000092556 | 0 | NA | NA | NA | NA | NA | Olfr755-ps1 | olfactory receptor 755, pseudogene 1 [So | 0 | 0 | 0 | 0 | 0 | 0 |
| 32188 | ENSMUSG00000092557 | 0 | NA | NA | NA | NA | NA | Gm8696 | predicted gene 8696 [Source:MGI Symbol | 0 | 0 | 0 | 0 | 0 | 0 |
| 32189 | ENSMUSG00000092558 | 258.6682283 | 0.170049949 | 0.170352921 | 0.998221505 | 0.318171961 | 0.764585353 | Med20 | mediator complex subunit 20 [Source:MG | 235.31 | 234.37 | 260.62 | 251.6 | 321.73 | 248.39 |
| 32190 | ENSMUSG00000092559 | 0 | NA | NA | NA | NA | NA | Gm20475 | predicted gene 20475 [Source:MGI Symbo | 0 | 0 | 0 | 0 | 0 | 0 |
| 32191 | ENSMUSG00000092560 | 0 | NA | NA | NA | NA | NA | Gm8750 | predicted pseudogene 8750 [Source:MGI | 0 | 0 | 0 | 0 | 0 | 0 |
| 32192 | ENSMUSG00000092561 | 0 | NA | NA | NA | NA | NA | Gm4011 | predicted gene 4011 [Source:MGI Symbol | 0 | 0 | 0 | 0 | 0 | 0 |
| 32193 | ENSMUSG00000092562 | 0 | NA | NA | NA | NA | NA | Gm20414 | predicted gene 20414 [Source:MGI Symbo | 0 | 0 | 0 | 0 | 0 | 0 |
| 32194 | ENSMUSG00000092563 | 0 | NA | NA | NA | NA | NA | Gm3617 | predicted gene 3617 [Source:MGI Symbol | 0 | 0 | 0 | 0 | 0 | 0 |
| 32195 | ENSMUSG00000092564 | 59.92493346 | 0.676334397 | 0.295144317 | 2.291537927 | 0.021932321 | 0.184542809 | BC051226 | cDNA sequence BC051226 [Source:MGI Sy | 41.761 | 54.48 | 42.262 | 65.55 | 73.071 | 82.424 |
| 32196 | ENSMUSG00000092565 | 0 | NA | NA | NA | NA | NA | Gm20477 | predicted gene 20477 [Source:MGI Symbo | 0 | 0 | 0 | 0 | 0 | 0 |

| | | | | | | | | | | | | | | | |
|---|---|---|---|---|---|---|---|---|---|---|---|---|---|---|---|
| 32197 | ENSMUSG00000092566 | 0 | NA | NA | NA | NA | NA | Gm20409 | predicted gene 20409 [Source:MGI Symbo | 0 | 0 | 0 | 0 | 0 | 0 |
| 32198 | ENSMUSG00000092567 | 0 | NA | NA | NA | NA | NA | Vmn2r-ps73 | vomeronasal 2, receptor, pseudogene 73 | 0 | 0 | 0 | 0 | 0 | 0 |
| 32199 | ENSMUSG00000092568 | 0 | NA | NA | NA | NA | NA | Olfr759-ps1 | olfactory receptor 759, pseudogene 1 [Sou | 0 | 0 | 0 | 0 | 0 | 0 |
| 32200 | ENSMUSG00000092569 | 18.99384771 | 0.539908159 | 0.531514678 | 1.015791627 | 0.309728624 | 0.758407791 | Gm20544 | predicted gene 20544 [Source:MGI Symbo | 16.062 | 11.307 | 19.119 | 29.883 | 13.087 | 24.505 |
| 32201 | ENSMUSG00000092570 | 0 | NA | NA | NA | NA | NA | Plut | PDX1 associated lncRNA, upregulator of tr | 0 | 0 | 0 | 0 | 0 | 0 |
| 32202 | ENSMUSG00000092571 | 0 | NA | NA | NA | NA | NA | Gm4134 | predicted gene 4134 [Source:MGI Symbol | 0 | 0 | 0 | 0 | 0 | 0 |
| 32203 | ENSMUSG00000092572 | 0 | NA | NA | NA | NA | NA | Serpinb10 | serine (or cysteine) peptidase inhibitor, cl | 0 | 0 | 0 | 0 | 0 | 0 |
| 32204 | ENSMUSG00000092573 | 0 | NA | NA | NA | NA | NA | Vmn2r-ps43 | vomeronasal 2, receptor, pseudogene 43 | 0 | 0 | 0 | 0 | 0 | 0 |
| 32205 | ENSMUSG00000092574 | 0.997622271 | -2.171875158 | 2.465204635 | -0.881012119 | 0.378311261 | NA | 2810047C21 | RIKEN cDNA 2810047C21 gene 1 [Source:I | 4.0155 | 0 | 1.0062 | 0.964 | 0 | 0 |
| 32206 | ENSMUSG00000092575 | 0 | NA | NA | NA | NA | NA | Gm5583 | predicted gene 5583 [Source:MGI Symbol | 0 | 0 | 0 | 0 | 0 | 0 |
| 32207 | ENSMUSG00000092576 | 0 | NA | NA | NA | NA | NA | Obox4-ps35 | oocyte specific homeobox 4, pseudogene | 0 | 0 | 0 | 0 | 0 | 0 |
| 32208 | ENSMUSG00000092577 | 0.524202404 | 2.600864256 | 3.301637836 | 0.787749713 | 0.430843121 | NA | Gm7393 | predicted gene 7393 [Source:MGI Symbol | 0 | 0 | 0 | 0.964 | 2.1812 | 0 |
| 32209 | ENSMUSG00000092578 | 0 | NA | NA | NA | NA | NA | 4930488B22 | RIKEN cDNA 4930488B22 gene [Source:M | 0 | 0 | 0 | 0 | 0 | 0 |
| 32210 | ENSMUSG00000092579 | 0 | NA | NA | NA | NA | NA | Vmn1r62 | vomeronasal 1 receptor 62 [Source:MGI S | 0 | 0 | 0 | 0 | 0 | 0 |
| 32211 | ENSMUSG00000092580 | 0 | NA | NA | NA | NA | NA | Vmn1r-ps45 | vomeronasal 1 receptor, pseudogene 45 [ | 0 | 0 | 0 | 0 | 0 | 0 |
| 32212 | ENSMUSG00000092581 | 0 | NA | NA | NA | NA | NA | Gm3211 | predicted gene 3211 [Source:MGI Symbol | 0 | 0 | 0 | 0 | 0 | 0 |
| 32213 | ENSMUSG00000092583 | 0 | NA | NA | NA | NA | NA | Vmn2r-ps53 | vomeronasal 2, receptor, pseudogene 53 | 0 | 0 | 0 | 0 | 0 | 0 |
| 32214 | ENSMUSG00000092584 | 0 | NA | NA | NA | NA | NA | Gm18199 | predicted gene, 18199 [Source:MGI Symb | 0 | 0 | 0 | 0 | 0 | 0 |
| 32215 | ENSMUSG00000092586 | 0 | NA | NA | NA | NA | NA | Ly6g6c | lymphocyte antigen 6 complex, locus G6C | 0 | 0 | 0 | 0 | 0 | 0 |
| 32216 | ENSMUSG00000092587 | 0 | NA | NA | NA | NA | NA | Vmn2r-ps85 | vomeronasal 2, receptor, pseudogene 85 | 0 | 0 | 0 | 0 | 0 | 0 |
| 32217 | ENSMUSG00000092588 | 0 | NA | NA | NA | NA | NA | Gm20534 | predicted gene 20534 [Source:MGI Symbo | 0 | 0 | 0 | 0 | 0 | 0 |
| 32218 | ENSMUSG00000092591 | 0 | NA | NA | NA | NA | NA | Gm20429 | predicted gene 20429 [Source:MGI Symbo | 0 | 0 | 0 | 0 | 0 | 0 |
| 32219 | ENSMUSG00000092592 | 0.8085244 | 0.747105454 | 2.417511592 | 0.309039037 | 0.757291832 | NA | Gm20449 | predicted gene 20449 [Source:MGI Symbo | 0.8031 | 0 | 1.0062 | 1.928 | 0 | 1.1138 |
| 32220 | ENSMUSG00000092593 | 7.322937267 | 0.840167444 | 0.949373574 | 0.884970329 | 0.376172634 | 0.809731241 | Gm20492 | predicted gene 20492 [Source:MGI Symbo | 1.6062 | 7.1954 | 7.0437 | 2.8919 | 19.631 | 5.5692 |
| 32221 | ENSMUSG00000092595 | 1.912126226 | -0.689350792 | 1.789147394 | -0.385295697 | 0.700018349 | NA | Gm20427 | predicted gene 20427 [Source:MGI Symbo | 0 | 2.0558 | 5.0312 | 0 | 3.2719 | 1.1138 |
| 32222 | ENSMUSG00000092596 | 0 | NA | NA | NA | NA | NA | Gm20490 | predicted gene 20490 [Source:MGI Symbo | 0 | 0 | 0 | 0 | 0 | 0 |
| 32223 | ENSMUSG00000092597 | 0 | NA | NA | NA | NA | NA | Gm20486 | predicted gene 20486 [Source:MGI Symbo | 0 | 0 | 0 | 0 | 0 | 0 |
| 32224 | ENSMUSG00000092599 | 0 | NA | NA | NA | NA | NA | 1700010K23 | RIKEN cDNA 1700010K23 gene [Source:M | 0 | 0 | 0 | 0 | 0 | 0 |
| 32225 | ENSMUSG00000092600 | 0 | NA | NA | NA | NA | NA | Gm20442 | predicted gene 20442 [Source:MGI Symbo | 0 | 0 | 0 | 0 | 0 | 0 |
| 32226 | ENSMUSG00000092601 | 0 | NA | NA | NA | NA | NA | Olfr760-ps1 | olfactory receptor 760, pseudogene 1 [Sou | 0 | 0 | 0 | 0 | 0 | 0 |
| 32227 | ENSMUSG00000092602 | 0.767390791 | -1.69152783 | 2.44773922 | -0.691057207 | 0.489529591 | NA | 4931413I07F | RIKEN cDNA 4931413I07 gene [Source:MG | 1.6062 | 1.0279 | 1.0062 | 0.964 | 0 | 0 |
| 32228 | ENSMUSG00000092603 | 0 | NA | NA | NA | NA | NA | Gm20386 | predicted gene 20386 [Source:MGI Symbo | 0 | 0 | 0 | 0 | 0 | 0 |
| 32229 | ENSMUSG00000092604 | 0 | NA | NA | NA | NA | NA | Gm20508 | predicted gene 20508 [Source:MGI Symbo | 0 | 0 | 0 | 0 | 0 | 0 |
| 32230 | ENSMUSG00000092606 | 0 | NA | NA | NA | NA | NA | Gm20450 | predicted gene 20450 [Source:MGI Symbo | 0 | 0 | 0 | 0 | 0 | 0 |
| 32231 | ENSMUSG00000092607 | 148.841876 | 0.131997671 | 0.201737417 | 0.654304358 | 0.512915739 | 0.884860213 | Scnm1 | sodium channel modifier 1 [Source:MGI S | 152.59 | 124.38 | 148.92 | 160.02 | 170.14 | 137 |
| 32232 | ENSMUSG00000092608 | 0 | NA | NA | NA | NA | NA | Vmn2r-ps81 | vomeronasal 2, receptor, pseudogene 81 | 0 | 0 | 0 | 0 | 0 | 0 |
| 32233 | ENSMUSG00000092609 | 0 | NA | NA | NA | NA | NA | Gm20481 | predicted gene 20481 [Source:MGI Symbo | 0 | 0 | 0 | 0 | 0 | 0 |
| 32234 | ENSMUSG00000092610 | 0 | NA | NA | NA | NA | NA | Vmn1r-ps67 | vomeronasal 1 receptor, pseudogene 67 [ | 0 | 0 | 0 | 0 | 0 | 0 |
| 32235 | ENSMUSG00000092611 | 0 | NA | NA | NA | NA | NA | Vmn2r-ps42 | vomeronasal 2, receptor, pseudogene 42 | 0 | 0 | 0 | 0 | 0 | 0 |
| 32236 | ENSMUSG00000092612 | 0 | NA | NA | NA | NA | NA | Gm18604 | predicted gene, 18604 [Source:MGI Symb | 0 | 0 | 0 | 0 | 0 | 0 |
| 32237 | ENSMUSG00000092614 | 0 | NA | NA | NA | NA | NA | Vmn2r-ps47 | vomeronasal 2, receptor, pseudogene 47 | 0 | 0 | 0 | 0 | 0 | 0 |
| 32238 | ENSMUSG00000092615 | 0 | NA | NA | NA | NA | NA | Olfr762-ps1 | olfactory receptor 762, pseudogene 1 [Sou | 0 | 0 | 0 | 0 | 0 | 0 |
| 32239 | ENSMUSG00000092616 | 0 | NA | NA | NA | NA | NA | Vmn1r-ps74 | vomeronasal 1 receptor, pseudogene 74 [ | 0 | 0 | 0 | 0 | 0 | 0 |
| 32240 | ENSMUSG00000092617 | 0 | NA | NA | NA | NA | NA | Gm8412 | predicted gene 8412 [Source:MGI Symbol | 0 | 0 | 0 | 0 | 0 | 0 |
| 32241 | ENSMUSG00000092618 | 0 | NA | NA | NA | NA | NA | Btnl6 | butyrophilin-like 6 [Source:MGI Symbol;A | 0 | 0 | 0 | 0 | 0 | 0 |
| 32242 | ENSMUSG00000092619 | 0 | NA | NA | NA | NA | NA | Gm20438 | predicted gene 20438 [Source:MGI Symbo | 0 | 0 | 0 | 0 | 0 | 0 |
| 32243 | ENSMUSG00000092620 | 0 | NA | NA | NA | NA | NA | Gm18201 | predicted gene, 18201 [Source:MGI Symb | 0 | 0 | 0 | 0 | 0 | 0 |
| 32244 | ENSMUSG00000092621 | 0 | NA | NA | NA | NA | NA | Vmn2r-ps66 | vomeronasal 2, receptor, pseudogene 66 | 0 | 0 | 0 | 0 | 0 | 0 |
| 32245 | ENSMUSG00000092622 | 3.929071179 | -0.295495807 | 1.042688717 | -0.283397914 | 0.776871827 | NA | Khdc3 | KH domain containing 3, subcortical mate | 4.8186 | 4.1117 | 4.025 | 2.8919 | 3.2719 | 4.4554 |
| 32246 | ENSMUSG00000092624 | 0.181769715 | 1.126924175 | 4.080472857 | 0.276174898 | 0.782413736 | NA | Gm3654 | predicted gene 3654 [Source:MGI Symbol | 0 | 0 | 0 | 0 | 1.0906 | 0 |
| 32247 | ENSMUSG00000092625 | 1.142077704 | 1.613415074 | 2.128062572 | 0.758161482 | 0.448354317 | NA | Gm20404 | predicted gene 20404 [Source:MGI Symbo | 1.6062 | 0 | 0 | 1.928 | 1.0906 | 2.2277 |
| 32248 | ENSMUSG00000092626 | 17.76109989 | 0.304572688 | 0.572007712 | 0.532462556 | 0.594405669 | 0.913168731 | 9130230N09 | RIKEN cDNA 9130230N09 gene [Source:M | 7.2279 | 18.503 | 22.137 | 13.496 | 21.812 | 23.391 |
| 32249 | ENSMUSG00000092627 | 0 | NA | NA | NA | NA | NA | D130058E05 | RIKEN cDNA D130058E05 gene [Source:M | 0 | 0 | 0 | 0 | 0 | 0 |
| 32250 | ENSMUSG00000092629 | 0 | NA | NA | NA | NA | NA | Vmn1r-ps85 | vomeronasal 1 receptor, pseudogene 85 [ | 0 | 0 | 0 | 0 | 0 | 0 |
| 32251 | ENSMUSG00000092633 | 0 | NA | NA | NA | NA | NA | Mir3962 | microRNA 3962 [Source:MGI Symbol;Acc: | 0 | 0 | 0 | 0 | 0 | 0 |
| 32252 | ENSMUSG00000092636 | 0 | NA | NA | NA | NA | NA | NA | NA | 0 | 0 | 0 | 0 | 0 | 0 |

| | | | | | | | | | | | | | | | |
|---|---|---|---|---|---|---|---|---|---|---|---|---|---|---|---|
| 32253 | ENSMUSG00000092638 | 0 | NA | NA | NA | NA | NA | Mir5112 | microRNA 5112 [Source:MGI Symbol;Acc: | 0 | 0 | 0 | 0 | 0 | 0 |
| 32254 | ENSMUSG00000092639 | 0 | NA | NA | NA | NA | NA | Mir3964 | microRNA 3964 [Source:MGI Symbol;Acc: | 0 | 0 | 0 | 0 | 0 | 0 |
| 32255 | ENSMUSG00000092641 | 0 | NA | NA | NA | NA | NA | NA | NA | 0 | 0 | 0 | 0 | 0 | 0 |
| 32256 | ENSMUSG00000092643 | 0 | NA | NA | NA | NA | NA | NA | NA | 0 | 0 | 0 | 0 | 0 | 0 |
| 32257 | ENSMUSG00000092644 | 0.439020386 | -2.189747911 | 3.52148864 | -0.621824499 | 0.534057275 | NA | NA | NA | 1.6062 | 1.0279 | 0 | 0 | 0 | 0 |
| 32258 | ENSMUSG00000092647 | 0 | NA | NA | NA | NA | NA | NA | NA | 0 | 0 | 0 | 0 | 0 | 0 |
| 32259 | ENSMUSG00000092648 | 0 | NA | NA | NA | NA | NA | NA | NA | 0 | 0 | 0 | 0 | 0 | 0 |
| 32260 | ENSMUSG00000092650 | 0 | NA | NA | NA | NA | NA | NA | NA | 0 | 0 | 0 | 0 | 0 | 0 |
| 32261 | ENSMUSG00000092652 | 0 | NA | NA | NA | NA | NA | Mir3104 | microRNA 3104 [Source:MGI Symbol;Acc: | 0 | 0 | 0 | 0 | 0 | 0 |
| 32262 | ENSMUSG00000092653 | 0 | NA | NA | NA | NA | NA | NA | NA | 0 | 0 | 0 | 0 | 0 | 0 |
| 32263 | ENSMUSG00000092655 | 0 | NA | NA | NA | NA | NA | NA | NA | 0 | 0 | 0 | 0 | 0 | 0 |
| 32264 | ENSMUSG00000092656 | 0 | NA | NA | NA | NA | NA | NA | NA | 0 | 0 | 0 | 0 | 0 | 0 |
| 32265 | ENSMUSG00000092657 | 0 | NA | NA | NA | NA | NA | NA | NA | 0 | 0 | 0 | 0 | 0 | 0 |
| 32266 | ENSMUSG00000092659 | 0.371281295 | 2.09253362 | 4.031852165 | 0.519000582 | 0.603760335 | NA | Mir3100 | microRNA 3100 [Source:MGI Symbol;Acc: | 0 | 0 | 0 | 0 | 0 | 2.2277 |
| 32267 | ENSMUSG00000092660 | 0 | NA | NA | NA | NA | NA | NA | NA | 0 | 0 | 0 | 0 | 0 | 0 |
| 32268 | ENSMUSG00000092661 | 0 | NA | NA | NA | NA | NA | NA | NA | 0 | 0 | 0 | 0 | 0 | 0 |
| 32269 | ENSMUSG00000092664 | 0 | NA | NA | NA | NA | NA | NA | NA | 0 | 0 | 0 | 0 | 0 | 0 |
| 32270 | ENSMUSG00000092666 | 0 | NA | NA | NA | NA | NA | NA | NA | 0 | 0 | 0 | 0 | 0 | 0 |
| 32271 | ENSMUSG00000092668 | 0 | NA | NA | NA | NA | NA | NA | NA | 0 | 0 | 0 | 0 | 0 | 0 |
| 32272 | ENSMUSG00000092671 | 0 | NA | NA | NA | NA | NA | NA | NA | 0 | 0 | 0 | 0 | 0 | 0 |
| 32273 | ENSMUSG00000092673 | 1.298844242 | -2.558331636 | 2.053310462 | -1.24595461 | 0.212781059 | NA | Mir3067 | microRNA 3067 [Source:MGI Symbol;Acc: | 1.6062 | 3.0838 | 2.0125 | 0 | 1.0906 | 0 |
| 32274 | ENSMUSG00000092674 | 0 | NA | NA | NA | NA | NA | Gm24105 | predicted gene, 24105 [Source:MGI Symb | 0 | 0 | 0 | 0 | 0 | 0 |
| 32275 | ENSMUSG00000092675 | 0.371281295 | 2.09253362 | 4.031852165 | 0.519000582 | 0.603760335 | NA | NA | NA | 0 | 0 | 0 | 0 | 0 | 2.2277 |
| 32276 | ENSMUSG00000092677 | 0 | NA | NA | NA | NA | NA | Mir5106 | microRNA 5106 [Source:MGI Symbol;Acc: | 0 | 0 | 0 | 0 | 0 | 0 |
| 32277 | ENSMUSG00000092679 | 0 | NA | NA | NA | NA | NA | Mir5129 | microRNA 5129 [Source:MGI Symbol;Acc: | 0 | 0 | 0 | 0 | 0 | 0 |
| 32278 | ENSMUSG00000092680 | 2.754946369 | 0.080625786 | 1.379051223 | 0.058464678 | 0.953378497 | NA | Snord55 | small nucleolar RNA, C/D box 55 [Source:I | 0 | 3.0838 | 5.0312 | 2.8919 | 2.1812 | 3.3415 |
| 32279 | ENSMUSG00000092683 | 0 | NA | NA | NA | NA | NA | NA | NA | 0 | 0 | 0 | 0 | 0 | 0 |
| 32280 | ENSMUSG00000092685 | 0 | NA | NA | NA | NA | NA | NA | NA | 0 | 0 | 0 | 0 | 0 | 0 |
| 32281 | ENSMUSG00000092686 | 0 | NA | NA | NA | NA | NA | NA | NA | 0 | 0 | 0 | 0 | 0 | 0 |
| 32282 | ENSMUSG00000092687 | 0 | NA | NA | NA | NA | NA | Gm25744 | predicted gene, 25744 [Source:MGI Symb | 0 | 0 | 0 | 0 | 0 | 0 |
| 32283 | ENSMUSG00000092693 | 0 | NA | NA | NA | NA | NA | NA | NA | 0 | 0 | 0 | 0 | 0 | 0 |
| 32284 | ENSMUSG00000092695 | 0.160662974 | 1.126924175 | 4.080472857 | 0.276174898 | 0.782413736 | NA | Mir3092 | microRNA 3092 [Source:MGI Symbol;Acc: | 0 | 0 | 0 | 0.964 | 0 | 0 |
| 32285 | ENSMUSG00000092696 | 0 | NA | NA | NA | NA | NA | Mir5133 | microRNA 5133 [Source:MGI Symbol;Acc: | 0 | 0 | 0 | 0 | 0 | 0 |
| 32286 | ENSMUSG00000092697 | 0 | NA | NA | NA | NA | NA | NA | NA | 0 | 0 | 0 | 0 | 0 | 0 |
| 32287 | ENSMUSG00000092698 | 0 | NA | NA | NA | NA | NA | NA | NA | 0 | 0 | 0 | 0 | 0 | 0 |
| 32288 | ENSMUSG00000092701 | 0 | NA | NA | NA | NA | NA | NA | NA | 0 | 0 | 0 | 0 | 0 | 0 |
| 32289 | ENSMUSG00000092702 | 0 | NA | NA | NA | NA | NA | Gm24514 | predicted gene, 24514 [Source:MGI Symb | 0 | 0 | 0 | 0 | 0 | 0 |
| 32290 | ENSMUSG00000092704 | 0 | NA | NA | NA | NA | NA | Gm24512 | predicted gene, 24512 [Source:MGI Symb | 0 | 0 | 0 | 0 | 0 | 0 |
| 32291 | ENSMUSG00000092705 | 0 | NA | NA | NA | NA | NA | NA | NA | 0 | 0 | 0 | 0 | 0 | 0 |
| 32292 | ENSMUSG00000092706 | 0 | NA | NA | NA | NA | NA | NA | NA | 0 | 0 | 0 | 0 | 0 | 0 |
| 32293 | ENSMUSG00000092710 | 0 | NA | NA | NA | NA | NA | Gm22857 | predicted gene, 22857 [Source:MGI Symb | 0 | 0 | 0 | 0 | 0 | 0 |
| 32294 | ENSMUSG00000092713 | 2.009720535 | 0.064883793 | 1.72239385 | 0.037670706 | 0.969950232 | NA | Gm22858 | predicted gene, 22858 [Source:MGI Symb | 4.8186 | 0 | 1.0062 | 2.8919 | 0 | 3.3415 |
| 32295 | ENSMUSG00000092714 | 0 | NA | NA | NA | NA | NA | NA | NA | 0 | 0 | 0 | 0 | 0 | 0 |
| 32296 | ENSMUSG00000092715 | 0 | NA | NA | NA | NA | NA | NA | NA | 0 | 0 | 0 | 0 | 0 | 0 |
| 32297 | ENSMUSG00000092718 | 0 | NA | NA | NA | NA | NA | NA | NA | 0 | 0 | 0 | 0 | 0 | 0 |
| 32298 | ENSMUSG00000092719 | 0 | NA | NA | NA | NA | NA | NA | NA | 0 | 0 | 0 | 0 | 0 | 0 |
| 32299 | ENSMUSG00000092720 | 0 | NA | NA | NA | NA | NA | NA | NA | 0 | 0 | 0 | 0 | 0 | 0 |
| 32300 | ENSMUSG00000092723 | 0 | NA | NA | NA | NA | NA | NA | NA | 0 | 0 | 0 | 0 | 0 | 0 |
| 32301 | ENSMUSG00000092724 | 0 | NA | NA | NA | NA | NA | NA | NA | 0 | 0 | 0 | 0 | 0 | 0 |
| 32302 | ENSMUSG00000092725 | 0 | NA | NA | NA | NA | NA | NA | NA | 0 | 0 | 0 | 0 | 0 | 0 |
| 32303 | ENSMUSG00000092727 | 0 | NA | NA | NA | NA | NA | Mir3062 | microRNA 3062 [Source:MGI Symbol;Acc: | 0 | 0 | 0 | 0 | 0 | 0 |
| 32304 | ENSMUSG00000092730 | 0 | NA | NA | NA | NA | NA | Snora24 | small nucleolar RNA, H/ACA box 24 [Sourc | 0 | 0 | 0 | 0 | 0 | 0 |
| 32305 | ENSMUSG00000092731 | 0 | NA | NA | NA | NA | NA | NA | NA | 0 | 0 | 0 | 0 | 0 | 0 |
| 32306 | ENSMUSG00000092732 | 0 | NA | NA | NA | NA | NA | NA | NA | 0 | 0 | 0 | 0 | 0 | 0 |
| 32307 | ENSMUSG00000092734 | 1.95270127 | 1.561935009 | 2.011044416 | 0.776678524 | 0.43734846 | NA | Mir5100 | microRNA 5100 [Source:MGI Symbol;Acc: | 0 | 1.0279 | 2.0125 | 8.6758 | 0 | 0 |
| 32308 | ENSMUSG00000092737 | 0 | NA | NA | NA | NA | NA | NA | NA | 0 | 0 | 0 | 0 | 0 | 0 |

| | | | | | | | | | | | | | | | |
|---|---|---|---|---|---|---|---|---|---|---|---|---|---|---|---|
| 32309 | ENSMUSG00000092738 | 0 | NA | NA | NA | NA | NA | Mir5135 | microRNA 5135 [Source:MGI Symbol;Acc: | 0 | 0 | 0 | 0 | 0 | 0 |
| 32310 | ENSMUSG00000092741 | 0 | NA | NA | NA | NA | NA | Mir3074-1 | microRNA 3074-1 [Source:MGI Symbol;Ac | 0 | 0 | 0 | 0 | 0 | 0 |
| 32311 | ENSMUSG00000092743 | 0 | NA | NA | NA | NA | NA | Gm22177 | predicted gene, 22177 [Source:MGI Symb | 0 | 0 | 0 | 0 | 0 | 0 |
| 32312 | ENSMUSG00000092745 | 0 | NA | NA | NA | NA | NA | Mir5122 | microRNA 5122 [Source:MGI Symbol;Acc: | 0 | 0 | 0 | 0 | 0 | 0 |
| 32313 | ENSMUSG00000092746 | 0.181769715 | 1.126924175 | 4.080472857 | 0.276174898 | 0.782413736 | NA | Rn7s6 | 7S RNA 6 [Source:MGI Symbol;Acc:MGI:9 | 0 | 0 | 0 | 0 | 1.0906 | 0 |
| 32314 | ENSMUSG00000092747 | 0 | NA | NA | NA | NA | NA | Gm22180 | predicted gene, 22180 [Source:MGI Symb | 0 | 0 | 0 | 0 | 0 | 0 |
| 32315 | ENSMUSG00000092748 | 0 | NA | NA | NA | NA | NA | NA | NA | 0 | 0 | 0 | 0 | 0 | 0 |
| 32316 | ENSMUSG00000092751 | 0 | NA | NA | NA | NA | NA | Mir3080 | microRNA 3080 [Source:MGI Symbol;Acc: | 0 | 0 | 0 | 0 | 0 | 0 |
| 32317 | ENSMUSG00000092752 | 0 | NA | NA | NA | NA | NA | Mir3473a | microRNA 3473a [Source:MGI Symbol;Acc | 0 | 0 | 0 | 0 | 0 | 0 |
| 32318 | ENSMUSG00000092753 | 0 | NA | NA | NA | NA | NA | NA | NA | 0 | 0 | 0 | 0 | 0 | 0 |
| 32319 | ENSMUSG00000092758 | 0.596071152 | -1.224343926 | 2.897583865 | -0.4225396 | 0.672631203 | NA | NA | NA | 1.6062 | 0 | 1.0062 | 0.964 | 0 | 0 |
| 32320 | ENSMUSG00000092759 | 0 | NA | NA | NA | NA | NA | NA | NA | 0 | 0 | 0 | 0 | 0 | 0 |
| 32321 | ENSMUSG00000092760 | 0 | NA | NA | NA | NA | NA | NA | NA | 0 | 0 | 0 | 0 | 0 | 0 |
| 32322 | ENSMUSG00000092761 | 0 | NA | NA | NA | NA | NA | Mir5120 | microRNA 5120 [Source:MGI Symbol;Acc: | 0 | 0 | 0 | 0 | 0 | 0 |
| 32323 | ENSMUSG00000092762 | 0 | NA | NA | NA | NA | NA | NA | NA | 0 | 0 | 0 | 0 | 0 | 0 |
| 32324 | ENSMUSG00000092763 | 0 | NA | NA | NA | NA | NA | NA | NA | 0 | 0 | 0 | 0 | 0 | 0 |
| 32325 | ENSMUSG00000092764 | 0 | NA | NA | NA | NA | NA | NA | NA | 0 | 0 | 0 | 0 | 0 | 0 |
| 32326 | ENSMUSG00000092765 | 0 | NA | NA | NA | NA | NA | Gm23363 | predicted gene, 23363 [Source:MGI Symb | 0 | 0 | 0 | 0 | 0 | 0 |
| 32327 | ENSMUSG00000092767 | 0 | NA | NA | NA | NA | NA | NA | NA | 0 | 0 | 0 | 0 | 0 | 0 |
| 32328 | ENSMUSG00000092770 | 0 | NA | NA | NA | NA | NA | Mirlet7a-2 | microRNA let7a-2 [Source:MGI Symbol;Ac | 0 | 0 | 0 | 0 | 0 | 0 |
| 32329 | ENSMUSG00000092774 | 0 | NA | NA | NA | NA | NA | NA | NA | 0 | 0 | 0 | 0 | 0 | 0 |
| 32330 | ENSMUSG00000092776 | 0 | NA | NA | NA | NA | NA | NA | NA | 0 | 0 | 0 | 0 | 0 | 0 |
| 32331 | ENSMUSG00000092777 | 0 | NA | NA | NA | NA | NA | NA | NA | 0 | 0 | 0 | 0 | 0 | 0 |
| 32332 | ENSMUSG00000092781 | 0 | NA | NA | NA | NA | NA | Mir3057 | microRNA 3057 [Source:MGI Symbol;Acc: | 0 | 0 | 0 | 0 | 0 | 0 |
| 32333 | ENSMUSG00000092784 | 0 | NA | NA | NA | NA | NA | Mir3077 | microRNA 3077 [Source:MGI Symbol;Acc: | 0 | 0 | 0 | 0 | 0 | 0 |
| 32334 | ENSMUSG00000092787 | 0 | NA | NA | NA | NA | NA | Mir5116 | microRNA 5116 [Source:MGI Symbol;Acc: | 0 | 0 | 0 | 0 | 0 | 0 |
| 32335 | ENSMUSG00000092790 | 0 | NA | NA | NA | NA | NA | NA | NA | 0 | 0 | 0 | 0 | 0 | 0 |
| 32336 | ENSMUSG00000092792 | 0 | NA | NA | NA | NA | NA | NA | NA | 0 | 0 | 0 | 0 | 0 | 0 |
| 32337 | ENSMUSG00000092793 | 0 | NA | NA | NA | NA | NA | Mir466m | microRNA 466m [Source:MGI Symbol;Acc | 0 | 0 | 0 | 0 | 0 | 0 |
| 32338 | ENSMUSG00000092794 | 0 | NA | NA | NA | NA | NA | Mir3110 | microRNA 3110 [Source:MGI Symbol;Acc: | 0 | 0 | 0 | 0 | 0 | 0 |
| 32339 | ENSMUSG00000092795 | 0.17131964 | -0.796629176 | 4.080472857 | -0.195229623 | 0.84521319 | NA | NA | NA | 0 | 1.0279 | 0 | 0 | 0 | 0 |
| 32340 | ENSMUSG00000092796 | 0 | NA | NA | NA | NA | NA | NA | NA | 0 | 0 | 0 | 0 | 0 | 0 |
| 32341 | ENSMUSG00000092799 | 0 | NA | NA | NA | NA | NA | NA | NA | 0 | 0 | 0 | 0 | 0 | 0 |
| 32342 | ENSMUSG00000092802 | 0 | NA | NA | NA | NA | NA | NA | NA | 0 | 0 | 0 | 0 | 0 | 0 |
| 32343 | ENSMUSG00000092805 | 0.545309144 | 2.651974629 | 3.666219369 | 0.72335405 | 0.469462394 | NA | Gm26461 | predicted gene, 26461 [Source:MGI Symb | 0 | 0 | 0 | 0 | 3.2719 | 0 |
| 32344 | ENSMUSG00000092806 | 0 | NA | NA | NA | NA | NA | NA | NA | 0 | 0 | 0 | 0 | 0 | 0 |
| 32345 | ENSMUSG00000092807 | 0 | NA | NA | NA | NA | NA | Mir199b | microRNA 199b [Source:MGI Symbol;Acc: | 0 | 0 | 0 | 0 | 0 | 0 |
| 32346 | ENSMUSG00000092811 | 0 | NA | NA | NA | NA | NA | NA | NA | 0 | 0 | 0 | 0 | 0 | 0 |
| 32347 | ENSMUSG00000092812 | 0 | NA | NA | NA | NA | NA | NA | NA | 0 | 0 | 0 | 0 | 0 | 0 |
| 32348 | ENSMUSG00000092813 | 0 | NA | NA | NA | NA | NA | Mir3086 | microRNA 3086 [Source:MGI Symbol;Acc: | 0 | 0 | 0 | 0 | 0 | 0 |
| 32349 | ENSMUSG00000092814 | 0.497389803 | 1.101799453 | 3.439268845 | 0.320358629 | 0.748696483 | NA | NA | NA | 0.8031 | 0 | 0 | 0 | 2.1812 | 0 |
| 32350 | ENSMUSG00000092815 | 0 | NA | NA | NA | NA | NA | Mir3099 | microRNA 3099 [Source:MGI Symbol;Acc: | 0 | 0 | 0 | 0 | 0 | 0 |
| 32351 | ENSMUSG00000092816 | 0 | NA | NA | NA | NA | NA | NA | NA | 0 | 0 | 0 | 0 | 0 | 0 |
| 32352 | ENSMUSG00000092818 | 0 | NA | NA | NA | NA | NA | NA | NA | 0 | 0 | 0 | 0 | 0 | 0 |
| 32353 | ENSMUSG00000092819 | 1.674228564 | 0.789871844 | 1.815541163 | 0.43506138 | 0.663517875 | NA | Gm23639 | predicted gene, 23639 [Source:MGI Symb | 1.6062 | 2.0558 | 0 | 1.928 | 0 | 4.4554 |
| 32354 | ENSMUSG00000092821 | 0 | NA | NA | NA | NA | NA | NA | NA | 0 | 0 | 0 | 0 | 0 | 0 |
| 32355 | ENSMUSG00000092822 | 0 | NA | NA | NA | NA | NA | NA | NA | 0 | 0 | 0 | 0 | 0 | 0 |
| 32356 | ENSMUSG00000092825 | 0 | NA | NA | NA | NA | NA | NA | NA | 0 | 0 | 0 | 0 | 0 | 0 |
| 32357 | ENSMUSG00000092826 | 0 | NA | NA | NA | NA | NA | NA | NA | 0 | 0 | 0 | 0 | 0 | 0 |
| 32358 | ENSMUSG00000092827 | 0.36353943 | 2.065989254 | 4.034137301 | 0.512126658 | 0.608562368 | NA | Mir3091 | microRNA 3091 [Source:MGI Symbol;Acc: | 0 | 0 | 0 | 0 | 2.1812 | 0 |
| 32359 | ENSMUSG00000092829 | 0 | NA | NA | NA | NA | NA | NA | NA | 0 | 0 | 0 | 0 | 0 | 0 |
| 32360 | ENSMUSG00000092830 | 0 | NA | NA | NA | NA | NA | Mir3963 | microRNA 3963 [Source:MGI Symbol;Acc: | 0 | 0 | 0 | 0 | 0 | 0 |
| 32361 | ENSMUSG00000092832 | 0 | NA | NA | NA | NA | NA | Mir3088 | microRNA 3088 [Source:MGI Symbol;Acc: | 0 | 0 | 0 | 0 | 0 | 0 |
| 32362 | ENSMUSG00000092833 | 0 | NA | NA | NA | NA | NA | NA | NA | 0 | 0 | 0 | 0 | 0 | 0 |
| 32363 | ENSMUSG00000092834 | 0 | NA | NA | NA | NA | NA | NA | NA | 0 | 0 | 0 | 0 | 0 | 0 |
| 32364 | ENSMUSG00000092835 | 0 | NA | NA | NA | NA | NA | NA | NA | 0 | 0 | 0 | 0 | 0 | 0 |

| | | | | | | | | | | | | | | | |
|---|---|---|---|---|---|---|---|---|---|---|---|---|---|---|---|
| 32365 | ENSMUSG00000092837 | 0 | NA | NA | NA | NA | NA | Rpph1 | ribonuclease P RNA component H1 [Sourc | 0 | 0 | 0 | 0 | 0 | 0 |
| 32366 | ENSMUSG00000092838 | 0 | NA | NA | NA | NA | NA | NA | NA | 0 | 0 | 0 | 0 | 0 | 0 |
| 32367 | ENSMUSG00000092840 | 0 | NA | NA | NA | NA | NA | NA | NA | 0 | 0 | 0 | 0 | 0 | 0 |
| 32368 | ENSMUSG00000092841 | 0.294513347 | 0.165143496 | 4.080472857 | 0.040471657 | 0.967717103 | NA | NA | NA | 0.8031 | 0 | 0 | 0.964 | 0 | 0 |
| 32369 | ENSMUSG00000092842 | 0 | NA | NA | NA | NA | NA | NA | NA | 0 | 0 | 0 | 0 | 0 | 0 |
| 32370 | ENSMUSG00000092843 | 0 | NA | NA | NA | NA | NA | NA | NA | 0 | 0 | 0 | 0 | 0 | 0 |
| 32371 | ENSMUSG00000092845 | 0.294513347 | 0.165143496 | 4.080472857 | 0.040471657 | 0.967717103 | NA | NA | NA | 0.8031 | 0 | 0 | 0.964 | 0 | 0 |
| 32372 | ENSMUSG00000092847 | 0 | NA | NA | NA | NA | NA | Mir344d-1 | microRNA 344d-1 [Source:MGI Symbol;Ac | 0 | 0 | 0 | 0 | 0 | 0 |
| 32373 | ENSMUSG00000092848 | 0 | NA | NA | NA | NA | NA | NA | NA | 0 | 0 | 0 | 0 | 0 | 0 |
| 32374 | ENSMUSG00000092849 | 0 | NA | NA | NA | NA | NA | NA | NA | 0 | 0 | 0 | 0 | 0 | 0 |
| 32375 | ENSMUSG00000092850 | 0 | NA | NA | NA | NA | NA | Mir344g | microRNA 344g [Source:MGI Symbol;Acc:I | 0 | 0 | 0 | 0 | 0 | 0 |
| 32376 | ENSMUSG00000092851 | 0.17131964 | -0.796629176 | 4.080472857 | -0.195229623 | 0.84521319 | NA | NA | NA | 0 | 1.0279 | 0 | 0 | 0 | 0 |
| 32377 | ENSMUSG00000092852 | 0 | NA | NA | NA | NA | NA | Mir5103 | microRNA 5103 [Source:MGI Symbol;Acc: | 0 | 0 | 0 | 0 | 0 | 0 |
| 32378 | ENSMUSG00000092853 | 0 | NA | NA | NA | NA | NA | NA | NA | 0 | 0 | 0 | 0 | 0 | 0 |
| 32379 | ENSMUSG00000092855 | 0 | NA | NA | NA | NA | NA | NA | NA | 0 | 0 | 0 | 0 | 0 | 0 |
| 32380 | ENSMUSG00000092856 | 0 | NA | NA | NA | NA | NA | NA | NA | 0 | 0 | 0 | 0 | 0 | 0 |
| 32381 | ENSMUSG00000092862 | 0.267700746 | -1.497481649 | 4.057001436 | -0.369110456 | 0.712045396 | NA | Mir3094 | microRNA 3094 [Source:MGI Symbol;Acc: | 1.6062 | 0 | 0 | 0 | 0 | 0 |
| 32382 | ENSMUSG00000092866 | 0 | NA | NA | NA | NA | NA | NA | NA | 0 | 0 | 0 | 0 | 0 | 0 |
| 32383 | ENSMUSG00000092867 | 2.581041133 | 2.157724218 | 1.484953633 | 1.453058311 | 0.146207565 | NA | Mir3058 | microRNA 3058 [Source:MGI Symbol;Acc: | 0.8031 | 2.0558 | 0 | 3.8559 | 6.5437 | 2.2277 |
| 32384 | ENSMUSG00000092868 | 0 | NA | NA | NA | NA | NA | Mir5113 | microRNA 5113 [Source:MGI Symbol;Acc: | 0 | 0 | 0 | 0 | 0 | 0 |
| 32385 | ENSMUSG00000092870 | 0 | NA | NA | NA | NA | NA | Mir3061 | microRNA 3061 [Source:MGI Symbol;Acc: | 0 | 0 | 0 | 0 | 0 | 0 |
| 32386 | ENSMUSG00000092871 | 0 | NA | NA | NA | NA | NA | NA | NA | 0 | 0 | 0 | 0 | 0 | 0 |
| 32387 | ENSMUSG00000092875 | 0 | NA | NA | NA | NA | NA | Gm22649 | predicted gene, 22649 [Source:MGI Symb | 0 | 0 | 0 | 0 | 0 | 0 |
| 32388 | ENSMUSG00000092876 | 0 | NA | NA | NA | NA | NA | NA | NA | 0 | 0 | 0 | 0 | 0 | 0 |
| 32389 | ENSMUSG00000092878 | 0 | NA | NA | NA | NA | NA | NA | NA | 0 | 0 | 0 | 0 | 0 | 0 |
| 32390 | ENSMUSG00000092879 | 0 | NA | NA | NA | NA | NA | Gm22650 | predicted gene, 22650 [Source:MGI Symb | 0 | 0 | 0 | 0 | 0 | 0 |
| 32391 | ENSMUSG00000092880 | 0 | NA | NA | NA | NA | NA | Mir3471-1 | microRNA 3471-1 [Source:MGI Symbol;Ac | 0 | 0 | 0 | 0 | 0 | 0 |
| 32392 | ENSMUSG00000092885 | 0 | NA | NA | NA | NA | NA | Gm22123 | predicted gene, 22123 [Source:MGI Symb | 0 | 0 | 0 | 0 | 0 | 0 |
| 32393 | ENSMUSG00000092887 | 1.336597737 | 1.762216136 | 1.911375842 | 0.921962127 | 0.356548331 | NA | Snord53 | small nucleolar RNA, C/D box 53 [Source:I | 0.8031 | 0 | 1.0062 | 2.8919 | 1.0906 | 2.2277 |
| 32394 | ENSMUSG00000092888 | 0 | NA | NA | NA | NA | NA | Gm22122 | predicted gene, 22122 [Source:MGI Symb | 0 | 0 | 0 | 0 | 0 | 0 |
| 32395 | ENSMUSG00000092889 | 3.734840888 | 0.816069641 | 1.112803308 | 0.733345808 | 0.463347543 | NA | NA | NA | 4.0155 | 2.0558 | 2.0125 | 7.7118 | 3.2719 | 3.3415 |
| 32396 | ENSMUSG00000092891 | 0 | NA | NA | NA | NA | NA | NA | NA | 0 | 0 | 0 | 0 | 0 | 0 |
| 32397 | ENSMUSG00000092894 | 0 | NA | NA | NA | NA | NA | NA | NA | 0 | 0 | 0 | 0 | 0 | 0 |
| 32398 | ENSMUSG00000092896 | 0 | NA | NA | NA | NA | NA | NA | NA | 0 | 0 | 0 | 0 | 0 | 0 |
| 32399 | ENSMUSG00000092897 | 0 | NA | NA | NA | NA | NA | NA | NA | 0 | 0 | 0 | 0 | 0 | 0 |
| 32400 | ENSMUSG00000092898 | 0 | NA | NA | NA | NA | NA | NA | NA | 0 | 0 | 0 | 0 | 0 | 0 |
| 32401 | ENSMUSG00000092901 | 0 | NA | NA | NA | NA | NA | Mir5130 | microRNA 5130 [Source:MGI Symbol;Acc: | 0 | 0 | 0 | 0 | 0 | 0 |
| 32402 | ENSMUSG00000092903 | 0 | NA | NA | NA | NA | NA | NA | NA | 0 | 0 | 0 | 0 | 0 | 0 |
| 32403 | ENSMUSG00000092905 | 0 | NA | NA | NA | NA | NA | Gm25735 | predicted gene, 25735 [Source:MGI Symb | 0 | 0 | 0 | 0 | 0 | 0 |
| 32404 | ENSMUSG00000092906 | 0 | NA | NA | NA | NA | NA | NA | NA | 0 | 0 | 0 | 0 | 0 | 0 |
| 32405 | ENSMUSG00000092907 | 0 | NA | NA | NA | NA | NA | Mir5132 | microRNA 5132 [Source:MGI Symbol;Acc: | 0 | 0 | 0 | 0 | 0 | 0 |
| 32406 | ENSMUSG00000092908 | 0 | NA | NA | NA | NA | NA | NA | NA | 0 | 0 | 0 | 0 | 0 | 0 |
| 32407 | ENSMUSG00000092909 | 0.656740811 | -0.013525372 | 3.112905782 | -0.004344934 | 0.996533255 | NA | NA | NA | 0 | 0 | 2.0125 | 1.928 | 0 | 0 |
| 32408 | ENSMUSG00000092912 | 0.267700746 | -1.497481649 | 4.057001436 | -0.369110456 | 0.712045396 | NA | Gm24086 | predicted gene, 24086 [Source:MGI Symb | 1.6062 | 0 | 0 | 0 | 0 | 0 |
| 32409 | ENSMUSG00000092913 | 0 | NA | NA | NA | NA | NA | NA | NA | 0 | 0 | 0 | 0 | 0 | 0 |
| 32410 | ENSMUSG00000092914 | 0 | NA | NA | NA | NA | NA | NA | NA | 0 | 0 | 0 | 0 | 0 | 0 |
| 32411 | ENSMUSG00000092915 | 0 | NA | NA | NA | NA | NA | NA | NA | 0 | 0 | 0 | 0 | 0 | 0 |
| 32412 | ENSMUSG00000092916 | 0 | NA | NA | NA | NA | NA | NA | NA | 0 | 0 | 0 | 0 | 0 | 0 |
| 32413 | ENSMUSG00000092917 | 0 | NA | NA | NA | NA | NA | NA | NA | 0 | 0 | 0 | 0 | 0 | 0 |
| 32414 | ENSMUSG00000092918 | 0 | NA | NA | NA | NA | NA | NA | NA | 0 | 0 | 0 | 0 | 0 | 0 |
| 32415 | ENSMUSG00000092919 | 0 | NA | NA | NA | NA | NA | NA | NA | 0 | 0 | 0 | 0 | 0 | 0 |
| 32416 | ENSMUSG00000092920 | 0 | NA | NA | NA | NA | NA | NA | NA | 0 | 0 | 0 | 0 | 0 | 0 |
| 32417 | ENSMUSG00000092921 | 0 | NA | NA | NA | NA | NA | Mir344b | microRNA 344b [Source:MGI Symbol;Acc: | 0 | 0 | 0 | 0 | 0 | 0 |
| 32418 | ENSMUSG00000092922 | 0 | NA | NA | NA | NA | NA | Mir5101 | microRNA 5101 [Source:MGI Symbol;Acc: | 0 | 0 | 0 | 0 | 0 | 0 |
| 32419 | ENSMUSG00000092923 | 0.480153995 | 1.043847617 | 3.151673025 | 0.331204287 | 0.740490182 | NA | NA | NA | 0.8031 | 0 | 0 | 0.964 | 0 | 1.1138 |
| 32420 | ENSMUSG00000092928 | 0 | NA | NA | NA | NA | NA | NA | NA | 0 | 0 | 0 | 0 | 0 | 0 |

| | | | | | | | | | | | | | | | |
|---|---|---|---|---|---|---|---|---|---|---|---|---|---|---|---|
| 32421 | ENSMUSG00000092931 | 0 | NA | NA | NA | NA | NA | NA | NA | 0 | 0 | 0 | 0 | 0 | 0 |
| 32422 | ENSMUSG00000092932 | 0 | NA | NA | NA | NA | NA | NA | NA | 0 | 0 | 0 | 0 | 0 | 0 |
| 32423 | ENSMUSG00000092934 | 0 | NA | NA | NA | NA | NA | NA | NA | 0 | 0 | 0 | 0 | 0 | 0 |
| 32424 | ENSMUSG00000092935 | 0 | NA | NA | NA | NA | NA | NA | NA | 0 | 0 | 0 | 0 | 0 | 0 |
| 32425 | ENSMUSG00000092939 | 0 | NA | NA | NA | NA | NA | NA | NA | 0 | 0 | 0 | 0 | 0 | 0 |
| 32426 | ENSMUSG00000092948 | 0 | NA | NA | NA | NA | NA | NA | NA | 0 | 0 | 0 | 0 | 0 | 0 |
| 32427 | ENSMUSG00000092949 | 0 | NA | NA | NA | NA | NA | NA | NA | 0 | 0 | 0 | 0 | 0 | 0 |
| 32428 | ENSMUSG00000092952 | 0 | NA | NA | NA | NA | NA | Mir3059 | microRNA 3059 [Source:MGI Symbol;Acc: | 0 | 0 | 0 | 0 | 0 | 0 |
| 32429 | ENSMUSG00000092953 | 0 | NA | NA | NA | NA | NA | NA | NA | 0 | 0 | 0 | 0 | 0 | 0 |
| 32430 | ENSMUSG00000092956 | 0 | NA | NA | NA | NA | NA | NA | NA | 0 | 0 | 0 | 0 | 0 | 0 |
| 32431 | ENSMUSG00000092957 | 0 | NA | NA | NA | NA | NA | NA | NA | 0 | 0 | 0 | 0 | 0 | 0 |
| 32432 | ENSMUSG00000092958 | 0 | NA | NA | NA | NA | NA | Mir5128 | microRNA 5128 [Source:MGI Symbol;Acc: | 0 | 0 | 0 | 0 | 0 | 0 |
| 32433 | ENSMUSG00000092961 | 0 | NA | NA | NA | NA | NA | NA | NA | 0 | 0 | 0 | 0 | 0 | 0 |
| 32434 | ENSMUSG00000092963 | 0 | NA | NA | NA | NA | NA | NA | NA | 0 | 0 | 0 | 0 | 0 | 0 |
| 32435 | ENSMUSG00000092965 | 0 | NA | NA | NA | NA | NA | NA | NA | 0 | 0 | 0 | 0 | 0 | 0 |
| 32436 | ENSMUSG00000092966 | 0 | NA | NA | NA | NA | NA | NA | NA | 0 | 0 | 0 | 0 | 0 | 0 |
| 32437 | ENSMUSG00000092972 | 0 | NA | NA | NA | NA | NA | Mir3078 | microRNA 3078 [Source:MGI Symbol;Acc: | 0 | 0 | 0 | 0 | 0 | 0 |
| 32438 | ENSMUSG00000092973 | 0 | NA | NA | NA | NA | NA | NA | NA | 0 | 0 | 0 | 0 | 0 | 0 |
| 32439 | ENSMUSG00000092974 | 0 | NA | NA | NA | NA | NA | Gm25072 | predicted gene, 25072 [Source:MGI Symb | 0 | 0 | 0 | 0 | 0 | 0 |
| 32440 | ENSMUSG00000092978 | 0 | NA | NA | NA | NA | NA | NA | NA | 0 | 0 | 0 | 0 | 0 | 0 |
| 32441 | ENSMUSG00000092979 | 0 | NA | NA | NA | NA | NA | NA | NA | 0 | 0 | 0 | 0 | 0 | 0 |
| 32442 | ENSMUSG00000092981 | 56.78126091 | 0.692735223 | 0.377511509 | 1.835004251 | 0.06650506 | 0.370971532 | Mir5125 | microRNA 5125 [Source:MGI Symbol;Acc: | 44.171 | 37.005 | 49.306 | 107.97 | 63.256 | 38.985 |
| 32443 | ENSMUSG00000092985 | 0.77082304 | -2.993021951 | 2.843236568 | -1.052681295 | 0.292487084 | NA | Mir378b | microRNA 378b [Source:MGI Symbol;Acc: | 1.6062 | 0 | 3.0187 | 0 | 0 | 0 |
| 32444 | ENSMUSG00000092986 | 0 | NA | NA | NA | NA | NA | Gm25574 | predicted gene, 25574 [Source:MGI Symb | 0 | 0 | 0 | 0 | 0 | 0 |
| 32445 | ENSMUSG00000092988 | 0 | NA | NA | NA | NA | NA | Mir3093 | microRNA 3093 [Source:MGI Symbol;Acc: | 0 | 0 | 0 | 0 | 0 | 0 |
| 32446 | ENSMUSG00000092990 | 0 | NA | NA | NA | NA | NA | Mir466q | microRNA 466q [Source:MGI Symbol;Acc: | 0 | 0 | 0 | 0 | 0 | 0 |
| 32447 | ENSMUSG00000092994 | 0.181769715 | 1.126924175 | 4.080472857 | 0.276174898 | 0.782413736 | NA | NA | NA | 0 | 0 | 0 | 0 | 1.0906 | 0 |
| 32448 | ENSMUSG00000092995 | 0 | NA | NA | NA | NA | NA | Mir16-1 | microRNA 16-1 [Source:MGI Symbol;Acc:I | 0 | 0 | 0 | 0 | 0 | 0 |
| 32449 | ENSMUSG00000092997 | 0 | NA | NA | NA | NA | NA | NA | NA | 0 | 0 | 0 | 0 | 0 | 0 |
| 32450 | ENSMUSG00000092998 | 0 | NA | NA | NA | NA | NA | Mir5099 | microRNA 5099 [Source:MGI Symbol;Acc: | 0 | 0 | 0 | 0 | 0 | 0 |
| 32451 | ENSMUSG00000092999 | 0 | NA | NA | NA | NA | NA | Gm23921 | predicted gene, 23921 [Source:MGI Symb | 0 | 0 | 0 | 0 | 0 | 0 |
| 32452 | ENSMUSG00000093000 | 0 | NA | NA | NA | NA | NA | NA | NA | 0 | 0 | 0 | 0 | 0 | 0 |
| 32453 | ENSMUSG00000093002 | 0 | NA | NA | NA | NA | NA | Gm24154 | predicted gene, 24154 [Source:MGI Symb | 0 | 0 | 0 | 0 | 0 | 0 |
| 32454 | ENSMUSG00000093003 | 0.160662974 | 1.126924175 | 4.080472857 | 0.276174898 | 0.782413736 | NA | Mir3473c | microRNA 3473c [Source:MGI Symbol;Acc | 0 | 0 | 0 | 0.964 | 0 | 0 |
| 32455 | ENSMUSG00000093004 | 0 | NA | NA | NA | NA | NA | NA | NA | 0 | 0 | 0 | 0 | 0 | 0 |
| 32456 | ENSMUSG00000093005 | 0 | NA | NA | NA | NA | NA | NA | NA | 0 | 0 | 0 | 0 | 0 | 0 |
| 32457 | ENSMUSG00000093006 | 0 | NA | NA | NA | NA | NA | NA | NA | 0 | 0 | 0 | 0 | 0 | 0 |
| 32458 | ENSMUSG00000093007 | 0 | NA | NA | NA | NA | NA | Mir15a | microRNA 15a [Source:MGI Symbol;Acc:M | 0 | 0 | 0 | 0 | 0 | 0 |
| 32459 | ENSMUSG00000093009 | 0 | NA | NA | NA | NA | NA | Snord52 | small nucleolar RNA, C/D box 52 [Source:I | 0 | 0 | 0 | 0 | 0 | 0 |
| 32460 | ENSMUSG00000093010 | 0 | NA | NA | NA | NA | NA | NA | NA | 0 | 0 | 0 | 0 | 0 | 0 |
| 32461 | ENSMUSG00000093011 | 0 | NA | NA | NA | NA | NA | Mir100 | microRNA 100 [Source:MGI Symbol;Acc:M | 0 | 0 | 0 | 0 | 0 | 0 |
| 32462 | ENSMUSG00000093013 | 0 | NA | NA | NA | NA | NA | NA | NA | 0 | 0 | 0 | 0 | 0 | 0 |
| 32463 | ENSMUSG00000093014 | 0 | NA | NA | NA | NA | NA | NA | NA | 0 | 0 | 0 | 0 | 0 | 0 |
| 32464 | ENSMUSG00000093015 | 0 | NA | NA | NA | NA | NA | NA | NA | 0 | 0 | 0 | 0 | 0 | 0 |
| 32465 | ENSMUSG00000093017 | 0 | NA | NA | NA | NA | NA | Mir5108 | microRNA 5108 [Source:MGI Symbol;Acc: | 0 | 0 | 0 | 0 | 0 | 0 |
| 32466 | ENSMUSG00000093019 | 0 | NA | NA | NA | NA | NA | NA | NA | 0 | 0 | 0 | 0 | 0 | 0 |
| 32467 | ENSMUSG00000093021 | 0 | NA | NA | NA | NA | NA | NA | NA | 0 | 0 | 0 | 0 | 0 | 0 |
| 32468 | ENSMUSG00000093023 | 0 | NA | NA | NA | NA | NA | NA | NA | 0 | 0 | 0 | 0 | 0 | 0 |
| 32469 | ENSMUSG00000093024 | 0 | NA | NA | NA | NA | NA | NA | NA | 0 | 0 | 0 | 0 | 0 | 0 |
| 32470 | ENSMUSG00000093025 | 0 | NA | NA | NA | NA | NA | NA | NA | 0 | 0 | 0 | 0 | 0 | 0 |
| 32471 | ENSMUSG00000093026 | 2.523395226 | -3.629669944 | 1.730335924 | -2.097667796 | 0.035934502 | NA | NA | NA | 4.8186 | 7.1954 | 2.0125 | 0 | 0 | 1.1138 |
| 32472 | ENSMUSG00000093029 | 0 | NA | NA | NA | NA | NA | NA | NA | 0 | 0 | 0 | 0 | 0 | 0 |
| 32473 | ENSMUSG00000093030 | 0.545309144 | 2.651974629 | 3.666219369 | 0.72335405 | 0.469462394 | NA | NA | NA | 0 | 0 | 0 | 0 | 3.2719 | 0 |
| 32474 | ENSMUSG00000093031 | 0.331982614 | 0.16514309 | 3.986751476 | 0.041422971 | 0.966958701 | NA | NA | NA | 0 | 1.0279 | 0 | 0.964 | 0 | 0 |
| 32475 | ENSMUSG00000093032 | 0 | NA | NA | NA | NA | NA | NA | NA | 0 | 0 | 0 | 0 | 0 | 0 |
| 32476 | ENSMUSG00000093033 | 0 | NA | NA | NA | NA | NA | NA | NA | 0 | 0 | 0 | 0 | 0 | 0 |

| | | | | | | | | | | | | | | | |
|---|---|---|---|---|---|---|---|---|---|---|---|---|---|---|---|
| 32477 | ENSMUSG00000093034 | 0 | NA | NA | NA | NA | NA | NA | NA | 0 | 0 | 0 | 0 | 0 | 0 |
| 32478 | ENSMUSG00000093036 | 0 | NA | NA | NA | NA | NA | Mir5127 | microRNA 5127 [Source:MGI Symbol;Acc: | 0 | 0 | 0 | 0 | 0 | 0 |
| 32479 | ENSMUSG00000093037 | 0 | NA | NA | NA | NA | NA | Mir3066 | microRNA 3066 [Source:MGI Symbol;Acc: | 0 | 0 | 0 | 0 | 0 | 0 |
| 32480 | ENSMUSG00000093039 | 0 | NA | NA | NA | NA | NA | Mir1912 | microRNA 1912 [Source:MGI Symbol;Acc: | 0 | 0 | 0 | 0 | 0 | 0 |
| 32481 | ENSMUSG00000093040 | 0 | NA | NA | NA | NA | NA | NA | NA | 0 | 0 | 0 | 0 | 0 | 0 |
| 32482 | ENSMUSG00000093041 | 0 | NA | NA | NA | NA | NA | NA | NA | 0 | 0 | 0 | 0 | 0 | 0 |
| 32483 | ENSMUSG00000093042 | 0 | NA | NA | NA | NA | NA | Mir3109 | microRNA 3109 [Source:MGI Symbol;Acc: | 0 | 0 | 0 | 0 | 0 | 0 |
| 32484 | ENSMUSG00000093044 | 1.099373405 | -1.223504443 | 2.147555505 | -0.56971959 | 0.568867901 | NA | Snord8 | small nucleolar RNA, C/D box 8 [Source:M | 1.6062 | 2.0558 | 1.0062 | 1.928 | 0 | 0 |
| 32485 | ENSMUSG00000093046 | 0 | NA | NA | NA | NA | NA | Mir3081 | microRNA 3081 [Source:MGI Symbol;Acc: | 0 | 0 | 0 | 0 | 0 | 0 |
| 32486 | ENSMUSG00000093051 | 0 | NA | NA | NA | NA | NA | NA | NA | 0 | 0 | 0 | 0 | 0 | 0 |
| 32487 | ENSMUSG00000093052 | 0 | NA | NA | NA | NA | NA | NA | NA | 0 | 0 | 0 | 0 | 0 | 0 |
| 32488 | ENSMUSG00000093054 | 0 | NA | NA | NA | NA | NA | NA | NA | 0 | 0 | 0 | 0 | 0 | 0 |
| 32489 | ENSMUSG00000093056 | 0 | NA | NA | NA | NA | NA | Gm24812 | predicted gene, 24812 [Source:MGI Symb | 0 | 0 | 0 | 0 | 0 | 0 |
| 32490 | ENSMUSG00000093058 | 0 | NA | NA | NA | NA | NA | NA | NA | 0 | 0 | 0 | 0 | 0 | 0 |
| 32491 | ENSMUSG00000093059 | 0 | NA | NA | NA | NA | NA | Gm24809 | predicted gene, 24809 [Source:MGI Symb | 0 | 0 | 0 | 0 | 0 | 0 |
| 32492 | ENSMUSG00000093060 | 0.36353943 | 2.065989254 | 4.034137301 | 0.512126658 | 0.608562368 | NA | Mir5104 | microRNA 5104 [Source:MGI Symbol;Acc: | 0 | 0 | 0 | 0 | 2.1812 | 0 |
| 32493 | ENSMUSG00000093061 | 0 | NA | NA | NA | NA | NA | Mir5098 | microRNA 5098 [Source:MGI Symbol;Acc: | 0 | 0 | 0 | 0 | 0 | 0 |
| 32494 | ENSMUSG00000093063 | 0 | NA | NA | NA | NA | NA | NA | NA | 0 | 0 | 0 | 0 | 0 | 0 |
| 32495 | ENSMUSG00000093064 | 0 | NA | NA | NA | NA | NA | Gm23153 | predicted gene, 23153 [Source:MGI Symb | 0 | 0 | 0 | 0 | 0 | 0 |
| 32496 | ENSMUSG00000093065 | 0 | NA | NA | NA | NA | NA | NA | NA | 0 | 0 | 0 | 0 | 0 | 0 |
| 32497 | ENSMUSG00000093067 | 0 | NA | NA | NA | NA | NA | NA | NA | 0 | 0 | 0 | 0 | 0 | 0 |
| 32498 | ENSMUSG00000093070 | 0 | NA | NA | NA | NA | NA | NA | NA | 0 | 0 | 0 | 0 | 0 | 0 |
| 32499 | ENSMUSG00000093071 | 0 | NA | NA | NA | NA | NA | NA | NA | 0 | 0 | 0 | 0 | 0 | 0 |
| 32500 | ENSMUSG00000093072 | 0.267700746 | -1.497481649 | 4.057001436 | -0.369110456 | 0.712045396 | NA | NA | NA | 1.6062 | 0 | 0 | 0 | 0 | 0 |
| 32501 | ENSMUSG00000093073 | 0 | NA | NA | NA | NA | NA | Mir124a-2 | microRNA 124a-2 [Source:MGI Symbol;Ac | 0 | 0 | 0 | 0 | 0 | 0 |
| 32502 | ENSMUSG00000093074 | 0 | NA | NA | NA | NA | NA | NA | NA | 0 | 0 | 0 | 0 | 0 | 0 |
| 32503 | ENSMUSG00000093075 | 0.535401493 | -2.48596965 | 3.684117979 | -0.674780141 | 0.499815459 | NA | NA | NA | 3.2124 | 0 | 0 | 0 | 0 | 0 |
| 32504 | ENSMUSG00000093076 | 0 | NA | NA | NA | NA | NA | Mir3082 | microRNA 3082 [Source:MGI Symbol;Acc: | 0 | 0 | 0 | 0 | 0 | 0 |
| 32505 | ENSMUSG00000093079 | 0 | NA | NA | NA | NA | NA | NA | NA | 0 | 0 | 0 | 0 | 0 | 0 |
| 32506 | ENSMUSG00000093080 | 3.056755729 | 0.121119702 | 1.256876939 | 0.096365601 | 0.923230211 | NA | Mir3060 | microRNA 3060 [Source:MGI Symbol;Acc: | 5.6217 | 1.0279 | 2.0125 | 1.928 | 2.1812 | 5.5692 |
| 32507 | ENSMUSG00000093081 | 0.167707431 | -0.796629176 | 4.080472857 | -0.195229623 | 0.84521319 | NA | Mir3075 | microRNA 3075 [Source:MGI Symbol;Acc: | 0 | 0 | 1.0062 | 0 | 0 | 0 |
| 32508 | ENSMUSG00000093082 | 0 | NA | NA | NA | NA | NA | NA | NA | 0 | 0 | 0 | 0 | 0 | 0 |
| 32509 | ENSMUSG00000093083 | 0 | NA | NA | NA | NA | NA | NA | NA | 0 | 0 | 0 | 0 | 0 | 0 |
| 32510 | ENSMUSG00000093085 | 0 | NA | NA | NA | NA | NA | NA | NA | 0 | 0 | 0 | 0 | 0 | 0 |
| 32511 | ENSMUSG00000093087 | 0 | NA | NA | NA | NA | NA | Mir3473d | microRNA 3473d [Source:MGI Symbol;Acc | 0 | 0 | 0 | 0 | 0 | 0 |
| 32512 | ENSMUSG00000093089 | 0 | NA | NA | NA | NA | NA | NA | NA | 0 | 0 | 0 | 0 | 0 | 0 |
| 32513 | ENSMUSG00000093090 | 0 | NA | NA | NA | NA | NA | Mir3073a | microRNA 3073a [Source:MGI Symbol;Acc | 0 | 0 | 0 | 0 | 0 | 0 |
| 32514 | ENSMUSG00000093091 | 0 | NA | NA | NA | NA | NA | Mir3089 | microRNA 3089 [Source:MGI Symbol;Acc: | 0 | 0 | 0 | 0 | 0 | 0 |
| 32515 | ENSMUSG00000093092 | 0 | NA | NA | NA | NA | NA | NA | NA | 0 | 0 | 0 | 0 | 0 | 0 |
| 32516 | ENSMUSG00000093093 | 0 | NA | NA | NA | NA | NA | Gm22642 | predicted gene, 22642 [Source:MGI Symb | 0 | 0 | 0 | 0 | 0 | 0 |
| 32517 | ENSMUSG00000093096 | 0 | NA | NA | NA | NA | NA | Mir3968 | microRNA 3968 [Source:MGI Symbol;Acc: | 0 | 0 | 0 | 0 | 0 | 0 |
| 32518 | ENSMUSG00000093097 | 0 | NA | NA | NA | NA | NA | Mir3097 | microRNA 3097 [Source:MGI Symbol;Acc: | 0 | 0 | 0 | 0 | 0 | 0 |
| 32519 | ENSMUSG00000093100 | 0 | NA | NA | NA | NA | NA | Mir3971 | microRNA 3971 [Source:MGI Symbol;Acc: | 0 | 0 | 0 | 0 | 0 | 0 |
| 32520 | ENSMUSG00000093101 | 0 | NA | NA | NA | NA | NA | NA | NA | 0 | 0 | 0 | 0 | 0 | 0 |
| 32521 | ENSMUSG00000093102 | 0 | NA | NA | NA | NA | NA | Mir3475 | microRNA 3475 [Source:MGI Symbol;Acc: | 0 | 0 | 0 | 0 | 0 | 0 |
| 32522 | ENSMUSG00000093105 | 0 | NA | NA | NA | NA | NA | NA | NA | 0 | 0 | 0 | 0 | 0 | 0 |
| 32523 | ENSMUSG00000093106 | 0 | NA | NA | NA | NA | NA | NA | NA | 0 | 0 | 0 | 0 | 0 | 0 |
| 32524 | ENSMUSG00000093107 | 0 | NA | NA | NA | NA | NA | NA | NA | 0 | 0 | 0 | 0 | 0 | 0 |
| 32525 | ENSMUSG00000093108 | 0 | NA | NA | NA | NA | NA | Mir344f | microRNA Mir344f [Source:MGI Symbol;A | 0 | 0 | 0 | 0 | 0 | 0 |
| 32526 | ENSMUSG00000093112 | 0 | NA | NA | NA | NA | NA | Gm22701 | predicted gene, 22701 [Source:MGI Symb | 0 | 0 | 0 | 0 | 0 | 0 |
| 32527 | ENSMUSG00000093113 | 0 | NA | NA | NA | NA | NA | NA | NA | 0 | 0 | 0 | 0 | 0 | 0 |
| 32528 | ENSMUSG00000093114 | 0 | NA | NA | NA | NA | NA | NA | NA | 0 | 0 | 0 | 0 | 0 | 0 |
| 32529 | ENSMUSG00000093115 | 0 | NA | NA | NA | NA | NA | NA | NA | 0 | 0 | 0 | 0 | 0 | 0 |
| 32530 | ENSMUSG00000093116 | 0 | NA | NA | NA | NA | NA | NA | NA | 0 | 0 | 0 | 0 | 0 | 0 |
| 32531 | ENSMUSG00000093117 | 0 | NA | NA | NA | NA | NA | Gm22699 | predicted gene, 22699 [Source:MGI Symb | 0 | 0 | 0 | 0 | 0 | 0 |
| 32532 | ENSMUSG00000093118 | 0 | NA | NA | NA | NA | NA | Mir3083 | microRNA 3083 [Source:MGI Symbol;Acc: | 0 | 0 | 0 | 0 | 0 | 0 |

| | | | | | | | | | | | | | | | |
|---|---|---|---|---|---|---|---|---|---|---|---|---|---|---|---|
| 32533 | ENSMUSG00000093119 | 0.17131964 | -0.796629176 | 4.080472857 | -0.195229623 | 0.84521319 | NA | NA | NA | 0 | 1.0279 | 0 | 0 | 0 | 0 |
| 32534 | ENSMUSG00000093120 | 0.17131964 | -0.796629176 | 4.080472857 | -0.195229623 | 0.84521319 | NA | Mir3474 | microRNA 3474 [Source:MGI Symbol;Acc: | 0 | 1.0279 | 0 | 0 | 0 | 0 |
| 32535 | ENSMUSG00000093122 | 0 | NA | NA | NA | NA | NA | Mir1264 | microRNA 1264 [Source:MGI Symbol;Acc: | 0 | 0 | 0 | 0 | 0 | 0 |
| 32536 | ENSMUSG00000093124 | 1.980329506 | 1.171502772 | 1.505034496 | 0.778389316 | 0.436339535 | NA | NA | NA | 1.6062 | 1.0279 | 1.0062 | 3.8559 | 3.2719 | 1.1138 |
| 32537 | ENSMUSG00000093125 | 0 | NA | NA | NA | NA | NA | NA | NA | 0 | 0 | 0 | 0 | 0 | 0 |
| 32538 | ENSMUSG00000093128 | 0 | NA | NA | NA | NA | NA | NA | NA | 0 | 0 | 0 | 0 | 0 | 0 |
| 32539 | ENSMUSG00000093129 | 0 | NA | NA | NA | NA | NA | NA | NA | 0 | 0 | 0 | 0 | 0 | 0 |
| 32540 | ENSMUSG00000093132 | 0 | NA | NA | NA | NA | NA | NA | NA | 0 | 0 | 0 | 0 | 0 | 0 |
| 32541 | ENSMUSG00000093133 | 0 | NA | NA | NA | NA | NA | NA | NA | 0 | 0 | 0 | 0 | 0 | 0 |
| 32542 | ENSMUSG00000093134 | 0 | NA | NA | NA | NA | NA | NA | NA | 0 | 0 | 0 | 0 | 0 | 0 |
| 32543 | ENSMUSG00000093137 | 0 | NA | NA | NA | NA | NA | Gm25995 | predicted gene, 25995 [Source:MGI Symb | 0 | 0 | 0 | 0 | 0 | 0 |
| 32544 | ENSMUSG00000093138 | 0 | NA | NA | NA | NA | NA | NA | NA | 0 | 0 | 0 | 0 | 0 | 0 |
| 32545 | ENSMUSG00000093139 | 0 | NA | NA | NA | NA | NA | Gm25999 | predicted gene, 25999 [Source:MGI Symb | 0 | 0 | 0 | 0 | 0 | 0 |
| 32546 | ENSMUSG00000093140 | 0.181769715 | 1.126924175 | 4.080472857 | 0.276174898 | 0.782413736 | NA | Mir28b | microRNA 28b [Source:MGI Symbol;Acc:M | 0 | 0 | 0 | 0 | 1.0906 | 0 |
| 32547 | ENSMUSG00000093142 | 0 | NA | NA | NA | NA | NA | Mir3106 | microRNA 3106 [Source:MGI Symbol;Acc: | 0 | 0 | 0 | 0 | 0 | 0 |
| 32548 | ENSMUSG00000093147 | 0 | NA | NA | NA | NA | NA | Gm23369 | predicted gene, 23369 [Source:MGI Symb | 0 | 0 | 0 | 0 | 0 | 0 |
| 32549 | ENSMUSG00000093148 | 0 | NA | NA | NA | NA | NA | NA | NA | 0 | 0 | 0 | 0 | 0 | 0 |
| 32550 | ENSMUSG00000093149 | 0 | NA | NA | NA | NA | NA | NA | NA | 0 | 0 | 0 | 0 | 0 | 0 |
| 32551 | ENSMUSG00000093152 | 0 | NA | NA | NA | NA | NA | NA | NA | 0 | 0 | 0 | 0 | 0 | 0 |
| 32552 | ENSMUSG00000093153 | 0 | NA | NA | NA | NA | NA | NA | NA | 0 | 0 | 0 | 0 | 0 | 0 |
| 32553 | ENSMUSG00000093155 | 0 | NA | NA | NA | NA | NA | NA | NA | 0 | 0 | 0 | 0 | 0 | 0 |
| 32554 | ENSMUSG00000093156 | 0 | NA | NA | NA | NA | NA | Mir3076 | microRNA 3076 [Source:MGI Symbol;Acc: | 0 | 0 | 0 | 0 | 0 | 0 |
| 32555 | ENSMUSG00000093157 | 0 | NA | NA | NA | NA | NA | Mir3098 | microRNA 3098 [Source:MGI Symbol;Acc: | 0 | 0 | 0 | 0 | 0 | 0 |
| 32556 | ENSMUSG00000093158 | 0 | NA | NA | NA | NA | NA | NA | NA | 0 | 0 | 0 | 0 | 0 | 0 |
| 32557 | ENSMUSG00000093159 | 0 | NA | NA | NA | NA | NA | NA | NA | 0 | 0 | 0 | 0 | 0 | 0 |
| 32558 | ENSMUSG00000093160 | 0 | NA | NA | NA | NA | NA | NA | NA | 0 | 0 | 0 | 0 | 0 | 0 |
| 32559 | ENSMUSG00000093161 | 0 | NA | NA | NA | NA | NA | NA | NA | 0 | 0 | 0 | 0 | 0 | 0 |
| 32560 | ENSMUSG00000093162 | 0 | NA | NA | NA | NA | NA | NA | NA | 0 | 0 | 0 | 0 | 0 | 0 |
| 32561 | ENSMUSG00000093163 | 0 | NA | NA | NA | NA | NA | NA | NA | 0 | 0 | 0 | 0 | 0 | 0 |
| 32562 | ENSMUSG00000093164 | 0 | NA | NA | NA | NA | NA | NA | NA | 0 | 0 | 0 | 0 | 0 | 0 |
| 32563 | ENSMUSG00000093167 | 0 | NA | NA | NA | NA | NA | Mir5131 | microRNA 5131 [Source:MGI Symbol;Acc: | 0 | 0 | 0 | 0 | 0 | 0 |
| 32564 | ENSMUSG00000093168 | 0.506734502 | -2.372847156 | 3.338060886 | -0.710845978 | 0.477179685 | NA | NA | NA | 0 | 1.0279 | 2.0125 | 0 | 0 | 0 |
| 32565 | ENSMUSG00000093169 | 0.133850373 | -0.796629176 | 4.080472857 | -0.195229623 | 0.84521319 | NA | Mir3079 | microRNA 3079 [Source:MGI Symbol;Acc: | 0.8031 | 0 | 0 | 0 | 0 | 0 |
| 32566 | ENSMUSG00000093170 | 0 | NA | NA | NA | NA | NA | NA | NA | 0 | 0 | 0 | 0 | 0 | 0 |
| 32567 | ENSMUSG00000093172 | 0 | NA | NA | NA | NA | NA | Mir3112 | microRNA 3112 [Source:MGI Symbol;Acc: | 0 | 0 | 0 | 0 | 0 | 0 |
| 32568 | ENSMUSG00000093173 | 0 | NA | NA | NA | NA | NA | NA | NA | 0 | 0 | 0 | 0 | 0 | 0 |
| 32569 | ENSMUSG00000093176 | 1.783638713 | 0.70073323 | 1.987117917 | 0.352637971 | 0.724359867 | NA | Mir5119 | microRNA 5119 [Source:MGI Symbol;Acc: | 0 | 4.1117 | 0 | 0 | 4.3625 | 2.2277 |
| 32570 | ENSMUSG00000093178 | 0.167707431 | -0.796629176 | 4.080472857 | -0.195229623 | 0.84521319 | NA | Snord87 | small nucleolar RNA, C/D box 87 [Source:I | 0 | 0 | 1.0062 | 0 | 0 | 0 |
| 32571 | ENSMUSG00000093180 | 0 | NA | NA | NA | NA | NA | NA | NA | 0 | 0 | 0 | 0 | 0 | 0 |
| 32572 | ENSMUSG00000093183 | 0 | NA | NA | NA | NA | NA | Gm25687 | predicted gene, 25687 [Source:MGI Symb | 0 | 0 | 0 | 0 | 0 | 0 |
| 32573 | ENSMUSG00000093184 | 0 | NA | NA | NA | NA | NA | NA | NA | 0 | 0 | 0 | 0 | 0 | 0 |
| 32574 | ENSMUSG00000093185 | 0 | NA | NA | NA | NA | NA | NA | NA | 0 | 0 | 0 | 0 | 0 | 0 |
| 32575 | ENSMUSG00000093187 | 0.133850373 | -0.796629176 | 4.080472857 | -0.195229623 | 0.84521319 | NA | Mir344-2 | microRNA 344-2 [Source:MGI Symbol;Acc | 0.8031 | 0 | 0 | 0 | 0 | 0 |
| 32576 | ENSMUSG00000093189 | 0 | NA | NA | NA | NA | NA | Gm25694 | predicted gene, 25694 [Source:MGI Symb | 0 | 0 | 0 | 0 | 0 | 0 |
| 32577 | ENSMUSG00000093190 | 0 | NA | NA | NA | NA | NA | NA | NA | 0 | 0 | 0 | 0 | 0 | 0 |
| 32578 | ENSMUSG00000093191 | 0 | NA | NA | NA | NA | NA | Gm22876 | predicted gene, 22876 [Source:MGI Symb | 0 | 0 | 0 | 0 | 0 | 0 |
| 32579 | ENSMUSG00000093192 | 0 | NA | NA | NA | NA | NA | NA | NA | 0 | 0 | 0 | 0 | 0 | 0 |
| 32580 | ENSMUSG00000093202 | 0 | NA | NA | NA | NA | NA | Mir3547 | microRNA 3547 [Source:MGI Symbol;Acc: | 0 | 0 | 0 | 0 | 0 | 0 |
| 32581 | ENSMUSG00000093203 | 0 | NA | NA | NA | NA | NA | NA | NA | 0 | 0 | 0 | 0 | 0 | 0 |
| 32582 | ENSMUSG00000093206 | 0 | NA | NA | NA | NA | NA | NA | NA | 0 | 0 | 0 | 0 | 0 | 0 |
| 32583 | ENSMUSG00000093207 | 0 | NA | NA | NA | NA | NA | NA | NA | 0 | 0 | 0 | 0 | 0 | 0 |
| 32584 | ENSMUSG00000093208 | 0 | NA | NA | NA | NA | NA | NA | NA | 0 | 0 | 0 | 0 | 0 | 0 |
| 32585 | ENSMUSG00000093211 | 0 | NA | NA | NA | NA | NA | NA | NA | 0 | 0 | 0 | 0 | 0 | 0 |
| 32586 | ENSMUSG00000093212 | 0 | NA | NA | NA | NA | NA | NA | NA | 0 | 0 | 0 | 0 | 0 | 0 |
| 32587 | ENSMUSG00000093214 | 0 | NA | NA | NA | NA | NA | NA | NA | 0 | 0 | 0 | 0 | 0 | 0 |
| 32588 | ENSMUSG00000093215 | 0 | NA | NA | NA | NA | NA | Mir3103 | microRNA 3103 [Source:MGI Symbol;Acc: | 0 | 0 | 0 | 0 | 0 | 0 |

| | | | | | | | | | | | | | | | |
|---|---|---|---|---|---|---|---|---|---|---|---|---|---|---|---|
| 32589 | ENSMUSG00000093216 | 0 | NA | NA | NA | NA | NA | NA | NA | 0 | 0 | 0 | 0 | 0 | 0 |
| 32590 | ENSMUSG00000093217 | 0 | NA | NA | NA | NA | NA | NA | NA | 0 | 0 | 0 | 0 | 0 | 0 |
| 32591 | ENSMUSG00000093218 | 0 | NA | NA | NA | NA | NA | NA | NA | 0 | 0 | 0 | 0 | 0 | 0 |
| 32592 | ENSMUSG00000093219 | 0 | NA | NA | NA | NA | NA | Mir3113 | microRNA 3113 [Source:MGI Symbol;Acc: | 0 | 0 | 0 | 0 | 0 | 0 |
| 32593 | ENSMUSG00000093221 | 0 | NA | NA | NA | NA | NA | Mir3105 | microRNA 3105 [Source:MGI Symbol;Acc: | 0 | 0 | 0 | 0 | 0 | 0 |
| 32594 | ENSMUSG00000093223 | 0 | NA | NA | NA | NA | NA | NA | NA | 0 | 0 | 0 | 0 | 0 | 0 |
| 32595 | ENSMUSG00000093224 | 0 | NA | NA | NA | NA | NA | NA | NA | 0 | 0 | 0 | 0 | 0 | 0 |
| 32596 | ENSMUSG00000093226 | 0 | NA | NA | NA | NA | NA | NA | NA | 0 | 0 | 0 | 0 | 0 | 0 |
| 32597 | ENSMUSG00000093227 | 0 | NA | NA | NA | NA | NA | Mir1251 | microRNA 1251 [Source:MGI Symbol;Acc: | 0 | 0 | 0 | 0 | 0 | 0 |
| 32598 | ENSMUSG00000093228 | 0 | NA | NA | NA | NA | NA | NA | NA | 0 | 0 | 0 | 0 | 0 | 0 |
| 32599 | ENSMUSG00000093230 | 0 | NA | NA | NA | NA | NA | NA | NA | 0 | 0 | 0 | 0 | 0 | 0 |
| 32600 | ENSMUSG00000093231 | 0 | NA | NA | NA | NA | NA | NA | NA | 0 | 0 | 0 | 0 | 0 | 0 |
| 32601 | ENSMUSG00000093232 | 0 | NA | NA | NA | NA | NA | NA | NA | 0 | 0 | 0 | 0 | 0 | 0 |
| 32602 | ENSMUSG00000093234 | 0 | NA | NA | NA | NA | NA | Mir3085 | microRNA 3085 [Source:MGI Symbol;Acc: | 0 | 0 | 0 | 0 | 0 | 0 |
| 32603 | ENSMUSG00000093236 | 0 | NA | NA | NA | NA | NA | NA | NA | 0 | 0 | 0 | 0 | 0 | 0 |
| 32604 | ENSMUSG00000093238 | 0 | NA | NA | NA | NA | NA | Mir9-3 | microRNA 9-3 [Source:MGI Symbol;Acc:M | 0 | 0 | 0 | 0 | 0 | 0 |
| 32605 | ENSMUSG00000093239 | 0.267700746 | -1.497481649 | 4.057001436 | -0.369110456 | 0.712045396 | NA | NA | NA | 1.6062 | 0 | 0 | 0 | 0 | 0 |
| 32606 | ENSMUSG00000093240 | 0 | NA | NA | NA | NA | NA | NA | NA | 0 | 0 | 0 | 0 | 0 | 0 |
| 32607 | ENSMUSG00000093241 | 0 | NA | NA | NA | NA | NA | Mir3108 | microRNA 3108 [Source:MGI Symbol;Acc: | 0 | 0 | 0 | 0 | 0 | 0 |
| 32608 | ENSMUSG00000093242 | 0 | NA | NA | NA | NA | NA | Gm24574 | predicted gene, 24574 [Source:MGI Symb | 0 | 0 | 0 | 0 | 0 | 0 |
| 32609 | ENSMUSG00000093243 | 0 | NA | NA | NA | NA | NA | NA | NA | 0 | 0 | 0 | 0 | 0 | 0 |
| 32610 | ENSMUSG00000093245 | 0 | NA | NA | NA | NA | NA | NA | NA | 0 | 0 | 0 | 0 | 0 | 0 |
| 32611 | ENSMUSG00000093246 | 0 | NA | NA | NA | NA | NA | Gm24572 | predicted gene, 24572 [Source:MGI Symb | 0 | 0 | 0 | 0 | 0 | 0 |
| 32612 | ENSMUSG00000093249 | 0.133850373 | -0.796629176 | 4.080472857 | -0.195229623 | 0.84521319 | NA | Gm24576 | predicted gene, 24576 [Source:MGI Symb | 0.8031 | 0 | 0 | 0 | 0 | 0 |
| 32613 | ENSMUSG00000093251 | 0 | NA | NA | NA | NA | NA | NA | NA | 0 | 0 | 0 | 0 | 0 | 0 |
| 32614 | ENSMUSG00000093254 | 0 | NA | NA | NA | NA | NA | Mir5126 | microRNA 5126 [Source:MGI Symbol;Acc: | 0 | 0 | 0 | 0 | 0 | 0 |
| 32615 | ENSMUSG00000093256 | 0 | NA | NA | NA | NA | NA | NA | NA | 0 | 0 | 0 | 0 | 0 | 0 |
| 32616 | ENSMUSG00000093258 | 0 | NA | NA | NA | NA | NA | Gm22929 | predicted gene, 22929 [Source:MGI Symb | 0 | 0 | 0 | 0 | 0 | 0 |
| 32617 | ENSMUSG00000093260 | 0 | NA | NA | NA | NA | NA | NA | NA | 0 | 0 | 0 | 0 | 0 | 0 |
| 32618 | ENSMUSG00000093261 | 0 | NA | NA | NA | NA | NA | NA | NA | 0 | 0 | 0 | 0 | 0 | 0 |
| 32619 | ENSMUSG00000093262 | 0.181769715 | 1.126924175 | 4.080472857 | 0.276174898 | 0.782413736 | NA | NA | NA | 0 | 0 | 0 | 0 | 1.0906 | 0 |
| 32620 | ENSMUSG00000093264 | 0 | NA | NA | NA | NA | NA | NA | NA | 0 | 0 | 0 | 0 | 0 | 0 |
| 32621 | ENSMUSG00000093266 | 0 | NA | NA | NA | NA | NA | NA | NA | 0 | 0 | 0 | 0 | 0 | 0 |
| 32622 | ENSMUSG00000093269 | 0 | NA | NA | NA | NA | NA | Mir5124a | microRNA 5124a [Source:MGI Symbol;Acc | 0 | 0 | 0 | 0 | 0 | 0 |
| 32623 | ENSMUSG00000093270 | 0 | NA | NA | NA | NA | NA | Mir5134 | microRNA 5134 [Source:MGI Symbol;Acc: | 0 | 0 | 0 | 0 | 0 | 0 |
| 32624 | ENSMUSG00000093271 | 0 | NA | NA | NA | NA | NA | NA | NA | 0 | 0 | 0 | 0 | 0 | 0 |
| 32625 | ENSMUSG00000093273 | 0 | NA | NA | NA | NA | NA | Mir3064 | microRNA 3064 [Source:MGI Symbol;Acc: | 0 | 0 | 0 | 0 | 0 | 0 |
| 32626 | ENSMUSG00000093274 | 0 | NA | NA | NA | NA | NA | NA | NA | 0 | 0 | 0 | 0 | 0 | 0 |
| 32627 | ENSMUSG00000093276 | 0 | NA | NA | NA | NA | NA | NA | NA | 0 | 0 | 0 | 0 | 0 | 0 |
| 32628 | ENSMUSG00000093277 | 0.428363721 | -0.543730382 | 3.624677344 | -0.15000794 | 0.880758351 | NA | Gm26236 | predicted gene, 26236 [Source:MGI Symb | 1.6062 | 0 | 0 | 0.964 | 0 | 0 |
| 32629 | ENSMUSG00000093278 | 0 | NA | NA | NA | NA | NA | Mir5110 | microRNA 5110 [Source:MGI Symbol;Acc: | 0 | 0 | 0 | 0 | 0 | 0 |
| 32630 | ENSMUSG00000093279 | 0 | NA | NA | NA | NA | NA | NA | NA | 0 | 0 | 0 | 0 | 0 | 0 |
| 32631 | ENSMUSG00000093280 | 0 | NA | NA | NA | NA | NA | Mir2861 | microRNA 2861 [Source:MGI Symbol;Acc: | 0 | 0 | 0 | 0 | 0 | 0 |
| 32632 | ENSMUSG00000093281 | 0 | NA | NA | NA | NA | NA | NA | NA | 0 | 0 | 0 | 0 | 0 | 0 |
| 32633 | ENSMUSG00000093282 | 0 | NA | NA | NA | NA | NA | NA | NA | 0 | 0 | 0 | 0 | 0 | 0 |
| 32634 | ENSMUSG00000093283 | 0 | NA | NA | NA | NA | NA | Gm22567 | predicted gene, 22567 [Source:MGI Symb | 0 | 0 | 0 | 0 | 0 | 0 |
| 32635 | ENSMUSG00000093286 | 0 | NA | NA | NA | NA | NA | NA | NA | 0 | 0 | 0 | 0 | 0 | 0 |
| 32636 | ENSMUSG00000093289 | 0 | NA | NA | NA | NA | NA | Snord92 | small nucleolar RNA, C/D box 92 [Source:I | 0 | 0 | 0 | 0 | 0 | 0 |
| 32637 | ENSMUSG00000093290 | 0 | NA | NA | NA | NA | NA | Mir3572 | microRNA 3572 [Source:MGI Symbol;Acc: | 0 | 0 | 0 | 0 | 0 | 0 |
| 32638 | ENSMUSG00000093291 | 0 | NA | NA | NA | NA | NA | Mir299b | microRNA 299b [Source:MGI Symbol;Acc: | 0 | 0 | 0 | 0 | 0 | 0 |
| 32639 | ENSMUSG00000093292 | 0 | NA | NA | NA | NA | NA | Mir5118 | microRNA 5118 [Source:MGI Symbol;Acc: | 0 | 0 | 0 | 0 | 0 | 0 |
| 32640 | ENSMUSG00000093293 | 0 | NA | NA | NA | NA | NA | Mir3095 | microRNA 3095 [Source:MGI Symbol;Acc: | 0 | 0 | 0 | 0 | 0 | 0 |
| 32641 | ENSMUSG00000093294 | 0 | NA | NA | NA | NA | NA | NA | NA | 0 | 0 | 0 | 0 | 0 | 0 |
| 32642 | ENSMUSG00000093296 | 0 | NA | NA | NA | NA | NA | Mir3102 | microRNA 3102 [Source:MGI Symbol;Acc: | 0 | 0 | 0 | 0 | 0 | 0 |
| 32643 | ENSMUSG00000093298 | 0 | NA | NA | NA | NA | NA | Mir3473b | microRNA 3473b [Source:MGI Symbol;Acc | 0 | 0 | 0 | 0 | 0 | 0 |
| 32644 | ENSMUSG00000093300 | 0 | NA | NA | NA | NA | NA | NA | NA | 0 | 0 | 0 | 0 | 0 | 0 |

| | | | | | | | | | | | | | | | |
|---|---|---|---|---|---|---|---|---|---|---|---|---|---|---|---|
| 32645 | ENSMUSG00000093301 | 0 | NA | NA | NA | NA | NA | NA | NA | 0 | 0 | 0 | 0 | 0 | 0 |
| 32646 | ENSMUSG00000093302 | 0 | NA | NA | NA | NA | NA | NA | NA | 0 | 0 | 0 | 0 | 0 | 0 |
| 32647 | ENSMUSG00000093303 | 0 | NA | NA | NA | NA | NA | NA | NA | 0 | 0 | 0 | 0 | 0 | 0 |
| 32648 | ENSMUSG00000093306 | 0 | NA | NA | NA | NA | NA | NA | NA | 0 | 0 | 0 | 0 | 0 | 0 |
| 32649 | ENSMUSG00000093311 | 0 | NA | NA | NA | NA | NA | Mir3965 | microRNA 3965 [Source:MGI Symbol;Acc: | 0 | 0 | 0 | 0 | 0 | 0 |
| 32650 | ENSMUSG00000093314 | 0 | NA | NA | NA | NA | NA | Mir5136 | microRNA 5136 [Source:MGI Symbol;Acc: | 0 | 0 | 0 | 0 | 0 | 0 |
| 32651 | ENSMUSG00000093315 | 0 | NA | NA | NA | NA | NA | Mir5114 | microRNA 5114 [Source:MGI Symbol;Acc: | 0 | 0 | 0 | 0 | 0 | 0 |
| 32652 | ENSMUSG00000093317 | 0 | NA | NA | NA | NA | NA | NA | NA | 0 | 0 | 0 | 0 | 0 | 0 |
| 32653 | ENSMUSG00000093319 | 0 | NA | NA | NA | NA | NA | NA | NA | 0 | 0 | 0 | 0 | 0 | 0 |
| 32654 | ENSMUSG00000093322 | 0 | NA | NA | NA | NA | NA | NA | NA | 0 | 0 | 0 | 0 | 0 | 0 |
| 32655 | ENSMUSG00000093323 | 0.685278559 | -2.809870862 | 3.376578027 | -0.832165239 | 0.405315682 | NA | NA | NA | 0 | 4.1117 | 0 | 0 | 0 | 0 |
| 32656 | ENSMUSG00000093325 | 0 | NA | NA | NA | NA | NA | Gm26061 | predicted gene, 26061 [Source:MGI Symb | 0 | 0 | 0 | 0 | 0 | 0 |
| 32657 | ENSMUSG00000093327 | 1.08321747 | -3.483110813 | 2.291372099 | -1.520098291 | 0.128486274 | NA | Mir5107 | microRNA 5107 [Source:MGI Symbol;Acc: | 2.4093 | 3.0838 | 1.0062 | 0 | 0 | 0 |
| 32658 | ENSMUSG00000093329 | 0 | NA | NA | NA | NA | NA | NA | NA | 0 | 0 | 0 | 0 | 0 | 0 |
| 32659 | ENSMUSG00000093331 | 0 | NA | NA | NA | NA | NA | NA | NA | 0 | 0 | 0 | 0 | 0 | 0 |
| 32660 | ENSMUSG00000093334 | 0 | NA | NA | NA | NA | NA | NA | NA | 0 | 0 | 0 | 0 | 0 | 0 |
| 32661 | ENSMUSG00000093335 | 0 | NA | NA | NA | NA | NA | NA | NA | 0 | 0 | 0 | 0 | 0 | 0 |
| 32662 | ENSMUSG00000093336 | 0.492645588 | 0.9565034 | 3.447648281 | 0.277436479 | 0.781444974 | NA | Mir1954 | microRNA 1954 [Source:MGI Symbol;Acc: | 0 | 1.0279 | 0 | 1.928 | 0 | 0 |
| 32663 | ENSMUSG00000093338 | 0 | NA | NA | NA | NA | NA | Mir3090 | microRNA 3090 [Source:MGI Symbol;Acc: | 0 | 0 | 0 | 0 | 0 | 0 |
| 32664 | ENSMUSG00000093340 | 0 | NA | NA | NA | NA | NA | NA | NA | 0 | 0 | 0 | 0 | 0 | 0 |
| 32665 | ENSMUSG00000093341 | 0 | NA | NA | NA | NA | NA | NA | NA | 0 | 0 | 0 | 0 | 0 | 0 |
| 32666 | ENSMUSG00000093342 | 0.353089354 | 0.165142706 | 3.896102244 | 0.042386646 | 0.966190474 | NA | NA | NA | 0 | 1.0279 | 0 | 0 | 1.0906 | 0 |
| 32667 | ENSMUSG00000093344 | 0 | NA | NA | NA | NA | NA | NA | NA | 0 | 0 | 0 | 0 | 0 | 0 |
| 32668 | ENSMUSG00000093345 | 0 | NA | NA | NA | NA | NA | Gm27956 | predicted gene, 27956 [Source:MGI Symb | 0 | 0 | 0 | 0 | 0 | 0 |
| 32669 | ENSMUSG00000093346 | 0.610340026 | -2.657561537 | 3.094276798 | -0.858863544 | 0.390415805 | NA | NA | NA | 1.6062 | 2.0558 | 0 | 0 | 0 | 0 |
| 32670 | ENSMUSG00000093348 | 0 | NA | NA | NA | NA | NA | Mir3101 | microRNA 3101 [Source:MGI Symbol;Acc: | 0 | 0 | 0 | 0 | 0 | 0 |
| 32671 | ENSMUSG00000093351 | 0 | NA | NA | NA | NA | NA | Mir3072 | microRNA 3072 [Source:MGI Symbol;Acc: | 0 | 0 | 0 | 0 | 0 | 0 |
| 32672 | ENSMUSG00000093352 | 0 | NA | NA | NA | NA | NA | NA | NA | 0 | 0 | 0 | 0 | 0 | 0 |
| 32673 | ENSMUSG00000093354 | 0 | NA | NA | NA | NA | NA | Mir125b-1 | microRNA 125b-1 [Source:MGI Symbol;Ac | 0 | 0 | 0 | 0 | 0 | 0 |
| 32674 | ENSMUSG00000093355 | 0 | NA | NA | NA | NA | NA | Snora26 | small nucleolar RNA, H/ACA box 26 [Sourc | 0 | 0 | 0 | 0 | 0 | 0 |
| 32675 | ENSMUSG00000093356 | 0 | NA | NA | NA | NA | NA | NA | NA | 0 | 0 | 0 | 0 | 0 | 0 |
| 32676 | ENSMUSG00000093358 | 0 | NA | NA | NA | NA | NA | Mir3063 | microRNA 3063 [Source:MGI Symbol;Acc: | 0 | 0 | 0 | 0 | 0 | 0 |
| 32677 | ENSMUSG00000093359 | 0 | NA | NA | NA | NA | NA | NA | NA | 0 | 0 | 0 | 0 | 0 | 0 |
| 32678 | ENSMUSG00000093362 | 0 | NA | NA | NA | NA | NA | NA | NA | 0 | 0 | 0 | 0 | 0 | 0 |
| 32679 | ENSMUSG00000093363 | 0 | NA | NA | NA | NA | NA | NA | NA | 0 | 0 | 0 | 0 | 0 | 0 |
| 32680 | ENSMUSG00000093365 | 0 | NA | NA | NA | NA | NA | NA | NA | 0 | 0 | 0 | 0 | 0 | 0 |
| 32681 | ENSMUSG00000093368 | 0 | NA | NA | NA | NA | NA | NA | NA | 0 | 0 | 0 | 0 | 0 | 0 |
| 32682 | ENSMUSG00000093373 | 0 | NA | NA | NA | NA | NA | NA | NA | 0 | 0 | 0 | 0 | 0 | 0 |
| 32683 | ENSMUSG00000093376 | 0 | NA | NA | NA | NA | NA | Vmn1r23 | vomeronasal 1 receptor 23 [Source:MGI S | 0 | 0 | 0 | 0 | 0 | 0 |
| 32684 | ENSMUSG00000093378 | 0 | NA | NA | NA | NA | NA | Gm20663 | predicted gene 20663 [Source:MGI Symbo | 0 | 0 | 0 | 0 | 0 | 0 |
| 32685 | ENSMUSG00000093380 | 0 | NA | NA | NA | NA | NA | Gm20685 | predicted gene 20685 [Source:MGI Symbo | 0 | 0 | 0 | 0 | 0 | 0 |
| 32686 | ENSMUSG00000093381 | 0 | NA | NA | NA | NA | NA | Gm20615 | predicted gene 20615 [Source:MGI Symbo | 0 | 0 | 0 | 0 | 0 | 0 |
| 32687 | ENSMUSG00000093382 | 0 | NA | NA | NA | NA | NA | Vmn1r-ps14 | vomeronasal 1 receptor, pseudogene 14 [ | 0 | 0 | 0 | 0 | 0 | 0 |
| 32688 | ENSMUSG00000093383 | 0 | NA | NA | NA | NA | NA | Gm20642 | predicted gene 20642 [Source:MGI Symbo | 0 | 0 | 0 | 0 | 0 | 0 |
| 32689 | ENSMUSG00000093384 | 0 | NA | NA | NA | NA | NA | Gm20689 | predicted gene 20689 [Source:MGI Symbo | 0 | 0 | 0 | 0 | 0 | 0 |
| 32690 | ENSMUSG00000093385 | 0 | NA | NA | NA | NA | NA | A330044P14 | RIKEN cDNA A330044P14 gene [Source:M | 0 | 0 | 0 | 0 | 0 | 0 |
| 32691 | ENSMUSG00000093386 | 0 | NA | NA | NA | NA | NA | 1700120O09 | RIKEN cDNA 1700120O09 gene [Source:M | 0 | 0 | 0 | 0 | 0 | 0 |
| 32692 | ENSMUSG00000093387 | 0.133850373 | -0.796629176 | 4.080472857 | -0.195229623 | 0.84521319 | NA | Gm20732 | predicted gene 20732 [Source:MGI Symbo | 0.8031 | 0 | 0 | 0 | 0 | 0 |
| 32693 | ENSMUSG00000093388 | 3.060637689 | -0.085992063 | 1.384759369 | -0.062098921 | 0.950484056 | NA | NA | NA | 2.4093 | 5.1396 | 2.0125 | 7.7118 | 1.0906 | 0 |
| 32694 | ENSMUSG00000093390 | 0.167707431 | -0.796629176 | 4.080472857 | -0.195229623 | 0.84521319 | NA | Gm20652 | predicted gene 20652 [Source:MGI Symbo | 0 | 0 | 1.0062 | 0 | 0 | 0 |
| 32695 | ENSMUSG00000093392 | 0 | NA | NA | NA | NA | NA | Gm6061 | predicted gene 6061 [Source:MGI Symbol | 0 | 0 | 0 | 0 | 0 | 0 |
| 32696 | ENSMUSG00000093394 | 5.702241242 | -0.594456276 | 0.902962615 | -0.658339854 | 0.510319774 | 0.883032087 | Gm20621 | predicted gene 20621 [Source:MGI Symbo | 6.4248 | 4.1117 | 10.062 | 4.8199 | 5.4531 | 3.3415 |
| 32697 | ENSMUSG00000093395 | 0 | NA | NA | NA | NA | NA | Vmn2r-ps6 | vomeronasal 2, receptor, pseudogene 6 [S | 0 | 0 | 0 | 0 | 0 | 0 |
| 32698 | ENSMUSG00000093397 | 0 | NA | NA | NA | NA | NA | Obox4-ps29 | oocyte specific homeobox 4, pseudogene | 0 | 0 | 0 | 0 | 0 | 0 |
| 32699 | ENSMUSG00000093398 | 0 | NA | NA | NA | NA | NA | Vmn1r-ps11 | vomeronasal 1 receptor, pseudogene 11 [ | 0 | 0 | 0 | 0 | 0 | 0 |
| 32700 | ENSMUSG00000093400 | 0.167707431 | -0.796629176 | 4.080472857 | -0.195229623 | 0.84521319 | NA | Gm20647 | predicted gene 20647 [Source:MGI Symbo | 0 | 0 | 1.0062 | 0 | 0 | 0 |

| | | | | | | | | | | | | | | | |
|---|---|---|---|---|---|---|---|---|---|---|---|---|---|---|---|
| 32701 | ENSMUSG00000093401 | 0 | NA | NA | NA | NA | NA | Vmn2r-ps12 | vomeronasal 2, receptor, pseudogene 126 | 0 | 0 | 0 | 0 | 0 | 0 |
| 32702 | ENSMUSG00000093402 | 15.9115995 | -0.423663841 | 0.593620262 | -0.713695047 | 0.47541577 | 0.86938592 | Gm18588 | predicted gene, 18588 [Source:MGI Symb | 12.85 | 27.754 | 14.087 | 7.7118 | 16.359 | 16.708 |
| 32703 | ENSMUSG00000093403 | 0 | NA | NA | NA | NA | NA | Mir5616 | microRNA 5616 [Source:MGI Symbol;Acc: | 0 | 0 | 0 | 0 | 0 | 0 |
| 32704 | ENSMUSG00000093404 | 0.503095663 | 2.549494177 | 3.348715477 | 0.761334964 | 0.446457022 | NA | NA | NA | 0 | 0 | 0 | 1.928 | 1.0906 | 0 |
| 32705 | ENSMUSG00000093405 | 105.4398724 | -0.222813842 | 0.249630593 | -0.892574259 | 0.372085209 | 0.807785387 | Gm20684 | predicted gene 20684 [Source:MGI Symbo | 104.4 | 90.457 | 145.91 | 94.47 | 92.703 | 104.7 |
| 32706 | ENSMUSG00000093406 | 0 | NA | NA | NA | NA | NA | Rpl48-ps1 | ribosomal protein L48, pseudogene 1 [Sou | 0 | 0 | 0 | 0 | 0 | 0 |
| 32707 | ENSMUSG00000093408 | 0 | NA | NA | NA | NA | NA | Mir5617 | microRNA 5617 [Source:MGI Symbol;Acc: | 0 | 0 | 0 | 0 | 0 | 0 |
| 32708 | ENSMUSG00000093409 | 0.663758638 | 2.950973782 | 3.025092005 | 0.975498853 | 0.32931287 | NA | NA | NA | 0 | 0 | 0 | 2.8919 | 1.0906 | 0 |
| 32709 | ENSMUSG00000093410 | 0 | NA | NA | NA | NA | NA | Gm20638 | predicted gene 20638 [Source:MGI Symbo | 0 | 0 | 0 | 0 | 0 | 0 |
| 32710 | ENSMUSG00000093413 | 0 | NA | NA | NA | NA | NA | Snora15 | small nucleolar RNA, H/ACA box 15 [Sourc | 0 | 0 | 0 | 0 | 0 | 0 |
| 32711 | ENSMUSG00000093416 | 0 | NA | NA | NA | NA | NA | Gm18294 | predicted gene, 18294 [Source:MGI Symb | 0 | 0 | 0 | 0 | 0 | 0 |
| 32712 | ENSMUSG00000093419 | 0 | NA | NA | NA | NA | NA | Gm19898 | predicted gene, 19898 [Source:MGI Symb | 0 | 0 | 0 | 0 | 0 | 0 |
| 32713 | ENSMUSG00000093420 | 0 | NA | NA | NA | NA | NA | Obox4-ps33 | oocyte specific homeobox 4, pseudogene | 0 | 0 | 0 | 0 | 0 | 0 |
| 32714 | ENSMUSG00000093421 | 0 | NA | NA | NA | NA | NA | Obox3-ps6 | oocyte specific homeobox 3, pseudogene | 0 | 0 | 0 | 0 | 0 | 0 |
| 32715 | ENSMUSG00000093424 | 4.949764599 | -0.843484692 | 0.957628744 | -0.880805529 | 0.378423087 | NA | 6330562C20 | RIKEN cDNA 6330562C20 gene [Source:M | 4.8186 | 8.2233 | 6.0375 | 2.8919 | 3.2719 | 4.4554 |
| 32716 | ENSMUSG00000093426 | 0 | NA | NA | NA | NA | NA | Vmn2r-ps12 | vomeronasal 2, receptor, pseudogene 123 | 0 | 0 | 0 | 0 | 0 | 0 |
| 32717 | ENSMUSG00000093427 | 0 | NA | NA | NA | NA | NA | Gm18513 | predicted gene, 18513 [Source:MGI Symb | 0 | 0 | 0 | 0 | 0 | 0 |
| 32718 | ENSMUSG00000093428 | 0 | NA | NA | NA | NA | NA | NA | NA | 0 | 0 | 0 | 0 | 0 | 0 |
| 32719 | ENSMUSG00000093429 | 0 | NA | NA | NA | NA | NA | Vmn1r-ps14 | vomeronasal 1 receptor, pseudogene 145 | 0 | 0 | 0 | 0 | 0 | 0 |
| 32720 | ENSMUSG00000093433 | 0 | NA | NA | NA | NA | NA | Gm18415 | predicted gene, 18415 [Source:MGI Symb | 0 | 0 | 0 | 0 | 0 | 0 |
| 32721 | ENSMUSG00000093434 | 0 | NA | NA | NA | NA | NA | Vmn2r-ps11 | vomeronasal 2, receptor, pseudogene 11 | 0 | 0 | 0 | 0 | 0 | 0 |
| 32722 | ENSMUSG00000093436 | 0 | NA | NA | NA | NA | NA | Gm20646 | predicted gene 20646 [Source:MGI Symbo | 0 | 0 | 0 | 0 | 0 | 0 |
| 32723 | ENSMUSG00000093437 | 0 | NA | NA | NA | NA | NA | Gm20702 | predicted gene 20702 [Source:MGI Symbo | 0 | 0 | 0 | 0 | 0 | 0 |
| 32724 | ENSMUSG00000093441 | 0 | NA | NA | NA | NA | NA | Mir5710 | microRNA 5710 [Source:MGI Symbol;Acc: | 0 | 0 | 0 | 0 | 0 | 0 |
| 32725 | ENSMUSG00000093443 | 0 | NA | NA | NA | NA | NA | Gm22871 | predicted gene, 22871 [Source:MGI Symb | 0 | 0 | 0 | 0 | 0 | 0 |
| 32726 | ENSMUSG00000093444 | 0 | NA | NA | NA | NA | NA | Vmn1r-ps15 | vomeronasal 1 receptor, pseudogene 152 | 0 | 0 | 0 | 0 | 0 | 0 |
| 32727 | ENSMUSG00000093445 | 1.000692056 | 0.147398412 | 2.220108818 | 0.066392427 | 0.9470654 | NA | Lrch4 | leucine-rich repeats and calponin homolo | 0.8031 | 2.0558 | 0 | 0.964 | 2.1812 | 0 |
| 32728 | ENSMUSG00000093446 | 0.160662974 | 1.126924175 | 4.080472857 | 0.276174898 | 0.782413736 | NA | Gm20694 | predicted gene 20694 [Source:MGI Symbo | 0 | 0 | 0 | 0.964 | 0 | 0 |
| 32729 | ENSMUSG00000093447 | 0 | NA | NA | NA | NA | NA | Gm5139 | predicted gene 5139 [Source:MGI Symbol | 0 | 0 | 0 | 0 | 0 | 0 |
| 32730 | ENSMUSG00000093449 | 0 | NA | NA | NA | NA | NA | Vmn1r-ps14 | vomeronasal 1 receptor, pseudogene 149 | 0 | 0 | 0 | 0 | 0 | 0 |
| 32731 | ENSMUSG00000093452 | 0 | NA | NA | NA | NA | NA | Zfhx2os | zinc finger homeobox 2, opposite strand [ | 0 | 0 | 0 | 0 | 0 | 0 |
| 32732 | ENSMUSG00000093453 | 0.17131964 | -0.796629176 | 4.080472857 | -0.195229623 | 0.84521319 | NA | Vmn1r-ps14 | vomeronasal 1 receptor, pseudogene 146 | 0 | 1.0279 | 0 | 0 | 0 | 0 |
| 32733 | ENSMUSG00000093454 | 0 | NA | NA | NA | NA | NA | Vmn1r-ps14 | vomeronasal 1 receptor, pseudogene 143 | 0 | 0 | 0 | 0 | 0 | 0 |
| 32734 | ENSMUSG00000093456 | 0 | NA | NA | NA | NA | NA | Gm20662 | predicted gene 20662 [Source:MGI Symbo | 0 | 0 | 0 | 0 | 0 | 0 |
| 32735 | ENSMUSG00000093458 | 9.982702671 | 0.550405466 | 0.787034212 | 0.69934122 | 0.484338812 | 0.872521682 | Gm20688 | predicted gene 20688 [Source:MGI Symbo | 3.2124 | 7.1954 | 14.087 | 13.496 | 17.45 | 4.4554 |
| 32736 | ENSMUSG00000093460 | 0 | NA | NA | NA | NA | NA | Six3os1 | SIX homeobox 3, opposite strand 1 [Sourc | 0 | 0 | 0 | 0 | 0 | 0 |
| 32737 | ENSMUSG00000093461 | 0 | NA | NA | NA | NA | NA | Vmn1r-ps21 | vomeronasal 1 receptor, pseudogene 21 [ | 0 | 0 | 0 | 0 | 0 | 0 |
| 32738 | ENSMUSG00000093465 | 0.531246861 | 1.101376188 | 3.36099124 | 0.327693858 | 0.743143147 | NA | Gm20682 | predicted gene 20682 [Source:MGI Symbo | 0 | 0 | 1.0062 | 0 | 2.1812 | 0 |
| 32739 | ENSMUSG00000093467 | 0 | NA | NA | NA | NA | NA | Gm20659 | predicted gene 20659 [Source:MGI Symbo | 0 | 0 | 0 | 0 | 0 | 0 |
| 32740 | ENSMUSG00000093469 | 0 | NA | NA | NA | NA | NA | Vmn2r-ps12 | vomeronasal 2, receptor, pseudogene 127 | 0 | 0 | 0 | 0 | 0 | 0 |
| 32741 | ENSMUSG00000093472 | 0 | NA | NA | NA | NA | NA | 4930438E09 | RIKEN cDNA 4930438E09 gene [Source:M | 0 | 0 | 0 | 0 | 0 | 0 |
| 32742 | ENSMUSG00000093473 | 0 | NA | NA | NA | NA | NA | Gm20620 | predicted gene 20620 [Source:MGI Symbo | 0 | 0 | 0 | 0 | 0 | 0 |
| 32743 | ENSMUSG00000093474 | 0 | NA | NA | NA | NA | NA | Vmn2r-ps12 | vomeronasal 2, receptor, pseudogene 12 | 0 | 0 | 0 | 0 | 0 | 0 |
| 32744 | ENSMUSG00000093476 | 0.401551119 | -2.072982726 | 4.008251979 | -0.517178744 | 0.605031388 | NA | NA | NA | 2.4093 | 0 | 0 | 0 | 0 | 0 |
| 32745 | ENSMUSG00000093479 | 0 | NA | NA | NA | NA | NA | Gm20629 | predicted gene 20629 [Source:MGI Symbo | 0 | 0 | 0 | 0 | 0 | 0 |
| 32746 | ENSMUSG00000093480 | 0 | NA | NA | NA | NA | NA | Mir5624 | microRNA 5624 [Source:MGI Symbol;Acc: | 0 | 0 | 0 | 0 | 0 | 0 |
| 32747 | ENSMUSG00000093482 | 0 | NA | NA | NA | NA | NA | Gm20619 | predicted gene 20619 [Source:MGI Symbo | 0 | 0 | 0 | 0 | 0 | 0 |
| 32748 | ENSMUSG00000093483 | 9.108809363 | 0.765689042 | 0.750929646 | 1.019654832 | 0.30789219 | 0.756343859 | AA465934 | expressed sequence AA465934 [Source:M | 8.031 | 4.1117 | 8.05 | 12.532 | 16.359 | 5.5692 |
| 32749 | ENSMUSG00000093484 | 0 | NA | NA | NA | NA | NA | Gm20657 | predicted gene 20657 [Source:MGI Symbo | 0 | 0 | 0 | 0 | 0 | 0 |
| 32750 | ENSMUSG00000093485 | 0 | NA | NA | NA | NA | NA | Gm20708 | predicted gene 20708 [Source:MGI Symbo | 0 | 0 | 0 | 0 | 0 | 0 |
| 32751 | ENSMUSG00000093487 | 0 | NA | NA | NA | NA | NA | Vmn1r-ps48 | vomeronasal 1 receptor, pseudogene 48 [ | 0 | 0 | 0 | 0 | 0 | 0 |
| 32752 | ENSMUSG00000093489 | 0 | NA | NA | NA | NA | NA | Gm20625 | predicted gene 20625 [Source:MGI Symbo | 0 | 0 | 0 | 0 | 0 | 0 |
| 32753 | ENSMUSG00000093490 | 0 | NA | NA | NA | NA | NA | Gm19932 | predicted gene, 19932 [Source:MGI Symb | 0 | 0 | 0 | 0 | 0 | 0 |
| 32754 | ENSMUSG00000093492 | 0.342432689 | 1.990653086 | 3.900281263 | 0.51038706 | 0.609780321 | NA | Gm20651 | predicted gene 20651 [Source:MGI Symbo | 0 | 0 | 0 | 0.964 | 1.0906 | 0 |
| 32755 | ENSMUSG00000093493 | 0 | NA | NA | NA | NA | NA | Vmn2r-ps13 | vomeronasal 2, receptor, pseudogene 133 | 0 | 0 | 0 | 0 | 0 | 0 |
| 32756 | ENSMUSG00000093495 | 0.817403785 | 0.568896211 | 2.872137803 | 0.198074135 | 0.842987061 | NA | NA | NA | 0 | 0 | 2.0125 | 2.8919 | 0 | 0 |

| | | | | | | | | | | | | | | | |
|---|---|---|---|---|---|---|---|---|---|---|---|---|---|---|---|
| 32757 | ENSMUSG00000093497 | 0 | NA | NA | NA | NA | NA | Gm20713 | predicted gene 20713 [Source:MGI Symbo | 0 | 0 | 0 | 0 | 0 | 0 |
| 32758 | ENSMUSG00000093498 | 0 | NA | NA | NA | NA | NA | Mir5615-2 | microRNA 5615-2 [Source:MGI Symbol;Ac | 0 | 0 | 0 | 0 | 0 | 0 |
| 32759 | ENSMUSG00000093499 | 0 | NA | NA | NA | NA | NA | NA | NA | 0 | 0 | 0 | 0 | 0 | 0 |
| 32760 | ENSMUSG00000093502 | 0 | NA | NA | NA | NA | NA | Gm20700 | predicted gene 20700 [Source:MGI Symbo | 0 | 0 | 0 | 0 | 0 | 0 |
| 32761 | ENSMUSG00000093503 | 0 | NA | NA | NA | NA | NA | Vmn2r-ps4 | vomeronasal 2, receptor, pseudogene 4 [S | 0 | 0 | 0 | 0 | 0 | 0 |
| 32762 | ENSMUSG00000093504 | 0 | NA | NA | NA | NA | NA | Vmn1r-ps24 | vomeronasal 1 receptor, pseudogene 24 [ | 0 | 0 | 0 | 0 | 0 | 0 |
| 32763 | ENSMUSG00000093505 | 0.267700746 | -1.497481649 | 4.057001436 | -0.369110456 | 0.712045396 | NA | Gm20691 | predicted gene 20691 [Source:MGI Symbo | 1.6062 | 0 | 0 | 0 | 0 | 0 |
| 32764 | ENSMUSG00000093507 | 1.257736137 | 0.234612575 | 2.011469531 | 0.116637399 | 0.9071474 | NA | Gm20627 | predicted gene 20627 [Source:MGI Symbo | 2.4093 | 1.0279 | 0 | 1.928 | 2.1812 | 0 |
| 32765 | ENSMUSG00000093508 | 0 | NA | NA | NA | NA | NA | Mir5709 | microRNA 5709 [Source:MGI Symbol;Acc: | 0 | 0 | 0 | 0 | 0 | 0 |
| 32766 | ENSMUSG00000093509 | 0 | NA | NA | NA | NA | NA | Obox3-ps3 | oocyte specific homeobox 3, pseudogene | 0 | 0 | 0 | 0 | 0 | 0 |
| 32767 | ENSMUSG00000093510 | 0 | NA | NA | NA | NA | NA | Vmn2r-ps21 | vomeronasal 2, receptor, pseudogene 21 | 0 | 0 | 0 | 0 | 0 | 0 |
| 32768 | ENSMUSG00000093513 | 0 | NA | NA | NA | NA | NA | Obox4-ps31 | oocyte specific homeobox 4, pseudogene | 0 | 0 | 0 | 0 | 0 | 0 |
| 32769 | ENSMUSG00000093515 | 0 | NA | NA | NA | NA | NA | Mir1231 | microRNA 1231 [Source:MGI Symbol;Acc: | 0 | 0 | 0 | 0 | 0 | 0 |
| 32770 | ENSMUSG00000093516 | 0 | NA | NA | NA | NA | NA | Gm19553 | predicted gene, 19553 [Source:MGI Symb | 0 | 0 | 0 | 0 | 0 | 0 |
| 32771 | ENSMUSG00000093518 | 0 | NA | NA | NA | NA | NA | Vmn1r-ps22 | vomeronasal 1 receptor, pseudogene 22 [ | 0 | 0 | 0 | 0 | 0 | 0 |
| 32772 | ENSMUSG00000093519 | 0 | NA | NA | NA | NA | NA | NA | NA | 0 | 0 | 0 | 0 | 0 | 0 |
| 32773 | ENSMUSG00000093520 | 0 | NA | NA | NA | NA | NA | Vmn1r-ps9 | vomeronasal 1 receptor, pseudogene 9 [S | 0 | 0 | 0 | 0 | 0 | 0 |
| 32774 | ENSMUSG00000093522 | 0 | NA | NA | NA | NA | NA | Vmn1r-ps19 | vomeronasal 1 receptor, pseudogene 19 [ | 0 | 0 | 0 | 0 | 0 | 0 |
| 32775 | ENSMUSG00000093525 | 0 | NA | NA | NA | NA | NA | Gm8974 | predicted gene 8974 [Source:MGI Symbol | 0 | 0 | 0 | 0 | 0 | 0 |
| 32776 | ENSMUSG00000093528 | 0 | NA | NA | NA | NA | NA | Nrg3os | neuregulin 3, opposite strand [Source:MG | 0 | 0 | 0 | 0 | 0 | 0 |
| 32777 | ENSMUSG00000093529 | 0 | NA | NA | NA | NA | NA | NA | NA | 0 | 0 | 0 | 0 | 0 | 0 |
| 32778 | ENSMUSG00000093530 | 0 | NA | NA | NA | NA | NA | Vmn1r-ps12 | vomeronasal 1 receptor, pseudogene 12 [ | 0 | 0 | 0 | 0 | 0 | 0 |
| 32779 | ENSMUSG00000093531 | 0 | NA | NA | NA | NA | NA | Vmn2r-ps11 | vomeronasal 2, receptor, pseudogene 11 | 0 | 0 | 0 | 0 | 0 | 0 |
| 32780 | ENSMUSG00000093533 | 0 | NA | NA | NA | NA | NA | Vmn2r127 | vomeronasal 2 receptor 127 [Source:MGI | 0 | 0 | 0 | 0 | 0 | 0 |
| 32781 | ENSMUSG00000093534 | 0 | NA | NA | NA | NA | NA | Vmn1r-ps51 | vomeronasal 1 receptor, pseudogene 51 [ | 0 | 0 | 0 | 0 | 0 | 0 |
| 32782 | ENSMUSG00000093535 | 0 | NA | NA | NA | NA | NA | 4930533I22R | RIKEN cDNA 4930533I22 gene [Source:MG | 0 | 0 | 0 | 0 | 0 | 0 |
| 32783 | ENSMUSG00000093536 | 0.455176322 | 0.955156269 | 3.546674842 | 0.269310357 | 0.787690862 | NA | Smim17 | small integral membrane protein 17 [Sour | 0.8031 | 0 | 0 | 1.928 | 0 | 0 |
| 32784 | ENSMUSG00000093537 | 0 | NA | NA | NA | NA | NA | Gm7584 | predicted gene 7584 [Source:MGI Symbol | 0 | 0 | 0 | 0 | 0 | 0 |
| 32785 | ENSMUSG00000093538 | 0 | NA | NA | NA | NA | NA | Gm20711 | predicted gene 20711 [Source:MGI Symbo | 0 | 0 | 0 | 0 | 0 | 0 |
| 32786 | ENSMUSG00000093539 | 0 | NA | NA | NA | NA | NA | Mir5621 | microRNA 5621 [Source:MGI Symbol;Acc: | 0 | 0 | 0 | 0 | 0 | 0 |
| 32787 | ENSMUSG00000093540 | 0 | NA | NA | NA | NA | NA | 4930505M18 | RIKEN cDNA 4930505M18 gene [Source:M | 0 | 0 | 0 | 0 | 0 | 0 |
| 32788 | ENSMUSG00000093541 | 0 | NA | NA | NA | NA | NA | Gm20660 | predicted gene 20660 [Source:MGI Symbo | 0 | 0 | 0 | 0 | 0 | 0 |
| 32789 | ENSMUSG00000093543 | 0 | NA | NA | NA | NA | NA | Gm8473 | predicted gene 8473 [Source:MGI Symbol | 0 | 0 | 0 | 0 | 0 | 0 |
| 32790 | ENSMUSG00000093544 | 0 | NA | NA | NA | NA | NA | Gm20613 | predicted gene 20613 [Source:MGI Symbo | 0 | 0 | 0 | 0 | 0 | 0 |
| 32791 | ENSMUSG00000093545 | 0 | NA | NA | NA | NA | NA | Gm5871 | predicted gene 5871 [Source:MGI Symbol | 0 | 0 | 0 | 0 | 0 | 0 |
| 32792 | ENSMUSG00000093546 | 0 | NA | NA | NA | NA | NA | Vmn2r-ps7 | vomeronasal 2, receptor, pseudogene 7 [S | 0 | 0 | 0 | 0 | 0 | 0 |
| 32793 | ENSMUSG00000093547 | 0 | NA | NA | NA | NA | NA | Gm20674 | predicted gene 20674 [Source:MGI Symbo | 0 | 0 | 0 | 0 | 0 | 0 |
| 32794 | ENSMUSG00000093548 | 0 | NA | NA | NA | NA | NA | Gm6407 | predicted gene 6407 [Source:MGI Symbol | 0 | 0 | 0 | 0 | 0 | 0 |
| 32795 | ENSMUSG00000093549 | 0 | NA | NA | NA | NA | NA | Obox4-ps36 | oocyte specific homeobox 4, pseudogene | 0 | 0 | 0 | 0 | 0 | 0 |
| 32796 | ENSMUSG00000093550 | 0 | NA | NA | NA | NA | NA | Higd1c | HIG1 domain family, member 1C [Source:I | 0 | 0 | 0 | 0 | 0 | 0 |
| 32797 | ENSMUSG00000093551 | 0 | NA | NA | NA | NA | NA | 4930505G20 | RIKEN cDNA 4930505G20 gene [Source:M | 0 | 0 | 0 | 0 | 0 | 0 |
| 32798 | ENSMUSG00000093552 | 0 | NA | NA | NA | NA | NA | Gm5777 | predicted gene 5777 [Source:MGI Symbol | 0 | 0 | 0 | 0 | 0 | 0 |
| 32799 | ENSMUSG00000093553 | 7.851252886 | 0.227982551 | 0.758456273 | 0.3005876 | 0.763728988 | 0.954980327 | Gm20633 | predicted gene 20633 [Source:MGI Symbo | 6.4248 | 9.2513 | 6.0375 | 7.7118 | 6.5437 | 11.138 |
| 32800 | ENSMUSG00000093554 | 0 | NA | NA | NA | NA | NA | Obox4-ps26 | oocyte specific homeobox 4, pseudogene | 0 | 0 | 0 | 0 | 0 | 0 |
| 32801 | ENSMUSG00000093556 | 0 | NA | NA | NA | NA | NA | Gm20641 | predicted gene 20641 [Source:MGI Symbo | 0 | 0 | 0 | 0 | 0 | 0 |
| 32802 | ENSMUSG00000093557 | 0 | NA | NA | NA | NA | NA | 4930555M17 | RIKEN cDNA 4930555M17 gene [Source:M | 0 | 0 | 0 | 0 | 0 | 0 |
| 32803 | ENSMUSG00000093559 | 0 | NA | NA | NA | NA | NA | Gm20705 | predicted gene 20705 [Source:MGI Symbo | 0 | 0 | 0 | 0 | 0 | 0 |
| 32804 | ENSMUSG00000093560 | 0 | NA | NA | NA | NA | NA | Gm6695 | predicted gene 6695 [Source:MGI Symbol | 0 | 0 | 0 | 0 | 0 | 0 |
| 32805 | ENSMUSG00000093562 | 0 | NA | NA | NA | NA | NA | Gm23090 | predicted gene, 23090 [Source:MGI Symb | 0 | 0 | 0 | 0 | 0 | 0 |
| 32806 | ENSMUSG00000093563 | 0 | NA | NA | NA | NA | NA | Gm20630 | predicted gene 20630 [Source:MGI Symbo | 0 | 0 | 0 | 0 | 0 | 0 |
| 32807 | ENSMUSG00000093564 | 0 | NA | NA | NA | NA | NA | Gm18396 | predicted gene, 18396 [Source:MGI Symb | 0 | 0 | 0 | 0 | 0 | 0 |
| 32808 | ENSMUSG00000093565 | 10.04951106 | 0.263674394 | 0.740399166 | 0.35612465 | 0.721747218 | 0.9447719 | Rab26os | RAB26, member RAS oncogene family, op | 3.2124 | 10.279 | 14.087 | 10.604 | 7.6343 | 14.48 |
| 32809 | ENSMUSG00000093567 | 0 | NA | NA | NA | NA | NA | Obox4-ps8 | oocyte specific homeobox 4, pseudogene | 0 | 0 | 0 | 0 | 0 | 0 |
| 32810 | ENSMUSG00000093568 | 0 | NA | NA | NA | NA | NA | Gm20611 | predicted gene 20611 [Source:MGI Symbo | 0 | 0 | 0 | 0 | 0 | 0 |
| 32811 | ENSMUSG00000093570 | 0 | NA | NA | NA | NA | NA | Gm20714 | predicted gene 20714 [Source:MGI Symbo | 0 | 0 | 0 | 0 | 0 | 0 |
| 32812 | ENSMUSG00000093571 | 0 | NA | NA | NA | NA | NA | NA | NA | 0 | 0 | 0 | 0 | 0 | 0 |

| | | | | | | | | | | | | | | | |
|---|---|---|---|---|---|---|---|---|---|---|---|---|---|---|---|
| 32813 | ENSMUSG00000093572 | 0 | NA | NA | NA | NA | NA | NA | NA | 0 | 0 | 0 | 0 | 0 | 0 |
| 32814 | ENSMUSG00000093573 | 0 | NA | NA | NA | NA | NA | Gm20610 | predicted gene 20610 [Source:MGI Symbo | 0 | 0 | 0 | 0 | 0 | 0 |
| 32815 | ENSMUSG00000093574 | 0 | NA | NA | NA | NA | NA | Gm20671 | predicted gene 20671 [Source:MGI Symbo | 0 | 0 | 0 | 0 | 0 | 0 |
| 32816 | ENSMUSG00000093575 | 1.023919079 | -2.141404304 | 2.462125713 | -0.869738004 | 0.384443598 | NA | Gm20695 | predicted gene 20695 [Source:MGI Symbo | 0 | 1.0279 | 4.025 | 0 | 1.0906 | 0 |
| 32817 | ENSMUSG00000093577 | 21.15169648 | 0.239898576 | 0.465978346 | 0.514827734 | 0.606673403 | 0.916376001 | Gm20632 | predicted gene 20632 [Source:MGI Symbo | 18.471 | 22.614 | 17.106 | 20.244 | 25.084 | 23.391 |
| 32818 | ENSMUSG00000093579 | 8.08540833 | -0.769973632 | 0.778694217 | -0.988801015 | 0.322760507 | 0.769926329 | Gm20036 | predicted gene, 20036 [Source:MGI Symb | 11.243 | 8.2233 | 11.069 | 4.8199 | 9.8156 | 3.3415 |
| 32819 | ENSMUSG00000093580 | 1.411970867 | 0.867051441 | 2.399641043 | 0.361325476 | 0.717856152 | NA | Gm20706 | predicted gene 20706 [Source:MGI Symbo | 0 | 0 | 3.0187 | 0 | 5.4531 | 0 |
| 32820 | ENSMUSG00000093581 | 0 | NA | NA | NA | NA | NA | Gm18006 | predicted gene, 18006 [Source:MGI Symb | 0 | 0 | 0 | 0 | 0 | 0 |
| 32821 | ENSMUSG00000093582 | 0 | NA | NA | NA | NA | NA | NA | NA | 0 | 0 | 0 | 0 | 0 | 0 |
| 32822 | ENSMUSG00000093584 | 0 | NA | NA | NA | NA | NA | Obox3-ps2 | oocyte specific homeobox 3, pseudogene | 0 | 0 | 0 | 0 | 0 | 0 |
| 32823 | ENSMUSG00000093587 | 0 | NA | NA | NA | NA | NA | Gm20554 | predicted gene, 20554 [Source:MGI Symb | 0 | 0 | 0 | 0 | 0 | 0 |
| 32824 | ENSMUSG00000093588 | 0 | NA | NA | NA | NA | NA | Vmn1r-ps23 | vomeronasal 1 receptor, pseudogene 23 [ | 0 | 0 | 0 | 0 | 0 | 0 |
| 32825 | ENSMUSG00000093589 | 0 | NA | NA | NA | NA | NA | Gm18367 | predicted gene, 18367 [Source:MGI Symb | 0 | 0 | 0 | 0 | 0 | 0 |
| 32826 | ENSMUSG00000093590 | 0 | NA | NA | NA | NA | NA | Gm20643 | predicted gene 20643 [Source:MGI Symbo | 0 | 0 | 0 | 0 | 0 | 0 |
| 32827 | ENSMUSG00000093592 | 0 | NA | NA | NA | NA | NA | Vmn2r-ps12 | vomeronasal 2, receptor, pseudogene 122 | 0 | 0 | 0 | 0 | 0 | 0 |
| 32828 | ENSMUSG00000093593 | 0 | NA | NA | NA | NA | NA | Gm20683 | predicted gene 20683 [Source:MGI Symbo | 0 | 0 | 0 | 0 | 0 | 0 |
| 32829 | ENSMUSG00000093594 | 2.71357954 | 1.25758433 | 1.304318322 | 0.964169796 | 0.334960805 | NA | Gm20707 | predicted gene 20707 [Source:MGI Symbo | 0.8031 | 2.0558 | 2.0125 | 4.8199 | 4.3625 | 2.2277 |
| 32830 | ENSMUSG00000093595 | 0 | NA | NA | NA | NA | NA | Vmn1r-ps49 | vomeronasal 1 receptor, pseudogene 49 [ | 0 | 0 | 0 | 0 | 0 | 0 |
| 32831 | ENSMUSG00000093598 | 0.181769715 | 1.126924175 | 4.080472857 | 0.276174898 | 0.782413736 | NA | A730085K08 | RIKEN cDNA A730085K08 gene [Source:M | 0 | 0 | 0 | 0 | 1.0906 | 0 |
| 32832 | ENSMUSG00000093599 | 0.852572809 | 2.030361656 | 2.559049725 | 0.793404534 | 0.42754217 | NA | Mir5620 | microRNA 5620 [Source:MGI Symbol;Acc: | 0 | 0 | 1.0062 | 1.928 | 2.1812 | 0 |
| 32833 | ENSMUSG00000093601 | 0 | NA | NA | NA | NA | NA | Gm19459 | predicted gene, 19459 [Source:MGI Symb | 0 | 0 | 0 | 0 | 0 | 0 |
| 32834 | ENSMUSG00000093602 | 0 | NA | NA | NA | NA | NA | Vmn2r-ps11 | vomeronasal 2, receptor, pseudogene 116 | 0 | 0 | 0 | 0 | 0 | 0 |
| 32835 | ENSMUSG00000093606 | 16.13964028 | 0.055388776 | 0.542697759 | 0.102061922 | 0.918707525 | 0.986374671 | B130034C11 | RIKEN cDNA B130034C11 gene [Source:M | 20.881 | 12.335 | 14.087 | 15.424 | 19.631 | 14.48 |
| 32836 | ENSMUSG00000093607 | 0 | NA | NA | NA | NA | NA | Vmn2r-ps11 | vomeronasal 2, receptor, pseudogene 118 | 0 | 0 | 0 | 0 | 0 | 0 |
| 32837 | ENSMUSG00000093610 | 3.676281022 | 0.856546582 | 1.222718674 | 0.700526294 | 0.48359869 | NA | Cyp2c53-ps | cytochrome P450, family 2, subfamily c, p | 0.8031 | 2.0558 | 5.0312 | 0.964 | 7.6343 | 5.5692 |
| 32838 | ENSMUSG00000093614 | 1.220137628 | -1.233564006 | 2.542691384 | -0.485141065 | 0.627576284 | NA | Gm20693 | predicted gene 20693 [Source:MGI Symbo | 0 | 5.1396 | 0 | 0 | 2.1812 | 0 |
| 32839 | ENSMUSG00000093615 | 0 | NA | NA | NA | NA | NA | Mir5618 | microRNA 5618 [Source:MGI Symbol;Acc: | 0 | 0 | 0 | 0 | 0 | 0 |
| 32840 | ENSMUSG00000093616 | 0 | NA | NA | NA | NA | NA | 1700055C04 | RIKEN cDNA 1700055C04 gene [Source:M | 0 | 0 | 0 | 0 | 0 | 0 |
| 32841 | ENSMUSG00000093617 | 0.133850373 | -0.796629176 | 4.080472857 | -0.195229623 | 0.84521319 | NA | 4933438K21 | RIKEN cDNA 4933438K21 gene [Source:M | 0.8031 | 0 | 0 | 0 | 0 | 0 |
| 32842 | ENSMUSG00000093618 | 0 | NA | NA | NA | NA | NA | Mir5627 | microRNA 5627 [Source:MGI Symbol;Acc: | 0 | 0 | 0 | 0 | 0 | 0 |
| 32843 | ENSMUSG00000093619 | 0.167707431 | -0.796629176 | 4.080472857 | -0.195229623 | 0.84521319 | NA | 4930535L15I | RIKEN cDNA 4930535L15 gene [Source:M | 0 | 0 | 1.0062 | 0 | 0 | 0 |
| 32844 | ENSMUSG00000093621 | 0 | NA | NA | NA | NA | NA | Vmn1r-ps10 | vomeronasal 1 receptor, pseudogene 10 [ | 0 | 0 | 0 | 0 | 0 | 0 |
| 32845 | ENSMUSG00000093622 | 6.516289 | 0.328936036 | 0.868058935 | 0.378932839 | 0.704737738 | 0.94295036 | Gm20703 | predicted gene 20703 [Source:MGI Symbo | 3.2124 | 9.2513 | 5.0312 | 10.604 | 6.5437 | 4.4554 |
| 32846 | ENSMUSG00000093623 | 0 | NA | NA | NA | NA | NA | Obox4-ps14 | oocyte specific homeobox 4, pseudogene | 0 | 0 | 0 | 0 | 0 | 0 |
| 32847 | ENSMUSG00000093624 | 0.167707431 | -0.796629176 | 4.080472857 | -0.195229623 | 0.84521319 | NA | Mipep-ps | mitochondrial intermediate peptidase, ps | 0 | 0 | 1.0062 | 0 | 0 | 0 |
| 32848 | ENSMUSG00000093625 | 0 | NA | NA | NA | NA | NA | Gm18953 | predicted gene, 18953 [Source:MGI Symb | 0 | 0 | 0 | 0 | 0 | 0 |
| 32849 | ENSMUSG00000093626 | 0.167707431 | -0.796629176 | 4.080472857 | -0.195229623 | 0.84521319 | NA | Ly6l | lymphocyte antigen 6 complex, locus L [Sc | 0 | 0 | 1.0062 | 0 | 0 | 0 |
| 32850 | ENSMUSG00000093627 | 0 | NA | NA | NA | NA | NA | Vmn2r-ps12 | vomeronasal 2, receptor, pseudogene 121 | 0 | 0 | 0 | 0 | 0 | 0 |
| 32851 | ENSMUSG00000093628 | 0.874583548 | 0.667972572 | 2.552079858 | 0.261736548 | 0.793524561 | NA | Gm20616 | predicted gene 20616 [Source:MGI Symbo | 0 | 2.0558 | 0 | 0.964 | 0 | 2.2277 |
| 32852 | ENSMUSG00000093629 | 2.282967182 | -0.197293231 | 1.508565883 | -0.130781979 | 0.89594778 | NA | Prox2os | prospero homeobox 2 opposite strand [Sc | 3.2124 | 2.0558 | 2.0125 | 0.964 | 5.4531 | 0 |
| 32853 | ENSMUSG00000093634 | 0 | NA | NA | NA | NA | NA | Gm10860 | predicted gene 10860 [Source:MGI Symbo | 0 | 0 | 0 | 0 | 0 | 0 |
| 32854 | ENSMUSG00000093637 | 7.921468092 | -0.316345407 | 0.75606997 | -0.418407581 | 0.675649145 | 0.932630311 | Gm20636 | predicted gene 20636 [Source:MGI Symbo | 8.031 | 10.279 | 8.05 | 5.7839 | 9.8156 | 5.5692 |
| 32855 | ENSMUSG00000093639 | 0 | NA | NA | NA | NA | NA | Gm52993 | predicted gene, 52993 [Source:MGI Symb | 0 | 0 | 0 | 0 | 0 | 0 |
| 32856 | ENSMUSG00000093642 | 0 | NA | NA | NA | NA | NA | Vmn2r-ps12 | vomeronasal 2, receptor, pseudogene 124 | 0 | 0 | 0 | 0 | 0 | 0 |
| 32857 | ENSMUSG00000093644 | 0 | NA | NA | NA | NA | NA | Gm19560 | predicted gene, 19560 [Source:MGI Symb | 0 | 0 | 0 | 0 | 0 | 0 |
| 32858 | ENSMUSG00000093645 | 0 | NA | NA | NA | NA | NA | Gm23187 | predicted gene, 23187 [Source:MGI Symb | 0 | 0 | 0 | 0 | 0 | 0 |
| 32859 | ENSMUSG00000093648 | 0 | NA | NA | NA | NA | NA | Vmn1r-ps14 | vomeronasal 1 receptor, pseudogene 148 | 0 | 0 | 0 | 0 | 0 | 0 |
| 32860 | ENSMUSG00000093649 | 0 | NA | NA | NA | NA | NA | Gm20614 | predicted gene 20614 [Source:MGI Symbo | 0 | 0 | 0 | 0 | 0 | 0 |
| 32861 | ENSMUSG00000093650 | 0 | NA | NA | NA | NA | NA | Gm20631 | predicted gene 20631 [Source:MGI Symbo | 0 | 0 | 0 | 0 | 0 | 0 |
| 32862 | ENSMUSG00000093651 | 0.133850373 | -0.796629176 | 4.080472857 | -0.195229623 | 0.84521319 | NA | Gm5873 | predicted gene 5873 [Source:MGI Symbol | 0.8031 | 0 | 0 | 0 | 0 | 0 |
| 32863 | ENSMUSG00000093652 | 0 | NA | NA | NA | NA | NA | Vmn2r-ps31 | vomeronasal 2, receptor, pseudogene 31 | 0 | 0 | 0 | 0 | 0 | 0 |
| 32864 | ENSMUSG00000093656 | 0.636946036 | 1.591416711 | 2.83781177 | 0.560790087 | 0.574940645 | NA | Gm20628 | predicted gene 20628 [Source:MGI Symbo | 0.8031 | 0 | 0 | 1.928 | 1.0906 | 0 |
| 32865 | ENSMUSG00000093658 | 0 | NA | NA | NA | NA | NA | Gm20617 | predicted gene 20617 [Source:MGI Symbo | 0 | 0 | 0 | 0 | 0 | 0 |
| 32866 | ENSMUSG00000093660 | 0 | NA | NA | NA | NA | NA | Gm26505 | predicted gene, 26505 [Source:MGI Symb | 0 | 0 | 0 | 0 | 0 | 0 |
| 32867 | ENSMUSG00000093661 | 178.662584 | -0.198004604 | 0.273120325 | -0.724972059 | 0.468469172 | 0.865461183 | Eif4e3 | eukaryotic translation initiation factor 4E | 160.62 | 200.44 | 211.31 | 90.614 | 199.58 | 209.4 |
| 32868 | ENSMUSG00000093662 | 0 | NA | NA | NA | NA | NA | Gm20649 | predicted gene 20649 [Source:MGI Symbo | 0 | 0 | 0 | 0 | 0 | 0 |

| | | | | | | | | | | | | | | | |
|---|---|---|---|---|---|---|---|---|---|---|---|---|---|---|---|
| 32869 | ENSMUSG00000093663 | 0 | NA | NA | NA | NA | NA | Gm7171 | predicted gene 7171 [Source:MGI Symbol | 0 | 0 | 0 | 0 | 0 | 0 |
| 32870 | ENSMUSG00000093664 | 0 | NA | NA | NA | NA | NA | Obox4-ps37 | oocyte specific homeobox 4, pseudogene | 0 | 0 | 0 | 0 | 0 | 0 |
| 32871 | ENSMUSG00000093665 | 0 | NA | NA | NA | NA | NA | Obox3-ps1 | oocyte specific homeobox 3, pseudogene | 0 | 0 | 0 | 0 | 0 | 0 |
| 32872 | ENSMUSG00000093666 | 0 | NA | NA | NA | NA | NA | Vmn1r-ps15 | vomeronasal 1 receptor, pseudogene 15 [ | 0 | 0 | 0 | 0 | 0 | 0 |
| 32873 | ENSMUSG00000093668 | 0.346303622 | 2.004339653 | 3.884003423 | 0.516049919 | 0.605819544 | NA | Pou5f2 | POU domain class 5, transcription factor 2 | 0 | 0 | 0 | 0.964 | 0 | 1.1138 |
| 32874 | ENSMUSG00000093669 | 0 | NA | NA | NA | NA | NA | Gm9399 | predicted gene 9399 [Source:MGI Symbol | 0 | 0 | 0 | 0 | 0 | 0 |
| 32875 | ENSMUSG00000093671 | 0 | NA | NA | NA | NA | NA | Gm20656 | predicted gene 20656 [Source:MGI Symbo | 0 | 0 | 0 | 0 | 0 | 0 |
| 32876 | ENSMUSG00000093672 | 0.294513347 | 0.165143496 | 4.080472857 | 0.040471657 | 0.967717103 | NA | Gm20655 | predicted gene 20655 [Source:MGI Symbo | 0.8031 | 0 | 0 | 0.964 | 0 | 0 |
| 32877 | ENSMUSG00000093673 | 0 | NA | NA | NA | NA | NA | Gm20644 | predicted gene 20644 [Source:MGI Symbo | 0 | 0 | 0 | 0 | 0 | 0 |
| 32878 | ENSMUSG00000093674 | 3289.606623 | 0.151171878 | 0.112523889 | 1.343464753 | 0.179121532 | 0.611235187 | Rpl41 | ribosomal protein L41 [Source:MGI Symbo | 3038.9 | 3232.8 | 3080.1 | 3174.4 | 3225 | 3986.4 |
| 32879 | ENSMUSG00000093675 | 0 | NA | NA | NA | NA | NA | Gm20618 | predicted gene 20618 [Source:MGI Symbo | 0 | 0 | 0 | 0 | 0 | 0 |
| 32880 | ENSMUSG00000093677 | 3.687744349 | -0.114221189 | 1.094521399 | -0.104357201 | 0.916885886 | NA | Gm20712 | predicted gene 20712 [Source:MGI Symbo | 2.4093 | 3.0838 | 6.0375 | 2.8919 | 4.3625 | 3.3415 |
| 32881 | ENSMUSG00000093681 | 0 | NA | NA | NA | NA | NA | Vmn2r-ps52 | vomeronasal 2, receptor, pseudogene 52 | 0 | 0 | 0 | 0 | 0 | 0 |
| 32882 | ENSMUSG00000093682 | 5.503772042 | 0.786414093 | 0.908556969 | 0.865563877 | 0.386729372 | 0.817307663 | Gm20681 | predicted gene 20681 [Source:MGI Symbo | 4.0155 | 3.0838 | 5.0312 | 7.7118 | 8.7249 | 4.4554 |
| 32883 | ENSMUSG00000093683 | 0 | NA | NA | NA | NA | NA | Obox4-ps34 | oocyte specific homeobox 4, pseudogene | 0 | 0 | 0 | 0 | 0 | 0 |
| 32884 | ENSMUSG00000093684 | 0.133850373 | -0.796629176 | 4.080472857 | -0.195229623 | 0.84521319 | NA | NA | NA | 0.8031 | 0 | 0 | 0 | 0 | 0 |
| 32885 | ENSMUSG00000093685 | 0 | NA | NA | NA | NA | NA | Gm18416 | predicted gene, 18416 [Source:MGI Symb | 0 | 0 | 0 | 0 | 0 | 0 |
| 32886 | ENSMUSG00000093686 | 1.581073115 | 1.933515655 | 1.930067172 | 1.001786716 | 0.316446614 | NA | Gm4705 | predicted gene 4705 [Source:MGI Symbol | 0 | 0 | 2.0125 | 1.928 | 1.0906 | 4.4554 |
| 32887 | ENSMUSG00000093687 | 0 | NA | NA | NA | NA | NA | Gm20637 | predicted gene 20637 [Source:MGI Symbo | 0 | 0 | 0 | 0 | 0 | 0 |
| 32888 | ENSMUSG00000093688 | 0 | NA | NA | NA | NA | NA | Gm8488 | predicted gene 8488 [Source:MGI Symbol | 0 | 0 | 0 | 0 | 0 | 0 |
| 32889 | ENSMUSG00000093689 | 0 | NA | NA | NA | NA | NA | Vmn1r-ps52 | vomeronasal 1 receptor, pseudogene 52 [ | 0 | 0 | 0 | 0 | 0 | 0 |
| 32890 | ENSMUSG00000093690 | 0.371281295 | 2.09253362 | 4.031852165 | 0.519000582 | 0.603760335 | NA | Mir5619 | microRNA 5619 [Source:MGI Symbol;Acc: | 0 | 0 | 0 | 0 | 0 | 2.2277 |
| 32891 | ENSMUSG00000093695 | 0.346303622 | 2.004339653 | 3.884003423 | 0.516049919 | 0.605819544 | NA | Gm20717 | predicted gene 20717 [Source:MGI Symbo | 0 | 0 | 0 | 0.964 | 0 | 1.1138 |
| 32892 | ENSMUSG00000093696 | 0 | NA | NA | NA | NA | NA | Vmn1r7 | vomeronasal 1 receptor 7 [Source:MGI Sy | 0 | 0 | 0 | 0 | 0 | 0 |
| 32893 | ENSMUSG00000093697 | 0 | NA | NA | NA | NA | NA | Gm20686 | predicted gene 20686 [Source:MGI Symbo | 0 | 0 | 0 | 0 | 0 | 0 |
| 32894 | ENSMUSG00000093699 | 0 | NA | NA | NA | NA | NA | Vmn2r-ps9 | vomeronasal 2, receptor, pseudogene 9 [S | 0 | 0 | 0 | 0 | 0 | 0 |
| 32895 | ENSMUSG00000093701 | 0 | NA | NA | NA | NA | NA | Gm21970 | predicted gene 21970 [Source:MGI Symbo | 0 | 0 | 0 | 0 | 0 | 0 |
| 32896 | ENSMUSG00000093702 | 0 | NA | NA | NA | NA | NA | Vmn1r-ps20 | vomeronasal 1 receptor, pseudogene 20 [ | 0 | 0 | 0 | 0 | 0 | 0 |
| 32897 | ENSMUSG00000093703 | 0 | NA | NA | NA | NA | NA | Gm18397 | predicted gene, 18397 [Source:MGI Symb | 0 | 0 | 0 | 0 | 0 | 0 |
| 32898 | ENSMUSG00000093705 | 0 | NA | NA | NA | NA | NA | Vmn1r-ps14 | vomeronasal 1 receptor, pseudogene 142 | 0 | 0 | 0 | 0 | 0 | 0 |
| 32899 | ENSMUSG00000093707 | 0 | NA | NA | NA | NA | NA | Obox4-ps9 | oocyte specific homeobox 4, pseudogene | 0 | 0 | 0 | 0 | 0 | 0 |
| 32900 | ENSMUSG00000093708 | 0 | NA | NA | NA | NA | NA | Gm20718 | predicted gene 20718 [Source:MGI Symbo | 0 | 0 | 0 | 0 | 0 | 0 |
| 32901 | ENSMUSG00000093710 | 0 | NA | NA | NA | NA | NA | Mir5625 | microRNA 5625 [Source:MGI Symbol;Acc: | 0 | 0 | 0 | 0 | 0 | 0 |
| 32902 | ENSMUSG00000093711 | 0 | NA | NA | NA | NA | NA | Vmn2r-ps12 | vomeronasal 2, receptor, pseudogene 125 | 0 | 0 | 0 | 0 | 0 | 0 |
| 32903 | ENSMUSG00000093714 | 0 | NA | NA | NA | NA | NA | Vmn2r-ps13 | vomeronasal 2, receptor, pseudogene 132 | 0 | 0 | 0 | 0 | 0 | 0 |
| 32904 | ENSMUSG00000093716 | 0 | NA | NA | NA | NA | NA | Gm19815 | predicted gene, 19815 [Source:MGI Symb | 0 | 0 | 0 | 0 | 0 | 0 |
| 32905 | ENSMUSG00000093719 | 0 | NA | NA | NA | NA | NA | Obox4-ps30 | oocyte specific homeobox 4, pseudogene | 0 | 0 | 0 | 0 | 0 | 0 |
| 32906 | ENSMUSG00000093720 | 1.295722852 | 0.191559112 | 1.971845392 | 0.097147126 | 0.922609557 | NA | Gm20635 | predicted gene 20635 [Source:MGI Symbo | 1.6062 | 0 | 2.0125 | 1.928 | 0 | 2.2277 |
| 32907 | ENSMUSG00000093721 | 0 | NA | NA | NA | NA | NA | Gm3896 | predicted gene 3896 [Source:MGI Symbol | 0 | 0 | 0 | 0 | 0 | 0 |
| 32908 | ENSMUSG00000093722 | 0.335414862 | -1.779047242 | 4.030804179 | -0.441362855 | 0.658950329 | NA | Gm20612 | predicted gene 20612 [Source:MGI Symbo | 0 | 0 | 2.0125 | 0 | 0 | 0 |
| 32909 | ENSMUSG00000093724 | 0 | NA | NA | NA | NA | NA | Vmn2r-ps32 | vomeronasal 2, receptor, pseudogene 32 | 0 | 0 | 0 | 0 | 0 | 0 |
| 32910 | ENSMUSG00000093725 | 0 | NA | NA | NA | NA | NA | Gm18595 | predicted gene, 18595 [Source:MGI Symb | 0 | 0 | 0 | 0 | 0 | 0 |
| 32911 | ENSMUSG00000093726 | 0 | NA | NA | NA | NA | NA | Gm20667 | predicted gene 20667 [Source:MGI Symbo | 0 | 0 | 0 | 0 | 0 | 0 |
| 32912 | ENSMUSG00000093727 | 0 | NA | NA | NA | NA | NA | Obox3-ps5 | oocyte specific homeobox 3, pseudogene | 0 | 0 | 0 | 0 | 0 | 0 |
| 32913 | ENSMUSG00000093728 | 0 | NA | NA | NA | NA | NA | Vmn1r-ps8 | vomeronasal 1 receptor, pseudogene 8 [S | 0 | 0 | 0 | 0 | 0 | 0 |
| 32914 | ENSMUSG00000093729 | 0 | NA | NA | NA | NA | NA | Vmn1r-ps14 | vomeronasal 1 receptor, pseudogene 141 | 0 | 0 | 0 | 0 | 0 | 0 |
| 32915 | ENSMUSG00000093730 | 0 | NA | NA | NA | NA | NA | Gm20690 | predicted gene 20690 [Source:MGI Symbo | 0 | 0 | 0 | 0 | 0 | 0 |
| 32916 | ENSMUSG00000093731 | 0 | NA | NA | NA | NA | NA | Vmn1r-ps18 | vomeronasal 1 receptor, pseudogene 18 [ | 0 | 0 | 0 | 0 | 0 | 0 |
| 32917 | ENSMUSG00000093732 | 0 | NA | NA | NA | NA | NA | Gm20609 | predicted gene 20609 [Source:MGI Symbo | 0 | 0 | 0 | 0 | 0 | 0 |
| 32918 | ENSMUSG00000093734 | 16.60847493 | 0.216233027 | 0.688831065 | 0.313913002 | 0.753587115 | 0.952989051 | NA | NA | 12.85 | 13.363 | 20.125 | 34.703 | 15.269 | 3.3415 |
| 32919 | ENSMUSG00000093736 | 0 | NA | NA | NA | NA | NA | Obox4-ps38 | oocyte specific homeobox 4, pseudogene | 0 | 0 | 0 | 0 | 0 | 0 |
| 32920 | ENSMUSG00000093738 | 0 | NA | NA | NA | NA | NA | AI606473 | expressed sequence AI606473 [Source:M( | 0 | 0 | 0 | 0 | 0 | 0 |
| 32921 | ENSMUSG00000093739 | 2.38495041 | 1.44767277 | 1.519219184 | 0.952905799 | 0.340637802 | NA | Gm20692 | predicted gene 20692 [Source:MGI Symbo | 0.8031 | 3.0838 | 0 | 3.8559 | 5.4531 | 1.1138 |
| 32922 | ENSMUSG00000093740 | 0 | NA | NA | NA | NA | NA | Gm18417 | predicted gene, 18417 [Source:MGI Symb | 0 | 0 | 0 | 0 | 0 | 0 |
| 32923 | ENSMUSG00000093741 | 0 | NA | NA | NA | NA | NA | Gm20672 | predicted gene 20672 [Source:MGI Symbo | 0 | 0 | 0 | 0 | 0 | 0 |
| 32924 | ENSMUSG00000093743 | 0 | NA | NA | NA | NA | NA | Vmn1r-ps17 | vomeronasal 1 receptor, pseudogene 17 [ | 0 | 0 | 0 | 0 | 0 | 0 |

| | | | | | | | | | | | | | | | |
|---|---|---|---|---|---|---|---|---|---|---|---|---|---|---|---|
| 32925 | ENSMUSG00000093744 | 0 | NA | NA | NA | NA | NA | Gm20675 | predicted gene 20675 [Source:MGI Symbo | 0 | 0 | 0 | 0 | 0 | 0 |
| 32926 | ENSMUSG00000093745 | 0 | NA | NA | NA | NA | NA | NA | NA | 0 | 0 | 0 | 0 | 0 | 0 |
| 32927 | ENSMUSG00000093746 | 0 | NA | NA | NA | NA | NA | Mir5622 | microRNA 5622 [Source:MGI Symbol;Acc: | 0 | 0 | 0 | 0 | 0 | 0 |
| 32928 | ENSMUSG00000093747 | 0 | NA | NA | NA | NA | NA | Gm20673 | predicted gene 20673 [Source:MGI Symbo | 0 | 0 | 0 | 0 | 0 | 0 |
| 32929 | ENSMUSG00000093748 | 0 | NA | NA | NA | NA | NA | Gm19031 | predicted gene, 19031 [Source:MGI Symb | 0 | 0 | 0 | 0 | 0 | 0 |
| 32930 | ENSMUSG00000093749 | 0 | NA | NA | NA | NA | NA | 4921523L03I | RIKEN cDNA 4921523L03 gene [Source:M( | 0 | 0 | 0 | 0 | 0 | 0 |
| 32931 | ENSMUSG00000093751 | 0 | NA | NA | NA | NA | NA | Gm3307 | predicted gene 3307 [Source:MGI Symbol | 0 | 0 | 0 | 0 | 0 | 0 |
| 32932 | ENSMUSG00000093752 | 0 | NA | NA | NA | NA | NA | Gm20716 | predicted gene 20716 [Source:MGI Symbo | 0 | 0 | 0 | 0 | 0 | 0 |
| 32933 | ENSMUSG00000093753 | 0 | NA | NA | NA | NA | NA | Gm28577 | predicted gene 28577 [Source:MGI Symbo | 0 | 0 | 0 | 0 | 0 | 0 |
| 32934 | ENSMUSG00000093755 | 0 | NA | NA | NA | NA | NA | Vmn1r39 | vomeronasal 1 receptor 39 [Source:MGI S | 0 | 0 | 0 | 0 | 0 | 0 |
| 32935 | ENSMUSG00000093756 | 0 | NA | NA | NA | NA | NA | Gm18594 | predicted gene, 18594 [Source:MGI Symb | 0 | 0 | 0 | 0 | 0 | 0 |
| 32936 | ENSMUSG00000093757 | 0 | NA | NA | NA | NA | NA | Mir5623 | microRNA 5623 [Source:MGI Symbol;Acc: | 0 | 0 | 0 | 0 | 0 | 0 |
| 32937 | ENSMUSG00000093758 | 0 | NA | NA | NA | NA | NA | Gm20701 | predicted gene 20701 [Source:MGI Symbo | 0 | 0 | 0 | 0 | 0 | 0 |
| 32938 | ENSMUSG00000093759 | 0.856470372 | -1.816219066 | 2.859255434 | -0.635207 | 0.525293418 | NA | Scarletltr | Scarletltr, erythroid developmental long ir | 0 | 0 | 4.025 | 0 | 0 | 1.1138 |
| 32939 | ENSMUSG00000093761 | 0 | NA | NA | NA | NA | NA | Obox6-ps1 | oocyte specific homeobox 6, pseudogene | 0 | 0 | 0 | 0 | 0 | 0 |
| 32940 | ENSMUSG00000093762 | 0 | NA | NA | NA | NA | NA | NA | NA | 0 | 0 | 0 | 0 | 0 | 0 |
| 32941 | ENSMUSG00000093763 | 0 | NA | NA | NA | NA | NA | Mir5626 | microRNA 5626 [Source:MGI Symbol;Acc: | 0 | 0 | 0 | 0 | 0 | 0 |
| 32942 | ENSMUSG00000093765 | 0 | NA | NA | NA | NA | NA | Gm20658 | predicted gene 20658 [Source:MGI Symbo | 0 | 0 | 0 | 0 | 0 | 0 |
| 32943 | ENSMUSG00000093766 | 0 | NA | NA | NA | NA | NA | Gm20679 | predicted gene 20679 [Source:MGI Symbo | 0 | 0 | 0 | 0 | 0 | 0 |
| 32944 | ENSMUSG00000093767 | 0 | NA | NA | NA | NA | NA | Vmn2r-ps13 | vomeronasal 2, receptor, pseudogene 13 | 0 | 0 | 0 | 0 | 0 | 0 |
| 32945 | ENSMUSG00000093768 | 0 | NA | NA | NA | NA | NA | Gm20650 | predicted gene 20650 [Source:MGI Symbo | 0 | 0 | 0 | 0 | 0 | 0 |
| 32946 | ENSMUSG00000093769 | 0 | NA | NA | NA | NA | NA | H3c14 | H3 clustered histone 14 [Source:MGI Sym | 0 | 0 | 0 | 0 | 0 | 0 |
| 32947 | ENSMUSG00000093770 | 0 | NA | NA | NA | NA | NA | Obox3-ps4 | oocyte specific homeobox, pseudogene 4 | 0 | 0 | 0 | 0 | 0 | 0 |
| 32948 | ENSMUSG00000093771 | 0 | NA | NA | NA | NA | NA | AU023070 | expressed sequence AU023070 [Source:M | 0 | 0 | 0 | 0 | 0 | 0 |
| 32949 | ENSMUSG00000093772 | 15.17444322 | 0.32154755 | 0.569491069 | 0.564622639 | 0.572330464 | 0.905173631 | 4931403E22 | RIKEN cDNA 4931403E22 gene [Source:M( | 16.062 | 13.363 | 11.069 | 24.099 | 13.087 | 13.366 |
| 32950 | ENSMUSG00000093773 | 0 | NA | NA | NA | NA | NA | Obox3-ps7 | oocyte specific homeobox 3, pseudogene | 0 | 0 | 0 | 0 | 0 | 0 |
| 32951 | ENSMUSG00000093774 | 0 | NA | NA | NA | NA | NA | Vmn1r-ps15 | vomeronasal 1 receptor, pseudogene 151 | 0 | 0 | 0 | 0 | 0 | 0 |
| 32952 | ENSMUSG00000093775 | 0 | NA | NA | NA | NA | NA | Vmn2r-ps11 | vomeronasal 2, receptor, pseudogene 114 | 0 | 0 | 0 | 0 | 0 | 0 |
| 32953 | ENSMUSG00000093776 | 0 | NA | NA | NA | NA | NA | Vmn2r-ps12 | vomeronasal 2, receptor, pseudogene 12( | 0 | 0 | 0 | 0 | 0 | 0 |
| 32954 | ENSMUSG00000093779 | 0.160662974 | 1.126924175 | 4.080472857 | 0.276174898 | 0.782413736 | NA | Gm20648 | predicted gene 20648 [Source:MGI Symbo | 0 | 0 | 0 | 0.964 | 0 | 0 |
| 32955 | ENSMUSG00000093780 | 0 | NA | NA | NA | NA | NA | NA | NA | 0 | 0 | 0 | 0 | 0 | 0 |
| 32956 | ENSMUSG00000093781 | 0 | NA | NA | NA | NA | NA | Vmn1r-ps47 | vomeronasal 1 receptor, pseudogene 47 [ | 0 | 0 | 0 | 0 | 0 | 0 |
| 32957 | ENSMUSG00000093782 | 0 | NA | NA | NA | NA | NA | NA | NA | 0 | 0 | 0 | 0 | 0 | 0 |
| 32958 | ENSMUSG00000093784 | 0 | NA | NA | NA | NA | NA | Gm22394 | predicted gene, 22394 [Source:MGI Symb | 0 | 0 | 0 | 0 | 0 | 0 |
| 32959 | ENSMUSG00000093788 | 0 | NA | NA | NA | NA | NA | Vmn1r-ps14 | vomeronasal 1 receptor, pseudogene 147 | 0 | 0 | 0 | 0 | 0 | 0 |
| 32960 | ENSMUSG00000093789 | 0.328370405 | 0.16514316 | 4.003149536 | 0.041253308 | 0.967093956 | NA | Methig1 | methyltransferase hypoxia inducible dom | 0 | 0 | 1.0062 | 0.964 | 0 | 0 |
| 32961 | ENSMUSG00000093791 | 0 | NA | NA | NA | NA | NA | NA | NA | 0 | 0 | 0 | 0 | 0 | 0 |
| 32962 | ENSMUSG00000093793 | 1.493337644 | 0.411943645 | 1.905771782 | 0.216155811 | 0.828866297 | NA | NA | NA | 0.8031 | 3.0838 | 0 | 2.8919 | 2.1812 | 0 |
| 32963 | ENSMUSG00000093796 | 0 | NA | NA | NA | NA | NA | Rps12-ps23 | ribosomal protein S12, pseudogene 23 [Sc | 0 | 0 | 0 | 0 | 0 | 0 |
| 32964 | ENSMUSG00000093798 | 10.28339367 | -0.179135847 | 0.724786841 | -0.247156594 | 0.804787036 | 0.963787787 | Gm8355 | predicted pseudogene 8355 [Source:MGI | 15.259 | 8.2233 | 9.0562 | 4.8199 | 7.6343 | 16.708 |
| 32965 | ENSMUSG00000093800 | 0 | NA | NA | NA | NA | NA | Gm20796 | predicted gene, 20796 [Source:MGI Symb | 0 | 0 | 0 | 0 | 0 | 0 |
| 32966 | ENSMUSG00000093801 | 0 | NA | NA | NA | NA | NA | Trav14n-2 | T cell receptor alpha variable 14N-2 [Sour | 0 | 0 | 0 | 0 | 0 | 0 |
| 32967 | ENSMUSG00000093802 | 0 | NA | NA | NA | NA | NA | NA | NA | 0 | 0 | 0 | 0 | 0 | 0 |
| 32968 | ENSMUSG00000093803 | 218.8671974 | 0.167793984 | 0.206352882 | 0.813140976 | 0.416137236 | 0.836616789 | Ppp2r3d | protein phosphatase 2 (formerly 2A), regu | 174.27 | 188.11 | 256.59 | 237.14 | 189.77 | 267.32 |
| 32969 | ENSMUSG00000093804 | 0 | NA | NA | NA | NA | NA | Olfr1303 | olfactory receptor 1303 [Source:MGI Syml | 0 | 0 | 0 | 0 | 0 | 0 |
| 32970 | ENSMUSG00000093805 | 0 | NA | NA | NA | NA | NA | Gal3st2b | galactose-3-O-sulfotransferase 2B [Source | 0 | 0 | 0 | 0 | 0 | 0 |
| 32971 | ENSMUSG00000093806 | 0 | NA | NA | NA | NA | NA | Asmt | acetylserotonin O-methyltransferase [Sou | 0 | 0 | 0 | 0 | 0 | 0 |
| 32972 | ENSMUSG00000093807 | 0 | NA | NA | NA | NA | NA | NA | NA | 0 | 0 | 0 | 0 | 0 | 0 |
| 32973 | ENSMUSG00000093812 | 0.328370405 | 0.16514316 | 4.003149536 | 0.041253308 | 0.967093956 | NA | Gm5627 | predicted gene 5627 [Source:MGI Symbol | 0 | 0 | 1.0062 | 0.964 | 0 | 0 |
| 32974 | ENSMUSG00000093813 | 0 | NA | NA | NA | NA | NA | Trav9n-2 | T cell receptor alpha variable 9N-2 [Sourc | 0 | 0 | 0 | 0 | 0 | 0 |
| 32975 | ENSMUSG00000093814 | 0 | NA | NA | NA | NA | NA | Gm9637 | predicted gene 9637 [Source:MGI Symbol | 0 | 0 | 0 | 0 | 0 | 0 |
| 32976 | ENSMUSG00000093815 | 0 | NA | NA | NA | NA | NA | NA | NA | 0 | 0 | 0 | 0 | 0 | 0 |
| 32977 | ENSMUSG00000093816 | 0 | NA | NA | NA | NA | NA | NA | NA | 0 | 0 | 0 | 0 | 0 | 0 |
| 32978 | ENSMUSG00000093817 | 0 | NA | NA | NA | NA | NA | Gm2101 | predicted gene 2101 [Source:MGI Symbol | 0 | 0 | 0 | 0 | 0 | 0 |
| 32979 | ENSMUSG00000093818 | 0 | NA | NA | NA | NA | NA | Ighd3-1 | immunoglobulin heavy diversity 3-1 [Sour | 0 | 0 | 0 | 0 | 0 | 0 |
| 32980 | ENSMUSG00000093820 | 0 | NA | NA | NA | NA | NA | Vmn2r21 | vomeronasal 2, receptor 21 [Source:MGI S | 0 | 0 | 0 | 0 | 0 | 0 |

| | | | | | | | | | | | | | | | |
|---|---|---|---|---|---|---|---|---|---|---|---|---|---|---|---|
| 32981 | ENSMUSG00000093822 | 0 | NA | NA | NA | NA | NA | NA | NA | 0 | 0 | 0 | 0 | 0 | 0 |
| 32982 | ENSMUSG00000093823 | 0 | NA | NA | NA | NA | NA | NA | NA | 0 | 0 | 0 | 0 | 0 | 0 |
| 32983 | ENSMUSG00000093824 | 0 | NA | NA | NA | NA | NA | NA | NA | 0 | 0 | 0 | 0 | 0 | 0 |
| 32984 | ENSMUSG00000093825 | 0 | NA | NA | NA | NA | NA | Olfr723 | olfactory receptor 723 [Source:MGI Symb | 0 | 0 | 0 | 0 | 0 | 0 |
| 32985 | ENSMUSG00000093826 | 4.960814745 | -0.720249416 | 0.980860957 | -0.734303279 | 0.46276392 | NA | Gm6900 | predicted gene 6900 [Source:MGI Symbol | 6.4248 | 5.1396 | 7.0437 | 6.7478 | 2.1812 | 2.2277 |
| 32986 | ENSMUSG00000093827 | 0 | NA | NA | NA | NA | NA | Gm10410 | predicted gene 10410 [Source:MGI Symbo | 0 | 0 | 0 | 0 | 0 | 0 |
| 32987 | ENSMUSG00000093828 | 0 | NA | NA | NA | NA | NA | NA | NA | 0 | 0 | 0 | 0 | 0 | 0 |
| 32988 | ENSMUSG00000093829 | 0 | NA | NA | NA | NA | NA | NA | NA | 0 | 0 | 0 | 0 | 0 | 0 |
| 32989 | ENSMUSG00000093830 | 0 | NA | NA | NA | NA | NA | NA | NA | 0 | 0 | 0 | 0 | 0 | 0 |
| 32990 | ENSMUSG00000093831 | 0 | NA | NA | NA | NA | NA | NA | NA | 0 | 0 | 0 | 0 | 0 | 0 |
| 32991 | ENSMUSG00000093833 | 0 | NA | NA | NA | NA | NA | Gm7970 | predicted gene 7970 [Source:MGI Symbol | 0 | 0 | 0 | 0 | 0 | 0 |
| 32992 | ENSMUSG00000093834 | 0 | NA | NA | NA | NA | NA | NA | NA | 0 | 0 | 0 | 0 | 0 | 0 |
| 32993 | ENSMUSG00000093835 | 0 | NA | NA | NA | NA | NA | NA | NA | 0 | 0 | 0 | 0 | 0 | 0 |
| 32994 | ENSMUSG00000093838 | 0 | NA | NA | NA | NA | NA | Ighv3-1 | immunoglobulin heavy variable 3-1 [Sourc | 0 | 0 | 0 | 0 | 0 | 0 |
| 32995 | ENSMUSG00000093839 | 0 | NA | NA | NA | NA | NA | Olfr462 | olfactory receptor 462 [Source:MGI Symb | 0 | 0 | 0 | 0 | 0 | 0 |
| 32996 | ENSMUSG00000093843 | 0.17131964 | -0.796629176 | 4.080472857 | -0.195229623 | 0.84521319 | NA | NA | NA | 0 | 1.0279 | 0 | 0 | 0 | 0 |
| 32997 | ENSMUSG00000093846 | 0 | NA | NA | NA | NA | NA | Gm4425 | predicted gene 4425 [Source:MGI Symbol | 0 | 0 | 0 | 0 | 0 | 0 |
| 32998 | ENSMUSG00000093847 | 0 | NA | NA | NA | NA | NA | Eif1ad15 | eukaryotic translation initiation factor 1A | 0 | 0 | 0 | 0 | 0 | 0 |
| 32999 | ENSMUSG00000093848 | 0 | NA | NA | NA | NA | NA | Gm20865 | predicted gene, 20865 [Source:MGI Symb | 0 | 0 | 0 | 0 | 0 | 0 |
| 33000 | ENSMUSG00000093849 | 0 | NA | NA | NA | NA | NA | Gm7083 | predicted gene 7083 [Source:MGI Symbol | 0 | 0 | 0 | 0 | 0 | 0 |
| 33001 | ENSMUSG00000093851 | 0 | NA | NA | NA | NA | NA | Gm15089 | predicted gene 15089 [Source:MGI Symbo | 0 | 0 | 0 | 0 | 0 | 0 |
| 33002 | ENSMUSG00000093852 | 0 | NA | NA | NA | NA | NA | NA | NA | 0 | 0 | 0 | 0 | 0 | 0 |
| 33003 | ENSMUSG00000093853 | 0 | NA | NA | NA | NA | NA | Vmn1r151 | vomeronasal 1 receptor 151 [Source:MGI | 0 | 0 | 0 | 0 | 0 | 0 |
| 33004 | ENSMUSG00000093854 | 0 | NA | NA | NA | NA | NA | NA | NA | 0 | 0 | 0 | 0 | 0 | 0 |
| 33005 | ENSMUSG00000093856 | 0 | NA | NA | NA | NA | NA | NA | NA | 0 | 0 | 0 | 0 | 0 | 0 |
| 33006 | ENSMUSG00000093857 | 0 | NA | NA | NA | NA | NA | NA | NA | 0 | 0 | 0 | 0 | 0 | 0 |
| 33007 | ENSMUSG00000093858 | 0.91169124 | -0.750330513 | 2.318519369 | -0.323624863 | 0.746222062 | NA | Gm19967 | predicted gene, 19967 [Source:MGI Symb | 2.4093 | 0 | 1.0062 | 0.964 | 1.0906 | 0 |
| 33008 | ENSMUSG00000093859 | 0 | NA | NA | NA | NA | NA | Gm7882 | predicted gene 7882 [Source:MGI Symbol | 0 | 0 | 0 | 0 | 0 | 0 |
| 33009 | ENSMUSG00000093860 | 0 | NA | NA | NA | NA | NA | NA | NA | 0 | 0 | 0 | 0 | 0 | 0 |
| 33010 | ENSMUSG00000093861 | 2.982003796 | -2.917602762 | 1.498852702 | -1.946557362 | 0.051587825 | NA | Igkv1-110 | immunoglobulin kappa variable 1-110 [So | 5.6217 | 6.1675 | 4.025 | 0.964 | 0 | 1.1138 |
| 33011 | ENSMUSG00000093862 | 0 | NA | NA | NA | NA | NA | Gm5117 | predicted gene 5117 [Source:MGI Symbol | 0 | 0 | 0 | 0 | 0 | 0 |
| 33012 | ENSMUSG00000093863 | 0 | NA | NA | NA | NA | NA | Gm7954 | predicted gene 7954 [Source:MGI Symbol | 0 | 0 | 0 | 0 | 0 | 0 |
| 33013 | ENSMUSG00000093864 | 0 | NA | NA | NA | NA | NA | NA | NA | 0 | 0 | 0 | 0 | 0 | 0 |
| 33014 | ENSMUSG00000093865 | 0 | NA | NA | NA | NA | NA | Lrit3 | leucine-rich repeat, immunoglobulin-like a | 0 | 0 | 0 | 0 | 0 | 0 |
| 33015 | ENSMUSG00000093866 | 0 | NA | NA | NA | NA | NA | Olfr802 | olfactory receptor 802 [Source:MGI Symb | 0 | 0 | 0 | 0 | 0 | 0 |
| 33016 | ENSMUSG00000093867 | 0 | NA | NA | NA | NA | NA | Gsdmcl2 | gasdermin C-like 2 [Source:MGI Symbol;A | 0 | 0 | 0 | 0 | 0 | 0 |
| 33017 | ENSMUSG00000093868 | 0 | NA | NA | NA | NA | NA | Gm20809 | predicted gene, 20809 [Source:MGI Symb | 0 | 0 | 0 | 0 | 0 | 0 |
| 33018 | ENSMUSG00000093869 | 0 | NA | NA | NA | NA | NA | NA | NA | 0 | 0 | 0 | 0 | 0 | 0 |
| 33019 | ENSMUSG00000093870 | 0 | NA | NA | NA | NA | NA | NA | NA | 0 | 0 | 0 | 0 | 0 | 0 |
| 33020 | ENSMUSG00000093871 | 0 | NA | NA | NA | NA | NA | Vmn1r253 | vomeronasal 1 receptor 253 [Source:MGI | 0 | 0 | 0 | 0 | 0 | 0 |
| 33021 | ENSMUSG00000093873 | 0 | NA | NA | NA | NA | NA | NA | NA | 0 | 0 | 0 | 0 | 0 | 0 |
| 33022 | ENSMUSG00000093874 | 0 | NA | NA | NA | NA | NA | NA | NA | 0 | 0 | 0 | 0 | 0 | 0 |
| 33023 | ENSMUSG00000093875 | 0 | NA | NA | NA | NA | NA | NA | NA | 0 | 0 | 0 | 0 | 0 | 0 |
| 33024 | ENSMUSG00000093876 | 0 | NA | NA | NA | NA | NA | Ighd5-3 | immunoglobulin heavy diversity 5-3 [Sour | 0 | 0 | 0 | 0 | 0 | 0 |
| 33025 | ENSMUSG00000093877 | 0 | NA | NA | NA | NA | NA | Olfr1348 | olfactory receptor 1348 [Source:MGI Sym | 0 | 0 | 0 | 0 | 0 | 0 |
| 33026 | ENSMUSG00000093878 | 0 | NA | NA | NA | NA | NA | NA | NA | 0 | 0 | 0 | 0 | 0 | 0 |
| 33027 | ENSMUSG00000093879 | 0 | NA | NA | NA | NA | NA | NA | NA | 0 | 0 | 0 | 0 | 0 | 0 |
| 33028 | ENSMUSG00000093880 | 0.17131964 | -0.796629176 | 4.080472857 | -0.195229623 | 0.84521319 | NA | Tmem181c-p | transmembrane protein 181C, pseudogen | 0 | 1.0279 | 0 | 0 | 0 | 0 |
| 33029 | ENSMUSG00000093883 | 0 | NA | NA | NA | NA | NA | Gm21866 | predicted gene, 21866 [Source:MGI Symb | 0 | 0 | 0 | 0 | 0 | 0 |
| 33030 | ENSMUSG00000093884 | 0 | NA | NA | NA | NA | NA | Olfr239 | olfactory receptor 239 [Source:MGI Symb | 0 | 0 | 0 | 0 | 0 | 0 |
| 33031 | ENSMUSG00000093885 | 0 | NA | NA | NA | NA | NA | Rpl35a-ps5 | ribosomal protein L35A, pseudogene 5 [Sc | 0 | 0 | 0 | 0 | 0 | 0 |
| 33032 | ENSMUSG00000093886 | 0.321325949 | 1.911575114 | 4.048267893 | 0.472195804 | 0.636787034 | NA | NA | NA | 0 | 0 | 0 | 1.928 | 0 | 0 |
| 33033 | ENSMUSG00000093887 | 0 | NA | NA | NA | NA | NA | Gm3033 | predicted gene 3033 [Source:MGI Symbol | 0 | 0 | 0 | 0 | 0 | 0 |
| 33034 | ENSMUSG00000093889 | 0 | NA | NA | NA | NA | NA | Ang-ps2 | angiogenin, ribonuclease A family, pseudo | 0 | 0 | 0 | 0 | 0 | 0 |
| 33035 | ENSMUSG00000093890 | 0 | NA | NA | NA | NA | NA | Vmn1r127 | vomeronasal 1 receptor 127 [Source:MGI | 0 | 0 | 0 | 0 | 0 | 0 |
| 33036 | ENSMUSG00000093891 | 0 | NA | NA | NA | NA | NA | Gm2866 | predicted gene 2866 [Source:MGI Symbol | 0 | 0 | 0 | 0 | 0 | 0 |

| | | | | | | | | | | | | | | | |
|---|---|---|---|---|---|---|---|---|---|---|---|---|---|---|---|
| 33037 | ENSMUSG00000093892 | 0 | NA | NA | NA | NA | NA | Gm3261 | predicted gene 3261 [Source:MGI Symbol | 0 | 0 | 0 | 0 | 0 | 0 |
| 33038 | ENSMUSG00000093893 | 0 | NA | NA | NA | NA | NA | NA | NA | 0 | 0 | 0 | 0 | 0 | 0 |
| 33039 | ENSMUSG00000093894 | 0.878040772 | -3.188971767 | 2.713192499 | -1.175357727 | 0.239851629 | NA | Ighv1-53 | immunoglobulin heavy variable 1-53 [Sou | 3.2124 | 2.0558 | 0 | 0 | 0 | 0 |
| 33040 | ENSMUSG00000093895 | 0 | NA | NA | NA | NA | NA | Gm21865 | predicted gene, 21865 [Source:MGI Symb | 0 | 0 | 0 | 0 | 0 | 0 |
| 33041 | ENSMUSG00000093896 | 0 | NA | NA | NA | NA | NA | Ighv1-76 | immunoglobulin heavy variable 1-76 [Sou | 0 | 0 | 0 | 0 | 0 | 0 |
| 33042 | ENSMUSG00000093897 | 2.06687675 | 0.170261953 | 1.78074797 | 0.095612606 | 0.923828253 | NA | NA | NA | 0.8031 | 0 | 5.0312 | 0 | 5.4531 | 1.1138 |
| 33043 | ENSMUSG00000093898 | 0 | NA | NA | NA | NA | NA | Gm33933 | predicted gene, 33933 [Source:MGI Symb | 0 | 0 | 0 | 0 | 0 | 0 |
| 33044 | ENSMUSG00000093899 | 0 | NA | NA | NA | NA | NA | Gm5645 | predicted pseudogene 5645 [Source:MGI | 0 | 0 | 0 | 0 | 0 | 0 |
| 33045 | ENSMUSG00000093900 | 0 | NA | NA | NA | NA | NA | NA | NA | 0 | 0 | 0 | 0 | 0 | 0 |
| 33046 | ENSMUSG00000093901 | 0 | NA | NA | NA | NA | NA | Olfr893 | olfactory receptor 893 [Source:MGI Symb | 0 | 0 | 0 | 0 | 0 | 0 |
| 33047 | ENSMUSG00000093902 | 0 | NA | NA | NA | NA | NA | NA | NA | 0 | 0 | 0 | 0 | 0 | 0 |
| 33048 | ENSMUSG00000093903 | 0 | NA | NA | NA | NA | NA | Gm21454 | predicted gene, 21454 [Source:MGI Symb | 0 | 0 | 0 | 0 | 0 | 0 |
| 33049 | ENSMUSG00000093904 | 1013.763984 | -0.008517577 | 0.112936143 | -0.075419404 | 0.939881021 | 0.988812319 | Tomm20 | translocase of outer mitochondrial memb | 938.02 | 1037.2 | 1075.7 | 963.98 | 1077.5 | 990.21 |
| 33050 | ENSMUSG00000093905 | 0 | NA | NA | NA | NA | NA | NA | NA | 0 | 0 | 0 | 0 | 0 | 0 |
| 33051 | ENSMUSG00000093906 | 0.267700746 | -1.497481649 | 4.057001436 | -0.369110456 | 0.712045396 | NA | Igkv9-129 | immunoglobulin kappa variable 9-129 [So | 1.6062 | 0 | 0 | 0 | 0 | 0 |
| 33052 | ENSMUSG00000093907 | 0 | NA | NA | NA | NA | NA | NA | NA | 0 | 0 | 0 | 0 | 0 | 0 |
| 33053 | ENSMUSG00000093908 | 0.649696354 | 1.537929172 | 3.118369374 | 0.493183773 | 0.621882743 | NA | Gm5784 | predicted gene 5784 [Source:MGI Symbol | 0 | 0 | 1.0062 | 2.8919 | 0 | 0 |
| 33054 | ENSMUSG00000093909 | 0 | NA | NA | NA | NA | NA | Gm3883 | predicted gene 3883 [Source:MGI Symbol | 0 | 0 | 0 | 0 | 0 | 0 |
| 33055 | ENSMUSG00000093910 | 0.301557804 | -1.64335916 | 4.042814367 | -0.406488899 | 0.684383412 | NA | Zfp853 | zinc finger protein 853 [Source:MGI Symb | 0.8031 | 0 | 1.0062 | 0 | 0 | 0 |
| 33056 | ENSMUSG00000093912 | 1.993027689 | 0.065074185 | 1.609977947 | 0.040419302 | 0.967758842 | NA | NA | NA | 0.8031 | 4.1117 | 1.0062 | 3.8559 | 2.1812 | 0 |
| 33057 | ENSMUSG00000093913 | 0 | NA | NA | NA | NA | NA | Obox4-ps28 | oocyte specific homeobox 4, pseudogene | 0 | 0 | 0 | 0 | 0 | 0 |
| 33058 | ENSMUSG00000093914 | 0 | NA | NA | NA | NA | NA | NA | NA | 0 | 0 | 0 | 0 | 0 | 0 |
| 33059 | ENSMUSG00000093915 | 0 | NA | NA | NA | NA | NA | Trav12n-3 | T cell receptor alpha variable 12N-3 [Sour | 0 | 0 | 0 | 0 | 0 | 0 |
| 33060 | ENSMUSG00000093916 | 4.993895261 | 3.865551805 | 1.404568338 | 2.752127967 | 0.005920938 | NA | Gm379 | predicted gene 379 [Source:MGI Symbol;A | 0 | 1.0279 | 1.0062 | 13.496 | 2.1812 | 12.252 |
| 33061 | ENSMUSG00000093917 | 0 | NA | NA | NA | NA | NA | Vmn1r254 | vomeronasal 1 receptor 254 [Source:MGI | 0 | 0 | 0 | 0 | 0 | 0 |
| 33062 | ENSMUSG00000093918 | 0 | NA | NA | NA | NA | NA | Gm21677 | predicted gene, 21677 [Source:MGI Symb | 0 | 0 | 0 | 0 | 0 | 0 |
| 33063 | ENSMUSG00000093919 | 0 | NA | NA | NA | NA | NA | Gm20760 | predicted gene, 20760 [Source:MGI Symb | 0 | 0 | 0 | 0 | 0 | 0 |
| 33064 | ENSMUSG00000093920 | 0 | NA | NA | NA | NA | NA | Olfr463 | olfactory receptor 463 [Source:MGI Symb | 0 | 0 | 0 | 0 | 0 | 0 |
| 33065 | ENSMUSG00000093921 | 0 | NA | NA | NA | NA | NA | NA | NA | 0 | 0 | 0 | 0 | 0 | 0 |
| 33066 | ENSMUSG00000093922 | 0 | NA | NA | NA | NA | NA | Rpl36-ps4 | ribosomal protein L36, pseudogene 4 [Sou | 0 | 0 | 0 | 0 | 0 | 0 |
| 33067 | ENSMUSG00000093923 | 0 | NA | NA | NA | NA | NA | Gm5935 | predicted gene 5935 [Source:MGI Symbol | 0 | 0 | 0 | 0 | 0 | 0 |
| 33068 | ENSMUSG00000093924 | 0 | NA | NA | NA | NA | NA | Gm12623 | predicted gene 12623 [Source:MGI Symbo | 0 | 0 | 0 | 0 | 0 | 0 |
| 33069 | ENSMUSG00000093926 | 0 | NA | NA | NA | NA | NA | Gm17027 | predicted gene 17027 [Source:MGI Symbo | 0 | 0 | 0 | 0 | 0 | 0 |
| 33070 | ENSMUSG00000093927 | 0 | NA | NA | NA | NA | NA | Gm20918 | predicted gene, 20918 [Source:MGI Symb | 0 | 0 | 0 | 0 | 0 | 0 |
| 33071 | ENSMUSG00000093928 | 0 | NA | NA | NA | NA | NA | NA | NA | 0 | 0 | 0 | 0 | 0 | 0 |
| 33072 | ENSMUSG00000093929 | 0 | NA | NA | NA | NA | NA | NA | NA | 0 | 0 | 0 | 0 | 0 | 0 |
| 33073 | ENSMUSG00000093930 | 1358.365247 | 0.149734097 | 0.184239415 | 0.812714788 | 0.4163816 | 0.836878476 | Hmgcs1 | 3-hydroxy-3-methylglutaryl-Coenzyme A s | 1436.7 | 1425.7 | 1000.2 | 1114.4 | 1316.4 | 1856.8 |
| 33074 | ENSMUSG00000093931 | 0 | NA | NA | NA | NA | NA | Amy2a3 | amylase 2a3 [Source:MGI Symbol;Acc:MG | 0 | 0 | 0 | 0 | 0 | 0 |
| 33075 | ENSMUSG00000093932 | 0 | NA | NA | NA | NA | NA | NA | NA | 0 | 0 | 0 | 0 | 0 | 0 |
| 33076 | ENSMUSG00000093933 | 0.17131964 | -0.796629176 | 4.080472857 | -0.195229623 | 0.84521319 | NA | NA | NA | 0 | 1.0279 | 0 | 0 | 0 | 0 |
| 33077 | ENSMUSG00000093934 | 0 | NA | NA | NA | NA | NA | Olfr971 | olfactory receptor 971 [Source:MGI Symb | 0 | 0 | 0 | 0 | 0 | 0 |
| 33078 | ENSMUSG00000093935 | 0 | NA | NA | NA | NA | NA | NA | NA | 0 | 0 | 0 | 0 | 0 | 0 |
| 33079 | ENSMUSG00000093937 | 0 | NA | NA | NA | NA | NA | NA | NA | 0 | 0 | 0 | 0 | 0 | 0 |
| 33080 | ENSMUSG00000093938 | 0 | NA | NA | NA | NA | NA | Evi2b | ecotropic viral integration site 2b [Source: | 0 | 0 | 0 | 0 | 0 | 0 |
| 33081 | ENSMUSG00000093939 | 0 | NA | NA | NA | NA | NA | Trav4d-3 | T cell receptor alpha variable 4D-3 [Source | 0 | 0 | 0 | 0 | 0 | 0 |
| 33082 | ENSMUSG00000093940 | 0 | NA | NA | NA | NA | NA | Obox4-ps19 | oocyte specific homeobox 4, pseudogene | 0 | 0 | 0 | 0 | 0 | 0 |
| 33083 | ENSMUSG00000093941 | 0 | NA | NA | NA | NA | NA | Vmn1r131 | vomeronasal 1 receptor 131 [Source:MGI | 0 | 0 | 0 | 0 | 0 | 0 |
| 33084 | ENSMUSG00000093942 | 0 | NA | NA | NA | NA | NA | Olfr46 | olfactory receptor 46 [Source:MGI Symbo | 0 | 0 | 0 | 0 | 0 | 0 |
| 33085 | ENSMUSG00000093943 | 0 | NA | NA | NA | NA | NA | NA | NA | 0 | 0 | 0 | 0 | 0 | 0 |
| 33086 | ENSMUSG00000093944 | 0 | NA | NA | NA | NA | NA | NA | NA | 0 | 0 | 0 | 0 | 0 | 0 |
| 33087 | ENSMUSG00000093945 | 0 | NA | NA | NA | NA | NA | Gm3543 | predicted gene 3543 [Source:MGI Symbol | 0 | 0 | 0 | 0 | 0 | 0 |
| 33088 | ENSMUSG00000093946 | 0 | NA | NA | NA | NA | NA | NA | NA | 0 | 0 | 0 | 0 | 0 | 0 |
| 33089 | ENSMUSG00000093947 | 0 | NA | NA | NA | NA | NA | NA | NA | 0 | 0 | 0 | 0 | 0 | 0 |
| 33090 | ENSMUSG00000093948 | 0 | NA | NA | NA | NA | NA | Gm10378 | predicted gene 10378 [Source:MGI Symbo | 0 | 0 | 0 | 0 | 0 | 0 |
| 33091 | ENSMUSG00000093949 | 0 | NA | NA | NA | NA | NA | Gm21754 | predicted gene, 21754 [Source:MGI Symb | 0 | 0 | 0 | 0 | 0 | 0 |
| 33092 | ENSMUSG00000093950 | 0 | NA | NA | NA | NA | NA | Gm20823 | predicted gene, 20823 [Source:MGI Symb | 0 | 0 | 0 | 0 | 0 | 0 |

| | | | | | | | | | | | | | | | |
|---|---|---|---|---|---|---|---|---|---|---|---|---|---|---|---|
| 33093 | ENSMUSG00000093952 | 0 | NA | NA | NA | NA | NA | NA | NA | 0 | 0 | 0 | 0 | 0 | 0 |
| 33094 | ENSMUSG00000093953 | 0 | NA | NA | NA | NA | NA | Trav3d-3 | T cell receptor alpha variable 3D-3 [Source | 0 | 0 | 0 | 0 | 0 | 0 |
| 33095 | ENSMUSG00000093954 | 2.649913884 | 0.71893828 | 1.791399732 | 0.40132767 | 0.688178894 | NA | Gm16867 | predicted gene, 16867 [Source:MGI Symb | 0 | 0 | 6.0375 | 0 | 7.6343 | 2.2277 |
| 33096 | ENSMUSG00000093955 | 0 | NA | NA | NA | NA | NA | Ighv1-34 | immunoglobulin heavy variable 1-34 [Sou | 0 | 0 | 0 | 0 | 0 | 0 |
| 33097 | ENSMUSG00000093956 | 0 | NA | NA | NA | NA | NA | NA | NA | 0 | 0 | 0 | 0 | 0 | 0 |
| 33098 | ENSMUSG00000093957 | 0 | NA | NA | NA | NA | NA | Esp3 | exocrine gland secreted peptide 3 [Source | 0 | 0 | 0 | 0 | 0 | 0 |
| 33099 | ENSMUSG00000093958 | 0 | NA | NA | NA | NA | NA | NA | NA | 0 | 0 | 0 | 0 | 0 | 0 |
| 33100 | ENSMUSG00000093959 | 0 | NA | NA | NA | NA | NA | NA | NA | 0 | 0 | 0 | 0 | 0 | 0 |
| 33101 | ENSMUSG00000093960 | 0 | NA | NA | NA | NA | NA | NA | NA | 0 | 0 | 0 | 0 | 0 | 0 |
| 33102 | ENSMUSG00000093961 | 0 | NA | NA | NA | NA | NA | Obox4-ps17 | oocyte specific homeobox 4, pseudogene | 0 | 0 | 0 | 0 | 0 | 0 |
| 33103 | ENSMUSG00000093962 | 0 | NA | NA | NA | NA | NA | Gm11756 | predicted gene 11756 [Source:MGI Symbo | 0 | 0 | 0 | 0 | 0 | 0 |
| 33104 | ENSMUSG00000093963 | 0 | NA | NA | NA | NA | NA | Gm21909 | predicted gene, 21909 [Source:MGI Symb | 0 | 0 | 0 | 0 | 0 | 0 |
| 33105 | ENSMUSG00000093964 | 0 | NA | NA | NA | NA | NA | NA | NA | 0 | 0 | 0 | 0 | 0 | 0 |
| 33106 | ENSMUSG00000093965 | 0 | NA | NA | NA | NA | NA | NA | NA | 0 | 0 | 0 | 0 | 0 | 0 |
| 33107 | ENSMUSG00000093966 | 0 | NA | NA | NA | NA | NA | Trav4-3 | T cell receptor alpha variable 4-3 [Source: | 0 | 0 | 0 | 0 | 0 | 0 |
| 33108 | ENSMUSG00000093967 | 1.706256207 | 0.038380257 | 1.836826358 | 0.020894875 | 0.983329515 | NA | NA | NA | 0 | 4.1117 | 1.0062 | 2.8919 | 0 | 2.2277 |
| 33109 | ENSMUSG00000093968 | 0 | NA | NA | NA | NA | NA | Gm8165 | predicted gene 8165 [Source:MGI Symbol | 0 | 0 | 0 | 0 | 0 | 0 |
| 33110 | ENSMUSG00000093969 | 0 | NA | NA | NA | NA | NA | Trav4n-4 | T cell receptor alpha variable 4N-4 [Source | 0 | 0 | 0 | 0 | 0 | 0 |
| 33111 | ENSMUSG00000093970 | 0 | NA | NA | NA | NA | NA | NA | NA | 0 | 0 | 0 | 0 | 0 | 0 |
| 33112 | ENSMUSG00000093971 | 0 | NA | NA | NA | NA | NA | NA | NA | 0 | 0 | 0 | 0 | 0 | 0 |
| 33113 | ENSMUSG00000093973 | 0 | NA | NA | NA | NA | NA | Mrgpra2a | MAS-related GPR, member A2A [Source:N | 0 | 0 | 0 | 0 | 0 | 0 |
| 33114 | ENSMUSG00000093974 | 0 | NA | NA | NA | NA | NA | NA | NA | 0 | 0 | 0 | 0 | 0 | 0 |
| 33115 | ENSMUSG00000093975 | 0.910352643 | 1.221794025 | 2.770215377 | 0.441046583 | 0.659179274 | NA | Vmn2r-ps19 | vomeronasal 2, receptor, pseudogene 19 | 1.6062 | 0 | 0 | 3.8559 | 0 | 0 |
| 33116 | ENSMUSG00000093976 | 0 | NA | NA | NA | NA | NA | Gm8674 | predicted gene 8674 [Source:MGI Symbol | 0 | 0 | 0 | 0 | 0 | 0 |
| 33117 | ENSMUSG00000093977 | 0 | NA | NA | NA | NA | NA | Gm14362 | predicted gene 14362 [Source:MGI Symbo | 0 | 0 | 0 | 0 | 0 | 0 |
| 33118 | ENSMUSG00000093978 | 0 | NA | NA | NA | NA | NA | Gm12612 | predicted gene 12612 [Source:MGI Symbo | 0 | 0 | 0 | 0 | 0 | 0 |
| 33119 | ENSMUSG00000093979 | 0.505131668 | 1.126696199 | 3.421159819 | 0.329331647 | 0.741905026 | NA | Gm2237 | predicted gene 2237 [Source:MGI Symbol | 0.8031 | 0 | 0 | 0 | 0 | 2.2277 |
| 33120 | ENSMUSG00000093980 | 0 | NA | NA | NA | NA | NA | Olfr493 | olfactory receptor 493 [Source:MGI Symb | 0 | 0 | 0 | 0 | 0 | 0 |
| 33121 | ENSMUSG00000093981 | 0 | NA | NA | NA | NA | NA | NA | NA | 0 | 0 | 0 | 0 | 0 | 0 |
| 33122 | ENSMUSG00000093983 | 0 | NA | NA | NA | NA | NA | NA | NA | 0 | 0 | 0 | 0 | 0 | 0 |
| 33123 | ENSMUSG00000093984 | 0 | NA | NA | NA | NA | NA | Gm7365 | predicted gene 7365 [Source:MGI Symbol | 0 | 0 | 0 | 0 | 0 | 0 |
| 33124 | ENSMUSG00000093985 | 0 | NA | NA | NA | NA | NA | Gm10406 | predicted gene 10406 [Source:MGI Symbo | 0 | 0 | 0 | 0 | 0 | 0 |
| 33125 | ENSMUSG00000093986 | 0 | NA | NA | NA | NA | NA | Vmn1r120 | vomeronasal 1 receptor 120 [Source:MGI | 0 | 0 | 0 | 0 | 0 | 0 |
| 33126 | ENSMUSG00000093987 | 0 | NA | NA | NA | NA | NA | Gm21704 | predicted gene, 21704 [Source:MGI Symb | 0 | 0 | 0 | 0 | 0 | 0 |
| 33127 | ENSMUSG00000093988 | 0 | NA | NA | NA | NA | NA | NA | NA | 0 | 0 | 0 | 0 | 0 | 0 |
| 33128 | ENSMUSG00000093989 | 6.255353986 | 0.257501174 | 0.946811629 | 0.271966636 | 0.785647673 | 0.960775321 | Rnasek | ribonuclease, RNase K [Source:MGI Symb | 4.0155 | 6.1675 | 7.0437 | 12.532 | 1.0906 | 6.6831 |
| 33129 | ENSMUSG00000093990 | 0 | NA | NA | NA | NA | NA | Rps12-ps12 | ribosomal protein S12, pseudogene 12 [Sc | 0 | 0 | 0 | 0 | 0 | 0 |
| 33130 | ENSMUSG00000093992 | 0 | NA | NA | NA | NA | NA | NA | NA | 0 | 0 | 0 | 0 | 0 | 0 |
| 33131 | ENSMUSG00000093993 | 0 | NA | NA | NA | NA | NA | Gm20736 | predicted gene, 20736 [Source:MGI Symb | 0 | 0 | 0 | 0 | 0 | 0 |
| 33132 | ENSMUSG00000093994 | 0 | NA | NA | NA | NA | NA | NA | NA | 0 | 0 | 0 | 0 | 0 | 0 |
| 33133 | ENSMUSG00000093995 | 0 | NA | NA | NA | NA | NA | NA | NA | 0 | 0 | 0 | 0 | 0 | 0 |
| 33134 | ENSMUSG00000093996 | 0 | NA | NA | NA | NA | NA | Fam205a3 | family with sequence similarity 205, meml | 0 | 0 | 0 | 0 | 0 | 0 |
| 33135 | ENSMUSG00000093997 | 0 | NA | NA | NA | NA | NA | NA | NA | 0 | 0 | 0 | 0 | 0 | 0 |
| 33136 | ENSMUSG00000093998 | 0 | NA | NA | NA | NA | NA | NA | NA | 0 | 0 | 0 | 0 | 0 | 0 |
| 33137 | ENSMUSG00000093999 | 0 | NA | NA | NA | NA | NA | NA | NA | 0 | 0 | 0 | 0 | 0 | 0 |
| 33138 | ENSMUSG00000094001 | 0 | NA | NA | NA | NA | NA | Vmn1r122 | vomeronasal 1 receptor 122 [Source:MGI | 0 | 0 | 0 | 0 | 0 | 0 |
| 33139 | ENSMUSG00000094002 | 0 | NA | NA | NA | NA | NA | Gm9866 | predicted gene 9866 [Source:MGI Symbol | 0 | 0 | 0 | 0 | 0 | 0 |
| 33140 | ENSMUSG00000094003 | 0 | NA | NA | NA | NA | NA | NA | NA | 0 | 0 | 0 | 0 | 0 | 0 |
| 33141 | ENSMUSG00000094004 | 0 | NA | NA | NA | NA | NA | Gm5128 | predicted gene 5128 [Source:MGI Symbol | 0 | 0 | 0 | 0 | 0 | 0 |
| 33142 | ENSMUSG00000094005 | 0 | NA | NA | NA | NA | NA | Gm12625 | predicted gene 12625 [Source:MGI Symbo | 0 | 0 | 0 | 0 | 0 | 0 |
| 33143 | ENSMUSG00000094006 | 0.17131964 | -0.796629176 | 4.080472857 | -0.195229623 | 0.84521319 | NA | Igkv4-59 | immunoglobulin kappa variable 4-59 [Sou | 0 | 1.0279 | 0 | 0 | 0 | 0 |
| 33144 | ENSMUSG00000094007 | 0 | NA | NA | NA | NA | NA | NA | NA | 0 | 0 | 0 | 0 | 0 | 0 |
| 33145 | ENSMUSG00000094008 | 0 | NA | NA | NA | NA | NA | NA | NA | 0 | 0 | 0 | 0 | 0 | 0 |
| 33146 | ENSMUSG00000094009 | 0 | NA | NA | NA | NA | NA | NA | NA | 0 | 0 | 0 | 0 | 0 | 0 |
| 33147 | ENSMUSG00000094010 | 0 | NA | NA | NA | NA | NA | Vmn1r119 | vomeronasal 1 receptor 119 [Source:MGI | 0 | 0 | 0 | 0 | 0 | 0 |
| 33148 | ENSMUSG00000094011 | 0 | NA | NA | NA | NA | NA | Vmn1r94 | vomeronasal 1 receptor 94 [Source:MGI S | 0 | 0 | 0 | 0 | 0 | 0 |

| | | | | | | | | | | | | | | | |
|---|---|---|---|---|---|---|---|---|---|---|---|---|---|---|---|
| 33149 | ENSMUSG00000094012 | 0 | NA | NA | NA | NA | NA | Gm10024 | predicted gene 10024 [Source:MGI Symbo | 0 | 0 | 0 | 0 | 0 | 0 |
| 33150 | ENSMUSG00000094013 | 0 | NA | NA | NA | NA | NA | NA | NA | 0 | 0 | 0 | 0 | 0 | 0 |
| 33151 | ENSMUSG00000094015 | 0 | NA | NA | NA | NA | NA | Rps12-ps20 | ribosomal protein S12, pseudogene 20 [So | 0 | 0 | 0 | 0 | 0 | 0 |
| 33152 | ENSMUSG00000094016 | 0 | NA | NA | NA | NA | NA | Trav15-1-dv6 | T cell receptor alpha variable 15-1-DV6-1 [ | 0 | 0 | 0 | 0 | 0 | 0 |
| 33153 | ENSMUSG00000094017 | 0.321325949 | 1.911575114 | 4.048267893 | 0.472195804 | 0.636787034 | NA | Gm13160 | predicted gene 13160 [Source:MGI Symbo | 0 | 0 | 0 | 1.928 | 0 | 0 |
| 33154 | ENSMUSG00000094018 | 0 | NA | NA | NA | NA | NA | S100a2 | S100 calcium binding protein A2 [Source:M | 0 | 0 | 0 | 0 | 0 | 0 |
| 33155 | ENSMUSG00000094020 | 0 | NA | NA | NA | NA | NA | NA | NA | 0 | 0 | 0 | 0 | 0 | 0 |
| 33156 | ENSMUSG00000094021 | 0 | NA | NA | NA | NA | NA | Gm3685 | predicted gene 3685 [Source:MGI Symbol | 0 | 0 | 0 | 0 | 0 | 0 |
| 33157 | ENSMUSG00000094023 | 0 | NA | NA | NA | NA | NA | Trav7d-4 | T cell receptor alpha variable 7D-4 [Source | 0 | 0 | 0 | 0 | 0 | 0 |
| 33158 | ENSMUSG00000094024 | 0 | NA | NA | NA | NA | NA | Esp18 | exocrine gland secreted peptide 18 [Sourc | 0 | 0 | 0 | 0 | 0 | 0 |
| 33159 | ENSMUSG00000094025 | 0 | NA | NA | NA | NA | NA | Gm8879 | predicted gene 8879 [Source:MGI Symbol | 0 | 0 | 0 | 0 | 0 | 0 |
| 33160 | ENSMUSG00000094026 | 0 | NA | NA | NA | NA | NA | NA | NA | 0 | 0 | 0 | 0 | 0 | 0 |
| 33161 | ENSMUSG00000094027 | 0.160662974 | 1.126924175 | 4.080472857 | 0.276174898 | 0.782413736 | NA | Gm21762 | predicted gene, 21762 [Source:MGI Symb | 0 | 0 | 0 | 0.964 | 0 | 0 |
| 33162 | ENSMUSG00000094028 | 0 | NA | NA | NA | NA | NA | Ighd4-1 | immunoglobulin heavy diversity 4-1 [Sour | 0 | 0 | 0 | 0 | 0 | 0 |
| 33163 | ENSMUSG00000094029 | 0 | NA | NA | NA | NA | NA | Ighv3-3 | immunoglobulin heavy variable V3-3 [Sou | 0 | 0 | 0 | 0 | 0 | 0 |
| 33164 | ENSMUSG00000094030 | 0 | NA | NA | NA | NA | NA | Gm21833 | predicted gene, 21833 [Source:MGI Symb | 0 | 0 | 0 | 0 | 0 | 0 |
| 33165 | ENSMUSG00000094031 | 0 | NA | NA | NA | NA | NA | NA | NA | 0 | 0 | 0 | 0 | 0 | 0 |
| 33166 | ENSMUSG00000094032 | 0 | NA | NA | NA | NA | NA | NA | NA | 0 | 0 | 0 | 0 | 0 | 0 |
| 33167 | ENSMUSG00000094033 | 0 | NA | NA | NA | NA | NA | NA | NA | 0 | 0 | 0 | 0 | 0 | 0 |
| 33168 | ENSMUSG00000094034 | 0 | NA | NA | NA | NA | NA | NA | NA | 0 | 0 | 0 | 0 | 0 | 0 |
| 33169 | ENSMUSG00000094035 | 0 | NA | NA | NA | NA | NA | Ldlrad2 | low density lipoprotein receptor class A de | 0 | 0 | 0 | 0 | 0 | 0 |
| 33170 | ENSMUSG00000094036 | 0 | NA | NA | NA | NA | NA | Gm6465 | predicted gene 6465 [Source:MGI Symbol | 0 | 0 | 0 | 0 | 0 | 0 |
| 33171 | ENSMUSG00000094037 | 0 | NA | NA | NA | NA | NA | NA | NA | 0 | 0 | 0 | 0 | 0 | 0 |
| 33172 | ENSMUSG00000094038 | 0 | NA | NA | NA | NA | NA | NA | NA | 0 | 0 | 0 | 0 | 0 | 0 |
| 33173 | ENSMUSG00000094039 | 0 | NA | NA | NA | NA | NA | NA | NA | 0 | 0 | 0 | 0 | 0 | 0 |
| 33174 | ENSMUSG00000094041 | 0.17131964 | -0.796629176 | 4.080472857 | -0.195229623 | 0.84521319 | NA | NA | NA | 0 | 1.0279 | 0 | 0 | 0 | 0 |
| 33175 | ENSMUSG00000094043 | 0 | NA | NA | NA | NA | NA | Pramel49 | PRAME like 49 [Source:MGI Symbol;Acc:M | 0 | 0 | 0 | 0 | 0 | 0 |
| 33176 | ENSMUSG00000094044 | 0 | NA | NA | NA | NA | NA | NA | NA | 0 | 0 | 0 | 0 | 0 | 0 |
| 33177 | ENSMUSG00000094048 | 0 | NA | NA | NA | NA | NA | NA | NA | 0 | 0 | 0 | 0 | 0 | 0 |
| 33178 | ENSMUSG00000094049 | 0 | NA | NA | NA | NA | NA | NA | NA | 0 | 0 | 0 | 0 | 0 | 0 |
| 33179 | ENSMUSG00000094050 | 0 | NA | NA | NA | NA | NA | NA | NA | 0 | 0 | 0 | 0 | 0 | 0 |
| 33180 | ENSMUSG00000094051 | 0 | NA | NA | NA | NA | NA | Ighv1-36 | immunoglobulin heavy variable 1-36 [Sour | 0 | 0 | 0 | 0 | 0 | 0 |
| 33181 | ENSMUSG00000094052 | 0 | NA | NA | NA | NA | NA | Gm21118 | predicted gene, 21118 [Source:MGI Symb | 0 | 0 | 0 | 0 | 0 | 0 |
| 33182 | ENSMUSG00000094053 | 0 | NA | NA | NA | NA | NA | Scgb2b7 | secretoglobin, family 2B, member 7 [Sour | 0 | 0 | 0 | 0 | 0 | 0 |
| 33183 | ENSMUSG00000094054 | 0.17131964 | -0.796629176 | 4.080472857 | -0.195229623 | 0.84521319 | NA | | | 0 | 1.0279 | 0 | 0 | 0 | 0 |
| 33184 | ENSMUSG00000094057 | 0 | NA | NA | NA | NA | NA | Ighd2-7 | immunoglobulin heavy diversity 2-7 [Sour | 0 | 0 | 0 | 0 | 0 | 0 |
| 33185 | ENSMUSG00000094059 | 0 | NA | NA | NA | NA | NA | NA | NA | 0 | 0 | 0 | 0 | 0 | 0 |
| 33186 | ENSMUSG00000094060 | 0 | NA | NA | NA | NA | NA | NA | NA | 0 | 0 | 0 | 0 | 0 | 0 |
| 33187 | ENSMUSG00000094063 | 0 | NA | NA | NA | NA | NA | Olfr638 | olfactory receptor 638 [Source:MGI Symb | 0 | 0 | 0 | 0 | 0 | 0 |
| 33188 | ENSMUSG00000094064 | 0 | NA | NA | NA | NA | NA | Gm21626 | predicted gene, 21626 [Source:MGI Symb | 0 | 0 | 0 | 0 | 0 | 0 |
| 33189 | ENSMUSG00000094065 | 0 | NA | NA | NA | NA | NA | Ccl21d | chemokine (C-C motif) ligand 21D [Source | 0 | 0 | 0 | 0 | 0 | 0 |
| 33190 | ENSMUSG00000094066 | 0 | NA | NA | NA | NA | NA | Fam205a2 | family with sequence similarity 205, meml | 0 | 0 | 0 | 0 | 0 | 0 |
| 33191 | ENSMUSG00000094067 | 0 | NA | NA | NA | NA | NA | NA | NA | 0 | 0 | 0 | 0 | 0 | 0 |
| 33192 | ENSMUSG00000094068 | 0 | NA | NA | NA | NA | NA | NA | NA | 0 | 0 | 0 | 0 | 0 | 0 |
| 33193 | ENSMUSG00000094069 | 0 | NA | NA | NA | NA | NA | NA | NA | 0 | 0 | 0 | 0 | 0 | 0 |
| 33194 | ENSMUSG00000094071 | 0 | NA | NA | NA | NA | NA | Gm21729 | predicted gene, 21729 [Source:MGI Symb | 0 | 0 | 0 | 0 | 0 | 0 |
| 33195 | ENSMUSG00000094072 | 0 | NA | NA | NA | NA | NA | Gm11105 | predicted gene 11105 [Source:MGI Symbo | 0 | 0 | 0 | 0 | 0 | 0 |
| 33196 | ENSMUSG00000094074 | 0 | NA | NA | NA | NA | NA | Olfr1292-ps1 | olfactory receptor 1292, pseudogene 1 [So | 0 | 0 | 0 | 0 | 0 | 0 |
| 33197 | ENSMUSG00000094075 | 0 | NA | NA | NA | NA | NA | Ighv1-80 | immunoglobulin heavy variable 1-80 [Sour | 0 | 0 | 0 | 0 | 0 | 0 |
| 33198 | ENSMUSG00000094076 | 0.185640647 | 1.126924175 | 4.080472857 | 0.276174898 | 0.782413736 | NA | Gm4767 | predicted gene 4767 [Source:MGI Symbol | 0 | 0 | 0 | 0 | 0 | 1.1138 |
| 33199 | ENSMUSG00000094077 | 0 | NA | NA | NA | NA | NA | NA | NA | 0 | 0 | 0 | 0 | 0 | 0 |
| 33200 | ENSMUSG00000094079 | 0 | NA | NA | NA | NA | NA | Rpl23a-ps12 | ribosomal protein L23A, pseudogene 12 [S | 0 | 0 | 0 | 0 | 0 | 0 |
| 33201 | ENSMUSG00000094080 | 0 | NA | NA | NA | NA | NA | Olfr8 | olfactory receptor 8 [Source:MGI Symbol; | 0 | 0 | 0 | 0 | 0 | 0 |
| 33202 | ENSMUSG00000094081 | 0 | NA | NA | NA | NA | NA | Gm20826 | predicted gene, 20826 [Source:MGI Symb | 0 | 0 | 0 | 0 | 0 | 0 |
| 33203 | ENSMUSG00000094082 | 0 | NA | NA | NA | NA | NA | Gm12452 | predicted gene 12452 [Source:MGI Symbo | 0 | 0 | 0 | 0 | 0 | 0 |
| 33204 | ENSMUSG00000094083 | 0 | NA | NA | NA | NA | NA | Gm1604b | predicted gene 1604b [Source:MGI Symbo | 0 | 0 | 0 | 0 | 0 | 0 |

| | | | | | | | | | | | | | | | |
|---|---|---|---|---|---|---|---|---|---|---|---|---|---|---|---|
| 33205 | ENSMUSG00000094084 | 0 | NA | NA | NA | NA | NA | Tdpoz1 | TD and POZ domain containing 1 [Source: | 0 | 0 | 0 | 0 | 0 | 0 |
| 33206 | ENSMUSG00000094085 | 0 | NA | NA | NA | NA | NA | Vmn1r114 | vomeronasal 1 receptor 114 [Source:MGI | 0 | 0 | 0 | 0 | 0 | 0 |
| 33207 | ENSMUSG00000094086 | 0 | NA | NA | NA | NA | NA | NA | NA | 0 | 0 | 0 | 0 | 0 | 0 |
| 33208 | ENSMUSG00000094087 | 0.167707431 | -0.796629176 | 4.080472857 | -0.195229623 | 0.84521319 | NA | Ighv1-61 | immunoglobulin heavy variable 1-61 [Sou | 0 | 0 | 1.0062 | 0 | 0 | 0 |
| 33209 | ENSMUSG00000094088 | 0.480153995 | 1.043847617 | 3.151673025 | 0.331204287 | 0.740490182 | NA | Ighv1-64 | immunoglobulin heavy variable 1-64 [Sou | 0.8031 | 0 | 0 | 0.964 | 0 | 1.1138 |
| 33210 | ENSMUSG00000094090 | 1.507505322 | -1.650924302 | 2.043553546 | -0.807869363 | 0.41916579 | NA | Gm8494 | predicted gene 8494 [Source:MGI Symbol | 0.8031 | 0 | 6.0375 | 0 | 1.0906 | 1.1138 |
| 33211 | ENSMUSG00000094091 | 21.62927789 | -0.633393097 | 0.604341053 | -1.048072266 | 0.294605313 | 0.745164683 | Gm21885 | predicted gene, 21885 [Source:MGI Symb | 48.989 | 16.447 | 13.081 | 10.604 | 26.175 | 14.48 |
| 33212 | ENSMUSG00000094092 | 1.714251055 | -0.542310995 | 1.715672439 | -0.316092386 | 0.751932386 | NA | Gm15921 | predicted gene 15921 [Source:MGI Symbo | 4.0155 | 1.0279 | 1.0062 | 0.964 | 3.2719 | 0 |
| 33213 | ENSMUSG00000094093 | 0 | NA | NA | NA | NA | NA | NA | NA | 0 | 0 | 0 | 0 | 0 | 0 |
| 33214 | ENSMUSG00000094094 | 0.36353943 | 2.065989254 | 4.034137301 | 0.512126658 | 0.608562368 | NA | Igkv5-45 | immunoglobulin kappa chain variable 5-4 | 0 | 0 | 0 | 0 | 2.1812 | 0 |
| 33215 | ENSMUSG00000094095 | 0 | NA | NA | NA | NA | NA | NA | NA | 0 | 0 | 0 | 0 | 0 | 0 |
| 33216 | ENSMUSG00000094096 | 0 | NA | NA | NA | NA | NA | NA | NA | 0 | 0 | 0 | 0 | 0 | 0 |
| 33217 | ENSMUSG00000094097 | 0 | NA | NA | NA | NA | NA | NA | NA | 0 | 0 | 0 | 0 | 0 | 0 |
| 33218 | ENSMUSG00000094098 | 0 | NA | NA | NA | NA | NA | Vmn2r44 | vomeronasal 2, receptor 44 [Source:MGI | 0 | 0 | 0 | 0 | 0 | 0 |
| 33219 | ENSMUSG00000094099 | 0 | NA | NA | NA | NA | NA | NA | NA | 0 | 0 | 0 | 0 | 0 | 0 |
| 33220 | ENSMUSG00000094101 | 0 | NA | NA | NA | NA | NA | NA | NA | 0 | 0 | 0 | 0 | 0 | 0 |
| 33221 | ENSMUSG00000094102 | 0 | NA | NA | NA | NA | NA | Ighv9-2 | immunoglobulin heavy variable V9-2 [Sou | 0 | 0 | 0 | 0 | 0 | 0 |
| 33222 | ENSMUSG00000094103 | 0 | NA | NA | NA | NA | NA | Fam177a2 | family with sequence similarity 177 memb | 0 | 0 | 0 | 0 | 0 | 0 |
| 33223 | ENSMUSG00000094105 | 0 | NA | NA | NA | NA | NA | Gm8922 | predicted gene 8922 [Source:MGI Symbol | 0 | 0 | 0 | 0 | 0 | 0 |
| 33224 | ENSMUSG00000094107 | 0 | NA | NA | NA | NA | NA | NA | NA | 0 | 0 | 0 | 0 | 0 | 0 |
| 33225 | ENSMUSG00000094108 | 0 | NA | NA | NA | NA | NA | Gm17258 | predicted gene, 17258 [Source:MGI Symb | 0 | 0 | 0 | 0 | 0 | 0 |
| 33226 | ENSMUSG00000094109 | 0 | NA | NA | NA | NA | NA | NA | NA | 0 | 0 | 0 | 0 | 0 | 0 |
| 33227 | ENSMUSG00000094111 | 0 | NA | NA | NA | NA | NA | Nutf2-ps2 | nuclear transport factor 2, pseudogene 2 | 0 | 0 | 0 | 0 | 0 | 0 |
| 33228 | ENSMUSG00000094112 | 0 | NA | NA | NA | NA | NA | 9330182O14 | RIKEN cDNA 9330182O14 gene [Source:M | 0 | 0 | 0 | 0 | 0 | 0 |
| 33229 | ENSMUSG00000094113 | 0 | NA | NA | NA | NA | NA | 5430401F13 | RIKEN cDNA 5430401F13 gene [Source:M | 0 | 0 | 0 | 0 | 0 | 0 |
| 33230 | ENSMUSG00000094115 | 0 | NA | NA | NA | NA | NA | Spin2-ps6 | spindlin family, member 2, pseudogene 6 | 0 | 0 | 0 | 0 | 0 | 0 |
| 33231 | ENSMUSG00000094116 | 0 | NA | NA | NA | NA | NA | Gm11237 | predicted gene 11237 [Source:MGI Symbo | 0 | 0 | 0 | 0 | 0 | 0 |
| 33232 | ENSMUSG00000094117 | 0 | NA | NA | NA | NA | NA | Igkv3-12 | immunoglobulin kappa variable 3-12 [Sou | 0 | 0 | 0 | 0 | 0 | 0 |
| 33233 | ENSMUSG00000094118 | 0 | NA | NA | NA | NA | NA | Gm16445 | predicted pseudogene 16445 [Source:MG | 0 | 0 | 0 | 0 | 0 | 0 |
| 33234 | ENSMUSG00000094119 | 0 | NA | NA | NA | NA | NA | Olfr612 | olfactory receptor 612 [Source:MGI Symb | 0 | 0 | 0 | 0 | 0 | 0 |
| 33235 | ENSMUSG00000094120 | 0 | NA | NA | NA | NA | NA | Gm3233 | predicted gene 3233 [Source:MGI Symbol | 0 | 0 | 0 | 0 | 0 | 0 |
| 33236 | ENSMUSG00000094121 | 0 | NA | NA | NA | NA | NA | NA | NA | 0 | 0 | 0 | 0 | 0 | 0 |
| 33237 | ENSMUSG00000094122 | 0.133850373 | -0.796629176 | 4.080472857 | -0.195229623 | 0.84521319 | NA | Rpl31-ps9 | ribosomal protein L31, pseudogene 9 [Sou | 0.8031 | 0 | 0 | 0 | 0 | 0 |
| 33238 | ENSMUSG00000094123 | 0 | NA | NA | NA | NA | NA | NA | NA | 0 | 0 | 0 | 0 | 0 | 0 |
| 33239 | ENSMUSG00000094124 | 0.315620088 | 0.165143421 | 4.063300985 | 0.040642675 | 0.967580763 | NA | Ighv1-74 | immunoglobulin heavy variable V1-74 [So | 0.8031 | 0 | 0 | 0 | 1.0906 | 0 |
| 33240 | ENSMUSG00000094125 | 0 | NA | NA | NA | NA | NA | Gm13698 | predicted gene 13698 [Source:MGI Symbo | 0 | 0 | 0 | 0 | 0 | 0 |
| 33241 | ENSMUSG00000094127 | 0 | NA | NA | NA | NA | NA | G530012D18 | RIKEN cDNA G530012D1 gene [Source:M | 0 | 0 | 0 | 0 | 0 | 0 |
| 33242 | ENSMUSG00000094128 | 0 | NA | NA | NA | NA | NA | NA | NA | 0 | 0 | 0 | 0 | 0 | 0 |
| 33243 | ENSMUSG00000094129 | 0 | NA | NA | NA | NA | NA | NA | NA | 0 | 0 | 0 | 0 | 0 | 0 |
| 33244 | ENSMUSG00000094131 | 0 | NA | NA | NA | NA | NA | NA | NA | 0 | 0 | 0 | 0 | 0 | 0 |
| 33245 | ENSMUSG00000094132 | 0 | NA | NA | NA | NA | NA | Gm3264 | predicted gene 3264 [Source:MGI Symbol | 0 | 0 | 0 | 0 | 0 | 0 |
| 33246 | ENSMUSG00000094133 | 0 | NA | NA | NA | NA | NA | Olfr1431 | olfactory receptor 1431 [Source:MGI Sym | 0 | 0 | 0 | 0 | 0 | 0 |
| 33247 | ENSMUSG00000094134 | 0 | NA | NA | NA | NA | NA | Ighv5-15 | immunoglobulin heavy variable 5-15 [Sou | 0 | 0 | 0 | 0 | 0 | 0 |
| 33248 | ENSMUSG00000094135 | 0 | NA | NA | NA | NA | NA | Gm21379 | predicted gene, 21379 [Source:MGI Symb | 0 | 0 | 0 | 0 | 0 | 0 |
| 33249 | ENSMUSG00000094136 | 0 | NA | NA | NA | NA | NA | Gm5973 | predicted gene 5973 [Source:MGI Symbol | 0 | 0 | 0 | 0 | 0 | 0 |
| 33250 | ENSMUSG00000094137 | 0 | NA | NA | NA | NA | NA | NA | NA | 0 | 0 | 0 | 0 | 0 | 0 |
| 33251 | ENSMUSG00000094139 | 0 | NA | NA | NA | NA | NA | NA | NA | 0 | 0 | 0 | 0 | 0 | 0 |
| 33252 | ENSMUSG00000094140 | 0 | NA | NA | NA | NA | NA | Olfr1512 | olfactory receptor 1512 [Source:MGI Sym | 0 | 0 | 0 | 0 | 0 | 0 |
| 33253 | ENSMUSG00000094144 | 0 | NA | NA | NA | NA | NA | Gm21312 | predicted gene, 21312 [Source:MGI Symb | 0 | 0 | 0 | 0 | 0 | 0 |
| 33254 | ENSMUSG00000094145 | 74.25923865 | 0.44809653 | 0.323787156 | 1.383923116 | 0.166381998 | 0.5925379 | Vmn2r20 | vomeronasal 2, receptor 20 [Source:MGI | 73.082 | 52.424 | 62.387 | 53.019 | 118.88 | 85.766 |
| 33255 | ENSMUSG00000094146 | 0 | NA | NA | NA | NA | NA | Gm10142 | predicted gene 10142 [Source:MGI Symbo | 0 | 0 | 0 | 0 | 0 | 0 |
| 33256 | ENSMUSG00000094147 | 0 | NA | NA | NA | NA | NA | NA | NA | 0 | 0 | 0 | 0 | 0 | 0 |
| 33257 | ENSMUSG00000094148 | 0 | NA | NA | NA | NA | NA | NA | NA | 0 | 0 | 0 | 0 | 0 | 0 |
| 33258 | ENSMUSG00000094149 | 0 | NA | NA | NA | NA | NA | Vmn1r252 | vomeronasal 1 receptor 252 [Source:MGI | 0 | 0 | 0 | 0 | 0 | 0 |
| 33259 | ENSMUSG00000094150 | 0 | NA | NA | NA | NA | NA | NA | NA | 0 | 0 | 0 | 0 | 0 | 0 |
| 33260 | ENSMUSG00000094151 | 0 | NA | NA | NA | NA | NA | Gm7233 | predicted gene 7233 [Source:MGI Symbol | 0 | 0 | 0 | 0 | 0 | 0 |

| | | | | | | | | | | | | | | | |
|---|---|---|---|---|---|---|---|---|---|---|---|---|---|---|---|
| 33261 | ENSMUSG00000094152 | 60.43458022 | 0.427474417 | 0.304734757 | 1.402775388 | 0.160683829 | 0.583512653 | Slc6a16 | solute carrier family 6, member 16 [Sourc | 57.823 | 41.117 | 55.343 | 59.767 | 85.068 | 63.489 |
| 33262 | ENSMUSG00000094153 | 0 | NA | NA | NA | NA | NA | Gm8807 | predicted gene 8807 [Source:MGI Symbol | 0 | 0 | 0 | 0 | 0 | 0 |
| 33263 | ENSMUSG00000094154 | 0 | NA | NA | NA | NA | NA | NA | NA | 0 | 0 | 0 | 0 | 0 | 0 |
| 33264 | ENSMUSG00000094155 | 0 | NA | NA | NA | NA | NA | NA | NA | 0 | 0 | 0 | 0 | 0 | 0 |
| 33265 | ENSMUSG00000094156 | 24.18474741 | -0.338599012 | 0.442526761 | -0.765149233 | 0.444182686 | 0.851512043 | Sult2a7 | sulfotransferase family 2A, dehydroepian | 28.109 | 27.754 | 25.156 | 21.208 | 26.175 | 16.708 |
| 33266 | ENSMUSG00000094157 | 0 | NA | NA | NA | NA | NA | Gm7995 | predicted gene 7995 [Source:MGI Symbol | 0 | 0 | 0 | 0 | 0 | 0 |
| 33267 | ENSMUSG00000094158 | 2.240417868 | -0.159192697 | 1.662364035 | -0.095762837 | 0.923708933 | NA | NA | NA | 0 | 5.1396 | 2.0125 | 1.928 | 4.3625 | 0 |
| 33268 | ENSMUSG00000094159 | 0 | NA | NA | NA | NA | NA | NA | NA | 0 | 0 | 0 | 0 | 0 | 0 |
| 33269 | ENSMUSG00000094160 | 0 | NA | NA | NA | NA | NA | Vmn1r-ps79 | vomeronasal 1 receptor, pseudogene 79 [ | 0 | 0 | 0 | 0 | 0 | 0 |
| 33270 | ENSMUSG00000094161 | 0 | NA | NA | NA | NA | NA | Gm21904 | predicted gene, 21904 [Source:MGI Symb | 0 | 0 | 0 | 0 | 0 | 0 |
| 33271 | ENSMUSG00000094162 | 0 | NA | NA | NA | NA | NA | Gm2457 | predicted gene, 2457 [Source:MGI Symbo | 0 | 0 | 0 | 0 | 0 | 0 |
| 33272 | ENSMUSG00000094163 | 0 | NA | NA | NA | NA | NA | Tdpoz5 | TD and POZ domain containing 5 [Source: | 0 | 0 | 0 | 0 | 0 | 0 |
| 33273 | ENSMUSG00000094164 | 0.301557804 | -1.64335916 | 4.042814367 | -0.406488899 | 0.684383412 | NA | Ighv2-3 | immunoglobulin heavy variable 2-3 [Sourc | 0.8031 | 0 | 1.0062 | 0 | 0 | 0 |
| 33274 | ENSMUSG00000094165 | 0 | NA | NA | NA | NA | NA | NA | NA | 0 | 0 | 0 | 0 | 0 | 0 |
| 33275 | ENSMUSG00000094167 | 0 | NA | NA | NA | NA | NA | Gm10580 | predicted gene 10580 [Source:MGI Symbo | 0 | 0 | 0 | 0 | 0 | 0 |
| 33276 | ENSMUSG00000094168 | 0 | NA | NA | NA | NA | NA | Gm21903 | predicted gene, 21903 [Source:MGI Symb | 0 | 0 | 0 | 0 | 0 | 0 |
| 33277 | ENSMUSG00000094170 | 0 | NA | NA | NA | NA | NA | NA | NA | 0 | 0 | 0 | 0 | 0 | 0 |
| 33278 | ENSMUSG00000094172 | 0 | NA | NA | NA | NA | NA | | | 0 | 0 | 0 | 0 | 0 | 0 |
| 33279 | ENSMUSG00000094173 | 0 | NA | NA | NA | NA | NA | NA | NA | 0 | 0 | 0 | 0 | 0 | 0 |
| 33280 | ENSMUSG00000094174 | 0.133850373 | -0.796629176 | 4.080472857 | -0.195229623 | 0.84521319 | NA | Ighv6-4 | immunoglobulin heavy variable V6-4 [Sou | 0.8031 | 0 | 0 | 0 | 0 | 0 |
| 33281 | ENSMUSG00000094175 | 0 | NA | NA | NA | NA | NA | NA | NA | 0 | 0 | 0 | 0 | 0 | 0 |
| 33282 | ENSMUSG00000094176 | 0 | NA | NA | NA | NA | NA | Trav6d-6 | T cell receptor alpha variable 6D-6 [Sourc | 0 | 0 | 0 | 0 | 0 | 0 |
| 33283 | ENSMUSG00000094177 | 0 | NA | NA | NA | NA | NA | Vmn2r-ps15 | vomeronasal 2, receptor, pseudogene 15 | 0 | 0 | 0 | 0 | 0 | 0 |
| 33284 | ENSMUSG00000094178 | 0 | NA | NA | NA | NA | NA | NA | NA | 0 | 0 | 0 | 0 | 0 | 0 |
| 33285 | ENSMUSG00000094179 | 0 | NA | NA | NA | NA | NA | NA | NA | 0 | 0 | 0 | 0 | 0 | 0 |
| 33286 | ENSMUSG00000094180 | 0 | NA | NA | NA | NA | NA | Defa-ps8 | defensin, alpha, pseudogene 8 [Source:M | 0 | 0 | 0 | 0 | 0 | 0 |
| 33287 | ENSMUSG00000094181 | 0 | NA | NA | NA | NA | NA | Gm21739 | predicted gene, 21739 [Source:MGI Symb | 0 | 0 | 0 | 0 | 0 | 0 |
| 33288 | ENSMUSG00000094182 | 0 | NA | NA | NA | NA | NA | Olfr937 | olfactory receptor 937 [Source:MGI Symb | 0 | 0 | 0 | 0 | 0 | 0 |
| 33289 | ENSMUSG00000094183 | 0 | NA | NA | NA | NA | NA | NA | NA | 0 | 0 | 0 | 0 | 0 | 0 |
| 33290 | ENSMUSG00000094184 | 0 | NA | NA | NA | NA | NA | NA | NA | 0 | 0 | 0 | 0 | 0 | 0 |
| 33291 | ENSMUSG00000094185 | 0 | NA | NA | NA | NA | NA | Gm5445 | predicted gene 5445 [Source:MGI Symbol | 0 | 0 | 0 | 0 | 0 | 0 |
| 33292 | ENSMUSG00000094186 | 0 | NA | NA | NA | NA | NA | Gm1553 | predicted gene 1553 [Source:MGI Symbol | 0 | 0 | 0 | 0 | 0 | 0 |
| 33293 | ENSMUSG00000094187 | 0 | NA | NA | NA | NA | NA | Vmn1r170 | vomeronasal 1 receptor 170 [Source:MGI | 0 | 0 | 0 | 0 | 0 | 0 |
| 33294 | ENSMUSG00000094188 | 0 | NA | NA | NA | NA | NA | NA | NA | 0 | 0 | 0 | 0 | 0 | 0 |
| 33295 | ENSMUSG00000094189 | 0 | NA | NA | NA | NA | NA | Gm6134 | predicted pseudogene 6134 [Source:MGI | 0 | 0 | 0 | 0 | 0 | 0 |
| 33296 | ENSMUSG00000094190 | 0 | NA | NA | NA | NA | NA | NA | NA | 0 | 0 | 0 | 0 | 0 | 0 |
| 33297 | ENSMUSG00000094191 | 0 | NA | NA | NA | NA | NA | Rpl31-ps14 | ribosomal protein L31, pseudogene 14 [Sc | 0 | 0 | 0 | 0 | 0 | 0 |
| 33298 | ENSMUSG00000094192 | 0 | NA | NA | NA | NA | NA | Olfr452 | olfactory receptor 452 [Source:MGI Symb | 0 | 0 | 0 | 0 | 0 | 0 |
| 33299 | ENSMUSG00000094194 | 0 | NA | NA | NA | NA | NA | Ighv5-16 | immunoglobulin heavy variable 5-16 [Sou | 0 | 0 | 0 | 0 | 0 | 0 |
| 33300 | ENSMUSG00000094195 | 0 | NA | NA | NA | NA | NA | Pramel44 | PRAME like 44 [Source:MGI Symbol;Acc:M | 0 | 0 | 0 | 0 | 0 | 0 |
| 33301 | ENSMUSG00000094196 | 0 | NA | NA | NA | NA | NA | Magea3 | MAGE family member A3 [Source:MGI Syr | 0 | 0 | 0 | 0 | 0 | 0 |
| 33302 | ENSMUSG00000094197 | 0 | NA | NA | NA | NA | NA | Olfr474 | olfactory receptor 474 [Source:MGI Symb | 0 | 0 | 0 | 0 | 0 | 0 |
| 33303 | ENSMUSG00000094198 | 0 | NA | NA | NA | NA | NA | Ighv1-50 | immunoglobulin heavy variable 1-50 [Sou | 0 | 0 | 0 | 0 | 0 | 0 |
| 33304 | ENSMUSG00000094199 | 0 | NA | NA | NA | NA | NA | NA | NA | 0 | 0 | 0 | 0 | 0 | 0 |
| 33305 | ENSMUSG00000094200 | 0 | NA | NA | NA | NA | NA | Olfr237 | olfactory receptor 237 [Source:MGI Symb | 0 | 0 | 0 | 0 | 0 | 0 |
| 33306 | ENSMUSG00000094201 | 0 | NA | NA | NA | NA | NA | NA | NA | 0 | 0 | 0 | 0 | 0 | 0 |
| 33307 | ENSMUSG00000094202 | 0 | NA | NA | NA | NA | NA | Mup-ps26 | major urinary protein, pseudogene 26 [So | 0 | 0 | 0 | 0 | 0 | 0 |
| 33308 | ENSMUSG00000094203 | 0 | NA | NA | NA | NA | NA | NA | NA | 0 | 0 | 0 | 0 | 0 | 0 |
| 33309 | ENSMUSG00000094204 | 0 | NA | NA | NA | NA | NA | Gm21915 | predicted gene, 21915 [Source:MGI Symb | 0 | 0 | 0 | 0 | 0 | 0 |
| 33310 | ENSMUSG00000094205 | 0 | NA | NA | NA | NA | NA | Gm8926 | predicted gene 8926 [Source:MGI Symbol | 0 | 0 | 0 | 0 | 0 | 0 |
| 33311 | ENSMUSG00000094208 | 0 | NA | NA | NA | NA | NA | Vmn1r130 | vomeronasal 1 receptor 130 [Source:MGI | 0 | 0 | 0 | 0 | 0 | 0 |
| 33312 | ENSMUSG00000094210 | 0 | NA | NA | NA | NA | NA | Gm23450 | predicted gene, 23450 [Source:MGI Symb | 0 | 0 | 0 | 0 | 0 | 0 |
| 33313 | ENSMUSG00000094211 | 0 | NA | NA | NA | NA | NA | NA | NA | 0 | 0 | 0 | 0 | 0 | 0 |
| 33314 | ENSMUSG00000094212 | 0 | NA | NA | NA | NA | NA | Trav14-3 | T cell receptor alpha variable 14-3 [Source | 0 | 0 | 0 | 0 | 0 | 0 |
| 33315 | ENSMUSG00000094213 | 0 | NA | NA | NA | NA | NA | NA | NA | 0 | 0 | 0 | 0 | 0 | 0 |
| 33316 | ENSMUSG00000094214 | 0 | NA | NA | NA | NA | NA | NA | NA | 0 | 0 | 0 | 0 | 0 | 0 |

| | | | | | | | | | | | | | | | |
|---|---|---|---|---|---|---|---|---|---|---|---|---|---|---|---|
| 33317 | ENSMUSG00000094215 | 0 | NA | NA | NA | NA | NA | Gm6489 | predicted gene 6489 [Source:MGI Symbol | 0 | 0 | 0 | 0 | 0 | 0 |
| 33318 | ENSMUSG00000094217 | 0 | NA | NA | NA | NA | NA | Gm20794 | predicted gene, 20794 [Source:MGI Symb | 0 | 0 | 0 | 0 | 0 | 0 |
| 33319 | ENSMUSG00000094218 | 0 | NA | NA | NA | NA | NA | NA | NA | 0 | 0 | 0 | 0 | 0 | 0 |
| 33320 | ENSMUSG00000094219 | 0 | NA | NA | NA | NA | NA | Calhm3 | calcium homeostasis modulator 3 [Source | 0 | 0 | 0 | 0 | 0 | 0 |
| 33321 | ENSMUSG00000094220 | 0 | NA | NA | NA | NA | NA | Trav6-3 | T cell receptor alpha variable 6-3 [Source: | 0 | 0 | 0 | 0 | 0 | 0 |
| 33322 | ENSMUSG00000094221 | 0 | NA | NA | NA | NA | NA | Vmn1r124 | vomeronasal 1 receptor 124 [Source:MGI | 0 | 0 | 0 | 0 | 0 | 0 |
| 33323 | ENSMUSG00000094222 | 7.710482624 | 1.960846466 | 0.887435323 | 2.209565492 | 0.027135332 | 0.21354341 | Mup-ps6 | major urinary protein, pseudogene 6 [Sou | 2.4093 | 3.0838 | 4.025 | 11.568 | 20.722 | 4.4554 |
| 33324 | ENSMUSG00000094226 | 0 | NA | NA | NA | NA | NA | Gm21759 | predicted gene, 21759 [Source:MGI Symb | 0 | 0 | 0 | 0 | 0 | 0 |
| 33325 | ENSMUSG00000094227 | 0 | NA | NA | NA | NA | NA | Gm21767 | predicted gene, 21767 [Source:MGI Symb | 0 | 0 | 0 | 0 | 0 | 0 |
| 33326 | ENSMUSG00000094229 | 0 | NA | NA | NA | NA | NA | NA | NA | 0 | 0 | 0 | 0 | 0 | 0 |
| 33327 | ENSMUSG00000094230 | 0 | NA | NA | NA | NA | NA | Gm21847 | predicted gene, 21847 [Source:MGI Symb | 0 | 0 | 0 | 0 | 0 | 0 |
| 33328 | ENSMUSG00000094231 | 0 | NA | NA | NA | NA | NA | Trav9n-4 | T cell receptor alpha variable 9N-4 [Source | 0 | 0 | 0 | 0 | 0 | 0 |
| 33329 | ENSMUSG00000094232 | 0 | NA | NA | NA | NA | NA | Gm21539 | predicted gene, 21539 [Source:MGI Symb | 0 | 0 | 0 | 0 | 0 | 0 |
| 33330 | ENSMUSG00000094233 | 0 | NA | NA | NA | NA | NA | Gm21276 | predicted gene, 21276 [Source:MGI Symb | 0 | 0 | 0 | 0 | 0 | 0 |
| 33331 | ENSMUSG00000094235 | 0 | NA | NA | NA | NA | NA | Mup-ps24 | major urinary protein, pseudogene 24 [So | 0 | 0 | 0 | 0 | 0 | 0 |
| 33332 | ENSMUSG00000094236 | 0 | NA | NA | NA | NA | NA | NA | NA | 0 | 0 | 0 | 0 | 0 | 0 |
| 33333 | ENSMUSG00000094237 | 0 | NA | NA | NA | NA | NA | AF067063 | cDNA sequence AF067063 [Source:MGI Sy | 0 | 0 | 0 | 0 | 0 | 0 |
| 33334 | ENSMUSG00000094238 | 0 | NA | NA | NA | NA | NA | Rps12-ps11 | ribosomal protein S12, pseudogene 11 [Sc | 0 | 0 | 0 | 0 | 0 | 0 |
| 33335 | ENSMUSG00000094239 | 0 | NA | NA | NA | NA | NA | NA | NA | 0 | 0 | 0 | 0 | 0 | 0 |
| 33336 | ENSMUSG00000094240 | 0 | NA | NA | NA | NA | NA | Gm14348 | predicted gene 14348 [Source:MGI Symbc | 0 | 0 | 0 | 0 | 0 | 0 |
| 33337 | ENSMUSG00000094241 | 0 | NA | NA | NA | NA | NA | Ighv1-24 | immunoglobulin heavy variable V1-24 [So | 0 | 0 | 0 | 0 | 0 | 0 |
| 33338 | ENSMUSG00000094242 | 0.856211648 | -1.81932939 | 2.591956202 | -0.701913631 | 0.482733027 | NA | Gm5456 | predicted gene 5456 [Source:MGI Symbol | 0 | 1.0279 | 3.0187 | 0 | 1.0906 | 0 |
| 33339 | ENSMUSG00000094243 | 0 | NA | NA | NA | NA | NA | Obox4-ps22 | oocyte specific homeobox 4, pseudogene | 0 | 0 | 0 | 0 | 0 | 0 |
| 33340 | ENSMUSG00000094244 | 0 | NA | NA | NA | NA | NA | NA | NA | 0 | 0 | 0 | 0 | 0 | 0 |
| 33341 | ENSMUSG00000094245 | 0.77650227 | 1.950354251 | 2.938137078 | 0.663806418 | 0.506814217 | NA | NA | NA | 0.8031 | 0 | 0 | 3.8559 | 0 | 0 |
| 33342 | ENSMUSG00000094247 | 0 | NA | NA | NA | NA | NA | NA | NA | 0 | 0 | 0 | 0 | 0 | 0 |
| 33343 | ENSMUSG00000094248 | 0 | NA | NA | NA | NA | NA | H2ac23 | H2A clustered histone 23 [Source:MGI Syr | 0 | 0 | 0 | 0 | 0 | 0 |
| 33344 | ENSMUSG00000094249 | 0 | NA | NA | NA | NA | NA | Gm20091 | predicted gene, 20091 [Source:MGI Symb | 0 | 0 | 0 | 0 | 0 | 0 |
| 33345 | ENSMUSG00000094250 | 0 | NA | NA | NA | NA | NA | Gm4479 | predicted gene 4479 [Source:MGI Symbol | 0 | 0 | 0 | 0 | 0 | 0 |
| 33346 | ENSMUSG00000094251 | 1.01026885 | 1.016270523 | 2.426385186 | 0.418841381 | 0.67533206 | NA | NA | NA | 0 | 2.0558 | 0 | 2.8919 | 0 | 1.1138 |
| 33347 | ENSMUSG00000094252 | 0 | NA | NA | NA | NA | NA | NA | NA | 0 | 0 | 0 | 0 | 0 | 0 |
| 33348 | ENSMUSG00000094253 | 0 | NA | NA | NA | NA | NA | Mir3471-2 | microRNA 3471-2 [Source:MGI Symbol;Ac | 0 | 0 | 0 | 0 | 0 | 0 |
| 33349 | ENSMUSG00000094254 | 0 | NA | NA | NA | NA | NA | Olfr969 | olfactory receptor 969 [Source:MGI Symb | 0 | 0 | 0 | 0 | 0 | 0 |
| 33350 | ENSMUSG00000094255 | 0 | NA | NA | NA | NA | NA | NA | NA | 0 | 0 | 0 | 0 | 0 | 0 |
| 33351 | ENSMUSG00000094256 | 0.492645588 | 0.9565034 | 3.447648281 | 0.277436479 | 0.781444974 | NA | Rpl21-ps7 | ribosomal protein L21, pseudogene 7 [Sou | 0 | 1.0279 | 0 | 1.928 | 0 | 0 |
| 33352 | ENSMUSG00000094257 | 2.000098776 | -2.240993541 | 1.726837083 | -1.297744624 | 0.194375104 | NA | Ap3s1-ps2 | adaptor-related protein complex 3, sigma | 0.8031 | 4.1117 | 5.0312 | 0.964 | 1.0906 | 0 |
| 33353 | ENSMUSG00000094258 | 0 | NA | NA | NA | NA | NA | Gm8159 | predicted gene 8159 [Source:MGI Symbol | 0 | 0 | 0 | 0 | 0 | 0 |
| 33354 | ENSMUSG00000094259 | 0 | NA | NA | NA | NA | NA | Gm14536 | predicted gene 14536 [Source:MGI Symbc | 0 | 0 | 0 | 0 | 0 | 0 |
| 33355 | ENSMUSG00000094261 | 0 | NA | NA | NA | NA | NA | NA | NA | 0 | 0 | 0 | 0 | 0 | 0 |
| 33356 | ENSMUSG00000094262 | 0 | NA | NA | NA | NA | NA | Igkv4-62 | immunoglobulin kappa variable 4-62 [Sou | 0 | 0 | 0 | 0 | 0 | 0 |
| 33357 | ENSMUSG00000094263 | 0 | NA | NA | NA | NA | NA | NA | NA | 0 | 0 | 0 | 0 | 0 | 0 |
| 33358 | ENSMUSG00000094264 | 0 | NA | NA | NA | NA | NA | Gm11320 | predicted gene 11320 [Source:MGI Symbc | 0 | 0 | 0 | 0 | 0 | 0 |
| 33359 | ENSMUSG00000094265 | 0 | NA | NA | NA | NA | NA | NA | NA | 0 | 0 | 0 | 0 | 0 | 0 |
| 33360 | ENSMUSG00000094266 | 0 | NA | NA | NA | NA | NA | Olfr340 | olfactory receptor 340 [Source:MGI Symb | 0 | 0 | 0 | 0 | 0 | 0 |
| 33361 | ENSMUSG00000094267 | 0 | NA | NA | NA | NA | NA | NA | NA | 0 | 0 | 0 | 0 | 0 | 0 |
| 33362 | ENSMUSG00000094268 | 0 | NA | NA | NA | NA | NA | Ighd5-8 | immunoglobulin heavy diversity 5-8 [Sour | 0 | 0 | 0 | 0 | 0 | 0 |
| 33363 | ENSMUSG00000094269 | 0 | NA | NA | NA | NA | NA | Olfr951 | olfactory receptor 951 [Source:MGI Symb | 0 | 0 | 0 | 0 | 0 | 0 |
| 33364 | ENSMUSG00000094270 | 0 | NA | NA | NA | NA | NA | NA | NA | 0 | 0 | 0 | 0 | 0 | 0 |
| 33365 | ENSMUSG00000094271 | 0 | NA | NA | NA | NA | NA | Gm13290 | predicted gene 13290 [Source:MGI Symbc | 0 | 0 | 0 | 0 | 0 | 0 |
| 33366 | ENSMUSG00000094273 | 0 | NA | NA | NA | NA | NA | Btbd35f17 | BTB domain containing 35, family membe | 0 | 0 | 0 | 0 | 0 | 0 |
| 33367 | ENSMUSG00000094274 | 0 | NA | NA | NA | NA | NA | NA | NA | 0 | 0 | 0 | 0 | 0 | 0 |
| 33368 | ENSMUSG00000094275 | 0 | NA | NA | NA | NA | NA | NA | NA | 0 | 0 | 0 | 0 | 0 | 0 |
| 33369 | ENSMUSG00000094276 | 0 | NA | NA | NA | NA | NA | Gm15068 | predicted gene 15068 [Source:MGI Symbc | 0 | 0 | 0 | 0 | 0 | 0 |
| 33370 | ENSMUSG00000094277 | 0 | NA | NA | NA | NA | NA | Gm21726 | predicted gene, 21726 [Source:MGI Symb | 0 | 0 | 0 | 0 | 0 | 0 |
| 33371 | ENSMUSG00000094278 | 0 | NA | NA | NA | NA | NA | Gm12626 | predicted gene 12626 [Source:MGI Symbc | 0 | 0 | 0 | 0 | 0 | 0 |
| 33372 | ENSMUSG00000094280 | 0 | NA | NA | NA | NA | NA | Vmn2r-ps51 | vomeronasal 2, receptor, pseudogene 51 | 0 | 0 | 0 | 0 | 0 | 0 |

| | | | | | | | | | | | | | | | |
|---|---|---|---|---|---|---|---|---|---|---|---|---|---|---|---|
| 33373 | ENSMUSG00000094281 | 8.87922209 | -0.302396935 | 0.748931633 | -0.403771081 | 0.686381064 | 0.936498039 | Gm8034 | predicted gene 8034 [Source:MGI Symbol | 16.062 | 7.1954 | 6.0375 | 9.6398 | 6.5437 | 7.7969 |
| 33374 | ENSMUSG00000094282 | 0 | NA | NA | NA | NA | NA | Cfap73 | cilia and flagella associated protein 73 [So | 0 | 0 | 0 | 0 | 0 | 0 |
| 33375 | ENSMUSG00000094283 | 0 | NA | NA | NA | NA | NA | NA | NA | 0 | 0 | 0 | 0 | 0 | 0 |
| 33376 | ENSMUSG00000094284 | 0 | NA | NA | NA | NA | NA | Vmn1r112 | vomeronasal 1 receptor 112 [Source:MGI | 0 | 0 | 0 | 0 | 0 | 0 |
| 33377 | ENSMUSG00000094285 | 0 | NA | NA | NA | NA | NA | Olfr743 | olfactory receptor 743 [Source:MGI Symb | 0 | 0 | 0 | 0 | 0 | 0 |
| 33378 | ENSMUSG00000094286 | 0 | NA | NA | NA | NA | NA | Gm12424 | predicted gene 12424 [Source:MGI Symbo | 0 | 0 | 0 | 0 | 0 | 0 |
| 33379 | ENSMUSG00000094287 | 0 | NA | NA | NA | NA | NA | NA | NA | 0 | 0 | 0 | 0 | 0 | 0 |
| 33380 | ENSMUSG00000094288 | 0 | NA | NA | NA | NA | NA | Astx2 | amplified spermatogenic transcripts X enc | 0 | 0 | 0 | 0 | 0 | 0 |
| 33381 | ENSMUSG00000094290 | 0 | NA | NA | NA | NA | NA | NA | NA | 0 | 0 | 0 | 0 | 0 | 0 |
| 33382 | ENSMUSG00000094292 | 0 | NA | NA | NA | NA | NA | NA | NA | 0 | 0 | 0 | 0 | 0 | 0 |
| 33383 | ENSMUSG00000094293 | 0 | NA | NA | NA | NA | NA | Gm3893 | predicted gene 3893 [Source:MGI Symbol | 0 | 0 | 0 | 0 | 0 | 0 |
| 33384 | ENSMUSG00000094294 | 0 | NA | NA | NA | NA | NA | Gm20909 | predicted gene, 20909 [Source:MGI Symb | 0 | 0 | 0 | 0 | 0 | 0 |
| 33385 | ENSMUSG00000094295 | 0 | NA | NA | NA | NA | NA | Olfr819 | olfactory receptor 819 [Source:MGI Symb | 0 | 0 | 0 | 0 | 0 | 0 |
| 33386 | ENSMUSG00000094296 | 0 | NA | NA | NA | NA | NA | Gm21798 | predicted gene, 21798 [Source:MGI Symb | 0 | 0 | 0 | 0 | 0 | 0 |
| 33387 | ENSMUSG00000094297 | 0 | NA | NA | NA | NA | NA | NA | NA | 0 | 0 | 0 | 0 | 0 | 0 |
| 33388 | ENSMUSG00000094298 | 0 | NA | NA | NA | NA | NA | Vmn1r241 | vomeronasal 1 receptor 241 [Source:MGI | 0 | 0 | 0 | 0 | 0 | 0 |
| 33389 | ENSMUSG00000094301 | 0 | NA | NA | NA | NA | NA | NA | NA | 0 | 0 | 0 | 0 | 0 | 0 |
| 33390 | ENSMUSG00000094303 | 0 | NA | NA | NA | NA | NA | | | 0 | 0 | 0 | 0 | 0 | 0 |
| 33391 | ENSMUSG00000094304 | 0 | NA | NA | NA | NA | NA | NA | NA | 0 | 0 | 0 | 0 | 0 | 0 |
| 33392 | ENSMUSG00000094305 | 0 | NA | NA | NA | NA | NA | Scgb2b20 | secretoglobin, family 2B, member 20 [Sou | 0 | 0 | 0 | 0 | 0 | 0 |
| 33393 | ENSMUSG00000094306 | 0.133850373 | -0.796629176 | 4.080472857 | -0.195229623 | 0.84521319 | NA | Gm24924 | predicted gene, 24924 [Source:MGI Symb | 0.8031 | 0 | 0 | 0 | 0 | 0 |
| 33394 | ENSMUSG00000094307 | 0 | NA | NA | NA | NA | NA | Btbd35f28 | BTB domain containing 35, family membe | 0 | 0 | 0 | 0 | 0 | 0 |
| 33395 | ENSMUSG00000094308 | 0 | NA | NA | NA | NA | NA | NA | NA | 0 | 0 | 0 | 0 | 0 | 0 |
| 33396 | ENSMUSG00000094309 | 0 | NA | NA | NA | NA | NA | NA | NA | 0 | 0 | 0 | 0 | 0 | 0 |
| 33397 | ENSMUSG00000094310 | 0 | NA | NA | NA | NA | NA | NA | NA | 0 | 0 | 0 | 0 | 0 | 0 |
| 33398 | ENSMUSG00000094311 | 0 | NA | NA | NA | NA | NA | Rfpl4b | ret finger protein-like 4B [Source:MGI Syn | 0 | 0 | 0 | 0 | 0 | 0 |
| 33399 | ENSMUSG00000094312 | 0 | NA | NA | NA | NA | NA | NA | NA | 0 | 0 | 0 | 0 | 0 | 0 |
| 33400 | ENSMUSG00000094313 | 0 | NA | NA | NA | NA | NA | NA | NA | 0 | 0 | 0 | 0 | 0 | 0 |
| 33401 | ENSMUSG00000094314 | 0 | NA | NA | NA | NA | NA | Gm4301 | predicted gene 4301 [Source:MGI Symbol | 0 | 0 | 0 | 0 | 0 | 0 |
| 33402 | ENSMUSG00000094315 | 0.167707431 | -0.796629176 | 4.080472857 | -0.195229623 | 0.84521319 | NA | Igkv4-78 | immunoglobulin kappa variable 4-78 [Sou | 0 | 0 | 1.0062 | 0 | 0 | 0 |
| 33403 | ENSMUSG00000094316 | 0 | NA | NA | NA | NA | NA | NA | NA | 0 | 0 | 0 | 0 | 0 | 0 |
| 33404 | ENSMUSG00000094317 | 0 | NA | NA | NA | NA | NA | Gm6660 | predicted gene 6660 [Source:MGI Symbol | 0 | 0 | 0 | 0 | 0 | 0 |
| 33405 | ENSMUSG00000094318 | 0 | NA | NA | NA | NA | NA | Gm13144 | predicted gene 13144 [Source:MGI Symbo | 0 | 0 | 0 | 0 | 0 | 0 |
| 33406 | ENSMUSG00000094319 | 0 | NA | NA | NA | NA | NA | Igkv4-54 | immunoglobulin kappa chain variable 4-5 | 0 | 0 | 0 | 0 | 0 | 0 |
| 33407 | ENSMUSG00000094320 | 3.295633612 | 0.274926673 | 1.274907924 | 0.215644336 | 0.829264993 | NA | Chchd2-ps | coiled-coil-helix-coiled-coil-helix domain c | 0.8031 | 7.1954 | 1.0062 | 1.928 | 3.2719 | 5.5692 |
| 33408 | ENSMUSG00000094321 | 0 | NA | NA | NA | NA | NA | Defa-ps5 | defensin, alpha, pseudogene 5 [Source:M | 0 | 0 | 0 | 0 | 0 | 0 |
| 33409 | ENSMUSG00000094322 | 0.371281295 | 2.09253362 | 4.031852165 | 0.519000582 | 0.603760335 | NA | Ighv9-4 | immunoglobulin heavy variable 9-4 [Sourc | 0 | 0 | 0 | 0 | 0 | 2.2277 |
| 33410 | ENSMUSG00000094323 | 0 | NA | NA | NA | NA | NA | Gm15256 | predicted gene 15256 [Source:MGI Symbo | 0 | 0 | 0 | 0 | 0 | 0 |
| 33411 | ENSMUSG00000094325 | 0 | NA | NA | NA | NA | NA | Gm20831 | predicted gene, 20831 [Source:MGI Symb | 0 | 0 | 0 | 0 | 0 | 0 |
| 33412 | ENSMUSG00000094326 | 0 | NA | NA | NA | NA | NA | Gm10045 | predicted pseudogene 10045 [Source:MG | 0 | 0 | 0 | 0 | 0 | 0 |
| 33413 | ENSMUSG00000094327 | 0 | NA | NA | NA | NA | NA | NA | NA | 0 | 0 | 0 | 0 | 0 | 0 |
| 33414 | ENSMUSG00000094328 | 0 | NA | NA | NA | NA | NA | Tdpoz9 | TD and POZ domain containing 9 [Source: | 0 | 0 | 0 | 0 | 0 | 0 |
| 33415 | ENSMUSG00000094330 | 0 | NA | NA | NA | NA | NA | NA | NA | 0 | 0 | 0 | 0 | 0 | 0 |
| 33416 | ENSMUSG00000094334 | 0 | NA | NA | NA | NA | NA | Fabp5l2 | fatty acid binding protein 5-like 2 [Source: | 0 | 0 | 0 | 0 | 0 | 0 |
| 33417 | ENSMUSG00000094335 | 1.129591625 | 1.441847052 | 2.185297448 | 0.659794415 | 0.509385768 | NA | Igkv1-117 | immunoglobulin kappa variable 1-117 [So | 0.8031 | 1.0279 | 0 | 3.8559 | 1.0906 | 0 |
| 33418 | ENSMUSG00000094336 | 0 | NA | NA | NA | NA | NA | Gm13693 | predicted gene 13693 [Source:MGI Symbo | 0 | 0 | 0 | 0 | 0 | 0 |
| 33419 | ENSMUSG00000094337 | 0 | NA | NA | NA | NA | NA | | | 0 | 0 | 0 | 0 | 0 | 0 |
| 33420 | ENSMUSG00000094338 | 0 | NA | NA | NA | NA | NA | H2bc13 | H2B clustered histone 13 [Source:MGI Syr | 0 | 0 | 0 | 0 | 0 | 0 |
| 33421 | ENSMUSG00000094342 | 0 | NA | NA | NA | NA | NA | Gm17522 | predicted gene, 17522 [Source:MGI Symb | 0 | 0 | 0 | 0 | 0 | 0 |
| 33422 | ENSMUSG00000094344 | 0.895690289 | 0.713065541 | 2.535541507 | 0.281228108 | 0.778535447 | NA | Gm11942 | predicted gene 11942 [Source:MGI Symbo | 0 | 2.0558 | 0 | 0 | 1.0906 | 2.2277 |
| 33423 | ENSMUSG00000094345 | 0.267700746 | -1.497481649 | 4.057001436 | -0.369110456 | 0.712045396 | NA | Igkv14-126 | immunoglobulin kappa variable 14-126 [S | 1.6062 | 0 | 0 | 0 | 0 | 0 |
| 33424 | ENSMUSG00000094346 | 0 | NA | NA | NA | NA | NA | NA | NA | 0 | 0 | 0 | 0 | 0 | 0 |
| 33425 | ENSMUSG00000094347 | 0.331982614 | 0.16514309 | 3.986751476 | 0.041422971 | 0.966958701 | NA | Olfr784 | olfactory receptor 784 [Source:MGI Symb | 0 | 1.0279 | 0 | 0.964 | 0 | 0 |
| 33426 | ENSMUSG00000094349 | 0 | NA | NA | NA | NA | NA | Gm8082 | predicted gene 8082 [Source:MGI Symbol | 0 | 0 | 0 | 0 | 0 | 0 |
| 33427 | ENSMUSG00000094350 | 0 | NA | NA | NA | NA | NA | NA | NA | 0 | 0 | 0 | 0 | 0 | 0 |
| 33428 | ENSMUSG00000094351 | 0 | NA | NA | NA | NA | NA | NA | NA | 0 | 0 | 0 | 0 | 0 | 0 |

| | | | | | | | | | | | | | | | |
|---|---|---|---|---|---|---|---|---|---|---|---|---|---|---|---|
| 33429 | ENSMUSG00000094352 | 0 | NA | NA | NA | NA | NA | NA | NA | 0 | 0 | 0 | 0 | 0 | 0 |
| 33430 | ENSMUSG00000094353 | 0 | NA | NA | NA | NA | NA | Olfr150 | olfactory receptor 150 [Source:MGI Symb | 0 | 0 | 0 | 0 | 0 | 0 |
| 33431 | ENSMUSG00000094354 | 0 | NA | NA | NA | NA | NA | Gm21882 | predicted gene, 21882 [Source:MGI Symb | 0 | 0 | 0 | 0 | 0 | 0 |
| 33432 | ENSMUSG00000094356 | 0.267700746 | -1.497481649 | 4.057001436 | -0.369110456 | 0.712045396 | NA | Igkv8-28 | immunoglobulin kappa variable 8-28 [Sou | 1.6062 | 0 | 0 | 0 | 0 | 0 |
| 33433 | ENSMUSG00000094357 | 0 | NA | NA | NA | NA | NA | NA | NA | 0 | 0 | 0 | 0 | 0 | 0 |
| 33434 | ENSMUSG00000094358 | 0 | NA | NA | NA | NA | NA | Gm14370 | predicted gene 14370 [Source:MGI Symbo | 0 | 0 | 0 | 0 | 0 | 0 |
| 33435 | ENSMUSG00000094359 | 0.321325949 | 1.911575114 | 4.048267893 | 0.472195804 | 0.636787034 | NA | NA | NA | 0 | 0 | 0 | 1.928 | 0 | 0 |
| 33436 | ENSMUSG00000094360 | 0 | NA | NA | NA | NA | NA | Gm14307 | predicted gene 14307 [Source:MGI Symbo | 0 | 0 | 0 | 0 | 0 | 0 |
| 33437 | ENSMUSG00000094361 | 0 | NA | NA | NA | NA | NA | NA | NA | 0 | 0 | 0 | 0 | 0 | 0 |
| 33438 | ENSMUSG00000094362 | 0 | NA | NA | NA | NA | NA | Defa33 | defensin, alpha, 33 [Source:MGI Symbol;A | 0 | 0 | 0 | 0 | 0 | 0 |
| 33439 | ENSMUSG00000094363 | 0 | NA | NA | NA | NA | NA | NA | NA | 0 | 0 | 0 | 0 | 0 | 0 |
| 33440 | ENSMUSG00000094364 | 0 | NA | NA | NA | NA | NA | NA | NA | 0 | 0 | 0 | 0 | 0 | 0 |
| 33441 | ENSMUSG00000094365 | 0.160662974 | 1.126924175 | 4.080472857 | 0.276174898 | 0.782413736 | NA | Gm21982 | predicted gene 21982 [Source:MGI Symbo | 0 | 0 | 0 | 0.964 | 0 | 0 |
| 33442 | ENSMUSG00000094366 | 0 | NA | NA | NA | NA | NA | NA | NA | 0 | 0 | 0 | 0 | 0 | 0 |
| 33443 | ENSMUSG00000094367 | 0 | NA | NA | NA | NA | NA | NA | NA | 0 | 0 | 0 | 0 | 0 | 0 |
| 33444 | ENSMUSG00000094369 | 0 | NA | NA | NA | NA | NA | NA | NA | 0 | 0 | 0 | 0 | 0 | 0 |
| 33445 | ENSMUSG00000094370 | 0 | NA | NA | NA | NA | NA | Gm3373 | predicted gene 3373 [Source:MGI Symbol | 0 | 0 | 0 | 0 | 0 | 0 |
| 33446 | ENSMUSG00000094371 | 0 | NA | NA | NA | NA | NA | Gm7905 | predicted pseudogene 7905 [Source:MGI | 0 | 0 | 0 | 0 | 0 | 0 |
| 33447 | ENSMUSG00000094372 | 0 | NA | NA | NA | NA | NA | Gm21839 | predicted gene, 21839 [Source:MGI Symb | 0 | 0 | 0 | 0 | 0 | 0 |
| 33448 | ENSMUSG00000094374 | 0.17131964 | -0.796629176 | 4.080472857 | -0.195229623 | 0.84521319 | NA | Gm5435 | predicted gene 5435 [Source:MGI Symbol | 0 | 1.0279 | 0 | 0 | 0 | 0 |
| 33449 | ENSMUSG00000094377 | 0 | NA | NA | NA | NA | NA | NA | NA | 0 | 0 | 0 | 0 | 0 | 0 |
| 33450 | ENSMUSG00000094378 | 0 | NA | NA | NA | NA | NA | Gm15097 | predicted gene 15097 [Source:MGI Symbo | 0 | 0 | 0 | 0 | 0 | 0 |
| 33451 | ENSMUSG00000094379 | 0 | NA | NA | NA | NA | NA | Vmn1r202 | vomeronasal 1 receptor 202 [Source:MGI | 0 | 0 | 0 | 0 | 0 | 0 |
| 33452 | ENSMUSG00000094380 | 0 | NA | NA | NA | NA | NA | Olfr904 | olfactory receptor 904 [Source:MGI Symb | 0 | 0 | 0 | 0 | 0 | 0 |
| 33453 | ENSMUSG00000094381 | 0 | NA | NA | NA | NA | NA | NA | NA | 0 | 0 | 0 | 0 | 0 | 0 |
| 33454 | ENSMUSG00000094382 | 0.813009891 | 0.960077285 | 2.883567268 | 0.332947768 | 0.739173711 | NA | NA | NA | 1.6062 | 0 | 0 | 0 | 3.2719 | 0 |
| 33455 | ENSMUSG00000094383 | 0 | NA | NA | NA | NA | NA | | | 0 | 0 | 0 | 0 | 0 | 0 |
| 33456 | ENSMUSG00000094384 | 0 | NA | NA | NA | NA | NA | NA | NA | 0 | 0 | 0 | 0 | 0 | 0 |
| 33457 | ENSMUSG00000094385 | 0 | NA | NA | NA | NA | NA | Vmn1r123 | vomeronasal 1 receptor 123 [Source:MGI | 0 | 0 | 0 | 0 | 0 | 0 |
| 33458 | ENSMUSG00000094386 | 0 | NA | NA | NA | NA | NA | NA | NA | 0 | 0 | 0 | 0 | 0 | 0 |
| 33459 | ENSMUSG00000094388 | 5.881644094 | -0.744827187 | 0.951668025 | -0.782654421 | 0.433830075 | 0.845264432 | Gm8783 | predicted pseudogene 8783 [Source:MGI | 8.031 | 4.1117 | 10.062 | 8.6758 | 2.1812 | 2.2277 |
| 33460 | ENSMUSG00000094389 | 0 | NA | NA | NA | NA | NA | NA | NA | 0 | 0 | 0 | 0 | 0 | 0 |
| 33461 | ENSMUSG00000094390 | 0 | NA | NA | NA | NA | NA | NA | NA | 0 | 0 | 0 | 0 | 0 | 0 |
| 33462 | ENSMUSG00000094391 | 0 | NA | NA | NA | NA | NA | Btbd35f29 | BTB domain containing 35, family membe | 0 | 0 | 0 | 0 | 0 | 0 |
| 33463 | ENSMUSG00000094392 | 5.194058363 | -0.741897342 | 0.928304012 | -0.799196526 | 0.424176466 | 0.840450466 | Gm3788 | predicted gene 3788 [Source:MGI Symbol | 7.2279 | 7.1954 | 5.0312 | 2.8919 | 4.3625 | 4.4554 |
| 33464 | ENSMUSG00000094394 | 0 | NA | NA | NA | NA | NA | NA | NA | 0 | 0 | 0 | 0 | 0 | 0 |
| 33465 | ENSMUSG00000094396 | 0 | NA | NA | NA | NA | NA | Vmn2r124 | vomeronasal 2, receptor 124 [Source:MGI | 0 | 0 | 0 | 0 | 0 | 0 |
| 33466 | ENSMUSG00000094399 | 0 | NA | NA | NA | NA | NA | Gm21477 | predicted gene, 21477 [Source:MGI Symb | 0 | 0 | 0 | 0 | 0 | 0 |
| 33467 | ENSMUSG00000094400 | 0 | NA | NA | NA | NA | NA | NA | NA | 0 | 0 | 0 | 0 | 0 | 0 |
| 33468 | ENSMUSG00000094401 | 0 | NA | NA | NA | NA | NA | Gm14365 | predicted gene 14365 [Source:MGI Symbo | 0 | 0 | 0 | 0 | 0 | 0 |
| 33469 | ENSMUSG00000094402 | 0 | NA | NA | NA | NA | NA | NA | NA | 0 | 0 | 0 | 0 | 0 | 0 |
| 33470 | ENSMUSG00000094403 | 0 | NA | NA | NA | NA | NA | NA | NA | 0 | 0 | 0 | 0 | 0 | 0 |
| 33471 | ENSMUSG00000094404 | 0 | NA | NA | NA | NA | NA | Gm21776 | predicted gene, 21776 [Source:MGI Symb | 0 | 0 | 0 | 0 | 0 | 0 |
| 33472 | ENSMUSG00000094405 | 0 | NA | NA | NA | NA | NA | NA | NA | 0 | 0 | 0 | 0 | 0 | 0 |
| 33473 | ENSMUSG00000094406 | 0 | NA | NA | NA | NA | NA | NA | NA | 0 | 0 | 0 | 0 | 0 | 0 |
| 33474 | ENSMUSG00000094407 | 0 | NA | NA | NA | NA | NA | NA | NA | 0 | 0 | 0 | 0 | 0 | 0 |
| 33475 | ENSMUSG00000094408 | 0 | NA | NA | NA | NA | NA | NA | NA | 0 | 0 | 0 | 0 | 0 | 0 |
| 33476 | ENSMUSG00000094409 | 0 | NA | NA | NA | NA | NA | Vmn2r-ps23 | vomeronasal 2, receptor, pseudogene 23 | 0 | 0 | 0 | 0 | 0 | 0 |
| 33477 | ENSMUSG00000094410 | 65.93948574 | 0.179515028 | 0.327798917 | 0.547637648 | 0.583940733 | 0.908955096 | Gm38394 | predicted gene, 38394 [Source:MGI Symb | 53.005 | 59.619 | 73.456 | 97.362 | 66.528 | 45.668 |
| 33478 | ENSMUSG00000094411 | 0 | NA | NA | NA | NA | NA | NA | NA | 0 | 0 | 0 | 0 | 0 | 0 |
| 33479 | ENSMUSG00000094412 | 0 | NA | NA | NA | NA | NA | Obox4-ps16 | oocyte specific homeobox 4, pseudogene | 0 | 0 | 0 | 0 | 0 | 0 |
| 33480 | ENSMUSG00000094413 | 0 | NA | NA | NA | NA | NA | Trav15n-1 | T cell receptor alpha variable 15N-1 [Sour | 0 | 0 | 0 | 0 | 0 | 0 |
| 33481 | ENSMUSG00000094415 | 0 | NA | NA | NA | NA | NA | NA | NA | 0 | 0 | 0 | 0 | 0 | 0 |
| 33482 | ENSMUSG00000094416 | 0 | NA | NA | NA | NA | NA | Obox4-ps11 | oocyte specific homeobox 4, pseudogene | 0 | 0 | 0 | 0 | 0 | 0 |
| 33483 | ENSMUSG00000094417 | 0.949367096 | -3.28842827 | 2.400393427 | -1.369953872 | 0.170701301 | NA | Gm13243 | predicted gene 13243 [Source:MGI Symbo | 1.6062 | 3.0838 | 1.0062 | 0 | 0 | 0 |
| 33484 | ENSMUSG00000094419 | 0 | NA | NA | NA | NA | NA | NA | NA | 0 | 0 | 0 | 0 | 0 | 0 |

| | | | | | | | | | | | | | | | |
|---|---|---|---|---|---|---|---|---|---|---|---|---|---|---|---|
| 33485 | ENSMUSG00000094420 | 1.127963287 | -1.166213631 | 2.155903317 | -0.540939671 | 0.588549166 | NA | Igkv10-96 | immunoglobulin kappa variable 10-96 [So | 1.6062 | 3.0838 | 0 | 0.964 | 0 | 1.1138 |
| 33486 | ENSMUSG00000094421 | 0.167707431 | -0.796629176 | 4.080472857 | -0.195229623 | 0.84521319 | NA | Gm20778 | predicted gene, 20778 [Source:MGI Symb | 0 | 0 | 1.0062 | 0 | 0 | 0 |
| 33487 | ENSMUSG00000094422 | 0 | NA | NA | NA | NA | NA | Olfr205 | olfactory receptor 205 [Source:MGI Symb | 0 | 0 | 0 | 0 | 0 | 0 |
| 33488 | ENSMUSG00000094423 | 0 | NA | NA | NA | NA | NA | Gm7950 | predicted pseudogene 7950 [Source:MGI | 0 | 0 | 0 | 0 | 0 | 0 |
| 33489 | ENSMUSG00000094424 | 0 | NA | NA | NA | NA | NA | Gm7386 | predicted pseudogene 7386 [Source:MGI | 0 | 0 | 0 | 0 | 0 | 0 |
| 33490 | ENSMUSG00000094426 | 0 | NA | NA | NA | NA | NA | Olfr478 | olfactory receptor 478 [Source:MGI Symb | 0 | 0 | 0 | 0 | 0 | 0 |
| 33491 | ENSMUSG00000094427 | 0 | NA | NA | NA | NA | NA | Gm15143 | predicted gene 15143 [Source:MGI Symbo | 0 | 0 | 0 | 0 | 0 | 0 |
| 33492 | ENSMUSG00000094428 | 0 | NA | NA | NA | NA | NA | NA | NA | 0 | 0 | 0 | 0 | 0 | 0 |
| 33493 | ENSMUSG00000094429 | 0 | NA | NA | NA | NA | NA | Gm19965 | predicted gene, 19965 [Source:MGI Symb | 0 | 0 | 0 | 0 | 0 | 0 |
| 33494 | ENSMUSG00000094430 | 0 | NA | NA | NA | NA | NA | Smbd1 | somatomedin B domain containing 1 [Sou | 0 | 0 | 0 | 0 | 0 | 0 |
| 33495 | ENSMUSG00000094431 | 0 | NA | NA | NA | NA | NA | NA | NA | 0 | 0 | 0 | 0 | 0 | 0 |
| 33496 | ENSMUSG00000094432 | 0 | NA | NA | NA | NA | NA | NA | NA | 0 | 0 | 0 | 0 | 0 | 0 |
| 33497 | ENSMUSG00000094433 | 0.462220778 | -0.685400455 | 3.191475091 | -0.21475977 | 0.82995462 | NA | Igkv5-43 | immunoglobulin kappa chain variable 5-4: | 0.8031 | 0 | 1.0062 | 0.964 | 0 | 0 |
| 33498 | ENSMUSG00000094435 | 0 | NA | NA | NA | NA | NA | NA | NA | 0 | 0 | 0 | 0 | 0 | 0 |
| 33499 | ENSMUSG00000094436 | 0 | NA | NA | NA | NA | NA | Gm16327 | predicted gene 16327 [Source:MGI Symbo | 0 | 0 | 0 | 0 | 0 | 0 |
| 33500 | ENSMUSG00000094437 | 0.160662974 | 1.126924175 | 4.080472857 | 0.276174898 | 0.782413736 | NA | Gm9830 | predicted gene 9830 [Source:MGI Symbol | 0 | 0 | 0 | 0.964 | 0 | 0 |
| 33501 | ENSMUSG00000094438 | 0 | NA | NA | NA | NA | NA | NA | NA | 0 | 0 | 0 | 0 | 0 | 0 |
| 33502 | ENSMUSG00000094439 | 1.167060891 | 1.325545444 | 2.321108505 | 0.571082929 | 0.567943429 | NA | Gm21969 | predicted gene 21969 [Source:MGI Symbo | 0 | 2.0558 | 0 | 3.8559 | 1.0906 | 0 |
| 33503 | ENSMUSG00000094440 | 0 | NA | NA | NA | NA | NA | NA | NA | 0 | 0 | 0 | 0 | 0 | 0 |
| 33504 | ENSMUSG00000094441 | 70.41885408 | 0.272462184 | 0.278451034 | 0.978492267 | 0.327830914 | 0.772937193 | Zfp955a | zinc finger protein 955A [Source:MGI Sym | 51.399 | 69.898 | 70.437 | 75.19 | 68.709 | 86.88 |
| 33505 | ENSMUSG00000094442 | 0 | NA | NA | NA | NA | NA | NA | NA | 0 | 0 | 0 | 0 | 0 | 0 |
| 33506 | ENSMUSG00000094443 | 0 | NA | NA | NA | NA | NA | Sgo2b | shugoshin 2B [Source:MGI Symbol;Acc:M | 0 | 0 | 0 | 0 | 0 | 0 |
| 33507 | ENSMUSG00000094444 | 0 | NA | NA | NA | NA | NA | Gm21562 | predicted gene, 21562 [Source:MGI Symb | 0 | 0 | 0 | 0 | 0 | 0 |
| 33508 | ENSMUSG00000094445 | 0 | NA | NA | NA | NA | NA | 1700015G11 | RIKEN cDNA 1700015G11 gene [Source:M | 0 | 0 | 0 | 0 | 0 | 0 |
| 33509 | ENSMUSG00000094446 | 0 | NA | NA | NA | NA | NA | Gm6344 | predicted gene 6344 [Source:MGI Symbol | 0 | 0 | 0 | 0 | 0 | 0 |
| 33510 | ENSMUSG00000094447 | 0 | NA | NA | NA | NA | NA | 9430069I07F | RIKEN cDNA 9430069I07 gene [Source:MC | 0 | 0 | 0 | 0 | 0 | 0 |
| 33511 | ENSMUSG00000094449 | 0 | NA | NA | NA | NA | NA | Olfr972 | olfactory receptor 972 [Source:MGI Symb | 0 | 0 | 0 | 0 | 0 | 0 |
| 33512 | ENSMUSG00000094452 | 0 | NA | NA | NA | NA | NA | NA | NA | 0 | 0 | 0 | 0 | 0 | 0 |
| 33513 | ENSMUSG00000094454 | 0 | NA | NA | NA | NA | NA | NA | NA | 0 | 0 | 0 | 0 | 0 | 0 |
| 33514 | ENSMUSG00000094455 | 0 | NA | NA | NA | NA | NA | NA | NA | 0 | 0 | 0 | 0 | 0 | 0 |
| 33515 | ENSMUSG00000094456 | 0 | NA | NA | NA | NA | NA | NA | NA | 0 | 0 | 0 | 0 | 0 | 0 |
| 33516 | ENSMUSG00000094457 | 0 | NA | NA | NA | NA | NA | NA | NA | 0 | 0 | 0 | 0 | 0 | 0 |
| 33517 | ENSMUSG00000094458 | 0 | NA | NA | NA | NA | NA | NA | NA | 0 | 0 | 0 | 0 | 0 | 0 |
| 33518 | ENSMUSG00000094460 | 0 | NA | NA | NA | NA | NA | Gm21560 | predicted gene, 21560 [Source:MGI Symb | 0 | 0 | 0 | 0 | 0 | 0 |
| 33519 | ENSMUSG00000094461 | 0 | NA | NA | NA | NA | NA | Olfr883 | olfactory receptor 883 [Source:MGI Symb | 0 | 0 | 0 | 0 | 0 | 0 |
| 33520 | ENSMUSG00000094462 | 3.28286472 | -0.425595763 | 1.163438815 | -0.365808462 | 0.714508005 | NA | Gm21028 | predicted gene, 21028 [Source:MGI Symb | 3.2124 | 2.0558 | 6.0375 | 2.8919 | 3.2719 | 2.2277 |
| 33521 | ENSMUSG00000094463 | 0.133850373 | -0.796629176 | 4.080472857 | -0.195229623 | 0.84521319 | NA | Gm8546 | predicted gene 8546 [Source:MGI Symbol | 0.8031 | 0 | 0 | 0 | 0 | 0 |
| 33522 | ENSMUSG00000094464 | 0 | NA | NA | NA | NA | NA | Olfr339 | olfactory receptor 339 [Source:MGI Symb | 0 | 0 | 0 | 0 | 0 | 0 |
| 33523 | ENSMUSG00000094465 | 0 | NA | NA | NA | NA | NA | NA | NA | 0 | 0 | 0 | 0 | 0 | 0 |
| 33524 | ENSMUSG00000094466 | 0 | NA | NA | NA | NA | NA | NA | NA | 0 | 0 | 0 | 0 | 0 | 0 |
| 33525 | ENSMUSG00000094467 | 0.17131964 | -0.796629176 | 4.080472857 | -0.195229623 | 0.84521319 | NA | NA | NA | 0 | 1.0279 | 0 | 0 | 0 | 0 |
| 33526 | ENSMUSG00000094468 | 0 | NA | NA | NA | NA | NA | Trav6-4 | T cell receptor alpha variable 6-4 [Source: | 0 | 0 | 0 | 0 | 0 | 0 |
| 33527 | ENSMUSG00000094470 | 0 | NA | NA | NA | NA | NA | NA | NA | 0 | 0 | 0 | 0 | 0 | 0 |
| 33528 | ENSMUSG00000094471 | 0 | NA | NA | NA | NA | NA | NA | NA | 0 | 0 | 0 | 0 | 0 | 0 |
| 33529 | ENSMUSG00000094472 | 0 | NA | NA | NA | NA | NA | Gm21897 | predicted gene, 21897 [Source:MGI Symb | 0 | 0 | 0 | 0 | 0 | 0 |
| 33530 | ENSMUSG00000094473 | 0 | NA | NA | NA | NA | NA | NA | NA | 0 | 0 | 0 | 0 | 0 | 0 |
| 33531 | ENSMUSG00000094474 | 0 | NA | NA | NA | NA | NA | | | 0 | 0 | 0 | 0 | 0 | 0 |
| 33532 | ENSMUSG00000094475 | 0 | NA | NA | NA | NA | NA | Gm11007 | predicted gene 11007 [Source:MGI Symbo | 0 | 0 | 0 | 0 | 0 | 0 |
| 33533 | ENSMUSG00000094476 | 0 | NA | NA | NA | NA | NA | NA | NA | 0 | 0 | 0 | 0 | 0 | 0 |
| 33534 | ENSMUSG00000094477 | 0 | NA | NA | NA | NA | NA | NA | NA | 0 | 0 | 0 | 0 | 0 | 0 |
| 33535 | ENSMUSG00000094478 | 0 | NA | NA | NA | NA | NA | Igkv3-3 | immunoglobulin kappa variable 3-3 [Sourc | 0 | 0 | 0 | 0 | 0 | 0 |
| 33536 | ENSMUSG00000094481 | 0 | NA | NA | NA | NA | NA | NA | NA | 0 | 0 | 0 | 0 | 0 | 0 |
| 33537 | ENSMUSG00000094482 | 0 | NA | NA | NA | NA | NA | Vmn2r-ps8 | vomeronasal 2, receptor, pseudogene 8 [S | 0 | 0 | 0 | 0 | 0 | 0 |
| 33538 | ENSMUSG00000094483 | 685.4800981 | -0.002772345 | 0.117778924 | -0.023538553 | 0.981220686 | 0.99820894 | Purb | purine rich element binding protein B [Sou | 705.93 | 696.93 | 655.07 | 670.93 | 650.01 | 734.02 |
| 33539 | ENSMUSG00000094484 | 0 | NA | NA | NA | NA | NA | Gm21244 | predicted gene, 21244 [Source:MGI Symb | 0 | 0 | 0 | 0 | 0 | 0 |
| 33540 | ENSMUSG00000094485 | 0 | NA | NA | NA | NA | NA | Gm11322 | predicted gene 11322 [Source:MGI Symbo | 0 | 0 | 0 | 0 | 0 | 0 |

| | | | | | | | | | | | | | | | |
|---|---|---|---|---|---|---|---|---|---|---|---|---|---|---|---|
| 33541 | ENSMUSG00000094486 | 0 | NA | NA | NA | NA | NA | NA | NA | 0 | 0 | 0 | 0 | 0 | 0 |
| 33542 | ENSMUSG00000094487 | 0 | NA | NA | NA | NA | NA | Gm21958 | predicted gene, 21958 [Source:MGI Symb | 0 | 0 | 0 | 0 | 0 | 0 |
| 33543 | ENSMUSG00000094488 | 0 | NA | NA | NA | NA | NA | Olfr393 | olfactory receptor 393 [Source:MGI Symb | 0 | 0 | 0 | 0 | 0 | 0 |
| 33544 | ENSMUSG00000094489 | 0 | NA | NA | NA | NA | NA | NA | NA | 0 | 0 | 0 | 0 | 0 | 0 |
| 33545 | ENSMUSG00000094491 | 0 | NA | NA | NA | NA | NA | Igkv1-133 | immunoglobulin kappa variable 1-133 [So | 0 | 0 | 0 | 0 | 0 | 0 |
| 33546 | ENSMUSG00000094493 | 0 | NA | NA | NA | NA | NA | Olfr700 | olfactory receptor 700 [Source:MGI Symb | 0 | 0 | 0 | 0 | 0 | 0 |
| 33547 | ENSMUSG00000094494 | 0.267700746 | -1.497481649 | 4.057001436 | -0.369110456 | 0.712045396 | NA | NA | NA | 1.6062 | 0 | 0 | 0 | 0 | 0 |
| 33548 | ENSMUSG00000094496 | 0 | NA | NA | NA | NA | NA | Olfr792 | olfactory receptor 792 [Source:MGI Symb | 0 | 0 | 0 | 0 | 0 | 0 |
| 33549 | ENSMUSG00000094497 | 0.439020386 | -2.189747911 | 3.52148864 | -0.621824499 | 0.534057275 | NA | Gm8210 | predicted pseudogene 8210 [Source:MGI | 1.6062 | 1.0279 | 0 | 0 | 0 | 0 |
| 33550 | ENSMUSG00000094498 | 0 | NA | NA | NA | NA | NA | Gm10582 | predicted gene 10582 [Source:MGI Symbo | 0 | 0 | 0 | 0 | 0 | 0 |
| 33551 | ENSMUSG00000094499 | 0 | NA | NA | NA | NA | NA | Vmn2r-ps10 | vomeronasal 2, receptor, pseudogene 10 | 0 | 0 | 0 | 0 | 0 | 0 |
| 33552 | ENSMUSG00000094500 | 0 | NA | NA | NA | NA | NA | Smim18 | small integral membrane protein 18 [Sour | 0 | 0 | 0 | 0 | 0 | 0 |
| 33553 | ENSMUSG00000094501 | 0 | NA | NA | NA | NA | NA | Vmn1r-ps73 | vomeronasal 1 receptor, pseudogene 73 [ | 0 | 0 | 0 | 0 | 0 | 0 |
| 33554 | ENSMUSG00000094502 | 0.17131964 | -0.796629176 | 4.080472857 | -0.195229623 | 0.84521319 | NA | Ighv1-69 | immunoglobulin heavy variable 1-69 [Sou | 0 | 1.0279 | 0 | 0 | 0 | 0 |
| 33555 | ENSMUSG00000094504 | 8.145491269 | 1.098259953 | 0.97845785 | 1.122439718 | 0.261675525 | 0.715612553 | Foxl3 | forkhead box L3 [Source:MGI Symbol;Acc: | 11.243 | 0 | 4.025 | 3.8559 | 15.269 | 14.48 |
| 33556 | ENSMUSG00000094505 | 0 | NA | NA | NA | NA | NA | Ighv8-6 | immunoglobulin heavy variable V8-6 [Sou | 0 | 0 | 0 | 0 | 0 | 0 |
| 33557 | ENSMUSG00000094506 | 0 | NA | NA | NA | NA | NA | NA | NA | 0 | 0 | 0 | 0 | 0 | 0 |
| 33558 | ENSMUSG00000094507 | 0 | NA | NA | NA | NA | NA | Gm21913 | predicted gene, 21913 [Source:MGI Symb | 0 | 0 | 0 | 0 | 0 | 0 |
| 33559 | ENSMUSG00000094508 | 0 | NA | NA | NA | NA | NA | NA | NA | 0 | 0 | 0 | 0 | 0 | 0 |
| 33560 | ENSMUSG00000094509 | 0 | NA | NA | NA | NA | NA | Ighv14-1 | immunoglobulin heavy variable 14-1 [Sou | 0 | 0 | 0 | 0 | 0 | 0 |
| 33561 | ENSMUSG00000094510 | 0 | NA | NA | NA | NA | NA | NA | NA | 0 | 0 | 0 | 0 | 0 | 0 |
| 33562 | ENSMUSG00000094511 | 0 | NA | NA | NA | NA | NA | Gm21693 | predicted gene, 21693 [Source:MGI Symb | 0 | 0 | 0 | 0 | 0 | 0 |
| 33563 | ENSMUSG00000094513 | 0 | NA | NA | NA | NA | NA | NA | NA | 0 | 0 | 0 | 0 | 0 | 0 |
| 33564 | ENSMUSG00000094514 | 0.267700746 | -1.497481649 | 4.057001436 | -0.369110456 | 0.712045396 | NA | | | 1.6062 | 0 | 0 | 0 | 0 | 0 |
| 33565 | ENSMUSG00000094515 | 0 | NA | NA | NA | NA | NA | NA | NA | 0 | 0 | 0 | 0 | 0 | 0 |
| 33566 | ENSMUSG00000094516 | 0 | NA | NA | NA | NA | NA | NA | NA | 0 | 0 | 0 | 0 | 0 | 0 |
| 33567 | ENSMUSG00000094520 | 0 | NA | NA | NA | NA | NA | Olfr635 | olfactory receptor 635 [Source:MGI Symb | 0 | 0 | 0 | 0 | 0 | 0 |
| 33568 | ENSMUSG00000094521 | 0 | NA | NA | NA | NA | NA | NA | NA | 0 | 0 | 0 | 0 | 0 | 0 |
| 33569 | ENSMUSG00000094522 | 0 | NA | NA | NA | NA | NA | NA | NA | 0 | 0 | 0 | 0 | 0 | 0 |
| 33570 | ENSMUSG00000094523 | 0 | NA | NA | NA | NA | NA | NA | NA | 0 | 0 | 0 | 0 | 0 | 0 |
| 33571 | ENSMUSG00000094524 | 0 | NA | NA | NA | NA | NA | NA | NA | 0 | 0 | 0 | 0 | 0 | 0 |
| 33572 | ENSMUSG00000094525 | 0.167707431 | -0.796629176 | 4.080472857 | -0.195229623 | 0.84521319 | NA | Trbv12-2 | T cell receptor beta, variable 12-2 [Source | 0 | 0 | 1.0062 | 0 | 0 | 0 |
| 33573 | ENSMUSG00000094526 | 1.191885234 | -2.403081538 | 2.596141673 | -0.925635747 | 0.354635309 | NA | Gm21451 | predicted gene, 21451 [Source:MGI Symb | 0 | 0 | 6.0375 | 0 | 0 | 1.1138 |
| 33574 | ENSMUSG00000094527 | 0 | NA | NA | NA | NA | NA | NA | NA | 0 | 0 | 0 | 0 | 0 | 0 |
| 33575 | ENSMUSG00000094528 | 0 | NA | NA | NA | NA | NA | NA | NA | 0 | 0 | 0 | 0 | 0 | 0 |
| 33576 | ENSMUSG00000094529 | 0 | NA | NA | NA | NA | NA | Gm21744 | predicted gene, 21744 [Source:MGI Symb | 0 | 0 | 0 | 0 | 0 | 0 |
| 33577 | ENSMUSG00000094530 | 5.956273423 | -1.294960734 | 0.969887418 | -1.33516603 | 0.181822012 | 0.615175107 | Gm21399 | predicted gene, 21399 [Source:MGI Symb | 3.2124 | 9.2513 | 13.081 | 5.7839 | 2.1812 | 2.2277 |
| 33578 | ENSMUSG00000094531 | 0 | NA | NA | NA | NA | NA | Olfr671 | olfactory receptor 671 [Source:MGI Symb | 0 | 0 | 0 | 0 | 0 | 0 |
| 33579 | ENSMUSG00000094532 | 0 | NA | NA | NA | NA | NA | Vmn1r259 | vomeronasal 1 receptor 259 [Source:MGI | 0 | 0 | 0 | 0 | 0 | 0 |
| 33580 | ENSMUSG00000094533 | 0 | NA | NA | NA | NA | NA | Ighv11-1 | immunoglobulin heavy variable 11-1 [Sou | 0 | 0 | 0 | 0 | 0 | 0 |
| 33581 | ENSMUSG00000094534 | 0 | NA | NA | NA | NA | NA | Gm20945 | predicted gene, 20945 [Source:MGI Symb | 0 | 0 | 0 | 0 | 0 | 0 |
| 33582 | ENSMUSG00000094535 | 0 | NA | NA | NA | NA | NA | Olfr850 | olfactory receptor 850 [Source:MGI Symb | 0 | 0 | 0 | 0 | 0 | 0 |
| 33583 | ENSMUSG00000094537 | 0 | NA | NA | NA | NA | NA | NA | NA | 0 | 0 | 0 | 0 | 0 | 0 |
| 33584 | ENSMUSG00000094538 | 0 | NA | NA | NA | NA | NA | NA | NA | 0 | 0 | 0 | 0 | 0 | 0 |
| 33585 | ENSMUSG00000094539 | 0 | NA | NA | NA | NA | NA | Olfr191 | olfactory receptor 191 [Source:MGI Symb | 0 | 0 | 0 | 0 | 0 | 0 |
| 33586 | ENSMUSG00000094540 | 0 | NA | NA | NA | NA | NA | Astx4c | amplified spermatogenic transcripts X enc | 0 | 0 | 0 | 0 | 0 | 0 |
| 33587 | ENSMUSG00000094541 | 0 | NA | NA | NA | NA | NA | NA | NA | 0 | 0 | 0 | 0 | 0 | 0 |
| 33588 | ENSMUSG00000094542 | 0 | NA | NA | NA | NA | NA | Vmn1r149 | vomeronasal 1 receptor 149 [Source:MGI | 0 | 0 | 0 | 0 | 0 | 0 |
| 33589 | ENSMUSG00000094543 | 0 | NA | NA | NA | NA | NA | Gm8212 | predicted gene 8212 [Source:MGI Symbol | 0 | 0 | 0 | 0 | 0 | 0 |
| 33590 | ENSMUSG00000094545 | 0 | NA | NA | NA | NA | NA | Vmn1r101 | vomeronasal 1 receptor 101 [Source:MGI | 0 | 0 | 0 | 0 | 0 | 0 |
| 33591 | ENSMUSG00000094546 | 0.181769715 | 1.126924175 | 4.080472857 | 0.276174898 | 0.782413736 | NA | Ighv1-26 | immunoglobulin heavy variable 1-26 [Sou | 0 | 0 | 0 | 0 | 1.0906 | 0 |
| 33592 | ENSMUSG00000094548 | 0 | NA | NA | NA | NA | NA | NA | NA | 0 | 0 | 0 | 0 | 0 | 0 |
| 33593 | ENSMUSG00000094550 | 0 | NA | NA | NA | NA | NA | NA | NA | 0 | 0 | 0 | 0 | 0 | 0 |
| 33594 | ENSMUSG00000094551 | 0 | NA | NA | NA | NA | NA | NA | NA | 0 | 0 | 0 | 0 | 0 | 0 |
| 33595 | ENSMUSG00000094552 | 0 | NA | NA | NA | NA | NA | Ighd3-2 | immunoglobulin heavy diversity 3-2 [Sour | 0 | 0 | 0 | 0 | 0 | 0 |
| 33596 | ENSMUSG00000094553 | 0 | NA | NA | NA | NA | NA | Vmn1r50 | vomeronasal 1 receptor 50 [Source:MGI S | 0 | 0 | 0 | 0 | 0 | 0 |

| | | | | | | | | | | | | | | | |
|---|---|---|---|---|---|---|---|---|---|---|---|---|---|---|---|
| 33597 | ENSMUSG00000094554 | 0 | NA | NA | NA | NA | NA | NA | NA | 0 | 0 | 0 | 0 | 0 | 0 |
| 33598 | ENSMUSG00000094555 | 0 | NA | NA | NA | NA | NA | Obox4-ps6 | oocyte specific homeobox 4, pseudogene | 0 | 0 | 0 | 0 | 0 | 0 |
| 33599 | ENSMUSG00000094556 | 0 | NA | NA | NA | NA | NA | Gm20821 | predicted gene, 20821 [Source:MGI Symb | 0 | 0 | 0 | 0 | 0 | 0 |
| 33600 | ENSMUSG00000094557 | 0 | NA | NA | NA | NA | NA | Rpl35a-ps7 | ribosomal protein L35A, pseudogene 7 [So | 0 | 0 | 0 | 0 | 0 | 0 |
| 33601 | ENSMUSG00000094558 | 0 | NA | NA | NA | NA | NA | Btbd35f11 | BTB domain containing 35, family membe | 0 | 0 | 0 | 0 | 0 | 0 |
| 33602 | ENSMUSG00000094559 | 0.133850373 | -0.796629176 | 4.080472857 | -0.195229623 | 0.84521319 | NA | Cyp2d34 | cytochrome P450, family 2, subfamily d, p | 0.8031 | 0 | 0 | 0 | 0 | 0 |
| 33603 | ENSMUSG00000094560 | 0 | NA | NA | NA | NA | NA | Pramel39 | PRAME like 39 [Source:MGI Symbol;Acc:M | 0 | 0 | 0 | 0 | 0 | 0 |
| 33604 | ENSMUSG00000094561 | 0.133850373 | -0.796629176 | 4.080472857 | -0.195229623 | 0.84521319 | NA | Ighv1-22 | immunoglobulin heavy variable 1-22 [Sou | 0.8031 | 0 | 0 | 0 | 0 | 0 |
| 33605 | ENSMUSG00000094562 | 0 | NA | NA | NA | NA | NA | Trav13-4-dv7 | T cell receptor alpha variable 13-4-DV7 [So | 0 | 0 | 0 | 0 | 0 | 0 |
| 33606 | ENSMUSG00000094563 | 0 | NA | NA | NA | NA | NA | NA | NA | 0 | 0 | 0 | 0 | 0 | 0 |
| 33607 | ENSMUSG00000094564 | 0 | NA | NA | NA | NA | NA | NA | NA | 0 | 0 | 0 | 0 | 0 | 0 |
| 33608 | ENSMUSG00000094565 | 0 | NA | NA | NA | NA | NA | NA | NA | 0 | 0 | 0 | 0 | 0 | 0 |
| 33609 | ENSMUSG00000094566 | 0 | NA | NA | NA | NA | NA | Gm12620 | predicted gene 12620 [Source:MGI Symbo | 0 | 0 | 0 | 0 | 0 | 0 |
| 33610 | ENSMUSG00000094567 | 0 | NA | NA | NA | NA | NA | NA | NA | 0 | 0 | 0 | 0 | 0 | 0 |
| 33611 | ENSMUSG00000094568 | 0.656740811 | -0.013525372 | 3.112905782 | -0.004344934 | 0.996533255 | NA | Smarce1-ps1 | SWI/SNF related, matrix associated, actin | 0 | 0 | 2.0125 | 1.928 | 0 | 0 |
| 33612 | ENSMUSG00000094569 | 0 | NA | NA | NA | NA | NA | Trbd2 | T cell receptor beta, D region 2 [Source:M | 0 | 0 | 0 | 0 | 0 | 0 |
| 33613 | ENSMUSG00000094570 | 0 | NA | NA | NA | NA | NA | Gm20931 | predicted gene, 20931 [Source:MGI Symb | 0 | 0 | 0 | 0 | 0 | 0 |
| 33614 | ENSMUSG00000094571 | 0 | NA | NA | NA | NA | NA | NA | NA | 0 | 0 | 0 | 0 | 0 | 0 |
| 33615 | ENSMUSG00000094572 | 0 | NA | NA | NA | NA | NA | NA | NA | 0 | 0 | 0 | 0 | 0 | 0 |
| 33616 | ENSMUSG00000094573 | 0 | NA | NA | NA | NA | NA | NA | NA | 0 | 0 | 0 | 0 | 0 | 0 |
| 33617 | ENSMUSG00000094574 | 0 | NA | NA | NA | NA | NA | Gm15027 | predicted gene 15027 [Source:MGI Symbo | 0 | 0 | 0 | 0 | 0 | 0 |
| 33618 | ENSMUSG00000094575 | 0 | NA | NA | NA | NA | NA | Gm20738 | predicted gene, 20738 [Source:MGI Symb | 0 | 0 | 0 | 0 | 0 | 0 |
| 33619 | ENSMUSG00000094576 | 0 | NA | NA | NA | NA | NA | NA | NA | 0 | 0 | 0 | 0 | 0 | 0 |
| 33620 | ENSMUSG00000094577 | 0 | NA | NA | NA | NA | NA | Samt1b | spermatogenesis associated multipass tra | 0 | 0 | 0 | 0 | 0 | 0 |
| 33621 | ENSMUSG00000094578 | 0 | NA | NA | NA | NA | NA | Gm3187 | predicted gene 3187 [Source:MGI Symbol | 0 | 0 | 0 | 0 | 0 | 0 |
| 33622 | ENSMUSG00000094579 | 0 | NA | NA | NA | NA | NA | NA | NA | 0 | 0 | 0 | 0 | 0 | 0 |
| 33623 | ENSMUSG00000094581 | 0 | NA | NA | NA | NA | NA | NA | NA | 0 | 0 | 0 | 0 | 0 | 0 |
| 33624 | ENSMUSG00000094582 | 0 | NA | NA | NA | NA | NA | Gm7736 | predicted gene 7736 [Source:MGI Symbol | 0 | 0 | 0 | 0 | 0 | 0 |
| 33625 | ENSMUSG00000094584 | 0 | NA | NA | NA | NA | NA | Ms4a18 | membrane-spanning 4-domains, subfamil | 0 | 0 | 0 | 0 | 0 | 0 |
| 33626 | ENSMUSG00000094585 | 0 | NA | NA | NA | NA | NA | NA | NA | 0 | 0 | 0 | 0 | 0 | 0 |
| 33627 | ENSMUSG00000094586 | 0 | NA | NA | NA | NA | NA | Vmn1r41 | vomeronasal 1 receptor 41 [Source:MGI S | 0 | 0 | 0 | 0 | 0 | 0 |
| 33628 | ENSMUSG00000094587 | 0 | NA | NA | NA | NA | NA | NA | NA | 0 | 0 | 0 | 0 | 0 | 0 |
| 33629 | ENSMUSG00000094588 | 0 | NA | NA | NA | NA | NA | Olfr898 | olfactory receptor 898 [Source:MGI Symb | 0 | 0 | 0 | 0 | 0 | 0 |
| 33630 | ENSMUSG00000094589 | 0 | NA | NA | NA | NA | NA | Vmn1r118 | vomeronasal 1 receptor 118 [Source:MGI | 0 | 0 | 0 | 0 | 0 | 0 |
| 33631 | ENSMUSG00000094590 | 0 | NA | NA | NA | NA | NA | Gm10251 | predicted gene 10251 [Source:MGI Symbo | 0 | 0 | 0 | 0 | 0 | 0 |
| 33632 | ENSMUSG00000094592 | 0 | NA | NA | NA | NA | NA | Samt1d | spermatogenesis associated multipass tra | 0 | 0 | 0 | 0 | 0 | 0 |
| 33633 | ENSMUSG00000094593 | 0 | NA | NA | NA | NA | NA | NA | NA | 0 | 0 | 0 | 0 | 0 | 0 |
| 33634 | ENSMUSG00000094594 | 0 | NA | NA | NA | NA | NA | Tpd52-ps | tumor protein D52, pseudogene [Source:M | 0 | 0 | 0 | 0 | 0 | 0 |
| 33635 | ENSMUSG00000094595 | 7.988563563 | -0.578745955 | 0.814073866 | -0.710925604 | 0.477130339 | 0.869671045 | Fsbp | fibrinogen silencer binding protein [Sourc | 6.4248 | 7.1954 | 15.094 | 3.8559 | 10.906 | 4.4554 |
| 33636 | ENSMUSG00000094596 | 0 | NA | NA | NA | NA | NA | Btbd35f20 | BTB domain containing 35, family membe | 0 | 0 | 0 | 0 | 0 | 0 |
| 33637 | ENSMUSG00000094597 | 0 | NA | NA | NA | NA | NA | Gm4810 | predicted gene 4810 [Source:MGI Symbol | 0 | 0 | 0 | 0 | 0 | 0 |
| 33638 | ENSMUSG00000094598 | 0 | NA | NA | NA | NA | NA | NA | NA | 0 | 0 | 0 | 0 | 0 | 0 |
| 33639 | ENSMUSG00000094599 | 0 | NA | NA | NA | NA | NA | NA | NA | 0 | 0 | 0 | 0 | 0 | 0 |
| 33640 | ENSMUSG00000094601 | 0 | NA | NA | NA | NA | NA | Gm5926 | predicted gene 5926 [Source:MGI Symbol | 0 | 0 | 0 | 0 | 0 | 0 |
| 33641 | ENSMUSG00000094602 | 0 | NA | NA | NA | NA | NA | Vmn1r169 | vomeronasal 1 receptor 169 [Source:MGI | 0 | 0 | 0 | 0 | 0 | 0 |
| 33642 | ENSMUSG00000094603 | 0 | NA | NA | NA | NA | NA | NA | NA | 0 | 0 | 0 | 0 | 0 | 0 |
| 33643 | ENSMUSG00000094604 | 0 | NA | NA | NA | NA | NA | NA | NA | 0 | 0 | 0 | 0 | 0 | 0 |
| 33644 | ENSMUSG00000094605 | 0.481988923 | 2.492401104 | 3.837077151 | 0.649557203 | 0.515978285 | NA | NA | NA | 0 | 0 | 0 | 2.8919 | 0 | 0 |
| 33645 | ENSMUSG00000094606 | 0 | NA | NA | NA | NA | NA | Vmn2r50 | vomeronasal 2, receptor 50 [Source:MGI S | 0 | 0 | 0 | 0 | 0 | 0 |
| 33646 | ENSMUSG00000094607 | 0 | NA | NA | NA | NA | NA | Trav12d-3 | T cell receptor alpha variable 12D-3 [Sourc | 0 | 0 | 0 | 0 | 0 | 0 |
| 33647 | ENSMUSG00000094608 | 0 | NA | NA | NA | NA | NA | NA | NA | 0 | 0 | 0 | 0 | 0 | 0 |
| 33648 | ENSMUSG00000094609 | 0 | NA | NA | NA | NA | NA | NA | NA | 0 | 0 | 0 | 0 | 0 | 0 |
| 33649 | ENSMUSG00000094610 | 0 | NA | NA | NA | NA | NA | NA | NA | 0 | 0 | 0 | 0 | 0 | 0 |
| 33650 | ENSMUSG00000094612 | 0 | NA | NA | NA | NA | NA | Olfr491 | olfactory receptor 491 [Source:MGI Symb | 0 | 0 | 0 | 0 | 0 | 0 |
| 33651 | ENSMUSG00000094613 | 15.54794357 | -0.103714965 | 0.636364235 | -0.162980506 | 0.870533784 | 0.978802475 | A630076J17Rik | RIKEN cDNA A630076J17 gene [Source:MG | 8.8341 | 22.614 | 17.106 | 17.352 | 21.812 | 5.5692 |
| 33652 | ENSMUSG00000094615 | 0 | NA | NA | NA | NA | NA | Gm4899 | predicted gene 4899 [Source:MGI Symbol | 0 | 0 | 0 | 0 | 0 | 0 |

| | | | | | | | | | | | | | | | |
|---|---|---|---|---|---|---|---|---|---|---|---|---|---|---|---|
| 33653 | ENSMUSG00000094616 | 0 | NA | NA | NA | NA | NA | Gm20888 | predicted gene, 20888 [Source:MGI Symb | 0 | 0 | 0 | 0 | 0 | 0 |
| 33654 | ENSMUSG00000094618 | 0 | NA | NA | NA | NA | NA | Gm13271 | predicted gene 13271 [Source:MGI Symbo | 0 | 0 | 0 | 0 | 0 | 0 |
| 33655 | ENSMUSG00000094619 | 0.185640647 | 1.126924175 | 4.080472857 | 0.276174898 | 0.782413736 | NA | Trav14d-3-d | T cell receptor alpha variable 14D-3-DV8 [ | 0 | 0 | 0 | 0 | 0 | 1.1138 |
| 33656 | ENSMUSG00000094620 | 0 | NA | NA | NA | NA | NA | NA | NA | 0 | 0 | 0 | 0 | 0 | 0 |
| 33657 | ENSMUSG00000094621 | 0 | NA | NA | NA | NA | NA | NA | NA | 0 | 0 | 0 | 0 | 0 | 0 |
| 33658 | ENSMUSG00000094622 | 7.321343854 | -0.560868973 | 0.859111655 | -0.652847589 | 0.51385454 | 0.885004045 | Gm3055 | predicted gene 3055 [Source:MGI Symbol | 16.865 | 4.1117 | 5.0312 | 5.7839 | 5.4531 | 6.6831 |
| 33659 | ENSMUSG00000094623 | 0 | NA | NA | NA | NA | NA | Gm21737 | predicted gene, 21737 [Source:MGI Symb | 0 | 0 | 0 | 0 | 0 | 0 |
| 33660 | ENSMUSG00000094624 | 0 | NA | NA | NA | NA | NA | Gm4836 | predicted gene 4836 [Source:MGI Symbol | 0 | 0 | 0 | 0 | 0 | 0 |
| 33661 | ENSMUSG00000094625 | 0 | NA | NA | NA | NA | NA | NA | NA | 0 | 0 | 0 | 0 | 0 | 0 |
| 33662 | ENSMUSG00000094626 | 0.335414862 | -1.779047242 | 4.030804179 | -0.441362855 | 0.658950329 | NA | Tmem121b | transmembrane protein 121B [Source:MG | 0 | 0 | 2.0125 | 0 | 0 | 0 |
| 33663 | ENSMUSG00000094627 | 0.335414862 | -1.779047242 | 4.030804179 | -0.441362855 | 0.658950329 | NA | Gm10182 | predicted pseudogene 10182 [Source:MG | 0 | 0 | 2.0125 | 0 | 0 | 0 |
| 33664 | ENSMUSG00000094628 | 0 | NA | NA | NA | NA | NA | Gm3252 | predicted gene 3252 [Source:MGI Symbol | 0 | 0 | 0 | 0 | 0 | 0 |
| 33665 | ENSMUSG00000094630 | 0 | NA | NA | NA | NA | NA | NA | NA | 0 | 0 | 0 | 0 | 0 | 0 |
| 33666 | ENSMUSG00000094631 | 0 | NA | NA | NA | NA | NA | NA | NA | 0 | 0 | 0 | 0 | 0 | 0 |
| 33667 | ENSMUSG00000094632 | 0 | NA | NA | NA | NA | NA | Gm10310 | predicted pseudogene 10310 [Source:MG | 0 | 0 | 0 | 0 | 0 | 0 |
| 33668 | ENSMUSG00000094634 | 0 | NA | NA | NA | NA | NA | Gm3468 | predicted gene 3468 [Source:MGI Symbol | 0 | 0 | 0 | 0 | 0 | 0 |
| 33669 | ENSMUSG00000094635 | 0 | NA | NA | NA | NA | NA | Gm7104 | predicted gene 7104 [Source:MGI Symbol | 0 | 0 | 0 | 0 | 0 | 0 |
| 33670 | ENSMUSG00000094636 | 0 | NA | NA | NA | NA | NA | NA | NA | 0 | 0 | 0 | 0 | 0 | 0 |
| 33671 | ENSMUSG00000094637 | 0 | NA | NA | NA | NA | NA | Vmn1r204 | vomeronasal 1 receptor 204 [Source:MGI | 0 | 0 | 0 | 0 | 0 | 0 |
| 33672 | ENSMUSG00000094638 | 0 | NA | NA | NA | NA | NA | Gm21972 | predicted gene 21972 [Source:MGI Symbo | 0 | 0 | 0 | 0 | 0 | 0 |
| 33673 | ENSMUSG00000094639 | 0 | NA | NA | NA | NA | NA | NA | NA | 0 | 0 | 0 | 0 | 0 | 0 |
| 33674 | ENSMUSG00000094641 | 0 | NA | NA | NA | NA | NA | NA | NA | 0 | 0 | 0 | 0 | 0 | 0 |
| 33675 | ENSMUSG00000094642 | 0 | NA | NA | NA | NA | NA | NA | NA | 0 | 0 | 0 | 0 | 0 | 0 |
| 33676 | ENSMUSG00000094643 | 0 | NA | NA | NA | NA | NA | Gm8303 | predicted gene 8303 [Source:MGI Symbol | 0 | 0 | 0 | 0 | 0 | 0 |
| 33677 | ENSMUSG00000094647 | 0 | NA | NA | NA | NA | NA | Gm21650 | predicted gene, 21650 [Source:MGI Symb | 0 | 0 | 0 | 0 | 0 | 0 |
| 33678 | ENSMUSG00000094648 | 0 | NA | NA | NA | NA | NA | Gm13287 | predicted gene 13287 [Source:MGI Symbo | 0 | 0 | 0 | 0 | 0 | 0 |
| 33679 | ENSMUSG00000094649 | 0 | NA | NA | NA | NA | NA | Gm7102 | predicted gene 7102 [Source:MGI Symbol | 0 | 0 | 0 | 0 | 0 | 0 |
| 33680 | ENSMUSG00000094650 | 1.775070186 | 2.254870257 | 1.908985045 | 1.181188016 | 0.237528041 | NA | NA | NA | 0.8031 | 0 | 1.0062 | 0 | 3.2719 | 5.5692 |
| 33681 | ENSMUSG00000094651 | 0.36353943 | 2.065989254 | 4.034137301 | 0.512126658 | 0.608562368 | NA | Gal3st2 | galactose-3-O-sulfotransferase 2 [Source: | 0 | 0 | 0 | 0 | 2.1812 | 0 |
| 33682 | ENSMUSG00000094652 | 0.133850373 | -0.796629176 | 4.080472857 | -0.195229623 | 0.84521319 | NA | Ighv1-42 | immunoglobulin heavy variable V1-42 [So | 0.8031 | 0 | 0 | 0 | 0 | 0 |
| 33683 | ENSMUSG00000094654 | 0 | NA | NA | NA | NA | NA | NA | NA | 0 | 0 | 0 | 0 | 0 | 0 |
| 33684 | ENSMUSG00000094655 | 0 | NA | NA | NA | NA | NA | NA | NA | 0 | 0 | 0 | 0 | 0 | 0 |
| 33685 | ENSMUSG00000094656 | 0 | NA | NA | NA | NA | NA | NA | NA | 0 | 0 | 0 | 0 | 0 | 0 |
| 33686 | ENSMUSG00000094657 | 0 | NA | NA | NA | NA | NA | NA | NA | 0 | 0 | 0 | 0 | 0 | 0 |
| 33687 | ENSMUSG00000094658 | 0 | NA | NA | NA | NA | NA | Rbmy | RNA binding motif protein, Y chromosome | 0 | 0 | 0 | 0 | 0 | 0 |
| 33688 | ENSMUSG00000094659 | 0 | NA | NA | NA | NA | NA | NA | NA | 0 | 0 | 0 | 0 | 0 | 0 |
| 33689 | ENSMUSG00000094660 | 0 | NA | NA | NA | NA | NA | Gm21394 | predicted gene, 21394 [Source:MGI Symb | 0 | 0 | 0 | 0 | 0 | 0 |
| 33690 | ENSMUSG00000094661 | 0 | NA | NA | NA | NA | NA | NA | NA | 0 | 0 | 0 | 0 | 0 | 0 |
| 33691 | ENSMUSG00000094662 | 0 | NA | NA | NA | NA | NA | Defa36 | defensin, alpha, 36 [Source:MGI Symbol;A | 0 | 0 | 0 | 0 | 0 | 0 |
| 33692 | ENSMUSG00000094663 | 0 | NA | NA | NA | NA | NA | NA | NA | 0 | 0 | 0 | 0 | 0 | 0 |
| 33693 | ENSMUSG00000094664 | 0.133850373 | -0.796629176 | 4.080472857 | -0.195229623 | 0.84521319 | NA | Rpl35a-ps6 | ribosomal protein L35A, pseudogene 6 [So | 0.8031 | 0 | 0 | 0 | 0 | 0 |
| 33694 | ENSMUSG00000094665 | 0 | NA | NA | NA | NA | NA | NA | NA | 0 | 0 | 0 | 0 | 0 | 0 |
| 33695 | ENSMUSG00000094667 | 0 | NA | NA | NA | NA | NA | NA | NA | 0 | 0 | 0 | 0 | 0 | 0 |
| 33696 | ENSMUSG00000094668 | 0 | NA | NA | NA | NA | NA | NA | NA | 0 | 0 | 0 | 0 | 0 | 0 |
| 33697 | ENSMUSG00000094669 | 0 | NA | NA | NA | NA | NA | Olfr441 | olfactory receptor 441 [Source:MGI Symb | 0 | 0 | 0 | 0 | 0 | 0 |
| 33698 | ENSMUSG00000094670 | 0 | NA | NA | NA | NA | NA | NA | NA | 0 | 0 | 0 | 0 | 0 | 0 |
| 33699 | ENSMUSG00000094672 | 0 | NA | NA | NA | NA | NA | Vmn2r25 | vomeronasal 2, receptor 25 [Source:MGI S | 0 | 0 | 0 | 0 | 0 | 0 |
| 33700 | ENSMUSG00000094673 | 0 | NA | NA | NA | NA | NA | Olfr1354 | olfactory receptor 1354 [Source:MGI Symb | 0 | 0 | 0 | 0 | 0 | 0 |
| 33701 | ENSMUSG00000094675 | 0 | NA | NA | NA | NA | NA | Astx4b | amplified spermatogenic transcripts X enc | 0 | 0 | 0 | 0 | 0 | 0 |
| 33702 | ENSMUSG00000094676 | 0 | NA | NA | NA | NA | NA | Gm10105 | predicted pseudogene 10105 [Source:MG | 0 | 0 | 0 | 0 | 0 | 0 |
| 33703 | ENSMUSG00000094677 | 0 | NA | NA | NA | NA | NA | NA | NA | 0 | 0 | 0 | 0 | 0 | 0 |
| 33704 | ENSMUSG00000094678 | 0 | NA | NA | NA | NA | NA | Olfr857 | olfactory receptor 857 [Source:MGI Symb | 0 | 0 | 0 | 0 | 0 | 0 |
| 33705 | ENSMUSG00000094679 | 0 | NA | NA | NA | NA | NA | Gm21721 | predicted gene, 21721 [Source:MGI Symb | 0 | 0 | 0 | 0 | 0 | 0 |
| 33706 | ENSMUSG00000094680 | 0 | NA | NA | NA | NA | NA | Vmn1r121 | vomeronasal 1 receptor 121 [Source:MGI | 0 | 0 | 0 | 0 | 0 | 0 |
| 33707 | ENSMUSG00000094681 | 0 | NA | NA | NA | NA | NA | NA | NA | 0 | 0 | 0 | 0 | 0 | 0 |
| 33708 | ENSMUSG00000094682 | 0 | NA | NA | NA | NA | NA | Gm6902 | predicted gene 6902 [Source:MGI Symbol | 0 | 0 | 0 | 0 | 0 | 0 |

| | | | | | | | | | | | | | | | |
|---|---|---|---|---|---|---|---|---|---|---|---|---|---|---|---|
| 33709 | ENSMUSG00000094683 | 3.927527118 | -1.434112498 | 1.123896383 | -1.27601843 | 0.201949004 | NA | Ap3m1-ps | adaptor-related protein complex 3, mu 1 s | 4.0155 | 5.1396 | 8.05 | 1.928 | 1.0906 | 3.3415 |
| 33710 | ENSMUSG00000094684 | 0 | NA | NA | NA | NA | NA | Gm16456 | predicted gene 16456 [Source:MGI Symbo | 0 | 0 | 0 | 0 | 0 | 0 |
| 33711 | ENSMUSG00000094685 | 2.162771125 | -0.545077627 | 1.415634139 | -0.385041313 | 0.700206807 | NA | Gm5900 | predicted pseudogene 5900 [Source:MGI | 1.6062 | 4.1117 | 2.0125 | 1.928 | 1.0906 | 2.2277 |
| 33712 | ENSMUSG00000094686 | 0 | NA | NA | NA | NA | NA | Ccl21a | chemokine (C-C motif) ligand 21A (serine) | 0 | 0 | 0 | 0 | 0 | 0 |
| 33713 | ENSMUSG00000094687 | 0 | NA | NA | NA | NA | NA | Defa25 | defensin, alpha, 25 [Source:MGI Symbol;A | 0 | 0 | 0 | 0 | 0 | 0 |
| 33714 | ENSMUSG00000094688 | 0 | NA | NA | NA | NA | NA | NA | NA | 0 | 0 | 0 | 0 | 0 | 0 |
| 33715 | ENSMUSG00000094689 | 1.038420047 | 0.084391034 | 2.062151212 | 0.040923786 | 0.967356655 | NA | Ighv1-81 | immunoglobulin heavy variable 1-81 [Sou | 0 | 2.0558 | 1.0062 | 0.964 | 1.0906 | 1.1138 |
| 33716 | ENSMUSG00000094690 | 0 | NA | NA | NA | NA | NA | 1600014C23 | RIKEN cDNA 1600014C23 gene [Source:M | 0 | 0 | 0 | 0 | 0 | 0 |
| 33717 | ENSMUSG00000094691 | 0 | NA | NA | NA | NA | NA | NA | NA | 0 | 0 | 0 | 0 | 0 | 0 |
| 33718 | ENSMUSG00000094692 | 0 | NA | NA | NA | NA | NA | Olfr738 | olfactory receptor 738 [Source:MGI Symb | 0 | 0 | 0 | 0 | 0 | 0 |
| 33719 | ENSMUSG00000094693 | 0 | NA | NA | NA | NA | NA | Gm4726 | predicted gene 4726 [Source:MGI Symbol | 0 | 0 | 0 | 0 | 0 | 0 |
| 33720 | ENSMUSG00000094694 | 0.17131964 | -0.796629176 | 4.080472857 | -0.195229623 | 0.84521319 | NA | Ighv1-9 | immunoglobulin heavy variable V1-9 [Sou | 0 | 1.0279 | 0 | 0 | 0 | 0 |
| 33721 | ENSMUSG00000094695 | 0 | NA | NA | NA | NA | NA | Gm21953 | predicted gene, 21953 [Source:MGI Symb | 0 | 0 | 0 | 0 | 0 | 0 |
| 33722 | ENSMUSG00000094696 | 0 | NA | NA | NA | NA | NA | Gm6632 | predicted pseudogene 6632 [Source:MGI | 0 | 0 | 0 | 0 | 0 | 0 |
| 33723 | ENSMUSG00000094698 | 0 | NA | NA | NA | NA | NA | Gm14596 | predicted gene 14596 [Source:MGI Symbo | 0 | 0 | 0 | 0 | 0 | 0 |
| 33724 | ENSMUSG00000094699 | 0 | NA | NA | NA | NA | NA | Gm28637 | predicted gene 28637 [Source:MGI Symbo | 0 | 0 | 0 | 0 | 0 | 0 |
| 33725 | ENSMUSG00000094700 | 0 | NA | NA | NA | NA | NA | Vmn1r158 | vomeronasal 1 receptor 158 [Source:MGI | 0 | 0 | 0 | 0 | 0 | 0 |
| 33726 | ENSMUSG00000094701 | 0 | NA | NA | NA | NA | NA | Olfr916 | olfactory receptor 916 [Source:MGI Symb | 0 | 0 | 0 | 0 | 0 | 0 |
| 33727 | ENSMUSG00000094702 | 0 | NA | NA | NA | NA | NA | NA | NA | 0 | 0 | 0 | 0 | 0 | 0 |
| 33728 | ENSMUSG00000094703 | 0 | NA | NA | NA | NA | NA | NA | NA | 0 | 0 | 0 | 0 | 0 | 0 |
| 33729 | ENSMUSG00000094704 | 0 | NA | NA | NA | NA | NA | NA | NA | 0 | 0 | 0 | 0 | 0 | 0 |
| 33730 | ENSMUSG00000094706 | 0 | NA | NA | NA | NA | NA | Gm10338 | predicted gene 10338 [Source:MGI Symbo | 0 | 0 | 0 | 0 | 0 | 0 |
| 33731 | ENSMUSG00000094707 | 2.525023564 | -1.453826053 | 1.446608176 | -1.004989518 | 0.314901897 | NA | A830019P07 | RIKEN cDNA A830019P07 gene [Source:M | 4.0155 | 5.1396 | 2.0125 | 2.8919 | 1.0906 | 0 |
| 33732 | ENSMUSG00000094708 | 3.445252841 | -1.14618085 | 1.169466309 | -0.980088816 | 0.32704228 | NA | Gm10359 | predicted gene 10359 [Source:MGI Symbo | 4.0155 | 6.1675 | 4.025 | 0.964 | 3.2719 | 2.2277 |
| 33733 | ENSMUSG00000094709 | 0 | NA | NA | NA | NA | NA | NA | NA | 0 | 0 | 0 | 0 | 0 | 0 |
| 33734 | ENSMUSG00000094710 | 0 | NA | NA | NA | NA | NA | NA | NA | 0 | 0 | 0 | 0 | 0 | 0 |
| 33735 | ENSMUSG00000094712 | 0 | NA | NA | NA | NA | NA | Gm15099 | predicted gene 15099 [Source:MGI Symbo | 0 | 0 | 0 | 0 | 0 | 0 |
| 33736 | ENSMUSG00000094713 | 0 | NA | NA | NA | NA | NA | NA | NA | 0 | 0 | 0 | 0 | 0 | 0 |
| 33737 | ENSMUSG00000094714 | 0 | NA | NA | NA | NA | NA | Gm16430 | predicted gene 16430 [Source:MGI Symbo | 0 | 0 | 0 | 0 | 0 | 0 |
| 33738 | ENSMUSG00000094715 | 0 | NA | NA | NA | NA | NA | Gm17078 | predicted gene 17078 [Source:MGI Symbo | 0 | 0 | 0 | 0 | 0 | 0 |
| 33739 | ENSMUSG00000094717 | 0 | NA | NA | NA | NA | NA | NA | NA | 0 | 0 | 0 | 0 | 0 | 0 |
| 33740 | ENSMUSG00000094718 | 0 | NA | NA | NA | NA | NA | NA | NA | 0 | 0 | 0 | 0 | 0 | 0 |
| 33741 | ENSMUSG00000094719 | 0 | NA | NA | NA | NA | NA | Gm5108 | predicted gene 5108 [Source:MGI Symbol | 0 | 0 | 0 | 0 | 0 | 0 |
| 33742 | ENSMUSG00000094720 | 0 | NA | NA | NA | NA | NA | NA | NA | 0 | 0 | 0 | 0 | 0 | 0 |
| 33743 | ENSMUSG00000094721 | 0 | NA | NA | NA | NA | NA | Olfr1462 | olfactory receptor 1462 [Source:MGI Sym | 0 | 0 | 0 | 0 | 0 | 0 |
| 33744 | ENSMUSG00000094722 | 0 | NA | NA | NA | NA | NA | | | 0 | 0 | 0 | 0 | 0 | 0 |
| 33745 | ENSMUSG00000094723 | 0 | NA | NA | NA | NA | NA | NA | NA | 0 | 0 | 0 | 0 | 0 | 0 |
| 33746 | ENSMUSG00000094724 | 9.828104939 | 0.556609539 | 0.692358962 | 0.80393202 | 0.421436236 | 0.838928396 | Rnaset2b | ribonuclease T2B [Source:MGI Symbol;Ac | 9.6372 | 6.1675 | 8.05 | 16.388 | 9.8156 | 8.9108 |
| 33747 | ENSMUSG00000094725 | 0 | NA | NA | NA | NA | NA | Gm15251 | predicted gene 15251 [Source:MGI Symbo | 0 | 0 | 0 | 0 | 0 | 0 |
| 33748 | ENSMUSG00000094726 | 0 | NA | NA | NA | NA | NA | NA | NA | 0 | 0 | 0 | 0 | 0 | 0 |
| 33749 | ENSMUSG00000094727 | 0 | NA | NA | NA | NA | NA | NA | NA | 0 | 0 | 0 | 0 | 0 | 0 |
| 33750 | ENSMUSG00000094728 | 0 | NA | NA | NA | NA | NA | | | 0 | 0 | 0 | 0 | 0 | 0 |
| 33751 | ENSMUSG00000094729 | 0 | NA | NA | NA | NA | NA | Gm20747 | predicted gene, 20747 [Source:MGI Symb | 0 | 0 | 0 | 0 | 0 | 0 |
| 33752 | ENSMUSG00000094730 | 0 | NA | NA | NA | NA | NA | NA | NA | 0 | 0 | 0 | 0 | 0 | 0 |
| 33753 | ENSMUSG00000094732 | 4.211602985 | -0.239860442 | 3.055935613 | -0.078490018 | 0.93743827 | NA | 1500015L24 | RIKEN cDNA 1500015L24 gene [Source:M | 13.653 | 0 | 0 | 2.8919 | 8.7249 | 0 |
| 33754 | ENSMUSG00000094733 | 0 | NA | NA | NA | NA | NA | Csta3 | cystatin A family member 3 [Source:MGI S | 0 | 0 | 0 | 0 | 0 | 0 |
| 33755 | ENSMUSG00000094734 | 0 | NA | NA | NA | NA | NA | Olfr799 | olfactory receptor 799 [Source:MGI Symb | 0 | 0 | 0 | 0 | 0 | 0 |
| 33756 | ENSMUSG00000094735 | 0 | NA | NA | NA | NA | NA | Vmn1r95 | vomeronasal 1 receptor, 95 [Source:MGI S | 0 | 0 | 0 | 0 | 0 | 0 |
| 33757 | ENSMUSG00000094736 | 0 | NA | NA | NA | NA | NA | Gm4073 | predicted gene 4073 [Source:MGI Symbol | 0 | 0 | 0 | 0 | 0 | 0 |
| 33758 | ENSMUSG00000094737 | 0 | NA | NA | NA | NA | NA | Gm17045 | predicted gene 17045 [Source:MGI Symbo | 0 | 0 | 0 | 0 | 0 | 0 |
| 33759 | ENSMUSG00000094738 | 0 | NA | NA | NA | NA | NA | NA | NA | 0 | 0 | 0 | 0 | 0 | 0 |
| 33760 | ENSMUSG00000094739 | 0 | NA | NA | NA | NA | NA | Gm20806 | predicted gene, 20806 [Source:MGI Symb | 0 | 0 | 0 | 0 | 0 | 0 |
| 33761 | ENSMUSG00000094741 | 0 | NA | NA | NA | NA | NA | | | 0 | 0 | 0 | 0 | 0 | 0 |
| 33762 | ENSMUSG00000094743 | 52.91479051 | 0.251679572 | 0.400189748 | 0.628900599 | 0.529414134 | 0.888334265 | Mup-ps7 | major urinary protein, pseudogene 7 [Sou | 66.657 | 19.53 | 58.362 | 61.695 | 58.893 | 52.351 |
| 33763 | ENSMUSG00000094745 | 0 | NA | NA | NA | NA | NA | Olfr954 | olfactory receptor 954 [Source:MGI Symb | 0 | 0 | 0 | 0 | 0 | 0 |
| 33764 | ENSMUSG00000094746 | 0 | NA | NA | NA | NA | NA | Gm20916 | predicted gene, 20916 [Source:MGI Symb | 0 | 0 | 0 | 0 | 0 | 0 |

| | | | | | | | | | | | | | | | |
|---|---|---|---|---|---|---|---|---|---|---|---|---|---|---|---|
| 33765 | ENSMUSG00000094747 | 0 | NA | NA | NA | NA | NA | Olfr1307 | olfactory receptor 1307 [Source:MGI Syml | 0 | 0 | 0 | 0 | 0 | 0 |
| 33766 | ENSMUSG00000094748 | 0 | NA | NA | NA | NA | NA | Vmn1r255 | vomeronasal 1 receptor 255 [Source:MGI | 0 | 0 | 0 | 0 | 0 | 0 |
| 33767 | ENSMUSG00000094750 | 0 | NA | NA | NA | NA | NA | NA | NA | 0 | 0 | 0 | 0 | 0 | 0 |
| 33768 | ENSMUSG00000094751 | 0 | NA | NA | NA | NA | NA | NA | NA | 0 | 0 | 0 | 0 | 0 | 0 |
| 33769 | ENSMUSG00000094752 | 0 | NA | NA | NA | NA | NA | Vmn2r-ps12 | vomeronasal 2, receptor, pseudogene 129 | 0 | 0 | 0 | 0 | 0 | 0 |
| 33770 | ENSMUSG00000094753 | 0 | NA | NA | NA | NA | NA | NA | NA | 0 | 0 | 0 | 0 | 0 | 0 |
| 33771 | ENSMUSG00000094754 | 0 | NA | NA | NA | NA | NA | NA | NA | 0 | 0 | 0 | 0 | 0 | 0 |
| 33772 | ENSMUSG00000094755 | 0 | NA | NA | NA | NA | NA | Olfr1453 | olfactory receptor 1453 [Source:MGI Syml | 0 | 0 | 0 | 0 | 0 | 0 |
| 33773 | ENSMUSG00000094757 | 0 | NA | NA | NA | NA | NA | Vmn1r242 | vomeronasal 1 receptor 242 [Source:MGI | 0 | 0 | 0 | 0 | 0 | 0 |
| 33774 | ENSMUSG00000094758 | 0 | NA | NA | NA | NA | NA | Trav13d-2 | T cell receptor alpha variable 13D-2 [Sour | 0 | 0 | 0 | 0 | 0 | 0 |
| 33775 | ENSMUSG00000094759 | 0 | NA | NA | NA | NA | NA | Gm10487 | predicted gene 10487 [Source:MGI Symbo | 0 | 0 | 0 | 0 | 0 | 0 |
| 33776 | ENSMUSG00000094760 | 0 | NA | NA | NA | NA | NA | NA | NA | 0 | 0 | 0 | 0 | 0 | 0 |
| 33777 | ENSMUSG00000094762 | 0 | NA | NA | NA | NA | NA | Gm10670 | predicted gene 10670 [Source:MGI Symbo | 0 | 0 | 0 | 0 | 0 | 0 |
| 33778 | ENSMUSG00000094763 | 0 | NA | NA | NA | NA | NA | Gm1647 | predicted gene 1647 [Source:MGI Symbol | 0 | 0 | 0 | 0 | 0 | 0 |
| 33779 | ENSMUSG00000094764 | 0 | NA | NA | NA | NA | NA | Olfr346 | olfactory receptor 346 [Source:MGI Symb | 0 | 0 | 0 | 0 | 0 | 0 |
| 33780 | ENSMUSG00000094765 | 0 | NA | NA | NA | NA | NA | NA | NA | 0 | 0 | 0 | 0 | 0 | 0 |
| 33781 | ENSMUSG00000094766 | 0 | NA | NA | NA | NA | NA | Trav7-4 | T cell receptor alpha variable 7-4 [Source: | 0 | 0 | 0 | 0 | 0 | 0 |
| 33782 | ENSMUSG00000094767 | 0 | NA | NA | NA | NA | NA | Gm8402 | predicted gene 8402 [Source:MGI Symbol | 0 | 0 | 0 | 0 | 0 | 0 |
| 33783 | ENSMUSG00000094768 | 0 | NA | NA | NA | NA | NA | NA | NA | 0 | 0 | 0 | 0 | 0 | 0 |
| 33784 | ENSMUSG00000094771 | 0 | NA | NA | NA | NA | NA | Gm23860 | predicted gene, 23860 [Source:MGI Symb | 0 | 0 | 0 | 0 | 0 | 0 |
| 33785 | ENSMUSG00000094772 | 11.16595315 | 0.161762808 | 0.673486962 | 0.240186993 | 0.810185297 | 0.966157155 | Rpl29-ps2 | ribosomal protein L29, pseudogene 2 [Sou | 7.2279 | 13.363 | 11.069 | 6.7478 | 17.45 | 11.138 |
| 33786 | ENSMUSG00000094773 | 0 | NA | NA | NA | NA | NA | Gm21812 | predicted gene, 21812 [Source:MGI Symb | 0 | 0 | 0 | 0 | 0 | 0 |
| 33787 | ENSMUSG00000094775 | 0 | NA | NA | NA | NA | NA | Gm2759 | predicted gene 2759 [Source:MGI Symbol | 0 | 0 | 0 | 0 | 0 | 0 |
| 33788 | ENSMUSG00000094777 | 0 | NA | NA | NA | NA | NA | H2ac24 | H2A clustered histone 24 [Source:MGI Syr | 0 | 0 | 0 | 0 | 0 | 0 |
| 33789 | ENSMUSG00000094778 | 0 | NA | NA | NA | NA | NA | Olfr143 | olfactory receptor 143 [Source:MGI Symb | 0 | 0 | 0 | 0 | 0 | 0 |
| 33790 | ENSMUSG00000094779 | 0 | NA | NA | NA | NA | NA | Trav13n-2 | T cell receptor alpha variable 13N-2 [Sour | 0 | 0 | 0 | 0 | 0 | 0 |
| 33791 | ENSMUSG00000094780 | 0 | NA | NA | NA | NA | NA | NA | NA | 0 | 0 | 0 | 0 | 0 | 0 |
| 33792 | ENSMUSG00000094782 | 0 | NA | NA | NA | NA | NA | Gm21256 | predicted gene, 21256 [Source:MGI Symb | 0 | 0 | 0 | 0 | 0 | 0 |
| 33793 | ENSMUSG00000094784 | 0 | NA | NA | NA | NA | NA | Gm8122 | predicted gene 8122 [Source:MGI Symbol | 0 | 0 | 0 | 0 | 0 | 0 |
| 33794 | ENSMUSG00000094785 | 0 | NA | NA | NA | NA | NA | NA | NA | 0 | 0 | 0 | 0 | 0 | 0 |
| 33795 | ENSMUSG00000094786 | 325.9979703 | -0.0610335 | 0.199396745 | -0.306090752 | 0.759535549 | 0.95402891 | Gm14403 | predicted gene 14403 [Source:MGI Symbo | 339.71 | 364.91 | 294.83 | 420.29 | 291.2 | 245.05 |
| 33796 | ENSMUSG00000094787 | 0 | NA | NA | NA | NA | NA | Ighv1-54 | immunoglobulin heavy variable V1-54 [So | 0 | 0 | 0 | 0 | 0 | 0 |
| 33797 | ENSMUSG00000094789 | 0 | NA | NA | NA | NA | NA | Gm28490 | predicted gene 28490 [Source:MGI Symbo | 0 | 0 | 0 | 0 | 0 | 0 |
| 33798 | ENSMUSG00000094790 | 0 | NA | NA | NA | NA | NA | Gm6625 | predicted gene 6625 [Source:MGI Symbol | 0 | 0 | 0 | 0 | 0 | 0 |
| 33799 | ENSMUSG00000094791 | 0 | NA | NA | NA | NA | NA | | | 0 | 0 | 0 | 0 | 0 | 0 |
| 33800 | ENSMUSG00000094792 | 0 | NA | NA | NA | NA | NA | Trav10d | T cell receptor alpha variable 10D [Source | 0 | 0 | 0 | 0 | 0 | 0 |
| 33801 | ENSMUSG00000094793 | 209.1586197 | 1.557200432 | 0.672419661 | 2.315816333 | 0.020568299 | 0.176557791 | Mup12 | major urinary protein 12 [Source:MGI Syn | 85.932 | 44.2 | 188.17 | 171.59 | 425.34 | 339.72 |
| 33802 | ENSMUSG00000094794 | 0 | NA | NA | NA | NA | NA | Gm6019 | predicted gene 6019 [Source:MGI Symbol | 0 | 0 | 0 | 0 | 0 | 0 |
| 33803 | ENSMUSG00000094796 | 1.843383844 | -3.159990241 | 1.853088091 | -1.705256354 | 0.088146624 | NA | BC147527 | cDNA sequence BC147527 [Source:MGI Sy | 4.0155 | 2.0558 | 4.025 | 0.964 | 0 | 0 |
| 33804 | ENSMUSG00000094797 | 0.635111108 | 0.379830552 | 2.868137728 | 0.132431071 | 0.89464334 | NA | Igkv6-15 | immunoglobulin kappa variable 6-15 [Sou | 1.6062 | 0 | 0 | 0 | 1.0906 | 1.1138 |
| 33805 | ENSMUSG00000094799 | 0 | NA | NA | NA | NA | NA | | | 0 | 0 | 0 | 0 | 0 | 0 |
| 33806 | ENSMUSG00000094800 | 0 | NA | NA | NA | NA | NA | NA | NA | 0 | 0 | 0 | 0 | 0 | 0 |
| 33807 | ENSMUSG00000094801 | 0 | NA | NA | NA | NA | NA | NA | NA | 0 | 0 | 0 | 0 | 0 | 0 |
| 33808 | ENSMUSG00000094802 | 0 | NA | NA | NA | NA | NA | Samt1 | spermatogenesis associated multipass tra | 0 | 0 | 0 | 0 | 0 | 0 |
| 33809 | ENSMUSG00000094803 | 0 | NA | NA | NA | NA | NA | NA | NA | 0 | 0 | 0 | 0 | 0 | 0 |
| 33810 | ENSMUSG00000094804 | 0 | NA | NA | NA | NA | NA | Gm21977 | predicted gene 21977 [Source:MGI Symbo | 0 | 0 | 0 | 0 | 0 | 0 |
| 33811 | ENSMUSG00000094805 | 0 | NA | NA | NA | NA | NA | Olfr311 | olfactory receptor 311 [Source:MGI Symb | 0 | 0 | 0 | 0 | 0 | 0 |
| 33812 | ENSMUSG00000094806 | 17091.31249 | -0.000425925 | 0.085682348 | -0.00497098 | 0.996033748 | 0.999003684 | Cyp2d10 | cytochrome P450, family 2, subfamily d, p | 16910 | 16556 | 17817 | 17982 | 16650 | 16633 |
| 33813 | ENSMUSG00000094807 | 0.181769715 | 1.126924175 | 4.080472857 | 0.276174898 | 0.782413736 | NA | NA | NA | 0 | 0 | 0 | 0 | 1.0906 | 0 |
| 33814 | ENSMUSG00000094808 | 0 | NA | NA | NA | NA | NA | 1810009J06I | RIKEN cDNA 1810009J06 gene [Source:MC | 0 | 0 | 0 | 0 | 0 | 0 |
| 33815 | ENSMUSG00000094810 | 0 | NA | NA | NA | NA | NA | Olfr907 | olfactory receptor 907 [Source:MGI Symb | 0 | 0 | 0 | 0 | 0 | 0 |
| 33816 | ENSMUSG00000094811 | 0 | NA | NA | NA | NA | NA | Gm21103 | predicted gene, 21103 [Source:MGI Symb | 0 | 0 | 0 | 0 | 0 | 0 |
| 33817 | ENSMUSG00000094812 | 0 | NA | NA | NA | NA | NA | NA | NA | 0 | 0 | 0 | 0 | 0 | 0 |
| 33818 | ENSMUSG00000094813 | 0 | NA | NA | NA | NA | NA | Gm20777 | predicted gene, 20777 [Source:MGI Symb | 0 | 0 | 0 | 0 | 0 | 0 |
| 33819 | ENSMUSG00000094814 | 0 | NA | NA | NA | NA | NA | Gm21973 | predicted gene 21973 [Source:MGI Symbo | 0 | 0 | 0 | 0 | 0 | 0 |
| 33820 | ENSMUSG00000094815 | 0 | NA | NA | NA | NA | NA | NA | NA | 0 | 0 | 0 | 0 | 0 | 0 |

| | | | | | | | | | | | | | | | |
|---|---|---|---|---|---|---|---|---|---|---|---|---|---|---|---|
| 33821 | ENSMUSG00000094817 | 0 | NA | NA | NA | NA | NA | NA | NA | 0 | 0 | 0 | 0 | 0 | 0 |
| 33822 | ENSMUSG00000094818 | 0 | NA | NA | NA | NA | NA | Defa32 | defensin, alpha, 32 [Source:MGI Symbol;A | 0 | 0 | 0 | 0 | 0 | 0 |
| 33823 | ENSMUSG00000094819 | 0.346303622 | 2.004339653 | 3.884003423 | 0.516049919 | 0.605819544 | NA | Olfr53 | olfactory receptor 53 [Source:MGI Symbo | 0 | 0 | 0 | 0.964 | 0 | 1.1138 |
| 33824 | ENSMUSG00000094820 | 0 | NA | NA | NA | NA | NA | NA | NA | 0 | 0 | 0 | 0 | 0 | 0 |
| 33825 | ENSMUSG00000094821 | 0 | NA | NA | NA | NA | NA | Gm21518 | predicted gene, 21518 [Source:MGI Symb | 0 | 0 | 0 | 0 | 0 | 0 |
| 33826 | ENSMUSG00000094822 | 0 | NA | NA | NA | NA | NA | Olfr243 | olfactory receptor 243 [Source:MGI Symb | 0 | 0 | 0 | 0 | 0 | 0 |
| 33827 | ENSMUSG00000094823 | 0 | NA | NA | NA | NA | NA | NA | NA | 0 | 0 | 0 | 0 | 0 | 0 |
| 33828 | ENSMUSG00000094824 | 0 | NA | NA | NA | NA | NA | NA | NA | 0 | 0 | 0 | 0 | 0 | 0 |
| 33829 | ENSMUSG00000094825 | 0 | NA | NA | NA | NA | NA | Gm3194 | predicted gene 3194 [Source:MGI Symbol | 0 | 0 | 0 | 0 | 0 | 0 |
| 33830 | ENSMUSG00000094826 | 0 | NA | NA | NA | NA | NA | NA | NA | 0 | 0 | 0 | 0 | 0 | 0 |
| 33831 | ENSMUSG00000094827 | 0 | NA | NA | NA | NA | NA | Gm5327 | predicted pseudogene 5327 [Source:MGI | 0 | 0 | 0 | 0 | 0 | 0 |
| 33832 | ENSMUSG00000094828 | 0 | NA | NA | NA | NA | NA | Trav3-3 | T cell receptor alpha variable 3-3 [Source: | 0 | 0 | 0 | 0 | 0 | 0 |
| 33833 | ENSMUSG00000094829 | 0 | NA | NA | NA | NA | NA | Gm21809 | predicted gene, 21809 [Source:MGI Symb | 0 | 0 | 0 | 0 | 0 | 0 |
| 33834 | ENSMUSG00000094830 | 0 | NA | NA | NA | NA | NA | NA | NA | 0 | 0 | 0 | 0 | 0 | 0 |
| 33835 | ENSMUSG00000094831 | 0 | NA | NA | NA | NA | NA | NA | NA | 0 | 0 | 0 | 0 | 0 | 0 |
| 33836 | ENSMUSG00000094832 | 0 | NA | NA | NA | NA | NA | Gm21846 | predicted gene, 21846 [Source:MGI Symb | 0 | 0 | 0 | 0 | 0 | 0 |
| 33837 | ENSMUSG00000094833 | 0 | NA | NA | NA | NA | NA | NA | NA | 0 | 0 | 0 | 0 | 0 | 0 |
| 33838 | ENSMUSG00000094834 | 0 | NA | NA | NA | NA | NA | NA | NA | 0 | 0 | 0 | 0 | 0 | 0 |
| 33839 | ENSMUSG00000094836 | 0 | NA | NA | NA | NA | NA | | | 0 | 0 | 0 | 0 | 0 | 0 |
| 33840 | ENSMUSG00000094837 | 0 | NA | NA | NA | NA | NA | NA | NA | 0 | 0 | 0 | 0 | 0 | 0 |
| 33841 | ENSMUSG00000094838 | 0 | NA | NA | NA | NA | NA | Gm20828 | predicted gene, 20828 [Source:MGI Symb | 0 | 0 | 0 | 0 | 0 | 0 |
| 33842 | ENSMUSG00000094840 | 0.17131964 | -0.796629176 | 4.080472857 | -0.195229623 | 0.84521319 | NA | Muc3a | mucin 3A, cell surface associated [Source: | 0 | 1.0279 | 0 | 0 | 0 | 0 |
| 33843 | ENSMUSG00000094841 | 0 | NA | NA | NA | NA | NA | Gm10610 | predicted gene 10610 [Source:MGI Symbo | 0 | 0 | 0 | 0 | 0 | 0 |
| 33844 | ENSMUSG00000094842 | 0.342639279 | -1.805976596 | 4.028547714 | -0.448294702 | 0.653940524 | NA | NA | NA | 0 | 2.0558 | 0 | 0 | 0 | 0 |
| 33845 | ENSMUSG00000094843 | 0 | NA | NA | NA | NA | NA | Ppp1r2-ps1 | protein phosphatase 1, regulatory inhibito | 0 | 0 | 0 | 0 | 0 | 0 |
| 33846 | ENSMUSG00000094845 | 0 | NA | NA | NA | NA | NA | Tmem95 | transmembrane protein 95 [Source:MGI S | 0 | 0 | 0 | 0 | 0 | 0 |
| 33847 | ENSMUSG00000094846 | 0 | NA | NA | NA | NA | NA | Olfr1487 | olfactory receptor 1487 [Source:MGI Sym | 0 | 0 | 0 | 0 | 0 | 0 |
| 33848 | ENSMUSG00000094847 | 0 | NA | NA | NA | NA | NA | NA | NA | 0 | 0 | 0 | 0 | 0 | 0 |
| 33849 | ENSMUSG00000094848 | 0 | NA | NA | NA | NA | NA | NA | NA | 0 | 0 | 0 | 0 | 0 | 0 |
| 33850 | ENSMUSG00000094849 | 0.160662974 | 1.126924175 | 4.080472857 | 0.276174898 | 0.782413736 | NA | Vmn2r-ps18 | vomeronasal 2, receptor, pseudogene 18 | 0 | 0 | 0 | 0.964 | 0 | 0 |
| 33851 | ENSMUSG00000094850 | 0 | NA | NA | NA | NA | NA | NA | NA | 0 | 0 | 0 | 0 | 0 | 0 |
| 33852 | ENSMUSG00000094852 | 0 | NA | NA | NA | NA | NA | Gm9034 | predicted pseudogene 9034 [Source:MGI | 0 | 0 | 0 | 0 | 0 | 0 |
| 33853 | ENSMUSG00000094854 | 0 | NA | NA | NA | NA | NA | Gm5334 | predicted pseudogene 5334 [Source:MGI | 0 | 0 | 0 | 0 | 0 | 0 |
| 33854 | ENSMUSG00000094855 | 0 | NA | NA | NA | NA | NA | | | 0 | 0 | 0 | 0 | 0 | 0 |
| 33855 | ENSMUSG00000094856 | 0 | NA | NA | NA | NA | NA | Gm21962 | predicted gene, 21962 [Source:MGI Symb | 0 | 0 | 0 | 0 | 0 | 0 |
| 33856 | ENSMUSG00000094858 | 0 | NA | NA | NA | NA | NA | Olfr1297 | olfactory receptor 1297 [Source:MGI Sym | 0 | 0 | 0 | 0 | 0 | 0 |
| 33857 | ENSMUSG00000094860 | 0 | NA | NA | NA | NA | NA | Btbd35f27 | BTB domain containing 35, family membe | 0 | 0 | 0 | 0 | 0 | 0 |
| 33858 | ENSMUSG00000094861 | 0 | NA | NA | NA | NA | NA | Mageb6b1 | MAGE family member B6B1 [Source:MGI S | 0 | 0 | 0 | 0 | 0 | 0 |
| 33859 | ENSMUSG00000094862 | 7.41084177 | 0.392585567 | 0.823801115 | 0.476553818 | 0.633679875 | 0.922736909 | Ighv1-56 | immunoglobulin heavy variable 1-56 [Sou | 4.0155 | 11.307 | 4.025 | 9.6398 | 5.4531 | 10.025 |
| 33860 | ENSMUSG00000094864 | 0 | NA | NA | NA | NA | NA | Gm17521 | predicted gene, 17521 [Source:MGI Symb | 0 | 0 | 0 | 0 | 0 | 0 |
| 33861 | ENSMUSG00000094865 | 0 | NA | NA | NA | NA | NA | Gm7426 | predicted gene 7426 [Source:MGI Symbol | 0 | 0 | 0 | 0 | 0 | 0 |
| 33862 | ENSMUSG00000094866 | 0 | NA | NA | NA | NA | NA | NA | NA | 0 | 0 | 0 | 0 | 0 | 0 |
| 33863 | ENSMUSG00000094867 | 0 | NA | NA | NA | NA | NA | NA | NA | 0 | 0 | 0 | 0 | 0 | 0 |
| 33864 | ENSMUSG00000094868 | 0 | NA | NA | NA | NA | NA | Dlx6os2 | distal-less homeobox 6, opposite strand 2 | 0 | 0 | 0 | 0 | 0 | 0 |
| 33865 | ENSMUSG00000094869 | 0 | NA | NA | NA | NA | NA | NA | NA | 0 | 0 | 0 | 0 | 0 | 0 |
| 33866 | ENSMUSG00000094870 | 211.4605795 | -0.088871492 | 0.173669419 | -0.511727928 | 0.608841438 | 0.916376001 | Zfp131 | zinc finger protein 131 [Source:MGI Symb | 241.73 | 200.44 | 211.31 | 214.97 | 188.68 | 211.63 |
| 33867 | ENSMUSG00000094872 | 0 | NA | NA | NA | NA | NA | Igkv9-120 | immunoglobulin kappa chain variable 9-1 | 0 | 0 | 0 | 0 | 0 | 0 |
| 33868 | ENSMUSG00000094873 | 0 | NA | NA | NA | NA | NA | Gm13812 | predicted gene 13812 [Source:MGI Symbo | 0 | 0 | 0 | 0 | 0 | 0 |
| 33869 | ENSMUSG00000094874 | 0 | NA | NA | NA | NA | NA | | | 0 | 0 | 0 | 0 | 0 | 0 |
| 33870 | ENSMUSG00000094875 | 0 | NA | NA | NA | NA | NA | NA | NA | 0 | 0 | 0 | 0 | 0 | 0 |
| 33871 | ENSMUSG00000094876 | 0 | NA | NA | NA | NA | NA | Btbd35f18 | BTB domain containing 35, family membe | 0 | 0 | 0 | 0 | 0 | 0 |
| 33872 | ENSMUSG00000094877 | 0 | NA | NA | NA | NA | NA | Trav3n-3 | T cell receptor alpha variable 3N-3 [Sourc | 0 | 0 | 0 | 0 | 0 | 0 |
| 33873 | ENSMUSG00000094878 | 0 | NA | NA | NA | NA | NA | Olfr130 | olfactory receptor 130 [Source:MGI Symb | 0 | 0 | 0 | 0 | 0 | 0 |
| 33874 | ENSMUSG00000094879 | 0 | NA | NA | NA | NA | NA | Vmn1r129 | vomeronasal 1 receptor 129 [Source:MGI | 0 | 0 | 0 | 0 | 0 | 0 |
| 33875 | ENSMUSG00000094880 | 0 | NA | NA | NA | NA | NA | Gm11109 | predicted gene 11109 [Source:MGI Symbo | 0 | 0 | 0 | 0 | 0 | 0 |
| 33876 | ENSMUSG00000094881 | 0 | NA | NA | NA | NA | NA | H2al2c | H2A histone family member L2C [Source:M | 0 | 0 | 0 | 0 | 0 | 0 |

| | | | | | | | | | | | | | | | |
|---|---|---|---|---|---|---|---|---|---|---|---|---|---|---|---|
| 33877 | ENSMUSG00000094883 | 0 | NA | NA | NA | NA | NA | NA | NA | 0 | 0 | 0 | 0 | 0 | 0 |
| 33878 | ENSMUSG00000094884 | 0 | NA | NA | NA | NA | NA | Olfr105-ps | olfactory receptor 105, pseudogene [Sour | 0 | 0 | 0 | 0 | 0 | 0 |
| 33879 | ENSMUSG00000094885 | 0 | NA | NA | NA | NA | NA | Ott | ovary testis transcribed [Source:MGI Syml | 0 | 0 | 0 | 0 | 0 | 0 |
| 33880 | ENSMUSG00000094886 | 0 | NA | NA | NA | NA | NA | Gm10784 | predicted pseudogene 10784 [Source:MG | 0 | 0 | 0 | 0 | 0 | 0 |
| 33881 | ENSMUSG00000094887 | 0 | NA | NA | NA | NA | NA | | | 0 | 0 | 0 | 0 | 0 | 0 |
| 33882 | ENSMUSG00000094888 | 0 | NA | NA | NA | NA | NA | NA | NA | 0 | 0 | 0 | 0 | 0 | 0 |
| 33883 | ENSMUSG00000094889 | 0 | NA | NA | NA | NA | NA | NA | NA | 0 | 0 | 0 | 0 | 0 | 0 |
| 33884 | ENSMUSG00000094890 | 0 | NA | NA | NA | NA | NA | NA | NA | 0 | 0 | 0 | 0 | 0 | 0 |
| 33885 | ENSMUSG00000094891 | 0 | NA | NA | NA | NA | NA | Olfr55 | olfactory receptor 55 [Source:MGI Symbo | 0 | 0 | 0 | 0 | 0 | 0 |
| 33886 | ENSMUSG00000094892 | 0 | NA | NA | NA | NA | NA | Vmn2r101 | vomeronasal 2, receptor 101 [Source:MGI | 0 | 0 | 0 | 0 | 0 | 0 |
| 33887 | ENSMUSG00000094893 | 0 | NA | NA | NA | NA | NA | Gm4791 | predicted gene 4791 [Source:MGI Symbol | 0 | 0 | 0 | 0 | 0 | 0 |
| 33888 | ENSMUSG00000094894 | 0 | NA | NA | NA | NA | NA | NA | NA | 0 | 0 | 0 | 0 | 0 | 0 |
| 33889 | ENSMUSG00000094895 | 0 | NA | NA | NA | NA | NA | NA | NA | 0 | 0 | 0 | 0 | 0 | 0 |
| 33890 | ENSMUSG00000094896 | 0.883540042 | 0.243447205 | 2.772289218 | 0.087814505 | 0.930024109 | NA | NA | NA | 2.4093 | 0 | 0 | 2.8919 | 0 | 0 |
| 33891 | ENSMUSG00000094897 | 0 | NA | NA | NA | NA | NA | NA | NA | 0 | 0 | 0 | 0 | 0 | 0 |
| 33892 | ENSMUSG00000094898 | 0 | NA | NA | NA | NA | NA | Vmn1r201 | vomeronasal 1 receptor 201 [Source:MGI | 0 | 0 | 0 | 0 | 0 | 0 |
| 33893 | ENSMUSG00000094901 | 0 | NA | NA | NA | NA | NA | NA | NA | 0 | 0 | 0 | 0 | 0 | 0 |
| 33894 | ENSMUSG00000094902 | 0 | NA | NA | NA | NA | NA | Igkv10-95 | immunoglobulin kappa variable 10-95 [So | 0 | 0 | 0 | 0 | 0 | 0 |
| 33895 | ENSMUSG00000094903 | 0 | NA | NA | NA | NA | NA | NA | NA | 0 | 0 | 0 | 0 | 0 | 0 |
| 33896 | ENSMUSG00000094904 | 0 | NA | NA | NA | NA | NA | H2al1d | H2A histone family member L1D [Source:I | 0 | 0 | 0 | 0 | 0 | 0 |
| 33897 | ENSMUSG00000094905 | 0 | NA | NA | NA | NA | NA | Vmn1r155 | vomeronasal 1 receptor 155 [Source:MGI | 0 | 0 | 0 | 0 | 0 | 0 |
| 33898 | ENSMUSG00000094906 | 0 | NA | NA | NA | NA | NA | NA | NA | 0 | 0 | 0 | 0 | 0 | 0 |
| 33899 | ENSMUSG00000094910 | 13.67043848 | -0.664754458 | 0.610401695 | -1.089044253 | 0.276134371 | 0.730038567 | D430019H16 | RIKEN cDNA D430019H16 gene [Source:M | 21.684 | 12.335 | 16.1 | 8.6758 | 7.6343 | 15.594 |
| 33900 | ENSMUSG00000094911 | 0 | NA | NA | NA | NA | NA | Gm21921 | predicted gene, 21921 [Source:MGI Symb | 0 | 0 | 0 | 0 | 0 | 0 |
| 33901 | ENSMUSG00000094912 | 0 | NA | NA | NA | NA | NA | NA | NA | 0 | 0 | 0 | 0 | 0 | 0 |
| 33902 | ENSMUSG00000094913 | 0 | NA | NA | NA | NA | NA | Gm9507 | predicted gene 9507 [Source:MGI Symbol | 0 | 0 | 0 | 0 | 0 | 0 |
| 33903 | ENSMUSG00000094914 | 0 | NA | NA | NA | NA | NA | Trav9d-2 | T cell receptor alpha variable 9D-2 [Sourc | 0 | 0 | 0 | 0 | 0 | 0 |
| 33904 | ENSMUSG00000094915 | 0 | NA | NA | NA | NA | NA | | | 0 | 0 | 0 | 0 | 0 | 0 |
| 33905 | ENSMUSG00000094916 | 0 | NA | NA | NA | NA | NA | NA | NA | 0 | 0 | 0 | 0 | 0 | 0 |
| 33906 | ENSMUSG00000094917 | 0 | NA | NA | NA | NA | NA | NA | NA | 0 | 0 | 0 | 0 | 0 | 0 |
| 33907 | ENSMUSG00000094918 | 0 | NA | NA | NA | NA | NA | Gm8765 | predicted gene 8765 [Source:MGI Symbol | 0 | 0 | 0 | 0 | 0 | 0 |
| 33908 | ENSMUSG00000094919 | 0 | NA | NA | NA | NA | NA | NA | NA | 0 | 0 | 0 | 0 | 0 | 0 |
| 33909 | ENSMUSG00000094920 | 0 | NA | NA | NA | NA | NA | Gm6605 | predicted gene 6605 [Source:MGI Symbol | 0 | 0 | 0 | 0 | 0 | 0 |
| 33910 | ENSMUSG00000094921 | 0 | NA | NA | NA | NA | NA | Vmn2r112 | vomeronasal 2, receptor 112 [Source:MGI | 0 | 0 | 0 | 0 | 0 | 0 |
| 33911 | ENSMUSG00000094922 | 0 | NA | NA | NA | NA | NA | NA | NA | 0 | 0 | 0 | 0 | 0 | 0 |
| 33912 | ENSMUSG00000094923 | 0 | NA | NA | NA | NA | NA | NA | NA | 0 | 0 | 0 | 0 | 0 | 0 |
| 33913 | ENSMUSG00000094925 | 0 | NA | NA | NA | NA | NA | Gm5798 | predicted gene 5798 [Source:MGI Symbol | 0 | 0 | 0 | 0 | 0 | 0 |
| 33914 | ENSMUSG00000094926 | 0 | NA | NA | NA | NA | NA | Gm16533 | predited gene 16533 [Source:MGI Symbol | 0 | 0 | 0 | 0 | 0 | 0 |
| 33915 | ENSMUSG00000094927 | 0 | NA | NA | NA | NA | NA | NA | NA | 0 | 0 | 0 | 0 | 0 | 0 |
| 33916 | ENSMUSG00000094928 | 0 | NA | NA | NA | NA | NA | 1700122O11 | RIKEN cDNA 1700122O11 gene [Source:M | 0 | 0 | 0 | 0 | 0 | 0 |
| 33917 | ENSMUSG00000094929 | 0 | NA | NA | NA | NA | NA | Gm3526 | predicted gene 3526 [Source:MGI Symbol | 0 | 0 | 0 | 0 | 0 | 0 |
| 33918 | ENSMUSG00000094930 | 0.167707431 | -0.796629176 | 4.080472857 | -0.195229623 | 0.84521319 | NA | Igkv6-25 | immunoglobulin kappa chain variable 6-2! | 0 | 0 | 1.0062 | 0 | 0 | 0 |
| 33919 | ENSMUSG00000094931 | 0 | NA | NA | NA | NA | NA | Vmn1r160 | vomeronasal 1 receptor 160 [Source:MGI | 0 | 0 | 0 | 0 | 0 | 0 |
| 33920 | ENSMUSG00000094932 | 0 | NA | NA | NA | NA | NA | Gm2007 | predicted gene 2007 [Source:MGI Symbol | 0 | 0 | 0 | 0 | 0 | 0 |
| 33921 | ENSMUSG00000094933 | 0 | NA | NA | NA | NA | NA | NA | NA | 0 | 0 | 0 | 0 | 0 | 0 |
| 33922 | ENSMUSG00000094934 | 0 | NA | NA | NA | NA | NA | Vmn1r163 | vomeronasal 1 receptor 163 [Source:MGI | 0 | 0 | 0 | 0 | 0 | 0 |
| 33923 | ENSMUSG00000094935 | 1.875095644 | -0.12049151 | 1.514722053 | -0.079546944 | 0.936597594 | NA | Gm9726 | predicted gene 9726 [Source:MGI Symbol | 0.8031 | 2.0558 | 3.0187 | 0.964 | 2.1812 | 2.2277 |
| 33924 | ENSMUSG00000094936 | 28.13385612 | 0.114087382 | 0.443064198 | 0.257496277 | 0.796795689 | 0.962236316 | Rbm4 | RNA binding motif protein 4 [Source:MGI | 19.274 | 37.005 | 25.156 | 35.667 | 30.537 | 21.163 |
| 33925 | ENSMUSG00000094937 | 0 | NA | NA | NA | NA | NA | NA | NA | 0 | 0 | 0 | 0 | 0 | 0 |
| 33926 | ENSMUSG00000094938 | 0.181769715 | 1.126924175 | 4.080472857 | 0.276174898 | 0.782413736 | NA | NA | NA | 0 | 0 | 0 | 0 | 1.0906 | 0 |
| 33927 | ENSMUSG00000094939 | 0 | NA | NA | NA | NA | NA | Gm21773 | predicted gene, 21773 [Source:MGI Symb | 0 | 0 | 0 | 0 | 0 | 0 |
| 33928 | ENSMUSG00000094940 | 0 | NA | NA | NA | NA | NA | Ighv1-84 | immunoglobulin heavy variable 1-84 [Sou | 0 | 0 | 0 | 0 | 0 | 0 |
| 33929 | ENSMUSG00000094941 | 0 | NA | NA | NA | NA | NA | Gm15093 | predicted gene 15093 [Source:MGI Symbo | 0 | 0 | 0 | 0 | 0 | 0 |
| 33930 | ENSMUSG00000094942 | 2.334167874 | -0.804328747 | 1.414626726 | -0.5685802 | 0.569641065 | NA | Gm3604 | predicted gene 3604 [Source:MGI Symbol | 4.8186 | 3.0838 | 1.0062 | 2.8919 | 1.0906 | 1.1138 |
| 33931 | ENSMUSG00000094943 | 0 | NA | NA | NA | NA | NA | NA | NA | 0 | 0 | 0 | 0 | 0 | 0 |
| 33932 | ENSMUSG00000094945 | 0 | NA | NA | NA | NA | NA | Rhox4a2 | reproductive homeobox 4A2 [Source:MGI | 0 | 0 | 0 | 0 | 0 | 0 |

| | | | | | | | | | | | | | | | |
|---|---|---|---|---|---|---|---|---|---|---|---|---|---|---|---|
| 33933 | ENSMUSG00000094946 | 0 | NA | NA | NA | NA | NA | NA | NA | 0 | 0 | 0 | 0 | 0 | 0 |
| 33934 | ENSMUSG00000094948 | 1.122573798 | -0.362418364 | 2.131244112 | -0.170050142 | 0.864970703 | NA | Gm21286 | predicted gene, 21286 [Source:MGI Symb | 0.8031 | 1.0279 | 2.0125 | 2.8919 | 0 | 0 |
| 33935 | ENSMUSG00000094950 | 0 | NA | NA | NA | NA | NA | Vmn2r66 | vomeronasal 2, receptor 66 [Source:MGI S | 0 | 0 | 0 | 0 | 0 | 0 |
| 33936 | ENSMUSG00000094951 | 0.181769715 | 1.126924175 | 4.080472857 | 0.276174898 | 0.782413736 | NA | Ighv5-6 | immunoglobulin heavy variable 5-6 [Sourc | 0 | 0 | 0 | 0 | 1.0906 | 0 |
| 33937 | ENSMUSG00000094952 | 0 | NA | NA | NA | NA | NA | NA | NA | 0 | 0 | 0 | 0 | 0 | 0 |
| 33938 | ENSMUSG00000094954 | 0 | NA | NA | NA | NA | NA | Gm8011 | predicted gene 8011 [Source:MGI Symbol | 0 | 0 | 0 | 0 | 0 | 0 |
| 33939 | ENSMUSG00000094955 | 2.578384528 | -0.53753913 | 1.31517956 | -0.408719194 | 0.682745746 | NA | Gm3699 | predicted gene 3699 [Source:MGI Symbol | 4.0155 | 4.1117 | 1.0062 | 1.928 | 2.1812 | 2.2277 |
| 33940 | ENSMUSG00000094956 | 0 | NA | NA | NA | NA | NA | NA | NA | 0 | 0 | 0 | 0 | 0 | 0 |
| 33941 | ENSMUSG00000094957 | 0 | NA | NA | NA | NA | NA | Ighd5-5 | immunoglobulin heavy diversity 5-5 [Sour | 0 | 0 | 0 | 0 | 0 | 0 |
| 33942 | ENSMUSG00000094958 | 8.860143448 | 0.534724625 | 0.793836106 | 0.673595748 | 0.500568357 | 0.880912608 | 3110021N24 | RIKEN cDNA 3110021N24 gene [Source:M | 5.6217 | 8.2233 | 8.05 | 20.244 | 5.4531 | 5.5692 |
| 33943 | ENSMUSG00000094959 | 0 | NA | NA | NA | NA | NA | NA | NA | 0 | 0 | 0 | 0 | 0 | 0 |
| 33944 | ENSMUSG00000094960 | 0 | NA | NA | NA | NA | NA | NA | NA | 0 | 0 | 0 | 0 | 0 | 0 |
| 33945 | ENSMUSG00000094962 | 0 | NA | NA | NA | NA | NA | Gm21954 | predicted gene, 21954 [Source:MGI Symb | 0 | 0 | 0 | 0 | 0 | 0 |
| 33946 | ENSMUSG00000094963 | 0 | NA | NA | NA | NA | NA | NA | NA | 0 | 0 | 0 | 0 | 0 | 0 |
| 33947 | ENSMUSG00000094964 | 0 | NA | NA | NA | NA | NA | Gm15101 | predicted gene 15101 [Source:MGI Symbo | 0 | 0 | 0 | 0 | 0 | 0 |
| 33948 | ENSMUSG00000094965 | 0 | NA | NA | NA | NA | NA | NA | NA | 0 | 0 | 0 | 0 | 0 | 0 |
| 33949 | ENSMUSG00000094966 | 0 | NA | NA | NA | NA | NA | Trav9-2 | T cell receptor alpha variable 9-2 [Source: | 0 | 0 | 0 | 0 | 0 | 0 |
| 33950 | ENSMUSG00000094967 | 0 | NA | NA | NA | NA | NA | Obox4-ps18 | oocyte specific homeobox 4, pseudogene | 0 | 0 | 0 | 0 | 0 | 0 |
| 33951 | ENSMUSG00000094968 | 0 | NA | NA | NA | NA | NA | NA | NA | 0 | 0 | 0 | 0 | 0 | 0 |
| 33952 | ENSMUSG00000094969 | 0 | NA | NA | NA | NA | NA | NA | NA | 0 | 0 | 0 | 0 | 0 | 0 |
| 33953 | ENSMUSG00000094970 | 0 | NA | NA | NA | NA | NA | Olfr943 | olfactory receptor 943 [Source:MGI Symb | 0 | 0 | 0 | 0 | 0 | 0 |
| 33954 | ENSMUSG00000094971 | 0 | NA | NA | NA | NA | NA | NA | NA | 0 | 0 | 0 | 0 | 0 | 0 |
| 33955 | ENSMUSG00000094972 | 0 | NA | NA | NA | NA | NA | NA | NA | 0 | 0 | 0 | 0 | 0 | 0 |
| 33956 | ENSMUSG00000094973 | 0 | NA | NA | NA | NA | NA | Eif4a3l1 | eukaryotic translation initiation factor 4A3 | 0 | 0 | 0 | 0 | 0 | 0 |
| 33957 | ENSMUSG00000094974 | 0 | NA | NA | NA | NA | NA | Rps19-ps2 | ribosomal protein S19, pseudogene 2 [Sou | 0 | 0 | 0 | 0 | 0 | 0 |
| 33958 | ENSMUSG00000094975 | 0 | NA | NA | NA | NA | NA | Gm14595 | predicted gene 14595 [Source:MGI Symbo | 0 | 0 | 0 | 0 | 0 | 0 |
| 33959 | ENSMUSG00000094976 | 0 | NA | NA | NA | NA | NA | NA | NA | 0 | 0 | 0 | 0 | 0 | 0 |
| 33960 | ENSMUSG00000094977 | 0 | NA | NA | NA | NA | NA | NA | NA | 0 | 0 | 0 | 0 | 0 | 0 |
| 33961 | ENSMUSG00000094978 | 0 | NA | NA | NA | NA | NA | Scgb1b20 | secretoglobin, family 1B, member 20 [Sou | 0 | 0 | 0 | 0 | 0 | 0 |
| 33962 | ENSMUSG00000094979 | 0 | NA | NA | NA | NA | NA | NA | NA | 0 | 0 | 0 | 0 | 0 | 0 |
| 33963 | ENSMUSG00000094980 | 0 | NA | NA | NA | NA | NA | NA | NA | 0 | 0 | 0 | 0 | 0 | 0 |
| 33964 | ENSMUSG00000094981 | 0 | NA | NA | NA | NA | NA | Vmn1r251 | vomeronasal 1 receptor 251 [Source:MGI | 0 | 0 | 0 | 0 | 0 | 0 |
| 33965 | ENSMUSG00000094982 | 0 | NA | NA | NA | NA | NA | NA | NA | 0 | 0 | 0 | 0 | 0 | 0 |
| 33966 | ENSMUSG00000094983 | 8.013080838 | 1.660485116 | 0.923102479 | 1.798809074 | 0.072048887 | 0.388994068 | NA | NA | 1.6062 | 1.0279 | 9.0562 | 14.46 | 16.359 | 5.5692 |
| 33967 | ENSMUSG00000094985 | 0.160662974 | 1.126924175 | 4.080472857 | 0.276174898 | 0.782413736 | NA | Topaz1 | testis and ovary specific PAZ domain cont | 0 | 0 | 0 | 0.964 | 0 | 0 |
| 33968 | ENSMUSG00000094986 | 0 | NA | NA | NA | NA | NA | Olfr1454 | olfactory receptor 1454 [Source:MGI Sym | 0 | 0 | 0 | 0 | 0 | 0 |
| 33969 | ENSMUSG00000094987 | 0 | NA | NA | NA | NA | NA | NA | NA | 0 | 0 | 0 | 0 | 0 | 0 |
| 33970 | ENSMUSG00000094989 | 2.172755886 | 0.844312688 | 1.429221469 | 0.590750074 | 0.554687892 | NA | Rpl9-ps4 | ribosomal protein L9, pseudogene 4 [Sour | 1.6062 | 2.0558 | 1.0062 | 2.8919 | 4.3625 | 1.1138 |
| 33971 | ENSMUSG00000094991 | 0 | NA | NA | NA | NA | NA | Gm21718 | predicted gene, 21718 [Source:MGI Symb | 0 | 0 | 0 | 0 | 0 | 0 |
| 33972 | ENSMUSG00000094992 | 0 | NA | NA | NA | NA | NA | Fbxw25 | F-box and WD-40 domain protein 25 [Sou | 0 | 0 | 0 | 0 | 0 | 0 |
| 33973 | ENSMUSG00000094993 | 0 | NA | NA | NA | NA | NA | Igkv4-51 | immunoglobulin kappa chain variable 4-51 | 0 | 0 | 0 | 0 | 0 | 0 |
| 33974 | ENSMUSG00000094994 | 0 | NA | NA | NA | NA | NA | NA | NA | 0 | 0 | 0 | 0 | 0 | 0 |
| 33975 | ENSMUSG00000094995 | 0 | NA | NA | NA | NA | NA | Pate3 | prostate and testis expressed 3 [Source:M | 0 | 0 | 0 | 0 | 0 | 0 |
| 33976 | ENSMUSG00000094997 | 0 | NA | NA | NA | NA | NA | NA | NA | 0 | 0 | 0 | 0 | 0 | 0 |
| 33977 | ENSMUSG00000094998 | 0 | NA | NA | NA | NA | NA | NA | NA | 0 | 0 | 0 | 0 | 0 | 0 |
| 33978 | ENSMUSG00000094999 | 0 | NA | NA | NA | NA | NA | Gm17412 | predicted gene, 17412 [Source:MGI Symb | 0 | 0 | 0 | 0 | 0 | 0 |
| 33979 | ENSMUSG00000095000 | 0 | NA | NA | NA | NA | NA | NA | NA | 0 | 0 | 0 | 0 | 0 | 0 |
| 33980 | ENSMUSG00000095002 | 0 | NA | NA | NA | NA | NA | Olfr798 | olfactory receptor 798 [Source:MGI Symb | 0 | 0 | 0 | 0 | 0 | 0 |
| 33981 | ENSMUSG00000095003 | 0 | NA | NA | NA | NA | NA | Gm25173 | predicted gene, 25173 [Source:MGI Symb | 0 | 0 | 0 | 0 | 0 | 0 |
| 33982 | ENSMUSG00000095005 | 0 | NA | NA | NA | NA | NA | Astx6 | amplified spermatogenic transcripts X enc | 0 | 0 | 0 | 0 | 0 | 0 |
| 33983 | ENSMUSG00000095006 | 0 | NA | NA | NA | NA | NA | NA | NA | 0 | 0 | 0 | 0 | 0 | 0 |
| 33984 | ENSMUSG00000095007 | 0.556921942 | 2.679043159 | 3.638243063 | 0.736356289 | 0.461513899 | NA | Igkv12-41 | immunoglobulin kappa chain variable 12-4 | 0 | 0 | 0 | 0 | 0 | 3.3415 |
| 33985 | ENSMUSG00000095008 | 0 | NA | NA | NA | NA | NA | Astx1c | amplified spermatogenic transcripts X enc | 0 | 0 | 0 | 0 | 0 | 0 |
| 33986 | ENSMUSG00000095009 | 0 | NA | NA | NA | NA | NA | NA | NA | 0 | 0 | 0 | 0 | 0 | 0 |
| 33987 | ENSMUSG00000095010 | 0 | NA | NA | NA | NA | NA | NA | NA | 0 | 0 | 0 | 0 | 0 | 0 |
| 33988 | ENSMUSG00000095011 | 0 | NA | NA | NA | NA | NA | Gm20838 | predicted gene, 20838 [Source:MGI Symb | 0 | 0 | 0 | 0 | 0 | 0 |

| | | | | | | | | | | | | | | | |
|---|---|---|---|---|---|---|---|---|---|---|---|---|---|---|---|
| 33989 | ENSMUSG00000095015 | 0 | NA | NA | NA | NA | NA | Gm16434 | predicted gene 16434 [Source:MGI Symbo | 0 | 0 | 0 | 0 | 0 | 0 |
| 33990 | ENSMUSG00000095016 | 0 | NA | NA | NA | NA | NA | NA | NA | 0 | 0 | 0 | 0 | 0 | 0 |
| 33991 | ENSMUSG00000095017 | 0 | NA | NA | NA | NA | NA | Rps12-ps15 | ribosomal protein S12, pseudogene 15 [Sc | 0 | 0 | 0 | 0 | 0 | 0 |
| 33992 | ENSMUSG00000095019 | 0 | NA | NA | NA | NA | NA | | | 0 | 0 | 0 | 0 | 0 | 0 |
| 33993 | ENSMUSG00000095020 | 0 | NA | NA | NA | NA | NA | NA | NA | 0 | 0 | 0 | 0 | 0 | 0 |
| 33994 | ENSMUSG00000095021 | 0 | NA | NA | NA | NA | NA | NA | NA | 0 | 0 | 0 | 0 | 0 | 0 |
| 33995 | ENSMUSG00000095022 | 0 | NA | NA | NA | NA | NA | Gm20765 | predicted gene, 20765 [Source:MGI Symb | 0 | 0 | 0 | 0 | 0 | 0 |
| 33996 | ENSMUSG00000095023 | 0 | NA | NA | NA | NA | NA | NA | NA | 0 | 0 | 0 | 0 | 0 | 0 |
| 33997 | ENSMUSG00000095024 | 0 | NA | NA | NA | NA | NA | Gm5458 | predicted gene 5458 [Source:MGI Symbol | 0 | 0 | 0 | 0 | 0 | 0 |
| 33998 | ENSMUSG00000095025 | 0 | NA | NA | NA | NA | NA | NA | NA | 0 | 0 | 0 | 0 | 0 | 0 |
| 33999 | ENSMUSG00000095026 | 37.51373938 | -0.048387955 | 0.393178515 | -0.123068665 | 0.902052725 | 0.983821605 | Gm3336 | predicted gene 3336 [Source:MGI Symbol | 38.549 | 30.838 | 45.281 | 52.055 | 31.628 | 26.732 |
| 34000 | ENSMUSG00000095027 | 0 | NA | NA | NA | NA | NA | NA | NA | 0 | 0 | 0 | 0 | 0 | 0 |
| 34001 | ENSMUSG00000095028 | 2.177118763 | -1.564570827 | 1.669143553 | -0.937349472 | 0.348578823 | NA | Sirpb1b | signal-regulatory protein beta 1B [Source: | 1.6062 | 3.0838 | 5.0312 | 0 | 0 | 3.3415 |
| 34002 | ENSMUSG00000095029 | 0 | NA | NA | NA | NA | NA | Gm36079 | predicted gene, 36079 [Source:MGI Symb | 0 | 0 | 0 | 0 | 0 | 0 |
| 34003 | ENSMUSG00000095030 | 0 | NA | NA | NA | NA | NA | Olfr1513 | olfactory receptor 1513 [Source:MGI Syml | 0 | 0 | 0 | 0 | 0 | 0 |
| 34004 | ENSMUSG00000095031 | 0 | NA | NA | NA | NA | NA | NA | NA | 0 | 0 | 0 | 0 | 0 | 0 |
| 34005 | ENSMUSG00000095032 | 0 | NA | NA | NA | NA | NA | Gm21310 | predicted gene, 21310 [Source:MGI Symb | 0 | 0 | 0 | 0 | 0 | 0 |
| 34006 | ENSMUSG00000095033 | 0 | NA | NA | NA | NA | NA | NA | NA | 0 | 0 | 0 | 0 | 0 | 0 |
| 34007 | ENSMUSG00000095034 | 0 | NA | NA | NA | NA | NA | Gm17577 | predicted gene, 17577 [Source:MGI Symb | 0 | 0 | 0 | 0 | 0 | 0 |
| 34008 | ENSMUSG00000095035 | 0 | NA | NA | NA | NA | NA | NA | NA | 0 | 0 | 0 | 0 | 0 | 0 |
| 34009 | ENSMUSG00000095037 | 0 | NA | NA | NA | NA | NA | NA | NA | 0 | 0 | 0 | 0 | 0 | 0 |
| 34010 | ENSMUSG00000095040 | 0.528073336 | 2.60973587 | 2.979250867 | 0.875970499 | 0.381046072 | NA | 1700001J03I | RIKEN cDNA 1700001J03 gene [Source:MG | 0 | 0 | 0 | 0.964 | 1.0906 | 1.1138 |
| 34011 | ENSMUSG00000095041 | 1716.995341 | -0.084360317 | 0.122795201 | -0.687000113 | 0.492082657 | 0.876368691 | | | 1905.8 | 1616.9 | 1779 | 1915.4 | 1541 | 1543.8 |
| 34012 | ENSMUSG00000095042 | 0.506734502 | -2.372847156 | 3.338060886 | -0.710845978 | 0.477179685 | NA | Gm12537 | predicted gene 12537 [Source:MGI Symbo | 0 | 1.0279 | 2.0125 | 0 | 0 | 0 |
| 34013 | ENSMUSG00000095044 | 0 | NA | NA | NA | NA | NA | Gm6401 | predicted gene 6401 [Source:MGI Symbol | 0 | 0 | 0 | 0 | 0 | 0 |
| 34014 | ENSMUSG00000095045 | 0 | NA | NA | NA | NA | NA | NA | NA | 0 | 0 | 0 | 0 | 0 | 0 |
| 34015 | ENSMUSG00000095047 | 0 | NA | NA | NA | NA | NA | NA | NA | 0 | 0 | 0 | 0 | 0 | 0 |
| 34016 | ENSMUSG00000095048 | 0 | NA | NA | NA | NA | NA | Gm11239 | predicted gene 11239 [Source:MGI Symbo | 0 | 0 | 0 | 0 | 0 | 0 |
| 34017 | ENSMUSG00000095049 | 0 | NA | NA | NA | NA | NA | Gm21756 | predicted gene, 21756 [Source:MGI Symb | 0 | 0 | 0 | 0 | 0 | 0 |
| 34018 | ENSMUSG00000095051 | 0 | NA | NA | NA | NA | NA | NA | NA | 0 | 0 | 0 | 0 | 0 | 0 |
| 34019 | ENSMUSG00000095052 | 0 | NA | NA | NA | NA | NA | NA | NA | 0 | 0 | 0 | 0 | 0 | 0 |
| 34020 | ENSMUSG00000095053 | 0 | NA | NA | NA | NA | NA | NA | NA | 0 | 0 | 0 | 0 | 0 | 0 |
| 34021 | ENSMUSG00000095054 | 0.17131964 | -0.796629176 | 4.080472857 | -0.195229623 | 0.84521319 | NA | Gm13146 | predicted gene 13146 [Source:MGI Symbo | 0 | 1.0279 | 0 | 0 | 0 | 0 |
| 34022 | ENSMUSG00000095056 | 0 | NA | NA | NA | NA | NA | Gm3159 | predicted gene 3159 [Source:MGI Symbol | 0 | 0 | 0 | 0 | 0 | 0 |
| 34023 | ENSMUSG00000095057 | 0 | NA | NA | NA | NA | NA | Zfp353-ps | zinc finger protein 353, pseudogene [Sour | 0 | 0 | 0 | 0 | 0 | 0 |
| 34024 | ENSMUSG00000095059 | 0 | NA | NA | NA | NA | NA | NA | NA | 0 | 0 | 0 | 0 | 0 | 0 |
| 34025 | ENSMUSG00000095060 | 0 | NA | NA | NA | NA | NA | NA | NA | 0 | 0 | 0 | 0 | 0 | 0 |
| 34026 | ENSMUSG00000095061 | 55.96351843 | 0.022884493 | 0.413239147 | 0.055378328 | 0.955837061 | 0.991678478 | E030018B13 | RIKEN cDNA E030018B13 gene [Source:M | 78.704 | 44.2 | 43.269 | 50.127 | 90.521 | 28.96 |
| 34027 | ENSMUSG00000095063 | 0 | NA | NA | NA | NA | NA | Slx | Sycp3 like X-linked [Source:MGI Symbol;A | 0 | 0 | 0 | 0 | 0 | 0 |
| 34028 | ENSMUSG00000095064 | 0 | NA | NA | NA | NA | NA | Vmn1r135 | vomeronasal 1 receptor 135 [Source:MGI | 0 | 0 | 0 | 0 | 0 | 0 |
| 34029 | ENSMUSG00000095065 | 0 | NA | NA | NA | NA | NA | Gm5947 | predicted pseudogene 5947 [Source:MGI | 0 | 0 | 0 | 0 | 0 | 0 |
| 34030 | ENSMUSG00000095066 | 0 | NA | NA | NA | NA | NA | Defa20 | defensin, alpha, 20 [Source:MGI Symbol;A | 0 | 0 | 0 | 0 | 0 | 0 |
| 34031 | ENSMUSG00000095068 | 0 | NA | NA | NA | NA | NA | NA | NA | 0 | 0 | 0 | 0 | 0 | 0 |
| 34032 | ENSMUSG00000095069 | 0 | NA | NA | NA | NA | NA | NA | NA | 0 | 0 | 0 | 0 | 0 | 0 |
| 34033 | ENSMUSG00000095070 | 0 | NA | NA | NA | NA | NA | NA | NA | 0 | 0 | 0 | 0 | 0 | 0 |
| 34034 | ENSMUSG00000095071 | 0 | NA | NA | NA | NA | NA | AF067061 | cDNA sequence AF067061 [Source:MGI Sy | 0 | 0 | 0 | 0 | 0 | 0 |
| 34035 | ENSMUSG00000095072 | 0 | NA | NA | NA | NA | NA | NA | NA | 0 | 0 | 0 | 0 | 0 | 0 |
| 34036 | ENSMUSG00000095073 | 0 | NA | NA | NA | NA | NA | NA | NA | 0 | 0 | 0 | 0 | 0 | 0 |
| 34037 | ENSMUSG00000095074 | 0 | NA | NA | NA | NA | NA | Pramel42 | PRAME like 42 [Source:MGI Symbol;Acc:M | 0 | 0 | 0 | 0 | 0 | 0 |
| 34038 | ENSMUSG00000095075 | 0 | NA | NA | NA | NA | NA | Olfr782 | olfactory receptor 782 [Source:MGI Symb | 0 | 0 | 0 | 0 | 0 | 0 |
| 34039 | ENSMUSG00000095076 | 0 | NA | NA | NA | NA | NA | | | 0 | 0 | 0 | 0 | 0 | 0 |
| 34040 | ENSMUSG00000095078 | 0 | NA | NA | NA | NA | NA | Gm5866 | predicted gene 5866 [Source:MGI Symbol | 0 | 0 | 0 | 0 | 0 | 0 |
| 34041 | ENSMUSG00000095079 | 19.15705818 | -1.324043482 | 0.512802759 | -2.58197418 | 0.009823693 | 0.107237662 | Igha | immunoglobulin heavy constant alpha [Sc | 24.093 | 31.865 | 26.162 | 9.6398 | 9.8156 | 13.366 |
| 34042 | ENSMUSG00000095081 | 0 | NA | NA | NA | NA | NA | Vmn1r257 | vomeronasal 1 receptor 257 [Source:MGI | 0 | 0 | 0 | 0 | 0 | 0 |
| 34043 | ENSMUSG00000095082 | 0 | NA | NA | NA | NA | NA | Gm15091 | predicted gene 15091 [Source:MGI Symbo | 0 | 0 | 0 | 0 | 0 | 0 |
| 34044 | ENSMUSG00000095085 | 0 | NA | NA | NA | NA | NA | NA | NA | 0 | 0 | 0 | 0 | 0 | 0 |

| | | | | | | | | | | | | | | | |
|---|---|---|---|---|---|---|---|---|---|---|---|---|---|---|---|
| 34045 | ENSMUSG00000095086 | 0 | NA | NA | NA | NA | NA | NA | NA | 0 | 0 | 0 | 0 | 0 | 0 |
| 34046 | ENSMUSG00000095087 | 1.308446539 | 2.811813438 | 2.250897484 | 1.24919658 | 0.211593183 | NA | NA | NA | 0.8031 | 0 | 0 | 4.8199 | 0 | 2.2277 |
| 34047 | ENSMUSG00000095088 | 0 | NA | NA | NA | NA | NA | Trav11d | T cell receptor alpha variable 11D [Source | 0 | 0 | 0 | 0 | 0 | 0 |
| 34048 | ENSMUSG00000095089 | 0 | NA | NA | NA | NA | NA | Gm7408 | predicted pseudogene 7408 [Source:MGI | 0 | 0 | 0 | 0 | 0 | 0 |
| 34049 | ENSMUSG00000095091 | 0 | NA | NA | NA | NA | NA | NA | NA | 0 | 0 | 0 | 0 | 0 | 0 |
| 34050 | ENSMUSG00000095092 | 0 | NA | NA | NA | NA | NA | | | 0 | 0 | 0 | 0 | 0 | 0 |
| 34051 | ENSMUSG00000095093 | 0 | NA | NA | NA | NA | NA | Vmn2r111 | vomeronasal 2, receptor 111 [Source:MGI | 0 | 0 | 0 | 0 | 0 | 0 |
| 34052 | ENSMUSG00000095094 | 0 | NA | NA | NA | NA | NA | NA | NA | 0 | 0 | 0 | 0 | 0 | 0 |
| 34053 | ENSMUSG00000095095 | 0 | NA | NA | NA | NA | NA | Olfr385 | olfactory receptor 385 [Source:MGI Symb | 0 | 0 | 0 | 0 | 0 | 0 |
| 34054 | ENSMUSG00000095096 | 0 | NA | NA | NA | NA | NA | Vmn2r-ps17 | vomeronasal 2, receptor, pseudogene 17 | 0 | 0 | 0 | 0 | 0 | 0 |
| 34055 | ENSMUSG00000095097 | 0 | NA | NA | NA | NA | NA | NA | NA | 0 | 0 | 0 | 0 | 0 | 0 |
| 34056 | ENSMUSG00000095098 | 352.9097641 | 0.445147462 | 0.14466738 | 3.07704101 | 0.002090665 | 0.036042174 | Ccdc85b | coiled-coil domain containing 85B [Source | 318.83 | 265.2 | 311.94 | 397.16 | 414.43 | 409.89 |
| 34057 | ENSMUSG00000095100 | 0 | NA | NA | NA | NA | NA | NA | NA | 0 | 0 | 0 | 0 | 0 | 0 |
| 34058 | ENSMUSG00000095101 | 0 | NA | NA | NA | NA | NA | Gm13285 | predicted gene 13285 [Source:MGI Symbo | 0 | 0 | 0 | 0 | 0 | 0 |
| 34059 | ENSMUSG00000095102 | 0 | NA | NA | NA | NA | NA | NA | NA | 0 | 0 | 0 | 0 | 0 | 0 |
| 34060 | ENSMUSG00000095103 | 0 | NA | NA | NA | NA | NA | Gm21171 | predicted gene, 21171 [Source:MGI Symb | 0 | 0 | 0 | 0 | 0 | 0 |
| 34061 | ENSMUSG00000095104 | 0 | NA | NA | NA | NA | NA | Esp15 | exocrine gland secreted peptide 15 [Sourc | 0 | 0 | 0 | 0 | 0 | 0 |
| 34062 | ENSMUSG00000095105 | 1.449109275 | -0.870864006 | 1.794516463 | -0.485291734 | 0.627469418 | NA | Edaradd | EDAR (ectodysplasin-A receptor)-associat | 1.6062 | 1.0279 | 3.0187 | 1.928 | 0 | 1.1138 |
| 34063 | ENSMUSG00000095107 | 0 | NA | NA | NA | NA | NA | Vmn2r-ps10 | vomeronasal 2, receptor, pseudogene 10 | 0 | 0 | 0 | 0 | 0 | 0 |
| 34064 | ENSMUSG00000095108 | 0 | NA | NA | NA | NA | NA | Gm17467 | predicted gene, 17467 [Source:MGI Symb | 0 | 0 | 0 | 0 | 0 | 0 |
| 34065 | ENSMUSG00000095110 | 0 | NA | NA | NA | NA | NA | 1700003E24 | RIKEN cDNA 1700003E24 gene [Source:M | 0 | 0 | 0 | 0 | 0 | 0 |
| 34066 | ENSMUSG00000095111 | 0 | NA | NA | NA | NA | NA | NA | NA | 0 | 0 | 0 | 0 | 0 | 0 |
| 34067 | ENSMUSG00000095112 | 0 | NA | NA | NA | NA | NA | NA | NA | 0 | 0 | 0 | 0 | 0 | 0 |
| 34068 | ENSMUSG00000095113 | 0 | NA | NA | NA | NA | NA | Gm3633 | predicted gene 3633 [Source:MGI Symbol | 0 | 0 | 0 | 0 | 0 | 0 |
| 34069 | ENSMUSG00000095115 | 89.46518086 | -0.663401821 | 0.27777942 | -2.388232432 | 0.016929631 | 0.1545157 | Itpripl2 | inositol 1,4,5-triphosphate receptor intera | 107.62 | 135.69 | 85.531 | 54.947 | 85.068 | 67.944 |
| 34070 | ENSMUSG00000095116 | 0 | NA | NA | NA | NA | NA | NA | NA | 0 | 0 | 0 | 0 | 0 | 0 |
| 34071 | ENSMUSG00000095117 | 0 | NA | NA | NA | NA | NA | Ighv8-9 | immunoglobulin heavy variable V8-9 [Sou | 0 | 0 | 0 | 0 | 0 | 0 |
| 34072 | ENSMUSG00000095118 | 0 | NA | NA | NA | NA | NA | NA | NA | 0 | 0 | 0 | 0 | 0 | 0 |
| 34073 | ENSMUSG00000095119 | 0 | NA | NA | NA | NA | NA | NA | NA | 0 | 0 | 0 | 0 | 0 | 0 |
| 34074 | ENSMUSG00000095121 | 0 | NA | NA | NA | NA | NA | NA | NA | 0 | 0 | 0 | 0 | 0 | 0 |
| 34075 | ENSMUSG00000095122 | 0 | NA | NA | NA | NA | NA | NA | NA | 0 | 0 | 0 | 0 | 0 | 0 |
| 34076 | ENSMUSG00000095123 | 69.19413057 | -0.007450939 | 0.369863706 | -0.020145093 | 0.983927629 | 0.998580735 | Gm21781 | predicted gene, 21781 [Source:MGI Symb | 83.523 | 47.284 | 77.481 | 108.93 | 55.622 | 42.326 |
| 34077 | ENSMUSG00000095125 | 0 | NA | NA | NA | NA | NA | Vmn1r198 | vomeronasal 1 receptor 198 [Source:MGI | 0 | 0 | 0 | 0 | 0 | 0 |
| 34078 | ENSMUSG00000095126 | 0 | NA | NA | NA | NA | NA | Trav6-7-dv9 | T cell receptor alpha variable 6-7-DV9 [So | 0 | 0 | 0 | 0 | 0 | 0 |
| 34079 | ENSMUSG00000095127 | 0.17131964 | -0.796629176 | 4.080472857 | -0.195229623 | 0.84521319 | NA | Ighv1-82 | immunoglobulin heavy variable 1-82 [Sou | 0 | 1.0279 | 0 | 0 | 0 | 0 |
| 34080 | ENSMUSG00000095128 | 0 | NA | NA | NA | NA | NA | Gm5709 | predicted gene 5709 [Source:MGI Symbol | 0 | 0 | 0 | 0 | 0 | 0 |
| 34081 | ENSMUSG00000095129 | 0 | NA | NA | NA | NA | NA | NA | NA | 0 | 0 | 0 | 0 | 0 | 0 |
| 34082 | ENSMUSG00000095130 | 0 | NA | NA | NA | NA | NA | Ighv1-39 | immunoglobulin heavy variable 1-39 [Sou | 0 | 0 | 0 | 0 | 0 | 0 |
| 34083 | ENSMUSG00000095131 | 0 | NA | NA | NA | NA | NA | NA | NA | 0 | 0 | 0 | 0 | 0 | 0 |
| 34084 | ENSMUSG00000095132 | 0.17131964 | -0.796629176 | 4.080472857 | -0.195229623 | 0.84521319 | NA | NA | NA | 0 | 1.0279 | 0 | 0 | 0 | 0 |
| 34085 | ENSMUSG00000095133 | 0 | NA | NA | NA | NA | NA | NA | NA | 0 | 0 | 0 | 0 | 0 | 0 |
| 34086 | ENSMUSG00000095134 | 0.876412963 | 2.129655479 | 2.836103289 | 0.750908998 | 0.452707425 | NA | Mid1-ps1 | midline 1, pseudogene 1 [Source:MGI Syn | 0.8031 | 0 | 0 | 0 | 0 | 4.4554 |
| 34087 | ENSMUSG00000095135 | 0 | NA | NA | NA | NA | NA | Gm21858 | predicted gene, 21858 [Source:MGI Symb | 0 | 0 | 0 | 0 | 0 | 0 |
| 34088 | ENSMUSG00000095136 | 0 | NA | NA | NA | NA | NA | Gm12616 | predicted gene 12616 [Source:MGI Symbo | 0 | 0 | 0 | 0 | 0 | 0 |
| 34089 | ENSMUSG00000095137 | 0 | NA | NA | NA | NA | NA | Rps12-ps17 | ribosomal protein S12, pseudogene 17 [So | 0 | 0 | 0 | 0 | 0 | 0 |
| 34090 | ENSMUSG00000095138 | 0 | NA | NA | NA | NA | NA | Olfr781 | olfactory receptor 781 [Source:MGI Symb | 0 | 0 | 0 | 0 | 0 | 0 |
| 34091 | ENSMUSG00000095139 | 0 | NA | NA | NA | NA | NA | Pou3f2 | POU domain, class 3, transcription factor | 0 | 0 | 0 | 0 | 0 | 0 |
| 34092 | ENSMUSG00000095140 | 0 | NA | NA | NA | NA | NA | NA | NA | 0 | 0 | 0 | 0 | 0 | 0 |
| 34093 | ENSMUSG00000095141 | 0 | NA | NA | NA | NA | NA | Gm28891 | predicted gene 28891 [Source:MGI Symbo | 0 | 0 | 0 | 0 | 0 | 0 |
| 34094 | ENSMUSG00000095142 | 0 | NA | NA | NA | NA | NA | NA | NA | 0 | 0 | 0 | 0 | 0 | 0 |
| 34095 | ENSMUSG00000095143 | 0 | NA | NA | NA | NA | NA | Hsd3b4 | hydroxy-delta-5-steroid dehydrogenase, 3 | 0 | 0 | 0 | 0 | 0 | 0 |
| 34096 | ENSMUSG00000095146 | 0 | NA | NA | NA | NA | NA | NA | NA | 0 | 0 | 0 | 0 | 0 | 0 |
| 34097 | ENSMUSG00000095147 | 0 | NA | NA | NA | NA | NA | NA | NA | 0 | 0 | 0 | 0 | 0 | 0 |
| 34098 | ENSMUSG00000095148 | 0 | NA | NA | NA | NA | NA | Gm20873 | predicted gene, 20873 [Source:MGI Symb | 0 | 0 | 0 | 0 | 0 | 0 |
| 34099 | ENSMUSG00000095149 | 0 | NA | NA | NA | NA | NA | NA | NA | 0 | 0 | 0 | 0 | 0 | 0 |
| 34100 | ENSMUSG00000095150 | 0 | NA | NA | NA | NA | NA | Rps19-ps13 | ribosomal protein S19, pseudogene 13 [Sc | 0 | 0 | 0 | 0 | 0 | 0 |

| | | | | | | | | | | | | | | | |
|---|---|---|---|---|---|---|---|---|---|---|---|---|---|---|---|
| 34101 | ENSMUSG00000095151 | 0 | NA | NA | NA | NA | NA | NA | NA | 0 | 0 | 0 | 0 | 0 | 0 |
| 34102 | ENSMUSG00000095152 | 0 | NA | NA | NA | NA | NA | Gm3657 | predicted gene 3657 [Source:MGI Symbol | 0 | 0 | 0 | 0 | 0 | 0 |
| 34103 | ENSMUSG00000095153 | 0 | NA | NA | NA | NA | NA | Gm21095 | predicted gene, 21095 [Source:MGI Symb | 0 | 0 | 0 | 0 | 0 | 0 |
| 34104 | ENSMUSG00000095154 | 0 | NA | NA | NA | NA | NA | NA | NA | 0 | 0 | 0 | 0 | 0 | 0 |
| 34105 | ENSMUSG00000095155 | 0 | NA | NA | NA | NA | NA | NA | NA | 0 | 0 | 0 | 0 | 0 | 0 |
| 34106 | ENSMUSG00000095156 | 0 | NA | NA | NA | NA | NA | Olfr1281 | olfactory receptor 1281 [Source:MGI Syml | 0 | 0 | 0 | 0 | 0 | 0 |
| 34107 | ENSMUSG00000095159 | 0.401551119 | -2.072982726 | 4.008251979 | -0.517178744 | 0.605031388 | NA | Tubb4b-ps1 | tubulin, beta 4B class IVB, pseudogene 1 [ | 2.4093 | 0 | 0 | 0 | 0 | 0 |
| 34108 | ENSMUSG00000095160 | 0.160662974 | 1.126924175 | 4.080472857 | 0.276174898 | 0.782413736 | NA | NA | NA | 0 | 0 | 0 | 0.964 | 0 | 0 |
| 34109 | ENSMUSG00000095163 | 0 | NA | NA | NA | NA | NA | Vmn1r138 | vomeronasal 1 receptor 138 [Source:MGI | 0 | 0 | 0 | 0 | 0 | 0 |
| 34110 | ENSMUSG00000095165 | 0 | NA | NA | NA | NA | NA | NA | NA | 0 | 0 | 0 | 0 | 0 | 0 |
| 34111 | ENSMUSG00000095168 | 0.531944269 | 2.618916564 | 3.285123409 | 0.797204926 | 0.425332026 | NA | Gm4795 | predicted pseudogene 4795 [Source:MGI | 0 | 0 | 0 | 0.964 | 0 | 2.2277 |
| 34112 | ENSMUSG00000095169 | 0 | NA | NA | NA | NA | NA | NA | NA | 0 | 0 | 0 | 0 | 0 | 0 |
| 34113 | ENSMUSG00000095170 | 0 | NA | NA | NA | NA | NA | Ighv8-11 | immunoglobulin heavy variable V8-11 [So | 0 | 0 | 0 | 0 | 0 | 0 |
| 34114 | ENSMUSG00000095171 | 0 | NA | NA | NA | NA | NA | Taar7b | trace amine-associated receptor 7B [Sour | 0 | 0 | 0 | 0 | 0 | 0 |
| 34115 | ENSMUSG00000095172 | 0 | NA | NA | NA | NA | NA | Gm20822 | predicted gene, 20822 [Source:MGI Symb | 0 | 0 | 0 | 0 | 0 | 0 |
| 34116 | ENSMUSG00000095174 | 0.160662974 | 1.126924175 | 4.080472857 | 0.276174898 | 0.782413736 | NA | Gm20069 | predicted gene, 20069 [Source:MGI Symb | 0 | 0 | 0 | 0.964 | 0 | 0 |
| 34117 | ENSMUSG00000095175 | 0 | NA | NA | NA | NA | NA | NA | NA | 0 | 0 | 0 | 0 | 0 | 0 |
| 34118 | ENSMUSG00000095176 | 0 | NA | NA | NA | NA | NA | NA | NA | 0 | 0 | 0 | 0 | 0 | 0 |
| 34119 | ENSMUSG00000095178 | 1.477647023 | -2.765922777 | 2.027309562 | -1.364331738 | 0.172463181 | NA | NA | NA | 1.6062 | 5.1396 | 1.0062 | 0 | 0 | 1.1138 |
| 34120 | ENSMUSG00000095179 | 0 | NA | NA | NA | NA | NA | Gm21638 | predicted gene, 21638 [Source:MGI Symb | 0 | 0 | 0 | 0 | 0 | 0 |
| 34121 | ENSMUSG00000095180 | 0.811904515 | -3.057764991 | 2.533063159 | -1.207141236 | 0.227377757 | NA | Rhox5 | reproductive homeobox 5 [Source:MGI Sy | 0.8031 | 2.0558 | 2.0125 | 0 | 0 | 0 |
| 34122 | ENSMUSG00000095182 | 0 | NA | NA | NA | NA | NA | NA | NA | 0 | 0 | 0 | 0 | 0 | 0 |
| 34123 | ENSMUSG00000095183 | 0 | NA | NA | NA | NA | NA | NA | NA | 0 | 0 | 0 | 0 | 0 | 0 |
| 34124 | ENSMUSG00000095184 | 0 | NA | NA | NA | NA | NA | NA | NA | 0 | 0 | 0 | 0 | 0 | 0 |
| 34125 | ENSMUSG00000095185 | 0 | NA | NA | NA | NA | NA | Vmn1r-ps75 | vomeronasal 1 receptor, pseudogene 75 [ | 0 | 0 | 0 | 0 | 0 | 0 |
| 34126 | ENSMUSG00000095186 | 0 | NA | NA | NA | NA | NA | Gm10718 | predicted gene 10718 [Source:MGI Symbo | 0 | 0 | 0 | 0 | 0 | 0 |
| 34127 | ENSMUSG00000095187 | 0 | NA | NA | NA | NA | NA | Olfr1381 | olfactory receptor 1381 [Source:MGI Syml | 0 | 0 | 0 | 0 | 0 | 0 |
| 34128 | ENSMUSG00000095188 | 0 | NA | NA | NA | NA | NA | NA | NA | 0 | 0 | 0 | 0 | 0 | 0 |
| 34129 | ENSMUSG00000095189 | 0 | NA | NA | NA | NA | NA | Olfr1472 | olfactory receptor 1472 [Source:MGI Syml | 0 | 0 | 0 | 0 | 0 | 0 |
| 34130 | ENSMUSG00000095190 | 0 | NA | NA | NA | NA | NA | Gm10668 | predicted gene 10668 [Source:MGI Symbo | 0 | 0 | 0 | 0 | 0 | 0 |
| 34131 | ENSMUSG00000095191 | 0 | NA | NA | NA | NA | NA | Vmn1r249 | vomeronasal 1 receptor 249 [Source:MGI | 0 | 0 | 0 | 0 | 0 | 0 |
| 34132 | ENSMUSG00000095192 | 0 | NA | NA | NA | NA | NA | Gm21529 | predicted gene, 21529 [Source:MGI Symb | 0 | 0 | 0 | 0 | 0 | 0 |
| 34133 | ENSMUSG00000095193 | 18.6577389 | 0.62256887 | 0.525514887 | 1.184683602 | 0.236142568 | 0.684756637 | Gm20939 | predicted gene, 20939 [Source:MGI Symb | 18.471 | 9.2513 | 16.1 | 17.352 | 21.812 | 28.96 |
| 34134 | ENSMUSG00000095194 | 0 | NA | NA | NA | NA | NA | Olfr936 | olfactory receptor 936 [Source:MGI Symb | 0 | 0 | 0 | 0 | 0 | 0 |
| 34135 | ENSMUSG00000095195 | 0 | NA | NA | NA | NA | NA | Gm3005 | predicted gene 3005 [Source:MGI Symbol | 0 | 0 | 0 | 0 | 0 | 0 |
| 34136 | ENSMUSG00000095196 | 0 | NA | NA | NA | NA | NA | Gm8305 | predicted gene 8305 [Source:MGI Symbol | 0 | 0 | 0 | 0 | 0 | 0 |
| 34137 | ENSMUSG00000095197 | 0 | NA | NA | NA | NA | NA | Ighv1-59 | immunoglobulin heavy variable V1-59 [So | 0 | 0 | 0 | 0 | 0 | 0 |
| 34138 | ENSMUSG00000095198 | 0 | NA | NA | NA | NA | NA | NA | NA | 0 | 0 | 0 | 0 | 0 | 0 |
| 34139 | ENSMUSG00000095199 | 0 | NA | NA | NA | NA | NA | Zfp967 | zinc finger protein 967 [Source:MGI Symb | 0 | 0 | 0 | 0 | 0 | 0 |
| 34140 | ENSMUSG00000095200 | 0 | NA | NA | NA | NA | NA | Ighv1-7 | immunoglobulin heavy variable V1-7 [Sou | 0 | 0 | 0 | 0 | 0 | 0 |
| 34141 | ENSMUSG00000095201 | 0 | NA | NA | NA | NA | NA | Vmn1r91 | vomeronasal 1 receptor 91 [Source:MGI S | 0 | 0 | 0 | 0 | 0 | 0 |
| 34142 | ENSMUSG00000095202 | 0 | NA | NA | NA | NA | NA | Gm6100 | predicted gene 6100 [Source:MGI Symbol | 0 | 0 | 0 | 0 | 0 | 0 |
| 34143 | ENSMUSG00000095203 | 0 | NA | NA | NA | NA | NA | Gm13810 | predicted gene 13810 [Source:MGI Symbo | 0 | 0 | 0 | 0 | 0 | 0 |
| 34144 | ENSMUSG00000095204 | 0 | NA | NA | NA | NA | NA | Ighv1-52 | immunoglobulin heavy variable 1-52 [Sou | 0 | 0 | 0 | 0 | 0 | 0 |
| 34145 | ENSMUSG00000095205 | 0 | NA | NA | NA | NA | NA | NA | NA | 0 | 0 | 0 | 0 | 0 | 0 |
| 34146 | ENSMUSG00000095207 | 0 | NA | NA | NA | NA | NA | NA | NA | 0 | 0 | 0 | 0 | 0 | 0 |
| 34147 | ENSMUSG00000095209 | 0 | NA | NA | NA | NA | NA | NA | NA | 0 | 0 | 0 | 0 | 0 | 0 |
| 34148 | ENSMUSG00000095210 | 0 | NA | NA | NA | NA | NA | Ighv5-9-1 | immunoglobulin heavy variable 5-9-1 [Sou | 0 | 0 | 0 | 0 | 0 | 0 |
| 34149 | ENSMUSG00000095212 | 0 | NA | NA | NA | NA | NA | Olfr473 | olfactory receptor 473 [Source:MGI Symb | 0 | 0 | 0 | 0 | 0 | 0 |
| 34150 | ENSMUSG00000095213 | 0 | NA | NA | NA | NA | NA | Gm9944 | predicted gene 9944 [Source:MGI Symbol | 0 | 0 | 0 | 0 | 0 | 0 |
| 34151 | ENSMUSG00000095214 | 0 | NA | NA | NA | NA | NA | NA | NA | 0 | 0 | 0 | 0 | 0 | 0 |
| 34152 | ENSMUSG00000095217 | 0.321325949 | 1.911575114 | 4.048267893 | 0.472195804 | 0.636787034 | NA | H2bc15 | H2B clustered histone 15 [Source:MGI Syr | 0 | 0 | 0 | 1.928 | 0 | 0 |
| 34153 | ENSMUSG00000095218 | 0 | NA | NA | NA | NA | NA | Olfr1338 | olfactory receptor 1338 [Source:MGI Syml | 0 | 0 | 0 | 0 | 0 | 0 |
| 34154 | ENSMUSG00000095220 | 0 | NA | NA | NA | NA | NA | NA | NA | 0 | 0 | 0 | 0 | 0 | 0 |
| 34155 | ENSMUSG00000095221 | 0 | NA | NA | NA | NA | NA | NA | NA | 0 | 0 | 0 | 0 | 0 | 0 |
| 34156 | ENSMUSG00000095222 | 0 | NA | NA | NA | NA | NA | NA | NA | 0 | 0 | 0 | 0 | 0 | 0 |

| | | | | | | | | | | | | | | | |
|---|---|---|---|---|---|---|---|---|---|---|---|---|---|---|---|
| 34157 | ENSMUSG00000095224 | 0 | NA | NA | NA | NA | NA | NA | NA | 0 | 0 | 0 | 0 | 0 | 0 |
| 34158 | ENSMUSG00000095226 | 0 | NA | NA | NA | NA | NA | Gm10377 | predicted gene 10377 [Source:MGI Symbo | 0 | 0 | 0 | 0 | 0 | 0 |
| 34159 | ENSMUSG00000095228 | 0 | NA | NA | NA | NA | NA | NA | NA | 0 | 0 | 0 | 0 | 0 | 0 |
| 34160 | ENSMUSG00000095229 | 0 | NA | NA | NA | NA | NA | NA | NA | 0 | 0 | 0 | 0 | 0 | 0 |
| 34161 | ENSMUSG00000095231 | 0 | NA | NA | NA | NA | NA | Gm14297 | predicted gene 14297 [Source:MGI Symbo | 0 | 0 | 0 | 0 | 0 | 0 |
| 34162 | ENSMUSG00000095232 | 0 | NA | NA | NA | NA | NA | Scgb1b19 | secretoglobin, family 1B, member 19 [Sou | 0 | 0 | 0 | 0 | 0 | 0 |
| 34163 | ENSMUSG00000095233 | 0 | NA | NA | NA | NA | NA | NA | NA | 0 | 0 | 0 | 0 | 0 | 0 |
| 34164 | ENSMUSG00000095234 | 0 | NA | NA | NA | NA | NA | Gm21586 | predicted gene, 21586 [Source:MGI Symb | 0 | 0 | 0 | 0 | 0 | 0 |
| 34165 | ENSMUSG00000095236 | 0 | NA | NA | NA | NA | NA | Olfr38 | olfactory receptor 38 [Source:MGI Symbo | 0 | 0 | 0 | 0 | 0 | 0 |
| 34166 | ENSMUSG00000095238 | 0 | NA | NA | NA | NA | NA | Gm4845 | predicted gene 4845 [Source:MGI Symbol | 0 | 0 | 0 | 0 | 0 | 0 |
| 34167 | ENSMUSG00000095239 | 0 | NA | NA | NA | NA | NA | Olfr497 | olfactory receptor 497 [Source:MGI Symb | 0 | 0 | 0 | 0 | 0 | 0 |
| 34168 | ENSMUSG00000095240 | 0 | NA | NA | NA | NA | NA | Cypt14 | cysteine-rich perinuclear theca 14 [Source | 0 | 0 | 0 | 0 | 0 | 0 |
| 34169 | ENSMUSG00000095241 | 0 | NA | NA | NA | NA | NA | Gm5478 | predicted pseudogene 5478 [Source:MGI | 0 | 0 | 0 | 0 | 0 | 0 |
| 34170 | ENSMUSG00000095242 | 0 | NA | NA | NA | NA | NA | Gm20834 | predicted gene, 20834 [Source:MGI Symb | 0 | 0 | 0 | 0 | 0 | 0 |
| 34171 | ENSMUSG00000095243 | 0 | NA | NA | NA | NA | NA | NA | NA | 0 | 0 | 0 | 0 | 0 | 0 |
| 34172 | ENSMUSG00000095245 | 0 | NA | NA | NA | NA | NA | NA | NA | 0 | 0 | 0 | 0 | 0 | 0 |
| 34173 | ENSMUSG00000095246 | 0 | NA | NA | NA | NA | NA | NA | NA | 0 | 0 | 0 | 0 | 0 | 0 |
| 34174 | ENSMUSG00000095247 | 0 | NA | NA | NA | NA | NA | NA | NA | 0 | 0 | 0 | 0 | 0 | 0 |
| 34175 | ENSMUSG00000095248 | 0 | NA | NA | NA | NA | NA | Olfr681 | olfactory receptor 681 [Source:MGI Symb | 0 | 0 | 0 | 0 | 0 | 0 |
| 34176 | ENSMUSG00000095249 | 0 | NA | NA | NA | NA | NA | NA | NA | 0 | 0 | 0 | 0 | 0 | 0 |
| 34177 | ENSMUSG00000095250 | 0 | NA | NA | NA | NA | NA | | | 0 | 0 | 0 | 0 | 0 | 0 |
| 34178 | ENSMUSG00000095251 | 0 | NA | NA | NA | NA | NA | Tdpoz7 | Td and POZ domain containing 7 [Source:I | 0 | 0 | 0 | 0 | 0 | 0 |
| 34179 | ENSMUSG00000095253 | 108.3078374 | 0.153812776 | 0.218188062 | 0.704955049 | 0.48083821 | 0.870278888 | Zfp799 | zinc finger protein 799 [Source:MGI Symb | 97.175 | 109.99 | 100.62 | 111.82 | 119.97 | 110.27 |
| 34180 | ENSMUSG00000095255 | 0 | NA | NA | NA | NA | NA | NA | NA | 0 | 0 | 0 | 0 | 0 | 0 |
| 34181 | ENSMUSG00000095256 | 0 | NA | NA | NA | NA | NA | NA | NA | 0 | 0 | 0 | 0 | 0 | 0 |
| 34182 | ENSMUSG00000095257 | 0 | NA | NA | NA | NA | NA | Scgb1b3 | secretoglobin, family 1B, member 3 [Sour | 0 | 0 | 0 | 0 | 0 | 0 |
| 34183 | ENSMUSG00000095258 | 0 | NA | NA | NA | NA | NA | Gm2163 | predicted gene 2163 [Source:MGI Symbol | 0 | 0 | 0 | 0 | 0 | 0 |
| 34184 | ENSMUSG00000095259 | 0 | NA | NA | NA | NA | NA | Gm14357 | predicted gene 14357 [Source:MGI Symbo | 0 | 0 | 0 | 0 | 0 | 0 |
| 34185 | ENSMUSG00000095260 | 0 | NA | NA | NA | NA | NA | NA | NA | 0 | 0 | 0 | 0 | 0 | 0 |
| 34186 | ENSMUSG00000095261 | 0 | NA | NA | NA | NA | NA | Gm17391 | predicted gene, 17391 [Source:MGI Symb | 0 | 0 | 0 | 0 | 0 | 0 |
| 34187 | ENSMUSG00000095262 | 0 | NA | NA | NA | NA | NA | Gm21893 | predicted gene, 21893 [Source:MGI Symb | 0 | 0 | 0 | 0 | 0 | 0 |
| 34188 | ENSMUSG00000095263 | 0 | NA | NA | NA | NA | NA | Gm21094 | predicted gene, 21094 [Source:MGI Symb | 0 | 0 | 0 | 0 | 0 | 0 |
| 34189 | ENSMUSG00000095266 | 0 | NA | NA | NA | NA | NA | Gm5225 | predicted gene 5225 [Source:MGI Symbol | 0 | 0 | 0 | 0 | 0 | 0 |
| 34190 | ENSMUSG00000095267 | 0 | NA | NA | NA | NA | NA | Ssty1 | spermiogenesis specific transcript on the ' | 0 | 0 | 0 | 0 | 0 | 0 |
| 34191 | ENSMUSG00000095269 | 0 | NA | NA | NA | NA | NA | Mir466b-8 | microRNA 466b-8 [Source:MGI Symbol;Ac | 0 | 0 | 0 | 0 | 0 | 0 |
| 34192 | ENSMUSG00000095270 | 0 | NA | NA | NA | NA | NA | Ifna9 | interferon alpha 9 [Source:MGI Symbol;A | 0 | 0 | 0 | 0 | 0 | 0 |
| 34193 | ENSMUSG00000095272 | 0 | NA | NA | NA | NA | NA | NA | NA | 0 | 0 | 0 | 0 | 0 | 0 |
| 34194 | ENSMUSG00000095273 | 0 | NA | NA | NA | NA | NA | Vmn1r142 | vomeronasal 1 receptor 142 [Source:MGI | 0 | 0 | 0 | 0 | 0 | 0 |
| 34195 | ENSMUSG00000095274 | 0 | NA | NA | NA | NA | NA | Obox4-ps27 | oocyte specific homeobox 4, pseudogene | 0 | 0 | 0 | 0 | 0 | 0 |
| 34196 | ENSMUSG00000095275 | 0 | NA | NA | NA | NA | NA | Vmn1r107 | vomeronasal 1 receptor 107 [Source:MGI | 0 | 0 | 0 | 0 | 0 | 0 |
| 34197 | ENSMUSG00000095276 | 0 | NA | NA | NA | NA | NA | Gfy | golgi-associated olfactory signaling regula | 0 | 0 | 0 | 0 | 0 | 0 |
| 34198 | ENSMUSG00000095277 | 0 | NA | NA | NA | NA | NA | Rps12-ps22 | ribosomal protein S12, pseudogene 22 [Sc | 0 | 0 | 0 | 0 | 0 | 0 |
| 34199 | ENSMUSG00000095278 | 0 | NA | NA | NA | NA | NA | NA | NA | 0 | 0 | 0 | 0 | 0 | 0 |
| 34200 | ENSMUSG00000095279 | 0 | NA | NA | NA | NA | NA | Gm10067 | predicted gene 10067 [Source:MGI Symbo | 0 | 0 | 0 | 0 | 0 | 0 |
| 34201 | ENSMUSG00000095280 | 0 | NA | NA | NA | NA | NA | NA | NA | 0 | 0 | 0 | 0 | 0 | 0 |
| 34202 | ENSMUSG00000095281 | 0.698954292 | 0.126053087 | 3.048972385 | 0.04134281 | 0.967022605 | NA | NA | NA | 0 | 0 | 2.0125 | 0 | 2.1812 | 0 |
| 34203 | ENSMUSG00000095282 | 0 | NA | NA | NA | NA | NA | NA | NA | 0 | 0 | 0 | 0 | 0 | 0 |
| 34204 | ENSMUSG00000095283 | 0 | NA | NA | NA | NA | NA | Astx1b | amplified spermatogenic transcripts X enc | 0 | 0 | 0 | 0 | 0 | 0 |
| 34205 | ENSMUSG00000095284 | 0 | NA | NA | NA | NA | NA | NA | NA | 0 | 0 | 0 | 0 | 0 | 0 |
| 34206 | ENSMUSG00000095285 | 0 | NA | NA | NA | NA | NA | Ighv5-9 | immunoglobulin heavy variable 5-9 [Sourc | 0 | 0 | 0 | 0 | 0 | 0 |
| 34207 | ENSMUSG00000095286 | 0 | NA | NA | NA | NA | NA | Olfr117 | olfactory receptor 117 [Source:MGI Symb | 0 | 0 | 0 | 0 | 0 | 0 |
| 34208 | ENSMUSG00000095287 | 0 | NA | NA | NA | NA | NA | Vmn1r-ps78 | vomeronasal 1 receptor, pseudogene 78 [ | 0 | 0 | 0 | 0 | 0 | 0 |
| 34209 | ENSMUSG00000095288 | 0 | NA | NA | NA | NA | NA | Gm7618 | predicted pseudogene 7618 [Source:MGI | 0 | 0 | 0 | 0 | 0 | 0 |
| 34210 | ENSMUSG00000095289 | 0 | NA | NA | NA | NA | NA | NA | NA | 0 | 0 | 0 | 0 | 0 | 0 |
| 34211 | ENSMUSG00000095290 | 0.439020386 | -2.189747911 | 3.52148864 | -0.621824499 | 0.534057275 | NA | NA | NA | 1.6062 | 1.0279 | 0 | 0 | 0 | 0 |
| 34212 | ENSMUSG00000095291 | 0 | NA | NA | NA | NA | NA | Gm12613 | predicted gene 12613 [Source:MGI Symbo | 0 | 0 | 0 | 0 | 0 | 0 |

| | | | | | | | | | | | | | | | |
|---|---|---|---|---|---|---|---|---|---|---|---|---|---|---|---|
| 34213 | ENSMUSG00000095292 | 0 | NA | NA | NA | NA | NA | NA | NA | 0 | 0 | 0 | 0 | 0 | 0 |
| 34214 | ENSMUSG00000095293 | 0 | NA | NA | NA | NA | NA | Gm10058 | predicted gene 10058 [Source:MGI Symbo | 0 | 0 | 0 | 0 | 0 | 0 |
| 34215 | ENSMUSG00000095294 | 0.999612184 | 2.312043103 | 2.464186403 | 0.938258201 | 0.348111735 | NA | Gm21119 | predicted gene, 21119 [Source:MGI Symb | 0 | 1.0279 | 0 | 3.8559 | 0 | 1.1138 |
| 34216 | ENSMUSG00000095295 | 0 | NA | NA | NA | NA | NA | Gm3488 | predicted gene, 3488 [Source:MGI Symbo | 0 | 0 | 0 | 0 | 0 | 0 |
| 34217 | ENSMUSG00000095296 | 0 | NA | NA | NA | NA | NA | Gm8906 | predicted gene 8906 [Source:MGI Symbol | 0 | 0 | 0 | 0 | 0 | 0 |
| 34218 | ENSMUSG00000095297 | 0 | NA | NA | NA | NA | NA | NA | NA | 0 | 0 | 0 | 0 | 0 | 0 |
| 34219 | ENSMUSG00000095298 | 1.570468055 | -0.984620644 | 1.876964216 | -0.524581468 | 0.599874172 | NA | Gm12407 | predicted gene 12407 [Source:MGI Symbo | 3.2124 | 0 | 3.0187 | 0.964 | 0 | 2.2277 |
| 34220 | ENSMUSG00000095299 | 0 | NA | NA | NA | NA | NA | NA | NA | 0 | 0 | 0 | 0 | 0 | 0 |
| 34221 | ENSMUSG00000095300 | 0 | NA | NA | NA | NA | NA | Gm906 | predicted gene 906 [Source:MGI Symbol;/ | 0 | 0 | 0 | 0 | 0 | 0 |
| 34222 | ENSMUSG00000095301 | 0 | NA | NA | NA | NA | NA | Olfr476 | olfactory receptor 476 [Source:MGI Symb | 0 | 0 | 0 | 0 | 0 | 0 |
| 34223 | ENSMUSG00000095302 | 0 | NA | NA | NA | NA | NA | Ssty2 | spermiogenesis specific transcript on the ' | 0 | 0 | 0 | 0 | 0 | 0 |
| 34224 | ENSMUSG00000095304 | 0 | NA | NA | NA | NA | NA | Plac9a | placenta specific 9a [Source:MGI Symbol;/ | 0 | 0 | 0 | 0 | 0 | 0 |
| 34225 | ENSMUSG00000095305 | 0 | NA | NA | NA | NA | NA | NA | NA | 0 | 0 | 0 | 0 | 0 | 0 |
| 34226 | ENSMUSG00000095307 | 0 | NA | NA | NA | NA | NA | Gm12573 | predicted gene 12573 [Source:MGI Symbo | 0 | 0 | 0 | 0 | 0 | 0 |
| 34227 | ENSMUSG00000095309 | 0 | NA | NA | NA | NA | NA | Vmn1r125 | vomeronasal 1 receptor 125 [Source:MGI | 0 | 0 | 0 | 0 | 0 | 0 |
| 34228 | ENSMUSG00000095310 | 0 | NA | NA | NA | NA | NA | NA | NA | 0 | 0 | 0 | 0 | 0 | 0 |
| 34229 | ENSMUSG00000095311 | 0.267700746 | -1.497481649 | 4.057001436 | -0.369110456 | 0.712045396 | NA | NA | NA | 1.6062 | 0 | 0 | 0 | 0 | 0 |
| 34230 | ENSMUSG00000095312 | 0 | NA | NA | NA | NA | NA | Olfr382 | olfactory receptor 382 [Source:MGI Symb | 0 | 0 | 0 | 0 | 0 | 0 |
| 34231 | ENSMUSG00000095314 | 0 | NA | NA | NA | NA | NA | NA | NA | 0 | 0 | 0 | 0 | 0 | 0 |
| 34232 | ENSMUSG00000095315 | 4.36690244 | 0.651367099 | 1.063851735 | 0.612272442 | 0.54035752 | NA | Gm10130 | predicted gene 10130 [Source:MGI Symbo | 4.0155 | 4.1117 | 2.0125 | 3.8559 | 2.1812 | 10.025 |
| 34233 | ENSMUSG00000095316 | 0 | NA | NA | NA | NA | NA | Gm21876 | predicted gene, 21876 [Source:MGI Symb | 0 | 0 | 0 | 0 | 0 | 0 |
| 34234 | ENSMUSG00000095318 | 0 | NA | NA | NA | NA | NA | Gm8024 | predicted gene 8024 [Source:MGI Symbol | 0 | 0 | 0 | 0 | 0 | 0 |
| 34235 | ENSMUSG00000095319 | 0 | NA | NA | NA | NA | NA | NA | NA | 0 | 0 | 0 | 0 | 0 | 0 |
| 34236 | ENSMUSG00000095320 | 0 | NA | NA | NA | NA | NA | NA | NA | 0 | 0 | 0 | 0 | 0 | 0 |
| 34237 | ENSMUSG00000095321 | 0 | NA | NA | NA | NA | NA | NA | NA | 0 | 0 | 0 | 0 | 0 | 0 |
| 34238 | ENSMUSG00000095322 | 0 | NA | NA | NA | NA | NA | Olfr957 | olfactory receptor 957 [Source:MGI Symb | 0 | 0 | 0 | 0 | 0 | 0 |
| 34239 | ENSMUSG00000095323 | 0 | NA | NA | NA | NA | NA | NA | NA | 0 | 0 | 0 | 0 | 0 | 0 |
| 34240 | ENSMUSG00000095324 | 0 | NA | NA | NA | NA | NA | Gm20825 | predicted gene, 20825 [Source:MGI Symb | 0 | 0 | 0 | 0 | 0 | 0 |
| 34241 | ENSMUSG00000095325 | 129.6546784 | 0.302175212 | 0.200326038 | 1.508417056 | 0.131447821 | 0.531226956 | Zfp870 | zinc finger protein 870 [Source:MGI Symb | 120.47 | 115.13 | 112.7 | 147.49 | 140.69 | 141.46 |
| 34242 | ENSMUSG00000095327 | 0 | NA | NA | NA | NA | NA | Gm14996 | predicted gene 14996 [Source:MGI Symbo | 0 | 0 | 0 | 0 | 0 | 0 |
| 34243 | ENSMUSG00000095328 | 0 | NA | NA | NA | NA | NA | Defa-ps6 | defensin, alpha, pseudogene 6 [Source:M | 0 | 0 | 0 | 0 | 0 | 0 |
| 34244 | ENSMUSG00000095330 | 0 | NA | NA | NA | NA | NA | Gm21834 | predicted gene, 21834 [Source:MGI Symb | 0 | 0 | 0 | 0 | 0 | 0 |
| 34245 | ENSMUSG00000095331 | 0 | NA | NA | NA | NA | NA | Ptma-ps1 | prothymosin alpha, pseudogene 1 [Source | 0 | 0 | 0 | 0 | 0 | 0 |
| 34246 | ENSMUSG00000095332 | 5.690638045 | -0.349914304 | 1.075129944 | -0.325462337 | 0.744831183 | 0.949997329 | Gm9821 | predicted gene 9821 [Source:MGI Symbol | 4.0155 | 9.2513 | 6.0375 | 11.568 | 3.2719 | 0 |
| 34247 | ENSMUSG00000095333 | 0 | NA | NA | NA | NA | NA | NA | NA | 0 | 0 | 0 | 0 | 0 | 0 |
| 34248 | ENSMUSG00000095334 | 0.77079641 | 0.856159616 | 2.652840961 | 0.322733111 | 0.746897374 | NA | Gm21984 | predicted gene 21984 [Source:MGI Symbo | 1.6062 | 0 | 0 | 1.928 | 1.0906 | 0 |
| 34249 | ENSMUSG00000095335 | 0 | NA | NA | NA | NA | NA | Igkv3-5 | immunoglobulin kappa chain variable 3-5 | 0 | 0 | 0 | 0 | 0 | 0 |
| 34250 | ENSMUSG00000095336 | 0 | NA | NA | NA | NA | NA | NA | NA | 0 | 0 | 0 | 0 | 0 | 0 |
| 34251 | ENSMUSG00000095338 | 0 | NA | NA | NA | NA | NA | Igkv3-9 | immunoglobulin kappa variable 3-9 [Sourc | 0 | 0 | 0 | 0 | 0 | 0 |
| 34252 | ENSMUSG00000095339 | 0 | NA | NA | NA | NA | NA | Zscan4b | zinc finger and SCAN domain containing 4 | 0 | 0 | 0 | 0 | 0 | 0 |
| 34253 | ENSMUSG00000095340 | 0 | NA | NA | NA | NA | NA | NA | NA | 0 | 0 | 0 | 0 | 0 | 0 |
| 34254 | ENSMUSG00000095341 | 0 | NA | NA | NA | NA | NA | Gm428 | predicted gene 428 [Source:MGI Symbol;/ | 0 | 0 | 0 | 0 | 0 | 0 |
| 34255 | ENSMUSG00000095342 | 0 | NA | NA | NA | NA | NA | NA | NA | 0 | 0 | 0 | 0 | 0 | 0 |
| 34256 | ENSMUSG00000095344 | 0 | NA | NA | NA | NA | NA | Gm6838 | predicted gene 6838 [Source:MGI Symbol | 0 | 0 | 0 | 0 | 0 | 0 |
| 34257 | ENSMUSG00000095345 | 0 | NA | NA | NA | NA | NA | Gm21821 | predicted gene, 21821 [Source:MGI Symb | 0 | 0 | 0 | 0 | 0 | 0 |
| 34258 | ENSMUSG00000095346 | 0 | NA | NA | NA | NA | NA | Gm21905 | predicted gene, 21905 [Source:MGI Symb | 0 | 0 | 0 | 0 | 0 | 0 |
| 34259 | ENSMUSG00000095347 | 0 | NA | NA | NA | NA | NA | NA | NA | 0 | 0 | 0 | 0 | 0 | 0 |
| 34260 | ENSMUSG00000095348 | 0 | NA | NA | NA | NA | NA | Gm3892 | predicted gene 3892 [Source:MGI Symbol | 0 | 0 | 0 | 0 | 0 | 0 |
| 34261 | ENSMUSG00000095349 | 0 | NA | NA | NA | NA | NA | Gm26097 | predicted gene, 26097 [Source:MGI Symb | 0 | 0 | 0 | 0 | 0 | 0 |
| 34262 | ENSMUSG00000095350 | 0 | NA | NA | NA | NA | NA | NA | NA | 0 | 0 | 0 | 0 | 0 | 0 |
| 34263 | ENSMUSG00000095351 | 0.305170013 | -1.657836477 | 4.041480335 | -0.410205257 | 0.681655385 | NA | Igkv3-2 | immunoglobulin kappa variable 3-2 [Sourc | 0.8031 | 1.0279 | 0 | 0 | 0 | 0 |
| 34264 | ENSMUSG00000095353 | 0 | NA | NA | NA | NA | NA | Gm21783 | predicted gene, 21783 [Source:MGI Symb | 0 | 0 | 0 | 0 | 0 | 0 |
| 34265 | ENSMUSG00000095356 | 0 | NA | NA | NA | NA | NA | Gm7398 | predicted gene 7398 [Source:MGI Symbol | 0 | 0 | 0 | 0 | 0 | 0 |
| 34266 | ENSMUSG00000095357 | 0 | NA | NA | NA | NA | NA | NA | NA | 0 | 0 | 0 | 0 | 0 | 0 |
| 34267 | ENSMUSG00000095358 | 0 | NA | NA | NA | NA | NA | Vmn1r165 | vomeronasal 1 receptor 165 [Source:MGI | 0 | 0 | 0 | 0 | 0 | 0 |
| 34268 | ENSMUSG00000095360 | 0 | NA | NA | NA | NA | NA | Gm3594 | predicted gene 3594 [Source:MGI Symbol | 0 | 0 | 0 | 0 | 0 | 0 |

| | | | | | | | | | | | | | | | |
|---|---|---|---|---|---|---|---|---|---|---|---|---|---|---|---|
| 34269 | ENSMUSG00000095361 | 0 | NA | NA | NA | NA | NA | NA | NA | 0 | 0 | 0 | 0 | 0 | 0 |
| 34270 | ENSMUSG00000095362 | 170.5811966 | 0.044939857 | 0.183598432 | 0.244772556 | 0.806632545 | 0.964718338 | Gm14325 | predicted gene 14325 [Source:MGI Symbo | 175.08 | 163.44 | 165.02 | 168.7 | 190.86 | 160.39 |
| 34271 | ENSMUSG00000095363 | 0 | NA | NA | NA | NA | NA | Gm4567 | predicted gene 4567 [Source:MGI Symbol | 0 | 0 | 0 | 0 | 0 | 0 |
| 34272 | ENSMUSG00000095365 | 0 | NA | NA | NA | NA | NA | Rbm31y | RNA binding motif 31, Y-linked [Source:M | 0 | 0 | 0 | 0 | 0 | 0 |
| 34273 | ENSMUSG00000095366 | 0 | NA | NA | NA | NA | NA | Gm21860 | predicted gene, 21860 [Source:MGI Symb | 0 | 0 | 0 | 0 | 0 | 0 |
| 34274 | ENSMUSG00000095368 | 0 | NA | NA | NA | NA | NA | Gm3012 | predicted gene 3012 [Source:MGI Symbol | 0 | 0 | 0 | 0 | 0 | 0 |
| 34275 | ENSMUSG00000095369 | 0 | NA | NA | NA | NA | NA | Gm21859 | predicted gene, 21859 [Source:MGI Symb | 0 | 0 | 0 | 0 | 0 | 0 |
| 34276 | ENSMUSG00000095370 | 0 | NA | NA | NA | NA | NA | Trav6n-7 | T cell receptor alpha variable 6N-7 [Sourc | 0 | 0 | 0 | 0 | 0 | 0 |
| 34277 | ENSMUSG00000095371 | 0 | NA | NA | NA | NA | NA | Gm9611 | predicted gene 9611 [Source:MGI Symbol | 0 | 0 | 0 | 0 | 0 | 0 |
| 34278 | ENSMUSG00000095372 | 0 | NA | NA | NA | NA | NA | NA | NA | 0 | 0 | 0 | 0 | 0 | 0 |
| 34279 | ENSMUSG00000095374 | 0 | NA | NA | NA | NA | NA | NA | NA | 0 | 0 | 0 | 0 | 0 | 0 |
| 34280 | ENSMUSG00000095376 | 0 | NA | NA | NA | NA | NA | NA | NA | 0 | 0 | 0 | 0 | 0 | 0 |
| 34281 | ENSMUSG00000095377 | 0 | NA | NA | NA | NA | NA | Olfr91 | olfactory receptor 91 [Source:MGI Symbo | 0 | 0 | 0 | 0 | 0 | 0 |
| 34282 | ENSMUSG00000095379 | 0 | NA | NA | NA | NA | NA | NA | NA | 0 | 0 | 0 | 0 | 0 | 0 |
| 34283 | ENSMUSG00000095380 | 0 | NA | NA | NA | NA | NA | NA | NA | 0 | 0 | 0 | 0 | 0 | 0 |
| 34284 | ENSMUSG00000095382 | 0 | NA | NA | NA | NA | NA | NA | NA | 0 | 0 | 0 | 0 | 0 | 0 |
| 34285 | ENSMUSG00000095383 | 0 | NA | NA | NA | NA | NA | Vmn1r152 | vomeronasal 1 receptor 152 [Source:MGI | 0 | 0 | 0 | 0 | 0 | 0 |
| 34286 | ENSMUSG00000095384 | 0 | NA | NA | NA | NA | NA | 1700001F09 | RIKEN cDNA 1700001F09 gene [Source:M | 0 | 0 | 0 | 0 | 0 | 0 |
| 34287 | ENSMUSG00000095385 | 24.93948243 | 0.041611772 | 0.472311377 | 0.088102413 | 0.929795278 | 0.988405116 | D630033O11 | RIKEN cDNA D630033O11 gene [Source:M | 16.865 | 28.782 | 28.175 | 16.388 | 33.809 | 25.618 |
| 34288 | ENSMUSG00000095386 | 13.71586716 | 0.762366178 | 0.619161909 | 1.231287272 | 0.218215441 | 0.663697152 | Gm17662 | predicted gene, 17662 [Source:MGI Symb | 11.243 | 8.2233 | 11.069 | 24.099 | 8.7249 | 18.935 |
| 34289 | ENSMUSG00000095387 | 0 | NA | NA | NA | NA | NA | Trav6d-4 | T cell receptor alpha variable 6D-4 [Sourc | 0 | 0 | 0 | 0 | 0 | 0 |
| 34290 | ENSMUSG00000095388 | 0 | NA | NA | NA | NA | NA | Gm10681 | predicted gene 10681 [Source:MGI Symbo | 0 | 0 | 0 | 0 | 0 | 0 |
| 34291 | ENSMUSG00000095389 | 0 | NA | NA | NA | NA | NA | NA | NA | 0 | 0 | 0 | 0 | 0 | 0 |
| 34292 | ENSMUSG00000095390 | 0 | NA | NA | NA | NA | NA | Olfr229 | olfactory receptor 229 [Source:MGI Symb | 0 | 0 | 0 | 0 | 0 | 0 |
| 34293 | ENSMUSG00000095392 | 0 | NA | NA | NA | NA | NA | Gm7350 | predicted gene 7350 [Source:MGI Symbol | 0 | 0 | 0 | 0 | 0 | 0 |
| 34294 | ENSMUSG00000095393 | 0 | NA | NA | NA | NA | NA | NA | NA | 0 | 0 | 0 | 0 | 0 | 0 |
| 34295 | ENSMUSG00000095394 | 0 | NA | NA | NA | NA | NA | Gm25485 | predicted gene, 25485 [Source:MGI Symb | 0 | 0 | 0 | 0 | 0 | 0 |
| 34296 | ENSMUSG00000095395 | 0 | NA | NA | NA | NA | NA | NA | NA | 0 | 0 | 0 | 0 | 0 | 0 |
| 34297 | ENSMUSG00000095396 | 0.36353943 | 2.065989254 | 4.034137301 | 0.512126658 | 0.608562368 | NA | NA | NA | 0 | 0 | 0 | 0 | 2.1812 | 0 |
| 34298 | ENSMUSG00000095398 | 0 | NA | NA | NA | NA | NA | Gm16404 | predicted gene 16404 [Source:MGI Symbo | 0 | 0 | 0 | 0 | 0 | 0 |
| 34299 | ENSMUSG00000095399 | 0 | NA | NA | NA | NA | NA | Ighd6-1 | immunoglobulin heavy diversity 6-1 [Sour | 0 | 0 | 0 | 0 | 0 | 0 |
| 34300 | ENSMUSG00000095401 | 0 | NA | NA | NA | NA | NA | Olfr804 | olfactory receptor 804 [Source:MGI Symb | 0 | 0 | 0 | 0 | 0 | 0 |
| 34301 | ENSMUSG00000095402 | 0 | NA | NA | NA | NA | NA | NA | NA | 0 | 0 | 0 | 0 | 0 | 0 |
| 34302 | ENSMUSG00000095403 | 12.56885813 | 0.28891934 | 0.624822321 | 0.462402399 | 0.643792779 | 0.925536106 | Gm21092 | predicted gene, 21092 [Source:MGI Symb | 9.6372 | 15.419 | 9.0562 | 19.28 | 11.997 | 10.025 |
| 34303 | ENSMUSG00000095407 | 0.572870759 | -2.577107685 | 3.182795569 | -0.809699407 | 0.418112959 | NA | Tmem200c | transmembrane protein 200C [Source:MG | 2.4093 | 1.0279 | 0 | 0 | 0 | 0 |
| 34304 | ENSMUSG00000095408 | 0 | NA | NA | NA | NA | NA | Gm8366 | predicted gene 8366 [Source:MGI Symbol | 0 | 0 | 0 | 0 | 0 | 0 |
| 34305 | ENSMUSG00000095409 | 0 | NA | NA | NA | NA | NA | Gm13043 | predicted gene 13043 [Source:MGI Symbo | 0 | 0 | 0 | 0 | 0 | 0 |
| 34306 | ENSMUSG00000095410 | 0 | NA | NA | NA | NA | NA | NA | NA | 0 | 0 | 0 | 0 | 0 | 0 |
| 34307 | ENSMUSG00000095411 | 0 | NA | NA | NA | NA | NA | Gm15260 | predicted gene 15260 [Source:MGI Symbo | 0 | 0 | 0 | 0 | 0 | 0 |
| 34308 | ENSMUSG00000095412 | 0 | NA | NA | NA | NA | NA | Gm5885 | predicted gene 5885 [Source:MGI Symbol | 0 | 0 | 0 | 0 | 0 | 0 |
| 34309 | ENSMUSG00000095413 | 0 | NA | NA | NA | NA | NA | H2al1e | H2A histone family member L1E [Source:N | 0 | 0 | 0 | 0 | 0 | 0 |
| 34310 | ENSMUSG00000095414 | 0 | NA | NA | NA | NA | NA | NA | NA | 0 | 0 | 0 | 0 | 0 | 0 |
| 34311 | ENSMUSG00000095416 | 0 | NA | NA | NA | NA | NA | Ighv1-12 | immunoglobulin heavy variable V1-12 [So | 0 | 0 | 0 | 0 | 0 | 0 |
| 34312 | ENSMUSG00000095418 | 0 | NA | NA | NA | NA | NA | NA | NA | 0 | 0 | 0 | 0 | 0 | 0 |
| 34313 | ENSMUSG00000095419 | 0.133850373 | -0.796629176 | 4.080472857 | -0.195229623 | 0.84521319 | NA | Gm14328 | predicted gene 14328 [Source:MGI Symbo | 0.8031 | 0 | 0 | 0 | 0 | 0 |
| 34314 | ENSMUSG00000095420 | 0 | NA | NA | NA | NA | NA | Astx1a | amplified spermatogenic transcripts X enc | 0 | 0 | 0 | 0 | 0 | 0 |
| 34315 | ENSMUSG00000095422 | 0 | NA | NA | NA | NA | NA | NA | NA | 0 | 0 | 0 | 0 | 0 | 0 |
| 34316 | ENSMUSG00000095423 | 0 | NA | NA | NA | NA | NA | Gm7375 | predicted pseudogene 7375 [Source:MGI | 0 | 0 | 0 | 0 | 0 | 0 |
| 34317 | ENSMUSG00000095424 | 0 | NA | NA | NA | NA | NA | NA | NA | 0 | 0 | 0 | 0 | 0 | 0 |
| 34318 | ENSMUSG00000095425 | 0 | NA | NA | NA | NA | NA | NA | NA | 0 | 0 | 0 | 0 | 0 | 0 |
| 34319 | ENSMUSG00000095426 | 0 | NA | NA | NA | NA | NA | Trav6d-5 | T cell receptor alpha variable 6D-5 [Sourc | 0 | 0 | 0 | 0 | 0 | 0 |
| 34320 | ENSMUSG00000095427 | 1.302947269 | 0.17976329 | 1.96941963 | 0.091277292 | 0.927272261 | NA | Rps2-ps6 | ribosomal protein S2, pseudogene 6 [Sour | 1.6062 | 2.0558 | 0 | 1.928 | 0 | 2.2277 |
| 34321 | ENSMUSG00000095428 | 0 | NA | NA | NA | NA | NA | NA | NA | 0 | 0 | 0 | 0 | 0 | 0 |
| 34322 | ENSMUSG00000095429 | 0 | NA | NA | NA | NA | NA | Ighv5-12 | immunoglobulin heavy variable 5-12 [Sour | 0 | 0 | 0 | 0 | 0 | 0 |
| 34323 | ENSMUSG00000095430 | 0 | NA | NA | NA | NA | NA | Vmn1r72 | vomeronasal 1 receptor 72 [Source:MGI S | 0 | 0 | 0 | 0 | 0 | 0 |
| 34324 | ENSMUSG00000095431 | 0 | NA | NA | NA | NA | NA | NA | NA | 0 | 0 | 0 | 0 | 0 | 0 |

| | | | | | | | | | | | | | | | |
|---|---|---|---|---|---|---|---|---|---|---|---|---|---|---|---|
| 34325 | ENSMUSG00000095432 | 95.06469886 | 0.291040854 | 0.241574034 | 1.20476878 | 0.228292575 | 0.674708584 | Zfp748 | zinc finger protein 748 [Source:MGI Symb | 74.689 | 92.513 | 89.556 | 97.362 | 95.974 | 120.3 |
| 34326 | ENSMUSG00000095433 | 0 | NA | NA | NA | NA | NA | Gm21914 | predicted gene, 21914 [Source:MGI Symb | 0 | 0 | 0 | 0 | 0 | 0 |
| 34327 | ENSMUSG00000095434 | 0 | NA | NA | NA | NA | NA | NA | NA | 0 | 0 | 0 | 0 | 0 | 0 |
| 34328 | ENSMUSG00000095435 | 0 | NA | NA | NA | NA | NA | NA | NA | 0 | 0 | 0 | 0 | 0 | 0 |
| 34329 | ENSMUSG00000095436 | 0 | NA | NA | NA | NA | NA | Gm15254 | predicted gene 15254 [Source:MGI Symbo | 0 | 0 | 0 | 0 | 0 | 0 |
| 34330 | ENSMUSG00000095437 | 0 | NA | NA | NA | NA | NA | NA | NA | 0 | 0 | 0 | 0 | 0 | 0 |
| 34331 | ENSMUSG00000095438 | 0 | NA | NA | NA | NA | NA | Mir133a-1hg | Mir133a-1, Mir1b and Mir1a-2 host gene | | 0 | 0 | 0 | 0 | 0 | 0 |
| 34332 | ENSMUSG00000095440 | 252.5891984 | 0.25754966 | 0.184382073 | 1.39682593 | 0.162465923 | 0.585865647 | Fignl2 | fidgetin-like 2 [Source:MGI Symbol;Acc:M | 183.91 | 245.67 | 261.62 | 273.77 | 246.48 | 304.08 |
| 34333 | ENSMUSG00000095442 | 0 | NA | NA | NA | NA | NA | Ighv1-4 | immunoglobulin heavy variable 1-4 [Sourc | 0 | 0 | 0 | 0 | 0 | 0 |
| 34334 | ENSMUSG00000095444 | 0 | NA | NA | NA | NA | NA | Ighd2-4 | immunoglobulin heavy diversity 2-4 [Sour | 0 | 0 | 0 | 0 | 0 | 0 |
| 34335 | ENSMUSG00000095445 | 0 | NA | NA | NA | NA | NA | H2al1i | H2A histone family member L1I [Source:N | 0 | 0 | 0 | 0 | 0 | 0 |
| 34336 | ENSMUSG00000095446 | 0 | NA | NA | NA | NA | NA | NA | NA | 0 | 0 | 0 | 0 | 0 | 0 |
| 34337 | ENSMUSG00000095447 | 0 | NA | NA | NA | NA | NA | NA | NA | 0 | 0 | 0 | 0 | 0 | 0 |
| 34338 | ENSMUSG00000095448 | 0 | NA | NA | NA | NA | NA | Olfr859 | olfactory receptor 859 [Source:MGI Symb | 0 | 0 | 0 | 0 | 0 | 0 |
| 34339 | ENSMUSG00000095449 | 0 | NA | NA | NA | NA | NA | NA | NA | 0 | 0 | 0 | 0 | 0 | 0 |
| 34340 | ENSMUSG00000095450 | 0 | NA | NA | NA | NA | NA | | | 0 | 0 | 0 | 0 | 0 | 0 |
| 34341 | ENSMUSG00000095452 | 0 | NA | NA | NA | NA | NA | Gm20795 | predicted gene, 20795 [Source:MGI Symb | 0 | 0 | 0 | 0 | 0 | 0 |
| 34342 | ENSMUSG00000095453 | 0 | NA | NA | NA | NA | NA | NA | NA | 0 | 0 | 0 | 0 | 0 | 0 |
| 34343 | ENSMUSG00000095455 | 0.181769715 | 1.126924175 | 4.080472857 | 0.276174898 | 0.782413736 | NA | Mup-ps3 | major urinary protein, pseudogene 3 [Sou | 0 | 0 | 0 | 0 | 1.0906 | 0 |
| 34344 | ENSMUSG00000095456 | 0 | NA | NA | NA | NA | NA | NA | NA | 0 | 0 | 0 | 0 | 0 | 0 |
| 34345 | ENSMUSG00000095457 | 0 | NA | NA | NA | NA | NA | Gvin-ps5 | GTPase, very large interferon inducible, ps | 0 | 0 | 0 | 0 | 0 | 0 |
| 34346 | ENSMUSG00000095458 | 0 | NA | NA | NA | NA | NA | NA | NA | 0 | 0 | 0 | 0 | 0 | 0 |
| 34347 | ENSMUSG00000095460 | 0 | NA | NA | NA | NA | NA | Vmn2r-ps16 | vomeronasal 2, receptor, pseudogene 16 | 0 | 0 | 0 | 0 | 0 | 0 |
| 34348 | ENSMUSG00000095462 | 0 | NA | NA | NA | NA | NA | Gm21725 | predicted gene, 21725 [Source:MGI Symb | 0 | 0 | 0 | 0 | 0 | 0 |
| 34349 | ENSMUSG00000095463 | 0 | NA | NA | NA | NA | NA | Entpd4 | ectonucleoside triphosphate diphosphohy | 0 | 0 | 0 | 0 | 0 | 0 |
| 34350 | ENSMUSG00000095464 | 0 | NA | NA | NA | NA | NA | Gm21987 | predicted gene 21987 [Source:MGI Symbo | 0 | 0 | 0 | 0 | 0 | 0 |
| 34351 | ENSMUSG00000095465 | 0 | NA | NA | NA | NA | NA | Gm6214 | predicted gene 6214 [Source:MGI Symbol | 0 | 0 | 0 | 0 | 0 | 0 |
| 34352 | ENSMUSG00000095466 | 0 | NA | NA | NA | NA | NA | Gm3015 | predicted gene 3015 [Source:MGI Symbol | 0 | 0 | 0 | 0 | 0 | 0 |
| 34353 | ENSMUSG00000095467 | 0 | NA | NA | NA | NA | NA | NA | NA | 0 | 0 | 0 | 0 | 0 | 0 |
| 34354 | ENSMUSG00000095469 | 0 | NA | NA | NA | NA | NA | Gm21965 | predicted gene, 21965 [Source:MGI Symb | 0 | 0 | 0 | 0 | 0 | 0 |
| 34355 | ENSMUSG00000095470 | 0 | NA | NA | NA | NA | NA | Gm21863 | predicted gene, 21863 [Source:MGI Symb | 0 | 0 | 0 | 0 | 0 | 0 |
| 34356 | ENSMUSG00000095472 | 0 | NA | NA | NA | NA | NA | NA | NA | 0 | 0 | 0 | 0 | 0 | 0 |
| 34357 | ENSMUSG00000095474 | 0 | NA | NA | NA | NA | NA | Cldn34c2 | claudin 34C2 [Source:MGI Symbol;Acc:MG | 0 | 0 | 0 | 0 | 0 | 0 |
| 34358 | ENSMUSG00000095475 | 0 | NA | NA | NA | NA | NA | | | 0 | 0 | 0 | 0 | 0 | 0 |
| 34359 | ENSMUSG00000095476 | 0 | NA | NA | NA | NA | NA | NA | NA | 0 | 0 | 0 | 0 | 0 | 0 |
| 34360 | ENSMUSG00000095477 | 0 | NA | NA | NA | NA | NA | NA | NA | 0 | 0 | 0 | 0 | 0 | 0 |
| 34361 | ENSMUSG00000095478 | 4.073991926 | 0.157851741 | 1.076037259 | 0.146697281 | 0.883370965 | NA | Gm9824 | predicted pseudogene 9824 [Source:MGI | 2.4093 | 5.1396 | 4.025 | 2.8919 | 2.1812 | 7.7969 |
| 34362 | ENSMUSG00000095479 | 0 | NA | NA | NA | NA | NA | NA | NA | 0 | 0 | 0 | 0 | 0 | 0 |
| 34363 | ENSMUSG00000095482 | 0 | NA | NA | NA | NA | NA | NA | NA | 0 | 0 | 0 | 0 | 0 | 0 |
| 34364 | ENSMUSG00000095483 | 0 | NA | NA | NA | NA | NA | Olfr776 | olfactory receptor 776 [Source:MGI Symb | 0 | 0 | 0 | 0 | 0 | 0 |
| 34365 | ENSMUSG00000095484 | 0 | NA | NA | NA | NA | NA | Olfr1480 | olfactory receptor 1480 [Source:MGI Sym | 0 | 0 | 0 | 0 | 0 | 0 |
| 34366 | ENSMUSG00000095486 | 0.185640647 | 1.126924175 | 4.080472857 | 0.276174898 | 0.782413736 | NA | Vmn2r22 | vomeronasal 2, receptor 22 [Source:MGI S | 0 | 0 | 0 | 0 | 0 | 1.1138 |
| 34367 | ENSMUSG00000095487 | 0 | NA | NA | NA | NA | NA | Scgb1b21 | secretoglobin, family 1B, member 21 [Sou | 0 | 0 | 0 | 0 | 0 | 0 |
| 34368 | ENSMUSG00000095489 | 0 | NA | NA | NA | NA | NA | NA | NA | 0 | 0 | 0 | 0 | 0 | 0 |
| 34369 | ENSMUSG00000095490 | 0 | NA | NA | NA | NA | NA | Gm7951 | predicted gene 7951 [Source:MGI Symbol | 0 | 0 | 0 | 0 | 0 | 0 |
| 34370 | ENSMUSG00000095491 | 0 | NA | NA | NA | NA | NA | NA | NA | 0 | 0 | 0 | 0 | 0 | 0 |
| 34371 | ENSMUSG00000095492 | 0 | NA | NA | NA | NA | NA | Gm14361 | predicted gene 14361 [Source:MGI Symbo | 0 | 0 | 0 | 0 | 0 | 0 |
| 34372 | ENSMUSG00000095493 | 0 | NA | NA | NA | NA | NA | A630023A22 | RIKEN cDNA A630023A22 gene [Source:M | 0 | 0 | 0 | 0 | 0 | 0 |
| 34373 | ENSMUSG00000095494 | 0 | NA | NA | NA | NA | NA | Gm7391 | predicted pseudogene 7391 [Source:MGI | 0 | 0 | 0 | 0 | 0 | 0 |
| 34374 | ENSMUSG00000095495 | 0 | NA | NA | NA | NA | NA | Trav9d-3 | T cell receptor alpha variable 9D-3 [Sourc | 0 | 0 | 0 | 0 | 0 | 0 |
| 34375 | ENSMUSG00000095497 | 0 | NA | NA | NA | NA | NA | Igkv1-122 | immunoglobulin kappa chain variable 1-1 | 0 | 0 | 0 | 0 | 0 | 0 |
| 34376 | ENSMUSG00000095498 | 0 | NA | NA | NA | NA | NA | Ifna1 | interferon alpha 1 [Source:MGI Symbol;Ac | 0 | 0 | 0 | 0 | 0 | 0 |
| 34377 | ENSMUSG00000095499 | 0 | NA | NA | NA | NA | NA | Rps12-ps14 | ribosomal protein S12, pseudogene 14 [Sc | 0 | 0 | 0 | 0 | 0 | 0 |
| 34378 | ENSMUSG00000095500 | 0 | NA | NA | NA | NA | NA | | | 0 | 0 | 0 | 0 | 0 | 0 |
| 34379 | ENSMUSG00000095501 | 0 | NA | NA | NA | NA | NA | NA | NA | 0 | 0 | 0 | 0 | 0 | 0 |
| 34380 | ENSMUSG00000095502 | 0 | NA | NA | NA | NA | NA | NA | NA | 0 | 0 | 0 | 0 | 0 | 0 |

| | | | | | | | | | | | | | | | | |
|---|---|---|---|---|---|---|---|---|---|---|---|---|---|---|---|---|
| 34381 | ENSMUSG00000095503 | 0 | NA | NA | NA | NA | NA | | Pramel43 | PRAME like 43 [Source:MGI Symbol;Acc:M | 0 | 0 | 0 | 0 | 0 | 0 |
| 34382 | ENSMUSG00000095504 | 0.506734502 | -2.372847156 | 3.338060886 | -0.710845978 | 0.477179685 | NA | | NA | NA | 0 | 1.0279 | 2.0125 | 0 | 0 | 0 |
| 34383 | ENSMUSG00000095505 | 0 | NA | NA | NA | NA | NA | | | | 0 | 0 | 0 | 0 | 0 | 0 |
| 34384 | ENSMUSG00000095506 | 0 | NA | NA | NA | NA | NA | | NA | NA | 0 | 0 | 0 | 0 | 0 | 0 |
| 34385 | ENSMUSG00000095507 | 0 | NA | NA | NA | NA | NA | | NA | NA | 0 | 0 | 0 | 0 | 0 | 0 |
| 34386 | ENSMUSG00000095508 | 0 | NA | NA | NA | NA | NA | | Gm20812 | predicted gene, 20812 [Source:MGI Symb | 0 | 0 | 0 | 0 | 0 | 0 |
| 34387 | ENSMUSG00000095509 | 0 | NA | NA | NA | NA | NA | | NA | NA | 0 | 0 | 0 | 0 | 0 | 0 |
| 34388 | ENSMUSG00000095510 | 0 | NA | NA | NA | NA | NA | | Rhox3f | reproductive homeobox 3F [Source:MGI S | 0 | 0 | 0 | 0 | 0 | 0 |
| 34389 | ENSMUSG00000095511 | 0 | NA | NA | NA | NA | NA | | NA | NA | 0 | 0 | 0 | 0 | 0 | 0 |
| 34390 | ENSMUSG00000095512 | 0 | NA | NA | NA | NA | NA | | Gm17222 | predicted gene 17222 [Source:MGI Symbo | 0 | 0 | 0 | 0 | 0 | 0 |
| 34391 | ENSMUSG00000095513 | 0 | NA | NA | NA | NA | NA | | NA | NA | 0 | 0 | 0 | 0 | 0 | 0 |
| 34392 | ENSMUSG00000095514 | 0 | NA | NA | NA | NA | NA | | D13Ertd608e | DNA segment, Chr 13, ERATO Doi 608, exp | 0 | 0 | 0 | 0 | 0 | 0 |
| 34393 | ENSMUSG00000095516 | 0 | NA | NA | NA | NA | NA | | NA | NA | 0 | 0 | 0 | 0 | 0 | 0 |
| 34394 | ENSMUSG00000095517 | 0 | NA | NA | NA | NA | NA | | Gm13249 | predicted gene 13249 [Source:MGI Symbo | 0 | 0 | 0 | 0 | 0 | 0 |
| 34395 | ENSMUSG00000095518 | 0 | NA | NA | NA | NA | NA | | Gm8068 | predicted gene 8068 [Source:MGI Symbol | 0 | 0 | 0 | 0 | 0 | 0 |
| 34396 | ENSMUSG00000095519 | 0 | NA | NA | NA | NA | NA | | Ighv1-66 | immunoglobulin heavy variable 1-66 [Sour | 0 | 0 | 0 | 0 | 0 | 0 |
| 34397 | ENSMUSG00000095520 | 0 | NA | NA | NA | NA | NA | | Gm20877 | predicted gene, 20877 [Source:MGI Symb | 0 | 0 | 0 | 0 | 0 | 0 |
| 34398 | ENSMUSG00000095521 | 0 | NA | NA | NA | NA | NA | | Astx5 | amplified spermatogenic transcripts X enc | 0 | 0 | 0 | 0 | 0 | 0 |
| 34399 | ENSMUSG00000095522 | 0 | NA | NA | NA | NA | NA | | Aadacl2fm3 | AADACL2 family member 3 [Source:MGI S | 0 | 0 | 0 | 0 | 0 | 0 |
| 34400 | ENSMUSG00000095523 | 0 | NA | NA | NA | NA | NA | | | | 0 | 0 | 0 | 0 | 0 | 0 |
| 34401 | ENSMUSG00000095524 | 1.877189296 | 0.333140416 | 1.646081549 | 0.202383907 | 0.839616608 | NA | | Gm15682 | predicted gene 15682 [Source:MGI Symbo | 0 | 1.0279 | 4.025 | 2.8919 | 1.0906 | 2.2277 |
| 34402 | ENSMUSG00000095525 | 0 | NA | NA | NA | NA | NA | | Olfr834 | olfactory receptor 834 [Source:MGI Symb | 0 | 0 | 0 | 0 | 0 | 0 |
| 34403 | ENSMUSG00000095526 | 3.028786147 | 0.509214093 | 1.341122824 | 0.379692362 | 0.70417379 | NA | | Gm10243 | predicted gene 10243 [Source:MGI Symbo | 2.4093 | 2.0558 | 3.0187 | 2.8919 | 0 | 7.7969 |
| 34404 | ENSMUSG00000095527 | 0 | NA | NA | NA | NA | NA | | Olfr888 | olfactory receptor 888 [Source:MGI Symb | 0 | 0 | 0 | 0 | 0 | 0 |
| 34405 | ENSMUSG00000095528 | 0 | NA | NA | NA | NA | NA | | Gm10375 | predicted gene 10375 [Source:MGI Symbo | 0 | 0 | 0 | 0 | 0 | 0 |
| 34406 | ENSMUSG00000095529 | 0 | NA | NA | NA | NA | NA | | NA | NA | 0 | 0 | 0 | 0 | 0 | 0 |
| 34407 | ENSMUSG00000095530 | 0 | NA | NA | NA | NA | NA | | NA | NA | 0 | 0 | 0 | 0 | 0 | 0 |
| 34408 | ENSMUSG00000095531 | 0 | NA | NA | NA | NA | NA | | NA | NA | 0 | 0 | 0 | 0 | 0 | 0 |
| 34409 | ENSMUSG00000095532 | 172.2502362 | 1.185303611 | 0.216196464 | 5.482530061 | 4.19E-08 | 5.23E-06 | | Mup-ps13 | major urinary protein, pseudogene 13 [So | 137.33 | 88.401 | 88.55 | 226.53 | 244.3 | 248.39 |
| 34410 | ENSMUSG00000095533 | 0 | NA | NA | NA | NA | NA | | Gm10413 | predicted gene 10413 [Source:MGI Symbo | 0 | 0 | 0 | 0 | 0 | 0 |
| 34411 | ENSMUSG00000095534 | 0 | NA | NA | NA | NA | NA | | Gm15102 | predicted gene 15102 [Source:MGI Symbo | 0 | 0 | 0 | 0 | 0 | 0 |
| 34412 | ENSMUSG00000095536 | 0 | NA | NA | NA | NA | NA | | NA | NA | 0 | 0 | 0 | 0 | 0 | 0 |
| 34413 | ENSMUSG00000095537 | 0 | NA | NA | NA | NA | NA | | NA | NA | 0 | 0 | 0 | 0 | 0 | 0 |
| 34414 | ENSMUSG00000095538 | 0 | NA | NA | NA | NA | NA | | Gm21983 | predicted gene 21983 [Source:MGI Symbo | 0 | 0 | 0 | 0 | 0 | 0 |
| 34415 | ENSMUSG00000095539 | 0.133850373 | -0.796629176 | 4.080472857 | -0.195229623 | 0.84521319 | NA | | NA | NA | 0.8031 | 0 | 0 | 0 | 0 | 0 |
| 34416 | ENSMUSG00000095540 | 0 | NA | NA | NA | NA | NA | | Esp4 | exocrine gland secreted peptide 4 [Source | 0 | 0 | 0 | 0 | 0 | 0 |
| 34417 | ENSMUSG00000095541 | 0 | NA | NA | NA | NA | NA | | NA | NA | 0 | 0 | 0 | 0 | 0 | 0 |
| 34418 | ENSMUSG00000095542 | 0 | NA | NA | NA | NA | NA | | NA | NA | 0 | 0 | 0 | 0 | 0 | 0 |
| 34419 | ENSMUSG00000095543 | 0 | NA | NA | NA | NA | NA | | Vmn1r148 | vomeronasal 1 receptor 148 [Source:MGI | 0 | 0 | 0 | 0 | 0 | 0 |
| 34420 | ENSMUSG00000095544 | 0 | NA | NA | NA | NA | NA | | Vmn2r-ps33 | vomeronasal 2, receptor, pseudogene 33 | 0 | 0 | 0 | 0 | 0 | 0 |
| 34421 | ENSMUSG00000095545 | 0 | NA | NA | NA | NA | NA | | Zfp969 | zinc finger protein 969 [Source:MGI Symb | 0 | 0 | 0 | 0 | 0 | 0 |
| 34422 | ENSMUSG00000095546 | 0 | NA | NA | NA | NA | NA | | Gm10230 | predicted gene 10230 [Source:MGI Symbo | 0 | 0 | 0 | 0 | 0 | 0 |
| 34423 | ENSMUSG00000095547 | 0 | NA | NA | NA | NA | NA | | Gm10719 | predicted gene 10719 [Source:MGI Symbo | 0 | 0 | 0 | 0 | 0 | 0 |
| 34424 | ENSMUSG00000095549 | 0 | NA | NA | NA | NA | NA | | Ighd5-1 | immunoglobulin heavy diversity 5-1 [Sour | 0 | 0 | 0 | 0 | 0 | 0 |
| 34425 | ENSMUSG00000095550 | 0 | NA | NA | NA | NA | NA | | Gm21671 | predicted gene, 21671 [Source:MGI Symb | 0 | 0 | 0 | 0 | 0 | 0 |
| 34426 | ENSMUSG00000095551 | 0 | NA | NA | NA | NA | NA | | Gm7980 | predicted gene 7980 [Source:MGI Symbol | 0 | 0 | 0 | 0 | 0 | 0 |
| 34427 | ENSMUSG00000095552 | 0 | NA | NA | NA | NA | NA | | NA | NA | 0 | 0 | 0 | 0 | 0 | 0 |
| 34428 | ENSMUSG00000095553 | 0 | NA | NA | NA | NA | NA | | Gm21916 | predicted gene, 21916 [Source:MGI Symb | 0 | 0 | 0 | 0 | 0 | 0 |
| 34429 | ENSMUSG00000095554 | 0 | NA | NA | NA | NA | NA | | Ighv1-16 | immunoglobulin heavy variable 1-16 [Sour | 0 | 0 | 0 | 0 | 0 | 0 |
| 34430 | ENSMUSG00000095555 | 0 | NA | NA | NA | NA | NA | | NA | NA | 0 | 0 | 0 | 0 | 0 | 0 |
| 34431 | ENSMUSG00000095557 | 0 | NA | NA | NA | NA | NA | | Gm3415 | predicted gene 3415 [Source:MGI Symbol | 0 | 0 | 0 | 0 | 0 | 0 |
| 34432 | ENSMUSG00000095558 | 0 | NA | NA | NA | NA | NA | | NA | NA | 0 | 0 | 0 | 0 | 0 | 0 |
| 34433 | ENSMUSG00000095560 | 0 | NA | NA | NA | NA | NA | | Trav14d-2 | T cell receptor alpha variable 14D-2 [Sour | 0 | 0 | 0 | 0 | 0 | 0 |
| 34434 | ENSMUSG00000095561 | 0 | NA | NA | NA | NA | NA | | Gm8383 | predicted gene 8383 [Source:MGI Symbol | 0 | 0 | 0 | 0 | 0 | 0 |
| 34435 | ENSMUSG00000095562 | 2.810857836 | -4.861280868 | 3.921628396 | -1.239607729 | 0.215120521 | NA | | | erythroid differentiation regulator 1 [Sour | 16.865 | 0 | 0 | 0 | 0 | 0 |
| 34436 | ENSMUSG00000095563 | 0 | NA | NA | NA | NA | NA | | Astx3 | amplified spermatogenic transcripts X enc | 0 | 0 | 0 | 0 | 0 | 0 |

| | | | | | | | | | | | | | | | |
|---|---|---|---|---|---|---|---|---|---|---|---|---|---|---|---|
| 34437 | ENSMUSG00000095564 | 0 | NA | NA | NA | NA | NA | Gm20474 | predicted gene 20474 [Source:MGI Symbo | 0 | 0 | 0 | 0 | 0 | 0 |
| 34438 | ENSMUSG00000095565 | 0.133850373 | -0.796629176 | 4.080472857 | -0.195229623 | 0.84521319 | NA | Ighv2-9-1 | immunoglobulin heavy variable 2-9-1 [Sou | 0.8031 | 0 | 0 | 0 | 0 | 0 |
| 34439 | ENSMUSG00000095567 | 481.2770167 | 0.07106762 | 0.128567769 | 0.55276389 | 0.580425098 | 0.908645447 | Noc2l | NOC2 like nucleolar associated transcripti | 454.56 | 487.23 | 466.9 | 495.48 | 524.59 | 458.9 |
| 34440 | ENSMUSG00000095569 | 0 | NA | NA | NA | NA | NA | Taar7d | trace amine-associated receptor 7D [Sour | 0 | 0 | 0 | 0 | 0 | 0 |
| 34441 | ENSMUSG00000095570 | 0 | NA | NA | NA | NA | NA | | | 0 | 0 | 0 | 0 | 0 | 0 |
| 34442 | ENSMUSG00000095571 | 0.17131964 | -0.796629176 | 4.080472857 | -0.195229623 | 0.84521319 | NA | Ighv5-17 | immunoglobulin heavy variable 5-17 [Sou | 0 | 1.0279 | 0 | 0 | 0 | 0 |
| 34443 | ENSMUSG00000095572 | 0 | NA | NA | NA | NA | NA | Trav4n-3 | T cell receptor alpha variable 4N-3 [Sourc | 0 | 0 | 0 | 0 | 0 | 0 |
| 34444 | ENSMUSG00000095573 | 0 | NA | NA | NA | NA | NA | H2al2b | H2A histone family member L2B [Source:M | 0 | 0 | 0 | 0 | 0 | 0 |
| 34445 | ENSMUSG00000095574 | 0 | NA | NA | NA | NA | NA | Trbv12-1 | T cell receptor beta, variable 12-1 [Source | 0 | 0 | 0 | 0 | 0 | 0 |
| 34446 | ENSMUSG00000095576 | 0 | NA | NA | NA | NA | NA | Fmo6 | flavin containing monooxygenase 6 [Sourc | 0 | 0 | 0 | 0 | 0 | 0 |
| 34447 | ENSMUSG00000095577 | 0 | NA | NA | NA | NA | NA | Gm6619 | predicted gene 6619 [Source:MGI Symbol | 0 | 0 | 0 | 0 | 0 | 0 |
| 34448 | ENSMUSG00000095578 | 0 | NA | NA | NA | NA | NA | Gm21760 | predicted gene, 21760 [Source:MGI Symb | 0 | 0 | 0 | 0 | 0 | 0 |
| 34449 | ENSMUSG00000095580 | 0 | NA | NA | NA | NA | NA | NA | NA | 0 | 0 | 0 | 0 | 0 | 0 |
| 34450 | ENSMUSG00000095581 | 0 | NA | NA | NA | NA | NA | NA | NA | 0 | 0 | 0 | 0 | 0 | 0 |
| 34451 | ENSMUSG00000095583 | 0 | NA | NA | NA | NA | NA | Ighv14-2 | immunoglobulin heavy variable 14-2 [Sou | 0 | 0 | 0 | 0 | 0 | 0 |
| 34452 | ENSMUSG00000095584 | 0 | NA | NA | NA | NA | NA | NA | NA | 0 | 0 | 0 | 0 | 0 | 0 |
| 34453 | ENSMUSG00000095585 | 0 | NA | NA | NA | NA | NA | NA | NA | 0 | 0 | 0 | 0 | 0 | 0 |
| 34454 | ENSMUSG00000095586 | 0 | NA | NA | NA | NA | NA | Olfr1287 | olfactory receptor 1287 [Source:MGI Sym | 0 | 0 | 0 | 0 | 0 | 0 |
| 34455 | ENSMUSG00000095587 | 0 | NA | NA | NA | NA | NA | NA | NA | 0 | 0 | 0 | 0 | 0 | 0 |
| 34456 | ENSMUSG00000095588 | 8.760779765 | -0.473207347 | 0.712487261 | -0.664162537 | 0.506586288 | 0.881669588 | Gm12350 | predicted gene 12350 [Source:MGI Symbo | 11.243 | 8.2233 | 11.069 | 7.7118 | 7.6343 | 6.6831 |
| 34457 | ENSMUSG00000095589 | 0 | NA | NA | NA | NA | NA | Ighv1-55 | immunoglobulin heavy variable 1-55 [Sou | 0 | 0 | 0 | 0 | 0 | 0 |
| 34458 | ENSMUSG00000095590 | 0 | NA | NA | NA | NA | NA | NA | NA | 0 | 0 | 0 | 0 | 0 | 0 |
| 34459 | ENSMUSG00000095591 | 0 | NA | NA | NA | NA | NA | Gm15045 | predicted gene 15045 [Source:MGI Symbo | 0 | 0 | 0 | 0 | 0 | 0 |
| 34460 | ENSMUSG00000095592 | 0 | NA | NA | NA | NA | NA | Ighd5-7 | immunoglobulin heavy diversity 5-7 [Sour | 0 | 0 | 0 | 0 | 0 | 0 |
| 34461 | ENSMUSG00000095593 | 0 | NA | NA | NA | NA | NA | Gm7138 | predicted gene 7138 [Source:MGI Symbol | 0 | 0 | 0 | 0 | 0 | 0 |
| 34462 | ENSMUSG00000095594 | 0 | NA | NA | NA | NA | NA | NA | NA | 0 | 0 | 0 | 0 | 0 | 0 |
| 34463 | ENSMUSG00000095595 | 0 | NA | NA | NA | NA | NA | Fam177a | family with sequence similarity 177, mem | 0 | 0 | 0 | 0 | 0 | 0 |
| 34464 | ENSMUSG00000095596 | 0.556921942 | 2.679043159 | 3.638243063 | 0.736356289 | 0.461513899 | NA | NA | NA | 0 | 0 | 0 | 0 | 0 | 3.3415 |
| 34465 | ENSMUSG00000095597 | 13.51899641 | 0.625218847 | 0.603913843 | 1.035278217 | 0.300538999 | 0.751875664 | Rps7-ps3 | ribosomal protein S7, pseudogene 3 [Sou | 9.6372 | 9.2513 | 13.081 | 19.28 | 9.8156 | 20.049 |
| 34466 | ENSMUSG00000095599 | 0 | NA | NA | NA | NA | NA | Gm6215 | predicted pseudogene 6215 [Source:MGI | 0 | 0 | 0 | 0 | 0 | 0 |
| 34467 | ENSMUSG00000095600 | 0.185640647 | 1.126924175 | 4.080472857 | 0.276174898 | 0.782413736 | NA | Gm9457 | predicted gene 9457 [Source:MGI Symbol | 0 | 0 | 0 | 0 | 0 | 1.1138 |
| 34468 | ENSMUSG00000095601 | 0 | NA | NA | NA | NA | NA | Rhox3a2 | reproductive homeobox 3A2 [Source:MGI | 0 | 0 | 0 | 0 | 0 | 0 |
| 34469 | ENSMUSG00000095602 | 0 | NA | NA | NA | NA | NA | NA | NA | 0 | 0 | 0 | 0 | 0 | 0 |
| 34470 | ENSMUSG00000095603 | 0 | NA | NA | NA | NA | NA | Olfr106-ps | olfactory receptor 106, pseudogene [Sour | 0 | 0 | 0 | 0 | 0 | 0 |
| 34471 | ENSMUSG00000095604 | 0 | NA | NA | NA | NA | NA | NA | NA | 0 | 0 | 0 | 0 | 0 | 0 |
| 34472 | ENSMUSG00000095605 | 0 | NA | NA | NA | NA | NA | Gm12624 | predicted gene 12624 [Source:MGI Symbo | 0 | 0 | 0 | 0 | 0 | 0 |
| 34473 | ENSMUSG00000095606 | 0 | NA | NA | NA | NA | NA | Gm21258 | predicted gene, 21258 [Source:MGI Symb | 0 | 0 | 0 | 0 | 0 | 0 |
| 34474 | ENSMUSG00000095607 | 0 | NA | NA | NA | NA | NA | Trav5-4 | T cell receptor alpha variable 5-4 [Source: | 0 | 0 | 0 | 0 | 0 | 0 |
| 34475 | ENSMUSG00000095608 | 0 | NA | NA | NA | NA | NA | Olfr247 | olfactory receptor 247 [Source:MGI Symb | 0 | 0 | 0 | 0 | 0 | 0 |
| 34476 | ENSMUSG00000095609 | 0 | NA | NA | NA | NA | NA | Gm21188 | predicted gene, 21188 [Source:MGI Symb | 0 | 0 | 0 | 0 | 0 | 0 |
| 34477 | ENSMUSG00000095610 | 0 | NA | NA | NA | NA | NA | NA | NA | 0 | 0 | 0 | 0 | 0 | 0 |
| 34478 | ENSMUSG00000095612 | 0 | NA | NA | NA | NA | NA | Ighv5-4 | immunoglobulin heavy variable 5-4 [Sourc | 0 | 0 | 0 | 0 | 0 | 0 |
| 34479 | ENSMUSG00000095614 | 0 | NA | NA | NA | NA | NA | Gm6291 | predicted gene 6291 [Source:MGI Symbol | 0 | 0 | 0 | 0 | 0 | 0 |
| 34480 | ENSMUSG00000095615 | 0.321325949 | 1.911575114 | 4.048267893 | 0.472195804 | 0.636787034 | NA | NA | NA | 0 | 0 | 0 | 1.928 | 0 | 0 |
| 34481 | ENSMUSG00000095616 | 0 | NA | NA | NA | NA | NA | Gm26244 | predicted gene, 26244 [Source:MGI Symb | 0 | 0 | 0 | 0 | 0 | 0 |
| 34482 | ENSMUSG00000095618 | 0 | NA | NA | NA | NA | NA | NA | NA | 0 | 0 | 0 | 0 | 0 | 0 |
| 34483 | ENSMUSG00000095619 | 0 | NA | NA | NA | NA | NA | Vmn1r157 | vomeronasal 1 receptor 157 [Source:MGI | 0 | 0 | 0 | 0 | 0 | 0 |
| 34484 | ENSMUSG00000095620 | 0.160662974 | 1.126924175 | 4.080472857 | 0.276174898 | 0.782413736 | NA | Csta2 | cystatin A family member 2 [Source:MGI S | 0 | 0 | 0 | 0.964 | 0 | 0 |
| 34485 | ENSMUSG00000095621 | 0 | NA | NA | NA | NA | NA | Gm15085 | predicted gene 15085 [Source:MGI Symbo | 0 | 0 | 0 | 0 | 0 | 0 |
| 34486 | ENSMUSG00000095623 | 0 | NA | NA | NA | NA | NA | NA | NA | 0 | 0 | 0 | 0 | 0 | 0 |
| 34487 | ENSMUSG00000095624 | 0 | NA | NA | NA | NA | NA | NA | NA | 0 | 0 | 0 | 0 | 0 | 0 |
| 34488 | ENSMUSG00000095625 | 0 | NA | NA | NA | NA | NA | Esp24 | exocrine gland secreted peptide 24 [Sourc | 0 | 0 | 0 | 0 | 0 | 0 |
| 34489 | ENSMUSG00000095626 | 0 | NA | NA | NA | NA | NA | Gm21572 | predicted gene, 21572 [Source:MGI Symb | 0 | 0 | 0 | 0 | 0 | 0 |
| 34490 | ENSMUSG00000095627 | 0 | NA | NA | NA | NA | NA | Gm8331 | predicted gene 8331 [Source:MGI Symbol | 0 | 0 | 0 | 0 | 0 | 0 |
| 34491 | ENSMUSG00000095628 | 0 | NA | NA | NA | NA | NA | NA | NA | 0 | 0 | 0 | 0 | 0 | 0 |
| 34492 | ENSMUSG00000095629 | 0 | NA | NA | NA | NA | NA | Vmn1r89 | vomeronasal 1 receptor 89 [Source:MGI S | 0 | 0 | 0 | 0 | 0 | 0 |

| | | | | | | | | | | | | | | | |
|---|---|---|---|---|---|---|---|---|---|---|---|---|---|---|---|
| 34493 | ENSMUSG00000095630 | 0 | NA | NA | NA | NA | NA | Igkv6-23 | immunoglobulin kappa variable 6-23 [Sou | 0 | 0 | 0 | 0 | 0 | 0 |
| 34494 | ENSMUSG00000095631 | 0 | NA | NA | NA | NA | NA | NA | NA | 0 | 0 | 0 | 0 | 0 | 0 |
| 34495 | ENSMUSG00000095632 | 0 | NA | NA | NA | NA | NA | Vmn1r175 | vomeronasal 1 receptor 175 [Source:MGI | 0 | 0 | 0 | 0 | 0 | 0 |
| 34496 | ENSMUSG00000095633 | 0 | NA | NA | NA | NA | NA | Igkv4-58 | immunoglobulin kappa variable 4-58 [Sou | 0 | 0 | 0 | 0 | 0 | 0 |
| 34497 | ENSMUSG00000095634 | 0 | NA | NA | NA | NA | NA | Gm20816 | predicted gene, 20816 [Source:MGI Symb | 0 | 0 | 0 | 0 | 0 | 0 |
| 34498 | ENSMUSG00000095635 | 0 | NA | NA | NA | NA | NA | NA | NA | 0 | 0 | 0 | 0 | 0 | 0 |
| 34499 | ENSMUSG00000095636 | 0 | NA | NA | NA | NA | NA | NA | NA | 0 | 0 | 0 | 0 | 0 | 0 |
| 34500 | ENSMUSG00000095637 | 0 | NA | NA | NA | NA | NA | NA | NA | 0 | 0 | 0 | 0 | 0 | 0 |
| 34501 | ENSMUSG00000095638 | 0 | NA | NA | NA | NA | NA | Gm8888 | predicted gene 8888 [Source:MGI Symbol | 0 | 0 | 0 | 0 | 0 | 0 |
| 34502 | ENSMUSG00000095639 | 0 | NA | NA | NA | NA | NA | NA | NA | 0 | 0 | 0 | 0 | 0 | 0 |
| 34503 | ENSMUSG00000095640 | 0 | NA | NA | NA | NA | NA | Olfr1434 | olfactory receptor 1434 [Source:MGI Sym | 0 | 0 | 0 | 0 | 0 | 0 |
| 34504 | ENSMUSG00000095641 | 0 | NA | NA | NA | NA | NA | NA | NA | 0 | 0 | 0 | 0 | 0 | 0 |
| 34505 | ENSMUSG00000095642 | 0.185640647 | 1.126924175 | 4.080472857 | 0.276174898 | 0.782413736 | NA | Ighv14-3 | immunoglobulin heavy variable V14-3 [So | 0 | 0 | 0 | 0 | 0 | 1.1138 |
| 34506 | ENSMUSG00000095643 | 0 | NA | NA | NA | NA | NA | Trav13d-3 | T cell receptor alpha variable 13D-3 [Sour | 0 | 0 | 0 | 0 | 0 | 0 |
| 34507 | ENSMUSG00000095645 | 0 | NA | NA | NA | NA | NA | NA | NA | 0 | 0 | 0 | 0 | 0 | 0 |
| 34508 | ENSMUSG00000095646 | 0 | NA | NA | NA | NA | NA | Trav10n | T cell receptor alpha variable 10N [Source | 0 | 0 | 0 | 0 | 0 | 0 |
| 34509 | ENSMUSG00000095647 | 0 | NA | NA | NA | NA | NA | Taar7a | trace amine-associated receptor 7A [Sour | 0 | 0 | 0 | 0 | 0 | 0 |
| 34510 | ENSMUSG00000095648 | 0.133850373 | -0.796629176 | 4.080472857 | -0.195229623 | 0.84521319 | NA | Gm2004 | predicted gene 2004 [Source:MGI Symbol | 0.8031 | 0 | 0 | 0 | 0 | 0 |
| 34511 | ENSMUSG00000095649 | 1.902450669 | -0.807948064 | 1.593070059 | -0.507164176 | 0.612039629 | NA | Gvin-ps3 | GTPase, very large interferon inducible, p | 3.2124 | 1.0279 | 3.0187 | 1.928 | 0 | 2.2277 |
| 34512 | ENSMUSG00000095650 | 0 | NA | NA | NA | NA | NA | Gm20854 | predicted gene, 20854 [Source:MGI Symb | 0 | 0 | 0 | 0 | 0 | 0 |
| 34513 | ENSMUSG00000095651 | 0 | NA | NA | NA | NA | NA | Gm21817 | predicted gene, 21817 [Source:MGI Symb | 0 | 0 | 0 | 0 | 0 | 0 |
| 34514 | ENSMUSG00000095652 | 0 | NA | NA | NA | NA | NA | NA | NA | 0 | 0 | 0 | 0 | 0 | 0 |
| 34515 | ENSMUSG00000095653 | 0 | NA | NA | NA | NA | NA | Gm21818 | predicted gene, 21818 [Source:MGI Symb | 0 | 0 | 0 | 0 | 0 | 0 |
| 34516 | ENSMUSG00000095654 | 1.656295348 | 0.173697343 | 1.678430807 | 0.103487938 | 0.917575722 | NA | Plscr5 | phospholipid scramblase family, member | 1.6062 | 2.0558 | 1.0062 | 1.928 | 0 | 3.3415 |
| 34517 | ENSMUSG00000095655 | 0 | NA | NA | NA | NA | NA | H2al1f | H2A histone family member L1F [Source:M | 0 | 0 | 0 | 0 | 0 | 0 |
| 34518 | ENSMUSG00000095656 | 0 | NA | NA | NA | NA | NA | Ighd2-8 | immunoglobulin heavy diversity 2-8 [Sour | 0 | 0 | 0 | 0 | 0 | 0 |
| 34519 | ENSMUSG00000095657 | 0 | NA | NA | NA | NA | NA | Gm15252 | predicted gene 15252 [Source:MGI Symbo | 0 | 0 | 0 | 0 | 0 | 0 |
| 34520 | ENSMUSG00000095658 | 0 | NA | NA | NA | NA | NA | Vmn2r130 | vomeronasal 2, receptor 130 [Source:MGI | 0 | 0 | 0 | 0 | 0 | 0 |
| 34521 | ENSMUSG00000095659 | 0 | NA | NA | NA | NA | NA | Spin2g | spindlin family, member 2G [Source:MGI S | 0 | 0 | 0 | 0 | 0 | 0 |
| 34522 | ENSMUSG00000095660 | 0 | NA | NA | NA | NA | NA | NA | NA | 0 | 0 | 0 | 0 | 0 | 0 |
| 34523 | ENSMUSG00000095661 | 0 | NA | NA | NA | NA | NA | Gm8935 | predicted pseudogene 8935 [Source:MGI | 0 | 0 | 0 | 0 | 0 | 0 |
| 34524 | ENSMUSG00000095662 | 0 | NA | NA | NA | NA | NA | H2al1g | H2A histone family member L1G [Source:I | 0 | 0 | 0 | 0 | 0 | 0 |
| 34525 | ENSMUSG00000095663 | 0 | NA | NA | NA | NA | NA | Gm2310 | predicted gene 2310 [Source:MGI Symbol | 0 | 0 | 0 | 0 | 0 | 0 |
| 34526 | ENSMUSG00000095664 | 0 | NA | NA | NA | NA | NA | Vmn2r67 | vomeronasal 2, receptor 67 [Source:MGI S | 0 | 0 | 0 | 0 | 0 | 0 |
| 34527 | ENSMUSG00000095666 | 0 | NA | NA | NA | NA | NA | | | 0 | 0 | 0 | 0 | 0 | 0 |
| 34528 | ENSMUSG00000095667 | 0 | NA | NA | NA | NA | NA | Olfr854 | olfactory receptor 854 [Source:MGI Symb | 0 | 0 | 0 | 0 | 0 | 0 |
| 34529 | ENSMUSG00000095668 | 0.17131964 | -0.796629176 | 4.080472857 | -0.195229623 | 0.84521319 | NA | Trbd1 | T cell receptor beta, D region 1 [Source:M | 0 | 1.0279 | 0 | 0 | 0 | 0 |
| 34530 | ENSMUSG00000095670 | 0 | NA | NA | NA | NA | NA | Vmn1r30 | vomeronasal 1 receptor 30 [Source:MGI S | 0 | 0 | 0 | 0 | 0 | 0 |
| 34531 | ENSMUSG00000095671 | 0 | NA | NA | NA | NA | NA | NA | NA | 0 | 0 | 0 | 0 | 0 | 0 |
| 34532 | ENSMUSG00000095672 | 0 | NA | NA | NA | NA | NA | | | 0 | 0 | 0 | 0 | 0 | 0 |
| 34533 | ENSMUSG00000095673 | 0 | NA | NA | NA | NA | NA | Gm4308 | predicted gene 4308 [Source:MGI Symbol | 0 | 0 | 0 | 0 | 0 | 0 |
| 34534 | ENSMUSG00000095674 | 0 | NA | NA | NA | NA | NA | NA | NA | 0 | 0 | 0 | 0 | 0 | 0 |
| 34535 | ENSMUSG00000095675 | 0 | NA | NA | NA | NA | NA | Ccl21b | chemokine (C-C motif) ligand 21B (leucine | 0 | 0 | 0 | 0 | 0 | 0 |
| 34536 | ENSMUSG00000095676 | 0 | NA | NA | NA | NA | NA | NA | NA | 0 | 0 | 0 | 0 | 0 | 0 |
| 34537 | ENSMUSG00000095677 | 6.519922325 | -0.070520725 | 0.824749998 | -0.085505578 | 0.931859461 | 0.988420305 | Dynlt1f | dynein light chain Tctex-type 1F [Source:N | 4.8186 | 8.2233 | 7.0437 | 5.7839 | 5.4531 | 7.7969 |
| 34538 | ENSMUSG00000095678 | 0 | NA | NA | NA | NA | NA | NA | NA | 0 | 0 | 0 | 0 | 0 | 0 |
| 34539 | ENSMUSG00000095680 | 0 | NA | NA | NA | NA | NA | Gm12627 | predicted gene 12627 [Source:MGI Symbo | 0 | 0 | 0 | 0 | 0 | 0 |
| 34540 | ENSMUSG00000095681 | 0 | NA | NA | NA | NA | NA | Gm8281 | predicted gene, 8281 [Source:MGI Symbo | 0 | 0 | 0 | 0 | 0 | 0 |
| 34541 | ENSMUSG00000095682 | 0 | NA | NA | NA | NA | NA | Igkv3-1 | immunoglobulin kappa variable 3-1 [Sour | 0 | 0 | 0 | 0 | 0 | 0 |
| 34542 | ENSMUSG00000095683 | 0 | NA | NA | NA | NA | NA | Gm10662 | predicted gene 10662 [Source:MGI Symbo | 0 | 0 | 0 | 0 | 0 | 0 |
| 34543 | ENSMUSG00000095684 | 0 | NA | NA | NA | NA | NA | NA | NA | 0 | 0 | 0 | 0 | 0 | 0 |
| 34544 | ENSMUSG00000095685 | 0 | NA | NA | NA | NA | NA | NA | NA | 0 | 0 | 0 | 0 | 0 | 0 |
| 34545 | ENSMUSG00000095686 | 0 | NA | NA | NA | NA | NA | Gm3099 | predicted gene 3099 [Source:MGI Symbol | 0 | 0 | 0 | 0 | 0 | 0 |
| 34546 | ENSMUSG00000095687 | 2.57925781 | -0.469410676 | 1.480624493 | -0.317035601 | 0.751216589 | NA | Rnaset2a | ribonuclease T2A [Source:MGI Symbol;Ac | 4.8186 | 3.0838 | 1.0062 | 0 | 5.4531 | 1.1138 |
| 34547 | ENSMUSG00000095689 | 0 | NA | NA | NA | NA | NA | Trav12d-1 | T cell receptor alpha variable 12D-1 [Sour | 0 | 0 | 0 | 0 | 0 | 0 |
| 34548 | ENSMUSG00000095690 | 6.154425126 | -1.54317364 | 0.898243237 | -1.717990826 | 0.085798289 | 0.431795087 | Rab11b-ps2 | RAB11B, member RAS oncogene family, p | 7.2279 | 9.2513 | 11.069 | 3.8559 | 2.1812 | 3.3415 |

| | | | | | | | | | | | | | | | |
|---|---|---|---|---|---|---|---|---|---|---|---|---|---|---|---|
| 34549 | ENSMUSG00000095691 | 0 | NA | NA | NA | NA | NA | NA | NA | 0 | 0 | 0 | 0 | 0 | 0 |
| 34550 | ENSMUSG00000095693 | 0 | NA | NA | NA | NA | NA | Gm21820 | predicted gene, 21820 [Source:MGI Symb | 0 | 0 | 0 | 0 | 0 | 0 |
| 34551 | ENSMUSG00000095696 | 0 | NA | NA | NA | NA | NA | Olfr786 | olfactory receptor 786 [Source:MGI Symb | 0 | 0 | 0 | 0 | 0 | 0 |
| 34552 | ENSMUSG00000095697 | 0 | NA | NA | NA | NA | NA | NA | NA | 0 | 0 | 0 | 0 | 0 | 0 |
| 34553 | ENSMUSG00000095698 | 0 | NA | NA | NA | NA | NA | Rhox2d | reproductive homeobox 2D [Source:MGI S | 0 | 0 | 0 | 0 | 0 | 0 |
| 34554 | ENSMUSG00000095699 | 0 | NA | NA | NA | NA | NA | NA | NA | 0 | 0 | 0 | 0 | 0 | 0 |
| 34555 | ENSMUSG00000095700 | 0 | NA | NA | NA | NA | NA | Ighv10-3 | immunoglobulin heavy variable V10-3 [So | 0 | 0 | 0 | 0 | 0 | 0 |
| 34556 | ENSMUSG00000095701 | 0 | NA | NA | NA | NA | NA | NA | NA | 0 | 0 | 0 | 0 | 0 | 0 |
| 34557 | ENSMUSG00000095702 | 0 | NA | NA | NA | NA | NA | Gm15005 | predicted gene 15005 [Source:MGI Symbo | 0 | 0 | 0 | 0 | 0 | 0 |
| 34558 | ENSMUSG00000095703 | 0 | NA | NA | NA | NA | NA | NA | NA | 0 | 0 | 0 | 0 | 0 | 0 |
| 34559 | ENSMUSG00000095704 | 0.181769715 | 1.126924175 | 4.080472857 | 0.276174898 | 0.782413736 | NA | Gm8221 | predicted gene 8221 [Source:MGI Symbol | 0 | 0 | 0 | 0 | 1.0906 | 0 |
| 34560 | ENSMUSG00000095705 | 0 | NA | NA | NA | NA | NA | NA | NA | 0 | 0 | 0 | 0 | 0 | 0 |
| 34561 | ENSMUSG00000095706 | 0 | NA | NA | NA | NA | NA | Olfr209 | olfactory receptor 209 [Source:MGI Symb | 0 | 0 | 0 | 0 | 0 | 0 |
| 34562 | ENSMUSG00000095710 | 0 | NA | NA | NA | NA | NA | Gm8871 | predicted pseudogene 8871 [Source:MGI | 0 | 0 | 0 | 0 | 0 | 0 |
| 34563 | ENSMUSG00000095711 | 0 | NA | NA | NA | NA | NA | Trav14-2 | T cell receptor alpha variable 14-2 [Source | 0 | 0 | 0 | 0 | 0 | 0 |
| 34564 | ENSMUSG00000095712 | 0 | NA | NA | NA | NA | NA | NA | NA | 0 | 0 | 0 | 0 | 0 | 0 |
| 34565 | ENSMUSG00000095713 | 0 | NA | NA | NA | NA | NA | NA | NA | 0 | 0 | 0 | 0 | 0 | 0 |
| 34566 | ENSMUSG00000095714 | 0 | NA | NA | NA | NA | NA | NA | NA | 0 | 0 | 0 | 0 | 0 | 0 |
| 34567 | ENSMUSG00000095716 | 0 | NA | NA | NA | NA | NA | Btbd35f10 | BTB domain containing 35, family membe | 0 | 0 | 0 | 0 | 0 | 0 |
| 34568 | ENSMUSG00000095717 | 0 | NA | NA | NA | NA | NA | Eif1ad11 | eukaryotic translation initiation factor 1A | 0 | 0 | 0 | 0 | 0 | 0 |
| 34569 | ENSMUSG00000095718 | 0 | NA | NA | NA | NA | NA | Pramel40 | PRAME like 40 [Source:MGI Symbol;Acc:M | 0 | 0 | 0 | 0 | 0 | 0 |
| 34570 | ENSMUSG00000095721 | 0 | NA | NA | NA | NA | NA | Gm7137 | predicted gene 7137 [Source:MGI Symbol | 0 | 0 | 0 | 0 | 0 | 0 |
| 34571 | ENSMUSG00000095722 | 0 | NA | NA | NA | NA | NA | NA | NA | 0 | 0 | 0 | 0 | 0 | 0 |
| 34572 | ENSMUSG00000095723 | 0 | NA | NA | NA | NA | NA | NA | NA | 0 | 0 | 0 | 0 | 0 | 0 |
| 34573 | ENSMUSG00000095724 | 0 | NA | NA | NA | NA | NA | Eif1ad19 | eukaryotic translation initiation factor 1A | 0 | 0 | 0 | 0 | 0 | 0 |
| 34574 | ENSMUSG00000095725 | 0 | NA | NA | NA | NA | NA | Gm21801 | predicted gene, 21801 [Source:MGI Symb | 0 | 0 | 0 | 0 | 0 | 0 |
| 34575 | ENSMUSG00000095727 | 0 | NA | NA | NA | NA | NA | NA | NA | 0 | 0 | 0 | 0 | 0 | 0 |
| 34576 | ENSMUSG00000095728 | 0 | NA | NA | NA | NA | NA | | | 0 | 0 | 0 | 0 | 0 | 0 |
| 34577 | ENSMUSG00000095729 | 0 | NA | NA | NA | NA | NA | NA | NA | 0 | 0 | 0 | 0 | 0 | 0 |
| 34578 | ENSMUSG00000095730 | 2.089435426 | -1.620483723 | 1.559094697 | -1.039374789 | 0.298630464 | NA | Vmn2r29 | vomeronasal 2, receptor 29 [Source:MGI S | 2.4093 | 3.0838 | 4.025 | 1.928 | 1.0906 | 0 |
| 34579 | ENSMUSG00000095731 | 0.267700746 | -1.497481649 | 4.057001436 | -0.369110456 | 0.712045396 | NA | NA | NA | 1.6062 | 0 | 0 | 0 | 0 | 0 |
| 34580 | ENSMUSG00000095732 | 0 | NA | NA | NA | NA | NA | NA | NA | 0 | 0 | 0 | 0 | 0 | 0 |
| 34581 | ENSMUSG00000095734 | 0 | NA | NA | NA | NA | NA | NA | NA | 0 | 0 | 0 | 0 | 0 | 0 |
| 34582 | ENSMUSG00000095736 | 0 | NA | NA | NA | NA | NA | Trav7n-4 | T cell receptor alpha variable 7N-4 [Source | 0 | 0 | 0 | 0 | 0 | 0 |
| 34583 | ENSMUSG00000095737 | 0 | NA | NA | NA | NA | NA | Igkv11-125 | immunoglobulin kappa variable 11-125 [S | 0 | 0 | 0 | 0 | 0 | 0 |
| 34584 | ENSMUSG00000095738 | 0 | NA | NA | NA | NA | NA | NA | NA | 0 | 0 | 0 | 0 | 0 | 0 |
| 34585 | ENSMUSG00000095739 | 0 | NA | NA | NA | NA | NA | NA | NA | 0 | 0 | 0 | 0 | 0 | 0 |
| 34586 | ENSMUSG00000095740 | 0 | NA | NA | NA | NA | NA | NA | NA | 0 | 0 | 0 | 0 | 0 | 0 |
| 34587 | ENSMUSG00000095741 | 0 | NA | NA | NA | NA | NA | Rhox2f | reproductive homeobox 2F [Source:MGI S | 0 | 0 | 0 | 0 | 0 | 0 |
| 34588 | ENSMUSG00000095742 | 67.16706639 | 0.178336832 | 0.347168507 | 0.513689544 | 0.607469064 | 0.916376001 | | | 50.595 | 81.206 | 57.356 | 57.839 | 49.078 | 106.93 |
| 34589 | ENSMUSG00000095743 | 0 | NA | NA | NA | NA | NA | Gm17654 | predicted gene, 17654 [Source:MGI Symb | 0 | 0 | 0 | 0 | 0 | 0 |
| 34590 | ENSMUSG00000095744 | 0 | NA | NA | NA | NA | NA | NA | NA | 0 | 0 | 0 | 0 | 0 | 0 |
| 34591 | ENSMUSG00000095745 | 0 | NA | NA | NA | NA | NA | Vmn1r243 | vomeronasal 1 receptor 243 [Source:MGI | 0 | 0 | 0 | 0 | 0 | 0 |
| 34592 | ENSMUSG00000095747 | 0 | NA | NA | NA | NA | NA | NA | NA | 0 | 0 | 0 | 0 | 0 | 0 |
| 34593 | ENSMUSG00000095749 | 0 | NA | NA | NA | NA | NA | NA | NA | 0 | 0 | 0 | 0 | 0 | 0 |
| 34594 | ENSMUSG00000095752 | 0 | NA | NA | NA | NA | NA | NA | NA | 0 | 0 | 0 | 0 | 0 | 0 |
| 34595 | ENSMUSG00000095753 | 0.17131964 | -0.796629176 | 4.080472857 | -0.195229623 | 0.84521319 | NA | Igkv4-53 | immunoglobulin kappa variable 4-53 [Sou | 0 | 1.0279 | 0 | 0 | 0 | 0 |
| 34596 | ENSMUSG00000095754 | 0 | NA | NA | NA | NA | NA | Spin2f | spindlin family, member 2F [Source:MGI S | 0 | 0 | 0 | 0 | 0 | 0 |
| 34597 | ENSMUSG00000095755 | 0 | NA | NA | NA | NA | NA | NA | NA | 0 | 0 | 0 | 0 | 0 | 0 |
| 34598 | ENSMUSG00000095756 | 0 | NA | NA | NA | NA | NA | Trav6d-7 | T cell receptor alpha variable 6D-7 [Source | 0 | 0 | 0 | 0 | 0 | 0 |
| 34599 | ENSMUSG00000095757 | 0 | NA | NA | NA | NA | NA | NA | NA | 0 | 0 | 0 | 0 | 0 | 0 |
| 34600 | ENSMUSG00000095758 | 0 | NA | NA | NA | NA | NA | Vmn1r128 | vomeronasal 1 receptor 128 [Source:MGI | 0 | 0 | 0 | 0 | 0 | 0 |
| 34601 | ENSMUSG00000095759 | 0 | NA | NA | NA | NA | NA | Gm21828 | predicted gene, 21828 [Source:MGI Symb | 0 | 0 | 0 | 0 | 0 | 0 |
| 34602 | ENSMUSG00000095761 | 0.133850373 | -0.796629176 | 4.080472857 | -0.195229623 | 0.84521319 | NA | Ighv1-20 | immunoglobulin heavy variable V1-20 [So | 0.8031 | 0 | 0 | 0 | 0 | 0 |
| 34603 | ENSMUSG00000095762 | 13.59623057 | -0.129798563 | 0.580185996 | -0.223718884 | 0.822976051 | 0.969382451 | Gm16378 | predicted gene 16378 [Source:MGI Symbo | 11.243 | 13.363 | 18.112 | 13.496 | 11.997 | 13.366 |
| 34604 | ENSMUSG00000095763 | 0 | NA | NA | NA | NA | NA | | | 0 | 0 | 0 | 0 | 0 | 0 |

| | | | | | | | | | | | | | | | |
|---|---|---|---|---|---|---|---|---|---|---|---|---|---|---|---|
| 34605 | ENSMUSG00000095764 | 0 | NA | NA | NA | NA | NA | NA | NA | 0 | 0 | 0 | 0 | 0 | 0 |
| 34606 | ENSMUSG00000095765 | 0 | NA | NA | NA | NA | NA | Olfr741 | olfactory receptor 741 [Source:MGI Symb | 0 | 0 | 0 | 0 | 0 | 0 |
| 34607 | ENSMUSG00000095766 | 0.931459384 | 1.251134958 | 2.489841943 | 0.502495735 | 0.615318852 | NA | Gm21182 | predicted gene, 21182 [Source:MGI Symb | 1.6062 | 0 | 0 | 2.8919 | 1.0906 | 0 |
| 34608 | ENSMUSG00000095768 | 0 | NA | NA | NA | NA | NA | Vmn1r111 | vomeronasal 1 receptor 111 [Source:MGI | 0 | 0 | 0 | 0 | 0 | 0 |
| 34609 | ENSMUSG00000095769 | 0 | NA | NA | NA | NA | NA | Gm21891 | predicted gene, 21891 [Source:MGI Symb | 0 | 0 | 0 | 0 | 0 | 0 |
| 34610 | ENSMUSG00000095770 | 0 | NA | NA | NA | NA | NA | Gm21637 | predicted gene, 21637 [Source:MGI Symb | 0 | 0 | 0 | 0 | 0 | 0 |
| 34611 | ENSMUSG00000095771 | 0.698954292 | 0.126053087 | 3.048972385 | 0.04134281 | 0.967022605 | NA | Igkv14-111 | immunoglobulin kappa variable 14-111 [S | 0 | 0 | 2.0125 | 0 | 2.1812 | 0 |
| 34612 | ENSMUSG00000095772 | 0 | NA | NA | NA | NA | NA | NA | NA | 0 | 0 | 0 | 0 | 0 | 0 |
| 34613 | ENSMUSG00000095773 | 0 | NA | NA | NA | NA | NA | Vmn2r38 | vomeronasal 2, receptor 38 [Source:MGI S | 0 | 0 | 0 | 0 | 0 | 0 |
| 34614 | ENSMUSG00000095774 | 0 | NA | NA | NA | NA | NA | Olfr970 | olfactory receptor 970 [Source:MGI Symb | 0 | 0 | 0 | 0 | 0 | 0 |
| 34615 | ENSMUSG00000095776 | 0 | NA | NA | NA | NA | NA | NA | NA | 0 | 0 | 0 | 0 | 0 | 0 |
| 34616 | ENSMUSG00000095778 | 0 | NA | NA | NA | NA | NA | NA | NA | 0 | 0 | 0 | 0 | 0 | 0 |
| 34617 | ENSMUSG00000095779 | 0 | NA | NA | NA | NA | NA | | novel pseudogene | 0 | 0 | 0 | 0 | 0 | 0 |
| 34618 | ENSMUSG00000095780 | 0 | NA | NA | NA | NA | NA | NA | NA | 0 | 0 | 0 | 0 | 0 | 0 |
| 34619 | ENSMUSG00000095782 | 0 | NA | NA | NA | NA | NA | NA | NA | 0 | 0 | 0 | 0 | 0 | 0 |
| 34620 | ENSMUSG00000095785 | 0 | NA | NA | NA | NA | NA | Gm20924 | predicted gene, 20924 [Source:MGI Symb | 0 | 0 | 0 | 0 | 0 | 0 |
| 34621 | ENSMUSG00000095786 | 0 | NA | NA | NA | NA | NA | NA | NA | 0 | 0 | 0 | 0 | 0 | 0 |
| 34622 | ENSMUSG00000095787 | 0 | NA | NA | NA | NA | NA | | | 0 | 0 | 0 | 0 | 0 | 0 |
| 34623 | ENSMUSG00000095788 | 2.958809446 | -0.251321354 | 1.292266411 | -0.19448107 | 0.845799217 | NA | Sirpb1a | signal-regulatory protein beta 1A [Source: | 1.6062 | 5.1396 | 3.0187 | 5.7839 | 1.0906 | 1.1138 |
| 34624 | ENSMUSG00000095789 | 21.92496003 | 0.128987609 | 0.468122134 | 0.27554264 | 0.782899371 | 0.960670212 | Nupr1l | nuclear protein transcriptional regulator 1 | 19.274 | 22.614 | 21.131 | 28.919 | 22.903 | 16.708 |
| 34625 | ENSMUSG00000095791 | 0 | NA | NA | NA | NA | NA | NA | NA | 0 | 0 | 0 | 0 | 0 | 0 |
| 34626 | ENSMUSG00000095792 | 0 | NA | NA | NA | NA | NA | NA | NA | 0 | 0 | 0 | 0 | 0 | 0 |
| 34627 | ENSMUSG00000095793 | 0 | NA | NA | NA | NA | NA | Gm20855 | predicted gene, 20855 [Source:MGI Symb | 0 | 0 | 0 | 0 | 0 | 0 |
| 34628 | ENSMUSG00000095794 | 0.483327519 | -0.685704547 | 3.145069396 | -0.218025252 | 0.827409436 | NA | Igkv6-17 | immunoglobulin kappa variable 6-17 [Sou | 0.8031 | 0 | 1.0062 | 0 | 1.0906 | 0 |
| 34629 | ENSMUSG00000095796 | 0 | NA | NA | NA | NA | NA | NA | NA | 0 | 0 | 0 | 0 | 0 | 0 |
| 34630 | ENSMUSG00000095797 | 0 | NA | NA | NA | NA | NA | Gm8246 | predicted gene 8246 [Source:MGI Symbol | 0 | 0 | 0 | 0 | 0 | 0 |
| 34631 | ENSMUSG00000095798 | 0 | NA | NA | NA | NA | NA | NA | NA | 0 | 0 | 0 | 0 | 0 | 0 |
| 34632 | ENSMUSG00000095799 | 0 | NA | NA | NA | NA | NA | Eif1ad10 | eukaryotic translation initiation factor 1A | 0 | 0 | 0 | 0 | 0 | 0 |
| 34633 | ENSMUSG00000095802 | 0 | NA | NA | NA | NA | NA | Gm7415 | predicted pseudogene 7415 [Source:MGI | 0 | 0 | 0 | 0 | 0 | 0 |
| 34634 | ENSMUSG00000095804 | 0 | NA | NA | NA | NA | NA | Olfr824 | olfactory receptor 824 [Source:MGI Symb | 0 | 0 | 0 | 0 | 0 | 0 |
| 34635 | ENSMUSG00000095805 | 0 | NA | NA | NA | NA | NA | NA | NA | 0 | 0 | 0 | 0 | 0 | 0 |
| 34636 | ENSMUSG00000095806 | 0 | NA | NA | NA | NA | NA | Vmn1r244 | vomeronasal 1 receptor 244 [Source:MGI | 0 | 0 | 0 | 0 | 0 | 0 |
| 34637 | ENSMUSG00000095807 | 0 | NA | NA | NA | NA | NA | Gm21405 | predicted gene, 21405 [Source:MGI Symb | 0 | 0 | 0 | 0 | 0 | 0 |
| 34638 | ENSMUSG00000095809 | 0 | NA | NA | NA | NA | NA | Olfr1290 | olfactory receptor 1290 [Source:MGI Sym | 0 | 0 | 0 | 0 | 0 | 0 |
| 34639 | ENSMUSG00000095812 | 0 | NA | NA | NA | NA | NA | Gm5473 | predicted pseudogene 5473 [Source:MGI | 0 | 0 | 0 | 0 | 0 | 0 |
| 34640 | ENSMUSG00000095813 | 0 | NA | NA | NA | NA | NA | Defa-ps1 | defensin, alpha, pseudogene 1 [Source:M | 0 | 0 | 0 | 0 | 0 | 0 |
| 34641 | ENSMUSG00000095814 | 0 | NA | NA | NA | NA | NA | Btbd35f7 | BTB domain containing 35, family membe | 0 | 0 | 0 | 0 | 0 | 0 |
| 34642 | ENSMUSG00000095816 | 0 | NA | NA | NA | NA | NA | Vmn1r-ps70 | vomeronasal 1 receptor, pseudogene 70 [ | 0 | 0 | 0 | 0 | 0 | 0 |
| 34643 | ENSMUSG00000095817 | 0 | NA | NA | NA | NA | NA | Gm3238 | predicted gene 3238 [Source:MGI Symbol | 0 | 0 | 0 | 0 | 0 | 0 |
| 34644 | ENSMUSG00000095818 | 0 | NA | NA | NA | NA | NA | NA | NA | 0 | 0 | 0 | 0 | 0 | 0 |
| 34645 | ENSMUSG00000095820 | 0 | NA | NA | NA | NA | NA | Gm15098 | predicted gene 15098 [Source:MGI Symbo | 0 | 0 | 0 | 0 | 0 | 0 |
| 34646 | ENSMUSG00000095821 | 0 | NA | NA | NA | NA | NA | Tcstv3 | 2-cell-stage, variable group, member 3 [So | 0 | 0 | 0 | 0 | 0 | 0 |
| 34647 | ENSMUSG00000095822 | 0 | NA | NA | NA | NA | NA | Tdpoz9-ps1 | TD and POZ domain containing 9, pseudog | 0 | 0 | 0 | 0 | 0 | 0 |
| 34648 | ENSMUSG00000095824 | 0 | NA | NA | NA | NA | NA | Gm13697 | predicted gene 13697 [Source:MGI Symbo | 0 | 0 | 0 | 0 | 0 | 0 |
| 34649 | ENSMUSG00000095825 | 0 | NA | NA | NA | NA | NA | NA | NA | 0 | 0 | 0 | 0 | 0 | 0 |
| 34650 | ENSMUSG00000095826 | 0 | NA | NA | NA | NA | NA | Gm9002 | predicted gene 9002 [Source:MGI Symbol | 0 | 0 | 0 | 0 | 0 | 0 |
| 34651 | ENSMUSG00000095827 | 0 | NA | NA | NA | NA | NA | Gm5283 | predicted gene 5283 [Source:MGI Symbol | 0 | 0 | 0 | 0 | 0 | 0 |
| 34652 | ENSMUSG00000095828 | 0 | NA | NA | NA | NA | NA | NA | NA | 0 | 0 | 0 | 0 | 0 | 0 |
| 34653 | ENSMUSG00000095829 | 0 | NA | NA | NA | NA | NA | Speer1 | spermatogenesis associated glutamate (E) | 0 | 0 | 0 | 0 | 0 | 0 |
| 34654 | ENSMUSG00000095831 | 0 | NA | NA | NA | NA | NA | Olfr453 | olfactory receptor 453 [Source:MGI Symb | 0 | 0 | 0 | 0 | 0 | 0 |
| 34655 | ENSMUSG00000095834 | 0 | NA | NA | NA | NA | NA | Tdgf1-ps1 | teratocarcinoma-derived growth factor, p | 0 | 0 | 0 | 0 | 0 | 0 |
| 34656 | ENSMUSG00000095836 | 4.813257024 | 1.108161472 | 1.023456034 | 1.08276412 | 0.278913135 | NA | Gm21960 | predicted gene, 21960 [Source:MGI Symb | 4.0155 | 3.0838 | 2.0125 | 8.6758 | 2.1812 | 8.9108 |
| 34657 | ENSMUSG00000095837 | 0 | NA | NA | NA | NA | NA | Vmn1r246 | vomeronasal 1 receptor 246 [Source:MGI | 0 | 0 | 0 | 0 | 0 | 0 |
| 34658 | ENSMUSG00000095839 | 0 | NA | NA | NA | NA | NA | Olfr965 | olfactory receptor 965 [Source:MGI Symb | 0 | 0 | 0 | 0 | 0 | 0 |
| 34659 | ENSMUSG00000095841 | 0 | NA | NA | NA | NA | NA | Gm8532 | predicted gene 8532 [Source:MGI Symbol | 0 | 0 | 0 | 0 | 0 | 0 |
| 34660 | ENSMUSG00000095842 | 0 | NA | NA | NA | NA | NA | NA | NA | 0 | 0 | 0 | 0 | 0 | 0 |

| | | | | | | | | | | | | | | | |
|---|---|---|---|---|---|---|---|---|---|---|---|---|---|---|---|
| 34661 | ENSMUSG00000095843 | 0 | NA | NA | NA | NA | NA | NA | NA | 0 | 0 | 0 | 0 | 0 | 0 |
| 34662 | ENSMUSG00000095845 | 3.659122324 | 0.76557672 | 1.199046083 | 0.638488153 | 0.523155963 | NA | Gm5741 | predicted gene 5741 [Source:MGI Symbol | 4.0155 | 0 | 4.025 | 2.8919 | 5.4531 | 5.5692 |
| 34663 | ENSMUSG00000095847 | 0.767390791 | -1.69152783 | 2.44773922 | -0.691057207 | 0.489529591 | NA | Gm5451 | predicted gene 5451 [Source:MGI Symbol | 1.6062 | 1.0279 | 1.0062 | 0.964 | 0 | 0 |
| 34664 | ENSMUSG00000095849 | 0 | NA | NA | NA | NA | NA | NA | NA | 0 | 0 | 0 | 0 | 0 | 0 |
| 34665 | ENSMUSG00000095850 | 0 | NA | NA | NA | NA | NA | NA | NA | 0 | 0 | 0 | 0 | 0 | 0 |
| 34666 | ENSMUSG00000095851 | 0 | NA | NA | NA | NA | NA | NA | NA | 0 | 0 | 0 | 0 | 0 | 0 |
| 34667 | ENSMUSG00000095852 | 0 | NA | NA | NA | NA | NA | Gm10256 | predicted gene 10256 [Source:MGI Symbo | 0 | 0 | 0 | 0 | 0 | 0 |
| 34668 | ENSMUSG00000095853 | 0 | NA | NA | NA | NA | NA | NA | NA | 0 | 0 | 0 | 0 | 0 | 0 |
| 34669 | ENSMUSG00000095854 | 0 | NA | NA | NA | NA | NA | Trav14d-1 | T cell receptor alpha variable 14D-1 [Sour | 0 | 0 | 0 | 0 | 0 | 0 |
| 34670 | ENSMUSG00000095856 | 0 | NA | NA | NA | NA | NA | NA | NA | 0 | 0 | 0 | 0 | 0 | 0 |
| 34671 | ENSMUSG00000095857 | 0 | NA | NA | NA | NA | NA | NA | NA | 0 | 0 | 0 | 0 | 0 | 0 |
| 34672 | ENSMUSG00000095858 | 0 | NA | NA | NA | NA | NA | Olfr104-ps | olfactory receptor 104, pseudogene [Sour | 0 | 0 | 0 | 0 | 0 | 0 |
| 34673 | ENSMUSG00000095859 | 0 | NA | NA | NA | NA | NA | Ighv1-43 | immunoglobulin heavy variable V1-43 [So | 0 | 0 | 0 | 0 | 0 | 0 |
| 34674 | ENSMUSG00000095860 | 0 | NA | NA | NA | NA | NA | NA | NA | 0 | 0 | 0 | 0 | 0 | 0 |
| 34675 | ENSMUSG00000095862 | 0 | NA | NA | NA | NA | NA | Trav6-6 | T cell receptor alpha variable 6-6 [Source: | 0 | 0 | 0 | 0 | 0 | 0 |
| 34676 | ENSMUSG00000095863 | 0 | NA | NA | NA | NA | NA | Ighv1-67 | immunoglobulin heavy variable V1-67 [So | 0 | 0 | 0 | 0 | 0 | 0 |
| 34677 | ENSMUSG00000095864 | 0 | NA | NA | NA | NA | NA | Vmn1r77 | vomeronasal 1 receptor 77 [Source:MGI S | 0 | 0 | 0 | 0 | 0 | 0 |
| 34678 | ENSMUSG00000095865 | 0.469265235 | -2.274672822 | 3.439794491 | -0.661281605 | 0.508431736 | NA | Gm13237 | predicted gene 13237 [Source:MGI Symbo | 0.8031 | 0 | 2.0125 | 0 | 0 | 0 |
| 34679 | ENSMUSG00000095866 | 0 | NA | NA | NA | NA | NA | Ighv2-4 | immunoglobulin heavy variable V2-4 [Sou | 0 | 0 | 0 | 0 | 0 | 0 |
| 34680 | ENSMUSG00000095867 | 0 | NA | NA | NA | NA | NA | Gm20917 | predicted gene, 20917 [Source:MGI Symb | 0 | 0 | 0 | 0 | 0 | 0 |
| 34681 | ENSMUSG00000095868 | 1.388594601 | 0.820146163 | 1.971024992 | 0.416101351 | 0.677335841 | NA | NA | NA | 0 | 2.0558 | 1.0062 | 1.928 | 0 | 3.3415 |
| 34682 | ENSMUSG00000095869 | 0.342432689 | 1.990653086 | 3.900281263 | 0.51038706 | 0.609780321 | NA | Ppp1r2-ps6 | protein phosphatase 1, regulatory (inhibit | 0 | 0 | 0 | 0.964 | 1.0906 | 0 |
| 34683 | ENSMUSG00000095870 | 0 | NA | NA | NA | NA | NA | Lce1k | late cornified envelope 1K [Source:MGI Sy | 0 | 0 | 0 | 0 | 0 | 0 |
| 34684 | ENSMUSG00000095871 | 0.167707431 | -0.796629176 | 4.080472857 | -0.195229623 | 0.84521319 | NA | NA | NA | 0 | 0 | 1.0062 | 0 | 0 | 0 |
| 34685 | ENSMUSG00000095872 | 0 | NA | NA | NA | NA | NA | Gm15128 | predicted gene 15128 [Source:MGI Symbo | 0 | 0 | 0 | 0 | 0 | 0 |
| 34686 | ENSMUSG00000095873 | 0 | NA | NA | NA | NA | NA | Ang-ps3 | angiogenin, ribonuclease A family, pseudo | 0 | 0 | 0 | 0 | 0 | 0 |
| 34687 | ENSMUSG00000095874 | 0 | NA | NA | NA | NA | NA | Gm9630 | predicted gene 9630 [Source:MGI Symbol | 0 | 0 | 0 | 0 | 0 | 0 |
| 34688 | ENSMUSG00000095876 | 0 | NA | NA | NA | NA | NA | NA | NA | 0 | 0 | 0 | 0 | 0 | 0 |
| 34689 | ENSMUSG00000095878 | 0 | NA | NA | NA | NA | NA | NA | NA | 0 | 0 | 0 | 0 | 0 | 0 |
| 34690 | ENSMUSG00000095879 | 0 | NA | NA | NA | NA | NA | Gm21764 | predicted gene, 21764 [Source:MGI Symb | 0 | 0 | 0 | 0 | 0 | 0 |
| 34691 | ENSMUSG00000095880 | 0 | NA | NA | NA | NA | NA | NA | NA | 0 | 0 | 0 | 0 | 0 | 0 |
| 34692 | ENSMUSG00000095883 | 0 | NA | NA | NA | NA | NA | NA | NA | 0 | 0 | 0 | 0 | 0 | 0 |
| 34693 | ENSMUSG00000095884 | 0 | NA | NA | NA | NA | NA | NA | NA | 0 | 0 | 0 | 0 | 0 | 0 |
| 34694 | ENSMUSG00000095885 | 0 | NA | NA | NA | NA | NA | NA | NA | 0 | 0 | 0 | 0 | 0 | 0 |
| 34695 | ENSMUSG00000095886 | 0 | NA | NA | NA | NA | NA | Esp38 | exocrine gland secreted peptide 38 [Sourc | 0 | 0 | 0 | 0 | 0 | 0 |
| 34696 | ENSMUSG00000095887 | 0 | NA | NA | NA | NA | NA | Gm10096 | predicted gene 10096 [Source:MGI Symbo | 0 | 0 | 0 | 0 | 0 | 0 |
| 34697 | ENSMUSG00000095889 | 0 | NA | NA | NA | NA | NA | Ighv1-58 | immunoglobulin heavy variable 1-58 [Sou | 0 | 0 | 0 | 0 | 0 | 0 |
| 34698 | ENSMUSG00000095890 | 0 | NA | NA | NA | NA | NA | Gm7958 | predicted gene 7958 [Source:MGI Symbol | 0 | 0 | 0 | 0 | 0 | 0 |
| 34699 | ENSMUSG00000095891 | 0 | NA | NA | NA | NA | NA | Gm10717 | predicted gene 10717 [Source:MGI Symbo | 0 | 0 | 0 | 0 | 0 | 0 |
| 34700 | ENSMUSG00000095892 | 0.958298615 | -0.940364986 | 2.473304036 | -0.380205981 | 0.70379252 | NA | NA | NA | 0.8031 | 0 | 3.0187 | 1.928 | 0 | 0 |
| 34701 | ENSMUSG00000095893 | 0 | NA | NA | NA | NA | NA | Olfr884 | olfactory receptor 884 [Source:MGI Symb | 0 | 0 | 0 | 0 | 0 | 0 |
| 34702 | ENSMUSG00000095894 | 0 | NA | NA | NA | NA | NA | NA | NA | 0 | 0 | 0 | 0 | 0 | 0 |
| 34703 | ENSMUSG00000095895 | 0 | NA | NA | NA | NA | NA | Gm21963 | predicted gene, 21963 [Source:MGI Symb | 0 | 0 | 0 | 0 | 0 | 0 |
| 34704 | ENSMUSG00000095896 | 0 | NA | NA | NA | NA | NA | Ifna14 | interferon alpha 14 [Source:MGI Symbol;A | 0 | 0 | 0 | 0 | 0 | 0 |
| 34705 | ENSMUSG00000095897 | 0 | NA | NA | NA | NA | NA | Ighd2-5 | immunoglobulin heavy diversity 2-5 [Sour | 0 | 0 | 0 | 0 | 0 | 0 |
| 34706 | ENSMUSG00000095898 | 0 | NA | NA | NA | NA | NA | NA | NA | 0 | 0 | 0 | 0 | 0 | 0 |
| 34707 | ENSMUSG00000095899 | 0 | NA | NA | NA | NA | NA | Gm2318 | predicted gene 2318 [Source:MGI Symbol | 0 | 0 | 0 | 0 | 0 | 0 |
| 34708 | ENSMUSG00000095900 | 0 | NA | NA | NA | NA | NA | Gm20773 | predicted gene, 20773 [Source:MGI Symb | 0 | 0 | 0 | 0 | 0 | 0 |
| 34709 | ENSMUSG00000095901 | 0 | NA | NA | NA | NA | NA | Olfr538 | olfactory receptor 538 [Source:MGI Symb | 0 | 0 | 0 | 0 | 0 | 0 |
| 34710 | ENSMUSG00000095902 | 0 | NA | NA | NA | NA | NA | NA | NA | 0 | 0 | 0 | 0 | 0 | 0 |
| 34711 | ENSMUSG00000095903 | 0 | NA | NA | NA | NA | NA | Olfr968 | olfactory receptor 968 [Source:MGI Symb | 0 | 0 | 0 | 0 | 0 | 0 |
| 34712 | ENSMUSG00000095905 | 0 | NA | NA | NA | NA | NA | Selenok-ps4 | selenoprotein K, pseudogene 4 [Source:M | 0 | 0 | 0 | 0 | 0 | 0 |
| 34713 | ENSMUSG00000095906 | 0 | NA | NA | NA | NA | NA | Gm7421 | predicted gene 7421 [Source:MGI Symbol | 0 | 0 | 0 | 0 | 0 | 0 |
| 34714 | ENSMUSG00000095907 | 0 | NA | NA | NA | NA | NA | Gm14333 | predicted gene 14333 [Source:MGI Symbo | 0 | 0 | 0 | 0 | 0 | 0 |
| 34715 | ENSMUSG00000095909 | 0 | NA | NA | NA | NA | NA | Zfp997 | zinc finger protein 997 [Source:MGI Symb | 0 | 0 | 0 | 0 | 0 | 0 |
| 34716 | ENSMUSG00000095910 | 0 | NA | NA | NA | NA | NA | Olfr469 | olfactory receptor 469 [Source:MGI Symb | 0 | 0 | 0 | 0 | 0 | 0 |

| | | | | | | | | | | | | | | | |
|---|---|---|---|---|---|---|---|---|---|---|---|---|---|---|---|
| 34717 | ENSMUSG00000095911 | 0 | NA | NA | NA | NA | NA | NA | NA | 0 | 0 | 0 | 0 | 0 | 0 |
| 34718 | ENSMUSG00000095912 | 0 | NA | NA | NA | NA | NA | Gm3317 | predicted gene 3317 [Source:MGI Symbol | 0 | 0 | 0 | 0 | 0 | 0 |
| 34719 | ENSMUSG00000095913 | 2.885705841 | 0.078571253 | 1.396848555 | 0.056248942 | 0.955143493 | NA | Gm21844 | predicted gene, 21844 [Source:MGI Symb | 2.4093 | 3.0838 | 3.0187 | 7.7118 | 1.0906 | 0 |
| 34720 | ENSMUSG00000095914 | 0 | NA | NA | NA | NA | NA | Vmn2r48 | vomeronasal 2, receptor 48 [Source:MGI S | 0 | 0 | 0 | 0 | 0 | 0 |
| 34721 | ENSMUSG00000095915 | 0 | NA | NA | NA | NA | NA | NA | NA | 0 | 0 | 0 | 0 | 0 | 0 |
| 34722 | ENSMUSG00000095916 | 0 | NA | NA | NA | NA | NA | Vmn1r191 | vomeronasal 1 receptor 191 [Source:MGI | 0 | 0 | 0 | 0 | 0 | 0 |
| 34723 | ENSMUSG00000095917 | 0 | NA | NA | NA | NA | NA | Olfr740 | olfactory receptor 740 [Source:MGI Symb | 0 | 0 | 0 | 0 | 0 | 0 |
| 34724 | ENSMUSG00000095918 | 0 | NA | NA | NA | NA | NA | Gm5861 | predicted gene 5861 [Source:MGI Symbol | 0 | 0 | 0 | 0 | 0 | 0 |
| 34725 | ENSMUSG00000095920 | 0 | NA | NA | NA | NA | NA | Gm6686 | predicted pseudogene 6686 [Source:MGI | 0 | 0 | 0 | 0 | 0 | 0 |
| 34726 | ENSMUSG00000095922 | 0 | NA | NA | NA | NA | NA | Obox4-ps21 | oocyte specific homeobox 4, pseudogene | 0 | 0 | 0 | 0 | 0 | 0 |
| 34727 | ENSMUSG00000095923 | 0 | NA | NA | NA | NA | NA | Ighv1-37 | immunoglobulin heavy variable 1-37 [Sou | 0 | 0 | 0 | 0 | 0 | 0 |
| 34728 | ENSMUSG00000095924 | 0 | NA | NA | NA | NA | NA | NA | NA | 0 | 0 | 0 | 0 | 0 | 0 |
| 34729 | ENSMUSG00000095925 | 0 | NA | NA | NA | NA | NA | Defa-ps17 | defensin, alpha, pseudogene 17 [Source:N | 0 | 0 | 0 | 0 | 0 | 0 |
| 34730 | ENSMUSG00000095926 | 0 | NA | NA | NA | NA | NA | NA | NA | 0 | 0 | 0 | 0 | 0 | 0 |
| 34731 | ENSMUSG00000095927 | 0 | NA | NA | NA | NA | NA | Gm21822 | predicted gene, 21822 [Source:MGI Symb | 0 | 0 | 0 | 0 | 0 | 0 |
| 34732 | ENSMUSG00000095928 | 0 | NA | NA | NA | NA | NA | Olfr204 | olfactory receptor 204 [Source:MGI Symb | 0 | 0 | 0 | 0 | 0 | 0 |
| 34733 | ENSMUSG00000095929 | 0 | NA | NA | NA | NA | NA | Olfr487 | olfactory receptor 487 [Source:MGI Symb | 0 | 0 | 0 | 0 | 0 | 0 |
| 34734 | ENSMUSG00000095930 | 74.02668534 | 0.167897458 | 0.317218417 | 0.529280299 | 0.59661102 | 0.914226274 | Nim1k | NIM1 serine/threonine protein kinase [So | 74.689 | 55.508 | 78.487 | 51.091 | 76.343 | 108.04 |
| 34735 | ENSMUSG00000095931 | 0 | NA | NA | NA | NA | NA | Vmn1r159 | vomeronasal 1 receptor 159 [Source:MGI | 0 | 0 | 0 | 0 | 0 | 0 |
| 34736 | ENSMUSG00000095932 | 0 | NA | NA | NA | NA | NA | Vmn1r49 | vomeronasal 1, receptor 49 [Source:MGI S | 0 | 0 | 0 | 0 | 0 | 0 |
| 34737 | ENSMUSG00000095933 | 0 | NA | NA | NA | NA | NA | NA | NA | 0 | 0 | 0 | 0 | 0 | 0 |
| 34738 | ENSMUSG00000095934 | 0 | NA | NA | NA | NA | NA | Btbd35f13 | BTB domain containing 35, family membe | 0 | 0 | 0 | 0 | 0 | 0 |
| 34739 | ENSMUSG00000095935 | 0 | NA | NA | NA | NA | NA | Gm11238 | predicted gene 11238 [Source:MGI Symbo | 0 | 0 | 0 | 0 | 0 | 0 |
| 34740 | ENSMUSG00000095936 | 0 | NA | NA | NA | NA | NA | Zscan4e | zinc finger and SCAN domain containing 4 | 0 | 0 | 0 | 0 | 0 | 0 |
| 34741 | ENSMUSG00000095937 | 0 | NA | NA | NA | NA | NA | Gm12671 | predicted gene 12671 [Source:MGI Symbo | 0 | 0 | 0 | 0 | 0 | 0 |
| 34742 | ENSMUSG00000095938 | 0 | NA | NA | NA | NA | NA | NA | NA | 0 | 0 | 0 | 0 | 0 | 0 |
| 34743 | ENSMUSG00000095939 | 0 | NA | NA | NA | NA | NA | NA | NA | 0 | 0 | 0 | 0 | 0 | 0 |
| 34744 | ENSMUSG00000095940 | 0 | NA | NA | NA | NA | NA | Gm16416 | predicted gene 16416 [Source:MGI Symbo | 0 | 0 | 0 | 0 | 0 | 0 |
| 34745 | ENSMUSG00000095941 | 0 | NA | NA | NA | NA | NA | Tle7 | TLE family member 7 [Source:MGI Symbol | 0 | 0 | 0 | 0 | 0 | 0 |
| 34746 | ENSMUSG00000095942 | 0 | NA | NA | NA | NA | NA | NA | NA | 0 | 0 | 0 | 0 | 0 | 0 |
| 34747 | ENSMUSG00000095943 | 0 | NA | NA | NA | NA | NA | NA | NA | 0 | 0 | 0 | 0 | 0 | 0 |
| 34748 | ENSMUSG00000095944 | 0 | NA | NA | NA | NA | NA | NA | NA | 0 | 0 | 0 | 0 | 0 | 0 |
| 34749 | ENSMUSG00000095945 | 0 | NA | NA | NA | NA | NA | NA | NA | 0 | 0 | 0 | 0 | 0 | 0 |
| 34750 | ENSMUSG00000095946 | 0 | NA | NA | NA | NA | NA | NA | NA | 0 | 0 | 0 | 0 | 0 | 0 |
| 34751 | ENSMUSG00000095947 | 0 | NA | NA | NA | NA | NA | NA | NA | 0 | 0 | 0 | 0 | 0 | 0 |
| 34752 | ENSMUSG00000095948 | 0 | NA | NA | NA | NA | NA | Gm21708 | predicted gene, 21708 [Source:MGI Symb | 0 | 0 | 0 | 0 | 0 | 0 |
| 34753 | ENSMUSG00000095950 | 0 | NA | NA | NA | NA | NA | Gm20737 | predicted gene, 20737 [Source:MGI Symb | 0 | 0 | 0 | 0 | 0 | 0 |
| 34754 | ENSMUSG00000095951 | 0 | NA | NA | NA | NA | NA | NA | NA | 0 | 0 | 0 | 0 | 0 | 0 |
| 34755 | ENSMUSG00000095952 | 0 | NA | NA | NA | NA | NA | Obox4-ps12 | oocyte specific homeobox 4, pseudogene | 0 | 0 | 0 | 0 | 0 | 0 |
| 34756 | ENSMUSG00000095953 | 0 | NA | NA | NA | NA | NA | 6030469F06 | RIKEN cDNA 6030469F06 gene [Source:M | 0 | 0 | 0 | 0 | 0 | 0 |
| 34757 | ENSMUSG00000095954 | 0 | NA | NA | NA | NA | NA | Gm3183 | predicted gene 3183 [Source:MGI Symbol | 0 | 0 | 0 | 0 | 0 | 0 |
| 34758 | ENSMUSG00000095955 | 0 | NA | NA | NA | NA | NA | NA | NA | 0 | 0 | 0 | 0 | 0 | 0 |
| 34759 | ENSMUSG00000095956 | 0 | NA | NA | NA | NA | NA | 1700036A12 | RIKEN cDNA 1700036A12 gene [Source:M | 0 | 0 | 0 | 0 | 0 | 0 |
| 34760 | ENSMUSG00000095957 | 0 | NA | NA | NA | NA | NA | Olfr832 | olfactory receptor 832 [Source:MGI Symb | 0 | 0 | 0 | 0 | 0 | 0 |
| 34761 | ENSMUSG00000095958 | 0 | NA | NA | NA | NA | NA | Trav12-3 | T cell receptor alpha variable 12-3 [Source | 0 | 0 | 0 | 0 | 0 | 0 |
| 34762 | ENSMUSG00000095959 | 0 | NA | NA | NA | NA | NA | Gm10845 | predicted gene 10845 [Source:MGI Symbo | 0 | 0 | 0 | 0 | 0 | 0 |
| 34763 | ENSMUSG00000095960 | 0 | NA | NA | NA | NA | NA | Gm13120 | predicted gene 13120 [Source:MGI Symbo | 0 | 0 | 0 | 0 | 0 | 0 |
| 34764 | ENSMUSG00000095961 | 0 | NA | NA | NA | NA | NA | Vmn2r102 | vomeronasal 2, receptor 102 [Source:MGI | 0 | 0 | 0 | 0 | 0 | 0 |
| 34765 | ENSMUSG00000095962 | 0 | NA | NA | NA | NA | NA | Vmn1r137 | vomeronasal 1 receptor 137 [Source:MGI | 0 | 0 | 0 | 0 | 0 | 0 |
| 34766 | ENSMUSG00000095963 | 0 | NA | NA | NA | NA | NA | Gm7742 | predicted gene 7742 [Source:MGI Symbol | 0 | 0 | 0 | 0 | 0 | 0 |
| 34767 | ENSMUSG00000095964 | 0 | NA | NA | NA | NA | NA | Gm6221 | predicted pseudogene 6221 [Source:MGI | 0 | 0 | 0 | 0 | 0 | 0 |
| 34768 | ENSMUSG00000095965 | 0 | NA | NA | NA | NA | NA | NA | NA | 0 | 0 | 0 | 0 | 0 | 0 |
| 34769 | ENSMUSG00000095966 | 0 | NA | NA | NA | NA | NA | NA | NA | 0 | 0 | 0 | 0 | 0 | 0 |
| 34770 | ENSMUSG00000095968 | 0 | NA | NA | NA | NA | NA | Gm21842 | predicted gene, 21842 [Source:MGI Symb | 0 | 0 | 0 | 0 | 0 | 0 |
| 34771 | ENSMUSG00000095969 | 0.767390791 | -1.69152783 | 2.44773922 | -0.691057207 | 0.489529591 | NA | NA | NA | 1.6062 | 1.0279 | 1.0062 | 0.964 | 0 | 0 |
| 34772 | ENSMUSG00000095970 | 0 | NA | NA | NA | NA | NA | Gm19402 | predicted gene, 19402 [Source:MGI Symb | 0 | 0 | 0 | 0 | 0 | 0 |

| | | | | | | | | | | | | | | | |
|---|---|---|---|---|---|---|---|---|---|---|---|---|---|---|---|
| 34773 | ENSMUSG00000095971 | 0 | NA | NA | NA | NA | NA | Gm26436 | predicted gene, 26436 [Source:MGI Symb | 0 | 0 | 0 | 0 | 0 | 0 |
| 34774 | ENSMUSG00000095972 | 0 | NA | NA | NA | NA | NA | NA | NA | 0 | 0 | 0 | 0 | 0 | 0 |
| 34775 | ENSMUSG00000095973 | 0 | NA | NA | NA | NA | NA | Vmn1r116 | vomeronasal 1 receptor 116 [Source:MGI | 0 | 0 | 0 | 0 | 0 | 0 |
| 34776 | ENSMUSG00000095974 | 0 | NA | NA | NA | NA | NA | Obox4-ps13 | oocyte specific homeobox 4, pseudogene | 0 | 0 | 0 | 0 | 0 | 0 |
| 34777 | ENSMUSG00000095975 | 0.335414862 | -1.779047242 | 4.030804179 | -0.441362855 | 0.658950329 | NA | Cphx1 | cytoplasmic polyadenylated homeobox 1 | 0 | 0 | 2.0125 | 0 | 0 | 0 |
| 34778 | ENSMUSG00000095976 | 0 | NA | NA | NA | NA | NA | Gm5891 | predicted gene 5891 [Source:MGI Symbol | 0 | 0 | 0 | 0 | 0 | 0 |
| 34779 | ENSMUSG00000095977 | 0 | NA | NA | NA | NA | NA | NA | NA | 0 | 0 | 0 | 0 | 0 | 0 |
| 34780 | ENSMUSG00000095978 | 1.207232854 | 2.667147173 | 2.163434337 | 1.232830194 | 0.217639124 | NA | Mup-ps11 | major urinary protein, pseudogene 11 [So | 0.8031 | 0 | 0 | 0.964 | 4.3625 | 1.1138 |
| 34781 | ENSMUSG00000095979 | 0 | NA | NA | NA | NA | NA | Gm21209 | predicted gene, 21209 [Source:MGI Symb | 0 | 0 | 0 | 0 | 0 | 0 |
| 34782 | ENSMUSG00000095980 | 0 | NA | NA | NA | NA | NA | NA | NA | 0 | 0 | 0 | 0 | 0 | 0 |
| 34783 | ENSMUSG00000095981 | 0.301557804 | -1.64335916 | 4.042814367 | -0.406488899 | 0.684383412 | NA | Ighv10-1 | immunoglobulin heavy variable 10-1 [Sou | 0.8031 | 0 | 1.0062 | 0 | 0 | 0 |
| 34784 | ENSMUSG00000095984 | 0 | NA | NA | NA | NA | NA | Vmn1r256 | vomeronasal 1 receptor 256 [Source:MGI | 0 | 0 | 0 | 0 | 0 | 0 |
| 34785 | ENSMUSG00000095985 | 0 | NA | NA | NA | NA | NA | NA | NA | 0 | 0 | 0 | 0 | 0 | 0 |
| 34786 | ENSMUSG00000095986 | 0 | NA | NA | NA | NA | NA | NA | NA | 0 | 0 | 0 | 0 | 0 | 0 |
| 34787 | ENSMUSG00000095987 | 0 | NA | NA | NA | NA | NA | Gm13104 | predicted gene 13104 [Source:MGI Symbo | 0 | 0 | 0 | 0 | 0 | 0 |
| 34788 | ENSMUSG00000095988 | 0 | NA | NA | NA | NA | NA | Gm15042 | predicted gene 15042 [Source:MGI Symbo | 0 | 0 | 0 | 0 | 0 | 0 |
| 34789 | ENSMUSG00000095989 | 0 | NA | NA | NA | NA | NA | NA | NA | 0 | 0 | 0 | 0 | 0 | 0 |
| 34790 | ENSMUSG00000095990 | 61.63758719 | 0.401637362 | 0.304277983 | 1.319968531 | 0.186845525 | 0.623134349 | Zfp97 | zinc finger protein 97 [Source:MGI Symbo | 63.445 | 54.48 | 41.256 | 82.902 | 68.709 | 59.034 |
| 34791 | ENSMUSG00000095991 | 0 | NA | NA | NA | NA | NA | Gm17048 | predicted gene 17048 [Source:MGI Symbo | 0 | 0 | 0 | 0 | 0 | 0 |
| 34792 | ENSMUSG00000095992 | 0 | NA | NA | NA | NA | NA | Krtap22-2 | keratin associated protein 22-2 [Source:M | 0 | 0 | 0 | 0 | 0 | 0 |
| 34793 | ENSMUSG00000095993 | 0 | NA | NA | NA | NA | NA | Gm21060 | predicted gene, 21060 [Source:MGI Symb | 0 | 0 | 0 | 0 | 0 | 0 |
| 34794 | ENSMUSG00000095994 | 0 | NA | NA | NA | NA | NA | Gm15084 | predicted gene 15084 [Source:MGI Symbo | 0 | 0 | 0 | 0 | 0 | 0 |
| 34795 | ENSMUSG00000095996 | 0 | NA | NA | NA | NA | NA | Gm16513 | predicted gene, Gm16513 [Source:MGI Sy | 0 | 0 | 0 | 0 | 0 | 0 |
| 34796 | ENSMUSG00000095997 | 0 | NA | NA | NA | NA | NA | NA | NA | 0 | 0 | 0 | 0 | 0 | 0 |
| 34797 | ENSMUSG00000095998 | 0 | NA | NA | NA | NA | NA | NA | NA | 0 | 0 | 0 | 0 | 0 | 0 |
| 34798 | ENSMUSG00000096000 | 0 | NA | NA | NA | NA | NA | Olfr773 | olfactory receptor 773 [Source:MGI Symb | 0 | 0 | 0 | 0 | 0 | 0 |
| 34799 | ENSMUSG00000096001 | 1.006604507 | -1.038622879 | 2.695419007 | -0.385328914 | 0.699993742 | NA | 2610528A11 | RIKEN cDNA 2610528A11 gene [Source:M | 0 | 4.1117 | 0 | 1.928 | 0 | 0 |
| 34800 | ENSMUSG00000096002 | 0 | NA | NA | NA | NA | NA | Vmn2r53 | vomeronasal 2, receptor 53 [Source:MGI S | 0 | 0 | 0 | 0 | 0 | 0 |
| 34801 | ENSMUSG00000096003 | 0 | NA | NA | NA | NA | NA | Gm3500 | predicted gene 3500 [Source:MGI Symbol | 0 | 0 | 0 | 0 | 0 | 0 |
| 34802 | ENSMUSG00000096005 | 0 | NA | NA | NA | NA | NA | NA | NA | 0 | 0 | 0 | 0 | 0 | 0 |
| 34803 | ENSMUSG00000096006 | 2.958493074 | -0.845024232 | 1.280526691 | -0.659903645 | 0.509315665 | NA | Gm21596 | predicted gene, 21596 [Source:MGI Symb | 3.2124 | 7.1954 | 1.0062 | 1.928 | 2.1812 | 2.2277 |
| 34804 | ENSMUSG00000096008 | 0 | NA | NA | NA | NA | NA | Gm15100 | predicted gene 15100 [Source:MGI Symbo | 0 | 0 | 0 | 0 | 0 | 0 |
| 34805 | ENSMUSG00000096009 | 0 | NA | NA | NA | NA | NA | Olfr134 | olfactory receptor 134 [Source:MGI Symb | 0 | 0 | 0 | 0 | 0 | 0 |
| 34806 | ENSMUSG00000096010 | 0.469265235 | -2.274672822 | 3.439794491 | -0.661281605 | 0.508431736 | NA | H4f16 | H4 histone 16 [Source:MGI Symbol;Acc:M | 0.8031 | 0 | 2.0125 | 0 | 0 | 0 |
| 34807 | ENSMUSG00000096011 | 0 | NA | NA | NA | NA | NA | Ifna15 | interferon alpha 15 [Source:MGI Symbol;/ | 0 | 0 | 0 | 0 | 0 | 0 |
| 34808 | ENSMUSG00000096014 | 0 | NA | NA | NA | NA | NA | Sox1 | SRY (sex determining region Y)-box 1 [Sou | 0 | 0 | 0 | 0 | 0 | 0 |
| 34809 | ENSMUSG00000096015 | 0 | NA | NA | NA | NA | NA | NA | NA | 0 | 0 | 0 | 0 | 0 | 0 |
| 34810 | ENSMUSG00000096016 | 0 | NA | NA | NA | NA | NA | Gm20937 | predicted gene, 20937 [Source:MGI Symb | 0 | 0 | 0 | 0 | 0 | 0 |
| 34811 | ENSMUSG00000096017 | 0 | NA | NA | NA | NA | NA | NA | NA | 0 | 0 | 0 | 0 | 0 | 0 |
| 34812 | ENSMUSG00000096018 | 0 | NA | NA | NA | NA | NA | NA | NA | 0 | 0 | 0 | 0 | 0 | 0 |
| 34813 | ENSMUSG00000096019 | 0 | NA | NA | NA | NA | NA | NA | NA | 0 | 0 | 0 | 0 | 0 | 0 |
| 34814 | ENSMUSG00000096020 | 0.267700746 | -1.497481649 | 4.057001436 | -0.369110456 | 0.712045396 | NA | Ighv1-75 | immunoglobulin heavy variable 1-75 [Sou | 1.6062 | 0 | 0 | 0 | 0 | 0 |
| 34815 | ENSMUSG00000096021 | 0 | NA | NA | NA | NA | NA | NA | NA | 0 | 0 | 0 | 0 | 0 | 0 |
| 34816 | ENSMUSG00000096022 | 0.133850373 | -0.796629176 | 4.080472857 | -0.195229623 | 0.84521319 | NA | NA | NA | 0.8031 | 0 | 0 | 0 | 0 | 0 |
| 34817 | ENSMUSG00000096023 | 0 | NA | NA | NA | NA | NA | Gm3542 | predicted gene 3542 [Source:MGI Symbol | 0 | 0 | 0 | 0 | 0 | 0 |
| 34818 | ENSMUSG00000096024 | 0 | NA | NA | NA | NA | NA | Gm17174 | predicted gene 17174 [Source:MGI Symbo | 0 | 0 | 0 | 0 | 0 | 0 |
| 34819 | ENSMUSG00000096027 | 0 | NA | NA | NA | NA | NA | NA | NA | 0 | 0 | 0 | 0 | 0 | 0 |
| 34820 | ENSMUSG00000096028 | 0 | NA | NA | NA | NA | NA | NA | NA | 0 | 0 | 0 | 0 | 0 | 0 |
| 34821 | ENSMUSG00000096029 | 0 | NA | NA | NA | NA | NA | Olfr679 | olfactory receptor 679 [Source:MGI Symb | 0 | 0 | 0 | 0 | 0 | 0 |
| 34822 | ENSMUSG00000096030 | 0 | NA | NA | NA | NA | NA | NA | NA | 0 | 0 | 0 | 0 | 0 | 0 |
| 34823 | ENSMUSG00000096031 | 0 | NA | NA | NA | NA | NA | NA | NA | 0 | 0 | 0 | 0 | 0 | 0 |
| 34824 | ENSMUSG00000096033 | 0 | NA | NA | NA | NA | NA | Gm5899 | predicted pseudogene 5899 [Source:MGI | 0 | 0 | 0 | 0 | 0 | 0 |
| 34825 | ENSMUSG00000096035 | 0.36353943 | 2.065989254 | 4.034137301 | 0.512126658 | 0.608562368 | NA | Odaph | odontogenesis associated phosphoprotei | 0 | 0 | 0 | 0 | 2.1812 | 0 |
| 34826 | ENSMUSG00000096036 | 0 | NA | NA | NA | NA | NA | Gm21778 | predicted gene, 21778 [Source:MGI Symb | 0 | 0 | 0 | 0 | 0 | 0 |
| 34827 | ENSMUSG00000096037 | 0 | NA | NA | NA | NA | NA | NA | NA | 0 | 0 | 0 | 0 | 0 | 0 |
| 34828 | ENSMUSG00000096038 | 0 | NA | NA | NA | NA | NA | Trav13n-3 | T cell receptor alpha variable 13N-3 [Sour | 0 | 0 | 0 | 0 | 0 | 0 |

| | | | | | | | | | | | | | | | |
|---|---|---|---|---|---|---|---|---|---|---|---|---|---|---|---|
| 34829 | ENSMUSG00000096039 | 1.855618368 | -4.248257702 | 2.175051688 | -1.953175515 | 0.050798801 | NA | D830030K20 | RIKEN cDNA D830030K20 gene [Source:M | 0 | 3.0838 | 8.05 | 0 | 0 | 0 |
| 34830 | ENSMUSG00000096041 | 0 | NA | NA | NA | NA | NA | NA | NA | 0 | 0 | 0 | 0 | 0 | 0 |
| 34831 | ENSMUSG00000096042 | 0 | NA | NA | NA | NA | NA | Vmn2r125 | vomeronasal 2, receptor 125 [Source:MGI | 0 | 0 | 0 | 0 | 0 | 0 |
| 34832 | ENSMUSG00000096043 | 0 | NA | NA | NA | NA | NA | NA | NA | 0 | 0 | 0 | 0 | 0 | 0 |
| 34833 | ENSMUSG00000096044 | 0 | NA | NA | NA | NA | NA | Pramel33 | PRAME like 33 [Source:MGI Symbol;Acc:M | 0 | 0 | 0 | 0 | 0 | 0 |
| 34834 | ENSMUSG00000096045 | 0 | NA | NA | NA | NA | NA | Gm21698 | predicted gene, 21698 [Source:MGI Symb | 0 | 0 | 0 | 0 | 0 | 0 |
| 34835 | ENSMUSG00000096046 | 0 | NA | NA | NA | NA | NA | NA | NA | 0 | 0 | 0 | 0 | 0 | 0 |
| 34836 | ENSMUSG00000096047 | 0 | NA | NA | NA | NA | NA | NA | NA | 0 | 0 | 0 | 0 | 0 | 0 |
| 34837 | ENSMUSG00000096048 | 0 | NA | NA | NA | NA | NA | NA | NA | 0 | 0 | 0 | 0 | 0 | 0 |
| 34838 | ENSMUSG00000096049 | 0 | NA | NA | NA | NA | NA | Eif1ad17 | eukaryotic translation initiation factor 1A | 0 | 0 | 0 | 0 | 0 | 0 |
| 34839 | ENSMUSG00000096051 | 0 | NA | NA | NA | NA | NA | Vmn1r40 | vomeronasal 1 receptor 40 [Source:MGI S | 0 | 0 | 0 | 0 | 0 | 0 |
| 34840 | ENSMUSG00000096052 | 0.945754888 | -3.283756462 | 2.374797396 | -1.382752258 | 0.166740842 | NA | 9930004E17 | RIKEN cDNA 9930004E17 gene [Source:M | 1.6062 | 2.0558 | 2.0125 | 0 | 0 | 0 |
| 34841 | ENSMUSG00000096053 | 0 | NA | NA | NA | NA | NA | NA | NA | 0 | 0 | 0 | 0 | 0 | 0 |
| 34842 | ENSMUSG00000096054 | 376.1237571 | -0.373222196 | 0.14573916 | -2.560891633 | 0.010440392 | 0.112025093 | Syne1 | spectrin repeat containing, nuclear envelo | 394.32 | 435.84 | 443.75 | 329.68 | 293.38 | 359.77 |
| 34843 | ENSMUSG00000096055 | 0 | NA | NA | NA | NA | NA | Trav12n-1 | T cell receptor alpha variable 12N-1 [Sour | 0 | 0 | 0 | 0 | 0 | 0 |
| 34844 | ENSMUSG00000096056 | 0.181769715 | 1.126924175 | 4.080472857 | 0.276174898 | 0.782413736 | NA | Gm21986 | predicted gene 21986 [Source:MGI Symbo | 0 | 0 | 0 | 0 | 1.0906 | 0 |
| 34845 | ENSMUSG00000096057 | 0 | NA | NA | NA | NA | NA | NA | NA | 0 | 0 | 0 | 0 | 0 | 0 |
| 34846 | ENSMUSG00000096061 | 0 | NA | NA | NA | NA | NA | NA | NA | 0 | 0 | 0 | 0 | 0 | 0 |
| 34847 | ENSMUSG00000096065 | 0 | NA | NA | NA | NA | NA | Gm13155 | predicted gene 13155 [Source:MGI Symbo | 0 | 0 | 0 | 0 | 0 | 0 |
| 34848 | ENSMUSG00000096066 | 0 | NA | NA | NA | NA | NA | Pramel46 | PRAME like 46 [Source:MGI Symbol;Acc:M | 0 | 0 | 0 | 0 | 0 | 0 |
| 34849 | ENSMUSG00000096067 | 0 | NA | NA | NA | NA | NA | NA | NA | 0 | 0 | 0 | 0 | 0 | 0 |
| 34850 | ENSMUSG00000096068 | 0 | NA | NA | NA | NA | NA | Olfr486 | olfactory receptor 486 [Source:MGI Symb | 0 | 0 | 0 | 0 | 0 | 0 |
| 34851 | ENSMUSG00000096069 | 0 | NA | NA | NA | NA | NA | Olfr61 | olfactory receptor 61 [Source:MGI Symbo | 0 | 0 | 0 | 0 | 0 | 0 |
| 34852 | ENSMUSG00000096070 | 0 | NA | NA | NA | NA | NA | NA | NA | 0 | 0 | 0 | 0 | 0 | 0 |
| 34853 | ENSMUSG00000096071 | 0 | NA | NA | NA | NA | NA | Vmn1r143 | vomeronasal 1 receptor 143 [Source:MGI | 0 | 0 | 0 | 0 | 0 | 0 |
| 34854 | ENSMUSG00000096072 | 0 | NA | NA | NA | NA | NA | Mageb6b2 | MAGE family member B6B2 [Source:MGI S | 0 | 0 | 0 | 0 | 0 | 0 |
| 34855 | ENSMUSG00000096073 | 0 | NA | NA | NA | NA | NA | Vmn1r166 | vomeronasal 1 receptor 166 [Source:MGI | 0 | 0 | 0 | 0 | 0 | 0 |
| 34856 | ENSMUSG00000096074 | 0 | NA | NA | NA | NA | NA | Ighv1-72 | immunoglobulin heavy variable 1-72 [Sou | 0 | 0 | 0 | 0 | 0 | 0 |
| 34857 | ENSMUSG00000096075 | 0 | NA | NA | NA | NA | NA | NA | NA | 0 | 0 | 0 | 0 | 0 | 0 |
| 34858 | ENSMUSG00000096076 | 0 | NA | NA | NA | NA | NA | NA | NA | 0 | 0 | 0 | 0 | 0 | 0 |
| 34859 | ENSMUSG00000096078 | 0.160662974 | 1.126924175 | 4.080472857 | 0.276174898 | 0.782413736 | NA | Ighv1-62-2 | immunoglobulin heavy variable 1-62-2 [Sc | 0 | 0 | 0 | 0.964 | 0 | 0 |
| 34860 | ENSMUSG00000096079 | 0 | NA | NA | NA | NA | NA | NA | NA | 0 | 0 | 0 | 0 | 0 | 0 |
| 34861 | ENSMUSG00000096080 | 0 | NA | NA | NA | NA | NA | NA | NA | 0 | 0 | 0 | 0 | 0 | 0 |
| 34862 | ENSMUSG00000096081 | 0 | NA | NA | NA | NA | NA | Gm21991 | predicted gene 21991 [Source:MGI Symbo | 0 | 0 | 0 | 0 | 0 | 0 |
| 34863 | ENSMUSG00000096082 | 0 | NA | NA | NA | NA | NA | NA | NA | 0 | 0 | 0 | 0 | 0 | 0 |
| 34864 | ENSMUSG00000096083 | 0 | NA | NA | NA | NA | NA | Gm10428 | predicted gene 10428 [Source:MGI Symbo | 0 | 0 | 0 | 0 | 0 | 0 |
| 34865 | ENSMUSG00000096084 | 0 | NA | NA | NA | NA | NA | Gm21850 | predicted gene, 21850 [Source:MGI Symb | 0 | 0 | 0 | 0 | 0 | 0 |
| 34866 | ENSMUSG00000096085 | 0 | NA | NA | NA | NA | NA | NA | NA | 0 | 0 | 0 | 0 | 0 | 0 |
| 34867 | ENSMUSG00000096086 | 0 | NA | NA | NA | NA | NA | Gm4761 | predicted gene 4761 [Source:MGI Symbol | 0 | 0 | 0 | 0 | 0 | 0 |
| 34868 | ENSMUSG00000096087 | 0 | NA | NA | NA | NA | NA | NA | NA | 0 | 0 | 0 | 0 | 0 | 0 |
| 34869 | ENSMUSG00000096088 | 0 | NA | NA | NA | NA | NA | Gm12614 | predicted gene 12614 [Source:MGI Symbo | 0 | 0 | 0 | 0 | 0 | 0 |
| 34870 | ENSMUSG00000096090 | 0 | NA | NA | NA | NA | NA | NA | NA | 0 | 0 | 0 | 0 | 0 | 0 |
| 34871 | ENSMUSG00000096091 | 0 | NA | NA | NA | NA | NA | NA | NA | 0 | 0 | 0 | 0 | 0 | 0 |
| 34872 | ENSMUSG00000096092 | 0 | NA | NA | NA | NA | NA | NA | NA | 0 | 0 | 0 | 0 | 0 | 0 |
| 34873 | ENSMUSG00000096094 | 2.922621656 | 1.066669672 | 1.274192983 | 0.837133532 | 0.402517514 | NA | A630095N17 | RIKEN cDNA A630095N17 gene [Source:M | 1.6062 | 1.0279 | 3.0187 | 1.928 | 3.2719 | 6.6831 |
| 34874 | ENSMUSG00000096095 | 0 | NA | NA | NA | NA | NA | NA | NA | 0 | 0 | 0 | 0 | 0 | 0 |
| 34875 | ENSMUSG00000096096 | 0 | NA | NA | NA | NA | NA | Trav4-4-dv10 | T cell receptor alpha variable 4-4-DV10 [Sc | 0 | 0 | 0 | 0 | 0 | 0 |
| 34876 | ENSMUSG00000096097 | 0 | NA | NA | NA | NA | NA | H2al1c | H2A histone family member L1C [Source:M | 0 | 0 | 0 | 0 | 0 | 0 |
| 34877 | ENSMUSG00000096098 | 0 | NA | NA | NA | NA | NA | NA | NA | 0 | 0 | 0 | 0 | 0 | 0 |
| 34878 | ENSMUSG00000096099 | 0 | NA | NA | NA | NA | NA | Vmn1r220 | vomeronasal 1 receptor 220 [Source:MGI | 0 | 0 | 0 | 0 | 0 | 0 |
| 34879 | ENSMUSG00000096100 | 0 | NA | NA | NA | NA | NA | | | 0 | 0 | 0 | 0 | 0 | 0 |
| 34880 | ENSMUSG00000096102 | 0 | NA | NA | NA | NA | NA | Gm9167 | predicted gene 9167 [Source:MGI Symbol | 0 | 0 | 0 | 0 | 0 | 0 |
| 34881 | ENSMUSG00000096105 | 48.80230862 | -0.246550302 | 0.378777966 | -0.650909832 | 0.515104694 | 0.885627279 | Bhmt-ps1 | betaine-homocysteine methyltransferase, | 42.564 | 80.178 | 36.225 | 35.667 | 44.715 | 53.465 |
| 34882 | ENSMUSG00000096106 | 0 | NA | NA | NA | NA | NA | Trav5d-4 | T cell receptor alpha variable 5D-4 [Source | 0 | 0 | 0 | 0 | 0 | 0 |
| 34883 | ENSMUSG00000096107 | 1.195418679 | -1.279722102 | 2.166649774 | -0.590645575 | 0.554757921 | NA | Gm16505 | predicted gene 16505 [Source:MGI Symbo | 0 | 4.1117 | 1.0062 | 0.964 | 1.0906 | 0 |
| 34884 | ENSMUSG00000096108 | 0 | NA | NA | NA | NA | NA | Ighv11-2 | immunoglobulin heavy variable V11-2 [So | 0 | 0 | 0 | 0 | 0 | 0 |

| | | | | | | | | | | | | | | | |
|---|---|---|---|---|---|---|---|---|---|---|---|---|---|---|---|
| 34885 | ENSMUSG00000096109 | 0 | NA | NA | NA | NA | NA | Olfr1537 | olfactory receptor 1537 [Source:MGI Syml | 0 | 0 | 0 | 0 | 0 | 0 |
| 34886 | ENSMUSG00000096110 | 0 | NA | NA | NA | NA | NA | NA | NA | 0 | 0 | 0 | 0 | 0 | 0 |
| 34887 | ENSMUSG00000096111 | 7.347155886 | -0.02298174 | 0.790891582 | -0.029058016 | 0.97681832 | 0.997226157 | Gm21917 | predicted gene, 21917 [Source:MGI Symb | 8.031 | 5.1396 | 9.0562 | 8.6758 | 8.7249 | 4.4554 |
| 34888 | ENSMUSG00000096112 | 0.185640647 | 1.126924175 | 4.080472857 | 0.276174898 | 0.782413736 | NA | NA | NA | 0 | 0 | 0 | 0 | 0 | 1.1138 |
| 34889 | ENSMUSG00000096113 | 0 | NA | NA | NA | NA | NA | NA | NA | 0 | 0 | 0 | 0 | 0 | 0 |
| 34890 | ENSMUSG00000096114 | 0 | NA | NA | NA | NA | NA | NA | NA | 0 | 0 | 0 | 0 | 0 | 0 |
| 34891 | ENSMUSG00000096117 | 0 | NA | NA | NA | NA | NA | NA | NA | 0 | 0 | 0 | 0 | 0 | 0 |
| 34892 | ENSMUSG00000096119 | 0 | NA | NA | NA | NA | NA | Gm6228 | predicted gene 6228 [Source:MGI Symbol | 0 | 0 | 0 | 0 | 0 | 0 |
| 34893 | ENSMUSG00000096120 | 0 | NA | NA | NA | NA | NA | Gm21800 | predicted gene, 21800 [Source:MGI Symb | 0 | 0 | 0 | 0 | 0 | 0 |
| 34894 | ENSMUSG00000096122 | 0 | NA | NA | NA | NA | NA | Gm21943 | predicted gene, 21943 [Source:MGI Symb | 0 | 0 | 0 | 0 | 0 | 0 |
| 34895 | ENSMUSG00000096124 | 0 | NA | NA | NA | NA | NA | NA | NA | 0 | 0 | 0 | 0 | 0 | 0 |
| 34896 | ENSMUSG00000096126 | 0.339027071 | -1.79174149 | 3.919585585 | -0.457125237 | 0.647581032 | NA | NA | NA | 0 | 1.0279 | 1.0062 | 0 | 0 | 0 |
| 34897 | ENSMUSG00000096127 | 0 | NA | NA | NA | NA | NA | Gm14625 | predicted gene 14625 [Source:MGI Symbo | 0 | 0 | 0 | 0 | 0 | 0 |
| 34898 | ENSMUSG00000096129 | 0 | NA | NA | NA | NA | NA | Trav12d-2 | T cell receptor alpha variable 12D-2 [Sour | 0 | 0 | 0 | 0 | 0 | 0 |
| 34899 | ENSMUSG00000096130 | 0 | NA | NA | NA | NA | NA | NA | NA | 0 | 0 | 0 | 0 | 0 | 0 |
| 34900 | ENSMUSG00000096131 | 0 | NA | NA | NA | NA | NA | Gm9639 | predicted gene 9639 [Source:MGI Symbol | 0 | 0 | 0 | 0 | 0 | 0 |
| 34901 | ENSMUSG00000096132 | 0 | NA | NA | NA | NA | NA | NA | NA | 0 | 0 | 0 | 0 | 0 | 0 |
| 34902 | ENSMUSG00000096133 | 0 | NA | NA | NA | NA | NA | NA | NA | 0 | 0 | 0 | 0 | 0 | 0 |
| 34903 | ENSMUSG00000096134 | 0 | NA | NA | NA | NA | NA | NA | NA | 0 | 0 | 0 | 0 | 0 | 0 |
| 34904 | ENSMUSG00000096135 | 0 | NA | NA | NA | NA | NA | Rps12-ps18 | ribosomal protein S12, pseudogene 18 [Sc | 0 | 0 | 0 | 0 | 0 | 0 |
| 34905 | ENSMUSG00000096136 | 0 | NA | NA | NA | NA | NA | NA | NA | 0 | 0 | 0 | 0 | 0 | 0 |
| 34906 | ENSMUSG00000096137 | 0 | NA | NA | NA | NA | NA | Gm14527 | predicted gene 14527 [Source:MGI Symbo | 0 | 0 | 0 | 0 | 0 | 0 |
| 34907 | ENSMUSG00000096138 | 0 | NA | NA | NA | NA | NA | Trav7-6 | T cell receptor alpha variable 7-6 [Source: | 0 | 0 | 0 | 0 | 0 | 0 |
| 34908 | ENSMUSG00000096139 | 0 | NA | NA | NA | NA | NA | Pramel50 | PRAME like 50 [Source:MGI Symbol;Acc:M | 0 | 0 | 0 | 0 | 0 | 0 |
| 34909 | ENSMUSG00000096140 | 0 | NA | NA | NA | NA | NA | Ankrd66 | ankyrin repeat domain 66 [Source:MGI Sy | 0 | 0 | 0 | 0 | 0 | 0 |
| 34910 | ENSMUSG00000096141 | 0 | NA | NA | NA | NA | NA | Dnah7a | dynein, axonemal, heavy chain 7A [Source | 0 | 0 | 0 | 0 | 0 | 0 |
| 34911 | ENSMUSG00000096142 | 0 | NA | NA | NA | NA | NA | NA | NA | 0 | 0 | 0 | 0 | 0 | 0 |
| 34912 | ENSMUSG00000096144 | 0 | NA | NA | NA | NA | NA | B020004C17 | RIKEN cDNA B020004C17 gene [Source:M | 0 | 0 | 0 | 0 | 0 | 0 |
| 34913 | ENSMUSG00000096145 | 209.2227817 | 0.080589696 | 0.178816628 | 0.450683453 | 0.65221771 | 0.925650807 | Vkorc1 | vitamin K epoxide reductase complex, sub | 175.88 | 213.81 | 221.37 | 234.25 | 202.86 | 207.17 |
| 34914 | ENSMUSG00000096146 | 0 | NA | NA | NA | NA | NA | Kcnj11 | potassium inwardly rectifying channel, sul | 0 | 0 | 0 | 0 | 0 | 0 |
| 34915 | ENSMUSG00000096147 | 0 | NA | NA | NA | NA | NA | NA | NA | 0 | 0 | 0 | 0 | 0 | 0 |
| 34916 | ENSMUSG00000096148 | 0 | NA | NA | NA | NA | NA | NA | NA | 0 | 0 | 0 | 0 | 0 | 0 |
| 34917 | ENSMUSG00000096149 | 0 | NA | NA | NA | NA | NA | Trav6-5 | T cell receptor alpha variable 6-5 [Source: | 0 | 0 | 0 | 0 | 0 | 0 |
| 34918 | ENSMUSG00000096150 | 0 | NA | NA | NA | NA | NA | Ighv1-85 | immunoglobulin heavy variable 1-85 [Sou | 0 | 0 | 0 | 0 | 0 | 0 |
| 34919 | ENSMUSG00000096151 | 0 | NA | NA | NA | NA | NA | Olfr477 | olfactory receptor 477 [Source:MGI Symb | 0 | 0 | 0 | 0 | 0 | 0 |
| 34920 | ENSMUSG00000096152 | 0 | NA | NA | NA | NA | NA | Vmn1r240 | vomeronasal 1 receptor 240 [Source:MGI | 0 | 0 | 0 | 0 | 0 | 0 |
| 34921 | ENSMUSG00000096153 | 0 | NA | NA | NA | NA | NA | BC061195 | cDNA sequence BC061195 [Source:MGI Sy | 0 | 0 | 0 | 0 | 0 | 0 |
| 34922 | ENSMUSG00000096154 | 0 | NA | NA | NA | NA | NA | Gm13057 | predicted gene 13057 [Source:MGI Symbo | 0 | 0 | 0 | 0 | 0 | 0 |
| 34923 | ENSMUSG00000096156 | 0.133850373 | -0.796629176 | 4.080472857 | -0.195229623 | 0.84521319 | NA | Gm9242 | predicted pseudogene 9242 [Source:MGI | 0.8031 | 0 | 0 | 0 | 0 | 0 |
| 34924 | ENSMUSG00000096157 | 0.167707431 | -0.796629176 | 4.080472857 | -0.195229623 | 0.84521319 | NA | Gm5472 | predicted gene 5472 [Source:MGI Symbol | 0 | 0 | 1.0062 | 0 | 0 | 0 |
| 34925 | ENSMUSG00000096158 | 0 | NA | NA | NA | NA | NA | Gm21827 | predicted gene, 21827 [Source:MGI Symb | 0 | 0 | 0 | 0 | 0 | 0 |
| 34926 | ENSMUSG00000096159 | 0 | NA | NA | NA | NA | NA | NA | NA | 0 | 0 | 0 | 0 | 0 | 0 |
| 34927 | ENSMUSG00000096160 | 1.015555487 | -0.983278458 | 2.45507575 | -0.400508399 | 0.688782098 | NA | Gm3436 | predicted pseudogene 3436 [Source:MGI | 4.0155 | 0 | 0 | 0.964 | 0 | 1.1138 |
| 34928 | ENSMUSG00000096161 | 0 | NA | NA | NA | NA | NA | NA | NA | 0 | 0 | 0 | 0 | 0 | 0 |
| 34929 | ENSMUSG00000096163 | 0 | NA | NA | NA | NA | NA | NA | NA | 0 | 0 | 0 | 0 | 0 | 0 |
| 34930 | ENSMUSG00000096164 | 0 | NA | NA | NA | NA | NA | Vmn1r93 | vomeronasal 1 receptor 93 [Source:MGI S | 0 | 0 | 0 | 0 | 0 | 0 |
| 34931 | ENSMUSG00000096166 | 0 | NA | NA | NA | NA | NA | NA | NA | 0 | 0 | 0 | 0 | 0 | 0 |
| 34932 | ENSMUSG00000096167 | 0 | NA | NA | NA | NA | NA | Olfr887 | olfactory receptor 887 [Source:MGI Symb | 0 | 0 | 0 | 0 | 0 | 0 |
| 34933 | ENSMUSG00000096168 | 0 | NA | NA | NA | NA | NA | Trav5n-4 | T cell receptor alpha variable 5N-4 [Source | 0 | 0 | 0 | 0 | 0 | 0 |
| 34934 | ENSMUSG00000096169 | 0.811697924 | -0.441734208 | 2.416070595 | -0.182831664 | 0.854930103 | NA | Olfr1564 | olfactory receptor 1564 [Source:MGI Syml | 0.8031 | 0 | 2.0125 | 0.964 | 1.0906 | 0 |
| 34935 | ENSMUSG00000096170 | 0 | NA | NA | NA | NA | NA | NA | NA | 0 | 0 | 0 | 0 | 0 | 0 |
| 34936 | ENSMUSG00000096171 | 3.295641668 | -1.458570822 | 1.27855402 | -1.140797181 | 0.253954334 | NA | Gm8055 | predicted pseudogene 8055 [Source:MGI | 2.4093 | 2.0558 | 10.062 | 1.928 | 1.0906 | 2.2277 |
| 34937 | ENSMUSG00000096172 | 0 | NA | NA | NA | NA | NA | NA | NA | 0 | 0 | 0 | 0 | 0 | 0 |
| 34938 | ENSMUSG00000096173 | 1.592018993 | -0.520167212 | 1.865096875 | -0.278895547 | 0.780324986 | NA | Gm3150 | predicted gene 3150 [Source:MGI Symbol | 1.6062 | 3.0838 | 1.0062 | 3.8559 | 0 | 0 |
| 34939 | ENSMUSG00000096174 | 0.17131964 | -0.796629176 | 4.080472857 | -0.195229623 | 0.84521319 | NA | NA | NA | 0 | 1.0279 | 0 | 0 | 0 | 0 |
| 34940 | ENSMUSG00000096175 | 0 | NA | NA | NA | NA | NA | Gm21761 | predicted gene, 21761 [Source:MGI Symb | 0 | 0 | 0 | 0 | 0 | 0 |

| | | | | | | | | | | | | | | | |
|---|---|---|---|---|---|---|---|---|---|---|---|---|---|---|---|
| 34941 | ENSMUSG00000096176 | 0 | NA | NA | NA | NA | NA | Trdd2 | T cell receptor delta diversity 2 [Source:M | 0 | 0 | 0 | 0 | 0 | 0 |
| 34942 | ENSMUSG00000096177 | 0 | NA | NA | NA | NA | NA | Gm5070 | predicted gene 5070 [Source:MGI Symbol | 0 | 0 | 0 | 0 | 0 | 0 |
| 34943 | ENSMUSG00000096178 | 0 | NA | NA | NA | NA | NA | Gm20837 | predicted gene, 20837 [Source:MGI Symb | 0 | 0 | 0 | 0 | 0 | 0 |
| 34944 | ENSMUSG00000096179 | 0 | NA | NA | NA | NA | NA | Gm15090 | predicted gene 15090 [Source:MGI Symbo | 0 | 0 | 0 | 0 | 0 | 0 |
| 34945 | ENSMUSG00000096180 | 0 | NA | NA | NA | NA | NA | Vmn2r49 | vomeronasal 2, receptor 49 [Source:MGI S | 0 | 0 | 0 | 0 | 0 | 0 |
| 34946 | ENSMUSG00000096181 | 0 | NA | NA | NA | NA | NA | NA | NA | 0 | 0 | 0 | 0 | 0 | 0 |
| 34947 | ENSMUSG00000096182 | 0 | NA | NA | NA | NA | NA | NA | NA | 0 | 0 | 0 | 0 | 0 | 0 |
| 34948 | ENSMUSG00000096183 | 0 | NA | NA | NA | NA | NA | Gm3727 | predicted gene 3727 [Source:MGI Symbol | 0 | 0 | 0 | 0 | 0 | 0 |
| 34949 | ENSMUSG00000096184 | 0 | NA | NA | NA | NA | NA | NA | NA | 0 | 0 | 0 | 0 | 0 | 0 |
| 34950 | ENSMUSG00000096185 | 0 | NA | NA | NA | NA | NA | NA | NA | 0 | 0 | 0 | 0 | 0 | 0 |
| 34951 | ENSMUSG00000096186 | 0 | NA | NA | NA | NA | NA | NA | NA | 0 | 0 | 0 | 0 | 0 | 0 |
| 34952 | ENSMUSG00000096188 | 1164.846437 | -0.106251121 | 0.119272813 | -0.890824301 | 0.373023436 | 0.808422382 | Cmtm4 | CKLF-like MARVEL transmembrane domai | 1132.4 | 1282.8 | 1209.5 | 1260.9 | 1073.2 | 1030.3 |
| 34953 | ENSMUSG00000096189 | 0 | NA | NA | NA | NA | NA | NA | NA | 0 | 0 | 0 | 0 | 0 | 0 |
| 34954 | ENSMUSG00000096190 | 0 | NA | NA | NA | NA | NA | NA | NA | 0 | 0 | 0 | 0 | 0 | 0 |
| 34955 | ENSMUSG00000096191 | 0 | NA | NA | NA | NA | NA | NA | NA | 0 | 0 | 0 | 0 | 0 | 0 |
| 34956 | ENSMUSG00000096192 | 1.836055687 | -0.708234262 | 1.55908564 | -0.454262578 | 0.649639847 | NA | Gm18856 | predicted gene, 18856 [Source:MGI Symb | 0.8031 | 2.0558 | 4.025 | 1.928 | 1.0906 | 1.1138 |
| 34957 | ENSMUSG00000096193 | 0 | NA | NA | NA | NA | NA | NA | NA | 0 | 0 | 0 | 0 | 0 | 0 |
| 34958 | ENSMUSG00000096194 | 0 | NA | NA | NA | NA | NA | Btbd35f2 | BTB domain containing 35, family membe | 0 | 0 | 0 | 0 | 0 | 0 |
| 34959 | ENSMUSG00000096195 | 0 | NA | NA | NA | NA | NA | Gm26138 | predicted gene, 26138 [Source:MGI Symb | 0 | 0 | 0 | 0 | 0 | 0 |
| 34960 | ENSMUSG00000096196 | 0 | NA | NA | NA | NA | NA | NA | NA | 0 | 0 | 0 | 0 | 0 | 0 |
| 34961 | ENSMUSG00000096197 | 0 | NA | NA | NA | NA | NA | 4930555G01 | RIKEN cDNA 4930555G01 gene [Source:M | 0 | 0 | 0 | 0 | 0 | 0 |
| 34962 | ENSMUSG00000096198 | 0 | NA | NA | NA | NA | NA | NA | NA | 0 | 0 | 0 | 0 | 0 | 0 |
| 34963 | ENSMUSG00000096199 | 121.0090444 | -0.131704442 | 0.22225331 | -0.592587091 | 0.55345752 | 0.896339668 | Ptrhd1 | peptidyl-tRNA hydrolase domain containi | 126.09 | 122.32 | 130.81 | 93.506 | 134.15 | 119.18 |
| 34964 | ENSMUSG00000096200 | 0 | NA | NA | NA | NA | NA | NA | NA | 0 | 0 | 0 | 0 | 0 | 0 |
| 34965 | ENSMUSG00000096201 | 0 | NA | NA | NA | NA | NA | Gm10715 | predicted gene 10715 [Source:MGI Symbo | 0 | 0 | 0 | 0 | 0 | 0 |
| 34966 | ENSMUSG00000096202 | 0 | NA | NA | NA | NA | NA | NA | NA | 0 | 0 | 0 | 0 | 0 | 0 |
| 34967 | ENSMUSG00000096205 | 0.167707431 | -0.796629176 | 4.080472857 | -0.195229623 | 0.84521319 | NA | NA | NA | 0 | 0 | 1.0062 | 0 | 0 | 0 |
| 34968 | ENSMUSG00000096206 | 0 | NA | NA | NA | NA | NA | NA | NA | 0 | 0 | 0 | 0 | 0 | 0 |
| 34969 | ENSMUSG00000096208 | 0 | NA | NA | NA | NA | NA | NA | NA | 0 | 0 | 0 | 0 | 0 | 0 |
| 34970 | ENSMUSG00000096209 | 0 | NA | NA | NA | NA | NA | Olfr510 | olfactory receptor 510 [Source:MGI Symb | 0 | 0 | 0 | 0 | 0 | 0 |
| 34971 | ENSMUSG00000096210 | 2947.371112 | 0.271269237 | 0.1063686 | 2.550275521 | 0.010763782 | 0.114475984 | H1f0 | H1.0 linker histone [Source:MGI Symbol;A | 2548.2 | 2543.1 | 2922.1 | 3104 | 3054.8 | 3511.9 |
| 34972 | ENSMUSG00000096211 | 0 | NA | NA | NA | NA | NA | NA | NA | 0 | 0 | 0 | 0 | 0 | 0 |
| 34973 | ENSMUSG00000096212 | 0 | NA | NA | NA | NA | NA | NA | NA | 0 | 0 | 0 | 0 | 0 | 0 |
| 34974 | ENSMUSG00000096213 | 0 | NA | NA | NA | NA | NA | Gm15088 | predicted gene 15088 [Source:MGI Symbo | 0 | 0 | 0 | 0 | 0 | 0 |
| 34975 | ENSMUSG00000096214 | 0 | NA | NA | NA | NA | NA | NA | NA | 0 | 0 | 0 | 0 | 0 | 0 |
| 34976 | ENSMUSG00000096215 | 31.36356486 | 1.112026942 | 0.413145355 | 2.69161187 | 0.007110765 | 0.084997751 | Smim22 | small integral membrane protein 22 [Sour | 20.881 | 13.363 | 25.156 | 40.487 | 38.172 | 50.123 |
| 34977 | ENSMUSG00000096216 | 0 | NA | NA | NA | NA | NA | Gm21959 | predicted gene, 21959 [Source:MGI Symb | 0 | 0 | 0 | 0 | 0 | 0 |
| 34978 | ENSMUSG00000096217 | 0 | NA | NA | NA | NA | NA | Gm4904 | predicted gene 4904 [Source:MGI Symbol | 0 | 0 | 0 | 0 | 0 | 0 |
| 34979 | ENSMUSG00000096218 | 0 | NA | NA | NA | NA | NA | Gm2916 | predicted gene 2916 [Source:MGI Symbol | 0 | 0 | 0 | 0 | 0 | 0 |
| 34980 | ENSMUSG00000096219 | 0 | NA | NA | NA | NA | NA | NA | NA | 0 | 0 | 0 | 0 | 0 | 0 |
| 34981 | ENSMUSG00000096220 | 0 | NA | NA | NA | NA | NA | Olfr775 | olfactory receptor 775 [Source:MGI Symb | 0 | 0 | 0 | 0 | 0 | 0 |
| 34982 | ENSMUSG00000096221 | 6.32666378 | -0.394015659 | 0.954745552 | -0.412691798 | 0.679832435 | 0.934260934 | 1500002C15 | RIKEN cDNA 1500002C15 gene [Source:M | 3.2124 | 10.279 | 8.05 | 0.964 | 6.5437 | 8.9108 |
| 34983 | ENSMUSG00000096222 | 0 | NA | NA | NA | NA | NA | NA | NA | 0 | 0 | 0 | 0 | 0 | 0 |
| 34984 | ENSMUSG00000096223 | 0 | NA | NA | NA | NA | NA | Gm21874 | predicted gene, 21874 [Source:MGI Symb | 0 | 0 | 0 | 0 | 0 | 0 |
| 34985 | ENSMUSG00000096225 | 0 | NA | NA | NA | NA | NA | Lhx8 | LIM homeobox protein 8 [Source:MGI Syn | 0 | 0 | 0 | 0 | 0 | 0 |
| 34986 | ENSMUSG00000096228 | 0 | NA | NA | NA | NA | NA | Olfr5 | olfactory receptor 5 [Source:MGI Symbol; | 0 | 0 | 0 | 0 | 0 | 0 |
| 34987 | ENSMUSG00000096229 | 0 | NA | NA | NA | NA | NA | Olfr774 | olfactory receptor 774 [Source:MGI Symb | 0 | 0 | 0 | 0 | 0 | 0 |
| 34988 | ENSMUSG00000096230 | 0 | NA | NA | NA | NA | NA | Pramel36 | PRAME like 36 [Source:MGI Symbol;Acc:M | 0 | 0 | 0 | 0 | 0 | 0 |
| 34989 | ENSMUSG00000096231 | 0 | NA | NA | NA | NA | NA | Gm22584 | predicted gene, 22584 [Source:MGI Symb | 0 | 0 | 0 | 0 | 0 | 0 |
| 34990 | ENSMUSG00000096233 | 0.695728634 | -1.425610641 | 3.05889997 | -0.46605337 | 0.641177299 | NA | Gm13238 | predicted gene 13238 [Source:MGI Symbo | 0 | 3.0838 | 0 | 0 | 1.0906 | 0 |
| 34991 | ENSMUSG00000096235 | 0 | NA | NA | NA | NA | NA | Gm5123 | predicted pseudogene 5123 [Source:MGI | 0 | 0 | 0 | 0 | 0 | 0 |
| 34992 | ENSMUSG00000096236 | 0 | NA | NA | NA | NA | NA | | | 0 | 0 | 0 | 0 | 0 | 0 |
| 34993 | ENSMUSG00000096237 | 0 | NA | NA | NA | NA | NA | NA | NA | 0 | 0 | 0 | 0 | 0 | 0 |
| 34994 | ENSMUSG00000096238 | 0.160662974 | 1.126924175 | 4.080472857 | 0.276174898 | 0.782413736 | NA | NA | NA | 0 | 0 | 0 | 0.964 | 0 | 0 |
| 34995 | ENSMUSG00000096239 | 0 | NA | NA | NA | NA | NA | NA | NA | 0 | 0 | 0 | 0 | 0 | 0 |
| 34996 | ENSMUSG00000096240 | 0 | NA | NA | NA | NA | NA | Gm21830 | predicted gene, 21830 [Source:MGI Symb | 0 | 0 | 0 | 0 | 0 | 0 |

| | | | | | | | | | | | | | | | |
|---|---|---|---|---|---|---|---|---|---|---|---|---|---|---|---|
| 34997 | ENSMUSG00000096241 | 0 | NA | NA | NA | NA | NA | NA | NA | 0 | 0 | 0 | 0 | 0 | 0 |
| 34998 | ENSMUSG00000096243 | 0 | NA | NA | NA | NA | NA | NA | NA | 0 | 0 | 0 | 0 | 0 | 0 |
| 34999 | ENSMUSG00000096244 | 0 | NA | NA | NA | NA | NA | | vomeronasal 1 receptor Vmn1r187 [Sourc | 0 | 0 | 0 | 0 | 0 | 0 |
| 35000 | ENSMUSG00000096245 | 0 | NA | NA | NA | NA | NA | NA | NA | 0 | 0 | 0 | 0 | 0 | 0 |
| 35001 | ENSMUSG00000096247 | 0 | NA | NA | NA | NA | NA | NA | NA | 0 | 0 | 0 | 0 | 0 | 0 |
| 35002 | ENSMUSG00000096248 | 0 | NA | NA | NA | NA | NA | NA | NA | 0 | 0 | 0 | 0 | 0 | 0 |
| 35003 | ENSMUSG00000096249 | 0 | NA | NA | NA | NA | NA | NA | NA | 0 | 0 | 0 | 0 | 0 | 0 |
| 35004 | ENSMUSG00000096250 | 0 | NA | NA | NA | NA | NA | Ighd2-6 | immunoglobulin heavy diversity 2-6 [Sour | 0 | 0 | 0 | 0 | 0 | 0 |
| 35005 | ENSMUSG00000096254 | 0 | NA | NA | NA | NA | NA | Olfr724 | olfactory receptor 724 [Source:MGI Symb | 0 | 0 | 0 | 0 | 0 | 0 |
| 35006 | ENSMUSG00000096255 | 24.53211186 | -1.58241238 | 0.474996624 | -3.331418164 | 0.000864047 | 0.018793347 | Dynlt1b | dynein light chain Tctex-type 1B [Source:N | 44.171 | 29.81 | 36.225 | 10.604 | 16.359 | 10.025 |
| 35007 | ENSMUSG00000096256 | 0 | NA | NA | NA | NA | NA | 4933409K07 | RIKEN cDNA 4933409K07 gene [Source:M | 0 | 0 | 0 | 0 | 0 | 0 |
| 35008 | ENSMUSG00000096257 | 0 | NA | NA | NA | NA | NA | Ccer2 | coiled-coil glutamate-rich protein 2 [Sourc | 0 | 0 | 0 | 0 | 0 | 0 |
| 35009 | ENSMUSG00000096258 | 0 | NA | NA | NA | NA | NA | Rps16-ps3 | ribosomal protein S16, pseudogene 3 [Sou | 0 | 0 | 0 | 0 | 0 | 0 |
| 35010 | ENSMUSG00000096259 | 0 | NA | NA | NA | NA | NA | Pramel38 | PRAME like 38 [Source:MGI Symbol;Acc:N | 0 | 0 | 0 | 0 | 0 | 0 |
| 35011 | ENSMUSG00000096263 | 0 | NA | NA | NA | NA | NA | NA | NA | 0 | 0 | 0 | 0 | 0 | 0 |
| 35012 | ENSMUSG00000096264 | 0 | NA | NA | NA | NA | NA | NA | NA | 0 | 0 | 0 | 0 | 0 | 0 |
| 35013 | ENSMUSG00000096265 | 0.160662974 | 1.126924175 | 4.080472857 | 0.276174898 | 0.782413736 | NA | 4930467E23 | RIKEN cDNA 4930467E23 gene [Source:M | 0 | 0 | 0 | 0.964 | 0 | 0 |
| 35014 | ENSMUSG00000096268 | 0 | NA | NA | NA | NA | NA | Gm20914 | predicted gene, 20914 [Source:MGI Symb | 0 | 0 | 0 | 0 | 0 | 0 |
| 35015 | ENSMUSG00000096269 | 0 | NA | NA | NA | NA | NA | Rpl31-ps21 | ribosomal protein L31, pseudogene 21 [Sc | 0 | 0 | 0 | 0 | 0 | 0 |
| 35016 | ENSMUSG00000096270 | 0 | NA | NA | NA | NA | NA | Vmn2r-ps13 | vomeronasal 2, receptor, pseudogene 134 | 0 | 0 | 0 | 0 | 0 | 0 |
| 35017 | ENSMUSG00000096271 | 0 | NA | NA | NA | NA | NA | NA | NA | 0 | 0 | 0 | 0 | 0 | 0 |
| 35018 | ENSMUSG00000096272 | 0 | NA | NA | NA | NA | NA | Gm21588 | predicted gene, 21588 [Source:MGI Symb | 0 | 0 | 0 | 0 | 0 | 0 |
| 35019 | ENSMUSG00000096273 | 0 | NA | NA | NA | NA | NA | Olfr1474 | olfactory receptor 1474 [Source:MGI Syml | 0 | 0 | 0 | 0 | 0 | 0 |
| 35020 | ENSMUSG00000096274 | 0 | NA | NA | NA | NA | NA | NA | NA | 0 | 0 | 0 | 0 | 0 | 0 |
| 35021 | ENSMUSG00000096275 | 0 | NA | NA | NA | NA | NA | Gm20901 | predicted gene, 20901 [Source:MGI Symb | 0 | 0 | 0 | 0 | 0 | 0 |
| 35022 | ENSMUSG00000096276 | 0 | NA | NA | NA | NA | NA | Gm2042 | predicted gene 2042 [Source:MGI Symbol | 0 | 0 | 0 | 0 | 0 | 0 |
| 35023 | ENSMUSG00000096277 | 0 | NA | NA | NA | NA | NA | Gm21142 | predicted gene, 21142 [Source:MGI Symb | 0 | 0 | 0 | 0 | 0 | 0 |
| 35024 | ENSMUSG00000096278 | 0 | NA | NA | NA | NA | NA | Dcpp2 | demilune cell and parotid protein 2 [Sourc | 0 | 0 | 0 | 0 | 0 | 0 |
| 35025 | ENSMUSG00000096280 | 0.185640647 | 1.126924175 | 4.080472857 | 0.276174898 | 0.782413736 | NA | NA | NA | 0 | 0 | 0 | 0 | 0 | 1.1138 |
| 35026 | ENSMUSG00000096281 | 0 | NA | NA | NA | NA | NA | NA | NA | 0 | 0 | 0 | 0 | 0 | 0 |
| 35027 | ENSMUSG00000096282 | 0 | NA | NA | NA | NA | NA | NA | NA | 0 | 0 | 0 | 0 | 0 | 0 |
| 35028 | ENSMUSG00000096283 | 0 | NA | NA | NA | NA | NA | Gm6176 | predicted gene 6176 [Source:MGI Symbol | 0 | 0 | 0 | 0 | 0 | 0 |
| 35029 | ENSMUSG00000096284 | 0 | NA | NA | NA | NA | NA | Tcstv1 | 2-cell-stage, variable group, member 1 [Sc | 0 | 0 | 0 | 0 | 0 | 0 |
| 35030 | ENSMUSG00000096285 | 0 | NA | NA | NA | NA | NA | NA | NA | 0 | 0 | 0 | 0 | 0 | 0 |
| 35031 | ENSMUSG00000096286 | 0 | NA | NA | NA | NA | NA | Gm8266 | predicted gene 8266 [Source:MGI Symbol | 0 | 0 | 0 | 0 | 0 | 0 |
| 35032 | ENSMUSG00000096287 | 0 | NA | NA | NA | NA | NA | NA | NA | 0 | 0 | 0 | 0 | 0 | 0 |
| 35033 | ENSMUSG00000096288 | 0 | NA | NA | NA | NA | NA | NA | NA | 0 | 0 | 0 | 0 | 0 | 0 |
| 35034 | ENSMUSG00000096289 | 0 | NA | NA | NA | NA | NA | Olfr1484 | olfactory receptor 1484 [Source:MGI Syml | 0 | 0 | 0 | 0 | 0 | 0 |
| 35035 | ENSMUSG00000096290 | 0 | NA | NA | NA | NA | NA | NA | NA | 0 | 0 | 0 | 0 | 0 | 0 |
| 35036 | ENSMUSG00000096291 | 0 | NA | NA | NA | NA | NA | NA | NA | 0 | 0 | 0 | 0 | 0 | 0 |
| 35037 | ENSMUSG00000096292 | 0.505131668 | 1.126696199 | 3.421159819 | 0.329331647 | 0.741905026 | NA | NA | NA | 0.8031 | 0 | 0 | 0 | 0 | 2.2277 |
| 35038 | ENSMUSG00000096293 | 0 | NA | NA | NA | NA | NA | NA | NA | 0 | 0 | 0 | 0 | 0 | 0 |
| 35039 | ENSMUSG00000096294 | 0 | NA | NA | NA | NA | NA | Gm10302 | predicted gene 10302 [Source:MGI Symbo | 0 | 0 | 0 | 0 | 0 | 0 |
| 35040 | ENSMUSG00000096295 | 0 | NA | NA | NA | NA | NA | Defa2 | defensin, alpha, 2 [Source:MGI Symbol;Ac | 0 | 0 | 0 | 0 | 0 | 0 |
| 35041 | ENSMUSG00000096296 | 0.36353943 | 2.065989254 | 4.034137301 | 0.512126658 | 0.608562368 | NA | NA | NA | 0 | 0 | 0 | 0 | 2.1812 | 0 |
| 35042 | ENSMUSG00000096297 | 0 | NA | NA | NA | NA | NA | NA | NA | 0 | 0 | 0 | 0 | 0 | 0 |
| 35043 | ENSMUSG00000096298 | 0 | NA | NA | NA | NA | NA | NA | NA | 0 | 0 | 0 | 0 | 0 | 0 |
| 35044 | ENSMUSG00000096299 | 0.17131964 | -0.796629176 | 4.080472857 | -0.195229623 | 0.84521319 | NA | Gm21814 | predicted gene, 21814 [Source:MGI Symb | 0 | 1.0279 | 0 | 0 | 0 | 0 |
| 35045 | ENSMUSG00000096300 | 0 | NA | NA | NA | NA | NA | NA | NA | 0 | 0 | 0 | 0 | 0 | 0 |
| 35046 | ENSMUSG00000096301 | 0 | NA | NA | NA | NA | NA | Ighd6-2 | immunoglobulin heavy diversity 6-2 [Sour | 0 | 0 | 0 | 0 | 0 | 0 |
| 35047 | ENSMUSG00000096302 | 0 | NA | NA | NA | NA | NA | NA | NA | 0 | 0 | 0 | 0 | 0 | 0 |
| 35048 | ENSMUSG00000096303 | 0.503122293 | -2.366675966 | 3.785596405 | -0.625179156 | 0.531853481 | NA | NA | NA | 0 | 0 | 3.0187 | 0 | 0 | 0 |
| 35049 | ENSMUSG00000096304 | 0 | NA | NA | NA | NA | NA | Gm16451 | predicted pseudogene 16451 [Source:MG | 0 | 0 | 0 | 0 | 0 | 0 |
| 35050 | ENSMUSG00000096305 | 0 | NA | NA | NA | NA | NA | NA | NA | 0 | 0 | 0 | 0 | 0 | 0 |
| 35051 | ENSMUSG00000096306 | 0 | NA | NA | NA | NA | NA | NA | NA | 0 | 0 | 0 | 0 | 0 | 0 |
| 35052 | ENSMUSG00000096307 | 0 | NA | NA | NA | NA | NA | NA | NA | 0 | 0 | 0 | 0 | 0 | 0 |

| | | | | | | | | | | | | | | | |
|---|---|---|---|---|---|---|---|---|---|---|---|---|---|---|---|
| 35053 | ENSMUSG00000096308 | 0 | NA | NA | NA | NA | NA | Gm14598 | predicted gene 14598 [Source:MGI Symbo | 0 | 0 | 0 | 0 | 0 | 0 |
| 35054 | ENSMUSG00000096309 | 0 | NA | NA | NA | NA | NA | NA | NA | 0 | 0 | 0 | 0 | 0 | 0 |
| 35055 | ENSMUSG00000096310 | 0 | NA | NA | NA | NA | NA | NA | NA | 0 | 0 | 0 | 0 | 0 | 0 |
| 35056 | ENSMUSG00000096311 | 0 | NA | NA | NA | NA | NA | Gm3681 | predicted gene 3681 [Source:MGI Symbol | 0 | 0 | 0 | 0 | 0 | 0 |
| 35057 | ENSMUSG00000096313 | 0 | NA | NA | NA | NA | NA | NA | NA | 0 | 0 | 0 | 0 | 0 | 0 |
| 35058 | ENSMUSG00000096314 | 0 | NA | NA | NA | NA | NA | NA | NA | 0 | 0 | 0 | 0 | 0 | 0 |
| 35059 | ENSMUSG00000096316 | 0 | NA | NA | NA | NA | NA | Faiml | Fas apoptotic inhibitory molecule like [Sou | 0 | 0 | 0 | 0 | 0 | 0 |
| 35060 | ENSMUSG00000096318 | 0 | NA | NA | NA | NA | NA | Astx4a | amplified spermatogenic transcripts X enc | 0 | 0 | 0 | 0 | 0 | 0 |
| 35061 | ENSMUSG00000096320 | 0 | NA | NA | NA | NA | NA | Olfr1471 | olfactory receptor 1471 [Source:MGI Sym | 0 | 0 | 0 | 0 | 0 | 0 |
| 35062 | ENSMUSG00000096322 | 0 | NA | NA | NA | NA | NA | NA | NA | 0 | 0 | 0 | 0 | 0 | 0 |
| 35063 | ENSMUSG00000096323 | 0 | NA | NA | NA | NA | NA | Gm20767 | predicted gene, 20767 [Source:MGI Symb | 0 | 0 | 0 | 0 | 0 | 0 |
| 35064 | ENSMUSG00000096324 | 0 | NA | NA | NA | NA | NA | NA | NA | 0 | 0 | 0 | 0 | 0 | 0 |
| 35065 | ENSMUSG00000096326 | 0 | NA | NA | NA | NA | NA | Ighv1-78 | immunoglobulin heavy variable 1-78 [Sou | 0 | 0 | 0 | 0 | 0 | 0 |
| 35066 | ENSMUSG00000096327 | 0 | NA | NA | NA | NA | NA | Gm21980 | predicted gene 21980 [Source:MGI Symbo | 0 | 0 | 0 | 0 | 0 | 0 |
| 35067 | ENSMUSG00000096328 | 0 | NA | NA | NA | NA | NA | NA | NA | 0 | 0 | 0 | 0 | 0 | 0 |
| 35068 | ENSMUSG00000096329 | 0 | NA | NA | NA | NA | NA | Trav13d-4 | T cell receptor alpha variable 13D-4 [Sour | 0 | 0 | 0 | 0 | 0 | 0 |
| 35069 | ENSMUSG00000096330 | 0 | NA | NA | NA | NA | NA | Gm21976 | predicted gene 21976 [Source:MGI Symbo | 0 | 0 | 0 | 0 | 0 | 0 |
| 35070 | ENSMUSG00000096331 | 0 | NA | NA | NA | NA | NA | NA | NA | 0 | 0 | 0 | 0 | 0 | 0 |
| 35071 | ENSMUSG00000096332 | 0 | NA | NA | NA | NA | NA | Gm21241 | predicted gene, 21241 [Source:MGI Symb | 0 | 0 | 0 | 0 | 0 | 0 |
| 35072 | ENSMUSG00000096333 | 0 | NA | NA | NA | NA | NA | Gm13871 | predicted gene 13871 [Source:MGI Symbo | 0 | 0 | 0 | 0 | 0 | 0 |
| 35073 | ENSMUSG00000096335 | 0 | NA | NA | NA | NA | NA | NA | NA | 0 | 0 | 0 | 0 | 0 | 0 |
| 35074 | ENSMUSG00000096336 | 1.614903015 | -1.044002942 | 1.762685125 | -0.592279884 | 0.553663184 | NA | Igkv1-135 | immunoglobulin kappa variable 1-135 [So | 2.4093 | 4.1117 | 0 | 0.964 | 1.0906 | 1.1138 |
| 35075 | ENSMUSG00000096337 | 0 | NA | NA | NA | NA | NA | Gm13694 | predicted gene 13694 [Source:MGI Symbo | 0 | 0 | 0 | 0 | 0 | 0 |
| 35076 | ENSMUSG00000096338 | 0 | NA | NA | NA | NA | NA | Gm15086 | predicted gene 15086 [Source:MGI Symbo | 0 | 0 | 0 | 0 | 0 | 0 |
| 35077 | ENSMUSG00000096339 | 0 | NA | NA | NA | NA | NA | NA | NA | 0 | 0 | 0 | 0 | 0 | 0 |
| 35078 | ENSMUSG00000096340 | 0 | NA | NA | NA | NA | NA | Defa-ps10 | defensin, alpha, pseudogene 10 [Source:N | 0 | 0 | 0 | 0 | 0 | 0 |
| 35079 | ENSMUSG00000096341 | 0 | NA | NA | NA | NA | NA | NA | NA | 0 | 0 | 0 | 0 | 0 | 0 |
| 35080 | ENSMUSG00000096342 | 0 | NA | NA | NA | NA | NA | NA | NA | 0 | 0 | 0 | 0 | 0 | 0 |
| 35081 | ENSMUSG00000096343 | 0 | NA | NA | NA | NA | NA | NA | NA | 0 | 0 | 0 | 0 | 0 | 0 |
| 35082 | ENSMUSG00000096344 | 0 | NA | NA | NA | NA | NA | Gm6408 | predicted gene 6408 [Source:MGI Symbol | 0 | 0 | 0 | 0 | 0 | 0 |
| 35083 | ENSMUSG00000096345 | 0 | NA | NA | NA | NA | NA | Esp16 | exocrine gland secreted peptide 16 [Sourc | 0 | 0 | 0 | 0 | 0 | 0 |
| 35084 | ENSMUSG00000096346 | 0.133850373 | -0.796629176 | 4.080472857 | -0.195229623 | 0.84521319 | NA | NA | NA | 0.8031 | 0 | 0 | 0 | 0 | 0 |
| 35085 | ENSMUSG00000096347 | 0 | NA | NA | NA | NA | NA | Gm21758 | predicted gene, 21758 [Source:MGI Symb | 0 | 0 | 0 | 0 | 0 | 0 |
| 35086 | ENSMUSG00000096348 | 0 | NA | NA | NA | NA | NA | Vmn1r250 | vomeronasal 1 receptor 250 [Source:MGI | 0 | 0 | 0 | 0 | 0 | 0 |
| 35087 | ENSMUSG00000096349 | 0 | NA | NA | NA | NA | NA | NA | NA | 0 | 0 | 0 | 0 | 0 | 0 |
| 35088 | ENSMUSG00000096350 | 0 | NA | NA | NA | NA | NA | NA | NA | 0 | 0 | 0 | 0 | 0 | 0 |
| 35089 | ENSMUSG00000096351 | 0.52105551 | -0.812028029 | 3.385364865 | -0.239864257 | 0.810435491 | NA | Samd11 | sterile alpha motif domain containing 11 [ | 0 | 0 | 2.0125 | 0 | 0 | 1.1138 |
| 35090 | ENSMUSG00000096352 | 0 | NA | NA | NA | NA | NA | NA | NA | 0 | 0 | 0 | 0 | 0 | 0 |
| 35091 | ENSMUSG00000096353 | 0 | NA | NA | NA | NA | NA | NA | NA | 0 | 0 | 0 | 0 | 0 | 0 |
| 35092 | ENSMUSG00000096354 | 0 | NA | NA | NA | NA | NA | Gm8178 | predicted gene 8178 [Source:MGI Symbol | 0 | 0 | 0 | 0 | 0 | 0 |
| 35093 | ENSMUSG00000096355 | 0 | NA | NA | NA | NA | NA | Ighv8-4 | immunoglobulin heavy variable V8-4 [Sou | 0 | 0 | 0 | 0 | 0 | 0 |
| 35094 | ENSMUSG00000096356 | 0 | NA | NA | NA | NA | NA | Olfr889 | olfactory receptor 889 [Source:MGI Symb | 0 | 0 | 0 | 0 | 0 | 0 |
| 35095 | ENSMUSG00000096357 | 0 | NA | NA | NA | NA | NA | NA | NA | 0 | 0 | 0 | 0 | 0 | 0 |
| 35096 | ENSMUSG00000096358 | 0 | NA | NA | NA | NA | NA | NA | NA | 0 | 0 | 0 | 0 | 0 | 0 |
| 35097 | ENSMUSG00000096359 | 0 | NA | NA | NA | NA | NA | NA | NA | 0 | 0 | 0 | 0 | 0 | 0 |
| 35098 | ENSMUSG00000096360 | 0 | NA | NA | NA | NA | NA | NA | NA | 0 | 0 | 0 | 0 | 0 | 0 |
| 35099 | ENSMUSG00000096361 | 1.538110092 | -0.26348682 | 1.761941206 | -0.14954348 | 0.881124803 | NA | Gm5814 | predicted pseudogene 5814 [Source:MGI | 0 | 3.0838 | 2.0125 | 1.928 | 1.0906 | 1.1138 |
| 35100 | ENSMUSG00000096362 | 0 | NA | NA | NA | NA | NA | Rps12-ps21 | ribosomal protein S12, pseudogene 22 [Sc | 0 | 0 | 0 | 0 | 0 | 0 |
| 35101 | ENSMUSG00000096363 | 0 | NA | NA | NA | NA | NA | Gm12621 | predicted gene 12621 [Source:MGI Symbo | 0 | 0 | 0 | 0 | 0 | 0 |
| 35102 | ENSMUSG00000096365 | 0 | NA | NA | NA | NA | NA | Olfr1463 | olfactory receptor 1463 [Source:MGI Sym | 0 | 0 | 0 | 0 | 0 | 0 |
| 35103 | ENSMUSG00000096366 | 0.916590439 | 3.39841529 | 2.684534563 | 1.265923463 | 0.205540485 | NA | Gm13240 | predicted gene 13240 [Source:MGI Symbo | 0 | 0 | 0 | 0 | 3.2719 | 2.2277 |
| 35104 | ENSMUSG00000096367 | 0 | NA | NA | NA | NA | NA | NA | NA | 0 | 0 | 0 | 0 | 0 | 0 |
| 35105 | ENSMUSG00000096370 | 8.928893509 | -0.195470931 | 0.813783411 | -0.240200191 | 0.810175065 | 0.966157155 | Gm21992 | predicted gene 21992 [Source:MGI Symbo | 15.259 | 7.1954 | 6.0375 | 8.6758 | 14.178 | 2.2277 |
| 35106 | ENSMUSG00000096371 | 0 | NA | NA | NA | NA | NA | NA | NA | 0 | 0 | 0 | 0 | 0 | 0 |
| 35107 | ENSMUSG00000096372 | 0 | NA | NA | NA | NA | NA | Gm8138 | predicted gene 8138 [Source:MGI Symbol | 0 | 0 | 0 | 0 | 0 | 0 |
| 35108 | ENSMUSG00000096373 | 0 | NA | NA | NA | NA | NA | Vmn2r31 | vomeronasal 2, receptor 31 [Source:MGI S | 0 | 0 | 0 | 0 | 0 | 0 |

| | | | | | | | | | | | | | | | |
|---|---|---|---|---|---|---|---|---|---|---|---|---|---|---|---|
| 35109 | ENSMUSG00000096374 | 0 | NA | NA | NA | NA | NA | Gm6664 | predicted gene 6664 [Source:MGI Symbol | 0 | 0 | 0 | 0 | 0 | 0 |
| 35110 | ENSMUSG00000096375 | 0.133850373 | -0.796629176 | 4.080472857 | -0.195229623 | 0.84521319 | NA | Gm7094 | predicted pseudogene 7094 [Source:MGI | 0.8031 | 0 | 0 | 0 | 0 | 0 |
| 35111 | ENSMUSG00000096376 | 0 | NA | NA | NA | NA | NA | Gm14590 | predicted gene 14590 [Source:MGI Symbo | 0 | 0 | 0 | 0 | 0 | 0 |
| 35112 | ENSMUSG00000096377 | 0 | NA | NA | NA | NA | NA | Mir5123 | microRNA 5123 [Source:MGI Symbol;Acc: | 0 | 0 | 0 | 0 | 0 | 0 |
| 35113 | ENSMUSG00000096379 | 0 | NA | NA | NA | NA | NA | NA | NA | 0 | 0 | 0 | 0 | 0 | 0 |
| 35114 | ENSMUSG00000096380 | 0 | NA | NA | NA | NA | NA | Gm19668 | predicted gene, 19668 [Source:MGI Symb | 0 | 0 | 0 | 0 | 0 | 0 |
| 35115 | ENSMUSG00000096381 | 0 | NA | NA | NA | NA | NA | NA | NA | 0 | 0 | 0 | 0 | 0 | 0 |
| 35116 | ENSMUSG00000096383 | 0 | NA | NA | NA | NA | NA | NA | NA | 0 | 0 | 0 | 0 | 0 | 0 |
| 35117 | ENSMUSG00000096384 | 0 | NA | NA | NA | NA | NA | NA | NA | 0 | 0 | 0 | 0 | 0 | 0 |
| 35118 | ENSMUSG00000096385 | 0 | NA | NA | NA | NA | NA | Gm11168 | predicted gene 11168 [Source:MGI Symbo | 0 | 0 | 0 | 0 | 0 | 0 |
| 35119 | ENSMUSG00000096386 | 0 | NA | NA | NA | NA | NA | Vmn1r100 | vomeronasal 1 receptor 100 [Source:MGI | 0 | 0 | 0 | 0 | 0 | 0 |
| 35120 | ENSMUSG00000096387 | 0 | NA | NA | NA | NA | NA | NA | NA | 0 | 0 | 0 | 0 | 0 | 0 |
| 35121 | ENSMUSG00000096388 | 0 | NA | NA | NA | NA | NA | NA | NA | 0 | 0 | 0 | 0 | 0 | 0 |
| 35122 | ENSMUSG00000096390 | 0 | NA | NA | NA | NA | NA | NA | NA | 0 | 0 | 0 | 0 | 0 | 0 |
| 35123 | ENSMUSG00000096391 | 0.371281295 | 2.09253362 | 4.031852165 | 0.519000582 | 0.603760335 | NA | NA | NA | 0 | 0 | 0 | 0 | 0 | 2.2277 |
| 35124 | ENSMUSG00000096392 | 0 | NA | NA | NA | NA | NA | NA | NA | 0 | 0 | 0 | 0 | 0 | 0 |
| 35125 | ENSMUSG00000096393 | 0 | NA | NA | NA | NA | NA | Fam240a | family with sequence similarity 240 memb | 0 | 0 | 0 | 0 | 0 | 0 |
| 35126 | ENSMUSG00000096396 | 0 | NA | NA | NA | NA | NA | Ighd5-2 | immunoglobulin heavy diversity 5-2 [Sour | 0 | 0 | 0 | 0 | 0 | 0 |
| 35127 | ENSMUSG00000096397 | 0 | NA | NA | NA | NA | NA | Trav17 | T cell receptor alpha variable 17 [Source:M | 0 | 0 | 0 | 0 | 0 | 0 |
| 35128 | ENSMUSG00000096398 | 0 | NA | NA | NA | NA | NA | NA | NA | 0 | 0 | 0 | 0 | 0 | 0 |
| 35129 | ENSMUSG00000096399 | 0 | NA | NA | NA | NA | NA | Vmn2r35 | vomeronasal 2, receptor 35 [Source:MGI S | 0 | 0 | 0 | 0 | 0 | 0 |
| 35130 | ENSMUSG00000096400 | 0 | NA | NA | NA | NA | NA | Gm6098 | predicted gene 6098 [Source:MGI Symbol | 0 | 0 | 0 | 0 | 0 | 0 |
| 35131 | ENSMUSG00000096401 | 49.51771549 | -0.272346275 | 0.324267862 | -0.839880564 | 0.400975356 | 0.827897196 | Gm21811 | predicted gene, 21811 [Source:MGI Symb | 48.989 | 57.563 | 56.35 | 55.911 | 37.081 | 41.212 |
| 35132 | ENSMUSG00000096402 | 0 | NA | NA | NA | NA | NA | NA | NA | 0 | 0 | 0 | 0 | 0 | 0 |
| 35133 | ENSMUSG00000096403 | 0.353348078 | 0.165142701 | 3.895042568 | 0.042398176 | 0.966181282 | NA | Rnps1-ps | RNA binding protein with serine rich doma | 0 | 0 | 1.0062 | 0 | 0 | 1.1138 |
| 35134 | ENSMUSG00000096405 | 0 | NA | NA | NA | NA | NA | 4930474N05 | RIKEN cDNA 4930474N05 gene [Source:M | 0 | 0 | 0 | 0 | 0 | 0 |
| 35135 | ENSMUSG00000096406 | 0 | NA | NA | NA | NA | NA | NA | NA | 0 | 0 | 0 | 0 | 0 | 0 |
| 35136 | ENSMUSG00000096407 | 0 | NA | NA | NA | NA | NA | Ighv6-5 | immunoglobulin heavy variable V6-5 [Sou | 0 | 0 | 0 | 0 | 0 | 0 |
| 35137 | ENSMUSG00000096409 | 0 | NA | NA | NA | NA | NA | Olfr890 | olfactory receptor 890 [Source:MGI Symb | 0 | 0 | 0 | 0 | 0 | 0 |
| 35138 | ENSMUSG00000096410 | 0 | NA | NA | NA | NA | NA | Ighv1-19 | immunoglobulin heavy variable V1-19 [So | 0 | 0 | 0 | 0 | 0 | 0 |
| 35139 | ENSMUSG00000096411 | 0 | NA | NA | NA | NA | NA | NA | NA | 0 | 0 | 0 | 0 | 0 | 0 |
| 35140 | ENSMUSG00000096412 | 0 | NA | NA | NA | NA | NA | NA | NA | 0 | 0 | 0 | 0 | 0 | 0 |
| 35141 | ENSMUSG00000096413 | 0 | NA | NA | NA | NA | NA | Gm3616 | predicted gene 3616 [Source:MGI Symbol | 0 | 0 | 0 | 0 | 0 | 0 |
| 35142 | ENSMUSG00000096414 | 0 | NA | NA | NA | NA | NA | Obox4-ps15 | oocyte specific homeobox 4, pseudogene | 0 | 0 | 0 | 0 | 0 | 0 |
| 35143 | ENSMUSG00000096415 | 0 | NA | NA | NA | NA | NA | NA | NA | 0 | 0 | 0 | 0 | 0 | 0 |
| 35144 | ENSMUSG00000096417 | 0 | NA | NA | NA | NA | NA | Trav7-2 | T cell receptor alpha variable 7-2 [Source: | 0 | 0 | 0 | 0 | 0 | 0 |
| 35145 | ENSMUSG00000096420 | 0 | NA | NA | NA | NA | NA | Ighd5-6 | immunoglobulin heavy diversity 5-6 [Sour | 0 | 0 | 0 | 0 | 0 | 0 |
| 35146 | ENSMUSG00000096421 | 0 | NA | NA | NA | NA | NA | Gm10100 | predicted gene 10100 [Source:MGI Symbo | 0 | 0 | 0 | 0 | 0 | 0 |
| 35147 | ENSMUSG00000096422 | 0.305170013 | -1.657836477 | 4.041480335 | -0.410205257 | 0.681655385 | NA | Igkv12-44 | immunoglobulin kappa variable 12-44 [So | 0.8031 | 1.0279 | 0 | 0 | 0 | 0 |
| 35148 | ENSMUSG00000096423 | 0 | NA | NA | NA | NA | NA | Gm24978 | predicted gene, 24978 [Source:MGI Symb | 0 | 0 | 0 | 0 | 0 | 0 |
| 35149 | ENSMUSG00000096424 | 0 | NA | NA | NA | NA | NA | Olfr885 | olfactory receptor 885 [Source:MGI Symb | 0 | 0 | 0 | 0 | 0 | 0 |
| 35150 | ENSMUSG00000096425 | 0 | NA | NA | NA | NA | NA | NA | NA | 0 | 0 | 0 | 0 | 0 | 0 |
| 35151 | ENSMUSG00000096426 | 0 | NA | NA | NA | NA | NA | Btbd35f24 | BTB domain containing 35, family membe | 0 | 0 | 0 | 0 | 0 | 0 |
| 35152 | ENSMUSG00000096427 | 0 | NA | NA | NA | NA | NA | Olfr895 | olfactory receptor 895 [Source:MGI Symb | 0 | 0 | 0 | 0 | 0 | 0 |
| 35153 | ENSMUSG00000096429 | 0 | NA | NA | NA | NA | NA | Vmn2r-ps13 | vomeronasal 2, receptor, pseudogene 131 | 0 | 0 | 0 | 0 | 0 | 0 |
| 35154 | ENSMUSG00000096430 | 0 | NA | NA | NA | NA | NA | NA | NA | 0 | 0 | 0 | 0 | 0 | 0 |
| 35155 | ENSMUSG00000096431 | 0 | NA | NA | NA | NA | NA | NA | NA | 0 | 0 | 0 | 0 | 0 | 0 |
| 35156 | ENSMUSG00000096432 | 0 | NA | NA | NA | NA | NA | NA | NA | 0 | 0 | 0 | 0 | 0 | 0 |
| 35157 | ENSMUSG00000096433 | 95.81622147 | 0.153386604 | 0.242476615 | 0.632583081 | 0.527005939 | 0.888334265 | Zfp994 | zinc finger protein 994 [Source:MGI Symb | 80.31 | 91.485 | 100.62 | 94.47 | 117.79 | 90.221 |
| 35158 | ENSMUSG00000096434 | 0 | NA | NA | NA | NA | NA | NA | NA | 0 | 0 | 0 | 0 | 0 | 0 |
| 35159 | ENSMUSG00000096436 | 0 | NA | NA | NA | NA | NA | Olfr1437 | olfactory receptor 1437 [Source:MGI Syml | 0 | 0 | 0 | 0 | 0 | 0 |
| 35160 | ENSMUSG00000096437 | 0 | NA | NA | NA | NA | NA | Rps12-ps16 | ribosomal protein S12, pseudogene 16 [Sc | 0 | 0 | 0 | 0 | 0 | 0 |
| 35161 | ENSMUSG00000096438 | 0 | NA | NA | NA | NA | NA | Gapdh-ps15 | glyceraldehyde-3-phosphate dehydrogena | 0 | 0 | 0 | 0 | 0 | 0 |
| 35162 | ENSMUSG00000096439 | 0 | NA | NA | NA | NA | NA | Obox4-ps23 | oocyte specific homeobox 4, pseudogene | 0 | 0 | 0 | 0 | 0 | 0 |
| 35163 | ENSMUSG00000096440 | 0 | NA | NA | NA | NA | NA | Vmn2r-ps20 | vomeronasal 2, receptor, pseudogene 20 | 0 | 0 | 0 | 0 | 0 | 0 |
| 35164 | ENSMUSG00000096441 | 0 | NA | NA | NA | NA | NA | NA | NA | 0 | 0 | 0 | 0 | 0 | 0 |

| | | | | | | | | | | | | | | | |
|---|---|---|---|---|---|---|---|---|---|---|---|---|---|---|---|
| 35165 | ENSMUSG00000096442 | 0 | NA | NA | NA | NA | NA | Taar8a | trace amine-associated receptor 8A [Sour | 0 | 0 | 0 | 0 | 0 | 0 |
| 35166 | ENSMUSG00000096444 | 0 | NA | NA | NA | NA | NA | NA | NA | 0 | 0 | 0 | 0 | 0 | 0 |
| 35167 | ENSMUSG00000096445 | 0 | NA | NA | NA | NA | NA | Dcpp1 | demilune cell and parotid protein 1 [Sourc | 0 | 0 | 0 | 0 | 0 | 0 |
| 35168 | ENSMUSG00000096446 | 0 | NA | NA | NA | NA | NA | Gm8104 | predicted gene 8104 [Source:MGI Symbol | 0 | 0 | 0 | 0 | 0 | 0 |
| 35169 | ENSMUSG00000096447 | 0 | NA | NA | NA | NA | NA | NA | NA | 0 | 0 | 0 | 0 | 0 | 0 |
| 35170 | ENSMUSG00000096448 | 0 | NA | NA | NA | NA | NA | Gm5808 | predicted gene 5808 [Source:MGI Symbol | 0 | 0 | 0 | 0 | 0 | 0 |
| 35171 | ENSMUSG00000096449 | 0.167707431 | -0.796629176 | 4.080472857 | -0.195229623 | 0.84521319 | NA | Gm4076 | predicted gene 4076 [Source:MGI Symbol | 0 | 0 | 1.0062 | 0 | 0 | 0 |
| 35172 | ENSMUSG00000096450 | 0 | NA | NA | NA | NA | NA | Gm2877 | predicted gene 2877 [Source:MGI Symbol | 0 | 0 | 0 | 0 | 0 | 0 |
| 35173 | ENSMUSG00000096451 | 0 | NA | NA | NA | NA | NA | NA | NA | 0 | 0 | 0 | 0 | 0 | 0 |
| 35174 | ENSMUSG00000096452 | 0 | NA | NA | NA | NA | NA | Ighv1-77 | immunoglobulin heavy variable 1-77 [Sou | 0 | 0 | 0 | 0 | 0 | 0 |
| 35175 | ENSMUSG00000096453 | 0 | NA | NA | NA | NA | NA | Gm8320 | predicted gene 8320 [Source:MGI Symbol | 0 | 0 | 0 | 0 | 0 | 0 |
| 35176 | ENSMUSG00000096454 | 0 | NA | NA | NA | NA | NA | NA | NA | 0 | 0 | 0 | 0 | 0 | 0 |
| 35177 | ENSMUSG00000096455 | 0 | NA | NA | NA | NA | NA | NA | NA | 0 | 0 | 0 | 0 | 0 | 0 |
| 35178 | ENSMUSG00000096457 | 0 | NA | NA | NA | NA | NA | Gm10147 | predicted gene 10147 [Source:MGI Symbo | 0 | 0 | 0 | 0 | 0 | 0 |
| 35179 | ENSMUSG00000096458 | 0 | NA | NA | NA | NA | NA | Moap1 | modulator of apoptosis 1 [Source:MGI Syr | 0 | 0 | 0 | 0 | 0 | 0 |
| 35180 | ENSMUSG00000096459 | 0.342639279 | -1.805976596 | 4.028547714 | -0.448294702 | 0.653940524 | NA | Ighv9-3 | immunoglobulin heavy variable V9-3 [Sou | 0 | 2.0558 | 0 | 0 | 0 | 0 |
| 35181 | ENSMUSG00000096461 | 0 | NA | NA | NA | NA | NA | Igkv14-130 | immunoglobulin kappa variable 14-130 [S | 0 | 0 | 0 | 0 | 0 | 0 |
| 35182 | ENSMUSG00000096462 | 0 | NA | NA | NA | NA | NA | NA | NA | 0 | 0 | 0 | 0 | 0 | 0 |
| 35183 | ENSMUSG00000096463 | 0 | NA | NA | NA | NA | NA | Gm21750 | predicted gene, 21750 [Source:MGI Symb | 0 | 0 | 0 | 0 | 0 | 0 |
| 35184 | ENSMUSG00000096464 | 0 | NA | NA | NA | NA | NA | Ighv2-2 | immunoglobulin heavy variable 2-2 [Sourc | 0 | 0 | 0 | 0 | 0 | 0 |
| 35185 | ENSMUSG00000096465 | 0 | NA | NA | NA | NA | NA | Olfr488 | olfactory receptor 488 [Source:MGI Symb | 0 | 0 | 0 | 0 | 0 | 0 |
| 35186 | ENSMUSG00000096467 | 0 | NA | NA | NA | NA | NA | Scgb2b19 | secretoglobin, family 2B, member 19 [Sou | 0 | 0 | 0 | 0 | 0 | 0 |
| 35187 | ENSMUSG00000096468 | 0 | NA | NA | NA | NA | NA | Gm16405 | predicted gene 16405 [Source:MGI Symbo | 0 | 0 | 0 | 0 | 0 | 0 |
| 35188 | ENSMUSG00000096470 | 0 | NA | NA | NA | NA | NA | Gm9603 | predicted gene 9603 [Source:MGI Symbol | 0 | 0 | 0 | 0 | 0 | 0 |
| 35189 | ENSMUSG00000096472 | 66.04397861 | 0.611736742 | 0.292058461 | 2.094569492 | 0.03620928 | 0.256479442 | Cdkn2d | cyclin dependent kinase inhibitor 2D [Sou | 44.171 | 60.647 | 52.325 | 86.758 | 63.256 | 89.108 |
| 35190 | ENSMUSG00000096473 | 0 | NA | NA | NA | NA | NA | NA | NA | 0 | 0 | 0 | 0 | 0 | 0 |
| 35191 | ENSMUSG00000096474 | 1.192245155 | -0.39688883 | 2.299030883 | -0.172633101 | 0.862939832 | NA | Gm5561 | predicted gene 5561 [Source:MGI Symbol | 0 | 4.1117 | 0 | 1.928 | 0 | 1.1138 |
| 35192 | ENSMUSG00000096475 | 0 | NA | NA | NA | NA | NA | NA | NA | 0 | 0 | 0 | 0 | 0 | 0 |
| 35193 | ENSMUSG00000096476 | 0 | NA | NA | NA | NA | NA | Gm12453 | predicted gene 12453 [Source:MGI Symbo | 0 | 0 | 0 | 0 | 0 | 0 |
| 35194 | ENSMUSG00000096477 | 0 | NA | NA | NA | NA | NA | Olfr92 | olfactory receptor 92 [Source:MGI Symbo | 0 | 0 | 0 | 0 | 0 | 0 |
| 35195 | ENSMUSG00000096478 | 0 | NA | NA | NA | NA | NA | Trav9d-1 | T cell receptor alpha variable 9D-1 [Sourc | 0 | 0 | 0 | 0 | 0 | 0 |
| 35196 | ENSMUSG00000096479 | 0 | NA | NA | NA | NA | NA | NA | NA | 0 | 0 | 0 | 0 | 0 | 0 |
| 35197 | ENSMUSG00000096480 | 0 | NA | NA | NA | NA | NA | NA | NA | 0 | 0 | 0 | 0 | 0 | 0 |
| 35198 | ENSMUSG00000096481 | 0 | NA | NA | NA | NA | NA | Gm3250 | predicted gene 3250 [Source:MGI Symbol | 0 | 0 | 0 | 0 | 0 | 0 |
| 35199 | ENSMUSG00000096482 | 0 | NA | NA | NA | NA | NA | NA | NA | 0 | 0 | 0 | 0 | 0 | 0 |
| 35200 | ENSMUSG00000096483 | 0 | NA | NA | NA | NA | NA | Gm12615 | predicted gene 12615 [Source:MGI Symbo | 0 | 0 | 0 | 0 | 0 | 0 |
| 35201 | ENSMUSG00000096484 | 0 | NA | NA | NA | NA | NA | Gm13696 | predicted gene 13696 [Source:MGI Symbo | 0 | 0 | 0 | 0 | 0 | 0 |
| 35202 | ENSMUSG00000096485 | 0 | NA | NA | NA | NA | NA | Olfr1466 | olfactory receptor 1466 [Source:MGI Sym | 0 | 0 | 0 | 0 | 0 | 0 |
| 35203 | ENSMUSG00000096486 | 0 | NA | NA | NA | NA | NA | Gm5426 | predicted pseudogene 5426 [Source:MGI | 0 | 0 | 0 | 0 | 0 | 0 |
| 35204 | ENSMUSG00000096488 | 0 | NA | NA | NA | NA | NA | Gm10409 | predicted gene 10409 [Source:MGI Symbo | 0 | 0 | 0 | 0 | 0 | 0 |
| 35205 | ENSMUSG00000096490 | 0.294513347 | 0.165143496 | 4.080472857 | 0.040471657 | 0.967717103 | NA | Igkv10-94 | immunoglobulin kappa variable 10-94 [So | 0.8031 | 0 | 0 | 0.964 | 0 | 0 |
| 35206 | ENSMUSG00000096492 | 0 | NA | NA | NA | NA | NA | NA | NA | 0 | 0 | 0 | 0 | 0 | 0 |
| 35207 | ENSMUSG00000096496 | 0 | NA | NA | NA | NA | NA | NA | NA | 0 | 0 | 0 | 0 | 0 | 0 |
| 35208 | ENSMUSG00000096497 | 0 | NA | NA | NA | NA | NA | Olfr787 | olfactory receptor 787 [Source:MGI Symb | 0 | 0 | 0 | 0 | 0 | 0 |
| 35209 | ENSMUSG00000096498 | 0 | NA | NA | NA | NA | NA | Ighv2-5 | immunoglobulin heavy variable 2-5 [Sourc | 0 | 0 | 0 | 0 | 0 | 0 |
| 35210 | ENSMUSG00000096499 | 0 | NA | NA | NA | NA | NA | Ighv1-5 | immunoglobulin heavy variable V1-5 [Sou | 0 | 0 | 0 | 0 | 0 | 0 |
| 35211 | ENSMUSG00000096500 | 0 | NA | NA | NA | NA | NA | NA | NA | 0 | 0 | 0 | 0 | 0 | 0 |
| 35212 | ENSMUSG00000096502 | 0 | NA | NA | NA | NA | NA | NA | NA | 0 | 0 | 0 | 0 | 0 | 0 |
| 35213 | ENSMUSG00000096504 | 0 | NA | NA | NA | NA | NA | NA | NA | 0 | 0 | 0 | 0 | 0 | 0 |
| 35214 | ENSMUSG00000096505 | 0 | NA | NA | NA | NA | NA | Trav4d-4 | T cell receptor alpha variable 4D-4 [Sourc | 0 | 0 | 0 | 0 | 0 | 0 |
| 35215 | ENSMUSG00000096506 | 1.411814455 | 0.09922145 | 2.380429545 | 0.041682162 | 0.966752074 | NA | NA | NA | 4.0155 | 0 | 0 | 0 | 0 | 4.4554 |
| 35216 | ENSMUSG00000096507 | 0 | NA | NA | NA | NA | NA | Gm21772 | predicted gene, 21772 [Source:MGI Symb | 0 | 0 | 0 | 0 | 0 | 0 |
| 35217 | ENSMUSG00000096508 | 0 | NA | NA | NA | NA | NA | Gm7903 | predicted gene 7903 [Source:MGI Symbol | 0 | 0 | 0 | 0 | 0 | 0 |
| 35218 | ENSMUSG00000096510 | 0 | NA | NA | NA | NA | NA | NA | NA | 0 | 0 | 0 | 0 | 0 | 0 |
| 35219 | ENSMUSG00000096512 | 0 | NA | NA | NA | NA | NA | NA | NA | 0 | 0 | 0 | 0 | 0 | 0 |
| 35220 | ENSMUSG00000096513 | 0 | NA | NA | NA | NA | NA | Vmn1r247 | vomeronasal 1 receptor 247 [Source:MGI | 0 | 0 | 0 | 0 | 0 | 0 |

| | | | | | | | | | | | | | | | |
|---|---|---|---|---|---|---|---|---|---|---|---|---|---|---|---|
| 35221 | ENSMUSG00000096514 | 0 | NA | NA | NA | NA | NA | NA | NA | 0 | 0 | 0 | 0 | 0 | 0 |
| 35222 | ENSMUSG00000096515 | 0.342639279 | -1.805976596 | 4.028547714 | -0.448294702 | 0.653940524 | NA | Igkv14-100 | immunoglobulin kappa chain variable 14- | 0 | 2.0558 | 0 | 0 | 0 | 0 |
| 35223 | ENSMUSG00000096516 | 0 | NA | NA | NA | NA | NA | Olfr628 | olfactory receptor 628 [Source:MGI Symb | 0 | 0 | 0 | 0 | 0 | 0 |
| 35224 | ENSMUSG00000096517 | 0 | NA | NA | NA | NA | NA | Gm6919 | predicted gene 6919 [Source:MGI Symbol | 0 | 0 | 0 | 0 | 0 | 0 |
| 35225 | ENSMUSG00000096518 | 0 | NA | NA | NA | NA | NA | Gm21894 | predicted gene, 21894 [Source:MGI Symb | 0 | 0 | 0 | 0 | 0 | 0 |
| 35226 | ENSMUSG00000096519 | 0 | NA | NA | NA | NA | NA | Gm10721 | predicted gene 10721 [Source:MGI Symbo | 0 | 0 | 0 | 0 | 0 | 0 |
| 35227 | ENSMUSG00000096520 | 0 | NA | NA | NA | NA | NA | Gm3376 | predicted gene 3376 [Source:MGI Symbol | 0 | 0 | 0 | 0 | 0 | 0 |
| 35228 | ENSMUSG00000096521 | 0 | NA | NA | NA | NA | NA | Gm13137 | predicted gene 13137 [Source:MGI Symbo | 0 | 0 | 0 | 0 | 0 | 0 |
| 35229 | ENSMUSG00000096522 | 0 | NA | NA | NA | NA | NA | NA | NA | 0 | 0 | 0 | 0 | 0 | 0 |
| 35230 | ENSMUSG00000096523 | 0 | NA | NA | NA | NA | NA | NA | NA | 0 | 0 | 0 | 0 | 0 | 0 |
| 35231 | ENSMUSG00000096524 | 0 | NA | NA | NA | NA | NA | Gm14355 | predicted gene 14355 [Source:MGI Symbo | 0 | 0 | 0 | 0 | 0 | 0 |
| 35232 | ENSMUSG00000096525 | 0 | NA | NA | NA | NA | NA | Gm2663 | predicted gene 2663 [Source:MGI Symbol | 0 | 0 | 0 | 0 | 0 | 0 |
| 35233 | ENSMUSG00000096527 | 0 | NA | NA | NA | NA | NA | Gm6370 | predicted gene 6370 [Source:MGI Symbol | 0 | 0 | 0 | 0 | 0 | 0 |
| 35234 | ENSMUSG00000096528 | 0 | NA | NA | NA | NA | NA | G430049J08 | RIKEN cDNA G430049J08 gene [Source:M | 0 | 0 | 0 | 0 | 0 | 0 |
| 35235 | ENSMUSG00000096530 | 0 | NA | NA | NA | NA | NA | Gm11758 | predicted gene 11758 [Source:MGI Symbo | 0 | 0 | 0 | 0 | 0 | 0 |
| 35236 | ENSMUSG00000096531 | 0 | NA | NA | NA | NA | NA | Trav12n-2 | T cell receptor alpha variable 12N-2 [Sour | 0 | 0 | 0 | 0 | 0 | 0 |
| 35237 | ENSMUSG00000096532 | 0 | NA | NA | NA | NA | NA | Gm21633 | predicted gene, 21633 [Source:MGI Symb | 0 | 0 | 0 | 0 | 0 | 0 |
| 35238 | ENSMUSG00000096533 | 0 | NA | NA | NA | NA | NA | NA | NA | 0 | 0 | 0 | 0 | 0 | 0 |
| 35239 | ENSMUSG00000096534 | 0 | NA | NA | NA | NA | NA | Krtap16-3 | keratin associated protein 16-3 [Source:M | 0 | 0 | 0 | 0 | 0 | 0 |
| 35240 | ENSMUSG00000096535 | 0 | NA | NA | NA | NA | NA | NA | NA | 0 | 0 | 0 | 0 | 0 | 0 |
| 35241 | ENSMUSG00000096537 | 0 | NA | NA | NA | NA | NA | Fam240b | family with sequence similarity 240 memb | 0 | 0 | 0 | 0 | 0 | 0 |
| 35242 | ENSMUSG00000096538 | 0 | NA | NA | NA | NA | NA | Gm21853 | predicted gene, 21853 [Source:MGI Symb | 0 | 0 | 0 | 0 | 0 | 0 |
| 35243 | ENSMUSG00000096539 | 0 | NA | NA | NA | NA | NA | Gm7438 | predicted pseudogene 7438 [Source:MGI | 0 | 0 | 0 | 0 | 0 | 0 |
| 35244 | ENSMUSG00000096541 | 0 | NA | NA | NA | NA | NA | Trav9n-3 | T cell receptor alpha variable 9N-3 [Sourc | 0 | 0 | 0 | 0 | 0 | 0 |
| 35245 | ENSMUSG00000096544 | 23.54138097 | -0.00257458 | 0.498636882 | -0.005163237 | 0.995880351 | 0.999003684 | Gm4617 | predicted pseudogene 4617 [Source:MGI | 25.699 | 23.642 | 21.131 | 12.532 | 37.081 | 21.163 |
| 35246 | ENSMUSG00000096546 | 519.5463559 | 0.077648203 | 0.138182299 | 0.561925832 | 0.574166553 | 0.906012546 | Smlr1 | small leucine-rich protein 1 [Source:MGI S | 544.5 | 483.12 | 489.03 | 600.56 | 478.78 | 521.28 |
| 35247 | ENSMUSG00000096547 | 0.185640647 | 1.126924175 | 4.080472857 | 0.276174898 | 0.782413736 | NA | NA | NA | 0 | 0 | 0 | 0 | 0 | 1.1138 |
| 35248 | ENSMUSG00000096549 | 3.973719658 | -0.184426408 | 1.112761915 | -0.165737527 | 0.868363512 | NA | Prickle4 | prickle planar cell polarity protein 4 [Sour | 5.6217 | 2.0558 | 5.0312 | 6.7478 | 3.2719 | 1.1138 |
| 35249 | ENSMUSG00000096550 | 0 | NA | NA | NA | NA | NA | | predicted gene 16367 [Source:NCBI gene | 0 | 0 | 0 | 0 | 0 | 0 |
| 35250 | ENSMUSG00000096551 | 0 | NA | NA | NA | NA | NA | Trav10 | T cell receptor alpha variable 10 [Source:N | 0 | 0 | 0 | 0 | 0 | 0 |
| 35251 | ENSMUSG00000096553 | 0 | NA | NA | NA | NA | NA | Gm10097 | predicted gene 10097 [Source:MGI Symbo | 0 | 0 | 0 | 0 | 0 | 0 |
| 35252 | ENSMUSG00000096554 | 0 | NA | NA | NA | NA | NA | Olfr1282 | olfactory receptor 1282 [Source:MGI Sym | 0 | 0 | 0 | 0 | 0 | 0 |
| 35253 | ENSMUSG00000096555 | 0 | NA | NA | NA | NA | NA | Olfr944 | olfactory receptor 944 [Source:MGI Symb | 0 | 0 | 0 | 0 | 0 | 0 |
| 35254 | ENSMUSG00000096557 | 0 | NA | NA | NA | NA | NA | Gm7258 | predicted gene 7258 [Source:MGI Symbol | 0 | 0 | 0 | 0 | 0 | 0 |
| 35255 | ENSMUSG00000096558 | 0 | NA | NA | NA | NA | NA | NA | NA | 0 | 0 | 0 | 0 | 0 | 0 |
| 35256 | ENSMUSG00000096559 | 0.545309144 | 2.651974629 | 3.666219369 | 0.72335405 | 0.469462394 | NA | NA | NA | 0 | 0 | 0 | 0 | 3.2719 | 0 |
| 35257 | ENSMUSG00000096560 | 0 | NA | NA | NA | NA | NA | NA | NA | 0 | 0 | 0 | 0 | 0 | 0 |
| 35258 | ENSMUSG00000096561 | 0.17131964 | -0.796629176 | 4.080472857 | -0.195229623 | 0.84521319 | NA | NA | NA | 0 | 1.0279 | 0 | 0 | 0 | 0 |
| 35259 | ENSMUSG00000096563 | 0 | NA | NA | NA | NA | NA | NA | NA | 0 | 0 | 0 | 0 | 0 | 0 |
| 35260 | ENSMUSG00000096564 | 0 | NA | NA | NA | NA | NA | NA | NA | 0 | 0 | 0 | 0 | 0 | 0 |
| 35261 | ENSMUSG00000096565 | 0 | NA | NA | NA | NA | NA | NA | NA | 0 | 0 | 0 | 0 | 0 | 0 |
| 35262 | ENSMUSG00000096566 | 0 | NA | NA | NA | NA | NA | Olfr1306 | olfactory receptor 1306 [Source:MGI Sym | 0 | 0 | 0 | 0 | 0 | 0 |
| 35263 | ENSMUSG00000096567 | 0 | NA | NA | NA | NA | NA | NA | NA | 0 | 0 | 0 | 0 | 0 | 0 |
| 35264 | ENSMUSG00000096568 | 0 | NA | NA | NA | NA | NA | Ighd2-3 | immunoglobulin heavy diversity 2-3 [Sour | 0 | 0 | 0 | 0 | 0 | 0 |
| 35265 | ENSMUSG00000096569 | 0 | NA | NA | NA | NA | NA | Amy2a2 | amylase 2a2 [Source:MGI Symbol;Acc:MG | 0 | 0 | 0 | 0 | 0 | 0 |
| 35266 | ENSMUSG00000096573 | 0 | NA | NA | NA | NA | NA | 1700009J07I | RIKEN cDNA 1700009J07 gene [Source:M | 0 | 0 | 0 | 0 | 0 | 0 |
| 35267 | ENSMUSG00000096574 | 0 | NA | NA | NA | NA | NA | Gm2956 | predicted gene 2956 [Source:MGI Symbol | 0 | 0 | 0 | 0 | 0 | 0 |
| 35268 | ENSMUSG00000096576 | 0 | NA | NA | NA | NA | NA | Oog1 | oogenesin 1 [Source:MGI Symbol;Acc:MG | 0 | 0 | 0 | 0 | 0 | 0 |
| 35269 | ENSMUSG00000096577 | 0 | NA | NA | NA | NA | NA | Ighv1-71 | immunoglobulin heavy variable 1-71 [Sou | 0 | 0 | 0 | 0 | 0 | 0 |
| 35270 | ENSMUSG00000096578 | 0 | NA | NA | NA | NA | NA | Gm9529 | predicted gene 9529 [Source:MGI Symbol | 0 | 0 | 0 | 0 | 0 | 0 |
| 35271 | ENSMUSG00000096579 | 0 | NA | NA | NA | NA | NA | Gm17121 | predicted gene 17121 [Source:MGI Symbo | 0 | 0 | 0 | 0 | 0 | 0 |
| 35272 | ENSMUSG00000096580 | 0 | NA | NA | NA | NA | NA | Igkv1-132 | immunoglobulin kappa variable 1-132 [So | 0 | 0 | 0 | 0 | 0 | 0 |
| 35273 | ENSMUSG00000096582 | 0 | NA | NA | NA | NA | NA | Gm13289 | predicted gene 13289 [Source:MGI Symbo | 0 | 0 | 0 | 0 | 0 | 0 |
| 35274 | ENSMUSG00000096583 | 0 | NA | NA | NA | NA | NA | NA | NA | 0 | 0 | 0 | 0 | 0 | 0 |
| 35275 | ENSMUSG00000096584 | 0 | NA | NA | NA | NA | NA | Olfr611 | olfactory receptor 611 [Source:MGI Symb | 0 | 0 | 0 | 0 | 0 | 0 |
| 35276 | ENSMUSG00000096585 | 0 | NA | NA | NA | NA | NA | NA | NA | 0 | 0 | 0 | 0 | 0 | 0 |

| | | | | | | | | | | | | | | | |
|---|---|---|---|---|---|---|---|---|---|---|---|---|---|---|---|
| 35277 | ENSMUSG00000096586 | 2.315692234 | -0.304096193 | 1.455674948 | -0.208903913 | 0.834523249 | NA | NA | NA | 1.6062 | 4.1117 | 2.0125 | 2.8919 | 3.2719 | 0 |
| 35278 | ENSMUSG00000096587 | 0 | NA | NA | NA | NA | NA | NA | NA | 0 | 0 | 0 | 0 | 0 | 0 |
| 35279 | ENSMUSG00000096589 | 0 | NA | NA | NA | NA | NA | Gm14992 | predicted gene 14992 [Source:MGI Symbo | 0 | 0 | 0 | 0 | 0 | 0 |
| 35280 | ENSMUSG00000096591 | 0 | NA | NA | NA | NA | NA | Gm13272 | predicted gene 13272 [Source:MGI Symbo | 0 | 0 | 0 | 0 | 0 | 0 |
| 35281 | ENSMUSG00000096592 | 1.62555968 | -1.802316758 | 1.942100038 | -0.928024676 | 0.35339477 | NA | Gm15801 | predicted gene 15801 [Source:MGI Symbo | 2.4093 | 5.1396 | 0 | 0 | 1.0906 | 1.1138 |
| 35282 | ENSMUSG00000096593 | 0 | NA | NA | NA | NA | NA | Vmn2r54 | vomeronasal 2, receptor 54 [Source:MGI S | 0 | 0 | 0 | 0 | 0 | 0 |
| 35283 | ENSMUSG00000096594 | 0.133850373 | -0.796629176 | 4.080472857 | -0.195229623 | 0.84521319 | NA | Igkv8-19 | immunoglobulin kappa variable 8-19 [Sou | 0.8031 | 0 | 0 | 0 | 0 | 0 |
| 35284 | ENSMUSG00000096595 | 0 | NA | NA | NA | NA | NA | NA | NA | 0 | 0 | 0 | 0 | 0 | 0 |
| 35285 | ENSMUSG00000096596 | 0 | NA | NA | NA | NA | NA | Gm10591 | predicted gene 10591 [Source:MGI Symbo | 0 | 0 | 0 | 0 | 0 | 0 |
| 35286 | ENSMUSG00000096597 | 0 | NA | NA | NA | NA | NA | Gm21886 | predicted gene, 21886 [Source:MGI Symb | 0 | 0 | 0 | 0 | 0 | 0 |
| 35287 | ENSMUSG00000096598 | 0 | NA | NA | NA | NA | NA | NA | NA | 0 | 0 | 0 | 0 | 0 | 0 |
| 35288 | ENSMUSG00000096599 | 0 | NA | NA | NA | NA | NA | NA | NA | 0 | 0 | 0 | 0 | 0 | 0 |
| 35289 | ENSMUSG00000096600 | 0 | NA | NA | NA | NA | NA | Trav6d-3 | T cell receptor alpha variable 6D-3 [Source | 0 | 0 | 0 | 0 | 0 | 0 |
| 35290 | ENSMUSG00000096601 | 0 | NA | NA | NA | NA | NA | Vmn1r260 | vomeronasal 1 receptor 260 [Source:MGI | 0 | 0 | 0 | 0 | 0 | 0 |
| 35291 | ENSMUSG00000096604 | 0 | NA | NA | NA | NA | NA | NA | NA | 0 | 0 | 0 | 0 | 0 | 0 |
| 35292 | ENSMUSG00000096605 | 0 | NA | NA | NA | NA | NA | NA | NA | 0 | 0 | 0 | 0 | 0 | 0 |
| 35293 | ENSMUSG00000096606 | 82.35056368 | -0.60456878 | 0.272276566 | -2.220421644 | 0.026390159 | 0.210307031 | Tpbgl | trophoblast glycoprotein-like [Source:MG | 99.585 | 103.82 | 94.587 | 64.587 | 47.987 | 83.538 |
| 35294 | ENSMUSG00000096607 | 0 | NA | NA | NA | NA | NA | 7530416G11 | RIKEN cDNA 7530416G11 gene [Source:M | 0 | 0 | 0 | 0 | 0 | 0 |
| 35295 | ENSMUSG00000096613 | 0 | NA | NA | NA | NA | NA | NA | NA | 0 | 0 | 0 | 0 | 0 | 0 |
| 35296 | ENSMUSG00000096614 | 0 | NA | NA | NA | NA | NA | Gm2871 | predicted gene 2871 [Source:MGI Symbol | 0 | 0 | 0 | 0 | 0 | 0 |
| 35297 | ENSMUSG00000096615 | 0 | NA | NA | NA | NA | NA | Trav11 | T cell receptor alpha variable 11 [Source:N | 0 | 0 | 0 | 0 | 0 | 0 |
| 35298 | ENSMUSG00000096617 | 11.38252104 | -0.398693908 | 0.633051747 | -0.62979671 | 0.528827598 | 0.888334265 | Gm5559 | predicted gene 5559 [Source:MGI Symbol | 10.44 | 14.391 | 14.087 | 10.604 | 7.6343 | 11.138 |
| 35299 | ENSMUSG00000096618 | 0 | NA | NA | NA | NA | NA | Gm9286 | predicted gene 9286 [Source:MGI Symbol | 0 | 0 | 0 | 0 | 0 | 0 |
| 35300 | ENSMUSG00000096619 | 0 | NA | NA | NA | NA | NA | Eif1ad14 | eukaryotic translation initiation factor 1A | 0 | 0 | 0 | 0 | 0 | 0 |
| 35301 | ENSMUSG00000096620 | 0 | NA | NA | NA | NA | NA | Gm5169 | predicted gene 5169 [Source:MGI Symbol | 0 | 0 | 0 | 0 | 0 | 0 |
| 35302 | ENSMUSG00000096621 | 0 | NA | NA | NA | NA | NA | Gm5849 | predicted gene 5849 [Source:MGI Symbol | 0 | 0 | 0 | 0 | 0 | 0 |
| 35303 | ENSMUSG00000096624 | 0 | NA | NA | NA | NA | NA | NA | NA | 0 | 0 | 0 | 0 | 0 | 0 |
| 35304 | ENSMUSG00000096626 | 0 | NA | NA | NA | NA | NA | Gm21861 | predicted gene, 21861 [Source:MGI Symb | 0 | 0 | 0 | 0 | 0 | 0 |
| 35305 | ENSMUSG00000096627 | 0 | NA | NA | NA | NA | NA | NA | NA | 0 | 0 | 0 | 0 | 0 | 0 |
| 35306 | ENSMUSG00000096628 | 0 | NA | NA | NA | NA | NA | NA | NA | 0 | 0 | 0 | 0 | 0 | 0 |
| 35307 | ENSMUSG00000096629 | 0 | NA | NA | NA | NA | NA | Gm3383 | predicted gene 3383 [Source:MGI Symbol | 0 | 0 | 0 | 0 | 0 | 0 |
| 35308 | ENSMUSG00000096630 | 0.439020386 | -2.189747911 | 3.52148864 | -0.621824499 | 0.534057275 | NA | Vmn2r26 | vomeronasal 2, receptor 26 [Source:MGI S | 1.6062 | 1.0279 | 0 | 0 | 0 | 0 |
| 35309 | ENSMUSG00000096631 | 0 | NA | NA | NA | NA | NA | NA | NA | 0 | 0 | 0 | 0 | 0 | 0 |
| 35310 | ENSMUSG00000096632 | 0.476489653 | -2.293616034 | 3.421506946 | -0.670352587 | 0.502633051 | NA | Igkv9-124 | immunoglobulin kappa chain variable 9-12 | 0.8031 | 2.0558 | 0 | 0 | 0 | 0 |
| 35311 | ENSMUSG00000096633 | 0 | NA | NA | NA | NA | NA | NA | NA | 0 | 0 | 0 | 0 | 0 | 0 |
| 35312 | ENSMUSG00000096634 | 0 | NA | NA | NA | NA | NA | NA | NA | 0 | 0 | 0 | 0 | 0 | 0 |
| 35313 | ENSMUSG00000096635 | 0 | NA | NA | NA | NA | NA | NA | NA | 0 | 0 | 0 | 0 | 0 | 0 |
| 35314 | ENSMUSG00000096638 | 0.133850373 | -0.796629176 | 4.080472857 | -0.195229623 | 0.84521319 | NA | Ighv2-9 | immunoglobulin heavy variable 2-9 [Sourc | 0.8031 | 0 | 0 | 0 | 0 | 0 |
| 35315 | ENSMUSG00000096639 | 0 | NA | NA | NA | NA | NA | NA | NA | 0 | 0 | 0 | 0 | 0 | 0 |
| 35316 | ENSMUSG00000096640 | 0 | NA | NA | NA | NA | NA | Gm4312 | predicted gene 4312 [Source:MGI Symbol | 0 | 0 | 0 | 0 | 0 | 0 |
| 35317 | ENSMUSG00000096641 | 0 | NA | NA | NA | NA | NA | Gm904 | predicted gene 904 [Source:MGI Symbol;A | 0 | 0 | 0 | 0 | 0 | 0 |
| 35318 | ENSMUSG00000096642 | 0 | NA | NA | NA | NA | NA | NA | NA | 0 | 0 | 0 | 0 | 0 | 0 |
| 35319 | ENSMUSG00000096644 | 0 | NA | NA | NA | NA | NA | Magea1 | MAGE family member A1 [Source:MGI Syr | 0 | 0 | 0 | 0 | 0 | 0 |
| 35320 | ENSMUSG00000096645 | 0 | NA | NA | NA | NA | NA | Spin2e | spindlin family, member 2E [Source:MGI S | 0 | 0 | 0 | 0 | 0 | 0 |
| 35321 | ENSMUSG00000096646 | 0 | NA | NA | NA | NA | NA | NA | NA | 0 | 0 | 0 | 0 | 0 | 0 |
| 35322 | ENSMUSG00000096647 | 0.983456248 | 0.112283097 | 2.230039339 | 0.050350276 | 0.95984326 | NA | Tmem41b-ps | transmembrane protein 41B, pseudogene | 0.8031 | 2.0558 | 0 | 1.928 | 0 | 1.1138 |
| 35323 | ENSMUSG00000096648 | 0 | NA | NA | NA | NA | NA | NA | NA | 0 | 0 | 0 | 0 | 0 | 0 |
| 35324 | ENSMUSG00000096649 | 0 | NA | NA | NA | NA | NA | Ighv1-31 | immunoglobulin heavy variable 1-31 [Sou | 0 | 0 | 0 | 0 | 0 | 0 |
| 35325 | ENSMUSG00000096650 | 0 | NA | NA | NA | NA | NA | Gm20896 | predicted gene, 20896 [Source:MGI Symb | 0 | 0 | 0 | 0 | 0 | 0 |
| 35326 | ENSMUSG00000096651 | 0 | NA | NA | NA | NA | NA | NA | NA | 0 | 0 | 0 | 0 | 0 | 0 |
| 35327 | ENSMUSG00000096652 | 1.461595355 | -0.730731747 | 2.092015268 | -0.349295609 | 0.726867395 | NA | Mup-ps9 | major urinary protein, pseudogene 9 [Sou | 2.4093 | 0 | 3.0187 | 0 | 0 | 3.3415 |
| 35328 | ENSMUSG00000096653 | 0 | NA | NA | NA | NA | NA | Trav15d-1-d | T cell receptor alpha variable 15D-1-DV6D | 0 | 0 | 0 | 0 | 0 | 0 |
| 35329 | ENSMUSG00000096654 | 0 | NA | NA | NA | NA | NA | Mup-ps29 | major urinary protein, pseudogene 29 [So | 0 | 0 | 0 | 0 | 0 | 0 |
| 35330 | ENSMUSG00000096655 | 0.335414862 | -1.779047242 | 4.030804179 | -0.441362855 | 0.658950329 | NA | 1700065D16 | RIKEN cDNA 1700065D16 gene [Source:M | 0 | 0 | 2.0125 | 0 | 0 | 0 |
| 35331 | ENSMUSG00000096656 | 0 | NA | NA | NA | NA | NA | Trav12-2 | T cell receptor alpha variable 12-2 [Source | 0 | 0 | 0 | 0 | 0 | 0 |
| 35332 | ENSMUSG00000096658 | 0 | NA | NA | NA | NA | NA | Vmn2r39 | vomeronasal 2, receptor 39 [Source:MGI S | 0 | 0 | 0 | 0 | 0 | 0 |

| | | | | | | | | | | | | | | | |
|---|---|---|---|---|---|---|---|---|---|---|---|---|---|---|---|
| 35333 | ENSMUSG00000096659 | 0 | NA | NA | NA | NA | NA | NA | NA | 0 | 0 | 0 | 0 | 0 | 0 |
| 35334 | ENSMUSG00000096660 | 0 | NA | NA | NA | NA | NA | NA | NA | 0 | 0 | 0 | 0 | 0 | 0 |
| 35335 | ENSMUSG00000096661 | 0 | NA | NA | NA | NA | NA | Gm15069 | predicted gene 15069 [Source:MGI Symbo | 0 | 0 | 0 | 0 | 0 | 0 |
| 35336 | ENSMUSG00000096663 | 0 | NA | NA | NA | NA | NA | Vmn1r103 | vomeronasal 1 receptor 103 [Source:MGI | 0 | 0 | 0 | 0 | 0 | 0 |
| 35337 | ENSMUSG00000096664 | 0 | NA | NA | NA | NA | NA | Gm3409 | predicted gene 3409 [Source:MGI Symbol | 0 | 0 | 0 | 0 | 0 | 0 |
| 35338 | ENSMUSG00000096666 | 0 | NA | NA | NA | NA | NA | Gm20852 | predicted gene, 20852 [Source:MGI Symb | 0 | 0 | 0 | 0 | 0 | 0 |
| 35339 | ENSMUSG00000096667 | 0 | NA | NA | NA | NA | NA | NA | NA | 0 | 0 | 0 | 0 | 0 | 0 |
| 35340 | ENSMUSG00000096668 | 0 | NA | NA | NA | NA | NA | Gm3665 | predicted gene 3665 [Source:MGI Symbol | 0 | 0 | 0 | 0 | 0 | 0 |
| 35341 | ENSMUSG00000096670 | 0 | NA | NA | NA | NA | NA | Ighv2-6 | immunoglobulin heavy variable 2-6 [Sourc | 0 | 0 | 0 | 0 | 0 | 0 |
| 35342 | ENSMUSG00000096672 | 0.342639279 | -1.805976596 | 4.028547714 | -0.448294702 | 0.653940524 | NA | Ighv1-63 | immunoglobulin heavy variable V1-63 [So | 0 | 2.0558 | 0 | 0 | 0 | 0 |
| 35343 | ENSMUSG00000096673 | 0 | NA | NA | NA | NA | NA | NA | NA | 0 | 0 | 0 | 0 | 0 | 0 |
| 35344 | ENSMUSG00000096674 | 6145.927253 | 2.42949612 | 0.888316202 | 2.734945186 | 0.006239066 | 0.07732807 | Mup15 | major urinary protein 15 [Source:MGI Syn | 2027.8 | 587.97 | 3157.6 | 2937.2 | 15312 | 12853 |
| 35345 | ENSMUSG00000096675 | 0 | NA | NA | NA | NA | NA | NA | NA | 0 | 0 | 0 | 0 | 0 | 0 |
| 35346 | ENSMUSG00000096676 | 0 | NA | NA | NA | NA | NA | NA | NA | 0 | 0 | 0 | 0 | 0 | 0 |
| 35347 | ENSMUSG00000096677 | 0 | NA | NA | NA | NA | NA | NA | NA | 0 | 0 | 0 | 0 | 0 | 0 |
| 35348 | ENSMUSG00000096678 | 0 | NA | NA | NA | NA | NA | Trav9-4 | T cell receptor alpha variable 9-4 [Source: | 0 | 0 | 0 | 0 | 0 | 0 |
| 35349 | ENSMUSG00000096679 | 0 | NA | NA | NA | NA | NA | Olfr498 | olfactory receptor 498 [Source:MGI Symb | 0 | 0 | 0 | 0 | 0 | 0 |
| 35350 | ENSMUSG00000096680 | 0 | NA | NA | NA | NA | NA | | | 0 | 0 | 0 | 0 | 0 | 0 |
| 35351 | ENSMUSG00000096681 | 0 | NA | NA | NA | NA | NA | Rps19-ps1 | ribosomal protein S19, pseudogene 1 [Sou | 0 | 0 | 0 | 0 | 0 | 0 |
| 35352 | ENSMUSG00000096682 | 0 | NA | NA | NA | NA | NA | Ifna5 | interferon alpha 5 [Source:MGI Symbol;Ac | 0 | 0 | 0 | 0 | 0 | 0 |
| 35353 | ENSMUSG00000096683 | 0 | NA | NA | NA | NA | NA | NA | NA | 0 | 0 | 0 | 0 | 0 | 0 |
| 35354 | ENSMUSG00000096684 | 0 | NA | NA | NA | NA | NA | NA | NA | 0 | 0 | 0 | 0 | 0 | 0 |
| 35355 | ENSMUSG00000096685 | 0 | NA | NA | NA | NA | NA | Gm3642 | predicted gene 3642 [Source:MGI Symbol | 0 | 0 | 0 | 0 | 0 | 0 |
| 35356 | ENSMUSG00000096686 | 0 | NA | NA | NA | NA | NA | Gm20830 | predicted gene, 20830 [Source:MGI Symb | 0 | 0 | 0 | 0 | 0 | 0 |
| 35357 | ENSMUSG00000096687 | 89.19159396 | 0.027490485 | 0.25762398 | 0.106707789 | 0.915020803 | 0.98625062 | Mfsd4b4 | major facilitator superfamily domain cont | 91.554 | 75.038 | 98.612 | 107 | 89.431 | 73.514 |
| 35358 | ENSMUSG00000096688 | 5271.208159 | 2.629081364 | 0.924512955 | 2.843747455 | 0.004458637 | 0.06187407 | Mup17 | major urinary protein 17 [Source:MGI Syn | 2289.6 | 432.75 | 1678.4 | 2458.1 | 16017 | 8751.5 |
| 35359 | ENSMUSG00000096689 | 0 | NA | NA | NA | NA | NA | NA | NA | 0 | 0 | 0 | 0 | 0 | 0 |
| 35360 | ENSMUSG00000096690 | 0 | NA | NA | NA | NA | NA | Olfr1296-ps1 | olfactory receptor 1296, pseudogene 1 [So | 0 | 0 | 0 | 0 | 0 | 0 |
| 35361 | ENSMUSG00000096691 | 0 | NA | NA | NA | NA | NA | Vmn2r33 | vomeronasal 2, receptor 33 [Source:MGI S | 0 | 0 | 0 | 0 | 0 | 0 |
| 35362 | ENSMUSG00000096692 | 0 | NA | NA | NA | NA | NA | NA | NA | 0 | 0 | 0 | 0 | 0 | 0 |
| 35363 | ENSMUSG00000096694 | 0 | NA | NA | NA | NA | NA | NA | NA | 0 | 0 | 0 | 0 | 0 | 0 |
| 35364 | ENSMUSG00000096695 | 0 | NA | NA | NA | NA | NA | Olfr196 | olfactory receptor 196 [Source:MGI Symb | 0 | 0 | 0 | 0 | 0 | 0 |
| 35365 | ENSMUSG00000096696 | 59.17173858 | 0.975641746 | 0.326967083 | 2.983914276 | 0.002845865 | 0.045040215 | Zfp960 | zinc finger protein 960 [Source:MGI Symb | 42.564 | 33.921 | 43.269 | 105.07 | 57.803 | 72.4 |
| 35366 | ENSMUSG00000096697 | 0 | NA | NA | NA | NA | NA | Esp23 | exocrine gland secreted peptide 23 [Sourc | 0 | 0 | 0 | 0 | 0 | 0 |
| 35367 | ENSMUSG00000096698 | 0 | NA | NA | NA | NA | NA | Gm21780 | predicted gene, 21780 [Source:MGI Symb | 0 | 0 | 0 | 0 | 0 | 0 |
| 35368 | ENSMUSG00000096699 | 0.17131964 | -0.796629176 | 4.080472857 | -0.195229623 | 0.84521319 | NA | Rps19-ps4 | ribosomal protein S19, pseudogene 4 [Sou | 0 | 1.0279 | 0 | 0 | 0 | 0 |
| 35369 | ENSMUSG00000096700 | 0 | NA | NA | NA | NA | NA | Gm11236 | predicted gene 11236 [Source:MGI Symbo | 0 | 0 | 0 | 0 | 0 | 0 |
| 35370 | ENSMUSG00000096701 | 0 | NA | NA | NA | NA | NA | Gm9285 | predicted gene 9285 [Source:MGI Symbol | 0 | 0 | 0 | 0 | 0 | 0 |
| 35371 | ENSMUSG00000096703 | 0 | NA | NA | NA | NA | NA | Olfr1286 | olfactory receptor 1286 [Source:MGI Syml | 0 | 0 | 0 | 0 | 0 | 0 |
| 35372 | ENSMUSG00000096704 | 0 | NA | NA | NA | NA | NA | NA | NA | 0 | 0 | 0 | 0 | 0 | 0 |
| 35373 | ENSMUSG00000096705 | 0 | NA | NA | NA | NA | NA | Olfr1329 | olfactory receptor 1329 [Source:MGI Syml | 0 | 0 | 0 | 0 | 0 | 0 |
| 35374 | ENSMUSG00000096706 | 0 | NA | NA | NA | NA | NA | Gm21854 | predicted gene, 21854 [Source:MGI Symb | 0 | 0 | 0 | 0 | 0 | 0 |
| 35375 | ENSMUSG00000096707 | 0 | NA | NA | NA | NA | NA | Vmn2r-ps44 | vomeronasal 2, receptor, pseudogene 44 | 0 | 0 | 0 | 0 | 0 | 0 |
| 35376 | ENSMUSG00000096708 | 0 | NA | NA | NA | NA | NA | Olfr1475 | olfactory receptor 1475 [Source:MGI Syml | 0 | 0 | 0 | 0 | 0 | 0 |
| 35377 | ENSMUSG00000096709 | 0 | NA | NA | NA | NA | NA | Gm7352 | predicted gene 7352 [Source:MGI Symbol | 0 | 0 | 0 | 0 | 0 | 0 |
| 35378 | ENSMUSG00000096710 | 0 | NA | NA | NA | NA | NA | NA | NA | 0 | 0 | 0 | 0 | 0 | 0 |
| 35379 | ENSMUSG00000096711 | 0 | NA | NA | NA | NA | NA | Gm21599 | predicted gene, 21599 [Source:MGI Symb | 0 | 0 | 0 | 0 | 0 | 0 |
| 35380 | ENSMUSG00000096712 | 0.160662974 | 1.126924175 | 4.080472857 | 0.276174898 | 0.782413736 | NA | Gm15454 | predicted gene 15454 [Source:MGI Symbo | 0 | 0 | 0 | 0.964 | 0 | 0 |
| 35381 | ENSMUSG00000096713 | 0 | NA | NA | NA | NA | NA | NA | NA | 0 | 0 | 0 | 0 | 0 | 0 |
| 35382 | ENSMUSG00000096714 | 0 | NA | NA | NA | NA | NA | Olfr699 | olfactory receptor 699 [Source:MGI Symb | 0 | 0 | 0 | 0 | 0 | 0 |
| 35383 | ENSMUSG00000096715 | 0.371281295 | 2.09253362 | 4.031852165 | 0.519000582 | 0.603760335 | NA | Igkv3-4 | immunoglobulin kappa variable 3-4 [Sourc | 0 | 0 | 0 | 0 | 0 | 2.2277 |
| 35384 | ENSMUSG00000096716 | 0 | NA | NA | NA | NA | NA | Gm5755 | predicted pseudogene 5755 [Source:MGI | 0 | 0 | 0 | 0 | 0 | 0 |
| 35385 | ENSMUSG00000096717 | 0 | NA | NA | NA | NA | NA | Vmn2r98 | vomeronasal 2, receptor 98 [Source:MGI S | 0 | 0 | 0 | 0 | 0 | 0 |
| 35386 | ENSMUSG00000096718 | 1.805578744 | -3.094561821 | 1.874781216 | -1.650625574 | 0.098815053 | NA | Zfp781 | zinc finger protein 781 [Source:MGI Symb | 1.6062 | 4.1117 | 4.025 | 0 | 1.0906 | 0 |
| 35387 | ENSMUSG00000096719 | 0.321325949 | 1.911575114 | 4.048267893 | 0.472195804 | 0.636787034 | NA | Mrgpra2b | MAS-related GPR, member A2B [Source:N | 0 | 0 | 0 | 1.928 | 0 | 0 |
| 35388 | ENSMUSG00000096721 | 0 | NA | NA | NA | NA | NA | NA | NA | 0 | 0 | 0 | 0 | 0 | 0 |

| | | | | | | | | | | | | | | | |
|---|---|---|---|---|---|---|---|---|---|---|---|---|---|---|---|---|
| 35389 | ENSMUSG00000096722 | 0 | NA | NA | NA | NA | NA | Hmgb1-ps9 | high mobility group box 1, pseudogene 9 [ | 0 | 0 | 0 | 0 | 0 | 0 |
| 35390 | ENSMUSG00000096724 | 0 | NA | NA | NA | NA | NA | Gm1989 | predicted gene 1989 [Source:MGI Symbol | 0 | 0 | 0 | 0 | 0 | 0 |
| 35391 | ENSMUSG00000096726 | 0.706721132 | -2.881789029 | 2.959064625 | -0.973885127 | 0.330113579 | NA | Gm5558 | predicted gene 5558 [Source:MGI Symbol | 3.2124 | 1.0279 | 0 | 0 | 0 | 0 |
| 35392 | ENSMUSG00000096727 | 550.2805535 | 0.187188444 | 0.184904779 | 1.012350495 | 0.3113705 | 0.759810187 | Psmb9 | proteasome (prosome, macropain) subun | 631.24 | 443.03 | 467.9 | 446.32 | 629.29 | 683.9 |
| 35393 | ENSMUSG00000096728 | 0 | NA | NA | NA | NA | NA | | | 0 | 0 | 0 | 0 | 0 | 0 |
| 35394 | ENSMUSG00000096729 | 0 | NA | NA | NA | NA | NA | Gm13695 | predicted gene 13695 [Source:MGI Symbo | 0 | 0 | 0 | 0 | 0 | 0 |
| 35395 | ENSMUSG00000096730 | 0 | NA | NA | NA | NA | NA | NA | NA | 0 | 0 | 0 | 0 | 0 | 0 |
| 35396 | ENSMUSG00000096731 | 0 | NA | NA | NA | NA | NA | NA | NA | 0 | 0 | 0 | 0 | 0 | 0 |
| 35397 | ENSMUSG00000096732 | 0 | NA | NA | NA | NA | NA | Gm8897 | predicted gene 8897 [Source:MGI Symbol | 0 | 0 | 0 | 0 | 0 | 0 |
| 35398 | ENSMUSG00000096733 | 0 | NA | NA | NA | NA | NA | Gm15257 | predicted gene 15257 [Source:MGI Symbo | 0 | 0 | 0 | 0 | 0 | 0 |
| 35399 | ENSMUSG00000096734 | 0 | NA | NA | NA | NA | NA | Gm21836 | predicted gene, 21836 [Source:MGI Symb | 0 | 0 | 0 | 0 | 0 | 0 |
| 35400 | ENSMUSG00000096735 | 0 | NA | NA | NA | NA | NA | Vmn1r79 | vomeronasal 1 receptor 79 [Source:MGI S | 0 | 0 | 0 | 0 | 0 | 0 |
| 35401 | ENSMUSG00000096736 | 0 | NA | NA | NA | NA | NA | Gm17535 | predicted gene, 17535 [Source:MGI Symb | 0 | 0 | 0 | 0 | 0 | 0 |
| 35402 | ENSMUSG00000096737 | 0 | NA | NA | NA | NA | NA | Vmn1r117 | vomeronasal 1 receptor 117 [Source:MGI | 0 | 0 | 0 | 0 | 0 | 0 |
| 35403 | ENSMUSG00000096738 | 0 | NA | NA | NA | NA | NA | Gm24317 | predicted gene, 24317 [Source:MGI Symb | 0 | 0 | 0 | 0 | 0 | 0 |
| 35404 | ENSMUSG00000096740 | 48.56875116 | -0.154354409 | 0.347562918 | -0.44410494 | 0.656966715 | 0.926969717 | Lbhd1 | LBH domain containing 1 [Source:MGI Syr | 44.974 | 55.508 | 53.331 | 53.019 | 55.622 | 28.96 |
| 35405 | ENSMUSG00000096741 | 0 | NA | NA | NA | NA | NA | Vmn1r-ps76 | vomeronasal 1 receptor, pseudogene 76 [ | 0 | 0 | 0 | 0 | 0 | 0 |
| 35406 | ENSMUSG00000096742 | 0 | NA | NA | NA | NA | NA | Gm6367 | predicted gene 6367 [Source:MGI Symbol | 0 | 0 | 0 | 0 | 0 | 0 |
| 35407 | ENSMUSG00000096743 | 0 | NA | NA | NA | NA | NA | Vmn2r32 | vomeronasal 2, receptor 32 [Source:MGI ! | 0 | 0 | 0 | 0 | 0 | 0 |
| 35408 | ENSMUSG00000096744 | 0 | NA | NA | NA | NA | NA | Gm4307 | predicted gene 4307 [Source:MGI Symbol | 0 | 0 | 0 | 0 | 0 | 0 |
| 35409 | ENSMUSG00000096745 | 0 | NA | NA | NA | NA | NA | Gm12622 | predicted gene 12622 [Source:MGI Symbo | 0 | 0 | 0 | 0 | 0 | 0 |
| 35410 | ENSMUSG00000096746 | 0 | NA | NA | NA | NA | NA | Trav7d-3 | T cell receptor alpha variable 7D-3 [Sourc | 0 | 0 | 0 | 0 | 0 | 0 |
| 35411 | ENSMUSG00000096747 | 0 | NA | NA | NA | NA | NA | Olfr823 | olfactory receptor 823 [Source:MGI Symb | 0 | 0 | 0 | 0 | 0 | 0 |
| 35412 | ENSMUSG00000096748 | 0 | NA | NA | NA | NA | NA | NA | NA | 0 | 0 | 0 | 0 | 0 | 0 |
| 35413 | ENSMUSG00000096749 | 0 | NA | NA | NA | NA | NA | Trdd1 | T cell receptor delta diversity 1 [Source:M | 0 | 0 | 0 | 0 | 0 | 0 |
| 35414 | ENSMUSG00000096750 | 0 | NA | NA | NA | NA | NA | Gm11757 | predicted gene 11757 [Source:MGI Symbo | 0 | 0 | 0 | 0 | 0 | 0 |
| 35415 | ENSMUSG00000096751 | 1.809423047 | -0.681912948 | 1.561604424 | -0.436674575 | 0.662347371 | NA | Gm28373 | predicted gene 28373 [Source:MGI Symbo | 1.6062 | 4.1117 | 1.0062 | 1.928 | 1.0906 | 1.1138 |
| 35416 | ENSMUSG00000096753 | 0.79190315 | 0.90205007 | 2.632846219 | 0.342614036 | 0.731888841 | NA | Fam181a | family with sequence similarity 181, meml | 1.6062 | 0 | 0 | 0.964 | 2.1812 | 0 |
| 35417 | ENSMUSG00000096755 | 1.756087586 | 0.920606104 | 1.829689879 | 0.503148711 | 0.614859722 | NA | Ftl1-ps2 | ferritin light polypeptide, pseudogene 2 [S | 1.6062 | 2.0558 | 0 | 5.7839 | 1.0906 | 0 |
| 35418 | ENSMUSG00000096756 | 0 | NA | NA | NA | NA | NA | | | 0 | 0 | 0 | 0 | 0 | 0 |
| 35419 | ENSMUSG00000096757 | 0 | NA | NA | NA | NA | NA | Olfr251 | olfactory receptor 251 [Source:MGI Symb | 0 | 0 | 0 | 0 | 0 | 0 |
| 35420 | ENSMUSG00000096758 | 0 | NA | NA | NA | NA | NA | NA | NA | 0 | 0 | 0 | 0 | 0 | 0 |
| 35421 | ENSMUSG00000096759 | 0 | NA | NA | NA | NA | NA | NA | NA | 0 | 0 | 0 | 0 | 0 | 0 |
| 35422 | ENSMUSG00000096760 | 0 | NA | NA | NA | NA | NA | Vmn1r248 | vomeronasal 1 receptor 248 [Source:MGI | 0 | 0 | 0 | 0 | 0 | 0 |
| 35423 | ENSMUSG00000096761 | 0 | NA | NA | NA | NA | NA | Vmn1r245 | vomeronasal 1 receptor 245 [Source:MGI | 0 | 0 | 0 | 0 | 0 | 0 |
| 35424 | ENSMUSG00000096763 | 0 | NA | NA | NA | NA | NA | Rps12-ps13 | ribosomal protein S12, pseudogene 13 [Sc | 0 | 0 | 0 | 0 | 0 | 0 |
| 35425 | ENSMUSG00000096764 | 0 | NA | NA | NA | NA | NA | Gm21985 | predicted gene 21985 [Source:MGI Symbo | 0 | 0 | 0 | 0 | 0 | 0 |
| 35426 | ENSMUSG00000096765 | 0 | NA | NA | NA | NA | NA | NA | NA | 0 | 0 | 0 | 0 | 0 | 0 |
| 35427 | ENSMUSG00000096766 | 0.17131964 | -0.796629176 | 4.080472857 | -0.195229623 | 0.84521319 | NA | Gm23793 | predicted gene, 23793 [Source:MGI Symb | 0 | 1.0279 | 0 | 0 | 0 | 0 |
| 35428 | ENSMUSG00000096767 | 0 | NA | NA | NA | NA | NA | Ighv1-62-3 | immunoglobulin heavy variable 1-62-3 [Sc | 0 | 0 | 0 | 0 | 0 | 0 |
| 35429 | ENSMUSG00000096768 | 53.10758167 | -1.375478851 | 1.416912452 | -0.970757826 | 0.331668889 | 0.774747658 | Gm47283 | predicted gene, 47283 [Source:MGI Symb | 95.569 | 122.32 | 12.075 | 24.099 | 1.0906 | 63.489 |
| 35430 | ENSMUSG00000096769 | 0 | NA | NA | NA | NA | NA | Gm21117 | predicted gene, 21117 [Source:MGI Symb | 0 | 0 | 0 | 0 | 0 | 0 |
| 35431 | ENSMUSG00000096770 | 0 | NA | NA | NA | NA | NA | Amy2a4 | amylase 2a4 [Source:MGI Symbol;Acc:MG | 0 | 0 | 0 | 0 | 0 | 0 |
| 35432 | ENSMUSG00000096771 | 0 | NA | NA | NA | NA | NA | NA | NA | 0 | 0 | 0 | 0 | 0 | 0 |
| 35433 | ENSMUSG00000096772 | 0 | NA | NA | NA | NA | NA | NA | NA | 0 | 0 | 0 | 0 | 0 | 0 |
| 35434 | ENSMUSG00000096773 | 0 | NA | NA | NA | NA | NA | Olfr675 | olfactory receptor 675 [Source:MGI Symb | 0 | 0 | 0 | 0 | 0 | 0 |
| 35435 | ENSMUSG00000096774 | 0 | NA | NA | NA | NA | NA | NA | NA | 0 | 0 | 0 | 0 | 0 | 0 |
| 35436 | ENSMUSG00000096775 | 0 | NA | NA | NA | NA | NA | Gm5796 | predicted gene 5796 [Source:MGI Symbol | 0 | 0 | 0 | 0 | 0 | 0 |
| 35437 | ENSMUSG00000096776 | 0 | NA | NA | NA | NA | NA | | vomeronasal 1 receptor Vmn1r186 [Sourc | 0 | 0 | 0 | 0 | 0 | 0 |
| 35438 | ENSMUSG00000096777 | 0 | NA | NA | NA | NA | NA | Gm21723 | predicted gene, 21723 [Source:MGI Symb | 0 | 0 | 0 | 0 | 0 | 0 |
| 35439 | ENSMUSG00000096779 | 0 | NA | NA | NA | NA | NA | NA | NA | 0 | 0 | 0 | 0 | 0 | 0 |
| 35440 | ENSMUSG00000096780 | 15.06808473 | -0.889473314 | 0.623269255 | -1.427109242 | 0.153548409 | 0.570925073 | Tmem181b-| | transmembrane protein 181B, pseudogen | 28.912 | 19.53 | 10.062 | 8.6758 | 7.6343 | 15.594 |
| 35441 | ENSMUSG00000096782 | 0 | NA | NA | NA | NA | NA | NA | NA | 0 | 0 | 0 | 0 | 0 | 0 |
| 35442 | ENSMUSG00000096784 | 0 | NA | NA | NA | NA | NA | NA | NA | 0 | 0 | 0 | 0 | 0 | 0 |
| 35443 | ENSMUSG00000096785 | 0 | NA | NA | NA | NA | NA | NA | NA | 0 | 0 | 0 | 0 | 0 | 0 |
| 35444 | ENSMUSG00000096786 | 0 | NA | NA | NA | NA | NA | NA | NA | 0 | 0 | 0 | 0 | 0 | 0 |

| | | | | | | | | | | | | | | | |
|---|---|---|---|---|---|---|---|---|---|---|---|---|---|---|---|
| 35445 | ENSMUSG00000096788 | 0 | NA | NA | NA | NA | NA | Rhox2e | reproductive homeobox 2E [Source:MGI S | 0 | 0 | 0 | 0 | 0 | 0 |
| 35446 | ENSMUSG00000096790 | 0 | NA | NA | NA | NA | NA | NA | NA | 0 | 0 | 0 | 0 | 0 | 0 |
| 35447 | ENSMUSG00000096792 | 0 | NA | NA | NA | NA | NA | Gm21956 | predicted gene, 21956 [Source:MGI Symb | 0 | 0 | 0 | 0 | 0 | 0 |
| 35448 | ENSMUSG00000096793 | 0 | NA | NA | NA | NA | NA | Gm3002 | predicted gene 3002 [Source:MGI Symbol | 0 | 0 | 0 | 0 | 0 | 0 |
| 35449 | ENSMUSG00000096794 | 0 | NA | NA | NA | NA | NA | Olfr905 | olfactory receptor 905 [Source:MGI Symb | 0 | 0 | 0 | 0 | 0 | 0 |
| 35450 | ENSMUSG00000096795 | 10.93381348 | 0.666706564 | 0.64490354 | 1.033808195 | 0.30122584 | 0.7521219 | Zfp433 | zinc finger protein 433 [Source:MGI Symb | 8.031 | 10.279 | 7.0437 | 11.568 | 13.087 | 15.594 |
| 35451 | ENSMUSG00000096796 | 0 | NA | NA | NA | NA | NA | 1700021P04 | RIKEN cDNA 1700021P04 gene [Source:M | 0 | 0 | 0 | 0 | 0 | 0 |
| 35452 | ENSMUSG00000096797 | 0 | NA | NA | NA | NA | NA | Obox4-ps20 | oocyte specific homeobox 4, pseudogene | 0 | 0 | 0 | 0 | 0 | 0 |
| 35453 | ENSMUSG00000096798 | 0 | NA | NA | NA | NA | NA | Gm6309 | predicted gene 6309 [Source:MGI Symbol | 0 | 0 | 0 | 0 | 0 | 0 |
| 35454 | ENSMUSG00000096799 | 0 | NA | NA | NA | NA | NA | NA | NA | 0 | 0 | 0 | 0 | 0 | 0 |
| 35455 | ENSMUSG00000096800 | 0 | NA | NA | NA | NA | NA | Gm4725 | predicted gene 4725 [Source:MGI Symbol | 0 | 0 | 0 | 0 | 0 | 0 |
| 35456 | ENSMUSG00000096801 | 0 | NA | NA | NA | NA | NA | NA | NA | 0 | 0 | 0 | 0 | 0 | 0 |
| 35457 | ENSMUSG00000096802 | 2.780651939 | -0.464684765 | 1.436595952 | -0.323462393 | 0.746345084 | NA | NA | NA | 1.6062 | 6.1675 | 2.0125 | 5.7839 | 0 | 1.1138 |
| 35458 | ENSMUSG00000096803 | 0 | NA | NA | NA | NA | NA | Eif1ad16 | eukaryotic translation initiation factor 1A | 0 | 0 | 0 | 0 | 0 | 0 |
| 35459 | ENSMUSG00000096804 | 0 | NA | NA | NA | NA | NA | NA | NA | 0 | 0 | 0 | 0 | 0 | 0 |
| 35460 | ENSMUSG00000096805 | 0 | NA | NA | NA | NA | NA | Ighv9-1 | immunoglobulin heavy variable 9-1 [Sourc | 0 | 0 | 0 | 0 | 0 | 0 |
| 35461 | ENSMUSG00000096806 | 0 | NA | NA | NA | NA | NA | Olfr312 | olfactory receptor 312 [Source:MGI Symb | 0 | 0 | 0 | 0 | 0 | 0 |
| 35462 | ENSMUSG00000096808 | 0 | NA | NA | NA | NA | NA | | | 0 | 0 | 0 | 0 | 0 | 0 |
| 35463 | ENSMUSG00000096809 | 0 | NA | NA | NA | NA | NA | NA | NA | 0 | 0 | 0 | 0 | 0 | 0 |
| 35464 | ENSMUSG00000096810 | 0.167707431 | -0.796629176 | 4.080472857 | -0.195229623 | 0.84521319 | NA | Gm10481 | predicted gene 10481 [Source:MGI Symbo | 0 | 0 | 1.0062 | 0 | 0 | 0 |
| 35465 | ENSMUSG00000096811 | 0 | NA | NA | NA | NA | NA | NA | NA | 0 | 0 | 0 | 0 | 0 | 0 |
| 35466 | ENSMUSG00000096812 | 0 | NA | NA | NA | NA | NA | NA | NA | 0 | 0 | 0 | 0 | 0 | 0 |
| 35467 | ENSMUSG00000096813 | 0 | NA | NA | NA | NA | NA | Vmn1r126 | vomeronasal 1 receptor 126 [Source:MGI | 0 | 0 | 0 | 0 | 0 | 0 |
| 35468 | ENSMUSG00000096814 | 0 | NA | NA | NA | NA | NA | NA | NA | 0 | 0 | 0 | 0 | 0 | 0 |
| 35469 | ENSMUSG00000096815 | 0 | NA | NA | NA | NA | NA | NA | NA | 0 | 0 | 0 | 0 | 0 | 0 |
| 35470 | ENSMUSG00000096817 | 0 | NA | NA | NA | NA | NA | NA | NA | 0 | 0 | 0 | 0 | 0 | 0 |
| 35471 | ENSMUSG00000096818 | 0 | NA | NA | NA | NA | NA | Gm29231 | predicted gene 29231 [Source:MGI Symbo | 0 | 0 | 0 | 0 | 0 | 0 |
| 35472 | ENSMUSG00000096819 | 0 | NA | NA | NA | NA | NA | Srsy | serine-rich, secreted, Y-linked [Source:MG | 0 | 0 | 0 | 0 | 0 | 0 |
| 35473 | ENSMUSG00000096820 | 0 | NA | NA | NA | NA | NA | Gm21425 | predicted gene, 21425 [Source:MGI Symb | 0 | 0 | 0 | 0 | 0 | 0 |
| 35474 | ENSMUSG00000096821 | 0 | NA | NA | NA | NA | NA | NA | NA | 0 | 0 | 0 | 0 | 0 | 0 |
| 35475 | ENSMUSG00000096822 | 0 | NA | NA | NA | NA | NA | Olfr344 | olfactory receptor 344 [Source:MGI Symb | 0 | 0 | 0 | 0 | 0 | 0 |
| 35476 | ENSMUSG00000096823 | 0 | NA | NA | NA | NA | NA | NA | NA | 0 | 0 | 0 | 0 | 0 | 0 |
| 35477 | ENSMUSG00000096824 | 0 | NA | NA | NA | NA | NA | Ighv2-7 | immunoglobulin heavy variable 2-7 [Sourc | 0 | 0 | 0 | 0 | 0 | 0 |
| 35478 | ENSMUSG00000096825 | 0 | NA | NA | NA | NA | NA | Trav12-1 | T cell receptor alpha variable 12-1 [Source | 0 | 0 | 0 | 0 | 0 | 0 |
| 35479 | ENSMUSG00000096826 | 0 | NA | NA | NA | NA | NA | Ccl27b | chemokine (C-C motif) ligand 27b [Source | 0 | 0 | 0 | 0 | 0 | 0 |
| 35480 | ENSMUSG00000096827 | 0 | NA | NA | NA | NA | NA | Trav14n-1 | T cell receptor alpha variable 14N-1 [Sour | 0 | 0 | 0 | 0 | 0 | 0 |
| 35481 | ENSMUSG00000096828 | 0 | NA | NA | NA | NA | NA | Gm3618 | predicted gene 3618 [Source:MGI Symbol | 0 | 0 | 0 | 0 | 0 | 0 |
| 35482 | ENSMUSG00000096829 | 0 | NA | NA | NA | NA | NA | NA | NA | 0 | 0 | 0 | 0 | 0 | 0 |
| 35483 | ENSMUSG00000096830 | 0 | NA | NA | NA | NA | NA | NA | NA | 0 | 0 | 0 | 0 | 0 | 0 |
| 35484 | ENSMUSG00000096831 | 0 | NA | NA | NA | NA | NA | Rps12-ps6 | ribosomal protein S12, pseudogene 6 [Sou | 0 | 0 | 0 | 0 | 0 | 0 |
| 35485 | ENSMUSG00000096832 | 0 | NA | NA | NA | NA | NA | NA | NA | 0 | 0 | 0 | 0 | 0 | 0 |
| 35486 | ENSMUSG00000096833 | 0 | NA | NA | NA | NA | NA | Igkv4-55 | immunoglobulin kappa variable 4-55 [Sou | 0 | 0 | 0 | 0 | 0 | 0 |
| 35487 | ENSMUSG00000096834 | 0 | NA | NA | NA | NA | NA | Gm15127 | predicted gene 15127 [Source:MGI Symbo | 0 | 0 | 0 | 0 | 0 | 0 |
| 35488 | ENSMUSG00000096835 | 0 | NA | NA | NA | NA | NA | NA | NA | 0 | 0 | 0 | 0 | 0 | 0 |
| 35489 | ENSMUSG00000096838 | 0 | NA | NA | NA | NA | NA | NA | NA | 0 | 0 | 0 | 0 | 0 | 0 |
| 35490 | ENSMUSG00000096839 | 0 | NA | NA | NA | NA | NA | Gm3460 | predicted gene 3460 [Source:MGI Symbol | 0 | 0 | 0 | 0 | 0 | 0 |
| 35491 | ENSMUSG00000096840 | 0 | NA | NA | NA | NA | NA | Olfr136 | olfactory receptor 136 [Source:MGI Symb | 0 | 0 | 0 | 0 | 0 | 0 |
| 35492 | ENSMUSG00000096841 | 4.944317463 | -1.034195947 | 1.002502607 | -1.031614222 | 0.302252875 | NA | NA | NA | 5.6217 | 7.1954 | 7.0437 | 0.964 | 3.2719 | 5.5692 |
| 35493 | ENSMUSG00000096842 | 73.6788067 | 0.331070704 | 0.350494565 | 0.944581563 | 0.344872544 | 0.786033695 | Gm10736 | predicted gene 10736 [Source:MGI Symbo | 38.549 | 92.513 | 65.406 | 96.398 | 54.531 | 94.677 |
| 35494 | ENSMUSG00000096844 | 0.513958919 | -2.394274821 | 3.756096013 | -0.637437066 | 0.523840191 | NA | Igkv6-14 | immunoglobulin kappa variable 6-14 [Sou | 0 | 3.0838 | 0 | 0 | 0 | 0 |
| 35495 | ENSMUSG00000096845 | 0 | NA | NA | NA | NA | NA | NA | NA | 0 | 0 | 0 | 0 | 0 | 0 |
| 35496 | ENSMUSG00000096846 | 0 | NA | NA | NA | NA | NA | Gm8890 | predicted gene 8890 [Source:MGI Symbol | 0 | 0 | 0 | 0 | 0 | 0 |
| 35497 | ENSMUSG00000096847 | 0 | NA | NA | NA | NA | NA | Tmem151b | transmembrane protein 151B [Source:MG | 0 | 0 | 0 | 0 | 0 | 0 |
| 35498 | ENSMUSG00000096848 | 0 | NA | NA | NA | NA | NA | NA | NA | 0 | 0 | 0 | 0 | 0 | 0 |
| 35499 | ENSMUSG00000096849 | 0 | NA | NA | NA | NA | NA | NA | NA | 0 | 0 | 0 | 0 | 0 | 0 |
| 35500 | ENSMUSG00000096850 | 0 | NA | NA | NA | NA | NA | Gm21748 | predicted gene, 21748 [Source:MGI Symb | 0 | 0 | 0 | 0 | 0 | 0 |

| | | | | | | | | | | | | | | | |
|---|---|---|---|---|---|---|---|---|---|---|---|---|---|---|---|
| 35501 | ENSMUSG00000096851 | 0 | NA | NA | NA | NA | NA | NA | NA | 0 | 0 | 0 | 0 | 0 | 0 |
| 35502 | ENSMUSG00000096852 | 9.217036187 | -0.440094683 | 0.758725148 | -0.58004494 | 0.561884312 | 0.900565925 | Cyp2d12 | cytochrome P450, family 2, subfamily d, p | 10.44 | 9.2513 | 12.075 | 5.7839 | 3.2719 | 14.48 |
| 35503 | ENSMUSG00000096854 | 0 | NA | NA | NA | NA | NA | Ifnz | interferon zeta [Source:MGI Symbol;Acc:N | 0 | 0 | 0 | 0 | 0 | 0 |
| 35504 | ENSMUSG00000096855 | 0 | NA | NA | NA | NA | NA | NA | NA | 0 | 0 | 0 | 0 | 0 | 0 |
| 35505 | ENSMUSG00000096856 | 4.308559425 | -0.800851061 | 1.057106347 | -0.757587979 | 0.448697679 | NA | Gm10778 | predicted gene 10778 [Source:MGI Symbo | 7.2279 | 5.1396 | 4.025 | 2.8919 | 5.4531 | 1.1138 |
| 35506 | ENSMUSG00000096857 | 0 | NA | NA | NA | NA | NA | Gm7732 | predicted gene 7732 [Source:MGI Symbol | 0 | 0 | 0 | 0 | 0 | 0 |
| 35507 | ENSMUSG00000096858 | 0 | NA | NA | NA | NA | NA | Olfr805 | olfactory receptor 805 [Source:MGI Symb | 0 | 0 | 0 | 0 | 0 | 0 |
| 35508 | ENSMUSG00000096859 | 0 | NA | NA | NA | NA | NA | Vmn1r176 | vomeronasal 1 receptor 176 [Source:MGI | 0 | 0 | 0 | 0 | 0 | 0 |
| 35509 | ENSMUSG00000096861 | 0 | NA | NA | NA | NA | NA | Vmn2r68 | vomeronasal 2, receptor 68 [Source:MGI S | 0 | 0 | 0 | 0 | 0 | 0 |
| 35510 | ENSMUSG00000096862 | 3.09435999 | -0.59293964 | 1.393454082 | -0.425517889 | 0.670459183 | NA | Gm13301 | predicted gene 13301 [Source:MGI Symbo | 0 | 8.2233 | 3.0187 | 2.8919 | 1.0906 | 3.3415 |
| 35511 | ENSMUSG00000096863 | 0 | NA | NA | NA | NA | NA | NA | NA | 0 | 0 | 0 | 0 | 0 | 0 |
| 35512 | ENSMUSG00000096864 | 0 | NA | NA | NA | NA | NA | Gm21978 | predicted gene 21978 [Source:MGI Symbo | 0 | 0 | 0 | 0 | 0 | 0 |
| 35513 | ENSMUSG00000096865 | 0 | NA | NA | NA | NA | NA | NA | NA | 0 | 0 | 0 | 0 | 0 | 0 |
| 35514 | ENSMUSG00000096866 | 0 | NA | NA | NA | NA | NA | NA | NA | 0 | 0 | 0 | 0 | 0 | 0 |
| 35515 | ENSMUSG00000096867 | 0 | NA | NA | NA | NA | NA | Samt1c | spermatogenesis associated multipass tra | 0 | 0 | 0 | 0 | 0 | 0 |
| 35516 | ENSMUSG00000096868 | 0 | NA | NA | NA | NA | NA | NA | NA | 0 | 0 | 0 | 0 | 0 | 0 |
| 35517 | ENSMUSG00000096869 | 0 | NA | NA | NA | NA | NA | Gm6676 | predicted gene 6676 [Source:MGI Symbol | 0 | 0 | 0 | 0 | 0 | 0 |
| 35518 | ENSMUSG00000096870 | 0 | NA | NA | NA | NA | NA | Gm21816 | predicted gene, 21816 [Source:MGI Symb | 0 | 0 | 0 | 0 | 0 | 0 |
| 35519 | ENSMUSG00000096871 | 0 | NA | NA | NA | NA | NA | Gm10665 | predicted gene 10665 [Source:MGI Symbo | 0 | 0 | 0 | 0 | 0 | 0 |
| 35520 | ENSMUSG00000096872 | 0 | NA | NA | NA | NA | NA | Scgb2a2 | secretoglobin, family 2A, member 2 [Sour | 0 | 0 | 0 | 0 | 0 | 0 |
| 35521 | ENSMUSG00000096873 | 0 | NA | NA | NA | NA | NA | NA | NA | 0 | 0 | 0 | 0 | 0 | 0 |
| 35522 | ENSMUSG00000096874 | 0 | NA | NA | NA | NA | NA | Gm10312 | predicted gene 10312 [Source:MGI Symbo | 0 | 0 | 0 | 0 | 0 | 0 |
| 35523 | ENSMUSG00000096875 | 0 | NA | NA | NA | NA | NA | Gm21815 | predicted gene, 21815 [Source:MGI Symb | 0 | 0 | 0 | 0 | 0 | 0 |
| 35524 | ENSMUSG00000096876 | 0 | NA | NA | NA | NA | NA | NA | NA | 0 | 0 | 0 | 0 | 0 | 0 |
| 35525 | ENSMUSG00000096878 | 0 | NA | NA | NA | NA | NA | Gm21083 | predicted gene, 21083 [Source:MGI Symb | 0 | 0 | 0 | 0 | 0 | 0 |
| 35526 | ENSMUSG00000096879 | 0 | NA | NA | NA | NA | NA | Tdpoz8 | TD and POZ domain containing 8 [Source: | 0 | 0 | 0 | 0 | 0 | 0 |
| 35527 | ENSMUSG00000096880 | 0 | NA | NA | NA | NA | NA | NA | NA | 0 | 0 | 0 | 0 | 0 | 0 |
| 35528 | ENSMUSG00000096881 | 0 | NA | NA | NA | NA | NA | Gm20874 | predicted gene, 20874 [Source:MGI Symb | 0 | 0 | 0 | 0 | 0 | 0 |
| 35529 | ENSMUSG00000096882 | 0 | NA | NA | NA | NA | NA | NA | NA | 0 | 0 | 0 | 0 | 0 | 0 |
| 35530 | ENSMUSG00000096883 | 49.39996139 | -2.532879886 | 0.467962072 | -5.412575156 | 6.21E-08 | 7.13E-06 | Shisa8 | shisa family member 8 [Source:MGI Symb | 47.383 | 94.568 | 110.69 | 8.6758 | 26.175 | 8.9108 |
| 35531 | ENSMUSG00000096884 | 0 | NA | NA | NA | NA | NA | Ighd5-4 | immunoglobulin heavy diversity 5-4 [Sour | 0 | 0 | 0 | 0 | 0 | 0 |
| 35532 | ENSMUSG00000096885 | 0 | NA | NA | NA | NA | NA | Gm21797 | predicted gene, 21797 [Source:MGI Symb | 0 | 0 | 0 | 0 | 0 | 0 |
| 35533 | ENSMUSG00000096887 | 0 | NA | NA | NA | NA | NA | Gm20594 | predicted gene, 20594 [Source:MGI Symb | 0 | 0 | 0 | 0 | 0 | 0 |
| 35534 | ENSMUSG00000096889 | 0 | NA | NA | NA | NA | NA | NA | NA | 0 | 0 | 0 | 0 | 0 | 0 |
| 35535 | ENSMUSG00000096890 | 0.267700746 | -1.497481649 | 4.057001436 | -0.369110456 | 0.712045396 | NA | Gm20797 | predicted gene, 20797 [Source:MGI Symb | 1.6062 | 0 | 0 | 0 | 0 | 0 |
| 35536 | ENSMUSG00000096891 | 10.53387997 | 1.880075919 | 0.802430632 | 2.342976256 | 0.019130601 | 0.168170501 | Mup-ps8 | major urinary protein, pseudogene 8 [Sou | 7.2279 | 0 | 6.0375 | 12.532 | 21.812 | 15.594 |
| 35537 | ENSMUSG00000096893 | 0 | NA | NA | NA | NA | NA | NA | NA | 0 | 0 | 0 | 0 | 0 | 0 |
| 35538 | ENSMUSG00000096894 | 0 | NA | NA | NA | NA | NA | NA | NA | 0 | 0 | 0 | 0 | 0 | 0 |
| 35539 | ENSMUSG00000096895 | 0 | NA | NA | NA | NA | NA | NA | NA | 0 | 0 | 0 | 0 | 0 | 0 |
| 35540 | ENSMUSG00000096896 | 0 | NA | NA | NA | NA | NA | Gm25244 | predicted gene, 25244 [Source:MGI Symb | 0 | 0 | 0 | 0 | 0 | 0 |
| 35541 | ENSMUSG00000096897 | 0 | NA | NA | NA | NA | NA | Astx4d | amplified spermatogenic transcripts X enc | 0 | 0 | 0 | 0 | 0 | 0 |
| 35542 | ENSMUSG00000096898 | 0 | NA | NA | NA | NA | NA | Gm20867 | predicted gene, 20867 [Source:MGI Symb | 0 | 0 | 0 | 0 | 0 | 0 |
| 35543 | ENSMUSG00000096899 | 0 | NA | NA | NA | NA | NA | Gm8346 | predicted gene 8346 [Source:MGI Symbol | 0 | 0 | 0 | 0 | 0 | 0 |
| 35544 | ENSMUSG00000096900 | 0 | NA | NA | NA | NA | NA | Trav9-1 | T cell receptor alpha variable 9-1 [Source: | 0 | 0 | 0 | 0 | 0 | 0 |
| 35545 | ENSMUSG00000096901 | 0 | NA | NA | NA | NA | NA | Gm3029 | predicted gene 3029 [Source:MGI Symbol | 0 | 0 | 0 | 0 | 0 | 0 |
| 35546 | ENSMUSG00000096902 | 0 | NA | NA | NA | NA | NA | Gm20843 | predicted gene, 20843 [Source:MGI Symb | 0 | 0 | 0 | 0 | 0 | 0 |
| 35547 | ENSMUSG00000096903 | 0 | NA | NA | NA | NA | NA | Vmn1r104 | vomeronasal 1 receptor 104 [Source:MGI | 0 | 0 | 0 | 0 | 0 | 0 |
| 35548 | ENSMUSG00000096904 | 0 | NA | NA | NA | NA | NA | Lamtor3-ps | late endosomal/lysosomal adaptor, MAPK | 0 | 0 | 0 | 0 | 0 | 0 |
| 35549 | ENSMUSG00000096905 | 0 | NA | NA | NA | NA | NA | Gm8324 | predicted gene 8324 [Source:MGI Symbol | 0 | 0 | 0 | 0 | 0 | 0 |
| 35550 | ENSMUSG00000096906 | 0 | NA | NA | NA | NA | NA | NA | NA | 0 | 0 | 0 | 0 | 0 | 0 |
| 35551 | ENSMUSG00000096907 | 0 | NA | NA | NA | NA | NA | Gm6113 | predicted gene 6113 [Source:MGI Symbol | 0 | 0 | 0 | 0 | 0 | 0 |
| 35552 | ENSMUSG00000096908 | 0 | NA | NA | NA | NA | NA | Trav7-3 | T cell receptor alpha variable 7-3 [Source: | 0 | 0 | 0 | 0 | 0 | 0 |
| 35553 | ENSMUSG00000096910 | 73.71018174 | 0.146287263 | 0.326858817 | 0.447554894 | 0.654474462 | 0.925797231 | Zfp955b | zinc finger protein 955B [Source:MGI Sym | 56.217 | 58.591 | 95.593 | 99.29 | 51.259 | 81.311 |
| 35554 | ENSMUSG00000096911 | 0 | NA | NA | NA | NA | NA | Gm2165 | predicted gene 2165 [Source:MGI Symbol | 0 | 0 | 0 | 0 | 0 | 0 |
| 35555 | ENSMUSG00000096913 | 0 | NA | NA | NA | NA | NA | NA | NA | 0 | 0 | 0 | 0 | 0 | 0 |
| 35556 | ENSMUSG00000096914 | 0 | NA | NA | NA | NA | NA | Galntl6 | UDP-N-acetyl-alpha-D-galactosamine:poly | 0 | 0 | 0 | 0 | 0 | 0 |

| | | | | | | | | | | | | | | | |
|---|---|---|---|---|---|---|---|---|---|---|---|---|---|---|---|
| 35557 | ENSMUSG00000096915 | 0 | NA | NA | NA | NA | NA | Btbd35f6 | BTB domain containing 35, family membe | 0 | 0 | 0 | 0 | 0 | 0 |
| 35558 | ENSMUSG00000096916 | 2.847946833 | -1.638391397 | 1.300013376 | -1.260288107 | 0.207565449 | NA | Zfp850 | zinc finger protein 850 [Source:MGI Symb | 4.8186 | 4.1117 | 4.025 | 1.928 | 1.0906 | 1.1138 |
| 35559 | ENSMUSG00000096917 | 35.65694991 | 0.261369093 | 0.398487389 | 0.655903049 | 0.511886506 | 0.884156487 | 2500002B13 | RIKEN cDNA 2500002B13 gene [Source:M | 26.502 | 26.726 | 44.275 | 42.415 | 28.356 | 45.668 |
| 35560 | ENSMUSG00000096918 | 0 | NA | NA | NA | NA | NA | Gm16863 | predicted gene, 16863 [Source:MGI Symb | 0 | 0 | 0 | 0 | 0 | 0 |
| 35561 | ENSMUSG00000096919 | 0 | NA | NA | NA | NA | NA | NA | NA | 0 | 0 | 0 | 0 | 0 | 0 |
| 35562 | ENSMUSG00000096920 | 0 | NA | NA | NA | NA | NA | Rps19-ps8 | ribosomal protein S19, pseudogene 8 [Sou | 0 | 0 | 0 | 0 | 0 | 0 |
| 35563 | ENSMUSG00000096921 | 0 | NA | NA | NA | NA | NA | Gm26822 | predicted gene, 26822 [Source:MGI Symb | 0 | 0 | 0 | 0 | 0 | 0 |
| 35564 | ENSMUSG00000096922 | 0 | NA | NA | NA | NA | NA | Gm17308 | predicted gene, 17308 [Source:MGI Symb | 0 | 0 | 0 | 0 | 0 | 0 |
| 35565 | ENSMUSG00000096923 | 0.824421612 | 3.264491806 | 2.808832384 | 1.162223786 | 0.245144575 | NA | A730071L15 | RIKEN cDNA A730071L15Rik gene [Source | 0 | 0 | 0 | 3.8559 | 1.0906 | 0 |
| 35566 | ENSMUSG00000096924 | 0.476489653 | -2.293616034 | 3.421506946 | -0.670352587 | 0.502633051 | NA | NA | NA | 0.8031 | 2.0558 | 0 | 0 | 0 | 0 |
| 35567 | ENSMUSG00000096926 | 2.457367624 | 2.825367521 | 1.682702426 | 1.679065459 | 0.093139287 | NA | NA | NA | 0.8031 | 0 | 1.0062 | 9.6398 | 2.1812 | 1.1138 |
| 35568 | ENSMUSG00000096927 | 0 | NA | NA | NA | NA | NA | Gm3466 | predicted gene 3466 [Source:MGI Symbol | 0 | 0 | 0 | 0 | 0 | 0 |
| 35569 | ENSMUSG00000096929 | 56.26363065 | -0.286556382 | 0.341214121 | -0.839813961 | 0.401012704 | 0.827897196 | A330023F24 | RIKEN cDNA A330023F24 gene [Source:M | 61.036 | 50.368 | 74.462 | 71.334 | 43.625 | 36.757 |
| 35570 | ENSMUSG00000096930 | 0 | NA | NA | NA | NA | NA | Gm26657 | predicted gene, 26657 [Source:MGI Symb | 0 | 0 | 0 | 0 | 0 | 0 |
| 35571 | ENSMUSG00000096931 | 0 | NA | NA | NA | NA | NA | NA | NA | 0 | 0 | 0 | 0 | 0 | 0 |
| 35572 | ENSMUSG00000096932 | 2.459146031 | -1.382079257 | 1.471123404 | -0.939472007 | 0.347488458 | NA | 5033403F01 | RIKEN cDNA 5033403F01 gene [Source:M | 1.6062 | 4.1117 | 5.0312 | 2.8919 | 0 | 1.1138 |
| 35573 | ENSMUSG00000096934 | 0 | NA | NA | NA | NA | NA | Gm26658 | predicted gene, 26658 [Source:MGI Symb | 0 | 0 | 0 | 0 | 0 | 0 |
| 35574 | ENSMUSG00000096935 | 36.25700237 | 0.083298624 | 0.427805236 | 0.194711557 | 0.845618764 | 0.972803041 | 1700113A16 | RIKEN cDNA 1700113A16 gene [Source:M | 42.564 | 25.698 | 37.231 | 38.559 | 53.44 | 20.049 |
| 35575 | ENSMUSG00000096936 | 0 | NA | NA | NA | NA | NA | Gm3510 | predicted gene 3510 [Source:MGI Symbol | 0 | 0 | 0 | 0 | 0 | 0 |
| 35576 | ENSMUSG00000096937 | 0 | NA | NA | NA | NA | NA | Gm10825 | predicted gene 10825 [Source:MGI Symbo | 0 | 0 | 0 | 0 | 0 | 0 |
| 35577 | ENSMUSG00000096938 | 1.716758416 | 2.093188109 | 1.869472255 | 1.119667919 | 0.262855301 | NA | 9530052E02 | RIKEN cDNA 9530052E02 gene [Source:M | 0 | 0 | 2.0125 | 3.8559 | 1.0906 | 3.3415 |
| 35578 | ENSMUSG00000096939 | 0 | NA | NA | NA | NA | NA | Gm8321 | predicted gene 8321 [Source:MGI Symbol | 0 | 0 | 0 | 0 | 0 | 0 |
| 35579 | ENSMUSG00000096940 | 7.681853577 | -0.798132932 | 0.834497199 | -0.956423739 | 0.338858196 | 0.780858357 | Gm4755 | predicted gene 4755 [Source:MGI Symbol | 4.0155 | 10.279 | 15.094 | 6.7478 | 3.2719 | 6.6831 |
| 35580 | ENSMUSG00000096941 | 0 | NA | NA | NA | NA | NA | NA | NA | 0 | 0 | 0 | 0 | 0 | 0 |
| 35581 | ENSMUSG00000096942 | 9.482711956 | -0.49276201 | 0.739078234 | -0.666725101 | 0.504947742 | 0.881592135 | Rps19-ps6 | ribosomal protein S19, pseudogene 6 [Sou | 8.031 | 13.363 | 12.075 | 13.496 | 4.3625 | 5.5692 |
| 35582 | ENSMUSG00000096943 | 0.181769715 | 1.126924175 | 4.080472857 | 0.276174898 | 0.782413736 | NA | NA | NA | 0 | 0 | 0 | 0 | 1.0906 | 0 |
| 35583 | ENSMUSG00000096944 | 1.391561535 | 1.697168055 | 1.972565864 | 0.860386001 | 0.3895763 | NA | NA | NA | 0 | 0 | 2.0125 | 1.928 | 2.1812 | 2.2277 |
| 35584 | ENSMUSG00000096945 | 0.185640647 | 1.126924175 | 4.080472857 | 0.276174898 | 0.782413736 | NA | NA | NA | 0 | 0 | 0 | 0 | 0 | 1.1138 |
| 35585 | ENSMUSG00000096946 | 0.185640647 | 1.126924175 | 4.080472857 | 0.276174898 | 0.782413736 | NA | NA | NA | 0 | 0 | 0 | 0 | 0 | 1.1138 |
| 35586 | ENSMUSG00000096947 | 0 | NA | NA | NA | NA | NA | NA | NA | 0 | 0 | 0 | 0 | 0 | 0 |
| 35587 | ENSMUSG00000096948 | 0.685278559 | -2.809870862 | 3.376578027 | -0.832165239 | 0.405315682 | NA | Gm4221 | predicted gene 4221 [Source:MGI Symbol | 0 | 4.1117 | 0 | 0 | 0 | 0 |
| 35588 | ENSMUSG00000096949 | 0 | NA | NA | NA | NA | NA | Gm26897 | predicted gene, 26897 [Source:MGI Symb | 0 | 0 | 0 | 0 | 0 | 0 |
| 35589 | ENSMUSG00000096950 | 7.27104769 | 0.462566788 | 0.895035125 | 0.516814117 | 0.605285924 | 0.916376001 | Gm9530 | predicted gene 9530 [Source:MGI Symbol | 12.047 | 1.0279 | 5.0312 | 6.7478 | 7.6343 | 11.138 |
| 35590 | ENSMUSG00000096953 | 0.96895528 | -2.08853317 | 2.292908757 | -0.910866237 | 0.362365857 | NA | Gm26571 | predicted gene, 26571 [Source:MGI Symb | 0.8031 | 1.0279 | 3.0187 | 0.964 | 0 | 0 |
| 35591 | ENSMUSG00000096954 | 50.85624991 | 0.26382543 | 0.396179374 | 0.665924195 | 0.505459555 | 0.881592135 | NA | NA | 39.352 | 56.535 | 43.269 | 88.686 | 31.628 | 45.668 |
| 35592 | ENSMUSG00000096955 | 0 | NA | NA | NA | NA | NA | Gm26570 | predicted gene, 26570 [Source:MGI Symb | 0 | 0 | 0 | 0 | 0 | 0 |
| 35593 | ENSMUSG00000096956 | 24.35728137 | -0.40498127 | 0.459745373 | -0.880881665 | 0.378381872 | 0.811764373 | Snhg18 | small nucleolar RNA host gene 18 [Source | 35.336 | 21.586 | 26.162 | 22.171 | 15.269 | 25.618 |
| 35594 | ENSMUSG00000096957 | 12.7273138 | 0.323850533 | 0.667832494 | 0.484927786 | 0.627727572 | 0.920328819 | E230013L22I | RIKEN cDNA E230013L22 gene [Source:M | 14.456 | 8.2233 | 11.069 | 11.568 | 6.5437 | 24.505 |
| 35595 | ENSMUSG00000096958 | 0.185640647 | 1.126924175 | 4.080472857 | 0.276174898 | 0.782413736 | NA | NA | NA | 0 | 0 | 0 | 0 | 0 | 1.1138 |
| 35596 | ENSMUSG00000096959 | 6.236867091 | 0.330931372 | 0.879156036 | 0.376419382 | 0.706605139 | 0.94295036 | 4930509G22 | RIKEN cDNA 4930509G22 gene [Source:M | 6.4248 | 3.0838 | 7.0437 | 7.7118 | 9.8156 | 3.3415 |
| 35597 | ENSMUSG00000096960 | 2.398708466 | -0.436431065 | 1.424707791 | -0.30633023 | 0.759353225 | NA | A230028O05 | RIKEN cDNA A230028O05 gene [Source:M | 3.2124 | 3.0838 | 2.0125 | 3.8559 | 0 | 2.2277 |
| 35598 | ENSMUSG00000096962 | 0.677847551 | 0.063310748 | 2.809814301 | 0.022532004 | 0.982023583 | NA | Gm26622 | predicted gene, 26622 [Source:MGI Symb | 0 | 0 | 2.0125 | 0.964 | 1.0906 | 0 |
| 35599 | ENSMUSG00000096963 | 0.185640647 | 1.126924175 | 4.080472857 | 0.276174898 | 0.782413736 | NA | B230364G03 | RIKEN cDNA B230364G03 gene [Source:M | 0 | 0 | 0 | 0 | 0 | 1.1138 |
| 35600 | ENSMUSG00000096964 | 5.016722533 | 0.192435711 | 0.956435245 | 0.201200983 | 0.840541423 | NA | Gm6345 | predicted gene 6345 [Source:MGI Symbol | 4.8186 | 6.1675 | 3.0187 | 2.8919 | 7.6343 | 5.5692 |
| 35601 | ENSMUSG00000096965 | 0.160662974 | 1.126924175 | 4.080472857 | 0.276174898 | 0.782413736 | NA | 3300005D01 | RIKEN cDNA 3300005D01 gene [Source:M | 0 | 0 | 0 | 0.964 | 0 | 0 |
| 35602 | ENSMUSG00000096966 | 0 | NA | NA | NA | NA | NA | Gm18336 | predicted gene, 18336 [Source:MGI Symb | 0 | 0 | 0 | 0 | 0 | 0 |
| 35603 | ENSMUSG00000096967 | 11.26830985 | 0.586134776 | 0.653563713 | 0.896828823 | 0.369810265 | 0.80653433 | Gm26621 | predicted gene, 26621 [Source:MGI Symb | 8.8341 | 6.1675 | 12.075 | 17.352 | 9.8156 | 13.366 |
| 35604 | ENSMUSG00000096968 | 1.666745423 | 0.787596987 | 1.596783862 | 0.493239571 | 0.621843321 | NA | Gm26620 | predicted gene, 26620 [Source:MGI Symb | 1.6062 | 1.0279 | 1.0062 | 1.928 | 1.0906 | 3.3415 |
| 35605 | ENSMUSG00000096969 | 0 | NA | NA | NA | NA | NA | NA | NA | 0 | 0 | 0 | 0 | 0 | 0 |
| 35606 | ENSMUSG00000096971 | 110.1155843 | -0.265260938 | 0.221711539 | -1.196423691 | 0.231531261 | 0.678732905 | 4930556M19 | RIKEN cDNA 4930556M19 gene [Source:M | 120.47 | 115.13 | 124.77 | 89.65 | 97.065 | 113.61 |
| 35607 | ENSMUSG00000096972 | 0 | NA | NA | NA | NA | NA | NA | NA | 0 | 0 | 0 | 0 | 0 | 0 |
| 35608 | ENSMUSG00000096973 | 0 | NA | NA | NA | NA | NA | Gm5974 | predicted gene 5974 [Source:MGI Symbol | 0 | 0 | 0 | 0 | 0 | 0 |
| 35609 | ENSMUSG00000096974 | 4.489889029 | -0.391408927 | 1.051742914 | -0.372152663 | 0.709779188 | NA | Gm26881 | predicted gene, 26881 [Source:MGI Symb | 8.031 | 5.1396 | 2.0125 | 2.8919 | 2.1812 | 6.6831 |
| 35610 | ENSMUSG00000096975 | 1.412409551 | 1.709645529 | 1.917510359 | 0.891596502 | 0.372609246 | NA | NA | NA | 0 | 1.0279 | 1.0062 | 0.964 | 4.3625 | 1.1138 |
| 35611 | ENSMUSG00000096976 | 0.321325949 | 1.911575114 | 4.048267893 | 0.472195804 | 0.636787034 | NA | NA | NA | 0 | 0 | 0 | 1.928 | 0 | 0 |
| 35612 | ENSMUSG00000096977 | 0.356960287 | 0.16514264 | 3.880370363 | 0.042558474 | 0.966053499 | NA | Gm10827 | predicted gene 10827 [Source:MGI Symbo | 0 | 1.0279 | 0 | 0 | 0 | 1.1138 |

| | | | | | | | | | | | | | | | |
|---|---|---|---|---|---|---|---|---|---|---|---|---|---|---|---|
| 35613 | ENSMUSG00000096978 | 0 | NA | NA | NA | NA | NA | NA | NA | 0 | 0 | 0 | 0 | 0 | 0 |
| 35614 | ENSMUSG00000096979 | 9.051941771 | -1.458201799 | 0.923377701 | -1.579204044 | 0.114289263 | 0.496868018 | NA | NA | 11.243 | 23.642 | 5.0312 | 7.7118 | 0 | 6.6831 |
| 35615 | ENSMUSG00000096980 | 14.09510844 | 0.038120115 | 0.595418553 | 0.064022384 | 0.948952404 | 0.990731019 | NA | NA | 11.243 | 12.335 | 18.112 | 8.6758 | 15.269 | 18.935 |
| 35616 | ENSMUSG00000096981 | 9.179263531 | -0.134647659 | 0.733115482 | -0.183665006 | 0.854276263 | 0.973825032 | Gm16845 | predicted gene, 16845 [Source:MGI Symb | 5.6217 | 9.2513 | 14.087 | 9.6398 | 10.906 | 5.5692 |
| 35617 | ENSMUSG00000096982 | 0 | NA | NA | NA | NA | NA | Redrum | Redrum, erythroid developmental long int | 0 | 0 | 0 | 0 | 0 | 0 |
| 35618 | ENSMUSG00000096983 | 3.782341007 | 0.293766581 | 1.23688676 | 0.237504831 | 0.812265171 | NA | NA | NA | 8.031 | 0 | 2.0125 | 3.8559 | 5.4531 | 3.3415 |
| 35619 | ENSMUSG00000096984 | 0 | NA | NA | NA | NA | NA | NA | NA | 0 | 0 | 0 | 0 | 0 | 0 |
| 35620 | ENSMUSG00000096986 | 3.00782865 | 1.243609028 | 1.253993599 | 0.991718801 | 0.321334717 | NA | 4930509E16 | RIKEN cDNA 4930509E16 gene [Source:M | 3.2124 | 1.0279 | 1.0062 | 2.8919 | 5.4531 | 4.4554 |
| 35621 | ENSMUSG00000096987 | 0 | NA | NA | NA | NA | NA | NA | NA | 0 | 0 | 0 | 0 | 0 | 0 |
| 35622 | ENSMUSG00000096988 | 4.221017115 | 1.558693498 | 1.28816622 | 1.210009604 | 0.226275207 | NA | A930029G22 | RIKEN cDNA A930029G22 gene [Source:M | 2.4093 | 0 | 4.025 | 14.46 | 1.0906 | 3.3415 |
| 35623 | ENSMUSG00000096990 | 1.565258526 | 0.169278159 | 1.865825703 | 0.090725602 | 0.927710627 | NA | NA | NA | 2.4093 | 0 | 2.0125 | 3.8559 | 0 | 1.1138 |
| 35624 | ENSMUSG00000096991 | 9.767245458 | 0.72408362 | 0.807320208 | 0.896897678 | 0.369773519 | 0.80653433 | Gm26789 | predicted gene, 26789 [Source:MGI Symb | 4.0155 | 10.279 | 8.05 | 24.099 | 4.3625 | 7.7969 |
| 35625 | ENSMUSG00000096992 | 0 | NA | NA | NA | NA | NA | Gm26788 | predicted gene, 26788 [Source:MGI Symb | 0 | 0 | 0 | 0 | 0 | 0 |
| 35626 | ENSMUSG00000096993 | 2.013747879 | 0.603296586 | 1.726388613 | 0.349455842 | 0.726747117 | NA | Gm26787 | predicted gene, 26787 [Source:MGI Symb | 0.8031 | 0 | 4.025 | 2.8919 | 4.3625 | 0 |
| 35627 | ENSMUSG00000096995 | 4.49959673 | -1.1119056 | 1.04370503 | -1.065344678 | 0.286719982 | NA | 2810029C07 | RIKEN cDNA 2810029C07 gene [Source:M | 3.2124 | 8.2233 | 7.0437 | 1.928 | 4.3625 | 2.2277 |
| 35628 | ENSMUSG00000096996 | 9.210969818 | 0.36011759 | 0.74903612 | 0.480774666 | 0.630676658 | 0.921020101 | NA | NA | 4.0155 | 7.1954 | 13.081 | 6.7478 | 13.087 | 11.138 |
| 35629 | ENSMUSG00000096997 | 0 | NA | NA | NA | NA | NA | Gm26791 | predicted gene, 26791 [Source:MGI Symb | 0 | 0 | 0 | 0 | 0 | 0 |
| 35630 | ENSMUSG00000096999 | 0 | NA | NA | NA | NA | NA | Gm26793 | predicted gene, 26793 [Source:MGI Symb | 0 | 0 | 0 | 0 | 0 | 0 |
| 35631 | ENSMUSG00000097000 | 16.24259215 | -0.57507578 | 0.633122126 | -0.9083173 | 0.363710605 | 0.802534854 | Gm17435 | predicted gene, 17435 [Source:MGI Symb | 28.912 | 8.2233 | 21.131 | 19.28 | 6.5437 | 13.366 |
| 35632 | ENSMUSG00000097001 | 0 | NA | NA | NA | NA | NA | 1700054K02 | RIKEN cDNA 1700054K02 gene [Source:M | 0 | 0 | 0 | 0 | 0 | 0 |
| 35633 | ENSMUSG00000097002 | 0.160662974 | 1.126924175 | 4.080472857 | 0.276174898 | 0.782413736 | NA | Gm2670 | predicted gene 2670 [Source:MGI Symbol | 0 | 0 | 0 | 0.964 | 0 | 0 |
| 35634 | ENSMUSG00000097003 | 0 | NA | NA | NA | NA | NA | D930007P13 | Riken cDNA D930007P13 gene [Source:M | 0 | 0 | 0 | 0 | 0 | 0 |
| 35635 | ENSMUSG00000097004 | 0 | NA | NA | NA | NA | NA | 4731419I09F | RIKEN cDNA 4731419I09 gene [Source:MG | 0 | 0 | 0 | 0 | 0 | 0 |
| 35636 | ENSMUSG00000097005 | 3.11554384 | -0.631310596 | 1.20019353 | -0.526007331 | 0.598883113 | NA | NA | NA | 3.2124 | 6.1675 | 2.0125 | 2.8919 | 2.1812 | 2.2277 |
| 35637 | ENSMUSG00000097006 | 41.74078956 | -0.172114567 | 0.36811483 | -0.467556734 | 0.640101611 | 0.923932929 | 9530082P21 | RIKEN cDNA 9530082P21 gene [Source:M | 55.414 | 29.81 | 47.293 | 46.271 | 34.9 | 36.757 |
| 35638 | ENSMUSG00000097007 | 0.462220778 | -0.685400455 | 3.191475091 | -0.21475977 | 0.82995462 | NA | 4930432B10 | RIKEN cDNA 4930432B10 gene [Source:M | 0.8031 | 0 | 1.0062 | 0.964 | 0 | 0 |
| 35639 | ENSMUSG00000097008 | 1.295464128 | 0.179168128 | 1.7554591 | 0.102063402 | 0.91870635 | NA | NA | NA | 1.6062 | 1.0279 | 1.0062 | 1.928 | 1.0906 | 1.1138 |
| 35640 | ENSMUSG00000097009 | 0.185640647 | 1.126924175 | 4.080472857 | 0.276174898 | 0.782413736 | NA | Gm26861 | predicted gene, 26861 [Source:MGI Symb | 0 | 0 | 0 | 0 | 0 | 1.1138 |
| 35641 | ENSMUSG00000097010 | 0 | NA | NA | NA | NA | NA | Gm26709 | predicted gene, 26709 [Source:MGI Symb | 0 | 0 | 0 | 0 | 0 | 0 |
| 35642 | ENSMUSG00000097011 | 0.492645588 | 0.9565034 | 3.447648281 | 0.277436479 | 0.781444974 | NA | Gm4651 | predicted gene 4651 [Source:MGI Symbol | 0 | 1.0279 | 0 | 1.928 | 0 | 0 |
| 35643 | ENSMUSG00000097012 | 0 | NA | NA | NA | NA | NA | Gm16531 | predicted gene, 16531 [Source:MGI Symb | 0 | 0 | 0 | 0 | 0 | 0 |
| 35644 | ENSMUSG00000097013 | 3.534019899 | -0.048076465 | 1.496567227 | -0.032124494 | 0.97437277 | NA | NA | NA | 9.6372 | 1.0279 | 0 | 3.8559 | 0 | 6.6831 |
| 35645 | ENSMUSG00000097014 | 0.963765213 | -2.11522655 | 2.746264995 | -0.770219391 | 0.441169764 | NA | NA | NA | 4.8186 | 0 | 0 | 0.964 | 0 | 0 |
| 35646 | ENSMUSG00000097015 | 0 | NA | NA | NA | NA | NA | Gm26705 | predicted gene, 26705 [Source:MGI Symb | 0 | 0 | 0 | 0 | 0 | 0 |
| 35647 | ENSMUSG00000097016 | 23.79509799 | 0.677290214 | 0.506616354 | 1.336889756 | 0.181258625 | 0.614562765 | Gm26708 | predicted gene, 26708 [Source:MGI Symb | 27.305 | 9.2513 | 18.112 | 38.559 | 27.265 | 22.277 |
| 35648 | ENSMUSG00000097017 | 0 | NA | NA | NA | NA | NA | Gm26707 | predicted gene, 26707 [Source:MGI Symb | 0 | 0 | 0 | 0 | 0 | 0 |
| 35649 | ENSMUSG00000097018 | 0 | NA | NA | NA | NA | NA | Gm26872 | predicted gene, 26872 [Source:MGI Symb | 0 | 0 | 0 | 0 | 0 | 0 |
| 35650 | ENSMUSG00000097019 | 0 | NA | NA | NA | NA | NA | NA | NA | 0 | 0 | 0 | 0 | 0 | 0 |
| 35651 | ENSMUSG00000097020 | 5.779478891 | 0.548935911 | 0.895416414 | 0.613050981 | 0.539842631 | 0.892687582 | NA | NA | 4.0155 | 4.1117 | 6.0375 | 10.604 | 5.4531 | 4.4554 |
| 35652 | ENSMUSG00000097021 | 2.193862626 | 0.860317317 | 1.444266372 | 0.595677732 | 0.551390537 | NA | 4933433G15 | RIKEN cDNA 4933433G15 gene [Source:M | 1.6062 | 2.0558 | 1.0062 | 1.928 | 5.4531 | 1.1138 |
| 35653 | ENSMUSG00000097022 | 0.133850373 | -0.796629176 | 4.080472857 | -0.195229623 | 0.84521319 | NA | BC001981 | cDNA sequence BC001981 [Source:MGI Sy | 0.8031 | 0 | 0 | 0 | 0 | 0 |
| 35654 | ENSMUSG00000097023 | 0 | NA | NA | NA | NA | NA | Mir9-3hg | Mir9-3 host gene [Source:MGI Symbol;Acc | 0 | 0 | 0 | 0 | 0 | 0 |
| 35655 | ENSMUSG00000097024 | 0 | NA | NA | NA | NA | NA | Gm26557 | predicted gene, 26557 [Source:MGI Symb | 0 | 0 | 0 | 0 | 0 | 0 |
| 35656 | ENSMUSG00000097025 | 2.119595998 | -0.943554282 | 1.511766015 | -0.624140424 | 0.532535368 | NA | Gm26558 | predicted gene, 26558 [Source:MGI Symb | 3.2124 | 3.0838 | 2.0125 | 0 | 2.1812 | 2.2277 |
| 35657 | ENSMUSG00000097026 | 0 | NA | NA | NA | NA | NA | NA | NA | 0 | 0 | 0 | 0 | 0 | 0 |
| 35658 | ENSMUSG00000097027 | 0.481988923 | 2.492401104 | 3.837077151 | 0.649557203 | 0.515978285 | NA | Gm26559 | predicted gene, 26559 [Source:MGI Symb | 0 | 0 | 0 | 2.8919 | 0 | 0 |
| 35659 | ENSMUSG00000097028 | 0 | NA | NA | NA | NA | NA | Ptgs2os | prostaglandin-endoperoxide synthase 2, c | 0 | 0 | 0 | 0 | 0 | 0 |
| 35660 | ENSMUSG00000097029 | 0 | NA | NA | NA | NA | NA | Gm26560 | predicted gene, 26560 [Source:MGI Symb | 0 | 0 | 0 | 0 | 0 | 0 |
| 35661 | ENSMUSG00000097030 | 0 | NA | NA | NA | NA | NA | Gm26693 | predicted gene, 26693 [Source:MGI Symb | 0 | 0 | 0 | 0 | 0 | 0 |
| 35662 | ENSMUSG00000097031 | 0 | NA | NA | NA | NA | NA | Gm26820 | predicted gene, 26820 [Source:MGI Symb | 0 | 0 | 0 | 0 | 0 | 0 |
| 35663 | ENSMUSG00000097032 | 9.683974851 | 0.450601067 | 0.710541488 | 0.634165739 | 0.525972665 | 0.888334265 | 4930539J05I | RIKEN cDNA 4930539J05 gene [Source:MG | 11.243 | 4.1117 | 9.0562 | 10.604 | 14.178 | 8.9108 |
| 35664 | ENSMUSG00000097033 | 4.301437859 | -0.522593172 | 1.030637895 | -0.507057983 | 0.612114136 | NA | NA | NA | 4.0155 | 7.1954 | 4.025 | 2.8919 | 5.4531 | 2.2277 |
| 35665 | ENSMUSG00000097035 | 0.167707431 | -0.796629176 | 4.080472857 | -0.195229623 | 0.84521319 | NA | Gm26692 | predicted gene, 26692 [Source:MGI Symb | 0 | 0 | 1.0062 | 0 | 0 | 0 |
| 35666 | ENSMUSG00000097036 | 0.881705114 | -0.66605326 | 2.559736336 | -0.260203854 | 0.794706531 | NA | NA | NA | 3.2124 | 0 | 0 | 0.964 | 0 | 1.1138 |
| 35667 | ENSMUSG00000097037 | 0 | NA | NA | NA | NA | NA | NA | NA | 0 | 0 | 0 | 0 | 0 | 0 |
| 35668 | ENSMUSG00000097038 | 0 | NA | NA | NA | NA | NA | NA | NA | 0 | 0 | 0 | 0 | 0 | 0 |

| | | | | | | | | | | | | | | | |
|---|---|---|---|---|---|---|---|---|---|---|---|---|---|---|---|
| 35669 | ENSMUSG00000097039 | 19.44031018 | 0.825058736 | 0.581136536 | 1.419733031 | 0.155685418 | 0.574732247 | Pvt1 | Pvt1 oncogene [Source:MGI Symbol;Acc:M | 23.29 | 11.307 | 7.0437 | 14.46 | 33.809 | 26.732 |
| 35670 | ENSMUSG00000097040 | 0 | NA | NA | NA | NA | NA | 2610316D01 | RIKEN cDNA 2610316D01 gene [Source:M | 0 | 0 | 0 | 0 | 0 | 0 |
| 35671 | ENSMUSG00000097041 | 0 | NA | NA | NA | NA | NA | Pafah1b1-ps | platelet-activating factor acetylhydrolase, | 0 | 0 | 0 | 0 | 0 | 0 |
| 35672 | ENSMUSG00000097042 | 140.0242687 | 0.085382039 | 0.216208169 | 0.394906629 | 0.692911797 | 0.938679052 | Gm17491 | predicted gene, 17491 [Source:MGI Symb | 140.54 | 125.41 | 141.88 | 169.66 | 114.51 | 148.14 |
| 35673 | ENSMUSG00000097043 | 4.454533943 | 1.630538123 | 1.05719465 | 1.542325364 | 0.122994548 | NA | Gm10556 | predicted gene 10556 [Source:MGI Symbo | 2.4093 | 3.0838 | 1.0062 | 4.8199 | 8.7249 | 6.6831 |
| 35674 | ENSMUSG00000097044 | 0 | NA | NA | NA | NA | NA | 4933433F19 | RIKEN cDNA 4933433F19 gene [Source:M | 0 | 0 | 0 | 0 | 0 | 0 |
| 35675 | ENSMUSG00000097045 | 0.972799583 | 1.135651363 | 2.280068491 | 0.498077741 | 0.618429249 | NA | NA | NA | 0.8031 | 1.0279 | 0 | 2.8919 | 0 | 1.1138 |
| 35676 | ENSMUSG00000097047 | 39.76848716 | -0.236532231 | 0.353813963 | -0.668521472 | 0.50380078 | 0.881510796 | 1110020A21 | RIKEN cDNA 1110020A21 gene [Source:M | 33.73 | 48.312 | 47.293 | 37.595 | 33.809 | 37.871 |
| 35677 | ENSMUSG00000097048 | 129.5515463 | -0.23956611 | 0.228601546 | -1.047963648 | 0.294655356 | 0.745164683 | 1600020E01 | RIKEN cDNA 1600020E01 gene [Source:M | 116.45 | 160.36 | 144.9 | 139.78 | 116.7 | 99.132 |
| 35678 | ENSMUSG00000097049 | 0 | NA | NA | NA | NA | NA | 6530411M01 | RIKEN cDNA 6530411M01 gene [Source:M | 0 | 0 | 0 | 0 | 0 | 0 |
| 35679 | ENSMUSG00000097050 | 0 | NA | NA | NA | NA | NA | Gm9918 | predicted gene 9918 [Source:MGI Symbol | 0 | 0 | 0 | 0 | 0 | 0 |
| 35680 | ENSMUSG00000097051 | 0.829837731 | -1.7608533 | 2.399900965 | -0.733719152 | 0.463119924 | NA | NA | NA | 0.8031 | 2.0558 | 1.0062 | 0 | 0 | 1.1138 |
| 35681 | ENSMUSG00000097052 | 0.596071152 | -1.224343926 | 2.897583865 | -0.4225396 | 0.672631203 | NA | Snora43 | small nucleolar RNA, H/ACA box 43 [Sourc | 1.6062 | 0 | 1.0062 | 0.964 | 0 | 0 |
| 35682 | ENSMUSG00000097053 | 0 | NA | NA | NA | NA | NA | Gm1972 | predicted pseudogene 1972 [Source:MGI | 0 | 0 | 0 | 0 | 0 | 0 |
| 35683 | ENSMUSG00000097054 | 0 | NA | NA | NA | NA | NA | Gm11395 | predicted gene 11395 [Source:MGI Symbo | 0 | 0 | 0 | 0 | 0 | 0 |
| 35684 | ENSMUSG00000097055 | 1.062493721 | 1.134258317 | 2.64304732 | 0.429147942 | 0.667815565 | NA | Gm4419 | predicted gene 4419 [Source:MGI Symbol | 0 | 0 | 2.0125 | 0 | 4.3625 | 0 |
| 35685 | ENSMUSG00000097056 | 0.760346335 | -0.431432104 | 2.662343224 | -0.162049769 | 0.871266662 | NA | NA | NA | 1.6062 | 1.0279 | 0 | 1.928 | 0 | 0 |
| 35686 | ENSMUSG00000097057 | 2.363171465 | 0.571056288 | 1.610014488 | 0.354690155 | 0.722821727 | NA | Gm17638 | predicted gene, 17638 [Source:MGI Symb | 1.6062 | 4.1117 | 0 | 2.8919 | 0 | 5.5692 |
| 35687 | ENSMUSG00000097059 | 245.1832275 | -0.252736897 | 0.156578866 | -1.614118834 | 0.106501653 | 0.481264827 | NA | NA | 256.19 | 269.31 | 274.7 | 232.32 | 222.49 | 216.09 |
| 35688 | ENSMUSG00000097060 | 0.160662974 | 1.126924175 | 4.080472857 | 0.276174898 | 0.782413736 | NA | Gm26759 | predicted gene, 26759 [Source:MGI Symb | 0 | 0 | 0 | 0.964 | 0 | 0 |
| 35689 | ENSMUSG00000097061 | 70.80466877 | 0.12301596 | 0.283664186 | 0.433667577 | 0.664529855 | 0.929301301 | 9330151L19 | RIKEN cDNA 9330151L19 gene [Source:M | 67.461 | 66.815 | 69.431 | 94.47 | 67.618 | 59.034 |
| 35690 | ENSMUSG00000097062 | 91.06094379 | 0.245521914 | 0.248821746 | 0.986738168 | 0.323771014 | 0.770868138 | NA | NA | 68.264 | 91.485 | 90.562 | 87.722 | 102.52 | 105.82 |
| 35691 | ENSMUSG00000097063 | 0 | NA | NA | NA | NA | NA | Pantr2 | POU domain, class 3, transcription factor | 0 | 0 | 0 | 0 | 0 | 0 |
| 35692 | ENSMUSG00000097065 | 0.401551119 | -2.072982726 | 4.008251979 | -0.517178744 | 0.605031388 | NA | NA | NA | 2.4093 | 0 | 0 | 0 | 0 | 0 |
| 35693 | ENSMUSG00000097066 | 0 | NA | NA | NA | NA | NA | NA | NA | 0 | 0 | 0 | 0 | 0 | 0 |
| 35694 | ENSMUSG00000097067 | 0 | NA | NA | NA | NA | NA | Gm26743 | predicted gene, 26743 [Source:MGI Symb | 0 | 0 | 0 | 0 | 0 | 0 |
| 35695 | ENSMUSG00000097068 | 0 | NA | NA | NA | NA | NA | Gm26760 | predicted gene, 26760 [Source:MGI Symb | 0 | 0 | 0 | 0 | 0 | 0 |
| 35696 | ENSMUSG00000097069 | 14.3305377 | 0.384791022 | 0.586476248 | 0.656106744 | 0.511755445 | 0.884035682 | Gm16998 | predicted gene, 16998 [Source:MGI Symb | 16.865 | 9.2513 | 11.069 | 21.208 | 11.997 | 15.594 |
| 35697 | ENSMUSG00000097071 | 0 | NA | NA | NA | NA | NA | 4930544I03I | RIKEN cDNA 4930544I03 gene [Source:M | 0 | 0 | 0 | 0 | 0 | 0 |
| 35698 | ENSMUSG00000097072 | 0 | NA | NA | NA | NA | NA | Foxl2os | forkhead box L2, opposite strand [Source: | 0 | 0 | 0 | 0 | 0 | 0 |
| 35699 | ENSMUSG00000097073 | 31.99471666 | -0.281011296 | 0.424535245 | -0.661926894 | 0.508018075 | 0.882639452 | 9430037G07 | RIKEN cDNA 9430037G07 gene [Source:M | 48.186 | 24.67 | 32.2 | 32.775 | 20.722 | 33.415 |
| 35700 | ENSMUSG00000097074 | 0 | NA | NA | NA | NA | NA | 4833428L15 | RIKEN cDNA 4833428L15 gene [Source:M | 0 | 0 | 0 | 0 | 0 | 0 |
| 35701 | ENSMUSG00000097075 | 2.829936507 | -2.112527371 | 1.405896165 | -1.502619769 | 0.132937122 | NA | Cdiptos | CDIP transferase, opposite strand [Source | 1.6062 | 6.1675 | 6.0375 | 0.964 | 1.0906 | 1.1138 |
| 35702 | ENSMUSG00000097076 | 0 | NA | NA | NA | NA | NA | Platr7 | pluripotency associated transcript 7 [Sour | 0 | 0 | 0 | 0 | 0 | 0 |
| 35703 | ENSMUSG00000097077 | 0.167707431 | -0.796629176 | 4.080472857 | -0.195229623 | 0.84521319 | NA | Gm16712 | predicted gene, 16712 [Source:MGI Symb | 0 | 0 | 1.0062 | 0 | 0 | 0 |
| 35704 | ENSMUSG00000097078 | 0 | NA | NA | NA | NA | NA | Gm26566 | predicted gene, 26566 [Source:MGI Symb | 0 | 0 | 0 | 0 | 0 | 0 |
| 35705 | ENSMUSG00000097079 | 0 | NA | NA | NA | NA | NA | Gm26604 | predicted gene, 26604 [Source:MGI Symb | 0 | 0 | 0 | 0 | 0 | 0 |
| 35706 | ENSMUSG00000097080 | 32.77406794 | -0.087268194 | 0.40961825 | -0.213047621 | 0.831289819 | 0.97052742 | 1700086O06 | RIKEN cDNA 1700086O06 gene [Source:M | 26.502 | 27.754 | 47.293 | 35.667 | 33.809 | 25.618 |
| 35707 | ENSMUSG00000097081 | 0.133850373 | -0.796629176 | 4.080472857 | -0.195229623 | 0.84521319 | NA | Gm10425 | predicted gene 10425 [Source:MGI Symbo | 0.8031 | 0 | 0 | 0 | 0 | 0 |
| 35708 | ENSMUSG00000097082 | 3.727972533 | 1.29121242 | 1.160166363 | 1.11295454 | 0.265727954 | NA | 4933440J02I | RIKEN cDNA 4933440J02 gene [Source:M | 2.4093 | 1.0279 | 3.0187 | 4.8199 | 2.1812 | 8.9108 |
| 35709 | ENSMUSG00000097083 | 0 | NA | NA | NA | NA | NA | D930019O06 | RIKEN cDNA D930019O06 [Source:MGI Sy | 0 | 0 | 0 | 0 | 0 | 0 |
| 35710 | ENSMUSG00000097084 | 0 | NA | NA | NA | NA | NA | Foxl1 | forkhead box L1 [Source:MGI Symbol;Acc: | 0 | 0 | 0 | 0 | 0 | 0 |
| 35711 | ENSMUSG00000097085 | 0 | NA | NA | NA | NA | NA | Gm26634 | predicted gene, 26634 [Source:MGI Symb | 0 | 0 | 0 | 0 | 0 | 0 |
| 35712 | ENSMUSG00000097086 | 0 | NA | NA | NA | NA | NA | Gm7672 | predicted gene 7672 [Source:MGI Symbol | 0 | 0 | 0 | 0 | 0 | 0 |
| 35713 | ENSMUSG00000097087 | 0 | NA | NA | NA | NA | NA | Gm26878 | predicted gene, 26878 [Source:MGI Symb | 0 | 0 | 0 | 0 | 0 | 0 |
| 35714 | ENSMUSG00000097088 | 0.506966596 | 2.558866737 | 3.339834942 | 0.766165628 | 0.443577757 | NA | NA | NA | 0 | 0 | 0 | 1.928 | 0 | 1.1138 |
| 35715 | ENSMUSG00000097089 | 0 | NA | NA | NA | NA | NA | Gm26879 | predicted gene, 26879 [Source:MGI Symb | 0 | 0 | 0 | 0 | 0 | 0 |
| 35716 | ENSMUSG00000097090 | 12.73690417 | -0.438784572 | 0.625677854 | -0.701294714 | 0.483119112 | 0.871953436 | Gm26724 | predicted gene, 26724 [Source:MGI Symb | 9.6372 | 14.391 | 20.125 | 13.496 | 7.6343 | 11.138 |
| 35717 | ENSMUSG00000097091 | 0 | NA | NA | NA | NA | NA | 4930526L06 | RIKEN cDNA 4930526L06 gene [Source:M | 0 | 0 | 0 | 0 | 0 | 0 |
| 35718 | ENSMUSG00000097092 | 8.934321412 | 0.255176528 | 0.708152157 | 0.360341384 | 0.718591855 | 0.9447719 | Gm26725 | predicted gene, 26725 [Source:MGI Symb | 7.2279 | 9.2513 | 8.05 | 12.532 | 7.6343 | 8.9108 |
| 35719 | ENSMUSG00000097093 | 40.37736673 | -0.005368609 | 0.350903431 | -0.015299392 | 0.987793327 | 0.99859176 | C330013E15 | RIKEN cDNA C330013E15 gene [Source:M | 40.958 | 43.173 | 37.231 | 39.523 | 49.078 | 32.301 |
| 35720 | ENSMUSG00000097094 | 0 | NA | NA | NA | NA | NA | NA | NA | 0 | 0 | 0 | 0 | 0 | 0 |
| 35721 | ENSMUSG00000097095 | 0 | NA | NA | NA | NA | NA | NA | NA | 0 | 0 | 0 | 0 | 0 | 0 |
| 35722 | ENSMUSG00000097097 | 0.401551119 | -2.072982726 | 4.008251979 | -0.517178744 | 0.605031388 | NA | 5033426O07 | RIKEN cDNA 5033426O07 gene [Source:M | 2.4093 | 0 | 0 | 0 | 0 | 0 |
| 35723 | ENSMUSG00000097098 | 2.088122334 | 1.896720279 | 1.677736094 | 1.130523618 | 0.258255652 | NA | 9330111N05 | RIKEN cDNA 9330111N05 gene [Source:M | 1.6062 | 0 | 1.0062 | 7.7118 | 1.0906 | 1.1138 |
| 35724 | ENSMUSG00000097099 | 47.8007541 | -0.222590453 | 0.35141663 | -0.633408991 | 0.526466595 | 0.888334265 | Gm9917 | predicted gene 9917 [Source:MGI Symbol | 57.02 | 49.34 | 48.3 | 61.695 | 39.262 | 31.188 |

| | | | | | | | | | | | | | | | |
|---|---|---|---|---|---|---|---|---|---|---|---|---|---|---|---|
| 35725 | ENSMUSG00000097100 | 0.160662974 | 1.126924175 | 4.080472857 | 0.276174898 | 0.782413736 | NA | 9230104M0( | RIKEN cDNA 9230104M06 gene [Source:N | 0 | 0 | 0 | 0.964 | 0 | 0 |
| 35726 | ENSMUSG00000097101 | 162.110365 | 0.513208357 | 0.217745202 | 2.356921541 | 0.01842714 | 0.163315042 | 1810034E14 | RIKEN cDNA 1810034E14 gene [Source:M( | 146.16 | 108.96 | 144.9 | 160.98 | 178.86 | 232.79 |
| 35727 | ENSMUSG00000097103 | 1.004769579 | -2.106122861 | 2.47240066 | -0.851853381 | 0.394295474 | NA | NA | NA | 0.8031 | 4.1117 | 0 | 0 | 0 | 1.1138 |
| 35728 | ENSMUSG00000097104 | 0 | NA | NA | NA | NA | NA | Gm26579 | predicted gene, 26579 [Source:MGI Symb | 0 | 0 | 0 | 0 | 0 | 0 |
| 35729 | ENSMUSG00000097106 | 0 | NA | NA | NA | NA | NA | Rpl31-ps23 | ribosomal protein L31, pseudogene 23 [Sc | 0 | 0 | 0 | 0 | 0 | 0 |
| 35730 | ENSMUSG00000097107 | 0 | NA | NA | NA | NA | NA | Platr6 | pluripotency associated transcript 6 [Sour | 0 | 0 | 0 | 0 | 0 | 0 |
| 35731 | ENSMUSG00000097108 | 0.133850373 | -0.796629176 | 4.080472857 | -0.195229623 | 0.84521319 | NA | Gm26581 | predicted gene, 26581 [Source:MGI Symb | 0.8031 | 0 | 0 | 0 | 0 | 0 |
| 35732 | ENSMUSG00000097109 | 0 | NA | NA | NA | NA | NA | NA | NA | 0 | 0 | 0 | 0 | 0 | 0 |
| 35733 | ENSMUSG00000097110 | 0 | NA | NA | NA | NA | NA | Gm26726 | predicted gene, 26726 [Source:MGI Symb | 0 | 0 | 0 | 0 | 0 | 0 |
| 35734 | ENSMUSG00000097111 | 0 | NA | NA | NA | NA | NA | Peak1os | pseudopodium-enriched atypical kinase 1 | 0 | 0 | 0 | 0 | 0 | 0 |
| 35735 | ENSMUSG00000097112 | 0 | NA | NA | NA | NA | NA | NA | NA | 0 | 0 | 0 | 0 | 0 | 0 |
| 35736 | ENSMUSG00000097113 | 8.096875743 | -0.988013461 | 0.845640925 | -1.168360507 | 0.242661378 | 0.692154638 | Gm19705 | predicted gene, 19705 [Source:MGI Symb | 12.047 | 12.335 | 8.05 | 10.604 | 1.0906 | 4.4554 |
| 35737 | ENSMUSG00000097115 | 1.120686736 | -3.525616607 | 2.284576285 | -1.5432256 | 0.122776048 | NA | 1810019N24 | RIKEN cDNA 1810019N24 gene [Source:M | 1.6062 | 4.1117 | 1.0062 | 0 | 0 | 0 |
| 35738 | ENSMUSG00000097116 | 0 | NA | NA | NA | NA | NA | Gm26729 | predicted gene, 26729 [Source:MGI Symb | 0 | 0 | 0 | 0 | 0 | 0 |
| 35739 | ENSMUSG00000097117 | 0 | NA | NA | NA | NA | NA | NA | NA | 0 | 0 | 0 | 0 | 0 | 0 |
| 35740 | ENSMUSG00000097118 | 0 | NA | NA | NA | NA | NA | NA | NA | 0 | 0 | 0 | 0 | 0 | 0 |
| 35741 | ENSMUSG00000097119 | 92.59600763 | 0.044851861 | 0.239085964 | 0.187597215 | 0.851192409 | 0.973568625 | B230354K17 | RIKEN cDNA B230354K17 gene [Source:M | 78.704 | 92.513 | 102.64 | 95.434 | 91.612 | 94.677 |
| 35742 | ENSMUSG00000097120 | 0 | NA | NA | NA | NA | NA | Gm26887 | predicted gene, 26887 [Source:MGI Symb | 0 | 0 | 0 | 0 | 0 | 0 |
| 35743 | ENSMUSG00000097121 | 266.9635393 | 0.107009701 | 0.169021112 | 0.633114402 | 0.526658938 | 0.888334265 | D130020L05 | RIKEN cDNA D130020L05 gene [Source:M | 294.74 | 221 | 254.58 | 270.88 | 298.83 | 261.75 |
| 35744 | ENSMUSG00000097122 | 0.880724006 | 0.715724162 | 2.808295084 | 0.25486074 | 0.79883066 | NA | NA | NA | 0 | 0 | 2.0125 | 0 | 3.2719 | 0 |
| 35745 | ENSMUSG00000097123 | 22.03249685 | 0.819220038 | 0.516491957 | 1.586123513 | 0.112711294 | 0.492822297 | NA | NA | 9.6372 | 13.363 | 25.156 | 36.631 | 22.903 | 24.505 |
| 35746 | ENSMUSG00000097124 | 66.87918211 | 0.12006624 | 0.36430916 | 0.329572387 | 0.74172309 | 0.948901713 | A530020G2( | RIKEN cDNA A530020G20 gene [Source:M | 92.357 | 53.452 | 46.287 | 102.18 | 49.078 | 57.92 |
| 35747 | ENSMUSG00000097125 | 1.22362201 | -0.289417703 | 2.087230194 | -0.138661133 | 0.889717933 | NA | Gm26885 | predicted gene, 26885 [Source:MGI Symb | 0 | 1.0279 | 3.0187 | 0 | 2.1812 | 1.1138 |
| 35748 | ENSMUSG00000097127 | 0 | NA | NA | NA | NA | NA | Gm26886 | predicted gene, 26886 [Source:MGI Symb | 0 | 0 | 0 | 0 | 0 | 0 |
| 35749 | ENSMUSG00000097128 | 1.685019986 | 1.059807323 | 2.040743151 | 0.519324209 | 0.603534674 | NA | NA | NA | 3.2124 | 0 | 0 | 5.7839 | 0 | 1.1138 |
| 35750 | ENSMUSG00000097129 | 0.133850373 | -0.796629176 | 4.080472857 | -0.195229623 | 0.84521319 | NA | 4930507D05 | RIKEN cDNA 4930507D05 gene [Source:M | 0.8031 | 0 | 0 | 0 | 0 | 0 |
| 35751 | ENSMUSG00000097130 | 0 | NA | NA | NA | NA | NA | Gm5262 | predicted gene 5262 [Source:MGI Symbol | 0 | 0 | 0 | 0 | 0 | 0 |
| 35752 | ENSMUSG00000097131 | 120.446382 | -0.492727043 | 0.287691492 | -1.712692437 | 0.086769137 | 0.434115808 | NA | NA | 96.372 | 141.85 | 185.15 | 133.99 | 82.887 | 82.424 |
| 35753 | ENSMUSG00000097133 | 5.581786461 | -0.745710531 | 0.919573676 | -0.810930707 | 0.417405465 | 0.837009933 | Gm26628 | predicted gene, 26628 [Source:MGI Symb | 4.8186 | 7.1954 | 9.0562 | 5.7839 | 2.1812 | 4.4554 |
| 35754 | ENSMUSG00000097134 | 12.66607764 | -0.11768317 | 0.62524178 | -0.188220259 | 0.850703992 | 0.973463808 | 1110002J07I | RIKEN cDNA 1110002J07 gene [Source:M( | 12.047 | 11.307 | 16.1 | 6.7478 | 13.087 | 16.708 |
| 35755 | ENSMUSG00000097135 | 0 | NA | NA | NA | NA | NA | NA | NA | 0 | 0 | 0 | 0 | 0 | 0 |
| 35756 | ENSMUSG00000097136 | 1.949645972 | 0.690204738 | 1.601659659 | 0.430930962 | 0.666518569 | NA | NA | NA | 2.4093 | 1.0279 | 1.0062 | 2.8919 | 4.3625 | 0 |
| 35757 | ENSMUSG00000097137 | 1.305682109 | -1.392582258 | 2.033790705 | -0.684722501 | 0.493519047 | NA | Gm26627 | predicted gene, 26627 [Source:MGI Symb | 1.6062 | 1.0279 | 3.0187 | 0 | 2.1812 | 0 |
| 35758 | ENSMUSG00000097138 | 0 | NA | NA | NA | NA | NA | NA | NA | 0 | 0 | 0 | 0 | 0 | 0 |
| 35759 | ENSMUSG00000097139 | 0 | NA | NA | NA | NA | NA | Gm26626 | predicted gene, 26626 [Source:MGI Symb | 0 | 0 | 0 | 0 | 0 | 0 |
| 35760 | ENSMUSG00000097140 | 5.234370824 | -0.011496773 | 0.977116345 | -0.011766023 | 0.990612289 | 0.99859176 | Gm26779 | predicted gene, 26779 [Source:MGI Symb | 5.6217 | 4.1117 | 6.0375 | 6.7478 | 1.0906 | 7.7969 |
| 35761 | ENSMUSG00000097141 | 19.94150332 | -0.093378129 | 0.507927838 | -0.18384133 | 0.854137932 | 0.973825032 | Gm10524 | predicted gene 10524 [Source:MGI Symbo | 20.078 | 20.558 | 21.131 | 20.244 | 10.906 | 26.732 |
| 35762 | ENSMUSG00000097142 | 0 | NA | NA | NA | NA | NA | NA | NA | 0 | 0 | 0 | 0 | 0 | 0 |
| 35763 | ENSMUSG00000097145 | 18.98348872 | -0.121693996 | 0.514648788 | -0.23646028 | 0.813075526 | 0.966802988 | 9230114K14 | RIKEN cDNA 9230114K14 gene [Source:M( | 17.668 | 25.698 | 16.1 | 19.28 | 22.903 | 12.252 |
| 35764 | ENSMUSG00000097146 | 1.197325202 | -0.391798983 | 2.118729513 | -0.184921662 | 0.85329048 | NA | Gm4211 | predicted gene 4211 [Source:MGI Symbol | 4.0155 | 0 | 0 | 0.964 | 1.0906 | 1.1138 |
| 35765 | ENSMUSG00000097147 | 0 | NA | NA | NA | NA | NA | NA | NA | 0 | 0 | 0 | 0 | 0 | 0 |
| 35766 | ENSMUSG00000097148 | 192.6462436 | 1.561954961 | 0.274148525 | 5.697477161 | 1.22E-08 | 1.87E-06 | Gm3839 | predicted pseudogene 3839 [Source:MGI | 92.357 | 82.233 | 117.73 | 184.12 | 418.8 | 260.64 |
| 35767 | ENSMUSG00000097149 | 2.055939944 | -0.95025093 | 1.732961487 | -0.548339324 | 0.583458931 | NA | G630030J09 | RIKEN cDNA G630030J09 gene [Source:M( | 0 | 6.1675 | 2.0125 | 1.928 | 0 | 2.2277 |
| 35768 | ENSMUSG00000097150 | 4.235668452 | 0.56369751 | 1.170517028 | 0.481579932 | 0.630104385 | NA | Gm26513 | predicted gene, 26513 [Source:MGI Symb | 3.2124 | 3.0838 | 4.025 | 9.6398 | 5.4531 | 0 |
| 35769 | ENSMUSG00000097151 | 4.846525926 | 0.133561767 | 1.00763225 | 0.132550111 | 0.894549189 | NA | NA | NA | 4.8186 | 3.0838 | 6.0375 | 9.6398 | 3.2719 | 2.2277 |
| 35770 | ENSMUSG00000097152 | 7.597206227 | 0.015539829 | 0.794634184 | 0.019555954 | 0.984397601 | 0.998580735 | NA | NA | 6.4248 | 11.307 | 5.0312 | 9.6398 | 8.7249 | 4.4554 |
| 35771 | ENSMUSG00000097154 | 1.850840355 | -0.816767705 | 1.764256038 | -0.462953045 | 0.643398024 | NA | Gm26510 | predicted gene, 26510 [Source:MGI Symb | 4.0155 | 3.0838 | 0 | 2.8919 | 0 | 1.1138 |
| 35772 | ENSMUSG00000097155 | 1.212965344 | 0.520942145 | 2.088455765 | 0.249438917 | 0.803021285 | NA | Gm26511 | predicted gene, 26511 [Source:MGI Symb | 0 | 0 | 3.0187 | 0.964 | 2.1812 | 1.1138 |
| 35773 | ENSMUSG00000097156 | 0 | NA | NA | NA | NA | NA | Gm3764 | predicted gene 3764 [Source:MGI Symbol | 0 | 0 | 0 | 0 | 0 | 0 |
| 35774 | ENSMUSG00000097157 | 0 | NA | NA | NA | NA | NA | Gm26512 | predicted gene, 26512 [Source:MGI Symb | 0 | 0 | 0 | 0 | 0 | 0 |
| 35775 | ENSMUSG00000097158 | 8.03906572 | 0.107782792 | 0.771481795 | 0.139708796 | 0.888890077 | 0.982382091 | NA | NA | 8.031 | 5.1396 | 10.062 | 9.6398 | 10.906 | 4.4554 |
| 35776 | ENSMUSG00000097159 | 1.37618563 | 0.485221616 | 2.169618527 | 0.223643747 | 0.823034521 | NA | NA | NA | 2.4093 | 1.0279 | 0 | 4.8199 | 0 | 0 |
| 35777 | ENSMUSG00000097160 | 0.649696354 | 1.537929172 | 3.118369374 | 0.493183773 | 0.621882743 | NA | A630072L19 | RIKEN cDNA A630072L19 gene [Source:M( | 0 | 0 | 1.0062 | 2.8919 | 0 | 0 |
| 35778 | ENSMUSG00000097162 | 104.8926313 | -0.361244231 | 0.227701675 | -1.586480337 | 0.112630389 | 0.492766469 | 2310010J17I | RIKEN cDNA 2310010J17 gene [Source:M( | 118.86 | 111.02 | 123.77 | 84.83 | 85.068 | 105.82 |
| 35779 | ENSMUSG00000097163 | 0 | NA | NA | NA | NA | NA | BC051077 | cDNA sequence BC051077 [Source:MGI Sy | 0 | 0 | 0 | 0 | 0 | 0 |
| 35780 | ENSMUSG00000097164 | 50.99794182 | 0.049538885 | 0.328681173 | 0.150720178 | 0.880196455 | 0.981359269 | Cep83os | centrosomal protein 83, opposite strand [ | 59.43 | 44.2 | 46.287 | 40.487 | 64.346 | 51.237 |

| | | | | | | | | | | | | | | | |
|---|---|---|---|---|---|---|---|---|---|---|---|---|---|---|---|
| 35781 | ENSMUSG00000097165 | 0.465832987 | -0.698295221 | 3.18348759 | -0.219349126 | 0.826378096 | NA | NA | NA | 0.8031 | 1.0279 | 0 | 0.964 | 0 | 0 |
| 35782 | ENSMUSG00000097166 | 0.342639279 | -1.805976596 | 4.028547714 | -0.448294702 | 0.653940524 | NA | 9330179D12 | RIKEN cDNA 9330179D12 gene [Source:M | 0 | 2.0558 | 0 | 0 | 0 | 0 |
| 35783 | ENSMUSG00000097167 | 6.03892144 | -0.098823102 | 1.064051345 | -0.092874374 | 0.926003364 | 0.987546253 | Gm16740 | predicted gene, 16740 [Source:MGI Symb | 9.6372 | 0 | 9.0562 | 8.6758 | 2.1812 | 6.6831 |
| 35784 | ENSMUSG00000097168 | 1.845801186 | -0.083026757 | 2.201183858 | -0.037719138 | 0.969911617 | NA | NA | NA | 5.6217 | 0 | 0 | 0 | 5.4531 | 0 |
| 35785 | ENSMUSG00000097169 | 0 | NA | NA | NA | NA | NA | Gm26661 | predicted gene, 26661 [Source:MGI Symb | 0 | 0 | 0 | 0 | 0 | 0 |
| 35786 | ENSMUSG00000097170 | 3.811015166 | -0.020046173 | 1.206432932 | -0.016616069 | 0.986742905 | NA | Gm16982 | predicted gene, 16982 [Source:MGI Symb | 6.4248 | 0 | 5.0312 | 4.8199 | 4.3625 | 2.2277 |
| 35787 | ENSMUSG00000097171 | 0 | NA | NA | NA | NA | NA | Gm17644 | predicted gene, 17644 [Source:MGI Symb | 0 | 0 | 0 | 0 | 0 | 0 |
| 35788 | ENSMUSG00000097172 | 0 | NA | NA | NA | NA | NA | NA | NA | 0 | 0 | 0 | 0 | 0 | 0 |
| 35789 | ENSMUSG00000097173 | 0 | NA | NA | NA | NA | NA | NA | NA | 0 | 0 | 0 | 0 | 0 | 0 |
| 35790 | ENSMUSG00000097174 | 16.0323493 | 0.115192053 | 0.553297929 | 0.208191729 | 0.835079266 | 0.971073158 | Gm4890 | predicted gene 4890 [Source:MGI Symbol | 17.668 | 15.419 | 13.081 | 19.28 | 20.722 | 10.025 |
| 35791 | ENSMUSG00000097175 | 0 | NA | NA | NA | NA | NA | Gm27018 | predicted gene, 27018 [Source:MGI Symb | 0 | 0 | 0 | 0 | 0 | 0 |
| 35792 | ENSMUSG00000097176 | 1.367229137 | 0.773135066 | 2.151465153 | 0.359352818 | 0.719331166 | NA | NA | NA | 0 | 3.0838 | 0 | 2.8919 | 0 | 2.2277 |
| 35793 | ENSMUSG00000097177 | 28.40597429 | 0.460427863 | 0.459622336 | 1.001752585 | 0.316463103 | 0.763155505 | 9330159M07 | RIKEN cDNA 9330159M07 gene [Source:M | 24.896 | 25.698 | 21.131 | 31.811 | 49.078 | 17.822 |
| 35794 | ENSMUSG00000097178 | 0 | NA | NA | NA | NA | NA | 2310002F09 | RIKEN cDNA 2310002F09 gene [Source:M | 0 | 0 | 0 | 0 | 0 | 0 |
| 35795 | ENSMUSG00000097180 | 1.275928078 | 1.046915345 | 2.012681056 | 0.520159586 | 0.60295235 | NA | 2700038G22 | RIKEN cDNA 2700038G22 gene [Source:M | 2.4093 | 0 | 0 | 1.928 | 1.0906 | 2.2277 |
| 35796 | ENSMUSG00000097181 | 0.160662974 | 1.126924175 | 4.080472857 | 0.276174898 | 0.782413736 | NA | 4930594M17 | RIKEN cDNA 4930594M17 gene [Source:M | 0 | 0 | 0 | 0.964 | 0 | 0 |
| 35797 | ENSMUSG00000097182 | 0 | NA | NA | NA | NA | NA | A830009L08 | RIKEN cDNA A830009L08 gene [Source:M | 0 | 0 | 0 | 0 | 0 | 0 |
| 35798 | ENSMUSG00000097183 | 3.785514531 | 0.045314667 | 1.249571254 | 0.036264172 | 0.971071718 | NA | Gm17501 | predicted gene, 17501 [Source:MGI Symb | 8.031 | 0 | 3.0187 | 2.8919 | 6.5437 | 2.2277 |
| 35799 | ENSMUSG00000097184 | 12.49981859 | 0.068089787 | 0.639196579 | 0.106524017 | 0.915166601 | 0.98625062 | 4632428C04 | RIKEN cDNA 4632428C04 gene [Source:M | 11.243 | 7.1954 | 18.112 | 8.6758 | 14.178 | 15.594 |
| 35800 | ENSMUSG00000097185 | 1.566493383 | 1.095339961 | 1.736922175 | 0.630621208 | 0.528288228 | NA | Gm26596 | predicted gene, 26596 [Source:MGI Symb | 0 | 2.0558 | 1.0062 | 1.928 | 2.1812 | 2.2277 |
| 35801 | ENSMUSG00000097186 | 0 | NA | NA | NA | NA | NA | Gm26595 | predicted gene, 26595 [Source:MGI Symb | 0 | 0 | 0 | 0 | 0 | 0 |
| 35802 | ENSMUSG00000097187 | 32.2650034 | 0.041236707 | 0.502092892 | 0.082129637 | 0.934543626 | 0.988420305 | Gm19426 | predicted gene, 19426 [Source:MGI Symb | 14.456 | 31.865 | 49.306 | 26.027 | 21.812 | 50.123 |
| 35803 | ENSMUSG00000097189 | 2.524765368 | 1.575986932 | 1.38172183 | 1.140596391 | 0.25403792 | NA | Gm26594 | predicted gene, 26594 [Source:MGI Symb | 0.8031 | 2.0558 | 1.0062 | 5.7839 | 3.2719 | 2.2277 |
| 35804 | ENSMUSG00000097190 | 0 | NA | NA | NA | NA | NA | Gm11394 | predicted gene 11394 [Source:MGI Symbo | 0 | 0 | 0 | 0 | 0 | 0 |
| 35805 | ENSMUSG00000097191 | 0.538988726 | 1.126003851 | 3.343747182 | 0.336749099 | 0.736306047 | NA | Mm2pr | macrophage M2 polarization regulator [S | 0 | 0 | 1.0062 | 0 | 0 | 2.2277 |
| 35806 | ENSMUSG00000097193 | 6.865765907 | 0.834640385 | 1.014261635 | 0.822904423 | 0.410562347 | 0.833241465 | NA | NA | 9.6372 | 0 | 5.0312 | 14.46 | 8.7249 | 3.3415 |
| 35807 | ENSMUSG00000097194 | 38.03066448 | 0.253537256 | 0.46470741 | 0.545584708 | 0.585351439 | 0.909202452 | 9330175E14 | RIKEN cDNA 9330175E14 gene [Source:M | 26.502 | 24.67 | 53.331 | 67.478 | 28.356 | 27.846 |
| 35808 | ENSMUSG00000097195 | 104.258822 | 0.441743458 | 0.221019646 | 1.998661505 | 0.045644991 | 0.295628575 | Snhg5 | small nucleolar RNA host gene 5 [Source: | 88.341 | 90.457 | 86.537 | 123.39 | 124.33 | 112.5 |
| 35809 | ENSMUSG00000097196 | 0 | NA | NA | NA | NA | NA | Gm26665 | predicted gene, 26665 [Source:MGI Symb | 0 | 0 | 0 | 0 | 0 | 0 |
| 35810 | ENSMUSG00000097197 | 1.946389825 | 1.632806767 | 1.69009784 | 0.966101919 | 0.333993184 | NA | NA | NA | 0.8031 | 1.0279 | 1.0062 | 0 | 3.2719 | 5.5692 |
| 35811 | ENSMUSG00000097199 | 0 | NA | NA | NA | NA | NA | Gm26991 | predicted gene, 26991 [Source:MGI Symb | 0 | 0 | 0 | 0 | 0 | 0 |
| 35812 | ENSMUSG00000097201 | 0 | NA | NA | NA | NA | NA | NA | NA | 0 | 0 | 0 | 0 | 0 | 0 |
| 35813 | ENSMUSG00000097202 | 0 | NA | NA | NA | NA | NA | Gm26701 | predicted gene, 26701 [Source:MGI Symb | 0 | 0 | 0 | 0 | 0 | 0 |
| 35814 | ENSMUSG00000097203 | 108.4311188 | 0.959779141 | 0.29414443 | 3.262951946 | 0.001102582 | 0.022493298 | 4732419C18 | RIKEN cDNA 4732419C18 gene [Source:M | 94.766 | 72.982 | 52.325 | 107.97 | 126.51 | 196.04 |
| 35815 | ENSMUSG00000097204 | 20.44190147 | 0.159043999 | 0.487490846 | 0.326250227 | 0.744235043 | 0.949997329 | Gm17690 | predicted gene, 17690 [Source:MGI Symb | 19.274 | 16.447 | 22.137 | 16.388 | 28.356 | 20.049 |
| 35816 | ENSMUSG00000097205 | 0 | NA | NA | NA | NA | NA | Gm17540 | predicted gene, 17540 [Source:MGI Symb | 0 | 0 | 0 | 0 | 0 | 0 |
| 35817 | ENSMUSG00000097206 | 0 | NA | NA | NA | NA | NA | NA | NA | 0 | 0 | 0 | 0 | 0 | 0 |
| 35818 | ENSMUSG00000097207 | 12.42898255 | -0.253300523 | 0.631584651 | -0.40105554 | 0.688379234 | 0.936908509 | 6030443J06 | RIKEN cDNA 6030443J06 gene [Source:M | 16.062 | 8.2233 | 16.1 | 7.7118 | 11.997 | 14.48 |
| 35819 | ENSMUSG00000097208 | 13.89247547 | -0.662890193 | 0.587162461 | -1.128972366 | 0.258909491 | 0.712239465 | NA | NA | 17.668 | 13.363 | 20.125 | 13.496 | 10.906 | 7.7969 |
| 35820 | ENSMUSG00000097209 | 0.549180077 | 2.659271622 | 3.249668912 | 0.818320787 | 0.41317404 | NA | AU022754 | expressed sequence AU022754 [Source:M | 0 | 0 | 0 | 0 | 2.1812 | 1.1138 |
| 35821 | ENSMUSG00000097211 | 0 | NA | NA | NA | NA | NA | BC065403 | cDNA sequence BC065403 [Source:MGI Sy | 0 | 0 | 0 | 0 | 0 | 0 |
| 35822 | ENSMUSG00000097213 | 0 | NA | NA | NA | NA | NA | NA | NA | 0 | 0 | 0 | 0 | 0 | 0 |
| 35823 | ENSMUSG00000097214 | 0.499690045 | -0.82080011 | 3.113553401 | -0.263621658 | 0.792071468 | NA | Gm19791 | predicted gene, 19791 [Source:MGI Symb | 0 | 1.0279 | 1.0062 | 0.964 | 0 | 0 |
| 35824 | ENSMUSG00000097215 | 0 | NA | NA | NA | NA | NA | Gm26550 | predicted gene, 26550 [Source:MGI Symb | 0 | 0 | 0 | 0 | 0 | 0 |
| 35825 | ENSMUSG00000097216 | 0.185640647 | 1.126924175 | 4.080472857 | 0.276174898 | 0.782413736 | NA | 4932441J04 | RIKEN cDNA 4932441J04 gene [Source:M | 0 | 0 | 0 | 0 | 0 | 1.1138 |
| 35826 | ENSMUSG00000097217 | 8.92848607 | 0.352996139 | 0.72656819 | 0.485840344 | 0.627080367 | 0.920273791 | Gm26549 | predicted gene, 26549 [Source:MGI Symb | 11.243 | 5.1396 | 7.0437 | 12.532 | 9.8156 | 7.7969 |
| 35827 | ENSMUSG00000097218 | 0 | NA | NA | NA | NA | NA | Gm26552 | predicted gene, 26552 [Source:MGI Symb | 0 | 0 | 0 | 0 | 0 | 0 |
| 35828 | ENSMUSG00000097219 | 0.952772715 | -0.827065859 | 2.269489396 | -0.364428166 | 0.715538306 | NA | NA | NA | 1.6062 | 2.0558 | 0 | 0.964 | 1.0906 | 0 |
| 35829 | ENSMUSG00000097220 | 0 | NA | NA | NA | NA | NA | NA | NA | 0 | 0 | 0 | 0 | 0 | 0 |
| 35830 | ENSMUSG00000097221 | 0 | NA | NA | NA | NA | NA | 1810049J17 | RIKEN cDNA 1810049J17 gene [Source:M | 0 | 0 | 0 | 0 | 0 | 0 |
| 35831 | ENSMUSG00000097222 | 46.46909839 | -0.643000135 | 0.323629401 | -1.986840918 | 0.046940029 | 0.300126868 | Gata6os | GATA binding protein 6, opposite strand [ | 60.233 | 52.424 | 57.356 | 40.487 | 34.9 | 33.415 |
| 35832 | ENSMUSG00000097223 | 0.538988726 | 1.126003851 | 3.343747182 | 0.336749099 | 0.736306047 | NA | Gm2449 | predicted gene 2449 [Source:MGI Symbol | 0 | 0 | 1.0062 | 0 | 0 | 2.2277 |
| 35833 | ENSMUSG00000097224 | 0 | NA | NA | NA | NA | NA | Gm26716 | predicted gene, 26716 [Source:MGI Symb | 0 | 0 | 0 | 0 | 0 | 0 |
| 35834 | ENSMUSG00000097226 | 0 | NA | NA | NA | NA | NA | Gm26600 | predicted gene, 26600 [Source:MGI Symb | 0 | 0 | 0 | 0 | 0 | 0 |
| 35835 | ENSMUSG00000097228 | 0 | NA | NA | NA | NA | NA | Gm26597 | predicted gene, 26597 [Source:MGI Symb | 0 | 0 | 0 | 0 | 0 | 0 |
| 35836 | ENSMUSG00000097229 | 0.301557804 | -1.64335916 | 4.042814367 | -0.406488899 | 0.684383412 | NA | Platr23 | pluripotency associated transcript 23 [Sou | 0.8031 | 0 | 1.0062 | 0 | 0 | 0 |

| | | | | | | | | | | | | | | | |
|---|---|---|---|---|---|---|---|---|---|---|---|---|---|---|---|
| 35837 | ENSMUSG00000097230 | 0.17131964 | -0.796629176 | 4.080472857 | -0.195229623 | 0.84521319 | NA | NA | NA | 0 | 1.0279 | 0 | 0 | 0 | 0 |
| 35838 | ENSMUSG00000097231 | 7.05351582 | -0.011611109 | 0.846481633 | -0.013716906 | 0.989055836 | 0.99859176 | Gm26852 | predicted gene, 26852 [Source:MGI Symb | 8.031 | 4.1117 | 9.0562 | 5.7839 | 11.997 | 3.3415 |
| 35839 | ENSMUSG00000097232 | 0.321325949 | 1.911575114 | 4.048267893 | 0.472195804 | 0.636787034 | NA | Gm26854 | predicted gene, 26854 [Source:MGI Symb | 0 | 0 | 0 | 1.928 | 0 | 0 |
| 35840 | ENSMUSG00000097233 | 0.294513347 | 0.165143496 | 4.080472857 | 0.040471657 | 0.967717103 | NA | Gm17552 | predicted gene, 17552 [Source:MGI Symb | 0.8031 | 0 | 0 | 0.964 | 0 | 0 |
| 35841 | ENSMUSG00000097234 | 5.497943568 | -0.291910474 | 0.95324011 | -0.306229743 | 0.759429728 | 0.95402891 | NA | NA | 4.0155 | 4.1117 | 10.062 | 4.8199 | 2.1812 | 7.7969 |
| 35842 | ENSMUSG00000097235 | 0 | NA | NA | NA | NA | NA | NA | NA | 0 | 0 | 0 | 0 | 0 | 0 |
| 35843 | ENSMUSG00000097236 | 14.80409286 | 0.380580276 | 0.608456268 | 0.625485011 | 0.531652783 | 0.888947136 | NA | NA | 15.259 | 11.307 | 12.075 | 26.027 | 16.359 | 7.7969 |
| 35844 | ENSMUSG00000097237 | 0 | NA | NA | NA | NA | NA | Gm26519 | predicted gene, 26519 [Source:MGI Symb | 0 | 0 | 0 | 0 | 0 | 0 |
| 35845 | ENSMUSG00000097239 | 1.006244586 | -3.367758763 | 2.959437494 | -1.137972595 | 0.255131926 | NA | Gm27029 | predicted gene, 27029 [Source:MGI Symb | 0 | 0 | 6.0375 | 0 | 0 | 0 |
| 35846 | ENSMUSG00000097240 | 0.17131964 | -0.796629176 | 4.080472857 | -0.195229623 | 0.84521319 | NA | Gm26614 | predicted gene, 26614 [Source:MGI Symb | 0 | 1.0279 | 0 | 0 | 0 | 0 |
| 35847 | ENSMUSG00000097241 | 0 | NA | NA | NA | NA | NA | Gm26907 | predicted gene, 26907 [Source:MGI Symb | 0 | 0 | 0 | 0 | 0 | 0 |
| 35848 | ENSMUSG00000097242 | 16.41238624 | 0.338888529 | 0.534791398 | 0.633683582 | 0.526287342 | 0.888334265 | NA | NA | 16.062 | 13.363 | 14.087 | 24.099 | 15.269 | 15.594 |
| 35849 | ENSMUSG00000097244 | 0 | NA | NA | NA | NA | NA | Gm7928 | predicted gene 7928 [Source:MGI Symbol | 0 | 0 | 0 | 0 | 0 | 0 |
| 35850 | ENSMUSG00000097245 | 0.820751756 | 0.96244451 | 2.606827123 | 0.36920151 | 0.71197753 | NA | Gm5421 | predicted gene 5421 [Source:MGI Symbol | 1.6062 | 0 | 0 | 0 | 1.0906 | 2.2277 |
| 35851 | ENSMUSG00000097246 | 1.853833919 | -2.13156676 | 1.740984718 | -1.224345475 | 0.220821929 | NA | NA | NA | 4.0155 | 1.0279 | 4.025 | 0.964 | 1.0906 | 0 |
| 35852 | ENSMUSG00000097247 | 0 | NA | NA | NA | NA | NA | 1500012K07 | RIKEN cDNA 1500012K07 gene [Source:M | 0 | 0 | 0 | 0 | 0 | 0 |
| 35853 | ENSMUSG00000097248 | 0 | NA | NA | NA | NA | NA | Gm2694 | predicted gene 2694 [Source:MGI Symbol | 0 | 0 | 0 | 0 | 0 | 0 |
| 35854 | ENSMUSG00000097249 | 0 | NA | NA | NA | NA | NA | NA | NA | 0 | 0 | 0 | 0 | 0 | 0 |
| 35855 | ENSMUSG00000097250 | 0 | NA | NA | NA | NA | NA | Gm26771 | predicted gene, 26771 [Source:MGI Symb | 0 | 0 | 0 | 0 | 0 | 0 |
| 35856 | ENSMUSG00000097251 | 17.45118821 | 0.038084372 | 0.538930827 | 0.070666531 | 0.943663158 | 0.990077979 | 5033417F24 | RIKEN cDNA 5033417F24 gene [Source:M | 24.093 | 11.307 | 16.1 | 21.208 | 14.178 | 17.822 |
| 35857 | ENSMUSG00000097252 | 1.677608678 | -1.029528125 | 1.854119612 | -0.555265215 | 0.578713268 | NA | NA | NA | 1.6062 | 4.1117 | 1.0062 | 0 | 0 | 3.3415 |
| 35858 | ENSMUSG00000097253 | 2.192286423 | 0.471737428 | 1.519677119 | 0.310419511 | 0.756241959 | NA | Gm26770 | predicted gene, 26770 [Source:MGI Symb | 2.4093 | 1.0279 | 2.0125 | 0 | 4.3625 | 3.3415 |
| 35859 | ENSMUSG00000097254 | 5.795929154 | 0.540711576 | 0.973593096 | 0.555377373 | 0.578636566 | 0.908137944 | C430042M1 | RIKEN cDNA C430042M11 gene [Source:M | 4.0155 | 2.0558 | 8.05 | 1.928 | 9.8156 | 8.9108 |
| 35860 | ENSMUSG00000097255 | 0 | NA | NA | NA | NA | NA | Rpl31-ps7 | ribosomal protein L31, pseudogene 7 [Sou | 0 | 0 | 0 | 0 | 0 | 0 |
| 35861 | ENSMUSG00000097256 | 0 | NA | NA | NA | NA | NA | Gm26769 | predicted gene, 26769 [Source:MGI Symb | 0 | 0 | 0 | 0 | 0 | 0 |
| 35862 | ENSMUSG00000097257 | 0.469265235 | -2.274672822 | 3.439794491 | -0.661281605 | 0.508431736 | NA | NA | NA | 0.8031 | 0 | 2.0125 | 0 | 0 | 0 |
| 35863 | ENSMUSG00000097258 | 0.331982614 | 0.16514309 | 3.986751476 | 0.041422971 | 0.966958701 | NA | NA | NA | 0 | 1.0279 | 0 | 0.964 | 0 | 0 |
| 35864 | ENSMUSG00000097259 | 0.17131964 | -0.796629176 | 4.080472857 | -0.195229623 | 0.84521319 | NA | Gm26766 | predicted gene, 26766 [Source:MGI Symb | 0 | 1.0279 | 0 | 0 | 0 | 0 |
| 35865 | ENSMUSG00000097260 | 0 | NA | NA | NA | NA | NA | Gm5706 | predicted gene 5706 [Source:MGI Symbol | 0 | 0 | 0 | 0 | 0 | 0 |
| 35866 | ENSMUSG00000097261 | 1.627705466 | 0.148859422 | 1.683640856 | 0.088415188 | 0.92954669 | NA | 4930532G15 | RIKEN cDNA 4930532G15 gene [Source:M | 1.6062 | 1.0279 | 2.0125 | 2.8919 | 0 | 2.2277 |
| 35867 | ENSMUSG00000097262 | 0 | NA | NA | NA | NA | NA | 4933416M0 | RIKEN cDNA 4933416M07 gene [Source:M | 0 | 0 | 0 | 0 | 0 | 0 |
| 35868 | ENSMUSG00000097263 | 0.859617266 | 0.673351556 | 2.564846152 | 0.262530973 | 0.792912109 | NA | NA | NA | 0 | 0 | 2.0125 | 0.964 | 2.1812 | 0 |
| 35869 | ENSMUSG00000097264 | 0 | NA | NA | NA | NA | NA | Rpl31-ps24 | ribosomal protein L31, pseudogene 24 [Sc | 0 | 0 | 0 | 0 | 0 | 0 |
| 35870 | ENSMUSG00000097265 | 0 | NA | NA | NA | NA | NA | Gm26803 | predicted gene, 26803 [Source:MGI Symb | 0 | 0 | 0 | 0 | 0 | 0 |
| 35871 | ENSMUSG00000097266 | 0 | NA | NA | NA | NA | NA | Gm26802 | predicted gene, 26802 [Source:MGI Symb | 0 | 0 | 0 | 0 | 0 | 0 |
| 35872 | ENSMUSG00000097267 | 6.283416819 | 0.034066942 | 0.849521634 | 0.040101323 | 0.968012347 | 0.994231936 | 1700109K24 | RIKEN cDNA 1700109K24 gene [Source:M | 6.4248 | 4.1117 | 8.05 | 4.8199 | 8.7249 | 5.5692 |
| 35873 | ENSMUSG00000097268 | 0 | NA | NA | NA | NA | NA | Gm26805 | predicted gene, 26805 [Source:MGI Symb | 0 | 0 | 0 | 0 | 0 | 0 |
| 35874 | ENSMUSG00000097271 | 2.017825402 | -0.375571829 | 1.480073579 | -0.253752134 | 0.799687055 | NA | Gm9903 | predicted gene 9903 [Source:MGI Symbol | 0.8031 | 2.0558 | 4.025 | 1.928 | 2.1812 | 1.1138 |
| 35875 | ENSMUSG00000097272 | 0 | NA | NA | NA | NA | NA | NA | NA | 0 | 0 | 0 | 0 | 0 | 0 |
| 35876 | ENSMUSG00000097274 | 1.163184445 | 1.640230721 | 2.119996406 | 0.773695048 | 0.439111149 | NA | Gm47512 | predicted gene, 47512 [Source:MGI Symb | 1.6062 | 0 | 0 | 0.964 | 2.1812 | 2.2277 |
| 35877 | ENSMUSG00000097275 | 11.1797011 | 0.390356358 | 0.669322516 | 0.583211155 | 0.559751163 | 0.899475387 | Gm26648 | predicted gene, 26648 [Source:MGI Symb | 8.8341 | 10.279 | 10.062 | 20.244 | 7.6343 | 10.025 |
| 35878 | ENSMUSG00000097276 | 4.239167021 | -0.393322734 | 1.15303278 | -0.341120167 | 0.73301312 | NA | 4930525G2C | RIKEN cDNA 4930525G20 gene [Source:M | 3.2124 | 5.1396 | 6.0375 | 0 | 4.3625 | 6.6831 |
| 35879 | ENSMUSG00000097277 | 41.75085162 | -0.53629458 | 0.342511725 | -1.565769989 | 0.117402474 | 0.502643331 | 2900076A07 | RIKEN cDNA 2900076A07 gene [Source:M | 42.564 | 57.563 | 48.3 | 33.739 | 33.809 | 34.529 |
| 35880 | ENSMUSG00000097278 | 0 | NA | NA | NA | NA | NA | Gm26650 | predicted gene, 26650 [Source:MGI Symb | 0 | 0 | 0 | 0 | 0 | 0 |
| 35881 | ENSMUSG00000097279 | 0.160662974 | 1.126924175 | 4.080472857 | 0.276174898 | 0.782413736 | NA | Gm5106 | predicted gene 5106 [Source:MGI Symbol | 0 | 0 | 0 | 0.964 | 0 | 0 |
| 35882 | ENSMUSG00000097280 | 0 | NA | NA | NA | NA | NA | AI849053 | expressed sequence AI849053 [Source:M | 0 | 0 | 0 | 0 | 0 | 0 |
| 35883 | ENSMUSG00000097281 | 0 | NA | NA | NA | NA | NA | Gm26685 | predicted gene, 26685 [Source:MGI Symb | 0 | 0 | 0 | 0 | 0 | 0 |
| 35884 | ENSMUSG00000097282 | 0.89236178 | -1.92645888 | 2.54470475 | -0.757046129 | 0.449022227 | NA | 5031415H12 | RIKEN cDNA 5031415H12 gene [Source:M | 3.2124 | 1.0279 | 0 | 0 | 0 | 1.1138 |
| 35885 | ENSMUSG00000097283 | 0.133850373 | -0.796629176 | 4.080472857 | -0.195229623 | 0.84521319 | NA | Gm26686 | predicted gene, 26686 [Source:MGI Symb | 0.8031 | 0 | 0 | 0 | 0 | 0 |
| 35886 | ENSMUSG00000097284 | 40.06918759 | -0.061031949 | 0.384378244 | -0.158780966 | 0.873841446 | 0.979925081 | NA | NA | 55.414 | 31.865 | 35.219 | 47.235 | 28.356 | 42.326 |
| 35887 | ENSMUSG00000097285 | 0.647809292 | -2.733819264 | 3.061311498 | -0.893022244 | 0.371845261 | NA | NA | NA | 0.8031 | 3.0838 | 0 | 0 | 0 | 0 |
| 35888 | ENSMUSG00000097286 | 176.6087087 | 0.154586248 | 0.206878349 | 0.747232608 | 0.454923164 | 0.857337687 | Gm26684 | predicted gene, 26684 [Source:MGI Symb | 205.59 | 139.8 | 154.96 | 164.84 | 201.76 | 192.69 |
| 35889 | ENSMUSG00000097287 | 26.74911153 | -0.163612366 | 0.431925066 | -0.378798034 | 0.704837848 | 0.94295036 | D130017N0 | RIKEN cDNA D130017N08 gene [Source:M | 25.699 | 29.81 | 29.181 | 17.352 | 27.265 | 31.188 |
| 35890 | ENSMUSG00000097288 | 0 | NA | NA | NA | NA | NA | Gm2155 | predicted gene 2155 [Source:MGI Symbol | 0 | 0 | 0 | 0 | 0 | 0 |
| 35891 | ENSMUSG00000097289 | 0 | NA | NA | NA | NA | NA | NA | NA | 0 | 0 | 0 | 0 | 0 | 0 |
| 35892 | ENSMUSG00000097290 | 142.3380589 | 0.259033823 | 0.20187747 | 1.283123981 | 0.199448643 | 0.641417519 | 1300002E11 | RIKEN cDNA 1300002E11 gene [Source:M | 135.72 | 109.99 | 142.89 | 159.06 | 153.78 | 152.6 |

| | | | | | | | | | | | | | | | |
|---|---|---|---|---|---|---|---|---|---|---|---|---|---|---|---|
| 35893 | ENSMUSG00000097291 | 0 | NA | NA | NA | NA | NA | Gm18828 | predicted gene, 18828 [Source:MGI Symb | 0 | 0 | 0 | 0 | 0 | 0 |
| 35894 | ENSMUSG00000097292 | 4.62216924 | 0.833069529 | 1.0313898 | 0.807715501 | 0.419254378 | NA | A230107N01 | RIKEN cDNA A230107N01 gene [Source:M | 4.8186 | 2.0558 | 3.0187 | 6.7478 | 2.1812 | 8.9108 |
| 35895 | ENSMUSG00000097293 | 0 | NA | NA | NA | NA | NA | D630002J18 | RIKEN cDNA D630002J18 gene [Source:M | 0 | 0 | 0 | 0 | 0 | 0 |
| 35896 | ENSMUSG00000097294 | 5.825931281 | -0.031405867 | 0.948697549 | -0.033104193 | 0.973591499 | 0.996075612 | NA | NA | 5.6217 | 3.0838 | 9.0562 | 10.604 | 4.3625 | 2.2277 |
| 35897 | ENSMUSG00000097295 | 1.152998607 | 0.387569402 | 2.313943582 | 0.167493021 | 0.866982138 | NA | Hmgb1-ps8 | high mobility group box 1, pseudogene 8 | | 0 | 2.0558 | 1.0062 | 3.8559 | 0 | 0 |
| 35898 | ENSMUSG00000097296 | 9.784037666 | -0.887177735 | 0.732665871 | -1.210889943 | 0.225937588 | 0.6725584 | Gm26532 | predicted gene, 26532 [Source:MGI Symb | 8.8341 | 14.391 | 15.094 | 11.568 | 4.3625 | 4.4554 |
| 35899 | ENSMUSG00000097299 | 0 | NA | NA | NA | NA | NA | Gm26654 | predicted gene, 26654 [Source:MGI Symb | 0 | 0 | 0 | 0 | 0 | 0 |
| 35900 | ENSMUSG00000097300 | 1.149405861 | -1.242804601 | 2.372406571 | -0.523858185 | 0.60037718 | NA | NA | NA | 4.8186 | 0 | 0 | 0.964 | 0 | 1.1138 |
| 35901 | ENSMUSG00000097301 | 0 | NA | NA | NA | NA | NA | NA | NA | 0 | 0 | 0 | 0 | 0 | 0 |
| 35902 | ENSMUSG00000097302 | 0 | NA | NA | NA | NA | NA | 1700034B16 | RIKEN cDNA 1700034B16 gene [Source:M | 0 | 0 | 0 | 0 | 0 | 0 |
| 35903 | ENSMUSG00000097303 | 0.496077836 | -0.812642266 | 3.444120572 | -0.235950586 | 0.813471014 | NA | 3110083C13 | RIKEN cDNA 3110083C13 gene [Source:M | 0 | 0 | 2.0125 | 0.964 | 0 | 0 |
| 35904 | ENSMUSG00000097304 | 0 | NA | NA | NA | NA | NA | Rpl31-ps15 | ribosomal protein L31, pseudogene 15 [Sc | 0 | 0 | 0 | 0 | 0 | 0 |
| 35905 | ENSMUSG00000097305 | 1.445470437 | 1.188662981 | 1.824723707 | 0.651420802 | 0.514774886 | NA | Gm17276 | predicted gene, 17276 [Source:MGI Symb | 1.6062 | 0 | 1.0062 | 3.8559 | 1.0906 | 1.1138 |
| 35906 | ENSMUSG00000097307 | 0.17131964 | -0.796629176 | 4.080472857 | -0.195229623 | 0.84521319 | NA | Gm26834 | predicted gene, 26834 [Source:MGI Symb | 0 | 1.0279 | 0 | 0 | 0 | 0 |
| 35907 | ENSMUSG00000097308 | 1.493874554 | -0.797315781 | 2.351259729 | -0.339101534 | 0.734533243 | NA | Gm6410 | predicted gene 6410 [Source:MGI Symbol | 5.6217 | 0 | 0 | 0 | 0 | 3.3415 |
| 35908 | ENSMUSG00000097309 | 21.61177586 | 0.432569363 | 0.494154143 | 0.87537334 | 0.381370801 | 0.8130053 | Gm17249 | predicted gene, 17249 [Source:MGI Symb | 18.471 | 25.698 | 11.069 | 26.027 | 28.356 | 20.049 |
| 35909 | ENSMUSG00000097310 | 83.6619167 | -0.064679661 | 0.27211393 | -0.237693311 | 0.812118972 | 0.96653525 | A930038B10 | RIKEN cDNA A930038B10 gene [Source:M | 82.72 | 81.206 | 92.575 | 68.442 | 107.97 | 69.058 |
| 35910 | ENSMUSG00000097311 | 0.167707431 | -0.796629176 | 4.080472857 | -0.195229623 | 0.84521319 | NA | Gm26871 | predicted gene, 26871 [Source:MGI Symb | 0 | 0 | 1.0062 | 0 | 0 | 0 |
| 35911 | ENSMUSG00000097312 | 0 | NA | NA | NA | NA | NA | NA | NA | 0 | 0 | 0 | 0 | 0 | 0 |
| 35912 | ENSMUSG00000097313 | 1.679340755 | -1.206138109 | 1.839629894 | -0.65564172 | 0.512054675 | NA | Gm26569 | predicted gene, 26569 [Source:MGI Symb | 4.0155 | 0 | 3.0187 | 1.928 | 0 | 1.1138 |
| 35913 | ENSMUSG00000097315 | 0 | NA | NA | NA | NA | NA | Gm26568 | predicted gene, 26568 [Source:MGI Symb | 0 | 0 | 0 | 0 | 0 | 0 |
| 35914 | ENSMUSG00000097316 | 40.55903713 | -0.022290727 | 0.352883489 | -0.063167384 | 0.949633217 | 0.990731019 | Gm10516 | predicted gene 10516 [Source:MGI Symbo | 35.336 | 37.005 | 50.312 | 34.703 | 41.443 | 44.554 |
| 35915 | ENSMUSG00000097317 | 8.270059275 | -0.151353789 | 0.803249157 | -0.18842695 | 0.850541975 | 0.973432436 | NA | NA | 13.653 | 8.2233 | 4.025 | 3.8559 | 8.7249 | 11.138 |
| 35916 | ENSMUSG00000097318 | 27.68465975 | 0.172158125 | 0.407774544 | 0.422189485 | 0.672886716 | 0.931454408 | 1700007L15 | RIKEN cDNA 1700007L15 gene [Source:M | 27.305 | 22.614 | 28.175 | 31.811 | 28.356 | 27.846 |
| 35917 | ENSMUSG00000097320 | 27.04278321 | 0.429094252 | 0.443840977 | 0.966774756 | 0.333656647 | 0.775487565 | Tmem147os | transmembrane protein 147, opposite str | 22.487 | 21.586 | 25.156 | 33.739 | 40.353 | 18.935 |
| 35918 | ENSMUSG00000097321 | 182.8467012 | 0.047338263 | 0.230456713 | 0.205410648 | 0.837251302 | 0.971073158 | 1700028E10 | RIKEN cDNA 1700028E10 gene [Source:M | 137.33 | 173.72 | 229.42 | 211.11 | 199.58 | 145.91 |
| 35919 | ENSMUSG00000097322 | 1.922456421 | 1.627737368 | 2.018057085 | 0.806586385 | 0.419904821 | NA | A530083I20 | RIKEN cDNA A530083I20 gene [Source:M | 0.8031 | 2.0558 | 0 | 8.6758 | 0 | 0 |
| 35920 | ENSMUSG00000097323 | 0.622883753 | 0.113321311 | 2.884614835 | 0.039284729 | 0.968663382 | NA | 4930426I24 | RIKEN cDNA 4930426I24 gene [Source:M | 0.8031 | 0 | 1.0062 | 1.928 | 0 | 0 |
| 35921 | ENSMUSG00000097324 | 0.17131964 | -0.796629176 | 4.080472857 | -0.195229623 | 0.84521319 | NA | Carmn | cardiac mesoderm enhancer-associated n | 0 | 1.0279 | 0 | 0 | 0 | 0 |
| 35922 | ENSMUSG00000097325 | 0.963481514 | 0.243735673 | 2.098497405 | 0.116147712 | 0.907535476 | NA | Gm16897 | predicted gene, 16897 [Source:MGI Symb | 1.6062 | 0 | 1.0062 | 0.964 | 1.0906 | 1.1138 |
| 35923 | ENSMUSG00000097326 | 0 | NA | NA | NA | NA | NA | A330048O09 | RIKEN cDNA A330048O09 gene [Source:M | 0 | 0 | 0 | 0 | 0 | 0 |
| 35924 | ENSMUSG00000097327 | 46.01680519 | 0.025768039 | 0.329953216 | 0.078096039 | 0.937751658 | 0.988420305 | E030030I06 | RIKEN cDNA E030030I06 gene [Source:M | 51.399 | 35.977 | 49.306 | 50.127 | 43.625 | 45.668 |
| 35925 | ENSMUSG00000097328 | 101.359299 | 0.023984835 | 0.235315024 | 0.101926494 | 0.91881502 | 0.986374671 | Tnfsf12 | tumor necrosis factor (ligand) superfamily | 106.01 | 91.485 | 103.64 | 86.758 | 117.79 | 102.47 |
| 35926 | ENSMUSG00000097329 | 0.562214094 | -1.122836648 | 3.276281221 | -0.34271681 | 0.731811515 | NA | NA | NA | 2.4093 | 0 | 0 | 0.964 | 0 | 0 |
| 35927 | ENSMUSG00000097330 | 0.342639279 | -1.805976596 | 4.028547714 | -0.448294702 | 0.653940524 | NA | Gm26672 | predicted gene, 26672 [Source:MGI Symb | 0 | 2.0558 | 0 | 0 | 0 | 0 |
| 35928 | ENSMUSG00000097331 | 4.473332594 | -0.224698003 | 1.033822693 | -0.217346751 | 0.827938126 | NA | F420014N23 | RIKEN cDNA F420014N23 gene [Source:M | 2.4093 | 5.1396 | 7.0437 | 6.7478 | 3.2719 | 2.2277 |
| 35929 | ENSMUSG00000097332 | 0 | NA | NA | NA | NA | NA | NA | NA | 0 | 0 | 0 | 0 | 0 | 0 |
| 35930 | ENSMUSG00000097333 | 63.94168839 | -0.234419641 | 0.286338844 | -0.818679149 | 0.412969496 | 0.835177683 | Zfp87 | zinc finger protein 87 [Source:MGI Symbo | 61.036 | 75.038 | 71.443 | 53.983 | 69.8 | 52.351 |
| 35931 | ENSMUSG00000097334 | 5.298238982 | -0.454337565 | 0.932059463 | -0.487455557 | 0.62593553 | 0.919638611 | NA | NA | 7.2279 | 5.1396 | 6.0375 | 6.7478 | 2.1812 | 4.4554 |
| 35932 | ENSMUSG00000097335 | 1.912409926 | -2.118131359 | 1.772367538 | -1.195085846 | 0.232053491 | NA | NA | NA | 3.2124 | 2.0558 | 4.025 | 0 | 2.1812 | 0 |
| 35933 | ENSMUSG00000097336 | 34.70859556 | -0.136395883 | 0.404738242 | -0.336997765 | 0.736118585 | 0.948109711 | Fendrr | Foxf1 adjacent non-coding developmenta | 40.958 | 39.061 | 29.181 | 47.235 | 25.084 | 26.732 |
| 35934 | ENSMUSG00000097337 | 2.534156159 | -1.361935617 | 1.44754191 | -0.940860922 | 0.346776136 | NA | NA | NA | 4.8186 | 2.0558 | 4.025 | 0.964 | 0 | 3.3415 |
| 35935 | ENSMUSG00000097338 | 0 | NA | NA | NA | NA | NA | NA | NA | 0 | 0 | 0 | 0 | 0 | 0 |
| 35936 | ENSMUSG00000097339 | 0 | NA | NA | NA | NA | NA | Gm26671 | predicted gene, 26671 [Source:MGI Symb | 0 | 0 | 0 | 0 | 0 | 0 |
| 35937 | ENSMUSG00000097340 | 26.2114809 | 0.070182672 | 0.446820323 | 0.157071352 | 0.875188615 | 0.980512892 | NA | NA | 20.881 | 35.977 | 20.125 | 31.811 | 25.084 | 23.391 |
| 35938 | ENSMUSG00000097342 | 0 | NA | NA | NA | NA | NA | Gm26618 | predicted gene, 26618 [Source:MGI Symb | 0 | 0 | 0 | 0 | 0 | 0 |
| 35939 | ENSMUSG00000097343 | 12.44357331 | -0.220553512 | 0.608753053 | -0.362303746 | 0.717125059 | 0.9447719 | 9030407P20 | RIKEN cDNA 9030407P20 gene [Source:M | 12.85 | 10.279 | 17.106 | 13.496 | 10.906 | 10.025 |
| 35940 | ENSMUSG00000097344 | 0 | NA | NA | NA | NA | NA | NA | NA | 0 | 0 | 0 | 0 | 0 | 0 |
| 35941 | ENSMUSG00000097345 | 8.728835812 | 0.681534393 | 0.799491529 | 0.852459805 | 0.393958939 | 0.822827574 | NA | NA | 4.0155 | 7.1954 | 9.0562 | 13.496 | 15.269 | 3.3415 |
| 35942 | ENSMUSG00000097346 | 0 | NA | NA | NA | NA | NA | Rps18-ps4 | ribosomal protein S18, pseudogene 4 [Sou | 0 | 0 | 0 | 0 | 0 | 0 |
| 35943 | ENSMUSG00000097347 | 19.57448621 | -0.142497008 | 0.506516965 | -0.281327217 | 0.778459437 | 0.959559604 | NA | NA | 24.093 | 16.447 | 21.131 | 26.027 | 15.269 | 14.48 |
| 35944 | ENSMUSG00000097349 | 0 | NA | NA | NA | NA | NA | Gm26731 | predicted gene, 26731 [Source:MGI Symb | 0 | 0 | 0 | 0 | 0 | 0 |
| 35945 | ENSMUSG00000097350 | 9.972534869 | 0.436221992 | 0.672252767 | 0.648895792 | 0.516405735 | 0.886716042 | 4732491K20 | RIKEN cDNA 4732491K20 gene [Source:M | 7.2279 | 9.2513 | 9.0562 | 14.46 | 9.8156 | 10.025 |
| 35946 | ENSMUSG00000097352 | 1.052482331 | 1.09488462 | 2.207898318 | 0.495894494 | 0.619968851 | NA | C920009B18 | RIKEN cDNA C920009B18 gene [Source:M | 0 | 2.0558 | 0 | 0.964 | 2.1812 | 1.1138 |
| 35947 | ENSMUSG00000097353 | 67.59210986 | -0.368100488 | 0.293003546 | -1.256300455 | 0.209007057 | 0.65350014 | NA | NA | 85.129 | 76.066 | 67.418 | 71.334 | 62.165 | 43.44 |
| 35948 | ENSMUSG00000097354 | 386.3229933 | 0.540184728 | 0.180765886 | 2.988311233 | 0.002805237 | 0.044538916 | 2310001H17 | RIKEN cDNA 2310001H17 gene [Source:M | 269.84 | 288.84 | 386.4 | 452.11 | 384.99 | 535.76 |

| | | | | | | | | | | | | | | | |
|---|---|---|---|---|---|---|---|---|---|---|---|---|---|---|---|
| 35949 | ENSMUSG00000097355 | 0 | NA | NA | NA | NA | NA | NA | NA | 0 | 0 | 0 | 0 | 0 | 0 |
| 35950 | ENSMUSG00000097356 | 0 | NA | NA | NA | NA | NA | Gm26774 | predicted gene, 26774 [Source:MGI Symb | 0 | 0 | 0 | 0 | 0 | 0 |
| 35951 | ENSMUSG00000097357 | 24.5032239 | 0.059725622 | 0.459424355 | 0.130000992 | 0.896565642 | 0.982798755 | Gm16793 | predicted gene, 16793 [Source:MGI Symb | 18.471 | 22.614 | 31.194 | 31.811 | 23.994 | 18.935 |
| 35952 | ENSMUSG00000097358 | 16.96134848 | 0.207498166 | 0.601755264 | 0.344821523 | 0.730228552 | 0.946534338 | NA | NA | 17.668 | 16.447 | 13.081 | 17.352 | 30.537 | 6.6831 |
| 35953 | ENSMUSG00000097360 | 5.258283112 | 0.472040338 | 0.946164773 | 0.498898661 | 0.617850777 | 0.917509899 | 9430065F17 | RIKEN cDNA 9430065F17 gene [Source:M | 4.0155 | 5.1396 | 4.025 | 2.8919 | 5.4531 | 10.025 |
| 35954 | ENSMUSG00000097361 | 0 | NA | NA | NA | NA | NA | 4930550C17 | RIKEN cDNA 4930550C17 gene [Source:M | 0 | 0 | 0 | 0 | 0 | 0 |
| 35955 | ENSMUSG00000097362 | 0 | NA | NA | NA | NA | NA | Gm26544 | predicted gene, 26544 [Source:MGI Symb | 0 | 0 | 0 | 0 | 0 | 0 |
| 35956 | ENSMUSG00000097363 | 0.822793274 | -0.447303882 | 2.407293551 | -0.185811939 | 0.852592242 | NA | Gm26717 | predicted gene, 26717 [Source:MGI Symb | 0.8031 | 2.0558 | 0 | 0.964 | 0 | 1.1138 |
| 35957 | ENSMUSG00000097364 | 1.4022182 | 0.853620496 | 1.827897972 | 0.466995702 | 0.640502953 | NA | NA | NA | 0 | 1.0279 | 2.0125 | 0.964 | 2.1812 | 2.2277 |
| 35958 | ENSMUSG00000097365 | 0 | NA | NA | NA | NA | NA | C030034L19 | RIKEN cDNA C030034L19 gene [Source:M | 0 | 0 | 0 | 0 | 0 | 0 |
| 35959 | ENSMUSG00000097366 | 0 | NA | NA | NA | NA | NA | Gm8177 | predicted gene 8177 [Source:MGI Symbol | 0 | 0 | 0 | 0 | 0 | 0 |
| 35960 | ENSMUSG00000097367 | 0 | NA | NA | NA | NA | NA | NA | NA | 0 | 0 | 0 | 0 | 0 | 0 |
| 35961 | ENSMUSG00000097368 | 2.375146498 | -1.358195619 | 1.549173659 | -0.87672264 | 0.380637307 | NA | Gm10390 | predicted gene 10390 [Source:MGI Symbo | 7.2279 | 1.0279 | 2.0125 | 2.8919 | 1.0906 | 0 |
| 35962 | ENSMUSG00000097369 | 0.371281295 | 2.09253362 | 4.031852165 | 0.519000582 | 0.603760335 | NA | NA | NA | 0 | 0 | 0 | 0 | 0 | 2.2277 |
| 35963 | ENSMUSG00000097370 | 0.649696354 | 1.537929172 | 3.118369374 | 0.493183773 | 0.621882743 | NA | 4930467K11 | RIKEN cDNA 4930467K11 gene [Source:M | 0 | 0 | 1.0062 | 2.8919 | 0 | 0 |
| 35964 | ENSMUSG00000097371 | 0 | NA | NA | NA | NA | NA | Gm26875 | predicted gene, 26875 [Source:MGI Symb | 0 | 0 | 0 | 0 | 0 | 0 |
| 35965 | ENSMUSG00000097372 | 4.96151707 | 3.325818315 | 1.302146319 | 2.554104916 | 0.010646117 | NA | Gm26876 | predicted gene, 26876 [Source:MGI Symb | 1.6062 | 0 | 1.0062 | 18.316 | 3.2719 | 5.5692 |
| 35966 | ENSMUSG00000097373 | 0 | NA | NA | NA | NA | NA | Gm26877 | predicted gene, 26877 [Source:MGI Symb | 0 | 0 | 0 | 0 | 0 | 0 |
| 35967 | ENSMUSG00000097374 | 0.185640647 | 1.126924175 | 4.080472857 | 0.276174898 | 0.782413736 | NA | NA | NA | 0 | 0 | 0 | 0 | 0 | 1.1138 |
| 35968 | ENSMUSG00000097375 | 65.33285025 | 0.109584845 | 0.301764712 | 0.363146652 | 0.716495337 | 0.9447719 | 6720427I07R | RIKEN cDNA 6720427I07 gene [Source:MC | 73.082 | 60.647 | 54.337 | 52.055 | 86.159 | 65.717 |
| 35969 | ENSMUSG00000097377 | 0.663758638 | 2.950973782 | 3.025092005 | 0.975498853 | 0.32931287 | NA | NA | NA | 0 | 0 | 0 | 2.8919 | 1.0906 | 0 |
| 35970 | ENSMUSG00000097378 | 21.35502708 | 0.055748832 | 0.508918175 | 0.109543802 | 0.912771182 | 0.98625062 | B230208H11 | RIKEN cDNA B230208H11 gene [Source:M | 28.109 | 19.53 | 15.094 | 28.919 | 13.087 | 23.391 |
| 35971 | ENSMUSG00000097379 | 27.9862756 | 0.160584138 | 0.504085292 | 0.318565411 | 0.750056086 | 0.951817093 | Gm26873 | predicted gene, 26873 [Source:MGI Symb | 25.699 | 26.726 | 27.169 | 52.055 | 22.903 | 13.366 |
| 35972 | ENSMUSG00000097380 | 7.418621053 | 0.380391298 | 0.8682243 | 0.438125606 | 0.661295234 | 0.928341045 | NA | NA | 7.2279 | 6.1675 | 6.0375 | 17.352 | 3.2719 | 4.4554 |
| 35973 | ENSMUSG00000097381 | 0 | NA | NA | NA | NA | NA | A230087F16 | RIKEN cDNA A230087F16 gene [Source:M | 0 | 0 | 0 | 0 | 0 | 0 |
| 35974 | ENSMUSG00000097382 | 0 | NA | NA | NA | NA | NA | Gm26817 | predicted gene, 26817 [Source:MGI Symb | 0 | 0 | 0 | 0 | 0 | 0 |
| 35975 | ENSMUSG00000097383 | 28.77839529 | -0.222284629 | 0.431846158 | -0.514731057 | 0.606740968 | 0.916376001 | 1500026H17 | RIKEN cDNA 1500026H17 gene [Source:M | 37.746 | 35.977 | 19.119 | 26.991 | 29.447 | 23.391 |
| 35976 | ENSMUSG00000097384 | 0 | NA | NA | NA | NA | NA | Gm26815 | predicted gene, 26815 [Source:MGI Symb | 0 | 0 | 0 | 0 | 0 | 0 |
| 35977 | ENSMUSG00000097385 | 1.17206383 | 1.510749169 | 1.9879861 | 0.759939503 | 0.447290747 | NA | Gm26814 | predicted gene, 26814 [Source:MGI Symb | 0.8031 | 0 | 1.0062 | 1.928 | 2.1812 | 1.1138 |
| 35978 | ENSMUSG00000097386 | 0.476489653 | -2.293616034 | 3.421506946 | -0.670352587 | 0.502633051 | NA | 9130213A22 | RIKEN cDNA 9130213A22 gene [Source:M | 0.8031 | 2.0558 | 0 | 0 | 0 | 0 |
| 35979 | ENSMUSG00000097387 | 0.651473635 | 0.179413857 | 2.624432884 | 0.068362905 | 0.94549675 | NA | 4930563E18 | RIKEN cDNA 4930563E18 gene [Source:M | 0.8031 | 1.0279 | 0 | 0.964 | 0 | 1.1138 |
| 35980 | ENSMUSG00000097388 | 0.797609011 | 1.992900111 | 2.636035165 | 0.756021823 | 0.449636112 | NA | Gm3200 | predicted pseudogene 3200 [Source:MGI | 0.8031 | 0 | 0 | 2.8919 | 1.0906 | 0 |
| 35981 | ENSMUSG00000097390 | 0.647602702 | 0.168575675 | 2.628959883 | 0.064122574 | 0.948872627 | NA | NA | NA | 0.8031 | 1.0279 | 0 | 0.964 | 1.0906 | 0 |
| 35982 | ENSMUSG00000097391 | 0.185640647 | 1.126924175 | 4.080472857 | 0.276174898 | 0.782413736 | NA | Mirg | miRNA containing gene [Source:MGI Syml | 0 | 0 | 0 | 0 | 0 | 1.1138 |
| 35983 | ENSMUSG00000097392 | 168.0240422 | 0.198413102 | 0.18762326 | 1.057508019 | 0.290279797 | 0.740288056 | Thoc2l | THO complex subunit 2-like [Source:MGI S | 140.54 | 161.38 | 168.04 | 180.26 | 191.95 | 165.96 |
| 35984 | ENSMUSG00000097393 | 0 | NA | NA | NA | NA | NA | D030068K23 | RIKEN cDNA D030068K23 gene [Source:M | 0 | 0 | 0 | 0 | 0 | 0 |
| 35985 | ENSMUSG00000097396 | 0 | NA | NA | NA | NA | NA | Gm26556 | predicted gene, 26556 [Source:MGI Symb | 0 | 0 | 0 | 0 | 0 | 0 |
| 35986 | ENSMUSG00000097397 | 7.861745005 | 0.867337074 | 0.954980833 | 0.908224588 | 0.363759576 | 0.802534854 | Gm16861 | predicted gene, 16861 [Source:MGI Symb | 1.6062 | 11.307 | 4.025 | 19.28 | 8.7249 | 2.2277 |
| 35987 | ENSMUSG00000097398 | 0.556921942 | 2.679043159 | 3.638243063 | 0.736356289 | 0.461513899 | NA | Platr5 | pluripotency associated transcript 5 [Sour | 0 | 0 | 0 | 0 | 0 | 3.3415 |
| 35988 | ENSMUSG00000097399 | 0.36353943 | 2.065989254 | 4.034137301 | 0.512126658 | 0.608562368 | NA | Gm26555 | predicted gene, 26555 [Source:MGI Symb | 0 | 0 | 0 | 0 | 2.1812 | 0 |
| 35989 | ENSMUSG00000097401 | 0.506734502 | -2.372847156 | 3.338060886 | -0.710845978 | 0.477179685 | NA | Gm26744 | predicted gene, 26744 [Source:MGI Symb | 0 | 1.0279 | 2.0125 | 0 | 0 | 0 |
| 35990 | ENSMUSG00000097402 | 1.706922899 | 1.389751209 | 1.637256165 | 0.848829425 | 0.395976214 | NA | Gm17399 | predicted gene, 17399 [Source:MGI Symb | 0.8031 | 1.0279 | 1.0062 | 1.928 | 4.3625 | 1.1138 |
| 35991 | ENSMUSG00000097403 | 5.407365431 | -0.058753142 | 0.942138857 | -0.062361447 | 0.950274996 | 0.990815588 | 9230116N13 | RIKEN cDNA 9230116N13 gene [Source:M | 2.4093 | 7.1954 | 7.0437 | 4.8199 | 7.6343 | 3.3415 |
| 35992 | ENSMUSG00000097404 | 4.152646707 | 0.750510827 | 1.10341437 | 0.680171337 | 0.496395979 | NA | Gm10814 | predicted gene 10814 [Source:MGI Symbo | 3.2124 | 2.0558 | 4.025 | 5.7839 | 8.7249 | 1.1138 |
| 35993 | ENSMUSG00000097405 | 1.258021491 | -2.509349807 | 2.263169021 | -1.108777022 | 0.267526385 | NA | D630044L22 | RIKEN cDNA gene D630044L22 gene [Sour | 2.4093 | 0 | 4.025 | 0 | 0 | 1.1138 |
| 35994 | ENSMUSG00000097406 | 0 | NA | NA | NA | NA | NA | Gm26833 | predicted gene, 26833 [Source:MGI Symb | 0 | 0 | 0 | 0 | 0 | 0 |
| 35995 | ENSMUSG00000097407 | 1.008227331 | 2.377072972 | 2.239511216 | 1.061424901 | 0.288496845 | NA | 4933408J17R | RIKEN cDNA 4933408J17 gene [Source:MC | 0.8031 | 0 | 0 | 1.928 | 1.0906 | 2.2277 |
| 35996 | ENSMUSG00000097408 | 0 | NA | NA | NA | NA | NA | Gm26831 | predicted gene, 26831 [Source:MGI Symb | 0 | 0 | 0 | 0 | 0 | 0 |
| 35997 | ENSMUSG00000097409 | 0 | NA | NA | NA | NA | NA | Gm26832 | predicted gene, 26832 [Source:MGI Symb | 0 | 0 | 0 | 0 | 0 | 0 |
| 35998 | ENSMUSG00000097410 | 1.707905661 | 0.166649478 | 1.871711491 | 0.089035879 | 0.929053396 | NA | NA | NA | 0.8031 | 0 | 4.025 | 0.964 | 0 | 4.4554 |
| 35999 | ENSMUSG00000097411 | 2.447941428 | 0.116096613 | 1.497590333 | 0.077522278 | 0.93820807 | NA | B430218F22 | RIKEN cDNA B430218F22 gene [Source:M | 0 | 2.0558 | 5.0312 | 0.964 | 2.1812 | 4.4554 |
| 36000 | ENSMUSG00000097412 | 60.13132393 | 0.18063055 | 0.336603952 | 0.536626351 | 0.591525738 | 0.912621683 | 1810014B01 | RIKEN cDNA 1810014B01 gene [Source:M | 53.808 | 63.731 | 51.318 | 37.595 | 75.253 | 79.083 |
| 36001 | ENSMUSG00000097413 | 15.74631097 | 0.557419759 | 0.686678773 | 0.811762037 | 0.416928189 | 0.836878476 | A830052D11 | RIKEN cDNA A830052D11 gene [Source:M | 12.047 | 2.0558 | 24.15 | 23.135 | 15.269 | 17.822 |
| 36002 | ENSMUSG00000097414 | 4.620010394 | -0.375775633 | 1.072303976 | -0.350437601 | 0.726010312 | NA | B130046B21 | RIKEN cDNA B130046B21 gene [Source:M | 5.6217 | 3.0838 | 7.0437 | 8.6758 | 2.1812 | 1.1138 |
| 36003 | ENSMUSG00000097415 | 60.63433858 | -0.753011068 | 0.334736156 | -2.249565975 | 0.02447651 | 0.199836083 | AU020206 | expressed sequence AU020206 [Source:M | 74.689 | 94.568 | 59.368 | 57.839 | 29.447 | 47.895 |
| 36004 | ENSMUSG00000097416 | 2.36384367 | 1.433643755 | 1.520606358 | 0.942810575 | 0.345777801 | NA | Gm26670 | predicted gene, 26670 [Source:MGI Symb | 0.8031 | 3.0838 | 0 | 4.8199 | 4.3625 | 1.1138 |

| | | | | | | | | | | | | | | | |
|---|---|---|---|---|---|---|---|---|---|---|---|---|---|---|---|
| 36005 | ENSMUSG00000097417 | 15.30054276 | 0.22375703 | 0.655088382 | 0.341567698 | 0.73267625 | 0.947722168 | NA | NA | 16.062 | 13.363 | 13.081 | 32.775 | 8.7249 | 7.7969 |
| 36006 | ENSMUSG00000097418 | 0.335414862 | -1.779047242 | 4.030804179 | -0.441362855 | 0.658950329 | NA | Mir155hg | Mir155 host gene (non-protein coding) [S | 0 | 0 | 2.0125 | 0 | 0 | 0 |
| 36007 | ENSMUSG00000097419 | 0 | NA | NA | NA | NA | NA | Gm26666 | predicted gene, 26666 [Source:MGI Symb | 0 | 0 | 0 | 0 | 0 | 0 |
| 36008 | ENSMUSG00000097420 | 0 | NA | NA | NA | NA | NA | NA | NA | 0 | 0 | 0 | 0 | 0 | 0 |
| 36009 | ENSMUSG00000097421 | 0 | NA | NA | NA | NA | NA | D630011A20 | RIKEN cDNA D630011A20 gene [Source:M | 0 | 0 | 0 | 0 | 0 | 0 |
| 36010 | ENSMUSG00000097422 | 0 | NA | NA | NA | NA | NA | NA | NA | 0 | 0 | 0 | 0 | 0 | 0 |
| 36011 | ENSMUSG00000097423 | 0.321325949 | 1.911575114 | 4.048267893 | 0.472195804 | 0.636787034 | NA | NA | NA | 0 | 0 | 0 | 1.928 | 0 | 0 |
| 36012 | ENSMUSG00000097424 | 0.167707431 | -0.796629176 | 4.080472857 | -0.195229623 | 0.84521319 | NA | 1700030M09 | RIKEN cDNA 1700030M09 gene [Source:M | 0 | 0 | 1.0062 | 0 | 0 | 0 |
| 36013 | ENSMUSG00000097425 | 0 | NA | NA | NA | NA | NA | Vmn1r181 | vomeronasal 1 receptor 181 [Source:MGI | 0 | 0 | 0 | 0 | 0 | 0 |
| 36014 | ENSMUSG00000097426 | 1016.512531 | -0.386652928 | 0.413739351 | -0.934532639 | 0.350029208 | 0.791509322 | Gm8941 | predicted gene 8941 [Source:MGI Symbol | 1113.1 | 1164.6 | 1178.3 | 1195.3 | 1007.7 | 439.97 |
| 36015 | ENSMUSG00000097427 | 0 | NA | NA | NA | NA | NA | Gm6763 | predicted gene 6763 [Source:MGI Symbol | 0 | 0 | 0 | 0 | 0 | 0 |
| 36016 | ENSMUSG00000097428 | 0 | NA | NA | NA | NA | NA | AW047730 | expressed sequence AW047730 [Source:N | 0 | 0 | 0 | 0 | 0 | 0 |
| 36017 | ENSMUSG00000097429 | 0 | NA | NA | NA | NA | NA | Gm26520 | predicted gene, 26520 [Source:MGI Symb | 0 | 0 | 0 | 0 | 0 | 0 |
| 36018 | ENSMUSG00000097430 | 0 | NA | NA | NA | NA | NA | Gm10544 | predicted gene 10544 [Source:MGI Symbo | 0 | 0 | 0 | 0 | 0 | 0 |
| 36019 | ENSMUSG00000097431 | 42.59814233 | 0.285558285 | 0.363016563 | 0.786626051 | 0.431500805 | 0.844934876 | Gm26782 | predicted gene, 26782 [Source:MGI Symb | 28.109 | 47.284 | 40.25 | 54.947 | 35.99 | 49.009 |
| 36020 | ENSMUSG00000097433 | 1.288187577 | -1.445076435 | 1.907466647 | -0.757589359 | 0.448696852 | NA | Gm26781 | predicted gene, 26781 [Source:MGI Symb | 1.6062 | 2.0558 | 2.0125 | 0.964 | 1.0906 | 0 |
| 36021 | ENSMUSG00000097434 | 0 | NA | NA | NA | NA | NA | Gm16630 | predicted gene, 16630 [Source:MGI Symb | 0 | 0 | 0 | 0 | 0 | 0 |
| 36022 | ENSMUSG00000097435 | 0 | NA | NA | NA | NA | NA | Gm26783 | predicted gene, 26783 [Source:MGI Symb | 0 | 0 | 0 | 0 | 0 | 0 |
| 36023 | ENSMUSG00000097438 | 0 | NA | NA | NA | NA | NA | Gm26784 | predicted gene, 26784 [Source:MGI Symb | 0 | 0 | 0 | 0 | 0 | 0 |
| 36024 | ENSMUSG00000097439 | 7.804625281 | 0.035058413 | 0.798066603 | 0.043929181 | 0.964960854 | 0.993480943 | Gm16754 | predicted gene, 16754 [Source:MGI Symb | 12.85 | 5.1396 | 5.0312 | 10.604 | 7.6343 | 5.5692 |
| 36025 | ENSMUSG00000097440 | 8.120460254 | -1.370884361 | 0.783164458 | -1.750442511 | 0.080041986 | 0.415341728 | Gm6277 | predicted gene 6277 [Source:MGI Symbol | 8.8341 | 11.307 | 15.094 | 5.7839 | 4.3625 | 3.3415 |
| 36026 | ENSMUSG00000097441 | 0 | NA | NA | NA | NA | NA | Gm26633 | predicted gene, 26633 [Source:MGI Symb | 0 | 0 | 0 | 0 | 0 | 0 |
| 36027 | ENSMUSG00000097442 | 0.587191767 | -1.12394062 | 3.226482889 | -0.348348545 | 0.72757844 | NA | Gm26632 | predicted gene, 26632 [Source:MGI Symb | 2.4093 | 0 | 0 | 0 | 0 | 1.1138 |
| 36028 | ENSMUSG00000097443 | 7.526120291 | -0.770840471 | 0.85440837 | -0.902192087 | 0.366954837 | 0.805239112 | Gm17529 | predicted gene, 17529 [Source:MGI Symb | 3.2124 | 13.363 | 12.075 | 7.7118 | 5.4531 | 3.3415 |
| 36029 | ENSMUSG00000097444 | 1.359307311 | 0.057869561 | 1.97828699 | 0.029252359 | 0.976663323 | NA | NA | NA | 0 | 1.0279 | 3.0187 | 1.928 | 2.1812 | 0 |
| 36030 | ENSMUSG00000097445 | 11.94412365 | -0.700056203 | 0.672175976 | -1.041477571 | 0.297653958 | 0.748091545 | Gm26631 | predicted gene, 26631 [Source:MGI Symb | 8.031 | 12.335 | 24.15 | 10.604 | 7.6343 | 8.9108 |
| 36031 | ENSMUSG00000097446 | 0 | NA | NA | NA | NA | NA | Gm26629 | predicted gene, 26629 [Source:MGI Symb | 0 | 0 | 0 | 0 | 0 | 0 |
| 36032 | ENSMUSG00000097447 | 0 | NA | NA | NA | NA | NA | Gm26630 | predicted gene, 26630 [Source:MGI Symb | 0 | 0 | 0 | 0 | 0 | 0 |
| 36033 | ENSMUSG00000097448 | 4.800016417 | 2.374242688 | 1.080857026 | 2.196629741 | 0.0280469 | NA | Platr22 | pluripotency associated transcript 22 [Sou | 1.6062 | 1.0279 | 2.0125 | 8.6758 | 5.4531 | 10.025 |
| 36034 | ENSMUSG00000097450 | 0 | NA | NA | NA | NA | NA | Gm26893 | predicted gene, 26893 [Source:MGI Symb | 0 | 0 | 0 | 0 | 0 | 0 |
| 36035 | ENSMUSG00000097451 | 6.35618844 | 0.7406548 | 0.875679699 | 0.845805608 | 0.397661197 | 0.825336712 | Rian | RNA imprinted and accumulated in nucleu | 8.031 | 3.0838 | 3.0187 | 9.6398 | 5.4531 | 8.9108 |
| 36036 | ENSMUSG00000097452 | 0 | NA | NA | NA | NA | NA | Gm2824 | predicted gene 2824 [Source:MGI Symbol | 0 | 0 | 0 | 0 | 0 | 0 |
| 36037 | ENSMUSG00000097453 | 0.94782191 | 1.090542322 | 2.508664729 | 0.43471027 | 0.663772743 | NA | NA | NA | 0.8031 | 1.0279 | 0 | 3.8559 | 0 | 0 |
| 36038 | ENSMUSG00000097454 | 0 | NA | NA | NA | NA | NA | NA | NA | 0 | 0 | 0 | 0 | 0 | 0 |
| 36039 | ENSMUSG00000097455 | 0.321325949 | 1.911575114 | 4.048267893 | 0.472195804 | 0.636787034 | NA | Gm26891 | predicted gene, 26891 [Source:MGI Symb | 0 | 0 | 0 | 1.928 | 0 | 0 |
| 36040 | ENSMUSG00000097456 | 0 | NA | NA | NA | NA | NA | Gm16958 | predicted gene, 16958 [Source:MGI Symb | 0 | 0 | 0 | 0 | 0 | 0 |
| 36041 | ENSMUSG00000097457 | 2.249782557 | 1.338689993 | 1.429550099 | 0.936441468 | 0.349045936 | NA | NA | NA | 0.8031 | 1.0279 | 2.0125 | 1.928 | 3.2719 | 4.4554 |
| 36042 | ENSMUSG00000097458 | 7.951421156 | -0.718409969 | 0.956894384 | -0.75077248 | 0.452789594 | 0.856769845 | NA | NA | 9.6372 | 3.0838 | 17.106 | 13.496 | 3.2719 | 1.1138 |
| 36043 | ENSMUSG00000097459 | 0.133850373 | -0.796629176 | 4.080472857 | -0.195229623 | 0.84521319 | NA | NA | NA | 0.8031 | 0 | 0 | 0 | 0 | 0 |
| 36044 | ENSMUSG00000097460 | 1.725838878 | 0.771628 | 1.873274568 | 0.411913989 | 0.680402467 | NA | NA | NA | 0.8031 | 0 | 3.0187 | 0.964 | 0 | 5.5692 |
| 36045 | ENSMUSG00000097461 | 0 | NA | NA | NA | NA | NA | NA | NA | 0 | 0 | 0 | 0 | 0 | 0 |
| 36046 | ENSMUSG00000097462 | 5.124773557 | -0.955402501 | 0.951014394 | -1.004614132 | 0.31508269 | 0.761215823 | 9530026P05 | RIKEN cDNA 9530026P05 gene [Source:M | 8.031 | 8.2233 | 4.025 | 3.8559 | 3.2719 | 3.3415 |
| 36047 | ENSMUSG00000097463 | 0.294513347 | 0.165143496 | 4.080472857 | 0.040471657 | 0.967717103 | NA | 4930448E22 | RIKEN cDNA 4930448E22 gene [Source:M | 0.8031 | 0 | 0 | 0.964 | 0 | 0 |
| 36048 | ENSMUSG00000097464 | 1.386553083 | 1.815488084 | 1.932603549 | 0.93940016 | 0.347525331 | NA | Gm26736 | predicted gene, 26736 [Source:MGI Symb | 0.8031 | 0 | 1.0062 | 0.964 | 1.0906 | 4.4554 |
| 36049 | ENSMUSG00000097466 | 0 | NA | NA | NA | NA | NA | D430036J16 | RIKEN cDNA D430036J16 gene [Source:M | 0 | 0 | 0 | 0 | 0 | 0 |
| 36050 | ENSMUSG00000097467 | 1.113694413 | -0.251399571 | 1.97572234 | -0.127244384 | 0.898746978 | NA | Gm26737 | predicted gene, 26737 [Source:MGI Symb | 1.6062 | 1.0279 | 1.0062 | 1.928 | 0 | 1.1138 |
| 36051 | ENSMUSG00000097468 | 0 | NA | NA | NA | NA | NA | NA | NA | 0 | 0 | 0 | 0 | 0 | 0 |
| 36052 | ENSMUSG00000097469 | 0 | NA | NA | NA | NA | NA | NA | NA | 0 | 0 | 0 | 0 | 0 | 0 |
| 36053 | ENSMUSG00000097470 | 0 | NA | NA | NA | NA | NA | Gm26587 | predicted gene, 26587 [Source:MGI Symb | 0 | 0 | 0 | 0 | 0 | 0 |
| 36054 | ENSMUSG00000097471 | 0.677847551 | 0.063310748 | 2.809814301 | 0.022532004 | 0.982023583 | NA | 5830432E09 | RIKEN cDNA 5830432E09 gene [Source:M | 0 | 0 | 2.0125 | 0.964 | 1.0906 | 0 |
| 36055 | ENSMUSG00000097472 | 2.134729709 | 1.741424721 | 1.945160249 | 0.895260286 | 0.370647965 | NA | Gm26586 | predicted gene, 26586 [Source:MGI Symb | 0 | 0 | 3.0187 | 8.6758 | 0 | 1.1138 |
| 36056 | ENSMUSG00000097473 | 0 | NA | NA | NA | NA | NA | Gm26585 | predicted gene, 26585 [Source:MGI Symb | 0 | 0 | 0 | 0 | 0 | 0 |
| 36057 | ENSMUSG00000097474 | 12.75004866 | 0.055764042 | 0.64101987 | 0.086992688 | 0.930677324 | 0.988420305 | NA | NA | 12.047 | 15.419 | 10.062 | 11.568 | 20.722 | 6.6831 |
| 36058 | ENSMUSG00000097476 | 0.503122293 | -2.366675966 | 3.785596405 | -0.625179156 | 0.531853481 | NA | Gm26583 | predicted gene, 26583 [Source:MGI Symb | 0 | 0 | 3.0187 | 0 | 0 | 0 |
| 36059 | ENSMUSG00000097479 | 3.340069459 | -1.094210721 | 1.207591874 | -0.906109708 | 0.364877787 | NA | NA | NA | 6.4248 | 5.1396 | 2.0125 | 0.964 | 3.2719 | 2.2277 |
| 36060 | ENSMUSG00000097480 | 0 | NA | NA | NA | NA | NA | Gm9499 | predicted gene 9499 [Source:MGI Symbol | 0 | 0 | 0 | 0 | 0 | 0 |

| | | | | | | | | | | | | | | | |
|---|---|---|---|---|---|---|---|---|---|---|---|---|---|---|---|
| 36061 | ENSMUSG00000097481 | 0 | NA | NA | NA | NA | NA | NA | NA | 0 | 0 | 0 | 0 | 0 | 0 |
| 36062 | ENSMUSG00000097482 | 52.28142156 | -0.049356699 | 0.368693563 | -0.133869164 | 0.893506034 | 0.982562372 | Gm17634 | predicted gene, 17634 [Source:MGI Symb | 47.383 | 61.675 | 50.312 | 28.919 | 74.162 | 51.237 |
| 36063 | ENSMUSG00000097483 | 0 | NA | NA | NA | NA | NA | Gm26806 | predicted gene, 26806 [Source:MGI Symb | 0 | 0 | 0 | 0 | 0 | 0 |
| 36064 | ENSMUSG00000097484 | 0 | NA | NA | NA | NA | NA | Gm26807 | predicted gene, 26807 [Source:MGI Symb | 0 | 0 | 0 | 0 | 0 | 0 |
| 36065 | ENSMUSG00000097486 | 0 | NA | NA | NA | NA | NA | Gm17733 | predicted gene, 17733 [Source:MGI Symb | 0 | 0 | 0 | 0 | 0 | 0 |
| 36066 | ENSMUSG00000097487 | 4.065221794 | -0.668387386 | 1.063201163 | -0.62865562 | 0.529574539 | NA | Ptges3l | prostaglandin E synthase 3 like [Source:M | 4.8186 | 5.1396 | 5.0312 | 3.8559 | 1.0906 | 4.4554 |
| 36067 | ENSMUSG00000097488 | 13.83083575 | 0.444756947 | 0.625472425 | 0.711073629 | 0.477038611 | 0.869671045 | 4732487G21 | RIKEN cDNA 4732487G21 gene [Source:M | 9.6372 | 19.53 | 6.0375 | 12.532 | 18.541 | 16.708 |
| 36068 | ENSMUSG00000097489 | 6.116044333 | 0.804569057 | 0.873614418 | 0.920965864 | 0.357068249 | 0.796166767 | NA | NA | 3.2124 | 7.1954 | 3.0187 | 6.7478 | 8.7249 | 7.7969 |
| 36069 | ENSMUSG00000097491 | 0.319491021 | 0.16514334 | 4.044653141 | 0.040830038 | 0.967431393 | NA | Gm2366 | predicted gene 2366 [Source:MGI Symbol | 0.8031 | 0 | 0 | 0 | 0 | 1.1138 |
| 36070 | ENSMUSG00000097493 | 3.833879725 | 0.264247937 | 1.099180357 | 0.24040453 | 0.810016667 | NA | 9930014A18 | RIKEN cDNA 9930014A18 gene [Source:M | 2.4093 | 2.0558 | 6.0375 | 4.8199 | 5.4531 | 2.2277 |
| 36071 | ENSMUSG00000097494 | 3.891142111 | 0.141079882 | 1.178930736 | 0.119667659 | 0.90474642 | NA | 4933406C10 | RIKEN cDNA 4933406C10 gene [Source:M | 4.0155 | 4.1117 | 3.0187 | 6.7478 | 5.4531 | 0 |
| 36072 | ENSMUSG00000097495 | 0 | NA | NA | NA | NA | NA | Gm26651 | predicted gene, 26651 [Source:MGI Symb | 0 | 0 | 0 | 0 | 0 | 0 |
| 36073 | ENSMUSG00000097496 | 0 | NA | NA | NA | NA | NA | Gm26653 | predicted gene, 26653 [Source:MGI Symb | 0 | 0 | 0 | 0 | 0 | 0 |
| 36074 | ENSMUSG00000097497 | 0 | NA | NA | NA | NA | NA | Gm26652 | predicted gene, 26652 [Source:MGI Symb | 0 | 0 | 0 | 0 | 0 | 0 |
| 36075 | ENSMUSG00000097500 | 0 | NA | NA | NA | NA | NA | Gm26537 | predicted gene, 26537 [Source:MGI Symb | 0 | 0 | 0 | 0 | 0 | 0 |
| 36076 | ENSMUSG00000097501 | 8.431988652 | 1.573003205 | 0.803765503 | 1.957042444 | 0.050342478 | 0.315279767 | NA | NA | 6.4248 | 5.1396 | 1.0062 | 11.568 | 13.087 | 13.366 |
| 36077 | ENSMUSG00000097502 | 0 | NA | NA | NA | NA | NA | 4930528D03 | RIKEN cDNA 4930528D03 gene [Source:M | 0 | 0 | 0 | 0 | 0 | 0 |
| 36078 | ENSMUSG00000097503 | 17.06382276 | 0.682332519 | 0.519254631 | 1.314061499 | 0.188825509 | 0.625984911 | 3110045C21 | RIKEN cDNA 3110045C21 gene [Source:M | 12.85 | 15.419 | 11.069 | 21.208 | 22.903 | 18.935 |
| 36079 | ENSMUSG00000097504 | 22.28729396 | 1.380932352 | 0.490293191 | 2.816544013 | 0.004854339 | 0.065497352 | 4930516B21 | RIKEN cDNA 4930516B21 gene [Source:M | 8.8341 | 13.363 | 15.094 | 27.955 | 27.265 | 41.212 |
| 36080 | ENSMUSG00000097505 | 0 | NA | NA | NA | NA | NA | NA | NA | 0 | 0 | 0 | 0 | 0 | 0 |
| 36081 | ENSMUSG00000097507 | 0 | NA | NA | NA | NA | NA | NA | NA | 0 | 0 | 0 | 0 | 0 | 0 |
| 36082 | ENSMUSG00000097509 | 3.10578596 | 1.376480403 | 1.342876084 | 1.02502414 | 0.305351797 | NA | B230322F03 | RIKEN cDNA B230322F03 gene [Source:M | 4.0155 | 1.0279 | 0 | 4.8199 | 6.5437 | 2.2277 |
| 36083 | ENSMUSG00000097511 | 0 | NA | NA | NA | NA | NA | Gm16677 | predicted gene, 16677 [Source:MGI Symb | 0 | 0 | 0 | 0 | 0 | 0 |
| 36084 | ENSMUSG00000097513 | 0 | NA | NA | NA | NA | NA | Gm26696 | predicted gene, 26696 [Source:MGI Symb | 0 | 0 | 0 | 0 | 0 | 0 |
| 36085 | ENSMUSG00000097514 | 0.301557804 | -1.64335916 | 4.042814367 | -0.406488899 | 0.684383412 | NA | NA | NA | 0.8031 | 0 | 1.0062 | 0 | 0 | 0 |
| 36086 | ENSMUSG00000097515 | 1.03140222 | -2.145901642 | 2.420890855 | -0.886409909 | 0.375396682 | NA | 1700040D17 | RIKEN cDNA 1700040D17 gene [Source:M | 0 | 2.0558 | 3.0187 | 0 | 0 | 1.1138 |
| 36087 | ENSMUSG00000097516 | 0 | NA | NA | NA | NA | NA | 4930467D21 | RIKEN cDNA 4930467D21 gene [Source:M | 0 | 0 | 0 | 0 | 0 | 0 |
| 36088 | ENSMUSG00000097517 | 0 | NA | NA | NA | NA | NA | NA | NA | 0 | 0 | 0 | 0 | 0 | 0 |
| 36089 | ENSMUSG00000097518 | 1.191858604 | 0.487587703 | 2.09676255 | 0.232543119 | 0.816116201 | NA | NA | NA | 0 | 0 | 3.0187 | 1.928 | 1.0906 | 1.1138 |
| 36090 | ENSMUSG00000097519 | 0.922606629 | -1.977583741 | 2.486779435 | -0.795238899 | 0.426474548 | NA | 4930558J18I | RIKEN cDNA 4930558J18 gene [Source:M( | 2.4093 | 0 | 2.0125 | 0 | 0 | 1.1138 |
| 36091 | ENSMUSG00000097520 | 0 | NA | NA | NA | NA | NA | 4930488L21I | RIKEN cDNA 4930488L21 gene [Source:M( | 0 | 0 | 0 | 0 | 0 | 0 |
| 36092 | ENSMUSG00000097522 | 0 | NA | NA | NA | NA | NA | Gm26844 | predicted gene, 26844 [Source:MGI Symb | 0 | 0 | 0 | 0 | 0 | 0 |
| 36093 | ENSMUSG00000097523 | 10.05995397 | 0.467311824 | 0.672247935 | 0.695148024 | 0.486962541 | 0.873874218 | 1700022N22 | RIKEN cDNA 1700022N22 gene [Source:M | 8.031 | 8.2233 | 9.0562 | 9.6398 | 9.8156 | 15.594 |
| 36094 | ENSMUSG00000097524 | 0 | NA | NA | NA | NA | NA | NA | NA | 0 | 0 | 0 | 0 | 0 | 0 |
| 36095 | ENSMUSG00000097525 | 0.77079641 | 0.856159616 | 2.652840961 | 0.322733111 | 0.746897374 | NA | Platr31 | pluripotency associated transcript 31 [Sou | 1.6062 | 0 | 0 | 1.928 | 1.0906 | 0 |
| 36096 | ENSMUSG00000097526 | 0.99938009 | -1.013527422 | 2.420827554 | -0.418669814 | 0.67545746 | NA | NA | NA | 0 | 2.0558 | 2.0125 | 1.928 | 0 | 0 |
| 36097 | ENSMUSG00000097527 | 2.176730558 | -0.604704241 | 1.528877306 | -0.395521758 | 0.692457867 | NA | 1700112J16I | RIKEN cDNA 1700112J16 gene [Source:M( | 4.8186 | 0 | 3.0187 | 1.928 | 2.1812 | 1.1138 |
| 36098 | ENSMUSG00000097529 | 0 | NA | NA | NA | NA | NA | Rp31-ps19 | ribosomal protein L31, pseudogene 19 [Sc | 0 | 0 | 0 | 0 | 0 | 0 |
| 36099 | ENSMUSG00000097530 | 3.092861424 | 1.518908553 | 1.320254285 | 1.150466672 | 0.249951713 | NA | Kansl2-ps | KAT8 regulatory NSL complex subunit 2, p | 0.8031 | 1.0279 | 3.0187 | 4.8199 | 1.0906 | 7.7969 |
| 36100 | ENSMUSG00000097532 | 0 | NA | NA | NA | NA | NA | Gm4349 | predicted gene 4349 [Source:MGI Symbol | 0 | 0 | 0 | 0 | 0 | 0 |
| 36101 | ENSMUSG00000097533 | 0.844158739 | -0.371321695 | 2.388630557 | -0.155453799 | 0.876463574 | NA | Gm26590 | predicted gene, 26590 [Source:MGI Symb | 0.8031 | 1.0279 | 1.0062 | 0 | 0 | 2.2277 |
| 36102 | ENSMUSG00000097534 | 23.99848639 | 0.503280074 | 0.464772134 | 1.082853376 | 0.278873509 | 0.731519729 | Gm16675 | predicted gene, 16675 [Source:MGI Symb | 24.896 | 14.391 | 20.125 | 31.811 | 32.719 | 20.049 |
| 36103 | ENSMUSG00000097535 | 0 | NA | NA | NA | NA | NA | Gm26592 | predicted gene, 26592 [Source:MGI Symb | 0 | 0 | 0 | 0 | 0 | 0 |
| 36104 | ENSMUSG00000097536 | 20.27126181 | -0.285100087 | 0.548330485 | -0.519942069 | 0.603103953 | 0.915859383 | 2610037D02 | RIKEN cDNA 2610037D02 gene [Source:M | 36.14 | 13.363 | 17.106 | 23.135 | 19.631 | 12.252 |
| 36105 | ENSMUSG00000097537 | 5.524490277 | 0.050759862 | 0.908248665 | 0.055887627 | 0.955431328 | 0.991678478 | 2610020C07 | RIKEN cDNA 2610020C07 gene [Source:M | 8.031 | 4.1117 | 4.025 | 4.8199 | 4.3625 | 7.7969 |
| 36106 | ENSMUSG00000097538 | 0 | NA | NA | NA | NA | NA | Gm26939 | predicted gene, 26939 [Source:MGI Symb | 0 | 0 | 0 | 0 | 0 | 0 |
| 36107 | ENSMUSG00000097539 | 0 | NA | NA | NA | NA | NA | 4930477E14 | RIKEN cDNA 4930477E14 gene [Source:M | 0 | 0 | 0 | 0 | 0 | 0 |
| 36108 | ENSMUSG00000097540 | 0 | NA | NA | NA | NA | NA | NA | NA | 0 | 0 | 0 | 0 | 0 | 0 |
| 36109 | ENSMUSG00000097541 | 0 | NA | NA | NA | NA | NA | NA | NA | 0 | 0 | 0 | 0 | 0 | 0 |
| 36110 | ENSMUSG00000097542 | 0 | NA | NA | NA | NA | NA | NA | NA | 0 | 0 | 0 | 0 | 0 | 0 |
| 36111 | ENSMUSG00000097543 | 0 | NA | NA | NA | NA | NA | Gm805 | predicted gene 805 [Source:MGI Symbol;/ | 0 | 0 | 0 | 0 | 0 | 0 |
| 36112 | ENSMUSG00000097544 | 0 | NA | NA | NA | NA | NA | NA | NA | 0 | 0 | 0 | 0 | 0 | 0 |
| 36113 | ENSMUSG00000097545 | 0 | NA | NA | NA | NA | NA | Mir124a-1hg | Mir124-1 host gene (non-protein coding) | 0 | 0 | 0 | 0 | 0 | 0 |
| 36114 | ENSMUSG00000097546 | 0.931665974 | -0.888641078 | 2.464021547 | -0.360646634 | 0.718363623 | NA | Gm26749 | predicted gene, 26749 [Source:MGI Symb | 1.6062 | 2.0558 | 0 | 1.928 | 0 | 0 |
| 36115 | ENSMUSG00000097547 | 0 | NA | NA | NA | NA | NA | B230110C06 | RIKEN cDNA B230110C06 gene [Source:M | 0 | 0 | 0 | 0 | 0 | 0 |
| 36116 | ENSMUSG00000097548 | 0 | NA | NA | NA | NA | NA | Gm26748 | predicted gene, 26748 [Source:MGI Symb | 0 | 0 | 0 | 0 | 0 | 0 |

| | | | | | | | | | | | | | | | |
|---|---|---|---|---|---|---|---|---|---|---|---|---|---|---|---|
| 36117 | ENSMUSG00000097550 | 0 | NA | NA | NA | NA | NA | Gm21304 | predicted gene, 21304 [Source:MGI Symb | 0 | 0 | 0 | 0 | 0 | 0 |
| 36118 | ENSMUSG00000097551 | 0 | NA | NA | NA | NA | NA | NA | NA | 0 | 0 | 0 | 0 | 0 | 0 |
| 36119 | ENSMUSG00000097552 | 0 | NA | NA | NA | NA | NA | Rps19-ps12 | ribosomal protein S19, pseudogene 12 [Sc | 0 | 0 | 0 | 0 | 0 | 0 |
| 36120 | ENSMUSG00000097553 | 0 | NA | NA | NA | NA | NA | NA | NA | 0 | 0 | 0 | 0 | 0 | 0 |
| 36121 | ENSMUSG00000097554 | 0.315620088 | 0.165143421 | 4.063300985 | 0.040642675 | 0.967580763 | NA | NA | NA | 0.8031 | 0 | 0 | 0 | 1.0906 | 0 |
| 36122 | ENSMUSG00000097557 | 0 | NA | NA | NA | NA | NA | C130060C02 | RIKEN cDNA C130060C02 gene [Source:M | 0 | 0 | 0 | 0 | 0 | 0 |
| 36123 | ENSMUSG00000097558 | 0.185640647 | 1.126924175 | 4.080472857 | 0.276174898 | 0.782413736 | NA | Gm26902 | predicted gene, 26902 [Source:MGI Symb | 0 | 0 | 0 | 0 | 0 | 1.1138 |
| 36124 | ENSMUSG00000097559 | 4.439578149 | 0.842227848 | 1.182438597 | 0.712280409 | 0.476291153 | NA | D430018E03 | RIKEN cDNA D430018E03 gene [Source:M | 6.4248 | 0 | 3.0187 | 10.604 | 4.3625 | 2.2277 |
| 36125 | ENSMUSG00000097560 | 0.328370405 | 0.16514316 | 4.003149536 | 0.041253308 | 0.967093956 | NA | NA | NA | 0 | 0 | 1.0062 | 0.964 | 0 | 0 |
| 36126 | ENSMUSG00000097561 | 1.378811218 | 1.806657107 | 1.901640051 | 0.950052091 | 0.342085785 | NA | 4632411P08 | RIKEN cDNA 4632411P08 gene [Source:M | 0.8031 | 0 | 1.0062 | 0.964 | 3.2719 | 2.2277 |
| 36127 | ENSMUSG00000097562 | 0 | NA | NA | NA | NA | NA | Gm26639 | predicted gene, 26639 [Source:MGI Symb | 0 | 0 | 0 | 0 | 0 | 0 |
| 36128 | ENSMUSG00000097563 | 0 | NA | NA | NA | NA | NA | Gm26638 | predicted gene, 26638 [Source:MGI Symb | 0 | 0 | 0 | 0 | 0 | 0 |
| 36129 | ENSMUSG00000097564 | 12.25968023 | 0.12657859 | 0.690918935 | 0.183203243 | 0.85463855 | 0.97395562 | C430014B12 | RIKEN cDNA C430014B12 gene [Source:M | 17.668 | 13.363 | 4.025 | 16.388 | 7.6343 | 14.48 |
| 36130 | ENSMUSG00000097565 | 0 | NA | NA | NA | NA | NA | Gm26965 | predicted gene, 26965 [Source:MGI Symb | 0 | 0 | 0 | 0 | 0 | 0 |
| 36131 | ENSMUSG00000097566 | 3.961355892 | -0.413637179 | 1.15251089 | -0.358900886 | 0.719669237 | NA | NA | NA | 6.4248 | 6.1675 | 1.0062 | 5.7839 | 3.2719 | 1.1138 |
| 36132 | ENSMUSG00000097567 | 3.287335724 | -0.575853998 | 1.492976899 | -0.385708579 | 0.699712509 | NA | NA | NA | 4.8186 | 0 | 7.0437 | 6.7478 | 0 | 1.1138 |
| 36133 | ENSMUSG00000097569 | 0 | NA | NA | NA | NA | NA | Gm26640 | predicted gene, 26640 [Source:MGI Symb | 0 | 0 | 0 | 0 | 0 | 0 |
| 36134 | ENSMUSG00000097571 | 56.87142447 | 0.117250072 | 0.3070522 | 0.381857129 | 0.702567341 | 0.942405009 | Jpx | Jpx transcript, Xist activator (non-protein | 62.642 | 48.312 | 52.325 | 49.163 | 71.981 | 56.806 |
| 36135 | ENSMUSG00000097572 | 0.481988923 | 2.492401104 | 3.837077151 | 0.649557203 | 0.515978285 | NA | Gm26797 | predicted gene, 26797 [Source:MGI Symb | 0 | 0 | 0 | 2.8919 | 0 | 0 |
| 36136 | ENSMUSG00000097573 | 6.213594036 | -1.400349992 | 0.915706716 | -1.529256002 | 0.126200992 | 0.521094669 | G730003C15 | RIKEN cDNA G730003C15 gene [Source:M | 12.85 | 7.1954 | 7.0437 | 5.7839 | 2.1812 | 2.2277 |
| 36137 | ENSMUSG00000097574 | 97.76893566 | 0.220004964 | 0.250127353 | 0.879571792 | 0.379091326 | 0.812043903 | C920006O11 | RIKEN cDNA C920006O11 gene [Source:M | 106.01 | 86.345 | 78.487 | 126.28 | 98.156 | 91.335 |
| 36138 | ENSMUSG00000097575 | 0 | NA | NA | NA | NA | NA | Gm26796 | predicted gene, 26796 [Source:MGI Symb | 0 | 0 | 0 | 0 | 0 | 0 |
| 36139 | ENSMUSG00000097576 | 1.357756082 | -2.652992791 | 2.055796826 | -1.29049367 | 0.196879308 | NA | D930030I03 | RIKEN cDNA D930030I03 gene [Source:M | 4.0155 | 1.0279 | 2.0125 | 0 | 1.0906 | 0 |
| 36140 | ENSMUSG00000097577 | 14.49419922 | -0.223881268 | 0.593215987 | -0.37740262 | 0.70587442 | 0.94295036 | 6230400D17 | RIKEN cDNA 6230400D17 gene [Source:M | 18.471 | 13.363 | 15.094 | 20.244 | 11.997 | 7.7969 |
| 36141 | ENSMUSG00000097578 | 3.382340587 | -0.518820597 | 1.238940824 | -0.418761402 | 0.675390516 | NA | NA | NA | 4.8186 | 4.1117 | 3.0187 | 2.8919 | 5.4531 | 0 |
| 36142 | ENSMUSG00000097579 | 5.03744681 | 0.766912822 | 0.96087996 | 0.798135931 | 0.424791612 | NA | NA | NA | 4.0155 | 5.1396 | 2.0125 | 5.7839 | 4.3625 | 8.9108 |
| 36143 | ENSMUSG00000097580 | 0 | NA | NA | NA | NA | NA | Gm5432 | predicted gene 5432 [Source:MGI Symbol | 0 | 0 | 0 | 0 | 0 | 0 |
| 36144 | ENSMUSG00000097582 | 0 | NA | NA | NA | NA | NA | NA | NA | 0 | 0 | 0 | 0 | 0 | 0 |
| 36145 | ENSMUSG00000097583 | 11.06249311 | -0.256294758 | 0.708381697 | -0.361803191 | 0.717499107 | 0.9447719 | 6430590A07 | RIKEN cDNA 6430590A07 gene [Source:M | 4.8186 | 17.475 | 14.087 | 13.496 | 9.8156 | 6.6831 |
| 36146 | ENSMUSG00000097584 | 0 | NA | NA | NA | NA | NA | 1700125G22 | RIKEN cDNA 1700125G22 gene [Source:M | 0 | 0 | 0 | 0 | 0 | 0 |
| 36147 | ENSMUSG00000097585 | 5.626217334 | -1.115651735 | 1.070884855 | -1.041803635 | 0.297502728 | 0.748091545 | E230029C05 | RIKEN cDNA E230029C05 gene [Source:M | 4.8186 | 8.2233 | 10.062 | 1.928 | 8.7249 | 0 |
| 36148 | ENSMUSG00000097586 | 0 | NA | NA | NA | NA | NA | Gm26528 | predicted gene, 26528 [Source:MGI Symb | 0 | 0 | 0 | 0 | 0 | 0 |
| 36149 | ENSMUSG00000097587 | 0.294513347 | 0.165143496 | 4.080472857 | 0.040471657 | 0.967717103 | NA | 4930578M01 | RIKEN cDNA 4930578M01 gene [Source:M | 0.8031 | 0 | 0 | 0.964 | 0 | 0 |
| 36150 | ENSMUSG00000097589 | 215.2984925 | 0.187436767 | 0.214120961 | 0.875377946 | 0.381368296 | 0.8130053 | Dleu2 | deleted in lymphocytic leukemia, 2 [Sourc | 146.97 | 194.28 | 263.64 | 229.43 | 223.58 | 233.91 |
| 36151 | ENSMUSG00000097590 | 0 | NA | NA | NA | NA | NA | NA | NA | 0 | 0 | 0 | 0 | 0 | 0 |
| 36152 | ENSMUSG00000097591 | 0.824622689 | 0.984749497 | 2.871267556 | 0.342966818 | 0.731623423 | NA | A330032B11 | RIKEN cDNA A330032B11 gene [Source:M | 1.6062 | 0 | 0 | 0 | 0 | 3.3415 |
| 36153 | ENSMUSG00000097592 | 0 | NA | NA | NA | NA | NA | 4930500F10 | RIKEN cDNA 4930500F10 gene [Source:M | 0 | 0 | 0 | 0 | 0 | 0 |
| 36154 | ENSMUSG00000097593 | 0 | NA | NA | NA | NA | NA | Gm26676 | predicted gene, 26676 [Source:MGI Symb | 0 | 0 | 0 | 0 | 0 | 0 |
| 36155 | ENSMUSG00000097594 | 0 | NA | NA | NA | NA | NA | Gm2907 | predicted gene 2907 [Source:MGI Symbol | 0 | 0 | 0 | 0 | 0 | 0 |
| 36156 | ENSMUSG00000097595 | 4.839526974 | 1.524413177 | 1.202958392 | 1.267220202 | 0.205076564 | NA | 1500002F19 | RIKEN cDNA 1500002F19 gene [Source:M | 2.4093 | 4.1117 | 1.0062 | 10.604 | 10.906 | 0 |
| 36157 | ENSMUSG00000097596 | 0 | NA | NA | NA | NA | NA | Gm26673 | predicted gene, 26673 [Source:MGI Symb | 0 | 0 | 0 | 0 | 0 | 0 |
| 36158 | ENSMUSG00000097597 | 2.322943281 | -1.763180233 | 1.54280878 | -1.142837827 | 0.25310593 | NA | Gm26674 | predicted gene, 26674 [Source:MGI Symb | 1.6062 | 6.1675 | 3.0187 | 0.964 | 2.1812 | 0 |
| 36159 | ENSMUSG00000097598 | 0 | NA | NA | NA | NA | NA | NA | NA | 0 | 0 | 0 | 0 | 0 | 0 |
| 36160 | ENSMUSG00000097599 | 0 | NA | NA | NA | NA | NA | NA | NA | 0 | 0 | 0 | 0 | 0 | 0 |
| 36161 | ENSMUSG00000097601 | 12.28759933 | -0.183739413 | 0.638482895 | -0.287774996 | 0.773518978 | 0.958177785 | Gm26660 | predicted gene, 26660 [Source:MGI Symb | 17.668 | 14.391 | 7.0437 | 12.532 | 8.7249 | 13.366 |
| 36162 | ENSMUSG00000097602 | 0.730949792 | 3.072456834 | 2.934263219 | 1.04709653 | 0.29505506 | NA | 4930519P11 | RIKEN cDNA 4930519P11 gene [Source:M | 0 | 0 | 0 | 0 | 3.2719 | 1.1138 |
| 36163 | ENSMUSG00000097603 | 0.335414862 | -1.779047242 | 4.030804179 | -0.441362855 | 0.658950329 | NA | A430010J10 | RIKEN cDNA A430010J10 gene [Source:M | 0 | 0 | 2.0125 | 0 | 0 | 0 |
| 36164 | ENSMUSG00000097604 | 0.17131964 | -0.796629176 | 4.080472857 | -0.195229623 | 0.84521319 | NA | Gm17322 | predicted gene, 17322 [Source:MGI Symb | 0 | 1.0279 | 0 | 0 | 0 | 0 |
| 36165 | ENSMUSG00000097605 | 0 | NA | NA | NA | NA | NA | NA | NA | 0 | 0 | 0 | 0 | 0 | 0 |
| 36166 | ENSMUSG00000097608 | 0 | NA | NA | NA | NA | NA | Gm7952 | predicted gene 7952 [Source:MGI Symbol | 0 | 0 | 0 | 0 | 0 | 0 |
| 36167 | ENSMUSG00000097609 | 0 | NA | NA | NA | NA | NA | NA | NA | 0 | 0 | 0 | 0 | 0 | 0 |
| 36168 | ENSMUSG00000097610 | 0.486939728 | -0.698472448 | 3.137608162 | -0.222613026 | 0.823836697 | NA | A930012L18 | RIKEN cDNA A930012L18 gene [Source:M | 0.8031 | 1.0279 | 0 | 0 | 1.0906 | 0 |
| 36169 | ENSMUSG00000097612 | 0 | NA | NA | NA | NA | NA | Gm26509 | predicted gene, 26509 [Source:MGI Symb | 0 | 0 | 0 | 0 | 0 | 0 |
| 36170 | ENSMUSG00000097613 | 6.866044332 | 0.043885107 | 0.798337946 | 0.054970588 | 0.956161895 | 0.991678478 | Gm17597 | predicted gene, 17597 [Source:MGI Symb | 8.031 | 7.1954 | 5.0312 | 7.7118 | 6.5437 | 6.6831 |
| 36171 | ENSMUSG00000097614 | 0 | NA | NA | NA | NA | NA | NA | NA | 0 | 0 | 0 | 0 | 0 | 0 |
| 36172 | ENSMUSG00000097615 | 18.82476076 | -1.084504374 | 0.57210837 | -1.895627527 | 0.058009315 | 0.342343159 | Gm2061 | predicted gene 2061 [Source:MGI Symbol | 19.274 | 18.503 | 39.244 | 18.316 | 9.8156 | 7.7969 |

| | | | | | | | | | | | | | | | |
|---|---|---|---|---|---|---|---|---|---|---|---|---|---|---|---|
| 36173 | ENSMUSG00000097616 | 46.35814164 | -0.043029059 | 0.362792015 | -0.11860531 | 0.905588059 | 0.983972587 | 1110019D14 | RIKEN cDNA 1110019D14 gene [Source:M | 56.217 | 47.284 | 37.231 | 31.811 | 62.165 | 43.44 |
| 36174 | ENSMUSG00000097617 | 9.056351568 | -1.032706544 | 0.739976336 | -1.395594013 | 0.162836786 | 0.586471932 | Gm10687 | predicted gene 10687 [Source:MGI Symbo | 12.047 | 15.419 | 9.0562 | 6.7478 | 3.2719 | 7.7969 |
| 36175 | ENSMUSG00000097618 | 0 | NA | NA | NA | NA | NA | NA | NA | 0 | 0 | 0 | 0 | 0 | 0 |
| 36176 | ENSMUSG00000097619 | 0 | NA | NA | NA | NA | NA | 4833422M21 | RIKEN cDNA 4833422M21 gene [Source:M | 0 | 0 | 0 | 0 | 0 | 0 |
| 36177 | ENSMUSG00000097620 | 5.891137148 | -0.012967573 | 1.011402843 | -0.012821373 | 0.989770305 | 0.99859176 | 4921514A10 | RIKEN cDNA 4921514A10 gene [Source:M | 1.6062 | 10.279 | 6.0375 | 8.6758 | 7.6343 | 1.1138 |
| 36178 | ENSMUSG00000097621 | 0 | NA | NA | NA | NA | NA | Gm26562 | predicted gene, 26562 [Source:MGI Symb | 0 | 0 | 0 | 0 | 0 | 0 |
| 36179 | ENSMUSG00000097622 | 0.160662974 | 1.126924175 | 4.080472857 | 0.276174898 | 0.782413736 | NA | A330033J07I | RIKEN cDNA A330033J07 gene [Source:M( | 0 | 0 | 0 | 0.964 | 0 | 0 |
| 36180 | ENSMUSG00000097623 | 0 | NA | NA | NA | NA | NA | B230323A14 | RIKEN cDNA B230323A14 gene [Source:M | 0 | 0 | 0 | 0 | 0 | 0 |
| 36181 | ENSMUSG00000097624 | 0 | NA | NA | NA | NA | NA | Gm5091 | predicted gene 5091 [Source:MGI Symbol | 0 | 0 | 0 | 0 | 0 | 0 |
| 36182 | ENSMUSG00000097625 | 0 | NA | NA | NA | NA | NA | NA | NA | 0 | 0 | 0 | 0 | 0 | 0 |
| 36183 | ENSMUSG00000097626 | 0.353348078 | 0.165142701 | 3.895042568 | 0.042398176 | 0.966181282 | NA | 4921504A21 | RIKEN cDNA 4921504A21 gene [Source:M | 0 | 0 | 1.0062 | 0 | 0 | 1.1138 |
| 36184 | ENSMUSG00000097627 | 0 | NA | NA | NA | NA | NA | 4930519A11 | RIKEN cDNA 4930519A11 gene [Source:M | 0 | 0 | 0 | 0 | 0 | 0 |
| 36185 | ENSMUSG00000097628 | 0 | NA | NA | NA | NA | NA | Gm2383 | predicted gene 2383 [Source:MGI Symbol | 0 | 0 | 0 | 0 | 0 | 0 |
| 36186 | ENSMUSG00000097630 | 0 | NA | NA | NA | NA | NA | Gm26828 | predicted gene, 26828 [Source:MGI Symb | 0 | 0 | 0 | 0 | 0 | 0 |
| 36187 | ENSMUSG00000097632 | 1.145251229 | 0.664618229 | 1.963396633 | 0.338504313 | 0.734983177 | NA | NA | NA | 1.6062 | 0 | 1.0062 | 0.964 | 2.1812 | 1.1138 |
| 36188 | ENSMUSG00000097633 | 0.349477146 | 0.165142768 | 3.911021834 | 0.042224967 | 0.966319359 | NA | Gm16617 | predicted gene, 16617 [Source:MGI Symb | 0 | 0 | 1.0062 | 0 | 1.0906 | 0 |
| 36189 | ENSMUSG00000097634 | 0 | NA | NA | NA | NA | NA | Gm26827 | predicted gene, 26827 [Source:MGI Symb | 0 | 0 | 0 | 0 | 0 | 0 |
| 36190 | ENSMUSG00000097635 | 0 | NA | NA | NA | NA | NA | Gm26826 | predicted gene, 26826 [Source:MGI Symb | 0 | 0 | 0 | 0 | 0 | 0 |
| 36191 | ENSMUSG00000097636 | 2.050517782 | -3.304200343 | 1.850952789 | -1.785134857 | 0.074239445 | NA | Mirt1 | myocardial infarction associated transcrip | 4.0155 | 6.1675 | 1.0062 | 0 | 0 | 1.1138 |
| 36192 | ENSMUSG00000097637 | 17.20185037 | -1.078219888 | 0.600505702 | -1.795519816 | 0.072570918 | 0.390758026 | 4933417D19 | RIKEN cDNA 4933417D19 gene [Source:M | 24.093 | 23.642 | 22.137 | 4.8199 | 20.722 | 7.7969 |
| 36193 | ENSMUSG00000097638 | 0 | NA | NA | NA | NA | NA | Carlr | cardiac and apoptosis-related long non-cc | 0 | 0 | 0 | 0 | 0 | 0 |
| 36194 | ENSMUSG00000097639 | 9.6215913 | 0.750156339 | 0.733102709 | 1.023262266 | 0.30618386 | 0.755354485 | Platr4 | pluripotency associated transcript 4 [Sour | 6.4248 | 4.1117 | 11.069 | 17.352 | 7.6343 | 11.138 |
| 36195 | ENSMUSG00000097640 | 13.57485877 | 0.904176027 | 0.623755978 | 1.449566912 | 0.147179329 | 0.558351692 | Gm20033 | predicted gene, 20033 [Source:MGI Symb | 12.047 | 5.1396 | 11.069 | 20.244 | 21.812 | 11.138 |
| 36196 | ENSMUSG00000097641 | 0.160662974 | 1.126924175 | 4.080472857 | 0.276174898 | 0.782413736 | NA | Gm26865 | predicted gene, 26865 [Source:MGI Symb | 0 | 0 | 0 | 0.964 | 0 | 0 |
| 36197 | ENSMUSG00000097642 | 0 | NA | NA | NA | NA | NA | Gm26866 | predicted gene, 26866 [Source:MGI Symb | 0 | 0 | 0 | 0 | 0 | 0 |
| 36198 | ENSMUSG00000097643 | 4.991259784 | -0.202611168 | 1.081563299 | -0.187331771 | 0.851400514 | NA | A130051J06I | RIKEN cDNA A130051J0 gene [Source:MG | 0.8031 | 10.279 | 5.0312 | 3.8559 | 2.1812 | 7.7969 |
| 36199 | ENSMUSG00000097644 | 0 | NA | NA | NA | NA | NA | Gm26862 | predicted gene, 26862 [Source:MGI Symb | 0 | 0 | 0 | 0 | 0 | 0 |
| 36200 | ENSMUSG00000097645 | 0 | NA | NA | NA | NA | NA | NA | NA | 0 | 0 | 0 | 0 | 0 | 0 |
| 36201 | ENSMUSG00000097646 | 0 | NA | NA | NA | NA | NA | Gm26864 | predicted gene, 26864 [Source:MGI Symb | 0 | 0 | 0 | 0 | 0 | 0 |
| 36202 | ENSMUSG00000097648 | 0 | NA | NA | NA | NA | NA | 9330185C12 | RIKEN cDNA 9330185C12 gene [Source:M | 0 | 0 | 0 | 0 | 0 | 0 |
| 36203 | ENSMUSG00000097649 | 0.840029082 | -0.398118545 | 2.584217466 | -0.154057679 | 0.877564257 | NA | Gm10561 | predicted gene 10561 [Source:MGI Symbo | 0.8031 | 2.0558 | 0 | 0 | 2.1812 | 0 |
| 36204 | ENSMUSG00000097650 | 0 | NA | NA | NA | NA | NA | 4921507G05 | RIKEN cDNA 4921507G05 gene [Source:M | 0 | 0 | 0 | 0 | 0 | 0 |
| 36205 | ENSMUSG00000097651 | 0 | NA | NA | NA | NA | NA | 4930461G14 | RIKEN cDNA 4930461G14 gene [Source:M | 0 | 0 | 0 | 0 | 0 | 0 |
| 36206 | ENSMUSG00000097652 | 0 | NA | NA | NA | NA | NA | Mhrt | myosin heavy chain associated RNA transc | 0 | 0 | 0 | 0 | 0 | 0 |
| 36207 | ENSMUSG00000097653 | 0 | NA | NA | NA | NA | NA | Gm16387 | predicted gene 16387 [Source:MGI Symbo | 0 | 0 | 0 | 0 | 0 | 0 |
| 36208 | ENSMUSG00000097654 | 21.05421156 | -2.1787205 | 0.591942499 | -3.680628613 | 0.00023266 | 0.007054905 | Gm26714 | predicted gene, 26714 [Source:MGI Symb | 47.383 | 38.033 | 18.112 | 9.6398 | 9.8156 | 3.3415 |
| 36209 | ENSMUSG00000097655 | 0 | NA | NA | NA | NA | NA | Gm26713 | predicted gene, 26713 [Source:MGI Symb | 0 | 0 | 0 | 0 | 0 | 0 |
| 36210 | ENSMUSG00000097656 | 0 | NA | NA | NA | NA | NA | Gm26712 | predicted gene, 26712 [Source:MGI Symb | 0 | 0 | 0 | 0 | 0 | 0 |
| 36211 | ENSMUSG00000097657 | 0.133850373 | -0.796629176 | 4.080472857 | -0.195229623 | 0.84521319 | NA | Gm7389 | predicted gene 7389 [Source:MGI Symbol | 0.8031 | 0 | 0 | 0 | 0 | 0 |
| 36212 | ENSMUSG00000097658 | 27.05485332 | -0.079299544 | 0.416645553 | -0.19032855 | 0.849051683 | 0.973071229 | NA | NA | 31.321 | 29.81 | 22.137 | 25.063 | 27.265 | 26.732 |
| 36213 | ENSMUSG00000097659 | 0 | NA | NA | NA | NA | NA | NA | NA | 0 | 0 | 0 | 0 | 0 | 0 |
| 36214 | ENSMUSG00000097660 | 2.231615592 | -0.187967291 | 1.625608897 | -0.115628852 | 0.907946695 | NA | NA | NA | 4.0155 | 3.0838 | 0 | 1.928 | 4.3625 | 0 |
| 36215 | ENSMUSG00000097661 | 0.684865378 | 2.990343487 | 2.966805827 | 1.00793367 | 0.313486306 | NA | NA | NA | 0 | 0 | 0 | 1.928 | 2.1812 | 0 |
| 36216 | ENSMUSG00000097662 | 1.230846427 | -0.30110149 | 2.084603758 | -0.144440635 | 0.885152531 | NA | Gm2093 | predicted gene 2093 [Source:MGI Symbol | 0 | 3.0838 | 1.0062 | 0 | 2.1812 | 1.1138 |
| 36217 | ENSMUSG00000097664 | 3.687226901 | -0.122929865 | 1.137822617 | -0.108039569 | 0.913964304 | NA | NA | NA | 2.4093 | 5.1396 | 4.025 | 2.8919 | 6.5437 | 1.1138 |
| 36218 | ENSMUSG00000097665 | 0.870919206 | -1.842640947 | 2.845777392 | -0.64750003 | 0.517308375 | NA | Gm16853 | predicted gene, 16853 [Source:MGI Symb | 0 | 4.1117 | 0 | 0 | 0 | 1.1138 |
| 36219 | ENSMUSG00000097666 | 0 | NA | NA | NA | NA | NA | A330094K24 | RIKEN cDNA A330094K24 gene [Source:M | 0 | 0 | 0 | 0 | 0 | 0 |
| 36220 | ENSMUSG00000097667 | 1.520403456 | 1.267293396 | 1.809164653 | 0.700485383 | 0.48362423 | NA | NA | NA | 1.6062 | 0 | 1.0062 | 0.964 | 1.0906 | 4.4554 |
| 36221 | ENSMUSG00000097668 | 0.181769715 | 1.126924175 | 4.080472857 | 0.276174898 | 0.782413736 | NA | Gm26761 | predicted gene, 26761 [Source:MGI Symb | 0 | 0 | 0 | 0 | 1.0906 | 0 |
| 36222 | ENSMUSG00000097669 | 0 | NA | NA | NA | NA | NA | Gm8649 | predicted gene 8649 [Source:MGI Symbol | 0 | 0 | 0 | 0 | 0 | 0 |
| 36223 | ENSMUSG00000097670 | 0 | NA | NA | NA | NA | NA | Gm26607 | predicted gene, 26607 [Source:MGI Symb | 0 | 0 | 0 | 0 | 0 | 0 |
| 36224 | ENSMUSG00000097671 | 1.053024754 | -1.051058068 | 2.223622046 | -0.472678381 | 0.636442653 | NA | 1700015O11 | RIKEN cDNA 1700015O11 gene [Source:M | 3.2124 | 1.0279 | 0 | 0.964 | 0 | 1.1138 |
| 36225 | ENSMUSG00000097672 | 0 | NA | NA | NA | NA | NA | NA | NA | 0 | 0 | 0 | 0 | 0 | 0 |
| 36226 | ENSMUSG00000097673 | 65.62506125 | 0.012083843 | 0.303055402 | 0.03987338 | 0.968194074 | 0.994231936 | Gm26608 | predicted gene, 26608 [Source:MGI Symb | 51.399 | 60.647 | 84.525 | 77.118 | 63.256 | 56.806 |
| 36227 | ENSMUSG00000097674 | 1.89724114 | 0.13631334 | 1.626813721 | 0.083791609 | 0.933222119 | NA | NA | NA | 2.4093 | 1.0279 | 2.0125 | 4.8199 | 0 | 1.1138 |
| 36228 | ENSMUSG00000097675 | 2.778449923 | -1.099274369 | 1.365413963 | -0.80508505 | 0.420770602 | NA | 1700101I11F | RIKEN cDNA 1700101I11 gene [Source:M( | 7.2279 | 1.0279 | 3.0187 | 0.964 | 1.0906 | 3.3415 |

| | | | | | | | | | | | | | | | |
|---|---|---|---|---|---|---|---|---|---|---|---|---|---|---|---|
| 36229 | ENSMUSG00000097676 | 0 | NA | NA | NA | NA | NA | Gm26612 | predicted gene, 26612 [Source:MGI Symb | 0 | 0 | 0 | 0 | 0 | 0 |
| 36230 | ENSMUSG00000097677 | 0 | NA | NA | NA | NA | NA | Gm26611 | predicted gene, 26611 [Source:MGI Symb | 0 | 0 | 0 | 0 | 0 | 0 |
| 36231 | ENSMUSG00000097679 | 0 | NA | NA | NA | NA | NA | Rps19-ps5 | ribosomal protein S19, pseudogene 5 [Sou | 0 | 0 | 0 | 0 | 0 | 0 |
| 36232 | ENSMUSG00000097680 | 25.17447154 | -0.134132261 | 0.5060496 | -0.265057537 | 0.790965138 | 0.961019028 | Gm26642 | predicted gene, 26642 [Source:MGI Symb | 30.518 | 14.391 | 34.212 | 36.631 | 16.359 | 18.935 |
| 36233 | ENSMUSG00000097681 | 0 | NA | NA | NA | NA | NA | NA | NA | 0 | 0 | 0 | 0 | 0 | 0 |
| 36234 | ENSMUSG00000097682 | 0 | NA | NA | NA | NA | NA | 4930554H23 | RIKEN cDNA 4930554H23 gene [Source:M | 0 | 0 | 0 | 0 | 0 | 0 |
| 36235 | ENSMUSG00000097683 | 0 | NA | NA | NA | NA | NA | Gm26644 | predicted gene, 26644 [Source:MGI Symb | 0 | 0 | 0 | 0 | 0 | 0 |
| 36236 | ENSMUSG00000097684 | 0.181769715 | 1.126924175 | 4.080472857 | 0.276174898 | 0.782413736 | NA | Gm26645 | predicted gene, 26645 [Source:MGI Symb | 0 | 0 | 0 | 0 | 1.0906 | 0 |
| 36237 | ENSMUSG00000097685 | 0.267700746 | -1.497481649 | 4.057001436 | -0.369110456 | 0.712045396 | NA | Gm26646 | predicted gene, 26646 [Source:MGI Symb | 1.6062 | 0 | 0 | 0 | 0 | 0 |
| 36238 | ENSMUSG00000097686 | 0 | NA | NA | NA | NA | NA | Gm26647 | predicted gene, 26647 [Source:MGI Symb | 0 | 0 | 0 | 0 | 0 | 0 |
| 36239 | ENSMUSG00000097689 | 0 | NA | NA | NA | NA | NA | 4930471D02 | RIKEN cDNA 4930471D02 gene [Source:M | 0 | 0 | 0 | 0 | 0 | 0 |
| 36240 | ENSMUSG00000097690 | 1.291619825 | -2.551117698 | 2.090478447 | -1.220351112 | 0.222331802 | NA | 4930505N22 | RIKEN cDNA 4930505N22 gene [Source:M | 1.6062 | 1.0279 | 4.025 | 0 | 1.0906 | 0 |
| 36241 | ENSMUSG00000097691 | 103.7175453 | 0.37547658 | 0.265203536 | 1.415805332 | 0.156832508 | 0.575786484 | 9030616G12 | RIKEN cDNA 9030616G12 gene [Source:M | 83.523 | 66.815 | 120.75 | 127.25 | 100.34 | 123.64 |
| 36242 | ENSMUSG00000097692 | 2.356929582 | -0.828513848 | 1.469660314 | -0.563745132 | 0.572927597 | NA | A230060F14 | RIKEN cDNA A230060F14 gene [Source:M | 4.0155 | 1.0279 | 4.025 | 2.8919 | 2.1812 | 0 |
| 36243 | ENSMUSG00000097693 | 0.698954292 | 0.126053087 | 3.048972385 | 0.04134281 | 0.967022605 | NA | | RIKEN cDNA 4930471M09 gene [Source:N | 0 | 0 | 2.0125 | 0 | 2.1812 | 0 |
| 36244 | ENSMUSG00000097694 | 1.192013061 | -3.607193727 | 2.478410404 | -1.455446492 | 0.145545693 | NA | G730013B05 | RIKEN cDNA G730013B05 gene [Source:M | 0 | 5.1396 | 2.0125 | 0 | 0 | 0 |
| 36245 | ENSMUSG00000097695 | 0.167707431 | -0.796629176 | 4.080472857 | -0.195229623 | 0.84521319 | NA | NA | NA | 0 | 0 | 1.0062 | 0 | 0 | 0 |
| 36246 | ENSMUSG00000097697 | 0.17131964 | -0.796629176 | 4.080472857 | -0.195229623 | 0.84521319 | NA | 4833412C05 | RIKEN cDNA 4833412C05 gene [Source:M | 0 | 1.0279 | 0 | 0 | 0 | 0 |
| 36247 | ENSMUSG00000097698 | 0 | NA | NA | NA | NA | NA | NA | NA | 0 | 0 | 0 | 0 | 0 | 0 |
| 36248 | ENSMUSG00000097699 | 6.071253012 | -0.386038677 | 0.920132183 | -0.419546978 | 0.674816429 | 0.932315978 | 5430400D12 | RIKEN cDNA 5430400D12 gene [Source:M | 6.4248 | 6.1675 | 8.05 | 5.7839 | 1.0906 | 8.9108 |
| 36249 | ENSMUSG00000097700 | 0 | NA | NA | NA | NA | NA | Gm26738 | predicted gene, 26738 [Source:MGI Symb | 0 | 0 | 0 | 0 | 0 | 0 |
| 36250 | ENSMUSG00000097701 | 0 | NA | NA | NA | NA | NA | Gm17508 | predicted gene, 17508 [Source:MGI Symb | 0 | 0 | 0 | 0 | 0 | 0 |
| 36251 | ENSMUSG00000097702 | 2.202179059 | -1.776793002 | 1.627275735 | -1.091881949 | 0.274884993 | NA | Gm26739 | predicted gene, 26739 [Source:MGI Symb | 3.2124 | 3.0838 | 4.025 | 2.8919 | 0 | 0 |
| 36252 | ENSMUSG00000097703 | 1.405650449 | 0.138001964 | 2.148685932 | 0.064226214 | 0.948790104 | NA | NA | NA | 0 | 0 | 4.025 | 0 | 2.1812 | 2.2277 |
| 36253 | ENSMUSG00000097704 | 0 | NA | NA | NA | NA | NA | Gm26741 | predicted gene, 26741 [Source:MGI Symb | 0 | 0 | 0 | 0 | 0 | 0 |
| 36254 | ENSMUSG00000097705 | 1.351125335 | 0.898520505 | 1.866150402 | 0.481483434 | 0.63017295 | NA | Gm26740 | predicted gene, 26740 [Source:MGI Symb | 0.8031 | 1.0279 | 1.0062 | 1.928 | 0 | 3.3415 |
| 36255 | ENSMUSG00000097706 | 21.9809023 | 0.609153322 | 0.486407066 | 1.252352945 | 0.210441284 | 0.654419768 | E030037K01 | RIKEN cDNA E030037K01 gene [Source:M | 18.471 | 20.558 | 13.081 | 19.28 | 35.99 | 24.505 |
| 36256 | ENSMUSG00000097707 | 0.469265235 | -2.274672822 | 3.439794491 | -0.661281605 | 0.508431736 | NA | NA | NA | 0.8031 | 0 | 2.0125 | 0 | 0 | 0 |
| 36257 | ENSMUSG00000097709 | 0 | NA | NA | NA | NA | NA | 2810429I04F | RIKEN cDNA 2810429I04 gene [Source:MC | 0 | 0 | 0 | 0 | 0 | 0 |
| 36258 | ENSMUSG00000097710 | 0 | NA | NA | NA | NA | NA | 2310026I22F | RIKEN cDNA 2310026I22 gene [Source:MC | 0 | 0 | 0 | 0 | 0 | 0 |
| 36259 | ENSMUSG00000097711 | 0.371281295 | 2.09253362 | 4.031852165 | 0.519000582 | 0.603760335 | NA | Gm5523 | predicted pseudogene 5523 [Source:MGI | 0 | 0 | 0 | 0 | 0 | 2.2277 |
| 36260 | ENSMUSG00000097712 | 0.428363721 | -0.543730382 | 3.624677344 | -0.15000794 | 0.880758351 | NA | Gm10532 | predicted gene 10532 [Source:MGI Symbo | 1.6062 | 0 | 0 | 0.964 | 0 | 0 |
| 36261 | ENSMUSG00000097713 | 0 | NA | NA | NA | NA | NA | Gm7741 | predicted gene 7741 [Source:MGI Symbol | 0 | 0 | 0 | 0 | 0 | 0 |
| 36262 | ENSMUSG00000097714 | 0.301557804 | -1.64335916 | 4.042814367 | -0.406488899 | 0.684383412 | NA | Gm20109 | predicted gene, 20109 [Source:MGI Symb | 0.8031 | 0 | 1.0062 | 0 | 0 | 0 |
| 36263 | ENSMUSG00000097715 | 22.83873035 | -1.026364685 | 0.487236541 | -2.10650187 | 0.035160781 | 0.25151549 | Gpr137b-ps | G protein-coupled receptor 137B, pseudo | 37.746 | 34.949 | 19.119 | 15.424 | 13.087 | 16.708 |
| 36264 | ENSMUSG00000097716 | 0.160662974 | 1.126924175 | 4.080472857 | 0.276174898 | 0.782413736 | NA | 1700026J14F | RIKEN cDNA 1700026J14 gene [Source:MC | 0 | 0 | 0 | 0.964 | 0 | 0 |
| 36265 | ENSMUSG00000097717 | 0.17131964 | -0.796629176 | 4.080472857 | -0.195229623 | 0.84521319 | NA | 1700066J03F | RIKEN cDNA 1700066J03 gene [Source:MC | 0 | 1.0279 | 0 | 0 | 0 | 0 |
| 36266 | ENSMUSG00000097718 | 5.30210494 | 0.024532095 | 0.973287996 | 0.025205382 | 0.979891144 | 0.99820894 | NA | NA | 1.6062 | 8.2233 | 6.0375 | 3.8559 | 7.6343 | 4.4554 |
| 36267 | ENSMUSG00000097719 | 0 | NA | NA | NA | NA | NA | Rpl31-ps18 | ribosomal protein L31, pseudogene 18 [Sc | 0 | 0 | 0 | 0 | 0 | 0 |
| 36268 | ENSMUSG00000097720 | 0 | NA | NA | NA | NA | NA | NA | NA | 0 | 0 | 0 | 0 | 0 | 0 |
| 36269 | ENSMUSG00000097722 | 1.205688793 | -1.211715284 | 2.313668566 | -0.523720338 | 0.600473068 | NA | NA | NA | 0 | 1.0279 | 4.025 | 0 | 2.1812 | 0 |
| 36270 | ENSMUSG00000097723 | 0.160662974 | 1.126924175 | 4.080472857 | 0.276174898 | 0.782413736 | NA | NA | NA | 0 | 0 | 0 | 0.964 | 0 | 0 |
| 36271 | ENSMUSG00000097724 | 3.319868672 | 0.144724835 | 1.186145341 | 0.122012733 | 0.902888934 | NA | Gm26850 | predicted gene, 26850 [Source:MGI Symb | 2.4093 | 2.0558 | 5.0312 | 3.8559 | 5.4531 | 1.1138 |
| 36272 | ENSMUSG00000097725 | 1.638155541 | 0.768922798 | 1.682838996 | 0.456920002 | 0.647728546 | NA | 4930593A02 | RIKEN cDNA 4930593A02 gene [Source:M | 1.6062 | 0 | 2.0125 | 2.8919 | 1.0906 | 2.2277 |
| 36273 | ENSMUSG00000097726 | 0 | NA | NA | NA | NA | NA | 9530036O11 | RIKEN cDNA 9530036O11Rik [Source:MGI | 0 | 0 | 0 | 0 | 0 | 0 |
| 36274 | ENSMUSG00000097727 | 0.349477146 | 0.165142768 | 3.911021834 | 0.042224967 | 0.966319359 | NA | F630040K05 | RIKEN cDNA F630040K05 gene [Source:M | 0 | 0 | 1.0062 | 0 | 1.0906 | 0 |
| 36275 | ENSMUSG00000097728 | 3.242423006 | -0.618768838 | 1.38578502 | -0.446511421 | 0.655227865 | NA | NA | NA | 5.6217 | 1.0279 | 5.0312 | 0 | 1.0906 | 6.6831 |
| 36276 | ENSMUSG00000097729 | 43.41528168 | -0.370856234 | 0.393496287 | -0.942464381 | 0.345954939 | 0.786899236 | 2310015A10 | RIKEN cDNA 2310015A10 gene [Source:M | 49.792 | 37.005 | 60.375 | 55.911 | 23.994 | 33.415 |
| 36277 | ENSMUSG00000097730 | 266.3720463 | -0.428763962 | 0.1742429 | -2.460725587 | 0.013865637 | 0.135783736 | Gm26588 | predicted gene, 26588 [Source:MGI Symb | 351.76 | 302.21 | 262.63 | 244.85 | 201.76 | 235.02 |
| 36278 | ENSMUSG00000097731 | 1.660838485 | 1.346556011 | 1.746883686 | 0.770833239 | 0.440805782 | NA | NA | NA | 0.8031 | 1.0279 | 1.0062 | 3.8559 | 3.2719 | 0 |
| 36279 | ENSMUSG00000097734 | 0 | NA | NA | NA | NA | NA | NA | NA | 0 | 0 | 0 | 0 | 0 | 0 |
| 36280 | ENSMUSG00000097735 | 0 | NA | NA | NA | NA | NA | D930032P07 | RIKEN cDNA D930032P07 gene [Source:M | 0 | 0 | 0 | 0 | 0 | 0 |
| 36281 | ENSMUSG00000097736 | 0 | NA | NA | NA | NA | NA | 9530059O14 | RIKEN cDNA 9530059O14 gene [Source:M | 0 | 0 | 0 | 0 | 0 | 0 |
| 36282 | ENSMUSG00000097737 | 7.523740747 | -0.781708146 | 0.812465551 | -0.962143128 | 0.335977713 | 0.777657487 | NA | NA | 11.243 | 10.279 | 7.0437 | 7.7118 | 2.1812 | 6.6831 |
| 36283 | ENSMUSG00000097738 | 0 | NA | NA | NA | NA | NA | 4930445N18 | RIKEN cDNA 4930445N18 gene [Source:M | 0 | 0 | 0 | 0 | 0 | 0 |
| 36284 | ENSMUSG00000097739 | 0 | NA | NA | NA | NA | NA | Gm2539 | predicted gene, 2539 [Source:MGI Symbo | 0 | 0 | 0 | 0 | 0 | 0 |

| | | | | | | | | | | | | | | | |
|---|---|---|---|---|---|---|---|---|---|---|---|---|---|---|---|
| 36285 | ENSMUSG00000097740 | 0.481988923 | 2.492401104 | 3.837077151 | 0.649557203 | 0.515978285 | NA | E030044B06 | RIKEN cDNA E030044B06 gene [Source:M | 0 | 0 | 0 | 2.8919 | 0 | 0 |
| 36286 | ENSMUSG00000097742 | 0 | NA | NA | NA | NA | NA | Gm26535 | predicted gene, 26535 [Source:MGI Symb | 0 | 0 | 0 | 0 | 0 | 0 |
| 36287 | ENSMUSG00000097743 | 35.62253407 | 0.15007347 | 0.392136783 | 0.382706945 | 0.701937064 | 0.942189108 | Gm16973 | predicted gene, 16973 [Source:MGI Symb | 26.502 | 35.977 | 39.244 | 48.199 | 37.081 | 26.732 |
| 36288 | ENSMUSG00000097744 | 0.181769715 | 1.126924175 | 4.080472857 | 0.276174898 | 0.782413736 | NA | D030040B21 | RIKEN cDNA D030040B21 gene [Source:M | 0 | 0 | 0 | 0 | 1.0906 | 0 |
| 36289 | ENSMUSG00000097745 | 0 | NA | NA | NA | NA | NA | AI115009 | expressed sequence AI115009 [Source:M | 0 | 0 | 0 | 0 | 0 | 0 |
| 36290 | ENSMUSG00000097746 | 4.141788665 | -0.009770639 | 1.056580028 | -0.009247419 | 0.992621732 | NA | Gm6225 | predicted gene 6225 [Source:MGI Symbol | 2.4093 | 5.1396 | 5.0312 | 6.7478 | 2.1812 | 3.3415 |
| 36291 | ENSMUSG00000097747 | 4.572136256 | -0.706980492 | 1.030089528 | -0.686329171 | 0.492505558 | NA | NA | NA | 4.8186 | 8.2233 | 4.025 | 4.8199 | 1.0906 | 4.4554 |
| 36292 | ENSMUSG00000097748 | 0.538730002 | 1.11098254 | 3.037664833 | 0.365735722 | 0.714562288 | NA | Gm26533 | predicted gene, 26533 [Source:MGI Symb | 0 | 1.0279 | 0 | 0 | 1.0906 | 1.1138 |
| 36293 | ENSMUSG00000097749 | 1.452798593 | -0.998986143 | 2.141750486 | -0.466434419 | 0.64090458 | NA | Gm16938 | predicted gene, 16938 [Source:MGI Symb | 4.8186 | 0 | 1.0062 | 2.8919 | 0 | 0 |
| 36294 | ENSMUSG00000097750 | 179.0668078 | 0.145214325 | 0.248415874 | 0.584561376 | 0.558842685 | 0.899349136 | Gm4673 | predicted gene 4673 [Source:MGI Symbol | 133.31 | 157.27 | 220.37 | 226.53 | 131.96 | 204.95 |
| 36295 | ENSMUSG00000097751 | 0 | NA | NA | NA | NA | NA | NA | NA | 0 | 0 | 0 | 0 | 0 | 0 |
| 36296 | ENSMUSG00000097752 | 0 | NA | NA | NA | NA | NA | Gm26688 | predicted gene, 26688 [Source:MGI Symb | 0 | 0 | 0 | 0 | 0 | 0 |
| 36297 | ENSMUSG00000097753 | 0 | NA | NA | NA | NA | NA | NA | NA | 0 | 0 | 0 | 0 | 0 | 0 |
| 36298 | ENSMUSG00000097754 | 0 | NA | NA | NA | NA | NA | Ptgs2os2 | prostaglandin-endoperoxide synthase 2, c | 0 | 0 | 0 | 0 | 0 | 0 |
| 36299 | ENSMUSG00000097755 | 1.775948981 | 2.133892326 | 1.825236827 | 1.169104356 | 0.242361589 | NA | 2010110K18 | RIKEN cDNA 2010110K18 gene [Source:M | 0 | 1.0279 | 1.0062 | 0.964 | 6.5437 | 1.1138 |
| 36300 | ENSMUSG00000097756 | 0 | NA | NA | NA | NA | NA | A730056A06 | RIKEN cDNA A730056A06 gene [Source:M | 0 | 0 | 0 | 0 | 0 | 0 |
| 36301 | ENSMUSG00000097757 | 10.38562666 | -0.392277393 | 0.705642359 | -0.555915314 | 0.578268749 | 0.908137944 | NA | NA | 8.031 | 17.475 | 10.062 | 13.496 | 5.4531 | 7.7969 |
| 36302 | ENSMUSG00000097759 | 1.618625004 | 1.304555113 | 1.813545066 | 0.719339782 | 0.47193159 | NA | NA | NA | 0.8031 | 1.0279 | 1.0062 | 5.7839 | 1.0906 | 0 |
| 36303 | ENSMUSG00000097760 | 2.873564567 | 0.284421175 | 1.353701455 | 0.210106279 | 0.833584725 | NA | 6030442K20 | RIKEN cDNA 6030442K20 gene [Source:M | 5.6217 | 1.0279 | 1.0062 | 1.928 | 6.5437 | 1.1138 |
| 36304 | ENSMUSG00000097761 | 1.959377222 | -0.930141411 | 1.770808787 | -0.525263607 | 0.599399953 | NA | 4930534D22 | RIKEN cDNA 4930534D22 gene [Source:M | 1.6062 | 6.1675 | 0 | 2.8919 | 1.0906 | 0 |
| 36305 | ENSMUSG00000097762 | 70.52952926 | 0.466152513 | 0.426664037 | 1.092551685 | 0.274590686 | 0.729487777 | 4732463B04 | RIKEN cDNA 4732463B04 gene [Source:M | 77.098 | 46.256 | 54.337 | 128.21 | 89.431 | 27.846 |
| 36306 | ENSMUSG00000097763 | 0 | NA | NA | NA | NA | NA | Gm26636 | predicted gene, 26636 [Source:MGI Symb | 0 | 0 | 0 | 0 | 0 | 0 |
| 36307 | ENSMUSG00000097764 | 0 | NA | NA | NA | NA | NA | NA | NA | 0 | 0 | 0 | 0 | 0 | 0 |
| 36308 | ENSMUSG00000097766 | 5.511215192 | -0.237698578 | 1.058660768 | -0.224527616 | 0.822346781 | 0.969197142 | 5730420D15 | RIKEN cDNA 5730420D15 gene [Source:M | 4.8186 | 5.1396 | 8.05 | 10.604 | 0 | 4.4554 |
| 36309 | ENSMUSG00000097767 | 0 | NA | NA | NA | NA | NA | Miat | myocardial infarction associated transcrip | 0 | 0 | 0 | 0 | 0 | 0 |
| 36310 | ENSMUSG00000097768 | 5.143870923 | -1.421742805 | 0.982898209 | -1.446480207 | 0.14804256 | 0.559667428 | 2310043M15 | RIKEN cDNA 2310043M15 gene [Source:M | 7.2279 | 8.2233 | 7.0437 | 2.8919 | 4.3625 | 1.1138 |
| 36311 | ENSMUSG00000097769 | 18.81303249 | 0.345630986 | 0.521007814 | 0.663389255 | 0.507081286 | 0.88185929 | NA | NA | 15.259 | 20.558 | 14.087 | 29.883 | 15.269 | 17.822 |
| 36312 | ENSMUSG00000097770 | 0 | NA | NA | NA | NA | NA | NA | NA | 0 | 0 | 0 | 0 | 0 | 0 |
| 36313 | ENSMUSG00000097772 | 29.0289399 | 0.111394417 | 0.436410873 | 0.255251242 | 0.798529056 | 0.962424862 | 5430416N02 | RIKEN cDNA 5430416N02 gene [Source:M | 24.093 | 24.67 | 35.219 | 40.487 | 19.631 | 30.074 |
| 36314 | ENSMUSG00000097773 | 13.47608032 | 0.637852611 | 0.642118236 | 0.993356947 | 0.320536035 | 0.767391055 | Gm10614 | predicted gene 10614 [Source:MGI Symbo | 6.4248 | 12.335 | 13.081 | 18.316 | 22.903 | 7.7969 |
| 36315 | ENSMUSG00000097774 | 0 | NA | NA | NA | NA | NA | NA | NA | 0 | 0 | 0 | 0 | 0 | 0 |
| 36316 | ENSMUSG00000097775 | 0.335414862 | -1.779047242 | 4.030804179 | -0.441362855 | 0.658950329 | NA | NA | NA | 0 | 0 | 2.0125 | 0 | 0 | 0 |
| 36317 | ENSMUSG00000097776 | 0 | NA | NA | NA | NA | NA | Gm26795 | predicted gene, 26795 [Source:MGI Symb | 0 | 0 | 0 | 0 | 0 | 0 |
| 36318 | ENSMUSG00000097777 | 5.405715439 | -0.505769105 | 0.969760634 | -0.521540149 | 0.601990545 | 0.915461118 | Gm16794 | predicted gene, 16794 [Source:MGI Symb | 8.8341 | 6.1675 | 4.025 | 6.7478 | 1.0906 | 5.5692 |
| 36319 | ENSMUSG00000097779 | 0.774435248 | -2.999089927 | 2.565713408 | -1.168910728 | 0.242439601 | NA | 4833407H14 | RIKEN cDNA 4833407H14 gene [Source:M | 1.6062 | 1.0279 | 2.0125 | 0 | 0 | 0 |
| 36320 | ENSMUSG00000097780 | 0 | NA | NA | NA | NA | NA | Gm26746 | predicted gene, 26746 [Source:MGI Symb | 0 | 0 | 0 | 0 | 0 | 0 |
| 36321 | ENSMUSG00000097781 | 2.82964564 | -0.667430925 | 1.350154936 | -0.494336544 | 0.621068521 | NA | 9330136K24 | RIKEN cDNA 9330136K24 gene [Source:M | 3.2124 | 5.1396 | 2.0125 | 0 | 3.2719 | 3.3415 |
| 36322 | ENSMUSG00000097783 | 2.67693165 | 1.130965457 | 1.524634976 | 0.741794249 | 0.458212005 | NA | Gm26747 | predicted gene, 26747 [Source:MGI Symb | 0 | 3.0838 | 2.0125 | 0.964 | 1.0906 | 8.9108 |
| 36323 | ENSMUSG00000097784 | 3.772181799 | -0.829326614 | 1.145299877 | -0.724113074 | 0.468996319 | NA | A430105J06 | RIKEN cDNA A430105J06 gene [Source:M | 6.4248 | 2.0558 | 6.0375 | 4.8199 | 2.1812 | 1.1138 |
| 36324 | ENSMUSG00000097785 | 24.63291468 | 0.215009566 | 0.499882478 | 0.430120229 | 0.667108185 | 0.929709374 | B230217O12 | RIKEN cDNA B230217O12 gene [Source:M | 25.699 | 25.698 | 17.106 | 37.595 | 29.447 | 12.252 |
| 36325 | ENSMUSG00000097786 | 4.291272012 | 0.390724063 | 0.998385361 | 0.391355961 | 0.69553414 | NA | 4933429H19 | RIKEN cDNA 4933429H19 gene [Source:M | 4.0155 | 4.1117 | 3.0187 | 5.7839 | 4.3625 | 4.4554 |
| 36326 | ENSMUSG00000097787 | 23.39433792 | -0.388565157 | 0.546927516 | -0.71045092 | 0.477424556 | 0.869706396 | 2700046G09 | RIKEN cDNA 2700046G09 gene [Source:M | 12.85 | 28.782 | 38.237 | 25.063 | 9.8156 | 25.618 |
| 36327 | ENSMUSG00000097788 | 0 | NA | NA | NA | NA | NA | Gm16596 | predicted gene, 16596 [Source:MGI Symb | 0 | 0 | 0 | 0 | 0 | 0 |
| 36328 | ENSMUSG00000097789 | 11.61047666 | 0.317257698 | 0.694176436 | 0.457027467 | 0.647651303 | 0.925643431 | Gm2115 | predicted gene 2115 [Source:MGI Symbol | 9.6372 | 15.419 | 6.0375 | 14.46 | 18.541 | 5.5692 |
| 36329 | ENSMUSG00000097790 | 0.545309144 | 2.651974629 | 3.666219369 | 0.72335405 | 0.469462394 | NA | Gm16399 | predicted pseudogene 16399 [Source:MG | 0 | 0 | 0 | 0 | 3.2719 | 0 |
| 36330 | ENSMUSG00000097791 | 0 | NA | NA | NA | NA | NA | NA | NA | 0 | 0 | 0 | 0 | 0 | 0 |
| 36331 | ENSMUSG00000097792 | 0.472877444 | -2.283655516 | 3.093270275 | -0.738265755 | 0.460352972 | NA | Gm27042 | predicted gene, 27042 [Source:MGI Symb | 0.8031 | 1.0279 | 1.0062 | 0 | 0 | 0 |
| 36332 | ENSMUSG00000097793 | 12.86447744 | -0.248181378 | 0.6253011 | -0.396898995 | 0.691441945 | 0.938005655 | Gm17259 | predicted gene, 17259 [Source:MGI Symb | 18.471 | 11.307 | 12.075 | 15.424 | 6.5437 | 13.366 |
| 36333 | ENSMUSG00000097794 | 0.371281295 | 2.09253362 | 4.031852165 | 0.519000582 | 0.603760335 | NA | Gm2990 | predicted gene 2990 [Source:MGI Symbol | 0 | 0 | 0 | 0 | 0 | 2.2277 |
| 36334 | ENSMUSG00000097795 | 0.845761573 | -3.112447952 | 2.7647732 | -1.125751636 | 0.260270661 | NA | Gm7678 | predicted gene 7678 [Source:MGI Symbol | 0 | 2.0558 | 3.0187 | 0 | 0 | 0 |
| 36335 | ENSMUSG00000097796 | 0 | NA | NA | NA | NA | NA | NA | NA | 0 | 0 | 0 | 0 | 0 | 0 |
| 36336 | ENSMUSG00000097797 | 1.493880068 | -1.047551152 | 2.081176639 | -0.503345623 | 0.614721295 | NA | Gm26901 | predicted gene, 26901 [Source:MGI Symb | 4.0155 | 2.0558 | 0 | 2.8919 | 0 | 0 |
| 36337 | ENSMUSG00000097798 | 0 | NA | NA | NA | NA | NA | Gm26898 | predicted gene, 26898 [Source:MGI Symb | 0 | 0 | 0 | 0 | 0 | 0 |
| 36338 | ENSMUSG00000097799 | 0.835284868 | -0.623834031 | 2.861223713 | -0.218030498 | 0.827405349 | NA | NA | NA | 0 | 3.0838 | 0 | 1.928 | 0 | 0 |
| 36339 | ENSMUSG00000097800 | 0 | NA | NA | NA | NA | NA | NA | NA | 0 | 0 | 0 | 0 | 0 | 0 |
| 36340 | ENSMUSG00000097801 | 0 | NA | NA | NA | NA | NA | Gm26777 | predicted gene, 26777 [Source:MGI Symb | 0 | 0 | 0 | 0 | 0 | 0 |

| | | | | | | | | | | | | | | | |
|---|---|---|---|---|---|---|---|---|---|---|---|---|---|---|---|
| 36341 | ENSMUSG00000097803 | 0.728241582 | 1.712995752 | 3.00895953 | 0.569298369 | 0.569153674 | NA | Rpl31-ps16 | ribosomal protein L31, pseudogene 16 [Sc | 0 | 1.0279 | 0 | 0 | 0 | 3.3415 |
| 36342 | ENSMUSG00000097804 | 0 | NA | NA | NA | NA | NA | Gm16685 | predicted gene, 16685 [Source:MGI Symb | 0 | 0 | 0 | 0 | 0 | 0 |
| 36343 | ENSMUSG00000097805 | 0 | NA | NA | NA | NA | NA | Gm17473 | predicted gene, 17473 [Source:MGI Symb | 0 | 0 | 0 | 0 | 0 | 0 |
| 36344 | ENSMUSG00000097806 | 0.181769715 | 1.126924175 | 4.080472857 | 0.276174898 | 0.782413736 | NA | Gm6556 | predicted gene 6556 [Source:MGI Symbol | 0 | 0 | 0 | 0 | 1.0906 | 0 |
| 36345 | ENSMUSG00000097807 | 0 | NA | NA | NA | NA | NA | Gm26553 | predicted gene, 26553 [Source:MGI Symb | 0 | 0 | 0 | 0 | 0 | 0 |
| 36346 | ENSMUSG00000097809 | 0 | NA | NA | NA | NA | NA | Gm26554 | predicted gene, 26554 [Source:MGI Symb | 0 | 0 | 0 | 0 | 0 | 0 |
| 36347 | ENSMUSG00000097811 | 50.59314586 | -0.119418438 | 0.376107806 | -0.317511193 | 0.750855748 | 0.952074518 | NA | NA | 45.777 | 37.005 | 75.468 | 56.875 | 31.628 | 56.806 |
| 36348 | ENSMUSG00000097812 | 0 | NA | NA | NA | NA | NA | Gm26812 | predicted gene, 26812 [Source:MGI Symb | 0 | 0 | 0 | 0 | 0 | 0 |
| 36349 | ENSMUSG00000097814 | 0.342432689 | 1.990653086 | 3.900281263 | 0.51038706 | 0.609780321 | NA | Panct2 | pluripotency-associated noncoding transc | 0 | 0 | 0 | 0.964 | 1.0906 | 0 |
| 36350 | ENSMUSG00000097815 | 86.7411213 | -0.074286124 | 0.254556222 | -0.291826001 | 0.770419664 | 0.956515747 | NA | NA | 94.766 | 100.74 | 71.443 | 91.578 | 85.068 | 76.855 |
| 36351 | ENSMUSG00000097816 | 0.972567489 | -2.093337667 | 2.266151568 | -0.923741243 | 0.355621053 | NA | Gm26811 | predicted gene, 26811 [Source:MGI Symb | 0.8031 | 2.0558 | 2.0125 | 0.964 | 0 | 0 |
| 36352 | ENSMUSG00000097817 | 0.339027071 | -1.79174149 | 3.919585585 | -0.457125237 | 0.647581032 | NA | Gm26810 | predicted gene, 26810 [Source:MGI Symb | 0 | 1.0279 | 1.0062 | 0 | 0 | 0 |
| 36353 | ENSMUSG00000097818 | 0.928203237 | 3.416716801 | 3.030141393 | 1.12757669 | 0.259498736 | NA | NA | NA | 0 | 0 | 0 | 0 | 0 | 5.5692 |
| 36354 | ENSMUSG00000097819 | 9.397331195 | 0.34702172 | 0.70030615 | 0.495528591 | 0.620227045 | 0.917977899 | Gm26813 | predicted gene, 26813 [Source:MGI Symb | 10.44 | 9.2513 | 5.0312 | 9.6398 | 11.997 | 10.025 |
| 36355 | ENSMUSG00000097820 | 0.678054141 | -2.792799308 | 2.990673428 | -0.933836266 | 0.350388359 | NA | E530011L22I | RIKEN cDNA E530011L22 gene [Source:M( | 0 | 2.0558 | 2.0125 | 0 | 0 | 0 |
| 36356 | ENSMUSG00000097821 | 0 | NA | NA | NA | NA | NA | C130032M1( | RIKEN cDNA C130032M10 gene [Source:N | 0 | 0 | 0 | 0 | 0 | 0 |
| 36357 | ENSMUSG00000097822 | 0 | NA | NA | NA | NA | NA | Gm26655 | predicted gene, 26655 [Source:MGI Symb | 0 | 0 | 0 | 0 | 0 | 0 |
| 36358 | ENSMUSG00000097823 | 0.520796785 | -0.820103173 | 3.072141206 | -0.266948398 | 0.789508892 | NA | Gm16701 | predicted gene, 16701 [Source:MGI Symb | 0 | 1.0279 | 1.0062 | 0 | 1.0906 | 0 |
| 36359 | ENSMUSG00000097824 | 0 | NA | NA | NA | NA | NA | NA | NA | 0 | 0 | 0 | 0 | 0 | 0 |
| 36360 | ENSMUSG00000097825 | 0 | NA | NA | NA | NA | NA | 9630001P10 | RIKEN cDNA 9630001P10 gene [Source:M | 0 | 0 | 0 | 0 | 0 | 0 |
| 36361 | ENSMUSG00000097827 | 7.031226594 | -0.430217681 | 0.832858558 | -0.516555514 | 0.605466477 | 0.916376001 | 5330439K02 | RIKEN cDNA 5330439K02 gene [Source:M | 4.0155 | 8.2233 | 12.075 | 5.7839 | 7.6343 | 4.4554 |
| 36362 | ENSMUSG00000097828 | 5.06915897 | -0.675640809 | 0.99469896 | -0.679241495 | 0.496984859 | NA | 6430562O15 | RIKEN cDNA 6430562O15 gene [Source:M | 10.44 | 3.0838 | 5.0312 | 1.928 | 4.3625 | 5.5692 |
| 36363 | ENSMUSG00000097831 | 0 | NA | NA | NA | NA | NA | 4930540M0! | RIKEN cDNA 4930540M05 gene [Source:N | 0 | 0 | 0 | 0 | 0 | 0 |
| 36364 | ENSMUSG00000097832 | 0.185640647 | 1.126924175 | 4.080472857 | 0.276174898 | 0.782413736 | NA | Gm26912 | predicted gene, 26912 [Source:MGI Symb | 0 | 0 | 0 | 0 | 0 | 1.1138 |
| 36365 | ENSMUSG00000097833 | 0.703108924 | -2.875459482 | 2.96384054 | -0.970180225 | 0.331956666 | NA | Gm5976 | predicted gene 5976 [Source:MGI Symbol | 3.2124 | 0 | 1.0062 | 0 | 0 | 0 |
| 36366 | ENSMUSG00000097834 | 0 | NA | NA | NA | NA | NA | Gm26911 | predicted gene, 26911 [Source:MGI Symb | 0 | 0 | 0 | 0 | 0 | 0 |
| 36367 | ENSMUSG00000097835 | 8.45509692 | 0.665334825 | 0.759092825 | 0.876486779 | 0.380765461 | 0.812628154 | Gm26910 | predicted gene, 26910 [Source:MGI Symb | 6.4248 | 8.2233 | 5.0312 | 14.46 | 5.4531 | 11.138 |
| 36368 | ENSMUSG00000097836 | 0.99233012 | 0.312367548 | 2.428448675 | 0.128628433 | 0.897651667 | NA | Gm26903 | predicted gene, 26903 [Source:MGI Symb | 1.6062 | 0 | 1.0062 | 0 | 0 | 3.3415 |
| 36369 | ENSMUSG00000097837 | 0 | NA | NA | NA | NA | NA | NA | NA | 0 | 0 | 0 | 0 | 0 | 0 |
| 36370 | ENSMUSG00000097838 | 0.514011053 | 1.044022102 | 3.082659575 | 0.338675769 | 0.734853997 | NA | NA | NA | 0 | 0 | 1.0062 | 0.964 | 0 | 1.1138 |
| 36371 | ENSMUSG00000097839 | 2.469331868 | -0.77727385 | 1.421884041 | -0.54665066 | 0.584618757 | NA | NA | NA | 3.2124 | 2.0558 | 4.025 | 0 | 2.1812 | 3.3415 |
| 36372 | ENSMUSG00000097840 | 0.17131964 | -0.796629176 | 4.080472857 | -0.195229623 | 0.84521319 | NA | Gm26756 | predicted gene, 26756 [Source:MGI Symb | 0 | 1.0279 | 0 | 0 | 0 | 0 |
| 36373 | ENSMUSG00000097841 | 0 | NA | NA | NA | NA | NA | 4930511A02 | RIKEN cDNA 4930511A02 gene [Source:M | 0 | 0 | 0 | 0 | 0 | 0 |
| 36374 | ENSMUSG00000097842 | 10.46545811 | -0.396987516 | 0.667848158 | -0.594427807 | 0.552226014 | 0.895950316 | 9330104G04 | RIKEN cDNA 9330104G04 gene [Source:M | 15.259 | 12.335 | 8.05 | 10.604 | 7.6343 | 8.9108 |
| 36375 | ENSMUSG00000097843 | 0 | NA | NA | NA | NA | NA | NA | NA | 0 | 0 | 0 | 0 | 0 | 0 |
| 36376 | ENSMUSG00000097845 | 0.503302254 | -0.837877584 | 3.426698643 | -0.244514523 | 0.806832356 | NA | A230020J21I | RIKEN cDNA A230020J21 gene [Source:M( | 0 | 2.0558 | 0 | 0.964 | 0 | 0 |
| 36377 | ENSMUSG00000097847 | 0.813009891 | 0.960077285 | 2.883567268 | 0.332947768 | 0.739173711 | NA | 4930478K11 | RIKEN cDNA 4930478K11 gene [Source:M | 1.6062 | 0 | 0 | 0 | 3.2719 | 0 |
| 36378 | ENSMUSG00000097848 | 0 | NA | NA | NA | NA | NA | Gm807 | predicted gene 807 [Source:MGI Symbol;/ | 0 | 0 | 0 | 0 | 0 | 0 |
| 36379 | ENSMUSG00000097850 | 0 | NA | NA | NA | NA | NA | Mir205hg | Mir205 host gene [Source:MGI Symbol;Ac | 0 | 0 | 0 | 0 | 0 | 0 |
| 36380 | ENSMUSG00000097851 | 0 | NA | NA | NA | NA | NA | Platr30 | pluripotency associated transcript 30 [Sou | 0 | 0 | 0 | 0 | 0 | 0 |
| 36381 | ENSMUSG00000097852 | 11.13852077 | 0.621800827 | 0.671524914 | 0.925953474 | 0.354470159 | 0.794633876 | 4933405D12 | RIKEN cDNA 4933405D12 gene [Source:M | 8.031 | 7.1954 | 11.069 | 8.6758 | 20.722 | 11.138 |
| 36382 | ENSMUSG00000097853 | 0 | NA | NA | NA | NA | NA | Gm3532 | predicted gene 3532 [Source:MGI Symbol | 0 | 0 | 0 | 0 | 0 | 0 |
| 36383 | ENSMUSG00000097854 | 0 | NA | NA | NA | NA | NA | Gm26602 | predicted gene, 26602 [Source:MGI Symb | 0 | 0 | 0 | 0 | 0 | 0 |
| 36384 | ENSMUSG00000097855 | 17.16580122 | 0.462514688 | 0.659827787 | 0.700962732 | 0.483326273 | 0.871960882 | A930007I19I | RIKEN cDNA A930007I19 gene [Source:M( | 24.896 | 8.2233 | 10.062 | 36.631 | 9.8156 | 13.366 |
| 36385 | ENSMUSG00000097857 | 0 | NA | NA | NA | NA | NA | NA | NA | 0 | 0 | 0 | 0 | 0 | 0 |
| 36386 | ENSMUSG00000097858 | 0.133850373 | -0.796629176 | 4.080472857 | -0.195229623 | 0.84521319 | NA | 9530052C20 | RIKEN cDNA 9530052C20 gene [Source:M | 0.8031 | 0 | 0 | 0 | 0 | 0 |
| 36387 | ENSMUSG00000097859 | 0.321325949 | 1.911575114 | 4.048267893 | 0.472195804 | 0.636787034 | NA | NA | NA | 0 | 0 | 0 | 1.928 | 0 | 0 |
| 36388 | ENSMUSG00000097860 | 1.907897805 | -0.359142985 | 1.587659436 | -0.226209083 | 0.821038806 | NA | NA | NA | 2.4093 | 2.0558 | 2.0125 | 3.8559 | 0 | 1.1138 |
| 36389 | ENSMUSG00000097861 | 0 | NA | NA | NA | NA | NA | NA | NA | 0 | 0 | 0 | 0 | 0 | 0 |
| 36390 | ENSMUSG00000097862 | 0 | NA | NA | NA | NA | NA | NA | NA | 0 | 0 | 0 | 0 | 0 | 0 |
| 36391 | ENSMUSG00000097863 | 10.12057149 | 0.34821427 | 0.67641467 | 0.514794083 | 0.60669692 | 0.916376001 | 1010001B22 | RIKEN cDNA 1010001B22 gene [Source:M | 6.4248 | 11.307 | 9.0562 | 9.6398 | 9.8156 | 14.48 |
| 36392 | ENSMUSG00000097864 | 0.845941533 | -1.843011426 | 2.87307209 | -0.641477613 | 0.521212422 | NA | NA | NA | 0 | 4.1117 | 0 | 0.964 | 0 | 0 |
| 36393 | ENSMUSG00000097867 | 180.3738975 | 0.1318221 | 0.195362479 | 0.674756489 | 0.499830488 | 0.880716237 | NA | NA | 182.3 | 158.3 | 175.09 | 155.2 | 193.04 | 218.31 |
| 36394 | ENSMUSG00000097868 | 9.788129078 | 0.532507749 | 0.695039931 | 0.766154181 | 0.443584568 | 0.851079912 | NA | NA | 4.8186 | 11.307 | 8.05 | 12.532 | 11.997 | 10.025 |
| 36395 | ENSMUSG00000097869 | 0 | NA | NA | NA | NA | NA | NA | NA | 0 | 0 | 0 | 0 | 0 | 0 |
| 36396 | ENSMUSG00000097870 | 0.181769715 | 1.126924175 | 4.080472857 | 0.276174898 | 0.782413736 | NA | Gm26868 | predicted gene, 26868 [Source:MGI Symb | 0 | 0 | 0 | 0 | 1.0906 | 0 |

| | | | | | | | | | | | | | | | |
|---|---|---|---|---|---|---|---|---|---|---|---|---|---|---|---|
| 36397 | ENSMUSG00000097871 | 0 | NA | NA | NA | NA | NA | B230104I21I | RIKEN cDNA B230104I21 gene [Source:MC | 0 | 0 | 0 | 0 | 0 | 0 |
| 36398 | ENSMUSG00000097872 | 0 | NA | NA | NA | NA | NA | NA | NA | 0 | 0 | 0 | 0 | 0 | 0 |
| 36399 | ENSMUSG00000097873 | 0 | NA | NA | NA | NA | NA | NA | NA | 0 | 0 | 0 | 0 | 0 | 0 |
| 36400 | ENSMUSG00000097874 | 2.128527517 | -0.030186256 | 1.503788624 | -0.02007347 | 0.983984764 | NA | C030010L15 | RIKEN cDNA C030010L15 gene [Source:MG | 2.4093 | 0 | 4.025 | 1.928 | 2.1812 | 2.2277 |
| 36401 | ENSMUSG00000097875 | 0.663758638 | 2.950973782 | 3.025092005 | 0.975498853 | 0.32931287 | NA | Gm26704 | predicted gene, 26704 [Source:MGI Symb | 0 | 0 | 0 | 2.8919 | 1.0906 | 0 |
| 36402 | ENSMUSG00000097876 | 10.08288613 | -0.250323218 | 0.735192586 | -0.340486591 | 0.733490119 | 0.947915087 | Gm16892 | predicted gene, 16892 [Source:MGI Symb | 9.6372 | 12.335 | 11.069 | 16.388 | 3.2719 | 7.7969 |
| 36403 | ENSMUSG00000097877 | 0 | NA | NA | NA | NA | NA | Gm26703 | predicted gene, 26703 [Source:MGI Symb | 0 | 0 | 0 | 0 | 0 | 0 |
| 36404 | ENSMUSG00000097878 | 0 | NA | NA | NA | NA | NA | Gm8764 | predicted gene 8764 [Source:MGI Symbol | 0 | 0 | 0 | 0 | 0 | 0 |
| 36405 | ENSMUSG00000097879 | 4.013846655 | -0.645709345 | 1.220140467 | -0.529209024 | 0.596660458 | NA | NA | NA | 1.6062 | 4.1117 | 9.0562 | 3.8559 | 5.4531 | 0 |
| 36406 | ENSMUSG00000097881 | 0 | NA | NA | NA | NA | NA | Celrr | cerebellum expressed regulatory RNA [So | 0 | 0 | 0 | 0 | 0 | 0 |
| 36407 | ENSMUSG00000097882 | 0.928053766 | -0.877348406 | 2.28763571 | -0.383517534 | 0.701336072 | NA | 0610038B21 | RIKEN cDNA 0610038B21 gene [Source:M | 1.6062 | 1.0279 | 1.0062 | 1.928 | 0 | 0 |
| 36408 | ENSMUSG00000097883 | 0 | NA | NA | NA | NA | NA | Gm8357 | predicted gene 8357 [Source:MGI Symbol | 0 | 0 | 0 | 0 | 0 | 0 |
| 36409 | ENSMUSG00000097884 | 0 | NA | NA | NA | NA | NA | Gm26543 | predicted gene, 26543 [Source:MGI Symb | 0 | 0 | 0 | 0 | 0 | 0 |
| 36410 | ENSMUSG00000097885 | 3.906110048 | -0.037009298 | 1.130837181 | -0.032727345 | 0.973892018 | NA | 5031434O11 | RIKEN cDNA 5031434O11 gene [Source:M | 0.8031 | 5.1396 | 6.0375 | 4.8199 | 2.1812 | 4.4554 |
| 36411 | ENSMUSG00000097886 | 0 | NA | NA | NA | NA | NA | Gsg1l2 | GSG1-like 2 [Source:MGI Symbol;Acc:MGI | 0 | 0 | 0 | 0 | 0 | 0 |
| 36412 | ENSMUSG00000097887 | 2.038905513 | 1.135895373 | 1.498518544 | 0.758012223 | 0.448443665 | NA | Gm26542 | predicted gene, 26542 [Source:MGI Symb | 0.8031 | 2.0558 | 1.0062 | 2.8919 | 4.3625 | 1.1138 |
| 36413 | ENSMUSG00000097888 | 0.517623261 | 1.044040381 | 3.075707576 | 0.339447218 | 0.734272853 | NA | Gm26682 | predicted gene, 26682 [Source:MGI Symb | 0 | 1.0279 | 0 | 0.964 | 0 | 1.1138 |
| 36414 | ENSMUSG00000097889 | 2.567212069 | -0.458186779 | 1.355205382 | -0.338093978 | 0.735292368 | NA | NA | NA | 0.8031 | 4.1117 | 4.025 | 0.964 | 3.2719 | 2.2277 |
| 36415 | ENSMUSG00000097890 | 2.74421094 | 0.791579324 | 1.423382257 | 0.556125609 | 0.57812499 | NA | 4930547M16 | RIKEN cDNA 4930547M16 gene [Source:M | 0 | 5.1396 | 1.0062 | 4.8199 | 3.2719 | 2.2277 |
| 36416 | ENSMUSG00000097891 | 0.133850373 | -0.796629176 | 4.080472857 | -0.195229623 | 0.84521319 | NA | Gm3650 | predicted gene 3650 [Source:MGI Symbol | 0.8031 | 0 | 0 | 0 | 0 | 0 |
| 36417 | ENSMUSG00000097892 | 0.167707431 | -0.796629176 | 4.080472857 | -0.195229623 | 0.84521319 | NA | Gm26801 | predicted gene, 26801 [Source:MGI Symb | 0 | 0 | 1.0062 | 0 | 0 | 0 |
| 36418 | ENSMUSG00000097893 | 32.48278871 | -0.257324171 | 0.412258987 | -0.624180865 | 0.532508812 | 0.889143097 | 1700034P13 | RIKEN cDNA 1700034P13 gene [Source:M | 44.974 | 28.782 | 32.2 | 31.811 | 37.081 | 20.049 |
| 36419 | ENSMUSG00000097895 | 0 | NA | NA | NA | NA | NA | Gm18709 | predicted gene, 18709 [Source:MGI Symb | 0 | 0 | 0 | 0 | 0 | 0 |
| 36420 | ENSMUSG00000097896 | 0 | NA | NA | NA | NA | NA | Gm26800 | predicted gene, 26800 [Source:MGI Symb | 0 | 0 | 0 | 0 | 0 | 0 |
| 36421 | ENSMUSG00000097897 | 0 | NA | NA | NA | NA | NA | Rpl31-ps25 | ribosomal protein L31. pseudogene 25 [Sc | 0 | 0 | 0 | 0 | 0 | 0 |
| 36422 | ENSMUSG00000097898 | 0 | NA | NA | NA | NA | NA | 4930455B14 | RIKEN cDNA 4930455B14 gene [Source:M | 0 | 0 | 0 | 0 | 0 | 0 |
| 36423 | ENSMUSG00000097899 | 0 | NA | NA | NA | NA | NA | Gm16894 | predicted gene, 16894 [Source:MGI Symb | 0 | 0 | 0 | 0 | 0 | 0 |
| 36424 | ENSMUSG00000097900 | 0 | NA | NA | NA | NA | NA | Gm8652 | predicted gene 8652 [Source:MGI Symbol | 0 | 0 | 0 | 0 | 0 | 0 |
| 36425 | ENSMUSG00000097901 | 0 | NA | NA | NA | NA | NA | Gm26680 | predicted gene, 26680 [Source:MGI Symb | 0 | 0 | 0 | 0 | 0 | 0 |
| 36426 | ENSMUSG00000097902 | 0 | NA | NA | NA | NA | NA | Gm26531 | predicted gene, 26531 [Source:MGI Symb | 0 | 0 | 0 | 0 | 0 | 0 |
| 36427 | ENSMUSG00000097903 | 0.48903338 | 0.95637552 | 3.456690196 | 0.276673773 | 0.782030612 | NA | Gm26679 | predicted gene, 26679 [Source:MGI Symb | 0 | 0 | 1.0062 | 1.928 | 0 | 0 |
| 36428 | ENSMUSG00000097904 | 4.368492213 | -0.464530783 | 1.181636153 | -0.39312506 | 0.694227116 | NA | NA | NA | 3.2124 | 2.0558 | 10.062 | 8.6758 | 1.0906 | 1.1138 |
| 36429 | ENSMUSG00000097906 | 3.561350487 | 0.837860097 | 1.119261093 | 0.748583242 | 0.454108436 | NA | Gm9625 | predicted gene 9625 [Source:MGI Symbol | 1.6062 | 4.1117 | 2.0125 | 4.8199 | 4.3625 | 4.4554 |
| 36430 | ENSMUSG00000097907 | 0 | NA | NA | NA | NA | NA | Raxos1 | retina and anterior neural fold homeobox | 0 | 0 | 0 | 0 | 0 | 0 |
| 36431 | ENSMUSG00000097908 | 111.1208721 | 0.461431111 | 0.241255855 | 1.912621402 | 0.055796538 | 0.333350319 | 4933404O12 | RIKEN cDNA 4933404O12 gene [Source:M | 91.554 | 75.038 | 113.71 | 111.82 | 128.69 | 145.91 |
| 36432 | ENSMUSG00000097910 | 2.863811372 | -2.815925423 | 1.7754918 | -1.585997425 | 0.112739894 | NA | Hdnr | Hand2 downstream lncRNA [Source:MGI S | 8.8341 | 6.1675 | 0 | 0 | 2.1812 | 0 |
| 36433 | ENSMUSG00000097911 | 0 | NA | NA | NA | NA | NA | Gm26691 | predicted gene, 26691 [Source:MGI Symb | 0 | 0 | 0 | 0 | 0 | 0 |
| 36434 | ENSMUSG00000097912 | 0 | NA | NA | NA | NA | NA | NA | NA | 0 | 0 | 0 | 0 | 0 | 0 |
| 36435 | ENSMUSG00000097913 | 0 | NA | NA | NA | NA | NA | NA | NA | 0 | 0 | 0 | 0 | 0 | 0 |
| 36436 | ENSMUSG00000097914 | 0 | NA | NA | NA | NA | NA | Gm26838 | predicted gene, 26838 [Source:MGI Symb | 0 | 0 | 0 | 0 | 0 | 0 |
| 36437 | ENSMUSG00000097915 | 1.375844284 | 0.917838236 | 1.859671904 | 0.493548477 | 0.621625096 | NA | A330009N23 | RIKEN cDNA A330009N23 gene [Source:M | 0.8031 | 2.0558 | 0 | 0.964 | 1.0906 | 3.3415 |
| 36438 | ENSMUSG00000097917 | 1.576995064 | -1.769238182 | 1.885676301 | -0.938251269 | 0.348115297 | NA | Gm26839 | predicted gene, 26839 [Source:MGI Symb | 3.2124 | 2.0558 | 2.0125 | 0 | 2.1812 | 0 |
| 36439 | ENSMUSG00000097918 | 0 | NA | NA | NA | NA | NA | Ascl5 | achaete-scute family bHLH transcription fa | 0 | 0 | 0 | 0 | 0 | 0 |
| 36440 | ENSMUSG00000097919 | 0 | NA | NA | NA | NA | NA | Gm27021 | predicted gene, 27021 [Source:MGI Symb | 0 | 0 | 0 | 0 | 0 | 0 |
| 36441 | ENSMUSG00000097920 | 0 | NA | NA | NA | NA | NA | Gm7784 | predicted gene 7784 [Source:MGI Symbol | 0 | 0 | 0 | 0 | 0 | 0 |
| 36442 | ENSMUSG00000097921 | 1.165046003 | -0.264508229 | 1.980197771 | -0.133576673 | 0.893737331 | NA | Gm26576 | predicted gene, 26576 [Source:MGI Symb | 0.8031 | 0 | 3.0187 | 0.964 | 1.0906 | 1.1138 |
| 36443 | ENSMUSG00000097923 | 0 | NA | NA | NA | NA | NA | Gm26577 | predicted gene, 26577 [Source:MGI Symb | 0 | 0 | 0 | 0 | 0 | 0 |
| 36444 | ENSMUSG00000097924 | 0 | NA | NA | NA | NA | NA | A730020E08 | RIKEN cDNA A730020E08 gene [Source:M | 0 | 0 | 0 | 0 | 0 | 0 |
| 36445 | ENSMUSG00000097925 | 0 | NA | NA | NA | NA | NA | Gm26574 | predicted gene, 26574 [Source:MGI Symb | 0 | 0 | 0 | 0 | 0 | 0 |
| 36446 | ENSMUSG00000097926 | 0.17131964 | -0.796629176 | 4.080472857 | -0.195229623 | 0.84521319 | NA | Gm26575 | predicted gene, 26575 [Source:MGI Symb | 0 | 1.0279 | 0 | 0 | 0 | 0 |
| 36447 | ENSMUSG00000097927 | 0 | NA | NA | NA | NA | NA | Gm6999 | predicted gene 6999 [Source:MGI Symbol | 0 | 0 | 0 | 0 | 0 | 0 |
| 36448 | ENSMUSG00000097928 | 0.335414862 | -1.779047242 | 4.030804179 | -0.441362855 | 0.658950329 | NA | NA | NA | 0 | 0 | 2.0125 | 0 | 0 | 0 |
| 36449 | ENSMUSG00000097929 | 0 | NA | NA | NA | NA | NA | Tunar | Tcl1 upstream neural differentiation assoc | 0 | 0 | 0 | 0 | 0 | 0 |
| 36450 | ENSMUSG00000097930 | 84.30194019 | 0.359616911 | 0.313618327 | 1.146670585 | 0.251517792 | 0.703994757 | C330002G04 | RIKEN cDNA C330002G04 gene [Source:M | 42.564 | 80.178 | 99.618 | 107 | 82.887 | 93.563 |
| 36451 | ENSMUSG00000097931 | 0 | NA | NA | NA | NA | NA | Gm11392 | predicted gene 11392 [Source:MGI Symbo | 0 | 0 | 0 | 0 | 0 | 0 |
| 36452 | ENSMUSG00000097932 | 0 | NA | NA | NA | NA | NA | C430039J16I | RIKEN cDNA C430039J16 gene [Source:MC | 0 | 0 | 0 | 0 | 0 | 0 |

| | | | | | | | | | | | | | | | |
|---|---|---|---|---|---|---|---|---|---|---|---|---|---|---|---|
| 36453 | ENSMUSG00000097933 | 0.133850373 | -0.796629176 | 4.080472857 | -0.195229623 | 0.84521319 | NA | 1700039M1( | RIKEN cDNA 1700039M10 gene [Source:M | 0.8031 | 0 | 0 | 0 | 0 | 0 |
| 36454 | ENSMUSG00000097934 | 0 | NA | NA | NA | NA | NA | 6720483E21 | RIKEN cDNA 6720483E21 gene [Source:M( | 0 | 0 | 0 | 0 | 0 | 0 |
| 36455 | ENSMUSG00000097935 | 0 | NA | NA | NA | NA | NA | Gm3809 | predicted gene 3809 [Source:MGI Symbol | 0 | 0 | 0 | 0 | 0 | 0 |
| 36456 | ENSMUSG00000097936 | 0 | NA | NA | NA | NA | NA | Gm4819 | predicted gene 4819 [Source:MGI Symbol | 0 | 0 | 0 | 0 | 0 | 0 |
| 36457 | ENSMUSG00000097937 | 2.22371897 | -1.035243667 | 1.650140711 | -0.627366903 | 0.530418758 | NA | Gm6967 | predicted gene 6967 [Source:MGI Symbol | 4.8186 | 2.0558 | 2.0125 | 0 | 0 | 4.4554 |
| 36458 | ENSMUSG00000097938 | 0 | NA | NA | NA | NA | NA | Gm26933 | predicted gene, 26933 [Source:MGI Symb | 0 | 0 | 0 | 0 | 0 | 0 |
| 36459 | ENSMUSG00000097939 | 0 | NA | NA | NA | NA | NA | Gm8601 | predicted gene 8601 [Source:MGI Symbol | 0 | 0 | 0 | 0 | 0 | 0 |
| 36460 | ENSMUSG00000097940 | 0 | NA | NA | NA | NA | NA | Gm21392 | predicted gene, 21392 [Source:MGI Symb | 0 | 0 | 0 | 0 | 0 | 0 |
| 36461 | ENSMUSG00000097941 | 0 | NA | NA | NA | NA | NA | Gm8984 | predicted gene 8984 [Source:MGI Symbol | 0 | 0 | 0 | 0 | 0 | 0 |
| 36462 | ENSMUSG00000097942 | 0.328370405 | 0.16514316 | 4.003149536 | 0.041253308 | 0.967093956 | NA | Gm26968 | predicted gene, 26968 [Source:MGI Symb | 0 | 0 | 1.0062 | 0.964 | 0 | 0 |
| 36463 | ENSMUSG00000097943 | 0 | NA | NA | NA | NA | NA | Gm7833 | predicted gene 7833 [Source:MGI Symbol | 0 | 0 | 0 | 0 | 0 | 0 |
| 36464 | ENSMUSG00000097944 | 40.67276993 | -0.193463215 | 0.365416773 | -0.529431677 | 0.596506028 | 0.914226274 | A130014A01 | RIKEN cDNA A130014A01 gene [Source:M | 40.958 | 31.865 | 57.356 | 36.631 | 34.9 | 42.326 |
| 36465 | ENSMUSG00000097945 | 0 | NA | NA | NA | NA | NA | Gm17799 | predicted gene, 17799 [Source:MGI Symb | 0 | 0 | 0 | 0 | 0 | 0 |
| 36466 | ENSMUSG00000097946 | 0 | NA | NA | NA | NA | NA | Gm8343 | predicted gene 8343 [Source:MGI Symbol | 0 | 0 | 0 | 0 | 0 | 0 |
| 36467 | ENSMUSG00000097947 | 0 | NA | NA | NA | NA | NA | Gm26970 | predicted gene, 26970 [Source:MGI Symb | 0 | 0 | 0 | 0 | 0 | 0 |
| 36468 | ENSMUSG00000097948 | 0.160662974 | 1.126924175 | 4.080472857 | 0.276174898 | 0.782413736 | NA | Gm21430 | predicted gene, 21430 [Source:MGI Symb | 0 | 0 | 0 | 0.964 | 0 | 0 |
| 36469 | ENSMUSG00000097949 | 0 | NA | NA | NA | NA | NA | Gm5354 | predicted gene 5354 [Source:MGI Symbol | 0 | 0 | 0 | 0 | 0 | 0 |
| 36470 | ENSMUSG00000097950 | 0 | NA | NA | NA | NA | NA | Gm17800 | predicted gene, 17800 [Source:MGI Symb | 0 | 0 | 0 | 0 | 0 | 0 |
| 36471 | ENSMUSG00000097951 | 0 | NA | NA | NA | NA | NA | Gm7611 | predicted gene 7611 [Source:MGI Symbol | 0 | 0 | 0 | 0 | 0 | 0 |
| 36472 | ENSMUSG00000097952 | 0 | NA | NA | NA | NA | NA | Gm4398 | predicted gene 4398 [Source:MGI Symbol | 0 | 0 | 0 | 0 | 0 | 0 |
| 36473 | ENSMUSG00000097953 | 0 | NA | NA | NA | NA | NA | Gm26952 | predicted gene, 26952 [Source:MGI Symb | 0 | 0 | 0 | 0 | 0 | 0 |
| 36474 | ENSMUSG00000097954 | 0 | NA | NA | NA | NA | NA | Gm4217 | predicted gene 4217 [Source:MGI Symbol | 0 | 0 | 0 | 0 | 0 | 0 |
| 36475 | ENSMUSG00000097955 | 0 | NA | NA | NA | NA | NA | Gm26950 | predicted gene, 26950 [Source:MGI Symb | 0 | 0 | 0 | 0 | 0 | 0 |
| 36476 | ENSMUSG00000097957 | 0 | NA | NA | NA | NA | NA | Gm8993 | predicted gene 8993 [Source:MGI Symbol | 0 | 0 | 0 | 0 | 0 | 0 |
| 36477 | ENSMUSG00000097958 | 0 | NA | NA | NA | NA | NA | Gm8492 | predicted gene 8492 [Source:MGI Symbol | 0 | 0 | 0 | 0 | 0 | 0 |
| 36478 | ENSMUSG00000097959 | 0 | NA | NA | NA | NA | NA | Gm6931 | predicted gene 6931 [Source:MGI Symbol | 0 | 0 | 0 | 0 | 0 | 0 |
| 36479 | ENSMUSG00000097960 | 0 | NA | NA | NA | NA | NA | A330074K22 | RIKEN cDNA A330074K22 gene [Source:M | 0 | 0 | 0 | 0 | 0 | 0 |
| 36480 | ENSMUSG00000097961 | 0 | NA | NA | NA | NA | NA | Gm27000 | predicted gene, 27000 [Source:MGI Symb | 0 | 0 | 0 | 0 | 0 | 0 |
| 36481 | ENSMUSG00000097962 | 0 | NA | NA | NA | NA | NA | 1700034G24 | RIKEN cDNA 1700034G24 gene [Source:M | 0 | 0 | 0 | 0 | 0 | 0 |
| 36482 | ENSMUSG00000097963 | 0 | NA | NA | NA | NA | NA | Gm27001 | predicted gene, 27001 [Source:MGI Symb | 0 | 0 | 0 | 0 | 0 | 0 |
| 36483 | ENSMUSG00000097964 | 0 | NA | NA | NA | NA | NA | Gm21045 | predicted gene, 21045 [Source:MGI Symb | 0 | 0 | 0 | 0 | 0 | 0 |
| 36484 | ENSMUSG00000097965 | 0 | NA | NA | NA | NA | NA | Gm5839 | predicted gene 5839 [Source:MGI Symbol | 0 | 0 | 0 | 0 | 0 | 0 |
| 36485 | ENSMUSG00000097966 | 0 | NA | NA | NA | NA | NA | Gm9423 | predicted gene 9423 [Source:MGI Symbol | 0 | 0 | 0 | 0 | 0 | 0 |
| 36486 | ENSMUSG00000097967 | 0 | NA | NA | NA | NA | NA | Gm27002 | predicted gene, 27002 [Source:MGI Symb | 0 | 0 | 0 | 0 | 0 | 0 |
| 36487 | ENSMUSG00000097968 | 0 | NA | NA | NA | NA | NA | Tpi-rs2 | triosephosphate isomerase related seque | 0 | 0 | 0 | 0 | 0 | 0 |
| 36488 | ENSMUSG00000097970 | 21.1391293 | -0.157837659 | 0.529805516 | -0.297916224 | 0.765767105 | 0.954996635 | Gm27028 | predicted gene, 27028 [Source:MGI Symb | 15.259 | 31.865 | 20.125 | 29.883 | 17.45 | 12.252 |
| 36489 | ENSMUSG00000097971 | 1795.936562 | -0.278924838 | 0.129743933 | -2.149810263 | 0.031570226 | 0.236342058 | NA | NA | 1872 | 1682.7 | 2352.6 | 1637.8 | 1605.4 | 1625.1 |
| 36490 | ENSMUSG00000097972 | 0 | NA | NA | NA | NA | NA | Scgb2b15 | secretoglobin, family 2B, member 15 [Sou | 0 | 0 | 0 | 0 | 0 | 0 |
| 36491 | ENSMUSG00000097973 | 0 | NA | NA | NA | NA | NA | Gm4929 | predicted gene 4929 [Source:MGI Symbol | 0 | 0 | 0 | 0 | 0 | 0 |
| 36492 | ENSMUSG00000097974 | 9.880897148 | -0.835380487 | 0.739050648 | -1.130342676 | 0.258331859 | 0.711962318 | Gm10605 | predicted gene 10605 [Source:MGI Symbo | 10.44 | 17.475 | 10.062 | 5.7839 | 3.2719 | 12.252 |
| 36493 | ENSMUSG00000097975 | 0 | NA | NA | NA | NA | NA | Gm8079 | predicted gene 8079 [Source:MGI Symbol | 0 | 0 | 0 | 0 | 0 | 0 |
| 36494 | ENSMUSG00000097976 | 0 | NA | NA | NA | NA | NA | Gm26918 | predicted gene, 26918 [Source:MGI Symb | 0 | 0 | 0 | 0 | 0 | 0 |
| 36495 | ENSMUSG00000097977 | 0 | NA | NA | NA | NA | NA | Gm5652 | predicted gene 5652 [Source:MGI Symbol | 0 | 0 | 0 | 0 | 0 | 0 |
| 36496 | ENSMUSG00000097979 | 1.3343491 | -1.459952214 | 2.234814003 | -0.653276833 | 0.513577825 | NA | Gm4691 | predicted gene 4691 [Source:MGI Symbol | 4.8186 | 0 | 1.0062 | 0 | 2.1812 | 0 |
| 36497 | ENSMUSG00000097980 | 0 | NA | NA | NA | NA | NA | Gm26972 | predicted gene, 26972 [Source:MGI Symb | 0 | 0 | 0 | 0 | 0 | 0 |
| 36498 | ENSMUSG00000097981 | 0 | NA | NA | NA | NA | NA | Gm7583 | predicted gene 7583 [Source:MGI Symbol | 0 | 0 | 0 | 0 | 0 | 0 |
| 36499 | ENSMUSG00000097982 | 0 | NA | NA | NA | NA | NA | Scgb2b12 | secretoglobin, family 2B, member 12 [Sou | 0 | 0 | 0 | 0 | 0 | 0 |
| 36500 | ENSMUSG00000097983 | 0.920461372 | 3.403685117 | 2.681240704 | 1.269444072 | 0.204282726 | NA | Gm26971 | predicted gene, 26971 [Source:MGI Symb | 0 | 0 | 0 | 0 | 2.1812 | 3.3415 |
| 36501 | ENSMUSG00000097984 | 0 | NA | NA | NA | NA | NA | Gm8167 | predicted gene 8167 [Source:MGI Symbol | 0 | 0 | 0 | 0 | 0 | 0 |
| 36502 | ENSMUSG00000097985 | 0 | NA | NA | NA | NA | NA | Gm5274 | predicted gene 5274 [Source:MGI Symbol | 0 | 0 | 0 | 0 | 0 | 0 |
| 36503 | ENSMUSG00000097986 | 0 | NA | NA | NA | NA | NA | Gm26953 | predicted gene, 26953 [Source:MGI Symb | 0 | 0 | 0 | 0 | 0 | 0 |
| 36504 | ENSMUSG00000097987 | 0 | NA | NA | NA | NA | NA | Gm17786 | predicted gene, 17786 [Source:MGI Symb | 0 | 0 | 0 | 0 | 0 | 0 |
| 36505 | ENSMUSG00000097988 | 0 | NA | NA | NA | NA | NA | Gm10535 | predicted gene 10535 [Source:MGI Symbo | 0 | 0 | 0 | 0 | 0 | 0 |
| 36506 | ENSMUSG00000097989 | 0.96342938 | -2.048320316 | 2.460986955 | -0.832316608 | 0.405230258 | NA | Gm4335 | predicted gene 4335 [Source:MGI Symbol | 1.6062 | 3.0838 | 0 | 0 | 1.0906 | 0 |
| 36507 | ENSMUSG00000097990 | 3.694995396 | -0.948686695 | 1.148603764 | -0.82594775 | 0.408833732 | NA | Gm19557 | predicted gene, 19557 [Source:MGI Symb | 2.4093 | 5.1396 | 7.0437 | 0.964 | 3.2719 | 3.3415 |
| 36508 | ENSMUSG00000097991 | 0 | NA | NA | NA | NA | NA | Gm7656 | predicted gene 7656 [Source:MGI Symbol | 0 | 0 | 0 | 0 | 0 | 0 |

| | | | | | | | | | | | | | | | |
|---|---|---|---|---|---|---|---|---|---|---|---|---|---|---|---|
| 36509 | ENSMUSG00000097992 | 0 | NA | NA | NA | NA | NA | Gm27016 | predicted gene, 27016 [Source:MGI Symb | 0 | 0 | 0 | 0 | 0 | 0 |
| 36510 | ENSMUSG00000097993 | 0.328370405 | 0.16514316 | 4.003149536 | 0.041253308 | 0.967093956 | NA | Ptprv | protein tyrosine phosphatase, receptor ty | 0 | 0 | 1.0062 | 0.964 | 0 | 0 |
| 36511 | ENSMUSG00000097994 | 31.19851402 | -0.216880398 | 0.421448262 | -0.514607409 | 0.606827387 | 0.916376001 | Gm26982 | predicted gene, 26982 [Source:MGI Symb | 21.684 | 33.921 | 45.281 | 28.919 | 25.084 | 32.301 |
| 36512 | ENSMUSG00000097995 | 0 | NA | NA | NA | NA | NA | Gm24362 | predicted gene, 24362 [Source:MGI Symb | 0 | 0 | 0 | 0 | 0 | 0 |
| 36513 | ENSMUSG00000097996 | 0 | NA | NA | NA | NA | NA | Gm5313 | predicted gene 5313 [Source:MGI Symbol | 0 | 0 | 0 | 0 | 0 | 0 |
| 36514 | ENSMUSG00000097997 | 0.781453075 | -0.369215669 | 2.439554878 | -0.151345507 | 0.879703174 | NA | Gm27017 | predicted gene, 27017 [Source:MGI Symb | 1.6062 | 1.0279 | 0 | 0.964 | 1.0906 | 0 |
| 36515 | ENSMUSG00000097998 | 0 | NA | NA | NA | NA | NA | Gm27019 | predicted gene, 27019 [Source:MGI Symb | 0 | 0 | 0 | 0 | 0 | 0 |
| 36516 | ENSMUSG00000097999 | 0 | NA | NA | NA | NA | NA | Gm9061 | predicted gene 9061 [Source:MGI Symbol | 0 | 0 | 0 | 0 | 0 | 0 |
| 36517 | ENSMUSG00000098000 | 0.160662974 | 1.126924175 | 4.080472857 | 0.276174898 | 0.782413736 | NA | Gm27026 | predicted gene, 27026 [Source:MGI Symb | 0 | 0 | 0 | 0.964 | 0 | 0 |
| 36518 | ENSMUSG00000098001 | 0.401551119 | -2.072982726 | 4.008251979 | -0.517178744 | 0.605031388 | NA | Gm27048 | predicted gene, 27048 [Source:MGI Symb | 2.4093 | 0 | 0 | 0 | 0 | 0 |
| 36519 | ENSMUSG00000098002 | 0 | NA | NA | NA | NA | NA | Gm9658 | predicted gene 9658 [Source:MGI Symbol | 0 | 0 | 0 | 0 | 0 | 0 |
| 36520 | ENSMUSG00000098003 | 0 | NA | NA | NA | NA | NA | Gm6317 | predicted gene 6317 [Source:MGI Symbol | 0 | 0 | 0 | 0 | 0 | 0 |
| 36521 | ENSMUSG00000098004 | 0.744190399 | -2.947713985 | 2.874421214 | -1.025498271 | 0.305128141 | NA | Gm27027 | predicted gene, 27027 [Source:MGI Symb | 2.4093 | 2.0558 | 0 | 0 | 0 | 0 |
| 36522 | ENSMUSG00000098005 | 0 | NA | NA | NA | NA | NA | Gm6114 | predicted gene 6114 [Source:MGI Symbol | 0 | 0 | 0 | 0 | 0 | 0 |
| 36523 | ENSMUSG00000098006 | 0.428363721 | -0.543730382 | 3.624677344 | -0.15000794 | 0.880758351 | NA | Gm8565 | predicted gene 8565 [Source:MGI Symbol | 1.6062 | 0 | 0 | 0.964 | 0 | 0 |
| 36524 | ENSMUSG00000098007 | 0 | NA | NA | NA | NA | NA | Gm8614 | predicted gene 8614 [Source:MGI Symbol | 0 | 0 | 0 | 0 | 0 | 0 |
| 36525 | ENSMUSG00000098008 | 0.133850373 | -0.796629176 | 4.080472857 | -0.195229623 | 0.84521319 | NA | A930001A2C | RIKEN cDNA A930001A20 gene [Source:M | 0.8031 | 0 | 0 | 0 | 0 | 0 |
| 36526 | ENSMUSG00000098009 | 0 | NA | NA | NA | NA | NA | Gm5597 | predicted gene 5597 [Source:MGI Symbol | 0 | 0 | 0 | 0 | 0 | 0 |
| 36527 | ENSMUSG00000098010 | 0 | NA | NA | NA | NA | NA | Gm21542 | predicted gene, 21542 [Source:MGI Symb | 0 | 0 | 0 | 0 | 0 | 0 |
| 36528 | ENSMUSG00000098011 | 0 | NA | NA | NA | NA | NA | Gm21232 | predicted gene, 21232 [Source:MGI Symb | 0 | 0 | 0 | 0 | 0 | 0 |
| 36529 | ENSMUSG00000098012 | 0 | NA | NA | NA | NA | NA | Gm8907 | predicted gene 8907 [Source:MGI Symbol | 0 | 0 | 0 | 0 | 0 | 0 |
| 36530 | ENSMUSG00000098013 | 0 | NA | NA | NA | NA | NA | Gm9135 | predicted gene 9135 [Source:MGI Symbol | 0 | 0 | 0 | 0 | 0 | 0 |
| 36531 | ENSMUSG00000098014 | 26.63545607 | 1.191553333 | 0.434795955 | 2.740488542 | 0.006134792 | 0.076494511 | Gm26967 | predicted gene, 26967 [Source:MGI Symb | 16.062 | 17.475 | 15.094 | 30.847 | 45.806 | 34.529 |
| 36532 | ENSMUSG00000098015 | 0 | NA | NA | NA | NA | NA | Gm44527 | predicted gene 44527 [Source:MGI Symbo | 0 | 0 | 0 | 0 | 0 | 0 |
| 36533 | ENSMUSG00000098016 | 0 | NA | NA | NA | NA | NA | Gm9211 | predicted gene 9211 [Source:MGI Symbol | 0 | 0 | 0 | 0 | 0 | 0 |
| 36534 | ENSMUSG00000098017 | 0 | NA | NA | NA | NA | NA | Gm5342 | predicted gene 5342 [Source:MGI Symbol | 0 | 0 | 0 | 0 | 0 | 0 |
| 36535 | ENSMUSG00000098019 | 1.14577419 | 0.394191739 | 2.55065941 | 0.154545031 | 0.877180008 | NA | Gm2546 | predicted gene 2546 [Source:MGI Symbol | 0 | 0 | 3.0187 | 3.8559 | 0 | 0 |
| 36536 | ENSMUSG00000098021 | 2.88584957 | 0.939228898 | 1.546727057 | 0.607236354 | 0.54369407 | NA | Gm9522 | predicted gene 9522 [Source:MGI Symbol | 4.8186 | 0 | 1.0062 | 3.8559 | 7.6343 | 0 |
| 36537 | ENSMUSG00000098022 | 0.534859069 | 1.101323472 | 3.351640271 | 0.328592386 | 0.742463804 | NA | Zfp82 | zinc finger protein 82 [Source:MGI Symbo | 0 | 1.0279 | 0 | 0 | 2.1812 | 0 |
| 36538 | ENSMUSG00000098023 | 0 | NA | NA | NA | NA | NA | Gm19200 | predicted gene, 19200 [Source:MGI Symb | 0 | 0 | 0 | 0 | 0 | 0 |
| 36539 | ENSMUSG00000098024 | 22.14891711 | 0.841393823 | 0.491563146 | 1.711669865 | 0.086957525 | 0.434457217 | Gm27003 | predicted gene, 27003 [Source:MGI Symb | 20.078 | 9.2513 | 18.112 | 34.703 | 22.903 | 27.846 |
| 36540 | ENSMUSG00000098025 | 0 | NA | NA | NA | NA | NA | Gm27013 | predicted gene, 27013 [Source:MGI Symb | 0 | 0 | 0 | 0 | 0 | 0 |
| 36541 | ENSMUSG00000098026 | 0.321325949 | 1.911575114 | 4.048267893 | 0.472195804 | 0.636787034 | NA | Gm27014 | predicted gene, 27014 [Source:MGI Symb | 0 | 0 | 0 | 1.928 | 0 | 0 |
| 36542 | ENSMUSG00000098027 | 0 | NA | NA | NA | NA | NA | Gm27004 | predicted gene, 27004 [Source:MGI Symb | 0 | 0 | 0 | 0 | 0 | 0 |
| 36543 | ENSMUSG00000098028 | 0 | NA | NA | NA | NA | NA | Gm8824 | predicted gene 8824 [Source:MGI Symbol | 0 | 0 | 0 | 0 | 0 | 0 |
| 36544 | ENSMUSG00000098029 | 0 | NA | NA | NA | NA | NA | Gm7745 | predicted gene 7745 [Source:MGI Symbol | 0 | 0 | 0 | 0 | 0 | 0 |
| 36545 | ENSMUSG00000098030 | 0 | NA | NA | NA | NA | NA | Gm6771 | predicted gene 6771 [Source:MGI Symbol | 0 | 0 | 0 | 0 | 0 | 0 |
| 36546 | ENSMUSG00000098031 | 0 | NA | NA | NA | NA | NA | Gm9341 | predicted gene 9341 [Source:MGI Symbol | 0 | 0 | 0 | 0 | 0 | 0 |
| 36547 | ENSMUSG00000098033 | 0 | NA | NA | NA | NA | NA | Gm9381 | predicted gene 9381 [Source:MGI Symbol | 0 | 0 | 0 | 0 | 0 | 0 |
| 36548 | ENSMUSG00000098035 | 0 | NA | NA | NA | NA | NA | Gm5335 | predicted gene 5335 [Source:MGI Symbol | 0 | 0 | 0 | 0 | 0 | 0 |
| 36549 | ENSMUSG00000098036 | 0 | NA | NA | NA | NA | NA | Gm8955 | predicted gene 8955 [Source:MGI Symbol | 0 | 0 | 0 | 0 | 0 | 0 |
| 36550 | ENSMUSG00000098037 | 0 | NA | NA | NA | NA | NA | Gm7765 | predicted gene 7765 [Source:MGI Symbol | 0 | 0 | 0 | 0 | 0 | 0 |
| 36551 | ENSMUSG00000098038 | 0 | NA | NA | NA | NA | NA | Gm8793 | predicted gene 8793 [Source:MGI Symbol | 0 | 0 | 0 | 0 | 0 | 0 |
| 36552 | ENSMUSG00000098039 | 0.160662974 | 1.126924175 | 4.080472857 | 0.276174898 | 0.782413736 | NA | Gm26925 | predicted gene, 26925 [Source:MGI Symb | 0 | 0 | 0 | 0.964 | 0 | 0 |
| 36553 | ENSMUSG00000098040 | 0 | NA | NA | NA | NA | NA | Gm20757 | predicted gene, 20757 [Source:MGI Symb | 0 | 0 | 0 | 0 | 0 | 0 |
| 36554 | ENSMUSG00000098041 | 11.60667001 | 0.179495299 | 0.667338485 | 0.268971899 | 0.787951307 | 0.960870081 | Gm26981 | predicted gene, 26981 [Source:MGI Symb | 10.44 | 10.279 | 12.075 | 19.28 | 11.997 | 5.5692 |
| 36555 | ENSMUSG00000098042 | 0 | NA | NA | NA | NA | NA | Gm5710 | predicted gene 5710 [Source:MGI Symbol | 0 | 0 | 0 | 0 | 0 | 0 |
| 36556 | ENSMUSG00000098043 | 0 | NA | NA | NA | NA | NA | Gm26980 | predicted gene, 26980 [Source:MGI Symb | 0 | 0 | 0 | 0 | 0 | 0 |
| 36557 | ENSMUSG00000098044 | 0 | NA | NA | NA | NA | NA | Gm9162 | predicted gene 9162 [Source:MGI Symbol | 0 | 0 | 0 | 0 | 0 | 0 |
| 36558 | ENSMUSG00000098045 | 0 | NA | NA | NA | NA | NA | Gm26979 | predicted gene, 26979 [Source:MGI Symb | 0 | 0 | 0 | 0 | 0 | 0 |
| 36559 | ENSMUSG00000098046 | 0 | NA | NA | NA | NA | NA | Gm8254 | predicted gene 8254 [Source:MGI Symbol | 0 | 0 | 0 | 0 | 0 | 0 |
| 36560 | ENSMUSG00000098047 | 0 | NA | NA | NA | NA | NA | Gm18822 | predicted gene, 18822 [Source:MGI Symb | 0 | 0 | 0 | 0 | 0 | 0 |
| 36561 | ENSMUSG00000098048 | 0 | NA | NA | NA | NA | NA | Gm8330 | predicted gene 8330 [Source:MGI Symbol | 0 | 0 | 0 | 0 | 0 | 0 |
| 36562 | ENSMUSG00000098050 | 121.3859548 | 0.069621563 | 0.229798695 | 0.302967618 | 0.76191454 | 0.954306538 | Gm5345 | predicted gene 5345 [Source:MGI Symbol | 101.99 | 142.88 | 110.69 | 106.04 | 133.06 | 133.66 |
| 36563 | ENSMUSG00000098051 | 0 | NA | NA | NA | NA | NA | Gm27032 | predicted gene, 27032 [Source:MGI Symb | 0 | 0 | 0 | 0 | 0 | 0 |
| 36564 | ENSMUSG00000098052 | 0 | NA | NA | NA | NA | NA | Gm5300 | predicted gene 5300 [Source:MGI Symbol | 0 | 0 | 0 | 0 | 0 | 0 |

| | | | | | | | | | | | | | | | |
|---|---|---|---|---|---|---|---|---|---|---|---|---|---|---|---|
| 36565 | ENSMUSG00000098053 | 0 | NA | NA | NA | NA | NA | Gm27033 | predicted gene, 27033 [Source:MGI Symb | 0 | 0 | 0 | 0 | 0 | 0 |
| 36566 | ENSMUSG00000098054 | 0 | NA | NA | NA | NA | NA | Gm8192 | predicted gene 8192 [Source:MGI Symbol | 0 | 0 | 0 | 0 | 0 | 0 |
| 36567 | ENSMUSG00000098055 | 2.986548588 | -2.217135383 | 1.455665898 | -1.523107319 | 0.127731858 | NA | Gm26947 | predicted gene, 26947 [Source:MGI Symb | 1.6062 | 4.1117 | 9.0562 | 0.964 | 2.1812 | 0 |
| 36568 | ENSMUSG00000098056 | 0 | NA | NA | NA | NA | NA | Gm8100 | predicted gene 8100 [Source:MGI Symbol | 0 | 0 | 0 | 0 | 0 | 0 |
| 36569 | ENSMUSG00000098057 | 0.335414862 | -1.779047242 | 4.030804179 | -0.441362855 | 0.658950329 | NA | Gm6283 | predicted gene 6283 [Source:MGI Symbol | 0 | 0 | 2.0125 | 0 | 0 | 0 |
| 36570 | ENSMUSG00000098058 | 0 | NA | NA | NA | NA | NA | Gm8529 | predicted gene 8529 [Source:MGI Symbol | 0 | 0 | 0 | 0 | 0 | 0 |
| 36571 | ENSMUSG00000098059 | 0 | NA | NA | NA | NA | NA | Gm27034 | predicted gene, 27034 [Source:MGI Symb | 0 | 0 | 0 | 0 | 0 | 0 |
| 36572 | ENSMUSG00000098060 | 0 | NA | NA | NA | NA | NA | Gm4842 | predicted pseudogene 4842 [Source:MGI | 0 | 0 | 0 | 0 | 0 | 0 |
| 36573 | ENSMUSG00000098061 | 0 | NA | NA | NA | NA | NA | NA | NA | 0 | 0 | 0 | 0 | 0 | 0 |
| 36574 | ENSMUSG00000098062 | 5.873042536 | 0.926911105 | 0.907811369 | 1.021039322 | 0.307235809 | 0.756237058 | Gm26931 | predicted gene, 26931 [Source:MGI Symb | 4.0155 | 4.1117 | 4.025 | 8.6758 | 3.2719 | 11.138 |
| 36575 | ENSMUSG00000098064 | 0 | NA | NA | NA | NA | NA | Gm9192 | predicted gene 9192 [Source:MGI Symbol | 0 | 0 | 0 | 0 | 0 | 0 |
| 36576 | ENSMUSG00000098065 | 0 | NA | NA | NA | NA | NA | Gm5177 | predicted pseudogene 5177 [Source:MGI | 0 | 0 | 0 | 0 | 0 | 0 |
| 36577 | ENSMUSG00000098066 | 17.77199784 | -0.347620647 | 0.512788229 | -0.677902937 | 0.497833238 | 0.879044893 | Gm26944 | predicted gene, 26944 [Source:MGI Symb | 24.093 | 16.447 | 19.119 | 18.316 | 14.178 | 14.48 |
| 36578 | ENSMUSG00000098067 | 0.596071152 | -1.224343926 | 2.897583865 | -0.4225396 | 0.672631203 | NA | Gm26943 | predicted gene, 26943 [Source:MGI Symb | 1.6062 | 0 | 1.0062 | 0.964 | 0 | 0 |
| 36579 | ENSMUSG00000098068 | 0.133850373 | -0.796629176 | 4.080472857 | -0.195229623 | 0.84521319 | NA | Gm7909 | predicted gene 7909 [Source:MGI Symbol | 0.8031 | 0 | 0 | 0 | 0 | 0 |
| 36580 | ENSMUSG00000098069 | 0 | NA | NA | NA | NA | NA | Gm26942 | predicted gene, 26942 [Source:MGI Symb | 0 | 0 | 0 | 0 | 0 | 0 |
| 36581 | ENSMUSG00000098070 | 0 | NA | NA | NA | NA | NA | Gm8609 | predicted gene 8609 [Source:MGI Symbol | 0 | 0 | 0 | 0 | 0 | 0 |
| 36582 | ENSMUSG00000098071 | 0 | NA | NA | NA | NA | NA | Gm44525 | predicted gene 44525 [Source:MGI Symbo | 0 | 0 | 0 | 0 | 0 | 0 |
| 36583 | ENSMUSG00000098072 | 0 | NA | NA | NA | NA | NA | Gm26995 | predicted gene, 26995 [Source:MGI Symb | 0 | 0 | 0 | 0 | 0 | 0 |
| 36584 | ENSMUSG00000098073 | 0 | NA | NA | NA | NA | NA | Gm8048 | predicted gene 8048 [Source:MGI Symbol | 0 | 0 | 0 | 0 | 0 | 0 |
| 36585 | ENSMUSG00000098074 | 0 | NA | NA | NA | NA | NA | Gm26993 | predicted gene, 26993 [Source:MGI Symb | 0 | 0 | 0 | 0 | 0 | 0 |
| 36586 | ENSMUSG00000098075 | 0 | NA | NA | NA | NA | NA | Gm26994 | predicted gene, 26994 [Source:MGI Symb | 0 | 0 | 0 | 0 | 0 | 0 |
| 36587 | ENSMUSG00000098076 | 8.182789647 | 0.223985497 | 0.782677502 | 0.28617853 | 0.774741387 | 0.95883113 | Gm6981 | predicted pseudogene 6981 [Source:MGI | 11.243 | 8.2233 | 3.0187 | 6.7478 | 8.7249 | 11.138 |
| 36588 | ENSMUSG00000098077 | 0 | NA | NA | NA | NA | NA | Gm4960 | predicted gene 4960 [Source:MGI Symbol | 0 | 0 | 0 | 0 | 0 | 0 |
| 36589 | ENSMUSG00000098078 | 0 | NA | NA | NA | NA | NA | Gm26992 | predicted gene, 26992 [Source:MGI Symb | 0 | 0 | 0 | 0 | 0 | 0 |
| 36590 | ENSMUSG00000098079 | 0 | NA | NA | NA | NA | NA | Gm6773 | predicted gene 6773 [Source:MGI Symbol | 0 | 0 | 0 | 0 | 0 | 0 |
| 36591 | ENSMUSG00000098080 | 0 | NA | NA | NA | NA | NA | Gm5732 | predicted gene 5732 [Source:MGI Symbol | 0 | 0 | 0 | 0 | 0 | 0 |
| 36592 | ENSMUSG00000098081 | 0 | NA | NA | NA | NA | NA | Gm26913 | predicted gene, 26913 [Source:MGI Symb | 0 | 0 | 0 | 0 | 0 | 0 |
| 36593 | ENSMUSG00000098082 | 0 | NA | NA | NA | NA | NA | Gm20574 | predicted gene, 20574 [Source:MGI Symb | 0 | 0 | 0 | 0 | 0 | 0 |
| 36594 | ENSMUSG00000098083 | 0 | NA | NA | NA | NA | NA | Gm8957 | predicted gene 8957 [Source:MGI Symbol | 0 | 0 | 0 | 0 | 0 | 0 |
| 36595 | ENSMUSG00000098084 | 0 | NA | NA | NA | NA | NA | Gm26914 | predicted gene, 26914 [Source:MGI Symb | 0 | 0 | 0 | 0 | 0 | 0 |
| 36596 | ENSMUSG00000098085 | 0 | NA | NA | NA | NA | NA | Gm8919 | predicted gene 8919 [Source:MGI Symbol | 0 | 0 | 0 | 0 | 0 | 0 |
| 36597 | ENSMUSG00000098086 | 0 | NA | NA | NA | NA | NA | 1700120C18 | RIKEN cDNA 1700120C18 gene [Source:M | 0 | 0 | 0 | 0 | 0 | 0 |
| 36598 | ENSMUSG00000098087 | 0 | NA | NA | NA | NA | NA | Gm17750 | predicted gene, 17750 [Source:MGI Symb | 0 | 0 | 0 | 0 | 0 | 0 |
| 36599 | ENSMUSG00000098088 | 0 | NA | NA | NA | NA | NA | Gm26916 | predicted gene, 26916 [Source:MGI Symb | 0 | 0 | 0 | 0 | 0 | 0 |
| 36600 | ENSMUSG00000098089 | 0 | NA | NA | NA | NA | NA | Gm26915 | predicted gene, 26915 [Source:MGI Symb | 0 | 0 | 0 | 0 | 0 | 0 |
| 36601 | ENSMUSG00000098090 | 5.46038837 | 0.079913659 | 0.987722255 | 0.080907015 | 0.935515901 | 0.988420305 | 2700099C18 | RIKEN cDNA 2700099C18 gene [Source:M | 9.6372 | 5.1396 | 1.0062 | 4.8199 | 4.3625 | 7.7969 |
| 36602 | ENSMUSG00000098091 | 0.371281295 | 2.09253362 | 4.031852165 | 0.519000582 | 0.603760335 | NA | Gm16470 | predicted pseudogene 16470 [Source:MG | 0 | 0 | 0 | 0 | 0 | 2.2277 |
| 36603 | ENSMUSG00000098092 | 0 | NA | NA | NA | NA | NA | Gm2076 | predicted gene 2076 [Source:MGI Symbol | 0 | 0 | 0 | 0 | 0 | 0 |
| 36604 | ENSMUSG00000098093 | 0 | NA | NA | NA | NA | NA | Gm8707 | predicted gene 8707 [Source:MGI Symbol | 0 | 0 | 0 | 0 | 0 | 0 |
| 36605 | ENSMUSG00000098094 | 0 | NA | NA | NA | NA | NA | Scgb2b17 | secretoglobin, family 2B, member 17 [Sou | 0 | 0 | 0 | 0 | 0 | 0 |
| 36606 | ENSMUSG00000098095 | 0 | NA | NA | NA | NA | NA | Gm9274 | predicted gene 9274 [Source:MGI Symbol | 0 | 0 | 0 | 0 | 0 | 0 |
| 36607 | ENSMUSG00000098096 | 0 | NA | NA | NA | NA | NA | Gm26969 | predicted gene, 26969 [Source:MGI Symb | 0 | 0 | 0 | 0 | 0 | 0 |
| 36608 | ENSMUSG00000098097 | 0 | NA | NA | NA | NA | NA | 6530403H02 | RIKEN cDNA 6530403H02 gene [Source:M | 0 | 0 | 0 | 0 | 0 | 0 |
| 36609 | ENSMUSG00000098098 | 11.89664543 | -0.484252103 | 0.75591466 | -0.640617424 | 0.521771276 | 0.888132458 | Bvht | braveheart long non-coding RNA [Source: | 11.243 | 17.475 | 13.081 | 17.352 | 1.0906 | 11.138 |
| 36610 | ENSMUSG00000098099 | 0 | NA | NA | NA | NA | NA | Gm7456 | predicted gene 7456 [Source:MGI Symbol | 0 | 0 | 0 | 0 | 0 | 0 |
| 36611 | ENSMUSG00000098100 | 0 | NA | NA | NA | NA | NA | Gm7572 | predicted gene 7572 [Source:MGI Symbol | 0 | 0 | 0 | 0 | 0 | 0 |
| 36612 | ENSMUSG00000098101 | 0.321325949 | 1.911575114 | 4.048267893 | 0.472195804 | 0.636787034 | NA | Gm27006 | predicted gene, 27006 [Source:MGI Symb | 0 | 0 | 0 | 1.928 | 0 | 0 |
| 36613 | ENSMUSG00000098102 | 0 | NA | NA | NA | NA | NA | Gm2467 | predicted gene 2467 [Source:MGI Symbol | 0 | 0 | 0 | 0 | 0 | 0 |
| 36614 | ENSMUSG00000098103 | 0 | NA | NA | NA | NA | NA | Gm27005 | predicted gene, 27005 [Source:MGI Symb | 0 | 0 | 0 | 0 | 0 | 0 |
| 36615 | ENSMUSG00000098104 | 0 | NA | NA | NA | NA | NA | Gm6085 | predicted gene 6085 [Source:MGI Symbol | 0 | 0 | 0 | 0 | 0 | 0 |
| 36616 | ENSMUSG00000098105 | 0 | NA | NA | NA | NA | NA | Gm8825 | predicted gene 8825 [Source:MGI Symbol | 0 | 0 | 0 | 0 | 0 | 0 |
| 36617 | ENSMUSG00000098106 | 0.17131964 | -0.796629176 | 4.080472857 | -0.195229623 | 0.84521319 | NA | Gm5350 | predicted gene 5350 [Source:MGI Symbol | 0 | 1.0279 | 0 | 0 | 0 | 0 |
| 36618 | ENSMUSG00000098107 | 0 | NA | NA | NA | NA | NA | Gm27007 | predicted gene, 27007 [Source:MGI Symb | 0 | 0 | 0 | 0 | 0 | 0 |
| 36619 | ENSMUSG00000098108 | 0 | NA | NA | NA | NA | NA | Gm27008 | predicted gene, 27008 [Source:MGI Symb | 0 | 0 | 0 | 0 | 0 | 0 |
| 36620 | ENSMUSG00000098109 | 0 | NA | NA | NA | NA | NA | 9530085P06 | RIKEN cDNA 9530085P06 gene [Source:M | 0 | 0 | 0 | 0 | 0 | 0 |

| | | | | | | | | | | | | | | | |
|---|---|---|---|---|---|---|---|---|---|---|---|---|---|---|---|
| 36621 | ENSMUSG00000098110 | 0 | NA | NA | NA | NA | NA | Gm26956 | predicted gene, 26956 [Source:MGI Symb | 0 | 0 | 0 | 0 | 0 | 0 |
| 36622 | ENSMUSG00000098111 | 2.777059193 | -1.172158244 | 1.409683232 | -0.831504708 | 0.40568857 | NA | Gm4654 | predicted gene 4654 [Source:MGI Symbol | 6.4248 | 4.1117 | 1.0062 | 2.8919 | 0 | 2.2277 |
| 36623 | ENSMUSG00000098112 | 21.74683633 | -1.696884297 | 0.577809294 | -2.936754934 | 0.003316661 | 0.050232652 | Bin2 | bridging integrator 2 [Source:MGI Symbol | 15.259 | 49.34 | 35.219 | 9.6398 | 6.5437 | 14.48 |
| 36624 | ENSMUSG00000098113 | 0.476489653 | -2.293616034 | 3.421506946 | -0.670352587 | 0.502633051 | NA | Gm2445 | predicted gene 2445 [Source:MGI Symbol | 0.8031 | 2.0558 | 0 | 0 | 0 | 0 |
| 36625 | ENSMUSG00000098114 | 0 | NA | NA | NA | NA | NA | Gm26957 | predicted gene, 26957 [Source:MGI Symb | 0 | 0 | 0 | 0 | 0 | 0 |
| 36626 | ENSMUSG00000098115 | 0 | NA | NA | NA | NA | NA | Gm8816 | predicted gene 8816 [Source:MGI Symbol | 0 | 0 | 0 | 0 | 0 | 0 |
| 36627 | ENSMUSG00000098116 | 0 | NA | NA | NA | NA | NA | Gm7191 | predicted gene 7191 [Source:MGI Symbol | 0 | 0 | 0 | 0 | 0 | 0 |
| 36628 | ENSMUSG00000098118 | 0 | NA | NA | NA | NA | NA | Gm26960 | predicted gene, 26960 [Source:MGI Symb | 0 | 0 | 0 | 0 | 0 | 0 |
| 36629 | ENSMUSG00000098119 | 0 | NA | NA | NA | NA | NA | Gm26959 | predicted gene, 26959 [Source:MGI Symb | 0 | 0 | 0 | 0 | 0 | 0 |
| 36630 | ENSMUSG00000098120 | 0.367410362 | 2.078255025 | 3.800070157 | 0.54689912 | 0.58444804 | NA | Gm5914 | predicted gene 5914 [Source:MGI Symbol | 0 | 0 | 0 | 0 | 1.0906 | 1.1138 |
| 36631 | ENSMUSG00000098121 | 0 | NA | NA | NA | NA | NA | Gm27038 | predicted gene, 27038 [Source:MGI Symb | 0 | 0 | 0 | 0 | 0 | 0 |
| 36632 | ENSMUSG00000098122 | 0 | NA | NA | NA | NA | NA | Gm27039 | predicted gene, 27039 [Source:MGI Symb | 0 | 0 | 0 | 0 | 0 | 0 |
| 36633 | ENSMUSG00000098123 | 0.487198452 | -0.685759572 | 3.136894395 | -0.218610985 | 0.826953095 | NA | Gm4518 | predicted gene 4518 [Source:MGI Symbol | 0.8031 | 0 | 1.0062 | 0 | 0 | 1.1138 |
| 36634 | ENSMUSG00000098124 | 0 | NA | NA | NA | NA | NA | Gm27036 | predicted gene, 27036 [Source:MGI Symb | 0 | 0 | 0 | 0 | 0 | 0 |
| 36635 | ENSMUSG00000098125 | 0 | NA | NA | NA | NA | NA | Gm9578 | predicted gene 9578 [Source:MGI Symbol | 0 | 0 | 0 | 0 | 0 | 0 |
| 36636 | ENSMUSG00000098126 | 0 | NA | NA | NA | NA | NA | Gm27037 | predicted gene, 27037 [Source:MGI Symb | 0 | 0 | 0 | 0 | 0 | 0 |
| 36637 | ENSMUSG00000098127 | 0 | NA | NA | NA | NA | NA | Gm4665 | predicted gene 4665 [Source:MGI Symbol | 0 | 0 | 0 | 0 | 0 | 0 |
| 36638 | ENSMUSG00000098128 | 0 | NA | NA | NA | NA | NA | Gm3693 | predicted gene 3693 [Source:MGI Symbol | 0 | 0 | 0 | 0 | 0 | 0 |
| 36639 | ENSMUSG00000098129 | 0 | NA | NA | NA | NA | NA | Gm27040 | predicted gene, 27040 [Source:MGI Symb | 0 | 0 | 0 | 0 | 0 | 0 |
| 36640 | ENSMUSG00000098130 | 0 | NA | NA | NA | NA | NA | Gm17797 | predicted gene, 17797 [Source:MGI Symb | 0 | 0 | 0 | 0 | 0 | 0 |
| 36641 | ENSMUSG00000098131 | 0 | NA | NA | NA | NA | NA | Gm26989 | predicted gene, 26989 [Source:MGI Symb | 0 | 0 | 0 | 0 | 0 | 0 |
| 36642 | ENSMUSG00000098132 | 1.730277748 | -0.515396549 | 1.877908765 | -0.27445239 | 0.783736986 | NA | Rassf10 | Ras association (RalGDS/AF-6) domain far | 0 | 5.1396 | 1.0062 | 0.964 | 3.2719 | 0 |
| 36643 | ENSMUSG00000098134 | 133.9725716 | 0.118524065 | 0.209010738 | 0.567071655 | 0.5706655 | 0.904651508 | Rnf113a2 | ring finger protein 113A2 [Source:MGI Syr | 129.3 | 119.24 | 136.85 | 143.63 | 118.88 | 155.94 |
| 36644 | ENSMUSG00000098135 | 0 | NA | NA | NA | NA | NA | Gm6946 | predicted gene 6946 [Source:MGI Symbol | 0 | 0 | 0 | 0 | 0 | 0 |
| 36645 | ENSMUSG00000098136 | 0 | NA | NA | NA | NA | NA | Gm8513 | predicted gene 8513 [Source:MGI Symbol | 0 | 0 | 0 | 0 | 0 | 0 |
| 36646 | ENSMUSG00000098137 | 0 | NA | NA | NA | NA | NA | Gm21940 | predicted gene, 21940 [Source:MGI Symb | 0 | 0 | 0 | 0 | 0 | 0 |
| 36647 | ENSMUSG00000098138 | 0 | NA | NA | NA | NA | NA | Gm9081 | predicted gene 9081 [Source:MGI Symbol | 0 | 0 | 0 | 0 | 0 | 0 |
| 36648 | ENSMUSG00000098139 | 0 | NA | NA | NA | NA | NA | Gm26988 | predicted gene, 26988 [Source:MGI Symb | 0 | 0 | 0 | 0 | 0 | 0 |
| 36649 | ENSMUSG00000098140 | 0 | NA | NA | NA | NA | NA | Gm26938 | predicted gene, 26938 [Source:MGI Symb | 0 | 0 | 0 | 0 | 0 | 0 |
| 36650 | ENSMUSG00000098141 | 0 | NA | NA | NA | NA | NA | Gm6944 | predicted gene 6944 [Source:MGI Symbol | 0 | 0 | 0 | 0 | 0 | 0 |
| 36651 | ENSMUSG00000098142 | 0.647809292 | -2.733819264 | 3.061311498 | -0.893022244 | 0.371845261 | NA | Gm7507 | predicted gene 7507 [Source:MGI Symbol | 0.8031 | 3.0838 | 0 | 0 | 0 | 0 |
| 36652 | ENSMUSG00000098143 | 0 | NA | NA | NA | NA | NA | Gm26937 | predicted gene, 26937 [Source:MGI Symb | 0 | 0 | 0 | 0 | 0 | 0 |
| 36653 | ENSMUSG00000098144 | 0 | NA | NA | NA | NA | NA | 1700029N11 | RIKEN cDNA 1700029N11 gene [Source:M | 0 | 0 | 0 | 0 | 0 | 0 |
| 36654 | ENSMUSG00000098145 | 0 | NA | NA | NA | NA | NA | Gm26936 | predicted gene, 26936 [Source:MGI Symb | 0 | 0 | 0 | 0 | 0 | 0 |
| 36655 | ENSMUSG00000098146 | 4.473145466 | -0.205755094 | 1.012708155 | -0.203173139 | 0.838999707 | NA | Gm26935 | predicted gene, 26935 [Source:MGI Symb | 7.2279 | 2.0558 | 5.0312 | 4.8199 | 4.3625 | 3.3415 |
| 36656 | ENSMUSG00000098147 | 0 | NA | NA | NA | NA | NA | Gm17798 | predicted gene, 17798 [Source:MGI Symb | 0 | 0 | 0 | 0 | 0 | 0 |
| 36657 | ENSMUSG00000098149 | 0 | NA | NA | NA | NA | NA | Gapdh-ps14 | glyceraldehyde-3-phosphate dehydrogena | 0 | 0 | 0 | 0 | 0 | 0 |
| 36658 | ENSMUSG00000098150 | 0 | NA | NA | NA | NA | NA | 4921515E04 | RIKEN cDNA 4921515E04 gene [Source:M | 0 | 0 | 0 | 0 | 0 | 0 |
| 36659 | ENSMUSG00000098152 | 0 | NA | NA | NA | NA | NA | Gm27025 | predicted gene, 27025 [Source:MGI Symb | 0 | 0 | 0 | 0 | 0 | 0 |
| 36660 | ENSMUSG00000098153 | 0 | NA | NA | NA | NA | NA | Gm27024 | predicted gene, 27024 [Source:MGI Symb | 0 | 0 | 0 | 0 | 0 | 0 |
| 36661 | ENSMUSG00000098154 | 0 | NA | NA | NA | NA | NA | Gm5787 | predicted gene 5787 [Source:MGI Symbol | 0 | 0 | 0 | 0 | 0 | 0 |
| 36662 | ENSMUSG00000098155 | 0 | NA | NA | NA | NA | NA | Gm27022 | predicted gene, 27022 [Source:MGI Symb | 0 | 0 | 0 | 0 | 0 | 0 |
| 36663 | ENSMUSG00000098156 | 0.513958919 | -2.394274821 | 3.756096013 | -0.637437066 | 0.523840191 | NA | Gm27023 | predicted gene, 27023 [Source:MGI Symb | 0 | 3.0838 | 0 | 0 | 0 | 0 |
| 36664 | ENSMUSG00000098157 | 0 | NA | NA | NA | NA | NA | Gm9022 | predicted gene 9022 [Source:MGI Symbol | 0 | 0 | 0 | 0 | 0 | 0 |
| 36665 | ENSMUSG00000098158 | 0.133850373 | -0.796629176 | 4.080472857 | -0.195229623 | 0.84521319 | NA | Gm4804 | predicted gene 4804 [Source:MGI Symbol | 0.8031 | 0 | 0 | 0 | 0 | 0 |
| 36666 | ENSMUSG00000098159 | 0 | NA | NA | NA | NA | NA | Gm21614 | predicted gene, 21614 [Source:MGI Symb | 0 | 0 | 0 | 0 | 0 | 0 |
| 36667 | ENSMUSG00000098160 | 0 | NA | NA | NA | NA | NA | Gm26974 | predicted gene, 26974 [Source:MGI Symb | 0 | 0 | 0 | 0 | 0 | 0 |
| 36668 | ENSMUSG00000098161 | 0 | NA | NA | NA | NA | NA | Platr11 | pluripotency associated transcript 11 [Sou | 0 | 0 | 0 | 0 | 0 | 0 |
| 36669 | ENSMUSG00000098162 | 0 | NA | NA | NA | NA | NA | Gm26976 | predicted gene, 26976 [Source:MGI Symb | 0 | 0 | 0 | 0 | 0 | 0 |
| 36670 | ENSMUSG00000098163 | 0 | NA | NA | NA | NA | NA | Gm6216 | predicted gene 6216 [Source:MGI Symbol | 0 | 0 | 0 | 0 | 0 | 0 |
| 36671 | ENSMUSG00000098164 | 0 | NA | NA | NA | NA | NA | Gm5493 | predicted gene 5493 [Source:MGI Symbol | 0 | 0 | 0 | 0 | 0 | 0 |
| 36672 | ENSMUSG00000098165 | 0 | NA | NA | NA | NA | NA | Gm26977 | predicted gene, 26977 [Source:MGI Symb | 0 | 0 | 0 | 0 | 0 | 0 |
| 36673 | ENSMUSG00000098166 | 0 | NA | NA | NA | NA | NA | Gm26978 | predicted gene, 26978 [Source:MGI Symb | 0 | 0 | 0 | 0 | 0 | 0 |
| 36674 | ENSMUSG00000098167 | 0 | NA | NA | NA | NA | NA | Gm18848 | predicted gene, 18848 [Source:MGI Symb | 0 | 0 | 0 | 0 | 0 | 0 |
| 36675 | ENSMUSG00000098169 | 0 | NA | NA | NA | NA | NA | Gm5205 | predicted gene 5205 [Source:MGI Symbol | 0 | 0 | 0 | 0 | 0 | 0 |
| 36676 | ENSMUSG00000098170 | 0 | NA | NA | NA | NA | NA | Gm3890 | predicted gene 3890 [Source:MGI Symbol | 0 | 0 | 0 | 0 | 0 | 0 |

| | | | | | | | | | | | | | | | |
|---|---|---|---|---|---|---|---|---|---|---|---|---|---|---|---|
| 36677 | ENSMUSG00000098171 | 0 | NA | NA | NA | NA | NA | Gm9621 | predicted gene 9621 [Source:MGI Symbol | 0 | 0 | 0 | 0 | 0 | 0 |
| 36678 | ENSMUSG00000098172 | 0 | NA | NA | NA | NA | NA | Gm26973 | predicted gene, 26973 [Source:MGI Symb | 0 | 0 | 0 | 0 | 0 | 0 |
| 36679 | ENSMUSG00000098173 | 0 | NA | NA | NA | NA | NA | Gm26923 | predicted gene, 26923 [Source:MGI Symb | 0 | 0 | 0 | 0 | 0 | 0 |
| 36680 | ENSMUSG00000098174 | 0 | NA | NA | NA | NA | NA | Gm8183 | predicted gene 8183 [Source:MGI Symbol | 0 | 0 | 0 | 0 | 0 | 0 |
| 36681 | ENSMUSG00000098175 | 0 | NA | NA | NA | NA | NA | Gm7545 | predicted gene 7545 [Source:MGI Symbol | 0 | 0 | 0 | 0 | 0 | 0 |
| 36682 | ENSMUSG00000098176 | 62.64486883 | 0.130144082 | 0.286580538 | 0.454127427 | 0.649737114 | 0.925643431 | Ccdc166 | coiled-coil domain containing 166 [Source | 59.43 | 56.535 | 63.393 | 57.839 | 78.525 | 60.148 |
| 36683 | ENSMUSG00000098177 | 0 | NA | NA | NA | NA | NA | Gm4823 | predicted gene 4823 [Source:MGI Symbol | 0 | 0 | 0 | 0 | 0 | 0 |
| 36684 | ENSMUSG00000098178 | 0 | NA | NA | NA | NA | NA | NA | NA | 0 | 0 | 0 | 0 | 0 | 0 |
| 36685 | ENSMUSG00000098179 | 0 | NA | NA | NA | NA | NA | Gm8709 | predicted gene 8709 [Source:MGI Symbol | 0 | 0 | 0 | 0 | 0 | 0 |
| 36686 | ENSMUSG00000098180 | 0 | NA | NA | NA | NA | NA | Gm5430 | predicted gene 5430 [Source:MGI Symbol | 0 | 0 | 0 | 0 | 0 | 0 |
| 36687 | ENSMUSG00000098181 | 0 | NA | NA | NA | NA | NA | Rps12-ps24 | ribosomal protein S12, pseudogene 24 [Sc | 0 | 0 | 0 | 0 | 0 | 0 |
| 36688 | ENSMUSG00000098183 | 10.74237217 | -0.05848909 | 0.796713587 | -0.073412944 | 0.941477518 | 0.989358531 | Gm27010 | predicted gene, 27010 [Source:MGI Symb | 13.653 | 1.0279 | 18.112 | 9.6398 | 11.997 | 10.025 |
| 36689 | ENSMUSG00000098184 | 1.497208577 | 0.420217169 | 1.790833422 | 0.234648943 | 0.814481219 | NA | Gm27011 | predicted gene, 27011 [Source:MGI Symb | 0.8031 | 3.0838 | 0 | 2.8919 | 1.0906 | 1.1138 |
| 36690 | ENSMUSG00000098185 | 0 | NA | NA | NA | NA | NA | Gm9146 | predicted gene 9146 [Source:MGI Symbol | 0 | 0 | 0 | 0 | 0 | 0 |
| 36691 | ENSMUSG00000098186 | 0 | NA | NA | NA | NA | NA | Gm7132 | predicted gene 7132 [Source:MGI Symbol | 0 | 0 | 0 | 0 | 0 | 0 |
| 36692 | ENSMUSG00000098187 | 0 | NA | NA | NA | NA | NA | Gm8927 | predicted gene 8927 [Source:MGI Symbol | 0 | 0 | 0 | 0 | 0 | 0 |
| 36693 | ENSMUSG00000098188 | 185.6585832 | -0.281208167 | 0.189173136 | -1.486512158 | 0.137143704 | 0.541092888 | Sowahc | sosondowah ankyrin repeat domain famil | 220.85 | 204.56 | 185.15 | 159.06 | 149.41 | 194.92 |
| 36694 | ENSMUSG00000098189 | 0 | NA | NA | NA | NA | NA | Gm8604 | predicted gene 8604 [Source:MGI Symbol | 0 | 0 | 0 | 0 | 0 | 0 |
| 36695 | ENSMUSG00000098190 | 0 | NA | NA | NA | NA | NA | Gm5210 | predicted gene 5210 [Source:MGI Symbol | 0 | 0 | 0 | 0 | 0 | 0 |
| 36696 | ENSMUSG00000098191 | 0 | NA | NA | NA | NA | NA | Gm26962 | predicted gene, 26962 [Source:MGI Symb | 0 | 0 | 0 | 0 | 0 | 0 |
| 36697 | ENSMUSG00000098192 | 1.575367254 | 1.224528453 | 1.902889801 | 0.643509915 | 0.519893284 | NA | Gm3534 | predicted pseudogene 3534 [Source:MGI | 0.8031 | 0 | 2.0125 | 0 | 2.1812 | 4.4554 |
| 36698 | ENSMUSG00000098193 | 0 | NA | NA | NA | NA | NA | Gm26963 | predicted gene, 26963 [Source:MGI Symb | 0 | 0 | 0 | 0 | 0 | 0 |
| 36699 | ENSMUSG00000098194 | 0 | NA | NA | NA | NA | NA | Gm7286 | predicted gene 7286 [Source:MGI Symbol | 0 | 0 | 0 | 0 | 0 | 0 |
| 36700 | ENSMUSG00000098195 | 0 | NA | NA | NA | NA | NA | Gm7693 | predicted gene 7693 [Source:MGI Symbol | 0 | 0 | 0 | 0 | 0 | 0 |
| 36701 | ENSMUSG00000098196 | 8.394004718 | 0.455039287 | 0.775691965 | 0.586623696 | 0.557456465 | 0.898590249 | Gm26964 | predicted gene, 26964 [Source:MGI Symb | 4.0155 | 8.2233 | 9.0562 | 4.8199 | 11.997 | 12.252 |
| 36702 | ENSMUSG00000098197 | 0 | NA | NA | NA | NA | NA | BC051537 | cDNA sequence BC051537 [Source:MGI Sy | 0 | 0 | 0 | 0 | 0 | 0 |
| 36703 | ENSMUSG00000098198 | 0 | NA | NA | NA | NA | NA | Gm9169 | predicted gene 9169 [Source:MGI Symbol | 0 | 0 | 0 | 0 | 0 | 0 |
| 36704 | ENSMUSG00000098199 | 0.496077836 | -0.812642266 | 3.444120572 | -0.235950586 | 0.813471014 | NA | Gm9358 | predicted gene 9358 [Source:MGI Symbol | 0 | 0 | 2.0125 | 0.964 | 0 | 0 |
| 36705 | ENSMUSG00000098200 | 0 | NA | NA | NA | NA | NA | Gm7763 | predicted gene 7763 [Source:MGI Symbol | 0 | 0 | 0 | 0 | 0 | 0 |
| 36706 | ENSMUSG00000098201 | 0 | NA | NA | NA | NA | NA | Gm26983 | predicted gene, 26983 [Source:MGI Symb | 0 | 0 | 0 | 0 | 0 | 0 |
| 36707 | ENSMUSG00000098202 | 0 | NA | NA | NA | NA | NA | B830012L14 | RIKEN cDNA B830012L14 gene [Source:M | 0 | 0 | 0 | 0 | 0 | 0 |
| 36708 | ENSMUSG00000098203 | 0 | NA | NA | NA | NA | NA | Gm9568 | predicted gene 9568 [Source:MGI Symbol | 0 | 0 | 0 | 0 | 0 | 0 |
| 36709 | ENSMUSG00000098204 | 0 | NA | NA | NA | NA | NA | Gm26984 | predicted gene, 26984 [Source:MGI Symb | 0 | 0 | 0 | 0 | 0 | 0 |
| 36710 | ENSMUSG00000098205 | 0 | NA | NA | NA | NA | NA | Gm9149 | predicted gene 9149 [Source:MGI Symbol | 0 | 0 | 0 | 0 | 0 | 0 |
| 36711 | ENSMUSG00000098206 | 0.321325949 | 1.911575114 | 4.048267893 | 0.472195804 | 0.636787034 | NA | Mir181a-1hg | Mir181a-1 and Mir181b-1 host gene [Sour | 0 | 0 | 0 | 1.928 | 0 | 0 |
| 36712 | ENSMUSG00000098207 | 0.401551119 | -2.072982726 | 4.008251979 | -0.517178744 | 0.605031388 | NA | Arl14 | ADP-ribosylation factor-like 14 [Source:M | 2.4093 | 0 | 0 | 0 | 0 | 0 |
| 36713 | ENSMUSG00000098208 | 0 | NA | NA | NA | NA | NA | Gm8349 | predicted gene 8349 [Source:MGI Symbol | 0 | 0 | 0 | 0 | 0 | 0 |
| 36714 | ENSMUSG00000098209 | 0 | NA | NA | NA | NA | NA | Gm8245 | predicted gene 8245 [Source:MGI Symbol | 0 | 0 | 0 | 0 | 0 | 0 |
| 36715 | ENSMUSG00000098210 | 0 | NA | NA | NA | NA | NA | Gm5224 | predicted pseudogene 5224 [Source:MGI | 0 | 0 | 0 | 0 | 0 | 0 |
| 36716 | ENSMUSG00000098211 | 0.644197084 | -2.725354879 | 2.762261801 | -0.986638876 | 0.323819705 | NA | Gm27045 | predicted gene, 27045 [Source:MGI Symb | 0.8031 | 2.0558 | 1.0062 | 0 | 0 | 0 |
| 36717 | ENSMUSG00000098212 | 0 | NA | NA | NA | NA | NA | Gm53052 | predicted gene, 53052 [Source:MGI Symb | 0 | 0 | 0 | 0 | 0 | 0 |
| 36718 | ENSMUSG00000098213 | 0 | NA | NA | NA | NA | NA | Gm27044 | predicted gene, 27044 [Source:MGI Symb | 0 | 0 | 0 | 0 | 0 | 0 |
| 36719 | ENSMUSG00000098214 | 0 | NA | NA | NA | NA | NA | Gm27047 | predicted gene, 27047 [Source:MGI Symb | 0 | 0 | 0 | 0 | 0 | 0 |
| 36720 | ENSMUSG00000098215 | 0 | NA | NA | NA | NA | NA | Gm7069 | predicted gene 7069 [Source:MGI Symbol | 0 | 0 | 0 | 0 | 0 | 0 |
| 36721 | ENSMUSG00000098216 | 0 | NA | NA | NA | NA | NA | Gm5181 | predicted gene 5181 [Source:MGI Symbol | 0 | 0 | 0 | 0 | 0 | 0 |
| 36722 | ENSMUSG00000098217 | 0 | NA | NA | NA | NA | NA | Gm27046 | predicted gene, 27046 [Source:MGI Symb | 0 | 0 | 0 | 0 | 0 | 0 |
| 36723 | ENSMUSG00000098218 | 0 | NA | NA | NA | NA | NA | Gm27043 | predicted gene, 27043 [Source:MGI Symb | 0 | 0 | 0 | 0 | 0 | 0 |
| 36724 | ENSMUSG00000098219 | 0 | NA | NA | NA | NA | NA | Gm7481 | predicted gene 7481 [Source:MGI Symbol | 0 | 0 | 0 | 0 | 0 | 0 |
| 36725 | ENSMUSG00000098221 | 0.692116425 | -1.412606127 | 2.773087916 | -0.509398248 | 0.610473104 | NA | Gm27030 | predicted gene, 27030 [Source:MGI Symb | 0 | 2.0558 | 1.0062 | 0 | 1.0906 | 0 |
| 36726 | ENSMUSG00000098222 | 0 | NA | NA | NA | NA | NA | Gm8318 | predicted gene 8318 [Source:MGI Symbol | 0 | 0 | 0 | 0 | 0 | 0 |
| 36727 | ENSMUSG00000098223 | 0 | NA | NA | NA | NA | NA | Gm4891 | predicted gene 4891 [Source:MGI Symbol | 0 | 0 | 0 | 0 | 0 | 0 |
| 36728 | ENSMUSG00000098224 | 0 | NA | NA | NA | NA | NA | Gm7985 | predicted gene 7985 [Source:MGI Symbol | 0 | 0 | 0 | 0 | 0 | 0 |
| 36729 | ENSMUSG00000098225 | 0 | NA | NA | NA | NA | NA | Gm9556 | predicted gene 9556 [Source:MGI Symbol | 0 | 0 | 0 | 0 | 0 | 0 |
| 36730 | ENSMUSG00000098226 | 0 | NA | NA | NA | NA | NA | Gm5272 | predicted gene 5272 [Source:MGI Symbol | 0 | 0 | 0 | 0 | 0 | 0 |
| 36731 | ENSMUSG00000098227 | 0 | NA | NA | NA | NA | NA | Gm4812 | predicted gene 4812 [Source:MGI Symbol | 0 | 0 | 0 | 0 | 0 | 0 |
| 36732 | ENSMUSG00000098228 | 0 | NA | NA | NA | NA | NA | Gm21070 | predicted gene, 21070 [Source:MGI Symb | 0 | 0 | 0 | 0 | 0 | 0 |

| | | | | | | | | | | | | | | | |
|---|---|---|---|---|---|---|---|---|---|---|---|---|---|---|---|
| 36733 | ENSMUSG00000098229 | 0 | NA | NA | NA | NA | NA | Gm7719 | predicted gene 7719 [Source:MGI Symbol | 0 | 0 | 0 | 0 | 0 | 0 |
| 36734 | ENSMUSG00000098230 | 0 | NA | NA | NA | NA | NA | 1700095B10 | RIKEN cDNA 1700095B10 gene [Source:M | 0 | 0 | 0 | 0 | 0 | 0 |
| 36735 | ENSMUSG00000098231 | 0 | NA | NA | NA | NA | NA | Gm6463 | predicted gene 6463 [Source:MGI Symbol | 0 | 0 | 0 | 0 | 0 | 0 |
| 36736 | ENSMUSG00000098232 | 0.181769715 | 1.126924175 | 4.080472857 | 0.276174898 | 0.782413736 | NA | Gm3671 | predicted gene 3671 [Source:MGI Symbol | 0 | 0 | 0 | 0 | 1.0906 | 0 |
| 36737 | ENSMUSG00000098233 | 0 | NA | NA | NA | NA | NA | Gm26954 | predicted gene, 26954 [Source:MGI Symb | 0 | 0 | 0 | 0 | 0 | 0 |
| 36738 | ENSMUSG00000098234 | 179.1017047 | 0.439395798 | 0.192597198 | 2.281423627 | 0.022523391 | 0.187876777 | Snhg6 | small nucleolar RNA host gene 6 [Source:I | 134.12 | 177.83 | 144.9 | 220.75 | 186.5 | 210.52 |
| 36739 | ENSMUSG00000098235 | 1.070519286 | -0.990795636 | 2.211649349 | -0.447989478 | 0.654160791 | NA | Gm7497 | predicted gene 7497 [Source:MGI Symbol | 3.2124 | 0 | 1.0062 | 0 | 1.0906 | 1.1138 |
| 36740 | ENSMUSG00000098236 | 0 | NA | NA | NA | NA | NA | Gm7734 | predicted gene 7734 [Source:MGI Symbol | 0 | 0 | 0 | 0 | 0 | 0 |
| 36741 | ENSMUSG00000098237 | 0 | NA | NA | NA | NA | NA | Gm7577 | predicted gene 7577 [Source:MGI Symbol | 0 | 0 | 0 | 0 | 0 | 0 |
| 36742 | ENSMUSG00000098238 | 0 | NA | NA | NA | NA | NA | Gm8756 | predicted gene 8756 [Source:MGI Symbol | 0 | 0 | 0 | 0 | 0 | 0 |
| 36743 | ENSMUSG00000098240 | 0.513958919 | -2.394274821 | 3.756096013 | -0.637437066 | 0.523840191 | NA | Gm4575 | predicted gene 4575 [Source:MGI Symbol | 0 | 3.0838 | 0 | 0 | 0 | 0 |
| 36744 | ENSMUSG00000098241 | 0 | NA | NA | NA | NA | NA | H2-Pa | histocompatibility 2, P region alpha locus | 0 | 0 | 0 | 0 | 0 | 0 |
| 36745 | ENSMUSG00000098242 | 0 | NA | NA | NA | NA | NA | Gm26929 | predicted gene, 26929 [Source:MGI Symb | 0 | 0 | 0 | 0 | 0 | 0 |
| 36746 | ENSMUSG00000098243 | 0 | NA | NA | NA | NA | NA | Gm4258 | predicted gene 4258 [Source:MGI Symbol | 0 | 0 | 0 | 0 | 0 | 0 |
| 36747 | ENSMUSG00000098244 | 0 | NA | NA | NA | NA | NA | Gm26928 | predicted gene, 26928 [Source:MGI Symb | 0 | 0 | 0 | 0 | 0 | 0 |
| 36748 | ENSMUSG00000098245 | 0 | NA | NA | NA | NA | NA | Gm26927 | predicted gene, 26927 [Source:MGI Symb | 0 | 0 | 0 | 0 | 0 | 0 |
| 36749 | ENSMUSG00000098246 | 0 | NA | NA | NA | NA | NA | Gm7627 | predicted gene 7627 [Source:MGI Symbol | 0 | 0 | 0 | 0 | 0 | 0 |
| 36750 | ENSMUSG00000098247 | 0 | NA | NA | NA | NA | NA | Mir7220 | microRNA 7220 [Source:MGI Symbol;Acc: | 0 | 0 | 0 | 0 | 0 | 0 |
| 36751 | ENSMUSG00000098248 | 0 | NA | NA | NA | NA | NA | Gm27222 | predicted gene 27222 [Source:MGI Symbo | 0 | 0 | 0 | 0 | 0 | 0 |
| 36752 | ENSMUSG00000098249 | 0 | NA | NA | NA | NA | NA | NA | NA | 0 | 0 | 0 | 0 | 0 | 0 |
| 36753 | ENSMUSG00000098251 | 0 | NA | NA | NA | NA | NA | Mir7656 | microRNA 7656 [Source:MGI Symbol;Acc: | 0 | 0 | 0 | 0 | 0 | 0 |
| 36754 | ENSMUSG00000098252 | 0.160662974 | 1.126924175 | 4.080472857 | 0.276174898 | 0.782413736 | NA | Mir6361 | microRNA 6361 [Source:MGI Symbol;Acc: | 0 | 0 | 0 | 0.964 | 0 | 0 |
| 36755 | ENSMUSG00000098253 | 0 | NA | NA | NA | NA | NA | Mir7219 | microRNA 7219 [Source:MGI Symbol;Acc: | 0 | 0 | 0 | 0 | 0 | 0 |
| 36756 | ENSMUSG00000098254 | 0 | NA | NA | NA | NA | NA | NA | NA | 0 | 0 | 0 | 0 | 0 | 0 |
| 36757 | ENSMUSG00000098255 | 0 | NA | NA | NA | NA | NA | Gm27223 | predicted gene 27223 [Source:MGI Symbo | 0 | 0 | 0 | 0 | 0 | 0 |
| 36758 | ENSMUSG00000098257 | 0 | NA | NA | NA | NA | NA | Gm27169 | predicted gene 27169 [Source:MGI Symbo | 0 | 0 | 0 | 0 | 0 | 0 |
| 36759 | ENSMUSG00000098258 | 0 | NA | NA | NA | NA | NA | Mir6369 | microRNA 6369 [Source:MGI Symbol;Acc: | 0 | 0 | 0 | 0 | 0 | 0 |
| 36760 | ENSMUSG00000098259 | 1.092148988 | -1.219782293 | 2.337692795 | -0.521788961 | 0.601817277 | NA | Gm27616 | predicted gene, 27616 [Source:MGI Symb | 1.6062 | 0 | 3.0187 | 1.928 | 0 | 0 |
| 36761 | ENSMUSG00000098260 | 0 | NA | NA | NA | NA | NA | NA | NA | 0 | 0 | 0 | 0 | 0 | 0 |
| 36762 | ENSMUSG00000098261 | 0 | NA | NA | NA | NA | NA | Gm27521 | predicted gene, 27521 [Source:MGI Symb | 0 | 0 | 0 | 0 | 0 | 0 |
| 36763 | ENSMUSG00000098262 | 0 | NA | NA | NA | NA | NA | NA | NA | 0 | 0 | 0 | 0 | 0 | 0 |
| 36764 | ENSMUSG00000098263 | 0.335414862 | -1.779047242 | 4.030804179 | -0.441362855 | 0.658950329 | NA | Mir6938 | microRNA 6938 [Source:MGI Symbol;Acc: | 0 | 0 | 2.0125 | 0 | 0 | 0 |
| 36765 | ENSMUSG00000098264 | 0 | NA | NA | NA | NA | NA | Mir6414 | microRNA 6414 [Source:MGI Symbol;Acc: | 0 | 0 | 0 | 0 | 0 | 0 |
| 36766 | ENSMUSG00000098265 | 0 | NA | NA | NA | NA | NA | NA | NA | 0 | 0 | 0 | 0 | 0 | 0 |
| 36767 | ENSMUSG00000098266 | 0 | NA | NA | NA | NA | NA | NA | NA | 0 | 0 | 0 | 0 | 0 | 0 |
| 36768 | ENSMUSG00000098267 | 0 | NA | NA | NA | NA | NA | Mir7060 | microRNA 7060 [Source:MGI Symbol;Acc: | 0 | 0 | 0 | 0 | 0 | 0 |
| 36769 | ENSMUSG00000098268 | 0.956643647 | -0.817327529 | 2.26666353 | -0.360586174 | 0.718408827 | NA | NA | NA | 1.6062 | 2.0558 | 0 | 0.964 | 0 | 1.1138 |
| 36770 | ENSMUSG00000098269 | 1.67134182 | -1.857553697 | 2.125926162 | -0.873762095 | 0.382247827 | NA | Mir8094 | microRNA 8094 [Source:MGI Symbol;Acc: | 0.8031 | 0 | 7.0437 | 0 | 2.1812 | 0 |
| 36771 | ENSMUSG00000098270 | 0 | NA | NA | NA | NA | NA | Gm27221 | predicted gene 27221 [Source:MGI Symbo | 0 | 0 | 0 | 0 | 0 | 0 |
| 36772 | ENSMUSG00000098271 | 0.831492699 | -1.816502146 | 2.887260166 | -0.629143909 | 0.529254847 | NA | NA | NA | 0 | 0 | 4.025 | 0.964 | 0 | 0 |
| 36773 | ENSMUSG00000098272 | 0.160662974 | 1.126924175 | 4.080472857 | 0.276174898 | 0.782413736 | NA | Gm27233 | predicted gene 27233 [Source:MGI Symbo | 0 | 0 | 0 | 0.964 | 0 | 0 |
| 36774 | ENSMUSG00000098273 | 0 | NA | NA | NA | NA | NA | Mir6346 | microRNA 6346 [Source:MGI Symbol;Acc: | 0 | 0 | 0 | 0 | 0 | 0 |
| 36775 | ENSMUSG00000098274 | 9.632200817 | -0.337660213 | 0.757576798 | -0.44571087 | 0.655806112 | 0.926416603 | Rpl24 | ribosomal protein L24 [Source:MGI Symbo | 8.031 | 14.391 | 10.062 | 15.424 | 6.5437 | 3.3415 |
| 36776 | ENSMUSG00000098275 | 0.346303622 | 2.004339653 | 3.884003423 | 0.516049919 | 0.605819544 | NA | Gm18014 | predicted gene, 18014 [Source:MGI Symb | 0 | 0 | 0 | 0.964 | 0 | 1.1138 |
| 36777 | ENSMUSG00000098276 | 0 | NA | NA | NA | NA | NA | Mir6358 | microRNA 6358 [Source:MGI Symbol;Acc: | 0 | 0 | 0 | 0 | 0 | 0 |
| 36778 | ENSMUSG00000098278 | 0 | NA | NA | NA | NA | NA | Ranbp2-ps8 | RAN binding protein 2, pseudogene 8 [Sou | 0 | 0 | 0 | 0 | 0 | 0 |
| 36779 | ENSMUSG00000098279 | 0.160662974 | 1.126924175 | 4.080472857 | 0.276174898 | 0.782413736 | NA | Mir6908 | microRNA 6908 [Source:MGI Symbol;Acc: | 0 | 0 | 0 | 0.964 | 0 | 0 |
| 36780 | ENSMUSG00000098280 | 0.321325949 | 1.911575114 | 4.048267893 | 0.472195804 | 0.636787034 | NA | Mir7687 | microRNA 7687 [Source:MGI Symbol;Acc: | 0 | 0 | 0 | 1.928 | 0 | 0 |
| 36781 | ENSMUSG00000098281 | 0 | NA | NA | NA | NA | NA | NA | NA | 0 | 0 | 0 | 0 | 0 | 0 |
| 36782 | ENSMUSG00000098282 | 0 | NA | NA | NA | NA | NA | Mir6936 | microRNA 6936 [Source:MGI Symbol;Acc: | 0 | 0 | 0 | 0 | 0 | 0 |
| 36783 | ENSMUSG00000098284 | 12.42921413 | -1.023206009 | 0.698863563 | -1.464099809 | 0.143166679 | 0.552906769 | A330093E20 | RIKEN cDNA A330093E20 gene [Source:M | 15.259 | 20.558 | 14.087 | 3.8559 | 16.359 | 4.4554 |
| 36784 | ENSMUSG00000098285 | 0 | NA | NA | NA | NA | NA | Mir7670 | microRNA 7670 [Source:MGI Symbol;Acc: | 0 | 0 | 0 | 0 | 0 | 0 |
| 36785 | ENSMUSG00000098286 | 0.674674027 | 1.600443604 | 2.78690385 | 0.574272989 | 0.565783085 | NA | Mir8091 | microRNA 8091 [Source:MGI Symbol;Acc: | 0 | 0 | 1.0062 | 1.928 | 0 | 1.1138 |
| 36786 | ENSMUSG00000098287 | 0 | NA | NA | NA | NA | NA | Mir6355 | microRNA 6355 [Source:MGI Symbol;Acc: | 0 | 0 | 0 | 0 | 0 | 0 |
| 36787 | ENSMUSG00000098288 | 0.321325949 | 1.911575114 | 4.048267893 | 0.472195804 | 0.636787034 | NA | Mir6900 | microRNA 6900 [Source:MGI Symbol;Acc: | 0 | 0 | 0 | 1.928 | 0 | 0 |
| 36788 | ENSMUSG00000098289 | 0 | NA | NA | NA | NA | NA | Mir6715 | microRNA 6715 [Source:MGI Symbol;Acc: | 0 | 0 | 0 | 0 | 0 | 0 |

| | | | | | | | | | | | | | | | |
|---|---|---|---|---|---|---|---|---|---|---|---|---|---|---|---|
| 36789 | ENSMUSG00000098290 | 15.75958029 | 0.673687488 | 0.661479711 | 1.018455255 | 0.308461655 | 0.757218183 | Obox4-ps1 | oocyte specific homeobox 4, pseudogene | 3.2124 | 19.53 | 14.087 | 27.955 | 14.178 | 15.594 |
| 36790 | ENSMUSG00000098291 | 0.133850373 | -0.796629176 | 4.080472857 | -0.195229623 | 0.84521319 | NA | Mir6973a | microRNA 6973a [Source:MGI Symbol;Acc | 0.8031 | 0 | 0 | 0 | 0 | 0 |
| 36791 | ENSMUSG00000098292 | 0 | NA | NA | NA | NA | NA | Gm27194 | predicted gene 27194 [Source:MGI Symbo | 0 | 0 | 0 | 0 | 0 | 0 |
| 36792 | ENSMUSG00000098293 | 0 | NA | NA | NA | NA | NA | Mir6991 | microRNA 6991 [Source:MGI Symbol;Acc: | 0 | 0 | 0 | 0 | 0 | 0 |
| 36793 | ENSMUSG00000098294 | 0.160662974 | 1.126924175 | 4.080472857 | 0.276174898 | 0.782413736 | NA | Gm27204 | predicted gene 27204 [Source:MGI Symbo | 0 | 0 | 0 | 0.964 | 0 | 0 |
| 36794 | ENSMUSG00000098295 | 0 | NA | NA | NA | NA | NA | Gm19872 | predicted gene, 19872 [Source:MGI Symb | 0 | 0 | 0 | 0 | 0 | 0 |
| 36795 | ENSMUSG00000098296 | 1.28093653 | 0.935754013 | 2.008329591 | 0.465936476 | 0.641260971 | NA | Mir6918 | microRNA 6918 [Source:MGI Symbol;Acc: | 1.6062 | 0 | 1.0062 | 2.8919 | 2.1812 | 0 |
| 36796 | ENSMUSG00000098297 | 0 | NA | NA | NA | NA | NA | Mir6980 | microRNA 6980 [Source:MGI Symbol;Acc: | 0 | 0 | 0 | 0 | 0 | 0 |
| 36797 | ENSMUSG00000098298 | 0 | NA | NA | NA | NA | NA | NA | NA | 0 | 0 | 0 | 0 | 0 | 0 |
| 36798 | ENSMUSG00000098299 | 0 | NA | NA | NA | NA | NA | Mir7224 | microRNA 7224 [Source:MGI Symbol;Acc: | 0 | 0 | 0 | 0 | 0 | 0 |
| 36799 | ENSMUSG00000098300 | 0 | NA | NA | NA | NA | NA | Mir6540 | microRNA 6540 [Source:MGI Symbol;Acc: | 0 | 0 | 0 | 0 | 0 | 0 |
| 36800 | ENSMUSG00000098302 | 0 | NA | NA | NA | NA | NA | Gm28039 | predicted gene, 28039 [Source:MGI Symb | 0 | 0 | 0 | 0 | 0 | 0 |
| 36801 | ENSMUSG00000098303 | 0 | NA | NA | NA | NA | NA | NA | NA | 0 | 0 | 0 | 0 | 0 | 0 |
| 36802 | ENSMUSG00000098304 | 0 | NA | NA | NA | NA | NA | 1700027I24F | RIKEN cDNA 1700027I24 gene [Source:MC | 0 | 0 | 0 | 0 | 0 | 0 |
| 36803 | ENSMUSG00000098305 | 0.535401493 | -2.48596965 | 3.684117979 | -0.674780141 | 0.499815459 | NA | NA | NA | 3.2124 | 0 | 0 | 0 | 0 | 0 |
| 36804 | ENSMUSG00000098306 | 0 | NA | NA | NA | NA | NA | Gm28040 | predicted gene, 28040 [Source:MGI Symb | 0 | 0 | 0 | 0 | 0 | 0 |
| 36805 | ENSMUSG00000098307 | 0 | NA | NA | NA | NA | NA | NA | NA | 0 | 0 | 0 | 0 | 0 | 0 |
| 36806 | ENSMUSG00000098308 | 0 | NA | NA | NA | NA | NA | NA | NA | 0 | 0 | 0 | 0 | 0 | 0 |
| 36807 | ENSMUSG00000098309 | 0 | NA | NA | NA | NA | NA | Gtpbp4-ps2 | GTP binding protein 4, pseudogene 2 [Sou | 0 | 0 | 0 | 0 | 0 | 0 |
| 36808 | ENSMUSG00000098310 | 0 | NA | NA | NA | NA | NA | Mir8118 | microRNA 8118 [Source:MGI Symbol;Acc: | 0 | 0 | 0 | 0 | 0 | 0 |
| 36809 | ENSMUSG00000098311 | 0 | NA | NA | NA | NA | NA | Gm27191 | predicted gene 27191 [Source:MGI Symbo | 0 | 0 | 0 | 0 | 0 | 0 |
| 36810 | ENSMUSG00000098312 | 0 | NA | NA | NA | NA | NA | NA | NA | 0 | 0 | 0 | 0 | 0 | 0 |
| 36811 | ENSMUSG00000098313 | 0 | NA | NA | NA | NA | NA | Nr1h2-ps | nuclear receptor subfamily 1, group H, me | 0 | 0 | 0 | 0 | 0 | 0 |
| 36812 | ENSMUSG00000098314 | 0 | NA | NA | NA | NA | NA | NA | NA | 0 | 0 | 0 | 0 | 0 | 0 |
| 36813 | ENSMUSG00000098315 | 0 | NA | NA | NA | NA | NA | Mir21b | microRNA 21b [Source:MGI Symbol;Acc:M | 0 | 0 | 0 | 0 | 0 | 0 |
| 36814 | ENSMUSG00000098316 | 0 | NA | NA | NA | NA | NA | Mir7683 | microRNA 7683 [Source:MGI Symbol;Acc: | 0 | 0 | 0 | 0 | 0 | 0 |
| 36815 | ENSMUSG00000098317 | 0 | NA | NA | NA | NA | NA | Mir6926 | microRNA 6926 [Source:MGI Symbol;Acc: | 0 | 0 | 0 | 0 | 0 | 0 |
| 36816 | ENSMUSG00000098318 | 1.665019747 | -2.012182987 | 1.886498819 | -1.066622977 | 0.28614212 | NA | Lockd | lncRNA downstream of Cdkn1b [Source:M | 4.0155 | 1.0279 | 3.0187 | 1.928 | 0 | 0 |
| 36817 | ENSMUSG00000098319 | 0 | NA | NA | NA | NA | NA | Mir7038 | microRNA 7038 [Source:MGI Symbol;Acc: | 0 | 0 | 0 | 0 | 0 | 0 |
| 36818 | ENSMUSG00000098320 | 0 | NA | NA | NA | NA | NA | NA | NA | 0 | 0 | 0 | 0 | 0 | 0 |
| 36819 | ENSMUSG00000098322 | 0 | NA | NA | NA | NA | NA | 5830468F06 | RIKEN cDNA 5830468F06 gene [Source:M | 0 | 0 | 0 | 0 | 0 | 0 |
| 36820 | ENSMUSG00000098323 | 0 | NA | NA | NA | NA | NA | NA | NA | 0 | 0 | 0 | 0 | 0 | 0 |
| 36821 | ENSMUSG00000098325 | 0.294513347 | 0.165143496 | 4.080472857 | 0.040471657 | 0.967717103 | NA | Mir7048 | microRNA 7048 [Source:MGI Symbol;Acc: | 0.8031 | 0 | 0 | 0.964 | 0 | 0 |
| 36822 | ENSMUSG00000098326 | 0 | NA | NA | NA | NA | NA | NA | NA | 0 | 0 | 0 | 0 | 0 | 0 |
| 36823 | ENSMUSG00000098327 | 1.562622451 | 1.090250591 | 1.761022148 | 0.61910101 | 0.535849818 | NA | Mir7087 | microRNA 7087 [Source:MGI Symbol;Acc: | 0 | 2.0558 | 1.0062 | 1.928 | 3.2719 | 1.1138 |
| 36824 | ENSMUSG00000098328 | 0 | NA | NA | NA | NA | NA | NA | NA | 0 | 0 | 0 | 0 | 0 | 0 |
| 36825 | ENSMUSG00000098329 | 0 | NA | NA | NA | NA | NA | Mir6966 | microRNA 6966 [Source:MGI Symbol;Acc: | 0 | 0 | 0 | 0 | 0 | 0 |
| 36826 | ENSMUSG00000098330 | 69.8343526 | -0.896365164 | 0.370208492 | -2.421244197 | 0.015467483 | 0.145412036 | Gtpbp4-ps1 | GTP binding protein 4, pseudogene 1 [Sou | 65.051 | 91.485 | 116.72 | 76.154 | 27.265 | 42.326 |
| 36827 | ENSMUSG00000098331 | 0 | NA | NA | NA | NA | NA | Mir7069 | microRNA 7069 [Source:MGI Symbol;Acc: | 0 | 0 | 0 | 0 | 0 | 0 |
| 36828 | ENSMUSG00000098332 | 96.49517331 | -0.073012708 | 0.257341959 | -0.283718631 | 0.776626016 | 0.959275721 | Pigbos1 | Pigb opposite strand 1 [Source:MGI Symb | 113.24 | 90.457 | 92.575 | 78.082 | 119.97 | 84.652 |
| 36829 | ENSMUSG00000098333 | 0 | NA | NA | NA | NA | NA | Gm27158 | predicted gene 27158 [Source:MGI Symbo | 0 | 0 | 0 | 0 | 0 | 0 |
| 36830 | ENSMUSG00000098334 | 0 | NA | NA | NA | NA | NA | Mir7043 | microRNA 7043 [Source:MGI Symbol;Acc: | 0 | 0 | 0 | 0 | 0 | 0 |
| 36831 | ENSMUSG00000098335 | 0 | NA | NA | NA | NA | NA | Gm27156 | predicted gene 27156 [Source:MGI Symbo | 0 | 0 | 0 | 0 | 0 | 0 |
| 36832 | ENSMUSG00000098336 | 0 | NA | NA | NA | NA | NA | Mir6388 | microRNA 6388 [Source:MGI Symbol;Acc: | 0 | 0 | 0 | 0 | 0 | 0 |
| 36833 | ENSMUSG00000098338 | 0 | NA | NA | NA | NA | NA | Mir6941 | microRNA 6941 [Source:MGI Symbol;Acc: | 0 | 0 | 0 | 0 | 0 | 0 |
| 36834 | ENSMUSG00000098339 | 0 | NA | NA | NA | NA | NA | Mir6977 | microRNA 6977 [Source:MGI Symbol;Acc: | 0 | 0 | 0 | 0 | 0 | 0 |
| 36835 | ENSMUSG00000098340 | 0.185640647 | 1.126924175 | 4.080472857 | 0.276174898 | 0.782413736 | NA | Gm27253 | predicted gene 27253 [Source:MGI Symbo | 0 | 0 | 0 | 0 | 0 | 1.1138 |
| 36836 | ENSMUSG00000098341 | 0 | NA | NA | NA | NA | NA | Mir6944 | microRNA 6944 [Source:MGI Symbol;Acc: | 0 | 0 | 0 | 0 | 0 | 0 |
| 36837 | ENSMUSG00000098342 | 0 | NA | NA | NA | NA | NA | NA | NA | 0 | 0 | 0 | 0 | 0 | 0 |
| 36838 | ENSMUSG00000098343 | 0 | NA | NA | NA | NA | NA | Mir6240 | microRNA 6240 [Source:MGI Symbol;Acc: | 0 | 0 | 0 | 0 | 0 | 0 |
| 36839 | ENSMUSG00000098344 | 0 | NA | NA | NA | NA | NA | Mir3473f | microRNA 3473f [Source:MGI Symbol;Acc | 0 | 0 | 0 | 0 | 0 | 0 |
| 36840 | ENSMUSG00000098345 | 0 | NA | NA | NA | NA | NA | NA | NA | 0 | 0 | 0 | 0 | 0 | 0 |
| 36841 | ENSMUSG00000098346 | 0 | NA | NA | NA | NA | NA | Mir6986 | microRNA 6986 [Source:MGI Symbol;Acc: | 0 | 0 | 0 | 0 | 0 | 0 |
| 36842 | ENSMUSG00000098347 | 0 | NA | NA | NA | NA | NA | Mir6339 | microRNA 6339 [Source:MGI Symbol;Acc: | 0 | 0 | 0 | 0 | 0 | 0 |
| 36843 | ENSMUSG00000098348 | 0 | NA | NA | NA | NA | NA | Btg1c | BTG anti-proliferation factor 1C [Source:M | 0 | 0 | 0 | 0 | 0 | 0 |
| 36844 | ENSMUSG00000098349 | 0.167707431 | -0.796629176 | 4.080472857 | -0.195229623 | 0.84521319 | NA | NA | NA | 0 | 0 | 1.0062 | 0 | 0 | 0 |

| | | | | | | | | | | | | | | | |
|---|---|---|---|---|---|---|---|---|---|---|---|---|---|---|---|
| 36845 | ENSMUSG00000098350 | 1.167267481 | -0.459895871 | 2.542377405 | -0.180892054 | 0.856452306 | NA | NA | NA | 0 | 4.1117 | 0 | 2.8919 | 0 | 0 |
| 36846 | ENSMUSG00000098351 | 0 | NA | NA | NA | NA | NA | NA | NA | 0 | 0 | 0 | 0 | 0 | 0 |
| 36847 | ENSMUSG00000098353 | 0 | NA | NA | NA | NA | NA | Mir7057 | microRNA 7057 [Source:MGI Symbol;Acc: | 0 | 0 | 0 | 0 | 0 | 0 |
| 36848 | ENSMUSG00000098354 | 0 | NA | NA | NA | NA | NA | Rpap3-ps2 | RNA polymerase II associated protein 3, p | 0 | 0 | 0 | 0 | 0 | 0 |
| 36849 | ENSMUSG00000098355 | 0 | NA | NA | NA | NA | NA | NA | NA | 0 | 0 | 0 | 0 | 0 | 0 |
| 36850 | ENSMUSG00000098356 | 0 | NA | NA | NA | NA | NA | NA | NA | 0 | 0 | 0 | 0 | 0 | 0 |
| 36851 | ENSMUSG00000098357 | 0 | NA | NA | NA | NA | NA | NA | NA | 0 | 0 | 0 | 0 | 0 | 0 |
| 36852 | ENSMUSG00000098358 | 0 | NA | NA | NA | NA | NA | Gm27164 | predicted gene 27164 [Source:MGI Symbo | 0 | 0 | 0 | 0 | 0 | 0 |
| 36853 | ENSMUSG00000098359 | 0 | NA | NA | NA | NA | NA | NA | NA | 0 | 0 | 0 | 0 | 0 | 0 |
| 36854 | ENSMUSG00000098360 | 0 | NA | NA | NA | NA | NA | 4930535O05 | RIKEN cDNA 4930535O05 gene [Source:M | 0 | 0 | 0 | 0 | 0 | 0 |
| 36855 | ENSMUSG00000098361 | 0 | NA | NA | NA | NA | NA | Mir7086 | microRNA 7086 [Source:MGI Symbol;Acc: | 0 | 0 | 0 | 0 | 0 | 0 |
| 36856 | ENSMUSG00000098362 | 0 | NA | NA | NA | NA | NA | Mir130c | microRNA 130c [Source:MGI Symbol;Acc:I | 0 | 0 | 0 | 0 | 0 | 0 |
| 36857 | ENSMUSG00000098363 | 0.799645015 | 0.918084596 | 2.625685367 | 0.34965522 | 0.726597464 | NA | Mir6993 | microRNA 6993 [Source:MGI Symbol;Acc: | 1.6062 | 0 | 0 | 0.964 | 0 | 2.2277 |
| 36858 | ENSMUSG00000098364 | 0 | NA | NA | NA | NA | NA | NA | NA | 0 | 0 | 0 | 0 | 0 | 0 |
| 36859 | ENSMUSG00000098365 | 0 | NA | NA | NA | NA | NA | NA | NA | 0 | 0 | 0 | 0 | 0 | 0 |
| 36860 | ENSMUSG00000098366 | 0 | NA | NA | NA | NA | NA | Mir145b | microRNA 145b [Source:MGI Symbol;Acc: | 0 | 0 | 0 | 0 | 0 | 0 |
| 36861 | ENSMUSG00000098367 | 0 | NA | NA | NA | NA | NA | NA | NA | 0 | 0 | 0 | 0 | 0 | 0 |
| 36862 | ENSMUSG00000098368 | 0 | NA | NA | NA | NA | NA | NA | NA | 0 | 0 | 0 | 0 | 0 | 0 |
| 36863 | ENSMUSG00000098369 | 0 | NA | NA | NA | NA | NA | NA | NA | 0 | 0 | 0 | 0 | 0 | 0 |
| 36864 | ENSMUSG00000098370 | 0 | NA | NA | NA | NA | NA | Gm19541 | predicted gene, 19541 [Source:MGI Symb | 0 | 0 | 0 | 0 | 0 | 0 |
| 36865 | ENSMUSG00000098371 | 0 | NA | NA | NA | NA | NA | Gm28037 | predicted gene, 28037 [Source:MGI Symb | 0 | 0 | 0 | 0 | 0 | 0 |
| 36866 | ENSMUSG00000098373 | 0 | NA | NA | NA | NA | NA | NA | NA | 0 | 0 | 0 | 0 | 0 | 0 |
| 36867 | ENSMUSG00000098374 | 25.20993288 | 0.308771474 | 0.437036314 | 0.706512169 | 0.479869688 | 0.870278888 | Gm28043 | predicted gene, 28043 [Source:MGI Symb | 24.896 | 16.447 | 26.162 | 30.847 | 26.175 | 26.732 |
| 36868 | ENSMUSG00000098376 | 0.503095663 | 2.549494177 | 3.348715477 | 0.761334964 | 0.446457022 | NA | Mir6907 | microRNA 6907 [Source:MGI Symbol;Acc: | 0 | 0 | 0 | 1.928 | 1.0906 | 0 |
| 36869 | ENSMUSG00000098378 | 0 | NA | NA | NA | NA | NA | NA | NA | 0 | 0 | 0 | 0 | 0 | 0 |
| 36870 | ENSMUSG00000098379 | 0 | NA | NA | NA | NA | NA | Mir6347 | microRNA 6347 [Source:MGI Symbol;Acc: | 0 | 0 | 0 | 0 | 0 | 0 |
| 36871 | ENSMUSG00000098380 | 0.503122293 | -2.366675966 | 3.785596405 | -0.625179156 | 0.531853481 | NA | Mir6939 | microRNA 6939 [Source:MGI Symbol;Acc: | 0 | 0 | 3.0187 | 0 | 0 | 0 |
| 36872 | ENSMUSG00000098381 | 0 | NA | NA | NA | NA | NA | Mir7225 | microRNA 7225 [Source:MGI Symbol;Acc: | 0 | 0 | 0 | 0 | 0 | 0 |
| 36873 | ENSMUSG00000098382 | 0.428363721 | -0.543730382 | 3.624677344 | -0.15000794 | 0.880758351 | NA | Gtpbp4-ps4 | GTP binding protein 4, pseudogene 4 [Sou | 1.6062 | 0 | 0 | 0.964 | 0 | 0 |
| 36874 | ENSMUSG00000098383 | 0 | NA | NA | NA | NA | NA | Gm27197 | predicted gene 27197 [Source:MGI Symbo | 0 | 0 | 0 | 0 | 0 | 0 |
| 36875 | ENSMUSG00000098385 | 0 | NA | NA | NA | NA | NA | NA | NA | 0 | 0 | 0 | 0 | 0 | 0 |
| 36876 | ENSMUSG00000098386 | 0 | NA | NA | NA | NA | NA | Ranbp2-ps1( | RAN binding protein 2, pseudogene 10 [Sc | 0 | 0 | 0 | 0 | 0 | 0 |
| 36877 | ENSMUSG00000098387 | 0 | NA | NA | NA | NA | NA | Pet117 | PET117 homolog [Source:MGI Symbol;Acc | 0 | 0 | 0 | 0 | 0 | 0 |
| 36878 | ENSMUSG00000098388 | 0 | NA | NA | NA | NA | NA | Mir8109 | microRNA 8109 [Source:MGI Symbol;Acc: | 0 | 0 | 0 | 0 | 0 | 0 |
| 36879 | ENSMUSG00000098389 | 0 | NA | NA | NA | NA | NA | Mir6983 | microRNA 6983 [Source:MGI Symbol;Acc: | 0 | 0 | 0 | 0 | 0 | 0 |
| 36880 | ENSMUSG00000098390 | 0 | NA | NA | NA | NA | NA | Gm3434 | predicted gene 3434 [Source:MGI Symbol | 0 | 0 | 0 | 0 | 0 | 0 |
| 36881 | ENSMUSG00000098391 | 0 | NA | NA | NA | NA | NA | Gm27208 | predicted gene 27208 [Source:MGI Symbo | 0 | 0 | 0 | 0 | 0 | 0 |
| 36882 | ENSMUSG00000098392 | 0 | NA | NA | NA | NA | NA | NA | NA | 0 | 0 | 0 | 0 | 0 | 0 |
| 36883 | ENSMUSG00000098393 | 0 | NA | NA | NA | NA | NA | NA | NA | 0 | 0 | 0 | 0 | 0 | 0 |
| 36884 | ENSMUSG00000098394 | 0 | NA | NA | NA | NA | NA | NA | NA | 0 | 0 | 0 | 0 | 0 | 0 |
| 36885 | ENSMUSG00000098395 | 0 | NA | NA | NA | NA | NA | NA | NA | 0 | 0 | 0 | 0 | 0 | 0 |
| 36886 | ENSMUSG00000098396 | 0 | NA | NA | NA | NA | NA | NA | NA | 0 | 0 | 0 | 0 | 0 | 0 |
| 36887 | ENSMUSG00000098397 | 0.17131964 | -0.796629176 | 4.080472857 | -0.195229623 | 0.84521319 | NA | Mir7669 | microRNA 7669 [Source:MGI Symbol;Acc: | 0 | 1.0279 | 0 | 0 | 0 | 0 |
| 36888 | ENSMUSG00000098399 | 0 | NA | NA | NA | NA | NA | NA | NA | 0 | 0 | 0 | 0 | 0 | 0 |
| 36889 | ENSMUSG00000098400 | 0 | NA | NA | NA | NA | NA | Mir7680 | microRNA 7680 [Source:MGI Symbol;Acc: | 0 | 0 | 0 | 0 | 0 | 0 |
| 36890 | ENSMUSG00000098401 | 0 | NA | NA | NA | NA | NA | Mir6377 | microRNA 6377 [Source:MGI Symbol;Acc: | 0 | 0 | 0 | 0 | 0 | 0 |
| 36891 | ENSMUSG00000098402 | 0 | NA | NA | NA | NA | NA | NA | NA | 0 | 0 | 0 | 0 | 0 | 0 |
| 36892 | ENSMUSG00000098403 | 0.68145976 | 0.058694068 | 2.590797836 | 0.022654824 | 0.981925612 | NA | NA | NA | 0 | 1.0279 | 1.0062 | 0.964 | 1.0906 | 0 |
| 36893 | ENSMUSG00000098404 | 2.050286217 | -0.348418392 | 1.48740903 | -0.234245177 | 0.814794644 | NA | Mrip-ps | Mom radiation induced polyposis, pseudc | 0.8031 | 3.0838 | 3.0187 | 0.964 | 1.0906 | 3.3415 |
| 36894 | ENSMUSG00000098405 | 0 | NA | NA | NA | NA | NA | Gm27414 | predicted gene, 27414 [Source:MGI Symb | 0 | 0 | 0 | 0 | 0 | 0 |
| 36895 | ENSMUSG00000098406 | 0 | NA | NA | NA | NA | NA | Mir7222 | microRNA 7222 [Source:MGI Symbol;Acc: | 0 | 0 | 0 | 0 | 0 | 0 |
| 36896 | ENSMUSG00000098407 | 0 | NA | NA | NA | NA | NA | Mir6348 | microRNA 6348 [Source:MGI Symbol;Acc: | 0 | 0 | 0 | 0 | 0 | 0 |
| 36897 | ENSMUSG00000098408 | 0 | NA | NA | NA | NA | NA | Mir6237 | microRNA 6237 [Source:MGI Symbol;Acc: | 0 | 0 | 0 | 0 | 0 | 0 |
| 36898 | ENSMUSG00000098409 | 0 | NA | NA | NA | NA | NA | NA | NA | 0 | 0 | 0 | 0 | 0 | 0 |
| 36899 | ENSMUSG00000098410 | 0 | NA | NA | NA | NA | NA | Mir1957b | microRNA 1957b [Source:MGI Symbol;Acc | 0 | 0 | 0 | 0 | 0 | 0 |
| 36900 | ENSMUSG00000098412 | 0 | NA | NA | NA | NA | NA | Gm7870 | predicted gene 7870 [Source:MGI Symbol | 0 | 0 | 0 | 0 | 0 | 0 |

| | | | | | | | | | | | | | | | |
|---|---|---|---|---|---|---|---|---|---|---|---|---|---|---|---|
| 36901 | ENSMUSG00000098413 | 0 | NA | NA | NA | NA | NA | Mir6990 | microRNA 6990 [Source:MGI Symbol;Acc: | 0 | 0 | 0 | 0 | 0 | 0 |
| 36902 | ENSMUSG00000098414 | 0.181769715 | 1.126924175 | 4.080472857 | 0.276174898 | 0.782413736 | NA | Mir6909 | microRNA 6909 [Source:MGI Symbol;Acc: | 0 | 0 | 0 | 0 | 1.0906 | 0 |
| 36903 | ENSMUSG00000098415 | 0 | NA | NA | NA | NA | NA | Mir6928 | microRNA 6928 [Source:MGI Symbol;Acc: | 0 | 0 | 0 | 0 | 0 | 0 |
| 36904 | ENSMUSG00000098416 | 0 | NA | NA | NA | NA | NA | Mir7117 | microRNA 7117 [Source:MGI Symbol;Acc: | 0 | 0 | 0 | 0 | 0 | 0 |
| 36905 | ENSMUSG00000098417 | 0 | NA | NA | NA | NA | NA | Mir6379 | microRNA 6379 [Source:MGI Symbol;Acc: | 0 | 0 | 0 | 0 | 0 | 0 |
| 36906 | ENSMUSG00000098418 | 0 | NA | NA | NA | NA | NA | NA | NA | 0 | 0 | 0 | 0 | 0 | 0 |
| 36907 | ENSMUSG00000098419 | 0 | NA | NA | NA | NA | NA | Mir6963 | microRNA 6963 [Source:MGI Symbol;Acc: | 0 | 0 | 0 | 0 | 0 | 0 |
| 36908 | ENSMUSG00000098420 | 0 | NA | NA | NA | NA | NA | Gm27249 | predicted gene 27249 [Source:MGI Symbo | 0 | 0 | 0 | 0 | 0 | 0 |
| 36909 | ENSMUSG00000098421 | 0 | NA | NA | NA | NA | NA | Gm27190 | predicted gene 27190 [Source:MGI Symbo | 0 | 0 | 0 | 0 | 0 | 0 |
| 36910 | ENSMUSG00000098422 | 0 | NA | NA | NA | NA | NA | Mir7016 | microRNA 7016 [Source:MGI Symbol;Acc: | 0 | 0 | 0 | 0 | 0 | 0 |
| 36911 | ENSMUSG00000098423 | 0 | NA | NA | NA | NA | NA | NA | NA | 0 | 0 | 0 | 0 | 0 | 0 |
| 36912 | ENSMUSG00000098424 | 0.267700746 | -1.497481649 | 4.057001436 | -0.369110456 | 0.712045396 | NA | Gm27202 | predicted gene 27202 [Source:MGI Symbo | 1.6062 | 0 | 0 | 0 | 0 | 0 |
| 36913 | ENSMUSG00000098425 | 0 | NA | NA | NA | NA | NA | NA | NA | 0 | 0 | 0 | 0 | 0 | 0 |
| 36914 | ENSMUSG00000098426 | 0 | NA | NA | NA | NA | NA | Gm27149 | predicted gene 27149 [Source:MGI Symbo | 0 | 0 | 0 | 0 | 0 | 0 |
| 36915 | ENSMUSG00000098428 | 0 | NA | NA | NA | NA | NA | NA | NA | 0 | 0 | 0 | 0 | 0 | 0 |
| 36916 | ENSMUSG00000098429 | 0 | NA | NA | NA | NA | NA | Gm27226 | predicted gene 27226 [Source:MGI Symbo | 0 | 0 | 0 | 0 | 0 | 0 |
| 36917 | ENSMUSG00000098430 | 0 | NA | NA | NA | NA | NA | NA | NA | 0 | 0 | 0 | 0 | 0 | 0 |
| 36918 | ENSMUSG00000098431 | 0 | NA | NA | NA | NA | NA | Mir7666 | microRNA 7666 [Source:MGI Symbol;Acc: | 0 | 0 | 0 | 0 | 0 | 0 |
| 36919 | ENSMUSG00000098432 | 0 | NA | NA | NA | NA | NA | NA | NA | 0 | 0 | 0 | 0 | 0 | 0 |
| 36920 | ENSMUSG00000098433 | 0 | NA | NA | NA | NA | NA | Mir6391 | microRNA 6391 [Source:MGI Symbol;Acc: | 0 | 0 | 0 | 0 | 0 | 0 |
| 36921 | ENSMUSG00000098434 | 0 | NA | NA | NA | NA | NA | 2010110E17 | RIKEN cDNA 2010110E17 gene [Source:M | 0 | 0 | 0 | 0 | 0 | 0 |
| 36922 | ENSMUSG00000098435 | 0 | NA | NA | NA | NA | NA | NA | NA | 0 | 0 | 0 | 0 | 0 | 0 |
| 36923 | ENSMUSG00000098436 | 0 | NA | NA | NA | NA | NA | NA | NA | 0 | 0 | 0 | 0 | 0 | 0 |
| 36924 | ENSMUSG00000098437 | 0 | NA | NA | NA | NA | NA | Mir195b | microRNA 195b [Source:MGI Symbol;Acc: | 0 | 0 | 0 | 0 | 0 | 0 |
| 36925 | ENSMUSG00000098438 | 0 | NA | NA | NA | NA | NA | Gm27178 | predicted gene 27178 [Source:MGI Symbo | 0 | 0 | 0 | 0 | 0 | 0 |
| 36926 | ENSMUSG00000098439 | 0.506734502 | -2.372847156 | 3.338060886 | -0.710845978 | 0.477179685 | NA | Hm629797 | cDNA sequence HM629797 [Source:MGI S | 0 | 1.0279 | 2.0125 | 0 | 0 | 0 |
| 36927 | ENSMUSG00000098441 | 0 | NA | NA | NA | NA | NA | Mir6537 | microRNA 6537 [Source:MGI Symbol;Acc: | 0 | 0 | 0 | 0 | 0 | 0 |
| 36928 | ENSMUSG00000098442 | 0 | NA | NA | NA | NA | NA | Mir6959 | microRNA 6959 [Source:MGI Symbol;Acc: | 0 | 0 | 0 | 0 | 0 | 0 |
| 36929 | ENSMUSG00000098443 | 0 | NA | NA | NA | NA | NA | Mir7681 | microRNA 7681 [Source:MGI Symbol;Acc: | 0 | 0 | 0 | 0 | 0 | 0 |
| 36930 | ENSMUSG00000098444 | 0 | NA | NA | NA | NA | NA | Gm27811 | predicted gene, 27811 [Source:MGI Symb | 0 | 0 | 0 | 0 | 0 | 0 |
| 36931 | ENSMUSG00000098445 | 0 | NA | NA | NA | NA | NA | Mir6972 | microRNA 6972 [Source:MGI Symbol;Acc: | 0 | 0 | 0 | 0 | 0 | 0 |
| 36932 | ENSMUSG00000098446 | 0 | NA | NA | NA | NA | NA | Mir7242 | microRNA 7242 [Source:MGI Symbol;Acc: | 0 | 0 | 0 | 0 | 0 | 0 |
| 36933 | ENSMUSG00000098447 | 7.556590595 | 0.72229379 | 0.777762797 | 0.928681331 | 0.353054259 | 0.793439527 | Gm27184 | predicted gene 27184 [Source:MGI Symbo | 4.0155 | 6.1675 | 7.0437 | 11.568 | 7.6343 | 8.9108 |
| 36934 | ENSMUSG00000098448 | 0 | NA | NA | NA | NA | NA | Mir6362 | microRNA 6362 [Source:MGI Symbol;Acc: | 0 | 0 | 0 | 0 | 0 | 0 |
| 36935 | ENSMUSG00000098449 | 5.338777505 | -1.510731464 | 0.996526959 | -1.515996581 | 0.129520217 | 0.526822389 | Gm7467 | predicted gene 7467 [Source:MGI Symbol | 6.4248 | 12.335 | 5.0312 | 3.8559 | 3.2719 | 1.1138 |
| 36936 | ENSMUSG00000098451 | 0 | NA | NA | NA | NA | NA | Mir6952 | microRNA 6952 [Source:MGI Symbol;Acc: | 0 | 0 | 0 | 0 | 0 | 0 |
| 36937 | ENSMUSG00000098452 | 0 | NA | NA | NA | NA | NA | NA | NA | 0 | 0 | 0 | 0 | 0 | 0 |
| 36938 | ENSMUSG00000098453 | 0 | NA | NA | NA | NA | NA | Mir6997 | microRNA 6997 [Source:MGI Symbol;Acc: | 0 | 0 | 0 | 0 | 0 | 0 |
| 36939 | ENSMUSG00000098454 | 0 | NA | NA | NA | NA | NA | Mir7032 | microRNA 7032 [Source:MGI Symbol;Acc: | 0 | 0 | 0 | 0 | 0 | 0 |
| 36940 | ENSMUSG00000098455 | 0 | NA | NA | NA | NA | NA | NA | NA | 0 | 0 | 0 | 0 | 0 | 0 |
| 36941 | ENSMUSG00000098456 | 0 | NA | NA | NA | NA | NA | Gm12117 | predicted gene 12117 [Source:MGI Symbo | 0 | 0 | 0 | 0 | 0 | 0 |
| 36942 | ENSMUSG00000098457 | 0 | NA | NA | NA | NA | NA | Mir8097 | microRNA 8097 [Source:MGI Symbol;Acc: | 0 | 0 | 0 | 0 | 0 | 0 |
| 36943 | ENSMUSG00000098458 | 0 | NA | NA | NA | NA | NA | Mir6971 | microRNA 6971 [Source:MGI Symbol;Acc: | 0 | 0 | 0 | 0 | 0 | 0 |
| 36944 | ENSMUSG00000098459 | 0 | NA | NA | NA | NA | NA | NA | NA | 0 | 0 | 0 | 0 | 0 | 0 |
| 36945 | ENSMUSG00000098460 | 0 | NA | NA | NA | NA | NA | NA | NA | 0 | 0 | 0 | 0 | 0 | 0 |
| 36946 | ENSMUSG00000098462 | 0 | NA | NA | NA | NA | NA | NA | NA | 0 | 0 | 0 | 0 | 0 | 0 |
| 36947 | ENSMUSG00000098463 | 0 | NA | NA | NA | NA | NA | NA | NA | 0 | 0 | 0 | 0 | 0 | 0 |
| 36948 | ENSMUSG00000098464 | 0 | NA | NA | NA | NA | NA | Mir7002 | microRNA 7002 [Source:MGI Symbol;Acc: | 0 | 0 | 0 | 0 | 0 | 0 |
| 36949 | ENSMUSG00000098465 | 0 | NA | NA | NA | NA | NA | Mir1668 | microRNA 1668 [Source:MGI Symbol;Acc: | 0 | 0 | 0 | 0 | 0 | 0 |
| 36950 | ENSMUSG00000098466 | 0 | NA | NA | NA | NA | NA | NA | NA | 0 | 0 | 0 | 0 | 0 | 0 |
| 36951 | ENSMUSG00000098467 | 0 | NA | NA | NA | NA | NA | NA | NA | 0 | 0 | 0 | 0 | 0 | 0 |
| 36952 | ENSMUSG00000098468 | 0 | NA | NA | NA | NA | NA | Gm27186 | predicted gene 27186 [Source:MGI Symbo | 0 | 0 | 0 | 0 | 0 | 0 |
| 36953 | ENSMUSG00000098469 | 0 | NA | NA | NA | NA | NA | Ranbp2-ps4 | RAN binding protein 2, pseudogene 4 [Sou | 0 | 0 | 0 | 0 | 0 | 0 |
| 36954 | ENSMUSG00000098470 | 0.695522043 | 1.639439662 | 2.762392557 | 0.593485404 | 0.552856349 | NA | C1rb | complement component 1, r subcompone | 0 | 1.0279 | 0 | 0.964 | 2.1812 | 0 |
| 36955 | ENSMUSG00000098471 | 0 | NA | NA | NA | NA | NA | Gm27203 | predicted gene 27203 [Source:MGI Symbo | 0 | 0 | 0 | 0 | 0 | 0 |
| 36956 | ENSMUSG00000098472 | 0 | NA | NA | NA | NA | NA | NA | NA | 0 | 0 | 0 | 0 | 0 | 0 |

| | | | | | | | | | | | | | | | |
|---|---|---|---|---|---|---|---|---|---|---|---|---|---|---|---|
| 36957 | ENSMUSG00000098473 | 0 | NA | NA | NA | NA | NA | Mir7083 | microRNA 7083 [Source:MGI Symbol;Acc: | 0 | 0 | 0 | 0 | 0 | 0 |
| 36958 | ENSMUSG00000098474 | 0 | NA | NA | NA | NA | NA | Mir6919 | microRNA 6919 [Source:MGI Symbol;Acc: | 0 | 0 | 0 | 0 | 0 | 0 |
| 36959 | ENSMUSG00000098475 | 0 | NA | NA | NA | NA | NA | Mir6982 | microRNA 6982 [Source:MGI Symbol;Acc: | 0 | 0 | 0 | 0 | 0 | 0 |
| 36960 | ENSMUSG00000098476 | 0 | NA | NA | NA | NA | NA | NA | NA | 0 | 0 | 0 | 0 | 0 | 0 |
| 36961 | ENSMUSG00000098477 | 0 | NA | NA | NA | NA | NA | NA | NA | 0 | 0 | 0 | 0 | 0 | 0 |
| 36962 | ENSMUSG00000098478 | 0 | NA | NA | NA | NA | NA | NA | NA | 0 | 0 | 0 | 0 | 0 | 0 |
| 36963 | ENSMUSG00000098479 | 0 | NA | NA | NA | NA | NA | NA | NA | 0 | 0 | 0 | 0 | 0 | 0 |
| 36964 | ENSMUSG00000098480 | 0 | NA | NA | NA | NA | NA | Mir6387 | microRNA 6387 [Source:MGI Symbol;Acc: | 0 | 0 | 0 | 0 | 0 | 0 |
| 36965 | ENSMUSG00000098481 | 0 | NA | NA | NA | NA | NA | NA | NA | 0 | 0 | 0 | 0 | 0 | 0 |
| 36966 | ENSMUSG00000098482 | 0 | NA | NA | NA | NA | NA | Mir6363 | microRNA 6363 [Source:MGI Symbol;Acc: | 0 | 0 | 0 | 0 | 0 | 0 |
| 36967 | ENSMUSG00000098483 | 0.696064467 | -1.534444742 | 3.043667477 | -0.504143358 | 0.61416064 | NA | Mir7078 | microRNA 7078 [Source:MGI Symbol;Acc: | 3.2124 | 0 | 0 | 0.964 | 0 | 0 |
| 36968 | ENSMUSG00000098484 | 0 | NA | NA | NA | NA | NA | NA | NA | 0 | 0 | 0 | 0 | 0 | 0 |
| 36969 | ENSMUSG00000098485 | 0 | NA | NA | NA | NA | NA | Gm6956 | predicted gene 6956 [Source:MGI Symbol | 0 | 0 | 0 | 0 | 0 | 0 |
| 36970 | ENSMUSG00000098486 | 0 | NA | NA | NA | NA | NA | NA | NA | 0 | 0 | 0 | 0 | 0 | 0 |
| 36971 | ENSMUSG00000098487 | 0 | NA | NA | NA | NA | NA | Mir6899 | microRNA 6899 [Source:MGI Symbol;Acc: | 0 | 0 | 0 | 0 | 0 | 0 |
| 36972 | ENSMUSG00000098488 | 5.625911979 | -0.603688092 | 0.911491057 | -0.6623083 | 0.507773659 | 0.882426793 | Pla2g4b | phospholipase A2, group IVB (cytosolic) [S | 7.2279 | 5.1396 | 8.05 | 6.7478 | 4.3625 | 2.2277 |
| 36973 | ENSMUSG00000098489 | 0 | NA | NA | NA | NA | NA | Mir7678 | microRNA 7678 [Source:MGI Symbol;Acc: | 0 | 0 | 0 | 0 | 0 | 0 |
| 36974 | ENSMUSG00000098490 | 0 | NA | NA | NA | NA | NA | Mir7089 | microRNA 7089 [Source:MGI Symbol;Acc: | 0 | 0 | 0 | 0 | 0 | 0 |
| 36975 | ENSMUSG00000098491 | 0 | NA | NA | NA | NA | NA | NA | NA | 0 | 0 | 0 | 0 | 0 | 0 |
| 36976 | ENSMUSG00000098492 | 0.428363721 | -0.543730382 | 3.624677344 | -0.15000794 | 0.880758351 | NA | Gm19353 | predicted gene, 19353 [Source:MGI Symb | 1.6062 | 0 | 0 | 0.964 | 0 | 0 |
| 36977 | ENSMUSG00000098493 | 0 | NA | NA | NA | NA | NA | Mir6383 | microRNA 6383 [Source:MGI Symbol;Acc: | 0 | 0 | 0 | 0 | 0 | 0 |
| 36978 | ENSMUSG00000098494 | 0 | NA | NA | NA | NA | NA | NA | NA | 0 | 0 | 0 | 0 | 0 | 0 |
| 36979 | ENSMUSG00000098497 | 0 | NA | NA | NA | NA | NA | Mir6932 | microRNA 6932 [Source:MGI Symbol;Acc: | 0 | 0 | 0 | 0 | 0 | 0 |
| 36980 | ENSMUSG00000098498 | 0 | NA | NA | NA | NA | NA | Mir8099-1 | microRNA 8099-1 [Source:MGI Symbol;Ac | 0 | 0 | 0 | 0 | 0 | 0 |
| 36981 | ENSMUSG00000098499 | 0 | NA | NA | NA | NA | NA | Mir6975 | microRNA 6975 [Source:MGI Symbol;Acc: | 0 | 0 | 0 | 0 | 0 | 0 |
| 36982 | ENSMUSG00000098500 | 0 | NA | NA | NA | NA | NA | Mir6396 | microRNA 6396 [Source:MGI Symbol;Acc: | 0 | 0 | 0 | 0 | 0 | 0 |
| 36983 | ENSMUSG00000098501 | 0 | NA | NA | NA | NA | NA | Gm19272 | predicted gene, 19272 [Source:MGI Symb | 0 | 0 | 0 | 0 | 0 | 0 |
| 36984 | ENSMUSG00000098502 | 0 | NA | NA | NA | NA | NA | NA | NA | 0 | 0 | 0 | 0 | 0 | 0 |
| 36985 | ENSMUSG00000098503 | 0 | NA | NA | NA | NA | NA | NA | NA | 0 | 0 | 0 | 0 | 0 | 0 |
| 36986 | ENSMUSG00000098504 | 0 | NA | NA | NA | NA | NA | Mir6351 | microRNA 6351 [Source:MGI Symbol;Acc: | 0 | 0 | 0 | 0 | 0 | 0 |
| 36987 | ENSMUSG00000098505 | 0 | NA | NA | NA | NA | NA | Gm14496 | predicted gene 14496 [Source:MGI Symbo | 0 | 0 | 0 | 0 | 0 | 0 |
| 36988 | ENSMUSG00000098506 | 0 | NA | NA | NA | NA | NA | Phf20-ps | PHD finger protein 20, pseudogene [Sourc | 0 | 0 | 0 | 0 | 0 | 0 |
| 36989 | ENSMUSG00000098507 | 0 | NA | NA | NA | NA | NA | NA | NA | 0 | 0 | 0 | 0 | 0 | 0 |
| 36990 | ENSMUSG00000098508 | 0 | NA | NA | NA | NA | NA | Ranbp2-ps2 | RAN binding protein 2, pseudogene 2 [Sou | 0 | 0 | 0 | 0 | 0 | 0 |
| 36991 | ENSMUSG00000098509 | 0 | NA | NA | NA | NA | NA | Rgs21 | regulator of G-protein signalling 21 [Sourc | 0 | 0 | 0 | 0 | 0 | 0 |
| 36992 | ENSMUSG00000098510 | 0 | NA | NA | NA | NA | NA | Mir6968 | microRNA 6968 [Source:MGI Symbol;Acc: | 0 | 0 | 0 | 0 | 0 | 0 |
| 36993 | ENSMUSG00000098511 | 0 | NA | NA | NA | NA | NA | Mir7072 | microRNA 7072 [Source:MGI Symbol;Acc: | 0 | 0 | 0 | 0 | 0 | 0 |
| 36994 | ENSMUSG00000098513 | 0 | NA | NA | NA | NA | NA | Mir6399 | microRNA 6399 [Source:MGI Symbol;Acc: | 0 | 0 | 0 | 0 | 0 | 0 |
| 36995 | ENSMUSG00000098514 | 0 | NA | NA | NA | NA | NA | NA | NA | 0 | 0 | 0 | 0 | 0 | 0 |
| 36996 | ENSMUSG00000098515 | 0 | NA | NA | NA | NA | NA | Gm27195 | predicted gene 27195 [Source:MGI Symbo | 0 | 0 | 0 | 0 | 0 | 0 |
| 36997 | ENSMUSG00000098516 | 1.20981845 | -1.185265591 | 2.547138868 | -0.465332144 | 0.641693619 | NA | Mir7651 | microRNA 7651 [Source:MGI Symbol;Acc: | 0 | 0 | 5.0312 | 0 | 0 | 2.2277 |
| 36998 | ENSMUSG00000098518 | 0 | NA | NA | NA | NA | NA | NA | NA | 0 | 0 | 0 | 0 | 0 | 0 |
| 36999 | ENSMUSG00000098519 | 0 | NA | NA | NA | NA | NA | Gm27166 | predicted gene 27166 [Source:MGI Symbo | 0 | 0 | 0 | 0 | 0 | 0 |
| 37000 | ENSMUSG00000098520 | 0 | NA | NA | NA | NA | NA | Ranbp2-ps1 | RAN binding protein 2, pseudogene 1 [Sou | 0 | 0 | 0 | 0 | 0 | 0 |
| 37001 | ENSMUSG00000098521 | 0 | NA | NA | NA | NA | NA | Mir7243 | microRNA 7243 [Source:MGI Symbol;Acc: | 0 | 0 | 0 | 0 | 0 | 0 |
| 37002 | ENSMUSG00000098522 | 0 | NA | NA | NA | NA | NA | Mir6412 | microRNA 6412 [Source:MGI Symbol;Acc: | 0 | 0 | 0 | 0 | 0 | 0 |
| 37003 | ENSMUSG00000098524 | 0 | NA | NA | NA | NA | NA | NA | NA | 0 | 0 | 0 | 0 | 0 | 0 |
| 37004 | ENSMUSG00000098525 | 0 | NA | NA | NA | NA | NA | NA | NA | 0 | 0 | 0 | 0 | 0 | 0 |
| 37005 | ENSMUSG00000098526 | 0 | NA | NA | NA | NA | NA | NA | NA | 0 | 0 | 0 | 0 | 0 | 0 |
| 37006 | ENSMUSG00000098527 | 0 | NA | NA | NA | NA | NA | Mir6961 | microRNA 6961 [Source:MGI Symbol;Acc: | 0 | 0 | 0 | 0 | 0 | 0 |
| 37007 | ENSMUSG00000098528 | 0 | NA | NA | NA | NA | NA | Zfp572 | zinc finger protein 572 [Source:MGI Symb | 0 | 0 | 0 | 0 | 0 | 0 |
| 37008 | ENSMUSG00000098529 | 0.481988923 | 2.492401104 | 3.837077151 | 0.649557203 | 0.515978285 | NA | NA | NA | 0 | 0 | 0 | 2.8919 | 0 | 0 |
| 37009 | ENSMUSG00000098530 | 0 | NA | NA | NA | NA | NA | Gm28051 | predicted gene, 28051 [Source:MGI Symb | 0 | 0 | 0 | 0 | 0 | 0 |
| 37010 | ENSMUSG00000098531 | 0 | NA | NA | NA | NA | NA | Mir7041 | microRNA 7041 [Source:MGI Symbol;Acc: | 0 | 0 | 0 | 0 | 0 | 0 |
| 37011 | ENSMUSG00000098532 | 0 | NA | NA | NA | NA | NA | Mir7065 | microRNA 7065 [Source:MGI Symbol;Acc: | 0 | 0 | 0 | 0 | 0 | 0 |
| 37012 | ENSMUSG00000098533 | 0 | NA | NA | NA | NA | NA | Mir6373 | microRNA 6373 [Source:MGI Symbol;Acc: | 0 | 0 | 0 | 0 | 0 | 0 |

| | | | | | | | | | | | | | | | |
|---|---|---|---|---|---|---|---|---|---|---|---|---|---|---|---|
| 37013 | ENSMUSG00000098534 | 0 | NA | NA | NA | NA | NA | Gm27167 | predicted gene 27167 [Source:MGI Symbc | 0 | 0 | 0 | 0 | 0 | 0 |
| 37014 | ENSMUSG00000098535 | 0 | NA | NA | NA | NA | NA | NA | NA | 0 | 0 | 0 | 0 | 0 | 0 |
| 37015 | ENSMUSG00000098536 | 0 | NA | NA | NA | NA | NA | NA | NA | 0 | 0 | 0 | 0 | 0 | 0 |
| 37016 | ENSMUSG00000098537 | 0 | NA | NA | NA | NA | NA | NA | NA | 0 | 0 | 0 | 0 | 0 | 0 |
| 37017 | ENSMUSG00000098538 | 0 | NA | NA | NA | NA | NA | NA | NA | 0 | 0 | 0 | 0 | 0 | 0 |
| 37018 | ENSMUSG00000098539 | 0 | NA | NA | NA | NA | NA | Mir133c | microRNA 133c [Source:MGI Symbol;Acc:I | 0 | 0 | 0 | 0 | 0 | 0 |
| 37019 | ENSMUSG00000098540 | 0.160662974 | 1.126924175 | 4.080472857 | 0.276174898 | 0.782413736 | NA | Mir7059 | microRNA 7059 [Source:MGI Symbol;Acc: | 0 | 0 | 0 | 0.964 | 0 | 0 |
| 37020 | ENSMUSG00000098541 | 0 | NA | NA | NA | NA | NA | NA | NA | 0 | 0 | 0 | 0 | 0 | 0 |
| 37021 | ENSMUSG00000098542 | 0 | NA | NA | NA | NA | NA | NA | NA | 0 | 0 | 0 | 0 | 0 | 0 |
| 37022 | ENSMUSG00000098543 | 0 | NA | NA | NA | NA | NA | NA | NA | 0 | 0 | 0 | 0 | 0 | 0 |
| 37023 | ENSMUSG00000098544 | 0.678054141 | -2.792799308 | 2.990673428 | -0.933836266 | 0.350388359 | NA | Mir7211 | microRNA 7211 [Source:MGI Symbol;Acc: | 0 | 2.0558 | 2.0125 | 0 | 0 | 0 |
| 37024 | ENSMUSG00000098547 | 0 | NA | NA | NA | NA | NA | Mir6921 | microRNA 6921 [Source:MGI Symbol;Acc: | 0 | 0 | 0 | 0 | 0 | 0 |
| 37025 | ENSMUSG00000098548 | 0 | NA | NA | NA | NA | NA | Mir6996 | microRNA 6996 [Source:MGI Symbol;Acc: | 0 | 0 | 0 | 0 | 0 | 0 |
| 37026 | ENSMUSG00000098549 | 0 | NA | NA | NA | NA | NA | Gm5269 | predicted gene 5269 [Source:MGI Symbol | 0 | 0 | 0 | 0 | 0 | 0 |
| 37027 | ENSMUSG00000098550 | 0 | NA | NA | NA | NA | NA | Mir7079 | microRNA 7079 [Source:MGI Symbol;Acc: | 0 | 0 | 0 | 0 | 0 | 0 |
| 37028 | ENSMUSG00000098552 | 0 | NA | NA | NA | NA | NA | Gm27217 | predicted gene 27217 [Source:MGI Symbc | 0 | 0 | 0 | 0 | 0 | 0 |
| 37029 | ENSMUSG00000098553 | 0.167707431 | -0.796629176 | 4.080472857 | -0.195229623 | 0.84521319 | NA | Mir3569 | microRNA 3569 [Source:MGI Symbol;Acc: | 0 | 0 | 1.0062 | 0 | 0 | 0 |
| 37030 | ENSMUSG00000098555 | 0 | NA | NA | NA | NA | NA | Mir6341 | microRNA 6341 [Source:MGI Symbol;Acc: | 0 | 0 | 0 | 0 | 0 | 0 |
| 37031 | ENSMUSG00000098557 | 79.97861933 | -1.064363058 | 0.325669832 | -3.268227372 | 0.001082234 | 0.022234986 | Kctd12 | potassium channel tetramerisation domai | 97.175 | 127.46 | 100.62 | 76.154 | 29.447 | 49.009 |
| 37032 | ENSMUSG00000098558 | 0 | NA | NA | NA | NA | NA | Mir7049 | microRNA 7049 [Source:MGI Symbol;Acc: | 0 | 0 | 0 | 0 | 0 | 0 |
| 37033 | ENSMUSG00000098559 | 0 | NA | NA | NA | NA | NA | Gm15013 | predicted gene 15013 [Source:MGI Symbc | 0 | 0 | 0 | 0 | 0 | 0 |
| 37034 | ENSMUSG00000098560 | 0 | NA | NA | NA | NA | NA | NA | NA | 0 | 0 | 0 | 0 | 0 | 0 |
| 37035 | ENSMUSG00000098561 | 0 | NA | NA | NA | NA | NA | Gtpbp4-ps5 | GTP binding protein 4, pseudogene 5 [Sou | 0 | 0 | 0 | 0 | 0 | 0 |
| 37036 | ENSMUSG00000098562 | 0 | NA | NA | NA | NA | NA | Mir6910 | microRNA 6910 [Source:MGI Symbol;Acc: | 0 | 0 | 0 | 0 | 0 | 0 |
| 37037 | ENSMUSG00000098563 | 0 | NA | NA | NA | NA | NA | Mir6402 | microRNA 6402 [Source:MGI Symbol;Acc: | 0 | 0 | 0 | 0 | 0 | 0 |
| 37038 | ENSMUSG00000098564 | 0 | NA | NA | NA | NA | NA | Ranbp2-ps3 | RAN binding protein 2, pseudogene 3 [Sou | 0 | 0 | 0 | 0 | 0 | 0 |
| 37039 | ENSMUSG00000098565 | 0 | NA | NA | NA | NA | NA | NA | NA | 0 | 0 | 0 | 0 | 0 | 0 |
| 37040 | ENSMUSG00000098566 | 0 | NA | NA | NA | NA | NA | NA | NA | 0 | 0 | 0 | 0 | 0 | 0 |
| 37041 | ENSMUSG00000098567 | 0 | NA | NA | NA | NA | NA | NA | NA | 0 | 0 | 0 | 0 | 0 | 0 |
| 37042 | ENSMUSG00000098568 | 0 | NA | NA | NA | NA | NA | Mir6987 | microRNA 6987 [Source:MGI Symbol;Acc: | 0 | 0 | 0 | 0 | 0 | 0 |
| 37043 | ENSMUSG00000098569 | 0.631240176 | 0.380643921 | 3.15447101 | 0.120668068 | 0.903953952 | NA | Mir7020 | microRNA 7020 [Source:MGI Symbol;Acc: | 1.6062 | 0 | 0 | 0 | 2.1812 | 0 |
| 37044 | ENSMUSG00000098570 | 0 | NA | NA | NA | NA | NA | NA | NA | 0 | 0 | 0 | 0 | 0 | 0 |
| 37045 | ENSMUSG00000098572 | 0 | NA | NA | NA | NA | NA | NA | NA | 0 | 0 | 0 | 0 | 0 | 0 |
| 37046 | ENSMUSG00000098573 | 0 | NA | NA | NA | NA | NA | Gm27232 | predicted gene 27232 [Source:MGI Symbc | 0 | 0 | 0 | 0 | 0 | 0 |
| 37047 | ENSMUSG00000098574 | 0 | NA | NA | NA | NA | NA | Mir7037 | microRNA 7037 [Source:MGI Symbol;Acc: | 0 | 0 | 0 | 0 | 0 | 0 |
| 37048 | ENSMUSG00000098575 | 0 | NA | NA | NA | NA | NA | NA | NA | 0 | 0 | 0 | 0 | 0 | 0 |
| 37049 | ENSMUSG00000098577 | 0 | NA | NA | NA | NA | NA | NA | NA | 0 | 0 | 0 | 0 | 0 | 0 |
| 37050 | ENSMUSG00000098578 | 0 | NA | NA | NA | NA | NA | Gm27449 | predicted gene, 27449 [Source:MGI Symb | 0 | 0 | 0 | 0 | 0 | 0 |
| 37051 | ENSMUSG00000098579 | 0.80281854 | -0.292476728 | 2.419692615 | -0.120873505 | 0.903791228 | NA | Mir6937 | microRNA 6937 [Source:MGI Symbol;Acc: | 1.6062 | 0 | 1.0062 | 0 | 1.0906 | 1.1138 |
| 37052 | ENSMUSG00000098580 | 0 | NA | NA | NA | NA | NA | Mir7023 | microRNA 7023 [Source:MGI Symbol;Acc: | 0 | 0 | 0 | 0 | 0 | 0 |
| 37053 | ENSMUSG00000098582 | 0 | NA | NA | NA | NA | NA | Mir7080 | microRNA 7080 [Source:MGI Symbol;Acc: | 0 | 0 | 0 | 0 | 0 | 0 |
| 37054 | ENSMUSG00000098583 | 0 | NA | NA | NA | NA | NA | Mir7044 | microRNA 7044 [Source:MGI Symbol;Acc: | 0 | 0 | 0 | 0 | 0 | 0 |
| 37055 | ENSMUSG00000098585 | 0 | NA | NA | NA | NA | NA | NA | NA | 0 | 0 | 0 | 0 | 0 | 0 |
| 37056 | ENSMUSG00000098586 | 0 | NA | NA | NA | NA | NA | NA | NA | 0 | 0 | 0 | 0 | 0 | 0 |
| 37057 | ENSMUSG00000098588 | 0 | NA | NA | NA | NA | NA | Mir6340 | microRNA 6340 [Source:MGI Symbol;Acc: | 0 | 0 | 0 | 0 | 0 | 0 |
| 37058 | ENSMUSG00000098589 | 0 | NA | NA | NA | NA | NA | NA | NA | 0 | 0 | 0 | 0 | 0 | 0 |
| 37059 | ENSMUSG00000098590 | 0 | NA | NA | NA | NA | NA | Gm1113 | predicted gene 1113 [Source:MGI Symbol | 0 | 0 | 0 | 0 | 0 | 0 |
| 37060 | ENSMUSG00000098591 | 0 | NA | NA | NA | NA | NA | NA | NA | 0 | 0 | 0 | 0 | 0 | 0 |
| 37061 | ENSMUSG00000098592 | 0.904646783 | 0.28560953 | 2.491718272 | 0.114623524 | 0.908743533 | NA | Mir6917 | microRNA 6917 [Source:MGI Symbol;Acc: | 2.4093 | 0 | 0 | 1.928 | 1.0906 | 0 |
| 37062 | ENSMUSG00000098593 | 0 | NA | NA | NA | NA | NA | Mir7064 | microRNA 7064 [Source:MGI Symbol;Acc: | 0 | 0 | 0 | 0 | 0 | 0 |
| 37063 | ENSMUSG00000098594 | 0 | NA | NA | NA | NA | NA | NA | NA | 0 | 0 | 0 | 0 | 0 | 0 |
| 37064 | ENSMUSG00000098595 | 0 | NA | NA | NA | NA | NA | NA | NA | 0 | 0 | 0 | 0 | 0 | 0 |
| 37065 | ENSMUSG00000098596 | 0 | NA | NA | NA | NA | NA | Mir7226 | microRNA 7226 [Source:MGI Symbol;Acc: | 0 | 0 | 0 | 0 | 0 | 0 |
| 37066 | ENSMUSG00000098597 | 0 | NA | NA | NA | NA | NA | Mir6418 | microRNA 6418 [Source:MGI Symbol;Acc: | 0 | 0 | 0 | 0 | 0 | 0 |
| 37067 | ENSMUSG00000098599 | 0 | NA | NA | NA | NA | NA | NA | NA | 0 | 0 | 0 | 0 | 0 | 0 |
| 37068 | ENSMUSG00000098600 | 0.928203237 | 3.416716801 | 3.030141393 | 1.12757669 | 0.259498736 | NA | Mir6929 | microRNA 6929 [Source:MGI Symbol;Acc: | 0 | 0 | 0 | 0 | 0 | 5.5692 |

| | | | | | | | | | | | | | | | |
|---|---|---|---|---|---|---|---|---|---|---|---|---|---|---|---|
| 37069 | ENSMUSG00000098601 | 0 | NA | NA | NA | NA | NA | NA | NA | 0 | 0 | 0 | 0 | 0 | 0 |
| 37070 | ENSMUSG00000098602 | 0.17131964 | -0.796629176 | 4.080472857 | -0.195229623 | 0.84521319 | NA | Mir6946 | microRNA 6946 [Source:MGI Symbol;Acc: | 0 | 1.0279 | 0 | 0 | 0 | 0 |
| 37071 | ENSMUSG00000098603 | 0 | NA | NA | NA | NA | NA | NA | NA | 0 | 0 | 0 | 0 | 0 | 0 |
| 37072 | ENSMUSG00000098604 | 0 | NA | NA | NA | NA | NA | NA | NA | 0 | 0 | 0 | 0 | 0 | 0 |
| 37073 | ENSMUSG00000098606 | 0 | NA | NA | NA | NA | NA | Mir6903 | microRNA 6903 [Source:MGI Symbol;Acc: | 0 | 0 | 0 | 0 | 0 | 0 |
| 37074 | ENSMUSG00000098607 | 0 | NA | NA | NA | NA | NA | Mir6360 | microRNA 6360 [Source:MGI Symbol;Acc: | 0 | 0 | 0 | 0 | 0 | 0 |
| 37075 | ENSMUSG00000098608 | 0 | NA | NA | NA | NA | NA | Mir6981 | microRNA 6981 [Source:MGI Symbol;Acc: | 0 | 0 | 0 | 0 | 0 | 0 |
| 37076 | ENSMUSG00000098609 | 1.696679414 | -0.461026215 | 1.627393494 | -0.283291175 | 0.776953642 | NA | Anxa11os | annexin A11, opposite strand [Source:MG | 0.8031 | 4.1117 | 1.0062 | 0.964 | 2.1812 | 1.1138 |
| 37077 | ENSMUSG00000098611 | 0 | NA | NA | NA | NA | NA | Mir6343 | microRNA 6343 [Source:MGI Symbol;Acc: | 0 | 0 | 0 | 0 | 0 | 0 |
| 37078 | ENSMUSG00000098612 | 0 | NA | NA | NA | NA | NA | Mir6241 | microRNA 6241 [Source:MGI Symbol;Acc: | 0 | 0 | 0 | 0 | 0 | 0 |
| 37079 | ENSMUSG00000098613 | 0 | NA | NA | NA | NA | NA | Mir6931 | microRNA 6931 [Source:MGI Symbol;Acc: | 0 | 0 | 0 | 0 | 0 | 0 |
| 37080 | ENSMUSG00000098614 | 0 | NA | NA | NA | NA | NA | NA | NA | 0 | 0 | 0 | 0 | 0 | 0 |
| 37081 | ENSMUSG00000098617 | 0 | NA | NA | NA | NA | NA | NA | NA | 0 | 0 | 0 | 0 | 0 | 0 |
| 37082 | ENSMUSG00000098618 | 0 | NA | NA | NA | NA | NA | NA | NA | 0 | 0 | 0 | 0 | 0 | 0 |
| 37083 | ENSMUSG00000098619 | 0 | NA | NA | NA | NA | NA | Mir6401 | microRNA 6401 [Source:MGI Symbol;Acc: | 0 | 0 | 0 | 0 | 0 | 0 |
| 37084 | ENSMUSG00000098620 | 2.038310417 | 0.087928797 | 1.729680023 | 0.050835297 | 0.959456764 | NA | Gm27209 | predicted gene 27209 [Source:MGI Symbo | 4.8186 | 1.0279 | 0 | 1.928 | 0 | 4.4554 |
| 37085 | ENSMUSG00000098621 | 0 | NA | NA | NA | NA | NA | NA | NA | 0 | 0 | 0 | 0 | 0 | 0 |
| 37086 | ENSMUSG00000098622 | 0 | NA | NA | NA | NA | NA | NA | NA | 0 | 0 | 0 | 0 | 0 | 0 |
| 37087 | ENSMUSG00000098623 | 0 | NA | NA | NA | NA | NA | Mir8112 | microRNA 8112 [Source:MGI Symbol;Acc: | 0 | 0 | 0 | 0 | 0 | 0 |
| 37088 | ENSMUSG00000098624 | 0 | NA | NA | NA | NA | NA | Mir7036 | microRNA 7036 [Source:MGI Symbol;Acc: | 0 | 0 | 0 | 0 | 0 | 0 |
| 37089 | ENSMUSG00000098625 | 0 | NA | NA | NA | NA | NA | Mir6953 | microRNA 6953 [Source:MGI Symbol;Acc: | 0 | 0 | 0 | 0 | 0 | 0 |
| 37090 | ENSMUSG00000098626 | 0 | NA | NA | NA | NA | NA | NA | NA | 0 | 0 | 0 | 0 | 0 | 0 |
| 37091 | ENSMUSG00000098627 | 0 | NA | NA | NA | NA | NA | 4930524O08 | RIKEN cDNA 4930524O08 gene [Source:M | 0 | 0 | 0 | 0 | 0 | 0 |
| 37092 | ENSMUSG00000098628 | 0 | NA | NA | NA | NA | NA | NA | NA | 0 | 0 | 0 | 0 | 0 | 0 |
| 37093 | ENSMUSG00000098629 | 0 | NA | NA | NA | NA | NA | NA | NA | 0 | 0 | 0 | 0 | 0 | 0 |
| 37094 | ENSMUSG00000098630 | 0 | NA | NA | NA | NA | NA | Gm27198 | predicted gene 27198 [Source:MGI Symbo | 0 | 0 | 0 | 0 | 0 | 0 |
| 37095 | ENSMUSG00000098631 | 0 | NA | NA | NA | NA | NA | Mir6957 | microRNA 6957 [Source:MGI Symbol;Acc: | 0 | 0 | 0 | 0 | 0 | 0 |
| 37096 | ENSMUSG00000098632 | 0 | NA | NA | NA | NA | NA | Mir7665 | microRNA 7665 [Source:MGI Symbol;Acc: | 0 | 0 | 0 | 0 | 0 | 0 |
| 37097 | ENSMUSG00000098634 | 0 | NA | NA | NA | NA | NA | Mir8096 | microRNA 8096 [Source:MGI Symbol;Acc: | 0 | 0 | 0 | 0 | 0 | 0 |
| 37098 | ENSMUSG00000098635 | 0 | NA | NA | NA | NA | NA | NA | NA | 0 | 0 | 0 | 0 | 0 | 0 |
| 37099 | ENSMUSG00000098636 | 0 | NA | NA | NA | NA | NA | NA | NA | 0 | 0 | 0 | 0 | 0 | 0 |
| 37100 | ENSMUSG00000098638 | 0 | NA | NA | NA | NA | NA | NA | NA | 0 | 0 | 0 | 0 | 0 | 0 |
| 37101 | ENSMUSG00000098640 | 0 | NA | NA | NA | NA | NA | Gm14214 | predicted gene 14214 [Source:MGI Symbo | 0 | 0 | 0 | 0 | 0 | 0 |
| 37102 | ENSMUSG00000098641 | 0 | NA | NA | NA | NA | NA | NA | NA | 0 | 0 | 0 | 0 | 0 | 0 |
| 37103 | ENSMUSG00000098642 | 0 | NA | NA | NA | NA | NA | NA | NA | 0 | 0 | 0 | 0 | 0 | 0 |
| 37104 | ENSMUSG00000098644 | 0 | NA | NA | NA | NA | NA | Bsph2-ps | binder of sperm protein homolog 2, pseu | 0 | 0 | 0 | 0 | 0 | 0 |
| 37105 | ENSMUSG00000098645 | 0 | NA | NA | NA | NA | NA | Gm27201 | predicted gene 27201 [Source:MGI Symbo | 0 | 0 | 0 | 0 | 0 | 0 |
| 37106 | ENSMUSG00000098646 | 0 | NA | NA | NA | NA | NA | Mir7084 | microRNA 7084 [Source:MGI Symbol;Acc: | 0 | 0 | 0 | 0 | 0 | 0 |
| 37107 | ENSMUSG00000098647 | 0 | NA | NA | NA | NA | NA | NA | NA | 0 | 0 | 0 | 0 | 0 | 0 |
| 37108 | ENSMUSG00000098648 | 0 | NA | NA | NA | NA | NA | Mir6239 | microRNA 6239 [Source:MGI Symbol;Acc: | 0 | 0 | 0 | 0 | 0 | 0 |
| 37109 | ENSMUSG00000098649 | 0 | NA | NA | NA | NA | NA | Gm27199 | predicted gene 27199 [Source:MGI Symbo | 0 | 0 | 0 | 0 | 0 | 0 |
| 37110 | ENSMUSG00000098650 | 0 | NA | NA | NA | NA | NA | Commd1b | COMM domain containing 1B [Source:MG | 0 | 0 | 0 | 0 | 0 | 0 |
| 37111 | ENSMUSG00000098651 | 0 | NA | NA | NA | NA | NA | Mir6999 | microRNA 6999 [Source:MGI Symbol;Acc: | 0 | 0 | 0 | 0 | 0 | 0 |
| 37112 | ENSMUSG00000098652 | 0 | NA | NA | NA | NA | NA | NA | NA | 0 | 0 | 0 | 0 | 0 | 0 |
| 37113 | ENSMUSG00000098653 | 0 | NA | NA | NA | NA | NA | NA | NA | 0 | 0 | 0 | 0 | 0 | 0 |
| 37114 | ENSMUSG00000098655 | 0 | NA | NA | NA | NA | NA | Mir7011 | microRNA 7011 [Source:MGI Symbol;Acc: | 0 | 0 | 0 | 0 | 0 | 0 |
| 37115 | ENSMUSG00000098656 | 0.294513347 | 0.165143496 | 4.080472857 | 0.040471657 | 0.967717103 | NA | Mir7028 | microRNA 7028 [Source:MGI Symbol;Acc: | 0.8031 | 0 | 0 | 0.964 | 0 | 0 |
| 37116 | ENSMUSG00000098657 | 0 | NA | NA | NA | NA | NA | NA | NA | 0 | 0 | 0 | 0 | 0 | 0 |
| 37117 | ENSMUSG00000098658 | 0.321325949 | 1.911575114 | 4.048267893 | 0.472195804 | 0.636787034 | NA | Mir6338 | microRNA 6338 [Source:MGI Symbol;Acc: | 0 | 0 | 0 | 1.928 | 0 | 0 |
| 37118 | ENSMUSG00000098659 | 0 | NA | NA | NA | NA | NA | 1110015O18 | RIKEN cDNA 1110015O18 gene [Source:M | 0 | 0 | 0 | 0 | 0 | 0 |
| 37119 | ENSMUSG00000098660 | 0 | NA | NA | NA | NA | NA | Mir6336 | microRNA 6336 [Source:MGI Symbol;Acc: | 0 | 0 | 0 | 0 | 0 | 0 |
| 37120 | ENSMUSG00000098661 | 3.782910827 | -0.103391 | 1.183611662 | -0.08735213 | 0.930391618 | NA | Mir7052 | microRNA 7052 [Source:MGI Symbol;Acc: | 1.6062 | 4.1117 | 6.0375 | 0.964 | 2.1812 | 7.7969 |
| 37121 | ENSMUSG00000098662 | 0 | NA | NA | NA | NA | NA | NA | NA | 0 | 0 | 0 | 0 | 0 | 0 |
| 37122 | ENSMUSG00000098663 | 0 | NA | NA | NA | NA | NA | Mir6958 | microRNA 6958 [Source:MGI Symbol;Acc: | 0 | 0 | 0 | 0 | 0 | 0 |
| 37123 | ENSMUSG00000098664 | 0 | NA | NA | NA | NA | NA | NA | NA | 0 | 0 | 0 | 0 | 0 | 0 |
| 37124 | ENSMUSG00000098665 | 0 | NA | NA | NA | NA | NA | NA | NA | 0 | 0 | 0 | 0 | 0 | 0 |

| | | | | | | | | | | | | | | | |
|---|---|---|---|---|---|---|---|---|---|---|---|---|---|---|---|
| 37125 | ENSMUSG00000098666 | 0 | NA | NA | NA | NA | NA | Mir6950 | microRNA 6950 [Source:MGI Symbol;Acc: | 0 | 0 | 0 | 0 | 0 | 0 |
| 37126 | ENSMUSG00000098667 | 0 | NA | NA | NA | NA | NA | NA | NA | 0 | 0 | 0 | 0 | 0 | 0 |
| 37127 | ENSMUSG00000098668 | 0 | NA | NA | NA | NA | NA | Gm27210 | predicted gene 27210 [Source:MGI Symbo | 0 | 0 | 0 | 0 | 0 | 0 |
| 37128 | ENSMUSG00000098669 | 0 | NA | NA | NA | NA | NA | NA | NA | 0 | 0 | 0 | 0 | 0 | 0 |
| 37129 | ENSMUSG00000098670 | 0.185640647 | 1.126924175 | 4.080472857 | 0.276174898 | 0.782413736 | NA | NA | NA | 0 | 0 | 0 | 0 | 0 | 1.1138 |
| 37130 | ENSMUSG00000098671 | 0 | NA | NA | NA | NA | NA | Mir7075 | microRNA 7075 [Source:MGI Symbol;Acc: | 0 | 0 | 0 | 0 | 0 | 0 |
| 37131 | ENSMUSG00000098672 | 0.781711799 | -0.350673833 | 2.438876442 | -0.143784994 | 0.885670253 | NA | Mir6911 | microRNA 6911 [Source:MGI Symbol;Acc: | 1.6062 | 0 | 1.0062 | 0.964 | 0 | 1.1138 |
| 37132 | ENSMUSG00000098674 | 0.524202404 | 2.600864256 | 3.301637836 | 0.787749713 | 0.430843121 | NA | NA | NA | 0 | 0 | 0 | 0.964 | 2.1812 | 0 |
| 37133 | ENSMUSG00000098675 | 0 | NA | NA | NA | NA | NA | Mir6364 | microRNA 6364 [Source:MGI Symbol;Acc: | 0 | 0 | 0 | 0 | 0 | 0 |
| 37134 | ENSMUSG00000098676 | 0 | NA | NA | NA | NA | NA | Mir6359 | microRNA 6359 [Source:MGI Symbol;Acc: | 0 | 0 | 0 | 0 | 0 | 0 |
| 37135 | ENSMUSG00000098677 | 0 | NA | NA | NA | NA | NA | NA | NA | 0 | 0 | 0 | 0 | 0 | 0 |
| 37136 | ENSMUSG00000098678 | 11.96026602 | -0.058848374 | 0.642383391 | -0.091609427 | 0.927008361 | 0.987712228 | Mroh6 | maestro heat-like repeat family member 6 | 7.2279 | 14.391 | 15.094 | 9.6398 | 9.8156 | 15.594 |
| 37137 | ENSMUSG00000098679 | 0.185640647 | 1.126924175 | 4.080472857 | 0.276174898 | 0.782413736 | NA | Mir7688 | microRNA 7688 [Source:MGI Symbol;Acc: | 0 | 0 | 0 | 0 | 0 | 1.1138 |
| 37138 | ENSMUSG00000098680 | 0 | NA | NA | NA | NA | NA | Mir6898 | microRNA 6898 [Source:MGI Symbol;Acc: | 0 | 0 | 0 | 0 | 0 | 0 |
| 37139 | ENSMUSG00000098682 | 0 | NA | NA | NA | NA | NA | Otx2os1 | orthodenticle homeobox 2 opposite stran | 0 | 0 | 0 | 0 | 0 | 0 |
| 37140 | ENSMUSG00000098683 | 0 | NA | NA | NA | NA | NA | NA | NA | 0 | 0 | 0 | 0 | 0 | 0 |
| 37141 | ENSMUSG00000098684 | 0 | NA | NA | NA | NA | NA | Gm27246 | predicted gene 27246 [Source:MGI Symbo | 0 | 0 | 0 | 0 | 0 | 0 |
| 37142 | ENSMUSG00000098685 | 0 | NA | NA | NA | NA | NA | Gm27224 | predicted gene 27224 [Source:MGI Symbo | 0 | 0 | 0 | 0 | 0 | 0 |
| 37143 | ENSMUSG00000098686 | 0 | NA | NA | NA | NA | NA | Mir6973b | microRNA 6973b [Source:MGI Symbol;Acc | 0 | 0 | 0 | 0 | 0 | 0 |
| 37144 | ENSMUSG00000098687 | 0 | NA | NA | NA | NA | NA | Mir7k | microRNA 7k [Source:MGI Symbol;Acc:MG | 0 | 0 | 0 | 0 | 0 | 0 |
| 37145 | ENSMUSG00000098688 | 0 | NA | NA | NA | NA | NA | NA | NA | 0 | 0 | 0 | 0 | 0 | 0 |
| 37146 | ENSMUSG00000098689 | 0 | NA | NA | NA | NA | NA | Mir6345 | microRNA 6345 [Source:MGI Symbol;Acc: | 0 | 0 | 0 | 0 | 0 | 0 |
| 37147 | ENSMUSG00000098690 | 0 | NA | NA | NA | NA | NA | NA | NA | 0 | 0 | 0 | 0 | 0 | 0 |
| 37148 | ENSMUSG00000098691 | 0 | NA | NA | NA | NA | NA | NA | NA | 0 | 0 | 0 | 0 | 0 | 0 |
| 37149 | ENSMUSG00000098692 | 0 | NA | NA | NA | NA | NA | Gm28041 | predicted gene, 28041 [Source:MGI Symb | 0 | 0 | 0 | 0 | 0 | 0 |
| 37150 | ENSMUSG00000098693 | 0 | NA | NA | NA | NA | NA | NA | NA | 0 | 0 | 0 | 0 | 0 | 0 |
| 37151 | ENSMUSG00000098694 | 0 | NA | NA | NA | NA | NA | NA | NA | 0 | 0 | 0 | 0 | 0 | 0 |
| 37152 | ENSMUSG00000098695 | 0 | NA | NA | NA | NA | NA | NA | NA | 0 | 0 | 0 | 0 | 0 | 0 |
| 37153 | ENSMUSG00000098696 | 0 | NA | NA | NA | NA | NA | NA | NA | 0 | 0 | 0 | 0 | 0 | 0 |
| 37154 | ENSMUSG00000098697 | 0 | NA | NA | NA | NA | NA | NA | NA | 0 | 0 | 0 | 0 | 0 | 0 |
| 37155 | ENSMUSG00000098698 | 0 | NA | NA | NA | NA | NA | Mir6960 | microRNA 6960 [Source:MGI Symbol;Acc: | 0 | 0 | 0 | 0 | 0 | 0 |
| 37156 | ENSMUSG00000098700 | 0 | NA | NA | NA | NA | NA | NA | NA | 0 | 0 | 0 | 0 | 0 | 0 |
| 37157 | ENSMUSG00000098701 | 0 | NA | NA | NA | NA | NA | Mir6914 | microRNA 6914 [Source:MGI Symbol;Acc: | 0 | 0 | 0 | 0 | 0 | 0 |
| 37158 | ENSMUSG00000098702 | 92.5467995 | 2.572436817 | 0.32545908 | 7.90402534 | 2.70E-15 | 1.86E-12 | 1500015A07 | RIKEN cDNA 1500015A07 gene [Source:M | 28.912 | 31.865 | 19.119 | 215.93 | 107.97 | 151.48 |
| 37159 | ENSMUSG00000098703 | 0 | NA | NA | NA | NA | NA | NA | NA | 0 | 0 | 0 | 0 | 0 | 0 |
| 37160 | ENSMUSG00000098704 | 0 | NA | NA | NA | NA | NA | Mir6934 | microRNA 6934 [Source:MGI Symbol;Acc: | 0 | 0 | 0 | 0 | 0 | 0 |
| 37161 | ENSMUSG00000098705 | 0 | NA | NA | NA | NA | NA | Gm3428 | predicted gene 3428 [Source:MGI Symbol | 0 | 0 | 0 | 0 | 0 | 0 |
| 37162 | ENSMUSG00000098706 | 0.667397476 | -1.404797968 | 2.804497115 | -0.500909044 | 0.616435137 | NA | Mir6979 | microRNA 6979 [Source:MGI Symbol;Acc: | 0 | 1.0279 | 2.0125 | 0.964 | 0 | 0 |
| 37163 | ENSMUSG00000098707 | 0 | NA | NA | NA | NA | NA | Gm27236 | predicted gene 27236 [Source:MGI Symbo | 0 | 0 | 0 | 0 | 0 | 0 |
| 37164 | ENSMUSG00000098708 | 86.86521499 | -0.407380509 | 0.267703075 | -1.521762529 | 0.128068593 | 0.524051388 | Gm27252 | predicted gene 27252 [Source:MGI Symbo | 83.523 | 119.24 | 94.587 | 66.514 | 91.612 | 65.717 |
| 37165 | ENSMUSG00000098709 | 0 | NA | NA | NA | NA | NA | NA | NA | 0 | 0 | 0 | 0 | 0 | 0 |
| 37166 | ENSMUSG00000098710 | 0 | NA | NA | NA | NA | NA | Mir7067 | microRNA 7067 [Source:MGI Symbol;Acc: | 0 | 0 | 0 | 0 | 0 | 0 |
| 37167 | ENSMUSG00000098711 | 0 | NA | NA | NA | NA | NA | NA | NA | 0 | 0 | 0 | 0 | 0 | 0 |
| 37168 | ENSMUSG00000098712 | 0 | NA | NA | NA | NA | NA | Mir7010 | microRNA 7010 [Source:MGI Symbol;Acc: | 0 | 0 | 0 | 0 | 0 | 0 |
| 37169 | ENSMUSG00000098713 | 0 | NA | NA | NA | NA | NA | Rps2-ps5 | ribosomal protein S2, pseudogene 5 [Sour | 0 | 0 | 0 | 0 | 0 | 0 |
| 37170 | ENSMUSG00000098714 | 0.133850373 | -0.796629176 | 4.080472857 | -0.195229623 | 0.84521319 | NA | Mir7663 | microRNA 7663 [Source:MGI Symbol;Acc: | 0.8031 | 0 | 0 | 0 | 0 | 0 |
| 37171 | ENSMUSG00000098715 | 0.556921942 | 2.679043159 | 3.638243063 | 0.736356289 | 0.461513899 | NA | Gm28053 | predicted gene, 28053 [Source:MGI Symb | 0 | 0 | 0 | 0 | 0 | 3.3415 |
| 37172 | ENSMUSG00000098716 | 0 | NA | NA | NA | NA | NA | NA | NA | 0 | 0 | 0 | 0 | 0 | 0 |
| 37173 | ENSMUSG00000098717 | 0 | NA | NA | NA | NA | NA | NA | NA | 0 | 0 | 0 | 0 | 0 | 0 |
| 37174 | ENSMUSG00000098718 | 0 | NA | NA | NA | NA | NA | Mir496b | microRNA 496b [Source:MGI Symbol;Acc: | 0 | 0 | 0 | 0 | 0 | 0 |
| 37175 | ENSMUSG00000098719 | 0.321325949 | 1.911575114 | 4.048267893 | 0.472195804 | 0.636787034 | NA | Mir7212 | microRNA 7212 [Source:MGI Symbol;Acc: | 0 | 0 | 0 | 1.928 | 0 | 0 |
| 37176 | ENSMUSG00000098720 | 0 | NA | NA | NA | NA | NA | Gm27239 | predicted gene 27239 [Source:MGI Symbo | 0 | 0 | 0 | 0 | 0 | 0 |
| 37177 | ENSMUSG00000098721 | 0 | NA | NA | NA | NA | NA | Mir6357 | microRNA 6357 [Source:MGI Symbol;Acc: | 0 | 0 | 0 | 0 | 0 | 0 |
| 37178 | ENSMUSG00000098722 | 0 | NA | NA | NA | NA | NA | NA | NA | 0 | 0 | 0 | 0 | 0 | 0 |
| 37179 | ENSMUSG00000098723 | 0 | NA | NA | NA | NA | NA | NA | NA | 0 | 0 | 0 | 0 | 0 | 0 |
| 37180 | ENSMUSG00000098725 | 0 | NA | NA | NA | NA | NA | NA | NA | 0 | 0 | 0 | 0 | 0 | 0 |

| | | | | | | | | | | | | | | | |
|---|---|---|---|---|---|---|---|---|---|---|---|---|---|---|---|
| 37181 | ENSMUSG00000098726 | 0 | NA | NA | NA | NA | NA | Mir6417 | microRNA 6417 [Source:MGI Symbol;Acc: | 0 | 0 | 0 | 0 | 0 | 0 |
| 37182 | ENSMUSG00000098727 | 0 | NA | NA | NA | NA | NA | 1700017I07F | RIKEN cDNA 1700017I07 gene [Source:MC | 0 | 0 | 0 | 0 | 0 | 0 |
| 37183 | ENSMUSG00000098728 | 0 | NA | NA | NA | NA | NA | NA | NA | 0 | 0 | 0 | 0 | 0 | 0 |
| 37184 | ENSMUSG00000098729 | 0.17131964 | -0.796629176 | 4.080472857 | -0.195229623 | 0.84521319 | NA | Gm27200 | predicted gene 27200 [Source:MGI Symbo | 0 | 1.0279 | 0 | 0 | 0 | 0 |
| 37185 | ENSMUSG00000098730 | 0.160662974 | 1.126924175 | 4.080472857 | 0.276174898 | 0.782413736 | NA | Mir6407 | microRNA 6407 [Source:MGI Symbol;Acc: | 0 | 0 | 0 | 0.964 | 0 | 0 |
| 37186 | ENSMUSG00000098731 | 1.470731809 | -0.289886275 | 2.127580163 | -0.136251634 | 0.891622357 | NA | Mir7027 | microRNA 7027 [Source:MGI Symbol;Acc: | 4.8186 | 0 | 0 | 2.8919 | 0 | 1.1138 |
| 37187 | ENSMUSG00000098732 | 0 | NA | NA | NA | NA | NA | NA | NA | 0 | 0 | 0 | 0 | 0 | 0 |
| 37188 | ENSMUSG00000098733 | 0.160662974 | 1.126924175 | 4.080472857 | 0.276174898 | 0.782413736 | NA | NA | NA | 0 | 0 | 0 | 0.964 | 0 | 0 |
| 37189 | ENSMUSG00000098734 | 0 | NA | NA | NA | NA | NA | Mir6335 | microRNA 6335 [Source:MGI Symbol;Acc: | 0 | 0 | 0 | 0 | 0 | 0 |
| 37190 | ENSMUSG00000098735 | 0 | NA | NA | NA | NA | NA | Mir7005 | microRNA 7005 [Source:MGI Symbol;Acc: | 0 | 0 | 0 | 0 | 0 | 0 |
| 37191 | ENSMUSG00000098736 | 0 | NA | NA | NA | NA | NA | NA | NA | 0 | 0 | 0 | 0 | 0 | 0 |
| 37192 | ENSMUSG00000098737 | 0 | NA | NA | NA | NA | NA | Mir7014 | microRNA 7014 [Source:MGI Symbol;Acc: | 0 | 0 | 0 | 0 | 0 | 0 |
| 37193 | ENSMUSG00000098739 | 0 | NA | NA | NA | NA | NA | Gm27151 | predicted gene 27151 [Source:MGI Symbo | 0 | 0 | 0 | 0 | 0 | 0 |
| 37194 | ENSMUSG00000098740 | 0 | NA | NA | NA | NA | NA | NA | NA | 0 | 0 | 0 | 0 | 0 | 0 |
| 37195 | ENSMUSG00000098741 | 18.77819025 | 1.497815278 | 0.6440446 | 2.325639058 | 0.020037814 | 0.173442151 | Obox4-ps2 | oocyte specific homeobox 4, pseudogene | 11.243 | 6.1675 | 12.075 | 51.091 | 9.8156 | 22.277 |
| 37196 | ENSMUSG00000098742 | 0 | NA | NA | NA | NA | NA | NA | NA | 0 | 0 | 0 | 0 | 0 | 0 |
| 37197 | ENSMUSG00000098743 | 0 | NA | NA | NA | NA | NA | NA | NA | 0 | 0 | 0 | 0 | 0 | 0 |
| 37198 | ENSMUSG00000098744 | 0 | NA | NA | NA | NA | NA | Ranbp2-ps7 | RAN binding protein 2, pseudogene 7 [Sou | 0 | 0 | 0 | 0 | 0 | 0 |
| 37199 | ENSMUSG00000098745 | 0 | NA | NA | NA | NA | NA | NA | NA | 0 | 0 | 0 | 0 | 0 | 0 |
| 37200 | ENSMUSG00000098747 | 231.2913931 | -0.075479873 | 0.175347498 | -0.430458798 | 0.666861932 | 0.929596279 | Gm27216 | predicted gene 27216 [Source:MGI Symbo | 274.66 | 221 | 215.34 | 208.22 | 227.94 | 240.59 |
| 37201 | ENSMUSG00000098748 | 0 | NA | NA | NA | NA | NA | Gm27189 | predicted gene 27189 [Source:MGI Symbo | 0 | 0 | 0 | 0 | 0 | 0 |
| 37202 | ENSMUSG00000098749 | 1.183004194 | -1.310594698 | 2.15604226 | -0.607870598 | 0.543273301 | NA | NA | NA | 4.0155 | 1.0279 | 0 | 0.964 | 1.0906 | 0 |
| 37203 | ENSMUSG00000098750 | 0 | NA | NA | NA | NA | NA | NA | NA | 0 | 0 | 0 | 0 | 0 | 0 |
| 37204 | ENSMUSG00000098751 | 0 | NA | NA | NA | NA | NA | Mir7672 | microRNA 7672 [Source:MGI Symbol;Acc: | 0 | 0 | 0 | 0 | 0 | 0 |
| 37205 | ENSMUSG00000098752 | 0 | NA | NA | NA | NA | NA | Mir6897 | microRNA 6897 [Source:MGI Symbol;Acc: | 0 | 0 | 0 | 0 | 0 | 0 |
| 37206 | ENSMUSG00000098754 | 0 | NA | NA | NA | NA | NA | Prn | prion protein readthrough transcript [Sou | 0 | 0 | 0 | 0 | 0 | 0 |
| 37207 | ENSMUSG00000098755 | 1.491322756 | -0.799032454 | 1.782155459 | -0.448351714 | 0.653899384 | NA | NA | NA | 1.6062 | 1.0279 | 3.0187 | 0 | 2.1812 | 1.1138 |
| 37208 | ENSMUSG00000098756 | 0 | NA | NA | NA | NA | NA | Mir378d | microRNA 378d [Source:MGI Symbol;Acc: | 0 | 0 | 0 | 0 | 0 | 0 |
| 37209 | ENSMUSG00000098757 | 0 | NA | NA | NA | NA | NA | NA | NA | 0 | 0 | 0 | 0 | 0 | 0 |
| 37210 | ENSMUSG00000098758 | 0 | NA | NA | NA | NA | NA | 7630403G23 | RIKEN cDNA 7630403G23 gene [Source:M | 0 | 0 | 0 | 0 | 0 | 0 |
| 37211 | ENSMUSG00000098759 | 0.331982614 | 0.16514309 | 3.986751476 | 0.041422971 | 0.966958701 | NA | Mir7063 | microRNA 7063 [Source:MGI Symbol;Acc: | 0 | 1.0279 | 0 | 0.964 | 0 | 0 |
| 37212 | ENSMUSG00000098760 | 0 | NA | NA | NA | NA | NA | Gm2164 | predicted gene 2164 [Source:MGI Symbol | 0 | 0 | 0 | 0 | 0 | 0 |
| 37213 | ENSMUSG00000098761 | 0 | NA | NA | NA | NA | NA | Gm18821 | predicted gene, 18821 [Source:MGI Symb | 0 | 0 | 0 | 0 | 0 | 0 |
| 37214 | ENSMUSG00000098762 | 0 | NA | NA | NA | NA | NA | Mir7045 | microRNA 7045 [Source:MGI Symbol;Acc: | 0 | 0 | 0 | 0 | 0 | 0 |
| 37215 | ENSMUSG00000098763 | 0 | NA | NA | NA | NA | NA | NA | NA | 0 | 0 | 0 | 0 | 0 | 0 |
| 37216 | ENSMUSG00000098764 | 0 | NA | NA | NA | NA | NA | Mir7667 | microRNA 7667 [Source:MGI Symbol;Acc: | 0 | 0 | 0 | 0 | 0 | 0 |
| 37217 | ENSMUSG00000098765 | 0 | NA | NA | NA | NA | NA | Mir7685 | microRNA 7685 [Source:MGI Symbol;Acc: | 0 | 0 | 0 | 0 | 0 | 0 |
| 37218 | ENSMUSG00000098766 | 1.28271381 | 0.267922475 | 1.87956843 | 0.142544677 | 0.886649792 | NA | Mir6386 | microRNA 6386 [Source:MGI Symbol;Acc: | 2.4093 | 1.0279 | 0 | 0.964 | 2.1812 | 1.1138 |
| 37219 | ENSMUSG00000098767 | 0 | NA | NA | NA | NA | NA | NA | NA | 0 | 0 | 0 | 0 | 0 | 0 |
| 37220 | ENSMUSG00000098768 | 0 | NA | NA | NA | NA | NA | Gm27207 | predicted gene 27207 [Source:MGI Symbo | 0 | 0 | 0 | 0 | 0 | 0 |
| 37221 | ENSMUSG00000098769 | 0.160662974 | 1.126924175 | 4.080472857 | 0.276174898 | 0.782413736 | NA | Mir8103 | microRNA 8103 [Source:MGI Symbol;Acc: | 0 | 0 | 0 | 0.964 | 0 | 0 |
| 37222 | ENSMUSG00000098770 | 0 | NA | NA | NA | NA | NA | NA | NA | 0 | 0 | 0 | 0 | 0 | 0 |
| 37223 | ENSMUSG00000098771 | 0 | NA | NA | NA | NA | NA | Mir6538 | microRNA 6538 [Source:MGI Symbol;Acc: | 0 | 0 | 0 | 0 | 0 | 0 |
| 37224 | ENSMUSG00000098772 | 0 | NA | NA | NA | NA | NA | NA | NA | 0 | 0 | 0 | 0 | 0 | 0 |
| 37225 | ENSMUSG00000098773 | 0 | NA | NA | NA | NA | NA | Gm27179 | predicted gene 27179 [Source:MGI Symbo | 0 | 0 | 0 | 0 | 0 | 0 |
| 37226 | ENSMUSG00000098774 | 0 | NA | NA | NA | NA | NA | Mir6930 | microRNA 6930 [Source:MGI Symbol;Acc: | 0 | 0 | 0 | 0 | 0 | 0 |
| 37227 | ENSMUSG00000098775 | 0 | NA | NA | NA | NA | NA | NA | NA | 0 | 0 | 0 | 0 | 0 | 0 |
| 37228 | ENSMUSG00000098776 | 0 | NA | NA | NA | NA | NA | NA | NA | 0 | 0 | 0 | 0 | 0 | 0 |
| 37229 | ENSMUSG00000098777 | 0 | NA | NA | NA | NA | NA | NA | NA | 0 | 0 | 0 | 0 | 0 | 0 |
| 37230 | ENSMUSG00000098778 | 0 | NA | NA | NA | NA | NA | Gm27231 | predicted gene 27231 [Source:MGI Symbo | 0 | 0 | 0 | 0 | 0 | 0 |
| 37231 | ENSMUSG00000098779 | 0 | NA | NA | NA | NA | NA | Gm8602 | predicted gene 8602 [Source:MGI Symbol | 0 | 0 | 0 | 0 | 0 | 0 |
| 37232 | ENSMUSG00000098780 | 0 | NA | NA | NA | NA | NA | Mir6390 | microRNA 6390 [Source:MGI Symbol;Acc: | 0 | 0 | 0 | 0 | 0 | 0 |
| 37233 | ENSMUSG00000098781 | 0.305170013 | -1.657836477 | 4.041480335 | -0.410205257 | 0.681655385 | NA | Gm28044 | predicted gene, 28044 [Source:MGI Symb | 0.8031 | 1.0279 | 0 | 0 | 0 | 0 |
| 37234 | ENSMUSG00000098782 | 0 | NA | NA | NA | NA | NA | Mrps36-ps3 | mitochondrial ribosomal protein S36, pseu | 0 | 0 | 0 | 0 | 0 | 0 |
| 37235 | ENSMUSG00000098783 | 0 | NA | NA | NA | NA | NA | Mir7050 | microRNA 7050 [Source:MGI Symbol;Acc: | 0 | 0 | 0 | 0 | 0 | 0 |
| 37236 | ENSMUSG00000098784 | 0 | NA | NA | NA | NA | NA | Gm27174 | predicted gene 27174 [Source:MGI Symbo | 0 | 0 | 0 | 0 | 0 | 0 |

| | | | | | | | | | | | | | | | |
|---|---|---|---|---|---|---|---|---|---|---|---|---|---|---|---|
| 37237 | ENSMUSG00000098785 | 0.825760208 | 0.783249908 | 2.404539989 | 0.325737942 | 0.744622635 | NA | Mir6395 | microRNA 6395 [Source:MGI Symbol;Acc: | 0.8031 | 0 | 1.0062 | 0.964 | 2.1812 | 0 |
| 37238 | ENSMUSG00000098786 | 0.167707431 | -0.796629176 | 4.080472857 | -0.195229623 | 0.84521319 | NA | Mir6995 | microRNA 6995 [Source:MGI Symbol;Acc: | 0 | 0 | 1.0062 | 0 | 0 | 0 |
| 37239 | ENSMUSG00000098787 | 0 | NA | NA | NA | NA | NA | Mir7088 | microRNA 7088 [Source:MGI Symbol;Acc: | 0 | 0 | 0 | 0 | 0 | 0 |
| 37240 | ENSMUSG00000098788 | 0 | NA | NA | NA | NA | NA | NA | NA | 0 | 0 | 0 | 0 | 0 | 0 |
| 37241 | ENSMUSG00000098789 | 0.517184577 | -0.812115922 | 3.393570547 | -0.239310163 | 0.810865085 | NA | Jmjd7 | jumonji domain containing 7 [Source:MGI | 0 | 0 | 2.0125 | 0 | 1.0906 | 0 |
| 37242 | ENSMUSG00000098790 | 2.540760277 | -1.960537831 | 1.525313484 | -1.285334361 | 0.198675468 | NA | Mir6942 | microRNA 6942 [Source:MGI Symbol;Acc: | 8.031 | 2.0558 | 2.0125 | 0.964 | 2.1812 | 0 |
| 37243 | ENSMUSG00000098791 | 0 | NA | NA | NA | NA | NA | NA | NA | 0 | 0 | 0 | 0 | 0 | 0 |
| 37244 | ENSMUSG00000098792 | 0 | NA | NA | NA | NA | NA | NA | NA | 0 | 0 | 0 | 0 | 0 | 0 |
| 37245 | ENSMUSG00000098793 | 0 | NA | NA | NA | NA | NA | NA | NA | 0 | 0 | 0 | 0 | 0 | 0 |
| 37246 | ENSMUSG00000098794 | 0 | NA | NA | NA | NA | NA | Gm28038 | predicted gene, 28038 [Source:MGI Symb | 0 | 0 | 0 | 0 | 0 | 0 |
| 37247 | ENSMUSG00000098795 | 0 | NA | NA | NA | NA | NA | NA | NA | 0 | 0 | 0 | 0 | 0 | 0 |
| 37248 | ENSMUSG00000098796 | 0 | NA | NA | NA | NA | NA | Gm27163 | predicted gene 27163 [Source:MGI Symbo | 0 | 0 | 0 | 0 | 0 | 0 |
| 37249 | ENSMUSG00000098797 | 0 | NA | NA | NA | NA | NA | Mir6344 | microRNA 6344 [Source:MGI Symbol;Acc: | 0 | 0 | 0 | 0 | 0 | 0 |
| 37250 | ENSMUSG00000098798 | 0 | NA | NA | NA | NA | NA | NA | NA | 0 | 0 | 0 | 0 | 0 | 0 |
| 37251 | ENSMUSG00000098799 | 0 | NA | NA | NA | NA | NA | Trgj4 | T cell receptor gamma joining 4 [Source:M | 0 | 0 | 0 | 0 | 0 | 0 |
| 37252 | ENSMUSG00000098800 | 0.160662974 | 1.126924175 | 4.080472857 | 0.276174898 | 0.782413736 | NA | Gm27244 | predicted gene 27244 [Source:MGI Symbo | 0 | 0 | 0 | 0.964 | 0 | 0 |
| 37253 | ENSMUSG00000098801 | 0 | NA | NA | NA | NA | NA | Gm27255 | predicted gene 27255 [Source:MGI Symbo | 0 | 0 | 0 | 0 | 0 | 0 |
| 37254 | ENSMUSG00000098802 | 0 | NA | NA | NA | NA | NA | Mir7046 | microRNA 7046 [Source:MGI Symbol;Acc: | 0 | 0 | 0 | 0 | 0 | 0 |
| 37255 | ENSMUSG00000098803 | 0 | NA | NA | NA | NA | NA | NA | NA | 0 | 0 | 0 | 0 | 0 | 0 |
| 37256 | ENSMUSG00000098805 | 0 | NA | NA | NA | NA | NA | NA | NA | 0 | 0 | 0 | 0 | 0 | 0 |
| 37257 | ENSMUSG00000098807 | 0 | NA | NA | NA | NA | NA | Mir300 | microRNA 300 [Source:MGI Symbol;Acc:M | 0 | 0 | 0 | 0 | 0 | 0 |
| 37258 | ENSMUSG00000098808 | 0 | NA | NA | NA | NA | NA | Mir7009 | microRNA 7009 [Source:MGI Symbol;Acc: | 0 | 0 | 0 | 0 | 0 | 0 |
| 37259 | ENSMUSG00000098809 | 0 | NA | NA | NA | NA | NA | NA | NA | 0 | 0 | 0 | 0 | 0 | 0 |
| 37260 | ENSMUSG00000098811 | 0.160662974 | 1.126924175 | 4.080472857 | 0.276174898 | 0.782413736 | NA | Mir6913 | microRNA 6913 [Source:MGI Symbol;Acc: | 0 | 0 | 0 | 0.964 | 0 | 0 |
| 37261 | ENSMUSG00000098812 | 0 | NA | NA | NA | NA | NA | Mir7578 | microRNA 7578 [Source:MGI Symbol;Acc: | 0 | 0 | 0 | 0 | 0 | 0 |
| 37262 | ENSMUSG00000098813 | 4.632128497 | 0.400740484 | 0.967916387 | 0.414023865 | 0.678856629 | NA | NA | NA | 4.8186 | 3.0838 | 4.025 | 4.8199 | 4.3625 | 6.6831 |
| 37263 | ENSMUSG00000098814 | 0 | NA | NA | NA | NA | NA | Igkv19-93 | immunoglobulin kappa chain variable 19-9 | 0 | 0 | 0 | 0 | 0 | 0 |
| 37264 | ENSMUSG00000098815 | 0 | NA | NA | NA | NA | NA | NA | NA | 0 | 0 | 0 | 0 | 0 | 0 |
| 37265 | ENSMUSG00000098816 | 0 | NA | NA | NA | NA | NA | NA | NA | 0 | 0 | 0 | 0 | 0 | 0 |
| 37266 | ENSMUSG00000098817 | 0 | NA | NA | NA | NA | NA | Mir6409 | microRNA 6409 [Source:MGI Symbol;Acc: | 0 | 0 | 0 | 0 | 0 | 0 |
| 37267 | ENSMUSG00000098818 | 0 | NA | NA | NA | NA | NA | NA | NA | 0 | 0 | 0 | 0 | 0 | 0 |
| 37268 | ENSMUSG00000098819 | 0 | NA | NA | NA | NA | NA | NA | NA | 0 | 0 | 0 | 0 | 0 | 0 |
| 37269 | ENSMUSG00000098820 | 0 | NA | NA | NA | NA | NA | NA | NA | 0 | 0 | 0 | 0 | 0 | 0 |
| 37270 | ENSMUSG00000098821 | 0 | NA | NA | NA | NA | NA | NA | NA | 0 | 0 | 0 | 0 | 0 | 0 |
| 37271 | ENSMUSG00000098822 | 0.185640647 | 1.126924175 | 4.080472857 | 0.276174898 | 0.782413736 | NA | Mir7039 | microRNA 7039 [Source:MGI Symbol;Acc: | 0 | 0 | 0 | 0 | 0 | 1.1138 |
| 37272 | ENSMUSG00000098823 | 0 | NA | NA | NA | NA | NA | Mir8111 | microRNA 8111 [Source:MGI Symbol;Acc: | 0 | 0 | 0 | 0 | 0 | 0 |
| 37273 | ENSMUSG00000098824 | 0 | NA | NA | NA | NA | NA | Mir7006 | microRNA 7006 [Source:MGI Symbol;Acc: | 0 | 0 | 0 | 0 | 0 | 0 |
| 37274 | ENSMUSG00000098825 | 0 | NA | NA | NA | NA | NA | NA | NA | 0 | 0 | 0 | 0 | 0 | 0 |
| 37275 | ENSMUSG00000098826 | 0 | NA | NA | NA | NA | NA | Mir7677 | microRNA 7677 [Source:MGI Symbol;Acc: | 0 | 0 | 0 | 0 | 0 | 0 |
| 37276 | ENSMUSG00000098827 | 0 | NA | NA | NA | NA | NA | NA | NA | 0 | 0 | 0 | 0 | 0 | 0 |
| 37277 | ENSMUSG00000098828 | 0 | NA | NA | NA | NA | NA | NA | NA | 0 | 0 | 0 | 0 | 0 | 0 |
| 37278 | ENSMUSG00000098829 | 0 | NA | NA | NA | NA | NA | Mir6354 | microRNA 6354 [Source:MGI Symbol;Acc: | 0 | 0 | 0 | 0 | 0 | 0 |
| 37279 | ENSMUSG00000098830 | 0 | NA | NA | NA | NA | NA | Mir7024 | microRNA 7024 [Source:MGI Symbol;Acc: | 0 | 0 | 0 | 0 | 0 | 0 |
| 37280 | ENSMUSG00000098831 | 0 | NA | NA | NA | NA | NA | Mir7119 | microRNA 7119 [Source:MGI Symbol;Acc: | 0 | 0 | 0 | 0 | 0 | 0 |
| 37281 | ENSMUSG00000098832 | 0 | NA | NA | NA | NA | NA | Kdm4dl | lysine (K)-specific demethylase 4D-like [So | 0 | 0 | 0 | 0 | 0 | 0 |
| 37282 | ENSMUSG00000098833 | 0 | NA | NA | NA | NA | NA | Mir6349 | microRNA 6349 [Source:MGI Symbol;Acc: | 0 | 0 | 0 | 0 | 0 | 0 |
| 37283 | ENSMUSG00000098836 | 0 | NA | NA | NA | NA | NA | NA | NA | 0 | 0 | 0 | 0 | 0 | 0 |
| 37284 | ENSMUSG00000098837 | 0 | NA | NA | NA | NA | NA | Gm20111 | predicted gene, 20111 [Source:MGI Symb | 0 | 0 | 0 | 0 | 0 | 0 |
| 37285 | ENSMUSG00000098838 | 0 | NA | NA | NA | NA | NA | Mir381 | microRNA 381 [Source:MGI Symbol;Acc:M | 0 | 0 | 0 | 0 | 0 | 0 |
| 37286 | ENSMUSG00000098839 | 0.321325949 | 1.911575114 | 4.048267893 | 0.472195804 | 0.636787034 | NA | Mir7054 | microRNA 7054 [Source:MGI Symbol;Acc: | 0 | 0 | 0 | 1.928 | 0 | 0 |
| 37287 | ENSMUSG00000098840 | 0 | NA | NA | NA | NA | NA | NA | NA | 0 | 0 | 0 | 0 | 0 | 0 |
| 37288 | ENSMUSG00000098842 | 0 | NA | NA | NA | NA | NA | NA | NA | 0 | 0 | 0 | 0 | 0 | 0 |
| 37289 | ENSMUSG00000098843 | 0 | NA | NA | NA | NA | NA | Gm27240 | predicted gene 27240 [Source:MGI Symbo | 0 | 0 | 0 | 0 | 0 | 0 |
| 37290 | ENSMUSG00000098844 | 0 | NA | NA | NA | NA | NA | Mir6992 | microRNA 6992 [Source:MGI Symbol;Acc: | 0 | 0 | 0 | 0 | 0 | 0 |
| 37291 | ENSMUSG00000098845 | 0 | NA | NA | NA | NA | NA | NA | NA | 0 | 0 | 0 | 0 | 0 | 0 |
| 37292 | ENSMUSG00000098846 | 0 | NA | NA | NA | NA | NA | NA | NA | 0 | 0 | 0 | 0 | 0 | 0 |

| | | | | | | | | | | | | | | | |
|---|---|---|---|---|---|---|---|---|---|---|---|---|---|---|---|
| 37293 | ENSMUSG00000098847 | 0 | NA | NA | NA | NA | NA | Gm27235 | predicted gene 27235 [Source:MGI Symbo | 0 | 0 | 0 | 0 | 0 | 0 |
| 37294 | ENSMUSG00000098848 | 0 | NA | NA | NA | NA | NA | NA | NA | 0 | 0 | 0 | 0 | 0 | 0 |
| 37295 | ENSMUSG00000098849 | 0 | NA | NA | NA | NA | NA | NA | NA | 0 | 0 | 0 | 0 | 0 | 0 |
| 37296 | ENSMUSG00000098850 | 0 | NA | NA | NA | NA | NA | NA | NA | 0 | 0 | 0 | 0 | 0 | 0 |
| 37297 | ENSMUSG00000098851 | 0 | NA | NA | NA | NA | NA | Mir6353 | microRNA 6353 [Source:MGI Symbol;Acc: | 0 | 0 | 0 | 0 | 0 | 0 |
| 37298 | ENSMUSG00000098852 | 0 | NA | NA | NA | NA | NA | NA | NA | 0 | 0 | 0 | 0 | 0 | 0 |
| 37299 | ENSMUSG00000098853 | 0 | NA | NA | NA | NA | NA | Gm27393 | predicted gene, 27393 [Source:MGI Symb | 0 | 0 | 0 | 0 | 0 | 0 |
| 37300 | ENSMUSG00000098854 | 0 | NA | NA | NA | NA | NA | Gm5118 | predicted gene 5118 [Source:MGI Symbol | 0 | 0 | 0 | 0 | 0 | 0 |
| 37301 | ENSMUSG00000098855 | 0 | NA | NA | NA | NA | NA | Mir6923 | microRNA 6923 [Source:MGI Symbol;Acc: | 0 | 0 | 0 | 0 | 0 | 0 |
| 37302 | ENSMUSG00000098856 | 0 | NA | NA | NA | NA | NA | Mir8113 | microRNA 8113 [Source:MGI Symbol;Acc: | 0 | 0 | 0 | 0 | 0 | 0 |
| 37303 | ENSMUSG00000098857 | 0 | NA | NA | NA | NA | NA | NA | NA | 0 | 0 | 0 | 0 | 0 | 0 |
| 37304 | ENSMUSG00000098858 | 0 | NA | NA | NA | NA | NA | Mir6541 | microRNA 6541 [Source:MGI Symbol;Acc: | 0 | 0 | 0 | 0 | 0 | 0 |
| 37305 | ENSMUSG00000098859 | 0 | NA | NA | NA | NA | NA | NA | NA | 0 | 0 | 0 | 0 | 0 | 0 |
| 37306 | ENSMUSG00000098860 | 0 | NA | NA | NA | NA | NA | NA | NA | 0 | 0 | 0 | 0 | 0 | 0 |
| 37307 | ENSMUSG00000098861 | 0 | NA | NA | NA | NA | NA | Ranbp2-ps9 | RAN binding protein 2, pseudogene 9 [Sou | 0 | 0 | 0 | 0 | 0 | 0 |
| 37308 | ENSMUSG00000098862 | 0 | NA | NA | NA | NA | NA | Mir6378 | microRNA 6378 [Source:MGI Symbol;Acc: | 0 | 0 | 0 | 0 | 0 | 0 |
| 37309 | ENSMUSG00000098863 | 0 | NA | NA | NA | NA | NA | Mir8101 | microRNA 8101 [Source:MGI Symbol;Acc: | 0 | 0 | 0 | 0 | 0 | 0 |
| 37310 | ENSMUSG00000098864 | 0.267700746 | -1.497481649 | 4.057001436 | -0.369110456 | 0.712045396 | NA | NA | NA | 1.6062 | 0 | 0 | 0 | 0 | 0 |
| 37311 | ENSMUSG00000098865 | 0 | NA | NA | NA | NA | NA | NA | NA | 0 | 0 | 0 | 0 | 0 | 0 |
| 37312 | ENSMUSG00000098866 | 0 | NA | NA | NA | NA | NA | Mir6539 | microRNA 6539 [Source:MGI Symbol;Acc: | 0 | 0 | 0 | 0 | 0 | 0 |
| 37313 | ENSMUSG00000098867 | 363.6917692 | 0.397693137 | 0.190100895 | 2.092010858 | 0.036437544 | 0.257340157 | NA | NA | 306.79 | 265.2 | 370.3 | 521.51 | 369.72 | 348.63 |
| 37314 | ENSMUSG00000098868 | 0 | NA | NA | NA | NA | NA | NA | NA | 0 | 0 | 0 | 0 | 0 | 0 |
| 37315 | ENSMUSG00000098869 | 0 | NA | NA | NA | NA | NA | NA | NA | 0 | 0 | 0 | 0 | 0 | 0 |
| 37316 | ENSMUSG00000098871 | 0 | NA | NA | NA | NA | NA | Mir6381 | microRNA 6381 [Source:MGI Symbol;Acc: | 0 | 0 | 0 | 0 | 0 | 0 |
| 37317 | ENSMUSG00000098873 | 0 | NA | NA | NA | NA | NA | Mir7668 | microRNA 7668 [Source:MGI Symbol;Acc: | 0 | 0 | 0 | 0 | 0 | 0 |
| 37318 | ENSMUSG00000098874 | 0.133850373 | -0.796629176 | 4.080472857 | -0.195229623 | 0.84521319 | NA | NA | NA | 0.8031 | 0 | 0 | 0 | 0 | 0 |
| 37319 | ENSMUSG00000098875 | 0 | NA | NA | NA | NA | NA | Gm27211 | predicted gene 27211 [Source:MGI Symbo | 0 | 0 | 0 | 0 | 0 | 0 |
| 37320 | ENSMUSG00000098876 | 0 | NA | NA | NA | NA | NA | Mir6350 | microRNA 6350 [Source:MGI Symbol;Acc: | 0 | 0 | 0 | 0 | 0 | 0 |
| 37321 | ENSMUSG00000098877 | 0.181769715 | 1.126924175 | 4.080472857 | 0.276174898 | 0.782413736 | NA | NA | NA | 0 | 0 | 0 | 0 | 1.0906 | 0 |
| 37322 | ENSMUSG00000098878 | 0 | NA | NA | NA | NA | NA | NA | NA | 0 | 0 | 0 | 0 | 0 | 0 |
| 37323 | ENSMUSG00000098879 | 0 | NA | NA | NA | NA | NA | Mir7675 | microRNA 7675 [Source:MGI Symbol;Acc: | 0 | 0 | 0 | 0 | 0 | 0 |
| 37324 | ENSMUSG00000098880 | 0 | NA | NA | NA | NA | NA | NA | NA | 0 | 0 | 0 | 0 | 0 | 0 |
| 37325 | ENSMUSG00000098881 | 0 | NA | NA | NA | NA | NA | Mir6927 | microRNA 6927 [Source:MGI Symbol;Acc: | 0 | 0 | 0 | 0 | 0 | 0 |
| 37326 | ENSMUSG00000098882 | 0.133850373 | -0.796629176 | 4.080472857 | -0.195229623 | 0.84521319 | NA | Mir6392 | microRNA 6392 [Source:MGI Symbol;Acc: | 0.8031 | 0 | 0 | 0 | 0 | 0 |
| 37327 | ENSMUSG00000098883 | 0 | NA | NA | NA | NA | NA | NA | NA | 0 | 0 | 0 | 0 | 0 | 0 |
| 37328 | ENSMUSG00000098884 | 0 | NA | NA | NA | NA | NA | Mir7689 | microRNA 7689 [Source:MGI Symbol;Acc: | 0 | 0 | 0 | 0 | 0 | 0 |
| 37329 | ENSMUSG00000098885 | 0.167707431 | -0.796629176 | 4.080472857 | -0.195229623 | 0.84521319 | NA | Mir7659 | microRNA 7659 [Source:MGI Symbol;Acc: | 0 | 0 | 1.0062 | 0 | 0 | 0 |
| 37330 | ENSMUSG00000098886 | 0 | NA | NA | NA | NA | NA | Mir6385 | microRNA 6385 [Source:MGI Symbol;Acc: | 0 | 0 | 0 | 0 | 0 | 0 |
| 37331 | ENSMUSG00000098887 | 0 | NA | NA | NA | NA | NA | Mir6413 | microRNA 6413 [Source:MGI Symbol;Acc: | 0 | 0 | 0 | 0 | 0 | 0 |
| 37332 | ENSMUSG00000098888 | 0 | NA | NA | NA | NA | NA | Gm18486 | predicted gene, 18486 [Source:MGI Symb | 0 | 0 | 0 | 0 | 0 | 0 |
| 37333 | ENSMUSG00000098889 | 4.527286461 | 0.339886972 | 1.110765093 | 0.305993566 | 0.759609545 | NA | Gm27206 | predicted gene 27206 [Source:MGI Symbo | 8.8341 | 1.0279 | 2.0125 | 8.6758 | 3.2719 | 3.3415 |
| 37334 | ENSMUSG00000098890 | 0 | NA | NA | NA | NA | NA | Mir7008 | microRNA 7008 [Source:MGI Symbol;Acc: | 0 | 0 | 0 | 0 | 0 | 0 |
| 37335 | ENSMUSG00000098891 | 0 | NA | NA | NA | NA | NA | NA | NA | 0 | 0 | 0 | 0 | 0 | 0 |
| 37336 | ENSMUSG00000098893 | 0.465832987 | -0.698295221 | 3.18348759 | -0.219349126 | 0.826378096 | NA | Gm17828 | predicted gene, 17828 [Source:MGI Symb | 0.8031 | 1.0279 | 0 | 0.964 | 0 | 0 |
| 37337 | ENSMUSG00000098894 | 0 | NA | NA | NA | NA | NA | Gm27234 | predicted gene 27234 [Source:MGI Symbo | 0 | 0 | 0 | 0 | 0 | 0 |
| 37338 | ENSMUSG00000098895 | 0.315620088 | 0.165143421 | 4.063300985 | 0.040642675 | 0.967580763 | NA | Gm21013 | predicted gene, 21013 [Source:MGI Symb | 0.8031 | 0 | 0 | 0 | 1.0906 | 0 |
| 37339 | ENSMUSG00000098896 | 1.376981803 | 0.805578381 | 1.97448793 | 0.407993571 | 0.683278395 | NA | Mir3473g | microRNA 3473g [Source:MGI Symbol;Acc | 0 | 2.0558 | 1.0062 | 1.928 | 3.2719 | 0 |
| 37340 | ENSMUSG00000098897 | 0 | NA | NA | NA | NA | NA | Gm19013 | predicted gene, 19013 [Source:MGI Symb | 0 | 0 | 0 | 0 | 0 | 0 |
| 37341 | ENSMUSG00000098898 | 0.167707431 | -0.796629176 | 4.080472857 | -0.195229623 | 0.84521319 | NA | NA | NA | 0 | 0 | 1.0062 | 0 | 0 | 0 |
| 37342 | ENSMUSG00000098899 | 0 | NA | NA | NA | NA | NA | NA | NA | 0 | 0 | 0 | 0 | 0 | 0 |
| 37343 | ENSMUSG00000098900 | 4.592681999 | -0.747284959 | 1.063259749 | -0.70282446 | 0.482165152 | NA | Gm18190 | predicted gene, 18190 [Source:MGI Symb | 3.2124 | 4.1117 | 10.062 | 5.7839 | 3.2719 | 1.1138 |
| 37344 | ENSMUSG00000098901 | 0 | NA | NA | NA | NA | NA | NA | NA | 0 | 0 | 0 | 0 | 0 | 0 |
| 37345 | ENSMUSG00000098903 | 0 | NA | NA | NA | NA | NA | NA | NA | 0 | 0 | 0 | 0 | 0 | 0 |
| 37346 | ENSMUSG00000098904 | 0.160662974 | 1.126924175 | 4.080472857 | 0.276174898 | 0.782413736 | NA | Mir6998 | microRNA 6998 [Source:MGI Symbol;Acc: | 0 | 0 | 0 | 0.964 | 0 | 0 |
| 37347 | ENSMUSG00000098905 | 80.68353889 | 0.289829367 | 0.258091888 | 1.12296969 | 0.261450368 | 0.71553796 | Zfp953 | zinc finger protein 953 [Source:MGI Symb | 63.445 | 77.094 | 77.481 | 82.902 | 80.706 | 102.47 |
| 37348 | ENSMUSG00000098906 | 0 | NA | NA | NA | NA | NA | Mir7652 | microRNA 7652 [Source:MGI Symbol;Acc: | 0 | 0 | 0 | 0 | 0 | 0 |

| | | | | | | | | | | | | | | | |
|---|---|---|---|---|---|---|---|---|---|---|---|---|---|---|---|
| 37349 | ENSMUSG00000098907 | 0 | NA | NA | NA | NA | NA | Mir7082 | microRNA 7082 [Source:MGI Symbol;Acc: | 0 | 0 | 0 | 0 | 0 | 0 |
| 37350 | ENSMUSG00000098908 | 0 | NA | NA | NA | NA | NA | NA | NA | 0 | 0 | 0 | 0 | 0 | 0 |
| 37351 | ENSMUSG00000098909 | 0 | NA | NA | NA | NA | NA | Gm27161 | predicted gene 27161 [Source:MGI Symbo | 0 | 0 | 0 | 0 | 0 | 0 |
| 37352 | ENSMUSG00000098910 | 0 | NA | NA | NA | NA | NA | Mir6416 | microRNA 6416 [Source:MGI Symbol;Acc: | 0 | 0 | 0 | 0 | 0 | 0 |
| 37353 | ENSMUSG00000098912 | 21.29215526 | -0.4170491 | 0.505123473 | -0.825637934 | 0.40900951 | 0.833065722 | 1500004A13 | RIKEN cDNA 1500004A13 gene [Source:M | 22.487 | 23.642 | 27.169 | 27.955 | 10.906 | 15.594 |
| 37354 | ENSMUSG00000098914 | 0 | NA | NA | NA | NA | NA | NA | NA | 0 | 0 | 0 | 0 | 0 | 0 |
| 37355 | ENSMUSG00000098915 | 0.68145976 | 0.058694068 | 2.590797836 | 0.022654824 | 0.981925612 | NA | Rpl15-ps2 | ribosomal protein L15, pseudogene 2 [Sou | 0 | 1.0279 | 1.0062 | 0.964 | 1.0906 | 0 |
| 37356 | ENSMUSG00000098916 | 0 | NA | NA | NA | NA | NA | NA | NA | 0 | 0 | 0 | 0 | 0 | 0 |
| 37357 | ENSMUSG00000098917 | 0 | NA | NA | NA | NA | NA | Mir7654 | microRNA 7654 [Source:MGI Symbol;Acc: | 0 | 0 | 0 | 0 | 0 | 0 |
| 37358 | ENSMUSG00000098918 | 0.695780767 | 1.647560897 | 2.554296336 | 0.645015566 | 0.518917098 | NA | Mir6481 | microRNA 6481 [Source:MGI Symbol;Acc: | 0 | 0 | 1.0062 | 0.964 | 1.0906 | 1.1138 |
| 37359 | ENSMUSG00000098919 | 0.349477146 | 0.165142768 | 3.911021834 | 0.042224967 | 0.966319359 | NA | Mir8102 | microRNA 8102 [Source:MGI Symbol;Acc: | 0 | 0 | 1.0062 | 0 | 1.0906 | 0 |
| 37360 | ENSMUSG00000098920 | 0 | NA | NA | NA | NA | NA | NA | NA | 0 | 0 | 0 | 0 | 0 | 0 |
| 37361 | ENSMUSG00000098922 | 0 | NA | NA | NA | NA | NA | Mir8106 | microRNA 8106 [Source:MGI Symbol;Acc: | 0 | 0 | 0 | 0 | 0 | 0 |
| 37362 | ENSMUSG00000098923 | 166.9807321 | -0.121310538 | 0.203675102 | -0.595608085 | 0.551437074 | 0.89589637 | Tmem185b | transmembrane protein 185B [Source:MG | 193.55 | 169.61 | 157.98 | 129.17 | 175.59 | 175.99 |
| 37363 | ENSMUSG00000098924 | 0 | NA | NA | NA | NA | NA | Gm27245 | predicted gene 27245 [Source:MGI Symbo | 0 | 0 | 0 | 0 | 0 | 0 |
| 37364 | ENSMUSG00000098925 | 0 | NA | NA | NA | NA | NA | NA | NA | 0 | 0 | 0 | 0 | 0 | 0 |
| 37365 | ENSMUSG00000098926 | 0 | NA | NA | NA | NA | NA | Mir6904 | microRNA 6904 [Source:MGI Symbol;Acc: | 0 | 0 | 0 | 0 | 0 | 0 |
| 37366 | ENSMUSG00000098928 | 0 | NA | NA | NA | NA | NA | Mir6988 | microRNA 6988 [Source:MGI Symbol;Acc: | 0 | 0 | 0 | 0 | 0 | 0 |
| 37367 | ENSMUSG00000098929 | 1.38268215 | 1.810549411 | 1.900767638 | 0.952535899 | 0.340825268 | NA | Mir6400 | microRNA 6400 [Source:MGI Symbol;Acc: | 0.8031 | 0 | 1.0062 | 0.964 | 2.1812 | 3.3415 |
| 37368 | ENSMUSG00000098931 | 0 | NA | NA | NA | NA | NA | Mir8099-2 | microRNA 8099-2 [Source:MGI Symbol;Ac | 0 | 0 | 0 | 0 | 0 | 0 |
| 37369 | ENSMUSG00000098932 | 0 | NA | NA | NA | NA | NA | Mir6410 | microRNA 6410 [Source:MGI Symbol;Acc: | 0 | 0 | 0 | 0 | 0 | 0 |
| 37370 | ENSMUSG00000098934 | 0.906682787 | -0.583665192 | 2.768799895 | -0.210800785 | 0.833042726 | NA | Gvin-ps4 | GTPase, very large interferon inducible, p: | 3.2124 | 0 | 0 | 0 | 0 | 2.2277 |
| 37371 | ENSMUSG00000098935 | 0 | NA | NA | NA | NA | NA | Gm27225 | predicted gene 27225 [Source:MGI Symbo | 0 | 0 | 0 | 0 | 0 | 0 |
| 37372 | ENSMUSG00000098937 | 0 | NA | NA | NA | NA | NA | Mir7241 | microRNA 7241 [Source:MGI Symbol;Acc: | 0 | 0 | 0 | 0 | 0 | 0 |
| 37373 | ENSMUSG00000098938 | 0 | NA | NA | NA | NA | NA | Mir6393 | microRNA 6393 [Source:MGI Symbol;Acc: | 0 | 0 | 0 | 0 | 0 | 0 |
| 37374 | ENSMUSG00000098939 | 0 | NA | NA | NA | NA | NA | NA | NA | 0 | 0 | 0 | 0 | 0 | 0 |
| 37375 | ENSMUSG00000098940 | 0 | NA | NA | NA | NA | NA | NA | NA | 0 | 0 | 0 | 0 | 0 | 0 |
| 37376 | ENSMUSG00000098942 | 0.160662974 | 1.126924175 | 4.080472857 | 0.276174898 | 0.782413736 | NA | Mir8100 | microRNA 8100 [Source:MGI Symbol;Acc: | 0 | 0 | 0 | 0.964 | 0 | 0 |
| 37377 | ENSMUSG00000098943 | 0 | NA | NA | NA | NA | NA | NA | NA | 0 | 0 | 0 | 0 | 0 | 0 |
| 37378 | ENSMUSG00000098944 | 0 | NA | NA | NA | NA | NA | Mir8120 | microRNA 8120 [Source:MGI Symbol;Acc: | 0 | 0 | 0 | 0 | 0 | 0 |
| 37379 | ENSMUSG00000098945 | 0 | NA | NA | NA | NA | NA | Gm27150 | predicted gene 27150 [Source:MGI Symbo | 0 | 0 | 0 | 0 | 0 | 0 |
| 37380 | ENSMUSG00000098946 | 0 | NA | NA | NA | NA | NA | NA | NA | 0 | 0 | 0 | 0 | 0 | 0 |
| 37381 | ENSMUSG00000098947 | 0 | NA | NA | NA | NA | NA | V1rg10 | vomeronasal 1 receptor, G10 [Source:MG | 0 | 0 | 0 | 0 | 0 | 0 |
| 37382 | ENSMUSG00000098948 | 0 | NA | NA | NA | NA | NA | NA | NA | 0 | 0 | 0 | 0 | 0 | 0 |
| 37383 | ENSMUSG00000098949 | 0 | NA | NA | NA | NA | NA | Gm27160 | predicted gene 27160 [Source:MGI Symbo | 0 | 0 | 0 | 0 | 0 | 0 |
| 37384 | ENSMUSG00000098950 | 0 | NA | NA | NA | NA | NA | Gm28036 | predicted gene, 28036 [Source:MGI Symb | 0 | 0 | 0 | 0 | 0 | 0 |
| 37385 | ENSMUSG00000098951 | 0.160662974 | 1.126924175 | 4.080472857 | 0.276174898 | 0.782413736 | NA | Mir7647 | microRNA 7647 [Source:MGI Symbol;Acc: | 0 | 0 | 0 | 0.964 | 0 | 0 |
| 37386 | ENSMUSG00000098952 | 0 | NA | NA | NA | NA | NA | Mir6405 | microRNA 6405 [Source:MGI Symbol;Acc: | 0 | 0 | 0 | 0 | 0 | 0 |
| 37387 | ENSMUSG00000098953 | 0 | NA | NA | NA | NA | NA | NA | NA | 0 | 0 | 0 | 0 | 0 | 0 |
| 37388 | ENSMUSG00000098954 | 0 | NA | NA | NA | NA | NA | Mir7658 | microRNA 7658 [Source:MGI Symbol;Acc: | 0 | 0 | 0 | 0 | 0 | 0 |
| 37389 | ENSMUSG00000098955 | 0 | NA | NA | NA | NA | NA | Mir6935 | microRNA 6935 [Source:MGI Symbol;Acc: | 0 | 0 | 0 | 0 | 0 | 0 |
| 37390 | ENSMUSG00000098956 | 0 | NA | NA | NA | NA | NA | NA | NA | 0 | 0 | 0 | 0 | 0 | 0 |
| 37391 | ENSMUSG00000098957 | 0.831492699 | -1.816502146 | 2.887260166 | -0.629143909 | 0.529254847 | NA | Mir8105 | microRNA 8105 [Source:MGI Symbol;Acc: | 0 | 0 | 4.025 | 0.964 | 0 | 0 |
| 37392 | ENSMUSG00000098958 | 0 | NA | NA | NA | NA | NA | Mir6933 | microRNA 6933 [Source:MGI Symbol;Acc: | 0 | 0 | 0 | 0 | 0 | 0 |
| 37393 | ENSMUSG00000098959 | 0 | NA | NA | NA | NA | NA | Gm27230 | predicted gene 27230 [Source:MGI Symbo | 0 | 0 | 0 | 0 | 0 | 0 |
| 37394 | ENSMUSG00000098960 | 0 | NA | NA | NA | NA | NA | Mir7056 | microRNA 7056 [Source:MGI Symbol;Acc: | 0 | 0 | 0 | 0 | 0 | 0 |
| 37395 | ENSMUSG00000098961 | 0 | NA | NA | NA | NA | NA | Mir6949 | microRNA 6949 [Source:MGI Symbol;Acc: | 0 | 0 | 0 | 0 | 0 | 0 |
| 37396 | ENSMUSG00000098962 | 0 | NA | NA | NA | NA | NA | NA | NA | 0 | 0 | 0 | 0 | 0 | 0 |
| 37397 | ENSMUSG00000098963 | 0 | NA | NA | NA | NA | NA | NA | NA | 0 | 0 | 0 | 0 | 0 | 0 |
| 37398 | ENSMUSG00000098964 | 0 | NA | NA | NA | NA | NA | NA | NA | 0 | 0 | 0 | 0 | 0 | 0 |
| 37399 | ENSMUSG00000098965 | 0 | NA | NA | NA | NA | NA | NA | NA | 0 | 0 | 0 | 0 | 0 | 0 |
| 37400 | ENSMUSG00000098966 | 0 | NA | NA | NA | NA | NA | NA | NA | 0 | 0 | 0 | 0 | 0 | 0 |
| 37401 | ENSMUSG00000098967 | 0 | NA | NA | NA | NA | NA | NA | NA | 0 | 0 | 0 | 0 | 0 | 0 |
| 37402 | ENSMUSG00000098968 | 0 | NA | NA | NA | NA | NA | NA | NA | 0 | 0 | 0 | 0 | 0 | 0 |
| 37403 | ENSMUSG00000098969 | 0 | NA | NA | NA | NA | NA | NA | NA | 0 | 0 | 0 | 0 | 0 | 0 |
| 37404 | ENSMUSG00000098970 | 0 | NA | NA | NA | NA | NA | Gm27549 | predicted gene, 27549 [Source:MGI Symb | 0 | 0 | 0 | 0 | 0 | 0 |

| | | | | | | | | | | | | | | | |
|---|---|---|---|---|---|---|---|---|---|---|---|---|---|---|---|
| 37405 | ENSMUSG00000098972 | 0 | NA | NA | NA | NA | NA | Mir8093 | microRNA 8093 [Source:MGI Symbol;Acc: | 0 | 0 | 0 | 0 | 0 | 0 |
| 37406 | ENSMUSG00000098973 | 0 | NA | NA | NA | NA | NA | Mir6236 | microRNA 6236 [Source:MGI Symbol;Acc: | 0 | 0 | 0 | 0 | 0 | 0 |
| 37407 | ENSMUSG00000098974 | 0 | NA | NA | NA | NA | NA | NA | NA | 0 | 0 | 0 | 0 | 0 | 0 |
| 37408 | ENSMUSG00000098975 | 0 | NA | NA | NA | NA | NA | Gm27177 | predicted gene 27177 [Source:MGI Symbo | 0 | 0 | 0 | 0 | 0 | 0 |
| 37409 | ENSMUSG00000098976 | 0 | NA | NA | NA | NA | NA | Mir6769b | microRNA 6769b [Source:MGI Symbol;Acc | 0 | 0 | 0 | 0 | 0 | 0 |
| 37410 | ENSMUSG00000098977 | 0 | NA | NA | NA | NA | NA | Gm27196 | predicted gene 27196 [Source:MGI Symbo | 0 | 0 | 0 | 0 | 0 | 0 |
| 37411 | ENSMUSG00000098978 | 0 | NA | NA | NA | NA | NA | NA | NA | 0 | 0 | 0 | 0 | 0 | 0 |
| 37412 | ENSMUSG00000098979 | 0 | NA | NA | NA | NA | NA | NA | NA | 0 | 0 | 0 | 0 | 0 | 0 |
| 37413 | ENSMUSG00000098980 | 8.851847006 | 1.961061404 | 0.793235217 | 2.472231894 | 0.013427239 | 0.132518282 | Mir6397 | microRNA 6397 [Source:MGI Symbol;Acc: | 5.6217 | 2.0558 | 3.0187 | 19.28 | 11.997 | 11.138 |
| 37414 | ENSMUSG00000098981 | 0 | NA | NA | NA | NA | NA | Mir7236 | microRNA 7236 [Source:MGI Symbol;Acc: | 0 | 0 | 0 | 0 | 0 | 0 |
| 37415 | ENSMUSG00000098982 | 0 | NA | NA | NA | NA | NA | NA | NA | 0 | 0 | 0 | 0 | 0 | 0 |
| 37416 | ENSMUSG00000098983 | 0 | NA | NA | NA | NA | NA | NA | NA | 0 | 0 | 0 | 0 | 0 | 0 |
| 37417 | ENSMUSG00000098984 | 0 | NA | NA | NA | NA | NA | Mir6420 | microRNA 6420 [Source:MGI Symbol;Acc: | 0 | 0 | 0 | 0 | 0 | 0 |
| 37418 | ENSMUSG00000098985 | 0.349477146 | 0.165142768 | 3.911021834 | 0.042224967 | 0.966319359 | NA | Gm27219 | predicted gene 27219 [Source:MGI Symbo | 0 | 0 | 1.0062 | 0 | 1.0906 | 0 |
| 37419 | ENSMUSG00000098986 | 0 | NA | NA | NA | NA | NA | NA | NA | 0 | 0 | 0 | 0 | 0 | 0 |
| 37420 | ENSMUSG00000098987 | 0 | NA | NA | NA | NA | NA | Gm8982 | predicted gene 8982 [Source:MGI Symbol | 0 | 0 | 0 | 0 | 0 | 0 |
| 37421 | ENSMUSG00000098988 | 0 | NA | NA | NA | NA | NA | NA | NA | 0 | 0 | 0 | 0 | 0 | 0 |
| 37422 | ENSMUSG00000098989 | 0 | NA | NA | NA | NA | NA | Mir7223 | microRNA 7223 [Source:MGI Symbol;Acc: | 0 | 0 | 0 | 0 | 0 | 0 |
| 37423 | ENSMUSG00000098990 | 0 | NA | NA | NA | NA | NA | NA | NA | 0 | 0 | 0 | 0 | 0 | 0 |
| 37424 | ENSMUSG00000098991 | 0 | NA | NA | NA | NA | NA | NA | NA | 0 | 0 | 0 | 0 | 0 | 0 |
| 37425 | ENSMUSG00000098992 | 0 | NA | NA | NA | NA | NA | NA | NA | 0 | 0 | 0 | 0 | 0 | 0 |
| 37426 | ENSMUSG00000098993 | 0 | NA | NA | NA | NA | NA | NA | NA | 0 | 0 | 0 | 0 | 0 | 0 |
| 37427 | ENSMUSG00000098994 | 0 | NA | NA | NA | NA | NA | NA | NA | 0 | 0 | 0 | 0 | 0 | 0 |
| 37428 | ENSMUSG00000098995 | 0 | NA | NA | NA | NA | NA | NA | NA | 0 | 0 | 0 | 0 | 0 | 0 |
| 37429 | ENSMUSG00000098996 | 0 | NA | NA | NA | NA | NA | NA | NA | 0 | 0 | 0 | 0 | 0 | 0 |
| 37430 | ENSMUSG00000098997 | 0 | NA | NA | NA | NA | NA | Mir6901 | microRNA 6901 [Source:MGI Symbol;Acc: | 0 | 0 | 0 | 0 | 0 | 0 |
| 37431 | ENSMUSG00000098998 | 0 | NA | NA | NA | NA | NA | NA | NA | 0 | 0 | 0 | 0 | 0 | 0 |
| 37432 | ENSMUSG00000098999 | 0 | NA | NA | NA | NA | NA | NA | NA | 0 | 0 | 0 | 0 | 0 | 0 |
| 37433 | ENSMUSG00000099000 | 0 | NA | NA | NA | NA | NA | NA | NA | 0 | 0 | 0 | 0 | 0 | 0 |
| 37434 | ENSMUSG00000099001 | 0 | NA | NA | NA | NA | NA | NA | NA | 0 | 0 | 0 | 0 | 0 | 0 |
| 37435 | ENSMUSG00000099002 | 0 | NA | NA | NA | NA | NA | NA | NA | 0 | 0 | 0 | 0 | 0 | 0 |
| 37436 | ENSMUSG00000099003 | 0 | NA | NA | NA | NA | NA | Mir7035 | microRNA 7035 [Source:MGI Symbol;Acc: | 0 | 0 | 0 | 0 | 0 | 0 |
| 37437 | ENSMUSG00000099004 | 0.17131964 | -0.796629176 | 4.080472857 | -0.195229623 | 0.84521319 | NA | NA | NA | 0 | 1.0279 | 0 | 0 | 0 | 0 |
| 37438 | ENSMUSG00000099005 | 0 | NA | NA | NA | NA | NA | Mir6398 | microRNA 6398 [Source:MGI Symbol;Acc: | 0 | 0 | 0 | 0 | 0 | 0 |
| 37439 | ENSMUSG00000099006 | 0 | NA | NA | NA | NA | NA | Mir6974 | microRNA 6974 [Source:MGI Symbol;Acc: | 0 | 0 | 0 | 0 | 0 | 0 |
| 37440 | ENSMUSG00000099008 | 0.36353943 | 2.065989254 | 4.034137301 | 0.512126658 | 0.608562368 | NA | Gm17976 | predicted gene, 17976 [Source:MGI Symb | 0 | 0 | 0 | 0 | 2.1812 | 0 |
| 37441 | ENSMUSG00000099009 | 0 | NA | NA | NA | NA | NA | Rdh16f1 | RDH16 family member 1 [Source:MGI Sym | 0 | 0 | 0 | 0 | 0 | 0 |
| 37442 | ENSMUSG00000099010 | 0 | NA | NA | NA | NA | NA | NA | NA | 0 | 0 | 0 | 0 | 0 | 0 |
| 37443 | ENSMUSG00000099011 | 0 | NA | NA | NA | NA | NA | Mir6380 | microRNA 6380 [Source:MGI Symbol;Acc: | 0 | 0 | 0 | 0 | 0 | 0 |
| 37444 | ENSMUSG00000099012 | 0 | NA | NA | NA | NA | NA | NA | NA | 0 | 0 | 0 | 0 | 0 | 0 |
| 37445 | ENSMUSG00000099013 | 0 | NA | NA | NA | NA | NA | Gm27153 | predicted gene 27153 [Source:MGI Symbo | 0 | 0 | 0 | 0 | 0 | 0 |
| 37446 | ENSMUSG00000099014 | 0 | NA | NA | NA | NA | NA | NA | NA | 0 | 0 | 0 | 0 | 0 | 0 |
| 37447 | ENSMUSG00000099015 | 0 | NA | NA | NA | NA | NA | Mir6372 | microRNA 6372 [Source:MGI Symbol;Acc: | 0 | 0 | 0 | 0 | 0 | 0 |
| 37448 | ENSMUSG00000099016 | 0 | NA | NA | NA | NA | NA | Gm27227 | predicted gene 27227 [Source:MGI Symbo | 0 | 0 | 0 | 0 | 0 | 0 |
| 37449 | ENSMUSG00000099017 | 0 | NA | NA | NA | NA | NA | Mir7058 | microRNA 7058 [Source:MGI Symbol;Acc: | 0 | 0 | 0 | 0 | 0 | 0 |
| 37450 | ENSMUSG00000099018 | 0 | NA | NA | NA | NA | NA | NA | NA | 0 | 0 | 0 | 0 | 0 | 0 |
| 37451 | ENSMUSG00000099019 | 0 | NA | NA | NA | NA | NA | Gm27254 | predicted gene 27254 [Source:MGI Symbo | 0 | 0 | 0 | 0 | 0 | 0 |
| 37452 | ENSMUSG00000099020 | 0 | NA | NA | NA | NA | NA | Gm27159 | predicted gene 27159 [Source:MGI Symbo | 0 | 0 | 0 | 0 | 0 | 0 |
| 37453 | ENSMUSG00000099021 | 0.315620088 | 0.165143421 | 4.063300985 | 0.040642675 | 0.967580763 | NA | NA | NA | 0.8031 | 0 | 0 | 0 | 1.0906 | 0 |
| 37454 | ENSMUSG00000099022 | 0 | NA | NA | NA | NA | NA | NA | NA | 0 | 0 | 0 | 0 | 0 | 0 |
| 37455 | ENSMUSG00000099024 | 0.528073336 | 2.60973587 | 2.979250867 | 0.875970499 | 0.381046072 | NA | NA | NA | 0 | 0 | 0 | 0.964 | 1.0906 | 1.1138 |
| 37456 | ENSMUSG00000099025 | 0.160662974 | 1.126924175 | 4.080472857 | 0.276174898 | 0.782413736 | NA | Gm27162 | predicted gene 27162 [Source:MGI Symbo | 0 | 0 | 0 | 0.964 | 0 | 0 |
| 37457 | ENSMUSG00000099027 | 0 | NA | NA | NA | NA | NA | Gm27176 | predicted gene 27176 [Source:MGI Symbo | 0 | 0 | 0 | 0 | 0 | 0 |
| 37458 | ENSMUSG00000099030 | 0 | NA | NA | NA | NA | NA | Mir6394 | microRNA 6394 [Source:MGI Symbol;Acc: | 0 | 0 | 0 | 0 | 0 | 0 |
| 37459 | ENSMUSG00000099031 | 5.811649964 | -0.239101466 | 0.876863417 | -0.272678118 | 0.785100657 | 0.960670212 | Mir7240 | microRNA 7240 [Source:MGI Symbol;Acc: | 5.6217 | 7.1954 | 6.0375 | 3.8559 | 4.3625 | 7.7969 |
| 37460 | ENSMUSG00000099032 | 2.717940762 | -0.343907492 | 1.30472299 | -0.263586597 | 0.792098488 | NA | Tcf24 | transcription factor 24 [Source:MGI Symb | 4.0155 | 4.1117 | 1.0062 | 3.8559 | 1.0906 | 2.2277 |

| | | | | | | | | | | | | | | | |
|---|---|---|---|---|---|---|---|---|---|---|---|---|---|---|---|
| 37461 | ENSMUSG00000099033 | 0.506734502 | -2.372847156 | 3.338060886 | -0.710845978 | 0.477179685 | NA | Mir7013 | microRNA 7013 [Source:MGI Symbol;Acc: | 0 | 1.0279 | 2.0125 | 0 | 0 | 0 |
| 37462 | ENSMUSG00000099034 | 58.21536863 | -0.030383765 | 0.367991097 | -0.082566577 | 0.934196178 | 0.988420305 | 2810039B14 | RIKEN cDNA 2810039B14 gene [Source:M | 42.564 | 83.261 | 51.318 | 82.902 | 45.806 | 43.44 |
| 37463 | ENSMUSG00000099035 | 0 | NA | NA | NA | NA | NA | Mir7040 | microRNA 7040 [Source:MGI Symbol;Acc: | 0 | 0 | 0 | 0 | 0 | 0 |
| 37464 | ENSMUSG00000099036 | 0 | NA | NA | NA | NA | NA | Mir378c | microRNA 378c [Source:MGI Symbol;Acc: | 0 | 0 | 0 | 0 | 0 | 0 |
| 37465 | ENSMUSG00000099037 | 0 | NA | NA | NA | NA | NA | NA | NA | 0 | 0 | 0 | 0 | 0 | 0 |
| 37466 | ENSMUSG00000099039 | 0 | NA | NA | NA | NA | NA | NA | NA | 0 | 0 | 0 | 0 | 0 | 0 |
| 37467 | ENSMUSG00000099041 | 0 | NA | NA | NA | NA | NA | Gm28035 | predicted gene, 28035 [Source:MGI Symb | 0 | 0 | 0 | 0 | 0 | 0 |
| 37468 | ENSMUSG00000099042 | 0 | NA | NA | NA | NA | NA | Mir7066 | microRNA 7066 [Source:MGI Symbol;Acc: | 0 | 0 | 0 | 0 | 0 | 0 |
| 37469 | ENSMUSG00000099043 | 0.160662974 | 1.126924175 | 4.080472857 | 0.276174898 | 0.782413736 | NA | Mir7022 | microRNA 7022 [Source:MGI Symbol;Acc: | 0 | 0 | 0 | 0.964 | 0 | 0 |
| 37470 | ENSMUSG00000099045 | 0 | NA | NA | NA | NA | NA | Mir6374 | microRNA 6374 [Source:MGI Symbol;Acc: | 0 | 0 | 0 | 0 | 0 | 0 |
| 37471 | ENSMUSG00000099047 | 0 | NA | NA | NA | NA | NA | Mir7029 | microRNA 7029 [Source:MGI Symbol;Acc: | 0 | 0 | 0 | 0 | 0 | 0 |
| 37472 | ENSMUSG00000099049 | 0 | NA | NA | NA | NA | NA | NA | NA | 0 | 0 | 0 | 0 | 0 | 0 |
| 37473 | ENSMUSG00000099050 | 0.160662974 | 1.126924175 | 4.080472857 | 0.276174898 | 0.782413736 | NA | Mir7655 | microRNA 7655 [Source:MGI Symbol;Acc: | 0 | 0 | 0 | 0.964 | 0 | 0 |
| 37474 | ENSMUSG00000099051 | 0.854892513 | -1.854622498 | 2.846973444 | -0.651436529 | 0.514764737 | NA | Gm27241 | predicted gene 27241 [Source:MGI Symbo | 4.0155 | 0 | 0 | 0 | 0 | 1.1138 |
| 37475 | ENSMUSG00000099052 | 0 | NA | NA | NA | NA | NA | NA | NA | 0 | 0 | 0 | 0 | 0 | 0 |
| 37476 | ENSMUSG00000099053 | 0 | NA | NA | NA | NA | NA | NA | NA | 0 | 0 | 0 | 0 | 0 | 0 |
| 37477 | ENSMUSG00000099054 | 0 | NA | NA | NA | NA | NA | Gm27242 | predicted gene 27242 [Source:MGI Symbo | 0 | 0 | 0 | 0 | 0 | 0 |
| 37478 | ENSMUSG00000099055 | 5.357552917 | 2.595386792 | 1.056452998 | 2.45669878 | 0.01402202 | 0.136450958 | Zfp-ps | zinc finger protein, pseudogene [Source:N | 2.4093 | 1.0279 | 1.0062 | 6.7478 | 9.8156 | 11.138 |
| 37479 | ENSMUSG00000099056 | 0 | NA | NA | NA | NA | NA | Gtpbp4-ps3 | GTP binding protein 4, pseudogene 3 [Sou | 0 | 0 | 0 | 0 | 0 | 0 |
| 37480 | ENSMUSG00000099057 | 0 | NA | NA | NA | NA | NA | Gm27185 | predicted gene 27185 [Source:MGI Symbo | 0 | 0 | 0 | 0 | 0 | 0 |
| 37481 | ENSMUSG00000099058 | 0 | NA | NA | NA | NA | NA | Mir6943 | microRNA 6943 [Source:MGI Symbol;Acc: | 0 | 0 | 0 | 0 | 0 | 0 |
| 37482 | ENSMUSG00000099059 | 0 | NA | NA | NA | NA | NA | NA | NA | 0 | 0 | 0 | 0 | 0 | 0 |
| 37483 | ENSMUSG00000099060 | 0 | NA | NA | NA | NA | NA | Mir7025 | microRNA 7025 [Source:MGI Symbol;Acc: | 0 | 0 | 0 | 0 | 0 | 0 |
| 37484 | ENSMUSG00000099061 | 0.777840866 | -0.360978154 | 2.442491672 | -0.147790945 | 0.882507755 | NA | Gm28050 | predicted gene, 28050 [Source:MGI Symb | 1.6062 | 0 | 1.0062 | 0.964 | 1.0906 | 0 |
| 37485 | ENSMUSG00000099062 | 0 | NA | NA | NA | NA | NA | Mir7077 | microRNA 7077 [Source:MGI Symbol;Acc: | 0 | 0 | 0 | 0 | 0 | 0 |
| 37486 | ENSMUSG00000099063 | 0.321325949 | 1.911575114 | 4.048267893 | 0.472195804 | 0.636787034 | NA | Mir7031 | microRNA 7031 [Source:MGI Symbol;Acc: | 0 | 0 | 0 | 1.928 | 0 | 0 |
| 37487 | ENSMUSG00000099064 | 0 | NA | NA | NA | NA | NA | Gm27148 | predicted gene 27148 [Source:MGI Symbo | 0 | 0 | 0 | 0 | 0 | 0 |
| 37488 | ENSMUSG00000099066 | 0 | NA | NA | NA | NA | NA | Mir6978 | microRNA 6978 [Source:MGI Symbol;Acc: | 0 | 0 | 0 | 0 | 0 | 0 |
| 37489 | ENSMUSG00000099068 | 0.670829724 | -2.782124174 | 3.401449285 | -0.817923168 | 0.41340106 | NA | NA | NA | 0 | 0 | 4.025 | 0 | 0 | 0 |
| 37490 | ENSMUSG00000099069 | 0 | NA | NA | NA | NA | NA | Mir8090 | microRNA 8090 [Source:MGI Symbol;Acc: | 0 | 0 | 0 | 0 | 0 | 0 |
| 37491 | ENSMUSG00000099070 | 0 | NA | NA | NA | NA | NA | Mir8119 | microRNA 8119 [Source:MGI Symbol;Acc: | 0 | 0 | 0 | 0 | 0 | 0 |
| 37492 | ENSMUSG00000099071 | 0 | NA | NA | NA | NA | NA | Mir8095 | microRNA 8095 [Source:MGI Symbol;Acc: | 0 | 0 | 0 | 0 | 0 | 0 |
| 37493 | ENSMUSG00000099072 | 0 | NA | NA | NA | NA | NA | Mir7673 | microRNA 7673 [Source:MGI Symbol;Acc: | 0 | 0 | 0 | 0 | 0 | 0 |
| 37494 | ENSMUSG00000099073 | 0 | NA | NA | NA | NA | NA | NA | NA | 0 | 0 | 0 | 0 | 0 | 0 |
| 37495 | ENSMUSG00000099074 | 0 | NA | NA | NA | NA | NA | NA | NA | 0 | 0 | 0 | 0 | 0 | 0 |
| 37496 | ENSMUSG00000099075 | 0 | NA | NA | NA | NA | NA | Mir7674 | microRNA 7674 [Source:MGI Symbol;Acc: | 0 | 0 | 0 | 0 | 0 | 0 |
| 37497 | ENSMUSG00000099076 | 0 | NA | NA | NA | NA | NA | Mir7070 | microRNA 7070 [Source:MGI Symbol;Acc: | 0 | 0 | 0 | 0 | 0 | 0 |
| 37498 | ENSMUSG00000099078 | 0 | NA | NA | NA | NA | NA | Mir8110 | microRNA 8110 [Source:MGI Symbol;Acc: | 0 | 0 | 0 | 0 | 0 | 0 |
| 37499 | ENSMUSG00000099079 | 0 | NA | NA | NA | NA | NA | Mir7068 | microRNA 7068 [Source:MGI Symbol;Acc: | 0 | 0 | 0 | 0 | 0 | 0 |
| 37500 | ENSMUSG00000099080 | 0 | NA | NA | NA | NA | NA | Mir7071 | microRNA 7071 [Source:MGI Symbol;Acc: | 0 | 0 | 0 | 0 | 0 | 0 |
| 37501 | ENSMUSG00000099081 | 0 | NA | NA | NA | NA | NA | NA | NA | 0 | 0 | 0 | 0 | 0 | 0 |
| 37502 | ENSMUSG00000099082 | 0 | NA | NA | NA | NA | NA | Mir7036b | microRNA 7036b [Source:MGI Symbol;Acc | 0 | 0 | 0 | 0 | 0 | 0 |
| 37503 | ENSMUSG00000099083 | 384.3980853 | 0.039081505 | 0.146397236 | 0.266955213 | 0.789503645 | 0.961009963 | Atf7 | activating transcription factor 7 [Source:N | 389.5 | 413.22 | 335.08 | 420.29 | 376.26 | 372.02 |
| 37504 | ENSMUSG00000099085 | 0 | NA | NA | NA | NA | NA | Gm8151 | predicted gene 8151 [Source:MGI Symbol | 0 | 0 | 0 | 0 | 0 | 0 |
| 37505 | ENSMUSG00000099086 | 0 | NA | NA | NA | NA | NA | NA | NA | 0 | 0 | 0 | 0 | 0 | 0 |
| 37506 | ENSMUSG00000099087 | 0 | NA | NA | NA | NA | NA | Ranbp2-ps5 | RAN binding protein 2, pseudogene 5 [Sou | 0 | 0 | 0 | 0 | 0 | 0 |
| 37507 | ENSMUSG00000099088 | 0 | NA | NA | NA | NA | NA | NA | NA | 0 | 0 | 0 | 0 | 0 | 0 |
| 37508 | ENSMUSG00000099089 | 0 | NA | NA | NA | NA | NA | NA | NA | 0 | 0 | 0 | 0 | 0 | 0 |
| 37509 | ENSMUSG00000099091 | 0 | NA | NA | NA | NA | NA | Mir7003 | microRNA 7003 [Source:MGI Symbol;Acc: | 0 | 0 | 0 | 0 | 0 | 0 |
| 37510 | ENSMUSG00000099092 | 0 | NA | NA | NA | NA | NA | Mir7062 | microRNA 7062 [Source:MGI Symbol;Acc: | 0 | 0 | 0 | 0 | 0 | 0 |
| 37511 | ENSMUSG00000099093 | 0 | NA | NA | NA | NA | NA | Mir7000 | microRNA 7000 [Source:MGI Symbol;Acc: | 0 | 0 | 0 | 0 | 0 | 0 |
| 37512 | ENSMUSG00000099094 | 0 | NA | NA | NA | NA | NA | Gm19569 | predicted gene, 19569 [Source:MGI Symb | 0 | 0 | 0 | 0 | 0 | 0 |
| 37513 | ENSMUSG00000099095 | 0 | NA | NA | NA | NA | NA | Mir6376 | microRNA 6376 [Source:MGI Symbol;Acc: | 0 | 0 | 0 | 0 | 0 | 0 |
| 37514 | ENSMUSG00000099096 | 0 | NA | NA | NA | NA | NA | Mir6945 | microRNA 6945 [Source:MGI Symbol;Acc: | 0 | 0 | 0 | 0 | 0 | 0 |
| 37515 | ENSMUSG00000099097 | 0 | NA | NA | NA | NA | NA | NA | NA | 0 | 0 | 0 | 0 | 0 | 0 |
| 37516 | ENSMUSG00000099098 | 2.49302255 | -2.567667522 | 1.632123451 | -1.573206683 | 0.115670968 | NA | 1110035H17 | RIKEN cDNA 1110035H17 gene [Source:M | 5.6217 | 3.0838 | 4.025 | 0 | 0 | 2.2277 |

| | | | | | | | | | | | | | | | |
|---|---|---|---|---|---|---|---|---|---|---|---|---|---|---|---|
| 37517 | ENSMUSG00000099099 | 0 | NA | NA | NA | NA | NA | NA | NA | 0 | 0 | 0 | 0 | 0 | 0 |
| 37518 | ENSMUSG00000099101 | 0 | NA | NA | NA | NA | NA | Mir7653 | microRNA 7653 [Source:MGI Symbol;Acc: | 0 | 0 | 0 | 0 | 0 | 0 |
| 37519 | ENSMUSG00000099102 | 0 | NA | NA | NA | NA | NA | Gm11983 | predicted gene 11983 [Source:MGI Symbo | 0 | 0 | 0 | 0 | 0 | 0 |
| 37520 | ENSMUSG00000099103 | 0 | NA | NA | NA | NA | NA | NA | NA | 0 | 0 | 0 | 0 | 0 | 0 |
| 37521 | ENSMUSG00000099104 | 0 | NA | NA | NA | NA | NA | Gm17087 | predicted gene 17087 [Source:MGI Symbo | 0 | 0 | 0 | 0 | 0 | 0 |
| 37522 | ENSMUSG00000099105 | 0 | NA | NA | NA | NA | NA | Mir7007 | microRNA 7007 [Source:MGI Symbol;Acc: | 0 | 0 | 0 | 0 | 0 | 0 |
| 37523 | ENSMUSG00000099106 | 0 | NA | NA | NA | NA | NA | NA | NA | 0 | 0 | 0 | 0 | 0 | 0 |
| 37524 | ENSMUSG00000099107 | 0 | NA | NA | NA | NA | NA | NA | NA | 0 | 0 | 0 | 0 | 0 | 0 |
| 37525 | ENSMUSG00000099108 | 0 | NA | NA | NA | NA | NA | NA | NA | 0 | 0 | 0 | 0 | 0 | 0 |
| 37526 | ENSMUSG00000099109 | 0 | NA | NA | NA | NA | NA | Mir7115 | microRNA 7115 [Source:MGI Symbol;Acc: | 0 | 0 | 0 | 0 | 0 | 0 |
| 37527 | ENSMUSG00000099110 | 1.013933192 | 3.558346573 | 2.586848119 | 1.375552955 | 0.16896008 | NA | Mir6951 | microRNA 6951 [Source:MGI Symbol;Acc: | 0 | 0 | 0 | 3.8559 | 0 | 2.2277 |
| 37528 | ENSMUSG00000099111 | 0.727078859 | 3.067238961 | 3.29680021 | 0.930368468 | 0.352180339 | NA | NA | NA | 0 | 0 | 0 | 0 | 4.3625 | 0 |
| 37529 | ENSMUSG00000099112 | 0 | NA | NA | NA | NA | NA | NA | NA | 0 | 0 | 0 | 0 | 0 | 0 |
| 37530 | ENSMUSG00000099114 | 0 | NA | NA | NA | NA | NA | NA | NA | 0 | 0 | 0 | 0 | 0 | 0 |
| 37531 | ENSMUSG00000099115 | 0 | NA | NA | NA | NA | NA | Gm17190 | predicted gene 17190 [Source:MGI Symbo | 0 | 0 | 0 | 0 | 0 | 0 |
| 37532 | ENSMUSG00000099117 | 0 | NA | NA | NA | NA | NA | NA | NA | 0 | 0 | 0 | 0 | 0 | 0 |
| 37533 | ENSMUSG00000099118 | 0 | NA | NA | NA | NA | NA | Mir8092 | microRNA 8092 [Source:MGI Symbol;Acc: | 0 | 0 | 0 | 0 | 0 | 0 |
| 37534 | ENSMUSG00000099119 | 0 | NA | NA | NA | NA | NA | NA | NA | 0 | 0 | 0 | 0 | 0 | 0 |
| 37535 | ENSMUSG00000099120 | 0 | NA | NA | NA | NA | NA | Gm27193 | predicted gene 27193 [Source:MGI Symbo | 0 | 0 | 0 | 0 | 0 | 0 |
| 37536 | ENSMUSG00000099123 | 0 | NA | NA | NA | NA | NA | Mir6965 | microRNA 6965 [Source:MGI Symbol;Acc: | 0 | 0 | 0 | 0 | 0 | 0 |
| 37537 | ENSMUSG00000099124 | 0 | NA | NA | NA | NA | NA | Mir8117 | microRNA 8117 [Source:MGI Symbol;Acc: | 0 | 0 | 0 | 0 | 0 | 0 |
| 37538 | ENSMUSG00000099125 | 0 | NA | NA | NA | NA | NA | NA | NA | 0 | 0 | 0 | 0 | 0 | 0 |
| 37539 | ENSMUSG00000099126 | 0 | NA | NA | NA | NA | NA | NA | NA | 0 | 0 | 0 | 0 | 0 | 0 |
| 37540 | ENSMUSG00000099127 | 0 | NA | NA | NA | NA | NA | NA | NA | 0 | 0 | 0 | 0 | 0 | 0 |
| 37541 | ENSMUSG00000099128 | 0 | NA | NA | NA | NA | NA | NA | NA | 0 | 0 | 0 | 0 | 0 | 0 |
| 37542 | ENSMUSG00000099129 | 0 | NA | NA | NA | NA | NA | Mir7076 | microRNA 7076 [Source:MGI Symbol;Acc: | 0 | 0 | 0 | 0 | 0 | 0 |
| 37543 | ENSMUSG00000099130 | 0 | NA | NA | NA | NA | NA | NA | NA | 0 | 0 | 0 | 0 | 0 | 0 |
| 37544 | ENSMUSG00000099131 | 0 | NA | NA | NA | NA | NA | Mir7074 | microRNA 7074 [Source:MGI Symbol;Acc: | 0 | 0 | 0 | 0 | 0 | 0 |
| 37545 | ENSMUSG00000099132 | 0 | NA | NA | NA | NA | NA | NA | NA | 0 | 0 | 0 | 0 | 0 | 0 |
| 37546 | ENSMUSG00000099133 | 0 | NA | NA | NA | NA | NA | NA | NA | 0 | 0 | 0 | 0 | 0 | 0 |
| 37547 | ENSMUSG00000099134 | 0 | NA | NA | NA | NA | NA | Mir7018 | microRNA 7018 [Source:MGI Symbol;Acc: | 0 | 0 | 0 | 0 | 0 | 0 |
| 37548 | ENSMUSG00000099135 | 0 | NA | NA | NA | NA | NA | NA | NA | 0 | 0 | 0 | 0 | 0 | 0 |
| 37549 | ENSMUSG00000099136 | 0 | NA | NA | NA | NA | NA | NA | NA | 0 | 0 | 0 | 0 | 0 | 0 |
| 37550 | ENSMUSG00000099137 | 6.930331713 | 0.113062943 | 0.853640866 | 0.132447904 | 0.894630027 | 0.982798755 | Gm10603 | predicted gene 10603 [Source:MGI Symbo | 4.8186 | 10.279 | 5.0312 | 11.568 | 6.5437 | 3.3415 |
| 37551 | ENSMUSG00000099138 | 0 | NA | NA | NA | NA | NA | Gm5746 | predicted gene 5746 [Source:MGI Symbol | 0 | 0 | 0 | 0 | 0 | 0 |
| 37552 | ENSMUSG00000099139 | 0 | NA | NA | NA | NA | NA | 1110028F18 | RIKEN cDNA 1110028F18 gene [Source:M | 0 | 0 | 0 | 0 | 0 | 0 |
| 37553 | ENSMUSG00000099140 | 0.51014012 | 1.031855138 | 3.09002894 | 0.333930574 | 0.738431948 | NA | Mir7237 | microRNA 7237 [Source:MGI Symbol;Acc: | 0 | 0 | 1.0062 | 0.964 | 1.0906 | 0 |
| 37554 | ENSMUSG00000099141 | 0 | NA | NA | NA | NA | NA | Mir6352 | microRNA 6352 [Source:MGI Symbol;Acc: | 0 | 0 | 0 | 0 | 0 | 0 |
| 37555 | ENSMUSG00000099142 | 0 | NA | NA | NA | NA | NA | Mir8108 | microRNA 8108 [Source:MGI Symbol;Acc: | 0 | 0 | 0 | 0 | 0 | 0 |
| 37556 | ENSMUSG00000099143 | 0 | NA | NA | NA | NA | NA | NA | NA | 0 | 0 | 0 | 0 | 0 | 0 |
| 37557 | ENSMUSG00000099144 | 0 | NA | NA | NA | NA | NA | NA | NA | 0 | 0 | 0 | 0 | 0 | 0 |
| 37558 | ENSMUSG00000099145 | 0.942142679 | -3.279214146 | 2.405018921 | -1.363487878 | 0.172728802 | NA | Mir8116 | microRNA 8116 [Source:MGI Symbol;Acc: | 1.6062 | 1.0279 | 3.0187 | 0 | 0 | 0 |
| 37559 | ENSMUSG00000099146 | 900.8170279 | 0.359683133 | 0.125899108 | 2.856915669 | 0.004277795 | 0.060247345 | 0610031O16 | RIKEN cDNA 0610031O16 gene [Source:M | 723.6 | 739.07 | 905.62 | 1045.9 | 1033.9 | 956.79 |
| 37560 | ENSMUSG00000099147 | 0 | NA | NA | NA | NA | NA | Mir7081 | microRNA 7081 [Source:MGI Symbol;Acc: | 0 | 0 | 0 | 0 | 0 | 0 |
| 37561 | ENSMUSG00000099148 | 0 | NA | NA | NA | NA | NA | Gm3331 | predicted gene 3331 [Source:MGI Symbol | 0 | 0 | 0 | 0 | 0 | 0 |
| 37562 | ENSMUSG00000099149 | 0 | NA | NA | NA | NA | NA | Mir6370 | microRNA 6370 [Source:MGI Symbol;Acc: | 0 | 0 | 0 | 0 | 0 | 0 |
| 37563 | ENSMUSG00000099150 | 0 | NA | NA | NA | NA | NA | Gm27243 | predicted gene 27243 [Source:MGI Symbo | 0 | 0 | 0 | 0 | 0 | 0 |
| 37564 | ENSMUSG00000099151 | 0 | NA | NA | NA | NA | NA | NA | NA | 0 | 0 | 0 | 0 | 0 | 0 |
| 37565 | ENSMUSG00000099152 | 0 | NA | NA | NA | NA | NA | Mir6969 | microRNA 6969 [Source:MGI Symbol;Acc: | 0 | 0 | 0 | 0 | 0 | 0 |
| 37566 | ENSMUSG00000099153 | 0 | NA | NA | NA | NA | NA | Mir7671 | microRNA 7671 [Source:MGI Symbol;Acc: | 0 | 0 | 0 | 0 | 0 | 0 |
| 37567 | ENSMUSG00000099154 | 0 | NA | NA | NA | NA | NA | NA | NA | 0 | 0 | 0 | 0 | 0 | 0 |
| 37568 | ENSMUSG00000099155 | 0 | NA | NA | NA | NA | NA | NA | NA | 0 | 0 | 0 | 0 | 0 | 0 |
| 37569 | ENSMUSG00000099157 | 0.497389803 | 1.101799453 | 3.439268845 | 0.320358629 | 0.748696483 | NA | NA | NA | 0.8031 | 0 | 0 | 0 | 2.1812 | 0 |
| 37570 | ENSMUSG00000099158 | 0 | NA | NA | NA | NA | NA | Mir6368 | microRNA 6368 [Source:MGI Symbol;Acc: | 0 | 0 | 0 | 0 | 0 | 0 |
| 37571 | ENSMUSG00000099159 | 0 | NA | NA | NA | NA | NA | NA | NA | 0 | 0 | 0 | 0 | 0 | 0 |
| 37572 | ENSMUSG00000099160 | 0.328370405 | 0.16514316 | 4.003149536 | 0.041253308 | 0.967093956 | NA | Mir6415 | microRNA 6415 [Source:MGI Symbol;Acc: | 0 | 0 | 1.0062 | 0.964 | 0 | 0 |

| | | | | | | | | | | | | | | | |
|---|---|---|---|---|---|---|---|---|---|---|---|---|---|---|---|
| 37573 | ENSMUSG00000099161 | 0 | NA | NA | NA | NA | NA | Mir6419 | microRNA 6419 [Source:MGI Symbol;Acc: | 0 | 0 | 0 | 0 | 0 | 0 |
| 37574 | ENSMUSG00000099162 | 0 | NA | NA | NA | NA | NA | Gm27168 | predicted gene 27168 [Source:MGI Symbo | 0 | 0 | 0 | 0 | 0 | 0 |
| 37575 | ENSMUSG00000099163 | 0 | NA | NA | NA | NA | NA | Mir7660 | microRNA 7660 [Source:MGI Symbol;Acc: | 0 | 0 | 0 | 0 | 0 | 0 |
| 37576 | ENSMUSG00000099165 | 0 | NA | NA | NA | NA | NA | Mir6940 | microRNA 6940 [Source:MGI Symbol;Acc: | 0 | 0 | 0 | 0 | 0 | 0 |
| 37577 | ENSMUSG00000099166 | 0 | NA | NA | NA | NA | NA | NA | NA | 0 | 0 | 0 | 0 | 0 | 0 |
| 37578 | ENSMUSG00000099168 | 0 | NA | NA | NA | NA | NA | NA | NA | 0 | 0 | 0 | 0 | 0 | 0 |
| 37579 | ENSMUSG00000099169 | 0 | NA | NA | NA | NA | NA | Mir7j | microRNA 7j [Source:MGI Symbol;Acc:MG | 0 | 0 | 0 | 0 | 0 | 0 |
| 37580 | ENSMUSG00000099170 | 0 | NA | NA | NA | NA | NA | 5730403I07F | RIKEN cDNA 5730403I07 gene [Source:MG | 0 | 0 | 0 | 0 | 0 | 0 |
| 37581 | ENSMUSG00000099172 | 0 | NA | NA | NA | NA | NA | Mir3620 | microRNA 3620 [Source:MGI Symbol;Acc: | 0 | 0 | 0 | 0 | 0 | 0 |
| 37582 | ENSMUSG00000099173 | 0 | NA | NA | NA | NA | NA | NA | NA | 0 | 0 | 0 | 0 | 0 | 0 |
| 37583 | ENSMUSG00000099174 | 0 | NA | NA | NA | NA | NA | Mir6406 | microRNA 6406 [Source:MGI Symbol;Acc: | 0 | 0 | 0 | 0 | 0 | 0 |
| 37584 | ENSMUSG00000099176 | 0 | NA | NA | NA | NA | NA | Mir6238 | microRNA 6238 [Source:MGI Symbol;Acc: | 0 | 0 | 0 | 0 | 0 | 0 |
| 37585 | ENSMUSG00000099178 | 0 | NA | NA | NA | NA | NA | Gm8669 | predicted gene 8669 [Source:MGI Symbol | 0 | 0 | 0 | 0 | 0 | 0 |
| 37586 | ENSMUSG00000099179 | 0.321325949 | 1.911575114 | 4.048267893 | 0.472195804 | 0.636787034 | NA | NA | NA | 0 | 0 | 0 | 1.928 | 0 | 0 |
| 37587 | ENSMUSG00000099180 | 0 | NA | NA | NA | NA | NA | Mir6956 | microRNA 6956 [Source:MGI Symbol;Acc: | 0 | 0 | 0 | 0 | 0 | 0 |
| 37588 | ENSMUSG00000099181 | 0.524202404 | 2.600864256 | 3.301637836 | 0.787749713 | 0.430843121 | NA | Mir6905 | microRNA 6905 [Source:MGI Symbol;Acc: | 0 | 0 | 0 | 0.964 | 2.1812 | 0 |
| 37589 | ENSMUSG00000099182 | 0 | NA | NA | NA | NA | NA | Gm8676 | predicted gene 8676 [Source:MGI Symbol | 0 | 0 | 0 | 0 | 0 | 0 |
| 37590 | ENSMUSG00000099183 | 0 | NA | NA | NA | NA | NA | NA | NA | 0 | 0 | 0 | 0 | 0 | 0 |
| 37591 | ENSMUSG00000099184 | 0 | NA | NA | NA | NA | NA | Mir6337 | microRNA 6337 [Source:MGI Symbol;Acc: | 0 | 0 | 0 | 0 | 0 | 0 |
| 37592 | ENSMUSG00000099185 | 0 | NA | NA | NA | NA | NA | Mir7662 | microRNA 7662 [Source:MGI Symbol;Acc: | 0 | 0 | 0 | 0 | 0 | 0 |
| 37593 | ENSMUSG00000099186 | 0 | NA | NA | NA | NA | NA | NA | NA | 0 | 0 | 0 | 0 | 0 | 0 |
| 37594 | ENSMUSG00000099188 | 0 | NA | NA | NA | NA | NA | Gm27248 | predicted gene 27248 [Source:MGI Symbo | 0 | 0 | 0 | 0 | 0 | 0 |
| 37595 | ENSMUSG00000099189 | 0 | NA | NA | NA | NA | NA | NA | NA | 0 | 0 | 0 | 0 | 0 | 0 |
| 37596 | ENSMUSG00000099190 | 0 | NA | NA | NA | NA | NA | Gm27188 | predicted gene 27188 [Source:MGI Symbo | 0 | 0 | 0 | 0 | 0 | 0 |
| 37597 | ENSMUSG00000099191 | 0 | NA | NA | NA | NA | NA | NA | NA | 0 | 0 | 0 | 0 | 0 | 0 |
| 37598 | ENSMUSG00000099192 | 0 | NA | NA | NA | NA | NA | NA | NA | 0 | 0 | 0 | 0 | 0 | 0 |
| 37599 | ENSMUSG00000099193 | 0 | NA | NA | NA | NA | NA | NA | NA | 0 | 0 | 0 | 0 | 0 | 0 |
| 37600 | ENSMUSG00000099196 | 1.042698947 | 2.426023578 | 2.689838701 | 0.901921583 | 0.367098525 | NA | Mir1258 | microRNA 1258 [Source:MGI Symbol;Acc: | 0.8031 | 0 | 0 | 0 | 5.4531 | 0 |
| 37601 | ENSMUSG00000099197 | 0 | NA | NA | NA | NA | NA | Mir7034 | microRNA 7034 [Source:MGI Symbol;Acc: | 0 | 0 | 0 | 0 | 0 | 0 |
| 37602 | ENSMUSG00000099198 | 0 | NA | NA | NA | NA | NA | Obox4-ps39 | oocyte specific homeobox 4, pseudogene | 0 | 0 | 0 | 0 | 0 | 0 |
| 37603 | ENSMUSG00000099199 | 0 | NA | NA | NA | NA | NA | Mir7012 | microRNA 7012 [Source:MGI Symbol;Acc: | 0 | 0 | 0 | 0 | 0 | 0 |
| 37604 | ENSMUSG00000099200 | 0 | NA | NA | NA | NA | NA | Mir7019 | microRNA 7019 [Source:MGI Symbol;Acc: | 0 | 0 | 0 | 0 | 0 | 0 |
| 37605 | ENSMUSG00000099201 | 0 | NA | NA | NA | NA | NA | Gm27187 | predicted gene 27187 [Source:MGI Symbo | 0 | 0 | 0 | 0 | 0 | 0 |
| 37606 | ENSMUSG00000099202 | 0 | NA | NA | NA | NA | NA | Mir7017 | microRNA 7017 [Source:MGI Symbol;Acc: | 0 | 0 | 0 | 0 | 0 | 0 |
| 37607 | ENSMUSG00000099203 | 0 | NA | NA | NA | NA | NA | Mir6389 | microRNA 6389 [Source:MGI Symbol;Acc: | 0 | 0 | 0 | 0 | 0 | 0 |
| 37608 | ENSMUSG00000099204 | 0 | NA | NA | NA | NA | NA | NA | NA | 0 | 0 | 0 | 0 | 0 | 0 |
| 37609 | ENSMUSG00000099205 | 0 | NA | NA | NA | NA | NA | NA | NA | 0 | 0 | 0 | 0 | 0 | 0 |
| 37610 | ENSMUSG00000099206 | 0.653308562 | 1.538013299 | 3.112409277 | 0.494155223 | 0.62119656 | NA | NA | NA | 0 | 1.0279 | 0 | 2.8919 | 0 | 0 |
| 37611 | ENSMUSG00000099207 | 0 | NA | NA | NA | NA | NA | Gm10637 | predicted gene 10637 [Source:MGI Symbo | 0 | 0 | 0 | 0 | 0 | 0 |
| 37612 | ENSMUSG00000099208 | 0 | NA | NA | NA | NA | NA | NA | NA | 0 | 0 | 0 | 0 | 0 | 0 |
| 37613 | ENSMUSG00000099209 | 0 | NA | NA | NA | NA | NA | NA | NA | 0 | 0 | 0 | 0 | 0 | 0 |
| 37614 | ENSMUSG00000099210 | 0.743983808 | -0.269067535 | 2.702716736 | -0.099554471 | 0.920698041 | NA | Mir7051 | microRNA 7051 [Source:MGI Symbol;Acc: | 2.4093 | 0 | 0 | 0.964 | 1.0906 | 0 |
| 37615 | ENSMUSG00000099211 | 0 | NA | NA | NA | NA | NA | NA | NA | 0 | 0 | 0 | 0 | 0 | 0 |
| 37616 | ENSMUSG00000099212 | 0 | NA | NA | NA | NA | NA | Mir7033 | microRNA 7033 [Source:MGI Symbol;Acc: | 0 | 0 | 0 | 0 | 0 | 0 |
| 37617 | ENSMUSG00000099213 | 0 | NA | NA | NA | NA | NA | NA | NA | 0 | 0 | 0 | 0 | 0 | 0 |
| 37618 | ENSMUSG00000099214 | 0 | NA | NA | NA | NA | NA | Mir6365 | microRNA 6365 [Source:MGI Symbol;Acc: | 0 | 0 | 0 | 0 | 0 | 0 |
| 37619 | ENSMUSG00000099215 | 0 | NA | NA | NA | NA | NA | NA | NA | 0 | 0 | 0 | 0 | 0 | 0 |
| 37620 | ENSMUSG00000099216 | 0.133850373 | -0.796629176 | 4.080472857 | -0.195229623 | 0.84521319 | NA | Obox8 | oocyte specific homeobox 8 [Source:MGI | 0.8031 | 0 | 0 | 0 | 0 | 0 |
| 37621 | ENSMUSG00000099218 | 0 | NA | NA | NA | NA | NA | Mir5124b | microRNA 5124b [Source:MGI Symbol;Acc | 0 | 0 | 0 | 0 | 0 | 0 |
| 37622 | ENSMUSG00000099219 | 0 | NA | NA | NA | NA | NA | NA | NA | 0 | 0 | 0 | 0 | 0 | 0 |
| 37623 | ENSMUSG00000099220 | 0 | NA | NA | NA | NA | NA | Gm27220 | predicted gene 27220 [Source:MGI Symbo | 0 | 0 | 0 | 0 | 0 | 0 |
| 37624 | ENSMUSG00000099221 | 0 | NA | NA | NA | NA | NA | Mir6371 | microRNA 6371 [Source:MGI Symbol;Acc: | 0 | 0 | 0 | 0 | 0 | 0 |
| 37625 | ENSMUSG00000099222 | 0 | NA | NA | NA | NA | NA | Mir7213 | microRNA 7213 [Source:MGI Symbol;Acc: | 0 | 0 | 0 | 0 | 0 | 0 |
| 37626 | ENSMUSG00000099223 | 0 | NA | NA | NA | NA | NA | Mir7221 | microRNA 7221 [Source:MGI Symbol;Acc: | 0 | 0 | 0 | 0 | 0 | 0 |
| 37627 | ENSMUSG00000099224 | 0 | NA | NA | NA | NA | NA | NA | NA | 0 | 0 | 0 | 0 | 0 | 0 |
| 37628 | ENSMUSG00000099225 | 0 | NA | NA | NA | NA | NA | Mir8104 | microRNA 8104 [Source:MGI Symbol;Acc: | 0 | 0 | 0 | 0 | 0 | 0 |

| | | | | | | | | | | | | | | | |
|---|---|---|---|---|---|---|---|---|---|---|---|---|---|---|---|
| 37629 | ENSMUSG00000099226 | 0 | NA | NA | NA | NA | NA | Mir7091 | microRNA 7091 [Source:MGI Symbol;Acc: | 0 | 0 | 0 | 0 | 0 | 0 |
| 37630 | ENSMUSG00000099227 | 0 | NA | NA | NA | NA | NA | Mir8114 | microRNA 8114 [Source:MGI Symbol;Acc: | 0 | 0 | 0 | 0 | 0 | 0 |
| 37631 | ENSMUSG00000099228 | 0 | NA | NA | NA | NA | NA | NA | NA | 0 | 0 | 0 | 0 | 0 | 0 |
| 37632 | ENSMUSG00000099229 | 0 | NA | NA | NA | NA | NA | NA | NA | 0 | 0 | 0 | 0 | 0 | 0 |
| 37633 | ENSMUSG00000099230 | 0 | NA | NA | NA | NA | NA | Mir6989 | microRNA 6989 [Source:MGI Symbol;Acc: | 0 | 0 | 0 | 0 | 0 | 0 |
| 37634 | ENSMUSG00000099232 | 0 | NA | NA | NA | NA | NA | NA | NA | 0 | 0 | 0 | 0 | 0 | 0 |
| 37635 | ENSMUSG00000099233 | 0 | NA | NA | NA | NA | NA | NA | NA | 0 | 0 | 0 | 0 | 0 | 0 |
| 37636 | ENSMUSG00000099235 | 0 | NA | NA | NA | NA | NA | NA | NA | 0 | 0 | 0 | 0 | 0 | 0 |
| 37637 | ENSMUSG00000099236 | 0 | NA | NA | NA | NA | NA | Mir668 | microRNA 668 [Source:MGI Symbol;Acc:M | 0 | 0 | 0 | 0 | 0 | 0 |
| 37638 | ENSMUSG00000099237 | 0 | NA | NA | NA | NA | NA | Mir7216 | microRNA 7216 [Source:MGI Symbol;Acc: | 0 | 0 | 0 | 0 | 0 | 0 |
| 37639 | ENSMUSG00000099238 | 1.152036961 | -0.177087519 | 2.115009693 | -0.08372894 | 0.933271947 | NA | Mir7001 | microRNA 7001 [Source:MGI Symbol;Acc: | 1.6062 | 1.0279 | 1.0062 | 0 | 3.2719 | 0 |
| 37640 | ENSMUSG00000099239 | 0 | NA | NA | NA | NA | NA | Gm27155 | predicted gene 27155 [Source:MGI Symbo | 0 | 0 | 0 | 0 | 0 | 0 |
| 37641 | ENSMUSG00000099240 | 0 | NA | NA | NA | NA | NA | NA | NA | 0 | 0 | 0 | 0 | 0 | 0 |
| 37642 | ENSMUSG00000099241 | 0.17131964 | -0.796629176 | 4.080472857 | -0.195229623 | 0.84521319 | NA | Gvin-ps2 | GTPase, very large interferon inducible, ps | 0 | 1.0279 | 0 | 0 | 0 | 0 |
| 37643 | ENSMUSG00000099242 | 23.25978947 | 1.229919013 | 0.533814349 | 2.304020143 | 0.021221513 | 0.180908987 | Obox4-ps3 | oocyte specific homeobox 4, pseudogene | 20.881 | 14.391 | 6.0375 | 23.135 | 29.447 | 45.668 |
| 37644 | ENSMUSG00000099243 | 0 | NA | NA | NA | NA | NA | NA | NA | 0 | 0 | 0 | 0 | 0 | 0 |
| 37645 | ENSMUSG00000099244 | 0 | NA | NA | NA | NA | NA | NA | NA | 0 | 0 | 0 | 0 | 0 | 0 |
| 37646 | ENSMUSG00000099245 | 0 | NA | NA | NA | NA | NA | Mir8098 | microRNA 8098 [Source:MGI Symbol;Acc: | 0 | 0 | 0 | 0 | 0 | 0 |
| 37647 | ENSMUSG00000099246 | 0 | NA | NA | NA | NA | NA | NA | NA | 0 | 0 | 0 | 0 | 0 | 0 |
| 37648 | ENSMUSG00000099247 | 0 | NA | NA | NA | NA | NA | NA | NA | 0 | 0 | 0 | 0 | 0 | 0 |
| 37649 | ENSMUSG00000099248 | 0 | NA | NA | NA | NA | NA | NA | NA | 0 | 0 | 0 | 0 | 0 | 0 |
| 37650 | ENSMUSG00000099250 | 0.870506025 | 3.333445469 | 2.471995426 | 1.348483672 | 0.177502868 | NA | NA | NA | 0 | 0 | 0 | 1.928 | 2.1812 | 1.1138 |
| 37651 | ENSMUSG00000099252 | 0 | NA | NA | NA | NA | NA | NA | NA | 0 | 0 | 0 | 0 | 0 | 0 |
| 37652 | ENSMUSG00000099253 | 0 | NA | NA | NA | NA | NA | Gm7863 | predicted gene 7863 [Source:MGI Symbol | 0 | 0 | 0 | 0 | 0 | 0 |
| 37653 | ENSMUSG00000099254 | 0 | NA | NA | NA | NA | NA | Mir7030 | microRNA 7030 [Source:MGI Symbol;Acc: | 0 | 0 | 0 | 0 | 0 | 0 |
| 37654 | ENSMUSG00000099255 | 0 | NA | NA | NA | NA | NA | NA | NA | 0 | 0 | 0 | 0 | 0 | 0 |
| 37655 | ENSMUSG00000099256 | 0 | NA | NA | NA | NA | NA | Gm27165 | predicted gene 27165 [Source:MGI Symbo | 0 | 0 | 0 | 0 | 0 | 0 |
| 37656 | ENSMUSG00000099257 | 0 | NA | NA | NA | NA | NA | Mir6342 | microRNA 6342 [Source:MGI Symbol;Acc: | 0 | 0 | 0 | 0 | 0 | 0 |
| 37657 | ENSMUSG00000099258 | 0 | NA | NA | NA | NA | NA | Mir7047 | microRNA 7047 [Source:MGI Symbol;Acc: | 0 | 0 | 0 | 0 | 0 | 0 |
| 37658 | ENSMUSG00000099259 | 0 | NA | NA | NA | NA | NA | Mir6922 | microRNA 6922 [Source:MGI Symbol;Acc: | 0 | 0 | 0 | 0 | 0 | 0 |
| 37659 | ENSMUSG00000099260 | 0 | NA | NA | NA | NA | NA | Mir6954 | microRNA 6954 [Source:MGI Symbol;Acc: | 0 | 0 | 0 | 0 | 0 | 0 |
| 37660 | ENSMUSG00000099261 | 0 | NA | NA | NA | NA | NA | Mir6984 | microRNA 6984 [Source:MGI Symbol;Acc: | 0 | 0 | 0 | 0 | 0 | 0 |
| 37661 | ENSMUSG00000099263 | 0 | NA | NA | NA | NA | NA | NA | NA | 0 | 0 | 0 | 0 | 0 | 0 |
| 37662 | ENSMUSG00000099264 | 0 | NA | NA | NA | NA | NA | Mir6411 | microRNA 6411 [Source:MGI Symbol;Acc: | 0 | 0 | 0 | 0 | 0 | 0 |
| 37663 | ENSMUSG00000099266 | 0 | NA | NA | NA | NA | NA | 1700063H06 | RIKEN cDNA 1700063H06 gene [Source:M | 0 | 0 | 0 | 0 | 0 | 0 |
| 37664 | ENSMUSG00000099267 | 0.455176322 | 0.955156269 | 3.546674842 | 0.269310357 | 0.787690862 | NA | Mir6955 | microRNA 6955 [Source:MGI Symbol;Acc: | 0.8031 | 0 | 0 | 1.928 | 0 | 0 |
| 37665 | ENSMUSG00000099268 | 0 | NA | NA | NA | NA | NA | NA | NA | 0 | 0 | 0 | 0 | 0 | 0 |
| 37666 | ENSMUSG00000099269 | 0 | NA | NA | NA | NA | NA | Calm5 | calmodulin 5 [Source:MGI Symbol;Acc:MC | 0 | 0 | 0 | 0 | 0 | 0 |
| 37667 | ENSMUSG00000099270 | 0 | NA | NA | NA | NA | NA | NA | NA | 0 | 0 | 0 | 0 | 0 | 0 |
| 37668 | ENSMUSG00000099272 | 0 | NA | NA | NA | NA | NA | Mir7093 | microRNA 7093 [Source:MGI Symbol;Acc: | 0 | 0 | 0 | 0 | 0 | 0 |
| 37669 | ENSMUSG00000099273 | 0 | NA | NA | NA | NA | NA | NA | NA | 0 | 0 | 0 | 0 | 0 | 0 |
| 37670 | ENSMUSG00000099274 | 0 | NA | NA | NA | NA | NA | Gm27218 | predicted gene 27218 [Source:MGI Symbo | 0 | 0 | 0 | 0 | 0 | 0 |
| 37671 | ENSMUSG00000099275 | 0 | NA | NA | NA | NA | NA | Mir8107 | microRNA 8107 [Source:MGI Symbol;Acc: | 0 | 0 | 0 | 0 | 0 | 0 |
| 37672 | ENSMUSG00000099276 | 0 | NA | NA | NA | NA | NA | Gm27228 | predicted gene 27228 [Source:MGI Symbo | 0 | 0 | 0 | 0 | 0 | 0 |
| 37673 | ENSMUSG00000099278 | 0 | NA | NA | NA | NA | NA | NA | NA | 0 | 0 | 0 | 0 | 0 | 0 |
| 37674 | ENSMUSG00000099279 | 0 | NA | NA | NA | NA | NA | Mir6408 | microRNA 6408 [Source:MGI Symbol;Acc: | 0 | 0 | 0 | 0 | 0 | 0 |
| 37675 | ENSMUSG00000099280 | 0 | NA | NA | NA | NA | NA | Gm27991 | predicted gene, 27991 [Source:MGI Symb | 0 | 0 | 0 | 0 | 0 | 0 |
| 37676 | ENSMUSG00000099283 | 0 | NA | NA | NA | NA | NA | Mir6912 | microRNA 6912 [Source:MGI Symbol;Acc: | 0 | 0 | 0 | 0 | 0 | 0 |
| 37677 | ENSMUSG00000099284 | 0.514011053 | 1.044022102 | 3.082659575 | 0.338675769 | 0.734853997 | NA | Mir7026 | microRNA 7026 [Source:MGI Symbol;Acc: | 0 | 0 | 1.0062 | 0.964 | 0 | 1.1138 |
| 37678 | ENSMUSG00000099285 | 0 | NA | NA | NA | NA | NA | NA | NA | 0 | 0 | 0 | 0 | 0 | 0 |
| 37679 | ENSMUSG00000099286 | 0 | NA | NA | NA | NA | NA | Mir7235 | microRNA 7235 [Source:MGI Symbol;Acc: | 0 | 0 | 0 | 0 | 0 | 0 |
| 37680 | ENSMUSG00000099287 | 0 | NA | NA | NA | NA | NA | NA | NA | 0 | 0 | 0 | 0 | 0 | 0 |
| 37681 | ENSMUSG00000099289 | 0 | NA | NA | NA | NA | NA | Mir6403 | microRNA 6403 [Source:MGI Symbol;Acc: | 0 | 0 | 0 | 0 | 0 | 0 |
| 37682 | ENSMUSG00000099290 | 0 | NA | NA | NA | NA | NA | Mir6985 | microRNA 6985 [Source:MGI Symbol;Acc: | 0 | 0 | 0 | 0 | 0 | 0 |
| 37683 | ENSMUSG00000099291 | 0 | NA | NA | NA | NA | NA | NA | NA | 0 | 0 | 0 | 0 | 0 | 0 |
| 37684 | ENSMUSG00000099292 | 0 | NA | NA | NA | NA | NA | Mir6916 | microRNA 6916 [Source:MGI Symbol;Acc: | 0 | 0 | 0 | 0 | 0 | 0 |

| | | | | | | | | | | | | | | | |
|---|---|---|---|---|---|---|---|---|---|---|---|---|---|---|---|
| 37685 | ENSMUSG00000099293 | 0 | NA | NA | NA | NA | NA | Mir6366 | microRNA 6366 [Source:MGI Symbol;Acc: | 0 | 0 | 0 | 0 | 0 | 0 |
| 37686 | ENSMUSG00000099294 | 0 | NA | NA | NA | NA | NA | Gm11214 | predicted gene 11214 [Source:MGI Symbo | 0 | 0 | 0 | 0 | 0 | 0 |
| 37687 | ENSMUSG00000099295 | 0 | NA | NA | NA | NA | NA | NA | NA | 0 | 0 | 0 | 0 | 0 | 0 |
| 37688 | ENSMUSG00000099298 | 0 | NA | NA | NA | NA | NA | Ranbp2-ps6 | RAN binding protein 2, pseudogene 6 [Sou | 0 | 0 | 0 | 0 | 0 | 0 |
| 37689 | ENSMUSG00000099299 | 0 | NA | NA | NA | NA | NA | NA | NA | 0 | 0 | 0 | 0 | 0 | 0 |
| 37690 | ENSMUSG00000099300 | 0 | NA | NA | NA | NA | NA | Mir6356 | microRNA 6356 [Source:MGI Symbol;Acc: | 0 | 0 | 0 | 0 | 0 | 0 |
| 37691 | ENSMUSG00000099301 | 0.339027071 | -1.79174149 | 3.919585585 | -0.457125237 | 0.647581032 | NA | Mir3473e | microRNA 3473e [Source:MGI Symbol;Acc | 0 | 1.0279 | 1.0062 | 0 | 0 | 0 |
| 37692 | ENSMUSG00000099302 | 0 | NA | NA | NA | NA | NA | NA | NA | 0 | 0 | 0 | 0 | 0 | 0 |
| 37693 | ENSMUSG00000099303 | 0 | NA | NA | NA | NA | NA | NA | NA | 0 | 0 | 0 | 0 | 0 | 0 |
| 37694 | ENSMUSG00000099304 | 0.267700746 | -1.497481649 | 4.057001436 | -0.369110456 | 0.712045396 | NA | NA | NA | 1.6062 | 0 | 0 | 0 | 0 | 0 |
| 37695 | ENSMUSG00000099307 | 0 | NA | NA | NA | NA | NA | NA | NA | 0 | 0 | 0 | 0 | 0 | 0 |
| 37696 | ENSMUSG00000099308 | 0 | NA | NA | NA | NA | NA | Mir6367 | microRNA 6367 [Source:MGI Symbol;Acc: | 0 | 0 | 0 | 0 | 0 | 0 |
| 37697 | ENSMUSG00000099309 | 3.975211585 | 0.325643837 | 1.234501928 | 0.263785604 | 0.791945128 | NA | Mir7085 | microRNA 7085 [Source:MGI Symbol;Acc: | 6.4248 | 4.1117 | 0 | 6.7478 | 5.4531 | 1.1138 |
| 37698 | ENSMUSG00000099310 | 0 | NA | NA | NA | NA | NA | Gm19427 | predicted gene, 19427 [Source:MGI Symb | 0 | 0 | 0 | 0 | 0 | 0 |
| 37699 | ENSMUSG00000099311 | 0 | NA | NA | NA | NA | NA | Mir6244 | microRNA 6244 [Source:MGI Symbol;Acc: | 0 | 0 | 0 | 0 | 0 | 0 |
| 37700 | ENSMUSG00000099312 | 0 | NA | NA | NA | NA | NA | NA | NA | 0 | 0 | 0 | 0 | 0 | 0 |
| 37701 | ENSMUSG00000099313 | 0.160662974 | 1.126924175 | 4.080472857 | 0.276174898 | 0.782413736 | NA | Gm27152 | predicted gene 27152 [Source:MGI Symbo | 0 | 0 | 0 | 0.964 | 0 | 0 |
| 37702 | ENSMUSG00000099314 | 0 | NA | NA | NA | NA | NA | NA | NA | 0 | 0 | 0 | 0 | 0 | 0 |
| 37703 | ENSMUSG00000099315 | 0 | NA | NA | NA | NA | NA | Gm27175 | predicted gene 27175 [Source:MGI Symbo | 0 | 0 | 0 | 0 | 0 | 0 |
| 37704 | ENSMUSG00000099316 | 0 | NA | NA | NA | NA | NA | NA | NA | 0 | 0 | 0 | 0 | 0 | 0 |
| 37705 | ENSMUSG00000099317 | 0 | NA | NA | NA | NA | NA | NA | NA | 0 | 0 | 0 | 0 | 0 | 0 |
| 37706 | ENSMUSG00000099318 | 0 | NA | NA | NA | NA | NA | NA | NA | 0 | 0 | 0 | 0 | 0 | 0 |
| 37707 | ENSMUSG00000099319 | 0 | NA | NA | NA | NA | NA | Mir7648 | microRNA 7648 [Source:MGI Symbol;Acc: | 0 | 0 | 0 | 0 | 0 | 0 |
| 37708 | ENSMUSG00000099320 | 0.672839099 | 0.267535205 | 2.816195809 | 0.094998794 | 0.924315785 | NA | Mir7664 | microRNA 7664 [Source:MGI Symbol;Acc: | 0.8031 | 0 | 1.0062 | 0 | 0 | 2.2277 |
| 37709 | ENSMUSG00000099321 | 0 | NA | NA | NA | NA | NA | Mir6404 | microRNA 6404 [Source:MGI Symbol;Acc: | 0 | 0 | 0 | 0 | 0 | 0 |
| 37710 | ENSMUSG00000099322 | 0 | NA | NA | NA | NA | NA | NA | NA | 0 | 0 | 0 | 0 | 0 | 0 |
| 37711 | ENSMUSG00000099323 | 0 | NA | NA | NA | NA | NA | NA | NA | 0 | 0 | 0 | 0 | 0 | 0 |
| 37712 | ENSMUSG00000099324 | 0 | NA | NA | NA | NA | NA | Mir8115 | microRNA 8115 [Source:MGI Symbol;Acc: | 0 | 0 | 0 | 0 | 0 | 0 |
| 37713 | ENSMUSG00000099325 | 0 | NA | NA | NA | NA | NA | NA | NA | 0 | 0 | 0 | 0 | 0 | 0 |
| 37714 | ENSMUSG00000099326 | 0 | NA | NA | NA | NA | NA | Mir21c | microRNA 21c [Source:MGI Symbol;Acc:M | 0 | 0 | 0 | 0 | 0 | 0 |
| 37715 | ENSMUSG00000099328 | 0 | NA | NA | NA | NA | NA | NA | NA | 0 | 0 | 0 | 0 | 0 | 0 |
| 37716 | ENSMUSG00000099329 | 0 | NA | NA | NA | NA | NA | Gm28052 | predicted gene, 28052 [Source:MGI Symb | 0 | 0 | 0 | 0 | 0 | 0 |
| 37717 | ENSMUSG00000099330 | 0 | NA | NA | NA | NA | NA | Mir7053 | microRNA 7053 [Source:MGI Symbol;Acc: | 0 | 0 | 0 | 0 | 0 | 0 |
| 37718 | ENSMUSG00000099331 | 0 | NA | NA | NA | NA | NA | Mir7239 | microRNA 7239 [Source:MGI Symbol;Acc: | 0 | 0 | 0 | 0 | 0 | 0 |
| 37719 | ENSMUSG00000099332 | 0 | NA | NA | NA | NA | NA | NA | NA | 0 | 0 | 0 | 0 | 0 | 0 |
| 37720 | ENSMUSG00000099334 | 0 | NA | NA | NA | NA | NA | NA | NA | 0 | 0 | 0 | 0 | 0 | 0 |
| 37721 | ENSMUSG00000099335 | 0 | NA | NA | NA | NA | NA | Gm29635 | predicted gene 29635 [Source:MGI Symbo | 0 | 0 | 0 | 0 | 0 | 0 |
| 37722 | ENSMUSG00000099336 | 0 | NA | NA | NA | NA | NA | Gm29348 | predicted gene 29348 [Source:MGI Symbo | 0 | 0 | 0 | 0 | 0 | 0 |
| 37723 | ENSMUSG00000099337 | 0 | NA | NA | NA | NA | NA | Gm29486 | predicted gene 29486 [Source:MGI Symbo | 0 | 0 | 0 | 0 | 0 | 0 |
| 37724 | ENSMUSG00000099338 | 67.11722984 | 0.61689239 | 0.317718226 | 1.941633621 | 0.05218147 | 0.321125242 | 2810030D12 | RIKEN cDNA 2810030D12 gene [Source:M | 73.082 | 37.005 | 48.3 | 95.434 | 69.8 | 79.083 |
| 37725 | ENSMUSG00000099339 | 0 | NA | NA | NA | NA | NA | Gm21751 | predicted gene, 21751 [Source:MGI Symb | 0 | 0 | 0 | 0 | 0 | 0 |
| 37726 | ENSMUSG00000099340 | 0 | NA | NA | NA | NA | NA | Gm29540 | predicted gene 29540 [Source:MGI Symbo | 0 | 0 | 0 | 0 | 0 | 0 |
| 37727 | ENSMUSG00000099341 | 0 | NA | NA | NA | NA | NA | Gm29090 | predicted gene 29090 [Source:MGI Symbo | 0 | 0 | 0 | 0 | 0 | 0 |
| 37728 | ENSMUSG00000099342 | 3.247257012 | -1.088534612 | 1.276572007 | -0.852701302 | 0.393824969 | NA | Gm18180 | predicted gene, 18180 [Source:MGI Symb | 3.2124 | 1.0279 | 9.0562 | 2.8919 | 2.1812 | 1.1138 |
| 37729 | ENSMUSG00000099343 | 33.96591509 | 0.315201267 | 0.446563488 | 0.705837525 | 0.480289183 | 0.870278888 | Gm28836 | predicted gene 28836 [Source:MGI Symbo | 46.58 | 25.698 | 18.112 | 37.595 | 49.078 | 26.732 |
| 37730 | ENSMUSG00000099344 | 0 | NA | NA | NA | NA | NA | Gm6669 | predicted pseudogene 6669 [Source:MGI | 0 | 0 | 0 | 0 | 0 | 0 |
| 37731 | ENSMUSG00000099346 | 0 | NA | NA | NA | NA | NA | Gm20895 | predicted gene, 20895 [Source:MGI Symb | 0 | 0 | 0 | 0 | 0 | 0 |
| 37732 | ENSMUSG00000099347 | 0 | NA | NA | NA | NA | NA | Gm28093 | predicted gene 28093 [Source:MGI Symbo | 0 | 0 | 0 | 0 | 0 | 0 |
| 37733 | ENSMUSG00000099348 | 0 | NA | NA | NA | NA | NA | Gm28132 | predicted gene 28132 [Source:MGI Symbo | 0 | 0 | 0 | 0 | 0 | 0 |
| 37734 | ENSMUSG00000099349 | 0 | NA | NA | NA | NA | NA | 1700047G03 | RIKEN cDNA 1700047G03 gene [Source:M | 0 | 0 | 0 | 0 | 0 | 0 |
| 37735 | ENSMUSG00000099350 | 0 | NA | NA | NA | NA | NA | Gm29623 | predicted gene 29623 [Source:MGI Symbo | 0 | 0 | 0 | 0 | 0 | 0 |
| 37736 | ENSMUSG00000099351 | 0 | NA | NA | NA | NA | NA | Gm28776 | predicted gene 28776 [Source:MGI Symbo | 0 | 0 | 0 | 0 | 0 | 0 |
| 37737 | ENSMUSG00000099352 | 0 | NA | NA | NA | NA | NA | Gm28920 | predicted gene 28920 [Source:MGI Symbo | 0 | 0 | 0 | 0 | 0 | 0 |
| 37738 | ENSMUSG00000099353 | 0 | NA | NA | NA | NA | NA | 1700031M1( | RIKEN cDNA 1700031M16 gene [Source:M | 0 | 0 | 0 | 0 | 0 | 0 |
| 37739 | ENSMUSG00000099354 | 0.17131964 | -0.796629176 | 4.080472857 | -0.195229623 | 0.84521319 | NA | 1700124L16 | RIKEN cDNA 1700124L16 gene [Source:M( | 0 | 1.0279 | 0 | 0 | 0 | 0 |
| 37740 | ENSMUSG00000099355 | 0 | NA | NA | NA | NA | NA | Gm29342 | predicted gene 29342 [Source:MGI Symbo | 0 | 0 | 0 | 0 | 0 | 0 |

| | | | | | | | | | | | | | | | |
|---|---|---|---|---|---|---|---|---|---|---|---|---|---|---|---|
| 37741 | ENSMUSG00000099357 | 0 | NA | NA | NA | NA | NA | Scgb1b1-ps | secretoglobin, family 1B, member 1, pseu | 0 | 0 | 0 | 0 | 0 | 0 |
| 37742 | ENSMUSG00000099359 | 0 | NA | NA | NA | NA | NA | Gm28934 | predicted gene 28934 [Source:MGI Symbo | 0 | 0 | 0 | 0 | 0 | 0 |
| 37743 | ENSMUSG00000099360 | 0 | NA | NA | NA | NA | NA | 1700022H16 | RIKEN cDNA 1700022H16 gene [Source:M | 0 | 0 | 0 | 0 | 0 | 0 |
| 37744 | ENSMUSG00000099361 | 0 | NA | NA | NA | NA | NA | Gm28492 | predicted gene 28492 [Source:MGI Symbo | 0 | 0 | 0 | 0 | 0 | 0 |
| 37745 | ENSMUSG00000099362 | 0 | NA | NA | NA | NA | NA | Gm28970 | predicted gene 28970 [Source:MGI Symbo | 0 | 0 | 0 | 0 | 0 | 0 |
| 37746 | ENSMUSG00000099363 | 0 | NA | NA | NA | NA | NA | Gm28478 | predicted gene 28478 [Source:MGI Symbo | 0 | 0 | 0 | 0 | 0 | 0 |
| 37747 | ENSMUSG00000099364 | 0 | NA | NA | NA | NA | NA | 5730419F03 | RIKEN cDNA 5730419F03 gene [Source:M | 0 | 0 | 0 | 0 | 0 | 0 |
| 37748 | ENSMUSG00000099366 | 0 | NA | NA | NA | NA | NA | Gm2497 | predicted gene 2497 [Source:MGI Symbol | 0 | 0 | 0 | 0 | 0 | 0 |
| 37749 | ENSMUSG00000099367 | 0 | NA | NA | NA | NA | NA | Gm29512 | predicted gene 29512 [Source:MGI Symbo | 0 | 0 | 0 | 0 | 0 | 0 |
| 37750 | ENSMUSG00000099368 | 0 | NA | NA | NA | NA | NA | Gm21910 | predicted gene, 21910 [Source:MGI Symb | 0 | 0 | 0 | 0 | 0 | 0 |
| 37751 | ENSMUSG00000099369 | 0 | NA | NA | NA | NA | NA | Gm28838 | predicted gene 28838 [Source:MGI Symbo | 0 | 0 | 0 | 0 | 0 | 0 |
| 37752 | ENSMUSG00000099370 | 0 | NA | NA | NA | NA | NA | Platr10 | pluripotency associated transcript 10 [Sou | 0 | 0 | 0 | 0 | 0 | 0 |
| 37753 | ENSMUSG00000099371 | 0 | NA | NA | NA | NA | NA | Gm20848 | predicted gene, 20848 [Source:MGI Symb | 0 | 0 | 0 | 0 | 0 | 0 |
| 37754 | ENSMUSG00000099372 | 0 | NA | NA | NA | NA | NA | Gm28509 | predicted gene 28509 [Source:MGI Symbo | 0 | 0 | 0 | 0 | 0 | 0 |
| 37755 | ENSMUSG00000099373 | 0.703263909 | 1.656064937 | 2.753745348 | 0.601386377 | 0.54758267 | NA | Gm19667 | predicted gene, 19667 [Source:MGI Symb | 0 | 1.0279 | 0 | 0.964 | 0 | 2.2277 |
| 37756 | ENSMUSG00000099374 | 0 | NA | NA | NA | NA | NA | Gm21344 | predicted gene, 21344 [Source:MGI Symb | 0 | 0 | 0 | 0 | 0 | 0 |
| 37757 | ENSMUSG00000099375 | 0 | NA | NA | NA | NA | NA | Gm28187 | predicted gene 28187 [Source:MGI Symbo | 0 | 0 | 0 | 0 | 0 | 0 |
| 37758 | ENSMUSG00000099376 | 0 | NA | NA | NA | NA | NA | Gm19124 | predicted gene, 19124 [Source:MGI Symb | 0 | 0 | 0 | 0 | 0 | 0 |
| 37759 | ENSMUSG00000099377 | 0.496077836 | -0.812642266 | 3.444120572 | -0.235950586 | 0.813471014 | NA | Gm6159 | predicted gene 6159 [Source:MGI Symbol | 0 | 0 | 2.0125 | 0.964 | 0 | 0 |
| 37760 | ENSMUSG00000099378 | 0 | NA | NA | NA | NA | NA | 1700067G17 | RIKEN cDNA 1700067G17 gene [Source:M | 0 | 0 | 0 | 0 | 0 | 0 |
| 37761 | ENSMUSG00000099379 | 0 | NA | NA | NA | NA | NA | Gm28993 | predicted gene 28993 [Source:MGI Symbo | 0 | 0 | 0 | 0 | 0 | 0 |
| 37762 | ENSMUSG00000099380 | 0 | NA | NA | NA | NA | NA | Gm28130 | predicted gene 28130 [Source:MGI Symbo | 0 | 0 | 0 | 0 | 0 | 0 |
| 37763 | ENSMUSG00000099381 | 0.133850373 | -0.796629176 | 4.080472857 | -0.195229623 | 0.84521319 | NA | Gm18303 | predicted gene, 18303 [Source:MGI Symb | 0.8031 | 0 | 0 | 0 | 0 | 0 |
| 37764 | ENSMUSG00000099382 | 0 | NA | NA | NA | NA | NA | Gm28631 | predicted gene 28631 [Source:MGI Symbo | 0 | 0 | 0 | 0 | 0 | 0 |
| 37765 | ENSMUSG00000099383 | 0 | NA | NA | NA | NA | NA | Gm28925 | predicted gene 28925 [Source:MGI Symbo | 0 | 0 | 0 | 0 | 0 | 0 |
| 37766 | ENSMUSG00000099384 | 0.301557804 | -1.64335916 | 4.042814367 | -0.406488899 | 0.684383412 | NA | 1700110C19 | RIKEN cDNA 1700110C19 gene [Source:M | 0.8031 | 0 | 1.0062 | 0 | 0 | 0 |
| 37767 | ENSMUSG00000099385 | 0 | NA | NA | NA | NA | NA | Gm9540 | predicted gene 9540 [Source:MGI Symbol | 0 | 0 | 0 | 0 | 0 | 0 |
| 37768 | ENSMUSG00000099386 | 0 | NA | NA | NA | NA | NA | Gm28255 | predicted gene 28255 [Source:MGI Symbo | 0 | 0 | 0 | 0 | 0 | 0 |
| 37769 | ENSMUSG00000099387 | 0 | NA | NA | NA | NA | NA | Gm29327 | predicted gene 29327 [Source:MGI Symbo | 0 | 0 | 0 | 0 | 0 | 0 |
| 37770 | ENSMUSG00000099388 | 0 | NA | NA | NA | NA | NA | Gm28966 | predicted gene 28966 [Source:MGI Symbo | 0 | 0 | 0 | 0 | 0 | 0 |
| 37771 | ENSMUSG00000099389 | 0 | NA | NA | NA | NA | NA | Gm29603 | predicted gene 29603 [Source:MGI Symbo | 0 | 0 | 0 | 0 | 0 | 0 |
| 37772 | ENSMUSG00000099390 | 0 | NA | NA | NA | NA | NA | Gm29225 | predicted gene 29225 [Source:MGI Symbo | 0 | 0 | 0 | 0 | 0 | 0 |
| 37773 | ENSMUSG00000099391 | 0 | NA | NA | NA | NA | NA | 1700003C15 | RIKEN cDNA 1700003C15 gene [Source:M | 0 | 0 | 0 | 0 | 0 | 0 |
| 37774 | ENSMUSG00000099392 | 0 | NA | NA | NA | NA | NA | Gm29414 | predicted gene 29414 [Source:MGI Symbo | 0 | 0 | 0 | 0 | 0 | 0 |
| 37775 | ENSMUSG00000099393 | 0 | NA | NA | NA | NA | NA | Gm28179 | predicted gene 28179 [Source:MGI Symbo | 0 | 0 | 0 | 0 | 0 | 0 |
| 37776 | ENSMUSG00000099394 | 0 | NA | NA | NA | NA | NA | Gm9142 | predicted gene 9142 [Source:MGI Symbol | 0 | 0 | 0 | 0 | 0 | 0 |
| 37777 | ENSMUSG00000099395 | 0 | NA | NA | NA | NA | NA | Gm28138 | predicted gene 28138 [Source:MGI Symbo | 0 | 0 | 0 | 0 | 0 | 0 |
| 37778 | ENSMUSG00000099396 | 0 | NA | NA | NA | NA | NA | Gm29263 | predicted gene 29263 [Source:MGI Symbo | 0 | 0 | 0 | 0 | 0 | 0 |
| 37779 | ENSMUSG00000099397 | 0 | NA | NA | NA | NA | NA | Gm7809 | predicted gene 7809 [Source:MGI Symbol | 0 | 0 | 0 | 0 | 0 | 0 |
| 37780 | ENSMUSG00000099398 | 1.927401184 | -4.32025846 | 2.113451212 | -2.044172316 | 0.040936527 | NA | Ms4a14 | membrane-spanning 4-domains, subfamil | 6.4248 | 5.1396 | 0 | 0 | 0 | 0 |
| 37781 | ENSMUSG00000099399 | 0 | NA | NA | NA | NA | NA | Gm28301 | predicted gene 28301 [Source:MGI Symbo | 0 | 0 | 0 | 0 | 0 | 0 |
| 37782 | ENSMUSG00000099400 | 0 | NA | NA | NA | NA | NA | Kat6b-ps1 | K(lysine) acetyltransferase 6B, pseudogen | 0 | 0 | 0 | 0 | 0 | 0 |
| 37783 | ENSMUSG00000099401 | 0.449470461 | -0.545136533 | 3.562105917 | -0.15303771 | 0.878368537 | NA | Gm28375 | predicted gene 28375 [Source:MGI Symbo | 1.6062 | 0 | 0 | 0 | 1.0906 | 0 |
| 37784 | ENSMUSG00000099403 | 0 | NA | NA | NA | NA | NA | Gm29599 | predicted gene 29599 [Source:MGI Symbo | 0 | 0 | 0 | 0 | 0 | 0 |
| 37785 | ENSMUSG00000099404 | 13.96541935 | -1.080043906 | 0.706502029 | -1.528720176 | 0.126333828 | 0.521167236 | Gm28172 | predicted gene 28172 [Source:MGI Symbo | 8.031 | 26.726 | 22.137 | 3.8559 | 16.359 | 6.6831 |
| 37786 | ENSMUSG00000099405 | 0 | NA | NA | NA | NA | NA | Gm8506 | predicted gene 8506 [Source:MGI Symbol | 0 | 0 | 0 | 0 | 0 | 0 |
| 37787 | ENSMUSG00000099406 | 0 | NA | NA | NA | NA | NA | Gm29408 | predicted gene 29408 [Source:MGI Symbo | 0 | 0 | 0 | 0 | 0 | 0 |
| 37788 | ENSMUSG00000099407 | 0 | NA | NA | NA | NA | NA | 1700060O08 | RIKEN cDNA 1700060O08 gene [Source:M | 0 | 0 | 0 | 0 | 0 | 0 |
| 37789 | ENSMUSG00000099408 | 0 | NA | NA | NA | NA | NA | Gm28311 | predicted gene 28311 [Source:MGI Symbo | 0 | 0 | 0 | 0 | 0 | 0 |
| 37790 | ENSMUSG00000099409 | 0 | NA | NA | NA | NA | NA | Gm28751 | predicted gene 28751 [Source:MGI Symbo | 0 | 0 | 0 | 0 | 0 | 0 |
| 37791 | ENSMUSG00000099410 | 0 | NA | NA | NA | NA | NA | Gm29355 | predicted gene 29355 [Source:MGI Symbo | 0 | 0 | 0 | 0 | 0 | 0 |
| 37792 | ENSMUSG00000099411 | 0 | NA | NA | NA | NA | NA | 2310015D24 | RIKEN cDNA 2310015D24 gene [Source:M | 0 | 0 | 0 | 0 | 0 | 0 |
| 37793 | ENSMUSG00000099412 | 0 | NA | NA | NA | NA | NA | Gm28223 | predicted gene 28223 [Source:MGI Symbo | 0 | 0 | 0 | 0 | 0 | 0 |
| 37794 | ENSMUSG00000099413 | 4.154771076 | 0.670565148 | 1.125792342 | 0.595638399 | 0.551416818 | NA | Gm17767 | predicted gene, 17767 [Source:MGI Symb | 1.6062 | 4.1117 | 4.025 | 9.6398 | 1.0906 | 4.4554 |
| 37795 | ENSMUSG00000099414 | 0 | NA | NA | NA | NA | NA | Gm28113 | predicted gene 28113 [Source:MGI Symbo | 0 | 0 | 0 | 0 | 0 | 0 |
| 37796 | ENSMUSG00000099415 | 0 | NA | NA | NA | NA | NA | Gm29285 | predicted gene 29285 [Source:MGI Symbo | 0 | 0 | 0 | 0 | 0 | 0 |

| | | | | | | | | | | | | | | | |
|---|---|---|---|---|---|---|---|---|---|---|---|---|---|---|---|
| 37797 | ENSMUSG00000099416 | 0 | NA | NA | NA | NA | NA | Gm28540 | predicted gene 28540 [Source:MGI Symbo | 0 | 0 | 0 | 0 | 0 | 0 |
| 37798 | ENSMUSG00000099417 | 0 | NA | NA | NA | NA | NA | 1700028B04 | RIKEN cDNA 1700028B04 gene [Source:M | 0 | 0 | 0 | 0 | 0 | 0 |
| 37799 | ENSMUSG00000099418 | 2.647385635 | 0.704870852 | 1.430294524 | 0.492815179 | 0.622143184 | NA | Gm6657 | predicted gene 6657 [Source:MGI Symbol | 0 | 3.0838 | 3.0187 | 0.964 | 4.3625 | 4.4554 |
| 37800 | ENSMUSG00000099419 | 0 | NA | NA | NA | NA | NA | 1700001D01 | RIKEN cDNA 1700001D01 gene [Source:M | 0 | 0 | 0 | 0 | 0 | 0 |
| 37801 | ENSMUSG00000099420 | 0 | NA | NA | NA | NA | NA | Gm13279 | predicted gene 13279 [Source:MGI Symbo | 0 | 0 | 0 | 0 | 0 | 0 |
| 37802 | ENSMUSG00000099421 | 0 | NA | NA | NA | NA | NA | Gm29067 | predicted gene 29067 [Source:MGI Symbo | 0 | 0 | 0 | 0 | 0 | 0 |
| 37803 | ENSMUSG00000099422 | 0.51034671 | -2.381743948 | 3.331081356 | -0.715006238 | 0.474605189 | NA | Gm4275 | predicted gene 4275 [Source:MGI Symbol | 0 | 2.0558 | 1.0062 | 0 | 0 | 0 |
| 37804 | ENSMUSG00000099423 | 0 | NA | NA | NA | NA | NA | Gm17826 | predicted gene, 17826 [Source:MGI Symb | 0 | 0 | 0 | 0 | 0 | 0 |
| 37805 | ENSMUSG00000099424 | 0 | NA | NA | NA | NA | NA | Gm28641 | predicted gene 28641 [Source:MGI Symbo | 0 | 0 | 0 | 0 | 0 | 0 |
| 37806 | ENSMUSG00000099425 | 0 | NA | NA | NA | NA | NA | Gm29573 | predicted gene 29573 [Source:MGI Symbo | 0 | 0 | 0 | 0 | 0 | 0 |
| 37807 | ENSMUSG00000099426 | 0 | NA | NA | NA | NA | NA | Gm28761 | predicted gene 28761 [Source:MGI Symbo | 0 | 0 | 0 | 0 | 0 | 0 |
| 37808 | ENSMUSG00000099427 | 0 | NA | NA | NA | NA | NA | Gm18700 | predicted gene, 18700 [Source:MGI Symb | 0 | 0 | 0 | 0 | 0 | 0 |
| 37809 | ENSMUSG00000099428 | 0 | NA | NA | NA | NA | NA | Gm28155 | predicted gene 28155 [Source:MGI Symbo | 0 | 0 | 0 | 0 | 0 | 0 |
| 37810 | ENSMUSG00000099429 | 0 | NA | NA | NA | NA | NA | 1600019K03 | RIKEN cDNA 1600019K03 gene [Source:M | 0 | 0 | 0 | 0 | 0 | 0 |
| 37811 | ENSMUSG00000099430 | 0 | NA | NA | NA | NA | NA | Gm28878 | predicted gene 28878 [Source:MGI Symbo | 0 | 0 | 0 | 0 | 0 | 0 |
| 37812 | ENSMUSG00000099431 | 0 | NA | NA | NA | NA | NA | Gm28985 | predicted gene 28985 [Source:MGI Symbo | 0 | 0 | 0 | 0 | 0 | 0 |
| 37813 | ENSMUSG00000099432 | 0 | NA | NA | NA | NA | NA | Gm18849 | predicted gene, 18849 [Source:MGI Symb | 0 | 0 | 0 | 0 | 0 | 0 |
| 37814 | ENSMUSG00000099433 | 0 | NA | NA | NA | NA | NA | Gm28344 | predicted gene 28344 [Source:MGI Symbo | 0 | 0 | 0 | 0 | 0 | 0 |
| 37815 | ENSMUSG00000099434 | 0 | NA | NA | NA | NA | NA | Gm20934 | predicted gene, 20934 [Source:MGI Symb | 0 | 0 | 0 | 0 | 0 | 0 |
| 37816 | ENSMUSG00000099435 | 0 | NA | NA | NA | NA | NA | 1700065I16F | RIKEN cDNA 1700065I16 gene [Source:M | 0 | 0 | 0 | 0 | 0 | 0 |
| 37817 | ENSMUSG00000099436 | 0 | NA | NA | NA | NA | NA | Gm28977 | predicted gene 28977 [Source:MGI Symbo | 0 | 0 | 0 | 0 | 0 | 0 |
| 37818 | ENSMUSG00000099437 | 0 | NA | NA | NA | NA | NA | Gm29135 | predicted gene 29135 [Source:MGI Symbo | 0 | 0 | 0 | 0 | 0 | 0 |
| 37819 | ENSMUSG00000099438 | 0 | NA | NA | NA | NA | NA | Gm21808 | predicted gene, 21808 [Source:MGI Symb | 0 | 0 | 0 | 0 | 0 | 0 |
| 37820 | ENSMUSG00000099439 | 0 | NA | NA | NA | NA | NA | Gm29142 | predicted gene 29142 [Source:MGI Symbo | 0 | 0 | 0 | 0 | 0 | 0 |
| 37821 | ENSMUSG00000099440 | 0 | NA | NA | NA | NA | NA | NA | NA | 0 | 0 | 0 | 0 | 0 | 0 |
| 37822 | ENSMUSG00000099441 | 0 | NA | NA | NA | NA | NA | Gm29268 | predicted gene 29268 [Source:MGI Symbo | 0 | 0 | 0 | 0 | 0 | 0 |
| 37823 | ENSMUSG00000099442 | 0 | NA | NA | NA | NA | NA | Gm28352 | predicted gene 28352 [Source:MGI Symbo | 0 | 0 | 0 | 0 | 0 | 0 |
| 37824 | ENSMUSG00000099443 | 0 | NA | NA | NA | NA | NA | H2al1h | H2A histone family member L1H [Source:I | 0 | 0 | 0 | 0 | 0 | 0 |
| 37825 | ENSMUSG00000099444 | 0 | NA | NA | NA | NA | NA | Gm29240 | predicted gene 29240 [Source:MGI Symbo | 0 | 0 | 0 | 0 | 0 | 0 |
| 37826 | ENSMUSG00000099445 | 0 | NA | NA | NA | NA | NA | Gm28629 | predicted gene 28629 [Source:MGI Symbo | 0 | 0 | 0 | 0 | 0 | 0 |
| 37827 | ENSMUSG00000099446 | 0 | NA | NA | NA | NA | NA | 2310005E17 | RIKEN cDNA 2310005E17 gene [Source:M | 0 | 0 | 0 | 0 | 0 | 0 |
| 37828 | ENSMUSG00000099447 | 0 | NA | NA | NA | NA | NA | 2310030A07 | RIKEN cDNA 2310030A07 gene [Source:M | 0 | 0 | 0 | 0 | 0 | 0 |
| 37829 | ENSMUSG00000099448 | 0 | NA | NA | NA | NA | NA | Gm18801 | predicted gene, 18801 [Source:MGI Symb | 0 | 0 | 0 | 0 | 0 | 0 |
| 37830 | ENSMUSG00000099449 | 0 | NA | NA | NA | NA | NA | Gm28401 | predicted gene 28401 [Source:MGI Symbo | 0 | 0 | 0 | 0 | 0 | 0 |
| 37831 | ENSMUSG00000099450 | 0 | NA | NA | NA | NA | NA | Gm6799 | predicted gene 6799 [Source:MGI Symbol | 0 | 0 | 0 | 0 | 0 | 0 |
| 37832 | ENSMUSG00000099451 | 0 | NA | NA | NA | NA | NA | Gm29192 | predicted gene 29192 [Source:MGI Symbo | 0 | 0 | 0 | 0 | 0 | 0 |
| 37833 | ENSMUSG00000099452 | 0 | NA | NA | NA | NA | NA | Gm28502 | predicted gene 28502 [Source:MGI Symbo | 0 | 0 | 0 | 0 | 0 | 0 |
| 37834 | ENSMUSG00000099453 | 0 | NA | NA | NA | NA | NA | Gm18248 | predicted gene, 18248 [Source:MGI Symb | 0 | 0 | 0 | 0 | 0 | 0 |
| 37835 | ENSMUSG00000099454 | 0 | NA | NA | NA | NA | NA | Gm8367 | predicted gene 8367 [Source:MGI Symbol | 0 | 0 | 0 | 0 | 0 | 0 |
| 37836 | ENSMUSG00000099455 | 0 | NA | NA | NA | NA | NA | Gm29288 | predicted gene 29288 [Source:MGI Symbo | 0 | 0 | 0 | 0 | 0 | 0 |
| 37837 | ENSMUSG00000099456 | 0 | NA | NA | NA | NA | NA | Gm28293 | predicted gene 28293 [Source:MGI Symbo | 0 | 0 | 0 | 0 | 0 | 0 |
| 37838 | ENSMUSG00000099457 | 0 | NA | NA | NA | NA | NA | Gm21794 | predicted gene, 21794 [Source:MGI Symb | 0 | 0 | 0 | 0 | 0 | 0 |
| 37839 | ENSMUSG00000099458 | 0 | NA | NA | NA | NA | NA | Gm19123 | predicted gene, 19123 [Source:MGI Symb | 0 | 0 | 0 | 0 | 0 | 0 |
| 37840 | ENSMUSG00000099459 | 0 | NA | NA | NA | NA | NA | Gm28556 | predicted gene 28556 [Source:MGI Symbo | 0 | 0 | 0 | 0 | 0 | 0 |
| 37841 | ENSMUSG00000099460 | 0 | NA | NA | NA | NA | NA | Gm21735 | predicted gene, 21735 [Source:MGI Symb | 0 | 0 | 0 | 0 | 0 | 0 |
| 37842 | ENSMUSG00000099461 | 0 | NA | NA | NA | NA | NA | Gm28521 | predicted gene 28521 [Source:MGI Symbo | 0 | 0 | 0 | 0 | 0 | 0 |
| 37843 | ENSMUSG00000099462 | 0 | NA | NA | NA | NA | NA | Gm28549 | predicted gene 28549 [Source:MGI Symbo | 0 | 0 | 0 | 0 | 0 | 0 |
| 37844 | ENSMUSG00000099463 | 0 | NA | NA | NA | NA | NA | Gm29081 | predicted gene 29081 [Source:MGI Symbo | 0 | 0 | 0 | 0 | 0 | 0 |
| 37845 | ENSMUSG00000099464 | 0 | NA | NA | NA | NA | NA | Gm5260 | predicted gene 5260 [Source:MGI Symbol | 0 | 0 | 0 | 0 | 0 | 0 |
| 37846 | ENSMUSG00000099465 | 0.428363721 | -0.543730382 | 3.624677344 | -0.15000794 | 0.880758351 | NA | Gm3830 | predicted gene 3830 [Source:MGI Symbol | 1.6062 | 0 | 0 | 0.964 | 0 | 0 |
| 37847 | ENSMUSG00000099466 | 0 | NA | NA | NA | NA | NA | Gm29304 | predicted gene 29304 [Source:MGI Symbo | 0 | 0 | 0 | 0 | 0 | 0 |
| 37848 | ENSMUSG00000099467 | 0 | NA | NA | NA | NA | NA | Gm28116 | predicted gene 28116 [Source:MGI Symbo | 0 | 0 | 0 | 0 | 0 | 0 |
| 37849 | ENSMUSG00000099468 | 0 | NA | NA | NA | NA | NA | Gm28271 | predicted gene 28271 [Source:MGI Symbo | 0 | 0 | 0 | 0 | 0 | 0 |
| 37850 | ENSMUSG00000099469 | 0 | NA | NA | NA | NA | NA | Gm29141 | predicted gene 29141 [Source:MGI Symbo | 0 | 0 | 0 | 0 | 0 | 0 |
| 37851 | ENSMUSG00000099470 | 0.167707431 | -0.796629176 | 4.080472857 | -0.195229623 | 0.84521319 | NA | Gm29340 | predicted gene 29340 [Source:MGI Symbo | 0 | 0 | 1.0062 | 0 | 0 | 0 |
| 37852 | ENSMUSG00000099471 | 4.141265703 | 0.057407323 | 1.015449353 | 0.05653391 | 0.954916482 | NA | Gm8451 | predicted gene 8451 [Source:MGI Symbol | 4.0155 | 5.1396 | 3.0187 | 3.8559 | 4.3625 | 4.4554 |

| | | | | | | | | | | | | | | | |
|---|---|---|---|---|---|---|---|---|---|---|---|---|---|---|---|
| 37853 | ENSMUSG00000099472 | 0 | NA | NA | NA | NA | NA | Gm29539 | predicted gene 29539 [Source:MGI Symbo | 0 | 0 | 0 | 0 | 0 | 0 |
| 37854 | ENSMUSG00000099473 | 0 | NA | NA | NA | NA | NA | Gm18775 | predicted gene, 18775 [Source:MGI Symb | 0 | 0 | 0 | 0 | 0 | 0 |
| 37855 | ENSMUSG00000099474 | 0.185640647 | 1.126924175 | 4.080472857 | 0.276174898 | 0.782413736 | NA | 1700097N02 | RIKEN cDNA 1700097N02 gene [Source:M | 0 | 0 | 0 | 0 | 0 | 1.1138 |
| 37856 | ENSMUSG00000099476 | 0 | NA | NA | NA | NA | NA | 4930488N24 | RIKEN cDNA 4930488N24 gene [Source:M | 0 | 0 | 0 | 0 | 0 | 0 |
| 37857 | ENSMUSG00000099477 | 0 | NA | NA | NA | NA | NA | Gm28349 | predicted gene 28349 [Source:MGI Symbo | 0 | 0 | 0 | 0 | 0 | 0 |
| 37858 | ENSMUSG00000099478 | 55.83543045 | 0.002778359 | 0.351309594 | 0.007908578 | 0.993689934 | 0.99859176 | Gm28370 | predicted gene 28370 [Source:MGI Symbo | 49.792 | 48.312 | 69.431 | 64.587 | 32.719 | 70.172 |
| 37859 | ENSMUSG00000099479 | 0 | NA | NA | NA | NA | NA | Gm29146 | predicted gene 29146 [Source:MGI Symbo | 0 | 0 | 0 | 0 | 0 | 0 |
| 37860 | ENSMUSG00000099480 | 0 | NA | NA | NA | NA | NA | Gm28124 | predicted gene 28124 [Source:MGI Symbo | 0 | 0 | 0 | 0 | 0 | 0 |
| 37861 | ENSMUSG00000099481 | 29.97536292 | 0.540816507 | 0.393317698 | 1.375011879 | 0.169127762 | 0.596414836 | Xndc1 | Xrcc1 N-terminal domain containing 1 [So | 22.487 | 24.67 | 26.162 | 33.739 | 33.809 | 38.985 |
| 37862 | ENSMUSG00000099482 | 0 | NA | NA | NA | NA | NA | Gm28955 | predicted gene 28955 [Source:MGI Symbo | 0 | 0 | 0 | 0 | 0 | 0 |
| 37863 | ENSMUSG00000099483 | 0 | NA | NA | NA | NA | NA | Gm28476 | predicted gene 28476 [Source:MGI Symbo | 0 | 0 | 0 | 0 | 0 | 0 |
| 37864 | ENSMUSG00000099484 | 0 | NA | NA | NA | NA | NA | Gm28282 | predicted gene 28282 [Source:MGI Symbo | 0 | 0 | 0 | 0 | 0 | 0 |
| 37865 | ENSMUSG00000099485 | 0 | NA | NA | NA | NA | NA | Gm29221 | predicted gene 29221 [Source:MGI Symbo | 0 | 0 | 0 | 0 | 0 | 0 |
| 37866 | ENSMUSG00000099486 | 0 | NA | NA | NA | NA | NA | Olfr1228 | olfactory receptor 1228 [Source:MGI Sym | 0 | 0 | 0 | 0 | 0 | 0 |
| 37867 | ENSMUSG00000099487 | 0 | NA | NA | NA | NA | NA | Gm21848 | predicted gene, 21848 [Source:MGI Symb | 0 | 0 | 0 | 0 | 0 | 0 |
| 37868 | ENSMUSG00000099488 | 1.006424547 | -2.146212304 | 2.438584396 | -0.880105814 | 0.37880199 | NA | Gm19774 | predicted gene, 19774 [Source:MGI Symb | 0 | 2.0558 | 3.0187 | 0.964 | 0 | 0 |
| 37869 | ENSMUSG00000099489 | 0 | NA | NA | NA | NA | NA | Gm21787 | predicted gene, 21787 [Source:MGI Symb | 0 | 0 | 0 | 0 | 0 | 0 |
| 37870 | ENSMUSG00000099490 | 0 | NA | NA | NA | NA | NA | Gm18802 | predicted gene, 18802 [Source:MGI Symb | 0 | 0 | 0 | 0 | 0 | 0 |
| 37871 | ENSMUSG00000099491 | 0 | NA | NA | NA | NA | NA | 4930533D04 | RIKEN cDNA 4930533D04 gene [Source:M | 0 | 0 | 0 | 0 | 0 | 0 |
| 37872 | ENSMUSG00000099492 | 1.237943018 | 0.557813438 | 2.236241869 | 0.249442355 | 0.803018626 | NA | Gm5525 | predicted gene 5525 [Source:MGI Symbol | 0 | 0 | 3.0187 | 0 | 2.1812 | 2.2277 |
| 37873 | ENSMUSG00000099493 | 0 | NA | NA | NA | NA | NA | Gm21753 | predicted gene, 21753 [Source:MGI Symb | 0 | 0 | 0 | 0 | 0 | 0 |
| 37874 | ENSMUSG00000099494 | 0 | NA | NA | NA | NA | NA | Gm28197 | predicted gene 28197 [Source:MGI Symbo | 0 | 0 | 0 | 0 | 0 | 0 |
| 37875 | ENSMUSG00000099495 | 0 | NA | NA | NA | NA | NA | Gm20800 | predicted gene, 20800 [Source:MGI Symb | 0 | 0 | 0 | 0 | 0 | 0 |
| 37876 | ENSMUSG00000099496 | 0 | NA | NA | NA | NA | NA | Gm28365 | predicted gene 28365 [Source:MGI Symbo | 0 | 0 | 0 | 0 | 0 | 0 |
| 37877 | ENSMUSG00000099497 | 0 | NA | NA | NA | NA | NA | Gm29147 | predicted gene 29147 [Source:MGI Symbo | 0 | 0 | 0 | 0 | 0 | 0 |
| 37878 | ENSMUSG00000099498 | 0 | NA | NA | NA | NA | NA | Gm29283 | predicted gene 29283 [Source:MGI Symbo | 0 | 0 | 0 | 0 | 0 | 0 |
| 37879 | ENSMUSG00000099499 | 0 | NA | NA | NA | NA | NA | Gm28787 | predicted gene 28787 [Source:MGI Symbo | 0 | 0 | 0 | 0 | 0 | 0 |
| 37880 | ENSMUSG00000099500 | 0 | NA | NA | NA | NA | NA | Gm29531 | predicted gene 29531 [Source:MGI Symbo | 0 | 0 | 0 | 0 | 0 | 0 |
| 37881 | ENSMUSG00000099501 | 0 | NA | NA | NA | NA | NA | Gm29662 | predicted gene 29662 [Source:MGI Symbo | 0 | 0 | 0 | 0 | 0 | 0 |
| 37882 | ENSMUSG00000099502 | 0 | NA | NA | NA | NA | NA | Gm28640 | predicted gene 28640 [Source:MGI Symbo | 0 | 0 | 0 | 0 | 0 | 0 |
| 37883 | ENSMUSG00000099503 | 0 | NA | NA | NA | NA | NA | Gm28822 | predicted gene 28822 [Source:MGI Symbo | 0 | 0 | 0 | 0 | 0 | 0 |
| 37884 | ENSMUSG00000099504 | 0 | NA | NA | NA | NA | NA | Gm28335 | predicted gene 28335 [Source:MGI Symbo | 0 | 0 | 0 | 0 | 0 | 0 |
| 37885 | ENSMUSG00000099505 | 0 | NA | NA | NA | NA | NA | Gm28442 | predicted gene 28442 [Source:MGI Symbo | 0 | 0 | 0 | 0 | 0 | 0 |
| 37886 | ENSMUSG00000099506 | 0 | NA | NA | NA | NA | NA | Gm28732 | predicted gene 28732 [Source:MGI Symbo | 0 | 0 | 0 | 0 | 0 | 0 |
| 37887 | ENSMUSG00000099507 | 0 | NA | NA | NA | NA | NA | Gm18970 | predicted gene, 18970 [Source:MGI Symb | 0 | 0 | 0 | 0 | 0 | 0 |
| 37888 | ENSMUSG00000099508 | 0 | NA | NA | NA | NA | NA | 1700030L20I | RIKEN cDNA 1700030L20 gene [Source:M( | 0 | 0 | 0 | 0 | 0 | 0 |
| 37889 | ENSMUSG00000099509 | 0 | NA | NA | NA | NA | NA | Gm29307 | predicted gene 29307 [Source:MGI Symbo | 0 | 0 | 0 | 0 | 0 | 0 |
| 37890 | ENSMUSG00000099510 | 0 | NA | NA | NA | NA | NA | Gm28501 | predicted gene 28501 [Source:MGI Symbo | 0 | 0 | 0 | 0 | 0 | 0 |
| 37891 | ENSMUSG00000099511 | 0 | NA | NA | NA | NA | NA | Topbp1-ps1 | topoisomerase (DNA) II binding protein 1, | 0 | 0 | 0 | 0 | 0 | 0 |
| 37892 | ENSMUSG00000099512 | 0 | NA | NA | NA | NA | NA | Gm28703 | predicted gene 28703 [Source:MGI Symbo | 0 | 0 | 0 | 0 | 0 | 0 |
| 37893 | ENSMUSG00000099513 | 0 | NA | NA | NA | NA | NA | Gm29119 | predicted gene 29119 [Source:MGI Symbo | 0 | 0 | 0 | 0 | 0 | 0 |
| 37894 | ENSMUSG00000099514 | 0 | NA | NA | NA | NA | NA | Gm28241 | predicted gene 28241 [Source:MGI Symbo | 0 | 0 | 0 | 0 | 0 | 0 |
| 37895 | ENSMUSG00000099516 | 0 | NA | NA | NA | NA | NA | Gm29066 | predicted gene 29066 [Source:MGI Symbo | 0 | 0 | 0 | 0 | 0 | 0 |
| 37896 | ENSMUSG00000099517 | 0.267700746 | -1.497481649 | 4.057001436 | -0.369110456 | 0.712045396 | NA | H3c8 | H3 clustered histone 8 [Source:MGI Symb | 1.6062 | 0 | 0 | 0 | 0 | 0 |
| 37897 | ENSMUSG00000099518 | 0 | NA | NA | NA | NA | NA | Gm13275 | predicted gene 13275 [Source:MGI Symbo | 0 | 0 | 0 | 0 | 0 | 0 |
| 37898 | ENSMUSG00000099519 | 0 | NA | NA | NA | NA | NA | Gm29253 | predicted gene 29253 [Source:MGI Symbo | 0 | 0 | 0 | 0 | 0 | 0 |
| 37899 | ENSMUSG00000099520 | 0 | NA | NA | NA | NA | NA | Gm29061 | predicted gene 29061 [Source:MGI Symbo | 0 | 0 | 0 | 0 | 0 | 0 |
| 37900 | ENSMUSG00000099521 | 0 | NA | NA | NA | NA | NA | Gm28309 | predicted gene 28309 [Source:MGI Symbo | 0 | 0 | 0 | 0 | 0 | 0 |
| 37901 | ENSMUSG00000099522 | 0 | NA | NA | NA | NA | NA | Gm28752 | predicted gene 28752 [Source:MGI Symbo | 0 | 0 | 0 | 0 | 0 | 0 |
| 37902 | ENSMUSG00000099523 | 0 | NA | NA | NA | NA | NA | Gm28327 | predicted gene 28327 [Source:MGI Symbo | 0 | 0 | 0 | 0 | 0 | 0 |
| 37903 | ENSMUSG00000099524 | 0 | NA | NA | NA | NA | NA | Gm20860 | predicted gene, 20860 [Source:MGI Symb | 0 | 0 | 0 | 0 | 0 | 0 |
| 37904 | ENSMUSG00000099525 | 0 | NA | NA | NA | NA | NA | Gm21872 | predicted gene, 21872 [Source:MGI Symb | 0 | 0 | 0 | 0 | 0 | 0 |
| 37905 | ENSMUSG00000099526 | 0 | NA | NA | NA | NA | NA | Gm29313 | predicted gene 29313 [Source:MGI Symbo | 0 | 0 | 0 | 0 | 0 | 0 |
| 37906 | ENSMUSG00000099527 | 0 | NA | NA | NA | NA | NA | Gm21819 | predicted gene, 21819 [Source:MGI Symb | 0 | 0 | 0 | 0 | 0 | 0 |
| 37907 | ENSMUSG00000099528 | 0 | NA | NA | NA | NA | NA | 2410012E07 | RIKEN cDNA 2410012E07 gene [Source:M( | 0 | 0 | 0 | 0 | 0 | 0 |
| 37908 | ENSMUSG00000099529 | 0 | NA | NA | NA | NA | NA | Gm29390 | predicted gene 29390 [Source:MGI Symbo | 0 | 0 | 0 | 0 | 0 | 0 |

| | | | | | | | | | | | | | | | |
|---|---|---|---|---|---|---|---|---|---|---|---|---|---|---|---|
| 37909 | ENSMUSG00000099530 | 0 | NA | NA | NA | NA | NA | Gm29644 | predicted gene 29644 [Source:MGI Symbc | 0 | 0 | 0 | 0 | 0 | 0 |
| 37910 | ENSMUSG00000099531 | 0 | NA | NA | NA | NA | NA | Gm20869 | predicted gene, 20869 [Source:MGI Symb | 0 | 0 | 0 | 0 | 0 | 0 |
| 37911 | ENSMUSG00000099532 | 0 | NA | NA | NA | NA | NA | Gm29098 | predicted gene 29098 [Source:MGI Symbc | 0 | 0 | 0 | 0 | 0 | 0 |
| 37912 | ENSMUSG00000099534 | 0 | NA | NA | NA | NA | NA | Gm28169 | predicted gene 28169 [Source:MGI Symbc | 0 | 0 | 0 | 0 | 0 | 0 |
| 37913 | ENSMUSG00000099535 | 0 | NA | NA | NA | NA | NA | Gm28542 | predicted gene 28542 [Source:MGI Symbc | 0 | 0 | 0 | 0 | 0 | 0 |
| 37914 | ENSMUSG00000099536 | 0 | NA | NA | NA | NA | NA | Gm9569 | predicted gene 9569 [Source:MGI Symbol | 0 | 0 | 0 | 0 | 0 | 0 |
| 37915 | ENSMUSG00000099537 | 0 | NA | NA | NA | NA | NA | Gm28938 | predicted gene 28938 [Source:MGI Symbc | 0 | 0 | 0 | 0 | 0 | 0 |
| 37916 | ENSMUSG00000099539 | 0 | NA | NA | NA | NA | NA | Gm8282 | predicted gene 8282 [Source:MGI Symbol | 0 | 0 | 0 | 0 | 0 | 0 |
| 37917 | ENSMUSG00000099541 | 0 | NA | NA | NA | NA | NA | Gm21488 | predicted gene, 21488 [Source:MGI Symb | 0 | 0 | 0 | 0 | 0 | 0 |
| 37918 | ENSMUSG00000099542 | 0 | NA | NA | NA | NA | NA | Gm28504 | predicted gene 28504 [Source:MGI Symbc | 0 | 0 | 0 | 0 | 0 | 0 |
| 37919 | ENSMUSG00000099543 | 0.928203237 | 3.416716801 | 3.030141393 | 1.12757669 | 0.259498736 | NA | Gm29322 | predicted gene 29322 [Source:MGI Symbc | 0 | 0 | 0 | 0 | 0 | 5.5692 |
| 37920 | ENSMUSG00000099544 | 0 | NA | NA | NA | NA | NA | Gm18699 | predicted gene, 18699 [Source:MGI Symb | 0 | 0 | 0 | 0 | 0 | 0 |
| 37921 | ENSMUSG00000099545 | 0 | NA | NA | NA | NA | NA | Gm13276 | predicted gene 13276 [Source:MGI Symbc | 0 | 0 | 0 | 0 | 0 | 0 |
| 37922 | ENSMUSG00000099546 | 0 | NA | NA | NA | NA | NA | Gm21802 | predicted gene, 21802 [Source:MGI Symb | 0 | 0 | 0 | 0 | 0 | 0 |
| 37923 | ENSMUSG00000099547 | 0 | NA | NA | NA | NA | NA | Gm28678 | predicted gene 28678 [Source:MGI Symbc | 0 | 0 | 0 | 0 | 0 | 0 |
| 37924 | ENSMUSG00000099548 | 0 | NA | NA | NA | NA | NA | Gm29220 | predicted gene 29220 [Source:MGI Symbc | 0 | 0 | 0 | 0 | 0 | 0 |
| 37925 | ENSMUSG00000099549 | 0 | NA | NA | NA | NA | NA | Gm29432 | predicted gene 29432 [Source:MGI Symbc | 0 | 0 | 0 | 0 | 0 | 0 |
| 37926 | ENSMUSG00000099550 | 0 | NA | NA | NA | NA | NA | Gm28079 | predicted gene 28079 [Source:MGI Symbc | 0 | 0 | 0 | 0 | 0 | 0 |
| 37927 | ENSMUSG00000099551 | 0 | NA | NA | NA | NA | NA | Gm28072 | predicted gene 28072 [Source:MGI Symbc | 0 | 0 | 0 | 0 | 0 | 0 |
| 37928 | ENSMUSG00000099552 | 0 | NA | NA | NA | NA | NA | 9830004L10I | RIKEN cDNA 9830004L10 gene [Source:M( | 0 | 0 | 0 | 0 | 0 | 0 |
| 37929 | ENSMUSG00000099553 | 0 | NA | NA | NA | NA | NA | Gm29538 | predicted gene 29538 [Source:MGI Symbc | 0 | 0 | 0 | 0 | 0 | 0 |
| 37930 | ENSMUSG00000099554 | 0 | NA | NA | NA | NA | NA | Gm28259 | predicted gene 28259 [Source:MGI Symbc | 0 | 0 | 0 | 0 | 0 | 0 |
| 37931 | ENSMUSG00000099555 | 0 | NA | NA | NA | NA | NA | Gm28081 | predicted gene 28081 [Source:MGI Symbc | 0 | 0 | 0 | 0 | 0 | 0 |
| 37932 | ENSMUSG00000099556 | 12.50957316 | -0.299786218 | 0.605474146 | -0.495126374 | 0.620510918 | 0.917977899 | Gm28857 | predicted gene 28857 [Source:MGI Symbc | 16.865 | 14.391 | 10.062 | 10.604 | 11.997 | 11.138 |
| 37933 | ENSMUSG00000099558 | 0 | NA | NA | NA | NA | NA | Gm29295 | predicted gene 29295 [Source:MGI Symbc | 0 | 0 | 0 | 0 | 0 | 0 |
| 37934 | ENSMUSG00000099559 | 0 | NA | NA | NA | NA | NA | Gm29063 | predicted gene 29063 [Source:MGI Symbc | 0 | 0 | 0 | 0 | 0 | 0 |
| 37935 | ENSMUSG00000099560 | 0 | NA | NA | NA | NA | NA | Mrgprx3-ps | MAS-related GPR, member X3, pseudoger | 0 | 0 | 0 | 0 | 0 | 0 |
| 37936 | ENSMUSG00000099561 | 0 | NA | NA | NA | NA | NA | Gm28410 | predicted gene 28410 [Source:MGI Symbc | 0 | 0 | 0 | 0 | 0 | 0 |
| 37937 | ENSMUSG00000099562 | 0 | NA | NA | NA | NA | NA | Gm19027 | predicted gene, 19027 [Source:MGI Symb | 0 | 0 | 0 | 0 | 0 | 0 |
| 37938 | ENSMUSG00000099563 | 0.160662974 | 1.126924175 | 4.080472857 | 0.276174898 | 0.782413736 | NA | Gm19125 | predicted gene, 19125 [Source:MGI Symb | 0 | 0 | 0 | 0.964 | 0 | 0 |
| 37939 | ENSMUSG00000099564 | 0 | NA | NA | NA | NA | NA | Gm28729 | predicted gene 28729 [Source:MGI Symbc | 0 | 0 | 0 | 0 | 0 | 0 |
| 37940 | ENSMUSG00000099565 | 0 | NA | NA | NA | NA | NA | Gm28738 | predicted gene 28738 [Source:MGI Symbc | 0 | 0 | 0 | 0 | 0 | 0 |
| 37941 | ENSMUSG00000099566 | 0 | NA | NA | NA | NA | NA | Gm28099 | predicted gene 28099 [Source:MGI Symbc | 0 | 0 | 0 | 0 | 0 | 0 |
| 37942 | ENSMUSG00000099568 | 19.71374149 | 0.135129564 | 0.601966301 | 0.224480281 | 0.822383609 | 0.969197142 | Gm28513 | predicted gene 28513 [Source:MGI Symbc | 20.881 | 21.586 | 14.087 | 37.595 | 17.45 | 6.6831 |
| 37943 | ENSMUSG00000099569 | 0 | NA | NA | NA | NA | NA | Gm18301 | predicted gene, 18301 [Source:MGI Symb | 0 | 0 | 0 | 0 | 0 | 0 |
| 37944 | ENSMUSG00000099570 | 0 | NA | NA | NA | NA | NA | Gm29000 | predicted gene 29000 [Source:MGI Symbc | 0 | 0 | 0 | 0 | 0 | 0 |
| 37945 | ENSMUSG00000099571 | 0 | NA | NA | NA | NA | NA | Gm18182 | predicted gene, 18182 [Source:MGI Symb | 0 | 0 | 0 | 0 | 0 | 0 |
| 37946 | ENSMUSG00000099572 | 0 | NA | NA | NA | NA | NA | Gm28951 | predicted gene 28951 [Source:MGI Symbc | 0 | 0 | 0 | 0 | 0 | 0 |
| 37947 | ENSMUSG00000099573 | 0 | NA | NA | NA | NA | NA | Gm28854 | predicted gene 28854 [Source:MGI Symbc | 0 | 0 | 0 | 0 | 0 | 0 |
| 37948 | ENSMUSG00000099574 | 0 | NA | NA | NA | NA | NA | 1700121L03I | RIKEN cDNA 1700121L03 gene [Source:M( | 0 | 0 | 0 | 0 | 0 | 0 |
| 37949 | ENSMUSG00000099575 | 0 | NA | NA | NA | NA | NA | Gm29350 | predicted gene 29350 [Source:MGI Symbc | 0 | 0 | 0 | 0 | 0 | 0 |
| 37950 | ENSMUSG00000099576 | 0 | NA | NA | NA | NA | NA | Gm29040 | predicted gene 29040 [Source:MGI Symbc | 0 | 0 | 0 | 0 | 0 | 0 |
| 37951 | ENSMUSG00000099577 | 0 | NA | NA | NA | NA | NA | Gm3605 | predicted gene 3605 [Source:MGI Symbol | 0 | 0 | 0 | 0 | 0 | 0 |
| 37952 | ENSMUSG00000099578 | 0 | NA | NA | NA | NA | NA | Gm5731 | predicted gene 5731 [Source:MGI Symbol | 0 | 0 | 0 | 0 | 0 | 0 |
| 37953 | ENSMUSG00000099579 | 0 | NA | NA | NA | NA | NA | 1700120K04 | RIKEN cDNA 1700120K04 gene [Source:M( | 0 | 0 | 0 | 0 | 0 | 0 |
| 37954 | ENSMUSG00000099581 | 0 | NA | NA | NA | NA | NA | Scgb1b10 | secretoglobin, family 1B, member 10 [Sou | 0 | 0 | 0 | 0 | 0 | 0 |
| 37955 | ENSMUSG00000099582 | 0 | NA | NA | NA | NA | NA | Gm29014 | predicted gene 29014 [Source:MGI Symbc | 0 | 0 | 0 | 0 | 0 | 0 |
| 37956 | ENSMUSG00000099583 | 0.679159517 | 1.687875387 | 3.079592583 | 0.548083989 | 0.583634234 | NA | H3c4 | H3 clustered histone 4 [Source:MGI Symb | 0.8031 | 0 | 0 | 0 | 3.2719 | 0 |
| 37957 | ENSMUSG00000099584 | 0 | NA | NA | NA | NA | NA | Gm28877 | predicted gene 28877 [Source:MGI Symbc | 0 | 0 | 0 | 0 | 0 | 0 |
| 37958 | ENSMUSG00000099585 | 0 | NA | NA | NA | NA | NA | Gm29544 | predicted gene 29544 [Source:MGI Symbc | 0 | 0 | 0 | 0 | 0 | 0 |
| 37959 | ENSMUSG00000099586 | 0 | NA | NA | NA | NA | NA | Gm28222 | predicted gene 28222 [Source:MGI Symbc | 0 | 0 | 0 | 0 | 0 | 0 |
| 37960 | ENSMUSG00000099587 | 0 | NA | NA | NA | NA | NA | Gm28967 | predicted gene 28967 [Source:MGI Symbc | 0 | 0 | 0 | 0 | 0 | 0 |
| 37961 | ENSMUSG00000099588 | 0 | NA | NA | NA | NA | NA | Gm29298 | predicted gene 29298 [Source:MGI Symbc | 0 | 0 | 0 | 0 | 0 | 0 |
| 37962 | ENSMUSG00000099589 | 0 | NA | NA | NA | NA | NA | Gm21898 | predicted gene, 21898 [Source:MGI Symb | 0 | 0 | 0 | 0 | 0 | 0 |
| 37963 | ENSMUSG00000099590 | 14.94131113 | 0.10304801 | 0.585696604 | 0.175940938 | 0.860340343 | 0.975844003 | C330022C24 | RIKEN cDNA C330022C24 gene [Source:M | 13.653 | 19.53 | 10.062 | 15.424 | 9.8156 | 21.163 |
| 37964 | ENSMUSG00000099591 | 0 | NA | NA | NA | NA | NA | Gm17969 | predicted gene, 17969 [Source:MGI Symb | 0 | 0 | 0 | 0 | 0 | 0 |

| | | | | | | | | | | | | | | | |
|---|---|---|---|---|---|---|---|---|---|---|---|---|---|---|---|
| 37965 | ENSMUSG00000099592 | 0 | NA | NA | NA | NA | NA | Gm5264 | predicted gene 5264 [Source:MGI Symbol | 0 | 0 | 0 | 0 | 0 | 0 |
| 37966 | ENSMUSG00000099593 | 0 | NA | NA | NA | NA | NA | Gm28566 | predicted gene 28566 [Source:MGI Symbo | 0 | 0 | 0 | 0 | 0 | 0 |
| 37967 | ENSMUSG00000099594 | 0 | NA | NA | NA | NA | NA | Gm21740 | predicted gene, 21740 [Source:MGI Symb | 0 | 0 | 0 | 0 | 0 | 0 |
| 37968 | ENSMUSG00000099595 | 0 | NA | NA | NA | NA | NA | Gm29406 | predicted gene 29406 [Source:MGI Symbo | 0 | 0 | 0 | 0 | 0 | 0 |
| 37969 | ENSMUSG00000099596 | 0 | NA | NA | NA | NA | NA | Zbed4-ps3 | zinc finger, BED type containing 4, pseudo | 0 | 0 | 0 | 0 | 0 | 0 |
| 37970 | ENSMUSG00000099598 | 0 | NA | NA | NA | NA | NA | Scgb2b28-ps | secretoglobin, family 2B, member 28, pse | 0 | 0 | 0 | 0 | 0 | 0 |
| 37971 | ENSMUSG00000099599 | 0 | NA | NA | NA | NA | NA | Gm28548 | predicted gene 28548 [Source:MGI Symbo | 0 | 0 | 0 | 0 | 0 | 0 |
| 37972 | ENSMUSG00000099600 | 0 | NA | NA | NA | NA | NA | Gm29126 | predicted gene 29126 [Source:MGI Symbo | 0 | 0 | 0 | 0 | 0 | 0 |
| 37973 | ENSMUSG00000099601 | 0 | NA | NA | NA | NA | NA | Gm29671 | predicted gene 29671 [Source:MGI Symbo | 0 | 0 | 0 | 0 | 0 | 0 |
| 37974 | ENSMUSG00000099603 | 0 | NA | NA | NA | NA | NA | Mrgprc3-ps | MAS-related GPR, member C3, pseudoger | 0 | 0 | 0 | 0 | 0 | 0 |
| 37975 | ENSMUSG00000099604 | 0 | NA | NA | NA | NA | NA | Gm6651 | predicted gene 6651 [Source:MGI Symbol | 0 | 0 | 0 | 0 | 0 | 0 |
| 37976 | ENSMUSG00000099605 | 0 | NA | NA | NA | NA | NA | 1700066N21 | RIKEN cDNA 1700066N21 gene [Source:M | 0 | 0 | 0 | 0 | 0 | 0 |
| 37977 | ENSMUSG00000099606 | 0 | NA | NA | NA | NA | NA | Gm28537 | predicted gene 28537 [Source:MGI Symbo | 0 | 0 | 0 | 0 | 0 | 0 |
| 37978 | ENSMUSG00000099607 | 0 | NA | NA | NA | NA | NA | Gm28205 | predicted gene 28205 [Source:MGI Symbo | 0 | 0 | 0 | 0 | 0 | 0 |
| 37979 | ENSMUSG00000099608 | 0 | NA | NA | NA | NA | NA | 4933411E06 | RIKEN cDNA 4933411E06 gene [Source:M | 0 | 0 | 0 | 0 | 0 | 0 |
| 37980 | ENSMUSG00000099609 | 0 | NA | NA | NA | NA | NA | Gm29008 | predicted gene 29008 [Source:MGI Symbo | 0 | 0 | 0 | 0 | 0 | 0 |
| 37981 | ENSMUSG00000099610 | 0 | NA | NA | NA | NA | NA | Gm37545 | predicted gene, 37545 [Source:MGI Symb | 0 | 0 | 0 | 0 | 0 | 0 |
| 37982 | ENSMUSG00000099611 | 0 | NA | NA | NA | NA | NA | Vmn1r189 | vomeronasal 1 receptor 189 [Source:MGI | 0 | 0 | 0 | 0 | 0 | 0 |
| 37983 | ENSMUSG00000099612 | 0 | NA | NA | NA | NA | NA | Scgb1b8-ps | secretoglobin, family 1B, member 8, pseu | 0 | 0 | 0 | 0 | 0 | 0 |
| 37984 | ENSMUSG00000099613 | 0 | NA | NA | NA | NA | NA | Gm28313 | predicted gene 28313 [Source:MGI Symbo | 0 | 0 | 0 | 0 | 0 | 0 |
| 37985 | ENSMUSG00000099614 | 0 | NA | NA | NA | NA | NA | Uba1y-ps2 | ubiquitin-activating enzyme, Chr Y, pseud | 0 | 0 | 0 | 0 | 0 | 0 |
| 37986 | ENSMUSG00000099615 | 0 | NA | NA | NA | NA | NA | Gm28362 | predicted gene 28362 [Source:MGI Symbo | 0 | 0 | 0 | 0 | 0 | 0 |
| 37987 | ENSMUSG00000099616 | 0 | NA | NA | NA | NA | NA | 4932413F04 | RIKEN cDNA 4932413F04 gene [Source:M | 0 | 0 | 0 | 0 | 0 | 0 |
| 37988 | ENSMUSG00000099617 | 0 | NA | NA | NA | NA | NA | Gm6550 | predicted gene 6550 [Source:MGI Symbol | 0 | 0 | 0 | 0 | 0 | 0 |
| 37989 | ENSMUSG00000099618 | 0.455176322 | 0.955156269 | 3.546674842 | 0.269310357 | 0.787690862 | NA | Gm28118 | predicted gene 28118 [Source:MGI Symbo | 0.8031 | 0 | 0 | 1.928 | 0 | 0 |
| 37990 | ENSMUSG00000099619 | 0 | NA | NA | NA | NA | NA | Gm28300 | predicted gene 28300 [Source:MGI Symbo | 0 | 0 | 0 | 0 | 0 | 0 |
| 37991 | ENSMUSG00000099620 | 0 | NA | NA | NA | NA | NA | Mrgprb13 | MAS-related GPR, member B13 [Source:M | 0 | 0 | 0 | 0 | 0 | 0 |
| 37992 | ENSMUSG00000099621 | 0 | NA | NA | NA | NA | NA | Gm20919 | predicted gene, 20919 [Source:MGI Symb | 0 | 0 | 0 | 0 | 0 | 0 |
| 37993 | ENSMUSG00000099622 | 0 | NA | NA | NA | NA | NA | Gm29188 | predicted gene 29188 [Source:MGI Symbo | 0 | 0 | 0 | 0 | 0 | 0 |
| 37994 | ENSMUSG00000099623 | 0 | NA | NA | NA | NA | NA | Gm4526 | predicted gene 4526 [Source:MGI Symbol | 0 | 0 | 0 | 0 | 0 | 0 |
| 37995 | ENSMUSG00000099624 | 0 | NA | NA | NA | NA | NA | Gm7889 | predicted gene 7889 [Source:MGI Symbol | 0 | 0 | 0 | 0 | 0 | 0 |
| 37996 | ENSMUSG00000099625 | 0 | NA | NA | NA | NA | NA | Gm29325 | predicted gene 29325 [Source:MGI Symbo | 0 | 0 | 0 | 0 | 0 | 0 |
| 37997 | ENSMUSG00000099626 | 0 | NA | NA | NA | NA | NA | Gm28185 | predicted gene 28185 [Source:MGI Symbo | 0 | 0 | 0 | 0 | 0 | 0 |
| 37998 | ENSMUSG00000099627 | 0 | NA | NA | NA | NA | NA | Gm6757 | predicted gene 6757 [Source:MGI Symbol | 0 | 0 | 0 | 0 | 0 | 0 |
| 37999 | ENSMUSG00000099628 | 0 | NA | NA | NA | NA | NA | Gm28427 | predicted gene 28427 [Source:MGI Symbo | 0 | 0 | 0 | 0 | 0 | 0 |
| 38000 | ENSMUSG00000099629 | 0 | NA | NA | NA | NA | NA | Gm29515 | predicted gene 29515 [Source:MGI Symbo | 0 | 0 | 0 | 0 | 0 | 0 |
| 38001 | ENSMUSG00000099630 | 0 | NA | NA | NA | NA | NA | 1700096J18I | RIKEN cDNA 1700096J18 gene [Source:M | 0 | 0 | 0 | 0 | 0 | 0 |
| 38002 | ENSMUSG00000099631 | 0.160662974 | 1.126924175 | 4.080472857 | 0.276174898 | 0.782413736 | NA | Gm8641 | predicted gene 8641 [Source:MGI Symbol | 0 | 0 | 0 | 0.964 | 0 | 0 |
| 38003 | ENSMUSG00000099632 | 18.56825823 | -0.117996323 | 0.558042546 | -0.211446821 | 0.832538624 | 0.97056812 | 2900093K20 | RIKEN cDNA 2900093K20 gene [Source:M | 11.243 | 26.726 | 20.125 | 13.496 | 13.087 | 26.732 |
| 38004 | ENSMUSG00000099633 | 0 | NA | NA | NA | NA | NA | Gm29071 | predicted gene 29071 [Source:MGI Symbo | 0 | 0 | 0 | 0 | 0 | 0 |
| 38005 | ENSMUSG00000099634 | 0 | NA | NA | NA | NA | NA | Gm29462 | predicted gene 29462 [Source:MGI Symbo | 0 | 0 | 0 | 0 | 0 | 0 |
| 38006 | ENSMUSG00000099635 | 0 | NA | NA | NA | NA | NA | Gm28528 | predicted gene 28528 [Source:MGI Symbo | 0 | 0 | 0 | 0 | 0 | 0 |
| 38007 | ENSMUSG00000099636 | 0 | NA | NA | NA | NA | NA | Gm28422 | predicted gene 28422 [Source:MGI Symbo | 0 | 0 | 0 | 0 | 0 | 0 |
| 38008 | ENSMUSG00000099637 | 0 | NA | NA | NA | NA | NA | Gm28737 | predicted gene 28737 [Source:MGI Symbo | 0 | 0 | 0 | 0 | 0 | 0 |
| 38009 | ENSMUSG00000099638 | 0 | NA | NA | NA | NA | NA | Gm28279 | predicted gene 28279 [Source:MGI Symbo | 0 | 0 | 0 | 0 | 0 | 0 |
| 38010 | ENSMUSG00000099639 | 0 | NA | NA | NA | NA | NA | 1700084F23 | RIKEN cDNA 1700084F23 gene [Source:M | 0 | 0 | 0 | 0 | 0 | 0 |
| 38011 | ENSMUSG00000099640 | 0 | NA | NA | NA | NA | NA | Gm28543 | predicted gene 28543 [Source:MGI Symbo | 0 | 0 | 0 | 0 | 0 | 0 |
| 38012 | ENSMUSG00000099641 | 0 | NA | NA | NA | NA | NA | Gm29448 | predicted gene 29448 [Source:MGI Symbo | 0 | 0 | 0 | 0 | 0 | 0 |
| 38013 | ENSMUSG00000099642 | 0 | NA | NA | NA | NA | NA | Gm28348 | predicted gene 28348 [Source:MGI Symbo | 0 | 0 | 0 | 0 | 0 | 0 |
| 38014 | ENSMUSG00000099643 | 0 | NA | NA | NA | NA | NA | Gm29162 | predicted gene 29162 [Source:MGI Symbo | 0 | 0 | 0 | 0 | 0 | 0 |
| 38015 | ENSMUSG00000099644 | 0 | NA | NA | NA | NA | NA | Gm28626 | predicted gene 28626 [Source:MGI Symbo | 0 | 0 | 0 | 0 | 0 | 0 |
| 38016 | ENSMUSG00000099645 | 0 | NA | NA | NA | NA | NA | Gm20832 | predicted gene, 20832 [Source:MGI Symb | 0 | 0 | 0 | 0 | 0 | 0 |
| 38017 | ENSMUSG00000099646 | 0 | NA | NA | NA | NA | NA | Gm28541 | predicted gene 28541 [Source:MGI Symbo | 0 | 0 | 0 | 0 | 0 | 0 |
| 38018 | ENSMUSG00000099647 | 2.38902242 | 0.665634086 | 1.595672099 | 0.417149667 | 0.676568938 | NA | Gm5776 | predicted gene 5776 [Source:MGI Symbol | 2.4093 | 3.0838 | 0 | 0 | 3.2719 | 5.5692 |
| 38019 | ENSMUSG00000099648 | 0 | NA | NA | NA | NA | NA | Gm5975 | predicted gene 5975 [Source:MGI Symbol | 0 | 0 | 0 | 0 | 0 | 0 |
| 38020 | ENSMUSG00000099649 | 0 | NA | NA | NA | NA | NA | Gm28406 | predicted gene 28406 [Source:MGI Symbo | 0 | 0 | 0 | 0 | 0 | 0 |

| | | | | | | | | | | | | | | | |
|---|---|---|---|---|---|---|---|---|---|---|---|---|---|---|---|
| 38021 | ENSMUSG00000099650 | 0 | NA | NA | NA | NA | NA | Gm28129 | predicted gene 28129 [Source:MGI Symbo | 0 | 0 | 0 | 0 | 0 | 0 |
| 38022 | ENSMUSG00000099651 | 0 | NA | NA | NA | NA | NA | Gm29078 | predicted gene 29078 [Source:MGI Symbo | 0 | 0 | 0 | 0 | 0 | 0 |
| 38023 | ENSMUSG00000099652 | 0 | NA | NA | NA | NA | NA | Gm28833 | predicted gene 28833 [Source:MGI Symbo | 0 | 0 | 0 | 0 | 0 | 0 |
| 38024 | ENSMUSG00000099653 | 0 | NA | NA | NA | NA | NA | Gm28984 | predicted gene 28984 [Source:MGI Symbo | 0 | 0 | 0 | 0 | 0 | 0 |
| 38025 | ENSMUSG00000099654 | 0 | NA | NA | NA | NA | NA | Gm28236 | predicted gene 28236 [Source:MGI Symbo | 0 | 0 | 0 | 0 | 0 | 0 |
| 38026 | ENSMUSG00000099655 | 0.935330317 | 1.257565026 | 2.487133718 | 0.505628232 | 0.613117658 | NA | 2310034G01 | RIKEN cDNA 2310034G01 gene [Source:M | 1.6062 | 0 | 0 | 2.8919 | 0 | 1.1138 |
| 38027 | ENSMUSG00000099656 | 0 | NA | NA | NA | NA | NA | Tent2-ps2 | terminal nucleotidyltransferase 2, pseudo | 0 | 0 | 0 | 0 | 0 | 0 |
| 38028 | ENSMUSG00000099657 | 0 | NA | NA | NA | NA | NA | Gm28667 | predicted gene 28667 [Source:MGI Symbo | 0 | 0 | 0 | 0 | 0 | 0 |
| 38029 | ENSMUSG00000099658 | 0 | NA | NA | NA | NA | NA | Gm28980 | predicted gene 28980 [Source:MGI Symbo | 0 | 0 | 0 | 0 | 0 | 0 |
| 38030 | ENSMUSG00000099659 | 0 | NA | NA | NA | NA | NA | Gm9322 | predicted gene 9322 [Source:MGI Symbol | 0 | 0 | 0 | 0 | 0 | 0 |
| 38031 | ENSMUSG00000099660 | 0 | NA | NA | NA | NA | NA | Gm2316 | predicted gene 2316 [Source:MGI Symbol | 0 | 0 | 0 | 0 | 0 | 0 |
| 38032 | ENSMUSG00000099661 | 0 | NA | NA | NA | NA | NA | Gm28781 | predicted gene 28781 [Source:MGI Symbo | 0 | 0 | 0 | 0 | 0 | 0 |
| 38033 | ENSMUSG00000099662 | 0.36353943 | 2.065989254 | 4.034137301 | 0.512126658 | 0.608562368 | NA | Ppp1r2-ps2 | protein phosphatase 1, regulatory (inhibit | 0 | 0 | 0 | 0 | 2.1812 | 0 |
| 38034 | ENSMUSG00000099663 | 0 | NA | NA | NA | NA | NA | Gm29201 | predicted gene 29201 [Source:MGI Symbo | 0 | 0 | 0 | 0 | 0 | 0 |
| 38035 | ENSMUSG00000099664 | 0 | NA | NA | NA | NA | NA | Gm28511 | predicted gene 28511 [Source:MGI Symbo | 0 | 0 | 0 | 0 | 0 | 0 |
| 38036 | ENSMUSG00000099665 | 0 | NA | NA | NA | NA | NA | Gm29482 | predicted gene 29482 [Source:MGI Symbo | 0 | 0 | 0 | 0 | 0 | 0 |
| 38037 | ENSMUSG00000099666 | 0 | NA | NA | NA | NA | NA | Gm29511 | predicted gene 29511 [Source:MGI Symbo | 0 | 0 | 0 | 0 | 0 | 0 |
| 38038 | ENSMUSG00000099667 | 0 | NA | NA | NA | NA | NA | Gm21901 | predicted gene, 21901 [Source:MGI Symb | 0 | 0 | 0 | 0 | 0 | 0 |
| 38039 | ENSMUSG00000099668 | 0 | NA | NA | NA | NA | NA | Gm29522 | predicted gene 29522 [Source:MGI Symbo | 0 | 0 | 0 | 0 | 0 | 0 |
| 38040 | ENSMUSG00000099669 | 9.999004828 | -0.492067837 | 0.706030353 | -0.696949975 | 0.485834101 | 0.872998689 | Gm29012 | predicted gene 29012 [Source:MGI Symbo | 17.668 | 9.2513 | 8.05 | 9.6398 | 9.8156 | 5.5692 |
| 38041 | ENSMUSG00000099670 | 0 | NA | NA | NA | NA | NA | Gm28423 | predicted gene 28423 [Source:MGI Symbo | 0 | 0 | 0 | 0 | 0 | 0 |
| 38042 | ENSMUSG00000099671 | 0 | NA | NA | NA | NA | NA | Gm18183 | predicted gene, 18183 [Source:MGI Symb | 0 | 0 | 0 | 0 | 0 | 0 |
| 38043 | ENSMUSG00000099672 | 1.177510966 | 2.58959483 | 2.314810155 | 1.118707219 | 0.263265065 | NA | Gm28564 | predicted gene 28564 [Source:MGI Symbo | 0 | 1.0279 | 0 | 3.8559 | 2.1812 | 0 |
| 38044 | ENSMUSG00000099673 | 0 | NA | NA | NA | NA | NA | Scgb2b30 | secretoglobin, family 2B, member 30 [Sou | 0 | 0 | 0 | 0 | 0 | 0 |
| 38045 | ENSMUSG00000099674 | 0 | NA | NA | NA | NA | NA | Gm29166 | predicted gene 29166 [Source:MGI Symbo | 0 | 0 | 0 | 0 | 0 | 0 |
| 38046 | ENSMUSG00000099675 | 0 | NA | NA | NA | NA | NA | Gm28717 | predicted gene 28717 [Source:MGI Symbo | 0 | 0 | 0 | 0 | 0 | 0 |
| 38047 | ENSMUSG00000099676 | 0 | NA | NA | NA | NA | NA | Gm29528 | predicted gene 29528 [Source:MGI Symbo | 0 | 0 | 0 | 0 | 0 | 0 |
| 38048 | ENSMUSG00000099677 | 0 | NA | NA | NA | NA | NA | Gm21722 | predicted gene, 21722 [Source:MGI Symb | 0 | 0 | 0 | 0 | 0 | 0 |
| 38049 | ENSMUSG00000099678 | 0 | NA | NA | NA | NA | NA | Gm28351 | predicted gene 28351 [Source:MGI Symbo | 0 | 0 | 0 | 0 | 0 | 0 |
| 38050 | ENSMUSG00000099679 | 0 | NA | NA | NA | NA | NA | Gm28899 | predicted gene 28899 [Source:MGI Symbo | 0 | 0 | 0 | 0 | 0 | 0 |
| 38051 | ENSMUSG00000099681 | 48.02675161 | -0.434915699 | 0.347812996 | -1.250429696 | 0.211142622 | 0.655691903 | 1700052K11 | RIKEN cDNA 1700052K11 gene [Source:M | 73.885 | 39.061 | 52.325 | 42.415 | 39.262 | 41.212 |
| 38052 | ENSMUSG00000099682 | 0 | NA | NA | NA | NA | NA | Gm28292 | predicted gene 28292 [Source:MGI Symbo | 0 | 0 | 0 | 0 | 0 | 0 |
| 38053 | ENSMUSG00000099683 | 0 | NA | NA | NA | NA | NA | Gm28214 | predicted gene 28214 [Source:MGI Symbo | 0 | 0 | 0 | 0 | 0 | 0 |
| 38054 | ENSMUSG00000099684 | 0 | NA | NA | NA | NA | NA | Gm28356 | predicted gene 28356 [Source:MGI Symbo | 0 | 0 | 0 | 0 | 0 | 0 |
| 38055 | ENSMUSG00000099685 | 0 | NA | NA | NA | NA | NA | Gm29617 | predicted gene 29617 [Source:MGI Symbo | 0 | 0 | 0 | 0 | 0 | 0 |
| 38056 | ENSMUSG00000099686 | 0 | NA | NA | NA | NA | NA | Gm29460 | predicted gene 29460 [Source:MGI Symbo | 0 | 0 | 0 | 0 | 0 | 0 |
| 38057 | ENSMUSG00000099687 | 0 | NA | NA | NA | NA | NA | Olfr623 | olfactory receptor 623 [Source:MGI Symb | 0 | 0 | 0 | 0 | 0 | 0 |
| 38058 | ENSMUSG00000099688 | 0 | NA | NA | NA | NA | NA | Gm29535 | predicted gene 29535 [Source:MGI Symbo | 0 | 0 | 0 | 0 | 0 | 0 |
| 38059 | ENSMUSG00000099689 | 78.32086323 | 0.223915301 | 0.256471195 | 0.873062183 | 0.382629185 | 0.814473662 | Zfp383 | zinc finger protein 383 [Source:MGI Symb | 63.445 | 77.094 | 76.475 | 81.938 | 78.525 | 92.449 |
| 38060 | ENSMUSG00000099690 | 0 | NA | NA | NA | NA | NA | Gm29513 | predicted gene 29513 [Source:MGI Symbo | 0 | 0 | 0 | 0 | 0 | 0 |
| 38061 | ENSMUSG00000099691 | 0 | NA | NA | NA | NA | NA | Gm29518 | predicted gene 29518 [Source:MGI Symbo | 0 | 0 | 0 | 0 | 0 | 0 |
| 38062 | ENSMUSG00000099692 | 0 | NA | NA | NA | NA | NA | Gm5696 | predicted gene 5696 [Source:MGI Symbol | 0 | 0 | 0 | 0 | 0 | 0 |
| 38063 | ENSMUSG00000099693 | 0 | NA | NA | NA | NA | NA | Gm29284 | predicted gene 29284 [Source:MGI Symbo | 0 | 0 | 0 | 0 | 0 | 0 |
| 38064 | ENSMUSG00000099694 | 0 | NA | NA | NA | NA | NA | Gm29610 | predicted gene 29610 [Source:MGI Symbo | 0 | 0 | 0 | 0 | 0 | 0 |
| 38065 | ENSMUSG00000099695 | 0 | NA | NA | NA | NA | NA | Gm28488 | predicted gene 28488 [Source:MGI Symbo | 0 | 0 | 0 | 0 | 0 | 0 |
| 38066 | ENSMUSG00000099696 | 0 | NA | NA | NA | NA | NA | 2900052N01 | RIKEN cDNA 2900052N01 gene [Source:M | 0 | 0 | 0 | 0 | 0 | 0 |
| 38067 | ENSMUSG00000099698 | 0 | NA | NA | NA | NA | NA | Gm28740 | predicted gene 28740 [Source:MGI Symbo | 0 | 0 | 0 | 0 | 0 | 0 |
| 38068 | ENSMUSG00000099699 | 0 | NA | NA | NA | NA | NA | Gm29421 | predicted gene 29421 [Source:MGI Symbo | 0 | 0 | 0 | 0 | 0 | 0 |
| 38069 | ENSMUSG00000099700 | 0 | NA | NA | NA | NA | NA | Gm29645 | predicted gene 29645 [Source:MGI Symbo | 0 | 0 | 0 | 0 | 0 | 0 |
| 38070 | ENSMUSG00000099701 | 0 | NA | NA | NA | NA | NA | Gm19136 | predicted gene, 19136 [Source:MGI Symb | 0 | 0 | 0 | 0 | 0 | 0 |
| 38071 | ENSMUSG00000099702 | 0 | NA | NA | NA | NA | NA | Gm29013 | predicted gene 29013 [Source:MGI Symbo | 0 | 0 | 0 | 0 | 0 | 0 |
| 38072 | ENSMUSG00000099703 | 0.321325949 | 1.911575114 | 4.048267893 | 0.472195804 | 0.636787034 | NA | NA | NA | 0 | 0 | 0 | 1.928 | 0 | 0 |
| 38073 | ENSMUSG00000099704 | 0 | NA | NA | NA | NA | NA | Gm28842 | predicted gene 28842 [Source:MGI Symbo | 0 | 0 | 0 | 0 | 0 | 0 |
| 38074 | ENSMUSG00000099706 | 0 | NA | NA | NA | NA | NA | Gm29536 | predicted gene 29536 [Source:MGI Symbo | 0 | 0 | 0 | 0 | 0 | 0 |
| 38075 | ENSMUSG00000099707 | 452.7895548 | 0.096305249 | 0.147145188 | 0.654491329 | 0.512795312 | 0.884860213 | Gm8883 | predicted gene 8883 [Source:MGI Symbol | 508.36 | 417.33 | 386.4 | 485.84 | 476.6 | 442.2 |
| 38076 | ENSMUSG00000099708 | 6.889710039 | -0.109129083 | 0.800929572 | -0.136253033 | 0.891621252 | 0.982489666 | Gm29361 | predicted gene 29361 [Source:MGI Symbo | 7.2279 | 7.1954 | 7.0437 | 7.7118 | 4.3625 | 7.7969 |

| | | | | | | | | | | | | | | | |
|---|---|---|---|---|---|---|---|---|---|---|---|---|---|---|---|
| 38077 | ENSMUSG00000099709 | 0 | NA | NA | NA | NA | NA | Gm21396 | predicted gene, 21396 [Source:MGI Symb | 0 | 0 | 0 | 0 | 0 | 0 |
| 38078 | ENSMUSG00000099711 | 0 | NA | NA | NA | NA | NA | Btbd35f21 | BTB domain containing 35, family membe | 0 | 0 | 0 | 0 | 0 | 0 |
| 38079 | ENSMUSG00000099712 | 0 | NA | NA | NA | NA | NA | Gm29411 | predicted gene 29411 [Source:MGI Symbo | 0 | 0 | 0 | 0 | 0 | 0 |
| 38080 | ENSMUSG00000099713 | 0 | NA | NA | NA | NA | NA | Gm21923 | predicted gene, 21923 [Source:MGI Symb | 0 | 0 | 0 | 0 | 0 | 0 |
| 38081 | ENSMUSG00000099714 | 0 | NA | NA | NA | NA | NA | Gm20587 | predicted gene, 20587 [Source:MGI Symb | 0 | 0 | 0 | 0 | 0 | 0 |
| 38082 | ENSMUSG00000099715 | 0 | NA | NA | NA | NA | NA | Gm28077 | predicted gene 28077 [Source:MGI Symbo | 0 | 0 | 0 | 0 | 0 | 0 |
| 38083 | ENSMUSG00000099716 | 0 | NA | NA | NA | NA | NA | Gm29082 | predicted gene 29082 [Source:MGI Symbo | 0 | 0 | 0 | 0 | 0 | 0 |
| 38084 | ENSMUSG00000099717 | 0.335414862 | -1.779047242 | 4.030804179 | -0.441362855 | 0.658950329 | NA | Gm9451 | predicted gene 9451 [Source:MGI Symbol | 0 | 0 | 2.0125 | 0 | 0 | 0 |
| 38085 | ENSMUSG00000099718 | 0 | NA | NA | NA | NA | NA | Gm2087 | predicted gene 2087 [Source:MGI Symbol | 0 | 0 | 0 | 0 | 0 | 0 |
| 38086 | ENSMUSG00000099719 | 0.339027071 | -1.79174149 | 3.919585585 | -0.457125237 | 0.647581032 | NA | Gm28802 | predicted gene 28802 [Source:MGI Symbo | 0 | 1.0279 | 1.0062 | 0 | 0 | 0 |
| 38087 | ENSMUSG00000099721 | 0 | NA | NA | NA | NA | NA | Gm28435 | predicted gene 28435 [Source:MGI Symbo | 0 | 0 | 0 | 0 | 0 | 0 |
| 38088 | ENSMUSG00000099722 | 0 | NA | NA | NA | NA | NA | Gm29154 | predicted gene 29154 [Source:MGI Symbo | 0 | 0 | 0 | 0 | 0 | 0 |
| 38089 | ENSMUSG00000099723 | 0 | NA | NA | NA | NA | NA | Gm28595 | predicted gene 28595 [Source:MGI Symbo | 0 | 0 | 0 | 0 | 0 | 0 |
| 38090 | ENSMUSG00000099724 | 0 | NA | NA | NA | NA | NA | 4930554C24 | RIKEN cDNA 4930554C24 gene [Source:M | 0 | 0 | 0 | 0 | 0 | 0 |
| 38091 | ENSMUSG00000099725 | 0 | NA | NA | NA | NA | NA | Gm28734 | predicted gene 28734 [Source:MGI Symbo | 0 | 0 | 0 | 0 | 0 | 0 |
| 38092 | ENSMUSG00000099726 | 0 | NA | NA | NA | NA | NA | Gm29160 | predicted gene 29160 [Source:MGI Symbo | 0 | 0 | 0 | 0 | 0 | 0 |
| 38093 | ENSMUSG00000099727 | 0 | NA | NA | NA | NA | NA | Gm29401 | predicted gene 29401 [Source:MGI Symbo | 0 | 0 | 0 | 0 | 0 | 0 |
| 38094 | ENSMUSG00000099728 | 0 | NA | NA | NA | NA | NA | Gm8510 | predicted gene 8510 [Source:MGI Symbol | 0 | 0 | 0 | 0 | 0 | 0 |
| 38095 | ENSMUSG00000099729 | 0 | NA | NA | NA | NA | NA | Scgb2b11 | secretoglobin, family 2B, member 11 [Sou | 0 | 0 | 0 | 0 | 0 | 0 |
| 38096 | ENSMUSG00000099730 | 0 | NA | NA | NA | NA | NA | Gm28547 | predicted gene 28547 [Source:MGI Symbo | 0 | 0 | 0 | 0 | 0 | 0 |
| 38097 | ENSMUSG00000099731 | 0 | NA | NA | NA | NA | NA | Gm29181 | predicted gene 29181 [Source:MGI Symbo | 0 | 0 | 0 | 0 | 0 | 0 |
| 38098 | ENSMUSG00000099732 | 0 | NA | NA | NA | NA | NA | Gm17752 | predicted gene, 17752 [Source:MGI Symb | 0 | 0 | 0 | 0 | 0 | 0 |
| 38099 | ENSMUSG00000099733 | 0 | NA | NA | NA | NA | NA | Gm21880 | predicted gene, 21880 [Source:MGI Symb | 0 | 0 | 0 | 0 | 0 | 0 |
| 38100 | ENSMUSG00000099734 | 0 | NA | NA | NA | NA | NA | Gm28159 | predicted gene 28159 [Source:MGI Symbo | 0 | 0 | 0 | 0 | 0 | 0 |
| 38101 | ENSMUSG00000099735 | 0 | NA | NA | NA | NA | NA | 1700122H20 | RIKEN cDNA 1700122H20 gene [Source:M | 0 | 0 | 0 | 0 | 0 | 0 |
| 38102 | ENSMUSG00000099736 | 0 | NA | NA | NA | NA | NA | Gm20863 | predicted gene, 20863 [Source:MGI Symb | 0 | 0 | 0 | 0 | 0 | 0 |
| 38103 | ENSMUSG00000099737 | 0 | NA | NA | NA | NA | NA | Gm29655 | predicted gene 29655 [Source:MGI Symbo | 0 | 0 | 0 | 0 | 0 | 0 |
| 38104 | ENSMUSG00000099738 | 0 | NA | NA | NA | NA | NA | Gm28483 | predicted gene 28483 [Source:MGI Symbo | 0 | 0 | 0 | 0 | 0 | 0 |
| 38105 | ENSMUSG00000099739 | 0 | NA | NA | NA | NA | NA | Gm20804 | predicted gene, 20804 [Source:MGI Symb | 0 | 0 | 0 | 0 | 0 | 0 |
| 38106 | ENSMUSG00000099740 | 0 | NA | NA | NA | NA | NA | Gm28897 | predicted gene 28897 [Source:MGI Symbo | 0 | 0 | 0 | 0 | 0 | 0 |
| 38107 | ENSMUSG00000099741 | 0 | NA | NA | NA | NA | NA | Gm5253 | predicted gene 5253 [Source:MGI Symbol | 0 | 0 | 0 | 0 | 0 | 0 |
| 38108 | ENSMUSG00000099742 | 0 | NA | NA | NA | NA | NA | Gm29148 | predicted gene 29148 [Source:MGI Symbo | 0 | 0 | 0 | 0 | 0 | 0 |
| 38109 | ENSMUSG00000099744 | 0 | NA | NA | NA | NA | NA | Tmem167-ps | transmembrane protein 167, pseudogene | 0 | 0 | 0 | 0 | 0 | 0 |
| 38110 | ENSMUSG00000099745 | 0 | NA | NA | NA | NA | NA | Gm28736 | predicted gene 28736 [Source:MGI Symbo | 0 | 0 | 0 | 0 | 0 | 0 |
| 38111 | ENSMUSG00000099746 | 0.684865378 | 2.990343487 | 2.966805827 | 1.00793367 | 0.313486306 | NA | Ppnr | per-pentamer repeat gene [Source:MGI S | 0 | 0 | 0 | 1.928 | 2.1812 | 0 |
| 38112 | ENSMUSG00000099747 | 0 | NA | NA | NA | NA | NA | 1700025K24 | RIKEN cDNA 1700025K24 gene [Source:M | 0 | 0 | 0 | 0 | 0 | 0 |
| 38113 | ENSMUSG00000099748 | 0 | NA | NA | NA | NA | NA | Uba1y-ps1 | ubiquitin-activating enzyme, Chr Y, pseud | 0 | 0 | 0 | 0 | 0 | 0 |
| 38114 | ENSMUSG00000099749 | 0 | NA | NA | NA | NA | NA | Gm29323 | predicted gene 29323 [Source:MGI Symbo | 0 | 0 | 0 | 0 | 0 | 0 |
| 38115 | ENSMUSG00000099750 | 0 | NA | NA | NA | NA | NA | Gm18827 | predicted gene, 18827 [Source:MGI Symb | 0 | 0 | 0 | 0 | 0 | 0 |
| 38116 | ENSMUSG00000099751 | 0 | NA | NA | NA | NA | NA | 1700022A22 | RIKEN cDNA 1700022A22 gene [Source:M | 0 | 0 | 0 | 0 | 0 | 0 |
| 38117 | ENSMUSG00000099752 | 0 | NA | NA | NA | NA | NA | Gm29018 | predicted gene 29018 [Source:MGI Symbo | 0 | 0 | 0 | 0 | 0 | 0 |
| 38118 | ENSMUSG00000099753 | 0 | NA | NA | NA | NA | NA | Gm28058 | predicted gene 28058 [Source:MGI Symbo | 0 | 0 | 0 | 0 | 0 | 0 |
| 38119 | ENSMUSG00000099754 | 0 | NA | NA | NA | NA | NA | Vmn1r-ps34 | vomeronasal 1 receptor, pseudogene 34 [ | 0 | 0 | 0 | 0 | 0 | 0 |
| 38120 | ENSMUSG00000099755 | 0 | NA | NA | NA | NA | NA | Gm21114 | predicted gene, 21114 [Source:MGI Symb | 0 | 0 | 0 | 0 | 0 | 0 |
| 38121 | ENSMUSG00000099756 | 0 | NA | NA | NA | NA | NA | Gm28763 | predicted gene 28763 [Source:MGI Symbo | 0 | 0 | 0 | 0 | 0 | 0 |
| 38122 | ENSMUSG00000099757 | 4.522072784 | -1.976893773 | 1.297614528 | -1.523483077 | 0.127637891 | NA | BE692007 | expressed sequence BE692007 [Source:M | 3.2124 | 11.307 | 7.0437 | 0 | 0 | 5.5692 |
| 38123 | ENSMUSG00000099758 | 0.663758638 | 2.950973782 | 3.025092005 | 0.975498853 | 0.32931287 | NA | Gm10830 | predicted gene 10830 [Source:MGI Symbo | 0 | 0 | 0 | 2.8919 | 1.0906 | 0 |
| 38124 | ENSMUSG00000099759 | 15.34728753 | -0.770539576 | 0.603534821 | -1.276711051 | 0.201704278 | 0.646025395 | 1700030C10 | RIKEN cDNA 1700030C10 gene [Source:M | 18.471 | 25.698 | 14.087 | 17.352 | 10.906 | 5.5692 |
| 38125 | ENSMUSG00000099760 | 1.440467498 | 1.046009396 | 2.168701849 | 0.482320516 | 0.629578275 | NA | Gm28800 | predicted gene 28800 [Source:MGI Symbo | 0.8031 | 2.0558 | 0 | 5.7839 | 0 | 0 |
| 38126 | ENSMUSG00000099761 | 0 | NA | NA | NA | NA | NA | Gm29158 | predicted gene 29158 [Source:MGI Symbo | 0 | 0 | 0 | 0 | 0 | 0 |
| 38127 | ENSMUSG00000099762 | 0 | NA | NA | NA | NA | NA | Gm21149 | predicted gene, 21149 [Source:MGI Symb | 0 | 0 | 0 | 0 | 0 | 0 |
| 38128 | ENSMUSG00000099764 | 10.18415133 | -0.062216594 | 0.729434284 | -0.08529431 | 0.932027415 | 0.988420305 | Rps10-ps2 | ribosomal protein S10, pseudogene 2 [Sou | 12.85 | 4.1117 | 14.087 | 5.7839 | 10.906 | 13.366 |
| 38129 | ENSMUSG00000099765 | 0 | NA | NA | NA | NA | NA | Gm4454 | predicted gene 4454 [Source:MGI Symbol | 0 | 0 | 0 | 0 | 0 | 0 |
| 38130 | ENSMUSG00000099766 | 0 | NA | NA | NA | NA | NA | Gm28767 | predicted gene 28767 [Source:MGI Symbo | 0 | 0 | 0 | 0 | 0 | 0 |
| 38131 | ENSMUSG00000099767 | 0 | NA | NA | NA | NA | NA | Gm28884 | predicted gene 28884 [Source:MGI Symbo | 0 | 0 | 0 | 0 | 0 | 0 |
| 38132 | ENSMUSG00000099768 | 0 | NA | NA | NA | NA | NA | Gm29293 | predicted gene 29293 [Source:MGI Symbo | 0 | 0 | 0 | 0 | 0 | 0 |

| | | | | | | | | | | | | | | | |
|---|---|---|---|---|---|---|---|---|---|---|---|---|---|---|---|
| 38133 | ENSMUSG00000099770 | 0 | NA | NA | NA | NA | NA | Gm28623 | predicted gene 28623 [Source:MGI Symbo | 0 | 0 | 0 | 0 | 0 | 0 |
| 38134 | ENSMUSG00000099771 | 0 | NA | NA | NA | NA | NA | Gm19257 | predicted gene, 19257 [Source:MGI Symb | 0 | 0 | 0 | 0 | 0 | 0 |
| 38135 | ENSMUSG00000099772 | 0 | NA | NA | NA | NA | NA | Gm29085 | predicted gene 29085 [Source:MGI Symbo | 0 | 0 | 0 | 0 | 0 | 0 |
| 38136 | ENSMUSG00000099773 | 0 | NA | NA | NA | NA | NA | Hmgb1-rs16 | high mobility group box 1, related sequen | 0 | 0 | 0 | 0 | 0 | 0 |
| 38137 | ENSMUSG00000099774 | 0 | NA | NA | NA | NA | NA | Gm29341 | predicted gene 29341 [Source:MGI Symbo | 0 | 0 | 0 | 0 | 0 | 0 |
| 38138 | ENSMUSG00000099775 | 0.267700746 | -1.497481649 | 4.057001436 | -0.369110456 | 0.712045396 | NA | Gm5960 | predicted gene 5960 [Source:MGI Symbol | 1.6062 | 0 | 0 | 0 | 0 | 0 |
| 38139 | ENSMUSG00000099776 | 0 | NA | NA | NA | NA | NA | Gm28954 | predicted gene 28954 [Source:MGI Symbo | 0 | 0 | 0 | 0 | 0 | 0 |
| 38140 | ENSMUSG00000099777 | 0 | NA | NA | NA | NA | NA | Gm21777 | predicted gene, 21777 [Source:MGI Symb | 0 | 0 | 0 | 0 | 0 | 0 |
| 38141 | ENSMUSG00000099778 | 0 | NA | NA | NA | NA | NA | Gm28134 | predicted gene 28134 [Source:MGI Symbo | 0 | 0 | 0 | 0 | 0 | 0 |
| 38142 | ENSMUSG00000099779 | 0 | NA | NA | NA | NA | NA | Gm8228 | predicted gene 8228 [Source:MGI Symbol | 0 | 0 | 0 | 0 | 0 | 0 |
| 38143 | ENSMUSG00000099780 | 0 | NA | NA | NA | NA | NA | Gm29537 | predicted gene 29537 [Source:MGI Symbo | 0 | 0 | 0 | 0 | 0 | 0 |
| 38144 | ENSMUSG00000099781 | 0 | NA | NA | NA | NA | NA | Gm28583 | predicted gene 28583 [Source:MGI Symbo | 0 | 0 | 0 | 0 | 0 | 0 |
| 38145 | ENSMUSG00000099782 | 0 | NA | NA | NA | NA | NA | Gm20903 | predicted gene, 20903 [Source:MGI Symb | 0 | 0 | 0 | 0 | 0 | 0 |
| 38146 | ENSMUSG00000099783 | 0 | NA | NA | NA | NA | NA | Gm28589 | predicted gene 28589 [Source:MGI Symbo | 0 | 0 | 0 | 0 | 0 | 0 |
| 38147 | ENSMUSG00000099784 | 0 | NA | NA | NA | NA | NA | Dalir | DNMT1 associated long intergenic non-co | 0 | 0 | 0 | 0 | 0 | 0 |
| 38148 | ENSMUSG00000099785 | 0 | NA | NA | NA | NA | NA | Gm28425 | predicted gene 28425 [Source:MGI Symbo | 0 | 0 | 0 | 0 | 0 | 0 |
| 38149 | ENSMUSG00000099786 | 0 | NA | NA | NA | NA | NA | Gm29215 | predicted gene 29215 [Source:MGI Symbo | 0 | 0 | 0 | 0 | 0 | 0 |
| 38150 | ENSMUSG00000099787 | 0 | NA | NA | NA | NA | NA | Vmn1r192 | vomeronasal 1 receptor 192 [Source:MGI | 0 | 0 | 0 | 0 | 0 | 0 |
| 38151 | ENSMUSG00000099789 | 0 | NA | NA | NA | NA | NA | Gm28565 | predicted gene 28565 [Source:MGI Symbo | 0 | 0 | 0 | 0 | 0 | 0 |
| 38152 | ENSMUSG00000099790 | 0 | NA | NA | NA | NA | NA | Gm28457 | predicted gene 28457 [Source:MGI Symbo | 0 | 0 | 0 | 0 | 0 | 0 |
| 38153 | ENSMUSG00000099791 | 0 | NA | NA | NA | NA | NA | Gm20891 | predicted gene, 20891 [Source:MGI Symb | 0 | 0 | 0 | 0 | 0 | 0 |
| 38154 | ENSMUSG00000099792 | 0 | NA | NA | NA | NA | NA | Gm29564 | predicted gene 29564 [Source:MGI Symbo | 0 | 0 | 0 | 0 | 0 | 0 |
| 38155 | ENSMUSG00000099793 | 0 | NA | NA | NA | NA | NA | Gm28775 | predicted gene 28775 [Source:MGI Symbo | 0 | 0 | 0 | 0 | 0 | 0 |
| 38156 | ENSMUSG00000099794 | 0 | NA | NA | NA | NA | NA | 1700128A07 | RIKEN cDNA 1700128A07 gene [Source:M | 0 | 0 | 0 | 0 | 0 | 0 |
| 38157 | ENSMUSG00000099795 | 0 | NA | NA | NA | NA | NA | Gm29492 | predicted gene 29492 [Source:MGI Symbo | 0 | 0 | 0 | 0 | 0 | 0 |
| 38158 | ENSMUSG00000099796 | 0 | NA | NA | NA | NA | NA | Gm7893 | predicted gene 7893 [Source:MGI Symbol | 0 | 0 | 0 | 0 | 0 | 0 |
| 38159 | ENSMUSG00000099797 | 0 | NA | NA | NA | NA | NA | Gm3145 | predicted gene 3145 [Source:MGI Symbol | 0 | 0 | 0 | 0 | 0 | 0 |
| 38160 | ENSMUSG00000099798 | 0 | NA | NA | NA | NA | NA | Gm29168 | predicted gene 29168 [Source:MGI Symbo | 0 | 0 | 0 | 0 | 0 | 0 |
| 38161 | ENSMUSG00000099799 | 0 | NA | NA | NA | NA | NA | Gm28260 | predicted gene 28260 [Source:MGI Symbo | 0 | 0 | 0 | 0 | 0 | 0 |
| 38162 | ENSMUSG00000099800 | 0 | NA | NA | NA | NA | NA | Gm28570 | predicted gene 28570 [Source:MGI Symbo | 0 | 0 | 0 | 0 | 0 | 0 |
| 38163 | ENSMUSG00000099801 | 0 | NA | NA | NA | NA | NA | Gm29011 | predicted gene 29011 [Source:MGI Symbo | 0 | 0 | 0 | 0 | 0 | 0 |
| 38164 | ENSMUSG00000099802 | 0 | NA | NA | NA | NA | NA | Gm28493 | predicted gene 28493 [Source:MGI Symbo | 0 | 0 | 0 | 0 | 0 | 0 |
| 38165 | ENSMUSG00000099803 | 1.954066496 | -0.978619262 | 1.808069458 | -0.541250922 | 0.588334643 | NA | Gm28863 | predicted gene 28863 [Source:MGI Symbo | 0.8031 | 1.0279 | 6.0375 | 3.8559 | 0 | 0 |
| 38166 | ENSMUSG00000099804 | 0 | NA | NA | NA | NA | NA | Gm28497 | predicted gene 28497 [Source:MGI Symbo | 0 | 0 | 0 | 0 | 0 | 0 |
| 38167 | ENSMUSG00000099805 | 0 | NA | NA | NA | NA | NA | Gm28161 | predicted gene 28161 [Source:MGI Symbo | 0 | 0 | 0 | 0 | 0 | 0 |
| 38168 | ENSMUSG00000099806 | 0 | NA | NA | NA | NA | NA | Gm28841 | predicted gene 28841 [Source:MGI Symbo | 0 | 0 | 0 | 0 | 0 | 0 |
| 38169 | ENSMUSG00000099807 | 0 | NA | NA | NA | NA | NA | Gm28771 | predicted gene 28771 [Source:MGI Symbo | 0 | 0 | 0 | 0 | 0 | 0 |
| 38170 | ENSMUSG00000099808 | 0 | NA | NA | NA | NA | NA | 1700019P21 | RIKEN cDNA 1700019P21 gene [Source:M | 0 | 0 | 0 | 0 | 0 | 0 |
| 38171 | ENSMUSG00000099809 | 0 | NA | NA | NA | NA | NA | Gm18665 | predicted gene, 18665 [Source:MGI Symb | 0 | 0 | 0 | 0 | 0 | 0 |
| 38172 | ENSMUSG00000099810 | 1.259829789 | 0.908327631 | 2.052575775 | 0.44253062 | 0.658105277 | NA | Gm29509 | predicted gene 29509 [Source:MGI Symbo | 1.6062 | 0 | 1.0062 | 3.8559 | 1.0906 | 0 |
| 38173 | ENSMUSG00000099811 | 0 | NA | NA | NA | NA | NA | Gm28411 | predicted gene 28411 [Source:MGI Symbo | 0 | 0 | 0 | 0 | 0 | 0 |
| 38174 | ENSMUSG00000099812 | 0 | NA | NA | NA | NA | NA | Gm28515 | predicted gene 28515 [Source:MGI Symbo | 0 | 0 | 0 | 0 | 0 | 0 |
| 38175 | ENSMUSG00000099814 | 0 | NA | NA | NA | NA | NA | Gm28338 | predicted gene 28338 [Source:MGI Symbo | 0 | 0 | 0 | 0 | 0 | 0 |
| 38176 | ENSMUSG00000099815 | 0 | NA | NA | NA | NA | NA | Gm28430 | predicted gene 28430 [Source:MGI Symbo | 0 | 0 | 0 | 0 | 0 | 0 |
| 38177 | ENSMUSG00000099816 | 0 | NA | NA | NA | NA | NA | Gm28552 | predicted gene 28552 [Source:MGI Symbo | 0 | 0 | 0 | 0 | 0 | 0 |
| 38178 | ENSMUSG00000099817 | 0 | NA | NA | NA | NA | NA | Gm29317 | predicted gene 29317 [Source:MGI Symbo | 0 | 0 | 0 | 0 | 0 | 0 |
| 38179 | ENSMUSG00000099818 | 0 | NA | NA | NA | NA | NA | Gm29652 | predicted gene 29652 [Source:MGI Symbo | 0 | 0 | 0 | 0 | 0 | 0 |
| 38180 | ENSMUSG00000099819 | 0 | NA | NA | NA | NA | NA | Gm28728 | predicted gene 28728 [Source:MGI Symbo | 0 | 0 | 0 | 0 | 0 | 0 |
| 38181 | ENSMUSG00000099820 | 0 | NA | NA | NA | NA | NA | Olfr1036 | olfactory receptor 1036 [Source:MGI Sym | 0 | 0 | 0 | 0 | 0 | 0 |
| 38182 | ENSMUSG00000099821 | 0 | NA | NA | NA | NA | NA | Gm28408 | predicted gene 28408 [Source:MGI Symbo | 0 | 0 | 0 | 0 | 0 | 0 |
| 38183 | ENSMUSG00000099822 | 0 | NA | NA | NA | NA | NA | Gm28332 | predicted gene 28332 [Source:MGI Symbo | 0 | 0 | 0 | 0 | 0 | 0 |
| 38184 | ENSMUSG00000099823 | 0 | NA | NA | NA | NA | NA | Gm28054 | predicted gene 28054 [Source:MGI Symbo | 0 | 0 | 0 | 0 | 0 | 0 |
| 38185 | ENSMUSG00000099824 | 0 | NA | NA | NA | NA | NA | Gm28218 | predicted gene 28218 [Source:MGI Symbo | 0 | 0 | 0 | 0 | 0 | 0 |
| 38186 | ENSMUSG00000099825 | 4.716772538 | -0.587470345 | 1.028528166 | -0.571175749 | 0.567880516 | NA | 1810012K16 | RIKEN cDNA 1810012K16 gene [Source:M | 8.8341 | 4.1117 | 4.025 | 5.7839 | 1.0906 | 4.4554 |
| 38187 | ENSMUSG00000099826 | 0 | NA | NA | NA | NA | NA | Scgb2b10 | secretoglobin, family 2B, member 10 [Sou | 0 | 0 | 0 | 0 | 0 | 0 |
| 38188 | ENSMUSG00000099827 | 0 | NA | NA | NA | NA | NA | Gm29520 | predicted gene 29520 [Source:MGI Symbo | 0 | 0 | 0 | 0 | 0 | 0 |

| | | | | | | | | | | | | | | | |
|---|---|---|---|---|---|---|---|---|---|---|---|---|---|---|---|
| 38189 | ENSMUSG00000099828 | 0 | NA | NA | NA | NA | NA | Gm28796 | predicted gene 28796 [Source:MGI Symbc | 0 | 0 | 0 | 0 | 0 | 0 |
| 38190 | ENSMUSG00000099829 | 0 | NA | NA | NA | NA | NA | Gm29380 | predicted gene 29380 [Source:MGI Symbc | 0 | 0 | 0 | 0 | 0 | 0 |
| 38191 | ENSMUSG00000099831 | 0 | NA | NA | NA | NA | NA | Gm28683 | predicted gene 28683 [Source:MGI Symbc | 0 | 0 | 0 | 0 | 0 | 0 |
| 38192 | ENSMUSG00000099832 | 0 | NA | NA | NA | NA | NA | Gm29025 | predicted gene 29025 [Source:MGI Symbc | 0 | 0 | 0 | 0 | 0 | 0 |
| 38193 | ENSMUSG00000099833 | 0 | NA | NA | NA | NA | NA | Gm29165 | predicted gene 29165 [Source:MGI Symbc | 0 | 0 | 0 | 0 | 0 | 0 |
| 38194 | ENSMUSG00000099834 | 0 | NA | NA | NA | NA | NA | Gm21728 | predicted gene, 21728 [Source:MGI Symb | 0 | 0 | 0 | 0 | 0 | 0 |
| 38195 | ENSMUSG00000099835 | 0 | NA | NA | NA | NA | NA | Gm29668 | predicted gene 29668 [Source:MGI Symbc | 0 | 0 | 0 | 0 | 0 | 0 |
| 38196 | ENSMUSG00000099836 | 0 | NA | NA | NA | NA | NA | Gm29204 | predicted gene 29204 [Source:MGI Symbc | 0 | 0 | 0 | 0 | 0 | 0 |
| 38197 | ENSMUSG00000099838 | 0 | NA | NA | NA | NA | NA | Gm8521 | predicted gene 8521 [Source:MGI Symbol | 0 | 0 | 0 | 0 | 0 | 0 |
| 38198 | ENSMUSG00000099839 | 0 | NA | NA | NA | NA | NA | Gm29374 | predicted gene 29374 [Source:MGI Symbc | 0 | 0 | 0 | 0 | 0 | 0 |
| 38199 | ENSMUSG00000099840 | 0 | NA | NA | NA | NA | NA | Gm21409 | predicted gene, 21409 [Source:MGI Symb | 0 | 0 | 0 | 0 | 0 | 0 |
| 38200 | ENSMUSG00000099841 | 0 | NA | NA | NA | NA | NA | Gm28855 | predicted gene 28855 [Source:MGI Symbc | 0 | 0 | 0 | 0 | 0 | 0 |
| 38201 | ENSMUSG00000099842 | 0 | NA | NA | NA | NA | NA | Gm4034 | predicted gene 4034 [Source:MGI Symbol | 0 | 0 | 0 | 0 | 0 | 0 |
| 38202 | ENSMUSG00000099843 | 73.59799318 | 0.547155622 | 0.299701432 | 1.825669032 | 0.067900131 | 0.375219813 | Gm7160 | predicted gene 7160 [Source:MGI Symbol | 53.808 | 60.647 | 65.406 | 113.75 | 59.984 | 87.994 |
| 38203 | ENSMUSG00000099845 | 0 | NA | NA | NA | NA | NA | Gm29273 | predicted gene 29273 [Source:MGI Symbc | 0 | 0 | 0 | 0 | 0 | 0 |
| 38204 | ENSMUSG00000099846 | 0 | NA | NA | NA | NA | NA | Gm6819 | predicted gene 6819 [Source:MGI Symbol | 0 | 0 | 0 | 0 | 0 | 0 |
| 38205 | ENSMUSG00000099847 | 2.42518002 | 3.746711532 | 1.914517244 | 1.957000672 | 0.050347389 | NA | Gm29458 | predicted gene 29458 [Source:MGI Symbc | 0 | 0 | 1.0062 | 4.8199 | 8.7249 | 0 |
| 38206 | ENSMUSG00000099848 | 0 | NA | NA | NA | NA | NA | Gm29337 | predicted gene 29337 [Source:MGI Symbc | 0 | 0 | 0 | 0 | 0 | 0 |
| 38207 | ENSMUSG00000099849 | 0 | NA | NA | NA | NA | NA | Gm29274 | predicted gene 29274 [Source:MGI Symbc | 0 | 0 | 0 | 0 | 0 | 0 |
| 38208 | ENSMUSG00000099850 | 0 | NA | NA | NA | NA | NA | Gm28906 | predicted gene 28906 [Source:MGI Symbc | 0 | 0 | 0 | 0 | 0 | 0 |
| 38209 | ENSMUSG00000099851 | 0 | NA | NA | NA | NA | NA | Gm29542 | predicted gene 29542 [Source:MGI Symbc | 0 | 0 | 0 | 0 | 0 | 0 |
| 38210 | ENSMUSG00000099852 | 0 | NA | NA | NA | NA | NA | Gm29002 | predicted gene 29002 [Source:MGI Symbc | 0 | 0 | 0 | 0 | 0 | 0 |
| 38211 | ENSMUSG00000099853 | 0 | NA | NA | NA | NA | NA | Gm29328 | predicted gene 29328 [Source:MGI Symbc | 0 | 0 | 0 | 0 | 0 | 0 |
| 38212 | ENSMUSG00000099854 | 0 | NA | NA | NA | NA | NA | Gm28710 | predicted gene 28710 [Source:MGI Symbc | 0 | 0 | 0 | 0 | 0 | 0 |
| 38213 | ENSMUSG00000099855 | 0 | NA | NA | NA | NA | NA | Gm28486 | predicted gene 28486 [Source:MGI Symbc | 0 | 0 | 0 | 0 | 0 | 0 |
| 38214 | ENSMUSG00000099856 | 0 | NA | NA | NA | NA | NA | Gm20906 | predicted gene, 20906 [Source:MGI Symb | 0 | 0 | 0 | 0 | 0 | 0 |
| 38215 | ENSMUSG00000099858 | 165.7100075 | -0.104062421 | 0.223415776 | -0.465779197 | 0.641373557 | 0.924019354 | Gm6652 | predicted gene 6652 [Source:MGI Symbol | 130.1 | 200.44 | 185.15 | 148.45 | 189.77 | 140.34 |
| 38216 | ENSMUSG00000099860 | 0 | NA | NA | NA | NA | NA | Gm29456 | predicted gene 29456 [Source:MGI Symbc | 0 | 0 | 0 | 0 | 0 | 0 |
| 38217 | ENSMUSG00000099861 | 0 | NA | NA | NA | NA | NA | Gm29049 | predicted gene 29049 [Source:MGI Symbc | 0 | 0 | 0 | 0 | 0 | 0 |
| 38218 | ENSMUSG00000099862 | 0 | NA | NA | NA | NA | NA | Gm28936 | predicted gene 28936 [Source:MGI Symbc | 0 | 0 | 0 | 0 | 0 | 0 |
| 38219 | ENSMUSG00000099863 | 0 | NA | NA | NA | NA | NA | 1700031L13I | RIKEN cDNA 1700031L13 gene [Source:M( | 0 | 0 | 0 | 0 | 0 | 0 |
| 38220 | ENSMUSG00000099864 | 0 | NA | NA | NA | NA | NA | Gm8391 | predicted gene 8391 [Source:MGI Symbol | 0 | 0 | 0 | 0 | 0 | 0 |
| 38221 | ENSMUSG00000099865 | 0 | NA | NA | NA | NA | NA | Gm29206 | predicted gene 29206 [Source:MGI Symbc | 0 | 0 | 0 | 0 | 0 | 0 |
| 38222 | ENSMUSG00000099866 | 0 | NA | NA | NA | NA | NA | Gm28889 | predicted gene 28889 [Source:MGI Symbc | 0 | 0 | 0 | 0 | 0 | 0 |
| 38223 | ENSMUSG00000099867 | 0 | NA | NA | NA | NA | NA | Gm29320 | predicted gene 29320 [Source:MGI Symbc | 0 | 0 | 0 | 0 | 0 | 0 |
| 38224 | ENSMUSG00000099868 | 0 | NA | NA | NA | NA | NA | Gm28792 | predicted gene 28792 [Source:MGI Symbc | 0 | 0 | 0 | 0 | 0 | 0 |
| 38225 | ENSMUSG00000099869 | 0 | NA | NA | NA | NA | NA | 1700030F04 | RIKEN cDNA 1700030F04 gene [Source:M( | 0 | 0 | 0 | 0 | 0 | 0 |
| 38226 | ENSMUSG00000099870 | 0 | NA | NA | NA | NA | NA | Gm7200 | predicted gene 7200 [Source:MGI Symbol | 0 | 0 | 0 | 0 | 0 | 0 |
| 38227 | ENSMUSG00000099871 | 0.267700746 | -1.497481649 | 4.057001436 | -0.369110456 | 0.712045396 | NA | Gm21742 | predicted gene, 21742 [Source:MGI Symb | 1.6062 | 0 | 0 | 0 | 0 | 0 |
| 38228 | ENSMUSG00000099872 | 0 | NA | NA | NA | NA | NA | 1700074H08 | RIKEN cDNA 1700074H08 gene [Source:M | 0 | 0 | 0 | 0 | 0 | 0 |
| 38229 | ENSMUSG00000099873 | 0 | NA | NA | NA | NA | NA | Gm28452 | predicted gene 28452 [Source:MGI Symbc | 0 | 0 | 0 | 0 | 0 | 0 |
| 38230 | ENSMUSG00000099874 | 2.105300494 | 0.937218682 | 1.526300158 | 0.614046115 | 0.539184855 | NA | Gm29629 | predicted gene 29629 [Source:MGI Symbc | 3.2124 | 1.0279 | 0 | 2.8919 | 3.2719 | 2.2277 |
| 38231 | ENSMUSG00000099875 | 0.476489653 | -2.293616034 | 3.421506946 | -0.670352587 | 0.502633051 | NA | Rbm3-ps | RNA binding motif (RNP1, RRM) protein 3 | 0.8031 | 2.0558 | 0 | 0 | 0 | 0 |
| 38232 | ENSMUSG00000099876 | 15.40384841 | -0.645288625 | 0.597969661 | -1.079132718 | 0.280528568 | 0.733467233 | Gm29650 | predicted gene 29650 [Source:MGI Symbc | 24.896 | 15.419 | 16.1 | 17.352 | 13.087 | 5.5692 |
| 38233 | ENSMUSG00000099877 | 0 | NA | NA | NA | NA | NA | Pinc | pregnancy induced noncoding RNA [Sourc | 0 | 0 | 0 | 0 | 0 | 0 |
| 38234 | ENSMUSG00000099878 | 0 | NA | NA | NA | NA | NA | Gm28178 | predicted gene 28178 [Source:MGI Symbc | 0 | 0 | 0 | 0 | 0 | 0 |
| 38235 | ENSMUSG00000099879 | 1.456333692 | -0.878911562 | 1.793036822 | -0.490180431 | 0.624006227 | NA | Gm29123 | predicted gene 29123 [Source:MGI Symbc | 1.6062 | 3.0838 | 1.0062 | 1.928 | 0 | 1.1138 |
| 38236 | ENSMUSG00000099880 | 0 | NA | NA | NA | NA | NA | Gm29316 | predicted gene 29316 [Source:MGI Symbc | 0 | 0 | 0 | 0 | 0 | 0 |
| 38237 | ENSMUSG00000099881 | 226.9942854 | 0.032515404 | 0.22841927 | 0.142349653 | 0.886803829 | 0.982382091 | 2810013P06 | RIKEN cDNA 2810013P06 gene [Source:M( | 273.86 | 185.03 | 214.33 | 295.94 | 226.85 | 165.96 |
| 38238 | ENSMUSG00000099882 | 0 | NA | NA | NA | NA | NA | Gm28388 | predicted gene 28388 [Source:MGI Symbc | 0 | 0 | 0 | 0 | 0 | 0 |
| 38239 | ENSMUSG00000099883 | 0 | NA | NA | NA | NA | NA | Gm21784 | predicted gene, 21784 [Source:MGI Symb | 0 | 0 | 0 | 0 | 0 | 0 |
| 38240 | ENSMUSG00000099884 | 0 | NA | NA | NA | NA | NA | Gm8204 | predicted gene 8204 [Source:MGI Symbol | 0 | 0 | 0 | 0 | 0 | 0 |
| 38241 | ENSMUSG00000099886 | 0 | NA | NA | NA | NA | NA | Gm2324 | predicted gene 2324 [Source:MGI Symbol | 0 | 0 | 0 | 0 | 0 | 0 |
| 38242 | ENSMUSG00000099888 | 0 | NA | NA | NA | NA | NA | Gm29552 | predicted gene 29552 [Source:MGI Symbc | 0 | 0 | 0 | 0 | 0 | 0 |
| 38243 | ENSMUSG00000099889 | 12.62574745 | -2.621939527 | 0.782368176 | -3.351286014 | 0.000804372 | 0.017762494 | Mrgprb11-p: | MAS-related GPR, member B11, pseudoge | 27.305 | 25.698 | 12.075 | 1.928 | 7.6343 | 1.1138 |
| 38244 | ENSMUSG00000099890 | 0 | NA | NA | NA | NA | NA | Gm28262 | predicted gene 28262 [Source:MGI Symbc | 0 | 0 | 0 | 0 | 0 | 0 |

| | | | | | | | | | | | | | | | |
|---|---|---|---|---|---|---|---|---|---|---|---|---|---|---|---|
| 38245 | ENSMUSG00000099891 | 0.342432689 | 1.990653086 | 3.900281263 | 0.51038706 | 0.609780321 | NA | Gm5575 | predicted gene 5575 [Source:MGI Symbol | 0 | 0 | 0 | 0.964 | 1.0906 | 0 |
| 38246 | ENSMUSG00000099892 | 0 | NA | NA | NA | NA | NA | Gm18185 | predicted gene, 18185 [Source:MGI Symb | 0 | 0 | 0 | 0 | 0 | 0 |
| 38247 | ENSMUSG00000099893 | 0 | NA | NA | NA | NA | NA | Gm29245 | predicted gene 29245 [Source:MGI Symbo | 0 | 0 | 0 | 0 | 0 | 0 |
| 38248 | ENSMUSG00000099894 | 0 | NA | NA | NA | NA | NA | Gm20911 | predicted gene, 20911 [Source:MGI Symb | 0 | 0 | 0 | 0 | 0 | 0 |
| 38249 | ENSMUSG00000099895 | 0 | NA | NA | NA | NA | NA | Gm28153 | predicted gene 28153 [Source:MGI Symbo | 0 | 0 | 0 | 0 | 0 | 0 |
| 38250 | ENSMUSG00000099896 | 0 | NA | NA | NA | NA | NA | Gm29278 | predicted gene 29278 [Source:MGI Symbo | 0 | 0 | 0 | 0 | 0 | 0 |
| 38251 | ENSMUSG00000099897 | 0 | NA | NA | NA | NA | NA | Gm28426 | predicted gene 28426 [Source:MGI Symbo | 0 | 0 | 0 | 0 | 0 | 0 |
| 38252 | ENSMUSG00000099898 | 0 | NA | NA | NA | NA | NA | Scgb1b17 | secretoglobin, family 1B, member 17 [Sou | 0 | 0 | 0 | 0 | 0 | 0 |
| 38253 | ENSMUSG00000099899 | 0 | NA | NA | NA | NA | NA | Gm28784 | predicted gene 28784 [Source:MGI Symbo | 0 | 0 | 0 | 0 | 0 | 0 |
| 38254 | ENSMUSG00000099900 | 0 | NA | NA | NA | NA | NA | Scgb2b21 | secretoglobin, family 2B, member 21 [Sou | 0 | 0 | 0 | 0 | 0 | 0 |
| 38255 | ENSMUSG00000099901 | 0 | NA | NA | NA | NA | NA | Zbed4-ps2 | zinc finger, BED type containing 4, pseudo | 0 | 0 | 0 | 0 | 0 | 0 |
| 38256 | ENSMUSG00000099902 | 1.065309757 | 0.682201659 | 2.353125625 | 0.289912979 | 0.771882812 | NA | Gm12115 | predicted gene 12115 [Source:MGI Symbo | 2.4093 | 0 | 0 | 2.8919 | 1.0906 | 0 |
| 38257 | ENSMUSG00000099903 | 0 | NA | NA | NA | NA | NA | Gm29070 | predicted gene 29070 [Source:MGI Symbo | 0 | 0 | 0 | 0 | 0 | 0 |
| 38258 | ENSMUSG00000099904 | 0 | NA | NA | NA | NA | NA | Gm28270 | predicted gene 28270 [Source:MGI Symbo | 0 | 0 | 0 | 0 | 0 | 0 |
| 38259 | ENSMUSG00000099905 | 0 | NA | NA | NA | NA | NA | Scgb1b14-ps | secretoglobin, family 1B, member 14, pse | 0 | 0 | 0 | 0 | 0 | 0 |
| 38260 | ENSMUSG00000099906 | 0 | NA | NA | NA | NA | NA | Gm28653 | predicted gene 28653 [Source:MGI Symbo | 0 | 0 | 0 | 0 | 0 | 0 |
| 38261 | ENSMUSG00000099907 | 0 | NA | NA | NA | NA | NA | Gm10421 | predicted gene 10421 [Source:MGI Symbo | 0 | 0 | 0 | 0 | 0 | 0 |
| 38262 | ENSMUSG00000099908 | 0 | NA | NA | NA | NA | NA | Gm28539 | predicted gene 28539 [Source:MGI Symbo | 0 | 0 | 0 | 0 | 0 | 0 |
| 38263 | ENSMUSG00000099910 | 0 | NA | NA | NA | NA | NA | Gm29651 | predicted gene 29651 [Source:MGI Symbo | 0 | 0 | 0 | 0 | 0 | 0 |
| 38264 | ENSMUSG00000099911 | 0 | NA | NA | NA | NA | NA | 1700044C05 | RIKEN cDNA 1700044C05 gene [Source:M | 0 | 0 | 0 | 0 | 0 | 0 |
| 38265 | ENSMUSG00000099912 | 0 | NA | NA | NA | NA | NA | Gm28368 | predicted gene 28368 [Source:MGI Symbo | 0 | 0 | 0 | 0 | 0 | 0 |
| 38266 | ENSMUSG00000099913 | 2.325935719 | 1.005442098 | 1.499826255 | 0.670372381 | 0.502620436 | NA | Gm28551 | predicted gene 28551 [Source:MGI Symbo | 1.6062 | 1.0279 | 2.0125 | 3.8559 | 5.4531 | 0 |
| 38267 | ENSMUSG00000099914 | 0 | NA | NA | NA | NA | NA | Gm28837 | predicted gene 28837 [Source:MGI Symbo | 0 | 0 | 0 | 0 | 0 | 0 |
| 38268 | ENSMUSG00000099915 | 0 | NA | NA | NA | NA | NA | Gm21191 | predicted gene, 21191 [Source:MGI Symb | 0 | 0 | 0 | 0 | 0 | 0 |
| 38269 | ENSMUSG00000099916 | 0 | NA | NA | NA | NA | NA | Gm28318 | predicted gene 28318 [Source:MGI Symbo | 0 | 0 | 0 | 0 | 0 | 0 |
| 38270 | ENSMUSG00000099917 | 0 | NA | NA | NA | NA | NA | Vmn1r215 | vomeronasal 1 receptor 215 [Source:MGI | 0 | 0 | 0 | 0 | 0 | 0 |
| 38271 | ENSMUSG00000099918 | 0 | NA | NA | NA | NA | NA | Gm28066 | predicted gene 28066 [Source:MGI Symbo | 0 | 0 | 0 | 0 | 0 | 0 |
| 38272 | ENSMUSG00000099920 | 0 | NA | NA | NA | NA | NA | Gm29223 | predicted gene 29223 [Source:MGI Symbo | 0 | 0 | 0 | 0 | 0 | 0 |
| 38273 | ENSMUSG00000099921 | 0 | NA | NA | NA | NA | NA | Gm28971 | predicted gene 28971 [Source:MGI Symbo | 0 | 0 | 0 | 0 | 0 | 0 |
| 38274 | ENSMUSG00000099922 | 1.274506859 | 3.875655733 | 2.465658267 | 1.571854374 | 0.115984328 | NA | Gm29016 | predicted gene 29016 [Source:MGI Symbo | 0 | 0 | 0 | 0.964 | 0 | 6.6831 |
| 38275 | ENSMUSG00000099923 | 0.305170013 | -1.657836477 | 4.041480335 | -0.410205257 | 0.681655385 | NA | 1700105P06 | RIKEN cDNA 1700105P06 gene [Source:M | 0.8031 | 1.0279 | 0 | 0 | 0 | 0 |
| 38276 | ENSMUSG00000099924 | 0 | NA | NA | NA | NA | NA | Gm28320 | predicted gene 28320 [Source:MGI Symbo | 0 | 0 | 0 | 0 | 0 | 0 |
| 38277 | ENSMUSG00000099925 | 0 | NA | NA | NA | NA | NA | Gm28827 | predicted gene 28827 [Source:MGI Symbo | 0 | 0 | 0 | 0 | 0 | 0 |
| 38278 | ENSMUSG00000099926 | 0 | NA | NA | NA | NA | NA | Gm6693 | predicted gene 6693 [Source:MGI Symbol | 0 | 0 | 0 | 0 | 0 | 0 |
| 38279 | ENSMUSG00000099927 | 0 | NA | NA | NA | NA | NA | Gm8226 | predicted gene 8226 [Source:MGI Symbol | 0 | 0 | 0 | 0 | 0 | 0 |
| 38280 | ENSMUSG00000099928 | 0 | NA | NA | NA | NA | NA | Gm29202 | predicted gene 29202 [Source:MGI Symbo | 0 | 0 | 0 | 0 | 0 | 0 |
| 38281 | ENSMUSG00000099929 | 0 | NA | NA | NA | NA | NA | Gm28098 | predicted gene 28098 [Source:MGI Symbo | 0 | 0 | 0 | 0 | 0 | 0 |
| 38282 | ENSMUSG00000099930 | 0 | NA | NA | NA | NA | NA | Gm2396 | predicted gene 2396 [Source:MGI Symbol | 0 | 0 | 0 | 0 | 0 | 0 |
| 38283 | ENSMUSG00000099931 | 0 | NA | NA | NA | NA | NA | Gm29358 | predicted gene 29358 [Source:MGI Symbo | 0 | 0 | 0 | 0 | 0 | 0 |
| 38284 | ENSMUSG00000099932 | 0 | NA | NA | NA | NA | NA | Scgb2b14-ps | secretoglobin, family 2B, member 14, pse | 0 | 0 | 0 | 0 | 0 | 0 |
| 38285 | ENSMUSG00000099933 | 0 | NA | NA | NA | NA | NA | 4930452N14 | RIKEN cDNA 4930452N14 gene [Source:M | 0 | 0 | 0 | 0 | 0 | 0 |
| 38286 | ENSMUSG00000099934 | 0 | NA | NA | NA | NA | NA | Gm17983 | predicted gene, 17983 [Source:MGI Symb | 0 | 0 | 0 | 0 | 0 | 0 |
| 38287 | ENSMUSG00000099935 | 0 | NA | NA | NA | NA | NA | Gm21918 | predicted gene, 21918 [Source:MGI Symb | 0 | 0 | 0 | 0 | 0 | 0 |
| 38288 | ENSMUSG00000099936 | 0 | NA | NA | NA | NA | NA | Gm28867 | predicted gene 28867 [Source:MGI Symbo | 0 | 0 | 0 | 0 | 0 | 0 |
| 38289 | ENSMUSG00000099937 | 0 | NA | NA | NA | NA | NA | Gm28872 | predicted gene 28872 [Source:MGI Symbo | 0 | 0 | 0 | 0 | 0 | 0 |
| 38290 | ENSMUSG00000099938 | 0 | NA | NA | NA | NA | NA | C030007H22 | RIKEN cDNA C030007H22 gene [Source:M | 0 | 0 | 0 | 0 | 0 | 0 |
| 38291 | ENSMUSG00000099939 | 0 | NA | NA | NA | NA | NA | Gm28330 | predicted gene 28330 [Source:MGI Symbo | 0 | 0 | 0 | 0 | 0 | 0 |
| 38292 | ENSMUSG00000099940 | 0 | NA | NA | NA | NA | NA | Gm29265 | predicted gene 29265 [Source:MGI Symbo | 0 | 0 | 0 | 0 | 0 | 0 |
| 38293 | ENSMUSG00000099941 | 0 | NA | NA | NA | NA | NA | Gm20118 | predicted gene, 20118 [Source:MGI Symb | 0 | 0 | 0 | 0 | 0 | 0 |
| 38294 | ENSMUSG00000099942 | 0 | NA | NA | NA | NA | NA | Gm28275 | predicted gene 28275 [Source:MGI Symbo | 0 | 0 | 0 | 0 | 0 | 0 |
| 38295 | ENSMUSG00000099943 | 0 | NA | NA | NA | NA | NA | Gm18181 | predicted gene, 18181 [Source:MGI Symb | 0 | 0 | 0 | 0 | 0 | 0 |
| 38296 | ENSMUSG00000099944 | 0 | NA | NA | NA | NA | NA | Gm21875 | predicted gene, 21875 [Source:MGI Symb | 0 | 0 | 0 | 0 | 0 | 0 |
| 38297 | ENSMUSG00000099945 | 0 | NA | NA | NA | NA | NA | 1700126A01 | RIKEN cDNA 1700126A01 gene [Source:M | 0 | 0 | 0 | 0 | 0 | 0 |
| 38298 | ENSMUSG00000099946 | 0 | NA | NA | NA | NA | NA | Gm9496 | predicted gene 9496 [Source:MGI Symbol | 0 | 0 | 0 | 0 | 0 | 0 |
| 38299 | ENSMUSG00000099947 | 0 | NA | NA | NA | NA | NA | Gm28660 | predicted gene 28660 [Source:MGI Symbo | 0 | 0 | 0 | 0 | 0 | 0 |
| 38300 | ENSMUSG00000099948 | 0 | NA | NA | NA | NA | NA | Gm29424 | predicted gene 29424 [Source:MGI Symbo | 0 | 0 | 0 | 0 | 0 | 0 |

| | | | | | | | | | | | | | | | |
|---|---|---|---|---|---|---|---|---|---|---|---|---|---|---|---|
| 38301 | ENSMUSG00000099949 | 0 | NA | NA | NA | NA | NA | Gm29124 | predicted gene 29124 [Source:MGI Symbo | 0 | 0 | 0 | 0 | 0 | 0 |
| 38302 | ENSMUSG00000099950 | 0 | NA | NA | NA | NA | NA | 9130227L01 | RIKEN cDNA 9130227L01 gene [Source:M | 0 | 0 | 0 | 0 | 0 | 0 |
| 38303 | ENSMUSG00000099951 | 0 | NA | NA | NA | NA | NA | Gm9226 | predicted gene 9226 [Source:MGI Symbol | 0 | 0 | 0 | 0 | 0 | 0 |
| 38304 | ENSMUSG00000099952 | 0 | NA | NA | NA | NA | NA | Gm18356 | predicted gene, 18356 [Source:MGI Symb | 0 | 0 | 0 | 0 | 0 | 0 |
| 38305 | ENSMUSG00000099953 | 0 | NA | NA | NA | NA | NA | Gm28699 | predicted gene 28699 [Source:MGI Symbo | 0 | 0 | 0 | 0 | 0 | 0 |
| 38306 | ENSMUSG00000099954 | 0 | NA | NA | NA | NA | NA | Gm28112 | predicted gene 28112 [Source:MGI Symbo | 0 | 0 | 0 | 0 | 0 | 0 |
| 38307 | ENSMUSG00000099955 | 0 | NA | NA | NA | NA | NA | Gm28880 | predicted gene 28880 [Source:MGI Symbo | 0 | 0 | 0 | 0 | 0 | 0 |
| 38308 | ENSMUSG00000099956 | 0 | NA | NA | NA | NA | NA | Gm29365 | predicted gene 29365 [Source:MGI Symbo | 0 | 0 | 0 | 0 | 0 | 0 |
| 38309 | ENSMUSG00000099957 | 0 | NA | NA | NA | NA | NA | 2610027F03 | RIKEN cDNA 2610027F03 gene [Source:M | 0 | 0 | 0 | 0 | 0 | 0 |
| 38310 | ENSMUSG00000099958 | 0 | NA | NA | NA | NA | NA | 1700010B13 | RIKEN cDNA 1700010B13 gene [Source:M | 0 | 0 | 0 | 0 | 0 | 0 |
| 38311 | ENSMUSG00000099959 | 0 | NA | NA | NA | NA | NA | Gm29487 | predicted gene 29487 [Source:MGI Symbo | 0 | 0 | 0 | 0 | 0 | 0 |
| 38312 | ENSMUSG00000099960 | 0.160662974 | 1.126924175 | 4.080472857 | 0.276174898 | 0.782413736 | NA | Gm28590 | predicted gene 28590 [Source:MGI Symbo | 0 | 0 | 0 | 0.964 | 0 | 0 |
| 38313 | ENSMUSG00000099961 | 0 | NA | NA | NA | NA | NA | Scgb2b8-ps | secretoglobin, family 2B, member 8, pseu | 0 | 0 | 0 | 0 | 0 | 0 |
| 38314 | ENSMUSG00000099962 | 0 | NA | NA | NA | NA | NA | Gm28613 | predicted gene 28613 [Source:MGI Symbo | 0 | 0 | 0 | 0 | 0 | 0 |
| 38315 | ENSMUSG00000099963 | 0 | NA | NA | NA | NA | NA | Gm28463 | predicted gene 28463 [Source:MGI Symbo | 0 | 0 | 0 | 0 | 0 | 0 |
| 38316 | ENSMUSG00000099964 | 0 | NA | NA | NA | NA | NA | Gm29514 | predicted gene 29514 [Source:MGI Symbo | 0 | 0 | 0 | 0 | 0 | 0 |
| 38317 | ENSMUSG00000099965 | 0 | NA | NA | NA | NA | NA | Gm28399 | predicted gene 28399 [Source:MGI Symbo | 0 | 0 | 0 | 0 | 0 | 0 |
| 38318 | ENSMUSG00000099966 | 85.74707923 | 0.335446965 | 0.266065481 | 1.260768453 | 0.207392283 | 0.652364421 | 2810402E24 | RIKEN cDNA 2810402E24 gene [Source:M | 65.854 | 84.289 | 77.481 | 80.01 | 119.97 | 86.88 |
| 38319 | ENSMUSG00000099967 | 0 | NA | NA | NA | NA | NA | Gm28212 | predicted gene 28212 [Source:MGI Symbo | 0 | 0 | 0 | 0 | 0 | 0 |
| 38320 | ENSMUSG00000099968 | 0 | NA | NA | NA | NA | NA | 1700029J03 | RIKEN cDNA 1700029J03 gene [Source:M | 0 | 0 | 0 | 0 | 0 | 0 |
| 38321 | ENSMUSG00000099969 | 0.133850373 | -0.796629176 | 4.080472857 | -0.195229623 | 0.84521319 | NA | Gm28960 | predicted gene 28960 [Source:MGI Symbo | 0.8031 | 0 | 0 | 0 | 0 | 0 |
| 38322 | ENSMUSG00000099970 | 0 | NA | NA | NA | NA | NA | Rhoay-ps2 | ras homolog A translocated to the Y, pseu | 0 | 0 | 0 | 0 | 0 | 0 |
| 38323 | ENSMUSG00000099971 | 0.185640647 | 1.126924175 | 4.080472857 | 0.276174898 | 0.782413736 | NA | Gm28287 | predicted gene 28287 [Source:MGI Symbo | 0 | 0 | 0 | 0 | 0 | 1.1138 |
| 38324 | ENSMUSG00000099972 | 0 | NA | NA | NA | NA | NA | Gm21248 | predicted gene, 21248 [Source:MGI Symb | 0 | 0 | 0 | 0 | 0 | 0 |
| 38325 | ENSMUSG00000099973 | 0 | NA | NA | NA | NA | NA | Gm18796 | predicted gene, 18796 [Source:MGI Symb | 0 | 0 | 0 | 0 | 0 | 0 |
| 38326 | ENSMUSG00000099974 | 2.144916673 | -1.599166571 | 1.634870878 | -0.978160779 | 0.327994811 | NA | Bcl2a1d | B cell leukemia/lymphoma 2 related prote | 1.6062 | 1.0279 | 7.0437 | 0.964 | 0 | 2.2277 |
| 38327 | ENSMUSG00000099975 | 0 | NA | NA | NA | NA | NA | Gm21617 | predicted gene, 21617 [Source:MGI Symb | 0 | 0 | 0 | 0 | 0 | 0 |
| 38328 | ENSMUSG00000099976 | 0 | NA | NA | NA | NA | NA | Gm20935 | predicted gene, 20935 [Source:MGI Symb | 0 | 0 | 0 | 0 | 0 | 0 |
| 38329 | ENSMUSG00000099977 | 0 | NA | NA | NA | NA | NA | Gm29286 | predicted gene 29286 [Source:MGI Symbo | 0 | 0 | 0 | 0 | 0 | 0 |
| 38330 | ENSMUSG00000099978 | 0 | NA | NA | NA | NA | NA | Gm29407 | predicted gene 29407 [Source:MGI Symbo | 0 | 0 | 0 | 0 | 0 | 0 |
| 38331 | ENSMUSG00000099979 | 0 | NA | NA | NA | NA | NA | Gm5896 | predicted gene 5896 [Source:MGI Symbol | 0 | 0 | 0 | 0 | 0 | 0 |
| 38332 | ENSMUSG00000099980 | 2.905565809 | 1.442006224 | 1.40358123 | 1.027376395 | 0.304243262 | NA | Gm5619 | predicted gene 5619 [Source:MGI Symbol | 1.6062 | 3.0838 | 0 | 3.8559 | 1.0906 | 7.7969 |
| 38333 | ENSMUSG00000099981 | 0 | NA | NA | NA | NA | NA | Gm21136 | predicted gene, 21136 [Source:MGI Symb | 0 | 0 | 0 | 0 | 0 | 0 |
| 38334 | ENSMUSG00000099982 | 0 | NA | NA | NA | NA | NA | Gm28670 | predicted gene 28670 [Source:MGI Symbo | 0 | 0 | 0 | 0 | 0 | 0 |
| 38335 | ENSMUSG00000099983 | 0 | NA | NA | NA | NA | NA | 1700022E09 | RIKEN cDNA 1700022E09 gene [Source:M | 0 | 0 | 0 | 0 | 0 | 0 |
| 38336 | ENSMUSG00000099985 | 0 | NA | NA | NA | NA | NA | Gm28317 | predicted gene 28317 [Source:MGI Symbo | 0 | 0 | 0 | 0 | 0 | 0 |
| 38337 | ENSMUSG00000099986 | 0 | NA | NA | NA | NA | NA | Gm29120 | predicted gene 29120 [Source:MGI Symbo | 0 | 0 | 0 | 0 | 0 | 0 |
| 38338 | ENSMUSG00000099988 | 0 | NA | NA | NA | NA | NA | 1700054A03 | RIKEN cDNA 1700054A03 gene [Source:M | 0 | 0 | 0 | 0 | 0 | 0 |
| 38339 | ENSMUSG00000099989 | 0 | NA | NA | NA | NA | NA | Gm20281 | predicted gene, 20281 [Source:MGI Symb | 0 | 0 | 0 | 0 | 0 | 0 |
| 38340 | ENSMUSG00000099990 | 0 | NA | NA | NA | NA | NA | Gm29578 | predicted gene 29578 [Source:MGI Symbo | 0 | 0 | 0 | 0 | 0 | 0 |
| 38341 | ENSMUSG00000099991 | 0 | NA | NA | NA | NA | NA | Gm29660 | predicted gene 29660 [Source:MGI Symbo | 0 | 0 | 0 | 0 | 0 | 0 |
| 38342 | ENSMUSG00000099992 | 0 | NA | NA | NA | NA | NA | Gm21435 | predicted gene, 21435 [Source:MGI Symb | 0 | 0 | 0 | 0 | 0 | 0 |
| 38343 | ENSMUSG00000099993 | 0 | NA | NA | NA | NA | NA | Gm20930 | predicted gene, 20930 [Source:MGI Symb | 0 | 0 | 0 | 0 | 0 | 0 |
| 38344 | ENSMUSG00000099994 | 0 | NA | NA | NA | NA | NA | 1700024B18 | RIKEN cDNA 1700024B18 gene [Source:M | 0 | 0 | 0 | 0 | 0 | 0 |
| 38345 | ENSMUSG00000099995 | 0 | NA | NA | NA | NA | NA | 1700036G14 | RIKEN cDNA 1700036G14 gene [Source:M | 0 | 0 | 0 | 0 | 0 | 0 |
| 38346 | ENSMUSG00000099996 | 0 | NA | NA | NA | NA | NA | Gm29065 | predicted gene 29065 [Source:MGI Symbo | 0 | 0 | 0 | 0 | 0 | 0 |
| 38347 | ENSMUSG00000099997 | 0.185640647 | 1.126924175 | 4.080472857 | 0.276174898 | 0.782413736 | NA | Tubb4b-ps2 | tubulin, beta 4B class IVB, pseudogene 2 [ | 0 | 0 | 0 | 0 | 0 | 1.1138 |
| 38348 | ENSMUSG00000099998 | 1.220009801 | -0.284706657 | 2.284449446 | -0.124628128 | 0.900817959 | NA | Gm6501 | predicted gene 6501 [Source:MGI Symbol | 0 | 0 | 4.025 | 0 | 2.1812 | 1.1138 |
| 38349 | ENSMUSG00000099999 | 0 | NA | NA | NA | NA | NA | Gm21795 | predicted gene, 21795 [Source:MGI Symb | 0 | 0 | 0 | 0 | 0 | 0 |
| 38350 | ENSMUSG00000100000 | 0 | NA | NA | NA | NA | NA | 1700023F02 | RIKEN cDNA 1700023F02 gene [Source:M | 0 | 0 | 0 | 0 | 0 | 0 |
| 38351 | ENSMUSG00000100001 | 0 | NA | NA | NA | NA | NA | 1810007D17 | RIKEN cDNA 1810007D17 gene [Source:M | 0 | 0 | 0 | 0 | 0 | 0 |
| 38352 | ENSMUSG00000100002 | 1.322276729 | 0.865032458 | 1.778635474 | 0.486346118 | 0.626721786 | NA | Gm29642 | predicted gene 29642 [Source:MGI Symbo | 0.8031 | 1.0279 | 1.0062 | 2.8919 | 1.0906 | 1.1138 |
| 38353 | ENSMUSG00000100003 | 0 | NA | NA | NA | NA | NA | Gm29646 | predicted gene 29646 [Source:MGI Symbo | 0 | 0 | 0 | 0 | 0 | 0 |
| 38354 | ENSMUSG00000100004 | 0 | NA | NA | NA | NA | NA | Taar8c | trace amine-associated receptor 8C [Sour | 0 | 0 | 0 | 0 | 0 | 0 |
| 38355 | ENSMUSG00000100005 | 12.56158602 | -0.6360452 | 0.689653596 | -0.922267648 | 0.356388985 | 0.795787977 | B130024G19 | RIKEN cDNA B130024G19 gene [Source:M | 18.471 | 17.475 | 10.062 | 18.316 | 4.3625 | 6.6831 |
| 38356 | ENSMUSG00000100006 | 0 | NA | NA | NA | NA | NA | Gm21971 | predicted gene 21971 [Source:MGI Symbo | 0 | 0 | 0 | 0 | 0 | 0 |

| | | | | | | | | | | | | | | | |
|---|---|---|---|---|---|---|---|---|---|---|---|---|---|---|---|
| 38357 | ENSMUSG00000100007 | 0.267700746 | -1.497481649 | 4.057001436 | -0.369110456 | 0.712045396 | NA | Gm5527 | predicted gene 5527 [Source:MGI Symbol | 1.6062 | 0 | 0 | 0 | 0 | 0 |
| 38358 | ENSMUSG00000100008 | 0 | NA | NA | NA | NA | NA | Gm28481 | predicted gene 28481 [Source:MGI Symbo | 0 | 0 | 0 | 0 | 0 | 0 |
| 38359 | ENSMUSG00000100009 | 1.912099596 | 0.902160609 | 1.618682754 | 0.557342448 | 0.577293479 | NA | Gm7967 | predicted gene 7967 [Source:MGI Symbol | 0 | 2.0558 | 2.0125 | 1.928 | 4.3625 | 1.1138 |
| 38360 | ENSMUSG00000100010 | 0 | NA | NA | NA | NA | NA | 1700010I02F | RIKEN cDNA 1700010I02 gene [Source:MG | 0 | 0 | 0 | 0 | 0 | 0 |
| 38361 | ENSMUSG00000100011 | 0 | NA | NA | NA | NA | NA | Gm28458 | predicted gene 28458 [Source:MGI Symbo | 0 | 0 | 0 | 0 | 0 | 0 |
| 38362 | ENSMUSG00000100012 | 0 | NA | NA | NA | NA | NA | Gm29207 | predicted gene 29207 [Source:MGI Symbo | 0 | 0 | 0 | 0 | 0 | 0 |
| 38363 | ENSMUSG00000100013 | 0 | NA | NA | NA | NA | NA | Gm19480 | predicted gene, 19480 [Source:MGI Symb | 0 | 0 | 0 | 0 | 0 | 0 |
| 38364 | ENSMUSG00000100014 | 0 | NA | NA | NA | NA | NA | Gm29290 | predicted gene 29290 [Source:MGI Symbo | 0 | 0 | 0 | 0 | 0 | 0 |
| 38365 | ENSMUSG00000100015 | 0 | NA | NA | NA | NA | NA | Gm28952 | predicted gene 28952 [Source:MGI Symbo | 0 | 0 | 0 | 0 | 0 | 0 |
| 38366 | ENSMUSG00000100016 | 0 | NA | NA | NA | NA | NA | Olfr1215 | olfactory receptor 1215 [Source:MGI Sym | 0 | 0 | 0 | 0 | 0 | 0 |
| 38367 | ENSMUSG00000100017 | 155.1625265 | 0.027865692 | 0.193626886 | 0.143914371 | 0.885568087 | 0.982048364 | 2410022M11 | RIKEN cDNA 2410022M11 gene [Source:M | 172.67 | 150.08 | 137.86 | 162.91 | 154.87 | 152.6 |
| 38368 | ENSMUSG00000100018 | 0 | NA | NA | NA | NA | NA | Gm7776 | predicted gene 7776 [Source:MGI Symbol | 0 | 0 | 0 | 0 | 0 | 0 |
| 38369 | ENSMUSG00000100019 | 0 | NA | NA | NA | NA | NA | Gm18766 | predicted gene, 18766 [Source:MGI Symb | 0 | 0 | 0 | 0 | 0 | 0 |
| 38370 | ENSMUSG00000100020 | 0 | NA | NA | NA | NA | NA | Gm19086 | predicted gene, 19086 [Source:MGI Symb | 0 | 0 | 0 | 0 | 0 | 0 |
| 38371 | ENSMUSG00000100021 | 0 | NA | NA | NA | NA | NA | Gm18697 | predicted gene, 18697 [Source:MGI Symb | 0 | 0 | 0 | 0 | 0 | 0 |
| 38372 | ENSMUSG00000100022 | 0.267700746 | -1.497481649 | 4.057001436 | -0.369110456 | 0.712045396 | NA | Gm29590 | predicted gene 29590 [Source:MGI Symbo | 1.6062 | 0 | 0 | 0 | 0 | 0 |
| 38373 | ENSMUSG00000100023 | 0 | NA | NA | NA | NA | NA | Gm28753 | predicted gene 28753 [Source:MGI Symbo | 0 | 0 | 0 | 0 | 0 | 0 |
| 38374 | ENSMUSG00000100024 | 0 | NA | NA | NA | NA | NA | Gm21782 | predicted gene, 21782 [Source:MGI Symb | 0 | 0 | 0 | 0 | 0 | 0 |
| 38375 | ENSMUSG00000100025 | 0 | NA | NA | NA | NA | NA | Gm8141 | predicted gene 8141 [Source:MGI Symbol | 0 | 0 | 0 | 0 | 0 | 0 |
| 38376 | ENSMUSG00000100026 | 0 | NA | NA | NA | NA | NA | Gm29105 | predicted gene 29105 [Source:MGI Symbo | 0 | 0 | 0 | 0 | 0 | 0 |
| 38377 | ENSMUSG00000100027 | 0 | NA | NA | NA | NA | NA | Gm28477 | predicted gene 28477 [Source:MGI Symbo | 0 | 0 | 0 | 0 | 0 | 0 |
| 38378 | ENSMUSG00000100028 | 0 | NA | NA | NA | NA | NA | Gm29092 | predicted gene 29092 [Source:MGI Symbo | 0 | 0 | 0 | 0 | 0 | 0 |
| 38379 | ENSMUSG00000100029 | 0.804680097 | -3.048447205 | 2.843448763 | -1.072095001 | 0.283677364 | NA | D7Ertd128e | DNA segment, Chr 7, ERATO Doi 128, expr | 0.8031 | 0 | 4.025 | 0 | 0 | 0 |
| 38380 | ENSMUSG00000100031 | 0 | NA | NA | NA | NA | NA | Gm18727 | predicted gene, 18727 [Source:MGI Symb | 0 | 0 | 0 | 0 | 0 | 0 |
| 38381 | ENSMUSG00000100032 | 0 | NA | NA | NA | NA | NA | Gm20817 | predicted gene, 20817 [Source:MGI Symb | 0 | 0 | 0 | 0 | 0 | 0 |
| 38382 | ENSMUSG00000100033 | 0 | NA | NA | NA | NA | NA | Gm8337 | predicted gene 8337 [Source:MGI Symbol | 0 | 0 | 0 | 0 | 0 | 0 |
| 38383 | ENSMUSG00000100034 | 0 | NA | NA | NA | NA | NA | Gm28469 | predicted gene 28469 [Source:MGI Symbo | 0 | 0 | 0 | 0 | 0 | 0 |
| 38384 | ENSMUSG00000100035 | 0 | NA | NA | NA | NA | NA | Gm29473 | predicted gene 29473 [Source:MGI Symbo | 0 | 0 | 0 | 0 | 0 | 0 |
| 38385 | ENSMUSG00000100036 | 0 | NA | NA | NA | NA | NA | Gm28671 | predicted gene 28671 [Source:MGI Symbo | 0 | 0 | 0 | 0 | 0 | 0 |
| 38386 | ENSMUSG00000100037 | 0 | NA | NA | NA | NA | NA | Gm29103 | predicted gene 29103 [Source:MGI Symbo | 0 | 0 | 0 | 0 | 0 | 0 |
| 38387 | ENSMUSG00000100038 | 0 | NA | NA | NA | NA | NA | Gm29580 | predicted gene 29580 [Source:MGI Symbo | 0 | 0 | 0 | 0 | 0 | 0 |
| 38388 | ENSMUSG00000100039 | 0 | NA | NA | NA | NA | NA | Gm28959 | predicted gene 28959 [Source:MGI Symbo | 0 | 0 | 0 | 0 | 0 | 0 |
| 38389 | ENSMUSG00000100040 | 0 | NA | NA | NA | NA | NA | Gm28467 | predicted gene 28467 [Source:MGI Symbo | 0 | 0 | 0 | 0 | 0 | 0 |
| 38390 | ENSMUSG00000100041 | 0 | NA | NA | NA | NA | NA | Gm5702 | predicted gene 5702 [Source:MGI Symbol | 0 | 0 | 0 | 0 | 0 | 0 |
| 38391 | ENSMUSG00000100042 | 0 | NA | NA | NA | NA | NA | Gm29658 | predicted gene 29658 [Source:MGI Symbo | 0 | 0 | 0 | 0 | 0 | 0 |
| 38392 | ENSMUSG00000100043 | 0 | NA | NA | NA | NA | NA | Gm29433 | predicted gene 29433 [Source:MGI Symbo | 0 | 0 | 0 | 0 | 0 | 0 |
| 38393 | ENSMUSG00000100044 | 0 | NA | NA | NA | NA | NA | Gm9553 | predicted gene 9553 [Source:MGI Symbol | 0 | 0 | 0 | 0 | 0 | 0 |
| 38394 | ENSMUSG00000100045 | 0 | NA | NA | NA | NA | NA | Gm20929 | predicted gene, 20929 [Source:MGI Symb | 0 | 0 | 0 | 0 | 0 | 0 |
| 38395 | ENSMUSG00000100046 | 0 | NA | NA | NA | NA | NA | Gm29232 | predicted gene 29232 [Source:MGI Symbo | 0 | 0 | 0 | 0 | 0 | 0 |
| 38396 | ENSMUSG00000100047 | 0 | NA | NA | NA | NA | NA | Gm28141 | predicted gene 28141 [Source:MGI Symbo | 0 | 0 | 0 | 0 | 0 | 0 |
| 38397 | ENSMUSG00000100048 | 0 | NA | NA | NA | NA | NA | Gm28603 | predicted gene 28603 [Source:MGI Symbo | 0 | 0 | 0 | 0 | 0 | 0 |
| 38398 | ENSMUSG00000100049 | 0 | NA | NA | NA | NA | NA | Gm28506 | predicted gene 28506 [Source:MGI Symbo | 0 | 0 | 0 | 0 | 0 | 0 |
| 38399 | ENSMUSG00000100050 | 0 | NA | NA | NA | NA | NA | Gm28121 | predicted gene 28121 [Source:MGI Symbo | 0 | 0 | 0 | 0 | 0 | 0 |
| 38400 | ENSMUSG00000100051 | 0 | NA | NA | NA | NA | NA | Gm28568 | predicted gene 28568 [Source:MGI Symbo | 0 | 0 | 0 | 0 | 0 | 0 |
| 38401 | ENSMUSG00000100052 | 0 | NA | NA | NA | NA | NA | Gm28353 | predicted gene 28353 [Source:MGI Symbo | 0 | 0 | 0 | 0 | 0 | 0 |
| 38402 | ENSMUSG00000100053 | 0 | NA | NA | NA | NA | NA | Gm28154 | predicted gene 28154 [Source:MGI Symbo | 0 | 0 | 0 | 0 | 0 | 0 |
| 38403 | ENSMUSG00000100054 | 0 | NA | NA | NA | NA | NA | Gm28202 | predicted gene 28202 [Source:MGI Symbo | 0 | 0 | 0 | 0 | 0 | 0 |
| 38404 | ENSMUSG00000100055 | 0 | NA | NA | NA | NA | NA | Gm20890 | predicted gene, 20890 [Source:MGI Symb | 0 | 0 | 0 | 0 | 0 | 0 |
| 38405 | ENSMUSG00000100056 | 0 | NA | NA | NA | NA | NA | Gm28173 | predicted gene 28173 [Source:MGI Symbo | 0 | 0 | 0 | 0 | 0 | 0 |
| 38406 | ENSMUSG00000100057 | 0 | NA | NA | NA | NA | NA | Gm28690 | predicted gene 28690 [Source:MGI Symbo | 0 | 0 | 0 | 0 | 0 | 0 |
| 38407 | ENSMUSG00000100058 | 0 | NA | NA | NA | NA | NA | Scgb2b18 | secretoglobin, family 2B, member 18 [Sou | 0 | 0 | 0 | 0 | 0 | 0 |
| 38408 | ENSMUSG00000100059 | 0 | NA | NA | NA | NA | NA | Gm29140 | predicted gene 29140 [Source:MGI Symbo | 0 | 0 | 0 | 0 | 0 | 0 |
| 38409 | ENSMUSG00000100060 | 0.181769715 | 1.126924175 | 4.080472857 | 0.276174898 | 0.782413736 | NA | Gm17944 | predicted gene, 17944 [Source:MGI Symb | 0 | 0 | 0 | 0 | 1.0906 | 0 |
| 38410 | ENSMUSG00000100061 | 0 | NA | NA | NA | NA | NA | Gm29212 | predicted gene 29212 [Source:MGI Symbo | 0 | 0 | 0 | 0 | 0 | 0 |
| 38411 | ENSMUSG00000100062 | 0 | NA | NA | NA | NA | NA | 1700063A18 | RIKEN cDNA 1700063A18 gene [Source:M | 0 | 0 | 0 | 0 | 0 | 0 |
| 38412 | ENSMUSG00000100063 | 0 | NA | NA | NA | NA | NA | Gm28538 | predicted gene 28538 [Source:MGI Symbo | 0 | 0 | 0 | 0 | 0 | 0 |

| | | | | | | | | | | | | | | | |
|---|---|---|---|---|---|---|---|---|---|---|---|---|---|---|---|
| 38413 | ENSMUSG00000100064 | 0 | NA | NA | NA | NA | NA | Gm28339 | predicted gene 28339 [Source:MGI Symbo | 0 | 0 | 0 | 0 | 0 | 0 |
| 38414 | ENSMUSG00000100065 | 0 | NA | NA | NA | NA | NA | Gm18969 | predicted gene, 18969 [Source:MGI Symb | 0 | 0 | 0 | 0 | 0 | 0 |
| 38415 | ENSMUSG00000100066 | 0 | NA | NA | NA | NA | NA | Gm21835 | predicted gene, 21835 [Source:MGI Symb | 0 | 0 | 0 | 0 | 0 | 0 |
| 38416 | ENSMUSG00000100067 | 0 | NA | NA | NA | NA | NA | Gm28407 | predicted gene 28407 [Source:MGI Symbo | 0 | 0 | 0 | 0 | 0 | 0 |
| 38417 | ENSMUSG00000100068 | 0 | NA | NA | NA | NA | NA | Gm18558 | predicted gene, 18558 [Source:MGI Symb | 0 | 0 | 0 | 0 | 0 | 0 |
| 38418 | ENSMUSG00000100069 | 0 | NA | NA | NA | NA | NA | Gm6801 | predicted gene 6801 [Source:MGI Symbol | 0 | 0 | 0 | 0 | 0 | 0 |
| 38419 | ENSMUSG00000100070 | 0 | NA | NA | NA | NA | NA | Gm28994 | predicted gene 28994 [Source:MGI Symbo | 0 | 0 | 0 | 0 | 0 | 0 |
| 38420 | ENSMUSG00000100071 | 1.563139899 | 1.105870765 | 1.892780217 | 0.584257356 | 0.559047178 | NA | Gm28707 | predicted gene 28707 [Source:MGI Symbo | 0 | 0 | 3.0187 | 1.928 | 1.0906 | 3.3415 |
| 38421 | ENSMUSG00000100072 | 0 | NA | NA | NA | NA | NA | Gm21275 | predicted gene, 21275 [Source:MGI Symb | 0 | 0 | 0 | 0 | 0 | 0 |
| 38422 | ENSMUSG00000100073 | 0 | NA | NA | NA | NA | NA | Gm29404 | predicted gene 29404 [Source:MGI Symbo | 0 | 0 | 0 | 0 | 0 | 0 |
| 38423 | ENSMUSG00000100074 | 0 | NA | NA | NA | NA | NA | Gm29524 | predicted gene 29524 [Source:MGI Symbo | 0 | 0 | 0 | 0 | 0 | 0 |
| 38424 | ENSMUSG00000100075 | 134.9369025 | 1.197345971 | 0.325244024 | 3.681377313 | 0.000231977 | 0.007054905 | 1700018L02I | RIKEN cDNA 1700018L02 gene [Source:MG | 49.792 | 80.178 | 116.72 | 240.03 | 110.15 | 212.74 |
| 38425 | ENSMUSG00000100076 | 0 | NA | NA | NA | NA | NA | Gm20963 | predicted gene, 20963 [Source:MGI Symb | 0 | 0 | 0 | 0 | 0 | 0 |
| 38426 | ENSMUSG00000100077 | 0 | NA | NA | NA | NA | NA | Gm21826 | predicted gene, 21826 [Source:MGI Symb | 0 | 0 | 0 | 0 | 0 | 0 |
| 38427 | ENSMUSG00000100078 | 6.070277417 | -0.240118823 | 0.937483413 | -0.256131276 | 0.797849472 | 0.962377049 | Gm6170 | predicted gene 6170 [Source:MGI Symbol | 1.6062 | 8.2233 | 10.062 | 7.7118 | 4.3625 | 4.4554 |
| 38428 | ENSMUSG00000100079 | 0 | NA | NA | NA | NA | NA | Ifnab | interferon alpha B [Source:MGI Symbol;A | 0 | 0 | 0 | 0 | 0 | 0 |
| 38429 | ENSMUSG00000100080 | 0 | NA | NA | NA | NA | NA | Gm29364 | predicted gene 29364 [Source:MGI Symbo | 0 | 0 | 0 | 0 | 0 | 0 |
| 38430 | ENSMUSG00000100081 | 0 | NA | NA | NA | NA | NA | Gm29434 | predicted gene 29434 [Source:MGI Symbo | 0 | 0 | 0 | 0 | 0 | 0 |
| 38431 | ENSMUSG00000100082 | 0 | NA | NA | NA | NA | NA | Gm28414 | predicted gene 28414 [Source:MGI Symbo | 0 | 0 | 0 | 0 | 0 | 0 |
| 38432 | ENSMUSG00000100083 | 0 | NA | NA | NA | NA | NA | Gm29467 | predicted gene 29467 [Source:MGI Symbo | 0 | 0 | 0 | 0 | 0 | 0 |
| 38433 | ENSMUSG00000100084 | 4.808140746 | 1.508940693 | 0.999115958 | 1.510275841 | 0.130973054 | NA | 2310068J16I | RIKEN cDNA 2310068J16 gene [Source:MG | 2.4093 | 2.0558 | 3.0187 | 4.8199 | 7.6343 | 8.9108 |
| 38434 | ENSMUSG00000100085 | 0 | NA | NA | NA | NA | NA | Gm21413 | predicted gene, 21413 [Source:MGI Symb | 0 | 0 | 0 | 0 | 0 | 0 |
| 38435 | ENSMUSG00000100086 | 0 | NA | NA | NA | NA | NA | Gm21271 | predicted gene, 21271 [Source:MGI Symb | 0 | 0 | 0 | 0 | 0 | 0 |
| 38436 | ENSMUSG00000100087 | 3.000609746 | 1.1393072 | 1.280755001 | 0.889559049 | 0.373702703 | NA | Gm29475 | predicted gene 29475 [Source:MGI Symbo | 1.6062 | 1.0279 | 3.0187 | 5.7839 | 5.4531 | 1.1138 |
| 38437 | ENSMUSG00000100088 | 0 | NA | NA | NA | NA | NA | Gm28801 | predicted gene 28801 [Source:MGI Symbo | 0 | 0 | 0 | 0 | 0 | 0 |
| 38438 | ENSMUSG00000100089 | 0 | NA | NA | NA | NA | NA | Gm29305 | predicted gene 29305 [Source:MGI Symbo | 0 | 0 | 0 | 0 | 0 | 0 |
| 38439 | ENSMUSG00000100090 | 0 | NA | NA | NA | NA | NA | Gm29199 | predicted gene 29199 [Source:MGI Symbo | 0 | 0 | 0 | 0 | 0 | 0 |
| 38440 | ENSMUSG00000100091 | 0 | NA | NA | NA | NA | NA | Gm28597 | predicted gene 28597 [Source:MGI Symbo | 0 | 0 | 0 | 0 | 0 | 0 |
| 38441 | ENSMUSG00000100092 | 0 | NA | NA | NA | NA | NA | Gm21088 | predicted gene, 21088 [Source:MGI Symb | 0 | 0 | 0 | 0 | 0 | 0 |
| 38442 | ENSMUSG00000100093 | 0 | NA | NA | NA | NA | NA | Gm29649 | predicted gene 29649 [Source:MGI Symbo | 0 | 0 | 0 | 0 | 0 | 0 |
| 38443 | ENSMUSG00000100094 | 647.6211275 | 0.840114274 | 0.193737347 | 4.336356865 | 1.45E-05 | 0.00072252 | 1810008I18I | RIKEN cDNA 1810008I18 gene [Source:MG | 621.6 | 392.66 | 376.34 | 696.96 | 816.87 | 981.3 |
| 38444 | ENSMUSG00000100095 | 0 | NA | NA | NA | NA | NA | Gm28234 | predicted gene 28234 [Source:MGI Symbo | 0 | 0 | 0 | 0 | 0 | 0 |
| 38445 | ENSMUSG00000100096 | 0 | NA | NA | NA | NA | NA | Gm28760 | predicted gene 28760 [Source:MGI Symbo | 0 | 0 | 0 | 0 | 0 | 0 |
| 38446 | ENSMUSG00000100097 | 0 | NA | NA | NA | NA | NA | Gm28464 | predicted gene 28464 [Source:MGI Symbo | 0 | 0 | 0 | 0 | 0 | 0 |
| 38447 | ENSMUSG00000100098 | 0 | NA | NA | NA | NA | NA | Gm21387 | predicted gene, 21387 [Source:MGI Symb | 0 | 0 | 0 | 0 | 0 | 0 |
| 38448 | ENSMUSG00000100099 | 0 | NA | NA | NA | NA | NA | Gm28879 | predicted gene 28879 [Source:MGI Symbo | 0 | 0 | 0 | 0 | 0 | 0 |
| 38449 | ENSMUSG00000100100 | 0 | NA | NA | NA | NA | NA | Gm29626 | predicted gene 29626 [Source:MGI Symbo | 0 | 0 | 0 | 0 | 0 | 0 |
| 38450 | ENSMUSG00000100101 | 0 | NA | NA | NA | NA | NA | Gm29591 | predicted gene 29591 [Source:MGI Symbo | 0 | 0 | 0 | 0 | 0 | 0 |
| 38451 | ENSMUSG00000100103 | 0 | NA | NA | NA | NA | NA | Gm29553 | predicted gene 29553 [Source:MGI Symbo | 0 | 0 | 0 | 0 | 0 | 0 |
| 38452 | ENSMUSG00000100104 | 0.654647159 | -1.320638211 | 2.599368548 | -0.508061164 | 0.61141045 | NA | Gm5644 | predicted gene 5644 [Source:MGI Symbol | 0.8031 | 1.0279 | 1.0062 | 0 | 1.0906 | 0 |
| 38453 | ENSMUSG00000100105 | 0 | NA | NA | NA | NA | NA | Gm29330 | predicted gene 29330 [Source:MGI Symbo | 0 | 0 | 0 | 0 | 0 | 0 |
| 38454 | ENSMUSG00000100106 | 3.549356352 | -0.622738249 | 1.212264085 | -0.513698506 | 0.607462797 | NA | Gm28856 | predicted gene 28856 [Source:MGI Symbo | 0.8031 | 6.1675 | 6.0375 | 3.8559 | 1.0906 | 3.3415 |
| 38455 | ENSMUSG00000100107 | 0 | NA | NA | NA | NA | NA | Gm28567 | predicted gene 28567 [Source:MGI Symbo | 0 | 0 | 0 | 0 | 0 | 0 |
| 38456 | ENSMUSG00000100108 | 0.17131964 | -0.796629176 | 4.080472857 | -0.195229623 | 0.84521319 | NA | Gm28100 | predicted gene 28100 [Source:MGI Symbo | 0 | 1.0279 | 0 | 0 | 0 | 0 |
| 38457 | ENSMUSG00000100109 | 0 | NA | NA | NA | NA | NA | Gm29366 | predicted gene 29366 [Source:MGI Symbo | 0 | 0 | 0 | 0 | 0 | 0 |
| 38458 | ENSMUSG00000100110 | 0 | NA | NA | NA | NA | NA | Gm28783 | predicted gene 28783 [Source:MGI Symbo | 0 | 0 | 0 | 0 | 0 | 0 |
| 38459 | ENSMUSG00000100111 | 0 | NA | NA | NA | NA | NA | A330087D11 | RIKEN cDNA A330087D11 gene [Source:M | 0 | 0 | 0 | 0 | 0 | 0 |
| 38460 | ENSMUSG00000100112 | 0 | NA | NA | NA | NA | NA | Gm29258 | predicted gene 29258 [Source:MGI Symbo | 0 | 0 | 0 | 0 | 0 | 0 |
| 38461 | ENSMUSG00000100113 | 0 | NA | NA | NA | NA | NA | Gm28592 | predicted gene 28592 [Source:MGI Symbo | 0 | 0 | 0 | 0 | 0 | 0 |
| 38462 | ENSMUSG00000100115 | 0 | NA | NA | NA | NA | NA | Gm28174 | predicted gene 28174 [Source:MGI Symbo | 0 | 0 | 0 | 0 | 0 | 0 |
| 38463 | ENSMUSG00000100116 | 0 | NA | NA | NA | NA | NA | Gm20861 | predicted gene, 20861 [Source:MGI Symb | 0 | 0 | 0 | 0 | 0 | 0 |
| 38464 | ENSMUSG00000100117 | 0 | NA | NA | NA | NA | NA | Gm28986 | predicted gene 28986 [Source:MGI Symbo | 0 | 0 | 0 | 0 | 0 | 0 |
| 38465 | ENSMUSG00000100118 | 0 | NA | NA | NA | NA | NA | Gm29117 | predicted gene 29117 [Source:MGI Symbo | 0 | 0 | 0 | 0 | 0 | 0 |
| 38466 | ENSMUSG00000100119 | 0 | NA | NA | NA | NA | NA | 2700089I24I | RIKEN cDNA 2700089I24 gene [Source:MG | 0 | 0 | 0 | 0 | 0 | 0 |
| 38467 | ENSMUSG00000100120 | 0 | NA | NA | NA | NA | NA | Gm53065 | predicted gene 53065 [Source:MGI Symbo | 0 | 0 | 0 | 0 | 0 | 0 |
| 38468 | ENSMUSG00000100121 | 0 | NA | NA | NA | NA | NA | 1700025N23 | RIKEN cDNA 1700025N23 gene [Source:M | 0 | 0 | 0 | 0 | 0 | 0 |

| | | | | | | | | | | | | | | | |
|---|---|---|---|---|---|---|---|---|---|---|---|---|---|---|---|
| 38469 | ENSMUSG00000100122 | 0 | NA | NA | NA | NA | NA | Gm28267 | predicted gene 28267 [Source:MGI Symbo | 0 | 0 | 0 | 0 | 0 | 0 |
| 38470 | ENSMUSG00000100123 | 0 | NA | NA | NA | NA | NA | Gm28087 | predicted gene 28087 [Source:MGI Symbo | 0 | 0 | 0 | 0 | 0 | 0 |
| 38471 | ENSMUSG00000100124 | 0 | NA | NA | NA | NA | NA | Gm28575 | predicted gene 28575 [Source:MGI Symbo | 0 | 0 | 0 | 0 | 0 | 0 |
| 38472 | ENSMUSG00000100125 | 0 | NA | NA | NA | NA | NA | Gm28739 | predicted gene 28739 [Source:MGI Symbo | 0 | 0 | 0 | 0 | 0 | 0 |
| 38473 | ENSMUSG00000100126 | 0 | NA | NA | NA | NA | NA | Gm29584 | predicted gene 29584 [Source:MGI Symbo | 0 | 0 | 0 | 0 | 0 | 0 |
| 38474 | ENSMUSG00000100127 | 0 | NA | NA | NA | NA | NA | Gsdmcl-ps | gasdermin C-like, pseudogene [Source:MG | 0 | 0 | 0 | 0 | 0 | 0 |
| 38475 | ENSMUSG00000100128 | 0 | NA | NA | NA | NA | NA | Vmn1r-ps25 | vomeronasal 1 receptor, pseudogene 25 [ | 0 | 0 | 0 | 0 | 0 | 0 |
| 38476 | ENSMUSG00000100129 | 0.133850373 | -0.796629176 | 4.080472857 | -0.195229623 | 0.84521319 | NA | Gm29064 | predicted gene 29064 [Source:MGI Symbo | 0.8031 | 0 | 0 | 0 | 0 | 0 |
| 38477 | ENSMUSG00000100130 | 0.160662974 | 1.126924175 | 4.080472857 | 0.276174898 | 0.782413736 | NA | Gm28379 | predicted gene 28379 [Source:MGI Symbo | 0 | 0 | 0 | 0.964 | 0 | 0 |
| 38478 | ENSMUSG00000100131 | 258.3791584 | 0.04094614 | 0.15834679 | 0.25858522 | 0.795955288 | 0.962236316 | Gm28439 | predicted gene 28439 [Source:MGI Symbo | 267.43 | 256.98 | 239.49 | 262.2 | 282.47 | 241.7 |
| 38479 | ENSMUSG00000100132 | 0 | NA | NA | NA | NA | NA | Gm18177 | predicted gene, 18177 [Source:MGI Symb | 0 | 0 | 0 | 0 | 0 | 0 |
| 38480 | ENSMUSG00000100133 | 0 | NA | NA | NA | NA | NA | Tgif2lx1 | TGFB-induced factor homeobox 2-like, X-l | 0 | 0 | 0 | 0 | 0 | 0 |
| 38481 | ENSMUSG00000100134 | 0 | NA | NA | NA | NA | NA | 1700065O2C | RIKEN cDNA 1700065O20 gene [Source:M | 0 | 0 | 0 | 0 | 0 | 0 |
| 38482 | ENSMUSG00000100135 | 0.133850373 | -0.796629176 | 4.080472857 | -0.195229623 | 0.84521319 | NA | Gm28643 | predicted gene 28643 [Source:MGI Symbo | 0.8031 | 0 | 0 | 0 | 0 | 0 |
| 38483 | ENSMUSG00000100136 | 0 | NA | NA | NA | NA | NA | Gm20933 | predicted gene, 20933 [Source:MGI Symb | 0 | 0 | 0 | 0 | 0 | 0 |
| 38484 | ENSMUSG00000100137 | 0 | NA | NA | NA | NA | NA | Gm29352 | predicted gene 29352 [Source:MGI Symbo | 0 | 0 | 0 | 0 | 0 | 0 |
| 38485 | ENSMUSG00000100138 | 0 | NA | NA | NA | NA | NA | Gm28745 | predicted gene 28745 [Source:MGI Symbo | 0 | 0 | 0 | 0 | 0 | 0 |
| 38486 | ENSMUSG00000100140 | 0 | NA | NA | NA | NA | NA | Gm28789 | predicted gene 28789 [Source:MGI Symbo | 0 | 0 | 0 | 0 | 0 | 0 |
| 38487 | ENSMUSG00000100141 | 0 | NA | NA | NA | NA | NA | Gm6445 | predicted gene 6445 [Source:MGI Symbol | 0 | 0 | 0 | 0 | 0 | 0 |
| 38488 | ENSMUSG00000100142 | 0 | NA | NA | NA | NA | NA | Gm21340 | predicted gene, 21340 [Source:MGI Symb | 0 | 0 | 0 | 0 | 0 | 0 |
| 38489 | ENSMUSG00000100144 | 0 | NA | NA | NA | NA | NA | Gm28208 | predicted gene 28208 [Source:MGI Symbo | 0 | 0 | 0 | 0 | 0 | 0 |
| 38490 | ENSMUSG00000100145 | 0 | NA | NA | NA | NA | NA | Gm8502 | predicted gene 8502 [Source:MGI Symbol | 0 | 0 | 0 | 0 | 0 | 0 |
| 38491 | ENSMUSG00000100146 | 0 | NA | NA | NA | NA | NA | 1700020M21 | RIKEN cDNA 1700020M21 gene [Source:M | 0 | 0 | 0 | 0 | 0 | 0 |
| 38492 | ENSMUSG00000100147 | 13.68681897 | -0.529945747 | 0.665882066 | -0.795855264 | 0.426116166 | 0.841641981 | 1700047M11 | RIKEN cDNA 1700047M11 gene [Source:M | 19.274 | 9.2513 | 20.125 | 20.244 | 6.5437 | 6.6831 |
| 38493 | ENSMUSG00000100148 | 0 | NA | NA | NA | NA | NA | Gm28158 | predicted gene 28158 [Source:MGI Symbo | 0 | 0 | 0 | 0 | 0 | 0 |
| 38494 | ENSMUSG00000100149 | 0 | NA | NA | NA | NA | NA | Gm28701 | predicted gene 28701 [Source:MGI Symbo | 0 | 0 | 0 | 0 | 0 | 0 |
| 38495 | ENSMUSG00000100150 | 0.724629373 | 1.713042495 | 3.014019035 | 0.56835822 | 0.569791754 | NA | Gm19585 | predicted gene, 19585 [Source:MGI Symb | 0 | 0 | 1.0062 | 0 | 0 | 3.3415 |
| 38496 | ENSMUSG00000100151 | 0 | NA | NA | NA | NA | NA | Gm28382 | predicted gene 28382 [Source:MGI Symbo | 0 | 0 | 0 | 0 | 0 | 0 |
| 38497 | ENSMUSG00000100152 | 0 | NA | NA | NA | NA | NA | Gm29598 | predicted gene 29598 [Source:MGI Symbo | 0 | 0 | 0 | 0 | 0 | 0 |
| 38498 | ENSMUSG00000100153 | 8.798630597 | 0.245533988 | 0.718461384 | 0.341749735 | 0.73253924 | 0.947714452 | Ppp1ccb | protein phosphatase 1 catalytic subunit ga | 8.8341 | 7.1954 | 8.05 | 7.7118 | 7.6343 | 13.366 |
| 38499 | ENSMUSG00000100154 | 0 | NA | NA | NA | NA | NA | Gm29052 | predicted gene 29052 [Source:MGI Symbo | 0 | 0 | 0 | 0 | 0 | 0 |
| 38500 | ENSMUSG00000100155 | 0 | NA | NA | NA | NA | NA | Gm2109 | predicted gene 2109 [Source:MGI Symbol | 0 | 0 | 0 | 0 | 0 | 0 |
| 38501 | ENSMUSG00000100156 | 0 | NA | NA | NA | NA | NA | Gm29053 | predicted gene 29053 [Source:MGI Symbo | 0 | 0 | 0 | 0 | 0 | 0 |
| 38502 | ENSMUSG00000100157 | 13.67155834 | -0.013108466 | 0.763226307 | -0.017175071 | 0.986296949 | 0.99859176 | 2310034O05 | RIKEN cDNA 2310034O05 gene [Source:M | 25.699 | 3.0838 | 12.075 | 5.7839 | 11.997 | 23.391 |
| 38503 | ENSMUSG00000100158 | 0 | NA | NA | NA | NA | NA | Gm28119 | predicted gene 28119 [Source:MGI Symbo | 0 | 0 | 0 | 0 | 0 | 0 |
| 38504 | ENSMUSG00000100159 | 0 | NA | NA | NA | NA | NA | Gm28808 | predicted gene 28808 [Source:MGI Symbo | 0 | 0 | 0 | 0 | 0 | 0 |
| 38505 | ENSMUSG00000100160 | 0 | NA | NA | NA | NA | NA | Gm28527 | predicted gene 28527 [Source:MGI Symbo | 0 | 0 | 0 | 0 | 0 | 0 |
| 38506 | ENSMUSG00000100161 | 0 | NA | NA | NA | NA | NA | Gm28445 | predicted gene 28445 [Source:MGI Symbo | 0 | 0 | 0 | 0 | 0 | 0 |
| 38507 | ENSMUSG00000100162 | 0 | NA | NA | NA | NA | NA | Gm20687 | predicted gene 20687 [Source:MGI Symbo | 0 | 0 | 0 | 0 | 0 | 0 |
| 38508 | ENSMUSG00000100164 | 25.62117753 | -0.085743795 | 0.48082728 | -0.178325561 | 0.858467305 | 0.975439379 | 2610306M01 | RIKEN cDNA 2610306M01 gene [Source:M | 24.093 | 30.838 | 24.15 | 19.28 | 15.269 | 40.098 |
| 38509 | ENSMUSG00000100165 | 0 | NA | NA | NA | NA | NA | Gm28253 | predicted gene 28253 [Source:MGI Symbo | 0 | 0 | 0 | 0 | 0 | 0 |
| 38510 | ENSMUSG00000100166 | 0 | NA | NA | NA | NA | NA | B230110G15 | RIKEN cDNA B230110G15 gene [Source:M | 0 | 0 | 0 | 0 | 0 | 0 |
| 38511 | ENSMUSG00000100167 | 0 | NA | NA | NA | NA | NA | Gm28207 | predicted gene 28207 [Source:MGI Symbo | 0 | 0 | 0 | 0 | 0 | 0 |
| 38512 | ENSMUSG00000100168 | 0 | NA | NA | NA | NA | NA | Gm28526 | predicted gene 28526 [Source:MGI Symbo | 0 | 0 | 0 | 0 | 0 | 0 |
| 38513 | ENSMUSG00000100169 | 0 | NA | NA | NA | NA | NA | Gm29087 | predicted gene 29087 [Source:MGI Symbo | 0 | 0 | 0 | 0 | 0 | 0 |
| 38514 | ENSMUSG00000100170 | 0 | NA | NA | NA | NA | NA | Gm28942 | predicted gene 28942 [Source:MGI Symbo | 0 | 0 | 0 | 0 | 0 | 0 |
| 38515 | ENSMUSG00000100171 | 0 | NA | NA | NA | NA | NA | Gm21201 | predicted gene, 21201 [Source:MGI Symb | 0 | 0 | 0 | 0 | 0 | 0 |
| 38516 | ENSMUSG00000100172 | 0 | NA | NA | NA | NA | NA | Gm4412 | predicted gene 4412 [Source:MGI Symbol | 0 | 0 | 0 | 0 | 0 | 0 |
| 38517 | ENSMUSG00000100174 | 10.23429819 | 0.681112433 | 0.672545085 | 1.012738697 | 0.311184989 | 0.759659017 | Gm28719 | predicted gene 28719 [Source:MGI Symbo | 6.4248 | 6.1675 | 11.069 | 13.496 | 11.997 | 12.252 |
| 38518 | ENSMUSG00000100175 | 0 | NA | NA | NA | NA | NA | 1700025M24 | RIKEN cDNA 1700025M24 gene [Source:M | 0 | 0 | 0 | 0 | 0 | 0 |
| 38519 | ENSMUSG00000100176 | 0.321325949 | 1.911575114 | 4.048267893 | 0.472195804 | 0.636787034 | NA | Gm28586 | predicted gene 28586 [Source:MGI Symbo | 0 | 0 | 0 | 1.928 | 0 | 0 |
| 38520 | ENSMUSG00000100177 | 0 | NA | NA | NA | NA | NA | Gm29489 | predicted gene 29489 [Source:MGI Symbo | 0 | 0 | 0 | 0 | 0 | 0 |
| 38521 | ENSMUSG00000100178 | 0 | NA | NA | NA | NA | NA | Gm5267 | predicted gene 5267 [Source:MGI Symbol | 0 | 0 | 0 | 0 | 0 | 0 |
| 38522 | ENSMUSG00000100179 | 0 | NA | NA | NA | NA | NA | Gm21715 | predicted gene, 21715 [Source:MGI Symb | 0 | 0 | 0 | 0 | 0 | 0 |
| 38523 | ENSMUSG00000100180 | 0 | NA | NA | NA | NA | NA | Gm28140 | predicted gene 28140 [Source:MGI Symbo | 0 | 0 | 0 | 0 | 0 | 0 |
| 38524 | ENSMUSG00000100182 | 0 | NA | NA | NA | NA | NA | 1810006J02F | RIKEN cDNA 1810006J02 gene [Source:MG | 0 | 0 | 0 | 0 | 0 | 0 |

| | | | | | | | | | | | | | | | |
|---|---|---|---|---|---|---|---|---|---|---|---|---|---|---|---|
| 38525 | ENSMUSG00000100183 | 0.59968336 | -1.233555632 | 2.892386901 | -0.426483618 | 0.669755482 | NA | Gm28512 | predicted gene 28512 [Source:MGI Symbo | 1.6062 | 1.0279 | 0 | 0.964 | 0 | 0 |
| 38526 | ENSMUSG00000100184 | 0 | NA | NA | NA | NA | NA | Gm29558 | predicted gene 29558 [Source:MGI Symbo | 0 | 0 | 0 | 0 | 0 | 0 |
| 38527 | ENSMUSG00000100185 | 0 | NA | NA | NA | NA | NA | Gm28634 | predicted gene 28634 [Source:MGI Symbo | 0 | 0 | 0 | 0 | 0 | 0 |
| 38528 | ENSMUSG00000100186 | 0 | NA | NA | NA | NA | NA | Taar8b | trace amine-associated receptor 8B [Sour | 0 | 0 | 0 | 0 | 0 | 0 |
| 38529 | ENSMUSG00000100187 | 0 | NA | NA | NA | NA | NA | Gm29324 | predicted gene 29324 [Source:MGI Symbo | 0 | 0 | 0 | 0 | 0 | 0 |
| 38530 | ENSMUSG00000100188 | 0 | NA | NA | NA | NA | NA | Gm29561 | predicted gene 29561 [Source:MGI Symbo | 0 | 0 | 0 | 0 | 0 | 0 |
| 38531 | ENSMUSG00000100189 | 0 | NA | NA | NA | NA | NA | Gm28096 | predicted gene 28096 [Source:MGI Symbo | 0 | 0 | 0 | 0 | 0 | 0 |
| 38532 | ENSMUSG00000100190 | 0 | NA | NA | NA | NA | NA | Krtap28-10 | keratin associated protein 28-10 [Source:I | 0 | 0 | 0 | 0 | 0 | 0 |
| 38533 | ENSMUSG00000100191 | 0 | NA | NA | NA | NA | NA | Gm7740 | predicted gene 7740 [Source:MGI Symbol | 0 | 0 | 0 | 0 | 0 | 0 |
| 38534 | ENSMUSG00000100192 | 0 | NA | NA | NA | NA | NA | Gm29587 | predicted gene 29587 [Source:MGI Symbo | 0 | 0 | 0 | 0 | 0 | 0 |
| 38535 | ENSMUSG00000100193 | 0 | NA | NA | NA | NA | NA | Gm29079 | predicted gene 29079 [Source:MGI Symbo | 0 | 0 | 0 | 0 | 0 | 0 |
| 38536 | ENSMUSG00000100194 | 0 | NA | NA | NA | NA | NA | Tgif2lx2 | TGFB-induced factor homeobox 2-like, X-l | 0 | 0 | 0 | 0 | 0 | 0 |
| 38537 | ENSMUSG00000100195 | 0 | NA | NA | NA | NA | NA | Gm29244 | predicted gene 29244 [Source:MGI Symbo | 0 | 0 | 0 | 0 | 0 | 0 |
| 38538 | ENSMUSG00000100196 | 0 | NA | NA | NA | NA | NA | Gm20827 | predicted gene, 20827 [Source:MGI Symb | 0 | 0 | 0 | 0 | 0 | 0 |
| 38539 | ENSMUSG00000100197 | 0 | NA | NA | NA | NA | NA | Gm28638 | predicted gene 28638 [Source:MGI Symbo | 0 | 0 | 0 | 0 | 0 | 0 |
| 38540 | ENSMUSG00000100198 | 0 | NA | NA | NA | NA | NA | 1700030O2C | RIKEN cDNA 1700030O20 gene [Source:M | 0 | 0 | 0 | 0 | 0 | 0 |
| 38541 | ENSMUSG00000100199 | 12.85095095 | -0.242806145 | 0.622320663 | -0.390162434 | 0.696416438 | 0.939992732 | Gm20324 | predicted gene, 20324 [Source:MGI Symb | 19.274 | 14.391 | 8.05 | 14.46 | 10.906 | 10.025 |
| 38542 | ENSMUSG00000100200 | 2.987296476 | -0.619427046 | 1.264843947 | -0.489726063 | 0.624327757 | NA | H2al1m | H2A histone family member L1M [Source: | 4.8186 | 2.0558 | 4.025 | 4.8199 | 1.0906 | 1.1138 |
| 38543 | ENSMUSG00000100201 | 10.3581638 | 0.33870726 | 0.667278319 | 0.507595182 | 0.61173727 | 0.916420248 | Gm29093 | predicted gene 29093 [Source:MGI Symbo | 7.2279 | 8.2233 | 12.075 | 12.532 | 8.7249 | 13.366 |
| 38544 | ENSMUSG00000100202 | 0 | NA | NA | NA | NA | NA | Gm29389 | predicted gene 29389 [Source:MGI Symbo | 0 | 0 | 0 | 0 | 0 | 0 |
| 38545 | ENSMUSG00000100203 | 0 | NA | NA | NA | NA | NA | Gm21924 | predicted gene, 21924 [Source:MGI Symb | 0 | 0 | 0 | 0 | 0 | 0 |
| 38546 | ENSMUSG00000100204 | 0 | NA | NA | NA | NA | NA | Gm4849 | predicted gene 4849 [Source:MGI Symbol | 0 | 0 | 0 | 0 | 0 | 0 |
| 38547 | ENSMUSG00000100205 | 0 | NA | NA | NA | NA | NA | Gm4850 | predicted pseudogene 4850 [Source:MGI | 0 | 0 | 0 | 0 | 0 | 0 |
| 38548 | ENSMUSG00000100206 | 0 | NA | NA | NA | NA | NA | Gm28681 | predicted gene 28681 [Source:MGI Symbo | 0 | 0 | 0 | 0 | 0 | 0 |
| 38549 | ENSMUSG00000100207 | 0 | NA | NA | NA | NA | NA | Gm28203 | predicted gene 28203 [Source:MGI Symbo | 0 | 0 | 0 | 0 | 0 | 0 |
| 38550 | ENSMUSG00000100209 | 0 | NA | NA | NA | NA | NA | Gm28793 | predicted gene 28793 [Source:MGI Symbo | 0 | 0 | 0 | 0 | 0 | 0 |
| 38551 | ENSMUSG00000100210 | 0.483327519 | -0.685704547 | 3.145069396 | -0.218025252 | 0.827409436 | NA | H3c7 | H3 clustered histone 7 [Source:MGI Symb | 0.8031 | 0 | 1.0062 | 0 | 1.0906 | 0 |
| 38552 | ENSMUSG00000100211 | 0 | NA | NA | NA | NA | NA | 1700064M1! | RIKEN cDNA 1700064M15 gene [Source:M | 0 | 0 | 0 | 0 | 0 | 0 |
| 38553 | ENSMUSG00000100212 | 0 | NA | NA | NA | NA | NA | Gm5522 | predicted gene 5522 [Source:MGI Symbol | 0 | 0 | 0 | 0 | 0 | 0 |
| 38554 | ENSMUSG00000100213 | 13.36833843 | 0.978457868 | 0.631730482 | 1.548853341 | 0.12141698 | 0.510744254 | Gm28151 | predicted gene 28151 [Source:MGI Symbo | 4.8186 | 11.307 | 11.069 | 15.424 | 13.087 | 24.505 |
| 38555 | ENSMUSG00000100215 | 0 | NA | NA | NA | NA | NA | Gm8292 | predicted gene 8292 [Source:MGI Symbol | 0 | 0 | 0 | 0 | 0 | 0 |
| 38556 | ENSMUSG00000100216 | 0 | NA | NA | NA | NA | NA | Gm21867 | predicted gene, 21867 [Source:MGI Symb | 0 | 0 | 0 | 0 | 0 | 0 |
| 38557 | ENSMUSG00000100217 | 0 | NA | NA | NA | NA | NA | 9930111H07 | RIKEN cDNA 9930111H07 gene [Source:M | 0 | 0 | 0 | 0 | 0 | 0 |
| 38558 | ENSMUSG00000100218 | 0 | NA | NA | NA | NA | NA | Gm8296 | predicted gene 8296 [Source:MGI Symbol | 0 | 0 | 0 | 0 | 0 | 0 |
| 38559 | ENSMUSG00000100219 | 0 | NA | NA | NA | NA | NA | 1700054K19 | RIKEN cDNA 1700054K19 gene [Source:M | 0 | 0 | 0 | 0 | 0 | 0 |
| 38560 | ENSMUSG00000100220 | 0 | NA | NA | NA | NA | NA | Gm29331 | predicted gene 29331 [Source:MGI Symbo | 0 | 0 | 0 | 0 | 0 | 0 |
| 38561 | ENSMUSG00000100221 | 0 | NA | NA | NA | NA | NA | 4930532M1{ | RIKEN cDNA 4930532M18 gene [Source:M | 0 | 0 | 0 | 0 | 0 | 0 |
| 38562 | ENSMUSG00000100222 | 0 | NA | NA | NA | NA | NA | Psmb6-ps2 | proteasome (prosome, macropain) subun | 0 | 0 | 0 | 0 | 0 | 0 |
| 38563 | ENSMUSG00000100223 | 0 | NA | NA | NA | NA | NA | Gm29403 | predicted gene 29403 [Source:MGI Symbo | 0 | 0 | 0 | 0 | 0 | 0 |
| 38564 | ENSMUSG00000100225 | 0 | NA | NA | NA | NA | NA | Gm29097 | predicted gene 29097 [Source:MGI Symbo | 0 | 0 | 0 | 0 | 0 | 0 |
| 38565 | ENSMUSG00000100226 | 0 | NA | NA | NA | NA | NA | Gm20904 | predicted gene, 20904 [Source:MGI Symb | 0 | 0 | 0 | 0 | 0 | 0 |
| 38566 | ENSMUSG00000100227 | 0 | NA | NA | NA | NA | NA | Gm28291 | predicted gene 28291 [Source:MGI Symbo | 0 | 0 | 0 | 0 | 0 | 0 |
| 38567 | ENSMUSG00000100228 | 0.305170013 | -1.657836477 | 4.041480335 | -0.410205257 | 0.681655385 | NA | Gm29257 | predicted gene 29257 [Source:MGI Symbo | 0.8031 | 1.0279 | 0 | 0 | 0 | 0 |
| 38568 | ENSMUSG00000100229 | 0 | NA | NA | NA | NA | NA | Gm21896 | predicted gene, 21896 [Source:MGI Symb | 0 | 0 | 0 | 0 | 0 | 0 |
| 38569 | ENSMUSG00000100230 | 0 | NA | NA | NA | NA | NA | Gm28650 | predicted gene 28650 [Source:MGI Symbo | 0 | 0 | 0 | 0 | 0 | 0 |
| 38570 | ENSMUSG00000100231 | 0 | NA | NA | NA | NA | NA | Gm29554 | predicted gene 29554 [Source:MGI Symbo | 0 | 0 | 0 | 0 | 0 | 0 |
| 38571 | ENSMUSG00000100232 | 0 | NA | NA | NA | NA | NA | Gm28947 | predicted gene 28947 [Source:MGI Symbo | 0 | 0 | 0 | 0 | 0 | 0 |
| 38572 | ENSMUSG00000100233 | 0 | NA | NA | NA | NA | NA | Gm20811 | predicted gene, 20811 [Source:MGI Symb | 0 | 0 | 0 | 0 | 0 | 0 |
| 38573 | ENSMUSG00000100234 | 0 | NA | NA | NA | NA | NA | Gm13277 | predicted gene 13277 [Source:MGI Symbo | 0 | 0 | 0 | 0 | 0 | 0 |
| 38574 | ENSMUSG00000100235 | 51.37583791 | 0.249715755 | 0.35572108 | 0.701998755 | 0.48267994 | 0.871710696 | Gm28557 | predicted gene 28557 [Source:MGI Symbo | 46.58 | 46.256 | 48.3 | 78.082 | 55.622 | 33.415 |
| 38575 | ENSMUSG00000100236 | 0 | NA | NA | NA | NA | NA | Gm3841 | predicted gene 3841 [Source:MGI Symbol | 0 | 0 | 0 | 0 | 0 | 0 |
| 38576 | ENSMUSG00000100237 | 6.905969713 | -0.292938226 | 0.839507965 | -0.348940377 | 0.727134071 | 0.945907265 | Gm28644 | predicted gene 28644 [Source:MGI Symbo | 9.6372 | 4.1117 | 9.0562 | 8.6758 | 3.2719 | 6.6831 |
| 38577 | ENSMUSG00000100238 | 0 | NA | NA | NA | NA | NA | Gm21450 | predicted gene, 21450 [Source:MGI Symb | 0 | 0 | 0 | 0 | 0 | 0 |
| 38578 | ENSMUSG00000100239 | 0 | NA | NA | NA | NA | NA | Gm28146 | predicted gene 28146 [Source:MGI Symbo | 0 | 0 | 0 | 0 | 0 | 0 |
| 38579 | ENSMUSG00000100240 | 0 | NA | NA | NA | NA | NA | Gm20820 | predicted gene, 20820 [Source:MGI Symb | 0 | 0 | 0 | 0 | 0 | 0 |
| 38580 | ENSMUSG00000100241 | 0 | NA | NA | NA | NA | NA | Slc18a3 | solute carrier family 18 (vesicular monoan | 0 | 0 | 0 | 0 | 0 | 0 |

| | | | | | | | | | | | | | | | |
|---|---|---|---|---|---|---|---|---|---|---|---|---|---|---|---|
| 38581 | ENSMUSG00000100242 | 0 | NA | NA | NA | NA | NA | Gm28206 | predicted gene 28206 [Source:MGI Symbo | 0 | 0 | 0 | 0 | 0 | 0 |
| 38582 | ENSMUSG00000100243 | 0 | NA | NA | NA | NA | NA | Gm28367 | predicted gene 28367 [Source:MGI Symbo | 0 | 0 | 0 | 0 | 0 | 0 |
| 38583 | ENSMUSG00000100244 | 0 | NA | NA | NA | NA | NA | Gm29279 | predicted gene 29279 [Source:MGI Symbo | 0 | 0 | 0 | 0 | 0 | 0 |
| 38584 | ENSMUSG00000100245 | 0 | NA | NA | NA | NA | NA | Gm21282 | predicted gene, 21282 [Source:MGI Symb | 0 | 0 | 0 | 0 | 0 | 0 |
| 38585 | ENSMUSG00000100246 | 0 | NA | NA | NA | NA | NA | Gm20879 | predicted gene, 20879 [Source:MGI Symb | 0 | 0 | 0 | 0 | 0 | 0 |
| 38586 | ENSMUSG00000100247 | 0 | NA | NA | NA | NA | NA | A530053M1 | RIKEN cDNA A530053M12 gene [Source:N | 0 | 0 | 0 | 0 | 0 | 0 |
| 38587 | ENSMUSG00000100248 | 0 | NA | NA | NA | NA | NA | Gm28950 | predicted gene 28950 [Source:MGI Symbo | 0 | 0 | 0 | 0 | 0 | 0 |
| 38588 | ENSMUSG00000100249 | 0 | NA | NA | NA | NA | NA | Btbd35f23 | BTB domain containing 35, family membe | 0 | 0 | 0 | 0 | 0 | 0 |
| 38589 | ENSMUSG00000100250 | 0 | NA | NA | NA | NA | NA | Gm18353 | predicted gene, 18353 [Source:MGI Symb | 0 | 0 | 0 | 0 | 0 | 0 |
| 38590 | ENSMUSG00000100251 | 0 | NA | NA | NA | NA | NA | Gm29615 | predicted gene 29615 [Source:MGI Symbo | 0 | 0 | 0 | 0 | 0 | 0 |
| 38591 | ENSMUSG00000100252 | 0 | NA | NA | NA | NA | NA | Mir124-2hg | Mir124-2 host gene (non-protein coding) | 0 | 0 | 0 | 0 | 0 | 0 |
| 38592 | ENSMUSG00000100253 | 0 | NA | NA | NA | NA | NA | 1700020N18 | RIKEN cDNA 1700020N18 gene [Source:M | 0 | 0 | 0 | 0 | 0 | 0 |
| 38593 | ENSMUSG00000100254 | 0 | NA | NA | NA | NA | NA | Trpc2 | transient receptor potential cation chann | 0 | 0 | 0 | 0 | 0 | 0 |
| 38594 | ENSMUSG00000100255 | 0 | NA | NA | NA | NA | NA | Gm28196 | predicted gene 28196 [Source:MGI Symbo | 0 | 0 | 0 | 0 | 0 | 0 |
| 38595 | ENSMUSG00000100256 | 0 | NA | NA | NA | NA | NA | Gm28851 | predicted gene 28851 [Source:MGI Symbo | 0 | 0 | 0 | 0 | 0 | 0 |
| 38596 | ENSMUSG00000100257 | 1.127524603 | -2.321202569 | 2.140371369 | -1.084485899 | 0.278149416 | NA | C4bp-ps1 | complement component 4 binding protei | 1.6062 | 2.0558 | 2.0125 | 0 | 1.0906 | 0 |
| 38597 | ENSMUSG00000100258 | 0 | NA | NA | NA | NA | NA | Gm29045 | predicted gene 29045 [Source:MGI Symbo | 0 | 0 | 0 | 0 | 0 | 0 |
| 38598 | ENSMUSG00000100260 | 0 | NA | NA | NA | NA | NA | Gm28608 | predicted gene 28608 [Source:MGI Symbo | 0 | 0 | 0 | 0 | 0 | 0 |
| 38599 | ENSMUSG00000100261 | 0 | NA | NA | NA | NA | NA | Gm6473 | predicted gene 6473 [Source:MGI Symbol | 0 | 0 | 0 | 0 | 0 | 0 |
| 38600 | ENSMUSG00000100262 | 0 | NA | NA | NA | NA | NA | Gm28398 | predicted gene 28398 [Source:MGI Symbo | 0 | 0 | 0 | 0 | 0 | 0 |
| 38601 | ENSMUSG00000100263 | 0 | NA | NA | NA | NA | NA | Gm28468 | predicted gene 28468 [Source:MGI Symbo | 0 | 0 | 0 | 0 | 0 | 0 |
| 38602 | ENSMUSG00000100264 | 0 | NA | NA | NA | NA | NA | Gm21419 | predicted gene, 21419 [Source:MGI Symb | 0 | 0 | 0 | 0 | 0 | 0 |
| 38603 | ENSMUSG00000100265 | 0 | NA | NA | NA | NA | NA | Gm28363 | predicted gene 28363 [Source:MGI Symbo | 0 | 0 | 0 | 0 | 0 | 0 |
| 38604 | ENSMUSG00000100266 | 0 | NA | NA | NA | NA | NA | Gm5372 | predicted gene 5372 [Source:MGI Symbol | 0 | 0 | 0 | 0 | 0 | 0 |
| 38605 | ENSMUSG00000100267 | 0 | NA | NA | NA | NA | NA | Gm28779 | predicted gene 28779 [Source:MGI Symbo | 0 | 0 | 0 | 0 | 0 | 0 |
| 38606 | ENSMUSG00000100268 | 0 | NA | NA | NA | NA | NA | Gm29468 | predicted gene 29468 [Source:MGI Symbo | 0 | 0 | 0 | 0 | 0 | 0 |
| 38607 | ENSMUSG00000100269 | 0 | NA | NA | NA | NA | NA | Gm28782 | predicted gene 28782 [Source:MGI Symbo | 0 | 0 | 0 | 0 | 0 | 0 |
| 38608 | ENSMUSG00000100270 | 0 | NA | NA | NA | NA | NA | Gm21803 | predicted gene, 21803 [Source:MGI Symb | 0 | 0 | 0 | 0 | 0 | 0 |
| 38609 | ENSMUSG00000100271 | 0 | NA | NA | NA | NA | NA | Gm21127 | predicted gene, 21127 [Source:MGI Symb | 0 | 0 | 0 | 0 | 0 | 0 |
| 38610 | ENSMUSG00000100272 | 0 | NA | NA | NA | NA | NA | Gm28785 | predicted gene 28785 [Source:MGI Symbo | 0 | 0 | 0 | 0 | 0 | 0 |
| 38611 | ENSMUSG00000100273 | 0 | NA | NA | NA | NA | NA | Gm29526 | predicted gene 29526 [Source:MGI Symbo | 0 | 0 | 0 | 0 | 0 | 0 |
| 38612 | ENSMUSG00000100274 | 0 | NA | NA | NA | NA | NA | 1700006F04 | RIKEN cDNA 1700006F04 gene [Source:M | 0 | 0 | 0 | 0 | 0 | 0 |
| 38613 | ENSMUSG00000100275 | 0 | NA | NA | NA | NA | NA | Gm28915 | predicted gene 28915 [Source:MGI Symbo | 0 | 0 | 0 | 0 | 0 | 0 |
| 38614 | ENSMUSG00000100276 | 0 | NA | NA | NA | NA | NA | Gm29618 | predicted gene 29618 [Source:MGI Symbo | 0 | 0 | 0 | 0 | 0 | 0 |
| 38615 | ENSMUSG00000100277 | 102.3522654 | -1.275250628 | 0.248944478 | -5.122630705 | 3.01E-07 | 2.81E-05 | 1810053B23 | RIKEN cDNA 1810053B23 gene [Source:M | 142.15 | 164.47 | 127.79 | 52.055 | 73.071 | 54.578 |
| 38616 | ENSMUSG00000100278 | 0 | NA | NA | NA | NA | NA | Gm28082 | predicted gene 28082 [Source:MGI Symbo | 0 | 0 | 0 | 0 | 0 | 0 |
| 38617 | ENSMUSG00000100279 | 0 | NA | NA | NA | NA | NA | Gm29387 | predicted gene 29387 [Source:MGI Symbo | 0 | 0 | 0 | 0 | 0 | 0 |
| 38618 | ENSMUSG00000100280 | 0.678054141 | -2.792799308 | 2.990673428 | -0.933836266 | 0.350388359 | NA | Gm28417 | predicted gene 28417 [Source:MGI Symbo | 0 | 2.0558 | 2.0125 | 0 | 0 | 0 |
| 38619 | ENSMUSG00000100281 | 2.833362116 | 0.759245692 | 1.34354519 | 0.565106182 | 0.572001544 | NA | 4930422M2 | RIKEN cDNA 4930422M22 gene [Source:N | 3.2124 | 0 | 3.0187 | 1.928 | 3.2719 | 5.5692 |
| 38620 | ENSMUSG00000100282 | 0 | NA | NA | NA | NA | NA | 1700041C23 | RIKEN cDNA 1700041C23 gene [Source:M | 0 | 0 | 0 | 0 | 0 | 0 |
| 38621 | ENSMUSG00000100283 | 0 | NA | NA | NA | NA | NA | Gm29024 | predicted gene 29024 [Source:MGI Symbo | 0 | 0 | 0 | 0 | 0 | 0 |
| 38622 | ENSMUSG00000100284 | 0 | NA | NA | NA | NA | NA | Gm29239 | predicted gene 29239 [Source:MGI Symbo | 0 | 0 | 0 | 0 | 0 | 0 |
| 38623 | ENSMUSG00000100285 | 0 | NA | NA | NA | NA | NA | Gm21147 | predicted gene, 21147 [Source:MGI Symb | 0 | 0 | 0 | 0 | 0 | 0 |
| 38624 | ENSMUSG00000100286 | 0 | NA | NA | NA | NA | NA | Gm28696 | predicted gene 28696 [Source:MGI Symbo | 0 | 0 | 0 | 0 | 0 | 0 |
| 38625 | ENSMUSG00000100287 | 2.050447603 | -1.094369729 | 1.878585219 | -0.582549952 | 0.560196305 | NA | Gm28068 | predicted gene 28068 [Source:MGI Symbo | 2.4093 | 0 | 6.0375 | 3.8559 | 0 | 0 |
| 38626 | ENSMUSG00000100288 | 0 | NA | NA | NA | NA | NA | Gm28345 | predicted gene 28345 [Source:MGI Symbo | 0 | 0 | 0 | 0 | 0 | 0 |
| 38627 | ENSMUSG00000100289 | 0 | NA | NA | NA | NA | NA | Gm28917 | predicted gene 28917 [Source:MGI Symbo | 0 | 0 | 0 | 0 | 0 | 0 |
| 38628 | ENSMUSG00000100290 | 0 | NA | NA | NA | NA | NA | Gm18552 | predicted gene, 18552 [Source:MGI Symb | 0 | 0 | 0 | 0 | 0 | 0 |
| 38629 | ENSMUSG00000100291 | 0 | NA | NA | NA | NA | NA | 2310069B03 | RIKEN cDNA 2310069B03 gene [Source:M | 0 | 0 | 0 | 0 | 0 | 0 |
| 38630 | ENSMUSG00000100292 | 0 | NA | NA | NA | NA | NA | Gm29198 | predicted gene 29198 [Source:MGI Symbo | 0 | 0 | 0 | 0 | 0 | 0 |
| 38631 | ENSMUSG00000100294 | 0 | NA | NA | NA | NA | NA | 1700034K08 | RIKEN cDNA 1700034K08 gene [Source:M | 0 | 0 | 0 | 0 | 0 | 0 |
| 38632 | ENSMUSG00000100295 | 0 | NA | NA | NA | NA | NA | Gm28165 | predicted gene 28165 [Source:MGI Symbo | 0 | 0 | 0 | 0 | 0 | 0 |
| 38633 | ENSMUSG00000100296 | 0 | NA | NA | NA | NA | NA | Vmn1r205 | vomeronasal 1 receptor 205 [Source:MGI | 0 | 0 | 0 | 0 | 0 | 0 |
| 38634 | ENSMUSG00000100297 | 0 | NA | NA | NA | NA | NA | 1700044K03 | RIKEN cDNA 1700044K03 gene [Source:M | 0 | 0 | 0 | 0 | 0 | 0 |
| 38635 | ENSMUSG00000100298 | 0 | NA | NA | NA | NA | NA | Gm28126 | predicted gene 28126 [Source:MGI Symbo | 0 | 0 | 0 | 0 | 0 | 0 |
| 38636 | ENSMUSG00000100299 | 0 | NA | NA | NA | NA | NA | Gm28584 | predicted gene 28584 [Source:MGI Symbo | 0 | 0 | 0 | 0 | 0 | 0 |

| | | | | | | | | | | | | | | | |
|---|---|---|---|---|---|---|---|---|---|---|---|---|---|---|---|
| 38637 | ENSMUSG00000100300 | 0 | NA | NA | NA | NA | NA | Gm28191 | predicted gene 28191 [Source:MGI Symbo | 0 | 0 | 0 | 0 | 0 | 0 |
| 38638 | ENSMUSG00000100301 | 2.629683985 | -1.396667371 | 1.443625289 | -0.967472225 | 0.333308019 | NA | 6030407O03 | RIKEN cDNA 6030407O03 gene [Source:M | 3.2124 | 6.1675 | 2.0125 | 0 | 3.2719 | 1.1138 |
| 38639 | ENSMUSG00000100302 | 0 | NA | NA | NA | NA | NA | Gm29670 | predicted gene 29670 [Source:MGI Symbo | 0 | 0 | 0 | 0 | 0 | 0 |
| 38640 | ENSMUSG00000100303 | 0 | NA | NA | NA | NA | NA | 2600014E21 | RIKEN cDNA 2600014E21 gene [Source:M | 0 | 0 | 0 | 0 | 0 | 0 |
| 38641 | ENSMUSG00000100304 | 0 | NA | NA | NA | NA | NA | Gm28190 | predicted gene 28190 [Source:MGI Symbo | 0 | 0 | 0 | 0 | 0 | 0 |
| 38642 | ENSMUSG00000100305 | 0 | NA | NA | NA | NA | NA | Gm28360 | predicted gene 28360 [Source:MGI Symbo | 0 | 0 | 0 | 0 | 0 | 0 |
| 38643 | ENSMUSG00000100306 | 0 | NA | NA | NA | NA | NA | Gm29362 | predicted gene 29362 [Source:MGI Symbo | 0 | 0 | 0 | 0 | 0 | 0 |
| 38644 | ENSMUSG00000100307 | 0 | NA | NA | NA | NA | NA | Gm29546 | predicted gene 29546 [Source:MGI Symbo | 0 | 0 | 0 | 0 | 0 | 0 |
| 38645 | ENSMUSG00000100308 | 0 | NA | NA | NA | NA | NA | Gm29410 | predicted gene 29410 [Source:MGI Symbo | 0 | 0 | 0 | 0 | 0 | 0 |
| 38646 | ENSMUSG00000100309 | 0 | NA | NA | NA | NA | NA | Gm6644 | predicted gene 6644 [Source:MGI Symbol | 0 | 0 | 0 | 0 | 0 | 0 |
| 38647 | ENSMUSG00000100310 | 0 | NA | NA | NA | NA | NA | Gm19641 | predicted gene, 19641 [Source:MGI Symb | 0 | 0 | 0 | 0 | 0 | 0 |
| 38648 | ENSMUSG00000100311 | 0 | NA | NA | NA | NA | NA | Gm29581 | predicted gene 29581 [Source:MGI Symbo | 0 | 0 | 0 | 0 | 0 | 0 |
| 38649 | ENSMUSG00000100312 | 0 | NA | NA | NA | NA | NA | Gm29048 | predicted gene 29048 [Source:MGI Symbo | 0 | 0 | 0 | 0 | 0 | 0 |
| 38650 | ENSMUSG00000100313 | 1.567093454 | 0.688414251 | 2.068504584 | 0.332807699 | 0.739279446 | NA | Gm28323 | predicted gene 28323 [Source:MGI Symbo | 1.6062 | 0 | 2.0125 | 5.7839 | 0 | 0 |
| 38651 | ENSMUSG00000100314 | 0 | NA | NA | NA | NA | NA | 1700010J16I | RIKEN cDNA 1700010J16 gene [Source:M | 0 | 0 | 0 | 0 | 0 | 0 |
| 38652 | ENSMUSG00000100315 | 0 | NA | NA | NA | NA | NA | 1700031P21 | RIKEN cDNA 1700031P21 gene [Source:M | 0 | 0 | 0 | 0 | 0 | 0 |
| 38653 | ENSMUSG00000100316 | 0 | NA | NA | NA | NA | NA | Gm28812 | predicted gene 28812 [Source:MGI Symbo | 0 | 0 | 0 | 0 | 0 | 0 |
| 38654 | ENSMUSG00000100317 | 0 | NA | NA | NA | NA | NA | Gm28224 | predicted gene 28224 [Source:MGI Symbo | 0 | 0 | 0 | 0 | 0 | 0 |
| 38655 | ENSMUSG00000100318 | 0 | NA | NA | NA | NA | NA | Gm18799 | predicted gene, 18799 [Source:MGI Symb | 0 | 0 | 0 | 0 | 0 | 0 |
| 38656 | ENSMUSG00000100319 | 0.346303622 | 2.004339653 | 3.884003423 | 0.516049919 | 0.605819544 | NA | Gm5256 | predicted gene 5256 [Source:MGI Symbol | 0 | 0 | 0 | 0.964 | 0 | 1.1138 |
| 38657 | ENSMUSG00000100320 | 2.248159733 | -0.146190544 | 1.513474436 | -0.096592675 | 0.923049874 | NA | Gm29417 | predicted gene 29417 [Source:MGI Symbo | 0 | 5.1396 | 2.0125 | 1.928 | 2.1812 | 2.2277 |
| 38658 | ENSMUSG00000100321 | 0 | NA | NA | NA | NA | NA | Gm29579 | predicted gene 29579 [Source:MGI Symbo | 0 | 0 | 0 | 0 | 0 | 0 |
| 38659 | ENSMUSG00000100322 | 0 | NA | NA | NA | NA | NA | Gm29624 | predicted gene 29624 [Source:MGI Symbo | 0 | 0 | 0 | 0 | 0 | 0 |
| 38660 | ENSMUSG00000100324 | 0 | NA | NA | NA | NA | NA | Gm9156 | predicted gene 9156 [Source:MGI Symbol | 0 | 0 | 0 | 0 | 0 | 0 |
| 38661 | ENSMUSG00000100325 | 0 | NA | NA | NA | NA | NA | Gm28604 | predicted gene 28604 [Source:MGI Symbo | 0 | 0 | 0 | 0 | 0 | 0 |
| 38662 | ENSMUSG00000100326 | 0 | NA | NA | NA | NA | NA | Gm28974 | predicted gene 28974 [Source:MGI Symbo | 0 | 0 | 0 | 0 | 0 | 0 |
| 38663 | ENSMUSG00000100328 | 0 | NA | NA | NA | NA | NA | Gm28816 | predicted gene 28816 [Source:MGI Symbo | 0 | 0 | 0 | 0 | 0 | 0 |
| 38664 | ENSMUSG00000100329 | 0 | NA | NA | NA | NA | NA | Gm29622 | predicted gene 29622 [Source:MGI Symbo | 0 | 0 | 0 | 0 | 0 | 0 |
| 38665 | ENSMUSG00000100330 | 0 | NA | NA | NA | NA | NA | Gm28531 | predicted gene 28531 [Source:MGI Symbo | 0 | 0 | 0 | 0 | 0 | 0 |
| 38666 | ENSMUSG00000100331 | 0 | NA | NA | NA | NA | NA | Gm20880 | predicted gene, 20880 [Source:MGI Symb | 0 | 0 | 0 | 0 | 0 | 0 |
| 38667 | ENSMUSG00000100332 | 0 | NA | NA | NA | NA | NA | Gm29250 | predicted gene 29250 [Source:MGI Symbo | 0 | 0 | 0 | 0 | 0 | 0 |
| 38668 | ENSMUSG00000100333 | 0 | NA | NA | NA | NA | NA | Gm28561 | predicted gene 28561 [Source:MGI Symbo | 0 | 0 | 0 | 0 | 0 | 0 |
| 38669 | ENSMUSG00000100334 | 8.598408923 | -1.211107967 | 0.83677344 | -1.447354695 | 0.147797608 | 0.559179858 | C230024C17 | RIKEN cDNA C230024C17 gene [Source:M | 21.684 | 8.2233 | 6.0375 | 5.7839 | 7.6343 | 2.2277 |
| 38670 | ENSMUSG00000100335 | 0 | NA | NA | NA | NA | NA | 2310008N11 | RIKEN cDNA 2310008N11 gene [Source:M | 0 | 0 | 0 | 0 | 0 | 0 |
| 38671 | ENSMUSG00000100336 | 0 | NA | NA | NA | NA | NA | Gm29601 | predicted gene 29601 [Source:MGI Symbo | 0 | 0 | 0 | 0 | 0 | 0 |
| 38672 | ENSMUSG00000100337 | 0 | NA | NA | NA | NA | NA | Gm5066 | predicted gene 5066 [Source:MGI Symbol | 0 | 0 | 0 | 0 | 0 | 0 |
| 38673 | ENSMUSG00000100338 | 0 | NA | NA | NA | NA | NA | Gm28870 | predicted gene 28870 [Source:MGI Symbo | 0 | 0 | 0 | 0 | 0 | 0 |
| 38674 | ENSMUSG00000100339 | 0 | NA | NA | NA | NA | NA | Gm21864 | predicted gene, 21864 [Source:MGI Symb | 0 | 0 | 0 | 0 | 0 | 0 |
| 38675 | ENSMUSG00000100340 | 0 | NA | NA | NA | NA | NA | Gm28923 | predicted gene 28923 [Source:MGI Symbo | 0 | 0 | 0 | 0 | 0 | 0 |
| 38676 | ENSMUSG00000100341 | 0 | NA | NA | NA | NA | NA | Gm29478 | predicted gene 29478 [Source:MGI Symbo | 0 | 0 | 0 | 0 | 0 | 0 |
| 38677 | ENSMUSG00000100343 | 0 | NA | NA | NA | NA | NA | Gm20923 | predicted gene, 20923 [Source:MGI Symb | 0 | 0 | 0 | 0 | 0 | 0 |
| 38678 | ENSMUSG00000100344 | 0 | NA | NA | NA | NA | NA | Gm29640 | predicted gene 29640 [Source:MGI Symbo | 0 | 0 | 0 | 0 | 0 | 0 |
| 38679 | ENSMUSG00000100345 | 0.342432689 | 1.990653086 | 3.900281263 | 0.51038706 | 0.609780321 | NA | Gm29418 | predicted gene 29418 [Source:MGI Symbo | 0 | 0 | 0 | 0.964 | 1.0906 | 0 |
| 38680 | ENSMUSG00000100346 | 0 | NA | NA | NA | NA | NA | Gm28996 | predicted gene 28996 [Source:MGI Symbo | 0 | 0 | 0 | 0 | 0 | 0 |
| 38681 | ENSMUSG00000100347 | 3.633575069 | 0.190186013 | 1.193699222 | 0.159324903 | 0.873412901 | NA | Gm7895 | predicted gene 7895 [Source:MGI Symbol | 4.0155 | 5.1396 | 1.0062 | 2.8919 | 7.6343 | 1.1138 |
| 38682 | ENSMUSG00000100349 | 0 | NA | NA | NA | NA | NA | Gm28188 | predicted gene 28188 [Source:MGI Symbo | 0 | 0 | 0 | 0 | 0 | 0 |
| 38683 | ENSMUSG00000100350 | 0 | NA | NA | NA | NA | NA | Gm29091 | predicted gene 29091 [Source:MGI Symbo | 0 | 0 | 0 | 0 | 0 | 0 |
| 38684 | ENSMUSG00000100351 | 0 | NA | NA | NA | NA | NA | Gm7867 | predicted gene 7867 [Source:MGI Symbol | 0 | 0 | 0 | 0 | 0 | 0 |
| 38685 | ENSMUSG00000100352 | 0 | NA | NA | NA | NA | NA | Gm28104 | predicted gene 28104 [Source:MGI Symbo | 0 | 0 | 0 | 0 | 0 | 0 |
| 38686 | ENSMUSG00000100353 | 0 | NA | NA | NA | NA | NA | Gm28859 | predicted gene 28859 [Source:MGI Symbo | 0 | 0 | 0 | 0 | 0 | 0 |
| 38687 | ENSMUSG00000100354 | 0 | NA | NA | NA | NA | NA | Gm29113 | predicted gene 29113 [Source:MGI Symbo | 0 | 0 | 0 | 0 | 0 | 0 |
| 38688 | ENSMUSG00000100355 | 0 | NA | NA | NA | NA | NA | Gm28107 | predicted gene 28107 [Source:MGI Symbo | 0 | 0 | 0 | 0 | 0 | 0 |
| 38689 | ENSMUSG00000100356 | 0 | NA | NA | NA | NA | NA | Gm18797 | predicted gene, 18797 [Source:MGI Symb | 0 | 0 | 0 | 0 | 0 | 0 |
| 38690 | ENSMUSG00000100357 | 0 | NA | NA | NA | NA | NA | Gm28181 | predicted gene 28181 [Source:MGI Symbo | 0 | 0 | 0 | 0 | 0 | 0 |
| 38691 | ENSMUSG00000100358 | 0 | NA | NA | NA | NA | NA | Scgb1b4-ps | secretoglobin, family 1B, member 4, pseu | 0 | 0 | 0 | 0 | 0 | 0 |
| 38692 | ENSMUSG00000100359 | 0 | NA | NA | NA | NA | NA | Gm21766 | predicted gene, 21766 [Source:MGI Symb | 0 | 0 | 0 | 0 | 0 | 0 |

| | | | | | | | | | | | | | | | |
|---|---|---|---|---|---|---|---|---|---|---|---|---|---|---|---|
| 38693 | ENSMUSG00000100360 | 0 | NA | NA | NA | NA | NA | Gm28649 | predicted gene 28649 [Source:MGI Symbo | 0 | 0 | 0 | 0 | 0 | 0 |
| 38694 | ENSMUSG00000100361 | 0 | NA | NA | NA | NA | NA | Gm7063 | predicted gene 7063 [Source:MGI Symbol | 0 | 0 | 0 | 0 | 0 | 0 |
| 38695 | ENSMUSG00000100362 | 0 | NA | NA | NA | NA | NA | Gm1833 | predicted gene 1833 [Source:MGI Symbol | 0 | 0 | 0 | 0 | 0 | 0 |
| 38696 | ENSMUSG00000100363 | 0 | NA | NA | NA | NA | NA | Gm20805 | predicted gene, 20805 [Source:MGI Symb | 0 | 0 | 0 | 0 | 0 | 0 |
| 38697 | ENSMUSG00000100364 | 0 | NA | NA | NA | NA | NA | 1700121L16I | RIKEN cDNA 1700121L16 gene [Source:M( | 0 | 0 | 0 | 0 | 0 | 0 |
| 38698 | ENSMUSG00000100365 | 0 | NA | NA | NA | NA | NA | Gm29319 | predicted gene 29319 [Source:MGI Symbo | 0 | 0 | 0 | 0 | 0 | 0 |
| 38699 | ENSMUSG00000100366 | 0 | NA | NA | NA | NA | NA | Gm29568 | predicted gene 29568 [Source:MGI Symbo | 0 | 0 | 0 | 0 | 0 | 0 |
| 38700 | ENSMUSG00000100367 | 0.17131964 | -0.796629176 | 4.080472857 | -0.195229623 | 0.84521319 | NA | Gm29336 | predicted gene 29336 [Source:MGI Symbo | 0 | 1.0279 | 0 | 0 | 0 | 0 |
| 38701 | ENSMUSG00000100368 | 0 | NA | NA | NA | NA | NA | Gm28656 | predicted gene 28656 [Source:MGI Symbo | 0 | 0 | 0 | 0 | 0 | 0 |
| 38702 | ENSMUSG00000100369 | 0 | NA | NA | NA | NA | NA | Gm29527 | predicted gene 29527 [Source:MGI Symbo | 0 | 0 | 0 | 0 | 0 | 0 |
| 38703 | ENSMUSG00000100372 | 0 | NA | NA | NA | NA | NA | 1700003I22F | RIKEN cDNA 1700003I22 gene [Source:MG | 0 | 0 | 0 | 0 | 0 | 0 |
| 38704 | ENSMUSG00000100373 | 0 | NA | NA | NA | NA | NA | Gm29003 | predicted gene 29003 [Source:MGI Symbo | 0 | 0 | 0 | 0 | 0 | 0 |
| 38705 | ENSMUSG00000100374 | 0 | NA | NA | NA | NA | NA | Gm4451 | predicted gene 4451 [Source:MGI Symbol | 0 | 0 | 0 | 0 | 0 | 0 |
| 38706 | ENSMUSG00000100375 | 0 | NA | NA | NA | NA | NA | Gm28114 | predicted gene 28114 [Source:MGI Symbo | 0 | 0 | 0 | 0 | 0 | 0 |
| 38707 | ENSMUSG00000100376 | 0 | NA | NA | NA | NA | NA | Gm28060 | predicted gene 28060 [Source:MGI Symbo | 0 | 0 | 0 | 0 | 0 | 0 |
| 38708 | ENSMUSG00000100377 | 0 | NA | NA | NA | NA | NA | Gm28369 | predicted gene 28369 [Source:MGI Symbo | 0 | 0 | 0 | 0 | 0 | 0 |
| 38709 | ENSMUSG00000100378 | 0 | NA | NA | NA | NA | NA | Rpap3-ps1 | RNA polymerase II associated protein 3, p | 0 | 0 | 0 | 0 | 0 | 0 |
| 38710 | ENSMUSG00000100379 | 0 | NA | NA | NA | NA | NA | Gm28677 | predicted gene 28677 [Source:MGI Symbo | 0 | 0 | 0 | 0 | 0 | 0 |
| 38711 | ENSMUSG00000100380 | 0 | NA | NA | NA | NA | NA | Gm17867 | predicted gene, 17867 [Source:MGI Symb | 0 | 0 | 0 | 0 | 0 | 0 |
| 38712 | ENSMUSG00000100381 | 0 | NA | NA | NA | NA | NA | Gm28094 | predicted gene 28094 [Source:MGI Symbo | 0 | 0 | 0 | 0 | 0 | 0 |
| 38713 | ENSMUSG00000100382 | 0 | NA | NA | NA | NA | NA | Gm28924 | predicted gene 28924 [Source:MGI Symbo | 0 | 0 | 0 | 0 | 0 | 0 |
| 38714 | ENSMUSG00000100383 | 0 | NA | NA | NA | NA | NA | Gm28688 | predicted gene 28688 [Source:MGI Symbo | 0 | 0 | 0 | 0 | 0 | 0 |
| 38715 | ENSMUSG00000100384 | 0 | NA | NA | NA | NA | NA | Gm28489 | predicted gene 28489 [Source:MGI Symbo | 0 | 0 | 0 | 0 | 0 | 0 |
| 38716 | ENSMUSG00000100385 | 0 | NA | NA | NA | NA | NA | Gm29227 | predicted gene 29227 [Source:MGI Symbo | 0 | 0 | 0 | 0 | 0 | 0 |
| 38717 | ENSMUSG00000100386 | 0 | NA | NA | NA | NA | NA | Scgb2b13-ps | secretoglobin, family 2B, member 13, pse | 0 | 0 | 0 | 0 | 0 | 0 |
| 38718 | ENSMUSG00000100387 | 0 | NA | NA | NA | NA | NA | Gm28514 | predicted gene 28514 [Source:MGI Symbo | 0 | 0 | 0 | 0 | 0 | 0 |
| 38719 | ENSMUSG00000100388 | 0 | NA | NA | NA | NA | NA | Gm29116 | predicted gene 29116 [Source:MGI Symbo | 0 | 0 | 0 | 0 | 0 | 0 |
| 38720 | ENSMUSG00000100389 | 0 | NA | NA | NA | NA | NA | Rbm6-ps2 | RNA binding motif protein 6, pseudogene | 0 | 0 | 0 | 0 | 0 | 0 |
| 38721 | ENSMUSG00000100390 | 0 | NA | NA | NA | NA | NA | Gm20881 | predicted gene, 20881 [Source:MGI Symb | 0 | 0 | 0 | 0 | 0 | 0 |
| 38722 | ENSMUSG00000100391 | 0 | NA | NA | NA | NA | NA | Gm28978 | predicted gene 28978 [Source:MGI Symbo | 0 | 0 | 0 | 0 | 0 | 0 |
| 38723 | ENSMUSG00000100392 | 0 | NA | NA | NA | NA | NA | Gm28935 | predicted gene 28935 [Source:MGI Symbo | 0 | 0 | 0 | 0 | 0 | 0 |
| 38724 | ENSMUSG00000100393 | 0 | NA | NA | NA | NA | NA | Gm29491 | predicted gene 29491 [Source:MGI Symbo | 0 | 0 | 0 | 0 | 0 | 0 |
| 38725 | ENSMUSG00000100394 | 0.985111216 | -0.018548963 | 2.684120522 | -0.00691063 | 0.994486159 | NA | Gm28791 | predicted gene 28791 [Source:MGI Symbo | 0 | 0 | 3.0187 | 2.8919 | 0 | 0 |
| 38726 | ENSMUSG00000100395 | 0 | NA | NA | NA | NA | NA | Gm28448 | predicted gene 28448 [Source:MGI Symbo | 0 | 0 | 0 | 0 | 0 | 0 |
| 38727 | ENSMUSG00000100396 | 0.59968336 | -1.233555632 | 2.892386901 | -0.426483618 | 0.669755482 | NA | Gm29367 | predicted gene 29367 [Source:MGI Symbo | 1.6062 | 1.0279 | 0 | 0.964 | 0 | 0 |
| 38728 | ENSMUSG00000100397 | 0 | NA | NA | NA | NA | NA | Gm28071 | predicted gene 28071 [Source:MGI Symbo | 0 | 0 | 0 | 0 | 0 | 0 |
| 38729 | ENSMUSG00000100398 | 0 | NA | NA | NA | NA | NA | Gm5251 | predicted gene 5251 [Source:MGI Symbol | 0 | 0 | 0 | 0 | 0 | 0 |
| 38730 | ENSMUSG00000100399 | 0 | NA | NA | NA | NA | NA | Gm28491 | predicted gene 28491 [Source:MGI Symbo | 0 | 0 | 0 | 0 | 0 | 0 |
| 38731 | ENSMUSG00000100400 | 0 | NA | NA | NA | NA | NA | Gm29326 | predicted gene 29326 [Source:MGI Symbo | 0 | 0 | 0 | 0 | 0 | 0 |
| 38732 | ENSMUSG00000100401 | 0 | NA | NA | NA | NA | NA | Gm29248 | predicted gene 29248 [Source:MGI Symbo | 0 | 0 | 0 | 0 | 0 | 0 |
| 38733 | ENSMUSG00000100402 | 0 | NA | NA | NA | NA | NA | Gm28505 | predicted gene 28505 [Source:MGI Symbo | 0 | 0 | 0 | 0 | 0 | 0 |
| 38734 | ENSMUSG00000100403 | 0 | NA | NA | NA | NA | NA | Gm21838 | predicted gene, 21838 [Source:MGI Symb | 0 | 0 | 0 | 0 | 0 | 0 |
| 38735 | ENSMUSG00000100404 | 0 | NA | NA | NA | NA | NA | Gm21577 | predicted gene, 21577 [Source:MGI Symb | 0 | 0 | 0 | 0 | 0 | 0 |
| 38736 | ENSMUSG00000100405 | 0.17131964 | -0.796629176 | 4.080472857 | -0.195229623 | 0.84521319 | NA | Gm28911 | predicted gene 28911 [Source:MGI Symbo | 0 | 1.0279 | 0 | 0 | 0 | 0 |
| 38737 | ENSMUSG00000100406 | 0 | NA | NA | NA | NA | NA | Gm28120 | predicted gene 28120 [Source:MGI Symbo | 0 | 0 | 0 | 0 | 0 | 0 |
| 38738 | ENSMUSG00000100407 | 0 | NA | NA | NA | NA | NA | Gm28472 | predicted gene 28472 [Source:MGI Symbo | 0 | 0 | 0 | 0 | 0 | 0 |
| 38739 | ENSMUSG00000100408 | 0 | NA | NA | NA | NA | NA | Gm28885 | predicted gene 28885 [Source:MGI Symbo | 0 | 0 | 0 | 0 | 0 | 0 |
| 38740 | ENSMUSG00000100410 | 0 | NA | NA | NA | NA | NA | 2310020H05 | RIKEN cDNA 2310020H05 gene [Source:M | 0 | 0 | 0 | 0 | 0 | 0 |
| 38741 | ENSMUSG00000100411 | 0 | NA | NA | NA | NA | NA | Gm29269 | predicted gene 29269 [Source:MGI Symbo | 0 | 0 | 0 | 0 | 0 | 0 |
| 38742 | ENSMUSG00000100412 | 0 | NA | NA | NA | NA | NA | Gm28572 | predicted gene 28572 [Source:MGI Symbo | 0 | 0 | 0 | 0 | 0 | 0 |
| 38743 | ENSMUSG00000100413 | 0 | NA | NA | NA | NA | NA | Gm21506 | predicted gene, 21506 [Source:MGI Symb | 0 | 0 | 0 | 0 | 0 | 0 |
| 38744 | ENSMUSG00000100415 | 0 | NA | NA | NA | NA | NA | Gm28089 | predicted gene 28089 [Source:MGI Symbo | 0 | 0 | 0 | 0 | 0 | 0 |
| 38745 | ENSMUSG00000100416 | 0 | NA | NA | NA | NA | NA | Gm29501 | predicted gene 29501 [Source:MGI Symbo | 0 | 0 | 0 | 0 | 0 | 0 |
| 38746 | ENSMUSG00000100417 | 0.339027071 | -1.79174149 | 3.919585585 | -0.457125237 | 0.647581032 | NA | Gm28578 | predicted gene 28578 [Source:MGI Symbo | 0 | 1.0279 | 1.0062 | 0 | 0 | 0 |
| 38747 | ENSMUSG00000100418 | 0 | NA | NA | NA | NA | NA | Gm28239 | predicted gene 28239 [Source:MGI Symbo | 0 | 0 | 0 | 0 | 0 | 0 |
| 38748 | ENSMUSG00000100420 | 0 | NA | NA | NA | NA | NA | Gm28616 | predicted gene 28616 [Source:MGI Symbo | 0 | 0 | 0 | 0 | 0 | 0 |

| | | | | | | | | | | | | | | | |
|---|---|---|---|---|---|---|---|---|---|---|---|---|---|---|---|
| 38749 | ENSMUSG00000100421 | 0.181769715 | 1.126924175 | 4.080472857 | 0.276174898 | 0.782413736 | NA | Gm29044 | predicted gene 29044 [Source:MGI Symbo | 0 | 0 | 0 | 0 | 1.0906 | 0 |
| 38750 | ENSMUSG00000100422 | 0 | NA | NA | NA | NA | NA | Gm29226 | predicted gene 29226 [Source:MGI Symbo | 0 | 0 | 0 | 0 | 0 | 0 |
| 38751 | ENSMUSG00000100423 | 0 | NA | NA | NA | NA | NA | Gm28754 | predicted gene 28754 [Source:MGI Symbo | 0 | 0 | 0 | 0 | 0 | 0 |
| 38752 | ENSMUSG00000100424 | 0 | NA | NA | NA | NA | NA | Gm29557 | predicted gene 29557 [Source:MGI Symbo | 0 | 0 | 0 | 0 | 0 | 0 |
| 38753 | ENSMUSG00000100425 | 0 | NA | NA | NA | NA | NA | Gm29131 | predicted gene 29131 [Source:MGI Symbo | 0 | 0 | 0 | 0 | 0 | 0 |
| 38754 | ENSMUSG00000100426 | 2.14148277 | -0.666277817 | 1.683866134 | -0.395683364 | 0.692338629 | NA | Gm4208 | predicted gene 4208 [Source:MGI Symbol | 4.8186 | 3.0838 | 0 | 3.8559 | 1.0906 | 0 |
| 38755 | ENSMUSG00000100428 | 0 | NA | NA | NA | NA | NA | Gm29413 | predicted gene 29413 [Source:MGI Symbo | 0 | 0 | 0 | 0 | 0 | 0 |
| 38756 | ENSMUSG00000100429 | 0 | NA | NA | NA | NA | NA | Gm29186 | predicted gene 29186 [Source:MGI Symbo | 0 | 0 | 0 | 0 | 0 | 0 |
| 38757 | ENSMUSG00000100430 | 0 | NA | NA | NA | NA | NA | 1700095A13 | RIKEN cDNA 1700095A13 gene [Source:M | 0 | 0 | 0 | 0 | 0 | 0 |
| 38758 | ENSMUSG00000100431 | 0 | NA | NA | NA | NA | NA | 4930558N11 | RIKEN cDNA 4930558N11 gene [Source:M | 0 | 0 | 0 | 0 | 0 | 0 |
| 38759 | ENSMUSG00000100432 | 0 | NA | NA | NA | NA | NA | Gm7539 | predicted gene 7539 [Source:MGI Symbol | 0 | 0 | 0 | 0 | 0 | 0 |
| 38760 | ENSMUSG00000100433 | 0 | NA | NA | NA | NA | NA | Gm21198 | predicted gene, 21198 [Source:MGI Symb | 0 | 0 | 0 | 0 | 0 | 0 |
| 38761 | ENSMUSG00000100434 | 0 | NA | NA | NA | NA | NA | Gm3489 | predicted gene 3489 [Source:MGI Symbol | 0 | 0 | 0 | 0 | 0 | 0 |
| 38762 | ENSMUSG00000100436 | 0 | NA | NA | NA | NA | NA | Gm29657 | predicted gene 29657 [Source:MGI Symbo | 0 | 0 | 0 | 0 | 0 | 0 |
| 38763 | ENSMUSG00000100437 | 0 | NA | NA | NA | NA | NA | Gm29152 | predicted gene 29152 [Source:MGI Symbo | 0 | 0 | 0 | 0 | 0 | 0 |
| 38764 | ENSMUSG00000100438 | 0 | NA | NA | NA | NA | NA | Gm21825 | predicted gene, 21825 [Source:MGI Symb | 0 | 0 | 0 | 0 | 0 | 0 |
| 38765 | ENSMUSG00000100439 | 0 | NA | NA | NA | NA | NA | Gm28922 | predicted gene 28922 [Source:MGI Symbo | 0 | 0 | 0 | 0 | 0 | 0 |
| 38766 | ENSMUSG00000100440 | 0 | NA | NA | NA | NA | NA | Gm28405 | predicted gene 28405 [Source:MGI Symbo | 0 | 0 | 0 | 0 | 0 | 0 |
| 38767 | ENSMUSG00000100441 | 0 | NA | NA | NA | NA | NA | Gm7266 | predicted gene 7266 [Source:MGI Symbol | 0 | 0 | 0 | 0 | 0 | 0 |
| 38768 | ENSMUSG00000100442 | 0 | NA | NA | NA | NA | NA | Gm20866 | predicted gene, 20866 [Source:MGI Symb | 0 | 0 | 0 | 0 | 0 | 0 |
| 38769 | ENSMUSG00000100443 | 0 | NA | NA | NA | NA | NA | Scgb2b16-ps | secretoglobin, family 2B, member 16, pse | 0 | 0 | 0 | 0 | 0 | 0 |
| 38770 | ENSMUSG00000100444 | 0 | NA | NA | NA | NA | NA | C530043A13 | RIKEN cDNA C530043A13 gene [Source:M | 0 | 0 | 0 | 0 | 0 | 0 |
| 38771 | ENSMUSG00000100445 | 0 | NA | NA | NA | NA | NA | Gm29550 | predicted gene 29550 [Source:MGI Symbo | 0 | 0 | 0 | 0 | 0 | 0 |
| 38772 | ENSMUSG00000100446 | 0 | NA | NA | NA | NA | NA | Gm28242 | predicted gene 28242 [Source:MGI Symbo | 0 | 0 | 0 | 0 | 0 | 0 |
| 38773 | ENSMUSG00000100447 | 0 | NA | NA | NA | NA | NA | Gm29378 | predicted gene 29378 [Source:MGI Symbo | 0 | 0 | 0 | 0 | 0 | 0 |
| 38774 | ENSMUSG00000100448 | 0 | NA | NA | NA | NA | NA | H2al1k | H2A histone family member L1K [Source:N | 0 | 0 | 0 | 0 | 0 | 0 |
| 38775 | ENSMUSG00000100449 | 0 | NA | NA | NA | NA | NA | Gm28757 | predicted gene 28757 [Source:MGI Symbo | 0 | 0 | 0 | 0 | 0 | 0 |
| 38776 | ENSMUSG00000100450 | 0 | NA | NA | NA | NA | NA | Gm9443 | predicted gene 9443 [Source:MGI Symbol | 0 | 0 | 0 | 0 | 0 | 0 |
| 38777 | ENSMUSG00000100451 | 0 | NA | NA | NA | NA | NA | Gm29665 | predicted gene 29665 [Source:MGI Symbo | 0 | 0 | 0 | 0 | 0 | 0 |
| 38778 | ENSMUSG00000100452 | 0 | NA | NA | NA | NA | NA | Gm29628 | predicted gene 29628 [Source:MGI Symbo | 0 | 0 | 0 | 0 | 0 | 0 |
| 38779 | ENSMUSG00000100453 | 0 | NA | NA | NA | NA | NA | Gm28397 | predicted gene 28397 [Source:MGI Symbo | 0 | 0 | 0 | 0 | 0 | 0 |
| 38780 | ENSMUSG00000100454 | 0.476283062 | 1.031501528 | 3.159856949 | 0.326439312 | 0.744091998 | NA | Gm28901 | predicted gene 28901 [Source:MGI Symbo | 0.8031 | 0 | 0 | 0.964 | 1.0906 | 0 |
| 38781 | ENSMUSG00000100455 | 4.274372037 | 0.331037114 | 1.13771945 | 0.290965505 | 0.771077702 | NA | Gm29170 | predicted gene 29170 [Source:MGI Symbo | 7.2279 | 1.0279 | 3.0187 | 7.7118 | 1.0906 | 5.5692 |
| 38782 | ENSMUSG00000100456 | 0 | NA | NA | NA | NA | NA | Gm21868 | predicted gene, 21868 [Source:MGI Symb | 0 | 0 | 0 | 0 | 0 | 0 |
| 38783 | ENSMUSG00000100457 | 1.932327542 | 1.00886694 | 1.656942162 | 0.608872755 | 0.542608784 | NA | D830032E09 | RIKEN cDNA D830032E09 gene [Source:M | 0.8031 | 1.0279 | 2.0125 | 0 | 2.1812 | 5.5692 |
| 38784 | ENSMUSG00000100458 | 0 | NA | NA | NA | NA | NA | Olfr1060-ps1 | olfactory receptor 1060, pseudogene 1 [S | 0 | 0 | 0 | 0 | 0 | 0 |
| 38785 | ENSMUSG00000100459 | 0 | NA | NA | NA | NA | NA | Gm5833 | predicted gene 5833 [Source:MGI Symbol | 0 | 0 | 0 | 0 | 0 | 0 |
| 38786 | ENSMUSG00000100460 | 0 | NA | NA | NA | NA | NA | Gm28145 | predicted gene 28145 [Source:MGI Symbo | 0 | 0 | 0 | 0 | 0 | 0 |
| 38787 | ENSMUSG00000100461 | 0 | NA | NA | NA | NA | NA | Gm28948 | predicted gene 28948 [Source:MGI Symbo | 0 | 0 | 0 | 0 | 0 | 0 |
| 38788 | ENSMUSG00000100462 | 0 | NA | NA | NA | NA | NA | Gm28115 | predicted gene 28115 [Source:MGI Symbo | 0 | 0 | 0 | 0 | 0 | 0 |
| 38789 | ENSMUSG00000100463 | 0 | NA | NA | NA | NA | NA | Gm28848 | predicted gene 28848 [Source:MGI Symbo | 0 | 0 | 0 | 0 | 0 | 0 |
| 38790 | ENSMUSG00000100464 | 0 | NA | NA | NA | NA | NA | Gm21771 | predicted gene, 21771 [Source:MGI Symb | 0 | 0 | 0 | 0 | 0 | 0 |
| 38791 | ENSMUSG00000100465 | 0.133850373 | -0.796629176 | 4.080472857 | -0.195229623 | 0.84521319 | NA | Gm29264 | predicted gene 29264 [Source:MGI Symbo | 0.8031 | 0 | 0 | 0 | 0 | 0 |
| 38792 | ENSMUSG00000100466 | 0 | NA | NA | NA | NA | NA | Gm29572 | predicted gene 29572 [Source:MGI Symbo | 0 | 0 | 0 | 0 | 0 | 0 |
| 38793 | ENSMUSG00000100467 | 0 | NA | NA | NA | NA | NA | Gm29582 | predicted gene 29582 [Source:MGI Symbo | 0 | 0 | 0 | 0 | 0 | 0 |
| 38794 | ENSMUSG00000100468 | 0 | NA | NA | NA | NA | NA | Tmem167-ps | transmembrane protein 167, pseudogene | 0 | 0 | 0 | 0 | 0 | 0 |
| 38795 | ENSMUSG00000100469 | 0 | NA | NA | NA | NA | NA | Gm28895 | predicted gene 28895 [Source:MGI Symbo | 0 | 0 | 0 | 0 | 0 | 0 |
| 38796 | ENSMUSG00000100470 | 0 | NA | NA | NA | NA | NA | Gm29479 | predicted gene 29479 [Source:MGI Symbo | 0 | 0 | 0 | 0 | 0 | 0 |
| 38797 | ENSMUSG00000100471 | 0 | NA | NA | NA | NA | NA | Gm28409 | predicted gene 28409 [Source:MGI Symbo | 0 | 0 | 0 | 0 | 0 | 0 |
| 38798 | ENSMUSG00000100472 | 0 | NA | NA | NA | NA | NA | Gm28918 | predicted gene 28918 [Source:MGI Symbo | 0 | 0 | 0 | 0 | 0 | 0 |
| 38799 | ENSMUSG00000100473 | 0 | NA | NA | NA | NA | NA | Gm21804 | predicted gene, 21804 [Source:MGI Symb | 0 | 0 | 0 | 0 | 0 | 0 |
| 38800 | ENSMUSG00000100474 | 0 | NA | NA | NA | NA | NA | Gm29614 | predicted gene 29614 [Source:MGI Symbo | 0 | 0 | 0 | 0 | 0 | 0 |
| 38801 | ENSMUSG00000100475 | 0 | NA | NA | NA | NA | NA | Gm29543 | predicted gene 29543 [Source:MGI Symbo | 0 | 0 | 0 | 0 | 0 | 0 |
| 38802 | ENSMUSG00000100476 | 0 | NA | NA | NA | NA | NA | Nanog-ps2 | Nanog pseudogene 2 [Source:MGI Symbo | 0 | 0 | 0 | 0 | 0 | 0 |
| 38803 | ENSMUSG00000100477 | 0 | NA | NA | NA | NA | NA | Gm28507 | predicted gene 28507 [Source:MGI Symbo | 0 | 0 | 0 | 0 | 0 | 0 |
| 38804 | ENSMUSG00000100478 | 0 | NA | NA | NA | NA | NA | Gm29632 | predicted gene 29632 [Source:MGI Symbo | 0 | 0 | 0 | 0 | 0 | 0 |

| | | | | | | | | | | | | | | | |
|---|---|---|---|---|---|---|---|---|---|---|---|---|---|---|---|
| 38805 | ENSMUSG00000100479 | 0 | NA | NA | NA | NA | NA | Gm28921 | predicted gene 28921 [Source:MGI Symbo | 0 | 0 | 0 | 0 | 0 | 0 |
| 38806 | ENSMUSG00000100480 | 2.206017848 | -0.135988269 | 1.558387311 | -0.087262177 | 0.930463118 | NA | Gm29156 | predicted gene 29156 [Source:MGI Symbo | 4.8186 | 1.0279 | 1.0062 | 1.928 | 0 | 4.4554 |
| 38807 | ENSMUSG00000100481 | 0.353089354 | 0.165142706 | 3.896102244 | 0.042386646 | 0.966190474 | NA | Gm7114 | predicted gene 7114 [Source:MGI Symbol | 0 | 1.0279 | 0 | 0 | 1.0906 | 0 |
| 38808 | ENSMUSG00000100483 | 0 | NA | NA | NA | NA | NA | Gm28237 | predicted gene 28237 [Source:MGI Symbo | 0 | 0 | 0 | 0 | 0 | 0 |
| 38809 | ENSMUSG00000100484 | 0 | NA | NA | NA | NA | NA | Gm7133 | predicted gene 7133 [Source:MGI Symbol | 0 | 0 | 0 | 0 | 0 | 0 |
| 38810 | ENSMUSG00000100485 | 0 | NA | NA | NA | NA | NA | Gm20920 | predicted gene, 20920 [Source:MGI Symb | 0 | 0 | 0 | 0 | 0 | 0 |
| 38811 | ENSMUSG00000100486 | 0.503302254 | -0.837877584 | 3.426698643 | -0.244514523 | 0.806832356 | NA | Gm4131 | predicted gene 4131 [Source:MGI Symbol | 0 | 2.0558 | 0 | 0.964 | 0 | 0 |
| 38812 | ENSMUSG00000100487 | 0 | NA | NA | NA | NA | NA | 1700072B07 | RIKEN cDNA 1700072B07 gene [Source:M | 0 | 0 | 0 | 0 | 0 | 0 |
| 38813 | ENSMUSG00000100488 | 0 | NA | NA | NA | NA | NA | Gm29559 | predicted gene 29559 [Source:MGI Symbo | 0 | 0 | 0 | 0 | 0 | 0 |
| 38814 | ENSMUSG00000100489 | 0 | NA | NA | NA | NA | NA | Gm28303 | predicted gene 28303 [Source:MGI Symbo | 0 | 0 | 0 | 0 | 0 | 0 |
| 38815 | ENSMUSG00000100490 | 2.712732888 | 0.040467299 | 1.535614522 | 0.026352511 | 0.978976171 | NA | Gm28455 | predicted gene 28455 [Source:MGI Symbo | 0 | 3.0838 | 5.0312 | 4.8199 | 0 | 3.3415 |
| 38816 | ENSMUSG00000100491 | 0 | NA | NA | NA | NA | NA | Gm28331 | predicted gene 28331 [Source:MGI Symbo | 0 | 0 | 0 | 0 | 0 | 0 |
| 38817 | ENSMUSG00000100492 | 0 | NA | NA | NA | NA | NA | Gm28672 | predicted gene 28672 [Source:MGI Symbo | 0 | 0 | 0 | 0 | 0 | 0 |
| 38818 | ENSMUSG00000100494 | 0 | NA | NA | NA | NA | NA | Gm29033 | predicted gene 29033 [Source:MGI Symbo | 0 | 0 | 0 | 0 | 0 | 0 |
| 38819 | ENSMUSG00000100495 | 0 | NA | NA | NA | NA | NA | Gm29369 | predicted gene 29369 [Source:MGI Symbo | 0 | 0 | 0 | 0 | 0 | 0 |
| 38820 | ENSMUSG00000100496 | 0 | NA | NA | NA | NA | NA | Gm28639 | predicted gene 28639 [Source:MGI Symbo | 0 | 0 | 0 | 0 | 0 | 0 |
| 38821 | ENSMUSG00000100497 | 0 | NA | NA | NA | NA | NA | Gm28624 | predicted gene 28624 [Source:MGI Symbo | 0 | 0 | 0 | 0 | 0 | 0 |
| 38822 | ENSMUSG00000100499 | 0 | NA | NA | NA | NA | NA | Gm28460 | predicted gene 28460 [Source:MGI Symbo | 0 | 0 | 0 | 0 | 0 | 0 |
| 38823 | ENSMUSG00000100500 | 0 | NA | NA | NA | NA | NA | Gm29080 | predicted gene 29080 [Source:MGI Symbo | 0 | 0 | 0 | 0 | 0 | 0 |
| 38824 | ENSMUSG00000100501 | 0 | NA | NA | NA | NA | NA | Gm29466 | predicted gene 29466 [Source:MGI Symbo | 0 | 0 | 0 | 0 | 0 | 0 |
| 38825 | ENSMUSG00000100502 | 0.545309144 | 2.651974629 | 3.666219369 | 0.72335405 | 0.469462394 | NA | Gm28286 | predicted gene 28286 [Source:MGI Symbo | 0 | 0 | 0 | 0 | 3.2719 | 0 |
| 38826 | ENSMUSG00000100503 | 0 | NA | NA | NA | NA | NA | Gm28927 | predicted gene 28927 [Source:MGI Symbo | 0 | 0 | 0 | 0 | 0 | 0 |
| 38827 | ENSMUSG00000100504 | 0 | NA | NA | NA | NA | NA | Gm28926 | predicted gene 28926 [Source:MGI Symbo | 0 | 0 | 0 | 0 | 0 | 0 |
| 38828 | ENSMUSG00000100505 | 0 | NA | NA | NA | NA | NA | Gm13283 | predicted gene 13283 [Source:MGI Symbo | 0 | 0 | 0 | 0 | 0 | 0 |
| 38829 | ENSMUSG00000100506 | 0 | NA | NA | NA | NA | NA | Nanog-ps1 | Nanog pseudogene 1 [Source:MGI Symbo | 0 | 0 | 0 | 0 | 0 | 0 |
| 38830 | ENSMUSG00000100507 | 0 | NA | NA | NA | NA | NA | Gm21906 | predicted gene, 21906 [Source:MGI Symb | 0 | 0 | 0 | 0 | 0 | 0 |
| 38831 | ENSMUSG00000100508 | 0 | NA | NA | NA | NA | NA | Gm28617 | predicted gene 28617 [Source:MGI Symbo | 0 | 0 | 0 | 0 | 0 | 0 |
| 38832 | ENSMUSG00000100509 | 0 | NA | NA | NA | NA | NA | Gm21810 | predicted gene, 21810 [Source:MGI Symb | 0 | 0 | 0 | 0 | 0 | 0 |
| 38833 | ENSMUSG00000100510 | 60.76673534 | -0.957637383 | 0.338907756 | -2.825657912 | 0.004718362 | 0.06420853 | Hand2os1 | Hand2, opposite strand 1 [Source:MGI Sy | 80.31 | 90.457 | 70.437 | 61.695 | 31.628 | 30.074 |
| 38834 | ENSMUSG00000100511 | 0 | NA | NA | NA | NA | NA | 1700111N16 | RIKEN cDNA 1700111N16 gene [Source:M | 0 | 0 | 0 | 0 | 0 | 0 |
| 38835 | ENSMUSG00000100512 | 7.844778021 | -0.016794074 | 0.828506085 | -0.020270309 | 0.983827741 | 0.998580735 | Ovol3 | ovo like zinc finger 3 [Source:MGI Symbol; | 2.4093 | 13.363 | 8.05 | 6.7478 | 9.8156 | 6.6831 |
| 38836 | ENSMUSG00000100513 | 0 | NA | NA | NA | NA | NA | Rpl23a-ps11 | ribosomal protein L23A, pseudogene 11 [S | 0 | 0 | 0 | 0 | 0 | 0 |
| 38837 | ENSMUSG00000100514 | 1.252811962 | -0.644181249 | 2.204770402 | -0.292176114 | 0.770151973 | NA | Gm12960 | predicted gene 12960 [Source:MGI Symbo | 1.6062 | 0 | 3.0187 | 2.8919 | 0 | 0 |
| 38838 | ENSMUSG00000100515 | 0 | NA | NA | NA | NA | NA | Gm17916 | predicted gene, 17916 [Source:MGI Symb | 0 | 0 | 0 | 0 | 0 | 0 |
| 38839 | ENSMUSG00000100516 | 0 | NA | NA | NA | NA | NA | Gm29332 | predicted gene 29332 [Source:MGI Symbo | 0 | 0 | 0 | 0 | 0 | 0 |
| 38840 | ENSMUSG00000100517 | 0 | NA | NA | NA | NA | NA | Gm29627 | predicted gene 29627 [Source:MGI Symbo | 0 | 0 | 0 | 0 | 0 | 0 |
| 38841 | ENSMUSG00000100518 | 0 | NA | NA | NA | NA | NA | 4930440C22 | RIKEN cDNA 4930440C22 gene [Source:M | 0 | 0 | 0 | 0 | 0 | 0 |
| 38842 | ENSMUSG00000100519 | 0 | NA | NA | NA | NA | NA | Gm20801 | predicted gene, 20801 [Source:MGI Symb | 0 | 0 | 0 | 0 | 0 | 0 |
| 38843 | ENSMUSG00000100520 | 0 | NA | NA | NA | NA | NA | Gm20885 | predicted gene, 20885 [Source:MGI Symb | 0 | 0 | 0 | 0 | 0 | 0 |
| 38844 | ENSMUSG00000100521 | 0 | NA | NA | NA | NA | NA | Gm29058 | predicted gene 29058 [Source:MGI Symbo | 0 | 0 | 0 | 0 | 0 | 0 |
| 38845 | ENSMUSG00000100522 | 0 | NA | NA | NA | NA | NA | Gm19280 | predicted gene, 19280 [Source:MGI Symb | 0 | 0 | 0 | 0 | 0 | 0 |
| 38846 | ENSMUSG00000100523 | 0 | NA | NA | NA | NA | NA | Gm29608 | predicted gene 29608 [Source:MGI Symbo | 0 | 0 | 0 | 0 | 0 | 0 |
| 38847 | ENSMUSG00000100524 | 0 | NA | NA | NA | NA | NA | Gm28418 | predicted gene 28418 [Source:MGI Symbo | 0 | 0 | 0 | 0 | 0 | 0 |
| 38848 | ENSMUSG00000100525 | 0 | NA | NA | NA | NA | NA | Gm20058 | predicted gene, 20058 [Source:MGI Symb | 0 | 0 | 0 | 0 | 0 | 0 |
| 38849 | ENSMUSG00000100526 | 0 | NA | NA | NA | NA | NA | Gm29309 | predicted gene 29309 [Source:MGI Symbo | 0 | 0 | 0 | 0 | 0 | 0 |
| 38850 | ENSMUSG00000100527 | 0 | NA | NA | NA | NA | NA | Gm28359 | predicted gene 28359 [Source:MGI Symbo | 0 | 0 | 0 | 0 | 0 | 0 |
| 38851 | ENSMUSG00000100528 | 0 | NA | NA | NA | NA | NA | Gm28881 | predicted gene 28881 [Source:MGI Symbo | 0 | 0 | 0 | 0 | 0 | 0 |
| 38852 | ENSMUSG00000100529 | 0 | NA | NA | NA | NA | NA | Gm19261 | predicted gene, 19261 [Source:MGI Symb | 0 | 0 | 0 | 0 | 0 | 0 |
| 38853 | ENSMUSG00000100530 | 0 | NA | NA | NA | NA | NA | Gm18117 | predicted gene, 18117 [Source:MGI Symb | 0 | 0 | 0 | 0 | 0 | 0 |
| 38854 | ENSMUSG00000100531 | 0 | NA | NA | NA | NA | NA | Gm28475 | predicted gene 28475 [Source:MGI Symbo | 0 | 0 | 0 | 0 | 0 | 0 |
| 38855 | ENSMUSG00000100532 | 0 | NA | NA | NA | NA | NA | Gm28225 | predicted gene 28225 [Source:MGI Symbo | 0 | 0 | 0 | 0 | 0 | 0 |
| 38856 | ENSMUSG00000100533 | 0 | NA | NA | NA | NA | NA | Gm29504 | predicted gene 29504 [Source:MGI Symbo | 0 | 0 | 0 | 0 | 0 | 0 |
| 38857 | ENSMUSG00000100534 | 0 | NA | NA | NA | NA | NA | Gm816 | predicted gene 816 [Source:MGI Symbol; | 0 | 0 | 0 | 0 | 0 | 0 |
| 38858 | ENSMUSG00000100535 | 0 | NA | NA | NA | NA | NA | Gm20870 | predicted gene, 20870 [Source:MGI Symb | 0 | 0 | 0 | 0 | 0 | 0 |
| 38859 | ENSMUSG00000100537 | 0 | NA | NA | NA | NA | NA | Gm28485 | predicted gene 28485 [Source:MGI Symbo | 0 | 0 | 0 | 0 | 0 | 0 |
| 38860 | ENSMUSG00000100538 | 0 | NA | NA | NA | NA | NA | Gm28340 | predicted gene 28340 [Source:MGI Symbo | 0 | 0 | 0 | 0 | 0 | 0 |

| | | | | | | | | | | | | | | | |
|---|---|---|---|---|---|---|---|---|---|---|---|---|---|---|---|
| 38861 | ENSMUSG00000100539 | 0 | NA | NA | NA | NA | NA | Gm20913 | predicted gene, 20913 [Source:MGI Symb | 0 | 0 | 0 | 0 | 0 | 0 |
| 38862 | ENSMUSG00000100540 | 0 | NA | NA | NA | NA | NA | Gm20799 | predicted gene, 20799 [Source:MGI Symb | 0 | 0 | 0 | 0 | 0 | 0 |
| 38863 | ENSMUSG00000100541 | 0 | NA | NA | NA | NA | NA | Gm28871 | predicted gene 28871 [Source:MGI Symbo | 0 | 0 | 0 | 0 | 0 | 0 |
| 38864 | ENSMUSG00000100542 | 0 | NA | NA | NA | NA | NA | Gm28097 | predicted gene 28097 [Source:MGI Symbo | 0 | 0 | 0 | 0 | 0 | 0 |
| 38865 | ENSMUSG00000100543 | 0 | NA | NA | NA | NA | NA | Gm29127 | predicted gene 29127 [Source:MGI Symbo | 0 | 0 | 0 | 0 | 0 | 0 |
| 38866 | ENSMUSG00000100544 | 0 | NA | NA | NA | NA | NA | Gm28606 | predicted gene 28606 [Source:MGI Symbo | 0 | 0 | 0 | 0 | 0 | 0 |
| 38867 | ENSMUSG00000100546 | 9.879145143 | -0.36750161 | 0.721414381 | -0.509418193 | 0.610459127 | 0.916376001 | Gm29483 | predicted gene 29483 [Source:MGI Symbo | 12.047 | 13.363 | 8.05 | 11.568 | 10.906 | 3.3415 |
| 38868 | ENSMUSG00000100547 | 0 | NA | NA | NA | NA | NA | Gm28611 | predicted gene 28611 [Source:MGI Symbo | 0 | 0 | 0 | 0 | 0 | 0 |
| 38869 | ENSMUSG00000100548 | 1.42753722 | 0.459331161 | 1.934778185 | 0.23740766 | 0.812340548 | NA | Gm29585 | predicted gene 29585 [Source:MGI Symbo | 1.6062 | 0 | 2.0125 | 3.8559 | 1.0906 | 0 |
| 38870 | ENSMUSG00000100549 | 0 | NA | NA | NA | NA | NA | Ifna11 | interferon alpha 11 [Source:MGI Symbol;/ | 0 | 0 | 0 | 0 | 0 | 0 |
| 38871 | ENSMUSG00000100550 | 0.469265235 | -2.274672822 | 3.439794491 | -0.661281605 | 0.508431736 | NA | 2310039L15I | RIKEN cDNA 2310039L15 gene [Source:M( | 0.8031 | 0 | 2.0125 | 0 | 0 | 0 |
| 38872 | ENSMUSG00000100551 | 0 | NA | NA | NA | NA | NA | Gm19503 | predicted gene, 19503 [Source:MGI Symb | 0 | 0 | 0 | 0 | 0 | 0 |
| 38873 | ENSMUSG00000100552 | 0 | NA | NA | NA | NA | NA | Gm29019 | predicted gene 29019 [Source:MGI Symbo | 0 | 0 | 0 | 0 | 0 | 0 |
| 38874 | ENSMUSG00000100553 | 0 | NA | NA | NA | NA | NA | Gm17751 | predicted gene, 17751 [Source:MGI Symb | 0 | 0 | 0 | 0 | 0 | 0 |
| 38875 | ENSMUSG00000100554 | 0 | NA | NA | NA | NA | NA | Gm7690 | predicted gene 7690 [Source:MGI Symbol | 0 | 0 | 0 | 0 | 0 | 0 |
| 38876 | ENSMUSG00000100555 | 0 | NA | NA | NA | NA | NA | Gm8173 | predicted gene 8173 [Source:MGI Symbol | 0 | 0 | 0 | 0 | 0 | 0 |
| 38877 | ENSMUSG00000100556 | 0 | NA | NA | NA | NA | NA | Gm28902 | predicted gene 28902 [Source:MGI Symbo | 0 | 0 | 0 | 0 | 0 | 0 |
| 38878 | ENSMUSG00000100557 | 0 | NA | NA | NA | NA | NA | 1700029E06 | RIKEN cDNA 1700029E06 gene [Source:M( | 0 | 0 | 0 | 0 | 0 | 0 |
| 38879 | ENSMUSG00000100558 | 0.133850373 | -0.796629176 | 4.080472857 | -0.195229623 | 0.84521319 | NA | 1700025F24 | RIKEN cDNA 1700025F24 gene [Source:M( | 0.8031 | 0 | 0 | 0 | 0 | 0 |
| 38880 | ENSMUSG00000100559 | 0 | NA | NA | NA | NA | NA | Gm20803 | predicted gene, 20803 [Source:MGI Symb | 0 | 0 | 0 | 0 | 0 | 0 |
| 38881 | ENSMUSG00000100560 | 0 | NA | NA | NA | NA | NA | Gm21180 | predicted gene, 21180 [Source:MGI Symb | 0 | 0 | 0 | 0 | 0 | 0 |
| 38882 | ENSMUSG00000100561 | 0 | NA | NA | NA | NA | NA | Gm7516 | predicted gene 7516 [Source:MGI Symbol | 0 | 0 | 0 | 0 | 0 | 0 |
| 38883 | ENSMUSG00000100562 | 0 | NA | NA | NA | NA | NA | Gm29611 | predicted gene 29611 [Source:MGI Symbo | 0 | 0 | 0 | 0 | 0 | 0 |
| 38884 | ENSMUSG00000100563 | 0 | NA | NA | NA | NA | NA | Gm3096 | predicted gene 3096 [Source:MGI Symbol | 0 | 0 | 0 | 0 | 0 | 0 |
| 38885 | ENSMUSG00000100564 | 0 | NA | NA | NA | NA | NA | Gm28682 | predicted gene 28682 [Source:MGI Symbo | 0 | 0 | 0 | 0 | 0 | 0 |
| 38886 | ENSMUSG00000100565 | 0 | NA | NA | NA | NA | NA | Gm28091 | predicted gene 28091 [Source:MGI Symbo | 0 | 0 | 0 | 0 | 0 | 0 |
| 38887 | ENSMUSG00000100566 | 0 | NA | NA | NA | NA | NA | Tmem167-ps | transmembrane protein 167, pseudogene | 0 | 0 | 0 | 0 | 0 | 0 |
| 38888 | ENSMUSG00000100567 | 0 | NA | NA | NA | NA | NA | Gm28958 | predicted gene 28958 [Source:MGI Symbo | 0 | 0 | 0 | 0 | 0 | 0 |
| 38889 | ENSMUSG00000100568 | 0 | NA | NA | NA | NA | NA | Gm29602 | predicted gene 29602 [Source:MGI Symbo | 0 | 0 | 0 | 0 | 0 | 0 |
| 38890 | ENSMUSG00000100569 | 0 | NA | NA | NA | NA | NA | Gm28646 | predicted gene 28646 [Source:MGI Symbo | 0 | 0 | 0 | 0 | 0 | 0 |
| 38891 | ENSMUSG00000100570 | 0 | NA | NA | NA | NA | NA | Gm29399 | predicted gene 29399 [Source:MGI Symbo | 0 | 0 | 0 | 0 | 0 | 0 |
| 38892 | ENSMUSG00000100571 | 0 | NA | NA | NA | NA | NA | Gm29474 | predicted gene 29474 [Source:MGI Symbo | 0 | 0 | 0 | 0 | 0 | 0 |
| 38893 | ENSMUSG00000100572 | 0 | NA | NA | NA | NA | NA | 2410021H03 | RIKEN cDNA 2410021H03 gene [Source:M | 0 | 0 | 0 | 0 | 0 | 0 |
| 38894 | ENSMUSG00000100573 | 0 | NA | NA | NA | NA | NA | 1700081H04 | RIKEN cDNA 1700081H04 gene [Source:M | 0 | 0 | 0 | 0 | 0 | 0 |
| 38895 | ENSMUSG00000100574 | 0 | NA | NA | NA | NA | NA | Gm28674 | predicted gene 28674 [Source:MGI Symbo | 0 | 0 | 0 | 0 | 0 | 0 |
| 38896 | ENSMUSG00000100575 | 0 | NA | NA | NA | NA | NA | Gm21489 | predicted gene, 21489 [Source:MGI Symb | 0 | 0 | 0 | 0 | 0 | 0 |
| 38897 | ENSMUSG00000100576 | 0 | NA | NA | NA | NA | NA | Gm28975 | predicted gene 28975 [Source:MGI Symbo | 0 | 0 | 0 | 0 | 0 | 0 |
| 38898 | ENSMUSG00000100577 | 0 | NA | NA | NA | NA | NA | Gm29429 | predicted gene 29429 [Source:MGI Symbo | 0 | 0 | 0 | 0 | 0 | 0 |
| 38899 | ENSMUSG00000100578 | 0 | NA | NA | NA | NA | NA | Gm28709 | predicted gene 28709 [Source:MGI Symbo | 0 | 0 | 0 | 0 | 0 | 0 |
| 38900 | ENSMUSG00000100579 | 0 | NA | NA | NA | NA | NA | Gm28692 | predicted gene 28692 [Source:MGI Symbo | 0 | 0 | 0 | 0 | 0 | 0 |
| 38901 | ENSMUSG00000100580 | 0 | NA | NA | NA | NA | NA | 4933436I20R | RIKEN cDNA 4933436I20 gene [Source:M( | 0 | 0 | 0 | 0 | 0 | 0 |
| 38902 | ENSMUSG00000100581 | 0 | NA | NA | NA | NA | NA | Gm21785 | predicted gene, 21785 [Source:MGI Symb | 0 | 0 | 0 | 0 | 0 | 0 |
| 38903 | ENSMUSG00000100582 | 0 | NA | NA | NA | NA | NA | Gm21428 | predicted gene, 21428 [Source:MGI Symb | 0 | 0 | 0 | 0 | 0 | 0 |
| 38904 | ENSMUSG00000100583 | 0.797815601 | -0.524044714 | 2.626768505 | -0.199501674 | 0.841870334 | NA | Gm29443 | predicted gene 29443 [Source:MGI Symbo | 0.8031 | 2.0558 | 0 | 1.928 | 0 | 0 |
| 38905 | ENSMUSG00000100584 | 0 | NA | NA | NA | NA | NA | Gm28733 | predicted gene 28733 [Source:MGI Symbo | 0 | 0 | 0 | 0 | 0 | 0 |
| 38906 | ENSMUSG00000100585 | 0 | NA | NA | NA | NA | NA | 1700108J01I | RIKEN cDNA 1700108J01 gene [Source:M( | 0 | 0 | 0 | 0 | 0 | 0 |
| 38907 | ENSMUSG00000100586 | 32.3245576 | 1.017025171 | 0.445664408 | 2.282042616 | 0.022486824 | 0.187679993 | Vmn1r90 | vomeronasal 1 receptor 90 [Source:MGI S | 21.684 | 15.419 | 27.169 | 64.587 | 29.447 | 35.643 |
| 38908 | ENSMUSG00000100587 | 0 | NA | NA | NA | NA | NA | Gm28797 | predicted gene 28797 [Source:MGI Symbo | 0 | 0 | 0 | 0 | 0 | 0 |
| 38909 | ENSMUSG00000100588 | 0 | NA | NA | NA | NA | NA | Gm29497 | predicted gene 29497 [Source:MGI Symbo | 0 | 0 | 0 | 0 | 0 | 0 |
| 38910 | ENSMUSG00000100589 | 0 | NA | NA | NA | NA | NA | 4930533P14 | RIKEN cDNA 4930533P14 gene [Source:M( | 0 | 0 | 0 | 0 | 0 | 0 |
| 38911 | ENSMUSG00000100590 | 0 | NA | NA | NA | NA | NA | Gm4785 | predicted gene 4785 [Source:MGI Symbol | 0 | 0 | 0 | 0 | 0 | 0 |
| 38912 | ENSMUSG00000100591 | 0 | NA | NA | NA | NA | NA | Gm21919 | predicted gene, 21919 [Source:MGI Symb | 0 | 0 | 0 | 0 | 0 | 0 |
| 38913 | ENSMUSG00000100592 | 0 | NA | NA | NA | NA | NA | Gm28645 | predicted gene 28645 [Source:MGI Symbo | 0 | 0 | 0 | 0 | 0 | 0 |
| 38914 | ENSMUSG00000100593 | 12.35567094 | -0.243450142 | 0.604659094 | -0.4026238 | 0.687224998 | 0.936854135 | 1700119H24 | RIKEN cDNA 1700119H24 gene [Source:M | 13.653 | 14.391 | 12.075 | 8.6758 | 13.087 | 12.252 |
| 38915 | ENSMUSG00000100594 | 84.95812136 | 0.27601156 | 0.280198523 | 0.985057153 | 0.324595998 | 0.771309263 | 2810414N06 | RIKEN cDNA 2810414N06 gene [Source:M | 75.492 | 87.373 | 67.418 | 66.514 | 94.884 | 118.07 |
| 38916 | ENSMUSG00000100595 | 0 | NA | NA | NA | NA | NA | Gm19087 | predicted gene, 19087 [Source:MGI Symb | 0 | 0 | 0 | 0 | 0 | 0 |

| | | | | | | | | | | | | | | | |
|---|---|---|---|---|---|---|---|---|---|---|---|---|---|---|---|
| 38917 | ENSMUSG00000100596 | 0 | NA | NA | NA | NA | NA | Gm29502 | predicted gene 29502 [Source:MGI Symbo | 0 | 0 | 0 | 0 | 0 | 0 |
| 38918 | ENSMUSG00000100597 | 0 | NA | NA | NA | NA | NA | Gm29349 | predicted gene 29349 [Source:MGI Symbo | 0 | 0 | 0 | 0 | 0 | 0 |
| 38919 | ENSMUSG00000100598 | 0 | NA | NA | NA | NA | NA | Gm28742 | predicted gene 28742 [Source:MGI Symbo | 0 | 0 | 0 | 0 | 0 | 0 |
| 38920 | ENSMUSG00000100599 | 3.419520641 | 0.19966284 | 1.26211398 | 0.158197154 | 0.874301446 | NA | 1700120C14 | RIKEN cDNA 1700120C14 gene [Source:M | 2.4093 | 3.0838 | 4.025 | 0 | 6.5437 | 4.4554 |
| 38921 | ENSMUSG00000100600 | 0 | NA | NA | NA | NA | NA | A230077H06 | RIKEN cDNA A230077H06 gene [Source:M | 0 | 0 | 0 | 0 | 0 | 0 |
| 38922 | ENSMUSG00000100602 | 0 | NA | NA | NA | NA | NA | Gm19498 | predicted gene, 19498 [Source:MGI Symb | 0 | 0 | 0 | 0 | 0 | 0 |
| 38923 | ENSMUSG00000100603 | 0 | NA | NA | NA | NA | NA | 1700129L04 | RIKEN cDNA 1700129L04 gene [Source:M | 0 | 0 | 0 | 0 | 0 | 0 |
| 38924 | ENSMUSG00000100604 | 0 | NA | NA | NA | NA | NA | Gm29256 | predicted gene 29256 [Source:MGI Symbo | 0 | 0 | 0 | 0 | 0 | 0 |
| 38925 | ENSMUSG00000100605 | 0 | NA | NA | NA | NA | NA | Gm29243 | predicted gene 29243 [Source:MGI Symbo | 0 | 0 | 0 | 0 | 0 | 0 |
| 38926 | ENSMUSG00000100606 | 0 | NA | NA | NA | NA | NA | Gm29612 | predicted gene 29612 [Source:MGI Symbo | 0 | 0 | 0 | 0 | 0 | 0 |
| 38927 | ENSMUSG00000100607 | 0 | NA | NA | NA | NA | NA | Gm29379 | predicted gene 29379 [Source:MGI Symbo | 0 | 0 | 0 | 0 | 0 | 0 |
| 38928 | ENSMUSG00000100608 | 0 | NA | NA | NA | NA | NA | Gm21996 | predicted gene 21996 [Source:MGI Symbo | 0 | 0 | 0 | 0 | 0 | 0 |
| 38929 | ENSMUSG00000100609 | 0 | NA | NA | NA | NA | NA | Gm28687 | predicted gene 28687 [Source:MGI Symbo | 0 | 0 | 0 | 0 | 0 | 0 |
| 38930 | ENSMUSG00000100610 | 0 | NA | NA | NA | NA | NA | Gm29031 | predicted gene 29031 [Source:MGI Symbo | 0 | 0 | 0 | 0 | 0 | 0 |
| 38931 | ENSMUSG00000100611 | 0 | NA | NA | NA | NA | NA | Gm21247 | predicted gene, 21247 [Source:MGI Symb | 0 | 0 | 0 | 0 | 0 | 0 |
| 38932 | ENSMUSG00000100613 | 0 | NA | NA | NA | NA | NA | Gm5058 | predicted gene 5058 [Source:MGI Symbol | 0 | 0 | 0 | 0 | 0 | 0 |
| 38933 | ENSMUSG00000100614 | 0 | NA | NA | NA | NA | NA | Gm28963 | predicted gene 28963 [Source:MGI Symbo | 0 | 0 | 0 | 0 | 0 | 0 |
| 38934 | ENSMUSG00000100615 | 0.181769715 | 1.126924175 | 4.080472857 | 0.276174898 | 0.782413736 | NA | Gm5511 | predicted gene 5511 [Source:MGI Symbol | 0 | 0 | 0 | 0 | 1.0906 | 0 |
| 38935 | ENSMUSG00000100616 | 0 | NA | NA | NA | NA | NA | Gm28076 | predicted gene 28076 [Source:MGI Symbo | 0 | 0 | 0 | 0 | 0 | 0 |
| 38936 | ENSMUSG00000100617 | 0 | NA | NA | NA | NA | NA | Gm7145 | predicted gene 7145 [Source:MGI Symbol | 0 | 0 | 0 | 0 | 0 | 0 |
| 38937 | ENSMUSG00000100618 | 0.435408177 | -2.179616019 | 3.531869654 | -0.617128103 | 0.537150234 | NA | NA | NA | 1.6062 | 0 | 1.0062 | 0 | 0 | 0 |
| 38938 | ENSMUSG00000100619 | 0 | NA | NA | NA | NA | NA | Gm8446 | predicted gene 8446 [Source:MGI Symbol | 0 | 0 | 0 | 0 | 0 | 0 |
| 38939 | ENSMUSG00000100620 | 0.48903338 | 0.95637552 | 3.456690196 | 0.276673773 | 0.782030612 | NA | Gm28277 | predicted gene 28277 [Source:MGI Symbo | 0 | 0 | 1.0062 | 1.928 | 0 | 0 |
| 38940 | ENSMUSG00000100621 | 0 | NA | NA | NA | NA | NA | Gm28374 | predicted gene 28374 [Source:MGI Symbo | 0 | 0 | 0 | 0 | 0 | 0 |
| 38941 | ENSMUSG00000100622 | 0.462220778 | -0.685400455 | 3.191475091 | -0.21475977 | 0.82995462 | NA | Gm20379 | predicted gene, 20379 [Source:MGI Symb | 0.8031 | 0 | 1.0062 | 0.964 | 0 | 0 |
| 38942 | ENSMUSG00000100623 | 0 | NA | NA | NA | NA | NA | Gm29428 | predicted gene 29428 [Source:MGI Symbo | 0 | 0 | 0 | 0 | 0 | 0 |
| 38943 | ENSMUSG00000100624 | 0 | NA | NA | NA | NA | NA | Gm28101 | predicted gene 28101 [Source:MGI Symbo | 0 | 0 | 0 | 0 | 0 | 0 |
| 38944 | ENSMUSG00000100625 | 0.181769715 | 1.126924175 | 4.080472857 | 0.276174898 | 0.782413736 | NA | 1700016G22 | RIKEN cDNA 1700016G22 gene [Source:M | 0 | 0 | 0 | 0 | 1.0906 | 0 |
| 38945 | ENSMUSG00000100626 | 0 | NA | NA | NA | NA | NA | H2al1a | H2A histone family member L1A [Source:I | 0 | 0 | 0 | 0 | 0 | 0 |
| 38946 | ENSMUSG00000100627 | 3.62202942 | 5.384156572 | 1.793497466 | 3.002043033 | 0.002681743 | NA | A830008E24 | RIKEN cDNA A830008E24 gene [Source:M | 0 | 0 | 0 | 0.964 | 18.541 | 2.2277 |
| 38947 | ENSMUSG00000100628 | 0 | NA | NA | NA | NA | NA | Gm29648 | predicted gene 29648 [Source:MGI Symbo | 0 | 0 | 0 | 0 | 0 | 0 |
| 38948 | ENSMUSG00000100629 | 19.12171352 | -0.060321335 | 0.543037165 | -0.111081412 | 0.911551788 | 0.98589875 | Gm28192 | predicted gene 28192 [Source:MGI Symbo | 12.047 | 26.726 | 20.125 | 25.063 | 19.631 | 11.138 |
| 38949 | ENSMUSG00000100630 | 0 | NA | NA | NA | NA | NA | Gm28450 | predicted gene 28450 [Source:MGI Symbo | 0 | 0 | 0 | 0 | 0 | 0 |
| 38950 | ENSMUSG00000100632 | 1.10120282 | -0.134983118 | 2.137435647 | -0.063151898 | 0.949645548 | NA | 9430060I03 | RIKEN cDNA 9430060I03 gene [Source:M | 2.4093 | 0 | 1.0062 | 0.964 | 0 | 2.2277 |
| 38951 | ENSMUSG00000100634 | 0 | NA | NA | NA | NA | NA | Gm28171 | predicted gene 28171 [Source:MGI Symbo | 0 | 0 | 0 | 0 | 0 | 0 |
| 38952 | ENSMUSG00000100635 | 0 | NA | NA | NA | NA | NA | Gm29157 | predicted gene 29157 [Source:MGI Symbo | 0 | 0 | 0 | 0 | 0 | 0 |
| 38953 | ENSMUSG00000100636 | 0 | NA | NA | NA | NA | NA | Gm3551 | predicted gene 3551 [Source:MGI Symbol | 0 | 0 | 0 | 0 | 0 | 0 |
| 38954 | ENSMUSG00000100637 | 0 | NA | NA | NA | NA | NA | Gm20819 | predicted gene, 20819 [Source:MGI Symb | 0 | 0 | 0 | 0 | 0 | 0 |
| 38955 | ENSMUSG00000100639 | 0 | NA | NA | NA | NA | NA | Scgb1b6-ps | secretoglobin, family 1B, member 6, pseu | 0 | 0 | 0 | 0 | 0 | 0 |
| 38956 | ENSMUSG00000100640 | 0 | NA | NA | NA | NA | NA | Gm29073 | predicted gene 29073 [Source:MGI Symbo | 0 | 0 | 0 | 0 | 0 | 0 |
| 38957 | ENSMUSG00000100641 | 0 | NA | NA | NA | NA | NA | Gm28413 | predicted gene 28413 [Source:MGI Symbo | 0 | 0 | 0 | 0 | 0 | 0 |
| 38958 | ENSMUSG00000100642 | 0 | NA | NA | NA | NA | NA | Gm28230 | predicted gene 28230 [Source:MGI Symbo | 0 | 0 | 0 | 0 | 0 | 0 |
| 38959 | ENSMUSG00000100643 | 0 | NA | NA | NA | NA | NA | Gm21163 | predicted gene, 21163 [Source:MGI Symb | 0 | 0 | 0 | 0 | 0 | 0 |
| 38960 | ENSMUSG00000100644 | 0 | NA | NA | NA | NA | NA | Gm20793 | predicted gene, 20793 [Source:MGI Symb | 0 | 0 | 0 | 0 | 0 | 0 |
| 38961 | ENSMUSG00000100645 | 0 | NA | NA | NA | NA | NA | Gm28576 | predicted gene 28576 [Source:MGI Symbo | 0 | 0 | 0 | 0 | 0 | 0 |
| 38962 | ENSMUSG00000100646 | 0 | NA | NA | NA | NA | NA | Gm28193 | predicted gene 28193 [Source:MGI Symbo | 0 | 0 | 0 | 0 | 0 | 0 |
| 38963 | ENSMUSG00000100647 | 0 | NA | NA | NA | NA | NA | Gm29027 | predicted gene 29027 [Source:MGI Symbo | 0 | 0 | 0 | 0 | 0 | 0 |
| 38964 | ENSMUSG00000100648 | 0 | NA | NA | NA | NA | NA | Gm28659 | predicted gene 28659 [Source:MGI Symbo | 0 | 0 | 0 | 0 | 0 | 0 |
| 38965 | ENSMUSG00000100649 | 0 | NA | NA | NA | NA | NA | Gm28428 | predicted gene 28428 [Source:MGI Symbo | 0 | 0 | 0 | 0 | 0 | 0 |
| 38966 | ENSMUSG00000100650 | 0 | NA | NA | NA | NA | NA | Gm28807 | predicted gene 28807 [Source:MGI Symbo | 0 | 0 | 0 | 0 | 0 | 0 |
| 38967 | ENSMUSG00000100651 | 0 | NA | NA | NA | NA | NA | Gm28810 | predicted gene 28810 [Source:MGI Symbo | 0 | 0 | 0 | 0 | 0 | 0 |
| 38968 | ENSMUSG00000100652 | 0 | NA | NA | NA | NA | NA | Gm7634 | predicted gene 7634 [Source:MGI Symbol | 0 | 0 | 0 | 0 | 0 | 0 |
| 38969 | ENSMUSG00000100653 | 0 | NA | NA | NA | NA | NA | Gm28238 | predicted gene 28238 [Source:MGI Symbo | 0 | 0 | 0 | 0 | 0 | 0 |
| 38970 | ENSMUSG00000100654 | 0 | NA | NA | NA | NA | NA | Gm4207 | predicted gene 4207 [Source:MGI Symbol | 0 | 0 | 0 | 0 | 0 | 0 |
| 38971 | ENSMUSG00000100655 | 0 | NA | NA | NA | NA | NA | Gm21462 | predicted gene, 21462 [Source:MGI Symb | 0 | 0 | 0 | 0 | 0 | 0 |
| 38972 | ENSMUSG00000100656 | 0 | NA | NA | NA | NA | NA | Gm8525 | predicted gene 8525 [Source:MGI Symbol | 0 | 0 | 0 | 0 | 0 | 0 |

| | | | | | | | | | | | | | | | |
|---|---|---|---|---|---|---|---|---|---|---|---|---|---|---|---|
| 38973 | ENSMUSG00000100657 | 0 | NA | NA | NA | NA | NA | Gm28862 | predicted gene 28862 [Source:MGI Symbo | 0 | 0 | 0 | 0 | 0 | 0 |
| 38974 | ENSMUSG00000100658 | 6.313354369 | -1.199629787 | 0.908423614 | -1.32056209 | 0.18664742 | 0.6230483 | F730311O21 | RIKEN cDNA F730311O21 gene [Source:M | 8.031 | 14.391 | 4.025 | 4.8199 | 3.2719 | 3.3415 |
| 38975 | ENSMUSG00000100659 | 0 | NA | NA | NA | NA | NA | Gm28636 | predicted gene 28636 [Source:MGI Symbo | 0 | 0 | 0 | 0 | 0 | 0 |
| 38976 | ENSMUSG00000100660 | 0 | NA | NA | NA | NA | NA | Kat6b-ps2 | k(lysine) acetyltransferase 6B, pseudogen | 0 | 0 | 0 | 0 | 0 | 0 |
| 38977 | ENSMUSG00000100661 | 0 | NA | NA | NA | NA | NA | Gm29471 | predicted gene 29471 [Source:MGI Symbo | 0 | 0 | 0 | 0 | 0 | 0 |
| 38978 | ENSMUSG00000100662 | 0 | NA | NA | NA | NA | NA | Gm19029 | predicted gene, 19029 [Source:MGI Symb | 0 | 0 | 0 | 0 | 0 | 0 |
| 38979 | ENSMUSG00000100663 | 0 | NA | NA | NA | NA | NA | Gm29604 | predicted gene 29604 [Source:MGI Symbo | 0 | 0 | 0 | 0 | 0 | 0 |
| 38980 | ENSMUSG00000100664 | 0 | NA | NA | NA | NA | NA | 6030442E23 | RIKEN cDNA 6030442E23 gene [Source:M | 0 | 0 | 0 | 0 | 0 | 0 |
| 38981 | ENSMUSG00000100665 | 0 | NA | NA | NA | NA | NA | Mrgprc1-ps | MAS-related GPR, member C1, pseudoger | 0 | 0 | 0 | 0 | 0 | 0 |
| 38982 | ENSMUSG00000100666 | 1.819407807 | 0.98233427 | 1.753384678 | 0.560250287 | 0.575308731 | NA | 1700007F19 | RIKEN cDNA 1700007F19 gene [Source:M | 1.6062 | 2.0558 | 0 | 2.8919 | 4.3625 | 0 |
| 38983 | ENSMUSG00000100667 | 0 | NA | NA | NA | NA | NA | 1700040F15 | RIKEN cDNA 1700040F15 gene [Source:M | 0 | 0 | 0 | 0 | 0 | 0 |
| 38984 | ENSMUSG00000100668 | 0 | NA | NA | NA | NA | NA | 1700083H02 | RIKEN cDNA 1700083H02 gene [Source:M | 0 | 0 | 0 | 0 | 0 | 0 |
| 38985 | ENSMUSG00000100669 | 0 | NA | NA | NA | NA | NA | Gm21111 | predicted gene, 21111 [Source:MGI Symb | 0 | 0 | 0 | 0 | 0 | 0 |
| 38986 | ENSMUSG00000100670 | 0 | NA | NA | NA | NA | NA | Gm28744 | predicted gene 28744 [Source:MGI Symbo | 0 | 0 | 0 | 0 | 0 | 0 |
| 38987 | ENSMUSG00000100671 | 0 | NA | NA | NA | NA | NA | Gm28322 | predicted gene 28322 [Source:MGI Symbo | 0 | 0 | 0 | 0 | 0 | 0 |
| 38988 | ENSMUSG00000100672 | 0 | NA | NA | NA | NA | NA | Gm28404 | predicted gene 28404 [Source:MGI Symbo | 0 | 0 | 0 | 0 | 0 | 0 |
| 38989 | ENSMUSG00000100673 | 0 | NA | NA | NA | NA | NA | Gm29574 | predicted gene 29574 [Source:MGI Symbo | 0 | 0 | 0 | 0 | 0 | 0 |
| 38990 | ENSMUSG00000100674 | 0 | NA | NA | NA | NA | NA | Gm28806 | predicted gene 28806 [Source:MGI Symbo | 0 | 0 | 0 | 0 | 0 | 0 |
| 38991 | ENSMUSG00000100675 | 0 | NA | NA | NA | NA | NA | Gm28607 | predicted gene 28607 [Source:MGI Symbo | 0 | 0 | 0 | 0 | 0 | 0 |
| 38992 | ENSMUSG00000100676 | 0 | NA | NA | NA | NA | NA | Gm28860 | predicted gene 28860 [Source:MGI Symbo | 0 | 0 | 0 | 0 | 0 | 0 |
| 38993 | ENSMUSG00000100677 | 0 | NA | NA | NA | NA | NA | Gm21184 | predicted gene, 21184 [Source:MGI Symb | 0 | 0 | 0 | 0 | 0 | 0 |
| 38994 | ENSMUSG00000100678 | 0 | NA | NA | NA | NA | NA | Gm29251 | predicted gene 29251 [Source:MGI Symbo | 0 | 0 | 0 | 0 | 0 | 0 |
| 38995 | ENSMUSG00000100679 | 0 | NA | NA | NA | NA | NA | Gm28778 | predicted gene 28778 [Source:MGI Symbo | 0 | 0 | 0 | 0 | 0 | 0 |
| 38996 | ENSMUSG00000100680 | 41.39427936 | 0.231761291 | 0.372004856 | 0.623006091 | 0.533280519 | 0.889143097 | 1810044D09 | RIKEN cDNA 1810044D09 gene [Source:M | 36.943 | 29.81 | 47.293 | 32.775 | 43.625 | 57.92 |
| 38997 | ENSMUSG00000100681 | 0 | NA | NA | NA | NA | NA | 1700064J06I | RIKEN cDNA 1700064J06 gene [Source:M( | 0 | 0 | 0 | 0 | 0 | 0 |
| 38998 | ENSMUSG00000100682 | 0 | NA | NA | NA | NA | NA | Gm28103 | predicted gene 28103 [Source:MGI Symbo | 0 | 0 | 0 | 0 | 0 | 0 |
| 38999 | ENSMUSG00000100683 | 0 | NA | NA | NA | NA | NA | Gm28186 | predicted gene 28186 [Source:MGI Symbo | 0 | 0 | 0 | 0 | 0 | 0 |
| 39000 | ENSMUSG00000100684 | 0 | NA | NA | NA | NA | NA | Gm20268 | predicted gene, 20268 [Source:MGI Symb | 0 | 0 | 0 | 0 | 0 | 0 |
| 39001 | ENSMUSG00000100685 | 0 | NA | NA | NA | NA | NA | Gm28610 | predicted gene 28610 [Source:MGI Symbo | 0 | 0 | 0 | 0 | 0 | 0 |
| 39002 | ENSMUSG00000100687 | 0 | NA | NA | NA | NA | NA | Gm18357 | predicted gene, 18357 [Source:MGI Symb | 0 | 0 | 0 | 0 | 0 | 0 |
| 39003 | ENSMUSG00000100689 | 0 | NA | NA | NA | NA | NA | Taar7e | trace amine-associated receptor 7E [Sourc | 0 | 0 | 0 | 0 | 0 | 0 |
| 39004 | ENSMUSG00000100690 | 0 | NA | NA | NA | NA | NA | Gm4322 | predicted gene 4322 [Source:MGI Symbol | 0 | 0 | 0 | 0 | 0 | 0 |
| 39005 | ENSMUSG00000100691 | 132.8647891 | 0.020247882 | 0.2070323 | 0.097800594 | 0.922090636 | 0.986685804 | 2010320M18 | RIKEN cDNA 2010320M18 gene [Source:M | 141.35 | 133.63 | 120.75 | 145.56 | 117.79 | 138.12 |
| 39006 | ENSMUSG00000100692 | 0 | NA | NA | NA | NA | NA | Gm21672 | predicted gene, 21672 [Source:MGI Symb | 0 | 0 | 0 | 0 | 0 | 0 |
| 39007 | ENSMUSG00000100693 | 0 | NA | NA | NA | NA | NA | Gm28278 | predicted gene 28278 [Source:MGI Symbo | 0 | 0 | 0 | 0 | 0 | 0 |
| 39008 | ENSMUSG00000100694 | 0 | NA | NA | NA | NA | NA | Gm29182 | predicted gene 29182 [Source:MGI Symbo | 0 | 0 | 0 | 0 | 0 | 0 |
| 39009 | ENSMUSG00000100695 | 0 | NA | NA | NA | NA | NA | Gm28110 | predicted gene 28110 [Source:MGI Symbo | 0 | 0 | 0 | 0 | 0 | 0 |
| 39010 | ENSMUSG00000100696 | 0 | NA | NA | NA | NA | NA | Gm8419 | predicted gene 8419 [Source:MGI Symbol | 0 | 0 | 0 | 0 | 0 | 0 |
| 39011 | ENSMUSG00000100697 | 0 | NA | NA | NA | NA | NA | Gm29184 | predicted gene 29184 [Source:MGI Symbo | 0 | 0 | 0 | 0 | 0 | 0 |
| 39012 | ENSMUSG00000100698 | 0 | NA | NA | NA | NA | NA | Gm20902 | predicted gene, 20902 [Source:MGI Symb | 0 | 0 | 0 | 0 | 0 | 0 |
| 39013 | ENSMUSG00000100699 | 0 | NA | NA | NA | NA | NA | Gm21765 | predicted gene, 21765 [Source:MGI Symb | 0 | 0 | 0 | 0 | 0 | 0 |
| 39014 | ENSMUSG00000100700 | 0 | NA | NA | NA | NA | NA | Gm6137 | predicted gene 6137 [Source:MGI Symbol | 0 | 0 | 0 | 0 | 0 | 0 |
| 39015 | ENSMUSG00000100701 | 0.492645588 | 0.9565034 | 3.447648281 | 0.277436479 | 0.781444974 | NA | Gm19587 | predicted gene, 19587 [Source:MGI Symb | 0 | 1.0279 | 0 | 1.928 | 0 | 0 |
| 39016 | ENSMUSG00000100702 | 0 | NA | NA | NA | NA | NA | Gm8976 | predicted gene 8976 [Source:MGI Symbol | 0 | 0 | 0 | 0 | 0 | 0 |
| 39017 | ENSMUSG00000100703 | 0 | NA | NA | NA | NA | NA | Gm28358 | predicted gene 28358 [Source:MGI Symbo | 0 | 0 | 0 | 0 | 0 | 0 |
| 39018 | ENSMUSG00000100704 | 0 | NA | NA | NA | NA | NA | Gm28434 | predicted gene 28434 [Source:MGI Symbo | 0 | 0 | 0 | 0 | 0 | 0 |
| 39019 | ENSMUSG00000100705 | 0 | NA | NA | NA | NA | NA | Gm28227 | predicted gene 28227 [Source:MGI Symbo | 0 | 0 | 0 | 0 | 0 | 0 |
| 39020 | ENSMUSG00000100706 | 9.19722957 | 0.700603809 | 0.772388069 | 0.907061926 | 0.364374051 | 0.802897466 | Gm19744 | predicted gene, 19744 [Source:MGI Symb | 9.6372 | 7.1954 | 4.025 | 5.7839 | 19.631 | 8.9108 |
| 39021 | ENSMUSG00000100707 | 0.850737881 | 0.833803351 | 2.386603902 | 0.349368133 | 0.726812954 | NA | Gm28523 | predicted gene 28523 [Source:MGI Symbo | 0.8031 | 0 | 1.0062 | 0 | 2.1812 | 1.1138 |
| 39022 | ENSMUSG00000100708 | 0 | NA | NA | NA | NA | NA | Gm21173 | predicted gene, 21173 [Source:MGI Symb | 0 | 0 | 0 | 0 | 0 | 0 |
| 39023 | ENSMUSG00000100709 | 0 | NA | NA | NA | NA | NA | Gm28295 | predicted gene 28295 [Source:MGI Symbo | 0 | 0 | 0 | 0 | 0 | 0 |
| 39024 | ENSMUSG00000100710 | 0 | NA | NA | NA | NA | NA | Gm29197 | predicted gene 29197 [Source:MGI Symbo | 0 | 0 | 0 | 0 | 0 | 0 |
| 39025 | ENSMUSG00000100711 | 0 | NA | NA | NA | NA | NA | Gm29461 | predicted gene 29461 [Source:MGI Symbo | 0 | 0 | 0 | 0 | 0 | 0 |
| 39026 | ENSMUSG00000100712 | 0 | NA | NA | NA | NA | NA | Gm28446 | predicted gene 28446 [Source:MGI Symbo | 0 | 0 | 0 | 0 | 0 | 0 |
| 39027 | ENSMUSG00000100713 | 0 | NA | NA | NA | NA | NA | Ifna7 | interferon alpha 7 [Source:MGI Symbol;A | 0 | 0 | 0 | 0 | 0 | 0 |
| 39028 | ENSMUSG00000100714 | 1.035091538 | -2.228354257 | 2.240983885 | -0.994364249 | 0.320045567 | NA | 2010308F09 | RIKEN cDNA 2010308F09 gene [Source:M | 3.2124 | 1.0279 | 1.0062 | 0.964 | 0 | 0 |

| | | | | | | | | | | | | | | | |
|---|---|---|---|---|---|---|---|---|---|---|---|---|---|---|---|
| 39029 | ENSMUSG00000100715 | 0 | NA | NA | NA | NA | NA | Gm28074 | predicted gene 28074 [Source:MGI Symbo | 0 | 0 | 0 | 0 | 0 | 0 |
| 39030 | ENSMUSG00000100716 | 0 | NA | NA | NA | NA | NA | Gm28550 | predicted gene 28550 [Source:MGI Symbo | 0 | 0 | 0 | 0 | 0 | 0 |
| 39031 | ENSMUSG00000100717 | 0.267700746 | -1.497481649 | 4.057001436 | -0.369110456 | 0.712045396 | NA | 1700120G07 | RIKEN cDNA 1700120G07 gene [Source:M | 1.6062 | 0 | 0 | 0 | 0 | 0 |
| 39032 | ENSMUSG00000100718 | 0 | NA | NA | NA | NA | NA | Gm21717 | predicted gene, 21717 [Source:MGI Symb | 0 | 0 | 0 | 0 | 0 | 0 |
| 39033 | ENSMUSG00000100719 | 0 | NA | NA | NA | NA | NA | Gm28786 | predicted gene 28786 [Source:MGI Symbo | 0 | 0 | 0 | 0 | 0 | 0 |
| 39034 | ENSMUSG00000100720 | 0.453341394 | -0.545391191 | 3.551113277 | -0.153583158 | 0.877938417 | NA | Gm10075 | predicted gene 10075 [Source:MGI Symbo | 1.6062 | 0 | 0 | 0 | 0 | 1.1138 |
| 39035 | ENSMUSG00000100721 | 0 | NA | NA | NA | NA | NA | Gm9271 | predicted gene 9271 [Source:MGI Symbol | 0 | 0 | 0 | 0 | 0 | 0 |
| 39036 | ENSMUSG00000100722 | 0 | NA | NA | NA | NA | NA | Gm29569 | predicted gene 29569 [Source:MGI Symbo | 0 | 0 | 0 | 0 | 0 | 0 |
| 39037 | ENSMUSG00000100723 | 0 | NA | NA | NA | NA | NA | Gm28160 | predicted gene 28160 [Source:MGI Symbo | 0 | 0 | 0 | 0 | 0 | 0 |
| 39038 | ENSMUSG00000100724 | 0 | NA | NA | NA | NA | NA | Gm28533 | predicted gene 28533 [Source:MGI Symbo | 0 | 0 | 0 | 0 | 0 | 0 |
| 39039 | ENSMUSG00000100726 | 0 | NA | NA | NA | NA | NA | Gm21497 | predicted gene, 21497 [Source:MGI Symb | 0 | 0 | 0 | 0 | 0 | 0 |
| 39040 | ENSMUSG00000100727 | 0 | NA | NA | NA | NA | NA | 1700003P14 | RIKEN cDNA 1700003P14 gene [Source:M | 0 | 0 | 0 | 0 | 0 | 0 |
| 39041 | ENSMUSG00000100728 | 0 | NA | NA | NA | NA | NA | Gm29431 | predicted gene 29431 [Source:MGI Symbo | 0 | 0 | 0 | 0 | 0 | 0 |
| 39042 | ENSMUSG00000100730 | 0 | NA | NA | NA | NA | NA | Gm20776 | predicted gene, 20776 [Source:MGI Symb | 0 | 0 | 0 | 0 | 0 | 0 |
| 39043 | ENSMUSG00000100731 | 0 | NA | NA | NA | NA | NA | Gm29523 | predicted gene 29523 [Source:MGI Symbo | 0 | 0 | 0 | 0 | 0 | 0 |
| 39044 | ENSMUSG00000100732 | 0 | NA | NA | NA | NA | NA | Topbp1-ps2 | topoisomerase (DNA) II binding protein 1, | 0 | 0 | 0 | 0 | 0 | 0 |
| 39045 | ENSMUSG00000100733 | 6.405420874 | -0.167495097 | 0.835910486 | -0.200374442 | 0.841187747 | 0.971854408 | 4932411K12 | RIKEN cDNA 4932411K12 gene [Source:M | 8.031 | 6.1675 | 6.0375 | 3.8559 | 6.5437 | 7.7969 |
| 39046 | ENSMUSG00000100734 | 0 | NA | NA | NA | NA | NA | Rpl23a-ps8 | ribosomal protein L23A, pseudogene 8 [So | 0 | 0 | 0 | 0 | 0 | 0 |
| 39047 | ENSMUSG00000100736 | 0.133850373 | -0.796629176 | 4.080472857 | -0.195229623 | 0.84521319 | NA | Gm5147 | predicted gene 5147 [Source:MGI Symbol | 0.8031 | 0 | 0 | 0 | 0 | 0 |
| 39048 | ENSMUSG00000100737 | 0 | NA | NA | NA | NA | NA | Gm21774 | predicted gene, 21774 [Source:MGI Symb | 0 | 0 | 0 | 0 | 0 | 0 |
| 39049 | ENSMUSG00000100738 | 0 | NA | NA | NA | NA | NA | 2010106C02 | RIKEN cDNA 2010106C02 gene [Source:M | 0 | 0 | 0 | 0 | 0 | 0 |
| 39050 | ENSMUSG00000100739 | 0 | NA | NA | NA | NA | NA | Gm28296 | predicted gene 28296 [Source:MGI Symbo | 0 | 0 | 0 | 0 | 0 | 0 |
| 39051 | ENSMUSG00000100740 | 0 | NA | NA | NA | NA | NA | Gm29476 | predicted gene 29476 [Source:MGI Symbo | 0 | 0 | 0 | 0 | 0 | 0 |
| 39052 | ENSMUSG00000100741 | 0 | NA | NA | NA | NA | NA | Gm28814 | predicted gene 28814 [Source:MGI Symbo | 0 | 0 | 0 | 0 | 0 | 0 |
| 39053 | ENSMUSG00000100742 | 0 | NA | NA | NA | NA | NA | Gm28953 | predicted gene 28953 [Source:MGI Symbo | 0 | 0 | 0 | 0 | 0 | 0 |
| 39054 | ENSMUSG00000100743 | 0.160662974 | 1.126924175 | 4.080472857 | 0.276174898 | 0.782413736 | NA | Scgb1b25-ps | secretoglobin, family 1B, member 25, pse | 0 | 0 | 0 | 0.964 | 0 | 0 |
| 39055 | ENSMUSG00000100744 | 0 | NA | NA | NA | NA | NA | Gm17764 | predicted gene, 17764 [Source:MGI Symb | 0 | 0 | 0 | 0 | 0 | 0 |
| 39056 | ENSMUSG00000100745 | 0 | NA | NA | NA | NA | NA | Gm29077 | predicted gene 29077 [Source:MGI Symbo | 0 | 0 | 0 | 0 | 0 | 0 |
| 39057 | ENSMUSG00000100746 | 0 | NA | NA | NA | NA | NA | Gm28716 | predicted gene 28716 [Source:MGI Symbo | 0 | 0 | 0 | 0 | 0 | 0 |
| 39058 | ENSMUSG00000100747 | 7.706327153 | 0.990665107 | 0.813441357 | 1.217869118 | 0.223273713 | 0.668757712 | 1700084E18 | RIKEN cDNA 1700084E18 gene [Source:M | 7.2279 | 2.0558 | 6.0375 | 7.7118 | 8.7249 | 14.48 |
| 39059 | ENSMUSG00000100749 | 0 | NA | NA | NA | NA | NA | Gm20798 | predicted gene, 20798 [Source:MGI Symb | 0 | 0 | 0 | 0 | 0 | 0 |
| 39060 | ENSMUSG00000100750 | 0 | NA | NA | NA | NA | NA | Gm29084 | predicted gene 29084 [Source:MGI Symbo | 0 | 0 | 0 | 0 | 0 | 0 |
| 39061 | ENSMUSG00000100751 | 0 | NA | NA | NA | NA | NA | Gm5116 | predicted gene 5116 [Source:MGI Symbol | 0 | 0 | 0 | 0 | 0 | 0 |
| 39062 | ENSMUSG00000100752 | 0 | NA | NA | NA | NA | NA | Gm28847 | predicted gene 28847 [Source:MGI Symbo | 0 | 0 | 0 | 0 | 0 | 0 |
| 39063 | ENSMUSG00000100753 | 0 | NA | NA | NA | NA | NA | Gm29435 | predicted gene 29435 [Source:MGI Symbo | 0 | 0 | 0 | 0 | 0 | 0 |
| 39064 | ENSMUSG00000100754 | 0 | NA | NA | NA | NA | NA | Scgb2b5-ps | secretoglobin, family 2B, member 5, pseu | 0 | 0 | 0 | 0 | 0 | 0 |
| 39065 | ENSMUSG00000100755 | 0 | NA | NA | NA | NA | NA | Rps23-ps1 | ribosomal protein S23, pseudogene 1 [Sou | 0 | 0 | 0 | 0 | 0 | 0 |
| 39066 | ENSMUSG00000100756 | 0 | NA | NA | NA | NA | NA | Gm21257 | predicted gene, 21257 [Source:MGI Symb | 0 | 0 | 0 | 0 | 0 | 0 |
| 39067 | ENSMUSG00000100757 | 0 | NA | NA | NA | NA | NA | Gm29613 | predicted gene 29613 [Source:MGI Symbo | 0 | 0 | 0 | 0 | 0 | 0 |
| 39068 | ENSMUSG00000100758 | 0 | NA | NA | NA | NA | NA | Gm3332 | predicted gene 3332 [Source:MGI Symbol | 0 | 0 | 0 | 0 | 0 | 0 |
| 39069 | ENSMUSG00000100759 | 0 | NA | NA | NA | NA | NA | 1700029B22 | RIKEN cDNA 1700029B22 gene [Source:M | 0 | 0 | 0 | 0 | 0 | 0 |
| 39070 | ENSMUSG00000100760 | 0 | NA | NA | NA | NA | NA | Gm4035 | predicted gene 4035 [Source:MGI Symbol | 0 | 0 | 0 | 0 | 0 | 0 |
| 39071 | ENSMUSG00000100761 | 0 | NA | NA | NA | NA | NA | Gm29344 | predicted gene 29344 [Source:MGI Symbo | 0 | 0 | 0 | 0 | 0 | 0 |
| 39072 | ENSMUSG00000100762 | 0 | NA | NA | NA | NA | NA | Gm8403 | predicted gene 8403 [Source:MGI Symbol | 0 | 0 | 0 | 0 | 0 | 0 |
| 39073 | ENSMUSG00000100763 | 0.328370405 | 0.16514316 | 4.003149536 | 0.041253308 | 0.967093956 | NA | Gm28777 | predicted gene 28777 [Source:MGI Symbo | 0 | 0 | 1.0062 | 0.964 | 0 | 0 |
| 39074 | ENSMUSG00000100764 | 0.462220778 | -0.685400455 | 3.191475091 | -0.21475977 | 0.82995462 | NA | Gm29155 | predicted gene 29155 [Source:MGI Symbo | 0.8031 | 0 | 1.0062 | 0.964 | 0 | 0 |
| 39075 | ENSMUSG00000100765 | 0 | NA | NA | NA | NA | NA | Gm28684 | predicted gene 28684 [Source:MGI Symbo | 0 | 0 | 0 | 0 | 0 | 0 |
| 39076 | ENSMUSG00000100767 | 0 | NA | NA | NA | NA | NA | Gm3508 | predicted gene 3508 [Source:MGI Symbol | 0 | 0 | 0 | 0 | 0 | 0 |
| 39077 | ENSMUSG00000100768 | 0 | NA | NA | NA | NA | NA | Gm29055 | predicted gene 29055 [Source:MGI Symbo | 0 | 0 | 0 | 0 | 0 | 0 |
| 39078 | ENSMUSG00000100769 | 0 | NA | NA | NA | NA | NA | Gm28893 | predicted gene 28893 [Source:MGI Symbo | 0 | 0 | 0 | 0 | 0 | 0 |
| 39079 | ENSMUSG00000100770 | 0 | NA | NA | NA | NA | NA | Gm28466 | predicted gene 28466 [Source:MGI Symbo | 0 | 0 | 0 | 0 | 0 | 0 |
| 39080 | ENSMUSG00000100771 | 0 | NA | NA | NA | NA | NA | Gm28618 | predicted gene 28618 [Source:MGI Symbo | 0 | 0 | 0 | 0 | 0 | 0 |
| 39081 | ENSMUSG00000100772 | 0 | NA | NA | NA | NA | NA | Gm28350 | predicted gene 28350 [Source:MGI Symbo | 0 | 0 | 0 | 0 | 0 | 0 |
| 39082 | ENSMUSG00000100773 | 0 | NA | NA | NA | NA | NA | Gm28366 | predicted gene 28366 [Source:MGI Symbo | 0 | 0 | 0 | 0 | 0 | 0 |
| 39083 | ENSMUSG00000100774 | 1.718587302 | -1.908667498 | 1.840722588 | -1.036912086 | 0.299776831 | NA | Gm7329 | predicted gene 7329 [Source:MGI Symbol | 4.0155 | 2.0558 | 2.0125 | 0 | 0 | 2.2277 |
| 39084 | ENSMUSG00000100775 | 2.696162117 | 1.104537299 | 1.453939638 | 0.759685801 | 0.447442418 | NA | Gm29107 | predicted gene 29107 [Source:MGI Symbo | 4.0155 | 1.0279 | 0 | 6.7478 | 3.2719 | 1.1138 |

| | | | | | | | | | | | | | | | |
|---|---|---|---|---|---|---|---|---|---|---|---|---|---|---|---|
| 39085 | ENSMUSG00000100776 | 0 | NA | NA | NA | NA | NA | Gm18254 | predicted gene, 18254 [Source:MGI Symb | 0 | 0 | 0 | 0 | 0 | 0 |
| 39086 | ENSMUSG00000100777 | 0 | NA | NA | NA | NA | NA | 4930598F16 | RIKEN cDNA 4930598F16 gene [Source:M | 0 | 0 | 0 | 0 | 0 | 0 |
| 39087 | ENSMUSG00000100778 | 0 | NA | NA | NA | NA | NA | Gm28908 | predicted gene 28908 [Source:MGI Symbo | 0 | 0 | 0 | 0 | 0 | 0 |
| 39088 | ENSMUSG00000100779 | 0 | NA | NA | NA | NA | NA | Gm29292 | predicted gene 29292 [Source:MGI Symbo | 0 | 0 | 0 | 0 | 0 | 0 |
| 39089 | ENSMUSG00000100780 | 0 | NA | NA | NA | NA | NA | Gm28579 | predicted gene 28579 [Source:MGI Symbo | 0 | 0 | 0 | 0 | 0 | 0 |
| 39090 | ENSMUSG00000100781 | 0.133850373 | -0.796629176 | 4.080472857 | -0.195229623 | 0.84521319 | NA | Gm29481 | predicted gene 29481 [Source:MGI Symbo | 0.8031 | 0 | 0 | 0 | 0 | 0 |
| 39091 | ENSMUSG00000100782 | 0 | NA | NA | NA | NA | NA | Gm28231 | predicted gene 28231 [Source:MGI Symbo | 0 | 0 | 0 | 0 | 0 | 0 |
| 39092 | ENSMUSG00000100783 | 0.346303622 | 2.004339653 | 3.884003423 | 0.516049919 | 0.605819544 | NA | 2310047D07 | RIKEN cDNA 2310047D07 gene [Source:M | 0 | 0 | 0 | 0.964 | 0 | 1.1138 |
| 39093 | ENSMUSG00000100784 | 0 | NA | NA | NA | NA | NA | Gm28421 | predicted gene 28421 [Source:MGI Symbo | 0 | 0 | 0 | 0 | 0 | 0 |
| 39094 | ENSMUSG00000100785 | 0 | NA | NA | NA | NA | NA | Gm29333 | predicted gene 29333 [Source:MGI Symbo | 0 | 0 | 0 | 0 | 0 | 0 |
| 39095 | ENSMUSG00000100786 | 0.160662974 | 1.126924175 | 4.080472857 | 0.276174898 | 0.782413736 | NA | Gm28940 | predicted gene 28940 [Source:MGI Symbo | 0 | 0 | 0 | 0.964 | 0 | 0 |
| 39096 | ENSMUSG00000100787 | 0 | NA | NA | NA | NA | NA | Gm28316 | predicted gene 28316 [Source:MGI Symbo | 0 | 0 | 0 | 0 | 0 | 0 |
| 39097 | ENSMUSG00000100788 | 0 | NA | NA | NA | NA | NA | Gm21122 | predicted gene, 21122 [Source:MGI Symb | 0 | 0 | 0 | 0 | 0 | 0 |
| 39098 | ENSMUSG00000100789 | 0.133850373 | -0.796629176 | 4.080472857 | -0.195229623 | 0.84521319 | NA | Gm29508 | predicted gene 29508 [Source:MGI Symbo | 0.8031 | 0 | 0 | 0 | 0 | 0 |
| 39099 | ENSMUSG00000100790 | 0 | NA | NA | NA | NA | NA | Gm28518 | predicted gene 28518 [Source:MGI Symbo | 0 | 0 | 0 | 0 | 0 | 0 |
| 39100 | ENSMUSG00000100791 | 0 | NA | NA | NA | NA | NA | Gm28199 | predicted gene 28199 [Source:MGI Symbo | 0 | 0 | 0 | 0 | 0 | 0 |
| 39101 | ENSMUSG00000100792 | 0 | NA | NA | NA | NA | NA | Gm29161 | predicted gene 29161 [Source:MGI Symbo | 0 | 0 | 0 | 0 | 0 | 0 |
| 39102 | ENSMUSG00000100794 | 0.36353943 | 2.065989254 | 4.034137301 | 0.512126658 | 0.608562368 | NA | Gm29667 | predicted gene 29667 [Source:MGI Symbo | 0 | 0 | 0 | 0 | 2.1812 | 0 |
| 39103 | ENSMUSG00000100795 | 0 | NA | NA | NA | NA | NA | Gm29208 | predicted gene 29208 [Source:MGI Symbo | 0 | 0 | 0 | 0 | 0 | 0 |
| 39104 | ENSMUSG00000100796 | 0 | NA | NA | NA | NA | NA | Gm28163 | predicted gene 28163 [Source:MGI Symbo | 0 | 0 | 0 | 0 | 0 | 0 |
| 39105 | ENSMUSG00000100797 | 0 | NA | NA | NA | NA | NA | Gm28596 | predicted gene 28596 [Source:MGI Symbo | 0 | 0 | 0 | 0 | 0 | 0 |
| 39106 | ENSMUSG00000100798 | 0.335414862 | -1.779047242 | 4.030804179 | -0.441362855 | 0.658950329 | NA | Gm19589 | predicted gene, 19589 [Source:MGI Symb | 0 | 0 | 2.0125 | 0 | 0 | 0 |
| 39107 | ENSMUSG00000100799 | 0 | NA | NA | NA | NA | NA | 1700027H10 | RIKEN cDNA 1700027H10 gene [Source:M | 0 | 0 | 0 | 0 | 0 | 0 |
| 39108 | ENSMUSG00000100800 | 0 | NA | NA | NA | NA | NA | Gm21301 | predicted gene, 21301 [Source:MGI Symb | 0 | 0 | 0 | 0 | 0 | 0 |
| 39109 | ENSMUSG00000100801 | 0 | NA | NA | NA | NA | NA | Gm15459 | predicted gene 15459 [Source:MGI Symbo | 0 | 0 | 0 | 0 | 0 | 0 |
| 39110 | ENSMUSG00000100802 | 0 | NA | NA | NA | NA | NA | Gm20889 | predicted gene, 20889 [Source:MGI Symb | 0 | 0 | 0 | 0 | 0 | 0 |
| 39111 | ENSMUSG00000100803 | 0 | NA | NA | NA | NA | NA | Gm28251 | predicted gene 28251 [Source:MGI Symbo | 0 | 0 | 0 | 0 | 0 | 0 |
| 39112 | ENSMUSG00000100804 | 0 | NA | NA | NA | NA | NA | Gm28150 | predicted gene 28150 [Source:MGI Symbo | 0 | 0 | 0 | 0 | 0 | 0 |
| 39113 | ENSMUSG00000100805 | 0 | NA | NA | NA | NA | NA | Gm29108 | predicted gene 29108 [Source:MGI Symbo | 0 | 0 | 0 | 0 | 0 | 0 |
| 39114 | ENSMUSG00000100806 | 0 | NA | NA | NA | NA | NA | Gm28396 | predicted gene 28396 [Source:MGI Symbo | 0 | 0 | 0 | 0 | 0 | 0 |
| 39115 | ENSMUSG00000100807 | 0 | NA | NA | NA | NA | NA | Gm29521 | predicted gene 29521 [Source:MGI Symbo | 0 | 0 | 0 | 0 | 0 | 0 |
| 39116 | ENSMUSG00000100808 | 0 | NA | NA | NA | NA | NA | Gm28594 | predicted gene 28594 [Source:MGI Symbo | 0 | 0 | 0 | 0 | 0 | 0 |
| 39117 | ENSMUSG00000100809 | 0 | NA | NA | NA | NA | NA | Gm28612 | predicted gene 28612 [Source:MGI Symbo | 0 | 0 | 0 | 0 | 0 | 0 |
| 39118 | ENSMUSG00000100810 | 0 | NA | NA | NA | NA | NA | Gm29086 | predicted gene 29086 [Source:MGI Symbo | 0 | 0 | 0 | 0 | 0 | 0 |
| 39119 | ENSMUSG00000100811 | 0 | NA | NA | NA | NA | NA | Gm29083 | predicted gene 29083 [Source:MGI Symbo | 0 | 0 | 0 | 0 | 0 | 0 |
| 39120 | ENSMUSG00000100812 | 0 | NA | NA | NA | NA | NA | A730098A19 | RIKEN cDNA A730098A19 gene [Source:M | 0 | 0 | 0 | 0 | 0 | 0 |
| 39121 | ENSMUSG00000100813 | 2.207309824 | -2.425896407 | 1.676805552 | -1.446736865 | 0.147970635 | NA | Gm28874 | predicted gene 28874 [Source:MGI Symbo | 4.0155 | 6.1675 | 1.0062 | 0.964 | 1.0906 | 0 |
| 39122 | ENSMUSG00000100814 | 0 | NA | NA | NA | NA | NA | Gm28669 | predicted gene 28669 [Source:MGI Symbo | 0 | 0 | 0 | 0 | 0 | 0 |
| 39123 | ENSMUSG00000100815 | 0 | NA | NA | NA | NA | NA | Gm29112 | predicted gene 29112 [Source:MGI Symbo | 0 | 0 | 0 | 0 | 0 | 0 |
| 39124 | ENSMUSG00000100816 | 0 | NA | NA | NA | NA | NA | Gm28321 | predicted gene 28321 [Source:MGI Symbo | 0 | 0 | 0 | 0 | 0 | 0 |
| 39125 | ENSMUSG00000100817 | 0 | NA | NA | NA | NA | NA | Gm29302 | predicted gene 29302 [Source:MGI Symbo | 0 | 0 | 0 | 0 | 0 | 0 |
| 39126 | ENSMUSG00000100818 | 0 | NA | NA | NA | NA | NA | Gm21287 | predicted gene, 21287 [Source:MGI Symb | 0 | 0 | 0 | 0 | 0 | 0 |
| 39127 | ENSMUSG00000100819 | 2.739564951 | -0.315669289 | 1.35233209 | -0.233425866 | 0.81543073 | NA | Gm2693 | predicted gene 2693 [Source:MGI Symbol | 4.0155 | 2.0558 | 3.0187 | 2.8919 | 0 | 4.4554 |
| 39128 | ENSMUSG00000100820 | 0 | NA | NA | NA | NA | NA | Gm28945 | predicted gene 28945 [Source:MGI Symbo | 0 | 0 | 0 | 0 | 0 | 0 |
| 39129 | ENSMUSG00000100821 | 0 | NA | NA | NA | NA | NA | Gm28894 | predicted gene 28894 [Source:MGI Symbo | 0 | 0 | 0 | 0 | 0 | 0 |
| 39130 | ENSMUSG00000100822 | 0 | NA | NA | NA | NA | NA | Gm29275 | predicted gene 29275 [Source:MGI Symbo | 0 | 0 | 0 | 0 | 0 | 0 |
| 39131 | ENSMUSG00000100823 | 0 | NA | NA | NA | NA | NA | Gm28704 | predicted gene 28704 [Source:MGI Symbo | 0 | 0 | 0 | 0 | 0 | 0 |
| 39132 | ENSMUSG00000100825 | 0 | NA | NA | NA | NA | NA | Itpa-ps2 | inosine triphosphatase (nucleoside tripho | 0 | 0 | 0 | 0 | 0 | 0 |
| 39133 | ENSMUSG00000100826 | 45.2019725 | -0.081341672 | 0.389873601 | -0.208636007 | 0.834732399 | 0.971073158 | Snhg14 | small nucleolar RNA host gene 14 [Source | 54.611 | 38.033 | 46.287 | 24.099 | 45.806 | 62.375 |
| 39134 | ENSMUSG00000100827 | 0 | NA | NA | NA | NA | NA | Gm29069 | predicted gene 29069 [Source:MGI Symbo | 0 | 0 | 0 | 0 | 0 | 0 |
| 39135 | ENSMUSG00000100830 | 0 | NA | NA | NA | NA | NA | Gm28441 | predicted gene 28441 [Source:MGI Symbo | 0 | 0 | 0 | 0 | 0 | 0 |
| 39136 | ENSMUSG00000100831 | 0 | NA | NA | NA | NA | NA | Gm17847 | predicted gene, 17847 [Source:MGI Symb | 0 | 0 | 0 | 0 | 0 | 0 |
| 39137 | ENSMUSG00000100832 | 0 | NA | NA | NA | NA | NA | Gm29260 | predicted gene 29260 [Source:MGI Symbo | 0 | 0 | 0 | 0 | 0 | 0 |
| 39138 | ENSMUSG00000100833 | 0.596071152 | -1.224343926 | 2.897583865 | -0.4225396 | 0.672631203 | NA | Gm28988 | predicted gene 28988 [Source:MGI Symbo | 1.6062 | 0 | 1.0062 | 0.964 | 0 | 0 |
| 39139 | ENSMUSG00000100834 | 0 | NA | NA | NA | NA | NA | Gm21285 | predicted gene, 21285 [Source:MGI Symb | 0 | 0 | 0 | 0 | 0 | 0 |
| 39140 | ENSMUSG00000100835 | 0 | NA | NA | NA | NA | NA | Gm28907 | predicted gene 28907 [Source:MGI Symbo | 0 | 0 | 0 | 0 | 0 | 0 |

| | | | | | | | | | | | | | | | |
|---|---|---|---|---|---|---|---|---|---|---|---|---|---|---|---|
| 39141 | ENSMUSG00000100836 | 0 | NA | NA | NA | NA | NA | Gm28215 | predicted gene 28215 [Source:MGI Symbo | 0 | 0 | 0 | 0 | 0 | 0 |
| 39142 | ENSMUSG00000100837 | 0.727078859 | 3.067238961 | 3.29680021 | 0.930368468 | 0.352180339 | NA | 1700063D05 | RIKEN cDNA 1700063D05 gene [Source:M | 0 | 0 | 0 | 0 | 4.3625 | 0 |
| 39143 | ENSMUSG00000100838 | 0 | NA | NA | NA | NA | NA | Gm29094 | predicted gene 29094 [Source:MGI Symbo | 0 | 0 | 0 | 0 | 0 | 0 |
| 39144 | ENSMUSG00000100839 | 0 | NA | NA | NA | NA | NA | Gm5920 | predicted gene 5920 [Source:MGI Symbol | 0 | 0 | 0 | 0 | 0 | 0 |
| 39145 | ENSMUSG00000100841 | 0 | NA | NA | NA | NA | NA | Gm21115 | predicted gene, 21115 [Source:MGI Symb | 0 | 0 | 0 | 0 | 0 | 0 |
| 39146 | ENSMUSG00000100842 | 0 | NA | NA | NA | NA | NA | Gm17790 | predicted gene, 17790 [Source:MGI Symb | 0 | 0 | 0 | 0 | 0 | 0 |
| 39147 | ENSMUSG00000100843 | 0 | NA | NA | NA | NA | NA | Gm8307 | predicted gene 8307 [Source:MGI Symbol | 0 | 0 | 0 | 0 | 0 | 0 |
| 39148 | ENSMUSG00000100844 | 0 | NA | NA | NA | NA | NA | 1810007C17 | RIKEN cDNA 1810007C17 gene [Source:M | 0 | 0 | 0 | 0 | 0 | 0 |
| 39149 | ENSMUSG00000100845 | 0 | NA | NA | NA | NA | NA | Gm28770 | predicted gene 28770 [Source:MGI Symbo | 0 | 0 | 0 | 0 | 0 | 0 |
| 39150 | ENSMUSG00000100846 | 0 | NA | NA | NA | NA | NA | Gm28845 | predicted gene 28845 [Source:MGI Symbo | 0 | 0 | 0 | 0 | 0 | 0 |
| 39151 | ENSMUSG00000100847 | 0 | NA | NA | NA | NA | NA | Gm28676 | predicted gene 28676 [Source:MGI Symbo | 0 | 0 | 0 | 0 | 0 | 0 |
| 39152 | ENSMUSG00000100848 | 0 | NA | NA | NA | NA | NA | Gm29281 | predicted gene 29281 [Source:MGI Symbo | 0 | 0 | 0 | 0 | 0 | 0 |
| 39153 | ENSMUSG00000100849 | 0 | NA | NA | NA | NA | NA | Gm28663 | predicted gene 28663 [Source:MGI Symbo | 0 | 0 | 0 | 0 | 0 | 0 |
| 39154 | ENSMUSG00000100850 | 0 | NA | NA | NA | NA | NA | 4930434B07 | RIKEN cDNA 4930434B07 gene [Source:M | 0 | 0 | 0 | 0 | 0 | 0 |
| 39155 | ENSMUSG00000100851 | 0 | NA | NA | NA | NA | NA | NA | NA | 0 | 0 | 0 | 0 | 0 | 0 |
| 39156 | ENSMUSG00000100852 | 0 | NA | NA | NA | NA | NA | Gm29056 | predicted gene 29056 [Source:MGI Symbo | 0 | 0 | 0 | 0 | 0 | 0 |
| 39157 | ENSMUSG00000100853 | 0 | NA | NA | NA | NA | NA | Gm28232 | predicted gene 28232 [Source:MGI Symbo | 0 | 0 | 0 | 0 | 0 | 0 |
| 39158 | ENSMUSG00000100854 | 0 | NA | NA | NA | NA | NA | Gm28057 | predicted gene 28057 [Source:MGI Symbo | 0 | 0 | 0 | 0 | 0 | 0 |
| 39159 | ENSMUSG00000100856 | 0 | NA | NA | NA | NA | NA | Gm21627 | predicted gene, 21627 [Source:MGI Symb | 0 | 0 | 0 | 0 | 0 | 0 |
| 39160 | ENSMUSG00000100857 | 0 | NA | NA | NA | NA | NA | 1700041M1 | RIKEN cDNA 1700041M19 gene [Source:M | 0 | 0 | 0 | 0 | 0 | 0 |
| 39161 | ENSMUSG00000100858 | 0 | NA | NA | NA | NA | NA | Gm28400 | predicted gene 28400 [Source:MGI Symbo | 0 | 0 | 0 | 0 | 0 | 0 |
| 39162 | ENSMUSG00000100859 | 0 | NA | NA | NA | NA | NA | Gm28725 | predicted gene 28725 [Source:MGI Symbo | 0 | 0 | 0 | 0 | 0 | 0 |
| 39163 | ENSMUSG00000100860 | 0.160662974 | 1.126924175 | 4.080472857 | 0.276174898 | 0.782413736 | NA | 2010009K17 | RIKEN cDNA 2010009K17 gene [Source:M | 0 | 0 | 0 | 0.964 | 0 | 0 |
| 39164 | ENSMUSG00000100861 | 0 | NA | NA | NA | NA | NA | Gm29382 | predicted gene 29382 [Source:MGI Symbo | 0 | 0 | 0 | 0 | 0 | 0 |
| 39165 | ENSMUSG00000100862 | 667.5251362 | 0.149970789 | 0.142232505 | 1.054405875 | 0.291697133 | 0.742016767 | Gm10925 | predicted gene 10925 [Source:MGI Symbo | 625.62 | 634.23 | 637.96 | 572.6 | 738.35 | 796.4 |
| 39166 | ENSMUSG00000100863 | 6.422656681 | -0.161168333 | 0.852672304 | -0.189015559 | 0.850080622 | 0.973301217 | Gm12669 | predicted gene 12669 [Source:MGI Symbo | 8.031 | 6.1675 | 6.0375 | 2.8919 | 8.7249 | 6.6831 |
| 39167 | ENSMUSG00000100864 | 0 | NA | NA | NA | NA | NA | Gm21302 | predicted gene, 21302 [Source:MGI Symb | 0 | 0 | 0 | 0 | 0 | 0 |
| 39168 | ENSMUSG00000100865 | 11.84916174 | 0.576083947 | 0.690996808 | 0.833699868 | 0.404450133 | 0.829271989 | Gm9320 | predicted gene 9320 [Source:MGI Symbol | 6.4248 | 7.1954 | 15.094 | 20.244 | 6.5437 | 15.594 |
| 39169 | ENSMUSG00000100866 | 0 | NA | NA | NA | NA | NA | Gm28830 | predicted gene 28830 [Source:MGI Symbo | 0 | 0 | 0 | 0 | 0 | 0 |
| 39170 | ENSMUSG00000100867 | 0 | NA | NA | NA | NA | NA | Gm18981 | predicted gene, 18981 [Source:MGI Symb | 0 | 0 | 0 | 0 | 0 | 0 |
| 39171 | ENSMUSG00000100868 | 0 | NA | NA | NA | NA | NA | Gm28095 | predicted gene 28095 [Source:MGI Symbo | 0 | 0 | 0 | 0 | 0 | 0 |
| 39172 | ENSMUSG00000100869 | 0 | NA | NA | NA | NA | NA | Gm20846 | predicted gene, 20846 [Source:MGI Symb | 0 | 0 | 0 | 0 | 0 | 0 |
| 39173 | ENSMUSG00000100870 | 0 | NA | NA | NA | NA | NA | Gm21720 | predicted gene, 21720 [Source:MGI Symb | 0 | 0 | 0 | 0 | 0 | 0 |
| 39174 | ENSMUSG00000100871 | 0 | NA | NA | NA | NA | NA | Gm29213 | predicted gene 29213 [Source:MGI Symbo | 0 | 0 | 0 | 0 | 0 | 0 |
| 39175 | ENSMUSG00000100872 | 0.133850373 | -0.796629176 | 4.080472857 | -0.195229623 | 0.84521319 | NA | 1700065J18 | RIKEN cDNA 1700065J18 gene [Source:M | 0.8031 | 0 | 0 | 0 | 0 | 0 |
| 39176 | ENSMUSG00000100873 | 0 | NA | NA | NA | NA | NA | Gm28243 | predicted gene 28243 [Source:MGI Symbo | 0 | 0 | 0 | 0 | 0 | 0 |
| 39177 | ENSMUSG00000100874 | 0 | NA | NA | NA | NA | NA | Gm28598 | predicted gene 28598 [Source:MGI Symbo | 0 | 0 | 0 | 0 | 0 | 0 |
| 39178 | ENSMUSG00000100875 | 0 | NA | NA | NA | NA | NA | Gm28987 | predicted gene 28987 [Source:MGI Symbo | 0 | 0 | 0 | 0 | 0 | 0 |
| 39179 | ENSMUSG00000100876 | 7.445966716 | -0.454317975 | 0.786398003 | -0.577720154 | 0.56345307 | 0.900821918 | 2810454H06 | RIKEN cDNA 2810454H06 gene [Source:M | 5.6217 | 9.2513 | 11.069 | 7.7118 | 5.4531 | 5.5692 |
| 39180 | ENSMUSG00000100877 | 0 | NA | NA | NA | NA | NA | Gm28534 | predicted gene 28534 [Source:MGI Symbo | 0 | 0 | 0 | 0 | 0 | 0 |
| 39181 | ENSMUSG00000100878 | 0 | NA | NA | NA | NA | NA | Gm29196 | predicted gene 29196 [Source:MGI Symbo | 0 | 0 | 0 | 0 | 0 | 0 |
| 39182 | ENSMUSG00000100879 | 0 | NA | NA | NA | NA | NA | Gm28743 | predicted gene 28743 [Source:MGI Symbo | 0 | 0 | 0 | 0 | 0 | 0 |
| 39183 | ENSMUSG00000100880 | 0.342639279 | -1.805976596 | 4.028547714 | -0.448294702 | 0.653940524 | NA | Gm6610 | predicted gene 6610 [Source:MGI Symbol | 0 | 2.0558 | 0 | 0 | 0 | 0 |
| 39184 | ENSMUSG00000100881 | 0 | NA | NA | NA | NA | NA | Gm21824 | predicted gene, 21824 [Source:MGI Symb | 0 | 0 | 0 | 0 | 0 | 0 |
| 39185 | ENSMUSG00000100882 | 0 | NA | NA | NA | NA | NA | Gm20849 | predicted gene, 20849 [Source:MGI Symb | 0 | 0 | 0 | 0 | 0 | 0 |
| 39186 | ENSMUSG00000100883 | 0 | NA | NA | NA | NA | NA | Gm28773 | predicted gene 28773 [Source:MGI Symbo | 0 | 0 | 0 | 0 | 0 | 0 |
| 39187 | ENSMUSG00000100884 | 0 | NA | NA | NA | NA | NA | Gm28281 | predicted gene 28281 [Source:MGI Symbo | 0 | 0 | 0 | 0 | 0 | 0 |
| 39188 | ENSMUSG00000100885 | 0 | NA | NA | NA | NA | NA | Gm29446 | predicted gene 29446 [Source:MGI Symbo | 0 | 0 | 0 | 0 | 0 | 0 |
| 39189 | ENSMUSG00000100886 | 0 | NA | NA | NA | NA | NA | Gm7195 | predicted gene 7195 [Source:MGI Symbol | 0 | 0 | 0 | 0 | 0 | 0 |
| 39190 | ENSMUSG00000100887 | 0 | NA | NA | NA | NA | NA | Gm29297 | predicted gene 29297 [Source:MGI Symbo | 0 | 0 | 0 | 0 | 0 | 0 |
| 39191 | ENSMUSG00000100888 | 0 | NA | NA | NA | NA | NA | Gm29567 | predicted gene 29567 [Source:MGI Symbo | 0 | 0 | 0 | 0 | 0 | 0 |
| 39192 | ENSMUSG00000100889 | 0 | NA | NA | NA | NA | NA | Gm28204 | predicted gene 28204 [Source:MGI Symbo | 0 | 0 | 0 | 0 | 0 | 0 |
| 39193 | ENSMUSG00000100890 | 0 | NA | NA | NA | NA | NA | 1700085C21 | RIKEN cDNA 1700085C21 gene [Source:M | 0 | 0 | 0 | 0 | 0 | 0 |
| 39194 | ENSMUSG00000100891 | 0 | NA | NA | NA | NA | NA | 2810049E08 | RIKEN cDNA 2810049E08 gene [Source:M | 0 | 0 | 0 | 0 | 0 | 0 |
| 39195 | ENSMUSG00000100892 | 0 | NA | NA | NA | NA | NA | Gm20897 | predicted gene, 20897 [Source:MGI Symb | 0 | 0 | 0 | 0 | 0 | 0 |
| 39196 | ENSMUSG00000100893 | 0 | NA | NA | NA | NA | NA | Gm28083 | predicted gene 28083 [Source:MGI Symbo | 0 | 0 | 0 | 0 | 0 | 0 |

| | | | | | | | | | | | | | | | |
|---|---|---|---|---|---|---|---|---|---|---|---|---|---|---|---|
| 39197 | ENSMUSG00000100894 | 0 | NA | NA | NA | NA | NA | Gm8304 | predicted gene 8304 [Source:MGI Symbol | 0 | 0 | 0 | 0 | 0 | 0 |
| 39198 | ENSMUSG00000100895 | 0 | NA | NA | NA | NA | NA | Gm28685 | predicted gene 28685 [Source:MGI Symbo | 0 | 0 | 0 | 0 | 0 | 0 |
| 39199 | ENSMUSG00000100896 | 0 | NA | NA | NA | NA | NA | Gm28217 | predicted gene 28217 [Source:MGI Symbo | 0 | 0 | 0 | 0 | 0 | 0 |
| 39200 | ENSMUSG00000100897 | 0 | NA | NA | NA | NA | NA | Gm29634 | predicted gene 29634 [Source:MGI Symbo | 0 | 0 | 0 | 0 | 0 | 0 |
| 39201 | ENSMUSG00000100898 | 0 | NA | NA | NA | NA | NA | Gm29178 | predicted gene 29178 [Source:MGI Symbo | 0 | 0 | 0 | 0 | 0 | 0 |
| 39202 | ENSMUSG00000100899 | 0 | NA | NA | NA | NA | NA | Olfr1167 | olfactory receptor 1167 [Source:MGI Sym | 0 | 0 | 0 | 0 | 0 | 0 |
| 39203 | ENSMUSG00000100900 | 0 | NA | NA | NA | NA | NA | Gm28850 | predicted gene 28850 [Source:MGI Symbo | 0 | 0 | 0 | 0 | 0 | 0 |
| 39204 | ENSMUSG00000100901 | 0 | NA | NA | NA | NA | NA | Gm18829 | predicted gene, 18829 [Source:MGI Symb | 0 | 0 | 0 | 0 | 0 | 0 |
| 39205 | ENSMUSG00000100902 | 0 | NA | NA | NA | NA | NA | Gm20850 | predicted gene, 20850 [Source:MGI Symb | 0 | 0 | 0 | 0 | 0 | 0 |
| 39206 | ENSMUSG00000100903 | 0 | NA | NA | NA | NA | NA | Scgb2b31-ps | secretoglobin, family 2B, member 31, pse | 0 | 0 | 0 | 0 | 0 | 0 |
| 39207 | ENSMUSG00000100904 | 0 | NA | NA | NA | NA | NA | Gm28559 | predicted gene 28559 [Source:MGI Symbo | 0 | 0 | 0 | 0 | 0 | 0 |
| 39208 | ENSMUSG00000100905 | 0.181769715 | 1.126924175 | 4.080472857 | 0.276174898 | 0.782413736 | NA | Gm28075 | predicted gene 28075 [Source:MGI Symbo | 0 | 0 | 0 | 0 | 1.0906 | 0 |
| 39209 | ENSMUSG00000100906 | 0 | NA | NA | NA | NA | NA | Gm29238 | predicted gene 29238 [Source:MGI Symbo | 0 | 0 | 0 | 0 | 0 | 0 |
| 39210 | ENSMUSG00000100907 | 0 | NA | NA | NA | NA | NA | Gm29310 | predicted gene 29310 [Source:MGI Symbo | 0 | 0 | 0 | 0 | 0 | 0 |
| 39211 | ENSMUSG00000100908 | 0 | NA | NA | NA | NA | NA | Gm28888 | predicted gene 28888 [Source:MGI Symbo | 0 | 0 | 0 | 0 | 0 | 0 |
| 39212 | ENSMUSG00000100910 | 0 | NA | NA | NA | NA | NA | Gm29128 | predicted gene 29128 [Source:MGI Symbo | 0 | 0 | 0 | 0 | 0 | 0 |
| 39213 | ENSMUSG00000100911 | 0 | NA | NA | NA | NA | NA | 1700027F09 | RIKEN cDNA 1700027F09 gene [Source:M | 0 | 0 | 0 | 0 | 0 | 0 |
| 39214 | ENSMUSG00000100912 | 0 | NA | NA | NA | NA | NA | Gm2021 | predicted gene 2021 [Source:MGI Symbol | 0 | 0 | 0 | 0 | 0 | 0 |
| 39215 | ENSMUSG00000100913 | 0 | NA | NA | NA | NA | NA | Gm8392 | predicted gene 8392 [Source:MGI Symbol | 0 | 0 | 0 | 0 | 0 | 0 |
| 39216 | ENSMUSG00000100914 | 0 | NA | NA | NA | NA | NA | Gm28679 | predicted gene 28679 [Source:MGI Symbo | 0 | 0 | 0 | 0 | 0 | 0 |
| 39217 | ENSMUSG00000100915 | 0 | NA | NA | NA | NA | NA | Gm29321 | predicted gene 29321 [Source:MGI Symbo | 0 | 0 | 0 | 0 | 0 | 0 |
| 39218 | ENSMUSG00000100916 | 0 | NA | NA | NA | NA | NA | Lhb | luteinizing hormone beta [Source:MGI Sy | 0 | 0 | 0 | 0 | 0 | 0 |
| 39219 | ENSMUSG00000100917 | 0 | NA | NA | NA | NA | NA | Gm18751 | predicted gene, 18751 [Source:MGI Symb | 0 | 0 | 0 | 0 | 0 | 0 |
| 39220 | ENSMUSG00000100918 | 0 | NA | NA | NA | NA | NA | Gm19224 | predicted gene, 19224 [Source:MGI Symb | 0 | 0 | 0 | 0 | 0 | 0 |
| 39221 | ENSMUSG00000100919 | 0 | NA | NA | NA | NA | NA | Gm28655 | predicted gene 28655 [Source:MGI Symbo | 0 | 0 | 0 | 0 | 0 | 0 |
| 39222 | ENSMUSG00000100920 | 0 | NA | NA | NA | NA | NA | Gm18348 | predicted gene, 18348 [Source:MGI Symb | 0 | 0 | 0 | 0 | 0 | 0 |
| 39223 | ENSMUSG00000100922 | 0 | NA | NA | NA | NA | NA | Gm8520 | predicted gene 8520 [Source:MGI Symbol | 0 | 0 | 0 | 0 | 0 | 0 |
| 39224 | ENSMUSG00000100923 | 0 | NA | NA | NA | NA | NA | Olfr10 | olfactory receptor 10 [Source:MGI Symbo | 0 | 0 | 0 | 0 | 0 | 0 |
| 39225 | ENSMUSG00000100925 | 0 | NA | NA | NA | NA | NA | Gm28898 | predicted gene 28898 [Source:MGI Symbo | 0 | 0 | 0 | 0 | 0 | 0 |
| 39226 | ENSMUSG00000100926 | 0 | NA | NA | NA | NA | NA | Gm5329 | predicted gene 5329 [Source:MGI Symbol | 0 | 0 | 0 | 0 | 0 | 0 |
| 39227 | ENSMUSG00000100927 | 0.847770947 | -0.376623742 | 2.576917887 | -0.146152791 | 0.883800772 | NA | Gm28536 | predicted gene 28536 [Source:MGI Symbo | 0.8031 | 2.0558 | 0 | 0 | 0 | 2.2277 |
| 39228 | ENSMUSG00000100928 | 0 | NA | NA | NA | NA | NA | Gm28487 | predicted gene 28487 [Source:MGI Symbo | 0 | 0 | 0 | 0 | 0 | 0 |
| 39229 | ENSMUSG00000100929 | 0 | NA | NA | NA | NA | NA | Gm28064 | predicted gene 28064 [Source:MGI Symbo | 0 | 0 | 0 | 0 | 0 | 0 |
| 39230 | ENSMUSG00000100930 | 0 | NA | NA | NA | NA | NA | Gm28866 | predicted gene 28866 [Source:MGI Symbo | 0 | 0 | 0 | 0 | 0 | 0 |
| 39231 | ENSMUSG00000100931 | 1.36157541 | 1.788493635 | 1.905524006 | 0.938583628 | 0.347944562 | NA | Gm28085 | predicted gene 28085 [Source:MGI Symbo | 0.8031 | 0 | 1.0062 | 1.928 | 1.0906 | 3.3415 |
| 39232 | ENSMUSG00000100932 | 0.979405355 | -0.876544108 | 2.280749483 | -0.384322835 | 0.700739185 | NA | Gm29459 | predicted gene 29459 [Source:MGI Symbo | 0.8031 | 0 | 3.0187 | 0.964 | 1.0906 | 0 |
| 39233 | ENSMUSG00000100933 | 0 | NA | NA | NA | NA | NA | Gm28143 | predicted gene 28143 [Source:MGI Symbo | 0 | 0 | 0 | 0 | 0 | 0 |
| 39234 | ENSMUSG00000100935 | 0 | NA | NA | NA | NA | NA | Gm29592 | predicted gene 29592 [Source:MGI Symbo | 0 | 0 | 0 | 0 | 0 | 0 |
| 39235 | ENSMUSG00000100936 | 0 | NA | NA | NA | NA | NA | Gm28865 | predicted gene 28865 [Source:MGI Symbo | 0 | 0 | 0 | 0 | 0 | 0 |
| 39236 | ENSMUSG00000100937 | 3.548013668 | -0.614402596 | 1.332548709 | -0.461073274 | 0.644746036 | NA | Nscme3l | NSE3 homolog, SMC5-SMC6 complex com | 0.8031 | 3.0838 | 9.0562 | 2.8919 | 5.4531 | 0 |
| 39237 | ENSMUSG00000100938 | 0 | NA | NA | NA | NA | NA | Gm28726 | predicted gene 28726 [Source:MGI Symbo | 0 | 0 | 0 | 0 | 0 | 0 |
| 39238 | ENSMUSG00000100939 | 0 | NA | NA | NA | NA | NA | Gm28998 | predicted gene 28998 [Source:MGI Symbo | 0 | 0 | 0 | 0 | 0 | 0 |
| 39239 | ENSMUSG00000100940 | 0 | NA | NA | NA | NA | NA | Gm29372 | predicted gene 29372 [Source:MGI Symbo | 0 | 0 | 0 | 0 | 0 | 0 |
| 39240 | ENSMUSG00000100941 | 0 | NA | NA | NA | NA | NA | Gm29597 | predicted gene 29597 [Source:MGI Symbo | 0 | 0 | 0 | 0 | 0 | 0 |
| 39241 | ENSMUSG00000100942 | 0 | NA | NA | NA | NA | NA | Gm28560 | predicted gene 28560 [Source:MGI Symbo | 0 | 0 | 0 | 0 | 0 | 0 |
| 39242 | ENSMUSG00000100943 | 0 | NA | NA | NA | NA | NA | Gm29229 | predicted gene 29229 [Source:MGI Symbo | 0 | 0 | 0 | 0 | 0 | 0 |
| 39243 | ENSMUSG00000100944 | 0 | NA | NA | NA | NA | NA | Gm28900 | predicted gene 28900 [Source:MGI Symbo | 0 | 0 | 0 | 0 | 0 | 0 |
| 39244 | ENSMUSG00000100945 | 0.51014012 | 1.031855138 | 3.09002894 | 0.333930574 | 0.738431948 | NA | Gm29529 | predicted gene 29529 [Source:MGI Symbo | 0 | 0 | 1.0062 | 0.964 | 1.0906 | 0 |
| 39245 | ENSMUSG00000100946 | 0 | NA | NA | NA | NA | NA | Gm28473 | predicted gene 28473 [Source:MGI Symbo | 0 | 0 | 0 | 0 | 0 | 0 |
| 39246 | ENSMUSG00000100947 | 0 | NA | NA | NA | NA | NA | Gm28949 | predicted gene 28949 [Source:MGI Symbo | 0 | 0 | 0 | 0 | 0 | 0 |
| 39247 | ENSMUSG00000100948 | 0 | NA | NA | NA | NA | NA | Gm28395 | predicted gene 28395 [Source:MGI Symbo | 0 | 0 | 0 | 0 | 0 | 0 |
| 39248 | ENSMUSG00000100949 | 0 | NA | NA | NA | NA | NA | Scgb1b13-ps | secretoglobin, family 1B, member 13, pse | 0 | 0 | 0 | 0 | 0 | 0 |
| 39249 | ENSMUSG00000100950 | 0 | NA | NA | NA | NA | NA | Taar7f | trace amine-associated receptor 7F [Sour | 0 | 0 | 0 | 0 | 0 | 0 |
| 39250 | ENSMUSG00000100951 | 0 | NA | NA | NA | NA | NA | Gm29339 | predicted gene 29339 [Source:MGI Symbo | 0 | 0 | 0 | 0 | 0 | 0 |
| 39251 | ENSMUSG00000100952 | 0 | NA | NA | NA | NA | NA | Gm28545 | predicted gene 28545 [Source:MGI Symbo | 0 | 0 | 0 | 0 | 0 | 0 |
| 39252 | ENSMUSG00000100953 | 0 | NA | NA | NA | NA | NA | Gm28280 | predicted gene 28280 [Source:MGI Symbo | 0 | 0 | 0 | 0 | 0 | 0 |

| | | | | | | | | | | | | | | | |
|---|---|---|---|---|---|---|---|---|---|---|---|---|---|---|---|
| 39253 | ENSMUSG00000100954 | 36.40463324 | -0.260639434 | 0.386669043 | -0.674063359 | 0.500271033 | 0.880716237 | Gm10138 | predicted gene 10138 [Source:MGI Symbo | 28.912 | 47.284 | 43.269 | 40.487 | 26.175 | 32.301 |
| 39254 | ENSMUSG00000100955 | 0 | NA | NA | NA | NA | NA | Gm18728 | predicted gene, 18728 [Source:MGI Symb | 0 | 0 | 0 | 0 | 0 | 0 |
| 39255 | ENSMUSG00000100956 | 0 | NA | NA | NA | NA | NA | Gm28558 | predicted gene 28558 [Source:MGI Symbo | 0 | 0 | 0 | 0 | 0 | 0 |
| 39256 | ENSMUSG00000100957 | 0 | NA | NA | NA | NA | NA | Gm28762 | predicted gene 28762 [Source:MGI Symbo | 0 | 0 | 0 | 0 | 0 | 0 |
| 39257 | ENSMUSG00000100958 | 0 | NA | NA | NA | NA | NA | Gm29351 | predicted gene 29351 [Source:MGI Symbo | 0 | 0 | 0 | 0 | 0 | 0 |
| 39258 | ENSMUSG00000100959 | 0.674621893 | -1.426050892 | 3.091195043 | -0.461326727 | 0.644564212 | NA | Idi1-ps5 | isopentenyl-diphosphate delta isomerase, | 0 | 3.0838 | 0 | 0.964 | 0 | 0 |
| 39259 | ENSMUSG00000100960 | 0 | NA | NA | NA | NA | NA | Gm21156 | predicted gene, 21156 [Source:MGI Symb | 0 | 0 | 0 | 0 | 0 | 0 |
| 39260 | ENSMUSG00000100961 | 0 | NA | NA | NA | NA | NA | Gm28394 | predicted gene 28394 [Source:MGI Symbo | 0 | 0 | 0 | 0 | 0 | 0 |
| 39261 | ENSMUSG00000100962 | 0 | NA | NA | NA | NA | NA | Gm29133 | predicted gene 29133 [Source:MGI Symbo | 0 | 0 | 0 | 0 | 0 | 0 |
| 39262 | ENSMUSG00000100963 | 0 | NA | NA | NA | NA | NA | Gm28372 | predicted gene 28372 [Source:MGI Symbo | 0 | 0 | 0 | 0 | 0 | 0 |
| 39263 | ENSMUSG00000100964 | 0 | NA | NA | NA | NA | NA | Gm29503 | predicted gene 29503 [Source:MGI Symbo | 0 | 0 | 0 | 0 | 0 | 0 |
| 39264 | ENSMUSG00000100965 | 0 | NA | NA | NA | NA | NA | Gm28449 | predicted gene 28449 [Source:MGI Symbo | 0 | 0 | 0 | 0 | 0 | 0 |
| 39265 | ENSMUSG00000100966 | 0 | NA | NA | NA | NA | NA | Gm5700 | predicted gene 5700 [Source:MGI Symbol | 0 | 0 | 0 | 0 | 0 | 0 |
| 39266 | ENSMUSG00000100967 | 6.100741679 | 0.450999485 | 0.872917315 | 0.516657738 | 0.605395103 | 0.916376001 | Gm29666 | predicted gene 29666 [Source:MGI Symbo | 7.2279 | 3.0838 | 5.0312 | 5.7839 | 5.4531 | 10.025 |
| 39267 | ENSMUSG00000100968 | 0 | NA | NA | NA | NA | NA | Gm2098 | predicted gene 2098 [Source:MGI Symbol | 0 | 0 | 0 | 0 | 0 | 0 |
| 39268 | ENSMUSG00000100969 | 0 | NA | NA | NA | NA | NA | 1700030N03 | RIKEN cDNA 1700030N03 gene [Source:M | 0 | 0 | 0 | 0 | 0 | 0 |
| 39269 | ENSMUSG00000100970 | 0 | NA | NA | NA | NA | NA | Gm29270 | predicted gene 29270 [Source:MGI Symbo | 0 | 0 | 0 | 0 | 0 | 0 |
| 39270 | ENSMUSG00000100971 | 0 | NA | NA | NA | NA | NA | Gm28824 | predicted gene 28824 [Source:MGI Symbo | 0 | 0 | 0 | 0 | 0 | 0 |
| 39271 | ENSMUSG00000100972 | 0 | NA | NA | NA | NA | NA | Gm28553 | predicted gene 28553 [Source:MGI Symbo | 0 | 0 | 0 | 0 | 0 | 0 |
| 39272 | ENSMUSG00000100973 | 0 | NA | NA | NA | NA | NA | Gm29299 | predicted gene 29299 [Source:MGI Symbo | 0 | 0 | 0 | 0 | 0 | 0 |
| 39273 | ENSMUSG00000100974 | 0 | NA | NA | NA | NA | NA | Gm28459 | predicted gene 28459 [Source:MGI Symbo | 0 | 0 | 0 | 0 | 0 | 0 |
| 39274 | ENSMUSG00000100975 | 32.62881511 | 0.897802108 | 0.475378585 | 1.888604442 | 0.058944851 | 0.345033723 | Gm28875 | predicted gene 28875 [Source:MGI Symbo | 31.321 | 27.754 | 9.0562 | 49.163 | 28.356 | 50.123 |
| 39275 | ENSMUSG00000100976 | 0 | NA | NA | NA | NA | NA | Gm29134 | predicted gene 29134 [Source:MGI Symbo | 0 | 0 | 0 | 0 | 0 | 0 |
| 39276 | ENSMUSG00000100977 | 0 | NA | NA | NA | NA | NA | Gm29334 | predicted gene 29334 [Source:MGI Symbo | 0 | 0 | 0 | 0 | 0 | 0 |
| 39277 | ENSMUSG00000100978 | 0 | NA | NA | NA | NA | NA | Gm20912 | predicted gene, 20912 [Source:MGI Symb | 0 | 0 | 0 | 0 | 0 | 0 |
| 39278 | ENSMUSG00000100979 | 0 | NA | NA | NA | NA | NA | Gm29347 | predicted gene 29347 [Source:MGI Symbo | 0 | 0 | 0 | 0 | 0 | 0 |
| 39279 | ENSMUSG00000100980 | 0.667397476 | -1.404797968 | 2.804497115 | -0.500909044 | 0.616435137 | NA | Gm29100 | predicted gene 29100 [Source:MGI Symbo | 0 | 1.0279 | 2.0125 | 0.964 | 0 | 0 |
| 39280 | ENSMUSG00000100981 | 0 | NA | NA | NA | NA | NA | Gm29072 | predicted gene 29072 [Source:MGI Symbo | 0 | 0 | 0 | 0 | 0 | 0 |
| 39281 | ENSMUSG00000100982 | 0 | NA | NA | NA | NA | NA | Gm21837 | predicted gene, 21837 [Source:MGI Symb | 0 | 0 | 0 | 0 | 0 | 0 |
| 39282 | ENSMUSG00000100983 | 0 | NA | NA | NA | NA | NA | Gm29030 | predicted gene 29030 [Source:MGI Symbo | 0 | 0 | 0 | 0 | 0 | 0 |
| 39283 | ENSMUSG00000100984 | 0 | NA | NA | NA | NA | NA | Gm28357 | predicted gene 28357 [Source:MGI Symbo | 0 | 0 | 0 | 0 | 0 | 0 |
| 39284 | ENSMUSG00000100985 | 0 | NA | NA | NA | NA | NA | Gm10640 | predicted gene 10640 [Source:MGI Symbo | 0 | 0 | 0 | 0 | 0 | 0 |
| 39285 | ENSMUSG00000100986 | 0 | NA | NA | NA | NA | NA | Gm21741 | predicted gene, 21741 [Source:MGI Symb | 0 | 0 | 0 | 0 | 0 | 0 |
| 39286 | ENSMUSG00000100987 | 0.556921942 | 2.679043159 | 3.638243063 | 0.736356289 | 0.461513899 | NA | Gm1627 | predicted gene 1627 [Source:MGI Symbol | 0 | 0 | 0 | 0 | 0 | 3.3415 |
| 39287 | ENSMUSG00000100988 | 0 | NA | NA | NA | NA | NA | Gm28136 | predicted gene 28136 [Source:MGI Symbo | 0 | 0 | 0 | 0 | 0 | 0 |
| 39288 | ENSMUSG00000100989 | 0 | NA | NA | NA | NA | NA | Gm29647 | predicted gene 29647 [Source:MGI Symbo | 0 | 0 | 0 | 0 | 0 | 0 |
| 39289 | ENSMUSG00000100990 | 0 | NA | NA | NA | NA | NA | Gm7891 | predicted gene 7891 [Source:MGI Symbol | 0 | 0 | 0 | 0 | 0 | 0 |
| 39290 | ENSMUSG00000100991 | 0 | NA | NA | NA | NA | NA | Gm21856 | predicted gene, 21856 [Source:MGI Symb | 0 | 0 | 0 | 0 | 0 | 0 |
| 39291 | ENSMUSG00000100992 | 0 | NA | NA | NA | NA | NA | Gm23925 | predicted gene, 23925 [Source:MGI Symb | 0 | 0 | 0 | 0 | 0 | 0 |
| 39292 | ENSMUSG00000100993 | 0 | NA | NA | NA | NA | NA | Gm21895 | predicted gene, 21895 [Source:MGI Symb | 0 | 0 | 0 | 0 | 0 | 0 |
| 39293 | ENSMUSG00000100994 | 0 | NA | NA | NA | NA | NA | Gm21333 | predicted gene, 21333 [Source:MGI Symb | 0 | 0 | 0 | 0 | 0 | 0 |
| 39294 | ENSMUSG00000100995 | 0 | NA | NA | NA | NA | NA | Gm29525 | predicted gene 29525 [Source:MGI Symbo | 0 | 0 | 0 | 0 | 0 | 0 |
| 39295 | ENSMUSG00000100996 | 0 | NA | NA | NA | NA | NA | Gm29169 | predicted gene 29169 [Source:MGI Symbo | 0 | 0 | 0 | 0 | 0 | 0 |
| 39296 | ENSMUSG00000100998 | 0 | NA | NA | NA | NA | NA | Gm28864 | predicted gene 28864 [Source:MGI Symbo | 0 | 0 | 0 | 0 | 0 | 0 |
| 39297 | ENSMUSG00000100999 | 0 | NA | NA | NA | NA | NA | Gm29145 | predicted gene 29145 [Source:MGI Symbo | 0 | 0 | 0 | 0 | 0 | 0 |
| 39298 | ENSMUSG00000101000 | 0 | NA | NA | NA | NA | NA | Gm5371 | predicted gene 5371 [Source:MGI Symbol | 0 | 0 | 0 | 0 | 0 | 0 |
| 39299 | ENSMUSG00000101001 | 0 | NA | NA | NA | NA | NA | Gm3121 | predicted gene 3121 [Source:MGI Symbol | 0 | 0 | 0 | 0 | 0 | 0 |
| 39300 | ENSMUSG00000101002 | 0 | NA | NA | NA | NA | NA | Gm28944 | predicted gene 28944 [Source:MGI Symbo | 0 | 0 | 0 | 0 | 0 | 0 |
| 39301 | ENSMUSG00000101003 | 0 | NA | NA | NA | NA | NA | Gm28456 | predicted gene 28456 [Source:MGI Symbo | 0 | 0 | 0 | 0 | 0 | 0 |
| 39302 | ENSMUSG00000101004 | 0 | NA | NA | NA | NA | NA | Gm8596 | predicted gene 8596 [Source:MGI Symbol | 0 | 0 | 0 | 0 | 0 | 0 |
| 39303 | ENSMUSG00000101005 | 0 | NA | NA | NA | NA | NA | Gm20871 | predicted gene, 20871 [Source:MGI Symb | 0 | 0 | 0 | 0 | 0 | 0 |
| 39304 | ENSMUSG00000101006 | 0 | NA | NA | NA | NA | NA | Gm28299 | predicted gene 28299 [Source:MGI Symbo | 0 | 0 | 0 | 0 | 0 | 0 |
| 39305 | ENSMUSG00000101007 | 0 | NA | NA | NA | NA | NA | Gm29541 | predicted gene 29541 [Source:MGI Symbo | 0 | 0 | 0 | 0 | 0 | 0 |
| 39306 | ENSMUSG00000101008 | 0 | NA | NA | NA | NA | NA | Gm29442 | predicted gene 29442 [Source:MGI Symbo | 0 | 0 | 0 | 0 | 0 | 0 |
| 39307 | ENSMUSG00000101009 | 1.038420047 | 0.084391034 | 2.062151212 | 0.040923786 | 0.967356655 | NA | 1700108F19 | RIKEN cDNA 1700108F19 gene [Source:M | 0 | 2.0558 | 1.0062 | 0.964 | 1.0906 | 1.1138 |
| 39308 | ENSMUSG00000101010 | 0 | NA | NA | NA | NA | NA | Gm21889 | predicted gene, 21889 [Source:MGI Symb | 0 | 0 | 0 | 0 | 0 | 0 |

| | | | | | | | | | | | | | | | |
|---|---|---|---|---|---|---|---|---|---|---|---|---|---|---|---|
| 39309 | ENSMUSG00000101011 | 0 | NA | NA | NA | NA | NA | Gm29398 | predicted gene 29398 [Source:MGI Symbo | 0 | 0 | 0 | 0 | 0 | 0 |
| 39310 | ENSMUSG00000101012 | 0 | NA | NA | NA | NA | NA | 1700008K24 | RIKEN cDNA 1700008K24 gene [Source:M | 0 | 0 | 0 | 0 | 0 | 0 |
| 39311 | ENSMUSG00000101013 | 4.172025818 | -1.604699705 | 1.143251027 | -1.403628483 | 0.160429509 | NA | A630072M1 | RIKEN cDNA A630072M18 gene [Source:N | 9.6372 | 7.1954 | 2.0125 | 2.8919 | 2.1812 | 1.1138 |
| 39312 | ENSMUSG00000101014 | 0 | NA | NA | NA | NA | NA | Gm28289 | predicted gene 28289 [Source:MGI Symbo | 0 | 0 | 0 | 0 | 0 | 0 |
| 39313 | ENSMUSG00000101015 | 0 | NA | NA | NA | NA | NA | Gm18444 | predicted gene, 18444 [Source:MGI Symb | 0 | 0 | 0 | 0 | 0 | 0 |
| 39314 | ENSMUSG00000101016 | 0 | NA | NA | NA | NA | NA | Gm2325 | predicted gene 2325 [Source:MGI Symbol | 0 | 0 | 0 | 0 | 0 | 0 |
| 39315 | ENSMUSG00000101017 | 0 | NA | NA | NA | NA | NA | Gm21658 | predicted gene, 21658 [Source:MGI Symb | 0 | 0 | 0 | 0 | 0 | 0 |
| 39316 | ENSMUSG00000101018 | 0 | NA | NA | NA | NA | NA | Gm28910 | predicted gene 28910 [Source:MGI Symbo | 0 | 0 | 0 | 0 | 0 | 0 |
| 39317 | ENSMUSG00000101019 | 0 | NA | NA | NA | NA | NA | Gm28943 | predicted gene 28943 [Source:MGI Symbo | 0 | 0 | 0 | 0 | 0 | 0 |
| 39318 | ENSMUSG00000101020 | 0 | NA | NA | NA | NA | NA | Gm28377 | predicted gene 28377 [Source:MGI Symbo | 0 | 0 | 0 | 0 | 0 | 0 |
| 39319 | ENSMUSG00000101021 | 0 | NA | NA | NA | NA | NA | Gm29222 | predicted gene 29222 [Source:MGI Symbo | 0 | 0 | 0 | 0 | 0 | 0 |
| 39320 | ENSMUSG00000101022 | 0 | NA | NA | NA | NA | NA | Gm20921 | predicted gene, 20921 [Source:MGI Symb | 0 | 0 | 0 | 0 | 0 | 0 |
| 39321 | ENSMUSG00000101023 | 0 | NA | NA | NA | NA | NA | Gm21152 | predicted gene, 21152 [Source:MGI Symb | 0 | 0 | 0 | 0 | 0 | 0 |
| 39322 | ENSMUSG00000101024 | 0.17131964 | -0.796629176 | 4.080472857 | -0.195229623 | 0.84521319 | NA | Gm5692 | predicted gene 5692 [Source:MGI Symbol | 0 | 1.0279 | 0 | 0 | 0 | 0 |
| 39323 | ENSMUSG00000101025 | 0 | NA | NA | NA | NA | NA | Gm29262 | predicted gene 29262 [Source:MGI Symbo | 0 | 0 | 0 | 0 | 0 | 0 |
| 39324 | ENSMUSG00000101026 | 0 | NA | NA | NA | NA | NA | Ly6g6g | lymphocyte antigen 6 complex, locus G6G | 0 | 0 | 0 | 0 | 0 | 0 |
| 39325 | ENSMUSG00000101027 | 0 | NA | NA | NA | NA | NA | Gm28794 | predicted gene 28794 [Source:MGI Symbo | 0 | 0 | 0 | 0 | 0 | 0 |
| 39326 | ENSMUSG00000101028 | 0 | NA | NA | NA | NA | NA | Gm28723 | predicted gene 28723 [Source:MGI Symbo | 0 | 0 | 0 | 0 | 0 | 0 |
| 39327 | ENSMUSG00000101029 | 0 | NA | NA | NA | NA | NA | Gm29439 | predicted gene 29439 [Source:MGI Symbo | 0 | 0 | 0 | 0 | 0 | 0 |
| 39328 | ENSMUSG00000101030 | 0 | NA | NA | NA | NA | NA | Gm28429 | predicted gene 28429 [Source:MGI Symbo | 0 | 0 | 0 | 0 | 0 | 0 |
| 39329 | ENSMUSG00000101031 | 0 | NA | NA | NA | NA | NA | Ms4a12 | membrane-spanning 4-domains, subfamil | 0 | 0 | 0 | 0 | 0 | 0 |
| 39330 | ENSMUSG00000101032 | 0 | NA | NA | NA | NA | NA | Gm29377 | predicted gene 29377 [Source:MGI Symbo | 0 | 0 | 0 | 0 | 0 | 0 |
| 39331 | ENSMUSG00000101033 | 0 | NA | NA | NA | NA | NA | Gm18776 | predicted gene, 18776 [Source:MGI Symb | 0 | 0 | 0 | 0 | 0 | 0 |
| 39332 | ENSMUSG00000101035 | 0 | NA | NA | NA | NA | NA | Gm28451 | predicted gene 28451 [Source:MGI Symbo | 0 | 0 | 0 | 0 | 0 | 0 |
| 39333 | ENSMUSG00000101036 | 0 | NA | NA | NA | NA | NA | Gm28588 | predicted gene 28588 [Source:MGI Symbo | 0 | 0 | 0 | 0 | 0 | 0 |
| 39334 | ENSMUSG00000101037 | 10.08754696 | -0.5312995 | 0.72459321 | -0.733238308 | 0.463413094 | 0.861543316 | Gm28424 | predicted gene 28424 [Source:MGI Symbo | 18.471 | 6.1675 | 11.069 | 11.568 | 5.4531 | 7.7969 |
| 39335 | ENSMUSG00000101038 | 0 | NA | NA | NA | NA | NA | Gm28272 | predicted gene 28272 [Source:MGI Symbo | 0 | 0 | 0 | 0 | 0 | 0 |
| 39336 | ENSMUSG00000101039 | 0 | NA | NA | NA | NA | NA | Scgb2b29-ps | secretoglobin, family 2B, member 29, pse | 0 | 0 | 0 | 0 | 0 | 0 |
| 39337 | ENSMUSG00000101040 | 0 | NA | NA | NA | NA | NA | Gm29498 | predicted gene 29498 [Source:MGI Symbo | 0 | 0 | 0 | 0 | 0 | 0 |
| 39338 | ENSMUSG00000101041 | 0 | NA | NA | NA | NA | NA | Gm29353 | predicted gene 29353 [Source:MGI Symbo | 0 | 0 | 0 | 0 | 0 | 0 |
| 39339 | ENSMUSG00000101042 | 0 | NA | NA | NA | NA | NA | Gm28167 | predicted gene 28167 [Source:MGI Symbo | 0 | 0 | 0 | 0 | 0 | 0 |
| 39340 | ENSMUSG00000101043 | 0 | NA | NA | NA | NA | NA | Gm29075 | predicted gene 29075 [Source:MGI Symbo | 0 | 0 | 0 | 0 | 0 | 0 |
| 39341 | ENSMUSG00000101044 | 0 | NA | NA | NA | NA | NA | Gm28420 | predicted gene 28420 [Source:MGI Symbo | 0 | 0 | 0 | 0 | 0 | 0 |
| 39342 | ENSMUSG00000101045 | 0 | NA | NA | NA | NA | NA | Gm29224 | predicted gene 29224 [Source:MGI Symbo | 0 | 0 | 0 | 0 | 0 | 0 |
| 39343 | ENSMUSG00000101046 | 0 | NA | NA | NA | NA | NA | Gm28803 | predicted gene 28803 [Source:MGI Symbo | 0 | 0 | 0 | 0 | 0 | 0 |
| 39344 | ENSMUSG00000101049 | 0 | NA | NA | NA | NA | NA | Gm8856 | predicted gene 8856 [Source:MGI Symbol | 0 | 0 | 0 | 0 | 0 | 0 |
| 39345 | ENSMUSG00000101050 | 0 | NA | NA | NA | NA | NA | Gm28774 | predicted gene 28774 [Source:MGI Symbo | 0 | 0 | 0 | 0 | 0 | 0 |
| 39346 | ENSMUSG00000101051 | 0 | NA | NA | NA | NA | NA | 1700100L14 | RIKEN cDNA 1700100L14 gene [Source:M | 0 | 0 | 0 | 0 | 0 | 0 |
| 39347 | ENSMUSG00000101052 | 0 | NA | NA | NA | NA | NA | Gm28471 | predicted gene 28471 [Source:MGI Symbo | 0 | 0 | 0 | 0 | 0 | 0 |
| 39348 | ENSMUSG00000101053 | 0 | NA | NA | NA | NA | NA | Gm20815 | predicted gene, 20815 [Source:MGI Symb | 0 | 0 | 0 | 0 | 0 | 0 |
| 39349 | ENSMUSG00000101054 | 0 | NA | NA | NA | NA | NA | Gm18406 | predicted gene, 18406 [Source:MGI Symb | 0 | 0 | 0 | 0 | 0 | 0 |
| 39350 | ENSMUSG00000101055 | 0 | NA | NA | NA | NA | NA | Gm28443 | predicted gene 28443 [Source:MGI Symbo | 0 | 0 | 0 | 0 | 0 | 0 |
| 39351 | ENSMUSG00000101056 | 0 | NA | NA | NA | NA | NA | Gm28329 | predicted gene 28329 [Source:MGI Symbo | 0 | 0 | 0 | 0 | 0 | 0 |
| 39352 | ENSMUSG00000101057 | 0 | NA | NA | NA | NA | NA | Gm29057 | predicted gene 29057 [Source:MGI Symbo | 0 | 0 | 0 | 0 | 0 | 0 |
| 39353 | ENSMUSG00000101058 | 0 | NA | NA | NA | NA | NA | Gm21829 | predicted gene, 21829 [Source:MGI Symb | 0 | 0 | 0 | 0 | 0 | 0 |
| 39354 | ENSMUSG00000101059 | 2.831739292 | -0.354338425 | 1.227496449 | -0.288667577 | 0.772835777 | NA | Gm4017 | predicted gene 4017 [Source:MGI Symbol | 2.4093 | 4.1117 | 3.0187 | 1.928 | 2.1812 | 3.3415 |
| 39355 | ENSMUSG00000101060 | 0 | NA | NA | NA | NA | NA | Gm29565 | predicted gene 29565 [Source:MGI Symbo | 0 | 0 | 0 | 0 | 0 | 0 |
| 39356 | ENSMUSG00000101061 | 0 | NA | NA | NA | NA | NA | Platr1 | pluripotency associated transcript 1 [Sour | 0 | 0 | 0 | 0 | 0 | 0 |
| 39357 | ENSMUSG00000101062 | 0 | NA | NA | NA | NA | NA | Gm28821 | predicted gene 28821 [Source:MGI Symbo | 0 | 0 | 0 | 0 | 0 | 0 |
| 39358 | ENSMUSG00000101063 | 0 | NA | NA | NA | NA | NA | Mrgprb7-ps | MAS-related GPR, member B7, pseudoger | 0 | 0 | 0 | 0 | 0 | 0 |
| 39359 | ENSMUSG00000101064 | 0 | NA | NA | NA | NA | NA | Gm18798 | predicted gene, 18798 [Source:MGI Symb | 0 | 0 | 0 | 0 | 0 | 0 |
| 39360 | ENSMUSG00000101065 | 0 | NA | NA | NA | NA | NA | Scgb1b5-ps | secretoglobin, family 1B, member 5, pseu | 0 | 0 | 0 | 0 | 0 | 0 |
| 39361 | ENSMUSG00000101066 | 0 | NA | NA | NA | NA | NA | Gm8498 | predicted gene 8498 [Source:MGI Symbol | 0 | 0 | 0 | 0 | 0 | 0 |
| 39362 | ENSMUSG00000101067 | 0 | NA | NA | NA | NA | NA | Gm29007 | predicted gene 29007 [Source:MGI Symbo | 0 | 0 | 0 | 0 | 0 | 0 |
| 39363 | ENSMUSG00000101068 | 0 | NA | NA | NA | NA | NA | Gm28759 | predicted gene 28759 [Source:MGI Symbo | 0 | 0 | 0 | 0 | 0 | 0 |
| 39364 | ENSMUSG00000101069 | 0 | NA | NA | NA | NA | NA | Gm28194 | predicted gene 28194 [Source:MGI Symbo | 0 | 0 | 0 | 0 | 0 | 0 |

| | | | | | | | | | | | | | | | |
|---|---|---|---|---|---|---|---|---|---|---|---|---|---|---|---|
| 39365 | ENSMUSG00000101070 | 0 | NA | NA | NA | NA | NA | Gm28695 | predicted gene 28695 [Source:MGI Symbo | 0 | 0 | 0 | 0 | 0 | 0 |
| 39366 | ENSMUSG00000101071 | 0 | NA | NA | NA | NA | NA | Gm29577 | predicted gene 29577 [Source:MGI Symbo | 0 | 0 | 0 | 0 | 0 | 0 |
| 39367 | ENSMUSG00000101072 | 0 | NA | NA | NA | NA | NA | Gm29470 | predicted gene 29470 [Source:MGI Symbo | 0 | 0 | 0 | 0 | 0 | 0 |
| 39368 | ENSMUSG00000101073 | 0 | NA | NA | NA | NA | NA | Vmn1r200 | vomeronasal 1 receptor 200 [Source:MGI | 0 | 0 | 0 | 0 | 0 | 0 |
| 39369 | ENSMUSG00000101074 | 0 | NA | NA | NA | NA | NA | Gm29022 | predicted gene 29022 [Source:MGI Symbo | 0 | 0 | 0 | 0 | 0 | 0 |
| 39370 | ENSMUSG00000101075 | 0 | NA | NA | NA | NA | NA | Gm29517 | predicted gene 29517 [Source:MGI Symbo | 0 | 0 | 0 | 0 | 0 | 0 |
| 39371 | ENSMUSG00000101076 | 0 | NA | NA | NA | NA | NA | Gm4545 | predicted gene 4545 [Source:MGI Symbol | 0 | 0 | 0 | 0 | 0 | 0 |
| 39372 | ENSMUSG00000101077 | 0.160662974 | 1.126924175 | 4.080472857 | 0.276174898 | 0.782413736 | NA | Gm29164 | predicted gene 29164 [Source:MGI Symbo | 0 | 0 | 0 | 0.964 | 0 | 0 |
| 39373 | ENSMUSG00000101078 | 0 | NA | NA | NA | NA | NA | Olfr1166 | olfactory receptor 1166 [Source:MGI Sym | 0 | 0 | 0 | 0 | 0 | 0 |
| 39374 | ENSMUSG00000101079 | 0 | NA | NA | NA | NA | NA | Gm28991 | predicted gene 28991 [Source:MGI Symbo | 0 | 0 | 0 | 0 | 0 | 0 |
| 39375 | ENSMUSG00000101080 | 0 | NA | NA | NA | NA | NA | Gm28633 | predicted gene 28633 [Source:MGI Symbo | 0 | 0 | 0 | 0 | 0 | 0 |
| 39376 | ENSMUSG00000101081 | 0 | NA | NA | NA | NA | NA | Gm29193 | predicted gene 29193 [Source:MGI Symbo | 0 | 0 | 0 | 0 | 0 | 0 |
| 39377 | ENSMUSG00000101082 | 0.294513347 | 0.165143496 | 4.080472857 | 0.040471657 | 0.967717103 | NA | Gm28069 | predicted gene 28069 [Source:MGI Symbo | 0.8031 | 0 | 0 | 0.964 | 0 | 0 |
| 39378 | ENSMUSG00000101083 | 0 | NA | NA | NA | NA | NA | Gm29631 | predicted gene 29631 [Source:MGI Symbo | 0 | 0 | 0 | 0 | 0 | 0 |
| 39379 | ENSMUSG00000101084 | 0 | NA | NA | NA | NA | NA | Gm28228 | predicted gene 28228 [Source:MGI Symbo | 0 | 0 | 0 | 0 | 0 | 0 |
| 39380 | ENSMUSG00000101085 | 0 | NA | NA | NA | NA | NA | Gm8531 | predicted gene 8531 [Source:MGI Symbol | 0 | 0 | 0 | 0 | 0 | 0 |
| 39381 | ENSMUSG00000101086 | 0 | NA | NA | NA | NA | NA | Gm28651 | predicted gene 28651 [Source:MGI Symbo | 0 | 0 | 0 | 0 | 0 | 0 |
| 39382 | ENSMUSG00000101087 | 0 | NA | NA | NA | NA | NA | Gm2707 | predicted gene 2707 [Source:MGI Symbol | 0 | 0 | 0 | 0 | 0 | 0 |
| 39383 | ENSMUSG00000101088 | 0 | NA | NA | NA | NA | NA | 1810018F18 | RIKEN cDNA 1810018F18 gene [Source:M | 0 | 0 | 0 | 0 | 0 | 0 |
| 39384 | ENSMUSG00000101089 | 0 | NA | NA | NA | NA | NA | 2610016A17 | RIKEN cDNA 2610016A17 gene [Source:M | 0 | 0 | 0 | 0 | 0 | 0 |
| 39385 | ENSMUSG00000101090 | 0 | NA | NA | NA | NA | NA | Gm28619 | predicted gene 28619 [Source:MGI Symbo | 0 | 0 | 0 | 0 | 0 | 0 |
| 39386 | ENSMUSG00000101091 | 0 | NA | NA | NA | NA | NA | Gm29384 | predicted gene 29384 [Source:MGI Symbo | 0 | 0 | 0 | 0 | 0 | 0 |
| 39387 | ENSMUSG00000101092 | 0 | NA | NA | NA | NA | NA | Gm21796 | predicted gene, 21796 [Source:MGI Symb | 0 | 0 | 0 | 0 | 0 | 0 |
| 39388 | ENSMUSG00000101093 | 0 | NA | NA | NA | NA | NA | Eef1a1-ps2 | eukaryotic translation elongation factor 1 | 0 | 0 | 0 | 0 | 0 | 0 |
| 39389 | ENSMUSG00000101094 | 0 | NA | NA | NA | NA | NA | Gm29533 | predicted gene 29533 [Source:MGI Symbo | 0 | 0 | 0 | 0 | 0 | 0 |
| 39390 | ENSMUSG00000101095 | 0 | NA | NA | NA | NA | NA | Gm29329 | predicted gene 29329 [Source:MGI Symbo | 0 | 0 | 0 | 0 | 0 | 0 |
| 39391 | ENSMUSG00000101096 | 0 | NA | NA | NA | NA | NA | Gm21524 | predicted gene, 21524 [Source:MGI Symb | 0 | 0 | 0 | 0 | 0 | 0 |
| 39392 | ENSMUSG00000101097 | 0 | NA | NA | NA | NA | NA | Gm6679 | predicted gene 6679 [Source:MGI Symbol | 0 | 0 | 0 | 0 | 0 | 0 |
| 39393 | ENSMUSG00000101098 | 0 | NA | NA | NA | NA | NA | Gm20808 | predicted gene, 20808 [Source:MGI Symb | 0 | 0 | 0 | 0 | 0 | 0 |
| 39394 | ENSMUSG00000101099 | 0 | NA | NA | NA | NA | NA | Gm29191 | predicted gene 29191 [Source:MGI Symbo | 0 | 0 | 0 | 0 | 0 | 0 |
| 39395 | ENSMUSG00000101100 | 0 | NA | NA | NA | NA | NA | Gm28326 | predicted gene 28326 [Source:MGI Symbo | 0 | 0 | 0 | 0 | 0 | 0 |
| 39396 | ENSMUSG00000101101 | 0 | NA | NA | NA | NA | NA | Gm18350 | predicted gene, 18350 [Source:MGI Symb | 0 | 0 | 0 | 0 | 0 | 0 |
| 39397 | ENSMUSG00000101102 | 0 | NA | NA | NA | NA | NA | Gm28609 | predicted gene 28609 [Source:MGI Symbo | 0 | 0 | 0 | 0 | 0 | 0 |
| 39398 | ENSMUSG00000101103 | 0 | NA | NA | NA | NA | NA | Gm21843 | predicted gene, 21843 [Source:MGI Symb | 0 | 0 | 0 | 0 | 0 | 0 |
| 39399 | ENSMUSG00000101104 | 0 | NA | NA | NA | NA | NA | Gm29247 | predicted gene 29247 [Source:MGI Symbo | 0 | 0 | 0 | 0 | 0 | 0 |
| 39400 | ENSMUSG00000101105 | 0 | NA | NA | NA | NA | NA | Gm21755 | predicted gene, 21755 [Source:MGI Symb | 0 | 0 | 0 | 0 | 0 | 0 |
| 39401 | ENSMUSG00000101106 | 0 | NA | NA | NA | NA | NA | Gm28819 | predicted gene 28819 [Source:MGI Symbo | 0 | 0 | 0 | 0 | 0 | 0 |
| 39402 | ENSMUSG00000101107 | 0 | NA | NA | NA | NA | NA | Gm28587 | predicted gene 28587 [Source:MGI Symbo | 0 | 0 | 0 | 0 | 0 | 0 |
| 39403 | ENSMUSG00000101108 | 0 | NA | NA | NA | NA | NA | Gm29038 | predicted gene 29038 [Source:MGI Symbo | 0 | 0 | 0 | 0 | 0 | 0 |
| 39404 | ENSMUSG00000101109 | 0 | NA | NA | NA | NA | NA | Gm28697 | predicted gene 28697 [Source:MGI Symbo | 0 | 0 | 0 | 0 | 0 | 0 |
| 39405 | ENSMUSG00000101110 | 0 | NA | NA | NA | NA | NA | Gm28890 | predicted gene 28890 [Source:MGI Symbo | 0 | 0 | 0 | 0 | 0 | 0 |
| 39406 | ENSMUSG00000101111 | 461.0721789 | 0.106465435 | 0.150111126 | 0.709244127 | 0.478172994 | 0.869794311 | Gm28437 | predicted gene 28437 [Source:MGI Symbo | 480.26 | 460.51 | 390.42 | 427.04 | 462.42 | 545.78 |
| 39407 | ENSMUSG00000101112 | 0 | NA | NA | NA | NA | NA | Gm29185 | predicted gene 29185 [Source:MGI Symbo | 0 | 0 | 0 | 0 | 0 | 0 |
| 39408 | ENSMUSG00000101113 | 0 | NA | NA | NA | NA | NA | 1700025B11 | RIKEN cDNA 1700025B11 gene [Source:M | 0 | 0 | 0 | 0 | 0 | 0 |
| 39409 | ENSMUSG00000101114 | 0 | NA | NA | NA | NA | NA | Gm28628 | predicted gene 28628 [Source:MGI Symbo | 0 | 0 | 0 | 0 | 0 | 0 |
| 39410 | ENSMUSG00000101115 | 0 | NA | NA | NA | NA | NA | Gm28664 | predicted gene 28664 [Source:MGI Symbo | 0 | 0 | 0 | 0 | 0 | 0 |
| 39411 | ENSMUSG00000101116 | 0 | NA | NA | NA | NA | NA | Gm28416 | predicted gene 28416 [Source:MGI Symbo | 0 | 0 | 0 | 0 | 0 | 0 |
| 39412 | ENSMUSG00000101117 | 0 | NA | NA | NA | NA | NA | Gm28341 | predicted gene 28341 [Source:MGI Symbo | 0 | 0 | 0 | 0 | 0 | 0 |
| 39413 | ENSMUSG00000101118 | 0 | NA | NA | NA | NA | NA | Gm29034 | predicted gene 29034 [Source:MGI Symbo | 0 | 0 | 0 | 0 | 0 | 0 |
| 39414 | ENSMUSG00000101119 | 0 | NA | NA | NA | NA | NA | Gm29060 | predicted gene 29060 [Source:MGI Symbo | 0 | 0 | 0 | 0 | 0 | 0 |
| 39415 | ENSMUSG00000101120 | 0 | NA | NA | NA | NA | NA | Gm29452 | predicted gene 29452 [Source:MGI Symbo | 0 | 0 | 0 | 0 | 0 | 0 |
| 39416 | ENSMUSG00000101122 | 15.51200479 | 0.86214876 | 0.59766588 | 1.442526315 | 0.149153954 | 0.561319344 | Gm17971 | predicted gene, 17971 [Source:MGI Symb | 14.456 | 14.391 | 4.025 | 18.316 | 20.722 | 21.163 |
| 39417 | ENSMUSG00000101123 | 0 | NA | NA | NA | NA | NA | Gm28654 | predicted gene 28654 [Source:MGI Symbo | 0 | 0 | 0 | 0 | 0 | 0 |
| 39418 | ENSMUSG00000101124 | 0 | NA | NA | NA | NA | NA | Gm28371 | predicted gene 28371 [Source:MGI Symbo | 0 | 0 | 0 | 0 | 0 | 0 |
| 39419 | ENSMUSG00000101125 | 0 | NA | NA | NA | NA | NA | Gm29625 | predicted gene 29625 [Source:MGI Symbo | 0 | 0 | 0 | 0 | 0 | 0 |
| 39420 | ENSMUSG00000101126 | 0 | NA | NA | NA | NA | NA | Gm10538 | predicted gene 10538 [Source:MGI Symbo | 0 | 0 | 0 | 0 | 0 | 0 |

| | | | | | | | | | | | | | | | |
|---|---|---|---|---|---|---|---|---|---|---|---|---|---|---|---|
| 39421 | ENSMUSG00000101127 | 0 | NA | NA | NA | NA | NA | Gm28585 | predicted gene 28585 [Source:MGI Symbo | 0 | 0 | 0 | 0 | 0 | 0 |
| 39422 | ENSMUSG00000101128 | 0 | NA | NA | NA | NA | NA | Gm21869 | predicted gene, 21869 [Source:MGI Symb | 0 | 0 | 0 | 0 | 0 | 0 |
| 39423 | ENSMUSG00000101129 | 0 | NA | NA | NA | NA | NA | Gm5254 | predicted gene 5254 [Source:MGI Symbol | 0 | 0 | 0 | 0 | 0 | 0 |
| 39424 | ENSMUSG00000101130 | 0 | NA | NA | NA | NA | NA | Gm19637 | predicted gene, 19637 [Source:MGI Symb | 0 | 0 | 0 | 0 | 0 | 0 |
| 39425 | ENSMUSG00000101131 | 0 | NA | NA | NA | NA | NA | Gm29370 | predicted gene 29370 [Source:MGI Symbo | 0 | 0 | 0 | 0 | 0 | 0 |
| 39426 | ENSMUSG00000101132 | 0.181769715 | 1.126924175 | 4.080472857 | 0.276174898 | 0.782413736 | NA | Gm8000 | predicted gene 8000 [Source:MGI Symbol | 0 | 0 | 0 | 0 | 1.0906 | 0 |
| 39427 | ENSMUSG00000101133 | 1.586106543 | 0.185815366 | 1.715039116 | 0.108344681 | 0.913722281 | NA | Gm29050 | predicted gene 29050 [Source:MGI Symbo | 2.4093 | 1.0279 | 1.0062 | 2.8919 | 2.1812 | 0 |
| 39428 | ENSMUSG00000101134 | 0 | NA | NA | NA | NA | NA | Gm28482 | predicted gene 28482 [Source:MGI Symbo | 0 | 0 | 0 | 0 | 0 | 0 |
| 39429 | ENSMUSG00000101135 | 0 | NA | NA | NA | NA | NA | Gm17981 | predicted gene, 17981 [Source:MGI Symb | 0 | 0 | 0 | 0 | 0 | 0 |
| 39430 | ENSMUSG00000101136 | 0 | NA | NA | NA | NA | NA | Gm8739 | predicted gene 8739 [Source:MGI Symbol | 0 | 0 | 0 | 0 | 0 | 0 |
| 39431 | ENSMUSG00000101137 | 0 | NA | NA | NA | NA | NA | Rpl23a-ps7 | ribosomal protein L23A, pseudogene 7 [Sc | 0 | 0 | 0 | 0 | 0 | 0 |
| 39432 | ENSMUSG00000101138 | 0 | NA | NA | NA | NA | NA | Gm29210 | predicted gene 29210 [Source:MGI Symbo | 0 | 0 | 0 | 0 | 0 | 0 |
| 39433 | ENSMUSG00000101139 | 0 | NA | NA | NA | NA | NA | Gm29639 | predicted gene 29639 [Source:MGI Symbo | 0 | 0 | 0 | 0 | 0 | 0 |
| 39434 | ENSMUSG00000101140 | 0.335414862 | -1.779047242 | 4.030804179 | -0.441362855 | 0.658950329 | NA | Gm28117 | predicted gene 28117 [Source:MGI Symbo | 0 | 0 | 2.0125 | 0 | 0 | 0 |
| 39435 | ENSMUSG00000101141 | 0 | NA | NA | NA | NA | NA | Gm21139 | predicted gene, 21139 [Source:MGI Symb | 0 | 0 | 0 | 0 | 0 | 0 |
| 39436 | ENSMUSG00000101142 | 0 | NA | NA | NA | NA | NA | Gm28142 | predicted gene 28142 [Source:MGI Symbo | 0 | 0 | 0 | 0 | 0 | 0 |
| 39437 | ENSMUSG00000101143 | 0 | NA | NA | NA | NA | NA | Gm21151 | predicted gene, 21151 [Source:MGI Symb | 0 | 0 | 0 | 0 | 0 | 0 |
| 39438 | ENSMUSG00000101144 | 0 | NA | NA | NA | NA | NA | Gm29054 | predicted gene 29054 [Source:MGI Symbo | 0 | 0 | 0 | 0 | 0 | 0 |
| 39439 | ENSMUSG00000101145 | 0 | NA | NA | NA | NA | NA | Gm28412 | predicted gene 28412 [Source:MGI Symbo | 0 | 0 | 0 | 0 | 0 | 0 |
| 39440 | ENSMUSG00000101146 | 0 | NA | NA | NA | NA | NA | Gm20814 | predicted gene, 20814 [Source:MGI Symb | 0 | 0 | 0 | 0 | 0 | 0 |
| 39441 | ENSMUSG00000101147 | 0 | NA | NA | NA | NA | NA | Gm20818 | predicted gene, 20818 [Source:MGI Symb | 0 | 0 | 0 | 0 | 0 | 0 |
| 39442 | ENSMUSG00000101148 | 0 | NA | NA | NA | NA | NA | Gm28200 | predicted gene 28200 [Source:MGI Symbo | 0 | 0 | 0 | 0 | 0 | 0 |
| 39443 | ENSMUSG00000101149 | 0 | NA | NA | NA | NA | NA | Gm20892 | predicted gene, 20892 [Source:MGI Symb | 0 | 0 | 0 | 0 | 0 | 0 |
| 39444 | ENSMUSG00000101150 | 0 | NA | NA | NA | NA | NA | Gm29405 | predicted gene 29405 [Source:MGI Symbo | 0 | 0 | 0 | 0 | 0 | 0 |
| 39445 | ENSMUSG00000101151 | 0 | NA | NA | NA | NA | NA | Gm17929 | predicted gene, 17929 [Source:MGI Symb | 0 | 0 | 0 | 0 | 0 | 0 |
| 39446 | ENSMUSG00000101152 | 0 | NA | NA | NA | NA | NA | Gm28305 | predicted gene 28305 [Source:MGI Symbo | 0 | 0 | 0 | 0 | 0 | 0 |
| 39447 | ENSMUSG00000101153 | 0 | NA | NA | NA | NA | NA | 4933404I11F | RIKEN cDNA 4933404I11 gene [Source:MC | 0 | 0 | 0 | 0 | 0 | 0 |
| 39448 | ENSMUSG00000101154 | 0 | NA | NA | NA | NA | NA | Gm17777 | predicted gene, 17777 [Source:MGI Symb | 0 | 0 | 0 | 0 | 0 | 0 |
| 39449 | ENSMUSG00000101155 | 0 | NA | NA | NA | NA | NA | Sly | Sycp3 like Y-linked [Source:MGI Symbol;A | 0 | 0 | 0 | 0 | 0 | 0 |
| 39450 | ENSMUSG00000101156 | 0 | NA | NA | NA | NA | NA | Gm29114 | predicted gene 29114 [Source:MGI Symbo | 0 | 0 | 0 | 0 | 0 | 0 |
| 39451 | ENSMUSG00000101157 | 0 | NA | NA | NA | NA | NA | Gm20905 | predicted gene, 20905 [Source:MGI Symb | 0 | 0 | 0 | 0 | 0 | 0 |
| 39452 | ENSMUSG00000101158 | 0 | NA | NA | NA | NA | NA | Gm20894 | predicted gene, 20894 [Source:MGI Symb | 0 | 0 | 0 | 0 | 0 | 0 |
| 39453 | ENSMUSG00000101159 | 0 | NA | NA | NA | NA | NA | Gm28184 | predicted gene 28184 [Source:MGI Symbo | 0 | 0 | 0 | 0 | 0 | 0 |
| 39454 | ENSMUSG00000101160 | 0 | NA | NA | NA | NA | NA | Gm29338 | predicted gene 29338 [Source:MGI Symbo | 0 | 0 | 0 | 0 | 0 | 0 |
| 39455 | ENSMUSG00000101161 | 0 | NA | NA | NA | NA | NA | Gm21799 | predicted gene, 21799 [Source:MGI Symb | 0 | 0 | 0 | 0 | 0 | 0 |
| 39456 | ENSMUSG00000101162 | 0.335414862 | -1.779047242 | 4.030804179 | -0.441362855 | 0.658950329 | NA | Gm26728 | predicted gene, 26728 [Source:MGI Symb | 0 | 0 | 2.0125 | 0 | 0 | 0 |
| 39457 | ENSMUSG00000101163 | 0 | NA | NA | NA | NA | NA | Gm13278 | predicted gene 13278 [Source:MGI Symbo | 0 | 0 | 0 | 0 | 0 | 0 |
| 39458 | ENSMUSG00000101164 | 0 | NA | NA | NA | NA | NA | Gm29218 | predicted gene 29218 [Source:MGI Symbo | 0 | 0 | 0 | 0 | 0 | 0 |
| 39459 | ENSMUSG00000101166 | 0 | NA | NA | NA | NA | NA | Gm28496 | predicted gene 28496 [Source:MGI Symbo | 0 | 0 | 0 | 0 | 0 | 0 |
| 39460 | ENSMUSG00000101167 | 0.305170013 | -1.657836477 | 4.041480335 | -0.410205257 | 0.681655385 | NA | Macroh2a3 | macroH2A.3 histone [Source:MGI Symbol | 0.8031 | 1.0279 | 0 | 0 | 0 | 0 |
| 39461 | ENSMUSG00000101168 | 0.181769715 | 1.126924175 | 4.080472857 | 0.276174898 | 0.782413736 | NA | Gm28892 | predicted gene 28892 [Source:MGI Symbo | 0 | 0 | 0 | 0 | 1.0906 | 0 |
| 39462 | ENSMUSG00000101170 | 0 | NA | NA | NA | NA | NA | Gm29343 | predicted gene 29343 [Source:MGI Symbo | 0 | 0 | 0 | 0 | 0 | 0 |
| 39463 | ENSMUSG00000101171 | 0 | NA | NA | NA | NA | NA | Gm7593 | predicted gene 7593 [Source:MGI Symbol | 0 | 0 | 0 | 0 | 0 | 0 |
| 39464 | ENSMUSG00000101172 | 0 | NA | NA | NA | NA | NA | Gm28535 | predicted gene 28535 [Source:MGI Symbo | 0 | 0 | 0 | 0 | 0 | 0 |
| 39465 | ENSMUSG00000101173 | 0 | NA | NA | NA | NA | NA | Gm29519 | predicted gene 29519 [Source:MGI Symbo | 0 | 0 | 0 | 0 | 0 | 0 |
| 39466 | ENSMUSG00000101174 | 0 | NA | NA | NA | NA | NA | Hoxd4 | homeobox D4 [Source:MGI Symbol;Acc:M | 0 | 0 | 0 | 0 | 0 | 0 |
| 39467 | ENSMUSG00000101177 | 0 | NA | NA | NA | NA | NA | Gm29586 | predicted gene 29586 [Source:MGI Symbo | 0 | 0 | 0 | 0 | 0 | 0 |
| 39468 | ENSMUSG00000101178 | 0 | NA | NA | NA | NA | NA | Gm28909 | predicted gene 28909 [Source:MGI Symbo | 0 | 0 | 0 | 0 | 0 | 0 |
| 39469 | ENSMUSG00000101179 | 0 | NA | NA | NA | NA | NA | Gm29455 | predicted gene 29455 [Source:MGI Symbo | 0 | 0 | 0 | 0 | 0 | 0 |
| 39470 | ENSMUSG00000101180 | 0 | NA | NA | NA | NA | NA | Gm2914 | predicted gene 2914 [Source:MGI Symbol | 0 | 0 | 0 | 0 | 0 | 0 |
| 39471 | ENSMUSG00000101181 | 0 | NA | NA | NA | NA | NA | Vmn2r-ps14 | vomeronasal 2, receptor, pseudogene 14 | 0 | 0 | 0 | 0 | 0 | 0 |
| 39472 | ENSMUSG00000101182 | 0 | NA | NA | NA | NA | NA | Gm29209 | predicted gene 29209 [Source:MGI Symbo | 0 | 0 | 0 | 0 | 0 | 0 |
| 39473 | ENSMUSG00000101183 | 0 | NA | NA | NA | NA | NA | Gm28614 | predicted gene 28614 [Source:MGI Symbo | 0 | 0 | 0 | 0 | 0 | 0 |
| 39474 | ENSMUSG00000101184 | 0 | NA | NA | NA | NA | NA | Gm28861 | predicted gene 28861 [Source:MGI Symbo | 0 | 0 | 0 | 0 | 0 | 0 |
| 39475 | ENSMUSG00000101185 | 0 | NA | NA | NA | NA | NA | Scgb1b16-ps | secretoglobin, family 1B, member 16, pse | 0 | 0 | 0 | 0 | 0 | 0 |
| 39476 | ENSMUSG00000101186 | 0 | NA | NA | NA | NA | NA | Gm29235 | predicted gene 29235 [Source:MGI Symbo | 0 | 0 | 0 | 0 | 0 | 0 |

| | | | | | | | | | | | | | | | |
|---|---|---|---|---|---|---|---|---|---|---|---|---|---|---|---|
| 39477 | ENSMUSG00000101188 | 30.25107037 | 0.002306916 | 0.395303701 | 0.005835807 | 0.995343726 | 0.998796424 | Eif4a-ps4 | eukaryotic translation initiation factor 4A, | 25.699 | 32.893 | 32.2 | 28.919 | 27.265 | 34.529 |
| 39478 | ENSMUSG00000101189 | 0 | NA | NA | NA | NA | NA | 1700029M03 | RIKEN cDNA 1700029M03 gene [Source:M | 0 | 0 | 0 | 0 | 0 | 0 |
| 39479 | ENSMUSG00000101190 | 0 | NA | NA | NA | NA | NA | Gm28274 | predicted gene 28274 [Source:MGI Symbo | 0 | 0 | 0 | 0 | 0 | 0 |
| 39480 | ENSMUSG00000101191 | 0.342639279 | -1.805976596 | 4.028547714 | -0.448294702 | 0.653940524 | NA | Gm28809 | predicted gene 28809 [Source:MGI Symbo | 0 | 2.0558 | 0 | 0 | 0 | 0 |
| 39481 | ENSMUSG00000101192 | 0 | NA | NA | NA | NA | NA | Gm29122 | predicted gene 29122 [Source:MGI Symbo | 0 | 0 | 0 | 0 | 0 | 0 |
| 39482 | ENSMUSG00000101193 | 0 | NA | NA | NA | NA | NA | Gm21849 | predicted gene, 21849 [Source:MGI Symb | 0 | 0 | 0 | 0 | 0 | 0 |
| 39483 | ENSMUSG00000101194 | 0 | NA | NA | NA | NA | NA | Gm29149 | predicted gene 29149 [Source:MGI Symbo | 0 | 0 | 0 | 0 | 0 | 0 |
| 39484 | ENSMUSG00000101195 | 0 | NA | NA | NA | NA | NA | Gm9239 | predicted gene 9239 [Source:MGI Symbol | 0 | 0 | 0 | 0 | 0 | 0 |
| 39485 | ENSMUSG00000101196 | 0 | NA | NA | NA | NA | NA | Gm6866 | predicted gene 6866 [Source:MGI Symbol | 0 | 0 | 0 | 0 | 0 | 0 |
| 39486 | ENSMUSG00000101197 | 0 | NA | NA | NA | NA | NA | Gm29032 | predicted gene 29032 [Source:MGI Symbo | 0 | 0 | 0 | 0 | 0 | 0 |
| 39487 | ENSMUSG00000101198 | 0 | NA | NA | NA | NA | NA | Gm29021 | predicted gene 29021 [Source:MGI Symbo | 0 | 0 | 0 | 0 | 0 | 0 |
| 39488 | ENSMUSG00000101199 | 0 | NA | NA | NA | NA | NA | Gm9687 | predicted gene 9687 [Source:MGI Symbol | 0 | 0 | 0 | 0 | 0 | 0 |
| 39489 | ENSMUSG00000101200 | 0 | NA | NA | NA | NA | NA | Gm20802 | predicted gene, 20802 [Source:MGI Symb | 0 | 0 | 0 | 0 | 0 | 0 |
| 39490 | ENSMUSG00000101201 | 0 | NA | NA | NA | NA | NA | Gm28125 | predicted gene 28125 [Source:MGI Symbo | 0 | 0 | 0 | 0 | 0 | 0 |
| 39491 | ENSMUSG00000101202 | 0 | NA | NA | NA | NA | NA | Gm28749 | predicted gene 28749 [Source:MGI Symbo | 0 | 0 | 0 | 0 | 0 | 0 |
| 39492 | ENSMUSG00000101203 | 0 | NA | NA | NA | NA | NA | Gm28615 | predicted gene 28615 [Source:MGI Symbo | 0 | 0 | 0 | 0 | 0 | 0 |
| 39493 | ENSMUSG00000101204 | 0 | NA | NA | NA | NA | NA | Gm29271 | predicted gene 29271 [Source:MGI Symbo | 0 | 0 | 0 | 0 | 0 | 0 |
| 39494 | ENSMUSG00000101205 | 0 | NA | NA | NA | NA | NA | Gm29308 | predicted gene 29308 [Source:MGI Symbo | 0 | 0 | 0 | 0 | 0 | 0 |
| 39495 | ENSMUSG00000101206 | 0 | NA | NA | NA | NA | NA | Gm5266 | predicted gene 5266 [Source:MGI Symbol | 0 | 0 | 0 | 0 | 0 | 0 |
| 39496 | ENSMUSG00000101207 | 0 | NA | NA | NA | NA | NA | Gm28310 | predicted gene 28310 [Source:MGI Symbo | 0 | 0 | 0 | 0 | 0 | 0 |
| 39497 | ENSMUSG00000101208 | 0 | NA | NA | NA | NA | NA | Gm19891 | predicted gene, 19891 [Source:MGI Symb | 0 | 0 | 0 | 0 | 0 | 0 |
| 39498 | ENSMUSG00000101209 | 0.185640647 | 1.126924175 | 4.080472857 | 0.276174898 | 0.782413736 | NA | Gm8524 | predicted gene 8524 [Source:MGI Symbol | 0 | 0 | 0 | 0 | 0 | 1.1138 |
| 39499 | ENSMUSG00000101210 | 23.38384016 | 0.202930418 | 0.512768686 | 0.395754311 | 0.692286285 | 0.938271067 | Gm28720 | predicted gene 28720 [Source:MGI Symbo | 16.865 | 18.503 | 30.187 | 39.523 | 19.631 | 15.594 |
| 39500 | ENSMUSG00000101211 | 30.99079992 | 0.184842771 | 0.439717775 | 0.42036684 | 0.674217489 | 0.932114677 | Gm28818 | predicted gene 28818 [Source:MGI Symbo | 36.943 | 17.475 | 32.2 | 22.171 | 38.172 | 38.985 |
| 39501 | ENSMUSG00000101212 | 0 | NA | NA | NA | NA | NA | Gm29354 | predicted gene 29354 [Source:MGI Symbo | 0 | 0 | 0 | 0 | 0 | 0 |
| 39502 | ENSMUSG00000101213 | 0 | NA | NA | NA | NA | NA | Gm28788 | predicted gene 28788 [Source:MGI Symbo | 0 | 0 | 0 | 0 | 0 | 0 |
| 39503 | ENSMUSG00000101214 | 0 | NA | NA | NA | NA | NA | Gm28403 | predicted gene 28403 [Source:MGI Symbo | 0 | 0 | 0 | 0 | 0 | 0 |
| 39504 | ENSMUSG00000101215 | 0 | NA | NA | NA | NA | NA | Gm28569 | predicted gene 28569 [Source:MGI Symbo | 0 | 0 | 0 | 0 | 0 | 0 |
| 39505 | ENSMUSG00000101216 | 0 | NA | NA | NA | NA | NA | Gm29451 | predicted gene 29451 [Source:MGI Symbo | 0 | 0 | 0 | 0 | 0 | 0 |
| 39506 | ENSMUSG00000101218 | 0 | NA | NA | NA | NA | NA | Gm29416 | predicted gene 29416 [Source:MGI Symbo | 0 | 0 | 0 | 0 | 0 | 0 |
| 39507 | ENSMUSG00000101219 | 0 | NA | NA | NA | NA | NA | Gm20302 | predicted gene, 20302 [Source:MGI Symb | 0 | 0 | 0 | 0 | 0 | 0 |
| 39508 | ENSMUSG00000101220 | 0 | NA | NA | NA | NA | NA | Gm6717 | predicted gene 6717 [Source:MGI Symbol | 0 | 0 | 0 | 0 | 0 | 0 |
| 39509 | ENSMUSG00000101221 | 0 | NA | NA | NA | NA | NA | Gm21881 | predicted gene, 21881 [Source:MGI Symb | 0 | 0 | 0 | 0 | 0 | 0 |
| 39510 | ENSMUSG00000101222 | 0 | NA | NA | NA | NA | NA | Gm28315 | predicted gene 28315 [Source:MGI Symbo | 0 | 0 | 0 | 0 | 0 | 0 |
| 39511 | ENSMUSG00000101223 | 0.160662974 | 1.126924175 | 4.080472857 | 0.276174898 | 0.782413736 | NA | 4930568A12 | RIKEN cDNA 4930568A12 gene [Source:M | 0 | 0 | 0 | 0.964 | 0 | 0 |
| 39512 | ENSMUSG00000101224 | 0 | NA | NA | NA | NA | NA | Gm28209 | predicted gene 28209 [Source:MGI Symbo | 0 | 0 | 0 | 0 | 0 | 0 |
| 39513 | ENSMUSG00000101225 | 23.6010257 | 0.484269034 | 0.468749637 | 1.033108074 | 0.301553325 | 0.752246851 | 1700008J07R | RIKEN cDNA 1700008J07 gene [Source:M( | 13.653 | 19.53 | 26.162 | 33.739 | 22.903 | 25.618 |
| 39514 | ENSMUSG00000101226 | 0 | NA | NA | NA | NA | NA | Gm20954 | predicted gene, 20954 [Source:MGI Symb | 0 | 0 | 0 | 0 | 0 | 0 |
| 39515 | ENSMUSG00000101227 | 0 | NA | NA | NA | NA | NA | Gm29506 | predicted gene 29506 [Source:MGI Symbo | 0 | 0 | 0 | 0 | 0 | 0 |
| 39516 | ENSMUSG00000101228 | 0 | NA | NA | NA | NA | NA | Gm21330 | predicted gene, 21330 [Source:MGI Symb | 0 | 0 | 0 | 0 | 0 | 0 |
| 39517 | ENSMUSG00000101229 | 0 | NA | NA | NA | NA | NA | Gm29046 | predicted gene 29046 [Source:MGI Symbo | 0 | 0 | 0 | 0 | 0 | 0 |
| 39518 | ENSMUSG00000101230 | 0 | NA | NA | NA | NA | NA | Gm21862 | predicted gene, 21862 [Source:MGI Symb | 0 | 0 | 0 | 0 | 0 | 0 |
| 39519 | ENSMUSG00000101231 | 0 | NA | NA | NA | NA | NA | Gm28283 | predicted gene 28283 [Source:MGI Symbo | 0 | 0 | 0 | 0 | 0 | 0 |
| 39520 | ENSMUSG00000101232 | 0 | NA | NA | NA | NA | NA | Scgb1b15 | secretoglobin, family 1B, member 15 [Sou | 0 | 0 | 0 | 0 | 0 | 0 |
| 39521 | ENSMUSG00000101233 | 0 | NA | NA | NA | NA | NA | Gm29242 | predicted gene 29242 [Source:MGI Symbo | 0 | 0 | 0 | 0 | 0 | 0 |
| 39522 | ENSMUSG00000101234 | 0 | NA | NA | NA | NA | NA | 1700007E05 | RIKEN cDNA 1700007E05 gene [Source:M( | 0 | 0 | 0 | 0 | 0 | 0 |
| 39523 | ENSMUSG00000101235 | 0 | NA | NA | NA | NA | NA | Gm18033 | predicted gene, 18033 [Source:MGI Symb | 0 | 0 | 0 | 0 | 0 | 0 |
| 39524 | ENSMUSG00000101236 | 0.665794642 | 1.656189793 | 2.801711575 | 0.591135007 | 0.554429965 | NA | Gm6750 | predicted gene 6750 [Source:MGI Symbol | 0.8031 | 0 | 0 | 0.964 | 0 | 2.2277 |
| 39525 | ENSMUSG00000101237 | 0 | NA | NA | NA | NA | NA | Gm29464 | predicted gene 29464 [Source:MGI Symbo | 0 | 0 | 0 | 0 | 0 | 0 |
| 39526 | ENSMUSG00000101238 | 0 | NA | NA | NA | NA | NA | Gm28123 | predicted gene 28123 [Source:MGI Symbo | 0 | 0 | 0 | 0 | 0 | 0 |
| 39527 | ENSMUSG00000101239 | 0 | NA | NA | NA | NA | NA | Gm28304 | predicted gene 28304 [Source:MGI Symbo | 0 | 0 | 0 | 0 | 0 | 0 |
| 39528 | ENSMUSG00000101240 | 0 | NA | NA | NA | NA | NA | Gm28266 | predicted gene 28266 [Source:MGI Symbo | 0 | 0 | 0 | 0 | 0 | 0 |
| 39529 | ENSMUSG00000101241 | 0 | NA | NA | NA | NA | NA | Gm29111 | predicted gene 29111 [Source:MGI Symbo | 0 | 0 | 0 | 0 | 0 | 0 |
| 39530 | ENSMUSG00000101242 | 0 | NA | NA | NA | NA | NA | Gm28319 | predicted gene 28319 [Source:MGI Symbo | 0 | 0 | 0 | 0 | 0 | 0 |
| 39531 | ENSMUSG00000101243 | 0 | NA | NA | NA | NA | NA | Gm28919 | predicted gene 28919 [Source:MGI Symbo | 0 | 0 | 0 | 0 | 0 | 0 |
| 39532 | ENSMUSG00000101244 | 0 | NA | NA | NA | NA | NA | Gm29368 | predicted gene 29368 [Source:MGI Symbo | 0 | 0 | 0 | 0 | 0 | 0 |

| | | | | | | | | | | | | | | | |
|---|---|---|---|---|---|---|---|---|---|---|---|---|---|---|---|
| 39533 | ENSMUSG00000101245 | 0 | NA | NA | NA | NA | NA | Gm28965 | predicted gene 28965 [Source:MGI Symbo | 0 | 0 | 0 | 0 | 0 | 0 |
| 39534 | ENSMUSG00000101246 | 0 | NA | NA | NA | NA | NA | Gm29026 | predicted gene 29026 [Source:MGI Symbo | 0 | 0 | 0 | 0 | 0 | 0 |
| 39535 | ENSMUSG00000101247 | 0 | NA | NA | NA | NA | NA | Gm29042 | predicted gene 29042 [Source:MGI Symbo | 0 | 0 | 0 | 0 | 0 | 0 |
| 39536 | ENSMUSG00000101248 | 0 | NA | NA | NA | NA | NA | Scgb1b9-ps | secretoglobin, family 1B, member 9, pseu | 0 | 0 | 0 | 0 | 0 | 0 |
| 39537 | ENSMUSG00000101249 | 379.3697523 | 0.200759032 | 0.16294514 | 1.232065169 | 0.217924741 | 0.663697152 | Gm29216 | predicted gene 29216 [Source:MGI Symbo | 338.91 | 402.94 | 316.97 | 345.1 | 411.16 | 461.13 |
| 39538 | ENSMUSG00000101250 | 0 | NA | NA | NA | NA | NA | Gm28105 | predicted gene 28105 [Source:MGI Symbo | 0 | 0 | 0 | 0 | 0 | 0 |
| 39539 | ENSMUSG00000101251 | 0 | NA | NA | NA | NA | NA | Gm3496 | predicted gene 3496 [Source:MGI Symbol | 0 | 0 | 0 | 0 | 0 | 0 |
| 39540 | ENSMUSG00000101252 | 0 | NA | NA | NA | NA | NA | Ifna6 | interferon alpha 6 [Source:MGI Symbol;Ac | 0 | 0 | 0 | 0 | 0 | 0 |
| 39541 | ENSMUSG00000101253 | 0 | NA | NA | NA | NA | NA | Gm29088 | predicted gene 29088 [Source:MGI Symbo | 0 | 0 | 0 | 0 | 0 | 0 |
| 39542 | ENSMUSG00000101254 | 0 | NA | NA | NA | NA | NA | Gm20932 | predicted gene, 20932 [Source:MGI Symb | 0 | 0 | 0 | 0 | 0 | 0 |
| 39543 | ENSMUSG00000101255 | 0 | NA | NA | NA | NA | NA | Gm21841 | predicted gene, 21841 [Source:MGI Symb | 0 | 0 | 0 | 0 | 0 | 0 |
| 39544 | ENSMUSG00000101256 | 0 | NA | NA | NA | NA | NA | Gm28133 | predicted gene 28133 [Source:MGI Symbo | 0 | 0 | 0 | 0 | 0 | 0 |
| 39545 | ENSMUSG00000101257 | 0 | NA | NA | NA | NA | NA | 2310015K22 | RIKEN cDNA 2310015K22 gene [Source:M | 0 | 0 | 0 | 0 | 0 | 0 |
| 39546 | ENSMUSG00000101258 | 0 | NA | NA | NA | NA | NA | Gm29477 | predicted gene 29477 [Source:MGI Symbo | 0 | 0 | 0 | 0 | 0 | 0 |
| 39547 | ENSMUSG00000101259 | 0 | NA | NA | NA | NA | NA | Gm21242 | predicted gene, 21242 [Source:MGI Symb | 0 | 0 | 0 | 0 | 0 | 0 |
| 39548 | ENSMUSG00000101260 | 0 | NA | NA | NA | NA | NA | Gm28780 | predicted gene 28780 [Source:MGI Symbo | 0 | 0 | 0 | 0 | 0 | 0 |
| 39549 | ENSMUSG00000101261 | 0 | NA | NA | NA | NA | NA | 1700109I08R | RIKEN cDNA 1700109I08 gene [Source:MG | 0 | 0 | 0 | 0 | 0 | 0 |
| 39550 | ENSMUSG00000101262 | 0 | NA | NA | NA | NA | NA | Gm8326 | predicted gene 8326 [Source:MGI Symbol | 0 | 0 | 0 | 0 | 0 | 0 |
| 39551 | ENSMUSG00000101263 | 0 | NA | NA | NA | NA | NA | Gm29121 | predicted gene 29121 [Source:MGI Symbo | 0 | 0 | 0 | 0 | 0 | 0 |
| 39552 | ENSMUSG00000101264 | 1.205662164 | 1.402791949 | 2.113435109 | 0.663749714 | 0.506850515 | NA | Gm28347 | predicted gene 28347 [Source:MGI Symbo | 0 | 1.0279 | 1.0062 | 1.928 | 3.2719 | 0 |
| 39553 | ENSMUSG00000101265 | 0 | NA | NA | NA | NA | NA | Gm29099 | predicted gene 29099 [Source:MGI Symbo | 0 | 0 | 0 | 0 | 0 | 0 |
| 39554 | ENSMUSG00000101266 | 0 | NA | NA | NA | NA | NA | Gm29234 | predicted gene 29234 [Source:MGI Symbo | 0 | 0 | 0 | 0 | 0 | 0 |
| 39555 | ENSMUSG00000101267 | 0 | NA | NA | NA | NA | NA | Gm21912 | predicted gene, 21912 [Source:MGI Symb | 0 | 0 | 0 | 0 | 0 | 0 |
| 39556 | ENSMUSG00000101268 | 1.944901758 | 0.603035859 | 1.595984096 | 0.377845782 | 0.705545161 | NA | 2010310C07 | RIKEN cDNA 2010310C07 gene [Source:M | 1.6062 | 2.0558 | 1.0062 | 4.8199 | 2.1812 | 0 |
| 39557 | ENSMUSG00000101270 | 0 | NA | NA | NA | NA | NA | Scgb2b9-ps | secretoglobin, family 2B, member 9, pseu | 0 | 0 | 0 | 0 | 0 | 0 |
| 39558 | ENSMUSG00000101271 | 0 | NA | NA | NA | NA | NA | Gm28334 | predicted gene 28334 [Source:MGI Symbo | 0 | 0 | 0 | 0 | 0 | 0 |
| 39559 | ENSMUSG00000101272 | 0 | NA | NA | NA | NA | NA | Gm36782 | predicted gene, 36782 [Source:MGI Symb | 0 | 0 | 0 | 0 | 0 | 0 |
| 39560 | ENSMUSG00000101273 | 0 | NA | NA | NA | NA | NA | Gm28831 | predicted gene 28831 [Source:MGI Symbo | 0 | 0 | 0 | 0 | 0 | 0 |
| 39561 | ENSMUSG00000101275 | 0 | NA | NA | NA | NA | NA | 1700012E03 | RIKEN cDNA 1700012E03 gene [Source:M | 0 | 0 | 0 | 0 | 0 | 0 |
| 39562 | ENSMUSG00000101276 | 0 | NA | NA | NA | NA | NA | Gm28657 | predicted gene 28657 [Source:MGI Symbo | 0 | 0 | 0 | 0 | 0 | 0 |
| 39563 | ENSMUSG00000101277 | 0 | NA | NA | NA | NA | NA | Gm20847 | predicted gene, 20847 [Source:MGI Symb | 0 | 0 | 0 | 0 | 0 | 0 |
| 39564 | ENSMUSG00000101278 | 0 | NA | NA | NA | NA | NA | Gm29381 | predicted gene 29381 [Source:MGI Symbo | 0 | 0 | 0 | 0 | 0 | 0 |
| 39565 | ENSMUSG00000101279 | 0 | NA | NA | NA | NA | NA | Gm18342 | predicted gene, 18342 [Source:MGI Symb | 0 | 0 | 0 | 0 | 0 | 0 |
| 39566 | ENSMUSG00000101280 | 0 | NA | NA | NA | NA | NA | Gm8600 | predicted gene 8600 [Source:MGI Symbol | 0 | 0 | 0 | 0 | 0 | 0 |
| 39567 | ENSMUSG00000101281 | 0 | NA | NA | NA | NA | NA | Gm8449 | predicted gene 8449 [Source:MGI Symbol | 0 | 0 | 0 | 0 | 0 | 0 |
| 39568 | ENSMUSG00000101282 | 0 | NA | NA | NA | NA | NA | Gm28152 | predicted gene 28152 [Source:MGI Symbo | 0 | 0 | 0 | 0 | 0 | 0 |
| 39569 | ENSMUSG00000101283 | 0 | NA | NA | NA | NA | NA | Gm29314 | predicted gene 29314 [Source:MGI Symbo | 0 | 0 | 0 | 0 | 0 | 0 |
| 39570 | ENSMUSG00000101284 | 0 | NA | NA | NA | NA | NA | Gm28811 | predicted gene 28811 [Source:MGI Symbo | 0 | 0 | 0 | 0 | 0 | 0 |
| 39571 | ENSMUSG00000101285 | 0 | NA | NA | NA | NA | NA | Gm28600 | predicted gene 28600 [Source:MGI Symbo | 0 | 0 | 0 | 0 | 0 | 0 |
| 39572 | ENSMUSG00000101286 | 0 | NA | NA | NA | NA | NA | Gm21317 | predicted gene, 21317 [Source:MGI Symb | 0 | 0 | 0 | 0 | 0 | 0 |
| 39573 | ENSMUSG00000101287 | 0 | NA | NA | NA | NA | NA | Gm28166 | predicted gene 28166 [Source:MGI Symbo | 0 | 0 | 0 | 0 | 0 | 0 |
| 39574 | ENSMUSG00000101288 | 0 | NA | NA | NA | NA | NA | Gm28342 | predicted gene 28342 [Source:MGI Symbo | 0 | 0 | 0 | 0 | 0 | 0 |
| 39575 | ENSMUSG00000101289 | 0.321325949 | 1.911575114 | 4.048267893 | 0.472195804 | 0.636787034 | NA | Gm3454 | predicted gene 3454 [Source:MGI Symbol | 0 | 0 | 0 | 1.928 | 0 | 0 |
| 39576 | ENSMUSG00000101290 | 0 | NA | NA | NA | NA | NA | Gm8480 | predicted gene 8480 [Source:MGI Symbol | 0 | 0 | 0 | 0 | 0 | 0 |
| 39577 | ENSMUSG00000101291 | 0 | NA | NA | NA | NA | NA | Gm29194 | predicted gene 29194 [Source:MGI Symbo | 0 | 0 | 0 | 0 | 0 | 0 |
| 39578 | ENSMUSG00000101292 | 0 | NA | NA | NA | NA | NA | Gm29496 | predicted gene 29496 [Source:MGI Symbo | 0 | 0 | 0 | 0 | 0 | 0 |
| 39579 | ENSMUSG00000101294 | 0 | NA | NA | NA | NA | NA | Btbd35f15 | BTB domain containing 35, family membe | 0 | 0 | 0 | 0 | 0 | 0 |
| 39580 | ENSMUSG00000101295 | 0 | NA | NA | NA | NA | NA | Gm28180 | predicted gene 28180 [Source:MGI Symbo | 0 | 0 | 0 | 0 | 0 | 0 |
| 39581 | ENSMUSG00000101296 | 0 | NA | NA | NA | NA | NA | Scgb2b32-ps | secretoglobin, family 2B, member 32, pse | 0 | 0 | 0 | 0 | 0 | 0 |
| 39582 | ENSMUSG00000101297 | 0 | NA | NA | NA | NA | NA | Gm18447 | predicted gene, 18447 [Source:MGI Symb | 0 | 0 | 0 | 0 | 0 | 0 |
| 39583 | ENSMUSG00000101298 | 0 | NA | NA | NA | NA | NA | Gm28308 | predicted gene 28308 [Source:MGI Symbo | 0 | 0 | 0 | 0 | 0 | 0 |
| 39584 | ENSMUSG00000101299 | 0 | NA | NA | NA | NA | NA | Gm28175 | predicted gene 28175 [Source:MGI Symbo | 0 | 0 | 0 | 0 | 0 | 0 |
| 39585 | ENSMUSG00000101300 | 0 | NA | NA | NA | NA | NA | Gm29130 | predicted gene 29130 [Source:MGI Symbo | 0 | 0 | 0 | 0 | 0 | 0 |
| 39586 | ENSMUSG00000101301 | 0 | NA | NA | NA | NA | NA | Gm29214 | predicted gene 29214 [Source:MGI Symbo | 0 | 0 | 0 | 0 | 0 | 0 |
| 39587 | ENSMUSG00000101302 | 0 | NA | NA | NA | NA | NA | Gm28519 | predicted gene 28519 [Source:MGI Symbo | 0 | 0 | 0 | 0 | 0 | 0 |
| 39588 | ENSMUSG00000101303 | 0 | NA | NA | NA | NA | NA | B020011L13 | RIKEN cDNA B020011L13 gene [Source:M | 0 | 0 | 0 | 0 | 0 | 0 |

| | | | | | | | | | | | | | | | |
|---|---|---|---|---|---|---|---|---|---|---|---|---|---|---|---|
| 39589 | ENSMUSG00000101304 | 10.92197191 | 0.216662522 | 0.703886112 | 0.307809059 | 0.758227628 | 0.953675669 | Plet1os | placenta expressed transcript 1, opposite | 16.865 | 9.2513 | 4.025 | 14.46 | 10.906 | 10.025 |
| 39590 | ENSMUSG00000101305 | 0 | NA | NA | NA | NA | NA | Gm29171 | predicted gene 29171 [Source:MGI Symbo | 0 | 0 | 0 | 0 | 0 | 0 |
| 39591 | ENSMUSG00000101306 | 0 | NA | NA | NA | NA | NA | Gm5252 | predicted gene 5252 [Source:MGI Symbol | 0 | 0 | 0 | 0 | 0 | 0 |
| 39592 | ENSMUSG00000101307 | 0 | NA | NA | NA | NA | NA | Gm7356 | predicted gene 7356 [Source:MGI Symbol | 0 | 0 | 0 | 0 | 0 | 0 |
| 39593 | ENSMUSG00000101308 | 0.319491021 | 0.16514334 | 4.044653141 | 0.040830038 | 0.967431393 | NA | Gm28989 | predicted gene 28989 [Source:MGI Symbo | 0.8031 | 0 | 0 | 0 | 0 | 1.1138 |
| 39594 | ENSMUSG00000101309 | 0 | NA | NA | NA | NA | NA | Gm29397 | predicted gene 29397 [Source:MGI Symbo | 0 | 0 | 0 | 0 | 0 | 0 |
| 39595 | ENSMUSG00000101310 | 0 | NA | NA | NA | NA | NA | Gm21281 | predicted gene, 21281 [Source:MGI Symb | 0 | 0 | 0 | 0 | 0 | 0 |
| 39596 | ENSMUSG00000101311 | 0 | NA | NA | NA | NA | NA | Gm28250 | predicted gene 28250 [Source:MGI Symbo | 0 | 0 | 0 | 0 | 0 | 0 |
| 39597 | ENSMUSG00000101312 | 0 | NA | NA | NA | NA | NA | Gm21135 | predicted gene, 21135 [Source:MGI Symb | 0 | 0 | 0 | 0 | 0 | 0 |
| 39598 | ENSMUSG00000101313 | 0 | NA | NA | NA | NA | NA | Gm29219 | predicted gene 29219 [Source:MGI Symbo | 0 | 0 | 0 | 0 | 0 | 0 |
| 39599 | ENSMUSG00000101314 | 0 | NA | NA | NA | NA | NA | Gm29663 | predicted gene 29663 [Source:MGI Symbo | 0 | 0 | 0 | 0 | 0 | 0 |
| 39600 | ENSMUSG00000101315 | 0 | NA | NA | NA | NA | NA | Krtap28-13 | keratin associated protein 28-13 [Source:I | 0 | 0 | 0 | 0 | 0 | 0 |
| 39601 | ENSMUSG00000101316 | 1.289758267 | -0.470778025 | 1.884798362 | -0.249776334 | 0.802760323 | NA | Gm12663 | predicted gene 12663 [Source:MGI Symbo | 2.4093 | 1.0279 | 1.0062 | 0 | 2.1812 | 1.1138 |
| 39602 | ENSMUSG00000101317 | 0 | NA | NA | NA | NA | NA | Gm28284 | predicted gene 28284 [Source:MGI Symbo | 0 | 0 | 0 | 0 | 0 | 0 |
| 39603 | ENSMUSG00000101318 | 0 | NA | NA | NA | NA | NA | Gm29109 | predicted gene 29109 [Source:MGI Symbo | 0 | 0 | 0 | 0 | 0 | 0 |
| 39604 | ENSMUSG00000101319 | 0 | NA | NA | NA | NA | NA | Gm28957 | predicted gene 28957 [Source:MGI Symbo | 0 | 0 | 0 | 0 | 0 | 0 |
| 39605 | ENSMUSG00000101320 | 17.24959965 | -0.330335211 | 0.542511169 | -0.608900295 | 0.542590529 | 0.894095573 | Gm28529 | predicted gene 28529 [Source:MGI Symbo | 24.093 | 13.363 | 20.125 | 17.352 | 18.541 | 10.025 |
| 39606 | ENSMUSG00000101321 | 0 | NA | NA | NA | NA | NA | Gm28817 | predicted gene 28817 [Source:MGI Symbo | 0 | 0 | 0 | 0 | 0 | 0 |
| 39607 | ENSMUSG00000101323 | 0 | NA | NA | NA | NA | NA | Gm21877 | predicted gene, 21877 [Source:MGI Symb | 0 | 0 | 0 | 0 | 0 | 0 |
| 39608 | ENSMUSG00000101324 | 0 | NA | NA | NA | NA | NA | Gm29187 | predicted gene 29187 [Source:MGI Symbo | 0 | 0 | 0 | 0 | 0 | 0 |
| 39609 | ENSMUSG00000101325 | 0 | NA | NA | NA | NA | NA | Scgb2b4-ps | secretoglobin, family 2B, member 4, pseu | 0 | 0 | 0 | 0 | 0 | 0 |
| 39610 | ENSMUSG00000101326 | 0 | NA | NA | NA | NA | NA | Gm28835 | predicted gene 28835 [Source:MGI Symbo | 0 | 0 | 0 | 0 | 0 | 0 |
| 39611 | ENSMUSG00000101327 | 0 | NA | NA | NA | NA | NA | Rpl23a-ps10 | ribosomal protein L23A, pseudogene 10 [S | 0 | 0 | 0 | 0 | 0 | 0 |
| 39612 | ENSMUSG00000101328 | 0 | NA | NA | NA | NA | NA | Rpl30-ps6 | ribosomal protein L30, pseudogene 6 [Sou | 0 | 0 | 0 | 0 | 0 | 0 |
| 39613 | ENSMUSG00000101329 | 0 | NA | NA | NA | NA | NA | Gm28226 | predicted gene 28226 [Source:MGI Symbo | 0 | 0 | 0 | 0 | 0 | 0 |
| 39614 | ENSMUSG00000101330 | 0.181769715 | 1.126924175 | 4.080472857 | 0.276174898 | 0.782413736 | NA | Gm10193 | predicted gene 10193 [Source:MGI Symbo | 0 | 0 | 0 | 0 | 1.0906 | 0 |
| 39615 | ENSMUSG00000101331 | 0 | NA | NA | NA | NA | NA | Gm28148 | predicted gene 28148 [Source:MGI Symbo | 0 | 0 | 0 | 0 | 0 | 0 |
| 39616 | ENSMUSG00000101332 | 0 | NA | NA | NA | NA | NA | Gm28240 | predicted gene 28240 [Source:MGI Symbo | 0 | 0 | 0 | 0 | 0 | 0 |
| 39617 | ENSMUSG00000101333 | 0 | NA | NA | NA | NA | NA | Gm4753 | predicted gene 4753 [Source:MGI Symbol | 0 | 0 | 0 | 0 | 0 | 0 |
| 39618 | ENSMUSG00000101334 | 0 | NA | NA | NA | NA | NA | 1700028M03 | RIKEN cDNA 1700028M03 gene [Source:M | 0 | 0 | 0 | 0 | 0 | 0 |
| 39619 | ENSMUSG00000101335 | 36.38671255 | 0.451715146 | 0.36596783 | 1.234302879 | 0.217090063 | 0.663070346 | Gm28229 | predicted gene 28229 [Source:MGI Symbo | 31.321 | 29.81 | 31.194 | 51.091 | 39.262 | 35.643 |
| 39620 | ENSMUSG00000101336 | 0 | NA | NA | NA | NA | NA | Gm28480 | predicted gene 28480 [Source:MGI Symbo | 0 | 0 | 0 | 0 | 0 | 0 |
| 39621 | ENSMUSG00000101337 | 0.501260735 | 1.111687294 | 3.109072805 | 0.357562323 | 0.720670879 | NA | Dnah7c | dynein, axonemal, heavy chain 7C [Source | 0.8031 | 0 | 0 | 0 | 1.0906 | 1.1138 |
| 39622 | ENSMUSG00000101338 | 0 | NA | NA | NA | NA | NA | Gm28139 | predicted gene 28139 [Source:MGI Symbo | 0 | 0 | 0 | 0 | 0 | 0 |
| 39623 | ENSMUSG00000101339 | 0 | NA | NA | NA | NA | NA | Gm29180 | predicted gene 29180 [Source:MGI Symbo | 0 | 0 | 0 | 0 | 0 | 0 |
| 39624 | ENSMUSG00000101340 | 0 | NA | NA | NA | NA | NA | Gm29607 | predicted gene 29607 [Source:MGI Symbo | 0 | 0 | 0 | 0 | 0 | 0 |
| 39625 | ENSMUSG00000101341 | 0 | NA | NA | NA | NA | NA | Gm29076 | predicted gene 29076 [Source:MGI Symbo | 0 | 0 | 0 | 0 | 0 | 0 |
| 39626 | ENSMUSG00000101342 | 0 | NA | NA | NA | NA | NA | Gm28078 | predicted gene 28078 [Source:MGI Symbo | 0 | 0 | 0 | 0 | 0 | 0 |
| 39627 | ENSMUSG00000101343 | 0 | NA | NA | NA | NA | NA | Gm29469 | predicted gene 29469 [Source:MGI Symbo | 0 | 0 | 0 | 0 | 0 | 0 |
| 39628 | ENSMUSG00000101344 | 0 | NA | NA | NA | NA | NA | Gm29183 | predicted gene 29183 [Source:MGI Symbo | 0 | 0 | 0 | 0 | 0 | 0 |
| 39629 | ENSMUSG00000101345 | 0 | NA | NA | NA | NA | NA | Gm28195 | predicted gene 28195 [Source:MGI Symbo | 0 | 0 | 0 | 0 | 0 | 0 |
| 39630 | ENSMUSG00000101346 | 0 | NA | NA | NA | NA | NA | Gm29505 | predicted gene 29505 [Source:MGI Symbo | 0 | 0 | 0 | 0 | 0 | 0 |
| 39631 | ENSMUSG00000101347 | 0 | NA | NA | NA | NA | NA | Gm28219 | predicted gene 28219 [Source:MGI Symbo | 0 | 0 | 0 | 0 | 0 | 0 |
| 39632 | ENSMUSG00000101348 | 0 | NA | NA | NA | NA | NA | Gm7135 | predicted gene 7135 [Source:MGI Symbol | 0 | 0 | 0 | 0 | 0 | 0 |
| 39633 | ENSMUSG00000101349 | 0 | NA | NA | NA | NA | NA | Gm28162 | predicted gene 28162 [Source:MGI Symbo | 0 | 0 | 0 | 0 | 0 | 0 |
| 39634 | ENSMUSG00000101350 | 0 | NA | NA | NA | NA | NA | Gm28962 | predicted gene 28962 [Source:MGI Symbo | 0 | 0 | 0 | 0 | 0 | 0 |
| 39635 | ENSMUSG00000101351 | 0 | NA | NA | NA | NA | NA | Gm28573 | predicted gene 28573 [Source:MGI Symbo | 0 | 0 | 0 | 0 | 0 | 0 |
| 39636 | ENSMUSG00000101352 | 0 | NA | NA | NA | NA | NA | Gm28665 | predicted gene 28665 [Source:MGI Symbo | 0 | 0 | 0 | 0 | 0 | 0 |
| 39637 | ENSMUSG00000101353 | 0 | NA | NA | NA | NA | NA | Gm29203 | predicted gene 29203 [Source:MGI Symbo | 0 | 0 | 0 | 0 | 0 | 0 |
| 39638 | ENSMUSG00000101354 | 0 | NA | NA | NA | NA | NA | Gm28973 | predicted gene 28973 [Source:MGI Symbo | 0 | 0 | 0 | 0 | 0 | 0 |
| 39639 | ENSMUSG00000101355 | 0 | NA | NA | NA | NA | NA | H3c10 | H3 clustered histone 10 [Source:MGI Sym | 0 | 0 | 0 | 0 | 0 | 0 |
| 39640 | ENSMUSG00000101356 | 6.39164886 | 1.650813229 | 0.975297053 | 1.692626081 | 0.090526664 | 0.444459491 | Gm28876 | predicted gene 28876 [Source:MGI Symbo | 3.2124 | 0 | 6.0375 | 12.532 | 6.5437 | 10.025 |
| 39641 | ENSMUSG00000101357 | 0 | NA | NA | NA | NA | NA | Gm2495 | predicted gene 2495 [Source:MGI Symbol | 0 | 0 | 0 | 0 | 0 | 0 |
| 39642 | ENSMUSG00000101358 | 0 | NA | NA | NA | NA | NA | Gm29661 | predicted gene 29661 [Source:MGI Symbo | 0 | 0 | 0 | 0 | 0 | 0 |
| 39643 | ENSMUSG00000101359 | 0 | NA | NA | NA | NA | NA | Gm28211 | predicted gene 28211 [Source:MGI Symbo | 0 | 0 | 0 | 0 | 0 | 0 |
| 39644 | ENSMUSG00000101360 | 0 | NA | NA | NA | NA | NA | 4933417O13 | RIKEN cDNA 4933417O13 gene [Source:M | 0 | 0 | 0 | 0 | 0 | 0 |

| | | | | | | | | | | | | | | | |
|---|---|---|---|---|---|---|---|---|---|---|---|---|---|---|---|
| 39645 | ENSMUSG00000101361 | 0 | NA | NA | NA | NA | NA | Gm7358 | predicted gene 7358 [Source:MGI Symbol | 0 | 0 | 0 | 0 | 0 | 0 |
| 39646 | ENSMUSG00000101362 | 0 | NA | NA | NA | NA | NA | Mrgprb9-ps | MAS-related GPR, member B9, pseudoger | 0 | 0 | 0 | 0 | 0 | 0 |
| 39647 | ENSMUSG00000101363 | 0.321325949 | 1.911575114 | 4.048267893 | 0.472195804 | 0.636787034 | NA | Gm29490 | predicted gene 29490 [Source:MGI Symbc | 0 | 0 | 0 | 1.928 | 0 | 0 |
| 39648 | ENSMUSG00000101364 | 0 | NA | NA | NA | NA | NA | Gm29300 | predicted gene 29300 [Source:MGI Symbc | 0 | 0 | 0 | 0 | 0 | 0 |
| 39649 | ENSMUSG00000101365 | 0.17131964 | -0.796629176 | 4.080472857 | -0.195229623 | 0.84521319 | NA | Gm19325 | predicted gene, 19325 [Source:MGI Symb | 0 | 1.0279 | 0 | 0 | 0 | 0 |
| 39650 | ENSMUSG00000101366 | 0 | NA | NA | NA | NA | NA | Gm28147 | predicted gene 28147 [Source:MGI Symbc | 0 | 0 | 0 | 0 | 0 | 0 |
| 39651 | ENSMUSG00000101367 | 0 | NA | NA | NA | NA | NA | Gm29484 | predicted gene 29484 [Source:MGI Symbc | 0 | 0 | 0 | 0 | 0 | 0 |
| 39652 | ENSMUSG00000101368 | 0 | NA | NA | NA | NA | NA | Gm28290 | predicted gene 28290 [Source:MGI Symbc | 0 | 0 | 0 | 0 | 0 | 0 |
| 39653 | ENSMUSG00000101369 | 0 | NA | NA | NA | NA | NA | Gm28080 | predicted gene 28080 [Source:MGI Symbc | 0 | 0 | 0 | 0 | 0 | 0 |
| 39654 | ENSMUSG00000101370 | 0 | NA | NA | NA | NA | NA | Gm29139 | predicted gene 29139 [Source:MGI Symbc | 0 | 0 | 0 | 0 | 0 | 0 |
| 39655 | ENSMUSG00000101371 | 0.160662974 | 1.126924175 | 4.080472857 | 0.276174898 | 0.782413736 | NA | Gm9114 | predicted gene 9114 [Source:MGI Symbol | 0 | 0 | 0 | 0.964 | 0 | 0 |
| 39656 | ENSMUSG00000101372 | 0 | NA | NA | NA | NA | NA | Gm19028 | predicted gene, 19028 [Source:MGI Symb | 0 | 0 | 0 | 0 | 0 | 0 |
| 39657 | ENSMUSG00000101373 | 0 | NA | NA | NA | NA | NA | Zfp813-ps | zinc finger protein 813, pseudogene [Sour | 0 | 0 | 0 | 0 | 0 | 0 |
| 39658 | ENSMUSG00000101374 | 0 | NA | NA | NA | NA | NA | Gm28844 | predicted gene 28844 [Source:MGI Symbc | 0 | 0 | 0 | 0 | 0 | 0 |
| 39659 | ENSMUSG00000101375 | 0 | NA | NA | NA | NA | NA | Gm28383 | predicted gene 28383 [Source:MGI Symbc | 0 | 0 | 0 | 0 | 0 | 0 |
| 39660 | ENSMUSG00000101376 | 0 | NA | NA | NA | NA | NA | Gm18291 | predicted gene, 18291 [Source:MGI Symb | 0 | 0 | 0 | 0 | 0 | 0 |
| 39661 | ENSMUSG00000101377 | 0 | NA | NA | NA | NA | NA | Gm28336 | predicted gene 28336 [Source:MGI Symbc | 0 | 0 | 0 | 0 | 0 | 0 |
| 39662 | ENSMUSG00000101378 | 1.220732185 | -1.349517554 | 1.980457861 | -0.681416949 | 0.495607701 | NA | Gm28086 | predicted gene 28086 [Source:MGI Symbc | 3.2124 | 1.0279 | 1.0062 | 0.964 | 0 | 1.1138 |
| 39663 | ENSMUSG00000101379 | 0 | NA | NA | NA | NA | NA | Gm28092 | predicted gene 28092 [Source:MGI Symbc | 0 | 0 | 0 | 0 | 0 | 0 |
| 39664 | ENSMUSG00000101380 | 0 | NA | NA | NA | NA | NA | Gm28769 | predicted gene 28769 [Source:MGI Symbc | 0 | 0 | 0 | 0 | 0 | 0 |
| 39665 | ENSMUSG00000101381 | 0 | NA | NA | NA | NA | NA | Btbd35f16 | BTB domain containing 35, family membe | 0 | 0 | 0 | 0 | 0 | 0 |
| 39666 | ENSMUSG00000101382 | 0 | NA | NA | NA | NA | NA | Gm28213 | predicted gene 28213 [Source:MGI Symbc | 0 | 0 | 0 | 0 | 0 | 0 |
| 39667 | ENSMUSG00000101383 | 0 | NA | NA | NA | NA | NA | Gm28843 | predicted gene 28843 [Source:MGI Symbc | 0 | 0 | 0 | 0 | 0 | 0 |
| 39668 | ENSMUSG00000101385 | 0 | NA | NA | NA | NA | NA | Gm5261 | predicted gene 5261 [Source:MGI Symbol | 0 | 0 | 0 | 0 | 0 | 0 |
| 39669 | ENSMUSG00000101386 | 0 | NA | NA | NA | NA | NA | Gm29346 | predicted gene 29346 [Source:MGI Symbc | 0 | 0 | 0 | 0 | 0 | 0 |
| 39670 | ENSMUSG00000101387 | 0 | NA | NA | NA | NA | NA | Pced1c-ps | PC-esterase domain containing 1C, pseud | 0 | 0 | 0 | 0 | 0 | 0 |
| 39671 | ENSMUSG00000101388 | 0 | NA | NA | NA | NA | NA | Gm28444 | predicted gene 28444 [Source:MGI Symbc | 0 | 0 | 0 | 0 | 0 | 0 |
| 39672 | ENSMUSG00000101389 | 1.552662665 | -2.883080985 | 2.134239277 | -1.35087055 | 0.176736904 | NA | Ms4a4a | membrane-spanning 4-domains, subfamil | 3.2124 | 5.1396 | 0 | 0.964 | 0 | 0 |
| 39673 | ENSMUSG00000101390 | 0 | NA | NA | NA | NA | NA | Gm29420 | predicted gene 29420 [Source:MGI Symbc | 0 | 0 | 0 | 0 | 0 | 0 |
| 39674 | ENSMUSG00000101391 | 0 | NA | NA | NA | NA | NA | Olfr1217 | olfactory receptor 1217 [Source:MGI Syml | 0 | 0 | 0 | 0 | 0 | 0 |
| 39675 | ENSMUSG00000101392 | 1.183004194 | -1.310594698 | 2.15604226 | -0.607870598 | 0.543273301 | NA | NA | NA | 4.0155 | 1.0279 | 0 | 0.964 | 1.0906 | 0 |
| 39676 | ENSMUSG00000101393 | 0 | NA | NA | NA | NA | NA | Gm29447 | predicted gene 29447 [Source:MGI Symbc | 0 | 0 | 0 | 0 | 0 | 0 |
| 39677 | ENSMUSG00000101394 | 0 | NA | NA | NA | NA | NA | Gm28183 | predicted gene 28183 [Source:MGI Symbc | 0 | 0 | 0 | 0 | 0 | 0 |
| 39678 | ENSMUSG00000101395 | 0 | NA | NA | NA | NA | NA | Gm8241 | predicted gene 8241 [Source:MGI Symbol | 0 | 0 | 0 | 0 | 0 | 0 |
| 39679 | ENSMUSG00000101396 | 0 | NA | NA | NA | NA | NA | Gm28510 | predicted gene 28510 [Source:MGI Symbc | 0 | 0 | 0 | 0 | 0 | 0 |
| 39680 | ENSMUSG00000101397 | 998.7565768 | 0.405279952 | 0.179556403 | 2.257117793 | 0.024000715 | 0.197242243 | Mug-ps1 | murinoglobulin, pseudogene 1 [Source:M | 967.74 | 738.05 | 872.41 | 1437.3 | 1151.7 | 825.36 |
| 39681 | ENSMUSG00000101398 | 0 | NA | NA | NA | NA | NA | Gm2617 | predicted gene 2617 [Source:MGI Symbol | 0 | 0 | 0 | 0 | 0 | 0 |
| 39682 | ENSMUSG00000101399 | 0 | NA | NA | NA | NA | NA | Gm28961 | predicted gene 28961 [Source:MGI Symbc | 0 | 0 | 0 | 0 | 0 | 0 |
| 39683 | ENSMUSG00000101400 | 0.979405355 | -0.876544108 | 2.280749483 | -0.384322835 | 0.700739185 | NA | Gm29596 | predicted gene 29596 [Source:MGI Symbc | 0.8031 | 0 | 3.0187 | 0.964 | 1.0906 | 0 |
| 39684 | ENSMUSG00000101401 | 0 | NA | NA | NA | NA | NA | Scgb1b18 | secretoglobin, family 1B, member 18 [Sou | 0 | 0 | 0 | 0 | 0 | 0 |
| 39685 | ENSMUSG00000101402 | 0 | NA | NA | NA | NA | NA | Gm28673 | predicted gene 28673 [Source:MGI Symbc | 0 | 0 | 0 | 0 | 0 | 0 |
| 39686 | ENSMUSG00000101403 | 0 | NA | NA | NA | NA | NA | Gm20882 | predicted gene, 20882 [Source:MGI Symb | 0 | 0 | 0 | 0 | 0 | 0 |
| 39687 | ENSMUSG00000101404 | 0 | NA | NA | NA | NA | NA | 1700009C05 | RIKEN cDNA 1700009C05 gene [Source:M | 0 | 0 | 0 | 0 | 0 | 0 |
| 39688 | ENSMUSG00000101405 | 0 | NA | NA | NA | NA | NA | Gm28758 | predicted gene 28758 [Source:MGI Symbc | 0 | 0 | 0 | 0 | 0 | 0 |
| 39689 | ENSMUSG00000101406 | 0.367410362 | 2.078255025 | 3.800070157 | 0.54689912 | 0.58444804 | NA | Gm29200 | predicted gene 29200 [Source:MGI Symbc | 0 | 0 | 0 | 0 | 1.0906 | 1.1138 |
| 39690 | ENSMUSG00000101407 | 0 | NA | NA | NA | NA | NA | Gm29532 | predicted gene 29532 [Source:MGI Symbc | 0 | 0 | 0 | 0 | 0 | 0 |
| 39691 | ENSMUSG00000101408 | 0 | NA | NA | NA | NA | NA | Gm21155 | predicted gene, 21155 [Source:MGI Symb | 0 | 0 | 0 | 0 | 0 | 0 |
| 39692 | ENSMUSG00000101409 | 0 | NA | NA | NA | NA | NA | 1700122D07 | RIKEN cDNA 1700122D07 gene [Source:M | 0 | 0 | 0 | 0 | 0 | 0 |
| 39693 | ENSMUSG00000101410 | 0.267700746 | -1.497481649 | 4.057001436 | -0.369110456 | 0.712045396 | NA | Gm28581 | predicted gene 28581 [Source:MGI Symbc | 1.6062 | 0 | 0 | 0 | 0 | 0 |
| 39694 | ENSMUSG00000101411 | 0 | NA | NA | NA | NA | NA | Gm28111 | predicted gene 28111 [Source:MGI Symbc | 0 | 0 | 0 | 0 | 0 | 0 |
| 39695 | ENSMUSG00000101412 | 0 | NA | NA | NA | NA | NA | Gm28131 | predicted gene 28131 [Source:MGI Symbc | 0 | 0 | 0 | 0 | 0 | 0 |
| 39696 | ENSMUSG00000101413 | 0 | NA | NA | NA | NA | NA | Gm29485 | predicted gene 29485 [Source:MGI Symbc | 0 | 0 | 0 | 0 | 0 | 0 |
| 39697 | ENSMUSG00000101414 | 0 | NA | NA | NA | NA | NA | Gm29101 | predicted gene 29101 [Source:MGI Symbc | 0 | 0 | 0 | 0 | 0 | 0 |
| 39698 | ENSMUSG00000101415 | 0 | NA | NA | NA | NA | NA | Gm28168 | predicted gene 28168 [Source:MGI Symbc | 0 | 0 | 0 | 0 | 0 | 0 |
| 39699 | ENSMUSG00000101416 | 0 | NA | NA | NA | NA | NA | Gm29318 | predicted gene 29318 [Source:MGI Symbc | 0 | 0 | 0 | 0 | 0 | 0 |
| 39700 | ENSMUSG00000101417 | 0 | NA | NA | NA | NA | NA | Gm21647 | predicted gene, 21647 [Source:MGI Symb | 0 | 0 | 0 | 0 | 0 | 0 |

| | | | | | | | | | | | | | | |
|---|---|---|---|---|---|---|---|---|---|---|---|---|---|---|
| 39701 | ENSMUSG00000101418 | 0 | NA | NA | NA | NA | NA | Gm28983 | predicted gene 28983 [Source:MGI Symbo | 0 | 0 | 0 | 0 | 0 | 0 |
| 39702 | ENSMUSG00000101419 | 0 | NA | NA | NA | NA | NA | Gm29600 | predicted gene 29600 [Source:MGI Symbo | 0 | 0 | 0 | 0 | 0 | 0 |
| 39703 | ENSMUSG00000101420 | 0 | NA | NA | NA | NA | NA | Gm21902 | predicted gene, 21902 [Source:MGI Symb | 0 | 0 | 0 | 0 | 0 | 0 |
| 39704 | ENSMUSG00000101421 | 0 | NA | NA | NA | NA | NA | Gm28433 | predicted gene 28433 [Source:MGI Symbo | 0 | 0 | 0 | 0 | 0 | 0 |
| 39705 | ENSMUSG00000101422 | 0 | NA | NA | NA | NA | NA | Gm29301 | predicted gene 29301 [Source:MGI Symbo | 0 | 0 | 0 | 0 | 0 | 0 |
| 39706 | ENSMUSG00000101423 | 0 | NA | NA | NA | NA | NA | Gm28298 | predicted gene 28298 [Source:MGI Symbo | 0 | 0 | 0 | 0 | 0 | 0 |
| 39707 | ENSMUSG00000101424 | 0 | NA | NA | NA | NA | NA | Gm28073 | predicted gene 28073 [Source:MGI Symbo | 0 | 0 | 0 | 0 | 0 | 0 |
| 39708 | ENSMUSG00000101425 | 0 | NA | NA | NA | NA | NA | Gm29255 | predicted gene 29255 [Source:MGI Symbo | 0 | 0 | 0 | 0 | 0 | 0 |
| 39709 | ENSMUSG00000101426 | 0 | NA | NA | NA | NA | NA | Gm20978 | predicted gene, 20978 [Source:MGI Symb | 0 | 0 | 0 | 0 | 0 | 0 |
| 39710 | ENSMUSG00000101427 | 0 | NA | NA | NA | NA | NA | Gm6003 | predicted gene 6003 [Source:MGI Symbol | 0 | 0 | 0 | 0 | 0 | 0 |
| 39711 | ENSMUSG00000101428 | 0 | NA | NA | NA | NA | NA | Gm29272 | predicted gene 29272 [Source:MGI Symbo | 0 | 0 | 0 | 0 | 0 | 0 |
| 39712 | ENSMUSG00000101429 | 0 | NA | NA | NA | NA | NA | BC055402 | cDNA sequence BC055402 [Source:MGI Sy | 0 | 0 | 0 | 0 | 0 | 0 |
| 39713 | ENSMUSG00000101431 | 0 | NA | NA | NA | NA | NA | Gm7901 | predicted gene 7901 [Source:MGI Symbol | 0 | 0 | 0 | 0 | 0 | 0 |
| 39714 | ENSMUSG00000101433 | 0 | NA | NA | NA | NA | NA | Gm29441 | predicted gene 29441 [Source:MGI Symbo | 0 | 0 | 0 | 0 | 0 | 0 |
| 39715 | ENSMUSG00000101434 | 0 | NA | NA | NA | NA | NA | Gm29211 | predicted gene 29211 [Source:MGI Symbo | 0 | 0 | 0 | 0 | 0 | 0 |
| 39716 | ENSMUSG00000101435 | 0 | NA | NA | NA | NA | NA | Gm28772 | predicted gene 28772 [Source:MGI Symbo | 0 | 0 | 0 | 0 | 0 | 0 |
| 39717 | ENSMUSG00000101436 | 0 | NA | NA | NA | NA | NA | Gm5157 | predicted gene 5157 [Source:MGI Symbol | 0 | 0 | 0 | 0 | 0 | 0 |
| 39718 | ENSMUSG00000101437 | 0.160662974 | 1.126924175 | 4.080472857 | 0.276174898 | 0.782413736 | NA | Gm29454 | predicted gene 29454 [Source:MGI Symbo | 0 | 0 | 0 | 0.964 | 0 | 0 |
| 39719 | ENSMUSG00000101438 | 0 | NA | NA | NA | NA | NA | Gm19412 | predicted gene, 19412 [Source:MGI Symb | 0 | 0 | 0 | 0 | 0 | 0 |
| 39720 | ENSMUSG00000101439 | 0.556921942 | 2.679043159 | 3.638243063 | 0.736356289 | 0.461513899 | NA | Gm336 | predicted pseudogene 336 [Source:MGI S | 0 | 0 | 0 | 0 | 0 | 3.3415 |
| 39721 | ENSMUSG00000101440 | 0 | NA | NA | NA | NA | NA | Gm29425 | predicted gene 29425 [Source:MGI Symbo | 0 | 0 | 0 | 0 | 0 | 0 |
| 39722 | ENSMUSG00000101441 | 0 | NA | NA | NA | NA | NA | Gm29363 | predicted gene 29363 [Source:MGI Symbo | 0 | 0 | 0 | 0 | 0 | 0 |
| 39723 | ENSMUSG00000101442 | 0 | NA | NA | NA | NA | NA | Gm28795 | predicted gene 28795 [Source:MGI Symbo | 0 | 0 | 0 | 0 | 0 | 0 |
| 39724 | ENSMUSG00000101443 | 0 | NA | NA | NA | NA | NA | Rhoay-ps1 | ras homolog A translocated to the Y, pseu | 0 | 0 | 0 | 0 | 0 | 0 |
| 39725 | ENSMUSG00000101444 | 0 | NA | NA | NA | NA | NA | Gm29179 | predicted gene 29179 [Source:MGI Symbo | 0 | 0 | 0 | 0 | 0 | 0 |
| 39726 | ENSMUSG00000101445 | 0 | NA | NA | NA | NA | NA | Gm28932 | predicted gene 28932 [Source:MGI Symbo | 0 | 0 | 0 | 0 | 0 | 0 |
| 39727 | ENSMUSG00000101446 | 0 | NA | NA | NA | NA | NA | Gm21749 | predicted gene, 21749 [Source:MGI Symb | 0 | 0 | 0 | 0 | 0 | 0 |
| 39728 | ENSMUSG00000101447 | 0 | NA | NA | NA | NA | NA | Gm28826 | predicted gene 28826 [Source:MGI Symbo | 0 | 0 | 0 | 0 | 0 | 0 |
| 39729 | ENSMUSG00000101448 | 0 | NA | NA | NA | NA | NA | Gm28494 | predicted gene 28494 [Source:MGI Symbo | 0 | 0 | 0 | 0 | 0 | 0 |
| 39730 | ENSMUSG00000101449 | 0 | NA | NA | NA | NA | NA | Gm17791 | predicted gene, 17791 [Source:MGI Symb | 0 | 0 | 0 | 0 | 0 | 0 |
| 39731 | ENSMUSG00000101450 | 3.589610937 | -0.409092614 | 1.200926096 | -0.340647618 | 0.733368877 | NA | Gm28941 | predicted gene 28941 [Source:MGI Symbo | 3.2124 | 4.1117 | 5.0312 | 4.8199 | 4.3625 | 0 |
| 39732 | ENSMUSG00000101451 | 0 | NA | NA | NA | NA | NA | Gm29430 | predicted gene 29430 [Source:MGI Symbo | 0 | 0 | 0 | 0 | 0 | 0 |
| 39733 | ENSMUSG00000101452 | 0.603115608 | -2.642822926 | 3.106511347 | -0.850736608 | 0.394915683 | NA | Gm28530 | predicted gene 28530 [Source:MGI Symbo | 1.6062 | 0 | 2.0125 | 0 | 0 | 0 |
| 39734 | ENSMUSG00000101453 | 0 | NA | NA | NA | NA | NA | Gm28189 | predicted gene 28189 [Source:MGI Symbo | 0 | 0 | 0 | 0 | 0 | 0 |
| 39735 | ENSMUSG00000101454 | 0 | NA | NA | NA | NA | NA | Gm1988 | predicted gene 1988 [Source:MGI Symbol | 0 | 0 | 0 | 0 | 0 | 0 |
| 39736 | ENSMUSG00000101455 | 0 | NA | NA | NA | NA | NA | Gm21805 | predicted gene, 21805 [Source:MGI Symb | 0 | 0 | 0 | 0 | 0 | 0 |
| 39737 | ENSMUSG00000101456 | 0 | NA | NA | NA | NA | NA | Gm29043 | predicted gene 29043 [Source:MGI Symbo | 0 | 0 | 0 | 0 | 0 | 0 |
| 39738 | ENSMUSG00000101457 | 0 | NA | NA | NA | NA | NA | Gm18701 | predicted gene, 18701 [Source:MGI Symb | 0 | 0 | 0 | 0 | 0 | 0 |
| 39739 | ENSMUSG00000101459 | 0 | NA | NA | NA | NA | NA | Gm19193 | predicted gene, 19193 [Source:MGI Symb | 0 | 0 | 0 | 0 | 0 | 0 |
| 39740 | ENSMUSG00000101462 | 0.367410362 | 2.078255025 | 3.800070157 | 0.54689912 | 0.58444804 | NA | Gm3052 | predicted gene 3052 [Source:MGI Symbol | 0 | 0 | 0 | 0 | 1.0906 | 1.1138 |
| 39741 | ENSMUSG00000101463 | 0 | NA | NA | NA | NA | NA | Gm28750 | predicted gene 28750 [Source:MGI Symbo | 0 | 0 | 0 | 0 | 0 | 0 |
| 39742 | ENSMUSG00000101464 | 0 | NA | NA | NA | NA | NA | Gm7436 | predicted gene 7436 [Source:MGI Symbol | 0 | 0 | 0 | 0 | 0 | 0 |
| 39743 | ENSMUSG00000101465 | 0 | NA | NA | NA | NA | NA | Gm29190 | predicted gene 29190 [Source:MGI Symbo | 0 | 0 | 0 | 0 | 0 | 0 |
| 39744 | ENSMUSG00000101466 | 0 | NA | NA | NA | NA | NA | Gm29548 | predicted gene 29548 [Source:MGI Symbo | 0 | 0 | 0 | 0 | 0 | 0 |
| 39745 | ENSMUSG00000101467 | 0 | NA | NA | NA | NA | NA | Gm28662 | predicted gene 28662 [Source:MGI Symbo | 0 | 0 | 0 | 0 | 0 | 0 |
| 39746 | ENSMUSG00000101468 | 0 | NA | NA | NA | NA | NA | Gm7761 | predicted gene 7761 [Source:MGI Symbol | 0 | 0 | 0 | 0 | 0 | 0 |
| 39747 | ENSMUSG00000101469 | 0 | NA | NA | NA | NA | NA | Gm29556 | predicted gene 29556 [Source:MGI Symbo | 0 | 0 | 0 | 0 | 0 | 0 |
| 39748 | ENSMUSG00000101470 | 0 | NA | NA | NA | NA | NA | Gm21129 | predicted gene, 21129 [Source:MGI Symb | 0 | 0 | 0 | 0 | 0 | 0 |
| 39749 | ENSMUSG00000101471 | 0 | NA | NA | NA | NA | NA | Gm29276 | predicted gene 29276 [Source:MGI Symbo | 0 | 0 | 0 | 0 | 0 | 0 |
| 39750 | ENSMUSG00000101472 | 0 | NA | NA | NA | NA | NA | Gm28517 | predicted gene 28517 [Source:MGI Symbo | 0 | 0 | 0 | 0 | 0 | 0 |
| 39751 | ENSMUSG00000101473 | 0 | NA | NA | NA | NA | NA | Gm20893 | predicted gene, 20893 [Source:MGI Symb | 0 | 0 | 0 | 0 | 0 | 0 |
| 39752 | ENSMUSG00000101474 | 0 | NA | NA | NA | NA | NA | Gm29656 | predicted gene 29656 [Source:MGI Symbo | 0 | 0 | 0 | 0 | 0 | 0 |
| 39753 | ENSMUSG00000101475 | 0 | NA | NA | NA | NA | NA | Gm21316 | predicted gene, 21316 [Source:MGI Symb | 0 | 0 | 0 | 0 | 0 | 0 |
| 39754 | ENSMUSG00000101476 | 0 | NA | NA | NA | NA | NA | Gm29570 | predicted gene 29570 [Source:MGI Symbo | 0 | 0 | 0 | 0 | 0 | 0 |
| 39755 | ENSMUSG00000101477 | 0 | NA | NA | NA | NA | NA | Gm18448 | predicted gene, 18448 [Source:MGI Symb | 0 | 0 | 0 | 0 | 0 | 0 |
| 39756 | ENSMUSG00000101478 | 0 | NA | NA | NA | NA | NA | Gm20859 | predicted gene, 20859 [Source:MGI Symb | 0 | 0 | 0 | 0 | 0 | 0 |

| | | | | | | | | | | | | | | | |
|---|---|---|---|---|---|---|---|---|---|---|---|---|---|---|---|
| 39757 | ENSMUSG00000101479 | 0 | NA | NA | NA | NA | NA | Gm28997 | predicted gene 28997 [Source:MGI Symbo | 0 | 0 | 0 | 0 | 0 | 0 |
| 39758 | ENSMUSG00000101480 | 0 | NA | NA | NA | NA | NA | Olfr1220 | olfactory receptor 1220 [Source:MGI Sym | 0 | 0 | 0 | 0 | 0 | 0 |
| 39759 | ENSMUSG00000101481 | 0 | NA | NA | NA | NA | NA | 1700123O21 | RIKEN cDNA 1700123O21 gene [Source:M | 0 | 0 | 0 | 0 | 0 | 0 |
| 39760 | ENSMUSG00000101482 | 0 | NA | NA | NA | NA | NA | Gm28883 | predicted gene 28883 [Source:MGI Symbo | 0 | 0 | 0 | 0 | 0 | 0 |
| 39761 | ENSMUSG00000101483 | 0 | NA | NA | NA | NA | NA | 1700016L21 | RIKEN cDNA 1700016L21 gene [Source:M | 0 | 0 | 0 | 0 | 0 | 0 |
| 39762 | ENSMUSG00000101484 | 0 | NA | NA | NA | NA | NA | Gm28798 | predicted gene 28798 [Source:MGI Symbo | 0 | 0 | 0 | 0 | 0 | 0 |
| 39763 | ENSMUSG00000101486 | 0 | NA | NA | NA | NA | NA | Gm4405 | predicted gene 4405 [Source:MGI Symbol | 0 | 0 | 0 | 0 | 0 | 0 |
| 39764 | ENSMUSG00000101487 | 0 | NA | NA | NA | NA | NA | Scgb2b33 | secretoglobin, family 2B, member 33 [Sou | 0 | 0 | 0 | 0 | 0 | 0 |
| 39765 | ENSMUSG00000101489 | 0 | NA | NA | NA | NA | NA | Gm28090 | predicted gene 28090 [Source:MGI Symbo | 0 | 0 | 0 | 0 | 0 | 0 |
| 39766 | ENSMUSG00000101490 | 0 | NA | NA | NA | NA | NA | Gm20927 | predicted gene, 20927 [Source:MGI Symb | 0 | 0 | 0 | 0 | 0 | 0 |
| 39767 | ENSMUSG00000101491 | 0 | NA | NA | NA | NA | NA | Gm28632 | predicted gene 28632 [Source:MGI Symbo | 0 | 0 | 0 | 0 | 0 | 0 |
| 39768 | ENSMUSG00000101492 | 0 | NA | NA | NA | NA | NA | Gm29089 | predicted gene 29089 [Source:MGI Symbo | 0 | 0 | 0 | 0 | 0 | 0 |
| 39769 | ENSMUSG00000101493 | 11.50259311 | 0.217760123 | 0.79369263 | 0.274363292 | 0.78380545 | 0.960670212 | 2810405F17 | RIKEN cDNA 2810405F17 gene [Source:M | 17.668 | 1.0279 | 13.081 | 17.352 | 7.6343 | 12.252 |
| 39770 | ENSMUSG00000101494 | 0 | NA | NA | NA | NA | NA | Gm21792 | predicted gene, 21792 [Source:MGI Symb | 0 | 0 | 0 | 0 | 0 | 0 |
| 39771 | ENSMUSG00000101495 | 0 | NA | NA | NA | NA | NA | Gm28912 | predicted gene 28912 [Source:MGI Symbo | 0 | 0 | 0 | 0 | 0 | 0 |
| 39772 | ENSMUSG00000101496 | 0 | NA | NA | NA | NA | NA | Gm8885 | predicted gene 8885 [Source:MGI Symbol | 0 | 0 | 0 | 0 | 0 | 0 |
| 39773 | ENSMUSG00000101497 | 0 | NA | NA | NA | NA | NA | Gm29391 | predicted gene 29391 [Source:MGI Symbo | 0 | 0 | 0 | 0 | 0 | 0 |
| 39774 | ENSMUSG00000101498 | 0 | NA | NA | NA | NA | NA | Gm2950 | predicted gene 2950 [Source:MGI Symbol | 0 | 0 | 0 | 0 | 0 | 0 |
| 39775 | ENSMUSG00000101499 | 0 | NA | NA | NA | NA | NA | Vmn2r-ps13 | vomeronasal 2, receptor, pseudogene 13 | 0 | 0 | 0 | 0 | 0 | 0 |
| 39776 | ENSMUSG00000101500 | 0 | NA | NA | NA | NA | NA | Btbd35f12 | BTB domain containing 35, family membe | 0 | 0 | 0 | 0 | 0 | 0 |
| 39777 | ENSMUSG00000101501 | 0.936952612 | -3.291460614 | 3.015666682 | -1.091453719 | 0.275073286 | NA | Rps27a-ps1 | ribosomal protein S27A, pseudogene 1 [S | 5.6217 | 0 | 0 | 0 | 0 | 0 |
| 39778 | ENSMUSG00000101502 | 0.167707431 | -0.796629176 | 4.080472857 | -0.195229623 | 0.84521319 | NA | Rpl7a-ps10 | ribosomal protein L7A, pseudogene 10 [S | 0 | 0 | 1.0062 | 0 | 0 | 0 |
| 39779 | ENSMUSG00000101503 | 0 | NA | NA | NA | NA | NA | Gm28364 | predicted gene 28364 [Source:MGI Symbo | 0 | 0 | 0 | 0 | 0 | 0 |
| 39780 | ENSMUSG00000101504 | 0 | NA | NA | NA | NA | NA | Gm28257 | predicted gene 28257 [Source:MGI Symbo | 0 | 0 | 0 | 0 | 0 | 0 |
| 39781 | ENSMUSG00000101505 | 0 | NA | NA | NA | NA | NA | 1700109G14 | RIKEN cDNA 1700109G14 gene [Source:M | 0 | 0 | 0 | 0 | 0 | 0 |
| 39782 | ENSMUSG00000101506 | 0 | NA | NA | NA | NA | NA | Gm28385 | predicted gene 28385 [Source:MGI Symbo | 0 | 0 | 0 | 0 | 0 | 0 |
| 39783 | ENSMUSG00000101507 | 0 | NA | NA | NA | NA | NA | Gm29457 | predicted gene 29457 [Source:MGI Symbo | 0 | 0 | 0 | 0 | 0 | 0 |
| 39784 | ENSMUSG00000101508 | 0 | NA | NA | NA | NA | NA | Gm28832 | predicted gene 28832 [Source:MGI Symbo | 0 | 0 | 0 | 0 | 0 | 0 |
| 39785 | ENSMUSG00000101509 | 0 | NA | NA | NA | NA | NA | Gm29641 | predicted gene 29641 [Source:MGI Symbo | 0 | 0 | 0 | 0 | 0 | 0 |
| 39786 | ENSMUSG00000101510 | 0 | NA | NA | NA | NA | NA | Gm21573 | predicted gene, 21573 [Source:MGI Symb | 0 | 0 | 0 | 0 | 0 | 0 |
| 39787 | ENSMUSG00000101511 | 0 | NA | NA | NA | NA | NA | Gm5681 | predicted gene 5681 [Source:MGI Symbol | 0 | 0 | 0 | 0 | 0 | 0 |
| 39788 | ENSMUSG00000101512 | 0 | NA | NA | NA | NA | NA | Gm28135 | predicted gene 28135 [Source:MGI Symbo | 0 | 0 | 0 | 0 | 0 | 0 |
| 39789 | ENSMUSG00000101513 | 0 | NA | NA | NA | NA | NA | Gm28853 | predicted gene 28853 [Source:MGI Symbo | 0 | 0 | 0 | 0 | 0 | 0 |
| 39790 | ENSMUSG00000101514 | 283.7059686 | -0.261514779 | 0.147974487 | -1.767296408 | 0.077178601 | 0.406818982 | Gm5524 | predicted gene 5524 [Source:MGI Symbol | 318.83 | 295.01 | 313.95 | 256.42 | 257.39 | 260.64 |
| 39791 | ENSMUSG00000101515 | 0 | NA | NA | NA | NA | NA | Gm21899 | predicted gene, 21899 [Source:MGI Symb | 0 | 0 | 0 | 0 | 0 | 0 |
| 39792 | ENSMUSG00000101516 | 0 | NA | NA | NA | NA | NA | Gm29074 | predicted gene 29074 [Source:MGI Symbo | 0 | 0 | 0 | 0 | 0 | 0 |
| 39793 | ENSMUSG00000101517 | 529.5995673 | -0.702319612 | 0.176564605 | -3.97769197 | 6.96E-05 | 0.00269121 | 4732465J04 | RIKEN cDNA 4732465J04 gene [Source:M | 632.04 | 551.99 | 784.87 | 490.66 | 332.64 | 385.39 |
| 39794 | ENSMUSG00000101518 | 0 | NA | NA | NA | NA | NA | Gm20868 | predicted gene, 20868 [Source:MGI Symb | 0 | 0 | 0 | 0 | 0 | 0 |
| 39795 | ENSMUSG00000101519 | 0 | NA | NA | NA | NA | NA | Ska2l-ps | spindle and kinetochore associated comp | 0 | 0 | 0 | 0 | 0 | 0 |
| 39796 | ENSMUSG00000101520 | 0 | NA | NA | NA | NA | NA | Scgb1b12 | secretoglobin, family 1B, member 12 [Sou | 0 | 0 | 0 | 0 | 0 | 0 |
| 39797 | ENSMUSG00000101521 | 0 | NA | NA | NA | NA | NA | Gm28393 | predicted gene 28393 [Source:MGI Symbo | 0 | 0 | 0 | 0 | 0 | 0 |
| 39798 | ENSMUSG00000101522 | 0 | NA | NA | NA | NA | NA | Gm7837 | predicted gene 7837 [Source:MGI Symbol | 0 | 0 | 0 | 0 | 0 | 0 |
| 39799 | ENSMUSG00000101523 | 10.85052404 | 0.640314694 | 0.650243205 | 0.984731081 | 0.324756181 | 0.771309263 | Csnk2a1-ps3 | casein kinase 2, alpha 1 polypeptide, pseu | 7.2279 | 10.279 | 8.05 | 16.388 | 10.906 | 12.252 |
| 39800 | ENSMUSG00000101525 | 0 | NA | NA | NA | NA | NA | Gm28593 | predicted gene 28593 [Source:MGI Symbo | 0 | 0 | 0 | 0 | 0 | 0 |
| 39801 | ENSMUSG00000101526 | 0 | NA | NA | NA | NA | NA | Gm28622 | predicted gene 28622 [Source:MGI Symbo | 0 | 0 | 0 | 0 | 0 | 0 |
| 39802 | ENSMUSG00000101527 | 0 | NA | NA | NA | NA | NA | Gm21908 | predicted gene, 21908 [Source:MGI Symb | 0 | 0 | 0 | 0 | 0 | 0 |
| 39803 | ENSMUSG00000101528 | 0 | NA | NA | NA | NA | NA | Gm29110 | predicted gene 29110 [Source:MGI Symbo | 0 | 0 | 0 | 0 | 0 | 0 |
| 39804 | ENSMUSG00000101529 | 0 | NA | NA | NA | NA | NA | Gm29636 | predicted gene 29636 [Source:MGI Symbo | 0 | 0 | 0 | 0 | 0 | 0 |
| 39805 | ENSMUSG00000101530 | 0 | NA | NA | NA | NA | NA | Gm29259 | predicted gene 29259 [Source:MGI Symbo | 0 | 0 | 0 | 0 | 0 | 0 |
| 39806 | ENSMUSG00000101531 | 0 | NA | NA | NA | NA | NA | Gm29672 | predicted gene 29672 [Source:MGI Symbo | 0 | 0 | 0 | 0 | 0 | 0 |
| 39807 | ENSMUSG00000101532 | 0 | NA | NA | NA | NA | NA | Zbed4-ps1 | zinc finger, BED type containing 4, pseudo | 0 | 0 | 0 | 0 | 0 | 0 |
| 39808 | ENSMUSG00000101533 | 0 | NA | NA | NA | NA | NA | Gm29393 | predicted gene 29393 [Source:MGI Symbo | 0 | 0 | 0 | 0 | 0 | 0 |
| 39809 | ENSMUSG00000101534 | 0 | NA | NA | NA | NA | NA | Gm29306 | predicted gene 29306 [Source:MGI Symbo | 0 | 0 | 0 | 0 | 0 | 0 |
| 39810 | ENSMUSG00000101535 | 0.705972119 | 3.02851815 | 2.966479847 | 1.020913104 | 0.30729561 | NA | Gm29383 | predicted gene 29383 [Source:MGI Symbo | 0 | 0 | 0 | 0.964 | 3.2719 | 0 |
| 39811 | ENSMUSG00000101536 | 0 | NA | NA | NA | NA | NA | Gm28834 | predicted gene 28834 [Source:MGI Symbo | 0 | 0 | 0 | 0 | 0 | 0 |
| 39812 | ENSMUSG00000101537 | 33.51503345 | 0.058673169 | 0.402849038 | 0.145645548 | 0.884201209 | 0.981769403 | Gm29571 | predicted gene 29571 [Source:MGI Symbo | 38.549 | 28.782 | 31.194 | 46.271 | 23.994 | 32.301 |

| | | | | | | | | | | | | | | | |
|---|---|---|---|---|---|---|---|---|---|---|---|---|---|---|---|
| 39813 | ENSMUSG00000101538 | 0 | NA | NA | NA | NA | NA | Gm29029 | predicted gene 29029 [Source:MGI Symbo | 0 | 0 | 0 | 0 | 0 | 0 |
| 39814 | ENSMUSG00000101539 | 0 | NA | NA | NA | NA | NA | Gm28252 | predicted gene 28252 [Source:MGI Symbo | 0 | 0 | 0 | 0 | 0 | 0 |
| 39815 | ENSMUSG00000101540 | 0 | NA | NA | NA | NA | NA | Gm28465 | predicted gene 28465 [Source:MGI Symbo | 0 | 0 | 0 | 0 | 0 | 0 |
| 39816 | ENSMUSG00000101541 | 0 | NA | NA | NA | NA | NA | Gm28554 | predicted gene 28554 [Source:MGI Symbo | 0 | 0 | 0 | 0 | 0 | 0 |
| 39817 | ENSMUSG00000101542 | 0 | NA | NA | NA | NA | NA | Gm29303 | predicted gene 29303 [Source:MGI Symbo | 0 | 0 | 0 | 0 | 0 | 0 |
| 39818 | ENSMUSG00000101543 | 0 | NA | NA | NA | NA | NA | Gm18352 | predicted gene, 18352 [Source:MGI Symb | 0 | 0 | 0 | 0 | 0 | 0 |
| 39819 | ENSMUSG00000101544 | 0 | NA | NA | NA | NA | NA | Gm29422 | predicted gene 29422 [Source:MGI Symbo | 0 | 0 | 0 | 0 | 0 | 0 |
| 39820 | ENSMUSG00000101545 | 0 | NA | NA | NA | NA | NA | Gm4373 | predicted gene 4373 [Source:MGI Symbol | 0 | 0 | 0 | 0 | 0 | 0 |
| 39821 | ENSMUSG00000101546 | 0 | NA | NA | NA | NA | NA | Gm28747 | predicted gene 28747 [Source:MGI Symbo | 0 | 0 | 0 | 0 | 0 | 0 |
| 39822 | ENSMUSG00000101547 | 0 | NA | NA | NA | NA | NA | Gm29252 | predicted gene 29252 [Source:MGI Symbo | 0 | 0 | 0 | 0 | 0 | 0 |
| 39823 | ENSMUSG00000101548 | 0 | NA | NA | NA | NA | NA | Gm9220 | predicted gene 9220 [Source:MGI Symbol | 0 | 0 | 0 | 0 | 0 | 0 |
| 39824 | ENSMUSG00000101549 | 0 | NA | NA | NA | NA | NA | Gm29630 | predicted gene 29630 [Source:MGI Symbo | 0 | 0 | 0 | 0 | 0 | 0 |
| 39825 | ENSMUSG00000101550 | 0 | NA | NA | NA | NA | NA | Gm29385 | predicted gene 29385 [Source:MGI Symbo | 0 | 0 | 0 | 0 | 0 | 0 |
| 39826 | ENSMUSG00000101551 | 0 | NA | NA | NA | NA | NA | Gm8870 | predicted gene 8870 [Source:MGI Symbol | 0 | 0 | 0 | 0 | 0 | 0 |
| 39827 | ENSMUSG00000101552 | 0 | NA | NA | NA | NA | NA | Gm29118 | predicted gene 29118 [Source:MGI Symbo | 0 | 0 | 0 | 0 | 0 | 0 |
| 39828 | ENSMUSG00000101553 | 0 | NA | NA | NA | NA | NA | 1700049E22 | RIKEN cDNA 1700049E22 gene [Source:M | 0 | 0 | 0 | 0 | 0 | 0 |
| 39829 | ENSMUSG00000101554 | 0 | NA | NA | NA | NA | NA | Scgb2b1 | secretoglobin, family 2B, member 1 [Sour | 0 | 0 | 0 | 0 | 0 | 0 |
| 39830 | ENSMUSG00000101555 | 0.706178709 | 0.10198885 | 3.03884937 | 0.033561667 | 0.97322669 | NA | Gm28731 | predicted gene 28731 [Source:MGI Symbo | 0 | 2.0558 | 0 | 0 | 2.1812 | 0 |
| 39831 | ENSMUSG00000101557 | 0 | NA | NA | NA | NA | NA | Gm28314 | predicted gene 28314 [Source:MGI Symbo | 0 | 0 | 0 | 0 | 0 | 0 |
| 39832 | ENSMUSG00000101558 | 0 | NA | NA | NA | NA | NA | Gm28647 | predicted gene 28647 [Source:MGI Symbo | 0 | 0 | 0 | 0 | 0 | 0 |
| 39833 | ENSMUSG00000101559 | 0.133850373 | -0.796629176 | 4.080472857 | -0.195229623 | 0.84521319 | NA | Gm28258 | predicted gene 28258 [Source:MGI Symbo | 0.8031 | 0 | 0 | 0 | 0 | 0 |
| 39834 | ENSMUSG00000101560 | 0 | NA | NA | NA | NA | NA | Gm28127 | predicted gene 28127 [Source:MGI Symbo | 0 | 0 | 0 | 0 | 0 | 0 |
| 39835 | ENSMUSG00000101562 | 0 | NA | NA | NA | NA | NA | Gm29144 | predicted gene 29144 [Source:MGI Symbo | 0 | 0 | 0 | 0 | 0 | 0 |
| 39836 | ENSMUSG00000101563 | 0 | NA | NA | NA | NA | NA | Gm7364 | predicted gene 7364 [Source:MGI Symbol | 0 | 0 | 0 | 0 | 0 | 0 |
| 39837 | ENSMUSG00000101564 | 0 | NA | NA | NA | NA | NA | Gm5703 | predicted gene 5703 [Source:MGI Symbol | 0 | 0 | 0 | 0 | 0 | 0 |
| 39838 | ENSMUSG00000101565 | 0 | NA | NA | NA | NA | NA | 1700003L19I | RIKEN cDNA 1700003L19 gene [Source:M | 0 | 0 | 0 | 0 | 0 | 0 |
| 39839 | ENSMUSG00000101566 | 0 | NA | NA | NA | NA | NA | Gm28276 | predicted gene 28276 [Source:MGI Symbo | 0 | 0 | 0 | 0 | 0 | 0 |
| 39840 | ENSMUSG00000101567 | 0 | NA | NA | NA | NA | NA | Txn-ps1 | thioredoxin, pseudogene 1 [Source:MGI S | 0 | 0 | 0 | 0 | 0 | 0 |
| 39841 | ENSMUSG00000101568 | 0 | NA | NA | NA | NA | NA | 1700040F17 | RIKEN cDNA 1700040F17 gene [Source:M | 0 | 0 | 0 | 0 | 0 | 0 |
| 39842 | ENSMUSG00000101569 | 0 | NA | NA | NA | NA | NA | Gm28254 | predicted gene 28254 [Source:MGI Symbo | 0 | 0 | 0 | 0 | 0 | 0 |
| 39843 | ENSMUSG00000101570 | 0 | NA | NA | NA | NA | NA | Gm28233 | predicted gene 28233 [Source:MGI Symbo | 0 | 0 | 0 | 0 | 0 | 0 |
| 39844 | ENSMUSG00000101571 | 0 | NA | NA | NA | NA | NA | Gm6187 | predicted gene 6187 [Source:MGI Symbol | 0 | 0 | 0 | 0 | 0 | 0 |
| 39845 | ENSMUSG00000101573 | 0 | NA | NA | NA | NA | NA | Gm28361 | predicted gene 28361 [Source:MGI Symbo | 0 | 0 | 0 | 0 | 0 | 0 |
| 39846 | ENSMUSG00000101574 | 0 | NA | NA | NA | NA | NA | 1700112H15 | RIKEN cDNA 1700112H15 gene [Source:M | 0 | 0 | 0 | 0 | 0 | 0 |
| 39847 | ENSMUSG00000101575 | 0 | NA | NA | NA | NA | NA | Gm20898 | predicted gene, 20898 [Source:MGI Symb | 0 | 0 | 0 | 0 | 0 | 0 |
| 39848 | ENSMUSG00000101576 | 0 | NA | NA | NA | NA | NA | Gm20915 | predicted gene, 20915 [Source:MGI Symb | 0 | 0 | 0 | 0 | 0 | 0 |
| 39849 | ENSMUSG00000101578 | 0 | NA | NA | NA | NA | NA | Vmn1r206 | vomeronasal 1 receptor 206 [Source:MGI | 0 | 0 | 0 | 0 | 0 | 0 |
| 39850 | ENSMUSG00000101579 | 0 | NA | NA | NA | NA | NA | Gm21181 | predicted gene, 21181 [Source:MGI Symb | 0 | 0 | 0 | 0 | 0 | 0 |
| 39851 | ENSMUSG00000101580 | 0 | NA | NA | NA | NA | NA | Gm28201 | predicted gene 28201 [Source:MGI Symbo | 0 | 0 | 0 | 0 | 0 | 0 |
| 39852 | ENSMUSG00000101581 | 0 | NA | NA | NA | NA | NA | C430002N11 | RIKEN cDNA C430002N11 gene [Source:M | 0 | 0 | 0 | 0 | 0 | 0 |
| 39853 | ENSMUSG00000101582 | 0 | NA | NA | NA | NA | NA | Gm28828 | predicted gene 28828 [Source:MGI Symbo | 0 | 0 | 0 | 0 | 0 | 0 |
| 39854 | ENSMUSG00000101583 | 2.83027234 | -2.169160603 | 1.420794942 | -1.526723202 | 0.126829855 | NA | Gm20753 | predicted gene, 20753 [Source:MGI Symb | 4.8186 | 3.0838 | 6.0375 | 1.928 | 0 | 1.1138 |
| 39855 | ENSMUSG00000101584 | 0 | NA | NA | NA | NA | NA | 4930519F24 | RIKEN cDNA 4930519F24 gene [Source:M | 0 | 0 | 0 | 0 | 0 | 0 |
| 39856 | ENSMUSG00000101585 | 1.914890129 | -2.237354827 | 1.789439998 | -1.250310058 | 0.211186306 | NA | 1600010M0 | RIKEN cDNA 1600010M07 gene [Source:M | 2.4093 | 5.1396 | 2.0125 | 1.928 | 0 | 0 |
| 39857 | ENSMUSG00000101586 | 11.18253825 | 0.123129295 | 0.717087017 | 0.171707606 | 0.863667406 | 0.976954351 | Gm29017 | predicted gene 29017 [Source:MGI Symbo | 12.85 | 8.2233 | 11.069 | 18.316 | 3.2719 | 13.366 |
| 39858 | ENSMUSG00000101587 | 1.027504657 | 1.047266995 | 2.389110177 | 0.43835023 | 0.661132419 | NA | Gm29036 | predicted gene 29036 [Source:MGI Symbo | 0 | 2.0558 | 0 | 1.928 | 2.1812 | 0 |
| 39859 | ENSMUSG00000101588 | 1.034549114 | 0.072428415 | 2.202180142 | 0.032889414 | 0.973762774 | NA | Gm28265 | predicted gene 28265 [Source:MGI Symbo | 0 | 2.0558 | 1.0062 | 0.964 | 2.1812 | 0 |
| 39860 | ENSMUSG00000101589 | 0.335414862 | -1.779047242 | 4.030804179 | -0.441362855 | 0.658950329 | NA | Rbm6-ps1 | RNA binding motif protein 6, pseudogene | 0 | 0 | 2.0125 | 0 | 0 | 0 |
| 39861 | ENSMUSG00000101590 | 0 | NA | NA | NA | NA | NA | Gm29104 | predicted gene 29104 [Source:MGI Symbo | 0 | 0 | 0 | 0 | 0 | 0 |
| 39862 | ENSMUSG00000101592 | 0 | NA | NA | NA | NA | NA | Gm28689 | predicted gene 28689 [Source:MGI Symbo | 0 | 0 | 0 | 0 | 0 | 0 |
| 39863 | ENSMUSG00000101593 | 0 | NA | NA | NA | NA | NA | Gm20842 | predicted gene, 20842 [Source:MGI Symb | 0 | 0 | 0 | 0 | 0 | 0 |
| 39864 | ENSMUSG00000101594 | 0 | NA | NA | NA | NA | NA | Gm21779 | predicted gene, 21779 [Source:MGI Symb | 0 | 0 | 0 | 0 | 0 | 0 |
| 39865 | ENSMUSG00000101595 | 0 | NA | NA | NA | NA | NA | Gm29395 | predicted gene 29395 [Source:MGI Symbo | 0 | 0 | 0 | 0 | 0 | 0 |
| 39866 | ENSMUSG00000101596 | 5.21883459 | 4.907841507 | 1.539626609 | 3.187682961 | 0.001434177 | 0.02740282 | Gm28182 | predicted gene 28182 [Source:MGI Symbo | 0 | 0 | 1.0062 | 12.532 | 2.1812 | 15.594 |
| 39867 | ENSMUSG00000101597 | 5.137959228 | 0.552181725 | 1.100445999 | 0.501779938 | 0.615822327 | 0.917509899 | Gm5621 | predicted gene 5621 [Source:MGI Symbol | 2.4093 | 2.0558 | 8.05 | 3.8559 | 1.0906 | 13.366 |
| 39868 | ENSMUSG00000101598 | 0 | NA | NA | NA | NA | NA | Gm29449 | predicted gene 29449 [Source:MGI Symbo | 0 | 0 | 0 | 0 | 0 | 0 |

| | | | | | | | | | | | | | | | |
|---|---|---|---|---|---|---|---|---|---|---|---|---|---|---|---|
| 39869 | ENSMUSG00000101599 | 62.15344122 | 0.67854077 | 0.286192394 | 2.370925241 | 0.01774362 | 0.15881418 | Gm20342 | predicted gene, 20342 [Source:MGI Symb | 49.792 | 42.145 | 51.318 | 75.19 | 68.709 | 85.766 |
| 39870 | ENSMUSG00000101600 | 0 | NA | NA | NA | NA | NA | 4930599A14 | RIKEN cDNA 4930599A14 gene [Source:M | 0 | 0 | 0 | 0 | 0 | 0 |
| 39871 | ENSMUSG00000101601 | 0 | NA | NA | NA | NA | NA | Gm28210 | predicted gene 28210 [Source:MGI Symbo | 0 | 0 | 0 | 0 | 0 | 0 |
| 39872 | ENSMUSG00000101602 | 0 | NA | NA | NA | NA | NA | Gm28700 | predicted gene 28700 [Source:MGI Symbo | 0 | 0 | 0 | 0 | 0 | 0 |
| 39873 | ENSMUSG00000101603 | 0 | NA | NA | NA | NA | NA | Gm28730 | predicted gene 28730 [Source:MGI Symbo | 0 | 0 | 0 | 0 | 0 | 0 |
| 39874 | ENSMUSG00000101605 | 0 | NA | NA | NA | NA | NA | Ace3 | angiotensin I converting enzyme (peptidyl | 0 | 0 | 0 | 0 | 0 | 0 |
| 39875 | ENSMUSG00000101606 | 0 | NA | NA | NA | NA | NA | Gm29051 | predicted gene 29051 [Source:MGI Symbo | 0 | 0 | 0 | 0 | 0 | 0 |
| 39876 | ENSMUSG00000101607 | 0 | NA | NA | NA | NA | NA | Gm28939 | predicted gene 28939 [Source:MGI Symbo | 0 | 0 | 0 | 0 | 0 | 0 |
| 39877 | ENSMUSG00000101608 | 0 | NA | NA | NA | NA | NA | Gm28652 | predicted gene 28652 [Source:MGI Symbo | 0 | 0 | 0 | 0 | 0 | 0 |
| 39878 | ENSMUSG00000101609 | 33.78948793 | 0.187778481 | 0.43738504 | 0.429320767 | 0.667689806 | 0.930033258 | Kcnq1ot1 | KCNQ1 overlapping transcript 1 [Source:M | 32.124 | 23.642 | 39.244 | 55.911 | 25.084 | 26.732 |
| 39879 | ENSMUSG00000101610 | 0.160662974 | 1.126924175 | 4.080472857 | 0.276174898 | 0.782413736 | NA | Gm7560 | predicted gene 7560 [Source:MGI Symbol | 0 | 0 | 0 | 0.964 | 0 | 0 |
| 39880 | ENSMUSG00000101611 | 0 | NA | NA | NA | NA | NA | Gm29357 | predicted gene 29357 [Source:MGI Symbo | 0 | 0 | 0 | 0 | 0 | 0 |
| 39881 | ENSMUSG00000101612 | 0 | NA | NA | NA | NA | NA | Gm7933 | predicted gene 7933 [Source:MGI Symbol | 0 | 0 | 0 | 0 | 0 | 0 |
| 39882 | ENSMUSG00000101613 | 0 | NA | NA | NA | NA | NA | Gm28516 | predicted gene 28516 [Source:MGI Symbo | 0 | 0 | 0 | 0 | 0 | 0 |
| 39883 | ENSMUSG00000101614 | 0 | NA | NA | NA | NA | NA | Gm29547 | predicted gene 29547 [Source:MGI Symbo | 0 | 0 | 0 | 0 | 0 | 0 |
| 39884 | ENSMUSG00000101615 | 0 | NA | NA | NA | NA | NA | Gm8495 | predicted gene 8495 [Source:MGI Symbol | 0 | 0 | 0 | 0 | 0 | 0 |
| 39885 | ENSMUSG00000101616 | 0 | NA | NA | NA | NA | NA | Gm29047 | predicted gene 29047 [Source:MGI Symbo | 0 | 0 | 0 | 0 | 0 | 0 |
| 39886 | ENSMUSG00000101617 | 0 | NA | NA | NA | NA | NA | Gm6947 | predicted gene 6947 [Source:MGI Symbol | 0 | 0 | 0 | 0 | 0 | 0 |
| 39887 | ENSMUSG00000101618 | 0 | NA | NA | NA | NA | NA | Gm18443 | predicted gene, 18443 [Source:MGI Symb | 0 | 0 | 0 | 0 | 0 | 0 |
| 39888 | ENSMUSG00000101619 | 0 | NA | NA | NA | NA | NA | Gm28436 | predicted gene 28436 [Source:MGI Symbo | 0 | 0 | 0 | 0 | 0 | 0 |
| 39889 | ENSMUSG00000101621 | 0 | NA | NA | NA | NA | NA | Gm28905 | predicted gene 28905 [Source:MGI Symbo | 0 | 0 | 0 | 0 | 0 | 0 |
| 39890 | ENSMUSG00000101622 | 0 | NA | NA | NA | NA | NA | Mrgpra18-ps | MAS-related GPR, member A18, pseudoge | 0 | 0 | 0 | 0 | 0 | 0 |
| 39891 | ENSMUSG00000101623 | 0 | NA | NA | NA | NA | NA | Gm21245 | predicted gene, 21245 [Source:MGI Symb | 0 | 0 | 0 | 0 | 0 | 0 |
| 39892 | ENSMUSG00000101624 | 0 | NA | NA | NA | NA | NA | Gm28882 | predicted gene 28882 [Source:MGI Symbo | 0 | 0 | 0 | 0 | 0 | 0 |
| 39893 | ENSMUSG00000101625 | 0.596071152 | -1.224343926 | 2.897583865 | -0.4225396 | 0.672631203 | NA | Gm29371 | predicted gene 29371 [Source:MGI Symbo | 1.6062 | 0 | 1.0062 | 0.964 | 0 | 0 |
| 39894 | ENSMUSG00000101626 | 0 | NA | NA | NA | NA | NA | Gm29386 | predicted gene 29386 [Source:MGI Symbo | 0 | 0 | 0 | 0 | 0 | 0 |
| 39895 | ENSMUSG00000101627 | 0 | NA | NA | NA | NA | NA | Gm28432 | predicted gene 28432 [Source:MGI Symbo | 0 | 0 | 0 | 0 | 0 | 0 |
| 39896 | ENSMUSG00000101628 | 0 | NA | NA | NA | NA | NA | Gm28177 | predicted gene 28177 [Source:MGI Symbo | 0 | 0 | 0 | 0 | 0 | 0 |
| 39897 | ENSMUSG00000101629 | 0 | NA | NA | NA | NA | NA | Gm29588 | predicted gene 29588 [Source:MGI Symbo | 0 | 0 | 0 | 0 | 0 | 0 |
| 39898 | ENSMUSG00000101630 | 0 | NA | NA | NA | NA | NA | Gm29261 | predicted gene 29261 [Source:MGI Symbo | 0 | 0 | 0 | 0 | 0 | 0 |
| 39899 | ENSMUSG00000101631 | 0 | NA | NA | NA | NA | NA | Gm20884 | predicted gene, 20884 [Source:MGI Symb | 0 | 0 | 0 | 0 | 0 | 0 |
| 39900 | ENSMUSG00000101632 | 0 | NA | NA | NA | NA | NA | Gm28264 | predicted gene 28264 [Source:MGI Symbo | 0 | 0 | 0 | 0 | 0 | 0 |
| 39901 | ENSMUSG00000101633 | 0 | NA | NA | NA | NA | NA | Gm29839 | predicted gene, 29839 [Source:MGI Symb | 0 | 0 | 0 | 0 | 0 | 0 |
| 39902 | ENSMUSG00000101634 | 0 | NA | NA | NA | NA | NA | 1700066B17 | RIKEN cDNA 1700066B17 gene [Source:M | 0 | 0 | 0 | 0 | 0 | 0 |
| 39903 | ENSMUSG00000101636 | 0 | NA | NA | NA | NA | NA | 4930579H20 | RIKEN cDNA 4930579H20 gene [Source:M | 0 | 0 | 0 | 0 | 0 | 0 |
| 39904 | ENSMUSG00000101637 | 0 | NA | NA | NA | NA | NA | Gm28896 | predicted gene 28896 [Source:MGI Symbo | 0 | 0 | 0 | 0 | 0 | 0 |
| 39905 | ENSMUSG00000101638 | 0 | NA | NA | NA | NA | NA | Scgb1b11 | secretoglobin, family 1B, member 11 [Sou | 0 | 0 | 0 | 0 | 0 | 0 |
| 39906 | ENSMUSG00000101639 | 0 | NA | NA | NA | NA | NA | Gm8597 | predicted gene 8597 [Source:MGI Symbol | 0 | 0 | 0 | 0 | 0 | 0 |
| 39907 | ENSMUSG00000101640 | 0 | NA | NA | NA | NA | NA | Gm28376 | predicted gene 28376 [Source:MGI Symbo | 0 | 0 | 0 | 0 | 0 | 0 |
| 39908 | ENSMUSG00000101641 | 0 | NA | NA | NA | NA | NA | Gm29560 | predicted gene 29560 [Source:MGI Symbo | 0 | 0 | 0 | 0 | 0 | 0 |
| 39909 | ENSMUSG00000101642 | 0 | NA | NA | NA | NA | NA | Gm29315 | predicted gene 29315 [Source:MGI Symbo | 0 | 0 | 0 | 0 | 0 | 0 |
| 39910 | ENSMUSG00000101643 | 0 | NA | NA | NA | NA | NA | Gm28307 | predicted gene 28307 [Source:MGI Symbo | 0 | 0 | 0 | 0 | 0 | 0 |
| 39911 | ENSMUSG00000101644 | 0 | NA | NA | NA | NA | NA | Gm28343 | predicted gene 28343 [Source:MGI Symbo | 0 | 0 | 0 | 0 | 0 | 0 |
| 39912 | ENSMUSG00000101645 | 0 | NA | NA | NA | NA | NA | Gm28635 | predicted gene 28635 [Source:MGI Symbo | 0 | 0 | 0 | 0 | 0 | 0 |
| 39913 | ENSMUSG00000101646 | 0 | NA | NA | NA | NA | NA | Gm28106 | predicted gene 28106 [Source:MGI Symbo | 0 | 0 | 0 | 0 | 0 | 0 |
| 39914 | ENSMUSG00000101647 | 0 | NA | NA | NA | NA | NA | Gm29534 | predicted gene 29534 [Source:MGI Symbo | 0 | 0 | 0 | 0 | 0 | 0 |
| 39915 | ENSMUSG00000101648 | 0 | NA | NA | NA | NA | NA | Gm29388 | predicted gene 29388 [Source:MGI Symbo | 0 | 0 | 0 | 0 | 0 | 0 |
| 39916 | ENSMUSG00000101649 | 0 | NA | NA | NA | NA | NA | Gm29444 | predicted gene 29444 [Source:MGI Symbo | 0 | 0 | 0 | 0 | 0 | 0 |
| 39917 | ENSMUSG00000101650 | 0 | NA | NA | NA | NA | NA | Gm6820 | predicted gene 6820 [Source:MGI Symbol | 0 | 0 | 0 | 0 | 0 | 0 |
| 39918 | ENSMUSG00000101651 | 0 | NA | NA | NA | NA | NA | Gm28402 | predicted gene 28402 [Source:MGI Symbo | 0 | 0 | 0 | 0 | 0 | 0 |
| 39919 | ENSMUSG00000101652 | 0 | NA | NA | NA | NA | NA | Gm6060 | predicted gene 6060 [Source:MGI Symbol | 0 | 0 | 0 | 0 | 0 | 0 |
| 39920 | ENSMUSG00000101653 | 0 | NA | NA | NA | NA | NA | Gm20792 | predicted gene, 20792 [Source:MGI Symb | 0 | 0 | 0 | 0 | 0 | 0 |
| 39921 | ENSMUSG00000101654 | 0 | NA | NA | NA | NA | NA | Gm29653 | predicted gene 29653 [Source:MGI Symbo | 0 | 0 | 0 | 0 | 0 | 0 |
| 39922 | ENSMUSG00000101655 | 34.7386304 | 0.161372348 | 0.377984928 | 0.426927997 | 0.669431773 | 0.930837335 | 2310040G24 | RIKEN cDNA 2310040G24 gene [Source:M | 31.321 | 28.782 | 38.237 | 30.847 | 44.715 | 34.529 |
| 39923 | ENSMUSG00000101656 | 0 | NA | NA | NA | NA | NA | Gm8004 | predicted gene 8004 [Source:MGI Symbol | 0 | 0 | 0 | 0 | 0 | 0 |
| 39924 | ENSMUSG00000101657 | 0 | NA | NA | NA | NA | NA | Gm20864 | predicted gene, 20864 [Source:MGI Symb | 0 | 0 | 0 | 0 | 0 | 0 |

| | | | | | | | | | | | | | | | |
|---|---|---|---|---|---|---|---|---|---|---|---|---|---|---|---|
| 39925 | ENSMUSG00000101658 | 0 | NA | NA | NA | NA | NA | Scgb1b28-ps | secretoglobin, family 1B, member 28, pseu | 0 | 0 | 0 | 0 | 0 | 0 |
| 39926 | ENSMUSG00000101659 | 0 | NA | NA | NA | NA | NA | Gm29009 | predicted gene 29009 [Source:MGI Symbo | 0 | 0 | 0 | 0 | 0 | 0 |
| 39927 | ENSMUSG00000101660 | 0 | NA | NA | NA | NA | NA | Gm18666 | predicted gene, 18666 [Source:MGI Symb | 0 | 0 | 0 | 0 | 0 | 0 |
| 39928 | ENSMUSG00000101661 | 0 | NA | NA | NA | NA | NA | Gm28392 | predicted gene 28392 [Source:MGI Symbo | 0 | 0 | 0 | 0 | 0 | 0 |
| 39929 | ENSMUSG00000101662 | 0.469265235 | -2.274672822 | 3.439794491 | -0.661281605 | 0.508431736 | NA | Gm28499 | predicted gene 28499 [Source:MGI Symbo | 0.8031 | 0 | 2.0125 | 0 | 0 | 0 |
| 39930 | ENSMUSG00000101664 | 0 | NA | NA | NA | NA | NA | Mrgpra14 | MAS-related GPR, member A14 [Source:M | 0 | 0 | 0 | 0 | 0 | 0 |
| 39931 | ENSMUSG00000101665 | 0 | NA | NA | NA | NA | NA | Gm21920 | predicted gene, 21920 [Source:MGI Symb | 0 | 0 | 0 | 0 | 0 | 0 |
| 39932 | ENSMUSG00000101666 | 0 | NA | NA | NA | NA | NA | Rhoay-ps3 | ras homolog A translocated to the Y, pseu | 0 | 0 | 0 | 0 | 0 | 0 |
| 39933 | ENSMUSG00000101667 | 0 | NA | NA | NA | NA | NA | Gm29289 | predicted gene 29289 [Source:MGI Symbo | 0 | 0 | 0 | 0 | 0 | 0 |
| 39934 | ENSMUSG00000101668 | 0 | NA | NA | NA | NA | NA | Gm29549 | predicted gene 29549 [Source:MGI Symbo | 0 | 0 | 0 | 0 | 0 | 0 |
| 39935 | ENSMUSG00000101669 | 0 | NA | NA | NA | NA | NA | Gm28937 | predicted gene 28937 [Source:MGI Symbo | 0 | 0 | 0 | 0 | 0 | 0 |
| 39936 | ENSMUSG00000101670 | 0 | NA | NA | NA | NA | NA | Gm21470 | predicted gene, 21470 [Source:MGI Symb | 0 | 0 | 0 | 0 | 0 | 0 |
| 39937 | ENSMUSG00000101671 | 0 | NA | NA | NA | NA | NA | Gm28221 | predicted gene 28221 [Source:MGI Symbo | 0 | 0 | 0 | 0 | 0 | 0 |
| 39938 | ENSMUSG00000101672 | 0 | NA | NA | NA | NA | NA | Gm28149 | predicted gene 28149 [Source:MGI Symbo | 0 | 0 | 0 | 0 | 0 | 0 |
| 39939 | ENSMUSG00000101673 | 0 | NA | NA | NA | NA | NA | Gm28070 | predicted gene 28070 [Source:MGI Symbo | 0 | 0 | 0 | 0 | 0 | 0 |
| 39940 | ENSMUSG00000101674 | 0 | NA | NA | NA | NA | NA | 4930444A19 | RIKEN cDNA 4930444A19 gene [Source:M | 0 | 0 | 0 | 0 | 0 | 0 |
| 39941 | ENSMUSG00000101675 | 0 | NA | NA | NA | NA | NA | Gm28461 | predicted gene 28461 [Source:MGI Symbo | 0 | 0 | 0 | 0 | 0 | 0 |
| 39942 | ENSMUSG00000101676 | 0 | NA | NA | NA | NA | NA | Gm21832 | predicted gene, 21832 [Source:MGI Symb | 0 | 0 | 0 | 0 | 0 | 0 |
| 39943 | ENSMUSG00000101678 | 2.794476615 | -1.08611908 | 1.296698072 | -0.837603682 | 0.402253324 | NA | Gm29609 | predicted gene 29609 [Source:MGI Symbo | 3.2124 | 5.1396 | 3.0187 | 0.964 | 1.0906 | 3.3415 |
| 39944 | ENSMUSG00000101679 | 0 | NA | NA | NA | NA | NA | Gm20936 | predicted gene, 20936 [Source:MGI Symb | 0 | 0 | 0 | 0 | 0 | 0 |
| 39945 | ENSMUSG00000101680 | 0 | NA | NA | NA | NA | NA | Gm29015 | predicted gene 29015 [Source:MGI Symbo | 0 | 0 | 0 | 0 | 0 | 0 |
| 39946 | ENSMUSG00000101681 | 0 | NA | NA | NA | NA | NA | Gm28522 | predicted gene 28522 [Source:MGI Symbo | 0 | 0 | 0 | 0 | 0 | 0 |
| 39947 | ENSMUSG00000101682 | 0 | NA | NA | NA | NA | NA | Gm29228 | predicted gene 29228 [Source:MGI Symbo | 0 | 0 | 0 | 0 | 0 | 0 |
| 39948 | ENSMUSG00000101683 | 0 | NA | NA | NA | NA | NA | 1700028D13 | RIKEN cDNA 1700028D13 gene [Source:M | 0 | 0 | 0 | 0 | 0 | 0 |
| 39949 | ENSMUSG00000101684 | 0 | NA | NA | NA | NA | NA | Gm28946 | predicted gene 28946 [Source:MGI Symbo | 0 | 0 | 0 | 0 | 0 | 0 |
| 39950 | ENSMUSG00000101685 | 0 | NA | NA | NA | NA | NA | Gm17837 | predicted gene, 17837 [Source:MGI Symb | 0 | 0 | 0 | 0 | 0 | 0 |
| 39951 | ENSMUSG00000101686 | 0 | NA | NA | NA | NA | NA | Gm29633 | predicted gene 29633 [Source:MGI Symbo | 0 | 0 | 0 | 0 | 0 | 0 |
| 39952 | ENSMUSG00000101687 | 0 | NA | NA | NA | NA | NA | Gm4854 | predicted gene 4854 [Source:MGI Symbol | 0 | 0 | 0 | 0 | 0 | 0 |
| 39953 | ENSMUSG00000101688 | 0 | NA | NA | NA | NA | NA | Gm20841 | predicted gene, 20841 [Source:MGI Symb | 0 | 0 | 0 | 0 | 0 | 0 |
| 39954 | ENSMUSG00000101689 | 0 | NA | NA | NA | NA | NA | Gm28599 | predicted gene 28599 [Source:MGI Symbo | 0 | 0 | 0 | 0 | 0 | 0 |
| 39955 | ENSMUSG00000101690 | 0 | NA | NA | NA | NA | NA | Gm28390 | predicted gene 28390 [Source:MGI Symbo | 0 | 0 | 0 | 0 | 0 | 0 |
| 39956 | ENSMUSG00000101691 | 0 | NA | NA | NA | NA | NA | Gm29280 | predicted gene 29280 [Source:MGI Symbo | 0 | 0 | 0 | 0 | 0 | 0 |
| 39957 | ENSMUSG00000101692 | 0 | NA | NA | NA | NA | NA | Gm28244 | predicted gene 28244 [Source:MGI Symbo | 0 | 0 | 0 | 0 | 0 | 0 |
| 39958 | ENSMUSG00000101693 | 7.341279665 | -0.069446833 | 0.816499936 | -0.085054304 | 0.932218219 | 0.988420305 | Gm19461 | predicted gene, 19461 [Source:MGI Symb | 6.4248 | 8.2233 | 8.05 | 12.532 | 4.3625 | 4.4554 |
| 39959 | ENSMUSG00000101694 | 0 | NA | NA | NA | NA | NA | Gm29392 | predicted gene 29392 [Source:MGI Symbo | 0 | 0 | 0 | 0 | 0 | 0 |
| 39960 | ENSMUSG00000101695 | 0 | NA | NA | NA | NA | NA | 1700094J05I | RIKEN cDNA 1700094J05 gene [Source:MG | 0 | 0 | 0 | 0 | 0 | 0 |
| 39961 | ENSMUSG00000101698 | 0.321325949 | 1.911575114 | 4.048267893 | 0.472195804 | 0.636787034 | NA | Gm29562 | predicted gene 29562 [Source:MGI Symbo | 0 | 0 | 0 | 1.928 | 0 | 0 |
| 39962 | ENSMUSG00000101699 | 0 | NA | NA | NA | NA | NA | Gm28508 | predicted gene 28508 [Source:MGI Symbo | 0 | 0 | 0 | 0 | 0 | 0 |
| 39963 | ENSMUSG00000101700 | 0 | NA | NA | NA | NA | NA | Gm29359 | predicted gene 29359 [Source:MGI Symbo | 0 | 0 | 0 | 0 | 0 | 0 |
| 39964 | ENSMUSG00000101701 | 0 | NA | NA | NA | NA | NA | 4930521E06 | RIKEN cDNA 4930521E06 gene [Source:MG | 0 | 0 | 0 | 0 | 0 | 0 |
| 39965 | ENSMUSG00000101702 | 0 | NA | NA | NA | NA | NA | 1700072G22 | RIKEN cDNA 1700072G22 gene [Source:M | 0 | 0 | 0 | 0 | 0 | 0 |
| 39966 | ENSMUSG00000101703 | 0 | NA | NA | NA | NA | NA | Gm1982 | predicted gene 1982 [Source:MGI Symbol | 0 | 0 | 0 | 0 | 0 | 0 |
| 39967 | ENSMUSG00000101704 | 0 | NA | NA | NA | NA | NA | Gm21878 | predicted gene, 21878 [Source:MGI Symb | 0 | 0 | 0 | 0 | 0 | 0 |
| 39968 | ENSMUSG00000101705 | 0 | NA | NA | NA | NA | NA | Gm28380 | predicted gene 28380 [Source:MGI Symbo | 0 | 0 | 0 | 0 | 0 | 0 |
| 39969 | ENSMUSG00000101706 | 0 | NA | NA | NA | NA | NA | Gm28431 | predicted gene 28431 [Source:MGI Symbo | 0 | 0 | 0 | 0 | 0 | 0 |
| 39970 | ENSMUSG00000101707 | 0 | NA | NA | NA | NA | NA | Gm28916 | predicted gene 28916 [Source:MGI Symbo | 0 | 0 | 0 | 0 | 0 | 0 |
| 39971 | ENSMUSG00000101708 | 0.562214094 | -1.122836648 | 3.276281221 | -0.34271681 | 0.731811515 | NA | Gm28453 | predicted gene 28453 [Source:MGI Symbo | 2.4093 | 0 | 0 | 0.964 | 0 | 0 |
| 39972 | ENSMUSG00000101709 | 0 | NA | NA | NA | NA | NA | Gm28337 | predicted gene 28337 [Source:MGI Symbo | 0 | 0 | 0 | 0 | 0 | 0 |
| 39973 | ENSMUSG00000101710 | 0 | NA | NA | NA | NA | NA | Gm20922 | predicted gene, 20922 [Source:MGI Symb | 0 | 0 | 0 | 0 | 0 | 0 |
| 39974 | ENSMUSG00000101711 | 0.637152627 | -1.330052897 | 2.84179579 | -0.468032538 | 0.639761321 | NA | Gm28306 | predicted gene 28306 [Source:MGI Symbo | 0.8031 | 2.0558 | 0 | 0.964 | 0 | 0 |
| 39975 | ENSMUSG00000101712 | 0 | NA | NA | NA | NA | NA | Gm28648 | predicted gene 28648 [Source:MGI Symbo | 0 | 0 | 0 | 0 | 0 | 0 |
| 39976 | ENSMUSG00000101713 | 0 | NA | NA | NA | NA | NA | Gm28668 | predicted gene 28668 [Source:MGI Symbo | 0 | 0 | 0 | 0 | 0 | 0 |
| 39977 | ENSMUSG00000101714 | 0 | NA | NA | NA | NA | NA | Gm28415 | predicted gene 28415 [Source:MGI Symbo | 0 | 0 | 0 | 0 | 0 | 0 |
| 39978 | ENSMUSG00000101715 | 0 | NA | NA | NA | NA | NA | Gm21855 | predicted gene, 21855 [Source:MGI Symb | 0 | 0 | 0 | 0 | 0 | 0 |
| 39979 | ENSMUSG00000101717 | 0 | NA | NA | NA | NA | NA | Gm45784 | predicted gene 45784 [Source:MGI Symbo | 0 | 0 | 0 | 0 | 0 | 0 |
| 39980 | ENSMUSG00000101718 | 0 | NA | NA | NA | NA | NA | Gm20862 | predicted gene, 20862 [Source:MGI Symb | 0 | 0 | 0 | 0 | 0 | 0 |

| | | | | | | | | | | | | | | | |
|---|---|---|---|---|---|---|---|---|---|---|---|---|---|---|---|
| 39981 | ENSMUSG00000101719 | 0 | NA | NA | NA | NA | NA | Gm28216 | predicted gene 28216 [Source:MGI Symbo | 0 | 0 | 0 | 0 | 0 | 0 |
| 39982 | ENSMUSG00000101720 | 0 | NA | NA | NA | NA | NA | Gm29005 | predicted gene 29005 [Source:MGI Symbo | 0 | 0 | 0 | 0 | 0 | 0 |
| 39983 | ENSMUSG00000101721 | 0 | NA | NA | NA | NA | NA | Gm8354 | predicted gene 8354 [Source:MGI Symbol | 0 | 0 | 0 | 0 | 0 | 0 |
| 39984 | ENSMUSG00000101722 | 0 | NA | NA | NA | NA | NA | Gm29125 | predicted gene 29125 [Source:MGI Symbo | 0 | 0 | 0 | 0 | 0 | 0 |
| 39985 | ENSMUSG00000101723 | 0 | NA | NA | NA | NA | NA | Gm28355 | predicted gene 28355 [Source:MGI Symbo | 0 | 0 | 0 | 0 | 0 | 0 |
| 39986 | ENSMUSG00000101724 | 0 | NA | NA | NA | NA | NA | Gm29453 | predicted gene 29453 [Source:MGI Symbo | 0 | 0 | 0 | 0 | 0 | 0 |
| 39987 | ENSMUSG00000101725 | 0 | NA | NA | NA | NA | NA | Gm29423 | predicted gene 29423 [Source:MGI Symbo | 0 | 0 | 0 | 0 | 0 | 0 |
| 39988 | ENSMUSG00000101726 | 0 | NA | NA | NA | NA | NA | Gm7858 | predicted gene 7858 [Source:MGI Symbol | 0 | 0 | 0 | 0 | 0 | 0 |
| 39989 | ENSMUSG00000101727 | 0 | NA | NA | NA | NA | NA | Gm29495 | predicted gene 29495 [Source:MGI Symbo | 0 | 0 | 0 | 0 | 0 | 0 |
| 39990 | ENSMUSG00000101728 | 0 | NA | NA | NA | NA | NA | Gm28765 | predicted gene 28765 [Source:MGI Symbo | 0 | 0 | 0 | 0 | 0 | 0 |
| 39991 | ENSMUSG00000101729 | 1.277763006 | 1.830198481 | 2.0837712 | 0.878310671 | 0.379775147 | NA | Gm17796 | predicted gene, 17796 [Source:MGI Symb | 1.6062 | 0 | 0 | 3.8559 | 1.0906 | 1.1138 |
| 39992 | ENSMUSG00000101730 | 0 | NA | NA | NA | NA | NA | Gm28580 | predicted gene 28580 [Source:MGI Symbo | 0 | 0 | 0 | 0 | 0 | 0 |
| 39993 | ENSMUSG00000101731 | 0.181769715 | 1.126924175 | 4.080472857 | 0.276174898 | 0.782413736 | NA | 3830432H09 | RIKEN cDNA 3830432H09 gene [Source:M | 0 | 0 | 0 | 0 | 1.0906 | 0 |
| 39994 | ENSMUSG00000101732 | 0 | NA | NA | NA | NA | NA | Gm28328 | predicted gene 28328 [Source:MGI Symbo | 0 | 0 | 0 | 0 | 0 | 0 |
| 39995 | ENSMUSG00000101733 | 0 | NA | NA | NA | NA | NA | Gm29189 | predicted gene 29189 [Source:MGI Symbo | 0 | 0 | 0 | 0 | 0 | 0 |
| 39996 | ENSMUSG00000101734 | 0 | NA | NA | NA | NA | NA | 4933400C23 | RIKEN cDNA 4933400C23 gene [Source:M | 0 | 0 | 0 | 0 | 0 | 0 |
| 39997 | ENSMUSG00000101735 | 0 | NA | NA | NA | NA | NA | Gm18505 | predicted gene, 18505 [Source:MGI Symb | 0 | 0 | 0 | 0 | 0 | 0 |
| 39998 | ENSMUSG00000101736 | 0 | NA | NA | NA | NA | NA | Gm29335 | predicted gene 29335 [Source:MGI Symbo | 0 | 0 | 0 | 0 | 0 | 0 |
| 39999 | ENSMUSG00000101737 | 0 | NA | NA | NA | NA | NA | Gm28849 | predicted gene 28849 [Source:MGI Symbo | 0 | 0 | 0 | 0 | 0 | 0 |
| 40000 | ENSMUSG00000101739 | 0 | NA | NA | NA | NA | NA | Gm5733 | predicted gene 5733 [Source:MGI Symbol | 0 | 0 | 0 | 0 | 0 | 0 |
| 40001 | ENSMUSG00000101740 | 0 | NA | NA | NA | NA | NA | Gm29360 | predicted gene 29360 [Source:MGI Symbo | 0 | 0 | 0 | 0 | 0 | 0 |
| 40002 | ENSMUSG00000101741 | 2.977669204 | -2.167104854 | 1.425856461 | -1.51986186 | 0.128545698 | NA | Gm28055 | predicted gene 28055 [Source:MGI Symbo | 2.4093 | 4.1117 | 8.05 | 0 | 2.1812 | 1.1138 |
| 40003 | ENSMUSG00000101742 | 0 | NA | NA | NA | NA | NA | Gm21380 | predicted gene, 21380 [Source:MGI Symb | 0 | 0 | 0 | 0 | 0 | 0 |
| 40004 | ENSMUSG00000101743 | 0 | NA | NA | NA | NA | NA | Gm28249 | predicted gene 28249 [Source:MGI Symbo | 0 | 0 | 0 | 0 | 0 | 0 |
| 40005 | ENSMUSG00000101744 | 0 | NA | NA | NA | NA | NA | Gm28325 | predicted gene 28325 [Source:MGI Symbo | 0 | 0 | 0 | 0 | 0 | 0 |
| 40006 | ENSMUSG00000101745 | 0 | NA | NA | NA | NA | NA | Gm28764 | predicted gene 28764 [Source:MGI Symbo | 0 | 0 | 0 | 0 | 0 | 0 |
| 40007 | ENSMUSG00000101746 | 0 | NA | NA | NA | NA | NA | 2310043L19I | RIKEN cDNA 2310043L19 gene [Source:M( | 0 | 0 | 0 | 0 | 0 | 0 |
| 40008 | ENSMUSG00000101747 | 0 | NA | NA | NA | NA | NA | Gm21683 | predicted gene, 21683 [Source:MGI Symb | 0 | 0 | 0 | 0 | 0 | 0 |
| 40009 | ENSMUSG00000101748 | 0 | NA | NA | NA | NA | NA | Gm21823 | predicted gene, 21823 [Source:MGI Symb | 0 | 0 | 0 | 0 | 0 | 0 |
| 40010 | ENSMUSG00000101750 | 0.17131964 | -0.796629176 | 4.080472857 | -0.195229623 | 0.84521319 | NA | Olfr1392 | olfactory receptor 1392 [Source:MGI Syml | 0 | 1.0279 | 0 | 0 | 0 | 0 |
| 40011 | ENSMUSG00000101751 | 0 | NA | NA | NA | NA | NA | Gm2427 | predicted gene 2427 [Source:MGI Symbol | 0 | 0 | 0 | 0 | 0 | 0 |
| 40012 | ENSMUSG00000101752 | 0 | NA | NA | NA | NA | NA | Gm20887 | predicted gene, 20887 [Source:MGI Symb | 0 | 0 | 0 | 0 | 0 | 0 |
| 40013 | ENSMUSG00000101753 | 0 | NA | NA | NA | NA | NA | Gm28799 | predicted gene 28799 [Source:MGI Symbo | 0 | 0 | 0 | 0 | 0 | 0 |
| 40014 | ENSMUSG00000101754 | 0 | NA | NA | NA | NA | NA | Gm2987 | predicted gene 2987 [Source:MGI Symbol | 0 | 0 | 0 | 0 | 0 | 0 |
| 40015 | ENSMUSG00000101755 | 0 | NA | NA | NA | NA | NA | Gm29507 | predicted gene 29507 [Source:MGI Symbo | 0 | 0 | 0 | 0 | 0 | 0 |
| 40016 | ENSMUSG00000101756 | 0 | NA | NA | NA | NA | NA | Gm29402 | predicted gene 29402 [Source:MGI Symbo | 0 | 0 | 0 | 0 | 0 | 0 |
| 40017 | ENSMUSG00000101757 | 0 | NA | NA | NA | NA | NA | Gm28273 | predicted gene 28273 [Source:MGI Symbo | 0 | 0 | 0 | 0 | 0 | 0 |
| 40018 | ENSMUSG00000101758 | 0 | NA | NA | NA | NA | NA | Gm28137 | predicted gene 28137 [Source:MGI Symbo | 0 | 0 | 0 | 0 | 0 | 0 |
| 40019 | ENSMUSG00000101759 | 0 | NA | NA | NA | NA | NA | Gm5734 | predicted gene 5734 [Source:MGI Symbol | 0 | 0 | 0 | 0 | 0 | 0 |
| 40020 | ENSMUSG00000101760 | 0 | NA | NA | NA | NA | NA | Gm8384 | predicted gene 8384 [Source:MGI Symbol | 0 | 0 | 0 | 0 | 0 | 0 |
| 40021 | ENSMUSG00000101761 | 0 | NA | NA | NA | NA | NA | Gm28297 | predicted gene 28297 [Source:MGI Symbo | 0 | 0 | 0 | 0 | 0 | 0 |
| 40022 | ENSMUSG00000101762 | 0 | NA | NA | NA | NA | NA | 1700047L14I | RIKEN cDNA 1700047L14 gene [Source:M( | 0 | 0 | 0 | 0 | 0 | 0 |
| 40023 | ENSMUSG00000101763 | 0 | NA | NA | NA | NA | NA | Gm29068 | predicted gene 29068 [Source:MGI Symbo | 0 | 0 | 0 | 0 | 0 | 0 |
| 40024 | ENSMUSG00000101764 | 0 | NA | NA | NA | NA | NA | 4930556G22 | RIKEN cDNA 4930556G22 gene [Source:M | 0 | 0 | 0 | 0 | 0 | 0 |
| 40025 | ENSMUSG00000101765 | 0 | NA | NA | NA | NA | NA | Gm21673 | predicted gene, 21673 [Source:MGI Symb | 0 | 0 | 0 | 0 | 0 | 0 |
| 40026 | ENSMUSG00000101766 | 0 | NA | NA | NA | NA | NA | Gm20908 | predicted gene, 20908 [Source:MGI Symb | 0 | 0 | 0 | 0 | 0 | 0 |
| 40027 | ENSMUSG00000101768 | 0 | NA | NA | NA | NA | NA | Gm28694 | predicted gene 28694 [Source:MGI Symbo | 0 | 0 | 0 | 0 | 0 | 0 |
| 40028 | ENSMUSG00000101769 | 0 | NA | NA | NA | NA | NA | Gm29356 | predicted gene 29356 [Source:MGI Symbo | 0 | 0 | 0 | 0 | 0 | 0 |
| 40029 | ENSMUSG00000101770 | 0 | NA | NA | NA | NA | NA | Gm28718 | predicted gene 28718 [Source:MGI Symbo | 0 | 0 | 0 | 0 | 0 | 0 |
| 40030 | ENSMUSG00000101771 | 0 | NA | NA | NA | NA | NA | Gm28176 | predicted gene 28176 [Source:MGI Symbo | 0 | 0 | 0 | 0 | 0 | 0 |
| 40031 | ENSMUSG00000101772 | 0 | NA | NA | NA | NA | NA | Gm28829 | predicted gene 28829 [Source:MGI Symbo | 0 | 0 | 0 | 0 | 0 | 0 |
| 40032 | ENSMUSG00000101773 | 0 | NA | NA | NA | NA | NA | 2210420H20 | RIKEN cDNA 2210420H20 gene [Source:M | 0 | 0 | 0 | 0 | 0 | 0 |
| 40033 | ENSMUSG00000101774 | 0 | NA | NA | NA | NA | NA | Gm29616 | predicted gene 29616 [Source:MGI Symbo | 0 | 0 | 0 | 0 | 0 | 0 |
| 40034 | ENSMUSG00000101775 | 0 | NA | NA | NA | NA | NA | Gm28088 | predicted gene 28088 [Source:MGI Symbo | 0 | 0 | 0 | 0 | 0 | 0 |
| 40035 | ENSMUSG00000101776 | 0 | NA | NA | NA | NA | NA | Gm28268 | predicted gene 28268 [Source:MGI Symbo | 0 | 0 | 0 | 0 | 0 | 0 |
| 40036 | ENSMUSG00000101777 | 0 | NA | NA | NA | NA | NA | Gm9127 | predicted gene 9127 [Source:MGI Symbol | 0 | 0 | 0 | 0 | 0 | 0 |

| | | | | | | | | | | | | | | | |
|---|---|---|---|---|---|---|---|---|---|---|---|---|---|---|---|
| 40037 | ENSMUSG00000101778 | 0.17131964 | -0.796629176 | 4.080472857 | -0.195229623 | 0.84521319 | NA | Gm29488 | predicted gene 29488 [Source:MGI Symbo | 0 | 1.0279 | 0 | 0 | 0 | 0 |
| 40038 | ENSMUSG00000101779 | 0 | NA | NA | NA | NA | NA | Gm28170 | predicted gene 28170 [Source:MGI Symbo | 0 | 0 | 0 | 0 | 0 | 0 |
| 40039 | ENSMUSG00000101780 | 0 | NA | NA | NA | NA | NA | Gm7255 | predicted gene 7255 [Source:MGI Symbol | 0 | 0 | 0 | 0 | 0 | 0 |
| 40040 | ENSMUSG00000101781 | 0 | NA | NA | NA | NA | NA | Gm28333 | predicted gene 28333 [Source:MGI Symbo | 0 | 0 | 0 | 0 | 0 | 0 |
| 40041 | ENSMUSG00000101782 | 0 | NA | NA | NA | NA | NA | Gm28964 | predicted gene 28964 [Source:MGI Symbo | 0 | 0 | 0 | 0 | 0 | 0 |
| 40042 | ENSMUSG00000101783 | 0 | NA | NA | NA | NA | NA | Gm28698 | predicted gene 28698 [Source:MGI Symbo | 0 | 0 | 0 | 0 | 0 | 0 |
| 40043 | ENSMUSG00000101784 | 0 | NA | NA | NA | NA | NA | Gm7553 | predicted gene 7553 [Source:MGI Symbol | 0 | 0 | 0 | 0 | 0 | 0 |
| 40044 | ENSMUSG00000101785 | 0 | NA | NA | NA | NA | NA | Gm28702 | predicted gene 28702 [Source:MGI Symbo | 0 | 0 | 0 | 0 | 0 | 0 |
| 40045 | ENSMUSG00000101786 | 0 | NA | NA | NA | NA | NA | Gm28840 | predicted gene 28840 [Source:MGI Symbo | 0 | 0 | 0 | 0 | 0 | 0 |
| 40046 | ENSMUSG00000101787 | 0 | NA | NA | NA | NA | NA | Gm20853 | predicted gene, 20853 [Source:MGI Symb | 0 | 0 | 0 | 0 | 0 | 0 |
| 40047 | ENSMUSG00000101788 | 0 | NA | NA | NA | NA | NA | Gm20813 | predicted gene, 20813 [Source:MGI Symb | 0 | 0 | 0 | 0 | 0 | 0 |
| 40048 | ENSMUSG00000101789 | 0 | NA | NA | NA | NA | NA | Gm28956 | predicted gene 28956 [Source:MGI Symbo | 0 | 0 | 0 | 0 | 0 | 0 |
| 40049 | ENSMUSG00000101790 | 0 | NA | NA | NA | NA | NA | Gm29059 | predicted gene 29059 [Source:MGI Symbo | 0 | 0 | 0 | 0 | 0 | 0 |
| 40050 | ENSMUSG00000101791 | 0 | NA | NA | NA | NA | NA | 2210011K15 | RIKEN cDNA 2210011K15 gene [Source:M | 0 | 0 | 0 | 0 | 0 | 0 |
| 40051 | ENSMUSG00000101792 | 0 | NA | NA | NA | NA | NA | Gm28447 | predicted gene 28447 [Source:MGI Symbo | 0 | 0 | 0 | 0 | 0 | 0 |
| 40052 | ENSMUSG00000101793 | 0 | NA | NA | NA | NA | NA | Gm28500 | predicted gene 28500 [Source:MGI Symbo | 0 | 0 | 0 | 0 | 0 | 0 |
| 40053 | ENSMUSG00000101794 | 0 | NA | NA | NA | NA | NA | Gm29217 | predicted gene 29217 [Source:MGI Symbo | 0 | 0 | 0 | 0 | 0 | 0 |
| 40054 | ENSMUSG00000101795 | 2.419169931 | 0.000856552 | 1.445075846 | 0.000592738 | 0.999527063 | NA | Gm5835 | predicted gene 5835 [Source:MGI Symbol | 3.2124 | 0 | 4.025 | 2.8919 | 3.2719 | 1.1138 |
| 40055 | ENSMUSG00000101796 | 0 | NA | NA | NA | NA | NA | Gm29277 | predicted gene 29277 [Source:MGI Symbo | 0 | 0 | 0 | 0 | 0 | 0 |
| 40056 | ENSMUSG00000101797 | 0.160662974 | 1.126924175 | 4.080472857 | 0.276174898 | 0.782413736 | NA | Gm29266 | predicted gene 29266 [Source:MGI Symbo | 0 | 0 | 0 | 0.964 | 0 | 0 |
| 40057 | ENSMUSG00000101798 | 0.160662974 | 1.126924175 | 4.080472857 | 0.276174898 | 0.782413736 | NA | Gm28995 | predicted gene 28995 [Source:MGI Symbo | 0 | 0 | 0 | 0.964 | 0 | 0 |
| 40058 | ENSMUSG00000101799 | 0 | NA | NA | NA | NA | NA | 9330175M2( | RIKEN cDNA 9330175M20 gene [Source:M | 0 | 0 | 0 | 0 | 0 | 0 |
| 40059 | ENSMUSG00000101800 | 0 | NA | NA | NA | NA | NA | Rnf170-ps | ring finger protein 170, pseudogene [Sour | 0 | 0 | 0 | 0 | 0 | 0 |
| 40060 | ENSMUSG00000101801 | 0 | NA | NA | NA | NA | NA | 1700020G17 | RIKEN cDNA 1700020G17 gene [Source:M | 0 | 0 | 0 | 0 | 0 | 0 |
| 40061 | ENSMUSG00000101802 | 0 | NA | NA | NA | NA | NA | Gm29445 | predicted gene 29445 [Source:MGI Symbo | 0 | 0 | 0 | 0 | 0 | 0 |
| 40062 | ENSMUSG00000101803 | 0 | NA | NA | NA | NA | NA | Gm28419 | predicted gene 28419 [Source:MGI Symbo | 0 | 0 | 0 | 0 | 0 | 0 |
| 40063 | ENSMUSG00000101804 | 0 | NA | NA | NA | NA | NA | Gm28813 | predicted gene 28813 [Source:MGI Symbo | 0 | 0 | 0 | 0 | 0 | 0 |
| 40064 | ENSMUSG00000101805 | 0 | NA | NA | NA | NA | NA | Gm28768 | predicted gene 28768 [Source:MGI Symbo | 0 | 0 | 0 | 0 | 0 | 0 |
| 40065 | ENSMUSG00000101806 | 0 | NA | NA | NA | NA | NA | Gm28261 | predicted gene 28261 [Source:MGI Symbo | 0 | 0 | 0 | 0 | 0 | 0 |
| 40066 | ENSMUSG00000101807 | 0 | NA | NA | NA | NA | NA | Gm9120 | predicted gene 9120 [Source:MGI Symbol | 0 | 0 | 0 | 0 | 0 | 0 |
| 40067 | ENSMUSG00000101808 | 0 | NA | NA | NA | NA | NA | Gm29654 | predicted gene 29654 [Source:MGI Symbo | 0 | 0 | 0 | 0 | 0 | 0 |
| 40068 | ENSMUSG00000101809 | 0 | NA | NA | NA | NA | NA | Gm28852 | predicted gene 28852 [Source:MGI Symbo | 0 | 0 | 0 | 0 | 0 | 0 |
| 40069 | ENSMUSG00000101810 | 0 | NA | NA | NA | NA | NA | 1700066C05 | RIKEN cDNA 1700066C05 gene [Source:M | 0 | 0 | 0 | 0 | 0 | 0 |
| 40070 | ENSMUSG00000101811 | 0 | NA | NA | NA | NA | NA | Gm28312 | predicted gene 28312 [Source:MGI Symbo | 0 | 0 | 0 | 0 | 0 | 0 |
| 40071 | ENSMUSG00000101812 | 0 | NA | NA | NA | NA | NA | D530049I02I | RIKEN cDNA D530049I02 gene [Source:M | 0 | 0 | 0 | 0 | 0 | 0 |
| 40072 | ENSMUSG00000101813 | 0 | NA | NA | NA | NA | NA | Gm28503 | predicted gene 28503 [Source:MGI Symbo | 0 | 0 | 0 | 0 | 0 | 0 |
| 40073 | ENSMUSG00000101814 | 0 | NA | NA | NA | NA | NA | Gm17807 | predicted gene, 17807 [Source:MGI Symb | 0 | 0 | 0 | 0 | 0 | 0 |
| 40074 | ENSMUSG00000101815 | 0 | NA | NA | NA | NA | NA | Gm29028 | predicted gene 29028 [Source:MGI Symbo | 0 | 0 | 0 | 0 | 0 | 0 |
| 40075 | ENSMUSG00000101816 | 0 | NA | NA | NA | NA | NA | Gm29659 | predicted gene 29659 [Source:MGI Symbo | 0 | 0 | 0 | 0 | 0 | 0 |
| 40076 | ENSMUSG00000101817 | 0 | NA | NA | NA | NA | NA | Gm28288 | predicted gene 28288 [Source:MGI Symbo | 0 | 0 | 0 | 0 | 0 | 0 |
| 40077 | ENSMUSG00000101818 | 0 | NA | NA | NA | NA | NA | Gm28269 | predicted gene 28269 [Source:MGI Symbo | 0 | 0 | 0 | 0 | 0 | 0 |
| 40078 | ENSMUSG00000101819 | 0 | NA | NA | NA | NA | NA | H2al1b | H2A histone family member L1B [Source:M | 0 | 0 | 0 | 0 | 0 | 0 |
| 40079 | ENSMUSG00000101820 | 0 | NA | NA | NA | NA | NA | Gm29023 | predicted gene 29023 [Source:MGI Symbo | 0 | 0 | 0 | 0 | 0 | 0 |
| 40080 | ENSMUSG00000101821 | 0 | NA | NA | NA | NA | NA | Gm21308 | predicted gene, 21308 [Source:MGI Symb | 0 | 0 | 0 | 0 | 0 | 0 |
| 40081 | ENSMUSG00000101822 | 0 | NA | NA | NA | NA | NA | Gm29396 | predicted gene 29396 [Source:MGI Symbo | 0 | 0 | 0 | 0 | 0 | 0 |
| 40082 | ENSMUSG00000101823 | 3.70258779 | -2.146687536 | 1.268428245 | -1.692399664 | 0.090569796 | NA | Gm29438 | predicted gene 29438 [Source:MGI Symbo | 4.0155 | 6.1675 | 8.05 | 2.8919 | 1.0906 | 0 |
| 40083 | ENSMUSG00000101824 | 0 | NA | NA | NA | NA | NA | Gm28256 | predicted gene 28256 [Source:MGI Symbo | 0 | 0 | 0 | 0 | 0 | 0 |
| 40084 | ENSMUSG00000101825 | 0 | NA | NA | NA | NA | NA | Gm21806 | predicted gene, 21806 [Source:MGI Symb | 0 | 0 | 0 | 0 | 0 | 0 |
| 40085 | ENSMUSG00000101826 | 0 | NA | NA | NA | NA | NA | Gm8581 | predicted gene 8581 [Source:MGI Symbol | 0 | 0 | 0 | 0 | 0 | 0 |
| 40086 | ENSMUSG00000101827 | 0 | NA | NA | NA | NA | NA | Gm28686 | predicted gene 28686 [Source:MGI Symbo | 0 | 0 | 0 | 0 | 0 | 0 |
| 40087 | ENSMUSG00000101829 | 0 | NA | NA | NA | NA | NA | Gm21752 | predicted gene, 21752 [Source:MGI Symb | 0 | 0 | 0 | 0 | 0 | 0 |
| 40088 | ENSMUSG00000101830 | 0 | NA | NA | NA | NA | NA | Gm20907 | predicted gene, 20907 [Source:MGI Symb | 0 | 0 | 0 | 0 | 0 | 0 |
| 40089 | ENSMUSG00000101831 | 0 | NA | NA | NA | NA | NA | Gm29412 | predicted gene 29412 [Source:MGI Symbo | 0 | 0 | 0 | 0 | 0 | 0 |
| 40090 | ENSMUSG00000101832 | 1.898475997 | 0.882175135 | 1.609412306 | 0.54813495 | 0.583599244 | NA | Gm28982 | predicted gene 28982 [Source:MGI Symbo | 0 | 3.0838 | 1.0062 | 2.8919 | 2.1812 | 2.2277 |
| 40091 | ENSMUSG00000101833 | 0 | NA | NA | NA | NA | NA | Gm20791 | predicted gene, 20791 [Source:MGI Symb | 0 | 0 | 0 | 0 | 0 | 0 |
| 40092 | ENSMUSG00000101834 | 0 | NA | NA | NA | NA | NA | Gm5829 | predicted gene 5829 [Source:MGI Symbol | 0 | 0 | 0 | 0 | 0 | 0 |

| | | | | | | | | | | | | | | | |
|---|---|---|---|---|---|---|---|---|---|---|---|---|---|---|---|
| 40093 | ENSMUSG00000101835 | 0 | NA | NA | NA | NA | NA | Gm21734 | predicted gene, 21734 [Source:MGI Symb | 0 | 0 | 0 | 0 | 0 | 0 |
| 40094 | ENSMUSG00000101836 | 0 | NA | NA | NA | NA | NA | Gm28968 | predicted gene 28968 [Source:MGI Symbo | 0 | 0 | 0 | 0 | 0 | 0 |
| 40095 | ENSMUSG00000101837 | 0 | NA | NA | NA | NA | NA | Gm20810 | predicted gene, 20810 [Source:MGI Symb | 0 | 0 | 0 | 0 | 0 | 0 |
| 40096 | ENSMUSG00000101838 | 0 | NA | NA | NA | NA | NA | Gm18800 | predicted gene, 18800 [Source:MGI Symb | 0 | 0 | 0 | 0 | 0 | 0 |
| 40097 | ENSMUSG00000101839 | 0 | NA | NA | NA | NA | NA | Gm28247 | predicted gene 28247 [Source:MGI Symbo | 0 | 0 | 0 | 0 | 0 | 0 |
| 40098 | ENSMUSG00000101840 | 0 | NA | NA | NA | NA | NA | Gm28294 | predicted gene 28294 [Source:MGI Symbo | 0 | 0 | 0 | 0 | 0 | 0 |
| 40099 | ENSMUSG00000101841 | 1.00095078 | 0.163345763 | 1.967377922 | 0.083027141 | 0.933829959 | NA | Gm11993 | predicted gene 11993 [Source:MGI Symbo | 0.8031 | 1.0279 | 1.0062 | 0.964 | 1.0906 | 1.1138 |
| 40100 | ENSMUSG00000101842 | 0 | NA | NA | NA | NA | NA | Gm28391 | predicted gene 28391 [Source:MGI Symbo | 0 | 0 | 0 | 0 | 0 | 0 |
| 40101 | ENSMUSG00000101843 | 0 | NA | NA | NA | NA | NA | Gm28746 | predicted gene 28746 [Source:MGI Symbo | 0 | 0 | 0 | 0 | 0 | 0 |
| 40102 | ENSMUSG00000101844 | 0 | NA | NA | NA | NA | NA | Gm20844 | predicted gene, 20844 [Source:MGI Symb | 0 | 0 | 0 | 0 | 0 | 0 |
| 40103 | ENSMUSG00000101845 | 1.477233843 | 0.510573446 | 1.660191398 | 0.307538906 | 0.758433214 | NA | Gm28198 | predicted gene 28198 [Source:MGI Symbo | 1.6062 | 1.0279 | 1.0062 | 1.928 | 2.1812 | 1.1138 |
| 40104 | ENSMUSG00000101846 | 0 | NA | NA | NA | NA | NA | Gm19927 | predicted gene, 19927 [Source:MGI Symb | 0 | 0 | 0 | 0 | 0 | 0 |
| 40105 | ENSMUSG00000101847 | 0 | NA | NA | NA | NA | NA | Gm20749 | predicted gene, 20749 [Source:MGI Symb | 0 | 0 | 0 | 0 | 0 | 0 |
| 40106 | ENSMUSG00000101848 | 0 | NA | NA | NA | NA | NA | 4933417E11 | RIKEN cDNA 4933417E11 gene [Source:M | 0 | 0 | 0 | 0 | 0 | 0 |
| 40107 | ENSMUSG00000101849 | 0 | NA | NA | NA | NA | NA | Gm28386 | predicted gene 28386 [Source:MGI Symbo | 0 | 0 | 0 | 0 | 0 | 0 |
| 40108 | ENSMUSG00000101850 | 0 | NA | NA | NA | NA | NA | Gm29437 | predicted gene 29437 [Source:MGI Symbo | 0 | 0 | 0 | 0 | 0 | 0 |
| 40109 | ENSMUSG00000101851 | 0 | NA | NA | NA | NA | NA | Gm28474 | predicted gene 28474 [Source:MGI Symbo | 0 | 0 | 0 | 0 | 0 | 0 |
| 40110 | ENSMUSG00000101852 | 0 | NA | NA | NA | NA | NA | Gm20886 | predicted gene, 20886 [Source:MGI Symb | 0 | 0 | 0 | 0 | 0 | 0 |
| 40111 | ENSMUSG00000101853 | 0 | NA | NA | NA | NA | NA | Gm28627 | predicted gene 28627 [Source:MGI Symbo | 0 | 0 | 0 | 0 | 0 | 0 |
| 40112 | ENSMUSG00000101854 | 0 | NA | NA | NA | NA | NA | 1700026F02 | RIKEN cDNA 1700026F02 gene [Source:M | 0 | 0 | 0 | 0 | 0 | 0 |
| 40113 | ENSMUSG00000101855 | 0 | NA | NA | NA | NA | NA | 1700010N08 | RIKEN cDNA 1700010N08 gene [Source:M | 0 | 0 | 0 | 0 | 0 | 0 |
| 40114 | ENSMUSG00000101856 | 56.59057925 | -0.071292771 | 0.307199291 | -0.232073359 | 0.816481036 | 0.967767553 | 1700096K18 | RIKEN cDNA 1700096K18 gene [Source:M | 69.87 | 56.535 | 47.293 | 56.875 | 61.075 | 47.895 |
| 40115 | ENSMUSG00000101857 | 0 | NA | NA | NA | NA | NA | Gm21757 | predicted gene, 21757 [Source:MGI Symb | 0 | 0 | 0 | 0 | 0 | 0 |
| 40116 | ENSMUSG00000101858 | 0 | NA | NA | NA | NA | NA | Gm28144 | predicted gene 28144 [Source:MGI Symbo | 0 | 0 | 0 | 0 | 0 | 0 |
| 40117 | ENSMUSG00000101859 | 0 | NA | NA | NA | NA | NA | Gm29233 | predicted gene 29233 [Source:MGI Symbo | 0 | 0 | 0 | 0 | 0 | 0 |
| 40118 | ENSMUSG00000101860 | 0.181769715 | 1.126924175 | 4.080472857 | 0.276174898 | 0.782413736 | NA | Mindy4b-ps | MINDY lysine 48 deubiquitinase 4B, pseu | 0 | 0 | 0 | 0 | 1.0906 | 0 |
| 40119 | ENSMUSG00000101861 | 0 | NA | NA | NA | NA | NA | Gm29450 | predicted gene 29450 [Source:MGI Symbo | 0 | 0 | 0 | 0 | 0 | 0 |
| 40120 | ENSMUSG00000101862 | 0 | NA | NA | NA | NA | NA | Gm8918 | predicted gene 8918 [Source:MGI Symbol | 0 | 0 | 0 | 0 | 0 | 0 |
| 40121 | ENSMUSG00000101863 | 0 | NA | NA | NA | NA | NA | Gm29400 | predicted gene 29400 [Source:MGI Symbo | 0 | 0 | 0 | 0 | 0 | 0 |
| 40122 | ENSMUSG00000101864 | 0 | NA | NA | NA | NA | NA | Gm21634 | predicted gene, 21634 [Source:MGI Symb | 0 | 0 | 0 | 0 | 0 | 0 |
| 40123 | ENSMUSG00000101865 | 0 | NA | NA | NA | NA | NA | Gm9775 | predicted gene 9775 [Source:MGI Symbol | 0 | 0 | 0 | 0 | 0 | 0 |
| 40124 | ENSMUSG00000101866 | 0 | NA | NA | NA | NA | NA | Gm28109 | predicted gene 28109 [Source:MGI Symbo | 0 | 0 | 0 | 0 | 0 | 0 |
| 40125 | ENSMUSG00000101867 | 0.167707431 | -0.796629176 | 4.080472857 | -0.195229623 | 0.84521319 | NA | C78334 | expressed sequence C78334 [Source:MGI | 0 | 0 | 1.0062 | 0 | 0 | 0 |
| 40126 | ENSMUSG00000101868 | 0 | NA | NA | NA | NA | NA | Gm28680 | predicted gene 28680 [Source:MGI Symbo | 0 | 0 | 0 | 0 | 0 | 0 |
| 40127 | ENSMUSG00000101869 | 0 | NA | NA | NA | NA | NA | Gm29500 | predicted gene 29500 [Source:MGI Symbo | 0 | 0 | 0 | 0 | 0 | 0 |
| 40128 | ENSMUSG00000101870 | 0 | NA | NA | NA | NA | NA | Gm21745 | predicted gene, 21745 [Source:MGI Symb | 0 | 0 | 0 | 0 | 0 | 0 |
| 40129 | ENSMUSG00000101871 | 0 | NA | NA | NA | NA | NA | Gm28389 | predicted gene 28389 [Source:MGI Symbo | 0 | 0 | 0 | 0 | 0 | 0 |
| 40130 | ENSMUSG00000101872 | 2.479755313 | 0.835538912 | 1.352075125 | 0.617967816 | 0.536596554 | NA | Gm29237 | predicted gene 29237 [Source:MGI Symbo | 3.2124 | 1.0279 | 1.0062 | 1.928 | 4.3625 | 3.3415 |
| 40131 | ENSMUSG00000101873 | 0 | NA | NA | NA | NA | NA | Gm28546 | predicted gene 28546 [Source:MGI Symbo | 0 | 0 | 0 | 0 | 0 | 0 |
| 40132 | ENSMUSG00000101874 | 0 | NA | NA | NA | NA | NA | Olfr1391 | olfactory receptor 1391 [Source:MGI Syml | 0 | 0 | 0 | 0 | 0 | 0 |
| 40133 | ENSMUSG00000101875 | 0 | NA | NA | NA | NA | NA | Gm6028 | predicted gene 6028 [Source:MGI Symbol | 0 | 0 | 0 | 0 | 0 | 0 |
| 40134 | ENSMUSG00000101877 | 0 | NA | NA | NA | NA | NA | Gm8176 | predicted gene 8176 [Source:MGI Symbol | 0 | 0 | 0 | 0 | 0 | 0 |
| 40135 | ENSMUSG00000101878 | 0 | NA | NA | NA | NA | NA | Gm8203 | predicted pseudogene 8203 [Source:MGI | 0 | 0 | 0 | 0 | 0 | 0 |
| 40136 | ENSMUSG00000101879 | 0 | NA | NA | NA | NA | NA | Gm28820 | predicted gene 28820 [Source:MGI Symbo | 0 | 0 | 0 | 0 | 0 | 0 |
| 40137 | ENSMUSG00000101880 | 0 | NA | NA | NA | NA | NA | Gm29282 | predicted gene 29282 [Source:MGI Symbo | 0 | 0 | 0 | 0 | 0 | 0 |
| 40138 | ENSMUSG00000101881 | 0 | NA | NA | NA | NA | NA | Gm21076 | predicted gene, 21076 [Source:MGI Symb | 0 | 0 | 0 | 0 | 0 | 0 |
| 40139 | ENSMUSG00000101882 | 0 | NA | NA | NA | NA | NA | Gm29545 | predicted gene 29545 [Source:MGI Symbo | 0 | 0 | 0 | 0 | 0 | 0 |
| 40140 | ENSMUSG00000101883 | 0 | NA | NA | NA | NA | NA | Gm21840 | predicted gene, 21840 [Source:MGI Symb | 0 | 0 | 0 | 0 | 0 | 0 |
| 40141 | ENSMUSG00000101884 | 0 | NA | NA | NA | NA | NA | Gm28658 | predicted gene 28658 [Source:MGI Symbo | 0 | 0 | 0 | 0 | 0 | 0 |
| 40142 | ENSMUSG00000101885 | 0 | NA | NA | NA | NA | NA | Gm28235 | predicted gene 28235 [Source:MGI Symbo | 0 | 0 | 0 | 0 | 0 | 0 |
| 40143 | ENSMUSG00000101886 | 0 | NA | NA | NA | NA | NA | Gm28324 | predicted gene 28324 [Source:MGI Symbo | 0 | 0 | 0 | 0 | 0 | 0 |
| 40144 | ENSMUSG00000101887 | 0 | NA | NA | NA | NA | NA | Gm21654 | predicted gene, 21654 [Source:MGI Symb | 0 | 0 | 0 | 0 | 0 | 0 |
| 40145 | ENSMUSG00000101888 | 0 | NA | NA | NA | NA | NA | 1700025H01 | RIKEN cDNA 1700025H01 gene [Source:M | 0 | 0 | 0 | 0 | 0 | 0 |
| 40146 | ENSMUSG00000101890 | 0 | NA | NA | NA | NA | NA | Gm53054 | predicted gene 53054 [Source:MGI Symbo | 0 | 0 | 0 | 0 | 0 | 0 |
| 40147 | ENSMUSG00000101891 | 0 | NA | NA | NA | NA | NA | Gm29132 | predicted gene 29132 [Source:MGI Symbo | 0 | 0 | 0 | 0 | 0 | 0 |
| 40148 | ENSMUSG00000101892 | 133.0688979 | -0.122000032 | 0.20023332 | -0.609289362 | 0.542332657 | 0.894095573 | 9130401M01 | RIKEN cDNA 9130401M01 gene [Source:M | 149.38 | 133.63 | 132.82 | 130.14 | 123.24 | 129.21 |

| | | | | | | | | | | | | | | | |
|---|---|---|---|---|---|---|---|---|---|---|---|---|---|---|---|
| 40149 | ENSMUSG00000101893 | 0 | NA | NA | NA | NA | NA | Gm28839 | predicted gene 28839 [Source:MGI Symbo | 0 | 0 | 0 | 0 | 0 | 0 |
| 40150 | ENSMUSG00000101894 | 0 | NA | NA | NA | NA | NA | 1700016P04 | RIKEN cDNA 1700016P04 gene [Source:M | 0 | 0 | 0 | 0 | 0 | 0 |
| 40151 | ENSMUSG00000101895 | 0 | NA | NA | NA | NA | NA | Gm28981 | predicted gene 28981 [Source:MGI Symbo | 0 | 0 | 0 | 0 | 0 | 0 |
| 40152 | ENSMUSG00000101896 | 0 | NA | NA | NA | NA | NA | Gm6244 | predicted gene 6244 [Source:MGI Symbol | 0 | 0 | 0 | 0 | 0 | 0 |
| 40153 | ENSMUSG00000101897 | 0 | NA | NA | NA | NA | NA | Gm29137 | predicted gene 29137 [Source:MGI Symbo | 0 | 0 | 0 | 0 | 0 | 0 |
| 40154 | ENSMUSG00000101898 | 0 | NA | NA | NA | NA | NA | Gm28122 | predicted gene 28122 [Source:MGI Symbo | 0 | 0 | 0 | 0 | 0 | 0 |
| 40155 | ENSMUSG00000101899 | 0 | NA | NA | NA | NA | NA | Gm29175 | predicted gene 29175 [Source:MGI Symbo | 0 | 0 | 0 | 0 | 0 | 0 |
| 40156 | ENSMUSG00000101900 | 0 | NA | NA | NA | NA | NA | Mrgprb12-ps | MAS-related GPR, member B12, pseudoge | 0 | 0 | 0 | 0 | 0 | 0 |
| 40157 | ENSMUSG00000101901 | 0 | NA | NA | NA | NA | NA | Gm9230 | predicted gene 9230 [Source:MGI Symbol | 0 | 0 | 0 | 0 | 0 | 0 |
| 40158 | ENSMUSG00000101902 | 0 | NA | NA | NA | NA | NA | Gm29115 | predicted gene 29115 [Source:MGI Symbo | 0 | 0 | 0 | 0 | 0 | 0 |
| 40159 | ENSMUSG00000101903 | 0 | NA | NA | NA | NA | NA | Gm29291 | predicted gene 29291 [Source:MGI Symbo | 0 | 0 | 0 | 0 | 0 | 0 |
| 40160 | ENSMUSG00000101904 | 5.972753049 | -0.339046945 | 0.899155742 | -0.377072547 | 0.706119693 | 0.94295036 | Gm29427 | predicted gene 29427 [Source:MGI Symbo | 4.8186 | 7.1954 | 8.05 | 4.8199 | 8.7249 | 2.2277 |
| 40161 | ENSMUSG00000101905 | 1.807485267 | -2.01902646 | 1.885314789 | -1.07092273 | 0.284204175 | NA | 4930439D14 | RIKEN cDNA 4930439D14 gene [Source:M | 5.6217 | 0 | 3.0187 | 0 | 1.0906 | 1.1138 |
| 40162 | ENSMUSG00000101906 | 0 | NA | NA | NA | NA | NA | Mrgprc2-ps | MAS-related GPR, member C2, pseudoger | 0 | 0 | 0 | 0 | 0 | 0 |
| 40163 | ENSMUSG00000101907 | 0 | NA | NA | NA | NA | NA | Gm29472 | predicted gene 29472 [Source:MGI Symbo | 0 | 0 | 0 | 0 | 0 | 0 |
| 40164 | ENSMUSG00000101908 | 0 | NA | NA | NA | NA | NA | Gm21642 | predicted gene, 21642 [Source:MGI Symb | 0 | 0 | 0 | 0 | 0 | 0 |
| 40165 | ENSMUSG00000101909 | 0 | NA | NA | NA | NA | NA | Gm28724 | predicted gene 28724 [Source:MGI Symbo | 0 | 0 | 0 | 0 | 0 | 0 |
| 40166 | ENSMUSG00000101910 | 0 | NA | NA | NA | NA | NA | Gm28691 | predicted gene 28691 [Source:MGI Symbo | 0 | 0 | 0 | 0 | 0 | 0 |
| 40167 | ENSMUSG00000101911 | 0 | NA | NA | NA | NA | NA | Gm28381 | predicted gene 28381 [Source:MGI Symbo | 0 | 0 | 0 | 0 | 0 | 0 |
| 40168 | ENSMUSG00000101912 | 0 | NA | NA | NA | NA | NA | 1700052I22R | RIKEN cDNA 1700052I22 gene [Source:MC | 0 | 0 | 0 | 0 | 0 | 0 |
| 40169 | ENSMUSG00000101913 | 0 | NA | NA | NA | NA | NA | Gm28999 | predicted gene 28999 [Source:MGI Symbo | 0 | 0 | 0 | 0 | 0 | 0 |
| 40170 | ENSMUSG00000101914 | 0 | NA | NA | NA | NA | NA | Gm21790 | predicted gene, 21790 [Source:MGI Symb | 0 | 0 | 0 | 0 | 0 | 0 |
| 40171 | ENSMUSG00000101915 | 0 | NA | NA | NA | NA | NA | Gm28102 | predicted gene 28102 [Source:MGI Symbo | 0 | 0 | 0 | 0 | 0 | 0 |
| 40172 | ENSMUSG00000101916 | 0 | NA | NA | NA | NA | NA | Gm18501 | predicted gene, 18501 [Source:MGI Symb | 0 | 0 | 0 | 0 | 0 | 0 |
| 40173 | ENSMUSG00000101917 | 0 | NA | NA | NA | NA | NA | Gm29312 | predicted gene 29312 [Source:MGI Symbo | 0 | 0 | 0 | 0 | 0 | 0 |
| 40174 | ENSMUSG00000101918 | 0 | NA | NA | NA | NA | NA | Olfr1225 | olfactory receptor 1225 [Source:MGI Syml | 0 | 0 | 0 | 0 | 0 | 0 |
| 40175 | ENSMUSG00000101919 | 0 | NA | NA | NA | NA | NA | Gm29664 | predicted gene 29664 [Source:MGI Symbo | 0 | 0 | 0 | 0 | 0 | 0 |
| 40176 | ENSMUSG00000101920 | 0 | NA | NA | NA | NA | NA | Gm28454 | predicted gene 28454 [Source:MGI Symbo | 0 | 0 | 0 | 0 | 0 | 0 |
| 40177 | ENSMUSG00000101921 | 0 | NA | NA | NA | NA | NA | Gm28825 | predicted gene 28825 [Source:MGI Symbo | 0 | 0 | 0 | 0 | 0 | 0 |
| 40178 | ENSMUSG00000101922 | 0 | NA | NA | NA | NA | NA | Gm29345 | predicted gene 29345 [Source:MGI Symbo | 0 | 0 | 0 | 0 | 0 | 0 |
| 40179 | ENSMUSG00000101923 | 0 | NA | NA | NA | NA | NA | Gm29041 | predicted gene 29041 [Source:MGI Symbo | 0 | 0 | 0 | 0 | 0 | 0 |
| 40180 | ENSMUSG00000101925 | 0 | NA | NA | NA | NA | NA | Gm28156 | predicted gene 28156 [Source:MGI Symbo | 0 | 0 | 0 | 0 | 0 | 0 |
| 40181 | ENSMUSG00000101927 | 0 | NA | NA | NA | NA | NA | Gm28992 | predicted gene 28992 [Source:MGI Symbo | 0 | 0 | 0 | 0 | 0 | 0 |
| 40182 | ENSMUSG00000101928 | 0 | NA | NA | NA | NA | NA | Gm20824 | predicted gene, 20824 [Source:MGI Symb | 0 | 0 | 0 | 0 | 0 | 0 |
| 40183 | ENSMUSG00000101929 | 0 | NA | NA | NA | NA | NA | Gm28245 | predicted gene 28245 [Source:MGI Symbo | 0 | 0 | 0 | 0 | 0 | 0 |
| 40184 | ENSMUSG00000101930 | 0 | NA | NA | NA | NA | NA | Gm5441 | predicted gene 5441 [Source:MGI Symbol | 0 | 0 | 0 | 0 | 0 | 0 |
| 40185 | ENSMUSG00000101931 | 0 | NA | NA | NA | NA | NA | Gm28108 | predicted gene 28108 [Source:MGI Symbo | 0 | 0 | 0 | 0 | 0 | 0 |
| 40186 | ENSMUSG00000101932 | 0.36353943 | 2.065989254 | 4.034137301 | 0.512126658 | 0.608562368 | NA | Gm10074 | predicted gene 10074 [Source:MGI Symbo | 0 | 0 | 0 | 0 | 2.1812 | 0 |
| 40187 | ENSMUSG00000101933 | 0 | NA | NA | NA | NA | NA | Gm20835 | predicted gene, 20835 [Source:MGI Symb | 0 | 0 | 0 | 0 | 0 | 0 |
| 40188 | ENSMUSG00000101934 | 0.503122293 | -2.366675966 | 3.785596405 | -0.625179156 | 0.531853481 | NA | Gm18304 | predicted gene, 18304 [Source:MGI Symb | 0 | 0 | 3.0187 | 0 | 0 | 0 |
| 40189 | ENSMUSG00000101935 | 0 | NA | NA | NA | NA | NA | Gm17748 | predicted gene, 17748 [Source:MGI Symb | 0 | 0 | 0 | 0 | 0 | 0 |
| 40190 | ENSMUSG00000101936 | 0 | NA | NA | NA | NA | NA | Gm28164 | predicted gene 28164 [Source:MGI Symbo | 0 | 0 | 0 | 0 | 0 | 0 |
| 40191 | ENSMUSG00000101937 | 0 | NA | NA | NA | NA | NA | Gm5268 | predicted gene 5268 [Source:MGI Symbol | 0 | 0 | 0 | 0 | 0 | 0 |
| 40192 | ENSMUSG00000101938 | 0 | NA | NA | NA | NA | NA | Gm29605 | predicted gene 29605 [Source:MGI Symbo | 0 | 0 | 0 | 0 | 0 | 0 |
| 40193 | ENSMUSG00000101939 | 66.35098583 | -0.035776532 | 0.30799328 | -0.116160106 | 0.907525653 | 0.984582287 | Gm28438 | predicted gene 28438 [Source:MGI Symbo | 73.885 | 67.843 | 59.368 | 46.271 | 87.249 | 63.489 |
| 40194 | ENSMUSG00000101940 | 0 | NA | NA | NA | NA | NA | Akp-ps1 | alkaline phosphatase pseudogene 1 [Sour | 0 | 0 | 0 | 0 | 0 | 0 |
| 40195 | ENSMUSG00000101941 | 0 | NA | NA | NA | NA | NA | Gm28979 | predicted gene 28979 [Source:MGI Symbo | 0 | 0 | 0 | 0 | 0 | 0 |
| 40196 | ENSMUSG00000101942 | 0.65103495 | -1.312578592 | 2.821398903 | -0.465222621 | 0.641772041 | NA | Gm19582 | predicted gene, 19582 [Source:MGI Symb | 0.8031 | 0 | 2.0125 | 0 | 1.0906 | 0 |
| 40197 | ENSMUSG00000101943 | 0 | NA | NA | NA | NA | NA | Gm28532 | predicted gene 28532 [Source:MGI Symbo | 0 | 0 | 0 | 0 | 0 | 0 |
| 40198 | ENSMUSG00000101944 | 0 | NA | NA | NA | NA | NA | Mrgprc4-ps | MAS-related GPR, member C4, pseudoger | 0 | 0 | 0 | 0 | 0 | 0 |
| 40199 | ENSMUSG00000101945 | 0 | NA | NA | NA | NA | NA | Btf3-ps17 | basic transcription factor 3, pseudogene 1 | 0 | 0 | 0 | 0 | 0 | 0 |
| 40200 | ENSMUSG00000101947 | 0 | NA | NA | NA | NA | NA | Gm29167 | predicted gene 29167 [Source:MGI Symbo | 0 | 0 | 0 | 0 | 0 | 0 |
| 40201 | ENSMUSG00000101948 | 0 | NA | NA | NA | NA | NA | Gm4557 | predicted gene 4557 [Source:MGI Symbol | 0 | 0 | 0 | 0 | 0 | 0 |
| 40202 | ENSMUSG00000101949 | 0 | NA | NA | NA | NA | NA | Gm5370 | predicted gene 5370 [Source:MGI Symbol | 0 | 0 | 0 | 0 | 0 | 0 |
| 40203 | ENSMUSG00000101950 | 0 | NA | NA | NA | NA | NA | Gm29267 | predicted gene 29267 [Source:MGI Symbo | 0 | 0 | 0 | 0 | 0 | 0 |
| 40204 | ENSMUSG00000101951 | 0 | NA | NA | NA | NA | NA | Gm28721 | predicted gene 28721 [Source:MGI Symbo | 0 | 0 | 0 | 0 | 0 | 0 |

| | | | | | | | | | | | | | | | |
|---|---|---|---|---|---|---|---|---|---|---|---|---|---|---|---|
| 40205 | ENSMUSG00000101952 | 0.342639279 | -1.805976596 | 4.028547714 | -0.448294702 | 0.653940524 | NA | Gm10550 | predicted gene 10550 [Source:MGI Symbo | 0 | 2.0558 | 0 | 0 | 0 | 0 |
| 40206 | ENSMUSG00000101953 | 0 | NA | NA | NA | NA | NA | Gm29241 | predicted gene 29241 [Source:MGI Symbo | 0 | 0 | 0 | 0 | 0 | 0 |
| 40207 | ENSMUSG00000101954 | 0 | NA | NA | NA | NA | NA | Gm28128 | predicted gene 28128 [Source:MGI Symbo | 0 | 0 | 0 | 0 | 0 | 0 |
| 40208 | ENSMUSG00000101955 | 0.556921942 | 2.679043159 | 3.638243063 | 0.736356289 | 0.461513899 | NA | Tuba5-ps | tubulin alpha 5, pseudogene [Source:MGI | 0 | 0 | 0 | 0 | 0 | 3.3415 |
| 40209 | ENSMUSG00000101956 | 0 | NA | NA | NA | NA | NA | Gm28574 | predicted gene 28574 [Source:MGI Symbo | 0 | 0 | 0 | 0 | 0 | 0 |
| 40210 | ENSMUSG00000101957 | 0 | NA | NA | NA | NA | NA | Gm29576 | predicted gene 29576 [Source:MGI Symbo | 0 | 0 | 0 | 0 | 0 | 0 |
| 40211 | ENSMUSG00000101958 | 0 | NA | NA | NA | NA | NA | Gm19085 | predicted gene, 19085 [Source:MGI Symb | 0 | 0 | 0 | 0 | 0 | 0 |
| 40212 | ENSMUSG00000101959 | 0 | NA | NA | NA | NA | NA | Ldhal6b | lactate dehydrogenase A-like 6B [Source:N | 0 | 0 | 0 | 0 | 0 | 0 |
| 40213 | ENSMUSG00000101960 | 0 | NA | NA | NA | NA | NA | Gm29311 | predicted gene 29311 [Source:MGI Symbo | 0 | 0 | 0 | 0 | 0 | 0 |
| 40214 | ENSMUSG00000101961 | 0 | NA | NA | NA | NA | NA | Gm28220 | predicted gene 28220 [Source:MGI Symbo | 0 | 0 | 0 | 0 | 0 | 0 |
| 40215 | ENSMUSG00000101962 | 0 | NA | NA | NA | NA | NA | Gm28741 | predicted gene 28741 [Source:MGI Symbo | 0 | 0 | 0 | 0 | 0 | 0 |
| 40216 | ENSMUSG00000101963 | 0 | NA | NA | NA | NA | NA | 1700001J11I | RIKEN cDNA 1700001J11 gene [Source:MG | 0 | 0 | 0 | 0 | 0 | 0 |
| 40217 | ENSMUSG00000101964 | 0.877291758 | 2.066311913 | 2.565974136 | 0.805273866 | 0.420661659 | NA | Gm28693 | predicted gene 28693 [Source:MGI Symbo | 0 | 1.0279 | 0 | 0.964 | 3.2719 | 0 |
| 40218 | ENSMUSG00000101965 | 0 | NA | NA | NA | NA | NA | Gm29129 | predicted gene 29129 [Source:MGI Symbo | 0 | 0 | 0 | 0 | 0 | 0 |
| 40219 | ENSMUSG00000101966 | 0 | NA | NA | NA | NA | NA | Gm5248 | predicted gene 5248 [Source:MGI Symbol | 0 | 0 | 0 | 0 | 0 | 0 |
| 40220 | ENSMUSG00000101967 | 0 | NA | NA | NA | NA | NA | Gm29555 | predicted gene 29555 [Source:MGI Symbo | 0 | 0 | 0 | 0 | 0 | 0 |
| 40221 | ENSMUSG00000101968 | 0 | NA | NA | NA | NA | NA | 1700027A15 | RIKEN cDNA 1700027A15 gene [Source:M | 0 | 0 | 0 | 0 | 0 | 0 |
| 40222 | ENSMUSG00000101969 | 0.703108924 | -2.875459482 | 2.96384054 | -0.970180225 | 0.331956666 | NA | Gm20125 | predicted gene, 20125 [Source:MGI Symb | 3.2124 | 0 | 1.0062 | 0 | 0 | 0 |
| 40223 | ENSMUSG00000101970 | 232.207126 | -0.152005456 | 0.162794676 | -0.933724981 | 0.350445776 | 0.791877747 | Chaserr | CHD2 adjacent suppressive regulatory RN | 257.8 | 245.67 | 229.42 | 215.93 | 209.4 | 235.02 |
| 40224 | ENSMUSG00000101971 | 0 | NA | NA | NA | NA | NA | Gm28246 | predicted gene 28246 [Source:MGI Symbo | 0 | 0 | 0 | 0 | 0 | 0 |
| 40225 | ENSMUSG00000101972 | 0 | NA | NA | NA | NA | NA | H3c11 | H3 clustered histone 11 [Source:MGI Sym | 0 | 0 | 0 | 0 | 0 | 0 |
| 40226 | ENSMUSG00000101973 | 0 | NA | NA | NA | NA | NA | Gm29516 | predicted gene 29516 [Source:MGI Symbo | 0 | 0 | 0 | 0 | 0 | 0 |
| 40227 | ENSMUSG00000101974 | 0 | NA | NA | NA | NA | NA | Gm28384 | predicted gene 28384 [Source:MGI Symbo | 0 | 0 | 0 | 0 | 0 | 0 |
| 40228 | ENSMUSG00000101975 | 0 | NA | NA | NA | NA | NA | Scgb1b34-ps | secretoglobin, family 1B, member 34, pse | 0 | 0 | 0 | 0 | 0 | 0 |
| 40229 | ENSMUSG00000101977 | 0 | NA | NA | NA | NA | NA | Gm28846 | predicted gene 28846 [Source:MGI Symbo | 0 | 0 | 0 | 0 | 0 | 0 |
| 40230 | ENSMUSG00000101978 | 0 | NA | NA | NA | NA | NA | 1700063O14 | RIKEN cDNA 1700063O14 gene [Source:M | 0 | 0 | 0 | 0 | 0 | 0 |
| 40231 | ENSMUSG00000101980 | 0 | NA | NA | NA | NA | NA | Gm21922 | predicted gene, 21922 [Source:MGI Symb | 0 | 0 | 0 | 0 | 0 | 0 |
| 40232 | ENSMUSG00000101981 | 0 | NA | NA | NA | NA | NA | Gm28157 | predicted gene 28157 [Source:MGI Symbo | 0 | 0 | 0 | 0 | 0 | 0 |
| 40233 | ENSMUSG00000101982 | 0 | NA | NA | NA | NA | NA | Gm28990 | predicted gene 28990 [Source:MGI Symbo | 0 | 0 | 0 | 0 | 0 | 0 |
| 40234 | ENSMUSG00000101983 | 0.160662974 | 1.126924175 | 4.080472857 | 0.276174898 | 0.782413736 | NA | Gm28976 | predicted gene 28976 [Source:MGI Symbo | 0 | 0 | 0 | 0.964 | 0 | 0 |
| 40235 | ENSMUSG00000101984 | 0 | NA | NA | NA | NA | NA | Gm29436 | predicted gene 29436 [Source:MGI Symbo | 0 | 0 | 0 | 0 | 0 | 0 |
| 40236 | ENSMUSG00000101985 | 0 | NA | NA | NA | NA | NA | Gm28929 | predicted gene 28929 [Source:MGI Symbo | 0 | 0 | 0 | 0 | 0 | 0 |
| 40237 | ENSMUSG00000101986 | 0 | NA | NA | NA | NA | NA | Gm6198 | predicted gene 6198 [Source:MGI Symbol | 0 | 0 | 0 | 0 | 0 | 0 |
| 40238 | ENSMUSG00000101987 | 0 | NA | NA | NA | NA | NA | Gm21268 | predicted gene, 21268 [Source:MGI Symb | 0 | 0 | 0 | 0 | 0 | 0 |
| 40239 | ENSMUSG00000101988 | 0 | NA | NA | NA | NA | NA | Gm28484 | predicted gene 28484 [Source:MGI Symbo | 0 | 0 | 0 | 0 | 0 | 0 |
| 40240 | ENSMUSG00000101989 | 0 | NA | NA | NA | NA | NA | Gm28346 | predicted gene 28346 [Source:MGI Symbo | 0 | 0 | 0 | 0 | 0 | 0 |
| 40241 | ENSMUSG00000101990 | 0 | NA | NA | NA | NA | NA | Gm20856 | predicted gene, 20856 [Source:MGI Symb | 0 | 0 | 0 | 0 | 0 | 0 |
| 40242 | ENSMUSG00000101991 | 0 | NA | NA | NA | NA | NA | Gm21724 | predicted gene, 21724 [Source:MGI Symb | 0 | 0 | 0 | 0 | 0 | 0 |
| 40243 | ENSMUSG00000101992 | 0 | NA | NA | NA | NA | NA | Gm29249 | predicted gene 29249 [Source:MGI Symbo | 0 | 0 | 0 | 0 | 0 | 0 |
| 40244 | ENSMUSG00000101993 | 0 | NA | NA | NA | NA | NA | Gm20987 | predicted gene, 20987 [Source:MGI Symb | 0 | 0 | 0 | 0 | 0 | 0 |
| 40245 | ENSMUSG00000101994 | 0 | NA | NA | NA | NA | NA | Gm18875 | predicted gene, 18875 [Source:MGI Symb | 0 | 0 | 0 | 0 | 0 | 0 |
| 40246 | ENSMUSG00000101995 | 1.05829193 | -1.166662428 | 2.355445425 | -0.495304377 | 0.620385282 | NA | Gm29480 | predicted gene 29480 [Source:MGI Symbo | 2.4093 | 0 | 2.0125 | 1.928 | 0 | 0 |
| 40247 | ENSMUSG00000101996 | 0 | NA | NA | NA | NA | NA | Gm19206 | predicted gene, 19206 [Source:MGI Symb | 0 | 0 | 0 | 0 | 0 | 0 |
| 40248 | ENSMUSG00000101997 | 0 | NA | NA | NA | NA | NA | Gm28722 | predicted gene 28722 [Source:MGI Symbo | 0 | 0 | 0 | 0 | 0 | 0 |
| 40249 | ENSMUSG00000101998 | 0 | NA | NA | NA | NA | NA | Gm21530 | predicted gene, 21530 [Source:MGI Symb | 0 | 0 | 0 | 0 | 0 | 0 |
| 40250 | ENSMUSG00000102000 | 0.160662974 | 1.126924175 | 4.080472857 | 0.276174898 | 0.782413736 | NA | 1700006H21 | RIKEN cDNA 1700006H21 gene [Source:M | 0 | 0 | 0 | 0.964 | 0 | 0 |
| 40251 | ENSMUSG00000102001 | 0 | NA | NA | NA | NA | NA | Gm28755 | predicted gene 28755 [Source:MGI Symbo | 0 | 0 | 0 | 0 | 0 | 0 |
| 40252 | ENSMUSG00000102002 | 0 | NA | NA | NA | NA | NA | Scgb2b6 | secretoglobin, family 2B, member 6 [Sour | 0 | 0 | 0 | 0 | 0 | 0 |
| 40253 | ENSMUSG00000102003 | 0 | NA | NA | NA | NA | NA | Gm28666 | predicted gene 28666 [Source:MGI Symbo | 0 | 0 | 0 | 0 | 0 | 0 |
| 40254 | ENSMUSG00000102004 | 0 | NA | NA | NA | NA | NA | 4933402C06 | RIKEN cDNA 4933402C06 gene [Source:M | 0 | 0 | 0 | 0 | 0 | 0 |
| 40255 | ENSMUSG00000102005 | 0 | NA | NA | NA | NA | NA | Gm29409 | predicted gene 29409 [Source:MGI Symbo | 0 | 0 | 0 | 0 | 0 | 0 |
| 40256 | ENSMUSG00000102006 | 0 | NA | NA | NA | NA | NA | Gm28591 | predicted gene 28591 [Source:MGI Symbo | 0 | 0 | 0 | 0 | 0 | 0 |
| 40257 | ENSMUSG00000102008 | 0 | NA | NA | NA | NA | NA | Gm28495 | predicted gene 28495 [Source:MGI Symbo | 0 | 0 | 0 | 0 | 0 | 0 |
| 40258 | ENSMUSG00000102009 | 0.17131964 | -0.796629176 | 4.080472857 | -0.195229623 | 0.84521319 | NA | 4933400F21 | RIKEN cDNA 4933400F21 gene [Source:M | 0 | 1.0279 | 0 | 0 | 0 | 0 |
| 40259 | ENSMUSG00000102010 | 0 | NA | NA | NA | NA | NA | Gm28248 | predicted gene 28248 [Source:MGI Symbo | 0 | 0 | 0 | 0 | 0 | 0 |
| 40260 | ENSMUSG00000102011 | 0 | NA | NA | NA | NA | NA | Gm28930 | predicted gene 28930 [Source:MGI Symbo | 0 | 0 | 0 | 0 | 0 | 0 |

| | | | | | | | | | | | | | | | |
|---|---|---|---|---|---|---|---|---|---|---|---|---|---|---|---|
| 40261 | ENSMUSG00000102013 | 0 | NA | NA | NA | NA | NA | E330023G01 | RIKEN cDNA E330023G01 gene [Source:M | 0 | 0 | 0 | 0 | 0 | 0 |
| 40262 | ENSMUSG00000102014 | 1.070235586 | 1.150616445 | 2.365541688 | 0.486407173 | 0.626678506 | NA | 2900009J06I | RIKEN cDNA 2900009J06 gene [Source:M( | 0 | 0 | 2.0125 | 0 | 2.1812 | 2.2277 |
| 40263 | ENSMUSG00000102015 | 0 | NA | NA | NA | NA | NA | Gm21476 | predicted gene, 21476 [Source:MGI Symb | 0 | 0 | 0 | 0 | 0 | 0 |
| 40264 | ENSMUSG00000102016 | 0 | NA | NA | NA | NA | NA | Gm29606 | predicted gene 29606 [Source:MGI Symbo | 0 | 0 | 0 | 0 | 0 | 0 |
| 40265 | ENSMUSG00000102017 | 0 | NA | NA | NA | NA | NA | Gm6124 | predicted gene 6124 [Source:MGI Symbol | 0 | 0 | 0 | 0 | 0 | 0 |
| 40266 | ENSMUSG00000102018 | 14.35229061 | -0.316266447 | 0.59384191 | -0.532576839 | 0.594326538 | 0.913144152 | Iqank1 | IQ motif and ankyrin repeat containing 1 [ | 19.274 | 15.419 | 13.081 | 16.388 | 15.269 | 6.6831 |
| 40267 | ENSMUSG00000102019 | 0 | NA | NA | NA | NA | NA | Gm21873 | predicted gene, 21873 [Source:MGI Symb | 0 | 0 | 0 | 0 | 0 | 0 |
| 40268 | ENSMUSG00000102020 | 0 | NA | NA | NA | NA | NA | Gm28620 | predicted gene 28620 [Source:MGI Symbo | 0 | 0 | 0 | 0 | 0 | 0 |
| 40269 | ENSMUSG00000102022 | 0 | NA | NA | NA | NA | NA | Gm28735 | predicted gene 28735 [Source:MGI Symbo | 0 | 0 | 0 | 0 | 0 | 0 |
| 40270 | ENSMUSG00000102023 | 0 | NA | NA | NA | NA | NA | Gm8708 | predicted gene 8708 [Source:MGI Symbol | 0 | 0 | 0 | 0 | 0 | 0 |
| 40271 | ENSMUSG00000102025 | 0 | NA | NA | NA | NA | NA | Gm29575 | predicted gene 29575 [Source:MGI Symbo | 0 | 0 | 0 | 0 | 0 | 0 |
| 40272 | ENSMUSG00000102026 | 0 | NA | NA | NA | NA | NA | Gm28571 | predicted gene 28571 [Source:MGI Symbo | 0 | 0 | 0 | 0 | 0 | 0 |
| 40273 | ENSMUSG00000102027 | 1.204865991 | 0.967559073 | 2.50299455 | 0.386560599 | 0.699081532 | NA | Gm28914 | predicted gene 28914 [Source:MGI Symbo | 2.4093 | 0 | 0 | 4.8199 | 0 | 0 |
| 40274 | ENSMUSG00000102028 | 0 | NA | NA | NA | NA | NA | Gm3142 | predicted gene 3142 [Source:MGI Symbol | 0 | 0 | 0 | 0 | 0 | 0 |
| 40275 | ENSMUSG00000102029 | 0 | NA | NA | NA | NA | NA | Gm29530 | predicted gene 29530 [Source:MGI Symbo | 0 | 0 | 0 | 0 | 0 | 0 |
| 40276 | ENSMUSG00000102030 | 0 | NA | NA | NA | NA | NA | Gm29106 | predicted gene 29106 [Source:MGI Symbo | 0 | 0 | 0 | 0 | 0 | 0 |
| 40277 | ENSMUSG00000102031 | 0 | NA | NA | NA | NA | NA | Gm29426 | predicted gene 29426 [Source:MGI Symbo | 0 | 0 | 0 | 0 | 0 | 0 |
| 40278 | ENSMUSG00000102032 | 0 | NA | NA | NA | NA | NA | Gm28705 | predicted gene 28705 [Source:MGI Symbo | 0 | 0 | 0 | 0 | 0 | 0 |
| 40279 | ENSMUSG00000102033 | 0 | NA | NA | NA | NA | NA | Gm28263 | predicted gene 28263 [Source:MGI Symbo | 0 | 0 | 0 | 0 | 0 | 0 |
| 40280 | ENSMUSG00000102035 | 0 | NA | NA | NA | NA | NA | Gm21733 | predicted gene, 21733 [Source:MGI Symb | 0 | 0 | 0 | 0 | 0 | 0 |
| 40281 | ENSMUSG00000102036 | 0 | NA | NA | NA | NA | NA | 1700063K16 | RIKEN cDNA 1700063K16 gene [Source:M | 0 | 0 | 0 | 0 | 0 | 0 |
| 40282 | ENSMUSG00000102037 | 2.410806579 | -1.187170365 | 1.609468948 | -0.737616197 | 0.46074771 | NA | Bcl2a1a | B cell leukemia/lymphoma 2 related prote | 0.8031 | 8.2233 | 1.0062 | 0 | 1.0906 | 3.3415 |
| 40283 | ENSMUSG00000102038 | 0 | NA | NA | NA | NA | NA | Gm12345 | predicted gene 12345 [Source:MGI Symbo | 0 | 0 | 0 | 0 | 0 | 0 |
| 40284 | ENSMUSG00000102039 | 0 | NA | NA | NA | NA | NA | Gm20788 | predicted gene, 20788 [Source:MGI Symb | 0 | 0 | 0 | 0 | 0 | 0 |
| 40285 | ENSMUSG00000102041 | 0 | NA | NA | NA | NA | NA | Rpl23a-ps9 | ribosomal protein L23A, pseudogene 9 [Sc | 0 | 0 | 0 | 0 | 0 | 0 |
| 40286 | ENSMUSG00000102042 | 0 | NA | NA | NA | NA | NA | Gm29195 | predicted gene 29195 [Source:MGI Symbo | 0 | 0 | 0 | 0 | 0 | 0 |
| 40287 | ENSMUSG00000102043 | 0 | NA | NA | NA | NA | NA | Gm9541 | predicted gene 9541 [Source:MGI Symbol | 0 | 0 | 0 | 0 | 0 | 0 |
| 40288 | ENSMUSG00000102044 | 0 | NA | NA | NA | NA | NA | Gm21249 | predicted gene, 21249 [Source:MGI Symb | 0 | 0 | 0 | 0 | 0 | 0 |
| 40289 | ENSMUSG00000102045 | 0 | NA | NA | NA | NA | NA | Gm21294 | predicted gene, 21294 [Source:MGI Symb | 0 | 0 | 0 | 0 | 0 | 0 |
| 40290 | ENSMUSG00000102046 | 0 | NA | NA | NA | NA | NA | 2810404M0: | RIKEN cDNA 2810404M03 gene [Source:M | 0 | 0 | 0 | 0 | 0 | 0 |
| 40291 | ENSMUSG00000102047 | 0 | NA | NA | NA | NA | NA | 2010010A06 | RIKEN cDNA 2010010A06 gene [Source:M | 0 | 0 | 0 | 0 | 0 | 0 |
| 40292 | ENSMUSG00000102048 | 0 | NA | NA | NA | NA | NA | Gm6075 | predicted gene 6075 [Source:MGI Symbol | 0 | 0 | 0 | 0 | 0 | 0 |
| 40293 | ENSMUSG00000102049 | 0 | NA | NA | NA | NA | NA | NA | NA | 0 | 0 | 0 | 0 | 0 | 0 |
| 40294 | ENSMUSG00000102050 | 0 | NA | NA | NA | NA | NA | Gm29499 | predicted gene 29499 [Source:MGI Symbo | 0 | 0 | 0 | 0 | 0 | 0 |
| 40295 | ENSMUSG00000102051 | 0 | NA | NA | NA | NA | NA | Ly6a2 | lymphocyte antigen 6 complex, locus A2 [ | 0 | 0 | 0 | 0 | 0 | 0 |
| 40296 | ENSMUSG00000102052 | 0 | NA | NA | NA | NA | NA | Gm28387 | predicted gene 28387 [Source:MGI Symbo | 0 | 0 | 0 | 0 | 0 | 0 |
| 40297 | ENSMUSG00000102053 | 0 | NA | NA | NA | NA | NA | Gm4064 | predicted gene 4064 [Source:MGI Symbol | 0 | 0 | 0 | 0 | 0 | 0 |
| 40298 | ENSMUSG00000102054 | 0 | NA | NA | NA | NA | NA | Gm29373 | predicted gene 29373 [Source:MGI Symbo | 0 | 0 | 0 | 0 | 0 | 0 |
| 40299 | ENSMUSG00000102055 | 4.304915301 | 0.290733341 | 1.308624731 | 0.222167085 | 0.824183816 | NA | Gm28913 | predicted gene 28913 [Source:MGI Symbo | 6.4248 | 2.0558 | 3.0187 | 0.964 | 0 | 13.366 |
| 40300 | ENSMUSG00000102056 | 0 | NA | NA | NA | NA | NA | Gm28354 | predicted gene 28354 [Source:MGI Symbo | 0 | 0 | 0 | 0 | 0 | 0 |
| 40301 | ENSMUSG00000102057 | 0 | NA | NA | NA | NA | NA | Gm29594 | predicted gene 29594 [Source:MGI Symbo | 0 | 0 | 0 | 0 | 0 | 0 |
| 40302 | ENSMUSG00000102058 | 0 | NA | NA | NA | NA | NA | Gm28470 | predicted gene 28470 [Source:MGI Symbo | 0 | 0 | 0 | 0 | 0 | 0 |
| 40303 | ENSMUSG00000102059 | 22.23520256 | 0.650002371 | 0.469797819 | 1.383578946 | 0.166487419 | 0.5925379 | Gm20257 | predicted gene, 20257 [Source:MGI Symb | 19.274 | 20.558 | 12.075 | 30.847 | 27.265 | 23.391 |
| 40304 | ENSMUSG00000102060 | 0.881162691 | 2.072029295 | 2.358945437 | 0.878371014 | 0.379742409 | NA | 1700061E17 | RIKEN cDNA 1700061E17 gene [Source:M | 0 | 1.0279 | 0 | 0.964 | 2.1812 | 1.1138 |
| 40305 | ENSMUSG00000102061 | 0 | NA | NA | NA | NA | NA | Gm28823 | predicted gene 28823 [Source:MGI Symbo | 0 | 0 | 0 | 0 | 0 | 0 |
| 40306 | ENSMUSG00000102062 | 0 | NA | NA | NA | NA | NA | Gm29246 | predicted gene 29246 [Source:MGI Symbo | 0 | 0 | 0 | 0 | 0 | 0 |
| 40307 | ENSMUSG00000102063 | 0 | NA | NA | NA | NA | NA | Gm28706 | predicted gene 28706 [Source:MGI Symbo | 0 | 0 | 0 | 0 | 0 | 0 |
| 40308 | ENSMUSG00000102064 | 0 | NA | NA | NA | NA | NA | Gm28625 | predicted gene 28625 [Source:MGI Symbo | 0 | 0 | 0 | 0 | 0 | 0 |
| 40309 | ENSMUSG00000102065 | 0 | NA | NA | NA | NA | NA | Gm28302 | predicted gene 28302 [Source:MGI Symbo | 0 | 0 | 0 | 0 | 0 | 0 |
| 40310 | ENSMUSG00000102067 | 0 | NA | NA | NA | NA | NA | Gm29230 | predicted gene 29230 [Source:MGI Symbo | 0 | 0 | 0 | 0 | 0 | 0 |
| 40311 | ENSMUSG00000102068 | 0 | NA | NA | NA | NA | NA | Mrgprc7-ps | MAS-related GPR, member C7, pseudoger | 0 | 0 | 0 | 0 | 0 | 0 |
| 40312 | ENSMUSG00000102069 | 0 | NA | NA | NA | NA | NA | 1700012I11I | RIKEN cDNA 1700012I11 gene [Source:M( | 0 | 0 | 0 | 0 | 0 | 0 |
| 40313 | ENSMUSG00000102070 | 566.6132549 | 0.175497088 | 0.126864264 | 1.383345339 | 0.166559002 | 0.592646841 | Gm28661 | predicted gene 28661 [Source:MGI Symbo | 544.5 | 519.1 | 532.3 | 562.96 | 575.85 | 664.96 |
| 40314 | ENSMUSG00000102071 | 0 | NA | NA | NA | NA | NA | Gm28815 | predicted gene 28815 [Source:MGI Symbo | 0 | 0 | 0 | 0 | 0 | 0 |
| 40315 | ENSMUSG00000102072 | 0 | NA | NA | NA | NA | NA | Gm29551 | predicted gene 29551 [Source:MGI Symbo | 0 | 0 | 0 | 0 | 0 | 0 |
| 40316 | ENSMUSG00000102073 | 0 | NA | NA | NA | NA | NA | Gm6818 | predicted gene 6818 [Source:MGI Symbol | 0 | 0 | 0 | 0 | 0 | 0 |

| | | | | | | | | | | | | | | | |
|---|---|---|---|---|---|---|---|---|---|---|---|---|---|---|---|
| 40317 | ENSMUSG00000102074 | 0 | NA | NA | NA | NA | NA | Gm29020 | predicted gene 29020 [Source:MGI Symbo | 0 | 0 | 0 | 0 | 0 | 0 |
| 40318 | ENSMUSG00000102075 | 0 | NA | NA | NA | NA | NA | Gm5368 | predicted gene 5368 [Source:MGI Symbol | 0 | 0 | 0 | 0 | 0 | 0 |
| 40319 | ENSMUSG00000102076 | 0 | NA | NA | NA | NA | NA | Gm20561 | predicted gene, 20561 [Source:MGI Symb | 0 | 0 | 0 | 0 | 0 | 0 |
| 40320 | ENSMUSG00000102077 | 0 | NA | NA | NA | NA | NA | Gm29001 | predicted gene 29001 [Source:MGI Symbo | 0 | 0 | 0 | 0 | 0 | 0 |
| 40321 | ENSMUSG00000102079 | 0 | NA | NA | NA | NA | NA | Gm21879 | predicted gene, 21879 [Source:MGI Symb | 0 | 0 | 0 | 0 | 0 | 0 |
| 40322 | ENSMUSG00000102080 | 0 | NA | NA | NA | NA | NA | Gm5092 | predicted gene 5092 [Source:MGI Symbol | 0 | 0 | 0 | 0 | 0 | 0 |
| 40323 | ENSMUSG00000102081 | 0 | NA | NA | NA | NA | NA | Gm28886 | predicted gene 28886 [Source:MGI Symbo | 0 | 0 | 0 | 0 | 0 | 0 |
| 40324 | ENSMUSG00000102082 | 0 | NA | NA | NA | NA | NA | Gm28675 | predicted gene 28675 [Source:MGI Symbo | 0 | 0 | 0 | 0 | 0 | 0 |
| 40325 | ENSMUSG00000102083 | 0 | NA | NA | NA | NA | NA | Gm5259 | predicted gene 5259 [Source:MGI Symbol | 0 | 0 | 0 | 0 | 0 | 0 |
| 40326 | ENSMUSG00000102084 | 0.181769715 | 1.126924175 | 4.080472857 | 0.276174898 | 0.782413736 | NA | Gm29638 | predicted gene 29638 [Source:MGI Symbo | 0 | 0 | 0 | 0 | 1.0906 | 0 |
| 40327 | ENSMUSG00000102085 | 0 | NA | NA | NA | NA | NA | 4930435C17 | RIKEN cDNA 4930435C17 gene [Source:M | 0 | 0 | 0 | 0 | 0 | 0 |
| 40328 | ENSMUSG00000102087 | 0 | NA | NA | NA | NA | NA | Gm28790 | predicted gene 28790 [Source:MGI Symbo | 0 | 0 | 0 | 0 | 0 | 0 |
| 40329 | ENSMUSG00000102088 | 1.038213457 | 2.372140906 | 2.221591093 | 1.067766662 | 0.28562578 | NA | Gm21064 | predicted gene, 21064 [Source:MGI Symb | 0 | 0 | 1.0062 | 1.928 | 2.1812 | 1.1138 |
| 40330 | ENSMUSG00000102090 | 0 | NA | NA | NA | NA | NA | Gm8964 | predicted gene 8964 [Source:MGI Symbol | 0 | 0 | 0 | 0 | 0 | 0 |
| 40331 | ENSMUSG00000102091 | 54.43506658 | -0.190213705 | 0.301422344 | -0.631053766 | 0.52800537 | 0.888334265 | Olfr1034 | olfactory receptor 1034 [Source:MGI Sym | 57.02 | 64.759 | 52.325 | 50.127 | 56.712 | 45.668 |
| 40332 | ENSMUSG00000102092 | 0 | NA | NA | NA | NA | NA | Gm21892 | predicted gene, 21892 [Source:MGI Symb | 0 | 0 | 0 | 0 | 0 | 0 |
| 40333 | ENSMUSG00000102093 | 0 | NA | NA | NA | NA | NA | Gm17813 | predicted gene, 17813 [Source:MGI Symb | 0 | 0 | 0 | 0 | 0 | 0 |
| 40334 | ENSMUSG00000102095 | 1254.265419 | -0.339240161 | 0.390600518 | -0.868509246 | 0.385115613 | 0.816437825 | C730036E19 | RIKEN cDNA C730036E19 gene [Source:M | 1343.6 | 1437 | 1422.8 | 1435.4 | 1310.9 | 575.86 |
| 40335 | ENSMUSG00000102096 | 0 | NA | NA | NA | NA | NA | 1700101O22 | RIKEN cDNA 1700101O22 gene [Source:M | 0 | 0 | 0 | 0 | 0 | 0 |
| 40336 | ENSMUSG00000102097 | 0 | NA | NA | NA | NA | NA | A930006L05 | RIKEN cDNA A930006L05 gene [Source:M | 0 | 0 | 0 | 0 | 0 | 0 |
| 40337 | ENSMUSG00000102098 | 0 | NA | NA | NA | NA | NA | 2310016D03 | RIKEN cDNA 2310016D03 gene [Source:M | 0 | 0 | 0 | 0 | 0 | 0 |
| 40338 | ENSMUSG00000102099 | 0 | NA | NA | NA | NA | NA | 1700011B04 | RIKEN cDNA 1700011B04 gene [Source:M | 0 | 0 | 0 | 0 | 0 | 0 |
| 40339 | ENSMUSG00000102100 | 0 | NA | NA | NA | NA | NA | Gm18351 | predicted gene, 18351 [Source:MGI Symb | 0 | 0 | 0 | 0 | 0 | 0 |
| 40340 | ENSMUSG00000102101 | 95.60429873 | -0.11158164 | 0.236405054 | -0.471993461 | 0.636931456 | 0.923478764 | Zbtb11os1 | zinc finger and BTB domain containing 11, | 101.19 | 107.93 | 88.55 | 81.938 | 94.884 | 99.132 |
| 40341 | ENSMUSG00000102102 | 0 | NA | NA | NA | NA | NA | Gm29159 | predicted gene 29159 [Source:MGI Symbo | 0 | 0 | 0 | 0 | 0 | 0 |
| 40342 | ENSMUSG00000102103 | 0 | NA | NA | NA | NA | NA | Gm21851 | predicted gene, 21851 [Source:MGI Symb | 0 | 0 | 0 | 0 | 0 | 0 |
| 40343 | ENSMUSG00000102104 | 0 | NA | NA | NA | NA | NA | Gm28858 | predicted gene 28858 [Source:MGI Symbo | 0 | 0 | 0 | 0 | 0 | 0 |
| 40344 | ENSMUSG00000102105 | 0 | NA | NA | NA | NA | NA | Gm17946 | predicted gene, 17946 [Source:MGI Symb | 0 | 0 | 0 | 0 | 0 | 0 |
| 40345 | ENSMUSG00000102106 | 0.36353943 | 2.065989254 | 4.034137301 | 0.512126658 | 0.608562368 | NA | 2310043O21 | RIKEN cDNA 2310043O21 gene [Source:M | 0 | 0 | 0 | 0 | 2.1812 | 0 |
| 40346 | ENSMUSG00000102107 | 0 | NA | NA | NA | NA | NA | Gm29062 | predicted gene 29062 [Source:MGI Symbo | 0 | 0 | 0 | 0 | 0 | 0 |
| 40347 | ENSMUSG00000102108 | 0 | NA | NA | NA | NA | NA | Gm29037 | predicted gene 29037 [Source:MGI Symbo | 0 | 0 | 0 | 0 | 0 | 0 |
| 40348 | ENSMUSG00000102109 | 0 | NA | NA | NA | NA | NA | Gm29563 | predicted gene 29563 [Source:MGI Symbo | 0 | 0 | 0 | 0 | 0 | 0 |
| 40349 | ENSMUSG00000102110 | 0 | NA | NA | NA | NA | NA | Gm28440 | predicted gene 28440 [Source:MGI Symbo | 0 | 0 | 0 | 0 | 0 | 0 |
| 40350 | ENSMUSG00000102111 | 0 | NA | NA | NA | NA | NA | Gm29566 | predicted gene 29566 [Source:MGI Symbo | 0 | 0 | 0 | 0 | 0 | 0 |
| 40351 | ENSMUSG00000102112 | 4.716226255 | -0.212078523 | 1.123617471 | -0.188746196 | 0.850291743 | NA | 1810041H14 | RIKEN cDNA 1810041H14 gene [Source:M | 4.0155 | 2.0558 | 9.0562 | 0.964 | 2.1812 | 10.025 |
| 40352 | ENSMUSG00000102113 | 0 | NA | NA | NA | NA | NA | Gm28061 | predicted gene 28061 [Source:MGI Symbo | 0 | 0 | 0 | 0 | 0 | 0 |
| 40353 | ENSMUSG00000102114 | 0.185640647 | 1.126924175 | 4.080472857 | 0.276174898 | 0.782413736 | NA | BC048559 | cDNA sequence BC048559 [Source:MGI Sy | 0 | 0 | 0 | 0 | 0 | 1.1138 |
| 40354 | ENSMUSG00000102115 | 0 | NA | NA | NA | NA | NA | Gm28766 | predicted gene 28766 [Source:MGI Symbo | 0 | 0 | 0 | 0 | 0 | 0 |
| 40355 | ENSMUSG00000102116 | 0 | NA | NA | NA | NA | NA | Gm28887 | predicted gene 28887 [Source:MGI Symbo | 0 | 0 | 0 | 0 | 0 | 0 |
| 40356 | ENSMUSG00000102117 | 0 | NA | NA | NA | NA | NA | Rpsa-ps1 | ribosomal protein SA, pseudogene 1 [Sou | 0 | 0 | 0 | 0 | 0 | 0 |
| 40357 | ENSMUSG00000102118 | 0 | NA | NA | NA | NA | NA | Gm29637 | predicted gene 29637 [Source:MGI Symbo | 0 | 0 | 0 | 0 | 0 | 0 |
| 40358 | ENSMUSG00000102119 | 0 | NA | NA | NA | NA | NA | Gm28462 | predicted gene 28462 [Source:MGI Symbo | 0 | 0 | 0 | 0 | 0 | 0 |
| 40359 | ENSMUSG00000102120 | 0 | NA | NA | NA | NA | NA | Gm28630 | predicted gene 28630 [Source:MGI Symbo | 0 | 0 | 0 | 0 | 0 | 0 |
| 40360 | ENSMUSG00000102121 | 0 | NA | NA | NA | NA | NA | Gm28065 | predicted gene 28065 [Source:MGI Symbo | 0 | 0 | 0 | 0 | 0 | 0 |
| 40361 | ENSMUSG00000102122 | 0 | NA | NA | NA | NA | NA | Gm20883 | predicted gene, 20883 [Source:MGI Symb | 0 | 0 | 0 | 0 | 0 | 0 |
| 40362 | ENSMUSG00000102123 | 0 | NA | NA | NA | NA | NA | Gm4319 | predicted gene 4319 [Source:MGI Symbol | 0 | 0 | 0 | 0 | 0 | 0 |
| 40363 | ENSMUSG00000102124 | 0 | NA | NA | NA | NA | NA | Gm29296 | predicted gene 29296 [Source:MGI Symbo | 0 | 0 | 0 | 0 | 0 | 0 |
| 40364 | ENSMUSG00000102125 | 0 | NA | NA | NA | NA | NA | Gm28520 | predicted gene 28520 [Source:MGI Symbo | 0 | 0 | 0 | 0 | 0 | 0 |
| 40365 | ENSMUSG00000102126 | 0 | NA | NA | NA | NA | NA | Gm29143 | predicted gene 29143 [Source:MGI Symbo | 0 | 0 | 0 | 0 | 0 | 0 |
| 40366 | ENSMUSG00000102127 | 0 | NA | NA | NA | NA | NA | Gm28972 | predicted gene 28972 [Source:MGI Symbo | 0 | 0 | 0 | 0 | 0 | 0 |
| 40367 | ENSMUSG00000102128 | 0 | NA | NA | NA | NA | NA | Gm18349 | predicted gene, 18349 [Source:MGI Symb | 0 | 0 | 0 | 0 | 0 | 0 |
| 40368 | ENSMUSG00000102129 | 0 | NA | NA | NA | NA | NA | Gm28969 | predicted gene 28969 [Source:MGI Symbo | 0 | 0 | 0 | 0 | 0 | 0 |
| 40369 | ENSMUSG00000102131 | 0 | NA | NA | NA | NA | NA | Gm21160 | predicted gene, 21160 [Source:MGI Symb | 0 | 0 | 0 | 0 | 0 | 0 |
| 40370 | ENSMUSG00000102132 | 0 | NA | NA | NA | NA | NA | Gm21813 | predicted gene, 21813 [Source:MGI Symb | 0 | 0 | 0 | 0 | 0 | 0 |
| 40371 | ENSMUSG00000102133 | 0 | NA | NA | NA | NA | NA | Gm37106 | predicted gene, 37106 [Source:MGI Symb | 0 | 0 | 0 | 0 | 0 | 0 |
| 40372 | ENSMUSG00000102134 | 0 | NA | NA | NA | NA | NA | Gm9931 | predicted gene 9931 [Source:MGI Symbol | 0 | 0 | 0 | 0 | 0 | 0 |

| | | | | | | | | | | | | | | | |
|---|---|---|---|---|---|---|---|---|---|---|---|---|---|---|---|
| 40373 | ENSMUSG00000102135 | 0 | NA | NA | NA | NA | NA | Gm37108 | predicted gene, 37108 [Source:MGI Symb | 0 | 0 | 0 | 0 | 0 | 0 |
| 40374 | ENSMUSG00000102136 | 0 | NA | NA | NA | NA | NA | Gm37107 | predicted gene, 37107 [Source:MGI Symb | 0 | 0 | 0 | 0 | 0 | 0 |
| 40375 | ENSMUSG00000102137 | 0 | NA | NA | NA | NA | NA | Rpl28-ps2 | ribosomal protein L28, pseudogene 2 [Sou | 0 | 0 | 0 | 0 | 0 | 0 |
| 40376 | ENSMUSG00000102138 | 0 | NA | NA | NA | NA | NA | Gm31942 | predicted gene, 31942 [Source:MGI Symb | 0 | 0 | 0 | 0 | 0 | 0 |
| 40377 | ENSMUSG00000102139 | 0.181769715 | 1.126924175 | 4.080472857 | 0.276174898 | 0.782413736 | NA | Gm37109 | predicted gene, 37109 [Source:MGI Symb | 0 | 0 | 0 | 0 | 1.0906 | 0 |
| 40378 | ENSMUSG00000102140 | 0.524202404 | 2.600864256 | 3.301637836 | 0.787749713 | 0.430843121 | NA | Gm38058 | predicted gene, 38058 [Source:MGI Symb | 0 | 0 | 0 | 0.964 | 2.1812 | 0 |
| 40379 | ENSMUSG00000102141 | 0 | NA | NA | NA | NA | NA | Gm3952 | predicted gene 3952 [Source:MGI Symbol | 0 | 0 | 0 | 0 | 0 | 0 |
| 40380 | ENSMUSG00000102142 | 0 | NA | NA | NA | NA | NA | Gm26930 | predicted gene, 26930 [Source:MGI Symb | 0 | 0 | 0 | 0 | 0 | 0 |
| 40381 | ENSMUSG00000102143 | 6.584209706 | -0.000377713 | 0.916500255 | -0.000412125 | 0.999671172 | 0.999897338 | Gm38057 | predicted gene, 38057 [Source:MGI Symb | 1.6062 | 11.307 | 7.0437 | 9.6398 | 5.4531 | 4.4554 |
| 40382 | ENSMUSG00000102144 | 0.181769715 | 1.126924175 | 4.080472857 | 0.276174898 | 0.782413736 | NA | 4930429P21 | RIKEN cDNA 4930429P21 gene [Source:M | 0 | 0 | 0 | 0 | 1.0906 | 0 |
| 40383 | ENSMUSG00000102145 | 0 | NA | NA | NA | NA | NA | Gm38056 | predicted gene, 38056 [Source:MGI Symb | 0 | 0 | 0 | 0 | 0 | 0 |
| 40384 | ENSMUSG00000102146 | 0 | NA | NA | NA | NA | NA | Gm38054 | predicted gene, 38054 [Source:MGI Symb | 0 | 0 | 0 | 0 | 0 | 0 |
| 40385 | ENSMUSG00000102147 | 0 | NA | NA | NA | NA | NA | Gm38055 | predicted gene, 38055 [Source:MGI Symb | 0 | 0 | 0 | 0 | 0 | 0 |
| 40386 | ENSMUSG00000102148 | 0 | NA | NA | NA | NA | NA | Gm38059 | predicted gene, 38059 [Source:MGI Symb | 0 | 0 | 0 | 0 | 0 | 0 |
| 40387 | ENSMUSG00000102149 | 0 | NA | NA | NA | NA | NA | Gm9723 | predicted gene 9723 [Source:MGI Symbol | 0 | 0 | 0 | 0 | 0 | 0 |
| 40388 | ENSMUSG00000102150 | 0 | NA | NA | NA | NA | NA | Gm37473 | predicted gene, 37473 [Source:MGI Symb | 0 | 0 | 0 | 0 | 0 | 0 |
| 40389 | ENSMUSG00000102151 | 0.695728634 | -1.425610641 | 3.05889997 | -0.46605337 | 0.641177299 | NA | Gm37472 | predicted gene, 37472 [Source:MGI Symb | 0 | 3.0838 | 0 | 0 | 1.0906 | 0 |
| 40390 | ENSMUSG00000102152 | 0.660353019 | -0.01500701 | 2.83315355 | -0.005296928 | 0.995773683 | NA | Gm37475 | predicted gene, 37475 [Source:MGI Symb | 0 | 1.0279 | 1.0062 | 1.928 | 0 | 0 |
| 40391 | ENSMUSG00000102153 | 0.542600935 | 1.125931187 | 3.335923981 | 0.33751704 | 0.735727169 | NA | Gm37474 | predicted gene, 37474 [Source:MGI Symb | 0 | 1.0279 | 0 | 0 | 0 | 2.2277 |
| 40392 | ENSMUSG00000102154 | 0.160662974 | 1.126924175 | 4.080472857 | 0.276174898 | 0.782413736 | NA | Gm37469 | predicted gene, 37469 [Source:MGI Symb | 0 | 0 | 0 | 0.964 | 0 | 0 |
| 40393 | ENSMUSG00000102155 | 0 | NA | NA | NA | NA | NA | Gm37468 | predicted gene, 37468 [Source:MGI Symb | 0 | 0 | 0 | 0 | 0 | 0 |
| 40394 | ENSMUSG00000102157 | 0.321325949 | 1.911575114 | 4.048267893 | 0.472195804 | 0.636787034 | NA | Gm37470 | predicted gene, 37470 [Source:MGI Symb | 0 | 0 | 0 | 1.928 | 0 | 0 |
| 40395 | ENSMUSG00000102158 | 0 | NA | NA | NA | NA | NA | Gm34342 | predicted gene, 34342 [Source:MGI Symb | 0 | 0 | 0 | 0 | 0 | 0 |
| 40396 | ENSMUSG00000102159 | 0 | NA | NA | NA | NA | NA | Gm37476 | predicted gene, 37476 [Source:MGI Symb | 0 | 0 | 0 | 0 | 0 | 0 |
| 40397 | ENSMUSG00000102160 | 0 | NA | NA | NA | NA | NA | Gm36944 | predicted gene, 36944 [Source:MGI Symb | 0 | 0 | 0 | 0 | 0 | 0 |
| 40398 | ENSMUSG00000102162 | 0 | NA | NA | NA | NA | NA | Gm2367 | predicted gene 2367 [Source:MGI Symbol | 0 | 0 | 0 | 0 | 0 | 0 |
| 40399 | ENSMUSG00000102163 | 0.353089354 | 0.165142706 | 3.896102244 | 0.042386646 | 0.966190474 | NA | Gm36945 | predicted gene, 36945 [Source:MGI Symb | 0 | 1.0279 | 0 | 0 | 1.0906 | 0 |
| 40400 | ENSMUSG00000102164 | 0 | NA | NA | NA | NA | NA | Gm36946 | predicted gene, 36946 [Source:MGI Symb | 0 | 0 | 0 | 0 | 0 | 0 |
| 40401 | ENSMUSG00000102166 | 0 | NA | NA | NA | NA | NA | Gm36947 | predicted gene, 36947 [Source:MGI Symb | 0 | 0 | 0 | 0 | 0 | 0 |
| 40402 | ENSMUSG00000102167 | 0 | NA | NA | NA | NA | NA | Gm2345 | predicted gene 2345 [Source:MGI Symbol | 0 | 0 | 0 | 0 | 0 | 0 |
| 40403 | ENSMUSG00000102168 | 0 | NA | NA | NA | NA | NA | Gm36943 | predicted gene, 36943 [Source:MGI Symb | 0 | 0 | 0 | 0 | 0 | 0 |
| 40404 | ENSMUSG00000102169 | 0 | NA | NA | NA | NA | NA | Gm9113 | predicted pseudogene 9113 [Source:MGI | 0 | 0 | 0 | 0 | 0 | 0 |
| 40405 | ENSMUSG00000102170 | 0 | NA | NA | NA | NA | NA | Gm20781 | predicted gene, 20781 [Source:MGI Symb | 0 | 0 | 0 | 0 | 0 | 0 |
| 40406 | ENSMUSG00000102172 | 0 | NA | NA | NA | NA | NA | Gm37873 | predicted gene, 37873 [Source:MGI Symb | 0 | 0 | 0 | 0 | 0 | 0 |
| 40407 | ENSMUSG00000102173 | 0 | NA | NA | NA | NA | NA | Gm37872 | predicted gene, 37872 [Source:MGI Symb | 0 | 0 | 0 | 0 | 0 | 0 |
| 40408 | ENSMUSG00000102174 | 0 | NA | NA | NA | NA | NA | Gm36398 | predicted gene, 36398 [Source:MGI Symb | 0 | 0 | 0 | 0 | 0 | 0 |
| 40409 | ENSMUSG00000102175 | 0 | NA | NA | NA | NA | NA | Gm6119 | predicted gene 6119 [Source:MGI Symbol | 0 | 0 | 0 | 0 | 0 | 0 |
| 40410 | ENSMUSG00000102176 | 0 | NA | NA | NA | NA | NA | Gm37871 | predicted gene, 37871 [Source:MGI Symb | 0 | 0 | 0 | 0 | 0 | 0 |
| 40411 | ENSMUSG00000102177 | 0 | NA | NA | NA | NA | NA | Gm37870 | predicted gene, 37870 [Source:MGI Symb | 0 | 0 | 0 | 0 | 0 | 0 |
| 40412 | ENSMUSG00000102178 | 0 | NA | NA | NA | NA | NA | Gm37869 | predicted gene, 37869 [Source:MGI Symb | 0 | 0 | 0 | 0 | 0 | 0 |
| 40413 | ENSMUSG00000102179 | 1.473931963 | -3.934826526 | 2.281835847 | -1.724412618 | 0.084633381 | NA | Gm37868 | predicted gene, 37868 [Source:MGI Symb | 4.8186 | 0 | 4.025 | 0 | 0 | 0 |
| 40414 | ENSMUSG00000102180 | 0 | NA | NA | NA | NA | NA | Gm37189 | predicted gene, 37189 [Source:MGI Symb | 0 | 0 | 0 | 0 | 0 | 0 |
| 40415 | ENSMUSG00000102181 | 0 | NA | NA | NA | NA | NA | Gm37190 | predicted gene, 37190 [Source:MGI Symb | 0 | 0 | 0 | 0 | 0 | 0 |
| 40416 | ENSMUSG00000102182 | 0 | NA | NA | NA | NA | NA | Gm37187 | predicted gene, 37187 [Source:MGI Symb | 0 | 0 | 0 | 0 | 0 | 0 |
| 40417 | ENSMUSG00000102184 | 0 | NA | NA | NA | NA | NA | Gm37192 | predicted gene, 37192 [Source:MGI Symb | 0 | 0 | 0 | 0 | 0 | 0 |
| 40418 | ENSMUSG00000102185 | 0 | NA | NA | NA | NA | NA | 5033404E19 | RIKEN cDNA 5033404E19 gene [Source:M | 0 | 0 | 0 | 0 | 0 | 0 |
| 40419 | ENSMUSG00000102186 | 0 | NA | NA | NA | NA | NA | Gm37191 | predicted gene, 37191 [Source:MGI Symb | 0 | 0 | 0 | 0 | 0 | 0 |
| 40420 | ENSMUSG00000102187 | 0 | NA | NA | NA | NA | NA | Gm7897 | predicted gene 7897 [Source:MGI Symbol | 0 | 0 | 0 | 0 | 0 | 0 |
| 40421 | ENSMUSG00000102188 | 0 | NA | NA | NA | NA | NA | Gm37193 | predicted gene, 37193 [Source:MGI Symb | 0 | 0 | 0 | 0 | 0 | 0 |
| 40422 | ENSMUSG00000102189 | 0 | NA | NA | NA | NA | NA | Gm37194 | predicted gene, 37194 [Source:MGI Symb | 0 | 0 | 0 | 0 | 0 | 0 |
| 40423 | ENSMUSG00000102190 | 0.160662974 | 1.126924175 | 4.080472857 | 0.276174898 | 0.782413736 | NA | Gm38166 | predicted gene, 38166 [Source:MGI Symb | 0 | 0 | 0 | 0.964 | 0 | 0 |
| 40424 | ENSMUSG00000102191 | 0 | NA | NA | NA | NA | NA | Gm36569 | predicted gene, 36569 [Source:MGI Symb | 0 | 0 | 0 | 0 | 0 | 0 |
| 40425 | ENSMUSG00000102192 | 0 | NA | NA | NA | NA | NA | Gm38167 | predicted gene, 38167 [Source:MGI Symb | 0 | 0 | 0 | 0 | 0 | 0 |
| 40426 | ENSMUSG00000102193 | 0.531246861 | 1.101376188 | 3.36099124 | 0.327693858 | 0.743143147 | NA | Gm7299 | predicted gene 7299 [Source:MGI Symbol | 0 | 0 | 1.0062 | 0 | 2.1812 | 0 |
| 40427 | ENSMUSG00000102194 | 0 | NA | NA | NA | NA | NA | Gm38169 | predicted gene, 38169 [Source:MGI Symb | 0 | 0 | 0 | 0 | 0 | 0 |
| 40428 | ENSMUSG00000102195 | 0 | NA | NA | NA | NA | NA | Gm38168 | predicted gene, 38168 [Source:MGI Symb | 0 | 0 | 0 | 0 | 0 | 0 |

| | | | | | | | | | | | | | | | |
|---|---|---|---|---|---|---|---|---|---|---|---|---|---|---|---|
| 40429 | ENSMUSG00000102196 | 0 | NA | NA | NA | NA | NA | Gm38171 | predicted gene, 38171 [Source:MGI Symb | 0 | 0 | 0 | 0 | 0 | 0 |
| 40430 | ENSMUSG00000102197 | 0 | NA | NA | NA | NA | NA | Gm38170 | predicted gene, 38170 [Source:MGI Symb | 0 | 0 | 0 | 0 | 0 | 0 |
| 40431 | ENSMUSG00000102198 | 0 | NA | NA | NA | NA | NA | Gm38173 | predicted gene, 38173 [Source:MGI Symb | 0 | 0 | 0 | 0 | 0 | 0 |
| 40432 | ENSMUSG00000102199 | 0 | NA | NA | NA | NA | NA | Gm38172 | predicted gene, 38172 [Source:MGI Symb | 0 | 0 | 0 | 0 | 0 | 0 |
| 40433 | ENSMUSG00000102200 | 0 | NA | NA | NA | NA | NA | Gm36957 | predicted gene, 36957 [Source:MGI Symb | 0 | 0 | 0 | 0 | 0 | 0 |
| 40434 | ENSMUSG00000102201 | 0 | NA | NA | NA | NA | NA | Gm36956 | predicted gene, 36956 [Source:MGI Symb | 0 | 0 | 0 | 0 | 0 | 0 |
| 40435 | ENSMUSG00000102202 | 6.551226517 | -1.643004432 | 0.965439344 | -1.701820464 | 0.088789029 | 0.43935837 | Gm33280 | predicted gene, 33280 [Source:MGI Symb | 10.44 | 8.2233 | 11.069 | 2.8919 | 0 | 6.6831 |
| 40436 | ENSMUSG00000102203 | 0 | NA | NA | NA | NA | NA | 4930412E21 | RIKEN cDNA 4930412E21 gene [Source:M | 0 | 0 | 0 | 0 | 0 | 0 |
| 40437 | ENSMUSG00000102204 | 0 | NA | NA | NA | NA | NA | Gm36958 | predicted gene, 36958 [Source:MGI Symb | 0 | 0 | 0 | 0 | 0 | 0 |
| 40438 | ENSMUSG00000102205 | 1.444663776 | 4.056420342 | 2.111175933 | 1.921403271 | 0.054680887 | NA | 9430092D12 | RIKEN cDNA 9430092D12 gene [Source:M | 0 | 0 | 0 | 0.964 | 4.3625 | 3.3415 |
| 40439 | ENSMUSG00000102206 | 0 | NA | NA | NA | NA | NA | Pcdha11 | protocadherin alpha 11 [Source:MGI Syml | 0 | 0 | 0 | 0 | 0 | 0 |
| 40440 | ENSMUSG00000102207 | 0 | NA | NA | NA | NA | NA | Gm10344 | predicted gene 10344 [Source:MGI Symbo | 0 | 0 | 0 | 0 | 0 | 0 |
| 40441 | ENSMUSG00000102208 | 0 | NA | NA | NA | NA | NA | Gm32033 | predicted gene, 32033 [Source:MGI Symb | 0 | 0 | 0 | 0 | 0 | 0 |
| 40442 | ENSMUSG00000102211 | 0.706437433 | 0.130348649 | 2.564119607 | 0.050835635 | 0.959456495 | NA | Gm37490 | predicted gene, 37490 [Source:MGI Symb | 0 | 1.0279 | 1.0062 | 0 | 1.0906 | 1.1138 |
| 40443 | ENSMUSG00000102212 | 0 | NA | NA | NA | NA | NA | C230085N15 | RIKEN cDNA C230085N15 gene [Source:M | 0 | 0 | 0 | 0 | 0 | 0 |
| 40444 | ENSMUSG00000102213 | 0 | NA | NA | NA | NA | NA | Gm37489 | predicted gene, 37489 [Source:MGI Symb | 0 | 0 | 0 | 0 | 0 | 0 |
| 40445 | ENSMUSG00000102214 | 0 | NA | NA | NA | NA | NA | Gm37492 | predicted gene, 37492 [Source:MGI Symb | 0 | 0 | 0 | 0 | 0 | 0 |
| 40446 | ENSMUSG00000102215 | 0 | NA | NA | NA | NA | NA | Gm2464 | predicted gene 2464 [Source:MGI Symbol | 0 | 0 | 0 | 0 | 0 | 0 |
| 40447 | ENSMUSG00000102216 | 0 | NA | NA | NA | NA | NA | Gm19231 | predicted gene, 19231 [Source:MGI Symb | 0 | 0 | 0 | 0 | 0 | 0 |
| 40448 | ENSMUSG00000102217 | 0 | NA | NA | NA | NA | NA | Gm37491 | predicted gene, 37491 [Source:MGI Symb | 0 | 0 | 0 | 0 | 0 | 0 |
| 40449 | ENSMUSG00000102218 | 0 | NA | NA | NA | NA | NA | Gm37493 | predicted gene, 37493 [Source:MGI Symb | 0 | 0 | 0 | 0 | 0 | 0 |
| 40450 | ENSMUSG00000102219 | 0 | NA | NA | NA | NA | NA | Gm6606 | predicted gene 6606 [Source:MGI Symbol | 0 | 0 | 0 | 0 | 0 | 0 |
| 40451 | ENSMUSG00000102220 | 0 | NA | NA | NA | NA | NA | Rpl23a-ps5 | ribosomal protein L23A, pseudogene 5 [Sc | 0 | 0 | 0 | 0 | 0 | 0 |
| 40452 | ENSMUSG00000102221 | 0 | NA | NA | NA | NA | NA | Gm37301 | predicted gene, 37301 [Source:MGI Symb | 0 | 0 | 0 | 0 | 0 | 0 |
| 40453 | ENSMUSG00000102222 | 0 | NA | NA | NA | NA | NA | Pcdhga10 | protocadherin gamma subfamily A, 10 [So | 0 | 0 | 0 | 0 | 0 | 0 |
| 40454 | ENSMUSG00000102223 | 0 | NA | NA | NA | NA | NA | Gm37300 | predicted gene, 37300 [Source:MGI Symb | 0 | 0 | 0 | 0 | 0 | 0 |
| 40455 | ENSMUSG00000102224 | 0 | NA | NA | NA | NA | NA | 4930447F24 | RIKEN cDNA 4930447F24 gene [Source:M | 0 | 0 | 0 | 0 | 0 | 0 |
| 40456 | ENSMUSG00000102225 | 0.319491021 | 0.16514334 | 4.044653141 | 0.040830038 | 0.967431393 | NA | Gm37983 | predicted gene, 37983 [Source:MGI Symb | 0.8031 | 0 | 0 | 0 | 0 | 1.1138 |
| 40457 | ENSMUSG00000102227 | 0 | NA | NA | NA | NA | NA | Gm37302 | predicted gene, 37302 [Source:MGI Symb | 0 | 0 | 0 | 0 | 0 | 0 |
| 40458 | ENSMUSG00000102229 | 0 | NA | NA | NA | NA | NA | Gm37299 | predicted gene, 37299 [Source:MGI Symb | 0 | 0 | 0 | 0 | 0 | 0 |
| 40459 | ENSMUSG00000102230 | 0 | NA | NA | NA | NA | NA | Gm31335 | predicted gene, 31335 [Source:MGI Symb | 0 | 0 | 0 | 0 | 0 | 0 |
| 40460 | ENSMUSG00000102231 | 0 | NA | NA | NA | NA | NA | 1700074A21 | RIKEN cDNA 1700074A21 gene [Source:M | 0 | 0 | 0 | 0 | 0 | 0 |
| 40461 | ENSMUSG00000102232 | 0 | NA | NA | NA | NA | NA | Gm19035 | predicted gene, 19035 [Source:MGI Symb | 0 | 0 | 0 | 0 | 0 | 0 |
| 40462 | ENSMUSG00000102233 | 0 | NA | NA | NA | NA | NA | Gm37889 | predicted gene, 37889 [Source:MGI Symb | 0 | 0 | 0 | 0 | 0 | 0 |
| 40463 | ENSMUSG00000102234 | 0 | NA | NA | NA | NA | NA | Gm37885 | predicted gene, 37885 [Source:MGI Symb | 0 | 0 | 0 | 0 | 0 | 0 |
| 40464 | ENSMUSG00000102235 | 0 | NA | NA | NA | NA | NA | Gm37886 | predicted gene, 37886 [Source:MGI Symb | 0 | 0 | 0 | 0 | 0 | 0 |
| 40465 | ENSMUSG00000102236 | 0 | NA | NA | NA | NA | NA | Gm37887 | predicted gene, 37887 [Source:MGI Symb | 0 | 0 | 0 | 0 | 0 | 0 |
| 40466 | ENSMUSG00000102238 | 0 | NA | NA | NA | NA | NA | Gm37890 | predicted gene, 37890 [Source:MGI Symb | 0 | 0 | 0 | 0 | 0 | 0 |
| 40467 | ENSMUSG00000102239 | 0 | NA | NA | NA | NA | NA | Gm38296 | predicted gene, 38296 [Source:MGI Symb | 0 | 0 | 0 | 0 | 0 | 0 |
| 40468 | ENSMUSG00000102240 | 0 | NA | NA | NA | NA | NA | Gm37717 | predicted gene, 37717 [Source:MGI Symb | 0 | 0 | 0 | 0 | 0 | 0 |
| 40469 | ENSMUSG00000102241 | 0.335414862 | -1.779047242 | 4.030804179 | -0.441362855 | 0.658950329 | NA | Gm37716 | predicted gene, 37716 [Source:MGI Symb | 0 | 0 | 2.0125 | 0 | 0 | 0 |
| 40470 | ENSMUSG00000102242 | 0 | NA | NA | NA | NA | NA | Gm37719 | predicted gene, 37719 [Source:MGI Symb | 0 | 0 | 0 | 0 | 0 | 0 |
| 40471 | ENSMUSG00000102243 | 0.321325949 | 1.911575114 | 4.048267893 | 0.472195804 | 0.636787034 | NA | Gm37718 | predicted gene, 37718 [Source:MGI Symb | 0 | 0 | 0 | 1.928 | 0 | 0 |
| 40472 | ENSMUSG00000102244 | 0 | NA | NA | NA | NA | NA | Gm37714 | predicted gene, 37714 [Source:MGI Symb | 0 | 0 | 0 | 0 | 0 | 0 |
| 40473 | ENSMUSG00000102245 | 0 | NA | NA | NA | NA | NA | Gm37713 | predicted gene, 37713 [Source:MGI Symb | 0 | 0 | 0 | 0 | 0 | 0 |
| 40474 | ENSMUSG00000102246 | 0 | NA | NA | NA | NA | NA | 9430037O13 | RIKEN cDNA 9430037O13 gene [Source:M | 0 | 0 | 0 | 0 | 0 | 0 |
| 40475 | ENSMUSG00000102247 | 0 | NA | NA | NA | NA | NA | Gm37715 | predicted gene, 37715 [Source:MGI Symb | 0 | 0 | 0 | 0 | 0 | 0 |
| 40476 | ENSMUSG00000102248 | 0 | NA | NA | NA | NA | NA | Gm30280 | predicted gene, 30280 [Source:MGI Symb | 0 | 0 | 0 | 0 | 0 | 0 |
| 40477 | ENSMUSG00000102249 | 0 | NA | NA | NA | NA | NA | 4930423C22 | RIKEN cDNA 4930423C22 gene [Source:M | 0 | 0 | 0 | 0 | 0 | 0 |
| 40478 | ENSMUSG00000102250 | 0 | NA | NA | NA | NA | NA | Gm38260 | predicted gene, 38260 [Source:MGI Symb | 0 | 0 | 0 | 0 | 0 | 0 |
| 40479 | ENSMUSG00000102251 | 0 | NA | NA | NA | NA | NA | Gm37654 | predicted gene, 37654 [Source:MGI Symb | 0 | 0 | 0 | 0 | 0 | 0 |
| 40480 | ENSMUSG00000102252 | 73.22060786 | 0.029717082 | 0.283324981 | 0.104886911 | 0.916465545 | 0.986374671 | Snrpn | small nuclear ribonucleoprotein N [Source | 73.885 | 60.647 | 82.512 | 56.875 | 78.525 | 86.88 |
| 40481 | ENSMUSG00000102253 | 0 | NA | NA | NA | NA | NA | Gm38259 | predicted gene, 38259 [Source:MGI Symb | 0 | 0 | 0 | 0 | 0 | 0 |
| 40482 | ENSMUSG00000102254 | 0.48903338 | 0.95637552 | 3.456690196 | 0.276673773 | 0.782030612 | NA | Gm7496 | predicted gene 7496 [Source:MGI Symbol | 0 | 0 | 1.0062 | 1.928 | 0 | 0 |
| 40483 | ENSMUSG00000102255 | 0 | NA | NA | NA | NA | NA | Gm36891 | predicted gene, 36891 [Source:MGI Symb | 0 | 0 | 0 | 0 | 0 | 0 |
| 40484 | ENSMUSG00000102256 | 0 | NA | NA | NA | NA | NA | Gm38258 | predicted gene, 38258 [Source:MGI Symb | 0 | 0 | 0 | 0 | 0 | 0 |

| | | | | | | | | | | | | | | | |
|---|---|---|---|---|---|---|---|---|---|---|---|---|---|---|---|
| 40485 | ENSMUSG00000102257 | 0 | NA | NA | NA | NA | NA | Gm37649 | predicted gene, 37649 [Source:MGI Symb | 0 | 0 | 0 | 0 | 0 | 0 |
| 40486 | ENSMUSG00000102258 | 1.568612011 | 0.154524965 | 1.865099064 | 0.082850809 | 0.933970169 | NA | Gm38257 | predicted gene, 38257 [Source:MGI Symb | 2.4093 | 2.0558 | 0 | 3.8559 | 1.0906 | 0 |
| 40487 | ENSMUSG00000102259 | 1.059061473 | 2.395592859 | 2.430477497 | 0.985647002 | 0.324306365 | NA | Gm38230 | predicted gene, 38230 [Source:MGI Symb | 0 | 1.0279 | 0 | 0.964 | 4.3625 | 0 |
| 40488 | ENSMUSG00000102260 | 0 | NA | NA | NA | NA | NA | D330025C20 | RIKEN cDNA D330025C20 gene [Source:M | 0 | 0 | 0 | 0 | 0 | 0 |
| 40489 | ENSMUSG00000102261 | 0 | NA | NA | NA | NA | NA | Gm18864 | predicted gene, 18864 [Source:MGI Symb | 0 | 0 | 0 | 0 | 0 | 0 |
| 40490 | ENSMUSG00000102262 | 0 | NA | NA | NA | NA | NA | Gm38107 | predicted gene, 38107 [Source:MGI Symb | 0 | 0 | 0 | 0 | 0 | 0 |
| 40491 | ENSMUSG00000102263 | 0 | NA | NA | NA | NA | NA | Gm31415 | predicted gene, 31415 [Source:MGI Symb | 0 | 0 | 0 | 0 | 0 | 0 |
| 40492 | ENSMUSG00000102264 | 0 | NA | NA | NA | NA | NA | Gm30638 | predicted gene, 30638 [Source:MGI Symb | 0 | 0 | 0 | 0 | 0 | 0 |
| 40493 | ENSMUSG00000102265 | 0 | NA | NA | NA | NA | NA | Gm32447 | predicted gene, 32447 [Source:MGI Symb | 0 | 0 | 0 | 0 | 0 | 0 |
| 40494 | ENSMUSG00000102266 | 0 | NA | NA | NA | NA | NA | Gm38106 | predicted gene, 38106 [Source:MGI Symb | 0 | 0 | 0 | 0 | 0 | 0 |
| 40495 | ENSMUSG00000102267 | 0 | NA | NA | NA | NA | NA | 9530003O04 | RIKEN cDNA 9530003O04 gene [Source:M | 0 | 0 | 0 | 0 | 0 | 0 |
| 40496 | ENSMUSG00000102268 | 0 | NA | NA | NA | NA | NA | Gm8826 | predicted gene 8826 [Source:MGI Symbol | 0 | 0 | 0 | 0 | 0 | 0 |
| 40497 | ENSMUSG00000102269 | 1.989932929 | 0.084282994 | 1.610165056 | 0.05234432 | 0.95825434 | NA | Gm7357 | predicted gene 7357 [Source:MGI Symbol | 0.8031 | 1.0279 | 4.025 | 3.8559 | 0 | 2.2277 |
| 40498 | ENSMUSG00000102270 | 0 | NA | NA | NA | NA | NA | Gm8388 | predicted gene 8388 [Source:MGI Symbol | 0 | 0 | 0 | 0 | 0 | 0 |
| 40499 | ENSMUSG00000102271 | 0 | NA | NA | NA | NA | NA | C630050I24I | RIKEN cDNA C630050I24 gene [Source:M | 0 | 0 | 0 | 0 | 0 | 0 |
| 40500 | ENSMUSG00000102272 | 0 | NA | NA | NA | NA | NA | Gm7512 | predicted gene 7512 [Source:MGI Symbol | 0 | 0 | 0 | 0 | 0 | 0 |
| 40501 | ENSMUSG00000102273 | 0 | NA | NA | NA | NA | NA | Gm34550 | predicted gene, 34550 [Source:MGI Symb | 0 | 0 | 0 | 0 | 0 | 0 |
| 40502 | ENSMUSG00000102274 | 0 | NA | NA | NA | NA | NA | Gm4855 | predicted gene 4855 [Source:MGI Symbol | 0 | 0 | 0 | 0 | 0 | 0 |
| 40503 | ENSMUSG00000102275 | 7.03654425 | -0.059746711 | 0.850036636 | -0.070287219 | 0.943965055 | 0.990077979 | Gm37144 | predicted gene, 37144 [Source:MGI Symb | 6.4248 | 5.1396 | 10.062 | 9.6398 | 8.7249 | 2.2277 |
| 40504 | ENSMUSG00000102276 | 0 | NA | NA | NA | NA | NA | 2900069G24 | RIKEN cDNA 2900069G24 gene [Source:M | 0 | 0 | 0 | 0 | 0 | 0 |
| 40505 | ENSMUSG00000102277 | 0 | NA | NA | NA | NA | NA | A130050O07 | RIKEN cDNA A130050O07 gene [Source:M | 0 | 0 | 0 | 0 | 0 | 0 |
| 40506 | ENSMUSG00000102278 | 0 | NA | NA | NA | NA | NA | Gm37145 | predicted gene, 37145 [Source:MGI Symb | 0 | 0 | 0 | 0 | 0 | 0 |
| 40507 | ENSMUSG00000102279 | 0 | NA | NA | NA | NA | NA | Gm37146 | predicted gene, 37146 [Source:MGI Symb | 0 | 0 | 0 | 0 | 0 | 0 |
| 40508 | ENSMUSG00000102280 | 0 | NA | NA | NA | NA | NA | Gm36999 | predicted gene, 36999 [Source:MGI Symb | 0 | 0 | 0 | 0 | 0 | 0 |
| 40509 | ENSMUSG00000102281 | 0 | NA | NA | NA | NA | NA | Ighv1-30 | immunoglobulin heavy variable 1-30 [Sou | 0 | 0 | 0 | 0 | 0 | 0 |
| 40510 | ENSMUSG00000102282 | 0.810301681 | -0.283528992 | 2.609307867 | -0.108660613 | 0.913471683 | NA | A930032L01 | RIKEN cDNA A930032L01 gene [Source:M | 1.6062 | 1.0279 | 0 | 0 | 0 | 2.2277 |
| 40511 | ENSMUSG00000102283 | 0 | NA | NA | NA | NA | NA | Gm37000 | predicted gene, 37000 [Source:MGI Symb | 0 | 0 | 0 | 0 | 0 | 0 |
| 40512 | ENSMUSG00000102284 | 0 | NA | NA | NA | NA | NA | Gm37002 | predicted gene, 37002 [Source:MGI Symb | 0 | 0 | 0 | 0 | 0 | 0 |
| 40513 | ENSMUSG00000102285 | 0 | NA | NA | NA | NA | NA | Gm37001 | predicted gene, 37001 [Source:MGI Symb | 0 | 0 | 0 | 0 | 0 | 0 |
| 40514 | ENSMUSG00000102286 | 0 | NA | NA | NA | NA | NA | Gm37004 | predicted gene, 37004 [Source:MGI Symb | 0 | 0 | 0 | 0 | 0 | 0 |
| 40515 | ENSMUSG00000102287 | 0 | NA | NA | NA | NA | NA | Gm37003 | predicted gene, 37003 [Source:MGI Symb | 0 | 0 | 0 | 0 | 0 | 0 |
| 40516 | ENSMUSG00000102288 | 0 | NA | NA | NA | NA | NA | Gm36998 | predicted gene, 36998 [Source:MGI Symb | 0 | 0 | 0 | 0 | 0 | 0 |
| 40517 | ENSMUSG00000102289 | 0 | NA | NA | NA | NA | NA | Gm31258 | predicted gene, 31258 [Source:MGI Symb | 0 | 0 | 0 | 0 | 0 | 0 |
| 40518 | ENSMUSG00000102290 | 0 | NA | NA | NA | NA | NA | Gm21678 | predicted gene, 21678 [Source:MGI Symb | 0 | 0 | 0 | 0 | 0 | 0 |
| 40519 | ENSMUSG00000102291 | 0 | NA | NA | NA | NA | NA | Gm37542 | predicted gene, 37542 [Source:MGI Symb | 0 | 0 | 0 | 0 | 0 | 0 |
| 40520 | ENSMUSG00000102293 | 0.160662974 | 1.126924175 | 4.080472857 | 0.276174898 | 0.782413736 | NA | Gm37541 | predicted gene, 37541 [Source:MGI Symb | 0 | 0 | 0 | 0.964 | 0 | 0 |
| 40521 | ENSMUSG00000102294 | 0 | NA | NA | NA | NA | NA | Gm37544 | predicted gene, 37544 [Source:MGI Symb | 0 | 0 | 0 | 0 | 0 | 0 |
| 40522 | ENSMUSG00000102295 | 0 | NA | NA | NA | NA | NA | Ighv1-57 | immunoglobulin heavy variable V1-57 [So | 0 | 0 | 0 | 0 | 0 | 0 |
| 40523 | ENSMUSG00000102296 | 0 | NA | NA | NA | NA | NA | Gm37543 | predicted gene, 37543 [Source:MGI Symb | 0 | 0 | 0 | 0 | 0 | 0 |
| 40524 | ENSMUSG00000102297 | 1.275928078 | 1.046915345 | 2.012681056 | 0.520159586 | 0.60295235 | NA | 5730585A16 | RIKEN cDNA 5730585A16 gene [Source:M | 2.4093 | 0 | 0 | 1.928 | 1.0906 | 2.2277 |
| 40525 | ENSMUSG00000102298 | 0 | NA | NA | NA | NA | NA | Gm33707 | predicted gene, 33707 [Source:MGI Symb | 0 | 0 | 0 | 0 | 0 | 0 |
| 40526 | ENSMUSG00000102299 | 3.546259937 | -0.397104638 | 1.211843136 | -0.327686502 | 0.74314871 | NA | Gm37539 | predicted gene, 37539 [Source:MGI Symb | 4.0155 | 4.1117 | 4.025 | 5.7839 | 0 | 3.3415 |
| 40527 | ENSMUSG00000102300 | 0 | NA | NA | NA | NA | NA | Gm34106 | predicted gene, 34106 [Source:MGI Symb | 0 | 0 | 0 | 0 | 0 | 0 |
| 40528 | ENSMUSG00000102301 | 0 | NA | NA | NA | NA | NA | Ighv8-2 | immunoglobulin heavy variable V8-2 [Sou | 0 | 0 | 0 | 0 | 0 | 0 |
| 40529 | ENSMUSG00000102302 | 14.37046256 | -0.613700258 | 0.613571555 | -1.00020976 | 0.317209007 | 0.763794086 | Gm38190 | predicted gene, 38190 [Source:MGI Symb | 17.668 | 19.53 | 15.094 | 16.388 | 13.087 | 4.4554 |
| 40530 | ENSMUSG00000102303 | 0 | NA | NA | NA | NA | NA | Gm38191 | predicted gene, 38191 [Source:MGI Symb | 0 | 0 | 0 | 0 | 0 | 0 |
| 40531 | ENSMUSG00000102304 | 0 | NA | NA | NA | NA | NA | Gm35843 | predicted gene, 35843 [Source:MGI Symb | 0 | 0 | 0 | 0 | 0 | 0 |
| 40532 | ENSMUSG00000102305 | 4.069267474 | 1.085070536 | 1.136400083 | 0.954831447 | 0.339662942 | NA | Gm38192 | predicted gene, 38192 [Source:MGI Symb | 5.6217 | 1.0279 | 1.0062 | 5.7839 | 7.6343 | 3.3415 |
| 40533 | ENSMUSG00000102307 | 0 | NA | NA | NA | NA | NA | Gm38194 | predicted gene, 38194 [Source:MGI Symb | 0 | 0 | 0 | 0 | 0 | 0 |
| 40534 | ENSMUSG00000102308 | 0 | NA | NA | NA | NA | NA | 2310046K23 | RIKEN cDNA 2310046K23 gene [Source:M | 0 | 0 | 0 | 0 | 0 | 0 |
| 40535 | ENSMUSG00000102309 | 0 | NA | NA | NA | NA | NA | Gm38189 | predicted gene, 38189 [Source:MGI Symb | 0 | 0 | 0 | 0 | 0 | 0 |
| 40536 | ENSMUSG00000102310 | 0 | NA | NA | NA | NA | NA | Gm37631 | predicted gene, 37631 [Source:MGI Symb | 0 | 0 | 0 | 0 | 0 | 0 |
| 40537 | ENSMUSG00000102311 | 0 | NA | NA | NA | NA | NA | Gm5543 | predicted gene 5543 [Source:MGI Symbol | 0 | 0 | 0 | 0 | 0 | 0 |
| 40538 | ENSMUSG00000102312 | 0 | NA | NA | NA | NA | NA | Pcdha3 | protocadherin alpha 3 [Source:MGI Symb | 0 | 0 | 0 | 0 | 0 | 0 |
| 40539 | ENSMUSG00000102313 | 0 | NA | NA | NA | NA | NA | Ighv1-62-1 | immunoglobulin heavy variable 1-62-1 [Sc | 0 | 0 | 0 | 0 | 0 | 0 |
| 40540 | ENSMUSG00000102314 | 0 | NA | NA | NA | NA | NA | Gm37630 | predicted gene, 37630 [Source:MGI Symb | 0 | 0 | 0 | 0 | 0 | 0 |

| | | | | | | | | | | | | | | | |
|---|---|---|---|---|---|---|---|---|---|---|---|---|---|---|---|
| 40541 | ENSMUSG00000102315 | 0 | NA | NA | NA | NA | NA | Ighv1-28 | immunoglobulin heavy variable 1-28 [Sou | 0 | 0 | 0 | 0 | 0 | 0 |
| 40542 | ENSMUSG00000102316 | 0 | NA | NA | NA | NA | NA | Gm37629 | predicted gene, 37629 [Source:MGI Symb | 0 | 0 | 0 | 0 | 0 | 0 |
| 40543 | ENSMUSG00000102317 | 0 | NA | NA | NA | NA | NA | Gm37628 | predicted gene, 37628 [Source:MGI Symb | 0 | 0 | 0 | 0 | 0 | 0 |
| 40544 | ENSMUSG00000102318 | 0 | NA | NA | NA | NA | NA | Gm37627 | predicted gene, 37627 [Source:MGI Symb | 0 | 0 | 0 | 0 | 0 | 0 |
| 40545 | ENSMUSG00000102319 | 0 | NA | NA | NA | NA | NA | Gm37626 | predicted gene, 37626 [Source:MGI Symb | 0 | 0 | 0 | 0 | 0 | 0 |
| 40546 | ENSMUSG00000102320 | 0 | NA | NA | NA | NA | NA | Gm37791 | predicted gene, 37791 [Source:MGI Symb | 0 | 0 | 0 | 0 | 0 | 0 |
| 40547 | ENSMUSG00000102321 | 0 | NA | NA | NA | NA | NA | Gm37792 | predicted gene, 37792 [Source:MGI Symb | 0 | 0 | 0 | 0 | 0 | 0 |
| 40548 | ENSMUSG00000102322 | 0 | NA | NA | NA | NA | NA | 4930566N20 | RIKEN cDNA 4930566N20 gene [Source:M | 0 | 0 | 0 | 0 | 0 | 0 |
| 40549 | ENSMUSG00000102323 | 0.68145976 | 0.058694068 | 2.590797836 | 0.022654824 | 0.981925612 | NA | Gm37790 | predicted gene, 37790 [Source:MGI Symb | 0 | 1.0279 | 1.0062 | 0.964 | 1.0906 | 0 |
| 40550 | ENSMUSG00000102324 | 1.659572611 | -2.975216116 | 2.091510308 | -1.422520417 | 0.154875229 | NA | Gm19721 | predicted gene, 19721 [Source:MGI Symb | 4.8186 | 0 | 4.025 | 0 | 0 | 1.1138 |
| 40551 | ENSMUSG00000102325 | 0 | NA | NA | NA | NA | NA | Gm37789 | predicted gene, 37789 [Source:MGI Symb | 0 | 0 | 0 | 0 | 0 | 0 |
| 40552 | ENSMUSG00000102326 | 0 | NA | NA | NA | NA | NA | Gm37788 | predicted gene, 37788 [Source:MGI Symb | 0 | 0 | 0 | 0 | 0 | 0 |
| 40553 | ENSMUSG00000102327 | 0 | NA | NA | NA | NA | NA | Trbv12-3 | T cell receptor beta, variable 12-3 [Source | 0 | 0 | 0 | 0 | 0 | 0 |
| 40554 | ENSMUSG00000102328 | 0 | NA | NA | NA | NA | NA | Gm37793 | predicted gene, 37793 [Source:MGI Symb | 0 | 0 | 0 | 0 | 0 | 0 |
| 40555 | ENSMUSG00000102329 | 2.886675932 | -2.123711391 | 1.5752035 | -1.348213987 | 0.177589567 | NA | Gm10851 | predicted gene 10851 [Source:MGI Symbo | 4.8186 | 8.2233 | 1.0062 | 0 | 3.2719 | 0 |
| 40556 | ENSMUSG00000102330 | 0 | NA | NA | NA | NA | NA | Gm37234 | predicted gene, 37234 [Source:MGI Symb | 0 | 0 | 0 | 0 | 0 | 0 |
| 40557 | ENSMUSG00000102331 | 0 | NA | NA | NA | NA | NA | Gm19938 | predicted gene, 19938 [Source:MGI Symb | 0 | 0 | 0 | 0 | 0 | 0 |
| 40558 | ENSMUSG00000102332 | 0 | NA | NA | NA | NA | NA | Gm19331 | predicted gene, 19331 [Source:MGI Symb | 0 | 0 | 0 | 0 | 0 | 0 |
| 40559 | ENSMUSG00000102333 | 0 | NA | NA | NA | NA | NA | Gm37235 | predicted gene, 37235 [Source:MGI Symb | 0 | 0 | 0 | 0 | 0 | 0 |
| 40560 | ENSMUSG00000102334 | 0 | NA | NA | NA | NA | NA | Gm18865 | predicted gene, 18865 [Source:MGI Symb | 0 | 0 | 0 | 0 | 0 | 0 |
| 40561 | ENSMUSG00000102335 | 0 | NA | NA | NA | NA | NA | Gm37231 | predicted gene, 37231 [Source:MGI Symb | 0 | 0 | 0 | 0 | 0 | 0 |
| 40562 | ENSMUSG00000102336 | 0.717171207 | -1.535127875 | 3.012718241 | -0.509549102 | 0.61036739 | NA | Gm37233 | predicted gene, 37233 [Source:MGI Symb | 3.2124 | 0 | 0 | 0 | 1.0906 | 0 |
| 40563 | ENSMUSG00000102337 | 0 | NA | NA | NA | NA | NA | Gm37232 | predicted gene, 37232 [Source:MGI Symb | 0 | 0 | 0 | 0 | 0 | 0 |
| 40564 | ENSMUSG00000102338 | 0 | NA | NA | NA | NA | NA | Gm37236 | predicted gene, 37236 [Source:MGI Symb | 0 | 0 | 0 | 0 | 0 | 0 |
| 40565 | ENSMUSG00000102339 | 0 | NA | NA | NA | NA | NA | Gm21443 | predicted gene, 21443 [Source:MGI Symb | 0 | 0 | 0 | 0 | 0 | 0 |
| 40566 | ENSMUSG00000102341 | 0 | NA | NA | NA | NA | NA | Gm18952 | predicted gene, 18952 [Source:MGI Symb | 0 | 0 | 0 | 0 | 0 | 0 |
| 40567 | ENSMUSG00000102342 | 0 | NA | NA | NA | NA | NA | Gm37380 | predicted gene, 37380 [Source:MGI Symb | 0 | 0 | 0 | 0 | 0 | 0 |
| 40568 | ENSMUSG00000102343 | 0 | NA | NA | NA | NA | NA | Gm37381 | predicted gene, 37381 [Source:MGI Symb | 0 | 0 | 0 | 0 | 0 | 0 |
| 40569 | ENSMUSG00000102344 | 2.232722623 | 0.90822192 | 1.666836553 | 0.544877612 | 0.585837695 | NA | 9430053O09 | RIKEN cDNA 9430053O09 gene [Source:M | 1.6062 | 0 | 3.0187 | 0 | 6.5437 | 2.2277 |
| 40570 | ENSMUSG00000102345 | 0 | NA | NA | NA | NA | NA | Gm37377 | predicted gene, 37377 [Source:MGI Symb | 0 | 0 | 0 | 0 | 0 | 0 |
| 40571 | ENSMUSG00000102346 | 0 | NA | NA | NA | NA | NA | Gm37378 | predicted gene, 37378 [Source:MGI Symb | 0 | 0 | 0 | 0 | 0 | 0 |
| 40572 | ENSMUSG00000102347 | 0 | NA | NA | NA | NA | NA | Gm21469 | predicted gene, 21469 [Source:MGI Symb | 0 | 0 | 0 | 0 | 0 | 0 |
| 40573 | ENSMUSG00000102348 | 0 | NA | NA | NA | NA | NA | Gm10568 | predicted gene 10568 [Source:MGI Symbo | 0 | 0 | 0 | 0 | 0 | 0 |
| 40574 | ENSMUSG00000102349 | 0 | NA | NA | NA | NA | NA | Gm37376 | predicted gene, 37376 [Source:MGI Symb | 0 | 0 | 0 | 0 | 0 | 0 |
| 40575 | ENSMUSG00000102350 | 0 | NA | NA | NA | NA | NA | Gm37434 | predicted gene, 37434 [Source:MGI Symb | 0 | 0 | 0 | 0 | 0 | 0 |
| 40576 | ENSMUSG00000102352 | 0.133850373 | -0.796629176 | 4.080472857 | -0.195229623 | 0.84521319 | NA | Gm38346 | predicted gene, 38346 [Source:MGI Symb | 0.8031 | 0 | 0 | 0 | 0 | 0 |
| 40577 | ENSMUSG00000102353 | 111.4549572 | 0.28072216 | 0.241271914 | 1.163509481 | 0.244622854 | 0.694577264 | Gm38345 | predicted gene, 38345 [Source:MGI Symb | 83.523 | 127.46 | 91.568 | 130.14 | 109.06 | 126.98 |
| 40578 | ENSMUSG00000102354 | 0 | NA | NA | NA | NA | NA | Gm38350 | predicted gene, 38350 [Source:MGI Symb | 0 | 0 | 0 | 0 | 0 | 0 |
| 40579 | ENSMUSG00000102355 | 0 | NA | NA | NA | NA | NA | Gm38349 | predicted gene, 38349 [Source:MGI Symb | 0 | 0 | 0 | 0 | 0 | 0 |
| 40580 | ENSMUSG00000102356 | 0.342639279 | -1.805976596 | 4.028547714 | -0.448294702 | 0.653940524 | NA | 1700047N06 | RIKEN cDNA 1700047N06 gene [Source:M | 0 | 2.0558 | 0 | 0 | 0 | 0 |
| 40581 | ENSMUSG00000102357 | 0 | NA | NA | NA | NA | NA | Gm38348 | predicted gene, 38348 [Source:MGI Symb | 0 | 0 | 0 | 0 | 0 | 0 |
| 40582 | ENSMUSG00000102358 | 0 | NA | NA | NA | NA | NA | Gm21412 | predicted gene, 21412 [Source:MGI Symb | 0 | 0 | 0 | 0 | 0 | 0 |
| 40583 | ENSMUSG00000102359 | 0 | NA | NA | NA | NA | NA | Gm38351 | predicted gene, 38351 [Source:MGI Symb | 0 | 0 | 0 | 0 | 0 | 0 |
| 40584 | ENSMUSG00000102360 | 0 | NA | NA | NA | NA | NA | Gm37043 | predicted gene, 37043 [Source:MGI Symb | 0 | 0 | 0 | 0 | 0 | 0 |
| 40585 | ENSMUSG00000102362 | 0 | NA | NA | NA | NA | NA | 4930509J09I | RIKEN cDNA 4930509J09 gene [Source:M( | 0 | 0 | 0 | 0 | 0 | 0 |
| 40586 | ENSMUSG00000102363 | 0.133850373 | -0.796629176 | 4.080472857 | -0.195229623 | 0.84521319 | NA | Gm32999 | predicted gene, 32999 [Source:MGI Symb | 0.8031 | 0 | 0 | 0 | 0 | 0 |
| 40587 | ENSMUSG00000102364 | 0 | NA | NA | NA | NA | NA | Ighv8-5 | immunoglobulin heavy variable V8-5 [Sou | 0 | 0 | 0 | 0 | 0 | 0 |
| 40588 | ENSMUSG00000102365 | 0.267700746 | -1.497481649 | 4.057001436 | -0.369110456 | 0.712045396 | NA | Gm37045 | predicted gene, 37045 [Source:MGI Symb | 1.6062 | 0 | 0 | 0 | 0 | 0 |
| 40589 | ENSMUSG00000102366 | 0 | NA | NA | NA | NA | NA | Gm37044 | predicted gene, 37044 [Source:MGI Symb | 0 | 0 | 0 | 0 | 0 | 0 |
| 40590 | ENSMUSG00000102368 | 0 | NA | NA | NA | NA | NA | 4930590L20I | RIKEN cDNA 4930590L20 gene [Source:M( | 0 | 0 | 0 | 0 | 0 | 0 |
| 40591 | ENSMUSG00000102369 | 0 | NA | NA | NA | NA | NA | 9530022L04I | RIKEN cDNA 9530022L04 gene [Source:M( | 0 | 0 | 0 | 0 | 0 | 0 |
| 40592 | ENSMUSG00000102370 | 7.029706912 | 0.823452845 | 0.855505491 | 0.962533675 | 0.335781596 | 0.777327945 | Gm37973 | predicted gene, 37973 [Source:MGI Symb | 3.2124 | 3.0838 | 9.0562 | 12.532 | 8.7249 | 5.5692 |
| 40593 | ENSMUSG00000102371 | 0 | NA | NA | NA | NA | NA | Olfr258-ps1 | olfactory receptor 258, pseudogene 1 [Sou | 0 | 0 | 0 | 0 | 0 | 0 |
| 40594 | ENSMUSG00000102372 | 0 | NA | NA | NA | NA | NA | A430050A11 | RIKEN cDNA A430050A11 gene [Source:M | 0 | 0 | 0 | 0 | 0 | 0 |
| 40595 | ENSMUSG00000102373 | 0 | NA | NA | NA | NA | NA | 9530018H14 | RIKEN cDNA 9530018H14 gene [Source:M | 0 | 0 | 0 | 0 | 0 | 0 |
| 40596 | ENSMUSG00000102374 | 8.37150316 | 0.213317064 | 0.809774601 | 0.263427704 | 0.79222094 | 0.961434599 | Gm38387 | predicted gene, 38387 [Source:MGI Symb | 4.0155 | 8.2233 | 11.069 | 3.8559 | 15.269 | 7.7969 |

| | | | | | | | | | | | | | | | |
|---|---|---|---|---|---|---|---|---|---|---|---|---|---|---|---|
| 40597 | ENSMUSG00000102375 | 20.1122754 | 1.670765073 | 0.627094854 | 2.664294027 | 0.00771501 | 0.089343394 | A930036K24 | RIKEN cDNA A930036K24 gene [Source:M | 20.078 | 5.1396 | 3.0187 | 24.099 | 33.809 | 34.529 |
| 40598 | ENSMUSG00000102376 | 2.397229371 | -0.732046825 | 1.786635002 | -0.409734962 | 0.68200038 | NA | Gm37975 | predicted gene, 37975 [Source:MGI Symb | 4.8186 | 4.1117 | 0 | 0 | 5.4531 | 0 |
| 40599 | ENSMUSG00000102377 | 0 | NA | NA | NA | NA | NA | Gm37974 | predicted gene, 37974 [Source:MGI Symb | 0 | 0 | 0 | 0 | 0 | 0 |
| 40600 | ENSMUSG00000102378 | 0 | NA | NA | NA | NA | NA | Gm37972 | predicted gene, 37972 [Source:MGI Symb | 0 | 0 | 0 | 0 | 0 | 0 |
| 40601 | ENSMUSG00000102379 | 0 | NA | NA | NA | NA | NA | Gm37971 | predicted gene, 37971 [Source:MGI Symb | 0 | 0 | 0 | 0 | 0 | 0 |
| 40602 | ENSMUSG00000102380 | 0.874428563 | -3.186291795 | 2.489702555 | -1.279788137 | 0.200619657 | NA | Gm38140 | predicted gene, 38140 [Source:MGI Symb | 3.2124 | 1.0279 | 1.0062 | 0 | 0 | 0 |
| 40603 | ENSMUSG00000102381 | 0 | NA | NA | NA | NA | NA | Ighv8-7 | immunoglobulin heavy variable V8-7 [Sou | 0 | 0 | 0 | 0 | 0 | 0 |
| 40604 | ENSMUSG00000102382 | 0 | NA | NA | NA | NA | NA | Gm38141 | predicted gene, 38141 [Source:MGI Symb | 0 | 0 | 0 | 0 | 0 | 0 |
| 40605 | ENSMUSG00000102383 | 0 | NA | NA | NA | NA | NA | Gm38142 | predicted gene, 38142 [Source:MGI Symb | 0 | 0 | 0 | 0 | 0 | 0 |
| 40606 | ENSMUSG00000102384 | 0 | NA | NA | NA | NA | NA | Gm38143 | predicted gene, 38143 [Source:MGI Symb | 0 | 0 | 0 | 0 | 0 | 0 |
| 40607 | ENSMUSG00000102385 | 0 | NA | NA | NA | NA | NA | Gm38144 | predicted gene, 38144 [Source:MGI Symb | 0 | 0 | 0 | 0 | 0 | 0 |
| 40608 | ENSMUSG00000102386 | 0 | NA | NA | NA | NA | NA | 2900022M0 | RIKEN cDNA 2900022M07 gene [Source:M | 0 | 0 | 0 | 0 | 0 | 0 |
| 40609 | ENSMUSG00000102387 | 0 | NA | NA | NA | NA | NA | Gm38145 | predicted gene, 38145 [Source:MGI Symb | 0 | 0 | 0 | 0 | 0 | 0 |
| 40610 | ENSMUSG00000102388 | 0 | NA | NA | NA | NA | NA | Gm33815 | predicted gene, 33815 [Source:MGI Symb | 0 | 0 | 0 | 0 | 0 | 0 |
| 40611 | ENSMUSG00000102389 | 0 | NA | NA | NA | NA | NA | D630023O1 | RIKEN cDNA D630023O14 gene [Source:M | 0 | 0 | 0 | 0 | 0 | 0 |
| 40612 | ENSMUSG00000102390 | 0 | NA | NA | NA | NA | NA | Gm37579 | predicted gene, 37579 [Source:MGI Symb | 0 | 0 | 0 | 0 | 0 | 0 |
| 40613 | ENSMUSG00000102391 | 0 | NA | NA | NA | NA | NA | Gm37578 | predicted gene, 37578 [Source:MGI Symb | 0 | 0 | 0 | 0 | 0 | 0 |
| 40614 | ENSMUSG00000102392 | 0 | NA | NA | NA | NA | NA | Gm37577 | predicted gene, 37577 [Source:MGI Symb | 0 | 0 | 0 | 0 | 0 | 0 |
| 40615 | ENSMUSG00000102393 | 0 | NA | NA | NA | NA | NA | Gm37576 | predicted gene, 37576 [Source:MGI Symb | 0 | 0 | 0 | 0 | 0 | 0 |
| 40616 | ENSMUSG00000102394 | 0.167707431 | -0.796629176 | 4.080472857 | -0.195229623 | 0.84521319 | NA | Gm37575 | predicted gene, 37575 [Source:MGI Symb | 0 | 0 | 1.0062 | 0 | 0 | 0 |
| 40617 | ENSMUSG00000102396 | 0 | NA | NA | NA | NA | NA | Gm37574 | predicted gene, 37574 [Source:MGI Symb | 0 | 0 | 0 | 0 | 0 | 0 |
| 40618 | ENSMUSG00000102397 | 0 | NA | NA | NA | NA | NA | Gm37573 | predicted gene, 37573 [Source:MGI Symb | 0 | 0 | 0 | 0 | 0 | 0 |
| 40619 | ENSMUSG00000102398 | 0.562214094 | -1.122836648 | 3.276281221 | -0.34271681 | 0.731811515 | NA | Gm37580 | predicted gene, 37580 [Source:MGI Symb | 2.4093 | 0 | 0 | 0.964 | 0 | 0 |
| 40620 | ENSMUSG00000102399 | 0 | NA | NA | NA | NA | NA | Gm5049 | predicted gene 5049 [Source:MGI Symbol | 0 | 0 | 0 | 0 | 0 | 0 |
| 40621 | ENSMUSG00000102400 | 0 | NA | NA | NA | NA | NA | Gm37865 | predicted gene, 37865 [Source:MGI Symb | 0 | 0 | 0 | 0 | 0 | 0 |
| 40622 | ENSMUSG00000102401 | 0.181769715 | 1.126924175 | 4.080472857 | 0.276174898 | 0.782413736 | NA | Gm37864 | predicted gene, 37864 [Source:MGI Symb | 0 | 0 | 0 | 0 | 1.0906 | 0 |
| 40623 | ENSMUSG00000102402 | 0 | NA | NA | NA | NA | NA | Gm20840 | predicted gene, 20840 [Source:MGI Symb | 0 | 0 | 0 | 0 | 0 | 0 |
| 40624 | ENSMUSG00000102403 | 0 | NA | NA | NA | NA | NA | Gm38187 | predicted gene, 38187 [Source:MGI Symb | 0 | 0 | 0 | 0 | 0 | 0 |
| 40625 | ENSMUSG00000102404 | 0 | NA | NA | NA | NA | NA | 5530400K19 | RIKEN cDNA 5530400K19 gene [Source:M | 0 | 0 | 0 | 0 | 0 | 0 |
| 40626 | ENSMUSG00000102405 | 0 | NA | NA | NA | NA | NA | Gm37863 | predicted gene, 37863 [Source:MGI Symb | 0 | 0 | 0 | 0 | 0 | 0 |
| 40627 | ENSMUSG00000102406 | 0 | NA | NA | NA | NA | NA | A730094K22 | RIKEN cDNA A730094K22 gene [Source:M | 0 | 0 | 0 | 0 | 0 | 0 |
| 40628 | ENSMUSG00000102407 | 0 | NA | NA | NA | NA | NA | Gm37862 | predicted gene, 37862 [Source:MGI Symb | 0 | 0 | 0 | 0 | 0 | 0 |
| 40629 | ENSMUSG00000102408 | 0 | NA | NA | NA | NA | NA | 4930568G15 | RIKEN cDNA 4930568G15 gene [Source:M | 0 | 0 | 0 | 0 | 0 | 0 |
| 40630 | ENSMUSG00000102409 | 0 | NA | NA | NA | NA | NA | Trbv18 | T cell receptor beta variable 18 [Source:M | 0 | 0 | 0 | 0 | 0 | 0 |
| 40631 | ENSMUSG00000102410 | 0 | NA | NA | NA | NA | NA | Gm36935 | predicted gene, 36935 [Source:MGI Symb | 0 | 0 | 0 | 0 | 0 | 0 |
| 40632 | ENSMUSG00000102411 | 26.64654733 | 0.250866281 | 0.498146596 | 0.503599308 | 0.614542979 | 0.917509899 | Gm36936 | predicted gene, 36936 [Source:MGI Symb | 22.487 | 18.503 | 32.2 | 42.415 | 13.087 | 31.188 |
| 40633 | ENSMUSG00000102413 | 0.160662974 | 1.126924175 | 4.080472857 | 0.276174898 | 0.782413736 | NA | Gm36937 | predicted gene, 36937 [Source:MGI Symb | 0 | 0 | 0 | 0.964 | 0 | 0 |
| 40634 | ENSMUSG00000102414 | 0 | NA | NA | NA | NA | NA | Gm36938 | predicted gene, 36938 [Source:MGI Symb | 0 | 0 | 0 | 0 | 0 | 0 |
| 40635 | ENSMUSG00000102415 | 1.20269523 | 0.466011887 | 2.092262081 | 0.222731125 | 0.823744776 | NA | A430110C17 | RIKEN cDNA A430110C17 gene [Source:M | 0 | 3.0838 | 0 | 1.928 | 1.0906 | 1.1138 |
| 40636 | ENSMUSG00000102416 | 0 | NA | NA | NA | NA | NA | 4933424G06 | RIKEN cDNA 4933424G06 gene [Source:M | 0 | 0 | 0 | 0 | 0 | 0 |
| 40637 | ENSMUSG00000102417 | 0 | NA | NA | NA | NA | NA | Gm36939 | predicted gene, 36939 [Source:MGI Symb | 0 | 0 | 0 | 0 | 0 | 0 |
| 40638 | ENSMUSG00000102418 | 0.658052777 | 1.6393978 | 2.811180186 | 0.583170658 | 0.559778421 | NA | Sh2d1b1 | SH2 domain containing 1B1 [Source:MGI S | 0.8031 | 0 | 0 | 0.964 | 2.1812 | 0 |
| 40639 | ENSMUSG00000102419 | 0 | NA | NA | NA | NA | NA | Gm36940 | predicted gene, 36940 [Source:MGI Symb | 0 | 0 | 0 | 0 | 0 | 0 |
| 40640 | ENSMUSG00000102420 | 0 | NA | NA | NA | NA | NA | Gm37464 | predicted gene, 37464 [Source:MGI Symb | 0 | 0 | 0 | 0 | 0 | 0 |
| 40641 | ENSMUSG00000102421 | 0 | NA | NA | NA | NA | NA | Gm37325 | predicted gene, 37325 [Source:MGI Symb | 0 | 0 | 0 | 0 | 0 | 0 |
| 40642 | ENSMUSG00000102422 | 0 | NA | NA | NA | NA | NA | Iqschfp | Iqcj and Schip1 fusion protein [Source:MG | 0 | 0 | 0 | 0 | 0 | 0 |
| 40643 | ENSMUSG00000102423 | 0.181769715 | 1.126924175 | 4.080472857 | 0.276174898 | 0.782413736 | NA | Gm37465 | predicted gene, 37465 [Source:MGI Symb | 0 | 0 | 0 | 0 | 1.0906 | 0 |
| 40644 | ENSMUSG00000102424 | 0 | NA | NA | NA | NA | NA | Paupar | Pax6 upstream antisense RNA [Source:MG | 0 | 0 | 0 | 0 | 0 | 0 |
| 40645 | ENSMUSG00000102425 | 29.73346628 | 0.453625959 | 0.412747424 | 1.099040073 | 0.271750587 | 0.727526428 | Gm26616 | predicted gene, 26616 [Source:MGI Symb | 22.487 | 28.782 | 24.15 | 33.739 | 43.625 | 25.618 |
| 40646 | ENSMUSG00000102427 | 0 | NA | NA | NA | NA | NA | Gm37463 | predicted gene, 37463 [Source:MGI Symb | 0 | 0 | 0 | 0 | 0 | 0 |
| 40647 | ENSMUSG00000102428 | 3.746087136 | 1.253463605 | 1.118057274 | 1.121108582 | 0.262241646 | NA | Pcdhga12 | protocadherin gamma subfamily A, 12 [So | 1.6062 | 2.0558 | 3.0187 | 4.8199 | 7.6343 | 3.3415 |
| 40648 | ENSMUSG00000102429 | 0 | NA | NA | NA | NA | NA | Gm37462 | predicted gene, 37462 [Source:MGI Symb | 0 | 0 | 0 | 0 | 0 | 0 |
| 40649 | ENSMUSG00000102430 | 2.835022597 | 0.597743413 | 1.450704192 | 0.412036731 | 0.680312501 | NA | Gm38051 | predicted gene, 38051 [Source:MGI Symb | 0.8031 | 0 | 6.0375 | 5.7839 | 3.2719 | 1.1138 |
| 40650 | ENSMUSG00000102432 | 8.211976808 | 1.361860018 | 0.853275008 | 1.596038798 | 0.110480131 | 0.488680312 | Gm29856 | predicted gene, 29856 [Source:MGI Symb | 0.8031 | 5.1396 | 8.05 | 7.7118 | 13.087 | 14.48 |
| 40651 | ENSMUSG00000102433 | 0 | NA | NA | NA | NA | NA | Gm38050 | predicted gene, 38050 [Source:MGI Symb | 0 | 0 | 0 | 0 | 0 | 0 |
| 40652 | ENSMUSG00000102434 | 0 | NA | NA | NA | NA | NA | Gm38049 | predicted gene, 38049 [Source:MGI Symb | 0 | 0 | 0 | 0 | 0 | 0 |

| | | | | | | | | | | | | | | | |
|---|---|---|---|---|---|---|---|---|---|---|---|---|---|---|---|
| 40653 | ENSMUSG00000102436 | 0 | NA | NA | NA | NA | NA | Gm38047 | predicted gene, 38047 [Source:MGI Symb | 0 | 0 | 0 | 0 | 0 | 0 |
| 40654 | ENSMUSG00000102437 | 0.167707431 | -0.796629176 | 4.080472857 | -0.195229623 | 0.84521319 | NA | NA | NA | 0 | 0 | 1.0062 | 0 | 0 | 0 |
| 40655 | ENSMUSG00000102438 | 0 | NA | NA | NA | NA | NA | Gm17953 | predicted gene, 17953 [Source:MGI Symb | 0 | 0 | 0 | 0 | 0 | 0 |
| 40656 | ENSMUSG00000102439 | 0 | NA | NA | NA | NA | NA | Flg | filaggrin [Source:MGI Symbol;Acc:MGI:95 | 0 | 0 | 0 | 0 | 0 | 0 |
| 40657 | ENSMUSG00000102440 | 0.898398499 | 2.103385297 | 2.811138121 | 0.74823264 | 0.454319847 | NA | Pcdhga9 | protocadherin gamma subfamily A, 9 [Sou | 0 | 1.0279 | 0 | 0 | 4.3625 | 0 |
| 40658 | ENSMUSG00000102441 | 0 | NA | NA | NA | NA | NA | Gm38032 | predicted gene, 38032 [Source:MGI Symb | 0 | 0 | 0 | 0 | 0 | 0 |
| 40659 | ENSMUSG00000102442 | 0 | NA | NA | NA | NA | NA | Gm37103 | predicted gene, 37103 [Source:MGI Symb | 0 | 0 | 0 | 0 | 0 | 0 |
| 40660 | ENSMUSG00000102443 | 0 | NA | NA | NA | NA | NA | Gm37102 | predicted gene, 37102 [Source:MGI Symb | 0 | 0 | 0 | 0 | 0 | 0 |
| 40661 | ENSMUSG00000102445 | 0 | NA | NA | NA | NA | NA | Gm37104 | predicted gene, 37104 [Source:MGI Symb | 0 | 0 | 0 | 0 | 0 | 0 |
| 40662 | ENSMUSG00000102446 | 0 | NA | NA | NA | NA | NA | Gm17965 | predicted gene, 17965 [Source:MGI Symb | 0 | 0 | 0 | 0 | 0 | 0 |
| 40663 | ENSMUSG00000102447 | 0 | NA | NA | NA | NA | NA | Gm6621 | predicted gene 6621 [Source:MGI Symbol | 0 | 0 | 0 | 0 | 0 | 0 |
| 40664 | ENSMUSG00000102448 | 0.481988923 | 2.492401104 | 3.837077151 | 0.649557203 | 0.515978285 | NA | Gm37101 | predicted gene, 37101 [Source:MGI Symb | 0 | 0 | 0 | 2.8919 | 0 | 0 |
| 40665 | ENSMUSG00000102449 | 0.17131964 | -0.796629176 | 4.080472857 | -0.195229623 | 0.84521319 | NA | Gm37100 | predicted gene, 37100 [Source:MGI Symb | 0 | 1.0279 | 0 | 0 | 0 | 0 |
| 40666 | ENSMUSG00000102450 | 0 | NA | NA | NA | NA | NA | Gm38343 | predicted gene, 38343 [Source:MGI Symb | 0 | 0 | 0 | 0 | 0 | 0 |
| 40667 | ENSMUSG00000102451 | 0 | NA | NA | NA | NA | NA | Gm37655 | predicted gene, 37655 [Source:MGI Symb | 0 | 0 | 0 | 0 | 0 | 0 |
| 40668 | ENSMUSG00000102452 | 0.524667718 | -0.819976748 | 3.064824146 | -0.267544469 | 0.78904998 | NA | Gm38344 | predicted gene, 38344 [Source:MGI Symb | 0 | 1.0279 | 1.0062 | 0 | 0 | 1.1138 |
| 40669 | ENSMUSG00000102453 | 0 | NA | NA | NA | NA | NA | Ighv1-48 | immunoglobulin heavy variable V1-48 [So | 0 | 0 | 0 | 0 | 0 | 0 |
| 40670 | ENSMUSG00000102456 | 0 | NA | NA | NA | NA | NA | Gm37508 | predicted gene, 37508 [Source:MGI Symb | 0 | 0 | 0 | 0 | 0 | 0 |
| 40671 | ENSMUSG00000102457 | 1.124428187 | -1.312889854 | 2.591707867 | -0.506573241 | 0.612454288 | NA | Gm38342 | predicted gene, 38342 [Source:MGI Symb | 4.8186 | 0 | 0 | 1.928 | 0 | 0 |
| 40672 | ENSMUSG00000102458 | 0 | NA | NA | NA | NA | NA | Pisrt1 | polled intersex syndrome regulated transc | 0 | 0 | 0 | 0 | 0 | 0 |
| 40673 | ENSMUSG00000102459 | 0 | NA | NA | NA | NA | NA | Gm38352 | predicted gene, 38352 [Source:MGI Symb | 0 | 0 | 0 | 0 | 0 | 0 |
| 40674 | ENSMUSG00000102460 | 0 | NA | NA | NA | NA | NA | Gm38197 | predicted gene, 38197 [Source:MGI Symb | 0 | 0 | 0 | 0 | 0 | 0 |
| 40675 | ENSMUSG00000102461 | 0 | NA | NA | NA | NA | NA | Gm37166 | predicted gene, 37166 [Source:MGI Symb | 0 | 0 | 0 | 0 | 0 | 0 |
| 40676 | ENSMUSG00000102462 | 0 | NA | NA | NA | NA | NA | Gm5542 | predicted gene 5542 [Source:MGI Symbol | 0 | 0 | 0 | 0 | 0 | 0 |
| 40677 | ENSMUSG00000102463 | 0.167707431 | -0.796629176 | 4.080472857 | -0.195229623 | 0.84521319 | NA | 1700003N22 | RIKEN cDNA 1700003N22 gene [Source:M | 0 | 0 | 1.0062 | 0 | 0 | 0 |
| 40678 | ENSMUSG00000102464 | 0 | NA | NA | NA | NA | NA | Gm38199 | predicted gene, 38199 [Source:MGI Symb | 0 | 0 | 0 | 0 | 0 | 0 |
| 40679 | ENSMUSG00000102465 | 0 | NA | NA | NA | NA | NA | Gm38198 | predicted gene, 38198 [Source:MGI Symb | 0 | 0 | 0 | 0 | 0 | 0 |
| 40680 | ENSMUSG00000102466 | 0.17131964 | -0.796629176 | 4.080472857 | -0.195229623 | 0.84521319 | NA | Gm38200 | predicted gene, 38200 [Source:MGI Symb | 0 | 1.0279 | 0 | 0 | 0 | 0 |
| 40681 | ENSMUSG00000102467 | 0.133850373 | -0.796629176 | 4.080472857 | -0.195229623 | 0.84521319 | NA | Gm37167 | predicted gene, 37167 [Source:MGI Symb | 0.8031 | 0 | 0 | 0 | 0 | 0 |
| 40682 | ENSMUSG00000102469 | 0 | NA | NA | NA | NA | NA | Gm38201 | predicted gene, 38201 [Source:MGI Symb | 0 | 0 | 0 | 0 | 0 | 0 |
| 40683 | ENSMUSG00000102470 | 0 | NA | NA | NA | NA | NA | Gm37244 | predicted gene, 37244 [Source:MGI Symb | 0 | 0 | 0 | 0 | 0 | 0 |
| 40684 | ENSMUSG00000102471 | 13.36436901 | 0.243398302 | 0.796022247 | 0.305768216 | 0.75978113 | 0.95402891 | Gm5100 | predicted gene 5100 [Source:MGI Symbol | 9.6372 | 4.1117 | 23.144 | 27.955 | 11.997 | 3.3415 |
| 40685 | ENSMUSG00000102473 | 0 | NA | NA | NA | NA | NA | Gm37243 | predicted gene, 37243 [Source:MGI Symb | 0 | 0 | 0 | 0 | 0 | 0 |
| 40686 | ENSMUSG00000102474 | 0 | NA | NA | NA | NA | NA | 2610012C04 | RIKEN cDNA 2610012C04 gene [Source:M | 0 | 0 | 0 | 0 | 0 | 0 |
| 40687 | ENSMUSG00000102475 | 0 | NA | NA | NA | NA | NA | Gm37246 | predicted gene, 37246 [Source:MGI Symb | 0 | 0 | 0 | 0 | 0 | 0 |
| 40688 | ENSMUSG00000102476 | 0 | NA | NA | NA | NA | NA | Gm37245 | predicted gene, 37245 [Source:MGI Symb | 0 | 0 | 0 | 0 | 0 | 0 |
| 40689 | ENSMUSG00000102478 | 0.181769715 | 1.126924175 | 4.080472857 | 0.276174898 | 0.782413736 | NA | BC085271 | cDNA sequence BC085271 [Source:MGI Sy | 0 | 0 | 0 | 0 | 1.0906 | 0 |
| 40690 | ENSMUSG00000102479 | 0 | NA | NA | NA | NA | NA | Gm18551 | predicted gene, 18551 [Source:MGI Symb | 0 | 0 | 0 | 0 | 0 | 0 |
| 40691 | ENSMUSG00000102480 | 0 | NA | NA | NA | NA | NA | Gm37926 | predicted gene, 37926 [Source:MGI Symb | 0 | 0 | 0 | 0 | 0 | 0 |
| 40692 | ENSMUSG00000102481 | 0 | NA | NA | NA | NA | NA | Gm37925 | predicted gene, 37925 [Source:MGI Symb | 0 | 0 | 0 | 0 | 0 | 0 |
| 40693 | ENSMUSG00000102482 | 0 | NA | NA | NA | NA | NA | Gm37924 | predicted gene, 37924 [Source:MGI Symb | 0 | 0 | 0 | 0 | 0 | 0 |
| 40694 | ENSMUSG00000102483 | 0 | NA | NA | NA | NA | NA | Catspere1 | cation channel sperm associated auxiliary | 0 | 0 | 0 | 0 | 0 | 0 |
| 40695 | ENSMUSG00000102484 | 0 | NA | NA | NA | NA | NA | Gm37923 | predicted gene, 37923 [Source:MGI Symb | 0 | 0 | 0 | 0 | 0 | 0 |
| 40696 | ENSMUSG00000102485 | 0 | NA | NA | NA | NA | NA | Gm37922 | predicted gene, 37922 [Source:MGI Symb | 0 | 0 | 0 | 0 | 0 | 0 |
| 40697 | ENSMUSG00000102486 | 0 | NA | NA | NA | NA | NA | Gm37921 | predicted gene, 37921 [Source:MGI Symb | 0 | 0 | 0 | 0 | 0 | 0 |
| 40698 | ENSMUSG00000102487 | 0 | NA | NA | NA | NA | NA | Gm21871 | predicted gene, 21871 [Source:MGI Symb | 0 | 0 | 0 | 0 | 0 | 0 |
| 40699 | ENSMUSG00000102488 | 0 | NA | NA | NA | NA | NA | Gm37920 | predicted gene, 37920 [Source:MGI Symb | 0 | 0 | 0 | 0 | 0 | 0 |
| 40700 | ENSMUSG00000102489 | 0 | NA | NA | NA | NA | NA | Dnmt3a-ps1 | DNA methyltransferase 3A, pseudogene 1 | 0 | 0 | 0 | 0 | 0 | 0 |
| 40701 | ENSMUSG00000102490 | 0 | NA | NA | NA | NA | NA | Gm36985 | predicted gene, 36985 [Source:MGI Symb | 0 | 0 | 0 | 0 | 0 | 0 |
| 40702 | ENSMUSG00000102491 | 0 | NA | NA | NA | NA | NA | Gm36986 | predicted gene, 36986 [Source:MGI Symb | 0 | 0 | 0 | 0 | 0 | 0 |
| 40703 | ENSMUSG00000102492 | 0 | NA | NA | NA | NA | NA | Gm36987 | predicted gene, 36987 [Source:MGI Symb | 0 | 0 | 0 | 0 | 0 | 0 |
| 40704 | ENSMUSG00000102493 | 0.167707431 | -0.796629176 | 4.080472857 | -0.195229623 | 0.84521319 | NA | Gm8010 | predicted gene 8010 [Source:MGI Symbol | 0 | 0 | 1.0062 | 0 | 0 | 0 |
| 40705 | ENSMUSG00000102494 | 0 | NA | NA | NA | NA | NA | Gm36988 | predicted gene, 36988 [Source:MGI Symb | 0 | 0 | 0 | 0 | 0 | 0 |
| 40706 | ENSMUSG00000102495 | 0.181769715 | 1.126924175 | 4.080472857 | 0.276174898 | 0.782413736 | NA | Gm26524 | predicted gene, 26524 [Source:MGI Symb | 0 | 0 | 0 | 0 | 1.0906 | 0 |
| 40707 | ENSMUSG00000102496 | 6.065031129 | -0.897445021 | 1.020402594 | -0.879500921 | 0.379129734 | 0.812043903 | Gm36989 | predicted gene, 36989 [Source:MGI Symb | 6.4248 | 7.1954 | 10.062 | 2.8919 | 9.8156 | 0 |
| 40708 | ENSMUSG00000102497 | 0 | NA | NA | NA | NA | NA | Ighv13-1 | immunoglobulin heavy variable 13-1 [Sou | 0 | 0 | 0 | 0 | 0 | 0 |

| | | | | | | | | | | | | | | | |
|---|---|---|---|---|---|---|---|---|---|---|---|---|---|---|---|
| 40709 | ENSMUSG00000102498 | 0 | NA | NA | NA | NA | NA | Gm19445 | predicted gene, 19445 [Source:MGI Symb | 0 | 0 | 0 | 0 | 0 | 0 |
| 40710 | ENSMUSG00000102499 | 0 | NA | NA | NA | NA | NA | Gm36984 | predicted gene, 36984 [Source:MGI Symb | 0 | 0 | 0 | 0 | 0 | 0 |
| 40711 | ENSMUSG00000102500 | 0 | NA | NA | NA | NA | NA | Gm37371 | predicted gene, 37371 [Source:MGI Symb | 0 | 0 | 0 | 0 | 0 | 0 |
| 40712 | ENSMUSG00000102501 | 0 | NA | NA | NA | NA | NA | Gm37372 | predicted gene, 37372 [Source:MGI Symb | 0 | 0 | 0 | 0 | 0 | 0 |
| 40713 | ENSMUSG00000102502 | 0 | NA | NA | NA | NA | NA | Ighv1-10 | immunoglobulin heavy variable V1-10 [So | 0 | 0 | 0 | 0 | 0 | 0 |
| 40714 | ENSMUSG00000102503 | 0 | NA | NA | NA | NA | NA | Gm37370 | predicted gene, 37370 [Source:MGI Symb | 0 | 0 | 0 | 0 | 0 | 0 |
| 40715 | ENSMUSG00000102504 | 0 | NA | NA | NA | NA | NA | Gm21955 | predicted gene, 21955 [Source:MGI Symb | 0 | 0 | 0 | 0 | 0 | 0 |
| 40716 | ENSMUSG00000102505 | 1.090618289 | 3.652571688 | 2.872889547 | 1.27139301 | 0.203588868 | NA | 1600017P15 | RIKEN cDNA 1600017P15 gene [Source:M | 0 | 0 | 0 | 0 | 6.5437 | 0 |
| 40717 | ENSMUSG00000102506 | 0 | NA | NA | NA | NA | NA | Gm37369 | predicted gene, 37369 [Source:MGI Symb | 0 | 0 | 0 | 0 | 0 | 0 |
| 40718 | ENSMUSG00000102507 | 0 | NA | NA | NA | NA | NA | Gm21350 | predicted gene, 21350 [Source:MGI Symb | 0 | 0 | 0 | 0 | 0 | 0 |
| 40719 | ENSMUSG00000102508 | 0 | NA | NA | NA | NA | NA | Gm37367 | predicted gene, 37367 [Source:MGI Symb | 0 | 0 | 0 | 0 | 0 | 0 |
| 40720 | ENSMUSG00000102509 | 0 | NA | NA | NA | NA | NA | Gm37368 | predicted gene, 37368 [Source:MGI Symb | 0 | 0 | 0 | 0 | 0 | 0 |
| 40721 | ENSMUSG00000102510 | 0 | NA | NA | NA | NA | NA | Olfr1398-ps1 | olfactory receptor 1398, pseudogene 1 [S | 0 | 0 | 0 | 0 | 0 | 0 |
| 40722 | ENSMUSG00000102511 | 0 | NA | NA | NA | NA | NA | Gm38086 | predicted gene, 38086 [Source:MGI Symb | 0 | 0 | 0 | 0 | 0 | 0 |
| 40723 | ENSMUSG00000102512 | 0 | NA | NA | NA | NA | NA | Ighv1-29 | immunoglobulin heavy variable V1-29 [So | 0 | 0 | 0 | 0 | 0 | 0 |
| 40724 | ENSMUSG00000102513 | 0 | NA | NA | NA | NA | NA | Gm32347 | predicted gene, 32347 [Source:MGI Symb | 0 | 0 | 0 | 0 | 0 | 0 |
| 40725 | ENSMUSG00000102514 | 0 | NA | NA | NA | NA | NA | Gm35323 | predicted gene, 35323 [Source:MGI Symb | 0 | 0 | 0 | 0 | 0 | 0 |
| 40726 | ENSMUSG00000102515 | 0 | NA | NA | NA | NA | NA | Ighv12-2 | immunoglobulin heavy variable V12-2 [So | 0 | 0 | 0 | 0 | 0 | 0 |
| 40727 | ENSMUSG00000102516 | 0.497389803 | 1.101799453 | 3.439268845 | 0.320358629 | 0.748696483 | NA | Gm38340 | predicted gene, 38340 [Source:MGI Symb | 0.8031 | 0 | 0 | 0 | 2.1812 | 0 |
| 40728 | ENSMUSG00000102517 | 0 | NA | NA | NA | NA | NA | Gm38085 | predicted gene, 38085 [Source:MGI Symb | 0 | 0 | 0 | 0 | 0 | 0 |
| 40729 | ENSMUSG00000102518 | 0 | NA | NA | NA | NA | NA | Gm38339 | predicted gene, 38339 [Source:MGI Symb | 0 | 0 | 0 | 0 | 0 | 0 |
| 40730 | ENSMUSG00000102520 | 0 | NA | NA | NA | NA | NA | Gm36941 | predicted gene, 36941 [Source:MGI Symb | 0 | 0 | 0 | 0 | 0 | 0 |
| 40731 | ENSMUSG00000102521 | 0 | NA | NA | NA | NA | NA | Gm37781 | predicted gene, 37781 [Source:MGI Symb | 0 | 0 | 0 | 0 | 0 | 0 |
| 40732 | ENSMUSG00000102522 | 0 | NA | NA | NA | NA | NA | Gm37782 | predicted gene, 37782 [Source:MGI Symb | 0 | 0 | 0 | 0 | 0 | 0 |
| 40733 | ENSMUSG00000102523 | 0.167707431 | -0.796629176 | 4.080472857 | -0.195229623 | 0.84521319 | NA | Gm37783 | predicted gene, 37783 [Source:MGI Symb | 0 | 0 | 1.0062 | 0 | 0 | 0 |
| 40734 | ENSMUSG00000102524 | 0 | NA | NA | NA | NA | NA | Ighv1-2 | immunoglobulin heavy variable 1-2 [Sourc | 0 | 0 | 0 | 0 | 0 | 0 |
| 40735 | ENSMUSG00000102525 | 0 | NA | NA | NA | NA | NA | Gm37784 | predicted gene, 37784 [Source:MGI Symb | 0 | 0 | 0 | 0 | 0 | 0 |
| 40736 | ENSMUSG00000102526 | 3.195981643 | 0.219654978 | 1.253541248 | 0.175227563 | 0.860900828 | NA | Gm37785 | predicted gene, 37785 [Source:MGI Symb | 0.8031 | 6.1675 | 2.0125 | 5.7839 | 2.1812 | 2.2277 |
| 40737 | ENSMUSG00000102528 | 0 | NA | NA | NA | NA | NA | Ighv1-45 | immunoglobulin heavy variable V1-45 [So | 0 | 0 | 0 | 0 | 0 | 0 |
| 40738 | ENSMUSG00000102529 | 0 | NA | NA | NA | NA | NA | Gm29740 | predicted gene, 29740 [Source:MGI Symb | 0 | 0 | 0 | 0 | 0 | 0 |
| 40739 | ENSMUSG00000102530 | 0 | NA | NA | NA | NA | NA | Gm37228 | predicted gene, 37228 [Source:MGI Symb | 0 | 0 | 0 | 0 | 0 | 0 |
| 40740 | ENSMUSG00000102531 | 0.133850373 | -0.796629176 | 4.080472857 | -0.195229623 | 0.84521319 | NA | 1110002O04 | RIKEN cDNA 1110002O04 gene [Source:M | 0.8031 | 0 | 0 | 0 | 0 | 0 |
| 40741 | ENSMUSG00000102532 | 0 | NA | NA | NA | NA | NA | Gm37227 | predicted gene, 37227 [Source:MGI Symb | 0 | 0 | 0 | 0 | 0 | 0 |
| 40742 | ENSMUSG00000102533 | 9.282066481 | 0.288180053 | 0.72417141 | 0.397944533 | 0.690671073 | 0.937606823 | Gm37226 | predicted gene, 37226 [Source:MGI Symb | 4.8186 | 13.363 | 7.0437 | 10.604 | 8.7249 | 11.138 |
| 40743 | ENSMUSG00000102534 | 0 | NA | NA | NA | NA | NA | Gm37225 | predicted gene, 37225 [Source:MGI Symb | 0 | 0 | 0 | 0 | 0 | 0 |
| 40744 | ENSMUSG00000102535 | 0 | NA | NA | NA | NA | NA | Ighv1-41 | immunoglobulin heavy variable 1-41 [Sou | 0 | 0 | 0 | 0 | 0 | 0 |
| 40745 | ENSMUSG00000102536 | 0 | NA | NA | NA | NA | NA | 1700039I01F | RIKEN cDNA 1700039I01 gene [Source:MG | 0 | 0 | 0 | 0 | 0 | 0 |
| 40746 | ENSMUSG00000102537 | 0 | NA | NA | NA | NA | NA | Gm37224 | predicted gene, 37224 [Source:MGI Symb | 0 | 0 | 0 | 0 | 0 | 0 |
| 40747 | ENSMUSG00000102538 | 0 | NA | NA | NA | NA | NA | Gm37230 | predicted gene, 37230 [Source:MGI Symb | 0 | 0 | 0 | 0 | 0 | 0 |
| 40748 | ENSMUSG00000102539 | 0 | NA | NA | NA | NA | NA | Gm37229 | predicted gene, 37229 [Source:MGI Symb | 0 | 0 | 0 | 0 | 0 | 0 |
| 40749 | ENSMUSG00000102540 | 0 | NA | NA | NA | NA | NA | Gm38134 | predicted gene, 38134 [Source:MGI Symb | 0 | 0 | 0 | 0 | 0 | 0 |
| 40750 | ENSMUSG00000102541 | 0 | NA | NA | NA | NA | NA | Gm38135 | predicted gene, 38135 [Source:MGI Symb | 0 | 0 | 0 | 0 | 0 | 0 |
| 40751 | ENSMUSG00000102542 | 0 | NA | NA | NA | NA | NA | Gm38133 | predicted gene, 38133 [Source:MGI Symb | 0 | 0 | 0 | 0 | 0 | 0 |
| 40752 | ENSMUSG00000102543 | 0.167707431 | -0.796629176 | 4.080472857 | -0.195229623 | 0.84521319 | NA | Pcdhgc5 | protocadherin gamma subfamily C, 5 [Sou | 0 | 0 | 1.0062 | 0 | 0 | 0 |
| 40753 | ENSMUSG00000102544 | 1.380956475 | -1.598416342 | 2.020048339 | -0.791276283 | 0.428782785 | NA | Gm5103 | predicted gene 5103 [Source:MGI Symbol | 3.2124 | 0 | 3.0187 | 0.964 | 1.0906 | 0 |
| 40754 | ENSMUSG00000102545 | 13.48034443 | 0.313354741 | 0.594276512 | 0.527287777 | 0.597993761 | 0.915029425 | 6430573P05 | RIKEN cDNA 6430573P05 gene [Source:M | 8.8341 | 12.335 | 15.094 | 19.28 | 13.087 | 12.252 |
| 40755 | ENSMUSG00000102546 | 0 | NA | NA | NA | NA | NA | Gm9722 | predicted gene 9722 [Source:MGI Symbol | 0 | 0 | 0 | 0 | 0 | 0 |
| 40756 | ENSMUSG00000102547 | 0 | NA | NA | NA | NA | NA | Gm38136 | predicted gene, 38136 [Source:MGI Symb | 0 | 0 | 0 | 0 | 0 | 0 |
| 40757 | ENSMUSG00000102548 | 0 | NA | NA | NA | NA | NA | Gm16701 | predicted gene, 16701 [Source:MGI Symb | 0 | 0 | 0 | 0 | 0 | 0 |
| 40758 | ENSMUSG00000102549 | 2.431293548 | 1.486807021 | 1.401263507 | 1.061047414 | 0.288668354 | NA | Gm38137 | predicted gene, 38137 [Source:MGI Symb | 0.8031 | 2.0558 | 1.0062 | 1.928 | 5.4531 | 3.3415 |
| 40759 | ENSMUSG00000102550 | 0 | NA | NA | NA | NA | NA | Gm8276 | predicted gene 8276 [Source:MGI Symbol | 0 | 0 | 0 | 0 | 0 | 0 |
| 40760 | ENSMUSG00000102551 | 0 | NA | NA | NA | NA | NA | Gm37927 | predicted gene, 37927 [Source:MGI Symb | 0 | 0 | 0 | 0 | 0 | 0 |
| 40761 | ENSMUSG00000102552 | 0 | NA | NA | NA | NA | NA | Gm21763 | predicted gene, 21763 [Source:MGI Symb | 0 | 0 | 0 | 0 | 0 | 0 |
| 40762 | ENSMUSG00000102553 | 6.090234257 | 0.978542254 | 0.952032972 | 1.027844921 | 0.304022782 | 0.752961647 | D930036K23 | RIKEN cDNA D930036K23 gene [Source:M | 8.031 | 1.0279 | 3.0187 | 5.7839 | 11.997 | 6.6831 |
| 40763 | ENSMUSG00000102554 | 0.713016575 | 1.687601183 | 3.028998059 | 0.557148321 | 0.577426095 | NA | Gm37568 | predicted gene, 37568 [Source:MGI Symb | 0 | 0 | 1.0062 | 0 | 3.2719 | 0 |
| 40764 | ENSMUSG00000102555 | 23.31017593 | 0.413398681 | 0.479882802 | 0.861457587 | 0.388986071 | 0.818727377 | 6430511E19 | RIKEN cDNA 6430511E19 gene [Source:M | 22.487 | 12.335 | 25.156 | 34.703 | 22.903 | 22.277 |

| | | | | | | | | | | | | | | | |
|---|---|---|---|---|---|---|---|---|---|---|---|---|---|---|---|
| 40765 | ENSMUSG00000102556 | 0.335414862 | -1.779047242 | 4.030804179 | -0.441362855 | 0.658950329 | NA | Gm37569 | predicted gene, 37569 [Source:MGI Symb | 0 | 0 | 2.0125 | 0 | 0 | 0 |
| 40766 | ENSMUSG00000102558 | 0.367410362 | 2.078255025 | 3.800070157 | 0.54689912 | 0.58444804 | NA | Gm37571 | predicted gene, 37571 [Source:MGI Symb | 0 | 0 | 0 | 0 | 1.0906 | 1.1138 |
| 40767 | ENSMUSG00000102559 | 9.31345217 | 0.064021841 | 0.759005275 | 0.084349666 | 0.932778425 | 0.988420305 | Gm37570 | predicted gene, 37570 [Source:MGI Symb | 12.047 | 10.279 | 5.0312 | 16.388 | 5.4531 | 6.6831 |
| 40768 | ENSMUSG00000102561 | 0 | NA | NA | NA | NA | NA | Gm37039 | predicted gene, 37039 [Source:MGI Symb | 0 | 0 | 0 | 0 | 0 | 0 |
| 40769 | ENSMUSG00000102562 | 5.269927825 | -0.402422892 | 0.969303465 | -0.415167082 | 0.678019594 | 0.93325821 | Gm37694 | predicted gene, 37694 [Source:MGI Symb | 4.8186 | 7.1954 | 6.0375 | 4.8199 | 7.6343 | 1.1138 |
| 40770 | ENSMUSG00000102563 | 0 | NA | NA | NA | NA | NA | Gm37695 | predicted gene, 37695 [Source:MGI Symb | 0 | 0 | 0 | 0 | 0 | 0 |
| 40771 | ENSMUSG00000102564 | 3.081273601 | 0.668099344 | 1.217864506 | 0.548582655 | 0.583291892 | NA | Gm37035 | predicted gene, 37035 [Source:MGI Symb | 4.0155 | 2.0558 | 1.0062 | 4.8199 | 4.3625 | 2.2277 |
| 40772 | ENSMUSG00000102565 | 0 | NA | NA | NA | NA | NA | Gm37036 | predicted gene, 37036 [Source:MGI Symb | 0 | 0 | 0 | 0 | 0 | 0 |
| 40773 | ENSMUSG00000102566 | 0 | NA | NA | NA | NA | NA | Gm37037 | predicted gene, 37037 [Source:MGI Symb | 0 | 0 | 0 | 0 | 0 | 0 |
| 40774 | ENSMUSG00000102568 | 0 | NA | NA | NA | NA | NA | Gm37040 | predicted gene, 37040 [Source:MGI Symb | 0 | 0 | 0 | 0 | 0 | 0 |
| 40775 | ENSMUSG00000102569 | 0 | NA | NA | NA | NA | NA | Gm37041 | predicted gene, 37041 [Source:MGI Symb | 0 | 0 | 0 | 0 | 0 | 0 |
| 40776 | ENSMUSG00000102570 | 1.70359439 | 0.018069916 | 1.820119283 | 0.009927875 | 0.992078832 | NA | Gm37968 | predicted gene, 37968 [Source:MGI Symb | 4.0155 | 0 | 1.0062 | 1.928 | 3.2719 | 0 |
| 40777 | ENSMUSG00000102571 | 0 | NA | NA | NA | NA | NA | Gm37967 | predicted gene, 37967 [Source:MGI Symb | 0 | 0 | 0 | 0 | 0 | 0 |
| 40778 | ENSMUSG00000102572 | 0 | NA | NA | NA | NA | NA | Gm37966 | predicted gene, 37966 [Source:MGI Symb | 0 | 0 | 0 | 0 | 0 | 0 |
| 40779 | ENSMUSG00000102573 | 2.023267025 | 0.18982517 | 1.464796042 | 0.129591537 | 0.896889599 | NA | Gm7265 | predicted gene 7265 [Source:MGI Symbol | 1.6062 | 1.0279 | 3.0187 | 0.964 | 2.1812 | 3.3415 |
| 40780 | ENSMUSG00000102574 | 0 | NA | NA | NA | NA | NA | Gm33206 | predicted gene, 33206 [Source:MGI Symb | 0 | 0 | 0 | 0 | 0 | 0 |
| 40781 | ENSMUSG00000102575 | 0 | NA | NA | NA | NA | NA | Gm37241 | predicted gene, 37241 [Source:MGI Symb | 0 | 0 | 0 | 0 | 0 | 0 |
| 40782 | ENSMUSG00000102576 | 0 | NA | NA | NA | NA | NA | Gm37875 | predicted gene, 37875 [Source:MGI Symb | 0 | 0 | 0 | 0 | 0 | 0 |
| 40783 | ENSMUSG00000102577 | 6.359291494 | -0.669930539 | 0.881365124 | -0.760105568 | 0.447191485 | 0.853566538 | Gm37969 | predicted gene, 37969 [Source:MGI Symb | 3.2124 | 10.279 | 10.062 | 5.7839 | 4.3625 | 4.4554 |
| 40784 | ENSMUSG00000102578 | 0 | NA | NA | NA | NA | NA | Gm10576 | predicted gene 10576 [Source:MGI Symbo | 0 | 0 | 0 | 0 | 0 | 0 |
| 40785 | ENSMUSG00000102579 | 0 | NA | NA | NA | NA | NA | Gm37965 | predicted gene, 37965 [Source:MGI Symb | 0 | 0 | 0 | 0 | 0 | 0 |
| 40786 | ENSMUSG00000102580 | 0 | NA | NA | NA | NA | NA | Gm37442 | predicted gene, 37442 [Source:MGI Symb | 0 | 0 | 0 | 0 | 0 | 0 |
| 40787 | ENSMUSG00000102581 | 0 | NA | NA | NA | NA | NA | Gm37443 | predicted gene, 37443 [Source:MGI Symb | 0 | 0 | 0 | 0 | 0 | 0 |
| 40788 | ENSMUSG00000102582 | 0 | NA | NA | NA | NA | NA | Gm37440 | predicted gene, 37440 [Source:MGI Symb | 0 | 0 | 0 | 0 | 0 | 0 |
| 40789 | ENSMUSG00000102583 | 0 | NA | NA | NA | NA | NA | Gm37441 | predicted gene, 37441 [Source:MGI Symb | 0 | 0 | 0 | 0 | 0 | 0 |
| 40790 | ENSMUSG00000102584 | 0 | NA | NA | NA | NA | NA | Gm37438 | predicted gene, 37438 [Source:MGI Symb | 0 | 0 | 0 | 0 | 0 | 0 |
| 40791 | ENSMUSG00000102585 | 0 | NA | NA | NA | NA | NA | Gm37439 | predicted gene, 37439 [Source:MGI Symb | 0 | 0 | 0 | 0 | 0 | 0 |
| 40792 | ENSMUSG00000102586 | 0 | NA | NA | NA | NA | NA | Gm38003 | predicted gene, 38003 [Source:MGI Symb | 0 | 0 | 0 | 0 | 0 | 0 |
| 40793 | ENSMUSG00000102587 | 0 | NA | NA | NA | NA | NA | Ighv1-13 | immunoglobulin heavy variable V1-13 [So | 0 | 0 | 0 | 0 | 0 | 0 |
| 40794 | ENSMUSG00000102588 | 0 | NA | NA | NA | NA | NA | Gm37444 | predicted gene, 37444 [Source:MGI Symb | 0 | 0 | 0 | 0 | 0 | 0 |
| 40795 | ENSMUSG00000102589 | 0 | NA | NA | NA | NA | NA | Gm37445 | predicted gene, 37445 [Source:MGI Symb | 0 | 0 | 0 | 0 | 0 | 0 |
| 40796 | ENSMUSG00000102590 | 0 | NA | NA | NA | NA | NA | Mannr | Mecom adjacent non-protein coding RNA | 0 | 0 | 0 | 0 | 0 | 0 |
| 40797 | ENSMUSG00000102591 | 0 | NA | NA | NA | NA | NA | Gm38383 | predicted gene, 38383 [Source:MGI Symb | 0 | 0 | 0 | 0 | 0 | 0 |
| 40798 | ENSMUSG00000102592 | 0 | NA | NA | NA | NA | NA | Gm38385 | predicted gene, 38385 [Source:MGI Symb | 0 | 0 | 0 | 0 | 0 | 0 |
| 40799 | ENSMUSG00000102593 | 0.810359328 | 1.950931649 | 2.89643804 | 0.673562363 | 0.500589588 | NA | Gm38384 | predicted gene, 38384 [Source:MGI Symb | 0 | 0 | 1.0062 | 3.8559 | 0 | 0 |
| 40800 | ENSMUSG00000102594 | 0.133850373 | -0.796629176 | 4.080472857 | -0.195229623 | 0.84521319 | NA | Gm38381 | predicted gene, 38381 [Source:MGI Symb | 0.8031 | 0 | 0 | 0 | 0 | 0 |
| 40801 | ENSMUSG00000102595 | 1.702153181 | -1.954371304 | 1.934234428 | -1.010410773 | 0.312298529 | NA | Gm30097 | predicted gene, 30097 [Source:MGI Symb | 0 | 5.1396 | 3.0187 | 0.964 | 1.0906 | 0 |
| 40802 | ENSMUSG00000102596 | 0 | NA | NA | NA | NA | NA | Gm20203 | predicted gene, 20203 [Source:MGI Symb | 0 | 0 | 0 | 0 | 0 | 0 |
| 40803 | ENSMUSG00000102597 | 0 | NA | NA | NA | NA | NA | Gm38382 | predicted gene, 38382 [Source:MGI Symb | 0 | 0 | 0 | 0 | 0 | 0 |
| 40804 | ENSMUSG00000102598 | 0 | NA | NA | NA | NA | NA | Gm35670 | predicted gene, 35670 [Source:MGI Symb | 0 | 0 | 0 | 0 | 0 | 0 |
| 40805 | ENSMUSG00000102599 | 0 | NA | NA | NA | NA | NA | Gm38386 | predicted gene, 38386 [Source:MGI Symb | 0 | 0 | 0 | 0 | 0 | 0 |
| 40806 | ENSMUSG00000102600 | 0 | NA | NA | NA | NA | NA | Gm37266 | predicted gene, 37266 [Source:MGI Symb | 0 | 0 | 0 | 0 | 0 | 0 |
| 40807 | ENSMUSG00000102601 | 0 | NA | NA | NA | NA | NA | Ighv3-7 | immunoglobulin heavy variable V3-7 [Sou | 0 | 0 | 0 | 0 | 0 | 0 |
| 40808 | ENSMUSG00000102602 | 71.35814169 | -0.808351796 | 0.356832427 | -2.265354083 | 0.02349096 | 0.193934498 | A930004J17I | RIKEN cDNA A930004J17 gene [Source:M( | 111.63 | 95.596 | 65.406 | 65.55 | 64.346 | 25.618 |
| 40809 | ENSMUSG00000102603 | 0 | NA | NA | NA | NA | NA | Olfr431-ps1 | olfactory receptor 431, pseudogene 1 [So | 0 | 0 | 0 | 0 | 0 | 0 |
| 40810 | ENSMUSG00000102605 | 0 | NA | NA | NA | NA | NA | Gm37264 | predicted gene, 37264 [Source:MGI Symb | 0 | 0 | 0 | 0 | 0 | 0 |
| 40811 | ENSMUSG00000102606 | 0 | NA | NA | NA | NA | NA | Rpl21-ps1 | ribosomal protein 21, pseudogene 1 [Sour | 0 | 0 | 0 | 0 | 0 | 0 |
| 40812 | ENSMUSG00000102607 | 0 | NA | NA | NA | NA | NA | Gm37265 | predicted gene, 37265 [Source:MGI Symb | 0 | 0 | 0 | 0 | 0 | 0 |
| 40813 | ENSMUSG00000102608 | 0 | NA | NA | NA | NA | NA | Gm37267 | predicted gene, 37267 [Source:MGI Symb | 0 | 0 | 0 | 0 | 0 | 0 |
| 40814 | ENSMUSG00000102609 | 0 | NA | NA | NA | NA | NA | Gm18432 | predicted gene, 18432 [Source:MGI Symb | 0 | 0 | 0 | 0 | 0 | 0 |
| 40815 | ENSMUSG00000102610 | 0 | NA | NA | NA | NA | NA | Gm37846 | predicted gene, 37846 [Source:MGI Symb | 0 | 0 | 0 | 0 | 0 | 0 |
| 40816 | ENSMUSG00000102611 | 0 | NA | NA | NA | NA | NA | Gm37847 | predicted gene, 37847 [Source:MGI Symb | 0 | 0 | 0 | 0 | 0 | 0 |
| 40817 | ENSMUSG00000102612 | 0 | NA | NA | NA | NA | NA | Gm37845 | predicted gene, 37845 [Source:MGI Symb | 0 | 0 | 0 | 0 | 0 | 0 |
| 40818 | ENSMUSG00000102613 | 0 | NA | NA | NA | NA | NA | Gm20836 | predicted gene, 20836 [Source:MGI Symb | 0 | 0 | 0 | 0 | 0 | 0 |
| 40819 | ENSMUSG00000102614 | 0 | NA | NA | NA | NA | NA | Gm37843 | predicted gene, 37843 [Source:MGI Symb | 0 | 0 | 0 | 0 | 0 | 0 |
| 40820 | ENSMUSG00000102615 | 2.951836056 | -2.971475162 | 1.652043594 | -1.798666315 | 0.07207148 | NA | Gm37844 | predicted gene, 37844 [Source:MGI Symb | 9.6372 | 5.1396 | 1.0062 | 1.928 | 0 | 0 |

| | | | | | | | | | | | | | | | |
|---|---|---|---|---|---|---|---|---|---|---|---|---|---|---|---|
| 40821 | ENSMUSG00000102616 | 0 | NA | NA | NA | NA | NA | Gm37841 | predicted gene, 37841 [Source:MGI Symb | 0 | 0 | 0 | 0 | 0 | 0 |
| 40822 | ENSMUSG00000102617 | 0 | NA | NA | NA | NA | NA | Gm37842 | predicted gene, 37842 [Source:MGI Symb | 0 | 0 | 0 | 0 | 0 | 0 |
| 40823 | ENSMUSG00000102618 | 0 | NA | NA | NA | NA | NA | Gm37848 | predicted gene, 37848 [Source:MGI Symb | 0 | 0 | 0 | 0 | 0 | 0 |
| 40824 | ENSMUSG00000102619 | 0.17131964 | -0.796629176 | 4.080472857 | -0.195229623 | 0.84521319 | NA | Gm37849 | predicted gene, 37849 [Source:MGI Symb | 0 | 1.0279 | 0 | 0 | 0 | 0 |
| 40825 | ENSMUSG00000102620 | 0 | NA | NA | NA | NA | NA | Gm37675 | predicted gene, 37675 [Source:MGI Symb | 0 | 0 | 0 | 0 | 0 | 0 |
| 40826 | ENSMUSG00000102621 | 0 | NA | NA | NA | NA | NA | Gm37674 | predicted gene, 37674 [Source:MGI Symb | 0 | 0 | 0 | 0 | 0 | 0 |
| 40827 | ENSMUSG00000102622 | 0 | NA | NA | NA | NA | NA | Trbv7 | T cell receptor beta variable 7 [Source:MG | 0 | 0 | 0 | 0 | 0 | 0 |
| 40828 | ENSMUSG00000102623 | 0 | NA | NA | NA | NA | NA | Gm37673 | predicted gene, 37673 [Source:MGI Symb | 0 | 0 | 0 | 0 | 0 | 0 |
| 40829 | ENSMUSG00000102624 | 0 | NA | NA | NA | NA | NA | Gm37059 | predicted gene, 37059 [Source:MGI Symb | 0 | 0 | 0 | 0 | 0 | 0 |
| 40830 | ENSMUSG00000102625 | 0 | NA | NA | NA | NA | NA | Gm37672 | predicted gene, 37672 [Source:MGI Symb | 0 | 0 | 0 | 0 | 0 | 0 |
| 40831 | ENSMUSG00000102626 | 0.160662974 | 1.126924175 | 4.080472857 | 0.276174898 | 0.782413736 | NA | Gm18553 | predicted gene, 18553 [Source:MGI Symb | 0 | 0 | 0 | 0.964 | 0 | 0 |
| 40832 | ENSMUSG00000102627 | 0 | NA | NA | NA | NA | NA | Snurf | SNRPN upstream reading frame [Source:N | 0 | 0 | 0 | 0 | 0 | 0 |
| 40833 | ENSMUSG00000102628 | 0 | NA | NA | NA | NA | NA | Gm37671 | predicted gene, 37671 [Source:MGI Symb | 0 | 0 | 0 | 0 | 0 | 0 |
| 40834 | ENSMUSG00000102629 | 0 | NA | NA | NA | NA | NA | Gm29999 | predicted gene, 29999 [Source:MGI Symb | 0 | 0 | 0 | 0 | 0 | 0 |
| 40835 | ENSMUSG00000102630 | 0.758511407 | -1.621915645 | 2.674939911 | -0.606337226 | 0.544290844 | NA | Gm37289 | predicted gene, 37289 [Source:MGI Symb | 2.4093 | 1.0279 | 0 | 0 | 0 | 1.1138 |
| 40836 | ENSMUSG00000102632 | 0.902269431 | 2.107869146 | 2.547671313 | 0.827370915 | 0.40802686 | NA | Gm37786 | predicted gene, 37786 [Source:MGI Symb | 0 | 1.0279 | 0 | 0 | 3.2719 | 1.1138 |
| 40837 | ENSMUSG00000102633 | 0 | NA | NA | NA | NA | NA | Gm38224 | predicted gene, 38224 [Source:MGI Symb | 0 | 0 | 0 | 0 | 0 | 0 |
| 40838 | ENSMUSG00000102634 | 0 | NA | NA | NA | NA | NA | Gm38225 | predicted gene, 38225 [Source:MGI Symb | 0 | 0 | 0 | 0 | 0 | 0 |
| 40839 | ENSMUSG00000102635 | 0.774667342 | 0.864398069 | 2.649064859 | 0.326303097 | 0.744195045 | NA | Gm36638 | predicted gene, 36638 [Source:MGI Symb | 1.6062 | 0 | 0 | 1.928 | 0 | 1.1138 |
| 40840 | ENSMUSG00000102636 | 0 | NA | NA | NA | NA | NA | Gm31466 | predicted gene, 31466 [Source:MGI Symb | 0 | 0 | 0 | 0 | 0 | 0 |
| 40841 | ENSMUSG00000102637 | 0 | NA | NA | NA | NA | NA | Gm8684 | predicted gene 8684 [Source:MGI Symbol | 0 | 0 | 0 | 0 | 0 | 0 |
| 40842 | ENSMUSG00000102639 | 0.181769715 | 1.126924175 | 4.080472857 | 0.276174898 | 0.782413736 | NA | Gm38223 | predicted gene, 38223 [Source:MGI Symb | 0 | 0 | 0 | 0 | 1.0906 | 0 |
| 40843 | ENSMUSG00000102641 | 0 | NA | NA | NA | NA | NA | Ighv1-3 | immunoglobulin heavy variable V1-3 [Sou | 0 | 0 | 0 | 0 | 0 | 0 |
| 40844 | ENSMUSG00000102642 | 0 | NA | NA | NA | NA | NA | A130048G24 | RIKEN cDNA A130048G24 gene [Source:M | 0 | 0 | 0 | 0 | 0 | 0 |
| 40845 | ENSMUSG00000102643 | 0 | NA | NA | NA | NA | NA | Gm37547 | predicted gene, 37547 [Source:MGI Symb | 0 | 0 | 0 | 0 | 0 | 0 |
| 40846 | ENSMUSG00000102644 | 216.1755236 | -0.008804555 | 0.175065182 | -0.050293008 | 0.959888895 | 0.992198455 | Thap6 | THAP domain containing 6 [Source:MGI Sy | 243.34 | 189.14 | 217.35 | 214.97 | 205.04 | 227.22 |
| 40847 | ENSMUSG00000102645 | 0 | NA | NA | NA | NA | NA | Ighv5-5 | immunoglobulin heavy variable V5-5 [Sou | 0 | 0 | 0 | 0 | 0 | 0 |
| 40848 | ENSMUSG00000102646 | 0 | NA | NA | NA | NA | NA | Gm19694 | predicted gene, 19694 [Source:MGI Symb | 0 | 0 | 0 | 0 | 0 | 0 |
| 40849 | ENSMUSG00000102647 | 0 | NA | NA | NA | NA | NA | Gm38024 | predicted gene, 38024 [Source:MGI Symb | 0 | 0 | 0 | 0 | 0 | 0 |
| 40850 | ENSMUSG00000102648 | 0 | NA | NA | NA | NA | NA | Gm38022 | predicted gene, 38022 [Source:MGI Symb | 0 | 0 | 0 | 0 | 0 | 0 |
| 40851 | ENSMUSG00000102649 | 0.481988923 | 2.492401104 | 3.837077151 | 0.649557203 | 0.515978285 | NA | Gm38021 | predicted gene, 38021 [Source:MGI Symb | 0 | 0 | 0 | 2.8919 | 0 | 0 |
| 40852 | ENSMUSG00000102650 | 0 | NA | NA | NA | NA | NA | Gm19863 | predicted gene, 19863 [Source:MGI Symb | 0 | 0 | 0 | 0 | 0 | 0 |
| 40853 | ENSMUSG00000102651 | 0.321325949 | 1.911575114 | 4.048267893 | 0.472195804 | 0.636787034 | NA | Gm37080 | predicted gene, 37080 [Source:MGI Symb | 0 | 0 | 0 | 1.928 | 0 | 0 |
| 40854 | ENSMUSG00000102652 | 0.674441933 | -2.786326546 | 3.022203109 | -0.921952114 | 0.356553554 | NA | Gm37078 | predicted gene, 37078 [Source:MGI Symb | 0 | 1.0279 | 3.0187 | 0 | 0 | 0 |
| 40855 | ENSMUSG00000102653 | 0 | NA | NA | NA | NA | NA | Gm37079 | predicted gene, 37079 [Source:MGI Symb | 0 | 0 | 0 | 0 | 0 | 0 |
| 40856 | ENSMUSG00000102654 | 0 | NA | NA | NA | NA | NA | Ighv1-21 | immunoglobulin heavy variable V1-21 [So | 0 | 0 | 0 | 0 | 0 | 0 |
| 40857 | ENSMUSG00000102655 | 0 | NA | NA | NA | NA | NA | Gm37081 | predicted gene, 37081 [Source:MGI Symb | 0 | 0 | 0 | 0 | 0 | 0 |
| 40858 | ENSMUSG00000102656 | 0 | NA | NA | NA | NA | NA | Ighv1-32 | immunoglobulin heavy variable V1-32 [So | 0 | 0 | 0 | 0 | 0 | 0 |
| 40859 | ENSMUSG00000102657 | 0 | NA | NA | NA | NA | NA | Gm37899 | predicted gene, 37899 [Source:MGI Symb | 0 | 0 | 0 | 0 | 0 | 0 |
| 40860 | ENSMUSG00000102658 | 0 | NA | NA | NA | NA | NA | Gm37076 | predicted gene, 37076 [Source:MGI Symb | 0 | 0 | 0 | 0 | 0 | 0 |
| 40861 | ENSMUSG00000102659 | 0 | NA | NA | NA | NA | NA | Gm37077 | predicted gene, 37077 [Source:MGI Symb | 0 | 0 | 0 | 0 | 0 | 0 |
| 40862 | ENSMUSG00000102660 | 0 | NA | NA | NA | NA | NA | Gm38378 | predicted gene, 38378 [Source:MGI Symb | 0 | 0 | 0 | 0 | 0 | 0 |
| 40863 | ENSMUSG00000102661 | 0 | NA | NA | NA | NA | NA | Gm34816 | predicted gene, 34816 [Source:MGI Symb | 0 | 0 | 0 | 0 | 0 | 0 |
| 40864 | ENSMUSG00000102662 | 1.347538101 | -0.743200427 | 2.011993948 | -0.369385021 | 0.711840761 | NA | Gm38377 | predicted gene, 38377 [Source:MGI Symb | 4.0155 | 1.0279 | 0 | 1.928 | 0 | 1.1138 |
| 40865 | ENSMUSG00000102663 | 0 | NA | NA | NA | NA | NA | Gm6631 | predicted gene 6631 [Source:MGI Symbol | 0 | 0 | 0 | 0 | 0 | 0 |
| 40866 | ENSMUSG00000102664 | 0.997338572 | 0.170071214 | 2.081193669 | 0.08171811 | 0.934870877 | NA | Gm38380 | predicted gene, 38380 [Source:MGI Symb | 0.8031 | 0 | 2.0125 | 0.964 | 1.0906 | 1.1138 |
| 40867 | ENSMUSG00000102665 | 0.660353019 | -0.01500701 | 2.83315355 | -0.005296928 | 0.995773683 | NA | Gm38379 | predicted gene, 38379 [Source:MGI Symb | 0 | 1.0279 | 1.0062 | 1.928 | 0 | 0 |
| 40868 | ENSMUSG00000102666 | 0 | NA | NA | NA | NA | NA | Gm8599 | predicted gene 8599 [Source:MGI Symbol | 0 | 0 | 0 | 0 | 0 | 0 |
| 40869 | ENSMUSG00000102667 | 0 | NA | NA | NA | NA | NA | Gm19901 | predicted gene, 19901 [Source:MGI Symb | 0 | 0 | 0 | 0 | 0 | 0 |
| 40870 | ENSMUSG00000102668 | 0 | NA | NA | NA | NA | NA | Gm29866 | predicted gene, 29866 [Source:MGI Symb | 0 | 0 | 0 | 0 | 0 | 0 |
| 40871 | ENSMUSG00000102670 | 0.160662974 | 1.126924175 | 4.080472857 | 0.276174898 | 0.782413736 | NA | Gm37436 | predicted gene, 37436 [Source:MGI Symb | 0 | 0 | 0 | 0.964 | 0 | 0 |
| 40872 | ENSMUSG00000102672 | 1.429082407 | -2.723467668 | 1.987237116 | -1.370479469 | 0.17053728 | NA | Gm37105 | predicted gene, 37105 [Source:MGI Symb | 2.4093 | 2.0558 | 3.0187 | 0 | 1.0906 | 0 |
| 40873 | ENSMUSG00000102673 | 0 | NA | NA | NA | NA | NA | Gm37437 | predicted gene, 37437 [Source:MGI Symb | 0 | 0 | 0 | 0 | 0 | 0 |
| 40874 | ENSMUSG00000102674 | 0 | NA | NA | NA | NA | NA | 8030442B05 | RIKEN cDNA 8030442B05 gene [Source:M | 0 | 0 | 0 | 0 | 0 | 0 |
| 40875 | ENSMUSG00000102676 | 0 | NA | NA | NA | NA | NA | Gm37435 | predicted gene, 37435 [Source:MGI Symb | 0 | 0 | 0 | 0 | 0 | 0 |
| 40876 | ENSMUSG00000102677 | 0 | NA | NA | NA | NA | NA | 4930535E02 | RIKEN cDNA 4930535E02 gene [Source:M | 0 | 0 | 0 | 0 | 0 | 0 |

| | | | | | | | | | | | | | | | |
|---|---|---|---|---|---|---|---|---|---|---|---|---|---|---|---|
| 40877 | ENSMUSG00000102678 | 0 | NA | NA | NA | NA | NA | Ighv1-21-1 | immunoglobulin heavy variable 1-21-1 [Sc | 0 | 0 | 0 | 0 | 0 | 0 |
| 40878 | ENSMUSG00000102679 | 0 | NA | NA | NA | NA | NA | Gm20557 | predicted gene, 20557 [Source:MGI Symb | 0 | 0 | 0 | 0 | 0 | 0 |
| 40879 | ENSMUSG00000102680 | 0 | NA | NA | NA | NA | NA | Gm37222 | predicted gene, 37222 [Source:MGI Symb | 0 | 0 | 0 | 0 | 0 | 0 |
| 40880 | ENSMUSG00000102681 | 0 | NA | NA | NA | NA | NA | Gm37221 | predicted gene, 37221 [Source:MGI Symb | 0 | 0 | 0 | 0 | 0 | 0 |
| 40881 | ENSMUSG00000102682 | 0 | NA | NA | NA | NA | NA | Gm37223 | predicted gene, 37223 [Source:MGI Symb | 0 | 0 | 0 | 0 | 0 | 0 |
| 40882 | ENSMUSG00000102683 | 0 | NA | NA | NA | NA | NA | Gm37375 | predicted gene, 37375 [Source:MGI Symb | 0 | 0 | 0 | 0 | 0 | 0 |
| 40883 | ENSMUSG00000102684 | 0 | NA | NA | NA | NA | NA | Gm37374 | predicted gene, 37374 [Source:MGI Symb | 0 | 0 | 0 | 0 | 0 | 0 |
| 40884 | ENSMUSG00000102685 | 0 | NA | NA | NA | NA | NA | Gm37373 | predicted gene, 37373 [Source:MGI Symb | 0 | 0 | 0 | 0 | 0 | 0 |
| 40885 | ENSMUSG00000102686 | 0 | NA | NA | NA | NA | NA | Gm37220 | predicted gene, 37220 [Source:MGI Symb | 0 | 0 | 0 | 0 | 0 | 0 |
| 40886 | ENSMUSG00000102687 | 0 | NA | NA | NA | NA | NA | Gm37219 | predicted gene, 37219 [Source:MGI Symb | 0 | 0 | 0 | 0 | 0 | 0 |
| 40887 | ENSMUSG00000102688 | 0 | NA | NA | NA | NA | NA | Gm37218 | predicted gene, 37218 [Source:MGI Symb | 0 | 0 | 0 | 0 | 0 | 0 |
| 40888 | ENSMUSG00000102689 | 0 | NA | NA | NA | NA | NA | Gm37217 | predicted gene, 37217 [Source:MGI Symb | 0 | 0 | 0 | 0 | 0 | 0 |
| 40889 | ENSMUSG00000102690 | 0 | NA | NA | NA | NA | NA | Gm37779 | predicted gene, 37779 [Source:MGI Symb | 0 | 0 | 0 | 0 | 0 | 0 |
| 40890 | ENSMUSG00000102691 | 2.153426426 | 0.423758361 | 1.488263802 | 0.284733366 | 0.775848426 | NA | Gm37780 | predicted gene, 37780 [Source:MGI Symb | 2.4093 | 3.0838 | 0 | 1.928 | 3.2719 | 2.2277 |
| 40891 | ENSMUSG00000102692 | 2.12150969 | -0.988174385 | 1.597071078 | -0.618741644 | 0.536086571 | NA | Dchs2 | dachsous cadherin related 2 [Source:MGI | 2.4093 | 0 | 6.0375 | 0.964 | 1.0906 | 2.2277 |
| 40892 | ENSMUSG00000102693 | 0 | NA | NA | NA | NA | NA | 4933401J01I | RIKEN cDNA 4933401J01 gene [Source:M( | 0 | 0 | 0 | 0 | 0 | 0 |
| 40893 | ENSMUSG00000102694 | 0.160662974 | 1.126924175 | 4.080472857 | 0.276174898 | 0.782413736 | NA | Gm37496 | predicted gene, 37496 [Source:MGI Symb | 0 | 0 | 0 | 0.964 | 0 | 0 |
| 40894 | ENSMUSG00000102696 | 0 | NA | NA | NA | NA | NA | Gm37778 | predicted gene, 37778 [Source:MGI Symb | 0 | 0 | 0 | 0 | 0 | 0 |
| 40895 | ENSMUSG00000102697 | 0.267700746 | -1.497481649 | 4.057001436 | -0.369110456 | 0.712045396 | NA | Pcdhac2 | protocadherin alpha subfamily C, 2 [Sourc | 1.6062 | 0 | 0 | 0 | 0 | 0 |
| 40896 | ENSMUSG00000102698 | 0 | NA | NA | NA | NA | NA | Gm37777 | predicted gene, 37777 [Source:MGI Symb | 0 | 0 | 0 | 0 | 0 | 0 |
| 40897 | ENSMUSG00000102699 | 0 | NA | NA | NA | NA | NA | 4930433B08 | RIKEN cDNA 4930433B08 gene [Source:M | 0 | 0 | 0 | 0 | 0 | 0 |
| 40898 | ENSMUSG00000102700 | 0 | NA | NA | NA | NA | NA | Gm38312 | predicted gene, 38312 [Source:MGI Symb | 0 | 0 | 0 | 0 | 0 | 0 |
| 40899 | ENSMUSG00000102701 | 9.393694909 | -0.393108365 | 0.787507718 | -0.499180334 | 0.617652348 | 0.917509899 | 9430083B18 | RIKEN cDNA 9430083B18 gene [Source:M | 9.6372 | 10.279 | 12.075 | 5.7839 | 16.359 | 2.2277 |
| 40900 | ENSMUSG00000102702 | 0 | NA | NA | NA | NA | NA | Gm37260 | predicted gene, 37260 [Source:MGI Symb | 0 | 0 | 0 | 0 | 0 | 0 |
| 40901 | ENSMUSG00000102703 | 1.905520454 | 0.348635131 | 1.584751361 | 0.219993584 | 0.825876152 | NA | Gm38313 | predicted gene, 38313 [Source:MGI Symb | 0 | 3.0838 | 2.0125 | 1.928 | 2.1812 | 2.2277 |
| 40902 | ENSMUSG00000102704 | 4.968804846 | -0.135473475 | 1.040273512 | -0.130228708 | 0.896385482 | NA | Gm38309 | predicted gene, 38309 [Source:MGI Symb | 2.4093 | 5.1396 | 8.05 | 0.964 | 5.4531 | 7.7969 |
| 40903 | ENSMUSG00000102705 | 0 | NA | NA | NA | NA | NA | 4632432E15 | RIKEN cDNA 4632432E15 gene [Source:M( | 0 | 0 | 0 | 0 | 0 | 0 |
| 40904 | ENSMUSG00000102706 | 0 | NA | NA | NA | NA | NA | Gm7417 | predicted gene 7417 [Source:MGI Symbol | 0 | 0 | 0 | 0 | 0 | 0 |
| 40905 | ENSMUSG00000102707 | 0 | NA | NA | NA | NA | NA | Gm38311 | predicted gene, 38311 [Source:MGI Symb | 0 | 0 | 0 | 0 | 0 | 0 |
| 40906 | ENSMUSG00000102709 | 0 | NA | NA | NA | NA | NA | Gm38308 | predicted gene, 38308 [Source:MGI Symb | 0 | 0 | 0 | 0 | 0 | 0 |
| 40907 | ENSMUSG00000102710 | 0 | NA | NA | NA | NA | NA | Gm4857 | predicted gene 4857 [Source:MGI Symbol | 0 | 0 | 0 | 0 | 0 | 0 |
| 40908 | ENSMUSG00000102712 | 0 | NA | NA | NA | NA | NA | Gm37758 | predicted gene, 37758 [Source:MGI Symb | 0 | 0 | 0 | 0 | 0 | 0 |
| 40909 | ENSMUSG00000102713 | 0 | NA | NA | NA | NA | NA | Gm29994 | predicted gene, 29994 [Source:MGI Symb | 0 | 0 | 0 | 0 | 0 | 0 |
| 40910 | ENSMUSG00000102714 | 0 | NA | NA | NA | NA | NA | Gm37618 | predicted gene, 37618 [Source:MGI Symb | 0 | 0 | 0 | 0 | 0 | 0 |
| 40911 | ENSMUSG00000102715 | 0 | NA | NA | NA | NA | NA | Gm6209 | predicted gene 6209 [Source:MGI Symbol | 0 | 0 | 0 | 0 | 0 | 0 |
| 40912 | ENSMUSG00000102716 | 0 | NA | NA | NA | NA | NA | Gm5099 | predicted gene 5099 [Source:MGI Symbol | 0 | 0 | 0 | 0 | 0 | 0 |
| 40913 | ENSMUSG00000102717 | 0 | NA | NA | NA | NA | NA | Gm37759 | predicted gene, 37759 [Source:MGI Symb | 0 | 0 | 0 | 0 | 0 | 0 |
| 40914 | ENSMUSG00000102718 | 0 | NA | NA | NA | NA | NA | Gm37761 | predicted gene, 37761 [Source:MGI Symb | 0 | 0 | 0 | 0 | 0 | 0 |
| 40915 | ENSMUSG00000102719 | 6.850903602 | -0.138947197 | 0.895221609 | -0.155209834 | 0.876655896 | 0.980632845 | Gm37760 | predicted gene, 37760 [Source:MGI Symb | 6.4248 | 5.1396 | 10.062 | 9.6398 | 8.7249 | 1.1138 |
| 40916 | ENSMUSG00000102721 | 0 | NA | NA | NA | NA | NA | A630081D01 | RIKEN cDNA A630081D01 gene [Source:M | 0 | 0 | 0 | 0 | 0 | 0 |
| 40917 | ENSMUSG00000102722 | 0.638982041 | 0.404325377 | 3.14176733 | 0.128693609 | 0.897600092 | NA | Gm37935 | predicted gene, 37935 [Source:MGI Symb | 1.6062 | 0 | 0 | 0 | 0 | 2.2277 |
| 40918 | ENSMUSG00000102723 | 0 | NA | NA | NA | NA | NA | Gm37936 | predicted gene, 37936 [Source:MGI Symb | 0 | 0 | 0 | 0 | 0 | 0 |
| 40919 | ENSMUSG00000102725 | 0 | NA | NA | NA | NA | NA | Gm37939 | predicted gene, 37939 [Source:MGI Symb | 0 | 0 | 0 | 0 | 0 | 0 |
| 40920 | ENSMUSG00000102726 | 0 | NA | NA | NA | NA | NA | Gm37937 | predicted gene, 37937 [Source:MGI Symb | 0 | 0 | 0 | 0 | 0 | 0 |
| 40921 | ENSMUSG00000102728 | 0 | NA | NA | NA | NA | NA | Gm37934 | predicted gene, 37934 [Source:MGI Symb | 0 | 0 | 0 | 0 | 0 | 0 |
| 40922 | ENSMUSG00000102729 | 0 | NA | NA | NA | NA | NA | Gm32444 | predicted gene, 32444 [Source:MGI Symb | 0 | 0 | 0 | 0 | 0 | 0 |
| 40923 | ENSMUSG00000102730 | 0 | NA | NA | NA | NA | NA | Gm38138 | predicted gene, 38138 [Source:MGI Symb | 0 | 0 | 0 | 0 | 0 | 0 |
| 40924 | ENSMUSG00000102731 | 0 | NA | NA | NA | NA | NA | Gm38261 | predicted gene, 38261 [Source:MGI Symb | 0 | 0 | 0 | 0 | 0 | 0 |
| 40925 | ENSMUSG00000102732 | 0 | NA | NA | NA | NA | NA | Gm37342 | predicted gene, 37342 [Source:MGI Symb | 0 | 0 | 0 | 0 | 0 | 0 |
| 40926 | ENSMUSG00000102733 | 0.160662974 | 1.126924175 | 4.080472857 | 0.276174898 | 0.782413736 | NA | Gm38262 | predicted gene, 38262 [Source:MGI Symb | 0 | 0 | 0 | 0.964 | 0 | 0 |
| 40927 | ENSMUSG00000102734 | 0 | NA | NA | NA | NA | NA | Gm37343 | predicted gene, 37343 [Source:MGI Symb | 0 | 0 | 0 | 0 | 0 | 0 |
| 40928 | ENSMUSG00000102735 | 0 | NA | NA | NA | NA | NA | Gm7369 | predicted gene 7369 [Source:MGI Symbol | 0 | 0 | 0 | 0 | 0 | 0 |
| 40929 | ENSMUSG00000102737 | 1.390010306 | -0.651990341 | 1.996396009 | -0.326583673 | 0.743982794 | NA | 1700016A09 | RIKEN cDNA 1700016A09 gene [Source:M | 4.0155 | 0 | 1.0062 | 0 | 1.0906 | 2.2277 |
| 40930 | ENSMUSG00000102738 | 0 | NA | NA | NA | NA | NA | 1700006P03 | RIKEN cDNA 1700006P03 gene [Source:M( | 0 | 0 | 0 | 0 | 0 | 0 |
| 40931 | ENSMUSG00000102739 | 0 | NA | NA | NA | NA | NA | Gm20772 | predicted gene, 20772 [Source:MGI Symb | 0 | 0 | 0 | 0 | 0 | 0 |
| 40932 | ENSMUSG00000102740 | 0 | NA | NA | NA | NA | NA | Gm37599 | predicted gene, 37599 [Source:MGI Symb | 0 | 0 | 0 | 0 | 0 | 0 |

| | | | | | | | | | | | | | | | |
|---|---|---|---|---|---|---|---|---|---|---|---|---|---|---|---|
| 40933 | ENSMUSG00000102741 | 0 | NA | NA | NA | NA | NA | Gm2979 | predicted gene 2979 [Source:MGI Symbol | 0 | 0 | 0 | 0 | 0 | 0 |
| 40934 | ENSMUSG00000102742 | 1.28134971 | -2.562434167 | 2.096087442 | -1.222484385 | 0.221524505 | NA | Pcdhga11 | protocadherin gamma subfamily A, 11 [So | 1.6062 | 4.1117 | 1.0062 | 0.964 | 0 | 0 |
| 40935 | ENSMUSG00000102743 | 0 | NA | NA | NA | NA | NA | Gm38108 | predicted gene, 38108 [Source:MGI Symb | 0 | 0 | 0 | 0 | 0 | 0 |
| 40936 | ENSMUSG00000102744 | 0 | NA | NA | NA | NA | NA | 5830444F18 | RIKEN cDNA 5830444F18 gene [Source:M | 0 | 0 | 0 | 0 | 0 | 0 |
| 40937 | ENSMUSG00000102745 | 0 | NA | NA | NA | NA | NA | Gm37600 | predicted gene, 37600 [Source:MGI Symb | 0 | 0 | 0 | 0 | 0 | 0 |
| 40938 | ENSMUSG00000102746 | 0 | NA | NA | NA | NA | NA | Gm37601 | predicted gene, 37601 [Source:MGI Symb | 0 | 0 | 0 | 0 | 0 | 0 |
| 40939 | ENSMUSG00000102747 | 0 | NA | NA | NA | NA | NA | Gm37602 | predicted gene, 37602 [Source:MGI Symb | 0 | 0 | 0 | 0 | 0 | 0 |
| 40940 | ENSMUSG00000102748 | 3.627106833 | -1.314815716 | 1.236299485 | -1.063509071 | 0.287551154 | NA | Pcdhgb2 | protocadherin gamma subfamily B, 2 [Sou | 2.4093 | 5.1396 | 8.05 | 2.8919 | 3.2719 | 0 |
| 40941 | ENSMUSG00000102749 | 0 | NA | NA | NA | NA | NA | Gm37598 | predicted gene, 37598 [Source:MGI Symb | 0 | 0 | 0 | 0 | 0 | 0 |
| 40942 | ENSMUSG00000102750 | 0 | NA | NA | NA | NA | NA | Gm37058 | predicted gene, 37058 [Source:MGI Symb | 0 | 0 | 0 | 0 | 0 | 0 |
| 40943 | ENSMUSG00000102751 | 0 | NA | NA | NA | NA | NA | Gm37057 | predicted gene, 37057 [Source:MGI Symb | 0 | 0 | 0 | 0 | 0 | 0 |
| 40944 | ENSMUSG00000102752 | 112.5229562 | 0.169871805 | 0.239955789 | 0.707929597 | 0.478988979 | 0.870169438 | Gm7694 | predicted gene 7694 [Source:MGI Symbol | 99.585 | 104.85 | 113.71 | 141.7 | 89.431 | 125.86 |
| 40945 | ENSMUSG00000102753 | 0 | NA | NA | NA | NA | NA | Gm37056 | predicted gene, 37056 [Source:MGI Symb | 0 | 0 | 0 | 0 | 0 | 0 |
| 40946 | ENSMUSG00000102754 | 0.954602129 | 0.390466883 | 2.454858538 | 0.159058812 | 0.873622538 | NA | Gm38183 | predicted gene, 38183 [Source:MGI Symb | 2.4093 | 0 | 0 | 0 | 1.0906 | 2.2277 |
| 40947 | ENSMUSG00000102755 | 0 | NA | NA | NA | NA | NA | Gm37055 | predicted gene, 37055 [Source:MGI Symb | 0 | 0 | 0 | 0 | 0 | 0 |
| 40948 | ENSMUSG00000102756 | 0 | NA | NA | NA | NA | NA | Gm37054 | predicted gene, 37054 [Source:MGI Symb | 0 | 0 | 0 | 0 | 0 | 0 |
| 40949 | ENSMUSG00000102757 | 0 | NA | NA | NA | NA | NA | Gm2756 | predicted gene 2756 [Source:MGI Symbol | 0 | 0 | 0 | 0 | 0 | 0 |
| 40950 | ENSMUSG00000102758 | 195.4840084 | -0.47478194 | 0.237970769 | -1.995127138 | 0.046029015 | 0.296818402 | Naaladl2 | N-acetylated alpha-linked acidic dipeptida | 164.64 | 301.18 | 217.35 | 195.69 | 157.05 | 137 |
| 40951 | ENSMUSG00000102759 | 0 | NA | NA | NA | NA | NA | Gm10463 | predicted gene 10463 [Source:MGI Symbo | 0 | 0 | 0 | 0 | 0 | 0 |
| 40952 | ENSMUSG00000102760 | 0 | NA | NA | NA | NA | NA | Gm37258 | predicted gene, 37258 [Source:MGI Symb | 0 | 0 | 0 | 0 | 0 | 0 |
| 40953 | ENSMUSG00000102761 | 0.185640647 | 1.126924175 | 4.080472857 | 0.276174898 | 0.782413736 | NA | Gm37184 | predicted gene, 37184 [Source:MGI Symb | 0 | 0 | 0 | 0 | 0 | 1.1138 |
| 40954 | ENSMUSG00000102762 | 0 | NA | NA | NA | NA | NA | 4930403P22 | RIKEN cDNA 4930403P22 gene [Source:M | 0 | 0 | 0 | 0 | 0 | 0 |
| 40955 | ENSMUSG00000102763 | 0 | NA | NA | NA | NA | NA | Gm37257 | predicted gene, 37257 [Source:MGI Symb | 0 | 0 | 0 | 0 | 0 | 0 |
| 40956 | ENSMUSG00000102764 | 0.328370405 | 0.16514316 | 4.003149536 | 0.041253308 | 0.967093956 | NA | Gm37183 | predicted gene, 37183 [Source:MGI Symb | 0 | 0 | 1.0062 | 0.964 | 0 | 0 |
| 40957 | ENSMUSG00000102765 | 0 | NA | NA | NA | NA | NA | Ighv1-62 | immunoglobulin heavy variable V1-62 [So | 0 | 0 | 0 | 0 | 0 | 0 |
| 40958 | ENSMUSG00000102766 | 0.305170013 | -1.657836477 | 4.041480335 | -0.410205257 | 0.681655385 | NA | Gm6420 | predicted gene 6420 [Source:MGI Symbol | 0.8031 | 1.0279 | 0 | 0 | 0 | 0 |
| 40959 | ENSMUSG00000102768 | 0 | NA | NA | NA | NA | NA | Gm19002 | predicted gene, 19002 [Source:MGI Symb | 0 | 0 | 0 | 0 | 0 | 0 |
| 40960 | ENSMUSG00000102769 | 0 | NA | NA | NA | NA | NA | Gm37185 | predicted gene, 37185 [Source:MGI Symb | 0 | 0 | 0 | 0 | 0 | 0 |
| 40961 | ENSMUSG00000102770 | 0 | NA | NA | NA | NA | NA | Gm38163 | predicted gene, 38163 [Source:MGI Symb | 0 | 0 | 0 | 0 | 0 | 0 |
| 40962 | ENSMUSG00000102771 | 0 | NA | NA | NA | NA | NA | Gm33320 | predicted gene, 33320 [Source:MGI Symb | 0 | 0 | 0 | 0 | 0 | 0 |
| 40963 | ENSMUSG00000102772 | 0 | NA | NA | NA | NA | NA | Gm32114 | predicted gene, 32114 [Source:MGI Symb | 0 | 0 | 0 | 0 | 0 | 0 |
| 40964 | ENSMUSG00000102773 | 0 | NA | NA | NA | NA | NA | Ighv1-17-1 | immunoglobulin heavy variable V1-17-1 [S | 0 | 0 | 0 | 0 | 0 | 0 |
| 40965 | ENSMUSG00000102774 | 0 | NA | NA | NA | NA | NA | Gm9748 | predicted gene 9748 [Source:MGI Symbol | 0 | 0 | 0 | 0 | 0 | 0 |
| 40966 | ENSMUSG00000102775 | 0.722877068 | -0.331317305 | 2.989821669 | -0.110815073 | 0.911762993 | NA | Gm38160 | predicted gene, 38160 [Source:MGI Symb | 2.4093 | 0 | 0 | 1.928 | 0 | 0 |
| 40967 | ENSMUSG00000102776 | 3.367094693 | 0.660864872 | 1.215526596 | 0.543686065 | 0.586657523 | NA | Gm38162 | predicted gene, 38162 [Source:MGI Symb | 0.8031 | 5.1396 | 2.0125 | 6.7478 | 3.2719 | 2.2277 |
| 40968 | ENSMUSG00000102777 | 0 | NA | NA | NA | NA | NA | Gm38161 | predicted gene, 38161 [Source:MGI Symb | 0 | 0 | 0 | 0 | 0 | 0 |
| 40969 | ENSMUSG00000102778 | 0 | NA | NA | NA | NA | NA | Gm38165 | predicted gene, 38165 [Source:MGI Symb | 0 | 0 | 0 | 0 | 0 | 0 |
| 40970 | ENSMUSG00000102779 | 0 | NA | NA | NA | NA | NA | Gm38164 | predicted gene, 38164 [Source:MGI Symb | 0 | 0 | 0 | 0 | 0 | 0 |
| 40971 | ENSMUSG00000102780 | 7.715217036 | -0.057809448 | 0.797373538 | -0.072499832 | 0.942204139 | 0.989798252 | Gm38253 | predicted gene, 38253 [Source:MGI Symb | 6.4248 | 10.279 | 7.0437 | 11.568 | 7.6343 | 3.3415 |
| 40972 | ENSMUSG00000102782 | 0.181769715 | 1.126924175 | 4.080472857 | 0.276174898 | 0.782413736 | NA | Gm37625 | predicted gene, 37625 [Source:MGI Symb | 0 | 0 | 0 | 0 | 1.0906 | 0 |
| 40973 | ENSMUSG00000102783 | 0 | NA | NA | NA | NA | NA | Gm38254 | predicted gene, 38254 [Source:MGI Symb | 0 | 0 | 0 | 0 | 0 | 0 |
| 40974 | ENSMUSG00000102784 | 0 | NA | NA | NA | NA | NA | Gm38251 | predicted gene, 38251 [Source:MGI Symb | 0 | 0 | 0 | 0 | 0 | 0 |
| 40975 | ENSMUSG00000102785 | 0 | NA | NA | NA | NA | NA | Gm2447 | predicted gene 2447 [Source:MGI Symbol | 0 | 0 | 0 | 0 | 0 | 0 |
| 40976 | ENSMUSG00000102786 | 0.181769715 | 1.126924175 | 4.080472857 | 0.276174898 | 0.782413736 | NA | Gm38252 | predicted gene, 38252 [Source:MGI Symb | 0 | 0 | 0 | 0 | 1.0906 | 0 |
| 40977 | ENSMUSG00000102787 | 0 | NA | NA | NA | NA | NA | Gm30174 | predicted gene, 30174 [Source:MGI Symb | 0 | 0 | 0 | 0 | 0 | 0 |
| 40978 | ENSMUSG00000102788 | 0 | NA | NA | NA | NA | NA | Gm38255 | predicted gene, 38255 [Source:MGI Symb | 0 | 0 | 0 | 0 | 0 | 0 |
| 40979 | ENSMUSG00000102789 | 0 | NA | NA | NA | NA | NA | Gm38256 | predicted gene, 38256 [Source:MGI Symb | 0 | 0 | 0 | 0 | 0 | 0 |
| 40980 | ENSMUSG00000102790 | 0 | NA | NA | NA | NA | NA | Gm37272 | predicted gene, 37272 [Source:MGI Symb | 0 | 0 | 0 | 0 | 0 | 0 |
| 40981 | ENSMUSG00000102791 | 0 | NA | NA | NA | NA | NA | Gm37709 | predicted gene, 37709 [Source:MGI Symb | 0 | 0 | 0 | 0 | 0 | 0 |
| 40982 | ENSMUSG00000102792 | 0 | NA | NA | NA | NA | NA | Gm32175 | predicted gene, 32175 [Source:MGI Symb | 0 | 0 | 0 | 0 | 0 | 0 |
| 40983 | ENSMUSG00000102793 | 0 | NA | NA | NA | NA | NA | Gm37708 | predicted gene, 37708 [Source:MGI Symb | 0 | 0 | 0 | 0 | 0 | 0 |
| 40984 | ENSMUSG00000102794 | 0 | NA | NA | NA | NA | NA | Gm37712 | predicted gene, 37712 [Source:MGI Symb | 0 | 0 | 0 | 0 | 0 | 0 |
| 40985 | ENSMUSG00000102795 | 0 | NA | NA | NA | NA | NA | Gm6236 | predicted gene 6236 [Source:MGI Symbol | 0 | 0 | 0 | 0 | 0 | 0 |
| 40986 | ENSMUSG00000102796 | 0 | NA | NA | NA | NA | NA | Gm37711 | predicted gene, 37711 [Source:MGI Symb | 0 | 0 | 0 | 0 | 0 | 0 |
| 40987 | ENSMUSG00000102797 | 0.321325949 | 1.911575114 | 4.048267893 | 0.472195804 | 0.636787034 | NA | Gm37710 | predicted gene, 37710 [Source:MGI Symb | 0 | 0 | 0 | 1.928 | 0 | 0 |
| 40988 | ENSMUSG00000102799 | 0 | NA | NA | NA | NA | NA | Gm32295 | predicted gene, 32295 [Source:MGI Symb | 0 | 0 | 0 | 0 | 0 | 0 |

| | | | | | | | | | | | | | | | |
|---|---|---|---|---|---|---|---|---|---|---|---|---|---|---|---|
| 40989 | ENSMUSG00000102800 | 0 | NA | NA | NA | NA | NA | Gm37479 | predicted gene, 37479 [Source:MGI Symb | 0 | 0 | 0 | 0 | 0 | 0 |
| 40990 | ENSMUSG00000102801 | 0 | NA | NA | NA | NA | NA | Gm37478 | predicted gene, 37478 [Source:MGI Symb | 0 | 0 | 0 | 0 | 0 | 0 |
| 40991 | ENSMUSG00000102802 | 0 | NA | NA | NA | NA | NA | Mgam2-ps | maltase-glucoamylase 2, pseudogene [Sou | 0 | 0 | 0 | 0 | 0 | 0 |
| 40992 | ENSMUSG00000102803 | 0 | NA | NA | NA | NA | NA | Gm37477 | predicted gene, 37477 [Source:MGI Symb | 0 | 0 | 0 | 0 | 0 | 0 |
| 40993 | ENSMUSG00000102804 | 0 | NA | NA | NA | NA | NA | Gm37481 | predicted gene, 37481 [Source:MGI Symb | 0 | 0 | 0 | 0 | 0 | 0 |
| 40994 | ENSMUSG00000102805 | 0.321325949 | 1.911575114 | 4.048267893 | 0.472195804 | 0.636787034 | NA | Gm37240 | predicted gene, 37240 [Source:MGI Symb | 0 | 0 | 0 | 1.928 | 0 | 0 |
| 40995 | ENSMUSG00000102806 | 0 | NA | NA | NA | NA | NA | Ighv1-19-1 | immunoglobulin heavy variable 1-19-1 [Sc | 0 | 0 | 0 | 0 | 0 | 0 |
| 40996 | ENSMUSG00000102807 | 0 | NA | NA | NA | NA | NA | Gm37480 | predicted gene, 37480 [Source:MGI Symb | 0 | 0 | 0 | 0 | 0 | 0 |
| 40997 | ENSMUSG00000102808 | 0 | NA | NA | NA | NA | NA | 5430420F09 | RIKEN cDNA 5430420F09 gene [Source:M | 0 | 0 | 0 | 0 | 0 | 0 |
| 40998 | ENSMUSG00000102809 | 0 | NA | NA | NA | NA | NA | Gm8643 | predicted gene 8643 [Source:MGI Symbol | 0 | 0 | 0 | 0 | 0 | 0 |
| 40999 | ENSMUSG00000102810 | 0 | NA | NA | NA | NA | NA | Gm37815 | predicted gene, 37815 [Source:MGI Symb | 0 | 0 | 0 | 0 | 0 | 0 |
| 41000 | ENSMUSG00000102813 | 12.44769941 | 0.209441014 | 0.734619941 | 0.285101183 | 0.775566624 | 0.95893551 | Gm37795 | predicted gene, 37795 [Source:MGI Symb | 10.44 | 2.0558 | 22.137 | 12.532 | 15.269 | 12.252 |
| 41001 | ENSMUSG00000102814 | 0 | NA | NA | NA | NA | NA | Gm37812 | predicted gene, 37812 [Source:MGI Symb | 0 | 0 | 0 | 0 | 0 | 0 |
| 41002 | ENSMUSG00000102815 | 0 | NA | NA | NA | NA | NA | Gm37813 | predicted gene, 37813 [Source:MGI Symb | 0 | 0 | 0 | 0 | 0 | 0 |
| 41003 | ENSMUSG00000102816 | 0 | NA | NA | NA | NA | NA | Gm37814 | predicted gene, 37814 [Source:MGI Symb | 0 | 0 | 0 | 0 | 0 | 0 |
| 41004 | ENSMUSG00000102817 | 4.183872664 | 0.421802932 | 1.08476126 | 0.388844022 | 0.697391534 | NA | Gm6185 | predicted gene 6185 [Source:MGI Symbol | 5.6217 | 1.0279 | 4.025 | 6.7478 | 5.4531 | 2.2277 |
| 41005 | ENSMUSG00000102818 | 0 | NA | NA | NA | NA | NA | Gm37796 | predicted gene, 37796 [Source:MGI Symb | 0 | 0 | 0 | 0 | 0 | 0 |
| 41006 | ENSMUSG00000102819 | 0.339027071 | -1.79174149 | 3.919585585 | -0.457125237 | 0.647581032 | NA | Gm37817 | predicted gene, 37817 [Source:MGI Symb | 0 | 1.0279 | 1.0062 | 0 | 0 | 0 |
| 41007 | ENSMUSG00000102820 | 0 | NA | NA | NA | NA | NA | Gm38353 | predicted gene, 38353 [Source:MGI Symb | 0 | 0 | 0 | 0 | 0 | 0 |
| 41008 | ENSMUSG00000102821 | 0 | NA | NA | NA | NA | NA | Trbv8 | T cell receptor beta variable 8 [Source:MG | 0 | 0 | 0 | 0 | 0 | 0 |
| 41009 | ENSMUSG00000102822 | 0 | NA | NA | NA | NA | NA | 1700113B19 | RIKEN cDNA 1700113B19 gene [Source:M | 0 | 0 | 0 | 0 | 0 | 0 |
| 41010 | ENSMUSG00000102824 | 6.38018999 | -0.226029921 | 0.871919147 | -0.259232662 | 0.795455732 | 0.962236316 | Pdcd5-ps | programmed cell death 5, pseudogene [Sc | 6.4248 | 9.2513 | 5.0312 | 7.7118 | 7.6343 | 2.2277 |
| 41011 | ENSMUSG00000102825 | 0 | NA | NA | NA | NA | NA | Gm36990 | predicted gene, 36990 [Source:MGI Symb | 0 | 0 | 0 | 0 | 0 | 0 |
| 41012 | ENSMUSG00000102826 | 0 | NA | NA | NA | NA | NA | Gm38181 | predicted gene, 38181 [Source:MGI Symb | 0 | 0 | 0 | 0 | 0 | 0 |
| 41013 | ENSMUSG00000102827 | 7.037979058 | 1.03835068 | 0.808759738 | 1.283880282 | 0.199183848 | 0.641237767 | Gm8242 | predicted gene 8242 [Source:MGI Symbol | 5.6217 | 4.1117 | 4.025 | 8.6758 | 11.997 | 7.7969 |
| 41014 | ENSMUSG00000102828 | 0 | NA | NA | NA | NA | NA | Gm38182 | predicted gene, 38182 [Source:MGI Symb | 0 | 0 | 0 | 0 | 0 | 0 |
| 41015 | ENSMUSG00000102830 | 0 | NA | NA | NA | NA | NA | Gm37012 | predicted gene, 37012 [Source:MGI Symb | 0 | 0 | 0 | 0 | 0 | 0 |
| 41016 | ENSMUSG00000102831 | 0 | NA | NA | NA | NA | NA | Gm37382 | predicted gene, 37382 [Source:MGI Symb | 0 | 0 | 0 | 0 | 0 | 0 |
| 41017 | ENSMUSG00000102832 | 0 | NA | NA | NA | NA | NA | Gm5190 | predicted gene 5190 [Source:MGI Symbol | 0 | 0 | 0 | 0 | 0 | 0 |
| 41018 | ENSMUSG00000102833 | 0.698954292 | 0.126053087 | 3.048972385 | 0.04134281 | 0.967022605 | NA | Gm9694 | predicted gene 9694 [Source:MGI Symbol | 0 | 0 | 2.0125 | 0 | 2.1812 | 0 |
| 41019 | ENSMUSG00000102834 | 0 | NA | NA | NA | NA | NA | 1700029B24 | RIKEN cDNA 1700029B24 gene [Source:M | 0 | 0 | 0 | 0 | 0 | 0 |
| 41020 | ENSMUSG00000102835 | 0 | NA | NA | NA | NA | NA | Gm37572 | predicted gene, 37572 [Source:MGI Symb | 0 | 0 | 0 | 0 | 0 | 0 |
| 41021 | ENSMUSG00000102836 | 0 | NA | NA | NA | NA | NA | Gm37013 | predicted gene, 37013 [Source:MGI Symb | 0 | 0 | 0 | 0 | 0 | 0 |
| 41022 | ENSMUSG00000102837 | 0.506966596 | 2.558866737 | 3.339834942 | 0.766165628 | 0.443577757 | NA | Gm37383 | predicted gene, 37383 [Source:MGI Symb | 0 | 0 | 0 | 1.928 | 0 | 1.1138 |
| 41023 | ENSMUSG00000102838 | 0 | NA | NA | NA | NA | NA | Gm37014 | predicted gene, 37014 [Source:MGI Symb | 0 | 0 | 0 | 0 | 0 | 0 |
| 41024 | ENSMUSG00000102839 | 0 | NA | NA | NA | NA | NA | Gm37385 | predicted gene, 37385 [Source:MGI Symb | 0 | 0 | 0 | 0 | 0 | 0 |
| 41025 | ENSMUSG00000102840 | 0.863926883 | 2.043289473 | 2.549701854 | 0.801383687 | 0.422909556 | NA | Gm38037 | predicted gene, 38037 [Source:MGI Symb | 0 | 1.0279 | 0 | 1.928 | 0 | 2.2277 |
| 41026 | ENSMUSG00000102841 | 0.328370405 | 0.16514316 | 4.003149536 | 0.041253308 | 0.967093956 | NA | Gm38036 | predicted gene, 38036 [Source:MGI Symb | 0 | 0 | 1.0062 | 0.964 | 0 | 0 |
| 41027 | ENSMUSG00000102842 | 0 | NA | NA | NA | NA | NA | Gm38035 | predicted gene, 38035 [Source:MGI Symb | 0 | 0 | 0 | 0 | 0 | 0 |
| 41028 | ENSMUSG00000102844 | 0.181769715 | 1.126924175 | 4.080472857 | 0.276174898 | 0.782413736 | NA | Gm38034 | predicted gene, 38034 [Source:MGI Symb | 0 | 0 | 0 | 0 | 1.0906 | 0 |
| 41029 | ENSMUSG00000102845 | 0 | NA | NA | NA | NA | NA | Gm38033 | predicted gene, 38033 [Source:MGI Symb | 0 | 0 | 0 | 0 | 0 | 0 |
| 41030 | ENSMUSG00000102846 | 0 | NA | NA | NA | NA | NA | Gm10292 | predicted gene 10292 [Source:MGI Symbo | 0 | 0 | 0 | 0 | 0 | 0 |
| 41031 | ENSMUSG00000102847 | 0 | NA | NA | NA | NA | NA | Gm34274 | predicted gene, 34274 [Source:MGI Symb | 0 | 0 | 0 | 0 | 0 | 0 |
| 41032 | ENSMUSG00000102848 | 0 | NA | NA | NA | NA | NA | Ighv5-7 | immunoglobulin heavy variable V5-7 [Sou | 0 | 0 | 0 | 0 | 0 | 0 |
| 41033 | ENSMUSG00000102849 | 0 | NA | NA | NA | NA | NA | Gm37658 | predicted gene, 37658 [Source:MGI Symb | 0 | 0 | 0 | 0 | 0 | 0 |
| 41034 | ENSMUSG00000102850 | 0 | NA | NA | NA | NA | NA | Gm37082 | predicted gene, 37082 [Source:MGI Symb | 0 | 0 | 0 | 0 | 0 | 0 |
| 41035 | ENSMUSG00000102851 | 0 | NA | NA | NA | NA | NA | Gm18956 | predicted gene, 18956 [Source:MGI Symb | 0 | 0 | 0 | 0 | 0 | 0 |
| 41036 | ENSMUSG00000102852 | 3.91297289 | -0.338477772 | 1.054068398 | -0.321115567 | 0.748122813 | NA | Gm37083 | predicted gene, 37083 [Source:MGI Symb | 4.0155 | 4.1117 | 5.0312 | 4.8199 | 3.2719 | 2.2277 |
| 41037 | ENSMUSG00000102853 | 0 | NA | NA | NA | NA | NA | Gm7270 | predicted gene 7270 [Source:MGI Symbol | 0 | 0 | 0 | 0 | 0 | 0 |
| 41038 | ENSMUSG00000102854 | 1.163448683 | 1.343870089 | 2.165028087 | 0.620717162 | 0.534785736 | NA | C130023A14 | RIKEN cDNA C130023A14 gene [Source:M | 0 | 1.0279 | 1.0062 | 3.8559 | 1.0906 | 0 |
| 41039 | ENSMUSG00000102855 | 0 | NA | NA | NA | NA | NA | Gm38002 | predicted gene, 38002 [Source:MGI Symb | 0 | 0 | 0 | 0 | 0 | 0 |
| 41040 | ENSMUSG00000102856 | 3.204758177 | -0.182326359 | 1.30689929 | -0.139510642 | 0.889046647 | NA | Gm37084 | predicted gene, 37084 [Source:MGI Symb | 3.2124 | 2.0558 | 5.0312 | 6.7478 | 2.1812 | 0 |
| 41041 | ENSMUSG00000102857 | 0 | NA | NA | NA | NA | NA | Gm37085 | predicted gene, 37085 [Source:MGI Symb | 0 | 0 | 0 | 0 | 0 | 0 |
| 41042 | ENSMUSG00000102858 | 0 | NA | NA | NA | NA | NA | Gm37086 | predicted gene, 37086 [Source:MGI Symb | 0 | 0 | 0 | 0 | 0 | 0 |
| 41043 | ENSMUSG00000102859 | 0 | NA | NA | NA | NA | NA | Gm8625 | predicted gene 8625 [Source:MGI Symbol | 0 | 0 | 0 | 0 | 0 | 0 |
| 41044 | ENSMUSG00000102860 | 0 | NA | NA | NA | NA | NA | Gm37638 | predicted gene, 37638 [Source:MGI Symb | 0 | 0 | 0 | 0 | 0 | 0 |

| | | | | | | | | | | | | | | | |
|---|---|---|---|---|---|---|---|---|---|---|---|---|---|---|---|
| 41045 | ENSMUSG00000102861 | 5.694969078 | 0.371862497 | 1.046853655 | 0.35521918 | 0.722425397 | 0.9447719 | Gm37637 | predicted gene, 37637 [Source:MGI Symb | 4.8186 | 4.1117 | 6.0375 | 11.568 | 7.6343 | 0 |
| 41046 | ENSMUSG00000102862 | 0 | NA | NA | NA | NA | NA | Gm37640 | predicted gene, 37640 [Source:MGI Symb | 0 | 0 | 0 | 0 | 0 | 0 |
| 41047 | ENSMUSG00000102863 | 1.068948596 | -2.273572368 | 2.191117856 | -1.037631254 | 0.299441762 | NA | Gm37639 | predicted gene, 37639 [Source:MGI Symb | 2.4093 | 1.0279 | 2.0125 | 0.964 | 0 | 0 |
| 41048 | ENSMUSG00000102864 | 0 | NA | NA | NA | NA | NA | Gm37634 | predicted gene, 37634 [Source:MGI Symb | 0 | 0 | 0 | 0 | 0 | 0 |
| 41049 | ENSMUSG00000102865 | 0 | NA | NA | NA | NA | NA | Gm9839 | predicted gene 9839 [Source:MGI Symbol | 0 | 0 | 0 | 0 | 0 | 0 |
| 41050 | ENSMUSG00000102866 | 0 | NA | NA | NA | NA | NA | Gm37636 | predicted gene, 37636 [Source:MGI Symb | 0 | 0 | 0 | 0 | 0 | 0 |
| 41051 | ENSMUSG00000102867 | 0 | NA | NA | NA | NA | NA | Gm37635 | predicted gene, 37635 [Source:MGI Symb | 0 | 0 | 0 | 0 | 0 | 0 |
| 41052 | ENSMUSG00000102868 | 0 | NA | NA | NA | NA | NA | Gm37633 | predicted gene, 37633 [Source:MGI Symb | 0 | 0 | 0 | 0 | 0 | 0 |
| 41053 | ENSMUSG00000102869 | 1740.02327 | 0.119553201 | 0.105195761 | 1.136483064 | 0.25575445 | 0.708918985 | Norad | non-coding RNA activated by DNA damag | 1634.3 | 1798.9 | 1570.7 | 1747.7 | 1750.4 | 1938.1 |
| 41054 | ENSMUSG00000102870 | 0 | NA | NA | NA | NA | NA | Gm17868 | predicted gene, 17868 [Source:MGI Symb | 0 | 0 | 0 | 0 | 0 | 0 |
| 41055 | ENSMUSG00000102871 | 0 | NA | NA | NA | NA | NA | Gm37143 | predicted gene, 37143 [Source:MGI Symb | 0 | 0 | 0 | 0 | 0 | 0 |
| 41056 | ENSMUSG00000102872 | 0 | NA | NA | NA | NA | NA | Gm38195 | predicted gene, 38195 [Source:MGI Symb | 0 | 0 | 0 | 0 | 0 | 0 |
| 41057 | ENSMUSG00000102874 | 0 | NA | NA | NA | NA | NA | Gm37137 | predicted gene, 37137 [Source:MGI Symb | 0 | 0 | 0 | 0 | 0 | 0 |
| 41058 | ENSMUSG00000102875 | 4.110846407 | -0.22993271 | 1.072650872 | -0.214359319 | 0.830266863 | NA | Gm37139 | predicted gene, 37139 [Source:MGI Symb | 3.2124 | 6.1675 | 4.025 | 5.7839 | 4.3625 | 1.1138 |
| 41059 | ENSMUSG00000102876 | 0 | NA | NA | NA | NA | NA | Gm37136 | predicted gene, 37136 [Source:MGI Symb | 0 | 0 | 0 | 0 | 0 | 0 |
| 41060 | ENSMUSG00000102877 | 0 | NA | NA | NA | NA | NA | Gm38314 | predicted gene, 38314 [Source:MGI Symb | 0 | 0 | 0 | 0 | 0 | 0 |
| 41061 | ENSMUSG00000102878 | 0 | NA | NA | NA | NA | NA | Ighv2-8 | immunoglobulin heavy variable V2-8 [Sou | 0 | 0 | 0 | 0 | 0 | 0 |
| 41062 | ENSMUSG00000102879 | 0.17131964 | -0.796629176 | 4.080472857 | -0.195229623 | 0.84521319 | NA | Gm37135 | predicted gene, 37135 [Source:MGI Symb | 0 | 1.0279 | 0 | 0 | 0 | 0 |
| 41063 | ENSMUSG00000102880 | 0 | NA | NA | NA | NA | NA | 4930517J16I | RIKEN cDNA 4930517J16 gene [Source:M( | 0 | 0 | 0 | 0 | 0 | 0 |
| 41064 | ENSMUSG00000102881 | 11.20790367 | 0.165975694 | 0.791956377 | 0.209576813 | 0.833997978 | 0.971005543 | Gm3807 | predicted gene 3807 [Source:MGI Symbol | 18.471 | 2.0558 | 11.069 | 20.244 | 8.7249 | 6.6831 |
| 41065 | ENSMUSG00000102882 | 0 | NA | NA | NA | NA | NA | Gm2065 | predicted gene 2065 [Source:MGI Symbol | 0 | 0 | 0 | 0 | 0 | 0 |
| 41066 | ENSMUSG00000102883 | 0 | NA | NA | NA | NA | NA | Gm37279 | predicted gene, 37279 [Source:MGI Symb | 0 | 0 | 0 | 0 | 0 | 0 |
| 41067 | ENSMUSG00000102884 | 0 | NA | NA | NA | NA | NA | Gm37281 | predicted gene, 37281 [Source:MGI Symb | 0 | 0 | 0 | 0 | 0 | 0 |
| 41068 | ENSMUSG00000102885 | 0 | NA | NA | NA | NA | NA | Gm37280 | predicted gene, 37280 [Source:MGI Symb | 0 | 0 | 0 | 0 | 0 | 0 |
| 41069 | ENSMUSG00000102886 | 0 | NA | NA | NA | NA | NA | Gm30667 | predicted gene, 30667 [Source:MGI Symb | 0 | 0 | 0 | 0 | 0 | 0 |
| 41070 | ENSMUSG00000102887 | 0 | NA | NA | NA | NA | NA | Gm10857 | predicted gene 10857 [Source:MGI Symbo | 0 | 0 | 0 | 0 | 0 | 0 |
| 41071 | ENSMUSG00000102888 | 0 | NA | NA | NA | NA | NA | Ighv1-11 | immunoglobulin heavy variable V1-11 [So | 0 | 0 | 0 | 0 | 0 | 0 |
| 41072 | ENSMUSG00000102889 | 0 | NA | NA | NA | NA | NA | Gm37282 | predicted gene, 37282 [Source:MGI Symb | 0 | 0 | 0 | 0 | 0 | 0 |
| 41073 | ENSMUSG00000102890 | 0 | NA | NA | NA | NA | NA | Gm7626 | predicted gene 7626 [Source:MGI Symbol | 0 | 0 | 0 | 0 | 0 | 0 |
| 41074 | ENSMUSG00000102891 | 0 | NA | NA | NA | NA | NA | Gm19114 | predicted gene, 19114 [Source:MGI Symb | 0 | 0 | 0 | 0 | 0 | 0 |
| 41075 | ENSMUSG00000102892 | 0 | NA | NA | NA | NA | NA | Gm37854 | predicted gene, 37854 [Source:MGI Symb | 0 | 0 | 0 | 0 | 0 | 0 |
| 41076 | ENSMUSG00000102893 | 0.160662974 | 1.126924175 | 4.080472857 | 0.276174898 | 0.782413736 | NA | Gm37855 | predicted gene, 37855 [Source:MGI Symb | 0 | 0 | 0 | 0.964 | 0 | 0 |
| 41077 | ENSMUSG00000102894 | 0.160662974 | 1.126924175 | 4.080472857 | 0.276174898 | 0.782413736 | NA | Gm37851 | predicted gene, 37851 [Source:MGI Symb | 0 | 0 | 0 | 0.964 | 0 | 0 |
| 41078 | ENSMUSG00000102895 | 0 | NA | NA | NA | NA | NA | 5830415G21 | RIKEN cDNA 5830415G21 gene [Source:M | 0 | 0 | 0 | 0 | 0 | 0 |
| 41079 | ENSMUSG00000102896 | 0 | NA | NA | NA | NA | NA | Gm37852 | predicted gene, 37852 [Source:MGI Symb | 0 | 0 | 0 | 0 | 0 | 0 |
| 41080 | ENSMUSG00000102897 | 0 | NA | NA | NA | NA | NA | Gm37853 | predicted gene, 37853 [Source:MGI Symb | 0 | 0 | 0 | 0 | 0 | 0 |
| 41081 | ENSMUSG00000102899 | 5.074328449 | 1.346945996 | 0.98974775 | 1.360898266 | 0.173545842 | NA | Gm37850 | predicted gene, 37850 [Source:MGI Symb | 1.6062 | 2.0558 | 5.0312 | 9.6398 | 6.5437 | 5.5692 |
| 41082 | ENSMUSG00000102900 | 0.185640647 | 1.126924175 | 4.080472857 | 0.276174898 | 0.782413736 | NA | Gm37811 | predicted gene, 37811 [Source:MGI Symb | 0 | 0 | 0 | 0 | 0 | 1.1138 |
| 41083 | ENSMUSG00000102901 | 0 | NA | NA | NA | NA | NA | Ighv5-1 | immunoglobulin heavy variable V5-1 [Sou | 0 | 0 | 0 | 0 | 0 | 0 |
| 41084 | ENSMUSG00000102902 | 0 | NA | NA | NA | NA | NA | Gm38263 | predicted gene, 38263 [Source:MGI Symb | 0 | 0 | 0 | 0 | 0 | 0 |
| 41085 | ENSMUSG00000102903 | 0 | NA | NA | NA | NA | NA | Trbv11 | T cell receptor beta variable 11 [Source:M | 0 | 0 | 0 | 0 | 0 | 0 |
| 41086 | ENSMUSG00000102904 | 0 | NA | NA | NA | NA | NA | Gm38265 | predicted gene, 38265 [Source:MGI Symb | 0 | 0 | 0 | 0 | 0 | 0 |
| 41087 | ENSMUSG00000102905 | 0.787158936 | 0.679629554 | 2.65851484 | 0.255642566 | 0.798226848 | NA | Gm38266 | predicted gene, 38266 [Source:MGI Symb | 0.8031 | 1.0279 | 0 | 2.8919 | 0 | 0 |
| 41088 | ENSMUSG00000102906 | 0 | NA | NA | NA | NA | NA | Gm20858 | predicted gene, 20858 [Source:MGI Symb | 0 | 0 | 0 | 0 | 0 | 0 |
| 41089 | ENSMUSG00000102907 | 0 | NA | NA | NA | NA | NA | Gm38264 | predicted gene, 38264 [Source:MGI Symb | 0 | 0 | 0 | 0 | 0 | 0 |
| 41090 | ENSMUSG00000102908 | 0 | NA | NA | NA | NA | NA | Gm7558 | predicted gene 7558 [Source:MGI Symbol | 0 | 0 | 0 | 0 | 0 | 0 |
| 41091 | ENSMUSG00000102909 | 0 | NA | NA | NA | NA | NA | Gm2453 | predicted gene 2453 [Source:MGI Symbol | 0 | 0 | 0 | 0 | 0 | 0 |
| 41092 | ENSMUSG00000102910 | 0 | NA | NA | NA | NA | NA | Gm37722 | predicted gene, 37722 [Source:MGI Symb | 0 | 0 | 0 | 0 | 0 | 0 |
| 41093 | ENSMUSG00000102911 | 0 | NA | NA | NA | NA | NA | Gm38017 | predicted gene, 38017 [Source:MGI Symb | 0 | 0 | 0 | 0 | 0 | 0 |
| 41094 | ENSMUSG00000102912 | 0.845528352 | 3.296539104 | 2.749625613 | 1.198904712 | 0.230564999 | NA | Gm20731 | predicted gene, 20731 [Source:MGI Symb | 0 | 0 | 0 | 2.8919 | 2.1812 | 0 |
| 41095 | ENSMUSG00000102913 | 0 | NA | NA | NA | NA | NA | Gm37723 | predicted gene, 37723 [Source:MGI Symb | 0 | 0 | 0 | 0 | 0 | 0 |
| 41096 | ENSMUSG00000102914 | 0.17131964 | -0.796629176 | 4.080472857 | -0.195229623 | 0.84521319 | NA | Gm35106 | predicted gene, 35106 [Source:MGI Symb | 0 | 1.0279 | 0 | 0 | 0 | 0 |
| 41097 | ENSMUSG00000102915 | 0 | NA | NA | NA | NA | NA | Gm32460 | predicted gene, 32460 [Source:MGI Symb | 0 | 0 | 0 | 0 | 0 | 0 |
| 41098 | ENSMUSG00000102916 | 0 | NA | NA | NA | NA | NA | Gm37725 | predicted gene, 37725 [Source:MGI Symb | 0 | 0 | 0 | 0 | 0 | 0 |
| 41099 | ENSMUSG00000102917 | 0 | NA | NA | NA | NA | NA | Gm37724 | predicted gene, 37724 [Source:MGI Symb | 0 | 0 | 0 | 0 | 0 | 0 |
| 41100 | ENSMUSG00000102918 | 13.88398186 | -1.000003882 | 0.648659832 | -1.541646072 | 0.123159623 | 0.513662281 | Pcdhgc3 | protocadherin gamma subfamily C, 3 [Sou | 28.109 | 12.335 | 15.094 | 13.496 | 9.8156 | 4.4554 |

| | | | | | | | | | | | | | | | |
|---|---|---|---|---|---|---|---|---|---|---|---|---|---|---|---|
| 41101 | ENSMUSG00000102919 | 0.59968336 | -1.233555632 | 2.892386901 | -0.426483618 | 0.669755482 | NA | Gm37726 | predicted gene, 37726 [Source:MGI Symb | 1.6062 | 1.0279 | 0 | 0.964 | 0 | 0 |
| 41102 | ENSMUSG00000102920 | 0 | NA | NA | NA | NA | NA | Olfr416-ps1 | olfactory receptor 416, pseudogene 1 [So | 0 | 0 | 0 | 0 | 0 | 0 |
| 41103 | ENSMUSG00000102921 | 0 | NA | NA | NA | NA | NA | Gm37153 | predicted gene, 37153 [Source:MGI Symb | 0 | 0 | 0 | 0 | 0 | 0 |
| 41104 | ENSMUSG00000102922 | 0 | NA | NA | NA | NA | NA | Gm34176 | predicted gene, 34176 [Source:MGI Symb | 0 | 0 | 0 | 0 | 0 | 0 |
| 41105 | ENSMUSG00000102923 | 10.82504452 | 0.416081386 | 0.64743686 | 0.642659403 | 0.520445129 | 0.887802551 | Gm31728 | predicted gene, 31728 [Source:MGI Symb | 10.44 | 8.2233 | 9.0562 | 9.6398 | 11.997 | 15.594 |
| 41106 | ENSMUSG00000102925 | 0 | NA | NA | NA | NA | NA | Gm37150 | predicted gene, 37150 [Source:MGI Symb | 0 | 0 | 0 | 0 | 0 | 0 |
| 41107 | ENSMUSG00000102926 | 0 | NA | NA | NA | NA | NA | Gm37151 | predicted gene, 37151 [Source:MGI Symb | 0 | 0 | 0 | 0 | 0 | 0 |
| 41108 | ENSMUSG00000102927 | 0.880724006 | 0.715724162 | 2.808295084 | 0.25486074 | 0.79883066 | NA | Gm37152 | predicted gene, 37152 [Source:MGI Symb | 0 | 0 | 2.0125 | 0 | 3.2719 | 0 |
| 41109 | ENSMUSG00000102928 | 0 | NA | NA | NA | NA | NA | Ighv1-46 | immunoglobulin heavy variable V1-46 [So | 0 | 0 | 0 | 0 | 0 | 0 |
| 41110 | ENSMUSG00000102929 | 9.985698128 | 0.374433111 | 0.730936545 | 0.512264866 | 0.608465651 | 0.916376001 | Gm37154 | predicted gene, 37154 [Source:MGI Symb | 12.85 | 4.1117 | 9.0562 | 17.352 | 7.6343 | 8.9108 |
| 41111 | ENSMUSG00000102930 | 0.133850373 | -0.796629176 | 4.080472857 | -0.195229623 | 0.84521319 | NA | Gm38115 | predicted gene, 38115 [Source:MGI Symb | 0.8031 | 0 | 0 | 0 | 0 | 0 |
| 41112 | ENSMUSG00000102931 | 4.696385749 | -1.492048055 | 1.116736684 | -1.336078662 | 0.181523564 | NA | Gm38114 | predicted gene, 38114 [Source:MGI Symb | 9.6372 | 1.0279 | 10.062 | 1.928 | 2.1812 | 3.3415 |
| 41113 | ENSMUSG00000102932 | 0 | NA | NA | NA | NA | NA | Gm38113 | predicted gene, 38113 [Source:MGI Symb | 0 | 0 | 0 | 0 | 0 | 0 |
| 41114 | ENSMUSG00000102933 | 0 | NA | NA | NA | NA | NA | Gm9116 | predicted gene 9116 [Source:MGI Symbol | 0 | 0 | 0 | 0 | 0 | 0 |
| 41115 | ENSMUSG00000102934 | 0 | NA | NA | NA | NA | NA | Gm38117 | predicted gene, 38117 [Source:MGI Symb | 0 | 0 | 0 | 0 | 0 | 0 |
| 41116 | ENSMUSG00000102935 | 1.88843389 | 1.576470303 | 1.813128491 | 0.869475225 | 0.384587253 | NA | Gm38355 | predicted gene, 38355 [Source:MGI Symb | 0.8031 | 2.0558 | 0 | 1.928 | 6.5437 | 0 |
| 41117 | ENSMUSG00000102936 | 0 | NA | NA | NA | NA | NA | Gm35584 | predicted gene, 35584 [Source:MGI Symb | 0 | 0 | 0 | 0 | 0 | 0 |
| 41118 | ENSMUSG00000102937 | 0 | NA | NA | NA | NA | NA | Gm38116 | predicted gene, 38116 [Source:MGI Symb | 0 | 0 | 0 | 0 | 0 | 0 |
| 41119 | ENSMUSG00000102939 | 0 | NA | NA | NA | NA | NA | Gm38111 | predicted gene, 38111 [Source:MGI Symb | 0 | 0 | 0 | 0 | 0 | 0 |
| 41120 | ENSMUSG00000102940 | 0 | NA | NA | NA | NA | NA | Gm37551 | predicted gene, 37551 [Source:MGI Symb | 0 | 0 | 0 | 0 | 0 | 0 |
| 41121 | ENSMUSG00000102941 | 0 | NA | NA | NA | NA | NA | Gm37552 | predicted gene, 37552 [Source:MGI Symb | 0 | 0 | 0 | 0 | 0 | 0 |
| 41122 | ENSMUSG00000102942 | 0 | NA | NA | NA | NA | NA | Ighv1-33 | immunoglobulin heavy variable 1-33 [Sou | 0 | 0 | 0 | 0 | 0 | 0 |
| 41123 | ENSMUSG00000102943 | 0 | NA | NA | NA | NA | NA | Gm37550 | predicted gene, 37550 [Source:MGI Symb | 0 | 0 | 0 | 0 | 0 | 0 |
| 41124 | ENSMUSG00000102945 | 0 | NA | NA | NA | NA | NA | Gm37884 | predicted gene, 37884 [Source:MGI Symb | 0 | 0 | 0 | 0 | 0 | 0 |
| 41125 | ENSMUSG00000102946 | 0 | NA | NA | NA | NA | NA | Gm37883 | predicted gene, 37883 [Source:MGI Symb | 0 | 0 | 0 | 0 | 0 | 0 |
| 41126 | ENSMUSG00000102948 | 0 | NA | NA | NA | NA | NA | Gm6101 | predicted gene 6101 [Source:MGI Symbol | 0 | 0 | 0 | 0 | 0 | 0 |
| 41127 | ENSMUSG00000102949 | 0 | NA | NA | NA | NA | NA | Gm37882 | predicted gene, 37882 [Source:MGI Symb | 0 | 0 | 0 | 0 | 0 | 0 |
| 41128 | ENSMUSG00000102950 | 0 | NA | NA | NA | NA | NA | Gm37018 | predicted gene, 37018 [Source:MGI Symb | 0 | 0 | 0 | 0 | 0 | 0 |
| 41129 | ENSMUSG00000102951 | 0 | NA | NA | NA | NA | NA | Gm37216 | predicted gene, 37216 [Source:MGI Symb | 0 | 0 | 0 | 0 | 0 | 0 |
| 41130 | ENSMUSG00000102952 | 0 | NA | NA | NA | NA | NA | Ighv1-25 | immunoglobulin heavy variable 1-25 [Sou | 0 | 0 | 0 | 0 | 0 | 0 |
| 41131 | ENSMUSG00000102953 | 0.771003 | -1.69715066 | 2.640552792 | -0.642725518 | 0.52040222 | NA | Gm37019 | predicted gene, 37019 [Source:MGI Symb | 1.6062 | 2.0558 | 0 | 0.964 | 0 | 0 |
| 41132 | ENSMUSG00000102955 | 0 | NA | NA | NA | NA | NA | Trbv24 | T cell receptor beta, variable 24 [Source:M | 0 | 0 | 0 | 0 | 0 | 0 |
| 41133 | ENSMUSG00000102956 | 0 | NA | NA | NA | NA | NA | Gm37017 | predicted gene, 37017 [Source:MGI Symb | 0 | 0 | 0 | 0 | 0 | 0 |
| 41134 | ENSMUSG00000102957 | 0 | NA | NA | NA | NA | NA | Gm21065 | predicted gene, 21065 [Source:MGI Symb | 0 | 0 | 0 | 0 | 0 | 0 |
| 41135 | ENSMUSG00000102958 | 0.319491021 | 0.16514334 | 4.044653141 | 0.040830038 | 0.967431393 | NA | Gm37016 | predicted gene, 37016 [Source:MGI Symb | 0.8031 | 0 | 0 | 0 | 0 | 1.1138 |
| 41136 | ENSMUSG00000102959 | 0 | NA | NA | NA | NA | NA | Gm37015 | predicted gene, 37015 [Source:MGI Symb | 0 | 0 | 0 | 0 | 0 | 0 |
| 41137 | ENSMUSG00000102960 | 1.03119563 | 0.094051261 | 2.229235409 | 0.042189919 | 0.966347299 | NA | Gm37943 | predicted gene, 37943 [Source:MGI Symb | 0 | 0 | 3.0187 | 0.964 | 1.0906 | 1.1138 |
| 41138 | ENSMUSG00000102961 | 0 | NA | NA | NA | NA | NA | Gm19918 | predicted gene, 19918 [Source:MGI Symb | 0 | 0 | 0 | 0 | 0 | 0 |
| 41139 | ENSMUSG00000102962 | 0 | NA | NA | NA | NA | NA | Gm37944 | predicted gene, 37944 [Source:MGI Symb | 0 | 0 | 0 | 0 | 0 | 0 |
| 41140 | ENSMUSG00000102963 | 0 | NA | NA | NA | NA | NA | Gm37945 | predicted gene, 37945 [Source:MGI Symb | 0 | 0 | 0 | 0 | 0 | 0 |
| 41141 | ENSMUSG00000102964 | 7.721150603 | -0.544557621 | 0.768043767 | -0.709018996 | 0.478312688 | 0.869794311 | 9430034N14 | RIKEN cDNA 9430034N14 gene [Source:M | 7.2279 | 10.279 | 10.062 | 7.7118 | 4.3625 | 6.6831 |
| 41142 | ENSMUSG00000102965 | 0 | NA | NA | NA | NA | NA | Gm20926 | predicted gene, 20926 [Source:MGI Symb | 0 | 0 | 0 | 0 | 0 | 0 |
| 41143 | ENSMUSG00000102966 | 0 | NA | NA | NA | NA | NA | Gm31123 | predicted gene, 31123 [Source:MGI Symb | 0 | 0 | 0 | 0 | 0 | 0 |
| 41144 | ENSMUSG00000102967 | 0.321325949 | 1.911575114 | 4.048267893 | 0.472195804 | 0.636787034 | NA | Gm37946 | predicted gene, 37946 [Source:MGI Symb | 0 | 0 | 0 | 1.928 | 0 | 0 |
| 41145 | ENSMUSG00000102968 | 0 | NA | NA | NA | NA | NA | Gm35070 | predicted gene, 35070 [Source:MGI Symb | 0 | 0 | 0 | 0 | 0 | 0 |
| 41146 | ENSMUSG00000102969 | 0 | NA | NA | NA | NA | NA | Gm33366 | predicted gene, 33366 [Source:MGI Symb | 0 | 0 | 0 | 0 | 0 | 0 |
| 41147 | ENSMUSG00000102970 | 1.204653358 | -3.653034181 | 2.782110532 | -1.31304423 | 0.189168045 | NA | Gm37349 | predicted gene, 37349 [Source:MGI Symb | 7.2279 | 0 | 0 | 0 | 0 | 0 |
| 41148 | ENSMUSG00000102971 | 0 | NA | NA | NA | NA | NA | A730062M1 | RIKEN cDNA A730062M13 gene [Source:N | 0 | 0 | 0 | 0 | 0 | 0 |
| 41149 | ENSMUSG00000102972 | 0 | NA | NA | NA | NA | NA | Gm37348 | predicted gene, 37348 [Source:MGI Symb | 0 | 0 | 0 | 0 | 0 | 0 |
| 41150 | ENSMUSG00000102973 | 0 | NA | NA | NA | NA | NA | E430014B02 | RIKEN cDNA E430014B02 gene [Source:M | 0 | 0 | 0 | 0 | 0 | 0 |
| 41151 | ENSMUSG00000102975 | 0 | NA | NA | NA | NA | NA | Gm37347 | predicted gene, 37347 [Source:MGI Symb | 0 | 0 | 0 | 0 | 0 | 0 |
| 41152 | ENSMUSG00000102976 | 121.2259634 | -0.154542342 | 0.212541236 | -0.727116984 | 0.467154296 | 0.864254244 | Zc3h11a | zinc finger CCCH type containing 11A [Sou | 134.12 | 123.35 | 125.78 | 128.21 | 113.42 | 102.47 |
| 41153 | ENSMUSG00000102977 | 0 | NA | NA | NA | NA | NA | Gm37346 | predicted gene, 37346 [Source:MGI Symb | 0 | 0 | 0 | 0 | 0 | 0 |
| 41154 | ENSMUSG00000102979 | 0 | NA | NA | NA | NA | NA | Gm37350 | predicted gene, 37350 [Source:MGI Symb | 0 | 0 | 0 | 0 | 0 | 0 |
| 41155 | ENSMUSG00000102980 | 0 | NA | NA | NA | NA | NA | Gm38321 | predicted gene, 38321 [Source:MGI Symb | 0 | 0 | 0 | 0 | 0 | 0 |
| 41156 | ENSMUSG00000102981 | 0 | NA | NA | NA | NA | NA | Gm38322 | predicted gene, 38322 [Source:MGI Symb | 0 | 0 | 0 | 0 | 0 | 0 |

| | | | | | | | | | | | | | | | |
|---|---|---|---|---|---|---|---|---|---|---|---|---|---|---|---|
| 41157 | ENSMUSG00000102982 | 24.6288614 | -0.413170603 | 0.506612944 | -0.81555477 | 0.414754827 | 0.836001897 | Gm38319 | predicted gene, 38319 [Source:MGI Symb | 37.746 | 12.335 | 34.212 | 26.027 | 19.631 | 17.822 |
| 41158 | ENSMUSG00000102983 | 2.491400255 | -0.466158338 | 1.40958632 | -0.330705776 | 0.740866739 | NA | Gm38320 | predicted gene, 38320 [Source:MGI Symb | 1.6062 | 3.0838 | 4.025 | 2.8919 | 0 | 3.3415 |
| 41159 | ENSMUSG00000102984 | 0 | NA | NA | NA | NA | NA | Gm34727 | predicted gene, 34727 [Source:MGI Symb | 0 | 0 | 0 | 0 | 0 | 0 |
| 41160 | ENSMUSG00000102985 | 0.371281295 | 2.09253362 | 4.031852165 | 0.519000582 | 0.603760335 | NA | 4930456G14 | RIKEN cDNA 4930456G14 gene [Source:M | 0 | 0 | 0 | 0 | 0 | 2.2277 |
| 41161 | ENSMUSG00000102986 | 0 | NA | NA | NA | NA | NA | Gm38323 | predicted gene, 38323 [Source:MGI Symb | 0 | 0 | 0 | 0 | 0 | 0 |
| 41162 | ENSMUSG00000102987 | 0 | NA | NA | NA | NA | NA | Gm38324 | predicted gene, 38324 [Source:MGI Symb | 0 | 0 | 0 | 0 | 0 | 0 |
| 41163 | ENSMUSG00000102988 | 0 | NA | NA | NA | NA | NA | Trbv22 | T cell receptor beta variable 22 [Source:M | 0 | 0 | 0 | 0 | 0 | 0 |
| 41164 | ENSMUSG00000102989 | 0 | NA | NA | NA | NA | NA | Gm38318 | predicted gene, 38318 [Source:MGI Symb | 0 | 0 | 0 | 0 | 0 | 0 |
| 41165 | ENSMUSG00000102990 | 0 | NA | NA | NA | NA | NA | Gm37764 | predicted gene, 37764 [Source:MGI Symb | 0 | 0 | 0 | 0 | 0 | 0 |
| 41166 | ENSMUSG00000102991 | 0 | NA | NA | NA | NA | NA | Ighv1-40 | immunoglobulin heavy variable V1-40 [So | 0 | 0 | 0 | 0 | 0 | 0 |
| 41167 | ENSMUSG00000102992 | 0 | NA | NA | NA | NA | NA | Gm37766 | predicted gene, 37766 [Source:MGI Symb | 0 | 0 | 0 | 0 | 0 | 0 |
| 41168 | ENSMUSG00000102993 | 0 | NA | NA | NA | NA | NA | Gm37765 | predicted gene, 37765 [Source:MGI Symb | 0 | 0 | 0 | 0 | 0 | 0 |
| 41169 | ENSMUSG00000102994 | 0.371281295 | 2.09253362 | 4.031852165 | 0.519000582 | 0.603760335 | NA | Gm37767 | predicted gene, 37767 [Source:MGI Symb | 0 | 0 | 0 | 0 | 0 | 2.2277 |
| 41170 | ENSMUSG00000102995 | 0 | NA | NA | NA | NA | NA | A330074H02 | RIKEN cDNA A330074H02 gene [Source:M | 0 | 0 | 0 | 0 | 0 | 0 |
| 41171 | ENSMUSG00000102997 | 0.267700746 | -1.497481649 | 4.057001436 | -0.369110456 | 0.712045396 | NA | Gm17244 | predicted gene, 17244 [Source:MGI Symb | 1.6062 | 0 | 0 | 0 | 0 | 0 |
| 41172 | ENSMUSG00000102998 | 0 | NA | NA | NA | NA | NA | Gm37763 | predicted gene, 37763 [Source:MGI Symb | 0 | 0 | 0 | 0 | 0 | 0 |
| 41173 | ENSMUSG00000102999 | 0.535401493 | -2.48596965 | 3.684117979 | -0.674780141 | 0.499815459 | NA | Gm37762 | predicted gene, 37762 [Source:MGI Symb | 3.2124 | 0 | 0 | 0 | 0 | 0 |
| 41174 | ENSMUSG00000103000 | 0.160662974 | 1.126924175 | 4.080472857 | 0.276174898 | 0.782413736 | NA | Gm37900 | predicted gene, 37900 [Source:MGI Symb | 0 | 0 | 0 | 0.964 | 0 | 0 |
| 41175 | ENSMUSG00000103001 | 0.481988923 | 2.492401104 | 3.837077151 | 0.649557203 | 0.515978285 | NA | A930005N03 | RIKEN cDNA A930005N03 gene [Source:M | 0 | 0 | 0 | 2.8919 | 0 | 0 |
| 41176 | ENSMUSG00000103002 | 0 | NA | NA | NA | NA | NA | Gm38075 | predicted gene, 38075 [Source:MGI Symb | 0 | 0 | 0 | 0 | 0 | 0 |
| 41177 | ENSMUSG00000103003 | 0 | NA | NA | NA | NA | NA | Gm38076 | predicted gene, 38076 [Source:MGI Symb | 0 | 0 | 0 | 0 | 0 | 0 |
| 41178 | ENSMUSG00000103004 | 0 | NA | NA | NA | NA | NA | 2900002M20 | RIKEN cDNA 2900002M20 gene [Source:M | 0 | 0 | 0 | 0 | 0 | 0 |
| 41179 | ENSMUSG00000103005 | 0 | NA | NA | NA | NA | NA | Gm37898 | predicted gene, 37898 [Source:MGI Symb | 0 | 0 | 0 | 0 | 0 | 0 |
| 41180 | ENSMUSG00000103006 | 2.356851347 | 3.728811294 | 1.757419374 | 2.121753834 | 0.033858415 | NA | 4933417C20 | RIKEN cDNA 4933417C20 gene [Source:M | 0.8031 | 0 | 0 | 6.7478 | 4.3625 | 2.2277 |
| 41181 | ENSMUSG00000103007 | 0 | NA | NA | NA | NA | NA | Gm20690 | predicted gene 20690 [Source:MGI Symbo | 0 | 0 | 0 | 0 | 0 | 0 |
| 41182 | ENSMUSG00000103008 | 0 | NA | NA | NA | NA | NA | Gm36468 | predicted gene, 36468 [Source:MGI Symb | 0 | 0 | 0 | 0 | 0 | 0 |
| 41183 | ENSMUSG00000103009 | 15.476353 | 0.242753255 | 0.540245507 | 0.449338777 | 0.653187289 | 0.925756399 | Gm4430 | predicted gene 4430 [Source:MGI Symbol | 16.062 | 11.307 | 15.094 | 17.352 | 17.45 | 15.594 |
| 41184 | ENSMUSG00000103010 | 0 | NA | NA | NA | NA | NA | Gm37504 | predicted gene, 37504 [Source:MGI Symb | 0 | 0 | 0 | 0 | 0 | 0 |
| 41185 | ENSMUSG00000103011 | 0 | NA | NA | NA | NA | NA | Gm8860 | predicted gene 8860 [Source:MGI Symbol | 0 | 0 | 0 | 0 | 0 | 0 |
| 41186 | ENSMUSG00000103012 | 0 | NA | NA | NA | NA | NA | Gm37548 | predicted gene, 37548 [Source:MGI Symb | 0 | 0 | 0 | 0 | 0 | 0 |
| 41187 | ENSMUSG00000103013 | 2.046931078 | 0.011531897 | 1.551731724 | 0.00743163 | 0.994070471 | NA | Gm37503 | predicted gene, 37503 [Source:MGI Symb | 4.0155 | 2.0558 | 0 | 2.8919 | 1.0906 | 2.2277 |
| 41188 | ENSMUSG00000103015 | 0.371281295 | 2.09253362 | 4.031852165 | 0.519000582 | 0.603760335 | NA | Gm37549 | predicted gene, 37549 [Source:MGI Symb | 0 | 0 | 0 | 0 | 0 | 2.2277 |
| 41189 | ENSMUSG00000103016 | 0.181769715 | 1.126924175 | 4.080472857 | 0.276174898 | 0.782413736 | NA | Gm37506 | predicted gene, 37506 [Source:MGI Symb | 0 | 0 | 0 | 0 | 1.0906 | 0 |
| 41190 | ENSMUSG00000103017 | 0 | NA | NA | NA | NA | NA | Gm37505 | predicted gene, 37505 [Source:MGI Symb | 0 | 0 | 0 | 0 | 0 | 0 |
| 41191 | ENSMUSG00000103018 | 0 | NA | NA | NA | NA | NA | Gm17968 | predicted gene, 17968 [Source:MGI Symb | 0 | 0 | 0 | 0 | 0 | 0 |
| 41192 | ENSMUSG00000103020 | 0 | NA | NA | NA | NA | NA | Gm37681 | predicted gene, 37681 [Source:MGI Symb | 0 | 0 | 0 | 0 | 0 | 0 |
| 41193 | ENSMUSG00000103021 | 0 | NA | NA | NA | NA | NA | Gm37682 | predicted gene, 37682 [Source:MGI Symb | 0 | 0 | 0 | 0 | 0 | 0 |
| 41194 | ENSMUSG00000103023 | 0 | NA | NA | NA | NA | NA | Gm20910 | predicted gene, 20910 [Source:MGI Symb | 0 | 0 | 0 | 0 | 0 | 0 |
| 41195 | ENSMUSG00000103024 | 1.10411762 | -1.060312722 | 2.162010164 | -0.49042911 | 0.623830282 | NA | Gm37685 | predicted gene, 37685 [Source:MGI Symb | 2.4093 | 1.0279 | 1.0062 | 0 | 2.1812 | 0 |
| 41196 | ENSMUSG00000103025 | 0 | NA | NA | NA | NA | NA | Gm37686 | predicted gene, 37686 [Source:MGI Symb | 0 | 0 | 0 | 0 | 0 | 0 |
| 41197 | ENSMUSG00000103026 | 0 | NA | NA | NA | NA | NA | Gm31326 | predicted gene, 31326 [Source:MGI Symb | 0 | 0 | 0 | 0 | 0 | 0 |
| 41198 | ENSMUSG00000103027 | 0 | NA | NA | NA | NA | NA | Gm37684 | predicted gene, 37684 [Source:MGI Symb | 0 | 0 | 0 | 0 | 0 | 0 |
| 41199 | ENSMUSG00000103028 | 0 | NA | NA | NA | NA | NA | Gm8747 | predicted gene 8747 [Source:MGI Symbol | 0 | 0 | 0 | 0 | 0 | 0 |
| 41200 | ENSMUSG00000103029 | 0 | NA | NA | NA | NA | NA | Gm37687 | predicted gene, 37687 [Source:MGI Symb | 0 | 0 | 0 | 0 | 0 | 0 |
| 41201 | ENSMUSG00000103030 | 9.250922608 | 0.125592002 | 0.724488441 | 0.173352665 | 0.86237423 | 0.976334255 | E330011M16 | RIKEN cDNA E330011M16 gene [Source:M | 11.243 | 9.2513 | 6.0375 | 13.496 | 5.4531 | 10.025 |
| 41202 | ENSMUSG00000103031 | 0 | NA | NA | NA | NA | NA | Gm29950 | predicted gene, 29950 [Source:MGI Symb | 0 | 0 | 0 | 0 | 0 | 0 |
| 41203 | ENSMUSG00000103032 | 7.142459518 | 0.89407011 | 0.841279877 | 1.062749906 | 0.287895382 | 0.73951397 | Gm37121 | predicted gene, 37121 [Source:MGI Symb | 4.8186 | 3.0838 | 7.0437 | 4.8199 | 14.178 | 8.9108 |
| 41204 | ENSMUSG00000103033 | 0 | NA | NA | NA | NA | NA | Ighv5-12-4 | immunoglobulin heavy variable 5-12-4 [Sc | 0 | 0 | 0 | 0 | 0 | 0 |
| 41205 | ENSMUSG00000103034 | 3.801877057 | 0.006604947 | 1.140696382 | 0.005790276 | 0.995380054 | NA | Gm8797 | predicted pseudogene 8797 [Source:MGI | 7.2279 | 1.0279 | 3.0187 | 3.8559 | 5.4531 | 2.2277 |
| 41206 | ENSMUSG00000103035 | 0 | NA | NA | NA | NA | NA | Gm37122 | predicted gene, 37122 [Source:MGI Symb | 0 | 0 | 0 | 0 | 0 | 0 |
| 41207 | ENSMUSG00000103037 | 0.778047457 | -3.004661392 | 2.562427823 | -1.172583815 | 0.240962743 | NA | Pcdhgb1 | protocadherin gamma subfamily B, 1 [Sou | 1.6062 | 2.0558 | 1.0062 | 0 | 0 | 0 |
| 41208 | ENSMUSG00000103038 | 0 | NA | NA | NA | NA | NA | Gm37124 | predicted gene, 37124 [Source:MGI Symb | 0 | 0 | 0 | 0 | 0 | 0 |
| 41209 | ENSMUSG00000103039 | 17.41619583 | 0.137890926 | 0.507410703 | 0.271754074 | 0.78581112 | 0.960775321 | Gm37123 | predicted gene, 37123 [Source:MGI Symb | 15.259 | 15.419 | 19.119 | 18.316 | 17.45 | 18.935 |
| 41210 | ENSMUSG00000103041 | 57.85803724 | 0.44053986 | 0.310430756 | 1.419124402 | 0.155862751 | 0.574797088 | Gm37305 | predicted gene, 37305 [Source:MGI Symb | 54.611 | 53.452 | 39.244 | 81.938 | 59.984 | 57.92 |
| 41211 | ENSMUSG00000103042 | 0 | NA | NA | NA | NA | NA | Gm2719 | predicted pseudogene 2719 [Source:MGI | 0 | 0 | 0 | 0 | 0 | 0 |
| 41212 | ENSMUSG00000103043 | 0 | NA | NA | NA | NA | NA | Gm37306 | predicted gene, 37306 [Source:MGI Symb | 0 | 0 | 0 | 0 | 0 | 0 |

| | | | | | | | | | | | | | | | |
|---|---|---|---|---|---|---|---|---|---|---|---|---|---|---|---|
| 41213 | ENSMUSG00000103044 | 0 | NA | NA | NA | NA | NA | Gm37307 | predicted gene, 37307 [Source:MGI Symb | 0 | 0 | 0 | 0 | 0 | 0 |
| 41214 | ENSMUSG00000103045 | 0 | NA | NA | NA | NA | NA | Gm37308 | predicted gene, 37308 [Source:MGI Symb | 0 | 0 | 0 | 0 | 0 | 0 |
| 41215 | ENSMUSG00000103046 | 5.896560197 | -0.226148991 | 1.04957704 | -0.215466786 | 0.829403405 | 0.969955221 | Gm37309 | predicted gene, 37309 [Source:MGI Symb | 4.0155 | 4.1117 | 11.069 | 9.6398 | 6.5437 | 0 |
| 41216 | ENSMUSG00000103047 | 0 | NA | NA | NA | NA | NA | Gm37310 | predicted gene, 37310 [Source:MGI Symb | 0 | 0 | 0 | 0 | 0 | 0 |
| 41217 | ENSMUSG00000103049 | 0.346303622 | 2.004339653 | 3.884003423 | 0.516049919 | 0.605819544 | NA | Gm37311 | predicted gene, 37311 [Source:MGI Symb | 0 | 0 | 0 | 0.964 | 0 | 1.1138 |
| 41218 | ENSMUSG00000103050 | 0 | NA | NA | NA | NA | NA | Gm38273 | predicted gene, 38273 [Source:MGI Symb | 0 | 0 | 0 | 0 | 0 | 0 |
| 41219 | ENSMUSG00000103051 | 0 | NA | NA | NA | NA | NA | Gm7871 | predicted gene 7871 [Source:MGI Symbol | 0 | 0 | 0 | 0 | 0 | 0 |
| 41220 | ENSMUSG00000103052 | 0 | NA | NA | NA | NA | NA | Gm38272 | predicted gene, 38272 [Source:MGI Symb | 0 | 0 | 0 | 0 | 0 | 0 |
| 41221 | ENSMUSG00000103053 | 0 | NA | NA | NA | NA | NA | Gm38271 | predicted gene, 38271 [Source:MGI Symb | 0 | 0 | 0 | 0 | 0 | 0 |
| 41222 | ENSMUSG00000103054 | 0 | NA | NA | NA | NA | NA | Gm38270 | predicted gene, 38270 [Source:MGI Symb | 0 | 0 | 0 | 0 | 0 | 0 |
| 41223 | ENSMUSG00000103055 | 0 | NA | NA | NA | NA | NA | Gm38269 | predicted gene, 38269 [Source:MGI Symb | 0 | 0 | 0 | 0 | 0 | 0 |
| 41224 | ENSMUSG00000103056 | 0.85143529 | 2.085519423 | 2.590856022 | 0.804953809 | 0.420846336 | NA | Gm38268 | predicted gene, 38268 [Source:MGI Symb | 0.8031 | 0 | 0 | 0.964 | 0 | 3.3415 |
| 41225 | ENSMUSG00000103058 | 0 | NA | NA | NA | NA | NA | Gm21845 | predicted gene, 21845 [Source:MGI Symb | 0 | 0 | 0 | 0 | 0 | 0 |
| 41226 | ENSMUSG00000103059 | 0.133850373 | -0.796629176 | 4.080472857 | -0.195229623 | 0.84521319 | NA | Gm18186 | predicted gene, 18186 [Source:MGI Symb | 0.8031 | 0 | 0 | 0 | 0 | 0 |
| 41227 | ENSMUSG00000103060 | 0 | NA | NA | NA | NA | NA | Gm36969 | predicted gene, 36969 [Source:MGI Symb | 0 | 0 | 0 | 0 | 0 | 0 |
| 41228 | ENSMUSG00000103061 | 0 | NA | NA | NA | NA | NA | Gm36970 | predicted gene, 36970 [Source:MGI Symb | 0 | 0 | 0 | 0 | 0 | 0 |
| 41229 | ENSMUSG00000103062 | 1.306410535 | 3.92918701 | 2.440139424 | 1.610230535 | 0.107347538 | NA | Gm37200 | predicted gene, 37200 [Source:MGI Symb | 0 | 0 | 0 | 6.7478 | 1.0906 | 0 |
| 41230 | ENSMUSG00000103063 | 0 | NA | NA | NA | NA | NA | Gm33163 | predicted gene, 33163 [Source:MGI Symb | 0 | 0 | 0 | 0 | 0 | 0 |
| 41231 | ENSMUSG00000103064 | 0 | NA | NA | NA | NA | NA | Gm36968 | predicted gene, 36968 [Source:MGI Symb | 0 | 0 | 0 | 0 | 0 | 0 |
| 41232 | ENSMUSG00000103065 | 0 | NA | NA | NA | NA | NA | Gm20236 | predicted gene, 20236 [Source:MGI Symb | 0 | 0 | 0 | 0 | 0 | 0 |
| 41233 | ENSMUSG00000103066 | 0 | NA | NA | NA | NA | NA | Gm36967 | predicted gene, 36967 [Source:MGI Symb | 0 | 0 | 0 | 0 | 0 | 0 |
| 41234 | ENSMUSG00000103067 | 0 | NA | NA | NA | NA | NA | Gm30414 | predicted gene, 30414 [Source:MGI Symb | 0 | 0 | 0 | 0 | 0 | 0 |
| 41235 | ENSMUSG00000103069 | 0 | NA | NA | NA | NA | NA | Gm36966 | predicted gene, 36966 [Source:MGI Symb | 0 | 0 | 0 | 0 | 0 | 0 |
| 41236 | ENSMUSG00000103070 | 0 | NA | NA | NA | NA | NA | Gm37903 | predicted gene, 37903 [Source:MGI Symb | 0 | 0 | 0 | 0 | 0 | 0 |
| 41237 | ENSMUSG00000103071 | 0.545309144 | 2.651974629 | 3.666219369 | 0.72335405 | 0.469462394 | NA | Gm38110 | predicted gene, 38110 [Source:MGI Symb | 0 | 0 | 0 | 0 | 3.2719 | 0 |
| 41238 | ENSMUSG00000103072 | 0 | NA | NA | NA | NA | NA | Gm37905 | predicted gene, 37905 [Source:MGI Symb | 0 | 0 | 0 | 0 | 0 | 0 |
| 41239 | ENSMUSG00000103073 | 0 | NA | NA | NA | NA | NA | Gm37904 | predicted gene, 37904 [Source:MGI Symb | 0 | 0 | 0 | 0 | 0 | 0 |
| 41240 | ENSMUSG00000103074 | 0 | NA | NA | NA | NA | NA | Gm20925 | predicted gene, 20925 [Source:MGI Symb | 0 | 0 | 0 | 0 | 0 | 0 |
| 41241 | ENSMUSG00000103075 | 0.181769715 | 1.126924175 | 4.080472857 | 0.276174898 | 0.782413736 | NA | Gm38109 | predicted gene, 38109 [Source:MGI Symb | 0 | 0 | 0 | 0 | 1.0906 | 0 |
| 41242 | ENSMUSG00000103076 | 3.482979706 | 1.196899101 | 1.215046508 | 0.985064435 | 0.324592422 | NA | Gm37902 | predicted gene, 37902 [Source:MGI Symb | 3.2124 | 3.0838 | 0 | 5.7839 | 4.3625 | 4.4554 |
| 41243 | ENSMUSG00000103077 | 0 | NA | NA | NA | NA | NA | Gm38371 | predicted gene, 38371 [Source:MGI Symb | 0 | 0 | 0 | 0 | 0 | 0 |
| 41244 | ENSMUSG00000103078 | 0 | NA | NA | NA | NA | NA | Gm37901 | predicted gene, 37901 [Source:MGI Symb | 0 | 0 | 0 | 0 | 0 | 0 |
| 41245 | ENSMUSG00000103079 | 0.401551119 | -2.072982726 | 4.008251979 | -0.517178744 | 0.605031388 | NA | Gm36307 | predicted gene, 36307 [Source:MGI Symb | 2.4093 | 0 | 0 | 0 | 0 | 0 |
| 41246 | ENSMUSG00000103080 | 0 | NA | NA | NA | NA | NA | Gm38123 | predicted gene, 38123 [Source:MGI Symb | 0 | 0 | 0 | 0 | 0 | 0 |
| 41247 | ENSMUSG00000103081 | 0.658052777 | 1.6393978 | 2.811180186 | 0.583170658 | 0.559778421 | NA | Pcdhgb8 | protocadherin gamma subfamily B, 8 [Sou | 0.8031 | 0 | 0 | 0.964 | 2.1812 | 0 |
| 41248 | ENSMUSG00000103082 | 0 | NA | NA | NA | NA | NA | Gm38124 | predicted gene, 38124 [Source:MGI Symb | 0 | 0 | 0 | 0 | 0 | 0 |
| 41249 | ENSMUSG00000103083 | 0 | NA | NA | NA | NA | NA | A630035G10 | RIKEN cDNA A630035G10 gene [Source:M | 0 | 0 | 0 | 0 | 0 | 0 |
| 41250 | ENSMUSG00000103084 | 0 | NA | NA | NA | NA | NA | Gm38119 | predicted gene, 38119 [Source:MGI Symb | 0 | 0 | 0 | 0 | 0 | 0 |
| 41251 | ENSMUSG00000103085 | 1.329269052 | -1.494954196 | 1.953578141 | -0.765239007 | 0.444129236 | NA | Gm38120 | predicted gene, 38120 [Source:MGI Symb | 0.8031 | 4.1117 | 1.0062 | 0.964 | 1.0906 | 0 |
| 41252 | ENSMUSG00000103086 | 0 | NA | NA | NA | NA | NA | Gm38121 | predicted gene, 38121 [Source:MGI Symb | 0 | 0 | 0 | 0 | 0 | 0 |
| 41253 | ENSMUSG00000103087 | 0 | NA | NA | NA | NA | NA | Gm38122 | predicted gene, 38122 [Source:MGI Symb | 0 | 0 | 0 | 0 | 0 | 0 |
| 41254 | ENSMUSG00000103088 | 3.740747825 | 0.590405771 | 1.151604649 | 0.512680955 | 0.608174514 | NA | Pcdhgb6 | protocadherin gamma subfamily B, 6 [Sou | 4.8186 | 2.0558 | 2.0125 | 5.7839 | 1.0906 | 6.6831 |
| 41255 | ENSMUSG00000103089 | 0.674674027 | 1.600443604 | 2.78690385 | 0.574272989 | 0.565783085 | NA | Gm38118 | predicted gene, 38118 [Source:MGI Symb | 0 | 0 | 1.0062 | 1.928 | 0 | 1.1138 |
| 41256 | ENSMUSG00000103090 | 0 | NA | NA | NA | NA | NA | Gm19214 | predicted gene, 19214 [Source:MGI Symb | 0 | 0 | 0 | 0 | 0 | 0 |
| 41257 | ENSMUSG00000103091 | 0 | NA | NA | NA | NA | NA | Gm37555 | predicted gene, 37555 [Source:MGI Symb | 0 | 0 | 0 | 0 | 0 | 0 |
| 41258 | ENSMUSG00000103092 | 0 | NA | NA | NA | NA | NA | Pcdha5 | protocadherin alpha 5 [Source:MGI Symb | 0 | 0 | 0 | 0 | 0 | 0 |
| 41259 | ENSMUSG00000103094 | 0.979791906 | -2.106318065 | 2.490808728 | -0.845636215 | 0.397755716 | NA | Gm37558 | predicted gene, 37558 [Source:MGI Symb | 0.8031 | 4.1117 | 0 | 0.964 | 0 | 0 |
| 41260 | ENSMUSG00000103095 | 0 | NA | NA | NA | NA | NA | Gm37557 | predicted gene, 37557 [Source:MGI Symb | 0 | 0 | 0 | 0 | 0 | 0 |
| 41261 | ENSMUSG00000103096 | 0 | NA | NA | NA | NA | NA | Gm8022 | predicted gene 8022 [Source:MGI Symbol | 0 | 0 | 0 | 0 | 0 | 0 |
| 41262 | ENSMUSG00000103097 | 0 | NA | NA | NA | NA | NA | Gm37556 | predicted gene, 37556 [Source:MGI Symb | 0 | 0 | 0 | 0 | 0 | 0 |
| 41263 | ENSMUSG00000103098 | 0 | NA | NA | NA | NA | NA | Gm37559 | predicted gene, 37559 [Source:MGI Symb | 0 | 0 | 0 | 0 | 0 | 0 |
| 41264 | ENSMUSG00000103099 | 0 | NA | NA | NA | NA | NA | Gm6074 | predicted gene 6074 [Source:MGI Symbol | 0 | 0 | 0 | 0 | 0 | 0 |
| 41265 | ENSMUSG00000103100 | 0 | NA | NA | NA | NA | NA | Gm37664 | predicted gene, 37664 [Source:MGI Symb | 0 | 0 | 0 | 0 | 0 | 0 |
| 41266 | ENSMUSG00000103101 | 0.367410362 | 2.078255025 | 3.800070157 | 0.54689912 | 0.58444804 | NA | Gm37048 | predicted gene, 37048 [Source:MGI Symb | 0 | 0 | 0 | 0 | 1.0906 | 1.1138 |
| 41267 | ENSMUSG00000103102 | 0 | NA | NA | NA | NA | NA | Gm37049 | predicted gene, 37049 [Source:MGI Symb | 0 | 0 | 0 | 0 | 0 | 0 |
| 41268 | ENSMUSG00000103103 | 7.822016074 | 0.438727402 | 0.775034718 | 0.566074515 | 0.571343127 | 0.904651508 | 4833445I07R | RIKEN cDNA 4833445I07 gene [Source:MC | 9.6372 | 6.1675 | 4.025 | 10.604 | 9.8156 | 6.6831 |

| | | | | | | | | | | | | | | | |
|---|---|---|---|---|---|---|---|---|---|---|---|---|---|---|---|
| 41269 | ENSMUSG00000103105 | 0.17131964 | -0.796629176 | 4.080472857 | -0.195229623 | 0.84521319 | NA | Gm18752 | predicted gene, 18752 [Source:MGI Symb | 0 | 1.0279 | 0 | 0 | 0 | 0 |
| 41270 | ENSMUSG00000103106 | 0 | NA | NA | NA | NA | NA | Gm37047 | predicted gene, 37047 [Source:MGI Symb | 0 | 0 | 0 | 0 | 0 | 0 |
| 41271 | ENSMUSG00000103107 | 0 | NA | NA | NA | NA | NA | Gm9867 | predicted gene 9867 [Source:MGI Symbol | 0 | 0 | 0 | 0 | 0 | 0 |
| 41272 | ENSMUSG00000103108 | 0 | NA | NA | NA | NA | NA | Rncr4 | retina expressed non-coding RNA 4 [Sourc | 0 | 0 | 0 | 0 | 0 | 0 |
| 41273 | ENSMUSG00000103110 | 0 | NA | NA | NA | NA | NA | Gm37586 | predicted gene, 37586 [Source:MGI Symb | 0 | 0 | 0 | 0 | 0 | 0 |
| 41274 | ENSMUSG00000103111 | 0 | NA | NA | NA | NA | NA | Ighv6-1 | immunoglobulin heavy variable V6-1 [Sou | 0 | 0 | 0 | 0 | 0 | 0 |
| 41275 | ENSMUSG00000103112 | 0 | NA | NA | NA | NA | NA | Gm38016 | predicted gene, 38016 [Source:MGI Symb | 0 | 0 | 0 | 0 | 0 | 0 |
| 41276 | ENSMUSG00000103113 | 0 | NA | NA | NA | NA | NA | Gm37585 | predicted gene, 37585 [Source:MGI Symb | 0 | 0 | 0 | 0 | 0 | 0 |
| 41277 | ENSMUSG00000103114 | 1.284548738 | 0.928524427 | 2.006876397 | 0.462671457 | 0.64359988 | NA | Gm32200 | predicted gene, 32200 [Source:MGI Symb | 1.6062 | 1.0279 | 0 | 2.8919 | 2.1812 | 0 |
| 41278 | ENSMUSG00000103115 | 0.804680097 | -3.048447205 | 2.843448763 | -1.072095001 | 0.283677364 | NA | Gm37584 | predicted gene, 37584 [Source:MGI Symb | 0.8031 | 0 | 4.025 | 0 | 0 | 0 |
| 41279 | ENSMUSG00000103116 | 0 | NA | NA | NA | NA | NA | 4930539M1 | RIKEN cDNA 4930539M17 gene [Source:N | 0 | 0 | 0 | 0 | 0 | 0 |
| 41280 | ENSMUSG00000103118 | 0 | NA | NA | NA | NA | NA | Gm37582 | predicted gene, 37582 [Source:MGI Symb | 0 | 0 | 0 | 0 | 0 | 0 |
| 41281 | ENSMUSG00000103119 | 0 | NA | NA | NA | NA | NA | Gm37583 | predicted gene, 37583 [Source:MGI Symb | 0 | 0 | 0 | 0 | 0 | 0 |
| 41282 | ENSMUSG00000103120 | 0 | NA | NA | NA | NA | NA | Gm37392 | predicted gene, 37392 [Source:MGI Symb | 0 | 0 | 0 | 0 | 0 | 0 |
| 41283 | ENSMUSG00000103121 | 0 | NA | NA | NA | NA | NA | Ighv15-1 | immunoglobulin heavy variable V15-1 [So | 0 | 0 | 0 | 0 | 0 | 0 |
| 41284 | ENSMUSG00000103122 | 0.481988923 | 2.492401104 | 3.837077151 | 0.649557203 | 0.515978285 | NA | Gm37391 | predicted gene, 37391 [Source:MGI Symb | 0 | 0 | 0 | 2.8919 | 0 | 0 |
| 41285 | ENSMUSG00000103123 | 0 | NA | NA | NA | NA | NA | Gm37390 | predicted gene, 37390 [Source:MGI Symb | 0 | 0 | 0 | 0 | 0 | 0 |
| 41286 | ENSMUSG00000103124 | 0 | NA | NA | NA | NA | NA | Gm37389 | predicted gene, 37389 [Source:MGI Symb | 0 | 0 | 0 | 0 | 0 | 0 |
| 41287 | ENSMUSG00000103125 | 0 | NA | NA | NA | NA | NA | Gm37388 | predicted gene, 37388 [Source:MGI Symb | 0 | 0 | 0 | 0 | 0 | 0 |
| 41288 | ENSMUSG00000103126 | 0.160662974 | 1.126924175 | 4.080472857 | 0.276174898 | 0.782413736 | NA | NA | NA | 0 | 0 | 0 | 0.964 | 0 | 0 |
| 41289 | ENSMUSG00000103127 | 0 | NA | NA | NA | NA | NA | Gm37386 | predicted gene, 37386 [Source:MGI Symb | 0 | 0 | 0 | 0 | 0 | 0 |
| 41290 | ENSMUSG00000103129 | 0 | NA | NA | NA | NA | NA | Gm37393 | predicted gene, 37393 [Source:MGI Symb | 0 | 0 | 0 | 0 | 0 | 0 |
| 41291 | ENSMUSG00000103130 | 0 | NA | NA | NA | NA | NA | F830212C03 | RIKEN cDNA F830212C03 gene [Source:M | 0 | 0 | 0 | 0 | 0 | 0 |
| 41292 | ENSMUSG00000103131 | 0 | NA | NA | NA | NA | NA | Gm37977 | predicted gene, 37977 [Source:MGI Symb | 0 | 0 | 0 | 0 | 0 | 0 |
| 41293 | ENSMUSG00000103132 | 0.331982614 | 0.16514309 | 3.986751476 | 0.041422971 | 0.966958701 | NA | Gm37978 | predicted gene, 37978 [Source:MGI Symb | 0 | 1.0279 | 0 | 0.964 | 0 | 0 |
| 41294 | ENSMUSG00000103133 | 4.795447547 | 0.279985776 | 0.966000036 | 0.289840337 | 0.771938387 | NA | Gm37303 | predicted gene, 37303 [Source:MGI Symb | 4.8186 | 5.1396 | 3.0187 | 4.8199 | 7.6343 | 3.3415 |
| 41295 | ENSMUSG00000103135 | 1.975326567 | 1.067103237 | 1.758261924 | 0.606908005 | 0.543911965 | NA | Gm37979 | predicted gene, 37979 [Source:MGI Symb | 0.8031 | 3.0838 | 0 | 5.7839 | 2.1812 | 0 |
| 41296 | ENSMUSG00000103136 | 0 | NA | NA | NA | NA | NA | Gm37980 | predicted gene, 37980 [Source:MGI Symb | 0 | 0 | 0 | 0 | 0 | 0 |
| 41297 | ENSMUSG00000103137 | 0 | NA | NA | NA | NA | NA | Gm37981 | predicted gene, 37981 [Source:MGI Symb | 0 | 0 | 0 | 0 | 0 | 0 |
| 41298 | ENSMUSG00000103138 | 2.366680822 | 0.070846062 | 1.510342815 | 0.046907273 | 0.962587132 | NA | Gm2238 | predicted gene 2238 [Source:MGI Symbol | 4.8186 | 1.0279 | 1.0062 | 2.8919 | 0 | 4.4554 |
| 41299 | ENSMUSG00000103139 | 0 | NA | NA | NA | NA | NA | Gm37982 | predicted gene, 37982 [Source:MGI Symb | 0 | 0 | 0 | 0 | 0 | 0 |
| 41300 | ENSMUSG00000103140 | 0 | NA | NA | NA | NA | NA | Gm37495 | predicted gene, 37495 [Source:MGI Symb | 0 | 0 | 0 | 0 | 0 | 0 |
| 41301 | ENSMUSG00000103141 | 0 | NA | NA | NA | NA | NA | Gm37741 | predicted gene, 37741 [Source:MGI Symb | 0 | 0 | 0 | 0 | 0 | 0 |
| 41302 | ENSMUSG00000103142 | 0 | NA | NA | NA | NA | NA | Gm37743 | predicted gene, 37743 [Source:MGI Symb | 0 | 0 | 0 | 0 | 0 | 0 |
| 41303 | ENSMUSG00000103143 | 0.542600935 | 1.125931187 | 3.335923981 | 0.33751704 | 0.735727169 | NA | Gm37742 | predicted gene, 37742 [Source:MGI Symb | 0 | 1.0279 | 0 | 0 | 0 | 2.2277 |
| 41304 | ENSMUSG00000103144 | 1.206359572 | 2.624312633 | 2.124387999 | 1.235326425 | 0.216709043 | NA | Pcdhga1 | protocadherin gamma subfamily A, 1 [Sou | 0 | 1.0279 | 0 | 2.8919 | 1.0906 | 2.2277 |
| 41305 | ENSMUSG00000103145 | 0 | NA | NA | NA | NA | NA | Gm37744 | predicted gene, 37744 [Source:MGI Symb | 0 | 0 | 0 | 0 | 0 | 0 |
| 41306 | ENSMUSG00000103146 | 0 | NA | NA | NA | NA | NA | Gm37745 | predicted gene, 37745 [Source:MGI Symb | 0 | 0 | 0 | 0 | 0 | 0 |
| 41307 | ENSMUSG00000103147 | 0 | NA | NA | NA | NA | NA | Gm7341 | predicted gene 7341 [Source:MGI Symbol | 0 | 0 | 0 | 0 | 0 | 0 |
| 41308 | ENSMUSG00000103148 | 0 | NA | NA | NA | NA | NA | Gm37746 | predicted gene, 37746 [Source:MGI Symb | 0 | 0 | 0 | 0 | 0 | 0 |
| 41309 | ENSMUSG00000103149 | 12.68091883 | 0.612426161 | 0.606764733 | 1.009330515 | 0.31281615 | 0.759921531 | AA914427 | EST AA914427 [Source:MGI Symbol;Acc:N | 13.653 | 8.2233 | 8.05 | 16.388 | 14.178 | 15.594 |
| 41310 | ENSMUSG00000103150 | 0 | NA | NA | NA | NA | NA | Gm38291 | predicted gene, 38291 [Source:MGI Symb | 0 | 0 | 0 | 0 | 0 | 0 |
| 41311 | ENSMUSG00000103151 | 0 | NA | NA | NA | NA | NA | Gm38292 | predicted gene, 38292 [Source:MGI Symb | 0 | 0 | 0 | 0 | 0 | 0 |
| 41312 | ENSMUSG00000103152 | 0 | NA | NA | NA | NA | NA | Gm8790 | predicted gene 8790 [Source:MGI Symbol | 0 | 0 | 0 | 0 | 0 | 0 |
| 41313 | ENSMUSG00000103153 | 0 | NA | NA | NA | NA | NA | Gm38290 | predicted gene, 38290 [Source:MGI Symb | 0 | 0 | 0 | 0 | 0 | 0 |
| 41314 | ENSMUSG00000103154 | 0 | NA | NA | NA | NA | NA | Gm38295 | predicted gene, 38295 [Source:MGI Symb | 0 | 0 | 0 | 0 | 0 | 0 |
| 41315 | ENSMUSG00000103155 | 0 | NA | NA | NA | NA | NA | Gm8109 | predicted gene 8109 [Source:MGI Symbol | 0 | 0 | 0 | 0 | 0 | 0 |
| 41316 | ENSMUSG00000103156 | 1.238329568 | -0.301762013 | 2.276277377 | -0.132568208 | 0.894534877 | NA | Gm38293 | predicted gene, 38293 [Source:MGI Symb | 0 | 4.1117 | 0 | 0 | 1.0906 | 2.2277 |
| 41317 | ENSMUSG00000103157 | 0 | NA | NA | NA | NA | NA | Gm38294 | predicted gene, 38294 [Source:MGI Symb | 0 | 0 | 0 | 0 | 0 | 0 |
| 41318 | ENSMUSG00000103158 | 0 | NA | NA | NA | NA | NA | Gm31925 | predicted gene, 31925 [Source:MGI Symb | 0 | 0 | 0 | 0 | 0 | 0 |
| 41319 | ENSMUSG00000103159 | 0 | NA | NA | NA | NA | NA | F830112A20 | RIKEN cDNA F830112A20 gene [Source:M | 0 | 0 | 0 | 0 | 0 | 0 |
| 41320 | ENSMUSG00000103160 | 1.777220956 | -0.71010784 | 1.634042197 | -0.434571299 | 0.663873632 | NA | C130012C08 | RIKEN cDNA C130012C08 gene [Source:M | 1.6062 | 2.0558 | 3.0187 | 2.8919 | 1.0906 | 0 |
| 41321 | ENSMUSG00000103161 | 0 | NA | NA | NA | NA | NA | Gm38148 | predicted gene, 38148 [Source:MGI Symb | 0 | 0 | 0 | 0 | 0 | 0 |
| 41322 | ENSMUSG00000103162 | 0.918735696 | -1.977544565 | 2.489980631 | -0.794200782 | 0.427078553 | NA | Gm38147 | predicted gene, 38147 [Source:MGI Symb | 2.4093 | 0 | 2.0125 | 0 | 1.0906 | 0 |
| 41323 | ENSMUSG00000103163 | 0 | NA | NA | NA | NA | NA | Gm38146 | predicted gene, 38146 [Source:MGI Symb | 0 | 0 | 0 | 0 | 0 | 0 |
| 41324 | ENSMUSG00000103164 | 0.453341394 | -0.545391191 | 3.551113277 | -0.153583158 | 0.877938417 | NA | Gm38150 | predicted gene, 38150 [Source:MGI Symb | 1.6062 | 0 | 0 | 0 | 0 | 1.1138 |

| | | | | | | | | | | | | | | | |
|---|---|---|---|---|---|---|---|---|---|---|---|---|---|---|---|
| 41325 | ENSMUSG00000103165 | 0 | NA | NA | NA | NA | NA | Gm37560 | predicted gene, 37560 [Source:MGI Symb | 0 | 0 | 0 | 0 | 0 | 0 |
| 41326 | ENSMUSG00000103166 | 0 | NA | NA | NA | NA | NA | Gm10537 | predicted gene 10537 [Source:MGI Symbc | 0 | 0 | 0 | 0 | 0 | 0 |
| 41327 | ENSMUSG00000103167 | 2.121993879 | 2.438194164 | 1.802586429 | 1.352608743 | 0.176180658 | NA | Gm38149 | predicted gene, 38149 [Source:MGI Symb | 0 | 2.0558 | 0 | 1.928 | 7.6343 | 1.1138 |
| 41328 | ENSMUSG00000103168 | 0 | NA | NA | NA | NA | NA | Gm30948 | predicted gene, 30948 [Source:MGI Symb | 0 | 0 | 0 | 0 | 0 | 0 |
| 41329 | ENSMUSG00000103170 | 0 | NA | NA | NA | NA | NA | Gm37170 | predicted gene, 37170 [Source:MGI Symb | 0 | 0 | 0 | 0 | 0 | 0 |
| 41330 | ENSMUSG00000103172 | 0 | NA | NA | NA | NA | NA | Gm37148 | predicted gene, 37148 [Source:MGI Symb | 0 | 0 | 0 | 0 | 0 | 0 |
| 41331 | ENSMUSG00000103174 | 0 | NA | NA | NA | NA | NA | Gm37168 | predicted gene, 37168 [Source:MGI Symb | 0 | 0 | 0 | 0 | 0 | 0 |
| 41332 | ENSMUSG00000103175 | 0.301557804 | -1.64335916 | 4.042814367 | -0.406488899 | 0.684383412 | NA | Gm37169 | predicted gene, 37169 [Source:MGI Symb | 0.8031 | 0 | 1.0062 | 0 | 0 | 0 |
| 41333 | ENSMUSG00000103176 | 0 | NA | NA | NA | NA | NA | Gm37147 | predicted gene, 37147 [Source:MGI Symb | 0 | 0 | 0 | 0 | 0 | 0 |
| 41334 | ENSMUSG00000103177 | 0 | NA | NA | NA | NA | NA | Gm18427 | predicted gene, 18427 [Source:MGI Symb | 0 | 0 | 0 | 0 | 0 | 0 |
| 41335 | ENSMUSG00000103178 | 0 | NA | NA | NA | NA | NA | Fgfr3-ps | fibroblast growth factor receptor 3, pseuc | 0 | 0 | 0 | 0 | 0 | 0 |
| 41336 | ENSMUSG00000103179 | 0 | NA | NA | NA | NA | NA | Gm37171 | predicted gene, 37171 [Source:MGI Symb | 0 | 0 | 0 | 0 | 0 | 0 |
| 41337 | ENSMUSG00000103180 | 0 | NA | NA | NA | NA | NA | Gm37089 | predicted gene, 37089 [Source:MGI Symb | 0 | 0 | 0 | 0 | 0 | 0 |
| 41338 | ENSMUSG00000103181 | 0 | NA | NA | NA | NA | NA | A330069K06 | RIKEN cDNA A330069K06 gene [Source:M | 0 | 0 | 0 | 0 | 0 | 0 |
| 41339 | ENSMUSG00000103182 | 0.167707431 | -0.796629176 | 4.080472857 | -0.195229623 | 0.84521319 | NA | Gm37091 | predicted gene, 37091 [Source:MGI Symb | 0 | 0 | 1.0062 | 0 | 0 | 0 |
| 41340 | ENSMUSG00000103183 | 33.27704132 | 0.4308796 | 0.395899663 | 1.088355563 | 0.27643817 | 0.730038567 | Gm37090 | predicted gene, 37090 [Source:MGI Symb | 36.943 | 26.726 | 21.131 | 44.343 | 35.99 | 34.529 |
| 41341 | ENSMUSG00000103184 | 0 | NA | NA | NA | NA | NA | Ighv5-13 | immunoglobulin heavy variable V5-13 [So | 0 | 0 | 0 | 0 | 0 | 0 |
| 41342 | ENSMUSG00000103185 | 0 | NA | NA | NA | NA | NA | Ighv5-11 | immunoglobulin heavy variable V5-11 [So | 0 | 0 | 0 | 0 | 0 | 0 |
| 41343 | ENSMUSG00000103186 | 0 | NA | NA | NA | NA | NA | Gm37088 | predicted gene, 37088 [Source:MGI Symb | 0 | 0 | 0 | 0 | 0 | 0 |
| 41344 | ENSMUSG00000103187 | 0 | NA | NA | NA | NA | NA | Gm37087 | predicted gene, 37087 [Source:MGI Symb | 0 | 0 | 0 | 0 | 0 | 0 |
| 41345 | ENSMUSG00000103188 | 0 | NA | NA | NA | NA | NA | Gm37093 | predicted gene, 37093 [Source:MGI Symb | 0 | 0 | 0 | 0 | 0 | 0 |
| 41346 | ENSMUSG00000103189 | 0.695522043 | 1.639439662 | 2.762392557 | 0.593485404 | 0.552856349 | NA | Gm37092 | predicted gene, 37092 [Source:MGI Symb | 0 | 1.0279 | 0 | 0.964 | 2.1812 | 0 |
| 41347 | ENSMUSG00000103190 | 0 | NA | NA | NA | NA | NA | Gm34716 | predicted gene, 34716 [Source:MGI Symb | 0 | 0 | 0 | 0 | 0 | 0 |
| 41348 | ENSMUSG00000103191 | 0.335414862 | -1.779047242 | 4.030804179 | -0.441362855 | 0.658950329 | NA | Gm37646 | predicted gene, 37646 [Source:MGI Symb | 0 | 0 | 2.0125 | 0 | 0 | 0 |
| 41349 | ENSMUSG00000103192 | 0 | NA | NA | NA | NA | NA | Gm37645 | predicted gene, 37645 [Source:MGI Symb | 0 | 0 | 0 | 0 | 0 | 0 |
| 41350 | ENSMUSG00000103193 | 0 | NA | NA | NA | NA | NA | Gm19066 | predicted gene, 19066 [Source:MGI Symb | 0 | 0 | 0 | 0 | 0 | 0 |
| 41351 | ENSMUSG00000103194 | 0 | NA | NA | NA | NA | NA | Gm37643 | predicted gene, 37643 [Source:MGI Symb | 0 | 0 | 0 | 0 | 0 | 0 |
| 41352 | ENSMUSG00000103195 | 0 | NA | NA | NA | NA | NA | Gm37644 | predicted gene, 37644 [Source:MGI Symb | 0 | 0 | 0 | 0 | 0 | 0 |
| 41353 | ENSMUSG00000103196 | 0 | NA | NA | NA | NA | NA | Gm37641 | predicted gene, 37641 [Source:MGI Symb | 0 | 0 | 0 | 0 | 0 | 0 |
| 41354 | ENSMUSG00000103197 | 0 | NA | NA | NA | NA | NA | Gm37642 | predicted gene, 37642 [Source:MGI Symb | 0 | 0 | 0 | 0 | 0 | 0 |
| 41355 | ENSMUSG00000103198 | 0 | NA | NA | NA | NA | NA | Gm9136 | predicted gene 9136 [Source:MGI Symbol | 0 | 0 | 0 | 0 | 0 | 0 |
| 41356 | ENSMUSG00000103199 | 0.181769715 | 1.126924175 | 4.080472857 | 0.276174898 | 0.782413736 | NA | Gm37648 | predicted gene, 37648 [Source:MGI Symb | 0 | 0 | 0 | 0 | 1.0906 | 0 |
| 41357 | ENSMUSG00000103200 | 1.705352209 | 0.640255105 | 1.797882078 | 0.356116295 | 0.721753474 | NA | Gm37328 | predicted gene, 37328 [Source:MGI Symb | 0 | 2.0558 | 2.0125 | 2.8919 | 3.2719 | 0 |
| 41358 | ENSMUSG00000103201 | 0 | NA | NA | NA | NA | NA | Gm37329 | predicted gene, 37329 [Source:MGI Symb | 0 | 0 | 0 | 0 | 0 | 0 |
| 41359 | ENSMUSG00000103202 | 0 | NA | NA | NA | NA | NA | Gm37326 | predicted gene, 37326 [Source:MGI Symb | 0 | 0 | 0 | 0 | 0 | 0 |
| 41360 | ENSMUSG00000103203 | 0 | NA | NA | NA | NA | NA | Gm37327 | predicted gene, 37327 [Source:MGI Symb | 0 | 0 | 0 | 0 | 0 | 0 |
| 41361 | ENSMUSG00000103204 | 0 | NA | NA | NA | NA | NA | Gm38139 | predicted gene, 38139 [Source:MGI Symb | 0 | 0 | 0 | 0 | 0 | 0 |
| 41362 | ENSMUSG00000103206 | 0 | NA | NA | NA | NA | NA | Gm8515 | predicted gene 8515 [Source:MGI Symbol | 0 | 0 | 0 | 0 | 0 | 0 |
| 41363 | ENSMUSG00000103207 | 0.321325949 | 1.911575114 | 4.048267893 | 0.472195804 | 0.636787034 | NA | Gm9874 | predicted gene 9874 [Source:MGI Symbol | 0 | 0 | 0 | 1.928 | 0 | 0 |
| 41364 | ENSMUSG00000103208 | 0 | NA | NA | NA | NA | NA | Gm37330 | predicted gene, 37330 [Source:MGI Symb | 0 | 0 | 0 | 0 | 0 | 0 |
| 41365 | ENSMUSG00000103209 | 0 | NA | NA | NA | NA | NA | Gm37331 | predicted gene, 37331 [Source:MGI Symb | 0 | 0 | 0 | 0 | 0 | 0 |
| 41366 | ENSMUSG00000103210 | 0 | NA | NA | NA | NA | NA | Gm5852 | predicted gene 5852 [Source:MGI Symbol | 0 | 0 | 0 | 0 | 0 | 0 |
| 41367 | ENSMUSG00000103211 | 0 | NA | NA | NA | NA | NA | Gm38288 | predicted gene, 38288 [Source:MGI Symb | 0 | 0 | 0 | 0 | 0 | 0 |
| 41368 | ENSMUSG00000103213 | 0 | NA | NA | NA | NA | NA | Gm38289 | predicted gene, 38289 [Source:MGI Symb | 0 | 0 | 0 | 0 | 0 | 0 |
| 41369 | ENSMUSG00000103214 | 0 | NA | NA | NA | NA | NA | Gm38286 | predicted gene, 38286 [Source:MGI Symb | 0 | 0 | 0 | 0 | 0 | 0 |
| 41370 | ENSMUSG00000103215 | 3.442131451 | -0.229840532 | 1.11662709 | -0.205834637 | 0.836920084 | NA | Gm38388 | predicted gene, 38388 [Source:MGI Symb | 4.0155 | 3.0838 | 4.025 | 2.8919 | 2.1812 | 4.4554 |
| 41371 | ENSMUSG00000103216 | 0 | NA | NA | NA | NA | NA | Gm37248 | predicted gene, 37248 [Source:MGI Symb | 0 | 0 | 0 | 0 | 0 | 0 |
| 41372 | ENSMUSG00000103217 | 0 | NA | NA | NA | NA | NA | Gm38287 | predicted gene, 38287 [Source:MGI Symb | 0 | 0 | 0 | 0 | 0 | 0 |
| 41373 | ENSMUSG00000103218 | 0 | NA | NA | NA | NA | NA | Gm7270 | predicted gene 7270 [Source:MGI Symbol | 0 | 0 | 0 | 0 | 0 | 0 |
| 41374 | ENSMUSG00000103219 | 0 | NA | NA | NA | NA | NA | Gm37787 | predicted gene, 37787 [Source:MGI Symb | 0 | 0 | 0 | 0 | 0 | 0 |
| 41375 | ENSMUSG00000103220 | 0.331982614 | 0.16514309 | 3.986751476 | 0.041422971 | 0.966958701 | NA | Gm37728 | predicted gene, 37728 [Source:MGI Symb | 0 | 1.0279 | 0 | 0.964 | 0 | 0 |
| 41376 | ENSMUSG00000103221 | 0 | NA | NA | NA | NA | NA | Sprr2c-ps | small proline-rich protein 2C, pseudogene | 0 | 0 | 0 | 0 | 0 | 0 |
| 41377 | ENSMUSG00000103222 | 78.2103875 | -0.118389223 | 0.279476282 | -0.423610986 | 0.671849542 | 0.931191575 | Gm37729 | predicted gene, 37729 [Source:MGI Symb | 98.782 | 66.815 | 78.487 | 90.614 | 62.165 | 72.4 |
| 41378 | ENSMUSG00000103223 | 0.181769715 | 1.126924175 | 4.080472857 | 0.276174898 | 0.782413736 | NA | Gm37730 | predicted gene, 37730 [Source:MGI Symb | 0 | 0 | 0 | 0 | 1.0906 | 0 |
| 41379 | ENSMUSG00000103224 | 0 | NA | NA | NA | NA | NA | Gm2077 | predicted gene 2077 [Source:MGI Symbol | 0 | 0 | 0 | 0 | 0 | 0 |
| 41380 | ENSMUSG00000103225 | 0 | NA | NA | NA | NA | NA | Gm37731 | predicted gene, 37731 [Source:MGI Symb | 0 | 0 | 0 | 0 | 0 | 0 |

| | | | | | | | | | | | | | | | |
|---|---|---|---|---|---|---|---|---|---|---|---|---|---|---|---|
| 41381 | ENSMUSG00000103226 | 0 | NA | NA | NA | NA | NA | Gm37732 | predicted gene, 37732 [Source:MGI Symb | 0 | 0 | 0 | 0 | 0 | 0 |
| 41382 | ENSMUSG00000103227 | 0.17131964 | -0.796629176 | 4.080472857 | -0.195229623 | 0.84521319 | NA | Gm37733 | predicted gene, 37733 [Source:MGI Symb | 0 | 1.0279 | 0 | 0 | 0 | 0 |
| 41383 | ENSMUSG00000103228 | 0 | NA | NA | NA | NA | NA | Gm37727 | predicted gene, 37727 [Source:MGI Symb | 0 | 0 | 0 | 0 | 0 | 0 |
| 41384 | ENSMUSG00000103230 | 0 | NA | NA | NA | NA | NA | Gm17970 | predicted gene, 17970 [Source:MGI Symb | 0 | 0 | 0 | 0 | 0 | 0 |
| 41385 | ENSMUSG00000103231 | 0 | NA | NA | NA | NA | NA | Gm21907 | predicted gene, 21907 [Source:MGI Symb | 0 | 0 | 0 | 0 | 0 | 0 |
| 41386 | ENSMUSG00000103232 | 0.17131964 | -0.796629176 | 4.080472857 | -0.195229623 | 0.84521319 | NA | Gm37160 | predicted gene, 37160 [Source:MGI Symb | 0 | 1.0279 | 0 | 0 | 0 | 0 |
| 41387 | ENSMUSG00000103233 | 0 | NA | NA | NA | NA | NA | Gm37159 | predicted gene, 37159 [Source:MGI Symb | 0 | 0 | 0 | 0 | 0 | 0 |
| 41388 | ENSMUSG00000103234 | 0 | NA | NA | NA | NA | NA | Gm37158 | predicted gene, 37158 [Source:MGI Symb | 0 | 0 | 0 | 0 | 0 | 0 |
| 41389 | ENSMUSG00000103236 | 0 | NA | NA | NA | NA | NA | Gm37157 | predicted gene, 37157 [Source:MGI Symb | 0 | 0 | 0 | 0 | 0 | 0 |
| 41390 | ENSMUSG00000103237 | 0 | NA | NA | NA | NA | NA | Gm18547 | predicted gene, 18547 [Source:MGI Symb | 0 | 0 | 0 | 0 | 0 | 0 |
| 41391 | ENSMUSG00000103238 | 1.332881261 | -1.503897601 | 2.106880042 | -0.713803146 | 0.475348914 | NA | Gm36527 | predicted gene, 36527 [Source:MGI Symb | 0.8031 | 5.1396 | 0 | 0.964 | 1.0906 | 0 |
| 41392 | ENSMUSG00000103239 | 0 | NA | NA | NA | NA | NA | Gm37155 | predicted gene, 37155 [Source:MGI Symb | 0 | 0 | 0 | 0 | 0 | 0 |
| 41393 | ENSMUSG00000103240 | 0 | NA | NA | NA | NA | NA | Gm37976 | predicted gene, 37976 [Source:MGI Symb | 0 | 0 | 0 | 0 | 0 | 0 |
| 41394 | ENSMUSG00000103241 | 0 | NA | NA | NA | NA | NA | Gm38089 | predicted gene, 38089 [Source:MGI Symb | 0 | 0 | 0 | 0 | 0 | 0 |
| 41395 | ENSMUSG00000103242 | 0 | NA | NA | NA | NA | NA | Gm34767 | predicted gene, 34767 [Source:MGI Symb | 0 | 0 | 0 | 0 | 0 | 0 |
| 41396 | ENSMUSG00000103243 | 0 | NA | NA | NA | NA | NA | Lce1d | late cornified envelope 1D [Source:MGI Sy | 0 | 0 | 0 | 0 | 0 | 0 |
| 41397 | ENSMUSG00000103244 | 0 | NA | NA | NA | NA | NA | Gm38092 | predicted gene, 38092 [Source:MGI Symb | 0 | 0 | 0 | 0 | 0 | 0 |
| 41398 | ENSMUSG00000103245 | 0 | NA | NA | NA | NA | NA | Pou5f1-rs3 | POU domain, class 5, transcription factor | 0 | 0 | 0 | 0 | 0 | 0 |
| 41399 | ENSMUSG00000103246 | 0 | NA | NA | NA | NA | NA | Gm38090 | predicted gene, 38090 [Source:MGI Symb | 0 | 0 | 0 | 0 | 0 | 0 |
| 41400 | ENSMUSG00000103247 | 3.643870387 | -0.02278111 | 1.170577227 | -0.019461432 | 0.984473004 | NA | Gm38091 | predicted gene, 38091 [Source:MGI Symb | 4.8186 | 3.0838 | 3.0187 | 0.964 | 2.1812 | 7.7969 |
| 41401 | ENSMUSG00000103248 | 0 | NA | NA | NA | NA | NA | Gm38087 | predicted gene, 38087 [Source:MGI Symb | 0 | 0 | 0 | 0 | 0 | 0 |
| 41402 | ENSMUSG00000103249 | 0 | NA | NA | NA | NA | NA | Gm38088 | predicted gene, 38088 [Source:MGI Symb | 0 | 0 | 0 | 0 | 0 | 0 |
| 41403 | ENSMUSG00000103250 | 0 | NA | NA | NA | NA | NA | Gm32180 | predicted gene, 32180 [Source:MGI Symb | 0 | 0 | 0 | 0 | 0 | 0 |
| 41404 | ENSMUSG00000103251 | 0.840287806 | -0.383646848 | 2.239474228 | -0.171311124 | 0.863979134 | NA | Gm37519 | predicted gene, 37519 [Source:MGI Symb | 0.8031 | 1.0279 | 1.0062 | 0 | 1.0906 | 1.1138 |
| 41405 | ENSMUSG00000103252 | 0 | NA | NA | NA | NA | NA | Gm37521 | predicted gene, 37521 [Source:MGI Symb | 0 | 0 | 0 | 0 | 0 | 0 |
| 41406 | ENSMUSG00000103253 | 0 | NA | NA | NA | NA | NA | Gm37520 | predicted gene, 37520 [Source:MGI Symb | 0 | 0 | 0 | 0 | 0 | 0 |
| 41407 | ENSMUSG00000103254 | 0 | NA | NA | NA | NA | NA | Ighv1-15 | immunoglobulin heavy variable 1-15 [Sou | 0 | 0 | 0 | 0 | 0 | 0 |
| 41408 | ENSMUSG00000103255 | 0 | NA | NA | NA | NA | NA | Pcdhac1 | protocadherin alpha subfamily C, 1 [Sourc | 0 | 0 | 0 | 0 | 0 | 0 |
| 41409 | ENSMUSG00000103256 | 0 | NA | NA | NA | NA | NA | Gm37647 | predicted gene, 37647 [Source:MGI Symb | 0 | 0 | 0 | 0 | 0 | 0 |
| 41410 | ENSMUSG00000103257 | 0 | NA | NA | NA | NA | NA | Gm7977 | predicted gene 7977 [Source:MGI Symbol | 0 | 0 | 0 | 0 | 0 | 0 |
| 41411 | ENSMUSG00000103258 | 1.468412105 | 1.945424723 | 2.004800999 | 0.970382958 | 0.33185564 | NA | Gm37518 | predicted gene, 37518 [Source:MGI Symb | 0.8031 | 0 | 1.0062 | 4.8199 | 2.1812 | 0 |
| 41412 | ENSMUSG00000103259 | 0 | NA | NA | NA | NA | NA | 2610105M2 | RIKEN cDNA 2610105M22 gene [Source:M | 0 | 0 | 0 | 0 | 0 | 0 |
| 41413 | ENSMUSG00000103260 | 0 | NA | NA | NA | NA | NA | Gm8146 | predicted gene 8146 [Source:MGI Symbol | 0 | 0 | 0 | 0 | 0 | 0 |
| 41414 | ENSMUSG00000103261 | 0.572870759 | -2.577107685 | 3.182795569 | -0.809699407 | 0.418112959 | NA | Gm36981 | predicted gene, 36981 [Source:MGI Symb | 2.4093 | 1.0279 | 0 | 0 | 0 | 0 |
| 41415 | ENSMUSG00000103262 | 0 | NA | NA | NA | NA | NA | Ighv5-8 | immunoglobulin heavy variable V5-8 [Sou | 0 | 0 | 0 | 0 | 0 | 0 |
| 41416 | ENSMUSG00000103263 | 0 | NA | NA | NA | NA | NA | Gm36982 | predicted gene, 36982 [Source:MGI Symb | 0 | 0 | 0 | 0 | 0 | 0 |
| 41417 | ENSMUSG00000103264 | 0 | NA | NA | NA | NA | NA | Gm30341 | predicted gene, 30341 [Source:MGI Symb | 0 | 0 | 0 | 0 | 0 | 0 |
| 41418 | ENSMUSG00000103265 | 0 | NA | NA | NA | NA | NA | Gm2053 | predicted gene 2053 [Source:MGI Symbol | 0 | 0 | 0 | 0 | 0 | 0 |
| 41419 | ENSMUSG00000103266 | 0 | NA | NA | NA | NA | NA | Gm36979 | predicted gene, 36979 [Source:MGI Symb | 0 | 0 | 0 | 0 | 0 | 0 |
| 41420 | ENSMUSG00000103267 | 0 | NA | NA | NA | NA | NA | Gm36980 | predicted gene, 36980 [Source:MGI Symb | 0 | 0 | 0 | 0 | 0 | 0 |
| 41421 | ENSMUSG00000103269 | 0 | NA | NA | NA | NA | NA | Gm36978 | predicted gene, 36978 [Source:MGI Symb | 0 | 0 | 0 | 0 | 0 | 0 |
| 41422 | ENSMUSG00000103270 | 1.47222539 | 0.602554691 | 1.7974176 | 0.335233554 | 0.737448912 | NA | Gm37915 | predicted gene, 37915 [Source:MGI Symb | 2.4093 | 1.0279 | 0 | 0.964 | 1.0906 | 3.3415 |
| 41423 | ENSMUSG00000103271 | 0 | NA | NA | NA | NA | NA | Ighv4-2 | immunoglobulin heavy variable V4-2 [Sou | 0 | 0 | 0 | 0 | 0 | 0 |
| 41424 | ENSMUSG00000103272 | 4.834570714 | -1.876289131 | 1.15031467 | -1.631109452 | 0.102867227 | NA | Gm37914 | predicted gene, 37914 [Source:MGI Symb | 13.653 | 3.0838 | 6.0375 | 2.8919 | 0 | 3.3415 |
| 41425 | ENSMUSG00000103273 | 0 | NA | NA | NA | NA | NA | Gm37913 | predicted gene, 37913 [Source:MGI Symb | 0 | 0 | 0 | 0 | 0 | 0 |
| 41426 | ENSMUSG00000103274 | 0 | NA | NA | NA | NA | NA | Gm37918 | predicted gene, 37918 [Source:MGI Symb | 0 | 0 | 0 | 0 | 0 | 0 |
| 41427 | ENSMUSG00000103275 | 0 | NA | NA | NA | NA | NA | Gm36983 | predicted gene, 36983 [Source:MGI Symb | 0 | 0 | 0 | 0 | 0 | 0 |
| 41428 | ENSMUSG00000103276 | 0 | NA | NA | NA | NA | NA | Gm37917 | predicted gene, 37917 [Source:MGI Symb | 0 | 0 | 0 | 0 | 0 | 0 |
| 41429 | ENSMUSG00000103277 | 0 | NA | NA | NA | NA | NA | Gm37916 | predicted gene, 37916 [Source:MGI Symb | 0 | 0 | 0 | 0 | 0 | 0 |
| 41430 | ENSMUSG00000103279 | 0 | NA | NA | NA | NA | NA | Gm3351 | predicted gene 3351 [Source:MGI Symbol | 0 | 0 | 0 | 0 | 0 | 0 |
| 41431 | ENSMUSG00000103280 | 1.692012309 | -0.682285877 | 1.980142714 | -0.344563991 | 0.73042218 | NA | Gm37277 | predicted gene, 37277 [Source:MGI Symb | 3.2124 | 3.0838 | 0 | 3.8559 | 0 | 0 |
| 41432 | ENSMUSG00000103281 | 0 | NA | NA | NA | NA | NA | Gm37278 | predicted gene, 37278 [Source:MGI Symb | 0 | 0 | 0 | 0 | 0 | 0 |
| 41433 | ENSMUSG00000103282 | 0 | NA | NA | NA | NA | NA | Gm37275 | predicted gene, 37275 [Source:MGI Symb | 0 | 0 | 0 | 0 | 0 | 0 |
| 41434 | ENSMUSG00000103284 | 0 | NA | NA | NA | NA | NA | 3110080O07 | RIKEN cDNA 3110080O07 gene [Source:M | 0 | 0 | 0 | 0 | 0 | 0 |
| 41435 | ENSMUSG00000103285 | 5.776068885 | -1.153275747 | 0.95588714 | -1.206497817 | 0.2276256 | 0.673700979 | Gm37274 | predicted gene, 37274 [Source:MGI Symb | 5.6217 | 6.1675 | 12.075 | 1.928 | 2.1812 | 6.6831 |
| 41436 | ENSMUSG00000103286 | 0 | NA | NA | NA | NA | NA | Gm5850 | predicted gene 5850 [Source:MGI Symbol | 0 | 0 | 0 | 0 | 0 | 0 |

| | | | | | | | | | | | | | | | |
|---|---|---|---|---|---|---|---|---|---|---|---|---|---|---|---|
| 41437 | ENSMUSG00000103287 | 0 | NA | NA | NA | NA | NA | 1700034J04I | RIKEN cDNA 1700034J04 gene [Source:M( | 0 | 0 | 0 | 0 | 0 | 0 |
| 41438 | ENSMUSG00000103288 | 0 | NA | NA | NA | NA | NA | Gm37273 | predicted gene, 37273 [Source:MGI Symb | 0 | 0 | 0 | 0 | 0 | 0 |
| 41439 | ENSMUSG00000103289 | 0 | NA | NA | NA | NA | NA | Gm31256 | predicted gene, 31256 [Source:MGI Symb | 0 | 0 | 0 | 0 | 0 | 0 |
| 41440 | ENSMUSG00000103290 | 0 | NA | NA | NA | NA | NA | Ighv1-23 | immunoglobulin heavy variable V1-23 [So | 0 | 0 | 0 | 0 | 0 | 0 |
| 41441 | ENSMUSG00000103291 | 0.356960287 | 0.16514264 | 3.880370363 | 0.042558474 | 0.966053499 | NA | Gm38235 | predicted gene, 38235 [Source:MGI Symb | 0 | 1.0279 | 0 | 0 | 0 | 1.1138 |
| 41442 | ENSMUSG00000103292 | 0 | NA | NA | NA | NA | NA | Gm35048 | predicted gene, 35048 [Source:MGI Symb | 0 | 0 | 0 | 0 | 0 | 0 |
| 41443 | ENSMUSG00000103293 | 0 | NA | NA | NA | NA | NA | Gm5842 | predicted gene 5842 [Source:MGI Symbol | 0 | 0 | 0 | 0 | 0 | 0 |
| 41444 | ENSMUSG00000103294 | 0 | NA | NA | NA | NA | NA | Gm38233 | predicted gene, 38233 [Source:MGI Symb | 0 | 0 | 0 | 0 | 0 | 0 |
| 41445 | ENSMUSG00000103295 | 0 | NA | NA | NA | NA | NA | Scp2-ps1 | sterol carrier protein 2, pseudogene 1 [So | 0 | 0 | 0 | 0 | 0 | 0 |
| 41446 | ENSMUSG00000103296 | 0 | NA | NA | NA | NA | NA | Gm38234 | predicted gene, 38234 [Source:MGI Symb | 0 | 0 | 0 | 0 | 0 | 0 |
| 41447 | ENSMUSG00000103297 | 0 | NA | NA | NA | NA | NA | Ighv3-2 | immunoglobulin heavy variable 3-2 [Sourc | 0 | 0 | 0 | 0 | 0 | 0 |
| 41448 | ENSMUSG00000103298 | 0 | NA | NA | NA | NA | NA | Gm38232 | predicted gene, 38232 [Source:MGI Symb | 0 | 0 | 0 | 0 | 0 | 0 |
| 41449 | ENSMUSG00000103299 | 0 | NA | NA | NA | NA | NA | Gm38231 | predicted gene, 38231 [Source:MGI Symb | 0 | 0 | 0 | 0 | 0 | 0 |
| 41450 | ENSMUSG00000103300 | 0 | NA | NA | NA | NA | NA | Ighv1-44 | immunoglobulin heavy variable V1-44 [So | 0 | 0 | 0 | 0 | 0 | 0 |
| 41451 | ENSMUSG00000103301 | 0 | NA | NA | NA | NA | NA | Gm37804 | predicted gene, 37804 [Source:MGI Symb | 0 | 0 | 0 | 0 | 0 | 0 |
| 41452 | ENSMUSG00000103302 | 0 | NA | NA | NA | NA | NA | Gm37803 | predicted gene, 37803 [Source:MGI Symb | 0 | 0 | 0 | 0 | 0 | 0 |
| 41453 | ENSMUSG00000103303 | 1.295205404 | 0.168345827 | 1.972095081 | 0.08536395 | 0.931972052 | NA | Gm37802 | predicted gene, 37802 [Source:MGI Symb | 1.6062 | 2.0558 | 0 | 1.928 | 2.1812 | 0 |
| 41454 | ENSMUSG00000103304 | 0 | NA | NA | NA | NA | NA | Gm5979 | predicted gene 5979 [Source:MGI Symbol | 0 | 0 | 0 | 0 | 0 | 0 |
| 41455 | ENSMUSG00000103305 | 0 | NA | NA | NA | NA | NA | Gm34068 | predicted gene, 34068 [Source:MGI Symb | 0 | 0 | 0 | 0 | 0 | 0 |
| 41456 | ENSMUSG00000103306 | 0 | NA | NA | NA | NA | NA | Gm37801 | predicted gene, 37801 [Source:MGI Symb | 0 | 0 | 0 | 0 | 0 | 0 |
| 41457 | ENSMUSG00000103307 | 0.642651897 | 2.906894826 | 3.440535777 | 0.844895974 | 0.398168923 | NA | Gm19335 | predicted gene, 19335 [Source:MGI Symb | 0 | 0 | 0 | 3.8559 | 0 | 0 |
| 41458 | ENSMUSG00000103308 | 0 | NA | NA | NA | NA | NA | Gm37800 | predicted gene, 37800 [Source:MGI Symb | 0 | 0 | 0 | 0 | 0 | 0 |
| 41459 | ENSMUSG00000103309 | 0 | NA | NA | NA | NA | NA | BC037039 | cDNA sequence BC037039 [Source:MGI Sy | 0 | 0 | 0 | 0 | 0 | 0 |
| 41460 | ENSMUSG00000103310 | 0 | NA | NA | NA | NA | NA | Pcdha12 | protocadherin alpha 12 [Source:MGI Syml | 0 | 0 | 0 | 0 | 0 | 0 |
| 41461 | ENSMUSG00000103311 | 0 | NA | NA | NA | NA | NA | Gm38229 | predicted gene, 38229 [Source:MGI Symb | 0 | 0 | 0 | 0 | 0 | 0 |
| 41462 | ENSMUSG00000103312 | 0 | NA | NA | NA | NA | NA | Gm6677 | predicted gene 6677 [Source:MGI Symbol | 0 | 0 | 0 | 0 | 0 | 0 |
| 41463 | ENSMUSG00000103313 | 18.02610737 | -0.78231699 | 0.527463954 | -1.483166733 | 0.138030117 | 0.542665838 | Gm38357 | predicted gene, 38357 [Source:MGI Symb | 17.668 | 28.782 | 22.137 | 16.388 | 9.8156 | 13.366 |
| 41464 | ENSMUSG00000103314 | 0 | NA | NA | NA | NA | NA | Gm38358 | predicted gene, 38358 [Source:MGI Symb | 0 | 0 | 0 | 0 | 0 | 0 |
| 41465 | ENSMUSG00000103315 | 0 | NA | NA | NA | NA | NA | Gm38341 | predicted gene, 38341 [Source:MGI Symb | 0 | 0 | 0 | 0 | 0 | 0 |
| 41466 | ENSMUSG00000103317 | 0 | NA | NA | NA | NA | NA | Gm38359 | predicted gene, 38359 [Source:MGI Symb | 0 | 0 | 0 | 0 | 0 | 0 |
| 41467 | ENSMUSG00000103318 | 0 | NA | NA | NA | NA | NA | Gm38356 | predicted gene, 38356 [Source:MGI Symb | 0 | 0 | 0 | 0 | 0 | 0 |
| 41468 | ENSMUSG00000103320 | 2.223544523 | -1.733553378 | 1.539916593 | -1.125744983 | 0.260273478 | NA | Gm37403 | predicted gene, 37403 [Source:MGI Symb | 3.2124 | 2.0558 | 5.0312 | 1.928 | 0 | 1.1138 |
| 41469 | ENSMUSG00000103321 | 0 | NA | NA | NA | NA | NA | 4933403L11I | RIKEN cDNA 4933403L11 gene [Source:M( | 0 | 0 | 0 | 0 | 0 | 0 |
| 41470 | ENSMUSG00000103322 | 0 | NA | NA | NA | NA | NA | Gm37404 | predicted gene, 37404 [Source:MGI Symb | 0 | 0 | 0 | 0 | 0 | 0 |
| 41471 | ENSMUSG00000103323 | 0 | NA | NA | NA | NA | NA | Gm21631 | predicted gene, 21631 [Source:MGI Symb | 0 | 0 | 0 | 0 | 0 | 0 |
| 41472 | ENSMUSG00000103324 | 1.188710055 | -0.543160199 | 2.270735428 | -0.239200125 | 0.810950405 | NA | Gm37402 | predicted gene, 37402 [Source:MGI Symb | 3.2124 | 1.0279 | 0 | 2.8919 | 0 | 0 |
| 41473 | ENSMUSG00000103325 | 0 | NA | NA | NA | NA | NA | Gm10531 | predicted gene 10531 [Source:MGI Symbo | 0 | 0 | 0 | 0 | 0 | 0 |
| 41474 | ENSMUSG00000103327 | 0 | NA | NA | NA | NA | NA | Gm36047 | predicted gene, 36047 [Source:MGI Symb | 0 | 0 | 0 | 0 | 0 | 0 |
| 41475 | ENSMUSG00000103328 | 0 | NA | NA | NA | NA | NA | Gm37406 | predicted gene, 37406 [Source:MGI Symb | 0 | 0 | 0 | 0 | 0 | 0 |
| 41476 | ENSMUSG00000103329 | 0 | NA | NA | NA | NA | NA | Gm42492 | predicted gene 42492 [Source:MGI Symbo | 0 | 0 | 0 | 0 | 0 | 0 |
| 41477 | ENSMUSG00000103330 | 0 | NA | NA | NA | NA | NA | Gm35108 | predicted gene, 35108 [Source:MGI Symb | 0 | 0 | 0 | 0 | 0 | 0 |
| 41478 | ENSMUSG00000103331 | 0 | NA | NA | NA | NA | NA | Gm37995 | predicted gene, 37995 [Source:MGI Symb | 0 | 0 | 0 | 0 | 0 | 0 |
| 41479 | ENSMUSG00000103332 | 0 | NA | NA | NA | NA | NA | Pcdhga2 | protocadherin gamma subfamily A, 2 [Sou | 0 | 0 | 0 | 0 | 0 | 0 |
| 41480 | ENSMUSG00000103333 | 0 | NA | NA | NA | NA | NA | Gm37994 | predicted gene, 37994 [Source:MGI Symb | 0 | 0 | 0 | 0 | 0 | 0 |
| 41481 | ENSMUSG00000103334 | 0 | NA | NA | NA | NA | NA | Gm29771 | predicted gene, 29771 [Source:MGI Symb | 0 | 0 | 0 | 0 | 0 | 0 |
| 41482 | ENSMUSG00000103336 | 0 | NA | NA | NA | NA | NA | Gm37993 | predicted gene, 37993 [Source:MGI Symb | 0 | 0 | 0 | 0 | 0 | 0 |
| 41483 | ENSMUSG00000103338 | 0 | NA | NA | NA | NA | NA | Olfr1409-ps1 | olfactory receptor 1409, pseudogene 1 [S( | 0 | 0 | 0 | 0 | 0 | 0 |
| 41484 | ENSMUSG00000103339 | 0 | NA | NA | NA | NA | NA | Gm37996 | predicted gene, 37996 [Source:MGI Symb | 0 | 0 | 0 | 0 | 0 | 0 |
| 41485 | ENSMUSG00000103340 | 0 | NA | NA | NA | NA | NA | Ighv1-1 | immunoglobulin heavy variable V1-1 [Sou | 0 | 0 | 0 | 0 | 0 | 0 |
| 41486 | ENSMUSG00000103341 | 0.267700746 | -1.497481649 | 4.057001436 | -0.369110456 | 0.712045396 | NA | 1700051O22 | RIKEN cDNA 1700051O22 Gene [Source:M | 1.6062 | 0 | 0 | 0 | 0 | 0 |
| 41487 | ENSMUSG00000103343 | 0.160662974 | 1.126924175 | 4.080472857 | 0.276174898 | 0.782413736 | NA | Gm37052 | predicted gene, 37052 [Source:MGI Symb | 0 | 0 | 0 | 0.964 | 0 | 0 |
| 41488 | ENSMUSG00000103345 | 0 | NA | NA | NA | NA | NA | Gm10723 | predicted gene 10723 [Source:MGI Symbo | 0 | 0 | 0 | 0 | 0 | 0 |
| 41489 | ENSMUSG00000103346 | 0 | NA | NA | NA | NA | NA | C230057A21 | RIKEN cDNA C230057A21 gene [Source:M | 0 | 0 | 0 | 0 | 0 | 0 |
| 41490 | ENSMUSG00000103347 | 0 | NA | NA | NA | NA | NA | Gm37734 | predicted gene, 37734 [Source:MGI Symb | 0 | 0 | 0 | 0 | 0 | 0 |
| 41491 | ENSMUSG00000103348 | 0 | NA | NA | NA | NA | NA | Gm37053 | predicted gene, 37053 [Source:MGI Symb | 0 | 0 | 0 | 0 | 0 | 0 |
| 41492 | ENSMUSG00000103349 | 0.267700746 | -1.497481649 | 4.057001436 | -0.369110456 | 0.712045396 | NA | Gm36888 | predicted gene, 36888 [Source:MGI Symb | 1.6062 | 0 | 0 | 0 | 0 | 0 |

| | | | | | | | | | | | | | | | |
|---|---|---|---|---|---|---|---|---|---|---|---|---|---|---|---|
| 41493 | ENSMUSG00000103350 | 0 | NA | NA | NA | NA | NA | Tdpoz6 | TD and POZ domain containing 6 [Source: | 0 | 0 | 0 | 0 | 0 | 0 |
| 41494 | ENSMUSG00000103351 | 0 | NA | NA | NA | NA | NA | Gm6350 | predicted gene 6350 [Source:MGI Symbol | 0 | 0 | 0 | 0 | 0 | 0 |
| 41495 | ENSMUSG00000103352 | 0 | NA | NA | NA | NA | NA | Gm38084 | predicted gene, 38084 [Source:MGI Symb | 0 | 0 | 0 | 0 | 0 | 0 |
| 41496 | ENSMUSG00000103353 | 0 | NA | NA | NA | NA | NA | Gm37597 | predicted gene, 37597 [Source:MGI Symb | 0 | 0 | 0 | 0 | 0 | 0 |
| 41497 | ENSMUSG00000103354 | 0 | NA | NA | NA | NA | NA | Gm38083 | predicted gene, 38083 [Source:MGI Symb | 0 | 0 | 0 | 0 | 0 | 0 |
| 41498 | ENSMUSG00000103355 | 0 | NA | NA | NA | NA | NA | Gm2147 | predicted gene 2147 [Source:MGI Symbol | 0 | 0 | 0 | 0 | 0 | 0 |
| 41499 | ENSMUSG00000103356 | 0 | NA | NA | NA | NA | NA | Gm37595 | predicted gene, 37595 [Source:MGI Symb | 0 | 0 | 0 | 0 | 0 | 0 |
| 41500 | ENSMUSG00000103357 | 0.133850373 | -0.796629176 | 4.080472857 | -0.195229623 | 0.84521319 | NA | Gm7804 | predicted gene 7804 [Source:MGI Symbol | 0.8031 | 0 | 0 | 0 | 0 | 0 |
| 41501 | ENSMUSG00000103358 | 0 | NA | NA | NA | NA | NA | Gm37593 | predicted gene, 37593 [Source:MGI Symb | 0 | 0 | 0 | 0 | 0 | 0 |
| 41502 | ENSMUSG00000103360 | 0.531944269 | 2.618916564 | 3.285123409 | 0.797204926 | 0.425332026 | NA | Gm38156 | predicted gene, 38156 [Source:MGI Symb | 0 | 0 | 0 | 0.964 | 0 | 2.2277 |
| 41503 | ENSMUSG00000103361 | 0 | NA | NA | NA | NA | NA | Gm36928 | predicted gene, 36928 [Source:MGI Symb | 0 | 0 | 0 | 0 | 0 | 0 |
| 41504 | ENSMUSG00000103362 | 0 | NA | NA | NA | NA | NA | Tdpoz2 | TD and POZ domain containing 2 [Source: | 0 | 0 | 0 | 0 | 0 | 0 |
| 41505 | ENSMUSG00000103364 | 0 | NA | NA | NA | NA | NA | Gm38157 | predicted gene, 38157 [Source:MGI Symb | 0 | 0 | 0 | 0 | 0 | 0 |
| 41506 | ENSMUSG00000103365 | 0 | NA | NA | NA | NA | NA | Gm36929 | predicted gene, 36929 [Source:MGI Symb | 0 | 0 | 0 | 0 | 0 | 0 |
| 41507 | ENSMUSG00000103366 | 0.133850373 | -0.796629176 | 4.080472857 | -0.195229623 | 0.84521319 | NA | C630004L07 | RIKEN cDNA C630004L07 gene [Source:M | 0.8031 | 0 | 0 | 0 | 0 | 0 |
| 41508 | ENSMUSG00000103367 | 2.592111867 | -1.942794979 | 1.620216197 | -1.199096134 | 0.230490567 | NA | Gm38158 | predicted gene, 38158 [Source:MGI Symb | 7.2279 | 1.0279 | 4.025 | 0 | 3.2719 | 0 |
| 41509 | ENSMUSG00000103368 | 0 | NA | NA | NA | NA | NA | Gm36927 | predicted gene, 36927 [Source:MGI Symb | 0 | 0 | 0 | 0 | 0 | 0 |
| 41510 | ENSMUSG00000103369 | 0 | NA | NA | NA | NA | NA | Gm6546 | predicted gene 6546 [Source:MGI Symbol | 0 | 0 | 0 | 0 | 0 | 0 |
| 41511 | ENSMUSG00000103370 | 0 | NA | NA | NA | NA | NA | Gm37178 | predicted gene, 37178 [Source:MGI Symb | 0 | 0 | 0 | 0 | 0 | 0 |
| 41512 | ENSMUSG00000103371 | 0 | NA | NA | NA | NA | NA | Gm35134 | predicted gene, 35134 [Source:MGI Symb | 0 | 0 | 0 | 0 | 0 | 0 |
| 41513 | ENSMUSG00000103372 | 0 | NA | NA | NA | NA | NA | Gm37237 | predicted gene, 37237 [Source:MGI Symb | 0 | 0 | 0 | 0 | 0 | 0 |
| 41514 | ENSMUSG00000103373 | 1.992821099 | 1.095504097 | 1.614970396 | 0.678343145 | 0.497554149 | NA | Gm37238 | predicted gene, 37238 [Source:MGI Symb | 0.8031 | 2.0558 | 1.0062 | 4.8199 | 3.2719 | 0 |
| 41515 | ENSMUSG00000103374 | 0 | NA | NA | NA | NA | NA | Gm37239 | predicted gene, 37239 [Source:MGI Symb | 0 | 0 | 0 | 0 | 0 | 0 |
| 41516 | ENSMUSG00000103375 | 0 | NA | NA | NA | NA | NA | Gm37181 | predicted gene, 37181 [Source:MGI Symb | 0 | 0 | 0 | 0 | 0 | 0 |
| 41517 | ENSMUSG00000103376 | 0 | NA | NA | NA | NA | NA | Gm37179 | predicted gene, 37179 [Source:MGI Symb | 0 | 0 | 0 | 0 | 0 | 0 |
| 41518 | ENSMUSG00000103377 | 0 | NA | NA | NA | NA | NA | Gm37180 | predicted gene, 37180 [Source:MGI Symb | 0 | 0 | 0 | 0 | 0 | 0 |
| 41519 | ENSMUSG00000103378 | 0 | NA | NA | NA | NA | NA | Gm37176 | predicted gene, 37176 [Source:MGI Symb | 0 | 0 | 0 | 0 | 0 | 0 |
| 41520 | ENSMUSG00000103379 | 0 | NA | NA | NA | NA | NA | Gm37177 | predicted gene, 37177 [Source:MGI Symb | 0 | 0 | 0 | 0 | 0 | 0 |
| 41521 | ENSMUSG00000103380 | 0.267700746 | -1.497481649 | 4.057001436 | -0.369110456 | 0.712045396 | NA | Gm37756 | predicted gene, 37756 [Source:MGI Symb | 1.6062 | 0 | 0 | 0 | 0 | 0 |
| 41522 | ENSMUSG00000103382 | 5.088125967 | 0.114271679 | 0.942914503 | 0.121189862 | 0.903540654 | 0.983821605 | Gm37755 | predicted gene, 37755 [Source:MGI Symb | 6.4248 | 5.1396 | 3.0187 | 3.8559 | 7.6343 | 4.4554 |
| 41523 | ENSMUSG00000103383 | 0 | NA | NA | NA | NA | NA | Gm37754 | predicted gene, 37754 [Source:MGI Symb | 0 | 0 | 0 | 0 | 0 | 0 |
| 41524 | ENSMUSG00000103384 | 0 | NA | NA | NA | NA | NA | Gm37753 | predicted gene, 37753 [Source:MGI Symb | 0 | 0 | 0 | 0 | 0 | 0 |
| 41525 | ENSMUSG00000103385 | 0 | NA | NA | NA | NA | NA | Gm37752 | predicted gene, 37752 [Source:MGI Symb | 0 | 0 | 0 | 0 | 0 | 0 |
| 41526 | ENSMUSG00000103386 | 0 | NA | NA | NA | NA | NA | Gm37751 | predicted gene, 37751 [Source:MGI Symb | 0 | 0 | 0 | 0 | 0 | 0 |
| 41527 | ENSMUSG00000103387 | 0 | NA | NA | NA | NA | NA | 1700030I03R | RIKEN cDNA 1700030I03 gene [Source:MC | 0 | 0 | 0 | 0 | 0 | 0 |
| 41528 | ENSMUSG00000103388 | 5.456293039 | 0.670912876 | 0.957616783 | 0.700606848 | 0.483548403 | 0.872075325 | Gm37581 | predicted gene, 37581 [Source:MGI Symb | 1.6062 | 3.0838 | 8.05 | 6.7478 | 5.4531 | 7.7969 |
| 41529 | ENSMUSG00000103389 | 0 | NA | NA | NA | NA | NA | Gm37757 | predicted gene, 37757 [Source:MGI Symb | 0 | 0 | 0 | 0 | 0 | 0 |
| 41530 | ENSMUSG00000103390 | 0 | NA | NA | NA | NA | NA | Gm38301 | predicted gene, 38301 [Source:MGI Symb | 0 | 0 | 0 | 0 | 0 | 0 |
| 41531 | ENSMUSG00000103391 | 0 | NA | NA | NA | NA | NA | Gm38302 | predicted gene, 38302 [Source:MGI Symb | 0 | 0 | 0 | 0 | 0 | 0 |
| 41532 | ENSMUSG00000103392 | 0 | NA | NA | NA | NA | NA | Gm38303 | predicted gene, 38303 [Source:MGI Symb | 0 | 0 | 0 | 0 | 0 | 0 |
| 41533 | ENSMUSG00000103393 | 0 | NA | NA | NA | NA | NA | Gm38304 | predicted gene, 38304 [Source:MGI Symb | 0 | 0 | 0 | 0 | 0 | 0 |
| 41534 | ENSMUSG00000103394 | 0 | NA | NA | NA | NA | NA | Gm10152 | predicted gene 10152 [Source:MGI Symbo | 0 | 0 | 0 | 0 | 0 | 0 |
| 41535 | ENSMUSG00000103395 | 0 | NA | NA | NA | NA | NA | Ighv1-27 | immunoglobulin heavy variable V1-27 [So | 0 | 0 | 0 | 0 | 0 | 0 |
| 41536 | ENSMUSG00000103396 | 0 | NA | NA | NA | NA | NA | Gm38305 | predicted gene, 38305 [Source:MGI Symb | 0 | 0 | 0 | 0 | 0 | 0 |
| 41537 | ENSMUSG00000103397 | 0 | NA | NA | NA | NA | NA | Gm38306 | predicted gene, 38306 [Source:MGI Symb | 0 | 0 | 0 | 0 | 0 | 0 |
| 41538 | ENSMUSG00000103398 | 0.328370405 | 0.16514316 | 4.003149536 | 0.041253308 | 0.967093956 | NA | Gm38307 | predicted gene, 38307 [Source:MGI Symb | 0 | 0 | 1.0062 | 0.964 | 0 | 0 |
| 41539 | ENSMUSG00000103399 | 0 | NA | NA | NA | NA | NA | Gm29747 | predicted gene, 29747 [Source:MGI Symb | 0 | 0 | 0 | 0 | 0 | 0 |
| 41540 | ENSMUSG00000103400 | 14.12624472 | -0.77638858 | 0.644016227 | -1.205541954 | 0.227994152 | 0.674239914 | Gm15853 | predicted gene 15853 [Source:MGI Symbo | 12.85 | 28.782 | 12.075 | 14.46 | 5.4531 | 11.138 |
| 41541 | ENSMUSG00000103401 | 0 | NA | NA | NA | NA | NA | Gm37929 | predicted gene, 37929 [Source:MGI Symb | 0 | 0 | 0 | 0 | 0 | 0 |
| 41542 | ENSMUSG00000103402 | 0 | NA | NA | NA | NA | NA | Gm37930 | predicted gene, 37930 [Source:MGI Symb | 0 | 0 | 0 | 0 | 0 | 0 |
| 41543 | ENSMUSG00000103403 | 0.160662974 | 1.126924175 | 4.080472857 | 0.276174898 | 0.782413736 | NA | Gm37931 | predicted gene, 37931 [Source:MGI Symb | 0 | 0 | 0 | 0.964 | 0 | 0 |
| 41544 | ENSMUSG00000103404 | 0 | NA | NA | NA | NA | NA | Gm37932 | predicted gene, 37932 [Source:MGI Symb | 0 | 0 | 0 | 0 | 0 | 0 |
| 41545 | ENSMUSG00000103406 | 0 | NA | NA | NA | NA | NA | Gm5541 | predicted gene 5541 [Source:MGI Symbol | 0 | 0 | 0 | 0 | 0 | 0 |
| 41546 | ENSMUSG00000103407 | 0 | NA | NA | NA | NA | NA | 4930540E01 | RIKEN cDNA 4930540E01 gene [Source:M | 0 | 0 | 0 | 0 | 0 | 0 |
| 41547 | ENSMUSG00000103408 | 0.181769715 | 1.126924175 | 4.080472857 | 0.276174898 | 0.782413736 | NA | Gm37933 | predicted gene, 37933 [Source:MGI Symb | 0 | 0 | 0 | 0 | 1.0906 | 0 |
| 41548 | ENSMUSG00000103409 | 1.117306622 | -0.258294525 | 2.105509541 | -0.122675542 | 0.902364032 | NA | Lsmem2 | leucine-rich single-pass membrane protei | 1.6062 | 2.0558 | 0 | 1.928 | 0 | 1.1138 |

| | | | | | | | | | | | | | | | |
|---|---|---|---|---|---|---|---|---|---|---|---|---|---|---|---|
| 41549 | ENSMUSG00000103410 | 0 | NA | NA | NA | NA | NA | Gm37341 | predicted gene, 37341 [Source:MGI Symb | 0 | 0 | 0 | 0 | 0 | 0 |
| 41550 | ENSMUSG00000103411 | 0.455176322 | 0.955156269 | 3.546674842 | 0.269310357 | 0.787690862 | NA | Gm18300 | predicted gene, 18300 [Source:MGI Symb | 0.8031 | 0 | 0 | 1.928 | 0 | 0 |
| 41551 | ENSMUSG00000103412 | 0 | NA | NA | NA | NA | NA | Gm37340 | predicted gene, 37340 [Source:MGI Symb | 0 | 0 | 0 | 0 | 0 | 0 |
| 41552 | ENSMUSG00000103413 | 0 | NA | NA | NA | NA | NA | Gm38389 | predicted gene, 38389 [Source:MGI Symb | 0 | 0 | 0 | 0 | 0 | 0 |
| 41553 | ENSMUSG00000103415 | 3.815887435 | 0.10957701 | 1.2691475 | 0.086339066 | 0.931196884 | NA | Gm37339 | predicted gene, 37339 [Source:MGI Symb | 4.8186 | 4.1117 | 2.0125 | 1.928 | 0 | 10.025 |
| 41554 | ENSMUSG00000103416 | 0 | NA | NA | NA | NA | NA | Gm37338 | predicted gene, 37338 [Source:MGI Symb | 0 | 0 | 0 | 0 | 0 | 0 |
| 41555 | ENSMUSG00000103417 | 0 | NA | NA | NA | NA | NA | Gm33112 | predicted gene, 33112 [Source:MGI Symb | 0 | 0 | 0 | 0 | 0 | 0 |
| 41556 | ENSMUSG00000103419 | 0 | NA | NA | NA | NA | NA | Gm29808 | predicted gene, 29808 [Source:MGI Symb | 0 | 0 | 0 | 0 | 0 | 0 |
| 41557 | ENSMUSG00000103420 | 0 | NA | NA | NA | NA | NA | Gm37537 | predicted gene, 37537 [Source:MGI Symb | 0 | 0 | 0 | 0 | 0 | 0 |
| 41558 | ENSMUSG00000103421 | 45.43198282 | 0.459661111 | 0.37669399 | 1.220250716 | 0.222369846 | 0.668407528 | Golt1a | golgi transport 1A [Source:MGI Symbol;Ac | 44.171 | 47.284 | 23.144 | 49.163 | 67.618 | 41.212 |
| 41559 | ENSMUSG00000103422 | 0 | NA | NA | NA | NA | NA | Gm37536 | predicted gene, 37536 [Source:MGI Symb | 0 | 0 | 0 | 0 | 0 | 0 |
| 41560 | ENSMUSG00000103423 | 0 | NA | NA | NA | NA | NA | Gm5845 | predicted gene 5845 [Source:MGI Symbol | 0 | 0 | 0 | 0 | 0 | 0 |
| 41561 | ENSMUSG00000103424 | 0 | NA | NA | NA | NA | NA | Gm37535 | predicted gene, 37535 [Source:MGI Symb | 0 | 0 | 0 | 0 | 0 | 0 |
| 41562 | ENSMUSG00000103425 | 0 | NA | NA | NA | NA | NA | Gm37075 | predicted gene, 37075 [Source:MGI Symb | 0 | 0 | 0 | 0 | 0 | 0 |
| 41563 | ENSMUSG00000103426 | 0 | NA | NA | NA | NA | NA | Gm37533 | predicted gene, 37533 [Source:MGI Symb | 0 | 0 | 0 | 0 | 0 | 0 |
| 41564 | ENSMUSG00000103427 | 0 | NA | NA | NA | NA | NA | Gm37534 | predicted gene, 37534 [Source:MGI Symb | 0 | 0 | 0 | 0 | 0 | 0 |
| 41565 | ENSMUSG00000103428 | 0.294513347 | 0.165143496 | 4.080472857 | 0.040471657 | 0.967717103 | NA | Gm20754 | predicted gene, 20754 [Source:MGI Symb | 0.8031 | 0 | 0 | 0.964 | 0 | 0 |
| 41566 | ENSMUSG00000103430 | 0 | NA | NA | NA | NA | NA | Gm36996 | predicted gene, 36996 [Source:MGI Symb | 0 | 0 | 0 | 0 | 0 | 0 |
| 41567 | ENSMUSG00000103431 | 0 | NA | NA | NA | NA | NA | Gm30686 | predicted gene, 30686 [Source:MGI Symb | 0 | 0 | 0 | 0 | 0 | 0 |
| 41568 | ENSMUSG00000103432 | 1.753840603 | -2.02343954 | 1.77638527 | -1.139076964 | 0.254671055 | NA | 6720464F23 | RIKEN cDNA 6720464F23 gene [Source:M | 2.4093 | 1.0279 | 5.0312 | 0.964 | 1.0906 | 0 |
| 41569 | ENSMUSG00000103433 | 0.133850373 | -0.796629176 | 4.080472857 | -0.195229623 | 0.84521319 | NA | Gm2272 | predicted gene 2272 [Source:MGI Symbol | 0.8031 | 0 | 0 | 0 | 0 | 0 |
| 41570 | ENSMUSG00000103434 | 0 | NA | NA | NA | NA | NA | Gm36993 | predicted gene, 36993 [Source:MGI Symb | 0 | 0 | 0 | 0 | 0 | 0 |
| 41571 | ENSMUSG00000103435 | 0 | NA | NA | NA | NA | NA | Gm36992 | predicted gene, 36992 [Source:MGI Symb | 0 | 0 | 0 | 0 | 0 | 0 |
| 41572 | ENSMUSG00000103436 | 2.727103846 | -1.229706386 | 1.565275663 | -0.785616499 | 0.432092196 | NA | Gm36995 | predicted gene, 36995 [Source:MGI Symb | 6.4248 | 4.1117 | 1.0062 | 4.8199 | 0 | 0 |
| 41573 | ENSMUSG00000103437 | 0 | NA | NA | NA | NA | NA | Gm36994 | predicted gene, 36994 [Source:MGI Symb | 0 | 0 | 0 | 0 | 0 | 0 |
| 41574 | ENSMUSG00000103438 | 0 | NA | NA | NA | NA | NA | Gm36991 | predicted gene, 36991 [Source:MGI Symb | 0 | 0 | 0 | 0 | 0 | 0 |
| 41575 | ENSMUSG00000103439 | 6.779358773 | 0.928477221 | 0.824756687 | 1.125758949 | 0.260267565 | 0.713728596 | Gm7019 | predicted pseudogene 7019 [Source:MGI | 4.8186 | 4.1117 | 5.0312 | 5.7839 | 10.906 | 10.025 |
| 41576 | ENSMUSG00000103440 | 0 | NA | NA | NA | NA | NA | Gm37131 | predicted gene, 37131 [Source:MGI Symb | 0 | 0 | 0 | 0 | 0 | 0 |
| 41577 | ENSMUSG00000103441 | 0 | NA | NA | NA | NA | NA | 7530428D23 | RIKEN cDNA 7530428D23 gene [Source:M | 0 | 0 | 0 | 0 | 0 | 0 |
| 41578 | ENSMUSG00000103442 | 0 | NA | NA | NA | NA | NA | Pcdha1 | protocadherin alpha 1 [Source:MGI Symb | 0 | 0 | 0 | 0 | 0 | 0 |
| 41579 | ENSMUSG00000103443 | 1.220266871 | 0.311146302 | 2.222383583 | 0.14000567 | 0.888655511 | NA | Gm37132 | predicted gene, 37132 [Source:MGI Symb | 3.2124 | 0 | 0 | 1.928 | 2.1812 | 0 |
| 41580 | ENSMUSG00000103444 | 0 | NA | NA | NA | NA | NA | Gm17836 | predicted gene, 17836 [Source:MGI Symb | 0 | 0 | 0 | 0 | 0 | 0 |
| 41581 | ENSMUSG00000103445 | 0 | NA | NA | NA | NA | NA | Gm36948 | predicted gene, 36948 [Source:MGI Symb | 0 | 0 | 0 | 0 | 0 | 0 |
| 41582 | ENSMUSG00000103446 | 0 | NA | NA | NA | NA | NA | Gm38203 | predicted gene, 38203 [Source:MGI Symb | 0 | 0 | 0 | 0 | 0 | 0 |
| 41583 | ENSMUSG00000103448 | 0 | NA | NA | NA | NA | NA | Gm37133 | predicted gene, 37133 [Source:MGI Symb | 0 | 0 | 0 | 0 | 0 | 0 |
| 41584 | ENSMUSG00000103449 | 0 | NA | NA | NA | NA | NA | Gm37134 | predicted gene, 37134 [Source:MGI Symb | 0 | 0 | 0 | 0 | 0 | 0 |
| 41585 | ENSMUSG00000103451 | 0 | NA | NA | NA | NA | NA | Gm33973 | predicted gene, 33973 [Source:MGI Symb | 0 | 0 | 0 | 0 | 0 | 0 |
| 41586 | ENSMUSG00000103452 | 0 | NA | NA | NA | NA | NA | 1700008A23 | RIKEN cDNA 1700008A23 gene [Source:M | 0 | 0 | 0 | 0 | 0 | 0 |
| 41587 | ENSMUSG00000103454 | 0 | NA | NA | NA | NA | NA | Gm34303 | predicted gene, 34303 [Source:MGI Symb | 0 | 0 | 0 | 0 | 0 | 0 |
| 41588 | ENSMUSG00000103456 | 0 | NA | NA | NA | NA | NA | Gm38097 | predicted gene, 38097 [Source:MGI Symb | 0 | 0 | 0 | 0 | 0 | 0 |
| 41589 | ENSMUSG00000103457 | 0 | NA | NA | NA | NA | NA | Gm37042 | predicted gene, 37042 [Source:MGI Symb | 0 | 0 | 0 | 0 | 0 | 0 |
| 41590 | ENSMUSG00000103458 | 0 | NA | NA | NA | NA | NA | Gm42416 | predicted gene, 42416 [Source:MGI Symb | 0 | 0 | 0 | 0 | 0 | 0 |
| 41591 | ENSMUSG00000103459 | 0 | NA | NA | NA | NA | NA | Gm38096 | predicted gene, 38096 [Source:MGI Symb | 0 | 0 | 0 | 0 | 0 | 0 |
| 41592 | ENSMUSG00000103460 | 0 | NA | NA | NA | NA | NA | Gm18491 | predicted gene, 18491 [Source:MGI Symb | 0 | 0 | 0 | 0 | 0 | 0 |
| 41593 | ENSMUSG00000103461 | 0 | NA | NA | NA | NA | NA | Gm37616 | predicted gene, 37616 [Source:MGI Symb | 0 | 0 | 0 | 0 | 0 | 0 |
| 41594 | ENSMUSG00000103462 | 0 | NA | NA | NA | NA | NA | Gm32181 | predicted gene, 32181 [Source:MGI Symb | 0 | 0 | 0 | 0 | 0 | 0 |
| 41595 | ENSMUSG00000103463 | 0 | NA | NA | NA | NA | NA | 1700063I16F | RIKEN cDNA 1700063I16 gene [Source:M | 0 | 0 | 0 | 0 | 0 | 0 |
| 41596 | ENSMUSG00000103464 | 0 | NA | NA | NA | NA | NA | Gm38246 | predicted gene, 38246 [Source:MGI Symb | 0 | 0 | 0 | 0 | 0 | 0 |
| 41597 | ENSMUSG00000103465 | 0 | NA | NA | NA | NA | NA | Gm37617 | predicted gene, 37617 [Source:MGI Symb | 0 | 0 | 0 | 0 | 0 | 0 |
| 41598 | ENSMUSG00000103466 | 0 | NA | NA | NA | NA | NA | Gm37247 | predicted gene, 37247 [Source:MGI Symb | 0 | 0 | 0 | 0 | 0 | 0 |
| 41599 | ENSMUSG00000103467 | 0.301557804 | -1.64335916 | 4.042814367 | -0.406488899 | 0.684383412 | NA | Gm38245 | predicted gene, 38245 [Source:MGI Symb | 0.8031 | 0 | 1.0062 | 0 | 0 | 0 |
| 41600 | ENSMUSG00000103468 | 0 | NA | NA | NA | NA | NA | Gm21440 | predicted gene, 21440 [Source:MGI Symb | 0 | 0 | 0 | 0 | 0 | 0 |
| 41601 | ENSMUSG00000103469 | 0.331982614 | 0.16514309 | 3.986751476 | 0.041422971 | 0.966958701 | NA | Gm9910 | predicted gene 9910 [Source:MGI Symbol | 0 | 1.0279 | 0 | 0.964 | 0 | 0 |
| 41602 | ENSMUSG00000103470 | 0 | NA | NA | NA | NA | NA | Gm38184 | predicted gene, 38184 [Source:MGI Symb | 0 | 0 | 0 | 0 | 0 | 0 |
| 41603 | ENSMUSG00000103471 | 0 | NA | NA | NA | NA | NA | Gm17966 | predicted gene, 17966 [Source:MGI Symb | 0 | 0 | 0 | 0 | 0 | 0 |
| 41604 | ENSMUSG00000103472 | 0.656740811 | -0.013525372 | 3.112905782 | -0.004344934 | 0.996533255 | NA | Pcdhga7 | protocadherin gamma subfamily A, 7 [Sou | 0 | 0 | 2.0125 | 1.928 | 0 | 0 |

| | | | | | | | | | | | | | | | |
|---|---|---|---|---|---|---|---|---|---|---|---|---|---|---|---|
| 41605 | ENSMUSG00000103473 | 0.305170013 | -1.657836477 | 4.041480335 | -0.410205257 | 0.681655385 | NA | Gm37696 | predicted gene, 37696 [Source:MGI Symb | 0.8031 | 1.0279 | 0 | 0 | 0 | 0 |
| 41606 | ENSMUSG00000103474 | 0 | NA | NA | NA | NA | NA | Gm37252 | predicted gene, 37252 [Source:MGI Symb | 0 | 0 | 0 | 0 | 0 | 0 |
| 41607 | ENSMUSG00000103475 | 0 | NA | NA | NA | NA | NA | Gm37697 | predicted gene, 37697 [Source:MGI Symb | 0 | 0 | 0 | 0 | 0 | 0 |
| 41608 | ENSMUSG00000103476 | 0 | NA | NA | NA | NA | NA | Strit1 | small transmembrane regulator of ion tra | 0 | 0 | 0 | 0 | 0 | 0 |
| 41609 | ENSMUSG00000103477 | 0 | NA | NA | NA | NA | NA | 5930409G06 | RIKEN cDNA 5930409G06 gene [Source:M | 0 | 0 | 0 | 0 | 0 | 0 |
| 41610 | ENSMUSG00000103478 | 0 | NA | NA | NA | NA | NA | 4930433J02I | RIKEN cDNA 4930433J02 gene [Source:M( | 0 | 0 | 0 | 0 | 0 | 0 |
| 41611 | ENSMUSG00000103479 | 0 | NA | NA | NA | NA | NA | Gm37698 | predicted gene, 37698 [Source:MGI Symb | 0 | 0 | 0 | 0 | 0 | 0 |
| 41612 | ENSMUSG00000103481 | 0 | NA | NA | NA | NA | NA | 1700016D08 | RIKEN cDNA 1700016D08 gene [Source:M | 0 | 0 | 0 | 0 | 0 | 0 |
| 41613 | ENSMUSG00000103482 | 0 | NA | NA | NA | NA | NA | Gm37999 | predicted gene, 37999 [Source:MGI Symb | 0 | 0 | 0 | 0 | 0 | 0 |
| 41614 | ENSMUSG00000103483 | 0 | NA | NA | NA | NA | NA | Gm20162 | predicted gene, 20162 [Source:MGI Symb | 0 | 0 | 0 | 0 | 0 | 0 |
| 41615 | ENSMUSG00000103484 | 0 | NA | NA | NA | NA | NA | Gm38000 | predicted gene, 38000 [Source:MGI Symb | 0 | 0 | 0 | 0 | 0 | 0 |
| 41616 | ENSMUSG00000103485 | 0 | NA | NA | NA | NA | NA | Gm38001 | predicted gene, 38001 [Source:MGI Symb | 0 | 0 | 0 | 0 | 0 | 0 |
| 41617 | ENSMUSG00000103486 | 0 | NA | NA | NA | NA | NA | Gm10657 | predicted gene 10657 [Source:MGI Symbo | 0 | 0 | 0 | 0 | 0 | 0 |
| 41618 | ENSMUSG00000103487 | 0 | NA | NA | NA | NA | NA | Gm30858 | predicted gene, 30858 [Source:MGI Symb | 0 | 0 | 0 | 0 | 0 | 0 |
| 41619 | ENSMUSG00000103488 | 0 | NA | NA | NA | NA | NA | Gm37997 | predicted gene, 37997 [Source:MGI Symb | 0 | 0 | 0 | 0 | 0 | 0 |
| 41620 | ENSMUSG00000103489 | 0 | NA | NA | NA | NA | NA | Gm37998 | predicted gene, 37998 [Source:MGI Symb | 0 | 0 | 0 | 0 | 0 | 0 |
| 41621 | ENSMUSG00000103491 | 0 | NA | NA | NA | NA | NA | Gm10730 | predicted gene 10730 [Source:MGI Symbo | 0 | 0 | 0 | 0 | 0 | 0 |
| 41622 | ENSMUSG00000103492 | 0.371281295 | 2.09253362 | 4.031852165 | 0.519000582 | 0.603760335 | NA | Gm37412 | predicted gene, 37412 [Source:MGI Symb | 0 | 0 | 0 | 0 | 0 | 2.2277 |
| 41623 | ENSMUSG00000103493 | 0.496077836 | -0.812642266 | 3.444120572 | -0.235950586 | 0.813471014 | NA | Gm37411 | predicted gene, 37411 [Source:MGI Symb | 0 | 0 | 2.0125 | 0.964 | 0 | 0 |
| 41624 | ENSMUSG00000103494 | 0 | NA | NA | NA | NA | NA | Gm37410 | predicted gene, 37410 [Source:MGI Symb | 0 | 0 | 0 | 0 | 0 | 0 |
| 41625 | ENSMUSG00000103495 | 0.696064467 | -1.534444742 | 3.043667477 | -0.504143358 | 0.61416064 | NA | Gm37409 | predicted gene, 37409 [Source:MGI Symb | 3.2124 | 0 | 0 | 0.964 | 0 | 0 |
| 41626 | ENSMUSG00000103497 | 0 | NA | NA | NA | NA | NA | Gm37407 | predicted gene, 37407 [Source:MGI Symb | 0 | 0 | 0 | 0 | 0 | 0 |
| 41627 | ENSMUSG00000103498 | 0 | NA | NA | NA | NA | NA | Gm18984 | predicted gene, 18984 [Source:MGI Symb | 0 | 0 | 0 | 0 | 0 | 0 |
| 41628 | ENSMUSG00000103499 | 0 | NA | NA | NA | NA | NA | Gm6558 | predicted gene 6558 [Source:MGI Symbol | 0 | 0 | 0 | 0 | 0 | 0 |
| 41629 | ENSMUSG00000103500 | 0 | NA | NA | NA | NA | NA | Gm38373 | predicted gene, 38373 [Source:MGI Symb | 0 | 0 | 0 | 0 | 0 | 0 |
| 41630 | ENSMUSG00000103501 | 0 | NA | NA | NA | NA | NA | Gm7412 | predicted gene 7412 [Source:MGI Symbol | 0 | 0 | 0 | 0 | 0 | 0 |
| 41631 | ENSMUSG00000103502 | 19.27695748 | -0.166164865 | 0.538603556 | -0.308510524 | 0.757693894 | 0.953466696 | 9330121J05I | RIKEN cDNA 9330121J05 gene [Source:M( | 23.29 | 19.53 | 18.112 | 11.568 | 13.087 | 30.074 |
| 41632 | ENSMUSG00000103503 | 0 | NA | NA | NA | NA | NA | 1700023G09 | RIKEN cDNA 1700023G09 gene [Source:M | 0 | 0 | 0 | 0 | 0 | 0 |
| 41633 | ENSMUSG00000103504 | 0 | NA | NA | NA | NA | NA | Gm38390 | predicted gene, 38390 [Source:MGI Symb | 0 | 0 | 0 | 0 | 0 | 0 |
| 41634 | ENSMUSG00000103505 | 0 | NA | NA | NA | NA | NA | Gm38374 | predicted gene, 38374 [Source:MGI Symb | 0 | 0 | 0 | 0 | 0 | 0 |
| 41635 | ENSMUSG00000103506 | 0.401551119 | -2.072982726 | 4.008251979 | -0.517178744 | 0.605031388 | NA | Gm38376 | predicted gene, 38376 [Source:MGI Symb | 2.4093 | 0 | 0 | 0 | 0 | 0 |
| 41636 | ENSMUSG00000103507 | 3.192597442 | 0.595181501 | 1.360843275 | 0.437362268 | 0.661848644 | NA | Gm38375 | predicted gene, 38375 [Source:MGI Symb | 1.6062 | 1.0279 | 5.0312 | 3.8559 | 7.6343 | 0 |
| 41637 | ENSMUSG00000103509 | 27.86929569 | -0.106811772 | 0.415167787 | -0.257273747 | 0.796967457 | 0.962236316 | Gm38372 | predicted gene, 38372 [Source:MGI Symb | 29.715 | 33.921 | 23.144 | 30.847 | 25.084 | 24.505 |
| 41638 | ENSMUSG00000103510 | 0 | NA | NA | NA | NA | NA | Gm30725 | predicted gene, 30725 [Source:MGI Symb | 0 | 0 | 0 | 0 | 0 | 0 |
| 41639 | ENSMUSG00000103511 | 0 | NA | NA | NA | NA | NA | Gm5285 | predicted gene 5285 [Source:MGI Symbol | 0 | 0 | 0 | 0 | 0 | 0 |
| 41640 | ENSMUSG00000103512 | 0 | NA | NA | NA | NA | NA | Gm37430 | predicted gene, 37430 [Source:MGI Symb | 0 | 0 | 0 | 0 | 0 | 0 |
| 41641 | ENSMUSG00000103513 | 0 | NA | NA | NA | NA | NA | Gm34780 | predicted gene, 34780 [Source:MGI Symb | 0 | 0 | 0 | 0 | 0 | 0 |
| 41642 | ENSMUSG00000103514 | 0 | NA | NA | NA | NA | NA | Gm37433 | predicted gene, 37433 [Source:MGI Symb | 0 | 0 | 0 | 0 | 0 | 0 |
| 41643 | ENSMUSG00000103516 | 0 | NA | NA | NA | NA | NA | Gm37431 | predicted gene, 37431 [Source:MGI Symb | 0 | 0 | 0 | 0 | 0 | 0 |
| 41644 | ENSMUSG00000103517 | 0.545309144 | 2.651974629 | 3.666219369 | 0.72335405 | 0.469462394 | NA | Gm37432 | predicted gene, 37432 [Source:MGI Symb | 0 | 0 | 0 | 0 | 3.2719 | 0 |
| 41645 | ENSMUSG00000103518 | 0 | NA | NA | NA | NA | NA | Gm37428 | predicted gene, 37428 [Source:MGI Symb | 0 | 0 | 0 | 0 | 0 | 0 |
| 41646 | ENSMUSG00000103519 | 0 | NA | NA | NA | NA | NA | Gm37429 | predicted gene, 37429 [Source:MGI Symb | 0 | 0 | 0 | 0 | 0 | 0 |
| 41647 | ENSMUSG00000103520 | 10.47101117 | -0.008122811 | 0.685612802 | -0.011847519 | 0.990547269 | 0.99859176 | 1110025M0! | RIKEN cDNA 1110025M09 gene [Source:N | 11.243 | 6.1675 | 14.087 | 12.532 | 6.5437 | 12.252 |
| 41648 | ENSMUSG00000103521 | 0 | NA | NA | NA | NA | NA | Gm29730 | predicted gene, 29730 [Source:MGI Symb | 0 | 0 | 0 | 0 | 0 | 0 |
| 41649 | ENSMUSG00000103522 | 0.670829724 | -2.782124174 | 3.401449285 | -0.817923168 | 0.41340106 | NA | 9430087J23I | RIKEN cDNA 9430087J23 gene [Source:M( | 0 | 0 | 4.025 | 0 | 0 | 0 |
| 41650 | ENSMUSG00000103523 | 0 | NA | NA | NA | NA | NA | 2210017I01F | RIKEN cDNA 2210017I01 gene [Source:M( | 0 | 0 | 0 | 0 | 0 | 0 |
| 41651 | ENSMUSG00000103524 | 0 | NA | NA | NA | NA | NA | Gm37263 | predicted gene, 37263 [Source:MGI Symb | 0 | 0 | 0 | 0 | 0 | 0 |
| 41652 | ENSMUSG00000103525 | 0.656740811 | -0.013525372 | 3.112905782 | -0.004344934 | 0.996533255 | NA | Gm37262 | predicted gene, 37262 [Source:MGI Symb | 0 | 0 | 2.0125 | 1.928 | 0 | 0 |
| 41653 | ENSMUSG00000103526 | 0 | NA | NA | NA | NA | NA | Gm18900 | predicted gene, 18900 [Source:MGI Symb | 0 | 0 | 0 | 0 | 0 | 0 |
| 41654 | ENSMUSG00000103527 | 0 | NA | NA | NA | NA | NA | Gm37261 | predicted gene, 37261 [Source:MGI Symb | 0 | 0 | 0 | 0 | 0 | 0 |
| 41655 | ENSMUSG00000103528 | 0 | NA | NA | NA | NA | NA | Gm31571 | predicted gene, 31571 [Source:MGI Symb | 0 | 0 | 0 | 0 | 0 | 0 |
| 41656 | ENSMUSG00000103529 | 0 | NA | NA | NA | NA | NA | A730089K16 | RIKEN cDNA A730089K16 gene [Source:M | 0 | 0 | 0 | 0 | 0 | 0 |
| 41657 | ENSMUSG00000103530 | 0 | NA | NA | NA | NA | NA | Gm37839 | predicted gene, 37839 [Source:MGI Symb | 0 | 0 | 0 | 0 | 0 | 0 |
| 41658 | ENSMUSG00000103531 | 0 | NA | NA | NA | NA | NA | 2610300A13 | RIKEN cDNA 2610300A13 gene [Source:M | 0 | 0 | 0 | 0 | 0 | 0 |
| 41659 | ENSMUSG00000103532 | 3.667509464 | 0.482779572 | 1.172584068 | 0.41172278 | 0.680542626 | NA | Gm4430 | predicted gene 4430 [Source:MGI Symbol | 4.0155 | 3.0838 | 2.0125 | 2.8919 | 1.0906 | 8.9108 |
| 41660 | ENSMUSG00000103533 | 0 | NA | NA | NA | NA | NA | Gm18493 | predicted gene, 18493 [Source:MGI Symb | 0 | 0 | 0 | 0 | 0 | 0 |

| | | | | | | | | | | | | | | | |
|---|---|---|---|---|---|---|---|---|---|---|---|---|---|---|---|
| 41661 | ENSMUSG00000103534 | 0 | NA | NA | NA | NA | NA | Gm37835 | predicted gene, 37835 [Source:MGI Symb | 0 | 0 | 0 | 0 | 0 | 0 |
| 41662 | ENSMUSG00000103535 | 0 | NA | NA | NA | NA | NA | Gm37836 | predicted gene, 37836 [Source:MGI Symb | 0 | 0 | 0 | 0 | 0 | 0 |
| 41663 | ENSMUSG00000103536 | 0 | NA | NA | NA | NA | NA | Gm37837 | predicted gene, 37837 [Source:MGI Symb | 0 | 0 | 0 | 0 | 0 | 0 |
| 41664 | ENSMUSG00000103537 | 0 | NA | NA | NA | NA | NA | Gm37838 | predicted gene, 37838 [Source:MGI Symb | 0 | 0 | 0 | 0 | 0 | 0 |
| 41665 | ENSMUSG00000103538 | 0 | NA | NA | NA | NA | NA | Olfr289-ps1 | olfactory receptor 289, pseudogene 1 [So | 0 | 0 | 0 | 0 | 0 | 0 |
| 41666 | ENSMUSG00000103539 | 0 | NA | NA | NA | NA | NA | Gm37834 | predicted gene, 37834 [Source:MGI Symb | 0 | 0 | 0 | 0 | 0 | 0 |
| 41667 | ENSMUSG00000103540 | 0 | NA | NA | NA | NA | NA | Gm37668 | predicted gene, 37668 [Source:MGI Symb | 0 | 0 | 0 | 0 | 0 | 0 |
| 41668 | ENSMUSG00000103541 | 0 | NA | NA | NA | NA | NA | Gm37667 | predicted gene, 37667 [Source:MGI Symb | 0 | 0 | 0 | 0 | 0 | 0 |
| 41669 | ENSMUSG00000103542 | 0 | NA | NA | NA | NA | NA | Gm34768 | predicted gene, 34768 [Source:MGI Symb | 0 | 0 | 0 | 0 | 0 | 0 |
| 41670 | ENSMUSG00000103543 | 0 | NA | NA | NA | NA | NA | Gm37669 | predicted gene, 37669 [Source:MGI Symb | 0 | 0 | 0 | 0 | 0 | 0 |
| 41671 | ENSMUSG00000103544 | 0 | NA | NA | NA | NA | NA | Gm10048 | predicted gene 10048 [Source:MGI Symbo | 0 | 0 | 0 | 0 | 0 | 0 |
| 41672 | ENSMUSG00000103545 | 0 | NA | NA | NA | NA | NA | 1700012J22I | RIKEN cDNA 1700012J22 gene [Source:MG | 0 | 0 | 0 | 0 | 0 | 0 |
| 41673 | ENSMUSG00000103546 | 0.877550482 | 2.072039088 | 2.361583201 | 0.877394066 | 0.380272635 | NA | Gm37666 | predicted gene, 37666 [Source:MGI Symb | 0 | 0 | 1.0062 | 0.964 | 2.1812 | 1.1138 |
| 41674 | ENSMUSG00000103547 | 0 | NA | NA | NA | NA | NA | Gm37665 | predicted gene, 37665 [Source:MGI Symb | 0 | 0 | 0 | 0 | 0 | 0 |
| 41675 | ENSMUSG00000103548 | 0.36353943 | 2.065989254 | 4.034137301 | 0.512126658 | 0.608562368 | NA | Gm37670 | predicted gene, 37670 [Source:MGI Symb | 0 | 0 | 0 | 0 | 2.1812 | 0 |
| 41676 | ENSMUSG00000103549 | 0 | NA | NA | NA | NA | NA | Gm17895 | predicted gene, 17895 [Source:MGI Symb | 0 | 0 | 0 | 0 | 0 | 0 |
| 41677 | ENSMUSG00000103552 | 0 | NA | NA | NA | NA | NA | Gm38217 | predicted gene, 38217 [Source:MGI Symb | 0 | 0 | 0 | 0 | 0 | 0 |
| 41678 | ENSMUSG00000103553 | 4.666148067 | -1.796701876 | 1.105913808 | -1.624631018 | 0.104241204 | NA | Gm38218 | predicted gene, 38218 [Source:MGI Symb | 5.6217 | 3.0838 | 13.081 | 2.8919 | 1.0906 | 2.2277 |
| 41679 | ENSMUSG00000103554 | 0 | NA | NA | NA | NA | NA | Ighv1-35 | immunoglobulin heavy variable 1-35 [Sou | 0 | 0 | 0 | 0 | 0 | 0 |
| 41680 | ENSMUSG00000103555 | 0 | NA | NA | NA | NA | NA | Gm37337 | predicted gene, 37337 [Source:MGI Symb | 0 | 0 | 0 | 0 | 0 | 0 |
| 41681 | ENSMUSG00000103556 | 0 | NA | NA | NA | NA | NA | Gm38215 | predicted gene, 38215 [Source:MGI Symb | 0 | 0 | 0 | 0 | 0 | 0 |
| 41682 | ENSMUSG00000103557 | 0 | NA | NA | NA | NA | NA | Gm38216 | predicted gene, 38216 [Source:MGI Symb | 0 | 0 | 0 | 0 | 0 | 0 |
| 41683 | ENSMUSG00000103558 | 0 | NA | NA | NA | NA | NA | Gm38220 | predicted gene, 38220 [Source:MGI Symb | 0 | 0 | 0 | 0 | 0 | 0 |
| 41684 | ENSMUSG00000103559 | 0 | NA | NA | NA | NA | NA | Gm38221 | predicted gene, 38221 [Source:MGI Symb | 0 | 0 | 0 | 0 | 0 | 0 |
| 41685 | ENSMUSG00000103560 | 12.05274879 | -0.110934609 | 0.81350375 | -0.136366439 | 0.891531604 | 0.982489666 | Gm38070 | predicted gene, 38070 [Source:MGI Symb | 3.2124 | 8.2233 | 26.162 | 3.8559 | 15.269 | 15.594 |
| 41686 | ENSMUSG00000103561 | 0 | NA | NA | NA | NA | NA | Gm38069 | predicted gene, 38069 [Source:MGI Symb | 0 | 0 | 0 | 0 | 0 | 0 |
| 41687 | ENSMUSG00000103562 | 0 | NA | NA | NA | NA | NA | 2810405F15 | RIKEN cDNA 2810405F15 gene [Source:M | 0 | 0 | 0 | 0 | 0 | 0 |
| 41688 | ENSMUSG00000103563 | 0 | NA | NA | NA | NA | NA | 8030445P17 | RIKEN cDNA 8030445P17 gene [Source:M | 0 | 0 | 0 | 0 | 0 | 0 |
| 41689 | ENSMUSG00000103564 | 0 | NA | NA | NA | NA | NA | Gm35067 | predicted gene, 35067 [Source:MGI Symb | 0 | 0 | 0 | 0 | 0 | 0 |
| 41690 | ENSMUSG00000103566 | 0 | NA | NA | NA | NA | NA | Gm38068 | predicted gene, 38068 [Source:MGI Symb | 0 | 0 | 0 | 0 | 0 | 0 |
| 41691 | ENSMUSG00000103567 | 0.610133435 | 0.304049584 | 2.904025954 | 0.104699334 | 0.91661439 | NA | Pcdhga5 | protocadherin gamma subfamily A, 5 [Sou | 1.6062 | 0 | 0 | 0.964 | 1.0906 | 0 |
| 41692 | ENSMUSG00000103568 | 0 | NA | NA | NA | NA | NA | Gm29741 | predicted gene, 29741 [Source:MGI Symb | 0 | 0 | 0 | 0 | 0 | 0 |
| 41693 | ENSMUSG00000103569 | 4.034745379 | 0.388775279 | 1.064698336 | 0.36515064 | 0.714998961 | NA | Gm38067 | predicted gene, 38067 [Source:MGI Symb | 2.4093 | 3.0838 | 5.0312 | 4.8199 | 2.1812 | 6.6831 |
| 41694 | ENSMUSG00000103570 | 1.456127102 | 0.486251466 | 1.686043828 | 0.28839788 | 0.77304219 | NA | C630004M2: | RIKEN cDNA C630004M23 gene [Source:N | 1.6062 | 1.0279 | 1.0062 | 2.8919 | 1.0906 | 1.1138 |
| 41695 | ENSMUSG00000103571 | 0 | NA | NA | NA | NA | NA | Gm37114 | predicted gene, 37114 [Source:MGI Symb | 0 | 0 | 0 | 0 | 0 | 0 |
| 41696 | ENSMUSG00000103572 | 0.356960287 | 0.16514264 | 3.880370363 | 0.042558474 | 0.966053499 | NA | Gm37115 | predicted gene, 37115 [Source:MGI Symb | 0 | 1.0279 | 0 | 0 | 0 | 1.1138 |
| 41697 | ENSMUSG00000103573 | 0 | NA | NA | NA | NA | NA | Gm18426 | predicted gene, 18426 [Source:MGI Symb | 0 | 0 | 0 | 0 | 0 | 0 |
| 41698 | ENSMUSG00000103574 | 3.655928809 | -0.407880774 | 1.099435488 | -0.370991093 | 0.710644166 | NA | Gm37116 | predicted gene, 37116 [Source:MGI Symb | 2.4093 | 5.1396 | 5.0312 | 3.8559 | 3.2719 | 2.2277 |
| 41699 | ENSMUSG00000103575 | 0 | NA | NA | NA | NA | NA | Gm37117 | predicted gene, 37117 [Source:MGI Symb | 0 | 0 | 0 | 0 | 0 | 0 |
| 41700 | ENSMUSG00000103576 | 0.267700746 | -1.497481649 | 4.057001436 | -0.369110456 | 0.712045396 | NA | Gm38188 | predicted gene, 38188 [Source:MGI Symb | 1.6062 | 0 | 0 | 0 | 0 | 0 |
| 41701 | ENSMUSG00000103577 | 0 | NA | NA | NA | NA | NA | 9330162B11 | RIKEN cDNA 9330162B11 gene [Source:M | 0 | 0 | 0 | 0 | 0 | 0 |
| 41702 | ENSMUSG00000103579 | 0 | NA | NA | NA | NA | NA | Gm37113 | predicted gene, 37113 [Source:MGI Symb | 0 | 0 | 0 | 0 | 0 | 0 |
| 41703 | ENSMUSG00000103580 | 0 | NA | NA | NA | NA | NA | Gm10417 | predicted gene 10417 [Source:MGI Symbo | 0 | 0 | 0 | 0 | 0 | 0 |
| 41704 | ENSMUSG00000103581 | 0 | NA | NA | NA | NA | NA | Gm37833 | predicted gene, 37833 [Source:MGI Symb | 0 | 0 | 0 | 0 | 0 | 0 |
| 41705 | ENSMUSG00000103583 | 0 | NA | NA | NA | NA | NA | Gm38325 | predicted gene, 38325 [Source:MGI Symb | 0 | 0 | 0 | 0 | 0 | 0 |
| 41706 | ENSMUSG00000103584 | 0 | NA | NA | NA | NA | NA | Gm38053 | predicted gene, 38053 [Source:MGI Symb | 0 | 0 | 0 | 0 | 0 | 0 |
| 41707 | ENSMUSG00000103585 | 2.073718175 | -2.306068798 | 1.664203538 | -1.385689157 | 0.165841841 | NA | Pcdhgb4 | protocadherin gamma subfamily B, 4 [Sou | 3.2124 | 5.1396 | 2.0125 | 0.964 | 0 | 1.1138 |
| 41708 | ENSMUSG00000103586 | 0.535401493 | -2.48596965 | 3.684117979 | -0.674780141 | 0.499815459 | NA | 5830405F06 | RIKEN cDNA 5830405F06 gene [Source:M | 3.2124 | 0 | 0 | 0 | 0 | 0 |
| 41709 | ENSMUSG00000103587 | 0 | NA | NA | NA | NA | NA | Gm38326 | predicted gene, 38326 [Source:MGI Symb | 0 | 0 | 0 | 0 | 0 | 0 |
| 41710 | ENSMUSG00000103588 | 0 | NA | NA | NA | NA | NA | Gm18445 | predicted gene, 18445 [Source:MGI Symb | 0 | 0 | 0 | 0 | 0 | 0 |
| 41711 | ENSMUSG00000103589 | 0 | NA | NA | NA | NA | NA | Gm37840 | predicted gene, 37840 [Source:MGI Symb | 0 | 0 | 0 | 0 | 0 | 0 |
| 41712 | ENSMUSG00000103590 | 2.332390593 | -0.434846526 | 1.487550736 | -0.292323828 | 0.770039042 | NA | Gm37353 | predicted gene, 37353 [Source:MGI Symb | 4.0155 | 2.0558 | 2.0125 | 4.8199 | 1.0906 | 0 |
| 41713 | ENSMUSG00000103591 | 1.004356399 | 2.372094719 | 2.241352889 | 1.058331658 | 0.289904265 | NA | Gm38365 | predicted gene, 38365 [Source:MGI Symb | 0.8031 | 0 | 0 | 1.928 | 2.1812 | 1.1138 |
| 41714 | ENSMUSG00000103592 | 1.839022621 | -0.150731079 | 1.797173305 | -0.083871198 | 0.933158839 | NA | Gm38364 | predicted gene, 38364 [Source:MGI Symb | 0.8031 | 0 | 5.0312 | 1.928 | 3.2719 | 0 |
| 41715 | ENSMUSG00000103593 | 1.152579385 | -2.386177204 | 2.369995502 | -1.006827735 | 0.314017564 | NA | Gm37352 | predicted gene, 37352 [Source:MGI Symb | 4.8186 | 0 | 1.0062 | 0 | 1.0906 | 0 |
| 41716 | ENSMUSG00000103594 | 0 | NA | NA | NA | NA | NA | A430034D21 | RIKEN cDNA A430034D21 gene [Source:M | 0 | 0 | 0 | 0 | 0 | 0 |

| | | | | | | | | | | | | | | | |
|---|---|---|---|---|---|---|---|---|---|---|---|---|---|---|---|
| 41717 | ENSMUSG00000103595 | 0 | NA | NA | NA | NA | NA | Gm37355 | predicted gene, 37355 [Source:MGI Symb | 0 | 0 | 0 | 0 | 0 | 0 |
| 41718 | ENSMUSG00000103596 | 0 | NA | NA | NA | NA | NA | Gm37354 | predicted gene, 37354 [Source:MGI Symb | 0 | 0 | 0 | 0 | 0 | 0 |
| 41719 | ENSMUSG00000103597 | 0 | NA | NA | NA | NA | NA | Gm19777 | predicted gene, 19777 [Source:MGI Symb | 0 | 0 | 0 | 0 | 0 | 0 |
| 41720 | ENSMUSG00000103598 | 0 | NA | NA | NA | NA | NA | Gm4856 | predicted gene 4856 [Source:MGI Symbol | 0 | 0 | 0 | 0 | 0 | 0 |
| 41721 | ENSMUSG00000103599 | 0.160662974 | 1.126924175 | 4.080472857 | 0.276174898 | 0.782413736 | NA | Gm37356 | predicted gene, 37356 [Source:MGI Symb | 0 | 0 | 0 | 0.964 | 0 | 0 |
| 41722 | ENSMUSG00000103600 | 0 | NA | NA | NA | NA | NA | Gm17952 | predicted gene, 17952 [Source:MGI Symb | 0 | 0 | 0 | 0 | 0 | 0 |
| 41723 | ENSMUSG00000103601 | 0 | NA | NA | NA | NA | NA | Gm18150 | predicted gene, 18150 [Source:MGI Symb | 0 | 0 | 0 | 0 | 0 | 0 |
| 41724 | ENSMUSG00000103602 | 0 | NA | NA | NA | NA | NA | Gm33191 | predicted gene, 33191 [Source:MGI Symb | 0 | 0 | 0 | 0 | 0 | 0 |
| 41725 | ENSMUSG00000103603 | 0 | NA | NA | NA | NA | NA | Gm37020 | predicted gene, 37020 [Source:MGI Symb | 0 | 0 | 0 | 0 | 0 | 0 |
| 41726 | ENSMUSG00000103604 | 0 | NA | NA | NA | NA | NA | Gm38205 | predicted gene, 38205 [Source:MGI Symb | 0 | 0 | 0 | 0 | 0 | 0 |
| 41727 | ENSMUSG00000103606 | 0 | NA | NA | NA | NA | NA | Gm412 | predicted gene 412 [Source:MGI Symbol;/ | 0 | 0 | 0 | 0 | 0 | 0 |
| 41728 | ENSMUSG00000103607 | 0 | NA | NA | NA | NA | NA | Gm37021 | predicted gene, 37021 [Source:MGI Symb | 0 | 0 | 0 | 0 | 0 | 0 |
| 41729 | ENSMUSG00000103608 | 0 | NA | NA | NA | NA | NA | 4930442P19 | RIKEN cDNA 4930442P19 gene [Source:M | 0 | 0 | 0 | 0 | 0 | 0 |
| 41730 | ENSMUSG00000103609 | 0 | NA | NA | NA | NA | NA | Gm37022 | predicted gene, 37022 [Source:MGI Symb | 0 | 0 | 0 | 0 | 0 | 0 |
| 41731 | ENSMUSG00000103610 | 0 | NA | NA | NA | NA | NA | Gm37947 | predicted gene, 37947 [Source:MGI Symb | 0 | 0 | 0 | 0 | 0 | 0 |
| 41732 | ENSMUSG00000103611 | 0.305170013 | -1.657836477 | 4.041480335 | -0.410205257 | 0.681655385 | NA | Gm37195 | predicted gene, 37195 [Source:MGI Symb | 0.8031 | 1.0279 | 0 | 0 | 0 | 0 |
| 41733 | ENSMUSG00000103612 | 0 | NA | NA | NA | NA | NA | Gm37948 | predicted gene, 37948 [Source:MGI Symb | 0 | 0 | 0 | 0 | 0 | 0 |
| 41734 | ENSMUSG00000103613 | 0.556921942 | 2.679043159 | 3.638243063 | 0.736356289 | 0.461513899 | NA | Gm34882 | predicted gene, 34882 [Source:MGI Symb | 0 | 0 | 0 | 0 | 0 | 3.3415 |
| 41735 | ENSMUSG00000103614 | 0.181769715 | 1.126924175 | 4.080472857 | 0.276174898 | 0.782413736 | NA | Gm37949 | predicted gene, 37949 [Source:MGI Symb | 0 | 0 | 0 | 0 | 1.0906 | 0 |
| 41736 | ENSMUSG00000103615 | 0 | NA | NA | NA | NA | NA | Gm30045 | predicted gene, 30045 [Source:MGI Symb | 0 | 0 | 0 | 0 | 0 | 0 |
| 41737 | ENSMUSG00000103616 | 0 | NA | NA | NA | NA | NA | 1700018A14 | RIKEN cDNA 1700018A14 gene [Source:M | 0 | 0 | 0 | 0 | 0 | 0 |
| 41738 | ENSMUSG00000103617 | 0 | NA | NA | NA | NA | NA | Gm37950 | predicted gene, 37950 [Source:MGI Symb | 0 | 0 | 0 | 0 | 0 | 0 |
| 41739 | ENSMUSG00000103618 | 0 | NA | NA | NA | NA | NA | Gm37951 | predicted gene, 37951 [Source:MGI Symb | 0 | 0 | 0 | 0 | 0 | 0 |
| 41740 | ENSMUSG00000103620 | 0 | NA | NA | NA | NA | NA | Gm37359 | predicted gene, 37359 [Source:MGI Symb | 0 | 0 | 0 | 0 | 0 | 0 |
| 41741 | ENSMUSG00000103621 | 4.972713197 | -0.139426352 | 1.1795972 | -0.118198273 | 0.905910558 | NA | Gm38366 | predicted gene, 38366 [Source:MGI Symb | 5.6217 | 0 | 10.062 | 8.6758 | 4.3625 | 1.1138 |
| 41742 | ENSMUSG00000103622 | 0.339027071 | -1.79174149 | 3.919585585 | -0.457125237 | 0.647581032 | NA | Gm38391 | predicted gene, 38391 [Source:MGI Symb | 0 | 1.0279 | 1.0062 | 0 | 0 | 0 |
| 41743 | ENSMUSG00000103624 | 0 | NA | NA | NA | NA | NA | Gm3081 | predicted gene 3081 [Source:MGI Symbol | 0 | 0 | 0 | 0 | 0 | 0 |
| 41744 | ENSMUSG00000103625 | 0.513958919 | -2.394274821 | 3.756096013 | -0.637437066 | 0.523840191 | NA | Gm37357 | predicted gene, 37357 [Source:MGI Symb | 0 | 3.0838 | 0 | 0 | 0 | 0 |
| 41745 | ENSMUSG00000103626 | 0 | NA | NA | NA | NA | NA | Gm37358 | predicted gene, 37358 [Source:MGI Symb | 0 | 0 | 0 | 0 | 0 | 0 |
| 41746 | ENSMUSG00000103628 | 0 | NA | NA | NA | NA | NA | Gm37361 | predicted gene, 37361 [Source:MGI Symb | 0 | 0 | 0 | 0 | 0 | 0 |
| 41747 | ENSMUSG00000103629 | 0 | NA | NA | NA | NA | NA | Gm5694 | predicted gene 5694 [Source:MGI Symbol | 0 | 0 | 0 | 0 | 0 | 0 |
| 41748 | ENSMUSG00000103630 | 0.556921942 | 2.679043159 | 3.638243063 | 0.736356289 | 0.461513899 | NA | Gm37242 | predicted gene, 37242 [Source:MGI Symb | 0 | 0 | 0 | 0 | 0 | 3.3415 |
| 41749 | ENSMUSG00000103631 | 0 | NA | NA | NA | NA | NA | Gm38060 | predicted gene, 38060 [Source:MGI Symb | 0 | 0 | 0 | 0 | 0 | 0 |
| 41750 | ENSMUSG00000103632 | 4.728999893 | -0.551400735 | 1.01575191 | -0.542849814 | 0.587233212 | NA | Gm38329 | predicted gene, 38329 [Source:MGI Symb | 9.6372 | 4.1117 | 3.0187 | 3.8559 | 2.1812 | 5.5692 |
| 41751 | ENSMUSG00000103633 | 0 | NA | NA | NA | NA | NA | Gm38328 | predicted gene, 38328 [Source:MGI Symb | 0 | 0 | 0 | 0 | 0 | 0 |
| 41752 | ENSMUSG00000103634 | 0.328370405 | 0.16514316 | 4.003149536 | 0.041253308 | 0.967093956 | NA | 3110062G12 | RIKEN cDNA 3110062G12 gene [Source:M | 0 | 0 | 1.0062 | 0.964 | 0 | 0 |
| 41753 | ENSMUSG00000103635 | 0 | NA | NA | NA | NA | NA | Gm38061 | predicted gene, 38061 [Source:MGI Symb | 0 | 0 | 0 | 0 | 0 | 0 |
| 41754 | ENSMUSG00000103636 | 0 | NA | NA | NA | NA | NA | Gm38331 | predicted gene, 38331 [Source:MGI Symb | 0 | 0 | 0 | 0 | 0 | 0 |
| 41755 | ENSMUSG00000103637 | 0 | NA | NA | NA | NA | NA | Gm38330 | predicted gene, 38330 [Source:MGI Symb | 0 | 0 | 0 | 0 | 0 | 0 |
| 41756 | ENSMUSG00000103638 | 0.167707431 | -0.796629176 | 4.080472857 | -0.195229623 | 0.84521319 | NA | Gm38327 | predicted gene, 38327 [Source:MGI Symb | 0 | 0 | 1.0062 | 0 | 0 | 0 |
| 41757 | ENSMUSG00000103639 | 0 | NA | NA | NA | NA | NA | Gm21791 | predicted gene, 21791 [Source:MGI Symb | 0 | 0 | 0 | 0 | 0 | 0 |
| 41758 | ENSMUSG00000103640 | 0 | NA | NA | NA | NA | NA | Gm31406 | predicted gene, 31406 [Source:MGI Symb | 0 | 0 | 0 | 0 | 0 | 0 |
| 41759 | ENSMUSG00000103641 | 0.133850373 | -0.796629176 | 4.080472857 | -0.195229623 | 0.84521319 | NA | Gm19024 | predicted gene, 19024 [Source:MGI Symb | 0.8031 | 0 | 0 | 0 | 0 | 0 |
| 41760 | ENSMUSG00000103642 | 0.17131964 | -0.796629176 | 4.080472857 | -0.195229623 | 0.84521319 | NA | Gm37769 | predicted gene, 37769 [Source:MGI Symb | 0 | 1.0279 | 0 | 0 | 0 | 0 |
| 41761 | ENSMUSG00000103643 | 0 | NA | NA | NA | NA | NA | Gm37770 | predicted gene, 37770 [Source:MGI Symb | 0 | 0 | 0 | 0 | 0 | 0 |
| 41762 | ENSMUSG00000103644 | 0 | NA | NA | NA | NA | NA | Gm29678 | predicted gene, 29678 [Source:MGI Symb | 0 | 0 | 0 | 0 | 0 | 0 |
| 41763 | ENSMUSG00000103646 | 0.133850373 | -0.796629176 | 4.080472857 | -0.195229623 | 0.84521319 | NA | Gm37706 | predicted gene, 37706 [Source:MGI Symb | 0.8031 | 0 | 0 | 0 | 0 | 0 |
| 41764 | ENSMUSG00000103647 | 0 | NA | NA | NA | NA | NA | Gm37707 | predicted gene, 37707 [Source:MGI Symb | 0 | 0 | 0 | 0 | 0 | 0 |
| 41765 | ENSMUSG00000103649 | 24.77831912 | -0.224597425 | 0.432581068 | -0.519203084 | 0.603619129 | 0.916106578 | Gm37768 | predicted gene, 37768 [Source:MGI Symb | 23.29 | 25.698 | 31.194 | 22.171 | 21.812 | 24.505 |
| 41766 | ENSMUSG00000103650 | 0 | NA | NA | NA | NA | NA | Gm37207 | predicted gene, 37207 [Source:MGI Symb | 0 | 0 | 0 | 0 | 0 | 0 |
| 41767 | ENSMUSG00000103651 | 0 | NA | NA | NA | NA | NA | Gm37206 | predicted gene, 37206 [Source:MGI Symb | 0 | 0 | 0 | 0 | 0 | 0 |
| 41768 | ENSMUSG00000103652 | 0 | NA | NA | NA | NA | NA | Gm37209 | predicted gene, 37209 [Source:MGI Symb | 0 | 0 | 0 | 0 | 0 | 0 |
| 41769 | ENSMUSG00000103653 | 38.16097419 | 1.16858168 | 0.372168573 | 3.139925733 | 0.001689907 | 0.030901151 | Gstp-ps | glutathione S-transferase, pi, pseudogene | 21.684 | 23.642 | 25.156 | 40.487 | 55.622 | 62.375 |
| 41770 | ENSMUSG00000103654 | 0 | NA | NA | NA | NA | NA | Gm37203 | predicted gene, 37203 [Source:MGI Symb | 0 | 0 | 0 | 0 | 0 | 0 |
| 41771 | ENSMUSG00000103655 | 0 | NA | NA | NA | NA | NA | Gm37202 | predicted gene, 37202 [Source:MGI Symb | 0 | 0 | 0 | 0 | 0 | 0 |
| 41772 | ENSMUSG00000103656 | 0 | NA | NA | NA | NA | NA | Gm37205 | predicted gene, 37205 [Source:MGI Symb | 0 | 0 | 0 | 0 | 0 | 0 |

| | | | | | | | | | | | | | | | |
|---|---|---|---|---|---|---|---|---|---|---|---|---|---|---|---|
| 41773 | ENSMUSG00000103657 | 0.589026695 | 0.241472922 | 3.228838524 | 0.074786311 | 0.940384733 | NA | Gm37204 | predicted gene, 37204 [Source:MGI Symb | 1.6062 | 0 | 0 | 1.928 | 0 | 0 |
| 41774 | ENSMUSG00000103658 | 0 | NA | NA | NA | NA | NA | Gm37201 | predicted gene, 37201 [Source:MGI Symb | 0 | 0 | 0 | 0 | 0 | 0 |
| 41775 | ENSMUSG00000103659 | 0.506734502 | -2.372847156 | 3.338060886 | -0.710845978 | 0.477179685 | NA | Gm32569 | predicted gene, 32569 [Source:MGI Symb | 0 | 1.0279 | 2.0125 | 0 | 0 | 0 |
| 41776 | ENSMUSG00000103660 | 0 | NA | NA | NA | NA | NA | Gm30705 | predicted gene, 30705 [Source:MGI Symb | 0 | 0 | 0 | 0 | 0 | 0 |
| 41777 | ENSMUSG00000103662 | 3.561429251 | 0.560974008 | 1.167922898 | 0.480317673 | 0.631001524 | NA | Gm34294 | predicted gene, 34294 [Source:MGI Symb | 1.6062 | 1.0279 | 6.0375 | 3.8559 | 3.2719 | 5.5692 |
| 41778 | ENSMUSG00000103663 | 0 | NA | NA | NA | NA | NA | 2900037B21 | RIKEN cDNA 2900037B21 gene [Source:M | 0 | 0 | 0 | 0 | 0 | 0 |
| 41779 | ENSMUSG00000103664 | 0 | NA | NA | NA | NA | NA | Gm36261 | predicted gene, 36261 [Source:MGI Symb | 0 | 0 | 0 | 0 | 0 | 0 |
| 41780 | ENSMUSG00000103665 | 0 | NA | NA | NA | NA | NA | Gm9227 | predicted gene 9227 [Source:MGI Symbol | 0 | 0 | 0 | 0 | 0 | 0 |
| 41781 | ENSMUSG00000103666 | 0 | NA | NA | NA | NA | NA | Gm38185 | predicted gene, 38185 [Source:MGI Symb | 0 | 0 | 0 | 0 | 0 | 0 |
| 41782 | ENSMUSG00000103667 | 0 | NA | NA | NA | NA | NA | Ighv1-8 | immunoglobulin heavy variable V1-8 [Sou | 0 | 0 | 0 | 0 | 0 | 0 |
| 41783 | ENSMUSG00000103668 | 0 | NA | NA | NA | NA | NA | Gm38186 | predicted gene, 38186 [Source:MGI Symb | 0 | 0 | 0 | 0 | 0 | 0 |
| 41784 | ENSMUSG00000103669 | 0 | NA | NA | NA | NA | NA | 2900042K21 | RIKEN cDNA 2900042K21 gene [Source:M | 0 | 0 | 0 | 0 | 0 | 0 |
| 41785 | ENSMUSG00000103670 | 0 | NA | NA | NA | NA | NA | Gm37622 | predicted gene, 37622 [Source:MGI Symb | 0 | 0 | 0 | 0 | 0 | 0 |
| 41786 | ENSMUSG00000103672 | 18.13009671 | 0.165755495 | 0.534173866 | 0.310302516 | 0.756330918 | 0.953085143 | Gm37621 | predicted gene, 37621 [Source:MGI Symb | 12.85 | 18.503 | 20.125 | 22.171 | 23.994 | 11.138 |
| 41787 | ENSMUSG00000103673 | 0 | NA | NA | NA | NA | NA | Gm37620 | predicted gene, 37620 [Source:MGI Symb | 0 | 0 | 0 | 0 | 0 | 0 |
| 41788 | ENSMUSG00000103675 | 0 | NA | NA | NA | NA | NA | Gm37991 | predicted gene, 37991 [Source:MGI Symb | 0 | 0 | 0 | 0 | 0 | 0 |
| 41789 | ENSMUSG00000103676 | 0.610133435 | 0.304049584 | 2.904025954 | 0.104699334 | 0.91661439 | NA | Gm38204 | predicted gene, 38204 [Source:MGI Symb | 1.6062 | 0 | 0 | 0.964 | 1.0906 | 0 |
| 41790 | ENSMUSG00000103677 | 0.506966596 | 2.558866737 | 3.339834942 | 0.766165628 | 0.443577757 | NA | Pcdhga4 | protocadherin gamma subfamily A, 4 [Sou | 0 | 0 | 0 | 1.928 | 0 | 1.1138 |
| 41791 | ENSMUSG00000103678 | 0 | NA | NA | NA | NA | NA | Gm37624 | predicted gene, 37624 [Source:MGI Symb | 0 | 0 | 0 | 0 | 0 | 0 |
| 41792 | ENSMUSG00000103679 | 0 | NA | NA | NA | NA | NA | Gm37623 | predicted gene, 37623 [Source:MGI Symb | 0 | 0 | 0 | 0 | 0 | 0 |
| 41793 | ENSMUSG00000103680 | 21.17852354 | -0.18598795 | 0.504552702 | -0.36861947 | 0.712411381 | 0.944239215 | Gm37072 | predicted gene, 37072 [Source:MGI Symb | 24.093 | 13.363 | 30.187 | 23.135 | 21.812 | 14.48 |
| 41794 | ENSMUSG00000103681 | 0 | NA | NA | NA | NA | NA | Gm37130 | predicted gene, 37130 [Source:MGI Symb | 0 | 0 | 0 | 0 | 0 | 0 |
| 41795 | ENSMUSG00000103684 | 0 | NA | NA | NA | NA | NA | Gm38180 | predicted gene, 38180 [Source:MGI Symb | 0 | 0 | 0 | 0 | 0 | 0 |
| 41796 | ENSMUSG00000103685 | 0 | NA | NA | NA | NA | NA | Gm37074 | predicted gene, 37074 [Source:MGI Symb | 0 | 0 | 0 | 0 | 0 | 0 |
| 41797 | ENSMUSG00000103686 | 0 | NA | NA | NA | NA | NA | Gm37073 | predicted gene, 37073 [Source:MGI Symb | 0 | 0 | 0 | 0 | 0 | 0 |
| 41798 | ENSMUSG00000103688 | 0 | NA | NA | NA | NA | NA | Gm6321 | predicted gene 6321 [Source:MGI Symbol | 0 | 0 | 0 | 0 | 0 | 0 |
| 41799 | ENSMUSG00000103689 | 0 | NA | NA | NA | NA | NA | Gm30598 | predicted gene, 30598 [Source:MGI Symb | 0 | 0 | 0 | 0 | 0 | 0 |
| 41800 | ENSMUSG00000103690 | 0 | NA | NA | NA | NA | NA | A730004F24 | RIKEN cDNA A730004F24 gene [Source:M | 0 | 0 | 0 | 0 | 0 | 0 |
| 41801 | ENSMUSG00000103691 | 0.339027071 | -1.79174149 | 3.919585585 | -0.457125237 | 0.647581032 | NA | Gm38019 | predicted gene, 38019 [Source:MGI Symb | 0 | 1.0279 | 1.0062 | 0 | 0 | 0 |
| 41802 | ENSMUSG00000103692 | 0 | NA | NA | NA | NA | NA | 4930503O07 | RIKEN cDNA 4930503O07 gene [Source:M | 0 | 0 | 0 | 0 | 0 | 0 |
| 41803 | ENSMUSG00000103693 | 1.513364513 | 1.841410709 | 1.985022185 | 0.927652458 | 0.353587877 | NA | Gm37529 | predicted gene, 37529 [Source:MGI Symb | 0 | 2.0558 | 0 | 4.8199 | 1.0906 | 1.1138 |
| 41804 | ENSMUSG00000103694 | 0 | NA | NA | NA | NA | NA | Gm37530 | predicted gene, 37530 [Source:MGI Symb | 0 | 0 | 0 | 0 | 0 | 0 |
| 41805 | ENSMUSG00000103695 | 0 | NA | NA | NA | NA | NA | A830029E22 | RIKEN cDNA A830029E22 gene [Source:M | 0 | 0 | 0 | 0 | 0 | 0 |
| 41806 | ENSMUSG00000103696 | 0 | NA | NA | NA | NA | NA | Gm37531 | predicted gene, 37531 [Source:MGI Symb | 0 | 0 | 0 | 0 | 0 | 0 |
| 41807 | ENSMUSG00000103697 | 2.472582501 | -1.287675456 | 1.629668007 | -0.790145876 | 0.42944258 | NA | Gm38020 | predicted gene, 38020 [Source:MGI Symb | 6.4248 | 0 | 4.025 | 0 | 3.2719 | 1.1138 |
| 41808 | ENSMUSG00000103698 | 0 | NA | NA | NA | NA | NA | Gm17877 | predicted gene, 17877 [Source:MGI Symb | 0 | 0 | 0 | 0 | 0 | 0 |
| 41809 | ENSMUSG00000103699 | 0 | NA | NA | NA | NA | NA | Gm38018 | predicted gene, 38018 [Source:MGI Symb | 0 | 0 | 0 | 0 | 0 | 0 |
| 41810 | ENSMUSG00000103700 | 0 | NA | NA | NA | NA | NA | Gm6555 | predicted gene 6555 [Source:MGI Symbol | 0 | 0 | 0 | 0 | 0 | 0 |
| 41811 | ENSMUSG00000103701 | 0.167707431 | -0.796629176 | 4.080472857 | -0.195229623 | 0.84521319 | NA | Gm10728 | predicted gene 10728 [Source:MGI Symbo | 0 | 0 | 1.0062 | 0 | 0 | 0 |
| 41812 | ENSMUSG00000103702 | 0 | NA | NA | NA | NA | NA | Gm37290 | predicted gene, 37290 [Source:MGI Symb | 0 | 0 | 0 | 0 | 0 | 0 |
| 41813 | ENSMUSG00000103703 | 18.85561853 | -0.177070614 | 0.494777095 | -0.35787957 | 0.72043344 | 0.9447719 | Gm42568 | predicted gene 42568 [Source:MGI Symbo | 19.274 | 20.558 | 20.125 | 13.496 | 19.631 | 20.049 |
| 41814 | ENSMUSG00000103704 | 2.121927258 | 0.720529199 | 1.632763489 | 0.441294286 | 0.658999962 | NA | Gm37291 | predicted gene, 37291 [Source:MGI Symb | 0.8031 | 3.0838 | 1.0062 | 6.7478 | 1.0906 | 0 |
| 41815 | ENSMUSG00000103705 | 0 | NA | NA | NA | NA | NA | 4930518J20 | RIKEN cDNA 4930518J20 gene [Source:M | 0 | 0 | 0 | 0 | 0 | 0 |
| 41816 | ENSMUSG00000103706 | 0.640816969 | 1.600078776 | 2.832778663 | 0.564844263 | 0.572179697 | NA | 6820402A03 | RIKEN cDNA 6820402A03 gene [Source:M | 0.8031 | 0 | 0 | 1.928 | 0 | 1.1138 |
| 41817 | ENSMUSG00000103707 | 0 | NA | NA | NA | NA | NA | Pcdha6 | protocadherin alpha 6 [Source:MGI Symb | 0 | 0 | 0 | 0 | 0 | 0 |
| 41818 | ENSMUSG00000103708 | 0 | NA | NA | NA | NA | NA | Gm33222 | predicted gene, 33222 [Source:MGI Symb | 0 | 0 | 0 | 0 | 0 | 0 |
| 41819 | ENSMUSG00000103709 | 0 | NA | NA | NA | NA | NA | Nras-ps2 | neuroblastoma ras oncogene, pseudogen | 0 | 0 | 0 | 0 | 0 | 0 |
| 41820 | ENSMUSG00000103710 | 0 | NA | NA | NA | NA | NA | D030062O11 | RIKEN cDNA D030062O11 gene [Source:M | 0 | 0 | 0 | 0 | 0 | 0 |
| 41821 | ENSMUSG00000103711 | 206.4508768 | 0.31011026 | 0.204630599 | 1.515463776 | 0.129654997 | 0.527222345 | Tstd1 | thiosulfate sulfurtransferase (rhodanese)- | 174.27 | 210.72 | 168.04 | 192.8 | 287.92 | 204.95 |
| 41822 | ENSMUSG00000103712 | 0 | NA | NA | NA | NA | NA | Gm37659 | predicted gene, 37659 [Source:MGI Symb | 0 | 0 | 0 | 0 | 0 | 0 |
| 41823 | ENSMUSG00000103713 | 0 | NA | NA | NA | NA | NA | Gm2136 | predicted gene 2136 [Source:MGI Symbol | 0 | 0 | 0 | 0 | 0 | 0 |
| 41824 | ENSMUSG00000103714 | 0 | NA | NA | NA | NA | NA | Gm20857 | predicted gene, 20857 [Source:MGI Symb | 0 | 0 | 0 | 0 | 0 | 0 |
| 41825 | ENSMUSG00000103715 | 29.08440777 | -0.72741151 | 0.437181441 | -1.663866399 | 0.096139131 | 0.457754528 | 4933431K14 | RIKEN cDNA 4933431K14 gene [Source:M | 45.777 | 25.698 | 37.231 | 26.027 | 15.269 | 24.505 |
| 41826 | ENSMUSG00000103716 | 0 | NA | NA | NA | NA | NA | Gm19430 | predicted gene, 19430 [Source:MGI Symb | 0 | 0 | 0 | 0 | 0 | 0 |
| 41827 | ENSMUSG00000103717 | 0.476283062 | 1.031501528 | 3.159856949 | 0.326439312 | 0.744091998 | NA | Gm38038 | predicted gene, 38038 [Source:MGI Symb | 0.8031 | 0 | 0 | 0.964 | 1.0906 | 0 |
| 41828 | ENSMUSG00000103718 | 0.328370405 | 0.16514316 | 4.003149536 | 0.041253308 | 0.967093956 | NA | Gm35299 | predicted gene, 35299 [Source:MGI Symb | 0 | 0 | 1.0062 | 0.964 | 0 | 0 |

| | | | | | | | | | | | | | | | |
|---|---|---|---|---|---|---|---|---|---|---|---|---|---|---|---|
| 41829 | ENSMUSG00000103719 | 0 | NA | NA | NA | NA | NA | Gm38039 | predicted gene, 38039 [Source:MGI Symb | 0 | 0 | 0 | 0 | 0 | 0 |
| 41830 | ENSMUSG00000103720 | 0 | NA | NA | NA | NA | NA | Gm37094 | predicted gene, 37094 [Source:MGI Symb | 0 | 0 | 0 | 0 | 0 | 0 |
| 41831 | ENSMUSG00000103722 | 0 | NA | NA | NA | NA | NA | 2700078F05 | RIKEN cDNA 2700078F05 gene [Source:M( | 0 | 0 | 0 | 0 | 0 | 0 |
| 41832 | ENSMUSG00000103723 | 0 | NA | NA | NA | NA | NA | Gm37095 | predicted gene, 37095 [Source:MGI Symb | 0 | 0 | 0 | 0 | 0 | 0 |
| 41833 | ENSMUSG00000103724 | 0 | NA | NA | NA | NA | NA | Gm37096 | predicted gene, 37096 [Source:MGI Symb | 0 | 0 | 0 | 0 | 0 | 0 |
| 41834 | ENSMUSG00000103725 | 0 | NA | NA | NA | NA | NA | Gm7103 | predicted gene 7103 [Source:MGI Symbol | 0 | 0 | 0 | 0 | 0 | 0 |
| 41835 | ENSMUSG00000103726 | 0 | NA | NA | NA | NA | NA | Gm30074 | predicted gene, 30074 [Source:MGI Symb | 0 | 0 | 0 | 0 | 0 | 0 |
| 41836 | ENSMUSG00000103727 | 0 | NA | NA | NA | NA | NA | Gm37097 | predicted gene, 37097 [Source:MGI Symb | 0 | 0 | 0 | 0 | 0 | 0 |
| 41837 | ENSMUSG00000103728 | 0 | NA | NA | NA | NA | NA | Gm37099 | predicted gene, 37099 [Source:MGI Symb | 0 | 0 | 0 | 0 | 0 | 0 |
| 41838 | ENSMUSG00000103729 | 0 | NA | NA | NA | NA | NA | Gm37098 | predicted gene, 37098 [Source:MGI Symb | 0 | 0 | 0 | 0 | 0 | 0 |
| 41839 | ENSMUSG00000103730 | 0 | NA | NA | NA | NA | NA | Gm38316 | predicted gene, 38316 [Source:MGI Symb | 0 | 0 | 0 | 0 | 0 | 0 |
| 41840 | ENSMUSG00000103731 | 11.93085997 | -0.538404714 | 0.666849904 | -0.807385157 | 0.419444616 | 0.837591673 | Gm17530 | predicted gene, 17530 [Source:MGI Symb | 8.031 | 17.475 | 17.106 | 13.496 | 5.4531 | 10.025 |
| 41841 | ENSMUSG00000103732 | 0.294513347 | 0.165143496 | 4.080472857 | 0.040471657 | 0.967717103 | NA | Gm38315 | predicted gene, 38315 [Source:MGI Symb | 0.8031 | 0 | 0 | 0.964 | 0 | 0 |
| 41842 | ENSMUSG00000103733 | 0 | NA | NA | NA | NA | NA | Gm6162 | predicted gene 6162 [Source:MGI Symbol | 0 | 0 | 0 | 0 | 0 | 0 |
| 41843 | ENSMUSG00000103734 | 0 | NA | NA | NA | NA | NA | Gm37651 | predicted gene, 37651 [Source:MGI Symb | 0 | 0 | 0 | 0 | 0 | 0 |
| 41844 | ENSMUSG00000103735 | 0.542600935 | 1.125931187 | 3.335923981 | 0.33751704 | 0.735727169 | NA | Gm38317 | predicted gene, 38317 [Source:MGI Symb | 0 | 1.0279 | 0 | 0 | 0 | 2.2277 |
| 41845 | ENSMUSG00000103736 | 0 | NA | NA | NA | NA | NA | Gm37650 | predicted gene, 37650 [Source:MGI Symb | 0 | 0 | 0 | 0 | 0 | 0 |
| 41846 | ENSMUSG00000103737 | 0 | NA | NA | NA | NA | NA | Gm7142 | predicted gene 7142 [Source:MGI Symbol | 0 | 0 | 0 | 0 | 0 | 0 |
| 41847 | ENSMUSG00000103738 | 0 | NA | NA | NA | NA | NA | Gm37652 | predicted gene, 37652 [Source:MGI Symb | 0 | 0 | 0 | 0 | 0 | 0 |
| 41848 | ENSMUSG00000103739 | 1.417087145 | -0.234962332 | 1.940873237 | -0.121060113 | 0.903643422 | NA | Gm37653 | predicted gene, 37653 [Source:MGI Symb | 1.6062 | 1.0279 | 2.0125 | 3.8559 | 0 | 0 |
| 41849 | ENSMUSG00000103740 | 0 | NA | NA | NA | NA | NA | Gm38242 | predicted gene, 38242 [Source:MGI Symb | 0 | 0 | 0 | 0 | 0 | 0 |
| 41850 | ENSMUSG00000103741 | 0 | NA | NA | NA | NA | NA | Gm38241 | predicted gene, 38241 [Source:MGI Symb | 0 | 0 | 0 | 0 | 0 | 0 |
| 41851 | ENSMUSG00000103742 | 0 | NA | NA | NA | NA | NA | Gm38240 | predicted gene, 38240 [Source:MGI Symb | 0 | 0 | 0 | 0 | 0 | 0 |
| 41852 | ENSMUSG00000103744 | 0 | NA | NA | NA | NA | NA | Gm38238 | predicted gene, 38238 [Source:MGI Symb | 0 | 0 | 0 | 0 | 0 | 0 |
| 41853 | ENSMUSG00000103745 | 0 | NA | NA | NA | NA | NA | Gm38237 | predicted gene, 38237 [Source:MGI Symb | 0 | 0 | 0 | 0 | 0 | 0 |
| 41854 | ENSMUSG00000103746 | 2.032662203 | 0.477921883 | 1.586158343 | 0.301307801 | 0.763179793 | NA | 1700001G17 | RIKEN cDNA 1700001G17 gene [Source:M | 4.0155 | 0 | 1.0062 | 3.8559 | 1.0906 | 2.2277 |
| 41855 | ENSMUSG00000103747 | 0 | NA | NA | NA | NA | NA | Gm38236 | predicted gene, 38236 [Source:MGI Symb | 0 | 0 | 0 | 0 | 0 | 0 |
| 41856 | ENSMUSG00000103748 | 0 | NA | NA | NA | NA | NA | Gm38243 | predicted gene, 38243 [Source:MGI Symb | 0 | 0 | 0 | 0 | 0 | 0 |
| 41857 | ENSMUSG00000103749 | 1.477388299 | -2.771040116 | 2.195717318 | -1.262020431 | 0.206941435 | NA | Pcdhgb5 | protocadherin gamma subfamily B, 5 [Sou | 1.6062 | 6.1675 | 0 | 0 | 1.0906 | 0 |
| 41858 | ENSMUSG00000103751 | 0 | NA | NA | NA | NA | NA | Gm37283 | predicted gene, 37283 [Source:MGI Symb | 0 | 0 | 0 | 0 | 0 | 0 |
| 41859 | ENSMUSG00000103752 | 0 | NA | NA | NA | NA | NA | Gm36467 | predicted gene, 36467 [Source:MGI Symb | 0 | 0 | 0 | 0 | 0 | 0 |
| 41860 | ENSMUSG00000103753 | 0 | NA | NA | NA | NA | NA | Gm6934 | predicted gene 6934 [Source:MGI Symbol | 0 | 0 | 0 | 0 | 0 | 0 |
| 41861 | ENSMUSG00000103754 | 0.535401493 | -2.48596965 | 3.684117979 | -0.674780141 | 0.499815459 | NA | Gm37284 | predicted gene, 37284 [Source:MGI Symb | 3.2124 | 0 | 0 | 0 | 0 | 0 |
| 41862 | ENSMUSG00000103755 | 0.133850373 | -0.796629176 | 4.080472857 | -0.195229623 | 0.84521319 | NA | Gm37805 | predicted gene, 37805 [Source:MGI Symb | 0.8031 | 0 | 0 | 0 | 0 | 0 |
| 41863 | ENSMUSG00000103756 | 8.05967252 | -0.75159707 | 0.867791667 | -0.86610312 | 0.386433614 | 0.817115886 | Gm37285 | predicted gene, 37285 [Source:MGI Symb | 16.865 | 11.307 | 2.0125 | 3.8559 | 7.6343 | 6.6831 |
| 41864 | ENSMUSG00000103757 | 0 | NA | NA | NA | NA | NA | Gm37286 | predicted gene, 37286 [Source:MGI Symb | 0 | 0 | 0 | 0 | 0 | 0 |
| 41865 | ENSMUSG00000103758 | 0 | NA | NA | NA | NA | NA | Gm37287 | predicted gene, 37287 [Source:MGI Symb | 0 | 0 | 0 | 0 | 0 | 0 |
| 41866 | ENSMUSG00000103759 | 0 | NA | NA | NA | NA | NA | Btf3-ps12 | basic transcription factor 3, pseudogene 1 | 0 | 0 | 0 | 0 | 0 | 0 |
| 41867 | ENSMUSG00000103760 | 0 | NA | NA | NA | NA | NA | Gm8009 | predicted gene 8009 [Source:MGI Symbol | 0 | 0 | 0 | 0 | 0 | 0 |
| 41868 | ENSMUSG00000103761 | 0 | NA | NA | NA | NA | NA | Gm37859 | predicted gene, 37859 [Source:MGI Symb | 0 | 0 | 0 | 0 | 0 | 0 |
| 41869 | ENSMUSG00000103762 | 0 | NA | NA | NA | NA | NA | Gm37861 | predicted gene, 37861 [Source:MGI Symb | 0 | 0 | 0 | 0 | 0 | 0 |
| 41870 | ENSMUSG00000103763 | 5.035958743 | 0.417151824 | 1.002010668 | 0.416314753 | 0.677179698 | NA | Gm37860 | predicted gene, 37860 [Source:MGI Symb | 4.8186 | 6.1675 | 2.0125 | 10.604 | 3.2719 | 3.3415 |
| 41871 | ENSMUSG00000103764 | 0 | NA | NA | NA | NA | NA | Gm18382 | predicted gene, 18382 [Source:MGI Symb | 0 | 0 | 0 | 0 | 0 | 0 |
| 41872 | ENSMUSG00000103765 | 0 | NA | NA | NA | NA | NA | Gm37857 | predicted gene, 37857 [Source:MGI Symb | 0 | 0 | 0 | 0 | 0 | 0 |
| 41873 | ENSMUSG00000103766 | 0.160662974 | 1.126924175 | 4.080472857 | 0.276174898 | 0.782413736 | NA | Gm38392 | predicted gene, 38392 [Source:MGI Symb | 0 | 0 | 0 | 0.964 | 0 | 0 |
| 41874 | ENSMUSG00000103767 | 0 | NA | NA | NA | NA | NA | Gm37858 | predicted gene, 37858 [Source:MGI Symb | 0 | 0 | 0 | 0 | 0 | 0 |
| 41875 | ENSMUSG00000103768 | 0.346303622 | 2.004339653 | 3.884003423 | 0.516049919 | 0.605819544 | NA | Gm37856 | predicted gene, 37856 [Source:MGI Symb | 0 | 0 | 0 | 0.964 | 0 | 1.1138 |
| 41876 | ENSMUSG00000103769 | 0 | NA | NA | NA | NA | NA | Gm7343 | predicted gene 7343 [Source:MGI Symbol | 0 | 0 | 0 | 0 | 0 | 0 |
| 41877 | ENSMUSG00000103770 | 0 | NA | NA | NA | NA | NA | Pcdha9 | protocadherin alpha 9 [Source:MGI Symb | 0 | 0 | 0 | 0 | 0 | 0 |
| 41878 | ENSMUSG00000103771 | 0.335414862 | -1.779047242 | 4.030804179 | -0.441362855 | 0.658950329 | NA | Gm36934 | predicted gene, 36934 [Source:MGI Symb | 0 | 0 | 2.0125 | 0 | 0 | 0 |
| 41879 | ENSMUSG00000103772 | 0 | NA | NA | NA | NA | NA | Gm36933 | predicted gene, 36933 [Source:MGI Symb | 0 | 0 | 0 | 0 | 0 | 0 |
| 41880 | ENSMUSG00000103773 | 0 | NA | NA | NA | NA | NA | Gm34217 | predicted gene, 34217 [Source:MGI Symb | 0 | 0 | 0 | 0 | 0 | 0 |
| 41881 | ENSMUSG00000103774 | 0 | NA | NA | NA | NA | NA | Gm36932 | predicted gene, 36932 [Source:MGI Symb | 0 | 0 | 0 | 0 | 0 | 0 |
| 41882 | ENSMUSG00000103775 | 0 | NA | NA | NA | NA | NA | Gm36997 | predicted gene, 36997 [Source:MGI Symb | 0 | 0 | 0 | 0 | 0 | 0 |
| 41883 | ENSMUSG00000103776 | 0 | NA | NA | NA | NA | NA | 2900035J10I | RIKEN cDNA 2900035J10 gene [Source:M( | 0 | 0 | 0 | 0 | 0 | 0 |
| 41884 | ENSMUSG00000103778 | 0 | NA | NA | NA | NA | NA | Gm36930 | predicted gene, 36930 [Source:MGI Symb | 0 | 0 | 0 | 0 | 0 | 0 |

| | | | | | | | | | | | | | | | |
|---|---|---|---|---|---|---|---|---|---|---|---|---|---|---|---|
| 41885 | ENSMUSG00000103779 | 0 | NA | NA | NA | NA | NA | Gm36931 | predicted gene, 36931 [Source:MGI Symb | 0 | 0 | 0 | 0 | 0 | 0 |
| 41886 | ENSMUSG00000103780 | 0 | NA | NA | NA | NA | NA | Gm37524 | predicted gene, 37524 [Source:MGI Symb | 0 | 0 | 0 | 0 | 0 | 0 |
| 41887 | ENSMUSG00000103781 | 0 | NA | NA | NA | NA | NA | Gm37523 | predicted gene, 37523 [Source:MGI Symb | 0 | 0 | 0 | 0 | 0 | 0 |
| 41888 | ENSMUSG00000103782 | 0 | NA | NA | NA | NA | NA | Gm37656 | predicted gene, 37656 [Source:MGI Symb | 0 | 0 | 0 | 0 | 0 | 0 |
| 41889 | ENSMUSG00000103783 | 0.674415303 | 1.59191567 | 2.786800062 | 0.57123426 | 0.567840858 | NA | Gm37522 | predicted gene, 37522 [Source:MGI Symb | 0 | 1.0279 | 0 | 1.928 | 1.0906 | 0 |
| 41890 | ENSMUSG00000103784 | 0 | NA | NA | NA | NA | NA | Gm37526 | predicted gene, 37526 [Source:MGI Symb | 0 | 0 | 0 | 0 | 0 | 0 |
| 41891 | ENSMUSG00000103785 | 0 | NA | NA | NA | NA | NA | Gm35025 | predicted gene, 35025 [Source:MGI Symb | 0 | 0 | 0 | 0 | 0 | 0 |
| 41892 | ENSMUSG00000103786 | 0 | NA | NA | NA | NA | NA | Gm37525 | predicted gene, 37525 [Source:MGI Symb | 0 | 0 | 0 | 0 | 0 | 0 |
| 41893 | ENSMUSG00000103787 | 0 | NA | NA | NA | NA | NA | Gm37657 | predicted gene, 37657 [Source:MGI Symb | 0 | 0 | 0 | 0 | 0 | 0 |
| 41894 | ENSMUSG00000103788 | 0 | NA | NA | NA | NA | NA | Gm37527 | predicted gene, 37527 [Source:MGI Symb | 0 | 0 | 0 | 0 | 0 | 0 |
| 41895 | ENSMUSG00000103789 | 1.68220395 | 1.376190897 | 1.709990903 | 0.80479428 | 0.420938403 | NA | Gm10129 | predicted gene 10129 [Source:MGI Symbo | 0.8031 | 0 | 2.0125 | 2.8919 | 3.2719 | 1.1138 |
| 41896 | ENSMUSG00000103790 | 0 | NA | NA | NA | NA | NA | Ighv1-17 | immunoglobulin heavy variable 1-17 [Sou | 0 | 0 | 0 | 0 | 0 | 0 |
| 41897 | ENSMUSG00000103791 | 0 | NA | NA | NA | NA | NA | Gm38095 | predicted gene, 38095 [Source:MGI Symb | 0 | 0 | 0 | 0 | 0 | 0 |
| 41898 | ENSMUSG00000103792 | 0 | NA | NA | NA | NA | NA | Gm3745 | predicted gene 3745 [Source:MGI Symbol | 0 | 0 | 0 | 0 | 0 | 0 |
| 41899 | ENSMUSG00000103793 | 1.487994247 | -2.804298781 | 2.158464401 | -1.299210114 | 0.193871831 | NA | Pcdhga6 | protocadherin gamma subfamily A, 6 [Sou | 4.8186 | 0 | 3.0187 | 0 | 1.0906 | 0 |
| 41900 | ENSMUSG00000103794 | 0 | NA | NA | NA | NA | NA | 4930590H14 | RIKEN cDNA 4930590H14 gene [Source:M | 0 | 0 | 0 | 0 | 0 | 0 |
| 41901 | ENSMUSG00000103795 | 0 | NA | NA | NA | NA | NA | Gm37990 | predicted gene, 37990 [Source:MGI Symb | 0 | 0 | 0 | 0 | 0 | 0 |
| 41902 | ENSMUSG00000103796 | 0 | NA | NA | NA | NA | NA | Gm38094 | predicted gene, 38094 [Source:MGI Symb | 0 | 0 | 0 | 0 | 0 | 0 |
| 41903 | ENSMUSG00000103797 | 0 | NA | NA | NA | NA | NA | Gm18964 | predicted gene, 18964 [Source:MGI Symb | 0 | 0 | 0 | 0 | 0 | 0 |
| 41904 | ENSMUSG00000103798 | 0 | NA | NA | NA | NA | NA | Gm2229 | predicted gene 2229 [Source:MGI Symbol | 0 | 0 | 0 | 0 | 0 | 0 |
| 41905 | ENSMUSG00000103799 | 0 | NA | NA | NA | NA | NA | Gm38093 | predicted gene, 38093 [Source:MGI Symb | 0 | 0 | 0 | 0 | 0 | 0 |
| 41906 | ENSMUSG00000103800 | 0 | NA | NA | NA | NA | NA | Pcdha8 | protocadherin alpha 8 [Source:MGI Symb | 0 | 0 | 0 | 0 | 0 | 0 |
| 41907 | ENSMUSG00000103801 | 0 | NA | NA | NA | NA | NA | Spin3-ps | spindlin family, member 3, pseudogene [S | 0 | 0 | 0 | 0 | 0 | 0 |
| 41908 | ENSMUSG00000103802 | 0 | NA | NA | NA | NA | NA | Gm21660 | predicted gene, 21660 [Source:MGI Symb | 0 | 0 | 0 | 0 | 0 | 0 |
| 41909 | ENSMUSG00000103803 | 0 | NA | NA | NA | NA | NA | Gm37064 | predicted gene, 37064 [Source:MGI Symb | 0 | 0 | 0 | 0 | 0 | 0 |
| 41910 | ENSMUSG00000103804 | 5.712406191 | 0.33481615 | 0.939505848 | 0.356374738 | 0.721559946 | 0.9447719 | Gm37062 | predicted gene, 37062 [Source:MGI Symb | 4.0155 | 4.1117 | 7.0437 | 5.7839 | 2.1812 | 11.138 |
| 41911 | ENSMUSG00000103805 | 0 | NA | NA | NA | NA | NA | Gm37063 | predicted gene, 37063 [Source:MGI Symb | 0 | 0 | 0 | 0 | 0 | 0 |
| 41912 | ENSMUSG00000103807 | 0 | NA | NA | NA | NA | NA | Trbv6 | T cell receptor beta variable 6 [Source:MG | 0 | 0 | 0 | 0 | 0 | 0 |
| 41913 | ENSMUSG00000103808 | 0 | NA | NA | NA | NA | NA | Gm37060 | predicted gene, 37060 [Source:MGI Symb | 0 | 0 | 0 | 0 | 0 | 0 |
| 41914 | ENSMUSG00000103809 | 0.167707431 | -0.796629176 | 4.080472857 | -0.195229623 | 0.84521319 | NA | Gm37061 | predicted gene, 37061 [Source:MGI Symb | 0 | 0 | 1.0062 | 0 | 0 | 0 |
| 41915 | ENSMUSG00000103810 | 0 | NA | NA | NA | NA | NA | Gm38005 | predicted gene, 38005 [Source:MGI Symb | 0 | 0 | 0 | 0 | 0 | 0 |
| 41916 | ENSMUSG00000103811 | 0.133850373 | -0.796629176 | 4.080472857 | -0.195229623 | 0.84521319 | NA | Gm38004 | predicted gene, 38004 [Source:MGI Symb | 0.8031 | 0 | 0 | 0 | 0 | 0 |
| 41917 | ENSMUSG00000103812 | 0 | NA | NA | NA | NA | NA | Gm33954 | predicted gene, 33954 [Source:MGI Symb | 0 | 0 | 0 | 0 | 0 | 0 |
| 41918 | ENSMUSG00000103813 | 0 | NA | NA | NA | NA | NA | Gm31422 | predicted gene, 31422 [Source:MGI Symb | 0 | 0 | 0 | 0 | 0 | 0 |
| 41919 | ENSMUSG00000103814 | 0 | NA | NA | NA | NA | NA | Gm38007 | predicted gene, 38007 [Source:MGI Symb | 0 | 0 | 0 | 0 | 0 | 0 |
| 41920 | ENSMUSG00000103815 | 0 | NA | NA | NA | NA | NA | Gm19194 | predicted gene, 19194 [Source:MGI Symb | 0 | 0 | 0 | 0 | 0 | 0 |
| 41921 | ENSMUSG00000103816 | 0 | NA | NA | NA | NA | NA | Gm38006 | predicted gene, 38006 [Source:MGI Symb | 0 | 0 | 0 | 0 | 0 | 0 |
| 41922 | ENSMUSG00000103817 | 0 | NA | NA | NA | NA | NA | Gm6680 | predicted gene 6680 [Source:MGI Symbol | 0 | 0 | 0 | 0 | 0 | 0 |
| 41923 | ENSMUSG00000103818 | 0 | NA | NA | NA | NA | NA | Gm38009 | predicted gene, 38009 [Source:MGI Symb | 0 | 0 | 0 | 0 | 0 | 0 |
| 41924 | ENSMUSG00000103819 | 0 | NA | NA | NA | NA | NA | Gm38008 | predicted gene, 38008 [Source:MGI Symb | 0 | 0 | 0 | 0 | 0 | 0 |
| 41925 | ENSMUSG00000103821 | 6.971148422 | -0.898121767 | 0.900506576 | -0.997351703 | 0.318593826 | 0.7652277 | D430013B06 | RIKEN cDNA D430013B06 gene [Source:M | 8.8341 | 15.419 | 3.0187 | 5.7839 | 6.5437 | 2.2277 |
| 41926 | ENSMUSG00000103822 | 0.331982614 | 0.16514309 | 3.986751476 | 0.041422971 | 0.966958701 | NA | 6030460B20 | RIKEN cDNA 6030460B20 gene [Source:M | 0 | 1.0279 | 0 | 0.964 | 0 | 0 |
| 41927 | ENSMUSG00000103823 | 0 | NA | NA | NA | NA | NA | Gm7899 | predicted gene 7899 [Source:MGI Symbol | 0 | 0 | 0 | 0 | 0 | 0 |
| 41928 | ENSMUSG00000103824 | 0 | NA | NA | NA | NA | NA | Gm38177 | predicted gene, 38177 [Source:MGI Symb | 0 | 0 | 0 | 0 | 0 | 0 |
| 41929 | ENSMUSG00000103825 | 0 | NA | NA | NA | NA | NA | Gm38178 | predicted gene, 38178 [Source:MGI Symb | 0 | 0 | 0 | 0 | 0 | 0 |
| 41930 | ENSMUSG00000103827 | 0.449470461 | -0.545136533 | 3.562105917 | -0.15303771 | 0.878368537 | NA | Gm32950 | predicted gene, 32950 [Source:MGI Symb | 1.6062 | 0 | 0 | 0 | 1.0906 | 0 |
| 41931 | ENSMUSG00000103828 | 0 | NA | NA | NA | NA | NA | 2610001A08 | RIKEN cDNA 2610001A08 gene [Source:M | 0 | 0 | 0 | 0 | 0 | 0 |
| 41932 | ENSMUSG00000103829 | 0 | NA | NA | NA | NA | NA | Gm38174 | predicted gene, 38174 [Source:MGI Symb | 0 | 0 | 0 | 0 | 0 | 0 |
| 41933 | ENSMUSG00000103830 | 0 | NA | NA | NA | NA | NA | Gm37609 | predicted gene, 37609 [Source:MGI Symb | 0 | 0 | 0 | 0 | 0 | 0 |
| 41934 | ENSMUSG00000103831 | 0 | NA | NA | NA | NA | NA | Gm37608 | predicted gene, 37608 [Source:MGI Symb | 0 | 0 | 0 | 0 | 0 | 0 |
| 41935 | ENSMUSG00000103832 | 0 | NA | NA | NA | NA | NA | Gm37611 | predicted gene, 37611 [Source:MGI Symb | 0 | 0 | 0 | 0 | 0 | 0 |
| 41936 | ENSMUSG00000103833 | 0 | NA | NA | NA | NA | NA | Gm49885 | predicted gene, 49885 [Source:MGI Symb | 0 | 0 | 0 | 0 | 0 | 0 |
| 41937 | ENSMUSG00000103834 | 0 | NA | NA | NA | NA | NA | Gm37613 | predicted gene, 37613 [Source:MGI Symb | 0 | 0 | 0 | 0 | 0 | 0 |
| 41938 | ENSMUSG00000103835 | 0.331982614 | 0.16514309 | 3.986751476 | 0.041422971 | 0.966958701 | NA | Gm37612 | predicted gene, 37612 [Source:MGI Symb | 0 | 1.0279 | 0 | 0.964 | 0 | 0 |
| 41939 | ENSMUSG00000103837 | 0 | NA | NA | NA | NA | NA | Gm37614 | predicted gene, 37614 [Source:MGI Symb | 0 | 0 | 0 | 0 | 0 | 0 |
| 41940 | ENSMUSG00000103839 | 5.558115241 | -0.491540165 | 0.902938755 | -0.54437819 | 0.586181251 | 0.910005969 | Gm37607 | predicted gene, 37607 [Source:MGI Symb | 7.2279 | 5.1396 | 7.0437 | 2.8919 | 4.3625 | 6.6831 |

| | | | | | | | | | | | | | | | |
|---|---|---|---|---|---|---|---|---|---|---|---|---|---|---|---|
| 41941 | ENSMUSG00000103840 | 0 | NA | NA | NA | NA | NA | 1700040K01 | RIKEN cDNA 1700040K01 gene [Source:M | 0 | 0 | 0 | 0 | 0 | 0 |
| 41942 | ENSMUSG00000103841 | 0 | NA | NA | NA | NA | NA | Gm37807 | predicted gene, 37807 [Source:MGI Symb | 0 | 0 | 0 | 0 | 0 | 0 |
| 41943 | ENSMUSG00000103842 | 0 | NA | NA | NA | NA | NA | Gm37808 | predicted gene, 37808 [Source:MGI Symb | 0 | 0 | 0 | 0 | 0 | 0 |
| 41944 | ENSMUSG00000103843 | 0.649696354 | 1.537929172 | 3.118369374 | 0.493183773 | 0.621882743 | NA | Gm31266 | predicted gene, 31266 [Source:MGI Symb | 0 | 0 | 1.0062 | 2.8919 | 0 | 0 |
| 41945 | ENSMUSG00000103844 | 0 | NA | NA | NA | NA | NA | 4933436E23 | RIKEN cDNA 4933436E23 gene [Source:M | 0 | 0 | 0 | 0 | 0 | 0 |
| 41946 | ENSMUSG00000103845 | 0 | NA | NA | NA | NA | NA | Gm19026 | predicted gene, 19026 [Source:MGI Symb | 0 | 0 | 0 | 0 | 0 | 0 |
| 41947 | ENSMUSG00000103846 | 0.315620088 | 0.165143421 | 4.063300985 | 0.040642675 | 0.967580763 | NA | Gm37809 | predicted gene, 37809 [Source:MGI Symb | 0.8031 | 0 | 0 | 0 | 1.0906 | 0 |
| 41948 | ENSMUSG00000103847 | 1.198850927 | -2.412458447 | 2.309606273 | -1.044532341 | 0.296239164 | NA | Gm20056 | predicted gene, 20056 [Source:MGI Symb | 0 | 3.0838 | 3.0187 | 0 | 1.0906 | 0 |
| 41949 | ENSMUSG00000103848 | 0 | NA | NA | NA | NA | NA | Gm17934 | predicted gene, 17934 [Source:MGI Symb | 0 | 0 | 0 | 0 | 0 | 0 |
| 41950 | ENSMUSG00000103849 | 0 | NA | NA | NA | NA | NA | Gm37806 | predicted gene, 37806 [Source:MGI Symb | 0 | 0 | 0 | 0 | 0 | 0 |
| 41951 | ENSMUSG00000103850 | 0 | NA | NA | NA | NA | NA | Gm3933 | predicted gene 3933 [Source:MGI Symbol | 0 | 0 | 0 | 0 | 0 | 0 |
| 41952 | ENSMUSG00000103851 | 0.849373781 | -3.117833363 | 2.761285217 | -1.129123983 | 0.258845535 | NA | Gm37606 | predicted gene, 37606 [Source:MGI Symb | 0 | 3.0838 | 2.0125 | 0 | 0 | 0 |
| 41953 | ENSMUSG00000103852 | 0 | NA | NA | NA | NA | NA | Gm18658 | predicted gene, 18658 [Source:MGI Symb | 0 | 0 | 0 | 0 | 0 | 0 |
| 41954 | ENSMUSG00000103854 | 0.167707431 | -0.796629176 | 4.080472857 | -0.195229623 | 0.84521319 | NA | Gm37250 | predicted gene, 37250 [Source:MGI Symb | 0 | 0 | 1.0062 | 0 | 0 | 0 |
| 41955 | ENSMUSG00000103856 | 0 | NA | NA | NA | NA | NA | Gm2474 | predicted gene 2474 [Source:MGI Symbol | 0 | 0 | 0 | 0 | 0 | 0 |
| 41956 | ENSMUSG00000103857 | 4.239275147 | 0.639200147 | 1.032594975 | 0.619023104 | 0.535901138 | NA | Gm37249 | predicted gene, 37249 [Source:MGI Symb | 4.8186 | 2.0558 | 3.0187 | 6.7478 | 5.4531 | 3.3415 |
| 41957 | ENSMUSG00000103858 | 0 | NA | NA | NA | NA | NA | Gm37604 | predicted gene, 37604 [Source:MGI Symb | 0 | 0 | 0 | 0 | 0 | 0 |
| 41958 | ENSMUSG00000103859 | 0 | NA | NA | NA | NA | NA | Gm37603 | predicted gene, 37603 [Source:MGI Symb | 0 | 0 | 0 | 0 | 0 | 0 |
| 41959 | ENSMUSG00000103860 | 0 | NA | NA | NA | NA | NA | Gm37418 | predicted gene, 37418 [Source:MGI Symb | 0 | 0 | 0 | 0 | 0 | 0 |
| 41960 | ENSMUSG00000103862 | 0 | NA | NA | NA | NA | NA | Gm10764 | predicted gene 10764 [Source:MGI Symbo | 0 | 0 | 0 | 0 | 0 | 0 |
| 41961 | ENSMUSG00000103863 | 0 | NA | NA | NA | NA | NA | Olfr1566-ps1 | olfactory receptor 1566, pseudogene 1 [S | 0 | 0 | 0 | 0 | 0 | 0 |
| 41962 | ENSMUSG00000103864 | 5.473537681 | 0.259219397 | 1.047504709 | 0.247463705 | 0.804549377 | 0.963787787 | Gm37415 | predicted gene, 37415 [Source:MGI Symb | 8.8341 | 3.0838 | 3.0187 | 13.496 | 2.1812 | 2.2277 |
| 41963 | ENSMUSG00000103865 | 0 | NA | NA | NA | NA | NA | Gm37416 | predicted gene, 37416 [Source:MGI Symb | 0 | 0 | 0 | 0 | 0 | 0 |
| 41964 | ENSMUSG00000103867 | 0.294513347 | 0.165143496 | 4.080472857 | 0.040471657 | 0.967717103 | NA | 9630010A21 | RIKEN cDNA 9630010A21 gene [Source:M | 0.8031 | 0 | 0 | 0.964 | 0 | 0 |
| 41965 | ENSMUSG00000103869 | 2.791795336 | -0.376541672 | 1.35193655 | -0.278520225 | 0.780613042 | NA | Gm37420 | predicted gene, 37420 [Source:MGI Symb | 2.4093 | 2.0558 | 5.0312 | 2.8919 | 4.3625 | 0 |
| 41966 | ENSMUSG00000103870 | 0 | NA | NA | NA | NA | NA | Gm38362 | predicted gene, 38362 [Source:MGI Symb | 0 | 0 | 0 | 0 | 0 | 0 |
| 41967 | ENSMUSG00000103871 | 0 | NA | NA | NA | NA | NA | Gm38361 | predicted gene, 38361 [Source:MGI Symb | 0 | 0 | 0 | 0 | 0 | 0 |
| 41968 | ENSMUSG00000103872 | 0 | NA | NA | NA | NA | NA | Gm38363 | predicted gene, 38363 [Source:MGI Symb | 0 | 0 | 0 | 0 | 0 | 0 |
| 41969 | ENSMUSG00000103873 | 0 | NA | NA | NA | NA | NA | Ighv1-38 | immunoglobulin heavy variable V1-38 [So | 0 | 0 | 0 | 0 | 0 | 0 |
| 41970 | ENSMUSG00000103874 | 0 | NA | NA | NA | NA | NA | Gm38360 | predicted gene, 38360 [Source:MGI Symb | 0 | 0 | 0 | 0 | 0 | 0 |
| 41971 | ENSMUSG00000103875 | 0 | NA | NA | NA | NA | NA | Ighv10-2 | immunoglobulin heavy variable V10-2 [So | 0 | 0 | 0 | 0 | 0 | 0 |
| 41972 | ENSMUSG00000103876 | 0 | NA | NA | NA | NA | NA | Gm17806 | predicted gene, 17806 [Source:MGI Symb | 0 | 0 | 0 | 0 | 0 | 0 |
| 41973 | ENSMUSG00000103877 | 0 | NA | NA | NA | NA | NA | Gm21358 | predicted gene, 21358 [Source:MGI Symb | 0 | 0 | 0 | 0 | 0 | 0 |
| 41974 | ENSMUSG00000103878 | 0 | NA | NA | NA | NA | NA | Gm8197 | predicted gene 8197 [Source:MGI Symbol | 0 | 0 | 0 | 0 | 0 | 0 |
| 41975 | ENSMUSG00000103880 | 0 | NA | NA | NA | NA | NA | Gm37009 | predicted gene, 37009 [Source:MGI Symb | 0 | 0 | 0 | 0 | 0 | 0 |
| 41976 | ENSMUSG00000103881 | 0.970525971 | -0.749213202 | 2.255458623 | -0.332177764 | 0.739755033 | NA | Gm10748 | predicted gene 10748 [Source:MGI Symbo | 1.6062 | 0 | 2.0125 | 0 | 1.0906 | 1.1138 |
| 41977 | ENSMUSG00000103882 | 0.353348078 | 0.165142701 | 3.895042568 | 0.042398176 | 0.966181282 | NA | Gm37452 | predicted gene, 37452 [Source:MGI Symb | 0 | 0 | 1.0062 | 0 | 0 | 1.1138 |
| 41978 | ENSMUSG00000103883 | 0 | NA | NA | NA | NA | NA | Gm37010 | predicted gene, 37010 [Source:MGI Symb | 0 | 0 | 0 | 0 | 0 | 0 |
| 41979 | ENSMUSG00000103884 | 0 | NA | NA | NA | NA | NA | Gm37005 | predicted gene, 37005 [Source:MGI Symb | 0 | 0 | 0 | 0 | 0 | 0 |
| 41980 | ENSMUSG00000103885 | 0 | NA | NA | NA | NA | NA | Gm37006 | predicted gene, 37006 [Source:MGI Symb | 0 | 0 | 0 | 0 | 0 | 0 |
| 41981 | ENSMUSG00000103886 | 0 | NA | NA | NA | NA | NA | Gm37007 | predicted gene, 37007 [Source:MGI Symb | 0 | 0 | 0 | 0 | 0 | 0 |
| 41982 | ENSMUSG00000103887 | 4.023881595 | -0.654079516 | 1.043773226 | -0.626649064 | 0.530889299 | NA | Gm37008 | predicted gene, 37008 [Source:MGI Symb | 5.6217 | 4.1117 | 5.0312 | 3.8559 | 2.1812 | 3.3415 |
| 41983 | ENSMUSG00000103889 | 0 | NA | NA | NA | NA | NA | Gm37011 | predicted gene, 37011 [Source:MGI Symb | 0 | 0 | 0 | 0 | 0 | 0 |
| 41984 | ENSMUSG00000103890 | 0 | NA | NA | NA | NA | NA | Gm28724 | predicted gene 28724 [Source:MGI Symbo | 0 | 0 | 0 | 0 | 0 | 0 |
| 41985 | ENSMUSG00000103891 | 0 | NA | NA | NA | NA | NA | Gm37941 | predicted gene, 37941 [Source:MGI Symb | 0 | 0 | 0 | 0 | 0 | 0 |
| 41986 | ENSMUSG00000103892 | 0 | NA | NA | NA | NA | NA | Gm31572 | predicted gene, 31572 [Source:MGI Symb | 0 | 0 | 0 | 0 | 0 | 0 |
| 41987 | ENSMUSG00000103893 | 0 | NA | NA | NA | NA | NA | Ighv5-18 | immunoglobulin heavy variable V5-18 [So | 0 | 0 | 0 | 0 | 0 | 0 |
| 41988 | ENSMUSG00000103894 | 0 | NA | NA | NA | NA | NA | Gm37129 | predicted gene, 37129 [Source:MGI Symb | 0 | 0 | 0 | 0 | 0 | 0 |
| 41989 | ENSMUSG00000103895 | 0 | NA | NA | NA | NA | NA | Gm37942 | predicted gene, 37942 [Source:MGI Symb | 0 | 0 | 0 | 0 | 0 | 0 |
| 41990 | ENSMUSG00000103896 | 0 | NA | NA | NA | NA | NA | Gm37128 | predicted gene, 37128 [Source:MGI Symb | 0 | 0 | 0 | 0 | 0 | 0 |
| 41991 | ENSMUSG00000103897 | 0.788497532 | -1.691662715 | 2.429158853 | -0.696398555 | 0.486179268 | NA | Pcdhga8 | protocadherin gamma subfamily A, 8 [Sou | 1.6062 | 1.0279 | 1.0062 | 0 | 1.0906 | 0 |
| 41992 | ENSMUSG00000103898 | 24.63005842 | 0.818848965 | 0.453364708 | 1.806159479 | 0.070893432 | 0.385556607 | Gm30238 | predicted gene, 30238 [Source:MGI Symb | 12.047 | 19.53 | 22.137 | 27.955 | 33.809 | 32.301 |
| 41993 | ENSMUSG00000103899 | 0.321325949 | 1.911575114 | 4.048267893 | 0.472195804 | 0.636787034 | NA | Gm37940 | predicted gene, 37940 [Source:MGI Symb | 0 | 0 | 0 | 1.928 | 0 | 0 |
| 41994 | ENSMUSG00000103900 | 0.831466069 | 1.993315099 | 2.60242648 | 0.765944827 | 0.443709132 | NA | Gm37500 | predicted gene, 37500 [Source:MGI Symb | 0 | 0 | 1.0062 | 2.8919 | 1.0906 | 0 |
| 41995 | ENSMUSG00000103901 | 7.084631109 | -0.103443934 | 0.88555699 | -0.116812284 | 0.907008809 | 0.984383096 | Gm37499 | predicted gene, 37499 [Source:MGI Symb | 5.6217 | 13.363 | 3.0187 | 2.8919 | 9.8156 | 7.7969 |
| 41996 | ENSMUSG00000103902 | 0 | NA | NA | NA | NA | NA | Gm37501 | predicted gene, 37501 [Source:MGI Symb | 0 | 0 | 0 | 0 | 0 | 0 |

| | | | | | | | | | | | | | | | |
|---|---|---|---|---|---|---|---|---|---|---|---|---|---|---|---|
| 41997 | ENSMUSG00000103903 | 0 | NA | NA | NA | NA | NA | Rps2-ps2 | ribosomal protein S2, pseudogene 2 [Sour | 0 | 0 | 0 | 0 | 0 | 0 |
| 41998 | ENSMUSG00000103904 | 0 | NA | NA | NA | NA | NA | Gm37497 | predicted gene, 37497 [Source:MGI Symb | 0 | 0 | 0 | 0 | 0 | 0 |
| 41999 | ENSMUSG00000103905 | 0 | NA | NA | NA | NA | NA | Gm34379 | predicted gene, 34379 [Source:MGI Symb | 0 | 0 | 0 | 0 | 0 | 0 |
| 42000 | ENSMUSG00000103906 | 99.31367334 | -0.368590829 | 0.240565717 | -1.53218353 | 0.125477152 | 0.51943375 | Tigd5 | tigger transposable element derived 5 [So | 97.175 | 106.9 | 131.82 | 78.082 | 88.34 | 93.563 |
| 42001 | ENSMUSG00000103907 | 0 | NA | NA | NA | NA | NA | Gm37498 | predicted gene, 37498 [Source:MGI Symb | 0 | 0 | 0 | 0 | 0 | 0 |
| 42002 | ENSMUSG00000103908 | 0 | NA | NA | NA | NA | NA | Gm37502 | predicted gene, 37502 [Source:MGI Symb | 0 | 0 | 0 | 0 | 0 | 0 |
| 42003 | ENSMUSG00000103909 | 0 | NA | NA | NA | NA | NA | Gm31511 | predicted gene, 31511 [Source:MGI Symb | 0 | 0 | 0 | 0 | 0 | 0 |
| 42004 | ENSMUSG00000103911 | 0 | NA | NA | NA | NA | NA | Gm38074 | predicted gene, 38074 [Source:MGI Symb | 0 | 0 | 0 | 0 | 0 | 0 |
| 42005 | ENSMUSG00000103912 | 0.342432689 | 1.990653086 | 3.900281263 | 0.51038706 | 0.609780321 | NA | Gm33819 | predicted gene, 33819 [Source:MGI Symb | 0 | 0 | 0 | 0.964 | 1.0906 | 0 |
| 42006 | ENSMUSG00000103913 | 0 | NA | NA | NA | NA | NA | Gm18403 | predicted gene, 18403 [Source:MGI Symb | 0 | 0 | 0 | 0 | 0 | 0 |
| 42007 | ENSMUSG00000103914 | 0 | NA | NA | NA | NA | NA | Gm38073 | predicted gene, 38073 [Source:MGI Symb | 0 | 0 | 0 | 0 | 0 | 0 |
| 42008 | ENSMUSG00000103915 | 0 | NA | NA | NA | NA | NA | Ighv1-6 | immunoglobulin heavy variable V1-6 [Sou | 0 | 0 | 0 | 0 | 0 | 0 |
| 42009 | ENSMUSG00000103916 | 0.181769715 | 1.126924175 | 4.080472857 | 0.276174898 | 0.782413736 | NA | Gm38071 | predicted gene, 38071 [Source:MGI Symb | 0 | 0 | 0 | 0 | 1.0906 | 0 |
| 42010 | ENSMUSG00000103917 | 0 | NA | NA | NA | NA | NA | Gm38072 | predicted gene, 38072 [Source:MGI Symb | 0 | 0 | 0 | 0 | 0 | 0 |
| 42011 | ENSMUSG00000103918 | 0 | NA | NA | NA | NA | NA | Ighv12-1 | immunoglobulin heavy variable V12-1 [So | 0 | 0 | 0 | 0 | 0 | 0 |
| 42012 | ENSMUSG00000103919 | 0 | NA | NA | NA | NA | NA | Gm21366 | predicted gene, 21366 [Source:MGI Symb | 0 | 0 | 0 | 0 | 0 | 0 |
| 42013 | ENSMUSG00000103920 | 0 | NA | NA | NA | NA | NA | Gm37897 | predicted gene, 37897 [Source:MGI Symb | 0 | 0 | 0 | 0 | 0 | 0 |
| 42014 | ENSMUSG00000103921 | 0 | NA | NA | NA | NA | NA | Gm5841 | predicted gene 5841 [Source:MGI Symbol | 0 | 0 | 0 | 0 | 0 | 0 |
| 42015 | ENSMUSG00000103922 | 1.074106519 | 1.156375985 | 2.39018303 | 0.483802274 | 0.628526201 | NA | Gm6123 | predicted gene 6123 [Source:MGI Symbol | 0 | 0 | 2.0125 | 0 | 1.0906 | 3.3415 |
| 42016 | ENSMUSG00000103923 | 0.181769715 | 1.126924175 | 4.080472857 | 0.276174898 | 0.782413736 | NA | Gm37896 | predicted gene, 37896 [Source:MGI Symb | 0 | 0 | 0 | 0 | 1.0906 | 0 |
| 42017 | ENSMUSG00000103924 | 0 | NA | NA | NA | NA | NA | Gm37895 | predicted gene, 37895 [Source:MGI Symb | 0 | 0 | 0 | 0 | 0 | 0 |
| 42018 | ENSMUSG00000103925 | 2.584174666 | -0.704065457 | 1.565413483 | -0.449763251 | 0.652881159 | NA | B230112G18 | RIKEN cDNA B230112G18 gene [Source:M | 1.6062 | 3.0838 | 5.0312 | 5.7839 | 0 | 0 |
| 42019 | ENSMUSG00000103926 | 0 | NA | NA | NA | NA | NA | Gm37894 | predicted gene, 37894 [Source:MGI Symb | 0 | 0 | 0 | 0 | 0 | 0 |
| 42020 | ENSMUSG00000103927 | 0 | NA | NA | NA | NA | NA | Gm9932 | predicted gene 9932 [Source:MGI Symbol | 0 | 0 | 0 | 0 | 0 | 0 |
| 42021 | ENSMUSG00000103928 | 31.83740765 | 0.304583584 | 0.38956474 | 0.781856141 | 0.434299123 | 0.845264432 | Gm37893 | predicted gene, 37893 [Source:MGI Symb | 30.518 | 29.81 | 25.156 | 39.523 | 38.172 | 27.846 |
| 42022 | ENSMUSG00000103929 | 0 | NA | NA | NA | NA | NA | Gm37892 | predicted gene, 37892 [Source:MGI Symb | 0 | 0 | 0 | 0 | 0 | 0 |
| 42023 | ENSMUSG00000103930 | 0.74256259 | 3.09446134 | 3.273353887 | 0.945348852 | 0.344480805 | NA | Gm36961 | predicted gene, 36961 [Source:MGI Symb | 0 | 0 | 0 | 0 | 0 | 4.4554 |
| 42024 | ENSMUSG00000103931 | 0 | NA | NA | NA | NA | NA | Gm36962 | predicted gene, 36962 [Source:MGI Symb | 0 | 0 | 0 | 0 | 0 | 0 |
| 42025 | ENSMUSG00000103932 | 0.160662974 | 1.126924175 | 4.080472857 | 0.276174898 | 0.782413736 | NA | Gm36963 | predicted gene, 36963 [Source:MGI Symb | 0 | 0 | 0 | 0.964 | 0 | 0 |
| 42026 | ENSMUSG00000103933 | 0.133850373 | -0.796629176 | 4.080472857 | -0.195229623 | 0.84521319 | NA | Gm36964 | predicted gene, 36964 [Source:MGI Symb | 0.8031 | 0 | 0 | 0 | 0 | 0 |
| 42027 | ENSMUSG00000103934 | 0 | NA | NA | NA | NA | NA | Ighv5-3 | immunoglobulin heavy variable V5-3 [Sou | 0 | 0 | 0 | 0 | 0 | 0 |
| 42028 | ENSMUSG00000103935 | 0 | NA | NA | NA | NA | NA | AY702102 | cDNA sequence AY702102 [Source:MGI Sy | 0 | 0 | 0 | 0 | 0 | 0 |
| 42029 | ENSMUSG00000103936 | 0 | NA | NA | NA | NA | NA | Gm36965 | predicted gene, 36965 [Source:MGI Symb | 0 | 0 | 0 | 0 | 0 | 0 |
| 42030 | ENSMUSG00000103937 | 0.956437057 | 1.282470111 | 2.271719443 | 0.56453719 | 0.572388597 | NA | Gm37186 | predicted gene, 37186 [Source:MGI Symb | 1.6062 | 0 | 0 | 1.928 | 1.0906 | 1.1138 |
| 42031 | ENSMUSG00000103938 | 0 | NA | NA | NA | NA | NA | Gm36960 | predicted gene, 36960 [Source:MGI Symb | 0 | 0 | 0 | 0 | 0 | 0 |
| 42032 | ENSMUSG00000103939 | 0 | NA | NA | NA | NA | NA | Ighv3-4 | immunoglobulin heavy variable V3-4 [Sou | 0 | 0 | 0 | 0 | 0 | 0 |
| 42033 | ENSMUSG00000103940 | 4.551380364 | 0.978238473 | 1.005746414 | 0.972649228 | 0.330727667 | NA | Gm38227 | predicted gene, 38227 [Source:MGI Symb | 4.0155 | 3.0838 | 2.0125 | 3.8559 | 6.5437 | 7.7969 |
| 42034 | ENSMUSG00000103942 | 0 | NA | NA | NA | NA | NA | Gm34732 | predicted gene, 34732 [Source:MGI Symb | 0 | 0 | 0 | 0 | 0 | 0 |
| 42035 | ENSMUSG00000103943 | 0 | NA | NA | NA | NA | NA | Gm37344 | predicted gene, 37344 [Source:MGI Symb | 0 | 0 | 0 | 0 | 0 | 0 |
| 42036 | ENSMUSG00000103944 | 0 | NA | NA | NA | NA | NA | Rpl26-ps3 | ribosomal protein L26, pseudogene 3 [Sou | 0 | 0 | 0 | 0 | 0 | 0 |
| 42037 | ENSMUSG00000103945 | 0.681718484 | 0.074918168 | 2.804856827 | 0.026710158 | 0.978690911 | NA | Gm38228 | predicted gene, 38228 [Source:MGI Symb | 0 | 0 | 2.0125 | 0.964 | 0 | 1.1138 |
| 42038 | ENSMUSG00000103947 | 0 | NA | NA | NA | NA | NA | Gm37345 | predicted gene, 37345 [Source:MGI Symb | 0 | 0 | 0 | 0 | 0 | 0 |
| 42039 | ENSMUSG00000103948 | 0 | NA | NA | NA | NA | NA | 4930594C11 | RIKEN cDNA 4930594C11 gene [Source:M | 0 | 0 | 0 | 0 | 0 | 0 |
| 42040 | ENSMUSG00000103949 | 0.305170013 | -1.657836477 | 4.041480335 | -0.410205257 | 0.681655385 | NA | Gm38226 | predicted gene, 38226 [Source:MGI Symb | 0.8031 | 1.0279 | 0 | 0 | 0 | 0 |
| 42041 | ENSMUSG00000103950 | 0 | NA | NA | NA | NA | NA | Gm8939 | predicted gene 8939 [Source:MGI Symbol | 0 | 0 | 0 | 0 | 0 | 0 |
| 42042 | ENSMUSG00000103951 | 1.341606189 | 1.631810917 | 2.190449601 | 0.744966201 | 0.456292153 | NA | Gm37269 | predicted gene, 37269 [Source:MGI Symb | 0 | 0 | 2.0125 | 3.8559 | 2.1812 | 0 |
| 42043 | ENSMUSG00000103952 | 0 | NA | NA | NA | NA | NA | Gm37268 | predicted gene, 37268 [Source:MGI Symb | 0 | 0 | 0 | 0 | 0 | 0 |
| 42044 | ENSMUSG00000103953 | 0.17131964 | -0.796629176 | 4.080472857 | -0.195229623 | 0.84521319 | NA | Gm29718 | predicted gene, 29718 [Source:MGI Symb | 0 | 1.0279 | 0 | 0 | 0 | 0 |
| 42045 | ENSMUSG00000103954 | 0 | NA | NA | NA | NA | NA | Gm37721 | predicted gene, 37721 [Source:MGI Symb | 0 | 0 | 0 | 0 | 0 | 0 |
| 42046 | ENSMUSG00000103955 | 0 | NA | NA | NA | NA | NA | Gm37270 | predicted gene, 37270 [Source:MGI Symb | 0 | 0 | 0 | 0 | 0 | 0 |
| 42047 | ENSMUSG00000103956 | 0 | NA | NA | NA | NA | NA | Gm37118 | predicted gene, 37118 [Source:MGI Symb | 0 | 0 | 0 | 0 | 0 | 0 |
| 42048 | ENSMUSG00000103957 | 5.653937782 | -0.138446373 | 0.997298264 | -0.138821432 | 0.889591258 | 0.982382091 | Gm10766 | predicted gene 10766 [Source:MGI Symbo | 9.6372 | 4.1117 | 4.025 | 10.604 | 1.0906 | 4.4554 |
| 42049 | ENSMUSG00000103958 | 0 | NA | NA | NA | NA | NA | Gm8023 | predicted gene 8023 [Source:MGI Symbol | 0 | 0 | 0 | 0 | 0 | 0 |
| 42050 | ENSMUSG00000103959 | 0 | NA | NA | NA | NA | NA | Gm37720 | predicted gene, 37720 [Source:MGI Symb | 0 | 0 | 0 | 0 | 0 | 0 |
| 42051 | ENSMUSG00000103960 | 0 | NA | NA | NA | NA | NA | Gm37119 | predicted gene, 37119 [Source:MGI Symb | 0 | 0 | 0 | 0 | 0 | 0 |
| 42052 | ENSMUSG00000103961 | 0 | NA | NA | NA | NA | NA | Gm36388 | predicted gene, 36388 [Source:MGI Symb | 0 | 0 | 0 | 0 | 0 | 0 |

| | | | | | | | | | | | | | | | |
|---|---|---|---|---|---|---|---|---|---|---|---|---|---|---|---|
| 42053 | ENSMUSG00000103962 | 0 | NA | NA | NA | NA | NA | Gm38213 | predicted gene, 38213 [Source:MGI Symb | 0 | 0 | 0 | 0 | 0 | 0 |
| 42054 | ENSMUSG00000103963 | 0 | NA | NA | NA | NA | NA | Gm30932 | predicted gene, 30932 [Source:MGI Symb | 0 | 0 | 0 | 0 | 0 | 0 |
| 42055 | ENSMUSG00000103964 | 0 | NA | NA | NA | NA | NA | Gm8407 | predicted gene 8407 [Source:MGI Symbol | 0 | 0 | 0 | 0 | 0 | 0 |
| 42056 | ENSMUSG00000103965 | 1.557717738 | -0.907768017 | 1.895587983 | -0.478884666 | 0.63202068 | NA | Gm30173 | predicted gene, 30173 [Source:MGI Symb | 4.0155 | 0 | 2.0125 | 0 | 1.0906 | 2.2277 |
| 42057 | ENSMUSG00000103966 | 0 | NA | NA | NA | NA | NA | Gm37120 | predicted gene, 37120 [Source:MGI Symb | 0 | 0 | 0 | 0 | 0 | 0 |
| 42058 | ENSMUSG00000103967 | 1.878242539 | 0.954312404 | 1.547105266 | 0.616837409 | 0.537341975 | NA | Gm38214 | predicted gene, 38214 [Source:MGI Symb | 0.8031 | 2.0558 | 1.0062 | 1.928 | 4.3625 | 1.1138 |
| 42059 | ENSMUSG00000103968 | 0 | NA | NA | NA | NA | NA | 2610509F24 | RIKEN cDNA 2610509F24 gene [Source:M | 0 | 0 | 0 | 0 | 0 | 0 |
| 42060 | ENSMUSG00000103969 | 0 | NA | NA | NA | NA | NA | Gm21890 | predicted gene, 21890 [Source:MGI Symb | 0 | 0 | 0 | 0 | 0 | 0 |
| 42061 | ENSMUSG00000103970 | 0.160662974 | 1.126924175 | 4.080472857 | 0.276174898 | 0.782413736 | NA | Gm37678 | predicted gene, 37678 [Source:MGI Symb | 0 | 0 | 0 | 0.964 | 0 | 0 |
| 42062 | ENSMUSG00000103971 | 0 | NA | NA | NA | NA | NA | Gm37679 | predicted gene, 37679 [Source:MGI Symb | 0 | 0 | 0 | 0 | 0 | 0 |
| 42063 | ENSMUSG00000103972 | 0 | NA | NA | NA | NA | NA | Gm37466 | predicted gene, 37466 [Source:MGI Symb | 0 | 0 | 0 | 0 | 0 | 0 |
| 42064 | ENSMUSG00000103973 | 2.981720097 | -1.668781586 | 1.303548568 | -1.280183667 | 0.200480549 | NA | BC055308 | cDNA sequence BC055308 [Source:MGI Sy | 2.4093 | 6.1675 | 5.0312 | 0.964 | 1.0906 | 2.2277 |
| 42065 | ENSMUSG00000103975 | 0 | NA | NA | NA | NA | NA | Gm37676 | predicted gene, 37676 [Source:MGI Symb | 0 | 0 | 0 | 0 | 0 | 0 |
| 42066 | ENSMUSG00000103976 | 0.951254158 | 0.070446961 | 2.457323444 | 0.028668168 | 0.977129244 | NA | Gm37677 | predicted gene, 37677 [Source:MGI Symb | 0.8031 | 0 | 2.0125 | 2.8919 | 0 | 0 |
| 42067 | ENSMUSG00000103977 | 0 | NA | NA | NA | NA | NA | Gm37071 | predicted gene, 37071 [Source:MGI Symb | 0 | 0 | 0 | 0 | 0 | 0 |
| 42068 | ENSMUSG00000103978 | 0 | NA | NA | NA | NA | NA | 4930474B08 | RIKEN cDNA 4930474B08 gene [Source:M | 0 | 0 | 0 | 0 | 0 | 0 |
| 42069 | ENSMUSG00000103979 | 0.315620088 | 0.165143421 | 4.063300985 | 0.040642675 | 0.967580763 | NA | Gm20305 | predicted gene, 20305 [Source:MGI Symb | 0.8031 | 0 | 0 | 0 | 1.0906 | 0 |
| 42070 | ENSMUSG00000103980 | 0 | NA | NA | NA | NA | NA | Gm37450 | predicted gene, 37450 [Source:MGI Symb | 0 | 0 | 0 | 0 | 0 | 0 |
| 42071 | ENSMUSG00000103981 | 0 | NA | NA | NA | NA | NA | Gm37449 | predicted gene, 37449 [Source:MGI Symb | 0 | 0 | 0 | 0 | 0 | 0 |
| 42072 | ENSMUSG00000103982 | 0 | NA | NA | NA | NA | NA | Gm37451 | predicted gene, 37451 [Source:MGI Symb | 0 | 0 | 0 | 0 | 0 | 0 |
| 42073 | ENSMUSG00000103983 | 25.4915472 | 0.319818418 | 0.431739992 | 0.740766258 | 0.458835177 | 0.859322431 | Gm20045 | predicted gene, 20045 [Source:MGI Symb | 18.471 | 22.614 | 27.169 | 31.811 | 27.265 | 25.618 |
| 42074 | ENSMUSG00000103984 | 2.251565352 | 0.754343294 | 1.514014095 | 0.498240602 | 0.618314468 | NA | Gm37447 | predicted gene, 37447 [Source:MGI Symb | 0 | 4.1117 | 1.0062 | 2.8919 | 3.2719 | 2.2277 |
| 42075 | ENSMUSG00000103985 | 0 | NA | NA | NA | NA | NA | Gm37446 | predicted gene, 37446 [Source:MGI Symb | 0 | 0 | 0 | 0 | 0 | 0 |
| 42076 | ENSMUSG00000103986 | 0 | NA | NA | NA | NA | NA | Gm5539 | predicted gene 5539 [Source:MGI Symbol | 0 | 0 | 0 | 0 | 0 | 0 |
| 42077 | ENSMUSG00000103987 | 0 | NA | NA | NA | NA | NA | Gm37448 | predicted gene, 37448 [Source:MGI Symb | 0 | 0 | 0 | 0 | 0 | 0 |
| 42078 | ENSMUSG00000103989 | 0 | NA | NA | NA | NA | NA | Ighv5-21 | immunoglobulin heavy variable V5-21 [So | 0 | 0 | 0 | 0 | 0 | 0 |
| 42079 | ENSMUSG00000103990 | 0 | NA | NA | NA | NA | NA | Ighv8-3 | immunoglobulin heavy variable V8-3 [Sou | 0 | 0 | 0 | 0 | 0 | 0 |
| 42080 | ENSMUSG00000103991 | 0 | NA | NA | NA | NA | NA | Gm38031 | predicted gene, 38031 [Source:MGI Symb | 0 | 0 | 0 | 0 | 0 | 0 |
| 42081 | ENSMUSG00000103992 | 0 | NA | NA | NA | NA | NA | Gm38029 | predicted gene, 38029 [Source:MGI Symb | 0 | 0 | 0 | 0 | 0 | 0 |
| 42082 | ENSMUSG00000103993 | 0 | NA | NA | NA | NA | NA | Gm38030 | predicted gene, 38030 [Source:MGI Symb | 0 | 0 | 0 | 0 | 0 | 0 |
| 42083 | ENSMUSG00000103994 | 0 | NA | NA | NA | NA | NA | Gm38028 | predicted gene, 38028 [Source:MGI Symb | 0 | 0 | 0 | 0 | 0 | 0 |
| 42084 | ENSMUSG00000103995 | 0 | NA | NA | NA | NA | NA | 5730488B01 | RIKEN cDNA 5730488B01 gene [Source:M | 0 | 0 | 0 | 0 | 0 | 0 |
| 42085 | ENSMUSG00000103996 | 0.181769715 | 1.126924175 | 4.080472857 | 0.276174898 | 0.782413736 | NA | 4833421G17 | RIKEN cDNA 4833421G17 gene [Source:M | 0 | 0 | 0 | 0 | 1.0906 | 0 |
| 42086 | ENSMUSG00000103997 | 0 | NA | NA | NA | NA | NA | Gm38027 | predicted gene, 38027 [Source:MGI Symb | 0 | 0 | 0 | 0 | 0 | 0 |
| 42087 | ENSMUSG00000103998 | 0 | NA | NA | NA | NA | NA | Gm38025 | predicted gene, 38025 [Source:MGI Symb | 0 | 0 | 0 | 0 | 0 | 0 |
| 42088 | ENSMUSG00000103999 | 0.319491021 | 0.16514334 | 4.044653141 | 0.040830038 | 0.967431393 | NA | Gm38026 | predicted gene, 38026 [Source:MGI Symb | 0.8031 | 0 | 0 | 0 | 0 | 1.1138 |
| 42089 | ENSMUSG00000104000 | 0 | NA | NA | NA | NA | NA | Gm38335 | predicted gene, 38335 [Source:MGI Symb | 0 | 0 | 0 | 0 | 0 | 0 |
| 42090 | ENSMUSG00000104001 | 0 | NA | NA | NA | NA | NA | Gm38334 | predicted gene, 38334 [Source:MGI Symb | 0 | 0 | 0 | 0 | 0 | 0 |
| 42091 | ENSMUSG00000104002 | 0.520796785 | -0.820103173 | 3.072141206 | -0.266948398 | 0.789508892 | NA | Gm38336 | predicted gene, 38336 [Source:MGI Symb | 0 | 1.0279 | 1.0062 | 0 | 1.0906 | 0 |
| 42092 | ENSMUSG00000104004 | 0 | NA | NA | NA | NA | NA | 4930565D16 | RIKEN cDNA 4930565D16 gene [Source:M | 0 | 0 | 0 | 0 | 0 | 0 |
| 42093 | ENSMUSG00000104006 | 0 | NA | NA | NA | NA | NA | Gm38333 | predicted gene, 38333 [Source:MGI Symb | 0 | 0 | 0 | 0 | 0 | 0 |
| 42094 | ENSMUSG00000104007 | 0 | NA | NA | NA | NA | NA | Gm38332 | predicted gene, 38332 [Source:MGI Symb | 0 | 0 | 0 | 0 | 0 | 0 |
| 42095 | ENSMUSG00000104009 | 0 | NA | NA | NA | NA | NA | Gm38337 | predicted gene, 38337 [Source:MGI Symb | 0 | 0 | 0 | 0 | 0 | 0 |
| 42096 | ENSMUSG00000104010 | 0.531246861 | 1.101376188 | 3.36099124 | 0.327693858 | 0.743143147 | NA | Gm37366 | predicted gene, 37366 [Source:MGI Symb | 0 | 0 | 1.0062 | 0 | 2.1812 | 0 |
| 42097 | ENSMUSG00000104011 | 33.71004919 | 0.134049441 | 0.384528856 | 0.348606974 | 0.727384392 | 0.945907265 | Gm32391 | predicted gene, 32391 [Source:MGI Symb | 39.352 | 23.642 | 33.206 | 36.631 | 34.9 | 34.529 |
| 42098 | ENSMUSG00000104012 | 0 | NA | NA | NA | NA | NA | Gm37364 | predicted gene, 37364 [Source:MGI Symb | 0 | 0 | 0 | 0 | 0 | 0 |
| 42099 | ENSMUSG00000104013 | 0 | NA | NA | NA | NA | NA | Gm37365 | predicted gene, 37365 [Source:MGI Symb | 0 | 0 | 0 | 0 | 0 | 0 |
| 42100 | ENSMUSG00000104014 | 0 | NA | NA | NA | NA | NA | 4930562F17 | RIKEN cDNA 4930562F17 gene [Source:M | 0 | 0 | 0 | 0 | 0 | 0 |
| 42101 | ENSMUSG00000104015 | 0 | NA | NA | NA | NA | NA | Gm10745 | predicted gene 10745 [Source:MGI Symbo | 0 | 0 | 0 | 0 | 0 | 0 |
| 42102 | ENSMUSG00000104016 | 7.889264876 | 0.670586487 | 0.776091739 | 0.864055695 | 0.387557296 | 0.817623793 | Gm37362 | predicted gene, 37362 [Source:MGI Symb | 8.031 | 5.1396 | 5.0312 | 11.568 | 11.997 | 5.5692 |
| 42103 | ENSMUSG00000104017 | 0 | NA | NA | NA | NA | NA | Gm37363 | predicted gene, 37363 [Source:MGI Symb | 0 | 0 | 0 | 0 | 0 | 0 |
| 42104 | ENSMUSG00000104018 | 1.309087727 | 0.212744688 | 1.993661878 | 0.106710516 | 0.915018639 | NA | 4833412K13 | RIKEN cDNA 4833412K13 gene [Source:M | 1.6062 | 0 | 2.0125 | 0.964 | 3.2719 | 0 |
| 42105 | ENSMUSG00000104020 | 0.17131964 | -0.796629176 | 4.080472857 | -0.195229623 | 0.84521319 | NA | Gm37215 | predicted gene, 37215 [Source:MGI Symb | 0 | 1.0279 | 0 | 0 | 0 | 0 |
| 42106 | ENSMUSG00000104021 | 0 | NA | NA | NA | NA | NA | Gm36536 | predicted gene, 36536 [Source:MGI Symb | 0 | 0 | 0 | 0 | 0 | 0 |
| 42107 | ENSMUSG00000104022 | 0.267700746 | -1.497481649 | 4.057001436 | -0.369110456 | 0.712045396 | NA | Gm37214 | predicted gene, 37214 [Source:MGI Symb | 1.6062 | 0 | 0 | 0 | 0 | 0 |
| 42108 | ENSMUSG00000104023 | 0 | NA | NA | NA | NA | NA | Gm37213 | predicted gene, 37213 [Source:MGI Symb | 0 | 0 | 0 | 0 | 0 | 0 |

| | | | | | | | | | | | | | | | |
|---|---|---|---|---|---|---|---|---|---|---|---|---|---|---|---|
| 42109 | ENSMUSG00000104024 | 0 | NA | NA | NA | NA | NA | Gm33571 | predicted gene, 33571 [Source:MGI Symb | 0 | 0 | 0 | 0 | 0 | 0 |
| 42110 | ENSMUSG00000104025 | 0 | NA | NA | NA | NA | NA | E330040D14 | RIKEN cDNA E330040D14 gene [Source:M | 0 | 0 | 0 | 0 | 0 | 0 |
| 42111 | ENSMUSG00000104026 | 0.674415303 | 1.59191567 | 2.786800062 | 0.57123426 | 0.567840858 | NA | Gm37212 | predicted gene, 37212 [Source:MGI Symb | 0 | 1.0279 | 0 | 1.928 | 1.0906 | 0 |
| 42112 | ENSMUSG00000104027 | 0 | NA | NA | NA | NA | NA | Gm37211 | predicted gene, 37211 [Source:MGI Symb | 0 | 0 | 0 | 0 | 0 | 0 |
| 42113 | ENSMUSG00000104028 | 0 | NA | NA | NA | NA | NA | Gm5851 | predicted gene 5851 [Source:MGI Symbol | 0 | 0 | 0 | 0 | 0 | 0 |
| 42114 | ENSMUSG00000104029 | 0 | NA | NA | NA | NA | NA | Gm37210 | predicted gene, 37210 [Source:MGI Symb | 0 | 0 | 0 | 0 | 0 | 0 |
| 42115 | ENSMUSG00000104030 | 2.171959713 | 0.637534248 | 1.598199525 | 0.398907795 | 0.689961146 | NA | 5330406M2: | RIKEN cDNA 5330406M23 gene [Source:M | 4.0155 | 1.0279 | 0 | 5.7839 | 1.0906 | 1.1138 |
| 42116 | ENSMUSG00000104031 | 1.779664929 | 0.032738715 | 1.974210357 | 0.016583195 | 0.986769131 | NA | Gm17771 | predicted gene, 17771 [Source:MGI Symb | 3.2124 | 0 | 2.0125 | 0 | 5.4531 | 0 |
| 42117 | ENSMUSG00000104032 | 0 | NA | NA | NA | NA | NA | Gm20833 | predicted gene, 20833 [Source:MGI Symb | 0 | 0 | 0 | 0 | 0 | 0 |
| 42118 | ENSMUSG00000104033 | 0 | NA | NA | NA | NA | NA | Gm37773 | predicted gene, 37773 [Source:MGI Symb | 0 | 0 | 0 | 0 | 0 | 0 |
| 42119 | ENSMUSG00000104034 | 0 | NA | NA | NA | NA | NA | 2900092N22 | RIKEN cDNA 2900092N22 gene [Source:M | 0 | 0 | 0 | 0 | 0 | 0 |
| 42120 | ENSMUSG00000104035 | 0 | NA | NA | NA | NA | NA | Gm37774 | predicted gene, 37774 [Source:MGI Symb | 0 | 0 | 0 | 0 | 0 | 0 |
| 42121 | ENSMUSG00000104036 | 0 | NA | NA | NA | NA | NA | Gm37775 | predicted gene, 37775 [Source:MGI Symb | 0 | 0 | 0 | 0 | 0 | 0 |
| 42122 | ENSMUSG00000104037 | 0 | NA | NA | NA | NA | NA | Gm37776 | predicted gene, 37776 [Source:MGI Symb | 0 | 0 | 0 | 0 | 0 | 0 |
| 42123 | ENSMUSG00000104038 | 0 | NA | NA | NA | NA | NA | Gm37771 | predicted gene, 37771 [Source:MGI Symb | 0 | 0 | 0 | 0 | 0 | 0 |
| 42124 | ENSMUSG00000104039 | 0 | NA | NA | NA | NA | NA | Gm37772 | predicted gene, 37772 [Source:MGI Symb | 0 | 0 | 0 | 0 | 0 | 0 |
| 42125 | ENSMUSG00000104040 | 0 | NA | NA | NA | NA | NA | Gm37563 | predicted gene, 37563 [Source:MGI Symb | 0 | 0 | 0 | 0 | 0 | 0 |
| 42126 | ENSMUSG00000104041 | 0 | NA | NA | NA | NA | NA | Gm37562 | predicted gene, 37562 [Source:MGI Symb | 0 | 0 | 0 | 0 | 0 | 0 |
| 42127 | ENSMUSG00000104042 | 0.881162691 | 2.072029295 | 2.358945437 | 0.878371014 | 0.379742409 | NA | Gm37564 | predicted gene, 37564 [Source:MGI Symb | 0 | 1.0279 | 0 | 0.964 | 2.1812 | 1.1138 |
| 42128 | ENSMUSG00000104043 | 2.854675447 | 0.069464958 | 1.349208603 | 0.051485706 | 0.958938491 | NA | Gm6525 | predicted pseudogene 6525 [Source:MGI | 3.2124 | 2.0558 | 3.0187 | 0 | 3.2719 | 5.5692 |
| 42129 | ENSMUSG00000104044 | 0 | NA | NA | NA | NA | NA | Gm37566 | predicted gene, 37566 [Source:MGI Symb | 0 | 0 | 0 | 0 | 0 | 0 |
| 42130 | ENSMUSG00000104045 | 0 | NA | NA | NA | NA | NA | Gm37565 | predicted gene, 37565 [Source:MGI Symb | 0 | 0 | 0 | 0 | 0 | 0 |
| 42131 | ENSMUSG00000104046 | 0 | NA | NA | NA | NA | NA | Gm37567 | predicted gene, 37567 [Source:MGI Symb | 0 | 0 | 0 | 0 | 0 | 0 |
| 42132 | ENSMUSG00000104048 | 0 | NA | NA | NA | NA | NA | Gm37561 | predicted gene, 37561 [Source:MGI Symb | 0 | 0 | 0 | 0 | 0 | 0 |
| 42133 | ENSMUSG00000104050 | 0 | NA | NA | NA | NA | NA | Gm38127 | predicted gene, 38127 [Source:MGI Symb | 0 | 0 | 0 | 0 | 0 | 0 |
| 42134 | ENSMUSG00000104051 | 0 | NA | NA | NA | NA | NA | Gm38128 | predicted gene, 38128 [Source:MGI Symb | 0 | 0 | 0 | 0 | 0 | 0 |
| 42135 | ENSMUSG00000104052 | 0 | NA | NA | NA | NA | NA | Gm38125 | predicted gene, 38125 [Source:MGI Symb | 0 | 0 | 0 | 0 | 0 | 0 |
| 42136 | ENSMUSG00000104053 | 0.79190315 | 0.90205007 | 2.632846219 | 0.342614036 | 0.731888841 | NA | Gm38126 | predicted gene, 38126 [Source:MGI Symb | 1.6062 | 0 | 0 | 0.964 | 2.1812 | 0 |
| 42137 | ENSMUSG00000104054 | 0 | NA | NA | NA | NA | NA | Gm38131 | predicted gene, 38131 [Source:MGI Symb | 0 | 0 | 0 | 0 | 0 | 0 |
| 42138 | ENSMUSG00000104055 | 0 | NA | NA | NA | NA | NA | Gm38132 | predicted gene, 38132 [Source:MGI Symb | 0 | 0 | 0 | 0 | 0 | 0 |
| 42139 | ENSMUSG00000104057 | 0 | NA | NA | NA | NA | NA | Gm38130 | predicted gene, 38130 [Source:MGI Symb | 0 | 0 | 0 | 0 | 0 | 0 |
| 42140 | ENSMUSG00000104058 | 0 | NA | NA | NA | NA | NA | NA | NA | 0 | 0 | 0 | 0 | 0 | 0 |
| 42141 | ENSMUSG00000104059 | 0 | NA | NA | NA | NA | NA | Gm9124 | predicted pseudogene 9124 [Source:MGI | 0 | 0 | 0 | 0 | 0 | 0 |
| 42142 | ENSMUSG00000104060 | 0 | NA | NA | NA | NA | NA | Gm37954 | predicted gene, 37954 [Source:MGI Symb | 0 | 0 | 0 | 0 | 0 | 0 |
| 42143 | ENSMUSG00000104061 | 1.673939123 | 1.598362957 | 2.284618494 | 0.699619197 | 0.48416515 | NA | Gm37953 | predicted gene, 37953 [Source:MGI Symb | 2.4093 | 0 | 0 | 0 | 7.6343 | 0 |
| 42144 | ENSMUSG00000104062 | 0 | NA | NA | NA | NA | NA | Gm37952 | predicted gene, 37952 [Source:MGI Symb | 0 | 0 | 0 | 0 | 0 | 0 |
| 42145 | ENSMUSG00000104063 | 1.136894805 | 0.527420275 | 2.120428083 | 0.248732923 | 0.80356738 | NA | Pcdhgb7 | protocadherin gamma subfamily B, 7 [Sou | 0.8031 | 0 | 2.0125 | 2.8919 | 0 | 1.1138 |
| 42146 | ENSMUSG00000104064 | 0.986629773 | -0.885187988 | 2.114354962 | -0.418656282 | 0.67546735 | NA | Gm37956 | predicted gene, 37956 [Source:MGI Symb | 0.8031 | 2.0558 | 1.0062 | 0.964 | 1.0906 | 0 |
| 42147 | ENSMUSG00000104065 | 0 | NA | NA | NA | NA | NA | Gm37197 | predicted gene, 37197 [Source:MGI Symb | 0 | 0 | 0 | 0 | 0 | 0 |
| 42148 | ENSMUSG00000104066 | 0 | NA | NA | NA | NA | NA | Gm37955 | predicted gene, 37955 [Source:MGI Symb | 0 | 0 | 0 | 0 | 0 | 0 |
| 42149 | ENSMUSG00000104067 | 0 | NA | NA | NA | NA | NA | Gm37196 | predicted gene, 37196 [Source:MGI Symb | 0 | 0 | 0 | 0 | 0 | 0 |
| 42150 | ENSMUSG00000104068 | 0 | NA | NA | NA | NA | NA | Gm37199 | predicted gene, 37199 [Source:MGI Symb | 0 | 0 | 0 | 0 | 0 | 0 |
| 42151 | ENSMUSG00000104069 | 0 | NA | NA | NA | NA | NA | Gm37198 | predicted gene, 37198 [Source:MGI Symb | 0 | 0 | 0 | 0 | 0 | 0 |
| 42152 | ENSMUSG00000104070 | 0 | NA | NA | NA | NA | NA | Gm36866 | predicted gene, 36866 [Source:MGI Symb | 0 | 0 | 0 | 0 | 0 | 0 |
| 42153 | ENSMUSG00000104071 | 0 | NA | NA | NA | NA | NA | Gm37028 | predicted gene, 37028 [Source:MGI Symb | 0 | 0 | 0 | 0 | 0 | 0 |
| 42154 | ENSMUSG00000104072 | 0 | NA | NA | NA | NA | NA | 4930448I06F | RIKEN cDNA 4930448I06 gene [Source:MC | 0 | 0 | 0 | 0 | 0 | 0 |
| 42155 | ENSMUSG00000104073 | 0 | NA | NA | NA | NA | NA | Gm37029 | predicted gene, 37029 [Source:MGI Symb | 0 | 0 | 0 | 0 | 0 | 0 |
| 42156 | ENSMUSG00000104074 | 0 | NA | NA | NA | NA | NA | Gm37025 | predicted gene, 37025 [Source:MGI Symb | 0 | 0 | 0 | 0 | 0 | 0 |
| 42157 | ENSMUSG00000104075 | 0 | NA | NA | NA | NA | NA | 5430433H01 | RIKEN cDNA 5430433H01 gene [Source:M | 0 | 0 | 0 | 0 | 0 | 0 |
| 42158 | ENSMUSG00000104076 | 0 | NA | NA | NA | NA | NA | Gm37026 | predicted gene, 37026 [Source:MGI Symb | 0 | 0 | 0 | 0 | 0 | 0 |
| 42159 | ENSMUSG00000104077 | 2.702101198 | -0.405983743 | 1.399422754 | -0.290108005 | 0.771733612 | NA | Gm37027 | predicted gene, 37027 [Source:MGI Symb | 3.2124 | 3.0838 | 3.0187 | 5.7839 | 0 | 1.1138 |
| 42160 | ENSMUSG00000104078 | 0 | NA | NA | NA | NA | NA | Gm37023 | predicted gene, 37023 [Source:MGI Symb | 0 | 0 | 0 | 0 | 0 | 0 |
| 42161 | ENSMUSG00000104079 | 0 | NA | NA | NA | NA | NA | Gm37024 | predicted gene, 37024 [Source:MGI Symb | 0 | 0 | 0 | 0 | 0 | 0 |
| 42162 | ENSMUSG00000104080 | 1.062932406 | 2.399885247 | 2.235174461 | 1.073690349 | 0.282961488 | NA | Gm38279 | predicted gene, 38279 [Source:MGI Symb | 0 | 1.0279 | 0 | 0.964 | 3.2719 | 1.1138 |
| 42163 | ENSMUSG00000104081 | 0 | NA | NA | NA | NA | NA | Gm38278 | predicted gene, 38278 [Source:MGI Symb | 0 | 0 | 0 | 0 | 0 | 0 |
| 42164 | ENSMUSG00000104082 | 0 | NA | NA | NA | NA | NA | Gm7115 | predicted gene 7115 [Source:MGI Symbol | 0 | 0 | 0 | 0 | 0 | 0 |

| | | | | | | | | | | | | | | | |
|---|---|---|---|---|---|---|---|---|---|---|---|---|---|---|---|
| 42165 | ENSMUSG00000104083 | 0 | NA | NA | NA | NA | NA | Gm29739 | predicted gene, 29739 [Source:MGI Symb | 0 | 0 | 0 | 0 | 0 | 0 |
| 42166 | ENSMUSG00000104084 | 0 | NA | NA | NA | NA | NA | Gm38276 | predicted gene, 38276 [Source:MGI Symb | 0 | 0 | 0 | 0 | 0 | 0 |
| 42167 | ENSMUSG00000104085 | 0 | NA | NA | NA | NA | NA | Gm21427 | predicted gene, 21427 [Source:MGI Symb | 0 | 0 | 0 | 0 | 0 | 0 |
| 42168 | ENSMUSG00000104086 | 0 | NA | NA | NA | NA | NA | 1700039M1! | RIKEN cDNA 1700039M15 gene [Source:M | 0 | 0 | 0 | 0 | 0 | 0 |
| 42169 | ENSMUSG00000104087 | 0 | NA | NA | NA | NA | NA | Gm38277 | predicted gene, 38277 [Source:MGI Symb | 0 | 0 | 0 | 0 | 0 | 0 |
| 42170 | ENSMUSG00000104088 | 0 | NA | NA | NA | NA | NA | Gm38275 | predicted gene, 38275 [Source:MGI Symb | 0 | 0 | 0 | 0 | 0 | 0 |
| 42171 | ENSMUSG00000104089 | 0 | NA | NA | NA | NA | NA | Gm38274 | predicted gene, 38274 [Source:MGI Symb | 0 | 0 | 0 | 0 | 0 | 0 |
| 42172 | ENSMUSG00000104091 | 7.433681115 | 1.399774361 | 0.816997886 | 1.713314545 | 0.086654688 | 0.433843339 | Gm37317 | predicted gene, 37317 [Source:MGI Symb | 3.2124 | 3.0838 | 6.0375 | 13.496 | 7.6343 | 11.138 |
| 42173 | ENSMUSG00000104092 | 0.185640647 | 1.126924175 | 4.080472857 | 0.276174898 | 0.782413736 | NA | A930014E01 | RIKEN cDNA A930014E01 gene [Source:M | 0 | 0 | 0 | 0 | 0 | 1.1138 |
| 42174 | ENSMUSG00000104093 | 0 | NA | NA | NA | NA | NA | A330015K06 | RIKEN cDNA A330015K06 gene [Source:M | 0 | 0 | 0 | 0 | 0 | 0 |
| 42175 | ENSMUSG00000104094 | 1.559164699 | -1.718751862 | 2.16488549 | -0.79392276 | 0.427240399 | NA | Gm37314 | predicted gene, 37314 [Source:MGI Symb | 0 | 6.1675 | 1.0062 | 0 | 2.1812 | 0 |
| 42176 | ENSMUSG00000104095 | 0.301557804 | -1.64335916 | 4.042814367 | -0.406488899 | 0.684383412 | NA | Gm37315 | predicted gene, 37315 [Source:MGI Symb | 0.8031 | 0 | 1.0062 | 0 | 0 | 0 |
| 42177 | ENSMUSG00000104096 | 0 | NA | NA | NA | NA | NA | Gm20356 | predicted gene, 20356 [Source:MGI Symb | 0 | 0 | 0 | 0 | 0 | 0 |
| 42178 | ENSMUSG00000104097 | 1.604562721 | 0.626214531 | 1.919543815 | 0.326230913 | 0.744249654 | NA | Gm37313 | predicted gene, 37313 [Source:MGI Symb | 0.8031 | 1.0279 | 2.0125 | 5.7839 | 0 | 0 |
| 42179 | ENSMUSG00000104098 | 0 | NA | NA | NA | NA | NA | AA619741 | expressed sequence AA619741 [Source:M | 0 | 0 | 0 | 0 | 0 | 0 |
| 42180 | ENSMUSG00000104099 | 0 | NA | NA | NA | NA | NA | Gm37312 | predicted gene, 37312 [Source:MGI Symb | 0 | 0 | 0 | 0 | 0 | 0 |
| 42181 | ENSMUSG00000104100 | 0 | NA | NA | NA | NA | NA | Gm37880 | predicted gene, 37880 [Source:MGI Symb | 0 | 0 | 0 | 0 | 0 | 0 |
| 42182 | ENSMUSG00000104101 | 18.41128682 | 0.704836862 | 0.538955966 | 1.307781909 | 0.190947292 | 0.628409928 | Gm37881 | predicted gene, 37881 [Source:MGI Symb | 13.653 | 7.1954 | 21.131 | 22.171 | 21.812 | 24.505 |
| 42183 | ENSMUSG00000104102 | 0 | NA | NA | NA | NA | NA | Gm32496 | predicted gene, 32496 [Source:MGI Symb | 0 | 0 | 0 | 0 | 0 | 0 |
| 42184 | ENSMUSG00000104103 | 0 | NA | NA | NA | NA | NA | Gm9517 | predicted gene 9517 [Source:MGI Symbol | 0 | 0 | 0 | 0 | 0 | 0 |
| 42185 | ENSMUSG00000104104 | 0 | NA | NA | NA | NA | NA | Gm37877 | predicted gene, 37877 [Source:MGI Symb | 0 | 0 | 0 | 0 | 0 | 0 |
| 42186 | ENSMUSG00000104105 | 0 | NA | NA | NA | NA | NA | Gm37878 | predicted gene, 37878 [Source:MGI Symb | 0 | 0 | 0 | 0 | 0 | 0 |
| 42187 | ENSMUSG00000104106 | 0 | NA | NA | NA | NA | NA | Gm10848 | predicted gene 10848 [Source:MGI Symbo | 0 | 0 | 0 | 0 | 0 | 0 |
| 42188 | ENSMUSG00000104107 | 0 | NA | NA | NA | NA | NA | Gm37879 | predicted gene, 37879 [Source:MGI Symb | 0 | 0 | 0 | 0 | 0 | 0 |
| 42189 | ENSMUSG00000104108 | 0 | NA | NA | NA | NA | NA | Gm37876 | predicted gene, 37876 [Source:MGI Symb | 0 | 0 | 0 | 0 | 0 | 0 |
| 42190 | ENSMUSG00000104109 | 0 | NA | NA | NA | NA | NA | Gm30292 | predicted gene, 30292 [Source:MGI Symb | 0 | 0 | 0 | 0 | 0 | 0 |
| 42191 | ENSMUSG00000104110 | 0 | NA | NA | NA | NA | NA | Gm37295 | predicted gene, 37295 [Source:MGI Symb | 0 | 0 | 0 | 0 | 0 | 0 |
| 42192 | ENSMUSG00000104111 | 0.838510525 | 0.626826968 | 2.582658213 | 0.242706126 | 0.808233062 | NA | Gm37294 | predicted gene, 37294 [Source:MGI Symb | 0 | 0 | 2.0125 | 1.928 | 1.0906 | 0 |
| 42193 | ENSMUSG00000104112 | 0 | NA | NA | NA | NA | NA | Gm37293 | predicted gene, 37293 [Source:MGI Symb | 0 | 0 | 0 | 0 | 0 | 0 |
| 42194 | ENSMUSG00000104113 | 3.054642615 | 0.472968952 | 1.185401969 | 0.398994573 | 0.689897203 | NA | Gm37292 | predicted gene, 37292 [Source:MGI Symb | 1.6062 | 3.0838 | 3.0187 | 2.8919 | 3.2719 | 4.4554 |
| 42195 | ENSMUSG00000104114 | 0 | NA | NA | NA | NA | NA | Gm37297 | predicted gene, 37297 [Source:MGI Symb | 0 | 0 | 0 | 0 | 0 | 0 |
| 42196 | ENSMUSG00000104115 | 0 | NA | NA | NA | NA | NA | Ighv8-1 | immunoglobulin heavy variable V8-1 [Sou | 0 | 0 | 0 | 0 | 0 | 0 |
| 42197 | ENSMUSG00000104116 | 4.081222745 | -0.996595959 | 1.099087229 | -0.906748739 | 0.364539683 | NA | Gm37296 | predicted gene, 37296 [Source:MGI Symb | 7.2279 | 3.0838 | 6.0375 | 4.8199 | 1.0906 | 2.2277 |
| 42198 | ENSMUSG00000104117 | 2.318270662 | -1.768019817 | 1.528304177 | -1.156850739 | 0.247333348 | NA | Gm20743 | predicted gene, 20743 [Source:MGI Symb | 5.6217 | 3.0838 | 2.0125 | 0.964 | 0 | 2.2277 |
| 42199 | ENSMUSG00000104118 | 0 | NA | NA | NA | NA | NA | Gm37298 | predicted gene, 37298 [Source:MGI Symb | 0 | 0 | 0 | 0 | 0 | 0 |
| 42200 | ENSMUSG00000104120 | 0 | NA | NA | NA | NA | NA | Gm37484 | predicted gene, 37484 [Source:MGI Symb | 0 | 0 | 0 | 0 | 0 | 0 |
| 42201 | ENSMUSG00000104121 | 0.301557804 | -1.64335916 | 4.042814367 | -0.406488899 | 0.684383412 | NA | Gm37485 | predicted gene, 37485 [Source:MGI Symb | 0.8031 | 0 | 1.0062 | 0 | 0 | 0 |
| 42202 | ENSMUSG00000104122 | 0.80281854 | -0.292476728 | 2.419692615 | -0.120873505 | 0.903791228 | NA | Gm37482 | predicted gene, 37482 [Source:MGI Symb | 1.6062 | 0 | 1.0062 | 0 | 1.0906 | 1.1138 |
| 42203 | ENSMUSG00000104123 | 0 | NA | NA | NA | NA | NA | Gm37483 | predicted gene, 37483 [Source:MGI Symb | 0 | 0 | 0 | 0 | 0 | 0 |
| 42204 | ENSMUSG00000104125 | 0.185640647 | 1.126924175 | 4.080472857 | 0.276174898 | 0.782413736 | NA | Gm37488 | predicted gene, 37488 [Source:MGI Symb | 0 | 0 | 0 | 0 | 0 | 1.1138 |
| 42205 | ENSMUSG00000104126 | 1.090055336 | -2.273671673 | 2.179733234 | -1.043096301 | 0.296903694 | NA | Gm37486 | predicted gene, 37486 [Source:MGI Symb | 2.4093 | 1.0279 | 2.0125 | 0 | 1.0906 | 0 |
| 42206 | ENSMUSG00000104127 | 0 | NA | NA | NA | NA | NA | Gm20928 | predicted gene, 20928 [Source:MGI Symb | 0 | 0 | 0 | 0 | 0 | 0 |
| 42207 | ENSMUSG00000104129 | 0 | NA | NA | NA | NA | NA | Ighv6-2 | immunoglobulin heavy variable V6-2 [Sou | 0 | 0 | 0 | 0 | 0 | 0 |
| 42208 | ENSMUSG00000104130 | 0 | NA | NA | NA | NA | NA | Gm36951 | predicted gene, 36951 [Source:MGI Symb | 0 | 0 | 0 | 0 | 0 | 0 |
| 42209 | ENSMUSG00000104131 | 0 | NA | NA | NA | NA | NA | Gm36950 | predicted gene, 36950 [Source:MGI Symb | 0 | 0 | 0 | 0 | 0 | 0 |
| 42210 | ENSMUSG00000104132 | 0 | NA | NA | NA | NA | NA | Gm36952 | predicted gene, 36952 [Source:MGI Symb | 0 | 0 | 0 | 0 | 0 | 0 |
| 42211 | ENSMUSG00000104133 | 0 | NA | NA | NA | NA | NA | 4921511E07 | RIKEN cDNA 4921511E07 gene [Source:M | 0 | 0 | 0 | 0 | 0 | 0 |
| 42212 | ENSMUSG00000104134 | 0 | NA | NA | NA | NA | NA | Gm2328 | predicted gene 2328 [Source:MGI Symbol | 0 | 0 | 0 | 0 | 0 | 0 |
| 42213 | ENSMUSG00000104135 | 0 | NA | NA | NA | NA | NA | Gm36953 | predicted gene, 36953 [Source:MGI Symb | 0 | 0 | 0 | 0 | 0 | 0 |
| 42214 | ENSMUSG00000104136 | 0.185640647 | 1.126924175 | 4.080472857 | 0.276174898 | 0.782413736 | NA | Gm36955 | predicted gene, 36955 [Source:MGI Symb | 0 | 0 | 0 | 0 | 0 | 1.1138 |
| 42215 | ENSMUSG00000104137 | 0 | NA | NA | NA | NA | NA | Gm36954 | predicted gene, 36954 [Source:MGI Symb | 0 | 0 | 0 | 0 | 0 | 0 |
| 42216 | ENSMUSG00000104138 | 0 | NA | NA | NA | NA | NA | Gm36949 | predicted gene, 36949 [Source:MGI Symb | 0 | 0 | 0 | 0 | 0 | 0 |
| 42217 | ENSMUSG00000104140 | 0 | NA | NA | NA | NA | NA | Gm37140 | predicted gene, 37140 [Source:MGI Symb | 0 | 0 | 0 | 0 | 0 | 0 |
| 42218 | ENSMUSG00000104141 | 0 | NA | NA | NA | NA | NA | Gm6140 | predicted gene 6140 [Source:MGI Symbol | 0 | 0 | 0 | 0 | 0 | 0 |
| 42219 | ENSMUSG00000104142 | 0 | NA | NA | NA | NA | NA | Gm37141 | predicted gene, 37141 [Source:MGI Symb | 0 | 0 | 0 | 0 | 0 | 0 |
| 42220 | ENSMUSG00000104144 | 0 | NA | NA | NA | NA | NA | Gm18433 | predicted gene, 18433 [Source:MGI Symb | 0 | 0 | 0 | 0 | 0 | 0 |

| | | | | | | | | | | | | | | | |
|---|---|---|---|---|---|---|---|---|---|---|---|---|---|---|---|
| 42221 | ENSMUSG00000104145 | 0.185640647 | 1.126924175 | 4.080472857 | 0.276174898 | 0.782413736 | NA | D130019J16 | RIKEN cDNA D130019J16 gene [Source:M | 0 | 0 | 0 | 0 | 0 | 1.1138 |
| 42222 | ENSMUSG00000104146 | 0 | NA | NA | NA | NA | NA | Gm30175 | predicted gene, 30175 [Source:MGI Symb | 0 | 0 | 0 | 0 | 0 | 0 |
| 42223 | ENSMUSG00000104147 | 0 | NA | NA | NA | NA | NA | Gm37142 | predicted gene, 37142 [Source:MGI Symb | 0 | 0 | 0 | 0 | 0 | 0 |
| 42224 | ENSMUSG00000104148 | 0 | NA | NA | NA | NA | NA | Pcdha2 | protocadherin alpha 2 [Source:MGI Symb | 0 | 0 | 0 | 0 | 0 | 0 |
| 42225 | ENSMUSG00000104149 | 6.645630184 | -0.190618067 | 1.100054468 | -0.173280571 | 0.862430895 | 0.976334255 | Gm37138 | predicted gene, 37138 [Source:MGI Symb | 8.8341 | 11.307 | 1.0062 | 0 | 9.8156 | 8.9108 |
| 42226 | ENSMUSG00000104150 | 2.131701041 | -0.485004904 | 1.570141498 | -0.308892482 | 0.757403316 | NA | Gm44573 | predicted gene 44573 [Source:MGI Symbo | 2.4093 | 0 | 5.0312 | 0.964 | 3.2719 | 1.1138 |
| 42227 | ENSMUSG00000104152 | 0 | NA | NA | NA | NA | NA | Ighv10-4 | immunoglobulin heavy variable V10-4 [So | 0 | 0 | 0 | 0 | 0 | 0 |
| 42228 | ENSMUSG00000104153 | 0 | NA | NA | NA | NA | NA | Gm38105 | predicted gene, 38105 [Source:MGI Symb | 0 | 0 | 0 | 0 | 0 | 0 |
| 42229 | ENSMUSG00000104154 | 0.843693424 | 2.072131851 | 2.387144628 | 0.868037833 | 0.385373622 | NA | Gm38104 | predicted gene, 38104 [Source:MGI Symb | 0.8031 | 0 | 0 | 0.964 | 2.1812 | 1.1138 |
| 42230 | ENSMUSG00000104155 | 0 | NA | NA | NA | NA | NA | Gm38103 | predicted gene, 38103 [Source:MGI Symb | 0 | 0 | 0 | 0 | 0 | 0 |
| 42231 | ENSMUSG00000104156 | 32.99616628 | -0.119877249 | 0.412466398 | -0.290635187 | 0.771330346 | 0.956965056 | Gm38102 | predicted gene, 38102 [Source:MGI Symb | 32.124 | 34.949 | 36.225 | 37.595 | 39.262 | 17.822 |
| 42232 | ENSMUSG00000104157 | 0 | NA | NA | NA | NA | NA | Gm38101 | predicted gene, 38101 [Source:MGI Symb | 0 | 0 | 0 | 0 | 0 | 0 |
| 42233 | ENSMUSG00000104158 | 0 | NA | NA | NA | NA | NA | Gm38100 | predicted gene, 38100 [Source:MGI Symb | 0 | 0 | 0 | 0 | 0 | 0 |
| 42234 | ENSMUSG00000104159 | 0 | NA | NA | NA | NA | NA | Gm38099 | predicted gene, 38099 [Source:MGI Symb | 0 | 0 | 0 | 0 | 0 | 0 |
| 42235 | ENSMUSG00000104161 | 0 | NA | NA | NA | NA | NA | Gm20089 | predicted gene, 20089 [Source:MGI Symb | 0 | 0 | 0 | 0 | 0 | 0 |
| 42236 | ENSMUSG00000104162 | 0 | NA | NA | NA | NA | NA | Gm37619 | predicted gene, 37619 [Source:MGI Symb | 0 | 0 | 0 | 0 | 0 | 0 |
| 42237 | ENSMUSG00000104163 | 0 | NA | NA | NA | NA | NA | Gm19057 | predicted gene, 19057 [Source:MGI Symb | 0 | 0 | 0 | 0 | 0 | 0 |
| 42238 | ENSMUSG00000104164 | 6.321615637 | -0.335615816 | 0.857056835 | -0.391591085 | 0.695360376 | 0.939992732 | Gm38248 | predicted gene, 38248 [Source:MGI Symb | 4.0155 | 7.1954 | 10.062 | 6.7478 | 5.4531 | 4.4554 |
| 42239 | ENSMUSG00000104165 | 0 | NA | NA | NA | NA | NA | Gm38249 | predicted gene, 38249 [Source:MGI Symb | 0 | 0 | 0 | 0 | 0 | 0 |
| 42240 | ENSMUSG00000104166 | 0 | NA | NA | NA | NA | NA | Gm38247 | predicted gene, 38247 [Source:MGI Symb | 0 | 0 | 0 | 0 | 0 | 0 |
| 42241 | ENSMUSG00000104167 | 0 | NA | NA | NA | NA | NA | A030012G06 | RIKEN cDNA A030012G06 gene [Source:M | 0 | 0 | 0 | 0 | 0 | 0 |
| 42242 | ENSMUSG00000104168 | 9.864766118 | 1.04248851 | 0.733032089 | 1.422159446 | 0.154979969 | 0.573306974 | Gm38250 | predicted gene, 38250 [Source:MGI Symb | 8.031 | 8.2233 | 3.0187 | 14.46 | 7.6343 | 17.822 |
| 42243 | ENSMUSG00000104169 | 0 | NA | NA | NA | NA | NA | Gm7430 | predicted gene 7430 [Source:MGI Symbol | 0 | 0 | 0 | 0 | 0 | 0 |
| 42244 | ENSMUSG00000104170 | 0 | NA | NA | NA | NA | NA | Gm37702 | predicted gene, 37702 [Source:MGI Symb | 0 | 0 | 0 | 0 | 0 | 0 |
| 42245 | ENSMUSG00000104171 | 0 | NA | NA | NA | NA | NA | Gm18296 | predicted gene, 18296 [Source:MGI Symb | 0 | 0 | 0 | 0 | 0 | 0 |
| 42246 | ENSMUSG00000104172 | 0 | NA | NA | NA | NA | NA | Gm37259 | predicted gene, 37259 [Source:MGI Symb | 0 | 0 | 0 | 0 | 0 | 0 |
| 42247 | ENSMUSG00000104173 | 0 | NA | NA | NA | NA | NA | Gm37703 | predicted gene, 37703 [Source:MGI Symb | 0 | 0 | 0 | 0 | 0 | 0 |
| 42248 | ENSMUSG00000104174 | 0 | NA | NA | NA | NA | NA | Gm37701 | predicted gene, 37701 [Source:MGI Symb | 0 | 0 | 0 | 0 | 0 | 0 |
| 42249 | ENSMUSG00000104175 | 0 | NA | NA | NA | NA | NA | Gm37700 | predicted gene, 37700 [Source:MGI Symb | 0 | 0 | 0 | 0 | 0 | 0 |
| 42250 | ENSMUSG00000104176 | 0 | NA | NA | NA | NA | NA | Ighv1-51 | immunoglobulin heavy variable V1-51 [So | 0 | 0 | 0 | 0 | 0 | 0 |
| 42251 | ENSMUSG00000104177 | 0 | NA | NA | NA | NA | NA | Gm2429 | predicted gene 2429 [Source:MGI Symbol | 0 | 0 | 0 | 0 | 0 | 0 |
| 42252 | ENSMUSG00000104178 | 0 | NA | NA | NA | NA | NA | Gm9916 | predicted gene 9916 [Source:MGI Symbol | 0 | 0 | 0 | 0 | 0 | 0 |
| 42253 | ENSMUSG00000104180 | 0 | NA | NA | NA | NA | NA | Gm37970 | predicted gene, 37970 [Source:MGI Symb | 0 | 0 | 0 | 0 | 0 | 0 |
| 42254 | ENSMUSG00000104181 | 0.9173971 | 0.153222854 | 2.48201849 | 0.061733164 | 0.950775329 | NA | Gm8850 | predicted gene 8850 [Source:MGI Symbol | 1.6062 | 0 | 1.0062 | 2.8919 | 0 | 0 |
| 42255 | ENSMUSG00000104182 | 0 | NA | NA | NA | NA | NA | Gm21362 | predicted gene, 21362 [Source:MGI Symb | 0 | 0 | 0 | 0 | 0 | 0 |
| 42256 | ENSMUSG00000104183 | 0.321325949 | 1.911575114 | 4.048267893 | 0.472195804 | 0.636787034 | NA | 1700054O19 | RIKEN cDNA 1700054O19 gene [Source:M | 0 | 0 | 0 | 1.928 | 0 | 0 |
| 42257 | ENSMUSG00000104184 | 0.589026695 | 0.241472922 | 3.228838524 | 0.074786311 | 0.940384733 | NA | Gm37818 | predicted gene, 37818 [Source:MGI Symb | 1.6062 | 0 | 0 | 1.928 | 0 | 0 |
| 42258 | ENSMUSG00000104185 | 0 | NA | NA | NA | NA | NA | Gm18692 | predicted gene, 18692 [Source:MGI Symb | 0 | 0 | 0 | 0 | 0 | 0 |
| 42259 | ENSMUSG00000104186 | 0 | NA | NA | NA | NA | NA | Gm37819 | predicted gene, 37819 [Source:MGI Symb | 0 | 0 | 0 | 0 | 0 | 0 |
| 42260 | ENSMUSG00000104187 | 0 | NA | NA | NA | NA | NA | Gm37820 | predicted gene, 37820 [Source:MGI Symb | 0 | 0 | 0 | 0 | 0 | 0 |
| 42261 | ENSMUSG00000104188 | 0 | NA | NA | NA | NA | NA | Gm37821 | predicted gene, 37821 [Source:MGI Symb | 0 | 0 | 0 | 0 | 0 | 0 |
| 42262 | ENSMUSG00000104189 | 0 | NA | NA | NA | NA | NA | Gm37822 | predicted gene, 37822 [Source:MGI Symb | 0 | 0 | 0 | 0 | 0 | 0 |
| 42263 | ENSMUSG00000104190 | 0.328370405 | 0.16514316 | 4.003149536 | 0.041253308 | 0.967093956 | NA | Gm10472 | predicted gene 10472 [Source:MGI Symbo | 0 | 0 | 1.0062 | 0.964 | 0 | 0 |
| 42264 | ENSMUSG00000104191 | 0 | NA | NA | NA | NA | NA | Gm30737 | predicted gene, 30737 [Source:MGI Symb | 0 | 0 | 0 | 0 | 0 | 0 |
| 42265 | ENSMUSG00000104192 | 0 | NA | NA | NA | NA | NA | Gm37253 | predicted gene, 37253 [Source:MGI Symb | 0 | 0 | 0 | 0 | 0 | 0 |
| 42266 | ENSMUSG00000104194 | 0 | NA | NA | NA | NA | NA | Gm33354 | predicted gene, 33354 [Source:MGI Symb | 0 | 0 | 0 | 0 | 0 | 0 |
| 42267 | ENSMUSG00000104195 | 3.495918039 | -0.141903066 | 1.362270476 | -0.104166587 | 0.91703715 | NA | B230377A18 | RIKEN cDNA B230377A18 gene [Source:M | 4.0155 | 0 | 7.0437 | 7.7118 | 1.0906 | 1.1138 |
| 42268 | ENSMUSG00000104197 | 0 | NA | NA | NA | NA | NA | Gm37632 | predicted gene, 37632 [Source:MGI Symb | 0 | 0 | 0 | 0 | 0 | 0 |
| 42269 | ENSMUSG00000104198 | 0 | NA | NA | NA | NA | NA | Trbv9 | T cell receptor beta variable 9 [Source:MG | 0 | 0 | 0 | 0 | 0 | 0 |
| 42270 | ENSMUSG00000104199 | 0 | NA | NA | NA | NA | NA | 4930420N18 | RIKEN cDNA 4930420N18 gene [Source:M | 0 | 0 | 0 | 0 | 0 | 0 |
| 42271 | ENSMUSG00000104200 | 0.167707431 | -0.796629176 | 4.080472857 | -0.195229623 | 0.84521319 | NA | Gm37399 | predicted gene, 37399 [Source:MGI Symb | 0 | 0 | 1.0062 | 0 | 0 | 0 |
| 42272 | ENSMUSG00000104201 | 0 | NA | NA | NA | NA | NA | 4930535G08 | RIKEN cDNA 4930535G08 gene [Source:M | 0 | 0 | 0 | 0 | 0 | 0 |
| 42273 | ENSMUSG00000104202 | 0.335414862 | -1.779047242 | 4.030804179 | -0.441362855 | 0.658950329 | NA | Gm8101 | predicted gene 8101 [Source:MGI Symbol | 0 | 0 | 2.0125 | 0 | 0 | 0 |
| 42274 | ENSMUSG00000104203 | 0 | NA | NA | NA | NA | NA | Gm37398 | predicted gene, 37398 [Source:MGI Symb | 0 | 0 | 0 | 0 | 0 | 0 |
| 42275 | ENSMUSG00000104204 | 9.933391532 | -0.499561632 | 0.703882594 | -0.709722951 | 0.477875957 | 0.869794311 | Gm37397 | predicted gene, 37397 [Source:MGI Symb | 13.653 | 9.2513 | 12.075 | 12.532 | 7.6343 | 4.4554 |
| 42276 | ENSMUSG00000104205 | 0 | NA | NA | NA | NA | NA | Gm37396 | predicted gene, 37396 [Source:MGI Symb | 0 | 0 | 0 | 0 | 0 | 0 |

| | | | | | | | | | | | | | | | |
|---|---|---|---|---|---|---|---|---|---|---|---|---|---|---|---|
| 42277 | ENSMUSG00000104206 | 0.133850373 | -0.796629176 | 4.080472857 | -0.195229623 | 0.84521319 | NA | Gm32250 | predicted gene, 32250 [Source:MGI Symb | 0.8031 | 0 | 0 | 0 | 0 | 0 |
| 42278 | ENSMUSG00000104207 | 0 | NA | NA | NA | NA | NA | Gm37395 | predicted gene, 37395 [Source:MGI Symb | 0 | 0 | 0 | 0 | 0 | 0 |
| 42279 | ENSMUSG00000104208 | 0 | NA | NA | NA | NA | NA | Gm37401 | predicted gene, 37401 [Source:MGI Symb | 0 | 0 | 0 | 0 | 0 | 0 |
| 42280 | ENSMUSG00000104209 | 0 | NA | NA | NA | NA | NA | Gm37400 | predicted gene, 37400 [Source:MGI Symb | 0 | 0 | 0 | 0 | 0 | 0 |
| 42281 | ENSMUSG00000104210 | 0 | NA | NA | NA | NA | NA | Gm37984 | predicted gene, 37984 [Source:MGI Symb | 0 | 0 | 0 | 0 | 0 | 0 |
| 42282 | ENSMUSG00000104211 | 0 | NA | NA | NA | NA | NA | Gm37985 | predicted gene, 37985 [Source:MGI Symb | 0 | 0 | 0 | 0 | 0 | 0 |
| 42283 | ENSMUSG00000104212 | 0.654905883 | -1.312574656 | 2.816093566 | -0.466097672 | 0.641145589 | NA | Gm37986 | predicted gene, 37986 [Source:MGI Symb | 0.8031 | 0 | 2.0125 | 0 | 0 | 1.1138 |
| 42284 | ENSMUSG00000104213 | 8.931225833 | -2.010711338 | 0.84242616 | -2.386810184 | 0.016995264 | 0.154824975 | Ighd | immunoglobulin heavy constant delta [So | 7.2279 | 23.642 | 12.075 | 2.8919 | 2.1812 | 5.5692 |
| 42285 | ENSMUSG00000104214 | 3.70472552 | 0.365708568 | 1.179077619 | 0.310164965 | 0.756435512 | NA | Gm36638 | predicted gene, 36638 [Source:MGI Symb | 1.6062 | 7.1954 | 1.0062 | 5.7839 | 2.1812 | 4.4554 |
| 42286 | ENSMUSG00000104215 | 0 | NA | NA | NA | NA | NA | Rpl13-ps4 | ribosomal protein L13, pseudogene 4 [Sou | 0 | 0 | 0 | 0 | 0 | 0 |
| 42287 | ENSMUSG00000104216 | 0 | NA | NA | NA | NA | NA | Gm37987 | predicted gene, 37987 [Source:MGI Symb | 0 | 0 | 0 | 0 | 0 | 0 |
| 42288 | ENSMUSG00000104217 | 0 | NA | NA | NA | NA | NA | Gm37988 | predicted gene, 37988 [Source:MGI Symb | 0 | 0 | 0 | 0 | 0 | 0 |
| 42289 | ENSMUSG00000104218 | 0.17131964 | -0.796629176 | 4.080472857 | -0.195229623 | 0.84521319 | NA | Gm9134 | predicted gene 9134 [Source:MGI Symbol | 0 | 1.0279 | 0 | 0 | 0 | 0 |
| 42290 | ENSMUSG00000104219 | 0 | NA | NA | NA | NA | NA | Gm37989 | predicted gene, 37989 [Source:MGI Symb | 0 | 0 | 0 | 0 | 0 | 0 |
| 42291 | ENSMUSG00000104220 | 0 | NA | NA | NA | NA | NA | 4930578I07F | RIKEN cDNA 4930578I07 gene [Source:MG | 0 | 0 | 0 | 0 | 0 | 0 |
| 42292 | ENSMUSG00000104221 | 0 | NA | NA | NA | NA | NA | Gm37051 | predicted gene, 37051 [Source:MGI Symb | 0 | 0 | 0 | 0 | 0 | 0 |
| 42293 | ENSMUSG00000104222 | 1.375844284 | 0.917838236 | 1.859671904 | 0.493548477 | 0.621625096 | NA | Gm7292 | predicted gene 7292 [Source:MGI Symbol | 0.8031 | 2.0558 | 0 | 0.964 | 1.0906 | 3.3415 |
| 42294 | ENSMUSG00000104223 | 0 | NA | NA | NA | NA | NA | Gm18428 | predicted gene, 18428 [Source:MGI Symb | 0 | 0 | 0 | 0 | 0 | 0 |
| 42295 | ENSMUSG00000104224 | 0 | NA | NA | NA | NA | NA | Gm37693 | predicted gene, 37693 [Source:MGI Symb | 0 | 0 | 0 | 0 | 0 | 0 |
| 42296 | ENSMUSG00000104225 | 0 | NA | NA | NA | NA | NA | 4930517L18I | RIKEN cDNA 4930517L18 gene [Source:MG | 0 | 0 | 0 | 0 | 0 | 0 |
| 42297 | ENSMUSG00000104226 | 0 | NA | NA | NA | NA | NA | Gm7470 | predicted gene 7470 [Source:MGI Symbol | 0 | 0 | 0 | 0 | 0 | 0 |
| 42298 | ENSMUSG00000104227 | 0 | NA | NA | NA | NA | NA | Gm37050 | predicted gene, 37050 [Source:MGI Symb | 0 | 0 | 0 | 0 | 0 | 0 |
| 42299 | ENSMUSG00000104228 | 0 | NA | NA | NA | NA | NA | Gm35955 | predicted gene, 35955 [Source:MGI Symb | 0 | 0 | 0 | 0 | 0 | 0 |
| 42300 | ENSMUSG00000104229 | 0 | NA | NA | NA | NA | NA | Gm18433 | predicted gene, 18433 [Source:MGI Symb | 0 | 0 | 0 | 0 | 0 | 0 |
| 42301 | ENSMUSG00000104230 | 0 | NA | NA | NA | NA | NA | Gm37591 | predicted gene, 37591 [Source:MGI Symb | 0 | 0 | 0 | 0 | 0 | 0 |
| 42302 | ENSMUSG00000104231 | 6.585011382 | 0.768484898 | 0.922057031 | 0.833446167 | 0.404593147 | 0.829271989 | Gm37592 | predicted gene, 37592 [Source:MGI Symb | 1.6062 | 5.1396 | 8.05 | 12.532 | 3.2719 | 8.9108 |
| 42303 | ENSMUSG00000104232 | 0 | NA | NA | NA | NA | NA | Gm37590 | predicted gene, 37590 [Source:MGI Symb | 0 | 0 | 0 | 0 | 0 | 0 |
| 42304 | ENSMUSG00000104233 | 0 | NA | NA | NA | NA | NA | Gm34023 | predicted gene, 34023 [Source:MGI Symb | 0 | 0 | 0 | 0 | 0 | 0 |
| 42305 | ENSMUSG00000104234 | 0.74256259 | 3.09446134 | 3.273353887 | 0.945348852 | 0.344480805 | NA | 5330439B14 | RIKEN cDNA 5330439B14 gene [Source:M | 0 | 0 | 0 | 0 | 0 | 4.4554 |
| 42306 | ENSMUSG00000104235 | 3.015912988 | -1.739330392 | 1.334483937 | -1.303373044 | 0.192447435 | NA | Gm37589 | predicted gene, 37589 [Source:MGI Symb | 4.8186 | 3.0838 | 6.0375 | 1.928 | 0 | 2.2277 |
| 42307 | ENSMUSG00000104236 | 0 | NA | NA | NA | NA | NA | 4930533L02I | RIKEN cDNA 4930533L02 gene [Source:MG | 0 | 0 | 0 | 0 | 0 | 0 |
| 42308 | ENSMUSG00000104237 | 0 | NA | NA | NA | NA | NA | Gm33533 | predicted gene, 33533 [Source:MGI Symb | 0 | 0 | 0 | 0 | 0 | 0 |
| 42309 | ENSMUSG00000104238 | 0.167707431 | -0.796629176 | 4.080472857 | -0.195229623 | 0.84521319 | NA | Gm37587 | predicted gene, 37587 [Source:MGI Symb | 0 | 0 | 1.0062 | 0 | 0 | 0 |
| 42310 | ENSMUSG00000104239 | 0.973879455 | -0.768123796 | 2.429069234 | -0.31622145 | 0.751834428 | NA | Gm37588 | predicted gene, 37588 [Source:MGI Symb | 1.6062 | 2.0558 | 0 | 0 | 2.1812 | 0 |
| 42311 | ENSMUSG00000104240 | 0 | NA | NA | NA | NA | NA | Gm36858 | predicted gene, 36858 [Source:MGI Symb | 0 | 0 | 0 | 0 | 0 | 0 |
| 42312 | ENSMUSG00000104241 | 1.120480146 | -1.168613372 | 2.009668602 | -0.581495562 | 0.560906509 | NA | Gm9442 | predicted gene 9442 [Source:MGI Symbol | 1.6062 | 2.0558 | 1.0062 | 0.964 | 1.0906 | 0 |
| 42313 | ENSMUSG00000104242 | 0 | NA | NA | NA | NA | NA | Gm38154 | predicted gene, 38154 [Source:MGI Symb | 0 | 0 | 0 | 0 | 0 | 0 |
| 42314 | ENSMUSG00000104243 | 0 | NA | NA | NA | NA | NA | Gm38153 | predicted gene, 38153 [Source:MGI Symb | 0 | 0 | 0 | 0 | 0 | 0 |
| 42315 | ENSMUSG00000104244 | 0 | NA | NA | NA | NA | NA | 2900018N21 | RIKEN cDNA 2900018N21 gene [Source:M | 0 | 0 | 0 | 0 | 0 | 0 |
| 42316 | ENSMUSG00000104245 | 15.00292251 | 0.467405127 | 0.597839677 | 0.781823531 | 0.43431829 | 0.845264432 | Gm38155 | predicted gene, 38155 [Source:MGI Symb | 14.456 | 12.335 | 11.069 | 27.955 | 14.178 | 10.025 |
| 42317 | ENSMUSG00000104247 | 0 | NA | NA | NA | NA | NA | Gm21768 | predicted gene, 21768 [Source:MGI Symb | 0 | 0 | 0 | 0 | 0 | 0 |
| 42318 | ENSMUSG00000104248 | 0 | NA | NA | NA | NA | NA | Gm38152 | predicted gene, 38152 [Source:MGI Symb | 0 | 0 | 0 | 0 | 0 | 0 |
| 42319 | ENSMUSG00000104249 | 0 | NA | NA | NA | NA | NA | Gm38151 | predicted gene, 38151 [Source:MGI Symb | 0 | 0 | 0 | 0 | 0 | 0 |
| 42320 | ENSMUSG00000104251 | 0 | NA | NA | NA | NA | NA | Gm18036 | predicted gene, 18036 [Source:MGI Symb | 0 | 0 | 0 | 0 | 0 | 0 |
| 42321 | ENSMUSG00000104252 | 0 | NA | NA | NA | NA | NA | Pcdha4 | protocadherin alpha 4 [Source:MGI Symb | 0 | 0 | 0 | 0 | 0 | 0 |
| 42322 | ENSMUSG00000104253 | 8.820860381 | -0.397547944 | 0.806200494 | -0.493112999 | 0.621932746 | 0.918578091 | Gm37174 | predicted gene, 37174 [Source:MGI Symb | 4.8186 | 14.391 | 11.069 | 10.604 | 9.8156 | 2.2277 |
| 42323 | ENSMUSG00000104254 | 0 | NA | NA | NA | NA | NA | Trbv27 | T cell receptor beta variable 27 [Source:M | 0 | 0 | 0 | 0 | 0 | 0 |
| 42324 | ENSMUSG00000104255 | 0 | NA | NA | NA | NA | NA | Gm37172 | predicted gene, 37172 [Source:MGI Symb | 0 | 0 | 0 | 0 | 0 | 0 |
| 42325 | ENSMUSG00000104256 | 0 | NA | NA | NA | NA | NA | Gm37173 | predicted gene, 37173 [Source:MGI Symb | 0 | 0 | 0 | 0 | 0 | 0 |
| 42326 | ENSMUSG00000104257 | 0 | NA | NA | NA | NA | NA | Gm20172 | predicted gene, 20172 [Source:MGI Symb | 0 | 0 | 0 | 0 | 0 | 0 |
| 42327 | ENSMUSG00000104258 | 0 | NA | NA | NA | NA | NA | Gm37175 | predicted gene, 37175 [Source:MGI Symb | 0 | 0 | 0 | 0 | 0 | 0 |
| 42328 | ENSMUSG00000104259 | 0 | NA | NA | NA | NA | NA | Gm37538 | predicted gene, 37538 [Source:MGI Symb | 0 | 0 | 0 | 0 | 0 | 0 |
| 42329 | ENSMUSG00000104260 | 0 | NA | NA | NA | NA | NA | Gm31186 | predicted gene, 31186 [Source:MGI Symb | 0 | 0 | 0 | 0 | 0 | 0 |
| 42330 | ENSMUSG00000104261 | 0 | NA | NA | NA | NA | NA | Gm17781 | predicted gene, 17781 [Source:MGI Symb | 0 | 0 | 0 | 0 | 0 | 0 |
| 42331 | ENSMUSG00000104262 | 0.401551119 | -2.072982726 | 4.008251979 | -0.517178744 | 0.605031388 | NA | Gm37747 | predicted gene, 37747 [Source:MGI Symb | 2.4093 | 0 | 0 | 0 | 0 | 0 |
| 42332 | ENSMUSG00000104263 | 3.878774486 | -0.192605222 | 1.102568166 | -0.174687813 | 0.861324944 | NA | 9430062P05 | RIKEN cDNA 9430062P05 gene [Source:M | 3.2124 | 5.1396 | 4.025 | 0.964 | 4.3625 | 5.5692 |

| | | | | | | | | | | | | | | | |
|---|---|---|---|---|---|---|---|---|---|---|---|---|---|---|---|
| 42333 | ENSMUSG00000104264 | 0 | NA | NA | NA | NA | NA | Gm18529 | predicted gene, 18529 [Source:MGI Symb | 0 | 0 | 0 | 0 | 0 | 0 |
| 42334 | ENSMUSG00000104265 | 0 | NA | NA | NA | NA | NA | Gm8869 | predicted gene 8869 [Source:MGI Symbol | 0 | 0 | 0 | 0 | 0 | 0 |
| 42335 | ENSMUSG00000104266 | 0.51014012 | 1.031855138 | 3.09002894 | 0.333930574 | 0.738431948 | NA | Gm37748 | predicted gene, 37748 [Source:MGI Symb | 0 | 0 | 1.0062 | 0.964 | 1.0906 | 0 |
| 42336 | ENSMUSG00000104267 | 0 | NA | NA | NA | NA | NA | Gm20807 | predicted gene, 20807 [Source:MGI Symb | 0 | 0 | 0 | 0 | 0 | 0 |
| 42337 | ENSMUSG00000104268 | 0.860262541 | -0.538502919 | 2.596000693 | -0.207435584 | 0.835669695 | NA | Gm37750 | predicted gene, 37750 [Source:MGI Symb | 0 | 3.0838 | 0 | 0.964 | 0 | 1.1138 |
| 42338 | ENSMUSG00000104269 | 0 | NA | NA | NA | NA | NA | Gm37749 | predicted gene, 37749 [Source:MGI Symb | 0 | 0 | 0 | 0 | 0 | 0 |
| 42339 | ENSMUSG00000104270 | 0 | NA | NA | NA | NA | NA | Gm38298 | predicted gene, 38298 [Source:MGI Symb | 0 | 0 | 0 | 0 | 0 | 0 |
| 42340 | ENSMUSG00000104271 | 0.267700746 | -1.497481649 | 4.057001436 | -0.369110456 | 0.712045396 | NA | Gm37891 | predicted gene, 37891 [Source:MGI Symb | 1.6062 | 0 | 0 | 0 | 0 | 0 |
| 42341 | ENSMUSG00000104272 | 0.346303622 | 2.004339653 | 3.884003423 | 0.516049919 | 0.605819544 | NA | Gm38297 | predicted gene, 38297 [Source:MGI Symb | 0 | 0 | 0 | 0.964 | 0 | 1.1138 |
| 42342 | ENSMUSG00000104273 | 0.439020386 | -2.189747911 | 3.52148864 | -0.621824499 | 0.534057275 | NA | A530064N14 | RIKEN cDNA A530064N14 gene [Source:M | 1.6062 | 1.0279 | 0 | 0 | 0 | 0 |
| 42343 | ENSMUSG00000104274 | 0 | NA | NA | NA | NA | NA | Gm8823 | predicted gene 8823 [Source:MGI Symbol | 0 | 0 | 0 | 0 | 0 | 0 |
| 42344 | ENSMUSG00000104275 | 0 | NA | NA | NA | NA | NA | Gm37867 | predicted gene, 37867 [Source:MGI Symb | 0 | 0 | 0 | 0 | 0 | 0 |
| 42345 | ENSMUSG00000104276 | 0 | NA | NA | NA | NA | NA | Gm37866 | predicted gene, 37866 [Source:MGI Symb | 0 | 0 | 0 | 0 | 0 | 0 |
| 42346 | ENSMUSG00000104277 | 0.503122293 | -2.366675966 | 3.785596405 | -0.625179156 | 0.531853481 | NA | Gm38299 | predicted gene, 38299 [Source:MGI Symb | 0 | 0 | 3.0187 | 0 | 0 | 0 |
| 42347 | ENSMUSG00000104278 | 0 | NA | NA | NA | NA | NA | Ighv5-10 | immunoglobulin heavy variable 5-10 [Sou | 0 | 0 | 0 | 0 | 0 | 0 |
| 42348 | ENSMUSG00000104279 | 8.123967597 | 0.065540836 | 0.764552084 | 0.085724488 | 0.931685435 | 0.988420305 | Gm38300 | predicted gene, 38300 [Source:MGI Symb | 5.6217 | 11.307 | 7.0437 | 11.568 | 7.6343 | 5.5692 |
| 42349 | ENSMUSG00000104280 | 0.513958919 | -2.394274821 | 3.756096013 | -0.637437066 | 0.523840191 | NA | Rps2-ps11 | ribosomal protein S2, pseudogene 11 [Sou | 0 | 3.0838 | 0 | 0 | 0 | 0 |
| 42350 | ENSMUSG00000104281 | 0 | NA | NA | NA | NA | NA | Gm37461 | predicted gene, 37461 [Source:MGI Symb | 0 | 0 | 0 | 0 | 0 | 0 |
| 42351 | ENSMUSG00000104282 | 0 | NA | NA | NA | NA | NA | Gm37460 | predicted gene, 37460 [Source:MGI Symb | 0 | 0 | 0 | 0 | 0 | 0 |
| 42352 | ENSMUSG00000104283 | 0 | NA | NA | NA | NA | NA | Gm37459 | predicted gene, 37459 [Source:MGI Symb | 0 | 0 | 0 | 0 | 0 | 0 |
| 42353 | ENSMUSG00000104284 | 0 | NA | NA | NA | NA | NA | Gm10862 | predicted gene 10862 [Source:MGI Symbo | 0 | 0 | 0 | 0 | 0 | 0 |
| 42354 | ENSMUSG00000104285 | 0 | NA | NA | NA | NA | NA | Gm37458 | predicted gene, 37458 [Source:MGI Symb | 0 | 0 | 0 | 0 | 0 | 0 |
| 42355 | ENSMUSG00000104286 | 0 | NA | NA | NA | NA | NA | Gm37457 | predicted gene, 37457 [Source:MGI Symb | 0 | 0 | 0 | 0 | 0 | 0 |
| 42356 | ENSMUSG00000104287 | 0 | NA | NA | NA | NA | NA | Gm37456 | predicted gene, 37456 [Source:MGI Symb | 0 | 0 | 0 | 0 | 0 | 0 |
| 42357 | ENSMUSG00000104288 | 0 | NA | NA | NA | NA | NA | Gm37455 | predicted gene, 37455 [Source:MGI Symb | 0 | 0 | 0 | 0 | 0 | 0 |
| 42358 | ENSMUSG00000104289 | 0 | NA | NA | NA | NA | NA | Gm37454 | predicted gene, 37454 [Source:MGI Symb | 0 | 0 | 0 | 0 | 0 | 0 |
| 42359 | ENSMUSG00000104290 | 0.401551119 | -2.072982726 | 4.008251979 | -0.517178744 | 0.605031388 | NA | Gm38041 | predicted gene, 38041 [Source:MGI Symb | 2.4093 | 0 | 0 | 0 | 0 | 0 |
| 42360 | ENSMUSG00000104291 | 0.684865378 | 2.990343487 | 2.966805827 | 1.00793367 | 0.313486306 | NA | A130071D04 | RIKEN cDNA A130071D04 gene [Source:M | 0 | 0 | 0 | 1.928 | 2.1812 | 0 |
| 42361 | ENSMUSG00000104292 | 0.267700746 | -1.497481649 | 4.057001436 | -0.369110456 | 0.712045396 | NA | Gm38042 | predicted gene, 38042 [Source:MGI Symb | 1.6062 | 0 | 0 | 0 | 0 | 0 |
| 42362 | ENSMUSG00000104293 | 4.380635292 | -0.136021843 | 0.995516037 | -0.136634507 | 0.8913197 | NA | Gm38043 | predicted gene, 38043 [Source:MGI Symb | 5.6217 | 3.0838 | 5.0312 | 4.8199 | 3.2719 | 4.4554 |
| 42363 | ENSMUSG00000104294 | 0 | NA | NA | NA | NA | NA | Gm38044 | predicted gene, 38044 [Source:MGI Symb | 0 | 0 | 0 | 0 | 0 | 0 |
| 42364 | ENSMUSG00000104295 | 0 | NA | NA | NA | NA | NA | Gm6197 | predicted gene 6197 [Source:MGI Symbol | 0 | 0 | 0 | 0 | 0 | 0 |
| 42365 | ENSMUSG00000104296 | 0 | NA | NA | NA | NA | NA | Gm38045 | predicted gene, 38045 [Source:MGI Symb | 0 | 0 | 0 | 0 | 0 | 0 |
| 42366 | ENSMUSG00000104297 | 0 | NA | NA | NA | NA | NA | Gm38046 | predicted gene, 38046 [Source:MGI Symb | 0 | 0 | 0 | 0 | 0 | 0 |
| 42367 | ENSMUSG00000104298 | 0 | NA | NA | NA | NA | NA | Gm38040 | predicted gene, 38040 [Source:MGI Symb | 0 | 0 | 0 | 0 | 0 | 0 |
| 42368 | ENSMUSG00000104299 | 0.503122293 | -2.366675966 | 3.785596405 | -0.625179156 | 0.531853481 | NA | Gm9924 | predicted gene 9924 [Source:MGI Symbol | 0 | 0 | 3.0187 | 0 | 0 | 0 |
| 42369 | ENSMUSG00000104301 | 0 | NA | NA | NA | NA | NA | Wdr49 | WD repeat domain 49 [Source:MGI Symbo | 0 | 0 | 0 | 0 | 0 | 0 |
| 42370 | ENSMUSG00000104302 | 0.328370405 | 0.16514316 | 4.003149536 | 0.041253308 | 0.967093956 | NA | Gm36975 | predicted gene, 36975 [Source:MGI Symb | 0 | 0 | 1.0062 | 0.964 | 0 | 0 |
| 42371 | ENSMUSG00000104303 | 0.185640647 | 1.126924175 | 4.080472857 | 0.276174898 | 0.782413736 | NA | Gm36976 | predicted gene, 36976 [Source:MGI Symb | 0 | 0 | 0 | 0 | 0 | 1.1138 |
| 42372 | ENSMUSG00000104304 | 0 | NA | NA | NA | NA | NA | Gm36974 | predicted gene, 36974 [Source:MGI Symb | 0 | 0 | 0 | 0 | 0 | 0 |
| 42373 | ENSMUSG00000104306 | 0.688942901 | 0.052211695 | 2.79596103 | 0.018673971 | 0.985101193 | NA | Gm36972 | predicted gene, 36972 [Source:MGI Symb | 0 | 2.0558 | 0 | 0.964 | 0 | 1.1138 |
| 42374 | ENSMUSG00000104307 | 0 | NA | NA | NA | NA | NA | Gm36973 | predicted gene, 36973 [Source:MGI Symb | 0 | 0 | 0 | 0 | 0 | 0 |
| 42375 | ENSMUSG00000104308 | 0 | NA | NA | NA | NA | NA | Gm36971 | predicted gene, 36971 [Source:MGI Symb | 0 | 0 | 0 | 0 | 0 | 0 |
| 42376 | ENSMUSG00000104309 | 0.17131964 | -0.796629176 | 4.080472857 | -0.195229623 | 0.84521319 | NA | Gm5846 | predicted gene 5846 [Source:MGI Symbol | 0 | 1.0279 | 0 | 0 | 0 | 0 |
| 42377 | ENSMUSG00000104310 | 0 | NA | NA | NA | NA | NA | Gm4081 | predicted gene 4081 [Source:MGI Symbol | 0 | 0 | 0 | 0 | 0 | 0 |
| 42378 | ENSMUSG00000104311 | 0 | NA | NA | NA | NA | NA | Gm37910 | predicted gene, 37910 [Source:MGI Symb | 0 | 0 | 0 | 0 | 0 | 0 |
| 42379 | ENSMUSG00000104312 | 0 | NA | NA | NA | NA | NA | Gm37912 | predicted gene, 37912 [Source:MGI Symb | 0 | 0 | 0 | 0 | 0 | 0 |
| 42380 | ENSMUSG00000104313 | 0 | NA | NA | NA | NA | NA | Gm37911 | predicted gene, 37911 [Source:MGI Symb | 0 | 0 | 0 | 0 | 0 | 0 |
| 42381 | ENSMUSG00000104314 | 0 | NA | NA | NA | NA | NA | Gm37907 | predicted gene, 37907 [Source:MGI Symb | 0 | 0 | 0 | 0 | 0 | 0 |
| 42382 | ENSMUSG00000104315 | 2.034703722 | -0.06993049 | 1.588290315 | -0.044028783 | 0.964881461 | NA | Gm37906 | predicted gene, 37906 [Source:MGI Symb | 3.2124 | 2.0558 | 1.0062 | 4.8199 | 0 | 1.1138 |
| 42383 | ENSMUSG00000104316 | 0.185640647 | 1.126924175 | 4.080472857 | 0.276174898 | 0.782413736 | NA | Gm37909 | predicted gene, 37909 [Source:MGI Symb | 0 | 0 | 0 | 0 | 0 | 1.1138 |
| 42384 | ENSMUSG00000104317 | 0 | NA | NA | NA | NA | NA | Gm37908 | predicted gene, 37908 [Source:MGI Symb | 0 | 0 | 0 | 0 | 0 | 0 |
| 42385 | ENSMUSG00000104318 | 0 | NA | NA | NA | NA | NA | Pcdha7 | protocadherin alpha 7 [Source:MGI Symb | 0 | 0 | 0 | 0 | 0 | 0 |
| 42386 | ENSMUSG00000104319 | 0 | NA | NA | NA | NA | NA | Ighv5-19 | immunoglobulin heavy variable V5-19 [So | 0 | 0 | 0 | 0 | 0 | 0 |
| 42387 | ENSMUSG00000104320 | 0 | NA | NA | NA | NA | NA | Ighv1-60 | immunoglobulin heavy variable V1-60 [So | 0 | 0 | 0 | 0 | 0 | 0 |
| 42388 | ENSMUSG00000104321 | 0 | NA | NA | NA | NA | NA | Gm37318 | predicted gene, 37318 [Source:MGI Symb | 0 | 0 | 0 | 0 | 0 | 0 |

| | | | | | | | | | | | | | | | |
|---|---|---|---|---|---|---|---|---|---|---|---|---|---|---|---|
| 42389 | ENSMUSG00000104323 | 0 | NA | NA | NA | NA | NA | Gm37319 | predicted gene, 37319 [Source:MGI Symb | 0 | 0 | 0 | 0 | 0 | 0 |
| 42390 | ENSMUSG00000104324 | 3.436611064 | -0.291610743 | 1.214622758 | -0.240083385 | 0.810265614 | NA | Gm37320 | predicted gene, 37320 [Source:MGI Symb | 3.2124 | 7.1954 | 1.0062 | 4.8199 | 3.2719 | 1.1138 |
| 42391 | ENSMUSG00000104325 | 0.342639279 | -1.805976596 | 4.028547714 | -0.448294702 | 0.653940524 | NA | Gm37321 | predicted gene, 37321 [Source:MGI Symb | 0 | 2.0558 | 0 | 0 | 0 | 0 |
| 42392 | ENSMUSG00000104326 | 0 | NA | NA | NA | NA | NA | Trbv28 | T cell receptor beta variable 28 [Source:M | 0 | 0 | 0 | 0 | 0 | 0 |
| 42393 | ENSMUSG00000104327 | 0 | NA | NA | NA | NA | NA | Gm37322 | predicted gene, 37322 [Source:MGI Symb | 0 | 0 | 0 | 0 | 0 | 0 |
| 42394 | ENSMUSG00000104328 | 0 | NA | NA | NA | NA | NA | Gm37323 | predicted gene, 37323 [Source:MGI Symb | 0 | 0 | 0 | 0 | 0 | 0 |
| 42395 | ENSMUSG00000104329 | 0 | NA | NA | NA | NA | NA | Gm37324 | predicted gene, 37324 [Source:MGI Symb | 0 | 0 | 0 | 0 | 0 | 0 |
| 42396 | ENSMUSG00000104330 | 0 | NA | NA | NA | NA | NA | Gm38285 | predicted gene, 38285 [Source:MGI Symb | 0 | 0 | 0 | 0 | 0 | 0 |
| 42397 | ENSMUSG00000104331 | 0 | NA | NA | NA | NA | NA | Gm38284 | predicted gene, 38284 [Source:MGI Symb | 0 | 0 | 0 | 0 | 0 | 0 |
| 42398 | ENSMUSG00000104332 | 1.721837935 | -3.047332428 | 2.107344196 | -1.446053489 | 0.1481622 | NA | Gm38283 | predicted gene, 38283 [Source:MGI Symb | 7.2279 | 0 | 2.0125 | 0 | 1.0906 | 0 |
| 42399 | ENSMUSG00000104333 | 0 | NA | NA | NA | NA | NA | Gm10253 | predicted gene 10253 [Source:MGI Symbo | 0 | 0 | 0 | 0 | 0 | 0 |
| 42400 | ENSMUSG00000104334 | 0 | NA | NA | NA | NA | NA | Gm38282 | predicted gene, 38282 [Source:MGI Symb | 0 | 0 | 0 | 0 | 0 | 0 |
| 42401 | ENSMUSG00000104335 | 0 | NA | NA | NA | NA | NA | Gm38281 | predicted gene, 38281 [Source:MGI Symb | 0 | 0 | 0 | 0 | 0 | 0 |
| 42402 | ENSMUSG00000104336 | 0 | NA | NA | NA | NA | NA | Gm34240 | predicted gene, 34240 [Source:MGI Symb | 0 | 0 | 0 | 0 | 0 | 0 |
| 42403 | ENSMUSG00000104337 | 0 | NA | NA | NA | NA | NA | Gm38280 | predicted gene, 38280 [Source:MGI Symb | 0 | 0 | 0 | 0 | 0 | 0 |
| 42404 | ENSMUSG00000104338 | 0 | NA | NA | NA | NA | NA | Gm9131 | predicted gene 9131 [Source:MGI Symbol | 0 | 0 | 0 | 0 | 0 | 0 |
| 42405 | ENSMUSG00000104339 | 0 | NA | NA | NA | NA | NA | C130089K02 | RIKEN cDNA C130089K02 gene [Source:M | 0 | 0 | 0 | 0 | 0 | 0 |
| 42406 | ENSMUSG00000104340 | 1.29933506 | 0.184946416 | 1.850499756 | 0.099944037 | 0.920388755 | NA | Gm10522 | predicted gene 10522 [Source:MGI Symbo | 1.6062 | 1.0279 | 1.0062 | 1.928 | 0 | 2.2277 |
| 42407 | ENSMUSG00000104341 | 0 | NA | NA | NA | NA | NA | Gm37688 | predicted gene, 37688 [Source:MGI Symb | 0 | 0 | 0 | 0 | 0 | 0 |
| 42408 | ENSMUSG00000104342 | 1.181149805 | -0.383612265 | 2.080148392 | -0.184415817 | 0.853687263 | NA | Gm36401 | predicted gene, 36401 [Source:MGI Symb | 0 | 2.0558 | 2.0125 | 1.928 | 1.0906 | 0 |
| 42409 | ENSMUSG00000104343 | 0 | NA | NA | NA | NA | NA | 5730408A14 | RIKEN cDNA 5730408A14 gene [Source:M | 0 | 0 | 0 | 0 | 0 | 0 |
| 42410 | ENSMUSG00000104344 | 19.66641136 | 1.074342553 | 0.541858004 | 1.982701269 | 0.047400808 | 0.302152946 | Gm38077 | predicted gene, 38077 [Source:MGI Symb | 13.653 | 13.363 | 11.069 | 42.415 | 17.45 | 20.049 |
| 42411 | ENSMUSG00000104345 | 0 | NA | NA | NA | NA | NA | Gm37690 | predicted gene, 37690 [Source:MGI Symb | 0 | 0 | 0 | 0 | 0 | 0 |
| 42412 | ENSMUSG00000104346 | 2.391244786 | -1.272982808 | 1.504044245 | -0.846373245 | 0.397344559 | NA | Pcdhga3 | protocadherin gamma subfamily A, 3 [Sou | 8.031 | 1.0279 | 1.0062 | 0.964 | 1.0906 | 2.2277 |
| 42413 | ENSMUSG00000104347 | 0.17131964 | -0.796629176 | 4.080472857 | -0.195229623 | 0.84521319 | NA | Gm37689 | predicted gene, 37689 [Source:MGI Symb | 0 | 1.0279 | 0 | 0 | 0 | 0 |
| 42414 | ENSMUSG00000104348 | 98.24271419 | -0.33020381 | 0.246594147 | -1.339057779 | 0.180551864 | 0.613893314 | Gm37691 | predicted gene, 37691 [Source:MGI Symb | 116.45 | 112.04 | 99.618 | 80.974 | 107.97 | 72.4 |
| 42415 | ENSMUSG00000104349 | 0 | NA | NA | NA | NA | NA | Gm37692 | predicted gene, 37692 [Source:MGI Symb | 0 | 0 | 0 | 0 | 0 | 0 |
| 42416 | ENSMUSG00000104350 | 0.160662974 | 1.126924175 | 4.080472857 | 0.276174898 | 0.782413736 | NA | Gm38244 | predicted gene, 38244 [Source:MGI Symb | 0 | 0 | 0 | 0.964 | 0 | 0 |
| 42417 | ENSMUSG00000104351 | 0.167707431 | -0.796629176 | 4.080472857 | -0.195229623 | 0.84521319 | NA | Gm37125 | predicted gene, 37125 [Source:MGI Symb | 0 | 0 | 1.0062 | 0 | 0 | 0 |
| 42418 | ENSMUSG00000104352 | 0 | NA | NA | NA | NA | NA | Gm7182 | predicted gene 7182 [Source:MGI Symbol | 0 | 0 | 0 | 0 | 0 | 0 |
| 42419 | ENSMUSG00000104354 | 0 | NA | NA | NA | NA | NA | Gm37126 | predicted gene, 37126 [Source:MGI Symb | 0 | 0 | 0 | 0 | 0 | 0 |
| 42420 | ENSMUSG00000104355 | 0 | NA | NA | NA | NA | NA | Gm7003 | predicted gene 7003 [Source:MGI Symbol | 0 | 0 | 0 | 0 | 0 | 0 |
| 42421 | ENSMUSG00000104356 | 0 | NA | NA | NA | NA | NA | Gm31260 | predicted gene, 31260 [Source:MGI Symb | 0 | 0 | 0 | 0 | 0 | 0 |
| 42422 | ENSMUSG00000104358 | 3.19355192 | 1.144326792 | 1.374513346 | 0.832532325 | 0.40510854 | NA | Gm37127 | predicted gene, 37127 [Source:MGI Symb | 4.8186 | 1.0279 | 0 | 6.7478 | 5.4531 | 1.1138 |
| 42423 | ENSMUSG00000104360 | 0.305170013 | -1.657836477 | 4.041480335 | -0.410205257 | 0.681655385 | NA | Mif-ps1 | macrophage migration inhibitory factor, p | 0.8031 | 1.0279 | 0 | 0 | 0 | 0 |
| 42424 | ENSMUSG00000104361 | 2.268031617 | -0.717475016 | 1.38584155 | -0.517717928 | 0.604655088 | NA | Gm38080 | predicted gene, 38080 [Source:MGI Symb | 2.4093 | 3.0838 | 3.0187 | 2.8919 | 1.0906 | 1.1138 |
| 42425 | ENSMUSG00000104362 | 1.482914199 | -2.794328659 | 2.013087349 | -1.388081178 | 0.165112327 | NA | Gm37928 | predicted gene, 37928 [Source:MGI Symb | 0.8031 | 4.1117 | 3.0187 | 0.964 | 0 | 0 |
| 42426 | ENSMUSG00000104363 | 0 | NA | NA | NA | NA | NA | Gm38081 | predicted gene, 38081 [Source:MGI Symb | 0 | 0 | 0 | 0 | 0 | 0 |
| 42427 | ENSMUSG00000104364 | 0 | NA | NA | NA | NA | NA | Gm38078 | predicted gene, 38078 [Source:MGI Symb | 0 | 0 | 0 | 0 | 0 | 0 |
| 42428 | ENSMUSG00000104365 | 0 | NA | NA | NA | NA | NA | 4930470B04 | RIKEN cDNA 4930470B04 gene [Source:M | 0 | 0 | 0 | 0 | 0 | 0 |
| 42429 | ENSMUSG00000104366 | 0 | NA | NA | NA | NA | NA | 4933409D19 | RIKEN cDNA 4933409D19 gene [Source:M | 0 | 0 | 0 | 0 | 0 | 0 |
| 42430 | ENSMUSG00000104367 | 12.21327536 | -0.158548657 | 0.655347596 | -0.24193063 | 0.808833916 | 0.965682062 | NA | NA | 20.078 | 12.335 | 6.0375 | 10.604 | 13.087 | 11.138 |
| 42431 | ENSMUSG00000104368 | 0.342432689 | 1.990653086 | 3.900281263 | 0.51038706 | 0.609780321 | NA | 2010103J01I | RIKEN cDNA 2010103J01 gene [Source:M( | 0 | 0 | 0 | 0.964 | 1.0906 | 0 |
| 42432 | ENSMUSG00000104369 | 0.802559816 | -0.30507865 | 2.617070748 | -0.116572565 | 0.90719878 | NA | Gm38082 | predicted gene, 38082 [Source:MGI Symb | 1.6062 | 1.0279 | 0 | 0 | 2.1812 | 0 |
| 42433 | ENSMUSG00000104370 | 0 | NA | NA | NA | NA | NA | Gm37514 | predicted gene, 37514 [Source:MGI Symb | 0 | 0 | 0 | 0 | 0 | 0 |
| 42434 | ENSMUSG00000104371 | 0 | NA | NA | NA | NA | NA | Gm37513 | predicted gene, 37513 [Source:MGI Symb | 0 | 0 | 0 | 0 | 0 | 0 |
| 42435 | ENSMUSG00000104373 | 0 | NA | NA | NA | NA | NA | Gm37511 | predicted gene, 37511 [Source:MGI Symb | 0 | 0 | 0 | 0 | 0 | 0 |
| 42436 | ENSMUSG00000104374 | 0 | NA | NA | NA | NA | NA | Gm37517 | predicted gene, 37517 [Source:MGI Symb | 0 | 0 | 0 | 0 | 0 | 0 |
| 42437 | ENSMUSG00000104375 | 0 | NA | NA | NA | NA | NA | Gm6300 | predicted gene 6300 [Source:MGI Symbol | 0 | 0 | 0 | 0 | 0 | 0 |
| 42438 | ENSMUSG00000104376 | 0.36353943 | 2.065989254 | 4.034137301 | 0.512126658 | 0.608562368 | NA | Gm37516 | predicted gene, 37516 [Source:MGI Symb | 0 | 0 | 0 | 0 | 2.1812 | 0 |
| 42439 | ENSMUSG00000104377 | 0 | NA | NA | NA | NA | NA | Gm37515 | predicted gene, 37515 [Source:MGI Symb | 0 | 0 | 0 | 0 | 0 | 0 |
| 42440 | ENSMUSG00000104378 | 0 | NA | NA | NA | NA | NA | Gm37510 | predicted gene, 37510 [Source:MGI Symb | 0 | 0 | 0 | 0 | 0 | 0 |
| 42441 | ENSMUSG00000104379 | 0.663785267 | -1.400021131 | 3.106754789 | -0.450637796 | 0.652250621 | NA | Gm37509 | predicted gene, 37509 [Source:MGI Symb | 0 | 0 | 3.0187 | 0.964 | 0 | 0 |
| 42442 | ENSMUSG00000104380 | 0 | NA | NA | NA | NA | NA | 4933436F18 | RIKEN cDNA 4933436F18 gene [Source:M( | 0 | 0 | 0 | 0 | 0 | 0 |
| 42443 | ENSMUSG00000104381 | 0 | NA | NA | NA | NA | NA | Gm37032 | predicted gene, 37032 [Source:MGI Symb | 0 | 0 | 0 | 0 | 0 | 0 |
| 42444 | ENSMUSG00000104382 | 0 | NA | NA | NA | NA | NA | 4930402C01 | RIKEN cDNA 4930402C01 gene [Source:M( | 0 | 0 | 0 | 0 | 0 | 0 |

| | | | | | | | | | | | | | | | |
|---|---|---|---|---|---|---|---|---|---|---|---|---|---|---|---|
| 42445 | ENSMUSG00000104383 | 1.472230904 | 0.383055431 | 1.948033802 | 0.196636953 | 0.844111651 | NA | Gm37553 | predicted gene, 37553 [Source:MGI Symb | 0.8031 | 3.0838 | 0 | 3.8559 | 1.0906 | 0 |
| 42446 | ENSMUSG00000104384 | 0 | NA | NA | NA | NA | NA | Gm37031 | predicted gene, 37031 [Source:MGI Symb | 0 | 0 | 0 | 0 | 0 | 0 |
| 42447 | ENSMUSG00000104385 | 0 | NA | NA | NA | NA | NA | Gm7449 | predicted gene 7449 [Source:MGI Symbol | 0 | 0 | 0 | 0 | 0 | 0 |
| 42448 | ENSMUSG00000104386 | 0 | NA | NA | NA | NA | NA | Gm37030 | predicted gene, 37030 [Source:MGI Symb | 0 | 0 | 0 | 0 | 0 | 0 |
| 42449 | ENSMUSG00000104388 | 1.733296815 | 1.318769158 | 2.005439485 | 0.657596087 | 0.51079771 | NA | Gm37033 | predicted gene, 37033 [Source:MGI Symb | 0 | 0 | 3.0187 | 1.928 | 5.4531 | 0 |
| 42450 | ENSMUSG00000104390 | 1.448120999 | 0.705574662 | 2.138746123 | 0.329901083 | 0.741474705 | NA | Gm17967 | predicted gene, 17967 [Source:MGI Symb | 3.2124 | 0 | 0 | 0 | 4.3625 | 1.1138 |
| 42451 | ENSMUSG00000104391 | 0 | NA | NA | NA | NA | NA | Gm37960 | predicted gene, 37960 [Source:MGI Symb | 0 | 0 | 0 | 0 | 0 | 0 |
| 42452 | ENSMUSG00000104392 | 11.02432448 | -0.810630313 | 0.687670851 | -1.178805692 | 0.238475557 | 0.687387498 | Gm37962 | predicted gene, 37962 [Source:MGI Symb | 9.6372 | 21.586 | 11.069 | 10.604 | 5.4531 | 7.7969 |
| 42453 | ENSMUSG00000104393 | 0 | NA | NA | NA | NA | NA | Gm37961 | predicted gene, 37961 [Source:MGI Symb | 0 | 0 | 0 | 0 | 0 | 0 |
| 42454 | ENSMUSG00000104394 | 70.72525081 | 0.065257652 | 0.293007267 | 0.222716837 | 0.823755896 | 0.96943211 | Gm37254 | predicted gene, 37254 [Source:MGI Symb | 65.051 | 80.178 | 62.387 | 81.938 | 51.259 | 83.538 |
| 42455 | ENSMUSG00000104395 | 0 | NA | NA | NA | NA | NA | Gm37957 | predicted gene, 37957 [Source:MGI Symb | 0 | 0 | 0 | 0 | 0 | 0 |
| 42456 | ENSMUSG00000104396 | 0 | NA | NA | NA | NA | NA | Gm37959 | predicted gene, 37959 [Source:MGI Symb | 0 | 0 | 0 | 0 | 0 | 0 |
| 42457 | ENSMUSG00000104397 | 0 | NA | NA | NA | NA | NA | Gm37958 | predicted gene, 37958 [Source:MGI Symb | 0 | 0 | 0 | 0 | 0 | 0 |
| 42458 | ENSMUSG00000104398 | 0 | NA | NA | NA | NA | NA | Gm37964 | predicted gene, 37964 [Source:MGI Symb | 0 | 0 | 0 | 0 | 0 | 0 |
| 42459 | ENSMUSG00000104399 | 16.18604275 | 1.107283682 | 0.636612507 | 1.73933699 | 0.081975504 | 0.421541345 | Gm37963 | predicted gene, 37963 [Source:MGI Symb | 9.6372 | 7.1954 | 14.087 | 38.559 | 9.8156 | 17.822 |
| 42460 | ENSMUSG00000104400 | 0.181769715 | 1.126924175 | 4.080472857 | 0.276174898 | 0.782413736 | NA | Gm38010 | predicted gene, 38010 [Source:MGI Symb | 0 | 0 | 0 | 0 | 1.0906 | 0 |
| 42461 | ENSMUSG00000104401 | 0 | NA | NA | NA | NA | NA | Gm20750 | predicted gene, 20750 [Source:MGI Symb | 0 | 0 | 0 | 0 | 0 | 0 |
| 42462 | ENSMUSG00000104402 | 0 | NA | NA | NA | NA | NA | Gm38012 | predicted gene, 38012 [Source:MGI Symb | 0 | 0 | 0 | 0 | 0 | 0 |
| 42463 | ENSMUSG00000104403 | 0 | NA | NA | NA | NA | NA | Gm38011 | predicted gene, 38011 [Source:MGI Symb | 0 | 0 | 0 | 0 | 0 | 0 |
| 42464 | ENSMUSG00000104404 | 0 | NA | NA | NA | NA | NA | Gm36728 | predicted gene, 36728 [Source:MGI Symb | 0 | 0 | 0 | 0 | 0 | 0 |
| 42465 | ENSMUSG00000104405 | 0 | NA | NA | NA | NA | NA | Gm31748 | predicted gene, 31748 [Source:MGI Symb | 0 | 0 | 0 | 0 | 0 | 0 |
| 42466 | ENSMUSG00000104406 | 0.74057819 | -2.943622841 | 2.624979752 | -1.12138878 | 0.26212241 | NA | Gm38014 | predicted gene, 38014 [Source:MGI Symb | 2.4093 | 1.0279 | 1.0062 | 0 | 0 | 0 |
| 42467 | ENSMUSG00000104407 | 0 | NA | NA | NA | NA | NA | Gm38013 | predicted gene, 38013 [Source:MGI Symb | 0 | 0 | 0 | 0 | 0 | 0 |
| 42468 | ENSMUSG00000104408 | 0 | NA | NA | NA | NA | NA | Gm38015 | predicted gene, 38015 [Source:MGI Symb | 0 | 0 | 0 | 0 | 0 | 0 |
| 42469 | ENSMUSG00000104409 | 0 | NA | NA | NA | NA | NA | Gm37467 | predicted gene, 37467 [Source:MGI Symb | 0 | 0 | 0 | 0 | 0 | 0 |
| 42470 | ENSMUSG00000104410 | 0 | NA | NA | NA | NA | NA | Gm37066 | predicted gene, 37066 [Source:MGI Symb | 0 | 0 | 0 | 0 | 0 | 0 |
| 42471 | ENSMUSG00000104411 | 0 | NA | NA | NA | NA | NA | Gm30340 | predicted gene, 30340 [Source:MGI Symb | 0 | 0 | 0 | 0 | 0 | 0 |
| 42472 | ENSMUSG00000104412 | 0 | NA | NA | NA | NA | NA | Gm37798 | predicted gene, 37798 [Source:MGI Symb | 0 | 0 | 0 | 0 | 0 | 0 |
| 42473 | ENSMUSG00000104413 | 0.17131964 | -0.796629176 | 4.080472857 | -0.195229623 | 0.84521319 | NA | Gm37065 | predicted gene, 37065 [Source:MGI Symb | 0 | 1.0279 | 0 | 0 | 0 | 0 |
| 42474 | ENSMUSG00000104414 | 0.671009685 | -1.413087452 | 2.800223337 | -0.504633839 | 0.613816038 | NA | Gm37799 | predicted gene, 37799 [Source:MGI Symb | 0 | 2.0558 | 1.0062 | 0.964 | 0 | 0 |
| 42475 | ENSMUSG00000104415 | 0.315620088 | 0.165143421 | 4.063300985 | 0.040642675 | 0.967580763 | NA | Gm37069 | predicted gene, 37069 [Source:MGI Symb | 0.8031 | 0 | 0 | 0 | 1.0906 | 0 |
| 42476 | ENSMUSG00000104416 | 0 | NA | NA | NA | NA | NA | Gm37067 | predicted gene, 37067 [Source:MGI Symb | 0 | 0 | 0 | 0 | 0 | 0 |
| 42477 | ENSMUSG00000104417 | 0 | NA | NA | NA | NA | NA | Gm37068 | predicted gene, 37068 [Source:MGI Symb | 0 | 0 | 0 | 0 | 0 | 0 |
| 42478 | ENSMUSG00000104418 | 0 | NA | NA | NA | NA | NA | Gm37070 | predicted gene, 37070 [Source:MGI Symb | 0 | 0 | 0 | 0 | 0 | 0 |
| 42479 | ENSMUSG00000104419 | 0 | NA | NA | NA | NA | NA | 2900092O11 | RIKEN cDNA 2900092O11 gene [Source:M | 0 | 0 | 0 | 0 | 0 | 0 |
| 42480 | ENSMUSG00000104420 | 0.321325949 | 1.911575114 | 4.048267893 | 0.472195804 | 0.636787034 | NA | E230020A03 | RIKEN cDNA E230020A03 gene [Source:M | 0 | 0 | 0 | 1.928 | 0 | 0 |
| 42481 | ENSMUSG00000104422 | 0 | NA | NA | NA | NA | NA | Ighv1-14 | immunoglobulin heavy variable 1-14 [Sou | 0 | 0 | 0 | 0 | 0 | 0 |
| 42482 | ENSMUSG00000104423 | 0 | NA | NA | NA | NA | NA | A030005K14 | RIKEN cDNA A030005K14 gene [Source:M | 0 | 0 | 0 | 0 | 0 | 0 |
| 42483 | ENSMUSG00000104424 | 0.133850373 | -0.796629176 | 4.080472857 | -0.195229623 | 0.84521319 | NA | Gm19058 | predicted gene, 19058 [Source:MGI Symb | 0.8031 | 0 | 0 | 0 | 0 | 0 |
| 42484 | ENSMUSG00000104425 | 0 | NA | NA | NA | NA | NA | Gm38370 | predicted gene, 38370 [Source:MGI Symb | 0 | 0 | 0 | 0 | 0 | 0 |
| 42485 | ENSMUSG00000104426 | 0 | NA | NA | NA | NA | NA | Gm38369 | predicted gene, 38369 [Source:MGI Symb | 0 | 0 | 0 | 0 | 0 | 0 |
| 42486 | ENSMUSG00000104427 | 0.36353943 | 2.065989254 | 4.034137301 | 0.512126658 | 0.608562368 | NA | Gm38368 | predicted gene, 38368 [Source:MGI Symb | 0 | 0 | 0 | 0 | 2.1812 | 0 |
| 42487 | ENSMUSG00000104428 | 0 | NA | NA | NA | NA | NA | Gm18299 | predicted gene, 18299 [Source:MGI Symb | 0 | 0 | 0 | 0 | 0 | 0 |
| 42488 | ENSMUSG00000104429 | 0 | NA | NA | NA | NA | NA | 2610034O05 | RIKEN cDNA 2610034O05 gene [Source:M | 0 | 0 | 0 | 0 | 0 | 0 |
| 42489 | ENSMUSG00000104430 | 0 | NA | NA | NA | NA | NA | Gm37425 | predicted gene, 37425 [Source:MGI Symb | 0 | 0 | 0 | 0 | 0 | 0 |
| 42490 | ENSMUSG00000104431 | 0 | NA | NA | NA | NA | NA | Gm37112 | predicted gene, 37112 [Source:MGI Symb | 0 | 0 | 0 | 0 | 0 | 0 |
| 42491 | ENSMUSG00000104432 | 0.331982614 | 0.16514309 | 3.986751476 | 0.041422971 | 0.966958701 | NA | A430027C01 | RIKEN cDNA A430027C01 gene [Source:M | 0 | 1.0279 | 0 | 0.964 | 0 | 0 |
| 42492 | ENSMUSG00000104433 | 0 | NA | NA | NA | NA | NA | Gm37426 | predicted gene, 37426 [Source:MGI Symb | 0 | 0 | 0 | 0 | 0 | 0 |
| 42493 | ENSMUSG00000104434 | 0 | NA | NA | NA | NA | NA | Gm37421 | predicted gene, 37421 [Source:MGI Symb | 0 | 0 | 0 | 0 | 0 | 0 |
| 42494 | ENSMUSG00000104435 | 33.80943986 | -0.024201414 | 0.414676957 | -0.058362089 | 0.953460212 | 0.9915542 | Gm37422 | predicted gene, 37422 [Source:MGI Symb | 43.368 | 21.586 | 37.231 | 43.379 | 29.447 | 27.846 |
| 42495 | ENSMUSG00000104436 | 1.605874687 | 1.392979365 | 1.881060613 | 0.740528697 | 0.458979255 | NA | Gm37423 | predicted gene, 37423 [Source:MGI Symb | 1.6062 | 1.0279 | 0 | 4.8199 | 2.1812 | 0 |
| 42496 | ENSMUSG00000104437 | 0 | NA | NA | NA | NA | NA | Gm37424 | predicted gene, 37424 [Source:MGI Symb | 0 | 0 | 0 | 0 | 0 | 0 |
| 42497 | ENSMUSG00000104438 | 0 | NA | NA | NA | NA | NA | Gm37427 | predicted gene, 37427 [Source:MGI Symb | 0 | 0 | 0 | 0 | 0 | 0 |
| 42498 | ENSMUSG00000104439 | 0 | NA | NA | NA | NA | NA | Gm20851 | predicted gene, 20851 [Source:MGI Symb | 0 | 0 | 0 | 0 | 0 | 0 |
| 42499 | ENSMUSG00000104440 | 0 | NA | NA | NA | NA | NA | Gm19030 | predicted gene, 19030 [Source:MGI Symb | 0 | 0 | 0 | 0 | 0 | 0 |
| 42500 | ENSMUSG00000104441 | 0 | NA | NA | NA | NA | NA | Gm35801 | predicted gene, 35801 [Source:MGI Symb | 0 | 0 | 0 | 0 | 0 | 0 |

| | | | | | | | | | | | | | | | |
|---|---|---|---|---|---|---|---|---|---|---|---|---|---|---|---|
| 42501 | ENSMUSG00000104442 | 0 | NA | NA | NA | NA | NA | 4932431L22I | RIKEN cDNA 4932431L22 gene [Source:M( | 0 | 0 | 0 | 0 | 0 | 0 |
| 42502 | ENSMUSG00000104443 | 11.30010464 | -0.100806174 | 0.731720806 | -0.137765898 | 0.890425437 | 0.982489666 | 4932442E05 | RIKEN cDNA 4932442E05 gene [Source:M( | 16.865 | 4.1117 | 14.087 | 17.352 | 9.8156 | 5.5692 |
| 42503 | ENSMUSG00000104444 | 0 | NA | NA | NA | NA | NA | Gm33051 | predicted gene, 33051 [Source:MGI Symb | 0 | 0 | 0 | 0 | 0 | 0 |
| 42504 | ENSMUSG00000104445 | 22.4387026 | 0.173079477 | 0.545107455 | 0.31751442 | 0.7508533 | 0.952074518 | Rhbg | Rhesus blood group-associated B glycopro | 35.336 | 13.363 | 14.087 | 14.46 | 25.084 | 32.301 |
| 42505 | ENSMUSG00000104446 | 0 | NA | NA | NA | NA | NA | Gm30312 | predicted gene, 30312 [Source:MGI Symb | 0 | 0 | 0 | 0 | 0 | 0 |
| 42506 | ENSMUSG00000104448 | 0 | NA | NA | NA | NA | NA | Gm37256 | predicted gene, 37256 [Source:MGI Symb | 0 | 0 | 0 | 0 | 0 | 0 |
| 42507 | ENSMUSG00000104449 | 2.701506641 | -0.356503604 | 1.58414865 | -0.225044288 | 0.821944821 | NA | Gm37255 | predicted gene, 37255 [Source:MGI Symb | 0 | 7.1954 | 2.0125 | 4.8199 | 2.1812 | 0 |
| 42508 | ENSMUSG00000104450 | 0 | NA | NA | NA | NA | NA | Gm37830 | predicted gene, 37830 [Source:MGI Symb | 0 | 0 | 0 | 0 | 0 | 0 |
| 42509 | ENSMUSG00000104451 | 0 | NA | NA | NA | NA | NA | Gm37831 | predicted gene, 37831 [Source:MGI Symb | 0 | 0 | 0 | 0 | 0 | 0 |
| 42510 | ENSMUSG00000104452 | 0 | NA | NA | NA | NA | NA | Ighv8-8 | immunoglobulin heavy variable 8-8 [Sourc | 0 | 0 | 0 | 0 | 0 | 0 |
| 42511 | ENSMUSG00000104453 | 12.90023628 | -0.261225512 | 0.617100409 | -0.423311196 | 0.672068227 | 0.931191575 | Gm37829 | predicted gene, 37829 [Source:MGI Symb | 9.6372 | 19.53 | 13.081 | 8.6758 | 11.997 | 14.48 |
| 42512 | ENSMUSG00000104454 | 0.503122293 | -2.366675966 | 3.785596405 | -0.625179156 | 0.531853481 | NA | Gm37827 | predicted gene, 37827 [Source:MGI Symb | 0 | 0 | 3.0187 | 0 | 0 | 0 |
| 42513 | ENSMUSG00000104455 | 0 | NA | NA | NA | NA | NA | Gm37828 | predicted gene, 37828 [Source:MGI Symb | 0 | 0 | 0 | 0 | 0 | 0 |
| 42514 | ENSMUSG00000104456 | 0 | NA | NA | NA | NA | NA | Gm37825 | predicted gene, 37825 [Source:MGI Symb | 0 | 0 | 0 | 0 | 0 | 0 |
| 42515 | ENSMUSG00000104457 | 0 | NA | NA | NA | NA | NA | Gm37826 | predicted gene, 37826 [Source:MGI Symb | 0 | 0 | 0 | 0 | 0 | 0 |
| 42516 | ENSMUSG00000104459 | 29.93036864 | 0.330418219 | 0.403969843 | 0.817927935 | 0.413398338 | 0.835385466 | Gm37824 | predicted gene, 37824 [Source:MGI Symb | 29.715 | 24.67 | 25.156 | 39.523 | 34.9 | 25.618 |
| 42517 | ENSMUSG00000104460 | 0 | NA | NA | NA | NA | NA | Gm9121 | predicted pseudogene 9121 [Source:MGI | 0 | 0 | 0 | 0 | 0 | 0 |
| 42518 | ENSMUSG00000104461 | 0 | NA | NA | NA | NA | NA | Gm37662 | predicted gene, 37662 [Source:MGI Symb | 0 | 0 | 0 | 0 | 0 | 0 |
| 42519 | ENSMUSG00000104462 | 0 | NA | NA | NA | NA | NA | Gm37661 | predicted gene, 37661 [Source:MGI Symb | 0 | 0 | 0 | 0 | 0 | 0 |
| 42520 | ENSMUSG00000104464 | 0 | NA | NA | NA | NA | NA | Gm18698 | predicted gene, 18698 [Source:MGI Symb | 0 | 0 | 0 | 0 | 0 | 0 |
| 42521 | ENSMUSG00000104465 | 0 | NA | NA | NA | NA | NA | BC002189 | cDNA sequence BC002189 [Source:MGI Sy | 0 | 0 | 0 | 0 | 0 | 0 |
| 42522 | ENSMUSG00000104466 | 0.301557804 | -1.64335916 | 4.042814367 | -0.406488899 | 0.684383412 | NA | Gm10513 | predicted gene 10513 [Source:MGI Symbo | 0.8031 | 0 | 1.0062 | 0 | 0 | 0 |
| 42523 | ENSMUSG00000104467 | 1.466731634 | -1.692377855 | 2.015245087 | -0.83978761 | 0.401027482 | NA | Gm37660 | predicted gene, 37660 [Source:MGI Symb | 1.6062 | 5.1396 | 0 | 0.964 | 1.0906 | 0 |
| 42524 | ENSMUSG00000104468 | 0 | NA | NA | NA | NA | NA | Gm21736 | predicted gene, 21736 [Source:MGI Symb | 0 | 0 | 0 | 0 | 0 | 0 |
| 42525 | ENSMUSG00000104469 | 0 | NA | NA | NA | NA | NA | Gm37663 | predicted gene, 37663 [Source:MGI Symb | 0 | 0 | 0 | 0 | 0 | 0 |
| 42526 | ENSMUSG00000104470 | 0 | NA | NA | NA | NA | NA | Gm38207 | predicted gene, 38207 [Source:MGI Symb | 0 | 0 | 0 | 0 | 0 | 0 |
| 42527 | ENSMUSG00000104471 | 0 | NA | NA | NA | NA | NA | Gm38208 | predicted gene, 38208 [Source:MGI Symb | 0 | 0 | 0 | 0 | 0 | 0 |
| 42528 | ENSMUSG00000104472 | 0 | NA | NA | NA | NA | NA | Gm38209 | predicted gene, 38209 [Source:MGI Symb | 0 | 0 | 0 | 0 | 0 | 0 |
| 42529 | ENSMUSG00000104473 | 0 | NA | NA | NA | NA | NA | Trbv25 | T cell receptor beta variable 25 [Source:M | 0 | 0 | 0 | 0 | 0 | 0 |
| 42530 | ENSMUSG00000104474 | 0 | NA | NA | NA | NA | NA | Gm38210 | predicted gene, 38210 [Source:MGI Symb | 0 | 0 | 0 | 0 | 0 | 0 |
| 42531 | ENSMUSG00000104476 | 0 | NA | NA | NA | NA | NA | Gm38211 | predicted gene, 38211 [Source:MGI Symb | 0 | 0 | 0 | 0 | 0 | 0 |
| 42532 | ENSMUSG00000104477 | 0 | NA | NA | NA | NA | NA | Gm18717 | predicted gene, 18717 [Source:MGI Symb | 0 | 0 | 0 | 0 | 0 | 0 |
| 42533 | ENSMUSG00000104478 | 0 | NA | NA | NA | NA | NA | Gm38212 | predicted gene, 38212 [Source:MGI Symb | 0 | 0 | 0 | 0 | 0 | 0 |
| 42534 | ENSMUSG00000104479 | 0.653308562 | 1.538013299 | 3.112409277 | 0.494155223 | 0.62119656 | NA | Gm10305 | predicted gene 10305 [Source:MGI Symbo | 0 | 1.0279 | 0 | 2.8919 | 0 | 0 |
| 42535 | ENSMUSG00000104480 | 0 | NA | NA | NA | NA | NA | Gm21661 | predicted gene, 21661 [Source:MGI Symb | 0 | 0 | 0 | 0 | 0 | 0 |
| 42536 | ENSMUSG00000104481 | 0 | NA | NA | NA | NA | NA | Gm38063 | predicted gene, 38063 [Source:MGI Symb | 0 | 0 | 0 | 0 | 0 | 0 |
| 42537 | ENSMUSG00000104482 | 0 | NA | NA | NA | NA | NA | Gm38064 | predicted gene, 38064 [Source:MGI Symb | 0 | 0 | 0 | 0 | 0 | 0 |
| 42538 | ENSMUSG00000104484 | 0 | NA | NA | NA | NA | NA | Gm33142 | predicted gene, 33142 [Source:MGI Symb | 0 | 0 | 0 | 0 | 0 | 0 |
| 42539 | ENSMUSG00000104485 | 0 | NA | NA | NA | NA | NA | Gm38065 | predicted gene, 38065 [Source:MGI Symb | 0 | 0 | 0 | 0 | 0 | 0 |
| 42540 | ENSMUSG00000104486 | 0 | NA | NA | NA | NA | NA | Gm38066 | predicted gene, 38066 [Source:MGI Symb | 0 | 0 | 0 | 0 | 0 | 0 |
| 42541 | ENSMUSG00000104487 | 0 | NA | NA | NA | NA | NA | Gm31373 | predicted gene, 31373 [Source:MGI Symb | 0 | 0 | 0 | 0 | 0 | 0 |
| 42542 | ENSMUSG00000104488 | 0 | NA | NA | NA | NA | NA | Gm38062 | predicted gene, 38062 [Source:MGI Symb | 0 | 0 | 0 | 0 | 0 | 0 |
| 42543 | ENSMUSG00000104489 | 0 | NA | NA | NA | NA | NA | Gm9700 | predicted gene 9700 [Source:MGI Symbol | 0 | 0 | 0 | 0 | 0 | 0 |
| 42544 | ENSMUSG00000104490 | 0 | NA | NA | NA | NA | NA | Gm20770 | predicted gene, 20770 [Source:MGI Symb | 0 | 0 | 0 | 0 | 0 | 0 |
| 42545 | ENSMUSG00000104491 | 0 | NA | NA | NA | NA | NA | Gm32340 | predicted gene, 32340 [Source:MGI Symb | 0 | 0 | 0 | 0 | 0 | 0 |
| 42546 | ENSMUSG00000104493 | 0 | NA | NA | NA | NA | NA | Gm19552 | predicted gene, 19552 [Source:MGI Symb | 0 | 0 | 0 | 0 | 0 | 0 |
| 42547 | ENSMUSG00000104494 | 0 | NA | NA | NA | NA | NA | Gm37111 | predicted gene, 37111 [Source:MGI Symb | 0 | 0 | 0 | 0 | 0 | 0 |
| 42548 | ENSMUSG00000104496 | 1.555836718 | 1.905225419 | 1.802547931 | 1.056962418 | 0.29052874 | NA | Gm5837 | predicted gene 5837 [Source:MGI Symbol | 0 | 1.0279 | 1.0062 | 2.8919 | 2.1812 | 2.2277 |
| 42549 | ENSMUSG00000104497 | 0.167707431 | -0.796629176 | 4.080472857 | -0.195229623 | 0.84521319 | NA | Gm10555 | predicted gene 10555 [Source:MGI Symbo | 0 | 0 | 1.0062 | 0 | 0 | 0 |
| 42550 | ENSMUSG00000104498 | 0 | NA | NA | NA | NA | NA | Gm37110 | predicted gene, 37110 [Source:MGI Symb | 0 | 0 | 0 | 0 | 0 | 0 |
| 42551 | ENSMUSG00000104499 | 0 | NA | NA | NA | NA | NA | Gm30353 | predicted gene, 30353 [Source:MGI Symb | 0 | 0 | 0 | 0 | 0 | 0 |
| 42552 | ENSMUSG00000104500 | 0 | NA | NA | NA | NA | NA | Gm36345 | predicted gene, 36345 [Source:MGI Symb | 0 | 0 | 0 | 0 | 0 | 0 |
| 42553 | ENSMUSG00000104501 | 18.80959207 | 1.37656456 | 0.576207518 | 2.389008328 | 0.016893919 | 0.154361118 | Gm37736 | predicted gene, 37736 [Source:MGI Symb | 17.668 | 8.2233 | 5.0312 | 21.208 | 25.084 | 35.643 |
| 42554 | ENSMUSG00000104502 | 0 | NA | NA | NA | NA | NA | Gm37737 | predicted gene, 37737 [Source:MGI Symb | 0 | 0 | 0 | 0 | 0 | 0 |
| 42555 | ENSMUSG00000104503 | 1.224344394 | -1.352088183 | 2.099147943 | -0.644112859 | 0.519502253 | NA | Gm37738 | predicted gene, 37738 [Source:MGI Symb | 3.2124 | 2.0558 | 0 | 0.964 | 0 | 1.1138 |
| 42556 | ENSMUSG00000104504 | 0 | NA | NA | NA | NA | NA | Gm7445 | predicted gene 7445 [Source:MGI Symbol | 0 | 0 | 0 | 0 | 0 | 0 |

| | | | | | | | | | | | | | | | |
|---|---|---|---|---|---|---|---|---|---|---|---|---|---|---|---|
| 42557 | ENSMUSG00000104505 | 0 | NA | NA | NA | NA | NA | Gm37739 | predicted gene, 37739 [Source:MGI Symb | 0 | 0 | 0 | 0 | 0 | 0 |
| 42558 | ENSMUSG00000104506 | 0 | NA | NA | NA | NA | NA | Gm37740 | predicted gene, 37740 [Source:MGI Symb | 0 | 0 | 0 | 0 | 0 | 0 |
| 42559 | ENSMUSG00000104507 | 0 | NA | NA | NA | NA | NA | A430027H14 | RIKEN cDNA A430027H14 gene [Source:M | 0 | 0 | 0 | 0 | 0 | 0 |
| 42560 | ENSMUSG00000104508 | 0.903225564 | 3.379710758 | 2.734057383 | 1.236152094 | 0.216402034 | NA | Gm37735 | predicted gene, 37735 [Source:MGI Symb | 0 | 0 | 0 | 0.964 | 0 | 4.4554 |
| 42561 | ENSMUSG00000104509 | 0 | NA | NA | NA | NA | NA | Gm33994 | predicted gene, 33994 [Source:MGI Symb | 0 | 0 | 0 | 0 | 0 | 0 |
| 42562 | ENSMUSG00000104510 | 0 | NA | NA | NA | NA | NA | Gm34015 | predicted gene, 34015 [Source:MGI Symb | 0 | 0 | 0 | 0 | 0 | 0 |
| 42563 | ENSMUSG00000104511 | 0 | NA | NA | NA | NA | NA | Gm18843 | predicted gene, 18843 [Source:MGI Symb | 0 | 0 | 0 | 0 | 0 | 0 |
| 42564 | ENSMUSG00000104512 | 0.160662974 | 1.126924175 | 4.080472857 | 0.276174898 | 0.782413736 | NA | Gm37165 | predicted gene, 37165 [Source:MGI Symb | 0 | 0 | 0 | 0.964 | 0 | 0 |
| 42565 | ENSMUSG00000104513 | 0 | NA | NA | NA | NA | NA | Gm29813 | predicted gene, 29813 [Source:MGI Symb | 0 | 0 | 0 | 0 | 0 | 0 |
| 42566 | ENSMUSG00000104514 | 0.17131964 | -0.796629176 | 4.080472857 | -0.195229623 | 0.84521319 | NA | Gm37164 | predicted gene, 37164 [Source:MGI Symb | 0 | 1.0279 | 0 | 0 | 0 | 0 |
| 42567 | ENSMUSG00000104515 | 16.10794931 | -1.070071997 | 0.594693748 | -1.799366481 | 0.071960728 | 0.388663171 | Gm37163 | predicted gene, 37163 [Source:MGI Symb | 22.487 | 23.642 | 19.119 | 3.8559 | 14.178 | 13.366 |
| 42568 | ENSMUSG00000104516 | 0 | NA | NA | NA | NA | NA | Gm9884 | predicted gene 9884 [Source:MGI Symbol | 0 | 0 | 0 | 0 | 0 | 0 |
| 42569 | ENSMUSG00000104517 | 0.531944269 | 2.618916564 | 3.285123409 | 0.797204926 | 0.425332026 | NA | Gm18407 | predicted gene, 18407 [Source:MGI Symb | 0 | 0 | 0 | 0.964 | 0 | 2.2277 |
| 42570 | ENSMUSG00000104518 | 0 | NA | NA | NA | NA | NA | Gm37162 | predicted gene, 37162 [Source:MGI Symb | 0 | 0 | 0 | 0 | 0 | 0 |
| 42571 | ENSMUSG00000104519 | 0.342639279 | -1.805976596 | 4.028547714 | -0.448294702 | 0.653940524 | NA | Gm37161 | predicted gene, 37161 [Source:MGI Symb | 0 | 2.0558 | 0 | 0 | 0 | 0 |
| 42572 | ENSMUSG00000104520 | 7.924415026 | -0.228993819 | 0.779376155 | -0.293816814 | 0.768897886 | 0.956016421 | Gm37336 | predicted gene, 37336 [Source:MGI Symb | 10.44 | 4.1117 | 11.069 | 8.6758 | 5.4531 | 7.7969 |
| 42573 | ENSMUSG00000104521 | 0 | NA | NA | NA | NA | NA | Gm34073 | predicted gene, 34073 [Source:MGI Symb | 0 | 0 | 0 | 0 | 0 | 0 |
| 42574 | ENSMUSG00000104522 | 0 | NA | NA | NA | NA | NA | Gm38159 | predicted gene, 38159 [Source:MGI Symb | 0 | 0 | 0 | 0 | 0 | 0 |
| 42575 | ENSMUSG00000104523 | 0 | NA | NA | NA | NA | NA | Gm37335 | predicted gene, 37335 [Source:MGI Symb | 0 | 0 | 0 | 0 | 0 | 0 |
| 42576 | ENSMUSG00000104524 | 0.692607243 | 3.004183657 | 2.957557172 | 1.015765202 | 0.309741211 | NA | Gm37333 | predicted gene, 37333 [Source:MGI Symb | 0 | 0 | 0 | 1.928 | 0 | 2.2277 |
| 42577 | ENSMUSG00000104525 | 0.367410362 | 2.078255025 | 3.800070157 | 0.54689912 | 0.58444804 | NA | Gm37334 | predicted gene, 37334 [Source:MGI Symb | 0 | 0 | 0 | 0 | 1.0906 | 1.1138 |
| 42578 | ENSMUSG00000104526 | 0 | NA | NA | NA | NA | NA | 1700045I11F | RIKEN cDNA 1700045I11 gene [Source:MG | 0 | 0 | 0 | 0 | 0 | 0 |
| 42579 | ENSMUSG00000104527 | 0 | NA | NA | NA | NA | NA | Gm37332 | predicted gene, 37332 [Source:MGI Symb | 0 | 0 | 0 | 0 | 0 | 0 |
| 42580 | ENSMUSG00000104528 | 53.00798424 | -1.031381066 | 0.434049587 | -2.376182575 | 0.017492805 | 0.157512348 | Gm43314 | predicted gene 43314 [Source:MGI Symbo | 41.761 | 85.317 | 86.537 | 22.171 | 59.984 | 22.277 |
| 42581 | ENSMUSG00000104529 | 0 | NA | NA | NA | NA | NA | Rbakdn | RB-associated KRAB zinc finger downstrea | 0 | 0 | 0 | 0 | 0 | 0 |
| 42582 | ENSMUSG00000104530 | 0 | NA | NA | NA | NA | NA | Gm6077 | predicted gene 6077 [Source:MGI Symbol | 0 | 0 | 0 | 0 | 0 | 0 |
| 42583 | ENSMUSG00000104531 | 0 | NA | NA | NA | NA | NA | Gm43731 | predicted gene 43731 [Source:MGI Symbo | 0 | 0 | 0 | 0 | 0 | 0 |
| 42584 | ENSMUSG00000104532 | 0 | NA | NA | NA | NA | NA | Gm43739 | predicted gene 43739 [Source:MGI Symbo | 0 | 0 | 0 | 0 | 0 | 0 |
| 42585 | ENSMUSG00000104533 | 0 | NA | NA | NA | NA | NA | Igkv5-40-1 | immunoglobulin kappa chain variable 5-4( | 0 | 0 | 0 | 0 | 0 | 0 |
| 42586 | ENSMUSG00000104534 | 0 | NA | NA | NA | NA | NA | Gm18660 | predicted gene, 18660 [Source:MGI Symb | 0 | 0 | 0 | 0 | 0 | 0 |
| 42587 | ENSMUSG00000104535 | 0.133850373 | -0.796629176 | 4.080472857 | -0.195229623 | 0.84521319 | NA | Gm42686 | predicted gene 42686 [Source:MGI Symbo | 0.8031 | 0 | 0 | 0 | 0 | 0 |
| 42588 | ENSMUSG00000104536 | 0 | NA | NA | NA | NA | NA | Gm23786 | predicted gene, 23786 [Source:MGI Symb | 0 | 0 | 0 | 0 | 0 | 0 |
| 42589 | ENSMUSG00000104537 | 0 | NA | NA | NA | NA | NA | NA | NA | 0 | 0 | 0 | 0 | 0 | 0 |
| 42590 | ENSMUSG00000104538 | 0 | NA | NA | NA | NA | NA | 1700016B01 | RIKEN cDNA 1700016B01 gene [Source:M | 0 | 0 | 0 | 0 | 0 | 0 |
| 42591 | ENSMUSG00000104539 | 0 | NA | NA | NA | NA | NA | Gm43035 | predicted gene 43035 [Source:MGI Symbo | 0 | 0 | 0 | 0 | 0 | 0 |
| 42592 | ENSMUSG00000104540 | 0 | NA | NA | NA | NA | NA | Gm3527 | predicted gene 3527 [Source:MGI Symbol | 0 | 0 | 0 | 0 | 0 | 0 |
| 42593 | ENSMUSG00000104541 | 0 | NA | NA | NA | NA | NA | Gm43114 | predicted gene 43114 [Source:MGI Symbo | 0 | 0 | 0 | 0 | 0 | 0 |
| 42594 | ENSMUSG00000104542 | 0 | NA | NA | NA | NA | NA | Ighv8-15 | immunoglobulin heavy variable V8-15 [So | 0 | 0 | 0 | 0 | 0 | 0 |
| 42595 | ENSMUSG00000104543 | 0 | NA | NA | NA | NA | NA | Gm6602 | predicted gene 6602 [Source:MGI Symbol | 0 | 0 | 0 | 0 | 0 | 0 |
| 42596 | ENSMUSG00000104544 | 0 | NA | NA | NA | NA | NA | NA | NA | 0 | 0 | 0 | 0 | 0 | 0 |
| 42597 | ENSMUSG00000104545 | 0 | NA | NA | NA | NA | NA | E030032P16 | RIKEN cDNA E030032P16 gene [Source:M | 0 | 0 | 0 | 0 | 0 | 0 |
| 42598 | ENSMUSG00000104546 | 17.04849226 | -0.265858905 | 0.560565374 | -0.474269225 | 0.635307935 | 0.923478764 | Gm43858 | predicted gene 43858 [Source:MGI Symbo | 14.456 | 19.53 | 22.137 | 24.099 | 9.8156 | 12.252 |
| 42599 | ENSMUSG00000104547 | 0 | NA | NA | NA | NA | NA | Gm43022 | predicted gene 43022 [Source:MGI Symbo | 0 | 0 | 0 | 0 | 0 | 0 |
| 42600 | ENSMUSG00000104548 | 8.310721677 | -0.140521897 | 0.809677503 | -0.173552923 | 0.862216832 | 0.976334255 | Gm43857 | predicted gene 43857 [Source:MGI Symbo | 8.031 | 4.1117 | 14.087 | 11.568 | 8.7249 | 3.3415 |
| 42601 | ENSMUSG00000104549 | 0 | NA | NA | NA | NA | NA | Gm43142 | predicted gene 43142 [Source:MGI Symbo | 0 | 0 | 0 | 0 | 0 | 0 |
| 42602 | ENSMUSG00000104550 | 0 | NA | NA | NA | NA | NA | Gm43589 | predicted gene 43589 [Source:MGI Symbo | 0 | 0 | 0 | 0 | 0 | 0 |
| 42603 | ENSMUSG00000104551 | 0 | NA | NA | NA | NA | NA | Igkv13-57-2 | immunoglobulin kappa variable 13-57-2 [S | 0 | 0 | 0 | 0 | 0 | 0 |
| 42604 | ENSMUSG00000104552 | 0 | NA | NA | NA | NA | NA | NA | NA | 0 | 0 | 0 | 0 | 0 | 0 |
| 42605 | ENSMUSG00000104553 | 0 | NA | NA | NA | NA | NA | 9330178D15 | RIKEN cDNA 9330178D15 gene [Source:M | 0 | 0 | 0 | 0 | 0 | 0 |
| 42606 | ENSMUSG00000104554 | 0 | NA | NA | NA | NA | NA | Gm4610 | predicted gene 4610 [Source:MGI Symbol | 0 | 0 | 0 | 0 | 0 | 0 |
| 42607 | ENSMUSG00000104555 | 0 | NA | NA | NA | NA | NA | Gm5983 | predicted gene 5983 [Source:MGI Symbol | 0 | 0 | 0 | 0 | 0 | 0 |
| 42608 | ENSMUSG00000104556 | 0 | NA | NA | NA | NA | NA | Gm43192 | predicted gene 43192 [Source:MGI Symbo | 0 | 0 | 0 | 0 | 0 | 0 |
| 42609 | ENSMUSG00000104557 | 0 | NA | NA | NA | NA | NA | Gm36520 | predicted gene, 36520 [Source:MGI Symb | 0 | 0 | 0 | 0 | 0 | 0 |
| 42610 | ENSMUSG00000104558 | 0 | NA | NA | NA | NA | NA | NA | NA | 0 | 0 | 0 | 0 | 0 | 0 |
| 42611 | ENSMUSG00000104559 | 0 | NA | NA | NA | NA | NA | Gm43118 | predicted gene 43118 [Source:MGI Symbo | 0 | 0 | 0 | 0 | 0 | 0 |
| 42612 | ENSMUSG00000104560 | 0 | NA | NA | NA | NA | NA | Gm8115 | predicted gene 8115 [Source:MGI Symbol | 0 | 0 | 0 | 0 | 0 | 0 |

| | | | | | | | | | | | | | | | |
|---|---|---|---|---|---|---|---|---|---|---|---|---|---|---|---|
| 42613 | ENSMUSG00000104561 | 0 | NA | NA | NA | NA | NA | NA | NA | 0 | 0 | 0 | 0 | 0 | 0 |
| 42614 | ENSMUSG00000104562 | 0 | NA | NA | NA | NA | NA | Gm42767 | predicted gene 42767 [Source:MGI Symbo | 0 | 0 | 0 | 0 | 0 | 0 |
| 42615 | ENSMUSG00000104563 | 0 | NA | NA | NA | NA | NA | Gm43041 | predicted gene 43041 [Source:MGI Symbo | 0 | 0 | 0 | 0 | 0 | 0 |
| 42616 | ENSMUSG00000104564 | 0 | NA | NA | NA | NA | NA | Gm17977 | predicted gene, 17977 [Source:MGI Symb | 0 | 0 | 0 | 0 | 0 | 0 |
| 42617 | ENSMUSG00000104566 | 0.160662974 | 1.126924175 | 4.080472857 | 0.276174898 | 0.782413736 | NA | Mir7227 | microRNA 7227 [Source:MGI Symbol;Acc: | 0 | 0 | 0 | 0.964 | 0 | 0 |
| 42618 | ENSMUSG00000104567 | 0 | NA | NA | NA | NA | NA | Gm43209 | predicted gene 43209 [Source:MGI Symbo | 0 | 0 | 0 | 0 | 0 | 0 |
| 42619 | ENSMUSG00000104568 | 0 | NA | NA | NA | NA | NA | Gm43255 | predicted gene 43255 [Source:MGI Symbo | 0 | 0 | 0 | 0 | 0 | 0 |
| 42620 | ENSMUSG00000104569 | 3.600389916 | -0.906582277 | 1.220845382 | -0.742585663 | 0.457732571 | NA | Gm43054 | predicted gene 43054 [Source:MGI Symbo | 4.8186 | 8.2233 | 1.0062 | 0.964 | 4.3625 | 2.2277 |
| 42621 | ENSMUSG00000104570 | 0 | NA | NA | NA | NA | NA | NA | NA | 0 | 0 | 0 | 0 | 0 | 0 |
| 42622 | ENSMUSG00000104571 | 0 | NA | NA | NA | NA | NA | Gm43010 | predicted gene 43010 [Source:MGI Symbo | 0 | 0 | 0 | 0 | 0 | 0 |
| 42623 | ENSMUSG00000104572 | 0 | NA | NA | NA | NA | NA | Fbxw23 | F-box and WD-40 domain protein 23 [Sou | 0 | 0 | 0 | 0 | 0 | 0 |
| 42624 | ENSMUSG00000104573 | 0 | NA | NA | NA | NA | NA | Gm42886 | predicted gene 42886 [Source:MGI Symbo | 0 | 0 | 0 | 0 | 0 | 0 |
| 42625 | ENSMUSG00000104574 | 0.698954292 | 0.126053087 | 3.048972385 | 0.04134281 | 0.967022605 | NA | Gm42836 | predicted gene 42836 [Source:MGI Symbo | 0 | 0 | 2.0125 | 0 | 2.1812 | 0 |
| 42626 | ENSMUSG00000104575 | 0 | NA | NA | NA | NA | NA | Igkv9-119 | immunoglobulin kappa chain variable 9-1 | 0 | 0 | 0 | 0 | 0 | 0 |
| 42627 | ENSMUSG00000104576 | 1.624914405 | -1.005835241 | 1.827637217 | -0.550347318 | 0.582081175 | NA | F830115B05 | RIKEN cDNA F830115B05 gene [Source:M | 2.4093 | 2.0558 | 2.0125 | 0 | 3.2719 | 0 |
| 42628 | ENSMUSG00000104577 | 0 | NA | NA | NA | NA | NA | Gm6641 | predicted gene 6641 [Source:MGI Symbol | 0 | 0 | 0 | 0 | 0 | 0 |
| 42629 | ENSMUSG00000104578 | 0 | NA | NA | NA | NA | NA | Gm8929 | predicted gene 8929 [Source:MGI Symbol | 0 | 0 | 0 | 0 | 0 | 0 |
| 42630 | ENSMUSG00000104579 | 0 | NA | NA | NA | NA | NA | 5830437K03 | RIKEN cDNA 5830437K03 gene [Source:M | 0 | 0 | 0 | 0 | 0 | 0 |
| 42631 | ENSMUSG00000104580 | 0 | NA | NA | NA | NA | NA | Gm42620 | predicted gene 42620 [Source:MGI Symbo | 0 | 0 | 0 | 0 | 0 | 0 |
| 42632 | ENSMUSG00000104581 | 0 | NA | NA | NA | NA | NA | Gm42497 | predicted gene 42497 [Source:MGI Symbo | 0 | 0 | 0 | 0 | 0 | 0 |
| 42633 | ENSMUSG00000104582 | 0 | NA | NA | NA | NA | NA | Mir380 | microRNA 380 [Source:MGI Symbol;Acc:M | 0 | 0 | 0 | 0 | 0 | 0 |
| 42634 | ENSMUSG00000104583 | 0 | NA | NA | NA | NA | NA | Gm42701 | predicted gene 42701 [Source:MGI Symbo | 0 | 0 | 0 | 0 | 0 | 0 |
| 42635 | ENSMUSG00000104584 | 0 | NA | NA | NA | NA | NA | Rpl18a-ps3 | ribosomal protein L18A, pseudogene 3 [S | 0 | 0 | 0 | 0 | 0 | 0 |
| 42636 | ENSMUSG00000104585 | 0.428363721 | -0.543730382 | 3.624677344 | -0.15000794 | 0.880758351 | NA | 4921511C10 | RIKEN cDNA 4921511C10 gene [Source:M | 1.6062 | 0 | 0 | 0.964 | 0 | 0 |
| 42637 | ENSMUSG00000104586 | 0.986423182 | 1.173929884 | 2.247995681 | 0.522211806 | 0.601522867 | NA | 4921539H07 | RIKEN cDNA 4921539H07 gene [Source:M | 0.8031 | 0 | 1.0062 | 1.928 | 2.1812 | 0 |
| 42638 | ENSMUSG00000104587 | 0 | NA | NA | NA | NA | NA | Gm43161 | predicted gene 43161 [Source:MGI Symbo | 0 | 0 | 0 | 0 | 0 | 0 |
| 42639 | ENSMUSG00000104588 | 0 | NA | NA | NA | NA | NA | Gm43666 | predicted gene 43666 [Source:MGI Symbo | 0 | 0 | 0 | 0 | 0 | 0 |
| 42640 | ENSMUSG00000104589 | 0 | NA | NA | NA | NA | NA | NA | NA | 0 | 0 | 0 | 0 | 0 | 0 |
| 42641 | ENSMUSG00000104590 | 0 | NA | NA | NA | NA | NA | Gm5034 | predicted gene 5034 [Source:MGI Symbol | 0 | 0 | 0 | 0 | 0 | 0 |
| 42642 | ENSMUSG00000104591 | 0 | NA | NA | NA | NA | NA | Gm43145 | predicted gene 43145 [Source:MGI Symbo | 0 | 0 | 0 | 0 | 0 | 0 |
| 42643 | ENSMUSG00000104592 | 0 | NA | NA | NA | NA | NA | Gm42721 | predicted gene 42721 [Source:MGI Symbo | 0 | 0 | 0 | 0 | 0 | 0 |
| 42644 | ENSMUSG00000104593 | 0 | NA | NA | NA | NA | NA | Ighv1-65 | immunoglobulin heavy variable V1-65 [So | 0 | 0 | 0 | 0 | 0 | 0 |
| 42645 | ENSMUSG00000104594 | 0 | NA | NA | NA | NA | NA | NA | NA | 0 | 0 | 0 | 0 | 0 | 0 |
| 42646 | ENSMUSG00000104595 | 0 | NA | NA | NA | NA | NA | Gm42593 | predicted gene 42593 [Source:MGI Symbo | 0 | 0 | 0 | 0 | 0 | 0 |
| 42647 | ENSMUSG00000104596 | 0 | NA | NA | NA | NA | NA | Gm19080 | predicted gene, 19080 [Source:MGI Symb | 0 | 0 | 0 | 0 | 0 | 0 |
| 42648 | ENSMUSG00000104597 | 0 | NA | NA | NA | NA | NA | B230334C09 | RIKEN cDNA B230334C09 gene [Source:M | 0 | 0 | 0 | 0 | 0 | 0 |
| 42649 | ENSMUSG00000104598 | 0 | NA | NA | NA | NA | NA | Gm30214 | predicted gene, 30214 [Source:MGI Symb | 0 | 0 | 0 | 0 | 0 | 0 |
| 42650 | ENSMUSG00000104599 | 0 | NA | NA | NA | NA | NA | NA | NA | 0 | 0 | 0 | 0 | 0 | 0 |
| 42651 | ENSMUSG00000104600 | 0 | NA | NA | NA | NA | NA | Dmrt1i | Dmrt1 interacting ncRNA [Source:MGI Syr | 0 | 0 | 0 | 0 | 0 | 0 |
| 42652 | ENSMUSG00000104601 | 0 | NA | NA | NA | NA | NA | Gm29745 | predicted gene, 29745 [Source:MGI Symb | 0 | 0 | 0 | 0 | 0 | 0 |
| 42653 | ENSMUSG00000104602 | 0 | NA | NA | NA | NA | NA | Gm42779 | predicted gene 42779 [Source:MGI Symbo | 0 | 0 | 0 | 0 | 0 | 0 |
| 42654 | ENSMUSG00000104603 | 0 | NA | NA | NA | NA | NA | NA | NA | 0 | 0 | 0 | 0 | 0 | 0 |
| 42655 | ENSMUSG00000104604 | 0 | NA | NA | NA | NA | NA | Gm42915 | predicted gene 42915 [Source:MGI Symbo | 0 | 0 | 0 | 0 | 0 | 0 |
| 42656 | ENSMUSG00000104605 | 0.476489653 | -2.293616034 | 3.421506946 | -0.670352587 | 0.502633051 | NA | Gm42922 | predicted gene 42922 [Source:MGI Symbo | 0.8031 | 2.0558 | 0 | 0 | 0 | 0 |
| 42657 | ENSMUSG00000104606 | 8.071712009 | 0.190381775 | 0.791716303 | 0.240467165 | 0.809968115 | 0.966157155 | Gm43409 | predicted gene 43409 [Source:MGI Symbo | 6.4248 | 10.279 | 6.0375 | 12.532 | 9.8156 | 3.3415 |
| 42658 | ENSMUSG00000104607 | 0 | NA | NA | NA | NA | NA | Gm31651 | predicted gene, 31651 [Source:MGI Symb | 0 | 0 | 0 | 0 | 0 | 0 |
| 42659 | ENSMUSG00000104608 | 0 | NA | NA | NA | NA | NA | NA | NA | 0 | 0 | 0 | 0 | 0 | 0 |
| 42660 | ENSMUSG00000104609 | 0 | NA | NA | NA | NA | NA | Igkv13-76 | immunoglobulin kappa chain variable 13- | 0 | 0 | 0 | 0 | 0 | 0 |
| 42661 | ENSMUSG00000104610 | 0 | NA | NA | NA | NA | NA | Gm567 | predicted gene 567 [Source:MGI Symbol;A | 0 | 0 | 0 | 0 | 0 | 0 |
| 42662 | ENSMUSG00000104612 | 0 | NA | NA | NA | NA | NA | Gm42449 | predicted gene 42449 [Source:MGI Symbo | 0 | 0 | 0 | 0 | 0 | 0 |
| 42663 | ENSMUSG00000104613 | 0 | NA | NA | NA | NA | NA | Gm8704 | predicted gene 8704 [Source:MGI Symbol | 0 | 0 | 0 | 0 | 0 | 0 |
| 42664 | ENSMUSG00000104614 | 0 | NA | NA | NA | NA | NA | Fbxw27 | F-box and WD-40 domain protein 27 [Sou | 0 | 0 | 0 | 0 | 0 | 0 |
| 42665 | ENSMUSG00000104615 | 0 | NA | NA | NA | NA | NA | Gm43804 | predicted gene 43804 [Source:MGI Symbo | 0 | 0 | 0 | 0 | 0 | 0 |
| 42666 | ENSMUSG00000104616 | 8.575609946 | -0.823483433 | 0.754216211 | -1.091840006 | 0.274903431 | 0.729578574 | Gm6069 | predicted gene 6069 [Source:MGI Symbol | 9.6372 | 11.307 | 12.075 | 9.6398 | 5.4531 | 3.3415 |
| 42667 | ENSMUSG00000104617 | 0 | NA | NA | NA | NA | NA | Gm43356 | predicted gene 43356 [Source:MGI Symbo | 0 | 0 | 0 | 0 | 0 | 0 |
| 42668 | ENSMUSG00000104618 | 0 | NA | NA | NA | NA | NA | Mir1839 | microRNA 1839 [Source:MGI Symbol;Acc: | 0 | 0 | 0 | 0 | 0 | 0 |

| | | | | | | | | | | | | | | | |
|---|---|---|---|---|---|---|---|---|---|---|---|---|---|---|---|
| 42669 | ENSMUSG00000104619 | 0 | NA | NA | NA | NA | NA | Gm42678 | predicted gene 42678 [Source:MGI Symbo | 0 | 0 | 0 | 0 | 0 | 0 |
| 42670 | ENSMUSG00000104620 | 0 | NA | NA | NA | NA | NA | Trav7-1 | T cell receptor alpha variable 7-1 [Source: | 0 | 0 | 0 | 0 | 0 | 0 |
| 42671 | ENSMUSG00000104621 | 0 | NA | NA | NA | NA | NA | Gm43185 | predicted gene 43185 [Source:MGI Symbo | 0 | 0 | 0 | 0 | 0 | 0 |
| 42672 | ENSMUSG00000104622 | 0 | NA | NA | NA | NA | NA | Gm42541 | predicted gene 42541 [Source:MGI Symbo | 0 | 0 | 0 | 0 | 0 | 0 |
| 42673 | ENSMUSG00000104623 | 0 | NA | NA | NA | NA | NA | Gm6394 | predicted gene 6394 [Source:MGI Symbol | 0 | 0 | 0 | 0 | 0 | 0 |
| 42674 | ENSMUSG00000104624 | 0 | NA | NA | NA | NA | NA | Gm43478 | predicted gene 43478 [Source:MGI Symbo | 0 | 0 | 0 | 0 | 0 | 0 |
| 42675 | ENSMUSG00000104625 | 0 | NA | NA | NA | NA | NA | Gm42448 | predicted gene 42448 [Source:MGI Symbo | 0 | 0 | 0 | 0 | 0 | 0 |
| 42676 | ENSMUSG00000104626 | 0 | NA | NA | NA | NA | NA | Gm42675 | predicted gene 42675 [Source:MGI Symbo | 0 | 0 | 0 | 0 | 0 | 0 |
| 42677 | ENSMUSG00000104627 | 0 | NA | NA | NA | NA | NA | Mir3535 | microRNA 3535 [Source:MGI Symbol;Acc: | 0 | 0 | 0 | 0 | 0 | 0 |
| 42678 | ENSMUSG00000104628 | 0 | NA | NA | NA | NA | NA | NA | NA | 0 | 0 | 0 | 0 | 0 | 0 |
| 42679 | ENSMUSG00000104629 | 0.167707431 | -0.796629176 | 4.080472857 | -0.195229623 | 0.84521319 | NA | Trav13n-1 | T cell receptor alpha variable 13N-1 [Sour | 0 | 0 | 1.0062 | 0 | 0 | 0 |
| 42680 | ENSMUSG00000104630 | 0 | NA | NA | NA | NA | NA | Trgj3 | T cell receptor gamma joining 3 [Source:M | 0 | 0 | 0 | 0 | 0 | 0 |
| 42681 | ENSMUSG00000104631 | 0 | NA | NA | NA | NA | NA | Gm43774 | predicted gene 43774 [Source:MGI Symbo | 0 | 0 | 0 | 0 | 0 | 0 |
| 42682 | ENSMUSG00000104632 | 0.342639279 | -1.805976596 | 4.028547714 | -0.448294702 | 0.653940524 | NA | Gm42909 | predicted gene 42909 [Source:MGI Symbo | 0 | 2.0558 | 0 | 0 | 0 | 0 |
| 42683 | ENSMUSG00000104633 | 0 | NA | NA | NA | NA | NA | Gm42421 | predicted gene, 42421 [Source:MGI Symb | 0 | 0 | 0 | 0 | 0 | 0 |
| 42684 | ENSMUSG00000104634 | 0 | NA | NA | NA | NA | NA | Gm42461 | predicted gene 42461 [Source:MGI Symbo | 0 | 0 | 0 | 0 | 0 | 0 |
| 42685 | ENSMUSG00000104635 | 0 | NA | NA | NA | NA | NA | NA | NA | 0 | 0 | 0 | 0 | 0 | 0 |
| 42686 | ENSMUSG00000104636 | 0.17131964 | -0.796629176 | 4.080472857 | -0.195229623 | 0.84521319 | NA | Gm43239 | predicted gene 43239 [Source:MGI Symbo | 0 | 1.0279 | 0 | 0 | 0 | 0 |
| 42687 | ENSMUSG00000104637 | 0 | NA | NA | NA | NA | NA | Gm42424 | predicted gene 42424 [Source:MGI Symbo | 0 | 0 | 0 | 0 | 0 | 0 |
| 42688 | ENSMUSG00000104638 | 0 | NA | NA | NA | NA | NA | 4930568A13 | RIKEN cDNA 4930568A13 gene [Source:M | 0 | 0 | 0 | 0 | 0 | 0 |
| 42689 | ENSMUSG00000104639 | 0 | NA | NA | NA | NA | NA | Gm42939 | predicted gene 42939 [Source:MGI Symbo | 0 | 0 | 0 | 0 | 0 | 0 |
| 42690 | ENSMUSG00000104640 | 0 | NA | NA | NA | NA | NA | Gm42623 | predicted gene 42623 [Source:MGI Symbo | 0 | 0 | 0 | 0 | 0 | 0 |
| 42691 | ENSMUSG00000104641 | 0.756734126 | -0.422689986 | 2.665959267 | -0.158550804 | 0.874022791 | NA | Gm43290 | predicted gene 43290 [Source:MGI Symbo | 1.6062 | 0 | 1.0062 | 1.928 | 0 | 0 |
| 42692 | ENSMUSG00000104642 | 0 | NA | NA | NA | NA | NA | Gm9207 | predicted gene 9207 [Source:MGI Symbol | 0 | 0 | 0 | 0 | 0 | 0 |
| 42693 | ENSMUSG00000104643 | 0 | NA | NA | NA | NA | NA | Gm8862 | predicted gene 8862 [Source:MGI Symbol | 0 | 0 | 0 | 0 | 0 | 0 |
| 42694 | ENSMUSG00000104644 | 0 | NA | NA | NA | NA | NA | Gm2314 | predicted gene 2314 [Source:MGI Symbol | 0 | 0 | 0 | 0 | 0 | 0 |
| 42695 | ENSMUSG00000104645 | 0 | NA | NA | NA | NA | NA | Gm30484 | predicted gene, 30484 [Source:MGI Symb | 0 | 0 | 0 | 0 | 0 | 0 |
| 42696 | ENSMUSG00000104646 | 0.346303622 | 2.004339653 | 3.884003423 | 0.516049919 | 0.605819544 | NA | Gm9173 | predicted gene 9173 [Source:MGI Symbol | 0 | 0 | 0 | 0.964 | 0 | 1.1138 |
| 42697 | ENSMUSG00000104647 | 0 | NA | NA | NA | NA | NA | Gm43485 | predicted gene 43485 [Source:MGI Symbo | 0 | 0 | 0 | 0 | 0 | 0 |
| 42698 | ENSMUSG00000104648 | 0 | NA | NA | NA | NA | NA | Gm42570 | predicted gene 42570 [Source:MGI Symbo | 0 | 0 | 0 | 0 | 0 | 0 |
| 42699 | ENSMUSG00000104649 | 1.160043064 | -0.43591056 | 2.278443107 | -0.191319484 | 0.848275297 | NA | Gm43712 | predicted gene 43712 [Source:MGI Symbo | 0 | 2.0558 | 2.0125 | 2.8919 | 0 | 0 |
| 42700 | ENSMUSG00000104650 | 0 | NA | NA | NA | NA | NA | Trav22 | T cell receptor alpha variable 22 [Source:N | 0 | 0 | 0 | 0 | 0 | 0 |
| 42701 | ENSMUSG00000104651 | 0 | NA | NA | NA | NA | NA | Gm21168 | predicted gene, 21168 [Source:MGI Symb | 0 | 0 | 0 | 0 | 0 | 0 |
| 42702 | ENSMUSG00000104652 | 0 | NA | NA | NA | NA | NA | NA | NA | 0 | 0 | 0 | 0 | 0 | 0 |
| 42703 | ENSMUSG00000104653 | 0.803102239 | -3.069617558 | 3.178309912 | -0.965801839 | 0.334143348 | NA | Gm43109 | predicted gene 43109 [Source:MGI Symbo | 4.8186 | 0 | 0 | 0 | 0 | 0 |
| 42704 | ENSMUSG00000104654 | 0.915510039 | -3.242382683 | 2.669421139 | -1.214638873 | 0.224503856 | NA | Gm43814 | predicted gene 43814 [Source:MGI Symbo | 2.4093 | 3.0838 | 0 | 0 | 0 | 0 |
| 42705 | ENSMUSG00000104655 | 0 | NA | NA | NA | NA | NA | Gm43317 | predicted gene 43317 [Source:MGI Symbo | 0 | 0 | 0 | 0 | 0 | 0 |
| 42706 | ENSMUSG00000104656 | 0 | NA | NA | NA | NA | NA | NA | NA | 0 | 0 | 0 | 0 | 0 | 0 |
| 42707 | ENSMUSG00000104657 | 0 | NA | NA | NA | NA | NA | NA | NA | 0 | 0 | 0 | 0 | 0 | 0 |
| 42708 | ENSMUSG00000104658 | 0 | NA | NA | NA | NA | NA | Gm30275 | predicted gene, 30275 [Source:MGI Symb | 0 | 0 | 0 | 0 | 0 | 0 |
| 42709 | ENSMUSG00000104659 | 0 | NA | NA | NA | NA | NA | Gm43619 | predicted gene 43619 [Source:MGI Symbo | 0 | 0 | 0 | 0 | 0 | 0 |
| 42710 | ENSMUSG00000104660 | 0 | NA | NA | NA | NA | NA | Gm43601 | predicted gene 43601 [Source:MGI Symbo | 0 | 0 | 0 | 0 | 0 | 0 |
| 42711 | ENSMUSG00000104661 | 0 | NA | NA | NA | NA | NA | Gm42768 | predicted gene 42768 [Source:MGI Symbo | 0 | 0 | 0 | 0 | 0 | 0 |
| 42712 | ENSMUSG00000104662 | 7.39349005 | -0.142027105 | 0.918947266 | -0.154554141 | 0.877172826 | 0.980632845 | Gm43267 | predicted gene 43267 [Source:MGI Symbo | 7.2279 | 4.1117 | 12.075 | 15.424 | 2.1812 | 3.3415 |
| 42713 | ENSMUSG00000104663 | 0 | NA | NA | NA | NA | NA | Gm3203 | predicted gene 3203 [Source:MGI Symbol | 0 | 0 | 0 | 0 | 0 | 0 |
| 42714 | ENSMUSG00000104664 | 75.71100236 | 0.756730062 | 0.382279104 | 1.979522435 | 0.047757216 | 0.303630934 | Idi1-ps2 | isopentenyl-diphosphate delta isomerase, | 72.279 | 48.312 | 48.3 | 161.95 | 62.165 | 61.261 |
| 42715 | ENSMUSG00000104665 | 0 | NA | NA | NA | NA | NA | Gm43366 | predicted gene 43366 [Source:MGI Symbo | 0 | 0 | 0 | 0 | 0 | 0 |
| 42716 | ENSMUSG00000104666 | 0 | NA | NA | NA | NA | NA | NA | NA | 0 | 0 | 0 | 0 | 0 | 0 |
| 42717 | ENSMUSG00000104667 | 0 | NA | NA | NA | NA | NA | Gm4961 | predicted pseudogene 4961 [Source:MGI | 0 | 0 | 0 | 0 | 0 | 0 |
| 42718 | ENSMUSG00000104668 | 0 | NA | NA | NA | NA | NA | Vmn2r-ps24 | vomeronasal 2, receptor, pseudogene 24 | 0 | 0 | 0 | 0 | 0 | 0 |
| 42719 | ENSMUSG00000104669 | 5.842158523 | -1.677001015 | 0.937159705 | -1.789450619 | 0.073542273 | 0.394114651 | Gm38411 | predicted gene, 38411 [Source:MGI Symb | 6.4248 | 11.307 | 9.0562 | 3.8559 | 2.1812 | 2.2277 |
| 42720 | ENSMUSG00000104670 | 0 | NA | NA | NA | NA | NA | Gm9369 | predicted pseudogene 9369 [Source:MGI | 0 | 0 | 0 | 0 | 0 | 0 |
| 42721 | ENSMUSG00000104671 | 0.849167191 | -0.527085716 | 2.389288686 | -0.220603613 | 0.825401088 | NA | Gm43062 | predicted gene 43062 [Source:MGI Symbo | 0 | 1.0279 | 2.0125 | 0.964 | 1.0906 | 0 |
| 42722 | ENSMUSG00000104672 | 0 | NA | NA | NA | NA | NA | Gm43665 | predicted gene 43665 [Source:MGI Symbo | 0 | 0 | 0 | 0 | 0 | 0 |
| 42723 | ENSMUSG00000104673 | 0 | NA | NA | NA | NA | NA | Gm43550 | predicted gene 43550 [Source:MGI Symbo | 0 | 0 | 0 | 0 | 0 | 0 |
| 42724 | ENSMUSG00000104674 | 0 | NA | NA | NA | NA | NA | Gm42756 | predicted gene 42756 [Source:MGI Symbo | 0 | 0 | 0 | 0 | 0 | 0 |

| | | | | | | | | | | | | | | | |
|---|---|---|---|---|---|---|---|---|---|---|---|---|---|---|---|
| 42725 | ENSMUSG00000104675 | 2.109379672 | -0.34802549 | 1.898895108 | -0.183277891 | 0.854579981 | NA | Gm43689 | predicted gene 43689 [Source:MGI Symbo | 0 | 2.0558 | 5.0312 | 0 | 0 | 5.5692 |
| 42726 | ENSMUSG00000104676 | 7.268904985 | -0.305352425 | 0.946911647 | -0.322471928 | 0.747095201 | 0.950561663 | Gm42777 | predicted gene 42777 [Source:MGI Symbo | 8.8341 | 3.0838 | 12.075 | 0.964 | 13.087 | 5.5692 |
| 42727 | ENSMUSG00000104677 | 0 | NA | NA | NA | NA | NA | Gm43376 | predicted gene 43376 [Source:MGI Symbo | 0 | 0 | 0 | 0 | 0 | 0 |
| 42728 | ENSMUSG00000104678 | 0 | NA | NA | NA | NA | NA | Gm4321 | predicted gene 4321 [Source:MGI Symbol | 0 | 0 | 0 | 0 | 0 | 0 |
| 42729 | ENSMUSG00000104679 | 0 | NA | NA | NA | NA | NA | Igkv4-60 | immunoglobulin kappa variable 4-60 [Sou | 0 | 0 | 0 | 0 | 0 | 0 |
| 42730 | ENSMUSG00000104680 | 0 | NA | NA | NA | NA | NA | Gm9415 | predicted gene 9415 [Source:MGI Symbol | 0 | 0 | 0 | 0 | 0 | 0 |
| 42731 | ENSMUSG00000104681 | 0 | NA | NA | NA | NA | NA | Gm43001 | predicted gene 43001 [Source:MGI Symbo | 0 | 0 | 0 | 0 | 0 | 0 |
| 42732 | ENSMUSG00000104682 | 2.976298464 | -0.128546906 | 1.2089111 | -0.106332803 | 0.915318306 | NA | Gm42636 | predicted gene 42636 [Source:MGI Symbo | 3.2124 | 2.0558 | 4.025 | 1.928 | 2.1812 | 4.4554 |
| 42733 | ENSMUSG00000104683 | 0 | NA | NA | NA | NA | NA | NA | NA | 0 | 0 | 0 | 0 | 0 | 0 |
| 42734 | ENSMUSG00000104684 | 1.840158714 | 0.925917867 | 1.651538692 | 0.560639524 | 0.575043302 | NA | 5430427N15 | RIKEN cDNA 5430427N15 gene [Source:M | 0.8031 | 1.0279 | 2.0125 | 3.8559 | 0 | 3.3415 |
| 42735 | ENSMUSG00000104685 | 0 | NA | NA | NA | NA | NA | Gm19202 | predicted gene, 19202 [Source:MGI Symb | 0 | 0 | 0 | 0 | 0 | 0 |
| 42736 | ENSMUSG00000104686 | 1.045903188 | 0.074846443 | 2.394919062 | 0.031252181 | 0.975068426 | NA | Gm42815 | predicted gene 42815 [Source:MGI Symbo | 0 | 3.0838 | 0 | 0.964 | 0 | 2.2277 |
| 42737 | ENSMUSG00000104687 | 0 | NA | NA | NA | NA | NA | Gm42899 | predicted gene 42899 [Source:MGI Symbo | 0 | 0 | 0 | 0 | 0 | 0 |
| 42738 | ENSMUSG00000104688 | 0 | NA | NA | NA | NA | NA | Gm18702 | predicted gene, 18702 [Source:MGI Symb | 0 | 0 | 0 | 0 | 0 | 0 |
| 42739 | ENSMUSG00000104689 | 3.133851075 | 0.759649165 | 1.565627573 | 0.485204258 | 0.627531462 | NA | NA | NA | 0 | 3.0838 | 4.025 | 10.604 | 1.0906 | 0 |
| 42740 | ENSMUSG00000104690 | 0 | NA | NA | NA | NA | NA | NA | NA | 0 | 0 | 0 | 0 | 0 | 0 |
| 42741 | ENSMUSG00000104691 | 0 | NA | NA | NA | NA | NA | Gm43021 | predicted gene 43021 [Source:MGI Symbo | 0 | 0 | 0 | 0 | 0 | 0 |
| 42742 | ENSMUSG00000104692 | 0 | NA | NA | NA | NA | NA | Gm42773 | predicted gene 42773 [Source:MGI Symbo | 0 | 0 | 0 | 0 | 0 | 0 |
| 42743 | ENSMUSG00000104693 | 0.476283062 | 1.031501528 | 3.159856949 | 0.326439312 | 0.744091998 | NA | Gm42941 | predicted gene 42941 [Source:MGI Symbo | 0.8031 | 0 | 0 | 0.964 | 1.0906 | 0 |
| 42744 | ENSMUSG00000104694 | 1.479611194 | -0.378768569 | 2.116041761 | -0.178998627 | 0.85793878 | NA | Gm43797 | predicted gene 43797 [Source:MGI Symbo | 4.0155 | 0 | 1.0062 | 3.8559 | 0 | 0 |
| 42745 | ENSMUSG00000104695 | 0 | NA | NA | NA | NA | NA | Gm30023 | predicted gene, 30023 [Source:MGI Symb | 0 | 0 | 0 | 0 | 0 | 0 |
| 42746 | ENSMUSG00000104696 | 0 | NA | NA | NA | NA | NA | Gm42946 | predicted gene 42946 [Source:MGI Symbo | 0 | 0 | 0 | 0 | 0 | 0 |
| 42747 | ENSMUSG00000104697 | 0 | NA | NA | NA | NA | NA | Igkv12-67 | immunoglobulin kappa chain variable 12-( | 0 | 0 | 0 | 0 | 0 | 0 |
| 42748 | ENSMUSG00000104698 | 0 | NA | NA | NA | NA | NA | Gm42602 | predicted gene 42602 [Source:MGI Symbo | 0 | 0 | 0 | 0 | 0 | 0 |
| 42749 | ENSMUSG00000104699 | 0.267700746 | -1.497481649 | 4.057001436 | -0.369110456 | 0.712045396 | NA | Rps4x-ps | Rps4x retrotransposed pseudogene [Sour | 1.6062 | 0 | 0 | 0 | 0 | 0 |
| 42750 | ENSMUSG00000104700 | 0 | NA | NA | NA | NA | NA | Gm26379 | predicted gene, 26379 [Source:MGI Symb | 0 | 0 | 0 | 0 | 0 | 0 |
| 42751 | ENSMUSG00000104701 | 0 | NA | NA | NA | NA | NA | Gm42555 | predicted gene 42555 [Source:MGI Symbo | 0 | 0 | 0 | 0 | 0 | 0 |
| 42752 | ENSMUSG00000104702 | 0 | NA | NA | NA | NA | NA | NA | NA | 0 | 0 | 0 | 0 | 0 | 0 |
| 42753 | ENSMUSG00000104703 | 0 | NA | NA | NA | NA | NA | 4921521D15 | RIKEN cDNA 4921521D15 gene [Source:M | 0 | 0 | 0 | 0 | 0 | 0 |
| 42754 | ENSMUSG00000104704 | 0.777840866 | -0.360978154 | 2.442491672 | -0.147790945 | 0.882507755 | NA | Gm43133 | predicted gene 43133 [Source:MGI Symbo | 1.6062 | 0 | 1.0062 | 0.964 | 1.0906 | 0 |
| 42755 | ENSMUSG00000104705 | 0 | NA | NA | NA | NA | NA | 4930405N21 | RIKEN cDNA 4930405N21 gene [Source:M | 0 | 0 | 0 | 0 | 0 | 0 |
| 42756 | ENSMUSG00000104706 | 1.117100031 | 1.582896982 | 2.163697584 | 0.731570342 | 0.464430856 | NA | Gm43421 | predicted gene 43421 [Source:MGI Symbo | 1.6062 | 0 | 0 | 2.8919 | 1.0906 | 1.1138 |
| 42757 | ENSMUSG00000104707 | 0 | NA | NA | NA | NA | NA | Gm43649 | predicted gene 43649 [Source:MGI Symbo | 0 | 0 | 0 | 0 | 0 | 0 |
| 42758 | ENSMUSG00000104708 | 0 | NA | NA | NA | NA | NA | Gm42503 | predicted gene 42503 [Source:MGI Symbo | 0 | 0 | 0 | 0 | 0 | 0 |
| 42759 | ENSMUSG00000104709 | 0 | NA | NA | NA | NA | NA | Gm42484 | predicted gene 42484 [Source:MGI Symbo | 0 | 0 | 0 | 0 | 0 | 0 |
| 42760 | ENSMUSG00000104710 | 0 | NA | NA | NA | NA | NA | Gm31243 | predicted gene, 31243 [Source:MGI Symb | 0 | 0 | 0 | 0 | 0 | 0 |
| 42761 | ENSMUSG00000104711 | 0 | NA | NA | NA | NA | NA | Gm43854 | predicted gene 43854 [Source:MGI Symbo | 0 | 0 | 0 | 0 | 0 | 0 |
| 42762 | ENSMUSG00000104712 | 0 | NA | NA | NA | NA | NA | Igkv4-75 | immunoglobulin kappa chain variable 4-7! | 0 | 0 | 0 | 0 | 0 | 0 |
| 42763 | ENSMUSG00000104713 | 71.96208431 | -0.046450842 | 0.316404718 | -0.146808311 | 0.883283325 | 0.981769403 | Gbp6 | guanylate binding protein 6 [Source:MGI S | 87.538 | 88.401 | 43.269 | 77.118 | 73.071 | 62.375 |
| 42764 | ENSMUSG00000104714 | 0 | NA | NA | NA | NA | NA | Trav5d-2 | T cell receptor alpha variable 5D-2 [Source | 0 | 0 | 0 | 0 | 0 | 0 |
| 42765 | ENSMUSG00000104715 | 0 | NA | NA | NA | NA | NA | Gm43515 | predicted gene 43515 [Source:MGI Symbo | 0 | 0 | 0 | 0 | 0 | 0 |
| 42766 | ENSMUSG00000104716 | 0 | NA | NA | NA | NA | NA | Gm30652 | predicted gene, 30652 [Source:MGI Symb | 0 | 0 | 0 | 0 | 0 | 0 |
| 42767 | ENSMUSG00000104717 | 0 | NA | NA | NA | NA | NA | Gm33969 | predicted gene, 33969 [Source:MGI Symb | 0 | 0 | 0 | 0 | 0 | 0 |
| 42768 | ENSMUSG00000104718 | 0 | NA | NA | NA | NA | NA | Gm31752 | predicted gene, 31752 [Source:MGI Symb | 0 | 0 | 0 | 0 | 0 | 0 |
| 42769 | ENSMUSG00000104719 | 0.945548297 | -0.814460806 | 2.273503937 | -0.358240332 | 0.720163467 | NA | 4933437G19 | RIKEN cDNA 4933437G19 gene [Source:M | 1.6062 | 0 | 2.0125 | 0.964 | 1.0906 | 0 |
| 42770 | ENSMUSG00000104720 | 2.953976867 | -0.485357762 | 1.320707933 | -0.36749818 | 0.713247449 | NA | Gm43138 | predicted gene 43138 [Source:MGI Symbo | 3.2124 | 4.1117 | 3.0187 | 1.928 | 5.4531 | 0 |
| 42771 | ENSMUSG00000104721 | 0.181769715 | 1.126924175 | 4.080472857 | 0.276174898 | 0.782413736 | NA | Gm42696 | predicted gene 42696 [Source:MGI Symbo | 0 | 0 | 0 | 0 | 1.0906 | 0 |
| 42772 | ENSMUSG00000104722 | 0 | NA | NA | NA | NA | NA | Gm43007 | predicted gene 43007 [Source:MGI Symbo | 0 | 0 | 0 | 0 | 0 | 0 |
| 42773 | ENSMUSG00000104723 | 0 | NA | NA | NA | NA | NA | Gm43106 | predicted gene 43106 [Source:MGI Symbo | 0 | 0 | 0 | 0 | 0 | 0 |
| 42774 | ENSMUSG00000104724 | 0 | NA | NA | NA | NA | NA | Gm43162 | predicted gene 43162 [Source:MGI Symbo | 0 | 0 | 0 | 0 | 0 | 0 |
| 42775 | ENSMUSG00000104725 | 0 | NA | NA | NA | NA | NA | Gm2860 | predicted gene 2860 [Source:MGI Symbol | 0 | 0 | 0 | 0 | 0 | 0 |
| 42776 | ENSMUSG00000104726 | 0 | NA | NA | NA | NA | NA | Gm18384 | predicted gene, 18384 [Source:MGI Symb | 0 | 0 | 0 | 0 | 0 | 0 |
| 42777 | ENSMUSG00000104727 | 0 | NA | NA | NA | NA | NA | Gm42500 | predicted gene 42500 [Source:MGI Symbo | 0 | 0 | 0 | 0 | 0 | 0 |
| 42778 | ENSMUSG00000104728 | 1.988019237 | 0.133606027 | 1.575387238 | 0.084808372 | 0.932413738 | NA | Gm42462 | predicted gene 42462 [Source:MGI Symbo | 1.6062 | 4.1117 | 0 | 2.8919 | 1.0906 | 2.2277 |
| 42779 | ENSMUSG00000104729 | 0 | NA | NA | NA | NA | NA | NA | NA | 0 | 0 | 0 | 0 | 0 | 0 |
| 42780 | ENSMUSG00000104730 | 0 | NA | NA | NA | NA | NA | Gm42511 | predicted gene 42511 [Source:MGI Symbo | 0 | 0 | 0 | 0 | 0 | 0 |

| | | | | | | | | | | | | | | | |
|---|---|---|---|---|---|---|---|---|---|---|---|---|---|---|---|
| 42781 | ENSMUSG00000104731 | 0 | NA | NA | NA | NA | NA | Gm43249 | predicted gene 43249 [Source:MGI Symbo | 0 | 0 | 0 | 0 | 0 | 0 |
| 42782 | ENSMUSG00000104732 | 0 | NA | NA | NA | NA | NA | Gm43346 | predicted gene 43346 [Source:MGI Symbo | 0 | 0 | 0 | 0 | 0 | 0 |
| 42783 | ENSMUSG00000104733 | 0 | NA | NA | NA | NA | NA | Gm29784 | predicted gene, 29784 [Source:MGI Symb | 0 | 0 | 0 | 0 | 0 | 0 |
| 42784 | ENSMUSG00000104734 | 0 | NA | NA | NA | NA | NA | NA | NA | 0 | 0 | 0 | 0 | 0 | 0 |
| 42785 | ENSMUSG00000104735 | 0 | NA | NA | NA | NA | NA | C130075A20 | RIKEN cDNA C130075A20 gene [Source:M | 0 | 0 | 0 | 0 | 0 | 0 |
| 42786 | ENSMUSG00000104736 | 0.160662974 | 1.126924175 | 4.080472857 | 0.276174898 | 0.782413736 | NA | Gm33609 | predicted gene, 33609 [Source:MGI Symb | 0 | 0 | 0 | 0.964 | 0 | 0 |
| 42787 | ENSMUSG00000104737 | 4.572033405 | -0.970454013 | 1.032454155 | -0.93994877 | 0.34724384 | NA | Gm42937 | predicted gene 42937 [Source:MGI Symbo | 8.031 | 4.1117 | 6.0375 | 4.8199 | 1.0906 | 3.3415 |
| 42788 | ENSMUSG00000104738 | 0 | NA | NA | NA | NA | NA | Trav3-2 | T cell receptor alpha variable 3-2 [Source: | 0 | 0 | 0 | 0 | 0 | 0 |
| 42789 | ENSMUSG00000104739 | 0 | NA | NA | NA | NA | NA | Gm3283 | predicted gene 3283 [Source:MGI Symbol | 0 | 0 | 0 | 0 | 0 | 0 |
| 42790 | ENSMUSG00000104740 | 0 | NA | NA | NA | NA | NA | Mir6516 | microRNA 6516 [Source:MGI Symbol;Acc: | 0 | 0 | 0 | 0 | 0 | 0 |
| 42791 | ENSMUSG00000104741 | 0 | NA | NA | NA | NA | NA | Mir3087 | microRNA 3087 [Source:MGI Symbol;Acc: | 0 | 0 | 0 | 0 | 0 | 0 |
| 42792 | ENSMUSG00000104742 | 0 | NA | NA | NA | NA | NA | Igkv9-128 | immunoglobulin kappa chain variable 9-1 | 0 | 0 | 0 | 0 | 0 | 0 |
| 42793 | ENSMUSG00000104743 | 0 | NA | NA | NA | NA | NA | NA | NA | 0 | 0 | 0 | 0 | 0 | 0 |
| 42794 | ENSMUSG00000104744 | 0 | NA | NA | NA | NA | NA | Gm18980 | predicted gene, 18980 [Source:MGI Symb | 0 | 0 | 0 | 0 | 0 | 0 |
| 42795 | ENSMUSG00000104745 | 0.335414862 | -1.779047242 | 4.030804179 | -0.441362855 | 0.658950329 | NA | Gm43073 | predicted gene 43073 [Source:MGI Symbo | 0 | 0 | 2.0125 | 0 | 0 | 0 |
| 42796 | ENSMUSG00000104746 | 0 | NA | NA | NA | NA | NA | NA | NA | 0 | 0 | 0 | 0 | 0 | 0 |
| 42797 | ENSMUSG00000104747 | 0 | NA | NA | NA | NA | NA | Mir292b | microRNA 292b [Source:MGI Symbol;Acc: | 0 | 0 | 0 | 0 | 0 | 0 |
| 42798 | ENSMUSG00000104748 | 3.77198827 | 0.427480979 | 1.282240645 | 0.333385921 | 0.738842989 | NA | Gm43592 | predicted gene 43592 [Source:MGI Symbo | 6.4248 | 3.0838 | 0 | 0.964 | 4.3625 | 7.7969 |
| 42799 | ENSMUSG00000104749 | 0 | NA | NA | NA | NA | NA | Gm42926 | predicted gene 42926 [Source:MGI Symbo | 0 | 0 | 0 | 0 | 0 | 0 |
| 42800 | ENSMUSG00000104750 | 0 | NA | NA | NA | NA | NA | NA | NA | 0 | 0 | 0 | 0 | 0 | 0 |
| 42801 | ENSMUSG00000104751 | 0 | NA | NA | NA | NA | NA | Gm43179 | predicted gene 43179 [Source:MGI Symbo | 0 | 0 | 0 | 0 | 0 | 0 |
| 42802 | ENSMUSG00000104752 | 0 | NA | NA | NA | NA | NA | Gm5565 | predicted gene 5565 [Source:MGI Symbol | 0 | 0 | 0 | 0 | 0 | 0 |
| 42803 | ENSMUSG00000104753 | 0 | NA | NA | NA | NA | NA | Gm35172 | predicted gene, 35172 [Source:MGI Symb | 0 | 0 | 0 | 0 | 0 | 0 |
| 42804 | ENSMUSG00000104754 | 0.535401493 | -2.48596965 | 3.684117979 | -0.674780141 | 0.499815459 | NA | Gm43116 | predicted gene 43116 [Source:MGI Symbo | 3.2124 | 0 | 0 | 0 | 0 | 0 |
| 42805 | ENSMUSG00000104755 | 0 | NA | NA | NA | NA | NA | Mir677 | microRNA 677 [Source:MGI Symbol;Acc:N | 0 | 0 | 0 | 0 | 0 | 0 |
| 42806 | ENSMUSG00000104756 | 0 | NA | NA | NA | NA | NA | Gm42464 | predicted gene 42464 [Source:MGI Symbo | 0 | 0 | 0 | 0 | 0 | 0 |
| 42807 | ENSMUSG00000104757 | 0 | NA | NA | NA | NA | NA | Gm44391 | predicted gene, 44391 [Source:MGI Symb | 0 | 0 | 0 | 0 | 0 | 0 |
| 42808 | ENSMUSG00000104758 | 0 | NA | NA | NA | NA | NA | Mir691 | microRNA 691 [Source:MGI Symbol;Acc:N | 0 | 0 | 0 | 0 | 0 | 0 |
| 42809 | ENSMUSG00000104759 | 0 | NA | NA | NA | NA | NA | Ranbp2-ps1 | RAN binding protein 2, pseudogene 12 [Sc | 0 | 0 | 0 | 0 | 0 | 0 |
| 42810 | ENSMUSG00000104760 | 0 | NA | NA | NA | NA | NA | Igkv3-8 | immunoglobulin kappa variable 3-8 [Sourc | 0 | 0 | 0 | 0 | 0 | 0 |
| 42811 | ENSMUSG00000104761 | 3.833676693 | -0.621176391 | 1.235314496 | -0.502848783 | 0.615070593 | NA | Gm43511 | predicted gene 43511 [Source:MGI Symbo | 4.8186 | 7.1954 | 2.0125 | 6.7478 | 0 | 2.2277 |
| 42812 | ENSMUSG00000104762 | 0 | NA | NA | NA | NA | NA | NA | NA | 0 | 0 | 0 | 0 | 0 | 0 |
| 42813 | ENSMUSG00000104763 | 0 | NA | NA | NA | NA | NA | 9830132P13 | RIKEN cDNA 9830132P13 gene [Source:M | 0 | 0 | 0 | 0 | 0 | 0 |
| 42814 | ENSMUSG00000104764 | 0 | NA | NA | NA | NA | NA | Gm43066 | predicted gene 43066 [Source:MGI Symbo | 0 | 0 | 0 | 0 | 0 | 0 |
| 42815 | ENSMUSG00000104765 | 0 | NA | NA | NA | NA | NA | Gm43058 | predicted gene 43058 [Source:MGI Symbo | 0 | 0 | 0 | 0 | 0 | 0 |
| 42816 | ENSMUSG00000104766 | 0 | NA | NA | NA | NA | NA | 4930573H18 | RIKEN cDNA 4930573H18 gene [Source:M | 0 | 0 | 0 | 0 | 0 | 0 |
| 42817 | ENSMUSG00000104767 | 0 | NA | NA | NA | NA | NA | NA | NA | 0 | 0 | 0 | 0 | 0 | 0 |
| 42818 | ENSMUSG00000104768 | 0 | NA | NA | NA | NA | NA | Gm38412 | predicted gene, 38412 [Source:MGI Symb | 0 | 0 | 0 | 0 | 0 | 0 |
| 42819 | ENSMUSG00000104769 | 0 | NA | NA | NA | NA | NA | Igkv8-34 | immunoglobulin kappa variable 8-34 [Sou | 0 | 0 | 0 | 0 | 0 | 0 |
| 42820 | ENSMUSG00000104770 | 0 | NA | NA | NA | NA | NA | Gm2939 | predicted gene 2939 [Source:MGI Symbol | 0 | 0 | 0 | 0 | 0 | 0 |
| 42821 | ENSMUSG00000104771 | 0 | NA | NA | NA | NA | NA | NA | NA | 0 | 0 | 0 | 0 | 0 | 0 |
| 42822 | ENSMUSG00000104772 | 0 | NA | NA | NA | NA | NA | Gm43347 | predicted gene 43347 [Source:MGI Symbo | 0 | 0 | 0 | 0 | 0 | 0 |
| 42823 | ENSMUSG00000104773 | 0 | NA | NA | NA | NA | NA | Mir7657 | microRNA 7657 [Source:MGI Symbol;Acc: | 0 | 0 | 0 | 0 | 0 | 0 |
| 42824 | ENSMUSG00000104774 | 0 | NA | NA | NA | NA | NA | Gm8920 | predicted gene 8920 [Source:MGI Symbol | 0 | 0 | 0 | 0 | 0 | 0 |
| 42825 | ENSMUSG00000104775 | 0 | NA | NA | NA | NA | NA | Gm8182 | predicted gene 8182 [Source:MGI Symbol | 0 | 0 | 0 | 0 | 0 | 0 |
| 42826 | ENSMUSG00000104776 | 2.042337761 | 0.625969637 | 1.494055127 | 0.418973588 | 0.675235435 | NA | Gm43691 | predicted gene 43691 [Source:MGI Symbo | 0.8031 | 1.0279 | 3.0187 | 1.928 | 4.3625 | 1.1138 |
| 42827 | ENSMUSG00000104777 | 0 | NA | NA | NA | NA | NA | Gm4865 | predicted gene 4865 [Source:MGI Symbol | 0 | 0 | 0 | 0 | 0 | 0 |
| 42828 | ENSMUSG00000104778 | 0 | NA | NA | NA | NA | NA | Gm43476 | predicted gene 43476 [Source:MGI Symbo | 0 | 0 | 0 | 0 | 0 | 0 |
| 42829 | ENSMUSG00000104779 | 0 | NA | NA | NA | NA | NA | Gm43559 | predicted gene 43559 [Source:MGI Symbo | 0 | 0 | 0 | 0 | 0 | 0 |
| 42830 | ENSMUSG00000104780 | 0 | NA | NA | NA | NA | NA | Gm19843 | predicted gene, 19843 [Source:MGI Symb | 0 | 0 | 0 | 0 | 0 | 0 |
| 42831 | ENSMUSG00000104781 | 0 | NA | NA | NA | NA | NA | Gm43303 | predicted gene 43303 [Source:MGI Symbo | 0 | 0 | 0 | 0 | 0 | 0 |
| 42832 | ENSMUSG00000104782 | 0 | NA | NA | NA | NA | NA | Gm43326 | predicted gene 43326 [Source:MGI Symbo | 0 | 0 | 0 | 0 | 0 | 0 |
| 42833 | ENSMUSG00000104783 | 0 | NA | NA | NA | NA | NA | NA | NA | 0 | 0 | 0 | 0 | 0 | 0 |
| 42834 | ENSMUSG00000104784 | 23.07184351 | -0.445924562 | 0.497179173 | -0.896909175 | 0.369767384 | 0.80653433 | Gm42984 | predicted gene 42984 [Source:MGI Symbo | 32.124 | 25.698 | 22.137 | 29.883 | 17.45 | 11.138 |
| 42835 | ENSMUSG00000104785 | 24.76952796 | 2.08491869 | 0.685991887 | 3.039276015 | 0.002371475 | 0.039642097 | Gm31121 | predicted gene, 31121 [Source:MGI Symb | 21.684 | 1.0279 | 5.0312 | 46.271 | 53.44 | 21.163 |
| 42836 | ENSMUSG00000104786 | 10.16100859 | -0.47370788 | 0.752676248 | -0.629364724 | 0.529110307 | 0.888334265 | Gm43573 | predicted gene 43573 [Source:MGI Symbo | 8.031 | 13.363 | 14.087 | 5.7839 | 16.359 | 3.3415 |

| | | | | | | | | | | | | | | | |
|---|---|---|---|---|---|---|---|---|---|---|---|---|---|---|---|
| 42837 | ENSMUSG00000104787 | 0 | NA | NA | NA | NA | NA | Gm19148 | predicted gene, 19148 [Source:MGI Symb | 0 | 0 | 0 | 0 | 0 | 0 |
| 42838 | ENSMUSG00000104788 | 0 | NA | NA | NA | NA | NA | Gm36448 | predicted gene, 36448 [Source:MGI Symb | 0 | 0 | 0 | 0 | 0 | 0 |
| 42839 | ENSMUSG00000104789 | 0 | NA | NA | NA | NA | NA | Gm43640 | predicted gene 43640 [Source:MGI Symbo | 0 | 0 | 0 | 0 | 0 | 0 |
| 42840 | ENSMUSG00000104790 | 0 | NA | NA | NA | NA | NA | Gm43276 | predicted gene 43276 [Source:MGI Symbo | 0 | 0 | 0 | 0 | 0 | 0 |
| 42841 | ENSMUSG00000104791 | 0 | NA | NA | NA | NA | NA | 4930511M1 | RIKEN cDNA 4930511M18 gene [Source:M | 0 | 0 | 0 | 0 | 0 | 0 |
| 42842 | ENSMUSG00000104792 | 0 | NA | NA | NA | NA | NA | Gm43308 | predicted gene 43308 [Source:MGI Symbo | 0 | 0 | 0 | 0 | 0 | 0 |
| 42843 | ENSMUSG00000104793 | 1.55300454 | 2.200654854 | 2.370535587 | 0.928336561 | 0.353233015 | NA | Gm43756 | predicted gene 43756 [Source:MGI Symbo | 1.6062 | 0 | 0 | 7.7118 | 0 | 0 |
| 42844 | ENSMUSG00000104794 | 0 | NA | NA | NA | NA | NA | Mir28c | microRNA 28c [Source:MGI Symbol;Acc:M | 0 | 0 | 0 | 0 | 0 | 0 |
| 42845 | ENSMUSG00000104795 | 2.693991356 | 0.881646034 | 1.377462325 | 0.640050924 | 0.522139493 | NA | Gm42783 | predicted gene 42783 [Source:MGI Symbo | 1.6062 | 4.1117 | 0 | 3.8559 | 4.3625 | 2.2277 |
| 42846 | ENSMUSG00000104796 | 0 | NA | NA | NA | NA | NA | NA | NA | 0 | 0 | 0 | 0 | 0 | 0 |
| 42847 | ENSMUSG00000104797 | 0 | NA | NA | NA | NA | NA | Mir1897 | microRNA 1897 [Source:MGI Symbol;Acc: | 0 | 0 | 0 | 0 | 0 | 0 |
| 42848 | ENSMUSG00000104798 | 3.706756539 | 2.77741419 | 1.354596553 | 2.050362659 | 0.040329054 | NA | Gm20121 | predicted gene, 20121 [Source:MGI Symb | 0.8031 | 1.0279 | 1.0062 | 3.8559 | 2.1812 | 13.366 |
| 42849 | ENSMUSG00000104799 | 0 | NA | NA | NA | NA | NA | Gm9361 | predicted gene 9361 [Source:MGI Symbol | 0 | 0 | 0 | 0 | 0 | 0 |
| 42850 | ENSMUSG00000104800 | 0 | NA | NA | NA | NA | NA | Gm36017 | predicted gene, 36017 [Source:MGI Symb | 0 | 0 | 0 | 0 | 0 | 0 |
| 42851 | ENSMUSG00000104801 | 0.133850373 | -0.796629176 | 4.080472857 | -0.195229623 | 0.84521319 | NA | Gm43834 | predicted gene 43834 [Source:MGI Symbo | 0.8031 | 0 | 0 | 0 | 0 | 0 |
| 42852 | ENSMUSG00000104802 | 8.363926768 | 0.24522186 | 0.754200231 | 0.325141587 | 0.745073916 | 0.949997329 | Gm5869 | predicted gene 5869 [Source:MGI Symbol | 4.8186 | 8.2233 | 10.062 | 10.604 | 10.906 | 5.5692 |
| 42853 | ENSMUSG00000104803 | 0 | NA | NA | NA | NA | NA | Gm43108 | predicted gene 43108 [Source:MGI Symbo | 0 | 0 | 0 | 0 | 0 | 0 |
| 42854 | ENSMUSG00000104804 | 0 | NA | NA | NA | NA | NA | Mir7233 | microRNA 7233 [Source:MGI Symbol;Acc: | 0 | 0 | 0 | 0 | 0 | 0 |
| 42855 | ENSMUSG00000104805 | 0 | NA | NA | NA | NA | NA | Gm43624 | predicted gene 43624 [Source:MGI Symbo | 0 | 0 | 0 | 0 | 0 | 0 |
| 42856 | ENSMUSG00000104806 | 0 | NA | NA | NA | NA | NA | Gm42566 | predicted gene 42566 [Source:MGI Symbo | 0 | 0 | 0 | 0 | 0 | 0 |
| 42857 | ENSMUSG00000104807 | 0 | NA | NA | NA | NA | NA | Gm42865 | predicted gene 42865 [Source:MGI Symbo | 0 | 0 | 0 | 0 | 0 | 0 |
| 42858 | ENSMUSG00000104808 | 0 | NA | NA | NA | NA | NA | Gm42619 | predicted gene 42619 [Source:MGI Symbo | 0 | 0 | 0 | 0 | 0 | 0 |
| 42859 | ENSMUSG00000104809 | 0 | NA | NA | NA | NA | NA | NA | NA | 0 | 0 | 0 | 0 | 0 | 0 |
| 42860 | ENSMUSG00000104810 | 0 | NA | NA | NA | NA | NA | Gm5684 | predicted gene 5684 [Source:MGI Symbol | 0 | 0 | 0 | 0 | 0 | 0 |
| 42861 | ENSMUSG00000104811 | 0 | NA | NA | NA | NA | NA | Gm33093 | predicted gene, 33093 [Source:MGI Symb | 0 | 0 | 0 | 0 | 0 | 0 |
| 42862 | ENSMUSG00000104812 | 0 | NA | NA | NA | NA | NA | NA | NA | 0 | 0 | 0 | 0 | 0 | 0 |
| 42863 | ENSMUSG00000104813 | 0 | NA | NA | NA | NA | NA | NA | NA | 0 | 0 | 0 | 0 | 0 | 0 |
| 42864 | ENSMUSG00000104814 | 13.55335689 | 0.8419597 | 0.630918237 | 1.33449891 | 0.182040404 | 0.615708221 | Gm42979 | predicted gene 42979 [Source:MGI Symbo | 13.653 | 10.279 | 5.0312 | 18.316 | 22.903 | 11.138 |
| 42865 | ENSMUSG00000104815 | 0 | NA | NA | NA | NA | NA | Trav5-2 | T cell receptor alpha variable 5-2 [Source: | 0 | 0 | 0 | 0 | 0 | 0 |
| 42866 | ENSMUSG00000104816 | 0 | NA | NA | NA | NA | NA | Gm43473 | predicted gene 43473 [Source:MGI Symbo | 0 | 0 | 0 | 0 | 0 | 0 |
| 42867 | ENSMUSG00000104817 | 0 | NA | NA | NA | NA | NA | NA | NA | 0 | 0 | 0 | 0 | 0 | 0 |
| 42868 | ENSMUSG00000104818 | 0.185640647 | 1.126924175 | 4.080472857 | 0.276174898 | 0.782413736 | NA | Gm43661 | predicted gene 43661 [Source:MGI Symbo | 0 | 0 | 0 | 0 | 0 | 1.1138 |
| 42869 | ENSMUSG00000104819 | 0 | NA | NA | NA | NA | NA | 4930515G16 | RIKEN cDNA 4930515G16 gene [Source:M | 0 | 0 | 0 | 0 | 0 | 0 |
| 42870 | ENSMUSG00000104820 | 0 | NA | NA | NA | NA | NA | NA | NA | 0 | 0 | 0 | 0 | 0 | 0 |
| 42871 | ENSMUSG00000104821 | 0 | NA | NA | NA | NA | NA | Gm42481 | predicted gene 42481 [Source:MGI Symbo | 0 | 0 | 0 | 0 | 0 | 0 |
| 42872 | ENSMUSG00000104822 | 2.324849805 | -1.212424709 | 1.425186761 | -0.850712863 | 0.394928876 | NA | Gm42967 | predicted gene 42967 [Source:MGI Symbo | 5.6217 | 2.0558 | 2.0125 | 0.964 | 2.1812 | 1.1138 |
| 42873 | ENSMUSG00000104823 | 0.305170013 | -1.657836477 | 4.041480335 | -0.410205257 | 0.681655385 | NA | Gm43182 | predicted gene 43182 [Source:MGI Symbo | 0.8031 | 1.0279 | 0 | 0 | 0 | 0 |
| 42874 | ENSMUSG00000104824 | 0 | NA | NA | NA | NA | NA | Gm21663 | predicted gene, 21663 [Source:MGI Symb | 0 | 0 | 0 | 0 | 0 | 0 |
| 42875 | ENSMUSG00000104825 | 0 | NA | NA | NA | NA | NA | Gm40330 | predicted gene, 40330 [Source:MGI Symb | 0 | 0 | 0 | 0 | 0 | 0 |
| 42876 | ENSMUSG00000104826 | 0 | NA | NA | NA | NA | NA | Gm42947 | predicted gene 42947 [Source:MGI Symbo | 0 | 0 | 0 | 0 | 0 | 0 |
| 42877 | ENSMUSG00000104827 | 0 | NA | NA | NA | NA | NA | NA | NA | 0 | 0 | 0 | 0 | 0 | 0 |
| 42878 | ENSMUSG00000104828 | 0 | NA | NA | NA | NA | NA | Gm42898 | predicted gene 42898 [Source:MGI Symbo | 0 | 0 | 0 | 0 | 0 | 0 |
| 42879 | ENSMUSG00000104829 | 0 | NA | NA | NA | NA | NA | Gm43685 | predicted gene 43685 [Source:MGI Symbo | 0 | 0 | 0 | 0 | 0 | 0 |
| 42880 | ENSMUSG00000104830 | 2.471140166 | 0.766218234 | 1.321822379 | 0.579668075 | 0.562138476 | NA | 5830487J09 | RIKEN cDNA 5830487J09 gene [Source:M | 2.4093 | 2.0558 | 1.0062 | 3.8559 | 3.2719 | 2.2277 |
| 42881 | ENSMUSG00000104831 | 0 | NA | NA | NA | NA | NA | B020017C02 | RIKEN cDNA B020017C02 gene [Source:M | 0 | 0 | 0 | 0 | 0 | 0 |
| 42882 | ENSMUSG00000104832 | 0 | NA | NA | NA | NA | NA | Igkv15-102 | immunoglobulin kappa chain variable 15- | 0 | 0 | 0 | 0 | 0 | 0 |
| 42883 | ENSMUSG00000104833 | 0 | NA | NA | NA | NA | NA | Gm42489 | predicted gene 42489 [Source:MGI Symbo | 0 | 0 | 0 | 0 | 0 | 0 |
| 42884 | ENSMUSG00000104834 | 0 | NA | NA | NA | NA | NA | Gm8861 | predicted gene 8861 [Source:MGI Symbol | 0 | 0 | 0 | 0 | 0 | 0 |
| 42885 | ENSMUSG00000104835 | 0.517623261 | 1.044040381 | 3.075707576 | 0.339447218 | 0.734272853 | NA | Gm5547 | predicted gene 5547 [Source:MGI Symbol | 0 | 1.0279 | 0 | 0.964 | 0 | 1.1138 |
| 42886 | ENSMUSG00000104836 | 0 | NA | NA | NA | NA | NA | NA | NA | 0 | 0 | 0 | 0 | 0 | 0 |
| 42887 | ENSMUSG00000104837 | 0 | NA | NA | NA | NA | NA | Gm42520 | predicted gene 42520 [Source:MGI Symbo | 0 | 0 | 0 | 0 | 0 | 0 |
| 42888 | ENSMUSG00000104838 | 0 | NA | NA | NA | NA | NA | Gm43710 | predicted gene 43710 [Source:MGI Symbo | 0 | 0 | 0 | 0 | 0 | 0 |
| 42889 | ENSMUSG00000104839 | 0 | NA | NA | NA | NA | NA | Gm42794 | predicted gene 42794 [Source:MGI Symbo | 0 | 0 | 0 | 0 | 0 | 0 |
| 42890 | ENSMUSG00000104840 | 0 | NA | NA | NA | NA | NA | Mir7230 | microRNA 7230 [Source:MGI Symbol;Acc: | 0 | 0 | 0 | 0 | 0 | 0 |
| 42891 | ENSMUSG00000104841 | 0 | NA | NA | NA | NA | NA | Gm17743 | predicted gene, 17743 [Source:MGI Symb | 0 | 0 | 0 | 0 | 0 | 0 |
| 42892 | ENSMUSG00000104842 | 0 | NA | NA | NA | NA | NA | Gm42821 | predicted gene 42821 [Source:MGI Symbo | 0 | 0 | 0 | 0 | 0 | 0 |

| | | | | | | | | | | | | | | | |
|---|---|---|---|---|---|---|---|---|---|---|---|---|---|---|---|
| 42893 | ENSMUSG00000104843 | 0 | NA | NA | NA | NA | NA | Gm5308 | predicted gene 5308 [Source:MGI Symbol | 0 | 0 | 0 | 0 | 0 | 0 |
| 42894 | ENSMUSG00000104844 | 0 | NA | NA | NA | NA | NA | NA | NA | 0 | 0 | 0 | 0 | 0 | 0 |
| 42895 | ENSMUSG00000104845 | 0 | NA | NA | NA | NA | NA | 1700041I07F | RIKEN cDNA 1700041I07 gene [Source:MC | 0 | 0 | 0 | 0 | 0 | 0 |
| 42896 | ENSMUSG00000104846 | 0.614004368 | 0.315622926 | 2.898299649 | 0.108899342 | 0.913282328 | NA | Gm43020 | predicted gene 43020 [Source:MGI Symbo | 1.6062 | 0 | 0 | 0.964 | 0 | 1.1138 |
| 42897 | ENSMUSG00000104847 | 0 | NA | NA | NA | NA | NA | Gm43177 | predicted gene 43177 [Source:MGI Symbo | 0 | 0 | 0 | 0 | 0 | 0 |
| 42898 | ENSMUSG00000104848 | 0 | NA | NA | NA | NA | NA | NA | NA | 0 | 0 | 0 | 0 | 0 | 0 |
| 42899 | ENSMUSG00000104849 | 0 | NA | NA | NA | NA | NA | Gm42924 | predicted gene 42924 [Source:MGI Symbo | 0 | 0 | 0 | 0 | 0 | 0 |
| 42900 | ENSMUSG00000104850 | 0 | NA | NA | NA | NA | NA | Gm42901 | predicted gene 42901 [Source:MGI Symbo | 0 | 0 | 0 | 0 | 0 | 0 |
| 42901 | ENSMUSG00000104851 | 0 | NA | NA | NA | NA | NA | E030026E10 | RIKEN cDNA E030026E10 gene [Source:M | 0 | 0 | 0 | 0 | 0 | 0 |
| 42902 | ENSMUSG00000104852 | 1.331414838 | 0.759462549 | 2.203025546 | 0.344736152 | 0.730292737 | NA | Gm43201 | predicted gene 43201 [Source:MGI Symbo | 0 | 0 | 3.0187 | 3.8559 | 0 | 1.1138 |
| 42903 | ENSMUSG00000104853 | 0 | NA | NA | NA | NA | NA | Olfr375-ps1 | olfactory receptor 375, pseudogene 1 [So | 0 | 0 | 0 | 0 | 0 | 0 |
| 42904 | ENSMUSG00000104854 | 0 | NA | NA | NA | NA | NA | Mir7116 | microRNA 7116 [Source:MGI Symbol;Acc: | 0 | 0 | 0 | 0 | 0 | 0 |
| 42905 | ENSMUSG00000104855 | 0 | NA | NA | NA | NA | NA | Gm43709 | predicted gene 43709 [Source:MGI Symbo | 0 | 0 | 0 | 0 | 0 | 0 |
| 42906 | ENSMUSG00000104856 | 0 | NA | NA | NA | NA | NA | NA | NA | 0 | 0 | 0 | 0 | 0 | 0 |
| 42907 | ENSMUSG00000104857 | 0 | NA | NA | NA | NA | NA | NA | NA | 0 | 0 | 0 | 0 | 0 | 0 |
| 42908 | ENSMUSG00000104858 | 0 | NA | NA | NA | NA | NA | Mir136 | microRNA 136 [Source:MGI Symbol;Acc:M | 0 | 0 | 0 | 0 | 0 | 0 |
| 42909 | ENSMUSG00000104859 | 0 | NA | NA | NA | NA | NA | A430072C10 | RIKEN cDNA A430072C10 gene [Source:M | 0 | 0 | 0 | 0 | 0 | 0 |
| 42910 | ENSMUSG00000104860 | 0 | NA | NA | NA | NA | NA | Gm42510 | predicted gene 42510 [Source:MGI Symbo | 0 | 0 | 0 | 0 | 0 | 0 |
| 42911 | ENSMUSG00000104861 | 0 | NA | NA | NA | NA | NA | 3110039M2( | RIKEN cDNA 3110039M20 gene [Source:M | 0 | 0 | 0 | 0 | 0 | 0 |
| 42912 | ENSMUSG00000104862 | 0.556921942 | 2.679043159 | 3.638243063 | 0.736356289 | 0.461513899 | NA | Gm6520 | predicted gene 6520 [Source:MGI Symbol | 0 | 0 | 0 | 0 | 0 | 3.3415 |
| 42913 | ENSMUSG00000104863 | 0 | NA | NA | NA | NA | NA | NA | NA | 0 | 0 | 0 | 0 | 0 | 0 |
| 42914 | ENSMUSG00000104864 | 0.17131964 | -0.796629176 | 4.080472857 | -0.195229623 | 0.84521319 | NA | Gm43655 | predicted gene 43655 [Source:MGI Symbo | 0 | 1.0279 | 0 | 0 | 0 | 0 |
| 42915 | ENSMUSG00000104865 | 0 | NA | NA | NA | NA | NA | Gm42708 | predicted gene 42708 [Source:MGI Symbo | 0 | 0 | 0 | 0 | 0 | 0 |
| 42916 | ENSMUSG00000104866 | 0 | NA | NA | NA | NA | NA | Gm42948 | predicted gene 42948 [Source:MGI Symbo | 0 | 0 | 0 | 0 | 0 | 0 |
| 42917 | ENSMUSG00000104867 | 0 | NA | NA | NA | NA | NA | Gm43728 | predicted gene 43728 [Source:MGI Symbo | 0 | 0 | 0 | 0 | 0 | 0 |
| 42918 | ENSMUSG00000104868 | 0 | NA | NA | NA | NA | NA | Gm9954 | predicted gene 9954 [Source:MGI Symbol | 0 | 0 | 0 | 0 | 0 | 0 |
| 42919 | ENSMUSG00000104869 | 0.339027071 | -1.79174149 | 3.919585585 | -0.457125237 | 0.647581032 | NA | Gm42544 | predicted gene 42544 [Source:MGI Symbo | 0 | 1.0279 | 1.0062 | 0 | 0 | 0 |
| 42920 | ENSMUSG00000104870 | 0 | NA | NA | NA | NA | NA | 4930448I18F | RIKEN cDNA 4930448I18 gene [Source:MC | 0 | 0 | 0 | 0 | 0 | 0 |
| 42921 | ENSMUSG00000104871 | 0 | NA | NA | NA | NA | NA | Gm42639 | predicted gene 42639 [Source:MGI Symbo | 0 | 0 | 0 | 0 | 0 | 0 |
| 42922 | ENSMUSG00000104872 | 3.095446792 | -0.352184691 | 1.357305951 | -0.259473327 | 0.79527006 | NA | 1700008B11 | RIKEN cDNA 1700008B11 gene [Source:M | 2.4093 | 1.0279 | 7.0437 | 4.8199 | 3.2719 | 0 |
| 42923 | ENSMUSG00000104873 | 1.365214248 | -0.551628455 | 2.008513699 | -0.274645104 | 0.78358891 | NA | Gm40332 | predicted gene, 40332 [Source:MGI Symb | 0.8031 | 1.0279 | 3.0187 | 0 | 0 | 3.3415 |
| 42924 | ENSMUSG00000104874 | 0.496077836 | -0.812642266 | 3.444120572 | -0.235950586 | 0.813471014 | NA | Gm43387 | predicted gene 43387 [Source:MGI Symbo | 0 | 0 | 2.0125 | 0.964 | 0 | 0 |
| 42925 | ENSMUSG00000104875 | 0 | NA | NA | NA | NA | NA | Gm43397 | predicted gene 43397 [Source:MGI Symbo | 0 | 0 | 0 | 0 | 0 | 0 |
| 42926 | ENSMUSG00000104876 | 0.816880823 | 0.95463775 | 2.610228283 | 0.365729602 | 0.714566855 | NA | Trdc | T cell receptor delta, constant region [Sou | 1.6062 | 0 | 0 | 0 | 2.1812 | 1.1138 |
| 42927 | ENSMUSG00000104877 | 0.637152627 | -1.330052897 | 2.84179579 | -0.468032538 | 0.639761321 | NA | Gm42651 | predicted gene 42651 [Source:MGI Symbo | 0.8031 | 2.0558 | 0 | 0.964 | 0 | 0 |
| 42928 | ENSMUSG00000104878 | 0 | NA | NA | NA | NA | NA | Gm43567 | predicted gene 43567 [Source:MGI Symbo | 0 | 0 | 0 | 0 | 0 | 0 |
| 42929 | ENSMUSG00000104879 | 0 | NA | NA | NA | NA | NA | NA | NA | 0 | 0 | 0 | 0 | 0 | 0 |
| 42930 | ENSMUSG00000104880 | 0 | NA | NA | NA | NA | NA | NA | NA | 0 | 0 | 0 | 0 | 0 | 0 |
| 42931 | ENSMUSG00000104881 | 0 | NA | NA | NA | NA | NA | Gm43711 | predicted gene 43711 [Source:MGI Symbo | 0 | 0 | 0 | 0 | 0 | 0 |
| 42932 | ENSMUSG00000104882 | 0.353348078 | 0.165142701 | 3.895042568 | 0.042398176 | 0.966181282 | NA | Gm43096 | predicted gene 43096 [Source:MGI Symbo | 0 | 0 | 1.0062 | 0 | 0 | 1.1138 |
| 42933 | ENSMUSG00000104883 | 0 | NA | NA | NA | NA | NA | Igkv14-134-1 | immunoglobulin kappa chain variable 14-1 | 0 | 0 | 0 | 0 | 0 | 0 |
| 42934 | ENSMUSG00000104884 | 0 | NA | NA | NA | NA | NA | Mir5046 | microRNA 5046 [Source:MGI Symbol;Acc: | 0 | 0 | 0 | 0 | 0 | 0 |
| 42935 | ENSMUSG00000104885 | 1.097615586 | -2.336177288 | 2.644223446 | -0.883502221 | 0.376964983 | NA | Gm43758 | predicted gene 43758 [Source:MGI Symbo | 5.6217 | 0 | 0 | 0.964 | 0 | 0 |
| 42936 | ENSMUSG00000104886 | 0 | NA | NA | NA | NA | NA | Gm43000 | predicted gene 43000 [Source:MGI Symbo | 0 | 0 | 0 | 0 | 0 | 0 |
| 42937 | ENSMUSG00000104887 | 0 | NA | NA | NA | NA | NA | Gm43158 | predicted gene 43158 [Source:MGI Symbo | 0 | 0 | 0 | 0 | 0 | 0 |
| 42938 | ENSMUSG00000104888 | 0 | NA | NA | NA | NA | NA | 1500005C15 | RIKEN cDNA 1500005C15 gene [Source:M | 0 | 0 | 0 | 0 | 0 | 0 |
| 42939 | ENSMUSG00000104889 | 0 | NA | NA | NA | NA | NA | Gm43043 | predicted gene 43043 [Source:MGI Symbo | 0 | 0 | 0 | 0 | 0 | 0 |
| 42940 | ENSMUSG00000104890 | 0 | NA | NA | NA | NA | NA | NA | NA | 0 | 0 | 0 | 0 | 0 | 0 |
| 42941 | ENSMUSG00000104891 | 5.208580748 | 0.252801824 | 1.010836446 | 0.250091719 | 0.80251642 | 0.9632861 | 2510017J16F | RIKEN cDNA 2510017J16 gene [Source:MC | 8.031 | 2.0558 | 4.025 | 2.8919 | 10.906 | 3.3415 |
| 42942 | ENSMUSG00000104892 | 0 | NA | NA | NA | NA | NA | Gm6615 | predicted gene 6615 [Source:MGI Symbol | 0 | 0 | 0 | 0 | 0 | 0 |
| 42943 | ENSMUSG00000104893 | 0 | NA | NA | NA | NA | NA | Gm2404 | predicted gene 2404 [Source:MGI Symbol | 0 | 0 | 0 | 0 | 0 | 0 |
| 42944 | ENSMUSG00000104894 | 0 | NA | NA | NA | NA | NA | Gm43507 | predicted gene 43507 [Source:MGI Symbo | 0 | 0 | 0 | 0 | 0 | 0 |
| 42945 | ENSMUSG00000104895 | 0.167707431 | -0.796629176 | 4.080472857 | -0.195229623 | 0.84521319 | NA | Gm8454 | predicted gene 8454 [Source:MGI Symbol | 0 | 0 | 1.0062 | 0 | 0 | 0 |
| 42946 | ENSMUSG00000104896 | 0 | NA | NA | NA | NA | NA | NA | NA | 0 | 0 | 0 | 0 | 0 | 0 |
| 42947 | ENSMUSG00000104897 | 2.33755403 | 0.856981405 | 1.645281228 | 0.520872293 | 0.602455738 | NA | Gm43203 | predicted gene 43203 [Source:MGI Symbo | 0 | 3.0838 | 2.0125 | 6.7478 | 2.1812 | 0 |
| 42948 | ENSMUSG00000104898 | 0 | NA | NA | NA | NA | NA | Gm30043 | predicted gene, 30043 [Source:MGI Symb | 0 | 0 | 0 | 0 | 0 | 0 |

| | | | | | | | | | | | | | | | |
|---|---|---|---|---|---|---|---|---|---|---|---|---|---|---|---|
| 42949 | ENSMUSG00000104899 | 0 | NA | NA | NA | NA | NA | Gm43855 | predicted gene 43855 [Source:MGI Symbo | 0 | 0 | 0 | 0 | 0 | 0 |
| 42950 | ENSMUSG00000104900 | 0.97984404 | 0.122644358 | 2.090083284 | 0.058679173 | 0.953207648 | NA | 4930596I21F | RIKEN cDNA 4930596I21 gene [Source:MC | 0.8031 | 1.0279 | 1.0062 | 1.928 | 0 | 1.1138 |
| 42951 | ENSMUSG00000104901 | 0 | NA | NA | NA | NA | NA | Gm42692 | predicted gene 42692 [Source:MGI Symbo | 0 | 0 | 0 | 0 | 0 | 0 |
| 42952 | ENSMUSG00000104902 | 0.315620088 | 0.165143421 | 4.063300985 | 0.040642675 | 0.967580763 | NA | Gm30517 | predicted gene, 30517 [Source:MGI Symb | 0.8031 | 0 | 0 | 0 | 1.0906 | 0 |
| 42953 | ENSMUSG00000104903 | 0 | NA | NA | NA | NA | NA | Gm43707 | predicted gene 43707 [Source:MGI Symbo | 0 | 0 | 0 | 0 | 0 | 0 |
| 42954 | ENSMUSG00000104904 | 0 | NA | NA | NA | NA | NA | 9330198I05F | RIKEN cDNA 9330198I05 gene [Source:MC | 0 | 0 | 0 | 0 | 0 | 0 |
| 42955 | ENSMUSG00000104905 | 0 | NA | NA | NA | NA | NA | Gm43718 | predicted gene 43718 [Source:MGI Symbo | 0 | 0 | 0 | 0 | 0 | 0 |
| 42956 | ENSMUSG00000104906 | 0 | NA | NA | NA | NA | NA | Gm43005 | predicted gene 43005 [Source:MGI Symbo | 0 | 0 | 0 | 0 | 0 | 0 |
| 42957 | ENSMUSG00000104908 | 0 | NA | NA | NA | NA | NA | 1700015C17 | RIKEN cDNA 1700015C17 gene [Source:M | 0 | 0 | 0 | 0 | 0 | 0 |
| 42958 | ENSMUSG00000104909 | 0 | NA | NA | NA | NA | NA | Gm43617 | predicted gene 43617 [Source:MGI Symbo | 0 | 0 | 0 | 0 | 0 | 0 |
| 42959 | ENSMUSG00000104910 | 0.339027071 | -1.79174149 | 3.919585585 | -0.457125237 | 0.647581032 | NA | Gm43331 | predicted gene 43331 [Source:MGI Symbo | 0 | 1.0279 | 1.0062 | 0 | 0 | 0 |
| 42960 | ENSMUSG00000104911 | 0 | NA | NA | NA | NA | NA | NA | NA | 0 | 0 | 0 | 0 | 0 | 0 |
| 42961 | ENSMUSG00000104912 | 0 | NA | NA | NA | NA | NA | Gm43023 | predicted gene 43023 [Source:MGI Symbo | 0 | 0 | 0 | 0 | 0 | 0 |
| 42962 | ENSMUSG00000104913 | 0 | NA | NA | NA | NA | NA | Gm6560 | predicted gene 6560 [Source:MGI Symbol | 0 | 0 | 0 | 0 | 0 | 0 |
| 42963 | ENSMUSG00000104914 | 0.692116425 | -1.412606127 | 2.773087916 | -0.509398248 | 0.610473104 | NA | Mir7061 | microRNA 7061 [Source:MGI Symbol;Acc: | 0 | 2.0558 | 1.0062 | 0 | 1.0906 | 0 |
| 42964 | ENSMUSG00000104915 | 0 | NA | NA | NA | NA | NA | Trav15n-2 | T cell receptor alpha variable 15N-2 [Sour | 0 | 0 | 0 | 0 | 0 | 0 |
| 42965 | ENSMUSG00000104916 | 0.335414862 | -1.779047242 | 4.030804179 | -0.441362855 | 0.658950329 | NA | Gm42830 | predicted gene 42830 [Source:MGI Symbo | 0 | 0 | 2.0125 | 0 | 0 | 0 |
| 42966 | ENSMUSG00000104917 | 0.877834182 | -0.676932822 | 2.563285057 | -0.264087999 | 0.791712111 | NA | Gm43289 | predicted gene 43289 [Source:MGI Symbo | 3.2124 | 0 | 0 | 0.964 | 1.0906 | 0 |
| 42967 | ENSMUSG00000104918 | 27.01544738 | 0.773952224 | 0.453100924 | 1.708123252 | 0.087613475 | 0.43652818 | Gm42944 | predicted gene 42944 [Source:MGI Symbo | 19.274 | 23.642 | 17.106 | 49.163 | 26.175 | 26.732 |
| 42968 | ENSMUSG00000104919 | 0 | NA | NA | NA | NA | NA | Gm42617 | predicted gene 42617 [Source:MGI Symbo | 0 | 0 | 0 | 0 | 0 | 0 |
| 42969 | ENSMUSG00000104920 | 1.341839409 | -2.630531392 | 2.253457836 | -1.16733109 | 0.243076688 | NA | Gm43215 | predicted gene 43215 [Source:MGI Symbo | 0 | 2.0558 | 5.0312 | 0.964 | 0 | 0 |
| 42970 | ENSMUSG00000104921 | 0 | NA | NA | NA | NA | NA | Snord3b-ps1 | small nucleolar RNA, C/D box 3B, pseudog | 0 | 0 | 0 | 0 | 0 | 0 |
| 42971 | ENSMUSG00000104922 | 0 | NA | NA | NA | NA | NA | Igkv13-74-1 | immunoglobulin kappa chain variable 13-7 | 0 | 0 | 0 | 0 | 0 | 0 |
| 42972 | ENSMUSG00000104923 | 0 | NA | NA | NA | NA | NA | Gm5865 | predicted gene 5865 [Source:MGI Symbol | 0 | 0 | 0 | 0 | 0 | 0 |
| 42973 | ENSMUSG00000104924 | 0 | NA | NA | NA | NA | NA | Gm43002 | predicted gene 43002 [Source:MGI Symbo | 0 | 0 | 0 | 0 | 0 | 0 |
| 42974 | ENSMUSG00000104925 | 1.206386202 | -0.322987335 | 2.094535446 | -0.15420476 | 0.877448289 | NA | Gm43061 | predicted gene 43061 [Source:MGI Symbo | 0 | 1.0279 | 3.0187 | 0.964 | 0 | 2.2277 |
| 42975 | ENSMUSG00000104926 | 0.979791906 | -2.106318065 | 2.490808728 | -0.845636215 | 0.397755716 | NA | Mir6947 | microRNA 6947 [Source:MGI Symbol;Acc: | 0.8031 | 4.1117 | 0 | 0.964 | 0 | 0 |
| 42976 | ENSMUSG00000104927 | 0.401551119 | -2.072982726 | 4.008251979 | -0.517178744 | 0.605031388 | NA | Gm43388 | predicted gene 43388 [Source:MGI Symbo | 2.4093 | 0 | 0 | 0 | 0 | 0 |
| 42977 | ENSMUSG00000104928 | 0 | NA | NA | NA | NA | NA | Gm42565 | predicted gene 42565 [Source:MGI Symbo | 0 | 0 | 0 | 0 | 0 | 0 |
| 42978 | ENSMUSG00000104929 | 0 | NA | NA | NA | NA | NA | Gm42766 | predicted gene 42766 [Source:MGI Symbo | 0 | 0 | 0 | 0 | 0 | 0 |
| 42979 | ENSMUSG00000104930 | 0 | NA | NA | NA | NA | NA | Gm43243 | predicted gene 43243 [Source:MGI Symbo | 0 | 0 | 0 | 0 | 0 | 0 |
| 42980 | ENSMUSG00000104931 | 0 | NA | NA | NA | NA | NA | Igkv1-136 | immunoglobulin kappa chain variable 1-13 | 0 | 0 | 0 | 0 | 0 | 0 |
| 42981 | ENSMUSG00000104932 | 0 | NA | NA | NA | NA | NA | Gm40323 | predicted gene, 40323 [Source:MGI Symb | 0 | 0 | 0 | 0 | 0 | 0 |
| 42982 | ENSMUSG00000104933 | 0 | NA | NA | NA | NA | NA | NA | NA | 0 | 0 | 0 | 0 | 0 | 0 |
| 42983 | ENSMUSG00000104934 | 0 | NA | NA | NA | NA | NA | Igkv11-118 | immunoglobulin kappa chain variable 11-1 | 0 | 0 | 0 | 0 | 0 | 0 |
| 42984 | ENSMUSG00000104935 | 0 | NA | NA | NA | NA | NA | Gm43414 | predicted gene 43414 [Source:MGI Symbo | 0 | 0 | 0 | 0 | 0 | 0 |
| 42985 | ENSMUSG00000104936 | 0.17131964 | -0.796629176 | 4.080472857 | -0.195229623 | 0.84521319 | NA | 2610011E03 | RIKEN cDNA 2610011E03 gene [Source:M | 0 | 1.0279 | 0 | 0 | 0 | 0 |
| 42986 | ENSMUSG00000104937 | 0 | NA | NA | NA | NA | NA | Gm43057 | predicted gene 43057 [Source:MGI Symbo | 0 | 0 | 0 | 0 | 0 | 0 |
| 42987 | ENSMUSG00000104938 | 0 | NA | NA | NA | NA | NA | Gm42840 | predicted gene 42840 [Source:MGI Symbo | 0 | 0 | 0 | 0 | 0 | 0 |
| 42988 | ENSMUSG00000104939 | 0 | NA | NA | NA | NA | NA | Gm33651 | predicted gene, 33651 [Source:MGI Symb | 0 | 0 | 0 | 0 | 0 | 0 |
| 42989 | ENSMUSG00000104940 | 0 | NA | NA | NA | NA | NA | Gm42908 | predicted gene 42908 [Source:MGI Symbo | 0 | 0 | 0 | 0 | 0 | 0 |
| 42990 | ENSMUSG00000104941 | 0 | NA | NA | NA | NA | NA | Gm8953 | predicted gene 8953 [Source:MGI Symbol | 0 | 0 | 0 | 0 | 0 | 0 |
| 42991 | ENSMUSG00000104942 | 0 | NA | NA | NA | NA | NA | Gm3321 | predicted gene 3321 [Source:MGI Symbol | 0 | 0 | 0 | 0 | 0 | 0 |
| 42992 | ENSMUSG00000104943 | 4.169241327 | 1.518260037 | 1.101158898 | 1.378783788 | 0.167961427 | NA | Gm42868 | predicted gene 42868 [Source:MGI Symbo | 2.4093 | 2.0558 | 2.0125 | 8.6758 | 7.6343 | 2.2277 |
| 42993 | ENSMUSG00000104944 | 0 | NA | NA | NA | NA | NA | Gm43405 | predicted gene 43405 [Source:MGI Symbo | 0 | 0 | 0 | 0 | 0 | 0 |
| 42994 | ENSMUSG00000104945 | 0.606727817 | -2.651277273 | 2.811524265 | -0.943003518 | 0.345679103 | NA | Gm35715 | predicted gene, 35715 [Source:MGI Symb | 1.6062 | 1.0279 | 1.0062 | 0 | 0 | 0 |
| 42995 | ENSMUSG00000104946 | 0.160662974 | 1.126924175 | 4.080472857 | 0.276174898 | 0.782413736 | NA | Gm19104 | predicted gene, 19104 [Source:MGI Symb | 0 | 0 | 0 | 0.964 | 0 | 0 |
| 42996 | ENSMUSG00000104947 | 0 | NA | NA | NA | NA | NA | Gm43721 | predicted gene 43721 [Source:MGI Symbo | 0 | 0 | 0 | 0 | 0 | 0 |
| 42997 | ENSMUSG00000104948 | 0 | NA | NA | NA | NA | NA | NA | NA | 0 | 0 | 0 | 0 | 0 | 0 |
| 42998 | ENSMUSG00000104949 | 0 | NA | NA | NA | NA | NA | Hcfc1r1-ps2 | host cell factor C1 regulator 1 (XPO1-depe | 0 | 0 | 0 | 0 | 0 | 0 |
| 42999 | ENSMUSG00000104950 | 42.70371086 | -0.316476908 | 0.375519313 | -0.842771324 | 0.399356349 | 0.8263661 | 4833413G10 | RIKEN cDNA 4833413G10 gene [Source:M | 37.746 | 49.34 | 55.343 | 42.415 | 47.987 | 23.391 |
| 43000 | ENSMUSG00000104951 | 3.276336057 | 0.088060374 | 1.229911659 | 0.071598943 | 0.942921082 | NA | Gm43413 | predicted gene 43413 [Source:MGI Symbo | 6.4248 | 1.0279 | 2.0125 | 5.7839 | 2.1812 | 2.2277 |
| 43001 | ENSMUSG00000104952 | 0 | NA | NA | NA | NA | NA | Gm38697 | predicted gene, 38697 [Source:MGI Symb | 0 | 0 | 0 | 0 | 0 | 0 |
| 43002 | ENSMUSG00000104953 | 0 | NA | NA | NA | NA | NA | NA | NA | 0 | 0 | 0 | 0 | 0 | 0 |
| 43003 | ENSMUSG00000104954 | 0 | NA | NA | NA | NA | NA | Gm42694 | predicted gene 42694 [Source:MGI Symbo | 0 | 0 | 0 | 0 | 0 | 0 |
| 43004 | ENSMUSG00000104955 | 0 | NA | NA | NA | NA | NA | 1700016F12 | RIKEN cDNA 1700016F12 gene [Source:M | 0 | 0 | 0 | 0 | 0 | 0 |

| | | | | | | | | | | | | | | | |
|---|---|---|---|---|---|---|---|---|---|---|---|---|---|---|---|
| 43005 | ENSMUSG00000104956 | 0 | NA | NA | NA | NA | NA | 4930429D17 | RIKEN cDNA 4930429D17 gene [Source:M | 0 | 0 | 0 | 0 | 0 | 0 |
| 43006 | ENSMUSG00000104957 | 0 | NA | NA | NA | NA | NA | Gm43354 | predicted gene 43354 [Source:MGI Symbo | 0 | 0 | 0 | 0 | 0 | 0 |
| 43007 | ENSMUSG00000104958 | 0 | NA | NA | NA | NA | NA | Mir7238 | microRNA 7238 [Source:MGI Symbol;Acc: | 0 | 0 | 0 | 0 | 0 | 0 |
| 43008 | ENSMUSG00000104959 | 0 | NA | NA | NA | NA | NA | Gm18131 | predicted gene, 18131 [Source:MGI Symb | 0 | 0 | 0 | 0 | 0 | 0 |
| 43009 | ENSMUSG00000104960 | 244.7487554 | -0.065623875 | 0.168535444 | -0.389377294 | 0.696997066 | 0.94023175 | Snhg8 | small nucleolar RNA host gene 8 [Source:I | 259.4 | 271.37 | 220.37 | 261.24 | 235.57 | 220.54 |
| 43010 | ENSMUSG00000104961 | 0 | NA | NA | NA | NA | NA | Gm43835 | predicted gene 43835 [Source:MGI Symbo | 0 | 0 | 0 | 0 | 0 | 0 |
| 43011 | ENSMUSG00000104962 | 0 | NA | NA | NA | NA | NA | Trgj1 | T cell receptor gamma joining 1 [Source:M | 0 | 0 | 0 | 0 | 0 | 0 |
| 43012 | ENSMUSG00000104963 | 0 | NA | NA | NA | NA | NA | Gm42974 | predicted gene 42974 [Source:MGI Symbo | 0 | 0 | 0 | 0 | 0 | 0 |
| 43013 | ENSMUSG00000104964 | 0 | NA | NA | NA | NA | NA | 4930597L12I | RIKEN cDNA 4930597L12 gene [Source:M( | 0 | 0 | 0 | 0 | 0 | 0 |
| 43014 | ENSMUSG00000104965 | 15.97504662 | -0.566104289 | 0.579045272 | -0.977651172 | 0.328246879 | 0.773161429 | Gm43437 | predicted gene 43437 [Source:MGI Symbo | 17.668 | 18.503 | 21.131 | 16.388 | 5.4531 | 16.708 |
| 43015 | ENSMUSG00000104966 | 0 | NA | NA | NA | NA | NA | Gm43273 | predicted gene 43273 [Source:MGI Symbo | 0 | 0 | 0 | 0 | 0 | 0 |
| 43016 | ENSMUSG00000104967 | 0 | NA | NA | NA | NA | NA | Gm43436 | predicted gene 43436 [Source:MGI Symbo | 0 | 0 | 0 | 0 | 0 | 0 |
| 43017 | ENSMUSG00000104968 | 0 | NA | NA | NA | NA | NA | Gm43803 | predicted gene 43803 [Source:MGI Symbo | 0 | 0 | 0 | 0 | 0 | 0 |
| 43018 | ENSMUSG00000104969 | 1.440467498 | 1.046009396 | 2.168701849 | 0.482320516 | 0.629578275 | NA | Gm43445 | predicted gene 43445 [Source:MGI Symbo | 0.8031 | 2.0558 | 0 | 5.7839 | 0 | 0 |
| 43019 | ENSMUSG00000104970 | 1.029799386 | 0.219173677 | 2.229624068 | 0.098300731 | 0.921693497 | NA | Gm43137 | predicted gene 43137 [Source:MGI Symbo | 0.8031 | 1.0279 | 1.0062 | 0 | 0 | 3.3415 |
| 43020 | ENSMUSG00000104971 | 0 | NA | NA | NA | NA | NA | 9430087B13 | RIKEN cDNA 9430087B13 gene [Source:M | 0 | 0 | 0 | 0 | 0 | 0 |
| 43021 | ENSMUSG00000104972 | 0 | NA | NA | NA | NA | NA | Gm19931 | predicted gene, 19931 [Source:MGI Symb | 0 | 0 | 0 | 0 | 0 | 0 |
| 43022 | ENSMUSG00000104973 | 2.839358844 | -0.095341426 | 1.458908818 | -0.065351189 | 0.947894387 | NA | A530041M0 | RIKEN cDNA A530041M06 gene [Source:M | 0.8031 | 1.0279 | 7.0437 | 4.8199 | 0 | 3.3415 |
| 43023 | ENSMUSG00000104974 | 0 | NA | NA | NA | NA | NA | Gm43686 | predicted gene 43686 [Source:MGI Symbo | 0 | 0 | 0 | 0 | 0 | 0 |
| 43024 | ENSMUSG00000104975 | 0 | NA | NA | NA | NA | NA | Iglj2 | immunoglobulin lambda joining 2 [Source | 0 | 0 | 0 | 0 | 0 | 0 |
| 43025 | ENSMUSG00000104976 | 0 | NA | NA | NA | NA | NA | Gm43060 | predicted gene 43060 [Source:MGI Symbo | 0 | 0 | 0 | 0 | 0 | 0 |
| 43026 | ENSMUSG00000104978 | 2.765996515 | 0.252397142 | 1.480264449 | 0.170508143 | 0.864610531 | NA | Gm31274 | predicted gene, 31274 [Source:MGI Symb | 1.6062 | 0 | 6.0375 | 6.7478 | 1.0906 | 1.1138 |
| 43027 | ENSMUSG00000104979 | 0 | NA | NA | NA | NA | NA | Mir6994 | microRNA 6994 [Source:MGI Symbol;Acc: | 0 | 0 | 0 | 0 | 0 | 0 |
| 43028 | ENSMUSG00000104980 | 0 | NA | NA | NA | NA | NA | 4930573I07F | RIKEN cDNA 4930573I07 gene [Source:MG | 0 | 0 | 0 | 0 | 0 | 0 |
| 43029 | ENSMUSG00000104981 | 0 | NA | NA | NA | NA | NA | Gm43736 | predicted gene 43736 [Source:MGI Symbo | 0 | 0 | 0 | 0 | 0 | 0 |
| 43030 | ENSMUSG00000104982 | 0 | NA | NA | NA | NA | NA | Gm32554 | predicted gene, 32554 [Source:MGI Symb | 0 | 0 | 0 | 0 | 0 | 0 |
| 43031 | ENSMUSG00000104983 | 1.526988112 | 1.86832238 | 1.965789709 | 0.950418232 | 0.341899784 | NA | Gm42763 | predicted gene 42763 [Source:MGI Symbo | 0 | 1.0279 | 1.0062 | 3.8559 | 3.2719 | 0 |
| 43032 | ENSMUSG00000104984 | 0 | NA | NA | NA | NA | NA | Gm5982 | predicted gene 5982 [Source:MGI Symbol | 0 | 0 | 0 | 0 | 0 | 0 |
| 43033 | ENSMUSG00000104985 | 0.133850373 | -0.796629176 | 4.080472857 | -0.195229623 | 0.84521319 | NA | Gm42917 | predicted gene 42917 [Source:MGI Symbo | 0.8031 | 0 | 0 | 0 | 0 | 0 |
| 43034 | ENSMUSG00000104986 | 2.683045478 | 2.853330156 | 1.570433638 | 1.816905909 | 0.069231522 | NA | 1600023N17 | RIKEN cDNA 1600023N17 gene [Source:M | 0 | 1.0279 | 1.0062 | 1.928 | 5.4531 | 6.6831 |
| 43035 | ENSMUSG00000104987 | 0 | NA | NA | NA | NA | NA | 2810432F15 | RIKEN cDNA 2810432F15 gene [Source:M( | 0 | 0 | 0 | 0 | 0 | 0 |
| 43036 | ENSMUSG00000104988 | 0.181769715 | 1.126924175 | 4.080472857 | 0.276174898 | 0.782413736 | NA | Gm43622 | predicted gene 43622 [Source:MGI Symbo | 0 | 0 | 0 | 0 | 1.0906 | 0 |
| 43037 | ENSMUSG00000104989 | 0 | NA | NA | NA | NA | NA | NA | NA | 0 | 0 | 0 | 0 | 0 | 0 |
| 43038 | ENSMUSG00000104990 | 0 | NA | NA | NA | NA | NA | Gm43504 | predicted gene 43504 [Source:MGI Symbo | 0 | 0 | 0 | 0 | 0 | 0 |
| 43039 | ENSMUSG00000104991 | 0 | NA | NA | NA | NA | NA | Gm44468 | predicted gene, 44468 [Source:MGI Symb | 0 | 0 | 0 | 0 | 0 | 0 |
| 43040 | ENSMUSG00000104992 | 0 | NA | NA | NA | NA | NA | Gm43163 | predicted gene 43163 [Source:MGI Symbo | 0 | 0 | 0 | 0 | 0 | 0 |
| 43041 | ENSMUSG00000104993 | 0 | NA | NA | NA | NA | NA | Gm43510 | predicted gene 43510 [Source:MGI Symbo | 0 | 0 | 0 | 0 | 0 | 0 |
| 43042 | ENSMUSG00000104994 | 0 | NA | NA | NA | NA | NA | Gm42989 | predicted gene 42989 [Source:MGI Symbo | 0 | 0 | 0 | 0 | 0 | 0 |
| 43043 | ENSMUSG00000104995 | 0 | NA | NA | NA | NA | NA | Gm42925 | predicted gene 42925 [Source:MGI Symbo | 0 | 0 | 0 | 0 | 0 | 0 |
| 43044 | ENSMUSG00000104996 | 0 | NA | NA | NA | NA | NA | Mir6924 | microRNA 6924 [Source:MGI Symbol;Acc: | 0 | 0 | 0 | 0 | 0 | 0 |
| 43045 | ENSMUSG00000104997 | 0 | NA | NA | NA | NA | NA | NA | NA | 0 | 0 | 0 | 0 | 0 | 0 |
| 43046 | ENSMUSG00000104998 | 0 | NA | NA | NA | NA | NA | Gm43822 | predicted gene 43822 [Source:MGI Symbo | 0 | 0 | 0 | 0 | 0 | 0 |
| 43047 | ENSMUSG00000104999 | 0.342639279 | -1.805976596 | 4.028547714 | -0.448294702 | 0.653940524 | NA | 4930477O15 | RIKEN cDNA 4930477O15 gene [Source:M | 0 | 2.0558 | 0 | 0 | 0 | 0 |
| 43048 | ENSMUSG00000105000 | 0 | NA | NA | NA | NA | NA | Gm43616 | predicted gene 43616 [Source:MGI Symbo | 0 | 0 | 0 | 0 | 0 | 0 |
| 43049 | ENSMUSG00000105001 | 0 | NA | NA | NA | NA | NA | NA | NA | 0 | 0 | 0 | 0 | 0 | 0 |
| 43050 | ENSMUSG00000105002 | 0.160662974 | 1.126924175 | 4.080472857 | 0.276174898 | 0.782413736 | NA | NA | NA | 0 | 0 | 0 | 0.964 | 0 | 0 |
| 43051 | ENSMUSG00000105003 | 0.684892008 | -1.399684857 | 3.073674502 | -0.455378361 | 0.64883706 | NA | Gm40055 | predicted gene, 40055 [Source:MGI Symb | 0 | 0 | 3.0187 | 0 | 1.0906 | 0 |
| 43052 | ENSMUSG00000105004 | 7.913084511 | -0.886670916 | 0.812680623 | -1.091044736 | 0.275253198 | 0.730038567 | Gm43075 | predicted gene 43075 [Source:MGI Symbo | 10.44 | 14.391 | 6.0375 | 6.7478 | 7.6343 | 2.2277 |
| 43053 | ENSMUSG00000105005 | 0 | NA | NA | NA | NA | NA | Gm9729 | predicted gene 9729 [Source:MGI Symbol | 0 | 0 | 0 | 0 | 0 | 0 |
| 43054 | ENSMUSG00000105006 | 1.817015969 | 1.42465994 | 1.870594821 | 0.761607979 | 0.446294012 | NA | Gm9484 | predicted gene 9484 [Source:MGI Symbol | 0 | 1.0279 | 2.0125 | 6.7478 | 0 | 1.1138 |
| 43055 | ENSMUSG00000105007 | 0 | NA | NA | NA | NA | NA | Gm43170 | predicted gene 43170 [Source:MGI Symbo | 0 | 0 | 0 | 0 | 0 | 0 |
| 43056 | ENSMUSG00000105008 | 0 | NA | NA | NA | NA | NA | Gm43652 | predicted gene 43652 [Source:MGI Symbo | 0 | 0 | 0 | 0 | 0 | 0 |
| 43057 | ENSMUSG00000105009 | 0 | NA | NA | NA | NA | NA | Gm42774 | predicted gene 42774 [Source:MGI Symbo | 0 | 0 | 0 | 0 | 0 | 0 |
| 43058 | ENSMUSG00000105010 | 0 | NA | NA | NA | NA | NA | Mir219b | microRNA 219b [Source:MGI Symbol;Acc: | 0 | 0 | 0 | 0 | 0 | 0 |
| 43059 | ENSMUSG00000105011 | 0 | NA | NA | NA | NA | NA | Gm43210 | predicted gene 43210 [Source:MGI Symbo | 0 | 0 | 0 | 0 | 0 | 0 |
| 43060 | ENSMUSG00000105012 | 0 | NA | NA | NA | NA | NA | Gm42813 | predicted gene 42813 [Source:MGI Symbo | 0 | 0 | 0 | 0 | 0 | 0 |

| | | | | | | | | | | | | | | | |
|---|---|---|---|---|---|---|---|---|---|---|---|---|---|---|---|
| 43061 | ENSMUSG00000105013 | 0 | NA | NA | NA | NA | NA | Gm7721 | predicted gene 7721 [Source:MGI Symbol | 0 | 0 | 0 | 0 | 0 | 0 |
| 43062 | ENSMUSG00000105014 | 0 | NA | NA | NA | NA | NA | NA | NA | 0 | 0 | 0 | 0 | 0 | 0 |
| 43063 | ENSMUSG00000105015 | 0 | NA | NA | NA | NA | NA | Gm6059 | predicted gene 6059 [Source:MGI Symbol | 0 | 0 | 0 | 0 | 0 | 0 |
| 43064 | ENSMUSG00000105016 | 0 | NA | NA | NA | NA | NA | Gm42569 | predicted gene 42569 [Source:MGI Symbo | 0 | 0 | 0 | 0 | 0 | 0 |
| 43065 | ENSMUSG00000105017 | 0 | NA | NA | NA | NA | NA | Gm43107 | predicted gene 43107 [Source:MGI Symbo | 0 | 0 | 0 | 0 | 0 | 0 |
| 43066 | ENSMUSG00000105018 | 0 | NA | NA | NA | NA | NA | Gm43546 | predicted gene 43546 [Source:MGI Symbo | 0 | 0 | 0 | 0 | 0 | 0 |
| 43067 | ENSMUSG00000105019 | 0 | NA | NA | NA | NA | NA | Gm43547 | predicted gene 43547 [Source:MGI Symbo | 0 | 0 | 0 | 0 | 0 | 0 |
| 43068 | ENSMUSG00000105020 | 0 | NA | NA | NA | NA | NA | Gm7205 | predicted gene 7205 [Source:MGI Symbol | 0 | 0 | 0 | 0 | 0 | 0 |
| 43069 | ENSMUSG00000105021 | 0 | NA | NA | NA | NA | NA | Gm8234 | predicted gene 8234 [Source:MGI Symbol | 0 | 0 | 0 | 0 | 0 | 0 |
| 43070 | ENSMUSG00000105022 | 0 | NA | NA | NA | NA | NA | NA | NA | 0 | 0 | 0 | 0 | 0 | 0 |
| 43071 | ENSMUSG00000105023 | 0 | NA | NA | NA | NA | NA | 1700012D16 | RIKEN cDNA 1700012D16 gene [Source:M | 0 | 0 | 0 | 0 | 0 | 0 |
| 43072 | ENSMUSG00000105024 | 0 | NA | NA | NA | NA | NA | Gm43561 | predicted gene 43561 [Source:MGI Symbo | 0 | 0 | 0 | 0 | 0 | 0 |
| 43073 | ENSMUSG00000105025 | 0 | NA | NA | NA | NA | NA | NA | NA | 0 | 0 | 0 | 0 | 0 | 0 |
| 43074 | ENSMUSG00000105026 | 0 | NA | NA | NA | NA | NA | Gm18719 | predicted gene, 18719 [Source:MGI Symb | 0 | 0 | 0 | 0 | 0 | 0 |
| 43075 | ENSMUSG00000105027 | 0 | NA | NA | NA | NA | NA | Gm43676 | predicted gene 43676 [Source:MGI Symbo | 0 | 0 | 0 | 0 | 0 | 0 |
| 43076 | ENSMUSG00000105028 | 0 | NA | NA | NA | NA | NA | Gm18747 | predicted gene, 18747 [Source:MGI Symb | 0 | 0 | 0 | 0 | 0 | 0 |
| 43077 | ENSMUSG00000105029 | 0 | NA | NA | NA | NA | NA | NA | NA | 0 | 0 | 0 | 0 | 0 | 0 |
| 43078 | ENSMUSG00000105030 | 0 | NA | NA | NA | NA | NA | Gm38671 | predicted gene, 38671 [Source:MGI Symb | 0 | 0 | 0 | 0 | 0 | 0 |
| 43079 | ENSMUSG00000105031 | 0.455176322 | 0.955156269 | 3.546674842 | 0.269310357 | 0.787690862 | NA | Gm3511 | predicted gene 3511 [Source:MGI Symbol | 0.8031 | 0 | 0 | 1.928 | 0 | 0 |
| 43080 | ENSMUSG00000105032 | 0 | NA | NA | NA | NA | NA | Gm43117 | predicted gene 43117 [Source:MGI Symbo | 0 | 0 | 0 | 0 | 0 | 0 |
| 43081 | ENSMUSG00000105033 | 0.160662974 | 1.126924175 | 4.080472857 | 0.276174898 | 0.782413736 | NA | Gm43423 | predicted gene 43423 [Source:MGI Symbo | 0 | 0 | 0 | 0.964 | 0 | 0 |
| 43082 | ENSMUSG00000105034 | 0 | NA | NA | NA | NA | NA | Gm43348 | predicted gene 43348 [Source:MGI Symbo | 0 | 0 | 0 | 0 | 0 | 0 |
| 43083 | ENSMUSG00000105035 | 0 | NA | NA | NA | NA | NA | Gm9682 | predicted pseudogene 9682 [Source:MGI | 0 | 0 | 0 | 0 | 0 | 0 |
| 43084 | ENSMUSG00000105036 | 0 | NA | NA | NA | NA | NA | 1110060G06 | RIKEN cDNA 1110060G06 gene [Source:M | 0 | 0 | 0 | 0 | 0 | 0 |
| 43085 | ENSMUSG00000105037 | 0 | NA | NA | NA | NA | NA | Gm25203 | predicted gene, 25203 [Source:MGI Symb | 0 | 0 | 0 | 0 | 0 | 0 |
| 43086 | ENSMUSG00000105038 | 0 | NA | NA | NA | NA | NA | Gm43044 | predicted gene 43044 [Source:MGI Symbo | 0 | 0 | 0 | 0 | 0 | 0 |
| 43087 | ENSMUSG00000105039 | 0 | NA | NA | NA | NA | NA | Gm32585 | predicted gene, 32585 [Source:MGI Symb | 0 | 0 | 0 | 0 | 0 | 0 |
| 43088 | ENSMUSG00000105040 | 0 | NA | NA | NA | NA | NA | Ube2l3-ps1 | ubiquitin-conjugating enzyme E2L 3, pseu | 0 | 0 | 0 | 0 | 0 | 0 |
| 43089 | ENSMUSG00000105041 | 0 | NA | NA | NA | NA | NA | Gm42676 | predicted gene 42676 [Source:MGI Symbo | 0 | 0 | 0 | 0 | 0 | 0 |
| 43090 | ENSMUSG00000105042 | 0.133850373 | -0.796629176 | 4.080472857 | -0.195229623 | 0.84521319 | NA | Gm42509 | predicted gene 42509 [Source:MGI Symbo | 0.8031 | 0 | 0 | 0 | 0 | 0 |
| 43091 | ENSMUSG00000105043 | 0 | NA | NA | NA | NA | NA | Igkv1-108 | immunoglobulin kappa variable 1-108 [So | 0 | 0 | 0 | 0 | 0 | 0 |
| 43092 | ENSMUSG00000105044 | 4.458102074 | -0.158821243 | 1.044604407 | -0.152039607 | 0.879155697 | NA | Gm10416 | predicted pseudogene 10416 [Source:MG | 4.0155 | 2.0558 | 8.05 | 3.8559 | 6.5437 | 2.2277 |
| 43093 | ENSMUSG00000105045 | 0 | NA | NA | NA | NA | NA | Rpl9-ps8 | ribosomal protein L9, pseudogene 8 [Sour | 0 | 0 | 0 | 0 | 0 | 0 |
| 43094 | ENSMUSG00000105046 | 0 | NA | NA | NA | NA | NA | NA | NA | 0 | 0 | 0 | 0 | 0 | 0 |
| 43095 | ENSMUSG00000105047 | 0 | NA | NA | NA | NA | NA | Gm42533 | predicted gene 42533 [Source:MGI Symbo | 0 | 0 | 0 | 0 | 0 | 0 |
| 43096 | ENSMUSG00000105048 | 0 | NA | NA | NA | NA | NA | Gm42966 | predicted gene 42966 [Source:MGI Symbo | 0 | 0 | 0 | 0 | 0 | 0 |
| 43097 | ENSMUSG00000105049 | 0 | NA | NA | NA | NA | NA | Gm42486 | predicted gene 42486 [Source:MGI Symbo | 0 | 0 | 0 | 0 | 0 | 0 |
| 43098 | ENSMUSG00000105050 | 0 | NA | NA | NA | NA | NA | Gm42482 | predicted gene 42482 [Source:MGI Symbo | 0 | 0 | 0 | 0 | 0 | 0 |
| 43099 | ENSMUSG00000105051 | 0 | NA | NA | NA | NA | NA | Gm4248 | predicted pseudogene 4248 [Source:MGI | 0 | 0 | 0 | 0 | 0 | 0 |
| 43100 | ENSMUSG00000105052 | 0 | NA | NA | NA | NA | NA | Gm21655 | predicted gene, 21655 [Source:MGI Symb | 0 | 0 | 0 | 0 | 0 | 0 |
| 43101 | ENSMUSG00000105053 | 0 | NA | NA | NA | NA | NA | Gm43064 | predicted gene 43064 [Source:MGI Symbo | 0 | 0 | 0 | 0 | 0 | 0 |
| 43102 | ENSMUSG00000105054 | 0 | NA | NA | NA | NA | NA | Gm43664 | predicted gene 43664 [Source:MGI Symbo | 0 | 0 | 0 | 0 | 0 | 0 |
| 43103 | ENSMUSG00000105055 | 0 | NA | NA | NA | NA | NA | Gm43079 | predicted gene 43079 [Source:MGI Symbo | 0 | 0 | 0 | 0 | 0 | 0 |
| 43104 | ENSMUSG00000105056 | 0 | NA | NA | NA | NA | NA | Gm42805 | predicted gene 42805 [Source:MGI Symbo | 0 | 0 | 0 | 0 | 0 | 0 |
| 43105 | ENSMUSG00000105057 | 0 | NA | NA | NA | NA | NA | Gm40190 | predicted gene, 40190 [Source:MGI Symb | 0 | 0 | 0 | 0 | 0 | 0 |
| 43106 | ENSMUSG00000105058 | 0 | NA | NA | NA | NA | NA | Gm43590 | predicted gene 43590 [Source:MGI Symbo | 0 | 0 | 0 | 0 | 0 | 0 |
| 43107 | ENSMUSG00000105059 | 0 | NA | NA | NA | NA | NA | Gm42498 | predicted gene 42498 [Source:MGI Symbo | 0 | 0 | 0 | 0 | 0 | 0 |
| 43108 | ENSMUSG00000105060 | 0 | NA | NA | NA | NA | NA | NA | NA | 0 | 0 | 0 | 0 | 0 | 0 |
| 43109 | ENSMUSG00000105061 | 0 | NA | NA | NA | NA | NA | Gm6740 | predicted gene 6740 [Source:MGI Symbol | 0 | 0 | 0 | 0 | 0 | 0 |
| 43110 | ENSMUSG00000105062 | 0 | NA | NA | NA | NA | NA | Gm43113 | predicted gene 43113 [Source:MGI Symbo | 0 | 0 | 0 | 0 | 0 | 0 |
| 43111 | ENSMUSG00000105063 | 0.17131964 | -0.796629176 | 4.080472857 | -0.195229623 | 0.84521319 | NA | Gm43488 | predicted gene 43488 [Source:MGI Symbo | 0 | 1.0279 | 0 | 0 | 0 | 0 |
| 43112 | ENSMUSG00000105064 | 0 | NA | NA | NA | NA | NA | NA | NA | 0 | 0 | 0 | 0 | 0 | 0 |
| 43113 | ENSMUSG00000105065 | 0.481988923 | 2.492401104 | 3.837077151 | 0.649557203 | 0.515978285 | NA | Gm42513 | predicted gene 42513 [Source:MGI Symbo | 0 | 0 | 0 | 2.8919 | 0 | 0 |
| 43114 | ENSMUSG00000105066 | 0 | NA | NA | NA | NA | NA | Iglj4 | immunoglobulin lambda joining 4 [Source | 0 | 0 | 0 | 0 | 0 | 0 |
| 43115 | ENSMUSG00000105067 | 0 | NA | NA | NA | NA | NA | Gm18018 | predicted gene, 18018 [Source:MGI Symb | 0 | 0 | 0 | 0 | 0 | 0 |
| 43116 | ENSMUSG00000105068 | 4.016520767 | -0.851778939 | 1.121621909 | -0.759417173 | 0.447603044 | NA | Gm30835 | predicted gene, 30835 [Source:MGI Symb | 7.2279 | 6.1675 | 2.0125 | 0.964 | 3.2719 | 4.4554 |

| | | | | | | | | | | | | | | | |
|---|---|---|---|---|---|---|---|---|---|---|---|---|---|---|---|
| 43117 | ENSMUSG00000105070 | 0 | NA | NA | NA | NA | NA | Trav3n-2 | T cell receptor alpha variable 3N-2 [Source | 0 | 0 | 0 | 0 | 0 | 0 |
| 43118 | ENSMUSG00000105071 | 0.353089354 | 0.165142706 | 3.896102244 | 0.042386646 | 0.966190474 | NA | Gm43336 | predicted gene 43336 [Source:MGI Symbo | 0 | 1.0279 | 0 | 0 | 1.0906 | 0 |
| 43119 | ENSMUSG00000105072 | 0 | NA | NA | NA | NA | NA | NA | NA | 0 | 0 | 0 | 0 | 0 | 0 |
| 43120 | ENSMUSG00000105073 | 0 | NA | NA | NA | NA | NA | Gm9317 | predicted gene 9317 [Source:MGI Symbol | 0 | 0 | 0 | 0 | 0 | 0 |
| 43121 | ENSMUSG00000105074 | 0 | NA | NA | NA | NA | NA | Gm43818 | predicted gene 43818 [Source:MGI Symbo | 0 | 0 | 0 | 0 | 0 | 0 |
| 43122 | ENSMUSG00000105075 | 0 | NA | NA | NA | NA | NA | Mark1-ps2 | MAP/microtubule affinity-regulating kinas | 0 | 0 | 0 | 0 | 0 | 0 |
| 43123 | ENSMUSG00000105076 | 2.58426649 | 0.00729686 | 1.597377404 | 0.004568025 | 0.996355256 | NA | A930003O13 | RIKEN cDNA A930003O13 gene [Source:M | 1.6062 | 3.0838 | 3.0187 | 0 | 0 | 7.7969 |
| 43124 | ENSMUSG00000105077 | 0 | NA | NA | NA | NA | NA | Gm4859 | predicted gene 4859 [Source:MGI Symbol | 0 | 0 | 0 | 0 | 0 | 0 |
| 43125 | ENSMUSG00000105078 | 0 | NA | NA | NA | NA | NA | Vamp9 | vesicle-associated membrane protein 9 [S | 0 | 0 | 0 | 0 | 0 | 0 |
| 43126 | ENSMUSG00000105079 | 0 | NA | NA | NA | NA | NA | Gm43715 | predicted gene 43715 [Source:MGI Symbo | 0 | 0 | 0 | 0 | 0 | 0 |
| 43127 | ENSMUSG00000105080 | 0 | NA | NA | NA | NA | NA | Trav7n-6 | T cell receptor alpha variable 7N-6 [Source | 0 | 0 | 0 | 0 | 0 | 0 |
| 43128 | ENSMUSG00000105081 | 0.267700746 | -1.497481649 | 4.057001436 | -0.369110456 | 0.712045396 | NA | Gm43110 | predicted gene 43110 [Source:MGI Symbo | 1.6062 | 0 | 0 | 0 | 0 | 0 |
| 43129 | ENSMUSG00000105082 | 0 | NA | NA | NA | NA | NA | Aqp10-ps | aquaporin 10, pseudogene [Source:MGI S | 0 | 0 | 0 | 0 | 0 | 0 |
| 43130 | ENSMUSG00000105083 | 0 | NA | NA | NA | NA | NA | Gm42699 | predicted gene 42699 [Source:MGI Symbo | 0 | 0 | 0 | 0 | 0 | 0 |
| 43131 | ENSMUSG00000105084 | 0 | NA | NA | NA | NA | NA | Gm43365 | predicted gene 43365 [Source:MGI Symbo | 0 | 0 | 0 | 0 | 0 | 0 |
| 43132 | ENSMUSG00000105085 | 0.160662974 | 1.126924175 | 4.080472857 | 0.276174898 | 0.782413736 | NA | Gm42567 | predicted gene 42567 [Source:MGI Symbo | 0 | 0 | 0 | 0.964 | 0 | 0 |
| 43133 | ENSMUSG00000105086 | 0 | NA | NA | NA | NA | NA | Gm42534 | predicted gene 42534 [Source:MGI Symbo | 0 | 0 | 0 | 0 | 0 | 0 |
| 43134 | ENSMUSG00000105087 | 0.335414862 | -1.779047242 | 4.030804179 | -0.441362855 | 0.658950329 | NA | Gm42552 | predicted gene 42552 [Source:MGI Symbo | 0 | 0 | 2.0125 | 0 | 0 | 0 |
| 43135 | ENSMUSG00000105088 | 0 | NA | NA | NA | NA | NA | Gm17978 | predicted gene, 17978 [Source:MGI Symb | 0 | 0 | 0 | 0 | 0 | 0 |
| 43136 | ENSMUSG00000105089 | 0 | NA | NA | NA | NA | NA | Gm43717 | predicted gene 43717 [Source:MGI Symbo | 0 | 0 | 0 | 0 | 0 | 0 |
| 43137 | ENSMUSG00000105090 | 0 | NA | NA | NA | NA | NA | Trav15-3 | T cell receptor alpha variable 15-3 [Source | 0 | 0 | 0 | 0 | 0 | 0 |
| 43138 | ENSMUSG00000105091 | 0 | NA | NA | NA | NA | NA | Gm43722 | predicted gene 43722 [Source:MGI Symbo | 0 | 0 | 0 | 0 | 0 | 0 |
| 43139 | ENSMUSG00000105092 | 0 | NA | NA | NA | NA | NA | Gm42468 | predicted gene 42468 [Source:MGI Symbo | 0 | 0 | 0 | 0 | 0 | 0 |
| 43140 | ENSMUSG00000105093 | 0.778047457 | -3.004661392 | 2.562427823 | -1.172583815 | 0.240962743 | NA | Gm17815 | predicted gene, 17815 [Source:MGI Symb | 1.6062 | 2.0558 | 1.0062 | 0 | 0 | 0 |
| 43141 | ENSMUSG00000105094 | 0 | NA | NA | NA | NA | NA | Obox4-ps10 | oocyte specific homeobox 4, pseudogene | 0 | 0 | 0 | 0 | 0 | 0 |
| 43142 | ENSMUSG00000105095 | 42.3686806 | 0.688053226 | 0.36254513 | 1.89784159 | 0.057716953 | 0.341829329 | 8430422M14 | RIKEN cDNA 8430422M14 gene [Source:M | 22.487 | 35.977 | 39.244 | 45.307 | 61.075 | 50.123 |
| 43143 | ENSMUSG00000105096 | 111.6771534 | 0.277024828 | 0.250306079 | 1.10674431 | 0.268404492 | 0.724039098 | Gbp10 | guanylate-binding protein 10 [Source:MG | 124.48 | 96.624 | 81.506 | 140.74 | 128.69 | 98.018 |
| 43144 | ENSMUSG00000105097 | 0 | NA | NA | NA | NA | NA | Gm42133 | predicted gene, 42133 [Source:MGI Symb | 0 | 0 | 0 | 0 | 0 | 0 |
| 43145 | ENSMUSG00000105098 | 0 | NA | NA | NA | NA | NA | NA | NA | 0 | 0 | 0 | 0 | 0 | 0 |
| 43146 | ENSMUSG00000105099 | 0 | NA | NA | NA | NA | NA | Gm42751 | predicted gene 42751 [Source:MGI Symbo | 0 | 0 | 0 | 0 | 0 | 0 |
| 43147 | ENSMUSG00000105100 | 20.33969331 | 0.02804157 | 0.553073075 | 0.050701384 | 0.959563474 | 0.992148164 | NA | NA | 20.881 | 8.2233 | 31.194 | 15.424 | 21.812 | 24.505 |
| 43148 | ENSMUSG00000105101 | 0 | NA | NA | NA | NA | NA | Gm43206 | predicted gene 43206 [Source:MGI Symbo | 0 | 0 | 0 | 0 | 0 | 0 |
| 43149 | ENSMUSG00000105102 | 0 | NA | NA | NA | NA | NA | Gm35507 | predicted gene, 35507 [Source:MGI Symb | 0 | 0 | 0 | 0 | 0 | 0 |
| 43150 | ENSMUSG00000105103 | 0 | NA | NA | NA | NA | NA | Gm43191 | predicted gene 43191 [Source:MGI Symbo | 0 | 0 | 0 | 0 | 0 | 0 |
| 43151 | ENSMUSG00000105104 | 0 | NA | NA | NA | NA | NA | Gm43729 | predicted gene 43729 [Source:MGI Symbo | 0 | 0 | 0 | 0 | 0 | 0 |
| 43152 | ENSMUSG00000105105 | 0 | NA | NA | NA | NA | NA | Gm31363 | predicted gene, 31363 [Source:MGI Symb | 0 | 0 | 0 | 0 | 0 | 0 |
| 43153 | ENSMUSG00000105106 | 0 | NA | NA | NA | NA | NA | Gm43657 | predicted gene 43657 [Source:MGI Symbo | 0 | 0 | 0 | 0 | 0 | 0 |
| 43154 | ENSMUSG00000105107 | 0.531944269 | 2.618916564 | 3.285123409 | 0.797204926 | 0.425332026 | NA | Gm43412 | predicted gene 43412 [Source:MGI Symbo | 0 | 0 | 0 | 0.964 | 0 | 2.2277 |
| 43155 | ENSMUSG00000105108 | 0 | NA | NA | NA | NA | NA | Gm42687 | predicted gene 42687 [Source:MGI Symbo | 0 | 0 | 0 | 0 | 0 | 0 |
| 43156 | ENSMUSG00000105109 | 0 | NA | NA | NA | NA | NA | NA | NA | 0 | 0 | 0 | 0 | 0 | 0 |
| 43157 | ENSMUSG00000105110 | 0 | NA | NA | NA | NA | NA | Igkv12-66 | immunoglobulin kappa chain variable 12-6 | 0 | 0 | 0 | 0 | 0 | 0 |
| 43158 | ENSMUSG00000105111 | 0 | NA | NA | NA | NA | NA | Gm43310 | predicted gene 43310 [Source:MGI Symbo | 0 | 0 | 0 | 0 | 0 | 0 |
| 43159 | ENSMUSG00000105112 | 0.481988923 | 2.492401104 | 3.837077151 | 0.649557203 | 0.515978285 | NA | Gm42778 | predicted gene 42778 [Source:MGI Symbo | 0 | 0 | 0 | 2.8919 | 0 | 0 |
| 43160 | ENSMUSG00000105113 | 0 | NA | NA | NA | NA | NA | Gm2622 | predicted gene 2622 [Source:MGI Symbol | 0 | 0 | 0 | 0 | 0 | 0 |
| 43161 | ENSMUSG00000105114 | 0 | NA | NA | NA | NA | NA | Amd-ps5 | S-adenosylmethionine decarboxylase, pse | 0 | 0 | 0 | 0 | 0 | 0 |
| 43162 | ENSMUSG00000105115 | 0 | NA | NA | NA | NA | NA | NA | NA | 0 | 0 | 0 | 0 | 0 | 0 |
| 43163 | ENSMUSG00000105116 | 0 | NA | NA | NA | NA | NA | NA | NA | 0 | 0 | 0 | 0 | 0 | 0 |
| 43164 | ENSMUSG00000105117 | 0 | NA | NA | NA | NA | NA | Gm42760 | predicted gene 42760 [Source:MGI Symbo | 0 | 0 | 0 | 0 | 0 | 0 |
| 43165 | ENSMUSG00000105118 | 0 | NA | NA | NA | NA | NA | NA | NA | 0 | 0 | 0 | 0 | 0 | 0 |
| 43166 | ENSMUSG00000105119 | 0 | NA | NA | NA | NA | NA | Gm43765 | predicted gene 43765 [Source:MGI Symbo | 0 | 0 | 0 | 0 | 0 | 0 |
| 43167 | ENSMUSG00000105120 | 0 | NA | NA | NA | NA | NA | Gm26413 | predicted gene, 26413 [Source:MGI Symb | 0 | 0 | 0 | 0 | 0 | 0 |
| 43168 | ENSMUSG00000105121 | 0 | NA | NA | NA | NA | NA | Gm43406 | predicted gene 43406 [Source:MGI Symbo | 0 | 0 | 0 | 0 | 0 | 0 |
| 43169 | ENSMUSG00000105122 | 0 | NA | NA | NA | NA | NA | NA | NA | 0 | 0 | 0 | 0 | 0 | 0 |
| 43170 | ENSMUSG00000105123 | 0 | NA | NA | NA | NA | NA | Ighv1-79 | immunoglobulin heavy variable 1-79 [Sou | 0 | 0 | 0 | 0 | 0 | 0 |
| 43171 | ENSMUSG00000105124 | 0 | NA | NA | NA | NA | NA | Gm5297 | predicted gene 5297 [Source:MGI Symbol | 0 | 0 | 0 | 0 | 0 | 0 |
| 43172 | ENSMUSG00000105125 | 0 | NA | NA | NA | NA | NA | Gm43468 | predicted gene 43468 [Source:MGI Symbo | 0 | 0 | 0 | 0 | 0 | 0 |

| | | | | | | | | | | | | | | | |
|---|---|---|---|---|---|---|---|---|---|---|---|---|---|---|---|
| 43173 | ENSMUSG00000105126 | 0 | NA | NA | NA | NA | NA | Gm43723 | predicted gene 43723 [Source:MGI Symbo | 0 | 0 | 0 | 0 | 0 | 0 |
| 43174 | ENSMUSG00000105127 | 0 | NA | NA | NA | NA | NA | Gm42470 | predicted gene 42470 [Source:MGI Symbo | 0 | 0 | 0 | 0 | 0 | 0 |
| 43175 | ENSMUSG00000105128 | 0.727078859 | 3.067238961 | 3.29680021 | 0.930368468 | 0.352180339 | NA | Gm42870 | predicted gene 42870 [Source:MGI Symbo | 0 | 0 | 0 | 0 | 4.3625 | 0 |
| 43176 | ENSMUSG00000105129 | 0 | NA | NA | NA | NA | NA | Gm19566 | predicted gene, 19566 [Source:MGI Symb | 0 | 0 | 0 | 0 | 0 | 0 |
| 43177 | ENSMUSG00000105130 | 1.99876018 | -1.024993246 | 1.895687002 | -0.540697512 | 0.588716093 | NA | Gm43825 | predicted gene 43825 [Source:MGI Symbo | 0 | 4.1117 | 4.025 | 3.8559 | 0 | 0 |
| 43178 | ENSMUSG00000105131 | 0 | NA | NA | NA | NA | NA | NA | NA | 0 | 0 | 0 | 0 | 0 | 0 |
| 43179 | ENSMUSG00000105132 | 0 | NA | NA | NA | NA | NA | Gm43250 | predicted gene 43250 [Source:MGI Symbo | 0 | 0 | 0 | 0 | 0 | 0 |
| 43180 | ENSMUSG00000105133 | 0 | NA | NA | NA | NA | NA | NA | NA | 0 | 0 | 0 | 0 | 0 | 0 |
| 43181 | ENSMUSG00000105134 | 0.160662974 | 1.126924175 | 4.080472857 | 0.276174898 | 0.782413736 | NA | Gm42923 | predicted gene 42923 [Source:MGI Symbo | 0 | 0 | 0 | 0.964 | 0 | 0 |
| 43182 | ENSMUSG00000105135 | 88.95789163 | -0.255938956 | 0.261752907 | -0.977788401 | 0.328178989 | 0.773161429 | Gm43667 | predicted gene 43667 [Source:MGI Symbo | 93.963 | 113.07 | 83.518 | 77.118 | 99.246 | 66.831 |
| 43183 | ENSMUSG00000105136 | 0 | NA | NA | NA | NA | NA | 8030487O14 | RIKEN cDNA 8030487O14 gene [Source:M | 0 | 0 | 0 | 0 | 0 | 0 |
| 43184 | ENSMUSG00000105137 | 1.634645655 | -4.067143069 | 2.074807359 | -1.960250936 | 0.049966468 | NA | Gm42869 | predicted gene 42869 [Source:MGI Symbo | 1.6062 | 7.1954 | 1.0062 | 0 | 0 | 0 |
| 43185 | ENSMUSG00000105138 | 0.51034671 | -2.381743948 | 3.331081356 | -0.715006238 | 0.474605189 | NA | Mir6964 | microRNA 6964 [Source:MGI Symbol;Acc: | 0 | 2.0558 | 1.0062 | 0 | 0 | 0 |
| 43186 | ENSMUSG00000105139 | 1.03140222 | -2.145901642 | 2.420890855 | -0.886409909 | 0.375396682 | NA | Gm19391 | predicted gene, 19391 [Source:MGI Symb | 0 | 2.0558 | 3.0187 | 0 | 0 | 1.1138 |
| 43187 | ENSMUSG00000105140 | 8.668100658 | -0.631445397 | 0.838596253 | -0.752979034 | 0.451462513 | 0.856162814 | Gm43127 | predicted gene 43127 [Source:MGI Symbo | 6.4248 | 11.307 | 14.087 | 12.532 | 6.5437 | 1.1138 |
| 43188 | ENSMUSG00000105141 | 0 | NA | NA | NA | NA | NA | Gm43828 | predicted gene 43828 [Source:MGI Symbo | 0 | 0 | 0 | 0 | 0 | 0 |
| 43189 | ENSMUSG00000105142 | 0 | NA | NA | NA | NA | NA | Mir7214 | microRNA 7214 [Source:MGI Symbol;Acc: | 0 | 0 | 0 | 0 | 0 | 0 |
| 43190 | ENSMUSG00000105143 | 0.638982041 | 0.404325377 | 3.14176733 | 0.128693609 | 0.897600092 | NA | Gm8069 | predicted pseudogene 8069 [Source:MGI | 1.6062 | 0 | 0 | 0 | 0 | 2.2277 |
| 43191 | ENSMUSG00000105144 | 1.140507014 | 0.523154827 | 1.987305671 | 0.263248294 | 0.79235921 | NA | Rpl21-ps11 | ribosomal protein L21, pseudogene 11 [Sc | 0.8031 | 1.0279 | 1.0062 | 2.8919 | 0 | 1.1138 |
| 43192 | ENSMUSG00000105145 | 0 | NA | NA | NA | NA | NA | NA | NA | 0 | 0 | 0 | 0 | 0 | 0 |
| 43193 | ENSMUSG00000105146 | 0 | NA | NA | NA | NA | NA | Gm35409 | predicted gene, 35409 [Source:MGI Symb | 0 | 0 | 0 | 0 | 0 | 0 |
| 43194 | ENSMUSG00000105147 | 0 | NA | NA | NA | NA | NA | Ckb-ps1 | creatine kinase, brain, pseudogene 1 [Sou | 0 | 0 | 0 | 0 | 0 | 0 |
| 43195 | ENSMUSG00000105148 | 0 | NA | NA | NA | NA | NA | Gm42700 | predicted gene 42700 [Source:MGI Symbo | 0 | 0 | 0 | 0 | 0 | 0 |
| 43196 | ENSMUSG00000105149 | 0 | NA | NA | NA | NA | NA | Gm42983 | predicted gene 42983 [Source:MGI Symbo | 0 | 0 | 0 | 0 | 0 | 0 |
| 43197 | ENSMUSG00000105150 | 0 | NA | NA | NA | NA | NA | Gm43615 | predicted gene 43615 [Source:MGI Symbo | 0 | 0 | 0 | 0 | 0 | 0 |
| 43198 | ENSMUSG00000105151 | 0 | NA | NA | NA | NA | NA | Gm43241 | predicted gene 43241 [Source:MGI Symbo | 0 | 0 | 0 | 0 | 0 | 0 |
| 43199 | ENSMUSG00000105152 | 0 | NA | NA | NA | NA | NA | Gm42864 | predicted gene 42864 [Source:MGI Symbo | 0 | 0 | 0 | 0 | 0 | 0 |
| 43200 | ENSMUSG00000105153 | 0 | NA | NA | NA | NA | NA | Gm3143 | predicted gene 3143 [Source:MGI Symbol | 0 | 0 | 0 | 0 | 0 | 0 |
| 43201 | ENSMUSG00000105154 | 0 | NA | NA | NA | NA | NA | Gm20128 | predicted gene, 20128 [Source:MGI Symb | 0 | 0 | 0 | 0 | 0 | 0 |
| 43202 | ENSMUSG00000105155 | 0 | NA | NA | NA | NA | NA | Gm42910 | predicted gene 42910 [Source:MGI Symbo | 0 | 0 | 0 | 0 | 0 | 0 |
| 43203 | ENSMUSG00000105156 | 3.006827013 | -0.893822576 | 1.277485332 | -0.699673455 | 0.484131258 | NA | Gm9057 | predicted gene 9057 [Source:MGI Symbol | 5.6217 | 1.0279 | 5.0312 | 1.928 | 1.0906 | 3.3415 |
| 43204 | ENSMUSG00000105157 | 0 | NA | NA | NA | NA | NA | Gm31831 | predicted gene, 31831 [Source:MGI Symb | 0 | 0 | 0 | 0 | 0 | 0 |
| 43205 | ENSMUSG00000105158 | 0 | NA | NA | NA | NA | NA | Gm42862 | predicted gene 42862 [Source:MGI Symbo | 0 | 0 | 0 | 0 | 0 | 0 |
| 43206 | ENSMUSG00000105159 | 0 | NA | NA | NA | NA | NA | Gm42988 | predicted gene 42988 [Source:MGI Symbo | 0 | 0 | 0 | 0 | 0 | 0 |
| 43207 | ENSMUSG00000105160 | 1.042032255 | 0.07171624 | 2.223606224 | 0.032252221 | 0.974270911 | NA | A530030E21 | RIKEN cDNA A530030E21 gene [Source:M | 0 | 3.0838 | 0 | 0.964 | 1.0906 | 1.1138 |
| 43208 | ENSMUSG00000105161 | 22.02673175 | -0.277721435 | 0.45790366 | -0.606506257 | 0.544178629 | 0.89481775 | Gm42595 | predicted gene 42595 [Source:MGI Symbo | 21.684 | 26.726 | 24.15 | 22.171 | 20.722 | 16.708 |
| 43209 | ENSMUSG00000105162 | 0 | NA | NA | NA | NA | NA | Gm42494 | predicted gene 42494 [Source:MGI Symbo | 0 | 0 | 0 | 0 | 0 | 0 |
| 43210 | ENSMUSG00000105163 | 0 | NA | NA | NA | NA | NA | Gm4873 | predicted gene 4873 [Source:MGI Symbol | 0 | 0 | 0 | 0 | 0 | 0 |
| 43211 | ENSMUSG00000105164 | 0 | NA | NA | NA | NA | NA | NA | NA | 0 | 0 | 0 | 0 | 0 | 0 |
| 43212 | ENSMUSG00000105165 | 0 | NA | NA | NA | NA | NA | Mir7686 | microRNA 7686 [Source:MGI Symbol;Acc: | 0 | 0 | 0 | 0 | 0 | 0 |
| 43213 | ENSMUSG00000105166 | 0 | NA | NA | NA | NA | NA | Gm42629 | predicted gene 42629 [Source:MGI Symbo | 0 | 0 | 0 | 0 | 0 | 0 |
| 43214 | ENSMUSG00000105167 | 0.812136609 | 0.736604728 | 2.414276032 | 0.305103774 | 0.760287117 | NA | Snord43 | small nucleolar RNA, C/D box 43 [Source:I | 0.8031 | 1.0279 | 0 | 1.928 | 0 | 1.1138 |
| 43215 | ENSMUSG00000105168 | 0 | NA | NA | NA | NA | NA | Gm30735 | predicted gene, 30735 [Source:MGI Symb | 0 | 0 | 0 | 0 | 0 | 0 |
| 43216 | ENSMUSG00000105169 | 0 | NA | NA | NA | NA | NA | NA | NA | 0 | 0 | 0 | 0 | 0 | 0 |
| 43217 | ENSMUSG00000105170 | 0 | NA | NA | NA | NA | NA | Gm19166 | predicted gene, 19166 [Source:MGI Symb | 0 | 0 | 0 | 0 | 0 | 0 |
| 43218 | ENSMUSG00000105171 | 0 | NA | NA | NA | NA | NA | Gm5288 | predicted gene 5288 [Source:MGI Symbol | 0 | 0 | 0 | 0 | 0 | 0 |
| 43219 | ENSMUSG00000105172 | 0 | NA | NA | NA | NA | NA | Gm42499 | predicted gene 42499 [Source:MGI Symbo | 0 | 0 | 0 | 0 | 0 | 0 |
| 43220 | ENSMUSG00000105173 | 0 | NA | NA | NA | NA | NA | Gm5715 | predicted gene 5715 [Source:MGI Symbol | 0 | 0 | 0 | 0 | 0 | 0 |
| 43221 | ENSMUSG00000105174 | 0 | NA | NA | NA | NA | NA | Gm43353 | predicted gene 43353 [Source:MGI Symbo | 0 | 0 | 0 | 0 | 0 | 0 |
| 43222 | ENSMUSG00000105175 | 0 | NA | NA | NA | NA | NA | Gm43097 | predicted gene 43097 [Source:MGI Symbo | 0 | 0 | 0 | 0 | 0 | 0 |
| 43223 | ENSMUSG00000105176 | 43.90321346 | -0.377251569 | 0.358432164 | -1.052504789 | 0.292568015 | 0.74357824 | Gm43668 | predicted gene 43668 [Source:MGI Symbo | 46.58 | 42.145 | 60.375 | 47.235 | 40.353 | 26.732 |
| 43224 | ENSMUSG00000105177 | 0 | NA | NA | NA | NA | NA | Gm43563 | predicted gene 43563 [Source:MGI Symbo | 0 | 0 | 0 | 0 | 0 | 0 |
| 43225 | ENSMUSG00000105178 | 0 | NA | NA | NA | NA | NA | Mir376a | microRNA 376a [Source:MGI Symbol;Acc: | 0 | 0 | 0 | 0 | 0 | 0 |
| 43226 | ENSMUSG00000105179 | 1.350427926 | 0.155292823 | 1.741700337 | 0.089161619 | 0.928953467 | NA | Gm42866 | predicted gene 42866 [Source:MGI Symbo | 0.8031 | 1.0279 | 2.0125 | 0.964 | 2.1812 | 1.1138 |
| 43227 | ENSMUSG00000105180 | 0 | NA | NA | NA | NA | NA | Gm4863 | predicted gene 4863 [Source:MGI Symbol | 0 | 0 | 0 | 0 | 0 | 0 |
| 43228 | ENSMUSG00000105181 | 0.185640647 | 1.126924175 | 4.080472857 | 0.276174898 | 0.782413736 | NA | Gm19620 | predicted gene, 19620 [Source:MGI Symb | 0 | 0 | 0 | 0 | 0 | 1.1138 |

| | | | | | | | | | | | | | | | |
|---|---|---|---|---|---|---|---|---|---|---|---|---|---|---|---|
| 43229 | ENSMUSG00000105182 | 0 | NA | NA | NA | NA | NA | NA | NA | 0 | 0 | 0 | 0 | 0 | 0 |
| 43230 | ENSMUSG00000105183 | 0 | NA | NA | NA | NA | NA | Gm42564 | predicted gene 42564 [Source:MGI Symbo | 0 | 0 | 0 | 0 | 0 | 0 |
| 43231 | ENSMUSG00000105184 | 0 | NA | NA | NA | NA | NA | Gm42537 | predicted gene 42537 [Source:MGI Symbo | 0 | 0 | 0 | 0 | 0 | 0 |
| 43232 | ENSMUSG00000105185 | 0 | NA | NA | NA | NA | NA | 4930519H02 | RIKEN cDNA 4930519H02 gene [Source:M | 0 | 0 | 0 | 0 | 0 | 0 |
| 43233 | ENSMUSG00000105186 | 0 | NA | NA | NA | NA | NA | Gm43778 | predicted gene 43778 [Source:MGI Symbo | 0 | 0 | 0 | 0 | 0 | 0 |
| 43234 | ENSMUSG00000105187 | 0 | NA | NA | NA | NA | NA | Gm23377 | predicted gene, 23377 [Source:MGI Symb | 0 | 0 | 0 | 0 | 0 | 0 |
| 43235 | ENSMUSG00000105188 | 0 | NA | NA | NA | NA | NA | Gm8190 | predicted gene 8190 [Source:MGI Symbol | 0 | 0 | 0 | 0 | 0 | 0 |
| 43236 | ENSMUSG00000105189 | 0.319491021 | 0.16514334 | 4.044653141 | 0.040830038 | 0.967431393 | NA | Gm7774 | predicted gene 7774 [Source:MGI Symbol | 0.8031 | 0 | 0 | 0 | 0 | 1.1138 |
| 43237 | ENSMUSG00000105190 | 0 | NA | NA | NA | NA | NA | Gm43470 | predicted gene 43470 [Source:MGI Symbo | 0 | 0 | 0 | 0 | 0 | 0 |
| 43238 | ENSMUSG00000105191 | 0 | NA | NA | NA | NA | NA | Gm4644 | predicted gene 4644 [Source:MGI Symbol | 0 | 0 | 0 | 0 | 0 | 0 |
| 43239 | ENSMUSG00000105192 | 37.20042842 | 2.320120649 | 0.563133183 | 4.120021197 | 3.79E-05 | 0.00168083 | Gm42796 | predicted gene 42796 [Source:MGI Symbo | 12.047 | 5.1396 | 20.125 | 111.82 | 26.175 | 47.895 |
| 43240 | ENSMUSG00000105193 | 0 | NA | NA | NA | NA | NA | Gm18971 | predicted gene, 18971 [Source:MGI Symb | 0 | 0 | 0 | 0 | 0 | 0 |
| 43241 | ENSMUSG00000105194 | 0 | NA | NA | NA | NA | NA | Gm6755 | predicted gene 6755 [Source:MGI Symbol | 0 | 0 | 0 | 0 | 0 | 0 |
| 43242 | ENSMUSG00000105195 | 0 | NA | NA | NA | NA | NA | Gm43584 | predicted gene 43584 [Source:MGI Symbo | 0 | 0 | 0 | 0 | 0 | 0 |
| 43243 | ENSMUSG00000105196 | 0 | NA | NA | NA | NA | NA | Mir142 | microRNA 142 [Source:MGI Symbol;Acc:M | 0 | 0 | 0 | 0 | 0 | 0 |
| 43244 | ENSMUSG00000105197 | 0 | NA | NA | NA | NA | NA | Gm43446 | predicted gene 43446 [Source:MGI Symbo | 0 | 0 | 0 | 0 | 0 | 0 |
| 43245 | ENSMUSG00000105198 | 2.018522811 | 0.107262899 | 1.561508707 | 0.068691835 | 0.945234917 | NA | Gm42502 | predicted gene 42502 [Source:MGI Symbo | 0.8031 | 2.0558 | 3.0187 | 2.8919 | 0 | 3.3415 |
| 43246 | ENSMUSG00000105199 | 0.684865378 | 2.990343487 | 2.966805827 | 1.00793367 | 0.313486306 | NA | Gm43581 | predicted gene 43581 [Source:MGI Symbo | 0 | 0 | 0 | 1.928 | 2.1812 | 0 |
| 43247 | ENSMUSG00000105200 | 0 | NA | NA | NA | NA | NA | Mir378a | microRNA 378a [Source:MGI Symbol;Acc: | 0 | 0 | 0 | 0 | 0 | 0 |
| 43248 | ENSMUSG00000105201 | 2.925101859 | 0.998479475 | 1.332746285 | 0.749189464 | 0.453743019 | NA | Gm43362 | predicted gene 43362 [Source:MGI Symbo | 0.8031 | 4.1117 | 1.0062 | 3.8559 | 1.0906 | 6.6831 |
| 43249 | ENSMUSG00000105202 | 0 | NA | NA | NA | NA | NA | C230031I18I | RIKEN cDNA C230031I18 gene [Source:MC | 0 | 0 | 0 | 0 | 0 | 0 |
| 43250 | ENSMUSG00000105203 | 0.17131964 | -0.796629176 | 4.080472857 | -0.195229623 | 0.84521319 | NA | 4933428P19 | RIKEN cDNA 4933428P19 gene [Source:M | 0 | 1.0279 | 0 | 0 | 0 | 0 |
| 43251 | ENSMUSG00000105204 | 0 | NA | NA | NA | NA | NA | Gm43738 | predicted gene 43738 [Source:MGI Symbo | 0 | 0 | 0 | 0 | 0 | 0 |
| 43252 | ENSMUSG00000105205 | 0 | NA | NA | NA | NA | NA | NA | NA | 0 | 0 | 0 | 0 | 0 | 0 |
| 43253 | ENSMUSG00000105206 | 0 | NA | NA | NA | NA | NA | Gm4870 | predicted gene 4870 [Source:MGI Symbol | 0 | 0 | 0 | 0 | 0 | 0 |
| 43254 | ENSMUSG00000105207 | 0 | NA | NA | NA | NA | NA | Gm42927 | predicted gene 42927 [Source:MGI Symbo | 0 | 0 | 0 | 0 | 0 | 0 |
| 43255 | ENSMUSG00000105208 | 0 | NA | NA | NA | NA | NA | Gm43525 | predicted gene 43525 [Source:MGI Symbo | 0 | 0 | 0 | 0 | 0 | 0 |
| 43256 | ENSMUSG00000105209 | 0 | NA | NA | NA | NA | NA | Gm43334 | predicted gene 43334 [Source:MGI Symbo | 0 | 0 | 0 | 0 | 0 | 0 |
| 43257 | ENSMUSG00000105210 | 0 | NA | NA | NA | NA | NA | Mir3544 | microRNA 3544 [Source:MGI Symbol;Acc: | 0 | 0 | 0 | 0 | 0 | 0 |
| 43258 | ENSMUSG00000105211 | 0 | NA | NA | NA | NA | NA | NA | NA | 0 | 0 | 0 | 0 | 0 | 0 |
| 43259 | ENSMUSG00000105212 | 0 | NA | NA | NA | NA | NA | Gm43557 | predicted gene 43557 [Source:MGI Symbo | 0 | 0 | 0 | 0 | 0 | 0 |
| 43260 | ENSMUSG00000105213 | 0 | NA | NA | NA | NA | NA | Gm43037 | predicted gene 43037 [Source:MGI Symbo | 0 | 0 | 0 | 0 | 0 | 0 |
| 43261 | ENSMUSG00000105214 | 0 | NA | NA | NA | NA | NA | NA | NA | 0 | 0 | 0 | 0 | 0 | 0 |
| 43262 | ENSMUSG00000105215 | 0 | NA | NA | NA | NA | NA | NA | NA | 0 | 0 | 0 | 0 | 0 | 0 |
| 43263 | ENSMUSG00000105216 | 0 | NA | NA | NA | NA | NA | Traj47 | T cell receptor alpha joining 47 [Source:M | 0 | 0 | 0 | 0 | 0 | 0 |
| 43264 | ENSMUSG00000105217 | 0 | NA | NA | NA | NA | NA | Gm42873 | predicted gene 42873 [Source:MGI Symbo | 0 | 0 | 0 | 0 | 0 | 0 |
| 43265 | ENSMUSG00000105218 | 0 | NA | NA | NA | NA | NA | Gm38413 | predicted gene, 38413 [Source:MGI Symb | 0 | 0 | 0 | 0 | 0 | 0 |
| 43266 | ENSMUSG00000105219 | 0 | NA | NA | NA | NA | NA | Gm43821 | predicted gene 43821 [Source:MGI Symbo | 0 | 0 | 0 | 0 | 0 | 0 |
| 43267 | ENSMUSG00000105220 | 0 | NA | NA | NA | NA | NA | Mir497 | microRNA 497 [Source:MGI Symbol;Acc:M | 0 | 0 | 0 | 0 | 0 | 0 |
| 43268 | ENSMUSG00000105221 | 0 | NA | NA | NA | NA | NA | Gm9332 | predicted gene 9332 [Source:MGI Symbol | 0 | 0 | 0 | 0 | 0 | 0 |
| 43269 | ENSMUSG00000105222 | 0 | NA | NA | NA | NA | NA | Gm42205 | predicted gene, 42205 [Source:MGI Symb | 0 | 0 | 0 | 0 | 0 | 0 |
| 43270 | ENSMUSG00000105223 | 0 | NA | NA | NA | NA | NA | Gm43433 | predicted gene 43433 [Source:MGI Symbo | 0 | 0 | 0 | 0 | 0 | 0 |
| 43271 | ENSMUSG00000105224 | 0.267700746 | -1.497481649 | 4.057001436 | -0.369110456 | 0.712045396 | NA | Gm3364 | predicted gene 3364 [Source:MGI Symbol | 1.6062 | 0 | 0 | 0 | 0 | 0 |
| 43272 | ENSMUSG00000105225 | 0 | NA | NA | NA | NA | NA | Gm19147 | predicted gene, 19147 [Source:MGI Symb | 0 | 0 | 0 | 0 | 0 | 0 |
| 43273 | ENSMUSG00000105226 | 0.946860264 | 0.386550567 | 2.714838108 | 0.142384389 | 0.886776392 | NA | A930028O11 | RIKEN cDNA A930028O11 gene [Source:M | 2.4093 | 0 | 0 | 0 | 3.2719 | 0 |
| 43274 | ENSMUSG00000105227 | 0 | NA | NA | NA | NA | NA | Gm36667 | predicted gene, 36667 [Source:MGI Symb | 0 | 0 | 0 | 0 | 0 | 0 |
| 43275 | ENSMUSG00000105228 | 0 | NA | NA | NA | NA | NA | Gm43340 | predicted gene 43340 [Source:MGI Symbo | 0 | 0 | 0 | 0 | 0 | 0 |
| 43276 | ENSMUSG00000105229 | 0.133850373 | -0.796629176 | 4.080472857 | -0.195229623 | 0.84521319 | NA | Gm43149 | predicted gene 43149 [Source:MGI Symbo | 0.8031 | 0 | 0 | 0 | 0 | 0 |
| 43277 | ENSMUSG00000105230 | 0.335414862 | -1.779047242 | 4.030804179 | -0.441362855 | 0.658950329 | NA | Gm42433 | predicted gene 42433 [Source:MGI Symbo | 0 | 0 | 2.0125 | 0 | 0 | 0 |
| 43278 | ENSMUSG00000105231 | 0 | NA | NA | NA | NA | NA | Iglj3 | immunoglobulin lambda joining 3 [Source | 0 | 0 | 0 | 0 | 0 | 0 |
| 43279 | ENSMUSG00000105232 | 0 | NA | NA | NA | NA | NA | NA | NA | 0 | 0 | 0 | 0 | 0 | 0 |
| 43280 | ENSMUSG00000105233 | 0 | NA | NA | NA | NA | NA | Gm20568 | predicted gene, 20568 [Source:MGI Symb | 0 | 0 | 0 | 0 | 0 | 0 |
| 43281 | ENSMUSG00000105234 | 0 | NA | NA | NA | NA | NA | 4930459L07I | RIKEN cDNA 4930459L07 gene [Source:MC | 0 | 0 | 0 | 0 | 0 | 0 |
| 43282 | ENSMUSG00000105235 | 0 | NA | NA | NA | NA | NA | Hcfc1r1-ps1 | host cell factor C1 regulator 1 (XPO1-depe | 0 | 0 | 0 | 0 | 0 | 0 |
| 43283 | ENSMUSG00000105236 | 0 | NA | NA | NA | NA | NA | NA | NA | 0 | 0 | 0 | 0 | 0 | 0 |
| 43284 | ENSMUSG00000105237 | 0 | NA | NA | NA | NA | NA | Gm42580 | predicted gene 42580 [Source:MGI Symbo | 0 | 0 | 0 | 0 | 0 | 0 |

| | | | | | | | | | | | | | | | |
|---|---|---|---|---|---|---|---|---|---|---|---|---|---|---|---|
| 43285 | ENSMUSG00000105238 | 0 | NA | NA | NA | NA | NA | NA | NA | 0 | 0 | 0 | 0 | 0 | 0 |
| 43286 | ENSMUSG00000105239 | 2.327103955 | 0.402769292 | 1.685765364 | 0.238923696 | 0.811164751 | NA | Gm43091 | predicted gene 43091 [Source:MGI Symbo | 0 | 4.1117 | 2.0125 | 6.7478 | 1.0906 | 0 |
| 43287 | ENSMUSG00000105240 | 0 | NA | NA | NA | NA | NA | Gm19418 | predicted gene, 19418 [Source:MGI Symb | 0 | 0 | 0 | 0 | 0 | 0 |
| 43288 | ENSMUSG00000105241 | 0.167707431 | -0.796629176 | 4.080472857 | -0.195229623 | 0.84521319 | NA | Gm43542 | predicted gene 43542 [Source:MGI Symbo | 0 | 0 | 1.0062 | 0 | 0 | 0 |
| 43289 | ENSMUSG00000105242 | 0 | NA | NA | NA | NA | NA | Gm9236 | predicted gene 9236 [Source:MGI Symbol | 0 | 0 | 0 | 0 | 0 | 0 |
| 43290 | ENSMUSG00000105243 | 2.13435569 | 0.023053468 | 1.560694533 | 0.014771287 | 0.988214646 | NA | Gm43444 | predicted gene 43444 [Source:MGI Symbo | 3.2124 | 3.0838 | 0 | 0.964 | 1.0906 | 4.4554 |
| 43291 | ENSMUSG00000105244 | 0 | NA | NA | NA | NA | NA | Gm5281 | predicted gene 5281 [Source:MGI Symbol | 0 | 0 | 0 | 0 | 0 | 0 |
| 43292 | ENSMUSG00000105245 | 3.39439207 | -0.226492332 | 1.199577771 | -0.188810044 | 0.8502417 | NA | Gm31305 | predicted gene, 31305 [Source:MGI Symb | 4.8186 | 5.1396 | 1.0062 | 3.8559 | 1.0906 | 4.4554 |
| 43293 | ENSMUSG00000105246 | 0.301557804 | -1.64335916 | 4.042814367 | -0.406488899 | 0.684383412 | NA | Gm21043 | predicted gene, 21043 [Source:MGI Symb | 0.8031 | 0 | 1.0062 | 0 | 0 | 0 |
| 43294 | ENSMUSG00000105247 | 0 | NA | NA | NA | NA | NA | Gm42519 | predicted gene 42519 [Source:MGI Symbo | 0 | 0 | 0 | 0 | 0 | 0 |
| 43295 | ENSMUSG00000105248 | 0 | NA | NA | NA | NA | NA | Gm5310 | predicted gene 5310 [Source:MGI Symbol | 0 | 0 | 0 | 0 | 0 | 0 |
| 43296 | ENSMUSG00000105249 | 0 | NA | NA | NA | NA | NA | NA | NA | 0 | 0 | 0 | 0 | 0 | 0 |
| 43297 | ENSMUSG00000105250 | 0 | NA | NA | NA | NA | NA | Gm29736 | predicted gene, 29736 [Source:MGI Symb | 0 | 0 | 0 | 0 | 0 | 0 |
| 43298 | ENSMUSG00000105251 | 0 | NA | NA | NA | NA | NA | 4930526M1( | RIKEN cDNA 4930526M16 gene [Source:M | 0 | 0 | 0 | 0 | 0 | 0 |
| 43299 | ENSMUSG00000105252 | 0 | NA | NA | NA | NA | NA | Gm42753 | predicted gene 42753 [Source:MGI Symbo | 0 | 0 | 0 | 0 | 0 | 0 |
| 43300 | ENSMUSG00000105253 | 0 | NA | NA | NA | NA | NA | Gm42522 | predicted gene 42522 [Source:MGI Symbo | 0 | 0 | 0 | 0 | 0 | 0 |
| 43301 | ENSMUSG00000105254 | 0 | NA | NA | NA | NA | NA | Gm42951 | predicted gene 42951 [Source:MGI Symbo | 0 | 0 | 0 | 0 | 0 | 0 |
| 43302 | ENSMUSG00000105255 | 0 | NA | NA | NA | NA | NA | Gm42413 | predicted gene, 42413 [Source:MGI Symb | 0 | 0 | 0 | 0 | 0 | 0 |
| 43303 | ENSMUSG00000105256 | 0 | NA | NA | NA | NA | NA | NA | NA | 0 | 0 | 0 | 0 | 0 | 0 |
| 43304 | ENSMUSG00000105257 | 0 | NA | NA | NA | NA | NA | Gm43144 | predicted gene 43144 [Source:MGI Symbo | 0 | 0 | 0 | 0 | 0 | 0 |
| 43305 | ENSMUSG00000105258 | 0 | NA | NA | NA | NA | NA | Gm40038 | predicted gene, 40038 [Source:MGI Symb | 0 | 0 | 0 | 0 | 0 | 0 |
| 43306 | ENSMUSG00000105259 | 0 | NA | NA | NA | NA | NA | Gm42874 | predicted gene 42874 [Source:MGI Symbo | 0 | 0 | 0 | 0 | 0 | 0 |
| 43307 | ENSMUSG00000105260 | 32.09002832 | 0.218648298 | 0.428154218 | 0.510676502 | 0.609577598 | 0.916376001 | Gm40040 | predicted gene, 40040 [Source:MGI Symb | 27.305 | 19.53 | 42.262 | 40.487 | 25.084 | 37.871 |
| 43308 | ENSMUSG00000105261 | 0 | NA | NA | NA | NA | NA | Gm43333 | predicted gene 43333 [Source:MGI Symbo | 0 | 0 | 0 | 0 | 0 | 0 |
| 43309 | ENSMUSG00000105262 | 0 | NA | NA | NA | NA | NA | Gm19175 | predicted gene, 19175 [Source:MGI Symb | 0 | 0 | 0 | 0 | 0 | 0 |
| 43310 | ENSMUSG00000105263 | 2.874966552 | 0.618431531 | 1.296668922 | 0.476938655 | 0.633405805 | NA | Gm42427 | predicted gene 42427 [Source:MGI Symbo | 0.8031 | 2.0558 | 4.025 | 4.8199 | 1.0906 | 4.4554 |
| 43311 | ENSMUSG00000105264 | 0 | NA | NA | NA | NA | NA | Gm42716 | predicted gene 42716 [Source:MGI Symbo | 0 | 0 | 0 | 0 | 0 | 0 |
| 43312 | ENSMUSG00000105265 | 0 | NA | NA | NA | NA | NA | Sox2ot | SOX2 overlapping transcript (non-protein | 0 | 0 | 0 | 0 | 0 | 0 |
| 43313 | ENSMUSG00000105266 | 0 | NA | NA | NA | NA | NA | Gm18546 | predicted gene, 18546 [Source:MGI Symb | 0 | 0 | 0 | 0 | 0 | 0 |
| 43314 | ENSMUSG00000105267 | 0 | NA | NA | NA | NA | NA | Gm42542 | predicted gene 42542 [Source:MGI Symbo | 0 | 0 | 0 | 0 | 0 | 0 |
| 43315 | ENSMUSG00000105269 | 0 | NA | NA | NA | NA | NA | Gm42999 | predicted gene 42999 [Source:MGI Symbo | 0 | 0 | 0 | 0 | 0 | 0 |
| 43316 | ENSMUSG00000105270 | 0 | NA | NA | NA | NA | NA | Gm42863 | predicted gene 42863 [Source:MGI Symbo | 0 | 0 | 0 | 0 | 0 | 0 |
| 43317 | ENSMUSG00000105271 | 0 | NA | NA | NA | NA | NA | Gm42875 | predicted gene 42875 [Source:MGI Symbo | 0 | 0 | 0 | 0 | 0 | 0 |
| 43318 | ENSMUSG00000105272 | 27.90270082 | -0.734038602 | 0.46989921 | -1.56211925 | 0.118259899 | 0.503832452 | Gm43071 | predicted gene 43071 [Source:MGI Symbo | 45.777 | 22.614 | 36.225 | 30.847 | 16.359 | 15.594 |
| 43319 | ENSMUSG00000105273 | 0 | NA | NA | NA | NA | NA | Gm43015 | predicted gene 43015 [Source:MGI Symbo | 0 | 0 | 0 | 0 | 0 | 0 |
| 43320 | ENSMUSG00000105274 | 0 | NA | NA | NA | NA | NA | 1700048F04 | RIKEN cDNA 1700048F04 gene [Source:M( | 0 | 0 | 0 | 0 | 0 | 0 |
| 43321 | ENSMUSG00000105275 | 1.138206772 | 1.608612704 | 2.129587282 | 0.755363595 | 0.450030851 | NA | Gm43214 | predicted gene 43214 [Source:MGI Symbo | 1.6062 | 0 | 0 | 1.928 | 2.1812 | 1.1138 |
| 43322 | ENSMUSG00000105277 | 0 | NA | NA | NA | NA | NA | Gm9504 | predicted gene 9504 [Source:MGI Symbol | 0 | 0 | 0 | 0 | 0 | 0 |
| 43323 | ENSMUSG00000105278 | 0 | NA | NA | NA | NA | NA | Gm42630 | predicted gene 42630 [Source:MGI Symbo | 0 | 0 | 0 | 0 | 0 | 0 |
| 43324 | ENSMUSG00000105279 | 0 | NA | NA | NA | NA | NA | Gm6260 | predicted gene 6260 [Source:MGI Symbol | 0 | 0 | 0 | 0 | 0 | 0 |
| 43325 | ENSMUSG00000105280 | 0 | NA | NA | NA | NA | NA | Pgk1-ps3 | phosphoglycerate kinase 1, pseudogene 3 | 0 | 0 | 0 | 0 | 0 | 0 |
| 43326 | ENSMUSG00000105281 | 0 | NA | NA | NA | NA | NA | Gm17835 | predicted gene, 17835 [Source:MGI Symb | 0 | 0 | 0 | 0 | 0 | 0 |
| 43327 | ENSMUSG00000105282 | 0 | NA | NA | NA | NA | NA | Gm42981 | predicted gene 42981 [Source:MGI Symbo | 0 | 0 | 0 | 0 | 0 | 0 |
| 43328 | ENSMUSG00000105283 | 1.088452502 | -0.017212433 | 2.349360385 | -0.007326434 | 0.994154404 | NA | Gm33370 | predicted gene, 33370 [Source:MGI Symb | 3.2124 | 0 | 0 | 0 | 1.0906 | 2.2277 |
| 43329 | ENSMUSG00000105284 | 0 | NA | NA | NA | NA | NA | Gm42693 | predicted gene 42693 [Source:MGI Symbo | 0 | 0 | 0 | 0 | 0 | 0 |
| 43330 | ENSMUSG00000105285 | 0 | NA | NA | NA | NA | NA | Gm43238 | predicted gene 43238 [Source:MGI Symbo | 0 | 0 | 0 | 0 | 0 | 0 |
| 43331 | ENSMUSG00000105286 | 0.160662974 | 1.126924175 | 4.080472857 | 0.276174898 | 0.782413736 | NA | Gm43140 | predicted gene 43140 [Source:MGI Symbo | 0 | 0 | 0 | 0.964 | 0 | 0 |
| 43332 | ENSMUSG00000105287 | 0 | NA | NA | NA | NA | NA | Gm43577 | predicted gene 43577 [Source:MGI Symbo | 0 | 0 | 0 | 0 | 0 | 0 |
| 43333 | ENSMUSG00000105288 | 0 | NA | NA | NA | NA | NA | NA | NA | 0 | 0 | 0 | 0 | 0 | 0 |
| 43334 | ENSMUSG00000105289 | 0 | NA | NA | NA | NA | NA | Gm43483 | predicted gene 43483 [Source:MGI Symbo | 0 | 0 | 0 | 0 | 0 | 0 |
| 43335 | ENSMUSG00000105290 | 0 | NA | NA | NA | NA | NA | Gm43528 | predicted gene 43528 [Source:MGI Symbo | 0 | 0 | 0 | 0 | 0 | 0 |
| 43336 | ENSMUSG00000105291 | 0 | NA | NA | NA | NA | NA | Gm18901 | predicted gene, 18901 [Source:MGI Symb | 0 | 0 | 0 | 0 | 0 | 0 |
| 43337 | ENSMUSG00000105292 | 0 | NA | NA | NA | NA | NA | NA | NA | 0 | 0 | 0 | 0 | 0 | 0 |
| 43338 | ENSMUSG00000105293 | 0 | NA | NA | NA | NA | NA | Gm42843 | predicted gene 42843 [Source:MGI Symbo | 0 | 0 | 0 | 0 | 0 | 0 |
| 43339 | ENSMUSG00000105294 | 0 | NA | NA | NA | NA | NA | Gm43304 | predicted gene 43304 [Source:MGI Symbo | 0 | 0 | 0 | 0 | 0 | 0 |
| 43340 | ENSMUSG00000105295 | 0 | NA | NA | NA | NA | NA | Gm42834 | predicted gene 42834 [Source:MGI Symbo | 0 | 0 | 0 | 0 | 0 | 0 |

| | | | | | | | | | | | | | | | |
|---|---|---|---|---|---|---|---|---|---|---|---|---|---|---|---|
| 43341 | ENSMUSG00000105296 | 0 | NA | NA | NA | NA | NA | Gm19708 | predicted gene, 19708 [Source:MGI Symb | 0 | 0 | 0 | 0 | 0 | 0 |
| 43342 | ENSMUSG00000105297 | 0 | NA | NA | NA | NA | NA | 1700052H01 | RIKEN cDNA 1700052H01 gene [Source:M | 0 | 0 | 0 | 0 | 0 | 0 |
| 43343 | ENSMUSG00000105298 | 0 | NA | NA | NA | NA | NA | Cldn25 | claudin 25 [Source:MGI Symbol;Acc:MGI:3 | 0 | 0 | 0 | 0 | 0 | 0 |
| 43344 | ENSMUSG00000105299 | 0 | NA | NA | NA | NA | NA | Gm33847 | predicted gene, 33847 [Source:MGI Symb | 0 | 0 | 0 | 0 | 0 | 0 |
| 43345 | ENSMUSG00000105300 | 0 | NA | NA | NA | NA | NA | Gm30613 | predicted gene, 30613 [Source:MGI Symb | 0 | 0 | 0 | 0 | 0 | 0 |
| 43346 | ENSMUSG00000105301 | 0 | NA | NA | NA | NA | NA | Igkv13-54-1 | immunoglobulin kappa chain variable 13-5 | 0 | 0 | 0 | 0 | 0 | 0 |
| 43347 | ENSMUSG00000105302 | 0 | NA | NA | NA | NA | NA | Gm19817 | predicted gene, 19817 [Source:MGI Symb | 0 | 0 | 0 | 0 | 0 | 0 |
| 43348 | ENSMUSG00000105303 | 0.524202404 | 2.600864256 | 3.301637836 | 0.787749713 | 0.430843121 | NA | Gm18555 | predicted gene, 18555 [Source:MGI Symb | 0 | 0 | 0 | 0.964 | 2.1812 | 0 |
| 43349 | ENSMUSG00000105304 | 0.133850373 | -0.796629176 | 4.080472857 | -0.195229623 | 0.84521319 | NA | Gm43696 | predicted gene 43696 [Source:MGI Symbo | 0.8031 | 0 | 0 | 0 | 0 | 0 |
| 43350 | ENSMUSG00000105305 | 0 | NA | NA | NA | NA | NA | Gm8872 | predicted gene 8872 [Source:MGI Symbol | 0 | 0 | 0 | 0 | 0 | 0 |
| 43351 | ENSMUSG00000105306 | 0 | NA | NA | NA | NA | NA | Gm43171 | predicted gene 43171 [Source:MGI Symbo | 0 | 0 | 0 | 0 | 0 | 0 |
| 43352 | ENSMUSG00000105307 | 0 | NA | NA | NA | NA | NA | Gm43839 | predicted gene 43839 [Source:MGI Symbo | 0 | 0 | 0 | 0 | 0 | 0 |
| 43353 | ENSMUSG00000105308 | 0 | NA | NA | NA | NA | NA | 4930572K03 | RIKEN cDNA 4930572K03 gene [Source:M | 0 | 0 | 0 | 0 | 0 | 0 |
| 43354 | ENSMUSG00000105309 | 0 | NA | NA | NA | NA | NA | Gm21461 | predicted gene, 21461 [Source:MGI Symb | 0 | 0 | 0 | 0 | 0 | 0 |
| 43355 | ENSMUSG00000105310 | 0 | NA | NA | NA | NA | NA | Gm42451 | predicted gene 42451 [Source:MGI Symbo | 0 | 0 | 0 | 0 | 0 | 0 |
| 43356 | ENSMUSG00000105311 | 0 | NA | NA | NA | NA | NA | Gm44574 | predicted gene 44574 [Source:MGI Symbo | 0 | 0 | 0 | 0 | 0 | 0 |
| 43357 | ENSMUSG00000105312 | 0 | NA | NA | NA | NA | NA | NA | NA | 0 | 0 | 0 | 0 | 0 | 0 |
| 43358 | ENSMUSG00000105313 | 0 | NA | NA | NA | NA | NA | Gm43583 | predicted gene 43583 [Source:MGI Symbo | 0 | 0 | 0 | 0 | 0 | 0 |
| 43359 | ENSMUSG00000105314 | 0 | NA | NA | NA | NA | NA | Ranbp2-ps1 | RAN binding protein 2, pseudogene 11 [Sc | 0 | 0 | 0 | 0 | 0 | 0 |
| 43360 | ENSMUSG00000105315 | 27.25356187 | 0.622939562 | 0.454720174 | 1.369940455 | 0.17070549 | 0.597556794 | Gm18635 | predicted gene, 18635 [Source:MGI Symb | 31.321 | 19.53 | 13.081 | 27.955 | 35.99 | 35.643 |
| 43361 | ENSMUSG00000105316 | 0 | NA | NA | NA | NA | NA | Gm42953 | predicted gene 42953 [Source:MGI Symbo | 0 | 0 | 0 | 0 | 0 | 0 |
| 43362 | ENSMUSG00000105317 | 0 | NA | NA | NA | NA | NA | Gbp2-ps | guanylate binding protein 2, pseudogene | 0 | 0 | 0 | 0 | 0 | 0 |
| 43363 | ENSMUSG00000105318 | 0 | NA | NA | NA | NA | NA | NA | NA | 0 | 0 | 0 | 0 | 0 | 0 |
| 43364 | ENSMUSG00000105319 | 0 | NA | NA | NA | NA | NA | Igkv15-101 | immunoglobulin kappa chain variable 15-1 | 0 | 0 | 0 | 0 | 0 | 0 |
| 43365 | ENSMUSG00000105320 | 0 | NA | NA | NA | NA | NA | Gm43016 | predicted gene 43016 [Source:MGI Symbo | 0 | 0 | 0 | 0 | 0 | 0 |
| 43366 | ENSMUSG00000105321 | 0 | NA | NA | NA | NA | NA | NA | NA | 0 | 0 | 0 | 0 | 0 | 0 |
| 43367 | ENSMUSG00000105322 | 0 | NA | NA | NA | NA | NA | Gm43751 | predicted gene 43751 [Source:MGI Symbo | 0 | 0 | 0 | 0 | 0 | 0 |
| 43368 | ENSMUSG00000105323 | 0 | NA | NA | NA | NA | NA | Igkv13-62-1 | immunoglobulin kappa chain variable 13-6 | 0 | 0 | 0 | 0 | 0 | 0 |
| 43369 | ENSMUSG00000105324 | 0 | NA | NA | NA | NA | NA | Gm42705 | predicted gene 42705 [Source:MGI Symbo | 0 | 0 | 0 | 0 | 0 | 0 |
| 43370 | ENSMUSG00000105325 | 0 | NA | NA | NA | NA | NA | Gm30946 | predicted gene, 30946 [Source:MGI Symb | 0 | 0 | 0 | 0 | 0 | 0 |
| 43371 | ENSMUSG00000105326 | 5.46347285 | -2.415809735 | 1.090833098 | -2.21464653 | 0.026784336 | 0.211649994 | 6030400A10 | RIKEN cDNA 6030400A10 gene [Source:M | 6.4248 | 7.1954 | 14.087 | 2.8919 | 2.1812 | 0 |
| 43372 | ENSMUSG00000105327 | 0.167707431 | -0.796629176 | 4.080472857 | -0.195229623 | 0.84521319 | NA | Gm9409 | predicted gene 9409 [Source:MGI Symbol | 0 | 0 | 1.0062 | 0 | 0 | 0 |
| 43373 | ENSMUSG00000105328 | 0.167707431 | -0.796629176 | 4.080472857 | -0.195229623 | 0.84521319 | NA | Gm43180 | predicted gene 43180 [Source:MGI Symbo | 0 | 0 | 1.0062 | 0 | 0 | 0 |
| 43374 | ENSMUSG00000105329 | 0 | NA | NA | NA | NA | NA | Mir7215 | microRNA 7215 [Source:MGI Symbol;Acc: | 0 | 0 | 0 | 0 | 0 | 0 |
| 43375 | ENSMUSG00000105330 | 0 | NA | NA | NA | NA | NA | NA | NA | 0 | 0 | 0 | 0 | 0 | 0 |
| 43376 | ENSMUSG00000105331 | 0 | NA | NA | NA | NA | NA | Gm29865 | predicted gene, 29865 [Source:MGI Symb | 0 | 0 | 0 | 0 | 0 | 0 |
| 43377 | ENSMUSG00000105332 | 0 | NA | NA | NA | NA | NA | Gm43316 | predicted gene 43316 [Source:MGI Symbo | 0 | 0 | 0 | 0 | 0 | 0 |
| 43378 | ENSMUSG00000105333 | 0 | NA | NA | NA | NA | NA | Mir684-2 | microRNA 684-2 [Source:MGI Symbol;Acc | 0 | 0 | 0 | 0 | 0 | 0 |
| 43379 | ENSMUSG00000105334 | 0 | NA | NA | NA | NA | NA | Gm42680 | predicted gene 42680 [Source:MGI Symbo | 0 | 0 | 0 | 0 | 0 | 0 |
| 43380 | ENSMUSG00000105335 | 0 | NA | NA | NA | NA | NA | Gm42423 | predicted gene 42423 [Source:MGI Symbo | 0 | 0 | 0 | 0 | 0 | 0 |
| 43381 | ENSMUSG00000105336 | 0 | NA | NA | NA | NA | NA | AU022133 | expressed sequence AU022133 [Source:M | 0 | 0 | 0 | 0 | 0 | 0 |
| 43382 | ENSMUSG00000105337 | 0 | NA | NA | NA | NA | NA | 1700094M2 | RIKEN cDNA 1700094M23 gene [Source:M | 0 | 0 | 0 | 0 | 0 | 0 |
| 43383 | ENSMUSG00000105338 | 0 | NA | NA | NA | NA | NA | Gm43802 | predicted gene 43802 [Source:MGI Symbo | 0 | 0 | 0 | 0 | 0 | 0 |
| 43384 | ENSMUSG00000105339 | 0 | NA | NA | NA | NA | NA | Gm42457 | predicted gene 42457 [Source:MGI Symbo | 0 | 0 | 0 | 0 | 0 | 0 |
| 43385 | ENSMUSG00000105340 | 0.321325949 | 1.911575114 | 4.048267893 | 0.472195804 | 0.636787034 | NA | Gm42878 | predicted gene 42878 [Source:MGI Symbo | 0 | 0 | 0 | 1.928 | 0 | 0 |
| 43386 | ENSMUSG00000105341 | 0.985291176 | 0.960545195 | 2.698810486 | 0.355914281 | 0.72190476 | NA | Gm43817 | predicted gene 43817 [Source:MGI Symbo | 0 | 2.0558 | 0 | 3.8559 | 0 | 0 |
| 43387 | ENSMUSG00000105342 | 0 | NA | NA | NA | NA | NA | Gm43244 | predicted gene 43244 [Source:MGI Symbo | 0 | 0 | 0 | 0 | 0 | 0 |
| 43388 | ENSMUSG00000105343 | 0 | NA | NA | NA | NA | NA | Gm43512 | predicted gene 43512 [Source:MGI Symbo | 0 | 0 | 0 | 0 | 0 | 0 |
| 43389 | ENSMUSG00000105344 | 0 | NA | NA | NA | NA | NA | Gm43556 | predicted gene 43556 [Source:MGI Symbo | 0 | 0 | 0 | 0 | 0 | 0 |
| 43390 | ENSMUSG00000105345 | 2.694508804 | 0.894478463 | 1.269537347 | 0.704570421 | 0.481077612 | NA | BC030343 | cDNA sequence BC030343 [Source:MGI Sy | 1.6062 | 2.0558 | 2.0125 | 3.8559 | 2.1812 | 4.4554 |
| 43391 | ENSMUSG00000105346 | 0 | NA | NA | NA | NA | NA | Gm18894 | predicted gene, 18894 [Source:MGI Symb | 0 | 0 | 0 | 0 | 0 | 0 |
| 43392 | ENSMUSG00000105347 | 0 | NA | NA | NA | NA | NA | Gm43503 | predicted gene 43503 [Source:MGI Symbo | 0 | 0 | 0 | 0 | 0 | 0 |
| 43393 | ENSMUSG00000105348 | 0 | NA | NA | NA | NA | NA | Gm35212 | predicted gene, 35212 [Source:MGI Symb | 0 | 0 | 0 | 0 | 0 | 0 |
| 43394 | ENSMUSG00000105349 | 3.077222708 | 0.360720623 | 1.328615182 | 0.271501205 | 0.786005573 | NA | Gm43604 | predicted gene 43604 [Source:MGI Symbo | 4.0155 | 0 | 4.025 | 3.8559 | 5.4531 | 1.1138 |
| 43395 | ENSMUSG00000105350 | 0.17131964 | -0.796629176 | 4.080472857 | -0.195229623 | 0.84521319 | NA | 4930537H20 | RIKEN cDNA 4930537H20 gene [Source:M | 0 | 1.0279 | 0 | 0 | 0 | 0 |
| 43396 | ENSMUSG00000105351 | 0 | NA | NA | NA | NA | NA | Gm43157 | predicted gene 43157 [Source:MGI Symbo | 0 | 0 | 0 | 0 | 0 | 0 |

| | | | | | | | | | | | | | | | |
|---|---|---|---|---|---|---|---|---|---|---|---|---|---|---|---|
| 43397 | ENSMUSG00000105352 | 0 | NA | NA | NA | NA | NA | C030018K13 | RIKEN cDNA C030018K13 gene [Source:M | 0 | 0 | 0 | 0 | 0 | 0 |
| 43398 | ENSMUSG00000105353 | 3.196130586 | -0.964106725 | 1.33139986 | -0.724130109 | 0.468985862 | NA | Gm42428 | predicted gene 42428 [Source:MGI Symbo | 2.4093 | 9.2513 | 1.0062 | 0.964 | 1.0906 | 4.4554 |
| 43399 | ENSMUSG00000105355 | 0.133850373 | -0.796629176 | 4.080472857 | -0.195229623 | 0.84521319 | NA | Gm40117 | predicted gene, 40117 [Source:MGI Symb | 0.8031 | 0 | 0 | 0 | 0 | 0 |
| 43400 | ENSMUSG00000105356 | 0 | NA | NA | NA | NA | NA | Gm42603 | predicted gene 42603 [Source:MGI Symbo | 0 | 0 | 0 | 0 | 0 | 0 |
| 43401 | ENSMUSG00000105357 | 0 | NA | NA | NA | NA | NA | Gm42647 | predicted gene 42647 [Source:MGI Symbo | 0 | 0 | 0 | 0 | 0 | 0 |
| 43402 | ENSMUSG00000105358 | 0.321325949 | 1.911575114 | 4.048267893 | 0.472195804 | 0.636787034 | NA | Gm43099 | predicted gene 43099 [Source:MGI Symbo | 0 | 0 | 0 | 1.928 | 0 | 0 |
| 43403 | ENSMUSG00000105359 | 0.185640647 | 1.126924175 | 4.080472857 | 0.276174898 | 0.782413736 | NA | Rpl21-ps10 | ribosomal protein L21, pseudogene 10 [Sc | 0 | 0 | 0 | 0 | 0 | 1.1138 |
| 43404 | ENSMUSG00000105360 | 0 | NA | NA | NA | NA | NA | Gm7824 | predicted gene 7824 [Source:MGI Symbol | 0 | 0 | 0 | 0 | 0 | 0 |
| 43405 | ENSMUSG00000105361 | 0 | NA | NA | NA | NA | NA | NA | NA | 0 | 0 | 0 | 0 | 0 | 0 |
| 43406 | ENSMUSG00000105362 | 1.486319818 | -0.924740954 | 1.787691823 | -0.517282085 | 0.604959258 | NA | Gm43474 | predicted gene 43474 [Source:MGI Symbo | 0.8031 | 3.0838 | 2.0125 | 1.928 | 1.0906 | 0 |
| 43407 | ENSMUSG00000105363 | 0 | NA | NA | NA | NA | NA | Igkv11-114 | immunoglobulin kappa chain variable [Sou | 0 | 0 | 0 | 0 | 0 | 0 |
| 43408 | ENSMUSG00000105364 | 0 | NA | NA | NA | NA | NA | Gm8924 | predicted gene 8924 [Source:MGI Symbol | 0 | 0 | 0 | 0 | 0 | 0 |
| 43409 | ENSMUSG00000105365 | 0 | NA | NA | NA | NA | NA | NA | NA | 0 | 0 | 0 | 0 | 0 | 0 |
| 43410 | ENSMUSG00000105366 | 0.167707431 | -0.796629176 | 4.080472857 | -0.195229623 | 0.84521319 | NA | Gm43719 | predicted gene 43719 [Source:MGI Symbo | 0 | 0 | 1.0062 | 0 | 0 | 0 |
| 43411 | ENSMUSG00000105367 | 0 | NA | NA | NA | NA | NA | NA | NA | 0 | 0 | 0 | 0 | 0 | 0 |
| 43412 | ENSMUSG00000105368 | 20.83048575 | 0.146884875 | 0.537219137 | 0.273417057 | 0.784532643 | 0.960670212 | Gm43759 | predicted gene 43759 [Source:MGI Symbo | 13.653 | 24.67 | 21.131 | 21.208 | 10.906 | 33.415 |
| 43413 | ENSMUSG00000105369 | 0 | NA | NA | NA | NA | NA | Igkv13-57-1 | immunoglobulin kappa chain variable 13-! | 0 | 0 | 0 | 0 | 0 | 0 |
| 43414 | ENSMUSG00000105370 | 0 | NA | NA | NA | NA | NA | Gm42718 | predicted gene 42718 [Source:MGI Symbo | 0 | 0 | 0 | 0 | 0 | 0 |
| 43415 | ENSMUSG00000105371 | 0 | NA | NA | NA | NA | NA | NA | NA | 0 | 0 | 0 | 0 | 0 | 0 |
| 43416 | ENSMUSG00000105372 | 0 | NA | NA | NA | NA | NA | Mir327 | microRNA 327 [Source:MGI Symbol;Acc:M | 0 | 0 | 0 | 0 | 0 | 0 |
| 43417 | ENSMUSG00000105373 | 0 | NA | NA | NA | NA | NA | Gm42429 | predicted gene 42429 [Source:MGI Symbo | 0 | 0 | 0 | 0 | 0 | 0 |
| 43418 | ENSMUSG00000105374 | 1.934394035 | -0.789005103 | 1.623000391 | -0.486139811 | 0.626868042 | NA | Gm42551 | predicted gene 42551 [Source:MGI Symbo | 3.2124 | 4.1117 | 0 | 0.964 | 1.0906 | 2.2277 |
| 43419 | ENSMUSG00000105375 | 0 | NA | NA | NA | NA | NA | Gm20239 | predicted gene, 20239 [Source:MGI Symb | 0 | 0 | 0 | 0 | 0 | 0 |
| 43420 | ENSMUSG00000105376 | 0.160662974 | 1.126924175 | 4.080472857 | 0.276174898 | 0.782413736 | NA | Gm36535 | predicted gene, 36535 [Source:MGI Symb | 0 | 0 | 0 | 0.964 | 0 | 0 |
| 43421 | ENSMUSG00000105377 | 0.160662974 | 1.126924175 | 4.080472857 | 0.276174898 | 0.782413736 | NA | Gm43148 | predicted gene 43148 [Source:MGI Symbo | 0 | 0 | 0 | 0.964 | 0 | 0 |
| 43422 | ENSMUSG00000105378 | 0.181769715 | 1.126924175 | 4.080472857 | 0.276174898 | 0.782413736 | NA | Gm42515 | predicted gene 42515 [Source:MGI Symbo | 0 | 0 | 0 | 0 | 1.0906 | 0 |
| 43423 | ENSMUSG00000105379 | 0.729921525 | -1.61423457 | 2.706831497 | -0.596355766 | 0.550937584 | NA | Mir1191b | microRNA 1191b [Source:MGI Symbol;Acc | 2.4093 | 0 | 1.0062 | 0.964 | 0 | 0 |
| 43424 | ENSMUSG00000105380 | 0 | NA | NA | NA | NA | NA | Trav12-4 | T cell receptor alpha variable 12-4 [Source | 0 | 0 | 0 | 0 | 0 | 0 |
| 43425 | ENSMUSG00000105381 | 0 | NA | NA | NA | NA | NA | Mir7234 | microRNA 7234 [Source:MGI Symbol;Acc: | 0 | 0 | 0 | 0 | 0 | 0 |
| 43426 | ENSMUSG00000105382 | 1.030936906 | 0.079807104 | 2.204048843 | 0.036209317 | 0.971115457 | NA | Gm43339 | predicted gene 43339 [Source:MGI Symbo | 0 | 1.0279 | 2.0125 | 0.964 | 2.1812 | 0 |
| 43427 | ENSMUSG00000105384 | 0 | NA | NA | NA | NA | NA | Gm43396 | predicted gene 43396 [Source:MGI Symbo | 0 | 0 | 0 | 0 | 0 | 0 |
| 43428 | ENSMUSG00000105385 | 0 | NA | NA | NA | NA | NA | 9230114J08I | RIKEN cDNA 9230114J08 gene [Source:MC | 0 | 0 | 0 | 0 | 0 | 0 |
| 43429 | ENSMUSG00000105386 | 0 | NA | NA | NA | NA | NA | Gm43104 | predicted gene 43104 [Source:MGI Symbo | 0 | 0 | 0 | 0 | 0 | 0 |
| 43430 | ENSMUSG00000105387 | 0 | NA | NA | NA | NA | NA | Gm29681 | predicted gene, 29681 [Source:MGI Symb | 0 | 0 | 0 | 0 | 0 | 0 |
| 43431 | ENSMUSG00000105388 | 8.871462418 | -0.174239593 | 0.722607433 | -0.241126211 | 0.809457298 | 0.966005805 | Rpl36a-ps2 | ribosomal protein L36A, pseudogene 2 [Sc | 8.031 | 7.1954 | 13.081 | 10.604 | 7.6343 | 6.6831 |
| 43432 | ENSMUSG00000105389 | 1.004562989 | 0.155751777 | 2.077846764 | 0.07495826 | 0.940247922 | NA | BB187690 | expressed sequence BB187690 [Source:M | 0.8031 | 2.0558 | 0 | 0.964 | 1.0906 | 1.1138 |
| 43433 | ENSMUSG00000105390 | 0 | NA | NA | NA | NA | NA | Gm8458 | predicted gene 8458 [Source:MGI Symbol | 0 | 0 | 0 | 0 | 0 | 0 |
| 43434 | ENSMUSG00000105391 | 6.399033606 | -0.961502443 | 0.996926055 | -0.964467162 | 0.334811764 | 0.776159726 | Gm43401 | predicted gene 43401 [Source:MGI Symbo | 11.243 | 7.1954 | 7.0437 | 9.6398 | 3.2719 | 0 |
| 43435 | ENSMUSG00000105392 | 0 | NA | NA | NA | NA | NA | Gm42684 | predicted gene 42684 [Source:MGI Symbo | 0 | 0 | 0 | 0 | 0 | 0 |
| 43436 | ENSMUSG00000105393 | 0 | NA | NA | NA | NA | NA | Trav7d-5 | T cell receptor alpha variable 7D-5 [Source | 0 | 0 | 0 | 0 | 0 | 0 |
| 43437 | ENSMUSG00000105394 | 0 | NA | NA | NA | NA | NA | Gm43115 | predicted gene 43115 [Source:MGI Symbo | 0 | 0 | 0 | 0 | 0 | 0 |
| 43438 | ENSMUSG00000105395 | 5.012925778 | 1.364544306 | 1.229102805 | 1.110195422 | 0.266914825 | NA | Gm42672 | predicted gene 42672 [Source:MGI Symbo | 2.4093 | 2.0558 | 4.025 | 18.316 | 3.2719 | 0 |
| 43439 | ENSMUSG00000105396 | 0 | NA | NA | NA | NA | NA | Gm6057 | predicted gene 6057 [Source:MGI Symbol | 0 | 0 | 0 | 0 | 0 | 0 |
| 43440 | ENSMUSG00000105397 | 0 | NA | NA | NA | NA | NA | Gm43471 | predicted gene 43471 [Source:MGI Symbo | 0 | 0 | 0 | 0 | 0 | 0 |
| 43441 | ENSMUSG00000105398 | 0 | NA | NA | NA | NA | NA | NA | NA | 0 | 0 | 0 | 0 | 0 | 0 |
| 43442 | ENSMUSG00000105399 | 0 | NA | NA | NA | NA | NA | Mir1843b | microRNA 1843b [Source:MGI Symbol;Acc | 0 | 0 | 0 | 0 | 0 | 0 |
| 43443 | ENSMUSG00000105400 | 0 | NA | NA | NA | NA | NA | Gm22897 | predicted gene, 22897 [Source:MGI Symb | 0 | 0 | 0 | 0 | 0 | 0 |
| 43444 | ENSMUSG00000105401 | 0 | NA | NA | NA | NA | NA | Gm42949 | predicted gene 42949 [Source:MGI Symbo | 0 | 0 | 0 | 0 | 0 | 0 |
| 43445 | ENSMUSG00000105402 | 0 | NA | NA | NA | NA | NA | Gm3716 | predicted gene 3716 [Source:MGI Symbol | 0 | 0 | 0 | 0 | 0 | 0 |
| 43446 | ENSMUSG00000105403 | 0.305170013 | -1.657836477 | 4.041480335 | -0.410205257 | 0.681655385 | NA | Gm43618 | predicted gene 43618 [Source:MGI Symbo | 0.8031 | 1.0279 | 0 | 0 | 0 | 0 |
| 43447 | ENSMUSG00000105404 | 1.184582053 | -1.265002395 | 2.170606727 | -0.582787467 | 0.560036383 | NA | Gm42807 | predicted gene 42807 [Source:MGI Symbo | 0 | 1.0279 | 4.025 | 0.964 | 1.0906 | 0 |
| 43448 | ENSMUSG00000105405 | 0 | NA | NA | NA | NA | NA | Gm42431 | predicted gene 42431 [Source:MGI Symbo | 0 | 0 | 0 | 0 | 0 | 0 |
| 43449 | ENSMUSG00000105406 | 0 | NA | NA | NA | NA | NA | Gm43247 | predicted gene 43247 [Source:MGI Symbo | 0 | 0 | 0 | 0 | 0 | 0 |
| 43450 | ENSMUSG00000105407 | 0 | NA | NA | NA | NA | NA | Gm36259 | predicted gene, 36259 [Source:MGI Symb | 0 | 0 | 0 | 0 | 0 | 0 |
| 43451 | ENSMUSG00000105408 | 6.466942159 | 0.043979377 | 0.921311826 | 0.047735605 | 0.961926958 | 0.992908678 | Gm7285 | predicted gene 7285 [Source:MGI Symbol | 12.85 | 2.0558 | 4.025 | 7.7118 | 4.3625 | 7.7969 |
| 43452 | ENSMUSG00000105409 | 0 | NA | NA | NA | NA | NA | NA | NA | 0 | 0 | 0 | 0 | 0 | 0 |

| | | | | | | | | | | | | | | | |
|---|---|---|---|---|---|---|---|---|---|---|---|---|---|---|---|
| 43453 | ENSMUSG00000105410 | 0 | NA | NA | NA | NA | NA | Gtpbp4-ps7 | GTP binding protein 4, pseudogene 7 [Sou | 0 | 0 | 0 | 0 | 0 | 0 |
| 43454 | ENSMUSG00000105411 | 0 | NA | NA | NA | NA | NA | NA | NA | 0 | 0 | 0 | 0 | 0 | 0 |
| 43455 | ENSMUSG00000105412 | 0.328370405 | 0.16514316 | 4.003149536 | 0.041253308 | 0.967093956 | NA | Gm43768 | predicted gene 43768 [Source:MGI Symbo | 0 | 0 | 1.0062 | 0.964 | 0 | 0 |
| 43456 | ENSMUSG00000105413 | 0 | NA | NA | NA | NA | NA | NA | NA | 0 | 0 | 0 | 0 | 0 | 0 |
| 43457 | ENSMUSG00000105415 | 0 | NA | NA | NA | NA | NA | NA | NA | 0 | 0 | 0 | 0 | 0 | 0 |
| 43458 | ENSMUSG00000105416 | 0.877498349 | -0.489540585 | 2.815048567 | -0.173901293 | 0.861943037 | NA | Gm42553 | predicted gene 42553 [Source:MGI Symbo | 0 | 3.0838 | 0 | 0 | 2.1812 | 0 |
| 43459 | ENSMUSG00000105418 | 0 | NA | NA | NA | NA | NA | Gm43650 | predicted gene 43650 [Source:MGI Symbo | 0 | 0 | 0 | 0 | 0 | 0 |
| 43460 | ENSMUSG00000105419 | 10.93193282 | -0.02329938 | 0.639638402 | -0.036425862 | 0.970942793 | 0.994966572 | Gm43205 | predicted gene 43205 [Source:MGI Symbo | 13.653 | 9.2513 | 10.062 | 10.604 | 11.997 | 10.025 |
| 43461 | ENSMUSG00000105420 | 0 | NA | NA | NA | NA | NA | NA | NA | 0 | 0 | 0 | 0 | 0 | 0 |
| 43462 | ENSMUSG00000105421 | 0 | NA | NA | NA | NA | NA | 5830411K02 | RIKEN cDNA 5830411K02 gene [Source:M | 0 | 0 | 0 | 0 | 0 | 0 |
| 43463 | ENSMUSG00000105422 | 0 | NA | NA | NA | NA | NA | Gm6089 | predicted gene 6089 [Source:MGI Symbol | 0 | 0 | 0 | 0 | 0 | 0 |
| 43464 | ENSMUSG00000105423 | 0 | NA | NA | NA | NA | NA | Gm36814 | predicted gene, 36814 [Source:MGI Symb | 0 | 0 | 0 | 0 | 0 | 0 |
| 43465 | ENSMUSG00000105424 | 0 | NA | NA | NA | NA | NA | NA | NA | 0 | 0 | 0 | 0 | 0 | 0 |
| 43466 | ENSMUSG00000105425 | 0 | NA | NA | NA | NA | NA | Gm43469 | predicted gene 43469 [Source:MGI Symbo | 0 | 0 | 0 | 0 | 0 | 0 |
| 43467 | ENSMUSG00000105426 | 0 | NA | NA | NA | NA | NA | 1700021F02 | RIKEN cDNA 1700021F02 gene [Source:M | 0 | 0 | 0 | 0 | 0 | 0 |
| 43468 | ENSMUSG00000105427 | 0 | NA | NA | NA | NA | NA | Gm31678 | predicted gene, 31678 [Source:MGI Symb | 0 | 0 | 0 | 0 | 0 | 0 |
| 43469 | ENSMUSG00000105428 | 0 | NA | NA | NA | NA | NA | Mir3068 | microRNA 3068 [Source:MGI Symbol;Acc: | 0 | 0 | 0 | 0 | 0 | 0 |
| 43470 | ENSMUSG00000105429 | 3.463811632 | 0.468037288 | 1.214267029 | 0.385448404 | 0.699905227 | NA | Gm43692 | predicted gene 43692 [Source:MGI Symbo | 5.6217 | 1.0279 | 2.0125 | 7.7118 | 2.1812 | 2.2277 |
| 43471 | ENSMUSG00000105430 | 0 | NA | NA | NA | NA | NA | Gm43301 | predicted gene 43301 [Source:MGI Symbo | 0 | 0 | 0 | 0 | 0 | 0 |
| 43472 | ENSMUSG00000105431 | 0 | NA | NA | NA | NA | NA | Gm42640 | predicted gene 42640 [Source:MGI Symbo | 0 | 0 | 0 | 0 | 0 | 0 |
| 43473 | ENSMUSG00000105432 | 0 | NA | NA | NA | NA | NA | Gm43218 | predicted gene 43218 [Source:MGI Symbo | 0 | 0 | 0 | 0 | 0 | 0 |
| 43474 | ENSMUSG00000105433 | 0 | NA | NA | NA | NA | NA | NA | NA | 0 | 0 | 0 | 0 | 0 | 0 |
| 43475 | ENSMUSG00000105434 | 1.830529787 | -0.654948551 | 1.558272847 | -0.420304154 | 0.674263276 | NA | Gm43359 | predicted gene 43359 [Source:MGI Symbo | 1.6062 | 4.1117 | 1.0062 | 0.964 | 2.1812 | 1.1138 |
| 43476 | ENSMUSG00000105435 | 0 | NA | NA | NA | NA | NA | NA | NA | 0 | 0 | 0 | 0 | 0 | 0 |
| 43477 | ENSMUSG00000105436 | 0 | NA | NA | NA | NA | NA | Gm43212 | predicted gene 43212 [Source:MGI Symbo | 0 | 0 | 0 | 0 | 0 | 0 |
| 43478 | ENSMUSG00000105437 | 0 | NA | NA | NA | NA | NA | Gm42450 | predicted gene 42450 [Source:MGI Symbo | 0 | 0 | 0 | 0 | 0 | 0 |
| 43479 | ENSMUSG00000105438 | 0 | NA | NA | NA | NA | NA | NA | NA | 0 | 0 | 0 | 0 | 0 | 0 |
| 43480 | ENSMUSG00000105439 | 0 | NA | NA | NA | NA | NA | A230098N1( | RIKEN cDNA A230098N10 gene [Source:M | 0 | 0 | 0 | 0 | 0 | 0 |
| 43481 | ENSMUSG00000105440 | 0 | NA | NA | NA | NA | NA | Gm31693 | predicted gene, 31693 [Source:MGI Symb | 0 | 0 | 0 | 0 | 0 | 0 |
| 43482 | ENSMUSG00000105441 | 0 | NA | NA | NA | NA | NA | Gm8041 | predicted gene 8041 [Source:MGI Symbol | 0 | 0 | 0 | 0 | 0 | 0 |
| 43483 | ENSMUSG00000105442 | 0.649696354 | 1.537929172 | 3.118369374 | 0.493183773 | 0.621882743 | NA | Gm42614 | predicted gene 42614 [Source:MGI Symbo | 0 | 0 | 1.0062 | 2.8919 | 0 | 0 |
| 43484 | ENSMUSG00000105443 | 0 | NA | NA | NA | NA | NA | Gm43837 | predicted gene 43837 [Source:MGI Symbo | 0 | 0 | 0 | 0 | 0 | 0 |
| 43485 | ENSMUSG00000105444 | 0 | NA | NA | NA | NA | NA | Gm10727 | predicted gene 10727 [Source:MGI Symbo | 0 | 0 | 0 | 0 | 0 | 0 |
| 43486 | ENSMUSG00000105445 | 0 | NA | NA | NA | NA | NA | Gm42972 | predicted gene 42972 [Source:MGI Symbo | 0 | 0 | 0 | 0 | 0 | 0 |
| 43487 | ENSMUSG00000105446 | 0.55305101 | 2.667977666 | 3.241982584 | 0.822946329 | 0.410538515 | NA | Gm42605 | predicted gene 42605 [Source:MGI Symbo | 0 | 0 | 0 | 0 | 1.0906 | 2.2277 |
| 43488 | ENSMUSG00000105447 | 5.36063983 | 0.43936064 | 0.987990492 | 0.444701284 | 0.656535642 | 0.926813539 | Gm43653 | predicted gene 43653 [Source:MGI Symbo | 1.6062 | 7.1954 | 5.0312 | 10.604 | 3.2719 | 4.4554 |
| 43489 | ENSMUSG00000105448 | 0 | NA | NA | NA | NA | NA | Gm25820 | predicted gene, 25820 [Source:MGI Symb | 0 | 0 | 0 | 0 | 0 | 0 |
| 43490 | ENSMUSG00000105449 | 18.75168512 | -0.035216455 | 0.527163091 | -0.066803719 | 0.946737963 | 0.990640085 | Gm43379 | predicted gene 43379 [Source:MGI Symbo | 15.259 | 28.782 | 13.081 | 21.208 | 16.359 | 17.822 |
| 43491 | ENSMUSG00000105450 | 0 | NA | NA | NA | NA | NA | NA | NA | 0 | 0 | 0 | 0 | 0 | 0 |
| 43492 | ENSMUSG00000105451 | 0 | NA | NA | NA | NA | NA | Gm43220 | predicted gene 43220 [Source:MGI Symbo | 0 | 0 | 0 | 0 | 0 | 0 |
| 43493 | ENSMUSG00000105452 | 3.60261281 | -0.149112888 | 1.147621782 | -0.129932083 | 0.896620161 | NA | Gm5276 | predicted gene 5276 [Source:MGI Symbol | 7.2279 | 2.0558 | 2.0125 | 4.8199 | 3.2719 | 2.2277 |
| 43494 | ENSMUSG00000105453 | 0.981678968 | 0.981156725 | 2.481402239 | 0.395404143 | 0.692544652 | NA | Gm43702 | predicted gene 43702 [Source:MGI Symbo | 0 | 1.0279 | 1.0062 | 3.8559 | 0 | 0 |
| 43495 | ENSMUSG00000105454 | 0 | NA | NA | NA | NA | NA | Gm43830 | predicted gene 43830 [Source:MGI Symbo | 0 | 0 | 0 | 0 | 0 | 0 |
| 43496 | ENSMUSG00000105455 | 0 | NA | NA | NA | NA | NA | NA | NA | 0 | 0 | 0 | 0 | 0 | 0 |
| 43497 | ENSMUSG00000105456 | 0 | NA | NA | NA | NA | NA | Gm43745 | predicted gene 43745 [Source:MGI Symbo | 0 | 0 | 0 | 0 | 0 | 0 |
| 43498 | ENSMUSG00000105457 | 0.653308562 | 1.538013299 | 3.112409277 | 0.494155223 | 0.62119656 | NA | Gm43200 | predicted gene 43200 [Source:MGI Symbo | 0 | 1.0279 | 0 | 2.8919 | 0 | 0 |
| 43499 | ENSMUSG00000105458 | 0 | NA | NA | NA | NA | NA | Mir3074-2 | microRNA 3074-2 [Source:MGI Symbol;Ac | 0 | 0 | 0 | 0 | 0 | 0 |
| 43500 | ENSMUSG00000105459 | 0 | NA | NA | NA | NA | NA | 4930449I04F | RIKEN cDNA 4930449I04 gene [Source:MG | 0 | 0 | 0 | 0 | 0 | 0 |
| 43501 | ENSMUSG00000105460 | 0 | NA | NA | NA | NA | NA | Gm42671 | predicted gene 42671 [Source:MGI Symbo | 0 | 0 | 0 | 0 | 0 | 0 |
| 43502 | ENSMUSG00000105461 | 0 | NA | NA | NA | NA | NA | Gm4742 | predicted gene 4742 [Source:MGI Symbol | 0 | 0 | 0 | 0 | 0 | 0 |
| 43503 | ENSMUSG00000105462 | 0.401551119 | -2.072982726 | 4.008251979 | -0.517178744 | 0.605031388 | NA | Igkv4-77 | immunoglobulin kappa variable 4-77 [Sou | 2.4093 | 0 | 0 | 0 | 0 | 0 |
| 43504 | ENSMUSG00000105463 | 0 | NA | NA | NA | NA | NA | Igkv2-113 | immunoglobulin kappa chain variable 2-1 | 0 | 0 | 0 | 0 | 0 | 0 |
| 43505 | ENSMUSG00000105464 | 0 | NA | NA | NA | NA | NA | Rpl18a-ps2 | ribosomal protein L18A, pseudogene 2 [Sc | 0 | 0 | 0 | 0 | 0 | 0 |
| 43506 | ENSMUSG00000105465 | 0 | NA | NA | NA | NA | NA | NA | NA | 0 | 0 | 0 | 0 | 0 | 0 |
| 43507 | ENSMUSG00000105466 | 0.17131964 | -0.796629176 | 4.080472857 | -0.195229623 | 0.84521319 | NA | Gm42998 | predicted gene 42998 [Source:MGI Symbo | 0 | 1.0279 | 0 | 0 | 0 | 0 |
| 43508 | ENSMUSG00000105467 | 0 | NA | NA | NA | NA | NA | Trav5n-3 | T cell receptor alpha variable 5N-3 [Sourc | 0 | 0 | 0 | 0 | 0 | 0 |

| | | | | | | | | | | | | | | | |
|---|---|---|---|---|---|---|---|---|---|---|---|---|---|---|---|
| 43509 | ENSMUSG00000105468 | 0 | NA | NA | NA | NA | NA | Mir6906 | microRNA 6906 [Source:MGI Symbol;Acc: | 0 | 0 | 0 | 0 | 0 | 0 |
| 43510 | ENSMUSG00000105469 | 0 | NA | NA | NA | NA | NA | A930036I15I | RIKEN cDNA A930036I15 gene [Source:M( | 0 | 0 | 0 | 0 | 0 | 0 |
| 43511 | ENSMUSG00000105470 | 0 | NA | NA | NA | NA | NA | Gm42589 | predicted gene 42589 [Source:MGI Symbo | 0 | 0 | 0 | 0 | 0 | 0 |
| 43512 | ENSMUSG00000105471 | 2.317785886 | 0.058646698 | 1.494774799 | 0.039234471 | 0.968703451 | NA | A430073D23 | RIKEN cDNA A430073D23 gene [Source:M | 0.8031 | 3.0838 | 3.0187 | 4.8199 | 2.1812 | 0 |
| 43513 | ENSMUSG00000105472 | 1.217274962 | 1.417673431 | 2.109212581 | 0.672133972 | 0.501498411 | NA | Gm43024 | predicted gene 43024 [Source:MGI Symbo | 0 | 1.0279 | 1.0062 | 1.928 | 0 | 3.3415 |
| 43514 | ENSMUSG00000105473 | 0 | NA | NA | NA | NA | NA | Mark1-ps1 | MAP/microtubule affinity regulating kinas | 0 | 0 | 0 | 0 | 0 | 0 |
| 43515 | ENSMUSG00000105474 | 0 | NA | NA | NA | NA | NA | Selenok-ps7 | selenoprotein K, pseudogene 7 [Source:M | 0 | 0 | 0 | 0 | 0 | 0 |
| 43516 | ENSMUSG00000105475 | 0 | NA | NA | NA | NA | NA | 4930431L21I | RIKEN cDNA 4930431L21 gene [Source:M( | 0 | 0 | 0 | 0 | 0 | 0 |
| 43517 | ENSMUSG00000105476 | 0 | NA | NA | NA | NA | NA | Gm35439 | predicted gene, 35439 [Source:MGI Symb | 0 | 0 | 0 | 0 | 0 | 0 |
| 43518 | ENSMUSG00000105477 | 0 | NA | NA | NA | NA | NA | NA | NA | 0 | 0 | 0 | 0 | 0 | 0 |
| 43519 | ENSMUSG00000105478 | 0 | NA | NA | NA | NA | NA | Gm43285 | predicted gene 43285 [Source:MGI Symbo | 0 | 0 | 0 | 0 | 0 | 0 |
| 43520 | ENSMUSG00000105479 | 0 | NA | NA | NA | NA | NA | Gm43132 | predicted gene 43132 [Source:MGI Symbo | 0 | 0 | 0 | 0 | 0 | 0 |
| 43521 | ENSMUSG00000105480 | 0 | NA | NA | NA | NA | NA | Igkv13-78-1 | immunoglobulin kappa variable 13-78-1 [S | 0 | 0 | 0 | 0 | 0 | 0 |
| 43522 | ENSMUSG00000105481 | 0 | NA | NA | NA | NA | NA | 6430500D05 | RIKEN cDNA 6430500D05 gene [Source:M | 0 | 0 | 0 | 0 | 0 | 0 |
| 43523 | ENSMUSG00000105482 | 0.531944269 | 2.618916564 | 3.285123409 | 0.797204926 | 0.425332026 | NA | Gm42596 | predicted gene 42596 [Source:MGI Symbo | 0 | 0 | 0 | 0.964 | 0 | 2.2277 |
| 43524 | ENSMUSG00000105483 | 0 | NA | NA | NA | NA | NA | 4930553I04I | RIKEN cDNA 4930553I04 gene [Source:M( | 0 | 0 | 0 | 0 | 0 | 0 |
| 43525 | ENSMUSG00000105484 | 0 | NA | NA | NA | NA | NA | Gm42775 | predicted gene 42775 [Source:MGI Symbo | 0 | 0 | 0 | 0 | 0 | 0 |
| 43526 | ENSMUSG00000105485 | 0.321325949 | 1.911575114 | 4.048267893 | 0.472195804 | 0.636787034 | NA | Gm42447 | predicted gene 42447 [Source:MGI Symbo | 0 | 0 | 0 | 1.928 | 0 | 0 |
| 43527 | ENSMUSG00000105486 | 0 | NA | NA | NA | NA | NA | Gm3372 | predicted gene 3372 [Source:MGI Symbol | 0 | 0 | 0 | 0 | 0 | 0 |
| 43528 | ENSMUSG00000105487 | 0 | NA | NA | NA | NA | NA | Gm43324 | predicted gene 43324 [Source:MGI Symbo | 0 | 0 | 0 | 0 | 0 | 0 |
| 43529 | ENSMUSG00000105488 | 0 | NA | NA | NA | NA | NA | NA | NA | 0 | 0 | 0 | 0 | 0 | 0 |
| 43530 | ENSMUSG00000105489 | 0 | NA | NA | NA | NA | NA | Gm38562 | predicted gene, 38562 [Source:MGI Symb | 0 | 0 | 0 | 0 | 0 | 0 |
| 43531 | ENSMUSG00000105490 | 0 | NA | NA | NA | NA | NA | 4933438B17 | RIKEN cDNA 4933438B17 gene [Source:M | 0 | 0 | 0 | 0 | 0 | 0 |
| 43532 | ENSMUSG00000105491 | 0 | NA | NA | NA | NA | NA | Gm42844 | predicted gene 42844 [Source:MGI Symbo | 0 | 0 | 0 | 0 | 0 | 0 |
| 43533 | ENSMUSG00000105492 | 0 | NA | NA | NA | NA | NA | Gm2505 | predicted gene 2505 [Source:MGI Symbol | 0 | 0 | 0 | 0 | 0 | 0 |
| 43534 | ENSMUSG00000105493 | 0 | NA | NA | NA | NA | NA | Gm43551 | predicted gene 43551 [Source:MGI Symbo | 0 | 0 | 0 | 0 | 0 | 0 |
| 43535 | ENSMUSG00000105494 | 0 | NA | NA | NA | NA | NA | NA | NA | 0 | 0 | 0 | 0 | 0 | 0 |
| 43536 | ENSMUSG00000105495 | 0 | NA | NA | NA | NA | NA | Gm42995 | predicted gene 42995 [Source:MGI Symbo | 0 | 0 | 0 | 0 | 0 | 0 |
| 43537 | ENSMUSG00000105496 | 0.160662974 | 1.126924175 | 4.080472857 | 0.276174898 | 0.782413736 | NA | Mir7004 | microRNA 7004 [Source:MGI Symbol;Acc: | 0 | 0 | 0 | 0.964 | 0 | 0 |
| 43538 | ENSMUSG00000105497 | 0 | NA | NA | NA | NA | NA | Mir191 | microRNA 191 [Source:MGI Symbol;Acc:M | 0 | 0 | 0 | 0 | 0 | 0 |
| 43539 | ENSMUSG00000105498 | 0 | NA | NA | NA | NA | NA | Gm42677 | predicted gene 42677 [Source:MGI Symbo | 0 | 0 | 0 | 0 | 0 | 0 |
| 43540 | ENSMUSG00000105499 | 0 | NA | NA | NA | NA | NA | Igkv3-11 | immunoglobulin kappa variable 3-11 [Sou | 0 | 0 | 0 | 0 | 0 | 0 |
| 43541 | ENSMUSG00000105500 | 0 | NA | NA | NA | NA | NA | Gm42591 | predicted gene 42591 [Source:MGI Symbo | 0 | 0 | 0 | 0 | 0 | 0 |
| 43542 | ENSMUSG00000105501 | 0 | NA | NA | NA | NA | NA | 5330426L24I | RIKEN cDNA 5330426L24 gene [Source:M( | 0 | 0 | 0 | 0 | 0 | 0 |
| 43543 | ENSMUSG00000105502 | 0 | NA | NA | NA | NA | NA | Gm42536 | predicted gene 42536 [Source:MGI Symbo | 0 | 0 | 0 | 0 | 0 | 0 |
| 43544 | ENSMUSG00000105503 | 0.342639279 | -1.805976596 | 4.028547714 | -0.448294702 | 0.653940524 | NA | Gm42982 | predicted gene 42982 [Source:MGI Symbo | 0 | 2.0558 | 0 | 0 | 0 | 0 |
| 43545 | ENSMUSG00000105504 | 18.15120618 | -0.855835752 | 0.542406348 | -1.577849807 | 0.114600118 | 0.497367836 | Gbp5 | guanylate binding protein 5 [Source:MGI S | 19.274 | 33.921 | 17.106 | 15.424 | 9.8156 | 13.366 |
| 43546 | ENSMUSG00000105505 | 0 | NA | NA | NA | NA | NA | Obox4-ps7 | oocyte specific homeobox 4, pseudogene | 0 | 0 | 0 | 0 | 0 | 0 |
| 43547 | ENSMUSG00000105506 | 5.76988467 | 1.426316342 | 1.049805894 | 1.358647679 | 0.174258259 | 0.604151393 | Gm43743 | predicted gene 43743 [Source:MGI Symbo | 3.2124 | 4.1117 | 2.0125 | 0.964 | 8.7249 | 15.594 |
| 43548 | ENSMUSG00000105507 | 0 | NA | NA | NA | NA | NA | Gm43853 | predicted gene 43853 [Source:MGI Symbo | 0 | 0 | 0 | 0 | 0 | 0 |
| 43549 | ENSMUSG00000105508 | 0 | NA | NA | NA | NA | NA | Gm32921 | predicted gene, 32921 [Source:MGI Symb | 0 | 0 | 0 | 0 | 0 | 0 |
| 43550 | ENSMUSG00000105509 | 34.63344482 | 0.124524559 | 0.391663753 | 0.317937409 | 0.750532415 | 0.952074518 | C130013H08 | RIKEN cDNA C130013H08 gene [Source:M | 44.171 | 23.642 | 31.194 | 32.775 | 39.262 | 36.757 |
| 43551 | ENSMUSG00000105510 | 0.133850373 | -0.796629176 | 4.080472857 | -0.195229623 | 0.84521319 | NA | Gm43815 | predicted gene 43815 [Source:MGI Symbo | 0.8031 | 0 | 0 | 0 | 0 | 0 |
| 43552 | ENSMUSG00000105511 | 0 | NA | NA | NA | NA | NA | Gm33758 | predicted gene, 33758 [Source:MGI Symb | 0 | 0 | 0 | 0 | 0 | 0 |
| 43553 | ENSMUSG00000105512 | 22.36480504 | 0.972069835 | 0.513695797 | 1.892306383 | 0.058450171 | 0.343085466 | Gm43714 | predicted gene 43714 [Source:MGI Symbo | 21.684 | 12.335 | 11.069 | 40.487 | 18.541 | 30.074 |
| 43554 | ENSMUSG00000105513 | 0 | NA | NA | NA | NA | NA | NA | NA | 0 | 0 | 0 | 0 | 0 | 0 |
| 43555 | ENSMUSG00000105514 | 4.395169291 | -1.483583879 | 1.384673516 | -1.071432263 | 0.283975113 | NA | Gm42913 | predicted gene 42913 [Source:MGI Symbo | 10.44 | 0 | 9.0562 | 5.7839 | 1.0906 | 0 |
| 43556 | ENSMUSG00000105515 | 0 | NA | NA | NA | NA | NA | Gm42454 | predicted gene 42454 [Source:MGI Symbo | 0 | 0 | 0 | 0 | 0 | 0 |
| 43557 | ENSMUSG00000105516 | 0 | NA | NA | NA | NA | NA | Gm36823 | predicted gene, 36823 [Source:MGI Symb | 0 | 0 | 0 | 0 | 0 | 0 |
| 43558 | ENSMUSG00000105517 | 4.009999272 | -0.669693799 | 1.086711528 | -0.616257196 | 0.537724785 | NA | Gm42436 | predicted gene 42436 [Source:MGI Symbo | 5.6217 | 6.1675 | 3.0187 | 4.8199 | 1.0906 | 3.3415 |
| 43559 | ENSMUSG00000105518 | 0 | NA | NA | NA | NA | NA | Gm42674 | predicted gene 42674 [Source:MGI Symbo | 0 | 0 | 0 | 0 | 0 | 0 |
| 43560 | ENSMUSG00000105519 | 1.681945226 | 1.3662437 | 1.743458822 | 0.783639787 | 0.433251503 | NA | E430021H15 | RIKEN cDNA E430021H15 gene [Source:M | 0.8031 | 1.0279 | 1.0062 | 2.8919 | 4.3625 | 0 |
| 43561 | ENSMUSG00000105520 | 0 | NA | NA | NA | NA | NA | Gm42837 | predicted gene 42837 [Source:MGI Symbo | 0 | 0 | 0 | 0 | 0 | 0 |
| 43562 | ENSMUSG00000105521 | 0 | NA | NA | NA | NA | NA | Gm35702 | predicted gene, 35702 [Source:MGI Symb | 0 | 0 | 0 | 0 | 0 | 0 |
| 43563 | ENSMUSG00000105522 | 0 | NA | NA | NA | NA | NA | Gm15235 | predicted gene 15235 [Source:MGI Symbo | 0 | 0 | 0 | 0 | 0 | 0 |
| 43564 | ENSMUSG00000105523 | 0 | NA | NA | NA | NA | NA | Gm43193 | predicted gene 43193 [Source:MGI Symbo | 0 | 0 | 0 | 0 | 0 | 0 |

| | | | | | | | | | | | | | | | |
|---|---|---|---|---|---|---|---|---|---|---|---|---|---|---|---|
| 43565 | ENSMUSG00000105524 | 0 | NA | NA | NA | NA | NA | Mir3065 | microRNA 3065 [Source:MGI Symbol;Acc: | 0 | 0 | 0 | 0 | 0 | 0 |
| 43566 | ENSMUSG00000105525 | 0 | NA | NA | NA | NA | NA | Gm42976 | predicted gene 42976 [Source:MGI Symbo | 0 | 0 | 0 | 0 | 0 | 0 |
| 43567 | ENSMUSG00000105526 | 0 | NA | NA | NA | NA | NA | Gm43490 | predicted gene 43490 [Source:MGI Symbo | 0 | 0 | 0 | 0 | 0 | 0 |
| 43568 | ENSMUSG00000105527 | 0 | NA | NA | NA | NA | NA | 4933424H11 | RIKEN cDNA 4933424H11 gene [Source:M | 0 | 0 | 0 | 0 | 0 | 0 |
| 43569 | ENSMUSG00000105528 | 0 | NA | NA | NA | NA | NA | Gm43519 | predicted gene 43519 [Source:MGI Symbo | 0 | 0 | 0 | 0 | 0 | 0 |
| 43570 | ENSMUSG00000105529 | 0 | NA | NA | NA | NA | NA | Gm9480 | predicted gene 9480 [Source:MGI Symbol | 0 | 0 | 0 | 0 | 0 | 0 |
| 43571 | ENSMUSG00000105530 | 0 | NA | NA | NA | NA | NA | Gm43829 | predicted gene 43829 [Source:MGI Symbo | 0 | 0 | 0 | 0 | 0 | 0 |
| 43572 | ENSMUSG00000105531 | 0 | NA | NA | NA | NA | NA | 4930458A03 | RIKEN cDNA 4930458A03 gene [Source:M | 0 | 0 | 0 | 0 | 0 | 0 |
| 43573 | ENSMUSG00000105532 | 0.267700746 | -1.497481649 | 4.057001436 | -0.369110456 | 0.712045396 | NA | Gm5279 | predicted gene 5279 [Source:MGI Symbol | 1.6062 | 0 | 0 | 0 | 0 | 0 |
| 43574 | ENSMUSG00000105533 | 0 | NA | NA | NA | NA | NA | Trav7d-2 | T cell receptor alpha variable 7D-2 [Source | 0 | 0 | 0 | 0 | 0 | 0 |
| 43575 | ENSMUSG00000105534 | 0.185640647 | 1.126924175 | 4.080472857 | 0.276174898 | 0.782413736 | NA | Gm42435 | predicted gene 42435 [Source:MGI Symbo | 0 | 0 | 0 | 0 | 0 | 1.1138 |
| 43576 | ENSMUSG00000105535 | 1.32957991 | 0.132730382 | 1.987140095 | 0.066794677 | 0.946745161 | NA | Gm43077 | predicted gene 43077 [Source:MGI Symbo | 0.8031 | 0 | 3.0187 | 1.928 | 0 | 2.2277 |
| 43577 | ENSMUSG00000105536 | 0 | NA | NA | NA | NA | NA | Gm4959 | predicted gene 4959 [Source:MGI Symbol | 0 | 0 | 0 | 0 | 0 | 0 |
| 43578 | ENSMUSG00000105537 | 0 | NA | NA | NA | NA | NA | NA | NA | 0 | 0 | 0 | 0 | 0 | 0 |
| 43579 | ENSMUSG00000105538 | 0 | NA | NA | NA | NA | NA | Gm2142 | predicted gene 2142 [Source:MGI Symbol | 0 | 0 | 0 | 0 | 0 | 0 |
| 43580 | ENSMUSG00000105539 | 0 | NA | NA | NA | NA | NA | Igkv13-80-1 | immunoglobulin kappa chain variable 13-8 | 0 | 0 | 0 | 0 | 0 | 0 |
| 43581 | ENSMUSG00000105540 | 0 | NA | NA | NA | NA | NA | Gm7988 | predicted gene 7988 [Source:MGI Symbol | 0 | 0 | 0 | 0 | 0 | 0 |
| 43582 | ENSMUSG00000105541 | 5.609884398 | 0.14654453 | 0.890383017 | 0.164585945 | 0.869269894 | 0.978605211 | Gm43136 | predicted gene 43136 [Source:MGI Symbo | 4.8186 | 7.1954 | 4.025 | 7.7118 | 5.4531 | 4.4554 |
| 43583 | ENSMUSG00000105542 | 0 | NA | NA | NA | NA | NA | Gm34248 | predicted gene, 34248 [Source:MGI Symb | 0 | 0 | 0 | 0 | 0 | 0 |
| 43584 | ENSMUSG00000105543 | 0 | NA | NA | NA | NA | NA | Gm43169 | predicted gene 43169 [Source:MGI Symbo | 0 | 0 | 0 | 0 | 0 | 0 |
| 43585 | ENSMUSG00000105544 | 0 | NA | NA | NA | NA | NA | NA | NA | 0 | 0 | 0 | 0 | 0 | 0 |
| 43586 | ENSMUSG00000105545 | 0.185640647 | 1.126924175 | 4.080472857 | 0.276174898 | 0.782413736 | NA | 4930599N24 | RIKEN cDNA 4930599N24 gene [Source:M | 0 | 0 | 0 | 0 | 0 | 1.1138 |
| 43587 | ENSMUSG00000105546 | 0 | NA | NA | NA | NA | NA | NA | NA | 0 | 0 | 0 | 0 | 0 | 0 |
| 43588 | ENSMUSG00000105547 | 0.640584875 | -2.718595279 | 2.766810251 | -0.982573806 | 0.325817242 | NA | Iglc3 | immunoglobulin lambda constant 3 [Sour | 0.8031 | 1.0279 | 2.0125 | 0 | 0 | 0 |
| 43589 | ENSMUSG00000105548 | 0 | NA | NA | NA | NA | NA | Trav15n-3 | T cell receptor alpha variable 15N-3 [Sour | 0 | 0 | 0 | 0 | 0 | 0 |
| 43590 | ENSMUSG00000105549 | 0 | NA | NA | NA | NA | NA | Gm43540 | predicted gene 43540 [Source:MGI Symbo | 0 | 0 | 0 | 0 | 0 | 0 |
| 43591 | ENSMUSG00000105550 | 0.684865378 | 2.990343487 | 2.966805827 | 1.00793367 | 0.313486306 | NA | Gm35585 | predicted gene, 35585 [Source:MGI Symb | 0 | 0 | 0 | 1.928 | 2.1812 | 0 |
| 43592 | ENSMUSG00000105551 | 0 | NA | NA | NA | NA | NA | Gm43199 | predicted gene 43199 [Source:MGI Symbo | 0 | 0 | 0 | 0 | 0 | 0 |
| 43593 | ENSMUSG00000105552 | 7.466401552 | 0.080793894 | 0.791837088 | 0.10203348 | 0.9187301 | 0.986374671 | Gm19710 | predicted gene, 19710 [Source:MGI Symb | 5.6217 | 6.1675 | 10.062 | 8.6758 | 9.8156 | 4.4554 |
| 43594 | ENSMUSG00000105553 | 0 | NA | NA | NA | NA | NA | Gm42514 | predicted gene 42514 [Source:MGI Symbo | 0 | 0 | 0 | 0 | 0 | 0 |
| 43595 | ENSMUSG00000105554 | 0 | NA | NA | NA | NA | NA | Gm7118 | predicted gene 7118 [Source:MGI Symbol | 0 | 0 | 0 | 0 | 0 | 0 |
| 43596 | ENSMUSG00000105555 | 0 | NA | NA | NA | NA | NA | Gm43856 | predicted gene 43856 [Source:MGI Symbo | 0 | 0 | 0 | 0 | 0 | 0 |
| 43597 | ENSMUSG00000105556 | 11.93617264 | 0.305062715 | 0.64989111 | 0.46940589 | 0.638779539 | 0.923932929 | Gm43080 | predicted gene 43080 [Source:MGI Symbo | 8.8341 | 8.2233 | 15.094 | 17.352 | 7.6343 | 14.48 |
| 43598 | ENSMUSG00000105557 | 0 | NA | NA | NA | NA | NA | Mir3966 | microRNA 3966 [Source:MGI Symbol;Acc: | 0 | 0 | 0 | 0 | 0 | 0 |
| 43599 | ENSMUSG00000105558 | 1.390864126 | 0.841680141 | 2.180887826 | 0.385934632 | 0.699545081 | NA | Gm5855 | predicted gene 5855 [Source:MGI Symbol | 0 | 0 | 3.0187 | 0.964 | 4.3625 | 0 |
| 43600 | ENSMUSG00000105559 | 0 | NA | NA | NA | NA | NA | Gm4860 | predicted gene 4860 [Source:MGI Symbol | 0 | 0 | 0 | 0 | 0 | 0 |
| 43601 | ENSMUSG00000105560 | 0 | NA | NA | NA | NA | NA | Gm42744 | predicted gene 42744 [Source:MGI Symbo | 0 | 0 | 0 | 0 | 0 | 0 |
| 43602 | ENSMUSG00000105561 | 1.812389981 | -0.115903558 | 1.598095237 | -0.072526064 | 0.942183264 | NA | Gm43462 | predicted gene 43462 [Source:MGI Symbo | 1.6062 | 2.0558 | 2.0125 | 1.928 | 3.2719 | 0 |
| 43603 | ENSMUSG00000105562 | 0 | NA | NA | NA | NA | NA | Gm43287 | predicted gene 43287 [Source:MGI Symbo | 0 | 0 | 0 | 0 | 0 | 0 |
| 43604 | ENSMUSG00000105563 | 0 | NA | NA | NA | NA | NA | NA | NA | 0 | 0 | 0 | 0 | 0 | 0 |
| 43605 | ENSMUSG00000105564 | 0 | NA | NA | NA | NA | NA | Gm8891 | predicted gene 8891 [Source:MGI Symbol | 0 | 0 | 0 | 0 | 0 | 0 |
| 43606 | ENSMUSG00000105565 | 1.938162117 | -1.43268062 | 1.796044462 | -0.797686611 | 0.425052376 | NA | Gm43566 | predicted gene 43566 [Source:MGI Symbo | 6.4248 | 0 | 2.0125 | 0.964 | 0 | 2.2277 |
| 43607 | ENSMUSG00000105566 | 0 | NA | NA | NA | NA | NA | Obox4-ps4 | oocyte specific homeobox 4, pseudogene | 0 | 0 | 0 | 0 | 0 | 0 |
| 43608 | ENSMUSG00000105567 | 0 | NA | NA | NA | NA | NA | NA | NA | 0 | 0 | 0 | 0 | 0 | 0 |
| 43609 | ENSMUSG00000105568 | 0 | NA | NA | NA | NA | NA | Gm42971 | predicted gene 42971 [Source:MGI Symbo | 0 | 0 | 0 | 0 | 0 | 0 |
| 43610 | ENSMUSG00000105569 | 0 | NA | NA | NA | NA | NA | NA | NA | 0 | 0 | 0 | 0 | 0 | 0 |
| 43611 | ENSMUSG00000105570 | 0 | NA | NA | NA | NA | NA | Gm42682 | predicted gene 42682 [Source:MGI Symbo | 0 | 0 | 0 | 0 | 0 | 0 |
| 43612 | ENSMUSG00000105571 | 0 | NA | NA | NA | NA | NA | Gm43662 | predicted gene 43662 [Source:MGI Symbo | 0 | 0 | 0 | 0 | 0 | 0 |
| 43613 | ENSMUSG00000105572 | 12.20209423 | 0.242865519 | 0.745626555 | 0.325720051 | 0.744636173 | 0.949997329 | Gm43300 | predicted gene 43300 [Source:MGI Symbo | 15.259 | 8.2233 | 10.062 | 22.171 | 15.269 | 2.2277 |
| 43614 | ENSMUSG00000105573 | 0 | NA | NA | NA | NA | NA | Gm43384 | predicted gene 43384 [Source:MGI Symbo | 0 | 0 | 0 | 0 | 0 | 0 |
| 43615 | ENSMUSG00000105574 | 0 | NA | NA | NA | NA | NA | Gm43648 | predicted gene 43648 [Source:MGI Symbo | 0 | 0 | 0 | 0 | 0 | 0 |
| 43616 | ENSMUSG00000105575 | 0 | NA | NA | NA | NA | NA | Gm43475 | predicted gene 43475 [Source:MGI Symbo | 0 | 0 | 0 | 0 | 0 | 0 |
| 43617 | ENSMUSG00000105576 | 0 | NA | NA | NA | NA | NA | Gm42164 | predicted gene, 42164 [Source:MGI Symb | 0 | 0 | 0 | 0 | 0 | 0 |
| 43618 | ENSMUSG00000105577 | 0 | NA | NA | NA | NA | NA | Gm34599 | predicted gene, 34599 [Source:MGI Symb | 0 | 0 | 0 | 0 | 0 | 0 |
| 43619 | ENSMUSG00000105578 | 0 | NA | NA | NA | NA | NA | Gm43663 | predicted gene 43663 [Source:MGI Symbo | 0 | 0 | 0 | 0 | 0 | 0 |
| 43620 | ENSMUSG00000105579 | 0 | NA | NA | NA | NA | NA | Gm43251 | predicted gene 43251 [Source:MGI Symbo | 0 | 0 | 0 | 0 | 0 | 0 |

| | | | | | | | | | | | | | | | |
|---|---|---|---|---|---|---|---|---|---|---|---|---|---|---|---|
| 43621 | ENSMUSG00000105580 | 0 | NA | NA | NA | NA | NA | NA | NA | 0 | 0 | 0 | 0 | 0 | 0 |
| 43622 | ENSMUSG00000105581 | 0 | NA | NA | NA | NA | NA | NA | NA | 0 | 0 | 0 | 0 | 0 | 0 |
| 43623 | ENSMUSG00000105582 | 0 | NA | NA | NA | NA | NA | Ighv1-68 | immunoglobulin heavy variable V1-68 [So | 0 | 0 | 0 | 0 | 0 | 0 |
| 43624 | ENSMUSG00000105583 | 0 | NA | NA | NA | NA | NA | Gm43777 | predicted gene 43777 [Source:MGI Symbo | 0 | 0 | 0 | 0 | 0 | 0 |
| 43625 | ENSMUSG00000105584 | 0 | NA | NA | NA | NA | NA | Gm42588 | predicted gene 42588 [Source:MGI Symbo | 0 | 0 | 0 | 0 | 0 | 0 |
| 43626 | ENSMUSG00000105585 | 0 | NA | NA | NA | NA | NA | NA | NA | 0 | 0 | 0 | 0 | 0 | 0 |
| 43627 | ENSMUSG00000105586 | 0.339027071 | -1.79174149 | 3.919585585 | -0.457125237 | 0.647581032 | NA | Gm43009 | predicted gene 43009 [Source:MGI Symbo | 0 | 1.0279 | 1.0062 | 0 | 0 | 0 |
| 43628 | ENSMUSG00000105588 | 0 | NA | NA | NA | NA | NA | Utp14b-ps1 | UTP14B small subunit processome compo | 0 | 0 | 0 | 0 | 0 | 0 |
| 43629 | ENSMUSG00000105589 | 0 | NA | NA | NA | NA | NA | Fbxw14 | F-box and WD-40 domain protein 14 [Sou | 0 | 0 | 0 | 0 | 0 | 0 |
| 43630 | ENSMUSG00000105590 | 0 | NA | NA | NA | NA | NA | Gm43364 | predicted gene 43364 [Source:MGI Symbo | 0 | 0 | 0 | 0 | 0 | 0 |
| 43631 | ENSMUSG00000105591 | 0 | NA | NA | NA | NA | NA | Gm43083 | predicted gene 43083 [Source:MGI Symbo | 0 | 0 | 0 | 0 | 0 | 0 |
| 43632 | ENSMUSG00000105592 | 0 | NA | NA | NA | NA | NA | Gm42527 | predicted gene 42527 [Source:MGI Symbo | 0 | 0 | 0 | 0 | 0 | 0 |
| 43633 | ENSMUSG00000105593 | 0 | NA | NA | NA | NA | NA | Gm42952 | predicted gene 42952 [Source:MGI Symbo | 0 | 0 | 0 | 0 | 0 | 0 |
| 43634 | ENSMUSG00000105594 | 0.321325949 | 1.911575114 | 4.048267893 | 0.472195804 | 0.636787034 | NA | Gm43398 | predicted gene 43398 [Source:MGI Symbo | 0 | 0 | 0 | 1.928 | 0 | 0 |
| 43635 | ENSMUSG00000105595 | 0 | NA | NA | NA | NA | NA | Gm49316 | predicted gene 49316 [Source:MGI Symbo | 0 | 0 | 0 | 0 | 0 | 0 |
| 43636 | ENSMUSG00000105596 | 0 | NA | NA | NA | NA | NA | Gm42650 | predicted gene 42650 [Source:MGI Symbo | 0 | 0 | 0 | 0 | 0 | 0 |
| 43637 | ENSMUSG00000105597 | 15.73733834 | 0.268426964 | 0.600266256 | 0.447179833 | 0.654745221 | 0.925797231 | 4633401B06 | RIKEN cDNA 4633401B06 gene [Source:M | 9.6372 | 12.335 | 21.131 | 26.027 | 15.269 | 10.025 |
| 43638 | ENSMUSG00000105598 | 0 | NA | NA | NA | NA | NA | Gm43186 | predicted gene 43186 [Source:MGI Symbo | 0 | 0 | 0 | 0 | 0 | 0 |
| 43639 | ENSMUSG00000105599 | 0 | NA | NA | NA | NA | NA | Gm9238 | predicted gene 9238 [Source:MGI Symbol | 0 | 0 | 0 | 0 | 0 | 0 |
| 43640 | ENSMUSG00000105600 | 0 | NA | NA | NA | NA | NA | Gm5987 | predicted gene 5987 [Source:MGI Symbol | 0 | 0 | 0 | 0 | 0 | 0 |
| 43641 | ENSMUSG00000105601 | 3.35691729 | -0.103452591 | 1.31797096 | -0.078493832 | 0.937435236 | NA | Gm42725 | predicted gene 42725 [Source:MGI Symbo | 7.2279 | 2.0558 | 1.0062 | 0.964 | 1.0906 | 7.7969 |
| 43642 | ENSMUSG00000105602 | 0 | NA | NA | NA | NA | NA | Gm10636 | predicted gene 10636 [Source:MGI Symbo | 0 | 0 | 0 | 0 | 0 | 0 |
| 43643 | ENSMUSG00000105603 | 0 | NA | NA | NA | NA | NA | Gm32996 | predicted gene, 32996 [Source:MGI Symb | 0 | 0 | 0 | 0 | 0 | 0 |
| 43644 | ENSMUSG00000105604 | 0 | NA | NA | NA | NA | NA | Gm42838 | predicted gene 42838 [Source:MGI Symbo | 0 | 0 | 0 | 0 | 0 | 0 |
| 43645 | ENSMUSG00000105605 | 0 | NA | NA | NA | NA | NA | Ighv1-86 | immunoglobulin heavy variable 1-86 [Sou | 0 | 0 | 0 | 0 | 0 | 0 |
| 43646 | ENSMUSG00000105606 | 0.17131964 | -0.796629176 | 4.080472857 | -0.195229623 | 0.84521319 | NA | Igkv2-109 | immunoglobulin kappa variable 2-109 [So | 0 | 1.0279 | 0 | 0 | 0 | 0 |
| 43647 | ENSMUSG00000105607 | 3.525710635 | 1.222631043 | 1.151786316 | 1.061508568 | 0.288458841 | NA | Gm43513 | predicted gene 43513 [Source:MGI Symbo | 3.2124 | 1.0279 | 2.0125 | 3.8559 | 4.3625 | 6.6831 |
| 43648 | ENSMUSG00000105608 | 0 | NA | NA | NA | NA | NA | Gm43408 | predicted gene 43408 [Source:MGI Symbo | 0 | 0 | 0 | 0 | 0 | 0 |
| 43649 | ENSMUSG00000105609 | 0.181769715 | 1.126924175 | 4.080472857 | 0.276174898 | 0.782413736 | NA | Gm42501 | predicted gene 42501 [Source:MGI Symbo | 0 | 0 | 0 | 0 | 1.0906 | 0 |
| 43650 | ENSMUSG00000105610 | 0 | NA | NA | NA | NA | NA | NA | NA | 0 | 0 | 0 | 0 | 0 | 0 |
| 43651 | ENSMUSG00000105611 | 0 | NA | NA | NA | NA | NA | Gm42938 | predicted gene 42938 [Source:MGI Symbo | 0 | 0 | 0 | 0 | 0 | 0 |
| 43652 | ENSMUSG00000105612 | 0 | NA | NA | NA | NA | NA | NA | NA | 0 | 0 | 0 | 0 | 0 | 0 |
| 43653 | ENSMUSG00000105613 | 0 | NA | NA | NA | NA | NA | Gm43684 | predicted gene 43684 [Source:MGI Symbo | 0 | 0 | 0 | 0 | 0 | 0 |
| 43654 | ENSMUSG00000105614 | 0 | NA | NA | NA | NA | NA | Dnd1-ps | DND microRNA-mediated repression inhib | 0 | 0 | 0 | 0 | 0 | 0 |
| 43655 | ENSMUSG00000105615 | 0 | NA | NA | NA | NA | NA | Gm31048 | predicted gene, 31048 [Source:MGI Symb | 0 | 0 | 0 | 0 | 0 | 0 |
| 43656 | ENSMUSG00000105616 | 0 | NA | NA | NA | NA | NA | Gm3010 | predicted gene 3010 [Source:MGI Symbol | 0 | 0 | 0 | 0 | 0 | 0 |
| 43657 | ENSMUSG00000105617 | 0 | NA | NA | NA | NA | NA | Gm43809 | predicted gene 43809 [Source:MGI Symbo | 0 | 0 | 0 | 0 | 0 | 0 |
| 43658 | ENSMUSG00000105618 | 0 | NA | NA | NA | NA | NA | Gm42916 | predicted gene 42916 [Source:MGI Symbo | 0 | 0 | 0 | 0 | 0 | 0 |
| 43659 | ENSMUSG00000105619 | 0 | NA | NA | NA | NA | NA | 9530034A14 | RIKEN cDNA 9530034A14 gene [Source:M | 0 | 0 | 0 | 0 | 0 | 0 |
| 43660 | ENSMUSG00000105620 | 0 | NA | NA | NA | NA | NA | Gm43047 | predicted gene 43047 [Source:MGI Symbo | 0 | 0 | 0 | 0 | 0 | 0 |
| 43661 | ENSMUSG00000105621 | 0 | NA | NA | NA | NA | NA | Mirlet7f-1 | microRNA let7f-1 [Source:MGI Symbol;Ac | 0 | 0 | 0 | 0 | 0 | 0 |
| 43662 | ENSMUSG00000105622 | 0.671009685 | -1.413087452 | 2.800223337 | -0.504633839 | 0.613816038 | NA | Gm42615 | predicted gene 42615 [Source:MGI Symbo | 0 | 2.0558 | 1.0062 | 0.964 | 0 | 0 |
| 43663 | ENSMUSG00000105624 | 0 | NA | NA | NA | NA | NA | Gm43187 | predicted gene 43187 [Source:MGI Symbo | 0 | 0 | 0 | 0 | 0 | 0 |
| 43664 | ENSMUSG00000105625 | 0 | NA | NA | NA | NA | NA | Gm31230 | predicted gene, 31230 [Source:MGI Symb | 0 | 0 | 0 | 0 | 0 | 0 |
| 43665 | ENSMUSG00000105626 | 0 | NA | NA | NA | NA | NA | NA | NA | 0 | 0 | 0 | 0 | 0 | 0 |
| 43666 | ENSMUSG00000105627 | 0 | NA | NA | NA | NA | NA | Gm43548 | predicted gene 43548 [Source:MGI Symbo | 0 | 0 | 0 | 0 | 0 | 0 |
| 43667 | ENSMUSG00000105628 | 0 | NA | NA | NA | NA | NA | Gm43350 | predicted gene 43350 [Source:MGI Symbo | 0 | 0 | 0 | 0 | 0 | 0 |
| 43668 | ENSMUSG00000105629 | 0 | NA | NA | NA | NA | NA | Trav7n-5 | T cell receptor alpha variable 7N-5 [Sourc | 0 | 0 | 0 | 0 | 0 | 0 |
| 43669 | ENSMUSG00000105630 | 0 | NA | NA | NA | NA | NA | Gm42543 | predicted gene 42543 [Source:MGI Symbo | 0 | 0 | 0 | 0 | 0 | 0 |
| 43670 | ENSMUSG00000105632 | 0.36353943 | 2.065989254 | 4.034137301 | 0.512126658 | 0.608562368 | NA | Gm43272 | predicted gene 43272 [Source:MGI Symbo | 0 | 0 | 0 | 0 | 2.1812 | 0 |
| 43671 | ENSMUSG00000105633 | 0 | NA | NA | NA | NA | NA | Gm32158 | predicted gene, 32158 [Source:MGI Symb | 0 | 0 | 0 | 0 | 0 | 0 |
| 43672 | ENSMUSG00000105634 | 0 | NA | NA | NA | NA | NA | Gm43367 | predicted gene 43367 [Source:MGI Symbo | 0 | 0 | 0 | 0 | 0 | 0 |
| 43673 | ENSMUSG00000105635 | 0 | NA | NA | NA | NA | NA | NA | NA | 0 | 0 | 0 | 0 | 0 | 0 |
| 43674 | ENSMUSG00000105636 | 1.074390219 | -0.973079277 | 2.37579194 | -0.409581016 | 0.682113325 | NA | Gm43625 | predicted gene 43625 [Source:MGI Symbo | 3.2124 | 0 | 1.0062 | 0 | 0 | 2.2277 |
| 43675 | ENSMUSG00000105637 | 0 | NA | NA | NA | NA | NA | Gm42480 | predicted gene 42480 [Source:MGI Symbo | 0 | 0 | 0 | 0 | 0 | 0 |
| 43676 | ENSMUSG00000105638 | 0 | NA | NA | NA | NA | NA | 4930408K08 | RIKEN cDNA 4930408K08 gene [Source:M | 0 | 0 | 0 | 0 | 0 | 0 |

| | | | | | | | | | | | | | | | |
|---|---|---|---|---|---|---|---|---|---|---|---|---|---|---|---|
| 43677 | ENSMUSG00000105639 | 0 | NA | NA | NA | NA | NA | Gm42516 | predicted gene 42516 [Source:MGI Symbo | 0 | 0 | 0 | 0 | 0 | 0 |
| 43678 | ENSMUSG00000105640 | 0 | NA | NA | NA | NA | NA | Gm42912 | predicted gene 42912 [Source:MGI Symbo | 0 | 0 | 0 | 0 | 0 | 0 |
| 43679 | ENSMUSG00000105641 | 0.321325949 | 1.911575114 | 4.048267893 | 0.472195804 | 0.636787034 | NA | Gm5853 | predicted gene 5853 [Source:MGI Symbol | 0 | 0 | 0 | 1.928 | 0 | 0 |
| 43680 | ENSMUSG00000105642 | 0 | NA | NA | NA | NA | NA | Gm43134 | predicted gene 43134 [Source:MGI Symbo | 0 | 0 | 0 | 0 | 0 | 0 |
| 43681 | ENSMUSG00000105643 | 2.74067584 | 0.689960775 | 1.309187542 | 0.527014467 | 0.598183542 | NA | Gm42611 | predicted gene 42611 [Source:MGI Symbo | 3.2124 | 2.0558 | 1.0062 | 5.7839 | 3.2719 | 1.1138 |
| 43682 | ENSMUSG00000105644 | 0 | NA | NA | NA | NA | NA | Gm19739 | predicted gene, 19739 [Source:MGI Symb | 0 | 0 | 0 | 0 | 0 | 0 |
| 43683 | ENSMUSG00000105645 | 0 | NA | NA | NA | NA | NA | Gm6649 | predicted gene 6649 [Source:MGI Symbol | 0 | 0 | 0 | 0 | 0 | 0 |
| 43684 | ENSMUSG00000105646 | 0.335414862 | -1.779047242 | 4.030804179 | -0.441362855 | 0.658950329 | NA | Gm30211 | predicted gene, 30211 [Source:MGI Symb | 0 | 0 | 2.0125 | 0 | 0 | 0 |
| 43685 | ENSMUSG00000105647 | 0 | NA | NA | NA | NA | NA | Gm43695 | predicted gene 43695 [Source:MGI Symbo | 0 | 0 | 0 | 0 | 0 | 0 |
| 43686 | ENSMUSG00000105648 | 0 | NA | NA | NA | NA | NA | Gm43505 | predicted gene 43505 [Source:MGI Symbo | 0 | 0 | 0 | 0 | 0 | 0 |
| 43687 | ENSMUSG00000105649 | 0 | NA | NA | NA | NA | NA | Gm43494 | predicted gene 43494 [Source:MGI Symbo | 0 | 0 | 0 | 0 | 0 | 0 |
| 43688 | ENSMUSG00000105650 | 0 | NA | NA | NA | NA | NA | Gm42889 | predicted gene 42889 [Source:MGI Symbo | 0 | 0 | 0 | 0 | 0 | 0 |
| 43689 | ENSMUSG00000105651 | 0 | NA | NA | NA | NA | NA | 1700017M07 | RIKEN cDNA 1700017M07 gene [Source:M | 0 | 0 | 0 | 0 | 0 | 0 |
| 43690 | ENSMUSG00000105652 | 0 | NA | NA | NA | NA | NA | 4930519L02I | RIKEN cDNA 4930519L02 gene [Source:M( | 0 | 0 | 0 | 0 | 0 | 0 |
| 43691 | ENSMUSG00000105653 | 0 | NA | NA | NA | NA | NA | Gm43831 | predicted gene 43831 [Source:MGI Symbo | 0 | 0 | 0 | 0 | 0 | 0 |
| 43692 | ENSMUSG00000105654 | 0 | NA | NA | NA | NA | NA | Gm29704 | predicted gene, 29704 [Source:MGI Symb | 0 | 0 | 0 | 0 | 0 | 0 |
| 43693 | ENSMUSG00000105655 | 3.743637122 | -1.016204445 | 1.09957163 | -0.924182124 | 0.355391501 | NA | Gm42659 | predicted gene 42659 [Source:MGI Symbo | 4.8186 | 6.1675 | 4.025 | 1.928 | 2.1812 | 3.3415 |
| 43694 | ENSMUSG00000105656 | 0 | NA | NA | NA | NA | NA | Gm43017 | predicted gene 43017 [Source:MGI Symbo | 0 | 0 | 0 | 0 | 0 | 0 |
| 43695 | ENSMUSG00000105657 | 0 | NA | NA | NA | NA | NA | Gm42833 | predicted gene 42833 [Source:MGI Symbo | 0 | 0 | 0 | 0 | 0 | 0 |
| 43696 | ENSMUSG00000105658 | 0 | NA | NA | NA | NA | NA | Gm43531 | predicted gene 43531 [Source:MGI Symbo | 0 | 0 | 0 | 0 | 0 | 0 |
| 43697 | ENSMUSG00000105659 | 0.520796785 | -0.820103173 | 3.072141206 | -0.266948398 | 0.789508892 | NA | 4930509H03 | RIKEN cDNA 4930509H03 gene [Source:M | 0 | 1.0279 | 1.0062 | 0 | 1.0906 | 0 |
| 43698 | ENSMUSG00000105660 | 0.736965982 | -2.935629304 | 2.883220453 | -1.018177192 | 0.308593757 | NA | Gm42975 | predicted gene 42975 [Source:MGI Symbo | 2.4093 | 0 | 2.0125 | 0 | 0 | 0 |
| 43699 | ENSMUSG00000105661 | 0 | NA | NA | NA | NA | NA | Gm43626 | predicted gene 43626 [Source:MGI Symbo | 0 | 0 | 0 | 0 | 0 | 0 |
| 43700 | ENSMUSG00000105662 | 0 | NA | NA | NA | NA | NA | Gm6639 | predicted gene 6639 [Source:MGI Symbol | 0 | 0 | 0 | 0 | 0 | 0 |
| 43701 | ENSMUSG00000105663 | 0 | NA | NA | NA | NA | NA | Gm43606 | predicted gene 43606 [Source:MGI Symbo | 0 | 0 | 0 | 0 | 0 | 0 |
| 43702 | ENSMUSG00000105664 | 0.185640647 | 1.126924175 | 4.080472857 | 0.276174898 | 0.782413736 | NA | Gm42797 | predicted gene 42797 [Source:MGI Symbo | 0 | 0 | 0 | 0 | 0 | 1.1138 |
| 43703 | ENSMUSG00000105665 | 0 | NA | NA | NA | NA | NA | Gm38395 | predicted gene, 38395 [Source:MGI Symb | 0 | 0 | 0 | 0 | 0 | 0 |
| 43704 | ENSMUSG00000105666 | 0 | NA | NA | NA | NA | NA | Gm18711 | predicted gene, 18711 [Source:MGI Symb | 0 | 0 | 0 | 0 | 0 | 0 |
| 43705 | ENSMUSG00000105667 | 0 | NA | NA | NA | NA | NA | Gm3945 | predicted gene 3945 [Source:MGI Symbol | 0 | 0 | 0 | 0 | 0 | 0 |
| 43706 | ENSMUSG00000105668 | 0 | NA | NA | NA | NA | NA | Igkv8-22 | immunoglobulin kappa chain variable 8-2: | 0 | 0 | 0 | 0 | 0 | 0 |
| 43707 | ENSMUSG00000105669 | 0 | NA | NA | NA | NA | NA | Gm18341 | predicted gene, 18341 [Source:MGI Symb | 0 | 0 | 0 | 0 | 0 | 0 |
| 43708 | ENSMUSG00000105670 | 0 | NA | NA | NA | NA | NA | NA | NA | 0 | 0 | 0 | 0 | 0 | 0 |
| 43709 | ENSMUSG00000105671 | 0 | NA | NA | NA | NA | NA | Gm31306 | predicted gene, 31306 [Source:MGI Symb | 0 | 0 | 0 | 0 | 0 | 0 |
| 43710 | ENSMUSG00000105672 | 0 | NA | NA | NA | NA | NA | Gm42816 | predicted gene 42816 [Source:MGI Symbo | 0 | 0 | 0 | 0 | 0 | 0 |
| 43711 | ENSMUSG00000105673 | 0 | NA | NA | NA | NA | NA | 1700081B01 | RIKEN cDNA 1700081B01 gene [Source:M | 0 | 0 | 0 | 0 | 0 | 0 |
| 43712 | ENSMUSG00000105674 | 1.006191327 | 3.548856365 | 2.592453122 | 1.36891824 | 0.171024834 | NA | Gm42437 | predicted gene 42437 [Source:MGI Symbo | 0 | 0 | 0 | 3.8559 | 2.1812 | 0 |
| 43713 | ENSMUSG00000105675 | 0 | NA | NA | NA | NA | NA | Gm43383 | predicted gene 43383 [Source:MGI Symbo | 0 | 0 | 0 | 0 | 0 | 0 |
| 43714 | ENSMUSG00000105676 | 0 | NA | NA | NA | NA | NA | Gm43403 | predicted gene 43403 [Source:MGI Symbo | 0 | 0 | 0 | 0 | 0 | 0 |
| 43715 | ENSMUSG00000105677 | 11.09494621 | -0.351913893 | 0.712001562 | -0.494260001 | 0.62112257 | 0.918069106 | Gm43328 | predicted gene 43328 [Source:MGI Symbo | 16.865 | 12.335 | 8.05 | 11.568 | 3.2719 | 14.48 |
| 43716 | ENSMUSG00000105678 | 0 | NA | NA | NA | NA | NA | Gm43705 | predicted gene 43705 [Source:MGI Symbo | 0 | 0 | 0 | 0 | 0 | 0 |
| 43717 | ENSMUSG00000105679 | 4.325743628 | 0.405519001 | 1.039134911 | 0.39024673 | 0.69635411 | NA | Gm43338 | predicted gene 43338 [Source:MGI Symbo | 4.0155 | 4.1117 | 3.0187 | 3.8559 | 8.7249 | 2.2277 |
| 43718 | ENSMUSG00000105680 | 0 | NA | NA | NA | NA | NA | Gm18340 | predicted gene, 18340 [Source:MGI Symb | 0 | 0 | 0 | 0 | 0 | 0 |
| 43719 | ENSMUSG00000105681 | 3.850495462 | 0.317949378 | 1.211314969 | 0.262482827 | 0.792949223 | NA | Gm43428 | predicted gene 43428 [Source:MGI Symbo | 3.2124 | 0 | 7.0437 | 2.8919 | 3.2719 | 6.6831 |
| 43720 | ENSMUSG00000105682 | 0 | NA | NA | NA | NA | NA | NA | NA | 0 | 0 | 0 | 0 | 0 | 0 |
| 43721 | ENSMUSG00000105683 | 0 | NA | NA | NA | NA | NA | Gm43487 | predicted gene 43487 [Source:MGI Symbo | 0 | 0 | 0 | 0 | 0 | 0 |
| 43722 | ENSMUSG00000105684 | 0 | NA | NA | NA | NA | NA | Gm42612 | predicted gene 42612 [Source:MGI Symbo | 0 | 0 | 0 | 0 | 0 | 0 |
| 43723 | ENSMUSG00000105685 | 0 | NA | NA | NA | NA | NA | Gm43315 | predicted gene 43315 [Source:MGI Symbo | 0 | 0 | 0 | 0 | 0 | 0 |
| 43724 | ENSMUSG00000105686 | 0.181769715 | 1.126924175 | 4.080472857 | 0.276174898 | 0.782413736 | NA | Gm35394 | predicted gene, 35394 [Source:MGI Symb | 0 | 0 | 0 | 0 | 1.0906 | 0 |
| 43725 | ENSMUSG00000105687 | 0 | NA | NA | NA | NA | NA | Gm6157 | predicted gene 6157 [Source:MGI Symbol | 0 | 0 | 0 | 0 | 0 | 0 |
| 43726 | ENSMUSG00000105688 | 0 | NA | NA | NA | NA | NA | 4930437M2: | RIKEN cDNA 4930437M23 gene [Source:M | 0 | 0 | 0 | 0 | 0 | 0 |
| 43727 | ENSMUSG00000105689 | 0 | NA | NA | NA | NA | NA | Gm43120 | predicted gene 43120 [Source:MGI Symbo | 0 | 0 | 0 | 0 | 0 | 0 |
| 43728 | ENSMUSG00000105690 | 0 | NA | NA | NA | NA | NA | Gm26265 | predicted gene, 26265 [Source:MGI Symb | 0 | 0 | 0 | 0 | 0 | 0 |
| 43729 | ENSMUSG00000105691 | 1.020693421 | -3.383091396 | 2.599252854 | -1.301563021 | 0.193065808 | NA | Gm42876 | predicted gene 42876 [Source:MGI Symbo | 0 | 4.1117 | 2.0125 | 0 | 0 | 0 |
| 43730 | ENSMUSG00000105692 | 36.49435446 | -0.105473851 | 0.374569829 | -0.281586618 | 0.778260503 | 0.959559604 | Gm43737 | predicted gene 43737 [Source:MGI Symbo | 36.14 | 42.145 | 35.219 | 31.811 | 45.806 | 27.846 |
| 43731 | ENSMUSG00000105693 | 0 | NA | NA | NA | NA | NA | Gm43706 | predicted gene 43706 [Source:MGI Symbo | 0 | 0 | 0 | 0 | 0 | 0 |
| 43732 | ENSMUSG00000105694 | 2.015032915 | -0.245136272 | 1.895974925 | -0.129292992 | 0.897125816 | NA | Gm43237 | predicted gene 43237 [Source:MGI Symbo | 2.4093 | 4.1117 | 0 | 0 | 0 | 5.5692 |

| | | | | | | | | | | | | | | | |
|---|---|---|---|---|---|---|---|---|---|---|---|---|---|---|---|
| 43733 | ENSMUSG00000105695 | 0 | NA | NA | NA | NA | NA | Gm43327 | predicted gene 43327 [Source:MGI Symbo | 0 | 0 | 0 | 0 | 0 | 0 |
| 43734 | ENSMUSG00000105696 | 0 | NA | NA | NA | NA | NA | Gm19211 | predicted gene, 19211 [Source:MGI Symb | 0 | 0 | 0 | 0 | 0 | 0 |
| 43735 | ENSMUSG00000105697 | 0 | NA | NA | NA | NA | NA | Mir7661 | microRNA 7661 [Source:MGI Symbol;Acc: | 0 | 0 | 0 | 0 | 0 | 0 |
| 43736 | ENSMUSG00000105698 | 0 | NA | NA | NA | NA | NA | Gm31881 | predicted gene, 31881 [Source:MGI Symb | 0 | 0 | 0 | 0 | 0 | 0 |
| 43737 | ENSMUSG00000105699 | 0 | NA | NA | NA | NA | NA | Gm43703 | predicted gene 43703 [Source:MGI Symbo | 0 | 0 | 0 | 0 | 0 | 0 |
| 43738 | ENSMUSG00000105700 | 0 | NA | NA | NA | NA | NA | Gm42772 | predicted gene 42772 [Source:MGI Symbo | 0 | 0 | 0 | 0 | 0 | 0 |
| 43739 | ENSMUSG00000105701 | 0 | NA | NA | NA | NA | NA | Gm42587 | predicted gene 42587 [Source:MGI Symbo | 0 | 0 | 0 | 0 | 0 | 0 |
| 43740 | ENSMUSG00000105702 | 0 | NA | NA | NA | NA | NA | Gm43780 | predicted gene 43780 [Source:MGI Symbo | 0 | 0 | 0 | 0 | 0 | 0 |
| 43741 | ENSMUSG00000105703 | 684.7027766 | -1.499878641 | 0.469518195 | -3.194505894 | 0.001400704 | 0.026905427 | Gm43305 | predicted gene 43305 [Source:MGI Symbo | 730.02 | 1660.1 | 645 | 328.72 | 297.74 | 446.65 |
| 43742 | ENSMUSG00000105704 | 0 | NA | NA | NA | NA | NA | Gm43055 | predicted gene 43055 [Source:MGI Symbo | 0 | 0 | 0 | 0 | 0 | 0 |
| 43743 | ENSMUSG00000105705 | 0 | NA | NA | NA | NA | NA | | anaphase-promoting complex subunit 5 p | 0 | 0 | 0 | 0 | 0 | 0 |
| 43744 | ENSMUSG00000105706 | 0 | NA | NA | NA | NA | NA | Gm42855 | predicted gene 42855 [Source:MGI Symbo | 0 | 0 | 0 | 0 | 0 | 0 |
| 43745 | ENSMUSG00000105707 | 0.167707431 | -0.796629176 | 4.080472857 | -0.195229623 | 0.84521319 | NA | Gm43753 | predicted gene 43753 [Source:MGI Symbo | 0 | 0 | 1.0062 | 0 | 0 | 0 |
| 43746 | ENSMUSG00000105708 | 4.347988116 | 0.718209999 | 1.079442943 | 0.665352443 | 0.505825095 | NA | Gm36070 | predicted gene, 36070 [Source:MGI Symb | 0.8031 | 4.1117 | 5.0312 | 2.8919 | 5.4531 | 7.7969 |
| 43747 | ENSMUSG00000105709 | 0 | NA | NA | NA | NA | NA | Gm42723 | predicted gene 42723 [Source:MGI Symbo | 0 | 0 | 0 | 0 | 0 | 0 |
| 43748 | ENSMUSG00000105710 | 8.984024076 | 1.064525525 | 0.763257991 | 1.394712585 | 0.163102528 | 0.586990831 | Gm43859 | predicted gene 43859 [Source:MGI Symbo | 2.4093 | 8.2233 | 7.0437 | 16.388 | 9.8156 | 10.025 |
| 43749 | ENSMUSG00000105711 | 0 | NA | NA | NA | NA | NA | Gm42598 | predicted gene 42598 [Source:MGI Symbo | 0 | 0 | 0 | 0 | 0 | 0 |
| 43750 | ENSMUSG00000105712 | 0.133850373 | -0.796629176 | 4.080472857 | -0.195229623 | 0.84521319 | NA | Gm5712 | predicted gene 5712 [Source:MGI Symbol | 0.8031 | 0 | 0 | 0 | 0 | 0 |
| 43751 | ENSMUSG00000105713 | 0 | NA | NA | NA | NA | NA | Gm43827 | predicted gene 43827 [Source:MGI Symbo | 0 | 0 | 0 | 0 | 0 | 0 |
| 43752 | ENSMUSG00000105714 | 0 | NA | NA | NA | NA | NA | Gm43380 | predicted gene 43380 [Source:MGI Symbo | 0 | 0 | 0 | 0 | 0 | 0 |
| 43753 | ENSMUSG00000105715 | 0 | NA | NA | NA | NA | NA | Gm31887 | predicted gene, 31887 [Source:MGI Symb | 0 | 0 | 0 | 0 | 0 | 0 |
| 43754 | ENSMUSG00000105716 | 0 | NA | NA | NA | NA | NA | Gm43522 | predicted gene 43522 [Source:MGI Symbo | 0 | 0 | 0 | 0 | 0 | 0 |
| 43755 | ENSMUSG00000105717 | 0 | NA | NA | NA | NA | NA | Gm43465 | predicted gene 43465 [Source:MGI Symbo | 0 | 0 | 0 | 0 | 0 | 0 |
| 43756 | ENSMUSG00000105718 | 0 | NA | NA | NA | NA | NA | Gm43576 | predicted gene 43576 [Source:MGI Symbo | 0 | 0 | 0 | 0 | 0 | 0 |
| 43757 | ENSMUSG00000105719 | 1.627034687 | -2.929784663 | 1.985254044 | -1.475773175 | 0.140004794 | NA | Gm42872 | predicted gene 42872 [Source:MGI Symbo | 1.6062 | 1.0279 | 6.0375 | 0 | 1.0906 | 0 |
| 43758 | ENSMUSG00000105720 | 0 | NA | NA | NA | NA | NA | Gm42440 | predicted gene 42440 [Source:MGI Symbo | 0 | 0 | 0 | 0 | 0 | 0 |
| 43759 | ENSMUSG00000105721 | 0 | NA | NA | NA | NA | NA | Trav7d-6 | T cell receptor alpha variable 7D-6 [Source | 0 | 0 | 0 | 0 | 0 | 0 |
| 43760 | ENSMUSG00000105722 | 0 | NA | NA | NA | NA | NA | Gm18187 | predicted gene, 18187 [Source:MGI Symb | 0 | 0 | 0 | 0 | 0 | 0 |
| 43761 | ENSMUSG00000105723 | 0 | NA | NA | NA | NA | NA | Mir7210 | microRNA 7210 [Source:MGI Symbol;Acc: | 0 | 0 | 0 | 0 | 0 | 0 |
| 43762 | ENSMUSG00000105724 | 0.77079641 | 0.856159616 | 2.652840961 | 0.322733111 | 0.746897374 | NA | NA | NA | 1.6062 | 0 | 0 | 1.928 | 1.0906 | 0 |
| 43763 | ENSMUSG00000105725 | 0 | NA | NA | NA | NA | NA | Igkv12-49 | immunoglobulin kappa chain variable 12-4 | 0 | 0 | 0 | 0 | 0 | 0 |
| 43764 | ENSMUSG00000105726 | 0 | NA | NA | NA | NA | NA | Gm42443 | predicted gene 42443 [Source:MGI Symbo | 0 | 0 | 0 | 0 | 0 | 0 |
| 43765 | ENSMUSG00000105727 | 0 | NA | NA | NA | NA | NA | Gm8828 | predicted gene 8828 [Source:MGI Symbol | 0 | 0 | 0 | 0 | 0 | 0 |
| 43766 | ENSMUSG00000105728 | 0.342639279 | -1.805976596 | 4.028547714 | -0.448294702 | 0.653940524 | NA | Gm42819 | predicted gene 42819 [Source:MGI Symbo | 0 | 2.0558 | 0 | 0 | 0 | 0 |
| 43767 | ENSMUSG00000105729 | 0 | NA | NA | NA | NA | NA | 5330425B07 | RIKEN cDNA 5330425B07 gene [Source:M | 0 | 0 | 0 | 0 | 0 | 0 |
| 43768 | ENSMUSG00000105730 | 0 | NA | NA | NA | NA | NA | Gm43838 | predicted gene 43838 [Source:MGI Symbo | 0 | 0 | 0 | 0 | 0 | 0 |
| 43769 | ENSMUSG00000105731 | 0 | NA | NA | NA | NA | NA | Gm43063 | predicted gene 43063 [Source:MGI Symbo | 0 | 0 | 0 | 0 | 0 | 0 |
| 43770 | ENSMUSG00000105732 | 0 | NA | NA | NA | NA | NA | Sult1c2-ps1 | sulfotransferase family, cytosolic, 1C, mer | 0 | 0 | 0 | 0 | 0 | 0 |
| 43771 | ENSMUSG00000105733 | 0 | NA | NA | NA | NA | NA | Gm42973 | predicted gene 42973 [Source:MGI Symbo | 0 | 0 | 0 | 0 | 0 | 0 |
| 43772 | ENSMUSG00000105734 | 0 | NA | NA | NA | NA | NA | 4930558C23 | RIKEN cDNA 4930558C23 gene [Source:M | 0 | 0 | 0 | 0 | 0 | 0 |
| 43773 | ENSMUSG00000105735 | 0 | NA | NA | NA | NA | NA | 4933415J04I | RIKEN cDNA 4933415J04 gene [Source:MG | 0 | 0 | 0 | 0 | 0 | 0 |
| 43774 | ENSMUSG00000105736 | 0 | NA | NA | NA | NA | NA | Gm42456 | predicted gene 42456 [Source:MGI Symbo | 0 | 0 | 0 | 0 | 0 | 0 |
| 43775 | ENSMUSG00000105737 | 0 | NA | NA | NA | NA | NA | Gm6105 | predicted gene 6105 [Source:MGI Symbol | 0 | 0 | 0 | 0 | 0 | 0 |
| 43776 | ENSMUSG00000105738 | 0 | NA | NA | NA | NA | NA | Mir6384 | microRNA 6384 [Source:MGI Symbol;Acc: | 0 | 0 | 0 | 0 | 0 | 0 |
| 43777 | ENSMUSG00000105739 | 0 | NA | NA | NA | NA | NA | Gm5708 | predicted gene 5708 [Source:MGI Symbol | 0 | 0 | 0 | 0 | 0 | 0 |
| 43778 | ENSMUSG00000105740 | 0 | NA | NA | NA | NA | NA | Gm42685 | predicted gene 42685 [Source:MGI Symbo | 0 | 0 | 0 | 0 | 0 | 0 |
| 43779 | ENSMUSG00000105741 | 9.384614676 | -0.056652524 | 0.699698284 | -0.080967076 | 0.935468136 | 0.988420305 | C79130 | expressed sequence C79130 [Source:MGI | 6.4248 | 10.279 | 12.075 | 7.7118 | 10.906 | 8.9108 |
| 43780 | ENSMUSG00000105742 | 10.03055397 | 0.583299282 | 0.741018754 | 0.787158595 | 0.431189032 | 0.84474037 | Gm42748 | predicted gene 42748 [Source:MGI Symbo | 4.0155 | 9.2513 | 11.069 | 19.28 | 6.5437 | 10.025 |
| 43781 | ENSMUSG00000105743 | 0 | NA | NA | NA | NA | NA | Mir1949 | microRNA 1949 [Source:MGI Symbol;Acc: | 0 | 0 | 0 | 0 | 0 | 0 |
| 43782 | ENSMUSG00000105744 | 0 | NA | NA | NA | NA | NA | Gm43141 | predicted gene 43141 [Source:MGI Symbo | 0 | 0 | 0 | 0 | 0 | 0 |
| 43783 | ENSMUSG00000105745 | 0.342639279 | -1.805976596 | 4.028547714 | -0.448294702 | 0.653940524 | NA | Gm42521 | predicted gene 42521 [Source:MGI Symbo | 0 | 2.0558 | 0 | 0 | 0 | 0 |
| 43784 | ENSMUSG00000105746 | 0.160662974 | 1.126924175 | 4.080472857 | 0.276174898 | 0.782413736 | NA | Gm43595 | predicted gene 43595 [Source:MGI Symbo | 0 | 0 | 0 | 0.964 | 0 | 0 |
| 43785 | ENSMUSG00000105747 | 0 | NA | NA | NA | NA | NA | Igkv13-64 | immunoglobulin kappa chain variable 13-6 | 0 | 0 | 0 | 0 | 0 | 0 |
| 43786 | ENSMUSG00000105748 | 3.192834749 | -0.398947682 | 1.342531153 | -0.297160838 | 0.766343719 | NA | Gm43088 | predicted gene 43088 [Source:MGI Symbo | 0.8031 | 6.1675 | 4.025 | 4.8199 | 0 | 3.3415 |
| 43787 | ENSMUSG00000105749 | 0 | NA | NA | NA | NA | NA | Gm9251 | predicted gene 9251 [Source:MGI Symbol | 0 | 0 | 0 | 0 | 0 | 0 |
| 43788 | ENSMUSG00000105750 | 0 | NA | NA | NA | NA | NA | Gm43130 | predicted gene 43130 [Source:MGI Symbo | 0 | 0 | 0 | 0 | 0 | 0 |

| | | | | | | | | | | | | | | | |
|---|---|---|---|---|---|---|---|---|---|---|---|---|---|---|---|
| 43789 | ENSMUSG00000105751 | 0 | NA | NA | NA | NA | NA | Gm43724 | predicted gene 43724 [Source:MGI Symbo | 0 | 0 | 0 | 0 | 0 | 0 |
| 43790 | ENSMUSG00000105752 | 0 | NA | NA | NA | NA | NA | Igkv13-82 | immunoglobulin kappa chain variable 13-8 | 0 | 0 | 0 | 0 | 0 | 0 |
| 43791 | ENSMUSG00000105753 | 0 | NA | NA | NA | NA | NA | Gm43847 | predicted gene 43847 [Source:MGI Symbo | 0 | 0 | 0 | 0 | 0 | 0 |
| 43792 | ENSMUSG00000105754 | 0 | NA | NA | NA | NA | NA | NA | NA | 0 | 0 | 0 | 0 | 0 | 0 |
| 43793 | ENSMUSG00000105755 | 0.335414862 | -1.779047242 | 4.030804179 | -0.441362855 | 0.658950329 | NA | Gm43514 | predicted gene 43514 [Source:MGI Symbo | 0 | 0 | 2.0125 | 0 | 0 | 0 |
| 43794 | ENSMUSG00000105756 | 0 | NA | NA | NA | NA | NA | NA | NA | 0 | 0 | 0 | 0 | 0 | 0 |
| 43795 | ENSMUSG00000105757 | 0 | NA | NA | NA | NA | NA | Ighv1-83 | immunoglobulin heavy variable 1-83 [Sou | 0 | 0 | 0 | 0 | 0 | 0 |
| 43796 | ENSMUSG00000105758 | 0 | NA | NA | NA | NA | NA | Mirlet7e | microRNA let7e [Source:MGI Symbol;Acc: | 0 | 0 | 0 | 0 | 0 | 0 |
| 43797 | ENSMUSG00000105759 | 0.294513347 | 0.165143496 | 4.080472857 | 0.040471657 | 0.967717103 | NA | Gm42425 | predicted gene 42425 [Source:MGI Symbo | 0.8031 | 0 | 0 | 0.964 | 0 | 0 |
| 43798 | ENSMUSG00000105760 | 0 | NA | NA | NA | NA | NA | Gm43211 | predicted gene 43211 [Source:MGI Symbo | 0 | 0 | 0 | 0 | 0 | 0 |
| 43799 | ENSMUSG00000105761 | 0 | NA | NA | NA | NA | NA | Gm43787 | predicted gene 43787 [Source:MGI Symbo | 0 | 0 | 0 | 0 | 0 | 0 |
| 43800 | ENSMUSG00000105762 | 0 | NA | NA | NA | NA | NA | Gm43605 | predicted gene 43605 [Source:MGI Symbo | 0 | 0 | 0 | 0 | 0 | 0 |
| 43801 | ENSMUSG00000105763 | 0 | NA | NA | NA | NA | NA | Gm19088 | predicted gene, 19088 [Source:MGI Symb | 0 | 0 | 0 | 0 | 0 | 0 |
| 43802 | ENSMUSG00000105764 | 0 | NA | NA | NA | NA | NA | Gm42859 | predicted gene 42859 [Source:MGI Symbo | 0 | 0 | 0 | 0 | 0 | 0 |
| 43803 | ENSMUSG00000105765 | 0 | NA | NA | NA | NA | NA | 4930539C22 | RIKEN cDNA 4930539C22 gene [Source:M | 0 | 0 | 0 | 0 | 0 | 0 |
| 43804 | ENSMUSG00000105766 | 0 | NA | NA | NA | NA | NA | Gm19666 | predicted gene, 19666 [Source:MGI Symb | 0 | 0 | 0 | 0 | 0 | 0 |
| 43805 | ENSMUSG00000105767 | 0 | NA | NA | NA | NA | NA | Gm43607 | predicted gene 43607 [Source:MGI Symbo | 0 | 0 | 0 | 0 | 0 | 0 |
| 43806 | ENSMUSG00000105768 | 0 | NA | NA | NA | NA | NA | NA | NA | 0 | 0 | 0 | 0 | 0 | 0 |
| 43807 | ENSMUSG00000105769 | 0.181769715 | 1.126924175 | 4.080472857 | 0.276174898 | 0.782413736 | NA | Gm43351 | predicted gene 43351 [Source:MGI Symbo | 0 | 0 | 0 | 0 | 1.0906 | 0 |
| 43808 | ENSMUSG00000105770 | 0 | NA | NA | NA | NA | NA | Mir664 | microRNA 664 [Source:MGI Symbol;Acc:M | 0 | 0 | 0 | 0 | 0 | 0 |
| 43809 | ENSMUSG00000105771 | 0 | NA | NA | NA | NA | NA | 2900064K03 | RIKEN cDNA 2900064K03 gene [Source:M | 0 | 0 | 0 | 0 | 0 | 0 |
| 43810 | ENSMUSG00000105772 | 0 | NA | NA | NA | NA | NA | Gm42496 | predicted gene 42496 [Source:MGI Symbo | 0 | 0 | 0 | 0 | 0 | 0 |
| 43811 | ENSMUSG00000105773 | 0 | NA | NA | NA | NA | NA | 4930563F08 | RIKEN cDNA 4930563F08 gene [Source:M | 0 | 0 | 0 | 0 | 0 | 0 |
| 43812 | ENSMUSG00000105774 | 0.181769715 | 1.126924175 | 4.080472857 | 0.276174898 | 0.782413736 | NA | Gm43006 | predicted gene 43006 [Source:MGI Symbo | 0 | 0 | 0 | 0 | 1.0906 | 0 |
| 43813 | ENSMUSG00000105775 | 0 | NA | NA | NA | NA | NA | Gm10290 | predicted pseudogene 10290 [Source:MG | 0 | 0 | 0 | 0 | 0 | 0 |
| 43814 | ENSMUSG00000105776 | 0.342639279 | -1.805976596 | 4.028547714 | -0.448294702 | 0.653940524 | NA | Gm43292 | predicted gene 43292 [Source:MGI Symbo | 0 | 2.0558 | 0 | 0 | 0 | 0 |
| 43815 | ENSMUSG00000105777 | 0 | NA | NA | NA | NA | NA | Gm43049 | predicted gene 43049 [Source:MGI Symbo | 0 | 0 | 0 | 0 | 0 | 0 |
| 43816 | ENSMUSG00000105778 | 0 | NA | NA | NA | NA | NA | Mir7229 | microRNA 7229 [Source:MGI Symbol;Acc: | 0 | 0 | 0 | 0 | 0 | 0 |
| 43817 | ENSMUSG00000105779 | 0 | NA | NA | NA | NA | NA | Gm6439 | predicted gene 6439 [Source:MGI Symbol | 0 | 0 | 0 | 0 | 0 | 0 |
| 43818 | ENSMUSG00000105780 | 0 | NA | NA | NA | NA | NA | NA | NA | 0 | 0 | 0 | 0 | 0 | 0 |
| 43819 | ENSMUSG00000105781 | 0 | NA | NA | NA | NA | NA | Igkv8-31 | immunoglobulin kappa chain variable 8-31 | 0 | 0 | 0 | 0 | 0 | 0 |
| 43820 | ENSMUSG00000105782 | 0 | NA | NA | NA | NA | NA | Gm42955 | predicted gene 42955 [Source:MGI Symbo | 0 | 0 | 0 | 0 | 0 | 0 |
| 43821 | ENSMUSG00000105783 | 0 | NA | NA | NA | NA | NA | NA | NA | 0 | 0 | 0 | 0 | 0 | 0 |
| 43822 | ENSMUSG00000105784 | 0 | NA | NA | NA | NA | NA | Gm35065 | predicted gene, 35065 [Source:MGI Symb | 0 | 0 | 0 | 0 | 0 | 0 |
| 43823 | ENSMUSG00000105785 | 0 | NA | NA | NA | NA | NA | Gm42526 | predicted gene 42526 [Source:MGI Symbo | 0 | 0 | 0 | 0 | 0 | 0 |
| 43824 | ENSMUSG00000105786 | 0 | NA | NA | NA | NA | NA | Gm18957 | predicted gene, 18957 [Source:MGI Symb | 0 | 0 | 0 | 0 | 0 | 0 |
| 43825 | ENSMUSG00000105787 | 0 | NA | NA | NA | NA | NA | Gm8099 | predicted gene 8099 [Source:MGI Symbol | 0 | 0 | 0 | 0 | 0 | 0 |
| 43826 | ENSMUSG00000105788 | 0 | NA | NA | NA | NA | NA | NA | NA | 0 | 0 | 0 | 0 | 0 | 0 |
| 43827 | ENSMUSG00000105789 | 0.36353943 | 2.065989254 | 4.034137301 | 0.512126658 | 0.608562368 | NA | Gm42997 | predicted gene 42997 [Source:MGI Symbo | 0 | 0 | 0 | 0 | 2.1812 | 0 |
| 43828 | ENSMUSG00000105790 | 0 | NA | NA | NA | NA | NA | NA | NA | 0 | 0 | 0 | 0 | 0 | 0 |
| 43829 | ENSMUSG00000105791 | 0 | NA | NA | NA | NA | NA | Gm43341 | predicted gene 43341 [Source:MGI Symbo | 0 | 0 | 0 | 0 | 0 | 0 |
| 43830 | ENSMUSG00000105792 | 0 | NA | NA | NA | NA | NA | Gm17887 | predicted gene, 17887 [Source:MGI Symb | 0 | 0 | 0 | 0 | 0 | 0 |
| 43831 | ENSMUSG00000105793 | 0 | NA | NA | NA | NA | NA | Gm17956 | predicted gene, 17956 [Source:MGI Symb | 0 | 0 | 0 | 0 | 0 | 0 |
| 43832 | ENSMUSG00000105794 | 0 | NA | NA | NA | NA | NA | Gm43516 | predicted gene 43516 [Source:MGI Symbo | 0 | 0 | 0 | 0 | 0 | 0 |
| 43833 | ENSMUSG00000105795 | 0 | NA | NA | NA | NA | NA | Gm3970 | predicted gene 3970 [Source:MGI Symbol | 0 | 0 | 0 | 0 | 0 | 0 |
| 43834 | ENSMUSG00000105796 | 0 | NA | NA | NA | NA | NA | Gm42845 | predicted gene 42845 [Source:MGI Symbo | 0 | 0 | 0 | 0 | 0 | 0 |
| 43835 | ENSMUSG00000105797 | 0 | NA | NA | NA | NA | NA | Gm5149 | predicted gene 5149 [Source:MGI Symbol | 0 | 0 | 0 | 0 | 0 | 0 |
| 43836 | ENSMUSG00000105798 | 0 | NA | NA | NA | NA | NA | Gm18982 | predicted gene, 18982 [Source:MGI Symb | 0 | 0 | 0 | 0 | 0 | 0 |
| 43837 | ENSMUSG00000105799 | 0 | NA | NA | NA | NA | NA | Mir126b | microRNA 126b [Source:MGI Symbol;Acc: | 0 | 0 | 0 | 0 | 0 | 0 |
| 43838 | ENSMUSG00000105800 | 0 | NA | NA | NA | NA | NA | Gm18985 | predicted gene, 18985 [Source:MGI Symb | 0 | 0 | 0 | 0 | 0 | 0 |
| 43839 | ENSMUSG00000105801 | 0.449470461 | -0.545136533 | 3.562105917 | -0.15303771 | 0.878368537 | NA | C430019N01 | RIKEN cDNA C430019N01 gene [Source:M | 1.6062 | 0 | 0 | 0 | 1.0906 | 0 |
| 43840 | ENSMUSG00000105802 | 42.91438374 | -0.081847699 | 0.340476232 | -0.240391815 | 0.810026523 | 0.966157155 | Gm43012 | predicted gene 43012 [Source:MGI Symbo | 44.171 | 41.117 | 47.293 | 51.091 | 38.172 | 35.643 |
| 43841 | ENSMUSG00000105803 | 0 | NA | NA | NA | NA | NA | Gm43526 | predicted gene 43526 [Source:MGI Symbo | 0 | 0 | 0 | 0 | 0 | 0 |
| 43842 | ENSMUSG00000105804 | 3.361261005 | 0.72362278 | 1.272276793 | 0.568762068 | 0.56951762 | NA | Gm43654 | predicted gene 43654 [Source:MGI Symbo | 1.6062 | 0 | 6.0375 | 4.8199 | 4.3625 | 3.3415 |
| 43843 | ENSMUSG00000105805 | 0 | NA | NA | NA | NA | NA | NA | NA | 0 | 0 | 0 | 0 | 0 | 0 |
| 43844 | ENSMUSG00000105806 | 0 | NA | NA | NA | NA | NA | Gm43345 | predicted gene 43345 [Source:MGI Symbo | 0 | 0 | 0 | 0 | 0 | 0 |

| | | | | | | | | | | | | | | | |
|---|---|---|---|---|---|---|---|---|---|---|---|---|---|---|---|
| 43845 | ENSMUSG00000105807 | 0 | NA | NA | NA | NA | NA | Gm42622 | predicted gene 42622 [Source:MGI Symbo | 0 | 0 | 0 | 0 | 0 | 0 |
| 43846 | ENSMUSG00000105808 | 0 | NA | NA | NA | NA | NA | Gm42627 | predicted gene 42627 [Source:MGI Symbo | 0 | 0 | 0 | 0 | 0 | 0 |
| 43847 | ENSMUSG00000105809 | 0 | NA | NA | NA | NA | NA | Gm42626 | predicted gene 42626 [Source:MGI Symbo | 0 | 0 | 0 | 0 | 0 | 0 |
| 43848 | ENSMUSG00000105810 | 0.321325949 | 1.911575114 | 4.048267893 | 0.472195804 | 0.636787034 | NA | Gm43435 | predicted gene 43435 [Source:MGI Symbo | 0 | 0 | 0 | 1.928 | 0 | 0 |
| 43849 | ENSMUSG00000105811 | 0 | NA | NA | NA | NA | NA | Gm42707 | predicted gene 42707 [Source:MGI Symbo | 0 | 0 | 0 | 0 | 0 | 0 |
| 43850 | ENSMUSG00000105813 | 8.991354246 | -0.682616366 | 0.7600845 | -0.898079577 | 0.369143126 | 0.80653433 | Gm42928 | predicted gene 42928 [Source:MGI Symbo | 12.047 | 5.1396 | 16.1 | 9.6398 | 5.4531 | 5.5692 |
| 43851 | ENSMUSG00000105814 | 4.586665809 | 0.19377482 | 1.084666367 | 0.178649238 | 0.85821313 | NA | Mir703 | microRNA 703 [Source:MGI Symbol;Acc:M | 0.8031 | 4.1117 | 8.05 | 5.7839 | 6.5437 | 2.2277 |
| 43852 | ENSMUSG00000105815 | 0 | NA | NA | NA | NA | NA | Gm43541 | predicted gene 43541 [Source:MGI Symbo | 0 | 0 | 0 | 0 | 0 | 0 |
| 43853 | ENSMUSG00000105816 | 0 | NA | NA | NA | NA | NA | D030025E07 | RIKEN cDNA D030025E07 gene [Source:M | 0 | 0 | 0 | 0 | 0 | 0 |
| 43854 | ENSMUSG00000105817 | 0 | NA | NA | NA | NA | NA | Igkv15-97 | immunoglobulin kappa chain variable 15-9 | 0 | 0 | 0 | 0 | 0 | 0 |
| 43855 | ENSMUSG00000105818 | 1.020693421 | -3.383091396 | 2.599252854 | -1.301563021 | 0.193065808 | NA | Gm43256 | predicted gene 43256 [Source:MGI Symbo | 0 | 4.1117 | 2.0125 | 0 | 0 | 0 |
| 43856 | ENSMUSG00000105819 | 0 | NA | NA | NA | NA | NA | Gm42579 | predicted gene 42579 [Source:MGI Symbo | 0 | 0 | 0 | 0 | 0 | 0 |
| 43857 | ENSMUSG00000105820 | 1.14024829 | 0.506576368 | 2.118787831 | 0.239087822 | 0.811037485 | NA | Gm43400 | predicted gene 43400 [Source:MGI Symbo | 0.8031 | 2.0558 | 0 | 2.8919 | 1.0906 | 0 |
| 43858 | ENSMUSG00000105821 | 0 | NA | NA | NA | NA | NA | NA | NA | 0 | 0 | 0 | 0 | 0 | 0 |
| 43859 | ENSMUSG00000105822 | 0 | NA | NA | NA | NA | NA | Gm42969 | predicted gene 42969 [Source:MGI Symbo | 0 | 0 | 0 | 0 | 0 | 0 |
| 43860 | ENSMUSG00000105823 | 0 | NA | NA | NA | NA | NA | Trdv3 | T cell receptor delta variable 3 [Source:MG | 0 | 0 | 0 | 0 | 0 | 0 |
| 43861 | ENSMUSG00000105824 | 0.983224154 | -3.332395149 | 2.379709572 | -1.40033691 | 0.161412453 | NA | Gm43466 | predicted gene 43466 [Source:MGI Symbo | 0.8031 | 3.0838 | 2.0125 | 0 | 0 | 0 |
| 43862 | ENSMUSG00000105825 | 0 | NA | NA | NA | NA | NA | NA | NA | 0 | 0 | 0 | 0 | 0 | 0 |
| 43863 | ENSMUSG00000105826 | 0.17131964 | -0.796629176 | 4.080472857 | -0.195229623 | 0.84521319 | NA | Gm42141 | predicted gene, 42141 [Source:MGI Symb | 0 | 1.0279 | 0 | 0 | 0 | 0 |
| 43864 | ENSMUSG00000105827 | 0 | NA | NA | NA | NA | NA | H2bc18 | H2B clustered histone 18 [Source:MGI Syr | 0 | 0 | 0 | 0 | 0 | 0 |
| 43865 | ENSMUSG00000105828 | 0 | NA | NA | NA | NA | NA | Gm43048 | predicted gene 43048 [Source:MGI Symbo | 0 | 0 | 0 | 0 | 0 | 0 |
| 43866 | ENSMUSG00000105829 | 0 | NA | NA | NA | NA | NA | Gm43669 | predicted gene 43669 [Source:MGI Symbo | 0 | 0 | 0 | 0 | 0 | 0 |
| 43867 | ENSMUSG00000105830 | 0 | NA | NA | NA | NA | NA | Trgj2 | T cell receptor gamma joining 2 [Source:M | 0 | 0 | 0 | 0 | 0 | 0 |
| 43868 | ENSMUSG00000105831 | 0 | NA | NA | NA | NA | NA | Gm43184 | predicted gene 43184 [Source:MGI Symbo | 0 | 0 | 0 | 0 | 0 | 0 |
| 43869 | ENSMUSG00000105832 | 0 | NA | NA | NA | NA | NA | Gm43841 | predicted gene 43841 [Source:MGI Symbo | 0 | 0 | 0 | 0 | 0 | 0 |
| 43870 | ENSMUSG00000105833 | 0 | NA | NA | NA | NA | NA | Gm4751 | predicted gene 4751 [Source:MGI Symbol | 0 | 0 | 0 | 0 | 0 | 0 |
| 43871 | ENSMUSG00000105834 | 4.141480349 | -1.931533634 | 1.200057325 | -1.609534473 | 0.107499523 | NA | 4930592C13 | RIKEN cDNA 4930592C13 gene [Source:M | 5.6217 | 2.0558 | 12.075 | 2.8919 | 1.0906 | 1.1138 |
| 43872 | ENSMUSG00000105835 | 0 | NA | NA | NA | NA | NA | Gm43552 | predicted gene 43552 [Source:MGI Symbo | 0 | 0 | 0 | 0 | 0 | 0 |
| 43873 | ENSMUSG00000105836 | 1.436617682 | -0.781446032 | 1.931604398 | -0.40455801 | 0.685802429 | NA | Gm17954 | predicted gene, 17954 [Source:MGI Symb | 2.4093 | 0 | 3.0187 | 0.964 | 0 | 2.2277 |
| 43874 | ENSMUSG00000105837 | 141.8385338 | -0.495669653 | 0.257714359 | -1.923329592 | 0.054438671 | 0.328575194 | Gm35986 | predicted gene, 35986 [Source:MGI Symb | 205.59 | 117.18 | 175.09 | 150.38 | 101.43 | 101.36 |
| 43875 | ENSMUSG00000105838 | 0 | NA | NA | NA | NA | NA | 4930425O10 | RIKEN cDNA 4930425O10 gene [Source:M | 0 | 0 | 0 | 0 | 0 | 0 |
| 43876 | ENSMUSG00000105839 | 0 | NA | NA | NA | NA | NA | NA | NA | 0 | 0 | 0 | 0 | 0 | 0 |
| 43877 | ENSMUSG00000105840 | 0 | NA | NA | NA | NA | NA | Gm6073 | predicted gene 6073 [Source:MGI Symbol | 0 | 0 | 0 | 0 | 0 | 0 |
| 43878 | ENSMUSG00000105841 | 0 | NA | NA | NA | NA | NA | Mir3960 | microRNA 3960 [Source:MGI Symbol;Acc: | 0 | 0 | 0 | 0 | 0 | 0 |
| 43879 | ENSMUSG00000105842 | 1.19918676 | -2.469682767 | 2.290014249 | -1.07845738 | 0.280829692 | NA | Gm43329 | predicted gene 43329 [Source:MGI Symbo | 3.2124 | 0 | 3.0187 | 0.964 | 0 | 0 |
| 43880 | ENSMUSG00000105843 | 1.9628361 | -4.3305005 | 1.900052792 | -2.279147463 | 0.022658301 | NA | Gm19439 | predicted gene, 19439 [Source:MGI Symb | 1.6062 | 5.1396 | 5.0312 | 0 | 0 | 0 |
| 43881 | ENSMUSG00000105844 | 0 | NA | NA | NA | NA | NA | NA | NA | 0 | 0 | 0 | 0 | 0 | 0 |
| 43882 | ENSMUSG00000105845 | 0 | NA | NA | NA | NA | NA | Gm42808 | predicted gene 42808 [Source:MGI Symbo | 0 | 0 | 0 | 0 | 0 | 0 |
| 43883 | ENSMUSG00000105846 | 0.349477146 | 0.165142768 | 3.911021834 | 0.042224967 | 0.966319359 | NA | Gm43258 | predicted gene 43258 [Source:MGI Symbo | 0 | 0 | 1.0062 | 0 | 1.0906 | 0 |
| 43884 | ENSMUSG00000105847 | 0 | NA | NA | NA | NA | NA | Mir7650 | microRNA 7650 [Source:MGI Symbol;Acc: | 0 | 0 | 0 | 0 | 0 | 0 |
| 43885 | ENSMUSG00000105848 | 0 | NA | NA | NA | NA | NA | Gm42683 | predicted gene 42683 [Source:MGI Symbo | 0 | 0 | 0 | 0 | 0 | 0 |
| 43886 | ENSMUSG00000105849 | 0 | NA | NA | NA | NA | NA | Mir409 | microRNA 409 [Source:MGI Symbol;Acc:M | 0 | 0 | 0 | 0 | 0 | 0 |
| 43887 | ENSMUSG00000105850 | 0 | NA | NA | NA | NA | NA | Gm2861 | predicted gene 2861 [Source:MGI Symbol | 0 | 0 | 0 | 0 | 0 | 0 |
| 43888 | ENSMUSG00000105851 | 9.844508582 | 0.315006058 | 0.675446897 | 0.466366875 | 0.640952918 | 0.923932929 | 9130604C24 | RIKEN cDNA 9130604C24 gene [Source:M | 8.031 | 9.2513 | 9.0562 | 9.6398 | 14.178 | 8.9108 |
| 43889 | ENSMUSG00000105852 | 0.945754888 | -3.283756462 | 2.374797396 | -1.382752258 | 0.166740842 | NA | Gm42890 | predicted gene 42890 [Source:MGI Symbo | 1.6062 | 2.0558 | 2.0125 | 0 | 0 | 0 |
| 43890 | ENSMUSG00000105853 | 0 | NA | NA | NA | NA | NA | Mir376b | microRNA 376b [Source:MGI Symbol;Acc: | 0 | 0 | 0 | 0 | 0 | 0 |
| 43891 | ENSMUSG00000105854 | 0 | NA | NA | NA | NA | NA | Gm8925 | predicted gene 8925 [Source:MGI Symbol | 0 | 0 | 0 | 0 | 0 | 0 |
| 43892 | ENSMUSG00000105855 | 0 | NA | NA | NA | NA | NA | Gm42681 | predicted gene 42681 [Source:MGI Symbo | 0 | 0 | 0 | 0 | 0 | 0 |
| 43893 | ENSMUSG00000105856 | 0 | NA | NA | NA | NA | NA | Gm43732 | predicted gene 43732 [Source:MGI Symbo | 0 | 0 | 0 | 0 | 0 | 0 |
| 43894 | ENSMUSG00000105857 | 0 | NA | NA | NA | NA | NA | Gm42817 | predicted gene 42817 [Source:MGI Symbo | 0 | 0 | 0 | 0 | 0 | 0 |
| 43895 | ENSMUSG00000105858 | 0.342639279 | -1.805976596 | 4.028547714 | -0.448294702 | 0.653940524 | NA | 2310074N15 | RIKEN cDNA 2310074N15 gene [Source:M | 0 | 2.0558 | 0 | 0 | 0 | 0 |
| 43896 | ENSMUSG00000105859 | 0 | NA | NA | NA | NA | NA | Gm19185 | predicted gene, 19185 [Source:MGI Symb | 0 | 0 | 0 | 0 | 0 | 0 |
| 43897 | ENSMUSG00000105860 | 0.842329324 | -1.832359871 | 2.605893773 | -0.703159848 | 0.481956138 | NA | Gm42578 | predicted gene 42578 [Source:MGI Symbo | 0 | 3.0838 | 1.0062 | 0.964 | 0 | 0 |
| 43898 | ENSMUSG00000105861 | 0.371281295 | 2.09253362 | 4.031852165 | 0.519000582 | 0.603760335 | NA | Gm43508 | predicted gene 43508 [Source:MGI Symbo | 0 | 0 | 0 | 0 | 0 | 2.2277 |
| 43899 | ENSMUSG00000105862 | 0 | NA | NA | NA | NA | NA | Igkv3-12-1 | immunoglobulin kappa chain variable 3-12 | 0 | 0 | 0 | 0 | 0 | 0 |
| 43900 | ENSMUSG00000105863 | 0 | NA | NA | NA | NA | NA | NA | NA | 0 | 0 | 0 | 0 | 0 | 0 |

| | | | | | | | | | | | | | | | |
|---|---|---|---|---|---|---|---|---|---|---|---|---|---|---|---|
| 43901 | ENSMUSG00000105864 | 0 | NA | NA | NA | NA | NA | Gm10484 | predicted gene 10484 [Source:MGI Symbo | 0 | 0 | 0 | 0 | 0 | 0 |
| 43902 | ENSMUSG00000105865 | 1.042111019 | -0.860289347 | 2.663009232 | -0.323051583 | 0.746656177 | NA | Gm34144 | predicted gene, 34144 [Source:MGI Symb | 0 | 0 | 4.025 | 0 | 0 | 2.2277 |
| 43903 | ENSMUSG00000105866 | 0.713920574 | 0.125792037 | 3.028037396 | 0.041542432 | 0.966863466 | NA | Gm6745 | predicted gene 6745 [Source:MGI Symbol | 0 | 2.0558 | 0 | 0 | 0 | 2.2277 |
| 43904 | ENSMUSG00000105867 | 0.160662974 | 1.126924175 | 4.080472857 | 0.276174898 | 0.782413736 | NA | Gm42517 | predicted gene 42517 [Source:MGI Symbo | 0 | 0 | 0 | 0.964 | 0 | 0 |
| 43905 | ENSMUSG00000105868 | 1.061904139 | -1.168441417 | 2.189134959 | -0.533745721 | 0.593517472 | NA | Gm43766 | predicted gene 43766 [Source:MGI Symbo | 2.4093 | 1.0279 | 1.0062 | 1.928 | 0 | 0 |
| 43906 | ENSMUSG00000105869 | 0 | NA | NA | NA | NA | NA | Gm9408 | predicted gene 9408 [Source:MGI Symbol | 0 | 0 | 0 | 0 | 0 | 0 |
| 43907 | ENSMUSG00000105870 | 0.335414862 | -1.779047242 | 4.030804179 | -0.441362855 | 0.658950329 | NA | 7330423F06 | RIKEN cDNA 7330423F06 gene [Source:M | 0 | 0 | 2.0125 | 0 | 0 | 0 |
| 43908 | ENSMUSG00000105871 | 0.160662974 | 1.126924175 | 4.080472857 | 0.276174898 | 0.782413736 | NA | Gm43147 | predicted gene 43147 [Source:MGI Symbo | 0 | 0 | 0 | 0.964 | 0 | 0 |
| 43909 | ENSMUSG00000105872 | 0 | NA | NA | NA | NA | NA | Gm9515 | predicted gene 9515 [Source:MGI Symbol | 0 | 0 | 0 | 0 | 0 | 0 |
| 43910 | ENSMUSG00000105873 | 0 | NA | NA | NA | NA | NA | Gm43708 | predicted gene 43708 [Source:MGI Symbo | 0 | 0 | 0 | 0 | 0 | 0 |
| 43911 | ENSMUSG00000105874 | 0 | NA | NA | NA | NA | NA | NA | NA | 0 | 0 | 0 | 0 | 0 | 0 |
| 43912 | ENSMUSG00000105875 | 1.271825051 | -1.34972375 | 2.043905257 | -0.660365125 | 0.509019547 | NA | Gm43518 | predicted gene 43518 [Source:MGI Symbo | 2.4093 | 1.0279 | 2.0125 | 0 | 2.1812 | 0 |
| 43913 | ENSMUSG00000105876 | 0.870712616 | 0.665767941 | 2.367602424 | 0.281199214 | 0.778557607 | NA | Gm43572 | predicted gene 43572 [Source:MGI Symbo | 0 | 2.0558 | 0 | 0.964 | 1.0906 | 1.1138 |
| 43914 | ENSMUSG00000105877 | 0 | NA | NA | NA | NA | NA | Gm42479 | predicted gene 42479 [Source:MGI Symbo | 0 | 0 | 0 | 0 | 0 | 0 |
| 43915 | ENSMUSG00000105878 | 0 | NA | NA | NA | NA | NA | NA | NA | 0 | 0 | 0 | 0 | 0 | 0 |
| 43916 | ENSMUSG00000105879 | 2.976505054 | -0.814771851 | 1.258362577 | -0.647485761 | 0.517317607 | NA | Gm6204 | predicted gene 6204 [Source:MGI Symbol | 3.2124 | 4.1117 | 4.025 | 0.964 | 1.0906 | 4.4554 |
| 43917 | ENSMUSG00000105880 | 0 | NA | NA | NA | NA | NA | Gm43678 | predicted gene 43678 [Source:MGI Symbo | 0 | 0 | 0 | 0 | 0 | 0 |
| 43918 | ENSMUSG00000105881 | 65.05856313 | 0.458160165 | 0.299929178 | 1.527561165 | 0.12662153 | 0.521187766 | 4932422M1 | RIKEN cDNA 4932422M17 gene [Source:M | 66.657 | 45.228 | 52.325 | 89.65 | 76.343 | 60.148 |
| 43919 | ENSMUSG00000105882 | 0 | NA | NA | NA | NA | NA | Gm42905 | predicted gene 42905 [Source:MGI Symbo | 0 | 0 | 0 | 0 | 0 | 0 |
| 43920 | ENSMUSG00000105883 | 0 | NA | NA | NA | NA | NA | Igkv4-65 | immunoglobulin kappa chain variable 4-6 | 0 | 0 | 0 | 0 | 0 | 0 |
| 43921 | ENSMUSG00000105884 | 0 | NA | NA | NA | NA | NA | NA | NA | 0 | 0 | 0 | 0 | 0 | 0 |
| 43922 | ENSMUSG00000105885 | 0 | NA | NA | NA | NA | NA | Sult1c2-ps2 | sulfotransferase family, cytosolic, 1C, mer | 0 | 0 | 0 | 0 | 0 | 0 |
| 43923 | ENSMUSG00000105886 | 0 | NA | NA | NA | NA | NA | Gm22813 | predicted gene, 22813 [Source:MGI Symb | 0 | 0 | 0 | 0 | 0 | 0 |
| 43924 | ENSMUSG00000105887 | 0 | NA | NA | NA | NA | NA | Trav5d-3 | T cell receptor alpha variable 5D-3 [Sourc | 0 | 0 | 0 | 0 | 0 | 0 |
| 43925 | ENSMUSG00000105888 | 0 | NA | NA | NA | NA | NA | BC037156 | cDNA sequence BC037156 [Source:MGI Sy | 0 | 0 | 0 | 0 | 0 | 0 |
| 43926 | ENSMUSG00000105889 | 0 | NA | NA | NA | NA | NA | Gm42776 | predicted gene 42776 [Source:MGI Symbo | 0 | 0 | 0 | 0 | 0 | 0 |
| 43927 | ENSMUSG00000105890 | 0 | NA | NA | NA | NA | NA | Gm42780 | predicted gene 42780 [Source:MGI Symbo | 0 | 0 | 0 | 0 | 0 | 0 |
| 43928 | ENSMUSG00000105891 | 2.760412967 | -0.3063309 | 1.244029674 | -0.24624083 | 0.80549581 | NA | A230001M1 | RIKEN cDNA A230001M10 gene [Source:M | 4.0155 | 3.0838 | 2.0125 | 1.928 | 2.1812 | 3.3415 |
| 43929 | ENSMUSG00000105892 | 0.36353943 | 2.065989254 | 4.034137301 | 0.512126658 | 0.608562368 | NA | Gm35013 | predicted gene, 35013 [Source:MGI Symb | 0 | 0 | 0 | 0 | 2.1812 | 0 |
| 43930 | ENSMUSG00000105893 | 0.353089354 | 0.165142706 | 3.896102244 | 0.042386646 | 0.966190474 | NA | Wdr46-ps | Wdr46 retrotransposed pseudogene [Sou | 0 | 1.0279 | 0 | 0 | 1.0906 | 0 |
| 43931 | ENSMUSG00000105894 | 0 | NA | NA | NA | NA | NA | Gm42710 | predicted gene 42710 [Source:MGI Symbo | 0 | 0 | 0 | 0 | 0 | 0 |
| 43932 | ENSMUSG00000105895 | 1.593796274 | -1.081739973 | 1.87424507 | -0.577160367 | 0.563831127 | NA | Gm42829 | predicted gene 42829 [Source:MGI Symbo | 2.4093 | 4.1117 | 0 | 1.928 | 0 | 1.1138 |
| 43933 | ENSMUSG00000105896 | 4.675226874 | -2.124692124 | 1.096180541 | -1.938268419 | 0.052590483 | NA | Gm43311 | predicted gene 43311 [Source:MGI Symbo | 9.6372 | 4.1117 | 9.0562 | 1.928 | 1.0906 | 2.2277 |
| 43934 | ENSMUSG00000105897 | 0 | NA | NA | NA | NA | NA | Gm7646 | predicted gene 7646 [Source:MGI Symbol | 0 | 0 | 0 | 0 | 0 | 0 |
| 43935 | ENSMUSG00000105898 | 0 | NA | NA | NA | NA | NA | Obox4-ps32 | oocyte specific homeobox 4, pseudogene | 0 | 0 | 0 | 0 | 0 | 0 |
| 43936 | ENSMUSG00000105899 | 0 | NA | NA | NA | NA | NA | NA | NA | 0 | 0 | 0 | 0 | 0 | 0 |
| 43937 | ENSMUSG00000105900 | 0 | NA | NA | NA | NA | NA | Gm43240 | predicted gene 43240 [Source:MGI Symbo | 0 | 0 | 0 | 0 | 0 | 0 |
| 43938 | ENSMUSG00000105901 | 0 | NA | NA | NA | NA | NA | Gm7291 | predicted gene 7291 [Source:MGI Symbol | 0 | 0 | 0 | 0 | 0 | 0 |
| 43939 | ENSMUSG00000105902 | 0 | NA | NA | NA | NA | NA | NA | NA | 0 | 0 | 0 | 0 | 0 | 0 |
| 43940 | ENSMUSG00000105903 | 0 | NA | NA | NA | NA | NA | 1700014F14 | RIKEN cDNA 1700014F14 gene [Source:M | 0 | 0 | 0 | 0 | 0 | 0 |
| 43941 | ENSMUSG00000105904 | 0 | NA | NA | NA | NA | NA | Mir24-1 | microRNA 24-1 [Source:MGI Symbol;Acc:I | 0 | 0 | 0 | 0 | 0 | 0 |
| 43942 | ENSMUSG00000105905 | 0 | NA | NA | NA | NA | NA | Gm7709 | predicted gene 7709 [Source:MGI Symbol | 0 | 0 | 0 | 0 | 0 | 0 |
| 43943 | ENSMUSG00000105906 | 4.099801764 | 0.764813317 | 1.196671607 | 0.639117126 | 0.52274674 | NA | Iglc1 | immunoglobulin lambda constant 1 [Sour | 4.0155 | 2.0558 | 3.0187 | 7.7118 | 0 | 7.7969 |
| 43944 | ENSMUSG00000105907 | 18.93679462 | 0.080946537 | 0.51449074 | 0.157333321 | 0.874982161 | 0.980503106 | Gm34333 | predicted gene, 34333 [Source:MGI Symb | 22.487 | 13.363 | 19.119 | 13.496 | 23.994 | 21.163 |
| 43945 | ENSMUSG00000105908 | 0.167707431 | -0.796629176 | 4.080472857 | -0.195229623 | 0.84521319 | NA | Mir6920 | microRNA 6920 [Source:MGI Symbol;Acc: | 0 | 0 | 1.0062 | 0 | 0 | 0 |
| 43946 | ENSMUSG00000105909 | 58.46519092 | -0.494974206 | 0.383522387 | -1.290600556 | 0.196842223 | 0.637822894 | Gm43189 | predicted gene 43189 [Source:MGI Symbo | 69.067 | 84.289 | 52.325 | 79.046 | 35.99 | 30.074 |
| 43947 | ENSMUSG00000105910 | 0.160662974 | 1.126924175 | 4.080472857 | 0.276174898 | 0.782413736 | NA | Gm43539 | predicted gene 43539 [Source:MGI Symbo | 0 | 0 | 0 | 0.964 | 0 | 0 |
| 43948 | ENSMUSG00000105911 | 0 | NA | NA | NA | NA | NA | NA | NA | 0 | 0 | 0 | 0 | 0 | 0 |
| 43949 | ENSMUSG00000105912 | 0.181769715 | 1.126924175 | 4.080472857 | 0.276174898 | 0.782413736 | NA | Gm10440 | predicted gene 10440 [Source:MGI Symbo | 0 | 0 | 0 | 0 | 1.0906 | 0 |
| 43950 | ENSMUSG00000105913 | 0 | NA | NA | NA | NA | NA | Mir5121 | microRNA 5121 [Source:MGI Symbol;Acc: | 0 | 0 | 0 | 0 | 0 | 0 |
| 43951 | ENSMUSG00000105914 | 0.185640647 | 1.126924175 | 4.080472857 | 0.276174898 | 0.782413736 | NA | Gm19243 | predicted gene, 19243 [Source:MGI Symb | 0 | 0 | 0 | 0 | 0 | 1.1138 |
| 43952 | ENSMUSG00000105915 | 0 | NA | NA | NA | NA | NA | Igkv12-42 | immunoglobulin kappa chain variable 12- | 0 | 0 | 0 | 0 | 0 | 0 |
| 43953 | ENSMUSG00000105916 | 0 | NA | NA | NA | NA | NA | Gm42453 | predicted gene 42453 [Source:MGI Symbo | 0 | 0 | 0 | 0 | 0 | 0 |
| 43954 | ENSMUSG00000105917 | 0 | NA | NA | NA | NA | NA | Gm43612 | predicted gene 43612 [Source:MGI Symbo | 0 | 0 | 0 | 0 | 0 | 0 |
| 43955 | ENSMUSG00000105918 | 0 | NA | NA | NA | NA | NA | NA | NA | 0 | 0 | 0 | 0 | 0 | 0 |
| 43956 | ENSMUSG00000105919 | 0 | NA | NA | NA | NA | NA | Gm7166 | predicted gene 7166 [Source:MGI Symbol | 0 | 0 | 0 | 0 | 0 | 0 |

| | | | | | | | | | | | | | | | |
|---|---|---|---|---|---|---|---|---|---|---|---|---|---|---|---|
| 43957 | ENSMUSG00000105920 | 0 | NA | NA | NA | NA | NA | Gm5408 | predicted pseudogene 5408 [Source:MGI | 0 | 0 | 0 | 0 | 0 | 0 |
| 43958 | ENSMUSG00000105921 | 0 | NA | NA | NA | NA | NA | Mark4-ps | MAP/microtubule affinity regulating kinas | 0 | 0 | 0 | 0 | 0 | 0 |
| 43959 | ENSMUSG00000105922 | 0 | NA | NA | NA | NA | NA | Gm42769 | predicted gene 42769 [Source:MGI Symbo | 0 | 0 | 0 | 0 | 0 | 0 |
| 43960 | ENSMUSG00000105923 | 0 | NA | NA | NA | NA | NA | A830019L24 | RIKEN cDNA A830019L24 gene [Source:M | 0 | 0 | 0 | 0 | 0 | 0 |
| 43961 | ENSMUSG00000105924 | 0 | NA | NA | NA | NA | NA | NA | NA | 0 | 0 | 0 | 0 | 0 | 0 |
| 43962 | ENSMUSG00000105925 | 0.305170013 | -1.657836477 | 4.041480335 | -0.410205257 | 0.681655385 | NA | Gm43846 | predicted gene 43846 [Source:MGI Symbo | 0.8031 | 1.0279 | 0 | 0 | 0 | 0 |
| 43963 | ENSMUSG00000105926 | 0 | NA | NA | NA | NA | NA | Gm43386 | predicted gene 43386 [Source:MGI Symbo | 0 | 0 | 0 | 0 | 0 | 0 |
| 43964 | ENSMUSG00000105927 | 0 | NA | NA | NA | NA | NA | NA | NA | 0 | 0 | 0 | 0 | 0 | 0 |
| 43965 | ENSMUSG00000105928 | 0.160662974 | 1.126924175 | 4.080472857 | 0.276174898 | 0.782413736 | NA | Gm9256 | predicted gene 9256 [Source:MGI Symbol | 0 | 0 | 0 | 0.964 | 0 | 0 |
| 43966 | ENSMUSG00000105929 | 2.388026825 | 0.910066903 | 1.641082091 | 0.554552943 | 0.579200484 | NA | Gm43693 | predicted gene 43693 [Source:MGI Symbo | 0 | 1.0279 | 4.025 | 4.8199 | 0 | 4.4554 |
| 43967 | ENSMUSG00000105930 | 0 | NA | NA | NA | NA | NA | Gm17889 | predicted gene, 17889 [Source:MGI Symb | 0 | 0 | 0 | 0 | 0 | 0 |
| 43968 | ENSMUSG00000105931 | 0 | NA | NA | NA | NA | NA | Gm43014 | predicted gene 43014 [Source:MGI Symbo | 0 | 0 | 0 | 0 | 0 | 0 |
| 43969 | ENSMUSG00000105932 | 0 | NA | NA | NA | NA | NA | Mir6925 | microRNA 6925 [Source:MGI Symbol;Acc: | 0 | 0 | 0 | 0 | 0 | 0 |
| 43970 | ENSMUSG00000105933 | 0.572870759 | -2.577107685 | 3.182795569 | -0.809699407 | 0.418112959 | NA | Gm43260 | predicted gene 43260 [Source:MGI Symbo | 2.4093 | 1.0279 | 0 | 0 | 0 | 0 |
| 43971 | ENSMUSG00000105934 | 0 | NA | NA | NA | NA | NA | Gm43575 | predicted gene 43575 [Source:MGI Symbo | 0 | 0 | 0 | 0 | 0 | 0 |
| 43972 | ENSMUSG00000105935 | 1.01026885 | 1.016270523 | 2.426385186 | 0.418841381 | 0.67533206 | NA | Gm43628 | predicted gene 43628 [Source:MGI Symbo | 0 | 2.0558 | 0 | 2.8919 | 0 | 1.1138 |
| 43973 | ENSMUSG00000105936 | 0 | NA | NA | NA | NA | NA | Gm43544 | predicted gene 43544 [Source:MGI Symbo | 0 | 0 | 0 | 0 | 0 | 0 |
| 43974 | ENSMUSG00000105937 | 0 | NA | NA | NA | NA | NA | Gm43602 | predicted gene 43602 [Source:MGI Symbo | 0 | 0 | 0 | 0 | 0 | 0 |
| 43975 | ENSMUSG00000105938 | 0 | NA | NA | NA | NA | NA | Gm33866 | predicted gene, 33866 [Source:MGI Symb | 0 | 0 | 0 | 0 | 0 | 0 |
| 43976 | ENSMUSG00000105939 | 0 | NA | NA | NA | NA | NA | Gm43322 | predicted gene 43322 [Source:MGI Symbo | 0 | 0 | 0 | 0 | 0 | 0 |
| 43977 | ENSMUSG00000105940 | 0 | NA | NA | NA | NA | NA | Gm42635 | predicted gene 42635 [Source:MGI Symbo | 0 | 0 | 0 | 0 | 0 | 0 |
| 43978 | ENSMUSG00000105941 | 6.655290488 | -0.798336037 | 0.904010012 | -0.883105304 | 0.377179379 | 0.81030298 | Gm42809 | predicted gene 42809 [Source:MGI Symbo | 16.062 | 5.1396 | 4.025 | 4.8199 | 6.5437 | 3.3415 |
| 43979 | ENSMUSG00000105942 | 0 | NA | NA | NA | NA | NA | Gm43175 | predicted gene 43175 [Source:MGI Symbo | 0 | 0 | 0 | 0 | 0 | 0 |
| 43980 | ENSMUSG00000105943 | 0 | NA | NA | NA | NA | NA | Gm2965 | predicted gene 2965 [Source:MGI Symbol | 0 | 0 | 0 | 0 | 0 | 0 |
| 43981 | ENSMUSG00000105944 | 0.133850373 | -0.796629176 | 4.080472857 | -0.195229623 | 0.84521319 | NA | Gm5075 | predicted gene 5075 [Source:MGI Symbol | 0.8031 | 0 | 0 | 0 | 0 | 0 |
| 43982 | ENSMUSG00000105945 | 0 | NA | NA | NA | NA | NA | Gm43570 | predicted gene 43570 [Source:MGI Symbo | 0 | 0 | 0 | 0 | 0 | 0 |
| 43983 | ENSMUSG00000105946 | 0 | NA | NA | NA | NA | NA | Gm43776 | predicted gene 43776 [Source:MGI Symbo | 0 | 0 | 0 | 0 | 0 | 0 |
| 43984 | ENSMUSG00000105947 | 0 | NA | NA | NA | NA | NA | Gm43789 | predicted gene 43789 [Source:MGI Symbo | 0 | 0 | 0 | 0 | 0 | 0 |
| 43985 | ENSMUSG00000105948 | 0 | NA | NA | NA | NA | NA | Gm43089 | predicted gene 43089 [Source:MGI Symbo | 0 | 0 | 0 | 0 | 0 | 0 |
| 43986 | ENSMUSG00000105949 | 0 | NA | NA | NA | NA | NA | Gm6510 | predicted gene 6510 [Source:MGI Symbol | 0 | 0 | 0 | 0 | 0 | 0 |
| 43987 | ENSMUSG00000105950 | 1.606287867 | -1.175097167 | 2.050416046 | -0.57310182 | 0.566575755 | NA | Gm43679 | predicted gene 43679 [Source:MGI Symbo | 1.6062 | 5.1396 | 0 | 2.8919 | 0 | 0 |
| 43988 | ENSMUSG00000105951 | 0 | NA | NA | NA | NA | NA | NA | NA | 0 | 0 | 0 | 0 | 0 | 0 |
| 43989 | ENSMUSG00000105952 | 0 | NA | NA | NA | NA | NA | Gm40153 | predicted gene, 40153 [Source:MGI Symb | 0 | 0 | 0 | 0 | 0 | 0 |
| 43990 | ENSMUSG00000105953 | 0 | NA | NA | NA | NA | NA | Trav15d-2-d | T cell receptor alpha variable 15D-2-DV6D | 0 | 0 | 0 | 0 | 0 | 0 |
| 43991 | ENSMUSG00000105954 | 0.267700746 | -1.497481649 | 4.057001436 | -0.369110456 | 0.712045396 | NA | Gm42793 | predicted gene 42793 [Source:MGI Symbo | 1.6062 | 0 | 0 | 0 | 0 | 0 |
| 43992 | ENSMUSG00000105955 | 0 | NA | NA | NA | NA | NA | Igkv12-40 | immunoglobulin kappa chain variable 12-4 | 0 | 0 | 0 | 0 | 0 | 0 |
| 43993 | ENSMUSG00000105956 | 0 | NA | NA | NA | NA | NA | Trav5n-2 | T cell receptor alpha variable 5N-2 [Sourc | 0 | 0 | 0 | 0 | 0 | 0 |
| 43994 | ENSMUSG00000105957 | 16.87042137 | 0.165135825 | 0.515681718 | 0.320228194 | 0.748795352 | 0.951074124 | Gm43254 | predicted gene 43254 [Source:MGI Symbo | 15.259 | 14.391 | 18.112 | 19.28 | 16.359 | 17.822 |
| 43995 | ENSMUSG00000105958 | 0 | NA | NA | NA | NA | NA | Gm43319 | predicted gene 43319 [Source:MGI Symbo | 0 | 0 | 0 | 0 | 0 | 0 |
| 43996 | ENSMUSG00000105959 | 0 | NA | NA | NA | NA | NA | D530037P16 | RIKEN cDNA D530037P16 gene [Source:M | 0 | 0 | 0 | 0 | 0 | 0 |
| 43997 | ENSMUSG00000105960 | 0 | NA | NA | NA | NA | NA | 6330410L21 | RIKEN cDNA 6330410L21 gene [Source:M | 0 | 0 | 0 | 0 | 0 | 0 |
| 43998 | ENSMUSG00000105961 | 0 | NA | NA | NA | NA | NA | Gm40123 | predicted gene, 40123 [Source:MGI Symb | 0 | 0 | 0 | 0 | 0 | 0 |
| 43999 | ENSMUSG00000105962 | 0 | NA | NA | NA | NA | NA | Gm42432 | predicted gene 42432 [Source:MGI Symbo | 0 | 0 | 0 | 0 | 0 | 0 |
| 44000 | ENSMUSG00000105963 | 0 | NA | NA | NA | NA | NA | Gm43647 | predicted gene 43647 [Source:MGI Symbo | 0 | 0 | 0 | 0 | 0 | 0 |
| 44001 | ENSMUSG00000105964 | 0 | NA | NA | NA | NA | NA | Mir219c | microRNA 219c [Source:MGI Symbol;Acc: | 0 | 0 | 0 | 0 | 0 | 0 |
| 44002 | ENSMUSG00000105965 | 0 | NA | NA | NA | NA | NA | Cycs-ps2 | cytochrome c, pseudogene 2 [Source:MGI | 0 | 0 | 0 | 0 | 0 | 0 |
| 44003 | ENSMUSG00000105966 | 0 | NA | NA | NA | NA | NA | Gm43242 | predicted gene 43242 [Source:MGI Symbo | 0 | 0 | 0 | 0 | 0 | 0 |
| 44004 | ENSMUSG00000105967 | 0 | NA | NA | NA | NA | NA | NA | NA | 0 | 0 | 0 | 0 | 0 | 0 |
| 44005 | ENSMUSG00000105968 | 0 | NA | NA | NA | NA | NA | Igkv13-71-1 | immunoglobulin kappa chain variable 13-7 | 0 | 0 | 0 | 0 | 0 | 0 |
| 44006 | ENSMUSG00000105969 | 0 | NA | NA | NA | NA | NA | Gm42525 | predicted gene 42525 [Source:MGI Symbo | 0 | 0 | 0 | 0 | 0 | 0 |
| 44007 | ENSMUSG00000105970 | 1.613384458 | -0.473440214 | 1.692060853 | -0.279800938 | 0.779630232 | NA | Gm43360 | predicted gene 43360 [Source:MGI Symbo | 1.6062 | 2.0558 | 2.0125 | 2.8919 | 0 | 1.1138 |
| 44008 | ENSMUSG00000105971 | 0 | NA | NA | NA | NA | NA | Gm43805 | predicted gene 43805 [Source:MGI Symbo | 0 | 0 | 0 | 0 | 0 | 0 |
| 44009 | ENSMUSG00000105972 | 0 | NA | NA | NA | NA | NA | Mir1843a | microRNA 1843a [Source:MGI Symbol;Acc | 0 | 0 | 0 | 0 | 0 | 0 |
| 44010 | ENSMUSG00000105973 | 0.181769715 | 1.126924175 | 4.080472857 | 0.276174898 | 0.782413736 | NA | Gm42881 | predicted gene 42881 [Source:MGI Symbo | 0 | 0 | 0 | 0 | 1.0906 | 0 |
| 44011 | ENSMUSG00000105974 | 0 | NA | NA | NA | NA | NA | Gm43318 | predicted gene 43318 [Source:MGI Symbo | 0 | 0 | 0 | 0 | 0 | 0 |
| 44012 | ENSMUSG00000105975 | 0 | NA | NA | NA | NA | NA | Gm9831 | predicted gene 9831 [Source:MGI Symbol | 0 | 0 | 0 | 0 | 0 | 0 |

| | | | | | | | | | | | | | | | |
|---|---|---|---|---|---|---|---|---|---|---|---|---|---|---|---|
| 44013 | ENSMUSG00000105976 | 1.197531793 | -2.437484061 | 2.303856673 | -1.058001607 | 0.29005471 | NA | Gm42782 | predicted gene 42782 [Source:MGI Symbc | 4.0155 | 2.0558 | 0 | 0 | 0 | 1.1138 |
| 44014 | ENSMUSG00000105977 | 0 | NA | NA | NA | NA | NA | Gm43555 | predicted gene 43555 [Source:MGI Symbc | 0 | 0 | 0 | 0 | 0 | 0 |
| 44015 | ENSMUSG00000105978 | 0 | NA | NA | NA | NA | NA | Gm43492 | predicted gene 43492 [Source:MGI Symbc | 0 | 0 | 0 | 0 | 0 | 0 |
| 44016 | ENSMUSG00000105979 | 0 | NA | NA | NA | NA | NA | Gm43411 | predicted gene 43411 [Source:MGI Symbc | 0 | 0 | 0 | 0 | 0 | 0 |
| 44017 | ENSMUSG00000105980 | 0 | NA | NA | NA | NA | NA | Gm42597 | predicted gene 42597 [Source:MGI Symbc | 0 | 0 | 0 | 0 | 0 | 0 |
| 44018 | ENSMUSG00000105981 | 3.087000578 | 0.102482924 | 1.319667895 | 0.077658117 | 0.938100011 | NA | 2810428J06I | RIKEN cDNA 2810428J06 gene [Source:M( | 4.8186 | 0 | 4.025 | 1.928 | 2.1812 | 5.5692 |
| 44019 | ENSMUSG00000105982 | 0 | NA | NA | NA | NA | NA | Mir6896 | microRNA 6896 [Source:MGI Symbol;Acc: | 0 | 0 | 0 | 0 | 0 | 0 |
| 44020 | ENSMUSG00000105983 | 0 | NA | NA | NA | NA | NA | Gm42770 | predicted gene 42770 [Source:MGI Symbc | 0 | 0 | 0 | 0 | 0 | 0 |
| 44021 | ENSMUSG00000105984 | 0 | NA | NA | NA | NA | NA | NA | NA | 0 | 0 | 0 | 0 | 0 | 0 |
| 44022 | ENSMUSG00000105985 | 0 | NA | NA | NA | NA | NA | Gm42993 | predicted gene 42993 [Source:MGI Symbc | 0 | 0 | 0 | 0 | 0 | 0 |
| 44023 | ENSMUSG00000105986 | 0.185640647 | 1.126924175 | 4.080472857 | 0.276174898 | 0.782413736 | NA | Gm43065 | predicted gene 43065 [Source:MGI Symbc | 0 | 0 | 0 | 0 | 0 | 1.1138 |
| 44024 | ENSMUSG00000105987 | 21.46276464 | -1.352412854 | 0.513892284 | -2.631704921 | 0.008495762 | 0.096227639 | AI506816 | expressed sequence AI506816 [Source:M( | 41.761 | 23.642 | 27.169 | 16.388 | 10.906 | 8.9108 |
| 44025 | ENSMUSG00000105988 | 0 | NA | NA | NA | NA | NA | NA | NA | 0 | 0 | 0 | 0 | 0 | 0 |
| 44026 | ENSMUSG00000105989 | 0 | NA | NA | NA | NA | NA | Igkv13-73-1 | immunoglobulin kappa chain variable 13-7 | 0 | 0 | 0 | 0 | 0 | 0 |
| 44027 | ENSMUSG00000105990 | 0 | NA | NA | NA | NA | NA | Gm43307 | predicted gene 43307 [Source:MGI Symbc | 0 | 0 | 0 | 0 | 0 | 0 |
| 44028 | ENSMUSG00000105991 | 0 | NA | NA | NA | NA | NA | 1700042D02 | RIKEN cDNA 1700042D02 gene [Source:M | 0 | 0 | 0 | 0 | 0 | 0 |
| 44029 | ENSMUSG00000105992 | 0.367410362 | 2.078255025 | 3.800070157 | 0.54689912 | 0.58444804 | NA | Gm43651 | predicted gene 43651 [Source:MGI Symbc | 0 | 0 | 0 | 0 | 1.0906 | 1.1138 |
| 44030 | ENSMUSG00000105993 | 0 | NA | NA | NA | NA | NA | Tex13c2 | TEX13 family member C2 [Source:MGI Syr | 0 | 0 | 0 | 0 | 0 | 0 |
| 44031 | ENSMUSG00000105994 | 0 | NA | NA | NA | NA | NA | Gm43427 | predicted gene 43427 [Source:MGI Symbc | 0 | 0 | 0 | 0 | 0 | 0 |
| 44032 | ENSMUSG00000105995 | 0 | NA | NA | NA | NA | NA | Gm43554 | predicted gene 43554 [Source:MGI Symbc | 0 | 0 | 0 | 0 | 0 | 0 |
| 44033 | ENSMUSG00000105996 | 0 | NA | NA | NA | NA | NA | Gm43219 | predicted gene 43219 [Source:MGI Symbc | 0 | 0 | 0 | 0 | 0 | 0 |
| 44034 | ENSMUSG00000105997 | 0 | NA | NA | NA | NA | NA | NA | NA | 0 | 0 | 0 | 0 | 0 | 0 |
| 44035 | ENSMUSG00000105998 | 0 | NA | NA | NA | NA | NA | NA | NA | 0 | 0 | 0 | 0 | 0 | 0 |
| 44036 | ENSMUSG00000105999 | 0 | NA | NA | NA | NA | NA | Gm2213 | predicted gene 2213 [Source:MGI Symbol | 0 | 0 | 0 | 0 | 0 | 0 |
| 44037 | ENSMUSG00000106000 | 0 | NA | NA | NA | NA | NA | Gm16508 | predicted gene 16508 [Source:MGI Symbc | 0 | 0 | 0 | 0 | 0 | 0 |
| 44038 | ENSMUSG00000106001 | 0 | NA | NA | NA | NA | NA | Gm42826 | predicted gene 42826 [Source:MGI Symbc | 0 | 0 | 0 | 0 | 0 | 0 |
| 44039 | ENSMUSG00000106002 | 0 | NA | NA | NA | NA | NA | 5430434I15I | RIKEN cDNA 5430434I15 gene [Source:M( | 0 | 0 | 0 | 0 | 0 | 0 |
| 44040 | ENSMUSG00000106003 | 0 | NA | NA | NA | NA | NA | Gm42465 | predicted gene 42465 [Source:MGI Symbc | 0 | 0 | 0 | 0 | 0 | 0 |
| 44041 | ENSMUSG00000106004 | 0 | NA | NA | NA | NA | NA | Gm43391 | predicted gene 43391 [Source:MGI Symbc | 0 | 0 | 0 | 0 | 0 | 0 |
| 44042 | ENSMUSG00000106005 | 0 | NA | NA | NA | NA | NA | Gm29151 | predicted gene 29151 [Source:MGI Symbc | 0 | 0 | 0 | 0 | 0 | 0 |
| 44043 | ENSMUSG00000106006 | 0 | NA | NA | NA | NA | NA | Gm17951 | predicted gene, 17951 [Source:MGI Symb | 0 | 0 | 0 | 0 | 0 | 0 |
| 44044 | ENSMUSG00000106007 | 0 | NA | NA | NA | NA | NA | NA | NA | 0 | 0 | 0 | 0 | 0 | 0 |
| 44045 | ENSMUSG00000106008 | 0 | NA | NA | NA | NA | NA | Gm42920 | predicted gene 42920 [Source:MGI Symbc | 0 | 0 | 0 | 0 | 0 | 0 |
| 44046 | ENSMUSG00000106009 | 0 | NA | NA | NA | NA | NA | Gm9379 | predicted gene 9379 [Source:MGI Symbol | 0 | 0 | 0 | 0 | 0 | 0 |
| 44047 | ENSMUSG00000106010 | 0 | NA | NA | NA | NA | NA | Gm42616 | predicted gene 42616 [Source:MGI Symbc | 0 | 0 | 0 | 0 | 0 | 0 |
| 44048 | ENSMUSG00000106011 | 0 | NA | NA | NA | NA | NA | Gm43833 | predicted gene 43833 [Source:MGI Symbc | 0 | 0 | 0 | 0 | 0 | 0 |
| 44049 | ENSMUSG00000106012 | 0 | NA | NA | NA | NA | NA | Gm43744 | predicted gene 43744 [Source:MGI Symbc | 0 | 0 | 0 | 0 | 0 | 0 |
| 44050 | ENSMUSG00000106013 | 0 | NA | NA | NA | NA | NA | 4933402J10I | RIKEN cDNA 4933402J10 gene [Source:M( | 0 | 0 | 0 | 0 | 0 | 0 |
| 44051 | ENSMUSG00000106014 | 0.160662974 | 1.126924175 | 4.080472857 | 0.276174898 | 0.782413736 | NA | NA | NA | 0 | 0 | 0 | 0.964 | 0 | 0 |
| 44052 | ENSMUSG00000106015 | 0 | NA | NA | NA | NA | NA | Gm42980 | predicted gene 42980 [Source:MGI Symbc | 0 | 0 | 0 | 0 | 0 | 0 |
| 44053 | ENSMUSG00000106016 | 0 | NA | NA | NA | NA | NA | Igkv4-56 | immunoglobulin kappa chain variable 4-5( | 0 | 0 | 0 | 0 | 0 | 0 |
| 44054 | ENSMUSG00000106017 | 0 | NA | NA | NA | NA | NA | NA | NA | 0 | 0 | 0 | 0 | 0 | 0 |
| 44055 | ENSMUSG00000106018 | 0 | NA | NA | NA | NA | NA | Gm43527 | predicted gene 43527 [Source:MGI Symbc | 0 | 0 | 0 | 0 | 0 | 0 |
| 44056 | ENSMUSG00000106019 | 187.9544573 | -0.236291705 | 0.17252094 | -1.36964072 | 0.170799082 | 0.597590208 | Gm43672 | predicted gene 43672 [Source:MGI Symbc | 211.22 | 198.39 | 200.24 | 177.37 | 172.32 | 168.19 |
| 44057 | ENSMUSG00000106020 | 0 | NA | NA | NA | NA | NA | 4930573C15 | RIKEN cDNA 4930573C15 gene [Source:M | 0 | 0 | 0 | 0 | 0 | 0 |
| 44058 | ENSMUSG00000106021 | 0 | NA | NA | NA | NA | NA | NA | NA | 0 | 0 | 0 | 0 | 0 | 0 |
| 44059 | ENSMUSG00000106022 | 7.837508099 | -0.697778813 | 0.833992685 | -0.836672582 | 0.402776635 | 0.828953151 | Gm42929 | predicted gene 42929 [Source:MGI Symbc | 4.8186 | 10.279 | 14.087 | 6.7478 | 2.1812 | 8.9108 |
| 44060 | ENSMUSG00000106023 | 0 | NA | NA | NA | NA | NA | Gm36831 | predicted gene, 36831 [Source:MGI Symb | 0 | 0 | 0 | 0 | 0 | 0 |
| 44061 | ENSMUSG00000106024 | 0 | NA | NA | NA | NA | NA | A530083M17 | RIKEN cDNA A530083M17 gene [Source:M | 0 | 0 | 0 | 0 | 0 | 0 |
| 44062 | ENSMUSG00000106025 | 5.394102641 | -0.510616548 | 0.939252371 | -0.543641479 | 0.586688211 | 0.910448513 | Gm42940 | predicted gene 42940 [Source:MGI Symbc | 8.8341 | 6.1675 | 4.025 | 6.7478 | 4.3625 | 2.2277 |
| 44063 | ENSMUSG00000106026 | 0 | NA | NA | NA | NA | NA | Gm43131 | predicted gene 43131 [Source:MGI Symbc | 0 | 0 | 0 | 0 | 0 | 0 |
| 44064 | ENSMUSG00000106027 | 0 | NA | NA | NA | NA | NA | NA | NA | 0 | 0 | 0 | 0 | 0 | 0 |
| 44065 | ENSMUSG00000106028 | 1.245767505 | 0.110185493 | 2.211495695 | 0.049823969 | 0.960262666 | NA | Gm8493 | predicted gene 8493 [Source:MGI Symbol | 1.6062 | 0 | 2.0125 | 3.8559 | 0 | 0 |
| 44066 | ENSMUSG00000106029 | 0 | NA | NA | NA | NA | NA | Gm4410 | predicted gene 4410 [Source:MGI Symbol | 0 | 0 | 0 | 0 | 0 | 0 |
| 44067 | ENSMUSG00000106030 | 59.42794415 | -0.730095212 | 0.460176106 | -1.586556108 | 0.112613215 | 0.492766469 | Gm43611 | predicted gene 43611 [Source:MGI Symbc | 85.932 | 49.34 | 87.543 | 80.01 | 39.262 | 14.48 |
| 44068 | ENSMUSG00000106031 | 0 | NA | NA | NA | NA | NA | Gm8848 | predicted gene 8848 [Source:MGI Symbol | 0 | 0 | 0 | 0 | 0 | 0 |

| | | | | | | | | | | | | | | | |
|---|---|---|---|---|---|---|---|---|---|---|---|---|---|---|---|
| 44069 | ENSMUSG00000106032 | 1.18868508 | 1.387790115 | 2.274389133 | 0.610181474 | 0.5417416 | NA | Gm42463 | predicted gene 42463 [Source:MGI Symbo | 0 | 0 | 2.0125 | 2.8919 | 0 | 2.2277 |
| 44070 | ENSMUSG00000106033 | 0 | NA | NA | NA | NA | NA | Gm43381 | predicted gene 43381 [Source:MGI Symbo | 0 | 0 | 0 | 0 | 0 | 0 |
| 44071 | ENSMUSG00000106034 | 0 | NA | NA | NA | NA | NA | 1700108N06 | RIKEN cDNA 1700108N06 gene [Source:M | 0 | 0 | 0 | 0 | 0 | 0 |
| 44072 | ENSMUSG00000106035 | 3.981041413 | 0.179537831 | 1.09188471 | 0.164429293 | 0.869393205 | NA | Gm43545 | predicted gene 43545 [Source:MGI Symbo | 4.0155 | 6.1675 | 1.0062 | 3.8559 | 3.2719 | 5.5692 |
| 44073 | ENSMUSG00000106036 | 0 | NA | NA | NA | NA | NA | Gm43608 | predicted gene 43608 [Source:MGI Symbo | 0 | 0 | 0 | 0 | 0 | 0 |
| 44074 | ENSMUSG00000106037 | 8.568358899 | -0.467187864 | 0.757458816 | -0.616783189 | 0.537377742 | 0.890758111 | Gm4332 | predicted gene 4332 [Source:MGI Symbol | 9.6372 | 9.2513 | 11.069 | 11.568 | 6.5437 | 3.3415 |
| 44075 | ENSMUSG00000106038 | 0 | NA | NA | NA | NA | NA | Gm4962 | predicted gene 4962 [Source:MGI Symbol | 0 | 0 | 0 | 0 | 0 | 0 |
| 44076 | ENSMUSG00000106039 | 0 | NA | NA | NA | NA | NA | Iglc4 | immunoglobulin lambda constant 4 [Sour | 0 | 0 | 0 | 0 | 0 | 0 |
| 44077 | ENSMUSG00000106040 | 0 | NA | NA | NA | NA | NA | Cyp3a63-ps | cytochrome P450, family 3, subfamily a, m | 0 | 0 | 0 | 0 | 0 | 0 |
| 44078 | ENSMUSG00000106041 | 0 | NA | NA | NA | NA | NA | Olfr241-ps1 | olfactory receptor 241, pseudogene 1 [Sou | 0 | 0 | 0 | 0 | 0 | 0 |
| 44079 | ENSMUSG00000106042 | 0 | NA | NA | NA | NA | NA | Gm42861 | predicted gene 42861 [Source:MGI Symbo | 0 | 0 | 0 | 0 | 0 | 0 |
| 44080 | ENSMUSG00000106043 | 0 | NA | NA | NA | NA | NA | Gm42968 | predicted gene 42968 [Source:MGI Symbo | 0 | 0 | 0 | 0 | 0 | 0 |
| 44081 | ENSMUSG00000106044 | 0.610133435 | 0.304049584 | 2.904025954 | 0.104699334 | 0.91661439 | NA | Gm42860 | predicted gene 42860 [Source:MGI Symbo | 1.6062 | 0 | 0 | 0.964 | 1.0906 | 0 |
| 44082 | ENSMUSG00000106045 | 0 | NA | NA | NA | NA | NA | Gm42996 | predicted gene 42996 [Source:MGI Symbo | 0 | 0 | 0 | 0 | 0 | 0 |
| 44083 | ENSMUSG00000106046 | 0 | NA | NA | NA | NA | NA | NA | NA | 0 | 0 | 0 | 0 | 0 | 0 |
| 44084 | ENSMUSG00000106047 | 0 | NA | NA | NA | NA | NA | Gm42942 | predicted gene 42942 [Source:MGI Symbo | 0 | 0 | 0 | 0 | 0 | 0 |
| 44085 | ENSMUSG00000106048 | 0.160662974 | 1.126924175 | 4.080472857 | 0.276174898 | 0.782413736 | NA | Gm42444 | predicted gene 42444 [Source:MGI Symbo | 0 | 0 | 0 | 0.964 | 0 | 0 |
| 44086 | ENSMUSG00000106049 | 0 | NA | NA | NA | NA | NA | Gm43019 | predicted gene 43019 [Source:MGI Symbo | 0 | 0 | 0 | 0 | 0 | 0 |
| 44087 | ENSMUSG00000106050 | 0 | NA | NA | NA | NA | NA | Ass-ps1 | argininosuccinate synthetase pseudogene | 0 | 0 | 0 | 0 | 0 | 0 |
| 44088 | ENSMUSG00000106051 | 0 | NA | NA | NA | NA | NA | Gm43160 | predicted gene 43160 [Source:MGI Symbo | 0 | 0 | 0 | 0 | 0 | 0 |
| 44089 | ENSMUSG00000106052 | 0.160662974 | 1.126924175 | 4.080472857 | 0.276174898 | 0.782413736 | NA | Gm43614 | predicted gene 43614 [Source:MGI Symbo | 0 | 0 | 0 | 0.964 | 0 | 0 |
| 44090 | ENSMUSG00000106053 | 0 | NA | NA | NA | NA | NA | Gm20752 | predicted gene, 20752 [Source:MGI Symb | 0 | 0 | 0 | 0 | 0 | 0 |
| 44091 | ENSMUSG00000106054 | 0 | NA | NA | NA | NA | NA | Gm43623 | predicted gene 43623 [Source:MGI Symbo | 0 | 0 | 0 | 0 | 0 | 0 |
| 44092 | ENSMUSG00000106056 | 0 | NA | NA | NA | NA | NA | 4930500L23I | RIKEN cDNA 4930500L23 gene [Source:M( | 0 | 0 | 0 | 0 | 0 | 0 |
| 44093 | ENSMUSG00000106057 | 0 | NA | NA | NA | NA | NA | Gm43560 | predicted gene 43560 [Source:MGI Symbo | 0 | 0 | 0 | 0 | 0 | 0 |
| 44094 | ENSMUSG00000106058 | 0 | NA | NA | NA | NA | NA | Gm43013 | predicted gene 43013 [Source:MGI Symbo | 0 | 0 | 0 | 0 | 0 | 0 |
| 44095 | ENSMUSG00000106059 | 0.838510525 | 0.626826968 | 2.582658213 | 0.242706126 | 0.808233062 | NA | Gm9364 | predicted gene 9364 [Source:MGI Symbol | 0 | 0 | 2.0125 | 1.928 | 1.0906 | 0 |
| 44096 | ENSMUSG00000106060 | 0 | NA | NA | NA | NA | NA | Gm42488 | predicted gene 42488 [Source:MGI Symbo | 0 | 0 | 0 | 0 | 0 | 0 |
| 44097 | ENSMUSG00000106061 | 0.36353943 | 2.065989254 | 4.034137301 | 0.512126658 | 0.608562368 | NA | Gm43773 | predicted gene 43773 [Source:MGI Symbo | 0 | 0 | 0 | 0 | 2.1812 | 0 |
| 44098 | ENSMUSG00000106062 | 2.291147732 | 0.223373015 | 1.453921431 | 0.15363486 | 0.877897649 | NA | Gm43820 | predicted gene 43820 [Source:MGI Symbo | 3.2124 | 3.0838 | 0 | 1.928 | 2.1812 | 3.3415 |
| 44099 | ENSMUSG00000106063 | 0 | NA | NA | NA | NA | NA | C030032O16 | RIKEN cDNA C030032O16 gene [Source:M | 0 | 0 | 0 | 0 | 0 | 0 |
| 44100 | ENSMUSG00000106064 | 0 | NA | NA | NA | NA | NA | Gm40263 | predicted gene, 40263 [Source:MGI Symb | 0 | 0 | 0 | 0 | 0 | 0 |
| 44101 | ENSMUSG00000106065 | 0 | NA | NA | NA | NA | NA | Gm42621 | predicted gene 42621 [Source:MGI Symbo | 0 | 0 | 0 | 0 | 0 | 0 |
| 44102 | ENSMUSG00000106066 | 0 | NA | NA | NA | NA | NA | Igkv2-105 | immunoglobulin kappa chain variable 2-1( | 0 | 0 | 0 | 0 | 0 | 0 |
| 44103 | ENSMUSG00000106067 | 0.685278559 | -2.809870862 | 3.376578027 | -0.832165239 | 0.405315682 | NA | Gm7902 | predicted gene 7902 [Source:MGI Symbol | 0 | 4.1117 | 0 | 0 | 0 | 0 |
| 44104 | ENSMUSG00000106068 | 0 | NA | NA | NA | NA | NA | Gm9497 | predicted gene 9497 [Source:MGI Symbol | 0 | 0 | 0 | 0 | 0 | 0 |
| 44105 | ENSMUSG00000106069 | 18.30975425 | -0.653903466 | 0.606777235 | -1.077666444 | 0.281182638 | 0.733710968 | Gm6135 | predicted gene 6135 [Source:MGI Symbo | 29.715 | 7.1954 | 30.187 | 17.352 | 9.8156 | 15.594 |
| 44106 | ENSMUSG00000106070 | 0 | NA | NA | NA | NA | NA | Gm43135 | predicted gene 43135 [Source:MGI Symbo | 0 | 0 | 0 | 0 | 0 | 0 |
| 44107 | ENSMUSG00000106071 | 0 | NA | NA | NA | NA | NA | Gm43826 | predicted gene 43826 [Source:MGI Symbo | 0 | 0 | 0 | 0 | 0 | 0 |
| 44108 | ENSMUSG00000106072 | 0 | NA | NA | NA | NA | NA | Gm42663 | predicted gene 42663 [Source:MGI Symbo | 0 | 0 | 0 | 0 | 0 | 0 |
| 44109 | ENSMUSG00000106073 | 19.31330509 | 0.644461894 | 0.601352538 | 1.071687327 | 0.283860495 | 0.736651636 | Gm42892 | predicted gene 42892 [Source:MGI Symbo | 8.031 | 15.419 | 22.137 | 39.523 | 19.631 | 11.138 |
| 44110 | ENSMUSG00000106074 | 0 | NA | NA | NA | NA | NA | Mir3969 | microRNA 3969 [Source:MGI Symbol;Acc: | 0 | 0 | 0 | 0 | 0 | 0 |
| 44111 | ENSMUSG00000106075 | 0 | NA | NA | NA | NA | NA | Gm42804 | predicted gene 42804 [Source:MGI Symbo | 0 | 0 | 0 | 0 | 0 | 0 |
| 44112 | ENSMUSG00000106076 | 0 | NA | NA | NA | NA | NA | Gm43840 | predicted gene 43840 [Source:MGI Symbo | 0 | 0 | 0 | 0 | 0 | 0 |
| 44113 | ENSMUSG00000106077 | 0 | NA | NA | NA | NA | NA | Gm5660 | predicted gene 5660 [Source:MGI Symbol | 0 | 0 | 0 | 0 | 0 | 0 |
| 44114 | ENSMUSG00000106078 | 0 | NA | NA | NA | NA | NA | Gm43697 | predicted gene 43697 [Source:MGI Symbo | 0 | 0 | 0 | 0 | 0 | 0 |
| 44115 | ENSMUSG00000106079 | 0 | NA | NA | NA | NA | NA | Gm43033 | predicted gene 43033 [Source:MGI Symbo | 0 | 0 | 0 | 0 | 0 | 0 |
| 44116 | ENSMUSG00000106080 | 0 | NA | NA | NA | NA | NA | Gm8836 | predicted gene 8836 [Source:MGI Symbol | 0 | 0 | 0 | 0 | 0 | 0 |
| 44117 | ENSMUSG00000106081 | 0 | NA | NA | NA | NA | NA | NA | NA | 0 | 0 | 0 | 0 | 0 | 0 |
| 44118 | ENSMUSG00000106082 | 0.321325949 | 1.911575114 | 4.048267893 | 0.472195804 | 0.636787034 | NA | 4930404A12 | RIKEN cDNA 4930404A12 gene [Source:M | 0 | 0 | 0 | 1.928 | 0 | 0 |
| 44119 | ENSMUSG00000106083 | 0 | NA | NA | NA | NA | NA | Gm42452 | predicted gene 42452 [Source:MGI Symbo | 0 | 0 | 0 | 0 | 0 | 0 |
| 44120 | ENSMUSG00000106084 | 0 | NA | NA | NA | NA | NA | Gm18163 | predicted gene, 18163 [Source:MGI Symb | 0 | 0 | 0 | 0 | 0 | 0 |
| 44121 | ENSMUSG00000106085 | 0 | NA | NA | NA | NA | NA | Gm43675 | predicted gene 43675 [Source:MGI Symbo | 0 | 0 | 0 | 0 | 0 | 0 |
| 44122 | ENSMUSG00000106086 | 0 | NA | NA | NA | NA | NA | Lef1os1 | LEF1 opposite strand RNA 1 [Source:MGI S | 0 | 0 | 0 | 0 | 0 | 0 |
| 44123 | ENSMUSG00000106087 | 52.43518022 | 0.332090172 | 0.327453146 | 1.014160885 | 0.31050599 | 0.759154604 | Gm43609 | predicted gene 43609 [Source:MGI Symbo | 36.14 | 49.34 | 54.337 | 72.298 | 51.259 | 51.237 |
| 44124 | ENSMUSG00000106088 | 0 | NA | NA | NA | NA | NA | NA | NA | 0 | 0 | 0 | 0 | 0 | 0 |

| | | | | | | | | | | | | | | | |
|---|---|---|---|---|---|---|---|---|---|---|---|---|---|---|---|
| 44125 | ENSMUSG00000106089 | 0 | NA | NA | NA | NA | NA | Gm9419 | predicted gene 9419 [Source:MGI Symbol | 0 | 0 | 0 | 0 | 0 | 0 |
| 44126 | ENSMUSG00000106090 | 0 | NA | NA | NA | NA | NA | Gm43704 | predicted gene 43704 [Source:MGI Symbc | 0 | 0 | 0 | 0 | 0 | 0 |
| 44127 | ENSMUSG00000106091 | 0 | NA | NA | NA | NA | NA | NA | NA | 0 | 0 | 0 | 0 | 0 | 0 |
| 44128 | ENSMUSG00000106092 | 0.17131964 | -0.796629176 | 4.080472857 | -0.195229623 | 0.84521319 | NA | Gm43072 | predicted gene 43072 [Source:MGI Symbc | 0 | 1.0279 | 0 | 0 | 0 | 0 |
| 44129 | ENSMUSG00000106093 | 0 | NA | NA | NA | NA | NA | Gm42722 | predicted gene 42722 [Source:MGI Symbc | 0 | 0 | 0 | 0 | 0 | 0 |
| 44130 | ENSMUSG00000106094 | 0 | NA | NA | NA | NA | NA | Gm18916 | predicted gene, 18916 [Source:MGI Symb | 0 | 0 | 0 | 0 | 0 | 0 |
| 44131 | ENSMUSG00000106095 | 0 | NA | NA | NA | NA | NA | 2010110G14 | RIKEN cDNA 2010110G14 gene [Source:M | 0 | 0 | 0 | 0 | 0 | 0 |
| 44132 | ENSMUSG00000106096 | 0 | NA | NA | NA | NA | NA | B230207O21 | RIKEN cDNA B230207O21 gene [Source:M | 0 | 0 | 0 | 0 | 0 | 0 |
| 44133 | ENSMUSG00000106097 | 0 | NA | NA | NA | NA | NA | Gm42625 | predicted gene 42625 [Source:MGI Symbc | 0 | 0 | 0 | 0 | 0 | 0 |
| 44134 | ENSMUSG00000106098 | 0 | NA | NA | NA | NA | NA | Igkv14-118-2 | immunoglobulin kappa chain variable 14-1 | 0 | 0 | 0 | 0 | 0 | 0 |
| 44135 | ENSMUSG00000106099 | 7.729207714 | 0.693463451 | 0.829530271 | 0.835971243 | 0.403171081 | 0.828953151 | Gm42664 | predicted gene 42664 [Source:MGI Symbc | 5.6217 | 5.1396 | 7.0437 | 16.388 | 3.2719 | 8.9108 |
| 44136 | ENSMUSG00000106100 | 0 | NA | NA | NA | NA | NA | Gm9314 | predicted gene 9314 [Source:MGI Symbol | 0 | 0 | 0 | 0 | 0 | 0 |
| 44137 | ENSMUSG00000106101 | 0 | NA | NA | NA | NA | NA | Gm43438 | predicted gene 43438 [Source:MGI Symbc | 0 | 0 | 0 | 0 | 0 | 0 |
| 44138 | ENSMUSG00000106102 | 0 | NA | NA | NA | NA | NA | Gm43493 | predicted gene 43493 [Source:MGI Symbc | 0 | 0 | 0 | 0 | 0 | 0 |
| 44139 | ENSMUSG00000106103 | 4.367657496 | 1.322111413 | 1.2019474 | 1.099974436 | 0.27134326 | NA | Gm43081 | predicted gene 43081 [Source:MGI Symbc | 2.4093 | 3.0838 | 2.0125 | 8.6758 | 0 | 10.025 |
| 44140 | ENSMUSG00000106104 | 0 | NA | NA | NA | NA | NA | Gm42660 | predicted gene 42660 [Source:MGI Symbc | 0 | 0 | 0 | 0 | 0 | 0 |
| 44141 | ENSMUSG00000106105 | 0 | NA | NA | NA | NA | NA | Gm4525 | predicted gene 4525 [Source:MGI Symbol | 0 | 0 | 0 | 0 | 0 | 0 |
| 44142 | ENSMUSG00000106106 | 56277.65496 | 0.013887431 | 0.62782129 | 0.022120039 | 0.982352202 | 0.998580735 | NA | NA | 53466 | 59918 | 54636 | 33669 | 110937 | 25040 |
| 44143 | ENSMUSG00000106107 | 0.17131964 | -0.796629176 | 4.080472857 | -0.195229623 | 0.84521319 | NA | Gm43190 | predicted gene 43190 [Source:MGI Symbc | 0 | 1.0279 | 0 | 0 | 0 | 0 |
| 44144 | ENSMUSG00000106108 | 0 | NA | NA | NA | NA | NA | Gm43221 | predicted gene 43221 [Source:MGI Symbc | 0 | 0 | 0 | 0 | 0 | 0 |
| 44145 | ENSMUSG00000106109 | 0 | NA | NA | NA | NA | NA | Gm42704 | predicted gene 42704 [Source:MGI Symbc | 0 | 0 | 0 | 0 | 0 | 0 |
| 44146 | ENSMUSG00000106110 | 1.900594625 | 0.122011041 | 1.751295284 | 0.069669028 | 0.944457094 | NA | Gm43571 | predicted gene 43571 [Source:MGI Symbc | 2.4093 | 3.0838 | 0 | 4.8199 | 1.0906 | 0 |
| 44147 | ENSMUSG00000106111 | 0 | NA | NA | NA | NA | NA | Gm5717 | predicted gene 5717 [Source:MGI Symbol | 0 | 0 | 0 | 0 | 0 | 0 |
| 44148 | ENSMUSG00000106112 | 0 | NA | NA | NA | NA | NA | Gm43434 | predicted gene 43434 [Source:MGI Symbc | 0 | 0 | 0 | 0 | 0 | 0 |
| 44149 | ENSMUSG00000106113 | 0 | NA | NA | NA | NA | NA | Gm43779 | predicted gene 43779 [Source:MGI Symbc | 0 | 0 | 0 | 0 | 0 | 0 |
| 44150 | ENSMUSG00000106114 | 0 | NA | NA | NA | NA | NA | Gm43286 | predicted gene 43286 [Source:MGI Symbc | 0 | 0 | 0 | 0 | 0 | 0 |
| 44151 | ENSMUSG00000106115 | 0.17131964 | -0.796629176 | 4.080472857 | -0.195229623 | 0.84521319 | NA | Gm43420 | predicted gene 43420 [Source:MGI Symbc | 0 | 1.0279 | 0 | 0 | 0 | 0 |
| 44152 | ENSMUSG00000106116 | 0 | NA | NA | NA | NA | NA | Gm18830 | predicted gene, 18830 [Source:MGI Symb | 0 | 0 | 0 | 0 | 0 | 0 |
| 44153 | ENSMUSG00000106117 | 0 | NA | NA | NA | NA | NA | Gm5284 | predicted gene 5284 [Source:MGI Symbol | 0 | 0 | 0 | 0 | 0 | 0 |
| 44154 | ENSMUSG00000106118 | 0 | NA | NA | NA | NA | NA | Gm18038 | predicted gene, 18038 [Source:MGI Symb | 0 | 0 | 0 | 0 | 0 | 0 |
| 44155 | ENSMUSG00000106119 | 0 | NA | NA | NA | NA | NA | Gm43656 | predicted gene 43656 [Source:MGI Symbc | 0 | 0 | 0 | 0 | 0 | 0 |
| 44156 | ENSMUSG00000106120 | 0 | NA | NA | NA | NA | NA | Gm42697 | predicted gene 42697 [Source:MGI Symbc | 0 | 0 | 0 | 0 | 0 | 0 |
| 44157 | ENSMUSG00000106121 | 0.133850373 | -0.796629176 | 4.080472857 | -0.195229623 | 0.84521319 | NA | Gm42679 | predicted gene 42679 [Source:MGI Symbc | 0.8031 | 0 | 0 | 0 | 0 | 0 |
| 44158 | ENSMUSG00000106122 | 0 | NA | NA | NA | NA | NA | Gm42610 | predicted gene 42610 [Source:MGI Symbc | 0 | 0 | 0 | 0 | 0 | 0 |
| 44159 | ENSMUSG00000106123 | 0 | NA | NA | NA | NA | NA | Gm42638 | predicted gene 42638 [Source:MGI Symbc | 0 | 0 | 0 | 0 | 0 | 0 |
| 44160 | ENSMUSG00000106124 | 0 | NA | NA | NA | NA | NA | Gm42539 | predicted gene 42539 [Source:MGI Symbc | 0 | 0 | 0 | 0 | 0 | 0 |
| 44161 | ENSMUSG00000106125 | 0.181769715 | 1.126924175 | 4.080472857 | 0.276174898 | 0.782413736 | NA | 4930512P04 | RIKEN cDNA 4930512P04 gene [Source:M | 0 | 0 | 0 | 0 | 1.0906 | 0 |
| 44162 | ENSMUSG00000106126 | 0 | NA | NA | NA | NA | NA | Gm43781 | predicted gene 43781 [Source:MGI Symbc | 0 | 0 | 0 | 0 | 0 | 0 |
| 44163 | ENSMUSG00000106127 | 0 | NA | NA | NA | NA | NA | 4930520M14 | RIKEN cDNA 4930520M14 gene [Source:M | 0 | 0 | 0 | 0 | 0 | 0 |
| 44164 | ENSMUSG00000106128 | 0 | NA | NA | NA | NA | NA | Gm43520 | predicted gene 43520 [Source:MGI Symbc | 0 | 0 | 0 | 0 | 0 | 0 |
| 44165 | ENSMUSG00000106129 | 0 | NA | NA | NA | NA | NA | Gm19210 | predicted gene, 19210 [Source:MGI Symb | 0 | 0 | 0 | 0 | 0 | 0 |
| 44166 | ENSMUSG00000106130 | 0 | NA | NA | NA | NA | NA | NA | NA | 0 | 0 | 0 | 0 | 0 | 0 |
| 44167 | ENSMUSG00000106131 | 0 | NA | NA | NA | NA | NA | Gm36382 | predicted gene, 36382 [Source:MGI Symb | 0 | 0 | 0 | 0 | 0 | 0 |
| 44168 | ENSMUSG00000106132 | 0 | NA | NA | NA | NA | NA | Gm43370 | predicted gene 43370 [Source:MGI Symbc | 0 | 0 | 0 | 0 | 0 | 0 |
| 44169 | ENSMUSG00000106133 | 2.859915993 | 1.082751376 | 1.248308293 | 0.867374976 | 0.385736588 | NA | Gm3724 | predicted gene 3724 [Source:MGI Symbol | 2.4093 | 1.0279 | 2.0125 | 2.8919 | 4.3625 | 4.4554 |
| 44170 | ENSMUSG00000106134 | 0 | NA | NA | NA | NA | NA | Gm42493 | predicted gene 42493 [Source:MGI Symbc | 0 | 0 | 0 | 0 | 0 | 0 |
| 44171 | ENSMUSG00000106135 | 0 | NA | NA | NA | NA | NA | Gm6543 | predicted gene 6543 [Source:MGI Symbol | 0 | 0 | 0 | 0 | 0 | 0 |
| 44172 | ENSMUSG00000106136 | 0 | NA | NA | NA | NA | NA | 4933408A14 | RIKEN cDNA 4933408A14 gene [Source:M | 0 | 0 | 0 | 0 | 0 | 0 |
| 44173 | ENSMUSG00000106137 | 0 | NA | NA | NA | NA | NA | Gm42538 | predicted gene 42538 [Source:MGI Symbc | 0 | 0 | 0 | 0 | 0 | 0 |
| 44174 | ENSMUSG00000106138 | 0 | NA | NA | NA | NA | NA | NA | NA | 0 | 0 | 0 | 0 | 0 | 0 |
| 44175 | ENSMUSG00000106139 | 3.944089066 | -1.775686927 | 1.245628615 | -1.425534791 | 0.154002671 | NA | Gm30648 | predicted gene, 30648 [Source:MGI Symb | 8.031 | 7.1954 | 3.0187 | 0.964 | 0 | 4.4554 |
| 44176 | ENSMUSG00000106140 | 0 | NA | NA | NA | NA | NA | Gm42208 | predicted gene, 42208 [Source:MGI Symb | 0 | 0 | 0 | 0 | 0 | 0 |
| 44177 | ENSMUSG00000106141 | 0 | NA | NA | NA | NA | NA | Gm18330 | predicted gene, 18330 [Source:MGI Symb | 0 | 0 | 0 | 0 | 0 | 0 |
| 44178 | ENSMUSG00000106142 | 0 | NA | NA | NA | NA | NA | Gm17937 | predicted gene, 17937 [Source:MGI Symb | 0 | 0 | 0 | 0 | 0 | 0 |
| 44179 | ENSMUSG00000106143 | 0 | NA | NA | NA | NA | NA | Gm43735 | predicted gene 43735 [Source:MGI Symbc | 0 | 0 | 0 | 0 | 0 | 0 |
| 44180 | ENSMUSG00000106144 | 0 | NA | NA | NA | NA | NA | Gm43972 | predicted gene, 43972 [Source:MGI Symb | 0 | 0 | 0 | 0 | 0 | 0 |

| | | | | | | | | | | | | | | | |
|---|---|---|---|---|---|---|---|---|---|---|---|---|---|---|---|
| 44181 | ENSMUSG00000106145 | 0 | NA | NA | NA | NA | NA | Mir7073 | microRNA 7073 [Source:MGI Symbol;Acc: | 0 | 0 | 0 | 0 | 0 | 0 |
| 44182 | ENSMUSG00000106146 | 0 | NA | NA | NA | NA | NA | Gm43045 | predicted gene 43045 [Source:MGI Symbo | 0 | 0 | 0 | 0 | 0 | 0 |
| 44183 | ENSMUSG00000106147 | 0 | NA | NA | NA | NA | NA | Snord3a | small nucleolar RNA, C/D box 3A [Source:I | 0 | 0 | 0 | 0 | 0 | 0 |
| 44184 | ENSMUSG00000106148 | 0 | NA | NA | NA | NA | NA | Gm32877 | predicted gene, 32877 [Source:MGI Symb | 0 | 0 | 0 | 0 | 0 | 0 |
| 44185 | ENSMUSG00000106149 | 0.517184577 | -0.812115922 | 3.393570547 | -0.239310163 | 0.810865085 | NA | Gm43430 | predicted gene 43430 [Source:MGI Symbo | 0 | 0 | 2.0125 | 0 | 1.0906 | 0 |
| 44186 | ENSMUSG00000106150 | 0 | NA | NA | NA | NA | NA | 4921527H02 | RIKEN cDNA 4921527H02 gene [Source:M | 0 | 0 | 0 | 0 | 0 | 0 |
| 44187 | ENSMUSG00000106151 | 0 | NA | NA | NA | NA | NA | NA | NA | 0 | 0 | 0 | 0 | 0 | 0 |
| 44188 | ENSMUSG00000106152 | 0 | NA | NA | NA | NA | NA | NA | NA | 0 | 0 | 0 | 0 | 0 | 0 |
| 44189 | ENSMUSG00000106153 | 0 | NA | NA | NA | NA | NA | Gm43677 | predicted gene 43677 [Source:MGI Symbo | 0 | 0 | 0 | 0 | 0 | 0 |
| 44190 | ENSMUSG00000106154 | 0 | NA | NA | NA | NA | NA | Mir7232 | microRNA 7232 [Source:MGI Symbol;Acc: | 0 | 0 | 0 | 0 | 0 | 0 |
| 44191 | ENSMUSG00000106155 | 0 | NA | NA | NA | NA | NA | Gm43495 | predicted gene 43495 [Source:MGI Symbo | 0 | 0 | 0 | 0 | 0 | 0 |
| 44192 | ENSMUSG00000106156 | 0 | NA | NA | NA | NA | NA | Gm21040 | predicted gene, 21040 [Source:MGI Symb | 0 | 0 | 0 | 0 | 0 | 0 |
| 44193 | ENSMUSG00000106157 | 0.401551119 | -2.072982726 | 4.008251979 | -0.517178744 | 0.605031388 | NA | 4930555A03 | RIKEN cDNA 4930555A03 gene [Source:M | 2.4093 | 0 | 0 | 0 | 0 | 0 |
| 44194 | ENSMUSG00000106158 | 0 | NA | NA | NA | NA | NA | Gm43216 | predicted gene 43216 [Source:MGI Symbo | 0 | 0 | 0 | 0 | 0 | 0 |
| 44195 | ENSMUSG00000106159 | 0 | NA | NA | NA | NA | NA | Gm31211 | predicted gene, 31211 [Source:MGI Symb | 0 | 0 | 0 | 0 | 0 | 0 |
| 44196 | ENSMUSG00000106160 | 0 | NA | NA | NA | NA | NA | Gm43610 | predicted gene 43610 [Source:MGI Symbo | 0 | 0 | 0 | 0 | 0 | 0 |
| 44197 | ENSMUSG00000106161 | 0.267700746 | -1.497481649 | 4.057001436 | -0.369110456 | 0.712045396 | NA | Pvrig-ps | poliovirus receptor related immunoglobul | 1.6062 | 0 | 0 | 0 | 0 | 0 |
| 44198 | ENSMUSG00000106162 | 0 | NA | NA | NA | NA | NA | Gm43613 | predicted gene 43613 [Source:MGI Symbo | 0 | 0 | 0 | 0 | 0 | 0 |
| 44199 | ENSMUSG00000106163 | 0 | NA | NA | NA | NA | NA | NA | NA | 0 | 0 | 0 | 0 | 0 | 0 |
| 44200 | ENSMUSG00000106164 | 2.548166838 | 1.188024063 | 1.33635087 | 0.889006091 | 0.373999806 | NA | 9430085M1 | RIKEN cDNA 9430085M18 gene [Source:M | 1.6062 | 1.0279 | 2.0125 | 2.8919 | 2.1812 | 5.5692 |
| 44201 | ENSMUSG00000106165 | 0 | NA | NA | NA | NA | NA | Gm3922 | predicted gene 3922 [Source:MGI Symbol | 0 | 0 | 0 | 0 | 0 | 0 |
| 44202 | ENSMUSG00000106166 | 0.83894921 | 2.000006572 | 2.595804237 | 0.770476657 | 0.441017196 | NA | Gm43484 | predicted gene 43484 [Source:MGI Symbo | 0 | 1.0279 | 0 | 2.8919 | 0 | 1.1138 |
| 44203 | ENSMUSG00000106168 | 0 | NA | NA | NA | NA | NA | Gm43538 | predicted gene 43538 [Source:MGI Symbo | 0 | 0 | 0 | 0 | 0 | 0 |
| 44204 | ENSMUSG00000106169 | 0.799645015 | 0.918084596 | 2.625685367 | 0.34965522 | 0.726597464 | NA | Mir6948 | microRNA 6948 [Source:MGI Symbol;Acc: | 1.6062 | 0 | 0 | 0.964 | 0 | 2.2277 |
| 44205 | ENSMUSG00000106170 | 0 | NA | NA | NA | NA | NA | Gm43393 | predicted gene 43393 [Source:MGI Symbo | 0 | 0 | 0 | 0 | 0 | 0 |
| 44206 | ENSMUSG00000106172 | 0 | NA | NA | NA | NA | NA | Gm17979 | predicted gene, 17979 [Source:MGI Symb | 0 | 0 | 0 | 0 | 0 | 0 |
| 44207 | ENSMUSG00000106173 | 0.181769715 | 1.126924175 | 4.080472857 | 0.276174898 | 0.782413736 | NA | Gm42764 | predicted gene 42764 [Source:MGI Symbo | 0 | 0 | 0 | 0 | 1.0906 | 0 |
| 44208 | ENSMUSG00000106174 | 0 | NA | NA | NA | NA | NA | Gm43235 | predicted gene 43235 [Source:MGI Symbo | 0 | 0 | 0 | 0 | 0 | 0 |
| 44209 | ENSMUSG00000106175 | 0 | NA | NA | NA | NA | NA | NA | NA | 0 | 0 | 0 | 0 | 0 | 0 |
| 44210 | ENSMUSG00000106176 | 0.267700746 | -1.497481649 | 4.057001436 | -0.369110456 | 0.712045396 | NA | Gm43730 | predicted gene 43730 [Source:MGI Symbo | 1.6062 | 0 | 0 | 0 | 0 | 0 |
| 44211 | ENSMUSG00000106177 | 0 | NA | NA | NA | NA | NA | Gm43491 | predicted gene 43491 [Source:MGI Symbo | 0 | 0 | 0 | 0 | 0 | 0 |
| 44212 | ENSMUSG00000106178 | 1.325992676 | -1.613440788 | 2.199576947 | -0.733523231 | 0.463239364 | NA | Gm42987 | predicted gene 42987 [Source:MGI Symbo | 4.0155 | 0 | 2.0125 | 1.928 | 0 | 0 |
| 44213 | ENSMUSG00000106179 | 0 | NA | NA | NA | NA | NA | NA | NA | 0 | 0 | 0 | 0 | 0 | 0 |
| 44214 | ENSMUSG00000106180 | 0 | NA | NA | NA | NA | NA | Pou5f1-rs11 | POU domain, class 5, transcription factor : | 0 | 0 | 0 | 0 | 0 | 0 |
| 44215 | ENSMUSG00000106181 | 0.17131964 | -0.796629176 | 4.080472857 | -0.195229623 | 0.84521319 | NA | Gm9523 | predicted gene 9523 [Source:MGI Symbol | 0 | 1.0279 | 0 | 0 | 0 | 0 |
| 44216 | ENSMUSG00000106182 | 0 | NA | NA | NA | NA | NA | Gm7442 | predicted gene 7442 [Source:MGI Symbol | 0 | 0 | 0 | 0 | 0 | 0 |
| 44217 | ENSMUSG00000106183 | 0 | NA | NA | NA | NA | NA | Mir101c | microRNA 101c [Source:MGI Symbol;Acc:I | 0 | 0 | 0 | 0 | 0 | 0 |
| 44218 | ENSMUSG00000106184 | 0.185640647 | 1.126924175 | 4.080472857 | 0.276174898 | 0.782413736 | NA | Gm43463 | predicted gene 43463 [Source:MGI Symbo | 0 | 0 | 0 | 0 | 0 | 1.1138 |
| 44219 | ENSMUSG00000106185 | 5.092953332 | 1.539363071 | 1.108391477 | 1.388826153 | 0.16488562 | 0.590031077 | Gm7631 | predicted gene 7631 [Source:MGI Symbol | 5.6217 | 0 | 2.0125 | 8.6758 | 10.906 | 3.3415 |
| 44220 | ENSMUSG00000106186 | 0.513958919 | -2.394274821 | 3.756096013 | -0.637437066 | 0.523840191 | NA | Gm43627 | predicted gene 43627 [Source:MGI Symbo | 0 | 3.0838 | 0 | 0 | 0 | 0 |
| 44221 | ENSMUSG00000106187 | 0 | NA | NA | NA | NA | NA | Igkv15-101-1 | immunoglobulin kappa chain variable 15-: | 0 | 0 | 0 | 0 | 0 | 0 |
| 44222 | ENSMUSG00000106188 | 0 | NA | NA | NA | NA | NA | Gm9710 | predicted gene 9710 [Source:MGI Symbol | 0 | 0 | 0 | 0 | 0 | 0 |
| 44223 | ENSMUSG00000106189 | 0 | NA | NA | NA | NA | NA | Gm42933 | predicted gene 42933 [Source:MGI Symbo | 0 | 0 | 0 | 0 | 0 | 0 |
| 44224 | ENSMUSG00000106190 | 0 | NA | NA | NA | NA | NA | Gm20768 | predicted gene, 20768 [Source:MGI Symb | 0 | 0 | 0 | 0 | 0 | 0 |
| 44225 | ENSMUSG00000106191 | 0.185640647 | 1.126924175 | 4.080472857 | 0.276174898 | 0.782413736 | NA | 4930557B06 | RIKEN cDNA 4930557B06 gene [Source:M | 0 | 0 | 0 | 0 | 0 | 1.1138 |
| 44226 | ENSMUSG00000106192 | 0 | NA | NA | NA | NA | NA | Gm42818 | predicted gene 42818 [Source:MGI Symbo | 0 | 0 | 0 | 0 | 0 | 0 |
| 44227 | ENSMUSG00000106193 | 0 | NA | NA | NA | NA | NA | Mir7090 | microRNA 7090 [Source:MGI Symbol;Acc: | 0 | 0 | 0 | 0 | 0 | 0 |
| 44228 | ENSMUSG00000106194 | 0.133850373 | -0.796629176 | 4.080472857 | -0.195229623 | 0.84521319 | NA | Gm43569 | predicted gene 43569 [Source:MGI Symbo | 0.8031 | 0 | 0 | 0 | 0 | 0 |
| 44229 | ENSMUSG00000106195 | 0.804680097 | -3.048447205 | 2.843448763 | -1.072095001 | 0.283677364 | NA | NA | NA | 0.8031 | 0 | 4.025 | 0 | 0 | 0 |
| 44230 | ENSMUSG00000106196 | 0 | NA | NA | NA | NA | NA | NA | NA | 0 | 0 | 0 | 0 | 0 | 0 |
| 44231 | ENSMUSG00000106197 | 3.425232015 | 0.347290532 | 1.306474188 | 0.265822728 | 0.790375739 | NA | Gm42717 | predicted gene 42717 [Source:MGI Symbo | 0 | 5.1396 | 4.025 | 4.8199 | 5.4531 | 1.1138 |
| 44232 | ENSMUSG00000106198 | 0 | NA | NA | NA | NA | NA | Gm42504 | predicted gene 42504 [Source:MGI Symbo | 0 | 0 | 0 | 0 | 0 | 0 |
| 44233 | ENSMUSG00000106199 | 0 | NA | NA | NA | NA | NA | Mir3967 | microRNA 3967 [Source:MGI Symbol;Acc: | 0 | 0 | 0 | 0 | 0 | 0 |
| 44234 | ENSMUSG00000106200 | 0 | NA | NA | NA | NA | NA | NA | NA | 0 | 0 | 0 | 0 | 0 | 0 |
| 44235 | ENSMUSG00000106201 | 0 | NA | NA | NA | NA | NA | Gm7815 | predicted gene 7815 [Source:MGI Symbol | 0 | 0 | 0 | 0 | 0 | 0 |
| 44236 | ENSMUSG00000106202 | 0 | NA | NA | NA | NA | NA | Gm43727 | predicted gene 43727 [Source:MGI Symbo | 0 | 0 | 0 | 0 | 0 | 0 |

| | | | | | | | | | | | | | | | |
|---|---|---|---|---|---|---|---|---|---|---|---|---|---|---|---|
| 44237 | ENSMUSG00000106203 | 0 | NA | NA | NA | NA | NA | NA | NA | 0 | 0 | 0 | 0 | 0 | 0 |
| 44238 | ENSMUSG00000106204 | 0 | NA | NA | NA | NA | NA | Gm21009 | predicted gene, 21009 [Source:MGI Symb | 0 | 0 | 0 | 0 | 0 | 0 |
| 44239 | ENSMUSG00000106205 | 10.2297859 | -0.879041077 | 0.713225955 | -1.232486102 | 0.217767555 | 0.663697152 | C230096K16 | RIKEN cDNA C230096K16 gene [Source:M | 18.471 | 8.2233 | 13.081 | 10.604 | 6.5437 | 4.4554 |
| 44240 | ENSMUSG00000106206 | 0.185640647 | 1.126924175 | 4.080472857 | 0.276174898 | 0.782413736 | NA | Gm43094 | predicted gene 43094 [Source:MGI Symbo | 0 | 0 | 0 | 0 | 0 | 1.1138 |
| 44241 | ENSMUSG00000106207 | 0 | NA | NA | NA | NA | NA | Gm19184 | predicted gene, 19184 [Source:MGI Symb | 0 | 0 | 0 | 0 | 0 | 0 |
| 44242 | ENSMUSG00000106208 | 0.572870759 | -2.577107685 | 3.182795569 | -0.809699407 | 0.418112959 | NA | Gm4734 | predicted gene 4734 [Source:MGI Symbol | 2.4093 | 1.0279 | 0 | 0 | 0 | 0 |
| 44243 | ENSMUSG00000106209 | 7.382478468 | 0.843784143 | 0.817268573 | 1.032444133 | 0.301864106 | 0.752248241 | Gm42918 | predicted gene 42918 [Source:MGI Symbo | 5.6217 | 7.1954 | 3.0187 | 9.6398 | 5.4531 | 13.366 |
| 44244 | ENSMUSG00000106210 | 0 | NA | NA | NA | NA | NA | 1700001N15 | RIKEN cDNA 1700001N15 gene [Source:M | 0 | 0 | 0 | 0 | 0 | 0 |
| 44245 | ENSMUSG00000106211 | 0 | NA | NA | NA | NA | NA | Gm42842 | predicted gene 42842 [Source:MGI Symbo | 0 | 0 | 0 | 0 | 0 | 0 |
| 44246 | ENSMUSG00000106212 | 0 | NA | NA | NA | NA | NA | Gm43112 | predicted gene 43112 [Source:MGI Symbo | 0 | 0 | 0 | 0 | 0 | 0 |
| 44247 | ENSMUSG00000106213 | 0 | NA | NA | NA | NA | NA | Igkv13-55-1 | immunoglobulin kappa chain variable 13-5 | 0 | 0 | 0 | 0 | 0 | 0 |
| 44248 | ENSMUSG00000106214 | 0 | NA | NA | NA | NA | NA | Gm43426 | predicted gene 43426 [Source:MGI Symbo | 0 | 0 | 0 | 0 | 0 | 0 |
| 44249 | ENSMUSG00000106215 | 0 | NA | NA | NA | NA | NA | Mir7218 | microRNA 7218 [Source:MGI Symbol;Acc: | 0 | 0 | 0 | 0 | 0 | 0 |
| 44250 | ENSMUSG00000106216 | 0 | NA | NA | NA | NA | NA | Igkv13-61-1 | immunoglobulin kappa chain variable 13-6 | 0 | 0 | 0 | 0 | 0 | 0 |
| 44251 | ENSMUSG00000106217 | 0 | NA | NA | NA | NA | NA | Gm43368 | predicted gene 43368 [Source:MGI Symbo | 0 | 0 | 0 | 0 | 0 | 0 |
| 44252 | ENSMUSG00000106218 | 0.649696354 | 1.537929172 | 3.118369374 | 0.493183773 | 0.621882743 | NA | Gm43713 | predicted gene 43713 [Source:MGI Symbo | 0 | 0 | 1.0062 | 2.8919 | 0 | 0 |
| 44253 | ENSMUSG00000106219 | 0 | NA | NA | NA | NA | NA | 5830416I19R | RIKEN cDNA 5830416I19 gene [Source:MG | 0 | 0 | 0 | 0 | 0 | 0 |
| 44254 | ENSMUSG00000106220 | 0 | NA | NA | NA | NA | NA | Gm43288 | predicted gene 43288 [Source:MGI Symbo | 0 | 0 | 0 | 0 | 0 | 0 |
| 44255 | ENSMUSG00000106221 | 0 | NA | NA | NA | NA | NA | NA | NA | 0 | 0 | 0 | 0 | 0 | 0 |
| 44256 | ENSMUSG00000106222 | 0 | NA | NA | NA | NA | NA | Rprl1 | ribonuclease P RNA-like 1 [Source:MGI Sy | 0 | 0 | 0 | 0 | 0 | 0 |
| 44257 | ENSMUSG00000106223 | 0 | NA | NA | NA | NA | NA | 2400006E01 | RIKEN cDNA 2400006E01 gene [Source:MG | 0 | 0 | 0 | 0 | 0 | 0 |
| 44258 | ENSMUSG00000106224 | 6.033784045 | 0.575795094 | 0.947376326 | 0.60777864 | 0.543334298 | 0.894449746 | Gm43823 | predicted gene 43823 [Source:MGI Symbo | 6.4248 | 1.0279 | 7.0437 | 9.6398 | 8.7249 | 3.3415 |
| 44259 | ENSMUSG00000106225 | 0 | NA | NA | NA | NA | NA | Gm43564 | predicted gene 43564 [Source:MGI Symbo | 0 | 0 | 0 | 0 | 0 | 0 |
| 44260 | ENSMUSG00000106226 | 0 | NA | NA | NA | NA | NA | Gm43586 | predicted gene 43586 [Source:MGI Symbo | 0 | 0 | 0 | 0 | 0 | 0 |
| 44261 | ENSMUSG00000106227 | 0.476283062 | 1.031501528 | 3.159856949 | 0.326439312 | 0.744091998 | NA | Gm43464 | predicted gene 43464 [Source:MGI Symbo | 0.8031 | 0 | 0 | 0.964 | 1.0906 | 0 |
| 44262 | ENSMUSG00000106228 | 0 | NA | NA | NA | NA | NA | Mir3073b | microRNA 3073b [Source:MGI Symbol;Acc | 0 | 0 | 0 | 0 | 0 | 0 |
| 44263 | ENSMUSG00000106229 | 3.48190096 | -0.581912691 | 1.228666892 | -0.473613064 | 0.635775858 | NA | Gm19409 | predicted gene, 19409 [Source:MGI Symb | 2.4093 | 5.1396 | 5.0312 | 3.8559 | 0 | 4.4554 |
| 44264 | ENSMUSG00000106230 | 0.603115608 | -2.642822926 | 3.106511347 | -0.850736608 | 0.394915683 | NA | Gm29811 | predicted gene, 29811 [Source:MGI Symb | 1.6062 | 0 | 2.0125 | 0 | 0 | 0 |
| 44265 | ENSMUSG00000106231 | 0 | NA | NA | NA | NA | NA | Gm18611 | predicted gene, 18611 [Source:MGI Symb | 0 | 0 | 0 | 0 | 0 | 0 |
| 44266 | ENSMUSG00000106232 | 0 | NA | NA | NA | NA | NA | Gm43236 | predicted gene 43236 [Source:MGI Symbo | 0 | 0 | 0 | 0 | 0 | 0 |
| 44267 | ENSMUSG00000106233 | 0 | NA | NA | NA | NA | NA | Mir509 | microRNA 509 [Source:MGI Symbol;Acc:M | 0 | 0 | 0 | 0 | 0 | 0 |
| 44268 | ENSMUSG00000106234 | 0 | NA | NA | NA | NA | NA | Gm43535 | predicted gene 43535 [Source:MGI Symbo | 0 | 0 | 0 | 0 | 0 | 0 |
| 44269 | ENSMUSG00000106235 | 0 | NA | NA | NA | NA | NA | Gm33347 | predicted gene, 33347 [Source:MGI Symb | 0 | 0 | 0 | 0 | 0 | 0 |
| 44270 | ENSMUSG00000106236 | 0 | NA | NA | NA | NA | NA | Gm38670 | predicted gene, 38670 [Source:MGI Symb | 0 | 0 | 0 | 0 | 0 | 0 |
| 44271 | ENSMUSG00000106237 | 10.9878388 | 0.207488009 | 0.667881531 | 0.310665887 | 0.756054633 | 0.953085143 | Gm8066 | predicted gene 8066 [Source:MGI Symbol | 6.4248 | 11.307 | 13.081 | 16.388 | 9.8156 | 8.9108 |
| 44272 | ENSMUSG00000106238 | 0 | NA | NA | NA | NA | NA | Gm42824 | predicted gene 42824 [Source:MGI Symbo | 0 | 0 | 0 | 0 | 0 | 0 |
| 44273 | ENSMUSG00000106239 | 0.453341394 | -0.545391191 | 3.551113277 | -0.153583158 | 0.877938417 | NA | Gm9260 | predicted gene 9260 [Source:MGI Symbol | 1.6062 | 0 | 0 | 0 | 0 | 1.1138 |
| 44274 | ENSMUSG00000106240 | 0 | NA | NA | NA | NA | NA | Gm3544 | predicted gene 3544 [Source:MGI Symbol | 0 | 0 | 0 | 0 | 0 | 0 |
| 44275 | ENSMUSG00000106241 | 1.304079275 | 0.316439097 | 1.995290604 | 0.158592987 | 0.873989554 | NA | Gm43143 | predicted gene 43143 [Source:MGI Symbo | 2.4093 | 0 | 1.0062 | 0 | 2.1812 | 2.2277 |
| 44276 | ENSMUSG00000106242 | 0 | NA | NA | NA | NA | NA | NA | NA | 0 | 0 | 0 | 0 | 0 | 0 |
| 44277 | ENSMUSG00000106243 | 0 | NA | NA | NA | NA | NA | Gm5050 | predicted gene 5050 [Source:MGI Symbol | 0 | 0 | 0 | 0 | 0 | 0 |
| 44278 | ENSMUSG00000106244 | 0 | NA | NA | NA | NA | NA | Gm43046 | predicted gene 43046 [Source:MGI Symbo | 0 | 0 | 0 | 0 | 0 | 0 |
| 44279 | ENSMUSG00000106245 | 0 | NA | NA | NA | NA | NA | Gm43824 | predicted gene 43824 [Source:MGI Symbo | 0 | 0 | 0 | 0 | 0 | 0 |
| 44280 | ENSMUSG00000106246 | 0 | NA | NA | NA | NA | NA | Gm18867 | predicted gene, 18867 [Source:MGI Symb | 0 | 0 | 0 | 0 | 0 | 0 |
| 44281 | ENSMUSG00000106247 | 0 | NA | NA | NA | NA | NA | Gm43720 | predicted gene 43720 [Source:MGI Symbo | 0 | 0 | 0 | 0 | 0 | 0 |
| 44282 | ENSMUSG00000106248 | 0 | NA | NA | NA | NA | NA | Gm9728 | predicted gene 9728 [Source:MGI Symbol | 0 | 0 | 0 | 0 | 0 | 0 |
| 44283 | ENSMUSG00000106249 | 0 | NA | NA | NA | NA | NA | Gm43349 | predicted gene 43349 [Source:MGI Symbo | 0 | 0 | 0 | 0 | 0 | 0 |
| 44284 | ENSMUSG00000106250 | 0 | NA | NA | NA | NA | NA | Gm43392 | predicted gene 43392 [Source:MGI Symbo | 0 | 0 | 0 | 0 | 0 | 0 |
| 44285 | ENSMUSG00000106251 | 1.270486455 | 0.142797745 | 1.88353307 | 0.075813771 | 0.93956726 | NA | Gm42658 | predicted gene 42658 [Source:MGI Symbo | 1.6062 | 1.0279 | 1.0062 | 2.8919 | 1.0906 | 0 |
| 44286 | ENSMUSG00000106252 | 0.301557804 | -1.64335916 | 4.042814367 | -0.406488899 | 0.684383412 | NA | Mir684-1 | microRNA 684-1 [Source:MGI Symbol;Acc | 0.8031 | 0 | 1.0062 | 0 | 0 | 0 |
| 44287 | ENSMUSG00000106253 | 0 | NA | NA | NA | NA | NA | Mir468 | microRNA 468 [Source:MGI Symbol;Acc:M | 0 | 0 | 0 | 0 | 0 | 0 |
| 44288 | ENSMUSG00000106254 | 0 | NA | NA | NA | NA | NA | Gm42907 | predicted gene 42907 [Source:MGI Symbo | 0 | 0 | 0 | 0 | 0 | 0 |
| 44289 | ENSMUSG00000106255 | 0 | NA | NA | NA | NA | NA | Gm36793 | predicted gene, 36793 [Source:MGI Symb | 0 | 0 | 0 | 0 | 0 | 0 |
| 44290 | ENSMUSG00000106256 | 0 | NA | NA | NA | NA | NA | Igkv12-47 | immunoglobulin kappa variable 12-47 [So | 0 | 0 | 0 | 0 | 0 | 0 |
| 44291 | ENSMUSG00000106257 | 0 | NA | NA | NA | NA | NA | Gm9023 | predicted gene 9023 [Source:MGI Symbol | 0 | 0 | 0 | 0 | 0 | 0 |
| 44292 | ENSMUSG00000106258 | 0 | NA | NA | NA | NA | NA | Rpl7a-ps7 | ribosomal protein L7A, pseudogene 7 [Sou | 0 | 0 | 0 | 0 | 0 | 0 |

| | | | | | | | | | | | | | | | |
|---|---|---|---|---|---|---|---|---|---|---|---|---|---|---|---|
| 44293 | ENSMUSG00000106259 | 0 | NA | NA | NA | NA | NA | Trav11n | T cell receptor alpha variable 11N [Source | 0 | 0 | 0 | 0 | 0 | 0 |
| 44294 | ENSMUSG00000106260 | 0 | NA | NA | NA | NA | NA | Gm43524 | predicted gene 43524 [Source:MGI Symbo | 0 | 0 | 0 | 0 | 0 | 0 |
| 44295 | ENSMUSG00000106261 | 0.17131964 | -0.796629176 | 4.080472857 | -0.195229623 | 0.84521319 | NA | Gm34086 | predicted gene, 34086 [Source:MGI Symb | 0 | 1.0279 | 0 | 0 | 0 | 0 |
| 44296 | ENSMUSG00000106262 | 1.927433856 | -0.948089143 | 1.710882067 | -0.554152248 | 0.579474655 | NA | Gm43375 | predicted gene 43375 [Source:MGI Symbo | 1.6062 | 3.0838 | 3.0187 | 3.8559 | 0 | 0 |
| 44297 | ENSMUSG00000106263 | 0 | NA | NA | NA | NA | NA | Gm33050 | predicted gene, 33050 [Source:MGI Symb | 0 | 0 | 0 | 0 | 0 | 0 |
| 44298 | ENSMUSG00000106264 | 0 | NA | NA | NA | NA | NA | Mir7646 | microRNA 7646 [Source:MGI Symbol;Acc: | 0 | 0 | 0 | 0 | 0 | 0 |
| 44299 | ENSMUSG00000106265 | 0 | NA | NA | NA | NA | NA | Gm42490 | predicted gene 42490 [Source:MGI Symbo | 0 | 0 | 0 | 0 | 0 | 0 |
| 44300 | ENSMUSG00000106266 | 0 | NA | NA | NA | NA | NA | Gm43447 | predicted gene 43447 [Source:MGI Symbo | 0 | 0 | 0 | 0 | 0 | 0 |
| 44301 | ENSMUSG00000106267 | 0 | NA | NA | NA | NA | NA | Gm40319 | predicted gene, 40319 [Source:MGI Symb | 0 | 0 | 0 | 0 | 0 | 0 |
| 44302 | ENSMUSG00000106268 | 0.160662974 | 1.126924175 | 4.080472857 | 0.276174898 | 0.782413736 | NA | Gm18492 | predicted gene, 18492 [Source:MGI Symb | 0 | 0 | 0 | 0.964 | 0 | 0 |
| 44303 | ENSMUSG00000106269 | 0 | NA | NA | NA | NA | NA | 4930580E04 | RIKEN cDNA 4930580E04 gene [Source:M | 0 | 0 | 0 | 0 | 0 | 0 |
| 44304 | ENSMUSG00000106270 | 0 | NA | NA | NA | NA | NA | C820005J03I | RIKEN cDNA C820005J03 gene [Source:M | 0 | 0 | 0 | 0 | 0 | 0 |
| 44305 | ENSMUSG00000106271 | 0 | NA | NA | NA | NA | NA | Gm43752 | predicted gene 43752 [Source:MGI Symbo | 0 | 0 | 0 | 0 | 0 | 0 |
| 44306 | ENSMUSG00000106272 | 0 | NA | NA | NA | NA | NA | NA | NA | 0 | 0 | 0 | 0 | 0 | 0 |
| 44307 | ENSMUSG00000106273 | 0 | NA | NA | NA | NA | NA | Gm43767 | predicted gene 43767 [Source:MGI Symbo | 0 | 0 | 0 | 0 | 0 | 0 |
| 44308 | ENSMUSG00000106274 | 0 | NA | NA | NA | NA | NA | Gm42828 | predicted gene 42828 [Source:MGI Symbo | 0 | 0 | 0 | 0 | 0 | 0 |
| 44309 | ENSMUSG00000106275 | 0 | NA | NA | NA | NA | NA | Gm42495 | predicted gene 42495 [Source:MGI Symbo | 0 | 0 | 0 | 0 | 0 | 0 |
| 44310 | ENSMUSG00000106276 | 0 | NA | NA | NA | NA | NA | Gm9397 | predicted gene 9397 [Source:MGI Symbol | 0 | 0 | 0 | 0 | 0 | 0 |
| 44311 | ENSMUSG00000106278 | 0 | NA | NA | NA | NA | NA | Mir3970 | microRNA 3970 [Source:MGI Symbol;Acc: | 0 | 0 | 0 | 0 | 0 | 0 |
| 44312 | ENSMUSG00000106279 | 0 | NA | NA | NA | NA | NA | NA | NA | 0 | 0 | 0 | 0 | 0 | 0 |
| 44313 | ENSMUSG00000106280 | 0 | NA | NA | NA | NA | NA | Gm42880 | predicted gene 42880 [Source:MGI Symbo | 0 | 0 | 0 | 0 | 0 | 0 |
| 44314 | ENSMUSG00000106281 | 0 | NA | NA | NA | NA | NA | NA | NA | 0 | 0 | 0 | 0 | 0 | 0 |
| 44315 | ENSMUSG00000106282 | 0.294513347 | 0.165143496 | 4.080472857 | 0.040471657 | 0.967717103 | NA | Gm2287 | predicted gene 2287 [Source:MGI Symbol | 0.8031 | 0 | 0 | 0.964 | 0 | 0 |
| 44316 | ENSMUSG00000106283 | 0 | NA | NA | NA | NA | NA | Rpl7a-ps2 | ribosomal protein L7A, pseudogene 2 [Sou | 0 | 0 | 0 | 0 | 0 | 0 |
| 44317 | ENSMUSG00000106284 | 0 | NA | NA | NA | NA | NA | Gm43139 | predicted gene 43139 [Source:MGI Symbo | 0 | 0 | 0 | 0 | 0 | 0 |
| 44318 | ENSMUSG00000106285 | 0 | NA | NA | NA | NA | NA | Gm43574 | predicted gene 43574 [Source:MGI Symbo | 0 | 0 | 0 | 0 | 0 | 0 |
| 44319 | ENSMUSG00000106286 | 0 | NA | NA | NA | NA | NA | Gm43424 | predicted gene 43424 [Source:MGI Symbo | 0 | 0 | 0 | 0 | 0 | 0 |
| 44320 | ENSMUSG00000106287 | 0.167707431 | -0.796629176 | 4.080472857 | -0.195229623 | 0.84521319 | NA | Gm43585 | predicted gene 43585 [Source:MGI Symbo | 0 | 0 | 1.0062 | 0 | 0 | 0 |
| 44321 | ENSMUSG00000106288 | 13.57612613 | -0.090391148 | 0.699088179 | -0.129298636 | 0.897121351 | 0.982798755 | Gm43593 | predicted gene 43593 [Source:MGI Symbo | 17.668 | 7.1954 | 17.106 | 15.424 | 20.722 | 3.3415 |
| 44322 | ENSMUSG00000106289 | 0 | NA | NA | NA | NA | NA | Trav23 | T cell receptor alpha variable 23 [Source:N | 0 | 0 | 0 | 0 | 0 | 0 |
| 44323 | ENSMUSG00000106290 | 24.03835226 | -0.134221498 | 0.492495402 | -0.272533506 | 0.785211832 | 0.960670212 | Gm43429 | predicted gene 43429 [Source:MGI Symbo | 16.865 | 26.726 | 32.2 | 30.847 | 13.087 | 24.505 |
| 44324 | ENSMUSG00000106291 | 2.820385218 | -0.357854748 | 1.256725058 | -0.284751821 | 0.775834286 | NA | 1110003F10 | RIKEN cDNA 1110003F10 gene [Source:M | 2.4093 | 3.0838 | 4.025 | 1.928 | 4.3625 | 1.1138 |
| 44325 | ENSMUSG00000106292 | 0 | NA | NA | NA | NA | NA | Gm43174 | predicted gene 43174 [Source:MGI Symbo | 0 | 0 | 0 | 0 | 0 | 0 |
| 44326 | ENSMUSG00000106293 | 0 | NA | NA | NA | NA | NA | Gm42476 | predicted gene 42476 [Source:MGI Symbo | 0 | 0 | 0 | 0 | 0 | 0 |
| 44327 | ENSMUSG00000106294 | 0.815516723 | -3.06336425 | 2.557025255 | -1.19801877 | 0.23090971 | NA | Gm8539 | predicted gene 8539 [Source:MGI Symbol | 0.8031 | 3.0838 | 1.0062 | 0 | 0 | 0 |
| 44328 | ENSMUSG00000106295 | 0 | NA | NA | NA | NA | NA | Gm42932 | predicted gene 42932 [Source:MGI Symbo | 0 | 0 | 0 | 0 | 0 | 0 |
| 44329 | ENSMUSG00000106296 | 0 | NA | NA | NA | NA | NA | 4632404M1( | RIKEN cDNA 4632404M16 gene [Source:M | 0 | 0 | 0 | 0 | 0 | 0 |
| 44330 | ENSMUSG00000106297 | 0 | NA | NA | NA | NA | NA | NA | NA | 0 | 0 | 0 | 0 | 0 | 0 |
| 44331 | ENSMUSG00000106298 | 0 | NA | NA | NA | NA | NA | Gm42958 | predicted gene 42958 [Source:MGI Symbo | 0 | 0 | 0 | 0 | 0 | 0 |
| 44332 | ENSMUSG00000106299 | 0 | NA | NA | NA | NA | NA | NA | NA | 0 | 0 | 0 | 0 | 0 | 0 |
| 44333 | ENSMUSG00000106300 | 0 | NA | NA | NA | NA | NA | Gm42765 | predicted gene 42765 [Source:MGI Symbo | 0 | 0 | 0 | 0 | 0 | 0 |
| 44334 | ENSMUSG00000106301 | 0 | NA | NA | NA | NA | NA | Gm8365 | predicted gene 8365 [Source:MGI Symbol | 0 | 0 | 0 | 0 | 0 | 0 |
| 44335 | ENSMUSG00000106302 | 0 | NA | NA | NA | NA | NA | Mir6902 | microRNA 6902 [Source:MGI Symbol;Acc: | 0 | 0 | 0 | 0 | 0 | 0 |
| 44336 | ENSMUSG00000106303 | 78.36654271 | 1.922700973 | 0.761042841 | 2.526403074 | 0.011523718 | 0.119817569 | Gm7652 | predicted gene 7652 [Source:MGI Symbol | 36.943 | 40.089 | 21.131 | 244.85 | 42.534 | 84.652 |
| 44337 | ENSMUSG00000106304 | 0 | NA | NA | NA | NA | NA | Gm42518 | predicted gene 42518 [Source:MGI Symbo | 0 | 0 | 0 | 0 | 0 | 0 |
| 44338 | ENSMUSG00000106305 | 0 | NA | NA | NA | NA | NA | Gm42911 | predicted gene 42911 [Source:MGI Symbo | 0 | 0 | 0 | 0 | 0 | 0 |
| 44339 | ENSMUSG00000106306 | 0.160662974 | 1.126924175 | 4.080472857 | 0.276174898 | 0.782413736 | NA | 4933401H06 | RIKEN cDNA 4933401H06 gene [Source:M | 0 | 0 | 0 | 0.964 | 0 | 0 |
| 44340 | ENSMUSG00000106307 | 0 | NA | NA | NA | NA | NA | Gm23370 | predicted gene, 23370 [Source:MGI Symb | 0 | 0 | 0 | 0 | 0 | 0 |
| 44341 | ENSMUSG00000106308 | 0 | NA | NA | NA | NA | NA | Gm42719 | predicted gene 42719 [Source:MGI Symbo | 0 | 0 | 0 | 0 | 0 | 0 |
| 44342 | ENSMUSG00000106309 | 0 | NA | NA | NA | NA | NA | Gm40414 | predicted gene, 40414 [Source:MGI Symb | 0 | 0 | 0 | 0 | 0 | 0 |
| 44343 | ENSMUSG00000106310 | 0 | NA | NA | NA | NA | NA | Gm43523 | predicted gene 43523 [Source:MGI Symbo | 0 | 0 | 0 | 0 | 0 | 0 |
| 44344 | ENSMUSG00000106311 | 0.342432689 | 1.990653086 | 3.900281263 | 0.51038706 | 0.609780321 | NA | Gm43755 | predicted gene 43755 [Source:MGI Symbo | 0 | 0 | 0 | 0.964 | 1.0906 | 0 |
| 44345 | ENSMUSG00000106312 | 0 | NA | NA | NA | NA | NA | NA | NA | 0 | 0 | 0 | 0 | 0 | 0 |
| 44346 | ENSMUSG00000106313 | 0 | NA | NA | NA | NA | NA | Gm9402 | predicted gene 9402 [Source:MGI Symbol | 0 | 0 | 0 | 0 | 0 | 0 |
| 44347 | ENSMUSG00000106314 | 0 | NA | NA | NA | NA | NA | Gm9632 | predicted gene 9632 [Source:MGI Symbol | 0 | 0 | 0 | 0 | 0 | 0 |
| 44348 | ENSMUSG00000106315 | 0 | NA | NA | NA | NA | NA | Gm42720 | predicted gene 42720 [Source:MGI Symbo | 0 | 0 | 0 | 0 | 0 | 0 |

| | | | | | | | | | | | | | | | |
|---|---|---|---|---|---|---|---|---|---|---|---|---|---|---|---|
| 44349 | ENSMUSG00000106316 | 0 | NA | NA | NA | NA | NA | Gm43467 | predicted gene 43467 [Source:MGI Symbo | 0 | 0 | 0 | 0 | 0 | 0 |
| 44350 | ENSMUSG00000106317 | 8.527330475 | 0.516384035 | 0.836131626 | 0.617587015 | 0.536847606 | 0.890579628 | Gm43848 | predicted gene 43848 [Source:MGI Symbo | 9.6372 | 9.2513 | 2.0125 | 3.8559 | 15.269 | 11.138 |
| 44351 | ENSMUSG00000106318 | 0 | NA | NA | NA | NA | NA | Igkv2-93-1 | immunoglobulin kappa chain variable 2-9: | 0 | 0 | 0 | 0 | 0 | 0 |
| 44352 | ENSMUSG00000106319 | 0 | NA | NA | NA | NA | NA | Gm42601 | predicted gene 42601 [Source:MGI Symbo | 0 | 0 | 0 | 0 | 0 | 0 |
| 44353 | ENSMUSG00000106320 | 0.185640647 | 1.126924175 | 4.080472857 | 0.276174898 | 0.782413736 | NA | Gm42992 | predicted gene 42992 [Source:MGI Symbo | 0 | 0 | 0 | 0 | 0 | 1.1138 |
| 44354 | ENSMUSG00000106321 | 0 | NA | NA | NA | NA | NA | Gm43674 | predicted gene 43674 [Source:MGI Symbo | 0 | 0 | 0 | 0 | 0 | 0 |
| 44355 | ENSMUSG00000106322 | 0.85435009 | 0.823372753 | 2.383559862 | 0.345438252 | 0.729764924 | NA | Gm43432 | predicted gene 43432 [Source:MGI Symbo | 0.8031 | 1.0279 | 0 | 0 | 2.1812 | 1.1138 |
| 44356 | ENSMUSG00000106323 | 0 | NA | NA | NA | NA | NA | Mir7228 | microRNA 7228 [Source:MGI Symbol;Acc: | 0 | 0 | 0 | 0 | 0 | 0 |
| 44357 | ENSMUSG00000106324 | 0 | NA | NA | NA | NA | NA | Gm43222 | predicted gene 43222 [Source:MGI Symbo | 0 | 0 | 0 | 0 | 0 | 0 |
| 44358 | ENSMUSG00000106325 | 0 | NA | NA | NA | NA | NA | Ighv8-10 | immunoglobulin heavy variable V8-10 [So | 0 | 0 | 0 | 0 | 0 | 0 |
| 44359 | ENSMUSG00000106326 | 0 | NA | NA | NA | NA | NA | Gm42977 | predicted gene 42977 [Source:MGI Symbo | 0 | 0 | 0 | 0 | 0 | 0 |
| 44360 | ENSMUSG00000106327 | 0 | NA | NA | NA | NA | NA | Ckb-ps2 | creatine kinase, brain, pseudogene 2 [Sou | 0 | 0 | 0 | 0 | 0 | 0 |
| 44361 | ENSMUSG00000106328 | 0 | NA | NA | NA | NA | NA | Mir7118 | microRNA 7118 [Source:MGI Symbol;Acc: | 0 | 0 | 0 | 0 | 0 | 0 |
| 44362 | ENSMUSG00000106329 | 2.567754493 | -1.39176861 | 1.347370332 | -1.0329518 | 0.301626456 | NA | Gm10652 | predicted gene 10652 [Source:MGI Symbo | 4.0155 | 3.0838 | 4.025 | 0.964 | 1.0906 | 2.2277 |
| 44363 | ENSMUSG00000106330 | 0 | NA | NA | NA | NA | NA | Gm43369 | predicted gene 43369 [Source:MGI Symbo | 0 | 0 | 0 | 0 | 0 | 0 |
| 44364 | ENSMUSG00000106331 | 0 | NA | NA | NA | NA | NA | Gm5309 | predicted gene 5309 [Source:MGI Symbol | 0 | 0 | 0 | 0 | 0 | 0 |
| 44365 | ENSMUSG00000106332 | 0 | NA | NA | NA | NA | NA | Gm43506 | predicted gene 43506 [Source:MGI Symbo | 0 | 0 | 0 | 0 | 0 | 0 |
| 44366 | ENSMUSG00000106333 | 0 | NA | NA | NA | NA | NA | 4930428O21 | RIKEN cDNA 4930428O21 gene [Source:M | 0 | 0 | 0 | 0 | 0 | 0 |
| 44367 | ENSMUSG00000106334 | 0 | NA | NA | NA | NA | NA | Gm43549 | predicted gene 43549 [Source:MGI Symbo | 0 | 0 | 0 | 0 | 0 | 0 |
| 44368 | ENSMUSG00000106335 | 0 | NA | NA | NA | NA | NA | Gm20223 | predicted gene, 20223 [Source:MGI Symb | 0 | 0 | 0 | 0 | 0 | 0 |
| 44369 | ENSMUSG00000106336 | 0 | NA | NA | NA | NA | NA | Mir129b | microRNA 129b [Source:MGI Symbol;Acc: | 0 | 0 | 0 | 0 | 0 | 0 |
| 44370 | ENSMUSG00000106337 | 0 | NA | NA | NA | NA | NA | Gm42478 | predicted gene 42478 [Source:MGI Symbo | 0 | 0 | 0 | 0 | 0 | 0 |
| 44371 | ENSMUSG00000106338 | 0 | NA | NA | NA | NA | NA | Gm43645 | predicted gene 43645 [Source:MGI Symbo | 0 | 0 | 0 | 0 | 0 | 0 |
| 44372 | ENSMUSG00000106339 | 1.106649956 | 0.595609412 | 2.134886147 | 0.278988841 | 0.780253388 | NA | Gm43489 | predicted gene 43489 [Source:MGI Symbo | 1.6062 | 1.0279 | 0 | 2.8919 | 0 | 1.1138 |
| 44373 | ENSMUSG00000106340 | 0 | NA | NA | NA | NA | NA | Mir290b | microRNA 290b [Source:MGI Symbol;Acc: | 0 | 0 | 0 | 0 | 0 | 0 |
| 44374 | ENSMUSG00000106341 | 0.335414862 | -1.779047242 | 4.030804179 | -0.441362855 | 0.658950329 | NA | Gm43330 | predicted gene 43330 [Source:MGI Symbo | 0 | 0 | 2.0125 | 0 | 0 | 0 |
| 44375 | ENSMUSG00000106342 | 0 | NA | NA | NA | NA | NA | Gm42891 | predicted gene 42891 [Source:MGI Symbo | 0 | 0 | 0 | 0 | 0 | 0 |
| 44376 | ENSMUSG00000106343 | 0 | NA | NA | NA | NA | NA | Gm18430 | predicted gene, 18430 [Source:MGI Symb | 0 | 0 | 0 | 0 | 0 | 0 |
| 44377 | ENSMUSG00000106344 | 0 | NA | NA | NA | NA | NA | Mir1892 | microRNA 1892 [Source:MGI Symbol;Acc: | 0 | 0 | 0 | 0 | 0 | 0 |
| 44378 | ENSMUSG00000106345 | 0 | NA | NA | NA | NA | NA | Gm42459 | predicted gene 42459 [Source:MGI Symbo | 0 | 0 | 0 | 0 | 0 | 0 |
| 44379 | ENSMUSG00000106346 | 0 | NA | NA | NA | NA | NA | Gtpbp4-ps6 | GTP binding protein 4, pseudogene 6 [Sou | 0 | 0 | 0 | 0 | 0 | 0 |
| 44380 | ENSMUSG00000106347 | 0 | NA | NA | NA | NA | NA | Mir5615-1 | microRNA 5615-1 [Source:MGI Symbol;Ac | 0 | 0 | 0 | 0 | 0 | 0 |
| 44381 | ENSMUSG00000106348 | 0 | NA | NA | NA | NA | NA | Mir1a-2 | microRNA 1a-2 [Source:MGI Symbol;Acc:I | 0 | 0 | 0 | 0 | 0 | 0 |
| 44382 | ENSMUSG00000106349 | 0 | NA | NA | NA | NA | NA | NA | NA | 0 | 0 | 0 | 0 | 0 | 0 |
| 44383 | ENSMUSG00000106350 | 0 | NA | NA | NA | NA | NA | Kif19b | kinesin family member 19B [Source:MGI S | 0 | 0 | 0 | 0 | 0 | 0 |
| 44384 | ENSMUSG00000106351 | 0 | NA | NA | NA | NA | NA | Gm8896 | predicted gene 8896 [Source:MGI Symbol | 0 | 0 | 0 | 0 | 0 | 0 |
| 44385 | ENSMUSG00000106352 | 369.3759751 | 0.020095275 | 0.144383003 | 0.139180334 | 0.889307649 | 0.982382091 | 5033403H07 | RIKEN cDNA 5033403H07 gene [Source:M | 360.59 | 344.35 | 395.45 | 370.17 | 341.36 | 404.33 |
| 44386 | ENSMUSG00000106353 | 0 | NA | NA | NA | NA | NA | Gm43726 | predicted gene 43726 [Source:MGI Symbo | 0 | 0 | 0 | 0 | 0 | 0 |
| 44387 | ENSMUSG00000106354 | 0 | NA | NA | NA | NA | NA | Gm42607 | predicted gene 42607 [Source:MGI Symbo | 0 | 0 | 0 | 0 | 0 | 0 |
| 44388 | ENSMUSG00000106355 | 0.545309144 | 2.651974629 | 3.666219369 | 0.72335405 | 0.469462394 | NA | NA | NA | 0 | 0 | 0 | 0 | 3.2719 | 0 |
| 44389 | ENSMUSG00000106356 | 0 | NA | NA | NA | NA | NA | Gm42438 | predicted gene 42438 [Source:MGI Symbo | 0 | 0 | 0 | 0 | 0 | 0 |
| 44390 | ENSMUSG00000106357 | 0.17131964 | -0.796629176 | 4.080472857 | -0.195229623 | 0.84521319 | NA | Gm8013 | predicted gene 8013 [Source:MGI Symbol | 0 | 1.0279 | 0 | 0 | 0 | 0 |
| 44391 | ENSMUSG00000106358 | 0 | NA | NA | NA | NA | NA | Gm7047 | predicted gene 7047 [Source:MGI Symbol | 0 | 0 | 0 | 0 | 0 | 0 |
| 44392 | ENSMUSG00000106359 | 0 | NA | NA | NA | NA | NA | 4930554G22 | RIKEN cDNA 4930554G22 gene [Source:M | 0 | 0 | 0 | 0 | 0 | 0 |
| 44393 | ENSMUSG00000106360 | 0 | NA | NA | NA | NA | NA | NA | NA | 0 | 0 | 0 | 0 | 0 | 0 |
| 44394 | ENSMUSG00000106361 | 0 | NA | NA | NA | NA | NA | Gm35066 | predicted gene, 35066 [Source:MGI Symb | 0 | 0 | 0 | 0 | 0 | 0 |
| 44395 | ENSMUSG00000106362 | 0 | NA | NA | NA | NA | NA | Gm47279 | predicted gene, 47279 [Source:MGI Symb | 0 | 0 | 0 | 0 | 0 | 0 |
| 44396 | ENSMUSG00000106363 | 0 | NA | NA | NA | NA | NA | Gm43402 | predicted gene 43402 [Source:MGI Symbo | 0 | 0 | 0 | 0 | 0 | 0 |
| 44397 | ENSMUSG00000106364 | 0.59968336 | -1.233555632 | 2.892386901 | -0.426483618 | 0.669755482 | NA | Gm42994 | predicted gene 42994 [Source:MGI Symbo | 1.6062 | 1.0279 | 0 | 0.964 | 0 | 0 |
| 44398 | ENSMUSG00000106365 | 11.94806671 | -0.082035858 | 0.723543311 | -0.113380716 | 0.909728728 | 0.985435587 | Gm43568 | predicted gene 43568 [Source:MGI Symbo | 9.6372 | 4.1117 | 23.144 | 11.568 | 7.6343 | 15.594 |
| 44399 | ENSMUSG00000106366 | 0 | NA | NA | NA | NA | NA | Mir3069 | microRNA 3069 [Source:MGI Symbol;Acc: | 0 | 0 | 0 | 0 | 0 | 0 |
| 44400 | ENSMUSG00000106367 | 0 | NA | NA | NA | NA | NA | Gm34078 | predicted gene, 34078 [Source:MGI Symb | 0 | 0 | 0 | 0 | 0 | 0 |
| 44401 | ENSMUSG00000106368 | 0.167707431 | -0.796629176 | 4.080472857 | -0.195229623 | 0.84521319 | NA | NA | NA | 0 | 0 | 1.0062 | 0 | 0 | 0 |
| 44402 | ENSMUSG00000106369 | 0.321325949 | 1.911575114 | 4.048267893 | 0.472195804 | 0.636787034 | NA | Gm43355 | predicted gene 43355 [Source:MGI Symbo | 0 | 0 | 0 | 1.928 | 0 | 0 |
| 44403 | ENSMUSG00000106370 | 0 | NA | NA | NA | NA | NA | Gm43382 | predicted gene 43382 [Source:MGI Symbo | 0 | 0 | 0 | 0 | 0 | 0 |
| 44404 | ENSMUSG00000106371 | 0 | NA | NA | NA | NA | NA | 9530097N15 | RIKEN cDNA 9530097N15 gene [Source:M | 0 | 0 | 0 | 0 | 0 | 0 |

| | | | | | | | | | | | | | | | |
|---|---|---|---|---|---|---|---|---|---|---|---|---|---|---|---|
| 44405 | ENSMUSG00000106372 | 0 | NA | NA | NA | NA | NA | Igkv14-126-1 | immunoglobulin kappa chain variable 14- | 0 | 0 | 0 | 0 | 0 | 0 |
| 44406 | ENSMUSG00000106373 | 0 | NA | NA | NA | NA | NA | Gm6522 | predicted gene 6522 [Source:MGI Symbol | 0 | 0 | 0 | 0 | 0 | 0 |
| 44407 | ENSMUSG00000106374 | 0.481988923 | 2.492401104 | 3.837077151 | 0.649557203 | 0.515978285 | NA | Gm27680 | predicted gene, 27680 [Source:MGI Symb | 0 | 0 | 0 | 2.8919 | 0 | 0 |
| 44408 | ENSMUSG00000106375 | 3.292626987 | 0.008840202 | 1.41415647 | 0.006251219 | 0.995012281 | NA | Gm43361 | predicted gene 43361 [Source:MGI Symbo | 0.8031 | 6.1675 | 3.0187 | 8.6758 | 1.0906 | 0 |
| 44409 | ENSMUSG00000106376 | 0 | NA | NA | NA | NA | NA | Gm34139 | predicted gene, 34139 [Source:MGI Symb | 0 | 0 | 0 | 0 | 0 | 0 |
| 44410 | ENSMUSG00000106377 | 0 | NA | NA | NA | NA | NA | NA | NA | 0 | 0 | 0 | 0 | 0 | 0 |
| 44411 | ENSMUSG00000106378 | 0 | NA | NA | NA | NA | NA | Gm42945 | predicted gene 42945 [Source:MGI Symbo | 0 | 0 | 0 | 0 | 0 | 0 |
| 44412 | ENSMUSG00000106379 | 0.185640647 | 1.126924175 | 4.080472857 | 0.276174898 | 0.782413736 | NA | Lhfpl3 | lipoma HMGIC fusion partner-like 3 [Sourc | 0 | 0 | 0 | 0 | 0 | 1.1138 |
| 44413 | ENSMUSG00000106380 | 0.906682787 | -0.583665192 | 2.768799895 | -0.210800785 | 0.833042726 | NA | Gm3519 | predicted gene 3519 [Source:MGI Symbol | 3.2124 | 0 | 0 | 0 | 0 | 2.2277 |
| 44414 | ENSMUSG00000106381 | 0 | NA | NA | NA | NA | NA | Igkv2-95-1 | immunoglobulin kappa chain variable [Sou | 0 | 0 | 0 | 0 | 0 | 0 |
| 44415 | ENSMUSG00000106382 | 0 | NA | NA | NA | NA | NA | Trav4-1 | T cell receptor alpha variable 4-1 [Source: | 0 | 0 | 0 | 0 | 0 | 0 |
| 44416 | ENSMUSG00000106383 | 3.332096028 | 0.194011915 | 1.171895823 | 0.165553892 | 0.868508035 | NA | E330034L11 | RIKEN cDNA E330034L11 gene [Source:M | 3.2124 | 2.0558 | 4.025 | 1.928 | 6.5437 | 2.2277 |
| 44417 | ENSMUSG00000106384 | 0 | NA | NA | NA | NA | NA | Gm43188 | predicted gene 43188 [Source:MGI Symbo | 0 | 0 | 0 | 0 | 0 | 0 |
| 44418 | ENSMUSG00000106385 | 0 | NA | NA | NA | NA | NA | Gm42943 | predicted gene 42943 [Source:MGI Symbo | 0 | 0 | 0 | 0 | 0 | 0 |
| 44419 | ENSMUSG00000106386 | 0 | NA | NA | NA | NA | NA | Gm42991 | predicted gene 42991 [Source:MGI Symbo | 0 | 0 | 0 | 0 | 0 | 0 |
| 44420 | ENSMUSG00000106387 | 0 | NA | NA | NA | NA | NA | Igkv4-73 | immunoglobulin kappa variable 4-73 [Sou | 0 | 0 | 0 | 0 | 0 | 0 |
| 44421 | ENSMUSG00000106388 | 0 | NA | NA | NA | NA | NA | Rhox7-ps2 | reproductive homeobox 7, pseudogene 2 | 0 | 0 | 0 | 0 | 0 | 0 |
| 44422 | ENSMUSG00000106389 | 0 | NA | NA | NA | NA | NA | Gm21006 | predicted gene, 21006 [Source:MGI Symb | 0 | 0 | 0 | 0 | 0 | 0 |
| 44423 | ENSMUSG00000106390 | 0 | NA | NA | NA | NA | NA | Gm5551 | predicted gene 5551 [Source:MGI Symbol | 0 | 0 | 0 | 0 | 0 | 0 |
| 44424 | ENSMUSG00000106391 | 6.484224826 | -0.136869661 | 0.935547117 | -0.146299057 | 0.88368531 | 0.981769403 | Gm42690 | predicted gene 42690 [Source:MGI Symbo | 8.031 | 6.1675 | 6.0375 | 0.964 | 5.4531 | 12.252 |
| 44425 | ENSMUSG00000106392 | 0 | NA | NA | NA | NA | NA | Gm5870 | predicted pseudogene 5870 [Source:MGI | 0 | 0 | 0 | 0 | 0 | 0 |
| 44426 | ENSMUSG00000106393 | 0 | NA | NA | NA | NA | NA | Gm43562 | predicted gene 43562 [Source:MGI Symbo | 0 | 0 | 0 | 0 | 0 | 0 |
| 44427 | ENSMUSG00000106394 | 0 | NA | NA | NA | NA | NA | Gm42698 | predicted gene 42698 [Source:MGI Symbo | 0 | 0 | 0 | 0 | 0 | 0 |
| 44428 | ENSMUSG00000106395 | 0 | NA | NA | NA | NA | NA | Gm42439 | predicted gene 42439 [Source:MGI Symbo | 0 | 0 | 0 | 0 | 0 | 0 |
| 44429 | ENSMUSG00000106396 | 0.803314871 | 3.228356836 | 3.175155518 | 1.0167555 | 0.309269756 | NA | Gm42902 | predicted gene 42902 [Source:MGI Symbo | 0 | 0 | 0 | 4.8199 | 0 | 0 |
| 44430 | ENSMUSG00000106397 | 51.16477326 | 0.803709474 | 0.463301987 | 1.734742127 | 0.082786491 | 0.424179628 | Gm21049 | predicted gene, 21049 [Source:MGI Symb | 45.777 | 34.949 | 31.194 | 123.39 | 33.809 | 37.871 |
| 44431 | ENSMUSG00000106398 | 0 | NA | NA | NA | NA | NA | Gm43357 | predicted gene 43357 [Source:MGI Symbo | 0 | 0 | 0 | 0 | 0 | 0 |
| 44432 | ENSMUSG00000106399 | 0 | NA | NA | NA | NA | NA | Gm43530 | predicted gene 43530 [Source:MGI Symbo | 0 | 0 | 0 | 0 | 0 | 0 |
| 44433 | ENSMUSG00000106400 | 0 | NA | NA | NA | NA | NA | Mir216c | microRNA 216c [Source:MGI Symbol;Acc:I | 0 | 0 | 0 | 0 | 0 | 0 |
| 44434 | ENSMUSG00000106401 | 0 | NA | NA | NA | NA | NA | Gm43284 | predicted gene 43284 [Source:MGI Symbo | 0 | 0 | 0 | 0 | 0 | 0 |
| 44435 | ENSMUSG00000106402 | 0 | NA | NA | NA | NA | NA | Gm17824 | predicted gene, 17824 [Source:MGI Symb | 0 | 0 | 0 | 0 | 0 | 0 |
| 44436 | ENSMUSG00000106403 | 0 | NA | NA | NA | NA | NA | Igkv20-101-2 | immunoglobulin kappa chain variable 20- | 0 | 0 | 0 | 0 | 0 | 0 |
| 44437 | ENSMUSG00000106404 | 0 | NA | NA | NA | NA | NA | Gm42919 | predicted gene 42919 [Source:MGI Symbo | 0 | 0 | 0 | 0 | 0 | 0 |
| 44438 | ENSMUSG00000106405 | 0 | NA | NA | NA | NA | NA | Iglj3p | immunoglobulin lambda joining 3 pseudo | 0 | 0 | 0 | 0 | 0 | 0 |
| 44439 | ENSMUSG00000106406 | 1.188014301 | -2.403107742 | 2.598729323 | -0.924724142 | 0.355109418 | NA | Mir7682 | microRNA 7682 [Source:MGI Symbol;Acc: | 0 | 0 | 6.0375 | 0 | 1.0906 | 0 |
| 44440 | ENSMUSG00000106407 | 0 | NA | NA | NA | NA | NA | Gm43439 | predicted gene 43439 [Source:MGI Symbo | 0 | 0 | 0 | 0 | 0 | 0 |
| 44441 | ENSMUSG00000106408 | 0 | NA | NA | NA | NA | NA | Gm43321 | predicted gene 43321 [Source:MGI Symbo | 0 | 0 | 0 | 0 | 0 | 0 |
| 44442 | ENSMUSG00000106409 | 0 | NA | NA | NA | NA | NA | Mir7092 | microRNA 7092 [Source:MGI Symbol;Acc: | 0 | 0 | 0 | 0 | 0 | 0 |
| 44443 | ENSMUSG00000106410 | 0 | NA | NA | NA | NA | NA | Gm29755 | predicted gene, 29755 [Source:MGI Symb | 0 | 0 | 0 | 0 | 0 | 0 |
| 44444 | ENSMUSG00000106411 | 0 | NA | NA | NA | NA | NA | Gm42445 | predicted gene 42445 [Source:MGI Symbo | 0 | 0 | 0 | 0 | 0 | 0 |
| 44445 | ENSMUSG00000106412 | 0 | NA | NA | NA | NA | NA | NA | NA | 0 | 0 | 0 | 0 | 0 | 0 |
| 44446 | ENSMUSG00000106413 | 0 | NA | NA | NA | NA | NA | Gm42823 | predicted gene 42823 [Source:MGI Symbo | 0 | 0 | 0 | 0 | 0 | 0 |
| 44447 | ENSMUSG00000106414 | 0 | NA | NA | NA | NA | NA | Gm43320 | predicted gene 43320 [Source:MGI Symbo | 0 | 0 | 0 | 0 | 0 | 0 |
| 44448 | ENSMUSG00000106415 | 0 | NA | NA | NA | NA | NA | Gm42893 | predicted gene 42893 [Source:MGI Symbo | 0 | 0 | 0 | 0 | 0 | 0 |
| 44449 | ENSMUSG00000106416 | 0.880724006 | 0.715724162 | 2.808295084 | 0.25486074 | 0.79883066 | NA | Gm5857 | predicted gene 5857 [Source:MGI Symbol | 0 | 0 | 2.0125 | 0 | 3.2719 | 0 |
| 44450 | ENSMUSG00000106417 | 0.921009309 | 0.144736979 | 2.479280948 | 0.058378611 | 0.953447051 | NA | Gm42628 | predicted gene 42628 [Source:MGI Symbo | 1.6062 | 1.0279 | 0 | 2.8919 | 0 | 0 |
| 44451 | ENSMUSG00000106418 | 0 | NA | NA | NA | NA | NA | NA | NA | 0 | 0 | 0 | 0 | 0 | 0 |
| 44452 | ENSMUSG00000106419 | 0 | NA | NA | NA | NA | NA | Gm5550 | predicted gene 5550 [Source:MGI Symbol | 0 | 0 | 0 | 0 | 0 | 0 |
| 44453 | ENSMUSG00000106420 | 0 | NA | NA | NA | NA | NA | Gm9245 | predicted gene 9245 [Source:MGI Symbol | 0 | 0 | 0 | 0 | 0 | 0 |
| 44454 | ENSMUSG00000106421 | 0.674621893 | -1.426050892 | 3.091195043 | -0.461326727 | 0.644564212 | NA | Gm9645 | predicted gene 9645 [Source:MGI Symbol | 0 | 3.0838 | 0 | 0.964 | 0 | 0 |
| 44455 | ENSMUSG00000106422 | 1.346789088 | 2.851836319 | 2.053945616 | 1.388467298 | 0.164994796 | NA | Gm42812 | predicted gene 42812 [Source:MGI Symbo | 0.8031 | 0 | 0 | 2.8919 | 3.2719 | 1.1138 |
| 44456 | ENSMUSG00000106423 | 0 | NA | NA | NA | NA | NA | Gm42825 | predicted gene 42825 [Source:MGI Symbo | 0 | 0 | 0 | 0 | 0 | 0 |
| 44457 | ENSMUSG00000106424 | 0.449470461 | -0.545136533 | 3.562105917 | -0.15303771 | 0.878368537 | NA | Gm34866 | predicted gene, 34866 [Source:MGI Symb | 1.6062 | 0 | 0 | 0 | 1.0906 | 0 |
| 44458 | ENSMUSG00000106425 | 0 | NA | NA | NA | NA | NA | Mir873b | microRNA 873b [Source:MGI Symbol;Acc: | 0 | 0 | 0 | 0 | 0 | 0 |
| 44459 | ENSMUSG00000106426 | 0 | NA | NA | NA | NA | NA | Gm36211 | predicted gene, 36211 [Source:MGI Symb | 0 | 0 | 0 | 0 | 0 | 0 |
| 44460 | ENSMUSG00000106427 | 1.79151646 | -4.199466609 | 1.946980263 | -2.156912779 | 0.031012461 | NA | Gm42820 | predicted gene 42820 [Source:MGI Symbo | 1.6062 | 4.1117 | 5.0312 | 0 | 0 | 0 |

| | | | | | | | | | | | | | | | |
|---|---|---|---|---|---|---|---|---|---|---|---|---|---|---|---|
| 44461 | ENSMUSG00000106428 | 0 | NA | NA | NA | NA | NA | Gm42667 | predicted gene 42667 [Source:MGI Symbo | 0 | 0 | 0 | 0 | 0 | 0 |
| 44462 | ENSMUSG00000106429 | 0 | NA | NA | NA | NA | NA | Gm10685 | predicted gene 10685 [Source:MGI Symbo | 0 | 0 | 0 | 0 | 0 | 0 |
| 44463 | ENSMUSG00000106430 | 2.018264087 | 0.103386347 | 1.465804285 | 0.070532163 | 0.943770102 | NA | Gm42460 | predicted gene 42460 [Source:MGI Symbo | 0.8031 | 3.0838 | 2.0125 | 2.8919 | 1.0906 | 2.2277 |
| 44464 | ENSMUSG00000106431 | 0 | NA | NA | NA | NA | NA | NA | NA | 0 | 0 | 0 | 0 | 0 | 0 |
| 44465 | ENSMUSG00000106432 | 0.17131964 | -0.796629176 | 4.080472857 | -0.195229623 | 0.84521319 | NA | Gm43259 | predicted gene 43259 [Source:MGI Symbo | 0 | 1.0279 | 0 | 0 | 0 | 0 |
| 44466 | ENSMUSG00000106433 | 0 | NA | NA | NA | NA | NA | Gm32102 | predicted gene, 32102 [Source:MGI Symb | 0 | 0 | 0 | 0 | 0 | 0 |
| 44467 | ENSMUSG00000106434 | 0 | NA | NA | NA | NA | NA | Gm42990 | predicted gene 42990 [Source:MGI Symbo | 0 | 0 | 0 | 0 | 0 | 0 |
| 44468 | ENSMUSG00000106435 | 0 | NA | NA | NA | NA | NA | Gm3103 | predicted gene 3103 [Source:MGI Symbol | 0 | 0 | 0 | 0 | 0 | 0 |
| 44469 | ENSMUSG00000106436 | 0 | NA | NA | NA | NA | NA | Mir7015 | microRNA 7015 [Source:MGI Symbol;Acc: | 0 | 0 | 0 | 0 | 0 | 0 |
| 44470 | ENSMUSG00000106437 | 0 | NA | NA | NA | NA | NA | Gm35667 | predicted gene, 35667 [Source:MGI Symb | 0 | 0 | 0 | 0 | 0 | 0 |
| 44471 | ENSMUSG00000106438 | 0 | NA | NA | NA | NA | NA | Gm32051 | predicted gene, 32051 [Source:MGI Symb | 0 | 0 | 0 | 0 | 0 | 0 |
| 44472 | ENSMUSG00000106439 | 0 | NA | NA | NA | NA | NA | NA | NA | 0 | 0 | 0 | 0 | 0 | 0 |
| 44473 | ENSMUSG00000106440 | 0 | NA | NA | NA | NA | NA | NA | NA | 0 | 0 | 0 | 0 | 0 | 0 |
| 44474 | ENSMUSG00000106441 | 0 | NA | NA | NA | NA | NA | Gm42921 | predicted gene 42921 [Source:MGI Symbo | 0 | 0 | 0 | 0 | 0 | 0 |
| 44475 | ENSMUSG00000106442 | 0 | NA | NA | NA | NA | NA | Gm42594 | predicted gene 42594 [Source:MGI Symbo | 0 | 0 | 0 | 0 | 0 | 0 |
| 44476 | ENSMUSG00000106443 | 0.335414862 | -1.779047242 | 4.030804179 | -0.441362855 | 0.658950329 | NA | Gm42702 | predicted gene 42702 [Source:MGI Symbo | 0 | 0 | 2.0125 | 0 | 0 | 0 |
| 44477 | ENSMUSG00000106444 | 0 | NA | NA | NA | NA | NA | Gm18748 | predicted gene, 18748 [Source:MGI Symb | 0 | 0 | 0 | 0 | 0 | 0 |
| 44478 | ENSMUSG00000106445 | 0 | NA | NA | NA | NA | NA | Gm21190 | predicted gene, 21190 [Source:MGI Symb | 0 | 0 | 0 | 0 | 0 | 0 |
| 44479 | ENSMUSG00000106446 | 0.17131964 | -0.796629176 | 4.080472857 | -0.195229623 | 0.84521319 | NA | Gm42970 | predicted gene 42970 [Source:MGI Symbo | 0 | 1.0279 | 0 | 0 | 0 | 0 |
| 44480 | ENSMUSG00000106447 | 0.349477146 | 0.165142768 | 3.911021834 | 0.042224967 | 0.966319359 | NA | Gm42957 | predicted gene 42957 [Source:MGI Symbo | 0 | 0 | 1.0062 | 0 | 1.0906 | 0 |
| 44481 | ENSMUSG00000106448 | 0 | NA | NA | NA | NA | NA | Gm43486 | predicted gene 43486 [Source:MGI Symbo | 0 | 0 | 0 | 0 | 0 | 0 |
| 44482 | ENSMUSG00000106450 | 0 | NA | NA | NA | NA | NA | Gm43018 | predicted gene 43018 [Source:MGI Symbo | 0 | 0 | 0 | 0 | 0 | 0 |
| 44483 | ENSMUSG00000106451 | 0.315620088 | 0.165143421 | 4.063300985 | 0.040642675 | 0.967580763 | NA | Gm42934 | predicted gene 42934 [Source:MGI Symbo | 0.8031 | 0 | 0 | 0 | 1.0906 | 0 |
| 44484 | ENSMUSG00000106452 | 0 | NA | NA | NA | NA | NA | Gm43749 | predicted gene 43749 [Source:MGI Symbo | 0 | 0 | 0 | 0 | 0 | 0 |
| 44485 | ENSMUSG00000106453 | 0 | NA | NA | NA | NA | NA | Gm9273 | predicted gene 9273 [Source:MGI Symbol | 0 | 0 | 0 | 0 | 0 | 0 |
| 44486 | ENSMUSG00000106454 | 0 | NA | NA | NA | NA | NA | Gm17775 | predicted gene, 17775 [Source:MGI Symb | 0 | 0 | 0 | 0 | 0 | 0 |
| 44487 | ENSMUSG00000106455 | 0.133850373 | -0.796629176 | 4.080472857 | -0.195229623 | 0.84521319 | NA | Gm5711 | predicted gene 5711 [Source:MGI Symbol | 0.8031 | 0 | 0 | 0 | 0 | 0 |
| 44488 | ENSMUSG00000106456 | 0.160662974 | 1.126924175 | 4.080472857 | 0.276174898 | 0.782413736 | NA | Gm9678 | predicted gene 9678 [Source:MGI Symbol | 0 | 0 | 0 | 0.964 | 0 | 0 |
| 44489 | ENSMUSG00000106457 | 2.226148994 | 0.281824621 | 1.631790615 | 0.172708813 | 0.862880316 | NA | Gm42585 | predicted gene 42585 [Source:MGI Symbo | 0 | 3.0838 | 3.0187 | 2.8919 | 4.3625 | 0 |
| 44490 | ENSMUSG00000106458 | 2.496221578 | -0.47453918 | 1.435739681 | -0.330518956 | 0.741007872 | NA | Gm42633 | predicted gene 42633 [Source:MGI Symbo | 5.6217 | 0 | 3.0187 | 1.928 | 2.1812 | 2.2277 |
| 44491 | ENSMUSG00000106459 | 0 | NA | NA | NA | NA | NA | Gm43245 | predicted gene 43245 [Source:MGI Symbo | 0 | 0 | 0 | 0 | 0 | 0 |
| 44492 | ENSMUSG00000106460 | 0 | NA | NA | NA | NA | NA | Gm17935 | predicted gene, 17935 [Source:MGI Symb | 0 | 0 | 0 | 0 | 0 | 0 |
| 44493 | ENSMUSG00000106461 | 0 | NA | NA | NA | NA | NA | Gm20755 | predicted gene, 20755 [Source:MGI Symb | 0 | 0 | 0 | 0 | 0 | 0 |
| 44494 | ENSMUSG00000106462 | 0 | NA | NA | NA | NA | NA | Gm43725 | predicted gene 43725 [Source:MGI Symbo | 0 | 0 | 0 | 0 | 0 | 0 |
| 44495 | ENSMUSG00000106463 | 0.181769715 | 1.126924175 | 4.080472857 | 0.276174898 | 0.782413736 | NA | Mir1291 | microRNA 1291 [Source:MGI Symbol;Acc: | 0 | 0 | 0 | 0 | 1.0906 | 0 |
| 44496 | ENSMUSG00000106464 | 40.05830033 | 0.46591956 | 0.384720332 | 1.211060402 | 0.225872256 | 0.6725584 | C130083M1 | RIKEN cDNA C130083M11 gene [Source:M | 41.761 | 19.53 | 39.244 | 39.523 | 50.168 | 50.123 |
| 44497 | ENSMUSG00000106465 | 0 | NA | NA | NA | NA | NA | Mir374c | microRNA 374c [Source:MGI Symbol;Acc: | 0 | 0 | 0 | 0 | 0 | 0 |
| 44498 | ENSMUSG00000106466 | 0 | NA | NA | NA | NA | NA | Gm40348 | predicted gene, 40348 [Source:MGI Symb | 0 | 0 | 0 | 0 | 0 | 0 |
| 44499 | ENSMUSG00000106467 | 0 | NA | NA | NA | NA | NA | Gm43395 | predicted gene 43395 [Source:MGI Symbo | 0 | 0 | 0 | 0 | 0 | 0 |
| 44500 | ENSMUSG00000106468 | 0.349477146 | 0.165142768 | 3.911021834 | 0.042224967 | 0.966319359 | NA | NA | NA | 0 | 0 | 1.0062 | 0 | 1.0906 | 0 |
| 44501 | ENSMUSG00000106469 | 0 | NA | NA | NA | NA | NA | C030015E24 | RIKEN cDNA C030015E24 gene [Source:M | 0 | 0 | 0 | 0 | 0 | 0 |
| 44502 | ENSMUSG00000106470 | 0 | NA | NA | NA | NA | NA | Mir7055 | microRNA 7055 [Source:MGI Symbol;Acc: | 0 | 0 | 0 | 0 | 0 | 0 |
| 44503 | ENSMUSG00000106471 | 0 | NA | NA | NA | NA | NA | Mir3107 | microRNA 3107 [Source:MGI Symbol;Acc: | 0 | 0 | 0 | 0 | 0 | 0 |
| 44504 | ENSMUSG00000106472 | 0 | NA | NA | NA | NA | NA | Gm43111 | predicted gene 43111 [Source:MGI Symbo | 0 | 0 | 0 | 0 | 0 | 0 |
| 44505 | ENSMUSG00000106473 | 0 | NA | NA | NA | NA | NA | Gm19089 | predicted gene, 19089 [Source:MGI Symb | 0 | 0 | 0 | 0 | 0 | 0 |
| 44506 | ENSMUSG00000106474 | 0 | NA | NA | NA | NA | NA | NA | NA | 0 | 0 | 0 | 0 | 0 | 0 |
| 44507 | ENSMUSG00000106475 | 1.117306622 | -0.258294525 | 2.105509541 | -0.122675542 | 0.902364032 | NA | Gm43011 | predicted gene 43011 [Source:MGI Symbo | 1.6062 | 2.0558 | 0 | 1.928 | 0 | 1.1138 |
| 44508 | ENSMUSG00000106476 | 0 | NA | NA | NA | NA | NA | Rpl36-ps9 | ribosomal protein L36, pseudogene 9 [Sou | 0 | 0 | 0 | 0 | 0 | 0 |
| 44509 | ENSMUSG00000106477 | 0 | NA | NA | NA | NA | NA | Gm8874 | predicted gene 8874 [Source:MGI Symbol | 0 | 0 | 0 | 0 | 0 | 0 |
| 44510 | ENSMUSG00000106478 | 0 | NA | NA | NA | NA | NA | Gm36551 | predicted gene, 36551 [Source:MGI Symb | 0 | 0 | 0 | 0 | 0 | 0 |
| 44511 | ENSMUSG00000106479 | 0.185640647 | 1.126924175 | 4.080472857 | 0.276174898 | 0.782413736 | NA | Mir721 | microRNA 721 [Source:MGI Symbol;Acc:M | 0 | 0 | 0 | 0 | 0 | 1.1138 |
| 44512 | ENSMUSG00000106480 | 0 | NA | NA | NA | NA | NA | Igkv14-118-1 | immunoglobulin kappa chain variable 14- | 0 | 0 | 0 | 0 | 0 | 0 |
| 44513 | ENSMUSG00000106481 | 0 | NA | NA | NA | NA | NA | Trav15d-3 | T cell receptor alpha variable 15D-3 [Sour | 0 | 0 | 0 | 0 | 0 | 0 |
| 44514 | ENSMUSG00000106482 | 0 | NA | NA | NA | NA | NA | Ighv8-16 | immunoglobulin heavy variable V8-16 [So | 0 | 0 | 0 | 0 | 0 | 0 |
| 44515 | ENSMUSG00000106483 | 0 | NA | NA | NA | NA | NA | Gm32736 | predicted gene, 32736 [Source:MGI Symb | 0 | 0 | 0 | 0 | 0 | 0 |
| 44516 | ENSMUSG00000106484 | 0.622883753 | 0.113321311 | 2.884614835 | 0.039284729 | 0.968663382 | NA | Gm17833 | predicted gene, 17833 [Source:MGI Symb | 0.8031 | 0 | 1.0062 | 1.928 | 0 | 0 |

| | | | | | | | | | | | | | | | |
|---|---|---|---|---|---|---|---|---|---|---|---|---|---|---|---|
| 44517 | ENSMUSG00000106485 | 0 | NA | NA | NA | NA | NA | 3830422I06F | RIKEN cDNA 3830422I06 gene [Source:MG | 0 | 0 | 0 | 0 | 0 | 0 |
| 44518 | ENSMUSG00000106486 | 0 | NA | NA | NA | NA | NA | Gm9771 | predicted gene 9771 [Source:MGI Symbol | 0 | 0 | 0 | 0 | 0 | 0 |
| 44519 | ENSMUSG00000106487 | 0 | NA | NA | NA | NA | NA | Gm43407 | predicted gene 43407 [Source:MGI Symbo | 0 | 0 | 0 | 0 | 0 | 0 |
| 44520 | ENSMUSG00000106488 | 0 | NA | NA | NA | NA | NA | NA | NA | 0 | 0 | 0 | 0 | 0 | 0 |
| 44521 | ENSMUSG00000106489 | 0 | NA | NA | NA | NA | NA | NA | NA | 0 | 0 | 0 | 0 | 0 | 0 |
| 44522 | ENSMUSG00000106490 | 0 | NA | NA | NA | NA | NA | Gm43283 | predicted gene 43283 [Source:MGI Symbo | 0 | 0 | 0 | 0 | 0 | 0 |
| 44523 | ENSMUSG00000106491 | 0 | NA | NA | NA | NA | NA | Gm42446 | predicted gene 42446 [Source:MGI Symbo | 0 | 0 | 0 | 0 | 0 | 0 |
| 44524 | ENSMUSG00000106492 | 2.423401911 | -3.561016223 | 1.736335004 | -2.050880857 | 0.040278551 | NA | Gm29707 | predicted gene, 29707 [Source:MGI Symb | 3.2124 | 7.1954 | 3.0187 | 0 | 0 | 1.1138 |
| 44525 | ENSMUSG00000106493 | 0 | NA | NA | NA | NA | NA | Mir9-1 | microRNA 9-1 [Source:MGI Symbol;Acc:M | 0 | 0 | 0 | 0 | 0 | 0 |
| 44526 | ENSMUSG00000106494 | 0 | NA | NA | NA | NA | NA | Igkv2-107 | immunoglobulin kappa variable 2-107 [So | 0 | 0 | 0 | 0 | 0 | 0 |
| 44527 | ENSMUSG00000106495 | 0 | NA | NA | NA | NA | NA | Gm42755 | predicted gene 42755 [Source:MGI Symbo | 0 | 0 | 0 | 0 | 0 | 0 |
| 44528 | ENSMUSG00000106496 | 0.160662974 | 1.126924175 | 4.080472857 | 0.276174898 | 0.782413736 | NA | Gm7733 | predicted gene 7733 [Source:MGI Symbol | 0 | 0 | 0 | 0.964 | 0 | 0 |
| 44529 | ENSMUSG00000106497 | 0 | NA | NA | NA | NA | NA | Gm43195 | predicted gene 43195 [Source:MGI Symbo | 0 | 0 | 0 | 0 | 0 | 0 |
| 44530 | ENSMUSG00000106498 | 0 | NA | NA | NA | NA | NA | Gm43032 | predicted gene 43032 [Source:MGI Symbo | 0 | 0 | 0 | 0 | 0 | 0 |
| 44531 | ENSMUSG00000106499 | 2.543370489 | 0.283423449 | 1.489043676 | 0.190339245 | 0.849043303 | NA | Gm9403 | predicted gene 9403 [Source:MGI Symbol | 0.8031 | 5.1396 | 1.0062 | 3.8559 | 0 | 4.4554 |
| 44532 | ENSMUSG00000106500 | 0 | NA | NA | NA | NA | NA | Gm42540 | predicted gene 42540 [Source:MGI Symbo | 0 | 0 | 0 | 0 | 0 | 0 |
| 44533 | ENSMUSG00000106501 | 1.921412152 | 1.612123667 | 1.581258082 | 1.019519638 | 0.307956335 | NA | Gm42711 | predicted gene 42711 [Source:MGI Symbo | 0.8031 | 1.0279 | 1.0062 | 0.964 | 3.2719 | 4.4554 |
| 44534 | ENSMUSG00000106502 | 0 | NA | NA | NA | NA | NA | Gm42795 | predicted gene 42795 [Source:MGI Symbo | 0 | 0 | 0 | 0 | 0 | 0 |
| 44535 | ENSMUSG00000106503 | 0 | NA | NA | NA | NA | NA | Gm42827 | predicted gene 42827 [Source:MGI Symbo | 0 | 0 | 0 | 0 | 0 | 0 |
| 44536 | ENSMUSG00000106504 | 0 | NA | NA | NA | NA | NA | Gm43529 | predicted gene 43529 [Source:MGI Symbo | 0 | 0 | 0 | 0 | 0 | 0 |
| 44537 | ENSMUSG00000106505 | 0 | NA | NA | NA | NA | NA | Gm32693 | predicted gene, 32693 [Source:MGI Symb | 0 | 0 | 0 | 0 | 0 | 0 |
| 44538 | ENSMUSG00000106506 | 0 | NA | NA | NA | NA | NA | 4930558F17 | RIKEN cDNA 4930558F17 gene [Source:M | 0 | 0 | 0 | 0 | 0 | 0 |
| 44539 | ENSMUSG00000106507 | 0 | NA | NA | NA | NA | NA | Gm43056 | predicted gene 43056 [Source:MGI Symbo | 0 | 0 | 0 | 0 | 0 | 0 |
| 44540 | ENSMUSG00000106508 | 0.36353943 | 2.065989254 | 4.034137301 | 0.512126658 | 0.608562368 | NA | 4933425M0 | RIKEN cDNA 4933425M03 gene [Source:M | 0 | 0 | 0 | 0 | 2.1812 | 0 |
| 44541 | ENSMUSG00000106509 | 0 | NA | NA | NA | NA | NA | Gm43394 | predicted gene 43394 [Source:MGI Symbo | 0 | 0 | 0 | 0 | 0 | 0 |
| 44542 | ENSMUSG00000106510 | 0 | NA | NA | NA | NA | NA | Gm18429 | predicted gene, 18429 [Source:MGI Symb | 0 | 0 | 0 | 0 | 0 | 0 |
| 44543 | ENSMUSG00000106511 | 0 | NA | NA | NA | NA | NA | Gm43521 | predicted gene 43521 [Source:MGI Symbo | 0 | 0 | 0 | 0 | 0 | 0 |
| 44544 | ENSMUSG00000106512 | 0 | NA | NA | NA | NA | NA | Gm43078 | predicted gene 43078 [Source:MGI Symbo | 0 | 0 | 0 | 0 | 0 | 0 |
| 44545 | ENSMUSG00000106513 | 0 | NA | NA | NA | NA | NA | NA | NA | 0 | 0 | 0 | 0 | 0 | 0 |
| 44546 | ENSMUSG00000106514 | 0.335414862 | -1.779047242 | 4.030804179 | -0.441362855 | 0.658950329 | NA | Mir6976 | microRNA 6976 [Source:MGI Symbol;Acc: | 0 | 0 | 2.0125 | 0 | 0 | 0 |
| 44547 | ENSMUSG00000106515 | 0 | NA | NA | NA | NA | NA | Gm30382 | predicted gene, 30382 [Source:MGI Symb | 0 | 0 | 0 | 0 | 0 | 0 |
| 44548 | ENSMUSG00000106516 | 0 | NA | NA | NA | NA | NA | Gm42441 | predicted gene 42441 [Source:MGI Symbo | 0 | 0 | 0 | 0 | 0 | 0 |
| 44549 | ENSMUSG00000106517 | 0 | NA | NA | NA | NA | NA | Gm42634 | predicted gene 42634 [Source:MGI Symbo | 0 | 0 | 0 | 0 | 0 | 0 |
| 44550 | ENSMUSG00000106518 | 1.31894822 | -0.787008657 | 2.20064999 | -0.357625547 | 0.720623557 | NA | Gm42759 | predicted gene 42759 [Source:MGI Symbo | 4.0155 | 0 | 1.0062 | 2.8919 | 0 | 0 |
| 44551 | ENSMUSG00000106519 | 0 | NA | NA | NA | NA | NA | Gm38509 | predicted gene, 38509 [Source:MGI Symb | 0 | 0 | 0 | 0 | 0 | 0 |
| 44552 | ENSMUSG00000106520 | 0 | NA | NA | NA | NA | NA | Gm8706 | predicted gene 8706 [Source:MGI Symbol | 0 | 0 | 0 | 0 | 0 | 0 |
| 44553 | ENSMUSG00000106521 | 0 | NA | NA | NA | NA | NA | NA | NA | 0 | 0 | 0 | 0 | 0 | 0 |
| 44554 | ENSMUSG00000106522 | 0 | NA | NA | NA | NA | NA | Gm42871 | predicted gene 42871 [Source:MGI Symbo | 0 | 0 | 0 | 0 | 0 | 0 |
| 44555 | ENSMUSG00000106523 | 0 | NA | NA | NA | NA | NA | NA | NA | 0 | 0 | 0 | 0 | 0 | 0 |
| 44556 | ENSMUSG00000106524 | 0 | NA | NA | NA | NA | NA | Gm42706 | predicted gene 42706 [Source:MGI Symbo | 0 | 0 | 0 | 0 | 0 | 0 |
| 44557 | ENSMUSG00000106525 | 0 | NA | NA | NA | NA | NA | Gm42695 | predicted gene 42695 [Source:MGI Symbo | 0 | 0 | 0 | 0 | 0 | 0 |
| 44558 | ENSMUSG00000106526 | 40.24395232 | -0.183699354 | 0.367745785 | -0.499528104 | 0.617407393 | 0.917509899 | Gm42604 | predicted gene 42604 [Source:MGI Symbo | 53.808 | 44.2 | 30.187 | 39.523 | 41.443 | 32.301 |
| 44559 | ENSMUSG00000106527 | 0 | NA | NA | NA | NA | NA | Gm43671 | predicted gene 43671 [Source:MGI Symbo | 0 | 0 | 0 | 0 | 0 | 0 |
| 44560 | ENSMUSG00000106528 | 0.17131964 | -0.796629176 | 4.080472857 | -0.195229623 | 0.84521319 | NA | Gm42841 | predicted gene 42841 [Source:MGI Symbo | 0 | 1.0279 | 0 | 0 | 0 | 0 |
| 44561 | ENSMUSG00000106529 | 0 | NA | NA | NA | NA | NA | Gm18451 | predicted gene, 18451 [Source:MGI Symb | 0 | 0 | 0 | 0 | 0 | 0 |
| 44562 | ENSMUSG00000106530 | 0 | NA | NA | NA | NA | NA | Gm42430 | predicted gene 42430 [Source:MGI Symbo | 0 | 0 | 0 | 0 | 0 | 0 |
| 44563 | ENSMUSG00000106531 | 0 | NA | NA | NA | NA | NA | Gm43233 | predicted gene 43233 [Source:MGI Symbo | 0 | 0 | 0 | 0 | 0 | 0 |
| 44564 | ENSMUSG00000106532 | 0 | NA | NA | NA | NA | NA | Gm5714 | predicted gene 5714 [Source:MGI Symbol | 0 | 0 | 0 | 0 | 0 | 0 |
| 44565 | ENSMUSG00000106533 | 0 | NA | NA | NA | NA | NA | Trav3d-2 | T cell receptor alpha variable 3D-2 [Source | 0 | 0 | 0 | 0 | 0 | 0 |
| 44566 | ENSMUSG00000106534 | 0.133850373 | -0.796629176 | 4.080472857 | -0.195229623 | 0.84521319 | NA | Gm6366 | predicted gene 6366 [Source:MGI Symbol | 0.8031 | 0 | 0 | 0 | 0 | 0 |
| 44567 | ENSMUSG00000106535 | 9.005407348 | 0.906647789 | 0.80316682 | 1.12884119 | 0.258964833 | 0.712239465 | Gm43031 | predicted gene 43031 [Source:MGI Symbo | 10.44 | 7.1954 | 1.0062 | 14.46 | 10.906 | 10.025 |
| 44568 | ENSMUSG00000106536 | 0.513958919 | -2.394274821 | 3.756096013 | -0.637437066 | 0.523840191 | NA | Gm42508 | predicted gene 42508 [Source:MGI Symbo | 0 | 3.0838 | 0 | 0 | 0 | 0 |
| 44569 | ENSMUSG00000106537 | 0 | NA | NA | NA | NA | NA | Gm43502 | predicted gene 43502 [Source:MGI Symbo | 0 | 0 | 0 | 0 | 0 | 0 |
| 44570 | ENSMUSG00000106538 | 43.20993196 | -0.438801419 | 0.41467423 | -1.058183478 | 0.289971802 | 0.740058153 | Gm30301 | predicted gene, 30301 [Source:MGI Symb | 48.186 | 34.949 | 66.412 | 56.875 | 29.447 | 23.391 |
| 44571 | ENSMUSG00000106539 | 0 | NA | NA | NA | NA | NA | Gm43670 | predicted gene 43670 [Source:MGI Symbo | 0 | 0 | 0 | 0 | 0 | 0 |
| 44572 | ENSMUSG00000106540 | 0 | NA | NA | NA | NA | NA | 4930590L14 | RIKEN cDNA 4930590L14 gene [Source:MG | 0 | 0 | 0 | 0 | 0 | 0 |

| | | | | | | | | | | | | | | | |
|---|---|---|---|---|---|---|---|---|---|---|---|---|---|---|---|
| 44573 | ENSMUSG00000106541 | 0 | NA | NA | NA | NA | NA | Gm43246 | predicted gene 43246 [Source:MGI Symbo | 0 | 0 | 0 | 0 | 0 | 0 |
| 44574 | ENSMUSG00000106542 | 0.465832987 | -0.698295221 | 3.18348759 | -0.219349126 | 0.826378096 | NA | Gm43410 | predicted gene 43410 [Source:MGI Symbo | 0.8031 | 1.0279 | 0 | 0.964 | 0 | 0 |
| 44575 | ENSMUSG00000106543 | 0 | NA | NA | NA | NA | NA | Gm43378 | predicted gene 43378 [Source:MGI Symbo | 0 | 0 | 0 | 0 | 0 | 0 |
| 44576 | ENSMUSG00000106544 | 0 | NA | NA | NA | NA | NA | NA | NA | 0 | 0 | 0 | 0 | 0 | 0 |
| 44577 | ENSMUSG00000106545 | 0 | NA | NA | NA | NA | NA | Gm33952 | predicted gene, 33952 [Source:MGI Symb | 0 | 0 | 0 | 0 | 0 | 0 |
| 44578 | ENSMUSG00000106546 | 0 | NA | NA | NA | NA | NA | Igkv13-56-1 | immunoglobulin kappa chain variable 13-! | 0 | 0 | 0 | 0 | 0 | 0 |
| 44579 | ENSMUSG00000106547 | 2.699511743 | 0.96458724 | 1.372517222 | 0.702786985 | 0.48218851 | NA | B230303O12 | RIKEN cDNA B230303O12 gene [Source:M | 2.4093 | 0 | 3.0187 | 1.928 | 3.2719 | 5.5692 |
| 44580 | ENSMUSG00000106548 | 0 | NA | NA | NA | NA | NA | Gm43390 | predicted gene 43390 [Source:MGI Symbo | 0 | 0 | 0 | 0 | 0 | 0 |
| 44581 | ENSMUSG00000106549 | 0 | NA | NA | NA | NA | NA | Gm42653 | predicted gene 42653 [Source:MGI Symbo | 0 | 0 | 0 | 0 | 0 | 0 |
| 44582 | ENSMUSG00000106550 | 0 | NA | NA | NA | NA | NA | Gm43646 | predicted gene 43646 [Source:MGI Symbo | 0 | 0 | 0 | 0 | 0 | 0 |
| 44583 | ENSMUSG00000106551 | 0 | NA | NA | NA | NA | NA | Gm1969 | predicted gene 1969 [Source:MGI Symbol | 0 | 0 | 0 | 0 | 0 | 0 |
| 44584 | ENSMUSG00000106552 | 0 | NA | NA | NA | NA | NA | Obox4-ps5 | oocyte specific homeobox 4, pseudogene | 0 | 0 | 0 | 0 | 0 | 0 |
| 44585 | ENSMUSG00000106553 | 0.435408177 | -2.179616019 | 3.531869654 | -0.617128103 | 0.537150234 | NA | 6720482G16 | RIKEN cDNA 6720482G16 gene [Source:M | 1.6062 | 0 | 1.0062 | 0 | 0 | 0 |
| 44586 | ENSMUSG00000106554 | 0 | NA | NA | NA | NA | NA | Gm33474 | predicted gene, 33474 [Source:MGI Symb | 0 | 0 | 0 | 0 | 0 | 0 |
| 44587 | ENSMUSG00000106555 | 0.465832987 | -0.698295221 | 3.18348759 | -0.219349126 | 0.826378096 | NA | Gm2479 | predicted gene 2479 [Source:MGI Symbol | 0.8031 | 1.0279 | 0 | 0.964 | 0 | 0 |
| 44588 | ENSMUSG00000106556 | 0 | NA | NA | NA | NA | NA | NA | NA | 0 | 0 | 0 | 0 | 0 | 0 |
| 44589 | ENSMUSG00000106557 | 0 | NA | NA | NA | NA | NA | Gm42442 | predicted gene 42442 [Source:MGI Symbo | 0 | 0 | 0 | 0 | 0 | 0 |
| 44590 | ENSMUSG00000106558 | 0.877291758 | 2.066311913 | 2.565974136 | 0.805273866 | 0.420661659 | NA | Gm5552 | predicted gene 5552 [Source:MGI Symbol | 0 | 1.0279 | 0 | 0.964 | 3.2719 | 0 |
| 44591 | ENSMUSG00000106559 | 0 | NA | NA | NA | NA | NA | Gm43680 | predicted gene 43680 [Source:MGI Symbo | 0 | 0 | 0 | 0 | 0 | 0 |
| 44592 | ENSMUSG00000106560 | 6.061235879 | -0.567703606 | 0.867976029 | -0.654054475 | 0.51307671 | 0.884860213 | Gm43119 | predicted gene 43119 [Source:MGI Symbo | 10.44 | 6.1675 | 5.0312 | 4.8199 | 5.4531 | 4.4554 |
| 44593 | ENSMUSG00000106561 | 0 | NA | NA | NA | NA | NA | NA | NA | 0 | 0 | 0 | 0 | 0 | 0 |
| 44594 | ENSMUSG00000106562 | 0.181769715 | 1.126924175 | 4.080472857 | 0.276174898 | 0.782413736 | NA | Gm5981 | predicted gene 5981 [Source:MGI Symbol | 0 | 0 | 0 | 0 | 1.0906 | 0 |
| 44595 | ENSMUSG00000106563 | 0 | NA | NA | NA | NA | NA | Mir6962 | microRNA 6962 [Source:MGI Symbol;Acc: | 0 | 0 | 0 | 0 | 0 | 0 |
| 44596 | ENSMUSG00000106564 | 0 | NA | NA | NA | NA | NA | Gm8969 | predicted gene 8969 [Source:MGI Symbol | 0 | 0 | 0 | 0 | 0 | 0 |
| 44597 | ENSMUSG00000106565 | 2.070105966 | -2.303509206 | 1.648333455 | -1.397477676 | 0.162269975 | NA | Gm43582 | predicted gene 43582 [Source:MGI Symbo | 3.2124 | 4.1117 | 3.0187 | 0.964 | 0 | 1.1138 |
| 44598 | ENSMUSG00000106566 | 0.133850373 | -0.796629176 | 4.080472857 | -0.195229623 | 0.84521319 | NA | Gm43836 | predicted gene 43836 [Source:MGI Symbo | 0.8031 | 0 | 0 | 0 | 0 | 0 |
| 44599 | ENSMUSG00000106567 | 0.160662974 | 1.126924175 | 4.080472857 | 0.276174898 | 0.782413736 | NA | 2010309G21 | RIKEN cDNA 2010309G21 gene [Source:M | 0 | 0 | 0 | 0.964 | 0 | 0 |
| 44600 | ENSMUSG00000106568 | 7.62697972 | -0.633181255 | 0.807297731 | -0.784321856 | 0.432851278 | 0.845264432 | Gm42814 | predicted gene 42814 [Source:MGI Symbo | 10.44 | 11.307 | 6.0375 | 4.8199 | 9.8156 | 3.3415 |
| 44601 | ENSMUSG00000106569 | 0 | NA | NA | NA | NA | NA | Gm43036 | predicted gene 43036 [Source:MGI Symbo | 0 | 0 | 0 | 0 | 0 | 0 |
| 44602 | ENSMUSG00000106570 | 0 | NA | NA | NA | NA | NA | C230071H17 | RIKEN cDNA C230071H17 gene [Source:M | 0 | 0 | 0 | 0 | 0 | 0 |
| 44603 | ENSMUSG00000106571 | 0 | NA | NA | NA | NA | NA | NA | NA | 0 | 0 | 0 | 0 | 0 | 0 |
| 44604 | ENSMUSG00000106572 | 0 | NA | NA | NA | NA | NA | 4933417G07 | RIKEN cDNA 4933417G07 gene [Source:M | 0 | 0 | 0 | 0 | 0 | 0 |
| 44605 | ENSMUSG00000106573 | 0 | NA | NA | NA | NA | NA | Gm43183 | predicted gene 43183 [Source:MGI Symbo | 0 | 0 | 0 | 0 | 0 | 0 |
| 44606 | ENSMUSG00000106574 | 2.184477936 | -0.746750096 | 1.632101807 | -0.457538919 | 0.647283735 | NA | Gm2451 | predicted pseudogene 2451 [Source:MGI | 3.2124 | 2.0558 | 3.0187 | 4.8199 | 0 | 0 |
| 44607 | ENSMUSG00000106575 | 0 | NA | NA | NA | NA | NA | Gm42613 | predicted gene 42613 [Source:MGI Symbo | 0 | 0 | 0 | 0 | 0 | 0 |
| 44608 | ENSMUSG00000106576 | 0 | NA | NA | NA | NA | NA | Gm4866 | predicted gene 4866 [Source:MGI Symbol | 0 | 0 | 0 | 0 | 0 | 0 |
| 44609 | ENSMUSG00000106577 | 0 | NA | NA | NA | NA | NA | Gm31026 | predicted gene, 31026 [Source:MGI Symb | 0 | 0 | 0 | 0 | 0 | 0 |
| 44610 | ENSMUSG00000106578 | 0 | NA | NA | NA | NA | NA | Mir7217 | microRNA 7217 [Source:MGI Symbol;Acc: | 0 | 0 | 0 | 0 | 0 | 0 |
| 44611 | ENSMUSG00000106579 | 0 | NA | NA | NA | NA | NA | Gm42771 | predicted gene 42771 [Source:MGI Symbo | 0 | 0 | 0 | 0 | 0 | 0 |
| 44612 | ENSMUSG00000106580 | 0 | NA | NA | NA | NA | NA | Mir7231 | microRNA 7231 [Source:MGI Symbol;Acc: | 0 | 0 | 0 | 0 | 0 | 0 |
| 44613 | ENSMUSG00000106581 | 0.167707431 | -0.796629176 | 4.080472857 | -0.195229623 | 0.84521319 | NA | Mir7042 | microRNA 7042 [Source:MGI Symbol;Acc: | 0 | 0 | 1.0062 | 0 | 0 | 0 |
| 44614 | ENSMUSG00000106582 | 0 | NA | NA | NA | NA | NA | Obox4-ps24 | oocyte specific homeobox 4, pseudogene | 0 | 0 | 0 | 0 | 0 | 0 |
| 44615 | ENSMUSG00000106583 | 0.181769715 | 1.126924175 | 4.080472857 | 0.276174898 | 0.782413736 | NA | 4930447N08 | RIKEN cDNA 4930447N08 gene [Source:M | 0 | 0 | 0 | 0 | 1.0906 | 0 |
| 44616 | ENSMUSG00000106584 | 0 | NA | NA | NA | NA | NA | 4930502C17 | RIKEN cDNA 4930502C17 gene [Source:M | 0 | 0 | 0 | 0 | 0 | 0 |
| 44617 | ENSMUSG00000106585 | 0 | NA | NA | NA | NA | NA | Gm25053 | predicted gene, 25053 [Source:MGI Symb | 0 | 0 | 0 | 0 | 0 | 0 |
| 44618 | ENSMUSG00000106586 | 0 | NA | NA | NA | NA | NA | 1700013M08 | RIKEN cDNA 1700013M08 gene [Source:M | 0 | 0 | 0 | 0 | 0 | 0 |
| 44619 | ENSMUSG00000106587 | 0 | NA | NA | NA | NA | NA | NA | NA | 0 | 0 | 0 | 0 | 0 | 0 |
| 44620 | ENSMUSG00000106588 | 0.133850373 | -0.796629176 | 4.080472857 | -0.195229623 | 0.84521319 | NA | Gm17590 | predicted gene, 17590 [Source:MGI Symb | 0.8031 | 0 | 0 | 0 | 0 | 0 |
| 44621 | ENSMUSG00000106589 | 0 | NA | NA | NA | NA | NA | 4931419H13 | RIKEN cDNA 4931419H13 gene [Source:M | 0 | 0 | 0 | 0 | 0 | 0 |
| 44622 | ENSMUSG00000106590 | 21.22366927 | 0.250674304 | 0.479948962 | 0.522293667 | 0.601465878 | 0.915461118 | Gm42879 | predicted gene 42879 [Source:MGI Symbo | 22.487 | 16.447 | 19.119 | 23.135 | 29.447 | 16.708 |
| 44623 | ENSMUSG00000106591 | 0 | NA | NA | NA | NA | NA | NA | NA | 0 | 0 | 0 | 0 | 0 | 0 |
| 44624 | ENSMUSG00000106592 | 0.569258551 | -2.569337407 | 3.18860282 | -0.805787849 | 0.420365185 | NA | Gm43070 | predicted gene 43070 [Source:MGI Symbo | 2.4093 | 0 | 1.0062 | 0 | 0 | 0 |
| 44625 | ENSMUSG00000106593 | 0 | NA | NA | NA | NA | NA | Gm7048 | predicted gene 7048 [Source:MGI Symbol | 0 | 0 | 0 | 0 | 0 | 0 |
| 44626 | ENSMUSG00000106594 | 0 | NA | NA | NA | NA | NA | 4930432H08 | RIKEN cDNA 4930432H08 gene [Source:M | 0 | 0 | 0 | 0 | 0 | 0 |
| 44627 | ENSMUSG00000106595 | 0 | NA | NA | NA | NA | NA | NA | NA | 0 | 0 | 0 | 0 | 0 | 0 |
| 44628 | ENSMUSG00000106596 | 0 | NA | NA | NA | NA | NA | Gm42822 | predicted gene 42822 [Source:MGI Symbo | 0 | 0 | 0 | 0 | 0 | 0 |

| | | | | | | | | | | | | | | | |
|---|---|---|---|---|---|---|---|---|---|---|---|---|---|---|---|
| 44629 | ENSMUSG00000106597 | 0 | NA | NA | NA | NA | NA | Mir7021 | microRNA 7021 [Source:MGI Symbol;Acc: | 0 | 0 | 0 | 0 | 0 | 0 |
| 44630 | ENSMUSG00000106598 | 0 | NA | NA | NA | NA | NA | Gm43532 | predicted gene 43532 [Source:MGI Symbo | 0 | 0 | 0 | 0 | 0 | 0 |
| 44631 | ENSMUSG00000106599 | 0 | NA | NA | NA | NA | NA | Ighv1-73 | immunoglobulin heavy variable 1-73 [Sou | 0 | 0 | 0 | 0 | 0 | 0 |
| 44632 | ENSMUSG00000106600 | 0 | NA | NA | NA | NA | NA | Mir6915 | microRNA 6915 [Source:MGI Symbol;Acc: | 0 | 0 | 0 | 0 | 0 | 0 |
| 44633 | ENSMUSG00000106601 | 0 | NA | NA | NA | NA | NA | Ighv1-70 | immunoglobulin heavy variable 1-70 [Sou | 0 | 0 | 0 | 0 | 0 | 0 |
| 44634 | ENSMUSG00000106602 | 0 | NA | NA | NA | NA | NA | 4930405L22I | RIKEN cDNA 4930405L22 gene [Source:M( | 0 | 0 | 0 | 0 | 0 | 0 |
| 44635 | ENSMUSG00000106603 | 0 | NA | NA | NA | NA | NA | Gm43509 | predicted gene 43509 [Source:MGI Symbo | 0 | 0 | 0 | 0 | 0 | 0 |
| 44636 | ENSMUSG00000106604 | 0 | NA | NA | NA | NA | NA | Gm42535 | predicted gene 42535 [Source:MGI Symbo | 0 | 0 | 0 | 0 | 0 | 0 |
| 44637 | ENSMUSG00000106605 | 0 | NA | NA | NA | NA | NA | NA | NA | 0 | 0 | 0 | 0 | 0 | 0 |
| 44638 | ENSMUSG00000106606 | 0 | NA | NA | NA | NA | NA | Gm43217 | predicted gene 43217 [Source:MGI Symbo | 0 | 0 | 0 | 0 | 0 | 0 |
| 44639 | ENSMUSG00000106607 | 0.435408177 | -2.179616019 | 3.531869654 | -0.617128103 | 0.537150234 | NA | Gm43178 | predicted gene 43178 [Source:MGI Symbo | 1.6062 | 0 | 1.0062 | 0 | 0 | 0 |
| 44640 | ENSMUSG00000106608 | 0 | NA | NA | NA | NA | NA | NA | NA | 0 | 0 | 0 | 0 | 0 | 0 |
| 44641 | ENSMUSG00000106609 | 0 | NA | NA | NA | NA | NA | Gm43181 | predicted gene 43181 [Source:MGI Symbo | 0 | 0 | 0 | 0 | 0 | 0 |
| 44642 | ENSMUSG00000106610 | 0 | NA | NA | NA | NA | NA | Gm43103 | predicted gene 43103 [Source:MGI Symbo | 0 | 0 | 0 | 0 | 0 | 0 |
| 44643 | ENSMUSG00000106611 | 0 | NA | NA | NA | NA | NA | NA | NA | 0 | 0 | 0 | 0 | 0 | 0 |
| 44644 | ENSMUSG00000106612 | 1.384568683 | -1.602668228 | 1.886518279 | -0.849537609 | 0.395582212 | NA | Gm43750 | predicted gene 43750 [Source:MGI Symbo | 3.2124 | 1.0279 | 2.0125 | 0.964 | 1.0906 | 0 |
| 44645 | ENSMUSG00000106613 | 0 | NA | NA | NA | NA | NA | Gm42652 | predicted gene 42652 [Source:MGI Symbo | 0 | 0 | 0 | 0 | 0 | 0 |
| 44646 | ENSMUSG00000106614 | 0 | NA | NA | NA | NA | NA | Gm43746 | predicted gene 43746 [Source:MGI Symbo | 0 | 0 | 0 | 0 | 0 | 0 |
| 44647 | ENSMUSG00000106615 | 0 | NA | NA | NA | NA | NA | Gm8968 | predicted gene 8968 [Source:MGI Symbol | 0 | 0 | 0 | 0 | 0 | 0 |
| 44648 | ENSMUSG00000106616 | 0 | NA | NA | NA | NA | NA | Obox4-ps25 | oocyte specific homeobox 4, pseudogene | 0 | 0 | 0 | 0 | 0 | 0 |
| 44649 | ENSMUSG00000106617 | 3.511389099 | -1.196857195 | 1.190341942 | -1.00547343 | 0.314668937 | NA | Gm36266 | predicted gene, 36266 [Source:MGI Symb | 6.4248 | 6.1675 | 2.0125 | 0.964 | 3.2719 | 2.2277 |
| 44650 | ENSMUSG00000106618 | 0 | NA | NA | NA | NA | NA | NA | NA | 0 | 0 | 0 | 0 | 0 | 0 |
| 44651 | ENSMUSG00000106619 | 2.54494118 | 0.746261381 | 1.451651546 | 0.514077488 | 0.607197816 | NA | Gm9353 | predicted gene 9353 [Source:MGI Symbol | 1.6062 | 4.1117 | 0 | 2.8919 | 1.0906 | 5.5692 |
| 44652 | ENSMUSG00000106620 | 0 | NA | NA | NA | NA | NA | Trav7-5 | T cell receptor alpha variable 7-5 [Source: | 0 | 0 | 0 | 0 | 0 | 0 |
| 44653 | ENSMUSG00000106621 | 0 | NA | NA | NA | NA | NA | Gm42752 | predicted gene 42752 [Source:MGI Symbo | 0 | 0 | 0 | 0 | 0 | 0 |
| 44654 | ENSMUSG00000106622 | 0 | NA | NA | NA | NA | NA | Gm5554 | predicted gene 5554 [Source:MGI Symbol | 0 | 0 | 0 | 0 | 0 | 0 |
| 44655 | ENSMUSG00000106623 | 0.531246861 | 1.101376188 | 3.36099124 | 0.327693858 | 0.743143147 | NA | Gm43335 | predicted gene 43335 [Source:MGI Symbo | 0 | 0 | 1.0062 | 0 | 2.1812 | 0 |
| 44656 | ENSMUSG00000106624 | 0 | NA | NA | NA | NA | NA | Gm8853 | predicted gene 8853 [Source:MGI Symbol | 0 | 0 | 0 | 0 | 0 | 0 |
| 44657 | ENSMUSG00000106625 | 0.185640647 | 1.126924175 | 4.080472857 | 0.276174898 | 0.782413736 | NA | Gm43344 | predicted gene 43344 [Source:MGI Symbo | 0 | 0 | 0 | 0 | 0 | 1.1138 |
| 44658 | ENSMUSG00000106626 | 0 | NA | NA | NA | NA | NA | NA | NA | 0 | 0 | 0 | 0 | 0 | 0 |
| 44659 | ENSMUSG00000106627 | 0 | NA | NA | NA | NA | NA | Gm21680 | predicted gene, 21680 [Source:MGI Symb | 0 | 0 | 0 | 0 | 0 | 0 |
| 44660 | ENSMUSG00000106628 | 0 | NA | NA | NA | NA | NA | Gm43558 | predicted gene 43558 [Source:MGI Symbo | 0 | 0 | 0 | 0 | 0 | 0 |
| 44661 | ENSMUSG00000106629 | 0.545309144 | 2.651974629 | 3.666219369 | 0.72335405 | 0.469462394 | NA | Gm40155 | predicted gene, 40155 [Source:MGI Symb | 0 | 0 | 0 | 0 | 3.2719 | 0 |
| 44662 | ENSMUSG00000106630 | 0.160662974 | 1.126924175 | 4.080472857 | 0.276174898 | 0.782413736 | NA | Igkv2-116 | immunoglobulin kappa variable 2-116 [So | 0 | 0 | 0 | 0.964 | 0 | 0 |
| 44663 | ENSMUSG00000106631 | 0.160662974 | 1.126924175 | 4.080472857 | 0.276174898 | 0.782413736 | NA | Gm42669 | predicted gene 42669 [Source:MGI Symbo | 0 | 0 | 0 | 0.964 | 0 | 0 |
| 44664 | ENSMUSG00000106632 | 0.513958919 | -2.394274821 | 3.756096013 | -0.637437066 | 0.523840191 | NA | Gm33466 | predicted gene, 33466 [Source:MGI Symb | 0 | 3.0838 | 0 | 0 | 0 | 0 |
| 44665 | ENSMUSG00000106633 | 0 | NA | NA | NA | NA | NA | Gm43431 | predicted gene 43431 [Source:MGI Symbo | 0 | 0 | 0 | 0 | 0 | 0 |
| 44666 | ENSMUSG00000106634 | 0 | NA | NA | NA | NA | NA | Gm43042 | predicted gene 43042 [Source:MGI Symbo | 0 | 0 | 0 | 0 | 0 | 0 |
| 44667 | ENSMUSG00000106635 | 0 | NA | NA | NA | NA | NA | Gm43098 | predicted gene 43098 [Source:MGI Symbo | 0 | 0 | 0 | 0 | 0 | 0 |
| 44668 | ENSMUSG00000106636 | 0.785324008 | -0.358914107 | 2.435960446 | -0.147339875 | 0.88286376 | NA | Gm43813 | predicted gene 43813 [Source:MGI Symbo | 1.6062 | 1.0279 | 0 | 0.964 | 0 | 1.1138 |
| 44669 | ENSMUSG00000106637 | 0 | NA | NA | NA | NA | NA | Gm42455 | predicted gene 42455 [Source:MGI Symbo | 0 | 0 | 0 | 0 | 0 | 0 |
| 44670 | ENSMUSG00000106639 | 0 | NA | NA | NA | NA | NA | Gm43121 | predicted gene 43121 [Source:MGI Symbo | 0 | 0 | 0 | 0 | 0 | 0 |
| 44671 | ENSMUSG00000106640 | 0 | NA | NA | NA | NA | NA | Gm20940 | predicted gene, 20940 [Source:MGI Symb | 0 | 0 | 0 | 0 | 0 | 0 |
| 44672 | ENSMUSG00000106641 | 0.867306998 | -1.831940351 | 2.582928097 | -0.709249457 | 0.478169687 | NA | Gm43534 | predicted gene 43534 [Source:MGI Symbo | 0 | 3.0838 | 1.0062 | 0 | 0 | 1.1138 |
| 44673 | ENSMUSG00000106642 | 0 | NA | NA | NA | NA | NA | Mir465d | microRNA 465d [Source:MGI Symbol;Acc: | 0 | 0 | 0 | 0 | 0 | 0 |
| 44674 | ENSMUSG00000106643 | 1.027298067 | 3.575098506 | 2.566323775 | 1.393081629 | 0.163595108 | NA | Gm43422 | predicted gene 43422 [Source:MGI Symbo | 0 | 0 | 0 | 2.8919 | 3.2719 | 0 |
| 44675 | ENSMUSG00000106644 | 0 | NA | NA | NA | NA | NA | Gm40331 | predicted gene, 40331 [Source:MGI Symb | 0 | 0 | 0 | 0 | 0 | 0 |
| 44676 | ENSMUSG00000106645 | 0 | NA | NA | NA | NA | NA | Gm42643 | predicted gene 42643 [Source:MGI Symbo | 0 | 0 | 0 | 0 | 0 | 0 |
| 44677 | ENSMUSG00000106646 | 0 | NA | NA | NA | NA | NA | Gm42483 | predicted gene 42483 [Source:MGI Symbo | 0 | 0 | 0 | 0 | 0 | 0 |
| 44678 | ENSMUSG00000106647 | 0 | NA | NA | NA | NA | NA | Gm42523 | predicted gene 42523 [Source:MGI Symbo | 0 | 0 | 0 | 0 | 0 | 0 |
| 44679 | ENSMUSG00000106648 | 0.499690045 | -0.82080011 | 3.113553401 | -0.263621658 | 0.792071468 | NA | Gm43176 | predicted gene 43176 [Source:MGI Symbo | 0 | 1.0279 | 1.0062 | 0.964 | 0 | 0 |
| 44680 | ENSMUSG00000106649 | 0 | NA | NA | NA | NA | NA | NA | NA | 0 | 0 | 0 | 0 | 0 | 0 |
| 44681 | ENSMUSG00000106650 | 0 | NA | NA | NA | NA | NA | Gm43404 | predicted gene 43404 [Source:MGI Symbo | 0 | 0 | 0 | 0 | 0 | 0 |
| 44682 | ENSMUSG00000106651 | 0 | NA | NA | NA | NA | NA | Gm42608 | predicted gene 42608 [Source:MGI Symbo | 0 | 0 | 0 | 0 | 0 | 0 |
| 44683 | ENSMUSG00000106652 | 0 | NA | NA | NA | NA | NA | Gm43358 | predicted gene 43358 [Source:MGI Symbo | 0 | 0 | 0 | 0 | 0 | 0 |
| 44684 | ENSMUSG00000106653 | 0 | NA | NA | NA | NA | NA | Mir299a | microRNA 299a [Source:MGI Symbol;Acc: | 0 | 0 | 0 | 0 | 0 | 0 |

| | | | | | | | | | | | | | | | |
|---|---|---|---|---|---|---|---|---|---|---|---|---|---|---|---|
| 44685 | ENSMUSG00000106654 | 0.976231831 | 0.127524978 | 2.234187259 | 0.057078912 | 0.954482335 | NA | Gm2623 | predicted gene 2623 [Source:MGI Symbol | 0.8031 | 0 | 2.0125 | 1.928 | 0 | 1.1138 |
| 44686 | ENSMUSG00000106655 | 0 | NA | NA | NA | NA | NA | Mir3961 | microRNA 3961 [Source:MGI Symbol;Acc: | 0 | 0 | 0 | 0 | 0 | 0 |
| 44687 | ENSMUSG00000106656 | 0 | NA | NA | NA | NA | NA | Gm42750 | predicted gene 42750 [Source:MGI Symbc | 0 | 0 | 0 | 0 | 0 | 0 |
| 44688 | ENSMUSG00000106657 | 0 | NA | NA | NA | NA | NA | NA | NA | 0 | 0 | 0 | 0 | 0 | 0 |
| 44689 | ENSMUSG00000106658 | 0 | NA | NA | NA | NA | NA | Gm36548 | predicted gene, 36548 [Source:MGI Symb | 0 | 0 | 0 | 0 | 0 | 0 |
| 44690 | ENSMUSG00000106659 | 1.822250473 | -0.849982706 | 1.781185644 | -0.477200514 | 0.633219345 | NA | Gm42161 | predicted gene, 42161 [Source:MGI Symb | 4.0155 | 2.0558 | 1.0062 | 3.8559 | 0 | 0 |
| 44691 | ENSMUSG00000106660 | 0 | NA | NA | NA | NA | NA | Gm43472 | predicted gene 43472 [Source:MGI Symbc | 0 | 0 | 0 | 0 | 0 | 0 |
| 44692 | ENSMUSG00000106661 | 0 | NA | NA | NA | NA | NA | Gm42832 | predicted gene 42832 [Source:MGI Symbc | 0 | 0 | 0 | 0 | 0 | 0 |
| 44693 | ENSMUSG00000106662 | 0.524408994 | -0.837196744 | 3.377823722 | -0.247850928 | 0.804249749 | NA | Gm43034 | predicted gene 43034 [Source:MGI Symbc | 0 | 2.0558 | 0 | 0 | 1.0906 | 0 |
| 44694 | ENSMUSG00000106663 | 0.684892008 | -1.399684857 | 3.073674502 | -0.455378361 | 0.64883706 | NA | Gm42839 | predicted gene 42839 [Source:MGI Symbc | 0 | 0 | 3.0187 | 0 | 1.0906 | 0 |
| 44695 | ENSMUSG00000106664 | 3.578825029 | -0.66276922 | 1.416036208 | -0.46804539 | 0.63975213 | NA | Gm17936 | predicted gene, 17936 [Source:MGI Symb | 0 | 8.2233 | 5.0312 | 3.8559 | 4.3625 | 0 |
| 44696 | ENSMUSG00000106665 | 2.997667787 | 0.021714714 | 1.464694191 | 0.014825425 | 0.988171455 | NA | Gm43389 | predicted gene 43389 [Source:MGI Symbc | 4.8186 | 4.1117 | 0 | 5.7839 | 3.2719 | 0 |
| 44697 | ENSMUSG00000106666 | 0 | NA | NA | NA | NA | NA | Gm43587 | predicted gene 43587 [Source:MGI Symbc | 0 | 0 | 0 | 0 | 0 | 0 |
| 44698 | ENSMUSG00000106667 | 0 | NA | NA | NA | NA | NA | Igkv3-6 | immunoglobulin kappa variable 3-6 [Sourc | 0 | 0 | 0 | 0 | 0 | 0 |
| 44699 | ENSMUSG00000106668 | 0 | NA | NA | NA | NA | NA | Iglj1 | immunoglobulin lambda joining 1 [Source | 0 | 0 | 0 | 0 | 0 | 0 |
| 44700 | ENSMUSG00000106669 | 0 | NA | NA | NA | NA | NA | Gm38433 | predicted gene, 38433 [Source:MGI Symb | 0 | 0 | 0 | 0 | 0 | 0 |
| 44701 | ENSMUSG00000106670 | 0 | NA | NA | NA | NA | NA | NA | NA | 0 | 0 | 0 | 0 | 0 | 0 |
| 44702 | ENSMUSG00000106671 | 0.480153995 | 1.043847617 | 3.151673025 | 0.331204287 | 0.740490182 | NA | Gm42900 | predicted gene 42900 [Source:MGI Symbc | 0.8031 | 0 | 0 | 0.964 | 0 | 1.1138 |
| 44703 | ENSMUSG00000106673 | 0 | NA | NA | NA | NA | NA | Gm43374 | predicted gene 43374 [Source:MGI Symbc | 0 | 0 | 0 | 0 | 0 | 0 |
| 44704 | ENSMUSG00000106674 | 0 | NA | NA | NA | NA | NA | Gm20005 | predicted gene, 20005 [Source:MGI Symb | 0 | 0 | 0 | 0 | 0 | 0 |
| 44705 | ENSMUSG00000106675 | 0 | NA | NA | NA | NA | NA | Gm43798 | predicted gene 43798 [Source:MGI Symbc | 0 | 0 | 0 | 0 | 0 | 0 |
| 44706 | ENSMUSG00000106676 | 0 | NA | NA | NA | NA | NA | Gm42895 | predicted gene 42895 [Source:MGI Symbc | 0 | 0 | 0 | 0 | 0 | 0 |
| 44707 | ENSMUSG00000106677 | 0 | NA | NA | NA | NA | NA | Ugt2a1 | UDP glucuronosyltransferase 2 family, pol | 0 | 0 | 0 | 0 | 0 | 0 |
| 44708 | ENSMUSG00000106678 | 3.7311782 | 0.093823523 | 1.161990526 | 0.080743794 | 0.935645708 | NA | Gm43457 | predicted gene 43457 [Source:MGI Symbc | 0.8031 | 3.0838 | 7.0437 | 4.8199 | 2.1812 | 4.4554 |
| 44709 | ENSMUSG00000106679 | 0 | NA | NA | NA | NA | NA | Gm42524 | predicted gene 42524 [Source:MGI Symbc | 0 | 0 | 0 | 0 | 0 | 0 |
| 44710 | ENSMUSG00000106680 | 3.159495439 | 1.571851919 | 1.517861037 | 1.035570373 | 0.30040262 | NA | Gm43852 | predicted gene 43852 [Source:MGI Symbc | 0.8031 | 0 | 4.025 | 8.6758 | 5.4531 | 0 |
| 44711 | ENSMUSG00000106681 | 0 | NA | NA | NA | NA | NA | Gm43004 | predicted gene 43004 [Source:MGI Symbc | 0 | 0 | 0 | 0 | 0 | 0 |
| 44712 | ENSMUSG00000106682 | 4.220153932 | -0.200627966 | 1.063570642 | -0.188636239 | 0.850377928 | NA | Gm42648 | predicted gene 42648 [Source:MGI Symbc | 2.4093 | 4.1117 | 7.0437 | 2.8919 | 2.1812 | 6.6831 |
| 44713 | ENSMUSG00000106683 | 0 | NA | NA | NA | NA | NA | Gm43263 | predicted gene 43263 [Source:MGI Symbc | 0 | 0 | 0 | 0 | 0 | 0 |
| 44714 | ENSMUSG00000106688 | 0.160662974 | 1.126924175 | 4.080472857 | 0.276174898 | 0.782413736 | NA | Gm42851 | predicted gene 42851 [Source:MGI Symbc | 0 | 0 | 0 | 0.964 | 0 | 0 |
| 44715 | ENSMUSG00000106689 | 0 | NA | NA | NA | NA | NA | Gm18345 | predicted gene, 18345 [Source:MGI Symb | 0 | 0 | 0 | 0 | 0 | 0 |
| 44716 | ENSMUSG00000106690 | 0 | NA | NA | NA | NA | NA | Gm6348 | predicted gene 6348 [Source:MGI Symbol | 0 | 0 | 0 | 0 | 0 | 0 |
| 44717 | ENSMUSG00000106691 | 0 | NA | NA | NA | NA | NA | 2700029L08I | RIKEN cDNA 2700029L08 gene [Source:M( | 0 | 0 | 0 | 0 | 0 | 0 |
| 44718 | ENSMUSG00000106692 | 0 | NA | NA | NA | NA | NA | Gm36447 | predicted gene, 36447 [Source:MGI Symb | 0 | 0 | 0 | 0 | 0 | 0 |
| 44719 | ENSMUSG00000106694 | 0 | NA | NA | NA | NA | NA | Gm42894 | predicted gene 42894 [Source:MGI Symbc | 0 | 0 | 0 | 0 | 0 | 0 |
| 44720 | ENSMUSG00000106696 | 0 | NA | NA | NA | NA | NA | Gm42729 | predicted gene 42729 [Source:MGI Symbc | 0 | 0 | 0 | 0 | 0 | 0 |
| 44721 | ENSMUSG00000106698 | 0 | NA | NA | NA | NA | NA | Gm43599 | predicted gene 43599 [Source:MGI Symbc | 0 | 0 | 0 | 0 | 0 | 0 |
| 44722 | ENSMUSG00000106699 | 0 | NA | NA | NA | NA | NA | Gm43224 | predicted gene 43224 [Source:MGI Symbc | 0 | 0 | 0 | 0 | 0 | 0 |
| 44723 | ENSMUSG00000106701 | 0 | NA | NA | NA | NA | NA | Gm6346 | predicted gene 6346 [Source:MGI Symbol | 0 | 0 | 0 | 0 | 0 | 0 |
| 44724 | ENSMUSG00000106702 | 0 | NA | NA | NA | NA | NA | Gm42846 | predicted gene 42846 [Source:MGI Symbc | 0 | 0 | 0 | 0 | 0 | 0 |
| 44725 | ENSMUSG00000106703 | 0 | NA | NA | NA | NA | NA | Vmn2r-ps26 | vomeronasal 2, receptor, pseudogene 26 | 0 | 0 | 0 | 0 | 0 | 0 |
| 44726 | ENSMUSG00000106704 | 0 | NA | NA | NA | NA | NA | Gm42799 | predicted gene 42799 [Source:MGI Symbc | 0 | 0 | 0 | 0 | 0 | 0 |
| 44727 | ENSMUSG00000106705 | 7.872256775 | 0.083202534 | 0.758109287 | 0.109750053 | 0.912607604 | 0.986248982 | Gm2602 | predicted gene 2602 [Source:MGI Symbol | 9.6372 | 5.1396 | 8.05 | 6.7478 | 7.6343 | 10.025 |
| 44728 | ENSMUSG00000106706 | 0.993674229 | -2.093174934 | 2.253073788 | -0.929030795 | 0.352873127 | NA | C530043K16 | RIKEN cDNA C530043K16 gene [Source:M | 0.8031 | 2.0558 | 2.0125 | 0 | 1.0906 | 0 |
| 44729 | ENSMUSG00000106707 | 0 | NA | NA | NA | NA | NA | Gm43748 | predicted gene 43748 [Source:MGI Symbc | 0 | 0 | 0 | 0 | 0 | 0 |
| 44730 | ENSMUSG00000106708 | 0 | NA | NA | NA | NA | NA | Gm43782 | predicted gene 43782 [Source:MGI Symbc | 0 | 0 | 0 | 0 | 0 | 0 |
| 44731 | ENSMUSG00000106709 | 6.727340263 | 0.389015281 | 0.915757741 | 0.424801521 | 0.670981369 | 0.93116767 | Gm30270 | predicted gene, 30270 [Source:MGI Symb | 7.2279 | 8.2233 | 2.0125 | 8.6758 | 11.997 | 2.2277 |
| 44732 | ENSMUSG00000106710 | 0 | NA | NA | NA | NA | NA | Gm43166 | predicted gene 43166 [Source:MGI Symbc | 0 | 0 | 0 | 0 | 0 | 0 |
| 44733 | ENSMUSG00000106712 | 0 | NA | NA | NA | NA | NA | Gm19649 | predicted gene, 19649 [Source:MGI Symb | 0 | 0 | 0 | 0 | 0 | 0 |
| 44734 | ENSMUSG00000106713 | 0 | NA | NA | NA | NA | NA | Gm42645 | predicted gene 42645 [Source:MGI Symbc | 0 | 0 | 0 | 0 | 0 | 0 |
| 44735 | ENSMUSG00000106714 | 0 | NA | NA | NA | NA | NA | Gm42546 | predicted gene 42546 [Source:MGI Symbc | 0 | 0 | 0 | 0 | 0 | 0 |
| 44736 | ENSMUSG00000106715 | 0 | NA | NA | NA | NA | NA | Tmem265 | transmembrane protein 265 [Source:MGI | 0 | 0 | 0 | 0 | 0 | 0 |
| 44737 | ENSMUSG00000106717 | 2.774532898 | 0.659513017 | 1.278792499 | 0.515731064 | 0.606042255 | NA | Gm42798 | predicted gene 42798 [Source:MGI Symbc | 2.4093 | 2.0558 | 2.0125 | 5.7839 | 3.2719 | 1.1138 |
| 44738 | ENSMUSG00000106718 | 0 | NA | NA | NA | NA | NA | Gm20052 | predicted gene, 20052 [Source:MGI Symb | 0 | 0 | 0 | 0 | 0 | 0 |
| 44739 | ENSMUSG00000106719 | 0 | NA | NA | NA | NA | NA | Gm42556 | predicted gene 42556 [Source:MGI Symbc | 0 | 0 | 0 | 0 | 0 | 0 |
| 44740 | ENSMUSG00000106720 | 0.160662974 | 1.126924175 | 4.080472857 | 0.276174898 | 0.782413736 | NA | Gm43795 | predicted gene 43795 [Source:MGI Symbc | 0 | 0 | 0 | 0.964 | 0 | 0 |

| | | | | | | | | | | | | | | | |
|---|---|---|---|---|---|---|---|---|---|---|---|---|---|---|---|
| 44741 | ENSMUSG00000106721 | 0 | NA | NA | NA | NA | NA | Gm5442 | predicted gene 5442 [Source:MGI Symbol | 0 | 0 | 0 | 0 | 0 | 0 |
| 44742 | ENSMUSG00000106725 | 0 | NA | NA | NA | NA | NA | Gm43363 | predicted gene 43363 [Source:MGI Symbo | 0 | 0 | 0 | 0 | 0 | 0 |
| 44743 | ENSMUSG00000106726 | 0 | NA | NA | NA | NA | NA | Gm43100 | predicted gene 43100 [Source:MGI Symbo | 0 | 0 | 0 | 0 | 0 | 0 |
| 44744 | ENSMUSG00000106729 | 0 | NA | NA | NA | NA | NA | Gm43165 | predicted gene 43165 [Source:MGI Symbo | 0 | 0 | 0 | 0 | 0 | 0 |
| 44745 | ENSMUSG00000106730 | 0 | NA | NA | NA | NA | NA | Gm42472 | predicted gene 42472 [Source:MGI Symbo | 0 | 0 | 0 | 0 | 0 | 0 |
| 44746 | ENSMUSG00000106732 | 0 | NA | NA | NA | NA | NA | Gm43792 | predicted gene 43792 [Source:MGI Symbo | 0 | 0 | 0 | 0 | 0 | 0 |
| 44747 | ENSMUSG00000106733 | 0 | NA | NA | NA | NA | NA | Gm36186 | predicted gene, 36186 [Source:MGI Symb | 0 | 0 | 0 | 0 | 0 | 0 |
| 44748 | ENSMUSG00000106734 | 58.26560077 | -0.58281457 | 0.311160862 | -1.873033026 | 0.061063825 | 0.352304411 | Gm20559 | predicted gene, 20559 [Source:MGI Symb | 65.051 | 86.345 | 58.362 | 48.199 | 38.172 | 53.465 |
| 44749 | ENSMUSG00000106735 | 0 | NA | NA | NA | NA | NA | A330058E17 | RIKEN cDNA A330058E17 gene [Source:M | 0 | 0 | 0 | 0 | 0 | 0 |
| 44750 | ENSMUSG00000106736 | 2.357781976 | 0.954706168 | 1.497603145 | 0.637489425 | 0.523806096 | NA | Gm42573 | predicted gene 42573 [Source:MGI Symbo | 0.8031 | 2.0558 | 2.0125 | 4.8199 | 0 | 4.4554 |
| 44751 | ENSMUSG00000106737 | 0 | NA | NA | NA | NA | NA | Gm42962 | predicted gene 42962 [Source:MGI Symbo | 0 | 0 | 0 | 0 | 0 | 0 |
| 44752 | ENSMUSG00000106738 | 0 | NA | NA | NA | NA | NA | Gm5 | predicted gene 5 [Source:MGI Symbol;Acc | 0 | 0 | 0 | 0 | 0 | 0 |
| 44753 | ENSMUSG00000106739 | 1.557981975 | -1.115189601 | 1.872883345 | -0.595439969 | 0.551549415 | NA | Gm43499 | predicted gene 43499 [Source:MGI Symbo | 2.4093 | 1.0279 | 3.0187 | 2.8919 | 0 | 0 |
| 44754 | ENSMUSG00000106740 | 0 | NA | NA | NA | NA | NA | Gm21221 | predicted gene, 21221 [Source:MGI Symb | 0 | 0 | 0 | 0 | 0 | 0 |
| 44755 | ENSMUSG00000106741 | 0 | NA | NA | NA | NA | NA | 4930513D17 | RIKEN cDNA 4930513D17 gene [Source:M | 0 | 0 | 0 | 0 | 0 | 0 |
| 44756 | ENSMUSG00000106742 | 0 | NA | NA | NA | NA | NA | Gm42474 | predicted gene 42474 [Source:MGI Symbo | 0 | 0 | 0 | 0 | 0 | 0 |
| 44757 | ENSMUSG00000106743 | 0 | NA | NA | NA | NA | NA | Gm42847 | predicted gene 42847 [Source:MGI Symbo | 0 | 0 | 0 | 0 | 0 | 0 |
| 44758 | ENSMUSG00000106745 | 0 | NA | NA | NA | NA | NA | Gm43342 | predicted gene 43342 [Source:MGI Symbo | 0 | 0 | 0 | 0 | 0 | 0 |
| 44759 | ENSMUSG00000106746 | 0 | NA | NA | NA | NA | NA | 2900064F13 | RIKEN cDNA 2900064F13 gene [Source:M | 0 | 0 | 0 | 0 | 0 | 0 |
| 44760 | ENSMUSG00000106747 | 0 | NA | NA | NA | NA | NA | Gm43025 | predicted gene 43025 [Source:MGI Symbo | 0 | 0 | 0 | 0 | 0 | 0 |
| 44761 | ENSMUSG00000106749 | 0 | NA | NA | NA | NA | NA | Gm43281 | predicted gene 43281 [Source:MGI Symbo | 0 | 0 | 0 | 0 | 0 | 0 |
| 44762 | ENSMUSG00000106750 | 0 | NA | NA | NA | NA | NA | Gm43151 | predicted gene 43151 [Source:MGI Symbo | 0 | 0 | 0 | 0 | 0 | 0 |
| 44763 | ENSMUSG00000106751 | 1.26003638 | -0.648637764 | 2.020901123 | -0.320964621 | 0.748237202 | NA | Gm42467 | predicted gene 42467 [Source:MGI Symbo | 1.6062 | 2.0558 | 1.0062 | 2.8919 | 0 | 0 |
| 44764 | ENSMUSG00000106752 | 0 | NA | NA | NA | NA | NA | Gm3348 | predicted gene 3348 [Source:MGI Symbol | 0 | 0 | 0 | 0 | 0 | 0 |
| 44765 | ENSMUSG00000106753 | 0 | NA | NA | NA | NA | NA | Gm43230 | predicted gene 43230 [Source:MGI Symbo | 0 | 0 | 0 | 0 | 0 | 0 |
| 44766 | ENSMUSG00000106754 | 0 | NA | NA | NA | NA | NA | Gm43101 | predicted gene 43101 [Source:MGI Symbo | 0 | 0 | 0 | 0 | 0 | 0 |
| 44767 | ENSMUSG00000106755 | 3.349977111 | -0.098564846 | 1.208147844 | -0.081583431 | 0.934977978 | NA | Tpi-rs11 | triosephosphate isomerase related seque | 3.2124 | 5.1396 | 2.0125 | 0.964 | 6.5437 | 2.2277 |
| 44768 | ENSMUSG00000106756 | 0 | NA | NA | NA | NA | NA | Gm32780 | predicted gene, 32780 [Source:MGI Symb | 0 | 0 | 0 | 0 | 0 | 0 |
| 44769 | ENSMUSG00000106757 | 2.364412723 | -0.826340771 | 1.372430673 | -0.602100191 | 0.547107447 | NA | Gm43482 | predicted gene 43482 [Source:MGI Symbo | 4.0155 | 2.0558 | 3.0187 | 2.8919 | 1.0906 | 1.1138 |
| 44770 | ENSMUSG00000106758 | 0 | NA | NA | NA | NA | NA | Gm42731 | predicted gene 42731 [Source:MGI Symbo | 0 | 0 | 0 | 0 | 0 | 0 |
| 44771 | ENSMUSG00000106759 | 0 | NA | NA | NA | NA | NA | | vomeronasal 2, receptor, pseudogene 28 | 0 | 0 | 0 | 0 | 0 | 0 |
| 44772 | ENSMUSG00000106760 | 0 | NA | NA | NA | NA | NA | Gm42802 | predicted gene 42802 [Source:MGI Symbo | 0 | 0 | 0 | 0 | 0 | 0 |
| 44773 | ENSMUSG00000106762 | 0 | NA | NA | NA | NA | NA | 4930478P22 | RIKEN cDNA 4930478P22 gene [Source:M | 0 | 0 | 0 | 0 | 0 | 0 |
| 44774 | ENSMUSG00000106763 | 0 | NA | NA | NA | NA | NA | Gm19102 | predicted gene, 19102 [Source:MGI Symb | 0 | 0 | 0 | 0 | 0 | 0 |
| 44775 | ENSMUSG00000106764 | 0.167707431 | -0.796629176 | 4.080472857 | -0.195229623 | 0.84521319 | NA | Gm42529 | predicted gene 42529 [Source:MGI Symbo | 0 | 0 | 1.0062 | 0 | 0 | 0 |
| 44776 | ENSMUSG00000106766 | 0 | NA | NA | NA | NA | NA | 4933424N20 | RIKEN cDNA 4933424N20 gene [Source:M | 0 | 0 | 0 | 0 | 0 | 0 |
| 44777 | ENSMUSG00000106767 | 3.559651442 | -1.276589756 | 1.156168813 | -1.104155156 | 0.269525839 | NA | Gm42727 | predicted gene 42727 [Source:MGI Symbo | 4.0155 | 4.1117 | 7.0437 | 2.8919 | 2.1812 | 1.1138 |
| 44778 | ENSMUSG00000106768 | 0 | NA | NA | NA | NA | NA | 4930480C01 | RIKEN cDNA 4930480C01 gene [Source:M | 0 | 0 | 0 | 0 | 0 | 0 |
| 44779 | ENSMUSG00000106771 | 0 | NA | NA | NA | NA | NA | Gm43459 | predicted gene 43459 [Source:MGI Symbo | 0 | 0 | 0 | 0 | 0 | 0 |
| 44780 | ENSMUSG00000106772 | 0 | NA | NA | NA | NA | NA | Gm43794 | predicted gene 43794 [Source:MGI Symbo | 0 | 0 | 0 | 0 | 0 | 0 |
| 44781 | ENSMUSG00000106773 | 0 | NA | NA | NA | NA | NA | Gm43799 | predicted gene 43799 [Source:MGI Symbo | 0 | 0 | 0 | 0 | 0 | 0 |
| 44782 | ENSMUSG00000106775 | 0 | NA | NA | NA | NA | NA | C130093G08 | RIKEN cDNA C130093G08 gene [Source:M | 0 | 0 | 0 | 0 | 0 | 0 |
| 44783 | ENSMUSG00000106776 | 0 | NA | NA | NA | NA | NA | Gm42646 | predicted gene 42646 [Source:MGI Symbo | 0 | 0 | 0 | 0 | 0 | 0 |
| 44784 | ENSMUSG00000106777 | 0 | NA | NA | NA | NA | NA | Gm43150 | predicted gene 43150 [Source:MGI Symbo | 0 | 0 | 0 | 0 | 0 | 0 |
| 44785 | ENSMUSG00000106779 | 0 | NA | NA | NA | NA | NA | Gm8579 | predicted gene 8579 [Source:MGI Symbol | 0 | 0 | 0 | 0 | 0 | 0 |
| 44786 | ENSMUSG00000106780 | 0 | NA | NA | NA | NA | NA | Gm7939 | predicted gene 7939 [Source:MGI Symbol | 0 | 0 | 0 | 0 | 0 | 0 |
| 44787 | ENSMUSG00000106783 | 2.474752375 | 0.76346293 | 1.402875896 | 0.544212737 | 0.586295088 | NA | Chaer1 | cardiac hypertrophy associated epigeneti | 2.4093 | 3.0838 | 0 | 3.8559 | 3.2719 | 2.2277 |
| 44788 | ENSMUSG00000106784 | 0 | NA | NA | NA | NA | NA | Gm43309 | predicted gene 43309 [Source:MGI Symbo | 0 | 0 | 0 | 0 | 0 | 0 |
| 44789 | ENSMUSG00000106785 | 0 | NA | NA | NA | NA | NA | Gm42713 | predicted gene 42713 [Source:MGI Symbo | 0 | 0 | 0 | 0 | 0 | 0 |
| 44790 | ENSMUSG00000106786 | 0 | NA | NA | NA | NA | NA | Gm42781 | predicted gene 42781 [Source:MGI Symbo | 0 | 0 | 0 | 0 | 0 | 0 |
| 44791 | ENSMUSG00000106788 | 18.18738235 | -0.181252715 | 0.544319327 | -0.332989674 | 0.739142078 | 0.948310621 | Gm43449 | predicted gene 43449 [Source:MGI Symbo | 19.274 | 26.726 | 12.075 | 21.208 | 10.906 | 18.935 |
| 44792 | ENSMUSG00000106791 | 0 | NA | NA | NA | NA | NA | 4930553P18 | RIKEN cDNA 4930553P18 gene [Source:M | 0 | 0 | 0 | 0 | 0 | 0 |
| 44793 | ENSMUSG00000106794 | 0 | NA | NA | NA | NA | NA | Gm40293 | predicted gene, 40293 [Source:MGI Symb | 0 | 0 | 0 | 0 | 0 | 0 |
| 44794 | ENSMUSG00000106795 | 0 | NA | NA | NA | NA | NA | Gm43050 | predicted gene 43050 [Source:MGI Symbo | 0 | 0 | 0 | 0 | 0 | 0 |
| 44795 | ENSMUSG00000106797 | 0 | NA | NA | NA | NA | NA | Gm42936 | predicted gene 42936 [Source:MGI Symbo | 0 | 0 | 0 | 0 | 0 | 0 |
| 44796 | ENSMUSG00000106798 | 0 | NA | NA | NA | NA | NA | Gm43275 | predicted gene 43275 [Source:MGI Symbo | 0 | 0 | 0 | 0 | 0 | 0 |

| | | | | | | | | | | | | | | | |
|---|---|---|---|---|---|---|---|---|---|---|---|---|---|---|---|
| 44797 | ENSMUSG00000106799 | 0 | NA | NA | NA | NA | NA | Gm20756 | predicted gene, 20756 [Source:MGI Symb | 0 | 0 | 0 | 0 | 0 | 0 |
| 44798 | ENSMUSG00000106800 | 0 | NA | NA | NA | NA | NA | 1700030N18 | RIKEN cDNA 1700030N18 gene [Source:M | 0 | 0 | 0 | 0 | 0 | 0 |
| 44799 | ENSMUSG00000106801 | 0 | NA | NA | NA | NA | NA | Gm42852 | predicted gene 42852 [Source:MGI Symbo | 0 | 0 | 0 | 0 | 0 | 0 |
| 44800 | ENSMUSG00000106802 | 0 | NA | NA | NA | NA | NA | Gm42885 | predicted gene 42885 [Source:MGI Symbo | 0 | 0 | 0 | 0 | 0 | 0 |
| 44801 | ENSMUSG00000106803 | 0.160662974 | 1.126924175 | 4.080472857 | 0.276174898 | 0.782413736 | NA | Gm8930 | predicted gene 8930 [Source:MGI Symbol | 0 | 0 | 0 | 0.964 | 0 | 0 |
| 44802 | ENSMUSG00000106805 | 0 | NA | NA | NA | NA | NA | Gm42714 | predicted gene 42714 [Source:MGI Symbo | 0 | 0 | 0 | 0 | 0 | 0 |
| 44803 | ENSMUSG00000106806 | 0 | NA | NA | NA | NA | NA | Gm17906 | predicted gene, 17906 [Source:MGI Symb | 0 | 0 | 0 | 0 | 0 | 0 |
| 44804 | ENSMUSG00000106807 | 0 | NA | NA | NA | NA | NA | Gm10441 | predicted gene 10441 [Source:MGI Symbo | 0 | 0 | 0 | 0 | 0 | 0 |
| 44805 | ENSMUSG00000106808 | 0 | NA | NA | NA | NA | NA | Gm42745 | predicted gene 42745 [Source:MGI Symbo | 0 | 0 | 0 | 0 | 0 | 0 |
| 44806 | ENSMUSG00000106810 | 0 | NA | NA | NA | NA | NA | Gm43600 | predicted gene 43600 [Source:MGI Symbo | 0 | 0 | 0 | 0 | 0 | 0 |
| 44807 | ENSMUSG00000106812 | 0 | NA | NA | NA | NA | NA | Gm28563 | predicted gene 28563 [Source:MGI Symbo | 0 | 0 | 0 | 0 | 0 | 0 |
| 44808 | ENSMUSG00000106813 | 0 | NA | NA | NA | NA | NA | Gm7181 | predicted gene 7181 [Source:MGI Symbol | 0 | 0 | 0 | 0 | 0 | 0 |
| 44809 | ENSMUSG00000106814 | 0 | NA | NA | NA | NA | NA | Gm43501 | predicted gene 43501 [Source:MGI Symbo | 0 | 0 | 0 | 0 | 0 | 0 |
| 44810 | ENSMUSG00000106815 | 1.240088275 | -3.678064142 | 2.41935062 | -1.520269163 | 0.128443341 | NA | Gm43658 | predicted gene 43658 [Source:MGI Symbo | 2.4093 | 0 | 5.0312 | 0 | 0 | 0 |
| 44811 | ENSMUSG00000106817 | 0 | NA | NA | NA | NA | NA | Gm43441 | predicted gene 43441 [Source:MGI Symbo | 0 | 0 | 0 | 0 | 0 | 0 |
| 44812 | ENSMUSG00000106818 | 0 | NA | NA | NA | NA | NA | Gm43790 | predicted gene 43790 [Source:MGI Symbo | 0 | 0 | 0 | 0 | 0 | 0 |
| 44813 | ENSMUSG00000106819 | 0 | NA | NA | NA | NA | NA | Gm43498 | predicted gene 43498 [Source:MGI Symbo | 0 | 0 | 0 | 0 | 0 | 0 |
| 44814 | ENSMUSG00000106820 | 1.452798593 | -0.998986143 | 2.141750486 | -0.466434419 | 0.64090458 | NA | D5Ertd605e | DNA segment, Chr 5, ERATO Doi 605, expr | 4.8186 | 0 | 1.0062 | 2.8919 | 0 | 0 |
| 44815 | ENSMUSG00000106821 | 0 | NA | NA | NA | NA | NA | Gm7961 | predicted gene 7961 [Source:MGI Symbol | 0 | 0 | 0 | 0 | 0 | 0 |
| 44816 | ENSMUSG00000106822 | 0.356960287 | 0.16514264 | 3.880370363 | 0.042558474 | 0.966053499 | NA | Gm43533 | predicted gene 43533 [Source:MGI Symbo | 0 | 1.0279 | 0 | 0 | 0 | 1.1138 |
| 44817 | ENSMUSG00000106824 | 0 | NA | NA | NA | NA | NA | Gm7697 | predicted gene 7697 [Source:MGI Symbol | 0 | 0 | 0 | 0 | 0 | 0 |
| 44818 | ENSMUSG00000106825 | 22.34861025 | 0.051184611 | 0.494628271 | 0.103480965 | 0.917581257 | 0.986374671 | 2510016D11 | RIKEN cDNA 2510016D11 gene [Source:M | 19.274 | 26.726 | 20.125 | 33.739 | 14.178 | 20.049 |
| 44819 | ENSMUSG00000106826 | 0 | NA | NA | NA | NA | NA | Gm42583 | predicted gene 42583 [Source:MGI Symbo | 0 | 0 | 0 | 0 | 0 | 0 |
| 44820 | ENSMUSG00000106827 | 0 | NA | NA | NA | NA | NA | Gm7556 | predicted gene 7556 [Source:MGI Symbol | 0 | 0 | 0 | 0 | 0 | 0 |
| 44821 | ENSMUSG00000106828 | 0 | NA | NA | NA | NA | NA | Gm34648 | predicted gene, 34648 [Source:MGI Symb | 0 | 0 | 0 | 0 | 0 | 0 |
| 44822 | ENSMUSG00000106829 | 0 | NA | NA | NA | NA | NA | Gm3786 | predicted gene 3786 [Source:MGI Symbol | 0 | 0 | 0 | 0 | 0 | 0 |
| 44823 | ENSMUSG00000106831 | 3.753400805 | 0.158725313 | 1.412772678 | 0.112350215 | 0.910545728 | NA | Ube2n-ps1 | ubiquitin-conjugating enzyme E2N, pseud | 5.6217 | 0 | 5.0312 | 9.6398 | 0 | 2.2277 |
| 44824 | ENSMUSG00000106832 | 0 | NA | NA | NA | NA | NA | Gm42632 | predicted gene 42632 [Source:MGI Symbo | 0 | 0 | 0 | 0 | 0 | 0 |
| 44825 | ENSMUSG00000106833 | 0 | NA | NA | NA | NA | NA | Gm42739 | predicted gene 42739 [Source:MGI Symbo | 0 | 0 | 0 | 0 | 0 | 0 |
| 44826 | ENSMUSG00000106834 | 0.185640647 | 1.126924175 | 4.080472857 | 0.276174898 | 0.782413736 | NA | Gm10459 | predicted gene 10459 [Source:MGI Symbo | 0 | 0 | 0 | 0 | 0 | 1.1138 |
| 44827 | ENSMUSG00000106835 | 0 | NA | NA | NA | NA | NA | Gm43812 | predicted gene 43812 [Source:MGI Symbo | 0 | 0 | 0 | 0 | 0 | 0 |
| 44828 | ENSMUSG00000106836 | 0 | NA | NA | NA | NA | NA | Gm43578 | predicted gene 43578 [Source:MGI Symbo | 0 | 0 | 0 | 0 | 0 | 0 |
| 44829 | ENSMUSG00000106837 | 0 | NA | NA | NA | NA | NA | D630030B08 | RIKEN cDNA D630030B08 gene [Source:M | 0 | 0 | 0 | 0 | 0 | 0 |
| 44830 | ENSMUSG00000106838 | 29.84539215 | -0.290836131 | 0.422899959 | -0.687718514 | 0.491630059 | 0.876368691 | 1810017P11 | RIKEN cDNA 1810017P11 gene [Source:M | 22.487 | 35.977 | 40.25 | 23.135 | 32.719 | 24.505 |
| 44831 | ENSMUSG00000106839 | 0 | NA | NA | NA | NA | NA | Gm43497 | predicted gene 43497 [Source:MGI Symbo | 0 | 0 | 0 | 0 | 0 | 0 |
| 44832 | ENSMUSG00000106840 | 0 | NA | NA | NA | NA | NA | Gm42790 | predicted gene 42790 [Source:MGI Symbo | 0 | 0 | 0 | 0 | 0 | 0 |
| 44833 | ENSMUSG00000106841 | 0 | NA | NA | NA | NA | NA | Gm42656 | predicted gene 42656 [Source:MGI Symbo | 0 | 0 | 0 | 0 | 0 | 0 |
| 44834 | ENSMUSG00000106843 | 0 | NA | NA | NA | NA | NA | Gm42458 | predicted gene 42458 [Source:MGI Symbo | 0 | 0 | 0 | 0 | 0 | 0 |
| 44835 | ENSMUSG00000106845 | 0 | NA | NA | NA | NA | NA | Gm43156 | predicted gene 43156 [Source:MGI Symbo | 0 | 0 | 0 | 0 | 0 | 0 |
| 44836 | ENSMUSG00000106846 | 0 | NA | NA | NA | NA | NA | Gm42786 | predicted gene 42786 [Source:MGI Symbo | 0 | 0 | 0 | 0 | 0 | 0 |
| 44837 | ENSMUSG00000106847 | 119.4499837 | -0.214957378 | 0.241976184 | -0.888341054 | 0.374357323 | 0.809332934 | Peg13 | paternally expressed 13 [Source:MGI Sym | 124.48 | 148.02 | 112.7 | 120.5 | 81.796 | 129.21 |
| 44838 | ENSMUSG00000106849 | 0 | NA | NA | NA | NA | NA | Gm43418 | predicted gene 43418 [Source:MGI Symbo | 0 | 0 | 0 | 0 | 0 | 0 |
| 44839 | ENSMUSG00000106851 | 0 | NA | NA | NA | NA | NA | 4930421C12 | RIKEN cDNA 4930421C12 gene [Source:M | 0 | 0 | 0 | 0 | 0 | 0 |
| 44840 | ENSMUSG00000106853 | 0 | NA | NA | NA | NA | NA | Gm7492 | predicted gene 7492 [Source:MGI Symbol | 0 | 0 | 0 | 0 | 0 | 0 |
| 44841 | ENSMUSG00000106854 | 0 | NA | NA | NA | NA | NA | Gm42850 | predicted gene 42850 [Source:MGI Symbo | 0 | 0 | 0 | 0 | 0 | 0 |
| 44842 | ENSMUSG00000106855 | 0 | NA | NA | NA | NA | NA | 5330437M03 | RIKEN cDNA 5330437M03 gene [Source:M | 0 | 0 | 0 | 0 | 0 | 0 |
| 44843 | ENSMUSG00000106856 | 0 | NA | NA | NA | NA | NA | Gm43761 | predicted gene 43761 [Source:MGI Symbo | 0 | 0 | 0 | 0 | 0 | 0 |
| 44844 | ENSMUSG00000106858 | 0.963765213 | -2.11522655 | 2.746264995 | -0.770219391 | 0.441169764 | NA | Gm43844 | predicted gene 43844 [Source:MGI Symbo | 4.8186 | 0 | 0 | 0.964 | 0 | 0 |
| 44845 | ENSMUSG00000106859 | 0 | NA | NA | NA | NA | NA | Gm42737 | predicted gene 42737 [Source:MGI Symbo | 0 | 0 | 0 | 0 | 0 | 0 |
| 44846 | ENSMUSG00000106860 | 0 | NA | NA | NA | NA | NA | 1700008H02 | RIKEN cDNA 1700008H02 gene [Source:M | 0 | 0 | 0 | 0 | 0 | 0 |
| 44847 | ENSMUSG00000106862 | 0 | NA | NA | NA | NA | NA | Gm42811 | predicted gene 42811 [Source:MGI Symbo | 0 | 0 | 0 | 0 | 0 | 0 |
| 44848 | ENSMUSG00000106863 | 8.894974974 | 1.36614167 | 0.877109194 | 1.557550279 | 0.119339893 | 0.505455992 | Gm42109 | predicted gene, 42109 [Source:MGI Symb | 9.6372 | 3.0838 | 2.0125 | 23.135 | 4.3625 | 11.138 |
| 44849 | ENSMUSG00000106864 | 422.6426186 | 0.160331377 | 0.153298285 | 1.045878483 | 0.29561714 | 0.746083809 | Gtf3c2 | general transcription factor IIIC, polypepti | 354.97 | 414.25 | 429.67 | 511.87 | 430.79 | 394.3 |
| 44850 | ENSMUSG00000106865 | 0 | NA | NA | NA | NA | NA | 4930519E07 | RIKEN cDNA 4930519E07 gene [Source:M | 0 | 0 | 0 | 0 | 0 | 0 |
| 44851 | ENSMUSG00000106866 | 0 | NA | NA | NA | NA | NA | Gm43659 | predicted gene 43659 [Source:MGI Symbo | 0 | 0 | 0 | 0 | 0 | 0 |
| 44852 | ENSMUSG00000106867 | 1.830581921 | 0.443049272 | 1.49628692 | 0.296099141 | 0.76715437 | NA | Gm43800 | predicted gene 43800 [Source:MGI Symbo | 1.6062 | 1.0279 | 2.0125 | 1.928 | 2.1812 | 2.2277 |

| | | | | | | | | | | | | | | | |
|---|---|---|---|---|---|---|---|---|---|---|---|---|---|---|---|
| 44853 | ENSMUSG00000106868 | 0 | NA | NA | NA | NA | NA | Gm19798 | predicted gene, 19798 [Source:MGI Symb | 0 | 0 | 0 | 0 | 0 | 0 |
| 44854 | ENSMUSG00000106869 | 0 | NA | NA | NA | NA | NA | Gm42473 | predicted gene 42473 [Source:MGI Symbc | 0 | 0 | 0 | 0 | 0 | 0 |
| 44855 | ENSMUSG00000106870 | 18.19795489 | -0.003768091 | 0.578960543 | -0.006508373 | 0.994807106 | 0.998676967 | Gm43681 | predicted gene 43681 [Source:MGI Symbc | 11.243 | 24.67 | 19.119 | 29.883 | 10.906 | 13.366 |
| 44856 | ENSMUSG00000106871 | 1.161355031 | 0.540929775 | 2.083844867 | 0.259582555 | 0.795185794 | NA | Gm3289 | predicted gene 3289 [Source:MGI Symbol | 0.8031 | 2.0558 | 0 | 1.928 | 2.1812 | 0 |
| 44857 | ENSMUSG00000106872 | 0 | NA | NA | NA | NA | NA | Gm2189 | predicted gene 2189 [Source:MGI Symbol | 0 | 0 | 0 | 0 | 0 | 0 |
| 44858 | ENSMUSG00000106873 | 0 | NA | NA | NA | NA | NA | BC049739 | cDNA sequence BC049739 [Source:MGI Sy | 0 | 0 | 0 | 0 | 0 | 0 |
| 44859 | ENSMUSG00000106874 | 2.228785069 | -0.309121998 | 1.536805687 | -0.201145793 | 0.840584576 | NA | Gm20186 | predicted gene, 20186 [Source:MGI Symb | 2.4093 | 1.0279 | 4.025 | 4.8199 | 1.0906 | 0 |
| 44860 | ENSMUSG00000106875 | 0 | NA | NA | NA | NA | NA | Gm43093 | predicted gene 43093 [Source:MGI Symbc | 0 | 0 | 0 | 0 | 0 | 0 |
| 44861 | ENSMUSG00000106876 | 0 | NA | NA | NA | NA | NA | Gm43456 | predicted gene 43456 [Source:MGI Symbc | 0 | 0 | 0 | 0 | 0 | 0 |
| 44862 | ENSMUSG00000106878 | 0 | NA | NA | NA | NA | NA | Gm17808 | predicted gene, 17808 [Source:MGI Symb | 0 | 0 | 0 | 0 | 0 | 0 |
| 44863 | ENSMUSG00000106879 | 0 | NA | NA | NA | NA | NA | Gm2566 | predicted gene 2566 [Source:MGI Symbol | 0 | 0 | 0 | 0 | 0 | 0 |
| 44864 | ENSMUSG00000106880 | 0 | NA | NA | NA | NA | NA | 4930425K10 | RIKEN cDNA 4930425K10 gene [Source:M | 0 | 0 | 0 | 0 | 0 | 0 |
| 44865 | ENSMUSG00000106883 | 0 | NA | NA | NA | NA | NA | Gm34091 | predicted gene, 34091 [Source:MGI Symb | 0 | 0 | 0 | 0 | 0 | 0 |
| 44866 | ENSMUSG00000106884 | 0 | NA | NA | NA | NA | NA | Gm20156 | predicted gene, 20156 [Source:MGI Symb | 0 | 0 | 0 | 0 | 0 | 0 |
| 44867 | ENSMUSG00000106885 | 0 | NA | NA | NA | NA | NA | Eif3s6-ps4 | eukaryotic translation initiation factor 3, s | 0 | 0 | 0 | 0 | 0 | 0 |
| 44868 | ENSMUSG00000106887 | 0 | NA | NA | NA | NA | NA | A930005G22 | RIKEN cDNA A930005G22 gene [Source:M | 0 | 0 | 0 | 0 | 0 | 0 |
| 44869 | ENSMUSG00000106888 | 0 | NA | NA | NA | NA | NA | Gm42848 | predicted gene 42848 [Source:MGI Symbc | 0 | 0 | 0 | 0 | 0 | 0 |
| 44870 | ENSMUSG00000106889 | 76.50295646 | 0.017334879 | 0.295037458 | 0.058754843 | 0.953147377 | 0.9915542 | Gm7463 | predicted gene 7463 [Source:MGI Symbol | 53.005 | 80.178 | 95.593 | 83.866 | 82.887 | 63.489 |
| 44871 | ENSMUSG00000106891 | 4.94547906 | 0.393449181 | 1.04281646 | 0.377294755 | 0.705954569 | NA | Gm43460 | predicted gene 43460 [Source:MGI Symbc | 5.6217 | 5.1396 | 2.0125 | 5.7839 | 1.0906 | 10.025 |
| 44872 | ENSMUSG00000106892 | 2.065123018 | 0.505285867 | 1.702516072 | 0.296787722 | 0.76662858 | NA | Gm42791 | predicted gene 42791 [Source:MGI Symbc | 4.0155 | 1.0279 | 0 | 2.8919 | 0 | 4.4554 |
| 44873 | ENSMUSG00000106893 | 0 | NA | NA | NA | NA | NA | Vmn1r261 | vomeronasal 1 receptor 261 [Source:MGI | 0 | 0 | 0 | 0 | 0 | 0 |
| 44874 | ENSMUSG00000106894 | 0.371281295 | 2.09253362 | 4.031852165 | 0.519000582 | 0.603760335 | NA | Gm17130 | predicted gene 17130 [Source:MGI Symbc | 0 | 0 | 0 | 0 | 0 | 2.2277 |
| 44875 | ENSMUSG00000106895 | 0.685071968 | 0.040605487 | 2.800847395 | 0.014497572 | 0.988433016 | NA | Gm4754 | predicted gene 4754 [Source:MGI Symbol | 0 | 2.0558 | 0 | 0.964 | 1.0906 | 0 |
| 44876 | ENSMUSG00000106896 | 2.003504394 | -0.91516881 | 1.583992422 | -0.577760851 | 0.56342559 | NA | G630022F23 | RIKEN cDNA G630022F23 gene [Source:M | 0.8031 | 3.0838 | 4.025 | 1.928 | 2.1812 | 0 |
| 44877 | ENSMUSG00000106897 | 0 | NA | NA | NA | NA | NA | Gm43027 | predicted gene 43027 [Source:MGI Symbc | 0 | 0 | 0 | 0 | 0 | 0 |
| 44878 | ENSMUSG00000106898 | 0 | NA | NA | NA | NA | NA | Gm43125 | predicted gene 43125 [Source:MGI Symbc | 0 | 0 | 0 | 0 | 0 | 0 |
| 44879 | ENSMUSG00000106899 | 0 | NA | NA | NA | NA | NA | 1700013A02 | RIKEN cDNA 1700013A02 gene [Source:M | 0 | 0 | 0 | 0 | 0 | 0 |
| 44880 | ENSMUSG00000106900 | 0 | NA | NA | NA | NA | NA | Gm2040 | predicted gene 2040 [Source:MGI Symbol | 0 | 0 | 0 | 0 | 0 | 0 |
| 44881 | ENSMUSG00000106902 | 0.371281295 | 2.09253362 | 4.031852165 | 0.519000582 | 0.603760335 | NA | 4930557J02I | RIKEN cDNA 4930557J02 gene [Source:M( | 0 | 0 | 0 | 0 | 0 | 2.2277 |
| 44882 | ENSMUSG00000106904 | 6.398020355 | -0.247534009 | 0.873593171 | -0.283351585 | 0.776907338 | 0.959295521 | Gm43029 | predicted gene 43029 [Source:MGI Symbc | 9.6372 | 5.1396 | 6.0375 | 7.7118 | 7.6343 | 2.2277 |
| 44883 | ENSMUSG00000106905 | 0 | NA | NA | NA | NA | NA | Gm42741 | predicted gene 42741 [Source:MGI Symbc | 0 | 0 | 0 | 0 | 0 | 0 |
| 44884 | ENSMUSG00000106906 | 0 | NA | NA | NA | NA | NA | Gm34728 | predicted gene, 34728 [Source:MGI Symb | 0 | 0 | 0 | 0 | 0 | 0 |
| 44885 | ENSMUSG00000106909 | 0 | NA | NA | NA | NA | NA | Gm43122 | predicted gene 43122 [Source:MGI Symbc | 0 | 0 | 0 | 0 | 0 | 0 |
| 44886 | ENSMUSG00000106910 | 0 | NA | NA | NA | NA | NA | Gm16273 | predicted gene 16273 [Source:MGI Symbc | 0 | 0 | 0 | 0 | 0 | 0 |
| 44887 | ENSMUSG00000106912 | 0 | NA | NA | NA | NA | NA | Gm43785 | predicted gene 43785 [Source:MGI Symbc | 0 | 0 | 0 | 0 | 0 | 0 |
| 44888 | ENSMUSG00000106913 | 0 | NA | NA | NA | NA | NA | 9430007M0! | RIKEN cDNA 9430007M09 gene [Source:M | 0 | 0 | 0 | 0 | 0 | 0 |
| 44889 | ENSMUSG00000106914 | 0 | NA | NA | NA | NA | NA | Gm42854 | predicted gene 42854 [Source:MGI Symbc | 0 | 0 | 0 | 0 | 0 | 0 |
| 44890 | ENSMUSG00000106915 | 0 | NA | NA | NA | NA | NA | Gm42655 | predicted gene 42655 [Source:MGI Symbc | 0 | 0 | 0 | 0 | 0 | 0 |
| 44891 | ENSMUSG00000106916 | 0 | NA | NA | NA | NA | NA | Gm3089 | predicted gene 3089 [Source:MGI Symbol | 0 | 0 | 0 | 0 | 0 | 0 |
| 44892 | ENSMUSG00000106917 | 0.133850373 | -0.796629176 | 4.080472857 | -0.195229623 | 0.84521319 | NA | Gm7832 | predicted gene 7832 [Source:MGI Symbol | 0.8031 | 0 | 0 | 0 | 0 | 0 |
| 44893 | ENSMUSG00000106918 | 296.4809389 | -0.029954633 | 0.150441258 | -0.199111823 | 0.842175273 | 0.971854408 | Mrpl33 | mitochondrial ribosomal protein L33 [Sou | 322.85 | 276.51 | 298.85 | 300.76 | 280.29 | 299.62 |
| 44894 | ENSMUSG00000106924 | 0 | NA | NA | NA | NA | NA | Gm42857 | predicted gene 42857 [Source:MGI Symbc | 0 | 0 | 0 | 0 | 0 | 0 |
| 44895 | ENSMUSG00000106925 | 0 | NA | NA | NA | NA | NA | Gm20072 | predicted gene, 20072 [Source:MGI Symb | 0 | 0 | 0 | 0 | 0 | 0 |
| 44896 | ENSMUSG00000106926 | 2.053975534 | -0.451523708 | 1.458954882 | -0.309484353 | 0.756953113 | NA | Rpl7-ps7 | ribosomal protein L7, pseudogene 7 [Sour | 4.0155 | 2.0558 | 1.0062 | 1.928 | 1.0906 | 2.2277 |
| 44897 | ENSMUSG00000106927 | 4.520862304 | 0.826819292 | 1.119090453 | 0.738831512 | 0.460009314 | NA | Gm43598 | predicted gene 43598 [Source:MGI Symbc | 5.6217 | 3.0838 | 1.0062 | 9.6398 | 1.0906 | 6.6831 |
| 44898 | ENSMUSG00000106928 | 8.489208843 | 0.613145592 | 0.848913119 | 0.722271312 | 0.470127687 | 0.867051954 | Gm43860 | predicted gene 43860 [Source:MGI Symbc | 4.0155 | 4.1117 | 12.075 | 9.6398 | 3.2719 | 17.822 |
| 44899 | ENSMUSG00000106930 | 0.99233012 | 0.312367548 | 2.428448675 | 0.128628433 | 0.897651667 | NA | Gm6450 | predicted gene 6450 [Source:MGI Symbol | 1.6062 | 0 | 1.0062 | 0 | 0 | 3.3415 |
| 44900 | ENSMUSG00000106932 | 1.266764993 | 3.866237933 | 2.205052686 | 1.753353993 | 0.079541263 | NA | 4930487D11 | RIKEN cDNA 4930487D11 gene [Source:M | 0 | 0 | 0 | 0.964 | 2.1812 | 4.4554 |
| 44901 | ENSMUSG00000106933 | 1.600788597 | -4.041370155 | 2.252751525 | -1.793970667 | 0.072817851 | NA | Gm43621 | predicted gene 43621 [Source:MGI Symbc | 2.4093 | 7.1954 | 0 | 0 | 0 | 0 |
| 44902 | ENSMUSG00000106934 | 0 | NA | NA | NA | NA | NA | Gm42506 | predicted gene 42506 [Source:MGI Symbc | 0 | 0 | 0 | 0 | 0 | 0 |
| 44903 | ENSMUSG00000106935 | 0 | NA | NA | NA | NA | NA | Gm19457 | predicted gene, 19457 [Source:MGI Symb | 0 | 0 | 0 | 0 | 0 | 0 |
| 44904 | ENSMUSG00000106936 | 0 | NA | NA | NA | NA | NA | Gm7554 | predicted gene 7554 [Source:MGI Symbol | 0 | 0 | 0 | 0 | 0 | 0 |
| 44905 | ENSMUSG00000106938 | 0.167707431 | -0.796629176 | 4.080472857 | -0.195229623 | 0.84521319 | NA | Gm31314 | predicted gene, 31314 [Source:MGI Symb | 0 | 0 | 1.0062 | 0 | 0 | 0 |
| 44906 | ENSMUSG00000106939 | 0.181769715 | 1.126924175 | 4.080472857 | 0.276174898 | 0.782413736 | NA | Gm29778 | predicted gene, 29778 [Source:MGI Symb | 0 | 0 | 0 | 0 | 1.0906 | 0 |
| 44907 | ENSMUSG00000106940 | 0.674621893 | -1.426050892 | 3.091195043 | -0.461326727 | 0.644564212 | NA | Gm42930 | predicted gene 42930 [Source:MGI Symbc | 0 | 3.0838 | 0 | 0.964 | 0 | 0 |
| 44908 | ENSMUSG00000106943 | 49.64765955 | 0.319060867 | 0.308070914 | 1.03567345 | 0.300354513 | 0.751875664 | Dancr | differentiation antagonizing non-protein c | 44.974 | 46.256 | 41.256 | 53.019 | 57.803 | 54.578 |

| | | | | | | | | | | | | | | | |
|---|---|---|---|---|---|---|---|---|---|---|---|---|---|---|---|
| 44909 | ENSMUSG00000106944 | 0 | NA | NA | NA | NA | NA | Gm43843 | predicted gene 43843 [Source:MGI Symbo | 0 | 0 | 0 | 0 | 0 | 0 |
| 44910 | ENSMUSG00000106945 | 0 | NA | NA | NA | NA | NA | Gm42961 | predicted gene 42961 [Source:MGI Symbo | 0 | 0 | 0 | 0 | 0 | 0 |
| 44911 | ENSMUSG00000106946 | 0.160662974 | 1.126924175 | 4.080472857 | 0.276174898 | 0.782413736 | NA | Gm42856 | predicted gene 42856 [Source:MGI Symbo | 0 | 0 | 0 | 0.964 | 0 | 0 |
| 44912 | ENSMUSG00000106947 | 0 | NA | NA | NA | NA | NA | 1700025A08 | RIKEN cDNA 1700025A08 gene [Source:M | 0 | 0 | 0 | 0 | 0 | 0 |
| 44913 | ENSMUSG00000106948 | 0 | NA | NA | NA | NA | NA | Gm42785 | predicted gene 42785 [Source:MGI Symbo | 0 | 0 | 0 | 0 | 0 | 0 |
| 44914 | ENSMUSG00000106950 | 0 | NA | NA | NA | NA | NA | 4930478M0 | RIKEN cDNA 4930478M09 gene [Source:M | 0 | 0 | 0 | 0 | 0 | 0 |
| 44915 | ENSMUSG00000106951 | 44.79512912 | 0.165098968 | 0.363257366 | 0.454495858 | 0.649471972 | 0.925643431 | 5930430L01 | RIKEN cDNA 5930430L01 gene [Source:M | 30.518 | 45.228 | 51.318 | 51.091 | 33.809 | 56.806 |
| 44916 | ENSMUSG00000106954 | 0 | NA | NA | NA | NA | NA | Gm43226 | predicted gene 43226 [Source:MGI Symbo | 0 | 0 | 0 | 0 | 0 | 0 |
| 44917 | ENSMUSG00000106955 | 0 | NA | NA | NA | NA | NA | Gm42960 | predicted gene 42960 [Source:MGI Symbo | 0 | 0 | 0 | 0 | 0 | 0 |
| 44918 | ENSMUSG00000106956 | 0 | NA | NA | NA | NA | NA | Gm19617 | predicted gene, 19617 [Source:MGI Symb | 0 | 0 | 0 | 0 | 0 | 0 |
| 44919 | ENSMUSG00000106957 | 0 | NA | NA | NA | NA | NA | Gm43085 | predicted gene 43085 [Source:MGI Symbo | 0 | 0 | 0 | 0 | 0 | 0 |
| 44920 | ENSMUSG00000106958 | 0 | NA | NA | NA | NA | NA | Gm35960 | predicted gene, 35960 [Source:MGI Symb | 0 | 0 | 0 | 0 | 0 | 0 |
| 44921 | ENSMUSG00000106959 | 1.379617879 | -0.182671712 | 2.126686134 | -0.085895003 | 0.931549884 | NA | Gm42548 | predicted gene 42548 [Source:MGI Symbo | 2.4093 | 0 | 2.0125 | 3.8559 | 0 | 0 |
| 44922 | ENSMUSG00000106960 | 0 | NA | NA | NA | NA | NA | Gm42673 | predicted gene 42673 [Source:MGI Symbo | 0 | 0 | 0 | 0 | 0 | 0 |
| 44923 | ENSMUSG00000106961 | 0 | NA | NA | NA | NA | NA | Gm43128 | predicted gene 43128 [Source:MGI Symbo | 0 | 0 | 0 | 0 | 0 | 0 |
| 44924 | ENSMUSG00000106962 | 0.716887508 | 1.693828953 | 2.739549946 | 0.618287305 | 0.536385969 | NA | Gm43633 | predicted gene 43633 [Source:MGI Symbo | 0 | 0 | 1.0062 | 0 | 2.1812 | 1.1138 |
| 44925 | ENSMUSG00000106963 | 0 | NA | NA | NA | NA | NA | Gm8675 | predicted gene 8675 [Source:MGI Symbol | 0 | 0 | 0 | 0 | 0 | 0 |
| 44926 | ENSMUSG00000106965 | 14.08531526 | 1.041272155 | 0.667400631 | 1.560190545 | 0.118714859 | 0.504308497 | Gm18290 | predicted gene, 18290 [Source:MGI Symb | 10.44 | 3.0838 | 14.087 | 25.063 | 21.812 | 10.025 |
| 44927 | ENSMUSG00000106967 | 0.838537155 | -3.104405377 | 3.144633814 | -0.987207275 | 0.323541036 | NA | Gm42477 | predicted gene 42477 [Source:MGI Symbo | 0 | 0 | 5.0312 | 0 | 0 | 0 |
| 44928 | ENSMUSG00000106968 | 0 | NA | NA | NA | NA | NA | C78283 | expressed sequence C78283 [Source:MGI | 0 | 0 | 0 | 0 | 0 | 0 |
| 44929 | ENSMUSG00000106969 | 0 | NA | NA | NA | NA | NA | Gm42882 | predicted gene 42882 [Source:MGI Symbo | 0 | 0 | 0 | 0 | 0 | 0 |
| 44930 | ENSMUSG00000106970 | 0 | NA | NA | NA | NA | NA | Gm43377 | predicted gene 43377 [Source:MGI Symbo | 0 | 0 | 0 | 0 | 0 | 0 |
| 44931 | ENSMUSG00000106971 | 0 | NA | NA | NA | NA | NA | Gm42965 | predicted gene 42965 [Source:MGI Symbo | 0 | 0 | 0 | 0 | 0 | 0 |
| 44932 | ENSMUSG00000106973 | 0.36353943 | 2.065989254 | 4.034137301 | 0.512126658 | 0.608562368 | NA | Gm43415 | predicted gene 43415 [Source:MGI Symbo | 0 | 0 | 0 | 0 | 2.1812 | 0 |
| 44933 | ENSMUSG00000106974 | 0 | NA | NA | NA | NA | NA | Vmn1r-ps5 | vomeronasal 1 receptor, pseudogene 5 [S | 0 | 0 | 0 | 0 | 0 | 0 |
| 44934 | ENSMUSG00000106975 | 0 | NA | NA | NA | NA | NA | Olfr428-ps1 | olfactory receptor 428, pseudogene 1 [So | 0 | 0 | 0 | 0 | 0 | 0 |
| 44935 | ENSMUSG00000106976 | 0.401551119 | -2.072982726 | 4.008251979 | -0.517178744 | 0.605031388 | NA | Gm42963 | predicted gene 42963 [Source:MGI Symbo | 2.4093 | 0 | 0 | 0 | 0 | 0 |
| 44936 | ENSMUSG00000106978 | 71.80503064 | -0.364351209 | 0.290220413 | -1.255429296 | 0.20932296 | 0.65350014 | N4bp2os | NEDD4 binding protein 2, opposite strand | 85.129 | 57.563 | 99.618 | 60.731 | 66.528 | 61.261 |
| 44937 | ENSMUSG00000106980 | 11.54934701 | -0.769207663 | 0.719735051 | -1.068737256 | 0.285188079 | 0.737769696 | Gm43690 | predicted gene 43690 [Source:MGI Symbo | 11.243 | 9.2513 | 23.144 | 4.8199 | 15.269 | 5.5692 |
| 44938 | ENSMUSG00000106981 | 18.86156484 | 0.044751322 | 0.545195451 | 0.082083081 | 0.934580647 | 0.988420305 | Gm45495 | predicted gene 45495 [Source:MGI Symbo | 19.274 | 10.279 | 26.162 | 21.208 | 23.994 | 12.252 |
| 44939 | ENSMUSG00000106982 | 0 | NA | NA | NA | NA | NA | Gm42665 | predicted gene 42665 [Source:MGI Symbo | 0 | 0 | 0 | 0 | 0 | 0 |
| 44940 | ENSMUSG00000106985 | 0 | NA | NA | NA | NA | NA | Gm42761 | predicted gene 42761 [Source:MGI Symbo | 0 | 0 | 0 | 0 | 0 | 0 |
| 44941 | ENSMUSG00000106986 | 0 | NA | NA | NA | NA | NA | Gm42887 | predicted gene 42887 [Source:MGI Symbo | 0 | 0 | 0 | 0 | 0 | 0 |
| 44942 | ENSMUSG00000106987 | 0 | NA | NA | NA | NA | NA | Gm42575 | predicted gene 42575 [Source:MGI Symbo | 0 | 0 | 0 | 0 | 0 | 0 |
| 44943 | ENSMUSG00000106988 | 0.949160506 | -0.820518204 | 2.131144287 | -0.385012976 | 0.700227802 | NA | Tsg101-ps | tumor susceptibility gene 101, pseudogen | 1.6062 | 1.0279 | 1.0062 | 0.964 | 1.0906 | 0 |
| 44944 | ENSMUSG00000106989 | 0 | NA | NA | NA | NA | NA | Gm5989 | predicted pseudogene 5989 [Source:MGI | 0 | 0 | 0 | 0 | 0 | 0 |
| 44945 | ENSMUSG00000106990 | 39.34425626 | 0.328723531 | 0.354257432 | 0.927922751 | 0.353447642 | 0.793953517 | Gm42547 | predicted gene 42547 [Source:MGI Symbo | 30.518 | 33.921 | 40.25 | 36.631 | 49.078 | 45.668 |
| 44946 | ENSMUSG00000106991 | 0 | NA | NA | NA | NA | NA | Gm43399 | predicted gene 43399 [Source:MGI Symbo | 0 | 0 | 0 | 0 | 0 | 0 |
| 44947 | ENSMUSG00000106992 | 0.160662974 | 1.126924175 | 4.080472857 | 0.276174898 | 0.782413736 | NA | Gm43167 | predicted gene 43167 [Source:MGI Symbo | 0 | 0 | 0 | 0.964 | 0 | 0 |
| 44948 | ENSMUSG00000106993 | 0.503122293 | -2.366675966 | 3.785596405 | -0.625179156 | 0.531853481 | NA | Gm43417 | predicted gene 43417 [Source:MGI Symbo | 0 | 0 | 3.0187 | 0 | 0 | 0 |
| 44949 | ENSMUSG00000106994 | 0 | NA | NA | NA | NA | NA | Gm10829 | predicted gene 10829 [Source:MGI Symbo | 0 | 0 | 0 | 0 | 0 | 0 |
| 44950 | ENSMUSG00000106995 | 0 | NA | NA | NA | NA | NA | Gm33167 | predicted gene, 33167 [Source:MGI Symb | 0 | 0 | 0 | 0 | 0 | 0 |
| 44951 | ENSMUSG00000106996 | 0 | NA | NA | NA | NA | NA | Gm31049 | predicted gene, 31049 [Source:MGI Symb | 0 | 0 | 0 | 0 | 0 | 0 |
| 44952 | ENSMUSG00000106999 | 0 | NA | NA | NA | NA | NA | Gm42734 | predicted gene 42734 [Source:MGI Symbo | 0 | 0 | 0 | 0 | 0 | 0 |
| 44953 | ENSMUSG00000107000 | 14.81176336 | 0.469583201 | 0.628951397 | 0.746612858 | 0.455297284 | 0.857339509 | Gm43481 | predicted gene 43481 [Source:MGI Symbo | 8.031 | 15.419 | 14.087 | 26.991 | 7.6343 | 16.708 |
| 44954 | ENSMUSG00000107002 | 832.5923575 | -0.094257327 | 0.122910502 | -0.766877731 | 0.443154215 | 0.850769625 | 0610012G03 | RIKEN cDNA 0610012G03 gene [Source:M | 949.27 | 832.61 | 795.94 | 741.3 | 832.14 | 844.29 |
| 44955 | ENSMUSG00000107004 | 0 | NA | NA | NA | NA | NA | Gm42657 | predicted gene 42657 [Source:MGI Symbo | 0 | 0 | 0 | 0 | 0 | 0 |
| 44956 | ENSMUSG00000107005 | 0.535117793 | 1.111049431 | 3.044213722 | 0.364970903 | 0.715133126 | NA | 9030607J07 | RIKEN cDNA 9030607J07 gene [Source:M | 0 | 0 | 1.0062 | 0 | 1.0906 | 1.1138 |
| 44957 | ENSMUSG00000107006 | 0.167707431 | -0.796629176 | 4.080472857 | -0.195229623 | 0.84521319 | NA | Gm5553 | predicted gene 5553 [Source:MGI Symbol | 0 | 0 | 1.0062 | 0 | 0 | 0 |
| 44958 | ENSMUSG00000107008 | 0 | NA | NA | NA | NA | NA | Gm2762 | predicted gene 2762 [Source:MGI Symbol | 0 | 0 | 0 | 0 | 0 | 0 |
| 44959 | ENSMUSG00000107009 | 0 | NA | NA | NA | NA | NA | 6720475M2 | RIKEN cDNA 6720475M21 gene [Source:M | 0 | 0 | 0 | 0 | 0 | 0 |
| 44960 | ENSMUSG00000107010 | 0 | NA | NA | NA | NA | NA | Gm43293 | predicted gene 43293 [Source:MGI Symbo | 0 | 0 | 0 | 0 | 0 | 0 |
| 44961 | ENSMUSG00000107011 | 0 | NA | NA | NA | NA | NA | Gm42906 | predicted gene 42906 [Source:MGI Symbo | 0 | 0 | 0 | 0 | 0 | 0 |
| 44962 | ENSMUSG00000107012 | 0 | NA | NA | NA | NA | NA | 4930528J11 | RIKEN cDNA 4930528J11 gene [Source:M | 0 | 0 | 0 | 0 | 0 | 0 |
| 44963 | ENSMUSG00000107013 | 0 | NA | NA | NA | NA | NA | Gm43635 | predicted gene 43635 [Source:MGI Symbo | 0 | 0 | 0 | 0 | 0 | 0 |
| 44964 | ENSMUSG00000107014 | 0.167707431 | -0.796629176 | 4.080472857 | -0.195229623 | 0.84521319 | NA | 1700126H18 | RIKEN cDNA 1700126H18 gene [Source:M | 0 | 0 | 1.0062 | 0 | 0 | 0 |

| | | | | | | | | | | | | | | | |
|---|---|---|---|---|---|---|---|---|---|---|---|---|---|---|---|
| 44965 | ENSMUSG00000107015 | 0.556921942 | 2.679043159 | 3.638243063 | 0.736356289 | 0.461513899 | NA | 2900076G11 | RIKEN cDNA 2900076G11 gene [Source:M | 0 | 0 | 0 | 0 | 0 | 3.3415 |
| 44966 | ENSMUSG00000107017 | 1.336752193 | -1.494756203 | 2.104889155 | -0.710135353 | 0.477620205 | NA | Gm43196 | predicted gene 43196 [Source:MGI Symbo | 0.8031 | 5.1396 | 0 | 0.964 | 0 | 1.1138 |
| 44967 | ENSMUSG00000107018 | 0 | NA | NA | NA | NA | NA | Gm43694 | predicted gene 43694 [Source:MGI Symbo | 0 | 0 | 0 | 0 | 0 | 0 |
| 44968 | ENSMUSG00000107019 | 0.847770947 | -0.376623742 | 2.576917887 | -0.146152791 | 0.883800772 | NA | Gm43682 | predicted gene 43682 [Source:MGI Symbo | 0.8031 | 2.0558 | 0 | 0 | 0 | 2.2277 |
| 44969 | ENSMUSG00000107020 | 0.503122293 | -2.366675966 | 3.785596405 | -0.625179156 | 0.531853481 | NA | Gm6051 | predicted gene 6051 [Source:MGI Symbol | 0 | 0 | 3.0187 | 0 | 0 | 0 |
| 44970 | ENSMUSG00000107021 | 0 | NA | NA | NA | NA | NA | Gm42853 | predicted gene 42853 [Source:MGI Symbo | 0 | 0 | 0 | 0 | 0 | 0 |
| 44971 | ENSMUSG00000107022 | 0 | NA | NA | NA | NA | NA | Gm36669 | predicted gene, 36669 [Source:MGI Symb | 0 | 0 | 0 | 0 | 0 | 0 |
| 44972 | ENSMUSG00000107023 | 15.0958617 | -0.198515247 | 0.559589422 | -0.354751608 | 0.722775685 | 0.9447719 | Gm42715 | predicted gene 42715 [Source:MGI Symbo | 12.85 | 20.558 | 15.094 | 14.46 | 10.906 | 16.708 |
| 44973 | ENSMUSG00000107026 | 0 | NA | NA | NA | NA | NA | Gm7504 | predicted gene 7504 [Source:MGI Symbol | 0 | 0 | 0 | 0 | 0 | 0 |
| 44974 | ENSMUSG00000107027 | 0 | NA | NA | NA | NA | NA | 1700019F05 | RIKEN cDNA 1700019F05 gene [Source:M | 0 | 0 | 0 | 0 | 0 | 0 |
| 44975 | ENSMUSG00000107028 | 0 | NA | NA | NA | NA | NA | Gm5043 | predicted gene 5043 [Source:MGI Symbol | 0 | 0 | 0 | 0 | 0 | 0 |
| 44976 | ENSMUSG00000107029 | 0.267700746 | -1.497481649 | 4.057001436 | -0.369110456 | 0.712045396 | NA | Gm43123 | predicted gene 43123 [Source:MGI Symbo | 1.6062 | 0 | 0 | 0 | 0 | 0 |
| 44977 | ENSMUSG00000107030 | 0 | NA | NA | NA | NA | NA | Gm5291 | predicted gene 5291 [Source:MGI Symbol | 0 | 0 | 0 | 0 | 0 | 0 |
| 44978 | ENSMUSG00000107031 | 0 | NA | NA | NA | NA | NA | Gm40292 | predicted gene, 40292 [Source:MGI Symb | 0 | 0 | 0 | 0 | 0 | 0 |
| 44979 | ENSMUSG00000107032 | 0 | NA | NA | NA | NA | NA | Gm5301 | predicted pseudogene 5301 [Source:MGI | 0 | 0 | 0 | 0 | 0 | 0 |
| 44980 | ENSMUSG00000107033 | 0 | NA | NA | NA | NA | NA | Gm43553 | predicted gene 43553 [Source:MGI Symbo | 0 | 0 | 0 | 0 | 0 | 0 |
| 44981 | ENSMUSG00000107034 | 0 | NA | NA | NA | NA | NA | Gm19060 | predicted gene, 19060 [Source:MGI Symb | 0 | 0 | 0 | 0 | 0 | 0 |
| 44982 | ENSMUSG00000107035 | 1.509255972 | -0.098793765 | 1.741928469 | -0.056715168 | 0.954772091 | NA | Ybx1-ps2 | Y box protein 1, pseudogene 2 [Source:M | 1.6062 | 1.0279 | 2.0125 | 0 | 2.1812 | 2.2277 |
| 44983 | ENSMUSG00000107036 | 0 | NA | NA | NA | NA | NA | Gm7596 | predicted gene 7596 [Source:MGI Symbol | 0 | 0 | 0 | 0 | 0 | 0 |
| 44984 | ENSMUSG00000107037 | 1.365214248 | -0.551628455 | 2.008513699 | -0.274645104 | 0.78358891 | NA | Gm29793 | predicted gene, 29793 [Source:MGI Symb | 0.8031 | 1.0279 | 3.0187 | 0 | 0 | 3.3415 |
| 44985 | ENSMUSG00000107040 | 0 | NA | NA | NA | NA | NA | Gm43791 | predicted gene 43791 [Source:MGI Symbo | 0 | 0 | 0 | 0 | 0 | 0 |
| 44986 | ENSMUSG00000107041 | 0.167707431 | -0.796629176 | 4.080472857 | -0.195229623 | 0.84521319 | NA | Gm42735 | predicted gene 42735 [Source:MGI Symbo | 0 | 0 | 1.0062 | 0 | 0 | 0 |
| 44987 | ENSMUSG00000107042 | 0 | NA | NA | NA | NA | NA | 4921513H07 | RIKEN cDNA 4921513H07 gene [Source:M | 0 | 0 | 0 | 0 | 0 | 0 |
| 44988 | ENSMUSG00000107043 | 0.36353943 | 2.065989254 | 4.034137301 | 0.512126658 | 0.608562368 | NA | Gm42849 | predicted gene 42849 [Source:MGI Symbo | 0 | 0 | 0 | 0 | 2.1812 | 0 |
| 44989 | ENSMUSG00000107044 | 0.160662974 | 1.126924175 | 4.080472857 | 0.276174898 | 0.782413736 | NA | Gm43086 | predicted gene 43086 [Source:MGI Symbo | 0 | 0 | 0 | 0.964 | 0 | 0 |
| 44990 | ENSMUSG00000107045 | 0 | NA | NA | NA | NA | NA | Gm43636 | predicted gene 43636 [Source:MGI Symbo | 0 | 0 | 0 | 0 | 0 | 0 |
| 44991 | ENSMUSG00000107046 | 0 | NA | NA | NA | NA | NA | Gm43849 | predicted gene 43849 [Source:MGI Symbo | 0 | 0 | 0 | 0 | 0 | 0 |
| 44992 | ENSMUSG00000107047 | 0 | NA | NA | NA | NA | NA | Gm43298 | predicted gene 43298 [Source:MGI Symbo | 0 | 0 | 0 | 0 | 0 | 0 |
| 44993 | ENSMUSG00000107048 | 0 | NA | NA | NA | NA | NA | Gm43807 | predicted gene 43807 [Source:MGI Symbo | 0 | 0 | 0 | 0 | 0 | 0 |
| 44994 | ENSMUSG00000107049 | 0 | NA | NA | NA | NA | NA | Gm43771 | predicted gene 43771 [Source:MGI Symbo | 0 | 0 | 0 | 0 | 0 | 0 |
| 44995 | ENSMUSG00000107050 | 1.031479329 | -2.220845979 | 2.403244684 | -0.924103148 | 0.355432614 | NA | C030017G13 | RIKEN cDNA C030017G13 gene [Source:M | 3.2124 | 0 | 2.0125 | 0.964 | 0 | 0 |
| 44996 | ENSMUSG00000107053 | 0 | NA | NA | NA | NA | NA | 1700021F13 | RIKEN cDNA 1700021F13 gene [Source:M | 0 | 0 | 0 | 0 | 0 | 0 |
| 44997 | ENSMUSG00000107054 | 0 | NA | NA | NA | NA | NA | C730045M1! | RIKEN cDNA C730045M19 gene [Source:M | 0 | 0 | 0 | 0 | 0 | 0 |
| 44998 | ENSMUSG00000107056 | 0 | NA | NA | NA | NA | NA | Gm21048 | predicted gene, 21048 [Source:MGI Symb | 0 | 0 | 0 | 0 | 0 | 0 |
| 44999 | ENSMUSG00000107058 | 0 | NA | NA | NA | NA | NA | Gm42545 | predicted gene 42545 [Source:MGI Symbo | 0 | 0 | 0 | 0 | 0 | 0 |
| 45000 | ENSMUSG00000107059 | 0 | NA | NA | NA | NA | NA | Gm43629 | predicted gene 43629 [Source:MGI Symbo | 0 | 0 | 0 | 0 | 0 | 0 |
| 45001 | ENSMUSG00000107060 | 0 | NA | NA | NA | NA | NA | Gm38901 | predicted gene, 38901 [Source:MGI Symb | 0 | 0 | 0 | 0 | 0 | 0 |
| 45002 | ENSMUSG00000107061 | 0.349477146 | 0.165142768 | 3.911021834 | 0.042224967 | 0.966319359 | NA | Gm19590 | predicted gene, 19590 [Source:MGI Symb | 0 | 0 | 1.0062 | 0 | 1.0906 | 0 |
| 45003 | ENSMUSG00000107062 | 0 | NA | NA | NA | NA | NA | Gm43580 | predicted gene 43580 [Source:MGI Symbo | 0 | 0 | 0 | 0 | 0 | 0 |
| 45004 | ENSMUSG00000107063 | 0 | NA | NA | NA | NA | NA | Gm38554 | predicted gene, 38554 [Source:MGI Symb | 0 | 0 | 0 | 0 | 0 | 0 |
| 45005 | ENSMUSG00000107064 | 0 | NA | NA | NA | NA | NA | Gm42561 | predicted gene 42561 [Source:MGI Symbo | 0 | 0 | 0 | 0 | 0 | 0 |
| 45006 | ENSMUSG00000107065 | 0 | NA | NA | NA | NA | NA | Gm43155 | predicted gene 43155 [Source:MGI Symbo | 0 | 0 | 0 | 0 | 0 | 0 |
| 45007 | ENSMUSG00000107066 | 1.597228522 | -1.842429345 | 1.781820152 | -1.034015326 | 0.301128998 | NA | Gm4482 | predicted gene 4482 [Source:MGI Symbol | 2.4093 | 3.0838 | 2.0125 | 0.964 | 0 | 1.1138 |
| 45008 | ENSMUSG00000107067 | 0 | NA | NA | NA | NA | NA | Gm42581 | predicted gene 42581 [Source:MGI Symbo | 0 | 0 | 0 | 0 | 0 | 0 |
| 45009 | ENSMUSG00000107068 | 0 | NA | NA | NA | NA | NA | Gm42742 | predicted gene 42742 [Source:MGI Symbo | 0 | 0 | 0 | 0 | 0 | 0 |
| 45010 | ENSMUSG00000107069 | 0 | NA | NA | NA | NA | NA | Gm43772 | predicted gene 43772 [Source:MGI Symbo | 0 | 0 | 0 | 0 | 0 | 0 |
| 45011 | ENSMUSG00000107070 | 0 | NA | NA | NA | NA | NA | Gm35191 | predicted gene, 35191 [Source:MGI Symb | 0 | 0 | 0 | 0 | 0 | 0 |
| 45012 | ENSMUSG00000107071 | 0 | NA | NA | NA | NA | NA | Gm42420 | predicted gene, 42420 [Source:MGI Symb | 0 | 0 | 0 | 0 | 0 | 0 |
| 45013 | ENSMUSG00000107072 | 0 | NA | NA | NA | NA | NA | Gm43090 | predicted gene 43090 [Source:MGI Symbo | 0 | 0 | 0 | 0 | 0 | 0 |
| 45014 | ENSMUSG00000107073 | 0 | NA | NA | NA | NA | NA | 4930428D20 | RIKEN cDNA 4930428D20 gene [Source:M | 0 | 0 | 0 | 0 | 0 | 0 |
| 45015 | ENSMUSG00000107074 | 0 | NA | NA | NA | NA | NA | Gm18100 | predicted gene, 18100 [Source:MGI Symb | 0 | 0 | 0 | 0 | 0 | 0 |
| 45016 | ENSMUSG00000107075 | 0.133850373 | -0.796629176 | 4.080472857 | -0.195229623 | 0.84521319 | NA | Gm43068 | predicted gene 43068 [Source:MGI Symbo | 0.8031 | 0 | 0 | 0 | 0 | 0 |
| 45017 | ENSMUSG00000107076 | 0.367410362 | 2.078255025 | 3.800070157 | 0.54689912 | 0.58444804 | NA | Gm43480 | predicted gene 43480 [Source:MGI Symbo | 0 | 0 | 0 | 0 | 1.0906 | 1.1138 |
| 45018 | ENSMUSG00000107078 | 0 | NA | NA | NA | NA | NA | Gm36840 | predicted gene, 36840 [Source:MGI Symb | 0 | 0 | 0 | 0 | 0 | 0 |
| 45019 | ENSMUSG00000107079 | 1.085104531 | -0.299366803 | 2.316360865 | -0.12924014 | 0.897167635 | NA | Gm35648 | predicted gene, 35648 [Source:MGI Symb | 1.6062 | 0 | 2.0125 | 2.8919 | 0 | 0 |
| 45020 | ENSMUSG00000107081 | 0.160662974 | 1.126924175 | 4.080472857 | 0.276174898 | 0.782413736 | NA | Gm43701 | predicted gene 43701 [Source:MGI Symbo | 0 | 0 | 0 | 0.964 | 0 | 0 |

| | | | | | | | | | | | | | | | |
|---|---|---|---|---|---|---|---|---|---|---|---|---|---|---|---|
| 45021 | ENSMUSG00000107082 | 0 | NA | NA | NA | NA | NA | Gm36378 | predicted gene, 36378 [Source:MGI Symb | 0 | 0 | 0 | 0 | 0 | 0 |
| 45022 | ENSMUSG00000107083 | 1.371973351 | 0.913315411 | 1.837007826 | 0.497175569 | 0.619065248 | NA | Gm43313 | predicted gene 43313 [Source:MGI Symbo | 0.8031 | 2.0558 | 0 | 0.964 | 2.1812 | 2.2277 |
| 45023 | ENSMUSG00000107084 | 0 | NA | NA | NA | NA | NA | Gm43269 | predicted gene 43269 [Source:MGI Symbo | 0 | 0 | 0 | 0 | 0 | 0 |
| 45024 | ENSMUSG00000107085 | 0.17131964 | -0.796629176 | 4.080472857 | -0.195229623 | 0.84521319 | NA | Gm42883 | predicted gene 42883 [Source:MGI Symbo | 0 | 1.0279 | 0 | 0 | 0 | 0 |
| 45025 | ENSMUSG00000107086 | 10.13426157 | 0.295801322 | 0.738877457 | 0.40033881 | 0.688906987 | 0.936908509 | Gm43808 | predicted gene 43808 [Source:MGI Symbo | 12.047 | 9.2513 | 6.0375 | 20.244 | 6.5437 | 6.6831 |
| 45026 | ENSMUSG00000107087 | 0 | NA | NA | NA | NA | NA | Gm34411 | predicted gene, 34411 [Source:MGI Symb | 0 | 0 | 0 | 0 | 0 | 0 |
| 45027 | ENSMUSG00000107089 | 0 | NA | NA | NA | NA | NA | Gm43443 | predicted gene 43443 [Source:MGI Symbo | 0 | 0 | 0 | 0 | 0 | 0 |
| 45028 | ENSMUSG00000107090 | 0 | NA | NA | NA | NA | NA | Gm42884 | predicted gene 42884 [Source:MGI Symbo | 0 | 0 | 0 | 0 | 0 | 0 |
| 45029 | ENSMUSG00000107091 | 0.160662974 | 1.126924175 | 4.080472857 | 0.276174898 | 0.782413736 | NA | Gm43343 | predicted gene 43343 [Source:MGI Symbo | 0 | 0 | 0 | 0.964 | 0 | 0 |
| 45030 | ENSMUSG00000107092 | 0.17131964 | -0.796629176 | 4.080472857 | -0.195229623 | 0.84521319 | NA | Gm7993 | predicted gene 7993 [Source:MGI Symbol | 0 | 1.0279 | 0 | 0 | 0 | 0 |
| 45031 | ENSMUSG00000107093 | 0 | NA | NA | NA | NA | NA | Gm43452 | predicted gene 43452 [Source:MGI Symbo | 0 | 0 | 0 | 0 | 0 | 0 |
| 45032 | ENSMUSG00000107094 | 2.044611373 | 1.64892965 | 1.619204935 | 1.018357599 | 0.308508045 | NA | Gm43294 | predicted gene 43294 [Source:MGI Symbo | 0 | 2.0558 | 1.0062 | 4.8199 | 3.2719 | 1.1138 |
| 45033 | ENSMUSG00000107096 | 50.5617815 | 0.026329635 | 0.323381673 | 0.081419689 | 0.935108192 | 0.988420305 | Gm43597 | predicted gene 43597 [Source:MGI Symbo | 49.792 | 54.48 | 46.287 | 64.587 | 41.443 | 46.781 |
| 45034 | ENSMUSG00000107097 | 0 | NA | NA | NA | NA | NA | Gm42584 | predicted gene 42584 [Source:MGI Symbo | 0 | 0 | 0 | 0 | 0 | 0 |
| 45035 | ENSMUSG00000107098 | 0 | NA | NA | NA | NA | NA | Gm29815 | predicted gene, 29815 [Source:MGI Symb | 0 | 0 | 0 | 0 | 0 | 0 |
| 45036 | ENSMUSG00000107100 | 0 | NA | NA | NA | NA | NA | Gm18482 | predicted gene, 18482 [Source:MGI Symb | 0 | 0 | 0 | 0 | 0 | 0 |
| 45037 | ENSMUSG00000107102 | 0.96895528 | -2.08853317 | 2.292908757 | -0.910866237 | 0.362365857 | NA | Gm42726 | predicted gene 42726 [Source:MGI Symbo | 0.8031 | 1.0279 | 3.0187 | 0.964 | 0 | 0 |
| 45038 | ENSMUSG00000107103 | 0 | NA | NA | NA | NA | NA | Gm43164 | predicted gene 43164 [Source:MGI Symbo | 0 | 0 | 0 | 0 | 0 | 0 |
| 45039 | ENSMUSG00000107105 | 0.133850373 | -0.796629176 | 4.080472857 | -0.195229623 | 0.84521319 | NA | Gm15820 | predicted gene 15820 [Source:MGI Symbo | 0.8031 | 0 | 0 | 0 | 0 | 0 |
| 45040 | ENSMUSG00000107106 | 0 | NA | NA | NA | NA | NA | Gm43760 | predicted gene 43760 [Source:MGI Symbo | 0 | 0 | 0 | 0 | 0 | 0 |
| 45041 | ENSMUSG00000107108 | 0 | NA | NA | NA | NA | NA | Gm9936 | predicted gene 9936 [Source:MGI Symbol | 0 | 0 | 0 | 0 | 0 | 0 |
| 45042 | ENSMUSG00000107109 | 0 | NA | NA | NA | NA | NA | Gm42571 | predicted gene 42571 [Source:MGI Symbo | 0 | 0 | 0 | 0 | 0 | 0 |
| 45043 | ENSMUSG00000107110 | 0 | NA | NA | NA | NA | NA | Gm42931 | predicted gene 42931 [Source:MGI Symbo | 0 | 0 | 0 | 0 | 0 | 0 |
| 45044 | ENSMUSG00000107111 | 0 | NA | NA | NA | NA | NA | Gm40304 | predicted gene, 40304 [Source:MGI Symb | 0 | 0 | 0 | 0 | 0 | 0 |
| 45045 | ENSMUSG00000107112 | 0 | NA | NA | NA | NA | NA | Gm36241 | predicted gene, 36241 [Source:MGI Symb | 0 | 0 | 0 | 0 | 0 | 0 |
| 45046 | ENSMUSG00000107113 | 0 | NA | NA | NA | NA | NA | Gm43092 | predicted gene 43092 [Source:MGI Symbo | 0 | 0 | 0 | 0 | 0 | 0 |
| 45047 | ENSMUSG00000107114 | 0 | NA | NA | NA | NA | NA | Gm43440 | predicted gene 43440 [Source:MGI Symbo | 0 | 0 | 0 | 0 | 0 | 0 |
| 45048 | ENSMUSG00000107115 | 0.133850373 | -0.796629176 | 4.080472857 | -0.195229623 | 0.84521319 | NA | Gm43052 | predicted gene 43052 [Source:MGI Symbo | 0.8031 | 0 | 0 | 0 | 0 | 0 |
| 45049 | ENSMUSG00000107116 | 0.267700746 | -1.497481649 | 4.057001436 | -0.369110456 | 0.712045396 | NA | Gm43274 | predicted gene 43274 [Source:MGI Symbo | 1.6062 | 0 | 0 | 0 | 0 | 0 |
| 45050 | ENSMUSG00000107117 | 0 | NA | NA | NA | NA | NA | Gm43842 | predicted gene 43842 [Source:MGI Symbo | 0 | 0 | 0 | 0 | 0 | 0 |
| 45051 | ENSMUSG00000107118 | 2.384464567 | -0.957608916 | 1.655342182 | -0.578496051 | 0.562929263 | NA | Gm42986 | predicted gene 42986 [Source:MGI Symbo | 6.4248 | 2.0558 | 1.0062 | 4.8199 | 0 | 0 |
| 45052 | ENSMUSG00000107120 | 2.851611175 | -0.812392464 | 1.257776155 | -0.645895902 | 0.518346774 | NA | Gm43059 | predicted gene 43059 [Source:MGI Symbo | 4.8186 | 2.0558 | 4.025 | 2.8919 | 1.0906 | 2.2277 |
| 45053 | ENSMUSG00000107121 | 15.58130399 | -0.329375921 | 0.563192011 | -0.584837702 | 0.558656851 | 0.899349136 | 1110006O24 | RIKEN cDNA 1110006O24 gene [Source:M | 14.456 | 17.475 | 20.125 | 11.568 | 9.8156 | 20.049 |
| 45054 | ENSMUSG00000107122 | 0 | NA | NA | NA | NA | NA | Gm32072 | predicted gene, 32072 [Source:MGI Symb | 0 | 0 | 0 | 0 | 0 | 0 |
| 45055 | ENSMUSG00000107123 | 0 | NA | NA | NA | NA | NA | Vmn2r-ps25 | vomeronasal 2, receptor, pseudogene 25 | 0 | 0 | 0 | 0 | 0 | 0 |
| 45056 | ENSMUSG00000107125 | 0 | NA | NA | NA | NA | NA | Gm3822 | predicted gene 3822 [Source:MGI Symbol | 0 | 0 | 0 | 0 | 0 | 0 |
| 45057 | ENSMUSG00000107126 | 0 | NA | NA | NA | NA | NA | Gm40289 | predicted gene, 40289 [Source:MGI Symb | 0 | 0 | 0 | 0 | 0 | 0 |
| 45058 | ENSMUSG00000107127 | 0 | NA | NA | NA | NA | NA | Gm6719 | predicted gene 6719 [Source:MGI Symbol | 0 | 0 | 0 | 0 | 0 | 0 |
| 45059 | ENSMUSG00000107128 | 0.160662974 | 1.126924175 | 4.080472857 | 0.276174898 | 0.782413736 | NA | Gm42867 | predicted gene 42867 [Source:MGI Symbo | 0 | 0 | 0 | 0.964 | 0 | 0 |
| 45060 | ENSMUSG00000107129 | 0 | NA | NA | NA | NA | NA | Gm43762 | predicted gene 43762 [Source:MGI Symbo | 0 | 0 | 0 | 0 | 0 | 0 |
| 45061 | ENSMUSG00000107130 | 0.133850373 | -0.796629176 | 4.080472857 | -0.195229623 | 0.84521319 | NA | Gm43264 | predicted gene 43264 [Source:MGI Symbo | 0.8031 | 0 | 0 | 0 | 0 | 0 |
| 45062 | ENSMUSG00000107131 | 0 | NA | NA | NA | NA | NA | Gm43763 | predicted gene 43763 [Source:MGI Symbo | 0 | 0 | 0 | 0 | 0 | 0 |
| 45063 | ENSMUSG00000107132 | 0 | NA | NA | NA | NA | NA | Gm15997 | predicted gene 15997 [Source:MGI Symbo | 0 | 0 | 0 | 0 | 0 | 0 |
| 45064 | ENSMUSG00000107133 | 0 | NA | NA | NA | NA | NA | Gm42787 | predicted gene 42787 [Source:MGI Symbo | 0 | 0 | 0 | 0 | 0 | 0 |
| 45065 | ENSMUSG00000107134 | 1.286848981 | 0.041445174 | 2.038986096 | 0.020326364 | 0.983783024 | NA | Gm42528 | predicted gene 42528 [Source:MGI Symbo | 0.8031 | 2.0558 | 1.0062 | 3.8559 | 0 | 0 |
| 45066 | ENSMUSG00000107135 | 0 | NA | NA | NA | NA | NA | Gm43806 | predicted gene 43806 [Source:MGI Symbo | 0 | 0 | 0 | 0 | 0 | 0 |
| 45067 | ENSMUSG00000107136 | 0 | NA | NA | NA | NA | NA | Gm42466 | predicted gene 42466 [Source:MGI Symbo | 0 | 0 | 0 | 0 | 0 | 0 |
| 45068 | ENSMUSG00000107139 | 0 | NA | NA | NA | NA | NA | Gm42475 | predicted gene 42475 [Source:MGI Symbo | 0 | 0 | 0 | 0 | 0 | 0 |
| 45069 | ENSMUSG00000107140 | 0 | NA | NA | NA | NA | NA | Gm43811 | predicted gene 43811 [Source:MGI Symbo | 0 | 0 | 0 | 0 | 0 | 0 |
| 45070 | ENSMUSG00000107141 | 0 | NA | NA | NA | NA | NA | Gm42654 | predicted gene 42654 [Source:MGI Symbo | 0 | 0 | 0 | 0 | 0 | 0 |
| 45071 | ENSMUSG00000107142 | 0 | NA | NA | NA | NA | NA | 4930566F21 | RIKEN cDNA 4930566F21 gene [Source:M | 0 | 0 | 0 | 0 | 0 | 0 |
| 45072 | ENSMUSG00000107143 | 22.81180088 | -0.050693992 | 0.47686161 | -0.106307555 | 0.915338338 | 0.98625062 | Gm6598 | predicted gene 6598 [Source:MGI Symbol | 28.912 | 22.614 | 18.112 | 30.847 | 17.45 | 18.935 |
| 45073 | ENSMUSG00000107144 | 0 | NA | NA | NA | NA | NA | 4930413E15 | RIKEN cDNA 4930413E15 gene [Source:M | 0 | 0 | 0 | 0 | 0 | 0 |
| 45074 | ENSMUSG00000107145 | 0.133850373 | -0.796629176 | 4.080472857 | -0.195229623 | 0.84521319 | NA | Gm43442 | predicted gene 43442 [Source:MGI Symbo | 0.8031 | 0 | 0 | 0 | 0 | 0 |
| 45075 | ENSMUSG00000107148 | 0 | NA | NA | NA | NA | NA | Gm42888 | predicted gene 42888 [Source:MGI Symbo | 0 | 0 | 0 | 0 | 0 | 0 |
| 45076 | ENSMUSG00000107149 | 0.294513347 | 0.165143496 | 4.080472857 | 0.040471657 | 0.967717103 | NA | Gm30708 | predicted gene, 30708 [Source:MGI Symb | 0.8031 | 0 | 0 | 0.964 | 0 | 0 |

| | | | | | | | | | | | | | | | |
|---|---|---|---|---|---|---|---|---|---|---|---|---|---|---|---|
| 45077 | ENSMUSG00000107151 | 0.17131964 | -0.796629176 | 4.080472857 | -0.195229623 | 0.84521319 | NA | Gm43810 | predicted gene 43810 [Source:MGI Symbo | 0 | 1.0279 | 0 | 0 | 0 | 0 |
| 45078 | ENSMUSG00000107152 | 0 | NA | NA | NA | NA | NA | Gm43152 | predicted gene 43152 [Source:MGI Symbo | 0 | 0 | 0 | 0 | 0 | 0 |
| 45079 | ENSMUSG00000107153 | 0 | NA | NA | NA | NA | NA | Gm38404 | predicted gene, 38404 [Source:MGI Symb | 0 | 0 | 0 | 0 | 0 | 0 |
| 45080 | ENSMUSG00000107154 | 0 | NA | NA | NA | NA | NA | Gm7041 | predicted gene 7041 [Source:MGI Symbol | 0 | 0 | 0 | 0 | 0 | 0 |
| 45081 | ENSMUSG00000107155 | 0.339027071 | -1.79174149 | 3.919585585 | -0.457125237 | 0.647581032 | NA | Gm43793 | predicted gene 43793 [Source:MGI Symbo | 0 | 1.0279 | 1.0062 | 0 | 0 | 0 |
| 45082 | ENSMUSG00000107157 | 0 | NA | NA | NA | NA | NA | Gm3170 | predicted gene 3170 [Source:MGI Symbol | 0 | 0 | 0 | 0 | 0 | 0 |
| 45083 | ENSMUSG00000107158 | 0.160662974 | 1.126924175 | 4.080472857 | 0.276174898 | 0.782413736 | NA | Gm42728 | predicted gene 42728 [Source:MGI Symbo | 0 | 0 | 0 | 0.964 | 0 | 0 |
| 45084 | ENSMUSG00000107159 | 0 | NA | NA | NA | NA | NA | Gm7919 | predicted gene 7919 [Source:MGI Symbol | 0 | 0 | 0 | 0 | 0 | 0 |
| 45085 | ENSMUSG00000107160 | 0 | NA | NA | NA | NA | NA | Gm42666 | predicted gene 42666 [Source:MGI Symbo | 0 | 0 | 0 | 0 | 0 | 0 |
| 45086 | ENSMUSG00000107161 | 3.544017341 | 0.38274353 | 1.193589506 | 0.320665964 | 0.748463543 | NA | Gm43850 | predicted gene 43850 [Source:MGI Symbo | 3.2124 | 2.0558 | 4.025 | 8.6758 | 2.1812 | 1.1138 |
| 45087 | ENSMUSG00000107162 | 0 | NA | NA | NA | NA | NA | Gm18593 | predicted gene, 18593 [Source:MGI Symb | 0 | 0 | 0 | 0 | 0 | 0 |
| 45088 | ENSMUSG00000107163 | 0 | NA | NA | NA | NA | NA | Gm43496 | predicted gene 43496 [Source:MGI Symbo | 0 | 0 | 0 | 0 | 0 | 0 |
| 45089 | ENSMUSG00000107164 | 0 | NA | NA | NA | NA | NA | 4933425D22 | RIKEN cDNA 4933425D22 gene [Source:M | 0 | 0 | 0 | 0 | 0 | 0 |
| 45090 | ENSMUSG00000107165 | 1.305682109 | -1.392582258 | 2.033790705 | -0.684722501 | 0.493519047 | NA | Gm43747 | predicted gene 43747 [Source:MGI Symbo | 1.6062 | 1.0279 | 3.0187 | 0 | 2.1812 | 0 |
| 45091 | ENSMUSG00000107167 | 0.949367096 | -3.28842827 | 2.400393427 | -1.369953872 | 0.170701301 | NA | Gm34653 | predicted gene, 34653 [Source:MGI Symb | 1.6062 | 3.0838 | 1.0062 | 0 | 0 | 0 |
| 45092 | ENSMUSG00000107168 | 0.321325949 | 1.911575114 | 4.048267893 | 0.472195804 | 0.636787034 | NA | Gm42507 | predicted gene 42507 [Source:MGI Symbo | 0 | 0 | 0 | 1.928 | 0 | 0 |
| 45093 | ENSMUSG00000107169 | 0 | NA | NA | NA | NA | NA | NA | NA | 0 | 0 | 0 | 0 | 0 | 0 |
| 45094 | ENSMUSG00000107171 | 1.759437212 | 0.708855825 | 1.720102069 | 0.412101024 | 0.680265378 | NA | Gm42572 | predicted gene 42572 [Source:MGI Symbo | 0 | 1.0279 | 3.0187 | 0.964 | 1.0906 | 4.4554 |
| 45095 | ENSMUSG00000107172 | 0 | NA | NA | NA | NA | NA | 4933430H06 | RIKEN cDNA 4933430H06 gene [Source:M | 0 | 0 | 0 | 0 | 0 | 0 |
| 45096 | ENSMUSG00000107173 | 0 | NA | NA | NA | NA | NA | Gm43266 | predicted gene 43266 [Source:MGI Symbo | 0 | 0 | 0 | 0 | 0 | 0 |
| 45097 | ENSMUSG00000107176 | 18.20315369 | 0.252380608 | 0.509381126 | 0.495465173 | 0.6202718 | 0.917977899 | Gm9794 | predicted pseudogene 9794 [Source:MGI | 14.456 | 16.447 | 19.119 | 25.063 | 18.541 | 15.594 |
| 45098 | ENSMUSG00000107177 | 0 | NA | NA | NA | NA | NA | Gm43845 | predicted gene 43845 [Source:MGI Symbo | 0 | 0 | 0 | 0 | 0 | 0 |
| 45099 | ENSMUSG00000107178 | 2.268801159 | 0.765722923 | 1.563530413 | 0.489739705 | 0.624318102 | NA | Gm42531 | predicted gene 42531 [Source:MGI Symbo | 0 | 4.1117 | 1.0062 | 1.928 | 5.4531 | 1.1138 |
| 45100 | ENSMUSG00000107179 | 0 | NA | NA | NA | NA | NA | Gm43069 | predicted gene 43069 [Source:MGI Symbo | 0 | 0 | 0 | 0 | 0 | 0 |
| 45101 | ENSMUSG00000107180 | 0 | NA | NA | NA | NA | NA | Gm43638 | predicted gene 43638 [Source:MGI Symbo | 0 | 0 | 0 | 0 | 0 | 0 |
| 45102 | ENSMUSG00000107181 | 0 | NA | NA | NA | NA | NA | Gm17768 | predicted gene, 17768 [Source:MGI Symb | 0 | 0 | 0 | 0 | 0 | 0 |
| 45103 | ENSMUSG00000107182 | 2.026911377 | -1.555317066 | 1.763476243 | -0.881960884 | 0.377797961 | NA | Gm43268 | predicted gene 43268 [Source:MGI Symbo | 0 | 4.1117 | 5.0312 | 1.928 | 1.0906 | 0 |
| 45104 | ENSMUSG00000107183 | 0 | NA | NA | NA | NA | NA | Gm18452 | predicted gene, 18452 [Source:MGI Symb | 0 | 0 | 0 | 0 | 0 | 0 |
| 45105 | ENSMUSG00000107184 | 0 | NA | NA | NA | NA | NA | Gm18222 | predicted gene, 18222 [Source:MGI Symb | 0 | 0 | 0 | 0 | 0 | 0 |
| 45106 | ENSMUSG00000107185 | 4.397017818 | -0.089677705 | 1.112197546 | -0.080631094 | 0.935735337 | NA | Gm42642 | predicted gene 42642 [Source:MGI Symbo | 2.4093 | 8.2233 | 3.0187 | 2.8919 | 8.7249 | 1.1138 |
| 45107 | ENSMUSG00000107187 | 0 | NA | NA | NA | NA | NA | Gm7663 | predicted gene 7663 [Source:MGI Symbol | 0 | 0 | 0 | 0 | 0 | 0 |
| 45108 | ENSMUSG00000107189 | 0 | NA | NA | NA | NA | NA | Gm42689 | predicted gene 42689 [Source:MGI Symbo | 0 | 0 | 0 | 0 | 0 | 0 |
| 45109 | ENSMUSG00000107191 | 0 | NA | NA | NA | NA | NA | Gm43579 | predicted gene 43579 [Source:MGI Symbo | 0 | 0 | 0 | 0 | 0 | 0 |
| 45110 | ENSMUSG00000107192 | 0 | NA | NA | NA | NA | NA | Gm19583 | predicted gene, 19583 [Source:MGI Symb | 0 | 0 | 0 | 0 | 0 | 0 |
| 45111 | ENSMUSG00000107193 | 0.589026695 | 0.241472922 | 3.228838524 | 0.074786311 | 0.940384733 | NA | Gm43700 | predicted gene 43700 [Source:MGI Symbo | 1.6062 | 0 | 0 | 1.928 | 0 | 0 |
| 45112 | ENSMUSG00000107194 | 0 | NA | NA | NA | NA | NA | Gm43173 | predicted gene 43173 [Source:MGI Symbo | 0 | 0 | 0 | 0 | 0 | 0 |
| 45113 | ENSMUSG00000107195 | 2.057149058 | -0.945565575 | 1.466912876 | -0.644595592 | 0.519189292 | NA | 4930589O11 | RIKEN cDNA 4930589O11 gene [Source:M | 4.0155 | 2.0558 | 2.0125 | 0.964 | 2.1812 | 1.1138 |
| 45114 | ENSMUSG00000107197 | 0 | NA | NA | NA | NA | NA | Gm43312 | predicted gene 43312 [Source:MGI Symbo | 0 | 0 | 0 | 0 | 0 | 0 |
| 45115 | ENSMUSG00000107198 | 89.44331084 | -1.203103332 | 0.36576715 | -3.289260205 | 0.001004511 | 0.020935517 | Gm19619 | predicted gene, 19619 [Source:MGI Symb | 202.38 | 108.96 | 62.387 | 55.911 | 47.987 | 59.034 |
| 45116 | ENSMUSG00000107199 | 0 | NA | NA | NA | NA | NA | Gm42550 | predicted gene 42550 [Source:MGI Symbo | 0 | 0 | 0 | 0 | 0 | 0 |
| 45117 | ENSMUSG00000107200 | 6.487507602 | -0.841356924 | 0.868533155 | -0.968710198 | 0.332689803 | 0.77523112 | Gm43039 | predicted gene 43039 [Source:MGI Symbo | 9.6372 | 6.1675 | 9.0562 | 1.928 | 5.4531 | 6.6831 |
| 45118 | ENSMUSG00000107201 | 4.897509011 | -0.625765478 | 0.960776526 | -0.651312206 | 0.514844971 | NA | 5930420M1 | RIKEN cDNA 5930420M18 gene [Source:N | 5.6217 | 7.1954 | 5.0312 | 3.8559 | 5.4531 | 2.2277 |
| 45119 | ENSMUSG00000107203 | 3.000377652 | 0.116435938 | 1.337660899 | 0.087044435 | 0.930636192 | NA | Gm7542 | predicted gene 7542 [Source:MGI Symbol | 1.6062 | 2.0558 | 5.0312 | 3.8559 | 5.4531 | 0 |
| 45120 | ENSMUSG00000107204 | 0 | NA | NA | NA | NA | NA | Gm43067 | predicted gene 43067 [Source:MGI Symbo | 0 | 0 | 0 | 0 | 0 | 0 |
| 45121 | ENSMUSG00000107205 | 2.912998243 | -0.061194428 | 1.327671968 | -0.046091527 | 0.963237299 | NA | Gm42576 | predicted gene 42576 [Source:MGI Symbo | 0.8031 | 6.1675 | 2.0125 | 1.928 | 5.4531 | 1.1138 |
| 45122 | ENSMUSG00000107206 | 0 | NA | NA | NA | NA | NA | Gm42736 | predicted gene 42736 [Source:MGI Symbo | 0 | 0 | 0 | 0 | 0 | 0 |
| 45123 | ENSMUSG00000107207 | 0 | NA | NA | NA | NA | NA | Gm42897 | predicted gene 42897 [Source:MGI Symbo | 0 | 0 | 0 | 0 | 0 | 0 |
| 45124 | ENSMUSG00000107208 | 0 | NA | NA | NA | NA | NA | D130004A15 | RIKEN cDNA D130004A15 gene [Source:M | 0 | 0 | 0 | 0 | 0 | 0 |
| 45125 | ENSMUSG00000107209 | 0 | NA | NA | NA | NA | NA | Gm42599 | predicted gene 42599 [Source:MGI Symbo | 0 | 0 | 0 | 0 | 0 | 0 |
| 45126 | ENSMUSG00000107211 | 1.14212931 | -3.574822843 | 2.305041306 | -1.550871489 | 0.120932484 | NA | Gm5864 | predicted gene 5864 [Source:MGI Symbol | 4.8186 | 1.0279 | 1.0062 | 0 | 0 | 0 |
| 45127 | ENSMUSG00000107212 | 0 | NA | NA | NA | NA | NA | Gm5302 | predicted gene 5302 [Source:MGI Symbol | 0 | 0 | 0 | 0 | 0 | 0 |
| 45128 | ENSMUSG00000107213 | 0 | NA | NA | NA | NA | NA | Gm3302 | predicted gene 3302 [Source:MGI Symbol | 0 | 0 | 0 | 0 | 0 | 0 |
| 45129 | ENSMUSG00000107214 | 0 | NA | NA | NA | NA | NA | 4930500F04 | RIKEN cDNA 4930500F04 gene [Source:M | 0 | 0 | 0 | 0 | 0 | 0 |
| 45130 | ENSMUSG00000107215 | 7.418733238 | -0.875139012 | 0.823048486 | -1.063289742 | 0.287650576 | 0.73951397 | Gm43197 | predicted gene 43197 [Source:MGI Symbo | 14.456 | 8.2233 | 6.0375 | 4.8199 | 7.6343 | 3.3415 |
| 45131 | ENSMUSG00000107216 | 3.26518306 | -0.405794263 | 1.313929508 | -0.308840209 | 0.757443081 | NA | 2210412B16 | RIKEN cDNA 2210412B16 gene [Source:M | 8.031 | 0 | 3.0187 | 1.928 | 3.2719 | 3.3415 |
| 45132 | ENSMUSG00000107217 | 0 | NA | NA | NA | NA | NA | 4930524B17 | RIKEN cDNA 4930524B17 gene [Source:M | 0 | 0 | 0 | 0 | 0 | 0 |

| | | | | | | | | | | | | | | | |
|---|---|---|---|---|---|---|---|---|---|---|---|---|---|---|---|
| 45133 | ENSMUSG00000107218 | 0 | NA | NA | NA | NA | NA | Gm42530 | predicted gene 42530 [Source:MGI Symbo | 0 | 0 | 0 | 0 | 0 | 0 |
| 45134 | ENSMUSG00000107219 | 0 | NA | NA | NA | NA | NA | Gm42738 | predicted gene 42738 [Source:MGI Symbo | 0 | 0 | 0 | 0 | 0 | 0 |
| 45135 | ENSMUSG00000107220 | 0 | NA | NA | NA | NA | NA | Gm9555 | predicted gene 9555 [Source:MGI Symbol | 0 | 0 | 0 | 0 | 0 | 0 |
| 45136 | ENSMUSG00000107222 | 0 | NA | NA | NA | NA | NA | Gm43198 | predicted gene 43198 [Source:MGI Symbo | 0 | 0 | 0 | 0 | 0 | 0 |
| 45137 | ENSMUSG00000107223 | 0 | NA | NA | NA | NA | NA | Gm18327 | predicted gene, 18327 [Source:MGI Symb | 0 | 0 | 0 | 0 | 0 | 0 |
| 45138 | ENSMUSG00000107225 | 2.246666152 | -0.750692007 | 1.497314672 | -0.50135888 | 0.616118574 | NA | Gm43637 | predicted gene 43637 [Source:MGI Symbo | 2.4093 | 4.1117 | 2.0125 | 3.8559 | 1.0906 | 0 |
| 45139 | ENSMUSG00000107226 | 0 | NA | NA | NA | NA | NA | Gm43594 | predicted gene 43594 [Source:MGI Symbo | 0 | 0 | 0 | 0 | 0 | 0 |
| 45140 | ENSMUSG00000107227 | 2.336004456 | -0.768794834 | 1.519301279 | -0.506018684 | 0.612843533 | NA | Gm42559 | predicted gene 42559 [Source:MGI Symbo | 0.8031 | 2.0558 | 6.0375 | 2.8919 | 0 | 2.2277 |
| 45141 | ENSMUSG00000107230 | 7.568145518 | 0.307649708 | 0.775916857 | 0.396498291 | 0.691737469 | 0.938106998 | Gm19265 | predicted gene, 19265 [Source:MGI Symb | 8.031 | 7.1954 | 5.0312 | 8.6758 | 10.906 | 5.5692 |
| 45142 | ENSMUSG00000107232 | 0 | NA | NA | NA | NA | NA | 1700017L05I | RIKEN cDNA 1700017L05 gene [Source:M( | 0 | 0 | 0 | 0 | 0 | 0 |
| 45143 | ENSMUSG00000107233 | 0 | NA | NA | NA | NA | NA | Gm42959 | predicted gene 42959 [Source:MGI Symbo | 0 | 0 | 0 | 0 | 0 | 0 |
| 45144 | ENSMUSG00000107234 | 0 | NA | NA | NA | NA | NA | Gm43371 | predicted gene 43371 [Source:MGI Symbo | 0 | 0 | 0 | 0 | 0 | 0 |
| 45145 | ENSMUSG00000107237 | 0 | NA | NA | NA | NA | NA | Gm29926 | predicted gene, 29926 [Source:MGI Symb | 0 | 0 | 0 | 0 | 0 | 0 |
| 45146 | ENSMUSG00000107238 | 0 | NA | NA | NA | NA | NA | Gm42758 | predicted gene 42758 [Source:MGI Symbo | 0 | 0 | 0 | 0 | 0 | 0 |
| 45147 | ENSMUSG00000107240 | 0 | NA | NA | NA | NA | NA | Gm43231 | predicted gene 43231 [Source:MGI Symbo | 0 | 0 | 0 | 0 | 0 | 0 |
| 45148 | ENSMUSG00000107241 | 0 | NA | NA | NA | NA | NA | Gm43129 | predicted gene 43129 [Source:MGI Symbo | 0 | 0 | 0 | 0 | 0 | 0 |
| 45149 | ENSMUSG00000107243 | 0 | NA | NA | NA | NA | NA | Gm43631 | predicted gene 43631 [Source:MGI Symbo | 0 | 0 | 0 | 0 | 0 | 0 |
| 45150 | ENSMUSG00000107244 | 0.301557804 | -1.64335916 | 4.042814367 | -0.406488899 | 0.684383412 | NA | Gm43699 | predicted gene 43699 [Source:MGI Symbo | 0.8031 | 0 | 1.0062 | 0 | 0 | 0 |
| 45151 | ENSMUSG00000107245 | 0 | NA | NA | NA | NA | NA | Gm42803 | predicted gene 42803 [Source:MGI Symbo | 0 | 0 | 0 | 0 | 0 | 0 |
| 45152 | ENSMUSG00000107246 | 0.62992821 | -1.312600165 | 2.851194982 | -0.460368433 | 0.645251789 | NA | Gm42560 | predicted gene 42560 [Source:MGI Symbo | 0.8031 | 0 | 2.0125 | 0.964 | 0 | 0 |
| 45153 | ENSMUSG00000107247 | 0 | NA | NA | NA | NA | NA | Gm43227 | predicted gene 43227 [Source:MGI Symbo | 0 | 0 | 0 | 0 | 0 | 0 |
| 45154 | ENSMUSG00000107248 | 0 | NA | NA | NA | NA | NA | Gm43851 | predicted gene 43851 [Source:MGI Symbo | 0 | 0 | 0 | 0 | 0 | 0 |
| 45155 | ENSMUSG00000107251 | 2.268238207 | -1.76290038 | 1.526648787 | -1.154751764 | 0.248192095 | NA | Gm43102 | predicted gene 43102 [Source:MGI Symbo | 2.4093 | 5.1396 | 3.0187 | 1.928 | 0 | 1.1138 |
| 45156 | ENSMUSG00000107252 | 2.615595599 | 2.808406407 | 1.605645758 | 1.749082195 | 0.080276813 | NA | Gm5767 | predicted gene 5767 [Source:MGI Symbol | 0 | 2.0558 | 0 | 4.8199 | 4.3625 | 4.4554 |
| 45157 | ENSMUSG00000107253 | 0 | NA | NA | NA | NA | NA | Gm43634 | predicted gene 43634 [Source:MGI Symbo | 0 | 0 | 0 | 0 | 0 | 0 |
| 45158 | ENSMUSG00000107254 | 0 | NA | NA | NA | NA | NA | Gm43769 | predicted gene 43769 [Source:MGI Symbo | 0 | 0 | 0 | 0 | 0 | 0 |
| 45159 | ENSMUSG00000107255 | 0.66192371 | 1.647602865 | 2.590567345 | 0.636000785 | 0.52477591 | NA | Gm43416 | predicted gene 43416 [Source:MGI Symbo | 0.8031 | 0 | 0 | 0.964 | 1.0906 | 1.1138 |
| 45160 | ENSMUSG00000107256 | 0 | NA | NA | NA | NA | NA | Gm13821 | predicted gene 13821 [Source:MGI Symbo | 0 | 0 | 0 | 0 | 0 | 0 |
| 45161 | ENSMUSG00000107257 | 0.181769715 | 1.126924175 | 4.080472857 | 0.276174898 | 0.782413736 | NA | Gm43028 | predicted gene 43028 [Source:MGI Symbo | 0 | 0 | 0 | 0 | 1.0906 | 0 |
| 45162 | ENSMUSG00000107258 | 0 | NA | NA | NA | NA | NA | Olfr1401-ps1 | olfactory receptor 1401, pseudogene 1 [S | 0 | 0 | 0 | 0 | 0 | 0 |
| 45163 | ENSMUSG00000107259 | 0.535401493 | -2.48596965 | 3.684117979 | -0.674780141 | 0.499815459 | NA | Gm5298 | predicted gene 5298 [Source:MGI Symbol | 3.2124 | 0 | 0 | 0 | 0 | 0 |
| 45164 | ENSMUSG00000107261 | 0.710308366 | 0.13927508 | 2.769785472 | 0.050283707 | 0.959896308 | NA | Trmt112-ps1 | tRNA methyltransferase 11-2, pseudogen | 0 | 1.0279 | 1.0062 | 0 | 0 | 2.2277 |
| 45165 | ENSMUSG00000107262 | 0 | NA | NA | NA | NA | NA | Gm18753 | predicted gene, 18753 [Source:MGI Symb | 0 | 0 | 0 | 0 | 0 | 0 |
| 45166 | ENSMUSG00000107263 | 0 | NA | NA | NA | NA | NA | Gm42471 | predicted gene 42471 [Source:MGI Symbo | 0 | 0 | 0 | 0 | 0 | 0 |
| 45167 | ENSMUSG00000107265 | 0 | NA | NA | NA | NA | NA | Gm15469 | predicted gene 15469 [Source:MGI Symbo | 0 | 0 | 0 | 0 | 0 | 0 |
| 45168 | ENSMUSG00000107266 | 0 | NA | NA | NA | NA | NA | Gm43698 | predicted gene 43698 [Source:MGI Symbo | 0 | 0 | 0 | 0 | 0 | 0 |
| 45169 | ENSMUSG00000107270 | 0 | NA | NA | NA | NA | NA | Tgif1-ps | TGFB-induced factor homeobox 1, pseudo | 0 | 0 | 0 | 0 | 0 | 0 |
| 45170 | ENSMUSG00000107271 | 0 | NA | NA | NA | NA | NA | BC028471 | cDNA sequence BC028471 [Source:MGI Sy | 0 | 0 | 0 | 0 | 0 | 0 |
| 45171 | ENSMUSG00000107272 | 0.167707431 | -0.796629176 | 4.080472857 | -0.195229623 | 0.84521319 | NA | Gm34583 | predicted gene, 34583 [Source:MGI Symb | 0 | 0 | 1.0062 | 0 | 0 | 0 |
| 45172 | ENSMUSG00000107276 | 1.576684734 | 1.906083427 | 1.929623865 | 0.987800504 | 0.32325036 | NA | Gm42858 | predicted gene 42858 [Source:MGI Symbo | 0 | 2.0558 | 0 | 1.928 | 4.3625 | 1.1138 |
| 45173 | ENSMUSG00000107277 | 24.85080572 | 0.938907909 | 0.518489687 | 1.810851659 | 0.070163817 | 0.383030326 | Gm10461 | predicted gene 10461 [Source:MGI Symbo | 17.668 | 19.53 | 14.087 | 55.911 | 19.631 | 22.277 |
| 45174 | ENSMUSG00000107278 | 5.712923411 | 0.098699479 | 0.912692566 | 0.108140992 | 0.913883852 | 0.98625062 | Gm42600 | predicted gene 42600 [Source:MGI Symbo | 6.4248 | 2.0558 | 8.05 | 6.7478 | 6.5437 | 4.4554 |
| 45175 | ENSMUSG00000107279 | 0 | NA | NA | NA | NA | NA | Gm43450 | predicted gene 43450 [Source:MGI Symbo | 0 | 0 | 0 | 0 | 0 | 0 |
| 45176 | ENSMUSG00000107280 | 0 | NA | NA | NA | NA | NA | Gm9495 | predicted gene 9495 [Source:MGI Symbol | 0 | 0 | 0 | 0 | 0 | 0 |
| 45177 | ENSMUSG00000107282 | 0 | NA | NA | NA | NA | NA | Gm43630 | predicted gene 43630 [Source:MGI Symbo | 0 | 0 | 0 | 0 | 0 | 0 |
| 45178 | ENSMUSG00000107283 | 438.1193504 | 0.089300143 | 0.142556566 | 0.626419011 | 0.531040143 | 0.888794797 | Mpv17 | MpV17 mitochondrial inner membrane pr | 469.81 | 414.25 | 388.41 | 409.69 | 480.96 | 465.59 |
| 45179 | ENSMUSG00000107284 | 0 | NA | NA | NA | NA | NA | Gm43280 | predicted gene 43280 [Source:MGI Symbo | 0 | 0 | 0 | 0 | 0 | 0 |
| 45180 | ENSMUSG00000107285 | 0 | NA | NA | NA | NA | NA | Gm3176 | predicted gene 3176 [Source:MGI Symbol | 0 | 0 | 0 | 0 | 0 | 0 |
| 45181 | ENSMUSG00000107286 | 0 | NA | NA | NA | NA | NA | Gm43788 | predicted gene 43788 [Source:MGI Symbo | 0 | 0 | 0 | 0 | 0 | 0 |
| 45182 | ENSMUSG00000107287 | 0 | NA | NA | NA | NA | NA | Gm43458 | predicted gene 43458 [Source:MGI Symbo | 0 | 0 | 0 | 0 | 0 | 0 |
| 45183 | ENSMUSG00000107288 | 0.808472266 | -1.769372749 | 2.633653003 | -0.671832146 | 0.501690561 | NA | Gm42903 | predicted gene 42903 [Source:MGI Symbo | 0.8031 | 3.0838 | 0 | 0.964 | 0 | 0 |
| 45184 | ENSMUSG00000107289 | 0 | NA | NA | NA | NA | NA | Gm43026 | predicted gene 43026 [Source:MGI Symbo | 0 | 0 | 0 | 0 | 0 | 0 |
| 45185 | ENSMUSG00000107290 | 0 | NA | NA | NA | NA | NA | Gm43282 | predicted gene 43282 [Source:MGI Symbo | 0 | 0 | 0 | 0 | 0 | 0 |
| 45186 | ENSMUSG00000107291 | 0.160662974 | 1.126924175 | 4.080472857 | 0.276174898 | 0.782413736 | NA | 3110031N09 | RIKEN cDNA 3110031N09 gene [Source:M | 0 | 0 | 0 | 0.964 | 0 | 0 |
| 45187 | ENSMUSG00000107292 | 0 | NA | NA | NA | NA | NA | Gm42733 | predicted gene 42733 [Source:MGI Symbo | 0 | 0 | 0 | 0 | 0 | 0 |
| 45188 | ENSMUSG00000107295 | 0 | NA | NA | NA | NA | NA | Gm42896 | predicted gene 42896 [Source:MGI Symbo | 0 | 0 | 0 | 0 | 0 | 0 |

| | | | | | | | | | | | | | | | |
|---|---|---|---|---|---|---|---|---|---|---|---|---|---|---|---|
| 45189 | ENSMUSG00000107296 | 12.47627866 | 0.187406196 | 0.710591152 | 0.263732803 | 0.791985817 | 0.961434599 | Gm43500 | predicted gene 43500 [Source:MGI Symbo | 12.85 | 4.1117 | 18.112 | 22.171 | 9.8156 | 7.7969 |
| 45190 | ENSMUSG00000107299 | 0 | NA | NA | NA | NA | NA | Gm43228 | predicted gene 43228 [Source:MGI Symbo | 0 | 0 | 0 | 0 | 0 | 0 |
| 45191 | ENSMUSG00000107300 | 0 | NA | NA | NA | NA | NA | Gm43279 | predicted gene 43279 [Source:MGI Symbo | 0 | 0 | 0 | 0 | 0 | 0 |
| 45192 | ENSMUSG00000107301 | 0 | NA | NA | NA | NA | NA | Gm43688 | predicted gene 43688 [Source:MGI Symbo | 0 | 0 | 0 | 0 | 0 | 0 |
| 45193 | ENSMUSG00000107303 | 0 | NA | NA | NA | NA | NA | Gm18861 | predicted gene, 18861 [Source:MGI Symb | 0 | 0 | 0 | 0 | 0 | 0 |
| 45194 | ENSMUSG00000107304 | 6.652247502 | 1.278514553 | 0.894038519 | 1.430044149 | 0.152704348 | 0.568523988 | Gm43775 | predicted gene 43775 [Source:MGI Symbo | 6.4248 | 1.0279 | 4.025 | 9.6398 | 6.5437 | 12.252 |
| 45195 | ENSMUSG00000107305 | 0 | NA | NA | NA | NA | NA | Gm43262 | predicted gene 43262 [Source:MGI Symbo | 0 | 0 | 0 | 0 | 0 | 0 |
| 45196 | ENSMUSG00000107306 | 0.671009685 | -1.413087452 | 2.800223337 | -0.504633839 | 0.613816038 | NA | Gm42577 | predicted gene 42577 [Source:MGI Symbo | 0 | 2.0558 | 1.0062 | 0.964 | 0 | 0 |
| 45197 | ENSMUSG00000107307 | 0 | NA | NA | NA | NA | NA | Gm42935 | predicted gene 42935 [Source:MGI Symbo | 0 | 0 | 0 | 0 | 0 | 0 |
| 45198 | ENSMUSG00000107308 | 0 | NA | NA | NA | NA | NA | Gm43234 | predicted gene 43234 [Source:MGI Symbo | 0 | 0 | 0 | 0 | 0 | 0 |
| 45199 | ENSMUSG00000107309 | 0 | NA | NA | NA | NA | NA | Gm43232 | predicted gene 43232 [Source:MGI Symbo | 0 | 0 | 0 | 0 | 0 | 0 |
| 45200 | ENSMUSG00000107310 | 0 | NA | NA | NA | NA | NA | Gm7452 | predicted pseudogene 7452 [Source:MGI | 0 | 0 | 0 | 0 | 0 | 0 |
| 45201 | ENSMUSG00000107312 | 0 | NA | NA | NA | NA | NA | Gm43229 | predicted gene 43229 [Source:MGI Symbo | 0 | 0 | 0 | 0 | 0 | 0 |
| 45202 | ENSMUSG00000107313 | 3.57523281 | 0.852461502 | 1.119341473 | 0.761574124 | 0.446314223 | NA | Gm43332 | predicted gene 43332 [Source:MGI Symbo | 1.6062 | 2.0558 | 4.025 | 3.8559 | 5.4531 | 4.4554 |
| 45203 | ENSMUSG00000107314 | 0 | NA | NA | NA | NA | NA | Gm20488 | predicted gene 20488 [Source:MGI Symbo | 0 | 0 | 0 | 0 | 0 | 0 |
| 45204 | ENSMUSG00000107315 | 0 | NA | NA | NA | NA | NA | BC080696 | cDNA sequence BC080696 [Source:MGI Sy | 0 | 0 | 0 | 0 | 0 | 0 |
| 45205 | ENSMUSG00000107316 | 0 | NA | NA | NA | NA | NA | Gm42964 | predicted gene 42964 [Source:MGI Symbo | 0 | 0 | 0 | 0 | 0 | 0 |
| 45206 | ENSMUSG00000107317 | 0 | NA | NA | NA | NA | NA | Gm19719 | predicted gene, 19719 [Source:MGI Symb | 0 | 0 | 0 | 0 | 0 | 0 |
| 45207 | ENSMUSG00000107318 | 0 | NA | NA | NA | NA | NA | Gm30003 | predicted gene, 30003 [Source:MGI Symb | 0 | 0 | 0 | 0 | 0 | 0 |
| 45208 | ENSMUSG00000107319 | 4.100184228 | 0.077762517 | 1.018194159 | 0.076372975 | 0.93912237 | NA | Gm42670 | predicted gene 42670 [Source:MGI Symbo | 4.8186 | 3.0838 | 4.025 | 3.8559 | 4.3625 | 4.4554 |
| 45209 | ENSMUSG00000107320 | 4.366645142 | 0.166195938 | 1.04352702 | 0.159263665 | 0.873461147 | NA | Gm42549 | predicted gene 42549 [Source:MGI Symbo | 3.2124 | 4.1117 | 5.0312 | 2.8919 | 8.7249 | 2.2277 |
| 45210 | ENSMUSG00000107322 | 0 | NA | NA | NA | NA | NA | 4933439J24I | RIKEN cDNA 4933439J24 gene [Source:MG | 0 | 0 | 0 | 0 | 0 | 0 |
| 45211 | ENSMUSG00000107325 | 3.939707854 | -2.220034361 | 1.251422744 | -1.774008321 | 0.076061774 | NA | Gm43425 | predicted gene 43425 [Source:MGI Symbo | 3.2124 | 10.279 | 6.0375 | 1.928 | 2.1812 | 0 |
| 45212 | ENSMUSG00000107326 | 0 | NA | NA | NA | NA | NA | Gm43038 | predicted gene 43038 [Source:MGI Symbo | 0 | 0 | 0 | 0 | 0 | 0 |
| 45213 | ENSMUSG00000107327 | 0 | NA | NA | NA | NA | NA | Gm7478 | predicted gene 7478 [Source:MGI Symbol | 0 | 0 | 0 | 0 | 0 | 0 |
| 45214 | ENSMUSG00000107331 | 40.54402129 | 0.182292681 | 0.386273773 | 0.471926115 | 0.636979527 | 0.923478764 | Gm42732 | predicted gene 42732 [Source:MGI Symbo | 32.124 | 28.782 | 53.331 | 53.983 | 32.719 | 42.326 |
| 45215 | ENSMUSG00000107333 | 0 | NA | NA | NA | NA | NA | Gm43008 | predicted gene 43008 [Source:MGI Symbo | 0 | 0 | 0 | 0 | 0 | 0 |
| 45216 | ENSMUSG00000107334 | 0 | NA | NA | NA | NA | NA | Gm7538 | predicted gene 7538 [Source:MGI Symbol | 0 | 0 | 0 | 0 | 0 | 0 |
| 45217 | ENSMUSG00000107335 | 2.378577858 | 0.09189951 | 1.512088228 | 0.060776553 | 0.951537164 | NA | Gm43372 | predicted gene 43372 [Source:MGI Symbo | 0.8031 | 6.1675 | 0 | 2.8919 | 2.1812 | 2.2277 |
| 45218 | ENSMUSG00000107336 | 1.456127102 | 0.486251466 | 1.686043828 | 0.28839788 | 0.77304219 | NA | Gm43461 | predicted gene 43461 [Source:MGI Symbo | 1.6062 | 1.0279 | 1.0062 | 2.8919 | 1.0906 | 1.1138 |
| 45219 | ENSMUSG00000107337 | 0 | NA | NA | NA | NA | NA | Gm43596 | predicted gene 43596 [Source:MGI Symbo | 0 | 0 | 0 | 0 | 0 | 0 |
| 45220 | ENSMUSG00000107340 | 0 | NA | NA | NA | NA | NA | Gm42789 | predicted gene 42789 [Source:MGI Symbo | 0 | 0 | 0 | 0 | 0 | 0 |
| 45221 | ENSMUSG00000107341 | 0 | NA | NA | NA | NA | NA | 1700065J11I | RIKEN cDNA 1700065J11 gene [Source:MG | 0 | 0 | 0 | 0 | 0 | 0 |
| 45222 | ENSMUSG00000107344 | 0 | NA | NA | NA | NA | NA | Gm2559 | predicted gene 2559 [Source:MGI Symbol | 0 | 0 | 0 | 0 | 0 | 0 |
| 45223 | ENSMUSG00000107346 | 0 | NA | NA | NA | NA | NA | Gm43225 | predicted gene 43225 [Source:MGI Symbo | 0 | 0 | 0 | 0 | 0 | 0 |
| 45224 | ENSMUSG00000107348 | 0 | NA | NA | NA | NA | NA | Gm6468 | predicted gene 6468 [Source:MGI Symbol | 0 | 0 | 0 | 0 | 0 | 0 |
| 45225 | ENSMUSG00000107349 | 0 | NA | NA | NA | NA | NA | Gm6655 | predicted gene 6655 [Source:MGI Symbol | 0 | 0 | 0 | 0 | 0 | 0 |
| 45226 | ENSMUSG00000107350 | 0.167707431 | -0.796629176 | 4.080472857 | -0.195229623 | 0.84521319 | NA | Gm19610 | predicted gene, 19610 [Source:MGI Symb | 0 | 0 | 1.0062 | 0 | 0 | 0 |
| 45227 | ENSMUSG00000107351 | 0.36353943 | 2.065989254 | 4.034137301 | 0.512126658 | 0.608562368 | NA | Gm43764 | predicted gene 43764 [Source:MGI Symbo | 0 | 0 | 0 | 0 | 2.1812 | 0 |
| 45228 | ENSMUSG00000107352 | 26.05462929 | 0.355692965 | 0.436568055 | 0.814748035 | 0.415216551 | 0.836001897 | Gm43660 | predicted gene 43660 [Source:MGI Symbo | 20.881 | 19.53 | 28.175 | 26.991 | 23.994 | 36.757 |
| 45229 | ENSMUSG00000107353 | 0 | NA | NA | NA | NA | NA | 4930430O22 | RIKEN cDNA 4930430O22 gene [Source:M | 0 | 0 | 0 | 0 | 0 | 0 |
| 45230 | ENSMUSG00000107354 | 0.462220778 | -0.685400455 | 3.191475091 | -0.21475977 | 0.82995462 | NA | Gm43040 | predicted gene 43040 [Source:MGI Symbo | 0.8031 | 0 | 1.0062 | 0.964 | 0 | 0 |
| 45231 | ENSMUSG00000107355 | 1.10982348 | -0.258217128 | 1.977413818 | -0.130583253 | 0.896104992 | NA | AI839979 | expressed sequence AI839979 [Source:MG | 1.6062 | 1.0279 | 1.0062 | 1.928 | 1.0906 | 0 |
| 45232 | ENSMUSG00000107356 | 0 | NA | NA | NA | NA | NA | Gm18118 | predicted gene, 18118 [Source:MGI Symb | 0 | 0 | 0 | 0 | 0 | 0 |
| 45233 | ENSMUSG00000107357 | 0 | NA | NA | NA | NA | NA | Gm42662 | predicted gene 42662 [Source:MGI Symbo | 0 | 0 | 0 | 0 | 0 | 0 |
| 45234 | ENSMUSG00000107358 | 0.181769715 | 1.126924175 | 4.080472857 | 0.276174898 | 0.782413736 | NA | Gm43168 | predicted gene 43168 [Source:MGI Symbo | 0 | 0 | 0 | 0 | 1.0906 | 0 |
| 45235 | ENSMUSG00000107359 | 0 | NA | NA | NA | NA | NA | Gm43295 | predicted gene 43295 [Source:MGI Symbo | 0 | 0 | 0 | 0 | 0 | 0 |
| 45236 | ENSMUSG00000107360 | 0 | NA | NA | NA | NA | NA | Gm42558 | predicted gene 42558 [Source:MGI Symbo | 0 | 0 | 0 | 0 | 0 | 0 |
| 45237 | ENSMUSG00000107362 | 0.992155673 | -1.012754205 | 2.706122858 | -0.374245464 | 0.708221702 | NA | Gm40309 | predicted gene, 40309 [Source:MGI Symb | 0 | 0 | 4.025 | 1.928 | 0 | 0 |
| 45238 | ENSMUSG00000107363 | 0 | NA | NA | NA | NA | NA | Gm43373 | predicted gene 43373 [Source:MGI Symbo | 0 | 0 | 0 | 0 | 0 | 0 |
| 45239 | ENSMUSG00000107364 | 0.321325949 | 1.911575114 | 4.048267893 | 0.472195804 | 0.636787034 | NA | Gm43051 | predicted gene 43051 [Source:MGI Symbo | 0 | 0 | 0 | 1.928 | 0 | 0 |
| 45240 | ENSMUSG00000107365 | 0 | NA | NA | NA | NA | NA | Gm43479 | predicted gene 43479 [Source:MGI Symbo | 0 | 0 | 0 | 0 | 0 | 0 |
| 45241 | ENSMUSG00000107368 | 1.479275361 | -0.255071331 | 1.798431719 | -0.141829867 | 0.887214392 | NA | 1700018A23 | RIKEN cDNA 1700018A23 gene [Source:M | 0.8031 | 3.0838 | 1.0062 | 2.8919 | 1.0906 | 0 |
| 45242 | ENSMUSG00000107369 | 2.889254888 | -0.382800725 | 1.335005565 | -0.286740921 | 0.774310702 | NA | Gstm2-ps1 | glutathione S-transferase mu 2 (muscle), p | 5.6217 | 3.0838 | 1.0062 | 0.964 | 1.0906 | 5.5692 |
| 45243 | ENSMUSG00000107370 | 0 | NA | NA | NA | NA | NA | Gm43588 | predicted gene 43588 [Source:MGI Symbo | 0 | 0 | 0 | 0 | 0 | 0 |
| 45244 | ENSMUSG00000107371 | 0 | NA | NA | NA | NA | NA | Gm43784 | predicted gene 43784 [Source:MGI Symbo | 0 | 0 | 0 | 0 | 0 | 0 |

| | | | | | | | | | | | | | | | |
|---|---|---|---|---|---|---|---|---|---|---|---|---|---|---|---|
| 45245 | ENSMUSG00000107373 | 0 | NA | NA | NA | NA | NA | Gm42668 | predicted gene 42668 [Source:MGI Symbo | 0 | 0 | 0 | 0 | 0 | 0 |
| 45246 | ENSMUSG00000107374 | 1.434968228 | -0.940318474 | 2.116368348 | -0.444307568 | 0.65682023 | NA | Gm43172 | predicted gene 43172 [Source:MGI Symbo | 1.6062 | 4.1117 | 0 | 2.8919 | 0 | 0 |
| 45247 | ENSMUSG00000107375 | 0 | NA | NA | NA | NA | NA | Gm43084 | predicted gene 43084 [Source:MGI Symbo | 0 | 0 | 0 | 0 | 0 | 0 |
| 45248 | ENSMUSG00000107376 | 0 | NA | NA | NA | NA | NA | Gm42661 | predicted gene 42661 [Source:MGI Symbo | 0 | 0 | 0 | 0 | 0 | 0 |
| 45249 | ENSMUSG00000107377 | 0 | NA | NA | NA | NA | NA | Gm42801 | predicted gene 42801 [Source:MGI Symbo | 0 | 0 | 0 | 0 | 0 | 0 |
| 45250 | ENSMUSG00000107378 | 0 | NA | NA | NA | NA | NA | Gm18995 | predicted gene, 18995 [Source:MGI Symb | 0 | 0 | 0 | 0 | 0 | 0 |
| 45251 | ENSMUSG00000107379 | 0 | NA | NA | NA | NA | NA | Gm43126 | predicted gene 43126 [Source:MGI Symbo | 0 | 0 | 0 | 0 | 0 | 0 |
| 45252 | ENSMUSG00000107380 | 0 | NA | NA | NA | NA | NA | Vmn1r-ps6 | vomeronasal 1 receptor, pseudogene 6 [S | 0 | 0 | 0 | 0 | 0 | 0 |
| 45253 | ENSMUSG00000107383 | 13.10491961 | 0.231194958 | 0.659871675 | 0.350363512 | 0.726065906 | 0.945513394 | Gm4366 | predicted gene 4366 [Source:MGI Symbol | 9.6372 | 20.558 | 6.0375 | 12.532 | 9.8156 | 20.049 |
| 45254 | ENSMUSG00000107384 | 0 | NA | NA | NA | NA | NA | Gm42557 | predicted gene 42557 [Source:MGI Symbo | 0 | 0 | 0 | 0 | 0 | 0 |
| 45255 | ENSMUSG00000107385 | 0 | NA | NA | NA | NA | NA | C330024D21 | RIKEN cDNA C330024D21 gene [Source:M | 0 | 0 | 0 | 0 | 0 | 0 |
| 45256 | ENSMUSG00000107386 | 0 | NA | NA | NA | NA | NA | Gm42800 | predicted gene 42800 [Source:MGI Symbo | 0 | 0 | 0 | 0 | 0 | 0 |
| 45257 | ENSMUSG00000107387 | 0 | NA | NA | NA | NA | NA | 5430435K18 | RIKEN cDNA 5430435K18 gene [Source:M | 0 | 0 | 0 | 0 | 0 | 0 |
| 45258 | ENSMUSG00000107388 | 0.321325949 | 1.911575114 | 4.048267893 | 0.472195804 | 0.636787034 | NA | Gm42788 | predicted gene 42788 [Source:MGI Symbo | 0 | 0 | 0 | 1.928 | 0 | 0 |
| 45259 | ENSMUSG00000107389 | 0 | NA | NA | NA | NA | NA | Gm43124 | predicted gene 43124 [Source:MGI Symbo | 0 | 0 | 0 | 0 | 0 | 0 |
| 45260 | ENSMUSG00000107390 | 7.112675397 | 1.659215178 | 0.904707829 | 1.83397902 | 0.066657108 | 0.371533109 | Gm43323 | predicted gene 43323 [Source:MGI Symbo | 3.2124 | 4.1117 | 3.0187 | 20.244 | 7.6343 | 4.4554 |
| 45261 | ENSMUSG00000107392 | 0 | NA | NA | NA | NA | NA | Pramel35 | PRAME like 35 [Source:MGI Symbol;Acc:M | 0 | 0 | 0 | 0 | 0 | 0 |
| 45262 | ENSMUSG00000107394 | 0 | NA | NA | NA | NA | NA | 1700012J15I | RIKEN cDNA 1700012J15 gene [Source:M | 0 | 0 | 0 | 0 | 0 | 0 |
| 45263 | ENSMUSG00000107395 | 0.506734502 | -2.372847156 | 3.338060886 | -0.710845978 | 0.477179685 | NA | Gm44331 | predicted gene, 44331 [Source:MGI Symb | 0 | 1.0279 | 2.0125 | 0 | 0 | 0 |
| 45264 | ENSMUSG00000107396 | 0 | NA | NA | NA | NA | NA | 4930425L21I | RIKEN cDNA 4930425L21 gene [Source:M | 0 | 0 | 0 | 0 | 0 | 0 |
| 45265 | ENSMUSG00000107397 | 0 | NA | NA | NA | NA | NA | 4930402H05 | RIKEN cDNA 4930402H05 gene [Source:M | 0 | 0 | 0 | 0 | 0 | 0 |
| 45266 | ENSMUSG00000107398 | 0 | NA | NA | NA | NA | NA | Gm44306 | predicted gene, 44306 [Source:MGI Symb | 0 | 0 | 0 | 0 | 0 | 0 |
| 45267 | ENSMUSG00000107399 | 0 | NA | NA | NA | NA | NA | | olfactory receptor 1544, pseudogene 1 | 0 | 0 | 0 | 0 | 0 | 0 |
| 45268 | ENSMUSG00000107400 | 0 | NA | NA | NA | NA | NA | Olfr797-ps1 | olfactory receptor 797, pseudogene 1 [So | 0 | 0 | 0 | 0 | 0 | 0 |
| 45269 | ENSMUSG00000107401 | 0 | NA | NA | NA | NA | NA | Gm6312 | predicted gene 6312 [Source:MGI Symbol | 0 | 0 | 0 | 0 | 0 | 0 |
| 45270 | ENSMUSG00000107402 | 10.20878254 | -0.063765014 | 0.681783858 | -0.09352673 | 0.925485116 | 0.987546253 | 4732416N19 | RIKEN cDNA 4732416N19 gene [Source:M | 12.047 | 8.2233 | 11.069 | 14.46 | 6.5437 | 8.9108 |
| 45271 | ENSMUSG00000107403 | 0 | NA | NA | NA | NA | NA | Gm8820 | predicted gene 8820 [Source:MGI Symbol | 0 | 0 | 0 | 0 | 0 | 0 |
| 45272 | ENSMUSG00000107404 | 0 | NA | NA | NA | NA | NA | Gm44125 | predicted gene, 44125 [Source:MGI Symb | 0 | 0 | 0 | 0 | 0 | 0 |
| 45273 | ENSMUSG00000107405 | 0 | NA | NA | NA | NA | NA | Gm44115 | predicted gene, 44115 [Source:MGI Symb | 0 | 0 | 0 | 0 | 0 | 0 |
| 45274 | ENSMUSG00000107406 | 0 | NA | NA | NA | NA | NA | 1700040L08I | RIKEN cDNA 1700040L08 gene [Source:M | 0 | 0 | 0 | 0 | 0 | 0 |
| 45275 | ENSMUSG00000107407 | 0 | NA | NA | NA | NA | NA | Gm44159 | predicted gene, 44159 [Source:MGI Symb | 0 | 0 | 0 | 0 | 0 | 0 |
| 45276 | ENSMUSG00000107408 | 0 | NA | NA | NA | NA | NA | 1700065L07I | RIKEN cDNA 1700065L07 gene [Source:M | 0 | 0 | 0 | 0 | 0 | 0 |
| 45277 | ENSMUSG00000107409 | 0 | NA | NA | NA | NA | NA | Gm43954 | predicted gene, 43954 [Source:MGI Symb | 0 | 0 | 0 | 0 | 0 | 0 |
| 45278 | ENSMUSG00000107410 | 0 | NA | NA | NA | NA | NA | Gm44023 | predicted gene, 44023 [Source:MGI Symb | 0 | 0 | 0 | 0 | 0 | 0 |
| 45279 | ENSMUSG00000107411 | 1.548870496 | -3.999065953 | 2.015830823 | -1.983830145 | 0.047274778 | NA | Gm19040 | predicted gene, 19040 [Source:MGI Symb | 3.2124 | 2.0558 | 4.025 | 0 | 0 | 0 |
| 45280 | ENSMUSG00000107412 | 0.654905883 | -1.312574656 | 2.816093566 | -0.466097672 | 0.641145589 | NA | Gm20383 | predicted gene, 20383 [Source:MGI Symb | 0.8031 | 0 | 2.0125 | 0 | 0 | 1.1138 |
| 45281 | ENSMUSG00000107413 | 0 | NA | NA | NA | NA | NA | Gm20553 | predicted gene, 20553 [Source:MGI Symb | 0 | 0 | 0 | 0 | 0 | 0 |
| 45282 | ENSMUSG00000107414 | 0 | NA | NA | NA | NA | NA | 6330419E04 | RIKEN cDNA 6330419E04 gene [Source:M | 0 | 0 | 0 | 0 | 0 | 0 |
| 45283 | ENSMUSG00000107415 | 0 | NA | NA | NA | NA | NA | Gm43929 | predicted gene, 43929 [Source:MGI Symb | 0 | 0 | 0 | 0 | 0 | 0 |
| 45284 | ENSMUSG00000107416 | 0 | NA | NA | NA | NA | NA | Gm44030 | predicted gene, 44030 [Source:MGI Symb | 0 | 0 | 0 | 0 | 0 | 0 |
| 45285 | ENSMUSG00000107417 | 0 | NA | NA | NA | NA | NA | Olfr1383 | olfactory receptor 1383 [Source:MGI Sym | 0 | 0 | 0 | 0 | 0 | 0 |
| 45286 | ENSMUSG00000107418 | 0 | NA | NA | NA | NA | NA | Gm4782 | predicted gene 4782 [Source:MGI Symbol | 0 | 0 | 0 | 0 | 0 | 0 |
| 45287 | ENSMUSG00000107419 | 0 | NA | NA | NA | NA | NA | Gm38839 | predicted gene, 38839 [Source:MGI Symb | 0 | 0 | 0 | 0 | 0 | 0 |
| 45288 | ENSMUSG00000107420 | 0 | NA | NA | NA | NA | NA | Gm43910 | predicted gene, 43910 [Source:MGI Symb | 0 | 0 | 0 | 0 | 0 | 0 |
| 45289 | ENSMUSG00000107421 | 0 | NA | NA | NA | NA | NA | Gm44177 | predicted gene, 44177 [Source:MGI Symb | 0 | 0 | 0 | 0 | 0 | 0 |
| 45290 | ENSMUSG00000107422 | 0 | NA | NA | NA | NA | NA | Gm44129 | predicted gene, 44129 [Source:MGI Symb | 0 | 0 | 0 | 0 | 0 | 0 |
| 45291 | ENSMUSG00000107423 | 0 | NA | NA | NA | NA | NA | Gm43977 | predicted gene, 43977 [Source:MGI Symb | 0 | 0 | 0 | 0 | 0 | 0 |
| 45292 | ENSMUSG00000107424 | 0 | NA | NA | NA | NA | NA | Gm18839 | predicted gene, 18839 [Source:MGI Symb | 0 | 0 | 0 | 0 | 0 | 0 |
| 45293 | ENSMUSG00000107425 | 0.17131964 | -0.796629176 | 4.080472857 | -0.195229623 | 0.84521319 | NA | Gm44216 | predicted gene, 44216 [Source:MGI Symb | 0 | 1.0279 | 0 | 0 | 0 | 0 |
| 45294 | ENSMUSG00000107427 | 0 | NA | NA | NA | NA | NA | Gm43948 | predicted gene, 43948 [Source:MGI Symb | 0 | 0 | 0 | 0 | 0 | 0 |
| 45295 | ENSMUSG00000107428 | 1.844850795 | -0.087524048 | 1.517453576 | -0.057678238 | 0.954004929 | NA | Gm44154 | predicted gene, 44154 [Source:MGI Symb | 1.6062 | 3.0838 | 1.0062 | 0.964 | 2.1812 | 2.2277 |
| 45296 | ENSMUSG00000107429 | 0 | NA | NA | NA | NA | NA | Gm44206 | predicted gene, 44206 [Source:MGI Symb | 0 | 0 | 0 | 0 | 0 | 0 |
| 45297 | ENSMUSG00000107430 | 0.965523032 | -0.944628791 | 2.267086424 | -0.416670834 | 0.67691919 | NA | Gm31532 | predicted gene, 31532 [Source:MGI Symb | 0.8031 | 2.0558 | 1.0062 | 1.928 | 0 | 0 |
| 45298 | ENSMUSG00000107431 | 0 | NA | NA | NA | NA | NA | Gm44114 | predicted gene, 44114 [Source:MGI Symb | 0 | 0 | 0 | 0 | 0 | 0 |
| 45299 | ENSMUSG00000107432 | 0.181769715 | 1.126924175 | 4.080472857 | 0.276174898 | 0.782413736 | NA | Gm36189 | predicted gene, 36189 [Source:MGI Symb | 0 | 0 | 0 | 0 | 1.0906 | 0 |
| 45300 | ENSMUSG00000107433 | 0 | NA | NA | NA | NA | NA | Gm43882 | predicted gene, 43882 [Source:MGI Symb | 0 | 0 | 0 | 0 | 0 | 0 |

| | | | | | | | | | | | | | | | |
|---|---|---|---|---|---|---|---|---|---|---|---|---|---|---|---|
| 45301 | ENSMUSG00000107434 | 0 | NA | NA | NA | NA | NA | Gm4640 | predicted gene 4640 [Source:MGI Symbol | 0 | 0 | 0 | 0 | 0 | 0 |
| 45302 | ENSMUSG00000107435 | 0 | NA | NA | NA | NA | NA | Gm19908 | predicted gene, 19908 [Source:MGI Symb | 0 | 0 | 0 | 0 | 0 | 0 |
| 45303 | ENSMUSG00000107436 | 0 | NA | NA | NA | NA | NA | Gm44416 | predicted gene, 44416 [Source:MGI Symb | 0 | 0 | 0 | 0 | 0 | 0 |
| 45304 | ENSMUSG00000107438 | 0 | NA | NA | NA | NA | NA | Gm43945 | predicted gene, 43945 [Source:MGI Symb | 0 | 0 | 0 | 0 | 0 | 0 |
| 45305 | ENSMUSG00000107439 | 0.342639279 | -1.805976596 | 4.028547714 | -0.448294702 | 0.653940524 | NA | Gm45060 | predicted gene 45060 [Source:MGI Symbo | 0 | 2.0558 | 0 | 0 | 0 | 0 |
| 45306 | ENSMUSG00000107440 | 1.497312316 | -1.827087666 | 1.954847722 | -0.934644497 | 0.349971539 | NA | Gm40377 | predicted gene, 40377 [Source:MGI Symb | 4.0155 | 1.0279 | 2.0125 | 1.928 | 0 | 0 |
| 45307 | ENSMUSG00000107441 | 0 | NA | NA | NA | NA | NA | Gm31747 | predicted gene, 31747 [Source:MGI Symb | 0 | 0 | 0 | 0 | 0 | 0 |
| 45308 | ENSMUSG00000107442 | 0 | NA | NA | NA | NA | NA | Gm43881 | predicted gene, 43881 [Source:MGI Symb | 0 | 0 | 0 | 0 | 0 | 0 |
| 45309 | ENSMUSG00000107443 | 0 | NA | NA | NA | NA | NA | Gm7468 | predicted gene 7468 [Source:MGI Symbol | 0 | 0 | 0 | 0 | 0 | 0 |
| 45310 | ENSMUSG00000107445 | 0 | NA | NA | NA | NA | NA | Gm44449 | predicted gene, 44449 [Source:MGI Symb | 0 | 0 | 0 | 0 | 0 | 0 |
| 45311 | ENSMUSG00000107446 | 0 | NA | NA | NA | NA | NA | Gm43906 | predicted gene, 43906 [Source:MGI Symb | 0 | 0 | 0 | 0 | 0 | 0 |
| 45312 | ENSMUSG00000107447 | 0 | NA | NA | NA | NA | NA | Gm44054 | predicted gene, 44054 [Source:MGI Symb | 0 | 0 | 0 | 0 | 0 | 0 |
| 45313 | ENSMUSG00000107448 | 0 | NA | NA | NA | NA | NA | Gm43947 | predicted gene, 43947 [Source:MGI Symb | 0 | 0 | 0 | 0 | 0 | 0 |
| 45314 | ENSMUSG00000107449 | 0 | NA | NA | NA | NA | NA | Gm19165 | predicted gene, 19165 [Source:MGI Symb | 0 | 0 | 0 | 0 | 0 | 0 |
| 45315 | ENSMUSG00000107450 | 0 | NA | NA | NA | NA | NA | Gm2522 | predicted gene 2522 [Source:MGI Symbol | 0 | 0 | 0 | 0 | 0 | 0 |
| 45316 | ENSMUSG00000107451 | 99.7292366 | -0.231843873 | 0.254243332 | -0.911897555 | 0.361822649 | 0.800840096 | Gm44421 | predicted gene, 44421 [Source:MGI Symb | 88.341 | 116.15 | 118.74 | 71.334 | 105.79 | 98.018 |
| 45317 | ENSMUSG00000107452 | 0 | NA | NA | NA | NA | NA | Gm18215 | predicted gene, 18215 [Source:MGI Symb | 0 | 0 | 0 | 0 | 0 | 0 |
| 45318 | ENSMUSG00000107453 | 0 | NA | NA | NA | NA | NA | Gm44145 | predicted gene, 44145 [Source:MGI Symb | 0 | 0 | 0 | 0 | 0 | 0 |
| 45319 | ENSMUSG00000107454 | 0 | NA | NA | NA | NA | NA | Gm43875 | predicted gene, 43875 [Source:MGI Symb | 0 | 0 | 0 | 0 | 0 | 0 |
| 45320 | ENSMUSG00000107455 | 2.40188199 | -0.852491143 | 1.354662631 | -0.629301439 | 0.529151729 | NA | Gm43868 | predicted gene, 43868 [Source:MGI Symb | 3.2124 | 3.0838 | 3.0187 | 2.8919 | 1.0906 | 1.1138 |
| 45321 | ENSMUSG00000107456 | 0 | NA | NA | NA | NA | NA | Gm10400 | predicted gene 10400 [Source:MGI Symbo | 0 | 0 | 0 | 0 | 0 | 0 |
| 45322 | ENSMUSG00000107457 | 0 | NA | NA | NA | NA | NA | Gm19173 | predicted gene, 19173 [Source:MGI Symb | 0 | 0 | 0 | 0 | 0 | 0 |
| 45323 | ENSMUSG00000107458 | 2.832339362 | -0.526339775 | 1.385999933 | -0.379754546 | 0.704127626 | NA | Gm44040 | predicted gene, 44040 [Source:MGI Symb | 4.0155 | 2.0558 | 4.025 | 5.7839 | 0 | 1.1138 |
| 45324 | ENSMUSG00000107459 | 0 | NA | NA | NA | NA | NA | Gm5110 | predicted gene 5110 [Source:MGI Symbol | 0 | 0 | 0 | 0 | 0 | 0 |
| 45325 | ENSMUSG00000107460 | 2.537716762 | 0.753869275 | 1.336038994 | 0.564256941 | 0.572579281 | NA | Gm44186 | predicted gene, 44186 [Source:MGI Symb | 1.6062 | 2.0558 | 2.0125 | 2.8919 | 1.0906 | 5.5692 |
| 45326 | ENSMUSG00000107461 | 0.528279927 | -0.837071592 | 3.369085264 | -0.248456636 | 0.803781118 | NA | Gm44045 | predicted gene, 44045 [Source:MGI Symb | 0 | 2.0558 | 0 | 0 | 0 | 1.1138 |
| 45327 | ENSMUSG00000107462 | 0 | NA | NA | NA | NA | NA | Gm7384 | predicted gene 7384 [Source:MGI Symbol | 0 | 0 | 0 | 0 | 0 | 0 |
| 45328 | ENSMUSG00000107463 | 0 | NA | NA | NA | NA | NA | Gm44294 | predicted gene, 44294 [Source:MGI Symb | 0 | 0 | 0 | 0 | 0 | 0 |
| 45329 | ENSMUSG00000107464 | 0 | NA | NA | NA | NA | NA | Gm44150 | predicted gene, 44150 [Source:MGI Symb | 0 | 0 | 0 | 0 | 0 | 0 |
| 45330 | ENSMUSG00000107465 | 0 | NA | NA | NA | NA | NA | Gm5990 | predicted gene 5990 [Source:MGI Symbol | 0 | 0 | 0 | 0 | 0 | 0 |
| 45331 | ENSMUSG00000107466 | 0 | NA | NA | NA | NA | NA | Gm44425 | predicted gene, 44425 [Source:MGI Symb | 0 | 0 | 0 | 0 | 0 | 0 |
| 45332 | ENSMUSG00000107467 | 0 | NA | NA | NA | NA | NA | Gm8992 | predicted gene 8992 [Source:MGI Symbol | 0 | 0 | 0 | 0 | 0 | 0 |
| 45333 | ENSMUSG00000107468 | 0.160662974 | 1.126924175 | 4.080472857 | 0.276174898 | 0.782413736 | NA | 5730507A11 | RIKEN cDNA 5730507A11 gene [Source:M | 0 | 0 | 0 | 0.964 | 0 | 0 |
| 45334 | ENSMUSG00000107469 | 0 | NA | NA | NA | NA | NA | Olfr436-ps1 | olfactory receptor 436, pseudogene 1 [Sou | 0 | 0 | 0 | 0 | 0 | 0 |
| 45335 | ENSMUSG00000107470 | 2.841176115 | 1.784788369 | 1.482724945 | 1.20372182 | 0.228697116 | NA | Gm3375 | predicted gene 3375 [Source:MGI Symbol | 0.8031 | 1.0279 | 2.0125 | 0 | 7.6343 | 5.5692 |
| 45336 | ENSMUSG00000107471 | 0 | NA | NA | NA | NA | NA | Gm44147 | predicted gene, 44147 [Source:MGI Symb | 0 | 0 | 0 | 0 | 0 | 0 |
| 45337 | ENSMUSG00000107472 | 0.331982614 | 0.16514309 | 3.986751476 | 0.041422971 | 0.966958701 | NA | Gm44265 | predicted gene, 44265 [Source:MGI Symb | 0 | 1.0279 | 0 | 0.964 | 0 | 0 |
| 45338 | ENSMUSG00000107473 | 0 | NA | NA | NA | NA | NA | Gm44255 | predicted gene, 44255 [Source:MGI Symb | 0 | 0 | 0 | 0 | 0 | 0 |
| 45339 | ENSMUSG00000107474 | 0 | NA | NA | NA | NA | NA | Gm43932 | predicted gene, 43932 [Source:MGI Symb | 0 | 0 | 0 | 0 | 0 | 0 |
| 45340 | ENSMUSG00000107475 | 0 | NA | NA | NA | NA | NA | Gm19177 | predicted gene, 19177 [Source:MGI Symb | 0 | 0 | 0 | 0 | 0 | 0 |
| 45341 | ENSMUSG00000107476 | 60.99965357 | 0.484747883 | 0.33570007 | 1.443991007 | 0.148741505 | 0.560697714 | Zfp862-ps | zinc finger protein 862, pseudogene [Sour | 57.823 | 32.893 | 61.381 | 54.947 | 67.618 | 91.335 |
| 45342 | ENSMUSG00000107477 | 1.356133787 | 0.795733596 | 1.862943989 | 0.427137692 | 0.669279041 | NA | Gm44366 | predicted gene, 44366 [Source:MGI Symb | 0 | 1.0279 | 2.0125 | 2.8919 | 1.0906 | 1.1138 |
| 45343 | ENSMUSG00000107478 | 0.676451308 | 0.255551166 | 2.811516859 | 0.09089441 | 0.927576492 | NA | Gm45234 | predicted gene 45234 [Source:MGI Symbo | 0.8031 | 1.0279 | 0 | 0 | 0 | 2.2277 |
| 45344 | ENSMUSG00000107479 | 0 | NA | NA | NA | NA | NA | 2610300M1: | RIKEN cDNA 2610300M13 gene [Source:M | 0 | 0 | 0 | 0 | 0 | 0 |
| 45345 | ENSMUSG00000107480 | 0 | NA | NA | NA | NA | NA | Gm44165 | predicted gene, 44165 [Source:MGI Symb | 0 | 0 | 0 | 0 | 0 | 0 |
| 45346 | ENSMUSG00000107481 | 1.192142303 | -1.40772128 | 2.268741164 | -0.620485625 | 0.534938115 | NA | 4833403J16I | RIKEN cDNA 4833403J16 gene [Source:MG | 3.2124 | 0 | 2.0125 | 1.928 | 0 | 0 |
| 45347 | ENSMUSG00000107482 | 0.371281295 | 2.09253362 | 4.031852165 | 0.519000582 | 0.603760335 | NA | Etfbl | electron transferring flavoprotein, beta po | 0 | 0 | 0 | 0 | 0 | 2.2277 |
| 45348 | ENSMUSG00000107483 | 0 | NA | NA | NA | NA | NA | Gm44288 | predicted gene, 44288 [Source:MGI Symb | 0 | 0 | 0 | 0 | 0 | 0 |
| 45349 | ENSMUSG00000107484 | 0 | NA | NA | NA | NA | NA | Nasp-ps1 | nuclear autoantigenic sperm protein (hist | 0 | 0 | 0 | 0 | 0 | 0 |
| 45350 | ENSMUSG00000107486 | 0 | NA | NA | NA | NA | NA | Trbv4 | T cell receptor beta, variable 10 [Source:M | 0 | 0 | 0 | 0 | 0 | 0 |
| 45351 | ENSMUSG00000107487 | 0 | NA | NA | NA | NA | NA | Gm44156 | predicted gene, 44156 [Source:MGI Symb | 0 | 0 | 0 | 0 | 0 | 0 |
| 45352 | ENSMUSG00000107488 | 0 | NA | NA | NA | NA | NA | Gm31579 | predicted gene, 31579 [Source:MGI Symb | 0 | 0 | 0 | 0 | 0 | 0 |
| 45353 | ENSMUSG00000107489 | 0 | NA | NA | NA | NA | NA | Gm18325 | predicted gene, 18325 [Source:MGI Symb | 0 | 0 | 0 | 0 | 0 | 0 |
| 45354 | ENSMUSG00000107490 | 0 | NA | NA | NA | NA | NA | Gm44058 | predicted gene, 44058 [Source:MGI Symb | 0 | 0 | 0 | 0 | 0 | 0 |
| 45355 | ENSMUSG00000107491 | 0 | NA | NA | NA | NA | NA | Gm17893 | predicted gene, 17893 [Source:MGI Symb | 0 | 0 | 0 | 0 | 0 | 0 |
| 45356 | ENSMUSG00000107492 | 0 | NA | NA | NA | NA | NA | Gm6590 | predicted gene 6590 [Source:MGI Symbol | 0 | 0 | 0 | 0 | 0 | 0 |

| | | | | | | | | | | | | | | | |
|---|---|---|---|---|---|---|---|---|---|---|---|---|---|---|---|
| 45357 | ENSMUSG00000107493 | 0 | NA | NA | NA | NA | NA | Gm44048 | predicted gene, 44048 [Source:MGI Symb | 0 | 0 | 0 | 0 | 0 | 0 |
| 45358 | ENSMUSG00000107494 | 0 | NA | NA | NA | NA | NA | NA | NA | 0 | 0 | 0 | 0 | 0 | 0 |
| 45359 | ENSMUSG00000107495 | 0 | NA | NA | NA | NA | NA | Gm44215 | predicted gene, 44215 [Source:MGI Symb | 0 | 0 | 0 | 0 | 0 | 0 |
| 45360 | ENSMUSG00000107496 | 0 | NA | NA | NA | NA | NA | 4933431M0 | RIKEN cDNA 4933431M02 gene [Source:N | 0 | 0 | 0 | 0 | 0 | 0 |
| 45361 | ENSMUSG00000107497 | 1.957387837 | 0.69362432 | 1.470808191 | 0.471594001 | 0.637216609 | NA | Gm44957 | predicted gene 44957 [Source:MGI Symbo | 2.4093 | 1.0279 | 1.0062 | 2.8919 | 2.1812 | 2.2277 |
| 45362 | ENSMUSG00000107498 | 0 | NA | NA | NA | NA | NA | Gm44240 | predicted gene, 44240 [Source:MGI Symb | 0 | 0 | 0 | 0 | 0 | 0 |
| 45363 | ENSMUSG00000107499 | 0.819181066 | -0.437943885 | 2.255670782 | -0.194152395 | 0.846056557 | NA | Ccdc142 | coiled-coil domain containing 142 [Source | 0.8031 | 1.0279 | 1.0062 | 0.964 | 0 | 1.1138 |
| 45364 | ENSMUSG00000107500 | 0 | NA | NA | NA | NA | NA | Gm2796 | predicted gene 2796 [Source:MGI Symbol | 0 | 0 | 0 | 0 | 0 | 0 |
| 45365 | ENSMUSG00000107501 | 0 | NA | NA | NA | NA | NA | Gm29781 | predicted gene, 29781 [Source:MGI Symb | 0 | 0 | 0 | 0 | 0 | 0 |
| 45366 | ENSMUSG00000107502 | 0.476489653 | -2.293616034 | 3.421506946 | -0.670352587 | 0.502633051 | NA | Gm44065 | predicted gene, 44065 [Source:MGI Symb | 0.8031 | 2.0558 | 0 | 0 | 0 | 0 |
| 45367 | ENSMUSG00000107503 | 0.181769715 | 1.126924175 | 4.080472857 | 0.276174898 | 0.782413736 | NA | Gm44009 | predicted gene, 44009 [Source:MGI Symb | 0 | 0 | 0 | 0 | 1.0906 | 0 |
| 45368 | ENSMUSG00000107504 | 17.51592167 | 0.136587269 | 0.603444474 | 0.226346043 | 0.82093229 | 0.969160361 | Gm35549 | predicted gene, 35549 [Source:MGI Symb | 13.653 | 16.447 | 20.125 | 26.027 | 5.4531 | 23.391 |
| 45369 | ENSMUSG00000107505 | 10.36477343 | 0.189373531 | 0.673521891 | 0.281169081 | 0.778580718 | 0.959559604 | Stambp-ps1 | STAM binding protein, pseudogene 1 [Sou | 8.8341 | 10.279 | 10.062 | 15.424 | 10.906 | 6.6831 |
| 45370 | ENSMUSG00000107506 | 0 | NA | NA | NA | NA | NA | Gm19254 | predicted gene, 19254 [Source:MGI Symb | 0 | 0 | 0 | 0 | 0 | 0 |
| 45371 | ENSMUSG00000107508 | 0 | NA | NA | NA | NA | NA | Gm44083 | predicted gene, 44083 [Source:MGI Symb | 0 | 0 | 0 | 0 | 0 | 0 |
| 45372 | ENSMUSG00000107509 | 3.236883158 | -0.107224614 | 1.320909034 | -0.081174866 | 0.935302888 | NA | Gm44941 | predicted gene 44941 [Source:MGI Symbo | 0 | 6.1675 | 4.025 | 4.8199 | 2.1812 | 2.2277 |
| 45373 | ENSMUSG00000107510 | 0 | NA | NA | NA | NA | NA | Gm44057 | predicted gene, 44057 [Source:MGI Symb | 0 | 0 | 0 | 0 | 0 | 0 |
| 45374 | ENSMUSG00000107511 | 0 | NA | NA | NA | NA | NA | Gm44440 | predicted gene, 44440 [Source:MGI Symb | 0 | 0 | 0 | 0 | 0 | 0 |
| 45375 | ENSMUSG00000107512 | 0 | NA | NA | NA | NA | NA | Gm44433 | predicted gene, 44433 [Source:MGI Symb | 0 | 0 | 0 | 0 | 0 | 0 |
| 45376 | ENSMUSG00000107513 | 0 | NA | NA | NA | NA | NA | Tgif2-ps1 | TGFB-induced factor homeobox 2, pseudo | 0 | 0 | 0 | 0 | 0 | 0 |
| 45377 | ENSMUSG00000107514 | 0 | NA | NA | NA | NA | NA | Gm32381 | predicted gene, 32381 [Source:MGI Symb | 0 | 0 | 0 | 0 | 0 | 0 |
| 45378 | ENSMUSG00000107515 | 0 | NA | NA | NA | NA | NA | Gm43886 | predicted gene, 43886 [Source:MGI Symb | 0 | 0 | 0 | 0 | 0 | 0 |
| 45379 | ENSMUSG00000107516 | 67.1898594 | -0.354747808 | 0.320435942 | -1.107078706 | 0.268259901 | 0.724039098 | Gm30784 | predicted gene, 30784 [Source:MGI Symb | 60.233 | 101.76 | 64.4 | 49.163 | 76.343 | 51.237 |
| 45380 | ENSMUSG00000107517 | 0 | NA | NA | NA | NA | NA | Gm44046 | predicted gene, 44046 [Source:MGI Symb | 0 | 0 | 0 | 0 | 0 | 0 |
| 45381 | ENSMUSG00000107518 | 0 | NA | NA | NA | NA | NA | Gm44016 | predicted gene, 44016 [Source:MGI Symb | 0 | 0 | 0 | 0 | 0 | 0 |
| 45382 | ENSMUSG00000107519 | 0 | NA | NA | NA | NA | NA | Gm26996 | predicted gene, 26996 [Source:MGI Symb | 0 | 0 | 0 | 0 | 0 | 0 |
| 45383 | ENSMUSG00000107520 | 0 | NA | NA | NA | NA | NA | Gm3726 | predicted gene 3726 [Source:MGI Symbol | 0 | 0 | 0 | 0 | 0 | 0 |
| 45384 | ENSMUSG00000107521 | 0 | NA | NA | NA | NA | NA | Gm43994 | predicted gene, 43994 [Source:MGI Symb | 0 | 0 | 0 | 0 | 0 | 0 |
| 45385 | ENSMUSG00000107522 | 0 | NA | NA | NA | NA | NA | Gm44988 | predicted gene 44988 [Source:MGI Symbo | 0 | 0 | 0 | 0 | 0 | 0 |
| 45386 | ENSMUSG00000107523 | 0 | NA | NA | NA | NA | NA | Gm44451 | predicted gene, 44451 [Source:MGI Symb | 0 | 0 | 0 | 0 | 0 | 0 |
| 45387 | ENSMUSG00000107524 | 4.272780231 | 0.667448806 | 1.236751379 | 0.539679047 | 0.589418392 | NA | Gm44136 | predicted gene, 44136 [Source:MGI Symb | 4.8186 | 4.1117 | 1.0062 | 13.496 | 1.0906 | 1.1138 |
| 45388 | ENSMUSG00000107525 | 0 | NA | NA | NA | NA | NA | Gm44071 | predicted gene, 44071 [Source:MGI Symb | 0 | 0 | 0 | 0 | 0 | 0 |
| 45389 | ENSMUSG00000107526 | 0.505131668 | 1.126696199 | 3.421159819 | 0.329331647 | 0.741905026 | NA | 1700003I16F | RIKEN cDNA 1700003I16 gene [Source:MG | 0.8031 | 0 | 0 | 0 | 0 | 2.2277 |
| 45390 | ENSMUSG00000107527 | 0 | NA | NA | NA | NA | NA | Gm44239 | predicted gene, 44239 [Source:MGI Symb | 0 | 0 | 0 | 0 | 0 | 0 |
| 45391 | ENSMUSG00000107528 | 0 | NA | NA | NA | NA | NA | Gm44442 | predicted gene, 44442 [Source:MGI Symb | 0 | 0 | 0 | 0 | 0 | 0 |
| 45392 | ENSMUSG00000107529 | 0 | NA | NA | NA | NA | NA | Gm44291 | predicted gene, 44291 [Source:MGI Symb | 0 | 0 | 0 | 0 | 0 | 0 |
| 45393 | ENSMUSG00000107530 | 0 | NA | NA | NA | NA | NA | Vmn1r-ps16 | vomeronasal 1 receptor, pseudogene 16 [ | 0 | 0 | 0 | 0 | 0 | 0 |
| 45394 | ENSMUSG00000107531 | 3.441717742 | -1.196439095 | 1.181562957 | -1.012590221 | 0.311255933 | NA | E330037G11 | RIKEN cDNA E330037G11 gene [Source:M | 7.2279 | 3.0838 | 4.025 | 1.928 | 3.2719 | 1.1138 |
| 45395 | ENSMUSG00000107532 | 0 | NA | NA | NA | NA | NA | Gm35931 | predicted gene, 35931 [Source:MGI Symb | 0 | 0 | 0 | 0 | 0 | 0 |
| 45396 | ENSMUSG00000107533 | 0 | NA | NA | NA | NA | NA | Gm44422 | predicted gene, 44422 [Source:MGI Symb | 0 | 0 | 0 | 0 | 0 | 0 |
| 45397 | ENSMUSG00000107534 | 0.301557804 | -1.64335916 | 4.042814367 | -0.406488899 | 0.684383412 | NA | Gm43920 | predicted gene, 43920 [Source:MGI Symb | 0.8031 | 0 | 1.0062 | 0 | 0 | 0 |
| 45398 | ENSMUSG00000107535 | 0.160662974 | 1.126924175 | 4.080472857 | 0.276174898 | 0.782413736 | NA | Gm44183 | predicted gene, 44183 [Source:MGI Symb | 0 | 0 | 0 | 0.964 | 0 | 0 |
| 45399 | ENSMUSG00000107536 | 0 | NA | NA | NA | NA | NA | Gm43955 | predicted gene, 43955 [Source:MGI Symb | 0 | 0 | 0 | 0 | 0 | 0 |
| 45400 | ENSMUSG00000107538 | 0.167707431 | -0.796629176 | 4.080472857 | -0.195229623 | 0.84521319 | NA | Gm44127 | predicted gene, 44127 [Source:MGI Symb | 0 | 0 | 1.0062 | 0 | 0 | 0 |
| 45401 | ENSMUSG00000107539 | 0 | NA | NA | NA | NA | NA | Gm44790 | predicted gene 44790 [Source:MGI Symbo | 0 | 0 | 0 | 0 | 0 | 0 |
| 45402 | ENSMUSG00000107540 | 0 | NA | NA | NA | NA | NA | Gm43911 | predicted gene, 43911 [Source:MGI Symb | 0 | 0 | 0 | 0 | 0 | 0 |
| 45403 | ENSMUSG00000107541 | 0 | NA | NA | NA | NA | NA | Gm44336 | predicted gene, 44336 [Source:MGI Symb | 0 | 0 | 0 | 0 | 0 | 0 |
| 45404 | ENSMUSG00000107542 | 0 | NA | NA | NA | NA | NA | Gm43969 | predicted gene, 43969 [Source:MGI Symb | 0 | 0 | 0 | 0 | 0 | 0 |
| 45405 | ENSMUSG00000107543 | 0 | NA | NA | NA | NA | NA | Gm34701 | predicted gene, 34701 [Source:MGI Symb | 0 | 0 | 0 | 0 | 0 | 0 |
| 45406 | ENSMUSG00000107544 | 0 | NA | NA | NA | NA | NA | Gm44426 | predicted gene, 44426 [Source:MGI Symb | 0 | 0 | 0 | 0 | 0 | 0 |
| 45407 | ENSMUSG00000107545 | 0 | NA | NA | NA | NA | NA | Gm44307 | predicted gene, 44307 [Source:MGI Symb | 0 | 0 | 0 | 0 | 0 | 0 |
| 45408 | ENSMUSG00000107546 | 0 | NA | NA | NA | NA | NA | Gm20560 | predicted gene, 20560 [Source:MGI Symb | 0 | 0 | 0 | 0 | 0 | 0 |
| 45409 | ENSMUSG00000107548 | 0 | NA | NA | NA | NA | NA | Gm43866 | predicted gene, 43866 [Source:MGI Symb | 0 | 0 | 0 | 0 | 0 | 0 |
| 45410 | ENSMUSG00000107549 | 2.49787513 | -1.896745059 | 1.545864975 | -1.226979775 | 0.219830193 | NA | Gm43961 | predicted gene, 43961 [Source:MGI Symb | 1.6062 | 8.2233 | 2.0125 | 0.964 | 2.1812 | 0 |
| 45411 | ENSMUSG00000107550 | 0 | NA | NA | NA | NA | NA | Gm44726 | predicted gene 44726 [Source:MGI Symbo | 0 | 0 | 0 | 0 | 0 | 0 |
| 45412 | ENSMUSG00000107551 | 0 | NA | NA | NA | NA | NA | Gm4605 | predicted gene 4605 [Source:MGI Symbol | 0 | 0 | 0 | 0 | 0 | 0 |

| | | | | | | | | | | | | | | | |
|---|---|---|---|---|---|---|---|---|---|---|---|---|---|---|---|
| 45413 | ENSMUSG00000107552 | 62.9779013 | -0.149766702 | 0.310326651 | -0.482609861 | 0.629372776 | 0.92057725 | Gm44096 | predicted gene, 44096 [Source:MGI Symb | 83.523 | 62.703 | 52.325 | 70.37 | 62.165 | 46.781 |
| 45414 | ENSMUSG00000107553 | 0 | NA | NA | NA | NA | NA | Gm44266 | predicted gene, 44266 [Source:MGI Symb | 0 | 0 | 0 | 0 | 0 | 0 |
| 45415 | ENSMUSG00000107555 | 0 | NA | NA | NA | NA | NA | Olfr795-ps1 | olfactory receptor 795, pseudogene 1 [So | 0 | 0 | 0 | 0 | 0 | 0 |
| 45416 | ENSMUSG00000107556 | 1.355668472 | 2.826061778 | 2.213380762 | 1.276807781 | 0.201670118 | NA | Gm44237 | predicted gene, 44237 [Source:MGI Symb | 0 | 0 | 1.0062 | 3.8559 | 3.2719 | 0 |
| 45417 | ENSMUSG00000107557 | 0 | NA | NA | NA | NA | NA | Gm21940 | predicted gene, 21940 [Source:MGI Symb | 0 | 0 | 0 | 0 | 0 | 0 |
| 45418 | ENSMUSG00000107558 | 0 | NA | NA | NA | NA | NA | 2010109P13 | RIKEN cDNA 2010109P13 gene [Source:M | 0 | 0 | 0 | 0 | 0 | 0 |
| 45419 | ENSMUSG00000107559 | 0 | NA | NA | NA | NA | NA | Gm44193 | predicted gene, 44193 [Source:MGI Symb | 0 | 0 | 0 | 0 | 0 | 0 |
| 45420 | ENSMUSG00000107560 | 6.813328642 | -0.405750356 | 0.85343655 | -0.475431192 | 0.634479666 | 0.923478764 | Gm44199 | predicted gene, 44199 [Source:MGI Symb | 8.031 | 11.307 | 4.025 | 8.6758 | 3.2719 | 5.5692 |
| 45421 | ENSMUSG00000107561 | 0 | NA | NA | NA | NA | NA | Gm44264 | predicted gene, 44264 [Source:MGI Symb | 0 | 0 | 0 | 0 | 0 | 0 |
| 45422 | ENSMUSG00000107562 | 0 | NA | NA | NA | NA | NA | Gm44412 | predicted gene, 44412 [Source:MGI Symb | 0 | 0 | 0 | 0 | 0 | 0 |
| 45423 | ENSMUSG00000107563 | 0 | NA | NA | NA | NA | NA | 4930587E11 | RIKEN cDNA 4930587E11 gene [Source:M | 0 | 0 | 0 | 0 | 0 | 0 |
| 45424 | ENSMUSG00000107564 | 0 | NA | NA | NA | NA | NA | Gm44379 | predicted gene, 44379 [Source:MGI Symb | 0 | 0 | 0 | 0 | 0 | 0 |
| 45425 | ENSMUSG00000107565 | 0 | NA | NA | NA | NA | NA | Gm34314 | predicted gene, 34314 [Source:MGI Symb | 0 | 0 | 0 | 0 | 0 | 0 |
| 45426 | ENSMUSG00000107566 | 0.160662974 | 1.126924175 | 4.080472857 | 0.276174898 | 0.782413736 | NA | Gm44332 | predicted gene, 44332 [Source:MGI Symb | 0 | 0 | 0 | 0.964 | 0 | 0 |
| 45427 | ENSMUSG00000107567 | 0.542600935 | 1.125931187 | 3.335923981 | 0.33751704 | 0.735727169 | NA | 4930480K02 | RIKEN cDNA 4930480K02 gene [Source:M | 0 | 1.0279 | 0 | 0 | 0 | 2.2277 |
| 45428 | ENSMUSG00000107568 | 0.17131964 | -0.796629176 | 4.080472857 | -0.195229623 | 0.84521319 | NA | Gm18840 | predicted gene, 18840 [Source:MGI Symb | 0 | 1.0279 | 0 | 0 | 0 | 0 |
| 45429 | ENSMUSG00000107569 | 0 | NA | NA | NA | NA | NA | Gm44268 | predicted gene, 44268 [Source:MGI Symb | 0 | 0 | 0 | 0 | 0 | 0 |
| 45430 | ENSMUSG00000107570 | 0.17131964 | -0.796629176 | 4.080472857 | -0.195229623 | 0.84521319 | NA | Gm44207 | predicted gene, 44207 [Source:MGI Symb | 0 | 1.0279 | 0 | 0 | 0 | 0 |
| 45431 | ENSMUSG00000107571 | 0 | NA | NA | NA | NA | NA | Gm43877 | predicted gene, 43877 [Source:MGI Symb | 0 | 0 | 0 | 0 | 0 | 0 |
| 45432 | ENSMUSG00000107572 | 0 | NA | NA | NA | NA | NA | Gm3961 | predicted gene 3961 [Source:MGI Symbol | 0 | 0 | 0 | 0 | 0 | 0 |
| 45433 | ENSMUSG00000107573 | 0 | NA | NA | NA | NA | NA | Olfr1389 | olfactory receptor 1389 [Source:MGI Sym | 0 | 0 | 0 | 0 | 0 | 0 |
| 45434 | ENSMUSG00000107574 | 0 | NA | NA | NA | NA | NA | Gm17834 | predicted gene, 17834 [Source:MGI Symb | 0 | 0 | 0 | 0 | 0 | 0 |
| 45435 | ENSMUSG00000107575 | 0 | NA | NA | NA | NA | NA | Gm44128 | predicted gene, 44128 [Source:MGI Symb | 0 | 0 | 0 | 0 | 0 | 0 |
| 45436 | ENSMUSG00000107576 | 0.160662974 | 1.126924175 | 4.080472857 | 0.276174898 | 0.782413736 | NA | Gm44093 | predicted gene, 44093 [Source:MGI Symb | 0 | 0 | 0 | 0.964 | 0 | 0 |
| 45437 | ENSMUSG00000107577 | 21.95504244 | -0.455779879 | 0.480390327 | -0.948769894 | 0.342737657 | 0.784392005 | Gm44103 | predicted gene, 44103 [Source:MGI Symb | 23.29 | 32.893 | 20.125 | 20.244 | 21.812 | 13.366 |
| 45438 | ENSMUSG00000107578 | 0 | NA | NA | NA | NA | NA | Gm32536 | predicted gene, 32536 [Source:MGI Symb | 0 | 0 | 0 | 0 | 0 | 0 |
| 45439 | ENSMUSG00000107579 | 0 | NA | NA | NA | NA | NA | Gm44176 | predicted gene, 44176 [Source:MGI Symb | 0 | 0 | 0 | 0 | 0 | 0 |
| 45440 | ENSMUSG00000107580 | 0 | NA | NA | NA | NA | NA | Gm44124 | predicted gene, 44124 [Source:MGI Symb | 0 | 0 | 0 | 0 | 0 | 0 |
| 45441 | ENSMUSG00000107581 | 0 | NA | NA | NA | NA | NA | Gm44828 | predicted gene 44828 [Source:MGI Symbo | 0 | 0 | 0 | 0 | 0 | 0 |
| 45442 | ENSMUSG00000107582 | 0 | NA | NA | NA | NA | NA | Gm44133 | predicted gene, 44133 [Source:MGI Symb | 0 | 0 | 0 | 0 | 0 | 0 |
| 45443 | ENSMUSG00000107583 | 0 | NA | NA | NA | NA | NA | Gm44104 | predicted gene, 44104 [Source:MGI Symb | 0 | 0 | 0 | 0 | 0 | 0 |
| 45444 | ENSMUSG00000107584 | 0 | NA | NA | NA | NA | NA | Gm18716 | predicted gene, 18716 [Source:MGI Symb | 0 | 0 | 0 | 0 | 0 | 0 |
| 45445 | ENSMUSG00000107585 | 0 | NA | NA | NA | NA | NA | 3300002P13 | RIKEN cDNA 3300002P13 gene [Source:M | 0 | 0 | 0 | 0 | 0 | 0 |
| 45446 | ENSMUSG00000107586 | 0.181769715 | 1.126924175 | 4.080472857 | 0.276174898 | 0.782413736 | NA | Gm44283 | predicted gene, 44283 [Source:MGI Symb | 0 | 0 | 0 | 0 | 1.0906 | 0 |
| 45447 | ENSMUSG00000107587 | 0 | NA | NA | NA | NA | NA | Gm44989 | predicted gene 44989 [Source:MGI Symbo | 0 | 0 | 0 | 0 | 0 | 0 |
| 45448 | ENSMUSG00000107588 | 0 | NA | NA | NA | NA | NA | Gm45021 | predicted gene 45021 [Source:MGI Symbo | 0 | 0 | 0 | 0 | 0 | 0 |
| 45449 | ENSMUSG00000107590 | 0 | NA | NA | NA | NA | NA | Gm43897 | predicted gene, 43897 [Source:MGI Symb | 0 | 0 | 0 | 0 | 0 | 0 |
| 45450 | ENSMUSG00000107591 | 0 | NA | NA | NA | NA | NA | Gm6794 | predicted gene 6794 [Source:MGI Symbol | 0 | 0 | 0 | 0 | 0 | 0 |
| 45451 | ENSMUSG00000107592 | 0 | NA | NA | NA | NA | NA | Tas2r145-ps | taste receptor, type 2, member 145, pseu | 0 | 0 | 0 | 0 | 0 | 0 |
| 45452 | ENSMUSG00000107593 | 0 | NA | NA | NA | NA | NA | Gm44097 | predicted gene, 44097 [Source:MGI Symb | 0 | 0 | 0 | 0 | 0 | 0 |
| 45453 | ENSMUSG00000107594 | 0 | NA | NA | NA | NA | NA | Tas2r142-ps | taste receptor, type 2, member 142, pseu | 0 | 0 | 0 | 0 | 0 | 0 |
| 45454 | ENSMUSG00000107595 | 0 | NA | NA | NA | NA | NA | Gm44441 | predicted gene, 44441 [Source:MGI Symb | 0 | 0 | 0 | 0 | 0 | 0 |
| 45455 | ENSMUSG00000107596 | 0 | NA | NA | NA | NA | NA | Gm18913 | predicted gene, 18913 [Source:MGI Symb | 0 | 0 | 0 | 0 | 0 | 0 |
| 45456 | ENSMUSG00000107597 | 0.17131964 | -0.796629176 | 4.080472857 | -0.195229623 | 0.84521319 | NA | Gm44764 | predicted gene 44764 [Source:MGI Symbo | 0 | 1.0279 | 0 | 0 | 0 | 0 |
| 45457 | ENSMUSG00000107598 | 0 | NA | NA | NA | NA | NA | Gm35216 | predicted gene, 35216 [Source:MGI Symb | 0 | 0 | 0 | 0 | 0 | 0 |
| 45458 | ENSMUSG00000107599 | 0 | NA | NA | NA | NA | NA | Gm43981 | predicted gene, 43981 [Source:MGI Symb | 0 | 0 | 0 | 0 | 0 | 0 |
| 45459 | ENSMUSG00000107600 | 0 | NA | NA | NA | NA | NA | Gm6308 | predicted gene 6308 [Source:MGI Symbol | 0 | 0 | 0 | 0 | 0 | 0 |
| 45460 | ENSMUSG00000107601 | 0 | NA | NA | NA | NA | NA | Gm2651 | predicted gene 2651 [Source:MGI Symbol | 0 | 0 | 0 | 0 | 0 | 0 |
| 45461 | ENSMUSG00000107602 | 0 | NA | NA | NA | NA | NA | Gm18691 | predicted gene, 18691 [Source:MGI Symb | 0 | 0 | 0 | 0 | 0 | 0 |
| 45462 | ENSMUSG00000107603 | 0 | NA | NA | NA | NA | NA | Gm43921 | predicted gene, 43921 [Source:MGI Symb | 0 | 0 | 0 | 0 | 0 | 0 |
| 45463 | ENSMUSG00000107604 | 0.637152627 | -1.330052897 | 2.84179579 | -0.468032538 | 0.639761321 | NA | Gm44041 | predicted gene, 44041 [Source:MGI Symb | 0.8031 | 2.0558 | 0 | 0.964 | 0 | 0 |
| 45464 | ENSMUSG00000107605 | 16.68875631 | 0.327314543 | 0.578572022 | 0.565728259 | 0.571578522 | 0.904651508 | Gm44117 | predicted gene, 44117 [Source:MGI Symb | 17.668 | 17.475 | 9.0562 | 9.6398 | 22.903 | 23.391 |
| 45465 | ENSMUSG00000107606 | 0 | NA | NA | NA | NA | NA | Gm43873 | predicted gene, 43873 [Source:MGI Symb | 0 | 0 | 0 | 0 | 0 | 0 |
| 45466 | ENSMUSG00000107608 | 13.59464131 | -0.392033184 | 0.638156594 | -0.614321292 | 0.539003035 | 0.89174748 | Gm43863 | predicted gene, 43863 [Source:MGI Symb | 12.047 | 9.2513 | 25.156 | 16.388 | 9.8156 | 8.9108 |
| 45467 | ENSMUSG00000107609 | 0 | NA | NA | NA | NA | NA | Gm20177 | predicted gene, 20177 [Source:MGI Symb | 0 | 0 | 0 | 0 | 0 | 0 |
| 45468 | ENSMUSG00000107610 | 5.053138858 | 1.143119026 | 1.065358338 | 1.072990171 | 0.283275526 | NA | D530018E20 | RIKEN cDNA D530018E20 gene [Source:M | 2.4093 | 2.0558 | 5.0312 | 7.7118 | 11.997 | 1.1138 |

| | | | | | | | | | | | | | | | |
|---|---|---|---|---|---|---|---|---|---|---|---|---|---|---|---|
| 45469 | ENSMUSG00000107611 | 0 | NA | NA | NA | NA | NA | Gm44131 | predicted gene, 44131 [Source:MGI Symb | 0 | 0 | 0 | 0 | 0 | 0 |
| 45470 | ENSMUSG00000107612 | 0 | NA | NA | NA | NA | NA | Gm44163 | predicted gene, 44163 [Source:MGI Symb | 0 | 0 | 0 | 0 | 0 | 0 |
| 45471 | ENSMUSG00000107613 | 0 | NA | NA | NA | NA | NA | Gm43927 | predicted gene, 43927 [Source:MGI Symb | 0 | 0 | 0 | 0 | 0 | 0 |
| 45472 | ENSMUSG00000107614 | 0 | NA | NA | NA | NA | NA | Gm44196 | predicted gene, 44196 [Source:MGI Symb | 0 | 0 | 0 | 0 | 0 | 0 |
| 45473 | ENSMUSG00000107615 | 0 | NA | NA | NA | NA | NA | Olfr793-ps1 | olfactory receptor 793, pseudogene 1 [So | 0 | 0 | 0 | 0 | 0 | 0 |
| 45474 | ENSMUSG00000107616 | 0 | NA | NA | NA | NA | NA | Gm7825 | predicted gene 7825 [Source:MGI Symbol | 0 | 0 | 0 | 0 | 0 | 0 |
| 45475 | ENSMUSG00000107618 | 0 | NA | NA | NA | NA | NA | Gm6261 | predicted gene 6261 [Source:MGI Symbol | 0 | 0 | 0 | 0 | 0 | 0 |
| 45476 | ENSMUSG00000107619 | 0 | NA | NA | NA | NA | NA | Gm32479 | predicted gene, 32479 [Source:MGI Symb | 0 | 0 | 0 | 0 | 0 | 0 |
| 45477 | ENSMUSG00000107620 | 0.342432689 | 1.990653086 | 3.900281263 | 0.51038706 | 0.609780321 | NA | Gm44256 | predicted gene, 44256 [Source:MGI Symb | 0 | 0 | 0 | 0.964 | 1.0906 | 0 |
| 45478 | ENSMUSG00000107622 | 0 | NA | NA | NA | NA | NA | 4930512J16I | RIKEN cDNA 4930512J16 gene [Source:M( | 0 | 0 | 0 | 0 | 0 | 0 |
| 45479 | ENSMUSG00000107623 | 0 | NA | NA | NA | NA | NA | Gm10415 | predicted gene 10415 [Source:MGI Symbo | 0 | 0 | 0 | 0 | 0 | 0 |
| 45480 | ENSMUSG00000107624 | 2.672007246 | 0.131907363 | 1.391856647 | 0.094770796 | 0.924496884 | NA | Gm44005 | predicted gene, 44005 [Source:MGI Symb | 1.6062 | 2.0558 | 4.025 | 2.8919 | 5.4531 | 0 |
| 45481 | ENSMUSG00000107625 | 0 | NA | NA | NA | NA | NA | Gm34149 | predicted gene, 34149 [Source:MGI Symb | 0 | 0 | 0 | 0 | 0 | 0 |
| 45482 | ENSMUSG00000107626 | 0 | NA | NA | NA | NA | NA | Gm44727 | predicted gene 44727 [Source:MGI Symbo | 0 | 0 | 0 | 0 | 0 | 0 |
| 45483 | ENSMUSG00000107627 | 0 | NA | NA | NA | NA | NA | Gm19244 | predicted gene, 19244 [Source:MGI Symb | 0 | 0 | 0 | 0 | 0 | 0 |
| 45484 | ENSMUSG00000107628 | 0 | NA | NA | NA | NA | NA | Gm44282 | predicted gene, 44282 [Source:MGI Symb | 0 | 0 | 0 | 0 | 0 | 0 |
| 45485 | ENSMUSG00000107629 | 0 | NA | NA | NA | NA | NA | Gm44011 | predicted gene, 44011 [Source:MGI Symb | 0 | 0 | 0 | 0 | 0 | 0 |
| 45486 | ENSMUSG00000107630 | 0 | NA | NA | NA | NA | NA | Gm44061 | predicted gene, 44061 [Source:MGI Symb | 0 | 0 | 0 | 0 | 0 | 0 |
| 45487 | ENSMUSG00000107631 | 0 | NA | NA | NA | NA | NA | Gm44233 | predicted gene, 44233 [Source:MGI Symb | 0 | 0 | 0 | 0 | 0 | 0 |
| 45488 | ENSMUSG00000107632 | 0 | NA | NA | NA | NA | NA | Gm44070 | predicted gene, 44070 [Source:MGI Symb | 0 | 0 | 0 | 0 | 0 | 0 |
| 45489 | ENSMUSG00000107633 | 0.160662974 | 1.126924175 | 4.080472857 | 0.276174898 | 0.782413736 | NA | Gm44078 | predicted gene, 44078 [Source:MGI Symb | 0 | 0 | 0 | 0.964 | 0 | 0 |
| 45490 | ENSMUSG00000107634 | 0 | NA | NA | NA | NA | NA | Gm36816 | predicted gene, 36816 [Source:MGI Symb | 0 | 0 | 0 | 0 | 0 | 0 |
| 45491 | ENSMUSG00000107635 | 0 | NA | NA | NA | NA | NA | Gm44247 | predicted gene, 44247 [Source:MGI Symb | 0 | 0 | 0 | 0 | 0 | 0 |
| 45492 | ENSMUSG00000107637 | 0 | NA | NA | NA | NA | NA | 1700069P05 | RIKEN cDNA 1700069P05 gene [Source:M | 0 | 0 | 0 | 0 | 0 | 0 |
| 45493 | ENSMUSG00000107638 | 0 | NA | NA | NA | NA | NA | Gm30524 | predicted gene, 30524 [Source:MGI Symb | 0 | 0 | 0 | 0 | 0 | 0 |
| 45494 | ENSMUSG00000107639 | 0 | NA | NA | NA | NA | NA | Gm4374 | predicted gene 4374 [Source:MGI Symbol | 0 | 0 | 0 | 0 | 0 | 0 |
| 45495 | ENSMUSG00000107640 | 0 | NA | NA | NA | NA | NA | Gm38804 | predicted gene, 38804 [Source:MGI Symb | 0 | 0 | 0 | 0 | 0 | 0 |
| 45496 | ENSMUSG00000107641 | 0 | NA | NA | NA | NA | NA | Gm32858 | predicted gene, 32858 [Source:MGI Symb | 0 | 0 | 0 | 0 | 0 | 0 |
| 45497 | ENSMUSG00000107642 | 0.17131964 | -0.796629176 | 4.080472857 | -0.195229623 | 0.84521319 | NA | Tas2r141-ps | taste receptor, type 2, member 141, pseu | 0 | 1.0279 | 0 | 0 | 0 | 0 |
| 45498 | ENSMUSG00000107643 | 0 | NA | NA | NA | NA | NA | Gm43885 | predicted gene, 43885 [Source:MGI Symb | 0 | 0 | 0 | 0 | 0 | 0 |
| 45499 | ENSMUSG00000107644 | 0 | NA | NA | NA | NA | NA | Gm6342 | predicted gene 6342 [Source:MGI Symbol | 0 | 0 | 0 | 0 | 0 | 0 |
| 45500 | ENSMUSG00000107645 | 0 | NA | NA | NA | NA | NA | Olfr1380 | olfactory receptor 1380 [Source:MGI Sym | 0 | 0 | 0 | 0 | 0 | 0 |
| 45501 | ENSMUSG00000107646 | 0 | NA | NA | NA | NA | NA | Gm44225 | predicted gene, 44225 [Source:MGI Symb | 0 | 0 | 0 | 0 | 0 | 0 |
| 45502 | ENSMUSG00000107647 | 1.472309668 | -0.240830353 | 1.951064645 | -0.123435353 | 0.901762364 | NA | Gm44445 | predicted gene, 44445 [Source:MGI Symb | 0.8031 | 0 | 4.025 | 2.8919 | 0 | 1.1138 |
| 45503 | ENSMUSG00000107648 | 0 | NA | NA | NA | NA | NA | Olfr764-ps1 | olfactory receptor 764, pseudogene 1 [So | 0 | 0 | 0 | 0 | 0 | 0 |
| 45504 | ENSMUSG00000107649 | 0 | NA | NA | NA | NA | NA | Gm44404 | predicted gene, 44404 [Source:MGI Symb | 0 | 0 | 0 | 0 | 0 | 0 |
| 45505 | ENSMUSG00000107650 | 0 | NA | NA | NA | NA | NA | Gm44032 | predicted gene, 44032 [Source:MGI Symb | 0 | 0 | 0 | 0 | 0 | 0 |
| 45506 | ENSMUSG00000107651 | 0 | NA | NA | NA | NA | NA | Gm44166 | predicted gene, 44166 [Source:MGI Symb | 0 | 0 | 0 | 0 | 0 | 0 |
| 45507 | ENSMUSG00000107652 | 0 | NA | NA | NA | NA | NA | Gm43894 | predicted gene, 43894 [Source:MGI Symb | 0 | 0 | 0 | 0 | 0 | 0 |
| 45508 | ENSMUSG00000107653 | 0 | NA | NA | NA | NA | NA | Gm31520 | predicted gene, 31520 [Source:MGI Symb | 0 | 0 | 0 | 0 | 0 | 0 |
| 45509 | ENSMUSG00000107655 | 9.446108803 | -0.35546083 | 0.712720947 | -0.498737734 | 0.617964158 | 0.917509899 | Gm44220 | predicted gene, 44220 [Source:MGI Symb | 14.456 | 10.279 | 7.0437 | 10.604 | 8.7249 | 5.5692 |
| 45510 | ENSMUSG00000107656 | 0 | NA | NA | NA | NA | NA | 4930563H07 | RIKEN cDNA 4930563H07 gene [Source:M | 0 | 0 | 0 | 0 | 0 | 0 |
| 45511 | ENSMUSG00000107657 | 0 | NA | NA | NA | NA | NA | Gm44446 | predicted gene, 44446 [Source:MGI Symb | 0 | 0 | 0 | 0 | 0 | 0 |
| 45512 | ENSMUSG00000107658 | 0.17131964 | -0.796629176 | 4.080472857 | -0.195229623 | 0.84521319 | NA | Gm44958 | predicted gene 44958 [Source:MGI Symbo | 0 | 1.0279 | 0 | 0 | 0 | 0 |
| 45513 | ENSMUSG00000107659 | 0 | NA | NA | NA | NA | NA | Gm44170 | predicted gene, 44170 [Source:MGI Symb | 0 | 0 | 0 | 0 | 0 | 0 |
| 45514 | ENSMUSG00000107660 | 0 | NA | NA | NA | NA | NA | Gm6559 | predicted gene 6559 [Source:MGI Symbol | 0 | 0 | 0 | 0 | 0 | 0 |
| 45515 | ENSMUSG00000107661 | 0 | NA | NA | NA | NA | NA | Gm18402 | predicted gene, 18402 [Source:MGI Symb | 0 | 0 | 0 | 0 | 0 | 0 |
| 45516 | ENSMUSG00000107662 | 0 | NA | NA | NA | NA | NA | Olfr772 | olfactory receptor 772 [Source:MGI Symb | 0 | 0 | 0 | 0 | 0 | 0 |
| 45517 | ENSMUSG00000107663 | 0 | NA | NA | NA | NA | NA | Gm44107 | predicted gene, 44107 [Source:MGI Symb | 0 | 0 | 0 | 0 | 0 | 0 |
| 45518 | ENSMUSG00000107664 | 0 | NA | NA | NA | NA | NA | Gm18007 | predicted gene, 18007 [Source:MGI Symb | 0 | 0 | 0 | 0 | 0 | 0 |
| 45519 | ENSMUSG00000107666 | 0 | NA | NA | NA | NA | NA | 4933406J09I | RIKEN cDNA 4933406J09 gene [Source:M( | 0 | 0 | 0 | 0 | 0 | 0 |
| 45520 | ENSMUSG00000107667 | 0 | NA | NA | NA | NA | NA | C530044C16 | RIKEN cDNA C530044C16 gene [Source:M | 0 | 0 | 0 | 0 | 0 | 0 |
| 45521 | ENSMUSG00000107668 | 0.906682787 | -0.583665192 | 2.768799895 | -0.210800785 | 0.833042726 | NA | Gm44424 | predicted gene, 44424 [Source:MGI Symb | 3.2124 | 0 | 0 | 0 | 0 | 2.2277 |
| 45522 | ENSMUSG00000107669 | 0 | NA | NA | NA | NA | NA | Gm44137 | predicted gene, 44137 [Source:MGI Symb | 0 | 0 | 0 | 0 | 0 | 0 |
| 45523 | ENSMUSG00000107671 | 0 | NA | NA | NA | NA | NA | Gm43976 | predicted gene, 43976 [Source:MGI Symb | 0 | 0 | 0 | 0 | 0 | 0 |
| 45524 | ENSMUSG00000107672 | 0 | NA | NA | NA | NA | NA | Gm3017 | predicted gene 3017 [Source:MGI Symbol | 0 | 0 | 0 | 0 | 0 | 0 |

| | | | | | | | | | | | | | | | |
|---|---|---|---|---|---|---|---|---|---|---|---|---|---|---|---|
| 45525 | ENSMUSG00000107673 | 0.167707431 | -0.796629176 | 4.080472857 | -0.195229623 | 0.84521319 | NA | Crs-ps | cone rod homeobox, pseudogene [Source | 0 | 0 | 1.0062 | 0 | 0 | 0 |
| 45526 | ENSMUSG00000107674 | 0 | NA | NA | NA | NA | NA | Gm44277 | predicted gene, 44277 [Source:MGI Symb | 0 | 0 | 0 | 0 | 0 | 0 |
| 45527 | ENSMUSG00000107675 | 0 | NA | NA | NA | NA | NA | Gm30332 | predicted gene, 30332 [Source:MGI Symb | 0 | 0 | 0 | 0 | 0 | 0 |
| 45528 | ENSMUSG00000107676 | 0 | NA | NA | NA | NA | NA | Gm44178 | predicted gene, 44178 [Source:MGI Symb | 0 | 0 | 0 | 0 | 0 | 0 |
| 45529 | ENSMUSG00000107677 | 0 | NA | NA | NA | NA | NA | Olfr316 | olfactory receptor 316 [Source:MGI Symb | 0 | 0 | 0 | 0 | 0 | 0 |
| 45530 | ENSMUSG00000107678 | 0 | NA | NA | NA | NA | NA | Vmn1r-ps13 | vomeronasal 1 receptor, pseudogene 13 [ | 0 | 0 | 0 | 0 | 0 | 0 |
| 45531 | ENSMUSG00000107679 | 0 | NA | NA | NA | NA | NA | 4930404I20F | RIKEN cDNA 4930404I20 gene [Source:MC | 0 | 0 | 0 | 0 | 0 | 0 |
| 45532 | ENSMUSG00000107680 | 0 | NA | NA | NA | NA | NA | Gm43903 | predicted gene, 43903 [Source:MGI Symb | 0 | 0 | 0 | 0 | 0 | 0 |
| 45533 | ENSMUSG00000107681 | 0 | NA | NA | NA | NA | NA | 4930528H21 | RIKEN cDNA 4930528H21 gene [Source:M | 0 | 0 | 0 | 0 | 0 | 0 |
| 45534 | ENSMUSG00000107682 | 0 | NA | NA | NA | NA | NA | Gm33024 | predicted gene, 33024 [Source:MGI Symb | 0 | 0 | 0 | 0 | 0 | 0 |
| 45535 | ENSMUSG00000107683 | 0 | NA | NA | NA | NA | NA | Gm44139 | predicted gene, 44139 [Source:MGI Symb | 0 | 0 | 0 | 0 | 0 | 0 |
| 45536 | ENSMUSG00000107684 | 0 | NA | NA | NA | NA | NA | Rab5a-ps | RAB5A, member RAS oncogene family, ps | 0 | 0 | 0 | 0 | 0 | 0 |
| 45537 | ENSMUSG00000107685 | 0 | NA | NA | NA | NA | NA | Gm5995 | predicted gene 5995 [Source:MGI Symbol | 0 | 0 | 0 | 0 | 0 | 0 |
| 45538 | ENSMUSG00000107686 | 0 | NA | NA | NA | NA | NA | Gm9038 | predicted gene 9038 [Source:MGI Symbol | 0 | 0 | 0 | 0 | 0 | 0 |
| 45539 | ENSMUSG00000107687 | 0 | NA | NA | NA | NA | NA | Gm44254 | predicted gene, 44254 [Source:MGI Symb | 0 | 0 | 0 | 0 | 0 | 0 |
| 45540 | ENSMUSG00000107688 | 0.462220778 | -0.685400455 | 3.191475091 | -0.21475977 | 0.82995462 | NA | Gm44091 | predicted gene, 44091 [Source:MGI Symb | 0.8031 | 0 | 1.0062 | 0.964 | 0 | 0 |
| 45541 | ENSMUSG00000107689 | 0.6621303 | -1.329905799 | 2.807279375 | -0.473734752 | 0.635689068 | NA | Gm44386 | predicted gene, 44386 [Source:MGI Symb | 0.8031 | 2.0558 | 0 | 0 | 0 | 1.1138 |
| 45542 | ENSMUSG00000107690 | 0 | NA | NA | NA | NA | NA | Gm44044 | predicted gene, 44044 [Source:MGI Symb | 0 | 0 | 0 | 0 | 0 | 0 |
| 45543 | ENSMUSG00000107691 | 0 | NA | NA | NA | NA | NA | Gm43901 | predicted gene, 43901 [Source:MGI Symb | 0 | 0 | 0 | 0 | 0 | 0 |
| 45544 | ENSMUSG00000107692 | 0 | NA | NA | NA | NA | NA | Gm19172 | predicted gene, 19172 [Source:MGI Symb | 0 | 0 | 0 | 0 | 0 | 0 |
| 45545 | ENSMUSG00000107693 | 0 | NA | NA | NA | NA | NA | Gm44312 | predicted gene, 44312 [Source:MGI Symb | 0 | 0 | 0 | 0 | 0 | 0 |
| 45546 | ENSMUSG00000107694 | 0 | NA | NA | NA | NA | NA | Gm8921 | predicted gene 8921 [Source:MGI Symbol | 0 | 0 | 0 | 0 | 0 | 0 |
| 45547 | ENSMUSG00000107695 | 0 | NA | NA | NA | NA | NA | Gm44289 | predicted gene, 44289 [Source:MGI Symb | 0 | 0 | 0 | 0 | 0 | 0 |
| 45548 | ENSMUSG00000107696 | 0 | NA | NA | NA | NA | NA | Gm44171 | predicted gene, 44171 [Source:MGI Symb | 0 | 0 | 0 | 0 | 0 | 0 |
| 45549 | ENSMUSG00000107697 | 0 | NA | NA | NA | NA | NA | Gm44399 | predicted gene, 44399 [Source:MGI Symb | 0 | 0 | 0 | 0 | 0 | 0 |
| 45550 | ENSMUSG00000107698 | 2.875555606 | -1.11228466 | 1.400145569 | -0.794406442 | 0.426958855 | NA | Gm44430 | predicted gene, 44430 [Source:MGI Symb | 1.6062 | 5.1396 | 5.0312 | 0 | 4.3625 | 1.1138 |
| 45551 | ENSMUSG00000107699 | 0 | NA | NA | NA | NA | NA | Gm43970 | predicted gene, 43970 [Source:MGI Symb | 0 | 0 | 0 | 0 | 0 | 0 |
| 45552 | ENSMUSG00000107700 | 0 | NA | NA | NA | NA | NA | Gm20144 | predicted gene, 20144 [Source:MGI Symb | 0 | 0 | 0 | 0 | 0 | 0 |
| 45553 | ENSMUSG00000107701 | 0 | NA | NA | NA | NA | NA | Gm19287 | predicted gene, 19287 [Source:MGI Symb | 0 | 0 | 0 | 0 | 0 | 0 |
| 45554 | ENSMUSG00000107702 | 0 | NA | NA | NA | NA | NA | Gm44140 | predicted gene, 44140 [Source:MGI Symb | 0 | 0 | 0 | 0 | 0 | 0 |
| 45555 | ENSMUSG00000107703 | 0 | NA | NA | NA | NA | NA | Gm44039 | predicted gene, 44039 [Source:MGI Symb | 0 | 0 | 0 | 0 | 0 | 0 |
| 45556 | ENSMUSG00000107704 | 2.355770946 | 0.325657609 | 1.648055523 | 0.197601115 | 0.843357162 | NA | Gm36355 | predicted gene, 36355 [Source:MGI Symb | 3.2124 | 3.0838 | 0 | 6.7478 | 1.0906 | 0 |
| 45557 | ENSMUSG00000107705 | 0 | NA | NA | NA | NA | NA | Gm45062 | predicted gene 45062 [Source:MGI Symbc | 0 | 0 | 0 | 0 | 0 | 0 |
| 45558 | ENSMUSG00000107706 | 0 | NA | NA | NA | NA | NA | Gm44022 | predicted gene, 44022 [Source:MGI Symb | 0 | 0 | 0 | 0 | 0 | 0 |
| 45559 | ENSMUSG00000107707 | 0 | NA | NA | NA | NA | NA | Gm44286 | predicted gene, 44286 [Source:MGI Symb | 0 | 0 | 0 | 0 | 0 | 0 |
| 45560 | ENSMUSG00000107708 | 0 | NA | NA | NA | NA | NA | Gm19131 | predicted gene, 19131 [Source:MGI Symb | 0 | 0 | 0 | 0 | 0 | 0 |
| 45561 | ENSMUSG00000107709 | 0 | NA | NA | NA | NA | NA | 9430018G01 | RIKEN cDNA 9430018G01 gene [Source:M | 0 | 0 | 0 | 0 | 0 | 0 |
| 45562 | ENSMUSG00000107710 | 0 | NA | NA | NA | NA | NA | Gm5307 | predicted gene 5307 [Source:MGI Symbol | 0 | 0 | 0 | 0 | 0 | 0 |
| 45563 | ENSMUSG00000107711 | 0 | NA | NA | NA | NA | NA | Olfr320 | olfactory receptor 320 [Source:MGI Symb | 0 | 0 | 0 | 0 | 0 | 0 |
| 45564 | ENSMUSG00000107712 | 0 | NA | NA | NA | NA | NA | Gm43870 | predicted gene, 43870 [Source:MGI Symb | 0 | 0 | 0 | 0 | 0 | 0 |
| 45565 | ENSMUSG00000107713 | 0 | NA | NA | NA | NA | NA | Gm6681 | predicted gene 6681 [Source:MGI Symbol | 0 | 0 | 0 | 0 | 0 | 0 |
| 45566 | ENSMUSG00000107714 | 0 | NA | NA | NA | NA | NA | Gm34933 | predicted gene, 34933 [Source:MGI Symb | 0 | 0 | 0 | 0 | 0 | 0 |
| 45567 | ENSMUSG00000107715 | 0 | NA | NA | NA | NA | NA | Gm44135 | predicted gene, 44135 [Source:MGI Symb | 0 | 0 | 0 | 0 | 0 | 0 |
| 45568 | ENSMUSG00000107716 | 0 | NA | NA | NA | NA | NA | Gm43997 | predicted gene, 43997 [Source:MGI Symb | 0 | 0 | 0 | 0 | 0 | 0 |
| 45569 | ENSMUSG00000107718 | 0 | NA | NA | NA | NA | NA | Vmn2r-ps34 | vomeronasal 2, receptor, pseudogene 34 | 0 | 0 | 0 | 0 | 0 | 0 |
| 45570 | ENSMUSG00000107719 | 1.833755445 | -0.084103654 | 1.53999718 | -0.054612862 | 0.956446891 | NA | Gm43937 | predicted gene, 43937 [Source:MGI Symb | 1.6062 | 1.0279 | 3.0187 | 0.964 | 3.2719 | 1.1138 |
| 45571 | ENSMUSG00000107720 | 0.439020386 | -2.189747911 | 3.52148864 | -0.621824499 | 0.534057275 | NA | Gm43946 | predicted gene, 43946 [Source:MGI Symb | 1.6062 | 1.0279 | 0 | 0 | 0 | 0 |
| 45572 | ENSMUSG00000107721 | 0 | NA | NA | NA | NA | NA | Gm44261 | predicted gene, 44261 [Source:MGI Symb | 0 | 0 | 0 | 0 | 0 | 0 |
| 45573 | ENSMUSG00000107722 | 0.133850373 | -0.796629176 | 4.080472857 | -0.195229623 | 0.84521319 | NA | 2900060B14 | RIKEN cDNA 2900060B14 gene [Source:M | 0.8031 | 0 | 0 | 0 | 0 | 0 |
| 45574 | ENSMUSG00000107723 | 0 | NA | NA | NA | NA | NA | Gm43964 | predicted gene, 43964 [Source:MGI Symb | 0 | 0 | 0 | 0 | 0 | 0 |
| 45575 | ENSMUSG00000107724 | 0 | NA | NA | NA | NA | NA | Gm16042 | predicted gene 16042 [Source:MGI Symbc | 0 | 0 | 0 | 0 | 0 | 0 |
| 45576 | ENSMUSG00000107725 | 0 | NA | NA | NA | NA | NA | Gm18484 | predicted gene, 18484 [Source:MGI Symb | 0 | 0 | 0 | 0 | 0 | 0 |
| 45577 | ENSMUSG00000107726 | 1.613152364 | -2.935044729 | 1.930291577 | -1.52051885 | 0.128380625 | NA | Gm44037 | predicted gene, 44037 [Source:MGI Symb | 1.6062 | 3.0838 | 4.025 | 0.964 | 0 | 0 |
| 45578 | ENSMUSG00000107727 | 0 | NA | NA | NA | NA | NA | Gm18010 | predicted gene, 18010 [Source:MGI Symb | 0 | 0 | 0 | 0 | 0 | 0 |
| 45579 | ENSMUSG00000107728 | 2.644347993 | -1.480610159 | 1.380814176 | -1.07227329 | 0.2835973 | NA | Gm38910 | predicted gene, 38910 [Source:MGI Symb | 1.6062 | 3.0838 | 7.0437 | 1.928 | 1.0906 | 1.1138 |
| 45580 | ENSMUSG00000107729 | 0 | NA | NA | NA | NA | NA | B230112I24F | RIKEN cDNA B230112I24 gene [Source:MC | 0 | 0 | 0 | 0 | 0 | 0 |

| | | | | | | | | | | | | | | | |
|---|---|---|---|---|---|---|---|---|---|---|---|---|---|---|---|
| 45581 | ENSMUSG00000107730 | 0 | NA | NA | NA | NA | NA | Gm44238 | predicted gene, 44238 [Source:MGI Symb | 0 | 0 | 0 | 0 | 0 | 0 |
| 45582 | ENSMUSG00000107731 | 0 | NA | NA | NA | NA | NA | Gm18326 | predicted gene, 18326 [Source:MGI Symb | 0 | 0 | 0 | 0 | 0 | 0 |
| 45583 | ENSMUSG00000107732 | 0 | NA | NA | NA | NA | NA | Gm44204 | predicted gene, 44204 [Source:MGI Symb | 0 | 0 | 0 | 0 | 0 | 0 |
| 45584 | ENSMUSG00000107733 | 0 | NA | NA | NA | NA | NA | Gm8774 | predicted gene 8774 [Source:MGI Symbol | 0 | 0 | 0 | 0 | 0 | 0 |
| 45585 | ENSMUSG00000107734 | 0 | NA | NA | NA | NA | NA | Gm30055 | predicted gene, 30055 [Source:MGI Symb | 0 | 0 | 0 | 0 | 0 | 0 |
| 45586 | ENSMUSG00000107735 | 0 | NA | NA | NA | NA | NA | Gm44173 | predicted gene, 44173 [Source:MGI Symb | 0 | 0 | 0 | 0 | 0 | 0 |
| 45587 | ENSMUSG00000107736 | 0.624661033 | -1.234026074 | 2.854658177 | -0.432285058 | 0.665534246 | NA | Gm44148 | predicted gene, 44148 [Source:MGI Symb | 1.6062 | 1.0279 | 0 | 0 | 0 | 1.1138 |
| 45588 | ENSMUSG00000107737 | 0 | NA | NA | NA | NA | NA | 1700027F06 | RIKEN cDNA 1700027F06 gene [Source:M( | 0 | 0 | 0 | 0 | 0 | 0 |
| 45589 | ENSMUSG00000107738 | 0 | NA | NA | NA | NA | NA | Gm32453 | predicted gene, 32453 [Source:MGI Symb | 0 | 0 | 0 | 0 | 0 | 0 |
| 45590 | ENSMUSG00000107739 | 0 | NA | NA | NA | NA | NA | Gm7812 | predicted gene 7812 [Source:MGI Symbol | 0 | 0 | 0 | 0 | 0 | 0 |
| 45591 | ENSMUSG00000107740 | 0 | NA | NA | NA | NA | NA | Gm18765 | predicted gene, 18765 [Source:MGI Symb | 0 | 0 | 0 | 0 | 0 | 0 |
| 45592 | ENSMUSG00000107741 | 59.58469353 | 0.279454174 | 0.314618495 | 0.888231872 | 0.374416038 | 0.809332934 | Gm2011 | predicted gene 2011 [Source:MGI Symbol | 70.673 | 41.117 | 49.306 | 75.19 | 61.075 | 60.148 |
| 45593 | ENSMUSG00000107742 | 6.841358055 | 0.611866988 | 0.858012032 | 0.713121687 | 0.475770461 | 0.86947987 | 9530085L11 | RIKEN cDNA 9530085L11 gene [Source:M( | 3.2124 | 4.1117 | 9.0562 | 12.532 | 5.4531 | 6.6831 |
| 45594 | ENSMUSG00000107743 | 0 | NA | NA | NA | NA | NA | Gm44087 | predicted gene, 44087 [Source:MGI Symb | 0 | 0 | 0 | 0 | 0 | 0 |
| 45595 | ENSMUSG00000107744 | 0 | NA | NA | NA | NA | NA | 1700072O05 | RIKEN cDNA 1700072O05 gene [Source:M | 0 | 0 | 0 | 0 | 0 | 0 |
| 45596 | ENSMUSG00000107745 | 1.184291185 | 1.680901049 | 2.289541188 | 0.734165019 | 0.462848171 | NA | Gm43917 | predicted gene, 43917 [Source:MGI Symb | 1.6062 | 0 | 0 | 0 | 3.2719 | 2.2277 |
| 45597 | ENSMUSG00000107746 | 0 | NA | NA | NA | NA | NA | Gm18690 | predicted gene, 18690 [Source:MGI Symb | 0 | 0 | 0 | 0 | 0 | 0 |
| 45598 | ENSMUSG00000107747 | 0 | NA | NA | NA | NA | NA | Gm5881 | predicted gene 5881 [Source:MGI Symbol | 0 | 0 | 0 | 0 | 0 | 0 |
| 45599 | ENSMUSG00000107748 | 0 | NA | NA | NA | NA | NA | Olfr766-ps1 | olfactory receptor 766, pseudogene 1 [So | 0 | 0 | 0 | 0 | 0 | 0 |
| 45600 | ENSMUSG00000107749 | 10.83979305 | -0.081209621 | 0.763008399 | -0.106433456 | 0.91523845 | 0.98625062 | Gm44321 | predicted gene, 44321 [Source:MGI Symb | 16.062 | 13.363 | 4.025 | 18.316 | 4.3625 | 8.9108 |
| 45601 | ENSMUSG00000107750 | 0.160662974 | 1.126924175 | 4.080472857 | 0.276174898 | 0.782413736 | NA | Gm44013 | predicted gene, 44013 [Source:MGI Symb | 0 | 0 | 0 | 0.964 | 0 | 0 |
| 45602 | ENSMUSG00000107751 | 0 | NA | NA | NA | NA | NA | Gm6749 | predicted pseudogene 6749 [Source:MGI | 0 | 0 | 0 | 0 | 0 | 0 |
| 45603 | ENSMUSG00000107752 | 0 | NA | NA | NA | NA | NA | Gm43934 | predicted gene, 43934 [Source:MGI Symb | 0 | 0 | 0 | 0 | 0 | 0 |
| 45604 | ENSMUSG00000107753 | 0 | NA | NA | NA | NA | NA | Gm44371 | predicted gene, 44371 [Source:MGI Symb | 0 | 0 | 0 | 0 | 0 | 0 |
| 45605 | ENSMUSG00000107754 | 0 | NA | NA | NA | NA | NA | Gm44990 | predicted gene 44990 [Source:MGI Symbo | 0 | 0 | 0 | 0 | 0 | 0 |
| 45606 | ENSMUSG00000107755 | 0 | NA | NA | NA | NA | NA | Mageb16-ps | MAGE family member B16, pseudogene 2 | 0 | 0 | 0 | 0 | 0 | 0 |
| 45607 | ENSMUSG00000107756 | 4.112391593 | -1.216816686 | 1.098863813 | -1.107340757 | 0.268146629 | NA | Gm44164 | predicted gene, 44164 [Source:MGI Symb | 4.0155 | 8.2233 | 5.0312 | 1.928 | 4.3625 | 1.1138 |
| 45608 | ENSMUSG00000107757 | 0 | NA | NA | NA | NA | NA | Gm43898 | predicted gene, 43898 [Source:MGI Symb | 0 | 0 | 0 | 0 | 0 | 0 |
| 45609 | ENSMUSG00000107758 | 0 | NA | NA | NA | NA | NA | Gm19253 | predicted gene, 19253 [Source:MGI Symb | 0 | 0 | 0 | 0 | 0 | 0 |
| 45610 | ENSMUSG00000107759 | 0 | NA | NA | NA | NA | NA | Gm44258 | predicted gene, 44258 [Source:MGI Symb | 0 | 0 | 0 | 0 | 0 | 0 |
| 45611 | ENSMUSG00000107760 | 3.267178486 | 0.70645119 | 1.21418131 | 0.581833359 | 0.560678932 | NA | Gm44401 | predicted gene, 44401 [Source:MGI Symb | 2.4093 | 3.0838 | 2.0125 | 7.7118 | 3.2719 | 1.1138 |
| 45612 | ENSMUSG00000107761 | 2.007632397 | -0.510260853 | 1.714441047 | -0.297625196 | 0.765989242 | NA | 2010008C14 | RIKEN cDNA 2010008C14 gene [Source:M( | 4.0155 | 3.0838 | 0 | 3.8559 | 1.0906 | 0 |
| 45613 | ENSMUSG00000107762 | 0 | NA | NA | NA | NA | NA | Gm18421 | predicted gene, 18421 [Source:MGI Symb | 0 | 0 | 0 | 0 | 0 | 0 |
| 45614 | ENSMUSG00000107763 | 1.191858604 | 0.487587703 | 2.09676255 | 0.232543119 | 0.816116201 | NA | Gm43962 | predicted gene, 43962 [Source:MGI Symb | 0 | 0 | 3.0187 | 1.928 | 1.0906 | 1.1138 |
| 45615 | ENSMUSG00000107764 | 0.549180077 | 2.659271622 | 3.249668912 | 0.818320787 | 0.41317404 | NA | Gm5318 | predicted gene 5318 [Source:MGI Symbol | 0 | 0 | 0 | 0 | 2.1812 | 1.1138 |
| 45616 | ENSMUSG00000107765 | 0 | NA | NA | NA | NA | NA | Gm34224 | predicted gene, 34224 [Source:MGI Symb | 0 | 0 | 0 | 0 | 0 | 0 |
| 45617 | ENSMUSG00000107766 | 0.185640647 | 1.126924175 | 4.080472857 | 0.276174898 | 0.782413736 | NA | Gm44073 | predicted gene, 44073 [Source:MGI Symb | 0 | 0 | 0 | 0 | 0 | 1.1138 |
| 45618 | ENSMUSG00000107767 | 0 | NA | NA | NA | NA | NA | Gm44448 | predicted gene, 44448 [Source:MGI Symb | 0 | 0 | 0 | 0 | 0 | 0 |
| 45619 | ENSMUSG00000107768 | 0 | NA | NA | NA | NA | NA | Gm44275 | predicted gene, 44275 [Source:MGI Symb | 0 | 0 | 0 | 0 | 0 | 0 |
| 45620 | ENSMUSG00000107769 | 0 | NA | NA | NA | NA | NA | Gm43959 | predicted gene, 43959 [Source:MGI Symb | 0 | 0 | 0 | 0 | 0 | 0 |
| 45621 | ENSMUSG00000107770 | 0.17131964 | -0.796629176 | 4.080472857 | -0.195229623 | 0.84521319 | NA | Gm44126 | predicted gene, 44126 [Source:MGI Symb | 0 | 1.0279 | 0 | 0 | 0 | 0 |
| 45622 | ENSMUSG00000107771 | 0 | NA | NA | NA | NA | NA | Gm8956 | predicted gene 8956 [Source:MGI Symbol | 0 | 0 | 0 | 0 | 0 | 0 |
| 45623 | ENSMUSG00000107772 | 0 | NA | NA | NA | NA | NA | Gm5311 | predicted gene 5311 [Source:MGI Symbol | 0 | 0 | 0 | 0 | 0 | 0 |
| 45624 | ENSMUSG00000107773 | 0.185640647 | 1.126924175 | 4.080472857 | 0.276174898 | 0.782413736 | NA | Gm43971 | predicted gene, 43971 [Source:MGI Symb | 0 | 0 | 0 | 0 | 0 | 1.1138 |
| 45625 | ENSMUSG00000107774 | 0 | NA | NA | NA | NA | NA | Gm43974 | predicted gene, 43974 [Source:MGI Symb | 0 | 0 | 0 | 0 | 0 | 0 |
| 45626 | ENSMUSG00000107775 | 0 | NA | NA | NA | NA | NA | Gm19764 | predicted gene, 19764 [Source:MGI Symb | 0 | 0 | 0 | 0 | 0 | 0 |
| 45627 | ENSMUSG00000107776 | 0 | NA | NA | NA | NA | NA | Gm44388 | predicted gene, 44388 [Source:MGI Symb | 0 | 0 | 0 | 0 | 0 | 0 |
| 45628 | ENSMUSG00000107777 | 0 | NA | NA | NA | NA | NA | Gm44687 | predicted gene 44687 [Source:MGI Symbo | 0 | 0 | 0 | 0 | 0 | 0 |
| 45629 | ENSMUSG00000107778 | 0 | NA | NA | NA | NA | NA | Gm44098 | predicted gene, 44098 [Source:MGI Symb | 0 | 0 | 0 | 0 | 0 | 0 |
| 45630 | ENSMUSG00000107779 | 0 | NA | NA | NA | NA | NA | Gm43874 | predicted gene, 43874 [Source:MGI Symb | 0 | 0 | 0 | 0 | 0 | 0 |
| 45631 | ENSMUSG00000107780 | 0 | NA | NA | NA | NA | NA | 9530013L04 | RIKEN cDNA 9530013L04 gene [Source:M( | 0 | 0 | 0 | 0 | 0 | 0 |
| 45632 | ENSMUSG00000107781 | 0 | NA | NA | NA | NA | NA | Gm44315 | predicted gene, 44315 [Source:MGI Symb | 0 | 0 | 0 | 0 | 0 | 0 |
| 45633 | ENSMUSG00000107782 | 0 | NA | NA | NA | NA | NA | Gm43888 | predicted gene, 43888 [Source:MGI Symb | 0 | 0 | 0 | 0 | 0 | 0 |
| 45634 | ENSMUSG00000107783 | 0 | NA | NA | NA | NA | NA | Gm44038 | predicted gene, 44038 [Source:MGI Symb | 0 | 0 | 0 | 0 | 0 | 0 |
| 45635 | ENSMUSG00000107784 | 0 | NA | NA | NA | NA | NA | Gm43943 | predicted gene, 43943 [Source:MGI Symb | 0 | 0 | 0 | 0 | 0 | 0 |
| 45636 | ENSMUSG00000107785 | 11.63228909 | 0.065736949 | 0.634250579 | 0.103645075 | 0.917451015 | 0.986374671 | Gm45083 | predicted gene 45083 [Source:MGI Symbo | 8.8341 | 10.279 | 15.094 | 13.496 | 8.7249 | 13.366 |

| | | | | | | | | | | | | | | | |
|---|---|---|---|---|---|---|---|---|---|---|---|---|---|---|---|
| 45637 | ENSMUSG00000107786 | 0 | NA | NA | NA | NA | NA | Gm44232 | predicted gene, 44232 [Source:MGI Symb | 0 | 0 | 0 | 0 | 0 | 0 |
| 45638 | ENSMUSG00000107787 | 0 | NA | NA | NA | NA | NA | Gm44526 | predicted gene 44526 [Source:MGI Symbo | 0 | 0 | 0 | 0 | 0 | 0 |
| 45639 | ENSMUSG00000107788 | 0 | NA | NA | NA | NA | NA | Gm6266 | predicted gene 6266 [Source:MGI Symbol | 0 | 0 | 0 | 0 | 0 | 0 |
| 45640 | ENSMUSG00000107789 | 0.371281295 | 2.09253362 | 4.031852165 | 0.519000582 | 0.603760335 | NA | Gm44965 | predicted gene 44965 [Source:MGI Symbo | 0 | 0 | 0 | 0 | 0 | 2.2277 |
| 45641 | ENSMUSG00000107790 | 0 | NA | NA | NA | NA | NA | Olfr783-ps1 | olfactory receptor 783, pseudogene 1 [So | 0 | 0 | 0 | 0 | 0 | 0 |
| 45642 | ENSMUSG00000107791 | 0 | NA | NA | NA | NA | NA | Gm30557 | predicted gene, 30557 [Source:MGI Symb | 0 | 0 | 0 | 0 | 0 | 0 |
| 45643 | ENSMUSG00000107792 | 2.36438722 | -4.605739352 | 1.767262036 | -2.606144 | 0.009156796 | NA | Gm43914 | predicted gene, 43914 [Source:MGI Symb | 4.0155 | 5.1396 | 5.0312 | 0 | 0 | 0 |
| 45644 | ENSMUSG00000107793 | 0 | NA | NA | NA | NA | NA | 4933406L23I | RIKEN cDNA 4933406L23 gene [Source:M( | 0 | 0 | 0 | 0 | 0 | 0 |
| 45645 | ENSMUSG00000107794 | 0 | NA | NA | NA | NA | NA | Gm44095 | predicted gene, 44095 [Source:MGI Symb | 0 | 0 | 0 | 0 | 0 | 0 |
| 45646 | ENSMUSG00000107795 | 0 | NA | NA | NA | NA | NA | Gm44105 | predicted gene, 44105 [Source:MGI Symb | 0 | 0 | 0 | 0 | 0 | 0 |
| 45647 | ENSMUSG00000107796 | 1.454137189 | -2.771385281 | 2.191258366 | -1.264746012 | 0.20596239 | NA | Gm44068 | predicted gene, 44068 [Source:MGI Symb | 5.6217 | 0 | 2.0125 | 0 | 1.0906 | 0 |
| 45648 | ENSMUSG00000107797 | 0 | NA | NA | NA | NA | NA | Gm10445 | predicted gene 10445 [Source:MGI Symbo | 0 | 0 | 0 | 0 | 0 | 0 |
| 45649 | ENSMUSG00000107798 | 0.160662974 | 1.126924175 | 4.080472857 | 0.276174898 | 0.782413736 | NA | Gm44064 | predicted gene, 44064 [Source:MGI Symb | 0 | 0 | 0 | 0.964 | 0 | 0 |
| 45650 | ENSMUSG00000107799 | 0 | NA | NA | NA | NA | NA | Gm38825 | predicted gene, 38825 [Source:MGI Symb | 0 | 0 | 0 | 0 | 0 | 0 |
| 45651 | ENSMUSG00000107800 | 4.934217338 | 2.863950046 | 1.272347399 | 2.2509183 | 0.024390712 | NA | Gm44478 | predicted gene, 44478 [Source:MGI Symb | 2.4093 | 0 | 1.0062 | 2.8919 | 4.3625 | 18.935 |
| 45652 | ENSMUSG00000107801 | 0.167707431 | -0.796629176 | 4.080472857 | -0.195229623 | 0.84521319 | NA | Gm44086 | predicted gene, 44086 [Source:MGI Symb | 0 | 0 | 1.0062 | 0 | 0 | 0 |
| 45653 | ENSMUSG00000107802 | 1.302767309 | -0.543952165 | 2.003182152 | -0.271544035 | 0.785972636 | NA | 1700126G02 | RIKEN cDNA 1700126G02 gene [Source:M | 1.6062 | 0 | 3.0187 | 0.964 | 0 | 2.2277 |
| 45654 | ENSMUSG00000107803 | 0 | NA | NA | NA | NA | NA | Olfr778-ps1 | olfactory receptor 778, pseudogene 1 [So | 0 | 0 | 0 | 0 | 0 | 0 |
| 45655 | ENSMUSG00000107804 | 0 | NA | NA | NA | NA | NA | Gm44439 | predicted gene, 44439 [Source:MGI Symb | 0 | 0 | 0 | 0 | 0 | 0 |
| 45656 | ENSMUSG00000107805 | 0 | NA | NA | NA | NA | NA | Gm44003 | predicted gene, 44003 [Source:MGI Symb | 0 | 0 | 0 | 0 | 0 | 0 |
| 45657 | ENSMUSG00000107806 | 0 | NA | NA | NA | NA | NA | Gm19726 | predicted gene, 19726 [Source:MGI Symb | 0 | 0 | 0 | 0 | 0 | 0 |
| 45658 | ENSMUSG00000107808 | 0 | NA | NA | NA | NA | NA | Gm18689 | predicted gene, 18689 [Source:MGI Symb | 0 | 0 | 0 | 0 | 0 | 0 |
| 45659 | ENSMUSG00000107809 | 0 | NA | NA | NA | NA | NA | Gm44209 | predicted gene, 44209 [Source:MGI Symb | 0 | 0 | 0 | 0 | 0 | 0 |
| 45660 | ENSMUSG00000107810 | 4.616004705 | -0.707229885 | 0.981573267 | -0.720506465 | 0.471213221 | NA | Gm18609 | predicted gene, 18609 [Source:MGI Symb | 4.0155 | 6.1675 | 7.0437 | 3.8559 | 3.2719 | 3.3415 |
| 45661 | ENSMUSG00000107811 | 0 | NA | NA | NA | NA | NA | Gm44000 | predicted gene, 44000 [Source:MGI Symb | 0 | 0 | 0 | 0 | 0 | 0 |
| 45662 | ENSMUSG00000107812 | 1.238381702 | 1.44940733 | 1.982388925 | 0.731141761 | 0.464692568 | NA | Gm43935 | predicted gene, 43935 [Source:MGI Symb | 0 | 1.0279 | 1.0062 | 0.964 | 1.0906 | 3.3415 |
| 45663 | ENSMUSG00000107813 | 3.64209862 | 0.121705552 | 1.102103901 | 0.110430198 | 0.912068205 | NA | Gm44434 | predicted gene, 44434 [Source:MGI Symb | 2.4093 | 4.1117 | 4.025 | 5.7839 | 2.1812 | 3.3415 |
| 45664 | ENSMUSG00000107814 | 0.965523032 | -0.944628791 | 2.267086424 | -0.416670834 | 0.67691919 | NA | Gm35876 | predicted gene, 35876 [Source:MGI Symb | 0.8031 | 2.0558 | 1.0062 | 1.928 | 0 | 0 |
| 45665 | ENSMUSG00000107815 | 0 | NA | NA | NA | NA | NA | Gm44257 | predicted gene, 44257 [Source:MGI Symb | 0 | 0 | 0 | 0 | 0 | 0 |
| 45666 | ENSMUSG00000107816 | 0 | NA | NA | NA | NA | NA | Gm29799 | predicted gene, 29799 [Source:MGI Symb | 0 | 0 | 0 | 0 | 0 | 0 |
| 45667 | ENSMUSG00000107817 | 0 | NA | NA | NA | NA | NA | Gm38840 | predicted gene, 38840 [Source:MGI Symb | 0 | 0 | 0 | 0 | 0 | 0 |
| 45668 | ENSMUSG00000107818 | 0 | NA | NA | NA | NA | NA | Gm44034 | predicted gene, 44034 [Source:MGI Symb | 0 | 0 | 0 | 0 | 0 | 0 |
| 45669 | ENSMUSG00000107819 | 0 | NA | NA | NA | NA | NA | Gm18862 | predicted gene, 18862 [Source:MGI Symb | 0 | 0 | 0 | 0 | 0 | 0 |
| 45670 | ENSMUSG00000107821 | 0 | NA | NA | NA | NA | NA | Gm8038 | predicted gene 8038 [Source:MGI Symbol | 0 | 0 | 0 | 0 | 0 | 0 |
| 45671 | ENSMUSG00000107822 | 0 | NA | NA | NA | NA | NA | Olfr1358 | olfactory receptor 1358 [Source:MGI Sym | 0 | 0 | 0 | 0 | 0 | 0 |
| 45672 | ENSMUSG00000107823 | 0 | NA | NA | NA | NA | NA | Gm44436 | predicted gene, 44436 [Source:MGI Symb | 0 | 0 | 0 | 0 | 0 | 0 |
| 45673 | ENSMUSG00000107824 | 0 | NA | NA | NA | NA | NA | Gm6443 | predicted gene 6443 [Source:MGI Symbol | 0 | 0 | 0 | 0 | 0 | 0 |
| 45674 | ENSMUSG00000107825 | 0 | NA | NA | NA | NA | NA | Gm44296 | predicted gene, 44296 [Source:MGI Symb | 0 | 0 | 0 | 0 | 0 | 0 |
| 45675 | ENSMUSG00000107826 | 0 | NA | NA | NA | NA | NA | Gm44242 | predicted gene, 44242 [Source:MGI Symb | 0 | 0 | 0 | 0 | 0 | 0 |
| 45676 | ENSMUSG00000107828 | 0 | NA | NA | NA | NA | NA | Gm18289 | predicted gene, 18289 [Source:MGI Symb | 0 | 0 | 0 | 0 | 0 | 0 |
| 45677 | ENSMUSG00000107829 | 0 | NA | NA | NA | NA | NA | Gm43985 | predicted gene, 43985 [Source:MGI Symb | 0 | 0 | 0 | 0 | 0 | 0 |
| 45678 | ENSMUSG00000107830 | 0 | NA | NA | NA | NA | NA | 4933439N06 | RIKEN cDNA 4933439N06 gene [Source:M | 0 | 0 | 0 | 0 | 0 | 0 |
| 45679 | ENSMUSG00000107831 | 0 | NA | NA | NA | NA | NA | Gm44420 | predicted gene, 44420 [Source:MGI Symb | 0 | 0 | 0 | 0 | 0 | 0 |
| 45680 | ENSMUSG00000107832 | 0 | NA | NA | NA | NA | NA | Gm18688 | predicted gene, 18688 [Source:MGI Symb | 0 | 0 | 0 | 0 | 0 | 0 |
| 45681 | ENSMUSG00000107833 | 0 | NA | NA | NA | NA | NA | Gm18791 | predicted gene, 18791 [Source:MGI Symb | 0 | 0 | 0 | 0 | 0 | 0 |
| 45682 | ENSMUSG00000107834 | 0 | NA | NA | NA | NA | NA | Gm43979 | predicted gene, 43979 [Source:MGI Symb | 0 | 0 | 0 | 0 | 0 | 0 |
| 45683 | ENSMUSG00000107835 | 1.487890508 | -0.138447183 | 1.659190934 | -0.083442586 | 0.933499628 | NA | Gm43999 | predicted gene, 43999 [Source:MGI Symb | 1.6062 | 2.0558 | 1.0062 | 0.964 | 2.1812 | 1.1138 |
| 45684 | ENSMUSG00000107836 | 0 | NA | NA | NA | NA | NA | Gm5315 | predicted gene 5315 [Source:MGI Symbol | 0 | 0 | 0 | 0 | 0 | 0 |
| 45685 | ENSMUSG00000107837 | 0 | NA | NA | NA | NA | NA | Gm44281 | predicted gene, 44281 [Source:MGI Symb | 0 | 0 | 0 | 0 | 0 | 0 |
| 45686 | ENSMUSG00000107838 | 231.0697814 | -0.094504322 | 0.188697454 | -0.500824576 | 0.616494588 | 0.917509899 | Gm45769 | predicted gene 45769 [Source:MGI Symbo | 244.14 | 244.64 | 227.41 | 246.78 | 253.02 | 170.42 |
| 45687 | ENSMUSG00000107839 | 0 | NA | NA | NA | NA | NA | 4930434O05 | RIKEN cDNA 4930434O05 gene [Source:M | 0 | 0 | 0 | 0 | 0 | 0 |
| 45688 | ENSMUSG00000107840 | 0 | NA | NA | NA | NA | NA | Gm44241 | predicted gene, 44241 [Source:MGI Symb | 0 | 0 | 0 | 0 | 0 | 0 |
| 45689 | ENSMUSG00000107841 | 0.185640647 | 1.126924175 | 4.080472857 | 0.276174898 | 0.782413736 | NA | Gm44185 | predicted gene, 44185 [Source:MGI Symb | 0 | 0 | 0 | 0 | 0 | 1.1138 |
| 45690 | ENSMUSG00000107842 | 0 | NA | NA | NA | NA | NA | Gm44089 | predicted gene, 44089 [Source:MGI Symb | 0 | 0 | 0 | 0 | 0 | 0 |
| 45691 | ENSMUSG00000107844 | 0 | NA | NA | NA | NA | NA | Gm8686 | predicted gene 8686 [Source:MGI Symbol | 0 | 0 | 0 | 0 | 0 | 0 |
| 45692 | ENSMUSG00000107845 | 0 | NA | NA | NA | NA | NA | Gm44212 | predicted gene, 44212 [Source:MGI Symb | 0 | 0 | 0 | 0 | 0 | 0 |

| | | | | | | | | | | | | | | | |
|---|---|---|---|---|---|---|---|---|---|---|---|---|---|---|---|
| 45693 | ENSMUSG00000107846 | 2.790895424 | 0.612357295 | 1.302494625 | 0.470141898 | 0.638253642 | NA | Gm43963 | predicted gene, 43963 [Source:MGI Symb | 1.6062 | 3.0838 | 2.0125 | 6.7478 | 2.1812 | 1.1138 |
| 45694 | ENSMUSG00000107847 | 0 | NA | NA | NA | NA | NA | Gm35822 | predicted gene, 35822 [Source:MGI Symb | 0 | 0 | 0 | 0 | 0 | 0 |
| 45695 | ENSMUSG00000107848 | 0 | NA | NA | NA | NA | NA | Gm44181 | predicted gene, 44181 [Source:MGI Symb | 0 | 0 | 0 | 0 | 0 | 0 |
| 45696 | ENSMUSG00000107849 | 0 | NA | NA | NA | NA | NA | Gm43904 | predicted gene, 43904 [Source:MGI Symb | 0 | 0 | 0 | 0 | 0 | 0 |
| 45697 | ENSMUSG00000107850 | 0 | NA | NA | NA | NA | NA | Gm4714 | predicted gene 4714 [Source:MGI Symbol | 0 | 0 | 0 | 0 | 0 | 0 |
| 45698 | ENSMUSG00000107851 | 0 | NA | NA | NA | NA | NA | Gm35736 | predicted gene, 35736 [Source:MGI Symb | 0 | 0 | 0 | 0 | 0 | 0 |
| 45699 | ENSMUSG00000107852 | 0 | NA | NA | NA | NA | NA | Gm6227 | predicted gene 6227 [Source:MGI Symbol | 0 | 0 | 0 | 0 | 0 | 0 |
| 45700 | ENSMUSG00000107853 | 0 | NA | NA | NA | NA | NA | Gm18584 | predicted gene, 18584 [Source:MGI Symb | 0 | 0 | 0 | 0 | 0 | 0 |
| 45701 | ENSMUSG00000107854 | 0 | NA | NA | NA | NA | NA | Gm5001 | predicted gene 5001 [Source:MGI Symbol | 0 | 0 | 0 | 0 | 0 | 0 |
| 45702 | ENSMUSG00000107855 | 0 | NA | NA | NA | NA | NA | Gm44110 | predicted gene, 44110 [Source:MGI Symb | 0 | 0 | 0 | 0 | 0 | 0 |
| 45703 | ENSMUSG00000107856 | 0 | NA | NA | NA | NA | NA | Gm7838 | predicted gene 7838 [Source:MGI Symbol | 0 | 0 | 0 | 0 | 0 | 0 |
| 45704 | ENSMUSG00000107857 | 0 | NA | NA | NA | NA | NA | Gm19596 | predicted gene, 19596 [Source:MGI Symb | 0 | 0 | 0 | 0 | 0 | 0 |
| 45705 | ENSMUSG00000107858 | 0 | NA | NA | NA | NA | NA | Gm43879 | predicted gene, 43879 [Source:MGI Symb | 0 | 0 | 0 | 0 | 0 | 0 |
| 45706 | ENSMUSG00000107859 | 0 | NA | NA | NA | NA | NA | Gm30731 | predicted gene, 30731 [Source:MGI Symb | 0 | 0 | 0 | 0 | 0 | 0 |
| 45707 | ENSMUSG00000107861 | 0.65103495 | -1.312578592 | 2.821398903 | -0.465222621 | 0.641772041 | NA | Gm44253 | predicted gene, 44253 [Source:MGI Symb | 0.8031 | 0 | 2.0125 | 0 | 1.0906 | 0 |
| 45708 | ENSMUSG00000107862 | 0 | NA | NA | NA | NA | NA | Gm44151 | predicted gene, 44151 [Source:MGI Symb | 0 | 0 | 0 | 0 | 0 | 0 |
| 45709 | ENSMUSG00000107863 | 0 | NA | NA | NA | NA | NA | Gm44389 | predicted gene, 44389 [Source:MGI Symb | 0 | 0 | 0 | 0 | 0 | 0 |
| 45710 | ENSMUSG00000107864 | 0 | NA | NA | NA | NA | NA | Gm44072 | predicted gene, 44072 [Source:MGI Symb | 0 | 0 | 0 | 0 | 0 | 0 |
| 45711 | ENSMUSG00000107865 | 0 | NA | NA | NA | NA | NA | Gm44417 | predicted gene, 44417 [Source:MGI Symb | 0 | 0 | 0 | 0 | 0 | 0 |
| 45712 | ENSMUSG00000107866 | 0 | NA | NA | NA | NA | NA | Gm43907 | predicted gene, 43907 [Source:MGI Symb | 0 | 0 | 0 | 0 | 0 | 0 |
| 45713 | ENSMUSG00000107867 | 0 | NA | NA | NA | NA | NA | Gm43992 | predicted gene, 43992 [Source:MGI Symb | 0 | 0 | 0 | 0 | 0 | 0 |
| 45714 | ENSMUSG00000107868 | 8.112614411 | -0.461592435 | 0.867482327 | -0.532105866 | 0.594652675 | 0.913409341 | Gm5112 | predicted gene 5112 [Source:MGI Symbol | 12.047 | 4.1117 | 12.075 | 10.604 | 8.7249 | 1.1138 |
| 45715 | ENSMUSG00000107869 | 0 | NA | NA | NA | NA | NA | Gm29848 | predicted gene, 29848 [Source:MGI Symb | 0 | 0 | 0 | 0 | 0 | 0 |
| 45716 | ENSMUSG00000107870 | 0 | NA | NA | NA | NA | NA | Gm43940 | predicted gene, 43940 [Source:MGI Symb | 0 | 0 | 0 | 0 | 0 | 0 |
| 45717 | ENSMUSG00000107872 | 0 | NA | NA | NA | NA | NA | Gm44511 | predicted gene 44511 [Source:MGI Symbc | 0 | 0 | 0 | 0 | 0 | 0 |
| 45718 | ENSMUSG00000107873 | 0 | NA | NA | NA | NA | NA | Gm19065 | predicted gene, 19065 [Source:MGI Symb | 0 | 0 | 0 | 0 | 0 | 0 |
| 45719 | ENSMUSG00000107874 | 0 | NA | NA | NA | NA | NA | Prpmp5 | proline-rich protein MP5 [Source:MGI Syn | 0 | 0 | 0 | 0 | 0 | 0 |
| 45720 | ENSMUSG00000107875 | 4.502535607 | 0.277232002 | 1.073557427 | 0.258236769 | 0.796224182 | NA | Gm44696 | predicted gene 44696 [Source:MGI Symbc | 4.0155 | 7.1954 | 1.0062 | 4.8199 | 2.1812 | 7.7969 |
| 45721 | ENSMUSG00000107876 | 7.713842738 | 1.367821388 | 0.85124094 | 1.606855737 | 0.108086018 | 0.484761602 | Gm43936 | predicted gene, 43936 [Source:MGI Symb | 4.8186 | 1.0279 | 7.0437 | 12.532 | 14.178 | 6.6831 |
| 45722 | ENSMUSG00000107877 | 0.965523032 | -0.944628791 | 2.267086424 | -0.416670834 | 0.67691919 | NA | Gm43951 | predicted gene, 43951 [Source:MGI Symb | 0.8031 | 2.0558 | 1.0062 | 1.928 | 0 | 0 |
| 45723 | ENSMUSG00000107878 | 0 | NA | NA | NA | NA | NA | Gm44106 | predicted gene, 44106 [Source:MGI Symb | 0 | 0 | 0 | 0 | 0 | 0 |
| 45724 | ENSMUSG00000107879 | 12.47149861 | 0.231906823 | 0.665480146 | 0.348480454 | 0.727479391 | 0.945907265 | 9330102E08 | RIKEN cDNA 9330102E08 gene [Source:M | 16.865 | 9.2513 | 8.05 | 7.7118 | 21.812 | 11.138 |
| 45725 | ENSMUSG00000107880 | 0 | NA | NA | NA | NA | NA | Gm44113 | predicted gene, 44113 [Source:MGI Symb | 0 | 0 | 0 | 0 | 0 | 0 |
| 45726 | ENSMUSG00000107881 | 1600.726249 | -0.178956476 | 0.102283428 | -1.7496136 | 0.080185011 | 0.415846541 | Gm44250 | predicted gene, 44250 [Source:MGI Symb | 1566 | 1742.3 | 1792.1 | 1476.8 | 1482.2 | 1544.9 |
| 45727 | ENSMUSG00000107882 | 0 | NA | NA | NA | NA | NA | Gm44278 | predicted gene, 44278 [Source:MGI Symb | 0 | 0 | 0 | 0 | 0 | 0 |
| 45728 | ENSMUSG00000107883 | 1.002521471 | 1.366653481 | 2.416683574 | 0.565507829 | 0.571728401 | NA | Gm44062 | predicted gene, 44062 [Source:MGI Symb | 1.6062 | 0 | 0 | 0 | 2.1812 | 2.2277 |
| 45729 | ENSMUSG00000107884 | 0 | NA | NA | NA | NA | NA | Gm44144 | predicted gene, 44144 [Source:MGI Symb | 0 | 0 | 0 | 0 | 0 | 0 |
| 45730 | ENSMUSG00000107885 | 0 | NA | NA | NA | NA | NA | Gm43993 | predicted gene, 43993 [Source:MGI Symb | 0 | 0 | 0 | 0 | 0 | 0 |
| 45731 | ENSMUSG00000107886 | 0 | NA | NA | NA | NA | NA | Gm43880 | predicted gene, 43880 [Source:MGI Symb | 0 | 0 | 0 | 0 | 0 | 0 |
| 45732 | ENSMUSG00000107887 | 0 | NA | NA | NA | NA | NA | Gm43950 | predicted gene, 43950 [Source:MGI Symb | 0 | 0 | 0 | 0 | 0 | 0 |
| 45733 | ENSMUSG00000107888 | 0 | NA | NA | NA | NA | NA | Gm44138 | predicted gene, 44138 [Source:MGI Symb | 0 | 0 | 0 | 0 | 0 | 0 |
| 45734 | ENSMUSG00000107889 | 0 | NA | NA | NA | NA | NA | Olfr440-ps1 | olfactory receptor 440, pseudogene 1 [Sou | 0 | 0 | 0 | 0 | 0 | 0 |
| 45735 | ENSMUSG00000107890 | 0 | NA | NA | NA | NA | NA | Gm43968 | predicted gene, 43968 [Source:MGI Symb | 0 | 0 | 0 | 0 | 0 | 0 |
| 45736 | ENSMUSG00000107891 | 0.160662974 | 1.126924175 | 4.080472857 | 0.276174898 | 0.782413736 | NA | Gm5051 | predicted gene 5051 [Source:MGI Symbol | 0 | 0 | 0 | 0.964 | 0 | 0 |
| 45737 | ENSMUSG00000107892 | 4.887686165 | -0.073284087 | 1.037260704 | -0.07065156 | 0.943675074 | NA | Gm44524 | predicted gene 44524 [Source:MGI Symbc | 4.0155 | 2.0558 | 9.0562 | 8.6758 | 2.1812 | 3.3415 |
| 45738 | ENSMUSG00000107893 | 0 | NA | NA | NA | NA | NA | Gm44192 | predicted gene, 44192 [Source:MGI Symb | 0 | 0 | 0 | 0 | 0 | 0 |
| 45739 | ENSMUSG00000107894 | 0.615839296 | 1.537131441 | 3.17677869 | 0.483864817 | 0.628481811 | NA | Gm10388 | predicted gene 10388 [Source:MGI Symbc | 0.8031 | 0 | 0 | 2.8919 | 0 | 0 |
| 45740 | ENSMUSG00000107895 | 0 | NA | NA | NA | NA | NA | 1700094M2 | RIKEN cDNA 1700094M24 gene [Source:M | 0 | 0 | 0 | 0 | 0 | 0 |
| 45741 | ENSMUSG00000107896 | 0 | NA | NA | NA | NA | NA | Gm8719 | predicted pseudogene 8719 [Source:MGI | 0 | 0 | 0 | 0 | 0 | 0 |
| 45742 | ENSMUSG00000107898 | 0 | NA | NA | NA | NA | NA | Gm53051 | predicted gene, 53051 [Source:MGI Symb | 0 | 0 | 0 | 0 | 0 | 0 |
| 45743 | ENSMUSG00000107899 | 0 | NA | NA | NA | NA | NA | Gm44069 | predicted gene, 44069 [Source:MGI Symb | 0 | 0 | 0 | 0 | 0 | 0 |
| 45744 | ENSMUSG00000107900 | 0.339027071 | -1.79174149 | 3.919585585 | -0.457125237 | 0.647581032 | NA | Gm5723 | predicted gene 5723 [Source:MGI Symbol | 0 | 1.0279 | 1.0062 | 0 | 0 | 0 |
| 45745 | ENSMUSG00000107901 | 0 | NA | NA | NA | NA | NA | Gm44074 | predicted gene, 44074 [Source:MGI Symb | 0 | 0 | 0 | 0 | 0 | 0 |
| 45746 | ENSMUSG00000107902 | 0 | NA | NA | NA | NA | NA | Gm32592 | predicted gene, 32592 [Source:MGI Symb | 0 | 0 | 0 | 0 | 0 | 0 |
| 45747 | ENSMUSG00000107903 | 0 | NA | NA | NA | NA | NA | Gm43930 | predicted gene, 43930 [Source:MGI Symb | 0 | 0 | 0 | 0 | 0 | 0 |
| 45748 | ENSMUSG00000107904 | 0 | NA | NA | NA | NA | NA | Gm44213 | predicted gene, 44213 [Source:MGI Symb | 0 | 0 | 0 | 0 | 0 | 0 |

| | | | | | | | | | | | | | | | |
|---|---|---|---|---|---|---|---|---|---|---|---|---|---|---|---|
| 45749 | ENSMUSG00000107905 | 0 | NA | NA | NA | NA | NA | Gm44230 | predicted gene, 44230 [Source:MGI Symb | 0 | 0 | 0 | 0 | 0 | 0 |
| 45750 | ENSMUSG00000107906 | 0 | NA | NA | NA | NA | NA | Eif4a3l2 | eukaryotic translation initiation factor 4A3 | 0 | 0 | 0 | 0 | 0 | 0 |
| 45751 | ENSMUSG00000107907 | 0 | NA | NA | NA | NA | NA | Gm8574 | predicted gene 8574 [Source:MGI Symbol | 0 | 0 | 0 | 0 | 0 | 0 |
| 45752 | ENSMUSG00000107908 | 0 | NA | NA | NA | NA | NA | Gm43895 | predicted gene, 43895 [Source:MGI Symb | 0 | 0 | 0 | 0 | 0 | 0 |
| 45753 | ENSMUSG00000107909 | 4.592188628 | 0.783147589 | 1.070949134 | 0.731264972 | 0.464617321 | NA | 2610017A05 | RIKEN cDNA 2610017A05 gene [Source:M | 4.0155 | 4.1117 | 2.0125 | 9.6398 | 1.0906 | 6.6831 |
| 45754 | ENSMUSG00000107911 | 0 | NA | NA | NA | NA | NA | Gm7565 | predicted gene 7565 [Source:MGI Symbol | 0 | 0 | 0 | 0 | 0 | 0 |
| 45755 | ENSMUSG00000107913 | 0 | NA | NA | NA | NA | NA | Gm43871 | predicted gene, 43871 [Source:MGI Symb | 0 | 0 | 0 | 0 | 0 | 0 |
| 45756 | ENSMUSG00000107914 | 0 | NA | NA | NA | NA | NA | Gm44076 | predicted gene, 44076 [Source:MGI Symb | 0 | 0 | 0 | 0 | 0 | 0 |
| 45757 | ENSMUSG00000107915 | 6.940233851 | 0.630424108 | 0.82155266 | 0.767356908 | 0.442869342 | 0.850769625 | Gm44014 | predicted gene, 44014 [Source:MGI Symb | 3.2124 | 8.2233 | 5.0312 | 8.6758 | 9.8156 | 6.6831 |
| 45758 | ENSMUSG00000107916 | 0 | NA | NA | NA | NA | NA | Gm18011 | predicted gene, 18011 [Source:MGI Symb | 0 | 0 | 0 | 0 | 0 | 0 |
| 45759 | ENSMUSG00000107917 | 0 | NA | NA | NA | NA | NA | Gm44235 | predicted gene, 44235 [Source:MGI Symb | 0 | 0 | 0 | 0 | 0 | 0 |
| 45760 | ENSMUSG00000107918 | 0 | NA | NA | NA | NA | NA | Gm8999 | predicted gene 8999 [Source:MGI Symbol | 0 | 0 | 0 | 0 | 0 | 0 |
| 45761 | ENSMUSG00000107919 | 0 | NA | NA | NA | NA | NA | Gm7946 | predicted gene 7946 [Source:MGI Symbol | 0 | 0 | 0 | 0 | 0 | 0 |
| 45762 | ENSMUSG00000107920 | 0 | NA | NA | NA | NA | NA | Gm7783 | predicted gene 7783 [Source:MGI Symbol | 0 | 0 | 0 | 0 | 0 | 0 |
| 45763 | ENSMUSG00000107921 | 0 | NA | NA | NA | NA | NA | Gm6423 | predicted gene 6423 [Source:MGI Symbol | 0 | 0 | 0 | 0 | 0 | 0 |
| 45764 | ENSMUSG00000107922 | 0.401551119 | -2.072982726 | 4.008251979 | -0.517178744 | 0.605031388 | NA | Gm44285 | predicted gene, 44285 [Source:MGI Symb | 2.4093 | 0 | 0 | 0 | 0 | 0 |
| 45765 | ENSMUSG00000107923 | 0 | NA | NA | NA | NA | NA | Gm43922 | predicted gene, 43922 [Source:MGI Symb | 0 | 0 | 0 | 0 | 0 | 0 |
| 45766 | ENSMUSG00000107924 | 0 | NA | NA | NA | NA | NA | 4930518I17F | RIKEN cDNA 4930518I17 gene [Source:MC | 0 | 0 | 0 | 0 | 0 | 0 |
| 45767 | ENSMUSG00000107925 | 0 | NA | NA | NA | NA | NA | Gm43957 | predicted gene, 43957 [Source:MGI Symb | 0 | 0 | 0 | 0 | 0 | 0 |
| 45768 | ENSMUSG00000107926 | 0 | NA | NA | NA | NA | NA | 4930504D19 | RIKEN cDNA 4930504D19 gene [Source:M | 0 | 0 | 0 | 0 | 0 | 0 |
| 45769 | ENSMUSG00000107927 | 0 | NA | NA | NA | NA | NA | Gm44090 | predicted gene, 44090 [Source:MGI Symb | 0 | 0 | 0 | 0 | 0 | 0 |
| 45770 | ENSMUSG00000107928 | 1.713377773 | -0.636291624 | 1.676800904 | -0.379467605 | 0.704340656 | NA | Gm45140 | predicted gene 45140 [Source:MGI Symbo | 3.2124 | 2.0558 | 1.0062 | 2.8919 | 0 | 1.1138 |
| 45771 | ENSMUSG00000107929 | 0.36353943 | 2.065989254 | 4.034137301 | 0.512126658 | 0.608562368 | NA | 4930595L18I | RIKEN cDNA 4930595L18 gene [Source:MC | 0 | 0 | 0 | 0 | 2.1812 | 0 |
| 45772 | ENSMUSG00000107930 | 0 | NA | NA | NA | NA | NA | Gm44162 | predicted gene, 44162 [Source:MGI Symb | 0 | 0 | 0 | 0 | 0 | 0 |
| 45773 | ENSMUSG00000107931 | 0 | NA | NA | NA | NA | NA | Gm44475 | predicted gene, 44475 [Source:MGI Symb | 0 | 0 | 0 | 0 | 0 | 0 |
| 45774 | ENSMUSG00000107932 | 3.694038137 | 0.96972966 | 1.164986936 | 0.832395308 | 0.40518585 | NA | Gm44432 | predicted gene, 44432 [Source:MGI Symb | 2.4093 | 3.0838 | 2.0125 | 4.8199 | 8.7249 | 1.1138 |
| 45775 | ENSMUSG00000107933 | 0 | NA | NA | NA | NA | NA | Gm30498 | predicted gene, 30498 [Source:MGI Symb | 0 | 0 | 0 | 0 | 0 | 0 |
| 45776 | ENSMUSG00000107934 | 0 | NA | NA | NA | NA | NA | Gm17822 | predicted gene, 17822 [Source:MGI Symb | 0 | 0 | 0 | 0 | 0 | 0 |
| 45777 | ENSMUSG00000107935 | 0 | NA | NA | NA | NA | NA | Gm4872 | predicted gene 4872 [Source:MGI Symbol | 0 | 0 | 0 | 0 | 0 | 0 |
| 45778 | ENSMUSG00000107936 | 0 | NA | NA | NA | NA | NA | Gm44155 | predicted gene, 44155 [Source:MGI Symb | 0 | 0 | 0 | 0 | 0 | 0 |
| 45779 | ENSMUSG00000107937 | 2.746149607 | -0.068969906 | 1.363941924 | -0.0505666 | 0.959670878 | NA | Gm44200 | predicted gene, 44200 [Source:MGI Symb | 2.4093 | 3.0838 | 3.0187 | 5.7839 | 2.1812 | 0 |
| 45780 | ENSMUSG00000107938 | 0 | NA | NA | NA | NA | NA | Gm32308 | predicted gene, 32308 [Source:MGI Symb | 0 | 0 | 0 | 0 | 0 | 0 |
| 45781 | ENSMUSG00000107940 | 0 | NA | NA | NA | NA | NA | Gm44100 | predicted gene, 44100 [Source:MGI Symb | 0 | 0 | 0 | 0 | 0 | 0 |
| 45782 | ENSMUSG00000107941 | 0 | NA | NA | NA | NA | NA | Gm44010 | predicted gene, 44010 [Source:MGI Symb | 0 | 0 | 0 | 0 | 0 | 0 |
| 45783 | ENSMUSG00000107942 | 0.449470461 | -0.545136533 | 3.562105917 | -0.15303771 | 0.878368537 | NA | Gm44411 | predicted gene, 44411 [Source:MGI Symb | 1.6062 | 0 | 0 | 0 | 1.0906 | 0 |
| 45784 | ENSMUSG00000107943 | 0 | NA | NA | NA | NA | NA | Gm18538 | predicted gene, 18538 [Source:MGI Symb | 0 | 0 | 0 | 0 | 0 | 0 |
| 45785 | ENSMUSG00000107944 | 6.369047122 | 0.871075003 | 0.945664453 | 0.921124824 | 0.356985261 | 0.796166767 | Gm44280 | predicted gene, 44280 [Source:MGI Symb | 2.4093 | 9.2513 | 2.0125 | 13.496 | 4.3625 | 6.6831 |
| 45786 | ENSMUSG00000107945 | 0 | NA | NA | NA | NA | NA | Rpl7a-ps14 | ribosomal protein L7A, pseudogene 14 [Sc | 0 | 0 | 0 | 0 | 0 | 0 |
| 45787 | ENSMUSG00000107946 | 1.788057582 | -0.727559766 | 1.810365742 | -0.401885514 | 0.687768284 | NA | Gm44066 | predicted gene, 44066 [Source:MGI Symb | 1.6062 | 5.1396 | 0 | 2.8919 | 1.0906 | 0 |
| 45788 | ENSMUSG00000107947 | 0 | NA | NA | NA | NA | NA | Gm44438 | predicted gene, 44438 [Source:MGI Symb | 0 | 0 | 0 | 0 | 0 | 0 |
| 45789 | ENSMUSG00000107948 | 0 | NA | NA | NA | NA | NA | AA545190 | EST AA545190 [Source:MGI Symbol;Acc:M | 0 | 0 | 0 | 0 | 0 | 0 |
| 45790 | ENSMUSG00000107949 | 0.17131964 | -0.796629176 | 4.080472857 | -0.195229623 | 0.84521319 | NA | Gm44157 | predicted gene, 44157 [Source:MGI Symb | 0 | 1.0279 | 0 | 0 | 0 | 0 |
| 45791 | ENSMUSG00000107950 | 0 | NA | NA | NA | NA | NA | Gm6375 | predicted gene 6375 [Source:MGI Symbol | 0 | 0 | 0 | 0 | 0 | 0 |
| 45792 | ENSMUSG00000107951 | 3.131203972 | 1.539296414 | 1.262070906 | 1.219659218 | 0.222594088 | NA | Gm6210 | predicted gene 6210 [Source:MGI Symbol | 0.8031 | 1.0279 | 3.0187 | 2.8919 | 4.3625 | 6.6831 |
| 45793 | ENSMUSG00000107952 | 0.185640647 | 1.126924175 | 4.080472857 | 0.276174898 | 0.782413736 | NA | Gm6352 | predicted gene 6352 [Source:MGI Symbol | 0 | 0 | 0 | 0 | 0 | 1.1138 |
| 45794 | ENSMUSG00000107953 | 0 | NA | NA | NA | NA | NA | Gm44211 | predicted gene, 44211 [Source:MGI Symb | 0 | 0 | 0 | 0 | 0 | 0 |
| 45795 | ENSMUSG00000107954 | 0 | NA | NA | NA | NA | NA | Gm43942 | predicted gene, 43942 [Source:MGI Symb | 0 | 0 | 0 | 0 | 0 | 0 |
| 45796 | ENSMUSG00000107955 | 0.335414862 | -1.779047242 | 4.030804179 | -0.441362855 | 0.658950329 | NA | Gm44027 | predicted gene, 44027 [Source:MGI Symb | 0 | 0 | 2.0125 | 0 | 0 | 0 |
| 45797 | ENSMUSG00000107956 | 0.181769715 | 1.126924175 | 4.080472857 | 0.276174898 | 0.782413736 | NA | Speer9-ps1 | spermatogenesis associated glutamate (E) | 0 | 0 | 0 | 0 | 1.0906 | 0 |
| 45798 | ENSMUSG00000107957 | 0 | NA | NA | NA | NA | NA | Gm44088 | predicted gene, 44088 [Source:MGI Symb | 0 | 0 | 0 | 0 | 0 | 0 |
| 45799 | ENSMUSG00000107959 | 0 | NA | NA | NA | NA | NA | Gm44111 | predicted gene, 44111 [Source:MGI Symb | 0 | 0 | 0 | 0 | 0 | 0 |
| 45800 | ENSMUSG00000107960 | 0 | NA | NA | NA | NA | NA | Gm35077 | predicted gene, 35077 [Source:MGI Symb | 0 | 0 | 0 | 0 | 0 | 0 |
| 45801 | ENSMUSG00000107961 | 2.152145477 | 2.469977755 | 1.941597629 | 1.27213678 | 0.203324525 | NA | Gm44036 | predicted gene, 44036 [Source:MGI Symb | 0 | 2.0558 | 0 | 8.6758 | 2.1812 | 0 |
| 45802 | ENSMUSG00000107962 | 18.8080506 | -1.240774069 | 0.577536776 | -2.148389714 | 0.031682809 | 0.236949406 | Gm43980 | predicted gene, 43980 [Source:MGI Symb | 29.715 | 11.307 | 38.237 | 11.568 | 11.997 | 10.025 |
| 45803 | ENSMUSG00000107964 | 0 | NA | NA | NA | NA | NA | 4930447C11 | RIKEN cDNA 4930447C11 gene [Source:M | 0 | 0 | 0 | 0 | 0 | 0 |
| 45804 | ENSMUSG00000107965 | 0 | NA | NA | NA | NA | NA | Gm44259 | predicted gene, 44259 [Source:MGI Symb | 0 | 0 | 0 | 0 | 0 | 0 |

| | | | | | | | | | | | | | | | |
|---|---|---|---|---|---|---|---|---|---|---|---|---|---|---|---|
| 45805 | ENSMUSG00000107966 | 0.301557804 | -1.64335916 | 4.042814367 | -0.406488899 | 0.684383412 | NA | Gm44001 | predicted gene, 44001 [Source:MGI Symb | 0.8031 | 0 | 1.0062 | 0 | 0 | 0 |
| 45806 | ENSMUSG00000107967 | 0 | NA | NA | NA | NA | NA | Gm44094 | predicted gene, 44094 [Source:MGI Symb | 0 | 0 | 0 | 0 | 0 | 0 |
| 45807 | ENSMUSG00000107968 | 0 | NA | NA | NA | NA | NA | Gm44419 | predicted gene, 44419 [Source:MGI Symb | 0 | 0 | 0 | 0 | 0 | 0 |
| 45808 | ENSMUSG00000107969 | 0.481988923 | 2.492401104 | 3.837077151 | 0.649557203 | 0.515978285 | NA | Gm44085 | predicted gene, 44085 [Source:MGI Symb | 0 | 0 | 0 | 2.8919 | 0 | 0 |
| 45809 | ENSMUSG00000107970 | 7.62117675 | -0.767107841 | 0.775259703 | -0.989484993 | 0.322425909 | 0.76988915 | Gm49058 | predicted gene, 49058 [Source:MGI Symb | 10.44 | 9.2513 | 9.0562 | 4.8199 | 4.3625 | 7.7969 |
| 45810 | ENSMUSG00000107971 | 2.986547462 | 0.775815027 | 1.277887444 | 0.607107481 | 0.543779586 | NA | Gm44260 | predicted gene, 44260 [Source:MGI Symb | 1.6062 | 1.0279 | 4.025 | 5.7839 | 4.3625 | 1.1138 |
| 45811 | ENSMUSG00000107972 | 0 | NA | NA | NA | NA | NA | Gm18198 | predicted gene, 18198 [Source:MGI Symb | 0 | 0 | 0 | 0 | 0 | 0 |
| 45812 | ENSMUSG00000107973 | 0 | NA | NA | NA | NA | NA | Gm43978 | predicted gene, 43978 [Source:MGI Symb | 0 | 0 | 0 | 0 | 0 | 0 |
| 45813 | ENSMUSG00000107975 | 0 | NA | NA | NA | NA | NA | Gm44118 | predicted gene, 44118 [Source:MGI Symb | 0 | 0 | 0 | 0 | 0 | 0 |
| 45814 | ENSMUSG00000107976 | 0 | NA | NA | NA | NA | NA | Gm44043 | predicted gene, 44043 [Source:MGI Symb | 0 | 0 | 0 | 0 | 0 | 0 |
| 45815 | ENSMUSG00000107977 | 0 | NA | NA | NA | NA | NA | Gm44121 | predicted gene, 44121 [Source:MGI Symb | 0 | 0 | 0 | 0 | 0 | 0 |
| 45816 | ENSMUSG00000107978 | 0 | NA | NA | NA | NA | NA | Gm44109 | predicted gene, 44109 [Source:MGI Symb | 0 | 0 | 0 | 0 | 0 | 0 |
| 45817 | ENSMUSG00000107979 | 0 | NA | NA | NA | NA | NA | Zfp813-ps | zinc finger protein 813, pseudogene [Sour | 0 | 0 | 0 | 0 | 0 | 0 |
| 45818 | ENSMUSG00000107980 | 0 | NA | NA | NA | NA | NA | Gm44142 | predicted gene, 44142 [Source:MGI Symb | 0 | 0 | 0 | 0 | 0 | 0 |
| 45819 | ENSMUSG00000107981 | 0 | NA | NA | NA | NA | NA | Gm6064 | predicted gene 6064 [Source:MGI Symbol | 0 | 0 | 0 | 0 | 0 | 0 |
| 45820 | ENSMUSG00000107982 | 0 | NA | NA | NA | NA | NA | Gm44219 | predicted gene, 44219 [Source:MGI Symb | 0 | 0 | 0 | 0 | 0 | 0 |
| 45821 | ENSMUSG00000107983 | 0 | NA | NA | NA | NA | NA | Gm43973 | predicted gene, 43973 [Source:MGI Symb | 0 | 0 | 0 | 0 | 0 | 0 |
| 45822 | ENSMUSG00000107984 | 0 | NA | NA | NA | NA | NA | Gm5306 | predicted gene 5306 [Source:MGI Symbol | 0 | 0 | 0 | 0 | 0 | 0 |
| 45823 | ENSMUSG00000107985 | 0.133850373 | -0.796629176 | 4.080472857 | -0.195229623 | 0.84521319 | NA | Gm35037 | predicted gene, 35037 [Source:MGI Symb | 0.8031 | 0 | 0 | 0 | 0 | 0 |
| 45824 | ENSMUSG00000107986 | 0 | NA | NA | NA | NA | NA | Olfr454-ps1 | olfactory receptor 454, pseudogene 1 [Sou | 0 | 0 | 0 | 0 | 0 | 0 |
| 45825 | ENSMUSG00000107987 | 0 | NA | NA | NA | NA | NA | Gm43905 | predicted gene, 43905 [Source:MGI Symb | 0 | 0 | 0 | 0 | 0 | 0 |
| 45826 | ENSMUSG00000107988 | 0 | NA | NA | NA | NA | NA | Gm44006 | predicted gene, 44006 [Source:MGI Symb | 0 | 0 | 0 | 0 | 0 | 0 |
| 45827 | ENSMUSG00000107989 | 0 | NA | NA | NA | NA | NA | Gm19692 | predicted gene, 19692 [Source:MGI Symb | 0 | 0 | 0 | 0 | 0 | 0 |
| 45828 | ENSMUSG00000107990 | 7.665740952 | -0.53201616 | 0.778514132 | -0.6833738 | 0.494370671 | 0.877473722 | Gm7890 | predicted gene 7890 [Source:MGI Symbol | 12.85 | 8.2233 | 6.0375 | 6.7478 | 5.4531 | 6.6831 |
| 45829 | ENSMUSG00000107991 | 0 | NA | NA | NA | NA | NA | Gm43919 | predicted gene, 43919 [Source:MGI Symb | 0 | 0 | 0 | 0 | 0 | 0 |
| 45830 | ENSMUSG00000107992 | 0 | NA | NA | NA | NA | NA | Gm44223 | predicted gene, 44223 [Source:MGI Symb | 0 | 0 | 0 | 0 | 0 | 0 |
| 45831 | ENSMUSG00000107993 | 0 | NA | NA | NA | NA | NA | G930045G22 | RIKEN cDNA G930045G22 gene [Source:M | 0 | 0 | 0 | 0 | 0 | 0 |
| 45832 | ENSMUSG00000107994 | 42.11594414 | -0.808270038 | 0.375065582 | -2.155009889 | 0.03116106 | 0.234334085 | D830050J10 | RIKEN cDNA D830050J10 gene [Source:M | 54.611 | 58.591 | 47.293 | 18.316 | 34.9 | 38.985 |
| 45833 | ENSMUSG00000107995 | 0 | NA | NA | NA | NA | NA | Gm44487 | predicted gene, 44487 [Source:MGI Symb | 0 | 0 | 0 | 0 | 0 | 0 |
| 45834 | ENSMUSG00000107996 | 0 | NA | NA | NA | NA | NA | Gm44228 | predicted gene, 44228 [Source:MGI Symb | 0 | 0 | 0 | 0 | 0 | 0 |
| 45835 | ENSMUSG00000107997 | 14.27859267 | -0.141065456 | 0.566005535 | -0.249229817 | 0.803183016 | 0.963596811 | Gm44243 | predicted gene, 44243 [Source:MGI Symb | 14.456 | 17.475 | 13.081 | 16.388 | 10.906 | 13.366 |
| 45836 | ENSMUSG00000107998 | 0 | NA | NA | NA | NA | NA | Gm44018 | predicted gene, 44018 [Source:MGI Symb | 0 | 0 | 0 | 0 | 0 | 0 |
| 45837 | ENSMUSG00000107999 | 0 | NA | NA | NA | NA | NA | Gm44123 | predicted gene, 44123 [Source:MGI Symb | 0 | 0 | 0 | 0 | 0 | 0 |
| 45838 | ENSMUSG00000108000 | 0 | NA | NA | NA | NA | NA | Gm44208 | predicted gene, 44208 [Source:MGI Symb | 0 | 0 | 0 | 0 | 0 | 0 |
| 45839 | ENSMUSG00000108002 | 0 | NA | NA | NA | NA | NA | Gm45063 | predicted gene 45063 [Source:MGI Symbo | 0 | 0 | 0 | 0 | 0 | 0 |
| 45840 | ENSMUSG00000108003 | 0 | NA | NA | NA | NA | NA | Gm35386 | predicted gene, 35386 [Source:MGI Symb | 0 | 0 | 0 | 0 | 0 | 0 |
| 45841 | ENSMUSG00000108004 | 0 | NA | NA | NA | NA | NA | Gm44080 | predicted gene, 44080 [Source:MGI Symb | 0 | 0 | 0 | 0 | 0 | 0 |
| 45842 | ENSMUSG00000108005 | 0 | NA | NA | NA | NA | NA | Gm44435 | predicted gene, 44435 [Source:MGI Symb | 0 | 0 | 0 | 0 | 0 | 0 |
| 45843 | ENSMUSG00000108006 | 0.167707431 | -0.796629176 | 4.080472857 | -0.195229623 | 0.84521319 | NA | Gm44418 | predicted gene, 44418 [Source:MGI Symb | 0 | 0 | 1.0062 | 0 | 0 | 0 |
| 45844 | ENSMUSG00000108007 | 0 | NA | NA | NA | NA | NA | Gm18540 | predicted gene, 18540 [Source:MGI Symb | 0 | 0 | 0 | 0 | 0 | 0 |
| 45845 | ENSMUSG00000108008 | 0 | NA | NA | NA | NA | NA | Gm43869 | predicted gene, 43869 [Source:MGI Symb | 0 | 0 | 0 | 0 | 0 | 0 |
| 45846 | ENSMUSG00000108009 | 0 | NA | NA | NA | NA | NA | Gm44279 | predicted gene, 44279 [Source:MGI Symb | 0 | 0 | 0 | 0 | 0 | 0 |
| 45847 | ENSMUSG00000108010 | 0 | NA | NA | NA | NA | NA | Gm38708 | predicted gene, 38708 [Source:MGI Symb | 0 | 0 | 0 | 0 | 0 | 0 |
| 45848 | ENSMUSG00000108011 | 0 | NA | NA | NA | NA | NA | Gm44596 | predicted gene 44596 [Source:MGI Symbo | 0 | 0 | 0 | 0 | 0 | 0 |
| 45849 | ENSMUSG00000108012 | 0 | NA | NA | NA | NA | NA | Gm44012 | predicted gene, 44012 [Source:MGI Symb | 0 | 0 | 0 | 0 | 0 | 0 |
| 45850 | ENSMUSG00000108013 | 0 | NA | NA | NA | NA | NA | Gm44052 | predicted gene, 44052 [Source:MGI Symb | 0 | 0 | 0 | 0 | 0 | 0 |
| 45851 | ENSMUSG00000108014 | 0.356960287 | 0.16514264 | 3.880370363 | 0.042558474 | 0.966053499 | NA | Gm44380 | predicted gene, 44380 [Source:MGI Symb | 0 | 1.0279 | 0 | 0 | 0 | 1.1138 |
| 45852 | ENSMUSG00000108015 | 0 | NA | NA | NA | NA | NA | Gm32591 | predicted gene, 32591 [Source:MGI Symb | 0 | 0 | 0 | 0 | 0 | 0 |
| 45853 | ENSMUSG00000108016 | 0 | NA | NA | NA | NA | NA | Gm8039 | predicted gene 8039 [Source:MGI Symbol | 0 | 0 | 0 | 0 | 0 | 0 |
| 45854 | ENSMUSG00000108017 | 0 | NA | NA | NA | NA | NA | Gm18539 | predicted gene, 18539 [Source:MGI Symb | 0 | 0 | 0 | 0 | 0 | 0 |
| 45855 | ENSMUSG00000108018 | 1.247492652 | -2.525342597 | 2.270614284 | -1.112184758 | 0.26605872 | NA | Gm43928 | predicted gene, 43928 [Source:MGI Symb | 2.4093 | 4.1117 | 0 | 0.964 | 0 | 0 |
| 45856 | ENSMUSG00000108019 | 0 | NA | NA | NA | NA | NA | Gm44108 | predicted gene, 44108 [Source:MGI Symb | 0 | 0 | 0 | 0 | 0 | 0 |
| 45857 | ENSMUSG00000108020 | 0 | NA | NA | NA | NA | NA | 6820426E19 | RIKEN cDNA 6820426E19 gene [Source:M | 0 | 0 | 0 | 0 | 0 | 0 |
| 45858 | ENSMUSG00000108021 | 0.133850373 | -0.796629176 | 4.080472857 | -0.195229623 | 0.84521319 | NA | Gm44049 | predicted gene, 44049 [Source:MGI Symb | 0.8031 | 0 | 0 | 0 | 0 | 0 |
| 45859 | ENSMUSG00000108022 | 0 | NA | NA | NA | NA | NA | Gm7298 | predicted gene 7298 [Source:MGI Symbol | 0 | 0 | 0 | 0 | 0 | 0 |
| 45860 | ENSMUSG00000108023 | 0 | NA | NA | NA | NA | NA | Gm44245 | predicted gene, 44245 [Source:MGI Symb | 0 | 0 | 0 | 0 | 0 | 0 |

| | | | | | | | | | | | | | | | |
|---|---|---|---|---|---|---|---|---|---|---|---|---|---|---|---|
| 45861 | ENSMUSG00000108024 | 5.746081634 | 0.275093947 | 0.903590733 | 0.304445295 | 0.760788664 | 0.95402891 | Gm43912 | predicted gene, 43912 [Source:MGI Symb | 6.4248 | 3.0838 | 6.0375 | 6.7478 | 3.2719 | 8.9108 |
| 45862 | ENSMUSG00000108025 | 0 | NA | NA | NA | NA | NA | Gm36503 | predicted gene, 36503 [Source:MGI Symb | 0 | 0 | 0 | 0 | 0 | 0 |
| 45863 | ENSMUSG00000108026 | 0 | NA | NA | NA | NA | NA | Gm44007 | predicted gene, 44007 [Source:MGI Symb | 0 | 0 | 0 | 0 | 0 | 0 |
| 45864 | ENSMUSG00000108027 | 0 | NA | NA | NA | NA | NA | Gm44271 | predicted gene, 44271 [Source:MGI Symb | 0 | 0 | 0 | 0 | 0 | 0 |
| 45865 | ENSMUSG00000108028 | 0 | NA | NA | NA | NA | NA | Gm8143 | predicted gene 8143 [Source:MGI Symbol | 0 | 0 | 0 | 0 | 0 | 0 |
| 45866 | ENSMUSG00000108029 | 0 | NA | NA | NA | NA | NA | Olfr1556-ps1 | olfactory receptor 1556, pseudogene 1 [So | 0 | 0 | 0 | 0 | 0 | 0 |
| 45867 | ENSMUSG00000108030 | 7.736156877 | 0.178349581 | 0.798980086 | 0.22322156 | 0.823363073 | 0.96943211 | 9530062K07 | RIKEN cDNA 9530062K07 gene [Source:M | 6.4248 | 11.307 | 4.025 | 4.8199 | 9.8156 | 10.025 |
| 45868 | ENSMUSG00000108032 | 0 | NA | NA | NA | NA | NA | Olfr210-ps1 | olfactory receptor 210, pseudogene 1 [So | 0 | 0 | 0 | 0 | 0 | 0 |
| 45869 | ENSMUSG00000108033 | 2.54987341 | -0.943537769 | 1.46071177 | -0.645943839 | 0.518315727 | NA | Gm44284 | predicted gene, 44284 [Source:MGI Symb | 4.0155 | 0 | 6.0375 | 1.928 | 1.0906 | 2.2277 |
| 45870 | ENSMUSG00000108034 | 8.621575007 | 0.393169498 | 0.812064085 | 0.484160679 | 0.628271841 | 0.920328819 | 4930557K07 | RIKEN cDNA 4930557K07 gene [Source:M | 7.2279 | 8.2233 | 7.0437 | 19.28 | 3.2719 | 6.6831 |
| 45871 | ENSMUSG00000108035 | 0 | NA | NA | NA | NA | NA | Gm44021 | predicted gene, 44021 [Source:MGI Symb | 0 | 0 | 0 | 0 | 0 | 0 |
| 45872 | ENSMUSG00000108036 | 0.17131964 | -0.796629176 | 4.080472857 | -0.195229623 | 0.84521319 | NA | Gm43967 | predicted gene, 43967 [Source:MGI Symb | 0 | 1.0279 | 0 | 0 | 0 | 0 |
| 45873 | ENSMUSG00000108037 | 3.909592776 | 0.405703117 | 1.068425634 | 0.379720501 | 0.7041529 | NA | Gm44597 | predicted gene 44597 [Source:MGI Symbo | 4.0155 | 2.0558 | 4.025 | 6.7478 | 3.2719 | 3.3415 |
| 45874 | ENSMUSG00000108038 | 0 | NA | NA | NA | NA | NA | D030044L04 | RIKEN cDNA D030044L04 gene [Source:M | 0 | 0 | 0 | 0 | 0 | 0 |
| 45875 | ENSMUSG00000108039 | 0 | NA | NA | NA | NA | NA | Gm43925 | predicted gene, 43925 [Source:MGI Symb | 0 | 0 | 0 | 0 | 0 | 0 |
| 45876 | ENSMUSG00000108040 | 0 | NA | NA | NA | NA | NA | Gm43900 | predicted gene, 43900 [Source:MGI Symb | 0 | 0 | 0 | 0 | 0 | 0 |
| 45877 | ENSMUSG00000108041 | 0 | NA | NA | NA | NA | NA | Gm5154 | predicted gene 5154 [Source:MGI Symbol | 0 | 0 | 0 | 0 | 0 | 0 |
| 45878 | ENSMUSG00000108042 | 1.631033447 | -4.063825688 | 2.026262421 | -2.005577188 | 0.044901378 | NA | B130021K23 | RIKEN cDNA B130021K23 gene [Source:M | 1.6062 | 6.1675 | 2.0125 | 0 | 0 | 0 |
| 45879 | ENSMUSG00000108043 | 0 | NA | NA | NA | NA | NA | Zim3 | zinc finger, imprinted 3 [Source:MGI Symb | 0 | 0 | 0 | 0 | 0 | 0 |
| 45880 | ENSMUSG00000108044 | 0 | NA | NA | NA | NA | NA | Gm44459 | predicted gene, 44459 [Source:MGI Symb | 0 | 0 | 0 | 0 | 0 | 0 |
| 45881 | ENSMUSG00000108045 | 0 | NA | NA | NA | NA | NA | Gm5576 | predicted pseudogene 5576 [Source:MGI | 0 | 0 | 0 | 0 | 0 | 0 |
| 45882 | ENSMUSG00000108046 | 2.710173922 | 0.468125033 | 1.280338554 | 0.365625976 | 0.71464419 | NA | Gm43924 | predicted gene, 43924 [Source:MGI Symb | 0.8031 | 3.0838 | 3.0187 | 3.8559 | 3.2719 | 2.2277 |
| 45883 | ENSMUSG00000108047 | 0 | NA | NA | NA | NA | NA | Gm44158 | predicted gene, 44158 [Source:MGI Symb | 0 | 0 | 0 | 0 | 0 | 0 |
| 45884 | ENSMUSG00000108048 | 0 | NA | NA | NA | NA | NA | Gm43990 | predicted gene, 43990 [Source:MGI Symb | 0 | 0 | 0 | 0 | 0 | 0 |
| 45885 | ENSMUSG00000108049 | 0 | NA | NA | NA | NA | NA | Gm44168 | predicted gene, 44168 [Source:MGI Symb | 0 | 0 | 0 | 0 | 0 | 0 |
| 45886 | ENSMUSG00000108050 | 0 | NA | NA | NA | NA | NA | Gm44472 | predicted gene, 44472 [Source:MGI Symb | 0 | 0 | 0 | 0 | 0 | 0 |
| 45887 | ENSMUSG00000108052 | 0 | NA | NA | NA | NA | NA | Gm8479 | predicted gene 8479 [Source:MGI Symbol | 0 | 0 | 0 | 0 | 0 | 0 |
| 45888 | ENSMUSG00000108053 | 0.967300313 | -2.048316886 | 2.458008888 | -0.833323629 | 0.404662234 | NA | Gm43890 | predicted gene, 43890 [Source:MGI Symb | 1.6062 | 3.0838 | 0 | 0 | 0 | 1.1138 |
| 45889 | ENSMUSG00000108054 | 0 | NA | NA | NA | NA | NA | Gm44149 | predicted gene, 44149 [Source:MGI Symb | 0 | 0 | 0 | 0 | 0 | 0 |
| 45890 | ENSMUSG00000108055 | 0 | NA | NA | NA | NA | NA | Gm43876 | predicted gene, 43876 [Source:MGI Symb | 0 | 0 | 0 | 0 | 0 | 0 |
| 45891 | ENSMUSG00000108056 | 0 | NA | NA | NA | NA | NA | Gm43941 | predicted gene, 43941 [Source:MGI Symb | 0 | 0 | 0 | 0 | 0 | 0 |
| 45892 | ENSMUSG00000108057 | 0 | NA | NA | NA | NA | NA | Gm44234 | predicted gene, 44234 [Source:MGI Symb | 0 | 0 | 0 | 0 | 0 | 0 |
| 45893 | ENSMUSG00000108058 | 0 | NA | NA | NA | NA | NA | Olfr451-ps1 | olfactory receptor 451, pseudogene 1 [So | 0 | 0 | 0 | 0 | 0 | 0 |
| 45894 | ENSMUSG00000108059 | 0.321325949 | 1.911575114 | 4.048267893 | 0.472195804 | 0.636787034 | NA | Gm44369 | predicted gene, 44369 [Source:MGI Symb | 0 | 0 | 0 | 1.928 | 0 | 0 |
| 45895 | ENSMUSG00000108060 | 0.36353943 | 2.065989254 | 4.034137301 | 0.512126658 | 0.608562368 | NA | 4921529L05 | RIKEN cDNA 4921529L05 gene [Source:M | 0 | 0 | 0 | 0 | 2.1812 | 0 |
| 45896 | ENSMUSG00000108062 | 0 | NA | NA | NA | NA | NA | Gm44028 | predicted gene, 44028 [Source:MGI Symb | 0 | 0 | 0 | 0 | 0 | 0 |
| 45897 | ENSMUSG00000108063 | 0 | NA | NA | NA | NA | NA | Gm3279 | predicted gene 3279 [Source:MGI Symbol | 0 | 0 | 0 | 0 | 0 | 0 |
| 45898 | ENSMUSG00000108064 | 28.12640383 | -0.380761691 | 0.489997064 | -0.777069333 | 0.437117866 | 0.846852749 | Gm44423 | predicted gene, 44423 [Source:MGI Symb | 30.518 | 37.005 | 28.175 | 34.703 | 29.447 | 8.9108 |
| 45899 | ENSMUSG00000108065 | 0 | NA | NA | NA | NA | NA | Gm18423 | predicted gene, 18423 [Source:MGI Symb | 0 | 0 | 0 | 0 | 0 | 0 |
| 45900 | ENSMUSG00000108066 | 0 | NA | NA | NA | NA | NA | Gm44134 | predicted gene, 44134 [Source:MGI Symb | 0 | 0 | 0 | 0 | 0 | 0 |
| 45901 | ENSMUSG00000108067 | 0.321325949 | 1.911575114 | 4.048267893 | 0.472195804 | 0.636787034 | NA | Gm43953 | predicted gene, 43953 [Source:MGI Symb | 0 | 0 | 0 | 1.928 | 0 | 0 |
| 45902 | ENSMUSG00000108068 | 0 | NA | NA | NA | NA | NA | Gm44075 | predicted gene, 44075 [Source:MGI Symb | 0 | 0 | 0 | 0 | 0 | 0 |
| 45903 | ENSMUSG00000108069 | 0 | NA | NA | NA | NA | NA | Hspd1-ps1 | heat shock protein 1 (chaperonin), pseudo | 0 | 0 | 0 | 0 | 0 | 0 |
| 45904 | ENSMUSG00000108070 | 0 | NA | NA | NA | NA | NA | Gm43872 | predicted gene, 43872 [Source:MGI Symb | 0 | 0 | 0 | 0 | 0 | 0 |
| 45905 | ENSMUSG00000108071 | 0 | NA | NA | NA | NA | NA | Gm3455 | predicted gene 3455 [Source:MGI Symbol | 0 | 0 | 0 | 0 | 0 | 0 |
| 45906 | ENSMUSG00000108072 | 0 | NA | NA | NA | NA | NA | Gm20362 | predicted gene, 20362 [Source:MGI Symb | 0 | 0 | 0 | 0 | 0 | 0 |
| 45907 | ENSMUSG00000108073 | 0 | NA | NA | NA | NA | NA | Gm6072 | predicted gene 6072 [Source:MGI Symbol | 0 | 0 | 0 | 0 | 0 | 0 |
| 45908 | ENSMUSG00000108074 | 0 | NA | NA | NA | NA | NA | 4930595O18 | RIKEN cDNA 4930595O18 gene [Source:M | 0 | 0 | 0 | 0 | 0 | 0 |
| 45909 | ENSMUSG00000108075 | 0.133850373 | -0.796629176 | 4.080472857 | -0.195229623 | 0.84521319 | NA | Gm45061 | predicted gene 45061 [Source:MGI Symbo | 0.8031 | 0 | 0 | 0 | 0 | 0 |
| 45910 | ENSMUSG00000108076 | 0 | NA | NA | NA | NA | NA | 9530086O07 | RIKEN cDNA 9530086O07 gene [Source:M | 0 | 0 | 0 | 0 | 0 | 0 |
| 45911 | ENSMUSG00000108077 | 0 | NA | NA | NA | NA | NA | 6330415B21 | RIKEN cDNA 6330415B21 gene [Source:M | 0 | 0 | 0 | 0 | 0 | 0 |
| 45912 | ENSMUSG00000108078 | 0 | NA | NA | NA | NA | NA | Gm8744 | predicted gene 8744 [Source:MGI Symbol | 0 | 0 | 0 | 0 | 0 | 0 |
| 45913 | ENSMUSG00000108079 | 0.52105551 | -0.812028029 | 3.385364865 | -0.239864257 | 0.810435491 | NA | Gm44210 | predicted gene, 44210 [Source:MGI Symb | 0 | 0 | 2.0125 | 0 | 0 | 1.1138 |
| 45914 | ENSMUSG00000108080 | 2.952844861 | -0.54670621 | 1.332654667 | -0.410238469 | 0.681631024 | NA | Gm10209 | predicted gene 10209 [Source:MGI Symbo | 2.4093 | 6.1675 | 2.0125 | 3.8559 | 3.2719 | 0 |
| 45915 | ENSMUSG00000108081 | 0.267700746 | -1.497481649 | 4.057001436 | -0.369110456 | 0.712045396 | NA | Gm44026 | predicted gene, 44026 [Source:MGI Symb | 1.6062 | 0 | 0 | 0 | 0 | 0 |
| 45916 | ENSMUSG00000108082 | 0 | NA | NA | NA | NA | NA | Btf3-ps6 | basic transcription factor 3, pseudogene 6 | 0 | 0 | 0 | 0 | 0 | 0 |

| | | | | | | | | | | | | | | | |
|---|---|---|---|---|---|---|---|---|---|---|---|---|---|---|---|
| 45917 | ENSMUSG00000108083 | 0.674674027 | 1.600443604 | 2.78690385 | 0.574272989 | 0.565783085 | NA | Mug4-ps | murinoglobulin 4, pseudogene [Source:M | 0 | 0 | 1.0062 | 1.928 | 0 | 1.1138 |
| 45918 | ENSMUSG00000108084 | 0.17131964 | -0.796629176 | 4.080472857 | -0.195229623 | 0.84521319 | NA | Gm7932 | predicted gene 7932 [Source:MGI Symbol | 0 | 1.0279 | 0 | 0 | 0 | 0 |
| 45919 | ENSMUSG00000108085 | 0 | NA | NA | NA | NA | NA | A930014E10 | RIKEN cDNA A930014E10 gene [Source:M | 0 | 0 | 0 | 0 | 0 | 0 |
| 45920 | ENSMUSG00000108086 | 0 | NA | NA | NA | NA | NA | Gm6786 | predicted gene 6786 [Source:MGI Symbol | 0 | 0 | 0 | 0 | 0 | 0 |
| 45921 | ENSMUSG00000108087 | 0 | NA | NA | NA | NA | NA | Gm43893 | predicted gene, 43893 [Source:MGI Symb | 0 | 0 | 0 | 0 | 0 | 0 |
| 45922 | ENSMUSG00000108088 | 0 | NA | NA | NA | NA | NA | Gm44229 | predicted gene, 44229 [Source:MGI Symb | 0 | 0 | 0 | 0 | 0 | 0 |
| 45923 | ENSMUSG00000108089 | 0 | NA | NA | NA | NA | NA | Gm3148 | predicted gene 3148 [Source:MGI Symbol | 0 | 0 | 0 | 0 | 0 | 0 |
| 45924 | ENSMUSG00000108090 | 0.167707431 | -0.796629176 | 4.080472857 | -0.195229623 | 0.84521319 | NA | Gm6637 | predicted gene 6637 [Source:MGI Symbol | 0 | 0 | 1.0062 | 0 | 0 | 0 |
| 45925 | ENSMUSG00000108091 | 0 | NA | NA | NA | NA | NA | Gm44221 | predicted gene, 44221 [Source:MGI Symb | 0 | 0 | 0 | 0 | 0 | 0 |
| 45926 | ENSMUSG00000108092 | 0 | NA | NA | NA | NA | NA | Gm44189 | predicted gene, 44189 [Source:MGI Symb | 0 | 0 | 0 | 0 | 0 | 0 |
| 45927 | ENSMUSG00000108093 | 0 | NA | NA | NA | NA | NA | Gm44102 | predicted gene, 44102 [Source:MGI Symb | 0 | 0 | 0 | 0 | 0 | 0 |
| 45928 | ENSMUSG00000108094 | 0 | NA | NA | NA | NA | NA | Gm31108 | predicted gene, 31108 [Source:MGI Symb | 0 | 0 | 0 | 0 | 0 | 0 |
| 45929 | ENSMUSG00000108095 | 0 | NA | NA | NA | NA | NA | Gm44067 | predicted gene, 44067 [Source:MGI Symb | 0 | 0 | 0 | 0 | 0 | 0 |
| 45930 | ENSMUSG00000108096 | 0 | NA | NA | NA | NA | NA | Minpp1-ps | multiple inositol polyphosphate histidine | 0 | 0 | 0 | 0 | 0 | 0 |
| 45931 | ENSMUSG00000108097 | 0 | NA | NA | NA | NA | NA | Gm44060 | predicted gene, 44060 [Source:MGI Symb | 0 | 0 | 0 | 0 | 0 | 0 |
| 45932 | ENSMUSG00000108098 | 0 | NA | NA | NA | NA | NA | Gm18838 | predicted gene, 18838 [Source:MGI Symb | 0 | 0 | 0 | 0 | 0 | 0 |
| 45933 | ENSMUSG00000108099 | 0 | NA | NA | NA | NA | NA | Gm43988 | predicted gene, 43988 [Source:MGI Symb | 0 | 0 | 0 | 0 | 0 | 0 |
| 45934 | ENSMUSG00000108100 | 0 | NA | NA | NA | NA | NA | Gm5876 | predicted gene 5876 [Source:MGI Symbol | 0 | 0 | 0 | 0 | 0 | 0 |
| 45935 | ENSMUSG00000108101 | 4.498334117 | 1.050441104 | 1.166747629 | 0.900315611 | 0.367952316 | NA | Gm45738 | predicted gene 45738 [Source:MGI Symbo | 5.6217 | 3.0838 | 0 | 10.604 | 5.4531 | 2.2277 |
| 45936 | ENSMUSG00000108102 | 0 | NA | NA | NA | NA | NA | Gm44437 | predicted gene, 44437 [Source:MGI Symb | 0 | 0 | 0 | 0 | 0 | 0 |
| 45937 | ENSMUSG00000108103 | 0 | NA | NA | NA | NA | NA | Gm44081 | predicted gene, 44081 [Source:MGI Symb | 0 | 0 | 0 | 0 | 0 | 0 |
| 45938 | ENSMUSG00000108104 | 0 | NA | NA | NA | NA | NA | Gm20997 | predicted gene, 20997 [Source:MGI Symb | 0 | 0 | 0 | 0 | 0 | 0 |
| 45939 | ENSMUSG00000108105 | 3.519524445 | -0.706890259 | 1.283537088 | -0.550736139 | 0.581814567 | NA | Gm5340 | predicted gene 5340 [Source:MGI Symbol | 8.031 | 2.0558 | 3.0187 | 5.7839 | 0 | 2.2277 |
| 45940 | ENSMUSG00000108106 | 0 | NA | NA | NA | NA | NA | Gm5886 | predicted gene 5886 [Source:MGI Symbol | 0 | 0 | 0 | 0 | 0 | 0 |
| 45941 | ENSMUSG00000108107 | 0 | NA | NA | NA | NA | NA | Gm5312 | predicted gene 5312 [Source:MGI Symbol | 0 | 0 | 0 | 0 | 0 | 0 |
| 45942 | ENSMUSG00000108108 | 0.542600935 | 1.125931187 | 3.335923981 | 0.33751704 | 0.735727169 | NA | Gm44270 | predicted gene, 44270 [Source:MGI Symb | 0 | 1.0279 | 0 | 0 | 0 | 2.2277 |
| 45943 | ENSMUSG00000108109 | 0.133850373 | -0.796629176 | 4.080472857 | -0.195229623 | 0.84521319 | NA | Gm44287 | predicted gene, 44287 [Source:MGI Symb | 0.8031 | 0 | 0 | 0 | 0 | 0 |
| 45944 | ENSMUSG00000108110 | 0 | NA | NA | NA | NA | NA | Gm18012 | predicted gene, 18012 [Source:MGI Symb | 0 | 0 | 0 | 0 | 0 | 0 |
| 45945 | ENSMUSG00000108111 | 0.435408177 | -2.179616019 | 3.531869654 | -0.617128103 | 0.537150234 | NA | Gm45901 | predicted gene 45901 [Source:MGI Symbo | 1.6062 | 0 | 1.0062 | 0 | 0 | 0 |
| 45946 | ENSMUSG00000108112 | 0 | NA | NA | NA | NA | NA | Gm45193 | predicted gene 45193 [Source:MGI Symbo | 0 | 0 | 0 | 0 | 0 | 0 |
| 45947 | ENSMUSG00000108113 | 0 | NA | NA | NA | NA | NA | Zim2 | zinc finger, imprinted 2 [Source:MGI Symt | 0 | 0 | 0 | 0 | 0 | 0 |
| 45948 | ENSMUSG00000108114 | 0 | NA | NA | NA | NA | NA | Olfr800 | olfactory receptor 800 [Source:MGI Symb | 0 | 0 | 0 | 0 | 0 | 0 |
| 45949 | ENSMUSG00000108115 | 1.82758427 | 0.539929346 | 1.971674318 | 0.273843069 | 0.784205224 | NA | Gm7039 | predicted gene 7039 [Source:MGI Symbol | 2.4093 | 0 | 2.0125 | 0 | 6.5437 | 0 |
| 45950 | ENSMUSG00000108117 | 0 | NA | NA | NA | NA | NA | Gm44262 | predicted gene, 44262 [Source:MGI Symb | 0 | 0 | 0 | 0 | 0 | 0 |
| 45951 | ENSMUSG00000108118 | 0 | NA | NA | NA | NA | NA | Gm34312 | predicted gene, 34312 [Source:MGI Symb | 0 | 0 | 0 | 0 | 0 | 0 |
| 45952 | ENSMUSG00000108119 | 1.727647176 | 4.323686 | 1.998996809 | 2.162927915 | 0.030546724 | NA | Gm43891 | predicted gene, 43891 [Source:MGI Symb | 0 | 0 | 0 | 4.8199 | 1.0906 | 4.4554 |
| 45953 | ENSMUSG00000108120 | 7.803935041 | -0.218357908 | 0.813814022 | -0.268314261 | 0.78845743 | 0.960872359 | 9930120I10F | RIKEN cDNA 9930120I10 gene [Source:MC | 8.031 | 6.1675 | 11.069 | 10.604 | 8.7249 | 2.2277 |
| 45954 | ENSMUSG00000108121 | 0 | NA | NA | NA | NA | NA | Gm43952 | predicted gene, 43952 [Source:MGI Symb | 0 | 0 | 0 | 0 | 0 | 0 |
| 45955 | ENSMUSG00000108122 | 1.719879807 | 0.071582039 | 1.668990294 | 0.042889428 | 0.965789676 | NA | Gm44116 | predicted gene, 44116 [Source:MGI Symb | 0 | 3.0838 | 2.0125 | 1.928 | 2.1812 | 1.1138 |
| 45956 | ENSMUSG00000108123 | 0 | NA | NA | NA | NA | NA | Gm43884 | predicted gene, 43884 [Source:MGI Symb | 0 | 0 | 0 | 0 | 0 | 0 |
| 45957 | ENSMUSG00000108124 | 0 | NA | NA | NA | NA | NA | Gm44146 | predicted gene, 44146 [Source:MGI Symb | 0 | 0 | 0 | 0 | 0 | 0 |
| 45958 | ENSMUSG00000108125 | 0.321325949 | 1.911575114 | 4.048267893 | 0.472195804 | 0.636787034 | NA | Gm17812 | predicted gene, 17812 [Source:MGI Symb | 0 | 0 | 0 | 1.928 | 0 | 0 |
| 45959 | ENSMUSG00000108126 | 0.556921942 | 2.679043159 | 3.638243063 | 0.736356289 | 0.461513899 | NA | Gm43909 | predicted gene, 43909 [Source:MGI Symb | 0 | 0 | 0 | 0 | 0 | 3.3415 |
| 45960 | ENSMUSG00000108127 | 0 | NA | NA | NA | NA | NA | Gm12845 | predicted gene 12845 [Source:MGI Symbo | 0 | 0 | 0 | 0 | 0 | 0 |
| 45961 | ENSMUSG00000108128 | 0 | NA | NA | NA | NA | NA | 4922502N22 | RIKEN cDNA 4922502N22 gene [Source:M | 0 | 0 | 0 | 0 | 0 | 0 |
| 45962 | ENSMUSG00000108129 | 0.758511407 | -1.621915645 | 2.674939911 | -0.606337226 | 0.544290844 | NA | 4930417O13 | RIKEN cDNA 4930417O13 gene [Source:M | 2.4093 | 1.0279 | 0 | 0 | 0 | 1.1138 |
| 45963 | ENSMUSG00000108130 | 0 | NA | NA | NA | NA | NA | Gm30781 | predicted gene, 30781 [Source:MGI Symb | 0 | 0 | 0 | 0 | 0 | 0 |
| 45964 | ENSMUSG00000108131 | 0 | NA | NA | NA | NA | NA | Gm44099 | predicted gene, 44099 [Source:MGI Symb | 0 | 0 | 0 | 0 | 0 | 0 |
| 45965 | ENSMUSG00000108132 | 0.401551119 | -2.072982726 | 4.008251979 | -0.517178744 | 0.605031388 | NA | Gm44175 | predicted gene, 44175 [Source:MGI Symb | 2.4093 | 0 | 0 | 0 | 0 | 0 |
| 45966 | ENSMUSG00000108134 | 0 | NA | NA | NA | NA | NA | Gm9560 | predicted gene 9560 [Source:MGI Symbol | 0 | 0 | 0 | 0 | 0 | 0 |
| 45967 | ENSMUSG00000108135 | 0 | NA | NA | NA | NA | NA | Gm8035 | predicted gene 8035 [Source:MGI Symbol | 0 | 0 | 0 | 0 | 0 | 0 |
| 45968 | ENSMUSG00000108137 | 4.087408935 | 0.662070786 | 1.062047307 | 0.62339105 | 0.533027578 | NA | Gm44053 | predicted gene, 44053 [Source:MGI Symb | 2.4093 | 2.0558 | 5.0312 | 2.8919 | 5.4531 | 6.6831 |
| 45969 | ENSMUSG00000108138 | 0 | NA | NA | NA | NA | NA | Gm18485 | predicted gene, 18485 [Source:MGI Symb | 0 | 0 | 0 | 0 | 0 | 0 |
| 45970 | ENSMUSG00000108139 | 0 | NA | NA | NA | NA | NA | Gm44055 | predicted gene, 44055 [Source:MGI Symb | 0 | 0 | 0 | 0 | 0 | 0 |
| 45971 | ENSMUSG00000108140 | 0 | NA | NA | NA | NA | NA | Gm44803 | predicted gene 44803 [Source:MGI Symbo | 0 | 0 | 0 | 0 | 0 | 0 |
| 45972 | ENSMUSG00000108141 | 0.481988923 | 2.492401104 | 3.837077151 | 0.649557203 | 0.515978285 | NA | Gm44079 | predicted gene, 44079 [Source:MGI Symb | 0 | 0 | 0 | 2.8919 | 0 | 0 |

| | | | | | | | | | | | | | | | |
|---|---|---|---|---|---|---|---|---|---|---|---|---|---|---|---|
| 45973 | ENSMUSG00000108142 | 0 | NA | NA | NA | NA | NA | Gm7498 | predicted gene 7498 [Source:MGI Symbol | 0 | 0 | 0 | 0 | 0 | 0 |
| 45974 | ENSMUSG00000108143 | 0 | NA | NA | NA | NA | NA | Gm44414 | predicted gene, 44414 [Source:MGI Symb | 0 | 0 | 0 | 0 | 0 | 0 |
| 45975 | ENSMUSG00000108144 | 0 | NA | NA | NA | NA | NA | Gm43975 | predicted gene, 43975 [Source:MGI Symb | 0 | 0 | 0 | 0 | 0 | 0 |
| 45976 | ENSMUSG00000108145 | 0.160662974 | 1.126924175 | 4.080472857 | 0.276174898 | 0.782413736 | NA | Gm38811 | predicted gene, 38811 [Source:MGI Symb | 0 | 0 | 0 | 0.964 | 0 | 0 |
| 45977 | ENSMUSG00000108146 | 0 | NA | NA | NA | NA | NA | Gm43889 | predicted gene, 43889 [Source:MGI Symb | 0 | 0 | 0 | 0 | 0 | 0 |
| 45978 | ENSMUSG00000108147 | 0 | NA | NA | NA | NA | NA | 4930443G03 | RIKEN cDNA 4930443G03 gene [Source:M | 0 | 0 | 0 | 0 | 0 | 0 |
| 45979 | ENSMUSG00000108148 | 0 | NA | NA | NA | NA | NA | Gm44020 | predicted gene, 44020 [Source:MGI Symb | 0 | 0 | 0 | 0 | 0 | 0 |
| 45980 | ENSMUSG00000108149 | 0 | NA | NA | NA | NA | NA | Olfr439-ps1 | olfactory receptor 439, pseudogene 1 [So | 0 | 0 | 0 | 0 | 0 | 0 |
| 45981 | ENSMUSG00000108150 | 0 | NA | NA | NA | NA | NA | Gm44470 | predicted gene, 44470 [Source:MGI Symb | 0 | 0 | 0 | 0 | 0 | 0 |
| 45982 | ENSMUSG00000108151 | 0 | NA | NA | NA | NA | NA | Gm33201 | predicted gene, 33201 [Source:MGI Symb | 0 | 0 | 0 | 0 | 0 | 0 |
| 45983 | ENSMUSG00000108152 | 0.133850373 | -0.796629176 | 4.080472857 | -0.195229623 | 0.84521319 | NA | Gm44152 | predicted gene, 44152 [Source:MGI Symb | 0.8031 | 0 | 0 | 0 | 0 | 0 |
| 45984 | ENSMUSG00000108153 | 0 | NA | NA | NA | NA | NA | Gm44402 | predicted gene, 44402 [Source:MGI Symb | 0 | 0 | 0 | 0 | 0 | 0 |
| 45985 | ENSMUSG00000108154 | 1.031143496 | -2.151070968 | 2.421593974 | -0.888287215 | 0.374386275 | NA | Gm44033 | predicted gene, 44033 [Source:MGI Symb | 0 | 3.0838 | 2.0125 | 0 | 1.0906 | 0 |
| 45986 | ENSMUSG00000108155 | 0 | NA | NA | NA | NA | NA | Gm44443 | predicted gene, 44443 [Source:MGI Symb | 0 | 0 | 0 | 0 | 0 | 0 |
| 45987 | ENSMUSG00000108156 | 0 | NA | NA | NA | NA | NA | Olfr326-ps1 | olfactory receptor 326, pseudogene 1 [So | 0 | 0 | 0 | 0 | 0 | 0 |
| 45988 | ENSMUSG00000108157 | 0 | NA | NA | NA | NA | NA | Olrf445-ps1 | olfactory receptor 445, pseudogene 1 [So | 0 | 0 | 0 | 0 | 0 | 0 |
| 45989 | ENSMUSG00000108158 | 0 | NA | NA | NA | NA | NA | Gm44002 | predicted gene, 44002 [Source:MGI Symb | 0 | 0 | 0 | 0 | 0 | 0 |
| 45990 | ENSMUSG00000108160 | 0 | NA | NA | NA | NA | NA | Gm19237 | predicted gene, 19237 [Source:MGI Symb | 0 | 0 | 0 | 0 | 0 | 0 |
| 45991 | ENSMUSG00000108161 | 0 | NA | NA | NA | NA | NA | Gm32914 | predicted gene, 32914 [Source:MGI Symb | 0 | 0 | 0 | 0 | 0 | 0 |
| 45992 | ENSMUSG00000108162 | 0 | NA | NA | NA | NA | NA | Gm20589 | predicted gene, 20589 [Source:MGI Symb | 0 | 0 | 0 | 0 | 0 | 0 |
| 45993 | ENSMUSG00000108163 | 0 | NA | NA | NA | NA | NA | Gm44431 | predicted gene, 44431 [Source:MGI Symb | 0 | 0 | 0 | 0 | 0 | 0 |
| 45994 | ENSMUSG00000108164 | 0 | NA | NA | NA | NA | NA | Gm43899 | predicted gene, 43899 [Source:MGI Symb | 0 | 0 | 0 | 0 | 0 | 0 |
| 45995 | ENSMUSG00000108166 | 2.166539207 | -1.097676044 | 1.657199367 | -0.662368129 | 0.507735324 | NA | Gm44017 | predicted gene, 44017 [Source:MGI Symb | 4.8186 | 0 | 4.025 | 1.928 | 0 | 2.2277 |
| 45996 | ENSMUSG00000108167 | 0 | NA | NA | NA | NA | NA | Olfr1386 | olfactory receptor 1386 [Source:MGI Sym | 0 | 0 | 0 | 0 | 0 | 0 |
| 45997 | ENSMUSG00000108168 | 0 | NA | NA | NA | NA | NA | Gm43864 | predicted gene, 43864 [Source:MGI Symb | 0 | 0 | 0 | 0 | 0 | 0 |
| 45998 | ENSMUSG00000108169 | 0 | NA | NA | NA | NA | NA | Gm43958 | predicted gene, 43958 [Source:MGI Symb | 0 | 0 | 0 | 0 | 0 | 0 |
| 45999 | ENSMUSG00000108170 | 8.129749441 | 1.133226737 | 0.875195981 | 1.294826258 | 0.195380172 | 0.635641153 | Gm43966 | predicted gene, 43966 [Source:MGI Symb | 8.031 | 6.1675 | 1.0062 | 19.28 | 8.7249 | 5.5692 |
| 46000 | ENSMUSG00000108171 | 1.399173391 | -2.736606948 | 2.259794983 | -1.210997886 | 0.225896215 | NA | Gm43915 | predicted gene, 43915 [Source:MGI Symb | 6.4248 | 0 | 1.0062 | 0.964 | 0 | 0 |
| 46001 | ENSMUSG00000108172 | 0 | NA | NA | NA | NA | NA | Pou5f1-rs5 | POU domain, class 5, transcription factor | 0 | 0 | 0 | 0 | 0 | 0 |
| 46002 | ENSMUSG00000108173 | 0 | NA | NA | NA | NA | NA | Gm44231 | predicted gene, 44231 [Source:MGI Symb | 0 | 0 | 0 | 0 | 0 | 0 |
| 46003 | ENSMUSG00000108174 | 0 | NA | NA | NA | NA | NA | Gm44092 | predicted gene, 44092 [Source:MGI Symb | 0 | 0 | 0 | 0 | 0 | 0 |
| 46004 | ENSMUSG00000108175 | 0 | NA | NA | NA | NA | NA | Gm16499 | predicted gene 16499 [Source:MGI Symbo | 0 | 0 | 0 | 0 | 0 | 0 |
| 46005 | ENSMUSG00000108176 | 0 | NA | NA | NA | NA | NA | Gm43965 | predicted gene, 43965 [Source:MGI Symb | 0 | 0 | 0 | 0 | 0 | 0 |
| 46006 | ENSMUSG00000108177 | 0 | NA | NA | NA | NA | NA | Gm7924 | predicted gene 7924 [Source:MGI Symbol | 0 | 0 | 0 | 0 | 0 | 0 |
| 46007 | ENSMUSG00000108178 | 0 | NA | NA | NA | NA | NA | Gm44267 | predicted gene, 44267 [Source:MGI Symb | 0 | 0 | 0 | 0 | 0 | 0 |
| 46008 | ENSMUSG00000108179 | 0 | NA | NA | NA | NA | NA | Gm44274 | predicted gene, 44274 [Source:MGI Symb | 0 | 0 | 0 | 0 | 0 | 0 |
| 46009 | ENSMUSG00000108180 | 0.944152054 | -0.673133018 | 2.474469057 | -0.272031293 | 0.785597958 | NA | Gm44141 | predicted gene, 44141 [Source:MGI Symb | 2.4093 | 1.0279 | 0 | 0 | 0 | 2.2277 |
| 46010 | ENSMUSG00000108181 | 32.04816298 | -0.119322251 | 0.384778753 | -0.310106132 | 0.756480249 | 0.953085143 | C030015A19 | RIKEN cDNA C030015A19 gene [Source:M | 32.124 | 32.893 | 35.219 | 33.739 | 33.809 | 24.505 |
| 46011 | ENSMUSG00000108183 | 0 | NA | NA | NA | NA | NA | Gm44227 | predicted gene, 44227 [Source:MGI Symb | 0 | 0 | 0 | 0 | 0 | 0 |
| 46012 | ENSMUSG00000108184 | 0 | NA | NA | NA | NA | NA | Gm44198 | predicted gene, 44198 [Source:MGI Symb | 0 | 0 | 0 | 0 | 0 | 0 |
| 46013 | ENSMUSG00000108185 | 0 | NA | NA | NA | NA | NA | Gm19653 | predicted gene, 19653 [Source:MGI Symb | 0 | 0 | 0 | 0 | 0 | 0 |
| 46014 | ENSMUSG00000108186 | 0 | NA | NA | NA | NA | NA | Gm8132 | predicted gene 8132 [Source:MGI Symbol | 0 | 0 | 0 | 0 | 0 | 0 |
| 46015 | ENSMUSG00000108187 | 0 | NA | NA | NA | NA | NA | 4930511E03 | RIKEN cDNA 4930511E03 gene [Source:M | 0 | 0 | 0 | 0 | 0 | 0 |
| 46016 | ENSMUSG00000108188 | 0 | NA | NA | NA | NA | NA | Gm43865 | predicted gene, 43865 [Source:MGI Symb | 0 | 0 | 0 | 0 | 0 | 0 |
| 46017 | ENSMUSG00000108189 | 0 | NA | NA | NA | NA | NA | Gm43887 | predicted gene, 43887 [Source:MGI Symb | 0 | 0 | 0 | 0 | 0 | 0 |
| 46018 | ENSMUSG00000108190 | 0.760346335 | -0.431432104 | 2.662343224 | -0.162049769 | 0.871266662 | NA | Gm44731 | predicted gene 44731 [Source:MGI Symbo | 1.6062 | 1.0279 | 0 | 1.928 | 0 | 0 |
| 46019 | ENSMUSG00000108191 | 0.36353943 | 2.065989254 | 4.034137301 | 0.512126658 | 0.608562368 | NA | Gm29803 | predicted gene, 29803 [Source:MGI Symb | 0 | 0 | 0 | 0 | 2.1812 | 0 |
| 46020 | ENSMUSG00000108192 | 0 | NA | NA | NA | NA | NA | Gm44410 | predicted gene, 44410 [Source:MGI Symb | 0 | 0 | 0 | 0 | 0 | 0 |
| 46021 | ENSMUSG00000108193 | 0 | NA | NA | NA | NA | NA | Olfr442-ps1 | olfactory receptor 442, pseudogene 1 [So | 0 | 0 | 0 | 0 | 0 | 0 |
| 46022 | ENSMUSG00000108194 | 0 | NA | NA | NA | NA | NA | 1700097M2: | RIKEN cDNA 1700097M23 gene [Source:M | 0 | 0 | 0 | 0 | 0 | 0 |
| 46023 | ENSMUSG00000108195 | 0 | NA | NA | NA | NA | NA | Gm44415 | predicted gene, 44415 [Source:MGI Symb | 0 | 0 | 0 | 0 | 0 | 0 |
| 46024 | ENSMUSG00000108196 | 1.059061473 | 2.395592859 | 2.430477497 | 0.985647002 | 0.324306365 | NA | Gm36328 | predicted gene, 36328 [Source:MGI Symb | 0 | 1.0279 | 0 | 0.964 | 4.3625 | 0 |
| 46025 | ENSMUSG00000108197 | 0.167707431 | -0.796629176 | 4.080472857 | -0.195229623 | 0.84521319 | NA | Gm44214 | predicted gene, 44214 [Source:MGI Symb | 0 | 0 | 1.0062 | 0 | 0 | 0 |
| 46026 | ENSMUSG00000108198 | 0 | NA | NA | NA | NA | NA | Gm44823 | predicted gene 44823 [Source:MGI Symbo | 0 | 0 | 0 | 0 | 0 | 0 |
| 46027 | ENSMUSG00000108199 | 1.133230463 | -1.272694062 | 2.134229246 | -0.596324909 | 0.550958194 | NA | Gm44249 | predicted gene, 44249 [Source:MGI Symb | 0.8031 | 2.0558 | 2.0125 | 1.928 | 0 | 0 |
| 46028 | ENSMUSG00000108200 | 0 | NA | NA | NA | NA | NA | Gm43931 | predicted gene, 43931 [Source:MGI Symb | 0 | 0 | 0 | 0 | 0 | 0 |

| | | | | | | | | | | | | | | | |
|---|---|---|---|---|---|---|---|---|---|---|---|---|---|---|---|
| 46029 | ENSMUSG00000108201 | 0 | NA | NA | NA | NA | NA | Gm43991 | predicted gene, 43991 [Source:MGI Symb | 0 | 0 | 0 | 0 | 0 | 0 |
| 46030 | ENSMUSG00000108202 | 0 | NA | NA | NA | NA | NA | Gm4874 | predicted gene 4874 [Source:MGI Symbol | 0 | 0 | 0 | 0 | 0 | 0 |
| 46031 | ENSMUSG00000108203 | 0 | NA | NA | NA | NA | NA | Gm43944 | predicted gene, 43944 [Source:MGI Symb | 0 | 0 | 0 | 0 | 0 | 0 |
| 46032 | ENSMUSG00000108204 | 0 | NA | NA | NA | NA | NA | Gm44180 | predicted gene, 44180 [Source:MGI Symb | 0 | 0 | 0 | 0 | 0 | 0 |
| 46033 | ENSMUSG00000108205 | 0 | NA | NA | NA | NA | NA | Gm19252 | predicted gene, 19252 [Source:MGI Symb | 0 | 0 | 0 | 0 | 0 | 0 |
| 46034 | ENSMUSG00000108206 | 5.39881494 | 0.375287653 | 1.10270972 | 0.340332226 | 0.733606351 | 0.947915087 | Gm44427 | predicted gene, 44427 [Source:MGI Symb | 8.8341 | 3.0838 | 2.0125 | 2.8919 | 1.0906 | 14.48 |
| 46035 | ENSMUSG00000108207 | 92.06331331 | 1.017321125 | 0.279776805 | 3.636188225 | 0.000276702 | 0.008134594 | 1810059H22 | RIKEN cDNA 1810059H22 gene [Source:M | 74.689 | 41.117 | 66.412 | 147.49 | 109.06 | 113.61 |
| 46036 | ENSMUSG00000108208 | 0 | NA | NA | NA | NA | NA | Gm44184 | predicted gene, 44184 [Source:MGI Symb | 0 | 0 | 0 | 0 | 0 | 0 |
| 46037 | ENSMUSG00000108209 | 0 | NA | NA | NA | NA | NA | Gm32222 | predicted gene, 32222 [Source:MGI Symb | 0 | 0 | 0 | 0 | 0 | 0 |
| 46038 | ENSMUSG00000108210 | 0.945754888 | -3.283756462 | 2.374797396 | -1.382752258 | 0.166740842 | NA | Gm35808 | predicted gene, 35808 [Source:MGI Symb | 1.6062 | 2.0558 | 2.0125 | 0 | 0 | 0 |
| 46039 | ENSMUSG00000108211 | 1.565026432 | -1.892347922 | 1.923521388 | -0.983793543 | 0.325217035 | NA | Gm44130 | predicted gene, 44130 [Source:MGI Symb | 2.4093 | 1.0279 | 4.025 | 1.928 | 0 | 0 |
| 46040 | ENSMUSG00000108212 | 0 | NA | NA | NA | NA | NA | Gm44132 | predicted gene, 44132 [Source:MGI Symb | 0 | 0 | 0 | 0 | 0 | 0 |
| 46041 | ENSMUSG00000108213 | 0.160662974 | 1.126924175 | 4.080472857 | 0.276174898 | 0.782413736 | NA | Gm44172 | predicted gene, 44172 [Source:MGI Symb | 0 | 0 | 0 | 0.964 | 0 | 0 |
| 46042 | ENSMUSG00000108214 | 0.331982614 | 0.16514309 | 3.986751476 | 0.041422971 | 0.966958701 | NA | Gm43982 | predicted gene, 43982 [Source:MGI Symb | 0 | 1.0279 | 0 | 0.964 | 0 | 0 |
| 46043 | ENSMUSG00000108215 | 0 | NA | NA | NA | NA | NA | Gm8083 | predicted gene 8083 [Source:MGI Symbol | 0 | 0 | 0 | 0 | 0 | 0 |
| 46044 | ENSMUSG00000108216 | 0 | NA | NA | NA | NA | NA | Gm44153 | predicted gene, 44153 [Source:MGI Symb | 0 | 0 | 0 | 0 | 0 | 0 |
| 46045 | ENSMUSG00000108217 | 0 | NA | NA | NA | NA | NA | Gm43883 | predicted gene, 43883 [Source:MGI Symb | 0 | 0 | 0 | 0 | 0 | 0 |
| 46046 | ENSMUSG00000108218 | 24.75536002 | -0.692330032 | 0.477467927 | -1.45000322 | 0.147057621 | 0.558278592 | Olfr1372-ps1 | olfactory receptor 1372, pseudogene 1 [S | 40.958 | 23.642 | 27.169 | 26.991 | 14.178 | 15.594 |
| 46047 | ENSMUSG00000108219 | 4.423131056 | -1.33745717 | 1.272488459 | -1.051056424 | 0.293232676 | NA | Gm44101 | predicted gene, 44101 [Source:MGI Symb | 5.6217 | 13.363 | 0 | 0.964 | 4.3625 | 2.2277 |
| 46048 | ENSMUSG00000108220 | 0 | NA | NA | NA | NA | NA | Gm44056 | predicted gene, 44056 [Source:MGI Symb | 0 | 0 | 0 | 0 | 0 | 0 |
| 46049 | ENSMUSG00000108221 | 0.328370405 | 0.16514316 | 4.003149536 | 0.041253308 | 0.967093956 | NA | Gm44203 | predicted gene, 44203 [Source:MGI Symb | 0 | 0 | 1.0062 | 0.964 | 0 | 0 |
| 46050 | ENSMUSG00000108222 | 0 | NA | NA | NA | NA | NA | Gm2691 | predicted gene 2691 [Source:MGI Symbol | 0 | 0 | 0 | 0 | 0 | 0 |
| 46051 | ENSMUSG00000108223 | 0 | NA | NA | NA | NA | NA | Gm43933 | predicted gene, 43933 [Source:MGI Symb | 0 | 0 | 0 | 0 | 0 | 0 |
| 46052 | ENSMUSG00000108224 | 0 | NA | NA | NA | NA | NA | Gm31861 | predicted gene, 31861 [Source:MGI Symb | 0 | 0 | 0 | 0 | 0 | 0 |
| 46053 | ENSMUSG00000108225 | 0 | NA | NA | NA | NA | NA | Gm43926 | predicted gene, 43926 [Source:MGI Symb | 0 | 0 | 0 | 0 | 0 | 0 |
| 46054 | ENSMUSG00000108226 | 0 | NA | NA | NA | NA | NA | Gm44063 | predicted gene, 44063 [Source:MGI Symb | 0 | 0 | 0 | 0 | 0 | 0 |
| 46055 | ENSMUSG00000108227 | 0 | NA | NA | NA | NA | NA | 4933412L11I | RIKEN cDNA 4933412L11 gene [Source:M | 0 | 0 | 0 | 0 | 0 | 0 |
| 46056 | ENSMUSG00000108228 | 0 | NA | NA | NA | NA | NA | 6430584L05I | RIKEN cDNA 6430584L05 gene [Source:M | 0 | 0 | 0 | 0 | 0 | 0 |
| 46057 | ENSMUSG00000108229 | 0 | NA | NA | NA | NA | NA | 1700102F20 | RIKEN cDNA 1700102F20 gene [Source:M | 0 | 0 | 0 | 0 | 0 | 0 |
| 46058 | ENSMUSG00000108230 | 0 | NA | NA | NA | NA | NA | Gm3549 | predicted gene 3549 [Source:MGI Symbol | 0 | 0 | 0 | 0 | 0 | 0 |
| 46059 | ENSMUSG00000108231 | 17.45715039 | 0.561654283 | 0.570055556 | 0.98526236 | 0.324495216 | 0.771309263 | Gm4045 | predicted gene 4045 [Source:MGI Symbol | 13.653 | 17.475 | 11.069 | 9.6398 | 26.175 | 26.732 |
| 46060 | ENSMUSG00000108232 | 0.803314871 | 3.228356836 | 3.175155518 | 1.0167555 | 0.309269756 | NA | Gm43916 | predicted gene, 43916 [Source:MGI Symb | 0 | 0 | 0 | 4.8199 | 0 | 0 |
| 46061 | ENSMUSG00000108233 | 0 | NA | NA | NA | NA | NA | Gm44084 | predicted gene, 44084 [Source:MGI Symb | 0 | 0 | 0 | 0 | 0 | 0 |
| 46062 | ENSMUSG00000108234 | 0.160662974 | 1.126924175 | 4.080472857 | 0.276174898 | 0.782413736 | NA | Gm44428 | predicted gene, 44428 [Source:MGI Symb | 0 | 0 | 0 | 0.964 | 0 | 0 |
| 46063 | ENSMUSG00000108235 | 0 | NA | NA | NA | NA | NA | Gm44248 | predicted gene, 44248 [Source:MGI Symb | 0 | 0 | 0 | 0 | 0 | 0 |
| 46064 | ENSMUSG00000108236 | 1.197041503 | 1.541473088 | 1.980336106 | 0.77838963 | 0.43633935 | NA | 0610033M10 | RIKEN cDNA 0610033M10 gene [Source:M | 0.8031 | 0 | 1.0062 | 0.964 | 2.1812 | 2.2277 |
| 46065 | ENSMUSG00000108237 | 0 | NA | NA | NA | NA | NA | Gm19757 | predicted gene, 19757 [Source:MGI Symb | 0 | 0 | 0 | 0 | 0 | 0 |
| 46066 | ENSMUSG00000108238 | 1.206231217 | -3.642380243 | 2.43040267 | -1.498673569 | 0.133958337 | NA | Gm43984 | predicted gene, 43984 [Source:MGI Symb | 3.2124 | 0 | 4.025 | 0 | 0 | 0 |
| 46067 | ENSMUSG00000108239 | 1.010036755 | -2.151347842 | 2.436507427 | -0.882963794 | 0.377255834 | NA | Gm19434 | predicted gene, 19434 [Source:MGI Symb | 0 | 3.0838 | 2.0125 | 0.964 | 0 | 0 |
| 46068 | ENSMUSG00000108240 | 0 | NA | NA | NA | NA | NA | Gm5317 | predicted gene 5317 [Source:MGI Symbol | 0 | 0 | 0 | 0 | 0 | 0 |
| 46069 | ENSMUSG00000108241 | 0 | NA | NA | NA | NA | NA | Gm44252 | predicted gene, 44252 [Source:MGI Symb | 0 | 0 | 0 | 0 | 0 | 0 |
| 46070 | ENSMUSG00000108242 | 0 | NA | NA | NA | NA | NA | 9330118I20F | RIKEN cDNA 9330118I20 gene [Source:M | 0 | 0 | 0 | 0 | 0 | 0 |
| 46071 | ENSMUSG00000108244 | 0 | NA | NA | NA | NA | NA | Gm44077 | predicted gene, 44077 [Source:MGI Symb | 0 | 0 | 0 | 0 | 0 | 0 |
| 46072 | ENSMUSG00000108245 | 0 | NA | NA | NA | NA | NA | Prb3-ps | proline-rich protein BstNI subfamily 3, pse | 0 | 0 | 0 | 0 | 0 | 0 |
| 46073 | ENSMUSG00000108246 | 0 | NA | NA | NA | NA | NA | Gm43896 | predicted gene, 43896 [Source:MGI Symb | 0 | 0 | 0 | 0 | 0 | 0 |
| 46074 | ENSMUSG00000108247 | 0 | NA | NA | NA | NA | NA | Gm19039 | predicted gene, 19039 [Source:MGI Symb | 0 | 0 | 0 | 0 | 0 | 0 |
| 46075 | ENSMUSG00000108248 | 0 | NA | NA | NA | NA | NA | Gm34391 | predicted gene, 34391 [Source:MGI Symb | 0 | 0 | 0 | 0 | 0 | 0 |
| 46076 | ENSMUSG00000108249 | 0 | NA | NA | NA | NA | NA | Gm43960 | predicted gene, 43960 [Source:MGI Symb | 0 | 0 | 0 | 0 | 0 | 0 |
| 46077 | ENSMUSG00000108250 | 0 | NA | NA | NA | NA | NA | Tas2r111-ps | taste receptor, type 2, member 111, pseu | 0 | 0 | 0 | 0 | 0 | 0 |
| 46078 | ENSMUSG00000108251 | 0 | NA | NA | NA | NA | NA | Gm44112 | predicted gene, 44112 [Source:MGI Symb | 0 | 0 | 0 | 0 | 0 | 0 |
| 46079 | ENSMUSG00000108252 | 0 | NA | NA | NA | NA | NA | Gm44059 | predicted gene, 44059 [Source:MGI Symb | 0 | 0 | 0 | 0 | 0 | 0 |
| 46080 | ENSMUSG00000108253 | 0 | NA | NA | NA | NA | NA | Olfr1542-ps1 | olfactory receptor 1542, pseudogene 1 [S | 0 | 0 | 0 | 0 | 0 | 0 |
| 46081 | ENSMUSG00000108254 | 0 | NA | NA | NA | NA | NA | Gm43908 | predicted gene, 43908 [Source:MGI Symb | 0 | 0 | 0 | 0 | 0 | 0 |
| 46082 | ENSMUSG00000108255 | 0 | NA | NA | NA | NA | NA | Gm16499 | predicted gene 16499 [Source:MGI Symbo | 0 | 0 | 0 | 0 | 0 | 0 |
| 46083 | ENSMUSG00000108256 | 0.988200463 | 0.281796504 | 2.227178257 | 0.126526246 | 0.899315375 | NA | Gm43923 | predicted gene, 43923 [Source:MGI Symb | 1.6062 | 1.0279 | 0 | 0 | 2.1812 | 1.1138 |
| 46084 | ENSMUSG00000108257 | 0 | NA | NA | NA | NA | NA | Gm44251 | predicted gene, 44251 [Source:MGI Symb | 0 | 0 | 0 | 0 | 0 | 0 |

| | | | | | | | | | | | | | | | |
|---|---|---|---|---|---|---|---|---|---|---|---|---|---|---|---|
| 46085 | ENSMUSG00000108258 | 0 | NA | NA | NA | NA | NA | Gm43913 | predicted gene, 43913 [Source:MGI Symb | 0 | 0 | 0 | 0 | 0 | 0 |
| 46086 | ENSMUSG00000108259 | 0 | NA | NA | NA | NA | NA | Gm6727 | predicted gene 6727 [Source:MGI Symbol | 0 | 0 | 0 | 0 | 0 | 0 |
| 46087 | ENSMUSG00000108260 | 0.660353019 | -0.01500701 | 2.83315355 | -0.005296928 | 0.995773683 | NA | Gm3793 | predicted gene 3793 [Source:MGI Symbol | 0 | 1.0279 | 1.0062 | 1.928 | 0 | 0 |
| 46088 | ENSMUSG00000108261 | 0 | NA | NA | NA | NA | NA | Tas2r146-ps | taste receptor, type 2, member 146, pseu | 0 | 0 | 0 | 0 | 0 | 0 |
| 46089 | ENSMUSG00000108262 | 0 | NA | NA | NA | NA | NA | Gm43867 | predicted gene, 43867 [Source:MGI Symb | 0 | 0 | 0 | 0 | 0 | 0 |
| 46090 | ENSMUSG00000108263 | 0 | NA | NA | NA | NA | NA | 1700051K13 | RIKEN cDNA 1700051K13 gene [Source:M | 0 | 0 | 0 | 0 | 0 | 0 |
| 46091 | ENSMUSG00000108264 | 0 | NA | NA | NA | NA | NA | Gm20371 | predicted gene, 20371 [Source:MGI Symb | 0 | 0 | 0 | 0 | 0 | 0 |
| 46092 | ENSMUSG00000108265 | 0 | NA | NA | NA | NA | NA | Olfr318 | olfactory receptor 318 [Source:MGI Symb | 0 | 0 | 0 | 0 | 0 | 0 |
| 46093 | ENSMUSG00000108266 | 0 | NA | NA | NA | NA | NA | Gm5305 | predicted gene 5305 [Source:MGI Symbol | 0 | 0 | 0 | 0 | 0 | 0 |
| 46094 | ENSMUSG00000108267 | 0 | NA | NA | NA | NA | NA | Gm44190 | predicted gene, 44190 [Source:MGI Symb | 0 | 0 | 0 | 0 | 0 | 0 |
| 46095 | ENSMUSG00000108268 | 0.653308562 | 1.538013299 | 3.112409277 | 0.494155223 | 0.62119656 | NA | Gm44187 | predicted gene, 44187 [Source:MGI Symb | 0 | 1.0279 | 0 | 2.8919 | 0 | 0 |
| 46096 | ENSMUSG00000108269 | 0 | NA | NA | NA | NA | NA | Gm18422 | predicted gene, 18422 [Source:MGI Symb | 0 | 0 | 0 | 0 | 0 | 0 |
| 46097 | ENSMUSG00000108270 | 0 | NA | NA | NA | NA | NA | Gm4602 | predicted gene 4602 [Source:MGI Symbol | 0 | 0 | 0 | 0 | 0 | 0 |
| 46098 | ENSMUSG00000108271 | 1.878242539 | 0.954312404 | 1.547105266 | 0.616837409 | 0.537341975 | NA | Gm44236 | predicted gene, 44236 [Source:MGI Symb | 0.8031 | 2.0558 | 1.0062 | 1.928 | 4.3625 | 1.1138 |
| 46099 | ENSMUSG00000108272 | 0 | NA | NA | NA | NA | NA | Gm44161 | predicted gene, 44161 [Source:MGI Symb | 0 | 0 | 0 | 0 | 0 | 0 |
| 46100 | ENSMUSG00000108274 | 0 | NA | NA | NA | NA | NA | Gm44169 | predicted gene, 44169 [Source:MGI Symb | 0 | 0 | 0 | 0 | 0 | 0 |
| 46101 | ENSMUSG00000108275 | 0 | NA | NA | NA | NA | NA | Gm7880 | predicted gene 7880 [Source:MGI Symbol | 0 | 0 | 0 | 0 | 0 | 0 |
| 46102 | ENSMUSG00000108276 | 0 | NA | NA | NA | NA | NA | Gm36640 | predicted gene, 36640 [Source:MGI Symb | 0 | 0 | 0 | 0 | 0 | 0 |
| 46103 | ENSMUSG00000108278 | 0.356960287 | 0.16514264 | 3.880370363 | 0.042558474 | 0.966053499 | NA | Gm44194 | predicted gene, 44194 [Source:MGI Symb | 0 | 1.0279 | 0 | 0 | 0 | 1.1138 |
| 46104 | ENSMUSG00000108279 | 0 | NA | NA | NA | NA | NA | Gm44024 | predicted gene, 44024 [Source:MGI Symb | 0 | 0 | 0 | 0 | 0 | 0 |
| 46105 | ENSMUSG00000108280 | 0.133850373 | -0.796629176 | 4.080472857 | -0.195229623 | 0.84521319 | NA | Gm8703 | predicted gene 8703 [Source:MGI Symbol | 0.8031 | 0 | 0 | 0 | 0 | 0 |
| 46106 | ENSMUSG00000108282 | 0.349477146 | 0.165142768 | 3.911021834 | 0.042224967 | 0.966319359 | NA | Gm44317 | predicted gene, 44317 [Source:MGI Symb | 0 | 0 | 1.0062 | 0 | 1.0906 | 0 |
| 46107 | ENSMUSG00000108283 | 0.720499717 | 1.693787846 | 2.735268864 | 0.619239984 | 0.535758275 | NA | Gm44008 | predicted gene, 44008 [Source:MGI Symb | 0 | 1.0279 | 0 | 0 | 2.1812 | 1.1138 |
| 46108 | ENSMUSG00000108284 | 0 | NA | NA | NA | NA | NA | Gm44182 | predicted gene, 44182 [Source:MGI Symb | 0 | 0 | 0 | 0 | 0 | 0 |
| 46109 | ENSMUSG00000108285 | 0 | NA | NA | NA | NA | NA | Gm34358 | predicted gene, 34358 [Source:MGI Symb | 0 | 0 | 0 | 0 | 0 | 0 |
| 46110 | ENSMUSG00000108286 | 0 | NA | NA | NA | NA | NA | Gm44224 | predicted gene, 44224 [Source:MGI Symb | 0 | 0 | 0 | 0 | 0 | 0 |
| 46111 | ENSMUSG00000108287 | 0 | NA | NA | NA | NA | NA | Gm44473 | predicted gene, 44473 [Source:MGI Symb | 0 | 0 | 0 | 0 | 0 | 0 |
| 46112 | ENSMUSG00000108288 | 11.33769593 | -1.165343548 | 0.666203428 | -1.749230787 | 0.080251134 | 0.41597095 | Gm44202 | predicted gene, 44202 [Source:MGI Symb | 14.456 | 21.586 | 11.069 | 7.7118 | 7.6343 | 5.5692 |
| 46113 | ENSMUSG00000108289 | 0 | NA | NA | NA | NA | NA | Gm44174 | predicted gene, 44174 [Source:MGI Symb | 0 | 0 | 0 | 0 | 0 | 0 |
| 46114 | ENSMUSG00000108290 | 0.859823856 | -1.825223727 | 2.563005556 | -0.712141931 | 0.476376891 | NA | Gm44120 | predicted gene, 44120 [Source:MGI Symb | 0 | 2.0558 | 2.0125 | 0 | 1.0906 | 0 |
| 46115 | ENSMUSG00000108291 | 0.476283062 | 1.031501528 | 3.159856949 | 0.326439312 | 0.744091998 | NA | Gm44292 | predicted gene, 44292 [Source:MGI Symb | 0.8031 | 0 | 0 | 0.964 | 1.0906 | 0 |
| 46116 | ENSMUSG00000108292 | 0 | NA | NA | NA | NA | NA | Gm44050 | predicted gene, 44050 [Source:MGI Symb | 0 | 0 | 0 | 0 | 0 | 0 |
| 46117 | ENSMUSG00000108293 | 0 | NA | NA | NA | NA | NA | Pla2g4c-ps | phospholipase A2, group IVC (cytosolic, ca | 0 | 0 | 0 | 0 | 0 | 0 |
| 46118 | ENSMUSG00000108294 | 0 | NA | NA | NA | NA | NA | Gm44429 | predicted gene, 44429 [Source:MGI Symb | 0 | 0 | 0 | 0 | 0 | 0 |
| 46119 | ENSMUSG00000108295 | 0 | NA | NA | NA | NA | NA | Gm36406 | predicted gene, 36406 [Source:MGI Symb | 0 | 0 | 0 | 0 | 0 | 0 |
| 46120 | ENSMUSG00000108296 | 0 | NA | NA | NA | NA | NA | Gm44295 | predicted gene, 44295 [Source:MGI Symb | 0 | 0 | 0 | 0 | 0 | 0 |
| 46121 | ENSMUSG00000108297 | 0 | NA | NA | NA | NA | NA | Gm44167 | predicted gene, 44167 [Source:MGI Symb | 0 | 0 | 0 | 0 | 0 | 0 |
| 46122 | ENSMUSG00000108298 | 2.003763119 | -0.905257485 | 1.527283951 | -0.59272376 | 0.553366038 | NA | Gm44226 | predicted gene, 44226 [Source:MGI Symb | 0.8031 | 2.0558 | 5.0312 | 1.928 | 1.0906 | 1.1138 |
| 46123 | ENSMUSG00000108299 | 0 | NA | NA | NA | NA | NA | Gm43892 | predicted gene, 43892 [Source:MGI Symb | 0 | 0 | 0 | 0 | 0 | 0 |
| 46124 | ENSMUSG00000108300 | 0 | NA | NA | NA | NA | NA | Gm44539 | predicted gene 44539 [Source:MGI Symbo | 0 | 0 | 0 | 0 | 0 | 0 |
| 46125 | ENSMUSG00000108301 | 0 | NA | NA | NA | NA | NA | Gm45127 | predicted gene 45127 [Source:MGI Symbo | 0 | 0 | 0 | 0 | 0 | 0 |
| 46126 | ENSMUSG00000108302 | 0 | NA | NA | NA | NA | NA | Gm18207 | predicted gene, 18207 [Source:MGI Symb | 0 | 0 | 0 | 0 | 0 | 0 |
| 46127 | ENSMUSG00000108303 | 0 | NA | NA | NA | NA | NA | Gm39075 | predicted gene, 39075 [Source:MGI Symb | 0 | 0 | 0 | 0 | 0 | 0 |
| 46128 | ENSMUSG00000108304 | 0 | NA | NA | NA | NA | NA | Gm4607 | predicted gene 4607 [Source:MGI Symbol | 0 | 0 | 0 | 0 | 0 | 0 |
| 46129 | ENSMUSG00000108305 | 0.321325949 | 1.911575114 | 4.048267893 | 0.472195804 | 0.636787034 | NA | Gm19078 | predicted gene, 19078 [Source:MGI Symb | 0 | 0 | 0 | 1.928 | 0 | 0 |
| 46130 | ENSMUSG00000108306 | 0 | NA | NA | NA | NA | NA | Gm44640 | predicted gene 44640 [Source:MGI Symbo | 0 | 0 | 0 | 0 | 0 | 0 |
| 46131 | ENSMUSG00000108307 | 0 | NA | NA | NA | NA | NA | Ceacam-ps1 | carcinoembryonic antigen-related cell adh | 0 | 0 | 0 | 0 | 0 | 0 |
| 46132 | ENSMUSG00000108308 | 0.692375149 | -1.404276945 | 2.772265464 | -0.506544905 | 0.612474175 | NA | Gm45218 | predicted gene 45218 [Source:MGI Symbo | 0 | 1.0279 | 2.0125 | 0 | 0 | 1.1138 |
| 46133 | ENSMUSG00000108309 | 0 | NA | NA | NA | NA | NA | Gm44703 | predicted gene 44703 [Source:MGI Symbo | 0 | 0 | 0 | 0 | 0 | 0 |
| 46134 | ENSMUSG00000108310 | 0.439020386 | -2.189747911 | 3.52148864 | -0.621824499 | 0.534057275 | NA | Gm44712 | predicted gene 44712 [Source:MGI Symbo | 1.6062 | 1.0279 | 0 | 0 | 0 | 0 |
| 46135 | ENSMUSG00000108311 | 0 | NA | NA | NA | NA | NA | Gm44737 | predicted gene 44737 [Source:MGI Symbo | 0 | 0 | 0 | 0 | 0 | 0 |
| 46136 | ENSMUSG00000108312 | 0 | NA | NA | NA | NA | NA | Gm44521 | predicted gene 44521 [Source:MGI Symbo | 0 | 0 | 0 | 0 | 0 | 0 |
| 46137 | ENSMUSG00000108313 | 0 | NA | NA | NA | NA | NA | Gm45052 | predicted gene 45052 [Source:MGI Symbo | 0 | 0 | 0 | 0 | 0 | 0 |
| 46138 | ENSMUSG00000108314 | 7.872494382 | 0.414280982 | 0.753112682 | 0.550091629 | 0.582256528 | 0.908645447 | Prkcz2 | protein kinase C, zeta 2 [Source:MGI Symb | 8.031 | 6.1675 | 6.0375 | 11.568 | 7.6343 | 7.7969 |
| 46139 | ENSMUSG00000108315 | 0 | NA | NA | NA | NA | NA | Gm44601 | predicted gene 44601 [Source:MGI Symbo | 0 | 0 | 0 | 0 | 0 | 0 |
| 46140 | ENSMUSG00000108316 | 0.160662974 | 1.126924175 | 4.080472857 | 0.276174898 | 0.782413736 | NA | Gm44589 | predicted gene 44589 [Source:MGI Symbo | 0 | 0 | 0 | 0.964 | 0 | 0 |

| | | | | | | | | | | | | | | | |
|---|---|---|---|---|---|---|---|---|---|---|---|---|---|---|---|
| 46141 | ENSMUSG00000108317 | 0 | NA | NA | NA | NA | NA | Gm44663 | predicted gene 44663 [Source:MGI Symbo | 0 | 0 | 0 | 0 | 0 | 0 |
| 46142 | ENSMUSG00000108318 | 0 | NA | NA | NA | NA | NA | Gm45170 | predicted gene 45170 [Source:MGI Symbo | 0 | 0 | 0 | 0 | 0 | 0 |
| 46143 | ENSMUSG00000108319 | 0.17131964 | -0.796629176 | 4.080472857 | -0.195229623 | 0.84521319 | NA | Gm44552 | predicted gene 44552 [Source:MGI Symbo | 0 | 1.0279 | 0 | 0 | 0 | 0 |
| 46144 | ENSMUSG00000108320 | 7.582804013 | 0.32154739 | 0.814626467 | 0.394717582 | 0.693051325 | 0.93867917 | Gm44877 | predicted gene 44877 [Source:MGI Symbo | 8.031 | 2.0558 | 10.062 | 7.7118 | 8.7249 | 8.9108 |
| 46145 | ENSMUSG00000108321 | 0 | NA | NA | NA | NA | NA | Gm44649 | predicted gene 44649 [Source:MGI Symbo | 0 | 0 | 0 | 0 | 0 | 0 |
| 46146 | ENSMUSG00000108322 | 0.690772315 | 1.713484714 | 3.063368077 | 0.559346664 | 0.575925154 | NA | 5430431A17 | RIKEN cDNA 5430431A17 gene [Source:M | 0.8031 | 0 | 0 | 0 | 0 | 3.3415 |
| 46147 | ENSMUSG00000108323 | 0 | NA | NA | NA | NA | NA | Gm44945 | predicted gene 44945 [Source:MGI Symbo | 0 | 0 | 0 | 0 | 0 | 0 |
| 46148 | ENSMUSG00000108324 | 0 | NA | NA | NA | NA | NA | Gm44608 | predicted gene 44608 [Source:MGI Symbo | 0 | 0 | 0 | 0 | 0 | 0 |
| 46149 | ENSMUSG00000108325 | 0 | NA | NA | NA | NA | NA | Gm44898 | predicted gene 44898 [Source:MGI Symbo | 0 | 0 | 0 | 0 | 0 | 0 |
| 46150 | ENSMUSG00000108326 | 0 | NA | NA | NA | NA | NA | Gm17838 | predicted gene, 17838 [Source:MGI Symb | 0 | 0 | 0 | 0 | 0 | 0 |
| 46151 | ENSMUSG00000108327 | 0 | NA | NA | NA | NA | NA | Gm44939 | predicted gene 44939 [Source:MGI Symbo | 0 | 0 | 0 | 0 | 0 | 0 |
| 46152 | ENSMUSG00000108328 | 0 | NA | NA | NA | NA | NA | Gm45196 | predicted gene 45196 [Source:MGI Symbo | 0 | 0 | 0 | 0 | 0 | 0 |
| 46153 | ENSMUSG00000108329 | 0 | NA | NA | NA | NA | NA | Gm44725 | predicted gene 44725 [Source:MGI Symbo | 0 | 0 | 0 | 0 | 0 | 0 |
| 46154 | ENSMUSG00000108330 | 0 | NA | NA | NA | NA | NA | Gm44882 | predicted gene 44882 [Source:MGI Symbo | 0 | 0 | 0 | 0 | 0 | 0 |
| 46155 | ENSMUSG00000108331 | 0 | NA | NA | NA | NA | NA | Gm5322 | predicted gene 5322 [Source:MGI Symbol | 0 | 0 | 0 | 0 | 0 | 0 |
| 46156 | ENSMUSG00000108332 | 1.445387814 | 1.750116901 | 2.197838421 | 0.796290066 | 0.425863458 | NA | D530033B14 | RIKEN cDNA D530033B14 gene [Source:M | 0 | 0 | 2.0125 | 0 | 1.0906 | 5.5692 |
| 46157 | ENSMUSG00000108333 | 2.583368005 | 0.271779605 | 1.402876759 | 0.193730207 | 0.846387138 | NA | Gm45095 | predicted gene 45095 [Source:MGI Symbo | 0 | 3.0838 | 4.025 | 2.8919 | 3.2719 | 2.2277 |
| 46158 | ENSMUSG00000108334 | 0 | NA | NA | NA | NA | NA | Gm30771 | predicted gene, 30771 [Source:MGI Symb | 0 | 0 | 0 | 0 | 0 | 0 |
| 46159 | ENSMUSG00000108335 | 0 | NA | NA | NA | NA | NA | Gm45215 | predicted gene 45215 [Source:MGI Symbo | 0 | 0 | 0 | 0 | 0 | 0 |
| 46160 | ENSMUSG00000108336 | 0 | NA | NA | NA | NA | NA | Olfr163-ps1 | olfactory receptor 163, pseudogene 1 [Sou | 0 | 0 | 0 | 0 | 0 | 0 |
| 46161 | ENSMUSG00000108337 | 0 | NA | NA | NA | NA | NA | Gm4971 | predicted gene 4971 [Source:MGI Symbol | 0 | 0 | 0 | 0 | 0 | 0 |
| 46162 | ENSMUSG00000108338 | 0.17131964 | -0.796629176 | 4.080472857 | -0.195229623 | 0.84521319 | NA | Gm44794 | predicted gene 44794 [Source:MGI Symbo | 0 | 1.0279 | 0 | 0 | 0 | 0 |
| 46163 | ENSMUSG00000108339 | 0 | NA | NA | NA | NA | NA | Klk1b28-ps | kallikrein 1-related peptidase b28, pseudo | 0 | 0 | 0 | 0 | 0 | 0 |
| 46164 | ENSMUSG00000108340 | 0 | NA | NA | NA | NA | NA | Gm44520 | predicted gene 44520 [Source:MGI Symbo | 0 | 0 | 0 | 0 | 0 | 0 |
| 46165 | ENSMUSG00000108341 | 0 | NA | NA | NA | NA | NA | Gm44746 | predicted gene 44746 [Source:MGI Symbo | 0 | 0 | 0 | 0 | 0 | 0 |
| 46166 | ENSMUSG00000108342 | 0 | NA | NA | NA | NA | NA | Gm29920 | predicted gene, 29920 [Source:MGI Symb | 0 | 0 | 0 | 0 | 0 | 0 |
| 46167 | ENSMUSG00000108343 | 0 | NA | NA | NA | NA | NA | Gm18307 | predicted gene, 18307 [Source:MGI Symb | 0 | 0 | 0 | 0 | 0 | 0 |
| 46168 | ENSMUSG00000108344 | 0.17131964 | -0.796629176 | 4.080472857 | -0.195229623 | 0.84521319 | NA | Gm8952 | predicted gene 8952 [Source:MGI Symbol | 0 | 1.0279 | 0 | 0 | 0 | 0 |
| 46169 | ENSMUSG00000108345 | 0 | NA | NA | NA | NA | NA | 4933435G04 | RIKEN cDNA 4933435G04 gene [Source:M | 0 | 0 | 0 | 0 | 0 | 0 |
| 46170 | ENSMUSG00000108346 | 0 | NA | NA | NA | NA | NA | Gm44980 | predicted gene 44980 [Source:MGI Symbo | 0 | 0 | 0 | 0 | 0 | 0 |
| 46171 | ENSMUSG00000108347 | 0.765090549 | -0.195763385 | 2.933488969 | -0.066733977 | 0.946793485 | NA | Gm19187 | predicted gene, 19187 [Source:MGI Symb | 2.4093 | 0 | 0 | 0 | 2.1812 | 0 |
| 46172 | ENSMUSG00000108348 | 0.401551119 | -2.072982726 | 4.008251979 | -0.517178744 | 0.605031388 | NA | Gm42372 | predicted gene, 42372 [Source:MGI Symb | 2.4093 | 0 | 0 | 0 | 0 | 0 |
| 46173 | ENSMUSG00000108349 | 0 | NA | NA | NA | NA | NA | Gm18599 | predicted gene, 18599 [Source:MGI Symb | 0 | 0 | 0 | 0 | 0 | 0 |
| 46174 | ENSMUSG00000108350 | 60.74826084 | -0.345743754 | 0.305192765 | -1.132870088 | 0.257268813 | 0.710872182 | Gm44950 | predicted gene 44950 [Source:MGI Symbo | 53.808 | 81.206 | 69.431 | 58.803 | 57.803 | 43.44 |
| 46175 | ENSMUSG00000108351 | 0 | NA | NA | NA | NA | NA | Gm36736 | predicted gene, 36736 [Source:MGI Symb | 0 | 0 | 0 | 0 | 0 | 0 |
| 46176 | ENSMUSG00000108352 | 0.339027071 | -1.79174149 | 3.919585585 | -0.457125237 | 0.647581032 | NA | Usf2-ps1 | upstream transcription factor 2, pseudoge | 0 | 1.0279 | 1.0062 | 0 | 0 | 0 |
| 46177 | ENSMUSG00000108353 | 0 | NA | NA | NA | NA | NA | Gm45205 | predicted gene 45205 [Source:MGI Symbo | 0 | 0 | 0 | 0 | 0 | 0 |
| 46178 | ENSMUSG00000108354 | 0 | NA | NA | NA | NA | NA | 4931431B13 | RIKEN cDNA 4931431B13 gene [Source:M | 0 | 0 | 0 | 0 | 0 | 0 |
| 46179 | ENSMUSG00000108355 | 0 | NA | NA | NA | NA | NA | Gm44604 | predicted gene 44604 [Source:MGI Symbo | 0 | 0 | 0 | 0 | 0 | 0 |
| 46180 | ENSMUSG00000108356 | 0 | NA | NA | NA | NA | NA | 9130221F21 | RIKEN cDNA 9130221F21 gene [Source:M | 0 | 0 | 0 | 0 | 0 | 0 |
| 46181 | ENSMUSG00000108357 | 0 | NA | NA | NA | NA | NA | Snrp1c-ps1 | U1 small nuclear ribonucleoprotein 1C, ps | 0 | 0 | 0 | 0 | 0 | 0 |
| 46182 | ENSMUSG00000108358 | 7.415948447 | -0.008467777 | 0.774285908 | -0.01093624 | 0.991274317 | 0.99859176 | Gm44509 | predicted gene 44509 [Source:MGI Symbo | 8.031 | 7.1954 | 7.0437 | 6.7478 | 5.4531 | 10.025 |
| 46183 | ENSMUSG00000108359 | 0.133850373 | -0.796629176 | 4.080472857 | -0.195229623 | 0.84521319 | NA | C430039J01I | RIKEN cDNA C430039J01 gene [Source:M | 0.8031 | 0 | 0 | 0 | 0 | 0 |
| 46184 | ENSMUSG00000108360 | 0 | NA | NA | NA | NA | NA | Gm36159 | predicted gene, 36159 [Source:MGI Symb | 0 | 0 | 0 | 0 | 0 | 0 |
| 46185 | ENSMUSG00000108361 | 0.321325949 | 1.911575114 | 4.048267893 | 0.472195804 | 0.636787034 | NA | Gm44796 | predicted gene 44796 [Source:MGI Symbo | 0 | 0 | 0 | 1.928 | 0 | 0 |
| 46186 | ENSMUSG00000108362 | 0 | NA | NA | NA | NA | NA | Gm18561 | predicted gene, 18561 [Source:MGI Symb | 0 | 0 | 0 | 0 | 0 | 0 |
| 46187 | ENSMUSG00000108363 | 0 | NA | NA | NA | NA | NA | Gm44615 | predicted gene 44615 [Source:MGI Symbo | 0 | 0 | 0 | 0 | 0 | 0 |
| 46188 | ENSMUSG00000108364 | 0 | NA | NA | NA | NA | NA | Gm38997 | predicted gene, 38997 [Source:MGI Symb | 0 | 0 | 0 | 0 | 0 | 0 |
| 46189 | ENSMUSG00000108365 | 1.106211272 | -0.253356961 | 2.11057745 | -0.120041537 | 0.904450244 | NA | Gm44951 | predicted gene 44951 [Source:MGI Symbo | 1.6062 | 0 | 2.0125 | 1.928 | 1.0906 | 0 |
| 46190 | ENSMUSG00000108366 | 1.181149805 | -0.383612265 | 2.080148392 | -0.184415817 | 0.853687263 | NA | Gm5586 | predicted gene 5586 [Source:MGI Symbol | 0 | 2.0558 | 2.0125 | 1.928 | 1.0906 | 0 |
| 46191 | ENSMUSG00000108367 | 0 | NA | NA | NA | NA | NA | Erfl | ETS repressor factor like [Source:MGI Sym | 0 | 0 | 0 | 0 | 0 | 0 |
| 46192 | ENSMUSG00000108368 | 49.11922872 | -0.122678896 | 0.478591244 | -0.256333349 | 0.797693449 | 0.962377049 | Gm45053 | predicted gene 45053 [Source:MGI Symbo | 43.368 | 49.34 | 61.381 | 85.794 | 40.353 | 14.48 |
| 46193 | ENSMUSG00000108370 | 0 | NA | NA | NA | NA | NA | Gm8209 | predicted gene 8209 [Source:MGI Symbol | 0 | 0 | 0 | 0 | 0 | 0 |
| 46194 | ENSMUSG00000108371 | 13.3811116 | -1.471891998 | 0.662320641 | -2.222325423 | 0.026261321 | 0.209626794 | Gm38832 | predicted gene, 38832 [Source:MGI Symb | 20.078 | 28.782 | 10.062 | 4.8199 | 7.6343 | 8.9108 |
| 46195 | ENSMUSG00000108372 | 0.349477146 | 0.165142768 | 3.911021834 | 0.042224967 | 0.966319359 | NA | Gm45124 | predicted gene 45124 [Source:MGI Symbo | 0 | 0 | 1.0062 | 0 | 1.0906 | 0 |
| 46196 | ENSMUSG00000108373 | 0 | NA | NA | NA | NA | NA | Gm45168 | predicted gene 45168 [Source:MGI Symbo | 0 | 0 | 0 | 0 | 0 | 0 |

| | | | | | | | | | | | | | | | |
|---|---|---|---|---|---|---|---|---|---|---|---|---|---|---|---|
| 46197 | ENSMUSG00000108374 | 0 | NA | NA | NA | NA | NA | Gm33626 | predicted gene, 33626 [Source:MGI Symb | 0 | 0 | 0 | 0 | 0 | 0 |
| 46198 | ENSMUSG00000108375 | 0 | NA | NA | NA | NA | NA | Gm45003 | predicted gene 45003 [Source:MGI Symbo | 0 | 0 | 0 | 0 | 0 | 0 |
| 46199 | ENSMUSG00000108376 | 0 | NA | NA | NA | NA | NA | Gm44874 | predicted gene 44874 [Source:MGI Symbo | 0 | 0 | 0 | 0 | 0 | 0 |
| 46200 | ENSMUSG00000108377 | 0 | NA | NA | NA | NA | NA | Gm18642 | predicted gene, 18642 [Source:MGI Symb | 0 | 0 | 0 | 0 | 0 | 0 |
| 46201 | ENSMUSG00000108378 | 0.160662974 | 1.126924175 | 4.080472857 | 0.276174898 | 0.782413736 | NA | Gm44641 | predicted gene 44641 [Source:MGI Symbo | 0 | 0 | 0 | 0.964 | 0 | 0 |
| 46202 | ENSMUSG00000108379 | 0 | NA | NA | NA | NA | NA | A730082K24 | RIKEN cDNA A730082K24 gene [Source:M | 0 | 0 | 0 | 0 | 0 | 0 |
| 46203 | ENSMUSG00000108380 | 0 | NA | NA | NA | NA | NA | Olfr737-ps1 | olfactory receptor 737, pseudogene 1 [Sou | 0 | 0 | 0 | 0 | 0 | 0 |
| 46204 | ENSMUSG00000108381 | 0 | NA | NA | NA | NA | NA | Gm9299 | predicted gene 9299 [Source:MGI Symbol | 0 | 0 | 0 | 0 | 0 | 0 |
| 46205 | ENSMUSG00000108382 | 0.36353943 | 2.065989254 | 4.034137301 | 0.512126658 | 0.608562368 | NA | 4930448A20 | RIKEN cDNA 4930448A20 gene [Source:M | 0 | 0 | 0 | 0 | 2.1812 | 0 |
| 46206 | ENSMUSG00000108383 | 0 | NA | NA | NA | NA | NA | Gm6567 | predicted gene 6567 [Source:MGI Symbol | 0 | 0 | 0 | 0 | 0 | 0 |
| 46207 | ENSMUSG00000108384 | 0 | NA | NA | NA | NA | NA | Gm45082 | predicted gene 45082 [Source:MGI Symbo | 0 | 0 | 0 | 0 | 0 | 0 |
| 46208 | ENSMUSG00000108385 | 0 | NA | NA | NA | NA | NA | Gm44730 | predicted gene 44730 [Source:MGI Symbo | 0 | 0 | 0 | 0 | 0 | 0 |
| 46209 | ENSMUSG00000108387 | 0 | NA | NA | NA | NA | NA | Gm31510 | predicted gene, 31510 [Source:MGI Symb | 0 | 0 | 0 | 0 | 0 | 0 |
| 46210 | ENSMUSG00000108388 | 0 | NA | NA | NA | NA | NA | Gm44673 | predicted gene 44673 [Source:MGI Symbo | 0 | 0 | 0 | 0 | 0 | 0 |
| 46211 | ENSMUSG00000108389 | 0.562214094 | -1.122836648 | 3.276281221 | -0.34271681 | 0.731811515 | NA | Gm45206 | predicted gene 45206 [Source:MGI Symbo | 2.4093 | 0 | 0 | 0.964 | 0 | 0 |
| 46212 | ENSMUSG00000108390 | 0 | NA | NA | NA | NA | NA | Gm39038 | predicted gene, 39038 [Source:MGI Symb | 0 | 0 | 0 | 0 | 0 | 0 |
| 46213 | ENSMUSG00000108391 | 3.778914272 | 0.456424425 | 1.376092997 | 0.33168138 | 0.740129862 | NA | Gm9768 | predicted gene 9768 [Source:MGI Symbol | 6.4248 | 3.0838 | 0 | 7.7118 | 5.4531 | 0 |
| 46214 | ENSMUSG00000108392 | 0 | NA | NA | NA | NA | NA | Gm9463 | predicted gene 9463 [Source:MGI Symbol | 0 | 0 | 0 | 0 | 0 | 0 |
| 46215 | ENSMUSG00000108393 | 0.346303622 | 2.004339653 | 3.884003423 | 0.516049919 | 0.605819544 | NA | Gm32633 | predicted gene, 32633 [Source:MGI Symb | 0 | 0 | 0 | 0.964 | 0 | 1.1138 |
| 46216 | ENSMUSG00000108394 | 1.184555423 | 1.374592059 | 2.121256755 | 0.648008336 | 0.516979559 | NA | Gm45477 | predicted gene 45477 [Source:MGI Symbo | 0 | 1.0279 | 1.0062 | 2.8919 | 2.1812 | 0 |
| 46217 | ENSMUSG00000108395 | 0 | NA | NA | NA | NA | NA | Gm34811 | predicted gene, 34811 [Source:MGI Symb | 0 | 0 | 0 | 0 | 0 | 0 |
| 46218 | ENSMUSG00000108396 | 0.181769715 | 1.126924175 | 4.080472857 | 0.276174898 | 0.782413736 | NA | Gm6905 | predicted gene 6905 [Source:MGI Symbol | 0 | 0 | 0 | 0 | 1.0906 | 0 |
| 46219 | ENSMUSG00000108397 | 0 | NA | NA | NA | NA | NA | Gm45171 | predicted gene 45171 [Source:MGI Symbo | 0 | 0 | 0 | 0 | 0 | 0 |
| 46220 | ENSMUSG00000108398 | 0 | NA | NA | NA | NA | NA | Gm30191 | predicted gene, 30191 [Source:MGI Symb | 0 | 0 | 0 | 0 | 0 | 0 |
| 46221 | ENSMUSG00000108400 | 0 | NA | NA | NA | NA | NA | Klk1b14-ps | kallikrein 1-related peptidase b14, pseudo | 0 | 0 | 0 | 0 | 0 | 0 |
| 46222 | ENSMUSG00000108401 | 0 | NA | NA | NA | NA | NA | Rpl19-ps10 | ribosomal protein L19, pseudogene 10 [So | 0 | 0 | 0 | 0 | 0 | 0 |
| 46223 | ENSMUSG00000108402 | 7.25753411 | 0.481597028 | 0.816094996 | 0.590123736 | 0.555107697 | 0.897628185 | 9430064I24R | RIKEN cDNA 9430064I24 gene [Source:MG | 4.0155 | 7.1954 | 7.0437 | 8.6758 | 4.3625 | 12.252 |
| 46224 | ENSMUSG00000108403 | 0 | NA | NA | NA | NA | NA | Gm45122 | predicted gene 45122 [Source:MGI Symbo | 0 | 0 | 0 | 0 | 0 | 0 |
| 46225 | ENSMUSG00000108404 | 0 | NA | NA | NA | NA | NA | Gm44972 | predicted gene 44972 [Source:MGI Symbo | 0 | 0 | 0 | 0 | 0 | 0 |
| 46226 | ENSMUSG00000108405 | 0 | NA | NA | NA | NA | NA | Ywhaq-ps1 | tyrosine 3-monooxygenase/tryptophan 5- | 0 | 0 | 0 | 0 | 0 | 0 |
| 46227 | ENSMUSG00000108406 | 0 | NA | NA | NA | NA | NA | Klk1b15-ps | kallikrein 1-related peptidase b15, pseudo | 0 | 0 | 0 | 0 | 0 | 0 |
| 46228 | ENSMUSG00000108407 | 0 | NA | NA | NA | NA | NA | Gm45093 | predicted gene 45093 [Source:MGI Symbo | 0 | 0 | 0 | 0 | 0 | 0 |
| 46229 | ENSMUSG00000108409 | 0 | NA | NA | NA | NA | NA | 1700025J12R | RIKEN cDNA 1700025J12 gene [Source:MG | 0 | 0 | 0 | 0 | 0 | 0 |
| 46230 | ENSMUSG00000108410 | 0 | NA | NA | NA | NA | NA | Gm44740 | predicted gene 44740 [Source:MGI Symbo | 0 | 0 | 0 | 0 | 0 | 0 |
| 46231 | ENSMUSG00000108411 | 0 | NA | NA | NA | NA | NA | Gm44818 | predicted gene 44818 [Source:MGI Symbo | 0 | 0 | 0 | 0 | 0 | 0 |
| 46232 | ENSMUSG00000108412 | 0.267700746 | -1.497481649 | 4.057001436 | -0.369110456 | 0.712045396 | NA | Gm45005 | predicted gene 45005 [Source:MGI Symbo | 1.6062 | 0 | 0 | 0 | 0 | 0 |
| 46233 | ENSMUSG00000108413 | 0.267700746 | -1.497481649 | 4.057001436 | -0.369110456 | 0.712045396 | NA | BC026762 | cDNA sequence BC026762 [Source:MGI Sy | 1.6062 | 0 | 0 | 0 | 0 | 0 |
| 46234 | ENSMUSG00000108414 | 159.8615914 | 0.703563065 | 0.196451774 | 3.581352573 | 0.00034182 | 0.009656415 | Snhg1 | small nucleolar RNA host gene 1 [Source:I | 127.69 | 124.38 | 112.7 | 201.47 | 221.4 | 171.53 |
| 46235 | ENSMUSG00000108415 | 0 | NA | NA | NA | NA | NA | Gm30146 | predicted gene, 30146 [Source:MGI Symb | 0 | 0 | 0 | 0 | 0 | 0 |
| 46236 | ENSMUSG00000108416 | 0 | NA | NA | NA | NA | NA | Gm44924 | predicted gene 44924 [Source:MGI Symbo | 0 | 0 | 0 | 0 | 0 | 0 |
| 46237 | ENSMUSG00000108417 | 0 | NA | NA | NA | NA | NA | Gm30928 | predicted gene, 30928 [Source:MGI Symb | 0 | 0 | 0 | 0 | 0 | 0 |
| 46238 | ENSMUSG00000108418 | 0.17131964 | -0.796629176 | 4.080472857 | -0.195229623 | 0.84521319 | NA | Gm44627 | predicted gene 44627 [Source:MGI Symbo | 0 | 1.0279 | 0 | 0 | 0 | 0 |
| 46239 | ENSMUSG00000108419 | 0 | NA | NA | NA | NA | NA | Gm45020 | predicted gene 45020 [Source:MGI Symbo | 0 | 0 | 0 | 0 | 0 | 0 |
| 46240 | ENSMUSG00000108420 | 0 | NA | NA | NA | NA | NA | Gm45141 | predicted gene 45141 [Source:MGI Symbo | 0 | 0 | 0 | 0 | 0 | 0 |
| 46241 | ENSMUSG00000108421 | 0 | NA | NA | NA | NA | NA | Gm44718 | predicted gene 44718 [Source:MGI Symbo | 0 | 0 | 0 | 0 | 0 | 0 |
| 46242 | ENSMUSG00000108422 | 0 | NA | NA | NA | NA | NA | Gm44765 | predicted gene 44765 [Source:MGI Symbo | 0 | 0 | 0 | 0 | 0 | 0 |
| 46243 | ENSMUSG00000108423 | 0 | NA | NA | NA | NA | NA | Gm44856 | predicted gene 44856 [Source:MGI Symbo | 0 | 0 | 0 | 0 | 0 | 0 |
| 46244 | ENSMUSG00000108424 | 0 | NA | NA | NA | NA | NA | Gm44590 | predicted gene 44590 [Source:MGI Symbo | 0 | 0 | 0 | 0 | 0 | 0 |
| 46245 | ENSMUSG00000108425 | 115.0092349 | -0.880697296 | 0.365718759 | -2.408127214 | 0.016034593 | 0.148963156 | Gm44706 | predicted gene 44706 [Source:MGI Symbo | 172.67 | 151.1 | 123.77 | 117.61 | 97.065 | 27.846 |
| 46246 | ENSMUSG00000108426 | 0 | NA | NA | NA | NA | NA | Olfr465-ps1 | olfactory receptor 465, pseudogene 1 [Sou | 0 | 0 | 0 | 0 | 0 | 0 |
| 46247 | ENSMUSG00000108427 | 0.160662974 | 1.126924175 | 4.080472857 | 0.276174898 | 0.782413736 | NA | Gm36696 | predicted gene, 36696 [Source:MGI Symb | 0 | 0 | 0 | 0.964 | 0 | 0 |
| 46248 | ENSMUSG00000108428 | 0 | NA | NA | NA | NA | NA | Gm45058 | predicted gene 45058 [Source:MGI Symbo | 0 | 0 | 0 | 0 | 0 | 0 |
| 46249 | ENSMUSG00000108429 | 0 | NA | NA | NA | NA | NA | Gm44603 | predicted gene 44603 [Source:MGI Symbo | 0 | 0 | 0 | 0 | 0 | 0 |
| 46250 | ENSMUSG00000108430 | 0 | NA | NA | NA | NA | NA | Gm36776 | predicted gene, 36776 [Source:MGI Symb | 0 | 0 | 0 | 0 | 0 | 0 |
| 46251 | ENSMUSG00000108431 | 0.453341394 | -0.545391191 | 3.551113277 | -0.153583158 | 0.877938417 | NA | Gm42375 | predicted gene, 42375 [Source:MGI Symb | 1.6062 | 0 | 0 | 0 | 0 | 1.1138 |
| 46252 | ENSMUSG00000108432 | 0 | NA | NA | NA | NA | NA | Gm44756 | predicted gene 44756 [Source:MGI Symbo | 0 | 0 | 0 | 0 | 0 | 0 |

| | | | | | | | | | | | | | | | |
|---|---|---|---|---|---|---|---|---|---|---|---|---|---|---|---|
| 46253 | ENSMUSG00000108433 | 0 | NA | NA | NA | NA | NA | Gm19761 | predicted gene, 19761 [Source:MGI Symb | 0 | 0 | 0 | 0 | 0 | 0 |
| 46254 | ENSMUSG00000108434 | 0 | NA | NA | NA | NA | NA | NA | NA | 0 | 0 | 0 | 0 | 0 | 0 |
| 46255 | ENSMUSG00000108435 | 39.74793115 | -0.527048539 | 0.368346796 | -1.430848714 | 0.152473578 | 0.568249894 | Gm45051 | predicted gene 45051 [Source:MGI Symbo | 47.383 | 51.396 | 42.262 | 43.379 | 23.994 | 30.074 |
| 46256 | ENSMUSG00000108436 | 44.78041295 | -0.091126735 | 0.413464963 | -0.220397718 | 0.825561422 | 0.96943211 | Gm44851 | predicted gene 44851 [Source:MGI Symbo | 48.186 | 45.228 | 45.281 | 51.091 | 61.075 | 17.822 |
| 46257 | ENSMUSG00000108437 | 0.328370405 | 0.16514316 | 4.003149536 | 0.041253308 | 0.967093956 | NA | Gm19366 | predicted gene, 19366 [Source:MGI Symb | 0 | 0 | 1.0062 | 0.964 | 0 | 0 |
| 46258 | ENSMUSG00000108438 | 0 | NA | NA | NA | NA | NA | Gm9278 | predicted gene 9278 [Source:MGI Symbol | 0 | 0 | 0 | 0 | 0 | 0 |
| 46259 | ENSMUSG00000108440 | 0 | NA | NA | NA | NA | NA | Klk1b19-ps | kallikrein 1-related peptidase b19, pseudo | 0 | 0 | 0 | 0 | 0 | 0 |
| 46260 | ENSMUSG00000108441 | 0 | NA | NA | NA | NA | NA | Gm32846 | predicted gene, 32846 [Source:MGI Symb | 0 | 0 | 0 | 0 | 0 | 0 |
| 46261 | ENSMUSG00000108442 | 0.342639279 | -1.805976596 | 4.028547714 | -0.448294702 | 0.653940524 | NA | Rpl15-ps5 | ribosomal protein L15, pseudogene 5 [Sou | 0 | 2.0558 | 0 | 0 | 0 | 0 |
| 46262 | ENSMUSG00000108443 | 0 | NA | NA | NA | NA | NA | Gm44510 | predicted gene 44510 [Source:MGI Symbo | 0 | 0 | 0 | 0 | 0 | 0 |
| 46263 | ENSMUSG00000108444 | 0 | NA | NA | NA | NA | NA | Klk2-ps | kallikrein-related peptidase 2, pseudogen | 0 | 0 | 0 | 0 | 0 | 0 |
| 46264 | ENSMUSG00000108445 | 0 | NA | NA | NA | NA | NA | Gm17918 | predicted gene, 17918 [Source:MGI Symb | 0 | 0 | 0 | 0 | 0 | 0 |
| 46265 | ENSMUSG00000108446 | 9.858725084 | 0.367410822 | 0.734922366 | 0.499931474 | 0.617123329 | 0.917509899 | Gm44997 | predicted gene 44997 [Source:MGI Symbo | 14.456 | 5.1396 | 6.0375 | 11.568 | 15.269 | 6.6831 |
| 46266 | ENSMUSG00000108447 | 0.958478575 | 0.054139297 | 2.45225727 | 0.022077331 | 0.982386269 | NA | Gm44567 | predicted gene 44567 [Source:MGI Symbo | 0.8031 | 2.0558 | 0 | 2.8919 | 0 | 0 |
| 46267 | ENSMUSG00000108448 | 0 | NA | NA | NA | NA | NA | Gm21269 | predicted gene, 21269 [Source:MGI Symb | 0 | 0 | 0 | 0 | 0 | 0 |
| 46268 | ENSMUSG00000108449 | 11.57721694 | -0.42860104 | 0.636638805 | -0.673224813 | 0.500804278 | 0.880974509 | Gm44507 | predicted gene 44507 [Source:MGI Symbo | 10.44 | 13.363 | 16.1 | 8.6758 | 13.087 | 7.7969 |
| 46269 | ENSMUSG00000108450 | 0 | NA | NA | NA | NA | NA | Gm44977 | predicted gene 44977 [Source:MGI Symbo | 0 | 0 | 0 | 0 | 0 | 0 |
| 46270 | ENSMUSG00000108451 | 0 | NA | NA | NA | NA | NA | Olfr185-ps1 | olfactory receptor 185, pseudogene 1 [So | 0 | 0 | 0 | 0 | 0 | 0 |
| 46271 | ENSMUSG00000108452 | 26.62433934 | -0.208685282 | 0.435924023 | -0.478719389 | 0.63213827 | 0.921791304 | 4930413G21 | RIKEN cDNA 4930413G21 gene [Source:M | 28.912 | 21.586 | 35.219 | 28.919 | 26.175 | 18.935 |
| 46272 | ENSMUSG00000108454 | 0 | NA | NA | NA | NA | NA | Gm44932 | predicted gene 44932 [Source:MGI Symbo | 0 | 0 | 0 | 0 | 0 | 0 |
| 46273 | ENSMUSG00000108455 | 0 | NA | NA | NA | NA | NA | Gm44745 | predicted gene 44745 [Source:MGI Symbo | 0 | 0 | 0 | 0 | 0 | 0 |
| 46274 | ENSMUSG00000108456 | 1.251746577 | 1.461824714 | 2.132479794 | 0.685504602 | 0.493025557 | NA | 4732496C06 | RIKEN cDNA 4732496C06 gene [Source:M | 0 | 1.0279 | 1.0062 | 0 | 4.3625 | 1.1138 |
| 46275 | ENSMUSG00000108457 | 0 | NA | NA | NA | NA | NA | Gm5027 | predicted gene 5027 [Source:MGI Symbol | 0 | 0 | 0 | 0 | 0 | 0 |
| 46276 | ENSMUSG00000108459 | 0 | NA | NA | NA | NA | NA | Gm44605 | predicted gene 44605 [Source:MGI Symbo | 0 | 0 | 0 | 0 | 0 | 0 |
| 46277 | ENSMUSG00000108460 | 2.162460268 | -1.631154901 | 1.769187347 | -0.921979746 | 0.35653914 | NA | Gm39041 | predicted gene, 39041 [Source:MGI Symb | 1.6062 | 8.2233 | 0 | 0.964 | 2.1812 | 0 |
| 46278 | ENSMUSG00000108461 | 18.07683806 | -0.005598689 | 0.507705368 | -0.011027437 | 0.991201556 | 0.99859176 | AV356131 | expressed sequence AV356131 [Source:M | 16.865 | 18.503 | 19.119 | 23.135 | 16.359 | 14.48 |
| 46279 | ENSMUSG00000108462 | 0 | NA | NA | NA | NA | NA | Gm45032 | predicted gene 45032 [Source:MGI Symbo | 0 | 0 | 0 | 0 | 0 | 0 |
| 46280 | ENSMUSG00000108464 | 0 | NA | NA | NA | NA | NA | Olfr192 | olfactory receptor 192 [Source:MGI Symb | 0 | 0 | 0 | 0 | 0 | 0 |
| 46281 | ENSMUSG00000108465 | 13.11855844 | -0.276611536 | 0.708720678 | -0.390296974 | 0.696316961 | 0.939992732 | Gm45110 | predicted gene 45110 [Source:MGI Symbo | 21.684 | 15.419 | 6.0375 | 21.208 | 5.4531 | 8.9108 |
| 46282 | ENSMUSG00000108466 | 0.331982614 | 0.16514309 | 3.986751476 | 0.041422971 | 0.966958701 | NA | Gm44771 | predicted gene 44771 [Source:MGI Symbo | 0 | 1.0279 | 0 | 0.964 | 0 | 0 |
| 46283 | ENSMUSG00000108467 | 0 | NA | NA | NA | NA | NA | Gm45211 | predicted gene 45211 [Source:MGI Symbo | 0 | 0 | 0 | 0 | 0 | 0 |
| 46284 | ENSMUSG00000108468 | 0 | NA | NA | NA | NA | NA | Gm30459 | predicted gene, 30459 [Source:MGI Symb | 0 | 0 | 0 | 0 | 0 | 0 |
| 46285 | ENSMUSG00000108469 | 0 | NA | NA | NA | NA | NA | Gm45054 | predicted gene 45054 [Source:MGI Symbo | 0 | 0 | 0 | 0 | 0 | 0 |
| 46286 | ENSMUSG00000108470 | 0 | NA | NA | NA | NA | NA | Gm4598 | predicted gene 4598 [Source:MGI Symbol | 0 | 0 | 0 | 0 | 0 | 0 |
| 46287 | ENSMUSG00000108471 | 0 | NA | NA | NA | NA | NA | Gm18855 | predicted gene, 18855 [Source:MGI Symb | 0 | 0 | 0 | 0 | 0 | 0 |
| 46288 | ENSMUSG00000108472 | 2.542596859 | -1.936107585 | 1.468880473 | -1.318083821 | 0.18747559 | NA | Gm45212 | predicted gene 45212 [Source:MGI Symbo | 4.0155 | 1.0279 | 7.0437 | 0.964 | 1.0906 | 1.1138 |
| 46289 | ENSMUSG00000108473 | 0 | NA | NA | NA | NA | NA | Gm44739 | predicted gene 44739 [Source:MGI Symbo | 0 | 0 | 0 | 0 | 0 | 0 |
| 46290 | ENSMUSG00000108474 | 0 | NA | NA | NA | NA | NA | Gm44895 | predicted gene 44895 [Source:MGI Symbo | 0 | 0 | 0 | 0 | 0 | 0 |
| 46291 | ENSMUSG00000108476 | 2.760600983 | -2.193661728 | 1.727724321 | -1.269682727 | 0.204197668 | NA | Gm44974 | predicted gene 44974 [Source:MGI Symbo | 5.6217 | 0 | 8.05 | 2.8919 | 0 | 0 |
| 46292 | ENSMUSG00000108477 | 16.05066054 | -1.101818167 | 0.672641614 | -1.638046389 | 0.101412017 | 0.46908328 | Gm44878 | predicted gene 44878 [Source:MGI Symbo | 29.715 | 30.838 | 5.0312 | 8.6758 | 10.906 | 11.138 |
| 46293 | ENSMUSG00000108478 | 0.371281295 | 2.09253362 | 4.031852165 | 0.519000582 | 0.603760335 | NA | Gm19514 | predicted gene, 19514 [Source:MGI Symb | 0 | 0 | 0 | 0 | 0 | 2.2277 |
| 46294 | ENSMUSG00000108479 | 0 | NA | NA | NA | NA | NA | 2200002A13 | RIKEN cDNA 2200002A13 gene [Source:M | 0 | 0 | 0 | 0 | 0 | 0 |
| 46295 | ENSMUSG00000108480 | 0 | NA | NA | NA | NA | NA | Gm36633 | predicted gene, 36633 [Source:MGI Symb | 0 | 0 | 0 | 0 | 0 | 0 |
| 46296 | ENSMUSG00000108481 | 0 | NA | NA | NA | NA | NA | Gm33248 | predicted gene, 33248 [Source:MGI Symb | 0 | 0 | 0 | 0 | 0 | 0 |
| 46297 | ENSMUSG00000108482 | 0 | NA | NA | NA | NA | NA | Gm45078 | predicted gene 45078 [Source:MGI Symbo | 0 | 0 | 0 | 0 | 0 | 0 |
| 46298 | ENSMUSG00000108483 | 0 | NA | NA | NA | NA | NA | Gm45184 | predicted gene 45184 [Source:MGI Symbo | 0 | 0 | 0 | 0 | 0 | 0 |
| 46299 | ENSMUSG00000108484 | 0.528073336 | 2.60973587 | 2.979250867 | 0.875970499 | 0.381046072 | NA | Gm18537 | predicted gene, 18537 [Source:MGI Symb | 0 | 0 | 0 | 0.964 | 1.0906 | 1.1138 |
| 46300 | ENSMUSG00000108485 | 0 | NA | NA | NA | NA | NA | Idi1-ps4 | isopentenyl-diphosphate delta isomerase, | 0 | 0 | 0 | 0 | 0 | 0 |
| 46301 | ENSMUSG00000108486 | 2.132610553 | -1.155689748 | 1.697561441 | -0.680794061 | 0.496001808 | NA | Gm44836 | predicted gene 44836 [Source:MGI Symbo | 0.8031 | 4.1117 | 4.025 | 3.8559 | 0 | 0 |
| 46302 | ENSMUSG00000108487 | 3.069480543 | -1.681951446 | 1.438591289 | -1.169165599 | 0.242336918 | NA | Cyp2a21-ps | cytochrome P450, family 2, subfamily a, p | 4.8186 | 4.1117 | 5.0312 | 0 | 0 | 4.4554 |
| 46303 | ENSMUSG00000108488 | 0 | NA | NA | NA | NA | NA | Gm44979 | predicted gene 44979 [Source:MGI Symbo | 0 | 0 | 0 | 0 | 0 | 0 |
| 46304 | ENSMUSG00000108489 | 0 | NA | NA | NA | NA | NA | Gm45025 | predicted gene 45025 [Source:MGI Symbo | 0 | 0 | 0 | 0 | 0 | 0 |
| 46305 | ENSMUSG00000108490 | 0 | NA | NA | NA | NA | NA | 1700110I07F | RIKEN cDNA 1700110I07 gene [Source:MG | 0 | 0 | 0 | 0 | 0 | 0 |
| 46306 | ENSMUSG00000108491 | 4.757112167 | 0.04306221 | 1.153269699 | 0.037339236 | 0.970214521 | NA | Gm44523 | predicted gene 44523 [Source:MGI Symbo | 0 | 6.1675 | 8.05 | 7.7118 | 3.2719 | 3.3415 |
| 46307 | ENSMUSG00000108492 | 0 | NA | NA | NA | NA | NA | Rpl18a-ps4 | ribosomal protein L18A, pseudogene 4 [So | 0 | 0 | 0 | 0 | 0 | 0 |
| 46308 | ENSMUSG00000108493 | 0.367410362 | 2.078255025 | 3.800070157 | 0.54689912 | 0.58444804 | NA | Gm45715 | predicted gene 45715 [Source:MGI Symbo | 0 | 0 | 0 | 0 | 1.0906 | 1.1138 |

| | | | | | | | | | | | | | | | |
|---|---|---|---|---|---|---|---|---|---|---|---|---|---|---|---|
| 46309 | ENSMUSG00000108494 | 0.514011053 | 1.044022102 | 3.082659575 | 0.338675769 | 0.734853997 | NA | Gm45203 | predicted gene 45203 [Source:MGI Symbo | 0 | 0 | 1.0062 | 0.964 | 0 | 1.1138 |
| 46310 | ENSMUSG00000108495 | 0 | NA | NA | NA | NA | NA | Gm45139 | predicted gene 45139 [Source:MGI Symbo | 0 | 0 | 0 | 0 | 0 | 0 |
| 46311 | ENSMUSG00000108496 | 0 | NA | NA | NA | NA | NA | Gm44768 | predicted gene 44768 [Source:MGI Symbo | 0 | 0 | 0 | 0 | 0 | 0 |
| 46312 | ENSMUSG00000108497 | 0 | NA | NA | NA | NA | NA | Gm44738 | predicted gene 44738 [Source:MGI Symbo | 0 | 0 | 0 | 0 | 0 | 0 |
| 46313 | ENSMUSG00000108498 | 0 | NA | NA | NA | NA | NA | Gm45026 | predicted gene 45026 [Source:MGI Symbo | 0 | 0 | 0 | 0 | 0 | 0 |
| 46314 | ENSMUSG00000108500 | 0.476489653 | -2.293616034 | 3.421506946 | -0.670352587 | 0.502633051 | NA | Gm45033 | predicted gene 45033 [Source:MGI Symbo | 0.8031 | 2.0558 | 0 | 0 | 0 | 0 |
| 46315 | ENSMUSG00000108502 | 0 | NA | NA | NA | NA | NA | Gm9240 | predicted gene 9240 [Source:MGI Symbol | 0 | 0 | 0 | 0 | 0 | 0 |
| 46316 | ENSMUSG00000108503 | 2.137903233 | 1.167777554 | 1.928773181 | 0.605450949 | 0.544879407 | NA | Gm20949 | predicted gene, 20949 [Source:MGI Symb | 0 | 0 | 4.025 | 7.7118 | 1.0906 | 0 |
| 46317 | ENSMUSG00000108504 | 1.565206392 | -1.1226443 | 1.871435756 | -0.599883964 | 0.54858357 | NA | Gm4265 | predicted gene 4265 [Source:MGI Symbol | 2.4093 | 3.0838 | 1.0062 | 2.8919 | 0 | 0 |
| 46318 | ENSMUSG00000108505 | 0 | NA | NA | NA | NA | NA | 1700095K22 | RIKEN cDNA 1700095K22 gene [Source:M | 0 | 0 | 0 | 0 | 0 | 0 |
| 46319 | ENSMUSG00000108506 | 1.390424016 | 1.823862407 | 2.081970201 | 0.876027143 | 0.381015278 | NA | Gm44705 | predicted gene 44705 [Source:MGI Symbo | 0.8031 | 0 | 1.0062 | 0.964 | 0 | 5.5692 |
| 46320 | ENSMUSG00000108507 | 0 | NA | NA | NA | NA | NA | 6720469O03 | RIKEN cDNA 6720469O03 gene [Source:M | 0 | 0 | 0 | 0 | 0 | 0 |
| 46321 | ENSMUSG00000108508 | 0 | NA | NA | NA | NA | NA | Gm44978 | predicted gene 44978 [Source:MGI Symbo | 0 | 0 | 0 | 0 | 0 | 0 |
| 46322 | ENSMUSG00000108509 | 0.167707431 | -0.796629176 | 4.080472857 | -0.195229623 | 0.84521319 | NA | Gm44780 | predicted gene 44780 [Source:MGI Symbo | 0 | 0 | 1.0062 | 0 | 0 | 0 |
| 46323 | ENSMUSG00000108510 | 0 | NA | NA | NA | NA | NA | Gm45192 | predicted gene 45192 [Source:MGI Symbo | 0 | 0 | 0 | 0 | 0 | 0 |
| 46324 | ENSMUSG00000108511 | 0 | NA | NA | NA | NA | NA | Gm44987 | predicted gene 44987 [Source:MGI Symbo | 0 | 0 | 0 | 0 | 0 | 0 |
| 46325 | ENSMUSG00000108512 | 0.185640647 | 1.126924175 | 4.080472857 | 0.276174898 | 0.782413736 | NA | Gm7238 | predicted gene 7238 [Source:MGI Symbol | 0 | 0 | 0 | 0 | 0 | 1.1138 |
| 46326 | ENSMUSG00000108513 | 0 | NA | NA | NA | NA | NA | Gm30075 | predicted gene, 30075 [Source:MGI Symb | 0 | 0 | 0 | 0 | 0 | 0 |
| 46327 | ENSMUSG00000108514 | 0 | NA | NA | NA | NA | NA | 2310001K20 | RIKEN cDNA 2310001K20 gene [Source:M | 0 | 0 | 0 | 0 | 0 | 0 |
| 46328 | ENSMUSG00000108515 | 0 | NA | NA | NA | NA | NA | Olfr717-ps1 | olfactory receptor 717, pseudogene 1 [Sou | 0 | 0 | 0 | 0 | 0 | 0 |
| 46329 | ENSMUSG00000108516 | 0 | NA | NA | NA | NA | NA | Gm44602 | predicted gene 44602 [Source:MGI Symbo | 0 | 0 | 0 | 0 | 0 | 0 |
| 46330 | ENSMUSG00000108517 | 0 | NA | NA | NA | NA | NA | Gm45125 | predicted gene 45125 [Source:MGI Symbo | 0 | 0 | 0 | 0 | 0 | 0 |
| 46331 | ENSMUSG00000108518 | 1.927485989 | 0.09716195 | 1.758808039 | 0.055243067 | 0.955944819 | NA | Gm5587 | predicted gene 5587 [Source:MGI Symbol | 1.6062 | 0 | 4.025 | 4.8199 | 0 | 1.1138 |
| 46332 | ENSMUSG00000108519 | 0 | NA | NA | NA | NA | NA | 4833421K07 | RIKEN cDNA 4833421K07 gene [Source:M | 0 | 0 | 0 | 0 | 0 | 0 |
| 46333 | ENSMUSG00000108520 | 0.72104214 | -1.535252325 | 3.007190864 | -0.510527065 | 0.609682258 | NA | Gm44684 | predicted gene 44684 [Source:MGI Symbo | 3.2124 | 0 | 0 | 0 | 0 | 1.1138 |
| 46334 | ENSMUSG00000108521 | 1.254537109 | -3.692333964 | 2.19368904 | -1.683161969 | 0.092343741 | NA | Gm44639 | predicted gene 44639 [Source:MGI Symbo | 2.4093 | 4.1117 | 1.0062 | 0 | 0 | 0 |
| 46335 | ENSMUSG00000108522 | 0 | NA | NA | NA | NA | NA | Gm44664 | predicted gene 44664 [Source:MGI Symbo | 0 | 0 | 0 | 0 | 0 | 0 |
| 46336 | ENSMUSG00000108523 | 0 | NA | NA | NA | NA | NA | 4933430L12 | RIKEN cDNA 4933430L12 gene [Source:M | 0 | 0 | 0 | 0 | 0 | 0 |
| 46337 | ENSMUSG00000108524 | 0.321325949 | 1.911575114 | 4.048267893 | 0.472195804 | 0.636787034 | NA | Gm44876 | predicted gene 44876 [Source:MGI Symbo | 0 | 0 | 0 | 1.928 | 0 | 0 |
| 46338 | ENSMUSG00000108525 | 0.335414862 | -1.779047242 | 4.030804179 | -0.441362855 | 0.658950329 | NA | Gm7443 | predicted gene 7443 [Source:MGI Symbol | 0 | 0 | 2.0125 | 0 | 0 | 0 |
| 46339 | ENSMUSG00000108526 | 0.6621303 | -1.329905799 | 2.807279375 | -0.473734752 | 0.635689068 | NA | Gm45828 | predicted gene 45828 [Source:MGI Symbo | 0.8031 | 2.0558 | 0 | 0 | 0 | 1.1138 |
| 46340 | ENSMUSG00000108527 | 0 | NA | NA | NA | NA | NA | Gm45173 | predicted gene 45173 [Source:MGI Symbo | 0 | 0 | 0 | 0 | 0 | 0 |
| 46341 | ENSMUSG00000108528 | 0 | NA | NA | NA | NA | NA | Gm8319 | predicted gene 8319 [Source:MGI Symbol | 0 | 0 | 0 | 0 | 0 | 0 |
| 46342 | ENSMUSG00000108529 | 0 | NA | NA | NA | NA | NA | Gm29759 | predicted gene, 29759 [Source:MGI Symb | 0 | 0 | 0 | 0 | 0 | 0 |
| 46343 | ENSMUSG00000108530 | 0 | NA | NA | NA | NA | NA | 4930429H19 | RIKEN cDNA 4930429H19 gene [Source:M | 0 | 0 | 0 | 0 | 0 | 0 |
| 46344 | ENSMUSG00000108531 | 0 | NA | NA | NA | NA | NA | Gm39091 | predicted gene, 39091 [Source:MGI Symb | 0 | 0 | 0 | 0 | 0 | 0 |
| 46345 | ENSMUSG00000108532 | 0 | NA | NA | NA | NA | NA | Gm32647 | predicted gene, 32647 [Source:MGI Symb | 0 | 0 | 0 | 0 | 0 | 0 |
| 46346 | ENSMUSG00000108533 | 0 | NA | NA | NA | NA | NA | Gm44792 | predicted gene 44792 [Source:MGI Symbo | 0 | 0 | 0 | 0 | 0 | 0 |
| 46347 | ENSMUSG00000108534 | 0 | NA | NA | NA | NA | NA | Olfr1360 | olfactory receptor 1360 [Source:MGI Syml | 0 | 0 | 0 | 0 | 0 | 0 |
| 46348 | ENSMUSG00000108535 | 0 | NA | NA | NA | NA | NA | Gm45071 | predicted gene 45071 [Source:MGI Symbo | 0 | 0 | 0 | 0 | 0 | 0 |
| 46349 | ENSMUSG00000108536 | 0.685330692 | 0.069534182 | 2.586556116 | 0.02688292 | 0.978553116 | NA | Gm45059 | predicted gene 45059 [Source:MGI Symbo | 0 | 1.0279 | 1.0062 | 0.964 | 0 | 1.1138 |
| 46350 | ENSMUSG00000108537 | 0 | NA | NA | NA | NA | NA | Olfr184-ps1 | olfactory receptor 184, pseudogene 1 [Sou | 0 | 0 | 0 | 0 | 0 | 0 |
| 46351 | ENSMUSG00000108538 | 0 | NA | NA | NA | NA | NA | Gm45197 | predicted gene 45197 [Source:MGI Symbo | 0 | 0 | 0 | 0 | 0 | 0 |
| 46352 | ENSMUSG00000108539 | 0 | NA | NA | NA | NA | NA | Klk1b12-ps | kallikrein 1-related peptidase b12, pseudo | 0 | 0 | 0 | 0 | 0 | 0 |
| 46353 | ENSMUSG00000108540 | 0 | NA | NA | NA | NA | NA | Gm4768 | predicted gene 4768 [Source:MGI Symbol | 0 | 0 | 0 | 0 | 0 | 0 |
| 46354 | ENSMUSG00000108541 | 0 | NA | NA | NA | NA | NA | Gm44795 | predicted gene 44795 [Source:MGI Symbo | 0 | 0 | 0 | 0 | 0 | 0 |
| 46355 | ENSMUSG00000108542 | 0.342639279 | -1.805976596 | 4.028547714 | -0.448294702 | 0.653940524 | NA | Gm44834 | predicted gene 44834 [Source:MGI Symbo | 0 | 2.0558 | 0 | 0 | 0 | 0 |
| 46356 | ENSMUSG00000108543 | 1.98657779 | 0.43989858 | 1.777258375 | 0.247515266 | 0.804509478 | NA | Gm44735 | predicted gene 44735 [Source:MGI Symbo | 4.0155 | 0 | 1.0062 | 5.7839 | 0 | 1.1138 |
| 46357 | ENSMUSG00000108544 | 0 | NA | NA | NA | NA | NA | Cyp2b26-ps | cytochrome P450, family 2, subfamily b, p | 0 | 0 | 0 | 0 | 0 | 0 |
| 46358 | ENSMUSG00000108545 | 0 | NA | NA | NA | NA | NA | Gm44613 | predicted gene 44613 [Source:MGI Symbo | 0 | 0 | 0 | 0 | 0 | 0 |
| 46359 | ENSMUSG00000108546 | 0 | NA | NA | NA | NA | NA | Gm44662 | predicted gene 44662 [Source:MGI Symbo | 0 | 0 | 0 | 0 | 0 | 0 |
| 46360 | ENSMUSG00000108547 | 1.017261173 | -2.166287499 | 2.717865413 | -0.797054736 | 0.425419243 | NA | Gm45186 | predicted gene 45186 [Source:MGI Symbo | 0 | 5.1396 | 0 | 0.964 | 0 | 0 |
| 46361 | ENSMUSG00000108548 | 0 | NA | NA | NA | NA | NA | Gm45006 | predicted gene 45006 [Source:MGI Symbo | 0 | 0 | 0 | 0 | 0 | 0 |
| 46362 | ENSMUSG00000108550 | 0 | NA | NA | NA | NA | NA | Gm44736 | predicted gene 44736 [Source:MGI Symbo | 0 | 0 | 0 | 0 | 0 | 0 |
| 46363 | ENSMUSG00000108551 | 7.569398031 | -0.027835162 | 0.832335735 | -0.033442229 | 0.973321935 | 0.995906088 | Gm20274 | predicted gene, 20274 [Source:MGI Symb | 4.8186 | 9.2513 | 9.0562 | 13.496 | 5.4531 | 3.3415 |
| 46364 | ENSMUSG00000108552 | 0 | NA | NA | NA | NA | NA | Gm45227 | predicted gene 45227 [Source:MGI Symbo | 0 | 0 | 0 | 0 | 0 | 0 |

| | | | | | | | | | | | | | | | |
|---|---|---|---|---|---|---|---|---|---|---|---|---|---|---|---|
| 46365 | ENSMUSG00000108553 | 0 | NA | NA | NA | NA | NA | Gm38569 | predicted gene, 38569 [Source:MGI Symb | 0 | 0 | 0 | 0 | 0 | 0 |
| 46366 | ENSMUSG00000108554 | 0.992155673 | -1.012754205 | 2.706122858 | -0.374245464 | 0.708221702 | NA | Gm44824 | predicted gene 44824 [Source:MGI Symbo | 0 | 0 | 4.025 | 1.928 | 0 | 0 |
| 46367 | ENSMUSG00000108555 | 1.937309364 | 3.388652841 | 1.82821598 | 1.853529822 | 0.063806456 | NA | NA | NA | 0 | 1.0279 | 0 | 2.8919 | 4.3625 | 3.3415 |
| 46368 | ENSMUSG00000108556 | 0 | NA | NA | NA | NA | NA | Gm17987 | predicted gene, 17987 [Source:MGI Symb | 0 | 0 | 0 | 0 | 0 | 0 |
| 46369 | ENSMUSG00000108557 | 0.133850373 | -0.796629176 | 4.080472857 | -0.195229623 | 0.84521319 | NA | Gm44835 | predicted gene 44835 [Source:MGI Symbo | 0.8031 | 0 | 0 | 0 | 0 | 0 |
| 46370 | ENSMUSG00000108558 | 0 | NA | NA | NA | NA | NA | Psg-ps1 | pregnancy specific glycoprotein, pseudog | 0 | 0 | 0 | 0 | 0 | 0 |
| 46371 | ENSMUSG00000108559 | 0 | NA | NA | NA | NA | NA | Gm44782 | predicted gene 44782 [Source:MGI Symbo | 0 | 0 | 0 | 0 | 0 | 0 |
| 46372 | ENSMUSG00000108560 | 0.181769715 | 1.126924175 | 4.080472857 | 0.276174898 | 0.782413736 | NA | Gm44695 | predicted gene 44695 [Source:MGI Symbo | 0 | 0 | 0 | 0 | 1.0906 | 0 |
| 46373 | ENSMUSG00000108561 | 0 | NA | NA | NA | NA | NA | Gm45219 | predicted gene 45219 [Source:MGI Symbo | 0 | 0 | 0 | 0 | 0 | 0 |
| 46374 | ENSMUSG00000108562 | 0 | NA | NA | NA | NA | NA | Gm44519 | predicted gene 44519 [Source:MGI Symbo | 0 | 0 | 0 | 0 | 0 | 0 |
| 46375 | ENSMUSG00000108563 | 12.02184034 | -0.547272919 | 0.632988258 | -0.86458621 | 0.387265944 | 0.817618864 | Gm44686 | predicted gene 44686 [Source:MGI Symbo | 14.456 | 10.279 | 18.112 | 10.604 | 11.997 | 6.6831 |
| 46376 | ENSMUSG00000108565 | 0 | NA | NA | NA | NA | NA | Olfr189-ps1 | olfactory receptor 189, pseudogene 1 [Sou | 0 | 0 | 0 | 0 | 0 | 0 |
| 46377 | ENSMUSG00000108566 | 0 | NA | NA | NA | NA | NA | Gm45031 | predicted gene 45031 [Source:MGI Symbo | 0 | 0 | 0 | 0 | 0 | 0 |
| 46378 | ENSMUSG00000108567 | 0 | NA | NA | NA | NA | NA | Gm44513 | predicted gene 44513 [Source:MGI Symbo | 0 | 0 | 0 | 0 | 0 | 0 |
| 46379 | ENSMUSG00000108568 | 0 | NA | NA | NA | NA | NA | Olfr277-ps1 | olfactory receptor 277, pseudogene 1 [Sou | 0 | 0 | 0 | 0 | 0 | 0 |
| 46380 | ENSMUSG00000108569 | 0 | NA | NA | NA | NA | NA | NA | NA | 0 | 0 | 0 | 0 | 0 | 0 |
| 46381 | ENSMUSG00000108570 | 0 | NA | NA | NA | NA | NA | Gm7726 | predicted gene 7726 [Source:MGI Symbol | 0 | 0 | 0 | 0 | 0 | 0 |
| 46382 | ENSMUSG00000108571 | 0.449470461 | -0.545136533 | 3.562105917 | -0.15303771 | 0.878368537 | NA | Gm6855 | predicted gene 6855 [Source:MGI Symbol | 1.6062 | 0 | 0 | 0 | 1.0906 | 0 |
| 46383 | ENSMUSG00000108572 | 0 | NA | NA | NA | NA | NA | Gm44883 | predicted gene 44883 [Source:MGI Symbo | 0 | 0 | 0 | 0 | 0 | 0 |
| 46384 | ENSMUSG00000108573 | 11.09467189 | -0.15229037 | 0.764620755 | -0.199171117 | 0.842128892 | 0.971854408 | Gm44986 | predicted gene 44986 [Source:MGI Symbo | 13.653 | 17.475 | 4.025 | 19.28 | 5.4531 | 6.6831 |
| 46385 | ENSMUSG00000108574 | 1.500899549 | -0.203928104 | 1.768908372 | -0.115284719 | 0.90821945 | NA | Gm44981 | predicted gene 44981 [Source:MGI Symbo | 0.8031 | 1.0279 | 3.0187 | 1.928 | 0 | 2.2277 |
| 46386 | ENSMUSG00000108575 | 19.01492819 | -0.160497978 | 0.511140028 | -0.314000019 | 0.753521024 | 0.952989051 | Gm35665 | predicted gene, 35665 [Source:MGI Symb | 21.684 | 18.503 | 20.125 | 24.099 | 18.541 | 11.138 |
| 46387 | ENSMUSG00000108576 | 0 | NA | NA | NA | NA | NA | Gm44778 | predicted gene 44778 [Source:MGI Symbo | 0 | 0 | 0 | 0 | 0 | 0 |
| 46388 | ENSMUSG00000108577 | 0 | NA | NA | NA | NA | NA | Gm17811 | predicted gene, 17811 [Source:MGI Symb | 0 | 0 | 0 | 0 | 0 | 0 |
| 46389 | ENSMUSG00000108578 | 0 | NA | NA | NA | NA | NA | Gm45700 | predicted gene 45700 [Source:MGI Symbo | 0 | 0 | 0 | 0 | 0 | 0 |
| 46390 | ENSMUSG00000108579 | 0 | NA | NA | NA | NA | NA | Gm29918 | predicted gene, 29918 [Source:MGI Symb | 0 | 0 | 0 | 0 | 0 | 0 |
| 46391 | ENSMUSG00000108580 | 0 | NA | NA | NA | NA | NA | Gm39094 | predicted gene, 39094 [Source:MGI Symb | 0 | 0 | 0 | 0 | 0 | 0 |
| 46392 | ENSMUSG00000108581 | 0 | NA | NA | NA | NA | NA | Gm44896 | predicted gene 44896 [Source:MGI Symbo | 0 | 0 | 0 | 0 | 0 | 0 |
| 46393 | ENSMUSG00000108582 | 0 | NA | NA | NA | NA | NA | Gm18973 | predicted gene, 18973 [Source:MGI Symb | 0 | 0 | 0 | 0 | 0 | 0 |
| 46394 | ENSMUSG00000108583 | 0 | NA | NA | NA | NA | NA | Gm44854 | predicted gene 44854 [Source:MGI Symbo | 0 | 0 | 0 | 0 | 0 | 0 |
| 46395 | ENSMUSG00000108584 | 0.36353943 | 2.065989254 | 4.034137301 | 0.512126658 | 0.608562368 | NA | Gm45216 | predicted gene 45216 [Source:MGI Symbo | 0 | 0 | 0 | 0 | 2.1812 | 0 |
| 46396 | ENSMUSG00000108586 | 0 | NA | NA | NA | NA | NA | Gm44976 | predicted gene 44976 [Source:MGI Symbo | 0 | 0 | 0 | 0 | 0 | 0 |
| 46397 | ENSMUSG00000108587 | 0 | NA | NA | NA | NA | NA | 1810049I09F | RIKEN cDNA 1810049I09 gene [Source:MG | 0 | 0 | 0 | 0 | 0 | 0 |
| 46398 | ENSMUSG00000108588 | 0 | NA | NA | NA | NA | NA | Gm44814 | predicted gene 44814 [Source:MGI Symbo | 0 | 0 | 0 | 0 | 0 | 0 |
| 46399 | ENSMUSG00000108589 | 0 | NA | NA | NA | NA | NA | Gm45225 | predicted gene 45225 [Source:MGI Symbo | 0 | 0 | 0 | 0 | 0 | 0 |
| 46400 | ENSMUSG00000108590 | 0 | NA | NA | NA | NA | NA | Gm44962 | predicted gene 44962 [Source:MGI Symbo | 0 | 0 | 0 | 0 | 0 | 0 |
| 46401 | ENSMUSG00000108591 | 32.4751694 | 0.238902081 | 0.465402995 | 0.513323041 | 0.607725369 | 0.916376001 | Particl | promoter of Mat2a antisense radiation in | 40.958 | 33.921 | 14.087 | 23.135 | 37.081 | 45.668 |
| 46402 | ENSMUSG00000108592 | 0 | NA | NA | NA | NA | NA | Gm38973 | predicted gene, 38973 [Source:MGI Symb | 0 | 0 | 0 | 0 | 0 | 0 |
| 46403 | ENSMUSG00000108593 | 0 | NA | NA | NA | NA | NA | Gm44581 | predicted gene 44581 [Source:MGI Symbo | 0 | 0 | 0 | 0 | 0 | 0 |
| 46404 | ENSMUSG00000108594 | 0 | NA | NA | NA | NA | NA | Gm44770 | predicted gene 44770 [Source:MGI Symbo | 0 | 0 | 0 | 0 | 0 | 0 |
| 46405 | ENSMUSG00000108596 | 0 | NA | NA | NA | NA | NA | Gm49368 | predicted gene, 49368 [Source:MGI Symb | 0 | 0 | 0 | 0 | 0 | 0 |
| 46406 | ENSMUSG00000108597 | 0 | NA | NA | NA | NA | NA | Gm44708 | predicted gene 44708 [Source:MGI Symbo | 0 | 0 | 0 | 0 | 0 | 0 |
| 46407 | ENSMUSG00000108598 | 0 | NA | NA | NA | NA | NA | Gm44665 | predicted gene 44665 [Source:MGI Symbo | 0 | 0 | 0 | 0 | 0 | 0 |
| 46408 | ENSMUSG00000108599 | 1.33249471 | -0.610195187 | 1.87565474 | -0.325323832 | 0.744935996 | NA | Gm10616 | predicted gene 10616 [Source:MGI Symbo | 0.8031 | 1.0279 | 3.0187 | 0.964 | 2.1812 | 0 |
| 46409 | ENSMUSG00000108600 | 0 | NA | NA | NA | NA | NA | Gm32884 | predicted gene, 32884 [Source:MGI Symb | 0 | 0 | 0 | 0 | 0 | 0 |
| 46410 | ENSMUSG00000108601 | 0.503122293 | -2.366675966 | 3.785596405 | -0.625179156 | 0.531853481 | NA | Gm44645 | predicted gene 44645 [Source:MGI Symbo | 0 | 0 | 3.0187 | 0 | 0 | 0 |
| 46411 | ENSMUSG00000108602 | 0 | NA | NA | NA | NA | NA | Klk1b10-ps | kallikrein 1-related peptidase b10, pseudo | 0 | 0 | 0 | 0 | 0 | 0 |
| 46412 | ENSMUSG00000108603 | 0.342639279 | -1.805976596 | 4.028547714 | -0.448294702 | 0.653940524 | NA | 5330411O13 | RIKEN cDNA 5330411O13 gene [Source:M | 0 | 2.0558 | 0 | 0 | 0 | 0 |
| 46413 | ENSMUSG00000108604 | 0 | NA | NA | NA | NA | NA | Gm44850 | predicted gene 44850 [Source:MGI Symbo | 0 | 0 | 0 | 0 | 0 | 0 |
| 46414 | ENSMUSG00000108605 | 0 | NA | NA | NA | NA | NA | Olfr396-ps1 | olfactory receptor 396, pseudogene 1 [Sou | 0 | 0 | 0 | 0 | 0 | 0 |
| 46415 | ENSMUSG00000108606 | 0 | NA | NA | NA | NA | NA | Cyp2b27-ps | cytochrome P450, family 2, subfamily b, p | 0 | 0 | 0 | 0 | 0 | 0 |
| 46416 | ENSMUSG00000108607 | 0 | NA | NA | NA | NA | NA | Gm44646 | predicted gene 44646 [Source:MGI Symbo | 0 | 0 | 0 | 0 | 0 | 0 |
| 46417 | ENSMUSG00000108608 | 0 | NA | NA | NA | NA | NA | Gm6916 | predicted pseudogene 6916 [Source:MGI | 0 | 0 | 0 | 0 | 0 | 0 |
| 46418 | ENSMUSG00000108609 | 0 | NA | NA | NA | NA | NA | Gm7424 | predicted gene 7424 [Source:MGI Symbol | 0 | 0 | 0 | 0 | 0 | 0 |
| 46419 | ENSMUSG00000108610 | 0 | NA | NA | NA | NA | NA | Gm29725 | predicted gene, 29725 [Source:MGI Symb | 0 | 0 | 0 | 0 | 0 | 0 |
| 46420 | ENSMUSG00000108611 | 0 | NA | NA | NA | NA | NA | Gm44879 | predicted gene 44879 [Source:MGI Symbo | 0 | 0 | 0 | 0 | 0 | 0 |

| | | | | | | | | | | | | | | | |
|---|---|---|---|---|---|---|---|---|---|---|---|---|---|---|---|
| 46421 | ENSMUSG00000108612 | 0.133850373 | -0.796629176 | 4.080472857 | -0.195229623 | 0.84521319 | NA | Gm44634 | predicted gene 44634 [Source:MGI Symbo | 0.8031 | 0 | 0 | 0 | 0 | 0 |
| 46422 | ENSMUSG00000108613 | 0 | NA | NA | NA | NA | NA | Gm36694 | predicted gene, 36694 [Source:MGI Symb | 0 | 0 | 0 | 0 | 0 | 0 |
| 46423 | ENSMUSG00000108614 | 2.951088697 | -1.207337712 | 1.259764256 | -0.958383845 | 0.337869238 | NA | 2610306O10 | RIKEN cDNA 2610306O10 gene [Source:M | 3.2124 | 3.0838 | 6.0375 | 0.964 | 2.1812 | 2.2277 |
| 46424 | ENSMUSG00000108615 | 0 | NA | NA | NA | NA | NA | Gm4973 | predicted gene 4973 [Source:MGI Symbol | 0 | 0 | 0 | 0 | 0 | 0 |
| 46425 | ENSMUSG00000108616 | 0 | NA | NA | NA | NA | NA | Gm35040 | predicted gene, 35040 [Source:MGI Symb | 0 | 0 | 0 | 0 | 0 | 0 |
| 46426 | ENSMUSG00000108617 | 0.335414862 | -1.779047242 | 4.030804179 | -0.441362855 | 0.658950329 | NA | Gm31749 | predicted gene, 31749 [Source:MGI Symb | 0 | 0 | 2.0125 | 0 | 0 | 0 |
| 46427 | ENSMUSG00000108618 | 5.133421376 | -0.137749094 | 0.922460873 | -0.149327844 | 0.881294945 | 0.981599421 | Gm44816 | predicted gene 44816 [Source:MGI Symbo | 4.0155 | 5.1396 | 7.0437 | 5.7839 | 4.3625 | 4.4554 |
| 46428 | ENSMUSG00000108619 | 0 | NA | NA | NA | NA | NA | Gm44839 | predicted gene 44839 [Source:MGI Symbo | 0 | 0 | 0 | 0 | 0 | 0 |
| 46429 | ENSMUSG00000108620 | 0 | NA | NA | NA | NA | NA | Gm20670 | predicted gene 20670 [Source:MGI Symbo | 0 | 0 | 0 | 0 | 0 | 0 |
| 46430 | ENSMUSG00000108621 | 823.8872403 | 0.2524104 | 0.12919336 | 1.953741279 | 0.050731822 | 0.316199857 | Gm33989 | predicted gene, 33989 [Source:MGI Symb | 726.81 | 664.03 | 865.37 | 893.61 | 931.39 | 862.12 |
| 46431 | ENSMUSG00000108622 | 0 | NA | NA | NA | NA | NA | Gm36864 | predicted gene, 36864 [Source:MGI Symb | 0 | 0 | 0 | 0 | 0 | 0 |
| 46432 | ENSMUSG00000108623 | 0 | NA | NA | NA | NA | NA | Gm45092 | predicted gene 45092 [Source:MGI Symbo | 0 | 0 | 0 | 0 | 0 | 0 |
| 46433 | ENSMUSG00000108624 | 45.26258487 | 0.146130912 | 0.334710943 | 0.436588392 | 0.662409882 | 0.9288455 | Gm45091 | predicted gene 45091 [Source:MGI Symbo | 39.352 | 51.396 | 38.237 | 42.415 | 55.622 | 44.554 |
| 46434 | ENSMUSG00000108625 | 0 | NA | NA | NA | NA | NA | Gm44711 | predicted gene 44711 [Source:MGI Symbo | 0 | 0 | 0 | 0 | 0 | 0 |
| 46435 | ENSMUSG00000108626 | 0 | NA | NA | NA | NA | NA | Gm40457 | predicted gene, 40457 [Source:MGI Symb | 0 | 0 | 0 | 0 | 0 | 0 |
| 46436 | ENSMUSG00000108627 | 0 | NA | NA | NA | NA | NA | Gm45079 | predicted gene 45079 [Source:MGI Symbo | 0 | 0 | 0 | 0 | 0 | 0 |
| 46437 | ENSMUSG00000108628 | 0.160662974 | 1.126924175 | 4.080472857 | 0.276174898 | 0.782413736 | NA | Gm5037 | predicted gene 5037 [Source:MGI Symbol | 0 | 0 | 0 | 0.964 | 0 | 0 |
| 46438 | ENSMUSG00000108629 | 0 | NA | NA | NA | NA | NA | Gm44720 | predicted gene 44720 [Source:MGI Symbo | 0 | 0 | 0 | 0 | 0 | 0 |
| 46439 | ENSMUSG00000108630 | 0 | NA | NA | NA | NA | NA | Gm44607 | predicted gene 44607 [Source:MGI Symbo | 0 | 0 | 0 | 0 | 0 | 0 |
| 46440 | ENSMUSG00000108633 | 0.267700746 | -1.497481649 | 4.057001436 | -0.369110456 | 0.712045396 | NA | Gm44694 | predicted gene 44694 [Source:MGI Symbo | 1.6062 | 0 | 0 | 0 | 0 | 0 |
| 46441 | ENSMUSG00000108634 | 0 | NA | NA | NA | NA | NA | Gm38534 | predicted gene, 38534 [Source:MGI Symb | 0 | 0 | 0 | 0 | 0 | 0 |
| 46442 | ENSMUSG00000108635 | 0 | NA | NA | NA | NA | NA | Gm44633 | predicted gene 44633 [Source:MGI Symbo | 0 | 0 | 0 | 0 | 0 | 0 |
| 46443 | ENSMUSG00000108636 | 0 | NA | NA | NA | NA | NA | Gm44650 | predicted gene 44650 [Source:MGI Symbo | 0 | 0 | 0 | 0 | 0 | 0 |
| 46444 | ENSMUSG00000108637 | 0.480153995 | 1.043847617 | 3.151673025 | 0.331204287 | 0.740490182 | NA | Gm4290 | predicted gene 4290 [Source:MGI Symbol | 0.8031 | 0 | 0 | 0.964 | 0 | 1.1138 |
| 46445 | ENSMUSG00000108638 | 0 | NA | NA | NA | NA | NA | Gm45150 | predicted gene 45150 [Source:MGI Symbo | 0 | 0 | 0 | 0 | 0 | 0 |
| 46446 | ENSMUSG00000108639 | 0 | NA | NA | NA | NA | NA | | olfactory receptor 1400, pseudogene 1 | 0 | 0 | 0 | 0 | 0 | 0 |
| 46447 | ENSMUSG00000108640 | 0 | NA | NA | NA | NA | NA | Gm44508 | predicted gene 44508 [Source:MGI Symbo | 0 | 0 | 0 | 0 | 0 | 0 |
| 46448 | ENSMUSG00000108641 | 0 | NA | NA | NA | NA | NA | Gm44807 | predicted gene 44807 [Source:MGI Symbo | 0 | 0 | 0 | 0 | 0 | 0 |
| 46449 | ENSMUSG00000108642 | 0 | NA | NA | NA | NA | NA | Gm44903 | predicted gene 44903 [Source:MGI Symbo | 0 | 0 | 0 | 0 | 0 | 0 |
| 46450 | ENSMUSG00000108643 | 0.267700746 | -1.497481649 | 4.057001436 | -0.369110456 | 0.712045396 | NA | Gm44931 | predicted gene 44931 [Source:MGI Symbo | 1.6062 | 0 | 0 | 0 | 0 | 0 |
| 46451 | ENSMUSG00000108644 | 0 | NA | NA | NA | NA | NA | Gm44920 | predicted gene 44920 [Source:MGI Symbo | 0 | 0 | 0 | 0 | 0 | 0 |
| 46452 | ENSMUSG00000108645 | 0 | NA | NA | NA | NA | NA | Gm44722 | predicted gene 44722 [Source:MGI Symbo | 0 | 0 | 0 | 0 | 0 | 0 |
| 46453 | ENSMUSG00000108646 | 0 | NA | NA | NA | NA | NA | Olfr162-ps1 | olfactory receptor 162, pseudogene 1 [Sou | 0 | 0 | 0 | 0 | 0 | 0 |
| 46454 | ENSMUSG00000108647 | 0 | NA | NA | NA | NA | NA | Gm17984 | predicted gene, 17984 [Source:MGI Symb | 0 | 0 | 0 | 0 | 0 | 0 |
| 46455 | ENSMUSG00000108648 | 0.133850373 | -0.796629176 | 4.080472857 | -0.195229623 | 0.84521319 | NA | Gm44674 | predicted gene 44674 [Source:MGI Symbo | 0.8031 | 0 | 0 | 0 | 0 | 0 |
| 46456 | ENSMUSG00000108649 | 0 | NA | NA | NA | NA | NA | Gm44855 | predicted gene 44855 [Source:MGI Symbo | 0 | 0 | 0 | 0 | 0 | 0 |
| 46457 | ENSMUSG00000108650 | 1.306617125 | 1.546876254 | 2.477123789 | 0.624464656 | 0.532322475 | NA | Gm44867 | predicted gene 44867 [Source:MGI Symbo | 0 | 2.0558 | 0 | 5.7839 | 0 | 0 |
| 46458 | ENSMUSG00000108651 | 0 | NA | NA | NA | NA | NA | Klk1b2-ps | kallikrein 1-related peptidase b2, pseudog | 0 | 0 | 0 | 0 | 0 | 0 |
| 46459 | ENSMUSG00000108652 | 0.483327519 | -0.685704547 | 3.145069396 | -0.218025252 | 0.827409436 | NA | 0610006L08I | RIKEN cDNA 0610006L08 gene [Source:MG | 0.8031 | 0 | 1.0062 | 0 | 1.0906 | 0 |
| 46460 | ENSMUSG00000108654 | 0 | NA | NA | NA | NA | NA | Gm9519 | predicted gene 9519 [Source:MGI Symbol | 0 | 0 | 0 | 0 | 0 | 0 |
| 46461 | ENSMUSG00000108655 | 7.444079654 | -0.856296152 | 0.79238455 | -1.080657305 | 0.279849582 | 0.732834426 | Gm44949 | predicted gene 44949 [Source:MGI Symbo | 6.4248 | 12.335 | 10.062 | 4.8199 | 5.4531 | 5.5692 |
| 46462 | ENSMUSG00000108657 | 6.505523441 | -0.75128398 | 0.846206798 | -0.887825509 | 0.374634619 | 0.809332934 | Gm35611 | predicted gene, 35611 [Source:MGI Symb | 11.243 | 6.1675 | 7.0437 | 5.7839 | 5.4531 | 3.3415 |
| 46463 | ENSMUSG00000108659 | 14.80802324 | 0.633689372 | 0.627458938 | 1.009929627 | 0.312529007 | 0.759830222 | Gm34121 | predicted gene, 34121 [Source:MGI Symb | 19.274 | 9.2513 | 6.0375 | 20.244 | 22.903 | 11.138 |
| 46464 | ENSMUSG00000108660 | 6.23998207 | 1.483779235 | 0.939389513 | 1.579514371 | 0.114218123 | 0.496868018 | Gm21284 | predicted gene, 21284 [Source:MGI Symb | 5.6217 | 1.0279 | 3.0187 | 6.7478 | 6.5437 | 14.48 |
| 46465 | ENSMUSG00000108661 | 0 | NA | NA | NA | NA | NA | Gm45172 | predicted gene 45172 [Source:MGI Symbo | 0 | 0 | 0 | 0 | 0 | 0 |
| 46466 | ENSMUSG00000108662 | 0 | NA | NA | NA | NA | NA | Olfr188-ps1 | olfactory receptor 188, pseudogene 1 [Sou | 0 | 0 | 0 | 0 | 0 | 0 |
| 46467 | ENSMUSG00000108663 | 0 | NA | NA | NA | NA | NA | Gm33080 | predicted gene, 33080 [Source:MGI Symb | 0 | 0 | 0 | 0 | 0 | 0 |
| 46468 | ENSMUSG00000108665 | 0 | NA | NA | NA | NA | NA | Gm44897 | predicted gene 44897 [Source:MGI Symbo | 0 | 0 | 0 | 0 | 0 | 0 |
| 46469 | ENSMUSG00000108666 | 0 | NA | NA | NA | NA | NA | Gm44538 | predicted gene 44538 [Source:MGI Symbo | 0 | 0 | 0 | 0 | 0 | 0 |
| 46470 | ENSMUSG00000108667 | 0 | NA | NA | NA | NA | NA | Gm45013 | predicted gene 45013 [Source:MGI Symbo | 0 | 0 | 0 | 0 | 0 | 0 |
| 46471 | ENSMUSG00000108668 | 0.267700746 | -1.497481649 | 4.057001436 | -0.369110456 | 0.712045396 | NA | Gm32816 | predicted gene, 32816 [Source:MGI Symb | 1.6062 | 0 | 0 | 0 | 0 | 0 |
| 46472 | ENSMUSG00000108669 | 0 | NA | NA | NA | NA | NA | 4930441H08 | RIKEN cDNA 4930441H08 gene [Source:M | 0 | 0 | 0 | 0 | 0 | 0 |
| 46473 | ENSMUSG00000108670 | 0 | NA | NA | NA | NA | NA | Gm44815 | predicted gene 44815 [Source:MGI Symbo | 0 | 0 | 0 | 0 | 0 | 0 |
| 46474 | ENSMUSG00000108671 | 0 | NA | NA | NA | NA | NA | 2700080J24I | RIKEN cDNA 2700080J24 gene [Source:MG | 0 | 0 | 0 | 0 | 0 | 0 |
| 46475 | ENSMUSG00000108672 | 0 | NA | NA | NA | NA | NA | Gm45008 | predicted gene 45008 [Source:MGI Symbo | 0 | 0 | 0 | 0 | 0 | 0 |
| 46476 | ENSMUSG00000108673 | 0 | NA | NA | NA | NA | NA | Gm44922 | predicted gene 44922 [Source:MGI Symbo | 0 | 0 | 0 | 0 | 0 | 0 |

| | | | | | | | | | | | | | | | |
|---|---|---|---|---|---|---|---|---|---|---|---|---|---|---|---|
| 46477 | ENSMUSG00000108674 | 0 | NA | NA | NA | NA | NA | Olfr1359 | olfactory receptor 1359 [Source:MGI Syml | 0 | 0 | 0 | 0 | 0 | 0 |
| 46478 | ENSMUSG00000108675 | 0 | NA | NA | NA | NA | NA | Gm44970 | predicted gene 44970 [Source:MGI Symbo | 0 | 0 | 0 | 0 | 0 | 0 |
| 46479 | ENSMUSG00000108676 | 0 | NA | NA | NA | NA | NA | 1700123J17I | RIKEN cDNA 1700123J17 gene [Source:M( | 0 | 0 | 0 | 0 | 0 | 0 |
| 46480 | ENSMUSG00000108677 | 0.133850373 | -0.796629176 | 4.080472857 | -0.195229623 | 0.84521319 | NA | Gm44759 | predicted gene 44759 [Source:MGI Symbo | 0.8031 | 0 | 0 | 0 | 0 | 0 |
| 46481 | ENSMUSG00000108678 | 0 | NA | NA | NA | NA | NA | Gm44758 | predicted gene 44758 [Source:MGI Symbo | 0 | 0 | 0 | 0 | 0 | 0 |
| 46482 | ENSMUSG00000108679 | 0 | NA | NA | NA | NA | NA | Gm45134 | predicted gene 45134 [Source:MGI Symbo | 0 | 0 | 0 | 0 | 0 | 0 |
| 46483 | ENSMUSG00000108681 | 7.118790351 | 0.848910727 | 0.939093696 | 0.903968081 | 0.366012327 | 0.804235439 | Gm17825 | predicted gene, 17825 [Source:MGI Symb | 3.2124 | 5.1396 | 7.0437 | 16.388 | 9.8156 | 1.1138 |
| 46484 | ENSMUSG00000108682 | 0 | NA | NA | NA | NA | NA | Gm29754 | predicted gene, 29754 [Source:MGI Symb | 0 | 0 | 0 | 0 | 0 | 0 |
| 46485 | ENSMUSG00000108683 | 0 | NA | NA | NA | NA | NA | Gm45017 | predicted gene 45017 [Source:MGI Symbo | 0 | 0 | 0 | 0 | 0 | 0 |
| 46486 | ENSMUSG00000108685 | 0 | NA | NA | NA | NA | NA | Gm45149 | predicted gene 45149 [Source:MGI Symbo | 0 | 0 | 0 | 0 | 0 | 0 |
| 46487 | ENSMUSG00000108686 | 0 | NA | NA | NA | NA | NA | Gm45080 | predicted gene 45080 [Source:MGI Symbo | 0 | 0 | 0 | 0 | 0 | 0 |
| 46488 | ENSMUSG00000108687 | 0 | NA | NA | NA | NA | NA | Gm45007 | predicted gene 45007 [Source:MGI Symbo | 0 | 0 | 0 | 0 | 0 | 0 |
| 46489 | ENSMUSG00000108688 | 0 | NA | NA | NA | NA | NA | Gm44985 | predicted gene 44985 [Source:MGI Symbo | 0 | 0 | 0 | 0 | 0 | 0 |
| 46490 | ENSMUSG00000108689 | 0.305170013 | -1.657836477 | 4.041480335 | -0.410205257 | 0.681655385 | NA | Gm45354 | predicted gene 45354 [Source:MGI Symbo | 0.8031 | 1.0279 | 0 | 0 | 0 | 0 |
| 46491 | ENSMUSG00000108691 | 0 | NA | NA | NA | NA | NA | Gm45226 | predicted gene 45226 [Source:MGI Symbo | 0 | 0 | 0 | 0 | 0 | 0 |
| 46492 | ENSMUSG00000108692 | 0 | NA | NA | NA | NA | NA | Gm45034 | predicted gene 45034 [Source:MGI Symbo | 0 | 0 | 0 | 0 | 0 | 0 |
| 46493 | ENSMUSG00000108693 | 0 | NA | NA | NA | NA | NA | Gm45153 | predicted gene 45153 [Source:MGI Symbo | 0 | 0 | 0 | 0 | 0 | 0 |
| 46494 | ENSMUSG00000108694 | 1.099522877 | 2.452134312 | 2.647755213 | 0.926118208 | 0.354384553 | NA | Gm7287 | predicted gene 7287 [Source:MGI Symbol | 0 | 1.0279 | 0 | 0 | 0 | 5.5692 |
| 46495 | ENSMUSG00000108695 | 0 | NA | NA | NA | NA | NA | Siglecl1 | Siglec family like 1 [Source:MGI Symbol;A( | 0 | 0 | 0 | 0 | 0 | 0 |
| 46496 | ENSMUSG00000108696 | 0 | NA | NA | NA | NA | NA | Gm5738 | predicted gene 5738 [Source:MGI Symbol | 0 | 0 | 0 | 0 | 0 | 0 |
| 46497 | ENSMUSG00000108697 | 0 | NA | NA | NA | NA | NA | 4930486N12 | RIKEN cDNA 4930486N12 gene [Source:M | 0 | 0 | 0 | 0 | 0 | 0 |
| 46498 | ENSMUSG00000108698 | 0 | NA | NA | NA | NA | NA | Gm44964 | predicted gene 44964 [Source:MGI Symbo | 0 | 0 | 0 | 0 | 0 | 0 |
| 46499 | ENSMUSG00000108699 | 0 | NA | NA | NA | NA | NA | Gm18988 | predicted gene, 18988 [Source:MGI Symb | 0 | 0 | 0 | 0 | 0 | 0 |
| 46500 | ENSMUSG00000108700 | 0 | NA | NA | NA | NA | NA | Gm7482 | predicted gene 7482 [Source:MGI Symbol | 0 | 0 | 0 | 0 | 0 | 0 |
| 46501 | ENSMUSG00000108701 | 6.53016581 | 0.386849222 | 0.81743396 | 0.473248288 | 0.636036051 | 0.923478764 | Gm44616 | predicted gene 44616 [Source:MGI Symbo | 4.8186 | 5.1396 | 7.0437 | 6.7478 | 7.6343 | 7.7969 |
| 46502 | ENSMUSG00000108702 | 0.538988726 | 1.126003851 | 3.343747182 | 0.336749099 | 0.736306047 | NA | Gm9333 | predicted gene 9333 [Source:MGI Symbol | 0 | 0 | 1.0062 | 0 | 0 | 2.2277 |
| 46503 | ENSMUSG00000108703 | 0.133850373 | -0.796629176 | 4.080472857 | -0.195229623 | 0.84521319 | NA | Gm44793 | predicted gene 44793 [Source:MGI Symbo | 0.8031 | 0 | 0 | 0 | 0 | 0 |
| 46504 | ENSMUSG00000108704 | 0.301557804 | -1.64335916 | 4.042814367 | -0.406488899 | 0.684383412 | NA | Gm44636 | predicted gene 44636 [Source:MGI Symbo | 0.8031 | 0 | 1.0062 | 0 | 0 | 0 |
| 46505 | ENSMUSG00000108705 | 0.439020386 | -2.189747911 | 3.52148864 | -0.621824499 | 0.534057275 | NA | Gm18905 | predicted gene, 18905 [Source:MGI Symb | 1.6062 | 1.0279 | 0 | 0 | 0 | 0 |
| 46506 | ENSMUSG00000108707 | 0 | NA | NA | NA | NA | NA | Gm44575 | predicted gene 44575 [Source:MGI Symbo | 0 | 0 | 0 | 0 | 0 | 0 |
| 46507 | ENSMUSG00000108708 | 0 | NA | NA | NA | NA | NA | Gm29977 | predicted gene, 29977 [Source:MGI Symb | 0 | 0 | 0 | 0 | 0 | 0 |
| 46508 | ENSMUSG00000108709 | 10.63794742 | -0.76528783 | 0.718042243 | -1.065797783 | 0.286515063 | 0.73880343 | 4933431G14 | RIKEN cDNA 4933431G14 gene [Source:M | 12.85 | 15.419 | 12.075 | 12.532 | 8.7249 | 2.2277 |
| 46509 | ENSMUSG00000108711 | 0 | NA | NA | NA | NA | NA | Gm38991 | predicted gene, 38991 [Source:MGI Symb | 0 | 0 | 0 | 0 | 0 | 0 |
| 46510 | ENSMUSG00000108712 | 0 | NA | NA | NA | NA | NA | Gm3189 | predicted gene 3189 [Source:MGI Symbol | 0 | 0 | 0 | 0 | 0 | 0 |
| 46511 | ENSMUSG00000108713 | 0 | NA | NA | NA | NA | NA | Gm33027 | predicted gene, 33027 [Source:MGI Symb | 0 | 0 | 0 | 0 | 0 | 0 |
| 46512 | ENSMUSG00000108714 | 0.534859069 | 1.101323472 | 3.351640271 | 0.328592386 | 0.742463804 | NA | Gm21057 | predicted gene, 21057 [Source:MGI Symb | 0 | 1.0279 | 0 | 0 | 2.1812 | 0 |
| 46513 | ENSMUSG00000108716 | 0 | NA | NA | NA | NA | NA | Gm44902 | predicted gene 44902 [Source:MGI Symbo | 0 | 0 | 0 | 0 | 0 | 0 |
| 46514 | ENSMUSG00000108717 | 0 | NA | NA | NA | NA | NA | Gm44940 | predicted gene 44940 [Source:MGI Symbo | 0 | 0 | 0 | 0 | 0 | 0 |
| 46515 | ENSMUSG00000108718 | 0 | NA | NA | NA | NA | NA | Gm44749 | predicted gene 44749 [Source:MGI Symbo | 0 | 0 | 0 | 0 | 0 | 0 |
| 46516 | ENSMUSG00000108719 | 0 | NA | NA | NA | NA | NA | Gm44757 | predicted gene 44757 [Source:MGI Symbo | 0 | 0 | 0 | 0 | 0 | 0 |
| 46517 | ENSMUSG00000108720 | 4.167975453 | -0.010060435 | 1.069914581 | -0.009403027 | 0.992497581 | NA | Gm44672 | predicted gene 44672 [Source:MGI Symbo | 6.4248 | 1.0279 | 5.0312 | 4.8199 | 4.3625 | 3.3415 |
| 46518 | ENSMUSG00000108721 | 0 | NA | NA | NA | NA | NA | Gm44721 | predicted gene 44721 [Source:MGI Symbo | 0 | 0 | 0 | 0 | 0 | 0 |
| 46519 | ENSMUSG00000108722 | 0 | NA | NA | NA | NA | NA | Gm34225 | predicted gene, 34225 [Source:MGI Symb | 0 | 0 | 0 | 0 | 0 | 0 |
| 46520 | ENSMUSG00000108723 | 0.301557804 | -1.64335916 | 4.042814367 | -0.406488899 | 0.684383412 | NA | Gm44837 | predicted gene 44837 [Source:MGI Symbo | 0.8031 | 0 | 1.0062 | 0 | 0 | 0 |
| 46521 | ENSMUSG00000108724 | 0 | NA | NA | NA | NA | NA | Gm44734 | predicted gene 44734 [Source:MGI Symbo | 0 | 0 | 0 | 0 | 0 | 0 |
| 46522 | ENSMUSG00000108725 | 0 | NA | NA | NA | NA | NA | Gm45004 | predicted gene 45004 [Source:MGI Symbo | 0 | 0 | 0 | 0 | 0 | 0 |
| 46523 | ENSMUSG00000108726 | 0 | NA | NA | NA | NA | NA | Gm30684 | predicted gene, 30684 [Source:MGI Symb | 0 | 0 | 0 | 0 | 0 | 0 |
| 46524 | ENSMUSG00000108727 | 0 | NA | NA | NA | NA | NA | Gm44779 | predicted gene 44779 [Source:MGI Symbo | 0 | 0 | 0 | 0 | 0 | 0 |
| 46525 | ENSMUSG00000108728 | 0 | NA | NA | NA | NA | NA | | olfactory receptor 240, pseudogene 1 | 0 | 0 | 0 | 0 | 0 | 0 |
| 46526 | ENSMUSG00000108729 | 0 | NA | NA | NA | NA | NA | Gm17831 | predicted gene, 17831 [Source:MGI Symb | 0 | 0 | 0 | 0 | 0 | 0 |
| 46527 | ENSMUSG00000108730 | 0 | NA | NA | NA | NA | NA | Gm45135 | predicted gene 45135 [Source:MGI Symbo | 0 | 0 | 0 | 0 | 0 | 0 |
| 46528 | ENSMUSG00000108731 | 0.535401493 | -2.48596965 | 3.684117979 | -0.674780141 | 0.499815459 | NA | Api5-ps | apoptosis inhibitor 5, pseudogene [Source | 3.2124 | 0 | 0 | 0 | 0 | 0 |
| 46529 | ENSMUSG00000108732 | 7.736710327 | -0.186344819 | 0.837153586 | -0.222593348 | 0.823852014 | 0.96943211 | 2310043P16 | RIKEN cDNA 2310043P16 gene [Source:M | 6.4248 | 14.391 | 4.025 | 10.604 | 7.6343 | 3.3415 |
| 46530 | ENSMUSG00000108734 | 0 | NA | NA | NA | NA | NA | Gm44817 | predicted gene 44817 [Source:MGI Symbo | 0 | 0 | 0 | 0 | 0 | 0 |
| 46531 | ENSMUSG00000108735 | 0 | NA | NA | NA | NA | NA | Gm44638 | predicted gene 44638 [Source:MGI Symbo | 0 | 0 | 0 | 0 | 0 | 0 |
| 46532 | ENSMUSG00000108736 | 0.758511407 | -1.621915645 | 2.674939911 | -0.606337226 | 0.544290844 | NA | Gm45151 | predicted gene 45151 [Source:MGI Symbo | 2.4093 | 1.0279 | 0 | 0 | 0 | 1.1138 |

| | | | | | | | | | | | | | | | |
|---|---|---|---|---|---|---|---|---|---|---|---|---|---|---|---|
| 46533 | ENSMUSG00000108737 | 0.486939728 | -0.698472448 | 3.137608162 | -0.222613026 | 0.823836697 | NA | BC060293 | cDNA sequence BC060293 [Source:MGI Sy | 0.8031 | 1.0279 | 0 | 0 | 1.0906 | 0 |
| 46534 | ENSMUSG00000108738 | 11.38756216 | -0.07165211 | 0.698925494 | -0.102517523 | 0.918345905 | 0.986374671 | Gm44777 | predicted gene 44777 [Source:MGI Symbo | 4.8186 | 12.335 | 18.112 | 15.424 | 8.7249 | 8.9108 |
| 46535 | ENSMUSG00000108739 | 0.569258551 | -2.569337407 | 3.18860282 | -0.805787849 | 0.420365185 | NA | Gm45096 | predicted gene 45096 [Source:MGI Symbo | 2.4093 | 0 | 1.0062 | 0 | 0 | 0 |
| 46536 | ENSMUSG00000108740 | 0 | NA | NA | NA | NA | NA | Gm6008 | predicted gene 6008 [Source:MGI Symbol | 0 | 0 | 0 | 0 | 0 | 0 |
| 46537 | ENSMUSG00000108741 | 1.009830165 | 0.037117158 | 2.215730638 | 0.016751656 | 0.986634737 | NA | Gm44553 | predicted gene 44553 [Source:MGI Symbo | 0 | 1.0279 | 2.0125 | 1.928 | 1.0906 | 0 |
| 46538 | ENSMUSG00000108742 | 0 | NA | NA | NA | NA | NA | Gm44923 | predicted gene 44923 [Source:MGI Symbo | 0 | 0 | 0 | 0 | 0 | 0 |
| 46539 | ENSMUSG00000108743 | 0 | NA | NA | NA | NA | NA | Gm44881 | predicted gene 44881 [Source:MGI Symbo | 0 | 0 | 0 | 0 | 0 | 0 |
| 46540 | ENSMUSG00000108744 | 0 | NA | NA | NA | NA | NA | Gm9367 | predicted gene 9367 [Source:MGI Symbol | 0 | 0 | 0 | 0 | 0 | 0 |
| 46541 | ENSMUSG00000108745 | 0 | NA | NA | NA | NA | NA | Gm17908 | predicted gene, 17908 [Source:MGI Symb | 0 | 0 | 0 | 0 | 0 | 0 |
| 46542 | ENSMUSG00000108746 | 0 | NA | NA | NA | NA | NA | Gm45198 | predicted gene 45198 [Source:MGI Symbo | 0 | 0 | 0 | 0 | 0 | 0 |
| 46543 | ENSMUSG00000108748 | 0 | NA | NA | NA | NA | NA | Gm44579 | predicted gene 44579 [Source:MGI Symbo | 0 | 0 | 0 | 0 | 0 | 0 |
| 46544 | ENSMUSG00000108749 | 0 | NA | NA | NA | NA | NA | Gm44767 | predicted gene 44767 [Source:MGI Symbo | 0 | 0 | 0 | 0 | 0 | 0 |
| 46545 | ENSMUSG00000108750 | 0.974396903 | -0.734194653 | 2.427679541 | -0.302426511 | 0.762326948 | NA | Gm44750 | predicted gene 44750 [Source:MGI Symbo | 1.6062 | 0 | 2.0125 | 0 | 0 | 2.2277 |
| 46546 | ENSMUSG00000108751 | 0 | NA | NA | NA | NA | NA | Gm45217 | predicted gene 45217 [Source:MGI Symbo | 0 | 0 | 0 | 0 | 0 | 0 |
| 46547 | ENSMUSG00000108752 | 0 | NA | NA | NA | NA | NA | Gm45191 | predicted gene 45191 [Source:MGI Symbo | 0 | 0 | 0 | 0 | 0 | 0 |
| 46548 | ENSMUSG00000108753 | 1.86080014 | 2.348210204 | 1.713620454 | 1.370321064 | 0.170586701 | NA | Gm45094 | predicted gene 45094 [Source:MGI Symbo | 0.8031 | 0 | 1.0062 | 3.8559 | 3.2719 | 2.2277 |
| 46549 | ENSMUSG00000108754 | 0 | NA | NA | NA | NA | NA | Gm44748 | predicted gene 44748 [Source:MGI Symbo | 0 | 0 | 0 | 0 | 0 | 0 |
| 46550 | ENSMUSG00000108756 | 0 | NA | NA | NA | NA | NA | Gm44894 | predicted gene 44894 [Source:MGI Symbo | 0 | 0 | 0 | 0 | 0 | 0 |
| 46551 | ENSMUSG00000108757 | 0 | NA | NA | NA | NA | NA | Gm45202 | predicted gene 45202 [Source:MGI Symbo | 0 | 0 | 0 | 0 | 0 | 0 |
| 46552 | ENSMUSG00000108758 | 0 | NA | NA | NA | NA | NA | Gm44921 | predicted gene 44921 [Source:MGI Symbo | 0 | 0 | 0 | 0 | 0 | 0 |
| 46553 | ENSMUSG00000108759 | 0 | NA | NA | NA | NA | NA | Klk1b18-ps | kallikrein 1-related peptidase b18, pseudo | 0 | 0 | 0 | 0 | 0 | 0 |
| 46554 | ENSMUSG00000108760 | 1.413649383 | 0.657908737 | 2.112220759 | 0.311477261 | 0.755437827 | NA | Gm31897 | predicted gene, 31897 [Source:MGI Symb | 3.2124 | 0 | 0 | 1.928 | 0 | 3.3415 |
| 46555 | ENSMUSG00000108761 | 0 | NA | NA | NA | NA | NA | Atp6v0c-ps1 | ATPase, H+ transporting, lysosomal V0 sul | 0 | 0 | 0 | 0 | 0 | 0 |
| 46556 | ENSMUSG00000108763 | 42.3747281 | 0.298613037 | 0.350192284 | 0.852711641 | 0.393819234 | 0.822674956 | Gm36028 | predicted gene, 36028 [Source:MGI Symb | 46.58 | 37.005 | 30.187 | 51.091 | 39.262 | 50.123 |
| 46557 | ENSMUSG00000108764 | 0 | NA | NA | NA | NA | NA | Gm2926 | predicted gene 2926 [Source:MGI Symbol | 0 | 0 | 0 | 0 | 0 | 0 |
| 46558 | ENSMUSG00000108765 | 0 | NA | NA | NA | NA | NA | Gm44580 | predicted gene 44580 [Source:MGI Symbo | 0 | 0 | 0 | 0 | 0 | 0 |
| 46559 | ENSMUSG00000108766 | 2.335693599 | -1.818243382 | 1.576329658 | -1.153466455 | 0.248718979 | NA | Gm44760 | predicted gene 44760 [Source:MGI Symbo | 0.8031 | 6.1675 | 4.025 | 1.928 | 1.0906 | 0 |
| 46560 | ENSMUSG00000108767 | 0 | NA | NA | NA | NA | NA | Gm29763 | predicted gene, 29763 [Source:MGI Symb | 0 | 0 | 0 | 0 | 0 | 0 |
| 46561 | ENSMUSG00000108768 | 0 | NA | NA | NA | NA | NA | Gm44971 | predicted gene 44971 [Source:MGI Symbo | 0 | 0 | 0 | 0 | 0 | 0 |
| 46562 | ENSMUSG00000108770 | 0 | NA | NA | NA | NA | NA | Gm44719 | predicted gene 44719 [Source:MGI Symbo | 0 | 0 | 0 | 0 | 0 | 0 |
| 46563 | ENSMUSG00000108771 | 0 | NA | NA | NA | NA | NA | Gm45210 | predicted gene 45210 [Source:MGI Symbo | 0 | 0 | 0 | 0 | 0 | 0 |
| 46564 | ENSMUSG00000108772 | 1.045955322 | 2.382148677 | 2.401164967 | 0.99208039 | 0.321158312 | NA | Gm6063 | predicted gene 6063 [Source:MGI Symbol | 0 | 0 | 1.0062 | 1.928 | 0 | 3.3415 |
| 46565 | ENSMUSG00000108773 | 0 | NA | NA | NA | NA | NA | Gm44506 | predicted gene 44506 [Source:MGI Symbo | 0 | 0 | 0 | 0 | 0 | 0 |
| 46566 | ENSMUSG00000108774 | 0 | NA | NA | NA | NA | NA | Gm45136 | predicted gene 45136 [Source:MGI Symbo | 0 | 0 | 0 | 0 | 0 | 0 |
| 46567 | ENSMUSG00000108775 | 0 | NA | NA | NA | NA | NA | Gm44540 | predicted gene 44540 [Source:MGI Symbo | 0 | 0 | 0 | 0 | 0 | 0 |
| 46568 | ENSMUSG00000108776 | 0 | NA | NA | NA | NA | NA | Gm45169 | predicted gene 45169 [Source:MGI Symbo | 0 | 0 | 0 | 0 | 0 | 0 |
| 46569 | ENSMUSG00000108777 | 0.17131964 | -0.796629176 | 4.080472857 | -0.195229623 | 0.84521319 | NA | Gm45081 | predicted gene 45081 [Source:MGI Symbo | 0 | 1.0279 | 0 | 0 | 0 | 0 |
| 46570 | ENSMUSG00000108778 | 0 | NA | NA | NA | NA | NA | Gm20083 | predicted gene, 20083 [Source:MGI Symb | 0 | 0 | 0 | 0 | 0 | 0 |
| 46571 | ENSMUSG00000108779 | 0 | NA | NA | NA | NA | NA | Gm45691 | predicted gene 45691 [Source:MGI Symbo | 0 | 0 | 0 | 0 | 0 | 0 |
| 46572 | ENSMUSG00000108780 | 0.727078859 | 3.067238961 | 3.29680021 | 0.930368468 | 0.352180339 | NA | 5430434F05 | RIKEN cDNA 5430434F05 gene [Source:M | 0 | 0 | 0 | 0 | 4.3625 | 0 |
| 46573 | ENSMUSG00000108781 | 0 | NA | NA | NA | NA | NA | Gm44707 | predicted gene 44707 [Source:MGI Symbo | 0 | 0 | 0 | 0 | 0 | 0 |
| 46574 | ENSMUSG00000108782 | 0 | NA | NA | NA | NA | NA | Gm32772 | predicted gene, 32772 [Source:MGI Symb | 0 | 0 | 0 | 0 | 0 | 0 |
| 46575 | ENSMUSG00000108783 | 0.643990493 | 0.179747576 | 2.633236711 | 0.068261078 | 0.945577807 | NA | Gm36546 | predicted gene, 36546 [Source:MGI Symb | 0.8031 | 0 | 1.0062 | 0.964 | 1.0906 | 0 |
| 46576 | ENSMUSG00000108784 | 0 | NA | NA | NA | NA | NA | 4930517G19 | RIKEN cDNA 4930517G19 gene [Source:M | 0 | 0 | 0 | 0 | 0 | 0 |
| 46577 | ENSMUSG00000108785 | 0 | NA | NA | NA | NA | NA | Gm4259 | predicted gene 4259 [Source:MGI Symbol | 0 | 0 | 0 | 0 | 0 | 0 |
| 46578 | ENSMUSG00000108786 | 0 | NA | NA | NA | NA | NA | Gm8309 | predicted gene 8309 [Source:MGI Symbol | 0 | 0 | 0 | 0 | 0 | 0 |
| 46579 | ENSMUSG00000108787 | 0 | NA | NA | NA | NA | NA | 2200007N16 | RIKEN cDNA 2200007N16 gene [Source:M | 0 | 0 | 0 | 0 | 0 | 0 |
| 46580 | ENSMUSG00000108788 | 0 | NA | NA | NA | NA | NA | 4921513I08R | RIKEN cDNA 4921513I08 gene [Source:M | 0 | 0 | 0 | 0 | 0 | 0 |
| 46581 | ENSMUSG00000108790 | 0 | NA | NA | NA | NA | NA | Gm44806 | predicted gene 44806 [Source:MGI Symbo | 0 | 0 | 0 | 0 | 0 | 0 |
| 46582 | ENSMUSG00000108791 | 0.133850373 | -0.796629176 | 4.080472857 | -0.195229623 | 0.84521319 | NA | 1700011D18 | RIKEN cDNA 1700011D18 gene [Source:M | 0.8031 | 0 | 0 | 0 | 0 | 0 |
| 46583 | ENSMUSG00000108792 | 0 | NA | NA | NA | NA | NA | Gm5418 | predicted gene 5418 [Source:MGI Symbol | 0 | 0 | 0 | 0 | 0 | 0 |
| 46584 | ENSMUSG00000108793 | 0 | NA | NA | NA | NA | NA | Gm45213 | predicted gene 45213 [Source:MGI Symbo | 0 | 0 | 0 | 0 | 0 | 0 |
| 46585 | ENSMUSG00000108794 | 0 | NA | NA | NA | NA | NA | Gm45195 | predicted gene 45195 [Source:MGI Symbo | 0 | 0 | 0 | 0 | 0 | 0 |
| 46586 | ENSMUSG00000108795 | 0 | NA | NA | NA | NA | NA | Gm45121 | predicted gene 45121 [Source:MGI Symbo | 0 | 0 | 0 | 0 | 0 | 0 |
| 46587 | ENSMUSG00000108796 | 0 | NA | NA | NA | NA | NA | Gm45126 | predicted gene 45126 [Source:MGI Symbo | 0 | 0 | 0 | 0 | 0 | 0 |
| 46588 | ENSMUSG00000108797 | 0 | NA | NA | NA | NA | NA | Klk1b23-ps | kallikrein 1-related peptidase b23, pseudo | 0 | 0 | 0 | 0 | 0 | 0 |

| | | | | | | | | | | | | | | | |
|---|---|---|---|---|---|---|---|---|---|---|---|---|---|---|---|
| 46589 | ENSMUSG00000108798 | 0 | NA | NA | NA | NA | NA | Gm9521 | predicted gene 9521 [Source:MGI Symbol | 0 | 0 | 0 | 0 | 0 | 0 |
| 46590 | ENSMUSG00000108799 | 4.239663353 | 0.23508232 | 1.039709365 | 0.226103878 | 0.821120628 | NA | Glud-ps | glutamate dehydrogenase, pseudogene [S | 1.6062 | 5.1396 | 5.0312 | 4.8199 | 3.2719 | 5.5692 |
| 46591 | ENSMUSG00000108800 | 0 | NA | NA | NA | NA | NA | 1700011C11 | RIKEN cDNA 1700011C11 gene [Source:M | 0 | 0 | 0 | 0 | 0 | 0 |
| 46592 | ENSMUSG00000108801 | 0 | NA | NA | NA | NA | NA | Gm39090 | predicted gene, 39090 [Source:MGI Symb | 0 | 0 | 0 | 0 | 0 | 0 |
| 46593 | ENSMUSG00000108802 | 0 | NA | NA | NA | NA | NA | Gm44769 | predicted gene 44769 [Source:MGI Symbo | 0 | 0 | 0 | 0 | 0 | 0 |
| 46594 | ENSMUSG00000108803 | 0 | NA | NA | NA | NA | NA | 4930533N22 | RIKEN cDNA 4930533N22 gene [Source:M | 0 | 0 | 0 | 0 | 0 | 0 |
| 46595 | ENSMUSG00000108804 | 0 | NA | NA | NA | NA | NA | Gm30437 | predicted gene, 30437 [Source:MGI Symb | 0 | 0 | 0 | 0 | 0 | 0 |
| 46596 | ENSMUSG00000108805 | 0 | NA | NA | NA | NA | NA | Olfr278-ps1 | olfactory receptor 278, pseudogene 1 [Sou | 0 | 0 | 0 | 0 | 0 | 0 |
| 46597 | ENSMUSG00000108806 | 2.308647777 | 0.107302592 | 1.488216637 | 0.07210146 | 0.942521164 | NA | Gm44729 | predicted gene 44729 [Source:MGI Symbo | 1.6062 | 4.1117 | 1.0062 | 3.8559 | 3.2719 | 0 |
| 46598 | ENSMUSG00000108807 | 0 | NA | NA | NA | NA | NA | 4833404L02 | RIKEN cDNA 4833404L02 gene [Source:M | 0 | 0 | 0 | 0 | 0 | 0 |
| 46599 | ENSMUSG00000108808 | 2.805028775 | -1.637192577 | 1.579109467 | -1.036782193 | 0.299837376 | NA | Gm45204 | predicted gene 45204 [Source:MGI Symbo | 9.6372 | 3.0838 | 0 | 1.928 | 2.1812 | 0 |
| 46600 | ENSMUSG00000108809 | 0 | NA | NA | NA | NA | NA | Gm45228 | predicted gene 45228 [Source:MGI Symbo | 0 | 0 | 0 | 0 | 0 | 0 |
| 46601 | ENSMUSG00000108810 | 0 | NA | NA | NA | NA | NA | Gm44629 | predicted gene 44629 [Source:MGI Symbo | 0 | 0 | 0 | 0 | 0 | 0 |
| 46602 | ENSMUSG00000108811 | 0 | NA | NA | NA | NA | NA | 4933432K03 | RIKEN cDNA 4933432K03 gene [Source:M | 0 | 0 | 0 | 0 | 0 | 0 |
| 46603 | ENSMUSG00000108812 | 0.503302254 | -0.837877584 | 3.426698643 | -0.244514523 | 0.806832356 | NA | C130083A15 | RIKEN cDNA C130083A15 gene [Source:M | 0 | 2.0558 | 0 | 0.964 | 0 | 0 |
| 46604 | ENSMUSG00000108813 | 1.494934965 | -0.803533485 | 1.757864854 | -0.457107657 | 0.647593667 | NA | Rpl19-ps9 | ribosomal protein L19, pseudogene 9 [Sou | 1.6062 | 2.0558 | 2.0125 | 0 | 2.1812 | 1.1138 |
| 46605 | ENSMUSG00000108814 | 0 | NA | NA | NA | NA | NA | Gm30717 | predicted gene, 30717 [Source:MGI Symb | 0 | 0 | 0 | 0 | 0 | 0 |
| 46606 | ENSMUSG00000108815 | 0 | NA | NA | NA | NA | NA | Gm49388 | predicted gene, 49388 [Source:MGI Symb | 0 | 0 | 0 | 0 | 0 | 0 |
| 46607 | ENSMUSG00000108816 | 0 | NA | NA | NA | NA | NA | Gm6230 | predicted gene 6230 [Source:MGI Symbol | 0 | 0 | 0 | 0 | 0 | 0 |
| 46608 | ENSMUSG00000108818 | 0.185640647 | 1.126924175 | 4.080472857 | 0.276174898 | 0.782413736 | NA | Gm44869 | predicted gene 44869 [Source:MGI Symbo | 0 | 0 | 0 | 0 | 0 | 1.1138 |
| 46609 | ENSMUSG00000108819 | 0 | NA | NA | NA | NA | NA | Gm45177 | predicted gene 45177 [Source:MGI Symbo | 0 | 0 | 0 | 0 | 0 | 0 |
| 46610 | ENSMUSG00000108820 | 34.00966122 | -2.117112744 | 0.461737254 | -4.585102726 | 4.54E-06 | 0.000282898 | Gm44620 | predicted gene 44620 [Source:MGI Symbo | 68.264 | 58.591 | 39.244 | 19.28 | 11.997 | 6.6831 |
| 46611 | ENSMUSG00000108821 | 0.679159517 | 1.687875387 | 3.079592583 | 0.548083989 | 0.583634234 | NA | 4933423L19 | RIKEN cDNA 4933423L19 gene [Source:M | 0.8031 | 0 | 0 | 0 | 3.2719 | 0 |
| 46612 | ENSMUSG00000108822 | 54.51550605 | -0.075700459 | 0.407168274 | -0.185919345 | 0.852508011 | 0.973781977 | Gm44787 | predicted gene 44787 [Source:MGI Symbo | 85.932 | 54.48 | 27.169 | 71.334 | 43.625 | 44.554 |
| 46613 | ENSMUSG00000108823 | 1.994186325 | -1.421337493 | 1.727478927 | -0.822781379 | 0.410632328 | NA | Gm18959 | predicted gene, 18959 [Source:MGI Symb | 1.6062 | 2.0558 | 5.0312 | 0 | 3.2719 | 0 |
| 46614 | ENSMUSG00000108825 | 114.5162163 | -0.127734126 | 0.310363318 | -0.411563217 | 0.680659597 | 0.934384556 | Gm45838 | predicted gene 45838 [Source:MGI Symbo | 144.56 | 97.652 | 116.72 | 160.02 | 106.88 | 61.261 |
| 46615 | ENSMUSG00000108826 | 0 | NA | NA | NA | NA | NA | 4931412I15 | RIKEN cDNA 4931412I15 gene [Source:M | 0 | 0 | 0 | 0 | 0 | 0 |
| 46616 | ENSMUSG00000108827 | 0 | NA | NA | NA | NA | NA | Olfr1310 | olfactory receptor 1310 [Source:MGI Sym | 0 | 0 | 0 | 0 | 0 | 0 |
| 46617 | ENSMUSG00000108828 | 0 | NA | NA | NA | NA | NA | Gm34549 | predicted gene, 34549 [Source:MGI Symb | 0 | 0 | 0 | 0 | 0 | 0 |
| 46618 | ENSMUSG00000108829 | 0 | NA | NA | NA | NA | NA | Vmn2r-ps89 | vomeronasal 2, receptor, pseudogene 89 | 0 | 0 | 0 | 0 | 0 | 0 |
| 46619 | ENSMUSG00000108830 | 0 | NA | NA | NA | NA | NA | Gm44661 | predicted gene 44661 [Source:MGI Symbo | 0 | 0 | 0 | 0 | 0 | 0 |
| 46620 | ENSMUSG00000108831 | 0 | NA | NA | NA | NA | NA | Gm7957 | predicted gene 7957 [Source:MGI Symbol | 0 | 0 | 0 | 0 | 0 | 0 |
| 46621 | ENSMUSG00000108832 | 0 | NA | NA | NA | NA | NA | Gm44832 | predicted gene 44832 [Source:MGI Symbo | 0 | 0 | 0 | 0 | 0 | 0 |
| 46622 | ENSMUSG00000108833 | 0 | NA | NA | NA | NA | NA | Gm17909 | predicted gene, 17909 [Source:MGI Symb | 0 | 0 | 0 | 0 | 0 | 0 |
| 46623 | ENSMUSG00000108834 | 0 | NA | NA | NA | NA | NA | Gm18458 | predicted gene, 18458 [Source:MGI Symb | 0 | 0 | 0 | 0 | 0 | 0 |
| 46624 | ENSMUSG00000108835 | 0 | NA | NA | NA | NA | NA | Gm44566 | predicted gene 44566 [Source:MGI Symbo | 0 | 0 | 0 | 0 | 0 | 0 |
| 46625 | ENSMUSG00000108836 | 0 | NA | NA | NA | NA | NA | Gm44813 | predicted gene 44813 [Source:MGI Symbo | 0 | 0 | 0 | 0 | 0 | 0 |
| 46626 | ENSMUSG00000108837 | 0 | NA | NA | NA | NA | NA | Gm44865 | predicted gene 44865 [Source:MGI Symbo | 0 | 0 | 0 | 0 | 0 | 0 |
| 46627 | ENSMUSG00000108838 | 0 | NA | NA | NA | NA | NA | Gm45189 | predicted gene 45189 [Source:MGI Symbo | 0 | 0 | 0 | 0 | 0 | 0 |
| 46628 | ENSMUSG00000108839 | 0 | NA | NA | NA | NA | NA | Gm44547 | predicted gene 44547 [Source:MGI Symbo | 0 | 0 | 0 | 0 | 0 | 0 |
| 46629 | ENSMUSG00000108840 | 0 | NA | NA | NA | NA | NA | Gm5320 | predicted gene 5320 [Source:MGI Symbol | 0 | 0 | 0 | 0 | 0 | 0 |
| 46630 | ENSMUSG00000108841 | 0 | NA | NA | NA | NA | NA | Frmpd2 | FERM and PDZ domain containing 2 [Sour | 0 | 0 | 0 | 0 | 0 | 0 |
| 46631 | ENSMUSG00000108842 | 9.722309551 | -0.4436378 | 0.742570391 | -0.597435349 | 0.550216759 | 0.895703112 | Gm45001 | predicted gene 45001 [Source:MGI Symbo | 10.44 | 7.1954 | 16.1 | 12.532 | 8.7249 | 3.3415 |
| 46632 | ENSMUSG00000108843 | 0 | NA | NA | NA | NA | NA | Gm4501 | predicted gene 4501 [Source:MGI Symbol | 0 | 0 | 0 | 0 | 0 | 0 |
| 46633 | ENSMUSG00000108844 | 0 | NA | NA | NA | NA | NA | Gm6702 | predicted gene 6702 [Source:MGI Symbol | 0 | 0 | 0 | 0 | 0 | 0 |
| 46634 | ENSMUSG00000108845 | 0 | NA | NA | NA | NA | NA | Gm45090 | predicted gene 45090 [Source:MGI Symbo | 0 | 0 | 0 | 0 | 0 | 0 |
| 46635 | ENSMUSG00000108846 | 0 | NA | NA | NA | NA | NA | Gm10623 | predicted gene 10623 [Source:MGI Symbo | 0 | 0 | 0 | 0 | 0 | 0 |
| 46636 | ENSMUSG00000108847 | 0 | NA | NA | NA | NA | NA | B830042I05 | RIKEN cDNA B830042I05 gene [Source:M | 0 | 0 | 0 | 0 | 0 | 0 |
| 46637 | ENSMUSG00000108848 | 0 | NA | NA | NA | NA | NA | 4933430N04 | RIKEN cDNA 4933430N04 gene [Source:M | 0 | 0 | 0 | 0 | 0 | 0 |
| 46638 | ENSMUSG00000108849 | 0 | NA | NA | NA | NA | NA | Gm35147 | predicted gene, 35147 [Source:MGI Symb | 0 | 0 | 0 | 0 | 0 | 0 |
| 46639 | ENSMUSG00000108850 | 0 | NA | NA | NA | NA | NA | Gm44914 | predicted gene 44914 [Source:MGI Symbo | 0 | 0 | 0 | 0 | 0 | 0 |
| 46640 | ENSMUSG00000108851 | 0 | NA | NA | NA | NA | NA | Gm44930 | predicted gene 44930 [Source:MGI Symbo | 0 | 0 | 0 | 0 | 0 | 0 |
| 46641 | ENSMUSG00000108852 | 1.042698947 | 2.426023578 | 2.689838701 | 0.901921583 | 0.367098525 | NA | Gm44911 | predicted gene 44911 [Source:MGI Symbo | 0.8031 | 0 | 0 | 0 | 5.4531 | 0 |
| 46642 | ENSMUSG00000108853 | 0 | NA | NA | NA | NA | NA | Gm18464 | predicted gene, 18464 [Source:MGI Symb | 0 | 0 | 0 | 0 | 0 | 0 |
| 46643 | ENSMUSG00000108854 | 0 | NA | NA | NA | NA | NA | D830036C21 | RIKEN cDNA D830036C21 gene [Source:M | 0 | 0 | 0 | 0 | 0 | 0 |
| 46644 | ENSMUSG00000108855 | 0 | NA | NA | NA | NA | NA | Gm44917 | predicted gene 44917 [Source:MGI Symbo | 0 | 0 | 0 | 0 | 0 | 0 |

| | | | | | | | | | | | | | | | |
|---|---|---|---|---|---|---|---|---|---|---|---|---|---|---|---|
| 46645 | ENSMUSG00000108856 | 0 | NA | NA | NA | NA | NA | Gm35309 | predicted gene, 35309 [Source:MGI Symb | 0 | 0 | 0 | 0 | 0 | 0 |
| 46646 | ENSMUSG00000108857 | 0 | NA | NA | NA | NA | NA | Gm44578 | predicted gene 44578 [Source:MGI Symbo | 0 | 0 | 0 | 0 | 0 | 0 |
| 46647 | ENSMUSG00000108858 | 0 | NA | NA | NA | NA | NA | Gm4587 | predicted gene 4587 [Source:MGI Symbol | 0 | 0 | 0 | 0 | 0 | 0 |
| 46648 | ENSMUSG00000108859 | 0 | NA | NA | NA | NA | NA | Gm44776 | predicted gene 44776 [Source:MGI Symbo | 0 | 0 | 0 | 0 | 0 | 0 |
| 46649 | ENSMUSG00000108860 | 0 | NA | NA | NA | NA | NA | Gm6070 | predicted gene 6070 [Source:MGI Symbol | 0 | 0 | 0 | 0 | 0 | 0 |
| 46650 | ENSMUSG00000108861 | 0 | NA | NA | NA | NA | NA | 5930435M05 | RIKEN cDNA 5930435M05 gene [Source:M | 0 | 0 | 0 | 0 | 0 | 0 |
| 46651 | ENSMUSG00000108862 | 0 | NA | NA | NA | NA | NA | Gm17991 | predicted gene, 17991 [Source:MGI Symb | 0 | 0 | 0 | 0 | 0 | 0 |
| 46652 | ENSMUSG00000108863 | 0 | NA | NA | NA | NA | NA | Gm5002 | predicted gene 5002 [Source:MGI Symbol | 0 | 0 | 0 | 0 | 0 | 0 |
| 46653 | ENSMUSG00000108864 | 0 | NA | NA | NA | NA | NA | Gm44933 | predicted gene 44933 [Source:MGI Symbo | 0 | 0 | 0 | 0 | 0 | 0 |
| 46654 | ENSMUSG00000108865 | 0 | NA | NA | NA | NA | NA | Gm44617 | predicted gene 44617 [Source:MGI Symbo | 0 | 0 | 0 | 0 | 0 | 0 |
| 46655 | ENSMUSG00000108866 | 0 | NA | NA | NA | NA | NA | Gm35082 | predicted gene, 35082 [Source:MGI Symb | 0 | 0 | 0 | 0 | 0 | 0 |
| 46656 | ENSMUSG00000108867 | 0 | NA | NA | NA | NA | NA | Gm44643 | predicted gene 44643 [Source:MGI Symbo | 0 | 0 | 0 | 0 | 0 | 0 |
| 46657 | ENSMUSG00000108868 | 0 | NA | NA | NA | NA | NA | Gm32850 | predicted gene, 32850 [Source:MGI Symb | 0 | 0 | 0 | 0 | 0 | 0 |
| 46658 | ENSMUSG00000108869 | 0.342639279 | -1.805976596 | 4.028547714 | -0.448294702 | 0.653940524 | NA | Gm44532 | predicted gene 44532 [Source:MGI Symbo | 0 | 2.0558 | 0 | 0 | 0 | 0 |
| 46659 | ENSMUSG00000108870 | 0 | NA | NA | NA | NA | NA | Gm44893 | predicted gene 44893 [Source:MGI Symbo | 0 | 0 | 0 | 0 | 0 | 0 |
| 46660 | ENSMUSG00000108871 | 0.556921942 | 2.679043159 | 3.638243063 | 0.736356289 | 0.461513899 | NA | Gm45102 | predicted gene 45102 [Source:MGI Symbo | 0 | 0 | 0 | 0 | 0 | 3.3415 |
| 46661 | ENSMUSG00000108873 | 0 | NA | NA | NA | NA | NA | Gm44609 | predicted gene 44609 [Source:MGI Symbo | 0 | 0 | 0 | 0 | 0 | 0 |
| 46662 | ENSMUSG00000108874 | 0 | NA | NA | NA | NA | NA | Gm9711 | predicted gene 9711 [Source:MGI Symbol | 0 | 0 | 0 | 0 | 0 | 0 |
| 46663 | ENSMUSG00000108875 | 0 | NA | NA | NA | NA | NA | Gm45064 | predicted gene 45064 [Source:MGI Symbo | 0 | 0 | 0 | 0 | 0 | 0 |
| 46664 | ENSMUSG00000108876 | 0 | NA | NA | NA | NA | NA | Gm44692 | predicted gene 44692 [Source:MGI Symbo | 0 | 0 | 0 | 0 | 0 | 0 |
| 46665 | ENSMUSG00000108877 | 0 | NA | NA | NA | NA | NA | Gm45157 | predicted gene 45157 [Source:MGI Symbo | 0 | 0 | 0 | 0 | 0 | 0 |
| 46666 | ENSMUSG00000108878 | 1.018883997 | 1.207572996 | 2.096893663 | 0.575886616 | 0.564691823 | NA | Gm49493 | predicted gene, 49493 [Source:MGI Symb | 0.8031 | 1.0279 | 0 | 0.964 | 1.0906 | 2.2277 |
| 46667 | ENSMUSG00000108879 | 0 | NA | NA | NA | NA | NA | Gm18056 | predicted gene, 18056 [Source:MGI Symb | 0 | 0 | 0 | 0 | 0 | 0 |
| 46668 | ENSMUSG00000108880 | 0 | NA | NA | NA | NA | NA | Gm44560 | predicted gene 44560 [Source:MGI Symbo | 0 | 0 | 0 | 0 | 0 | 0 |
| 46669 | ENSMUSG00000108881 | 0 | NA | NA | NA | NA | NA | Gm44849 | predicted gene 44849 [Source:MGI Symbo | 0 | 0 | 0 | 0 | 0 | 0 |
| 46670 | ENSMUSG00000108882 | 0 | NA | NA | NA | NA | NA | Gm44563 | predicted gene 44563 [Source:MGI Symbo | 0 | 0 | 0 | 0 | 0 | 0 |
| 46671 | ENSMUSG00000108883 | 7.240860498 | 0.029083408 | 0.904879052 | 0.032140657 | 0.97435988 | 0.996336532 | Gm44623 | predicted gene 44623 [Source:MGI Symbo | 7.2279 | 10.279 | 4.025 | 7.7118 | 13.087 | 1.1138 |
| 46672 | ENSMUSG00000108884 | 13.52491242 | 2.564603617 | 0.728452869 | 3.520617087 | 0.000430544 | 0.011200335 | Gm45792 | predicted gene 45792 [Source:MGI Symbo | 6.4248 | 3.0838 | 2.0125 | 36.631 | 19.631 | 13.366 |
| 46673 | ENSMUSG00000108885 | 0 | NA | NA | NA | NA | NA | 4930402F11 | RIKEN cDNA 4930402F11 gene [Source:M | 0 | 0 | 0 | 0 | 0 | 0 |
| 46674 | ENSMUSG00000108886 | 0.644197084 | -2.725354879 | 2.762261801 | -0.986638876 | 0.323819705 | NA | Gm44830 | predicted gene 44830 [Source:MGI Symbo | 0.8031 | 2.0558 | 1.0062 | 0 | 0 | 0 |
| 46675 | ENSMUSG00000108887 | 0 | NA | NA | NA | NA | NA | Olfr1479-ps1 | olfactory receptor 1479, pseudogene 1 [S | 0 | 0 | 0 | 0 | 0 | 0 |
| 46676 | ENSMUSG00000108888 | 2.852174127 | 1.079403548 | 1.275404856 | 0.846322282 | 0.397372981 | NA | Gm45231 | predicted gene 45231 [Source:MGI Symbo | 2.4093 | 1.0279 | 2.0125 | 2.8919 | 6.5437 | 2.2277 |
| 46677 | ENSMUSG00000108889 | 0 | NA | NA | NA | NA | NA | Olfr1555-ps1 | olfactory receptor 1555, pseudogene 1 [S | 0 | 0 | 0 | 0 | 0 | 0 |
| 46678 | ENSMUSG00000108890 | 0 | NA | NA | NA | NA | NA | Gm44847 | predicted gene 44847 [Source:MGI Symbo | 0 | 0 | 0 | 0 | 0 | 0 |
| 46679 | ENSMUSG00000108891 | 0 | NA | NA | NA | NA | NA | Olfr1291-ps1 | olfactory receptor 1291, pseudogene 1 [S | 0 | 0 | 0 | 0 | 0 | 0 |
| 46680 | ENSMUSG00000108892 | 0 | NA | NA | NA | NA | NA | Gm9449 | predicted gene 9449 [Source:MGI Symbol | 0 | 0 | 0 | 0 | 0 | 0 |
| 46681 | ENSMUSG00000108893 | 0 | NA | NA | NA | NA | NA | Gm44927 | predicted gene 44927 [Source:MGI Symbo | 0 | 0 | 0 | 0 | 0 | 0 |
| 46682 | ENSMUSG00000108894 | 0 | NA | NA | NA | NA | NA | Gm45841 | predicted gene 45841 [Source:MGI Symbo | 0 | 0 | 0 | 0 | 0 | 0 |
| 46683 | ENSMUSG00000108895 | 0.587191767 | -1.12394062 | 3.226482889 | -0.348348545 | 0.72757844 | NA | Gm45070 | predicted gene 45070 [Source:MGI Symbo | 2.4093 | 0 | 0 | 0 | 0 | 1.1138 |
| 46684 | ENSMUSG00000108896 | 0 | NA | NA | NA | NA | NA | Gm20136 | predicted gene, 20136 [Source:MGI Symb | 0 | 0 | 0 | 0 | 0 | 0 |
| 46685 | ENSMUSG00000108897 | 0.615839296 | 1.537131441 | 3.17677869 | 0.483864817 | 0.628481811 | NA | Gm44861 | predicted gene 44861 [Source:MGI Symbo | 0.8031 | 0 | 0 | 2.8919 | 0 | 0 |
| 46686 | ENSMUSG00000108899 | 0 | NA | NA | NA | NA | NA | Gm44811 | predicted gene 44811 [Source:MGI Symbo | 0 | 0 | 0 | 0 | 0 | 0 |
| 46687 | ENSMUSG00000108900 | 0 | NA | NA | NA | NA | NA | Ccdc194 | coiled-coil domain containing 194 [Source | 0 | 0 | 0 | 0 | 0 | 0 |
| 46688 | ENSMUSG00000108902 | 0 | NA | NA | NA | NA | NA | Gm9377 | predicted gene 9377 [Source:MGI Symbol | 0 | 0 | 0 | 0 | 0 | 0 |
| 46689 | ENSMUSG00000108903 | 0 | NA | NA | NA | NA | NA | Gm45029 | predicted gene 45029 [Source:MGI Symbo | 0 | 0 | 0 | 0 | 0 | 0 |
| 46690 | ENSMUSG00000108904 | 0 | NA | NA | NA | NA | NA | Gm45147 | predicted gene 45147 [Source:MGI Symbo | 0 | 0 | 0 | 0 | 0 | 0 |
| 46691 | ENSMUSG00000108905 | 0 | NA | NA | NA | NA | NA | Gm44618 | predicted gene 44618 [Source:MGI Symbo | 0 | 0 | 0 | 0 | 0 | 0 |
| 46692 | ENSMUSG00000108906 | 0 | NA | NA | NA | NA | NA | Gm8314 | predicted gene 8314 [Source:MGI Symbol | 0 | 0 | 0 | 0 | 0 | 0 |
| 46693 | ENSMUSG00000108907 | 0 | NA | NA | NA | NA | NA | Gm4541 | predicted gene 4541 [Source:MGI Symbol | 0 | 0 | 0 | 0 | 0 | 0 |
| 46694 | ENSMUSG00000108908 | 0 | NA | NA | NA | NA | NA | Olfr1293-ps | olfactory receptor 1293, pseudogene [Sou | 0 | 0 | 0 | 0 | 0 | 0 |
| 46695 | ENSMUSG00000108909 | 0 | NA | NA | NA | NA | NA | Gm9373 | predicted gene 9373 [Source:MGI Symbol | 0 | 0 | 0 | 0 | 0 | 0 |
| 46696 | ENSMUSG00000108910 | 0.17131964 | -0.796629176 | 4.080472857 | -0.195229623 | 0.84521319 | NA | Gm44625 | predicted gene 44625 [Source:MGI Symbo | 0 | 1.0279 | 0 | 0 | 0 | 0 |
| 46697 | ENSMUSG00000108911 | 0 | NA | NA | NA | NA | NA | Gm45113 | predicted gene 45113 [Source:MGI Symbo | 0 | 0 | 0 | 0 | 0 | 0 |
| 46698 | ENSMUSG00000108912 | 0.160662974 | 1.126924175 | 4.080472857 | 0.276174898 | 0.782413736 | NA | E230020D15 | RIKEN cDNA E230020D15 gene [Source:M | 0 | 0 | 0 | 0.964 | 0 | 0 |
| 46699 | ENSMUSG00000108913 | 0 | NA | NA | NA | NA | NA | Gm2448 | predicted gene 2448 [Source:MGI Symbol | 0 | 0 | 0 | 0 | 0 | 0 |
| 46700 | ENSMUSG00000108914 | 0 | NA | NA | NA | NA | NA | Gm44908 | predicted gene 44908 [Source:MGI Symbo | 0 | 0 | 0 | 0 | 0 | 0 |

| | | | | | | | | | | | | | | | |
|---|---|---|---|---|---|---|---|---|---|---|---|---|---|---|---|
| 46701 | ENSMUSG00000108915 | 5.228540108 | 0.952271547 | 1.047657832 | 0.908952826 | 0.363375027 | 0.802199995 | Gm44966 | predicted gene 44966 [Source:MGI Symbo | 1.6062 | 8.2233 | 1.0062 | 10.604 | 4.3625 | 5.5692 |
| 46702 | ENSMUSG00000108916 | 0 | NA | NA | NA | NA | NA | Gm18944 | predicted gene, 18944 [Source:MGI Symb | 0 | 0 | 0 | 0 | 0 | 0 |
| 46703 | ENSMUSG00000108917 | 0 | NA | NA | NA | NA | NA | Gm38451 | predicted gene, 38451 [Source:MGI Symb | 0 | 0 | 0 | 0 | 0 | 0 |
| 46704 | ENSMUSG00000108918 | 0 | NA | NA | NA | NA | NA | Gm44802 | predicted gene 44802 [Source:MGI Symbo | 0 | 0 | 0 | 0 | 0 | 0 |
| 46705 | ENSMUSG00000108919 | 0 | NA | NA | NA | NA | NA | Olfr1295 | olfactory receptor 1295 [Source:MGI Sym | 0 | 0 | 0 | 0 | 0 | 0 |
| 46706 | ENSMUSG00000108920 | 0 | NA | NA | NA | NA | NA | Gm44678 | predicted gene 44678 [Source:MGI Symbo | 0 | 0 | 0 | 0 | 0 | 0 |
| 46707 | ENSMUSG00000108921 | 0 | NA | NA | NA | NA | NA | Gm4083 | predicted gene 4083 [Source:MGI Symbol | 0 | 0 | 0 | 0 | 0 | 0 |
| 46708 | ENSMUSG00000108922 | 0 | NA | NA | NA | NA | NA | Gm44825 | predicted gene 44825 [Source:MGI Symbo | 0 | 0 | 0 | 0 | 0 | 0 |
| 46709 | ENSMUSG00000108923 | 15.03502197 | 0.483300129 | 0.60992947 | 0.792386912 | 0.428135108 | 0.842753542 | 4933431C10 | RIKEN cDNA 4933431C10 gene [Source:M | 15.259 | 10.279 | 12.075 | 26.027 | 7.6343 | 18.935 |
| 46710 | ENSMUSG00000108924 | 0 | NA | NA | NA | NA | NA | 4933436H12 | RIKEN cDNA 4933436H12 gene [Source:M | 0 | 0 | 0 | 0 | 0 | 0 |
| 46711 | ENSMUSG00000108925 | 0 | NA | NA | NA | NA | NA | Gm19880 | predicted gene, 19880 [Source:MGI Symb | 0 | 0 | 0 | 0 | 0 | 0 |
| 46712 | ENSMUSG00000108926 | 0.133850373 | -0.796629176 | 4.080472857 | -0.195229623 | 0.84521319 | NA | Gm10988 | predicted gene 10988 [Source:MGI Symbo | 0.8031 | 0 | 0 | 0 | 0 | 0 |
| 46713 | ENSMUSG00000108928 | 0 | NA | NA | NA | NA | NA | Vmn2r-ps61 | vomeronasal 2, receptor, pseudogene 61 | 0 | 0 | 0 | 0 | 0 | 0 |
| 46714 | ENSMUSG00000108929 | 10.7750906 | 0.215664239 | 0.681994279 | 0.316225877 | 0.751831068 | 0.952398368 | Cc2d2b | coiled-coil and C2 domain containing 2B [ | 9.6372 | 7.1954 | 13.081 | 10.604 | 17.45 | 6.6831 |
| 46715 | ENSMUSG00000108930 | 0 | NA | NA | NA | NA | NA | Gm10648 | predicted gene 10648 [Source:MGI Symbo | 0 | 0 | 0 | 0 | 0 | 0 |
| 46716 | ENSMUSG00000108931 | 0 | NA | NA | NA | NA | NA | Olfr1284 | olfactory receptor 1284 [Source:MGI Sym | 0 | 0 | 0 | 0 | 0 | 0 |
| 46717 | ENSMUSG00000108932 | 0 | NA | NA | NA | NA | NA | Gm31463 | predicted gene, 31463 [Source:MGI Symb | 0 | 0 | 0 | 0 | 0 | 0 |
| 46718 | ENSMUSG00000108933 | 0 | NA | NA | NA | NA | NA | Gm44582 | predicted gene 44582 [Source:MGI Symbo | 0 | 0 | 0 | 0 | 0 | 0 |
| 46719 | ENSMUSG00000108934 | 0 | NA | NA | NA | NA | NA | Gm44732 | predicted gene 44732 [Source:MGI Symbo | 0 | 0 | 0 | 0 | 0 | 0 |
| 46720 | ENSMUSG00000108935 | 1.163707407 | 1.351240998 | 2.323660064 | 0.58151406 | 0.560894045 | NA | 4930588G17 | RIKEN cDNA 4930588G17 gene [Source:M | 0 | 0 | 2.0125 | 3.8559 | 0 | 1.1138 |
| 46721 | ENSMUSG00000108936 | 0 | NA | NA | NA | NA | NA | Gm8745 | predicted gene 8745 [Source:MGI Symbol | 0 | 0 | 0 | 0 | 0 | 0 |
| 46722 | ENSMUSG00000108937 | 0 | NA | NA | NA | NA | NA | Gm18908 | predicted gene, 18908 [Source:MGI Symb | 0 | 0 | 0 | 0 | 0 | 0 |
| 46723 | ENSMUSG00000108938 | 0 | NA | NA | NA | NA | NA | Gm44583 | predicted gene 44583 [Source:MGI Symbo | 0 | 0 | 0 | 0 | 0 | 0 |
| 46724 | ENSMUSG00000108939 | 0 | NA | NA | NA | NA | NA | Olfr500-ps1 | olfactory receptor 500, pseudogene 1 [So | 0 | 0 | 0 | 0 | 0 | 0 |
| 46725 | ENSMUSG00000108940 | 0.462220778 | -0.685400455 | 3.191475091 | -0.21475977 | 0.82995462 | NA | Gm45719 | predicted gene 45719 [Source:MGI Symbo | 0.8031 | 0 | 1.0062 | 0.964 | 0 | 0 |
| 46726 | ENSMUSG00000108941 | 0 | NA | NA | NA | NA | NA | D530014G21 | RIKEN cDNA D530014G21 gene [Source:M | 0 | 0 | 0 | 0 | 0 | 0 |
| 46727 | ENSMUSG00000108942 | 0 | NA | NA | NA | NA | NA | Gm44660 | predicted gene 44660 [Source:MGI Symbo | 0 | 0 | 0 | 0 | 0 | 0 |
| 46728 | ENSMUSG00000108943 | 0.476283062 | 1.031501528 | 3.159856949 | 0.326439312 | 0.744091998 | NA | Gm44559 | predicted gene 44559 [Source:MGI Symbo | 0.8031 | 0 | 0 | 0.964 | 1.0906 | 0 |
| 46729 | ENSMUSG00000108944 | 0 | NA | NA | NA | NA | NA | Gm4558 | predicted gene 4558 [Source:MGI Symbol | 0 | 0 | 0 | 0 | 0 | 0 |
| 46730 | ENSMUSG00000108945 | 0 | NA | NA | NA | NA | NA | Olfr1481-ps1 | olfactory receptor 1481, pseudogene 1 [So | 0 | 0 | 0 | 0 | 0 | 0 |
| 46731 | ENSMUSG00000108946 | 0 | NA | NA | NA | NA | NA | Gm18598 | predicted gene, 18598 [Source:MGI Symb | 0 | 0 | 0 | 0 | 0 | 0 |
| 46732 | ENSMUSG00000108947 | 0 | NA | NA | NA | NA | NA | Olfr1422-ps1 | olfactory receptor 1422, pseudogene 1 [So | 0 | 0 | 0 | 0 | 0 | 0 |
| 46733 | ENSMUSG00000108948 | 0.160662974 | 1.126924175 | 4.080472857 | 0.276174898 | 0.782413736 | NA | Olfr695 | olfactory receptor 695 [Source:MGI Symb | 0 | 0 | 0 | 0.964 | 0 | 0 |
| 46734 | ENSMUSG00000108949 | 1.599295544 | 0.707867103 | 1.87485889 | 0.377557536 | 0.705759315 | NA | Gm45069 | predicted gene 45069 [Source:MGI Symbo | 1.6062 | 2.0558 | 0 | 4.8199 | 0 | 1.1138 |
| 46735 | ENSMUSG00000108950 | 0 | NA | NA | NA | NA | NA | 9130015G15 | RIKEN cDNA 9130015G15 gene [Source:M | 0 | 0 | 0 | 0 | 0 | 0 |
| 46736 | ENSMUSG00000108952 | 0 | NA | NA | NA | NA | NA | Gm39117 | predicted gene, 39117 [Source:MGI Symb | 0 | 0 | 0 | 0 | 0 | 0 |
| 46737 | ENSMUSG00000108953 | 0 | NA | NA | NA | NA | NA | Gm45129 | predicted gene 45129 [Source:MGI Symbo | 0 | 0 | 0 | 0 | 0 | 0 |
| 46738 | ENSMUSG00000108954 | 0.674674027 | 1.600443604 | 2.78690385 | 0.574272989 | 0.565783085 | NA | Gm44901 | predicted gene 44901 [Source:MGI Symbo | 0 | 0 | 1.0062 | 1.928 | 0 | 1.1138 |
| 46739 | ENSMUSG00000108955 | 0.919174381 | -0.746991182 | 2.31347519 | -0.322887051 | 0.746780783 | NA | Gm44775 | predicted gene 44775 [Source:MGI Symbo | 2.4093 | 1.0279 | 0 | 0.964 | 0 | 1.1138 |
| 46740 | ENSMUSG00000108956 | 0 | NA | NA | NA | NA | NA | Gm45179 | predicted gene 45179 [Source:MGI Symbo | 0 | 0 | 0 | 0 | 0 | 0 |
| 46741 | ENSMUSG00000108957 | 17.25336118 | -0.290919979 | 0.586660593 | -0.495891461 | 0.619970991 | 0.917977899 | Gm45235 | predicted gene 45235 [Source:MGI Symbo | 13.653 | 13.363 | 30.187 | 23.135 | 9.8156 | 13.366 |
| 46742 | ENSMUSG00000108958 | 0 | NA | NA | NA | NA | NA | Gm21032 | predicted gene, 21032 [Source:MGI Symb | 0 | 0 | 0 | 0 | 0 | 0 |
| 46743 | ENSMUSG00000108959 | 0.55305101 | 2.667977666 | 3.241982584 | 0.822946329 | 0.410538515 | NA | Gm44697 | predicted gene 44697 [Source:MGI Symbo | 0 | 0 | 0 | 0 | 1.0906 | 2.2277 |
| 46744 | ENSMUSG00000108960 | 0 | NA | NA | NA | NA | NA | Gm44943 | predicted gene 44943 [Source:MGI Symbo | 0 | 0 | 0 | 0 | 0 | 0 |
| 46745 | ENSMUSG00000108961 | 6.931200238 | 1.543442024 | 0.923535023 | 1.671232803 | 0.094675703 | 0.453643796 | Gm32540 | predicted gene, 32540 [Source:MGI Symb | 1.6062 | 3.0838 | 6.0375 | 8.6758 | 4.3625 | 17.822 |
| 46746 | ENSMUSG00000108962 | 0 | NA | NA | NA | NA | NA | Gm44959 | predicted gene 44959 [Source:MGI Symbo | 0 | 0 | 0 | 0 | 0 | 0 |
| 46747 | ENSMUSG00000108963 | 0 | NA | NA | NA | NA | NA | Gm16478 | predicted pseudogene 16478 [Source:MG | 0 | 0 | 0 | 0 | 0 | 0 |
| 46748 | ENSMUSG00000108964 | 0.656740811 | -0.013525372 | 3.112905782 | -0.004344934 | 0.996533255 | NA | Gm18600 | predicted gene, 18600 [Source:MGI Symb | 0 | 0 | 2.0125 | 1.928 | 0 | 0 |
| 46749 | ENSMUSG00000108965 | 0 | NA | NA | NA | NA | NA | Gm44551 | predicted gene 44551 [Source:MGI Symbo | 0 | 0 | 0 | 0 | 0 | 0 |
| 46750 | ENSMUSG00000108966 | 0 | NA | NA | NA | NA | NA | Gm45119 | predicted gene 45119 [Source:MGI Symbo | 0 | 0 | 0 | 0 | 0 | 0 |
| 46751 | ENSMUSG00000108967 | 0 | NA | NA | NA | NA | NA | Gm45181 | predicted gene 45181 [Source:MGI Symbo | 0 | 0 | 0 | 0 | 0 | 0 |
| 46752 | ENSMUSG00000108968 | 0 | NA | NA | NA | NA | NA | Gm45011 | predicted gene 45011 [Source:MGI Symbo | 0 | 0 | 0 | 0 | 0 | 0 |
| 46753 | ENSMUSG00000108969 | 23.84836797 | 0.805879662 | 0.512877103 | 1.571291949 | 0.11611485 | 0.500730933 | Gm45044 | predicted gene 45044 [Source:MGI Symbo | 9.6372 | 17.475 | 25.156 | 21.208 | 26.175 | 43.44 |
| 46754 | ENSMUSG00000108970 | 0 | NA | NA | NA | NA | NA | NA | NA | 0 | 0 | 0 | 0 | 0 | 0 |
| 46755 | ENSMUSG00000108971 | 0 | NA | NA | NA | NA | NA | Gm18210 | predicted gene, 18210 [Source:MGI Symb | 0 | 0 | 0 | 0 | 0 | 0 |
| 46756 | ENSMUSG00000108972 | 0 | NA | NA | NA | NA | NA | Gm7675 | predicted gene 7675 [Source:MGI Symbol | 0 | 0 | 0 | 0 | 0 | 0 |

| | | | | | | | | | | | | | | | |
|---|---|---|---|---|---|---|---|---|---|---|---|---|---|---|---|
| 46757 | ENSMUSG00000108973 | 0 | NA | NA | NA | NA | NA | Gm44555 | predicted gene 44555 [Source:MGI Symbo | 0 | 0 | 0 | 0 | 0 | 0 |
| 46758 | ENSMUSG00000108974 | 0 | NA | NA | NA | NA | NA | Olfr1483-ps1 | olfactory receptor 1483, pseudogene 1 [S | 0 | 0 | 0 | 0 | 0 | 0 |
| 46759 | ENSMUSG00000108975 | 0 | NA | NA | NA | NA | NA | Mlnr-ps | motilin receptor, pseudogene [Source:MG | 0 | 0 | 0 | 0 | 0 | 0 |
| 46760 | ENSMUSG00000108976 | 1.26553565 | 1.680641925 | 2.314608575 | 0.726102004 | 0.467776241 | NA | Gm29797 | predicted gene, 29797 [Source:MGI Symb | 0.8031 | 0 | 1.0062 | 5.7839 | 0 | 0 |
| 46761 | ENSMUSG00000108977 | 0 | NA | NA | NA | NA | NA | Gm39119 | predicted gene, 39119 [Source:MGI Symb | 0 | 0 | 0 | 0 | 0 | 0 |
| 46762 | ENSMUSG00000108978 | 0 | NA | NA | NA | NA | NA | Gm44689 | predicted gene 44689 [Source:MGI Symbo | 0 | 0 | 0 | 0 | 0 | 0 |
| 46763 | ENSMUSG00000108979 | 0.267700746 | -1.497481649 | 4.057001436 | -0.369110456 | 0.712045396 | NA | 1700016B15 | RIKEN cDNA 1700016B15 gene [Source:M | 1.6062 | 0 | 0 | 0 | 0 | 0 |
| 46764 | ENSMUSG00000108980 | 1.144113709 | 0.823834277 | 2.553104362 | 0.322679437 | 0.746938027 | NA | Gm45155 | predicted gene 45155 [Source:MGI Symbo | 2.4093 | 0 | 0 | 0 | 0 | 4.4554 |
| 46765 | ENSMUSG00000108981 | 0 | NA | NA | NA | NA | NA | Gm15262 | predicted gene 15262 [Source:MGI Symbo | 0 | 0 | 0 | 0 | 0 | 0 |
| 46766 | ENSMUSG00000108982 | 0 | NA | NA | NA | NA | NA | Gm17988 | predicted gene, 17988 [Source:MGI Symb | 0 | 0 | 0 | 0 | 0 | 0 |
| 46767 | ENSMUSG00000108983 | 0 | NA | NA | NA | NA | NA | Gm44926 | predicted gene 44926 [Source:MGI Symbo | 0 | 0 | 0 | 0 | 0 | 0 |
| 46768 | ENSMUSG00000108984 | 0.160662974 | 1.126924175 | 4.080472857 | 0.276174898 | 0.782413736 | NA | Gm44916 | predicted gene 44916 [Source:MGI Symbo | 0 | 0 | 0 | 0.964 | 0 | 0 |
| 46769 | ENSMUSG00000108985 | 0 | NA | NA | NA | NA | NA | A930030B08 | RIKEN cDNA A930030B08 gene [Source:M | 0 | 0 | 0 | 0 | 0 | 0 |
| 46770 | ENSMUSG00000108986 | 0 | NA | NA | NA | NA | NA | Gm32061 | predicted gene, 32061 [Source:MGI Symb | 0 | 0 | 0 | 0 | 0 | 0 |
| 46771 | ENSMUSG00000108987 | 0 | NA | NA | NA | NA | NA | Gm45056 | predicted gene 45056 [Source:MGI Symbo | 0 | 0 | 0 | 0 | 0 | 0 |
| 46772 | ENSMUSG00000108988 | 0 | NA | NA | NA | NA | NA | Gm44934 | predicted gene 44934 [Source:MGI Symbo | 0 | 0 | 0 | 0 | 0 | 0 |
| 46773 | ENSMUSG00000108989 | 0.160662974 | 1.126924175 | 4.080472857 | 0.276174898 | 0.782413736 | NA | Gm44984 | predicted gene 44984 [Source:MGI Symbo | 0 | 0 | 0 | 0.964 | 0 | 0 |
| 46774 | ENSMUSG00000108990 | 0.133850373 | -0.796629176 | 4.080472857 | -0.195229623 | 0.84521319 | NA | Gm44648 | predicted gene 44648 [Source:MGI Symbo | 0.8031 | 0 | 0 | 0 | 0 | 0 |
| 46775 | ENSMUSG00000108991 | 0.98432953 | 0.280713436 | 2.433774132 | 0.115340792 | 0.908175007 | NA | Gm45807 | predicted gene 45807 [Source:MGI Symbo | 1.6062 | 1.0279 | 0 | 0 | 3.2719 | 0 |
| 46776 | ENSMUSG00000108992 | 0 | NA | NA | NA | NA | NA | Gm19092 | predicted gene, 19092 [Source:MGI Symb | 0 | 0 | 0 | 0 | 0 | 0 |
| 46777 | ENSMUSG00000108993 | 0 | NA | NA | NA | NA | NA | Gm44846 | predicted gene 44846 [Source:MGI Symbo | 0 | 0 | 0 | 0 | 0 | 0 |
| 46778 | ENSMUSG00000108994 | 0 | NA | NA | NA | NA | NA | Gm31663 | predicted gene, 31663 [Source:MGI Symb | 0 | 0 | 0 | 0 | 0 | 0 |
| 46779 | ENSMUSG00000108995 | 0 | NA | NA | NA | NA | NA | Olfr485 | olfactory receptor 485 [Source:MGI Symb | 0 | 0 | 0 | 0 | 0 | 0 |
| 46780 | ENSMUSG00000108997 | 0 | NA | NA | NA | NA | NA | Gm45045 | predicted gene 45045 [Source:MGI Symbo | 0 | 0 | 0 | 0 | 0 | 0 |
| 46781 | ENSMUSG00000108998 | 0 | NA | NA | NA | NA | NA | Olfr505-ps1 | olfactory receptor 505, pseudogene 1 [So | 0 | 0 | 0 | 0 | 0 | 0 |
| 46782 | ENSMUSG00000109000 | 0 | NA | NA | NA | NA | NA | Gm18460 | predicted gene, 18460 [Source:MGI Symb | 0 | 0 | 0 | 0 | 0 | 0 |
| 46783 | ENSMUSG00000109001 | 0 | NA | NA | NA | NA | NA | Gm45024 | predicted gene 45024 [Source:MGI Symbo | 0 | 0 | 0 | 0 | 0 | 0 |
| 46784 | ENSMUSG00000109002 | 0 | NA | NA | NA | NA | NA | Gm38405 | predicted gene, 38405 [Source:MGI Symb | 0 | 0 | 0 | 0 | 0 | 0 |
| 46785 | ENSMUSG00000109003 | 1.89331256 | -0.754280296 | 1.534512313 | -0.491543984 | 0.623041751 | NA | Gm36011 | predicted gene, 36011 [Source:MGI Symb | 4.0155 | 2.0558 | 1.0062 | 0.964 | 1.0906 | 2.2277 |
| 46786 | ENSMUSG00000109004 | 0 | NA | NA | NA | NA | NA | Gm44594 | predicted gene 44594 [Source:MGI Symbo | 0 | 0 | 0 | 0 | 0 | 0 |
| 46787 | ENSMUSG00000109005 | 6.701123874 | 0.125211817 | 0.834660213 | 0.150015318 | 0.88075253 | 0.981448736 | Gm45221 | predicted gene 45221 [Source:MGI Symbo | 8.031 | 3.0838 | 8.05 | 6.7478 | 8.7249 | 5.5692 |
| 46788 | ENSMUSG00000109006 | 0 | NA | NA | NA | NA | NA | B230209E15 | RIKEN cDNA B230209E15 gene [Source:M | 0 | 0 | 0 | 0 | 0 | 0 |
| 46789 | ENSMUSG00000109007 | 0 | NA | NA | NA | NA | NA | Olfr1439-ps1 | olfactory receptor 1439, pseudogene 1 [S | 0 | 0 | 0 | 0 | 0 | 0 |
| 46790 | ENSMUSG00000109008 | 6.552364565 | -0.430872938 | 0.865661789 | -0.497738197 | 0.618668582 | 0.917509899 | Gm44899 | predicted gene 44899 [Source:MGI Symbo | 6.4248 | 4.1117 | 12.075 | 6.7478 | 3.2719 | 6.6831 |
| 46791 | ENSMUSG00000109009 | 0.167707431 | -0.796629176 | 4.080472857 | -0.195229623 | 0.84521319 | NA | Gm44929 | predicted gene 44929 [Source:MGI Symbo | 0 | 0 | 1.0062 | 0 | 0 | 0 |
| 46792 | ENSMUSG00000109010 | 0 | NA | NA | NA | NA | NA | Gm45027 | predicted gene 45027 [Source:MGI Symbo | 0 | 0 | 0 | 0 | 0 | 0 |
| 46793 | ENSMUSG00000109011 | 0 | NA | NA | NA | NA | NA | Gm44857 | predicted gene 44857 [Source:MGI Symbo | 0 | 0 | 0 | 0 | 0 | 0 |
| 46794 | ENSMUSG00000109012 | 0 | NA | NA | NA | NA | NA | Gm44820 | predicted gene 44820 [Source:MGI Symbo | 0 | 0 | 0 | 0 | 0 | 0 |
| 46795 | ENSMUSG00000109013 | 0 | NA | NA | NA | NA | NA | Gm44723 | predicted gene 44723 [Source:MGI Symbo | 0 | 0 | 0 | 0 | 0 | 0 |
| 46796 | ENSMUSG00000109014 | 0 | NA | NA | NA | NA | NA | Gm44546 | predicted gene 44546 [Source:MGI Symbo | 0 | 0 | 0 | 0 | 0 | 0 |
| 46797 | ENSMUSG00000109015 | 8.82322505 | -0.38999153 | 0.734658427 | -0.530847419 | 0.595524517 | 0.913808971 | Gm31024 | predicted gene, 31024 [Source:MGI Symb | 13.653 | 10.279 | 6.0375 | 8.6758 | 8.7249 | 5.5692 |
| 46798 | ENSMUSG00000109016 | 2.922885894 | 0.953385049 | 1.349921094 | 0.706252427 | 0.480031172 | NA | Gm44647 | predicted gene 44647 [Source:MGI Symbo | 0 | 2.0558 | 4.025 | 4.8199 | 2.1812 | 4.4554 |
| 46799 | ENSMUSG00000109017 | 0 | NA | NA | NA | NA | NA | Gm38979 | predicted gene, 38979 [Source:MGI Symb | 0 | 0 | 0 | 0 | 0 | 0 |
| 46800 | ENSMUSG00000109018 | 0 | NA | NA | NA | NA | NA | Gm44809 | predicted gene 44809 [Source:MGI Symbo | 0 | 0 | 0 | 0 | 0 | 0 |
| 46801 | ENSMUSG00000109019 | 0 | NA | NA | NA | NA | NA | Gm18455 | predicted gene, 18455 [Source:MGI Symb | 0 | 0 | 0 | 0 | 0 | 0 |
| 46802 | ENSMUSG00000109020 | 0 | NA | NA | NA | NA | NA | Olfr197 | olfactory receptor 197 [Source:MGI Symb | 0 | 0 | 0 | 0 | 0 | 0 |
| 46803 | ENSMUSG00000109021 | 0.181769715 | 1.126924175 | 4.080472857 | 0.276174898 | 0.782413736 | NA | Gm44812 | predicted gene 44812 [Source:MGI Symbo | 0 | 0 | 0 | 0 | 1.0906 | 0 |
| 46804 | ENSMUSG00000109022 | 0 | NA | NA | NA | NA | NA | Olfr1432 | olfactory receptor 1432 [Source:MGI Sym | 0 | 0 | 0 | 0 | 0 | 0 |
| 46805 | ENSMUSG00000109023 | 0 | NA | NA | NA | NA | NA | Gm44565 | predicted gene 44565 [Source:MGI Symbo | 0 | 0 | 0 | 0 | 0 | 0 |
| 46806 | ENSMUSG00000109024 | 0 | NA | NA | NA | NA | NA | Gm18063 | predicted gene, 18063 [Source:MGI Symb | 0 | 0 | 0 | 0 | 0 | 0 |
| 46807 | ENSMUSG00000109025 | 0 | NA | NA | NA | NA | NA | Gm6858 | predicted gene 6858 [Source:MGI Symbol | 0 | 0 | 0 | 0 | 0 | 0 |
| 46808 | ENSMUSG00000109026 | 0 | NA | NA | NA | NA | NA | Gm44683 | predicted gene 44683 [Source:MGI Symbo | 0 | 0 | 0 | 0 | 0 | 0 |
| 46809 | ENSMUSG00000109027 | 0 | NA | NA | NA | NA | NA | Olfr752-ps1 | olfactory receptor 752, pseudogene 1 [So | 0 | 0 | 0 | 0 | 0 | 0 |
| 46810 | ENSMUSG00000109028 | 0 | NA | NA | NA | NA | NA | Gm35842 | predicted gene, 35842 [Source:MGI Symb | 0 | 0 | 0 | 0 | 0 | 0 |
| 46811 | ENSMUSG00000109029 | 0 | NA | NA | NA | NA | NA | Gm44717 | predicted gene 44717 [Source:MGI Symbo | 0 | 0 | 0 | 0 | 0 | 0 |
| 46812 | ENSMUSG00000109030 | 0 | NA | NA | NA | NA | NA | Olfr1421-ps1 | olfactory receptor 1421, pseudogene 1 [S | 0 | 0 | 0 | 0 | 0 | 0 |

| | | | | | | | | | | | | | | | |
|---|---|---|---|---|---|---|---|---|---|---|---|---|---|---|---|
| 46813 | ENSMUSG00000109031 | 0 | NA | NA | NA | NA | NA | Gm45201 | predicted gene 45201 [Source:MGI Symbo | 0 | 0 | 0 | 0 | 0 | 0 |
| 46814 | ENSMUSG00000109032 | 3.839759504 | -1.082279052 | 1.101160474 | -0.982853161 | 0.325679713 | NA | Gm7972 | predicted gene 7972 [Source:MGI Symbol | 6.4248 | 6.1675 | 3.0187 | 1.928 | 3.2719 | 2.2277 |
| 46815 | ENSMUSG00000109033 | 0 | NA | NA | NA | NA | NA | Olfr1315-ps1 | olfactory receptor 1315, pseudogene 1 [So | 0 | 0 | 0 | 0 | 0 | 0 |
| 46816 | ENSMUSG00000109034 | 0 | NA | NA | NA | NA | NA | Gm44960 | predicted gene 44960 [Source:MGI Symbo | 0 | 0 | 0 | 0 | 0 | 0 |
| 46817 | ENSMUSG00000109035 | 0 | NA | NA | NA | NA | NA | Gm45209 | predicted gene 45209 [Source:MGI Symbo | 0 | 0 | 0 | 0 | 0 | 0 |
| 46818 | ENSMUSG00000109036 | 3.753341742 | -1.076204966 | 1.408060507 | -0.764317272 | 0.444678195 | NA | 2210406H18 | RIKEN cDNA 2210406H18 gene [Source:M | 4.0155 | 11.307 | 0 | 3.8559 | 0 | 3.3415 |
| 46819 | ENSMUSG00000109037 | 0 | NA | NA | NA | NA | NA | C030038I04R | RIKEN cDNA C030038I04 gene [Source:MC | 0 | 0 | 0 | 0 | 0 | 0 |
| 46820 | ENSMUSG00000109038 | 0.924441557 | -0.875430577 | 2.469021705 | -0.354565768 | 0.722914925 | NA | Gm45120 | predicted gene 45120 [Source:MGI Symbo | 1.6062 | 0 | 2.0125 | 1.928 | 0 | 0 |
| 46821 | ENSMUSG00000109039 | 0.538988726 | 1.126003851 | 3.343747182 | 0.336749099 | 0.736306047 | NA | Gm33570 | predicted gene, 33570 [Source:MGI Symb | 0 | 0 | 1.0062 | 0 | 0 | 2.2277 |
| 46822 | ENSMUSG00000109040 | 0 | NA | NA | NA | NA | NA | Gm44542 | predicted gene 44542 [Source:MGI Symbo | 0 | 0 | 0 | 0 | 0 | 0 |
| 46823 | ENSMUSG00000109041 | 0 | NA | NA | NA | NA | NA | Gm18465 | predicted gene, 18465 [Source:MGI Symb | 0 | 0 | 0 | 0 | 0 | 0 |
| 46824 | ENSMUSG00000109042 | 0 | NA | NA | NA | NA | NA | Gm44991 | predicted gene 44991 [Source:MGI Symbo | 0 | 0 | 0 | 0 | 0 | 0 |
| 46825 | ENSMUSG00000109043 | 0 | NA | NA | NA | NA | NA | Gm18060 | predicted gene, 18060 [Source:MGI Symb | 0 | 0 | 0 | 0 | 0 | 0 |
| 46826 | ENSMUSG00000109044 | 0 | NA | NA | NA | NA | NA | Gm44680 | predicted gene 44680 [Source:MGI Symbo | 0 | 0 | 0 | 0 | 0 | 0 |
| 46827 | ENSMUSG00000109045 | 2.126698102 | -0.572297305 | 1.507968309 | -0.379515472 | 0.704305117 | NA | Gm45084 | predicted gene 45084 [Source:MGI Symbo | 1.6062 | 2.0558 | 4.025 | 2.8919 | 2.1812 | 0 |
| 46828 | ENSMUSG00000109046 | 0.321325949 | 1.911575114 | 4.048267893 | 0.472195804 | 0.636787034 | NA | Gm44571 | predicted gene 44571 [Source:MGI Symbo | 0 | 0 | 0 | 1.928 | 0 | 0 |
| 46829 | ENSMUSG00000109047 | 0 | NA | NA | NA | NA | NA | Gm45115 | predicted gene 45115 [Source:MGI Symbo | 0 | 0 | 0 | 0 | 0 | 0 |
| 46830 | ENSMUSG00000109049 | 0 | NA | NA | NA | NA | NA | 4930479H17 | RIKEN cDNA 4930479H17 gene [Source:M | 0 | 0 | 0 | 0 | 0 | 0 |
| 46831 | ENSMUSG00000109050 | 0 | NA | NA | NA | NA | NA | Gm6290 | predicted gene 6290 [Source:MGI Symbol | 0 | 0 | 0 | 0 | 0 | 0 |
| 46832 | ENSMUSG00000109051 | 1.434937739 | 0.887225091 | 2.181643624 | 0.40667737 | 0.684244964 | NA | Gm44913 | predicted gene 44913 [Source:MGI Symbo | 0 | 1.0279 | 2.0125 | 0 | 0 | 5.5692 |
| 46833 | ENSMUSG00000109052 | 1.206075344 | -2.42209007 | 2.358860796 | -1.026805005 | 0.304512292 | NA | Gm45012 | predicted gene 45012 [Source:MGI Symbo | 0 | 5.1396 | 1.0062 | 0 | 1.0906 | 0 |
| 46834 | ENSMUSG00000109053 | 0 | NA | NA | NA | NA | NA | Gm44656 | predicted gene 44656 [Source:MGI Symbo | 0 | 0 | 0 | 0 | 0 | 0 |
| 46835 | ENSMUSG00000109054 | 0.371281295 | 2.09253362 | 4.031852165 | 0.519000582 | 0.603760335 | NA | Gm44789 | predicted gene 44789 [Source:MGI Symbo | 0 | 0 | 0 | 0 | 0 | 2.2277 |
| 46836 | ENSMUSG00000109055 | 0 | NA | NA | NA | NA | NA | Gm2676 | predicted gene 2676 [Source:MGI Symbol | 0 | 0 | 0 | 0 | 0 | 0 |
| 46837 | ENSMUSG00000109056 | 0.133850373 | -0.796629176 | 4.080472857 | -0.195229623 | 0.84521319 | NA | Leat1 | lncRNA Efnb2 associated transcript 1 [Sou | 0.8031 | 0 | 0 | 0 | 0 | 0 |
| 46838 | ENSMUSG00000109057 | 0.481988923 | 2.492401104 | 3.837077151 | 0.649557203 | 0.515978285 | NA | Gm44741 | predicted gene 44741 [Source:MGI Symbo | 0 | 0 | 0 | 2.8919 | 0 | 0 |
| 46839 | ENSMUSG00000109058 | 0 | NA | NA | NA | NA | NA | Olfr705 | olfactory receptor 705 [Source:MGI Symb | 0 | 0 | 0 | 0 | 0 | 0 |
| 46840 | ENSMUSG00000109059 | 0 | NA | NA | NA | NA | NA | Gm9354 | predicted gene 9354 [Source:MGI Symbol | 0 | 0 | 0 | 0 | 0 | 0 |
| 46841 | ENSMUSG00000109060 | 0 | NA | NA | NA | NA | NA | Gm4513 | predicted gene 4513 [Source:MGI Symbol | 0 | 0 | 0 | 0 | 0 | 0 |
| 46842 | ENSMUSG00000109061 | 0 | NA | NA | NA | NA | NA | Gm49320 | predicted gene, 49320 [Source:MGI Symb | 0 | 0 | 0 | 0 | 0 | 0 |
| 46843 | ENSMUSG00000109062 | 0 | NA | NA | NA | NA | NA | Gm44622 | predicted gene 44622 [Source:MGI Symbo | 0 | 0 | 0 | 0 | 0 | 0 |
| 46844 | ENSMUSG00000109063 | 0 | NA | NA | NA | NA | NA | Gm44961 | predicted gene 44961 [Source:MGI Symbo | 0 | 0 | 0 | 0 | 0 | 0 |
| 46845 | ENSMUSG00000109064 | 0 | NA | NA | NA | NA | NA | Gm39128 | predicted gene, 39128 [Source:MGI Symb | 0 | 0 | 0 | 0 | 0 | 0 |
| 46846 | ENSMUSG00000109065 | 0 | NA | NA | NA | NA | NA | Gm5324 | predicted gene 5324 [Source:MGI Symbol | 0 | 0 | 0 | 0 | 0 | 0 |
| 46847 | ENSMUSG00000109066 | 0 | NA | NA | NA | NA | NA | Gm4585 | predicted gene 4585 [Source:MGI Symbol | 0 | 0 | 0 | 0 | 0 | 0 |
| 46848 | ENSMUSG00000109067 | 0 | NA | NA | NA | NA | NA | Gm44784 | predicted gene 44784 [Source:MGI Symbo | 0 | 0 | 0 | 0 | 0 | 0 |
| 46849 | ENSMUSG00000109068 | 0 | NA | NA | NA | NA | NA | Gm45207 | predicted gene 45207 [Source:MGI Symbo | 0 | 0 | 0 | 0 | 0 | 0 |
| 46850 | ENSMUSG00000109069 | 0 | NA | NA | NA | NA | NA | Gm2734 | predicted gene 2734 [Source:MGI Symbol | 0 | 0 | 0 | 0 | 0 | 0 |
| 46851 | ENSMUSG00000109070 | 0 | NA | NA | NA | NA | NA | Olfr511-ps1 | olfactory receptor 511, pseudogene 1 [So | 0 | 0 | 0 | 0 | 0 | 0 |
| 46852 | ENSMUSG00000109071 | 0 | NA | NA | NA | NA | NA | Gm44840 | predicted gene 44840 [Source:MGI Symbo | 0 | 0 | 0 | 0 | 0 | 0 |
| 46853 | ENSMUSG00000109072 | 0 | NA | NA | NA | NA | NA | Gm18469 | predicted gene, 18469 [Source:MGI Symb | 0 | 0 | 0 | 0 | 0 | 0 |
| 46854 | ENSMUSG00000109073 | 0 | NA | NA | NA | NA | NA | Gm44975 | predicted gene 44975 [Source:MGI Symbo | 0 | 0 | 0 | 0 | 0 | 0 |
| 46855 | ENSMUSG00000109074 | 0 | NA | NA | NA | NA | NA | Gm45162 | predicted gene 45162 [Source:MGI Symbo | 0 | 0 | 0 | 0 | 0 | 0 |
| 46856 | ENSMUSG00000109076 | 0 | NA | NA | NA | NA | NA | Gm45145 | predicted gene 45145 [Source:MGI Symbo | 0 | 0 | 0 | 0 | 0 | 0 |
| 46857 | ENSMUSG00000109077 | 0 | NA | NA | NA | NA | NA | Gm8285 | predicted gene 8285 [Source:MGI Symbol | 0 | 0 | 0 | 0 | 0 | 0 |
| 46858 | ENSMUSG00000109078 | 0 | NA | NA | NA | NA | NA | Gm19033 | predicted gene, 19033 [Source:MGI Symb | 0 | 0 | 0 | 0 | 0 | 0 |
| 46859 | ENSMUSG00000109079 | 0 | NA | NA | NA | NA | NA | Gm44755 | predicted gene 44755 [Source:MGI Symbo | 0 | 0 | 0 | 0 | 0 | 0 |
| 46860 | ENSMUSG00000109080 | 0 | NA | NA | NA | NA | NA | Gm38944 | predicted gene, 38944 [Source:MGI Symb | 0 | 0 | 0 | 0 | 0 | 0 |
| 46861 | ENSMUSG00000109081 | 0 | NA | NA | NA | NA | NA | Gm44611 | predicted gene 44611 [Source:MGI Symbo | 0 | 0 | 0 | 0 | 0 | 0 |
| 46862 | ENSMUSG00000109082 | 0 | NA | NA | NA | NA | NA | Gm44586 | predicted gene 44586 [Source:MGI Symbo | 0 | 0 | 0 | 0 | 0 | 0 |
| 46863 | ENSMUSG00000109083 | 0 | NA | NA | NA | NA | NA | Gm44992 | predicted gene 44992 [Source:MGI Symbo | 0 | 0 | 0 | 0 | 0 | 0 |
| 46864 | ENSMUSG00000109084 | 0 | NA | NA | NA | NA | NA | Gm44955 | predicted gene 44955 [Source:MGI Symbo | 0 | 0 | 0 | 0 | 0 | 0 |
| 46865 | ENSMUSG00000109085 | 0 | NA | NA | NA | NA | NA | Gm18470 | predicted gene, 18470 [Source:MGI Symb | 0 | 0 | 0 | 0 | 0 | 0 |
| 46866 | ENSMUSG00000109086 | 0 | NA | NA | NA | NA | NA | Gm5596 | predicted gene 5596 [Source:MGI Symbol | 0 | 0 | 0 | 0 | 0 | 0 |
| 46867 | ENSMUSG00000109087 | 0 | NA | NA | NA | NA | NA | Gm45105 | predicted gene 45105 [Source:MGI Symbo | 0 | 0 | 0 | 0 | 0 | 0 |
| 46868 | ENSMUSG00000109088 | 0 | NA | NA | NA | NA | NA | Gm44593 | predicted gene 44593 [Source:MGI Symbo | 0 | 0 | 0 | 0 | 0 | 0 |

| | | | | | | | | | | | | | | | |
|---|---|---|---|---|---|---|---|---|---|---|---|---|---|---|---|
| 46869 | ENSMUSG00000109089 | 256.00945 | 0.188015339 | 0.198742594 | 0.946024378 | 0.344136152 | 0.785727752 | 4833411C07 | RIKEN cDNA 4833411C07 gene [Source:M | 231.29 | 209.7 | 277.72 | 343.18 | 224.67 | 249.5 |
| 46870 | ENSMUSG00000109090 | 0 | NA | NA | NA | NA | NA | Gm45088 | predicted gene 45088 [Source:MGI Symbo | 0 | 0 | 0 | 0 | 0 | 0 |
| 46871 | ENSMUSG00000109091 | 0 | NA | NA | NA | NA | NA | Gm44827 | predicted gene 44827 [Source:MGI Symbo | 0 | 0 | 0 | 0 | 0 | 0 |
| 46872 | ENSMUSG00000109092 | 0 | NA | NA | NA | NA | NA | Gm18057 | predicted gene, 18057 [Source:MGI Symb | 0 | 0 | 0 | 0 | 0 | 0 |
| 46873 | ENSMUSG00000109093 | 373.0835854 | -0.044987189 | 0.212526803 | -0.211677721 | 0.83235847 | 0.970533397 | Gm19950 | predicted gene, 19950 [Source:MGI Symb | 451.34 | 392.66 | 292.82 | 483.92 | 333.73 | 284.03 |
| 46874 | ENSMUSG00000109094 | 0 | NA | NA | NA | NA | NA | Gm44587 | predicted gene 44587 [Source:MGI Symbo | 0 | 0 | 0 | 0 | 0 | 0 |
| 46875 | ENSMUSG00000109095 | 0 | NA | NA | NA | NA | NA | Gm44799 | predicted gene 44799 [Source:MGI Symbo | 0 | 0 | 0 | 0 | 0 | 0 |
| 46876 | ENSMUSG00000109096 | 3.133446339 | 0.073707815 | 1.382373657 | 0.053319748 | 0.957477146 | NA | Gm44888 | predicted gene 44888 [Source:MGI Symbo | 4.0155 | 3.0838 | 2.0125 | 0.964 | 8.7249 | 0 |
| 46877 | ENSMUSG00000109097 | 1.697888529 | -0.455662931 | 2.041687872 | -0.223179526 | 0.823395786 | NA | Gm29683 | predicted gene, 29683 [Source:MGI Symb | 4.8186 | 0 | 1.0062 | 0 | 4.3625 | 0 |
| 46878 | ENSMUSG00000109098 | 0 | NA | NA | NA | NA | NA | Gm18062 | predicted gene, 18062 [Source:MGI Symb | 0 | 0 | 0 | 0 | 0 | 0 |
| 46879 | ENSMUSG00000109099 | 0 | NA | NA | NA | NA | NA | Gm20074 | predicted gene, 20074 [Source:MGI Symb | 0 | 0 | 0 | 0 | 0 | 0 |
| 46880 | ENSMUSG00000109100 | 0 | NA | NA | NA | NA | NA | Olfr1464-ps1 | olfactory receptor 1464, pseudogene 1 [So | 0 | 0 | 0 | 0 | 0 | 0 |
| 46881 | ENSMUSG00000109101 | 0 | NA | NA | NA | NA | NA | Gm45132 | predicted gene 45132 [Source:MGI Symbo | 0 | 0 | 0 | 0 | 0 | 0 |
| 46882 | ENSMUSG00000109102 | 0.545309144 | 2.651974629 | 3.666219369 | 0.72335405 | 0.469462394 | NA | 1700019A23 | RIKEN cDNA 1700019A23 gene [Source:M | 0 | 0 | 0 | 0 | 3.2719 | 0 |
| 46883 | ENSMUSG00000109103 | 0 | NA | NA | NA | NA | NA | Gm18877 | predicted gene, 18877 [Source:MGI Symb | 0 | 0 | 0 | 0 | 0 | 0 |
| 46884 | ENSMUSG00000109104 | 0.534859069 | 1.101323472 | 3.351640271 | 0.328592386 | 0.742463804 | NA | Gm6108 | predicted gene 6108 [Source:MGI Symbol | 0 | 1.0279 | 0 | 0 | 2.1812 | 0 |
| 46885 | ENSMUSG00000109105 | 0.506966596 | 2.558866737 | 3.339834942 | 0.766165628 | 0.443577757 | NA | Gm44873 | predicted gene 44873 [Source:MGI Symbo | 0 | 0 | 0 | 1.928 | 0 | 1.1138 |
| 46886 | ENSMUSG00000109106 | 0 | NA | NA | NA | NA | NA | 4933430H16 | RIKEN cDNA 4933430H16 gene [Source:M | 0 | 0 | 0 | 0 | 0 | 0 |
| 46887 | ENSMUSG00000109107 | 0 | NA | NA | NA | NA | NA | Gm44781 | predicted gene 44781 [Source:MGI Symbo | 0 | 0 | 0 | 0 | 0 | 0 |
| 46888 | ENSMUSG00000109108 | 11.1005555 | -0.092621155 | 0.743763715 | -0.124530349 | 0.900895372 | 0.983000175 | Gm36371 | predicted gene, 36371 [Source:MGI Symb | 8.031 | 15.419 | 11.069 | 15.424 | 2.1812 | 14.48 |
| 46889 | ENSMUSG00000109109 | 0.321325949 | 1.911575114 | 4.048267893 | 0.472195804 | 0.636787034 | NA | Gm44953 | predicted gene 44953 [Source:MGI Symbo | 0 | 0 | 0 | 1.928 | 0 | 0 |
| 46890 | ENSMUSG00000109110 | 0 | NA | NA | NA | NA | NA | Gm45014 | predicted gene 45014 [Source:MGI Symbo | 0 | 0 | 0 | 0 | 0 | 0 |
| 46891 | ENSMUSG00000109111 | 0 | NA | NA | NA | NA | NA | 6530437J22I | RIKEN cDNA 6530437J22 gene [Source:MC | 0 | 0 | 0 | 0 | 0 | 0 |
| 46892 | ENSMUSG00000109112 | 0.884878638 | -1.920491624 | 2.550705096 | -0.75292578 | 0.451494515 | NA | Gm44544 | predicted gene 44544 [Source:MGI Symbo | 3.2124 | 0 | 1.0062 | 0 | 1.0906 | 0 |
| 46893 | ENSMUSG00000109113 | 0 | NA | NA | NA | NA | NA | Gm32916 | predicted gene, 32916 [Source:MGI Symb | 0 | 0 | 0 | 0 | 0 | 0 |
| 46894 | ENSMUSG00000109114 | 0 | NA | NA | NA | NA | NA | Gm44753 | predicted gene 44753 [Source:MGI Symbo | 0 | 0 | 0 | 0 | 0 | 0 |
| 46895 | ENSMUSG00000109115 | 16.77230548 | -0.604871634 | 0.564855976 | -1.070842231 | 0.284240375 | 0.736651636 | Gm44669 | predicted gene 44669 [Source:MGI Symbo | 20.881 | 28.782 | 11.069 | 13.496 | 15.269 | 11.138 |
| 46896 | ENSMUSG00000109116 | 0 | NA | NA | NA | NA | NA | Sycp1-ps1 | synaptonemal complex protein 1, pseudo | 0 | 0 | 0 | 0 | 0 | 0 |
| 46897 | ENSMUSG00000109117 | 0 | NA | NA | NA | NA | NA | Gm18393 | predicted gene, 18393 [Source:MGI Symb | 0 | 0 | 0 | 0 | 0 | 0 |
| 46898 | ENSMUSG00000109118 | 11.94452688 | -0.214156826 | 0.629955927 | -0.339955252 | 0.733890226 | 0.948020741 | Gm32031 | predicted gene, 32031 [Source:MGI Symb | 16.062 | 13.363 | 9.0562 | 14.46 | 9.8156 | 8.9108 |
| 46899 | ENSMUSG00000109119 | 0.321325949 | 1.911575114 | 4.048267893 | 0.472195804 | 0.636787034 | NA | Gm44752 | predicted gene 44752 [Source:MGI Symbo | 0 | 0 | 0 | 1.928 | 0 | 0 |
| 46900 | ENSMUSG00000109121 | 0 | NA | NA | NA | NA | NA | Gm44644 | predicted gene 44644 [Source:MGI Symbo | 0 | 0 | 0 | 0 | 0 | 0 |
| 46901 | ENSMUSG00000109122 | 0.342639279 | -1.805976596 | 4.028547714 | -0.448294702 | 0.653940524 | NA | A230103L15 | RIKEN cDNA A230103L15 gene [Source:M | 0 | 2.0558 | 0 | 0 | 0 | 0 |
| 46902 | ENSMUSG00000109123 | 0 | NA | NA | NA | NA | NA | Gm39059 | predicted gene, 39059 [Source:MGI Symb | 0 | 0 | 0 | 0 | 0 | 0 |
| 46903 | ENSMUSG00000109124 | 0 | NA | NA | NA | NA | NA | Gm44516 | predicted gene 44516 [Source:MGI Symbo | 0 | 0 | 0 | 0 | 0 | 0 |
| 46904 | ENSMUSG00000109125 | 0 | NA | NA | NA | NA | NA | Gm45159 | predicted gene 45159 [Source:MGI Symbo | 0 | 0 | 0 | 0 | 0 | 0 |
| 46905 | ENSMUSG00000109126 | 0 | NA | NA | NA | NA | NA | Olfr1455-ps1 | olfactory receptor 1455, pseudogene 1 [So | 0 | 0 | 0 | 0 | 0 | 0 |
| 46906 | ENSMUSG00000109127 | 0 | NA | NA | NA | NA | NA | Gm31135 | predicted gene, 31135 [Source:MGI Symb | 0 | 0 | 0 | 0 | 0 | 0 |
| 46907 | ENSMUSG00000109128 | 0.481988923 | 2.492401104 | 3.837077151 | 0.649557203 | 0.515978285 | NA | Gm45022 | predicted gene 45022 [Source:MGI Symbo | 0 | 0 | 0 | 2.8919 | 0 | 0 |
| 46908 | ENSMUSG00000109129 | 0 | NA | NA | NA | NA | NA | Gm44973 | predicted gene 44973 [Source:MGI Symbo | 0 | 0 | 0 | 0 | 0 | 0 |
| 46909 | ENSMUSG00000109130 | 5.813783079 | -0.357834865 | 1.06268916 | -0.336725807 | 0.736323607 | 0.948109711 | Gm45187 | predicted gene 45187 [Source:MGI Symbo | 7.2279 | 9.2513 | 3.0187 | 0 | 9.8156 | 5.5692 |
| 46910 | ENSMUSG00000109131 | 0 | NA | NA | NA | NA | NA | Gm44862 | predicted gene 44862 [Source:MGI Symbo | 0 | 0 | 0 | 0 | 0 | 0 |
| 46911 | ENSMUSG00000109132 | 0 | NA | NA | NA | NA | NA | Gm44545 | predicted gene 44545 [Source:MGI Symbo | 0 | 0 | 0 | 0 | 0 | 0 |
| 46912 | ENSMUSG00000109133 | 0 | NA | NA | NA | NA | NA | Gm45098 | predicted gene 45098 [Source:MGI Symbo | 0 | 0 | 0 | 0 | 0 | 0 |
| 46913 | ENSMUSG00000109134 | 0 | NA | NA | NA | NA | NA | Gm45076 | predicted gene 45076 [Source:MGI Symbo | 0 | 0 | 0 | 0 | 0 | 0 |
| 46914 | ENSMUSG00000109135 | 0 | NA | NA | NA | NA | NA | Gm6240 | predicted gene 6240 [Source:MGI Symbol | 0 | 0 | 0 | 0 | 0 | 0 |
| 46915 | ENSMUSG00000109136 | 0 | NA | NA | NA | NA | NA | Gm45114 | predicted gene 45114 [Source:MGI Symbo | 0 | 0 | 0 | 0 | 0 | 0 |
| 46916 | ENSMUSG00000109137 | 0 | NA | NA | NA | NA | NA | Gm34908 | predicted gene, 34908 [Source:MGI Symb | 0 | 0 | 0 | 0 | 0 | 0 |
| 46917 | ENSMUSG00000109138 | 0 | NA | NA | NA | NA | NA | Gm18052 | predicted gene, 18052 [Source:MGI Symb | 0 | 0 | 0 | 0 | 0 | 0 |
| 46918 | ENSMUSG00000109139 | 0 | NA | NA | NA | NA | NA | Gm44798 | predicted gene 44798 [Source:MGI Symbo | 0 | 0 | 0 | 0 | 0 | 0 |
| 46919 | ENSMUSG00000109140 | 0 | NA | NA | NA | NA | NA | Gm45089 | predicted gene 45089 [Source:MGI Symbo | 0 | 0 | 0 | 0 | 0 | 0 |
| 46920 | ENSMUSG00000109141 | 13.84901107 | -1.295449603 | 0.749753135 | -1.727834859 | 0.084017836 | 0.427147267 | Gm30692 | predicted gene, 30692 [Source:MGI Symb | 27.305 | 23.642 | 8.05 | 9.6398 | 1.0906 | 13.366 |
| 46921 | ENSMUSG00000109142 | 0 | NA | NA | NA | NA | NA | Gm44600 | predicted gene 44600 [Source:MGI Symbo | 0 | 0 | 0 | 0 | 0 | 0 |
| 46922 | ENSMUSG00000109143 | 0 | NA | NA | NA | NA | NA | Gm10046 | predicted gene 10046 [Source:MGI Symbo | 0 | 0 | 0 | 0 | 0 | 0 |
| 46923 | ENSMUSG00000109145 | 0 | NA | NA | NA | NA | NA | Gm20744 | predicted gene, 20744 [Source:MGI Symb | 0 | 0 | 0 | 0 | 0 | 0 |
| 46924 | ENSMUSG00000109147 | 12.56900849 | 0.221237629 | 0.609605195 | 0.362919527 | 0.716665 | 0.9447719 | 4930431P19 | RIKEN cDNA 4930431P19 gene [Source:M | 14.456 | 8.2233 | 12.075 | 16.388 | 10.906 | 13.366 |

| | | | | | | | | | | | | | | | |
|---|---|---|---|---|---|---|---|---|---|---|---|---|---|---|---|
| 46925 | ENSMUSG00000109148 | 0 | NA | NA | NA | NA | NA | Olfr1452-ps1 | olfactory receptor 1452, pseudogene 1 [S | 0 | 0 | 0 | 0 | 0 | 0 |
| 46926 | ENSMUSG00000109149 | 0.133850373 | -0.796629176 | 4.080472857 | -0.195229623 | 0.84521319 | NA | Gm19656 | predicted gene, 19656 [Source:MGI Symb | 0.8031 | 0 | 0 | 0 | 0 | 0 |
| 46927 | ENSMUSG00000109150 | 0 | NA | NA | NA | NA | NA | Gm45793 | predicted gene 45793 [Source:MGI Symbo | 0 | 0 | 0 | 0 | 0 | 0 |
| 46928 | ENSMUSG00000109151 | 0 | NA | NA | NA | NA | NA | Gm44714 | predicted gene 44714 [Source:MGI Symbo | 0 | 0 | 0 | 0 | 0 | 0 |
| 46929 | ENSMUSG00000109152 | 0 | NA | NA | NA | NA | NA | Gm44666 | predicted gene 44666 [Source:MGI Symbo | 0 | 0 | 0 | 0 | 0 | 0 |
| 46930 | ENSMUSG00000109153 | 0 | NA | NA | NA | NA | NA | Gm38584 | predicted gene, 38584 [Source:MGI Symb | 0 | 0 | 0 | 0 | 0 | 0 |
| 46931 | ENSMUSG00000109154 | 0.160662974 | 1.126924175 | 4.080472857 | 0.276174898 | 0.782413736 | NA | Gm44822 | predicted gene 44822 [Source:MGI Symbo | 0 | 0 | 0 | 0.964 | 0 | 0 |
| 46932 | ENSMUSG00000109155 | 0 | NA | NA | NA | NA | NA | Gm18757 | predicted gene, 18757 [Source:MGI Symb | 0 | 0 | 0 | 0 | 0 | 0 |
| 46933 | ENSMUSG00000109156 | 0 | NA | NA | NA | NA | NA | Gm45194 | predicted gene 45194 [Source:MGI Symbo | 0 | 0 | 0 | 0 | 0 | 0 |
| 46934 | ENSMUSG00000109157 | 2.177965415 | 0.003893439 | 1.499743732 | 0.00259607 | 0.997928638 | NA | Gm44829 | predicted gene 44829 [Source:MGI Symbo | 2.4093 | 2.0558 | 2.0125 | 0 | 4.3625 | 2.2277 |
| 46935 | ENSMUSG00000109158 | 0 | NA | NA | NA | NA | NA | Gm45166 | predicted gene 45166 [Source:MGI Symbo | 0 | 0 | 0 | 0 | 0 | 0 |
| 46936 | ENSMUSG00000109159 | 0 | NA | NA | NA | NA | NA | Gm44657 | predicted gene 44657 [Source:MGI Symbo | 0 | 0 | 0 | 0 | 0 | 0 |
| 46937 | ENSMUSG00000109160 | 0 | NA | NA | NA | NA | NA | 4930598N05 | RIKEN cDNA 4930598N05 gene [Source:M | 0 | 0 | 0 | 0 | 0 | 0 |
| 46938 | ENSMUSG00000109161 | 0.499690045 | -0.82080011 | 3.113553401 | -0.263621658 | 0.792071468 | NA | Gm39033 | predicted gene, 39033 [Source:MGI Symb | 0 | 1.0279 | 1.0062 | 0.964 | 0 | 0 |
| 46939 | ENSMUSG00000109162 | 0 | NA | NA | NA | NA | NA | 2900027M19 | RIKEN cDNA 2900027M19 gene [Source:M | 0 | 0 | 0 | 0 | 0 | 0 |
| 46940 | ENSMUSG00000109163 | 0.181769715 | 1.126924175 | 4.080472857 | 0.276174898 | 0.782413736 | NA | 3110009M11 | RIKEN cDNA 3110009M11 gene [Source:M | 0 | 0 | 0 | 0 | 1.0906 | 0 |
| 46941 | ENSMUSG00000109164 | 0 | NA | NA | NA | NA | NA | Gm45085 | predicted gene 45085 [Source:MGI Symbo | 0 | 0 | 0 | 0 | 0 | 0 |
| 46942 | ENSMUSG00000109165 | 9.011144824 | -0.16159843 | 0.799923881 | -0.202017259 | 0.839903231 | 0.971838962 | Gm45148 | predicted gene 45148 [Source:MGI Symbo | 11.243 | 13.363 | 4.025 | 14.46 | 7.6343 | 3.3415 |
| 46943 | ENSMUSG00000109166 | 0 | NA | NA | NA | NA | NA | Gm5890 | predicted gene 5890 [Source:MGI Symbol | 0 | 0 | 0 | 0 | 0 | 0 |
| 46944 | ENSMUSG00000109167 | 0 | NA | NA | NA | NA | NA | Gm44652 | predicted gene 44652 [Source:MGI Symbo | 0 | 0 | 0 | 0 | 0 | 0 |
| 46945 | ENSMUSG00000109168 | 0 | NA | NA | NA | NA | NA | Gm44709 | predicted gene 44709 [Source:MGI Symbo | 0 | 0 | 0 | 0 | 0 | 0 |
| 46946 | ENSMUSG00000109169 | 0 | NA | NA | NA | NA | NA | Gm18061 | predicted gene, 18061 [Source:MGI Symb | 0 | 0 | 0 | 0 | 0 | 0 |
| 46947 | ENSMUSG00000109170 | 6.185146468 | -1.513880815 | 1.15247053 | -1.313596118 | 0.188982156 | 0.626143323 | Gm45086 | predicted gene 45086 [Source:MGI Symbo | 12.047 | 13.363 | 2.0125 | 0.964 | 8.7249 | 0 |
| 46948 | ENSMUSG00000109171 | 0.181769715 | 1.126924175 | 4.080472857 | 0.276174898 | 0.782413736 | NA | Gm5735 | predicted gene 5735 [Source:MGI Symbol | 0 | 0 | 0 | 0 | 1.0906 | 0 |
| 46949 | ENSMUSG00000109172 | 0 | NA | NA | NA | NA | NA | Gm21035 | predicted gene, 21035 [Source:MGI Symb | 0 | 0 | 0 | 0 | 0 | 0 |
| 46950 | ENSMUSG00000109173 | 0 | NA | NA | NA | NA | NA | Gm21036 | predicted gene, 21036 [Source:MGI Symb | 0 | 0 | 0 | 0 | 0 | 0 |
| 46951 | ENSMUSG00000109174 | 0 | NA | NA | NA | NA | NA | Gm6007 | predicted gene 6007 [Source:MGI Symbol | 0 | 0 | 0 | 0 | 0 | 0 |
| 46952 | ENSMUSG00000109175 | 0 | NA | NA | NA | NA | NA | Gm44947 | predicted gene 44947 [Source:MGI Symbo | 0 | 0 | 0 | 0 | 0 | 0 |
| 46953 | ENSMUSG00000109176 | 0 | NA | NA | NA | NA | NA | Zfp264 | zinc finger protein 264 [Source:MGI Symb | 0 | 0 | 0 | 0 | 0 | 0 |
| 46954 | ENSMUSG00000109177 | 0 | NA | NA | NA | NA | NA | Pafah1b1-ps | platelet-activating factor acetylhydrolase, | 0 | 0 | 0 | 0 | 0 | 0 |
| 46955 | ENSMUSG00000109178 | 0 | NA | NA | NA | NA | NA | Gm3257 | predicted gene 3257 [Source:MGI Symbol | 0 | 0 | 0 | 0 | 0 | 0 |
| 46956 | ENSMUSG00000109179 | 28.53558725 | -0.323204452 | 0.434511524 | -0.743834018 | 0.456976897 | 0.858627415 | Gm35339 | predicted gene, 35339 [Source:MGI Symb | 36.14 | 31.865 | 27.169 | 31.811 | 15.269 | 28.96 |
| 46957 | ENSMUSG00000109180 | 0 | NA | NA | NA | NA | NA | Gm44654 | predicted gene 44654 [Source:MGI Symbo | 0 | 0 | 0 | 0 | 0 | 0 |
| 46958 | ENSMUSG00000109181 | 0 | NA | NA | NA | NA | NA | Gm5605 | predicted gene 5605 [Source:MGI Symbol | 0 | 0 | 0 | 0 | 0 | 0 |
| 46959 | ENSMUSG00000109182 | 0 | NA | NA | NA | NA | NA | Gm18195 | predicted gene, 18195 [Source:MGI Symb | 0 | 0 | 0 | 0 | 0 | 0 |
| 46960 | ENSMUSG00000109183 | 0 | NA | NA | NA | NA | NA | Gm44761 | predicted gene 44761 [Source:MGI Symbo | 0 | 0 | 0 | 0 | 0 | 0 |
| 46961 | ENSMUSG00000109184 | 0 | NA | NA | NA | NA | NA | Olfr1478-ps1 | olfactory receptor 1478, pseudogene 1 [S | 0 | 0 | 0 | 0 | 0 | 0 |
| 46962 | ENSMUSG00000109185 | 0 | NA | NA | NA | NA | NA | Gm18268 | predicted gene, 18268 [Source:MGI Symb | 0 | 0 | 0 | 0 | 0 | 0 |
| 46963 | ENSMUSG00000109186 | 0 | NA | NA | NA | NA | NA | Gm34821 | predicted gene, 34821 [Source:MGI Symb | 0 | 0 | 0 | 0 | 0 | 0 |
| 46964 | ENSMUSG00000109188 | 0 | NA | NA | NA | NA | NA | Gm18213 | predicted gene, 18213 [Source:MGI Symb | 0 | 0 | 0 | 0 | 0 | 0 |
| 46965 | ENSMUSG00000109189 | 0 | NA | NA | NA | NA | NA | Olfr1476-ps1 | olfactory receptor 1476, pseudogene 1 [S | 0 | 0 | 0 | 0 | 0 | 0 |
| 46966 | ENSMUSG00000109190 | 0 | NA | NA | NA | NA | NA | Gm44621 | predicted gene 44621 [Source:MGI Symbo | 0 | 0 | 0 | 0 | 0 | 0 |
| 46967 | ENSMUSG00000109191 | 0 | NA | NA | NA | NA | NA | Gm45131 | predicted gene 45131 [Source:MGI Symbo | 0 | 0 | 0 | 0 | 0 | 0 |
| 46968 | ENSMUSG00000109192 | 0 | NA | NA | NA | NA | NA | Olfr468-ps1 | olfactory receptor 468, pseudogene 1 [Sou | 0 | 0 | 0 | 0 | 0 | 0 |
| 46969 | ENSMUSG00000109193 | 0 | NA | NA | NA | NA | NA | Gm39027 | predicted gene, 39027 [Source:MGI Symb | 0 | 0 | 0 | 0 | 0 | 0 |
| 46970 | ENSMUSG00000109194 | 2.260728436 | -0.295939842 | 1.49348948 | -0.198153282 | 0.842925138 | NA | Gm44570 | predicted gene 44570 [Source:MGI Symbo | 2.4093 | 4.1117 | 1.0062 | 3.8559 | 2.1812 | 0 |
| 46971 | ENSMUSG00000109195 | 0 | NA | NA | NA | NA | NA | Gm45038 | predicted gene 45038 [Source:MGI Symbo | 0 | 0 | 0 | 0 | 0 | 0 |
| 46972 | ENSMUSG00000109196 | 0 | NA | NA | NA | NA | NA | Gm44715 | predicted gene 44715 [Source:MGI Symbo | 0 | 0 | 0 | 0 | 0 | 0 |
| 46973 | ENSMUSG00000109197 | 0 | NA | NA | NA | NA | NA | 4933402C05 | RIKEN cDNA 4933402C05 gene [Source:M | 0 | 0 | 0 | 0 | 0 | 0 |
| 46974 | ENSMUSG00000109198 | 0.160662974 | 1.126924175 | 4.080472857 | 0.276174898 | 0.782413736 | NA | D7Bwg0826 | DNA segment, Chr 7, Brigham & Women's | 0 | 0 | 0 | 0.964 | 0 | 0 |
| 46975 | ENSMUSG00000109199 | 3.90194772 | -1.083864784 | 1.151983311 | -0.940868477 | 0.346772264 | NA | Gm44996 | predicted gene 44996 [Source:MGI Symbo | 5.6217 | 8.2233 | 2.0125 | 0.964 | 4.3625 | 2.2277 |
| 46976 | ENSMUSG00000109200 | 0 | NA | NA | NA | NA | NA | Gm44969 | predicted gene 44969 [Source:MGI Symbo | 0 | 0 | 0 | 0 | 0 | 0 |
| 46977 | ENSMUSG00000109201 | 0 | NA | NA | NA | NA | NA | Gm6518 | predicted gene 6518 [Source:MGI Symbol | 0 | 0 | 0 | 0 | 0 | 0 |
| 46978 | ENSMUSG00000109202 | 0 | NA | NA | NA | NA | NA | Gm21037 | predicted gene, 21037 [Source:MGI Symb | 0 | 0 | 0 | 0 | 0 | 0 |
| 46979 | ENSMUSG00000109203 | 0 | NA | NA | NA | NA | NA | Gm32849 | predicted gene, 32849 [Source:MGI Symb | 0 | 0 | 0 | 0 | 0 | 0 |
| 46980 | ENSMUSG00000109204 | 0 | NA | NA | NA | NA | NA | Gm9445 | predicted gene 9445 [Source:MGI Symbol | 0 | 0 | 0 | 0 | 0 | 0 |

| | | | | | | | | | | | | | | | |
|---|---|---|---|---|---|---|---|---|---|---|---|---|---|---|---|
| 46981 | ENSMUSG00000109205 | 2.628449656 | 0.919799588 | 1.374952982 | 0.66896803 | 0.503515872 | NA | Gm44954 | predicted gene 44954 [Source:MGI Symbo | 2.4093 | 0 | 3.0187 | 4.8199 | 2.1812 | 3.3415 |
| 46982 | ENSMUSG00000109206 | 2.314689471 | 0.369271876 | 1.576410481 | 0.234248554 | 0.814792023 | NA | Gm45137 | predicted gene 45137 [Source:MGI Symbo | 4.0155 | 1.0279 | 1.0062 | 6.7478 | 1.0906 | 0 |
| 46983 | ENSMUSG00000109207 | 0 | NA | NA | NA | NA | NA | Gm45232 | predicted gene 45232 [Source:MGI Symbo | 0 | 0 | 0 | 0 | 0 | 0 |
| 46984 | ENSMUSG00000109209 | 15.65195095 | 1.023417375 | 0.59063492 | 1.732741054 | 0.083141704 | 0.425083636 | Gm45104 | predicted gene 45104 [Source:MGI Symbo | 9.6372 | 12.335 | 9.0562 | 21.208 | 30.537 | 11.138 |
| 46985 | ENSMUSG00000109210 | 0 | NA | NA | NA | NA | NA | Gm18206 | predicted gene, 18206 [Source:MGI Symb | 0 | 0 | 0 | 0 | 0 | 0 |
| 46986 | ENSMUSG00000109211 | 0 | NA | NA | NA | NA | NA | Gm45146 | predicted gene 45146 [Source:MGI Symbo | 0 | 0 | 0 | 0 | 0 | 0 |
| 46987 | ENSMUSG00000109212 | 0 | NA | NA | NA | NA | NA | Olfr126 | olfactory receptor 126 [Source:MGI Symb | 0 | 0 | 0 | 0 | 0 | 0 |
| 46988 | ENSMUSG00000109213 | 0 | NA | NA | NA | NA | NA | Olfr499-ps1 | olfactory receptor 499, pseudogene 1 [So | 0 | 0 | 0 | 0 | 0 | 0 |
| 46989 | ENSMUSG00000109214 | 0 | NA | NA | NA | NA | NA | Gm44659 | predicted gene 44659 [Source:MGI Symbo | 0 | 0 | 0 | 0 | 0 | 0 |
| 46990 | ENSMUSG00000109215 | 0 | NA | NA | NA | NA | NA | 4930453L07I | RIKEN cDNA 4930453L07 gene [Source:M | 0 | 0 | 0 | 0 | 0 | 0 |
| 46991 | ENSMUSG00000109216 | 1.38559695 | 0.941357517 | 1.891340479 | 0.497719753 | 0.618681583 | NA | Gm44907 | predicted gene 44907 [Source:MGI Symbo | 0.8031 | 1.0279 | 1.0062 | 0 | 4.3625 | 1.1138 |
| 46992 | ENSMUSG00000109217 | 0 | NA | NA | NA | NA | NA | Gm44676 | predicted gene 44676 [Source:MGI Symbo | 0 | 0 | 0 | 0 | 0 | 0 |
| 46993 | ENSMUSG00000109218 | 0 | NA | NA | NA | NA | NA | Gm44872 | predicted gene 44872 [Source:MGI Symbo | 0 | 0 | 0 | 0 | 0 | 0 |
| 46994 | ENSMUSG00000109219 | 0 | NA | NA | NA | NA | NA | Olfr1299 | olfactory receptor 1299 [Source:MGI Sym | 0 | 0 | 0 | 0 | 0 | 0 |
| 46995 | ENSMUSG00000109220 | 0 | NA | NA | NA | NA | NA | Gm30196 | predicted gene, 30196 [Source:MGI Symb | 0 | 0 | 0 | 0 | 0 | 0 |
| 46996 | ENSMUSG00000109221 | 0 | NA | NA | NA | NA | NA | Gm9375 | predicted gene 9375 [Source:MGI Symbol | 0 | 0 | 0 | 0 | 0 | 0 |
| 46997 | ENSMUSG00000109222 | 10.74463364 | 0.436659505 | 0.703874553 | 0.620365523 | 0.535017165 | 0.889858671 | Gm10297 | predicted pseudogene 10297 [Source:MG | 12.047 | 4.1117 | 11.069 | 8.6758 | 18.541 | 10.025 |
| 46998 | ENSMUSG00000109223 | 0 | NA | NA | NA | NA | NA | Gm45116 | predicted gene 45116 [Source:MGI Symbo | 0 | 0 | 0 | 0 | 0 | 0 |
| 46999 | ENSMUSG00000109224 | 0 | NA | NA | NA | NA | NA | Tmem147os | transmembrane protein 147, opposite str | 0 | 0 | 0 | 0 | 0 | 0 |
| 47000 | ENSMUSG00000109225 | 0.353089354 | 0.165142706 | 3.896102244 | 0.042386646 | 0.966190474 | NA | 4930513N20 | RIKEN cDNA 4930513N20 gene [Source:M | 0 | 1.0279 | 0 | 0 | 1.0906 | 0 |
| 47001 | ENSMUSG00000109226 | 0 | NA | NA | NA | NA | NA | Gm45039 | predicted gene 45039 [Source:MGI Symbo | 0 | 0 | 0 | 0 | 0 | 0 |
| 47002 | ENSMUSG00000109227 | 0 | NA | NA | NA | NA | NA | Gm29791 | predicted gene, 29791 [Source:MGI Symb | 0 | 0 | 0 | 0 | 0 | 0 |
| 47003 | ENSMUSG00000109228 | 0 | NA | NA | NA | NA | NA | Fam81b | family with sequence similarity 81, memb | 0 | 0 | 0 | 0 | 0 | 0 |
| 47004 | ENSMUSG00000109229 | 0 | NA | NA | NA | NA | NA | Gm44858 | predicted gene 44858 [Source:MGI Symbo | 0 | 0 | 0 | 0 | 0 | 0 |
| 47005 | ENSMUSG00000109230 | 0 | NA | NA | NA | NA | NA | Olfr1470-ps1 | olfactory receptor 1470, pseudogene 1 [S | 0 | 0 | 0 | 0 | 0 | 0 |
| 47006 | ENSMUSG00000109231 | 4.104164413 | -0.761128528 | 1.095434657 | -0.694818739 | 0.487168903 | NA | Gm45737 | predicted gene 45737 [Source:MGI Symbo | 6.4248 | 3.0838 | 6.0375 | 5.7839 | 2.1812 | 1.1138 |
| 47007 | ENSMUSG00000109232 | 0 | NA | NA | NA | NA | NA | Gm44577 | predicted gene 44577 [Source:MGI Symbo | 0 | 0 | 0 | 0 | 0 | 0 |
| 47008 | ENSMUSG00000109233 | 0.267700746 | -1.497481649 | 4.057001436 | -0.369110456 | 0.712045396 | NA | Gm44866 | predicted gene 44866 [Source:MGI Symbo | 1.6062 | 0 | 0 | 0 | 0 | 0 |
| 47009 | ENSMUSG00000109234 | 0 | NA | NA | NA | NA | NA | Gm44561 | predicted gene 44561 [Source:MGI Symbo | 0 | 0 | 0 | 0 | 0 | 0 |
| 47010 | ENSMUSG00000109235 | 0 | NA | NA | NA | NA | NA | Gm5997 | predicted gene 5997 [Source:MGI Symbol | 0 | 0 | 0 | 0 | 0 | 0 |
| 47011 | ENSMUSG00000109236 | 0.167707431 | -0.796629176 | 4.080472857 | -0.195229623 | 0.84521319 | NA | Gm44833 | predicted gene 44833 [Source:MGI Symbo | 0 | 0 | 1.0062 | 0 | 0 | 0 |
| 47012 | ENSMUSG00000109237 | 0 | NA | NA | NA | NA | NA | 9130214F15 | RIKEN cDNA 9130214F15 gene [Source:M | 0 | 0 | 0 | 0 | 0 | 0 |
| 47013 | ENSMUSG00000109238 | 0 | NA | NA | NA | NA | NA | Gm35060 | predicted gene, 35060 [Source:MGI Symb | 0 | 0 | 0 | 0 | 0 | 0 |
| 47014 | ENSMUSG00000109239 | 0 | NA | NA | NA | NA | NA | Gm45176 | predicted gene 45176 [Source:MGI Symbo | 0 | 0 | 0 | 0 | 0 | 0 |
| 47015 | ENSMUSG00000109240 | 0 | NA | NA | NA | NA | NA | Gm44598 | predicted gene 44598 [Source:MGI Symbo | 0 | 0 | 0 | 0 | 0 | 0 |
| 47016 | ENSMUSG00000109241 | 0 | NA | NA | NA | NA | NA | Gm7434 | predicted gene 7434 [Source:MGI Symbol | 0 | 0 | 0 | 0 | 0 | 0 |
| 47017 | ENSMUSG00000109242 | 0 | NA | NA | NA | NA | NA | A930002H02 | RIKEN cDNA A930002H02 gene [Source:M | 0 | 0 | 0 | 0 | 0 | 0 |
| 47018 | ENSMUSG00000109243 | 1.503607759 | 0.456200103 | 1.903153515 | 0.239707464 | 0.810557048 | NA | Gm45867 | predicted gene 45867 [Source:MGI Symbo | 0.8031 | 0 | 3.0187 | 1.928 | 3.2719 | 0 |
| 47019 | ENSMUSG00000109244 | 13.29904003 | 0.130690756 | 0.605965442 | 0.215673613 | 0.829242171 | 0.969955221 | Gm44751 | predicted gene 44751 [Source:MGI Symbo | 8.8341 | 13.363 | 16.1 | 18.316 | 9.8156 | 13.366 |
| 47020 | ENSMUSG00000109245 | 0.695987358 | -1.412518496 | 2.768247659 | -0.510257271 | 0.609871234 | NA | Gm44860 | predicted gene 44860 [Source:MGI Symbo | 0 | 2.0558 | 1.0062 | 0 | 0 | 1.1138 |
| 47021 | ENSMUSG00000109246 | 0 | NA | NA | NA | NA | NA | Olfr712-ps1 | olfactory receptor 712, pseudogene 1 [So | 0 | 0 | 0 | 0 | 0 | 0 |
| 47022 | ENSMUSG00000109247 | 0 | NA | NA | NA | NA | NA | Gm8685 | predicted gene 8685 [Source:MGI Symbol | 0 | 0 | 0 | 0 | 0 | 0 |
| 47023 | ENSMUSG00000109248 | 0 | NA | NA | NA | NA | NA | Gm44993 | predicted gene 44993 [Source:MGI Symbo | 0 | 0 | 0 | 0 | 0 | 0 |
| 47024 | ENSMUSG00000109249 | 0 | NA | NA | NA | NA | NA | Gm44610 | predicted gene 44610 [Source:MGI Symbo | 0 | 0 | 0 | 0 | 0 | 0 |
| 47025 | ENSMUSG00000109250 | 0 | NA | NA | NA | NA | NA | Gm5053 | predicted gene 5053 [Source:MGI Symbol | 0 | 0 | 0 | 0 | 0 | 0 |
| 47026 | ENSMUSG00000109251 | 0.904853373 | -1.988674067 | 2.502700767 | -0.794611203 | 0.4268397 | NA | E230032D23 | RIKEN cDNA E230032D23 gene [Source:M | 2.4093 | 2.0558 | 0 | 0.964 | 0 | 0 |
| 47027 | ENSMUSG00000109252 | 0 | NA | NA | NA | NA | NA | Gm18453 | predicted gene, 18453 [Source:MGI Symb | 0 | 0 | 0 | 0 | 0 | 0 |
| 47028 | ENSMUSG00000109253 | 0 | NA | NA | NA | NA | NA | Gm4536 | predicted gene 4536 [Source:MGI Symbol | 0 | 0 | 0 | 0 | 0 | 0 |
| 47029 | ENSMUSG00000109254 | 0 | NA | NA | NA | NA | NA | Gm44588 | predicted gene 44588 [Source:MGI Symbo | 0 | 0 | 0 | 0 | 0 | 0 |
| 47030 | ENSMUSG00000109255 | 0 | NA | NA | NA | NA | NA | Gm7407 | predicted gene 7407 [Source:MGI Symbol | 0 | 0 | 0 | 0 | 0 | 0 |
| 47031 | ENSMUSG00000109256 | 0 | NA | NA | NA | NA | NA | Gm44681 | predicted gene 44681 [Source:MGI Symbo | 0 | 0 | 0 | 0 | 0 | 0 |
| 47032 | ENSMUSG00000109257 | 0 | NA | NA | NA | NA | NA | Gm45200 | predicted gene 45200 [Source:MGI Symbo | 0 | 0 | 0 | 0 | 0 | 0 |
| 47033 | ENSMUSG00000109258 | 0 | NA | NA | NA | NA | NA | 5330413D20 | RIKEN cDNA 5330413D20 gene [Source:M | 0 | 0 | 0 | 0 | 0 | 0 |
| 47034 | ENSMUSG00000109259 | 0 | NA | NA | NA | NA | NA | Gm18055 | predicted gene, 18055 [Source:MGI Symb | 0 | 0 | 0 | 0 | 0 | 0 |
| 47035 | ENSMUSG00000109260 | 0 | NA | NA | NA | NA | NA | Gm30954 | predicted gene, 30954 [Source:MGI Symb | 0 | 0 | 0 | 0 | 0 | 0 |
| 47036 | ENSMUSG00000109261 | 0 | NA | NA | NA | NA | NA | Gm44938 | predicted gene 44938 [Source:MGI Symbo | 0 | 0 | 0 | 0 | 0 | 0 |

| | | | | | | | | | | | | | | | |
|---|---|---|---|---|---|---|---|---|---|---|---|---|---|---|---|
| 47037 | ENSMUSG00000109262 | 25.05691891 | 0.271342259 | 0.460567071 | 0.589148197 | 0.555761864 | 0.897628185 | Gm44744 | predicted gene 44744 [Source:MGI Symbo | 28.109 | 26.726 | 13.081 | 25.063 | 26.175 | 31.188 |
| 47038 | ENSMUSG00000109263 | 0 | NA | NA | NA | NA | NA | Gm18058 | predicted gene, 18058 [Source:MGI Symb | 0 | 0 | 0 | 0 | 0 | 0 |
| 47039 | ENSMUSG00000109264 | 0 | NA | NA | NA | NA | NA | Gm44871 | predicted gene 44871 [Source:MGI Symbo | 0 | 0 | 0 | 0 | 0 | 0 |
| 47040 | ENSMUSG00000109266 | 0 | NA | NA | NA | NA | NA | Gm6426 | predicted gene 6426 [Source:MGI Symbol | 0 | 0 | 0 | 0 | 0 | 0 |
| 47041 | ENSMUSG00000109267 | 0 | NA | NA | NA | NA | NA | Gm45048 | predicted gene 45048 [Source:MGI Symbo | 0 | 0 | 0 | 0 | 0 | 0 |
| 47042 | ENSMUSG00000109268 | 0 | NA | NA | NA | NA | NA | 9530078K11 | RIKEN cDNA 9530078K11 gene [Source:M | 0 | 0 | 0 | 0 | 0 | 0 |
| 47043 | ENSMUSG00000109269 | 0 | NA | NA | NA | NA | NA | Gm18054 | predicted gene, 18054 [Source:MGI Symb | 0 | 0 | 0 | 0 | 0 | 0 |
| 47044 | ENSMUSG00000109270 | 0 | NA | NA | NA | NA | NA | Gm10584 | predicted gene 10584 [Source:MGI Symbo | 0 | 0 | 0 | 0 | 0 | 0 |
| 47045 | ENSMUSG00000109271 | 0 | NA | NA | NA | NA | NA | Gm39129 | predicted gene, 39129 [Source:MGI Symb | 0 | 0 | 0 | 0 | 0 | 0 |
| 47046 | ENSMUSG00000109272 | 0 | NA | NA | NA | NA | NA | Gm4528 | predicted gene 4528 [Source:MGI Symbol | 0 | 0 | 0 | 0 | 0 | 0 |
| 47047 | ENSMUSG00000109273 | 0 | NA | NA | NA | NA | NA | Gm45101 | predicted gene 45101 [Source:MGI Symbo | 0 | 0 | 0 | 0 | 0 | 0 |
| 47048 | ENSMUSG00000109274 | 55.18764838 | 0.501957288 | 0.369552671 | 1.358283479 | 0.17437375 | 0.604151393 | Gm45133 | predicted gene 45133 [Source:MGI Symbo | 31.321 | 63.731 | 42.262 | 50.127 | 52.35 | 91.335 |
| 47049 | ENSMUSG00000109275 | 0 | NA | NA | NA | NA | NA | Gm19182 | predicted gene, 19182 [Source:MGI Symb | 0 | 0 | 0 | 0 | 0 | 0 |
| 47050 | ENSMUSG00000109276 | 0 | NA | NA | NA | NA | NA | Gm4548 | predicted gene 4548 [Source:MGI Symbol | 0 | 0 | 0 | 0 | 0 | 0 |
| 47051 | ENSMUSG00000109277 | 0.133850373 | -0.796629176 | 4.080472857 | -0.195229623 | 0.84521319 | NA | Gm39121 | predicted gene, 39121 [Source:MGI Symb | 0.8031 | 0 | 0 | 0 | 0 | 0 |
| 47052 | ENSMUSG00000109278 | 0 | NA | NA | NA | NA | NA | Gm45144 | predicted gene 45144 [Source:MGI Symbo | 0 | 0 | 0 | 0 | 0 | 0 |
| 47053 | ENSMUSG00000109279 | 4.012759615 | 0.073976577 | 1.065527558 | 0.069427184 | 0.944649592 | NA | Gm45220 | predicted gene 45220 [Source:MGI Symbo | 5.6217 | 2.0558 | 4.025 | 5.7839 | 4.3625 | 2.2277 |
| 47054 | ENSMUSG00000109280 | 0 | NA | NA | NA | NA | NA | Gm35325 | predicted gene, 35325 [Source:MGI Symb | 0 | 0 | 0 | 0 | 0 | 0 |
| 47055 | ENSMUSG00000109281 | 0 | NA | NA | NA | NA | NA | Gm44651 | predicted gene 44651 [Source:MGI Symbo | 0 | 0 | 0 | 0 | 0 | 0 |
| 47056 | ENSMUSG00000109282 | 6.644212435 | 0.326359007 | 0.917037465 | 0.35588405 | 0.721927401 | 0.9447719 | Gm45188 | predicted gene 45188 [Source:MGI Symbo | 5.6217 | 5.1396 | 7.0437 | 15.424 | 2.1812 | 4.4554 |
| 47057 | ENSMUSG00000109283 | 0 | NA | NA | NA | NA | NA | Gm44733 | predicted gene 44733 [Source:MGI Symbo | 0 | 0 | 0 | 0 | 0 | 0 |
| 47058 | ENSMUSG00000109284 | 0 | NA | NA | NA | NA | NA | B230311B06 | RIKEN cDNA B230311B06 gene [Source:M | 0 | 0 | 0 | 0 | 0 | 0 |
| 47059 | ENSMUSG00000109285 | 0 | NA | NA | NA | NA | NA | Olfr1488-ps1 | olfactory receptor 1488, pseudogene 1 [S | 0 | 0 | 0 | 0 | 0 | 0 |
| 47060 | ENSMUSG00000109286 | 0 | NA | NA | NA | NA | NA | Gm44541 | predicted gene 44541 [Source:MGI Symbo | 0 | 0 | 0 | 0 | 0 | 0 |
| 47061 | ENSMUSG00000109287 | 0 | NA | NA | NA | NA | NA | Gm44713 | predicted gene 44713 [Source:MGI Symbo | 0 | 0 | 0 | 0 | 0 | 0 |
| 47062 | ENSMUSG00000109289 | 0 | NA | NA | NA | NA | NA | Gm32111 | predicted gene, 32111 [Source:MGI Symb | 0 | 0 | 0 | 0 | 0 | 0 |
| 47063 | ENSMUSG00000109290 | 0 | NA | NA | NA | NA | NA | Phgdh-ps1 | 3-phosphoglycerate dehydrogenase, pseu | 0 | 0 | 0 | 0 | 0 | 0 |
| 47064 | ENSMUSG00000109291 | 50.55039892 | 0.477753411 | 0.360870682 | 1.323890896 | 0.185539282 | 0.621183449 | Gm2814 | predicted gene 2814 [Source:MGI Symbol | 37.746 | 43.173 | 46.287 | 86.758 | 41.443 | 47.895 |
| 47065 | ENSMUSG00000109292 | 0 | NA | NA | NA | NA | NA | Gm44564 | predicted gene 44564 [Source:MGI Symbo | 0 | 0 | 0 | 0 | 0 | 0 |
| 47066 | ENSMUSG00000109293 | 4.526977258 | -0.248089665 | 0.981480737 | -0.252770794 | 0.800445338 | NA | Dcst2 | DC-STAMP domain containing 2 [Source:N | 5.6217 | 4.1117 | 5.0312 | 5.7839 | 3.2719 | 3.3415 |
| 47067 | ENSMUSG00000109294 | 0 | NA | NA | NA | NA | NA | Gm44808 | predicted gene 44808 [Source:MGI Symbo | 0 | 0 | 0 | 0 | 0 | 0 |
| 47068 | ENSMUSG00000109295 | 0.160662974 | 1.126924175 | 4.080472857 | 0.276174898 | 0.782413736 | NA | Gm5739 | predicted gene 5739 [Source:MGI Symbol | 0 | 0 | 0 | 0.964 | 0 | 0 |
| 47069 | ENSMUSG00000109297 | 13.10326477 | -0.740643201 | 0.678105285 | -1.092224493 | 0.274734439 | 0.729487777 | Gm31522 | predicted gene, 31522 [Source:MGI Symb | 22.487 | 20.558 | 6.0375 | 8.6758 | 14.178 | 6.6831 |
| 47070 | ENSMUSG00000109298 | 0 | NA | NA | NA | NA | NA | Gm44886 | predicted gene 44886 [Source:MGI Symbo | 0 | 0 | 0 | 0 | 0 | 0 |
| 47071 | ENSMUSG00000109299 | 0 | NA | NA | NA | NA | NA | Gm45164 | predicted gene 45164 [Source:MGI Symbo | 0 | 0 | 0 | 0 | 0 | 0 |
| 47072 | ENSMUSG00000109300 | 0 | NA | NA | NA | NA | NA | Gm18462 | predicted gene, 18462 [Source:MGI Symb | 0 | 0 | 0 | 0 | 0 | 0 |
| 47073 | ENSMUSG00000109301 | 0 | NA | NA | NA | NA | NA | Gm45167 | predicted gene 45167 [Source:MGI Symbo | 0 | 0 | 0 | 0 | 0 | 0 |
| 47074 | ENSMUSG00000109302 | 0.945548297 | -0.814460806 | 2.273503937 | -0.358240332 | 0.720163467 | NA | Gm44967 | predicted gene 44967 [Source:MGI Symbo | 1.6062 | 0 | 2.0125 | 0.964 | 1.0906 | 0 |
| 47075 | ENSMUSG00000109303 | 0.342639279 | -1.805976596 | 4.028547714 | -0.448294702 | 0.653940524 | NA | Gm44549 | predicted gene 44549 [Source:MGI Symbo | 0 | 2.0558 | 0 | 0 | 0 | 0 |
| 47076 | ENSMUSG00000109304 | 10.09087458 | 0.105766721 | 0.680905339 | 0.155332489 | 0.876559204 | 0.980632845 | Gm45112 | predicted gene 45112 [Source:MGI Symbo | 8.8341 | 13.363 | 7.0437 | 12.532 | 7.6343 | 11.138 |
| 47077 | ENSMUSG00000109305 | 1.119136036 | 0.770837187 | 2.32360793 | 0.331741503 | 0.740084459 | NA | Smim38 | small integral membrane protein 38 [Sour | 2.4093 | 0 | 0 | 0.964 | 0 | 3.3415 |
| 47078 | ENSMUSG00000109306 | 0 | NA | NA | NA | NA | NA | Gm18410 | predicted gene, 18410 [Source:MGI Symb | 0 | 0 | 0 | 0 | 0 | 0 |
| 47079 | ENSMUSG00000109307 | 0 | NA | NA | NA | NA | NA | Gm44700 | predicted gene 44700 [Source:MGI Symbo | 0 | 0 | 0 | 0 | 0 | 0 |
| 47080 | ENSMUSG00000109308 | 0 | NA | NA | NA | NA | NA | Gm44743 | predicted gene 44743 [Source:MGI Symbo | 0 | 0 | 0 | 0 | 0 | 0 |
| 47081 | ENSMUSG00000109309 | 0 | NA | NA | NA | NA | NA | Gm49772 | predicted gene, 49772 [Source:MGI Symb | 0 | 0 | 0 | 0 | 0 | 0 |
| 47082 | ENSMUSG00000109310 | 0 | NA | NA | NA | NA | NA | Gm44576 | predicted gene 44576 [Source:MGI Symbo | 0 | 0 | 0 | 0 | 0 | 0 |
| 47083 | ENSMUSG00000109311 | 0.703108924 | -2.875459482 | 2.96384054 | -0.970180225 | 0.331956666 | NA | AI314278 | expressed sequence AI314278 [Source:M | 3.2124 | 0 | 1.0062 | 0 | 0 | 0 |
| 47084 | ENSMUSG00000109312 | 0 | NA | NA | NA | NA | NA | Gm44682 | predicted gene 44682 [Source:MGI Symbo | 0 | 0 | 0 | 0 | 0 | 0 |
| 47085 | ENSMUSG00000109313 | 0 | NA | NA | NA | NA | NA | Gm44693 | predicted gene 44693 [Source:MGI Symbo | 0 | 0 | 0 | 0 | 0 | 0 |
| 47086 | ENSMUSG00000109314 | 3.979574462 | -0.909033651 | 1.082342282 | -0.839876319 | 0.400977736 | NA | Gm44699 | predicted gene 44699 [Source:MGI Symbo | 6.4248 | 5.1396 | 4.025 | 3.8559 | 1.0906 | 3.3415 |
| 47087 | ENSMUSG00000109315 | 0 | NA | NA | NA | NA | NA | Gm45037 | predicted gene 45037 [Source:MGI Symbo | 0 | 0 | 0 | 0 | 0 | 0 |
| 47088 | ENSMUSG00000109316 | 0 | NA | NA | NA | NA | NA | Gm45160 | predicted gene 45160 [Source:MGI Symbo | 0 | 0 | 0 | 0 | 0 | 0 |
| 47089 | ENSMUSG00000109317 | 0 | NA | NA | NA | NA | NA | Gm45041 | predicted gene 45041 [Source:MGI Symbo | 0 | 0 | 0 | 0 | 0 | 0 |
| 47090 | ENSMUSG00000109318 | 0 | NA | NA | NA | NA | NA | Gm17780 | predicted gene, 17780 [Source:MGI Symb | 0 | 0 | 0 | 0 | 0 | 0 |
| 47091 | ENSMUSG00000109319 | 0 | NA | NA | NA | NA | NA | Olfr1482-ps1 | olfactory receptor 1482, pseudogene 1 [S | 0 | 0 | 0 | 0 | 0 | 0 |
| 47092 | ENSMUSG00000109320 | 0 | NA | NA | NA | NA | NA | Gm44679 | predicted gene 44679 [Source:MGI Symbo | 0 | 0 | 0 | 0 | 0 | 0 |

| | | | | | | | | | | | | | | | |
|---|---|---|---|---|---|---|---|---|---|---|---|---|---|---|---|
| 47093 | ENSMUSG00000109321 | 0 | NA | NA | NA | NA | NA | A330076H08 | RIKEN cDNA A330076H08 gene [Source:M | 0 | 0 | 0 | 0 | 0 | 0 |
| 47094 | ENSMUSG00000109322 | 0 | NA | NA | NA | NA | NA | Olfr1283 | olfactory receptor 1283 [Source:MGI Syml | 0 | 0 | 0 | 0 | 0 | 0 |
| 47095 | ENSMUSG00000109323 | 0.305170013 | -1.657836477 | 4.041480335 | -0.410205257 | 0.681655385 | NA | Gm44968 | predicted gene 44968 [Source:MGI Symbo | 0.8031 | 1.0279 | 0 | 0 | 0 | 0 |
| 47096 | ENSMUSG00000109324 | 237.545811 | -0.052719931 | 0.164786505 | -0.319928691 | 0.749022387 | 0.951074124 | Prmt1 | protein arginine N-methyltransferase 1 [S | 232.1 | 244.64 | 249.55 | 260.27 | 215.94 | 222.77 |
| 47097 | ENSMUSG00000109325 | 2.901572035 | -0.8776905 | 1.260361065 | -0.696380208 | 0.486190755 | NA | Krt8-ps | keratin 8, pseudogene [Source:MGI Symb | 3.2124 | 4.1117 | 4.025 | 3.8559 | 1.0906 | 1.1138 |
| 47098 | ENSMUSG00000109326 | 0 | NA | NA | NA | NA | NA | Gm9165 | predicted gene 9165 [Source:MGI Symbol | 0 | 0 | 0 | 0 | 0 | 0 |
| 47099 | ENSMUSG00000109327 | 0 | NA | NA | NA | NA | NA | Gm44995 | predicted gene 44995 [Source:MGI Symbo | 0 | 0 | 0 | 0 | 0 | 0 |
| 47100 | ENSMUSG00000109328 | 0 | NA | NA | NA | NA | NA | Olfr1438-ps1 | olfactory receptor 1438, pseudogene 1 [S | 0 | 0 | 0 | 0 | 0 | 0 |
| 47101 | ENSMUSG00000109329 | 0 | NA | NA | NA | NA | NA | Gm33926 | predicted gene, 33926 [Source:MGI Symb | 0 | 0 | 0 | 0 | 0 | 0 |
| 47102 | ENSMUSG00000109330 | 0.160662974 | 1.126924175 | 4.080472857 | 0.276174898 | 0.782413736 | NA | Gm44754 | predicted gene 44754 [Source:MGI Symbo | 0 | 0 | 0 | 0.964 | 0 | 0 |
| 47103 | ENSMUSG00000109331 | 0 | NA | NA | NA | NA | NA | Gm42402 | predicted gene, 42402 [Source:MGI Symb | 0 | 0 | 0 | 0 | 0 | 0 |
| 47104 | ENSMUSG00000109332 | 0 | NA | NA | NA | NA | NA | Gm44742 | predicted gene 44742 [Source:MGI Symbo | 0 | 0 | 0 | 0 | 0 | 0 |
| 47105 | ENSMUSG00000109333 | 0.401551119 | -2.072982726 | 4.008251979 | -0.517178744 | 0.605031388 | NA | Gm30790 | predicted gene, 30790 [Source:MGI Symb | 2.4093 | 0 | 0 | 0 | 0 | 0 |
| 47106 | ENSMUSG00000109334 | 0 | NA | NA | NA | NA | NA | Gm44925 | predicted gene 44925 [Source:MGI Symbo | 0 | 0 | 0 | 0 | 0 | 0 |
| 47107 | ENSMUSG00000109335 | 0 | NA | NA | NA | NA | NA | Gm44845 | predicted gene 44845 [Source:MGI Symbo | 0 | 0 | 0 | 0 | 0 | 0 |
| 47108 | ENSMUSG00000109336 | 387.0651941 | -0.019987856 | 0.14403172 | -0.13877399 | 0.889628748 | 0.982382091 | Samd4b | sterile alpha motif domain containing 4B [ | 391.11 | 351.55 | 426.65 | 401.01 | 355.54 | 396.53 |
| 47109 | ENSMUSG00000109337 | 0.328370405 | 0.16514316 | 4.003149536 | 0.041253308 | 0.967093956 | NA | Gm45047 | predicted gene 45047 [Source:MGI Symbo | 0 | 0 | 1.0062 | 0.964 | 0 | 0 |
| 47110 | ENSMUSG00000109338 | 0 | NA | NA | NA | NA | NA | Gm45030 | predicted gene 45030 [Source:MGI Symbo | 0 | 0 | 0 | 0 | 0 | 0 |
| 47111 | ENSMUSG00000109339 | 0 | NA | NA | NA | NA | NA | Gm18127 | predicted gene, 18127 [Source:MGI Symb | 0 | 0 | 0 | 0 | 0 | 0 |
| 47112 | ENSMUSG00000109340 | 0 | NA | NA | NA | NA | NA | Gm33146 | predicted gene, 33146 [Source:MGI Symb | 0 | 0 | 0 | 0 | 0 | 0 |
| 47113 | ENSMUSG00000109341 | 0.185640647 | 1.126924175 | 4.080472857 | 0.276174898 | 0.782413736 | NA | Gm30873 | predicted gene, 30873 [Source:MGI Symb | 0 | 0 | 0 | 0 | 0 | 1.1138 |
| 47114 | ENSMUSG00000109342 | 3.105095419 | -1.283245464 | 1.541385547 | -0.832527246 | 0.405111406 | NA | Gm32983 | predicted gene, 32983 [Source:MGI Symb | 0 | 6.1675 | 7.0437 | 0.964 | 0 | 4.4554 |
| 47115 | ENSMUSG00000109343 | 0 | NA | NA | NA | NA | NA | Olfr1486-ps1 | olfactory receptor 1486, pseudogene 1 [S | 0 | 0 | 0 | 0 | 0 | 0 |
| 47116 | ENSMUSG00000109344 | 0 | NA | NA | NA | NA | NA | A26c2 | ANKRD26-like family C, member 2 [Source | 0 | 0 | 0 | 0 | 0 | 0 |
| 47117 | ENSMUSG00000109345 | 0 | NA | NA | NA | NA | NA | Gm4574 | predicted gene 4574 [Source:MGI Symbol | 0 | 0 | 0 | 0 | 0 | 0 |
| 47118 | ENSMUSG00000109346 | 0 | NA | NA | NA | NA | NA | Gm44515 | predicted gene 44515 [Source:MGI Symbo | 0 | 0 | 0 | 0 | 0 | 0 |
| 47119 | ENSMUSG00000109347 | 0 | NA | NA | NA | NA | NA | Gm45175 | predicted gene 45175 [Source:MGI Symbo | 0 | 0 | 0 | 0 | 0 | 0 |
| 47120 | ENSMUSG00000109348 | 0 | NA | NA | NA | NA | NA | Gm44852 | predicted gene 44852 [Source:MGI Symbo | 0 | 0 | 0 | 0 | 0 | 0 |
| 47121 | ENSMUSG00000109350 | 1.727647176 | 4.323686 | 1.998996809 | 2.162927915 | 0.030546724 | NA | Gm44805 | predicted gene 44805 [Source:MGI Symbo | 0 | 0 | 0 | 4.8199 | 1.0906 | 4.4554 |
| 47122 | ENSMUSG00000109351 | 0 | NA | NA | NA | NA | NA | Gm44632 | predicted gene 44632 [Source:MGI Symbo | 0 | 0 | 0 | 0 | 0 | 0 |
| 47123 | ENSMUSG00000109352 | 0.167707431 | -0.796629176 | 4.080472857 | -0.195229623 | 0.84521319 | NA | Gm18256 | predicted gene, 18256 [Source:MGI Symb | 0 | 0 | 1.0062 | 0 | 0 | 0 |
| 47124 | ENSMUSG00000109353 | 0 | NA | NA | NA | NA | NA | Gm45183 | predicted gene 45183 [Source:MGI Symbo | 0 | 0 | 0 | 0 | 0 | 0 |
| 47125 | ENSMUSG00000109354 | 0 | NA | NA | NA | NA | NA | Olfr706 | olfactory receptor 706 [Source:MGI Symb | 0 | 0 | 0 | 0 | 0 | 0 |
| 47126 | ENSMUSG00000109355 | 0 | NA | NA | NA | NA | NA | Gm4880 | predicted pseudogene 4880 [Source:MGI | 0 | 0 | 0 | 0 | 0 | 0 |
| 47127 | ENSMUSG00000109356 | 0.342432689 | 1.990653086 | 3.900281263 | 0.51038706 | 0.609780321 | NA | Gm45118 | predicted gene 45118 [Source:MGI Symbo | 0 | 0 | 0 | 0.964 | 1.0906 | 0 |
| 47128 | ENSMUSG00000109357 | 0 | NA | NA | NA | NA | NA | Gm45236 | predicted gene 45236 [Source:MGI Symbo | 0 | 0 | 0 | 0 | 0 | 0 |
| 47129 | ENSMUSG00000109358 | 0 | NA | NA | NA | NA | NA | Gm44716 | predicted gene 44716 [Source:MGI Symbo | 0 | 0 | 0 | 0 | 0 | 0 |
| 47130 | ENSMUSG00000109359 | 0 | NA | NA | NA | NA | NA | Gm44797 | predicted gene 44797 [Source:MGI Symbo | 0 | 0 | 0 | 0 | 0 | 0 |
| 47131 | ENSMUSG00000109360 | 0 | NA | NA | NA | NA | NA | Gm18049 | predicted gene, 18049 [Source:MGI Symb | 0 | 0 | 0 | 0 | 0 | 0 |
| 47132 | ENSMUSG00000109361 | 0.315620088 | 0.165143421 | 4.063300985 | 0.040642675 | 0.967580763 | NA | Gm44946 | predicted gene 44946 [Source:MGI Symbo | 0.8031 | 0 | 0 | 0 | 1.0906 | 0 |
| 47133 | ENSMUSG00000109362 | 0.685278559 | -2.809870862 | 3.376578027 | -0.832165239 | 0.405315682 | NA | Gm44562 | predicted gene 44562 [Source:MGI Symbo | 0 | 4.1117 | 0 | 0 | 0 | 0 |
| 47134 | ENSMUSG00000109363 | 0.860929232 | 2.10360141 | 2.850627967 | 0.73794316 | 0.46054899 | NA | Gm44668 | predicted gene 44668 [Source:MGI Symbo | 0.8031 | 0 | 0 | 0 | 4.3625 | 0 |
| 47135 | ENSMUSG00000109364 | 0.514011053 | 1.044022102 | 3.082659575 | 0.338675769 | 0.734853997 | NA | Gm44948 | predicted gene 44948 [Source:MGI Symbo | 0 | 0 | 1.0062 | 0.964 | 0 | 1.1138 |
| 47136 | ENSMUSG00000109365 | 0 | NA | NA | NA | NA | NA | Olfr696-ps1 | olfactory receptor 696, pseudogene 1 [So | 0 | 0 | 0 | 0 | 0 | 0 |
| 47137 | ENSMUSG00000109366 | 0.877291758 | 2.066311913 | 2.565974136 | 0.805273866 | 0.420661659 | NA | Gm44698 | predicted gene 44698 [Source:MGI Symbo | 0 | 1.0279 | 0 | 0.964 | 3.2719 | 0 |
| 47138 | ENSMUSG00000109367 | 2.934007874 | -0.092069376 | 1.50025865 | -0.061369002 | 0.951065339 | NA | Gm18782 | predicted gene, 18782 [Source:MGI Symb | 0 | 4.1117 | 5.0312 | 2.8919 | 0 | 5.5692 |
| 47139 | ENSMUSG00000109368 | 0 | NA | NA | NA | NA | NA | Gm45015 | predicted gene 45015 [Source:MGI Symbo | 0 | 0 | 0 | 0 | 0 | 0 |
| 47140 | ENSMUSG00000109369 | 0.505131668 | 1.126696199 | 3.421159819 | 0.329331647 | 0.741905026 | NA | Vmn2r-ps88 | vomeronasal 2, receptor, pseudogene 88 | 0.8031 | 0 | 0 | 0 | 0 | 2.2277 |
| 47141 | ENSMUSG00000109370 | 0.185640647 | 1.126924175 | 4.080472857 | 0.276174898 | 0.782413736 | NA | Gm44675 | predicted gene 44675 [Source:MGI Symbo | 0 | 0 | 0 | 0 | 0 | 1.1138 |
| 47142 | ENSMUSG00000109371 | 0 | NA | NA | NA | NA | NA | Gm44626 | predicted gene 44626 [Source:MGI Symbo | 0 | 0 | 0 | 0 | 0 | 0 |
| 47143 | ENSMUSG00000109372 | 0 | NA | NA | NA | NA | NA | Gm19410 | predicted gene, 19410 [Source:MGI Symb | 0 | 0 | 0 | 0 | 0 | 0 |
| 47144 | ENSMUSG00000109373 | 0 | NA | NA | NA | NA | NA | Gm44690 | predicted gene 44690 [Source:MGI Symbo | 0 | 0 | 0 | 0 | 0 | 0 |
| 47145 | ENSMUSG00000109374 | 1.755566279 | 0.705950551 | 1.6866093 | 0.418561994 | 0.675536271 | NA | Gm44773 | predicted gene 44773 [Source:MGI Symbo | 0 | 1.0279 | 3.0187 | 0.964 | 2.1812 | 3.3415 |
| 47146 | ENSMUSG00000109375 | 0 | NA | NA | NA | NA | NA | Gm44887 | predicted gene 44887 [Source:MGI Symbo | 0 | 0 | 0 | 0 | 0 | 0 |
| 47147 | ENSMUSG00000109376 | 0 | NA | NA | NA | NA | NA | Olfr761 | olfactory receptor 761 [Source:MGI Symb | 0 | 0 | 0 | 0 | 0 | 0 |
| 47148 | ENSMUSG00000109377 | 0 | NA | NA | NA | NA | NA | Gm7397 | predicted gene 7397 [Source:MGI Symbol | 0 | 0 | 0 | 0 | 0 | 0 |

| | | | | | | | | | | | | | | | |
|---|---|---|---|---|---|---|---|---|---|---|---|---|---|---|---|
| 47149 | ENSMUSG00000109378 | 16.40841501 | -0.331504351 | 0.608282918 | -0.544983824 | 0.585764643 | 0.90955446 | Gm49396 | predicted gene, 49396 [Source:MGI Symb | 18.471 | 11.307 | 25.156 | 18.316 | 19.631 | 5.5692 |
| 47150 | ENSMUSG00000109379 | 4.011782066 | -0.979673521 | 1.17609984 | -0.832984996 | 0.404853191 | NA | Gm44550 | predicted gene 44550 [Source:MGI Symbo | 4.8186 | 9.2513 | 2.0125 | 5.7839 | 1.0906 | 1.1138 |
| 47151 | ENSMUSG00000109380 | 0 | NA | NA | NA | NA | NA | Gm18461 | predicted gene, 18461 [Source:MGI Symb | 0 | 0 | 0 | 0 | 0 | 0 |
| 47152 | ENSMUSG00000109381 | 0.160662974 | 1.126924175 | 4.080472857 | 0.276174898 | 0.782413736 | NA | Gm44826 | predicted gene 44826 [Source:MGI Symbo | 0 | 0 | 0 | 0.964 | 0 | 0 |
| 47153 | ENSMUSG00000109382 | 0 | NA | NA | NA | NA | NA | 4930567K12 | RIKEN cDNA 4930567K12 gene [Source:M | 0 | 0 | 0 | 0 | 0 | 0 |
| 47154 | ENSMUSG00000109383 | 1.520512709 | -1.078635927 | 2.055777676 | -0.524685106 | 0.599802113 | NA | Gm36584 | predicted gene, 36584 [Source:MGI Symb | 3.2124 | 0 | 3.0187 | 2.8919 | 0 | 0 |
| 47155 | ENSMUSG00000109384 | 0 | NA | NA | NA | NA | NA | Gm17931 | predicted gene, 17931 [Source:MGI Symb | 0 | 0 | 0 | 0 | 0 | 0 |
| 47156 | ENSMUSG00000109385 | 0 | NA | NA | NA | NA | NA | Gm44518 | predicted gene 44518 [Source:MGI Symbo | 0 | 0 | 0 | 0 | 0 | 0 |
| 47157 | ENSMUSG00000109387 | 0 | NA | NA | NA | NA | NA | Gm45182 | predicted gene 45182 [Source:MGI Symbo | 0 | 0 | 0 | 0 | 0 | 0 |
| 47158 | ENSMUSG00000109388 | 0 | NA | NA | NA | NA | NA | Gm44688 | predicted gene 44688 [Source:MGI Symbo | 0 | 0 | 0 | 0 | 0 | 0 |
| 47159 | ENSMUSG00000109389 | 0 | NA | NA | NA | NA | NA | Olfr1300-ps1 | olfactory receptor 1300, pseudogene 1 [So | 0 | 0 | 0 | 0 | 0 | 0 |
| 47160 | ENSMUSG00000109390 | 5.425557372 | 0.131683155 | 0.920783823 | 0.14301202 | 0.886280688 | 0.98229112 | Gm45718 | predicted gene 45718 [Source:MGI Symbo | 2.4093 | 6.1675 | 7.0437 | 4.8199 | 6.5437 | 5.5692 |
| 47161 | ENSMUSG00000109391 | 0 | NA | NA | NA | NA | NA | Gm45023 | predicted gene 45023 [Source:MGI Symbo | 0 | 0 | 0 | 0 | 0 | 0 |
| 47162 | ENSMUSG00000109392 | 0 | NA | NA | NA | NA | NA | Gm5737 | predicted gene 5737 [Source:MGI Symbol | 0 | 0 | 0 | 0 | 0 | 0 |
| 47163 | ENSMUSG00000109393 | 0 | NA | NA | NA | NA | NA | Gm44936 | predicted gene 44936 [Source:MGI Symbo | 0 | 0 | 0 | 0 | 0 | 0 |
| 47164 | ENSMUSG00000109394 | 0 | NA | NA | NA | NA | NA | A230057D06 | RIKEN cDNA A230057D06 gene [Source:M | 0 | 0 | 0 | 0 | 0 | 0 |
| 47165 | ENSMUSG00000109395 | 0 | NA | NA | NA | NA | NA | Gm45199 | predicted gene 45199 [Source:MGI Symbo | 0 | 0 | 0 | 0 | 0 | 0 |
| 47166 | ENSMUSG00000109396 | 0 | NA | NA | NA | NA | NA | Gm4565 | predicted gene 4565 [Source:MGI Symbol | 0 | 0 | 0 | 0 | 0 | 0 |
| 47167 | ENSMUSG00000109397 | 0 | NA | NA | NA | NA | NA | Gm45117 | predicted gene 45117 [Source:MGI Symbo | 0 | 0 | 0 | 0 | 0 | 0 |
| 47168 | ENSMUSG00000109398 | 0.181769715 | 1.126924175 | 4.080472857 | 0.276174898 | 0.782413736 | NA | Gm3854 | predicted gene 3854 [Source:MGI Symbol | 0 | 0 | 0 | 0 | 1.0906 | 0 |
| 47169 | ENSMUSG00000109399 | 1.420151417 | 1.718951162 | 1.883250026 | 0.912757806 | 0.361369934 | NA | Gm44982 | predicted gene 44982 [Source:MGI Symbo | 0 | 1.0279 | 1.0062 | 0.964 | 2.1812 | 3.3415 |
| 47170 | ENSMUSG00000109400 | 0 | NA | NA | NA | NA | NA | Gm44533 | predicted gene 44533 [Source:MGI Symbo | 0 | 0 | 0 | 0 | 0 | 0 |
| 47171 | ENSMUSG00000109401 | 1.118438627 | -0.101338974 | 2.129153173 | -0.047595906 | 0.962038295 | NA | Cpeb1os1 | cytoplasmic polyadenylation element binc | 2.4093 | 0 | 1.0062 | 0 | 2.1812 | 1.1138 |
| 47172 | ENSMUSG00000109402 | 0 | NA | NA | NA | NA | NA | Gm45040 | predicted gene 45040 [Source:MGI Symbo | 0 | 0 | 0 | 0 | 0 | 0 |
| 47173 | ENSMUSG00000109403 | 0 | NA | NA | NA | NA | NA | Olfr1311 | olfactory receptor 1311 [Source:MGI Syml | 0 | 0 | 0 | 0 | 0 | 0 |
| 47174 | ENSMUSG00000109404 | 1.048611398 | 1.082000883 | 2.403683562 | 0.450142814 | 0.652607467 | NA | Gm19963 | predicted gene, 19963 [Source:MGI Symb | 0 | 2.0558 | 0 | 0.964 | 3.2719 | 0 |
| 47175 | ENSMUSG00000109405 | 0 | NA | NA | NA | NA | NA | Gm16443 | predicted gene 16443 [Source:MGI Symbo | 0 | 0 | 0 | 0 | 0 | 0 |
| 47176 | ENSMUSG00000109406 | 0 | NA | NA | NA | NA | NA | Olfr515-ps1 | olfactory receptor 515, pseudogene 1 [Sou | 0 | 0 | 0 | 0 | 0 | 0 |
| 47177 | ENSMUSG00000109407 | 0.160662974 | 1.126924175 | 4.080472857 | 0.276174898 | 0.782413736 | NA | Gm34664 | predicted gene, 34664 [Source:MGI Symb | 0 | 0 | 0 | 0.964 | 0 | 0 |
| 47178 | ENSMUSG00000109408 | 13.75099324 | -0.271289666 | 0.638517919 | -0.424874006 | 0.670928525 | 0.93116767 | A930037H05 | RIKEN cDNA A930037H05 gene [Source:M | 12.85 | 9.2513 | 23.144 | 17.352 | 6.5437 | 13.366 |
| 47179 | ENSMUSG00000109409 | 0 | NA | NA | NA | NA | NA | Gm9343 | predicted gene 9343 [Source:MGI Symbol | 0 | 0 | 0 | 0 | 0 | 0 |
| 47180 | ENSMUSG00000109410 | 0 | NA | NA | NA | NA | NA | Olfr1430-ps1 | olfactory receptor 1430, pseudogene 1 [So | 0 | 0 | 0 | 0 | 0 | 0 |
| 47181 | ENSMUSG00000109411 | 0 | NA | NA | NA | NA | NA | Gm18059 | predicted gene, 18059 [Source:MGI Symb | 0 | 0 | 0 | 0 | 0 | 0 |
| 47182 | ENSMUSG00000109412 | 0 | NA | NA | NA | NA | NA | Gm44535 | predicted gene 44535 [Source:MGI Symbo | 0 | 0 | 0 | 0 | 0 | 0 |
| 47183 | ENSMUSG00000109413 | 0 | NA | NA | NA | NA | NA | Gm45165 | predicted gene 45165 [Source:MGI Symbo | 0 | 0 | 0 | 0 | 0 | 0 |
| 47184 | ENSMUSG00000109414 | 0 | NA | NA | NA | NA | NA | 4930505K13 | RIKEN cDNA 4930505K13 gene [Source:M | 0 | 0 | 0 | 0 | 0 | 0 |
| 47185 | ENSMUSG00000109415 | 0 | NA | NA | NA | NA | NA | Olfr1460-ps1 | olfactory receptor 1460, pseudogene 1 [So | 0 | 0 | 0 | 0 | 0 | 0 |
| 47186 | ENSMUSG00000109416 | 0 | NA | NA | NA | NA | NA | Gm44863 | predicted gene 44863 [Source:MGI Symbo | 0 | 0 | 0 | 0 | 0 | 0 |
| 47187 | ENSMUSG00000109417 | 0 | NA | NA | NA | NA | NA | Gm44859 | predicted gene 44859 [Source:MGI Symbo | 0 | 0 | 0 | 0 | 0 | 0 |
| 47188 | ENSMUSG00000109418 | 0 | NA | NA | NA | NA | NA | Gm44800 | predicted gene 44800 [Source:MGI Symbo | 0 | 0 | 0 | 0 | 0 | 0 |
| 47189 | ENSMUSG00000109419 | 0 | NA | NA | NA | NA | NA | Gm45163 | predicted gene 45163 [Source:MGI Symbo | 0 | 0 | 0 | 0 | 0 | 0 |
| 47190 | ENSMUSG00000109420 | 4.157448269 | -0.198119078 | 1.021432258 | -0.193962034 | 0.84620561 | NA | Gm44702 | predicted gene 44702 [Source:MGI Symbo | 3.2124 | 4.1117 | 6.0375 | 3.8559 | 3.2719 | 4.4554 |
| 47191 | ENSMUSG00000109421 | 0 | NA | NA | NA | NA | NA | Gm44998 | predicted gene 44998 [Source:MGI Symbo | 0 | 0 | 0 | 0 | 0 | 0 |
| 47192 | ENSMUSG00000109422 | 0 | NA | NA | NA | NA | NA | Gm45174 | predicted gene 45174 [Source:MGI Symbo | 0 | 0 | 0 | 0 | 0 | 0 |
| 47193 | ENSMUSG00000109423 | 0 | NA | NA | NA | NA | NA | Olfr1304-ps1 | olfactory receptor 1304, pseudogene 1 [So | 0 | 0 | 0 | 0 | 0 | 0 |
| 47194 | ENSMUSG00000109424 | 2.838757119 | 0.338515413 | 1.24266835 | 0.272410103 | 0.785306706 | NA | Gm44658 | predicted gene 44658 [Source:MGI Symbo | 2.4093 | 4.1117 | 1.0062 | 2.8919 | 3.2719 | 3.3415 |
| 47195 | ENSMUSG00000109425 | 0 | NA | NA | NA | NA | NA | Gm18471 | predicted gene, 18471 [Source:MGI Symb | 0 | 0 | 0 | 0 | 0 | 0 |
| 47196 | ENSMUSG00000109426 | 0 | NA | NA | NA | NA | NA | Olfr496-ps1 | olfactory receptor 496, pseudogene 1 [Sou | 0 | 0 | 0 | 0 | 0 | 0 |
| 47197 | ENSMUSG00000109427 | 0.367410362 | 2.078255025 | 3.800070157 | 0.54689912 | 0.58444804 | NA | Gm45845 | predicted gene 45845 [Source:MGI Symbo | 0 | 0 | 0 | 0 | 1.0906 | 1.1138 |
| 47198 | ENSMUSG00000109428 | 9.920484803 | -0.595559253 | 0.744420788 | -0.800030389 | 0.42369319 | 0.84018348 | Gm45698 | predicted gene 45698 [Source:MGI Symbo | 18.471 | 9.2513 | 8.05 | 11.568 | 3.2719 | 8.9108 |
| 47199 | ENSMUSG00000109429 | 2.29871515 | -0.314337043 | 1.456788096 | -0.215774034 | 0.82916389 | NA | Gm45223 | predicted gene 45223 [Source:MGI Symbo | 1.6062 | 3.0838 | 3.0187 | 3.8559 | 0 | 2.2277 |
| 47200 | ENSMUSG00000109430 | 0 | NA | NA | NA | NA | NA | Gm44585 | predicted gene 44585 [Source:MGI Symbo | 0 | 0 | 0 | 0 | 0 | 0 |
| 47201 | ENSMUSG00000109431 | 0 | NA | NA | NA | NA | NA | Gm45161 | predicted gene 45161 [Source:MGI Symbo | 0 | 0 | 0 | 0 | 0 | 0 |
| 47202 | ENSMUSG00000109432 | 0 | NA | NA | NA | NA | NA | Gm44785 | predicted gene 44785 [Source:MGI Symbo | 0 | 0 | 0 | 0 | 0 | 0 |
| 47203 | ENSMUSG00000109433 | 0 | NA | NA | NA | NA | NA | Olfr1435-ps1 | olfactory receptor 1435, pseudogene 1 [So | 0 | 0 | 0 | 0 | 0 | 0 |
| 47204 | ENSMUSG00000109434 | 0 | NA | NA | NA | NA | NA | Gm44937 | predicted gene 44937 [Source:MGI Symbo | 0 | 0 | 0 | 0 | 0 | 0 |

| | | | | | | | | | | | | | | | |
|---|---|---|---|---|---|---|---|---|---|---|---|---|---|---|---|
| 47205 | ENSMUSG00000109435 | 0 | NA | NA | NA | NA | NA | Gm7067 | predicted gene 7067 [Source:MGI Symbol | 0 | 0 | 0 | 0 | 0 | 0 |
| 47206 | ENSMUSG00000109436 | 0 | NA | NA | NA | NA | NA | Gm45075 | predicted gene 45075 [Source:MGI Symbo | 0 | 0 | 0 | 0 | 0 | 0 |
| 47207 | ENSMUSG00000109437 | 0 | NA | NA | NA | NA | NA | Gm45072 | predicted gene 45072 [Source:MGI Symbo | 0 | 0 | 0 | 0 | 0 | 0 |
| 47208 | ENSMUSG00000109438 | 0.503122293 | -2.366675966 | 3.785596405 | -0.625179156 | 0.531853481 | NA | Gm45073 | predicted gene 45073 [Source:MGI Symbo | 0 | 0 | 3.0187 | 0 | 0 | 0 |
| 47209 | ENSMUSG00000109439 | 0 | NA | NA | NA | NA | NA | Gm44853 | predicted gene 44853 [Source:MGI Symbo | 0 | 0 | 0 | 0 | 0 | 0 |
| 47210 | ENSMUSG00000109440 | 0 | NA | NA | NA | NA | NA | Bc1-ps1 | brain cytoplasmic RNA 1, pseudogene 1 [S | 0 | 0 | 0 | 0 | 0 | 0 |
| 47211 | ENSMUSG00000109441 | 0 | NA | NA | NA | NA | NA | Gm44983 | predicted gene 44983 [Source:MGI Symbo | 0 | 0 | 0 | 0 | 0 | 0 |
| 47212 | ENSMUSG00000109442 | 0 | NA | NA | NA | NA | NA | Gm34964 | predicted gene, 34964 [Source:MGI Symb | 0 | 0 | 0 | 0 | 0 | 0 |
| 47213 | ENSMUSG00000109443 | 0 | NA | NA | NA | NA | NA | Gm45002 | predicted gene 45002 [Source:MGI Symbo | 0 | 0 | 0 | 0 | 0 | 0 |
| 47214 | ENSMUSG00000109444 | 0 | NA | NA | NA | NA | NA | Gm18466 | predicted gene, 18466 [Source:MGI Symb | 0 | 0 | 0 | 0 | 0 | 0 |
| 47215 | ENSMUSG00000109445 | 0.160662974 | 1.126924175 | 4.080472857 | 0.276174898 | 0.782413736 | NA | 4930560O18 | RIKEN cDNA 4930560O18 gene [Source:M | 0 | 0 | 0 | 0.964 | 0 | 0 |
| 47216 | ENSMUSG00000109446 | 0 | NA | NA | NA | NA | NA | Gm9195 | predicted gene 9195 [Source:MGI Symbol | 0 | 0 | 0 | 0 | 0 | 0 |
| 47217 | ENSMUSG00000109447 | 0 | NA | NA | NA | NA | NA | Gm3198 | predicted gene 3198 [Source:MGI Symbol | 0 | 0 | 0 | 0 | 0 | 0 |
| 47218 | ENSMUSG00000109448 | 0.17131964 | -0.796629176 | 4.080472857 | -0.195229623 | 0.84521319 | NA | Gm45224 | predicted gene 45224 [Source:MGI Symbo | 0 | 1.0279 | 0 | 0 | 0 | 0 |
| 47219 | ENSMUSG00000109449 | 0 | NA | NA | NA | NA | NA | Olfr1280 | olfactory receptor 1280 [Source:MGI Sym | 0 | 0 | 0 | 0 | 0 | 0 |
| 47220 | ENSMUSG00000109450 | 0 | NA | NA | NA | NA | NA | Gm39043 | predicted gene, 39043 [Source:MGI Symb | 0 | 0 | 0 | 0 | 0 | 0 |
| 47221 | ENSMUSG00000109451 | 0 | NA | NA | NA | NA | NA | Gm18129 | predicted gene, 18129 [Source:MGI Symb | 0 | 0 | 0 | 0 | 0 | 0 |
| 47222 | ENSMUSG00000109452 | 0.481988923 | 2.492401104 | 3.837077151 | 0.649557203 | 0.515978285 | NA | Gm44530 | predicted gene 44530 [Source:MGI Symbo | 0 | 0 | 0 | 2.8919 | 0 | 0 |
| 47223 | ENSMUSG00000109453 | 0.331982614 | 0.16514309 | 3.986751476 | 0.041422971 | 0.966958701 | NA | 4930405G09 | RIKEN cDNA 4930405G09 gene [Source:M | 0 | 1.0279 | 0 | 0.964 | 0 | 0 |
| 47224 | ENSMUSG00000109454 | 0 | NA | NA | NA | NA | NA | Gm34783 | predicted gene, 34783 [Source:MGI Symb | 0 | 0 | 0 | 0 | 0 | 0 |
| 47225 | ENSMUSG00000109455 | 1.748263098 | 1.3256548 | 1.673821253 | 0.79199305 | 0.428364728 | NA | Gm44710 | predicted gene 44710 [Source:MGI Symbo | 0 | 2.0558 | 1.0062 | 1.928 | 3.2719 | 2.2277 |
| 47226 | ENSMUSG00000109456 | 0 | NA | NA | NA | NA | NA | Gm44584 | predicted gene 44584 [Source:MGI Symbo | 0 | 0 | 0 | 0 | 0 | 0 |
| 47227 | ENSMUSG00000109457 | 0 | NA | NA | NA | NA | NA | Gm44909 | predicted gene 44909 [Source:MGI Symbo | 0 | 0 | 0 | 0 | 0 | 0 |
| 47228 | ENSMUSG00000109458 | 0 | NA | NA | NA | NA | NA | Gm44889 | predicted gene 44889 [Source:MGI Symbo | 0 | 0 | 0 | 0 | 0 | 0 |
| 47229 | ENSMUSG00000109459 | 4.569428347 | 0.338566884 | 1.045625795 | 0.323793546 | 0.746094343 | NA | Gm39526 | predicted gene, 39526 [Source:MGI Symb | 4.0155 | 5.1396 | 3.0187 | 8.6758 | 5.4531 | 1.1138 |
| 47230 | ENSMUSG00000109460 | 0 | NA | NA | NA | NA | NA | Gm45591 | predicted gene 45591 [Source:MGI Symbo | 0 | 0 | 0 | 0 | 0 | 0 |
| 47231 | ENSMUSG00000109461 | 0 | NA | NA | NA | NA | NA | Gm44848 | predicted gene 44848 [Source:MGI Symbo | 0 | 0 | 0 | 0 | 0 | 0 |
| 47232 | ENSMUSG00000109462 | 0 | NA | NA | NA | NA | NA | Gm6334 | predicted gene 6334 [Source:MGI Symbol | 0 | 0 | 0 | 0 | 0 | 0 |
| 47233 | ENSMUSG00000109463 | 0 | NA | NA | NA | NA | NA | Gm7506 | predicted gene 7506 [Source:MGI Symbol | 0 | 0 | 0 | 0 | 0 | 0 |
| 47234 | ENSMUSG00000109464 | 0.160662974 | 1.126924175 | 4.080472857 | 0.276174898 | 0.782413736 | NA | Gm34350 | predicted gene, 34350 [Source:MGI Symb | 0 | 0 | 0 | 0.964 | 0 | 0 |
| 47235 | ENSMUSG00000109465 | 0 | NA | NA | NA | NA | NA | Gm19032 | predicted gene, 19032 [Source:MGI Symb | 0 | 0 | 0 | 0 | 0 | 0 |
| 47236 | ENSMUSG00000109466 | 0 | NA | NA | NA | NA | NA | Gm7587 | predicted gene 7587 [Source:MGI Symbol | 0 | 0 | 0 | 0 | 0 | 0 |
| 47237 | ENSMUSG00000109467 | 0 | NA | NA | NA | NA | NA | Gm30646 | predicted gene, 30646 [Source:MGI Symb | 0 | 0 | 0 | 0 | 0 | 0 |
| 47238 | ENSMUSG00000109468 | 0 | NA | NA | NA | NA | NA | Gm45185 | predicted gene 45185 [Source:MGI Symbo | 0 | 0 | 0 | 0 | 0 | 0 |
| 47239 | ENSMUSG00000109469 | 0 | NA | NA | NA | NA | NA | Gm8486 | predicted gene 8486 [Source:MGI Symbol | 0 | 0 | 0 | 0 | 0 | 0 |
| 47240 | ENSMUSG00000109471 | 0 | NA | NA | NA | NA | NA | Gm19127 | predicted gene, 19127 [Source:MGI Symb | 0 | 0 | 0 | 0 | 0 | 0 |
| 47241 | ENSMUSG00000109472 | 0 | NA | NA | NA | NA | NA | Gm18943 | predicted gene, 18943 [Source:MGI Symb | 0 | 0 | 0 | 0 | 0 | 0 |
| 47242 | ENSMUSG00000109473 | 71.97989548 | 0.810996772 | 0.299656171 | 2.706424399 | 0.006801207 | 0.082389156 | B930025P03 | RIKEN cDNA B930025P03 gene [Source:M | 39.352 | 49.34 | 68.425 | 74.226 | 102.52 | 98.018 |
| 47243 | ENSMUSG00000109475 | 0 | NA | NA | NA | NA | NA | Gm34272 | predicted gene, 34272 [Source:MGI Symb | 0 | 0 | 0 | 0 | 0 | 0 |
| 47244 | ENSMUSG00000109476 | 0 | NA | NA | NA | NA | NA | Gm21034 | predicted gene, 21034 [Source:MGI Symb | 0 | 0 | 0 | 0 | 0 | 0 |
| 47245 | ENSMUSG00000109477 | 0 | NA | NA | NA | NA | NA | Gm18306 | predicted gene, 18306 [Source:MGI Symb | 0 | 0 | 0 | 0 | 0 | 0 |
| 47246 | ENSMUSG00000109478 | 0.77079641 | 0.856159616 | 2.652840961 | 0.322733111 | 0.746897374 | NA | Gm39078 | predicted gene, 39078 [Source:MGI Symb | 1.6062 | 0 | 0 | 1.928 | 1.0906 | 0 |
| 47247 | ENSMUSG00000109479 | 0 | NA | NA | NA | NA | NA | Gm44919 | predicted gene 44919 [Source:MGI Symbo | 0 | 0 | 0 | 0 | 0 | 0 |
| 47248 | ENSMUSG00000109480 | 0.185640647 | 1.126924175 | 4.080472857 | 0.276174898 | 0.782413736 | NA | Gm45042 | predicted gene 45042 [Source:MGI Symbo | 0 | 0 | 0 | 0 | 0 | 1.1138 |
| 47249 | ENSMUSG00000109481 | 0 | NA | NA | NA | NA | NA | Gm45130 | predicted gene 45130 [Source:MGI Symbo | 0 | 0 | 0 | 0 | 0 | 0 |
| 47250 | ENSMUSG00000109482 | 1700.197312 | 0.463464088 | 0.130988657 | 3.538200182 | 0.000402865 | 0.010687243 | Gm4756 | predicted gene 4756 [Source:MGI Symbol | 1278.5 | 1357.9 | 1652.3 | 1725.5 | 2142 | 2045 |
| 47251 | ENSMUSG00000109484 | 0 | NA | NA | NA | NA | NA | Gm4199 | predicted gene 4199 [Source:MGI Symbol | 0 | 0 | 0 | 0 | 0 | 0 |
| 47252 | ENSMUSG00000109485 | 1.129798215 | -0.382394472 | 2.319957523 | -0.164828221 | 0.86907919 | NA | Gm45016 | predicted gene 45016 [Source:MGI Symbo | 0.8031 | 3.0838 | 0 | 2.8919 | 0 | 0 |
| 47253 | ENSMUSG00000109487 | 0 | NA | NA | NA | NA | NA | Olfr1298 | olfactory receptor 1298 [Source:MGI Sym | 0 | 0 | 0 | 0 | 0 | 0 |
| 47254 | ENSMUSG00000109488 | 0 | NA | NA | NA | NA | NA | Gm6828 | predicted gene 6828 [Source:MGI Symbol | 0 | 0 | 0 | 0 | 0 | 0 |
| 47255 | ENSMUSG00000109489 | 1.133230463 | -1.272694062 | 2.134229246 | -0.596324909 | 0.550958194 | NA | Gm7451 | predicted gene 7451 [Source:MGI Symbol | 0.8031 | 2.0558 | 2.0125 | 1.928 | 0 | 0 |
| 47256 | ENSMUSG00000109490 | 0 | NA | NA | NA | NA | NA | Gm19126 | predicted gene, 19126 [Source:MGI Symb | 0 | 0 | 0 | 0 | 0 | 0 |
| 47257 | ENSMUSG00000109491 | 0 | NA | NA | NA | NA | NA | Gm15396 | predicted gene 15396 [Source:MGI Symbo | 0 | 0 | 0 | 0 | 0 | 0 |
| 47258 | ENSMUSG00000109492 | 0 | NA | NA | NA | NA | NA | Gm33882 | predicted gene, 33882 [Source:MGI Symb | 0 | 0 | 0 | 0 | 0 | 0 |
| 47259 | ENSMUSG00000109493 | 0 | NA | NA | NA | NA | NA | Gm45208 | predicted gene 45208 [Source:MGI Symbo | 0 | 0 | 0 | 0 | 0 | 0 |
| 47260 | ENSMUSG00000109494 | 0 | NA | NA | NA | NA | NA | Gm21033 | predicted gene, 21033 [Source:MGI Symb | 0 | 0 | 0 | 0 | 0 | 0 |

| | | | | | | | | | | | | | | | |
|---|---|---|---|---|---|---|---|---|---|---|---|---|---|---|---|
| 47261 | ENSMUSG00000109495 | 0 | NA | NA | NA | NA | NA | Gm47692 | predicted gene, 47692 [Source:MGI Symb | 0 | 0 | 0 | 0 | 0 | 0 |
| 47262 | ENSMUSG00000109496 | 0 | NA | NA | NA | NA | NA | Olfr1485-ps1 | olfactory receptor 1485, pseudogene 1 [S | 0 | 0 | 0 | 0 | 0 | 0 |
| 47263 | ENSMUSG00000109497 | 0 | NA | NA | NA | NA | NA | Olfr492 | olfactory receptor 492 [Source:MGI Symb | 0 | 0 | 0 | 0 | 0 | 0 |
| 47264 | ENSMUSG00000109498 | 4.062971503 | -1.565159086 | 1.378888478 | -1.135087508 | 0.256338655 | NA | Gm45222 | predicted gene 45222 [Source:MGI Symbo | 12.047 | 6.1675 | 0 | 2.8919 | 3.2719 | 0 |
| 47265 | ENSMUSG00000109499 | 0.160662974 | 1.126924175 | 4.080472857 | 0.276174898 | 0.782413736 | NA | Gm44864 | predicted gene 44864 [Source:MGI Symbo | 0 | 0 | 0 | 0.964 | 0 | 0 |
| 47266 | ENSMUSG00000109500 | 0.133850373 | -0.796629176 | 4.080472857 | -0.195229623 | 0.84521319 | NA | Gm44999 | predicted gene 44999 [Source:MGI Symbo | 0.8031 | 0 | 0 | 0 | 0 | 0 |
| 47267 | ENSMUSG00000109501 | 0 | NA | NA | NA | NA | NA | Gm14377 | predicted gene 14377 [Source:MGI Symbo | 0 | 0 | 0 | 0 | 0 | 0 |
| 47268 | ENSMUSG00000109502 | 0 | NA | NA | NA | NA | NA | Gm18672 | predicted gene, 18672 [Source:MGI Symb | 0 | 0 | 0 | 0 | 0 | 0 |
| 47269 | ENSMUSG00000109503 | 0 | NA | NA | NA | NA | NA | Gm44786 | predicted gene 44786 [Source:MGI Symbo | 0 | 0 | 0 | 0 | 0 | 0 |
| 47270 | ENSMUSG00000109504 | 0 | NA | NA | NA | NA | NA | Gm45156 | predicted gene 45156 [Source:MGI Symbo | 0 | 0 | 0 | 0 | 0 | 0 |
| 47271 | ENSMUSG00000109505 | 0 | NA | NA | NA | NA | NA | Gm44677 | predicted gene 44677 [Source:MGI Symbo | 0 | 0 | 0 | 0 | 0 | 0 |
| 47272 | ENSMUSG00000109506 | 0 | NA | NA | NA | NA | NA | Gm44724 | predicted gene 44724 [Source:MGI Symbo | 0 | 0 | 0 | 0 | 0 | 0 |
| 47273 | ENSMUSG00000109507 | 0 | NA | NA | NA | NA | NA | Gm18559 | predicted gene, 18559 [Source:MGI Symb | 0 | 0 | 0 | 0 | 0 | 0 |
| 47274 | ENSMUSG00000109508 | 0.185640647 | 1.126924175 | 4.080472857 | 0.276174898 | 0.782413736 | NA | Gm44956 | predicted gene 44956 [Source:MGI Symbo | 0 | 0 | 0 | 0 | 0 | 1.1138 |
| 47275 | ENSMUSG00000109509 | 0.839822492 | 2.066357436 | 2.600192809 | 0.794693928 | 0.426791566 | NA | Rps12-ps4 | ribosomal protein S12, pseudogene 4 [Sou | 0.8031 | 0 | 0 | 0.964 | 3.2719 | 0 |
| 47276 | ENSMUSG00000109510 | 0 | NA | NA | NA | NA | NA | Gm42417 | predicted gene, 42417 [Source:MGI Symb | 0 | 0 | 0 | 0 | 0 | 0 |
| 47277 | ENSMUSG00000109511 | 268.563702 | -0.551479041 | 0.185568298 | -2.97183866 | 0.002960222 | 0.046110081 | Nup62 | nucleoporin 62 [Source:MGI Symbol;Acc:I | 304.38 | 338.18 | 315.96 | 244.85 | 245.39 | 162.62 |
| 47278 | ENSMUSG00000109512 | 0 | NA | NA | NA | NA | NA | Gm45049 | predicted gene 45049 [Source:MGI Symbo | 0 | 0 | 0 | 0 | 0 | 0 |
| 47279 | ENSMUSG00000109513 | 0 | NA | NA | NA | NA | NA | Gm44928 | predicted gene 44928 [Source:MGI Symbo | 0 | 0 | 0 | 0 | 0 | 0 |
| 47280 | ENSMUSG00000109514 | 0 | NA | NA | NA | NA | NA | Gm19900 | predicted gene, 19900 [Source:MGI Symb | 0 | 0 | 0 | 0 | 0 | 0 |
| 47281 | ENSMUSG00000109515 | 0 | NA | NA | NA | NA | NA | Gm31343 | predicted gene, 31343 [Source:MGI Symb | 0 | 0 | 0 | 0 | 0 | 0 |
| 47282 | ENSMUSG00000109516 | 0 | NA | NA | NA | NA | NA | Gm6882 | predicted gene 6882 [Source:MGI Symbol | 0 | 0 | 0 | 0 | 0 | 0 |
| 47283 | ENSMUSG00000109517 | 0.160662974 | 1.126924175 | 4.080472857 | 0.276174898 | 0.782413736 | NA | Gm44763 | predicted gene 44763 [Source:MGI Symbo | 0 | 0 | 0 | 0.964 | 0 | 0 |
| 47284 | ENSMUSG00000109518 | 0 | NA | NA | NA | NA | NA | Gm44843 | predicted gene 44843 [Source:MGI Symbo | 0 | 0 | 0 | 0 | 0 | 0 |
| 47285 | ENSMUSG00000109519 | 0 | NA | NA | NA | NA | NA | Gm18806 | predicted gene, 18806 [Source:MGI Symb | 0 | 0 | 0 | 0 | 0 | 0 |
| 47286 | ENSMUSG00000109520 | 0 | NA | NA | NA | NA | NA | Olfr1493-ps1 | olfactory receptor 1493, pseudogene 1 [S | 0 | 0 | 0 | 0 | 0 | 0 |
| 47287 | ENSMUSG00000109521 | 0 | NA | NA | NA | NA | NA | Olfr1285 | olfactory receptor 1285 [Source:MGI Sym | 0 | 0 | 0 | 0 | 0 | 0 |
| 47288 | ENSMUSG00000109522 | 0 | NA | NA | NA | NA | NA | Olfr1319-ps1 | olfactory receptor 1319, pseudogene 1 [S | 0 | 0 | 0 | 0 | 0 | 0 |
| 47289 | ENSMUSG00000109523 | 0 | NA | NA | NA | NA | NA | Gdf1 | growth differentiation factor 1 [Source:M | 0 | 0 | 0 | 0 | 0 | 0 |
| 47290 | ENSMUSG00000109524 | 0 | NA | NA | NA | NA | NA | Gm44691 | predicted gene 44691 [Source:MGI Symbo | 0 | 0 | 0 | 0 | 0 | 0 |
| 47291 | ENSMUSG00000109525 | 0 | NA | NA | NA | NA | NA | Gm19091 | predicted gene, 19091 [Source:MGI Symb | 0 | 0 | 0 | 0 | 0 | 0 |
| 47292 | ENSMUSG00000109526 | 0.342639279 | -1.805976596 | 4.028547714 | -0.448294702 | 0.653940524 | NA | Gm9357 | predicted gene 9357 [Source:MGI Symbol | 0 | 2.0558 | 0 | 0 | 0 | 0 |
| 47293 | ENSMUSG00000109527 | 0 | NA | NA | NA | NA | NA | Olfr1492-ps1 | olfactory receptor 1492, pseudogene 1 [S | 0 | 0 | 0 | 0 | 0 | 0 |
| 47294 | ENSMUSG00000109528 | 0 | NA | NA | NA | NA | NA | Olfr1309 | olfactory receptor 1309 [Source:MGI Sym | 0 | 0 | 0 | 0 | 0 | 0 |
| 47295 | ENSMUSG00000109529 | 0 | NA | NA | NA | NA | NA | Gm44801 | predicted gene 44801 [Source:MGI Symbo | 0 | 0 | 0 | 0 | 0 | 0 |
| 47296 | ENSMUSG00000109530 | 0 | NA | NA | NA | NA | NA | Olfr1498-ps1 | olfactory receptor 1498, pseudogene 1 [S | 0 | 0 | 0 | 0 | 0 | 0 |
| 47297 | ENSMUSG00000109532 | 0 | NA | NA | NA | NA | NA | Gm42397 | predicted gene, 42397 [Source:MGI Symb | 0 | 0 | 0 | 0 | 0 | 0 |
| 47298 | ENSMUSG00000109533 | 0 | NA | NA | NA | NA | NA | Gm7394 | predicted gene 7394 [Source:MGI Symbol | 0 | 0 | 0 | 0 | 0 | 0 |
| 47299 | ENSMUSG00000109534 | 0 | NA | NA | NA | NA | NA | Gm44842 | predicted gene 44842 [Source:MGI Symbo | 0 | 0 | 0 | 0 | 0 | 0 |
| 47300 | ENSMUSG00000109535 | 0 | NA | NA | NA | NA | NA | Gm8728 | predicted gene 8728 [Source:MGI Symbol | 0 | 0 | 0 | 0 | 0 | 0 |
| 47301 | ENSMUSG00000109536 | 18.67881645 | 0.212584837 | 0.518923403 | 0.409665156 | 0.682051593 | 0.934933546 | 9330162G02 | RIKEN cDNA 9330162G02 gene [Source:M | 18.471 | 14.391 | 19.119 | 26.027 | 21.812 | 12.252 |
| 47302 | ENSMUSG00000109537 | 0 | NA | NA | NA | NA | NA | Olfr200-ps1 | olfactory receptor 200, pseudogene 1 [So | 0 | 0 | 0 | 0 | 0 | 0 |
| 47303 | ENSMUSG00000109538 | 0 | NA | NA | NA | NA | NA | Gm44910 | predicted gene 44910 [Source:MGI Symbo | 0 | 0 | 0 | 0 | 0 | 0 |
| 47304 | ENSMUSG00000109539 | 5.08875178 | 0.290233946 | 0.999370243 | 0.290416838 | 0.771497364 | 0.956965056 | Gm44667 | predicted gene 44667 [Source:MGI Symbo | 1.6062 | 9.2513 | 3.0187 | 6.7478 | 5.4531 | 4.4554 |
| 47305 | ENSMUSG00000109540 | 0 | NA | NA | NA | NA | NA | Gm44821 | predicted gene 44821 [Source:MGI Symbo | 0 | 0 | 0 | 0 | 0 | 0 |
| 47306 | ENSMUSG00000109541 | 0 | NA | NA | NA | NA | NA | Gm6112 | predicted gene 6112 [Source:MGI Symbol | 0 | 0 | 0 | 0 | 0 | 0 |
| 47307 | ENSMUSG00000109542 | 0 | NA | NA | NA | NA | NA | Olfr470 | olfactory receptor 470 [Source:MGI Symb | 0 | 0 | 0 | 0 | 0 | 0 |
| 47308 | ENSMUSG00000109543 | 0 | NA | NA | NA | NA | NA | Gm44844 | predicted gene 44844 [Source:MGI Symbo | 0 | 0 | 0 | 0 | 0 | 0 |
| 47309 | ENSMUSG00000109544 | 0 | NA | NA | NA | NA | NA | Olfr1468-ps1 | olfactory receptor 1468, pseudogene 1 [S | 0 | 0 | 0 | 0 | 0 | 0 |
| 47310 | ENSMUSG00000109545 | 0 | NA | NA | NA | NA | NA | Gm44788 | predicted gene 44788 [Source:MGI Symbo | 0 | 0 | 0 | 0 | 0 | 0 |
| 47311 | ENSMUSG00000109546 | 0 | NA | NA | NA | NA | NA | Gm18468 | predicted gene, 18468 [Source:MGI Symb | 0 | 0 | 0 | 0 | 0 | 0 |
| 47312 | ENSMUSG00000109547 | 0 | NA | NA | NA | NA | NA | Olfr1294 | olfactory receptor 1294 [Source:MGI Sym | 0 | 0 | 0 | 0 | 0 | 0 |
| 47313 | ENSMUSG00000109548 | 0 | NA | NA | NA | NA | NA | Gm45066 | predicted gene 45066 [Source:MGI Symbo | 0 | 0 | 0 | 0 | 0 | 0 |
| 47314 | ENSMUSG00000109549 | 0 | NA | NA | NA | NA | NA | Gm38941 | predicted gene, 38941 [Source:MGI Symb | 0 | 0 | 0 | 0 | 0 | 0 |
| 47315 | ENSMUSG00000109550 | 0 | NA | NA | NA | NA | NA | Gm44892 | predicted gene 44892 [Source:MGI Symbo | 0 | 0 | 0 | 0 | 0 | 0 |
| 47316 | ENSMUSG00000109551 | 0 | NA | NA | NA | NA | NA | 4930435N07 | RIKEN cDNA 4930435N07 gene [Source:M | 0 | 0 | 0 | 0 | 0 | 0 |

| | | | | | | | | | | | | | | | |
|---|---|---|---|---|---|---|---|---|---|---|---|---|---|---|---|
| 47317 | ENSMUSG00000109552 | 0 | NA | NA | NA | NA | NA | Gm44531 | predicted gene 44531 [Source:MGI Symbo | 0 | 0 | 0 | 0 | 0 | 0 |
| 47318 | ENSMUSG00000109553 | 0 | NA | NA | NA | NA | NA | Gm17992 | predicted gene, 17992 [Source:MGI Symb | 0 | 0 | 0 | 0 | 0 | 0 |
| 47319 | ENSMUSG00000109555 | 2.50362761 | -0.397726636 | 1.3461046 | -0.295464881 | 0.767638776 | NA | Gm44891 | predicted gene 44891 [Source:MGI Symbo | 2.4093 | 3.0838 | 3.0187 | 0.964 | 1.0906 | 4.4554 |
| 47320 | ENSMUSG00000109556 | 0 | NA | NA | NA | NA | NA | Gm38843 | predicted gene, 38843 [Source:MGI Symb | 0 | 0 | 0 | 0 | 0 | 0 |
| 47321 | ENSMUSG00000109557 | 0 | NA | NA | NA | NA | NA | Gm18463 | predicted gene, 18463 [Source:MGI Symb | 0 | 0 | 0 | 0 | 0 | 0 |
| 47322 | ENSMUSG00000109558 | 0 | NA | NA | NA | NA | NA | Gm45100 | predicted gene 45100 [Source:MGI Symbo | 0 | 0 | 0 | 0 | 0 | 0 |
| 47323 | ENSMUSG00000109559 | 8.948821852 | -0.318908248 | 0.744386932 | -0.428417312 | 0.668347323 | 0.930232456 | Gm34280 | predicted gene, 34280 [Source:MGI Symb | 10.44 | 14.391 | 5.0312 | 10.604 | 6.5437 | 6.6831 |
| 47324 | ENSMUSG00000109560 | 0 | NA | NA | NA | NA | NA | Gm8463 | predicted gene 8463 [Source:MGI Symbol | 0 | 0 | 0 | 0 | 0 | 0 |
| 47325 | ENSMUSG00000109561 | 0.346303622 | 2.004339653 | 3.884003423 | 0.516049919 | 0.605819544 | NA | Ankrd31 | ankyrin repeat domain 31 [Source:MGI Sy | 0 | 0 | 0 | 0.964 | 0 | 1.1138 |
| 47326 | ENSMUSG00000109562 | 0 | NA | NA | NA | NA | NA | Gm45178 | predicted gene 45178 [Source:MGI Symbo | 0 | 0 | 0 | 0 | 0 | 0 |
| 47327 | ENSMUSG00000109563 | 0 | NA | NA | NA | NA | NA | NA | NA | 0 | 0 | 0 | 0 | 0 | 0 |
| 47328 | ENSMUSG00000109564 | 16.62078279 | -0.25554368 | 0.531208838 | -0.481060671 | 0.630473379 | 0.921020101 | Muc16 | mucin 16 [Source:MGI Symbol;Acc:MGI:1 | 13.653 | 22.614 | 18.112 | 14.46 | 14.178 | 16.708 |
| 47329 | ENSMUSG00000109565 | 0 | NA | NA | NA | NA | NA | Gm6579 | predicted gene 6579 [Source:MGI Symbol | 0 | 0 | 0 | 0 | 0 | 0 |
| 47330 | ENSMUSG00000109566 | 0 | NA | NA | NA | NA | NA | Gm31401 | predicted gene, 31401 [Source:MGI Symb | 0 | 0 | 0 | 0 | 0 | 0 |
| 47331 | ENSMUSG00000109567 | 0 | NA | NA | NA | NA | NA | Gm18467 | predicted gene, 18467 [Source:MGI Symb | 0 | 0 | 0 | 0 | 0 | 0 |
| 47332 | ENSMUSG00000109568 | 0.160662974 | 1.126924175 | 4.080472857 | 0.276174898 | 0.782413736 | NA | Gm45074 | predicted gene 45074 [Source:MGI Symbo | 0 | 0 | 0 | 0.964 | 0 | 0 |
| 47333 | ENSMUSG00000109569 | 0 | NA | NA | NA | NA | NA | Gm44554 | predicted gene 44554 [Source:MGI Symbo | 0 | 0 | 0 | 0 | 0 | 0 |
| 47334 | ENSMUSG00000109570 | 1.782403855 | -0.073807118 | 1.524472841 | -0.048414846 | 0.961385628 | NA | Gm44624 | predicted gene 44624 [Source:MGI Symbo | 2.4093 | 2.0558 | 1.0062 | 1.928 | 2.1812 | 1.1138 |
| 47335 | ENSMUSG00000109571 | 0 | NA | NA | NA | NA | NA | Gm44655 | predicted gene 44655 [Source:MGI Symbo | 0 | 0 | 0 | 0 | 0 | 0 |
| 47336 | ENSMUSG00000109572 | 0 | NA | NA | NA | NA | NA | Cfap99 | cilia and flagella associated protein 99 [So | 0 | 0 | 0 | 0 | 0 | 0 |
| 47337 | ENSMUSG00000109573 | 0 | NA | NA | NA | NA | NA | Olfr1473-ps1 | olfactory receptor 1473, pseudogene 1 [S | 0 | 0 | 0 | 0 | 0 | 0 |
| 47338 | ENSMUSG00000109574 | 0 | NA | NA | NA | NA | NA | B230206I08I | RIKEN cDNA B230206I08 gene [Source:MG | 0 | 0 | 0 | 0 | 0 | 0 |
| 47339 | ENSMUSG00000109575 | 0 | NA | NA | NA | NA | NA | Gm44831 | predicted gene 44831 [Source:MGI Symbo | 0 | 0 | 0 | 0 | 0 | 0 |
| 47340 | ENSMUSG00000109576 | 0.743983808 | -0.269067535 | 2.702716736 | -0.099554471 | 0.920698041 | NA | Gm44704 | predicted gene 44704 [Source:MGI Symbo | 2.4093 | 0 | 0 | 0.964 | 1.0906 | 0 |
| 47341 | ENSMUSG00000109577 | 0 | NA | NA | NA | NA | NA | Gm44543 | predicted gene 44543 [Source:MGI Symbo | 0 | 0 | 0 | 0 | 0 | 0 |
| 47342 | ENSMUSG00000109578 | 0 | NA | NA | NA | NA | NA | Gm16475 | predicted pseudogene 16475 [Source:MG | 0 | 0 | 0 | 0 | 0 | 0 |
| 47343 | ENSMUSG00000109579 | 0 | NA | NA | NA | NA | NA | Gm35521 | predicted gene, 35521 [Source:MGI Symb | 0 | 0 | 0 | 0 | 0 | 0 |
| 47344 | ENSMUSG00000109580 | 0 | NA | NA | NA | NA | NA | Gm8268 | predicted gene 8268 [Source:MGI Symbol | 0 | 0 | 0 | 0 | 0 | 0 |
| 47345 | ENSMUSG00000109581 | 0 | NA | NA | NA | NA | NA | Gm19094 | predicted gene, 19094 [Source:MGI Symb | 0 | 0 | 0 | 0 | 0 | 0 |
| 47346 | ENSMUSG00000109582 | 0 | NA | NA | NA | NA | NA | 4933439N14 | RIKEN cDNA 4933439N14 gene [Source:M | 0 | 0 | 0 | 0 | 0 | 0 |
| 47347 | ENSMUSG00000109583 | 0 | NA | NA | NA | NA | NA | Gm39204 | predicted gene, 39204 [Source:MGI Symb | 0 | 0 | 0 | 0 | 0 | 0 |
| 47348 | ENSMUSG00000109584 | 1.313656068 | 0.977573871 | 1.997599366 | 0.48937434 | 0.624576699 | NA | Gm7997 | predicted gene 7997 [Source:MGI Symbol | 1.6062 | 0 | 1.0062 | 1.928 | 0 | 3.3415 |
| 47349 | ENSMUSG00000109585 | 0 | NA | NA | NA | NA | NA | Gm45358 | predicted gene 45358 [Source:MGI Symbo | 0 | 0 | 0 | 0 | 0 | 0 |
| 47350 | ENSMUSG00000109586 | 0 | NA | NA | NA | NA | NA | Gm6853 | predicted gene 6853 [Source:MGI Symbol | 0 | 0 | 0 | 0 | 0 | 0 |
| 47351 | ENSMUSG00000109587 | 39.07962168 | -0.226695608 | 0.392002485 | -0.578301458 | 0.56306061 | 0.900821918 | Gm31105 | predicted gene, 31105 [Source:MGI Symb | 32.124 | 49.34 | 45.281 | 38.559 | 46.897 | 22.277 |
| 47352 | ENSMUSG00000109588 | 0 | NA | NA | NA | NA | NA | Lnp1 | leukemia NUP98 fusion partner 1 [Source: | 0 | 0 | 0 | 0 | 0 | 0 |
| 47353 | ENSMUSG00000109589 | 0 | NA | NA | NA | NA | NA | Gm45267 | predicted gene 45267 [Source:MGI Symbo | 0 | 0 | 0 | 0 | 0 | 0 |
| 47354 | ENSMUSG00000109590 | 0 | NA | NA | NA | NA | NA | Gm32710 | predicted gene, 32710 [Source:MGI Symb | 0 | 0 | 0 | 0 | 0 | 0 |
| 47355 | ENSMUSG00000109591 | 0 | NA | NA | NA | NA | NA | Gm8077 | predicted gene 8077 [Source:MGI Symbol | 0 | 0 | 0 | 0 | 0 | 0 |
| 47356 | ENSMUSG00000109592 | 0 | NA | NA | NA | NA | NA | Gm45417 | predicted gene 45417 [Source:MGI Symbo | 0 | 0 | 0 | 0 | 0 | 0 |
| 47357 | ENSMUSG00000109593 | 0 | NA | NA | NA | NA | NA | Gm45431 | predicted gene 45431 [Source:MGI Symbo | 0 | 0 | 0 | 0 | 0 | 0 |
| 47358 | ENSMUSG00000109594 | 0 | NA | NA | NA | NA | NA | 1700047O18 | RIKEN cDNA 1700047O18 gene [Source:M | 0 | 0 | 0 | 0 | 0 | 0 |
| 47359 | ENSMUSG00000109595 | 0 | NA | NA | NA | NA | NA | Gm18204 | predicted gene, 18204 [Source:MGI Symb | 0 | 0 | 0 | 0 | 0 | 0 |
| 47360 | ENSMUSG00000109596 | 0 | NA | NA | NA | NA | NA | Olfr565-ps1 | olfactory receptor 565, pseudogene 1 [So | 0 | 0 | 0 | 0 | 0 | 0 |
| 47361 | ENSMUSG00000109597 | 0 | NA | NA | NA | NA | NA | Gm45492 | predicted gene 45492 [Source:MGI Symbo | 0 | 0 | 0 | 0 | 0 | 0 |
| 47362 | ENSMUSG00000109598 | 0 | NA | NA | NA | NA | NA | Gm36356 | predicted gene, 36356 [Source:MGI Symb | 0 | 0 | 0 | 0 | 0 | 0 |
| 47363 | ENSMUSG00000109599 | 16.7771418 | 0.15223199 | 0.582360382 | 0.261405127 | 0.793780104 | 0.961915077 | Gm31812 | predicted gene, 31812 [Source:MGI Symb | 24.896 | 14.391 | 8.05 | 12.532 | 19.631 | 21.163 |
| 47364 | ENSMUSG00000109600 | 0 | NA | NA | NA | NA | NA | Gm18212 | predicted gene, 18212 [Source:MGI Symb | 0 | 0 | 0 | 0 | 0 | 0 |
| 47365 | ENSMUSG00000109601 | 0 | NA | NA | NA | NA | NA | Zfp148-ps1 | zinc finger protein 148, pseudogene 1 [So | 0 | 0 | 0 | 0 | 0 | 0 |
| 47366 | ENSMUSG00000109602 | 0 | NA | NA | NA | NA | NA | Gm31597 | predicted gene, 31597 [Source:MGI Symb | 0 | 0 | 0 | 0 | 0 | 0 |
| 47367 | ENSMUSG00000109603 | 0.294513347 | 0.165143496 | 4.080472857 | 0.040471657 | 0.967717103 | NA | Gm32389 | predicted gene, 32389 [Source:MGI Symb | 0.8031 | 0 | 0 | 0.964 | 0 | 0 |
| 47368 | ENSMUSG00000109604 | 0 | NA | NA | NA | NA | NA | Gm45346 | predicted gene 45346 [Source:MGI Symbo | 0 | 0 | 0 | 0 | 0 | 0 |
| 47369 | ENSMUSG00000109605 | 0 | NA | NA | NA | NA | NA | Gm45303 | predicted gene 45303 [Source:MGI Symbo | 0 | 0 | 0 | 0 | 0 | 0 |
| 47370 | ENSMUSG00000109606 | 0 | NA | NA | NA | NA | NA | Gm45537 | predicted gene 45537 [Source:MGI Symbo | 0 | 0 | 0 | 0 | 0 | 0 |
| 47371 | ENSMUSG00000109607 | 0.465832987 | -0.698295221 | 3.18348759 | -0.219349126 | 0.826378096 | NA | Gm34533 | predicted gene, 34533 [Source:MGI Symb | 0.8031 | 1.0279 | 0 | 0.964 | 0 | 0 |
| 47372 | ENSMUSG00000109608 | 0.335414862 | -1.779047242 | 4.030804179 | -0.441362855 | 0.658950329 | NA | Gm45597 | predicted gene 45597 [Source:MGI Symbo | 0 | 0 | 2.0125 | 0 | 0 | 0 |

| | | | | | | | | | | | | | | | |
|---|---|---|---|---|---|---|---|---|---|---|---|---|---|---|---|
| 47373 | ENSMUSG00000109609 | 0 | NA | NA | NA | NA | NA | Gm4972 | predicted gene 4972 [Source:MGI Symbol | 0 | 0 | 0 | 0 | 0 | 0 |
| 47374 | ENSMUSG00000109610 | 0.535117793 | 1.111049431 | 3.044213722 | 0.364970903 | 0.715133126 | NA | Gm7432 | predicted gene 7432 [Source:MGI Symbol | 0 | 0 | 1.0062 | 0 | 1.0906 | 1.1138 |
| 47375 | ENSMUSG00000109611 | 0 | NA | NA | NA | NA | NA | Gm45473 | predicted gene 45473 [Source:MGI Symbo | 0 | 0 | 0 | 0 | 0 | 0 |
| 47376 | ENSMUSG00000109612 | 20.25051312 | 0.539948637 | 0.600820374 | 0.898685631 | 0.368820133 | 0.806506847 | Gm45253 | predicted gene 45253 [Source:MGI Symbo | 20.078 | 16.447 | 13.081 | 43.379 | 20.722 | 7.7969 |
| 47377 | ENSMUSG00000109613 | 0 | NA | NA | NA | NA | NA | Gm45405 | predicted gene 45405 [Source:MGI Symbo | 0 | 0 | 0 | 0 | 0 | 0 |
| 47378 | ENSMUSG00000109614 | 0 | NA | NA | NA | NA | NA | Gm45252 | predicted gene 45252 [Source:MGI Symbo | 0 | 0 | 0 | 0 | 0 | 0 |
| 47379 | ENSMUSG00000109615 | 0 | NA | NA | NA | NA | NA | Gm39231 | predicted gene, 39231 [Source:MGI Symb | 0 | 0 | 0 | 0 | 0 | 0 |
| 47380 | ENSMUSG00000109616 | 0 | NA | NA | NA | NA | NA | Gm45393 | predicted gene 45393 [Source:MGI Symbo | 0 | 0 | 0 | 0 | 0 | 0 |
| 47381 | ENSMUSG00000109617 | 0 | NA | NA | NA | NA | NA | Gm7669 | predicted gene 7669 [Source:MGI Symbol | 0 | 0 | 0 | 0 | 0 | 0 |
| 47382 | ENSMUSG00000109618 | 0.349477146 | 0.165142768 | 3.911021834 | 0.042224967 | 0.966319359 | NA | Gm45527 | predicted gene 45527 [Source:MGI Symbo | 0 | 0 | 1.0062 | 0 | 1.0906 | 0 |
| 47383 | ENSMUSG00000109619 | 0 | NA | NA | NA | NA | NA | Gm45672 | predicted gene 45672 [Source:MGI Symbo | 0 | 0 | 0 | 0 | 0 | 0 |
| 47384 | ENSMUSG00000109620 | 0 | NA | NA | NA | NA | NA | Gm48810 | predicted gene, 48810 [Source:MGI Symb | 0 | 0 | 0 | 0 | 0 | 0 |
| 47385 | ENSMUSG00000109621 | 0 | NA | NA | NA | NA | NA | Gm45370 | predicted gene 45370 [Source:MGI Symbo | 0 | 0 | 0 | 0 | 0 | 0 |
| 47386 | ENSMUSG00000109622 | 0 | NA | NA | NA | NA | NA | Gm45517 | predicted gene 45517 [Source:MGI Symbo | 0 | 0 | 0 | 0 | 0 | 0 |
| 47387 | ENSMUSG00000109623 | 0 | NA | NA | NA | NA | NA | Gm7562 | predicted gene 7562 [Source:MGI Symbol | 0 | 0 | 0 | 0 | 0 | 0 |
| 47388 | ENSMUSG00000109624 | 0 | NA | NA | NA | NA | NA | Saa-ps | serum amyloid A, pseudogene [Source:M | 0 | 0 | 0 | 0 | 0 | 0 |
| 47389 | ENSMUSG00000109625 | 0 | NA | NA | NA | NA | NA | 3110080E11 | RIKEN cDNA 3110080E11 gene [Source:M | 0 | 0 | 0 | 0 | 0 | 0 |
| 47390 | ENSMUSG00000109626 | 0 | NA | NA | NA | NA | NA | Gm18641 | predicted gene, 18641 [Source:MGI Symb | 0 | 0 | 0 | 0 | 0 | 0 |
| 47391 | ENSMUSG00000109627 | 0 | NA | NA | NA | NA | NA | Gm45590 | predicted gene 45590 [Source:MGI Symbo | 0 | 0 | 0 | 0 | 0 | 0 |
| 47392 | ENSMUSG00000109628 | 4693.719322 | 0.476034242 | 0.133050127 | 3.577856365 | 0.000346424 | 0.009767393 | BC024386 | cDNA sequence BC023486 [Source:MGI Sy | 3662.9 | 3574.1 | 4543.2 | 6388.3 | 5158.6 | 4835.2 |
| 47393 | ENSMUSG00000109629 | 0 | NA | NA | NA | NA | NA | Gm34099 | predicted gene, 34099 [Source:MGI Symb | 0 | 0 | 0 | 0 | 0 | 0 |
| 47394 | ENSMUSG00000109630 | 0 | NA | NA | NA | NA | NA | Gm45364 | predicted gene 45364 [Source:MGI Symbo | 0 | 0 | 0 | 0 | 0 | 0 |
| 47395 | ENSMUSG00000109631 | 0 | NA | NA | NA | NA | NA | Olfr494 | olfactory receptor 494 [Source:MGI Symb | 0 | 0 | 0 | 0 | 0 | 0 |
| 47396 | ENSMUSG00000109632 | 1.78578397 | -1.98897573 | 1.814938555 | -1.095891497 | 0.273126261 | NA | Gm31479 | predicted gene, 31479 [Source:MGI Symb | 2.4093 | 4.1117 | 2.0125 | 0 | 2.1812 | 0 |
| 47397 | ENSMUSG00000109633 | 0 | NA | NA | NA | NA | NA | C230079O03 | RIKEN cDNA C230079O03 gene [Source:M | 0 | 0 | 0 | 0 | 0 | 0 |
| 47398 | ENSMUSG00000109634 | 0.963977846 | 3.490969125 | 2.983271141 | 1.170181643 | 0.241927879 | NA | Gm45474 | predicted gene 45474 [Source:MGI Symbo | 0 | 0 | 0 | 5.7839 | 0 | 0 |
| 47399 | ENSMUSG00000109635 | 0 | NA | NA | NA | NA | NA | Gm45670 | predicted gene 45670 [Source:MGI Symbo | 0 | 0 | 0 | 0 | 0 | 0 |
| 47400 | ENSMUSG00000109636 | 0 | NA | NA | NA | NA | NA | Gm45644 | predicted gene 45644 [Source:MGI Symbo | 0 | 0 | 0 | 0 | 0 | 0 |
| 47401 | ENSMUSG00000109637 | 0 | NA | NA | NA | NA | NA | Gm31941 | predicted gene, 31941 [Source:MGI Symb | 0 | 0 | 0 | 0 | 0 | 0 |
| 47402 | ENSMUSG00000109638 | 1.455661788 | 2.064756333 | 2.132205591 | 0.968366438 | 0.332861395 | NA | Gm45630 | predicted gene 45630 [Source:MGI Symbo | 1.6062 | 0 | 0 | 3.8559 | 3.2719 | 0 |
| 47403 | ENSMUSG00000109639 | 0 | NA | NA | NA | NA | NA | Olfr489-ps1 | olfactory receptor 489, pseudogene 1 [So | 0 | 0 | 0 | 0 | 0 | 0 |
| 47404 | ENSMUSG00000109640 | 2.376587945 | -1.768006013 | 1.548310409 | -1.141893772 | 0.253498178 | NA | Gm45413 | predicted gene 45413 [Source:MGI Symbo | 4.8186 | 5.1396 | 1.0062 | 0 | 2.1812 | 1.1138 |
| 47405 | ENSMUSG00000109641 | 0 | NA | NA | NA | NA | NA | Gm45556 | predicted gene 45556 [Source:MGI Symbo | 0 | 0 | 0 | 0 | 0 | 0 |
| 47406 | ENSMUSG00000109642 | 0.638982041 | 0.404325377 | 3.14176733 | 0.128693609 | 0.897600092 | NA | Gm45289 | predicted gene 45289 [Source:MGI Symbo | 1.6062 | 0 | 0 | 0 | 0 | 2.2277 |
| 47407 | ENSMUSG00000109643 | 0 | NA | NA | NA | NA | NA | Gm31545 | predicted gene, 31545 [Source:MGI Symb | 0 | 0 | 0 | 0 | 0 | 0 |
| 47408 | ENSMUSG00000109644 | 2845.790341 | 0.17860856 | 0.094654118 | 1.886960283 | 0.059165668 | 0.346046187 | 0610005C13 | RIKEN cDNA 0610005C13 gene [Source:M | 2768.3 | 2559.5 | 2680.6 | 2873.6 | 3204.2 | 2988.4 |
| 47409 | ENSMUSG00000109645 | 0 | NA | NA | NA | NA | NA | Gm45246 | predicted gene 45246 [Source:MGI Symbo | 0 | 0 | 0 | 0 | 0 | 0 |
| 47410 | ENSMUSG00000109646 | 0 | NA | NA | NA | NA | NA | Gm45523 | predicted gene 45523 [Source:MGI Symbo | 0 | 0 | 0 | 0 | 0 | 0 |
| 47411 | ENSMUSG00000109647 | 0.506734502 | -2.372847156 | 3.338060886 | -0.710845978 | 0.477179685 | NA | Gm45510 | predicted gene 45510 [Source:MGI Symbo | 0 | 1.0279 | 2.0125 | 0 | 0 | 0 |
| 47412 | ENSMUSG00000109648 | 0 | NA | NA | NA | NA | NA | Svet1 | subventricular expressed transcript 1 [Sou | 0 | 0 | 0 | 0 | 0 | 0 |
| 47413 | ENSMUSG00000109649 | 0 | NA | NA | NA | NA | NA | Gm45521 | predicted gene 45521 [Source:MGI Symbo | 0 | 0 | 0 | 0 | 0 | 0 |
| 47414 | ENSMUSG00000109650 | 0 | NA | NA | NA | NA | NA | Gm45448 | predicted gene 45448 [Source:MGI Symbo | 0 | 0 | 0 | 0 | 0 | 0 |
| 47415 | ENSMUSG00000109651 | 0 | NA | NA | NA | NA | NA | Gm45463 | predicted gene 45463 [Source:MGI Symbo | 0 | 0 | 0 | 0 | 0 | 0 |
| 47416 | ENSMUSG00000109652 | 0.51034671 | -2.381743948 | 3.331081356 | -0.715006238 | 0.474605189 | NA | Gm45555 | predicted gene 45555 [Source:MGI Symbo | 0 | 2.0558 | 1.0062 | 0 | 0 | 0 |
| 47417 | ENSMUSG00000109653 | 0 | NA | NA | NA | NA | NA | Gm45608 | predicted gene 45608 [Source:MGI Symbo | 0 | 0 | 0 | 0 | 0 | 0 |
| 47418 | ENSMUSG00000109654 | 0 | NA | NA | NA | NA | NA | Gm3912 | predicted gene 3912 [Source:MGI Symbol | 0 | 0 | 0 | 0 | 0 | 0 |
| 47419 | ENSMUSG00000109655 | 0 | NA | NA | NA | NA | NA | Gm29735 | predicted gene, 29735 [Source:MGI Symb | 0 | 0 | 0 | 0 | 0 | 0 |
| 47420 | ENSMUSG00000109656 | 21.46611338 | -1.149795773 | 0.47839811 | -2.403428752 | 0.016242132 | 0.150497241 | Gm45548 | predicted gene 45548 [Source:MGI Symbo | 33.73 | 28.782 | 26.162 | 12.532 | 11.997 | 15.594 |
| 47421 | ENSMUSG00000109657 | 0 | NA | NA | NA | NA | NA | 1700008N11 | RIKEN cDNA 1700008N11 gene [Source:M | 0 | 0 | 0 | 0 | 0 | 0 |
| 47422 | ENSMUSG00000109658 | 0 | NA | NA | NA | NA | NA | Gm45660 | predicted gene 45660 [Source:MGI Symbo | 0 | 0 | 0 | 0 | 0 | 0 |
| 47423 | ENSMUSG00000109659 | 0 | NA | NA | NA | NA | NA | Olfr605 | olfactory receptor 605 [Source:MGI Symb | 0 | 0 | 0 | 0 | 0 | 0 |
| 47424 | ENSMUSG00000109661 | 0 | NA | NA | NA | NA | NA | NA | NA | 0 | 0 | 0 | 0 | 0 | 0 |
| 47425 | ENSMUSG00000109662 | 0 | NA | NA | NA | NA | NA | Gm45366 | predicted gene 45366 [Source:MGI Symbo | 0 | 0 | 0 | 0 | 0 | 0 |
| 47426 | ENSMUSG00000109663 | 0.17131964 | -0.796629176 | 4.080472857 | -0.195229623 | 0.84521319 | NA | Gm5331 | predicted gene 5331 [Source:MGI Symbol | 0 | 1.0279 | 0 | 0 | 0 | 0 |
| 47427 | ENSMUSG00000109664 | 0 | NA | NA | NA | NA | NA | Gm9520 | predicted gene 9520 [Source:MGI Symbol | 0 | 0 | 0 | 0 | 0 | 0 |
| 47428 | ENSMUSG00000109665 | 0 | NA | NA | NA | NA | NA | Gm45458 | predicted gene 45458 [Source:MGI Symbo | 0 | 0 | 0 | 0 | 0 | 0 |

| | | | | | | | | | | | | | | | |
|---|---|---|---|---|---|---|---|---|---|---|---|---|---|---|---|
| 47429 | ENSMUSG00000109666 | 0 | NA | NA | NA | NA | NA | Gm39154 | predicted gene, 39154 [Source:MGI Symb | 0 | 0 | 0 | 0 | 0 | 0 |
| 47430 | ENSMUSG00000109667 | 2.566260912 | -2.062535023 | 1.574975762 | -1.309566199 | 0.190342629 | NA | Gm36231 | predicted gene, 36231 [Source:MGI Symb | 6.4248 | 2.0558 | 4.025 | 2.8919 | 0 | 0 |
| 47431 | ENSMUSG00000109668 | 0 | NA | NA | NA | NA | NA | Gm8342 | predicted gene 8342 [Source:MGI Symbol | 0 | 0 | 0 | 0 | 0 | 0 |
| 47432 | ENSMUSG00000109669 | 0 | NA | NA | NA | NA | NA | Gm45472 | predicted gene 45472 [Source:MGI Symbo | 0 | 0 | 0 | 0 | 0 | 0 |
| 47433 | ENSMUSG00000109670 | 0 | NA | NA | NA | NA | NA | Gm45375 | predicted gene 45375 [Source:MGI Symbo | 0 | 0 | 0 | 0 | 0 | 0 |
| 47434 | ENSMUSG00000109671 | 0 | NA | NA | NA | NA | NA | Olfr674-ps1 | olfactory receptor 674, pseudogene 1 [So | 0 | 0 | 0 | 0 | 0 | 0 |
| 47435 | ENSMUSG00000109672 | 0 | NA | NA | NA | NA | NA | Gm45639 | predicted gene 45639 [Source:MGI Symbo | 0 | 0 | 0 | 0 | 0 | 0 |
| 47436 | ENSMUSG00000109673 | 0 | NA | NA | NA | NA | NA | Gm9712 | predicted gene 9712 [Source:MGI Symbol | 0 | 0 | 0 | 0 | 0 | 0 |
| 47437 | ENSMUSG00000109674 | 111.5069727 | 0.569136209 | 0.326004704 | 1.745791404 | 0.080847205 | 0.417633563 | Gm45470 | predicted gene 45470 [Source:MGI Symbo | 69.87 | 75.038 | 124.77 | 119.53 | 197.4 | 82.424 |
| 47438 | ENSMUSG00000109675 | 1.959371708 | -0.751859498 | 1.740144918 | -0.432067175 | 0.665692591 | NA | Nxpe1-ps | neurexophilin and PC-esterase domain fa | 3.2124 | 4.1117 | 0 | 0 | 1.0906 | 3.3415 |
| 47439 | ENSMUSG00000109676 | 0 | NA | NA | NA | NA | NA | Gm8913 | predicted gene 8913 [Source:MGI Symbol | 0 | 0 | 0 | 0 | 0 | 0 |
| 47440 | ENSMUSG00000109677 | 0 | NA | NA | NA | NA | NA | Gm45514 | predicted gene 45514 [Source:MGI Symbo | 0 | 0 | 0 | 0 | 0 | 0 |
| 47441 | ENSMUSG00000109678 | 0 | NA | NA | NA | NA | NA | Gm45626 | predicted gene 45626 [Source:MGI Symbo | 0 | 0 | 0 | 0 | 0 | 0 |
| 47442 | ENSMUSG00000109679 | 1.487425194 | 1.22639787 | 1.925715731 | 0.636853015 | 0.52422059 | NA | Gm45342 | predicted gene 45342 [Source:MGI Symbo | 1.6062 | 1.0279 | 0 | 1.928 | 4.3625 | 0 |
| 47443 | ENSMUSG00000109680 | 1.366997043 | -1.530520783 | 2.056209344 | -0.744340934 | 0.456670243 | NA | Gm32098 | predicted gene, 32098 [Source:MGI Symb | 0 | 4.1117 | 2.0125 | 0.964 | 0 | 1.1138 |
| 47444 | ENSMUSG00000109681 | 0 | NA | NA | NA | NA | NA | Gm17997 | predicted gene, 17997 [Source:MGI Symb | 0 | 0 | 0 | 0 | 0 | 0 |
| 47445 | ENSMUSG00000109682 | 0 | NA | NA | NA | NA | NA | Gm45307 | predicted gene 45307 [Source:MGI Symbo | 0 | 0 | 0 | 0 | 0 | 0 |
| 47446 | ENSMUSG00000109683 | 0 | NA | NA | NA | NA | NA | Gm7836 | predicted gene 7836 [Source:MGI Symbol | 0 | 0 | 0 | 0 | 0 | 0 |
| 47447 | ENSMUSG00000109684 | 0.160662974 | 1.126924175 | 4.080472857 | 0.276174898 | 0.782413736 | NA | Gm45745 | predicted gene 45745 [Source:MGI Symbo | 0 | 0 | 0 | 0.964 | 0 | 0 |
| 47448 | ENSMUSG00000109685 | 0.185640647 | 1.126924175 | 4.080472857 | 0.276174898 | 0.782413736 | NA | Gvin-ps1 | GTPase, very large interferon inducible, p | 0 | 0 | 0 | 0 | 0 | 1.1138 |
| 47449 | ENSMUSG00000109686 | 0 | NA | NA | NA | NA | NA | Gm45562 | predicted gene 45562 [Source:MGI Symbo | 0 | 0 | 0 | 0 | 0 | 0 |
| 47450 | ENSMUSG00000109687 | 0 | NA | NA | NA | NA | NA | Hspd1-ps2 | heat shock protein 1 (chaperonin), pseud | 0 | 0 | 0 | 0 | 0 | 0 |
| 47451 | ENSMUSG00000109688 | 0 | NA | NA | NA | NA | NA | Gm34597 | predicted gene, 34597 [Source:MGI Symb | 0 | 0 | 0 | 0 | 0 | 0 |
| 47452 | ENSMUSG00000109689 | 0 | NA | NA | NA | NA | NA | Gm45646 | predicted gene 45646 [Source:MGI Symbo | 0 | 0 | 0 | 0 | 0 | 0 |
| 47453 | ENSMUSG00000109690 | 0 | NA | NA | NA | NA | NA | Gm6376 | predicted gene 6376 [Source:MGI Symbol | 0 | 0 | 0 | 0 | 0 | 0 |
| 47454 | ENSMUSG00000109691 | 0 | NA | NA | NA | NA | NA | Gm8679 | predicted gene 8679 [Source:MGI Symbol | 0 | 0 | 0 | 0 | 0 | 0 |
| 47455 | ENSMUSG00000109692 | 0 | NA | NA | NA | NA | NA | Gm8575 | predicted gene 8575 [Source:MGI Symbol | 0 | 0 | 0 | 0 | 0 | 0 |
| 47456 | ENSMUSG00000109693 | 0.301557804 | -1.64335916 | 4.042814367 | -0.406488899 | 0.684383412 | NA | Gm45483 | predicted gene 45483 [Source:MGI Symbo | 0.8031 | 0 | 1.0062 | 0 | 0 | 0 |
| 47457 | ENSMUSG00000109694 | 0 | NA | NA | NA | NA | NA | Gm45500 | predicted gene 45500 [Source:MGI Symbo | 0 | 0 | 0 | 0 | 0 | 0 |
| 47458 | ENSMUSG00000109695 | 3.312559978 | 0.52431993 | 1.193417215 | 0.439343361 | 0.66041276 | NA | Gm31166 | predicted gene, 31166 [Source:MGI Symb | 4.0155 | 1.0279 | 3.0187 | 1.928 | 6.5437 | 3.3415 |
| 47459 | ENSMUSG00000109696 | 0 | NA | NA | NA | NA | NA | Gm36670 | predicted gene, 36670 [Source:MGI Symb | 0 | 0 | 0 | 0 | 0 | 0 |
| 47460 | ENSMUSG00000109697 | 2.589384195 | -1.486989513 | 1.437688838 | -1.034291617 | 0.300999855 | NA | Gm34030 | predicted gene, 34030 [Source:MGI Symb | 2.4093 | 3.0838 | 6.0375 | 2.8919 | 0 | 1.1138 |
| 47461 | ENSMUSG00000109698 | 23.73963431 | -0.529236603 | 0.445124443 | -1.188963249 | 0.234454129 | 0.682322842 | Gm45469 | predicted gene 45469 [Source:MGI Symbo | 28.109 | 26.726 | 29.181 | 15.424 | 20.722 | 22.277 |
| 47462 | ENSMUSG00000109699 | 0 | NA | NA | NA | NA | NA | Gm45277 | predicted gene 45277 [Source:MGI Symbo | 0 | 0 | 0 | 0 | 0 | 0 |
| 47463 | ENSMUSG00000109700 | 0 | NA | NA | NA | NA | NA | Gm21123 | predicted gene, 21123 [Source:MGI Symb | 0 | 0 | 0 | 0 | 0 | 0 |
| 47464 | ENSMUSG00000109701 | 0 | NA | NA | NA | NA | NA | Gm32531 | predicted gene, 32531 [Source:MGI Symb | 0 | 0 | 0 | 0 | 0 | 0 |
| 47465 | ENSMUSG00000109702 | 36.21735344 | 0.709607769 | 0.47219922 | 1.502772005 | 0.132897847 | 0.533805738 | Gm45674 | predicted gene 45674 [Source:MGI Symbo | 33.73 | 11.307 | 37.231 | 52.055 | 26.175 | 56.806 |
| 47466 | ENSMUSG00000109703 | 34.73846061 | -0.711069822 | 0.434665701 | -1.635900465 | 0.101860421 | 0.469655443 | Gm45299 | predicted gene 45299 [Source:MGI Symbo | 35.336 | 37.005 | 57.356 | 33.739 | 31.628 | 13.366 |
| 47467 | ENSMUSG00000109704 | 0 | NA | NA | NA | NA | NA | Gm38414 | predicted gene, 38414 [Source:MGI Symb | 0 | 0 | 0 | 0 | 0 | 0 |
| 47468 | ENSMUSG00000109705 | 0 | NA | NA | NA | NA | NA | Gm19096 | predicted gene, 19096 [Source:MGI Symb | 0 | 0 | 0 | 0 | 0 | 0 |
| 47469 | ENSMUSG00000109706 | 0 | NA | NA | NA | NA | NA | Fth-ps3 | ferritin heavy chain, pseudogene 3 [Sourc | 0 | 0 | 0 | 0 | 0 | 0 |
| 47470 | ENSMUSG00000109707 | 0 | NA | NA | NA | NA | NA | Gm4270 | predicted gene 4270 [Source:MGI Symbol | 0 | 0 | 0 | 0 | 0 | 0 |
| 47471 | ENSMUSG00000109708 | 2.935785455 | 1.79166172 | 1.44072103 | 1.243586845 | 0.21365166 | NA | Gm45809 | predicted gene 45809 [Source:MGI Symbo | 0 | 1.0279 | 3.0187 | 4.8199 | 7.6343 | 1.1138 |
| 47472 | ENSMUSG00000109709 | 0.371281295 | 2.09253362 | 4.031852165 | 0.519000582 | 0.603760335 | NA | Gm45290 | predicted gene 45290 [Source:MGI Symbo | 0 | 0 | 0 | 0 | 0 | 2.2277 |
| 47473 | ENSMUSG00000109710 | 0 | NA | NA | NA | NA | NA | Gm4974 | predicted gene 4974 [Source:MGI Symbol | 0 | 0 | 0 | 0 | 0 | 0 |
| 47474 | ENSMUSG00000109711 | 0 | NA | NA | NA | NA | NA | Gm45445 | predicted gene 45445 [Source:MGI Symbo | 0 | 0 | 0 | 0 | 0 | 0 |
| 47475 | ENSMUSG00000109712 | 0 | NA | NA | NA | NA | NA | Gm8667 | predicted gene 8667 [Source:MGI Symbol | 0 | 0 | 0 | 0 | 0 | 0 |
| 47476 | ENSMUSG00000109713 | 0.49081066 | -0.698504554 | 3.1295234 | -0.22319838 | 0.823381113 | NA | Pvrig | poliovirus receptor related immunoglobul | 0.8031 | 1.0279 | 0 | 0 | 0 | 1.1138 |
| 47477 | ENSMUSG00000109714 | 1.928824585 | -0.840932307 | 1.631154232 | -0.515544325 | 0.606172704 | NA | Gm35021 | predicted gene, 35021 [Source:MGI Symb | 2.4093 | 0 | 5.0312 | 1.928 | 1.0906 | 1.1138 |
| 47478 | ENSMUSG00000109715 | 4.970948449 | -0.870842223 | 0.953547377 | -0.913265815 | 0.361102757 | NA | Gm45606 | predicted gene 45606 [Source:MGI Symbo | 8.031 | 6.1675 | 5.0312 | 2.8919 | 4.3625 | 3.3415 |
| 47479 | ENSMUSG00000109716 | 0 | NA | NA | NA | NA | NA | Gm42196 | predicted gene, 42196 [Source:MGI Symb | 0 | 0 | 0 | 0 | 0 | 0 |
| 47480 | ENSMUSG00000109717 | 0.476489653 | -2.293616034 | 3.421506946 | -0.670352587 | 0.502633051 | NA | Gm34368 | predicted gene, 34368 [Source:MGI Symb | 0.8031 | 2.0558 | 0 | 0 | 0 | 0 |
| 47481 | ENSMUSG00000109718 | 0 | NA | NA | NA | NA | NA | Trim61 | tripartite motif-containing 61 [Source:MG | 0 | 0 | 0 | 0 | 0 | 0 |
| 47482 | ENSMUSG00000109719 | 0 | NA | NA | NA | NA | NA | Gm45266 | predicted gene 45266 [Source:MGI Symbo | 0 | 0 | 0 | 0 | 0 | 0 |
| 47483 | ENSMUSG00000109720 | 0 | NA | NA | NA | NA | NA | Gm18639 | predicted gene, 18639 [Source:MGI Symb | 0 | 0 | 0 | 0 | 0 | 0 |
| 47484 | ENSMUSG00000109721 | 0.267700746 | -1.497481649 | 4.057001436 | -0.369110456 | 0.712045396 | NA | Gm45280 | predicted gene 45280 [Source:MGI Symbo | 1.6062 | 0 | 0 | 0 | 0 | 0 |

| | | | | | | | | | | | | | | | |
|---|---|---|---|---|---|---|---|---|---|---|---|---|---|---|---|
| 47485 | ENSMUSG00000109722 | 0 | NA | NA | NA | NA | NA | Gm10578 | predicted gene 10578 [Source:MGI Symbo | 0 | 0 | 0 | 0 | 0 | 0 |
| 47486 | ENSMUSG00000109723 | 0 | NA | NA | NA | NA | NA | Gm5347 | predicted gene 5347 [Source:MGI Symbol | 0 | 0 | 0 | 0 | 0 | 0 |
| 47487 | ENSMUSG00000109724 | 8.888273449 | -2.615255732 | 1.117906407 | -2.339422796 | 0.019313563 | 0.169201316 | Gm18194 | predicted gene, 18194 [Source:MGI Symb | 20.078 | 25.698 | 0 | 0.964 | 4.3625 | 2.2277 |
| 47488 | ENSMUSG00000109725 | 0 | NA | NA | NA | NA | NA | Gm45265 | predicted gene 45265 [Source:MGI Symbo | 0 | 0 | 0 | 0 | 0 | 0 |
| 47489 | ENSMUSG00000109726 | 0 | NA | NA | NA | NA | NA | Gm45638 | predicted gene 45638 [Source:MGI Symbo | 0 | 0 | 0 | 0 | 0 | 0 |
| 47490 | ENSMUSG00000109727 | 3.960627994 | -0.028708327 | 1.059277488 | -0.027101801 | 0.978378539 | NA | Gm45464 | predicted gene 45464 [Source:MGI Symbo | 4.8186 | 3.0838 | 4.025 | 1.928 | 5.4531 | 4.4554 |
| 47491 | ENSMUSG00000109728 | 0.545309144 | 2.651974629 | 3.666219369 | 0.72335405 | 0.469462394 | NA | Gm45359 | predicted gene 45359 [Source:MGI Symbo | 0 | 0 | 0 | 0 | 3.2719 | 0 |
| 47492 | ENSMUSG00000109729 | 0.545309144 | 2.651974629 | 3.666219369 | 0.72335405 | 0.469462394 | NA | Gm45418 | predicted gene 45418 [Source:MGI Symbo | 0 | 0 | 0 | 0 | 3.2719 | 0 |
| 47493 | ENSMUSG00000109730 | 0 | NA | NA | NA | NA | NA | Gm7151 | predicted gene 7151 [Source:MGI Symbol | 0 | 0 | 0 | 0 | 0 | 0 |
| 47494 | ENSMUSG00000109731 | 19.1358343 | -0.698954304 | 0.578475503 | -1.208269495 | 0.226943618 | 0.67343689 | Gm45425 | predicted gene 45425 [Source:MGI Symbo | 27.305 | 12.335 | 31.194 | 7.7118 | 22.903 | 13.366 |
| 47495 | ENSMUSG00000109732 | 0 | NA | NA | NA | NA | NA | Gm45285 | predicted gene 45285 [Source:MGI Symbo | 0 | 0 | 0 | 0 | 0 | 0 |
| 47496 | ENSMUSG00000109733 | 0 | NA | NA | NA | NA | NA | Gm33940 | predicted gene, 33940 [Source:MGI Symb | 0 | 0 | 0 | 0 | 0 | 0 |
| 47497 | ENSMUSG00000109734 | 0 | NA | NA | NA | NA | NA | Gm45272 | predicted gene 45272 [Source:MGI Symbo | 0 | 0 | 0 | 0 | 0 | 0 |
| 47498 | ENSMUSG00000109735 | 0 | NA | NA | NA | NA | NA | Gm45667 | predicted gene 45667 [Source:MGI Symbo | 0 | 0 | 0 | 0 | 0 | 0 |
| 47499 | ENSMUSG00000109736 | 0 | NA | NA | NA | NA | NA | Gm8734 | predicted gene 8734 [Source:MGI Symbol | 0 | 0 | 0 | 0 | 0 | 0 |
| 47500 | ENSMUSG00000109737 | 0.401551119 | -2.072982726 | 4.008251979 | -0.517178744 | 0.605031388 | NA | Gm39566 | predicted gene, 39566 [Source:MGI Symb | 2.4093 | 0 | 0 | 0 | 0 | 0 |
| 47501 | ENSMUSG00000109738 | 0 | NA | NA | NA | NA | NA | Gm33096 | predicted gene, 33096 [Source:MGI Symb | 0 | 0 | 0 | 0 | 0 | 0 |
| 47502 | ENSMUSG00000109739 | 0.321325949 | 1.911575114 | 4.048267893 | 0.472195804 | 0.636787034 | NA | Gm17949 | predicted gene, 17949 [Source:MGI Symb | 0 | 0 | 0 | 1.928 | 0 | 0 |
| 47503 | ENSMUSG00000109740 | 0 | NA | NA | NA | NA | NA | Gm7561 | predicted gene 7561 [Source:MGI Symbol | 0 | 0 | 0 | 0 | 0 | 0 |
| 47504 | ENSMUSG00000109741 | 0 | NA | NA | NA | NA | NA | Gm45455 | predicted gene 45455 [Source:MGI Symbo | 0 | 0 | 0 | 0 | 0 | 0 |
| 47505 | ENSMUSG00000109742 | 0 | NA | NA | NA | NA | NA | Gm45558 | predicted gene 45558 [Source:MGI Symbo | 0 | 0 | 0 | 0 | 0 | 0 |
| 47506 | ENSMUSG00000109743 | 0 | NA | NA | NA | NA | NA | Gm45255 | predicted gene 45255 [Source:MGI Symbo | 0 | 0 | 0 | 0 | 0 | 0 |
| 47507 | ENSMUSG00000109744 | 0 | NA | NA | NA | NA | NA | Gm45461 | predicted gene 45461 [Source:MGI Symbo | 0 | 0 | 0 | 0 | 0 | 0 |
| 47508 | ENSMUSG00000109745 | 0 | NA | NA | NA | NA | NA | 1700018G05 | RIKEN cDNA 1700018G05 gene [Source:M | 0 | 0 | 0 | 0 | 0 | 0 |
| 47509 | ENSMUSG00000109746 | 0 | NA | NA | NA | NA | NA | Gm7499 | predicted gene 7499 [Source:MGI Symbol | 0 | 0 | 0 | 0 | 0 | 0 |
| 47510 | ENSMUSG00000109747 | 0 | NA | NA | NA | NA | NA | Gm45438 | predicted gene 45438 [Source:MGI Symbo | 0 | 0 | 0 | 0 | 0 | 0 |
| 47511 | ENSMUSG00000109748 | 0 | NA | NA | NA | NA | NA | Gm35392 | predicted gene, 35392 [Source:MGI Symb | 0 | 0 | 0 | 0 | 0 | 0 |
| 47512 | ENSMUSG00000109749 | 0 | NA | NA | NA | NA | NA | Gm18907 | predicted gene, 18907 [Source:MGI Symb | 0 | 0 | 0 | 0 | 0 | 0 |
| 47513 | ENSMUSG00000109750 | 0 | NA | NA | NA | NA | NA | Hmgb1-rs17 | high mobility group box 1, related sequen | 0 | 0 | 0 | 0 | 0 | 0 |
| 47514 | ENSMUSG00000109751 | 1.20979182 | 1.422832619 | 2.087243498 | 0.681680226 | 0.495441174 | NA | Gm45479 | predicted gene 45479 [Source:MGI Symbo | 0 | 0 | 2.0125 | 1.928 | 1.0906 | 2.2277 |
| 47515 | ENSMUSG00000109752 | 0 | NA | NA | NA | NA | NA | Gm18597 | predicted gene, 18597 [Source:MGI Symb | 0 | 0 | 0 | 0 | 0 | 0 |
| 47516 | ENSMUSG00000109753 | 0 | NA | NA | NA | NA | NA | Gm45633 | predicted gene 45633 [Source:MGI Symbo | 0 | 0 | 0 | 0 | 0 | 0 |
| 47517 | ENSMUSG00000109754 | 0.705972119 | 3.02851815 | 2.966479847 | 1.020913104 | 0.30729561 | NA | Gm39214 | predicted gene, 39214 [Source:MGI Symb | 0 | 0 | 0 | 0.964 | 3.2719 | 0 |
| 47518 | ENSMUSG00000109755 | 0 | NA | NA | NA | NA | NA | Gm8489 | predicted gene 8489 [Source:MGI Symbol | 0 | 0 | 0 | 0 | 0 | 0 |
| 47519 | ENSMUSG00000109756 | 0 | NA | NA | NA | NA | NA | Gm5339 | predicted gene 5339 [Source:MGI Symbol | 0 | 0 | 0 | 0 | 0 | 0 |
| 47520 | ENSMUSG00000109757 | 0 | NA | NA | NA | NA | NA | Gm7128 | predicted gene 7128 [Source:MGI Symbol | 0 | 0 | 0 | 0 | 0 | 0 |
| 47521 | ENSMUSG00000109758 | 0 | NA | NA | NA | NA | NA | Gm18854 | predicted gene, 18854 [Source:MGI Symb | 0 | 0 | 0 | 0 | 0 | 0 |
| 47522 | ENSMUSG00000109759 | 1.506366148 | -0.908924023 | 1.954963621 | -0.464931426 | 0.641980565 | NA | 1700011L03I | RIKEN cDNA 1700011L03 gene [Source:M( | 4.8186 | 1.0279 | 0 | 0.964 | 0 | 2.2277 |
| 47523 | ENSMUSG00000109760 | 0 | NA | NA | NA | NA | NA | Gm6503 | predicted gene 6503 [Source:MGI Symbol | 0 | 0 | 0 | 0 | 0 | 0 |
| 47524 | ENSMUSG00000109761 | 0 | NA | NA | NA | NA | NA | 5430403N17 | RIKEN cDNA 5430403N17 gene [Source:M | 0 | 0 | 0 | 0 | 0 | 0 |
| 47525 | ENSMUSG00000109762 | 0 | NA | NA | NA | NA | NA | Gm34779 | predicted gene, 34779 [Source:MGI Symb | 0 | 0 | 0 | 0 | 0 | 0 |
| 47526 | ENSMUSG00000109763 | 0 | NA | NA | NA | NA | NA | Gm18124 | predicted gene, 18124 [Source:MGI Symb | 0 | 0 | 0 | 0 | 0 | 0 |
| 47527 | ENSMUSG00000109764 | 3650.707743 | 0.407964354 | 0.097023823 | 4.204785392 | 2.61E-05 | 0.001207628 | Klkb1 | kallikrein B, plasma 1 [Source:MGI Symbo | 3136.9 | 3064.2 | 3211.9 | 3801.9 | 4438.8 | 4250.4 |
| 47528 | ENSMUSG00000109765 | 0 | NA | NA | NA | NA | NA | Gm18227 | predicted gene, 18227 [Source:MGI Symb | 0 | 0 | 0 | 0 | 0 | 0 |
| 47529 | ENSMUSG00000109766 | 0 | NA | NA | NA | NA | NA | Gm45415 | predicted gene 45415 [Source:MGI Symbo | 0 | 0 | 0 | 0 | 0 | 0 |
| 47530 | ENSMUSG00000109767 | 0 | NA | NA | NA | NA | NA | Gm45460 | predicted gene 45460 [Source:MGI Symbo | 0 | 0 | 0 | 0 | 0 | 0 |
| 47531 | ENSMUSG00000109768 | 0.305170013 | -1.657836477 | 4.041480335 | -0.410205257 | 0.681655385 | NA | Gm45446 | predicted gene 45446 [Source:MGI Symbo | 0.8031 | 1.0279 | 0 | 0 | 0 | 0 |
| 47532 | ENSMUSG00000109769 | 0 | NA | NA | NA | NA | NA | Gm39232 | predicted gene, 39232 [Source:MGI Symb | 0 | 0 | 0 | 0 | 0 | 0 |
| 47533 | ENSMUSG00000109770 | 0 | NA | NA | NA | NA | NA | Gm30085 | predicted gene, 30085 [Source:MGI Symb | 0 | 0 | 0 | 0 | 0 | 0 |
| 47534 | ENSMUSG00000109771 | 69.68738638 | 0.588375985 | 0.293822848 | 2.002485472 | 0.045232544 | 0.294174243 | Gm35315 | predicted gene, 35315 [Source:MGI Symb | 51.399 | 56.535 | 59.368 | 92.542 | 99.246 | 59.034 |
| 47535 | ENSMUSG00000109772 | 0 | NA | NA | NA | NA | NA | Gm39132 | predicted gene, 39132 [Source:MGI Symb | 0 | 0 | 0 | 0 | 0 | 0 |
| 47536 | ENSMUSG00000109773 | 6.707572347 | 0.322227349 | 0.836110747 | 0.385388359 | 0.699949706 | 0.941372511 | Gm34474 | predicted gene, 34474 [Source:MGI Symb | 5.6217 | 8.2233 | 4.025 | 5.7839 | 5.4531 | 11.138 |
| 47537 | ENSMUSG00000109774 | 0 | NA | NA | NA | NA | NA | Gm45602 | predicted gene 45602 [Source:MGI Symbo | 0 | 0 | 0 | 0 | 0 | 0 |
| 47538 | ENSMUSG00000109775 | 0 | NA | NA | NA | NA | NA | Gm45297 | predicted gene 45297 [Source:MGI Symbo | 0 | 0 | 0 | 0 | 0 | 0 |
| 47539 | ENSMUSG00000109776 | 0.36353943 | 2.065989254 | 4.034137301 | 0.512126658 | 0.608562368 | NA | Gm30400 | predicted gene, 30400 [Source:MGI Symb | 0 | 0 | 0 | 0 | 2.1812 | 0 |
| 47540 | ENSMUSG00000109777 | 0 | NA | NA | NA | NA | NA | Gm45384 | predicted gene 45384 [Source:MGI Symbo | 0 | 0 | 0 | 0 | 0 | 0 |

| | | | | | | | | | | | | | | | |
|---|---|---|---|---|---|---|---|---|---|---|---|---|---|---|---|
| 47541 | ENSMUSG00000109778 | 1.010263336 | 1.395395317 | 2.689636066 | 0.518804508 | 0.603897074 | NA | Gm45504 | predicted gene 45504 [Source:MGI Symbo | 1.6062 | 0 | 0 | 0 | 0 | 4.4554 |
| 47542 | ENSMUSG00000109779 | 0 | NA | NA | NA | NA | NA | Gm18780 | predicted gene, 18780 [Source:MGI Symb | 0 | 0 | 0 | 0 | 0 | 0 |
| 47543 | ENSMUSG00000109780 | 0.17131964 | -0.796629176 | 4.080472857 | -0.195229623 | 0.84521319 | NA | Gm45447 | predicted gene 45447 [Source:MGI Symbo | 0 | 1.0279 | 0 | 0 | 0 | 0 |
| 47544 | ENSMUSG00000109781 | 0.524408994 | -0.837196744 | 3.377823722 | -0.247850928 | 0.804249749 | NA | Gm45509 | predicted gene 45509 [Source:MGI Symbo | 0 | 2.0558 | 0 | 0 | 1.0906 | 0 |
| 47545 | ENSMUSG00000109782 | 0 | NA | NA | NA | NA | NA | NA | NA | 0 | 0 | 0 | 0 | 0 | 0 |
| 47546 | ENSMUSG00000109783 | 93.41351073 | 0.561170078 | 0.259836708 | 2.159702845 | 0.030795681 | 0.232721383 | Gm45338 | predicted gene 45338 [Source:MGI Symbo | 74.689 | 75.038 | 76.475 | 93.506 | 95.974 | 144.8 |
| 47547 | ENSMUSG00000109784 | 2.327020804 | -3.491706213 | 1.792615175 | -1.947828102 | 0.051435535 | NA | Gm45493 | predicted gene 45493 [Source:MGI Symbo | 1.6062 | 8.2233 | 3.0187 | 0 | 0 | 1.1138 |
| 47548 | ENSMUSG00000109785 | 0 | NA | NA | NA | NA | NA | Gm45542 | predicted gene 45542 [Source:MGI Symbo | 0 | 0 | 0 | 0 | 0 | 0 |
| 47549 | ENSMUSG00000109786 | 0 | NA | NA | NA | NA | NA | Gm7839 | predicted gene 7839 [Source:MGI Symbol | 0 | 0 | 0 | 0 | 0 | 0 |
| 47550 | ENSMUSG00000109787 | 5.29012914 | 0.18697027 | 0.913078614 | 0.204769083 | 0.837752544 | 0.971073158 | Gm45286 | predicted gene 45286 [Source:MGI Symbo | 5.6217 | 6.1675 | 3.0187 | 4.8199 | 6.5437 | 5.5692 |
| 47551 | ENSMUSG00000109788 | 0 | NA | NA | NA | NA | NA | Olfr471-ps1 | olfactory receptor 471, pseudogene 1 [So | 0 | 0 | 0 | 0 | 0 | 0 |
| 47552 | ENSMUSG00000109789 | 0 | NA | NA | NA | NA | NA | Gm2033 | predicted gene 2033 [Source:MGI Symbol | 0 | 0 | 0 | 0 | 0 | 0 |
| 47553 | ENSMUSG00000109790 | 0.319491021 | 0.16514334 | 4.044653141 | 0.040830038 | 0.967431393 | NA | Gm45594 | predicted gene 45594 [Source:MGI Symbo | 0.8031 | 0 | 0 | 0 | 0 | 1.1138 |
| 47554 | ENSMUSG00000109791 | 0 | NA | NA | NA | NA | NA | Gm45636 | predicted gene 45636 [Source:MGI Symbo | 0 | 0 | 0 | 0 | 0 | 0 |
| 47555 | ENSMUSG00000109792 | 0 | NA | NA | NA | NA | NA | Gm45426 | predicted gene 45426 [Source:MGI Symbo | 0 | 0 | 0 | 0 | 0 | 0 |
| 47556 | ENSMUSG00000109793 | 0.17131964 | -0.796629176 | 4.080472857 | -0.195229623 | 0.84521319 | NA | Gm10043 | predicted pseudogene 10043 [Source:MG | 0 | 1.0279 | 0 | 0 | 0 | 0 |
| 47557 | ENSMUSG00000109794 | 0 | NA | NA | NA | NA | NA | Gm45571 | predicted gene 45571 [Source:MGI Symbo | 0 | 0 | 0 | 0 | 0 | 0 |
| 47558 | ENSMUSG00000109795 | 0 | NA | NA | NA | NA | NA | Gm45522 | predicted gene 45522 [Source:MGI Symbo | 0 | 0 | 0 | 0 | 0 | 0 |
| 47559 | ENSMUSG00000109796 | 0 | NA | NA | NA | NA | NA | Gm6850 | predicted gene 6850 [Source:MGI Symbol | 0 | 0 | 0 | 0 | 0 | 0 |
| 47560 | ENSMUSG00000109797 | 0 | NA | NA | NA | NA | NA | Stambp-ps2 | STAM binding protein, pseudogene 2 [Sou | 0 | 0 | 0 | 0 | 0 | 0 |
| 47561 | ENSMUSG00000109798 | 0 | NA | NA | NA | NA | NA | Gm5996 | predicted gene 5996 [Source:MGI Symbol | 0 | 0 | 0 | 0 | 0 | 0 |
| 47562 | ENSMUSG00000109799 | 1.977240259 | 1.000347539 | 1.966054522 | 0.508809663 | 0.610885645 | NA | Gm45515 | predicted gene 45515 [Source:MGI Symbo | 0 | 0 | 4.025 | 6.7478 | 1.0906 | 0 |
| 47563 | ENSMUSG00000109800 | 0 | NA | NA | NA | NA | NA | Gm5130 | predicted gene 5130 [Source:MGI Symbol | 0 | 0 | 0 | 0 | 0 | 0 |
| 47564 | ENSMUSG00000109801 | 0 | NA | NA | NA | NA | NA | Olfr1316 | olfactory receptor 1316 [Source:MGI Sym | 0 | 0 | 0 | 0 | 0 | 0 |
| 47565 | ENSMUSG00000109802 | 0 | NA | NA | NA | NA | NA | Gm30808 | predicted gene, 30808 [Source:MGI Symb | 0 | 0 | 0 | 0 | 0 | 0 |
| 47566 | ENSMUSG00000109803 | 0 | NA | NA | NA | NA | NA | Gm45604 | predicted gene 45604 [Source:MGI Symbo | 0 | 0 | 0 | 0 | 0 | 0 |
| 47567 | ENSMUSG00000109805 | 0 | NA | NA | NA | NA | NA | Gm45886 | predicted gene 45886 [Source:MGI Symbo | 0 | 0 | 0 | 0 | 0 | 0 |
| 47568 | ENSMUSG00000109806 | 0 | NA | NA | NA | NA | NA | Olfr663 | olfactory receptor 663 [Source:MGI Symb | 0 | 0 | 0 | 0 | 0 | 0 |
| 47569 | ENSMUSG00000109807 | 6.635158604 | 0.153368884 | 0.971857562 | 0.157810044 | 0.874606484 | 0.980188144 | Gm45244 | predicted gene 45244 [Source:MGI Symbo | 4.8186 | 5.1396 | 9.0562 | 16.388 | 2.1812 | 2.2277 |
| 47570 | ENSMUSG00000109808 | 0 | NA | NA | NA | NA | NA | 5430402P08 | RIKEN cDNA 5430402P08 gene [Source:M | 0 | 0 | 0 | 0 | 0 | 0 |
| 47571 | ENSMUSG00000109809 | 0 | NA | NA | NA | NA | NA | Gm34066 | predicted gene, 34066 [Source:MGI Symb | 0 | 0 | 0 | 0 | 0 | 0 |
| 47572 | ENSMUSG00000109810 | 0.476283062 | 1.031501528 | 3.159856949 | 0.326439312 | 0.744091998 | NA | Gm45592 | predicted gene 45592 [Source:MGI Symbo | 0.8031 | 0 | 0 | 0.964 | 1.0906 | 0 |
| 47573 | ENSMUSG00000109811 | 0 | NA | NA | NA | NA | NA | Tpt1-ps4 | tumor protein, translationally-controlled, | 0 | 0 | 0 | 0 | 0 | 0 |
| 47574 | ENSMUSG00000109812 | 42.06291559 | -0.635300418 | 0.409437774 | -1.551640953 | 0.120748158 | 0.509184071 | Gm45640 | predicted gene 45640 [Source:MGI Symbo | 44.974 | 65.787 | 43.269 | 52.055 | 22.903 | 23.391 |
| 47575 | ENSMUSG00000109813 | 0.371281295 | 2.09253362 | 4.031852165 | 0.519000582 | 0.603760335 | NA | Olfr1279 | olfactory receptor 1279 [Source:MGI Sym | 0 | 0 | 0 | 0 | 0 | 2.2277 |
| 47576 | ENSMUSG00000109814 | 0.349477146 | 0.165142768 | 3.911021834 | 0.042224967 | 0.966319359 | NA | Gm45847 | predicted gene 45847 [Source:MGI Symbo | 0 | 0 | 1.0062 | 0 | 1.0906 | 0 |
| 47577 | ENSMUSG00000109815 | 0 | NA | NA | NA | NA | NA | Gm7082 | predicted gene 7082 [Source:MGI Symbol | 0 | 0 | 0 | 0 | 0 | 0 |
| 47578 | ENSMUSG00000109816 | 0.133850373 | -0.796629176 | 4.080472857 | -0.195229623 | 0.84521319 | NA | 4930483O08 | RIKEN cDNA 4930483O08 gene [Source:M | 0.8031 | 0 | 0 | 0 | 0 | 0 |
| 47579 | ENSMUSG00000109817 | 0 | NA | NA | NA | NA | NA | Gm10683 | predicted gene 10683 [Source:MGI Symbo | 0 | 0 | 0 | 0 | 0 | 0 |
| 47580 | ENSMUSG00000109818 | 0 | NA | NA | NA | NA | NA | Gm35713 | predicted gene, 35713 [Source:MGI Symb | 0 | 0 | 0 | 0 | 0 | 0 |
| 47581 | ENSMUSG00000109819 | 0 | NA | NA | NA | NA | NA | B930018H19 | RIKEN cDNA B930018H19 gene [Source:M | 0 | 0 | 0 | 0 | 0 | 0 |
| 47582 | ENSMUSG00000109820 | 0 | NA | NA | NA | NA | NA | Gm45654 | predicted gene 45654 [Source:MGI Symbo | 0 | 0 | 0 | 0 | 0 | 0 |
| 47583 | ENSMUSG00000109821 | 0 | NA | NA | NA | NA | NA | Gm6593 | predicted gene 6593 [Source:MGI Symbol | 0 | 0 | 0 | 0 | 0 | 0 |
| 47584 | ENSMUSG00000109822 | 0.353348078 | 0.165142701 | 3.895042568 | 0.042398176 | 0.966181282 | NA | Gm8556 | predicted gene 8556 [Source:MGI Symbol | 0 | 0 | 1.0062 | 0 | 0 | 1.1138 |
| 47585 | ENSMUSG00000109823 | 0 | NA | NA | NA | NA | NA | Gm45388 | predicted gene 45388 [Source:MGI Symbo | 0 | 0 | 0 | 0 | 0 | 0 |
| 47586 | ENSMUSG00000109824 | 0 | NA | NA | NA | NA | NA | Olfr643 | olfactory receptor 643 [Source:MGI Symb | 0 | 0 | 0 | 0 | 0 | 0 |
| 47587 | ENSMUSG00000109825 | 3.482895788 | -1.952432453 | 1.574381212 | -1.240126875 | 0.21492847 | NA | Gm45589 | predicted gene 45589 [Source:MGI Symbo | 14.456 | 2.0558 | 0 | 0 | 3.2719 | 1.1138 |
| 47588 | ENSMUSG00000109826 | 0 | NA | NA | NA | NA | NA | Gm45658 | predicted gene 45658 [Source:MGI Symbo | 0 | 0 | 0 | 0 | 0 | 0 |
| 47589 | ENSMUSG00000109827 | 0 | NA | NA | NA | NA | NA | Gm6213 | predicted gene 6213 [Source:MGI Symbol | 0 | 0 | 0 | 0 | 0 | 0 |
| 47590 | ENSMUSG00000109828 | 0 | NA | NA | NA | NA | NA | Gm45662 | predicted gene 45662 [Source:MGI Symbo | 0 | 0 | 0 | 0 | 0 | 0 |
| 47591 | ENSMUSG00000109829 | 4.464072172 | -0.047711041 | 1.205065562 | -0.039592071 | 0.968418349 | NA | Gm45605 | predicted gene 45605 [Source:MGI Symbo | 1.6062 | 3.0838 | 9.0562 | 8.6758 | 4.3625 | 0 |
| 47592 | ENSMUSG00000109830 | 0 | NA | NA | NA | NA | NA | Gm7965 | predicted gene 7965 [Source:MGI Symbol | 0 | 0 | 0 | 0 | 0 | 0 |
| 47593 | ENSMUSG00000109831 | 0 | NA | NA | NA | NA | NA | Gm36431 | predicted gene, 36431 [Source:MGI Symb | 0 | 0 | 0 | 0 | 0 | 0 |
| 47594 | ENSMUSG00000109832 | 0 | NA | NA | NA | NA | NA | Olfr662-ps1 | olfactory receptor 662, pseudogene 1 [So | 0 | 0 | 0 | 0 | 0 | 0 |
| 47595 | ENSMUSG00000109833 | 0 | NA | NA | NA | NA | NA | Gm40193 | predicted gene, 40193 [Source:MGI Symb | 0 | 0 | 0 | 0 | 0 | 0 |
| 47596 | ENSMUSG00000109834 | 0 | NA | NA | NA | NA | NA | Gm5650 | predicted gene 5650 [Source:MGI Symbol | 0 | 0 | 0 | 0 | 0 | 0 |

| | | | | | | | | | | | | | | | |
|---|---|---|---|---|---|---|---|---|---|---|---|---|---|---|---|
| 47597 | ENSMUSG00000109835 | 0 | NA | NA | NA | NA | NA | Olfr730 | olfactory receptor 730 [Source:MGI Symb | 0 | 0 | 0 | 0 | 0 | 0 |
| 47598 | ENSMUSG00000109836 | 86.92102847 | 0.083910609 | 0.2541551 | 0.330155125 | 0.741282752 | 0.948792385 | NA | NA | 70.673 | 93.541 | 89.556 | 99.29 | 91.612 | 76.855 |
| 47599 | ENSMUSG00000109837 | 0 | NA | NA | NA | NA | NA | Gm45655 | predicted gene 45655 [Source:MGI Symbo | 0 | 0 | 0 | 0 | 0 | 0 |
| 47600 | ENSMUSG00000109838 | 0 | NA | NA | NA | NA | NA | Gm45341 | predicted gene 45341 [Source:MGI Symbo | 0 | 0 | 0 | 0 | 0 | 0 |
| 47601 | ENSMUSG00000109839 | 0 | NA | NA | NA | NA | NA | Gm45677 | predicted gene 45677 [Source:MGI Symbo | 0 | 0 | 0 | 0 | 0 | 0 |
| 47602 | ENSMUSG00000109840 | 0 | NA | NA | NA | NA | NA | Gm45386 | predicted gene 45386 [Source:MGI Symbo | 0 | 0 | 0 | 0 | 0 | 0 |
| 47603 | ENSMUSG00000109841 | 53.24607317 | -0.653073161 | 0.342094105 | -1.909045351 | 0.056256242 | 0.335404072 | E330011O21 | RIKEN cDNA E330011O21 gene [Source:M | 63.445 | 59.619 | 72.45 | 48.199 | 51.259 | 24.505 |
| 47604 | ENSMUSG00000109842 | 0.36353943 | 2.065989254 | 4.034137301 | 0.512126658 | 0.608562368 | NA | Gm45502 | predicted gene 45502 [Source:MGI Symbo | 0 | 0 | 0 | 0 | 2.1812 | 0 |
| 47605 | ENSMUSG00000109843 | 0 | NA | NA | NA | NA | NA | Gm17841 | predicted gene, 17841 [Source:MGI Symb | 0 | 0 | 0 | 0 | 0 | 0 |
| 47606 | ENSMUSG00000109844 | 0 | NA | NA | NA | NA | NA | Gm31816 | predicted gene, 31816 [Source:MGI Symb | 0 | 0 | 0 | 0 | 0 | 0 |
| 47607 | ENSMUSG00000109845 | 0 | NA | NA | NA | NA | NA | Olfr581-ps1 | olfactory receptor 581, pseudogene 1 [So | 0 | 0 | 0 | 0 | 0 | 0 |
| 47608 | ENSMUSG00000109846 | 0 | NA | NA | NA | NA | NA | Gm45627 | predicted gene 45627 [Source:MGI Symbo | 0 | 0 | 0 | 0 | 0 | 0 |
| 47609 | ENSMUSG00000109847 | 0.17131964 | -0.796629176 | 4.080472857 | -0.195229623 | 0.84521319 | NA | Gm45278 | predicted gene 45278 [Source:MGI Symbo | 0 | 1.0279 | 0 | 0 | 0 | 0 |
| 47610 | ENSMUSG00000109848 | 0 | NA | NA | NA | NA | NA | Gm45391 | predicted gene 45391 [Source:MGI Symbo | 0 | 0 | 0 | 0 | 0 | 0 |
| 47611 | ENSMUSG00000109849 | 0.321325949 | 1.911575114 | 4.048267893 | 0.472195804 | 0.636787034 | NA | Gm45645 | predicted gene 45645 [Source:MGI Symbo | 0 | 0 | 0 | 1.928 | 0 | 0 |
| 47612 | ENSMUSG00000109850 | 0 | NA | NA | NA | NA | NA | Gm45692 | predicted gene 45692 [Source:MGI Symbo | 0 | 0 | 0 | 0 | 0 | 0 |
| 47613 | ENSMUSG00000109851 | 0 | NA | NA | NA | NA | NA | Gm39288 | predicted gene, 39288 [Source:MGI Symb | 0 | 0 | 0 | 0 | 0 | 0 |
| 47614 | ENSMUSG00000109852 | 0 | NA | NA | NA | NA | NA | Gm45360 | predicted gene 45360 [Source:MGI Symbo | 0 | 0 | 0 | 0 | 0 | 0 |
| 47615 | ENSMUSG00000109853 | 0 | NA | NA | NA | NA | NA | Gm33045 | predicted gene, 33045 [Source:MGI Symb | 0 | 0 | 0 | 0 | 0 | 0 |
| 47616 | ENSMUSG00000109854 | 0.17131964 | -0.796629176 | 4.080472857 | -0.195229623 | 0.84521319 | NA | Gm45345 | predicted gene 45345 [Source:MGI Symbo | 0 | 1.0279 | 0 | 0 | 0 | 0 |
| 47617 | ENSMUSG00000109855 | 0.321325949 | 1.911575114 | 4.048267893 | 0.472195804 | 0.636787034 | NA | Gm45613 | predicted gene 45613 [Source:MGI Symbo | 0 | 0 | 0 | 1.928 | 0 | 0 |
| 47618 | ENSMUSG00000109856 | 0 | NA | NA | NA | NA | NA | Gm19855 | predicted gene, 19855 [Source:MGI Symb | 0 | 0 | 0 | 0 | 0 | 0 |
| 47619 | ENSMUSG00000109857 | 0 | NA | NA | NA | NA | NA | Gm53058 | predicted gene 53058 [Source:MGI Symbo | 0 | 0 | 0 | 0 | 0 | 0 |
| 47620 | ENSMUSG00000109858 | 0 | NA | NA | NA | NA | NA | Pfn5 | profilin 5 [Source:MGI Symbol;Acc:MGI:5 | 0 | 0 | 0 | 0 | 0 | 0 |
| 47621 | ENSMUSG00000109859 | 0 | NA | NA | NA | NA | NA | Gm45618 | predicted gene 45618 [Source:MGI Symbo | 0 | 0 | 0 | 0 | 0 | 0 |
| 47622 | ENSMUSG00000109860 | 0 | NA | NA | NA | NA | NA | Gm17737 | predicted gene, 17737 [Source:MGI Symb | 0 | 0 | 0 | 0 | 0 | 0 |
| 47623 | ENSMUSG00000109861 | 0 | NA | NA | NA | NA | NA | 4930458B22 | RIKEN cDNA 4930458B22 gene [Source:M | 0 | 0 | 0 | 0 | 0 | 0 |
| 47624 | ENSMUSG00000109862 | 0 | NA | NA | NA | NA | NA | NA | NA | 0 | 0 | 0 | 0 | 0 | 0 |
| 47625 | ENSMUSG00000109863 | 2.000350332 | -1.437966513 | 1.645124265 | -0.874077748 | 0.382075914 | NA | Gm45643 | predicted gene 45643 [Source:MGI Symbo | 5.6217 | 2.0558 | 1.0062 | 0 | 1.0906 | 2.2277 |
| 47626 | ENSMUSG00000109864 | 3.686017547 | 0.84560308 | 1.26591201 | 0.667979349 | 0.504146775 | NA | Eid3 | EP300 interacting inhibitor of differentiati | 4.8186 | 0 | 3.0187 | 7.7118 | 5.4531 | 1.1138 |
| 47627 | ENSMUSG00000109865 | 289.8000574 | -0.028429275 | 0.160481061 | -0.177150346 | 0.859390297 | 0.975502162 | Hspa14 | heat shock protein 14 [Source:MGI Symbo | 317.23 | 269.31 | 290.8 | 284.37 | 256.3 | 320.79 |
| 47628 | ENSMUSG00000109866 | 4.394078402 | -2.362460419 | 1.290069304 | -1.831266283 | 0.067060808 | NA | Gm45714 | predicted gene 45714 [Source:MGI Symbo | 8.8341 | 4.1117 | 9.0562 | 0 | 4.3625 | 0 |
| 47629 | ENSMUSG00000109867 | 0 | NA | NA | NA | NA | NA | Gm5338 | predicted gene 5338 [Source:MGI Symbol | 0 | 0 | 0 | 0 | 0 | 0 |
| 47630 | ENSMUSG00000109868 | 0 | NA | NA | NA | NA | NA | Gm39244 | predicted gene, 39244 [Source:MGI Symb | 0 | 0 | 0 | 0 | 0 | 0 |
| 47631 | ENSMUSG00000109869 | 0 | NA | NA | NA | NA | NA | Gm39158 | predicted gene, 39158 [Source:MGI Symb | 0 | 0 | 0 | 0 | 0 | 0 |
| 47632 | ENSMUSG00000109870 | 0 | NA | NA | NA | NA | NA | Gm35850 | predicted gene, 35850 [Source:MGI Symb | 0 | 0 | 0 | 0 | 0 | 0 |
| 47633 | ENSMUSG00000109871 | 0 | NA | NA | NA | NA | NA | Gm45276 | predicted gene 45276 [Source:MGI Symbo | 0 | 0 | 0 | 0 | 0 | 0 |
| 47634 | ENSMUSG00000109872 | 0 | NA | NA | NA | NA | NA | Gm45312 | predicted gene 45312 [Source:MGI Symbo | 0 | 0 | 0 | 0 | 0 | 0 |
| 47635 | ENSMUSG00000109873 | 0.160662974 | 1.126924175 | 4.080472857 | 0.276174898 | 0.782413736 | NA | Gm45407 | predicted gene 45407 [Source:MGI Symbo | 0 | 0 | 0 | 0.964 | 0 | 0 |
| 47636 | ENSMUSG00000109874 | 0 | NA | NA | NA | NA | NA | Gm9911 | predicted gene 9911 [Source:MGI Symbol | 0 | 0 | 0 | 0 | 0 | 0 |
| 47637 | ENSMUSG00000109875 | 0 | NA | NA | NA | NA | NA | Gm6314 | predicted pseudogene 6314 [Source:MGI | 0 | 0 | 0 | 0 | 0 | 0 |
| 47638 | ENSMUSG00000109876 | 0.167707431 | -0.796629176 | 4.080472857 | -0.195229623 | 0.84521319 | NA | Gm45449 | predicted gene 45449 [Source:MGI Symbo | 0 | 0 | 1.0062 | 0 | 0 | 0 |
| 47639 | ENSMUSG00000109877 | 40.87250004 | -0.294702834 | 0.386997468 | -0.761511013 | 0.446351904 | 0.852777619 | Gm45609 | predicted gene 45609 [Source:MGI Symbo | 29.715 | 46.256 | 59.368 | 31.811 | 46.897 | 31.188 |
| 47640 | ENSMUSG00000109878 | 0 | NA | NA | NA | NA | NA | Olfr567-ps1 | olfactory receptor 567, pseudogene 1 [So | 0 | 0 | 0 | 0 | 0 | 0 |
| 47641 | ENSMUSG00000109879 | 0.328370405 | 0.16514316 | 4.003149536 | 0.041253308 | 0.967093956 | NA | Gm10997 | predicted gene 10997 [Source:MGI Symbo | 0 | 0 | 1.0062 | 0.964 | 0 | 0 |
| 47642 | ENSMUSG00000109880 | 0 | NA | NA | NA | NA | NA | Gm34096 | predicted gene, 34096 [Source:MGI Symb | 0 | 0 | 0 | 0 | 0 | 0 |
| 47643 | ENSMUSG00000109881 | 0 | NA | NA | NA | NA | NA | Gm45507 | predicted gene 45507 [Source:MGI Symbo | 0 | 0 | 0 | 0 | 0 | 0 |
| 47644 | ENSMUSG00000109882 | 0 | NA | NA | NA | NA | NA | Gm18706 | predicted gene, 18706 [Source:MGI Symb | 0 | 0 | 0 | 0 | 0 | 0 |
| 47645 | ENSMUSG00000109883 | 0 | NA | NA | NA | NA | NA | Gm18850 | predicted gene, 18850 [Source:MGI Symb | 0 | 0 | 0 | 0 | 0 | 0 |
| 47646 | ENSMUSG00000109884 | 0 | NA | NA | NA | NA | NA | Olfr490 | olfactory receptor 490 [Source:MGI Symb | 0 | 0 | 0 | 0 | 0 | 0 |
| 47647 | ENSMUSG00000109885 | 0 | NA | NA | NA | NA | NA | Gm6842 | predicted gene 6842 [Source:MGI Symbol | 0 | 0 | 0 | 0 | 0 | 0 |
| 47648 | ENSMUSG00000109886 | 0 | NA | NA | NA | NA | NA | Gm45271 | predicted gene 45271 [Source:MGI Symbo | 0 | 0 | 0 | 0 | 0 | 0 |
| 47649 | ENSMUSG00000109887 | 0 | NA | NA | NA | NA | NA | Gm28756 | predicted gene 28756 [Source:MGI Symbo | 0 | 0 | 0 | 0 | 0 | 0 |
| 47650 | ENSMUSG00000109888 | 0 | NA | NA | NA | NA | NA | Gm45600 | predicted gene 45600 [Source:MGI Symbo | 0 | 0 | 0 | 0 | 0 | 0 |
| 47651 | ENSMUSG00000109889 | 0 | NA | NA | NA | NA | NA | Gm3774 | predicted gene 3774 [Source:MGI Symbol | 0 | 0 | 0 | 0 | 0 | 0 |
| 47652 | ENSMUSG00000109890 | 0 | NA | NA | NA | NA | NA | AU023762 | expressed sequence AU023762 [Source:M | 0 | 0 | 0 | 0 | 0 | 0 |

| | | | | | | | | | | | | | | | |
|---|---|---|---|---|---|---|---|---|---|---|---|---|---|---|---|
| 47653 | ENSMUSG00000109891 | 0 | NA | NA | NA | NA | NA | Gm45339 | predicted gene 45339 [Source:MGI Symbo | 0 | 0 | 0 | 0 | 0 | 0 |
| 47654 | ENSMUSG00000109892 | 0 | NA | NA | NA | NA | NA | Olfr540-ps1 | olfactory receptor 540, pseudogene 1 [Sou | 0 | 0 | 0 | 0 | 0 | 0 |
| 47655 | ENSMUSG00000109893 | 0.17131964 | -0.796629176 | 4.080472857 | -0.195229623 | 0.84521319 | NA | Gm9655 | predicted gene 9655 [Source:MGI Symbol | 0 | 1.0279 | 0 | 0 | 0 | 0 |
| 47656 | ENSMUSG00000109894 | 0.514011053 | 1.044022102 | 3.082659575 | 0.338675769 | 0.734853997 | NA | Gm5904 | predicted gene 5904 [Source:MGI Symbol | 0 | 0 | 1.0062 | 0.964 | 0 | 1.1138 |
| 47657 | ENSMUSG00000109895 | 0 | NA | NA | NA | NA | NA | Gm2975 | predicted gene 2975 [Source:MGI Symbol | 0 | 0 | 0 | 0 | 0 | 0 |
| 47658 | ENSMUSG00000109896 | 0 | NA | NA | NA | NA | NA | Gm31409 | predicted gene, 31409 [Source:MGI Symb | 0 | 0 | 0 | 0 | 0 | 0 |
| 47659 | ENSMUSG00000109897 | 0 | NA | NA | NA | NA | NA | Gm6458 | predicted gene 6458 [Source:MGI Symbol | 0 | 0 | 0 | 0 | 0 | 0 |
| 47660 | ENSMUSG00000109898 | 0 | NA | NA | NA | NA | NA | 6330420H09 | RIKEN cDNA 6330420H09 gene [Source:M | 0 | 0 | 0 | 0 | 0 | 0 |
| 47661 | ENSMUSG00000109899 | 0 | NA | NA | NA | NA | NA | Gm6012 | predicted gene 6012 [Source:MGI Symbol | 0 | 0 | 0 | 0 | 0 | 0 |
| 47662 | ENSMUSG00000109900 | 0 | NA | NA | NA | NA | NA | Gm18638 | predicted gene, 18638 [Source:MGI Symb | 0 | 0 | 0 | 0 | 0 | 0 |
| 47663 | ENSMUSG00000109901 | 708.3171806 | -0.175525396 | 0.128480749 | -1.366161056 | 0.171888418 | 0.599537638 | Chmp1b | charged multivesicular body protein 1B [S | 774.19 | 717.49 | 762.73 | 724.91 | 694.72 | 575.86 |
| 47664 | ENSMUSG00000109902 | 0.803314871 | 3.228356836 | 3.175155518 | 1.0167555 | 0.309269756 | NA | Gm45693 | predicted gene 45693 [Source:MGI Symbo | 0 | 0 | 0 | 4.8199 | 0 | 0 |
| 47665 | ENSMUSG00000109903 | 0 | NA | NA | NA | NA | NA | Gm28710 | predicted gene 28710 [Source:MGI Symbo | 0 | 0 | 0 | 0 | 0 | 0 |
| 47666 | ENSMUSG00000109904 | 105.2450388 | -0.300118271 | 0.336659531 | -0.891459303 | 0.372682815 | 0.808168552 | Gm45819 | predicted gene 45819 [Source:MGI Symbo | 120.47 | 125.41 | 102.64 | 94.47 | 145.05 | 43.44 |
| 47667 | ENSMUSG00000109905 | 0 | NA | NA | NA | NA | NA | Gm7380 | predicted gene 7380 [Source:MGI Symbol | 0 | 0 | 0 | 0 | 0 | 0 |
| 47668 | ENSMUSG00000109906 | 0 | NA | NA | NA | NA | NA | Olfr546-ps1 | olfactory receptor 546, pseudogene 1 [Sou | 0 | 0 | 0 | 0 | 0 | 0 |
| 47669 | ENSMUSG00000109907 | 0 | NA | NA | NA | NA | NA | Gm45321 | predicted gene 45321 [Source:MGI Symbo | 0 | 0 | 0 | 0 | 0 | 0 |
| 47670 | ENSMUSG00000109909 | 0.17131964 | -0.796629176 | 4.080472857 | -0.195229623 | 0.84521319 | NA | Gm45663 | predicted gene 45663 [Source:MGI Symbo | 0 | 1.0279 | 0 | 0 | 0 | 0 |
| 47671 | ENSMUSG00000109910 | 0 | NA | NA | NA | NA | NA | Gm45240 | predicted gene 45240 [Source:MGI Symbo | 0 | 0 | 0 | 0 | 0 | 0 |
| 47672 | ENSMUSG00000109911 | 0 | NA | NA | NA | NA | NA | Gm45525 | predicted gene 45525 [Source:MGI Symbo | 0 | 0 | 0 | 0 | 0 | 0 |
| 47673 | ENSMUSG00000109912 | 0 | NA | NA | NA | NA | NA | Gm45245 | predicted gene 45245 [Source:MGI Symbo | 0 | 0 | 0 | 0 | 0 | 0 |
| 47674 | ENSMUSG00000109913 | 0 | NA | NA | NA | NA | NA | Gm30694 | predicted gene, 30694 [Source:MGI Symb | 0 | 0 | 0 | 0 | 0 | 0 |
| 47675 | ENSMUSG00000109914 | 0 | NA | NA | NA | NA | NA | Gm45498 | predicted gene 45498 [Source:MGI Symbo | 0 | 0 | 0 | 0 | 0 | 0 |
| 47676 | ENSMUSG00000109915 | 0 | NA | NA | NA | NA | NA | 7420700N18 | RIKEN cDNA 7420700N18 gene [Source:M | 0 | 0 | 0 | 0 | 0 | 0 |
| 47677 | ENSMUSG00000109916 | 0 | NA | NA | NA | NA | NA | Gm45551 | predicted gene 45551 [Source:MGI Symbo | 0 | 0 | 0 | 0 | 0 | 0 |
| 47678 | ENSMUSG00000109917 | 0 | NA | NA | NA | NA | NA | Gm45671 | predicted gene 45671 [Source:MGI Symbo | 0 | 0 | 0 | 0 | 0 | 0 |
| 47679 | ENSMUSG00000109918 | 0 | NA | NA | NA | NA | NA | Gm45374 | predicted gene 45374 [Source:MGI Symbo | 0 | 0 | 0 | 0 | 0 | 0 |
| 47680 | ENSMUSG00000109919 | 0 | NA | NA | NA | NA | NA | Gm45673 | predicted gene 45673 [Source:MGI Symbo | 0 | 0 | 0 | 0 | 0 | 0 |
| 47681 | ENSMUSG00000109920 | 0 | NA | NA | NA | NA | NA | Gm9604 | predicted gene 9604 [Source:MGI Symbol | 0 | 0 | 0 | 0 | 0 | 0 |
| 47682 | ENSMUSG00000109921 | 0 | NA | NA | NA | NA | NA | Gm33326 | predicted gene, 33326 [Source:MGI Symb | 0 | 0 | 0 | 0 | 0 | 0 |
| 47683 | ENSMUSG00000109922 | 0 | NA | NA | NA | NA | NA | Gm20948 | predicted gene, 20948 [Source:MGI Symb | 0 | 0 | 0 | 0 | 0 | 0 |
| 47684 | ENSMUSG00000109923 | 0 | NA | NA | NA | NA | NA | Gm20039 | predicted gene, 20039 [Source:MGI Symb | 0 | 0 | 0 | 0 | 0 | 0 |
| 47685 | ENSMUSG00000109924 | 0 | NA | NA | NA | NA | NA | Gm18781 | predicted gene, 18781 [Source:MGI Symb | 0 | 0 | 0 | 0 | 0 | 0 |
| 47686 | ENSMUSG00000109925 | 0 | NA | NA | NA | NA | NA | Gm45315 | predicted gene 45315 [Source:MGI Symbo | 0 | 0 | 0 | 0 | 0 | 0 |
| 47687 | ENSMUSG00000109926 | 0 | NA | NA | NA | NA | NA | Gm45808 | predicted gene 45808 [Source:MGI Symbo | 0 | 0 | 0 | 0 | 0 | 0 |
| 47688 | ENSMUSG00000109927 | 0 | NA | NA | NA | NA | NA | Gm45264 | predicted gene 45264 [Source:MGI Symbo | 0 | 0 | 0 | 0 | 0 | 0 |
| 47689 | ENSMUSG00000109928 | 0 | NA | NA | NA | NA | NA | Gm45539 | predicted gene 45539 [Source:MGI Symbo | 0 | 0 | 0 | 0 | 0 | 0 |
| 47690 | ENSMUSG00000109929 | 0 | NA | NA | NA | NA | NA | Gm45268 | predicted gene 45268 [Source:MGI Symbo | 0 | 0 | 0 | 0 | 0 | 0 |
| 47691 | ENSMUSG00000109930 | 0.167707431 | -0.796629176 | 4.080472857 | -0.195229623 | 0.84521319 | NA | Gm34730 | predicted gene, 34730 [Source:MGI Symb | 0 | 0 | 1.0062 | 0 | 0 | 0 |
| 47692 | ENSMUSG00000109931 | 9.537649863 | -0.966409232 | 0.874544235 | -1.105043283 | 0.269140835 | 0.724979259 | Gm39929 | predicted gene, 39929 [Source:MGI Symb | 23.29 | 11.307 | 3.0187 | 1.928 | 6.5437 | 11.138 |
| 47693 | ENSMUSG00000109932 | 0 | NA | NA | NA | NA | NA | Gm7319 | predicted gene 7319 [Source:MGI Symbol | 0 | 0 | 0 | 0 | 0 | 0 |
| 47694 | ENSMUSG00000109933 | 2.8375028 | 0.526046877 | 1.463527234 | 0.35943771 | 0.719267669 | NA | Gm7600 | predicted gene 7600 [Source:MGI Symbol | 0 | 3.0838 | 4.025 | 7.7118 | 1.0906 | 1.1138 |
| 47695 | ENSMUSG00000109934 | 0 | NA | NA | NA | NA | NA | Gm45392 | predicted gene 45392 [Source:MGI Symbo | 0 | 0 | 0 | 0 | 0 | 0 |
| 47696 | ENSMUSG00000109935 | 0 | NA | NA | NA | NA | NA | Gm7619 | predicted gene 7619 [Source:MGI Symbol | 0 | 0 | 0 | 0 | 0 | 0 |
| 47697 | ENSMUSG00000109936 | 0.49081066 | -0.698504554 | 3.1295234 | -0.22319838 | 0.823381113 | NA | Gm45889 | predicted gene 45889 [Source:MGI Symbo | 0.8031 | 1.0279 | 0 | 0 | 0 | 1.1138 |
| 47698 | ENSMUSG00000109937 | 0 | NA | NA | NA | NA | NA | Gm45247 | predicted gene 45247 [Source:MGI Symbo | 0 | 0 | 0 | 0 | 0 | 0 |
| 47699 | ENSMUSG00000109938 | 0 | NA | NA | NA | NA | NA | Gm6009 | predicted gene 6009 [Source:MGI Symbol | 0 | 0 | 0 | 0 | 0 | 0 |
| 47700 | ENSMUSG00000109939 | 0 | NA | NA | NA | NA | NA | Gm19232 | predicted gene, 19232 [Source:MGI Symb | 0 | 0 | 0 | 0 | 0 | 0 |
| 47701 | ENSMUSG00000109940 | 0.803314871 | 3.228356836 | 3.175155518 | 1.0167555 | 0.309269756 | NA | Gm45399 | predicted gene 45399 [Source:MGI Symbo | 0 | 0 | 0 | 4.8199 | 0 | 0 |
| 47702 | ENSMUSG00000109941 | 0 | NA | NA | NA | NA | NA | Exosc6 | exosome component 6 [Source:MGI Symb | 0 | 0 | 0 | 0 | 0 | 0 |
| 47703 | ENSMUSG00000109942 | 0 | NA | NA | NA | NA | NA | Gm45262 | predicted gene 45262 [Source:MGI Symbo | 0 | 0 | 0 | 0 | 0 | 0 |
| 47704 | ENSMUSG00000109943 | 0 | NA | NA | NA | NA | NA | Gm45610 | predicted gene 45610 [Source:MGI Symbo | 0 | 0 | 0 | 0 | 0 | 0 |
| 47705 | ENSMUSG00000109944 | 0 | NA | NA | NA | NA | NA | Gm32975 | predicted gene, 32975 [Source:MGI Symb | 0 | 0 | 0 | 0 | 0 | 0 |
| 47706 | ENSMUSG00000109945 | 0.656740811 | -0.013525372 | 3.112905782 | -0.004344934 | 0.996533255 | NA | Gm45601 | predicted gene 45601 [Source:MGI Symbo | 0 | 0 | 2.0125 | 1.928 | 0 | 0 |
| 47707 | ENSMUSG00000109946 | 15.14114254 | 0.085029914 | 0.632615054 | 0.134410198 | 0.893078218 | 0.982562372 | Brd3os | bromodomain containing 3, opposite stra | 26.502 | 9.2513 | 8.05 | 18.316 | 10.906 | 17.822 |
| 47708 | ENSMUSG00000109947 | 1.447506441 | 0.587345229 | 1.918165023 | 0.306201616 | 0.759451142 | NA | Gm45669 | predicted gene 45669 [Source:MGI Symbo | 2.4093 | 0 | 1.0062 | 1.928 | 0 | 3.3415 |

| | | | | | | | | | | | | | | | |
|---|---|---|---|---|---|---|---|---|---|---|---|---|---|---|---|
| 47709 | ENSMUSG00000109948 | 0 | NA | NA | NA | NA | NA | Gm45288 | predicted gene 45288 [Source:MGI Symbc | 0 | 0 | 0 | 0 | 0 | 0 |
| 47710 | ENSMUSG00000109949 | 0 | NA | NA | NA | NA | NA | Gm45402 | predicted gene 45402 [Source:MGI Symbc | 0 | 0 | 0 | 0 | 0 | 0 |
| 47711 | ENSMUSG00000109950 | 0 | NA | NA | NA | NA | NA | Gm45550 | predicted gene 45550 [Source:MGI Symbc | 0 | 0 | 0 | 0 | 0 | 0 |
| 47712 | ENSMUSG00000109951 | 0 | NA | NA | NA | NA | NA | Olfr601 | olfactory receptor 601 [Source:MGI Symb | 0 | 0 | 0 | 0 | 0 | 0 |
| 47713 | ENSMUSG00000109952 | 0 | NA | NA | NA | NA | NA | Gm45395 | predicted gene 45395 [Source:MGI Symbc | 0 | 0 | 0 | 0 | 0 | 0 |
| 47714 | ENSMUSG00000109953 | 3.370887749 | -0.337565752 | 1.320089764 | -0.255714241 | 0.798171498 | NA | 5430430B14 | RIKEN cDNA 5430430B14 gene [Source:M | 7.2279 | 2.0558 | 2.0125 | 6.7478 | 2.1812 | 0 |
| 47715 | ENSMUSG00000109954 | 0 | NA | NA | NA | NA | NA | Gm45371 | predicted gene 45371 [Source:MGI Symbc | 0 | 0 | 0 | 0 | 0 | 0 |
| 47716 | ENSMUSG00000109955 | 0 | NA | NA | NA | NA | NA | Gm18644 | predicted gene, 18644 [Source:MGI Symb | 0 | 0 | 0 | 0 | 0 | 0 |
| 47717 | ENSMUSG00000109956 | 1.83855588 | 2.452158623 | 1.99619886 | 1.228413999 | 0.219291596 | NA | Gm45428 | predicted gene 45428 [Source:MGI Symbc | 1.6062 | 0 | 0 | 3.8559 | 0 | 5.5692 |
| 47718 | ENSMUSG00000109957 | 1.74622158 | 2.227061855 | 1.782917768 | 1.249110809 | 0.211624548 | NA | Gm45353 | predicted gene 45353 [Source:MGI Symbc | 0.8031 | 0 | 1.0062 | 0.964 | 4.3625 | 3.3415 |
| 47719 | ENSMUSG00000109958 | 0 | NA | NA | NA | NA | NA | Gm32786 | predicted gene, 32786 [Source:MGI Symb | 0 | 0 | 0 | 0 | 0 | 0 |
| 47720 | ENSMUSG00000109959 | 0.17131964 | -0.796629176 | 4.080472857 | -0.195229623 | 0.84521319 | NA | Gm45355 | predicted gene 45355 [Source:MGI Symbc | 0 | 1.0279 | 0 | 0 | 0 | 0 |
| 47721 | ENSMUSG00000109960 | 0 | NA | NA | NA | NA | NA | Gm45575 | predicted gene 45575 [Source:MGI Symbc | 0 | 0 | 0 | 0 | 0 | 0 |
| 47722 | ENSMUSG00000109961 | 0 | NA | NA | NA | NA | NA | Gm31983 | predicted gene, 31983 [Source:MGI Symb | 0 | 0 | 0 | 0 | 0 | 0 |
| 47723 | ENSMUSG00000109962 | 0 | NA | NA | NA | NA | NA | 1700025L06I | RIKEN cDNA 1700025L06 gene [Source:M( | 0 | 0 | 0 | 0 | 0 | 0 |
| 47724 | ENSMUSG00000109963 | 0 | NA | NA | NA | NA | NA | Gm45635 | predicted gene 45635 [Source:MGI Symbc | 0 | 0 | 0 | 0 | 0 | 0 |
| 47725 | ENSMUSG00000109964 | 0 | NA | NA | NA | NA | NA | Gm45350 | predicted gene 45350 [Source:MGI Symbc | 0 | 0 | 0 | 0 | 0 | 0 |
| 47726 | ENSMUSG00000109965 | 0 | NA | NA | NA | NA | NA | Gm45274 | predicted gene 45274 [Source:MGI Symbc | 0 | 0 | 0 | 0 | 0 | 0 |
| 47727 | ENSMUSG00000109966 | 0 | NA | NA | NA | NA | NA | Gm34623 | predicted gene, 34623 [Source:MGI Symb | 0 | 0 | 0 | 0 | 0 | 0 |
| 47728 | ENSMUSG00000109967 | 0 | NA | NA | NA | NA | NA | Gm45883 | predicted gene 45883 [Source:MGI Symbc | 0 | 0 | 0 | 0 | 0 | 0 |
| 47729 | ENSMUSG00000109968 | 0 | NA | NA | NA | NA | NA | Gm33828 | predicted gene, 33828 [Source:MGI Symb | 0 | 0 | 0 | 0 | 0 | 0 |
| 47730 | ENSMUSG00000109969 | 0 | NA | NA | NA | NA | NA | Zscan4-ps3 | zinc finger and SCAN domain containing 4 | 0 | 0 | 0 | 0 | 0 | 0 |
| 47731 | ENSMUSG00000109970 | 0 | NA | NA | NA | NA | NA | Gm17995 | predicted gene, 17995 [Source:MGI Symb | 0 | 0 | 0 | 0 | 0 | 0 |
| 47732 | ENSMUSG00000109971 | 0 | NA | NA | NA | NA | NA | Gm45279 | predicted gene 45279 [Source:MGI Symbc | 0 | 0 | 0 | 0 | 0 | 0 |
| 47733 | ENSMUSG00000109972 | 0 | NA | NA | NA | NA | NA | Gm17989 | predicted gene, 17989 [Source:MGI Symb | 0 | 0 | 0 | 0 | 0 | 0 |
| 47734 | ENSMUSG00000109973 | 0 | NA | NA | NA | NA | NA | Gm45397 | predicted gene 45397 [Source:MGI Symbc | 0 | 0 | 0 | 0 | 0 | 0 |
| 47735 | ENSMUSG00000109974 | 0 | NA | NA | NA | NA | NA | Gm2221 | predicted gene 2221 [Source:MGI Symbol | 0 | 0 | 0 | 0 | 0 | 0 |
| 47736 | ENSMUSG00000109975 | 4.812373613 | -0.747462054 | 1.045142698 | -0.715177033 | 0.474499659 | NA | Gm19196 | predicted gene, 19196 [Source:MGI Symb | 8.8341 | 3.0838 | 6.0375 | 0.964 | 3.2719 | 6.6831 |
| 47737 | ENSMUSG00000109976 | 0 | NA | NA | NA | NA | NA | Gm19683 | predicted gene, 19683 [Source:MGI Symb | 0 | 0 | 0 | 0 | 0 | 0 |
| 47738 | ENSMUSG00000109977 | 0 | NA | NA | NA | NA | NA | Gm19140 | predicted gene, 19140 [Source:MGI Symb | 0 | 0 | 0 | 0 | 0 | 0 |
| 47739 | ENSMUSG00000109978 | 0 | NA | NA | NA | NA | NA | Gm45530 | predicted gene 45530 [Source:MGI Symbc | 0 | 0 | 0 | 0 | 0 | 0 |
| 47740 | ENSMUSG00000109979 | 0 | NA | NA | NA | NA | NA | Gm45596 | predicted gene 45596 [Source:MGI Symbc | 0 | 0 | 0 | 0 | 0 | 0 |
| 47741 | ENSMUSG00000109980 | 1.348883866 | -3.787569084 | 2.390514453 | -1.58441589 | 0.113099108 | NA | Gm45538 | predicted gene 45538 [Source:MGI Symbc | 0 | 2.0558 | 6.0375 | 0 | 0 | 0 |
| 47742 | ENSMUSG00000109981 | 0 | NA | NA | NA | NA | NA | Gm36849 | predicted gene, 36849 [Source:MGI Symb | 0 | 0 | 0 | 0 | 0 | 0 |
| 47743 | ENSMUSG00000109982 | 0.160662974 | 1.126924175 | 4.080472857 | 0.276174898 | 0.782413736 | NA | Gm45520 | predicted gene 45520 [Source:MGI Symbc | 0 | 0 | 0 | 0.964 | 0 | 0 |
| 47744 | ENSMUSG00000109983 | 0 | NA | NA | NA | NA | NA | Gm9685 | predicted gene 9685 [Source:MGI Symbol | 0 | 0 | 0 | 0 | 0 | 0 |
| 47745 | ENSMUSG00000109984 | 2.10486181 | 0.471133866 | 1.538103076 | 0.306308383 | 0.759369857 | NA | Gm38948 | predicted gene, 38948 [Source:MGI Symb | 3.2124 | 0 | 2.0125 | 1.928 | 4.3625 | 1.1138 |
| 47746 | ENSMUSG00000109985 | 0 | NA | NA | NA | NA | NA | Gm18640 | predicted gene, 18640 [Source:MGI Symb | 0 | 0 | 0 | 0 | 0 | 0 |
| 47747 | ENSMUSG00000109986 | 0 | NA | NA | NA | NA | NA | 4930465I24F | RIKEN cDNA 4930465I24 gene [Source:M( | 0 | 0 | 0 | 0 | 0 | 0 |
| 47748 | ENSMUSG00000109987 | 0 | NA | NA | NA | NA | NA | Gm18414 | predicted gene, 18414 [Source:MGI Symb | 0 | 0 | 0 | 0 | 0 | 0 |
| 47749 | ENSMUSG00000109988 | 0 | NA | NA | NA | NA | NA | Gm45651 | predicted gene 45651 [Source:MGI Symbc | 0 | 0 | 0 | 0 | 0 | 0 |
| 47750 | ENSMUSG00000109990 | 0 | NA | NA | NA | NA | NA | Gm45543 | predicted gene 45543 [Source:MGI Symbc | 0 | 0 | 0 | 0 | 0 | 0 |
| 47751 | ENSMUSG00000109991 | 0 | NA | NA | NA | NA | NA | Gm6284 | predicted gene 6284 [Source:MGI Symbol | 0 | 0 | 0 | 0 | 0 | 0 |
| 47752 | ENSMUSG00000109992 | 0 | NA | NA | NA | NA | NA | Gm45650 | predicted gene 45650 [Source:MGI Symbc | 0 | 0 | 0 | 0 | 0 | 0 |
| 47753 | ENSMUSG00000109993 | 0 | NA | NA | NA | NA | NA | Gm45659 | predicted gene 45659 [Source:MGI Symbc | 0 | 0 | 0 | 0 | 0 | 0 |
| 47754 | ENSMUSG00000109994 | 0 | NA | NA | NA | NA | NA | Gm45459 | predicted gene 45459 [Source:MGI Symbc | 0 | 0 | 0 | 0 | 0 | 0 |
| 47755 | ENSMUSG00000109995 | 0 | NA | NA | NA | NA | NA | B020031H02 | RIKEN cDNA B020031H02 gene [Source:M | 0 | 0 | 0 | 0 | 0 | 0 |
| 47756 | ENSMUSG00000109996 | 0 | NA | NA | NA | NA | NA | Gm31898 | predicted gene, 31898 [Source:MGI Symb | 0 | 0 | 0 | 0 | 0 | 0 |
| 47757 | ENSMUSG00000109997 | 0 | NA | NA | NA | NA | NA | 9430099M0( | RIKEN cDNA 9430099M06 gene [Source:N | 0 | 0 | 0 | 0 | 0 | 0 |
| 47758 | ENSMUSG00000109998 | 0.831672659 | -0.604339699 | 2.59720557 | -0.232688435 | 0.81600335 | NA | Gm45437 | predicted gene 45437 [Source:MGI Symbc | 0 | 2.0558 | 1.0062 | 1.928 | 0 | 0 |
| 47759 | ENSMUSG00000109999 | 0 | NA | NA | NA | NA | NA | Gm35368 | predicted gene, 35368 [Source:MGI Symb | 0 | 0 | 0 | 0 | 0 | 0 |
| 47760 | ENSMUSG00000110000 | 0 | NA | NA | NA | NA | NA | Gm33501 | predicted gene, 33501 [Source:MGI Symb | 0 | 0 | 0 | 0 | 0 | 0 |
| 47761 | ENSMUSG00000110001 | 0 | NA | NA | NA | NA | NA | C86187 | expressed sequence C86187 [Source:MGI | 0 | 0 | 0 | 0 | 0 | 0 |
| 47762 | ENSMUSG00000110002 | 0 | NA | NA | NA | NA | NA | Gm45680 | predicted gene 45680 [Source:MGI Symbc | 0 | 0 | 0 | 0 | 0 | 0 |
| 47763 | ENSMUSG00000110003 | 0 | NA | NA | NA | NA | NA | Gm45403 | predicted gene 45403 [Source:MGI Symbc | 0 | 0 | 0 | 0 | 0 | 0 |
| 47764 | ENSMUSG00000110004 | 0 | NA | NA | NA | NA | NA | Gm45440 | predicted gene 45440 [Source:MGI Symbc | 0 | 0 | 0 | 0 | 0 | 0 |

| | | | | | | | | | | | | | | | |
|---|---|---|---|---|---|---|---|---|---|---|---|---|---|---|---|
| 47765 | ENSMUSG00000110005 | 0 | NA | NA | NA | NA | NA | Gm18123 | predicted gene, 18123 [Source:MGI Symb | 0 | 0 | 0 | 0 | 0 | 0 |
| 47766 | ENSMUSG00000110006 | 0 | NA | NA | NA | NA | NA | Gm45490 | predicted gene 45490 [Source:MGI Symbo | 0 | 0 | 0 | 0 | 0 | 0 |
| 47767 | ENSMUSG00000110007 | 0 | NA | NA | NA | NA | NA | Gm45352 | predicted gene 45352 [Source:MGI Symbo | 0 | 0 | 0 | 0 | 0 | 0 |
| 47768 | ENSMUSG00000110008 | 0 | NA | NA | NA | NA | NA | Olfr560 | olfactory receptor 560 [Source:MGI Symb | 0 | 0 | 0 | 0 | 0 | 0 |
| 47769 | ENSMUSG00000110009 | 0 | NA | NA | NA | NA | NA | 4930543E12 | RIKEN cDNA 4930543E12 gene [Source:M | 0 | 0 | 0 | 0 | 0 | 0 |
| 47770 | ENSMUSG00000110010 | 2.457278959 | -0.313928103 | 1.479036305 | -0.21225179 | 0.831910605 | NA | Gm45629 | predicted gene 45629 [Source:MGI Symbo | 4.0155 | 2.0558 | 2.0125 | 0 | 1.0906 | 5.5692 |
| 47771 | ENSMUSG00000110011 | 0 | NA | NA | NA | NA | NA | Gm29817 | predicted gene, 29817 [Source:MGI Symb | 0 | 0 | 0 | 0 | 0 | 0 |
| 47772 | ENSMUSG00000110012 | 0 | NA | NA | NA | NA | NA | Olfr644 | olfactory receptor 644 [Source:MGI Symb | 0 | 0 | 0 | 0 | 0 | 0 |
| 47773 | ENSMUSG00000110013 | 0 | NA | NA | NA | NA | NA | 1190028D05 | RIKEN cDNA 1190028D05 gene [Source:M | 0 | 0 | 0 | 0 | 0 | 0 |
| 47774 | ENSMUSG00000110014 | 0 | NA | NA | NA | NA | NA | Gm5361 | predicted gene 5361 [Source:MGI Symbol | 0 | 0 | 0 | 0 | 0 | 0 |
| 47775 | ENSMUSG00000110015 | 0 | NA | NA | NA | NA | NA | Gm33968 | predicted gene, 33968 [Source:MGI Symb | 0 | 0 | 0 | 0 | 0 | 0 |
| 47776 | ENSMUSG00000110016 | 0 | NA | NA | NA | NA | NA | Gm20751 | predicted gene, 20751 [Source:MGI Symb | 0 | 0 | 0 | 0 | 0 | 0 |
| 47777 | ENSMUSG00000110017 | 0 | NA | NA | NA | NA | NA | Gm45237 | predicted gene 45237 [Source:MGI Symbo | 0 | 0 | 0 | 0 | 0 | 0 |
| 47778 | ENSMUSG00000110018 | 0.528279927 | -0.837071592 | 3.369085264 | -0.248456636 | 0.803781118 | NA | 5430437J10I | RIKEN cDNA 5430437J10 gene [Source:M | 0 | 2.0558 | 0 | 0 | 0 | 1.1138 |
| 47779 | ENSMUSG00000110019 | 0 | NA | NA | NA | NA | NA | Gm8748 | predicted gene 8748 [Source:MGI Symbol | 0 | 0 | 0 | 0 | 0 | 0 |
| 47780 | ENSMUSG00000110020 | 0.167707431 | -0.796629176 | 4.080472857 | -0.195229623 | 0.84521319 | NA | Gm45441 | predicted gene 45441 [Source:MGI Symbo | 0 | 0 | 1.0062 | 0 | 0 | 0 |
| 47781 | ENSMUSG00000110022 | 0 | NA | NA | NA | NA | NA | Gm45475 | predicted gene 45475 [Source:MGI Symbo | 0 | 0 | 0 | 0 | 0 | 0 |
| 47782 | ENSMUSG00000110023 | 0 | NA | NA | NA | NA | NA | Gm45505 | predicted gene 45505 [Source:MGI Symbo | 0 | 0 | 0 | 0 | 0 | 0 |
| 47783 | ENSMUSG00000110024 | 0 | NA | NA | NA | NA | NA | Gm7210 | predicted gene 7210 [Source:MGI Symbol | 0 | 0 | 0 | 0 | 0 | 0 |
| 47784 | ENSMUSG00000110025 | 0 | NA | NA | NA | NA | NA | Gm20586 | predicted gene, 20586 [Source:MGI Symb | 0 | 0 | 0 | 0 | 0 | 0 |
| 47785 | ENSMUSG00000110026 | 0 | NA | NA | NA | NA | NA | Gm35625 | predicted gene, 35625 [Source:MGI Symb | 0 | 0 | 0 | 0 | 0 | 0 |
| 47786 | ENSMUSG00000110027 | 0 | NA | NA | NA | NA | NA | C030029H02 | RIKEN cDNA C030029H02 gene [Source:M | 0 | 0 | 0 | 0 | 0 | 0 |
| 47787 | ENSMUSG00000110028 | 0 | NA | NA | NA | NA | NA | Gm34838 | predicted gene, 34838 [Source:MGI Symb | 0 | 0 | 0 | 0 | 0 | 0 |
| 47788 | ENSMUSG00000110030 | 0.371281295 | 2.09253362 | 4.031852165 | 0.519000582 | 0.603760335 | NA | Gm45546 | predicted gene 45546 [Source:MGI Symbo | 0 | 0 | 0 | 0 | 0 | 2.2277 |
| 47789 | ENSMUSG00000110031 | 0 | NA | NA | NA | NA | NA | Gm18205 | predicted gene, 18205 [Source:MGI Symb | 0 | 0 | 0 | 0 | 0 | 0 |
| 47790 | ENSMUSG00000110032 | 0 | NA | NA | NA | NA | NA | Gm8436 | predicted gene 8436 [Source:MGI Symbol | 0 | 0 | 0 | 0 | 0 | 0 |
| 47791 | ENSMUSG00000110033 | 0 | NA | NA | NA | NA | NA | | olfactory receptor 1527, pseudogene 1 | 0 | 0 | 0 | 0 | 0 | 0 |
| 47792 | ENSMUSG00000110034 | 0 | NA | NA | NA | NA | NA | Gm45379 | predicted gene 45379 [Source:MGI Symbo | 0 | 0 | 0 | 0 | 0 | 0 |
| 47793 | ENSMUSG00000110035 | 37.09336703 | -0.098841894 | 0.359068609 | -0.275273003 | 0.783106504 | 0.960670212 | Gm30931 | predicted gene, 30931 [Source:MGI Symb | 41.761 | 33.921 | 39.244 | 32.775 | 41.443 | 33.415 |
| 47794 | ENSMUSG00000110036 | 0 | NA | NA | NA | NA | NA | Gm6303 | predicted gene 6303 [Source:MGI Symbol | 0 | 0 | 0 | 0 | 0 | 0 |
| 47795 | ENSMUSG00000110037 | 0 | NA | NA | NA | NA | NA | 4930543N07 | RIKEN cDNA 4930543N07 gene [Source:M | 0 | 0 | 0 | 0 | 0 | 0 |
| 47796 | ENSMUSG00000110038 | 0.878040772 | -3.188971767 | 2.713192499 | -1.175357727 | 0.239851629 | NA | Gm45570 | predicted gene 45570 [Source:MGI Symbo | 3.2124 | 2.0558 | 0 | 0 | 0 | 0 |
| 47797 | ENSMUSG00000110039 | 0 | NA | NA | NA | NA | NA | Gm45541 | predicted gene 45541 [Source:MGI Symbo | 0 | 0 | 0 | 0 | 0 | 0 |
| 47798 | ENSMUSG00000110040 | 0 | NA | NA | NA | NA | NA | Gm49369 | predicted gene, 49369 [Source:MGI Symb | 0 | 0 | 0 | 0 | 0 | 0 |
| 47799 | ENSMUSG00000110041 | 0 | NA | NA | NA | NA | NA | Olfr673-ps1 | olfactory receptor 673, pseudogene 1 [So | 0 | 0 | 0 | 0 | 0 | 0 |
| 47800 | ENSMUSG00000110042 | 0 | NA | NA | NA | NA | NA | Gm4224 | predicted gene 4224 [Source:MGI Symbol | 0 | 0 | 0 | 0 | 0 | 0 |
| 47801 | ENSMUSG00000110043 | 0 | NA | NA | NA | NA | NA | Gm7328 | predicted gene 7328 [Source:MGI Symbol | 0 | 0 | 0 | 0 | 0 | 0 |
| 47802 | ENSMUSG00000110044 | 0.503302254 | -0.837877584 | 3.426698643 | -0.244514523 | 0.806832356 | NA | Gm45434 | predicted gene 45434 [Source:MGI Symbo | 0 | 2.0558 | 0 | 0.964 | 0 | 0 |
| 47803 | ENSMUSG00000110045 | 0 | NA | NA | NA | NA | NA | Gm38590 | predicted gene, 38590 [Source:MGI Symb | 0 | 0 | 0 | 0 | 0 | 0 |
| 47804 | ENSMUSG00000110046 | 0.17131964 | -0.796629176 | 4.080472857 | -0.195229623 | 0.84521319 | NA | Gm45484 | predicted gene 45484 [Source:MGI Symbo | 0 | 1.0279 | 0 | 0 | 0 | 0 |
| 47805 | ENSMUSG00000110047 | 0 | NA | NA | NA | NA | NA | A230085B16 | RIKEN cDNA A230085B16 gene [Source:M | 0 | 0 | 0 | 0 | 0 | 0 |
| 47806 | ENSMUSG00000110048 | 0 | NA | NA | NA | NA | NA | Gm45433 | predicted gene 45433 [Source:MGI Symbo | 0 | 0 | 0 | 0 | 0 | 0 |
| 47807 | ENSMUSG00000110049 | 0 | NA | NA | NA | NA | NA | Gm3836 | predicted gene 3836 [Source:MGI Symbol | 0 | 0 | 0 | 0 | 0 | 0 |
| 47808 | ENSMUSG00000110050 | 0 | NA | NA | NA | NA | NA | Gm45259 | predicted gene 45259 [Source:MGI Symbo | 0 | 0 | 0 | 0 | 0 | 0 |
| 47809 | ENSMUSG00000110051 | 0 | NA | NA | NA | NA | NA | Gm45322 | predicted gene 45322 [Source:MGI Symbo | 0 | 0 | 0 | 0 | 0 | 0 |
| 47810 | ENSMUSG00000110052 | 0.481988923 | 2.492401104 | 3.837077151 | 0.649557203 | 0.515978285 | NA | 1700020G03 | RIKEN cDNA 1700020G03 gene [Source:M | 0 | 0 | 0 | 2.8919 | 0 | 0 |
| 47811 | ENSMUSG00000110053 | 0 | NA | NA | NA | NA | NA | Gm29682 | predicted gene, 29682 [Source:MGI Symb | 0 | 0 | 0 | 0 | 0 | 0 |
| 47812 | ENSMUSG00000110054 | 0 | NA | NA | NA | NA | NA | Gm18851 | predicted gene, 18851 [Source:MGI Symb | 0 | 0 | 0 | 0 | 0 | 0 |
| 47813 | ENSMUSG00000110055 | 6.894480883 | -0.51288013 | 0.816282916 | -0.628311729 | 0.52979975 | 0.888334265 | Gm18857 | predicted gene, 18857 [Source:MGI Symb | 8.031 | 6.1675 | 10.062 | 3.8559 | 5.4531 | 7.7969 |
| 47814 | ENSMUSG00000110056 | 0 | NA | NA | NA | NA | NA | 1700014L14I | RIKEN cDNA 1700014L14 gene [Source:M | 0 | 0 | 0 | 0 | 0 | 0 |
| 47815 | ENSMUSG00000110057 | 3.128671636 | 0.857246329 | 1.371196114 | 0.625181416 | 0.531851997 | NA | Gm2225 | predicted gene 2225 [Source:MGI Symbol | 1.6062 | 1.0279 | 4.025 | 0 | 6.5437 | 5.5692 |
| 47816 | ENSMUSG00000110058 | 0.349477146 | 0.165142768 | 3.911021834 | 0.042224967 | 0.966319359 | NA | Gm5907 | predicted gene 5907 [Source:MGI Symbol | 0 | 0 | 1.0062 | 0 | 1.0906 | 0 |
| 47817 | ENSMUSG00000110059 | 0 | NA | NA | NA | NA | NA | Olfr528-ps1 | olfactory receptor 528, pseudogene 1 [So | 0 | 0 | 0 | 0 | 0 | 0 |
| 47818 | ENSMUSG00000110060 | 0 | NA | NA | NA | NA | NA | Gm9860 | predicted gene 9860 [Source:MGI Symbol | 0 | 0 | 0 | 0 | 0 | 0 |
| 47819 | ENSMUSG00000110061 | 0 | NA | NA | NA | NA | NA | Gm45337 | predicted gene 45337 [Source:MGI Symbo | 0 | 0 | 0 | 0 | 0 | 0 |
| 47820 | ENSMUSG00000110062 | 0.335414862 | -1.779047242 | 4.030804179 | -0.441362855 | 0.658950329 | NA | B430319F04 | RIKEN cDNA B430319F04 gene [Source:M | 0 | 0 | 2.0125 | 0 | 0 | 0 |

| | | | | | | | | | | | | | | | |
|---|---|---|---|---|---|---|---|---|---|---|---|---|---|---|---|
| 47821 | ENSMUSG00000110063 | 0 | NA | NA | NA | NA | NA | Gm31045 | predicted gene, 31045 [Source:MGI Symb | 0 | 0 | 0 | 0 | 0 | 0 |
| 47822 | ENSMUSG00000110064 | 0 | NA | NA | NA | NA | NA | Gm18320 | predicted gene, 18320 [Source:MGI Symb | 0 | 0 | 0 | 0 | 0 | 0 |
| 47823 | ENSMUSG00000110065 | 0 | NA | NA | NA | NA | NA | Gm45588 | predicted gene 45588 [Source:MGI Symbo | 0 | 0 | 0 | 0 | 0 | 0 |
| 47824 | ENSMUSG00000110066 | 0 | NA | NA | NA | NA | NA | Gm38430 | predicted gene, 38430 [Source:MGI Symb | 0 | 0 | 0 | 0 | 0 | 0 |
| 47825 | ENSMUSG00000110067 | 0 | NA | NA | NA | NA | NA | 6330411D24 | RIKEN cDNA 6330411D24 gene [Source:M | 0 | 0 | 0 | 0 | 0 | 0 |
| 47826 | ENSMUSG00000110068 | 0.89236178 | -1.92645888 | 2.54470475 | -0.757046129 | 0.449022227 | NA | Gm45294 | predicted gene 45294 [Source:MGI Symbo | 3.2124 | 1.0279 | 0 | 0 | 0 | 1.1138 |
| 47827 | ENSMUSG00000110069 | 0 | NA | NA | NA | NA | NA | Gm45422 | predicted gene 45422 [Source:MGI Symbo | 0 | 0 | 0 | 0 | 0 | 0 |
| 47828 | ENSMUSG00000110070 | 0 | NA | NA | NA | NA | NA | Gm45619 | predicted gene 45619 [Source:MGI Symbo | 0 | 0 | 0 | 0 | 0 | 0 |
| 47829 | ENSMUSG00000110071 | 0 | NA | NA | NA | NA | NA | Gm45512 | predicted gene 45512 [Source:MGI Symbo | 0 | 0 | 0 | 0 | 0 | 0 |
| 47830 | ENSMUSG00000110072 | 0 | NA | NA | NA | NA | NA | Gm45327 | predicted gene 45327 [Source:MGI Symbo | 0 | 0 | 0 | 0 | 0 | 0 |
| 47831 | ENSMUSG00000110073 | 0 | NA | NA | NA | NA | NA | Gm2875 | predicted gene 2875 [Source:MGI Symbol | 0 | 0 | 0 | 0 | 0 | 0 |
| 47832 | ENSMUSG00000110074 | 0 | NA | NA | NA | NA | NA | Olfr563-ps1 | olfactory receptor 563, pseudogene 1 [Sou | 0 | 0 | 0 | 0 | 0 | 0 |
| 47833 | ENSMUSG00000110075 | 0 | NA | NA | NA | NA | NA | Gm45305 | predicted gene 45305 [Source:MGI Symbo | 0 | 0 | 0 | 0 | 0 | 0 |
| 47834 | ENSMUSG00000110076 | 0 | NA | NA | NA | NA | NA | Gm45549 | predicted gene 45549 [Source:MGI Symbo | 0 | 0 | 0 | 0 | 0 | 0 |
| 47835 | ENSMUSG00000110077 | 2.596529849 | -1.069884915 | 1.621561495 | -0.659786828 | 0.509390637 | NA | Gm45632 | predicted gene 45632 [Source:MGI Symbo | 2.4093 | 8.2233 | 0 | 3.8559 | 1.0906 | 0 |
| 47836 | ENSMUSG00000110078 | 0 | NA | NA | NA | NA | NA | Gm45241 | predicted gene 45241 [Source:MGI Symbo | 0 | 0 | 0 | 0 | 0 | 0 |
| 47837 | ENSMUSG00000110079 | 0 | NA | NA | NA | NA | NA | Gm45292 | predicted gene 45292 [Source:MGI Symbo | 0 | 0 | 0 | 0 | 0 | 0 |
| 47838 | ENSMUSG00000110080 | 0 | NA | NA | NA | NA | NA | Gm6145 | predicted gene 6145 [Source:MGI Symbol | 0 | 0 | 0 | 0 | 0 | 0 |
| 47839 | ENSMUSG00000110081 | 0 | NA | NA | NA | NA | NA | Gm19164 | predicted gene, 19164 [Source:MGI Symb | 0 | 0 | 0 | 0 | 0 | 0 |
| 47840 | ENSMUSG00000110082 | 0 | NA | NA | NA | NA | NA | Gm19201 | predicted gene, 19201 [Source:MGI Symb | 0 | 0 | 0 | 0 | 0 | 0 |
| 47841 | ENSMUSG00000110083 | 0 | NA | NA | NA | NA | NA | Gm45382 | predicted gene 45382 [Source:MGI Symbo | 0 | 0 | 0 | 0 | 0 | 0 |
| 47842 | ENSMUSG00000110084 | 0 | NA | NA | NA | NA | NA | Gm45257 | predicted gene 45257 [Source:MGI Symbo | 0 | 0 | 0 | 0 | 0 | 0 |
| 47843 | ENSMUSG00000110085 | 0 | NA | NA | NA | NA | NA | Gm17911 | predicted gene, 17911 [Source:MGI Symb | 0 | 0 | 0 | 0 | 0 | 0 |
| 47844 | ENSMUSG00000110086 | 0.185640647 | 1.126924175 | 4.080472857 | 0.276174898 | 0.782413736 | NA | Gm45623 | predicted gene 45623 [Source:MGI Symbo | 0 | 0 | 0 | 0 | 0 | 1.1138 |
| 47845 | ENSMUSG00000110087 | 1.072354214 | -0.191149323 | 2.152025997 | -0.088822962 | 0.929222609 | NA | Gm45309 | predicted gene 45309 [Source:MGI Symbo | 2.4093 | 0 | 1.0062 | 1.928 | 1.0906 | 0 |
| 47846 | ENSMUSG00000110088 | 1.05230237 | 0.134697323 | 2.390801759 | 0.056339813 | 0.955071103 | NA | Gm45343 | predicted gene 45343 [Source:MGI Symbo | 0 | 0 | 3.0187 | 0 | 2.1812 | 1.1138 |
| 47847 | ENSMUSG00000110089 | 0.335414862 | -1.779047242 | 4.030804179 | -0.441362855 | 0.658950329 | NA | Gm36243 | predicted gene, 36243 [Source:MGI Symb | 0 | 0 | 2.0125 | 0 | 0 | 0 |
| 47848 | ENSMUSG00000110090 | 0 | NA | NA | NA | NA | NA | Gm45682 | predicted gene 45682 [Source:MGI Symbo | 0 | 0 | 0 | 0 | 0 | 0 |
| 47849 | ENSMUSG00000110091 | 0 | NA | NA | NA | NA | NA | Gm39115 | predicted gene, 39115 [Source:MGI Symb | 0 | 0 | 0 | 0 | 0 | 0 |
| 47850 | ENSMUSG00000110092 | 0 | NA | NA | NA | NA | NA | Olfr537-ps1 | olfactory receptor 537, pseudogene 1 [Sou | 0 | 0 | 0 | 0 | 0 | 0 |
| 47851 | ENSMUSG00000110093 | 4.622257376 | 0.627751302 | 1.136632667 | 0.552290393 | 0.580749412 | NA | Gm45282 | predicted gene 45282 [Source:MGI Symbo | 4.8186 | 4.1117 | 2.0125 | 13.496 | 2.1812 | 1.1138 |
| 47852 | ENSMUSG00000110094 | 0 | NA | NA | NA | NA | NA | Gm45326 | predicted gene 45326 [Source:MGI Symbo | 0 | 0 | 0 | 0 | 0 | 0 |
| 47853 | ENSMUSG00000110095 | 0 | NA | NA | NA | NA | NA | Gm7207 | predicted gene 7207 [Source:MGI Symbol | 0 | 0 | 0 | 0 | 0 | 0 |
| 47854 | ENSMUSG00000110096 | 0 | NA | NA | NA | NA | NA | Gm45544 | predicted gene 45544 [Source:MGI Symbo | 0 | 0 | 0 | 0 | 0 | 0 |
| 47855 | ENSMUSG00000110097 | 0 | NA | NA | NA | NA | NA | Zscan4a | zinc finger and SCAN domain containing 4 | 0 | 0 | 0 | 0 | 0 | 0 |
| 47856 | ENSMUSG00000110098 | 0 | NA | NA | NA | NA | NA | Gm45401 | predicted gene 45401 [Source:MGI Symbo | 0 | 0 | 0 | 0 | 0 | 0 |
| 47857 | ENSMUSG00000110099 | 7.481542053 | 0.066822024 | 0.899370838 | 0.074298633 | 0.940772765 | 0.989001277 | Gm45344 | predicted gene 45344 [Source:MGI Symbo | 9.6372 | 8.2233 | 4.025 | 7.7118 | 14.178 | 1.1138 |
| 47858 | ENSMUSG00000110100 | 0 | NA | NA | NA | NA | NA | Gm45261 | predicted gene 45261 [Source:MGI Symbo | 0 | 0 | 0 | 0 | 0 | 0 |
| 47859 | ENSMUSG00000110101 | 0 | NA | NA | NA | NA | NA | Gm6249 | predicted gene 6249 [Source:MGI Symbol | 0 | 0 | 0 | 0 | 0 | 0 |
| 47860 | ENSMUSG00000110102 | 0 | NA | NA | NA | NA | NA | Gm9132 | predicted gene 9132 [Source:MGI Symbol | 0 | 0 | 0 | 0 | 0 | 0 |
| 47861 | ENSMUSG00000110103 | 0 | NA | NA | NA | NA | NA | Zscan4-ps1 | zinc finger and SCAN domain containing 4 | 0 | 0 | 0 | 0 | 0 | 0 |
| 47862 | ENSMUSG00000110104 | 0 | NA | NA | NA | NA | NA | Gm45717 | predicted gene 45717 [Source:MGI Symbo | 0 | 0 | 0 | 0 | 0 | 0 |
| 47863 | ENSMUSG00000110105 | 15.10134776 | -0.393541006 | 0.585744184 | -0.671864982 | 0.501669655 | 0.880974509 | Gm45844 | predicted gene 45844 [Source:MGI Symbo | 21.684 | 19.53 | 10.062 | 10.604 | 10.906 | 17.822 |
| 47864 | ENSMUSG00000110106 | 0 | NA | NA | NA | NA | NA | Gm45666 | predicted gene 45666 [Source:MGI Symbo | 0 | 0 | 0 | 0 | 0 | 0 |
| 47865 | ENSMUSG00000110108 | 0 | NA | NA | NA | NA | NA | Gm3702 | predicted gene 3702 [Source:MGI Symbol | 0 | 0 | 0 | 0 | 0 | 0 |
| 47866 | ENSMUSG00000110109 | 0 | NA | NA | NA | NA | NA | Gm5599 | predicted gene 5599 [Source:MGI Symbol | 0 | 0 | 0 | 0 | 0 | 0 |
| 47867 | ENSMUSG00000110110 | 0.342639279 | -1.805976596 | 4.028547714 | -0.448294702 | 0.653940524 | NA | Gm30978 | predicted gene, 30978 [Source:MGI Symb | 0 | 2.0558 | 0 | 0 | 0 | 0 |
| 47868 | ENSMUSG00000110111 | 0 | NA | NA | NA | NA | NA | Gm7786 | predicted gene 7786 [Source:MGI Symbol | 0 | 0 | 0 | 0 | 0 | 0 |
| 47869 | ENSMUSG00000110112 | 0 | NA | NA | NA | NA | NA | Gm39481 | predicted gene, 39481 [Source:MGI Symb | 0 | 0 | 0 | 0 | 0 | 0 |
| 47870 | ENSMUSG00000110113 | 0 | NA | NA | NA | NA | NA | Gm45439 | predicted gene 45439 [Source:MGI Symbo | 0 | 0 | 0 | 0 | 0 | 0 |
| 47871 | ENSMUSG00000110114 | 0.48903338 | 0.95637552 | 3.456690196 | 0.276673773 | 0.782030612 | NA | Gm45471 | predicted gene 45471 [Source:MGI Symbo | 0 | 0 | 1.0062 | 1.928 | 0 | 0 |
| 47872 | ENSMUSG00000110115 | 0 | NA | NA | NA | NA | NA | Gm32486 | predicted gene, 32486 [Source:MGI Symb | 0 | 0 | 0 | 0 | 0 | 0 |
| 47873 | ENSMUSG00000110116 | 0 | NA | NA | NA | NA | NA | 4933430A20 | RIKEN cDNA 4933430A20 gene [Source:M | 0 | 0 | 0 | 0 | 0 | 0 |
| 47874 | ENSMUSG00000110117 | 0 | NA | NA | NA | NA | NA | Gm17990 | predicted gene, 17990 [Source:MGI Symb | 0 | 0 | 0 | 0 | 0 | 0 |
| 47875 | ENSMUSG00000110118 | 0 | NA | NA | NA | NA | NA | Gm5609 | predicted gene 5609 [Source:MGI Symbol | 0 | 0 | 0 | 0 | 0 | 0 |
| 47876 | ENSMUSG00000110119 | 0 | NA | NA | NA | NA | NA | Gm45593 | predicted gene 45593 [Source:MGI Symbo | 0 | 0 | 0 | 0 | 0 | 0 |

| | | | | | | | | | | | | | | | |
|---|---|---|---|---|---|---|---|---|---|---|---|---|---|---|---|
| 47877 | ENSMUSG00000110120 | 4.567030466 | -1.0150163 | 1.119862768 | -0.90637561 | 0.364737078 | NA | Gm45251 | predicted gene 45251 [Source:MGI Symbo | 7.2279 | 6.1675 | 5.0312 | 6.7478 | 0 | 2.2277 |
| 47878 | ENSMUSG00000110121 | 0 | NA | NA | NA | NA | NA | Gm45367 | predicted gene 45367 [Source:MGI Symbo | 0 | 0 | 0 | 0 | 0 | 0 |
| 47879 | ENSMUSG00000110122 | 0 | NA | NA | NA | NA | NA | Olfr562-ps1 | olfactory receptor 562, pseudogene 1 [Sou | 0 | 0 | 0 | 0 | 0 | 0 |
| 47880 | ENSMUSG00000110123 | 0.167707431 | -0.796629176 | 4.080472857 | -0.195229623 | 0.84521319 | NA | Gm18562 | predicted gene, 18562 [Source:MGI Symb | 0 | 0 | 1.0062 | 0 | 0 | 0 |
| 47881 | ENSMUSG00000110124 | 0.181769715 | 1.126924175 | 4.080472857 | 0.276174898 | 0.782413736 | NA | Gm4247 | predicted gene 4247 [Source:MGI Symbol | 0 | 0 | 0 | 0 | 1.0906 | 0 |
| 47882 | ENSMUSG00000110125 | 0 | NA | NA | NA | NA | NA | Gm33148 | predicted gene, 33148 [Source:MGI Symb | 0 | 0 | 0 | 0 | 0 | 0 |
| 47883 | ENSMUSG00000110126 | 0 | NA | NA | NA | NA | NA | Gm9347 | predicted gene 9347 [Source:MGI Symbol | 0 | 0 | 0 | 0 | 0 | 0 |
| 47884 | ENSMUSG00000110127 | 0.831672659 | -0.604339699 | 2.59720557 | -0.232688435 | 0.81600335 | NA | 6430710M23 | RIKEN cDNA 6430710M23 gene [Source:M | 0 | 2.0558 | 1.0062 | 1.928 | 0 | 0 |
| 47885 | ENSMUSG00000110128 | 0 | NA | NA | NA | NA | NA | Gm45398 | predicted gene 45398 [Source:MGI Symbo | 0 | 0 | 0 | 0 | 0 | 0 |
| 47886 | ENSMUSG00000110129 | 14.89179128 | -0.578103975 | 0.589315448 | -0.98097543 | 0.326604858 | 0.772276061 | Gm45453 | predicted gene 45453 [Source:MGI Symbo | 16.865 | 20.558 | 16.1 | 10.604 | 18.541 | 6.6831 |
| 47887 | ENSMUSG00000110130 | 0 | NA | NA | NA | NA | NA | Gm34419 | predicted gene, 34419 [Source:MGI Symb | 0 | 0 | 0 | 0 | 0 | 0 |
| 47888 | ENSMUSG00000110131 | 0 | NA | NA | NA | NA | NA | Gm18066 | predicted gene, 18066 [Source:MGI Symb | 0 | 0 | 0 | 0 | 0 | 0 |
| 47889 | ENSMUSG00000110132 | 0 | NA | NA | NA | NA | NA | Gm45394 | predicted gene 45394 [Source:MGI Symbo | 0 | 0 | 0 | 0 | 0 | 0 |
| 47890 | ENSMUSG00000110133 | 0 | NA | NA | NA | NA | NA | Gm8168 | predicted gene 8168 [Source:MGI Symbol | 0 | 0 | 0 | 0 | 0 | 0 |
| 47891 | ENSMUSG00000110134 | 0 | NA | NA | NA | NA | NA | Gm45334 | predicted gene 45334 [Source:MGI Symbo | 0 | 0 | 0 | 0 | 0 | 0 |
| 47892 | ENSMUSG00000110135 | 0 | NA | NA | NA | NA | NA | Gm45839 | predicted gene 45839 [Source:MGI Symbo | 0 | 0 | 0 | 0 | 0 | 0 |
| 47893 | ENSMUSG00000110136 | 0 | NA | NA | NA | NA | NA | Gm45785 | predicted gene 45785 [Source:MGI Symbo | 0 | 0 | 0 | 0 | 0 | 0 |
| 47894 | ENSMUSG00000110137 | 0 | NA | NA | NA | NA | NA | Gm45577 | predicted gene 45577 [Source:MGI Symbo | 0 | 0 | 0 | 0 | 0 | 0 |
| 47895 | ENSMUSG00000110138 | 0 | NA | NA | NA | NA | NA | 4831440D22 | RIKEN cDNA 4831440D22 gene [Source:M | 0 | 0 | 0 | 0 | 0 | 0 |
| 47896 | ENSMUSG00000110139 | 0 | NA | NA | NA | NA | NA | Gm45783 | predicted gene 45783 [Source:MGI Symbo | 0 | 0 | 0 | 0 | 0 | 0 |
| 47897 | ENSMUSG00000110140 | 0 | NA | NA | NA | NA | NA | 5430421F17 | RIKEN cDNA 5430421F17 gene [Source:M | 0 | 0 | 0 | 0 | 0 | 0 |
| 47898 | ENSMUSG00000110141 | 0 | NA | NA | NA | NA | NA | Gm45684 | predicted gene 45684 [Source:MGI Symbo | 0 | 0 | 0 | 0 | 0 | 0 |
| 47899 | ENSMUSG00000110142 | 0 | NA | NA | NA | NA | NA | Gm45872 | predicted gene 45872 [Source:MGI Symbo | 0 | 0 | 0 | 0 | 0 | 0 |
| 47900 | ENSMUSG00000110143 | 0 | NA | NA | NA | NA | NA | Olfr580-ps1 | olfactory receptor 580, pseudogene 1 [Sou | 0 | 0 | 0 | 0 | 0 | 0 |
| 47901 | ENSMUSG00000110144 | 0.633540418 | -1.320719359 | 2.623870149 | -0.503347835 | 0.61471974 | NA | Gm45564 | predicted gene 45564 [Source:MGI Symbo | 0.8031 | 1.0279 | 1.0062 | 0.964 | 0 | 0 |
| 47902 | ENSMUSG00000110145 | 0 | NA | NA | NA | NA | NA | Gm45683 | predicted gene 45683 [Source:MGI Symbo | 0 | 0 | 0 | 0 | 0 | 0 |
| 47903 | ENSMUSG00000110146 | 0 | NA | NA | NA | NA | NA | Gm45487 | predicted gene 45487 [Source:MGI Symbo | 0 | 0 | 0 | 0 | 0 | 0 |
| 47904 | ENSMUSG00000110147 | 0 | NA | NA | NA | NA | NA | Gm45324 | predicted gene 45324 [Source:MGI Symbo | 0 | 0 | 0 | 0 | 0 | 0 |
| 47905 | ENSMUSG00000110148 | 0.492645588 | 0.9565034 | 3.447648281 | 0.277436479 | 0.781444974 | NA | 5830408C22 | RIKEN cDNA 5830408C22 gene [Source:M | 0 | 1.0279 | 0 | 1.928 | 0 | 0 |
| 47906 | ENSMUSG00000110149 | 0 | NA | NA | NA | NA | NA | Gm7848 | predicted gene 7848 [Source:MGI Symbol | 0 | 0 | 0 | 0 | 0 | 0 |
| 47907 | ENSMUSG00000110150 | 0 | NA | NA | NA | NA | NA | Gm8523 | predicted gene 8523 [Source:MGI Symbol | 0 | 0 | 0 | 0 | 0 | 0 |
| 47908 | ENSMUSG00000110151 | 53.13281794 | 1.558394424 | 0.350628639 | 4.44457255 | 8.81E-06 | 0.000477078 | Gm38416 | predicted gene, 38416 [Source:MGI Symb | 29.715 | 25.698 | 25.156 | 64.587 | 110.15 | 63.489 |
| 47909 | ENSMUSG00000110152 | 0 | NA | NA | NA | NA | NA | Gm45689 | predicted gene 45689 [Source:MGI Symbo | 0 | 0 | 0 | 0 | 0 | 0 |
| 47910 | ENSMUSG00000110153 | 0 | NA | NA | NA | NA | NA | Gm17766 | predicted gene, 17766 [Source:MGI Symb | 0 | 0 | 0 | 0 | 0 | 0 |
| 47911 | ENSMUSG00000110154 | 0 | NA | NA | NA | NA | NA | Gm9731 | predicted gene 9731 [Source:MGI Symbol | 0 | 0 | 0 | 0 | 0 | 0 |
| 47912 | ENSMUSG00000110155 | 0 | NA | NA | NA | NA | NA | Gm35629 | predicted gene, 35629 [Source:MGI Symb | 0 | 0 | 0 | 0 | 0 | 0 |
| 47913 | ENSMUSG00000110156 | 46.96291537 | -0.075786596 | 0.342779568 | -0.221094264 | 0.825019037 | 0.96943211 | Gm42067 | predicted gene, 42067 [Source:MGI Symb | 61.036 | 33.921 | 49.306 | 48.199 | 42.534 | 46.781 |
| 47914 | ENSMUSG00000110157 | 0 | NA | NA | NA | NA | NA | Gm32507 | predicted gene, 32507 [Source:MGI Symb | 0 | 0 | 0 | 0 | 0 | 0 |
| 47915 | ENSMUSG00000110158 | 0 | NA | NA | NA | NA | NA | Gm45690 | predicted gene 45690 [Source:MGI Symbo | 0 | 0 | 0 | 0 | 0 | 0 |
| 47916 | ENSMUSG00000110159 | 1.946937762 | 0.179315356 | 1.471494763 | 0.121859323 | 0.903010431 | NA | Gm45501 | predicted gene 45501 [Source:MGI Symbo | 2.4093 | 2.0558 | 1.0062 | 2.8919 | 1.0906 | 2.2277 |
| 47917 | ENSMUSG00000110160 | 0 | NA | NA | NA | NA | NA | Gm45560 | predicted gene 45560 [Source:MGI Symbo | 0 | 0 | 0 | 0 | 0 | 0 |
| 47918 | ENSMUSG00000110161 | 9.822595409 | 0.72205749 | 0.744710793 | 0.969581074 | 0.332255351 | 0.775138789 | Gm45494 | predicted gene 45494 [Source:MGI Symbo | 4.0155 | 9.2513 | 9.0562 | 6.7478 | 9.8156 | 20.049 |
| 47919 | ENSMUSG00000110162 | 0 | NA | NA | NA | NA | NA | Gm45356 | predicted gene 45356 [Source:MGI Symbo | 0 | 0 | 0 | 0 | 0 | 0 |
| 47920 | ENSMUSG00000110163 | 0 | NA | NA | NA | NA | NA | Gm45389 | predicted gene 45389 [Source:MGI Symbo | 0 | 0 | 0 | 0 | 0 | 0 |
| 47921 | ENSMUSG00000110164 | 0 | NA | NA | NA | NA | NA | Gm7351 | predicted gene 7351 [Source:MGI Symbol | 0 | 0 | 0 | 0 | 0 | 0 |
| 47922 | ENSMUSG00000110165 | 33.10405911 | 0.08049452 | 0.399783365 | 0.201345346 | 0.840428547 | 0.971838962 | Btf3-ps18 | basic transcription factor 3, pseudogene 1 | 30.518 | 40.089 | 26.162 | 43.379 | 26.175 | 32.301 |
| 47923 | ENSMUSG00000110166 | 0.185640647 | 1.126924175 | 4.080472857 | 0.276174898 | 0.782413736 | NA | Gm45641 | predicted gene 45641 [Source:MGI Symbo | 0 | 0 | 0 | 0 | 0 | 1.1138 |
| 47924 | ENSMUSG00000110167 | 0.17131964 | -0.796629176 | 4.080472857 | -0.195229623 | 0.84521319 | NA | Gm45532 | predicted gene 45532 [Source:MGI Symbo | 0 | 1.0279 | 0 | 0 | 0 | 0 |
| 47925 | ENSMUSG00000110168 | 0 | NA | NA | NA | NA | NA | Gm45488 | predicted gene 45488 [Source:MGI Symbo | 0 | 0 | 0 | 0 | 0 | 0 |
| 47926 | ENSMUSG00000110169 | 0 | NA | NA | NA | NA | NA | Olfr534-ps1 | olfactory receptor 534, pseudogene 1 [Sou | 0 | 0 | 0 | 0 | 0 | 0 |
| 47927 | ENSMUSG00000110170 | 0.294513347 | 0.165143496 | 4.080472857 | 0.040471657 | 0.967717103 | NA | St6galnac2 | ST6 (alpha-N-acetyl-neuraminyl-2,3-beta-g | 0.8031 | 0 | 0 | 0.964 | 0 | 0 |
| 47928 | ENSMUSG00000110171 | 0 | NA | NA | NA | NA | NA | Olfr484 | olfactory receptor 484 [Source:MGI Symb | 0 | 0 | 0 | 0 | 0 | 0 |
| 47929 | ENSMUSG00000110172 | 0 | NA | NA | NA | NA | NA | Gm39147 | predicted gene, 39147 [Source:MGI Symb | 0 | 0 | 0 | 0 | 0 | 0 |
| 47930 | ENSMUSG00000110173 | 0 | NA | NA | NA | NA | NA | Gm6929 | predicted gene 6929 [Source:MGI Symbol | 0 | 0 | 0 | 0 | 0 | 0 |
| 47931 | ENSMUSG00000110174 | 0 | NA | NA | NA | NA | NA | Gm45499 | predicted gene 45499 [Source:MGI Symbo | 0 | 0 | 0 | 0 | 0 | 0 |
| 47932 | ENSMUSG00000110175 | 0 | NA | NA | NA | NA | NA | Gm18255 | predicted gene, 18255 [Source:MGI Symb | 0 | 0 | 0 | 0 | 0 | 0 |

| | | | | | | | | | | | | | | | |
|---|---|---|---|---|---|---|---|---|---|---|---|---|---|---|---|
| 47933 | ENSMUSG00000110176 | 0 | NA | NA | NA | NA | NA | Gm35520 | predicted gene, 35520 [Source:MGI Symb | 0 | 0 | 0 | 0 | 0 | 0 |
| 47934 | ENSMUSG00000110177 | 0 | NA | NA | NA | NA | NA | Gm6524 | predicted gene 6524 [Source:MGI Symbol | 0 | 0 | 0 | 0 | 0 | 0 |
| 47935 | ENSMUSG00000110178 | 0 | NA | NA | NA | NA | NA | Gm33831 | predicted gene, 33831 [Source:MGI Symb | 0 | 0 | 0 | 0 | 0 | 0 |
| 47936 | ENSMUSG00000110179 | 0 | NA | NA | NA | NA | NA | Gm39168 | predicted gene, 39168 [Source:MGI Symb | 0 | 0 | 0 | 0 | 0 | 0 |
| 47937 | ENSMUSG00000110180 | 0 | NA | NA | NA | NA | NA | Gm32567 | predicted gene, 32567 [Source:MGI Symb | 0 | 0 | 0 | 0 | 0 | 0 |
| 47938 | ENSMUSG00000110181 | 0.167707431 | -0.796629176 | 4.080472857 | -0.195229623 | 0.84521319 | NA | Gm45508 | predicted gene 45508 [Source:MGI Symbo | 0 | 0 | 1.0062 | 0 | 0 | 0 |
| 47939 | ENSMUSG00000110182 | 0 | NA | NA | NA | NA | NA | Gm45617 | predicted gene 45617 [Source:MGI Symbo | 0 | 0 | 0 | 0 | 0 | 0 |
| 47940 | ENSMUSG00000110183 | 0.77650227 | 1.950354251 | 2.938137078 | 0.663806418 | 0.506814217 | NA | Gm45411 | predicted gene 45411 [Source:MGI Symbo | 0.8031 | 0 | 0 | 3.8559 | 0 | 0 |
| 47941 | ENSMUSG00000110184 | 0 | NA | NA | NA | NA | NA | Gm45429 | predicted gene 45429 [Source:MGI Symbo | 0 | 0 | 0 | 0 | 0 | 0 |
| 47942 | ENSMUSG00000110185 | 186.8520793 | 0.177028005 | 0.193134089 | 0.916606724 | 0.35934876 | 0.799142137 | Igip | IgA inducing protein [Source:MGI Symbol; | 191.94 | 160.36 | 173.07 | 163.88 | 223.58 | 208.29 |
| 47943 | ENSMUSG00000110186 | 0 | NA | NA | NA | NA | NA | Gm7789 | predicted pseudogene 7789 [Source:MGI | 0 | 0 | 0 | 0 | 0 | 0 |
| 47944 | ENSMUSG00000110187 | 0 | NA | NA | NA | NA | NA | Gm45496 | predicted gene 45496 [Source:MGI Symbo | 0 | 0 | 0 | 0 | 0 | 0 |
| 47945 | ENSMUSG00000110188 | 0 | NA | NA | NA | NA | NA | Gm45432 | predicted gene 45432 [Source:MGI Symbo | 0 | 0 | 0 | 0 | 0 | 0 |
| 47946 | ENSMUSG00000110189 | 0 | NA | NA | NA | NA | NA | Gm45665 | predicted gene 45665 [Source:MGI Symbo | 0 | 0 | 0 | 0 | 0 | 0 |
| 47947 | ENSMUSG00000110190 | 0 | NA | NA | NA | NA | NA | Zscan4-ps2 | zinc finger and SCAN domain containing 4 | 0 | 0 | 0 | 0 | 0 | 0 |
| 47948 | ENSMUSG00000110192 | 0 | NA | NA | NA | NA | NA | Gm9652 | predicted gene 9652 [Source:MGI Symbol | 0 | 0 | 0 | 0 | 0 | 0 |
| 47949 | ENSMUSG00000110193 | 1.439532482 | -1.591016417 | 1.964358454 | -0.809942001 | 0.417973511 | NA | Gm45452 | predicted gene 45452 [Source:MGI Symbo | 2.4093 | 1.0279 | 3.0187 | 0 | 2.1812 | 0 |
| 47950 | ENSMUSG00000110194 | 0.428363721 | -0.543730382 | 3.624677344 | -0.15000794 | 0.880758351 | NA | Gm7192 | predicted gene 7192 [Source:MGI Symbol | 1.6062 | 0 | 0 | 0.964 | 0 | 0 |
| 47951 | ENSMUSG00000110195 | 0.663785267 | -1.400021131 | 3.106754789 | -0.450637796 | 0.652250621 | NA | Pde2a | phosphodiesterase 2A, cGMP-stimulated | 0 | 0 | 3.0187 | 0.964 | 0 | 0 |
| 47952 | ENSMUSG00000110196 | 0 | NA | NA | NA | NA | NA | Gm33601 | predicted gene, 33601 [Source:MGI Symb | 0 | 0 | 0 | 0 | 0 | 0 |
| 47953 | ENSMUSG00000110197 | 0.17131964 | -0.796629176 | 4.080472857 | -0.195229623 | 0.84521319 | NA | Gm45444 | predicted gene 45444 [Source:MGI Symbo | 0 | 1.0279 | 0 | 0 | 0 | 0 |
| 47954 | ENSMUSG00000110198 | 0 | NA | NA | NA | NA | NA | Gm45335 | predicted gene 45335 [Source:MGI Symbo | 0 | 0 | 0 | 0 | 0 | 0 |
| 47955 | ENSMUSG00000110199 | 7.737078304 | -0.638641164 | 0.940369577 | -0.679138479 | 0.497050124 | 0.878676728 | Olfr542-ps1 | olfactory receptor 542, pseudogene 1 [So | 8.031 | 13.363 | 7.0437 | 12.532 | 5.4531 | 0 |
| 47956 | ENSMUSG00000110200 | 0 | NA | NA | NA | NA | NA | Gm45316 | predicted gene 45316 [Source:MGI Symbo | 0 | 0 | 0 | 0 | 0 | 0 |
| 47957 | ENSMUSG00000110201 | 0 | NA | NA | NA | NA | NA | Gm45675 | predicted gene 45675 [Source:MGI Symbo | 0 | 0 | 0 | 0 | 0 | 0 |
| 47958 | ENSMUSG00000110202 | 0.167707431 | -0.796629176 | 4.080472857 | -0.195229623 | 0.84521319 | NA | Gm45435 | predicted gene 45435 [Source:MGI Symbo | 0 | 0 | 1.0062 | 0 | 0 | 0 |
| 47959 | ENSMUSG00000110203 | 0 | NA | NA | NA | NA | NA | Gm45365 | predicted gene 45365 [Source:MGI Symbo | 0 | 0 | 0 | 0 | 0 | 0 |
| 47960 | ENSMUSG00000110204 | 0 | NA | NA | NA | NA | NA | Olfr501-ps1 | olfactory receptor 501, pseudogene 1 [So | 0 | 0 | 0 | 0 | 0 | 0 |
| 47961 | ENSMUSG00000110205 | 0 | NA | NA | NA | NA | NA | Gm33594 | predicted gene, 33594 [Source:MGI Symb | 0 | 0 | 0 | 0 | 0 | 0 |
| 47962 | ENSMUSG00000110206 | 43.03691165 | -0.331170452 | 0.353876903 | -0.935835171 | 0.34935806 | 0.790730887 | Flt3l | FMS-like tyrosine kinase 3 ligand [Source:I | 46.58 | 46.256 | 51.318 | 49.163 | 38.172 | 26.732 |
| 47963 | ENSMUSG00000110207 | 0 | NA | NA | NA | NA | NA | Gm33586 | predicted gene, 33586 [Source:MGI Symb | 0 | 0 | 0 | 0 | 0 | 0 |
| 47964 | ENSMUSG00000110208 | 0 | NA | NA | NA | NA | NA | Gm45442 | predicted gene 45442 [Source:MGI Symbo | 0 | 0 | 0 | 0 | 0 | 0 |
| 47965 | ENSMUSG00000110209 | 0 | NA | NA | NA | NA | NA | Gm45868 | predicted gene 45868 [Source:MGI Symbo | 0 | 0 | 0 | 0 | 0 | 0 |
| 47966 | ENSMUSG00000110210 | 0 | NA | NA | NA | NA | NA | Gm33756 | predicted gene, 33756 [Source:MGI Symb | 0 | 0 | 0 | 0 | 0 | 0 |
| 47967 | ENSMUSG00000110211 | 0 | NA | NA | NA | NA | NA | Gm31294 | predicted gene, 31294 [Source:MGI Symb | 0 | 0 | 0 | 0 | 0 | 0 |
| 47968 | ENSMUSG00000110212 | 0 | NA | NA | NA | NA | NA | Gm35963 | predicted gene, 35963 [Source:MGI Symb | 0 | 0 | 0 | 0 | 0 | 0 |
| 47969 | ENSMUSG00000110213 | 0 | NA | NA | NA | NA | NA | Gm45628 | predicted gene 45628 [Source:MGI Symbo | 0 | 0 | 0 | 0 | 0 | 0 |
| 47970 | ENSMUSG00000110214 | 0 | NA | NA | NA | NA | NA | Gm45587 | predicted gene 45587 [Source:MGI Symbo | 0 | 0 | 0 | 0 | 0 | 0 |
| 47971 | ENSMUSG00000110215 | 0 | NA | NA | NA | NA | NA | Gm45598 | predicted gene 45598 [Source:MGI Symbo | 0 | 0 | 0 | 0 | 0 | 0 |
| 47972 | ENSMUSG00000110216 | 0.842329324 | -1.832359871 | 2.605893773 | -0.703159848 | 0.481956138 | NA | Gm36325 | predicted gene, 36325 [Source:MGI Symb | 0 | 3.0838 | 1.0062 | 0.964 | 0 | 0 |
| 47973 | ENSMUSG00000110217 | 0 | NA | NA | NA | NA | NA | Gm45408 | predicted gene 45408 [Source:MGI Symbo | 0 | 0 | 0 | 0 | 0 | 0 |
| 47974 | ENSMUSG00000110218 | 1.190410227 | -1.156645952 | 2.318925491 | -0.498785302 | 0.617930643 | NA | Tincr | TINCR ubiquitin domain containing [Sourc | 0.8031 | 4.1117 | 0 | 0 | 0 | 2.2277 |
| 47975 | ENSMUSG00000110219 | 0 | NA | NA | NA | NA | NA | Gm45317 | predicted gene 45317 [Source:MGI Symbo | 0 | 0 | 0 | 0 | 0 | 0 |
| 47976 | ENSMUSG00000110220 | 0 | NA | NA | NA | NA | NA | Gm45296 | predicted gene 45296 [Source:MGI Symbo | 0 | 0 | 0 | 0 | 0 | 0 |
| 47977 | ENSMUSG00000110221 | 11.14510022 | 1.290338945 | 0.818209137 | 1.577028276 | 0.114789019 | 0.497545211 | Gm36210 | predicted gene, 36210 [Source:MGI Symb | 7.2279 | 2.0558 | 10.062 | 12.532 | 30.537 | 4.4554 |
| 47978 | ENSMUSG00000110222 | 0 | NA | NA | NA | NA | NA | Gm39149 | predicted gene, 39149 [Source:MGI Symb | 0 | 0 | 0 | 0 | 0 | 0 |
| 47979 | ENSMUSG00000110223 | 0 | NA | NA | NA | NA | NA | Gm45489 | predicted gene 45489 [Source:MGI Symbo | 0 | 0 | 0 | 0 | 0 | 0 |
| 47980 | ENSMUSG00000110224 | 0 | NA | NA | NA | NA | NA | Gm6011 | predicted gene 6011 [Source:MGI Symbol | 0 | 0 | 0 | 0 | 0 | 0 |
| 47981 | ENSMUSG00000110225 | 0 | NA | NA | NA | NA | NA | Gm45528 | predicted gene 45528 [Source:MGI Symbo | 0 | 0 | 0 | 0 | 0 | 0 |
| 47982 | ENSMUSG00000110226 | 0 | NA | NA | NA | NA | NA | Gm7624 | predicted gene 7624 [Source:MGI Symbol | 0 | 0 | 0 | 0 | 0 | 0 |
| 47983 | ENSMUSG00000110227 | 0 | NA | NA | NA | NA | NA | Gm45634 | predicted gene 45634 [Source:MGI Symbo | 0 | 0 | 0 | 0 | 0 | 0 |
| 47984 | ENSMUSG00000110228 | 0 | NA | NA | NA | NA | NA | Gm2080 | predicted gene 2080 [Source:MGI Symbol | 0 | 0 | 0 | 0 | 0 | 0 |
| 47985 | ENSMUSG00000110229 | 0 | NA | NA | NA | NA | NA | Gm31784 | predicted gene, 31784 [Source:MGI Symb | 0 | 0 | 0 | 0 | 0 | 0 |
| 47986 | ENSMUSG00000110230 | 0 | NA | NA | NA | NA | NA | Gm45846 | predicted gene 45846 [Source:MGI Symbo | 0 | 0 | 0 | 0 | 0 | 0 |
| 47987 | ENSMUSG00000110231 | 0 | NA | NA | NA | NA | NA | Gm6329 | predicted gene 6329 [Source:MGI Symbol | 0 | 0 | 0 | 0 | 0 | 0 |
| 47988 | ENSMUSG00000110232 | 0 | NA | NA | NA | NA | NA | Gm18193 | predicted gene, 18193 [Source:MGI Symb | 0 | 0 | 0 | 0 | 0 | 0 |

| | | | | | | | | | | | | | | | |
|---|---|---|---|---|---|---|---|---|---|---|---|---|---|---|---|
| 47989 | ENSMUSG00000110233 | 0 | NA | NA | NA | NA | NA | Gm45427 | predicted gene 45427 [Source:MGI Symbo | 0 | 0 | 0 | 0 | 0 | 0 |
| 47990 | ENSMUSG00000110234 | 1.605119631 | -0.299293217 | 2.034218938 | -0.147129304 | 0.88302996 | NA | Gm45799 | predicted gene 45799 [Source:MGI Symbo | 3.2124 | 2.0558 | 0 | 0 | 4.3625 | 0 |
| 47991 | ENSMUSG00000110235 | 1.332933394 | 0.12098469 | 1.746119635 | 0.069287744 | 0.944760581 | NA | Gm5086 | predicted gene 5086 [Source:MGI Symbol | 0.8031 | 2.0558 | 1.0062 | 1.928 | 1.0906 | 1.1138 |
| 47992 | ENSMUSG00000110236 | 0 | NA | NA | NA | NA | NA | Gm40493 | predicted gene, 40493 [Source:MGI Symb | 0 | 0 | 0 | 0 | 0 | 0 |
| 47993 | ENSMUSG00000110237 | 7.835954857 | 0.480971984 | 0.87079599 | 0.552336011 | 0.580718162 | 0.908645447 | Gm18258 | predicted gene, 18258 [Source:MGI Symb | 2.4093 | 13.363 | 4.025 | 10.604 | 4.3625 | 12.252 |
| 47994 | ENSMUSG00000110238 | 0 | NA | NA | NA | NA | NA | Gm19269 | predicted gene, 19269 [Source:MGI Symb | 0 | 0 | 0 | 0 | 0 | 0 |
| 47995 | ENSMUSG00000110239 | 7.319980821 | 0.282301561 | 0.79610973 | 0.35460132 | 0.722888287 | 0.9447719 | Gm45620 | predicted gene 45620 [Source:MGI Symbo | 5.6217 | 8.2233 | 6.0375 | 8.6758 | 10.906 | 4.4554 |
| 47996 | ENSMUSG00000110240 | 0 | NA | NA | NA | NA | NA | Gm45486 | predicted gene 45486 [Source:MGI Symbo | 0 | 0 | 0 | 0 | 0 | 0 |
| 47997 | ENSMUSG00000110241 | 0 | NA | NA | NA | NA | NA | Gm45506 | predicted gene 45506 [Source:MGI Symbo | 0 | 0 | 0 | 0 | 0 | 0 |
| 47998 | ENSMUSG00000110242 | 0 | NA | NA | NA | NA | NA | G630064G18 | RIKEN cDNA G630064G18 gene [Source:M | 0 | 0 | 0 | 0 | 0 | 0 |
| 47999 | ENSMUSG00000110243 | 0.439020386 | -2.189747911 | 3.52148864 | -0.621824499 | 0.534057275 | NA | Gm45631 | predicted gene 45631 [Source:MGI Symbo | 1.6062 | 1.0279 | 0 | 0 | 0 | 0 |
| 48000 | ENSMUSG00000110244 | 0.730949792 | 3.072456834 | 2.934263219 | 1.04709653 | 0.29505506 | NA | Gm2253 | predicted gene 2253 [Source:MGI Symbol | 0 | 0 | 0 | 0 | 3.2719 | 1.1138 |
| 48001 | ENSMUSG00000110245 | 3.346039557 | -0.90544729 | 1.298776754 | -0.697153908 | 0.485706481 | NA | Gm20100 | predicted gene, 20100 [Source:MGI Symb | 4.0155 | 6.1675 | 3.0187 | 5.7839 | 1.0906 | 0 |
| 48002 | ENSMUSG00000110246 | 0 | NA | NA | NA | NA | NA | C130073E24 | RIKEN cDNA C130073E24 gene [Source:M | 0 | 0 | 0 | 0 | 0 | 0 |
| 48003 | ENSMUSG00000110247 | 0.503122293 | -2.366675966 | 3.785596405 | -0.625179156 | 0.531853481 | NA | Gm45269 | predicted gene 45269 [Source:MGI Symbo | 0 | 0 | 3.0187 | 0 | 0 | 0 |
| 48004 | ENSMUSG00000110248 | 0.160662974 | 1.126924175 | 4.080472857 | 0.276174898 | 0.782413736 | NA | Gm45615 | predicted gene 45615 [Source:MGI Symbo | 0 | 0 | 0 | 0.964 | 0 | 0 |
| 48005 | ENSMUSG00000110249 | 0 | NA | NA | NA | NA | NA | 9330121K16 | RIKEN cDNA 9330121K16 gene [Source:M | 0 | 0 | 0 | 0 | 0 | 0 |
| 48006 | ENSMUSG00000110250 | 0 | NA | NA | NA | NA | NA | Gm9077 | predicted gene 9077 [Source:MGI Symbol | 0 | 0 | 0 | 0 | 0 | 0 |
| 48007 | ENSMUSG00000110251 | 0.48903338 | 0.95637552 | 3.456690196 | 0.276673773 | 0.782030612 | NA | Gm2996 | predicted gene 2996 [Source:MGI Symbol | 0 | 0 | 1.0062 | 1.928 | 0 | 0 |
| 48008 | ENSMUSG00000110252 | 0 | NA | NA | NA | NA | NA | Gm2961 | predicted gene 2961 [Source:MGI Symbol | 0 | 0 | 0 | 0 | 0 | 0 |
| 48009 | ENSMUSG00000110253 | 0 | NA | NA | NA | NA | NA | Olfr495 | olfactory receptor 495 [Source:MGI Symb | 0 | 0 | 0 | 0 | 0 | 0 |
| 48010 | ENSMUSG00000110254 | 0 | NA | NA | NA | NA | NA | Gm47762 | predicted gene, 47762 [Source:MGI Symb | 0 | 0 | 0 | 0 | 0 | 0 |
| 48011 | ENSMUSG00000110255 | 0 | NA | NA | NA | NA | NA | Gm45400 | predicted gene 45400 [Source:MGI Symbo | 0 | 0 | 0 | 0 | 0 | 0 |
| 48012 | ENSMUSG00000110256 | 5.721671899 | -0.106971792 | 0.951498173 | -0.11242459 | 0.910486759 | 0.985650811 | Gm45412 | predicted gene 45412 [Source:MGI Symbo | 5.6217 | 8.2233 | 4.025 | 7.7118 | 7.6343 | 1.1138 |
| 48013 | ENSMUSG00000110257 | 0 | NA | NA | NA | NA | NA | Olfr529-ps1 | olfactory receptor 529, pseudogene 1 [So | 0 | 0 | 0 | 0 | 0 | 0 |
| 48014 | ENSMUSG00000110258 | 0 | NA | NA | NA | NA | NA | Gm45406 | predicted gene 45406 [Source:MGI Symbo | 0 | 0 | 0 | 0 | 0 | 0 |
| 48015 | ENSMUSG00000110259 | 0 | NA | NA | NA | NA | NA | Olfr65 | olfactory receptor 65 [Source:MGI Symbo | 0 | 0 | 0 | 0 | 0 | 0 |
| 48016 | ENSMUSG00000110260 | 0 | NA | NA | NA | NA | NA | Gm30504 | predicted gene, 30504 [Source:MGI Symb | 0 | 0 | 0 | 0 | 0 | 0 |
| 48017 | ENSMUSG00000110261 | 0 | NA | NA | NA | NA | NA | Gm45485 | predicted gene 45485 [Source:MGI Symbo | 0 | 0 | 0 | 0 | 0 | 0 |
| 48018 | ENSMUSG00000110262 | 0 | NA | NA | NA | NA | NA | Gm45614 | predicted gene 45614 [Source:MGI Symbo | 0 | 0 | 0 | 0 | 0 | 0 |
| 48019 | ENSMUSG00000110263 | 0 | NA | NA | NA | NA | NA | Gm45652 | predicted gene 45652 [Source:MGI Symbo | 0 | 0 | 0 | 0 | 0 | 0 |
| 48020 | ENSMUSG00000110264 | 0 | NA | NA | NA | NA | NA | E330018M18 | RIKEN cDNA E330018M18 gene [Source:M | 0 | 0 | 0 | 0 | 0 | 0 |
| 48021 | ENSMUSG00000110265 | 0 | NA | NA | NA | NA | NA | Gm45331 | predicted gene 45331 [Source:MGI Symbo | 0 | 0 | 0 | 0 | 0 | 0 |
| 48022 | ENSMUSG00000110266 | 0 | NA | NA | NA | NA | NA | Gm32742 | predicted gene, 32742 [Source:MGI Symb | 0 | 0 | 0 | 0 | 0 | 0 |
| 48023 | ENSMUSG00000110267 | 0 | NA | NA | NA | NA | NA | Gm45283 | predicted gene 45283 [Source:MGI Symbo | 0 | 0 | 0 | 0 | 0 | 0 |
| 48024 | ENSMUSG00000110268 | 0 | NA | NA | NA | NA | NA | Gm45436 | predicted gene 45436 [Source:MGI Symbo | 0 | 0 | 0 | 0 | 0 | 0 |
| 48025 | ENSMUSG00000110269 | 0 | NA | NA | NA | NA | NA | Gm45258 | predicted gene 45258 [Source:MGI Symbo | 0 | 0 | 0 | 0 | 0 | 0 |
| 48026 | ENSMUSG00000110270 | 0 | NA | NA | NA | NA | NA | Gm45607 | predicted gene 45607 [Source:MGI Symbo | 0 | 0 | 0 | 0 | 0 | 0 |
| 48027 | ENSMUSG00000110271 | 0 | NA | NA | NA | NA | NA | Gm45387 | predicted gene 45387 [Source:MGI Symbo | 0 | 0 | 0 | 0 | 0 | 0 |
| 48028 | ENSMUSG00000110272 | 0 | NA | NA | NA | NA | NA | Gm45310 | predicted gene 45310 [Source:MGI Symbo | 0 | 0 | 0 | 0 | 0 | 0 |
| 48029 | ENSMUSG00000110273 | 16.85502834 | 0.743844543 | 0.525826895 | 1.41461867 | 0.157180332 | 0.576436451 | Gm41231 | predicted gene, 41231 [Source:MGI Symb | 15.259 | 11.307 | 11.069 | 18.316 | 22.903 | 22.277 |
| 48030 | ENSMUSG00000110274 | 0 | NA | NA | NA | NA | NA | Gm36843 | predicted gene, 36843 [Source:MGI Symb | 0 | 0 | 0 | 0 | 0 | 0 |
| 48031 | ENSMUSG00000110275 | 4.108695636 | 0.917646831 | 1.06197291 | 0.864096271 | 0.387535007 | NA | Gm5905 | predicted gene 5905 [Source:MGI Symbol | 2.4093 | 4.1117 | 2.0125 | 2.8919 | 6.5437 | 6.6831 |
| 48032 | ENSMUSG00000110276 | 1.941904335 | 1.535280249 | 1.803191949 | 0.851423638 | 0.394534066 | NA | Gm45330 | predicted gene 45330 [Source:MGI Symbo | 0 | 1.0279 | 2.0125 | 1.928 | 0 | 6.6831 |
| 48033 | ENSMUSG00000110277 | 111.1803579 | 0.811391725 | 0.272992342 | 2.972214238 | 0.002956603 | 0.046110081 | Gm45871 | predicted gene 45871 [Source:MGI Symbo | 68.264 | 95.596 | 78.487 | 118.57 | 111.24 | 194.92 |
| 48034 | ENSMUSG00000110278 | 0 | NA | NA | NA | NA | NA | Gm5608 | predicted gene 5608 [Source:MGI Symbol | 0 | 0 | 0 | 0 | 0 | 0 |
| 48035 | ENSMUSG00000110279 | 2.826932217 | 1.189630683 | 1.448323426 | 0.821384686 | 0.411427183 | NA | Gm45552 | predicted gene 45552 [Source:MGI Symbo | 4.0155 | 0 | 1.0062 | 0.964 | 7.6343 | 3.3415 |
| 48036 | ENSMUSG00000110280 | 0 | NA | NA | NA | NA | NA | Gm45497 | predicted gene 45497 [Source:MGI Symbo | 0 | 0 | 0 | 0 | 0 | 0 |
| 48037 | ENSMUSG00000110281 | 0 | NA | NA | NA | NA | NA | Gm2516 | predicted gene 2516 [Source:MGI Symbol | 0 | 0 | 0 | 0 | 0 | 0 |
| 48038 | ENSMUSG00000110282 | 0 | NA | NA | NA | NA | NA | B930086L07 | RIKEN cDNA B930086L07 gene [Source:M | 0 | 0 | 0 | 0 | 0 | 0 |
| 48039 | ENSMUSG00000110283 | 0 | NA | NA | NA | NA | NA | Gm45249 | predicted gene 45249 [Source:MGI Symbo | 0 | 0 | 0 | 0 | 0 | 0 |
| 48040 | ENSMUSG00000110284 | 0 | NA | NA | NA | NA | NA | Gm45273 | predicted gene 45273 [Source:MGI Symbo | 0 | 0 | 0 | 0 | 0 | 0 |
| 48041 | ENSMUSG00000110285 | 0 | NA | NA | NA | NA | NA | Gm45524 | predicted gene 45524 [Source:MGI Symbo | 0 | 0 | 0 | 0 | 0 | 0 |
| 48042 | ENSMUSG00000110286 | 0 | NA | NA | NA | NA | NA | Gm7706 | predicted gene 7706 [Source:MGI Symbol | 0 | 0 | 0 | 0 | 0 | 0 |
| 48043 | ENSMUSG00000110287 | 0.267700746 | -1.497481649 | 4.057001436 | -0.369110456 | 0.712045396 | NA | Gm45313 | predicted gene 45313 [Source:MGI Symbo | 1.6062 | 0 | 0 | 0 | 0 | 0 |
| 48044 | ENSMUSG00000110288 | 0 | NA | NA | NA | NA | NA | Gm45678 | predicted gene 45678 [Source:MGI Symbo | 0 | 0 | 0 | 0 | 0 | 0 |

| | | | | | | | | | | | | | | | |
|---|---|---|---|---|---|---|---|---|---|---|---|---|---|---|---|
| 48045 | ENSMUSG00000110289 | 0.82991484 | -1.853997749 | 2.875131703 | -0.644839242 | 0.519031369 | NA | 4930412F12 | RIKEN cDNA 4930412F12 gene [Source:M( | 4.0155 | 0 | 0 | 0.964 | 0 | 0 |
| 48046 | ENSMUSG00000110290 | 0 | NA | NA | NA | NA | NA | Gm45336 | predicted gene 45336 [Source:MGI Symbo | 0 | 0 | 0 | 0 | 0 | 0 |
| 48047 | ENSMUSG00000110291 | 4.42517399 | -0.882090442 | 1.017920561 | -0.866561179 | 0.386182492 | NA | Gm37419 | predicted gene, 37419 [Source:MGI Symb | 8.031 | 5.1396 | 4.025 | 3.8559 | 3.2719 | 2.2277 |
| 48048 | ENSMUSG00000110292 | 0 | NA | NA | NA | NA | NA | Gm17850 | predicted gene, 17850 [Source:MGI Symb | 0 | 0 | 0 | 0 | 0 | 0 |
| 48049 | ENSMUSG00000110293 | 0 | NA | NA | NA | NA | NA | Gm45557 | predicted gene 45557 [Source:MGI Symbo | 0 | 0 | 0 | 0 | 0 | 0 |
| 48050 | ENSMUSG00000110294 | 0 | NA | NA | NA | NA | NA | Gm45430 | predicted gene 45430 [Source:MGI Symbo | 0 | 0 | 0 | 0 | 0 | 0 |
| 48051 | ENSMUSG00000110295 | 0 | NA | NA | NA | NA | NA | Gm45319 | predicted gene 45319 [Source:MGI Symbo | 0 | 0 | 0 | 0 | 0 | 0 |
| 48052 | ENSMUSG00000110296 | 0 | NA | NA | NA | NA | NA | Gm45533 | predicted gene 45533 [Source:MGI Symbo | 0 | 0 | 0 | 0 | 0 | 0 |
| 48053 | ENSMUSG00000110297 | 0 | NA | NA | NA | NA | NA | Gm45647 | predicted gene 45647 [Source:MGI Symbo | 0 | 0 | 0 | 0 | 0 | 0 |
| 48054 | ENSMUSG00000110298 | 2.367534642 | 0.972245575 | 1.382440448 | 0.703282066 | 0.481879985 | NA | Gm8189 | predicted gene 8189 [Source:MGI Symbol | 0.8031 | 1.0279 | 3.0187 | 3.8559 | 3.2719 | 2.2277 |
| 48055 | ENSMUSG00000110299 | 0.958478575 | 0.054139297 | 2.45225727 | 0.022077331 | 0.982386269 | NA | Gm17910 | predicted gene, 17910 [Source:MGI Symb | 0.8031 | 2.0558 | 0 | 2.8919 | 0 | 0 |
| 48056 | ENSMUSG00000110300 | 0 | NA | NA | NA | NA | NA | Gm18645 | predicted gene, 18645 [Source:MGI Symb | 0 | 0 | 0 | 0 | 0 | 0 |
| 48057 | ENSMUSG00000110301 | 0 | NA | NA | NA | NA | NA | Gm35363 | predicted gene, 35363 [Source:MGI Symb | 0 | 0 | 0 | 0 | 0 | 0 |
| 48058 | ENSMUSG00000110302 | 0.401551119 | -2.072982726 | 4.008251979 | -0.517178744 | 0.605031388 | NA | Gm45295 | predicted gene 45295 [Source:MGI Symbo | 2.4093 | 0 | 0 | 0 | 0 | 0 |
| 48059 | ENSMUSG00000110303 | 0 | NA | NA | NA | NA | NA | Gm45328 | predicted gene 45328 [Source:MGI Symbo | 0 | 0 | 0 | 0 | 0 | 0 |
| 48060 | ENSMUSG00000110304 | 0 | NA | NA | NA | NA | NA | Gm36380 | predicted gene, 36380 [Source:MGI Symb | 0 | 0 | 0 | 0 | 0 | 0 |
| 48061 | ENSMUSG00000110305 | 0 | NA | NA | NA | NA | NA | Gm45513 | predicted gene 45513 [Source:MGI Symbo | 0 | 0 | 0 | 0 | 0 | 0 |
| 48062 | ENSMUSG00000110306 | 0 | NA | NA | NA | NA | NA | NA | NA | 0 | 0 | 0 | 0 | 0 | 0 |
| 48063 | ENSMUSG00000110307 | 0 | NA | NA | NA | NA | NA | Gm3816 | predicted gene 3816 [Source:MGI Symbol | 0 | 0 | 0 | 0 | 0 | 0 |
| 48064 | ENSMUSG00000110308 | 0 | NA | NA | NA | NA | NA | Gm45516 | predicted gene 45516 [Source:MGI Symbo | 0 | 0 | 0 | 0 | 0 | 0 |
| 48065 | ENSMUSG00000110309 | 0 | NA | NA | NA | NA | NA | Gm7419 | predicted gene 7419 [Source:MGI Symbol | 0 | 0 | 0 | 0 | 0 | 0 |
| 48066 | ENSMUSG00000110310 | 0 | NA | NA | NA | NA | NA | 4930518J21I | RIKEN cDNA 4930518J21 gene [Source:M( | 0 | 0 | 0 | 0 | 0 | 0 |
| 48067 | ENSMUSG00000110311 | 0 | NA | NA | NA | NA | NA | Gm45410 | predicted gene 45410 [Source:MGI Symbo | 0 | 0 | 0 | 0 | 0 | 0 |
| 48068 | ENSMUSG00000110312 | 0 | NA | NA | NA | NA | NA | Gm45368 | predicted gene 45368 [Source:MGI Symbo | 0 | 0 | 0 | 0 | 0 | 0 |
| 48069 | ENSMUSG00000110313 | 0 | NA | NA | NA | NA | NA | Gm45420 | predicted gene 45420 [Source:MGI Symbo | 0 | 0 | 0 | 0 | 0 | 0 |
| 48070 | ENSMUSG00000110314 | 0 | NA | NA | NA | NA | NA | Gm45625 | predicted gene 45625 [Source:MGI Symbo | 0 | 0 | 0 | 0 | 0 | 0 |
| 48071 | ENSMUSG00000110315 | 0 | NA | NA | NA | NA | NA | Gm8423 | predicted gene 8423 [Source:MGI Symbol | 0 | 0 | 0 | 0 | 0 | 0 |
| 48072 | ENSMUSG00000110316 | 0 | NA | NA | NA | NA | NA | Gm45311 | predicted gene 45311 [Source:MGI Symbo | 0 | 0 | 0 | 0 | 0 | 0 |
| 48073 | ENSMUSG00000110317 | 51.58288192 | 0.619866118 | 0.32585576 | 1.902271477 | 0.057135673 | 0.339527678 | Gm45250 | predicted gene 45250 [Source:MGI Symbo | 41.761 | 42.145 | 38.237 | 78.082 | 46.897 | 62.375 |
| 48074 | ENSMUSG00000110318 | 1.010036755 | -2.151347842 | 2.436507427 | -0.882963794 | 0.377255834 | NA | Gm2767 | predicted gene 2767 [Source:MGI Symbol | 0 | 3.0838 | 2.0125 | 0.964 | 0 | 0 |
| 48075 | ENSMUSG00000110319 | 0 | NA | NA | NA | NA | NA | Gm45668 | predicted gene 45668 [Source:MGI Symbo | 0 | 0 | 0 | 0 | 0 | 0 |
| 48076 | ENSMUSG00000110320 | 0 | NA | NA | NA | NA | NA | Gm31152 | predicted gene, 31152 [Source:MGI Symb | 0 | 0 | 0 | 0 | 0 | 0 |
| 48077 | ENSMUSG00000110321 | 0 | NA | NA | NA | NA | NA | Gm39164 | predicted gene, 39164 [Source:MGI Symb | 0 | 0 | 0 | 0 | 0 | 0 |
| 48078 | ENSMUSG00000110322 | 0 | NA | NA | NA | NA | NA | Gm8110 | predicted gene 8110 [Source:MGI Symbol | 0 | 0 | 0 | 0 | 0 | 0 |
| 48079 | ENSMUSG00000110323 | 0 | NA | NA | NA | NA | NA | Olfr1345-ps1 | olfactory receptor 1345, pseudogene 1 [S( | 0 | 0 | 0 | 0 | 0 | 0 |
| 48080 | ENSMUSG00000110324 | 0 | NA | NA | NA | NA | NA | Gm40460 | predicted gene, 40460 [Source:MGI Symb | 0 | 0 | 0 | 0 | 0 | 0 |
| 48081 | ENSMUSG00000110325 | 0.36353943 | 2.065989254 | 4.034137301 | 0.512126658 | 0.608562368 | NA | Gm45293 | predicted gene 45293 [Source:MGI Symbo | 0 | 0 | 0 | 0 | 2.1812 | 0 |
| 48082 | ENSMUSG00000110326 | 0.8085244 | 0.747105454 | 2.417511592 | 0.309039037 | 0.757291832 | NA | Gm45378 | predicted gene 45378 [Source:MGI Symbo | 0.8031 | 0 | 1.0062 | 1.928 | 0 | 1.1138 |
| 48083 | ENSMUSG00000110327 | 0 | NA | NA | NA | NA | NA | Gm17994 | predicted gene, 17994 [Source:MGI Symb | 0 | 0 | 0 | 0 | 0 | 0 |
| 48084 | ENSMUSG00000110328 | 0 | NA | NA | NA | NA | NA | Gm45574 | predicted gene 45574 [Source:MGI Symbo | 0 | 0 | 0 | 0 | 0 | 0 |
| 48085 | ENSMUSG00000110329 | 0 | NA | NA | NA | NA | NA | Gm45304 | predicted gene 45304 [Source:MGI Symbo | 0 | 0 | 0 | 0 | 0 | 0 |
| 48086 | ENSMUSG00000110330 | 0 | NA | NA | NA | NA | NA | Gm45275 | predicted gene 45275 [Source:MGI Symbo | 0 | 0 | 0 | 0 | 0 | 0 |
| 48087 | ENSMUSG00000110331 | 4.632200631 | 0.939698466 | 1.0315103 | 0.910992809 | 0.362299162 | NA | Nudc-ps1 | nuclear distribution gene C homolog (Asp | 2.4093 | 4.1117 | 3.0187 | 2.8919 | 10.906 | 4.4554 |
| 48088 | ENSMUSG00000110332 | 1.565206392 | -1.1226443 | 1.871435756 | -0.599883964 | 0.54858357 | NA | Gm19935 | predicted gene, 19935 [Source:MGI Symb | 2.4093 | 3.0838 | 1.0062 | 2.8919 | 0 | 0 |
| 48089 | ENSMUSG00000110333 | 0 | NA | NA | NA | NA | NA | Gm45861 | predicted gene 45861 [Source:MGI Symbo | 0 | 0 | 0 | 0 | 0 | 0 |
| 48090 | ENSMUSG00000110334 | 0 | NA | NA | NA | NA | NA | Gm19921 | predicted gene, 19921 [Source:MGI Symb | 0 | 0 | 0 | 0 | 0 | 0 |
| 48091 | ENSMUSG00000110335 | 0 | NA | NA | NA | NA | NA | 1700015I17F | RIKEN cDNA 1700015I17 gene [Source:M( | 0 | 0 | 0 | 0 | 0 | 0 |
| 48092 | ENSMUSG00000110336 | 0 | NA | NA | NA | NA | NA | 4933400L20I | RIKEN cDNA 4933400L20 gene [Source:M( | 0 | 0 | 0 | 0 | 0 | 0 |
| 48093 | ENSMUSG00000110337 | 0 | NA | NA | NA | NA | NA | Gm3433 | predicted gene 3433 [Source:MGI Symbol | 0 | 0 | 0 | 0 | 0 | 0 |
| 48094 | ENSMUSG00000110338 | 0 | NA | NA | NA | NA | NA | Gm45584 | predicted gene 45584 [Source:MGI Symbo | 0 | 0 | 0 | 0 | 0 | 0 |
| 48095 | ENSMUSG00000110339 | 0 | NA | NA | NA | NA | NA | Vmn2r-ps35 | vomeronasal 2, receptor, pseudogene 35 | 0 | 0 | 0 | 0 | 0 | 0 |
| 48096 | ENSMUSG00000110340 | 0 | NA | NA | NA | NA | NA | 1600027J07F | RIKEN cDNA 1600027J07 gene [Source:M( | 0 | 0 | 0 | 0 | 0 | 0 |
| 48097 | ENSMUSG00000110341 | 9.291470992 | -0.77767474 | 0.835943869 | -0.930295405 | 0.352218156 | 0.792945339 | Gm45424 | predicted gene 45424 [Source:MGI Symbo | 17.668 | 10.279 | 7.0437 | 0.964 | 11.997 | 7.7969 |
| 48098 | ENSMUSG00000110342 | 0 | NA | NA | NA | NA | NA | Gm45263 | predicted gene 45263 [Source:MGI Symbo | 0 | 0 | 0 | 0 | 0 | 0 |
| 48099 | ENSMUSG00000110343 | 0 | NA | NA | NA | NA | NA | Gm45751 | predicted gene 45751 [Source:MGI Symbo | 0 | 0 | 0 | 0 | 0 | 0 |
| 48100 | ENSMUSG00000110344 | 102.5732651 | 0.040256053 | 0.228397505 | 0.176254346 | 0.860094127 | 0.97570916 | Gm45716 | predicted gene 45716 [Source:MGI Symbo | 88.341 | 106.9 | 108.67 | 109.89 | 103.61 | 98.018 |

| | | | | | | | | | | | | | | | |
|---|---|---|---|---|---|---|---|---|---|---|---|---|---|---|---|
| 48101 | ENSMUSG00000110345 | 0.524202404 | 2.600864256 | 3.301637836 | 0.787749713 | 0.430843121 | NA | Gm45572 | predicted gene 45572 [Source:MGI Symbo | 0 | 0 | 0 | 0.964 | 2.1812 | 0 |
| 48102 | ENSMUSG00000110346 | 0 | NA | NA | NA | NA | NA | Gm18122 | predicted gene, 18122 [Source:MGI Symb | 0 | 0 | 0 | 0 | 0 | 0 |
| 48103 | ENSMUSG00000110347 | 0 | NA | NA | NA | NA | NA | Gm45657 | predicted gene 45657 [Source:MGI Symbo | 0 | 0 | 0 | 0 | 0 | 0 |
| 48104 | ENSMUSG00000110348 | 0 | NA | NA | NA | NA | NA | Gm4157 | predicted gene 4157 [Source:MGI Symbol | 0 | 0 | 0 | 0 | 0 | 0 |
| 48105 | ENSMUSG00000110349 | 0 | NA | NA | NA | NA | NA | Gm18602 | predicted gene, 18602 [Source:MGI Symb | 0 | 0 | 0 | 0 | 0 | 0 |
| 48106 | ENSMUSG00000110350 | 1.134335839 | 1.615183059 | 2.312890309 | 0.698339672 | 0.484964795 | NA | Gm10252 | predicted gene 10252 [Source:MGI Symbo | 1.6062 | 0 | 0 | 1.928 | 3.2719 | 0 |
| 48107 | ENSMUSG00000110351 | 0 | NA | NA | NA | NA | NA | Gm32050 | predicted gene, 32050 [Source:MGI Symb | 0 | 0 | 0 | 0 | 0 | 0 |
| 48108 | ENSMUSG00000110352 | 0 | NA | NA | NA | NA | NA | Gm45687 | predicted gene 45687 [Source:MGI Symbo | 0 | 0 | 0 | 0 | 0 | 0 |
| 48109 | ENSMUSG00000110353 | 230.2741087 | -0.481505148 | 0.200043945 | -2.406996865 | 0.016084308 | 0.149271467 | Gm33543 | predicted gene, 33543 [Source:MGI Symb | 241.73 | 263.15 | 299.86 | 155.2 | 177.77 | 243.93 |
| 48110 | ENSMUSG00000110354 | 0 | NA | NA | NA | NA | NA | Gm45686 | predicted gene 45686 [Source:MGI Symbo | 0 | 0 | 0 | 0 | 0 | 0 |
| 48111 | ENSMUSG00000110355 | 0 | NA | NA | NA | NA | NA | Gm7514 | predicted gene 7514 [Source:MGI Symbol | 0 | 0 | 0 | 0 | 0 | 0 |
| 48112 | ENSMUSG00000110356 | 0 | NA | NA | NA | NA | NA | Gm35453 | predicted gene, 35453 [Source:MGI Symb | 0 | 0 | 0 | 0 | 0 | 0 |
| 48113 | ENSMUSG00000110357 | 3.016150595 | -0.711916583 | 1.458695499 | -0.488050168 | 0.625514307 | NA | A030001D2( | RIKEN cDNA A030001D20 gene [Source:M | 3.2124 | 4.1117 | 4.025 | 6.7478 | 0 | 0 |
| 48114 | ENSMUSG00000110358 | 0 | NA | NA | NA | NA | NA | A730009L09 | RIKEN cDNA A730009L09 gene [Source:M | 0 | 0 | 0 | 0 | 0 | 0 |
| 48115 | ENSMUSG00000110359 | 0 | NA | NA | NA | NA | NA | Gm45566 | predicted gene 45566 [Source:MGI Symbo | 0 | 0 | 0 | 0 | 0 | 0 |
| 48116 | ENSMUSG00000110360 | 4.288511969 | 1.066431494 | 1.221030855 | 0.873386196 | 0.382452612 | NA | Gm45380 | predicted gene 45380 [Source:MGI Symbo | 3.2124 | 4.1117 | 1.0062 | 8.6758 | 8.7249 | 0 |
| 48117 | ENSMUSG00000110361 | 0 | NA | NA | NA | NA | NA | Gm8291 | predicted gene 8291 [Source:MGI Symbol | 0 | 0 | 0 | 0 | 0 | 0 |
| 48118 | ENSMUSG00000110362 | 0 | NA | NA | NA | NA | NA | Gm45554 | predicted gene 45554 [Source:MGI Symbo | 0 | 0 | 0 | 0 | 0 | 0 |
| 48119 | ENSMUSG00000110363 | 0 | NA | NA | NA | NA | NA | Gm45576 | predicted gene 45576 [Source:MGI Symbo | 0 | 0 | 0 | 0 | 0 | 0 |
| 48120 | ENSMUSG00000110364 | 0 | NA | NA | NA | NA | NA | Gm45462 | predicted gene 45462 [Source:MGI Symbo | 0 | 0 | 0 | 0 | 0 | 0 |
| 48121 | ENSMUSG00000110365 | 0 | NA | NA | NA | NA | NA | Gm38947 | predicted gene, 38947 [Source:MGI Symb | 0 | 0 | 0 | 0 | 0 | 0 |
| 48122 | ENSMUSG00000110366 | 0.181769715 | 1.126924175 | 4.080472857 | 0.276174898 | 0.782413736 | NA | Gm45885 | predicted gene 45885 [Source:MGI Symbo | 0 | 0 | 0 | 0 | 1.0906 | 0 |
| 48123 | ENSMUSG00000110367 | 0 | NA | NA | NA | NA | NA | Gm34853 | predicted gene, 34853 [Source:MGI Symb | 0 | 0 | 0 | 0 | 0 | 0 |
| 48124 | ENSMUSG00000110368 | 0.321325949 | 1.911575114 | 4.048267893 | 0.472195804 | 0.636787034 | NA | Gm45518 | predicted gene 45518 [Source:MGI Symbo | 0 | 0 | 0 | 1.928 | 0 | 0 |
| 48125 | ENSMUSG00000110369 | 2.264130195 | 0.407327067 | 1.698705627 | 0.23978673 | 0.810495595 | NA | Gm45349 | predicted gene 45349 [Source:MGI Symbo | 0.8031 | 0 | 5.0312 | 0 | 2.1812 | 5.5692 |
| 48126 | ENSMUSG00000110370 | 0 | NA | NA | NA | NA | NA | Gm5749 | predicted gene 5749 [Source:MGI Symbol | 0 | 0 | 0 | 0 | 0 | 0 |
| 48127 | ENSMUSG00000110371 | 0 | NA | NA | NA | NA | NA | Gm9619 | predicted gene 9619 [Source:MGI Symbol | 0 | 0 | 0 | 0 | 0 | 0 |
| 48128 | ENSMUSG00000110372 | 0 | NA | NA | NA | NA | NA | Gm8096 | predicted gene 8096 [Source:MGI Symbol | 0 | 0 | 0 | 0 | 0 | 0 |
| 48129 | ENSMUSG00000110373 | 0.66192371 | 1.647602865 | 2.590567345 | 0.636000785 | 0.52477591 | NA | Gm42303 | predicted gene, 42303 [Source:MGI Symb | 0.8031 | 0 | 0 | 0.964 | 1.0906 | 1.1138 |
| 48130 | ENSMUSG00000110374 | 0 | NA | NA | NA | NA | NA | Gm31172 | predicted gene, 31172 [Source:MGI Symb | 0 | 0 | 0 | 0 | 0 | 0 |
| 48131 | ENSMUSG00000110375 | 0 | NA | NA | NA | NA | NA | Gm45323 | predicted gene 45323 [Source:MGI Symbo | 0 | 0 | 0 | 0 | 0 | 0 |
| 48132 | ENSMUSG00000110376 | 0 | NA | NA | NA | NA | NA | Gm45688 | predicted gene 45688 [Source:MGI Symbo | 0 | 0 | 0 | 0 | 0 | 0 |
| 48133 | ENSMUSG00000110377 | 2.237787296 | 3.623834178 | 1.776046449 | 2.040393808 | 0.041311118 | NA | Gm45363 | predicted gene 45363 [Source:MGI Symbo | 0 | 0 | 1.0062 | 5.7839 | 2.1812 | 4.4554 |
| 48134 | ENSMUSG00000110378 | 4.003649791 | -0.82753875 | 1.154465256 | -0.716815639 | 0.47348786 | NA | Gm45242 | predicted gene 45242 [Source:MGI Symbo | 3.2124 | 2.0558 | 10.062 | 0.964 | 3.2719 | 4.4554 |
| 48135 | ENSMUSG00000110379 | 0 | NA | NA | NA | NA | NA | Gm6909 | predicted gene 6909 [Source:MGI Symbol | 0 | 0 | 0 | 0 | 0 | 0 |
| 48136 | ENSMUSG00000110380 | 0 | NA | NA | NA | NA | NA | Gm45332 | predicted gene 45332 [Source:MGI Symbo | 0 | 0 | 0 | 0 | 0 | 0 |
| 48137 | ENSMUSG00000110381 | 0 | NA | NA | NA | NA | NA | Gm34370 | predicted gene, 34370 [Source:MGI Symb | 0 | 0 | 0 | 0 | 0 | 0 |
| 48138 | ENSMUSG00000110382 | 0 | NA | NA | NA | NA | NA | Gm29895 | predicted gene, 29895 [Source:MGI Symb | 0 | 0 | 0 | 0 | 0 | 0 |
| 48139 | ENSMUSG00000110383 | 0 | NA | NA | NA | NA | NA | Gm45580 | predicted gene 45580 [Source:MGI Symbo | 0 | 0 | 0 | 0 | 0 | 0 |
| 48140 | ENSMUSG00000110384 | 21.2789158 | -3.158650114 | 0.677375801 | -4.663069022 | 3.12E-06 | 0.000204816 | Gm45301 | predicted gene 45301 [Source:MGI Symbo | 16.865 | 58.591 | 39.244 | 1.928 | 4.3625 | 6.6831 |
| 48141 | ENSMUSG00000110385 | 0 | NA | NA | NA | NA | NA | Gm19093 | predicted gene, 19093 [Source:MGI Symb | 0 | 0 | 0 | 0 | 0 | 0 |
| 48142 | ENSMUSG00000110386 | 0 | NA | NA | NA | NA | NA | Gm42031 | predicted gene, 42031 [Source:MGI Symb | 0 | 0 | 0 | 0 | 0 | 0 |
| 48143 | ENSMUSG00000110387 | 0 | NA | NA | NA | NA | NA | Gm45478 | predicted gene 45478 [Source:MGI Symbo | 0 | 0 | 0 | 0 | 0 | 0 |
| 48144 | ENSMUSG00000110388 | 0 | NA | NA | NA | NA | NA | Gm30329 | predicted gene, 30329 [Source:MGI Symb | 0 | 0 | 0 | 0 | 0 | 0 |
| 48145 | ENSMUSG00000110389 | 0 | NA | NA | NA | NA | NA | Gm45579 | predicted gene 45579 [Source:MGI Symbo | 0 | 0 | 0 | 0 | 0 | 0 |
| 48146 | ENSMUSG00000110390 | 0 | NA | NA | NA | NA | NA | Gm45869 | predicted gene 45869 [Source:MGI Symbo | 0 | 0 | 0 | 0 | 0 | 0 |
| 48147 | ENSMUSG00000110391 | 0 | NA | NA | NA | NA | NA | Gm7269 | predicted gene 7269 [Source:MGI Symbol | 0 | 0 | 0 | 0 | 0 | 0 |
| 48148 | ENSMUSG00000110392 | 0 | NA | NA | NA | NA | NA | Gm45676 | predicted gene 45676 [Source:MGI Symbo | 0 | 0 | 0 | 0 | 0 | 0 |
| 48149 | ENSMUSG00000110393 | 12.65627118 | -0.221340926 | 0.619089845 | -0.357526339 | 0.720697812 | 0.9447719 | Gm36445 | predicted gene, 36445 [Source:MGI Symb | 15.259 | 16.447 | 9.0562 | 8.6758 | 10.906 | 15.594 |
| 48150 | ENSMUSG00000110394 | 0 | NA | NA | NA | NA | NA | Gm18991 | predicted gene, 18991 [Source:MGI Symb | 0 | 0 | 0 | 0 | 0 | 0 |
| 48151 | ENSMUSG00000110395 | 0 | NA | NA | NA | NA | NA | Gm4903 | predicted gene 4903 [Source:MGI Symbol | 0 | 0 | 0 | 0 | 0 | 0 |
| 48152 | ENSMUSG00000110396 | 0 | NA | NA | NA | NA | NA | Gm36737 | predicted gene, 36737 [Source:MGI Symb | 0 | 0 | 0 | 0 | 0 | 0 |
| 48153 | ENSMUSG00000110397 | 22.66191372 | 0.063374389 | 0.522331211 | 0.121329891 | 0.903429745 | 0.983821605 | Gm45540 | predicted gene 45540 [Source:MGI Symbo | 12.047 | 24.67 | 30.187 | 32.775 | 21.812 | 14.48 |
| 48154 | ENSMUSG00000110398 | 0 | NA | NA | NA | NA | NA | Gm45616 | predicted gene 45616 [Source:MGI Symbo | 0 | 0 | 0 | 0 | 0 | 0 |
| 48155 | ENSMUSG00000110399 | 0 | NA | NA | NA | NA | NA | Gm45238 | predicted gene 45238 [Source:MGI Symbo | 0 | 0 | 0 | 0 | 0 | 0 |
| 48156 | ENSMUSG00000110400 | 0.670803094 | 1.59186814 | 2.791527899 | 0.570249769 | 0.568508305 | NA | Gm45416 | predicted gene 45416 [Source:MGI Symbo | 0 | 0 | 1.0062 | 1.928 | 1.0906 | 0 |

| | | | | | | | | | | | | | | | |
|---|---|---|---|---|---|---|---|---|---|---|---|---|---|---|---|
| 48157 | ENSMUSG00000110401 | 0 | NA | NA | NA | NA | NA | Gm5740 | predicted gene 5740 [Source:MGI Symbol | 0 | 0 | 0 | 0 | 0 | 0 |
| 48158 | ENSMUSG00000110402 | 0 | NA | NA | NA | NA | NA | Gm7203 | predicted gene 7203 [Source:MGI Symbol | 0 | 0 | 0 | 0 | 0 | 0 |
| 48159 | ENSMUSG00000110403 | 0 | NA | NA | NA | NA | NA | Gm45553 | predicted gene 45553 [Source:MGI Symbo | 0 | 0 | 0 | 0 | 0 | 0 |
| 48160 | ENSMUSG00000110404 | 0.6621303 | -1.329905799 | 2.807279375 | -0.473734752 | 0.635689068 | NA | Rnf223 | ring finger 223 [Source:MGI Symbol;Acc:N | 0.8031 | 2.0558 | 0 | 0 | 0 | 1.1138 |
| 48161 | ENSMUSG00000110405 | 0 | NA | NA | NA | NA | NA | Gm45534 | predicted gene 45534 [Source:MGI Symbo | 0 | 0 | 0 | 0 | 0 | 0 |
| 48162 | ENSMUSG00000110406 | 0 | NA | NA | NA | NA | NA | 4930470O06 | RIKEN cDNA 4930470O06 gene [Source:M | 0 | 0 | 0 | 0 | 0 | 0 |
| 48163 | ENSMUSG00000110407 | 0 | NA | NA | NA | NA | NA | Gm45454 | predicted gene 45454 [Source:MGI Symbo | 0 | 0 | 0 | 0 | 0 | 0 |
| 48164 | ENSMUSG00000110408 | 0.684865378 | 2.990343487 | 2.966805827 | 1.00793367 | 0.313486306 | NA | NA | NA | 0 | 0 | 0 | 1.928 | 2.1812 | 0 |
| 48165 | ENSMUSG00000110409 | 0 | NA | NA | NA | NA | NA | Gm9908 | predicted gene 9908 [Source:MGI Symbol | 0 | 0 | 0 | 0 | 0 | 0 |
| 48166 | ENSMUSG00000110410 | 0 | NA | NA | NA | NA | NA | 4933421D24 | RIKEN cDNA 4933421D24 gene [Source:M | 0 | 0 | 0 | 0 | 0 | 0 |
| 48167 | ENSMUSG00000110411 | 10.89578269 | -0.006490453 | 0.642523866 | -0.010101497 | 0.991940309 | 0.99859176 | Gm45457 | predicted gene 45457 [Source:MGI Symbo | 10.44 | 9.2513 | 13.081 | 10.604 | 13.087 | 8.9108 |
| 48168 | ENSMUSG00000110412 | 0 | NA | NA | NA | NA | NA | Gm33175 | predicted gene, 33175 [Source:MGI Symb | 0 | 0 | 0 | 0 | 0 | 0 |
| 48169 | ENSMUSG00000110413 | 0 | NA | NA | NA | NA | NA | Gm45409 | predicted gene 45409 [Source:MGI Symbo | 0 | 0 | 0 | 0 | 0 | 0 |
| 48170 | ENSMUSG00000110414 | 1.468412105 | 1.945424723 | 2.004800999 | 0.970382958 | 0.33185564 | NA | Gm45481 | predicted gene 45481 [Source:MGI Symbo | 0.8031 | 0 | 1.0062 | 4.8199 | 2.1812 | 0 |
| 48171 | ENSMUSG00000110415 | 0 | NA | NA | NA | NA | NA | Gm6203 | predicted gene 6203 [Source:MGI Symbol | 0 | 0 | 0 | 0 | 0 | 0 |
| 48172 | ENSMUSG00000110416 | 0 | NA | NA | NA | NA | NA | Gm19248 | predicted gene, 19248 [Source:MGI Symb | 0 | 0 | 0 | 0 | 0 | 0 |
| 48173 | ENSMUSG00000110417 | 0 | NA | NA | NA | NA | NA | Gm31727 | predicted gene, 31727 [Source:MGI Symb | 0 | 0 | 0 | 0 | 0 | 0 |
| 48174 | ENSMUSG00000110419 | 1.386635706 | 1.244513227 | 2.170905042 | 0.573269306 | 0.566462364 | NA | AW046200 | expressed sequence AW046200 [Source:N | 2.4093 | 0 | 0 | 4.8199 | 1.0906 | 0 |
| 48175 | ENSMUSG00000110420 | 0 | NA | NA | NA | NA | NA | Gm45870 | predicted gene 45870 [Source:MGI Symbo | 0 | 0 | 0 | 0 | 0 | 0 |
| 48176 | ENSMUSG00000110421 | 0 | NA | NA | NA | NA | NA | Gm9689 | predicted gene 9689 [Source:MGI Symbol | 0 | 0 | 0 | 0 | 0 | 0 |
| 48177 | ENSMUSG00000110422 | 0.674441933 | -2.786326546 | 3.022203109 | -0.921952114 | 0.356553554 | NA | Gm17999 | predicted gene, 17999 [Source:MGI Symb | 0 | 1.0279 | 3.0187 | 0 | 0 | 0 |
| 48178 | ENSMUSG00000110423 | 0 | NA | NA | NA | NA | NA | NA | NA | 0 | 0 | 0 | 0 | 0 | 0 |
| 48179 | ENSMUSG00000110424 | 21.87174912 | -0.263735838 | 0.475036372 | -0.555190831 | 0.57876414 | 0.908239614 | 1700012D14 | RIKEN cDNA 1700012D14 gene [Source:M | 24.896 | 20.558 | 26.162 | 23.135 | 13.087 | 23.391 |
| 48180 | ENSMUSG00000110425 | 0 | NA | NA | NA | NA | NA | Gm32390 | predicted gene, 32390 [Source:MGI Symb | 0 | 0 | 0 | 0 | 0 | 0 |
| 48181 | ENSMUSG00000110426 | 0 | NA | NA | NA | NA | NA | Gm45757 | predicted gene 45757 [Source:MGI Symbo | 0 | 0 | 0 | 0 | 0 | 0 |
| 48182 | ENSMUSG00000110427 | 0 | NA | NA | NA | NA | NA | 4933406B17 | RIKEN cDNA 4933406B17 gene [Source:M | 0 | 0 | 0 | 0 | 0 | 0 |
| 48183 | ENSMUSG00000110428 | 0 | NA | NA | NA | NA | NA | Gm45812 | predicted gene 45812 [Source:MGI Symbo | 0 | 0 | 0 | 0 | 0 | 0 |
| 48184 | ENSMUSG00000110429 | 0 | NA | NA | NA | NA | NA | Gm2059 | predicted gene 2059 [Source:MGI Symbol | 0 | 0 | 0 | 0 | 0 | 0 |
| 48185 | ENSMUSG00000110430 | 0 | NA | NA | NA | NA | NA | Cmtm1 | CKLF-like MARVEL transmembrane domai | 0 | 0 | 0 | 0 | 0 | 0 |
| 48186 | ENSMUSG00000110431 | 0 | NA | NA | NA | NA | NA | Olfr892-ps1 | olfactory receptor 892, pseudogene 1 [Sou | 0 | 0 | 0 | 0 | 0 | 0 |
| 48187 | ENSMUSG00000110432 | 0 | NA | NA | NA | NA | NA | Gm45907 | predicted gene 45907 [Source:MGI Symbo | 0 | 0 | 0 | 0 | 0 | 0 |
| 48188 | ENSMUSG00000110433 | 4.463096039 | -0.346326957 | 1.424541403 | -0.243114701 | 0.807916543 | NA | Gm45820 | predicted gene 45820 [Source:MGI Symbo | 4.0155 | 0 | 11.069 | 10.604 | 1.0906 | 0 |
| 48189 | ENSMUSG00000110434 | 0 | NA | NA | NA | NA | NA | Rps7-ps1 | ribosomal protein S7, pseudogene 1 [Sour | 0 | 0 | 0 | 0 | 0 | 0 |
| 48190 | ENSMUSG00000110435 | 0 | NA | NA | NA | NA | NA | Gm3635 | predicted gene 3635 [Source:MGI Symbol | 0 | 0 | 0 | 0 | 0 | 0 |
| 48191 | ENSMUSG00000110437 | 0 | NA | NA | NA | NA | NA | Gm45876 | predicted gene 45876 [Source:MGI Symbo | 0 | 0 | 0 | 0 | 0 | 0 |
| 48192 | ENSMUSG00000110438 | 1.060069211 | -2.228626096 | 2.226826576 | -1.00080811 | 0.316919588 | NA | Gm45756 | predicted gene 45756 [Source:MGI Symbo | 3.2124 | 1.0279 | 1.0062 | 0 | 0 | 1.1138 |
| 48193 | ENSMUSG00000110439 | 830.2760931 | 0.401809814 | 0.354294531 | 1.13411238 | 0.256747411 | 0.710192112 | Mup22 | major urinary protein 22 [Source:MGI Syn | 624.01 | 528.35 | 994.17 | 1241.6 | 855.04 | 738.48 |
| 48194 | ENSMUSG00000110440 | 0 | NA | NA | NA | NA | NA | Gm45894 | predicted gene 45894 [Source:MGI Symbo | 0 | 0 | 0 | 0 | 0 | 0 |
| 48195 | ENSMUSG00000110441 | 0 | NA | NA | NA | NA | NA | Gm45790 | predicted gene 45790 [Source:MGI Symbo | 0 | 0 | 0 | 0 | 0 | 0 |
| 48196 | ENSMUSG00000110442 | 0 | NA | NA | NA | NA | NA | Gm45804 | predicted gene 45804 [Source:MGI Symbo | 0 | 0 | 0 | 0 | 0 | 0 |
| 48197 | ENSMUSG00000110443 | 0 | NA | NA | NA | NA | NA | Gm39197 | predicted gene, 39197 [Source:MGI Symb | 0 | 0 | 0 | 0 | 0 | 0 |
| 48198 | ENSMUSG00000110444 | 170.4894447 | 0.213011665 | 0.211692681 | 1.006230656 | 0.314304629 | 0.760853916 | Gm10033 | predicted gene 10033 [Source:MGI Symbo | 123.68 | 165.49 | 185.15 | 165.8 | 173.41 | 209.4 |
| 48199 | ENSMUSG00000110445 | 0.133850373 | -0.796629176 | 4.080472857 | -0.195229623 | 0.84521319 | NA | Gm45747 | predicted gene 45747 [Source:MGI Symbo | 0.8031 | 0 | 0 | 0 | 0 | 0 |
| 48200 | ENSMUSG00000110446 | 0 | NA | NA | NA | NA | NA | Gm18646 | predicted gene, 18646 [Source:MGI Symb | 0 | 0 | 0 | 0 | 0 | 0 |
| 48201 | ENSMUSG00000110447 | 0 | NA | NA | NA | NA | NA | Gm31518 | predicted gene, 31518 [Source:MGI Symb | 0 | 0 | 0 | 0 | 0 | 0 |
| 48202 | ENSMUSG00000110448 | 0 | NA | NA | NA | NA | NA | Gm33242 | predicted gene, 33242 [Source:MGI Symb | 0 | 0 | 0 | 0 | 0 | 0 |
| 48203 | ENSMUSG00000110449 | 0 | NA | NA | NA | NA | NA | Gm45780 | predicted gene 45780 [Source:MGI Symbo | 0 | 0 | 0 | 0 | 0 | 0 |
| 48204 | ENSMUSG00000110450 | 0 | NA | NA | NA | NA | NA | Gm7943 | predicted gene 7943 [Source:MGI Symbol | 0 | 0 | 0 | 0 | 0 | 0 |
| 48205 | ENSMUSG00000110452 | 0 | NA | NA | NA | NA | NA | Gm36247 | predicted gene, 36247 [Source:MGI Symb | 0 | 0 | 0 | 0 | 0 | 0 |
| 48206 | ENSMUSG00000110453 | 0.717171207 | -1.535127875 | 3.012718241 | -0.509549102 | 0.61036739 | NA | Gm32352 | predicted gene, 32352 [Source:MGI Symb | 3.2124 | 0 | 0 | 0 | 1.0906 | 0 |
| 48207 | ENSMUSG00000110454 | 0 | NA | NA | NA | NA | NA | Gm45911 | predicted gene 45911 [Source:MGI Symbo | 0 | 0 | 0 | 0 | 0 | 0 |
| 48208 | ENSMUSG00000110455 | 0 | NA | NA | NA | NA | NA | Gm45904 | predicted gene 45904 [Source:MGI Symbo | 0 | 0 | 0 | 0 | 0 | 0 |
| 48209 | ENSMUSG00000110456 | 2.461279673 | 1.435864274 | 1.480301167 | 0.969981181 | 0.332055873 | NA | Gm45865 | predicted gene 45865 [Source:MGI Symbo | 0 | 2.0558 | 2.0125 | 1.928 | 6.5437 | 2.2277 |
| 48210 | ENSMUSG00000110457 | 0 | NA | NA | NA | NA | NA | Gm45752 | predicted gene 45752 [Source:MGI Symbo | 0 | 0 | 0 | 0 | 0 | 0 |
| 48211 | ENSMUSG00000110458 | 7.448209611 | -0.05689898 | 0.831541594 | -0.068425897 | 0.945446607 | 0.990309554 | Gm21769 | predicted gene, 21769 [Source:MGI Symb | 5.6217 | 7.1954 | 10.062 | 8.6758 | 10.906 | 2.2277 |
| 48212 | ENSMUSG00000110459 | 14.46657461 | 1.257429619 | 0.577687411 | 2.176660933 | 0.029505875 | 0.226164801 | Gm31717 | predicted gene, 31717 [Source:MGI Symb | 6.4248 | 9.2513 | 10.062 | 20.244 | 18.541 | 22.277 |

| | | | | | | | | | | | | | | | |
|---|---|---|---|---|---|---|---|---|---|---|---|---|---|---|---|
| 48213 | ENSMUSG00000110460 | 0 | NA | NA | NA | NA | NA | Gm20786 | predicted gene, 20786 [Source:MGI Symb | 0 | 0 | 0 | 0 | 0 | 0 |
| 48214 | ENSMUSG00000110461 | 0 | NA | NA | NA | NA | NA | Gm45860 | predicted gene 45860 [Source:MGI Symbo | 0 | 0 | 0 | 0 | 0 | 0 |
| 48215 | ENSMUSG00000110462 | 0.401551119 | -2.072982726 | 4.008251979 | -0.517178744 | 0.605031388 | NA | Gm35572 | predicted gene, 35572 [Source:MGI Symb | 2.4093 | 0 | 0 | 0 | 0 | 0 |
| 48216 | ENSMUSG00000110463 | 0 | NA | NA | NA | NA | NA | Gm35998 | predicted gene, 35998 [Source:MGI Symb | 0 | 0 | 0 | 0 | 0 | 0 |
| 48217 | ENSMUSG00000110464 | 0 | NA | NA | NA | NA | NA | 1700061N14 | RIKEN cDNA 1700061N14 gene [Source:M | 0 | 0 | 0 | 0 | 0 | 0 |
| 48218 | ENSMUSG00000110465 | 0 | NA | NA | NA | NA | NA | Gm45832 | predicted gene 45832 [Source:MGI Symbo | 0 | 0 | 0 | 0 | 0 | 0 |
| 48219 | ENSMUSG00000110466 | 0 | NA | NA | NA | NA | NA | Gm45815 | predicted gene 45815 [Source:MGI Symbo | 0 | 0 | 0 | 0 | 0 | 0 |
| 48220 | ENSMUSG00000110467 | 0 | NA | NA | NA | NA | NA | Gm45781 | predicted gene 45781 [Source:MGI Symbo | 0 | 0 | 0 | 0 | 0 | 0 |
| 48221 | ENSMUSG00000110468 | 0 | NA | NA | NA | NA | NA | Gm39185 | predicted gene, 39185 [Source:MGI Symb | 0 | 0 | 0 | 0 | 0 | 0 |
| 48222 | ENSMUSG00000110469 | 0 | NA | NA | NA | NA | NA | Gm10358 | predicted gene 10358 [Source:MGI Symbo | 0 | 0 | 0 | 0 | 0 | 0 |
| 48223 | ENSMUSG00000110470 | 0 | NA | NA | NA | NA | NA | Gm45723 | predicted gene 45723 [Source:MGI Symbo | 0 | 0 | 0 | 0 | 0 | 0 |
| 48224 | ENSMUSG00000110471 | 0 | NA | NA | NA | NA | NA | Gm45823 | predicted gene 45823 [Source:MGI Symbo | 0 | 0 | 0 | 0 | 0 | 0 |
| 48225 | ENSMUSG00000110472 | 0 | NA | NA | NA | NA | NA | Gm45739 | predicted gene 45739 [Source:MGI Symbo | 0 | 0 | 0 | 0 | 0 | 0 |
| 48226 | ENSMUSG00000110473 | 0 | NA | NA | NA | NA | NA | Gm29705 | predicted gene, 29705 [Source:MGI Symb | 0 | 0 | 0 | 0 | 0 | 0 |
| 48227 | ENSMUSG00000110474 | 0 | NA | NA | NA | NA | NA | Gm45848 | predicted gene 45848 [Source:MGI Symbo | 0 | 0 | 0 | 0 | 0 | 0 |
| 48228 | ENSMUSG00000110475 | 7.572997558 | 0.147038102 | 0.806295271 | 0.1823626 | 0.855298174 | 0.974158771 | Gm31036 | predicted gene, 31036 [Source:MGI Symb | 11.243 | 7.1954 | 3.0187 | 9.6398 | 6.5437 | 7.7969 |
| 48229 | ENSMUSG00000110476 | 0 | NA | NA | NA | NA | NA | 4930422C21 | RIKEN cDNA 4930422C21 gene [Source:M | 0 | 0 | 0 | 0 | 0 | 0 |
| 48230 | ENSMUSG00000110477 | 0.160662974 | 1.126924175 | 4.080472857 | 0.276174898 | 0.782413736 | NA | Gm3045 | predicted gene 3045 [Source:MGI Symbol | 0 | 0 | 0 | 0.964 | 0 | 0 |
| 48231 | ENSMUSG00000110478 | 0 | NA | NA | NA | NA | NA | Gm18000 | predicted gene, 18000 [Source:MGI Symb | 0 | 0 | 0 | 0 | 0 | 0 |
| 48232 | ENSMUSG00000110479 | 0.167707431 | -0.796629176 | 4.080472857 | -0.195229623 | 0.84521319 | NA | Gm5743 | predicted gene 5743 [Source:MGI Symbol | 0 | 0 | 1.0062 | 0 | 0 | 0 |
| 48233 | ENSMUSG00000110480 | 0 | NA | NA | NA | NA | NA | Gm32842 | predicted gene, 32842 [Source:MGI Symb | 0 | 0 | 0 | 0 | 0 | 0 |
| 48234 | ENSMUSG00000110481 | 0 | NA | NA | NA | NA | NA | Gm45705 | predicted gene 45705 [Source:MGI Symbo | 0 | 0 | 0 | 0 | 0 | 0 |
| 48235 | ENSMUSG00000110482 | 0 | NA | NA | NA | NA | NA | Gm3889 | predicted gene 3889 [Source:MGI Symbol | 0 | 0 | 0 | 0 | 0 | 0 |
| 48236 | ENSMUSG00000110483 | 0 | NA | NA | NA | NA | NA | Gm45702 | predicted gene 45702 [Source:MGI Symbo | 0 | 0 | 0 | 0 | 0 | 0 |
| 48237 | ENSMUSG00000110484 | 0.167707431 | -0.796629176 | 4.080472857 | -0.195229623 | 0.84521319 | NA | Gm45712 | predicted gene 45712 [Source:MGI Symbo | 0 | 0 | 1.0062 | 0 | 0 | 0 |
| 48238 | ENSMUSG00000110485 | 0 | NA | NA | NA | NA | NA | Gm45740 | predicted gene 45740 [Source:MGI Symbo | 0 | 0 | 0 | 0 | 0 | 0 |
| 48239 | ENSMUSG00000110486 | 0.524202404 | 2.600864256 | 3.301637836 | 0.787749713 | 0.430843121 | NA | Gm45877 | predicted gene 45877 [Source:MGI Symbo | 0 | 0 | 0 | 0.964 | 2.1812 | 0 |
| 48240 | ENSMUSG00000110487 | 0 | NA | NA | NA | NA | NA | Gm18035 | predicted gene, 18035 [Source:MGI Symb | 0 | 0 | 0 | 0 | 0 | 0 |
| 48241 | ENSMUSG00000110488 | 62.00842679 | 0.866974844 | 0.391289036 | 2.21568908 | 0.026712803 | 0.211364326 | Gm45724 | predicted gene 45724 [Source:MGI Symbo | 32.927 | 35.977 | 63.393 | 126.28 | 58.893 | 54.578 |
| 48242 | ENSMUSG00000110489 | 0 | NA | NA | NA | NA | NA | Gm31659 | predicted gene, 31659 [Source:MGI Symb | 0 | 0 | 0 | 0 | 0 | 0 |
| 48243 | ENSMUSG00000110490 | 0 | NA | NA | NA | NA | NA | Vmn2r-ps91 | vomeronasal 2, receptor, pseudogene 91 | 0 | 0 | 0 | 0 | 0 | 0 |
| 48244 | ENSMUSG00000110491 | 0.328370405 | 0.16514316 | 4.003149536 | 0.041253308 | 0.967093956 | NA | Gm31138 | predicted gene, 31138 [Source:MGI Symb | 0 | 0 | 1.0062 | 0.964 | 0 | 0 |
| 48245 | ENSMUSG00000110492 | 0 | NA | NA | NA | NA | NA | Gm5358 | predicted gene 5358 [Source:MGI Symbol | 0 | 0 | 0 | 0 | 0 | 0 |
| 48246 | ENSMUSG00000110493 | 0 | NA | NA | NA | NA | NA | | olfactory receptor 841, pseudogene 1 | 0 | 0 | 0 | 0 | 0 | 0 |
| 48247 | ENSMUSG00000110494 | 22.86999399 | 0.635676029 | 0.584979721 | 1.08666336 | 0.277185612 | 0.730041765 | Gm31774 | predicted gene, 31774 [Source:MGI Symb | 10.44 | 20.558 | 23.144 | 50.127 | 21.812 | 11.138 |
| 48248 | ENSMUSG00000110495 | 0 | NA | NA | NA | NA | NA | Gm45879 | predicted gene 45879 [Source:MGI Symbo | 0 | 0 | 0 | 0 | 0 | 0 |
| 48249 | ENSMUSG00000110496 | 68.27916688 | 0.073949331 | 0.364757879 | 0.202735391 | 0.839341859 | 0.971685662 | Gm45909 | predicted gene 45909 [Source:MGI Symbo | 90.751 | 46.256 | 62.387 | 90.614 | 81.796 | 37.871 |
| 48250 | ENSMUSG00000110497 | 0 | NA | NA | NA | NA | NA | 9230110F11 | RIKEN cDNA 9230110F11 gene [Source:M | 0 | 0 | 0 | 0 | 0 | 0 |
| 48251 | ENSMUSG00000110498 | 1.319744392 | -0.523326266 | 1.997208713 | -0.262028832 | 0.793299214 | NA | A630001O12 | RIKEN cDNA A630001O12 gene [Source:M | 1.6062 | 1.0279 | 2.0125 | 0 | 3.2719 | 0 |
| 48252 | ENSMUSG00000110499 | 1.588008091 | 3.105077357 | 2.40190538 | 1.292755902 | 0.196095497 | NA | Gm45873 | predicted gene 45873 [Source:MGI Symbo | 0.8031 | 0 | 0 | 0 | 8.7249 | 0 |
| 48253 | ENSMUSG00000110500 | 0 | NA | NA | NA | NA | NA | Gm32568 | predicted gene, 32568 [Source:MGI Symb | 0 | 0 | 0 | 0 | 0 | 0 |
| 48254 | ENSMUSG00000110501 | 0.663758638 | 2.950973782 | 3.025092005 | 0.975498853 | 0.32931287 | NA | Gm45900 | predicted gene 45900 [Source:MGI Symbo | 0 | 0 | 0 | 2.8919 | 1.0906 | 0 |
| 48255 | ENSMUSG00000110502 | 0 | NA | NA | NA | NA | NA | Gm8804 | predicted gene 8804 [Source:MGI Symbol | 0 | 0 | 0 | 0 | 0 | 0 |
| 48256 | ENSMUSG00000110503 | 0 | NA | NA | NA | NA | NA | Gm45830 | predicted gene 45830 [Source:MGI Symbo | 0 | 0 | 0 | 0 | 0 | 0 |
| 48257 | ENSMUSG00000110504 | 5.353491994 | -0.943236568 | 1.069434853 | -0.88199535 | 0.377779322 | 0.810712182 | Gm45797 | predicted gene 45797 [Source:MGI Symbo | 8.031 | 7.1954 | 6.0375 | 8.6758 | 2.1812 | 0 |
| 48258 | ENSMUSG00000110505 | 0 | NA | NA | NA | NA | NA | Gm45842 | predicted gene 45842 [Source:MGI Symbo | 0 | 0 | 0 | 0 | 0 | 0 |
| 48259 | ENSMUSG00000110506 | 0.181769715 | 1.126924175 | 4.080472857 | 0.276174898 | 0.782413736 | NA | Gm8637 | predicted gene 8637 [Source:MGI Symbol | 0 | 0 | 0 | 0 | 1.0906 | 0 |
| 48260 | ENSMUSG00000110507 | 0 | NA | NA | NA | NA | NA | 8030455M16 | RIKEN cDNA 8030455M16 gene [Source:M | 0 | 0 | 0 | 0 | 0 | 0 |
| 48261 | ENSMUSG00000110508 | 0 | NA | NA | NA | NA | NA | Gm8940 | predicted gene 8940 [Source:MGI Symbol | 0 | 0 | 0 | 0 | 0 | 0 |
| 48262 | ENSMUSG00000110509 | 0 | NA | NA | NA | NA | NA | Gm45772 | predicted gene 45772 [Source:MGI Symbo | 0 | 0 | 0 | 0 | 0 | 0 |
| 48263 | ENSMUSG00000110510 | 0 | NA | NA | NA | NA | NA | Gm45908 | predicted gene 45908 [Source:MGI Symbo | 0 | 0 | 0 | 0 | 0 | 0 |
| 48264 | ENSMUSG00000110511 | 0 | NA | NA | NA | NA | NA | Gm45699 | predicted gene 45699 [Source:MGI Symbo | 0 | 0 | 0 | 0 | 0 | 0 |
| 48265 | ENSMUSG00000110512 | 0 | NA | NA | NA | NA | NA | Gm45827 | predicted gene 45827 [Source:MGI Symbo | 0 | 0 | 0 | 0 | 0 | 0 |
| 48266 | ENSMUSG00000110513 | 0 | NA | NA | NA | NA | NA | Olfr882-ps1 | olfactory receptor 882, pseudogene 1 [So | 0 | 0 | 0 | 0 | 0 | 0 |
| 48267 | ENSMUSG00000110514 | 0 | NA | NA | NA | NA | NA | Gm7688 | predicted gene 7688 [Source:MGI Symbol | 0 | 0 | 0 | 0 | 0 | 0 |
| 48268 | ENSMUSG00000110515 | 0 | NA | NA | NA | NA | NA | Gm9644 | predicted gene 9644 [Source:MGI Symbol | 0 | 0 | 0 | 0 | 0 | 0 |

| | | | | | | | | | | | | | | | |
|---|---|---|---|---|---|---|---|---|---|---|---|---|---|---|---|
| 48269 | ENSMUSG00000110516 | 0 | NA | NA | NA | NA | NA | Gm45758 | predicted gene 45758 [Source:MGI Symbo | 0 | 0 | 0 | 0 | 0 | 0 |
| 48270 | ENSMUSG00000110517 | 1.687852163 | 0.808204742 | 1.593943988 | 0.507047141 | 0.612121742 | NA | Gm31224 | predicted gene, 31224 [Source:MGI Symb | 1.6062 | 1.0279 | 1.0062 | 0.964 | 2.1812 | 3.3415 |
| 48271 | ENSMUSG00000110518 | 0 | NA | NA | NA | NA | NA | Olfr861-ps1 | olfactory receptor 861, pseudogene 1 [So | 0 | 0 | 0 | 0 | 0 | 0 |
| 48272 | ENSMUSG00000110519 | 0 | NA | NA | NA | NA | NA | Olfr839-ps1 | olfactory receptor 839, pseudogene 1 [So | 0 | 0 | 0 | 0 | 0 | 0 |
| 48273 | ENSMUSG00000110520 | 3.855683874 | 0.530880728 | 1.069787138 | 0.496248936 | 0.619718789 | NA | Gm45776 | predicted gene 45776 [Source:MGI Symbo | 2.4093 | 2.0558 | 5.0312 | 4.8199 | 4.3625 | 4.4554 |
| 48274 | ENSMUSG00000110521 | 1.216836277 | 0.526797018 | 2.086953666 | 0.252423917 | 0.800713416 | NA | Olfr856-ps1 | olfactory receptor 856, pseudogene 1 [So | 0 | 0 | 3.0187 | 0.964 | 1.0906 | 2.2277 |
| 48275 | ENSMUSG00000110522 | 0 | NA | NA | NA | NA | NA | Gm45735 | predicted gene 45735 [Source:MGI Symbo | 0 | 0 | 0 | 0 | 0 | 0 |
| 48276 | ENSMUSG00000110523 | 0 | NA | NA | NA | NA | NA | C230057M0 | RIKEN cDNA C230057M02 gene [Source:M | 0 | 0 | 0 | 0 | 0 | 0 |
| 48277 | ENSMUSG00000110524 | 0.160662974 | 1.126924175 | 4.080472857 | 0.276174898 | 0.782413736 | NA | Gm39228 | predicted gene, 39228 [Source:MGI Symb | 0 | 0 | 0 | 0.964 | 0 | 0 |
| 48278 | ENSMUSG00000110525 | 0 | NA | NA | NA | NA | NA | Gm45893 | predicted gene 45893 [Source:MGI Symbo | 0 | 0 | 0 | 0 | 0 | 0 |
| 48279 | ENSMUSG00000110526 | 0 | NA | NA | NA | NA | NA | Olfr151 | olfactory receptor 151 [Source:MGI Symb | 0 | 0 | 0 | 0 | 0 | 0 |
| 48280 | ENSMUSG00000110527 | 0 | NA | NA | NA | NA | NA | Gm7751 | predicted gene 7751 [Source:MGI Symbol | 0 | 0 | 0 | 0 | 0 | 0 |
| 48281 | ENSMUSG00000110528 | 0.505131668 | 1.126696199 | 3.421159819 | 0.329331647 | 0.741905026 | NA | Gm1943 | predicted gene 1943 [Source:MGI Symbol | 0.8031 | 0 | 0 | 0 | 0 | 2.2277 |
| 48282 | ENSMUSG00000110529 | 1.284961919 | -2.567677342 | 2.273365472 | -1.129460869 | 0.258703468 | NA | Gm45694 | predicted gene 45694 [Source:MGI Symbo | 1.6062 | 5.1396 | 0 | 0.964 | 0 | 0 |
| 48283 | ENSMUSG00000110530 | 0 | NA | NA | NA | NA | NA | Gm35934 | predicted gene, 35934 [Source:MGI Symb | 0 | 0 | 0 | 0 | 0 | 0 |
| 48284 | ENSMUSG00000110531 | 0.513752329 | 1.031892132 | 3.082998058 | 0.334704114 | 0.737848298 | NA | Gm7760 | predicted gene 7760 [Source:MGI Symbol | 0 | 1.0279 | 0 | 0.964 | 1.0906 | 0 |
| 48285 | ENSMUSG00000110532 | 0.305170013 | -1.657836477 | 4.041480335 | -0.410205257 | 0.681655385 | NA | Gm35857 | predicted gene, 35857 [Source:MGI Symb | 0.8031 | 1.0279 | 0 | 0 | 0 | 0 |
| 48286 | ENSMUSG00000110533 | 0 | NA | NA | NA | NA | NA | LOC665443 | RNA binding motif protein 7 pseudogene | 0 | 0 | 0 | 0 | 0 | 0 |
| 48287 | ENSMUSG00000110534 | 0 | NA | NA | NA | NA | NA | Gm45708 | predicted gene 45708 [Source:MGI Symbo | 0 | 0 | 0 | 0 | 0 | 0 |
| 48288 | ENSMUSG00000110535 | 0 | NA | NA | NA | NA | NA | Olfr880-ps1 | olfactory receptor 880, pseudogene 1 [So | 0 | 0 | 0 | 0 | 0 | 0 |
| 48289 | ENSMUSG00000110536 | 0 | NA | NA | NA | NA | NA | Gm45850 | predicted gene 45850 [Source:MGI Symbo | 0 | 0 | 0 | 0 | 0 | 0 |
| 48290 | ENSMUSG00000110537 | 3.216696319 | 0.662284063 | 1.290934209 | 0.513026968 | 0.607932457 | NA | Gm4316 | predicted gene 4316 [Source:MGI Symbol | 2.4093 | 0 | 5.0312 | 1.928 | 4.3625 | 5.5692 |
| 48291 | ENSMUSG00000110538 | 0.760346335 | -0.431432104 | 2.662343224 | -0.162049769 | 0.871266662 | NA | Gm33178 | predicted gene, 33178 [Source:MGI Symb | 1.6062 | 1.0279 | 0 | 1.928 | 0 | 0 |
| 48292 | ENSMUSG00000110539 | 0 | NA | NA | NA | NA | NA | Gm45864 | predicted gene 45864 [Source:MGI Symbo | 0 | 0 | 0 | 0 | 0 | 0 |
| 48293 | ENSMUSG00000110540 | 0.401551119 | -2.072982726 | 4.008251979 | -0.517178744 | 0.605031388 | NA | Gm45743 | predicted gene 45743 [Source:MGI Symbo | 2.4093 | 0 | 0 | 0 | 0 | 0 |
| 48294 | ENSMUSG00000110541 | 0 | NA | NA | NA | NA | NA | Gm10280 | predicted gene 10280 [Source:MGI Symbo | 0 | 0 | 0 | 0 | 0 | 0 |
| 48295 | ENSMUSG00000110542 | 0 | NA | NA | NA | NA | NA | Gm39139 | predicted gene, 39139 [Source:MGI Symb | 0 | 0 | 0 | 0 | 0 | 0 |
| 48296 | ENSMUSG00000110543 | 0 | NA | NA | NA | NA | NA | Umpk-ps | uridine monophosphate kinase, pseudoge | 0 | 0 | 0 | 0 | 0 | 0 |
| 48297 | ENSMUSG00000110544 | 0 | NA | NA | NA | NA | NA | Gm2607 | predicted gene 2607 [Source:MGI Symbol | 0 | 0 | 0 | 0 | 0 | 0 |
| 48298 | ENSMUSG00000110545 | 4.359837993 | -0.336490808 | 1.09488446 | -0.30732997 | 0.758592226 | NA | Gm7730 | predicted gene 7730 [Source:MGI Symbol | 2.4093 | 9.2513 | 3.0187 | 4.8199 | 1.0906 | 5.5692 |
| 48299 | ENSMUSG00000110546 | 0 | NA | NA | NA | NA | NA | Gm45734 | predicted gene 45734 [Source:MGI Symbo | 0 | 0 | 0 | 0 | 0 | 0 |
| 48300 | ENSMUSG00000110547 | 0.640816969 | 1.600078776 | 2.832778663 | 0.564844263 | 0.572179697 | NA | Gm29773 | predicted gene, 29773 [Source:MGI Symb | 0.8031 | 0 | 0 | 1.928 | 0 | 1.1138 |
| 48301 | ENSMUSG00000110548 | 0 | NA | NA | NA | NA | NA | Gm45721 | predicted gene 45721 [Source:MGI Symbo | 0 | 0 | 0 | 0 | 0 | 0 |
| 48302 | ENSMUSG00000110549 | 0.562214094 | -1.122836648 | 3.276281221 | -0.34271681 | 0.731811515 | NA | Gm45725 | predicted gene 45725 [Source:MGI Symbo | 2.4093 | 0 | 0 | 0.964 | 0 | 0 |
| 48303 | ENSMUSG00000110550 | 0 | NA | NA | NA | NA | NA | Gm45856 | predicted gene 45856 [Source:MGI Symbo | 0 | 0 | 0 | 0 | 0 | 0 |
| 48304 | ENSMUSG00000110551 | 0 | NA | NA | NA | NA | NA | Gm45750 | predicted gene 45750 [Source:MGI Symbo | 0 | 0 | 0 | 0 | 0 | 0 |
| 48305 | ENSMUSG00000110552 | 0 | NA | NA | NA | NA | NA | Gm45788 | predicted gene 45788 [Source:MGI Symbo | 0 | 0 | 0 | 0 | 0 | 0 |
| 48306 | ENSMUSG00000110553 | 0 | NA | NA | NA | NA | NA | Gm45742 | predicted gene 45742 [Source:MGI Symbo | 0 | 0 | 0 | 0 | 0 | 0 |
| 48307 | ENSMUSG00000110554 | 0 | NA | NA | NA | NA | NA | Gm35256 | predicted gene, 35256 [Source:MGI Symb | 0 | 0 | 0 | 0 | 0 | 0 |
| 48308 | ENSMUSG00000110555 | 0 | NA | NA | NA | NA | NA | Olfr863-ps1 | olfactory receptor 863, pseudogene 1 [So | 0 | 0 | 0 | 0 | 0 | 0 |
| 48309 | ENSMUSG00000110556 | 0.660353019 | -0.01500701 | 2.83315355 | -0.005296928 | 0.995773683 | NA | Gm45762 | predicted gene 45762 [Source:MGI Symbo | 0 | 1.0279 | 1.0062 | 1.928 | 0 | 0 |
| 48310 | ENSMUSG00000110557 | 0 | NA | NA | NA | NA | NA | Gm5159 | predicted gene 5159 [Source:MGI Symbol | 0 | 0 | 0 | 0 | 0 | 0 |
| 48311 | ENSMUSG00000110558 | 0.506734502 | -2.372847156 | 3.338060886 | -0.710845978 | 0.477179685 | NA | Gm45779 | predicted gene 45779 [Source:MGI Symbo | 0 | 1.0279 | 2.0125 | 0 | 0 | 0 |
| 48312 | ENSMUSG00000110559 | 0 | NA | NA | NA | NA | NA | Gm26843 | predicted gene, 26843 [Source:MGI Symb | 0 | 0 | 0 | 0 | 0 | 0 |
| 48313 | ENSMUSG00000110560 | 0 | NA | NA | NA | NA | NA | Olfr864-ps1 | olfactory receptor 864, pseudogene 1 [So | 0 | 0 | 0 | 0 | 0 | 0 |
| 48314 | ENSMUSG00000110561 | 0 | NA | NA | NA | NA | NA | Gm45888 | predicted gene 45888 [Source:MGI Symbo | 0 | 0 | 0 | 0 | 0 | 0 |
| 48315 | ENSMUSG00000110562 | 0 | NA | NA | NA | NA | NA | Gm9204 | predicted gene 9204 [Source:MGI Symbol | 0 | 0 | 0 | 0 | 0 | 0 |
| 48316 | ENSMUSG00000110563 | 0 | NA | NA | NA | NA | NA | Gm45791 | predicted gene 45791 [Source:MGI Symbo | 0 | 0 | 0 | 0 | 0 | 0 |
| 48317 | ENSMUSG00000110564 | 0 | NA | NA | NA | NA | NA | Gm45697 | predicted gene 45697 [Source:MGI Symbo | 0 | 0 | 0 | 0 | 0 | 0 |
| 48318 | ENSMUSG00000110565 | 0 | NA | NA | NA | NA | NA | Gm45814 | predicted gene 45814 [Source:MGI Symbo | 0 | 0 | 0 | 0 | 0 | 0 |
| 48319 | ENSMUSG00000110566 | 0 | NA | NA | NA | NA | NA | Olfr865-ps1 | olfactory receptor 865, pseudogene 1 [So | 0 | 0 | 0 | 0 | 0 | 0 |
| 48320 | ENSMUSG00000110567 | 0 | NA | NA | NA | NA | NA | Gm45703 | predicted gene 45703 [Source:MGI Symbo | 0 | 0 | 0 | 0 | 0 | 0 |
| 48321 | ENSMUSG00000110568 | 0.845941533 | -1.843011426 | 2.87307209 | -0.641477613 | 0.521212422 | NA | Gm7208 | predicted gene 7208 [Source:MGI Symbol | 0 | 4.1117 | 0 | 0.964 | 0 | 0 |
| 48322 | ENSMUSG00000110569 | 0.294513347 | 0.165143496 | 4.080472857 | 0.040471657 | 0.967717103 | NA | Gm18860 | predicted gene, 18860 [Source:MGI Symb | 0.8031 | 0 | 0 | 0.964 | 0 | 0 |
| 48323 | ENSMUSG00000110570 | 0.133850373 | -0.796629176 | 4.080472857 | -0.195229623 | 0.84521319 | NA | Gm45778 | predicted gene 45778 [Source:MGI Symbo | 0.8031 | 0 | 0 | 0 | 0 | 0 |
| 48324 | ENSMUSG00000110571 | 0 | NA | NA | NA | NA | NA | Gm45730 | predicted gene 45730 [Source:MGI Symbo | 0 | 0 | 0 | 0 | 0 | 0 |

| | | | | | | | | | | | | | | | |
|---|---|---|---|---|---|---|---|---|---|---|---|---|---|---|---|
| 48325 | ENSMUSG00000110572 | 0 | NA | NA | NA | NA | NA | Olfr852-ps1 | olfactory receptor 852, pseudogene 1 [Sou | 0 | 0 | 0 | 0 | 0 | 0 |
| 48326 | ENSMUSG00000110573 | 0 | NA | NA | NA | NA | NA | Gm5485 | predicted gene 5485 [Source:MGI Symbol | 0 | 0 | 0 | 0 | 0 | 0 |
| 48327 | ENSMUSG00000110574 | 2.781029418 | 1.30064829 | 1.322001435 | 0.983847865 | 0.325190321 | NA | Gm21112 | predicted gene, 21112 [Source:MGI Symb | 0.8031 | 1.0279 | 3.0187 | 1.928 | 5.4531 | 4.4554 |
| 48328 | ENSMUSG00000110575 | 2.592757141 | -2.65025359 | 1.581375747 | -1.675916426 | 0.093754564 | NA | 6330537M0 | RIKEN cDNA 6330537M06 gene [Source:M | 7.2279 | 4.1117 | 2.0125 | 0 | 1.0906 | 1.1138 |
| 48329 | ENSMUSG00000110576 | 0 | NA | NA | NA | NA | NA | Gm36368 | predicted gene, 36368 [Source:MGI Symb | 0 | 0 | 0 | 0 | 0 | 0 |
| 48330 | ENSMUSG00000110577 | 0 | NA | NA | NA | NA | NA | 1700112L15 | RIKEN cDNA 1700112L15 gene [Source:M | 0 | 0 | 0 | 0 | 0 | 0 |
| 48331 | ENSMUSG00000110578 | 0 | NA | NA | NA | NA | NA | Olfr858-ps1 | olfactory receptor 858, pseudogene 1 [Sou | 0 | 0 | 0 | 0 | 0 | 0 |
| 48332 | ENSMUSG00000110580 | 2.470701482 | 0.370804756 | 1.343402043 | 0.276019199 | 0.78253332 | NA | D830024N0 | RIKEN cDNA D830024N08 gene [Source:M | 2.4093 | 1.0279 | 3.0187 | 2.8919 | 4.3625 | 1.1138 |
| 48333 | ENSMUSG00000110581 | 0 | NA | NA | NA | NA | NA | Gm45789 | predicted gene 45789 [Source:MGI Symbo | 0 | 0 | 0 | 0 | 0 | 0 |
| 48334 | ENSMUSG00000110582 | 0 | NA | NA | NA | NA | NA | Gm30052 | predicted gene, 30052 [Source:MGI Symb | 0 | 0 | 0 | 0 | 0 | 0 |
| 48335 | ENSMUSG00000110583 | 0 | NA | NA | NA | NA | NA | Gm7639 | predicted gene 7639 [Source:MGI Symbol | 0 | 0 | 0 | 0 | 0 | 0 |
| 48336 | ENSMUSG00000110585 | 0 | NA | NA | NA | NA | NA | Gm45704 | predicted gene 45704 [Source:MGI Symbo | 0 | 0 | 0 | 0 | 0 | 0 |
| 48337 | ENSMUSG00000110586 | 0.685278559 | -2.809870862 | 3.376578027 | -0.832165239 | 0.405315682 | NA | 4930578M0 | RIKEN cDNA 4930578M07 gene [Source:M | 0 | 4.1117 | 0 | 0 | 0 | 0 |
| 48338 | ENSMUSG00000110587 | 0 | NA | NA | NA | NA | NA | Gm7344 | predicted gene 7344 [Source:MGI Symbol | 0 | 0 | 0 | 0 | 0 | 0 |
| 48339 | ENSMUSG00000110588 | 5.081020403 | -0.283081583 | 1.198916924 | -0.236114427 | 0.813343879 | NA | Gm45774 | predicted gene 45774 [Source:MGI Symbo | 5.6217 | 8.2233 | 3.0187 | 12.532 | 1.0906 | 0 |
| 48340 | ENSMUSG00000110589 | 1.481104775 | 0.514806832 | 1.659456273 | 0.310226211 | 0.75638894 | NA | Gm45754 | predicted gene 45754 [Source:MGI Symbo | 1.6062 | 1.0279 | 1.0062 | 1.928 | 1.0906 | 2.2277 |
| 48341 | ENSMUSG00000110590 | 0 | NA | NA | NA | NA | NA | Olfr848-ps1 | olfactory receptor 848, pseudogene 1 [Sou | 0 | 0 | 0 | 0 | 0 | 0 |
| 48342 | ENSMUSG00000110591 | 0 | NA | NA | NA | NA | NA | Gm31371 | predicted gene, 31371 [Source:MGI Symb | 0 | 0 | 0 | 0 | 0 | 0 |
| 48343 | ENSMUSG00000110592 | 0 | NA | NA | NA | NA | NA | 4930488N1 | RIKEN cDNA 4930488N15 gene [Source:M | 0 | 0 | 0 | 0 | 0 | 0 |
| 48344 | ENSMUSG00000110593 | 1.081124346 | 3.638382224 | 2.309068382 | 1.575692713 | 0.115096638 | NA | Gm45822 | predicted gene 45822 [Source:MGI Symbo | 0 | 0 | 0 | 0.964 | 2.1812 | 3.3415 |
| 48345 | ENSMUSG00000110594 | 0.185640647 | 1.126924175 | 4.080472857 | 0.276174898 | 0.782413736 | NA | Gm33103 | predicted gene, 33103 [Source:MGI Symb | 0 | 0 | 0 | 0 | 0 | 1.1138 |
| 48346 | ENSMUSG00000110595 | 0 | NA | NA | NA | NA | NA | Gm7990 | predicted gene 7990 [Source:MGI Symbol | 0 | 0 | 0 | 0 | 0 | 0 |
| 48347 | ENSMUSG00000110596 | 0 | NA | NA | NA | NA | NA | Gm45859 | predicted gene 45859 [Source:MGI Symbo | 0 | 0 | 0 | 0 | 0 | 0 |
| 48348 | ENSMUSG00000110597 | 16.6520916 | 1.383028813 | 0.576997323 | 2.396941473 | 0.016532564 | 0.152621202 | Gm8798 | predicted gene 8798 [Source:MGI Symbol | 10.44 | 6.1675 | 11.069 | 33.739 | 22.903 | 15.594 |
| 48349 | ENSMUSG00000110598 | 5.168119044 | -0.893111675 | 1.076872992 | -0.829356556 | 0.406902677 | 0.830690236 | Gm45854 | predicted gene 45854 [Source:MGI Symbo | 8.8341 | 6.1675 | 5.0312 | 0 | 7.6343 | 3.3415 |
| 48350 | ENSMUSG00000110599 | 0.538730002 | 1.11098254 | 3.037664833 | 0.365735722 | 0.714562288 | NA | 6030466F02 | RIKEN cDNA 6030466F02 gene [Source:M | 0 | 1.0279 | 0 | 0 | 1.0906 | 1.1138 |
| 48351 | ENSMUSG00000110600 | 0 | NA | NA | NA | NA | NA | Gm45878 | predicted gene 45878 [Source:MGI Symbo | 0 | 0 | 0 | 0 | 0 | 0 |
| 48352 | ENSMUSG00000110601 | 1.814690223 | -0.748633839 | 1.653030911 | -0.452885566 | 0.650631145 | NA | Gm45834 | predicted gene 45834 [Source:MGI Symbo | 0.8031 | 3.0838 | 3.0187 | 2.8919 | 1.0906 | 0 |
| 48353 | ENSMUSG00000110602 | 0 | NA | NA | NA | NA | NA | Olfr886-ps1 | olfactory receptor 886, pseudogene 1 [Sou | 0 | 0 | 0 | 0 | 0 | 0 |
| 48354 | ENSMUSG00000110603 | 0 | NA | NA | NA | NA | NA | Gm45741 | predicted gene 45741 [Source:MGI Symbo | 0 | 0 | 0 | 0 | 0 | 0 |
| 48355 | ENSMUSG00000110604 | 0 | NA | NA | NA | NA | NA | Gm45824 | predicted gene 45824 [Source:MGI Symbo | 0 | 0 | 0 | 0 | 0 | 0 |
| 48356 | ENSMUSG00000110605 | 10.3788047 | -0.154323533 | 0.649730933 | -0.237519141 | 0.812254072 | 0.96653525 | Gm32856 | predicted gene, 32856 [Source:MGI Symb | 10.44 | 11.307 | 11.069 | 9.6398 | 10.906 | 8.9108 |
| 48357 | ENSMUSG00000110606 | 0 | NA | NA | NA | NA | NA | Gm45829 | predicted gene 45829 [Source:MGI Symbo | 0 | 0 | 0 | 0 | 0 | 0 |
| 48358 | ENSMUSG00000110607 | 0 | NA | NA | NA | NA | NA | Gm45843 | predicted gene 45843 [Source:MGI Symbo | 0 | 0 | 0 | 0 | 0 | 0 |
| 48359 | ENSMUSG00000110608 | 0 | NA | NA | NA | NA | NA | Gm8838 | predicted gene 8838 [Source:MGI Symbol | 0 | 0 | 0 | 0 | 0 | 0 |
| 48360 | ENSMUSG00000110609 | 0 | NA | NA | NA | NA | NA | Gm7948 | predicted gene 7948 [Source:MGI Symbol | 0 | 0 | 0 | 0 | 0 | 0 |
| 48361 | ENSMUSG00000110610 | 0.878247891 | 3.344090894 | 2.718259224 | 1.230232519 | 0.218610046 | NA | Gm7769 | predicted gene 7769 [Source:MGI Symbol | 0 | 0 | 0 | 1.928 | 0 | 3.3415 |
| 48362 | ENSMUSG00000110611 | 3.150274708 | 1.92352149 | 1.343592905 | 1.431625221 | 0.152251108 | NA | Gm20163 | predicted gene, 20163 [Source:MGI Symb | 0 | 1.0279 | 3.0187 | 3.8559 | 6.5437 | 4.4554 |
| 48363 | ENSMUSG00000110612 | 0.503095663 | 2.549494177 | 3.348715477 | 0.761334964 | 0.446457022 | NA | Gm17827 | predicted gene, 17827 [Source:MGI Symb | 0 | 0 | 0 | 1.928 | 1.0906 | 0 |
| 48364 | ENSMUSG00000110613 | 21.38679605 | 0.450678773 | 0.521734684 | 0.863808344 | 0.387693184 | 0.817734468 | Lncbate1 | brown adipose tissue enriched long non-c | 16.865 | 9.2513 | 28.175 | 28.919 | 26.175 | 18.935 |
| 48365 | ENSMUSG00000110614 | 0.62992821 | -1.312600165 | 2.851194982 | -0.460368433 | 0.645251789 | NA | Gm45817 | predicted gene 45817 [Source:MGI Symbo | 0.8031 | 0 | 2.0125 | 0.964 | 0 | 0 |
| 48366 | ENSMUSG00000110615 | 5.155581359 | 0.083144596 | 0.943126617 | 0.088158466 | 0.929750728 | 0.988405116 | Gm45890 | predicted gene 45890 [Source:MGI Symbo | 4.8186 | 6.1675 | 4.025 | 3.8559 | 8.7249 | 3.3415 |
| 48367 | ENSMUSG00000110616 | 0 | NA | NA | NA | NA | NA | Gm36879 | predicted gene, 36879 [Source:MGI Symb | 0 | 0 | 0 | 0 | 0 | 0 |
| 48368 | ENSMUSG00000110617 | 0.167707431 | -0.796629176 | 4.080472857 | -0.195229623 | 0.84521319 | NA | Gm5910 | predicted gene 5910 [Source:MGI Symbol | 0 | 0 | 1.0062 | 0 | 0 | 0 |
| 48369 | ENSMUSG00000110618 | 0.339027071 | -1.79174149 | 3.919585585 | -0.457125237 | 0.647581032 | NA | Gm39822 | predicted gene, 39822 [Source:MGI Symb | 0 | 1.0279 | 1.0062 | 0 | 0 | 0 |
| 48370 | ENSMUSG00000110619 | 0 | NA | NA | NA | NA | NA | Gm7850 | predicted gene 7850 [Source:MGI Symbol | 0 | 0 | 0 | 0 | 0 | 0 |
| 48371 | ENSMUSG00000110620 | 0.133850373 | -0.796629176 | 4.080472857 | -0.195229623 | 0.84521319 | NA | Gm45733 | predicted gene 45733 [Source:MGI Symbo | 0.8031 | 0 | 0 | 0 | 0 | 0 |
| 48372 | ENSMUSG00000110621 | 0 | NA | NA | NA | NA | NA | Olfr837 | olfactory receptor 837 [Source:MGI Symb | 0 | 0 | 0 | 0 | 0 | 0 |
| 48373 | ENSMUSG00000110622 | 3.893262393 | -0.387299003 | 1.19920595 | -0.322962876 | 0.746723357 | NA | Iqcn | IQ motif containing N [Source:MGI Symbo | 3.2124 | 3.0838 | 7.0437 | 6.7478 | 3.2719 | 0 |
| 48374 | ENSMUSG00000110623 | 0 | NA | NA | NA | NA | NA | Gm32005 | predicted gene, 32005 [Source:MGI Symb | 0 | 0 | 0 | 0 | 0 | 0 |
| 48375 | ENSMUSG00000110624 | 0 | NA | NA | NA | NA | NA | Gm32052 | predicted gene, 32052 [Source:MGI Symb | 0 | 0 | 0 | 0 | 0 | 0 |
| 48376 | ENSMUSG00000110625 | 0 | NA | NA | NA | NA | NA | 5033426E14 | RIKEN cDNA 5033426E14 gene [Source:M | 0 | 0 | 0 | 0 | 0 | 0 |
| 48377 | ENSMUSG00000110626 | 0 | NA | NA | NA | NA | NA | Gm45805 | predicted gene 45805 [Source:MGI Symbo | 0 | 0 | 0 | 0 | 0 | 0 |
| 48378 | ENSMUSG00000110627 | 2.048740502 | -2.355112831 | 1.748190437 | -1.347171784 | 0.177924913 | NA | Gm18935 | predicted gene, 18935 [Source:MGI Symb | 3.2124 | 5.1396 | 2.0125 | 1.928 | 0 | 0 |
| 48379 | ENSMUSG00000110628 | 0 | NA | NA | NA | NA | NA | Gm37797 | predicted gene, 37797 [Source:MGI Symb | 0 | 0 | 0 | 0 | 0 | 0 |
| 48380 | ENSMUSG00000110629 | 0 | NA | NA | NA | NA | NA | 4930567H12 | RIKEN cDNA 4930567H12 gene [Source:M | 0 | 0 | 0 | 0 | 0 | 0 |

| | | | | | | | | | | | | | | | |
|---|---|---|---|---|---|---|---|---|---|---|---|---|---|---|---|
| 48381 | ENSMUSG00000110630 | 7.69183117 | -0.254795077 | 0.752792303 | -0.338466634 | 0.735011567 | 0.948109711 | K230015D01 | RIKEN cDNA K230015D01 gene [Source:M | 8.8341 | 7.1954 | 9.0562 | 6.7478 | 7.6343 | 6.6831 |
| 48382 | ENSMUSG00000110631 | 1581.456987 | -0.032900719 | 0.568640608 | -0.057858546 | 0.953861303 | 0.9915542 | Gm42047 | predicted gene, 42047 [Source:MGI Symb | 2161.1 | 1241.7 | 1395.7 | 2757.9 | 1057.9 | 874.37 |
| 48383 | ENSMUSG00000110632 | 0 | NA | NA | NA | NA | NA | Gm45880 | predicted gene 45880 [Source:MGI Symbo | 0 | 0 | 0 | 0 | 0 | 0 |
| 48384 | ENSMUSG00000110633 | 0 | NA | NA | NA | NA | NA | Gm32122 | predicted gene, 32122 [Source:MGI Symb | 0 | 0 | 0 | 0 | 0 | 0 |
| 48385 | ENSMUSG00000110634 | 3.436740307 | 0.215444622 | 1.23519596 | 0.174421411 | 0.861534289 | NA | Gm45895 | predicted gene 45895 [Source:MGI Symbo | 6.4248 | 2.0558 | 1.0062 | 6.7478 | 3.2719 | 1.1138 |
| 48386 | ENSMUSG00000110635 | 1.950607618 | 1.057150656 | 1.686681085 | 0.626763806 | 0.530814072 | NA | Gm45853 | predicted gene 45853 [Source:MGI Symbo | 0.8031 | 2.0558 | 1.0062 | 6.7478 | 1.0906 | 0 |
| 48387 | ENSMUSG00000110636 | 0 | NA | NA | NA | NA | NA | Gm39397 | predicted gene, 39397 [Source:MGI Symb | 0 | 0 | 0 | 0 | 0 | 0 |
| 48388 | ENSMUSG00000110637 | 1.586365267 | 0.194917051 | 1.714960808 | 0.113656854 | 0.909509816 | NA | Gm7807 | predicted gene 7807 [Source:MGI Symbol | 2.4093 | 0 | 2.0125 | 2.8919 | 1.0906 | 1.1138 |
| 48389 | ENSMUSG00000110638 | 0 | NA | NA | NA | NA | NA | Gm45887 | predicted gene 45887 [Source:MGI Symbo | 0 | 0 | 0 | 0 | 0 | 0 |
| 48390 | ENSMUSG00000110639 | 2.527553468 | -1.870791048 | 1.739252543 | -1.07562933 | 0.282093065 | NA | Gm6831 | predicted gene 6831 [Source:MGI Symbol | 0.8031 | 1.0279 | 10.062 | 0 | 3.2719 | 0 |
| 48391 | ENSMUSG00000110640 | 0 | NA | NA | NA | NA | NA | Gm45903 | predicted gene 45903 [Source:MGI Symbo | 0 | 0 | 0 | 0 | 0 | 0 |
| 48392 | ENSMUSG00000110641 | 0 | NA | NA | NA | NA | NA | Gm45826 | predicted gene 45826 [Source:MGI Symbo | 0 | 0 | 0 | 0 | 0 | 0 |
| 48393 | ENSMUSG00000110642 | 0 | NA | NA | NA | NA | NA | Gm45710 | predicted gene 45710 [Source:MGI Symbo | 0 | 0 | 0 | 0 | 0 | 0 |
| 48394 | ENSMUSG00000110643 | 0 | NA | NA | NA | NA | NA | Gm18924 | predicted gene, 18924 [Source:MGI Symb | 0 | 0 | 0 | 0 | 0 | 0 |
| 48395 | ENSMUSG00000110644 | 0.496077836 | -0.812642266 | 3.444120572 | -0.235950586 | 0.813471014 | NA | Gm7390 | predicted gene 7390 [Source:MGI Symbol | 0 | 0 | 2.0125 | 0.964 | 0 | 0 |
| 48396 | ENSMUSG00000110645 | 0 | NA | NA | NA | NA | NA | Olfr838-ps1 | olfactory receptor 838, pseudogene 1 [So | 0 | 0 | 0 | 0 | 0 | 0 |
| 48397 | ENSMUSG00000110646 | 0 | NA | NA | NA | NA | NA | Olfr369-ps1 | olfactory receptor 369, pseudogene 1 [So | 0 | 0 | 0 | 0 | 0 | 0 |
| 48398 | ENSMUSG00000110647 | 0.328370405 | 0.16514316 | 4.003149536 | 0.041253308 | 0.967093956 | NA | Gm17745 | predicted gene, 17745 [Source:MGI Symb | 0 | 0 | 1.0062 | 0.964 | 0 | 0 |
| 48399 | ENSMUSG00000110648 | 0.342432689 | 1.990653086 | 3.900281263 | 0.51038706 | 0.609780321 | NA | Gm38574 | predicted gene, 38574 [Source:MGI Symb | 0 | 0 | 0 | 0.964 | 1.0906 | 0 |
| 48400 | ENSMUSG00000110649 | 18.12099828 | 0.431706933 | 0.507872518 | 0.85003011 | 0.395308346 | 0.822933206 | Gm40466 | predicted gene, 40466 [Source:MGI Symb | 17.668 | 13.363 | 15.094 | 17.352 | 19.631 | 25.618 |
| 48401 | ENSMUSG00000110650 | 0.185640647 | 1.126924175 | 4.080472857 | 0.276174898 | 0.782413736 | NA | Gm19170 | predicted gene, 19170 [Source:MGI Symb | 0 | 0 | 0 | 0 | 0 | 1.1138 |
| 48402 | ENSMUSG00000110651 | 0 | NA | NA | NA | NA | NA | Gm45770 | predicted gene 45770 [Source:MGI Symbo | 0 | 0 | 0 | 0 | 0 | 0 |
| 48403 | ENSMUSG00000110652 | 0 | NA | NA | NA | NA | NA | Gm33023 | predicted gene, 33023 [Source:MGI Symb | 0 | 0 | 0 | 0 | 0 | 0 |
| 48404 | ENSMUSG00000110653 | 0 | NA | NA | NA | NA | NA | Gm34418 | predicted gene, 34418 [Source:MGI Symb | 0 | 0 | 0 | 0 | 0 | 0 |
| 48405 | ENSMUSG00000110654 | 0 | NA | NA | NA | NA | NA | Gm45746 | predicted gene 45746 [Source:MGI Symbo | 0 | 0 | 0 | 0 | 0 | 0 |
| 48406 | ENSMUSG00000110655 | 0 | NA | NA | NA | NA | NA | Gm45798 | predicted gene 45798 [Source:MGI Symbo | 0 | 0 | 0 | 0 | 0 | 0 |
| 48407 | ENSMUSG00000110656 | 4.322445835 | -0.364567545 | 1.075618859 | -0.338937479 | 0.734656829 | NA | Gm20735 | predicted gene, 20735 [Source:MGI Symb | 6.4248 | 6.1675 | 2.0125 | 5.7839 | 1.0906 | 4.4554 |
| 48408 | ENSMUSG00000110657 | 0.305170013 | -1.657836477 | 4.041480335 | -0.410205257 | 0.681655385 | NA | Gm7684 | predicted gene 7684 [Source:MGI Symbol | 0.8031 | 1.0279 | 0 | 0 | 0 | 0 |
| 48409 | ENSMUSG00000110658 | 0 | NA | NA | NA | NA | NA | Gm45857 | predicted gene 45857 [Source:MGI Symbo | 0 | 0 | 0 | 0 | 0 | 0 |
| 48410 | ENSMUSG00000110659 | 20.87280548 | 0.32725704 | 0.493087481 | 0.663689615 | 0.506888988 | 0.88185929 | Gm45727 | predicted gene 45727 [Source:MGI Symbo | 20.881 | 22.614 | 12.075 | 28.919 | 21.812 | 18.935 |
| 48411 | ENSMUSG00000110660 | 0 | NA | NA | NA | NA | NA | Gm5362 | predicted gene 5362 [Source:MGI Symbol | 0 | 0 | 0 | 0 | 0 | 0 |
| 48412 | ENSMUSG00000110661 | 1.113694413 | -0.251399571 | 1.97572234 | -0.127244384 | 0.898746978 | NA | Gm31805 | predicted gene, 31805 [Source:MGI Symb | 1.6062 | 1.0279 | 1.0062 | 1.928 | 0 | 1.1138 |
| 48413 | ENSMUSG00000110662 | 0 | NA | NA | NA | NA | NA | Gm45722 | predicted gene 45722 [Source:MGI Symbo | 0 | 0 | 0 | 0 | 0 | 0 |
| 48414 | ENSMUSG00000110663 | 19.49305479 | 0.813447768 | 0.491101866 | 1.656372789 | 0.097646348 | 0.460719048 | Gm39157 | predicted gene, 39157 [Source:MGI Symb | 16.865 | 11.307 | 14.087 | 25.063 | 22.903 | 26.732 |
| 48415 | ENSMUSG00000110664 | 0 | NA | NA | NA | NA | NA | Aprt-ps | adenine phosphoribosyl transferase, pseu | 0 | 0 | 0 | 0 | 0 | 0 |
| 48416 | ENSMUSG00000110665 | 0 | NA | NA | NA | NA | NA | Gm31786 | predicted gene, 31786 [Source:MGI Symb | 0 | 0 | 0 | 0 | 0 | 0 |
| 48417 | ENSMUSG00000110666 | 0 | NA | NA | NA | NA | NA | Gm9172 | predicted gene 9172 [Source:MGI Symbol | 0 | 0 | 0 | 0 | 0 | 0 |
| 48418 | ENSMUSG00000110667 | 0 | NA | NA | NA | NA | NA | Gm45720 | predicted gene 45720 [Source:MGI Symbo | 0 | 0 | 0 | 0 | 0 | 0 |
| 48419 | ENSMUSG00000110668 | 0 | NA | NA | NA | NA | NA | Gm19885 | predicted gene, 19885 [Source:MGI Symb | 0 | 0 | 0 | 0 | 0 | 0 |
| 48420 | ENSMUSG00000110669 | 6.640046248 | -0.310817756 | 0.835571105 | -0.371982414 | 0.709905943 | 0.944239215 | Gm10060 | predicted gene 10060 [Source:MGI Symbo | 8.8341 | 5.1396 | 8.05 | 6.7478 | 3.2719 | 7.7969 |
| 48421 | ENSMUSG00000110670 | 0 | NA | NA | NA | NA | NA | Gm30132 | predicted gene, 30132 [Source:MGI Symb | 0 | 0 | 0 | 0 | 0 | 0 |
| 48422 | ENSMUSG00000110671 | 0 | NA | NA | NA | NA | NA | Olfr840-ps1 | olfactory receptor 840, pseudogene 1 [So | 0 | 0 | 0 | 0 | 0 | 0 |
| 48423 | ENSMUSG00000110672 | 0 | NA | NA | NA | NA | NA | Gm45786 | predicted gene 45786 [Source:MGI Symbo | 0 | 0 | 0 | 0 | 0 | 0 |
| 48424 | ENSMUSG00000110673 | 0 | NA | NA | NA | NA | NA | Gm45833 | predicted gene 45833 [Source:MGI Symbo | 0 | 0 | 0 | 0 | 0 | 0 |
| 48425 | ENSMUSG00000110674 | 0 | NA | NA | NA | NA | NA | Gm45892 | predicted gene 45892 [Source:MGI Symbo | 0 | 0 | 0 | 0 | 0 | 0 |
| 48426 | ENSMUSG00000110675 | 0 | NA | NA | NA | NA | NA | Olfr879-ps1 | olfactory receptor 879, pseudogene 1 [So | 0 | 0 | 0 | 0 | 0 | 0 |
| 48427 | ENSMUSG00000110676 | 0 | NA | NA | NA | NA | NA | Gm45897 | predicted gene 45897 [Source:MGI Symbo | 0 | 0 | 0 | 0 | 0 | 0 |
| 48428 | ENSMUSG00000110677 | 0 | NA | NA | NA | NA | NA | Gm45777 | predicted gene 45777 [Source:MGI Symbo | 0 | 0 | 0 | 0 | 0 | 0 |
| 48429 | ENSMUSG00000110678 | 0 | NA | NA | NA | NA | NA | Gm18645 | predicted gene, 18645 [Source:MGI Symb | 0 | 0 | 0 | 0 | 0 | 0 |
| 48430 | ENSMUSG00000110679 | 2.132371819 | 1.406922304 | 1.527191594 | 0.921248067 | 0.356920928 | NA | Rpl10-ps5 | ribosomal protein L10, pseudogene 5 [Sou | 2.4093 | 0 | 1.0062 | 3.8559 | 2.1812 | 3.3415 |
| 48431 | ENSMUSG00000110680 | 0 | NA | NA | NA | NA | NA | Gm6796 | predicted pseudogene 6796 [Source:MGI | 0 | 0 | 0 | 0 | 0 | 0 |
| 48432 | ENSMUSG00000110681 | 0 | NA | NA | NA | NA | NA | Gm45796 | predicted gene 45796 [Source:MGI Symbo | 0 | 0 | 0 | 0 | 0 | 0 |
| 48433 | ENSMUSG00000110682 | 1.355875063 | 0.77795787 | 1.980777249 | 0.39275384 | 0.694501301 | NA | A530010L16 | RIKEN cDNA A530010L16 gene [Source:M | 0 | 2.0558 | 1.0062 | 2.8919 | 2.1812 | 0 |
| 48434 | ENSMUSG00000110683 | 1.171805106 | 1.498519501 | 2.132819898 | 0.702600113 | 0.482304992 | NA | Gm45874 | predicted gene 45874 [Source:MGI Symbo | 0.8031 | 1.0279 | 0 | 1.928 | 3.2719 | 0 |
| 48435 | ENSMUSG00000110684 | 0.321325949 | 1.911575114 | 4.048267893 | 0.472195804 | 0.636787034 | NA | Gm45866 | predicted gene 45866 [Source:MGI Symbo | 0 | 0 | 0 | 1.928 | 0 | 0 |
| 48436 | ENSMUSG00000110685 | 0 | NA | NA | NA | NA | NA | Gm5373 | predicted gene 5373 [Source:MGI Symbol | 0 | 0 | 0 | 0 | 0 | 0 |

| | | | | | | | | | | | | | | | |
|---|---|---|---|---|---|---|---|---|---|---|---|---|---|---|---|
| 48437 | ENSMUSG00000110686 | 3.928839613 | 0.790771408 | 1.125050904 | 0.702876115 | 0.482132957 | NA | Gm45855 | predicted gene 45855 [Source:MGI Symbo | 1.6062 | 1.0279 | 6.0375 | 3.8559 | 4.3625 | 6.6831 |
| 48438 | ENSMUSG00000110687 | 0 | NA | NA | NA | NA | NA | Gm45764 | predicted gene 45764 [Source:MGI Symbo | 0 | 0 | 0 | 0 | 0 | 0 |
| 48439 | ENSMUSG00000110688 | 0 | NA | NA | NA | NA | NA | Gm45696 | predicted gene 45696 [Source:MGI Symbo | 0 | 0 | 0 | 0 | 0 | 0 |
| 48440 | ENSMUSG00000110689 | 0.339027071 | -1.79174149 | 3.919585585 | -0.457125237 | 0.647581032 | NA | Gm45731 | predicted gene 45731 [Source:MGI Symbo | 0 | 1.0279 | 1.0062 | 0 | 0 | 0 |
| 48441 | ENSMUSG00000110690 | 2.551836165 | -1.031286913 | 1.433688834 | -0.719324088 | 0.471941258 | NA | Gm18001 | predicted gene, 18001 [Source:MGI Symb | 3.2124 | 5.1396 | 2.0125 | 3.8559 | 1.0906 | 0 |
| 48442 | ENSMUSG00000110691 | 0 | NA | NA | NA | NA | NA | Gm5363 | predicted gene 5363 [Source:MGI Symbol | 0 | 0 | 0 | 0 | 0 | 0 |
| 48443 | ENSMUSG00000110692 | 0 | NA | NA | NA | NA | NA | Gm33460 | predicted gene, 33460 [Source:MGI Symb | 0 | 0 | 0 | 0 | 0 | 0 |
| 48444 | ENSMUSG00000110693 | 0 | NA | NA | NA | NA | NA | Gm45899 | predicted gene 45899 [Source:MGI Symbo | 0 | 0 | 0 | 0 | 0 | 0 |
| 48445 | ENSMUSG00000110694 | 0 | NA | NA | NA | NA | NA | Gm45782 | predicted gene 45782 [Source:MGI Symbo | 0 | 0 | 0 | 0 | 0 | 0 |
| 48446 | ENSMUSG00000110695 | 0 | NA | NA | NA | NA | NA | Gm45766 | predicted gene 45766 [Source:MGI Symbo | 0 | 0 | 0 | 0 | 0 | 0 |
| 48447 | ENSMUSG00000110696 | 0 | NA | NA | NA | NA | NA | Gm45706 | predicted gene 45706 [Source:MGI Symbo | 0 | 0 | 0 | 0 | 0 | 0 |
| 48448 | ENSMUSG00000110697 | 9.557691067 | -1.008661041 | 0.72204663 | -1.396947231 | 0.16242944 | 0.585865647 | Gm31718 | predicted gene, 31718 [Source:MGI Symb | 12.85 | 12.335 | 13.081 | 5.7839 | 3.2719 | 10.025 |
| 48449 | ENSMUSG00000110698 | 0 | NA | NA | NA | NA | NA | Gm45875 | predicted gene 45875 [Source:MGI Symbo | 0 | 0 | 0 | 0 | 0 | 0 |
| 48450 | ENSMUSG00000110699 | 0 | NA | NA | NA | NA | NA | Gm20946 | predicted gene, 20946 [Source:MGI Symb | 0 | 0 | 0 | 0 | 0 | 0 |
| 48451 | ENSMUSG00000110700 | 0 | NA | NA | NA | NA | NA | Gm5357 | predicted gene 5357 [Source:MGI Symbol | 0 | 0 | 0 | 0 | 0 | 0 |
| 48452 | ENSMUSG00000110701 | 0 | NA | NA | NA | NA | NA | Gm6013 | predicted gene 6013 [Source:MGI Symbol | 0 | 0 | 0 | 0 | 0 | 0 |
| 48453 | ENSMUSG00000110702 | 14.43823816 | -0.115312706 | 0.688022018 | -0.167600314 | 0.866897724 | 0.977930657 | Gm45767 | predicted gene 45767 [Source:MGI Symbo | 4.8186 | 12.335 | 28.175 | 19.28 | 11.997 | 10.025 |
| 48454 | ENSMUSG00000110703 | 0 | NA | NA | NA | NA | NA | Gm45821 | predicted gene 45821 [Source:MGI Symbo | 0 | 0 | 0 | 0 | 0 | 0 |
| 48455 | ENSMUSG00000110704 | 0 | NA | NA | NA | NA | NA | Gm45732 | predicted gene 45732 [Source:MGI Symbo | 0 | 0 | 0 | 0 | 0 | 0 |
| 48456 | ENSMUSG00000110705 | 0 | NA | NA | NA | NA | NA | Gm45701 | predicted gene 45701 [Source:MGI Symbo | 0 | 0 | 0 | 0 | 0 | 0 |
| 48457 | ENSMUSG00000110706 | 2.78505404 | -1.586348506 | 1.553946904 | -1.020851164 | 0.307324959 | NA | Gm45709 | predicted gene 45709 [Source:MGI Symbo | 10.44 | 1.0279 | 1.0062 | 0.964 | 3.2719 | 0 |
| 48458 | ENSMUSG00000110707 | 0.342432689 | 1.990653086 | 3.900281263 | 0.51038706 | 0.609780321 | NA | Gm45795 | predicted gene 45795 [Source:MGI Symbo | 0 | 0 | 0 | 0.964 | 1.0906 | 0 |
| 48459 | ENSMUSG00000110708 | 0 | NA | NA | NA | NA | NA | Gnb1-ps3 | guanine nucleotide binding protein (G pro | 0 | 0 | 0 | 0 | 0 | 0 |
| 48460 | ENSMUSG00000110709 | 0 | NA | NA | NA | NA | NA | 1700018P08 | RIKEN cDNA 1700018P08 gene [Source:M | 0 | 0 | 0 | 0 | 0 | 0 |
| 48461 | ENSMUSG00000110710 | 1.073589071 | 1.123304884 | 2.389581492 | 0.470084359 | 0.638294748 | NA | C78859 | expressed sequence C78859 [Source:MGI | 0 | 2.0558 | 0 | 0 | 3.2719 | 1.1138 |
| 48462 | ENSMUSG00000110711 | 1.171831736 | -1.161928617 | 2.135676568 | -0.544056453 | 0.586402626 | NA | Gm45760 | predicted gene 45760 [Source:MGI Symbo | 0.8031 | 1.0279 | 3.0187 | 0 | 2.1812 | 0 |
| 48463 | ENSMUSG00000110712 | 0 | NA | NA | NA | NA | NA | Gm39662 | predicted gene, 39662 [Source:MGI Symb | 0 | 0 | 0 | 0 | 0 | 0 |
| 48464 | ENSMUSG00000110713 | 0 | NA | NA | NA | NA | NA | Gm30606 | predicted gene, 30606 [Source:MGI Symb | 0 | 0 | 0 | 0 | 0 | 0 |
| 48465 | ENSMUSG00000110714 | 0 | NA | NA | NA | NA | NA | Gm45910 | predicted gene 45910 [Source:MGI Symbo | 0 | 0 | 0 | 0 | 0 | 0 |
| 48466 | ENSMUSG00000110715 | 0.321325949 | 1.911575114 | 4.048267893 | 0.472195804 | 0.636787034 | NA | Gm5131 | predicted gene 5131 [Source:MGI Symbol | 0 | 0 | 0 | 1.928 | 0 | 0 |
| 48467 | ENSMUSG00000110716 | 0.133850373 | -0.796629176 | 4.080472857 | -0.195229623 | 0.84521319 | NA | Gm39271 | predicted gene, 39271 [Source:MGI Symb | 0.8031 | 0 | 0 | 0 | 0 | 0 |
| 48468 | ENSMUSG00000110717 | 0 | NA | NA | NA | NA | NA | 1700047G07 | RIKEN cDNA 1700047G07 gene [Source:M | 0 | 0 | 0 | 0 | 0 | 0 |
| 48469 | ENSMUSG00000110718 | 0 | NA | NA | NA | NA | NA | Gm45765 | predicted gene 45765 [Source:MGI Symbo | 0 | 0 | 0 | 0 | 0 | 0 |
| 48470 | ENSMUSG00000110719 | 0 | NA | NA | NA | NA | NA | Gm2716 | predicted gene 2716 [Source:MGI Symbol | 0 | 0 | 0 | 0 | 0 | 0 |
| 48471 | ENSMUSG00000110720 | 0 | NA | NA | NA | NA | NA | Gm46223 | predicted gene, 46223 [Source:MGI Symb | 0 | 0 | 0 | 0 | 0 | 0 |
| 48472 | ENSMUSG00000110721 | 0 | NA | NA | NA | NA | NA | Gm47231 | predicted gene, 47231 [Source:MGI Symb | 0 | 0 | 0 | 0 | 0 | 0 |
| 48473 | ENSMUSG00000110722 | 0 | NA | NA | NA | NA | NA | Gm48716 | predicted gene, 48716 [Source:MGI Symb | 0 | 0 | 0 | 0 | 0 | 0 |
| 48474 | ENSMUSG00000110723 | 0 | NA | NA | NA | NA | NA | Gm49353 | predicted gene, 49353 [Source:MGI Symb | 0 | 0 | 0 | 0 | 0 | 0 |
| 48475 | ENSMUSG00000110724 | 0 | NA | NA | NA | NA | NA | Gm47824 | predicted gene, 47824 [Source:MGI Symb | 0 | 0 | 0 | 0 | 0 | 0 |
| 48476 | ENSMUSG00000110725 | 0 | NA | NA | NA | NA | NA | Gm47129 | predicted gene, 47129 [Source:MGI Symb | 0 | 0 | 0 | 0 | 0 | 0 |
| 48477 | ENSMUSG00000110726 | 0.349477146 | 0.165142768 | 3.911021834 | 0.042224967 | 0.966319359 | NA | Gm18101 | predicted gene, 18101 [Source:MGI Symb | 0 | 0 | 1.0062 | 0 | 1.0906 | 0 |
| 48478 | ENSMUSG00000110727 | 0 | NA | NA | NA | NA | NA | Gm48784 | predicted gene, 48784 [Source:MGI Symb | 0 | 0 | 0 | 0 | 0 | 0 |
| 48479 | ENSMUSG00000110729 | 0.167707431 | -0.796629176 | 4.080472857 | -0.195229623 | 0.84521319 | NA | Gm47117 | predicted gene, 47117 [Source:MGI Symb | 0 | 0 | 1.0062 | 0 | 0 | 0 |
| 48480 | ENSMUSG00000110730 | 0 | NA | NA | NA | NA | NA | Gm19385 | predicted gene, 19385 [Source:MGI Symb | 0 | 0 | 0 | 0 | 0 | 0 |
| 48481 | ENSMUSG00000110731 | 0 | NA | NA | NA | NA | NA | Gm47049 | predicted gene, 47049 [Source:MGI Symb | 0 | 0 | 0 | 0 | 0 | 0 |
| 48482 | ENSMUSG00000110732 | 0 | NA | NA | NA | NA | NA | Gm48674 | predicted gene, 48674 [Source:MGI Symb | 0 | 0 | 0 | 0 | 0 | 0 |
| 48483 | ENSMUSG00000110734 | 0 | NA | NA | NA | NA | NA | Gm18934 | predicted gene, 18934 [Source:MGI Symb | 0 | 0 | 0 | 0 | 0 | 0 |
| 48484 | ENSMUSG00000110735 | 0 | NA | NA | NA | NA | NA | Gm47173 | predicted gene, 47173 [Source:MGI Symb | 0 | 0 | 0 | 0 | 0 | 0 |
| 48485 | ENSMUSG00000110736 | 0 | NA | NA | NA | NA | NA | Olfr942-ps1 | olfactory receptor 942, pseudogene 1 [So | 0 | 0 | 0 | 0 | 0 | 0 |
| 48486 | ENSMUSG00000110737 | 0 | NA | NA | NA | NA | NA | 4930444F02 | RIKEN cDNA 4930444F02 gene [Source:M | 0 | 0 | 0 | 0 | 0 | 0 |
| 48487 | ENSMUSG00000110738 | 0.513958919 | -2.394274821 | 3.756096013 | -0.637437066 | 0.523840191 | NA | Gm47092 | predicted gene, 47092 [Source:MGI Symb | 0 | 3.0838 | 0 | 0 | 0 | 0 |
| 48488 | ENSMUSG00000110739 | 0 | NA | NA | NA | NA | NA | Gm17950 | predicted gene, 17950 [Source:MGI Symb | 0 | 0 | 0 | 0 | 0 | 0 |
| 48489 | ENSMUSG00000110740 | 0.562214094 | -1.122836648 | 3.276281221 | -0.34271681 | 0.731811515 | NA | Gm48853 | predicted gene, 48853 [Source:MGI Symb | 2.4093 | 0 | 0 | 0.964 | 0 | 0 |
| 48490 | ENSMUSG00000110741 | 0 | NA | NA | NA | NA | NA | Gm48091 | predicted gene, 48091 [Source:MGI Symb | 0 | 0 | 0 | 0 | 0 | 0 |
| 48491 | ENSMUSG00000110744 | 1.004562989 | 0.155751777 | 2.077846764 | 0.07495826 | 0.940247922 | NA | Gm5171 | predicted gene 5171 [Source:MGI Symbol | 0.8031 | 2.0558 | 0 | 0.964 | 1.0906 | 1.1138 |
| 48492 | ENSMUSG00000110745 | 0 | NA | NA | NA | NA | NA | Gm8106 | predicted gene 8106 [Source:MGI Symbol | 0 | 0 | 0 | 0 | 0 | 0 |

| | | | | | | | | | | | | | | | |
|---|---|---|---|---|---|---|---|---|---|---|---|---|---|---|---|
| 48493 | ENSMUSG00000110747 | 0 | NA | NA | NA | NA | NA | Gm35657 | predicted gene, 35657 [Source:MGI Symb | 0 | 0 | 0 | 0 | 0 | 0 |
| 48494 | ENSMUSG00000110748 | 0 | NA | NA | NA | NA | NA | Gm19228 | predicted gene, 19228 [Source:MGI Symb | 0 | 0 | 0 | 0 | 0 | 0 |
| 48495 | ENSMUSG00000110749 | 0 | NA | NA | NA | NA | NA | Gm47931 | predicted gene, 47931 [Source:MGI Symb | 0 | 0 | 0 | 0 | 0 | 0 |
| 48496 | ENSMUSG00000110750 | 12.52572963 | 0.023898148 | 0.655066823 | 0.036482001 | 0.97089803 | 0.994966572 | Gm48702 | predicted gene, 48702 [Source:MGI Symb | 8.8341 | 18.503 | 10.062 | 12.532 | 18.541 | 6.6831 |
| 48497 | ENSMUSG00000110751 | 0.614004368 | 0.315622926 | 2.898299649 | 0.108899342 | 0.913282328 | NA | C230053D17 | RIKEN cDNA C230053D17 gene [Source:M | 1.6062 | 0 | 0 | 0.964 | 0 | 1.1138 |
| 48498 | ENSMUSG00000110752 | 0 | NA | NA | NA | NA | NA | D730003K21 | RIKEN cDNA D730003K21 gene [Source:M | 0 | 0 | 0 | 0 | 0 | 0 |
| 48499 | ENSMUSG00000110753 | 0 | NA | NA | NA | NA | NA | Gm18391 | predicted gene, 18391 [Source:MGI Symb | 0 | 0 | 0 | 0 | 0 | 0 |
| 48500 | ENSMUSG00000110754 | 0.160662974 | 1.126924175 | 4.080472857 | 0.276174898 | 0.782413736 | NA | 4930405J17I | RIKEN cDNA 4930405J17 gene [Source:MG | 0 | 0 | 0 | 0.964 | 0 | 0 |
| 48501 | ENSMUSG00000110755 | 260.0760391 | 0.197515732 | 0.599594644 | 0.329415438 | 0.7418417 | 0.948939213 | BC049987 | cDNA sequence BC049987 [Source:MGI Sy | 447.33 | 164.47 | 114.71 | 242.92 | 333.73 | 257.3 |
| 48502 | ENSMUSG00000110756 | 0 | NA | NA | NA | NA | NA | Gm47786 | predicted gene, 47786 [Source:MGI Symb | 0 | 0 | 0 | 0 | 0 | 0 |
| 48503 | ENSMUSG00000110757 | 2.773194302 | 1.37521595 | 1.371418786 | 1.002768785 | 0.315972433 | NA | Gm18228 | predicted gene, 18228 [Source:MGI Symb | 1.6062 | 2.0558 | 1.0062 | 8.6758 | 2.1812 | 1.1138 |
| 48504 | ENSMUSG00000110758 | 0 | NA | NA | NA | NA | NA | Gm48092 | predicted gene, 48092 [Source:MGI Symb | 0 | 0 | 0 | 0 | 0 | 0 |
| 48505 | ENSMUSG00000110759 | 0 | NA | NA | NA | NA | NA | Gm6607 | predicted gene 6607 [Source:MGI Symbol | 0 | 0 | 0 | 0 | 0 | 0 |
| 48506 | ENSMUSG00000110760 | 0 | NA | NA | NA | NA | NA | Gm39456 | predicted gene, 39456 [Source:MGI Symb | 0 | 0 | 0 | 0 | 0 | 0 |
| 48507 | ENSMUSG00000110761 | 0 | NA | NA | NA | NA | NA | Gm29761 | predicted gene, 29761 [Source:MGI Symb | 0 | 0 | 0 | 0 | 0 | 0 |
| 48508 | ENSMUSG00000110762 | 0 | NA | NA | NA | NA | NA | Gm48229 | predicted gene, 48229 [Source:MGI Symb | 0 | 0 | 0 | 0 | 0 | 0 |
| 48509 | ENSMUSG00000110765 | 0.636946036 | 1.591416711 | 2.83781177 | 0.560790087 | 0.574940645 | NA | Gm47468 | predicted gene, 47468 [Source:MGI Symb | 0.8031 | 0 | 0 | 1.928 | 1.0906 | 0 |
| 48510 | ENSMUSG00000110766 | 0 | NA | NA | NA | NA | NA | Gm47214 | predicted gene, 47214 [Source:MGI Symb | 0 | 0 | 0 | 0 | 0 | 0 |
| 48511 | ENSMUSG00000110767 | 0 | NA | NA | NA | NA | NA | Gm31432 | predicted gene, 31432 [Source:MGI Symb | 0 | 0 | 0 | 0 | 0 | 0 |
| 48512 | ENSMUSG00000110768 | 0.167707431 | -0.796629176 | 4.080472857 | -0.195229623 | 0.84521319 | NA | Gm18541 | predicted gene, 18541 [Source:MGI Symb | 0 | 0 | 1.0062 | 0 | 0 | 0 |
| 48513 | ENSMUSG00000110769 | 0 | NA | NA | NA | NA | NA | Gm47783 | predicted gene, 47783 [Source:MGI Symb | 0 | 0 | 0 | 0 | 0 | 0 |
| 48514 | ENSMUSG00000110770 | 0 | NA | NA | NA | NA | NA | Tpi-rs5 | triosephosphate isomerase related seque | 0 | 0 | 0 | 0 | 0 | 0 |
| 48515 | ENSMUSG00000110771 | 0 | NA | NA | NA | NA | NA | Gm47785 | predicted gene, 47785 [Source:MGI Symb | 0 | 0 | 0 | 0 | 0 | 0 |
| 48516 | ENSMUSG00000110772 | 1.461336631 | -0.75227185 | 1.787655625 | -0.420814747 | 0.673890362 | NA | Gm47856 | predicted gene, 47856 [Source:MGI Symb | 2.4093 | 1.0279 | 2.0125 | 0 | 1.0906 | 2.2277 |
| 48517 | ENSMUSG00000110773 | 0.520796785 | -0.820103173 | 3.072141206 | -0.266948398 | 0.789508892 | NA | Gm39326 | predicted gene, 39326 [Source:MGI Symb | 0 | 1.0279 | 1.0062 | 0 | 1.0906 | 0 |
| 48518 | ENSMUSG00000110774 | 3.685195871 | -0.580921737 | 1.299716806 | -0.446960241 | 0.654903767 | NA | Gm39318 | predicted gene, 39318 [Source:MGI Symb | 7.2279 | 2.0558 | 4.025 | 7.7118 | 1.0906 | 0 |
| 48519 | ENSMUSG00000110776 | 0 | NA | NA | NA | NA | NA | Gm2829 | predicted gene 2829 [Source:MGI Symbol | 0 | 0 | 0 | 0 | 0 | 0 |
| 48520 | ENSMUSG00000110778 | 0 | NA | NA | NA | NA | NA | Gm8171 | predicted gene 8171 [Source:MGI Symbol | 0 | 0 | 0 | 0 | 0 | 0 |
| 48521 | ENSMUSG00000110779 | 0 | NA | NA | NA | NA | NA | Gm48271 | predicted gene, 48271 [Source:MGI Symb | 0 | 0 | 0 | 0 | 0 | 0 |
| 48522 | ENSMUSG00000110780 | 0 | NA | NA | NA | NA | NA | Olfr949-ps1 | olfactory receptor 949, pseudogene 1 [So | 0 | 0 | 0 | 0 | 0 | 0 |
| 48523 | ENSMUSG00000110781 | 52.56258089 | 0.072585625 | 0.313243536 | 0.231722658 | 0.816753432 | 0.967845457 | Gm31992 | predicted gene, 31992 [Source:MGI Symb | 44.171 | 47.284 | 62.387 | 50.127 | 51.259 | 60.148 |
| 48524 | ENSMUSG00000110782 | 0 | NA | NA | NA | NA | NA | Gm48198 | predicted gene, 48198 [Source:MGI Symb | 0 | 0 | 0 | 0 | 0 | 0 |
| 48525 | ENSMUSG00000110783 | 0 | NA | NA | NA | NA | NA | Gm48821 | predicted gene, 48821 [Source:MGI Symb | 0 | 0 | 0 | 0 | 0 | 0 |
| 48526 | ENSMUSG00000110784 | 0 | NA | NA | NA | NA | NA | Gm39436 | predicted gene, 39436 [Source:MGI Symb | 0 | 0 | 0 | 0 | 0 | 0 |
| 48527 | ENSMUSG00000110786 | 0 | NA | NA | NA | NA | NA | Gm5119 | predicted gene 5119 [Source:MGI Symbol | 0 | 0 | 0 | 0 | 0 | 0 |
| 48528 | ENSMUSG00000110787 | 0 | NA | NA | NA | NA | NA | Gm6667 | predicted gene 6667 [Source:MGI Symbol | 0 | 0 | 0 | 0 | 0 | 0 |
| 48529 | ENSMUSG00000110788 | 0 | NA | NA | NA | NA | NA | Gm33700 | predicted gene, 33700 [Source:MGI Symb | 0 | 0 | 0 | 0 | 0 | 0 |
| 48530 | ENSMUSG00000110789 | 0 | NA | NA | NA | NA | NA | Gm33053 | predicted gene, 33053 [Source:MGI Symb | 0 | 0 | 0 | 0 | 0 | 0 |
| 48531 | ENSMUSG00000110790 | 0 | NA | NA | NA | NA | NA | Gm47079 | predicted gene, 47079 [Source:MGI Symb | 0 | 0 | 0 | 0 | 0 | 0 |
| 48532 | ENSMUSG00000110791 | 7.457707641 | 0.178787461 | 0.847298461 | 0.211008835 | 0.832880377 | 0.970611245 | Gm47508 | predicted gene, 47508 [Source:MGI Symb | 4.8186 | 10.279 | 6.0375 | 11.568 | 9.8156 | 2.2277 |
| 48533 | ENSMUSG00000110792 | 0 | NA | NA | NA | NA | NA | Olfr931-ps1 | olfactory receptor 931, pseudogene 1 [So | 0 | 0 | 0 | 0 | 0 | 0 |
| 48534 | ENSMUSG00000110793 | 0 | NA | NA | NA | NA | NA | Gm7395 | predicted gene 7395 [Source:MGI Symbol | 0 | 0 | 0 | 0 | 0 | 0 |
| 48535 | ENSMUSG00000110794 | 0 | NA | NA | NA | NA | NA | 4930401A07 | RIKEN cDNA 4930401A07 gene [Source:M | 0 | 0 | 0 | 0 | 0 | 0 |
| 48536 | ENSMUSG00000110795 | 0.17131964 | -0.796629176 | 4.080472857 | -0.195229623 | 0.84521319 | NA | Gm47179 | predicted gene, 47179 [Source:MGI Symb | 0 | 1.0279 | 0 | 0 | 0 | 0 |
| 48537 | ENSMUSG00000110799 | 0 | NA | NA | NA | NA | NA | 4930543K20 | RIKEN cDNA 4930543K20 gene [Source:M | 0 | 0 | 0 | 0 | 0 | 0 |
| 48538 | ENSMUSG00000110801 | 0 | NA | NA | NA | NA | NA | Gm48736 | predicted gene, 48736 [Source:MGI Symb | 0 | 0 | 0 | 0 | 0 | 0 |
| 48539 | ENSMUSG00000110802 | 0 | NA | NA | NA | NA | NA | Gm47141 | predicted gene, 47141 [Source:MGI Symb | 0 | 0 | 0 | 0 | 0 | 0 |
| 48540 | ENSMUSG00000110803 | 0.267700746 | -1.497481649 | 4.057001436 | -0.369110456 | 0.712045396 | NA | Gm20275 | predicted gene, 20275 [Source:MGI Symb | 1.6062 | 0 | 0 | 0 | 0 | 0 |
| 48541 | ENSMUSG00000110804 | 0 | NA | NA | NA | NA | NA | Olfr1084 | olfactory receptor 1084 [Source:MGI Sym | 0 | 0 | 0 | 0 | 0 | 0 |
| 48542 | ENSMUSG00000110808 | 0 | NA | NA | NA | NA | NA | Gm5364 | predicted gene 5364 [Source:MGI Symbol | 0 | 0 | 0 | 0 | 0 | 0 |
| 48543 | ENSMUSG00000110809 | 0 | NA | NA | NA | NA | NA | Gm46138 | predicted gene, 46138 [Source:MGI Symb | 0 | 0 | 0 | 0 | 0 | 0 |
| 48544 | ENSMUSG00000110810 | 0 | NA | NA | NA | NA | NA | Olfr1005-ps1 | olfactory receptor 1005, pseudogene 1 [S | 0 | 0 | 0 | 0 | 0 | 0 |
| 48545 | ENSMUSG00000110811 | 0 | NA | NA | NA | NA | NA | Gm48729 | predicted gene, 48729 [Source:MGI Symb | 0 | 0 | 0 | 0 | 0 | 0 |
| 48546 | ENSMUSG00000110812 | 0 | NA | NA | NA | NA | NA | Gm48407 | predicted gene, 48407 [Source:MGI Symb | 0 | 0 | 0 | 0 | 0 | 0 |
| 48547 | ENSMUSG00000110813 | 0 | NA | NA | NA | NA | NA | Gm31497 | predicted gene, 31497 [Source:MGI Symb | 0 | 0 | 0 | 0 | 0 | 0 |
| 48548 | ENSMUSG00000110814 | 0 | NA | NA | NA | NA | NA | Gm38661 | predicted gene, 38661 [Source:MGI Symb | 0 | 0 | 0 | 0 | 0 | 0 |

| | | | | | | | | | | | | | | | |
|---|---|---|---|---|---|---|---|---|---|---|---|---|---|---|---|
| 48549 | ENSMUSG00000110815 | 0 | NA | NA | NA | NA | NA | Gm47345 | predicted gene, 47345 [Source:MGI Symb | 0 | 0 | 0 | 0 | 0 | 0 |
| 48550 | ENSMUSG00000110816 | 1.191858604 | 0.487587703 | 2.09676255 | 0.232543119 | 0.816116201 | NA | Gm19178 | predicted gene, 19178 [Source:MGI Symb | 0 | 0 | 3.0187 | 1.928 | 1.0906 | 1.1138 |
| 48551 | ENSMUSG00000110817 | 0 | NA | NA | NA | NA | NA | Gm47723 | predicted gene, 47723 [Source:MGI Symb | 0 | 0 | 0 | 0 | 0 | 0 |
| 48552 | ENSMUSG00000110819 | 0 | NA | NA | NA | NA | NA | Olfr1252 | olfactory receptor 1252 [Source:MGI Symb | 0 | 0 | 0 | 0 | 0 | 0 |
| 48553 | ENSMUSG00000110821 | 0 | NA | NA | NA | NA | NA | Gm47176 | predicted gene, 47176 [Source:MGI Symb | 0 | 0 | 0 | 0 | 0 | 0 |
| 48554 | ENSMUSG00000110822 | 0 | NA | NA | NA | NA | NA | Gm18305 | predicted gene, 18305 [Source:MGI Symb | 0 | 0 | 0 | 0 | 0 | 0 |
| 48555 | ENSMUSG00000110823 | 0 | NA | NA | NA | NA | NA | Gm36799 | predicted gene, 36799 [Source:MGI Symb | 0 | 0 | 0 | 0 | 0 | 0 |
| 48556 | ENSMUSG00000110824 | 0 | NA | NA | NA | NA | NA | Gm48677 | predicted gene, 48677 [Source:MGI Symb | 0 | 0 | 0 | 0 | 0 | 0 |
| 48557 | ENSMUSG00000110825 | 0 | NA | NA | NA | NA | NA | Gm8959 | predicted gene 8959 [Source:MGI Symbol | 0 | 0 | 0 | 0 | 0 | 0 |
| 48558 | ENSMUSG00000110826 | 0 | NA | NA | NA | NA | NA | Gm47262 | predicted gene, 47262 [Source:MGI Symb | 0 | 0 | 0 | 0 | 0 | 0 |
| 48559 | ENSMUSG00000110827 | 36.12082168 | -0.13145344 | 0.37646234 | -0.349180849 | 0.726953542 | 0.945902846 | Gm32281 | predicted gene, 32281 [Source:MGI Symb | 41.761 | 44.2 | 27.169 | 30.847 | 35.99 | 36.757 |
| 48560 | ENSMUSG00000110828 | 0 | NA | NA | NA | NA | NA | Gm47517 | predicted gene, 47517 [Source:MGI Symb | 0 | 0 | 0 | 0 | 0 | 0 |
| 48561 | ENSMUSG00000110829 | 0 | NA | NA | NA | NA | NA | Gm47638 | predicted gene, 47638 [Source:MGI Symb | 0 | 0 | 0 | 0 | 0 | 0 |
| 48562 | ENSMUSG00000110830 | 3.999168159 | -0.751651148 | 1.225153394 | -0.61351595 | 0.53953524 | NA | Gm46218 | predicted gene, 46218 [Source:MGI Symb | 4.0155 | 5.1396 | 6.0375 | 7.7118 | 1.0906 | 0 |
| 48563 | ENSMUSG00000110831 | 0 | NA | NA | NA | NA | NA | Gm48159 | predicted gene, 48159 [Source:MGI Symb | 0 | 0 | 0 | 0 | 0 | 0 |
| 48564 | ENSMUSG00000110832 | 0 | NA | NA | NA | NA | NA | Gm48125 | predicted gene, 48125 [Source:MGI Symb | 0 | 0 | 0 | 0 | 0 | 0 |
| 48565 | ENSMUSG00000110833 | 0 | NA | NA | NA | NA | NA | Gm47981 | predicted gene, 47981 [Source:MGI Symb | 0 | 0 | 0 | 0 | 0 | 0 |
| 48566 | ENSMUSG00000110834 | 54.70396327 | -0.042719308 | 0.330524611 | -0.129246982 | 0.897162222 | 0.982798755 | Gm39469 | predicted gene, 39469 [Source:MGI Symb | 66.657 | 39.061 | 60.375 | 44.343 | 65.437 | 52.351 |
| 48567 | ENSMUSG00000110835 | 0 | NA | NA | NA | NA | NA | Gm48563 | predicted gene, 48563 [Source:MGI Symb | 0 | 0 | 0 | 0 | 0 | 0 |
| 48568 | ENSMUSG00000110836 | 0 | NA | NA | NA | NA | NA | Gm48801 | predicted gene, 48801 [Source:MGI Symb | 0 | 0 | 0 | 0 | 0 | 0 |
| 48569 | ENSMUSG00000110837 | 0 | NA | NA | NA | NA | NA | Gm18729 | predicted gene, 18729 [Source:MGI Symb | 0 | 0 | 0 | 0 | 0 | 0 |
| 48570 | ENSMUSG00000110838 | 7.226384733 | -0.357911013 | 0.784494799 | -0.456231212 | 0.648223723 | 0.925643431 | Gm47789 | predicted gene, 47789 [Source:MGI Symb | 8.031 | 10.279 | 6.0375 | 5.7839 | 6.5437 | 6.6831 |
| 48571 | ENSMUSG00000110839 | 0 | NA | NA | NA | NA | NA | Gm47800 | predicted gene, 47800 [Source:MGI Symb | 0 | 0 | 0 | 0 | 0 | 0 |
| 48572 | ENSMUSG00000110840 | 0.513958919 | -2.394274821 | 3.756096013 | -0.637437066 | 0.523840191 | NA | Gm47496 | predicted gene, 47496 [Source:MGI Symb | 0 | 3.0838 | 0 | 0 | 0 | 0 |
| 48573 | ENSMUSG00000110841 | 187.6778978 | -0.119225956 | 0.177493043 | -0.671721855 | 0.501760786 | 0.880974509 | Gpx4-ps2 | glutathione peroxidase 4, pseudogene 2 [! | 205.59 | 200.44 | 180.12 | 187.01 | 165.77 | 187.13 |
| 48574 | ENSMUSG00000110843 | 0 | NA | NA | NA | NA | NA | Gm35552 | predicted gene, 35552 [Source:MGI Symb | 0 | 0 | 0 | 0 | 0 | 0 |
| 48575 | ENSMUSG00000110844 | 0 | NA | NA | NA | NA | NA | Gm47680 | predicted gene, 47680 [Source:MGI Symb | 0 | 0 | 0 | 0 | 0 | 0 |
| 48576 | ENSMUSG00000110845 | 0.185640647 | 1.126924175 | 4.080472857 | 0.276174898 | 0.782413736 | NA | Gm47481 | predicted gene, 47481 [Source:MGI Symb | 0 | 0 | 0 | 0 | 0 | 1.1138 |
| 48577 | ENSMUSG00000110846 | 0 | NA | NA | NA | NA | NA | Gm18508 | predicted gene, 18508 [Source:MGI Symb | 0 | 0 | 0 | 0 | 0 | 0 |
| 48578 | ENSMUSG00000110847 | 0 | NA | NA | NA | NA | NA | Gm18705 | predicted gene, 18705 [Source:MGI Symb | 0 | 0 | 0 | 0 | 0 | 0 |
| 48579 | ENSMUSG00000110848 | 0 | NA | NA | NA | NA | NA | Gm47663 | predicted gene, 47663 [Source:MGI Symb | 0 | 0 | 0 | 0 | 0 | 0 |
| 48580 | ENSMUSG00000110849 | 0 | NA | NA | NA | NA | NA | | olfactory receptor 1545, pseudogene 1 | 0 | 0 | 0 | 0 | 0 | 0 |
| 48581 | ENSMUSG00000110851 | 0 | NA | NA | NA | NA | NA | Gm48902 | predicted gene, 48902 [Source:MGI Symb | 0 | 0 | 0 | 0 | 0 | 0 |
| 48582 | ENSMUSG00000110853 | 0 | NA | NA | NA | NA | NA | Gm47436 | predicted gene, 47436 [Source:MGI Symb | 0 | 0 | 0 | 0 | 0 | 0 |
| 48583 | ENSMUSG00000110854 | 0 | NA | NA | NA | NA | NA | Gm48345 | predicted gene, 48345 [Source:MGI Symb | 0 | 0 | 0 | 0 | 0 | 0 |
| 48584 | ENSMUSG00000110855 | 0 | NA | NA | NA | NA | NA | Olfr1007-ps1 | olfactory receptor 1007, pseudogene 1 [S | 0 | 0 | 0 | 0 | 0 | 0 |
| 48585 | ENSMUSG00000110856 | 0 | NA | NA | NA | NA | NA | Timm29-ps1 | translocase of inner mitochondrial memb | 0 | 0 | 0 | 0 | 0 | 0 |
| 48586 | ENSMUSG00000110857 | 0 | NA | NA | NA | NA | NA | Gm48330 | predicted gene, 48330 [Source:MGI Symb | 0 | 0 | 0 | 0 | 0 | 0 |
| 48587 | ENSMUSG00000110858 | 0 | NA | NA | NA | NA | NA | Gm18160 | predicted gene, 18160 [Source:MGI Symb | 0 | 0 | 0 | 0 | 0 | 0 |
| 48588 | ENSMUSG00000110862 | 0 | NA | NA | NA | NA | NA | Gm35835 | predicted gene, 35835 [Source:MGI Symb | 0 | 0 | 0 | 0 | 0 | 0 |
| 48589 | ENSMUSG00000110863 | 3.349031995 | 0.876119465 | 1.333900947 | 0.656809988 | 0.511303101 | NA | Gm34655 | predicted gene, 34655 [Source:MGI Symb | 4.0155 | 1.0279 | 2.0125 | 8.6758 | 4.3625 | 0 |
| 48590 | ENSMUSG00000110864 | 0 | NA | NA | NA | NA | NA | Gm31374 | predicted gene, 31374 [Source:MGI Symb | 0 | 0 | 0 | 0 | 0 | 0 |
| 48591 | ENSMUSG00000110865 | 0 | NA | NA | NA | NA | NA | Gm47046 | predicted gene, 47046 [Source:MGI Symb | 0 | 0 | 0 | 0 | 0 | 0 |
| 48592 | ENSMUSG00000110866 | 0 | NA | NA | NA | NA | NA | Gm48362 | predicted gene, 48362 [Source:MGI Symb | 0 | 0 | 0 | 0 | 0 | 0 |
| 48593 | ENSMUSG00000110867 | 0.642651897 | 2.906894826 | 3.440535777 | 0.844895974 | 0.398168923 | NA | Gm7368 | predicted gene 7368 [Source:MGI Symbol | 0 | 0 | 0 | 3.8559 | 0 | 0 |
| 48594 | ENSMUSG00000110869 | 0.82991484 | -1.853997749 | 2.875131703 | -0.644839242 | 0.519031369 | NA | Gm49628 | predicted gene, 49628 [Source:MGI Symb | 4.0155 | 0 | 0 | 0.964 | 0 | 0 |
| 48595 | ENSMUSG00000110870 | 0 | NA | NA | NA | NA | NA | Gm47081 | predicted gene, 47081 [Source:MGI Symb | 0 | 0 | 0 | 0 | 0 | 0 |
| 48596 | ENSMUSG00000110872 | 0 | NA | NA | NA | NA | NA | Gm48255 | predicted gene, 48255 [Source:MGI Symb | 0 | 0 | 0 | 0 | 0 | 0 |
| 48597 | ENSMUSG00000110873 | 0 | NA | NA | NA | NA | NA | Gm48652 | predicted gene, 48652 [Source:MGI Symb | 0 | 0 | 0 | 0 | 0 | 0 |
| 48598 | ENSMUSG00000110874 | 0 | NA | NA | NA | NA | NA | D030045P18 | RIKEN cDNA D030045P18 gene [Source:M | 0 | 0 | 0 | 0 | 0 | 0 |
| 48599 | ENSMUSG00000110875 | 0 | NA | NA | NA | NA | NA | Gm47100 | predicted gene, 47100 [Source:MGI Symb | 0 | 0 | 0 | 0 | 0 | 0 |
| 48600 | ENSMUSG00000110876 | 0 | NA | NA | NA | NA | NA | 4933411E02 | RIKEN cDNA 4933411E02 gene [Source:M | 0 | 0 | 0 | 0 | 0 | 0 |
| 48601 | ENSMUSG00000110877 | 0.48903338 | 0.95637552 | 3.456690196 | 0.276673773 | 0.782030612 | NA | Gm47427 | predicted gene, 47427 [Source:MGI Symb | 0 | 0 | 1.0062 | 1.928 | 0 | 0 |
| 48602 | ENSMUSG00000110878 | 0 | NA | NA | NA | NA | NA | Olfr1064-ps1 | olfactory receptor 1064, pseudogene 1 [S | 0 | 0 | 0 | 0 | 0 | 0 |
| 48603 | ENSMUSG00000110879 | 0 | NA | NA | NA | NA | NA | Olfr1010 | olfactory receptor 1010 [Source:MGI Symb | 0 | 0 | 0 | 0 | 0 | 0 |
| 48604 | ENSMUSG00000110880 | 0 | NA | NA | NA | NA | NA | Gm48431 | predicted gene, 48431 [Source:MGI Symb | 0 | 0 | 0 | 0 | 0 | 0 |

| | | | | | | | | | | | | | | | |
|---|---|---|---|---|---|---|---|---|---|---|---|---|---|---|---|
| 48605 | ENSMUSG00000110881 | 0 | NA | NA | NA | NA | NA | Olfr897-ps1 | olfactory receptor 897, pseudogene 1 [So | 0 | 0 | 0 | 0 | 0 | 0 |
| 48606 | ENSMUSG00000110882 | 0 | NA | NA | NA | NA | NA | Gm7642 | predicted gene 7642 [Source:MGI Symbol | 0 | 0 | 0 | 0 | 0 | 0 |
| 48607 | ENSMUSG00000110884 | 0 | NA | NA | NA | NA | NA | Gm47416 | predicted gene, 47416 [Source:MGI Symb | 0 | 0 | 0 | 0 | 0 | 0 |
| 48608 | ENSMUSG00000110885 | 0 | NA | NA | NA | NA | NA | Gm48739 | predicted gene, 48739 [Source:MGI Symb | 0 | 0 | 0 | 0 | 0 | 0 |
| 48609 | ENSMUSG00000110886 | 0 | NA | NA | NA | NA | NA | Olfr1108-ps1 | olfactory receptor 1108, pseudogene 1 [S | 0 | 0 | 0 | 0 | 0 | 0 |
| 48610 | ENSMUSG00000110887 | 0.133850373 | -0.796629176 | 4.080472857 | -0.195229623 | 0.84521319 | NA | Cbx3-ps8 | chromobox 3, pseudogene 8 [Source:MGI | 0.8031 | 0 | 0 | 0 | 0 | 0 |
| 48611 | ENSMUSG00000110888 | 0 | NA | NA | NA | NA | NA | Gm47237 | predicted gene, 47237 [Source:MGI Symb | 0 | 0 | 0 | 0 | 0 | 0 |
| 48612 | ENSMUSG00000110889 | 0 | NA | NA | NA | NA | NA | Gm19611 | predicted gene, 19611 [Source:MGI Symb | 0 | 0 | 0 | 0 | 0 | 0 |
| 48613 | ENSMUSG00000110891 | 0 | NA | NA | NA | NA | NA | Gm40639 | predicted gene, 40639 [Source:MGI Symb | 0 | 0 | 0 | 0 | 0 | 0 |
| 48614 | ENSMUSG00000110892 | 0 | NA | NA | NA | NA | NA | Gm18585 | predicted gene, 18585 [Source:MGI Symb | 0 | 0 | 0 | 0 | 0 | 0 |
| 48615 | ENSMUSG00000110893 | 0 | NA | NA | NA | NA | NA | Gm47392 | predicted gene, 47392 [Source:MGI Symb | 0 | 0 | 0 | 0 | 0 | 0 |
| 48616 | ENSMUSG00000110894 | 0 | NA | NA | NA | NA | NA | Gm48335 | predicted gene, 48335 [Source:MGI Symb | 0 | 0 | 0 | 0 | 0 | 0 |
| 48617 | ENSMUSG00000110895 | 0 | NA | NA | NA | NA | NA | Gm48822 | predicted gene, 48822 [Source:MGI Symb | 0 | 0 | 0 | 0 | 0 | 0 |
| 48618 | ENSMUSG00000110896 | 0 | NA | NA | NA | NA | NA | Gm48273 | predicted gene, 48273 [Source:MGI Symb | 0 | 0 | 0 | 0 | 0 | 0 |
| 48619 | ENSMUSG00000110897 | 0 | NA | NA | NA | NA | NA | Gm47085 | predicted gene, 47085 [Source:MGI Symb | 0 | 0 | 0 | 0 | 0 | 0 |
| 48620 | ENSMUSG00000110898 | 0 | NA | NA | NA | NA | NA | Gm40638 | predicted gene, 40638 [Source:MGI Symb | 0 | 0 | 0 | 0 | 0 | 0 |
| 48621 | ENSMUSG00000110899 | 6.817046544 | -0.128219739 | 0.891649562 | -0.143800597 | 0.885657932 | 0.982072702 | Gm48840 | predicted gene, 48840 [Source:MGI Symb | 7.2279 | 5.1396 | 9.0562 | 9.6398 | 8.7249 | 1.1138 |
| 48622 | ENSMUSG00000110901 | 0 | NA | NA | NA | NA | NA | Gm48393 | predicted gene, 48393 [Source:MGI Symb | 0 | 0 | 0 | 0 | 0 | 0 |
| 48623 | ENSMUSG00000110902 | 0 | NA | NA | NA | NA | NA | Gm33104 | predicted gene, 33104 [Source:MGI Symb | 0 | 0 | 0 | 0 | 0 | 0 |
| 48624 | ENSMUSG00000110903 | 0.167707431 | -0.796629176 | 4.080472857 | -0.195229623 | 0.84521319 | NA | Gm47857 | predicted gene, 47857 [Source:MGI Symb | 0 | 0 | 1.0062 | 0 | 0 | 0 |
| 48625 | ENSMUSG00000110905 | 0 | NA | NA | NA | NA | NA | Gm48007 | predicted gene, 48007 [Source:MGI Symb | 0 | 0 | 0 | 0 | 0 | 0 |
| 48626 | ENSMUSG00000110908 | 0 | NA | NA | NA | NA | NA | Gm48434 | predicted gene, 48434 [Source:MGI Symb | 0 | 0 | 0 | 0 | 0 | 0 |
| 48627 | ENSMUSG00000110909 | 0 | NA | NA | NA | NA | NA | B020014A21 | RIKEN cDNA B020014A21 gene [Source:M | 0 | 0 | 0 | 0 | 0 | 0 |
| 48628 | ENSMUSG00000110910 | 4.569241218 | 0.354420175 | 1.104383012 | 0.32092143 | 0.748269934 | NA | 4921534A09 | RIKEN cDNA 4921534A09 gene [Source:M | 8.8341 | 2.0558 | 1.0062 | 6.7478 | 6.5437 | 2.2277 |
| 48629 | ENSMUSG00000110911 | 0 | NA | NA | NA | NA | NA | Gm47068 | predicted gene, 47068 [Source:MGI Symb | 0 | 0 | 0 | 0 | 0 | 0 |
| 48630 | ENSMUSG00000110912 | 0 | NA | NA | NA | NA | NA | Olfr1080 | olfactory receptor 1080 [Source:MGI Sym | 0 | 0 | 0 | 0 | 0 | 0 |
| 48631 | ENSMUSG00000110913 | 0 | NA | NA | NA | NA | NA | Gm47192 | predicted gene, 47192 [Source:MGI Symb | 0 | 0 | 0 | 0 | 0 | 0 |
| 48632 | ENSMUSG00000110914 | 0 | NA | NA | NA | NA | NA | Gm48611 | predicted gene, 48611 [Source:MGI Symb | 0 | 0 | 0 | 0 | 0 | 0 |
| 48633 | ENSMUSG00000110915 | 0.503095663 | 2.549494177 | 3.348715477 | 0.761334964 | 0.446457022 | NA | Gm36198 | predicted gene, 36198 [Source:MGI Symb | 0 | 0 | 0 | 1.928 | 1.0906 | 0 |
| 48634 | ENSMUSG00000110916 | 0 | NA | NA | NA | NA | NA | 2310003N18 | RIKEN cDNA 2310003N18 gene [Source:M | 0 | 0 | 0 | 0 | 0 | 0 |
| 48635 | ENSMUSG00000110917 | 0 | NA | NA | NA | NA | NA | Gm33056 | predicted gene, 33056 [Source:MGI Symb | 0 | 0 | 0 | 0 | 0 | 0 |
| 48636 | ENSMUSG00000110918 | 0 | NA | NA | NA | NA | NA | Gm48862 | predicted gene, 48862 [Source:MGI Symb | 0 | 0 | 0 | 0 | 0 | 0 |
| 48637 | ENSMUSG00000110919 | 0 | NA | NA | NA | NA | NA | Dppa5b | developmental pluripotency associated 5I | 0 | 0 | 0 | 0 | 0 | 0 |
| 48638 | ENSMUSG00000110920 | 0 | NA | NA | NA | NA | NA | Gm48858 | predicted gene, 48858 [Source:MGI Symb | 0 | 0 | 0 | 0 | 0 | 0 |
| 48639 | ENSMUSG00000110922 | 0 | NA | NA | NA | NA | NA | Gm35177 | predicted gene, 35177 [Source:MGI Symb | 0 | 0 | 0 | 0 | 0 | 0 |
| 48640 | ENSMUSG00000110923 | 0 | NA | NA | NA | NA | NA | Gm47499 | predicted gene, 47499 [Source:MGI Symb | 0 | 0 | 0 | 0 | 0 | 0 |
| 48641 | ENSMUSG00000110924 | 0 | NA | NA | NA | NA | NA | Gm40518 | predicted gene, 40518 [Source:MGI Symb | 0 | 0 | 0 | 0 | 0 | 0 |
| 48642 | ENSMUSG00000110925 | 0 | NA | NA | NA | NA | NA | Gm47858 | predicted gene, 47858 [Source:MGI Symb | 0 | 0 | 0 | 0 | 0 | 0 |
| 48643 | ENSMUSG00000110926 | 0.342639279 | -1.805976596 | 4.028547714 | -0.448294702 | 0.653940524 | NA | Gm5917 | predicted gene 5917 [Source:MGI Symbol | 0 | 2.0558 | 0 | 0 | 0 | 0 |
| 48644 | ENSMUSG00000110927 | 0 | NA | NA | NA | NA | NA | Gm48506 | predicted gene, 48506 [Source:MGI Symb | 0 | 0 | 0 | 0 | 0 | 0 |
| 48645 | ENSMUSG00000110928 | 0.342432689 | 1.990653086 | 3.900281263 | 0.51038706 | 0.609780321 | NA | Gm48114 | predicted gene, 48114 [Source:MGI Symb | 0 | 0 | 0 | 0.964 | 1.0906 | 0 |
| 48646 | ENSMUSG00000110929 | 0 | NA | NA | NA | NA | NA | Gm47757 | predicted gene, 47757 [Source:MGI Symb | 0 | 0 | 0 | 0 | 0 | 0 |
| 48647 | ENSMUSG00000110930 | 1.323847419 | 1.876094015 | 2.007969798 | 0.934323822 | 0.35013688 | NA | Gm19936 | predicted gene, 19936 [Source:MGI Symb | 1.6062 | 0 | 0 | 1.928 | 2.1812 | 2.2277 |
| 48648 | ENSMUSG00000110932 | 0 | NA | NA | NA | NA | NA | Gm48640 | predicted gene, 48640 [Source:MGI Symb | 0 | 0 | 0 | 0 | 0 | 0 |
| 48649 | ENSMUSG00000110935 | 0 | NA | NA | NA | NA | NA | Gm8834 | predicted gene 8834 [Source:MGI Symbol | 0 | 0 | 0 | 0 | 0 | 0 |
| 48650 | ENSMUSG00000110936 | 0 | NA | NA | NA | NA | NA | Gm19212 | predicted gene, 19212 [Source:MGI Symb | 0 | 0 | 0 | 0 | 0 | 0 |
| 48651 | ENSMUSG00000110937 | 0 | NA | NA | NA | NA | NA | Gm47629 | predicted gene, 47629 [Source:MGI Symb | 0 | 0 | 0 | 0 | 0 | 0 |
| 48652 | ENSMUSG00000110938 | 0 | NA | NA | NA | NA | NA | 1700124M09 | RIKEN cDNA 1700124M09 gene [Source:M | 0 | 0 | 0 | 0 | 0 | 0 |
| 48653 | ENSMUSG00000110940 | 0 | NA | NA | NA | NA | NA | Olfr274-ps1 | olfactory receptor 274, pseudogene 1 [So | 0 | 0 | 0 | 0 | 0 | 0 |
| 48654 | ENSMUSG00000110941 | 0 | NA | NA | NA | NA | NA | Gm48477 | predicted gene, 48477 [Source:MGI Symb | 0 | 0 | 0 | 0 | 0 | 0 |
| 48655 | ENSMUSG00000110942 | 0 | NA | NA | NA | NA | NA | Gm47029 | predicted gene, 47029 [Source:MGI Symb | 0 | 0 | 0 | 0 | 0 | 0 |
| 48656 | ENSMUSG00000110943 | 0 | NA | NA | NA | NA | NA | Gm30674 | predicted gene, 30674 [Source:MGI Symb | 0 | 0 | 0 | 0 | 0 | 0 |
| 48657 | ENSMUSG00000110944 | 0 | NA | NA | NA | NA | NA | Gm19174 | predicted gene, 19174 [Source:MGI Symb | 0 | 0 | 0 | 0 | 0 | 0 |
| 48658 | ENSMUSG00000110945 | 20.07362333 | 0.327621963 | 0.505569438 | 0.648025648 | 0.516968362 | 0.886792029 | Gm9856 | predicted gene 9856 [Source:MGI Symbol | 20.881 | 10.279 | 22.137 | 24.099 | 18.541 | 24.505 |
| 48659 | ENSMUSG00000110946 | 0 | NA | NA | NA | NA | NA | Gm47604 | predicted gene, 47604 [Source:MGI Symb | 0 | 0 | 0 | 0 | 0 | 0 |
| 48660 | ENSMUSG00000110947 | 0 | NA | NA | NA | NA | NA | Olfr1333 | olfactory receptor 1333 [Source:MGI Sym | 0 | 0 | 0 | 0 | 0 | 0 |

| | | | | | | | | | | | | | | | |
|---|---|---|---|---|---|---|---|---|---|---|---|---|---|---|---|
| 48661 | ENSMUSG00000110949 | 5.247368023 | 1.983792332 | 1.052250611 | 1.885285038 | 0.059391365 | 0.346665337 | Nudt8 | nudix (nucleoside diphosphate linked moi | 3.2124 | 1.0279 | 2.0125 | 7.7118 | 14.178 | 3.3415 |
| 48662 | ENSMUSG00000110950 | 0 | NA | NA | NA | NA | NA | Gm39384 | predicted gene, 39384 [Source:MGI Symb | 0 | 0 | 0 | 0 | 0 | 0 |
| 48663 | ENSMUSG00000110951 | 0 | NA | NA | NA | NA | NA | Gm47001 | predicted gene, 47001 [Source:MGI Symb | 0 | 0 | 0 | 0 | 0 | 0 |
| 48664 | ENSMUSG00000110952 | 0 | NA | NA | NA | NA | NA | Gm18226 | predicted gene, 18226 [Source:MGI Symb | 0 | 0 | 0 | 0 | 0 | 0 |
| 48665 | ENSMUSG00000110953 | 0 | NA | NA | NA | NA | NA | Olfr1088-ps1 | olfactory receptor 1088, pseudogene 1 [S | 0 | 0 | 0 | 0 | 0 | 0 |
| 48666 | ENSMUSG00000110956 | 0 | NA | NA | NA | NA | NA | Gm48834 | predicted gene, 48834 [Source:MGI Symb | 0 | 0 | 0 | 0 | 0 | 0 |
| 48667 | ENSMUSG00000110957 | 0 | NA | NA | NA | NA | NA | Gm7439 | predicted gene 7439 [Source:MGI Symbol | 0 | 0 | 0 | 0 | 0 | 0 |
| 48668 | ENSMUSG00000110958 | 0 | NA | NA | NA | NA | NA | Gm33332 | predicted gene, 33332 [Source:MGI Symb | 0 | 0 | 0 | 0 | 0 | 0 |
| 48669 | ENSMUSG00000110959 | 0 | NA | NA | NA | NA | NA | Gm47198 | predicted gene, 47198 [Source:MGI Symb | 0 | 0 | 0 | 0 | 0 | 0 |
| 48670 | ENSMUSG00000110960 | 0.480153995 | 1.043847617 | 3.151673025 | 0.331204287 | 0.740490182 | NA | Supt4b | SPT4B, DSIF elongation factor subunit [So | 0.8031 | 0 | 0 | 0.964 | 0 | 1.1138 |
| 48671 | ENSMUSG00000110962 | 0.185640647 | 1.126924175 | 4.080472857 | 0.276174898 | 0.782413736 | NA | Gm47062 | predicted gene, 47062 [Source:MGI Symb | 0 | 0 | 0 | 0 | 0 | 1.1138 |
| 48672 | ENSMUSG00000110964 | 0 | NA | NA | NA | NA | NA | Gm20953 | predicted gene, 20953 [Source:MGI Symb | 0 | 0 | 0 | 0 | 0 | 0 |
| 48673 | ENSMUSG00000110966 | 0 | NA | NA | NA | NA | NA | Gm47255 | predicted gene, 47255 [Source:MGI Symb | 0 | 0 | 0 | 0 | 0 | 0 |
| 48674 | ENSMUSG00000110967 | 0 | NA | NA | NA | NA | NA | Gm40608 | predicted gene, 40608 [Source:MGI Symb | 0 | 0 | 0 | 0 | 0 | 0 |
| 48675 | ENSMUSG00000110968 | 0 | NA | NA | NA | NA | NA | Gm19208 | predicted gene, 19208 [Source:MGI Symb | 0 | 0 | 0 | 0 | 0 | 0 |
| 48676 | ENSMUSG00000110969 | 0 | NA | NA | NA | NA | NA | Gm47174 | predicted gene, 47174 [Source:MGI Symb | 0 | 0 | 0 | 0 | 0 | 0 |
| 48677 | ENSMUSG00000110970 | 0 | NA | NA | NA | NA | NA | Olfr156 | olfactory receptor 156 [Source:MGI Symb | 0 | 0 | 0 | 0 | 0 | 0 |
| 48678 | ENSMUSG00000110972 | 0 | NA | NA | NA | NA | NA | 5830462O15 | RIKEN cDNA 5830462O15 gene [Source:M | 0 | 0 | 0 | 0 | 0 | 0 |
| 48679 | ENSMUSG00000110973 | 0 | NA | NA | NA | NA | NA | Gm48130 | predicted gene, 48130 [Source:MGI Symb | 0 | 0 | 0 | 0 | 0 | 0 |
| 48680 | ENSMUSG00000110976 | 0 | NA | NA | NA | NA | NA | Gm48093 | predicted gene, 48093 [Source:MGI Symb | 0 | 0 | 0 | 0 | 0 | 0 |
| 48681 | ENSMUSG00000110977 | 0 | NA | NA | NA | NA | NA | Gm48598 | predicted gene, 48598 [Source:MGI Symb | 0 | 0 | 0 | 0 | 0 | 0 |
| 48682 | ENSMUSG00000110978 | 0 | NA | NA | NA | NA | NA | Olfr991-ps1 | olfactory receptor 991, pseudogene 1 [So | 0 | 0 | 0 | 0 | 0 | 0 |
| 48683 | ENSMUSG00000110980 | 0 | NA | NA | NA | NA | NA | Gm47204 | predicted gene, 47204 [Source:MGI Symb | 0 | 0 | 0 | 0 | 0 | 0 |
| 48684 | ENSMUSG00000110981 | 0 | NA | NA | NA | NA | NA | Gm35940 | predicted gene, 35940 [Source:MGI Symb | 0 | 0 | 0 | 0 | 0 | 0 |
| 48685 | ENSMUSG00000110982 | 0 | NA | NA | NA | NA | NA | Gm48565 | predicted gene, 48565 [Source:MGI Symb | 0 | 0 | 0 | 0 | 0 | 0 |
| 48686 | ENSMUSG00000110983 | 0 | NA | NA | NA | NA | NA | Gm47679 | predicted gene, 47679 [Source:MGI Symb | 0 | 0 | 0 | 0 | 0 | 0 |
| 48687 | ENSMUSG00000110984 | 0 | NA | NA | NA | NA | NA | Gm47131 | predicted gene, 47131 [Source:MGI Symb | 0 | 0 | 0 | 0 | 0 | 0 |
| 48688 | ENSMUSG00000110985 | 0 | NA | NA | NA | NA | NA | Olfr953-ps1 | olfactory receptor 953, pseudogene 1 [So | 0 | 0 | 0 | 0 | 0 | 0 |
| 48689 | ENSMUSG00000110986 | 0.267700746 | -1.497481649 | 4.057001436 | -0.369110456 | 0.712045396 | NA | Gm20276 | predicted gene, 20276 [Source:MGI Symb | 1.6062 | 0 | 0 | 0 | 0 | 0 |
| 48690 | ENSMUSG00000110987 | 0 | NA | NA | NA | NA | NA | Gm47048 | predicted gene, 47048 [Source:MGI Symb | 0 | 0 | 0 | 0 | 0 | 0 |
| 48691 | ENSMUSG00000110988 | 0 | NA | NA | NA | NA | NA | Gm29909 | predicted gene, 29909 [Source:MGI Symb | 0 | 0 | 0 | 0 | 0 | 0 |
| 48692 | ENSMUSG00000110989 | 0 | NA | NA | NA | NA | NA | Gm7444 | predicted gene 7444 [Source:MGI Symbol | 0 | 0 | 0 | 0 | 0 | 0 |
| 48693 | ENSMUSG00000110991 | 0 | NA | NA | NA | NA | NA | Olfr908 | olfactory receptor 908 [Source:MGI Symb | 0 | 0 | 0 | 0 | 0 | 0 |
| 48694 | ENSMUSG00000110992 | 0 | NA | NA | NA | NA | NA | Gm35371 | predicted gene, 35371 [Source:MGI Symb | 0 | 0 | 0 | 0 | 0 | 0 |
| 48695 | ENSMUSG00000110993 | 1.342303597 | 2.811990234 | 2.234743945 | 1.258305338 | 0.208281351 | NA | Gm47963 | predicted gene, 47963 [Source:MGI Symb | 0 | 0 | 1.0062 | 4.8199 | 0 | 2.2277 |
| 48696 | ENSMUSG00000110995 | 0 | NA | NA | NA | NA | NA | Gm18789 | predicted gene, 18789 [Source:MGI Symb | 0 | 0 | 0 | 0 | 0 | 0 |
| 48697 | ENSMUSG00000110996 | 54.05379716 | -0.031856307 | 0.348985899 | -0.091282504 | 0.927268119 | 0.987712228 | Gm36251 | predicted gene, 36251 [Source:MGI Symb | 40.155 | 70.926 | 53.331 | 62.659 | 35.99 | 61.261 |
| 48698 | ENSMUSG00000110997 | 1.755384664 | 0.536143138 | 1.662682837 | 0.32245665 | 0.747106773 | NA | 2610028D06 | RIKEN cDNA 2610028D06 gene [Source:M | 3.2124 | 0 | 1.0062 | 1.928 | 3.2719 | 1.1138 |
| 48699 | ENSMUSG00000110998 | 0 | NA | NA | NA | NA | NA | I730028E13F | RIKEN cDNA I730028E13 gene [Source:MG | 0 | 0 | 0 | 0 | 0 | 0 |
| 48700 | ENSMUSG00000110999 | 0 | NA | NA | NA | NA | NA | Olfr1075-ps1 | olfactory receptor 1075, pseudogene 1 [S | 0 | 0 | 0 | 0 | 0 | 0 |
| 48701 | ENSMUSG00000111000 | 0 | NA | NA | NA | NA | NA | Gm47108 | predicted gene, 47108 [Source:MGI Symb | 0 | 0 | 0 | 0 | 0 | 0 |
| 48702 | ENSMUSG00000111001 | 0 | NA | NA | NA | NA | NA | Gm8767 | predicted gene 8767 [Source:MGI Symbol | 0 | 0 | 0 | 0 | 0 | 0 |
| 48703 | ENSMUSG00000111002 | 0 | NA | NA | NA | NA | NA | Gm47674 | predicted gene, 47674 [Source:MGI Symb | 0 | 0 | 0 | 0 | 0 | 0 |
| 48704 | ENSMUSG00000111003 | 1.175676038 | 1.502328677 | 1.986024232 | 0.756450326 | 0.449379244 | NA | Gm47122 | predicted gene, 47122 [Source:MGI Symb | 0.8031 | 1.0279 | 0 | 1.928 | 2.1812 | 1.1138 |
| 48705 | ENSMUSG00000111004 | 0 | NA | NA | NA | NA | NA | Gm48272 | predicted gene, 48272 [Source:MGI Symb | 0 | 0 | 0 | 0 | 0 | 0 |
| 48706 | ENSMUSG00000111005 | 0 | NA | NA | NA | NA | NA | Gm33619 | predicted gene, 33619 [Source:MGI Symb | 0 | 0 | 0 | 0 | 0 | 0 |
| 48707 | ENSMUSG00000111006 | 0.349477146 | 0.165142768 | 3.911021834 | 0.042224967 | 0.966319359 | NA | Gm10826 | predicted gene 10826 [Source:MGI Symbo | 0 | 0 | 1.0062 | 0 | 1.0906 | 0 |
| 48708 | ENSMUSG00000111008 | 0 | NA | NA | NA | NA | NA | Gm31562 | predicted gene, 31562 [Source:MGI Symb | 0 | 0 | 0 | 0 | 0 | 0 |
| 48709 | ENSMUSG00000111010 | 0 | NA | NA | NA | NA | NA | Gm20734 | predicted gene, 20734 [Source:MGI Symb | 0 | 0 | 0 | 0 | 0 | 0 |
| 48710 | ENSMUSG00000111012 | 0 | NA | NA | NA | NA | NA | Gm7588 | predicted gene 7588 [Source:MGI Symbol | 0 | 0 | 0 | 0 | 0 | 0 |
| 48711 | ENSMUSG00000111013 | 52.3724255 | -0.454621922 | 0.962834421 | -0.472170409 | 0.636805159 | 0.923478764 | Gm32468 | predicted gene, 32468 [Source:MGI Symb | 102.8 | 63.731 | 15.094 | 54.947 | 66.528 | 11.138 |
| 48712 | ENSMUSG00000111014 | 0.654905883 | -1.312574656 | 2.816093566 | -0.466097672 | 0.641145589 | NA | Gm47795 | predicted gene, 47795 [Source:MGI Symb | 0.8031 | 0 | 2.0125 | 0 | 0 | 1.1138 |
| 48713 | ENSMUSG00000111015 | 0 | NA | NA | NA | NA | NA | Gm3126 | predicted gene 3126 [Source:MGI Symbol | 0 | 0 | 0 | 0 | 0 | 0 |
| 48714 | ENSMUSG00000111016 | 0 | NA | NA | NA | NA | NA | Gm48462 | predicted gene, 48462 [Source:MGI Symb | 0 | 0 | 0 | 0 | 0 | 0 |
| 48715 | ENSMUSG00000111017 | 0.531944269 | 2.618916564 | 3.285123409 | 0.797204926 | 0.425332026 | NA | Gm48314 | predicted gene, 48314 [Source:MGI Symb | 0 | 0 | 0 | 0.964 | 0 | 2.2277 |
| 48716 | ENSMUSG00000111018 | 0 | NA | NA | NA | NA | NA | Olfr941-ps1 | olfactory receptor 941, pseudogene 1 [So | 0 | 0 | 0 | 0 | 0 | 0 |

| | | | | | | | | | | | | | | | |
|---|---|---|---|---|---|---|---|---|---|---|---|---|---|---|---|
| 48717 | ENSMUSG00000111019 | 0 | NA | NA | NA | NA | NA | Gm39404 | predicted gene, 39404 [Source:MGI Symb | 0 | 0 | 0 | 0 | 0 | 0 |
| 48718 | ENSMUSG00000111020 | 0 | NA | NA | NA | NA | NA | Gm1715 | predicted gene 1715 [Source:MGI Symbol | 0 | 0 | 0 | 0 | 0 | 0 |
| 48719 | ENSMUSG00000111021 | 0 | NA | NA | NA | NA | NA | Olfr50 | olfactory receptor 50 [Source:MGI Symbo | 0 | 0 | 0 | 0 | 0 | 0 |
| 48720 | ENSMUSG00000111022 | 0 | NA | NA | NA | NA | NA | Gm36367 | predicted gene, 36367 [Source:MGI Symb | 0 | 0 | 0 | 0 | 0 | 0 |
| 48721 | ENSMUSG00000111023 | 0 | NA | NA | NA | NA | NA | Gm48391 | predicted gene, 48391 [Source:MGI Symb | 0 | 0 | 0 | 0 | 0 | 0 |
| 48722 | ENSMUSG00000111025 | 0 | NA | NA | NA | NA | NA | Gm47185 | predicted gene, 47185 [Source:MGI Symb | 0 | 0 | 0 | 0 | 0 | 0 |
| 48723 | ENSMUSG00000111027 | 1.401080681 | 0.9619189 | 2.04626315 | 0.470085629 | 0.638293841 | NA | Gm2981 | predicted gene 2981 [Source:MGI Symbol | 0.8031 | 1.0279 | 1.0062 | 0 | 0 | 5.5692 |
| 48724 | ENSMUSG00000111028 | 0.294513347 | 0.165143496 | 4.080472857 | 0.040471657 | 0.967717103 | NA | Gm5922 | predicted gene 5922 [Source:MGI Symbol | 0.8031 | 0 | 0 | 0.964 | 0 | 0 |
| 48725 | ENSMUSG00000111029 | 0 | NA | NA | NA | NA | NA | Olfr1244-ps1 | olfactory receptor 1244, pseudogene 1 [S | 0 | 0 | 0 | 0 | 0 | 0 |
| 48726 | ENSMUSG00000111030 | 0.787158936 | 0.679629554 | 2.65851484 | 0.255642566 | 0.798226848 | NA | Gm47234 | predicted gene, 47234 [Source:MGI Symb | 0.8031 | 1.0279 | 0 | 2.8919 | 0 | 0 |
| 48727 | ENSMUSG00000111031 | 0 | NA | NA | NA | NA | NA | Gm47458 | predicted gene, 47458 [Source:MGI Symb | 0 | 0 | 0 | 0 | 0 | 0 |
| 48728 | ENSMUSG00000111032 | 0 | NA | NA | NA | NA | NA | Olfr1543-ps1 | olfactory receptor 1543, pseudogene 1 [S | 0 | 0 | 0 | 0 | 0 | 0 |
| 48729 | ENSMUSG00000111033 | 0 | NA | NA | NA | NA | NA | Gm34322 | predicted gene, 34322 [Source:MGI Symb | 0 | 0 | 0 | 0 | 0 | 0 |
| 48730 | ENSMUSG00000111034 | 0.353348078 | 0.165142701 | 3.895042568 | 0.042398176 | 0.966181282 | NA | Gm48066 | predicted gene, 48066 [Source:MGI Symb | 0 | 0 | 1.0062 | 0 | 0 | 1.1138 |
| 48731 | ENSMUSG00000111035 | 0 | NA | NA | NA | NA | NA | Gm33054 | predicted gene, 33054 [Source:MGI Symb | 0 | 0 | 0 | 0 | 0 | 0 |
| 48732 | ENSMUSG00000111036 | 0 | NA | NA | NA | NA | NA | Gm47502 | predicted gene, 47502 [Source:MGI Symb | 0 | 0 | 0 | 0 | 0 | 0 |
| 48733 | ENSMUSG00000111037 | 0 | NA | NA | NA | NA | NA | Gm47235 | predicted gene, 47235 [Source:MGI Symb | 0 | 0 | 0 | 0 | 0 | 0 |
| 48734 | ENSMUSG00000111039 | 1.090055336 | -2.273671673 | 2.179733234 | -1.043096301 | 0.296903694 | NA | Gm47000 | predicted gene, 47000 [Source:MGI Symb | 2.4093 | 1.0279 | 2.0125 | 0 | 1.0906 | 0 |
| 48735 | ENSMUSG00000111041 | 0 | NA | NA | NA | NA | NA | Gm3011 | predicted gene 3011 [Source:MGI Symbol | 0 | 0 | 0 | 0 | 0 | 0 |
| 48736 | ENSMUSG00000111042 | 0 | NA | NA | NA | NA | NA | Gm18252 | predicted gene, 18252 [Source:MGI Symb | 0 | 0 | 0 | 0 | 0 | 0 |
| 48737 | ENSMUSG00000111043 | 0 | NA | NA | NA | NA | NA | 2210414F02 | RIKEN cDNA 2210414F02 gene [Source:M | 0 | 0 | 0 | 0 | 0 | 0 |
| 48738 | ENSMUSG00000111044 | 0 | NA | NA | NA | NA | NA | Gm47106 | predicted gene, 47106 [Source:MGI Symb | 0 | 0 | 0 | 0 | 0 | 0 |
| 48739 | ENSMUSG00000111045 | 0.167707431 | -0.796629176 | 4.080472857 | -0.195229623 | 0.84521319 | NA | Gm47598 | predicted gene, 47598 [Source:MGI Symb | 0 | 0 | 1.0062 | 0 | 0 | 0 |
| 48740 | ENSMUSG00000111046 | 0 | NA | NA | NA | NA | NA | Gm46197 | predicted gene, 46197 [Source:MGI Symb | 0 | 0 | 0 | 0 | 0 | 0 |
| 48741 | ENSMUSG00000111047 | 0 | NA | NA | NA | NA | NA | Gm4780 | predicted gene 4780 [Source:MGI Symbol | 0 | 0 | 0 | 0 | 0 | 0 |
| 48742 | ENSMUSG00000111048 | 0 | NA | NA | NA | NA | NA | Gm48173 | predicted gene, 48173 [Source:MGI Symb | 0 | 0 | 0 | 0 | 0 | 0 |
| 48743 | ENSMUSG00000111049 | 0 | NA | NA | NA | NA | NA | Gm34184 | predicted gene, 34184 [Source:MGI Symb | 0 | 0 | 0 | 0 | 0 | 0 |
| 48744 | ENSMUSG00000111050 | 0 | NA | NA | NA | NA | NA | Gm47877 | predicted gene, 47877 [Source:MGI Symb | 0 | 0 | 0 | 0 | 0 | 0 |
| 48745 | ENSMUSG00000111051 | 0 | NA | NA | NA | NA | NA | Gm47084 | predicted gene, 47084 [Source:MGI Symb | 0 | 0 | 0 | 0 | 0 | 0 |
| 48746 | ENSMUSG00000111052 | 0 | NA | NA | NA | NA | NA | Gm20139 | predicted gene, 20139 [Source:MGI Symb | 0 | 0 | 0 | 0 | 0 | 0 |
| 48747 | ENSMUSG00000111053 | 0 | NA | NA | NA | NA | NA | Gm49343 | predicted gene, 49343 [Source:MGI Symb | 0 | 0 | 0 | 0 | 0 | 0 |
| 48748 | ENSMUSG00000111054 | 0 | NA | NA | NA | NA | NA | 4930545L08 | RIKEN cDNA 4930545L08 gene [Source:M | 0 | 0 | 0 | 0 | 0 | 0 |
| 48749 | ENSMUSG00000111055 | 3.191102434 | 0.812623966 | 1.282727873 | 0.633512363 | 0.52639911 | NA | D030034A15 | RIKEN cDNA D030034A15 gene [Source:M | 4.8186 | 1.0279 | 1.0062 | 6.7478 | 1.0906 | 4.4554 |
| 48750 | ENSMUSG00000111056 | 0 | NA | NA | NA | NA | NA | Gm39317 | predicted gene, 39317 [Source:MGI Symb | 0 | 0 | 0 | 0 | 0 | 0 |
| 48751 | ENSMUSG00000111057 | 0 | NA | NA | NA | NA | NA | Gm3442 | predicted gene 3442 [Source:MGI Symbol | 0 | 0 | 0 | 0 | 0 | 0 |
| 48752 | ENSMUSG00000111058 | 0 | NA | NA | NA | NA | NA | Gm36033 | predicted gene, 36033 [Source:MGI Symb | 0 | 0 | 0 | 0 | 0 | 0 |
| 48753 | ENSMUSG00000111059 | 0 | NA | NA | NA | NA | NA | Gm33838 | predicted gene, 33838 [Source:MGI Symb | 0 | 0 | 0 | 0 | 0 | 0 |
| 48754 | ENSMUSG00000111060 | 0.36353943 | 2.065989254 | 4.034137301 | 0.512126658 | 0.608562368 | NA | Gm47238 | predicted gene, 47238 [Source:MGI Symb | 0 | 0 | 0 | 0 | 2.1812 | 0 |
| 48755 | ENSMUSG00000111061 | 0 | NA | NA | NA | NA | NA | Gm46102 | predicted gene, 46102 [Source:MGI Symb | 0 | 0 | 0 | 0 | 0 | 0 |
| 48756 | ENSMUSG00000111062 | 0 | NA | NA | NA | NA | NA | Gm47253 | predicted gene, 47253 [Source:MGI Symb | 0 | 0 | 0 | 0 | 0 | 0 |
| 48757 | ENSMUSG00000111063 | 1.144119223 | 0.512621048 | 2.117101743 | 0.242133402 | 0.808676797 | NA | Zfp660 | zinc finger protein 660 [Source:MGI Symb | 0.8031 | 2.0558 | 0 | 2.8919 | 0 | 1.1138 |
| 48758 | ENSMUSG00000111064 | 0 | NA | NA | NA | NA | NA | Gm30285 | predicted gene, 30285 [Source:MGI Symb | 0 | 0 | 0 | 0 | 0 | 0 |
| 48759 | ENSMUSG00000111065 | 0 | NA | NA | NA | NA | NA | Gm47676 | predicted gene, 47676 [Source:MGI Symb | 0 | 0 | 0 | 0 | 0 | 0 |
| 48760 | ENSMUSG00000111066 | 0 | NA | NA | NA | NA | NA | Gm30313 | predicted gene, 30313 [Source:MGI Symb | 0 | 0 | 0 | 0 | 0 | 0 |
| 48761 | ENSMUSG00000111067 | 7.160569235 | 0.982691733 | 0.897683518 | 1.09469731 | 0.273649267 | 0.728923893 | Gm48714 | predicted gene, 48714 [Source:MGI Symb | 2.4093 | 7.1954 | 5.0312 | 17.352 | 7.6343 | 3.3415 |
| 48762 | ENSMUSG00000111068 | 0 | NA | NA | NA | NA | NA | Gm35501 | predicted gene, 35501 [Source:MGI Symb | 0 | 0 | 0 | 0 | 0 | 0 |
| 48763 | ENSMUSG00000111069 | 0 | NA | NA | NA | NA | NA | Gm48646 | predicted gene, 48646 [Source:MGI Symb | 0 | 0 | 0 | 0 | 0 | 0 |
| 48764 | ENSMUSG00000111070 | 0 | NA | NA | NA | NA | NA | Gm47596 | predicted gene, 47596 [Source:MGI Symb | 0 | 0 | 0 | 0 | 0 | 0 |
| 48765 | ENSMUSG00000111071 | 0 | NA | NA | NA | NA | NA | Olfr1266-ps1 | olfactory receptor 1266, pseudogene 1 [S | 0 | 0 | 0 | 0 | 0 | 0 |
| 48766 | ENSMUSG00000111072 | 0 | NA | NA | NA | NA | NA | Gm9068 | predicted gene 9068 [Source:MGI Symbol | 0 | 0 | 0 | 0 | 0 | 0 |
| 48767 | ENSMUSG00000111073 | 0 | NA | NA | NA | NA | NA | Olfr977-ps1 | olfactory receptor 977, pseudogene 1 [So | 0 | 0 | 0 | 0 | 0 | 0 |
| 48768 | ENSMUSG00000111074 | 10.27601942 | 0.599343876 | 0.670446487 | 0.893947374 | 0.371350049 | 0.807214774 | Gm36435 | predicted gene, 36435 [Source:MGI Symb | 7.2279 | 9.2513 | 8.05 | 10.604 | 9.8156 | 16.708 |
| 48769 | ENSMUSG00000111075 | 0 | NA | NA | NA | NA | NA | Gm47964 | predicted gene, 47964 [Source:MGI Symb | 0 | 0 | 0 | 0 | 0 | 0 |
| 48770 | ENSMUSG00000111077 | 0 | NA | NA | NA | NA | NA | Gm48743 | predicted gene, 48743 [Source:MGI Symb | 0 | 0 | 0 | 0 | 0 | 0 |
| 48771 | ENSMUSG00000111078 | 0.453341394 | -0.545391191 | 3.551113277 | -0.153583158 | 0.877938417 | NA | Gm35028 | predicted gene, 35028 [Source:MGI Symb | 1.6062 | 0 | 0 | 0 | 0 | 1.1138 |
| 48772 | ENSMUSG00000111079 | 2.533949569 | -0.496436836 | 1.442758527 | -0.344088652 | 0.730779617 | NA | Gm47112 | predicted gene, 47112 [Source:MGI Symb | 4.8186 | 0 | 4.025 | 1.928 | 1.0906 | 3.3415 |

| | | | | | | | | | | | | | | | |
|---|---|---|---|---|---|---|---|---|---|---|---|---|---|---|---|
| 48773 | ENSMUSG00000111080 | 488.7377603 | -0.098368623 | 0.138994102 | -0.707717965 | 0.479120419 | 0.870169438 | Gm20300 | predicted gene, 20300 [Source:MGI Symb | 493.1 | 497.51 | 525.26 | 422.22 | 538.77 | 455.56 |
| 48774 | ENSMUSG00000111081 | 0 | NA | NA | NA | NA | NA | Gm47498 | predicted gene, 47498 [Source:MGI Symb | 0 | 0 | 0 | 0 | 0 | 0 |
| 48775 | ENSMUSG00000111082 | 11.63643271 | -0.493372892 | 0.653764262 | -0.754664824 | 0.450450117 | 0.855701405 | Gm30934 | predicted gene, 30934 [Source:MGI Symb | 11.243 | 15.419 | 14.087 | 4.8199 | 11.997 | 12.252 |
| 48776 | ENSMUSG00000111083 | 0 | NA | NA | NA | NA | NA | Gm48274 | predicted gene, 48274 [Source:MGI Symb | 0 | 0 | 0 | 0 | 0 | 0 |
| 48777 | ENSMUSG00000111084 | 0 | NA | NA | NA | NA | NA | Gm48292 | predicted gene, 48292 [Source:MGI Symb | 0 | 0 | 0 | 0 | 0 | 0 |
| 48778 | ENSMUSG00000111085 | 0 | NA | NA | NA | NA | NA | Gm7607 | predicted gene 7607 [Source:MGI Symbol | 0 | 0 | 0 | 0 | 0 | 0 |
| 48779 | ENSMUSG00000111086 | 0 | NA | NA | NA | NA | NA | Gm48671 | predicted gene, 48671 [Source:MGI Symb | 0 | 0 | 0 | 0 | 0 | 0 |
| 48780 | ENSMUSG00000111087 | 0 | NA | NA | NA | NA | NA | Gm47168 | predicted gene, 47168 [Source:MGI Symb | 0 | 0 | 0 | 0 | 0 | 0 |
| 48781 | ENSMUSG00000111088 | 7.952202105 | 0.133768403 | 0.775844019 | 0.172416619 | 0.863110007 | 0.976764193 | 9430081H08 | RIKEN cDNA 9430081H08 gene [Source:M | 10.44 | 8.2233 | 4.025 | 9.6398 | 9.8156 | 5.5692 |
| 48782 | ENSMUSG00000111089 | 0 | NA | NA | NA | NA | NA | Gm49427 | predicted gene, 49427 [Source:MGI Symb | 0 | 0 | 0 | 0 | 0 | 0 |
| 48783 | ENSMUSG00000111090 | 4.376143172 | 0.576598119 | 1.098464298 | 0.524912935 | 0.599643717 | NA | Gm30698 | predicted gene, 30698 [Source:MGI Symb | 2.4093 | 7.1954 | 1.0062 | 5.7839 | 7.6343 | 2.2277 |
| 48784 | ENSMUSG00000111091 | 0 | NA | NA | NA | NA | NA | Gm48651 | predicted gene, 48651 [Source:MGI Symb | 0 | 0 | 0 | 0 | 0 | 0 |
| 48785 | ENSMUSG00000111092 | 0 | NA | NA | NA | NA | NA | Gm17875 | predicted gene, 17875 [Source:MGI Symb | 0 | 0 | 0 | 0 | 0 | 0 |
| 48786 | ENSMUSG00000111093 | 0 | NA | NA | NA | NA | NA | Olfr1119-ps1 | olfactory receptor 1119, pseudogene 1 [S | 0 | 0 | 0 | 0 | 0 | 0 |
| 48787 | ENSMUSG00000111094 | 0 | NA | NA | NA | NA | NA | Gm34425 | predicted gene, 34425 [Source:MGI Symb | 0 | 0 | 0 | 0 | 0 | 0 |
| 48788 | ENSMUSG00000111095 | 0 | NA | NA | NA | NA | NA | Gm18165 | predicted gene, 18165 [Source:MGI Symb | 0 | 0 | 0 | 0 | 0 | 0 |
| 48789 | ENSMUSG00000111096 | 0 | NA | NA | NA | NA | NA | Gm47203 | predicted gene, 47203 [Source:MGI Symb | 0 | 0 | 0 | 0 | 0 | 0 |
| 48790 | ENSMUSG00000111097 | 0 | NA | NA | NA | NA | NA | Gm34069 | predicted gene, 34069 [Source:MGI Symb | 0 | 0 | 0 | 0 | 0 | 0 |
| 48791 | ENSMUSG00000111098 | 0 | NA | NA | NA | NA | NA | Gm48057 | predicted gene, 48057 [Source:MGI Symb | 0 | 0 | 0 | 0 | 0 | 0 |
| 48792 | ENSMUSG00000111100 | 0 | NA | NA | NA | NA | NA | Gm47535 | predicted gene, 47535 [Source:MGI Symb | 0 | 0 | 0 | 0 | 0 | 0 |
| 48793 | ENSMUSG00000111102 | 0 | NA | NA | NA | NA | NA | Gm48189 | predicted gene, 48189 [Source:MGI Symb | 0 | 0 | 0 | 0 | 0 | 0 |
| 48794 | ENSMUSG00000111103 | 0 | NA | NA | NA | NA | NA | 1700016J18I | RIKEN cDNA 1700016J18 gene [Source:M | 0 | 0 | 0 | 0 | 0 | 0 |
| 48795 | ENSMUSG00000111104 | 0 | NA | NA | NA | NA | NA | Gm8512 | predicted gene 8512 [Source:MGI Symbol | 0 | 0 | 0 | 0 | 0 | 0 |
| 48796 | ENSMUSG00000111105 | 0 | NA | NA | NA | NA | NA | Gm5023 | predicted gene 5023 [Source:MGI Symbol | 0 | 0 | 0 | 0 | 0 | 0 |
| 48797 | ENSMUSG00000111106 | 0 | NA | NA | NA | NA | NA | Gm3318 | predicted gene 3318 [Source:MGI Symbol | 0 | 0 | 0 | 0 | 0 | 0 |
| 48798 | ENSMUSG00000111107 | 0 | NA | NA | NA | NA | NA | Gm48737 | predicted gene, 48737 [Source:MGI Symb | 0 | 0 | 0 | 0 | 0 | 0 |
| 48799 | ENSMUSG00000111108 | 0.17131964 | -0.796629176 | 4.080472857 | -0.195229623 | 0.84521319 | NA | Gm17795 | predicted gene, 17795 [Source:MGI Symb | 0 | 1.0279 | 0 | 0 | 0 | 0 |
| 48800 | ENSMUSG00000111109 | 0 | NA | NA | NA | NA | NA | Gm30933 | predicted gene, 30933 [Source:MGI Symb | 0 | 0 | 0 | 0 | 0 | 0 |
| 48801 | ENSMUSG00000111110 | 0.799645015 | 0.918084596 | 2.625685367 | 0.34965522 | 0.726597464 | NA | Gm36855 | predicted gene, 36855 [Source:MGI Symb | 1.6062 | 0 | 0 | 0.964 | 0 | 2.2277 |
| 48802 | ENSMUSG00000111111 | 0 | NA | NA | NA | NA | NA | Gm47037 | predicted gene, 47037 [Source:MGI Symb | 0 | 0 | 0 | 0 | 0 | 0 |
| 48803 | ENSMUSG00000111112 | 0 | NA | NA | NA | NA | NA | Btf3-ps15 | basic transcription factor 3, pseudogene 1 | 0 | 0 | 0 | 0 | 0 | 0 |
| 48804 | ENSMUSG00000111114 | 0 | NA | NA | NA | NA | NA | Gm18703 | predicted gene, 18703 [Source:MGI Symb | 0 | 0 | 0 | 0 | 0 | 0 |
| 48805 | ENSMUSG00000111115 | 0 | NA | NA | NA | NA | NA | Gm47144 | predicted gene, 47144 [Source:MGI Symb | 0 | 0 | 0 | 0 | 0 | 0 |
| 48806 | ENSMUSG00000111116 | 0 | NA | NA | NA | NA | NA | Gm48065 | predicted gene, 48065 [Source:MGI Symb | 0 | 0 | 0 | 0 | 0 | 0 |
| 48807 | ENSMUSG00000111117 | 0 | NA | NA | NA | NA | NA | A730065G17 | RIKEN cDNA A730065G17 gene [Source:M | 0 | 0 | 0 | 0 | 0 | 0 |
| 48808 | ENSMUSG00000111118 | 10.37016345 | 0.81063969 | 0.665150651 | 1.218730956 | 0.222946324 | 0.668407528 | Gm6545 | predicted gene 6545 [Source:MGI Symbol | 6.4248 | 8.2233 | 8.05 | 16.388 | 11.997 | 11.138 |
| 48809 | ENSMUSG00000111119 | 0 | NA | NA | NA | NA | NA | Gm34005 | predicted gene, 34005 [Source:MGI Symb | 0 | 0 | 0 | 0 | 0 | 0 |
| 48810 | ENSMUSG00000111120 | 0 | NA | NA | NA | NA | NA | Gm49332 | predicted gene, 49332 [Source:MGI Symb | 0 | 0 | 0 | 0 | 0 | 0 |
| 48811 | ENSMUSG00000111121 | 0 | NA | NA | NA | NA | NA | Gm3523 | predicted gene 3523 [Source:MGI Symbol | 0 | 0 | 0 | 0 | 0 | 0 |
| 48812 | ENSMUSG00000111122 | 0 | NA | NA | NA | NA | NA | Gm5949 | predicted gene 5949 [Source:MGI Symbol | 0 | 0 | 0 | 0 | 0 | 0 |
| 48813 | ENSMUSG00000111124 | 0 | NA | NA | NA | NA | NA | Gm48373 | predicted gene, 48373 [Source:MGI Symb | 0 | 0 | 0 | 0 | 0 | 0 |
| 48814 | ENSMUSG00000111126 | 0 | NA | NA | NA | NA | NA | Gm39329 | predicted gene, 39329 [Source:MGI Symb | 0 | 0 | 0 | 0 | 0 | 0 |
| 48815 | ENSMUSG00000111127 | 0 | NA | NA | NA | NA | NA | Gm5010 | predicted gene 5010 [Source:MGI Symbol | 0 | 0 | 0 | 0 | 0 | 0 |
| 48816 | ENSMUSG00000111128 | 85.15594017 | 0.244147707 | 0.261328284 | 0.934256726 | 0.350171481 | 0.791589971 | Gm49338 | predicted gene, 49338 [Source:MGI Symb | 79.507 | 75.038 | 79.493 | 107.97 | 69.8 | 99.132 |
| 48817 | ENSMUSG00000111129 | 0.181769715 | 1.126924175 | 4.080472857 | 0.276174898 | 0.782413736 | NA | Gm47542 | predicted gene, 47542 [Source:MGI Symb | 0 | 0 | 0 | 0 | 1.0906 | 0 |
| 48818 | ENSMUSG00000111131 | 0 | NA | NA | NA | NA | NA | 9930024M1! | RIKEN cDNA 9930024M15 gene [Source:N | 0 | 0 | 0 | 0 | 0 | 0 |
| 48819 | ENSMUSG00000111132 | 0 | NA | NA | NA | NA | NA | Gm48142 | predicted gene, 48142 [Source:MGI Symb | 0 | 0 | 0 | 0 | 0 | 0 |
| 48820 | ENSMUSG00000111133 | 0 | NA | NA | NA | NA | NA | Gm5831 | predicted gene 5831 [Source:MGI Symbol | 0 | 0 | 0 | 0 | 0 | 0 |
| 48821 | ENSMUSG00000111135 | 0 | NA | NA | NA | NA | NA | Gm47670 | predicted gene, 47670 [Source:MGI Symb | 0 | 0 | 0 | 0 | 0 | 0 |
| 48822 | ENSMUSG00000111137 | 0 | NA | NA | NA | NA | NA | Gm2553 | predicted gene 2553 [Source:MGI Symbol | 0 | 0 | 0 | 0 | 0 | 0 |
| 48823 | ENSMUSG00000111139 | 0 | NA | NA | NA | NA | NA | Gm47497 | predicted gene, 47497 [Source:MGI Symb | 0 | 0 | 0 | 0 | 0 | 0 |
| 48824 | ENSMUSG00000111141 | 0 | NA | NA | NA | NA | NA | Gm47115 | predicted gene, 47115 [Source:MGI Symb | 0 | 0 | 0 | 0 | 0 | 0 |
| 48825 | ENSMUSG00000111142 | 0 | NA | NA | NA | NA | NA | Gm48131 | predicted gene, 48131 [Source:MGI Symb | 0 | 0 | 0 | 0 | 0 | 0 |
| 48826 | ENSMUSG00000111143 | 0 | NA | NA | NA | NA | NA | Gm47134 | predicted gene, 47134 [Source:MGI Symb | 0 | 0 | 0 | 0 | 0 | 0 |
| 48827 | ENSMUSG00000111144 | 0 | NA | NA | NA | NA | NA | Gm48193 | predicted gene, 48193 [Source:MGI Symb | 0 | 0 | 0 | 0 | 0 | 0 |
| 48828 | ENSMUSG00000111146 | 0 | NA | NA | NA | NA | NA | Gm35454 | predicted gene, 35454 [Source:MGI Symb | 0 | 0 | 0 | 0 | 0 | 0 |

| | | | | | | | | | | | | | | | |
|---|---|---|---|---|---|---|---|---|---|---|---|---|---|---|---|
| 48829 | ENSMUSG00000111147 | 2.644029968 | -1.969146122 | 1.747797197 | -1.126644513 | 0.259892811 | NA | Gm33699 | predicted gene, 33699 [Source:MGI Symb | 6.4248 | 6.1675 | 0 | 0 | 3.2719 | 0 |
| 48830 | ENSMUSG00000111148 | 0 | NA | NA | NA | NA | NA | Gm2774 | predicted gene 2774 [Source:MGI Symbol | 0 | 0 | 0 | 0 | 0 | 0 |
| 48831 | ENSMUSG00000111149 | 0 | NA | NA | NA | NA | NA | Gm26564 | predicted gene, 26564 [Source:MGI Symb | 0 | 0 | 0 | 0 | 0 | 0 |
| 48832 | ENSMUSG00000111151 | 0 | NA | NA | NA | NA | NA | Gm5120 | predicted gene 5120 [Source:MGI Symbol | 0 | 0 | 0 | 0 | 0 | 0 |
| 48833 | ENSMUSG00000111152 | 0 | NA | NA | NA | NA | NA | Gm48490 | predicted gene, 48490 [Source:MGI Symb | 0 | 0 | 0 | 0 | 0 | 0 |
| 48834 | ENSMUSG00000111153 | 0 | NA | NA | NA | NA | NA | Gm18820 | predicted gene, 18820 [Source:MGI Symb | 0 | 0 | 0 | 0 | 0 | 0 |
| 48835 | ENSMUSG00000111154 | 0 | NA | NA | NA | NA | NA | Gm47229 | predicted gene, 47229 [Source:MGI Symb | 0 | 0 | 0 | 0 | 0 | 0 |
| 48836 | ENSMUSG00000111155 | 0 | NA | NA | NA | NA | NA | Gm47197 | predicted gene, 47197 [Source:MGI Symb | 0 | 0 | 0 | 0 | 0 | 0 |
| 48837 | ENSMUSG00000111156 | 0 | NA | NA | NA | NA | NA | Gm48814 | predicted gene, 48814 [Source:MGI Symb | 0 | 0 | 0 | 0 | 0 | 0 |
| 48838 | ENSMUSG00000111157 | 0 | NA | NA | NA | NA | NA | Gm47887 | predicted gene, 47887 [Source:MGI Symb | 0 | 0 | 0 | 0 | 0 | 0 |
| 48839 | ENSMUSG00000111159 | 0 | NA | NA | NA | NA | NA | Olfr1337 | olfactory receptor 1337 [Source:MGI Sym | 0 | 0 | 0 | 0 | 0 | 0 |
| 48840 | ENSMUSG00000111160 | 0.17131964 | -0.796629176 | 4.080472857 | -0.195229623 | 0.84521319 | NA | Gm48855 | predicted gene, 48855 [Source:MGI Symb | 0 | 1.0279 | 0 | 0 | 0 | 0 |
| 48841 | ENSMUSG00000111161 | 0 | NA | NA | NA | NA | NA | Gm48849 | predicted gene, 48849 [Source:MGI Symb | 0 | 0 | 0 | 0 | 0 | 0 |
| 48842 | ENSMUSG00000111162 | 0 | NA | NA | NA | NA | NA | Gm7293 | predicted gene 7293 [Source:MGI Symbol | 0 | 0 | 0 | 0 | 0 | 0 |
| 48843 | ENSMUSG00000111163 | 1.067268653 | 0.134458805 | 2.381868772 | 0.056450971 | 0.954982553 | NA | Gm46136 | predicted gene, 46136 [Source:MGI Symb | 0 | 2.0558 | 1.0062 | 0 | 0 | 3.3415 |
| 48844 | ENSMUSG00000111164 | 0 | NA | NA | NA | NA | NA | Gm47803 | predicted gene, 47803 [Source:MGI Symb | 0 | 0 | 0 | 0 | 0 | 0 |
| 48845 | ENSMUSG00000111165 | 0.160662974 | 1.126924175 | 4.080472857 | 0.276174898 | 0.782413736 | NA | 4930553I21F | RIKEN cDNA 4930553I21 gene [Source:MC | 0 | 0 | 0 | 0.964 | 0 | 0 |
| 48846 | ENSMUSG00000111166 | 2.092395192 | -0.476011494 | 1.937122471 | -0.245731233 | 0.805890293 | NA | Tpi-rs4 | triosephosphate isomerase related seque | 7.2279 | 0 | 0 | 0.964 | 4.3625 | 0 |
| 48847 | ENSMUSG00000111167 | 0.17131964 | -0.796629176 | 4.080472857 | -0.195229623 | 0.84521319 | NA | Gm39321 | predicted gene, 39321 [Source:MGI Symb | 0 | 1.0279 | 0 | 0 | 0 | 0 |
| 48848 | ENSMUSG00000111168 | 0 | NA | NA | NA | NA | NA | Gm48468 | predicted gene, 48468 [Source:MGI Symb | 0 | 0 | 0 | 0 | 0 | 0 |
| 48849 | ENSMUSG00000111169 | 0 | NA | NA | NA | NA | NA | Gm34004 | predicted gene, 34004 [Source:MGI Symb | 0 | 0 | 0 | 0 | 0 | 0 |
| 48850 | ENSMUSG00000111171 | 0.678054141 | -2.792799308 | 2.990673428 | -0.933836266 | 0.350388359 | NA | Gm47815 | predicted gene, 47815 [Source:MGI Symb | 0 | 2.0558 | 2.0125 | 0 | 0 | 0 |
| 48851 | ENSMUSG00000111172 | 0 | NA | NA | NA | NA | NA | Gm39302 | predicted gene, 39302 [Source:MGI Symb | 0 | 0 | 0 | 0 | 0 | 0 |
| 48852 | ENSMUSG00000111173 | 0 | NA | NA | NA | NA | NA | Gm48297 | predicted gene, 48297 [Source:MGI Symb | 0 | 0 | 0 | 0 | 0 | 0 |
| 48853 | ENSMUSG00000111174 | 0 | NA | NA | NA | NA | NA | Olfr1240 | olfactory receptor 1240 [Source:MGI Sym | 0 | 0 | 0 | 0 | 0 | 0 |
| 48854 | ENSMUSG00000111175 | 0 | NA | NA | NA | NA | NA | Gm19640 | predicted gene, 19640 [Source:MGI Symb | 0 | 0 | 0 | 0 | 0 | 0 |
| 48855 | ENSMUSG00000111176 | 0 | NA | NA | NA | NA | NA | Gm47233 | predicted gene, 47233 [Source:MGI Symb | 0 | 0 | 0 | 0 | 0 | 0 |
| 48856 | ENSMUSG00000111177 | 0.342432689 | 1.990653086 | 3.900281263 | 0.51038706 | 0.609780321 | NA | Rps2-ps3 | ribosomal protein S2, pseudogene 3 [Sour | 0 | 0 | 0 | 0.964 | 1.0906 | 0 |
| 48857 | ENSMUSG00000111179 | 0 | NA | NA | NA | NA | NA | Olfr996 | olfactory receptor 996 [Source:MGI Symb | 0 | 0 | 0 | 0 | 0 | 0 |
| 48858 | ENSMUSG00000111180 | 0.17131964 | -0.796629176 | 4.080472857 | -0.195229623 | 0.84521319 | NA | Gm39348 | predicted gene, 39348 [Source:MGI Symb | 0 | 1.0279 | 0 | 0 | 0 | 0 |
| 48859 | ENSMUSG00000111181 | 0 | NA | NA | NA | NA | NA | Gm48124 | predicted gene, 48124 [Source:MGI Symb | 0 | 0 | 0 | 0 | 0 | 0 |
| 48860 | ENSMUSG00000111182 | 0 | NA | NA | NA | NA | NA | Gm35430 | predicted gene, 35430 [Source:MGI Symb | 0 | 0 | 0 | 0 | 0 | 0 |
| 48861 | ENSMUSG00000111183 | 0 | NA | NA | NA | NA | NA | Gm48486 | predicted gene, 48486 [Source:MGI Symb | 0 | 0 | 0 | 0 | 0 | 0 |
| 48862 | ENSMUSG00000111184 | 0 | NA | NA | NA | NA | NA | Gm49661 | predicted gene, 49661 [Source:MGI Symb | 0 | 0 | 0 | 0 | 0 | 0 |
| 48863 | ENSMUSG00000111186 | 0 | NA | NA | NA | NA | NA | Gm48135 | predicted gene, 48135 [Source:MGI Symb | 0 | 0 | 0 | 0 | 0 | 0 |
| 48864 | ENSMUSG00000111188 | 0 | NA | NA | NA | NA | NA | Gm19709 | predicted gene, 19709 [Source:MGI Symb | 0 | 0 | 0 | 0 | 0 | 0 |
| 48865 | ENSMUSG00000111189 | 1.602289108 | -0.475427307 | 1.86206979 | -0.255321959 | 0.798474441 | NA | Gm48293 | predicted gene, 48293 [Source:MGI Symb | 1.6062 | 0 | 4.025 | 2.8919 | 1.0906 | 0 |
| 48866 | ENSMUSG00000111190 | 0 | NA | NA | NA | NA | NA | Olfr947-ps1 | olfactory receptor 947, pseudogene 1 [So | 0 | 0 | 0 | 0 | 0 | 0 |
| 48867 | ENSMUSG00000111191 | 0 | NA | NA | NA | NA | NA | Gm6015 | predicted gene 6015 [Source:MGI Symbol | 0 | 0 | 0 | 0 | 0 | 0 |
| 48868 | ENSMUSG00000111192 | 0 | NA | NA | NA | NA | NA | Gm47128 | predicted gene, 47128 [Source:MGI Symb | 0 | 0 | 0 | 0 | 0 | 0 |
| 48869 | ENSMUSG00000111193 | 0 | NA | NA | NA | NA | NA | Gm35288 | predicted gene, 35288 [Source:MGI Symb | 0 | 0 | 0 | 0 | 0 | 0 |
| 48870 | ENSMUSG00000111194 | 0 | NA | NA | NA | NA | NA | Gm47180 | predicted gene, 47180 [Source:MGI Symb | 0 | 0 | 0 | 0 | 0 | 0 |
| 48871 | ENSMUSG00000111195 | 0 | NA | NA | NA | NA | NA | Gm48215 | predicted gene, 48215 [Source:MGI Symb | 0 | 0 | 0 | 0 | 0 | 0 |
| 48872 | ENSMUSG00000111196 | 0 | NA | NA | NA | NA | NA | Gm47581 | predicted gene, 47581 [Source:MGI Symb | 0 | 0 | 0 | 0 | 0 | 0 |
| 48873 | ENSMUSG00000111197 | 3.021354611 | -1.25381027 | 1.36395844 | -0.919243749 | 0.357968093 | NA | Gm10608 | predicted gene 10608 [Source:MGI Symbo | 5.6217 | 2.0558 | 5.0312 | 0.964 | 0 | 4.4554 |
| 48874 | ENSMUSG00000111198 | 0 | NA | NA | NA | NA | NA | Gm34263 | predicted gene, 34263 [Source:MGI Symb | 0 | 0 | 0 | 0 | 0 | 0 |
| 48875 | ENSMUSG00000111199 | 0 | NA | NA | NA | NA | NA | Gm47875 | predicted gene, 47875 [Source:MGI Symb | 0 | 0 | 0 | 0 | 0 | 0 |
| 48876 | ENSMUSG00000111200 | 0 | NA | NA | NA | NA | NA | Gm33728 | predicted gene, 33728 [Source:MGI Symb | 0 | 0 | 0 | 0 | 0 | 0 |
| 48877 | ENSMUSG00000111201 | 0 | NA | NA | NA | NA | NA | Gm47178 | predicted gene, 47178 [Source:MGI Symb | 0 | 0 | 0 | 0 | 0 | 0 |
| 48878 | ENSMUSG00000111202 | 0 | NA | NA | NA | NA | NA | Gm48275 | predicted gene, 48275 [Source:MGI Symb | 0 | 0 | 0 | 0 | 0 | 0 |
| 48879 | ENSMUSG00000111203 | 0 | NA | NA | NA | NA | NA | Gm48719 | predicted gene, 48719 [Source:MGI Symb | 0 | 0 | 0 | 0 | 0 | 0 |
| 48880 | ENSMUSG00000111204 | 0.167707431 | -0.796629176 | 4.080472857 | -0.195229623 | 0.84521319 | NA | Gm47320 | predicted gene, 47320 [Source:MGI Symb | 0 | 0 | 1.0062 | 0 | 0 | 0 |
| 48881 | ENSMUSG00000111205 | 0 | NA | NA | NA | NA | NA | Gm39307 | predicted gene, 39307 [Source:MGI Symb | 0 | 0 | 0 | 0 | 0 | 0 |
| 48882 | ENSMUSG00000111207 | 0 | NA | NA | NA | NA | NA | Gm8125 | predicted gene 8125 [Source:MGI Symbol | 0 | 0 | 0 | 0 | 0 | 0 |
| 48883 | ENSMUSG00000111208 | 0 | NA | NA | NA | NA | NA | Gm48197 | predicted gene, 48197 [Source:MGI Symb | 0 | 0 | 0 | 0 | 0 | 0 |
| 48884 | ENSMUSG00000111209 | 0 | NA | NA | NA | NA | NA | Gm7529 | predicted gene 7529 [Source:MGI Symbol | 0 | 0 | 0 | 0 | 0 | 0 |

| | | | | | | | | | | | | | | | |
|---|---|---|---|---|---|---|---|---|---|---|---|---|---|---|---|
| 48885 | ENSMUSG00000111210 | 0 | NA | NA | NA | NA | NA | Gm18378 | predicted gene, 18378 [Source:MGI Symb | 0 | 0 | 0 | 0 | 0 | 0 |
| 48886 | ENSMUSG00000111211 | 0.321325949 | 1.911575114 | 4.048267893 | 0.472195804 | 0.636787034 | NA | Gm48233 | predicted gene, 48233 [Source:MGI Symb | 0 | 0 | 0 | 1.928 | 0 | 0 |
| 48887 | ENSMUSG00000111212 | 0 | NA | NA | NA | NA | NA | 5330432J10I | RIKEN cDNA 5330432J10 gene [Source:M( | 0 | 0 | 0 | 0 | 0 | 0 |
| 48888 | ENSMUSG00000111213 | 0.167707431 | -0.796629176 | 4.080472857 | -0.195229623 | 0.84521319 | NA | Gm47867 | predicted gene, 47867 [Source:MGI Symb | 0 | 0 | 1.0062 | 0 | 0 | 0 |
| 48889 | ENSMUSG00000111215 | 0 | NA | NA | NA | NA | NA | Gm48654 | predicted gene, 48654 [Source:MGI Symb | 0 | 0 | 0 | 0 | 0 | 0 |
| 48890 | ENSMUSG00000111216 | 0 | NA | NA | NA | NA | NA | Gm47044 | predicted gene, 47044 [Source:MGI Symb | 0 | 0 | 0 | 0 | 0 | 0 |
| 48891 | ENSMUSG00000111217 | 0 | NA | NA | NA | NA | NA | Gm48711 | predicted gene, 48711 [Source:MGI Symb | 0 | 0 | 0 | 0 | 0 | 0 |
| 48892 | ENSMUSG00000111218 | 2.884367244 | 0.690900108 | 1.579585989 | 0.437393161 | 0.661826243 | NA | Gm46999 | predicted gene, 46999 [Source:MGI Symb | 1.6062 | 3.0838 | 2.0125 | 10.604 | 0 | 0 |
| 48893 | ENSMUSG00000111219 | 0 | NA | NA | NA | NA | NA | Gm38642 | predicted gene, 38642 [Source:MGI Symb | 0 | 0 | 0 | 0 | 0 | 0 |
| 48894 | ENSMUSG00000111221 | 0 | NA | NA | NA | NA | NA | Gm47133 | predicted gene, 47133 [Source:MGI Symb | 0 | 0 | 0 | 0 | 0 | 0 |
| 48895 | ENSMUSG00000111222 | 0 | NA | NA | NA | NA | NA | Gm34158 | predicted gene, 34158 [Source:MGI Symb | 0 | 0 | 0 | 0 | 0 | 0 |
| 48896 | ENSMUSG00000111223 | 0 | NA | NA | NA | NA | NA | Gm47089 | predicted gene, 47089 [Source:MGI Symb | 0 | 0 | 0 | 0 | 0 | 0 |
| 48897 | ENSMUSG00000111224 | 0 | NA | NA | NA | NA | NA | Gm18669 | predicted gene, 18669 [Source:MGI Symb | 0 | 0 | 0 | 0 | 0 | 0 |
| 48898 | ENSMUSG00000111225 | 3.416197645 | -0.537867167 | 1.22767006 | -0.438120293 | 0.661299085 | NA | Gm47790 | predicted gene, 47790 [Source:MGI Symb | 4.0155 | 4.1117 | 4.025 | 2.8919 | 5.4531 | 0 |
| 48899 | ENSMUSG00000111227 | 0 | NA | NA | NA | NA | NA | Gm20149 | predicted gene, 20149 [Source:MGI Symb | 0 | 0 | 0 | 0 | 0 | 0 |
| 48900 | ENSMUSG00000111228 | 0 | NA | NA | NA | NA | NA | Gm48789 | predicted gene, 48789 [Source:MGI Symb | 0 | 0 | 0 | 0 | 0 | 0 |
| 48901 | ENSMUSG00000111229 | 0 | NA | NA | NA | NA | NA | Gm39323 | predicted gene, 39323 [Source:MGI Symb | 0 | 0 | 0 | 0 | 0 | 0 |
| 48902 | ENSMUSG00000111231 | 0 | NA | NA | NA | NA | NA | Gm47054 | predicted gene, 47054 [Source:MGI Symb | 0 | 0 | 0 | 0 | 0 | 0 |
| 48903 | ENSMUSG00000111232 | 0 | NA | NA | NA | NA | NA | A730043L09 | RIKEN cDNA A730043L09 gene [Source:M( | 0 | 0 | 0 | 0 | 0 | 0 |
| 48904 | ENSMUSG00000111233 | 0 | NA | NA | NA | NA | NA | Gm47069 | predicted gene, 47069 [Source:MGI Symb | 0 | 0 | 0 | 0 | 0 | 0 |
| 48905 | ENSMUSG00000111234 | 0 | NA | NA | NA | NA | NA | 4930591E09 | RIKEN cDNA 4930591E09 gene [Source:M( | 0 | 0 | 0 | 0 | 0 | 0 |
| 48906 | ENSMUSG00000111235 | 0 | NA | NA | NA | NA | NA | Gm4657 | predicted gene 4657 [Source:MGI Symbol | 0 | 0 | 0 | 0 | 0 | 0 |
| 48907 | ENSMUSG00000111236 | 0 | NA | NA | NA | NA | NA | Gm31614 | predicted gene, 31614 [Source:MGI Symb | 0 | 0 | 0 | 0 | 0 | 0 |
| 48908 | ENSMUSG00000111238 | 0 | NA | NA | NA | NA | NA | Gm19113 | predicted gene, 19113 [Source:MGI Symb | 0 | 0 | 0 | 0 | 0 | 0 |
| 48909 | ENSMUSG00000111239 | 0 | NA | NA | NA | NA | NA | Olfr1248 | olfactory receptor 1248 [Source:MGI Syml | 0 | 0 | 0 | 0 | 0 | 0 |
| 48910 | ENSMUSG00000111240 | 0 | NA | NA | NA | NA | NA | Gm39312 | predicted gene, 39312 [Source:MGI Symb | 0 | 0 | 0 | 0 | 0 | 0 |
| 48911 | ENSMUSG00000111241 | 0.160662974 | 1.126924175 | 4.080472857 | 0.276174898 | 0.782413736 | NA | Gm19121 | predicted gene, 19121 [Source:MGI Symb | 0 | 0 | 0 | 0.964 | 0 | 0 |
| 48912 | ENSMUSG00000111242 | 0 | NA | NA | NA | NA | NA | Gm8155 | predicted gene 8155 [Source:MGI Symbol | 0 | 0 | 0 | 0 | 0 | 0 |
| 48913 | ENSMUSG00000111243 | 0 | NA | NA | NA | NA | NA | Gm4668 | predicted gene 4668 [Source:MGI Symbol | 0 | 0 | 0 | 0 | 0 | 0 |
| 48914 | ENSMUSG00000111244 | 0 | NA | NA | NA | NA | NA | Gm47130 | predicted gene, 47130 [Source:MGI Symb | 0 | 0 | 0 | 0 | 0 | 0 |
| 48915 | ENSMUSG00000111245 | 0 | NA | NA | NA | NA | NA | Gm39388 | predicted gene, 39388 [Source:MGI Symb | 0 | 0 | 0 | 0 | 0 | 0 |
| 48916 | ENSMUSG00000111246 | 6.053442409 | 0.272086377 | 0.863307992 | 0.315167216 | 0.752634696 | 0.952989051 | 4930517E14 | Riken cDNA 4930517E14 gene [Source:M( | 7.2279 | 5.1396 | 4.025 | 6.7478 | 8.7249 | 4.4554 |
| 48917 | ENSMUSG00000111248 | 0 | NA | NA | NA | NA | NA | Olfr1114-ps1 | olfactory receptor 1114, pseudogene 1 [S( | 0 | 0 | 0 | 0 | 0 | 0 |
| 48918 | ENSMUSG00000111249 | 0 | NA | NA | NA | NA | NA | Gm47717 | predicted gene, 47717 [Source:MGI Symb | 0 | 0 | 0 | 0 | 0 | 0 |
| 48919 | ENSMUSG00000111250 | 2.943709294 | -2.856886042 | 1.761269944 | -1.622060293 | 0.104790439 | NA | Gm47818 | predicted gene, 47818 [Source:MGI Symb | 6.4248 | 0 | 9.0562 | 0 | 2.1812 | 0 |
| 48920 | ENSMUSG00000111251 | 0 | NA | NA | NA | NA | NA | Gm18671 | predicted gene, 18671 [Source:MGI Symb | 0 | 0 | 0 | 0 | 0 | 0 |
| 48921 | ENSMUSG00000111252 | 0 | NA | NA | NA | NA | NA | Gm47317 | predicted gene, 47317 [Source:MGI Symb | 0 | 0 | 0 | 0 | 0 | 0 |
| 48922 | ENSMUSG00000111253 | 18.36391958 | 0.523421054 | 0.660728681 | 0.792187578 | 0.42825131 | 0.842753542 | Gm34654 | predicted gene, 34654 [Source:MGI Symb | 22.487 | 15.419 | 7.0437 | 16.388 | 7.6343 | 41.212 |
| 48923 | ENSMUSG00000111254 | 0 | NA | NA | NA | NA | NA | Gm48717 | predicted gene, 48717 [Source:MGI Symb | 0 | 0 | 0 | 0 | 0 | 0 |
| 48924 | ENSMUSG00000111256 | 0 | NA | NA | NA | NA | NA | Gm48094 | predicted gene, 48094 [Source:MGI Symb | 0 | 0 | 0 | 0 | 0 | 0 |
| 48925 | ENSMUSG00000111257 | 0 | NA | NA | NA | NA | NA | Gm21089 | predicted gene, 21089 [Source:MGI Symb | 0 | 0 | 0 | 0 | 0 | 0 |
| 48926 | ENSMUSG00000111258 | 0 | NA | NA | NA | NA | NA | Gm34397 | predicted gene, 34397 [Source:MGI Symb | 0 | 0 | 0 | 0 | 0 | 0 |
| 48927 | ENSMUSG00000111259 | 0 | NA | NA | NA | NA | NA | Olfr1328 | olfactory receptor 1328 [Source:MGI Syml | 0 | 0 | 0 | 0 | 0 | 0 |
| 48928 | ENSMUSG00000111260 | 0 | NA | NA | NA | NA | NA | Gm47714 | predicted gene, 47714 [Source:MGI Symb | 0 | 0 | 0 | 0 | 0 | 0 |
| 48929 | ENSMUSG00000111262 | 0 | NA | NA | NA | NA | NA | Gm47354 | predicted gene, 47354 [Source:MGI Symb | 0 | 0 | 0 | 0 | 0 | 0 |
| 48930 | ENSMUSG00000111263 | 0 | NA | NA | NA | NA | NA | Gm7680 | predicted gene 7680 [Source:MGI Symbol | 0 | 0 | 0 | 0 | 0 | 0 |
| 48931 | ENSMUSG00000111264 | 5.231848388 | 1.038338912 | 1.066534992 | 0.973562911 | 0.330273609 | 0.774747658 | E030022I16I | RIKEN cDNA E030022I16 gene [Source:M( | 3.2124 | 5.1396 | 2.0125 | 14.46 | 5.4531 | 1.1138 |
| 48932 | ENSMUSG00000111265 | 0 | NA | NA | NA | NA | NA | Gm47784 | predicted gene, 47784 [Source:MGI Symb | 0 | 0 | 0 | 0 | 0 | 0 |
| 48933 | ENSMUSG00000111266 | 0 | NA | NA | NA | NA | NA | Gm30570 | predicted gene, 30570 [Source:MGI Symb | 0 | 0 | 0 | 0 | 0 | 0 |
| 48934 | ENSMUSG00000111267 | 0 | NA | NA | NA | NA | NA | Gm18292 | predicted gene, 18292 [Source:MGI Symb | 0 | 0 | 0 | 0 | 0 | 0 |
| 48935 | ENSMUSG00000111268 | 0 | NA | NA | NA | NA | NA | Gm47126 | predicted gene, 47126 [Source:MGI Symb | 0 | 0 | 0 | 0 | 0 | 0 |
| 48936 | ENSMUSG00000111269 | 0 | NA | NA | NA | NA | NA | Gm47933 | predicted gene, 47933 [Source:MGI Symb | 0 | 0 | 0 | 0 | 0 | 0 |
| 48937 | ENSMUSG00000111270 | 0 | NA | NA | NA | NA | NA | Gm47066 | predicted gene, 47066 [Source:MGI Symb | 0 | 0 | 0 | 0 | 0 | 0 |
| 48938 | ENSMUSG00000111271 | 0 | NA | NA | NA | NA | NA | Gm48127 | predicted gene, 48127 [Source:MGI Symb | 0 | 0 | 0 | 0 | 0 | 0 |
| 48939 | ENSMUSG00000111272 | 0 | NA | NA | NA | NA | NA | Gm7787 | predicted gene 7787 [Source:MGI Symbol | 0 | 0 | 0 | 0 | 0 | 0 |
| 48940 | ENSMUSG00000111273 | 0 | NA | NA | NA | NA | NA | Olfr789 | olfactory receptor 789 [Source:MGI Symb | 0 | 0 | 0 | 0 | 0 | 0 |

| | | | | | | | | | | | | | | | |
|---|---|---|---|---|---|---|---|---|---|---|---|---|---|---|---|
| 48941 | ENSMUSG00000111274 | 0 | NA | NA | NA | NA | NA | Gm47409 | predicted gene, 47409 [Source:MGI Symb | 0 | 0 | 0 | 0 | 0 | 0 |
| 48942 | ENSMUSG00000111275 | 0 | NA | NA | NA | NA | NA | Gm29824 | predicted gene, 29824 [Source:MGI Symb | 0 | 0 | 0 | 0 | 0 | 0 |
| 48943 | ENSMUSG00000111276 | 0 | NA | NA | NA | NA | NA | Gm47778 | predicted gene, 47778 [Source:MGI Symb | 0 | 0 | 0 | 0 | 0 | 0 |
| 48944 | ENSMUSG00000111278 | 0 | NA | NA | NA | NA | NA | Gm32743 | predicted gene, 32743 [Source:MGI Symb | 0 | 0 | 0 | 0 | 0 | 0 |
| 48945 | ENSMUSG00000111280 | 0 | NA | NA | NA | NA | NA | Gm47239 | predicted gene, 47239 [Source:MGI Symb | 0 | 0 | 0 | 0 | 0 | 0 |
| 48946 | ENSMUSG00000111281 | 0 | NA | NA | NA | NA | NA | Gm48830 | predicted gene, 48830 [Source:MGI Symb | 0 | 0 | 0 | 0 | 0 | 0 |
| 48947 | ENSMUSG00000111282 | 44.9474281 | -1.841089952 | 0.54879878 | -3.354763203 | 0.000794329 | 0.017594459 | Gm47528 | predicted gene, 47528 [Source:MGI Symb | 106.01 | 70.926 | 34.212 | 36.631 | 17.45 | 4.4554 |
| 48948 | ENSMUSG00000111283 | 0 | NA | NA | NA | NA | NA | E230034D01 | RIKEN cDNA E230034D01 gene [Source:M | 0 | 0 | 0 | 0 | 0 | 0 |
| 48949 | ENSMUSG00000111284 | 0.185640647 | 1.126924175 | 4.080472857 | 0.276174898 | 0.782413736 | NA | Gm47334 | predicted gene, 47334 [Source:MGI Symb | 0 | 0 | 0 | 0 | 0 | 1.1138 |
| 48950 | ENSMUSG00000111285 | 0 | NA | NA | NA | NA | NA | Gm47051 | predicted gene, 47051 [Source:MGI Symb | 0 | 0 | 0 | 0 | 0 | 0 |
| 48951 | ENSMUSG00000111286 | 0 | NA | NA | NA | NA | NA | Gm18745 | predicted gene, 18745 [Source:MGI Symb | 0 | 0 | 0 | 0 | 0 | 0 |
| 48952 | ENSMUSG00000111290 | 0.59968336 | -1.233555632 | 2.892386901 | -0.426483618 | 0.669755482 | NA | Gm5122 | predicted gene 5122 [Source:MGI Symbol | 1.6062 | 1.0279 | 0 | 0.964 | 0 | 0 |
| 48953 | ENSMUSG00000111291 | 28.70732247 | 0.00712249 | 0.431715664 | 0.016498104 | 0.986837015 | 0.99859176 | Gm48604 | predicted gene, 48604 [Source:MGI Symb | 24.896 | 27.754 | 33.206 | 19.28 | 39.262 | 27.846 |
| 48954 | ENSMUSG00000111292 | 0.339027071 | -1.79174149 | 3.919585585 | -0.457125237 | 0.647581032 | NA | Gm48627 | predicted gene, 48627 [Source:MGI Symb | 0 | 1.0279 | 1.0062 | 0 | 0 | 0 |
| 48955 | ENSMUSG00000111293 | 0.328370405 | 0.16514316 | 4.003149536 | 0.041253308 | 0.967093956 | NA | Gm34006 | predicted gene, 34006 [Source:MGI Symb | 0 | 0 | 1.0062 | 0.964 | 0 | 0 |
| 48956 | ENSMUSG00000111294 | 0 | NA | NA | NA | NA | NA | Gm47215 | predicted gene, 47215 [Source:MGI Symb | 0 | 0 | 0 | 0 | 0 | 0 |
| 48957 | ENSMUSG00000111295 | 0 | NA | NA | NA | NA | NA | Olfr1092-ps1 | olfactory receptor 1092, pseudogene 1 [S | 0 | 0 | 0 | 0 | 0 | 0 |
| 48958 | ENSMUSG00000111296 | 0 | NA | NA | NA | NA | NA | 2010001M07 | RIKEN cDNA 2010001M07 gene [Source:M | 0 | 0 | 0 | 0 | 0 | 0 |
| 48959 | ENSMUSG00000111297 | 0 | NA | NA | NA | NA | NA | Gm6082 | predicted gene 6082 [Source:MGI Symbol | 0 | 0 | 0 | 0 | 0 | 0 |
| 48960 | ENSMUSG00000111298 | 0 | NA | NA | NA | NA | NA | Gm3473 | predicted gene 3473 [Source:MGI Symbol | 0 | 0 | 0 | 0 | 0 | 0 |
| 48961 | ENSMUSG00000111299 | 0 | NA | NA | NA | NA | NA | 4930540M03 | RIKEN cDNA 4930540M03 gene [Source:M | 0 | 0 | 0 | 0 | 0 | 0 |
| 48962 | ENSMUSG00000111300 | 0 | NA | NA | NA | NA | NA | Rpl26-ps1 | ribosomal protein L26, pseudogene 1 [Sou | 0 | 0 | 0 | 0 | 0 | 0 |
| 48963 | ENSMUSG00000111301 | 0.353089354 | 0.165142706 | 3.896102244 | 0.042386646 | 0.966190474 | NA | Gm30671 | predicted gene, 30671 [Source:MGI Symb | 0 | 1.0279 | 0 | 0 | 1.0906 | 0 |
| 48964 | ENSMUSG00000111302 | 0 | NA | NA | NA | NA | NA | | olfactory receptor 1236, pseudogene 1 | 0 | 0 | 0 | 0 | 0 | 0 |
| 48965 | ENSMUSG00000111303 | 0 | NA | NA | NA | NA | NA | 4930502A04 | RIKEN cDNA 4930502A04 gene [Source:M | 0 | 0 | 0 | 0 | 0 | 0 |
| 48966 | ENSMUSG00000111304 | 0 | NA | NA | NA | NA | NA | Gm47677 | predicted gene, 47677 [Source:MGI Symb | 0 | 0 | 0 | 0 | 0 | 0 |
| 48967 | ENSMUSG00000111305 | 0 | NA | NA | NA | NA | NA | Olfr966-ps1 | olfactory receptor 966, pseudogene 1 [Sou | 0 | 0 | 0 | 0 | 0 | 0 |
| 48968 | ENSMUSG00000111306 | 0 | NA | NA | NA | NA | NA | Olfr1065 | olfactory receptor 1065 [Source:MGI Sym | 0 | 0 | 0 | 0 | 0 | 0 |
| 48969 | ENSMUSG00000111308 | 0 | NA | NA | NA | NA | NA | Gm48710 | predicted gene, 48710 [Source:MGI Symb | 0 | 0 | 0 | 0 | 0 | 0 |
| 48970 | ENSMUSG00000111309 | 0 | NA | NA | NA | NA | NA | Gm18863 | predicted gene, 18863 [Source:MGI Symb | 0 | 0 | 0 | 0 | 0 | 0 |
| 48971 | ENSMUSG00000111310 | 0 | NA | NA | NA | NA | NA | Gm47005 | predicted gene, 47005 [Source:MGI Symb | 0 | 0 | 0 | 0 | 0 | 0 |
| 48972 | ENSMUSG00000111312 | 3.171515905 | 0.314449551 | 1.299976652 | 0.241888614 | 0.808866473 | NA | Gm47205 | predicted gene, 47205 [Source:MGI Symb | 2.4093 | 2.0558 | 4.025 | 3.8559 | 0 | 6.6831 |
| 48973 | ENSMUSG00000111313 | 0.160662974 | 1.126924175 | 4.080472857 | 0.276174898 | 0.782413736 | NA | Gm18942 | predicted gene, 18942 [Source:MGI Symb | 0 | 0 | 0 | 0.964 | 0 | 0 |
| 48974 | ENSMUSG00000111315 | 0 | NA | NA | NA | NA | NA | Gm47473 | predicted gene, 47473 [Source:MGI Symb | 0 | 0 | 0 | 0 | 0 | 0 |
| 48975 | ENSMUSG00000111316 | 0 | NA | NA | NA | NA | NA | Gm47344 | predicted gene, 47344 [Source:MGI Symb | 0 | 0 | 0 | 0 | 0 | 0 |
| 48976 | ENSMUSG00000111317 | 0 | NA | NA | NA | NA | NA | Gm47333 | predicted gene, 47333 [Source:MGI Symb | 0 | 0 | 0 | 0 | 0 | 0 |
| 48977 | ENSMUSG00000111318 | 0 | NA | NA | NA | NA | NA | Gm48483 | predicted gene, 48483 [Source:MGI Symb | 0 | 0 | 0 | 0 | 0 | 0 |
| 48978 | ENSMUSG00000111320 | 0 | NA | NA | NA | NA | NA | Gm48546 | predicted gene, 48546 [Source:MGI Symb | 0 | 0 | 0 | 0 | 0 | 0 |
| 48979 | ENSMUSG00000111322 | 0 | NA | NA | NA | NA | NA | Gm48002 | predicted gene, 48002 [Source:MGI Symb | 0 | 0 | 0 | 0 | 0 | 0 |
| 48980 | ENSMUSG00000111323 | 0 | NA | NA | NA | NA | NA | Gm48129 | predicted gene, 48129 [Source:MGI Symb | 0 | 0 | 0 | 0 | 0 | 0 |
| 48981 | ENSMUSG00000111324 | 0 | NA | NA | NA | NA | NA | Gm31410 | predicted gene, 31410 [Source:MGI Symb | 0 | 0 | 0 | 0 | 0 | 0 |
| 48982 | ENSMUSG00000111325 | 34.89222049 | 0.210446329 | 0.545726486 | 0.385626014 | 0.699773664 | 0.941362191 | Gm47140 | predicted gene, 47140 [Source:MGI Symb | 22.487 | 25.698 | 49.306 | 72.298 | 25.084 | 14.48 |
| 48983 | ENSMUSG00000111326 | 1.979651559 | -0.273344208 | 2.161110692 | -0.126483206 | 0.899349442 | NA | Gm3953 | predicted gene 3953 [Source:MGI Symbol | 6.4248 | 0 | 0 | 0 | 5.4531 | 0 |
| 48984 | ENSMUSG00000111327 | 0 | NA | NA | NA | NA | NA | Gm48401 | predicted gene, 48401 [Source:MGI Symb | 0 | 0 | 0 | 0 | 0 | 0 |
| 48985 | ENSMUSG00000111329 | 11.72076272 | -0.254219947 | 0.76114524 | -0.333996632 | 0.7383821 | 0.948137969 | A830035O19 | RIKEN cDNA A830035O19 gene [Source:M | 16.062 | 9.2513 | 13.081 | 23.135 | 5.4531 | 3.3415 |
| 48986 | ENSMUSG00000111330 | 0 | NA | NA | NA | NA | NA | Gm19790 | predicted gene, 19790 [Source:MGI Symb | 0 | 0 | 0 | 0 | 0 | 0 |
| 48987 | ENSMUSG00000111333 | 0 | NA | NA | NA | NA | NA | Gm48670 | predicted gene, 48670 [Source:MGI Symb | 0 | 0 | 0 | 0 | 0 | 0 |
| 48988 | ENSMUSG00000111334 | 8.047768703 | -0.693599753 | 0.784371064 | -0.884275039 | 0.376547759 | 0.809930558 | Gm30015 | predicted gene, 30015 [Source:MGI Symb | 9.6372 | 10.279 | 10.062 | 10.604 | 4.3625 | 3.3415 |
| 48989 | ENSMUSG00000111335 | 0 | NA | NA | NA | NA | NA | Gm39375 | predicted gene, 39375 [Source:MGI Symb | 0 | 0 | 0 | 0 | 0 | 0 |
| 48990 | ENSMUSG00000111336 | 0 | NA | NA | NA | NA | NA | Aldoa-ps4 | aldolase A, pseudogene 4 [Source:MGI Sy | 0 | 0 | 0 | 0 | 0 | 0 |
| 48991 | ENSMUSG00000111337 | 0 | NA | NA | NA | NA | NA | Gm6885 | predicted gene 6885 [Source:MGI Symbol | 0 | 0 | 0 | 0 | 0 | 0 |
| 48992 | ENSMUSG00000111338 | 0 | NA | NA | NA | NA | NA | Olfr29-ps1 | olfactory receptor 29, pseudogene 1 [Sou | 0 | 0 | 0 | 0 | 0 | 0 |
| 48993 | ENSMUSG00000111339 | 0 | NA | NA | NA | NA | NA | | post-GPI attachment to proteins 2 | 0 | 0 | 0 | 0 | 0 | 0 |
| 48994 | ENSMUSG00000111340 | 0.17131964 | -0.796629176 | 4.080472857 | -0.195229623 | 0.84521319 | NA | Gm47171 | predicted gene, 47171 [Source:MGI Symb | 0 | 1.0279 | 0 | 0 | 0 | 0 |
| 48995 | ENSMUSG00000111341 | 0 | NA | NA | NA | NA | NA | Gm47111 | predicted gene, 47111 [Source:MGI Symb | 0 | 0 | 0 | 0 | 0 | 0 |
| 48996 | ENSMUSG00000111342 | 0.133850373 | -0.796629176 | 4.080472857 | -0.195229623 | 0.84521319 | NA | 4933404K13 | RIKEN cDNA 4933404K13 gene [Source:M | 0.8031 | 0 | 0 | 0 | 0 | 0 |

| | | | | | | | | | | | | | | | |
|---|---|---|---|---|---|---|---|---|---|---|---|---|---|---|---|
| 48997 | ENSMUSG00000111343 | 0 | NA | NA | NA | NA | NA | Gm47175 | predicted gene, 47175 [Source:MGI Symb | 0 | 0 | 0 | 0 | 0 | 0 |
| 48998 | ENSMUSG00000111344 | 0 | NA | NA | NA | NA | NA | Gm48467 | predicted gene, 48467 [Source:MGI Symb | 0 | 0 | 0 | 0 | 0 | 0 |
| 48999 | ENSMUSG00000111345 | 2.656316625 | -1.413784008 | 1.420925064 | -0.994974361 | 0.319748734 | NA | Gm48562 | predicted gene, 48562 [Source:MGI Symb | 2.4093 | 4.1117 | 5.0312 | 0 | 3.2719 | 1.1138 |
| 49000 | ENSMUSG00000111346 | 0 | NA | NA | NA | NA | NA | Gm18586 | predicted gene, 18586 [Source:MGI Symb | 0 | 0 | 0 | 0 | 0 | 0 |
| 49001 | ENSMUSG00000111348 | 0.331982614 | 0.16514309 | 3.986751476 | 0.041422971 | 0.966958701 | NA | Gm19531 | predicted gene, 19531 [Source:MGI Symb | 0 | 1.0279 | 0 | 0.964 | 0 | 0 |
| 49002 | ENSMUSG00000111349 | 0 | NA | NA | NA | NA | NA | Gm48191 | predicted gene, 48191 [Source:MGI Symb | 0 | 0 | 0 | 0 | 0 | 0 |
| 49003 | ENSMUSG00000111350 | 0 | NA | NA | NA | NA | NA | Gm47182 | predicted gene, 47182 [Source:MGI Symb | 0 | 0 | 0 | 0 | 0 | 0 |
| 49004 | ENSMUSG00000111351 | 0 | NA | NA | NA | NA | NA | Gm48537 | predicted gene, 48537 [Source:MGI Symb | 0 | 0 | 0 | 0 | 0 | 0 |
| 49005 | ENSMUSG00000111352 | 0 | NA | NA | NA | NA | NA | Gm5369 | predicted gene 5369 [Source:MGI Symbol | 0 | 0 | 0 | 0 | 0 | 0 |
| 49006 | ENSMUSG00000111353 | 0 | NA | NA | NA | NA | NA | Gm19324 | predicted gene, 19324 [Source:MGI Symb | 0 | 0 | 0 | 0 | 0 | 0 |
| 49007 | ENSMUSG00000111354 | 0 | NA | NA | NA | NA | NA | Gm33914 | predicted gene, 33914 [Source:MGI Symb | 0 | 0 | 0 | 0 | 0 | 0 |
| 49008 | ENSMUSG00000111355 | 0 | NA | NA | NA | NA | NA | Idi1-ps3 | isopentenyl-diphosphate delta isomerase, | 0 | 0 | 0 | 0 | 0 | 0 |
| 49009 | ENSMUSG00000111356 | 1.017313306 | 0.03731332 | 2.211598506 | 0.016871652 | 0.986539008 | NA | Rpl32l | ribosomal protein L32-like [Source:MGI Sy | 0 | 2.0558 | 1.0062 | 1.928 | 0 | 1.1138 |
| 49010 | ENSMUSG00000111357 | 0 | NA | NA | NA | NA | NA | Gm6980 | predicted gene 6980 [Source:MGI Symbol | 0 | 0 | 0 | 0 | 0 | 0 |
| 49011 | ENSMUSG00000111358 | 0 | NA | NA | NA | NA | NA | 2900078I11F | RIKEN cDNA 2900078I11 gene [Source:MC | 0 | 0 | 0 | 0 | 0 | 0 |
| 49012 | ENSMUSG00000111359 | 0 | NA | NA | NA | NA | NA | Gm39336 | predicted gene, 39336 [Source:MGI Symb | 0 | 0 | 0 | 0 | 0 | 0 |
| 49013 | ENSMUSG00000111360 | 0 | NA | NA | NA | NA | NA | Gm38642 | predicted gene, 38642 [Source:MGI Symb | 0 | 0 | 0 | 0 | 0 | 0 |
| 49014 | ENSMUSG00000111361 | 0.720758441 | 1.701909892 | 2.735431843 | 0.622172289 | 0.533828585 | NA | Gm47445 | predicted gene, 47445 [Source:MGI Symb | 0 | 0 | 1.0062 | 0 | 1.0906 | 2.2277 |
| 49015 | ENSMUSG00000111362 | 0 | NA | NA | NA | NA | NA | Gm47389 | predicted gene, 47389 [Source:MGI Symb | 0 | 0 | 0 | 0 | 0 | 0 |
| 49016 | ENSMUSG00000111363 | 0 | NA | NA | NA | NA | NA | Gm35024 | predicted gene, 35024 [Source:MGI Symb | 0 | 0 | 0 | 0 | 0 | 0 |
| 49017 | ENSMUSG00000111364 | 0 | NA | NA | NA | NA | NA | Gm36745 | predicted gene, 36745 [Source:MGI Symb | 0 | 0 | 0 | 0 | 0 | 0 |
| 49018 | ENSMUSG00000111365 | 7.797221142 | -0.540051894 | 0.762955772 | -0.707841678 | 0.479043581 | 0.870169438 | Gm47772 | predicted gene, 47772 [Source:MGI Symb | 6.4248 | 10.279 | 11.069 | 5.7839 | 6.5437 | 6.6831 |
| 49019 | ENSMUSG00000111366 | 0 | NA | NA | NA | NA | NA | Gm47156 | predicted gene, 47156 [Source:MGI Symb | 0 | 0 | 0 | 0 | 0 | 0 |
| 49020 | ENSMUSG00000111368 | 13.94441356 | -0.287748912 | 0.56931092 | -0.505433678 | 0.61325427 | 0.916813074 | Gm47889 | predicted gene, 47889 [Source:MGI Symb | 18.471 | 13.363 | 14.087 | 13.496 | 11.997 | 12.252 |
| 49021 | ENSMUSG00000111369 | 0 | NA | NA | NA | NA | NA | Gm47043 | predicted gene, 47043 [Source:MGI Symb | 0 | 0 | 0 | 0 | 0 | 0 |
| 49022 | ENSMUSG00000111371 | 0 | NA | NA | NA | NA | NA | Gm47162 | predicted gene, 47162 [Source:MGI Symb | 0 | 0 | 0 | 0 | 0 | 0 |
| 49023 | ENSMUSG00000111373 | 0 | NA | NA | NA | NA | NA | Gm48818 | predicted gene, 48818 [Source:MGI Symb | 0 | 0 | 0 | 0 | 0 | 0 |
| 49024 | ENSMUSG00000111374 | 0 | NA | NA | NA | NA | NA | Gm50490 | predicted gene, 50490 [Source:MGI Symb | 0 | 0 | 0 | 0 | 0 | 0 |
| 49025 | ENSMUSG00000111375 | 31.12811345 | 0.534838434 | 0.464203581 | 1.152163523 | 0.249253884 | 0.701402369 | Btbd8 | BTB (POZ) domain containing 8 [Source:M | 16.865 | 22.614 | 37.231 | 54.947 | 27.265 | 27.846 |
| 49026 | ENSMUSG00000111376 | 0 | NA | NA | NA | NA | NA | Gm47099 | predicted gene, 47099 [Source:MGI Symb | 0 | 0 | 0 | 0 | 0 | 0 |
| 49027 | ENSMUSG00000111377 | 5.630726373 | -1.100067238 | 1.001981849 | -1.097891383 | 0.272251921 | 0.727618887 | Gm40634 | predicted gene, 40634 [Source:MGI Symb | 9.6372 | 10.279 | 3.0187 | 0.964 | 6.5437 | 3.3415 |
| 49028 | ENSMUSG00000111378 | 0 | NA | NA | NA | NA | NA | Gm47230 | predicted gene, 47230 [Source:MGI Symb | 0 | 0 | 0 | 0 | 0 | 0 |
| 49029 | ENSMUSG00000111379 | 0 | NA | NA | NA | NA | NA | Gm48346 | predicted gene, 48346 [Source:MGI Symb | 0 | 0 | 0 | 0 | 0 | 0 |
| 49030 | ENSMUSG00000111380 | 0 | NA | NA | NA | NA | NA | D9Wsu149 | DNA segment, Chr 9, Wayne State Univers | 0 | 0 | 0 | 0 | 0 | 0 |
| 49031 | ENSMUSG00000111381 | 0 | NA | NA | NA | NA | NA | Gm34424 | predicted gene, 34424 [Source:MGI Symb | 0 | 0 | 0 | 0 | 0 | 0 |
| 49032 | ENSMUSG00000111382 | 0.160662974 | 1.126924175 | 4.080472857 | 0.276174898 | 0.782413736 | NA | Gm47536 | predicted gene, 47536 [Source:MGI Symb | 0 | 0 | 0 | 0.964 | 0 | 0 |
| 49033 | ENSMUSG00000111383 | 0.167707431 | -0.796629176 | 4.080472857 | -0.195229623 | 0.84521319 | NA | Gm18914 | predicted gene, 18914 [Source:MGI Symb | 0 | 0 | 1.0062 | 0 | 0 | 0 |
| 49034 | ENSMUSG00000111384 | 0 | NA | NA | NA | NA | NA | Olfr1059-ps1 | olfactory receptor 1059, pseudogene 1 [S | 0 | 0 | 0 | 0 | 0 | 0 |
| 49035 | ENSMUSG00000111386 | 0 | NA | NA | NA | NA | NA | Gm48842 | predicted gene, 48842 [Source:MGI Symb | 0 | 0 | 0 | 0 | 0 | 0 |
| 49036 | ENSMUSG00000111387 | 0 | NA | NA | NA | NA | NA | Gm48803 | predicted gene, 48803 [Source:MGI Symb | 0 | 0 | 0 | 0 | 0 | 0 |
| 49037 | ENSMUSG00000111388 | 0 | NA | NA | NA | NA | NA | Gm7183 | predicted gene 7183 [Source:MGI Symbol | 0 | 0 | 0 | 0 | 0 | 0 |
| 49038 | ENSMUSG00000111389 | 0.185640647 | 1.126924175 | 4.080472857 | 0.276174898 | 0.782413736 | NA | Gm39465 | predicted gene, 39465 [Source:MGI Symb | 0 | 0 | 0 | 0 | 0 | 1.1138 |
| 49039 | ENSMUSG00000111390 | 1.528792023 | -0.842850825 | 1.775242115 | -0.474780774 | 0.634943238 | NA | Gm48796 | predicted gene, 48796 [Source:MGI Symb | 0.8031 | 2.0558 | 3.0187 | 0 | 2.1812 | 1.1138 |
| 49040 | ENSMUSG00000111394 | 8.240473269 | -0.977352325 | 0.758657295 | -1.288265902 | 0.197653418 | 0.639708893 | Gm49759 | predicted gene, 49759 [Source:MGI Symb | 14.456 | 9.2513 | 9.0562 | 6.7478 | 4.3625 | 5.5692 |
| 49041 | ENSMUSG00000111395 | 0 | NA | NA | NA | NA | NA | Gm18655 | predicted gene, 18655 [Source:MGI Symb | 0 | 0 | 0 | 0 | 0 | 0 |
| 49042 | ENSMUSG00000111396 | 0 | NA | NA | NA | NA | NA | Gm7333 | predicted gene 7333 [Source:MGI Symbol | 0 | 0 | 0 | 0 | 0 | 0 |
| 49043 | ENSMUSG00000111397 | 0 | NA | NA | NA | NA | NA | Gm48232 | predicted gene, 48232 [Source:MGI Symb | 0 | 0 | 0 | 0 | 0 | 0 |
| 49044 | ENSMUSG00000111398 | 0 | NA | NA | NA | NA | NA | D330037F02 | RIKEN cDNA D330037F02 gene [Source:M | 0 | 0 | 0 | 0 | 0 | 0 |
| 49045 | ENSMUSG00000111399 | 0 | NA | NA | NA | NA | NA | Gm47328 | predicted gene, 47328 [Source:MGI Symb | 0 | 0 | 0 | 0 | 0 | 0 |
| 49046 | ENSMUSG00000111400 | 0 | NA | NA | NA | NA | NA | Gm29961 | predicted gene, 29961 [Source:MGI Symb | 0 | 0 | 0 | 0 | 0 | 0 |
| 49047 | ENSMUSG00000111403 | 0 | NA | NA | NA | NA | NA | Gm47544 | predicted gene, 47544 [Source:MGI Symb | 0 | 0 | 0 | 0 | 0 | 0 |
| 49048 | ENSMUSG00000111404 | 0 | NA | NA | NA | NA | NA | Gm47169 | predicted gene, 47169 [Source:MGI Symb | 0 | 0 | 0 | 0 | 0 | 0 |
| 49049 | ENSMUSG00000111405 | 0 | NA | NA | NA | NA | NA | Gm2594 | predicted gene 2594 [Source:MGI Symbol | 0 | 0 | 0 | 0 | 0 | 0 |
| 49050 | ENSMUSG00000111406 | 0 | NA | NA | NA | NA | NA | Gm47449 | predicted gene, 47449 [Source:MGI Symb | 0 | 0 | 0 | 0 | 0 | 0 |
| 49051 | ENSMUSG00000111409 | 0 | NA | NA | NA | NA | NA | Gm49380 | predicted gene, 49380 [Source:MGI Symb | 0 | 0 | 0 | 0 | 0 | 0 |
| 49052 | ENSMUSG00000111410 | 0 | NA | NA | NA | NA | NA | Gm49337 | predicted gene, 49337 [Source:MGI Symb | 0 | 0 | 0 | 0 | 0 | 0 |

| | | | | | | | | | | | | | | | |
|---|---|---|---|---|---|---|---|---|---|---|---|---|---|---|---|
| 49053 | ENSMUSG00000111411 | 0 | NA | NA | NA | NA | NA | Gm33778 | predicted gene, 33778 [Source:MGI Symb | 0 | 0 | 0 | 0 | 0 | 0 |
| 49054 | ENSMUSG00000111412 | 0 | NA | NA | NA | NA | NA | Gm47047 | predicted gene, 47047 [Source:MGI Symb | 0 | 0 | 0 | 0 | 0 | 0 |
| 49055 | ENSMUSG00000111413 | 0 | NA | NA | NA | NA | NA | Gm47716 | predicted gene, 47716 [Source:MGI Symb | 0 | 0 | 0 | 0 | 0 | 0 |
| 49056 | ENSMUSG00000111414 | 0 | NA | NA | NA | NA | NA | Gm18103 | predicted gene, 18103 [Source:MGI Symb | 0 | 0 | 0 | 0 | 0 | 0 |
| 49057 | ENSMUSG00000111415 | 0.17131964 | -0.796629176 | 4.080472857 | -0.195229623 | 0.84521319 | NA | Gm39325 | predicted gene, 39325 [Source:MGI Symb | 0 | 1.0279 | 0 | 0 | 0 | 0 |
| 49058 | ENSMUSG00000111416 | 0 | NA | NA | NA | NA | NA | Gm48331 | predicted gene, 48331 [Source:MGI Symb | 0 | 0 | 0 | 0 | 0 | 0 |
| 49059 | ENSMUSG00000111417 | 3.310085517 | 0.936919526 | 1.390636377 | 0.673734372 | 0.500480205 | NA | Gm47430 | predicted gene, 47430 [Source:MGI Symb | 0.8031 | 0 | 6.0375 | 1.928 | 2.1812 | 8.9108 |
| 49060 | ENSMUSG00000111418 | 0 | NA | NA | NA | NA | NA | Gm18016 | predicted gene, 18016 [Source:MGI Symb | 0 | 0 | 0 | 0 | 0 | 0 |
| 49061 | ENSMUSG00000111419 | 0 | NA | NA | NA | NA | NA | Gm47568 | predicted gene, 47568 [Source:MGI Symb | 0 | 0 | 0 | 0 | 0 | 0 |
| 49062 | ENSMUSG00000111420 | 0 | NA | NA | NA | NA | NA | Olfr927-ps1 | olfactory receptor 927, pseudogene 1 [Sou | 0 | 0 | 0 | 0 | 0 | 0 |
| 49063 | ENSMUSG00000111421 | 0.48903338 | 0.95637552 | 3.456690196 | 0.276673773 | 0.782030612 | NA | Gm47643 | predicted gene, 47643 [Source:MGI Symb | 0 | 0 | 1.0062 | 1.928 | 0 | 0 |
| 49064 | ENSMUSG00000111422 | 2.911782848 | -1.210579513 | 1.354367071 | -0.89383413 | 0.371410645 | NA | 1700027J07I | RIKEN cDNA 1700027J07 gene [Source:MG | 8.031 | 3.0838 | 1.0062 | 0.964 | 3.2719 | 1.1138 |
| 49065 | ENSMUSG00000111423 | 0 | NA | NA | NA | NA | NA | Dppa5c | developmental pluripotency associated 5( | 0 | 0 | 0 | 0 | 0 | 0 |
| 49066 | ENSMUSG00000111424 | 6.531332392 | 0.290266472 | 0.896630286 | 0.32373039 | 0.746142161 | 0.949997329 | Gm48478 | predicted gene, 48478 [Source:MGI Symb | 6.4248 | 9.2513 | 2.0125 | 11.568 | 4.3625 | 5.5692 |
| 49067 | ENSMUSG00000111425 | 0.315620088 | 0.165143421 | 4.063300985 | 0.040642675 | 0.967580763 | NA | Gm47324 | predicted gene, 47324 [Source:MGI Symb | 0.8031 | 0 | 0 | 0 | 1.0906 | 0 |
| 49068 | ENSMUSG00000111426 | 0 | NA | NA | NA | NA | NA | Gm47318 | predicted gene, 47318 [Source:MGI Symb | 0 | 0 | 0 | 0 | 0 | 0 |
| 49069 | ENSMUSG00000111427 | 0 | NA | NA | NA | NA | NA | Gm6390 | predicted gene 6390 [Source:MGI Symbol | 0 | 0 | 0 | 0 | 0 | 0 |
| 49070 | ENSMUSG00000111428 | 0 | NA | NA | NA | NA | NA | Gm47146 | predicted gene, 47146 [Source:MGI Symb | 0 | 0 | 0 | 0 | 0 | 0 |
| 49071 | ENSMUSG00000111429 | 66.55571797 | 0.116956054 | 0.309606815 | 0.377756718 | 0.705611329 | 0.94295036 | Gm32511 | predicted gene, 32511 [Source:MGI Symb | 65.051 | 67.843 | 58.362 | 46.271 | 90.521 | 71.286 |
| 49072 | ENSMUSG00000111430 | 0 | NA | NA | NA | NA | NA | Olfr950-ps1 | olfactory receptor 950, pseudogene 1 [Sou | 0 | 0 | 0 | 0 | 0 | 0 |
| 49073 | ENSMUSG00000111431 | 0 | NA | NA | NA | NA | NA | Gm17779 | predicted gene, 17779 [Source:MGI Symb | 0 | 0 | 0 | 0 | 0 | 0 |
| 49074 | ENSMUSG00000111432 | 0.649696354 | 1.537929172 | 3.118369374 | 0.493183773 | 0.621882743 | NA | Gm47993 | predicted gene, 47993 [Source:MGI Symb | 0 | 0 | 1.0062 | 2.8919 | 0 | 0 |
| 49075 | ENSMUSG00000111433 | 0 | NA | NA | NA | NA | NA | Gm47378 | predicted gene, 47378 [Source:MGI Symb | 0 | 0 | 0 | 0 | 0 | 0 |
| 49076 | ENSMUSG00000111434 | 0.160662974 | 1.126924175 | 4.080472857 | 0.276174898 | 0.782413736 | NA | Gm47121 | predicted gene, 47121 [Source:MGI Symb | 0 | 0 | 0 | 0.964 | 0 | 0 |
| 49077 | ENSMUSG00000111435 | 0 | NA | NA | NA | NA | NA | Gm47358 | predicted gene, 47358 [Source:MGI Symb | 0 | 0 | 0 | 0 | 0 | 0 |
| 49078 | ENSMUSG00000111436 | 0 | NA | NA | NA | NA | NA | Gm47356 | predicted gene, 47356 [Source:MGI Symb | 0 | 0 | 0 | 0 | 0 | 0 |
| 49079 | ENSMUSG00000111437 | 0 | NA | NA | NA | NA | NA | Gm47236 | predicted gene, 47236 [Source:MGI Symb | 0 | 0 | 0 | 0 | 0 | 0 |
| 49080 | ENSMUSG00000111438 | 0 | NA | NA | NA | NA | NA | Rps2-ps12 | ribosomal protein S2, pseudogene 12 [Sou | 0 | 0 | 0 | 0 | 0 | 0 |
| 49081 | ENSMUSG00000111439 | 0 | NA | NA | NA | NA | NA | Gm47633 | predicted gene, 47633 [Source:MGI Symb | 0 | 0 | 0 | 0 | 0 | 0 |
| 49082 | ENSMUSG00000111440 | 0 | NA | NA | NA | NA | NA | Vmn1r-ps50 | vomeronasal 1 receptor, pseudogene 50 [ | 0 | 0 | 0 | 0 | 0 | 0 |
| 49083 | ENSMUSG00000111441 | 0 | NA | NA | NA | NA | NA | Gm47137 | predicted gene, 47137 [Source:MGI Symb | 0 | 0 | 0 | 0 | 0 | 0 |
| 49084 | ENSMUSG00000111442 | 0.367410362 | 2.078255025 | 3.800070157 | 0.54689912 | 0.58444804 | NA | Gm48334 | predicted gene, 48334 [Source:MGI Symb | 0 | 0 | 0 | 0 | 1.0906 | 1.1138 |
| 49085 | ENSMUSG00000111443 | 0 | NA | NA | NA | NA | NA | Gm46123 | predicted gene, 46123 [Source:MGI Symb | 0 | 0 | 0 | 0 | 0 | 0 |
| 49086 | ENSMUSG00000111444 | 0 | NA | NA | NA | NA | NA | Gm47923 | predicted gene, 47923 [Source:MGI Symb | 0 | 0 | 0 | 0 | 0 | 0 |
| 49087 | ENSMUSG00000111445 | 0 | NA | NA | NA | NA | NA | Gm47070 | predicted gene, 47070 [Source:MGI Symb | 0 | 0 | 0 | 0 | 0 | 0 |
| 49088 | ENSMUSG00000111446 | 0.185640647 | 1.126924175 | 4.080472857 | 0.276174898 | 0.782413736 | NA | Gm6581 | predicted gene 6581 [Source:MGI Symbol | 0 | 0 | 0 | 0 | 0 | 1.1138 |
| 49089 | ENSMUSG00000111447 | 4.027778031 | 0.563326978 | 1.10768989 | 0.508560188 | 0.611060541 | NA | Gm48249 | predicted gene, 48249 [Source:MGI Symb | 5.6217 | 1.0279 | 3.0187 | 6.7478 | 2.1812 | 5.5692 |
| 49090 | ENSMUSG00000111448 | 0 | NA | NA | NA | NA | NA | Olfr912 | olfactory receptor 912 [Source:MGI Symb | 0 | 0 | 0 | 0 | 0 | 0 |
| 49091 | ENSMUSG00000111449 | 0 | NA | NA | NA | NA | NA | Gm34105 | predicted gene, 34105 [Source:MGI Symb | 0 | 0 | 0 | 0 | 0 | 0 |
| 49092 | ENSMUSG00000111450 | 0 | NA | NA | NA | NA | NA | Gm33351 | predicted gene, 33351 [Source:MGI Symb | 0 | 0 | 0 | 0 | 0 | 0 |
| 49093 | ENSMUSG00000111451 | 0.335414862 | -1.779047242 | 4.030804179 | -0.441362855 | 0.658950329 | NA | Gm47321 | predicted gene, 47321 [Source:MGI Symb | 0 | 0 | 2.0125 | 0 | 0 | 0 |
| 49094 | ENSMUSG00000111453 | 0 | NA | NA | NA | NA | NA | Olfr925-ps1 | olfactory receptor 925, pseudogene 1 [Sou | 0 | 0 | 0 | 0 | 0 | 0 |
| 49095 | ENSMUSG00000111454 | 0 | NA | NA | NA | NA | NA | Olfr998 | olfactory receptor 998 [Source:MGI Symb | 0 | 0 | 0 | 0 | 0 | 0 |
| 49096 | ENSMUSG00000111455 | 0 | NA | NA | NA | NA | NA | Olfr1142-ps1 | olfactory receptor 1142, pseudogene 1 [So | 0 | 0 | 0 | 0 | 0 | 0 |
| 49097 | ENSMUSG00000111456 | 0 | NA | NA | NA | NA | NA | Olfr1245 | olfactory receptor 1245 [Source:MGI Syml | 0 | 0 | 0 | 0 | 0 | 0 |
| 49098 | ENSMUSG00000111457 | 0 | NA | NA | NA | NA | NA | Gm18996 | predicted gene, 18996 [Source:MGI Symb | 0 | 0 | 0 | 0 | 0 | 0 |
| 49099 | ENSMUSG00000111458 | 0 | NA | NA | NA | NA | NA | Gm47035 | predicted gene, 47035 [Source:MGI Symb | 0 | 0 | 0 | 0 | 0 | 0 |
| 49100 | ENSMUSG00000111459 | 0 | NA | NA | NA | NA | NA | Gm8031 | predicted gene 8031 [Source:MGI Symbol | 0 | 0 | 0 | 0 | 0 | 0 |
| 49101 | ENSMUSG00000111460 | 0 | NA | NA | NA | NA | NA | Olfr1227-ps1 | olfactory receptor 1227, pseudogene 1 [So | 0 | 0 | 0 | 0 | 0 | 0 |
| 49102 | ENSMUSG00000111462 | 0 | NA | NA | NA | NA | NA | Gm49317 | predicted gene, 49317 [Source:MGI Symb | 0 | 0 | 0 | 0 | 0 | 0 |
| 49103 | ENSMUSG00000111463 | 0 | NA | NA | NA | NA | NA | Gm48061 | predicted gene, 48061 [Source:MGI Symb | 0 | 0 | 0 | 0 | 0 | 0 |
| 49104 | ENSMUSG00000111464 | 0 | NA | NA | NA | NA | NA | Gm48675 | predicted gene, 48675 [Source:MGI Symb | 0 | 0 | 0 | 0 | 0 | 0 |
| 49105 | ENSMUSG00000111465 | 1.402760624 | -0.737730338 | 1.9516106 | -0.378011033 | 0.705422398 | NA | Gm47113 | predicted gene, 47113 [Source:MGI Symb | 3.2124 | 0 | 2.0125 | 0.964 | 0 | 2.2277 |
| 49106 | ENSMUSG00000111466 | 0 | NA | NA | NA | NA | NA | Gm47504 | predicted gene, 47504 [Source:MGI Symb | 0 | 0 | 0 | 0 | 0 | 0 |
| 49107 | ENSMUSG00000111467 | 0 | NA | NA | NA | NA | NA | Gm47775 | predicted gene, 47775 [Source:MGI Symb | 0 | 0 | 0 | 0 | 0 | 0 |
| 49108 | ENSMUSG00000111468 | 0 | NA | NA | NA | NA | NA | 5033425B01 | RIKEN cDNA 5033425B01 gene [Source:M | 0 | 0 | 0 | 0 | 0 | 0 |

| | | | | | | | | | | | | | | | |
|---|---|---|---|---|---|---|---|---|---|---|---|---|---|---|---|
| 49109 | ENSMUSG00000111470 | 0 | NA | NA | NA | NA | NA | Gm31822 | predicted gene, 31822 [Source:MGI Symb | 0 | 0 | 0 | 0 | 0 | 0 |
| 49110 | ENSMUSG00000111471 | 1.000898647 | -2.10615299 | 2.475207832 | -0.850899453 | 0.394825209 | NA | Gm47350 | predicted gene, 47350 [Source:MGI Symb | 0.8031 | 4.1117 | 0 | 0 | 1.0906 | 0 |
| 49111 | ENSMUSG00000111472 | 0 | NA | NA | NA | NA | NA | Gm47103 | predicted gene, 47103 [Source:MGI Symb | 0 | 0 | 0 | 0 | 0 | 0 |
| 49112 | ENSMUSG00000111473 | 0 | NA | NA | NA | NA | NA | LOC1026370 | immunoglobulin superfamily member 22- | 0 | 0 | 0 | 0 | 0 | 0 |
| 49113 | ENSMUSG00000111474 | 0 | NA | NA | NA | NA | NA | Gm32926 | predicted gene, 32926 [Source:MGI Symb | 0 | 0 | 0 | 0 | 0 | 0 |
| 49114 | ENSMUSG00000111477 | 0 | NA | NA | NA | NA | NA | Gm47668 | predicted gene, 47668 [Source:MGI Symb | 0 | 0 | 0 | 0 | 0 | 0 |
| 49115 | ENSMUSG00000111478 | 0 | NA | NA | NA | NA | NA | Gm18251 | predicted gene, 18251 [Source:MGI Symb | 0 | 0 | 0 | 0 | 0 | 0 |
| 49116 | ENSMUSG00000111479 | 0 | NA | NA | NA | NA | NA | Gm47559 | predicted gene, 47559 [Source:MGI Symb | 0 | 0 | 0 | 0 | 0 | 0 |
| 49117 | ENSMUSG00000111480 | 0 | NA | NA | NA | NA | NA | Gm48469 | predicted gene, 48469 [Source:MGI Symb | 0 | 0 | 0 | 0 | 0 | 0 |
| 49118 | ENSMUSG00000111481 | 1.559293941 | -0.364951707 | 1.735206242 | -0.210321804 | 0.833416519 | NA | Gm47652 | predicted gene, 47652 [Source:MGI Symb | 3.2124 | 1.0279 | 1.0062 | 1.928 | 2.1812 | 0 |
| 49119 | ENSMUSG00000111482 | 0 | NA | NA | NA | NA | NA | Gm47153 | predicted gene, 47153 [Source:MGI Symb | 0 | 0 | 0 | 0 | 0 | 0 |
| 49120 | ENSMUSG00000111483 | 0 | NA | NA | NA | NA | NA | Gm18997 | predicted gene, 18997 [Source:MGI Symb | 0 | 0 | 0 | 0 | 0 | 0 |
| 49121 | ENSMUSG00000111484 | 0 | NA | NA | NA | NA | NA | Gm18587 | predicted gene, 18587 [Source:MGI Symb | 0 | 0 | 0 | 0 | 0 | 0 |
| 49122 | ENSMUSG00000111485 | 0 | NA | NA | NA | NA | NA | Gm48443 | predicted gene, 48443 [Source:MGI Symb | 0 | 0 | 0 | 0 | 0 | 0 |
| 49123 | ENSMUSG00000111486 | 0 | NA | NA | NA | NA | NA | Gm47380 | predicted gene, 47380 [Source:MGI Symb | 0 | 0 | 0 | 0 | 0 | 0 |
| 49124 | ENSMUSG00000111487 | 0 | NA | NA | NA | NA | NA | 1700092E16 | RIKEN cDNA 1700092E16 gene [Source:M | 0 | 0 | 0 | 0 | 0 | 0 |
| 49125 | ENSMUSG00000111488 | 0 | NA | NA | NA | NA | NA | Gm31378 | predicted gene, 31378 [Source:MGI Symb | 0 | 0 | 0 | 0 | 0 | 0 |
| 49126 | ENSMUSG00000111489 | 0 | NA | NA | NA | NA | NA | 1700019J19I | RIKEN cDNA 1700019J19 gene [Source:M | 0 | 0 | 0 | 0 | 0 | 0 |
| 49127 | ENSMUSG00000111490 | 0 | NA | NA | NA | NA | NA | Gm47159 | predicted gene, 47159 [Source:MGI Symb | 0 | 0 | 0 | 0 | 0 | 0 |
| 49128 | ENSMUSG00000111491 | 0.167707431 | -0.796629176 | 4.080472857 | -0.195229623 | 0.84521319 | NA | Gm47503 | predicted gene, 47503 [Source:MGI Symb | 0 | 0 | 1.0062 | 0 | 0 | 0 |
| 49129 | ENSMUSG00000111492 | 0 | NA | NA | NA | NA | NA | Aldoa-ps3 | aldolase A, pseudogene 3 [Source:MGI Sy | 0 | 0 | 0 | 0 | 0 | 0 |
| 49130 | ENSMUSG00000111493 | 0 | NA | NA | NA | NA | NA | Olfr1091-ps1 | olfactory receptor 1091, pseudogene 1 [S | 0 | 0 | 0 | 0 | 0 | 0 |
| 49131 | ENSMUSG00000111494 | 0 | NA | NA | NA | NA | NA | 4930431F10 | RIKEN cDNA 4930431F10 gene [Source:M | 0 | 0 | 0 | 0 | 0 | 0 |
| 49132 | ENSMUSG00000111495 | 0.181769715 | 1.126924175 | 4.080472857 | 0.276174898 | 0.782413736 | NA | Gm5366 | predicted gene 5366 [Source:MGI Symbol | 0 | 0 | 0 | 0 | 1.0906 | 0 |
| 49133 | ENSMUSG00000111496 | 36.39733365 | -0.552969229 | 0.372520968 | -1.484397595 | 0.137703472 | 0.541918695 | Gm47483 | predicted gene, 47483 [Source:MGI Symb | 43.368 | 44.2 | 42.262 | 26.991 | 38.172 | 23.391 |
| 49134 | ENSMUSG00000111497 | 1.572250849 | -1.897026774 | 1.8859058 | -1.005896888 | 0.314465174 | NA | Gm38431 | predicted gene, 38431 [Source:MGI Symb | 2.4093 | 3.0838 | 2.0125 | 1.928 | 0 | 0 |
| 49135 | ENSMUSG00000111498 | 0 | NA | NA | NA | NA | NA | Gm47059 | predicted gene, 47059 [Source:MGI Symb | 0 | 0 | 0 | 0 | 0 | 0 |
| 49136 | ENSMUSG00000111499 | 0 | NA | NA | NA | NA | NA | Gm47562 | predicted gene, 47562 [Source:MGI Symb | 0 | 0 | 0 | 0 | 0 | 0 |
| 49137 | ENSMUSG00000111500 | 0 | NA | NA | NA | NA | NA | Gm48820 | predicted gene, 48820 [Source:MGI Symb | 0 | 0 | 0 | 0 | 0 | 0 |
| 49138 | ENSMUSG00000111503 | 0 | NA | NA | NA | NA | NA | Gm47034 | predicted gene, 47034 [Source:MGI Symb | 0 | 0 | 0 | 0 | 0 | 0 |
| 49139 | ENSMUSG00000111504 | 0 | NA | NA | NA | NA | NA | Gm48284 | predicted gene, 48284 [Source:MGI Symb | 0 | 0 | 0 | 0 | 0 | 0 |
| 49140 | ENSMUSG00000111505 | 0.678054141 | -2.792799308 | 2.990673428 | -0.933836266 | 0.350388359 | NA | Gm48673 | predicted gene, 48673 [Source:MGI Symb | 0 | 2.0558 | 2.0125 | 0 | 0 | 0 |
| 49141 | ENSMUSG00000111506 | 0 | NA | NA | NA | NA | NA | Gm48848 | predicted gene, 48848 [Source:MGI Symb | 0 | 0 | 0 | 0 | 0 | 0 |
| 49142 | ENSMUSG00000111507 | 0 | NA | NA | NA | NA | NA | Amd-ps6 | S-adenosylmethionine decarboxylase, pse | 0 | 0 | 0 | 0 | 0 | 0 |
| 49143 | ENSMUSG00000111508 | 0 | NA | NA | NA | NA | NA | Gm48634 | predicted gene, 48634 [Source:MGI Symb | 0 | 0 | 0 | 0 | 0 | 0 |
| 49144 | ENSMUSG00000111509 | 0 | NA | NA | NA | NA | NA | Gm47266 | predicted gene, 47266 [Source:MGI Symb | 0 | 0 | 0 | 0 | 0 | 0 |
| 49145 | ENSMUSG00000111510 | 0 | NA | NA | NA | NA | NA | Gm48387 | predicted gene, 48387 [Source:MGI Symb | 0 | 0 | 0 | 0 | 0 | 0 |
| 49146 | ENSMUSG00000111511 | 0 | NA | NA | NA | NA | NA | 6720432D03 | RIKEN cDNA 6720432D03 gene [Source:M | 0 | 0 | 0 | 0 | 0 | 0 |
| 49147 | ENSMUSG00000111512 | 0 | NA | NA | NA | NA | NA | Gm17853 | predicted gene, 17853 [Source:MGI Symb | 0 | 0 | 0 | 0 | 0 | 0 |
| 49148 | ENSMUSG00000111514 | 0.133850373 | -0.796629176 | 4.080472857 | -0.195229623 | 0.84521319 | NA | E230014E18 | RIKEN cDNA E230014E18 gene [Source:M | 0.8031 | 0 | 0 | 0 | 0 | 0 |
| 49149 | ENSMUSG00000111515 | 0 | NA | NA | NA | NA | NA | Gm5055 | predicted gene 5055 [Source:MGI Symbol | 0 | 0 | 0 | 0 | 0 | 0 |
| 49150 | ENSMUSG00000111517 | 0 | NA | NA | NA | NA | NA | Olfr1238 | olfactory receptor 1238 [Source:MGI Sym | 0 | 0 | 0 | 0 | 0 | 0 |
| 49151 | ENSMUSG00000111518 | 0 | NA | NA | NA | NA | NA | Olfr1073-ps1 | olfactory receptor 1073, pseudogene 1 [S | 0 | 0 | 0 | 0 | 0 | 0 |
| 49152 | ENSMUSG00000111520 | 0 | NA | NA | NA | NA | NA | Gm33125 | predicted gene, 33125 [Source:MGI Symb | 0 | 0 | 0 | 0 | 0 | 0 |
| 49153 | ENSMUSG00000111521 | 26.23288837 | -0.17621247 | 0.422505208 | -0.417065794 | 0.676630284 | 0.932891189 | Gm48529 | predicted gene, 48529 [Source:MGI Symb | 26.502 | 30.838 | 26.162 | 22.171 | 29.447 | 22.277 |
| 49154 | ENSMUSG00000111523 | 0 | NA | NA | NA | NA | NA | Gm47183 | predicted gene, 47183 [Source:MGI Symb | 0 | 0 | 0 | 0 | 0 | 0 |
| 49155 | ENSMUSG00000111524 | 0 | NA | NA | NA | NA | NA | Gm34113 | predicted gene, 34113 [Source:MGI Symb | 0 | 0 | 0 | 0 | 0 | 0 |
| 49156 | ENSMUSG00000111525 | 0 | NA | NA | NA | NA | NA | Gm47082 | predicted gene, 47082 [Source:MGI Symb | 0 | 0 | 0 | 0 | 0 | 0 |
| 49157 | ENSMUSG00000111526 | 0 | NA | NA | NA | NA | NA | Gm47433 | predicted gene, 47433 [Source:MGI Symb | 0 | 0 | 0 | 0 | 0 | 0 |
| 49158 | ENSMUSG00000111528 | 0.133850373 | -0.796629176 | 4.080472857 | -0.195229623 | 0.84521319 | NA | Gm39460 | predicted gene, 39460 [Source:MGI Symb | 0.8031 | 0 | 0 | 0 | 0 | 0 |
| 49159 | ENSMUSG00000111529 | 0 | NA | NA | NA | NA | NA | Gm48141 | predicted gene, 48141 [Source:MGI Symb | 0 | 0 | 0 | 0 | 0 | 0 |
| 49160 | ENSMUSG00000111530 | 0 | NA | NA | NA | NA | NA | Gm9487 | predicted gene 9487 [Source:MGI Symbol | 0 | 0 | 0 | 0 | 0 | 0 |
| 49161 | ENSMUSG00000111531 | 0.772832414 | -0.17263844 | 2.923794788 | -0.059046018 | 0.952915455 | NA | Gm32688 | predicted gene, 32688 [Source:MGI Symb | 2.4093 | 0 | 0 | 0 | 0 | 2.2277 |
| 49162 | ENSMUSG00000111532 | 0 | NA | NA | NA | NA | NA | Gm36485 | predicted gene, 36485 [Source:MGI Symb | 0 | 0 | 0 | 0 | 0 | 0 |
| 49163 | ENSMUSG00000111533 | 0 | NA | NA | NA | NA | NA | Gm39458 | predicted gene, 39458 [Source:MGI Symb | 0 | 0 | 0 | 0 | 0 | 0 |
| 49164 | ENSMUSG00000111534 | 0 | NA | NA | NA | NA | NA | Gm48846 | predicted gene, 48846 [Source:MGI Symb | 0 | 0 | 0 | 0 | 0 | 0 |

| | | | | | | | | | | | | | | | |
|---|---|---|---|---|---|---|---|---|---|---|---|---|---|---|---|
| 49165 | ENSMUSG00000111535 | 0.469265235 | -2.274672822 | 3.439794491 | -0.661281605 | 0.508431736 | NA | Gm35154 | predicted gene, 35154 [Source:MGI Symb | 0.8031 | 0 | 2.0125 | 0 | 0 | 0 |
| 49166 | ENSMUSG00000111537 | 0 | NA | NA | NA | NA | NA | Gm7653 | predicted gene 7653 [Source:MGI Symbol | 0 | 0 | 0 | 0 | 0 | 0 |
| 49167 | ENSMUSG00000111539 | 3.483885131 | 0.266726755 | 1.239648916 | 0.215163142 | 0.829640127 | NA | Gm48372 | predicted gene, 48372 [Source:MGI Symb | 2.4093 | 4.1117 | 3.0187 | 4.8199 | 6.5437 | 0 |
| 49168 | ENSMUSG00000111540 | 0.17131964 | -0.796629176 | 4.080472857 | -0.195229623 | 0.84521319 | NA | 4930584E12 | RIKEN cDNA 4930584E12 gene [Source:M | 0 | 1.0279 | 0 | 0 | 0 | 0 |
| 49169 | ENSMUSG00000111541 | 0 | NA | NA | NA | NA | NA | Gm47219 | predicted gene, 47219 [Source:MGI Symb | 0 | 0 | 0 | 0 | 0 | 0 |
| 49170 | ENSMUSG00000111542 | 0 | NA | NA | NA | NA | NA | Gm30619 | predicted gene, 30619 [Source:MGI Symb | 0 | 0 | 0 | 0 | 0 | 0 |
| 49171 | ENSMUSG00000111543 | 0 | NA | NA | NA | NA | NA | Olfr1139-ps1 | olfactory receptor 1139, pseudogene 1 [S | 0 | 0 | 0 | 0 | 0 | 0 |
| 49172 | ENSMUSG00000111544 | 0 | NA | NA | NA | NA | NA | 4930534H03 | RIKEN cDNA 4930534H03 gene [Source:M | 0 | 0 | 0 | 0 | 0 | 0 |
| 49173 | ENSMUSG00000111545 | 0 | NA | NA | NA | NA | NA | Gm18015 | predicted gene, 18015 [Source:MGI Symb | 0 | 0 | 0 | 0 | 0 | 0 |
| 49174 | ENSMUSG00000111546 | 0 | NA | NA | NA | NA | NA | Gm47050 | predicted gene, 47050 [Source:MGI Symb | 0 | 0 | 0 | 0 | 0 | 0 |
| 49175 | ENSMUSG00000111547 | 3.214199087 | 2.544587397 | 1.40714747 | 1.808330293 | 0.07055511 | NA | Gm48069 | predicted gene, 48069 [Source:MGI Symb | 0.8031 | 2.0558 | 0 | 8.6758 | 2.1812 | 5.5692 |
| 49176 | ENSMUSG00000111548 | 0 | NA | NA | NA | NA | NA | Gm8162 | predicted gene 8162 [Source:MGI Symbol | 0 | 0 | 0 | 0 | 0 | 0 |
| 49177 | ENSMUSG00000111549 | 0 | NA | NA | NA | NA | NA | Gm48172 | predicted gene, 48172 [Source:MGI Symb | 0 | 0 | 0 | 0 | 0 | 0 |
| 49178 | ENSMUSG00000111550 | 0 | NA | NA | NA | NA | NA | Gm40623 | predicted gene, 40623 [Source:MGI Symb | 0 | 0 | 0 | 0 | 0 | 0 |
| 49179 | ENSMUSG00000111551 | 0 | NA | NA | NA | NA | NA | Gm48128 | predicted gene, 48128 [Source:MGI Symb | 0 | 0 | 0 | 0 | 0 | 0 |
| 49180 | ENSMUSG00000111552 | 0 | NA | NA | NA | NA | NA | Gm7570 | predicted gene 7570 [Source:MGI Symbol | 0 | 0 | 0 | 0 | 0 | 0 |
| 49181 | ENSMUSG00000111553 | 0 | NA | NA | NA | NA | NA | Gm18892 | predicted gene, 18892 [Source:MGI Symb | 0 | 0 | 0 | 0 | 0 | 0 |
| 49182 | ENSMUSG00000111554 | 0 | NA | NA | NA | NA | NA | Gm232 | predicted gene 232 [Source:MGI Symbol;/ | 0 | 0 | 0 | 0 | 0 | 0 |
| 49183 | ENSMUSG00000111555 | 0 | NA | NA | NA | NA | NA | Gm47326 | predicted gene, 47326 [Source:MGI Symb | 0 | 0 | 0 | 0 | 0 | 0 |
| 49184 | ENSMUSG00000111556 | 0 | NA | NA | NA | NA | NA | Gm19299 | predicted gene, 19299 [Source:MGI Symb | 0 | 0 | 0 | 0 | 0 | 0 |
| 49185 | ENSMUSG00000111557 | 0 | NA | NA | NA | NA | NA | Gm18164 | predicted gene, 18164 [Source:MGI Symb | 0 | 0 | 0 | 0 | 0 | 0 |
| 49186 | ENSMUSG00000111558 | 0 | NA | NA | NA | NA | NA | Gm18891 | predicted gene, 18891 [Source:MGI Symb | 0 | 0 | 0 | 0 | 0 | 0 |
| 49187 | ENSMUSG00000111560 | 0 | NA | NA | NA | NA | NA | Gm35262 | predicted gene, 35262 [Source:MGI Symb | 0 | 0 | 0 | 0 | 0 | 0 |
| 49188 | ENSMUSG00000111561 | 0 | NA | NA | NA | NA | NA | Gm47957 | predicted gene, 47957 [Source:MGI Symb | 0 | 0 | 0 | 0 | 0 | 0 |
| 49189 | ENSMUSG00000111562 | 0 | NA | NA | NA | NA | NA | Gm19987 | predicted gene, 19987 [Source:MGI Symb | 0 | 0 | 0 | 0 | 0 | 0 |
| 49190 | ENSMUSG00000111563 | 0 | NA | NA | NA | NA | NA | Gm47485 | predicted gene, 47485 [Source:MGI Symb | 0 | 0 | 0 | 0 | 0 | 0 |
| 49191 | ENSMUSG00000111564 | 0 | NA | NA | NA | NA | NA | Stmn1-rs1 | stathmin 1, related sequence 1 [Source:M | 0 | 0 | 0 | 0 | 0 | 0 |
| 49192 | ENSMUSG00000111565 | 0.133850373 | -0.796629176 | 4.080472857 | -0.195229623 | 0.84521319 | NA | Gm34481 | predicted gene, 34481 [Source:MGI Symb | 0.8031 | 0 | 0 | 0 | 0 | 0 |
| 49193 | ENSMUSG00000111566 | 0 | NA | NA | NA | NA | NA | Olfr1001-ps1 | olfactory receptor 1001, pseudogene 1 [S | 0 | 0 | 0 | 0 | 0 | 0 |
| 49194 | ENSMUSG00000111567 | 0 | NA | NA | NA | NA | NA | Olfr1251 | olfactory receptor 1251 [Source:MGI Syml | 0 | 0 | 0 | 0 | 0 | 0 |
| 49195 | ENSMUSG00000111568 | 0 | NA | NA | NA | NA | NA | Gm18488 | predicted gene, 18488 [Source:MGI Symb | 0 | 0 | 0 | 0 | 0 | 0 |
| 49196 | ENSMUSG00000111569 | 0 | NA | NA | NA | NA | NA | Gm47410 | predicted gene, 47410 [Source:MGI Symb | 0 | 0 | 0 | 0 | 0 | 0 |
| 49197 | ENSMUSG00000111570 | 0 | NA | NA | NA | NA | NA | Gm32335 | predicted gene, 32335 [Source:MGI Symb | 0 | 0 | 0 | 0 | 0 | 0 |
| 49198 | ENSMUSG00000111572 | 0 | NA | NA | NA | NA | NA | Gm3028 | predicted gene 3028 [Source:MGI Symbol | 0 | 0 | 0 | 0 | 0 | 0 |
| 49199 | ENSMUSG00000111573 | 0 | NA | NA | NA | NA | NA | Olfr962-ps1 | olfactory receptor 962, pseudogene 1 [So | 0 | 0 | 0 | 0 | 0 | 0 |
| 49200 | ENSMUSG00000111574 | 0 | NA | NA | NA | NA | NA | Gm46146 | predicted gene, 46146 [Source:MGI Symb | 0 | 0 | 0 | 0 | 0 | 0 |
| 49201 | ENSMUSG00000111575 | 0 | NA | NA | NA | NA | NA | Gm47926 | predicted gene, 47926 [Source:MGI Symb | 0 | 0 | 0 | 0 | 0 | 0 |
| 49202 | ENSMUSG00000111576 | 0 | NA | NA | NA | NA | NA | Gm32284 | predicted gene, 32284 [Source:MGI Symb | 0 | 0 | 0 | 0 | 0 | 0 |
| 49203 | ENSMUSG00000111577 | 0 | NA | NA | NA | NA | NA | Gm47136 | predicted gene, 47136 [Source:MGI Symb | 0 | 0 | 0 | 0 | 0 | 0 |
| 49204 | ENSMUSG00000111578 | 0 | NA | NA | NA | NA | NA | Gm18166 | predicted gene, 18166 [Source:MGI Symb | 0 | 0 | 0 | 0 | 0 | 0 |
| 49205 | ENSMUSG00000111580 | 33.96251136 | 1.068025614 | 0.57209628 | 1.866863415 | 0.061920664 | 0.355884025 | Gm19572 | predicted gene, 19572 [Source:MGI Symb | 45.777 | 9.2513 | 10.062 | 34.703 | 33.809 | 70.172 |
| 49206 | ENSMUSG00000111581 | 0 | NA | NA | NA | NA | NA | Gm47364 | predicted gene, 47364 [Source:MGI Symb | 0 | 0 | 0 | 0 | 0 | 0 |
| 49207 | ENSMUSG00000111582 | 0 | NA | NA | NA | NA | NA | Gm47347 | predicted gene, 47347 [Source:MGI Symb | 0 | 0 | 0 | 0 | 0 | 0 |
| 49208 | ENSMUSG00000111583 | 2.55193439 | -2.612226528 | 1.695980423 | -1.540245684 | 0.123500478 | NA | Gm47962 | predicted gene, 47962 [Source:MGI Symb | 8.031 | 1.0279 | 4.025 | 0 | 0 | 2.2277 |
| 49209 | ENSMUSG00000111585 | 0 | NA | NA | NA | NA | NA | Gm47355 | predicted gene, 47355 [Source:MGI Symb | 0 | 0 | 0 | 0 | 0 | 0 |
| 49210 | ENSMUSG00000111587 | 0 | NA | NA | NA | NA | NA | Gm32338 | predicted gene, 32338 [Source:MGI Symb | 0 | 0 | 0 | 0 | 0 | 0 |
| 49211 | ENSMUSG00000111589 | 0 | NA | NA | NA | NA | NA | Gm4042 | predicted gene 4042 [Source:MGI Symbol | 0 | 0 | 0 | 0 | 0 | 0 |
| 49212 | ENSMUSG00000111590 | 0 | NA | NA | NA | NA | NA | Olfr1192-ps1 | olfactory receptor 1192, pseudogene 1 [S | 0 | 0 | 0 | 0 | 0 | 0 |
| 49213 | ENSMUSG00000111591 | 0 | NA | NA | NA | NA | NA | Gm47083 | predicted gene, 47083 [Source:MGI Symb | 0 | 0 | 0 | 0 | 0 | 0 |
| 49214 | ENSMUSG00000111593 | 0 | NA | NA | NA | NA | NA | Gm48742 | predicted gene, 48742 [Source:MGI Symb | 0 | 0 | 0 | 0 | 0 | 0 |
| 49215 | ENSMUSG00000111594 | 5.686129384 | 0.36908374 | 0.920705813 | 0.400870435 | 0.688515518 | 0.936908509 | Gm3365 | predicted gene 3365 [Source:MGI Symbol | 5.6217 | 7.1954 | 2.0125 | 3.8559 | 7.6343 | 7.7969 |
| 49216 | ENSMUSG00000111596 | 0 | NA | NA | NA | NA | NA | Gm8058 | predicted gene 8058 [Source:MGI Symbol | 0 | 0 | 0 | 0 | 0 | 0 |
| 49217 | ENSMUSG00000111597 | 0 | NA | NA | NA | NA | NA | Olfr989-ps1 | olfactory receptor 989, pseudogene 1 [So | 0 | 0 | 0 | 0 | 0 | 0 |
| 49218 | ENSMUSG00000111598 | 0 | NA | NA | NA | NA | NA | Gm47095 | predicted gene, 47095 [Source:MGI Symb | 0 | 0 | 0 | 0 | 0 | 0 |
| 49219 | ENSMUSG00000111600 | 0 | NA | NA | NA | NA | NA | Gm8049 | predicted gene 8049 [Source:MGI Symbol | 0 | 0 | 0 | 0 | 0 | 0 |
| 49220 | ENSMUSG00000111601 | 0 | NA | NA | NA | NA | NA | Gm18986 | predicted gene, 18986 [Source:MGI Symb | 0 | 0 | 0 | 0 | 0 | 0 |

| | | | | | | | | | | | | | | | |
|---|---|---|---|---|---|---|---|---|---|---|---|---|---|---|---|
| 49221 | ENSMUSG00000111602 | 0 | NA | NA | NA | NA | NA | Gm32014 | predicted gene, 32014 [Source:MGI Symb | 0 | 0 | 0 | 0 | 0 | 0 |
| 49222 | ENSMUSG00000111604 | 0 | NA | NA | NA | NA | NA | Gm18043 | predicted gene, 18043 [Source:MGI Symb | 0 | 0 | 0 | 0 | 0 | 0 |
| 49223 | ENSMUSG00000111605 | 0.931486014 | -2.031896103 | 2.484683032 | -0.817768736 | 0.413489253 | NA | 4632418H02 | RIKEN cDNA 4632418H02 gene [Source:M | 1.6062 | 0 | 3.0187 | 0.964 | 0 | 0 |
| 49224 | ENSMUSG00000111606 | 0 | NA | NA | NA | NA | NA | Gm7213 | predicted gene 7213 [Source:MGI Symbol | 0 | 0 | 0 | 0 | 0 | 0 |
| 49225 | ENSMUSG00000111608 | 5.849305592 | -0.890033923 | 0.908178854 | -0.980020531 | 0.327075984 | 0.772772049 | Gm48839 | predicted gene, 48839 [Source:MGI Symb | 9.6372 | 6.1675 | 7.0437 | 6.7478 | 3.2719 | 2.2277 |
| 49226 | ENSMUSG00000111609 | 0 | NA | NA | NA | NA | NA | Gm6713 | predicted gene 6713 [Source:MGI Symbol | 0 | 0 | 0 | 0 | 0 | 0 |
| 49227 | ENSMUSG00000111610 | 0 | NA | NA | NA | NA | NA | Gm34829 | predicted gene, 34829 [Source:MGI Symb | 0 | 0 | 0 | 0 | 0 | 0 |
| 49228 | ENSMUSG00000111611 | 0 | NA | NA | NA | NA | NA | Olfr157 | olfactory receptor 157 [Source:MGI Symb | 0 | 0 | 0 | 0 | 0 | 0 |
| 49229 | ENSMUSG00000111612 | 0 | NA | NA | NA | NA | NA | Gm47090 | predicted gene, 47090 [Source:MGI Symb | 0 | 0 | 0 | 0 | 0 | 0 |
| 49230 | ENSMUSG00000111613 | 0.133850373 | -0.796629176 | 4.080472857 | -0.195229623 | 0.84521319 | NA | Gm4977 | predicted gene 4977 [Source:MGI Symbol | 0.8031 | 0 | 0 | 0 | 0 | 0 |
| 49231 | ENSMUSG00000111615 | 0 | NA | NA | NA | NA | NA | Gm47741 | predicted gene, 47741 [Source:MGI Symb | 0 | 0 | 0 | 0 | 0 | 0 |
| 49232 | ENSMUSG00000111616 | 0 | NA | NA | NA | NA | NA | Gm18487 | predicted gene, 18487 [Source:MGI Symb | 0 | 0 | 0 | 0 | 0 | 0 |
| 49233 | ENSMUSG00000111617 | 0.133850373 | -0.796629176 | 4.080472857 | -0.195229623 | 0.84521319 | NA | Gm47148 | predicted gene, 47148 [Source:MGI Symb | 0.8031 | 0 | 0 | 0 | 0 | 0 |
| 49234 | ENSMUSG00000111618 | 9.858287754 | -0.381334025 | 0.71758071 | -0.531416214 | 0.595130388 | 0.913525602 | Gm47268 | predicted gene, 47268 [Source:MGI Symb | 12.047 | 7.1954 | 14.087 | 4.8199 | 7.6343 | 13.366 |
| 49235 | ENSMUSG00000111619 | 0 | NA | NA | NA | NA | NA | Gm48348 | predicted gene, 48348 [Source:MGI Symb | 0 | 0 | 0 | 0 | 0 | 0 |
| 49236 | ENSMUSG00000111620 | 2.319971362 | 1.305002039 | 1.593033703 | 0.819192988 | 0.412676315 | NA | Gm31848 | predicted gene, 31848 [Source:MGI Symb | 0 | 2.0558 | 2.0125 | 0.964 | 1.0906 | 7.7969 |
| 49237 | ENSMUSG00000111621 | 0 | NA | NA | NA | NA | NA | Gm7802 | predicted gene 7802 [Source:MGI Symbol | 0 | 0 | 0 | 0 | 0 | 0 |
| 49238 | ENSMUSG00000111622 | 0 | NA | NA | NA | NA | NA | Gm21041 | predicted gene, 21041 [Source:MGI Symb | 0 | 0 | 0 | 0 | 0 | 0 |
| 49239 | ENSMUSG00000111623 | 1.545645367 | 1.075941953 | 1.896485735 | 0.567334588 | 0.570486882 | NA | Gm48303 | predicted gene, 48303 [Source:MGI Symb | 0 | 1.0279 | 2.0125 | 2.8919 | 0 | 3.3415 |
| 49240 | ENSMUSG00000111625 | 0 | NA | NA | NA | NA | NA | Gm48097 | predicted gene, 48097 [Source:MGI Symb | 0 | 0 | 0 | 0 | 0 | 0 |
| 49241 | ENSMUSG00000111626 | 0 | NA | NA | NA | NA | NA | Gm48738 | predicted gene, 48738 [Source:MGI Symb | 0 | 0 | 0 | 0 | 0 | 0 |
| 49242 | ENSMUSG00000111627 | 0 | NA | NA | NA | NA | NA | Gm48749 | predicted gene, 48749 [Source:MGI Symb | 0 | 0 | 0 | 0 | 0 | 0 |
| 49243 | ENSMUSG00000111629 | 0.620790101 | -1.233953725 | 2.860353486 | -0.431399032 | 0.666178254 | NA | 4930442G15 | RIKEN cDNA 4930442G15 gene [Source:M | 1.6062 | 1.0279 | 0 | 0 | 1.0906 | 0 |
| 49244 | ENSMUSG00000111630 | 0 | NA | NA | NA | NA | NA | Gm47165 | predicted gene, 47165 [Source:MGI Symb | 0 | 0 | 0 | 0 | 0 | 0 |
| 49245 | ENSMUSG00000111631 | 1.243673853 | -0.5525115 | 1.901396787 | -0.290581905 | 0.771371101 | NA | Gm32017 | predicted gene, 32017 [Source:MGI Symb | 2.4093 | 1.0279 | 1.0062 | 1.928 | 1.0906 | 0 |
| 49246 | ENSMUSG00000111632 | 0 | NA | NA | NA | NA | NA | Gm47327 | predicted gene, 47327 [Source:MGI Symb | 0 | 0 | 0 | 0 | 0 | 0 |
| 49247 | ENSMUSG00000111633 | 0 | NA | NA | NA | NA | NA | Gm39377 | predicted gene, 39377 [Source:MGI Symb | 0 | 0 | 0 | 0 | 0 | 0 |
| 49248 | ENSMUSG00000111635 | 0 | NA | NA | NA | NA | NA | Gm47240 | predicted gene, 47240 [Source:MGI Symb | 0 | 0 | 0 | 0 | 0 | 0 |
| 49249 | ENSMUSG00000111637 | 0 | NA | NA | NA | NA | NA | Gm10688 | predicted gene 10688 [Source:MGI Symbo | 0 | 0 | 0 | 0 | 0 | 0 |
| 49250 | ENSMUSG00000111638 | 0 | NA | NA | NA | NA | NA | Gm47501 | predicted gene, 47501 [Source:MGI Symb | 0 | 0 | 0 | 0 | 0 | 0 |
| 49251 | ENSMUSG00000111639 | 0.710308366 | 0.13927508 | 2.769785472 | 0.050283707 | 0.959896308 | NA | 1700019L13I | RIKEN cDNA 1700019L13 gene [Source:M | 0 | 1.0279 | 1.0062 | 0 | 0 | 2.2277 |
| 49252 | ENSMUSG00000111641 | 0 | NA | NA | NA | NA | NA | Gm47388 | predicted gene, 47388 [Source:MGI Symb | 0 | 0 | 0 | 0 | 0 | 0 |
| 49253 | ENSMUSG00000111643 | 0 | NA | NA | NA | NA | NA | Gm48628 | predicted gene, 48628 [Source:MGI Symb | 0 | 0 | 0 | 0 | 0 | 0 |
| 49254 | ENSMUSG00000111645 | 0 | NA | NA | NA | NA | NA | Gm32171 | predicted gene, 32171 [Source:MGI Symb | 0 | 0 | 0 | 0 | 0 | 0 |
| 49255 | ENSMUSG00000111646 | 0 | NA | NA | NA | NA | NA | Gm40621 | predicted gene, 40621 [Source:MGI Symb | 0 | 0 | 0 | 0 | 0 | 0 |
| 49256 | ENSMUSG00000111648 | 0 | NA | NA | NA | NA | NA | Gm46103 | predicted gene, 46103 [Source:MGI Symb | 0 | 0 | 0 | 0 | 0 | 0 |
| 49257 | ENSMUSG00000111649 | 0 | NA | NA | NA | NA | NA | Gm48945 | predicted gene, 48945 [Source:MGI Symb | 0 | 0 | 0 | 0 | 0 | 0 |
| 49258 | ENSMUSG00000111651 | 0 | NA | NA | NA | NA | NA | Gm47199 | predicted gene, 47199 [Source:MGI Symb | 0 | 0 | 0 | 0 | 0 | 0 |
| 49259 | ENSMUSG00000111652 | 0 | NA | NA | NA | NA | NA | Gm48726 | predicted gene, 48726 [Source:MGI Symb | 0 | 0 | 0 | 0 | 0 | 0 |
| 49260 | ENSMUSG00000111654 | 4.447910723 | -0.383647081 | 1.0355283 | -0.3704844 | 0.711021598 | NA | Gm47077 | predicted gene, 47077 [Source:MGI Symb | 4.0155 | 2.0558 | 9.0562 | 3.8559 | 4.3625 | 3.3415 |
| 49261 | ENSMUSG00000111655 | 0 | NA | NA | NA | NA | NA | 4930434F21 | RIKEN cDNA 4930434F21 gene [Source:M | 0 | 0 | 0 | 0 | 0 | 0 |
| 49262 | ENSMUSG00000111656 | 1.714923488 | 1.405597587 | 1.715616761 | 0.819295789 | 0.412617675 | NA | Gm47232 | predicted gene, 47232 [Source:MGI Symb | 0.8031 | 0 | 2.0125 | 1.928 | 1.0906 | 4.4554 |
| 49263 | ENSMUSG00000111657 | 0 | NA | NA | NA | NA | NA | Gm31592 | predicted gene, 31592 [Source:MGI Symb | 0 | 0 | 0 | 0 | 0 | 0 |
| 49264 | ENSMUSG00000111658 | 0 | NA | NA | NA | NA | NA | B930092H01 | RIKEN cDNA B930092H01 gene [Source:M | 0 | 0 | 0 | 0 | 0 | 0 |
| 49265 | ENSMUSG00000111660 | 0 | NA | NA | NA | NA | NA | Olfr964-ps1 | olfactory receptor 964, pseudogene 1 [Sou | 0 | 0 | 0 | 0 | 0 | 0 |
| 49266 | ENSMUSG00000111661 | 0 | NA | NA | NA | NA | NA | Gm46142 | predicted gene, 46142 [Source:MGI Symb | 0 | 0 | 0 | 0 | 0 | 0 |
| 49267 | ENSMUSG00000111662 | 1.604298483 | 0.823063026 | 1.71362948 | 0.480303961 | 0.631011273 | NA | Gm7435 | predicted gene 7435 [Source:MGI Symbol | 2.4093 | 0 | 1.0062 | 2.8919 | 1.0906 | 2.2277 |
| 49268 | ENSMUSG00000111663 | 0 | NA | NA | NA | NA | NA | Gm48833 | predicted gene, 48833 [Source:MGI Symb | 0 | 0 | 0 | 0 | 0 | 0 |
| 49269 | ENSMUSG00000111664 | 0.185640647 | 1.126924175 | 4.080472857 | 0.276174898 | 0.782413736 | NA | Gm48865 | predicted gene, 48865 [Source:MGI Symb | 0 | 0 | 0 | 0 | 0 | 1.1138 |
| 49270 | ENSMUSG00000111665 | 0.837114282 | 0.78811204 | 2.395742738 | 0.328963552 | 0.742183238 | NA | A930033M1 | RIKEN cDNA A930033M14 gene [Source:N | 0.8031 | 1.0279 | 0 | 0.964 | 0 | 2.2277 |
| 49271 | ENSMUSG00000111666 | 7.289214665 | 0.236661763 | 0.815797981 | 0.290098491 | 0.77174089 | 0.957083104 | Gm30849 | predicted gene, 30849 [Source:MGI Symb | 4.8186 | 8.2233 | 7.0437 | 3.8559 | 11.997 | 7.7969 |
| 49272 | ENSMUSG00000111667 | 0 | NA | NA | NA | NA | NA | Gm47740 | predicted gene, 47740 [Source:MGI Symb | 0 | 0 | 0 | 0 | 0 | 0 |
| 49273 | ENSMUSG00000111668 | 0 | NA | NA | NA | NA | NA | Gm47213 | predicted gene, 47213 [Source:MGI Symb | 0 | 0 | 0 | 0 | 0 | 0 |
| 49274 | ENSMUSG00000111669 | 0 | NA | NA | NA | NA | NA | Gm31698 | predicted gene, 31698 [Source:MGI Symb | 0 | 0 | 0 | 0 | 0 | 0 |
| 49275 | ENSMUSG00000111670 | 0 | NA | NA | NA | NA | NA | Gm47500 | predicted gene, 47500 [Source:MGI Symb | 0 | 0 | 0 | 0 | 0 | 0 |
| 49276 | ENSMUSG00000111671 | 0 | NA | NA | NA | NA | NA | Gm6983 | predicted gene 6983 [Source:MGI Symbol | 0 | 0 | 0 | 0 | 0 | 0 |

| | | | | | | | | | | | | | | | |
|---|---|---|---|---|---|---|---|---|---|---|---|---|---|---|---|
| 49277 | ENSMUSG00000111672 | 0 | NA | NA | NA | NA | NA | Gm47934 | predicted gene, 47934 [Source:MGI Symb | 0 | 0 | 0 | 0 | 0 | 0 |
| 49278 | ENSMUSG00000111673 | 5.544413406 | 0.982748524 | 0.930409372 | 1.056253896 | 0.290852233 | 0.740649067 | Gm29724 | predicted gene, 29724 [Source:MGI Symb | 4.0155 | 5.1396 | 2.0125 | 7.7118 | 4.3625 | 10.025 |
| 49279 | ENSMUSG00000111674 | 2.555081285 | -1.411535129 | 1.443134453 | -0.978103686 | 0.328023045 | NA | Gm48728 | predicted gene, 48728 [Source:MGI Symb | 8.031 | 1.0279 | 2.0125 | 0.964 | 2.1812 | 1.1138 |
| 49280 | ENSMUSG00000111678 | 0 | NA | NA | NA | NA | NA | Olfr1077-ps1 | olfactory receptor 1077, pseudogene 1 [S | 0 | 0 | 0 | 0 | 0 | 0 |
| 49281 | ENSMUSG00000111679 | 0 | NA | NA | NA | NA | NA | Gm47063 | predicted gene, 47063 [Source:MGI Symb | 0 | 0 | 0 | 0 | 0 | 0 |
| 49282 | ENSMUSG00000111680 | 0 | NA | NA | NA | NA | NA | Gm18377 | predicted gene, 18377 [Source:MGI Symb | 0 | 0 | 0 | 0 | 0 | 0 |
| 49283 | ENSMUSG00000111681 | 21.80809654 | 0.799176177 | 0.508692453 | 1.571039972 | 0.116173364 | 0.500730933 | Gm47640 | predicted gene, 47640 [Source:MGI Symb | 15.259 | 15.419 | 17.106 | 26.027 | 41.443 | 15.594 |
| 49284 | ENSMUSG00000111682 | 0 | NA | NA | NA | NA | NA | Gm47543 | predicted gene, 47543 [Source:MGI Symb | 0 | 0 | 0 | 0 | 0 | 0 |
| 49285 | ENSMUSG00000111683 | 0 | NA | NA | NA | NA | NA | Gm49367 | predicted gene, 49367 [Source:MGI Symb | 0 | 0 | 0 | 0 | 0 | 0 |
| 49286 | ENSMUSG00000111684 | 0 | NA | NA | NA | NA | NA | Gm8543 | predicted gene 8543 [Source:MGI Symbol | 0 | 0 | 0 | 0 | 0 | 0 |
| 49287 | ENSMUSG00000111685 | 0.890584499 | -0.80930233 | 2.520154863 | -0.321131983 | 0.748110374 | NA | Gm10686 | predicted gene 10686 [Source:MGI Symbo | 2.4093 | 0 | 1.0062 | 1.928 | 0 | 0 |
| 49288 | ENSMUSG00000111686 | 0.904673413 | -3.227537609 | 2.678727401 | -1.204877214 | 0.228250706 | NA | Rab7-ps1 | RAB7, member RAS oncogene family, pseu | 2.4093 | 0 | 3.0187 | 0 | 0 | 0 |
| 49289 | ENSMUSG00000111687 | 0 | NA | NA | NA | NA | NA | Olfr956-ps1 | olfactory receptor 956, pseudogene 1 [So | 0 | 0 | 0 | 0 | 0 | 0 |
| 49290 | ENSMUSG00000111689 | 0 | NA | NA | NA | NA | NA | Olfr1082 | olfactory receptor 1082 [Source:MGI Sym | 0 | 0 | 0 | 0 | 0 | 0 |
| 49291 | ENSMUSG00000111690 | 0 | NA | NA | NA | NA | NA | Gm48720 | predicted gene, 48720 [Source:MGI Symb | 0 | 0 | 0 | 0 | 0 | 0 |
| 49292 | ENSMUSG00000111691 | 0 | NA | NA | NA | NA | NA | Gm47002 | predicted gene, 47002 [Source:MGI Symb | 0 | 0 | 0 | 0 | 0 | 0 |
| 49293 | ENSMUSG00000111692 | 0.185640647 | 1.126924175 | 4.080472857 | 0.276174898 | 0.782413736 | NA | Gm49373 | predicted gene, 49373 [Source:MGI Symb | 0 | 0 | 0 | 0 | 0 | 1.1138 |
| 49294 | ENSMUSG00000111693 | 0 | NA | NA | NA | NA | NA | Gm47986 | predicted gene, 47986 [Source:MGI Symb | 0 | 0 | 0 | 0 | 0 | 0 |
| 49295 | ENSMUSG00000111694 | 0 | NA | NA | NA | NA | NA | Gm48901 | predicted gene, 48901 [Source:MGI Symb | 0 | 0 | 0 | 0 | 0 | 0 |
| 49296 | ENSMUSG00000111695 | 0 | NA | NA | NA | NA | NA | Gm47565 | predicted gene, 47565 [Source:MGI Symb | 0 | 0 | 0 | 0 | 0 | 0 |
| 49297 | ENSMUSG00000111696 | 0 | NA | NA | NA | NA | NA | Gm47932 | predicted gene, 47932 [Source:MGI Symb | 0 | 0 | 0 | 0 | 0 | 0 |
| 49298 | ENSMUSG00000111697 | 0 | NA | NA | NA | NA | NA | Gm35548 | predicted gene, 35548 [Source:MGI Symb | 0 | 0 | 0 | 0 | 0 | 0 |
| 49299 | ENSMUSG00000111698 | 0 | NA | NA | NA | NA | NA | Gm47105 | predicted gene, 47105 [Source:MGI Symb | 0 | 0 | 0 | 0 | 0 | 0 |
| 49300 | ENSMUSG00000111699 | 0 | NA | NA | NA | NA | NA | C230099D08 | RIKEN cDNA C230099D08 gene [Source:M | 0 | 0 | 0 | 0 | 0 | 0 |
| 49301 | ENSMUSG00000111700 | 0 | NA | NA | NA | NA | NA | Gm49385 | predicted gene, 49385 [Source:MGI Symb | 0 | 0 | 0 | 0 | 0 | 0 |
| 49302 | ENSMUSG00000111702 | 0 | NA | NA | NA | NA | NA | Gm47945 | predicted gene, 47945 [Source:MGI Symb | 0 | 0 | 0 | 0 | 0 | 0 |
| 49303 | ENSMUSG00000111703 | 0 | NA | NA | NA | NA | NA | Gm47161 | predicted gene, 47161 [Source:MGI Symb | 0 | 0 | 0 | 0 | 0 | 0 |
| 49304 | ENSMUSG00000111704 | 0 | NA | NA | NA | NA | NA | Gm47379 | predicted gene, 47379 [Source:MGI Symb | 0 | 0 | 0 | 0 | 0 | 0 |
| 49305 | ENSMUSG00000111707 | 0 | NA | NA | NA | NA | NA | Gm9662 | predicted gene 9662 [Source:MGI Symbol | 0 | 0 | 0 | 0 | 0 | 0 |
| 49306 | ENSMUSG00000111708 | 0 | NA | NA | NA | NA | NA | Olfr929-ps1 | olfactory receptor 929, pseudogene 1 [So | 0 | 0 | 0 | 0 | 0 | 0 |
| 49307 | ENSMUSG00000111709 | 6.984572948 | -1.837849172 | 1.009490719 | -1.820570647 | 0.068672148 | 0.377670704 | Gm3776 | predicted gene 3776 [Source:MGI Symbol | 17.668 | 2.0558 | 13.081 | 5.7839 | 1.0906 | 2.2277 |
| 49308 | ENSMUSG00000111710 | 0 | NA | NA | NA | NA | NA | Gm47145 | predicted gene, 47145 [Source:MGI Symb | 0 | 0 | 0 | 0 | 0 | 0 |
| 49309 | ENSMUSG00000111711 | 0 | NA | NA | NA | NA | NA | Olfr1089 | olfactory receptor 1089 [Source:MGI Sym | 0 | 0 | 0 | 0 | 0 | 0 |
| 49310 | ENSMUSG00000111712 | 0.449470461 | -0.545136533 | 3.562105917 | -0.15303771 | 0.878368537 | NA | Gm39363 | predicted gene, 39363 [Source:MGI Symb | 1.6062 | 0 | 0 | 0 | 1.0906 | 0 |
| 49311 | ENSMUSG00000111713 | 0 | NA | NA | NA | NA | NA | Gm20234 | predicted gene, 20234 [Source:MGI Symb | 0 | 0 | 0 | 0 | 0 | 0 |
| 49312 | ENSMUSG00000111715 | 0 | NA | NA | NA | NA | NA | Olfr1246 | olfactory receptor 1246 [Source:MGI Sym | 0 | 0 | 0 | 0 | 0 | 0 |
| 49313 | ENSMUSG00000111716 | 0 | NA | NA | NA | NA | NA | Gm48554 | predicted gene, 48554 [Source:MGI Symb | 0 | 0 | 0 | 0 | 0 | 0 |
| 49314 | ENSMUSG00000111717 | 0 | NA | NA | NA | NA | NA | Gm39380 | predicted gene, 39380 [Source:MGI Symb | 0 | 0 | 0 | 0 | 0 | 0 |
| 49315 | ENSMUSG00000111719 | 0 | NA | NA | NA | NA | NA | Gm19137 | predicted gene, 19137 [Source:MGI Symb | 0 | 0 | 0 | 0 | 0 | 0 |
| 49316 | ENSMUSG00000111720 | 0 | NA | NA | NA | NA | NA | Gm39459 | predicted gene, 39459 [Source:MGI Symb | 0 | 0 | 0 | 0 | 0 | 0 |
| 49317 | ENSMUSG00000111722 | 1.55108699 | 2.025326771 | 1.985529864 | 1.02004347 | 0.307707845 | NA | Gm47270 | predicted gene, 47270 [Source:MGI Symb | 0.8031 | 0 | 1.0062 | 1.928 | 0 | 5.5692 |
| 49318 | ENSMUSG00000111725 | 0 | NA | NA | NA | NA | NA | Gm19219 | predicted gene, 19219 [Source:MGI Symb | 0 | 0 | 0 | 0 | 0 | 0 |
| 49319 | ENSMUSG00000111726 | 0 | NA | NA | NA | NA | NA | Gm40513 | predicted gene, 40513 [Source:MGI Symb | 0 | 0 | 0 | 0 | 0 | 0 |
| 49320 | ENSMUSG00000111728 | 0 | NA | NA | NA | NA | NA | Gm48850 | predicted gene, 48850 [Source:MGI Symb | 0 | 0 | 0 | 0 | 0 | 0 |
| 49321 | ENSMUSG00000111729 | 0 | NA | NA | NA | NA | NA | Gm48038 | predicted gene, 48038 [Source:MGI Symb | 0 | 0 | 0 | 0 | 0 | 0 |
| 49322 | ENSMUSG00000111730 | 0 | NA | NA | NA | NA | NA | Gm5950 | predicted gene 5950 [Source:MGI Symbol | 0 | 0 | 0 | 0 | 0 | 0 |
| 49323 | ENSMUSG00000111731 | 1.987807134 | 1.355970449 | 1.74858645 | 0.775466634 | 0.438063974 | NA | Gm48727 | predicted gene, 48727 [Source:MGI Symb | 3.2124 | 0 | 0 | 0.964 | 2.1812 | 5.5692 |
| 49324 | ENSMUSG00000111732 | 0 | NA | NA | NA | NA | NA | Olfr785 | olfactory receptor 785 [Source:MGI Symb | 0 | 0 | 0 | 0 | 0 | 0 |
| 49325 | ENSMUSG00000111733 | 0.367410362 | 2.078255025 | 3.800070157 | 0.54689912 | 0.58444804 | NA | Gm47181 | predicted gene, 47181 [Source:MGI Symb | 0 | 0 | 0 | 0 | 1.0906 | 1.1138 |
| 49326 | ENSMUSG00000111734 | 0 | NA | NA | NA | NA | NA | Gm29825 | predicted gene, 29825 [Source:MGI Symb | 0 | 0 | 0 | 0 | 0 | 0 |
| 49327 | ENSMUSG00000111736 | 0 | NA | NA | NA | NA | NA | Olfr1070-ps1 | olfactory receptor 1070, pseudogene 1 [S | 0 | 0 | 0 | 0 | 0 | 0 |
| 49328 | ENSMUSG00000111737 | 0 | NA | NA | NA | NA | NA | Gm47248 | predicted gene, 47248 [Source:MGI Symb | 0 | 0 | 0 | 0 | 0 | 0 |
| 49329 | ENSMUSG00000111738 | 2.673898395 | 2.144176803 | 1.651774688 | 1.298104892 | 0.194251294 | NA | Gm18329 | predicted gene, 18329 [Source:MGI Symb | 0 | 2.0558 | 1.0062 | 9.6398 | 0 | 3.3415 |
| 49330 | ENSMUSG00000111739 | 0 | NA | NA | NA | NA | NA | Gm47241 | predicted gene, 47241 [Source:MGI Symb | 0 | 0 | 0 | 0 | 0 | 0 |
| 49331 | ENSMUSG00000111740 | 9.910815826 | 0.278928874 | 0.73945916 | 0.377206598 | 0.706020077 | 0.94295036 | Gm49783 | predicted gene, 49783 [Source:MGI Symb | 9.6372 | 11.307 | 6.0375 | 19.28 | 7.6343 | 5.5692 |
| 49332 | ENSMUSG00000111741 | 0 | NA | NA | NA | NA | NA | Gm18903 | predicted gene, 18903 [Source:MGI Symb | 0 | 0 | 0 | 0 | 0 | 0 |

| | | | | | | | | | | | | | | | |
|---|---|---|---|---|---|---|---|---|---|---|---|---|---|---|---|
| 49333 | ENSMUSG00000111743 | 0 | NA | NA | NA | NA | NA | Gm48856 | predicted gene, 48856 [Source:MGI Symb | 0 | 0 | 0 | 0 | 0 | 0 |
| 49334 | ENSMUSG00000111744 | 0 | NA | NA | NA | NA | NA | Gm7780 | predicted gene 7780 [Source:MGI Symbol | 0 | 0 | 0 | 0 | 0 | 0 |
| 49335 | ENSMUSG00000111746 | 0 | NA | NA | NA | NA | NA | 4933422A05 | RIKEN cDNA 4933422A05 gene [Source:M | 0 | 0 | 0 | 0 | 0 | 0 |
| 49336 | ENSMUSG00000111747 | 0 | NA | NA | NA | NA | NA | Olfr1134 | olfactory receptor 1134 [Source:MGI Syml | 0 | 0 | 0 | 0 | 0 | 0 |
| 49337 | ENSMUSG00000111748 | 0.167707431 | -0.796629176 | 4.080472857 | -0.195229623 | 0.84521319 | NA | Gm34885 | predicted gene, 34885 [Source:MGI Symb | 0 | 0 | 1.0062 | 0 | 0 | 0 |
| 49338 | ENSMUSG00000111749 | 0 | NA | NA | NA | NA | NA | Gm47927 | predicted gene, 47927 [Source:MGI Symb | 0 | 0 | 0 | 0 | 0 | 0 |
| 49339 | ENSMUSG00000111750 | 0 | NA | NA | NA | NA | NA | Gm47886 | predicted gene, 47886 [Source:MGI Symb | 0 | 0 | 0 | 0 | 0 | 0 |
| 49340 | ENSMUSG00000111751 | 0 | NA | NA | NA | NA | NA | Gm48256 | predicted gene, 48256 [Source:MGI Symb | 0 | 0 | 0 | 0 | 0 | 0 |
| 49341 | ENSMUSG00000111752 | 0 | NA | NA | NA | NA | NA | Gm38575 | predicted gene, 38575 [Source:MGI Symb | 0 | 0 | 0 | 0 | 0 | 0 |
| 49342 | ENSMUSG00000111753 | 0 | NA | NA | NA | NA | NA | 1700054O05 | RIKEN cDNA 1700054O05 gene [Source:M | 0 | 0 | 0 | 0 | 0 | 0 |
| 49343 | ENSMUSG00000111754 | 0 | NA | NA | NA | NA | NA | Gm19994 | predicted gene, 19994 [Source:MGI Symb | 0 | 0 | 0 | 0 | 0 | 0 |
| 49344 | ENSMUSG00000111755 | 0 | NA | NA | NA | NA | NA | Gm47160 | predicted gene, 47160 [Source:MGI Symb | 0 | 0 | 0 | 0 | 0 | 0 |
| 49345 | ENSMUSG00000111756 | 0.133850373 | -0.796629176 | 4.080472857 | -0.195229623 | 0.84521319 | NA | Gm48183 | predicted gene, 48183 [Source:MGI Symb | 0.8031 | 0 | 0 | 0 | 0 | 0 |
| 49346 | ENSMUSG00000111757 | 0.294513347 | 0.165143496 | 4.080472857 | 0.040471657 | 0.967717103 | NA | Gm46124 | predicted gene, 46124 [Source:MGI Symb | 0.8031 | 0 | 0 | 0.964 | 0 | 0 |
| 49347 | ENSMUSG00000111758 | 0 | NA | NA | NA | NA | NA | Gm30373 | predicted gene, 30373 [Source:MGI Symb | 0 | 0 | 0 | 0 | 0 | 0 |
| 49348 | ENSMUSG00000111759 | 0 | NA | NA | NA | NA | NA | Gm4800 | predicted gene 4800 [Source:MGI Symbol | 0 | 0 | 0 | 0 | 0 | 0 |
| 49349 | ENSMUSG00000111760 | 0 | NA | NA | NA | NA | NA | Gm48337 | predicted gene, 48337 [Source:MGI Symb | 0 | 0 | 0 | 0 | 0 | 0 |
| 49350 | ENSMUSG00000111761 | 0.160662974 | 1.126924175 | 4.080472857 | 0.276174898 | 0.782413736 | NA | 9230112J17I | RIKEN cDNA 9230112J17 gene [Source:M( | 0 | 0 | 0 | 0.964 | 0 | 0 |
| 49351 | ENSMUSG00000111762 | 0 | NA | NA | NA | NA | NA | Olfr946-ps1 | olfactory receptor 946, pseudogene 1 [So | 0 | 0 | 0 | 0 | 0 | 0 |
| 49352 | ENSMUSG00000111763 | 0 | NA | NA | NA | NA | NA | Gm34973 | predicted gene, 34973 [Source:MGI Symb | 0 | 0 | 0 | 0 | 0 | 0 |
| 49353 | ENSMUSG00000111764 | 0 | NA | NA | NA | NA | NA | Gm47616 | predicted gene, 47616 [Source:MGI Symb | 0 | 0 | 0 | 0 | 0 | 0 |
| 49354 | ENSMUSG00000111765 | 0 | NA | NA | NA | NA | NA | Gm10635 | predicted gene 10635 [Source:MGI Symbo | 0 | 0 | 0 | 0 | 0 | 0 |
| 49355 | ENSMUSG00000111766 | 0 | NA | NA | NA | NA | NA | Gm47873 | predicted gene, 47873 [Source:MGI Symb | 0 | 0 | 0 | 0 | 0 | 0 |
| 49356 | ENSMUSG00000111767 | 0.906140364 | 2.113800522 | 2.518615882 | 0.839270703 | 0.401317419 | NA | 4930546K05 | RIKEN cDNA 4930546K05 gene [Source:M | 0 | 1.0279 | 0 | 0 | 2.1812 | 2.2277 |
| 49357 | ENSMUSG00000111769 | 1.357472383 | -0.566420676 | 1.86742493 | -0.303316437 | 0.761648721 | NA | Gm36278 | predicted gene, 36278 [Source:MGI Symb | 0.8031 | 1.0279 | 3.0187 | 0 | 2.1812 | 1.1138 |
| 49358 | ENSMUSG00000111770 | 0 | NA | NA | NA | NA | NA | Gm48811 | predicted gene, 48811 [Source:MGI Symb | 0 | 0 | 0 | 0 | 0 | 0 |
| 49359 | ENSMUSG00000111771 | 63.96324856 | -0.695052784 | 0.318415792 | -2.182846459 | 0.029047119 | 0.223715404 | Gm47465 | predicted gene, 47465 [Source:MGI Symb | 70.673 | 98.68 | 68.425 | 65.55 | 40.353 | 40.098 |
| 49360 | ENSMUSG00000111772 | 0 | NA | NA | NA | NA | NA | Olfr228 | olfactory receptor 228 [Source:MGI Symb | 0 | 0 | 0 | 0 | 0 | 0 |
| 49361 | ENSMUSG00000111774 | 0 | NA | NA | NA | NA | NA | Gm38398 | predicted gene, 38398 [Source:MGI Symb | 0 | 0 | 0 | 0 | 0 | 0 |
| 49362 | ENSMUSG00000111776 | 0 | NA | NA | NA | NA | NA | Gm47124 | predicted gene, 47124 [Source:MGI Symb | 0 | 0 | 0 | 0 | 0 | 0 |
| 49363 | ENSMUSG00000111777 | 0.744190399 | -2.947713985 | 2.874421214 | -1.025498271 | 0.305128141 | NA | Gm47147 | predicted gene, 47147 [Source:MGI Symb | 2.4093 | 2.0558 | 0 | 0 | 0 | 0 |
| 49364 | ENSMUSG00000111778 | 0 | NA | NA | NA | NA | NA | Gm30676 | predicted gene, 30676 [Source:MGI Symb | 0 | 0 | 0 | 0 | 0 | 0 |
| 49365 | ENSMUSG00000111780 | 0 | NA | NA | NA | NA | NA | Gm47960 | predicted gene, 47960 [Source:MGI Symb | 0 | 0 | 0 | 0 | 0 | 0 |
| 49366 | ENSMUSG00000111781 | 0 | NA | NA | NA | NA | NA | | olfactory receptor 939, pseudogene 1 | 0 | 0 | 0 | 0 | 0 | 0 |
| 49367 | ENSMUSG00000111782 | 0 | NA | NA | NA | NA | NA | Gm47091 | predicted gene, 47091 [Source:MGI Symb | 0 | 0 | 0 | 0 | 0 | 0 |
| 49368 | ENSMUSG00000111783 | 0 | NA | NA | NA | NA | NA | Gm8863 | predicted gene 8863 [Source:MGI Symbol | 0 | 0 | 0 | 0 | 0 | 0 |
| 49369 | ENSMUSG00000111785 | 0 | NA | NA | NA | NA | NA | Gm39464 | predicted gene, 39464 [Source:MGI Symb | 0 | 0 | 0 | 0 | 0 | 0 |
| 49370 | ENSMUSG00000111786 | 0 | NA | NA | NA | NA | NA | Gm47319 | predicted gene, 47319 [Source:MGI Symb | 0 | 0 | 0 | 0 | 0 | 0 |
| 49371 | ENSMUSG00000111787 | 0 | NA | NA | NA | NA | NA | Gm5035 | predicted gene 5035 [Source:MGI Symbol | 0 | 0 | 0 | 0 | 0 | 0 |
| 49372 | ENSMUSG00000111788 | 0 | NA | NA | NA | NA | NA | Gm8948 | predicted gene 8948 [Source:MGI Symbol | 0 | 0 | 0 | 0 | 0 | 0 |
| 49373 | ENSMUSG00000111789 | 0 | NA | NA | NA | NA | NA | Gm49323 | predicted gene, 49323 [Source:MGI Symb | 0 | 0 | 0 | 0 | 0 | 0 |
| 49374 | ENSMUSG00000111792 | 0 | NA | NA | NA | NA | NA | Gm33858 | predicted gene, 33858 [Source:MGI Symb | 0 | 0 | 0 | 0 | 0 | 0 |
| 49375 | ENSMUSG00000111793 | 4.29519455 | -0.809171007 | 1.025063648 | -0.789386112 | 0.42988637 | NA | Gm47950 | predicted gene, 47950 [Source:MGI Symb | 7.2279 | 5.1396 | 4.025 | 3.8559 | 2.1812 | 3.3415 |
| 49376 | ENSMUSG00000111794 | 0 | NA | NA | NA | NA | NA | Gm48294 | predicted gene, 48294 [Source:MGI Symb | 0 | 0 | 0 | 0 | 0 | 0 |
| 49377 | ENSMUSG00000111795 | 0 | NA | NA | NA | NA | NA | Gm47620 | predicted gene, 47620 [Source:MGI Symb | 0 | 0 | 0 | 0 | 0 | 0 |
| 49378 | ENSMUSG00000111796 | 0 | NA | NA | NA | NA | NA | Gm47575 | predicted gene, 47575 [Source:MGI Symb | 0 | 0 | 0 | 0 | 0 | 0 |
| 49379 | ENSMUSG00000111797 | 0 | NA | NA | NA | NA | NA | Gm48081 | predicted gene, 48081 [Source:MGI Symb | 0 | 0 | 0 | 0 | 0 | 0 |
| 49380 | ENSMUSG00000111799 | 0 | NA | NA | NA | NA | NA | Olfr928-ps1 | olfactory receptor 928, pseudogene 1 [So | 0 | 0 | 0 | 0 | 0 | 0 |
| 49381 | ENSMUSG00000111800 | 0 | NA | NA | NA | NA | NA | Gm31557 | predicted gene, 31557 [Source:MGI Symb | 0 | 0 | 0 | 0 | 0 | 0 |
| 49382 | ENSMUSG00000111801 | 0 | NA | NA | NA | NA | NA | Gm48288 | predicted gene, 48288 [Source:MGI Symb | 0 | 0 | 0 | 0 | 0 | 0 |
| 49383 | ENSMUSG00000111802 | 0 | NA | NA | NA | NA | NA | Olfr1041-ps1 | olfactory receptor 1041, pseudogene 1 [S( | 0 | 0 | 0 | 0 | 0 | 0 |
| 49384 | ENSMUSG00000111803 | 0 | NA | NA | NA | NA | NA | Gm47564 | predicted gene, 47564 [Source:MGI Symb | 0 | 0 | 0 | 0 | 0 | 0 |
| 49385 | ENSMUSG00000111805 | 0 | NA | NA | NA | NA | NA | Gm47274 | predicted gene, 47274 [Source:MGI Symb | 0 | 0 | 0 | 0 | 0 | 0 |
| 49386 | ENSMUSG00000111806 | 0.496077836 | -0.812642266 | 3.444120572 | -0.235950586 | 0.813471014 | NA | Gm47403 | predicted gene, 47403 [Source:MGI Symb | 0 | 0 | 2.0125 | 0.964 | 0 | 0 |
| 49387 | ENSMUSG00000111807 | 0 | NA | NA | NA | NA | NA | 4933433G08 | RIKEN cDNA 4933433G08 gene [Source:M | 0 | 0 | 0 | 0 | 0 | 0 |
| 49388 | ENSMUSG00000111808 | 0.133850373 | -0.796629176 | 4.080472857 | -0.195229623 | 0.84521319 | NA | Gm18225 | predicted gene, 18225 [Source:MGI Symb | 0.8031 | 0 | 0 | 0 | 0 | 0 |

| | | | | | | | | | | | | | | | |
|---|---|---|---|---|---|---|---|---|---|---|---|---|---|---|---|
| 49389 | ENSMUSG00000111809 | 0 | NA | NA | NA | NA | NA | Gm49334 | predicted gene, 49334 [Source:MGI Symb | 0 | 0 | 0 | 0 | 0 | 0 |
| 49390 | ENSMUSG00000111810 | 0 | NA | NA | NA | NA | NA | Gm29785 | predicted gene, 29785 [Source:MGI Symb | 0 | 0 | 0 | 0 | 0 | 0 |
| 49391 | ENSMUSG00000111811 | 0 | NA | NA | NA | NA | NA | Gm39327 | predicted gene, 39327 [Source:MGI Symb | 0 | 0 | 0 | 0 | 0 | 0 |
| 49392 | ENSMUSG00000111812 | 0.497389803 | 1.101799453 | 3.439268845 | 0.320358629 | 0.748696483 | NA | Gm7904 | predicted gene 7904 [Source:MGI Symbol | 0.8031 | 0 | 0 | 0 | 2.1812 | 0 |
| 49393 | ENSMUSG00000111814 | 0 | NA | NA | NA | NA | NA | Olfr896-ps1 | olfactory receptor 896, pseudogene 1 [Sou | 0 | 0 | 0 | 0 | 0 | 0 |
| 49394 | ENSMUSG00000111815 | 0 | NA | NA | NA | NA | NA | Gm6018 | predicted gene 6018 [Source:MGI Symbol | 0 | 0 | 0 | 0 | 0 | 0 |
| 49395 | ENSMUSG00000111816 | 0 | NA | NA | NA | NA | NA | Gm7229 | predicted gene 7229 [Source:MGI Symbol | 0 | 0 | 0 | 0 | 0 | 0 |
| 49396 | ENSMUSG00000111817 | 0.556921942 | 2.679043159 | 3.638243063 | 0.736356289 | 0.461513899 | NA | Gm47154 | predicted gene, 47154 [Source:MGI Symb | 0 | 0 | 0 | 0 | 0 | 3.3415 |
| 49397 | ENSMUSG00000111818 | 0 | NA | NA | NA | NA | NA | Gm17749 | predicted gene, 17749 [Source:MGI Symb | 0 | 0 | 0 | 0 | 0 | 0 |
| 49398 | ENSMUSG00000111819 | 0 | NA | NA | NA | NA | NA | Gm47110 | predicted gene, 47110 [Source:MGI Symb | 0 | 0 | 0 | 0 | 0 | 0 |
| 49399 | ENSMUSG00000111820 | 0 | NA | NA | NA | NA | NA | Gm36120 | predicted gene, 36120 [Source:MGI Symb | 0 | 0 | 0 | 0 | 0 | 0 |
| 49400 | ENSMUSG00000111821 | 0 | NA | NA | NA | NA | NA | Gm48545 | predicted gene, 48545 [Source:MGI Symb | 0 | 0 | 0 | 0 | 0 | 0 |
| 49401 | ENSMUSG00000111822 | 0 | NA | NA | NA | NA | NA | Gm48831 | predicted gene, 48831 [Source:MGI Symb | 0 | 0 | 0 | 0 | 0 | 0 |
| 49402 | ENSMUSG00000111823 | 0 | NA | NA | NA | NA | NA | Gm2517 | predicted gene 2517 [Source:MGI Symbol | 0 | 0 | 0 | 0 | 0 | 0 |
| 49403 | ENSMUSG00000111824 | 0 | NA | NA | NA | NA | NA | Gm47102 | predicted gene, 47102 [Source:MGI Symb | 0 | 0 | 0 | 0 | 0 | 0 |
| 49404 | ENSMUSG00000111825 | 0 | NA | NA | NA | NA | NA | Gm10646 | predicted gene 10646 [Source:MGI Symbo | 0 | 0 | 0 | 0 | 0 | 0 |
| 49405 | ENSMUSG00000111827 | 0 | NA | NA | NA | NA | NA | Gm8131 | predicted gene 8131 [Source:MGI Symbol | 0 | 0 | 0 | 0 | 0 | 0 |
| 49406 | ENSMUSG00000111828 | 12.7004686 | 0.295557246 | 0.701798448 | 0.421142632 | 0.673650932 | 0.931915649 | D830035M0 | RIKEN cDNA D830035M03 gene [Source:N | 21.684 | 6.1675 | 6.0375 | 11.568 | 20.722 | 10.025 |
| 49407 | ENSMUSG00000111829 | 0 | NA | NA | NA | NA | NA | Gm47475 | predicted gene, 47475 [Source:MGI Symb | 0 | 0 | 0 | 0 | 0 | 0 |
| 49408 | ENSMUSG00000111830 | 0 | NA | NA | NA | NA | NA | 4930568E12 | RIKEN cDNA 4930568E12 gene [Source:M | 0 | 0 | 0 | 0 | 0 | 0 |
| 49409 | ENSMUSG00000111831 | 0 | NA | NA | NA | NA | NA | Gm47064 | predicted gene, 47064 [Source:MGI Symb | 0 | 0 | 0 | 0 | 0 | 0 |
| 49410 | ENSMUSG00000111833 | 0 | NA | NA | NA | NA | NA | Gm48805 | predicted gene, 48805 [Source:MGI Symb | 0 | 0 | 0 | 0 | 0 | 0 |
| 49411 | ENSMUSG00000111834 | 0 | NA | NA | NA | NA | NA | Gm40020 | predicted gene, 40020 [Source:MGI Symb | 0 | 0 | 0 | 0 | 0 | 0 |
| 49412 | ENSMUSG00000111836 | 0 | NA | NA | NA | NA | NA | 4933407I05F | RIKEN cDNA 4933407I05 gene [Source:MG | 0 | 0 | 0 | 0 | 0 | 0 |
| 49413 | ENSMUSG00000111837 | 0 | NA | NA | NA | NA | NA | Gm47135 | predicted gene, 47135 [Source:MGI Symb | 0 | 0 | 0 | 0 | 0 | 0 |
| 49414 | ENSMUSG00000111839 | 0.167707431 | -0.796629176 | 4.080472857 | -0.195229623 | 0.84521319 | NA | Gm39383 | predicted gene, 39383 [Source:MGI Symb | 0 | 0 | 1.0062 | 0 | 0 | 0 |
| 49415 | ENSMUSG00000111840 | 0.428363721 | -0.543730382 | 3.624677344 | -0.15000794 | 0.880758351 | NA | Gm48832 | predicted gene, 48832 [Source:MGI Symb | 1.6062 | 0 | 0 | 0.964 | 0 | 0 |
| 49416 | ENSMUSG00000111841 | 0 | NA | NA | NA | NA | NA | Gm20745 | predicted gene, 20745 [Source:MGI Symb | 0 | 0 | 0 | 0 | 0 | 0 |
| 49417 | ENSMUSG00000111842 | 0 | NA | NA | NA | NA | NA | Gm49318 | predicted gene, 49318 [Source:MGI Symb | 0 | 0 | 0 | 0 | 0 | 0 |
| 49418 | ENSMUSG00000111843 | 0 | NA | NA | NA | NA | NA | 4933432G23 | RIKEN cDNA 4933432G23 gene [Source:M | 0 | 0 | 0 | 0 | 0 | 0 |
| 49419 | ENSMUSG00000111844 | 0.663965228 | -0.037580459 | 3.10181903 | -0.012115619 | 0.990333371 | NA | Gm47097 | predicted gene, 47097 [Source:MGI Symb | 0 | 2.0558 | 0 | 1.928 | 0 | 0 |
| 49420 | ENSMUSG00000111845 | 0 | NA | NA | NA | NA | NA | Gm7495 | predicted gene 7495 [Source:MGI Symbol | 0 | 0 | 0 | 0 | 0 | 0 |
| 49421 | ENSMUSG00000111846 | 0 | NA | NA | NA | NA | NA | Gm47271 | predicted gene, 47271 [Source:MGI Symb | 0 | 0 | 0 | 0 | 0 | 0 |
| 49422 | ENSMUSG00000111847 | 0 | NA | NA | NA | NA | NA | Gm8899 | predicted gene 8899 [Source:MGI Symbol | 0 | 0 | 0 | 0 | 0 | 0 |
| 49423 | ENSMUSG00000111849 | 0 | NA | NA | NA | NA | NA | Gm47186 | predicted gene, 47186 [Source:MGI Symb | 0 | 0 | 0 | 0 | 0 | 0 |
| 49424 | ENSMUSG00000111850 | 0 | NA | NA | NA | NA | NA | Gm47829 | predicted gene, 47829 [Source:MGI Symb | 0 | 0 | 0 | 0 | 0 | 0 |
| 49425 | ENSMUSG00000111852 | 0 | NA | NA | NA | NA | NA | Gm33180 | predicted gene, 33180 [Source:MGI Symb | 0 | 0 | 0 | 0 | 0 | 0 |
| 49426 | ENSMUSG00000111853 | 0 | NA | NA | NA | NA | NA | Gm31408 | predicted gene, 31408 [Source:MGI Symb | 0 | 0 | 0 | 0 | 0 | 0 |
| 49427 | ENSMUSG00000111854 | 0 | NA | NA | NA | NA | NA | Gm48740 | predicted gene, 48740 [Source:MGI Symb | 0 | 0 | 0 | 0 | 0 | 0 |
| 49428 | ENSMUSG00000111856 | 2.767664543 | 1.292731348 | 1.33256486 | 0.970107637 | 0.331992843 | NA | 6530413G14 | RIKEN cDNA 6530413G14 gene [Source:M | 0.8031 | 1.0279 | 3.0187 | 2.8919 | 2.1812 | 6.6831 |
| 49429 | ENSMUSG00000111857 | 2.321682323 | 2.728186846 | 1.654282123 | 1.649166613 | 0.099113506 | NA | 1190001M18 | RIKEN cDNA 1190001M18 gene [Source:N | 0.8031 | 0 | 1.0062 | 7.7118 | 2.1812 | 2.2277 |
| 49430 | ENSMUSG00000111858 | 0 | NA | NA | NA | NA | NA | Gm48277 | predicted gene, 48277 [Source:MGI Symb | 0 | 0 | 0 | 0 | 0 | 0 |
| 49431 | ENSMUSG00000111859 | 0 | NA | NA | NA | NA | NA | Gm47983 | predicted gene, 47983 [Source:MGI Symb | 0 | 0 | 0 | 0 | 0 | 0 |
| 49432 | ENSMUSG00000111862 | 0 | NA | NA | NA | NA | NA | Gm47828 | predicted gene, 47828 [Source:MGI Symb | 0 | 0 | 0 | 0 | 0 | 0 |
| 49433 | ENSMUSG00000111863 | 0 | NA | NA | NA | NA | NA | Olfr347 | olfactory receptor 347 [Source:MGI Symb | 0 | 0 | 0 | 0 | 0 | 0 |
| 49434 | ENSMUSG00000111864 | 9.539836784 | 0.776908512 | 0.944918432 | 0.822196377 | 0.41096514 | 0.833663203 | Rpl26-ps6 | ribosomal protein L26, pseudogene 6 [Sou | 0 | 7.1954 | 14.087 | 11.568 | 5.4531 | 18.935 |
| 49435 | ENSMUSG00000111865 | 0.315620088 | 0.165143421 | 4.063300985 | 0.040642675 | 0.967580763 | NA | Gm47687 | predicted gene, 47687 [Source:MGI Symb | 0.8031 | 0 | 0 | 0 | 1.0906 | 0 |
| 49436 | ENSMUSG00000111866 | 0 | NA | NA | NA | NA | NA | E130101E03 | RIKEN cDNA E130101E03 gene [Source:M | 0 | 0 | 0 | 0 | 0 | 0 |
| 49437 | ENSMUSG00000111867 | 0 | NA | NA | NA | NA | NA | Gm31013 | predicted gene, 31013 [Source:MGI Symb | 0 | 0 | 0 | 0 | 0 | 0 |
| 49438 | ENSMUSG00000111868 | 0 | NA | NA | NA | NA | NA | Gm47398 | predicted gene, 47398 [Source:MGI Symb | 0 | 0 | 0 | 0 | 0 | 0 |
| 49439 | ENSMUSG00000111869 | 0 | NA | NA | NA | NA | NA | Olfr342 | olfactory receptor 342 [Source:MGI Symb | 0 | 0 | 0 | 0 | 0 | 0 |
| 49440 | ENSMUSG00000111870 | 0.133850373 | -0.796629176 | 4.080472857 | -0.195229623 | 0.84521319 | NA | Gm7692 | predicted gene 7692 [Source:MGI Symbol | 0.8031 | 0 | 0 | 0 | 0 | 0 |
| 49441 | ENSMUSG00000111871 | 0 | NA | NA | NA | NA | NA | Gm47557 | predicted gene, 47557 [Source:MGI Symb | 0 | 0 | 0 | 0 | 0 | 0 |
| 49442 | ENSMUSG00000111873 | 0.535401493 | -2.48596965 | 3.684117979 | -0.674780141 | 0.499815459 | NA | Gm39463 | predicted gene, 39463 [Source:MGI Symb | 3.2124 | 0 | 0 | 0 | 0 | 0 |
| 49443 | ENSMUSG00000111874 | 0 | NA | NA | NA | NA | NA | Gm36539 | predicted gene, 36539 [Source:MGI Symb | 0 | 0 | 0 | 0 | 0 | 0 |
| 49444 | ENSMUSG00000111875 | 0.847564357 | 2.078206845 | 2.384595547 | 0.871513346 | 0.383473922 | NA | Gm5425 | predicted gene 5425 [Source:MGI Symbol | 0.8031 | 0 | 0 | 0.964 | 1.0906 | 2.2277 |

| | | | | | | | | | | | | | | | |
|---|---|---|---|---|---|---|---|---|---|---|---|---|---|---|---|
| 49445 | ENSMUSG00000111876 | 0 | NA | NA | NA | NA | NA | Gm18346 | predicted gene, 18346 [Source:MGI Symb | 0 | 0 | 0 | 0 | 0 | 0 |
| 49446 | ENSMUSG00000111877 | 4.662294583 | 0.588683794 | 0.976425911 | 0.60289653 | 0.546577524 | NA | Gm6477 | predicted gene 6477 [Source:MGI Symbol | 4.0155 | 5.1396 | 2.0125 | 5.7839 | 5.4531 | 5.5692 |
| 49447 | ENSMUSG00000111878 | 0 | NA | NA | NA | NA | NA | Gm47777 | predicted gene, 47777 [Source:MGI Symb | 0 | 0 | 0 | 0 | 0 | 0 |
| 49448 | ENSMUSG00000111880 | 0 | NA | NA | NA | NA | NA | Gm5040 | predicted gene 5040 [Source:MGI Symbol | 0 | 0 | 0 | 0 | 0 | 0 |
| 49449 | ENSMUSG00000111881 | 0 | NA | NA | NA | NA | NA | Vmn2r-ps92 | vomeronasal 2, receptor, pseudogene 92 | 0 | 0 | 0 | 0 | 0 | 0 |
| 49450 | ENSMUSG00000111882 | 0 | NA | NA | NA | NA | NA | Gm48864 | predicted gene, 48864 [Source:MGI Symb | 0 | 0 | 0 | 0 | 0 | 0 |
| 49451 | ENSMUSG00000111883 | 0.481988923 | 2.492401104 | 3.837077151 | 0.649557203 | 0.515978285 | NA | Gm31182 | predicted gene, 31182 [Source:MGI Symb | 0 | 0 | 0 | 2.8919 | 0 | 0 |
| 49452 | ENSMUSG00000111884 | 0 | NA | NA | NA | NA | NA | Gm19612 | predicted gene, 19612 [Source:MGI Symb | 0 | 0 | 0 | 0 | 0 | 0 |
| 49453 | ENSMUSG00000111885 | 0 | NA | NA | NA | NA | NA | Gm35326 | predicted gene, 35326 [Source:MGI Symb | 0 | 0 | 0 | 0 | 0 | 0 |
| 49454 | ENSMUSG00000111886 | 0.160662974 | 1.126924175 | 4.080472857 | 0.276174898 | 0.782413736 | NA | Gm47617 | predicted gene, 47617 [Source:MGI Symb | 0 | 0 | 0 | 0.964 | 0 | 0 |
| 49455 | ENSMUSG00000111887 | 0 | NA | NA | NA | NA | NA | Gm47610 | predicted gene, 47610 [Source:MGI Symb | 0 | 0 | 0 | 0 | 0 | 0 |
| 49456 | ENSMUSG00000111888 | 0 | NA | NA | NA | NA | NA | Gm46212 | predicted gene, 46212 [Source:MGI Symb | 0 | 0 | 0 | 0 | 0 | 0 |
| 49457 | ENSMUSG00000111889 | 0 | NA | NA | NA | NA | NA | Gm48071 | predicted gene, 48071 [Source:MGI Symb | 0 | 0 | 0 | 0 | 0 | 0 |
| 49458 | ENSMUSG00000111890 | 0 | NA | NA | NA | NA | NA | Gm48836 | predicted gene, 48836 [Source:MGI Symb | 0 | 0 | 0 | 0 | 0 | 0 |
| 49459 | ENSMUSG00000111891 | 0 | NA | NA | NA | NA | NA | Gm47366 | predicted gene, 47366 [Source:MGI Symb | 0 | 0 | 0 | 0 | 0 | 0 |
| 49460 | ENSMUSG00000111892 | 1.114567695 | -0.100982579 | 2.324776265 | -0.043437547 | 0.965352748 | NA | Gm47641 | predicted gene, 47641 [Source:MGI Symb | 2.4093 | 0 | 1.0062 | 0 | 3.2719 | 0 |
| 49461 | ENSMUSG00000111893 | 0 | NA | NA | NA | NA | NA | Gm47928 | predicted gene, 47928 [Source:MGI Symb | 0 | 0 | 0 | 0 | 0 | 0 |
| 49462 | ENSMUSG00000111894 | 0 | NA | NA | NA | NA | NA | Gm47843 | predicted gene, 47843 [Source:MGI Symb | 0 | 0 | 0 | 0 | 0 | 0 |
| 49463 | ENSMUSG00000111895 | 0 | NA | NA | NA | NA | NA | Gm47252 | predicted gene, 47252 [Source:MGI Symb | 0 | 0 | 0 | 0 | 0 | 0 |
| 49464 | ENSMUSG00000111896 | 0 | NA | NA | NA | NA | NA | Gm48196 | predicted gene, 48196 [Source:MGI Symb | 0 | 0 | 0 | 0 | 0 | 0 |
| 49465 | ENSMUSG00000111897 | 0 | NA | NA | NA | NA | NA | Gm19810 | predicted gene, 19810 [Source:MGI Symb | 0 | 0 | 0 | 0 | 0 | 0 |
| 49466 | ENSMUSG00000111898 | 0 | NA | NA | NA | NA | NA | Gm48608 | predicted gene, 48608 [Source:MGI Symb | 0 | 0 | 0 | 0 | 0 | 0 |
| 49467 | ENSMUSG00000111899 | 0.335414862 | -1.779047242 | 4.030804179 | -0.441362855 | 0.658950329 | NA | Gm18636 | predicted gene, 18636 [Source:MGI Symb | 0 | 0 | 2.0125 | 0 | 0 | 0 |
| 49468 | ENSMUSG00000111900 | 0 | NA | NA | NA | NA | NA | Gm47377 | predicted gene, 47377 [Source:MGI Symb | 0 | 0 | 0 | 0 | 0 | 0 |
| 49469 | ENSMUSG00000111901 | 0 | NA | NA | NA | NA | NA | Gm6150 | predicted gene 6150 [Source:MGI Symbol | 0 | 0 | 0 | 0 | 0 | 0 |
| 49470 | ENSMUSG00000111902 | 0 | NA | NA | NA | NA | NA | Gm4921 | predicted gene 4921 [Source:MGI Symbol | 0 | 0 | 0 | 0 | 0 | 0 |
| 49471 | ENSMUSG00000111903 | 0 | NA | NA | NA | NA | NA | 4930458K08 | RIKEN cDNA 4930458K08 gene [Source:M | 0 | 0 | 0 | 0 | 0 | 0 |
| 49472 | ENSMUSG00000111904 | 0 | NA | NA | NA | NA | NA | Gm2395 | predicted gene 2395 [Source:MGI Symbol | 0 | 0 | 0 | 0 | 0 | 0 |
| 49473 | ENSMUSG00000111905 | 2.153246466 | -0.014818016 | 1.421475055 | -0.010424394 | 0.991682687 | NA | Gm48045 | predicted gene, 48045 [Source:MGI Symb | 2.4093 | 1.0279 | 3.0187 | 0.964 | 3.2719 | 2.2277 |
| 49474 | ENSMUSG00000111906 | 0 | NA | NA | NA | NA | NA | Gm19233 | predicted gene, 19233 [Source:MGI Symb | 0 | 0 | 0 | 0 | 0 | 0 |
| 49475 | ENSMUSG00000111907 | 0 | NA | NA | NA | NA | NA | Gm48202 | predicted gene, 48202 [Source:MGI Symb | 0 | 0 | 0 | 0 | 0 | 0 |
| 49476 | ENSMUSG00000111908 | 0 | NA | NA | NA | NA | NA | Gm47516 | predicted gene, 47516 [Source:MGI Symb | 0 | 0 | 0 | 0 | 0 | 0 |
| 49477 | ENSMUSG00000111909 | 0 | NA | NA | NA | NA | NA | Gm18670 | predicted gene, 18670 [Source:MGI Symb | 0 | 0 | 0 | 0 | 0 | 0 |
| 49478 | ENSMUSG00000111910 | 0 | NA | NA | NA | NA | NA | Gm47472 | predicted gene, 47472 [Source:MGI Symb | 0 | 0 | 0 | 0 | 0 | 0 |
| 49479 | ENSMUSG00000111911 | 0 | NA | NA | NA | NA | NA | Gm47332 | predicted gene, 47332 [Source:MGI Symb | 0 | 0 | 0 | 0 | 0 | 0 |
| 49480 | ENSMUSG00000111912 | 0 | NA | NA | NA | NA | NA | Gm48521 | predicted gene, 48521 [Source:MGI Symb | 0 | 0 | 0 | 0 | 0 | 0 |
| 49481 | ENSMUSG00000111913 | 0 | NA | NA | NA | NA | NA | Gm49751 | predicted gene, 49751 [Source:MGI Symb | 0 | 0 | 0 | 0 | 0 | 0 |
| 49482 | ENSMUSG00000111914 | 0 | NA | NA | NA | NA | NA | Gm8796 | predicted gene 8796 [Source:MGI Symbol | 0 | 0 | 0 | 0 | 0 | 0 |
| 49483 | ENSMUSG00000111915 | 0 | NA | NA | NA | NA | NA | Gm3285 | predicted gene 3285 [Source:MGI Symbol | 0 | 0 | 0 | 0 | 0 | 0 |
| 49484 | ENSMUSG00000111916 | 0.160662974 | 1.126924175 | 4.080472857 | 0.276174898 | 0.782413736 | NA | 4921501I09F | RIKEN cDNA 4921501I09 gene [Source:MG | 0 | 0 | 0 | 0.964 | 0 | 0 |
| 49485 | ENSMUSG00000111917 | 0 | NA | NA | NA | NA | NA | Gm48793 | predicted gene, 48793 [Source:MGI Symb | 0 | 0 | 0 | 0 | 0 | 0 |
| 49486 | ENSMUSG00000111918 | 0 | NA | NA | NA | NA | NA | Gm47624 | predicted gene, 47624 [Source:MGI Symb | 0 | 0 | 0 | 0 | 0 | 0 |
| 49487 | ENSMUSG00000111919 | 0 | NA | NA | NA | NA | NA | Gm46192 | predicted gene, 46192 [Source:MGI Symb | 0 | 0 | 0 | 0 | 0 | 0 |
| 49488 | ENSMUSG00000111920 | 0 | NA | NA | NA | NA | NA | Gm7370 | predicted gene 7370 [Source:MGI Symbol | 0 | 0 | 0 | 0 | 0 | 0 |
| 49489 | ENSMUSG00000111921 | 0 | NA | NA | NA | NA | NA | Gm48406 | predicted gene, 48406 [Source:MGI Symb | 0 | 0 | 0 | 0 | 0 | 0 |
| 49490 | ENSMUSG00000111922 | 0 | NA | NA | NA | NA | NA | Gm47760 | predicted gene, 47760 [Source:MGI Symb | 0 | 0 | 0 | 0 | 0 | 0 |
| 49491 | ENSMUSG00000111923 | 36.25215136 | -0.518830791 | 0.471831694 | -1.099609877 | 0.271502136 | 0.727358196 | Gm34777 | predicted gene, 34777 [Source:MGI Symb | 60.233 | 21.586 | 46.287 | 44.343 | 28.356 | 16.708 |
| 49492 | ENSMUSG00000111924 | 0 | NA | NA | NA | NA | NA | 4930556N09 | RIKEN cDNA 4930556N09 gene [Source:M | 0 | 0 | 0 | 0 | 0 | 0 |
| 49493 | ENSMUSG00000111925 | 0 | NA | NA | NA | NA | NA | Gm48791 | predicted gene, 48791 [Source:MGI Symb | 0 | 0 | 0 | 0 | 0 | 0 |
| 49494 | ENSMUSG00000111926 | 2.369777237 | -0.246937615 | 1.386306992 | -0.178126213 | 0.858623856 | NA | Gm46204 | predicted gene, 46204 [Source:MGI Symb | 1.6062 | 3.0838 | 3.0187 | 0.964 | 1.0906 | 4.4554 |
| 49495 | ENSMUSG00000111927 | 0.36353943 | 2.065989254 | 4.034137301 | 0.512126658 | 0.608562368 | NA | Gm47770 | predicted gene, 47770 [Source:MGI Symb | 0 | 0 | 0 | 0 | 2.1812 | 0 |
| 49496 | ENSMUSG00000111928 | 0 | NA | NA | NA | NA | NA | Gm48082 | predicted gene, 48082 [Source:MGI Symb | 0 | 0 | 0 | 0 | 0 | 0 |
| 49497 | ENSMUSG00000111929 | 0.167707431 | -0.796629176 | 4.080472857 | -0.195229623 | 0.84521319 | NA | Gm48780 | predicted gene, 48780 [Source:MGI Symb | 0 | 0 | 1.0062 | 0 | 0 | 0 |
| 49498 | ENSMUSG00000111930 | 0 | NA | NA | NA | NA | NA | Gm47382 | predicted gene, 47382 [Source:MGI Symb | 0 | 0 | 0 | 0 | 0 | 0 |
| 49499 | ENSMUSG00000111931 | 0 | NA | NA | NA | NA | NA | Gm47032 | predicted gene, 47032 [Source:MGI Symb | 0 | 0 | 0 | 0 | 0 | 0 |
| 49500 | ENSMUSG00000111932 | 0.181769715 | 1.126924175 | 4.080472857 | 0.276174898 | 0.782413736 | NA | Gm18899 | predicted gene, 18899 [Source:MGI Symb | 0 | 0 | 0 | 0 | 1.0906 | 0 |

| | | | | | | | | | | | | | | | |
|---|---|---|---|---|---|---|---|---|---|---|---|---|---|---|---|
| 49501 | ENSMUSG00000111933 | 1.361316686 | 1.776258021 | 1.88191201 | 0.943858167 | 0.345242132 | NA | Gm36660 | predicted gene, 36660 [Source:MGI Symb | 0.8031 | 1.0279 | 0 | 1.928 | 2.1812 | 2.2277 |
| 49502 | ENSMUSG00000111934 | 0 | NA | NA | NA | NA | NA | Gm48123 | predicted gene, 48123 [Source:MGI Symb | 0 | 0 | 0 | 0 | 0 | 0 |
| 49503 | ENSMUSG00000111935 | 0 | NA | NA | NA | NA | NA | Gm47023 | predicted gene, 47023 [Source:MGI Symb | 0 | 0 | 0 | 0 | 0 | 0 |
| 49504 | ENSMUSG00000111936 | 0.481988923 | 2.492401104 | 3.837077151 | 0.649557203 | 0.515978285 | NA | Gm47540 | predicted gene, 47540 [Source:MGI Symb | 0 | 0 | 0 | 2.8919 | 0 | 0 |
| 49505 | ENSMUSG00000111937 | 0 | NA | NA | NA | NA | NA | Gm46203 | predicted gene, 46203 [Source:MGI Symb | 0 | 0 | 0 | 0 | 0 | 0 |
| 49506 | ENSMUSG00000111938 | 0 | NA | NA | NA | NA | NA | 2900045O2C | RIKEN cDNA 2900045O20 gene [Source:M | 0 | 0 | 0 | 0 | 0 | 0 |
| 49507 | ENSMUSG00000111939 | 0 | NA | NA | NA | NA | NA | Gm48177 | predicted gene, 48177 [Source:MGI Symb | 0 | 0 | 0 | 0 | 0 | 0 |
| 49508 | ENSMUSG00000111940 | 0.181769715 | 1.126924175 | 4.080472857 | 0.276174898 | 0.782413736 | NA | Gm4478 | predicted gene 4478 [Source:MGI Symbol | 0 | 0 | 0 | 0 | 1.0906 | 0 |
| 49509 | ENSMUSG00000111941 | 2.692183058 | -0.543549384 | 1.363468631 | -0.398651917 | 0.690149702 | NA | Gm33263 | predicted gene, 33263 [Source:MGI Symb | 2.4093 | 4.1117 | 3.0187 | 0 | 3.2719 | 3.3415 |
| 49510 | ENSMUSG00000111942 | 4.469693756 | 0.427781834 | 1.032563825 | 0.414290936 | 0.678661051 | NA | Gm5182 | predicted gene 5182 [Source:MGI Symbol | 2.4093 | 4.1117 | 5.0312 | 8.6758 | 4.3625 | 2.2277 |
| 49511 | ENSMUSG00000111943 | 0.542600935 | 1.125931187 | 3.335923981 | 0.33751704 | 0.735727169 | NA | Gm47862 | predicted gene, 47862 [Source:MGI Symb | 0 | 1.0279 | 0 | 0 | 0 | 2.2277 |
| 49512 | ENSMUSG00000111944 | 0 | NA | NA | NA | NA | NA | Gm18024 | predicted gene, 18024 [Source:MGI Symb | 0 | 0 | 0 | 0 | 0 | 0 |
| 49513 | ENSMUSG00000111945 | 0 | NA | NA | NA | NA | NA | Gm19169 | predicted gene, 19169 [Source:MGI Symb | 0 | 0 | 0 | 0 | 0 | 0 |
| 49514 | ENSMUSG00000111946 | 0 | NA | NA | NA | NA | NA | Gm47495 | predicted gene, 47495 [Source:MGI Symb | 0 | 0 | 0 | 0 | 0 | 0 |
| 49515 | ENSMUSG00000111947 | 0.824628202 | 0.545567773 | 2.864022412 | 0.19049005 | 0.848925139 | NA | Gm47438 | predicted gene, 47438 [Source:MGI Symb | 0 | 2.0558 | 0 | 2.8919 | 0 | 0 |
| 49516 | ENSMUSG00000111948 | 0 | NA | NA | NA | NA | NA | Gm47055 | predicted gene, 47055 [Source:MGI Symb | 0 | 0 | 0 | 0 | 0 | 0 |
| 49517 | ENSMUSG00000111949 | 0 | NA | NA | NA | NA | NA | Gm48679 | predicted gene, 48679 [Source:MGI Symb | 0 | 0 | 0 | 0 | 0 | 0 |
| 49518 | ENSMUSG00000111950 | 0 | NA | NA | NA | NA | NA | Gm48893 | predicted gene, 48893 [Source:MGI Symb | 0 | 0 | 0 | 0 | 0 | 0 |
| 49519 | ENSMUSG00000111951 | 0 | NA | NA | NA | NA | NA | Gm48755 | predicted gene, 48755 [Source:MGI Symb | 0 | 0 | 0 | 0 | 0 | 0 |
| 49520 | ENSMUSG00000111952 | 0 | NA | NA | NA | NA | NA | Gm4489 | predicted gene 4489 [Source:MGI Symbol | 0 | 0 | 0 | 0 | 0 | 0 |
| 49521 | ENSMUSG00000111953 | 0 | NA | NA | NA | NA | NA | Gm18637 | predicted gene, 18637 [Source:MGI Symb | 0 | 0 | 0 | 0 | 0 | 0 |
| 49522 | ENSMUSG00000111954 | 0 | NA | NA | NA | NA | NA | Gm7001 | predicted gene 7001 [Source:MGI Symbol | 0 | 0 | 0 | 0 | 0 | 0 |
| 49523 | ENSMUSG00000111955 | 0 | NA | NA | NA | NA | NA | Gm48148 | predicted gene, 48148 [Source:MGI Symb | 0 | 0 | 0 | 0 | 0 | 0 |
| 49524 | ENSMUSG00000111956 | 0 | NA | NA | NA | NA | NA | Gm29774 | predicted gene, 29774 [Source:MGI Symb | 0 | 0 | 0 | 0 | 0 | 0 |
| 49525 | ENSMUSG00000111957 | 0 | NA | NA | NA | NA | NA | Gm47582 | predicted gene, 47582 [Source:MGI Symb | 0 | 0 | 0 | 0 | 0 | 0 |
| 49526 | ENSMUSG00000111958 | 0 | NA | NA | NA | NA | NA | Gm47844 | predicted gene, 47844 [Source:MGI Symb | 0 | 0 | 0 | 0 | 0 | 0 |
| 49527 | ENSMUSG00000111959 | 0 | NA | NA | NA | NA | NA | Gm18503 | predicted gene, 18503 [Source:MGI Symb | 0 | 0 | 0 | 0 | 0 | 0 |
| 49528 | ENSMUSG00000111960 | 0.17131964 | -0.796629176 | 4.080472857 | -0.195229623 | 0.84521319 | NA | Gm46349 | predicted gene, 46349 [Source:MGI Symb | 0 | 1.0279 | 0 | 0 | 0 | 0 |
| 49529 | ENSMUSG00000111961 | 0 | NA | NA | NA | NA | NA | Gm47966 | predicted gene, 47966 [Source:MGI Symb | 0 | 0 | 0 | 0 | 0 | 0 |
| 49530 | ENSMUSG00000111962 | 0 | NA | NA | NA | NA | NA | Gm20110 | predicted gene, 20110 [Source:MGI Symb | 0 | 0 | 0 | 0 | 0 | 0 |
| 49531 | ENSMUSG00000111963 | 1.791716411 | 0.566500375 | 1.691822132 | 0.334846296 | 0.737741035 | NA | Gm32283 | predicted gene, 32283 [Source:MGI Symb | 3.2124 | 1.0279 | 0 | 0.964 | 1.0906 | 4.4554 |
| 49532 | ENSMUSG00000111964 | 0.621048825 | -1.224891493 | 2.859518283 | -0.428355888 | 0.668392035 | NA | Gm8942 | predicted gene 8942 [Source:MGI Symbol | 1.6062 | 0 | 1.0062 | 0 | 0 | 1.1138 |
| 49533 | ENSMUSG00000111965 | 0 | NA | NA | NA | NA | NA | Gm4797 | predicted gene 4797 [Source:MGI Symbol | 0 | 0 | 0 | 0 | 0 | 0 |
| 49534 | ENSMUSG00000111966 | 0 | NA | NA | NA | NA | NA | Gm5781 | predicted gene 5781 [Source:MGI Symbol | 0 | 0 | 0 | 0 | 0 | 0 |
| 49535 | ENSMUSG00000111967 | 0 | NA | NA | NA | NA | NA | Gm3678 | predicted gene 3678 [Source:MGI Symbol | 0 | 0 | 0 | 0 | 0 | 0 |
| 49536 | ENSMUSG00000111968 | 0.167707431 | -0.796629176 | 4.080472857 | -0.195229623 | 0.84521319 | NA | Gm8873 | predicted gene 8873 [Source:MGI Symbol | 0 | 0 | 1.0062 | 0 | 0 | 0 |
| 49537 | ENSMUSG00000111969 | 0 | NA | NA | NA | NA | NA | Gm9129 | predicted gene 9129 [Source:MGI Symbol | 0 | 0 | 0 | 0 | 0 | 0 |
| 49538 | ENSMUSG00000111970 | 0 | NA | NA | NA | NA | NA | Gm47715 | predicted gene, 47715 [Source:MGI Symb | 0 | 0 | 0 | 0 | 0 | 0 |
| 49539 | ENSMUSG00000111971 | 1.340468669 | 1.76797944 | 2.044838259 | 0.864606006 | 0.387255075 | NA | Gm48678 | predicted gene, 48678 [Source:MGI Symb | 0.8031 | 0 | 1.0062 | 2.8919 | 0 | 3.3415 |
| 49540 | ENSMUSG00000111972 | 0 | NA | NA | NA | NA | NA | Gm40778 | predicted gene, 40778 [Source:MGI Symb | 0 | 0 | 0 | 0 | 0 | 0 |
| 49541 | ENSMUSG00000111973 | 0.160662974 | 1.126924175 | 4.080472857 | 0.276174898 | 0.782413736 | NA | Gm48827 | predicted gene, 48827 [Source:MGI Symb | 0 | 0 | 0 | 0.964 | 0 | 0 |
| 49542 | ENSMUSG00000111974 | 0 | NA | NA | NA | NA | NA | Gm4803 | predicted gene 4803 [Source:MGI Symbol | 0 | 0 | 0 | 0 | 0 | 0 |
| 49543 | ENSMUSG00000111975 | 0 | NA | NA | NA | NA | NA | Gm46311 | predicted gene, 46311 [Source:MGI Symb | 0 | 0 | 0 | 0 | 0 | 0 |
| 49544 | ENSMUSG00000111976 | 0 | NA | NA | NA | NA | NA | Gm48178 | predicted gene, 48178 [Source:MGI Symb | 0 | 0 | 0 | 0 | 0 | 0 |
| 49545 | ENSMUSG00000111977 | 0.267700746 | -1.497481649 | 4.057001436 | -0.369110456 | 0.712045396 | NA | Gm47163 | predicted gene, 47163 [Source:MGI Symb | 1.6062 | 0 | 0 | 0 | 0 | 0 |
| 49546 | ENSMUSG00000111978 | 0 | NA | NA | NA | NA | NA | Gm48268 | predicted gene, 48268 [Source:MGI Symb | 0 | 0 | 0 | 0 | 0 | 0 |
| 49547 | ENSMUSG00000111979 | 0 | NA | NA | NA | NA | NA | Gm47522 | predicted gene, 47522 [Source:MGI Symb | 0 | 0 | 0 | 0 | 0 | 0 |
| 49548 | ENSMUSG00000111980 | 0 | NA | NA | NA | NA | NA | Gm20304 | predicted gene, 20304 [Source:MGI Symb | 0 | 0 | 0 | 0 | 0 | 0 |
| 49549 | ENSMUSG00000111981 | 0 | NA | NA | NA | NA | NA | Gm18515 | predicted gene, 18515 [Source:MGI Symb | 0 | 0 | 0 | 0 | 0 | 0 |
| 49550 | ENSMUSG00000111982 | 0.499690045 | -0.82080011 | 3.113553401 | -0.263621658 | 0.792071468 | NA | Gm35135 | predicted gene, 35135 [Source:MGI Symb | 0 | 1.0279 | 1.0062 | 0.964 | 0 | 0 |
| 49551 | ENSMUSG00000111983 | 0 | NA | NA | NA | NA | NA | Gm48666 | predicted gene, 48666 [Source:MGI Symb | 0 | 0 | 0 | 0 | 0 | 0 |
| 49552 | ENSMUSG00000111984 | 0 | NA | NA | NA | NA | NA | Gm46327 | predicted gene, 46327 [Source:MGI Symb | 0 | 0 | 0 | 0 | 0 | 0 |
| 49553 | ENSMUSG00000111985 | 0 | NA | NA | NA | NA | NA | Gm8290 | predicted gene 8290 [Source:MGI Symbol | 0 | 0 | 0 | 0 | 0 | 0 |
| 49554 | ENSMUSG00000111986 | 0 | NA | NA | NA | NA | NA | 1700008C04 | RIKEN cDNA 1700008C04 gene [Source:M | 0 | 0 | 0 | 0 | 0 | 0 |
| 49555 | ENSMUSG00000111987 | 0.321325949 | 1.911575114 | 4.048267893 | 0.472195804 | 0.636787034 | NA | Gm48809 | predicted gene, 48809 [Source:MGI Symb | 0 | 0 | 0 | 1.928 | 0 | 0 |
| 49556 | ENSMUSG00000111988 | 0 | NA | NA | NA | NA | NA | Gm19256 | predicted gene, 19256 [Source:MGI Symb | 0 | 0 | 0 | 0 | 0 | 0 |

| | | | | | | | | | | | | | | | |
|---|---|---|---|---|---|---|---|---|---|---|---|---|---|---|---|
| 49557 | ENSMUSG00000111989 | 0 | NA | NA | NA | NA | NA | Gm18696 | predicted gene, 18696 [Source:MGI Symb | 0 | 0 | 0 | 0 | 0 | 0 |
| 49558 | ENSMUSG00000111990 | 0 | NA | NA | NA | NA | NA | Gm47033 | predicted gene, 47033 [Source:MGI Symb | 0 | 0 | 0 | 0 | 0 | 0 |
| 49559 | ENSMUSG00000111991 | 0 | NA | NA | NA | NA | NA | Gm47835 | predicted gene, 47835 [Source:MGI Symb | 0 | 0 | 0 | 0 | 0 | 0 |
| 49560 | ENSMUSG00000111992 | 0 | NA | NA | NA | NA | NA | Gm47724 | predicted gene, 47724 [Source:MGI Symb | 0 | 0 | 0 | 0 | 0 | 0 |
| 49561 | ENSMUSG00000111993 | 0 | NA | NA | NA | NA | NA | Gm48018 | predicted gene, 48018 [Source:MGI Symb | 0 | 0 | 0 | 0 | 0 | 0 |
| 49562 | ENSMUSG00000111994 | 0 | NA | NA | NA | NA | NA | A330049N07 | RIKEN cDNA A330049N07 gene [Source:M | 0 | 0 | 0 | 0 | 0 | 0 |
| 49563 | ENSMUSG00000111995 | 0 | NA | NA | NA | NA | NA | Gm48697 | predicted gene, 48697 [Source:MGI Symb | 0 | 0 | 0 | 0 | 0 | 0 |
| 49564 | ENSMUSG00000111996 | 0 | NA | NA | NA | NA | NA | Gm47365 | predicted gene, 47365 [Source:MGI Symb | 0 | 0 | 0 | 0 | 0 | 0 |
| 49565 | ENSMUSG00000111997 | 0.686901383 | 1.702364833 | 2.776876575 | 0.613050233 | 0.539843125 | NA | Gm5176 | predicted gene 5176 [Source:MGI Symbol | 0.8031 | 0 | 0 | 0 | 1.0906 | 2.2277 |
| 49566 | ENSMUSG00000111998 | 0 | NA | NA | NA | NA | NA | Gm48465 | predicted gene, 48465 [Source:MGI Symb | 0 | 0 | 0 | 0 | 0 | 0 |
| 49567 | ENSMUSG00000111999 | 0 | NA | NA | NA | NA | NA | Gm5080 | predicted gene 5080 [Source:MGI Symbol | 0 | 0 | 0 | 0 | 0 | 0 |
| 49568 | ENSMUSG00000112000 | 1.827620501 | -0.303292172 | 2.008952629 | -0.150970296 | 0.879999148 | NA | Gm47956 | predicted gene, 47956 [Source:MGI Symb | 0 | 5.1396 | 1.0062 | 4.8199 | 0 | 0 |
| 49569 | ENSMUSG00000112001 | 0 | NA | NA | NA | NA | NA | Gm8112 | predicted gene 8112 [Source:MGI Symbol | 0 | 0 | 0 | 0 | 0 | 0 |
| 49570 | ENSMUSG00000112002 | 0 | NA | NA | NA | NA | NA | Gm9585 | predicted gene 9585 [Source:MGI Symbol | 0 | 0 | 0 | 0 | 0 | 0 |
| 49571 | ENSMUSG00000112003 | 0 | NA | NA | NA | NA | NA | Gm4864 | predicted gene 4864 [Source:MGI Symbol | 0 | 0 | 0 | 0 | 0 | 0 |
| 49572 | ENSMUSG00000112004 | 0 | NA | NA | NA | NA | NA | Gm8785 | predicted gene 8785 [Source:MGI Symbol | 0 | 0 | 0 | 0 | 0 | 0 |
| 49573 | ENSMUSG00000112005 | 0 | NA | NA | NA | NA | NA | Gm29818 | predicted gene, 29818 [Source:MGI Symb | 0 | 0 | 0 | 0 | 0 | 0 |
| 49574 | ENSMUSG00000112006 | 723.861419 | 0.082128399 | 0.129975019 | 0.63187834 | 0.527466378 | 0.888334265 | Gm48633 | predicted gene, 48633 [Source:MGI Symb | 628.03 | 716.46 | 765.75 | 673.82 | 797.24 | 761.87 |
| 49575 | ENSMUSG00000112007 | 0 | NA | NA | NA | NA | NA | Gm18119 | predicted gene, 18119 [Source:MGI Symb | 0 | 0 | 0 | 0 | 0 | 0 |
| 49576 | ENSMUSG00000112008 | 4.544736028 | -0.302766152 | 1.048062464 | -0.288881782 | 0.772671845 | NA | Gm47218 | predicted gene, 47218 [Source:MGI Symb | 4.0155 | 4.1117 | 7.0437 | 7.7118 | 3.2719 | 1.1138 |
| 49577 | ENSMUSG00000112009 | 1.134801153 | -0.214097227 | 1.862719624 | -0.114937978 | 0.908494283 | NA | Gm48591 | predicted gene, 48591 [Source:MGI Symb | 1.6062 | 1.0279 | 1.0062 | 0.964 | 1.0906 | 1.1138 |
| 49578 | ENSMUSG00000112010 | 0 | NA | NA | NA | NA | NA | Gm48461 | predicted gene, 48461 [Source:MGI Symb | 0 | 0 | 0 | 0 | 0 | 0 |
| 49579 | ENSMUSG00000112011 | 0 | NA | NA | NA | NA | NA | Gm9535 | predicted gene 9535 [Source:MGI Symbol | 0 | 0 | 0 | 0 | 0 | 0 |
| 49580 | ENSMUSG00000112012 | 0 | NA | NA | NA | NA | NA | Gm47025 | predicted gene, 47025 [Source:MGI Symb | 0 | 0 | 0 | 0 | 0 | 0 |
| 49581 | ENSMUSG00000112013 | 0 | NA | NA | NA | NA | NA | Gm47967 | predicted gene, 47967 [Source:MGI Symb | 0 | 0 | 0 | 0 | 0 | 0 |
| 49582 | ENSMUSG00000112014 | 5.154449881 | 1.573129313 | 0.997188139 | 1.577565207 | 0.114665531 | 0.497415802 | Gm48435 | predicted gene, 48435 [Source:MGI Symb | 0.8031 | 4.1117 | 3.0187 | 8.6758 | 7.6343 | 6.6831 |
| 49583 | ENSMUSG00000112015 | 0 | NA | NA | NA | NA | NA | Gm47592 | predicted gene, 47592 [Source:MGI Symb | 0 | 0 | 0 | 0 | 0 | 0 |
| 49584 | ENSMUSG00000112016 | 0 | NA | NA | NA | NA | NA | Gm31267 | predicted gene, 31267 [Source:MGI Symb | 0 | 0 | 0 | 0 | 0 | 0 |
| 49585 | ENSMUSG00000112017 | 0 | NA | NA | NA | NA | NA | Gm7842 | predicted gene 7842 [Source:MGI Symbol | 0 | 0 | 0 | 0 | 0 | 0 |
| 49586 | ENSMUSG00000112018 | 0 | NA | NA | NA | NA | NA | Gm47919 | predicted gene, 47919 [Source:MGI Symb | 0 | 0 | 0 | 0 | 0 | 0 |
| 49587 | ENSMUSG00000112019 | 0 | NA | NA | NA | NA | NA | Gm48748 | predicted gene, 48748 [Source:MGI Symb | 0 | 0 | 0 | 0 | 0 | 0 |
| 49588 | ENSMUSG00000112020 | 0 | NA | NA | NA | NA | NA | Gm48532 | predicted gene, 48532 [Source:MGI Symb | 0 | 0 | 0 | 0 | 0 | 0 |
| 49589 | ENSMUSG00000112021 | 0 | NA | NA | NA | NA | NA | Gm30346 | predicted gene, 30346 [Source:MGI Symb | 0 | 0 | 0 | 0 | 0 | 0 |
| 49590 | ENSMUSG00000112022 | 0 | NA | NA | NA | NA | NA | Gm2436 | predicted gene 2436 [Source:MGI Symbol | 0 | 0 | 0 | 0 | 0 | 0 |
| 49591 | ENSMUSG00000112023 | 30.92287998 | -1.216776733 | 0.401996655 | -3.026832981 | 0.002471305 | 0.040944961 | Lilr4b | leukocyte immunoglobulin-like receptor, s | 48.186 | 43.173 | 38.237 | 17.352 | 18.541 | 20.049 |
| 49592 | ENSMUSG00000112024 | 0 | NA | NA | NA | NA | NA | Gm47394 | predicted gene, 47394 [Source:MGI Symb | 0 | 0 | 0 | 0 | 0 | 0 |
| 49593 | ENSMUSG00000112025 | 0 | NA | NA | NA | NA | NA | Gm47669 | predicted gene, 47669 [Source:MGI Symb | 0 | 0 | 0 | 0 | 0 | 0 |
| 49594 | ENSMUSG00000112026 | 0 | NA | NA | NA | NA | NA | Gm6653 | predicted gene 6653 [Source:MGI Symbol | 0 | 0 | 0 | 0 | 0 | 0 |
| 49595 | ENSMUSG00000112027 | 0 | NA | NA | NA | NA | NA | Gm9045 | predicted gene 9045 [Source:MGI Symbol | 0 | 0 | 0 | 0 | 0 | 0 |
| 49596 | ENSMUSG00000112028 | 5.879788578 | 0.771396974 | 0.971031372 | 0.79440994 | 0.426956819 | 0.842220356 | 1700113B09 | RIKEN cDNA 1700113B09 gene [Source:M | 4.0155 | 2.0558 | 7.0437 | 14.46 | 4.3625 | 3.3415 |
| 49597 | ENSMUSG00000112029 | 0 | NA | NA | NA | NA | NA | Gm33337 | predicted gene, 33337 [Source:MGI Symb | 0 | 0 | 0 | 0 | 0 | 0 |
| 49598 | ENSMUSG00000112030 | 0.455176322 | 0.955156269 | 3.546674842 | 0.269310357 | 0.787690862 | NA | Gm35189 | predicted gene, 35189 [Source:MGI Symb | 0.8031 | 0 | 0 | 1.928 | 0 | 0 |
| 49599 | ENSMUSG00000112031 | 0 | NA | NA | NA | NA | NA | Gm48484 | predicted gene, 48484 [Source:MGI Symb | 0 | 0 | 0 | 0 | 0 | 0 |
| 49600 | ENSMUSG00000112032 | 0 | NA | NA | NA | NA | NA | Gm48036 | predicted gene, 48036 [Source:MGI Symb | 0 | 0 | 0 | 0 | 0 | 0 |
| 49601 | ENSMUSG00000112033 | 0 | NA | NA | NA | NA | NA | Gm48808 | predicted gene, 48808 [Source:MGI Symb | 0 | 0 | 0 | 0 | 0 | 0 |
| 49602 | ENSMUSG00000112034 | 0 | NA | NA | NA | NA | NA | Gm48363 | predicted gene, 48363 [Source:MGI Symb | 0 | 0 | 0 | 0 | 0 | 0 |
| 49603 | ENSMUSG00000112035 | 0 | NA | NA | NA | NA | NA | Gm49335 | predicted gene, 49335 [Source:MGI Symb | 0 | 0 | 0 | 0 | 0 | 0 |
| 49604 | ENSMUSG00000112036 | 0 | NA | NA | NA | NA | NA | Gm47801 | predicted gene, 47801 [Source:MGI Symb | 0 | 0 | 0 | 0 | 0 | 0 |
| 49605 | ENSMUSG00000112037 | 0 | NA | NA | NA | NA | NA | Gm48179 | predicted gene, 48179 [Source:MGI Symb | 0 | 0 | 0 | 0 | 0 | 0 |
| 49606 | ENSMUSG00000112038 | 0 | NA | NA | NA | NA | NA | Gm47056 | predicted gene, 47056 [Source:MGI Symb | 0 | 0 | 0 | 0 | 0 | 0 |
| 49607 | ENSMUSG00000112039 | 0 | NA | NA | NA | NA | NA | Gm5136 | predicted gene 5136 [Source:MGI Symbol | 0 | 0 | 0 | 0 | 0 | 0 |
| 49608 | ENSMUSG00000112040 | 0 | NA | NA | NA | NA | NA | Gm47396 | predicted gene, 47396 [Source:MGI Symb | 0 | 0 | 0 | 0 | 0 | 0 |
| 49609 | ENSMUSG00000112041 | 1.527685521 | 3.018588606 | 2.000675249 | 1.5087849 | 0.131353761 | NA | 9530020I12F | RIKEN cDNA 9530020I12 gene [Source:MG | 0 | 1.0279 | 0 | 4.8199 | 1.0906 | 2.2277 |
| 49610 | ENSMUSG00000112042 | 0 | NA | NA | NA | NA | NA | Gm47402 | predicted gene, 47402 [Source:MGI Symb | 0 | 0 | 0 | 0 | 0 | 0 |
| 49611 | ENSMUSG00000112043 | 0 | NA | NA | NA | NA | NA | Gm48774 | predicted gene, 48774 [Source:MGI Symb | 0 | 0 | 0 | 0 | 0 | 0 |
| 49612 | ENSMUSG00000112044 | 0 | NA | NA | NA | NA | NA | Gm48184 | predicted gene, 48184 [Source:MGI Symb | 0 | 0 | 0 | 0 | 0 | 0 |

| | | | | | | | | | | | | | | | |
|---|---|---|---|---|---|---|---|---|---|---|---|---|---|---|---|
| 49613 | ENSMUSG00000112045 | 0 | NA | NA | NA | NA | NA | Gm5649 | predicted gene 5649 [Source:MGI Symbol | 0 | 0 | 0 | 0 | 0 | 0 |
| 49614 | ENSMUSG00000112046 | 0 | NA | NA | NA | NA | NA | Gm5175 | predicted gene 5175 [Source:MGI Symbol | 0 | 0 | 0 | 0 | 0 | 0 |
| 49615 | ENSMUSG00000112047 | 0 | NA | NA | NA | NA | NA | Gm47852 | predicted gene, 47852 [Source:MGI Symb | 0 | 0 | 0 | 0 | 0 | 0 |
| 49616 | ENSMUSG00000112048 | 0 | NA | NA | NA | NA | NA | Gm48750 | predicted gene, 48750 [Source:MGI Symb | 0 | 0 | 0 | 0 | 0 | 0 |
| 49617 | ENSMUSG00000112049 | 0 | NA | NA | NA | NA | NA | Gm48269 | predicted gene, 48269 [Source:MGI Symb | 0 | 0 | 0 | 0 | 0 | 0 |
| 49618 | ENSMUSG00000112050 | 0 | NA | NA | NA | NA | NA | Gm29809 | predicted gene, 29809 [Source:MGI Symb | 0 | 0 | 0 | 0 | 0 | 0 |
| 49619 | ENSMUSG00000112051 | 0 | NA | NA | NA | NA | NA | Gm8158 | predicted gene 8158 [Source:MGI Symbol | 0 | 0 | 0 | 0 | 0 | 0 |
| 49620 | ENSMUSG00000112052 | 0 | NA | NA | NA | NA | NA | Gm17917 | predicted gene, 17917 [Source:MGI Symb | 0 | 0 | 0 | 0 | 0 | 0 |
| 49621 | ENSMUSG00000112053 | 0 | NA | NA | NA | NA | NA | Gm46328 | predicted gene, 46328 [Source:MGI Symb | 0 | 0 | 0 | 0 | 0 | 0 |
| 49622 | ENSMUSG00000112054 | 0 | NA | NA | NA | NA | NA | Gm33680 | predicted gene, 33680 [Source:MGI Symb | 0 | 0 | 0 | 0 | 0 | 0 |
| 49623 | ENSMUSG00000112055 | 0 | NA | NA | NA | NA | NA | Gm48773 | predicted gene, 48773 [Source:MGI Symb | 0 | 0 | 0 | 0 | 0 | 0 |
| 49624 | ENSMUSG00000112056 | 0 | NA | NA | NA | NA | NA | Gm48181 | predicted gene, 48181 [Source:MGI Symb | 0 | 0 | 0 | 0 | 0 | 0 |
| 49625 | ENSMUSG00000112057 | 0 | NA | NA | NA | NA | NA | Gm21321 | predicted gene, 21321 [Source:MGI Symb | 0 | 0 | 0 | 0 | 0 | 0 |
| 49626 | ENSMUSG00000112058 | 1.300492042 | -1.407579939 | 2.481546929 | -0.567218747 | 0.570565573 | NA | Gm47593 | predicted gene, 47593 [Source:MGI Symb | 5.6217 | 0 | 0 | 0 | 2.1812 | 0 |
| 49627 | ENSMUSG00000112059 | 0 | NA | NA | NA | NA | NA | Gm48282 | predicted gene, 48282 [Source:MGI Symb | 0 | 0 | 0 | 0 | 0 | 0 |
| 49628 | ENSMUSG00000112060 | 0 | NA | NA | NA | NA | NA | Gm48279 | predicted gene, 48279 [Source:MGI Symb | 0 | 0 | 0 | 0 | 0 | 0 |
| 49629 | ENSMUSG00000112061 | 1.902508316 | 0.06433555 | 1.936658998 | 0.033219865 | 0.973499257 | NA | Gm8249 | predicted gene 8249 [Source:MGI Symbol | 1.6062 | 0 | 4.025 | 5.7839 | 0 | 0 |
| 49630 | ENSMUSG00000112062 | 0 | NA | NA | NA | NA | NA | Gm47353 | predicted gene, 47353 [Source:MGI Symb | 0 | 0 | 0 | 0 | 0 | 0 |
| 49631 | ENSMUSG00000112063 | 0 | NA | NA | NA | NA | NA | Gm32220 | predicted gene, 32220 [Source:MGI Symb | 0 | 0 | 0 | 0 | 0 | 0 |
| 49632 | ENSMUSG00000112064 | 2.827920493 | 0.346456282 | 1.242776229 | 0.278776077 | 0.780416674 | NA | Gm47399 | predicted gene, 47399 [Source:MGI Symb | 2.4093 | 1.0279 | 4.025 | 2.8919 | 3.2719 | 3.3415 |
| 49633 | ENSMUSG00000112065 | 0 | NA | NA | NA | NA | NA | Gm47466 | predicted gene, 47466 [Source:MGI Symb | 0 | 0 | 0 | 0 | 0 | 0 |
| 49634 | ENSMUSG00000112066 | 0 | NA | NA | NA | NA | NA | Gm48380 | predicted gene, 48380 [Source:MGI Symb | 0 | 0 | 0 | 0 | 0 | 0 |
| 49635 | ENSMUSG00000112067 | 0 | NA | NA | NA | NA | NA | Gm48015 | predicted gene, 48015 [Source:MGI Symb | 0 | 0 | 0 | 0 | 0 | 0 |
| 49636 | ENSMUSG00000112068 | 0 | NA | NA | NA | NA | NA | Gm9182 | predicted gene 9182 [Source:MGI Symbol | 0 | 0 | 0 | 0 | 0 | 0 |
| 49637 | ENSMUSG00000112069 | 0 | NA | NA | NA | NA | NA | Gm9476 | predicted gene 9476 [Source:MGI Symbol | 0 | 0 | 0 | 0 | 0 | 0 |
| 49638 | ENSMUSG00000112070 | 0 | NA | NA | NA | NA | NA | Gm47809 | predicted gene, 47809 [Source:MGI Symb | 0 | 0 | 0 | 0 | 0 | 0 |
| 49639 | ENSMUSG00000112071 | 0.160662974 | 1.126924175 | 4.080472857 | 0.276174898 | 0.782413736 | NA | 5430401H09 | RIKEN cDNA 5430401H09 gene [Source:M | 0 | 0 | 0 | 0.964 | 0 | 0 |
| 49640 | ENSMUSG00000112072 | 0 | NA | NA | NA | NA | NA | Gm47244 | predicted gene, 47244 [Source:MGI Symb | 0 | 0 | 0 | 0 | 0 | 0 |
| 49641 | ENSMUSG00000112073 | 0 | NA | NA | NA | NA | NA | Gm47101 | predicted gene, 47101 [Source:MGI Symb | 0 | 0 | 0 | 0 | 0 | 0 |
| 49642 | ENSMUSG00000112074 | 0 | NA | NA | NA | NA | NA | Gm48278 | predicted gene, 48278 [Source:MGI Symb | 0 | 0 | 0 | 0 | 0 | 0 |
| 49643 | ENSMUSG00000112075 | 0 | NA | NA | NA | NA | NA | Gm47700 | predicted gene, 47700 [Source:MGI Symb | 0 | 0 | 0 | 0 | 0 | 0 |
| 49644 | ENSMUSG00000112076 | 0 | NA | NA | NA | NA | NA | Gm9368 | predicted gene 9368 [Source:MGI Symbol | 0 | 0 | 0 | 0 | 0 | 0 |
| 49645 | ENSMUSG00000112077 | 0 | NA | NA | NA | NA | NA | Gm47245 | predicted gene, 47245 [Source:MGI Symb | 0 | 0 | 0 | 0 | 0 | 0 |
| 49646 | ENSMUSG00000112078 | 0 | NA | NA | NA | NA | NA | Gm48466 | predicted gene, 48466 [Source:MGI Symb | 0 | 0 | 0 | 0 | 0 | 0 |
| 49647 | ENSMUSG00000112079 | 0 | NA | NA | NA | NA | NA | Gm19599 | predicted gene, 19599 [Source:MGI Symb | 0 | 0 | 0 | 0 | 0 | 0 |
| 49648 | ENSMUSG00000112080 | 0.481988923 | 2.492401104 | 3.837077151 | 0.649557203 | 0.515978285 | NA | Gm47595 | predicted gene, 47595 [Source:MGI Symb | 0 | 0 | 0 | 2.8919 | 0 | 0 |
| 49649 | ENSMUSG00000112081 | 0 | NA | NA | NA | NA | NA | Gm9004 | predicted gene 9004 [Source:MGI Symbol | 0 | 0 | 0 | 0 | 0 | 0 |
| 49650 | ENSMUSG00000112082 | 9.996506517 | -0.96850334 | 0.743243334 | -1.303077061 | 0.192548454 | 0.63052317 | Gm35035 | predicted gene, 35035 [Source:MGI Symb | 11.243 | 10.279 | 18.112 | 4.8199 | 3.2719 | 12.252 |
| 49651 | ENSMUSG00000112083 | 0 | NA | NA | NA | NA | NA | Gm47107 | predicted gene, 47107 [Source:MGI Symb | 0 | 0 | 0 | 0 | 0 | 0 |
| 49652 | ENSMUSG00000112084 | 0 | NA | NA | NA | NA | NA | Gm10773 | predicted gene 10773 [Source:MGI Symbo | 0 | 0 | 0 | 0 | 0 | 0 |
| 49653 | ENSMUSG00000112085 | 0 | NA | NA | NA | NA | NA | Gm48587 | predicted gene, 48587 [Source:MGI Symb | 0 | 0 | 0 | 0 | 0 | 0 |
| 49654 | ENSMUSG00000112086 | 0 | NA | NA | NA | NA | NA | Gm35412 | predicted gene, 35412 [Source:MGI Symb | 0 | 0 | 0 | 0 | 0 | 0 |
| 49655 | ENSMUSG00000112087 | 0 | NA | NA | NA | NA | NA | Gm40765 | predicted gene, 40765 [Source:MGI Symb | 0 | 0 | 0 | 0 | 0 | 0 |
| 49656 | ENSMUSG00000112088 | 0 | NA | NA | NA | NA | NA | Gm48531 | predicted gene, 48531 [Source:MGI Symb | 0 | 0 | 0 | 0 | 0 | 0 |
| 49657 | ENSMUSG00000112089 | 0 | NA | NA | NA | NA | NA | Gm47712 | predicted gene, 47712 [Source:MGI Symb | 0 | 0 | 0 | 0 | 0 | 0 |
| 49658 | ENSMUSG00000112090 | 0.167707431 | -0.796629176 | 4.080472857 | -0.195229623 | 0.84521319 | NA | Gm48231 | predicted gene, 48231 [Source:MGI Symb | 0 | 0 | 1.0062 | 0 | 0 | 0 |
| 49659 | ENSMUSG00000112091 | 0 | NA | NA | NA | NA | NA | Gm47787 | predicted gene, 47787 [Source:MGI Symb | 0 | 0 | 0 | 0 | 0 | 0 |
| 49660 | ENSMUSG00000112092 | 0 | NA | NA | NA | NA | NA | Gm32936 | predicted gene, 32936 [Source:MGI Symb | 0 | 0 | 0 | 0 | 0 | 0 |
| 49661 | ENSMUSG00000112093 | 0 | NA | NA | NA | NA | NA | Gm47941 | predicted gene, 47941 [Source:MGI Symb | 0 | 0 | 0 | 0 | 0 | 0 |
| 49662 | ENSMUSG00000112094 | 0 | NA | NA | NA | NA | NA | Gm47506 | predicted gene, 47506 [Source:MGI Symb | 0 | 0 | 0 | 0 | 0 | 0 |
| 49663 | ENSMUSG00000112095 | 2.135776909 | -1.088812939 | 1.674597338 | -0.650193879 | 0.515566994 | NA | A130077B15 | RIKEN cDNA A130077B15 gene [Source:M | 5.6217 | 3.0838 | 0 | 1.928 | 2.1812 | 0 |
| 49664 | ENSMUSG00000112096 | 3.223507028 | -0.336001956 | 1.220682214 | -0.275257518 | 0.7831184 | NA | A430103D13 | RIKEN cDNA A430103D13 gene [Source:M | 5.6217 | 2.0558 | 3.0187 | 0.964 | 5.4531 | 2.2277 |
| 49665 | ENSMUSG00000112097 | 0 | NA | NA | NA | NA | NA | Gm32515 | predicted gene, 32515 [Source:MGI Symb | 0 | 0 | 0 | 0 | 0 | 0 |
| 49666 | ENSMUSG00000112098 | 0 | NA | NA | NA | NA | NA | Gm18876 | predicted gene, 18876 [Source:MGI Symb | 0 | 0 | 0 | 0 | 0 | 0 |
| 49667 | ENSMUSG00000112099 | 0 | NA | NA | NA | NA | NA | Gm8960 | predicted gene 8960 [Source:MGI Symbol | 0 | 0 | 0 | 0 | 0 | 0 |
| 49668 | ENSMUSG00000112100 | 0 | NA | NA | NA | NA | NA | Gm9102 | predicted gene 9102 [Source:MGI Symbol | 0 | 0 | 0 | 0 | 0 | 0 |

| | | | | | | | | | | | | | | | |
|---|---|---|---|---|---|---|---|---|---|---|---|---|---|---|---|
| 49669 | ENSMUSG00000112101 | 0 | NA | NA | NA | NA | NA | Gm47924 | predicted gene, 47924 [Source:MGI Symb | 0 | 0 | 0 | 0 | 0 | 0 |
| 49670 | ENSMUSG00000112102 | 0 | NA | NA | NA | NA | NA | Gm32105 | predicted gene, 32105 [Source:MGI Symb | 0 | 0 | 0 | 0 | 0 | 0 |
| 49671 | ENSMUSG00000112103 | 0 | NA | NA | NA | NA | NA | Gm47018 | predicted gene, 47018 [Source:MGI Symb | 0 | 0 | 0 | 0 | 0 | 0 |
| 49672 | ENSMUSG00000112104 | 0 | NA | NA | NA | NA | NA | Gm35722 | predicted gene, 35722 [Source:MGI Symb | 0 | 0 | 0 | 0 | 0 | 0 |
| 49673 | ENSMUSG00000112105 | 1.41313359 | 0.139131381 | 1.987109017 | 0.070016984 | 0.944180141 | NA | Rps7-ps2 | ribosomal protein S7, pseudogene 2 [Sour | 0 | 1.0279 | 3.0187 | 0 | 1.0906 | 3.3415 |
| 49674 | ENSMUSG00000112106 | 0 | NA | NA | NA | NA | NA | Gm47026 | predicted gene, 47026 [Source:MGI Symb | 0 | 0 | 0 | 0 | 0 | 0 |
| 49675 | ENSMUSG00000112107 | 0 | NA | NA | NA | NA | NA | Gm47560 | predicted gene, 47560 [Source:MGI Symb | 0 | 0 | 0 | 0 | 0 | 0 |
| 49676 | ENSMUSG00000112108 | 0 | NA | NA | NA | NA | NA | 4930551I15F | RIKEN cDNA 4930551I15 gene [Source:MG | 0 | 0 | 0 | 0 | 0 | 0 |
| 49677 | ENSMUSG00000112109 | 10.07390931 | 0.278373249 | 0.824477426 | 0.337635986 | 0.73563752 | 0.948109711 | Gm30906 | predicted gene, 30906 [Source:MGI Symb | 12.047 | 1.0279 | 14.087 | 5.7839 | 16.359 | 11.138 |
| 49678 | ENSMUSG00000112110 | 15.30174787 | -0.310946027 | 0.605851157 | -0.513238315 | 0.607784627 | 0.916376001 | Gm15608 | predicted gene 15608 [Source:MGI Symbo | 24.093 | 19.53 | 7.0437 | 12.532 | 16.359 | 12.252 |
| 49679 | ENSMUSG00000112111 | 0 | NA | NA | NA | NA | NA | Gm47834 | predicted gene, 47834 [Source:MGI Symb | 0 | 0 | 0 | 0 | 0 | 0 |
| 49680 | ENSMUSG00000112112 | 0.133850373 | -0.796629176 | 4.080472857 | -0.195229623 | 0.84521319 | NA | Gm48508 | predicted gene, 48508 [Source:MGI Symb | 0.8031 | 0 | 0 | 0 | 0 | 0 |
| 49681 | ENSMUSG00000112113 | 0 | NA | NA | NA | NA | NA | Gm47698 | predicted gene, 47698 [Source:MGI Symb | 0 | 0 | 0 | 0 | 0 | 0 |
| 49682 | ENSMUSG00000112114 | 0 | NA | NA | NA | NA | NA | Gm7540 | predicted gene 7540 [Source:MGI Symbol | 0 | 0 | 0 | 0 | 0 | 0 |
| 49683 | ENSMUSG00000112115 | 0 | NA | NA | NA | NA | NA | Gm21027 | predicted gene, 21027 [Source:MGI Symb | 0 | 0 | 0 | 0 | 0 | 0 |
| 49684 | ENSMUSG00000112116 | 0.160662974 | 1.126924175 | 4.080472857 | 0.276174898 | 0.782413736 | NA | Gm46234 | predicted gene, 46234 [Source:MGI Symb | 0 | 0 | 0 | 0.964 | 0 | 0 |
| 49685 | ENSMUSG00000112117 | 0 | NA | NA | NA | NA | NA | Rmst | rhabdomyosarcoma 2 associated transcri | 0 | 0 | 0 | 0 | 0 | 0 |
| 49686 | ENSMUSG00000112118 | 0 | NA | NA | NA | NA | NA | Gm18039 | predicted gene, 18039 [Source:MGI Symb | 0 | 0 | 0 | 0 | 0 | 0 |
| 49687 | ENSMUSG00000112119 | 0 | NA | NA | NA | NA | NA | Gm47362 | predicted gene, 47362 [Source:MGI Symb | 0 | 0 | 0 | 0 | 0 | 0 |
| 49688 | ENSMUSG00000112120 | 0 | NA | NA | NA | NA | NA | Gm32255 | predicted gene, 32255 [Source:MGI Symb | 0 | 0 | 0 | 0 | 0 | 0 |
| 49689 | ENSMUSG00000112121 | 0 | NA | NA | NA | NA | NA | C230072F16 | RIKEN cDNA C230072F16 gene [Source:M | 0 | 0 | 0 | 0 | 0 | 0 |
| 49690 | ENSMUSG00000112122 | 0 | NA | NA | NA | NA | NA | Gm47628 | predicted gene, 47628 [Source:MGI Symb | 0 | 0 | 0 | 0 | 0 | 0 |
| 49691 | ENSMUSG00000112123 | 0 | NA | NA | NA | NA | NA | Gm5178 | predicted gene 5178 [Source:MGI Symbol | 0 | 0 | 0 | 0 | 0 | 0 |
| 49692 | ENSMUSG00000112124 | 0.610340026 | -2.657561537 | 3.094276798 | -0.858863544 | 0.390415805 | NA | Gm19056 | predicted gene, 19056 [Source:MGI Symb | 1.6062 | 2.0558 | 0 | 0 | 0 | 0 |
| 49693 | ENSMUSG00000112125 | 0 | NA | NA | NA | NA | NA | Gm48024 | predicted gene, 48024 [Source:MGI Symb | 0 | 0 | 0 | 0 | 0 | 0 |
| 49694 | ENSMUSG00000112126 | 0 | NA | NA | NA | NA | NA | Gm34544 | predicted gene, 34544 [Source:MGI Symb | 0 | 0 | 0 | 0 | 0 | 0 |
| 49695 | ENSMUSG00000112127 | 0 | NA | NA | NA | NA | NA | Gm48886 | predicted gene, 48886 [Source:MGI Symb | 0 | 0 | 0 | 0 | 0 | 0 |
| 49696 | ENSMUSG00000112128 | 0 | NA | NA | NA | NA | NA | Gm19217 | predicted gene, 19217 [Source:MGI Symb | 0 | 0 | 0 | 0 | 0 | 0 |
| 49697 | ENSMUSG00000112129 | 5757.03303 | 0.048767822 | 0.089213415 | 0.546642254 | 0.584624534 | 0.908955096 | Pbld1 | phenazine biosynthesis-like protein doma | 5580 | 5940.3 | 5458.9 | 5629.6 | 6134.7 | 5798.7 |
| 49698 | ENSMUSG00000112130 | 0 | NA | NA | NA | NA | NA | Gm47839 | predicted gene, 47839 [Source:MGI Symb | 0 | 0 | 0 | 0 | 0 | 0 |
| 49699 | ENSMUSG00000112131 | 0 | NA | NA | NA | NA | NA | Gm48724 | predicted gene, 48724 [Source:MGI Symb | 0 | 0 | 0 | 0 | 0 | 0 |
| 49700 | ENSMUSG00000112132 | 1.225999361 | -1.452919977 | 2.243334418 | -0.647660895 | 0.517204302 | NA | Gm36229 | predicted gene, 36229 [Source:MGI Symb | 2.4093 | 0 | 3.0187 | 1.928 | 0 | 0 |
| 49701 | ENSMUSG00000112133 | 0 | NA | NA | NA | NA | NA | Gm47247 | predicted gene, 47247 [Source:MGI Symb | 0 | 0 | 0 | 0 | 0 | 0 |
| 49702 | ENSMUSG00000112134 | 0 | NA | NA | NA | NA | NA | Gm40723 | predicted gene, 40723 [Source:MGI Symb | 0 | 0 | 0 | 0 | 0 | 0 |
| 49703 | ENSMUSG00000112135 | 0 | NA | NA | NA | NA | NA | Gm9029 | predicted gene 9029 [Source:MGI Symbol | 0 | 0 | 0 | 0 | 0 | 0 |
| 49704 | ENSMUSG00000112136 | 0 | NA | NA | NA | NA | NA | Gm40692 | predicted gene, 40692 [Source:MGI Symb | 0 | 0 | 0 | 0 | 0 | 0 |
| 49705 | ENSMUSG00000112137 | 0 | NA | NA | NA | NA | NA | Gm47865 | predicted gene, 47865 [Source:MGI Symb | 0 | 0 | 0 | 0 | 0 | 0 |
| 49706 | ENSMUSG00000112138 | 0.328370405 | 0.16514316 | 4.003149536 | 0.041253308 | 0.967093956 | NA | Gm47526 | predicted gene, 47526 [Source:MGI Symb | 0 | 0 | 1.0062 | 0.964 | 0 | 0 |
| 49707 | ENSMUSG00000112139 | 15.5480215 | -0.681995519 | 0.572201267 | -1.191880476 | 0.233308115 | 0.680673483 | Gm47431 | predicted gene, 47431 [Source:MGI Symb | 16.865 | 18.503 | 22.137 | 12.532 | 6.5437 | 16.708 |
| 49708 | ENSMUSG00000112140 | 0 | NA | NA | NA | NA | NA | Gm5780 | predicted gene 5780 [Source:MGI Symbol | 0 | 0 | 0 | 0 | 0 | 0 |
| 49709 | ENSMUSG00000112141 | 0 | NA | NA | NA | NA | NA | 4930563J15F | RIKEN cDNA 4930563J15 gene [Source:MG | 0 | 0 | 0 | 0 | 0 | 0 |
| 49710 | ENSMUSG00000112142 | 0 | NA | NA | NA | NA | NA | Gm47022 | predicted gene, 47022 [Source:MGI Symb | 0 | 0 | 0 | 0 | 0 | 0 |
| 49711 | ENSMUSG00000112143 | 0.160662974 | 1.126924175 | 4.080472857 | 0.276174898 | 0.782413736 | NA | Gm35721 | predicted gene, 35721 [Source:MGI Symb | 0 | 0 | 0 | 0.964 | 0 | 0 |
| 49712 | ENSMUSG00000112144 | 0.678054141 | -2.792799308 | 2.990673428 | -0.933836266 | 0.350388359 | NA | Gm47861 | predicted gene, 47861 [Source:MGI Symb | 0 | 2.0558 | 2.0125 | 0 | 0 | 0 |
| 49713 | ENSMUSG00000112145 | 0.48903338 | 0.95637552 | 3.456690196 | 0.276673773 | 0.782030612 | NA | Gm6218 | predicted gene 6218 [Source:MGI Symbol | 0 | 0 | 1.0062 | 1.928 | 0 | 0 |
| 49714 | ENSMUSG00000112146 | 1.138619952 | -2.330991236 | 2.345811896 | -0.993682077 | 0.320377671 | NA | Gm46210 | predicted gene, 46210 [Source:MGI Symb | 1.6062 | 4.1117 | 0 | 0 | 0 | 1.1138 |
| 49715 | ENSMUSG00000112147 | 0 | NA | NA | NA | NA | NA | Gm18244 | predicted gene, 18244 [Source:MGI Symb | 0 | 0 | 0 | 0 | 0 | 0 |
| 49716 | ENSMUSG00000112148 | 21.23639853 | -2.30447376 | 0.574791164 | -4.009236581 | 6.09E-05 | 0.002440664 | Lilrb4a | leukocyte immunoglobulin-like receptor, s | 31.321 | 45.228 | 29.181 | 2.8919 | 6.5437 | 12.252 |
| 49717 | ENSMUSG00000112149 | 0 | NA | NA | NA | NA | NA | Gm47611 | predicted gene, 47611 [Source:MGI Symb | 0 | 0 | 0 | 0 | 0 | 0 |
| 49718 | ENSMUSG00000112150 | 2.351610801 | 1.508606288 | 1.390522649 | 1.084920328 | 0.277956944 | NA | 9530018F02 | RIKEN cDNA 9530018F02 gene [Source:M | 1.6062 | 1.0279 | 1.0062 | 3.8559 | 3.2719 | 3.3415 |
| 49719 | ENSMUSG00000112151 | 0 | NA | NA | NA | NA | NA | Gm48680 | predicted gene, 48680 [Source:MGI Symb | 0 | 0 | 0 | 0 | 0 | 0 |
| 49720 | ENSMUSG00000112152 | 0 | NA | NA | NA | NA | NA | Gm47045 | predicted gene, 47045 [Source:MGI Symb | 0 | 0 | 0 | 0 | 0 | 0 |
| 49721 | ENSMUSG00000112153 | 0 | NA | NA | NA | NA | NA | Gm47591 | predicted gene, 47591 [Source:MGI Symb | 0 | 0 | 0 | 0 | 0 | 0 |
| 49722 | ENSMUSG00000112154 | 0 | NA | NA | NA | NA | NA | Gm48777 | predicted gene, 48777 [Source:MGI Symb | 0 | 0 | 0 | 0 | 0 | 0 |
| 49723 | ENSMUSG00000112155 | 1.520667694 | 1.047693729 | 1.913505628 | 0.547525815 | 0.58401754 | NA | 4930477N07 | RIKEN cDNA 4930477N07 gene [Source:M | 0 | 1.0279 | 2.0125 | 3.8559 | 0 | 2.2277 |
| 49724 | ENSMUSG00000112156 | 0 | NA | NA | NA | NA | NA | Gm48723 | predicted gene, 48723 [Source:MGI Symb | 0 | 0 | 0 | 0 | 0 | 0 |

| | | | | | | | | | | | | | | | |
|---|---|---|---|---|---|---|---|---|---|---|---|---|---|---|---|
| 49725 | ENSMUSG00000112157 | 0 | NA | NA | NA | NA | NA | 4930545H06 | RIKEN cDNA 4930545H06 gene [Source:M | 0 | 0 | 0 | 0 | 0 | 0 |
| 49726 | ENSMUSG00000112158 | 0 | NA | NA | NA | NA | NA | Gm47822 | predicted gene, 47822 [Source:MGI Symb | 0 | 0 | 0 | 0 | 0 | 0 |
| 49727 | ENSMUSG00000112159 | 0 | NA | NA | NA | NA | NA | Gm18777 | predicted gene, 18777 [Source:MGI Symb | 0 | 0 | 0 | 0 | 0 | 0 |
| 49728 | ENSMUSG00000112160 | 12.07006965 | 0.05625746 | 0.628895392 | 0.0894544 | 0.928720791 | 0.988375949 | BC024063 | cDNA sequence BC024063 [Source:MGI Sy | 7.2279 | 12.335 | 16.1 | 12.532 | 13.087 | 11.138 |
| 49729 | ENSMUSG00000112161 | 0 | NA | NA | NA | NA | NA | Gm34045 | predicted gene, 34045 [Source:MGI Symb | 0 | 0 | 0 | 0 | 0 | 0 |
| 49730 | ENSMUSG00000112162 | 0.17131964 | -0.796629176 | 4.080472857 | -0.195229623 | 0.84521319 | NA | Gm48758 | predicted gene, 48758 [Source:MGI Symb | 0 | 1.0279 | 0 | 0 | 0 | 0 |
| 49731 | ENSMUSG00000112163 | 0 | NA | NA | NA | NA | NA | Gm8188 | predicted gene 8188 [Source:MGI Symbol | 0 | 0 | 0 | 0 | 0 | 0 |
| 49732 | ENSMUSG00000112164 | 0 | NA | NA | NA | NA | NA | Gm19990 | predicted gene, 19990 [Source:MGI Symb | 0 | 0 | 0 | 0 | 0 | 0 |
| 49733 | ENSMUSG00000112165 | 0 | NA | NA | NA | NA | NA | Gm48851 | predicted gene, 48851 [Source:MGI Symb | 0 | 0 | 0 | 0 | 0 | 0 |
| 49734 | ENSMUSG00000112166 | 0 | NA | NA | NA | NA | NA | Gm47948 | predicted gene, 47948 [Source:MGI Symb | 0 | 0 | 0 | 0 | 0 | 0 |
| 49735 | ENSMUSG00000112167 | 0 | NA | NA | NA | NA | NA | Gm47053 | predicted gene, 47053 [Source:MGI Symb | 0 | 0 | 0 | 0 | 0 | 0 |
| 49736 | ENSMUSG00000112168 | 0 | NA | NA | NA | NA | NA | Gm34776 | predicted gene, 34776 [Source:MGI Symb | 0 | 0 | 0 | 0 | 0 | 0 |
| 49737 | ENSMUSG00000112169 | 0 | NA | NA | NA | NA | NA | Gm48536 | predicted gene, 48536 [Source:MGI Symb | 0 | 0 | 0 | 0 | 0 | 0 |
| 49738 | ENSMUSG00000112170 | 0 | NA | NA | NA | NA | NA | Gm9508 | predicted gene 9508 [Source:MGI Symbol | 0 | 0 | 0 | 0 | 0 | 0 |
| 49739 | ENSMUSG00000112171 | 0 | NA | NA | NA | NA | NA | Gm17820 | predicted gene, 17820 [Source:MGI Symb | 0 | 0 | 0 | 0 | 0 | 0 |
| 49740 | ENSMUSG00000112172 | 0 | NA | NA | NA | NA | NA | Gm32172 | predicted gene, 32172 [Source:MGI Symb | 0 | 0 | 0 | 0 | 0 | 0 |
| 49741 | ENSMUSG00000112173 | 0 | NA | NA | NA | NA | NA | Gm48145 | predicted gene, 48145 [Source:MGI Symb | 0 | 0 | 0 | 0 | 0 | 0 |
| 49742 | ENSMUSG00000112174 | 0 | NA | NA | NA | NA | NA | Gm48535 | predicted gene, 48535 [Source:MGI Symb | 0 | 0 | 0 | 0 | 0 | 0 |
| 49743 | ENSMUSG00000112175 | 0 | NA | NA | NA | NA | NA | 4930452L12I | RIKEN cDNA 4930452L12 gene [Source:MG | 0 | 0 | 0 | 0 | 0 | 0 |
| 49744 | ENSMUSG00000112176 | 0 | NA | NA | NA | NA | NA | Gm48205 | predicted gene, 48205 [Source:MGI Symb | 0 | 0 | 0 | 0 | 0 | 0 |
| 49745 | ENSMUSG00000112177 | 0 | NA | NA | NA | NA | NA | Gm18188 | predicted gene, 18188 [Source:MGI Symb | 0 | 0 | 0 | 0 | 0 | 0 |
| 49746 | ENSMUSG00000112178 | 0 | NA | NA | NA | NA | NA | Gm47630 | predicted gene, 47630 [Source:MGI Symb | 0 | 0 | 0 | 0 | 0 | 0 |
| 49747 | ENSMUSG00000112179 | 0 | NA | NA | NA | NA | NA | Gm3213 | predicted gene 3213 [Source:MGI Symbol | 0 | 0 | 0 | 0 | 0 | 0 |
| 49748 | ENSMUSG00000112180 | 0 | NA | NA | NA | NA | NA | Gm48085 | predicted gene, 48085 [Source:MGI Symb | 0 | 0 | 0 | 0 | 0 | 0 |
| 49749 | ENSMUSG00000112181 | 0 | NA | NA | NA | NA | NA | Gm47538 | predicted gene, 47538 [Source:MGI Symb | 0 | 0 | 0 | 0 | 0 | 0 |
| 49750 | ENSMUSG00000112182 | 0 | NA | NA | NA | NA | NA | Gm47845 | predicted gene, 47845 [Source:MGI Symb | 0 | 0 | 0 | 0 | 0 | 0 |
| 49751 | ENSMUSG00000112183 | 0 | NA | NA | NA | NA | NA | Gm48306 | predicted gene, 48306 [Source:MGI Symb | 0 | 0 | 0 | 0 | 0 | 0 |
| 49752 | ENSMUSG00000112184 | 0 | NA | NA | NA | NA | NA | Gm48826 | predicted gene, 48826 [Source:MGI Symb | 0 | 0 | 0 | 0 | 0 | 0 |
| 49753 | ENSMUSG00000112185 | 0 | NA | NA | NA | NA | NA | Gm47060 | predicted gene, 47060 [Source:MGI Symb | 0 | 0 | 0 | 0 | 0 | 0 |
| 49754 | ENSMUSG00000112186 | 0 | NA | NA | NA | NA | NA | Gm2412 | predicted gene 2412 [Source:MGI Symbol | 0 | 0 | 0 | 0 | 0 | 0 |
| 49755 | ENSMUSG00000112187 | 0 | NA | NA | NA | NA | NA | Gm6993 | predicted gene 6993 [Source:MGI Symbol | 0 | 0 | 0 | 0 | 0 | 0 |
| 49756 | ENSMUSG00000112188 | 0.995947842 | -0.043351441 | 2.675688395 | -0.016201977 | 0.987073258 | NA | Gm47708 | predicted gene, 47708 [Source:MGI Symb | 0 | 3.0838 | 0 | 2.8919 | 0 | 0 |
| 49757 | ENSMUSG00000112189 | 33.04860765 | 0.663438567 | 0.392612573 | 1.689804691 | 0.091065326 | 0.445203913 | C730027H18 | RIKEN cDNA C730027H18 gene [Source:M | 29.715 | 28.782 | 18.112 | 42.415 | 43.625 | 35.643 |
| 49758 | ENSMUSG00000112190 | 0 | NA | NA | NA | NA | NA | Gm47209 | predicted gene, 47209 [Source:MGI Symb | 0 | 0 | 0 | 0 | 0 | 0 |
| 49759 | ENSMUSG00000112191 | 0 | NA | NA | NA | NA | NA | Gm31931 | predicted gene, 31931 [Source:MGI Symb | 0 | 0 | 0 | 0 | 0 | 0 |
| 49760 | ENSMUSG00000112192 | 0 | NA | NA | NA | NA | NA | Gm48341 | predicted gene, 48341 [Source:MGI Symb | 0 | 0 | 0 | 0 | 0 | 0 |
| 49761 | ENSMUSG00000112193 | 0 | NA | NA | NA | NA | NA | Gm48206 | predicted gene, 48206 [Source:MGI Symb | 0 | 0 | 0 | 0 | 0 | 0 |
| 49762 | ENSMUSG00000112194 | 0 | NA | NA | NA | NA | NA | Gm48029 | predicted gene, 48029 [Source:MGI Symb | 0 | 0 | 0 | 0 | 0 | 0 |
| 49763 | ENSMUSG00000112195 | 0 | NA | NA | NA | NA | NA | Gm35405 | predicted gene, 35405 [Source:MGI Symb | 0 | 0 | 0 | 0 | 0 | 0 |
| 49764 | ENSMUSG00000112196 | 2.679852851 | -1.060843873 | 1.438804368 | -0.737309322 | 0.460934264 | NA | Gm8274 | predicted gene 8274 [Source:MGI Symbol | 4.8186 | 0 | 6.0375 | 1.928 | 2.1812 | 1.1138 |
| 49765 | ENSMUSG00000112197 | 0 | NA | NA | NA | NA | NA | Gm47337 | predicted gene, 47337 [Source:MGI Symb | 0 | 0 | 0 | 0 | 0 | 0 |
| 49766 | ENSMUSG00000112198 | 0.294513347 | 0.165143496 | 4.080472857 | 0.040471657 | 0.967717103 | NA | Gm4065 | predicted gene 4065 [Source:MGI Symbol | 0.8031 | 0 | 0 | 0.964 | 0 | 0 |
| 49767 | ENSMUSG00000112199 | 0 | NA | NA | NA | NA | NA | Gm19090 | predicted gene, 19090 [Source:MGI Symb | 0 | 0 | 0 | 0 | 0 | 0 |
| 49768 | ENSMUSG00000112200 | 0 | NA | NA | NA | NA | NA | Gm19161 | predicted gene, 19161 [Source:MGI Symb | 0 | 0 | 0 | 0 | 0 | 0 |
| 49769 | ENSMUSG00000112201 | 0.181769715 | 1.126924175 | 4.080472857 | 0.276174898 | 0.782413736 | NA | 4930532I03F | RIKEN cDNA 4930532I03 gene [Source:MG | 0 | 0 | 0 | 0 | 1.0906 | 0 |
| 49770 | ENSMUSG00000112202 | 0 | NA | NA | NA | NA | NA | Gm31181 | predicted gene, 31181 [Source:MGI Symb | 0 | 0 | 0 | 0 | 0 | 0 |
| 49771 | ENSMUSG00000112203 | 2.502696982 | 1.219710379 | 1.358182581 | 0.898045959 | 0.369161048 | NA | Gm49167 | predicted gene, 49167 [Source:MGI Symb | 2.4093 | 1.0279 | 1.0062 | 2.8919 | 5.4531 | 2.2277 |
| 49772 | ENSMUSG00000112204 | 0.74256259 | 3.09446134 | 3.273353887 | 0.945348852 | 0.344480805 | NA | Gm47553 | predicted gene, 47553 [Source:MGI Symb | 0 | 0 | 0 | 0 | 0 | 4.4554 |
| 49773 | ENSMUSG00000112205 | 0 | NA | NA | NA | NA | NA | Gm47546 | predicted gene, 47546 [Source:MGI Symb | 0 | 0 | 0 | 0 | 0 | 0 |
| 49774 | ENSMUSG00000112206 | 0 | NA | NA | NA | NA | NA | Gm30228 | predicted gene, 30228 [Source:MGI Symb | 0 | 0 | 0 | 0 | 0 | 0 |
| 49775 | ENSMUSG00000112207 | 0 | NA | NA | NA | NA | NA | Gm48448 | predicted gene, 48448 [Source:MGI Symb | 0 | 0 | 0 | 0 | 0 | 0 |
| 49776 | ENSMUSG00000112208 | 0 | NA | NA | NA | NA | NA | Gm40797 | predicted gene, 40797 [Source:MGI Symb | 0 | 0 | 0 | 0 | 0 | 0 |
| 49777 | ENSMUSG00000112209 | 0 | NA | NA | NA | NA | NA | Gm18730 | predicted gene, 18730 [Source:MGI Symb | 0 | 0 | 0 | 0 | 0 | 0 |
| 49778 | ENSMUSG00000112210 | 0 | NA | NA | NA | NA | NA | 4930407I19F | RIKEN cDNA 4930407I19 gene [Source:MG | 0 | 0 | 0 | 0 | 0 | 0 |
| 49779 | ENSMUSG00000112211 | 0 | NA | NA | NA | NA | NA | Gm47682 | predicted gene, 47682 [Source:MGI Symb | 0 | 0 | 0 | 0 | 0 | 0 |
| 49780 | ENSMUSG00000112212 | 0 | NA | NA | NA | NA | NA | Gm46189 | predicted gene, 46189 [Source:MGI Symb | 0 | 0 | 0 | 0 | 0 | 0 |

| | | | | | | | | | | | | | | | |
|---|---|---|---|---|---|---|---|---|---|---|---|---|---|---|---|
| 49781 | ENSMUSG00000112213 | 0 | NA | NA | NA | NA | NA | 4930555G07 | RIKEN cDNA 4930555G07 gene [Source:M | 0 | 0 | 0 | 0 | 0 | 0 |
| 49782 | ENSMUSG00000112214 | 0 | NA | NA | NA | NA | NA | Cyp3a61-ps1 | cytochrome P450, family 3, subfamily a, m | 0 | 0 | 0 | 0 | 0 | 0 |
| 49783 | ENSMUSG00000112215 | 0 | NA | NA | NA | NA | NA | Gm2089 | predicted gene 2089 [Source:MGI Symbol | 0 | 0 | 0 | 0 | 0 | 0 |
| 49784 | ENSMUSG00000112216 | 0 | NA | NA | NA | NA | NA | Gm32717 | predicted gene, 32717 [Source:MGI Symb | 0 | 0 | 0 | 0 | 0 | 0 |
| 49785 | ENSMUSG00000112217 | 0 | NA | NA | NA | NA | NA | Gm48055 | predicted gene, 48055 [Source:MGI Symb | 0 | 0 | 0 | 0 | 0 | 0 |
| 49786 | ENSMUSG00000112218 | 0 | NA | NA | NA | NA | NA | Gm18726 | predicted gene, 18726 [Source:MGI Symb | 0 | 0 | 0 | 0 | 0 | 0 |
| 49787 | ENSMUSG00000112219 | 0 | NA | NA | NA | NA | NA | Gm31904 | predicted gene, 31904 [Source:MGI Symb | 0 | 0 | 0 | 0 | 0 | 0 |
| 49788 | ENSMUSG00000112220 | 0 | NA | NA | NA | NA | NA | Gm36208 | predicted gene, 36208 [Source:MGI Symb | 0 | 0 | 0 | 0 | 0 | 0 |
| 49789 | ENSMUSG00000112221 | 0 | NA | NA | NA | NA | NA | Gm47142 | predicted gene, 47142 [Source:MGI Symb | 0 | 0 | 0 | 0 | 0 | 0 |
| 49790 | ENSMUSG00000112222 | 0 | NA | NA | NA | NA | NA | Tyms-ps | thymidylate synthase, pseudogene [Sourc | 0 | 0 | 0 | 0 | 0 | 0 |
| 49791 | ENSMUSG00000112223 | 0 | NA | NA | NA | NA | NA | Krtap10-10 | keratin associated protein 10-10 [Source:I | 0 | 0 | 0 | 0 | 0 | 0 |
| 49792 | ENSMUSG00000112224 | 0 | NA | NA | NA | NA | NA | Gm47944 | predicted gene, 47944 [Source:MGI Symb | 0 | 0 | 0 | 0 | 0 | 0 |
| 49793 | ENSMUSG00000112225 | 0 | NA | NA | NA | NA | NA | Gm48854 | predicted gene, 48854 [Source:MGI Symb | 0 | 0 | 0 | 0 | 0 | 0 |
| 49794 | ENSMUSG00000112226 | 0 | NA | NA | NA | NA | NA | Gm48786 | predicted gene, 48786 [Source:MGI Symb | 0 | 0 | 0 | 0 | 0 | 0 |
| 49795 | ENSMUSG00000112227 | 0 | NA | NA | NA | NA | NA | C330004P14 | RIKEN cDNA C330004P14 gene [Source:M | 0 | 0 | 0 | 0 | 0 | 0 |
| 49796 | ENSMUSG00000112228 | 0 | NA | NA | NA | NA | NA | Gm47524 | predicted gene, 47524 [Source:MGI Symb | 0 | 0 | 0 | 0 | 0 | 0 |
| 49797 | ENSMUSG00000112229 | 5.715920534 | -1.222270233 | 0.940165369 | -1.300058769 | 0.193580827 | 0.631833452 | Gm48086 | predicted gene, 48086 [Source:MGI Symb | 8.8341 | 7.1954 | 8.05 | 5.7839 | 1.0906 | 3.3415 |
| 49798 | ENSMUSG00000112230 | 0 | NA | NA | NA | NA | NA | Ifngas1 | Ifng antisense RNA 1 [Source:MGI Symbol | 0 | 0 | 0 | 0 | 0 | 0 |
| 49799 | ENSMUSG00000112231 | 0 | NA | NA | NA | NA | NA | Gm8435 | predicted gene 8435 [Source:MGI Symbol | 0 | 0 | 0 | 0 | 0 | 0 |
| 49800 | ENSMUSG00000112232 | 0 | NA | NA | NA | NA | NA | Gm48578 | predicted gene, 48578 [Source:MGI Symb | 0 | 0 | 0 | 0 | 0 | 0 |
| 49801 | ENSMUSG00000112233 | 0 | NA | NA | NA | NA | NA | Gm48698 | predicted gene, 48698 [Source:MGI Symb | 0 | 0 | 0 | 0 | 0 | 0 |
| 49802 | ENSMUSG00000112234 | 0.160662974 | 1.126924175 | 4.080472857 | 0.276174898 | 0.782413736 | NA | Gm48427 | predicted gene, 48427 [Source:MGI Symb | 0 | 0 | 0 | 0.964 | 0 | 0 |
| 49803 | ENSMUSG00000112235 | 0 | NA | NA | NA | NA | NA | Gm48307 | predicted gene, 48307 [Source:MGI Symb | 0 | 0 | 0 | 0 | 0 | 0 |
| 49804 | ENSMUSG00000112236 | 0 | NA | NA | NA | NA | NA | Gm48180 | predicted gene, 48180 [Source:MGI Symb | 0 | 0 | 0 | 0 | 0 | 0 |
| 49805 | ENSMUSG00000112237 | 7.175960214 | -0.156601453 | 0.806040271 | -0.1942849 | 0.845952808 | 0.972803041 | D630033A02 | RIKEN cDNA D630033A02 gene [Source:M | 6.4248 | 6.1675 | 10.062 | 3.8559 | 7.6343 | 8.9108 |
| 49806 | ENSMUSG00000112238 | 0 | NA | NA | NA | NA | NA | Gm48844 | predicted gene, 48844 [Source:MGI Symb | 0 | 0 | 0 | 0 | 0 | 0 |
| 49807 | ENSMUSG00000112239 | 3.289417233 | 0.771873184 | 1.21882505 | 0.633292845 | 0.526542425 | NA | Gm17823 | predicted gene, 17823 [Source:MGI Symb | 3.2124 | 1.0279 | 3.0187 | 4.8199 | 6.5437 | 1.1138 |
| 49808 | ENSMUSG00000112240 | 0 | NA | NA | NA | NA | NA | Gm48310 | predicted gene, 48310 [Source:MGI Symb | 0 | 0 | 0 | 0 | 0 | 0 |
| 49809 | ENSMUSG00000112241 | 0 | NA | NA | NA | NA | NA | Gm49325 | predicted gene, 49325 [Source:MGI Symb | 0 | 0 | 0 | 0 | 0 | 0 |
| 49810 | ENSMUSG00000112242 | 0 | NA | NA | NA | NA | NA | Gm10747 | predicted gene 10747 [Source:MGI Symbo | 0 | 0 | 0 | 0 | 0 | 0 |
| 49811 | ENSMUSG00000112243 | 0.331982614 | 0.16514309 | 3.986751476 | 0.041422971 | 0.966958701 | NA | Gm47342 | predicted gene, 47342 [Source:MGI Symb | 0 | 1.0279 | 0 | 0.964 | 0 | 0 |
| 49812 | ENSMUSG00000112244 | 0 | NA | NA | NA | NA | NA | Gm48788 | predicted gene, 48788 [Source:MGI Symb | 0 | 0 | 0 | 0 | 0 | 0 |
| 49813 | ENSMUSG00000112245 | 0 | NA | NA | NA | NA | NA | Gm8689 | predicted gene 8689 [Source:MGI Symbol | 0 | 0 | 0 | 0 | 0 | 0 |
| 49814 | ENSMUSG00000112246 | 0 | NA | NA | NA | NA | NA | Gm40761 | predicted gene, 40761 [Source:MGI Symb | 0 | 0 | 0 | 0 | 0 | 0 |
| 49815 | ENSMUSG00000112247 | 0 | NA | NA | NA | NA | NA | Gm47894 | predicted gene, 47894 [Source:MGI Symb | 0 | 0 | 0 | 0 | 0 | 0 |
| 49816 | ENSMUSG00000112248 | 0 | NA | NA | NA | NA | NA | Gm47910 | predicted gene, 47910 [Source:MGI Symb | 0 | 0 | 0 | 0 | 0 | 0 |
| 49817 | ENSMUSG00000112249 | 497.0023018 | 0.531357755 | 0.137182285 | 3.87337007 | 0.000107341 | 0.003833519 | Gm30262 | predicted gene, 30262 [Source:MGI Symb | 390.31 | 433.78 | 395.45 | 531.15 | 655.46 | 575.86 |
| 49818 | ENSMUSG00000112250 | 0 | NA | NA | NA | NA | NA | Gm47837 | predicted gene, 47837 [Source:MGI Symb | 0 | 0 | 0 | 0 | 0 | 0 |
| 49819 | ENSMUSG00000112251 | 0 | NA | NA | NA | NA | NA | Gm47488 | predicted gene, 47488 [Source:MGI Symb | 0 | 0 | 0 | 0 | 0 | 0 |
| 49820 | ENSMUSG00000112252 | 0 | NA | NA | NA | NA | NA | Gm32802 | predicted gene, 32802 [Source:MGI Symb | 0 | 0 | 0 | 0 | 0 | 0 |
| 49821 | ENSMUSG00000112253 | 0 | NA | NA | NA | NA | NA | Gm47329 | predicted gene, 47329 [Source:MGI Symb | 0 | 0 | 0 | 0 | 0 | 0 |
| 49822 | ENSMUSG00000112254 | 2.116041437 | -0.116651766 | 1.540260386 | -0.075735094 | 0.939629855 | NA | Gm47690 | predicted gene, 47690 [Source:MGI Symb | 1.6062 | 1.0279 | 4.025 | 3.8559 | 2.1812 | 0 |
| 49823 | ENSMUSG00000112255 | 1.037954732 | 2.366557412 | 2.405105848 | 0.98397225 | 0.325129157 | NA | Gm47594 | predicted gene, 47594 [Source:MGI Symb | 0 | 1.0279 | 0 | 1.928 | 3.2719 | 0 |
| 49824 | ENSMUSG00000112256 | 0 | NA | NA | NA | NA | NA | Gm48077 | predicted gene, 48077 [Source:MGI Symb | 0 | 0 | 0 | 0 | 0 | 0 |
| 49825 | ENSMUSG00000112257 | 0 | NA | NA | NA | NA | NA | Gm47623 | predicted gene, 47623 [Source:MGI Symb | 0 | 0 | 0 | 0 | 0 | 0 |
| 49826 | ENSMUSG00000112258 | 0 | NA | NA | NA | NA | NA | Gm10165 | predicted gene 10165 [Source:MGI Symbo | 0 | 0 | 0 | 0 | 0 | 0 |
| 49827 | ENSMUSG00000112259 | 0 | NA | NA | NA | NA | NA | Gm17980 | predicted gene, 17980 [Source:MGI Symb | 0 | 0 | 0 | 0 | 0 | 0 |
| 49828 | ENSMUSG00000112260 | 0 | NA | NA | NA | NA | NA | 3110043J17I | RIKEN cDNA 3110043J17 gene [Source:MC | 0 | 0 | 0 | 0 | 0 | 0 |
| 49829 | ENSMUSG00000112261 | 0 | NA | NA | NA | NA | NA | Gm47599 | predicted gene, 47599 [Source:MGI Symb | 0 | 0 | 0 | 0 | 0 | 0 |
| 49830 | ENSMUSG00000112262 | 0 | NA | NA | NA | NA | NA | Gm48013 | predicted gene, 48013 [Source:MGI Symb | 0 | 0 | 0 | 0 | 0 | 0 |
| 49831 | ENSMUSG00000112263 | 1.600168826 | 0.814054987 | 1.846189732 | 0.44093788 | 0.659257969 | NA | Gm47874 | predicted gene, 47874 [Source:MGI Symb | 2.4093 | 1.0279 | 0 | 2.8919 | 3.2719 | 0 |
| 49832 | ENSMUSG00000112264 | 0 | NA | NA | NA | NA | NA | Gm48505 | predicted gene, 48505 [Source:MGI Symb | 0 | 0 | 0 | 0 | 0 | 0 |
| 49833 | ENSMUSG00000112265 | 13.51817404 | -1.8605623 | 0.696695221 | -2.670554132 | 0.007572616 | 0.088616762 | Gm34983 | predicted gene, 34983 [Source:MGI Symb | 28.109 | 17.475 | 18.112 | 9.6398 | 1.0906 | 6.6831 |
| 49834 | ENSMUSG00000112266 | 0.160662974 | 1.126924175 | 4.080472857 | 0.276174898 | 0.782413736 | NA | Gm48843 | predicted gene, 48843 [Source:MGI Symb | 0 | 0 | 0 | 0.964 | 0 | 0 |
| 49835 | ENSMUSG00000112267 | 0 | NA | NA | NA | NA | NA | Gm47903 | predicted gene, 47903 [Source:MGI Symb | 0 | 0 | 0 | 0 | 0 | 0 |
| 49836 | ENSMUSG00000112268 | 0 | NA | NA | NA | NA | NA | Gm48460 | predicted gene, 48460 [Source:MGI Symb | 0 | 0 | 0 | 0 | 0 | 0 |

| | | | | | | | | | | | | | | | |
|---|---|---|---|---|---|---|---|---|---|---|---|---|---|---|---|
| 49837 | ENSMUSG00000112269 | 0.513752329 | 1.031892132 | 3.082998058 | 0.334704114 | 0.737848298 | NA | Gm6906 | predicted gene 6906 [Source:MGI Symbol | 0 | 1.0279 | 0 | 0.964 | 1.0906 | 0 |
| 49838 | ENSMUSG00000112270 | 0 | NA | NA | NA | NA | NA | Gm47613 | predicted gene, 47613 [Source:MGI Symb | 0 | 0 | 0 | 0 | 0 | 0 |
| 49839 | ENSMUSG00000112271 | 0 | NA | NA | NA | NA | NA | Gm9088 | predicted gene 9088 [Source:MGI Symbol | 0 | 0 | 0 | 0 | 0 | 0 |
| 49840 | ENSMUSG00000112272 | 0 | NA | NA | NA | NA | NA | Gm48087 | predicted gene, 48087 [Source:MGI Symb | 0 | 0 | 0 | 0 | 0 | 0 |
| 49841 | ENSMUSG00000112273 | 0 | NA | NA | NA | NA | NA | Gm29783 | predicted gene, 29783 [Source:MGI Symb | 0 | 0 | 0 | 0 | 0 | 0 |
| 49842 | ENSMUSG00000112274 | 2.706556199 | 0.589281804 | 1.40951586 | 0.418073908 | 0.675893081 | NA | Gm47525 | predicted gene, 47525 [Source:MGI Symb | 2.4093 | 0 | 4.025 | 0.964 | 3.2719 | 5.5692 |
| 49843 | ENSMUSG00000112275 | 8.677090203 | 0.251124765 | 0.718125137 | 0.349694994 | 0.726567611 | 0.945777495 | Gm47078 | predicted gene, 47078 [Source:MGI Symb | 10.44 | 7.1954 | 6.0375 | 9.6398 | 8.7249 | 10.025 |
| 49844 | ENSMUSG00000112276 | 7.759507041 | 1.634462949 | 0.822463259 | 1.987277766 | 0.046891624 | 0.300126868 | 5033421B08 | RIKEN cDNA 5033421B08 gene [Source:M | 3.2124 | 5.1396 | 3.0187 | 7.7118 | 17.45 | 10.025 |
| 49845 | ENSMUSG00000112277 | 0 | NA | NA | NA | NA | NA | Gm48867 | predicted gene, 48867 [Source:MGI Symb | 0 | 0 | 0 | 0 | 0 | 0 |
| 49846 | ENSMUSG00000112278 | 72.1232114 | -0.2480101 | 0.294997556 | -0.840719169 | 0.400505279 | 0.827266295 | Gm30025 | predicted gene, 30025 [Source:MGI Symb | 105.21 | 67.843 | 61.381 | 69.406 | 66.528 | 62.375 |
| 49847 | ENSMUSG00000112279 | 0 | NA | NA | NA | NA | NA | Gm18774 | predicted gene, 18774 [Source:MGI Symb | 0 | 0 | 0 | 0 | 0 | 0 |
| 49848 | ENSMUSG00000112280 | 7.863667131 | 0.687026676 | 0.7810834 | 0.879581714 | 0.379085948 | 0.812043903 | A830082N09 | RIKEN cDNA A830082N09 gene [Source:M | 8.8341 | 3.0838 | 6.0375 | 11.568 | 7.6343 | 10.025 |
| 49849 | ENSMUSG00000112281 | 0 | NA | NA | NA | NA | NA | Gm47226 | predicted gene, 47226 [Source:MGI Symb | 0 | 0 | 0 | 0 | 0 | 0 |
| 49850 | ENSMUSG00000112282 | 0 | NA | NA | NA | NA | NA | Gm19989 | predicted gene, 19989 [Source:MGI Symb | 0 | 0 | 0 | 0 | 0 | 0 |
| 49851 | ENSMUSG00000112283 | 0 | NA | NA | NA | NA | NA | Gm36719 | predicted gene, 36719 [Source:MGI Symb | 0 | 0 | 0 | 0 | 0 | 0 |
| 49852 | ENSMUSG00000112284 | 0 | NA | NA | NA | NA | NA | Gm48845 | predicted gene, 48845 [Source:MGI Symb | 0 | 0 | 0 | 0 | 0 | 0 |
| 49853 | ENSMUSG00000112285 | 0 | NA | NA | NA | NA | NA | Gm50278 | predicted gene, 50278 [Source:MGI Symb | 0 | 0 | 0 | 0 | 0 | 0 |
| 49854 | ENSMUSG00000112286 | 0.342432689 | 1.990653086 | 3.900281263 | 0.51038706 | 0.609780321 | NA | Gm47561 | predicted gene, 47561 [Source:MGI Symb | 0 | 0 | 0 | 0.964 | 1.0906 | 0 |
| 49855 | ENSMUSG00000112287 | 0 | NA | NA | NA | NA | NA | Gm47024 | predicted gene, 47024 [Source:MGI Symb | 0 | 0 | 0 | 0 | 0 | 0 |
| 49856 | ENSMUSG00000112288 | 2.797960997 | -0.674240774 | 1.344284192 | -0.501561186 | 0.615976228 | NA | Gm47879 | predicted gene, 47879 [Source:MGI Symb | 3.2124 | 1.0279 | 6.0375 | 0.964 | 1.0906 | 4.4554 |
| 49857 | ENSMUSG00000112289 | 0.321325949 | 1.911575114 | 4.048267893 | 0.472195804 | 0.636787034 | NA | 1110019B22 | RIKEN cDNA 1110019B22 gene [Source:M | 0 | 0 | 0 | 1.928 | 0 | 0 |
| 49858 | ENSMUSG00000112290 | 0.167707431 | -0.796629176 | 4.080472857 | -0.195229623 | 0.84521319 | NA | Gm33782 | predicted gene, 33782 [Source:MGI Symb | 0 | 0 | 1.0062 | 0 | 0 | 0 |
| 49859 | ENSMUSG00000112291 | 0 | NA | NA | NA | NA | NA | Gm48276 | predicted gene, 48276 [Source:MGI Symb | 0 | 0 | 0 | 0 | 0 | 0 |
| 49860 | ENSMUSG00000112292 | 0.503122293 | -2.366675966 | 3.785596405 | -0.625179156 | 0.531853481 | NA | Gm8275 | predicted gene 8275 [Source:MGI Symbol | 0 | 0 | 3.0187 | 0 | 0 | 0 |
| 49861 | ENSMUSG00000112293 | 0 | NA | NA | NA | NA | NA | Gm48509 | predicted gene, 48509 [Source:MGI Symb | 0 | 0 | 0 | 0 | 0 | 0 |
| 49862 | ENSMUSG00000112294 | 24.81070117 | -0.184835609 | 0.43113675 | -0.428716895 | 0.668129264 | 0.930136128 | Gm4129 | predicted gene 4129 [Source:MGI Symbol | 23.29 | 29.81 | 26.162 | 22.171 | 21.812 | 25.618 |
| 49863 | ENSMUSG00000112295 | 0 | NA | NA | NA | NA | NA | Gm47017 | predicted gene, 47017 [Source:MGI Symb | 0 | 0 | 0 | 0 | 0 | 0 |
| 49864 | ENSMUSG00000112296 | 0 | NA | NA | NA | NA | NA | Gm47768 | predicted gene, 47768 [Source:MGI Symb | 0 | 0 | 0 | 0 | 0 | 0 |
| 49865 | ENSMUSG00000112297 | 3.220883095 | -1.068439304 | 1.220955948 | -0.875084237 | 0.381528073 | NA | Gm17816 | predicted gene, 17816 [Source:MGI Symb | 4.0155 | 2.0558 | 7.0437 | 2.8919 | 1.0906 | 2.2277 |
| 49866 | ENSMUSG00000112298 | 0 | NA | NA | NA | NA | NA | Gm48088 | predicted gene, 48088 [Source:MGI Symb | 0 | 0 | 0 | 0 | 0 | 0 |
| 49867 | ENSMUSG00000112299 | 0 | NA | NA | NA | NA | NA | Gm17870 | predicted gene, 17870 [Source:MGI Symb | 0 | 0 | 0 | 0 | 0 | 0 |
| 49868 | ENSMUSG00000112300 | 30.13213736 | -0.571342871 | 0.42172635 | -1.354771574 | 0.175490346 | 0.605810563 | Gm47976 | predicted gene, 47976 [Source:MGI Symb | 32.927 | 33.921 | 41.256 | 22.171 | 33.809 | 16.708 |
| 49869 | ENSMUSG00000112301 | 1.996512071 | 0.589990934 | 1.60743875 | 0.367037894 | 0.713590752 | NA | Gm10752 | predicted gene 10752 [Source:MGI Symbo | 0.8031 | 0 | 4.025 | 3.8559 | 2.1812 | 1.1138 |
| 49870 | ENSMUSG00000112302 | 98.33113837 | 0.059184011 | 0.244768418 | 0.241795947 | 0.80893828 | 0.965705123 | Gm48226 | predicted gene, 48226 [Source:MGI Symb | 107.62 | 79.15 | 101.63 | 88.686 | 97.065 | 115.84 |
| 49871 | ENSMUSG00000112303 | 0 | NA | NA | NA | NA | NA | Gm48759 | predicted gene, 48759 [Source:MGI Symb | 0 | 0 | 0 | 0 | 0 | 0 |
| 49872 | ENSMUSG00000112304 | 0 | NA | NA | NA | NA | NA | Gm35206 | predicted gene, 35206 [Source:MGI Symb | 0 | 0 | 0 | 0 | 0 | 0 |
| 49873 | ENSMUSG00000112305 | 0 | NA | NA | NA | NA | NA | Gm48875 | predicted gene, 48875 [Source:MGI Symb | 0 | 0 | 0 | 0 | 0 | 0 |
| 49874 | ENSMUSG00000112306 | 0 | NA | NA | NA | NA | NA | Gm47744 | predicted gene, 47744 [Source:MGI Symb | 0 | 0 | 0 | 0 | 0 | 0 |
| 49875 | ENSMUSG00000112307 | 0 | NA | NA | NA | NA | NA | Gm48751 | predicted gene, 48751 [Source:MGI Symb | 0 | 0 | 0 | 0 | 0 | 0 |
| 49876 | ENSMUSG00000112308 | 0 | NA | NA | NA | NA | NA | Gm48512 | predicted gene, 48512 [Source:MGI Symb | 0 | 0 | 0 | 0 | 0 | 0 |
| 49877 | ENSMUSG00000112309 | 0 | NA | NA | NA | NA | NA | Gm48353 | predicted gene, 48353 [Source:MGI Symb | 0 | 0 | 0 | 0 | 0 | 0 |
| 49878 | ENSMUSG00000112310 | 0 | NA | NA | NA | NA | NA | NA | NA | 0 | 0 | 0 | 0 | 0 | 0 |
| 49879 | ENSMUSG00000112311 | 0.738691657 | 3.085611452 | 2.924644825 | 1.055038008 | 0.291407942 | NA | Gm8153 | predicted gene 8153 [Source:MGI Symbol | 0 | 0 | 0 | 0 | 1.0906 | 3.3415 |
| 49880 | ENSMUSG00000112312 | 0 | NA | NA | NA | NA | NA | Gm47755 | predicted gene, 47755 [Source:MGI Symb | 0 | 0 | 0 | 0 | 0 | 0 |
| 49881 | ENSMUSG00000112313 | 0 | NA | NA | NA | NA | NA | Gm5951 | predicted gene 5951 [Source:MGI Symbol | 0 | 0 | 0 | 0 | 0 | 0 |
| 49882 | ENSMUSG00000112314 | 0 | NA | NA | NA | NA | NA | Gm49454 | predicted gene, 49454 [Source:MGI Symb | 0 | 0 | 0 | 0 | 0 | 0 |
| 49883 | ENSMUSG00000112315 | 0 | NA | NA | NA | NA | NA | Gm5183 | predicted gene 5183 [Source:MGI Symbol | 0 | 0 | 0 | 0 | 0 | 0 |
| 49884 | ENSMUSG00000112316 | 0 | NA | NA | NA | NA | NA | Gm48425 | predicted gene, 48425 [Source:MGI Symb | 0 | 0 | 0 | 0 | 0 | 0 |
| 49885 | ENSMUSG00000112317 | 0.615839296 | 1.537131441 | 3.17677869 | 0.483864817 | 0.628481811 | NA | Gm47580 | predicted gene, 47580 [Source:MGI Symb | 0.8031 | 0 | 0 | 2.8919 | 0 | 0 |
| 49886 | ENSMUSG00000112318 | 0 | NA | NA | NA | NA | NA | Gm6721 | predicted gene 6721 [Source:MGI Symbol | 0 | 0 | 0 | 0 | 0 | 0 |
| 49887 | ENSMUSG00000112319 | 0.167707431 | -0.796629176 | 4.080472857 | -0.195229623 | 0.84521319 | NA | Gm47221 | predicted gene, 47221 [Source:MGI Symb | 0 | 0 | 1.0062 | 0 | 0 | 0 |
| 49888 | ENSMUSG00000112320 | 0 | NA | NA | NA | NA | NA | Gm48022 | predicted gene, 48022 [Source:MGI Symb | 0 | 0 | 0 | 0 | 0 | 0 |
| 49889 | ENSMUSG00000112321 | 0 | NA | NA | NA | NA | NA | Gm47866 | predicted gene, 47866 [Source:MGI Symb | 0 | 0 | 0 | 0 | 0 | 0 |
| 49890 | ENSMUSG00000112322 | 0 | NA | NA | NA | NA | NA | Gm48115 | predicted gene, 48115 [Source:MGI Symb | 0 | 0 | 0 | 0 | 0 | 0 |
| 49891 | ENSMUSG00000112323 | 0 | NA | NA | NA | NA | NA | Gm32828 | predicted gene, 32828 [Source:MGI Symb | 0 | 0 | 0 | 0 | 0 | 0 |
| 49892 | ENSMUSG00000112324 | 0 | NA | NA | NA | NA | NA | Gm47939 | predicted gene, 47939 [Source:MGI Symb | 0 | 0 | 0 | 0 | 0 | 0 |

| | | | | | | | | | | | | | | | |
|---|---|---|---|---|---|---|---|---|---|---|---|---|---|---|---|
| 49893 | ENSMUSG00000112325 | 0 | NA | NA | NA | NA | NA | Gm48454 | predicted gene, 48454 [Source:MGI Symb | 0 | 0 | 0 | 0 | 0 | 0 |
| 49894 | ENSMUSG00000112326 | 0 | NA | NA | NA | NA | NA | Gm19624 | predicted gene, 19624 [Source:MGI Symb | 0 | 0 | 0 | 0 | 0 | 0 |
| 49895 | ENSMUSG00000112327 | 0 | NA | NA | NA | NA | NA | Gm36827 | predicted gene, 36827 [Source:MGI Symb | 0 | 0 | 0 | 0 | 0 | 0 |
| 49896 | ENSMUSG00000112328 | 0 | NA | NA | NA | NA | NA | Gm47325 | predicted gene, 47325 [Source:MGI Symb | 0 | 0 | 0 | 0 | 0 | 0 |
| 49897 | ENSMUSG00000112329 | 0 | NA | NA | NA | NA | NA | Gm47908 | predicted gene, 47908 [Source:MGI Symb | 0 | 0 | 0 | 0 | 0 | 0 |
| 49898 | ENSMUSG00000112330 | 0 | NA | NA | NA | NA | NA | Gm3266 | predicted gene 3266 [Source:MGI Symbol | 0 | 0 | 0 | 0 | 0 | 0 |
| 49899 | ENSMUSG00000112331 | 0 | NA | NA | NA | NA | NA | Gm33035 | predicted gene, 33035 [Source:MGI Symb | 0 | 0 | 0 | 0 | 0 | 0 |
| 49900 | ENSMUSG00000112332 | 15.33396127 | 0.369510678 | 0.572946128 | 0.644930927 | 0.518971949 | 0.887592015 | 4930466K18 | RIKEN cDNA 4930466K18 gene [Source:M | 8.8341 | 12.335 | 19.119 | 15.424 | 21.812 | 14.48 |
| 49901 | ENSMUSG00000112333 | 0 | NA | NA | NA | NA | NA | 1700048P04 | RIKEN cDNA 1700048P04 gene [Source:M | 0 | 0 | 0 | 0 | 0 | 0 |
| 49902 | ENSMUSG00000112334 | 0 | NA | NA | NA | NA | NA | Gm48047 | predicted gene, 48047 [Source:MGI Symb | 0 | 0 | 0 | 0 | 0 | 0 |
| 49903 | ENSMUSG00000112335 | 0 | NA | NA | NA | NA | NA | Gm46188 | predicted gene, 46188 [Source:MGI Symb | 0 | 0 | 0 | 0 | 0 | 0 |
| 49904 | ENSMUSG00000112336 | 0.319491021 | 0.16514334 | 4.044653141 | 0.040830038 | 0.967431393 | NA | Gm47766 | predicted gene, 47766 [Source:MGI Symb | 0.8031 | 0 | 0 | 0 | 0 | 1.1138 |
| 49905 | ENSMUSG00000112337 | 0 | NA | NA | NA | NA | NA | Gm47614 | predicted gene, 47614 [Source:MGI Symb | 0 | 0 | 0 | 0 | 0 | 0 |
| 49906 | ENSMUSG00000112338 | 0.331982614 | 0.16514309 | 3.986751476 | 0.041422971 | 0.966958701 | NA | Gm47693 | predicted gene, 47693 [Source:MGI Symb | 0 | 1.0279 | 0 | 0.964 | 0 | 0 |
| 49907 | ENSMUSG00000112339 | 0.185640647 | 1.126924175 | 4.080472857 | 0.276174898 | 0.782413736 | NA | Gm47529 | predicted gene, 47529 [Source:MGI Symb | 0 | 0 | 0 | 0 | 0 | 1.1138 |
| 49908 | ENSMUSG00000112340 | 0 | NA | NA | NA | NA | NA | Gm47016 | predicted gene, 47016 [Source:MGI Symb | 0 | 0 | 0 | 0 | 0 | 0 |
| 49909 | ENSMUSG00000112341 | 0 | NA | NA | NA | NA | NA | Gm4796 | predicted gene 4796 [Source:MGI Symbol | 0 | 0 | 0 | 0 | 0 | 0 |
| 49910 | ENSMUSG00000112342 | 0 | NA | NA | NA | NA | NA | Gm47031 | predicted gene, 47031 [Source:MGI Symb | 0 | 0 | 0 | 0 | 0 | 0 |
| 49911 | ENSMUSG00000112343 | 0 | NA | NA | NA | NA | NA | Sfta3-ps | surfactant associated 3, pseudogene [Sou | 0 | 0 | 0 | 0 | 0 | 0 |
| 49912 | ENSMUSG00000112344 | 0 | NA | NA | NA | NA | NA | Gm32141 | predicted gene, 32141 [Source:MGI Symb | 0 | 0 | 0 | 0 | 0 | 0 |
| 49913 | ENSMUSG00000112345 | 0 | NA | NA | NA | NA | NA | Gm40652 | predicted gene, 40652 [Source:MGI Symb | 0 | 0 | 0 | 0 | 0 | 0 |
| 49914 | ENSMUSG00000112346 | 18.60737946 | 0.257542329 | 0.519251922 | 0.495987243 | 0.619903411 | 0.917977899 | Gm48768 | predicted gene, 48768 [Source:MGI Symb | 15.259 | 23.642 | 12.075 | 22.171 | 22.903 | 15.594 |
| 49915 | ENSMUSG00000112347 | 2.196215002 | 1.307128173 | 1.519414629 | 0.860284052 | 0.389632482 | NA | Gm47257 | predicted gene, 47257 [Source:MGI Symb | 0.8031 | 0 | 3.0187 | 3.8559 | 3.2719 | 2.2277 |
| 49916 | ENSMUSG00000112348 | 0.17131964 | -0.796629176 | 4.080472857 | -0.195229623 | 0.84521319 | NA | Gm36634 | predicted gene, 36634 [Source:MGI Symb | 0 | 1.0279 | 0 | 0 | 0 | 0 |
| 49917 | ENSMUSG00000112349 | 0.133850373 | -0.796629176 | 4.080472857 | -0.195229623 | 0.84521319 | NA | Gm48132 | predicted gene, 48132 [Source:MGI Symb | 0.8031 | 0 | 0 | 0 | 0 | 0 |
| 49918 | ENSMUSG00000112350 | 0 | NA | NA | NA | NA | NA | Gm47697 | predicted gene, 47697 [Source:MGI Symb | 0 | 0 | 0 | 0 | 0 | 0 |
| 49919 | ENSMUSG00000112351 | 0 | NA | NA | NA | NA | NA | Gm48326 | predicted gene, 48326 [Source:MGI Symb | 0 | 0 | 0 | 0 | 0 | 0 |
| 49920 | ENSMUSG00000112352 | 0.492645588 | 0.9565034 | 3.447648281 | 0.277436479 | 0.781444974 | NA | Gm40617 | predicted gene, 40617 [Source:MGI Symb | 0 | 1.0279 | 0 | 1.928 | 0 | 0 |
| 49921 | ENSMUSG00000112353 | 0 | NA | NA | NA | NA | NA | Gm40418 | predicted gene, 40418 [Source:MGI Symb | 0 | 0 | 0 | 0 | 0 | 0 |
| 49922 | ENSMUSG00000112354 | 0 | NA | NA | NA | NA | NA | Gm33843 | predicted gene, 33843 [Source:MGI Symb | 0 | 0 | 0 | 0 | 0 | 0 |
| 49923 | ENSMUSG00000112355 | 0 | NA | NA | NA | NA | NA | Gm48285 | predicted gene, 48285 [Source:MGI Symb | 0 | 0 | 0 | 0 | 0 | 0 |
| 49924 | ENSMUSG00000112357 | 0 | NA | NA | NA | NA | NA | Gm47864 | predicted gene, 47864 [Source:MGI Symb | 0 | 0 | 0 | 0 | 0 | 0 |
| 49925 | ENSMUSG00000112358 | 0 | NA | NA | NA | NA | NA | Gm33869 | predicted gene, 33869 [Source:MGI Symb | 0 | 0 | 0 | 0 | 0 | 0 |
| 49926 | ENSMUSG00000112359 | 0 | NA | NA | NA | NA | NA | Gm48475 | predicted gene, 48475 [Source:MGI Symb | 0 | 0 | 0 | 0 | 0 | 0 |
| 49927 | ENSMUSG00000112360 | 0.167707431 | -0.796629176 | 4.080472857 | -0.195229623 | 0.84521319 | NA | Gm48614 | predicted gene, 48614 [Source:MGI Symb | 0 | 0 | 1.0062 | 0 | 0 | 0 |
| 49928 | ENSMUSG00000112361 | 0 | NA | NA | NA | NA | NA | Gm47607 | predicted gene, 47607 [Source:MGI Symb | 0 | 0 | 0 | 0 | 0 | 0 |
| 49929 | ENSMUSG00000112362 | 0 | NA | NA | NA | NA | NA | Gm48507 | predicted gene, 48507 [Source:MGI Symb | 0 | 0 | 0 | 0 | 0 | 0 |
| 49930 | ENSMUSG00000112363 | 0 | NA | NA | NA | NA | NA | Gm4178 | predicted gene 4178 [Source:MGI Symbol | 0 | 0 | 0 | 0 | 0 | 0 |
| 49931 | ENSMUSG00000112364 | 0 | NA | NA | NA | NA | NA | 4930503E24 | RIKEN cDNA 4930503E24 gene [Source:M | 0 | 0 | 0 | 0 | 0 | 0 |
| 49932 | ENSMUSG00000112365 | 2.328540777 | -1.722551047 | 1.679027116 | -1.025922113 | 0.304928298 | NA | NA | NA | 5.6217 | 0 | 5.0312 | 0 | 1.0906 | 2.2277 |
| 49933 | ENSMUSG00000112366 | 131.1365292 | 0.437463533 | 0.237407994 | 1.842665553 | 0.065377856 | 0.367233132 | C730027H18 | RIKEN cDNA C730027H18 gene [Source:M | 122.07 | 88.401 | 123.77 | 186.05 | 141.78 | 124.75 |
| 49934 | ENSMUSG00000112367 | 0 | NA | NA | NA | NA | NA | A230081H15 | Riken cDNA A230081H15 gene [Source:M | 0 | 0 | 0 | 0 | 0 | 0 |
| 49935 | ENSMUSG00000112368 | 2.15368515 | 0.428694213 | 1.399175977 | 0.30639049 | 0.759307348 | NA | Gm49376 | predicted gene, 49376 [Source:MGI Symb | 2.4093 | 2.0558 | 1.0062 | 1.928 | 2.1812 | 3.3415 |
| 49936 | ENSMUSG00000112369 | 0 | NA | NA | NA | NA | NA | Gm47911 | predicted gene, 47911 [Source:MGI Symb | 0 | 0 | 0 | 0 | 0 | 0 |
| 49937 | ENSMUSG00000112370 | 0 | NA | NA | NA | NA | NA | Gm48385 | predicted gene, 48385 [Source:MGI Symb | 0 | 0 | 0 | 0 | 0 | 0 |
| 49938 | ENSMUSG00000112371 | 0 | NA | NA | NA | NA | NA | Gm7791 | predicted gene 7791 [Source:MGI Symbol | 0 | 0 | 0 | 0 | 0 | 0 |
| 49939 | ENSMUSG00000112372 | 0 | NA | NA | NA | NA | NA | Gm48779 | predicted gene, 48779 [Source:MGI Symb | 0 | 0 | 0 | 0 | 0 | 0 |
| 49940 | ENSMUSG00000112373 | 0 | NA | NA | NA | NA | NA | Gm18556 | predicted gene, 18556 [Source:MGI Symb | 0 | 0 | 0 | 0 | 0 | 0 |
| 49941 | ENSMUSG00000112374 | 10.43245795 | 0.006399804 | 0.801530635 | 0.007984479 | 0.993629375 | 0.99859176 | Gm48072 | predicted gene, 48072 [Source:MGI Symb | 12.047 | 1.0279 | 18.112 | 11.568 | 9.8156 | 10.025 |
| 49942 | ENSMUSG00000112375 | 8.80901495 | -1.008257191 | 0.759549152 | -1.327441664 | 0.184362625 | 0.619137675 | Gm38560 | predicted gene, 38560 [Source:MGI Symb | 12.047 | 10.279 | 13.081 | 8.6758 | 6.5437 | 2.2277 |
| 49943 | ENSMUSG00000112376 | 0 | NA | NA | NA | NA | NA | Gm40649 | predicted gene, 40649 [Source:MGI Symb | 0 | 0 | 0 | 0 | 0 | 0 |
| 49944 | ENSMUSG00000112377 | 0 | NA | NA | NA | NA | NA | Gm48794 | predicted gene, 48794 [Source:MGI Symb | 0 | 0 | 0 | 0 | 0 | 0 |
| 49945 | ENSMUSG00000112378 | 0 | NA | NA | NA | NA | NA | Gm47261 | predicted gene, 47261 [Source:MGI Symb | 0 | 0 | 0 | 0 | 0 | 0 |
| 49946 | ENSMUSG00000112379 | 0 | NA | NA | NA | NA | NA | Gm47900 | predicted gene, 47900 [Source:MGI Symb | 0 | 0 | 0 | 0 | 0 | 0 |
| 49947 | ENSMUSG00000112380 | 0 | NA | NA | NA | NA | NA | Gm9736 | predicted gene 9736 [Source:MGI Symbol | 0 | 0 | 0 | 0 | 0 | 0 |
| 49948 | ENSMUSG00000112381 | 0.849167191 | -0.527085716 | 2.389288686 | -0.220603613 | 0.825401088 | NA | Gm8170 | predicted gene 8170 [Source:MGI Symbol | 0 | 1.0279 | 2.0125 | 0.964 | 1.0906 | 0 |

| | | | | | | | | | | | | | | | |
|---|---|---|---|---|---|---|---|---|---|---|---|---|---|---|---|
| 49949 | ENSMUSG00000112382 | 0 | NA | NA | NA | NA | NA | Gm47228 | predicted gene, 47228 [Source:MGI Symb | 0 | 0 | 0 | 0 | 0 | 0 |
| 49950 | ENSMUSG00000112383 | 0 | NA | NA | NA | NA | NA | Gm47895 | predicted gene, 47895 [Source:MGI Symb | 0 | 0 | 0 | 0 | 0 | 0 |
| 49951 | ENSMUSG00000112384 | 4.551715669 | -0.655878082 | 1.063498291 | -0.61671757 | 0.53742103 | NA | Gm34921 | predicted gene, 34921 [Source:MGI Symb | 10.44 | 4.1117 | 2.0125 | 1.928 | 4.3625 | 4.4554 |
| 49952 | ENSMUSG00000112385 | 0 | NA | NA | NA | NA | NA | Gm48258 | predicted gene, 48258 [Source:MGI Symb | 0 | 0 | 0 | 0 | 0 | 0 |
| 49953 | ENSMUSG00000112386 | 0 | NA | NA | NA | NA | NA | Gm47752 | predicted gene, 47752 [Source:MGI Symb | 0 | 0 | 0 | 0 | 0 | 0 |
| 49954 | ENSMUSG00000112387 | 0 | NA | NA | NA | NA | NA | 1700021G15 | RIKEN cDNA 1700021G15 gene [Source:M | 0 | 0 | 0 | 0 | 0 | 0 |
| 49955 | ENSMUSG00000112388 | 21.25546043 | -0.29586564 | 0.488638303 | -0.605490069 | 0.544853422 | 0.8954353 | AI463170 | expressed sequence AI463170 [Source:M | 22.487 | 20.558 | 27.169 | 14.46 | 27.265 | 15.594 |
| 49956 | ENSMUSG00000112389 | 0 | NA | NA | NA | NA | NA | Gm48891 | predicted gene, 48891 [Source:MGI Symb | 0 | 0 | 0 | 0 | 0 | 0 |
| 49957 | ENSMUSG00000112390 | 0 | NA | NA | NA | NA | NA | Gm20208 | predicted gene, 20208 [Source:MGI Symb | 0 | 0 | 0 | 0 | 0 | 0 |
| 49958 | ENSMUSG00000112391 | 0 | NA | NA | NA | NA | NA | Gm48230 | predicted gene, 48230 [Source:MGI Symb | 0 | 0 | 0 | 0 | 0 | 0 |
| 49959 | ENSMUSG00000112392 | 0 | NA | NA | NA | NA | NA | Gm35240 | predicted gene, 35240 [Source:MGI Symb | 0 | 0 | 0 | 0 | 0 | 0 |
| 49960 | ENSMUSG00000112393 | 0.167707431 | -0.796629176 | 4.080472857 | -0.195229623 | 0.84521319 | NA | Gm48655 | predicted gene, 48655 [Source:MGI Symb | 0 | 0 | 1.0062 | 0 | 0 | 0 |
| 49961 | ENSMUSG00000112394 | 0 | NA | NA | NA | NA | NA | Gm48121 | predicted gene, 48121 [Source:MGI Symb | 0 | 0 | 0 | 0 | 0 | 0 |
| 49962 | ENSMUSG00000112395 | 0 | NA | NA | NA | NA | NA | Gm7198 | predicted gene 7198 [Source:MGI Symbol | 0 | 0 | 0 | 0 | 0 | 0 |
| 49963 | ENSMUSG00000112396 | 0 | NA | NA | NA | NA | NA | Gm47574 | predicted gene, 47574 [Source:MGI Symb | 0 | 0 | 0 | 0 | 0 | 0 |
| 49964 | ENSMUSG00000112397 | 1.133023873 | 0.521379031 | 2.122130646 | 0.245686585 | 0.805924858 | NA | Gm10824 | predicted gene 10824 [Source:MGI Symbo | 0.8031 | 0 | 2.0125 | 2.8919 | 1.0906 | 0 |
| 49965 | ENSMUSG00000112398 | 0 | NA | NA | NA | NA | NA | 9230102K24 | RIKEN cDNA 9230102K24 gene [Source:M | 0 | 0 | 0 | 0 | 0 | 0 |
| 49966 | ENSMUSG00000112399 | 0 | NA | NA | NA | NA | NA | Gm32235 | predicted gene, 32235 [Source:MGI Symb | 0 | 0 | 0 | 0 | 0 | 0 |
| 49967 | ENSMUSG00000112400 | 6.412375689 | 0.398756024 | 1.018784334 | 0.391403765 | 0.69549881 | 0.939992732 | Gm47725 | predicted gene, 47725 [Source:MGI Symb | 1.6062 | 1.0279 | 14.087 | 9.6398 | 6.5437 | 5.5692 |
| 49968 | ENSMUSG00000112401 | 0 | NA | NA | NA | NA | NA | Gm47999 | predicted gene, 47999 [Source:MGI Symb | 0 | 0 | 0 | 0 | 0 | 0 |
| 49969 | ENSMUSG00000112402 | 0 | NA | NA | NA | NA | NA | Gm10744 | predicted gene 10744 [Source:MGI Symbo | 0 | 0 | 0 | 0 | 0 | 0 |
| 49970 | ENSMUSG00000112403 | 0 | NA | NA | NA | NA | NA | Gm47227 | predicted gene, 47227 [Source:MGI Symb | 0 | 0 | 0 | 0 | 0 | 0 |
| 49971 | ENSMUSG00000112404 | 0 | NA | NA | NA | NA | NA | Gm32442 | predicted gene, 32442 [Source:MGI Symb | 0 | 0 | 0 | 0 | 0 | 0 |
| 49972 | ENSMUSG00000112405 | 0 | NA | NA | NA | NA | NA | Fkbp1a-ps2 | FK506 binding protein 1a, pseudogene 2 [ | 0 | 0 | 0 | 0 | 0 | 0 |
| 49973 | ENSMUSG00000112406 | 0 | NA | NA | NA | NA | NA | 4921515L22 | RIKEN cDNA 4921515L22 gene [Source:M | 0 | 0 | 0 | 0 | 0 | 0 |
| 49974 | ENSMUSG00000112407 | 0 | NA | NA | NA | NA | NA | Gm33340 | predicted gene, 33340 [Source:MGI Symb | 0 | 0 | 0 | 0 | 0 | 0 |
| 49975 | ENSMUSG00000112408 | 0 | NA | NA | NA | NA | NA | Gm8701 | predicted gene 8701 [Source:MGI Symbol | 0 | 0 | 0 | 0 | 0 | 0 |
| 49976 | ENSMUSG00000112409 | 0 | NA | NA | NA | NA | NA | C030006N10 | RIKEN cDNA C030006N10 gene [Source:M | 0 | 0 | 0 | 0 | 0 | 0 |
| 49977 | ENSMUSG00000112410 | 0 | NA | NA | NA | NA | NA | B230217J21 | RIKEN cDNA B230217J21 gene [Source:M | 0 | 0 | 0 | 0 | 0 | 0 |
| 49978 | ENSMUSG00000112411 | 0.670803094 | 1.59186814 | 2.791527899 | 0.570249769 | 0.568508305 | NA | Gm19337 | predicted gene, 19337 [Source:MGI Symb | 0 | 0 | 1.0062 | 1.928 | 1.0906 | 0 |
| 49979 | ENSMUSG00000112412 | 0 | NA | NA | NA | NA | NA | Gm35239 | predicted gene, 35239 [Source:MGI Symb | 0 | 0 | 0 | 0 | 0 | 0 |
| 49980 | ENSMUSG00000112413 | 0 | NA | NA | NA | NA | NA | Gm6867 | predicted gene 6867 [Source:MGI Symbol | 0 | 0 | 0 | 0 | 0 | 0 |
| 49981 | ENSMUSG00000112414 | 0 | NA | NA | NA | NA | NA | Gm34609 | predicted gene, 34609 [Source:MGI Symb | 0 | 0 | 0 | 0 | 0 | 0 |
| 49982 | ENSMUSG00000112415 | 0 | NA | NA | NA | NA | NA | Gm48426 | predicted gene, 48426 [Source:MGI Symb | 0 | 0 | 0 | 0 | 0 | 0 |
| 49983 | ENSMUSG00000112416 | 0 | NA | NA | NA | NA | NA | Gm47028 | predicted gene, 47028 [Source:MGI Symb | 0 | 0 | 0 | 0 | 0 | 0 |
| 49984 | ENSMUSG00000112417 | 0 | NA | NA | NA | NA | NA | A430028G04 | RIKEN cDNA A430028G04 gene [Source:M | 0 | 0 | 0 | 0 | 0 | 0 |
| 49985 | ENSMUSG00000112418 | 0 | NA | NA | NA | NA | NA | Gm9466 | predicted gene 9466 [Source:MGI Symbol | 0 | 0 | 0 | 0 | 0 | 0 |
| 49986 | ENSMUSG00000112419 | 0 | NA | NA | NA | NA | NA | Gm9044 | predicted gene 9044 [Source:MGI Symbol | 0 | 0 | 0 | 0 | 0 | 0 |
| 49987 | ENSMUSG00000112420 | 0 | NA | NA | NA | NA | NA | Gm48543 | predicted gene, 48543 [Source:MGI Symb | 0 | 0 | 0 | 0 | 0 | 0 |
| 49988 | ENSMUSG00000112421 | 0 | NA | NA | NA | NA | NA | 4933406B15 | RIKEN cDNA 4933406B15 gene [Source:M | 0 | 0 | 0 | 0 | 0 | 0 |
| 49989 | ENSMUSG00000112422 | 0 | NA | NA | NA | NA | NA | Gm31938 | predicted gene, 31938 [Source:MGI Symb | 0 | 0 | 0 | 0 | 0 | 0 |
| 49990 | ENSMUSG00000112423 | 0 | NA | NA | NA | NA | NA | Gm6333 | predicted gene 6333 [Source:MGI Symbol | 0 | 0 | 0 | 0 | 0 | 0 |
| 49991 | ENSMUSG00000112424 | 0 | NA | NA | NA | NA | NA | Gm6494 | predicted gene 6494 [Source:MGI Symbol | 0 | 0 | 0 | 0 | 0 | 0 |
| 49992 | ENSMUSG00000112425 | 0 | NA | NA | NA | NA | NA | Gm47896 | predicted gene, 47896 [Source:MGI Symb | 0 | 0 | 0 | 0 | 0 | 0 |
| 49993 | ENSMUSG00000112426 | 2.696654061 | -0.718227998 | 1.365122347 | -0.5261272 | 0.598799831 | NA | Gm47200 | predicted gene, 47200 [Source:MGI Symb | 4.0155 | 2.0558 | 4.025 | 3.8559 | 0 | 2.2277 |
| 49994 | ENSMUSG00000112427 | 0 | NA | NA | NA | NA | NA | Gm19168 | predicted gene, 19168 [Source:MGI Symb | 0 | 0 | 0 | 0 | 0 | 0 |
| 49995 | ENSMUSG00000112428 | 0 | NA | NA | NA | NA | NA | Gm48106 | predicted gene, 48106 [Source:MGI Symb | 0 | 0 | 0 | 0 | 0 | 0 |
| 49996 | ENSMUSG00000112429 | 0.995947842 | -0.043351441 | 2.675688395 | -0.016201977 | 0.987073258 | NA | Gm47093 | predicted gene, 47093 [Source:MGI Symb | 0 | 3.0838 | 0 | 2.8919 | 0 | 0 |
| 49997 | ENSMUSG00000112430 | 0 | NA | NA | NA | NA | NA | Gm30624 | predicted gene, 30624 [Source:MGI Symb | 0 | 0 | 0 | 0 | 0 | 0 |
| 49998 | ENSMUSG00000112431 | 0 | NA | NA | NA | NA | NA | Gm10741 | predicted gene 10741 [Source:MGI Symbo | 0 | 0 | 0 | 0 | 0 | 0 |
| 49999 | ENSMUSG00000112432 | 0 | NA | NA | NA | NA | NA | Gm18507 | predicted gene, 18507 [Source:MGI Symb | 0 | 0 | 0 | 0 | 0 | 0 |
| 50000 | ENSMUSG00000112433 | 17.49116237 | -0.719594977 | 0.553538157 | -1.29999164 | 0.193603834 | 0.631833452 | Gm30122 | predicted gene, 30122 [Source:MGI Symb | 22.487 | 29.81 | 13.081 | 16.388 | 9.8156 | 13.366 |
| 50001 | ENSMUSG00000112434 | 0 | NA | NA | NA | NA | NA | Gm7594 | predicted gene 7594 [Source:MGI Symbol | 0 | 0 | 0 | 0 | 0 | 0 |
| 50002 | ENSMUSG00000112435 | 0 | NA | NA | NA | NA | NA | Gm47965 | predicted gene, 47965 [Source:MGI Symb | 0 | 0 | 0 | 0 | 0 | 0 |
| 50003 | ENSMUSG00000112436 | 0 | NA | NA | NA | NA | NA | Gm48403 | predicted gene, 48403 [Source:MGI Symb | 0 | 0 | 0 | 0 | 0 | 0 |
| 50004 | ENSMUSG00000112437 | 0.642651897 | 2.906894826 | 3.440535777 | 0.844895974 | 0.398168923 | NA | Gm35188 | predicted gene, 35188 [Source:MGI Symb | 0 | 0 | 0 | 3.8559 | 0 | 0 |

| | | | | | | | | | | | | | | | |
|---|---|---|---|---|---|---|---|---|---|---|---|---|---|---|---|
| 50005 | ENSMUSG00000112438 | 0 | NA | NA | NA | NA | NA | Gm40810 | predicted gene, 40810 [Source:MGI Symb | 0 | 0 | 0 | 0 | 0 | 0 |
| 50006 | ENSMUSG00000112439 | 0 | NA | NA | NA | NA | NA | Gm33336 | predicted gene, 33336 [Source:MGI Symb | 0 | 0 | 0 | 0 | 0 | 0 |
| 50007 | ENSMUSG00000112440 | 0 | NA | NA | NA | NA | NA | Gm47722 | predicted gene, 47722 [Source:MGI Symb | 0 | 0 | 0 | 0 | 0 | 0 |
| 50008 | ENSMUSG00000112441 | 1.984122801 | -3.243759051 | 1.867811062 | -1.736663369 | 0.082446607 | NA | Gm48898 | predicted gene, 48898 [Source:MGI Symb | 1.6062 | 7.1954 | 2.0125 | 0 | 1.0906 | 0 |
| 50009 | ENSMUSG00000112442 | 1.621302197 | -1.001523998 | 1.854428868 | -0.540071402 | 0.589147792 | NA | Gm47992 | predicted gene, 47992 [Source:MGI Symb | 2.4093 | 1.0279 | 3.0187 | 0 | 3.2719 | 0 |
| 50010 | ENSMUSG00000112443 | 0.835104907 | -1.8196403 | 2.6116807 | -0.696731534 | 0.48597082 | NA | Gm19007 | predicted gene, 19007 [Source:MGI Symb | 0 | 1.0279 | 3.0187 | 0.964 | 0 | 0 |
| 50011 | ENSMUSG00000112444 | 0.185640647 | 1.126924175 | 4.080472857 | 0.276174898 | 0.782413736 | NA | 4930401C15 | RIKEN cDNA 4930401C15 gene [Source:M | 0 | 0 | 0 | 0 | 0 | 1.1138 |
| 50012 | ENSMUSG00000112445 | 0 | NA | NA | NA | NA | NA | Gm47830 | predicted gene, 47830 [Source:MGI Symb | 0 | 0 | 0 | 0 | 0 | 0 |
| 50013 | ENSMUSG00000112446 | 0.133850373 | -0.796629176 | 4.080472857 | -0.195229623 | 0.84521319 | NA | Gm47949 | predicted gene, 47949 [Source:MGI Symb | 0.8031 | 0 | 0 | 0 | 0 | 0 |
| 50014 | ENSMUSG00000112447 | 0 | NA | NA | NA | NA | NA | Gm47935 | predicted gene, 47935 [Source:MGI Symb | 0 | 0 | 0 | 0 | 0 | 0 |
| 50015 | ENSMUSG00000112448 | 0 | NA | NA | NA | NA | NA | Gm48182 | predicted gene, 48182 [Source:MGI Symb | 0 | 0 | 0 | 0 | 0 | 0 |
| 50016 | ENSMUSG00000112449 | 0.820751756 | 0.96244451 | 2.606827123 | 0.36920151 | 0.71197753 | NA | Srp54b | signal recognition particle 54B [Source:M( | 1.6062 | 0 | 0 | 0 | 1.0906 | 2.2277 |
| 50017 | ENSMUSG00000112450 | 0.167707431 | -0.796629176 | 4.080472857 | -0.195229623 | 0.84521319 | NA | Gm3619 | predicted gene 3619 [Source:MGI Symbol | 0 | 0 | 1.0062 | 0 | 0 | 0 |
| 50018 | ENSMUSG00000112451 | 0 | NA | NA | NA | NA | NA | Gm6714 | predicted gene 6714 [Source:MGI Symbol | 0 | 0 | 0 | 0 | 0 | 0 |
| 50019 | ENSMUSG00000112452 | 0 | NA | NA | NA | NA | NA | Gm48075 | predicted gene, 48075 [Source:MGI Symb | 0 | 0 | 0 | 0 | 0 | 0 |
| 50020 | ENSMUSG00000112453 | 0 | NA | NA | NA | NA | NA | Gm4781 | predicted gene 4781 [Source:MGI Symbol | 0 | 0 | 0 | 0 | 0 | 0 |
| 50021 | ENSMUSG00000112454 | 0 | NA | NA | NA | NA | NA | Gm48619 | predicted gene, 48619 [Source:MGI Symb | 0 | 0 | 0 | 0 | 0 | 0 |
| 50022 | ENSMUSG00000112455 | 0 | NA | NA | NA | NA | NA | Gm48289 | predicted gene, 48289 [Source:MGI Symb | 0 | 0 | 0 | 0 | 0 | 0 |
| 50023 | ENSMUSG00000112456 | 0 | NA | NA | NA | NA | NA | Gm6991 | predicted gene 6991 [Source:MGI Symbol | 0 | 0 | 0 | 0 | 0 | 0 |
| 50024 | ENSMUSG00000112457 | 0 | NA | NA | NA | NA | NA | Gm48687 | predicted gene, 48687 [Source:MGI Symb | 0 | 0 | 0 | 0 | 0 | 0 |
| 50025 | ENSMUSG00000112458 | 0.513958919 | -2.394274821 | 3.756096013 | -0.637437066 | 0.523840191 | NA | Gm40773 | predicted gene, 40773 [Source:MGI Symb | 0 | 3.0838 | 0 | 0 | 0 | 0 |
| 50026 | ENSMUSG00000112459 | 0.339027071 | -1.79174149 | 3.919585585 | -0.457125237 | 0.647581032 | NA | Gm47689 | predicted gene, 47689 [Source:MGI Symb | 0 | 1.0279 | 1.0062 | 0 | 0 | 0 |
| 50027 | ENSMUSG00000112460 | 0 | NA | NA | NA | NA | NA | Gm48120 | predicted gene, 48120 [Source:MGI Symb | 0 | 0 | 0 | 0 | 0 | 0 |
| 50028 | ENSMUSG00000112461 | 0.401551119 | -2.072982726 | 4.008251979 | -0.517178744 | 0.605031388 | NA | Gm47625 | predicted gene, 47625 [Source:MGI Symb | 2.4093 | 0 | 0 | 0 | 0 | 0 |
| 50029 | ENSMUSG00000112462 | 0.401551119 | -2.072982726 | 4.008251979 | -0.517178744 | 0.605031388 | NA | Gm48772 | predicted gene, 48772 [Source:MGI Symb | 2.4093 | 0 | 0 | 0 | 0 | 0 |
| 50030 | ENSMUSG00000112463 | 0 | NA | NA | NA | NA | NA | Gm47579 | predicted gene, 47579 [Source:MGI Symb | 0 | 0 | 0 | 0 | 0 | 0 |
| 50031 | ENSMUSG00000112464 | 0 | NA | NA | NA | NA | NA | Gm47609 | predicted gene, 47609 [Source:MGI Symb | 0 | 0 | 0 | 0 | 0 | 0 |
| 50032 | ENSMUSG00000112465 | 0.51014012 | 1.031855138 | 3.09002894 | 0.333930574 | 0.738431948 | NA | Gm18387 | predicted gene, 18387 [Source:MGI Symb | 0 | 0 | 1.0062 | 0.964 | 1.0906 | 0 |
| 50033 | ENSMUSG00000112466 | 0.315620088 | 0.165143421 | 4.063300985 | 0.040642675 | 0.967580763 | NA | Gm29674 | predicted gene, 29674 [Source:MGI Symb | 0.8031 | 0 | 0 | 0 | 1.0906 | 0 |
| 50034 | ENSMUSG00000112467 | 0 | NA | NA | NA | NA | NA | Gm48144 | predicted gene, 48144 [Source:MGI Symb | 0 | 0 | 0 | 0 | 0 | 0 |
| 50035 | ENSMUSG00000112468 | 0 | NA | NA | NA | NA | NA | Gm47720 | predicted gene, 47720 [Source:MGI Symb | 0 | 0 | 0 | 0 | 0 | 0 |
| 50036 | ENSMUSG00000112469 | 0 | NA | NA | NA | NA | NA | Gm47823 | predicted gene, 47823 [Source:MGI Symb | 0 | 0 | 0 | 0 | 0 | 0 |
| 50037 | ENSMUSG00000112470 | 0 | NA | NA | NA | NA | NA | 5330439M1( | RIKEN cDNA 5330439M10 gene [Source:M | 0 | 0 | 0 | 0 | 0 | 0 |
| 50038 | ENSMUSG00000112471 | 0 | NA | NA | NA | NA | NA | Gm5779 | predicted gene 5779 [Source:MGI Symbol | 0 | 0 | 0 | 0 | 0 | 0 |
| 50039 | ENSMUSG00000112472 | 0 | NA | NA | NA | NA | NA | Gm48083 | predicted gene, 48083 [Source:MGI Symb | 0 | 0 | 0 | 0 | 0 | 0 |
| 50040 | ENSMUSG00000112473 | 0 | NA | NA | NA | NA | NA | Gm48658 | predicted gene, 48658 [Source:MGI Symb | 0 | 0 | 0 | 0 | 0 | 0 |
| 50041 | ENSMUSG00000112474 | 2.448322766 | 2.799556596 | 1.646845771 | 1.699950684 | 0.089140202 | NA | Gm46209 | predicted gene, 46209 [Source:MGI Symb | 0.8031 | 0 | 1.0062 | 1.928 | 8.7249 | 2.2277 |
| 50042 | ENSMUSG00000112475 | 0 | NA | NA | NA | NA | NA | Gm47703 | predicted gene, 47703 [Source:MGI Symb | 0 | 0 | 0 | 0 | 0 | 0 |
| 50043 | ENSMUSG00000112476 | 0 | NA | NA | NA | NA | NA | Gm32365 | predicted gene, 32365 [Source:MGI Symb | 0 | 0 | 0 | 0 | 0 | 0 |
| 50044 | ENSMUSG00000112477 | 0 | NA | NA | NA | NA | NA | Gm48657 | predicted gene, 48657 [Source:MGI Symb | 0 | 0 | 0 | 0 | 0 | 0 |
| 50045 | ENSMUSG00000112478 | 2.034497131 | 0.891314147 | 1.629686971 | 0.546923528 | 0.584431271 | NA | Gm47761 | predicted gene, 47761 [Source:MGI Symb | 3.2124 | 0 | 1.0062 | 5.7839 | 1.0906 | 1.1138 |
| 50046 | ENSMUSG00000112479 | 0 | NA | NA | NA | NA | NA | Gm21026 | predicted gene, 21026 [Source:MGI Symb | 0 | 0 | 0 | 0 | 0 | 0 |
| 50047 | ENSMUSG00000112480 | 0 | NA | NA | NA | NA | NA | Gm4310 | predicted gene 4310 [Source:MGI Symbol | 0 | 0 | 0 | 0 | 0 | 0 |
| 50048 | ENSMUSG00000112481 | 0 | NA | NA | NA | NA | NA | Gm29684 | predicted gene, 29684 [Source:MGI Symb | 0 | 0 | 0 | 0 | 0 | 0 |
| 50049 | ENSMUSG00000112482 | 0 | NA | NA | NA | NA | NA | 1700025K04 | RIKEN cDNA 1700025K04 gene [Source:M( | 0 | 0 | 0 | 0 | 0 | 0 |
| 50050 | ENSMUSG00000112483 | 0 | NA | NA | NA | NA | NA | Gm48542 | predicted gene, 48542 [Source:MGI Symb | 0 | 0 | 0 | 0 | 0 | 0 |
| 50051 | ENSMUSG00000112484 | 0 | NA | NA | NA | NA | NA | 4930473O22 | RIKEN cDNA 4930473O22 gene [Source:M | 0 | 0 | 0 | 0 | 0 | 0 |
| 50052 | ENSMUSG00000112485 | 0 | NA | NA | NA | NA | NA | Gm47631 | predicted gene, 47631 [Source:MGI Symb | 0 | 0 | 0 | 0 | 0 | 0 |
| 50053 | ENSMUSG00000112486 | 0 | NA | NA | NA | NA | NA | Gm48659 | predicted gene, 48659 [Source:MGI Symb | 0 | 0 | 0 | 0 | 0 | 0 |
| 50054 | ENSMUSG00000112487 | 0 | NA | NA | NA | NA | NA | Gm18873 | predicted gene, 18873 [Source:MGI Symb | 0 | 0 | 0 | 0 | 0 | 0 |
| 50055 | ENSMUSG00000112488 | 0 | NA | NA | NA | NA | NA | Gm33981 | predicted gene, 33981 [Source:MGI Symb | 0 | 0 | 0 | 0 | 0 | 0 |
| 50056 | ENSMUSG00000112489 | 0.517184577 | -0.812115922 | 3.393570547 | -0.239310163 | 0.810865085 | NA | 9230116L04 | RIKEN cDNA 9230116L04 gene [Source:M( | 0 | 0 | 2.0125 | 0 | 1.0906 | 0 |
| 50057 | ENSMUSG00000112490 | 0 | NA | NA | NA | NA | NA | Gm48068 | predicted gene, 48068 [Source:MGI Symb | 0 | 0 | 0 | 0 | 0 | 0 |
| 50058 | ENSMUSG00000112491 | 1.659210148 | -0.419146134 | 1.600599804 | -0.261868165 | 0.793423084 | NA | Gm19801 | predicted gene, 19801 [Source:MGI Symb | 1.6062 | 3.0838 | 1.0062 | 0.964 | 2.1812 | 1.1138 |
| 50059 | ENSMUSG00000112492 | 0 | NA | NA | NA | NA | NA | Gm9111 | predicted gene 9111 [Source:MGI Symbol | 0 | 0 | 0 | 0 | 0 | 0 |
| 50060 | ENSMUSG00000112493 | 0 | NA | NA | NA | NA | NA | 4930486F22 | RIKEN cDNA 4930486F22 gene [Source:M( | 0 | 0 | 0 | 0 | 0 | 0 |

| | | | | | | | | | | | | | | | |
|---|---|---|---|---|---|---|---|---|---|---|---|---|---|---|---|
| 50061 | ENSMUSG00000112494 | 0 | NA | NA | NA | NA | NA | 4930444K16 | RIKEN cDNA 4930444K16 gene [Source:M | 0 | 0 | 0 | 0 | 0 | 0 |
| 50062 | ENSMUSG00000112495 | 0 | NA | NA | NA | NA | NA | Gm9048 | predicted gene 9048 [Source:MGI Symbol | 0 | 0 | 0 | 0 | 0 | 0 |
| 50063 | ENSMUSG00000112496 | 0 | NA | NA | NA | NA | NA | Gm48764 | predicted gene, 48764 [Source:MGI Symb | 0 | 0 | 0 | 0 | 0 | 0 |
| 50064 | ENSMUSG00000112497 | 0 | NA | NA | NA | NA | NA | 4930525C09 | RIKEN cDNA 4930525C09 gene [Source:M | 0 | 0 | 0 | 0 | 0 | 0 |
| 50065 | ENSMUSG00000112498 | 0 | NA | NA | NA | NA | NA | Gm47691 | predicted gene, 47691 [Source:MGI Symb | 0 | 0 | 0 | 0 | 0 | 0 |
| 50066 | ENSMUSG00000112499 | 0 | NA | NA | NA | NA | NA | Gm47736 | predicted gene, 47736 [Source:MGI Symb | 0 | 0 | 0 | 0 | 0 | 0 |
| 50067 | ENSMUSG00000112500 | 0 | NA | NA | NA | NA | NA | 4933411E08 | RIKEN cDNA 4933411E08 gene [Source:M | 0 | 0 | 0 | 0 | 0 | 0 |
| 50068 | ENSMUSG00000112501 | 0.181769715 | 1.126924175 | 4.080472857 | 0.276174898 | 0.782413736 | NA | Gm33091 | predicted gene, 33091 [Source:MGI Symb | 0 | 0 | 0 | 0 | 1.0906 | 0 |
| 50069 | ENSMUSG00000112502 | 0.335414862 | -1.779047242 | 4.030804179 | -0.441362855 | 0.658950329 | NA | Gm46194 | predicted gene, 46194 [Source:MGI Symb | 0 | 0 | 2.0125 | 0 | 0 | 0 |
| 50070 | ENSMUSG00000112503 | 0 | NA | NA | NA | NA | NA | Gm18409 | predicted gene, 18409 [Source:MGI Symb | 0 | 0 | 0 | 0 | 0 | 0 |
| 50071 | ENSMUSG00000112504 | 0 | NA | NA | NA | NA | NA | Gm47367 | predicted gene, 47367 [Source:MGI Symb | 0 | 0 | 0 | 0 | 0 | 0 |
| 50072 | ENSMUSG00000112505 | 0.181769715 | 1.126924175 | 4.080472857 | 0.276174898 | 0.782413736 | NA | Gm48610 | predicted gene, 48610 [Source:MGI Symb | 0 | 0 | 0 | 0 | 1.0906 | 0 |
| 50073 | ENSMUSG00000112506 | 0 | NA | NA | NA | NA | NA | Gm48224 | predicted gene, 48224 [Source:MGI Symb | 0 | 0 | 0 | 0 | 0 | 0 |
| 50074 | ENSMUSG00000112507 | 0 | NA | NA | NA | NA | NA | Gm47589 | predicted gene, 47589 [Source:MGI Symb | 0 | 0 | 0 | 0 | 0 | 0 |
| 50075 | ENSMUSG00000112509 | 0 | NA | NA | NA | NA | NA | Gm47211 | predicted gene, 47211 [Source:MGI Symb | 0 | 0 | 0 | 0 | 0 | 0 |
| 50076 | ENSMUSG00000112510 | 0 | NA | NA | NA | NA | NA | Gm47566 | predicted gene, 47566 [Source:MGI Symb | 0 | 0 | 0 | 0 | 0 | 0 |
| 50077 | ENSMUSG00000112511 | 0 | NA | NA | NA | NA | NA | Gm18704 | predicted gene, 18704 [Source:MGI Symb | 0 | 0 | 0 | 0 | 0 | 0 |
| 50078 | ENSMUSG00000112512 | 0.160662974 | 1.126924175 | 4.080472857 | 0.276174898 | 0.782413736 | NA | Gm48203 | predicted gene, 48203 [Source:MGI Symb | 0 | 0 | 0 | 0.964 | 0 | 0 |
| 50079 | ENSMUSG00000112513 | 0 | NA | NA | NA | NA | NA | 4930422I22F | RIKEN cDNA 4930422I22 gene [Source:MG | 0 | 0 | 0 | 0 | 0 | 0 |
| 50080 | ENSMUSG00000112514 | 0 | NA | NA | NA | NA | NA | Gm47897 | predicted gene, 47897 [Source:MGI Symb | 0 | 0 | 0 | 0 | 0 | 0 |
| 50081 | ENSMUSG00000112515 | 0 | NA | NA | NA | NA | NA | Gm4928 | predicted gene 4928 [Source:MGI Symbol | 0 | 0 | 0 | 0 | 0 | 0 |
| 50082 | ENSMUSG00000112516 | 0 | NA | NA | NA | NA | NA | Gm47968 | predicted gene, 47968 [Source:MGI Symb | 0 | 0 | 0 | 0 | 0 | 0 |
| 50083 | ENSMUSG00000112517 | 0 | NA | NA | NA | NA | NA | Gm48001 | predicted gene, 48001 [Source:MGI Symb | 0 | 0 | 0 | 0 | 0 | 0 |
| 50084 | ENSMUSG00000112518 | 0 | NA | NA | NA | NA | NA | Gm48117 | predicted gene, 48117 [Source:MGI Symb | 0 | 0 | 0 | 0 | 0 | 0 |
| 50085 | ENSMUSG00000112519 | 0 | NA | NA | NA | NA | NA | Gm47671 | predicted gene, 47671 [Source:MGI Symb | 0 | 0 | 0 | 0 | 0 | 0 |
| 50086 | ENSMUSG00000112520 | 0 | NA | NA | NA | NA | NA | Gm48379 | predicted gene, 48379 [Source:MGI Symb | 0 | 0 | 0 | 0 | 0 | 0 |
| 50087 | ENSMUSG00000112521 | 0 | NA | NA | NA | NA | NA | Gm29884 | predicted gene, 29884 [Source:MGI Symb | 0 | 0 | 0 | 0 | 0 | 0 |
| 50088 | ENSMUSG00000112522 | 0 | NA | NA | NA | NA | NA | Gm47632 | predicted gene, 47632 [Source:MGI Symb | 0 | 0 | 0 | 0 | 0 | 0 |
| 50089 | ENSMUSG00000112523 | 0 | NA | NA | NA | NA | NA | Gm48377 | predicted gene, 48377 [Source:MGI Symb | 0 | 0 | 0 | 0 | 0 | 0 |
| 50090 | ENSMUSG00000112524 | 0 | NA | NA | NA | NA | NA | Gm35274 | predicted gene, 35274 [Source:MGI Symb | 0 | 0 | 0 | 0 | 0 | 0 |
| 50091 | ENSMUSG00000112525 | 0 | NA | NA | NA | NA | NA | Gm47899 | predicted gene, 47899 [Source:MGI Symb | 0 | 0 | 0 | 0 | 0 | 0 |
| 50092 | ENSMUSG00000112526 | 0 | NA | NA | NA | NA | NA | Gm35865 | predicted gene, 35865 [Source:MGI Symb | 0 | 0 | 0 | 0 | 0 | 0 |
| 50093 | ENSMUSG00000112527 | 110.241944 | 0.93387816 | 0.306851251 | 3.043423017 | 0.002339033 | 0.039247993 | Gm35696 | predicted gene, 35696 [Source:MGI Symb | 56.217 | 58.591 | 112.7 | 104.11 | 150.51 | 179.33 |
| 50094 | ENSMUSG00000112528 | 0 | NA | NA | NA | NA | NA | Gm29794 | predicted gene, 29794 [Source:MGI Symb | 0 | 0 | 0 | 0 | 0 | 0 |
| 50095 | ENSMUSG00000112529 | 0 | NA | NA | NA | NA | NA | Gm46199 | predicted gene, 46199 [Source:MGI Symb | 0 | 0 | 0 | 0 | 0 | 0 |
| 50096 | ENSMUSG00000112530 | 0 | NA | NA | NA | NA | NA | Gm47738 | predicted gene, 47738 [Source:MGI Symb | 0 | 0 | 0 | 0 | 0 | 0 |
| 50097 | ENSMUSG00000112531 | 0 | NA | NA | NA | NA | NA | Gm2649 | predicted gene 2649 [Source:MGI Symbol | 0 | 0 | 0 | 0 | 0 | 0 |
| 50098 | ENSMUSG00000112532 | 10.2965485 | -0.111185209 | 0.70373937 | -0.157992025 | 0.874463083 | 0.980179326 | Gm36283 | predicted gene, 36283 [Source:MGI Symb | 12.85 | 8.2233 | 11.069 | 16.388 | 5.4531 | 7.7969 |
| 50099 | ENSMUSG00000112533 | 0 | NA | NA | NA | NA | NA | Gm20758 | predicted gene, 20758 [Source:MGI Symb | 0 | 0 | 0 | 0 | 0 | 0 |
| 50100 | ENSMUSG00000112534 | 0 | NA | NA | NA | NA | NA | Gm47940 | predicted gene, 47940 [Source:MGI Symb | 0 | 0 | 0 | 0 | 0 | 0 |
| 50101 | ENSMUSG00000112535 | 0.79190315 | 0.90205007 | 2.632846219 | 0.342614036 | 0.731888841 | NA | Gm47425 | predicted gene, 47425 [Source:MGI Symb | 1.6062 | 0 | 0 | 0.964 | 2.1812 | 0 |
| 50102 | ENSMUSG00000112536 | 0 | NA | NA | NA | NA | NA | Gm48195 | predicted gene, 48195 [Source:MGI Symb | 0 | 0 | 0 | 0 | 0 | 0 |
| 50103 | ENSMUSG00000112537 | 0 | NA | NA | NA | NA | NA | 4930543I11F | RIKEN cDNA 4930543I11 gene [Source:MG | 0 | 0 | 0 | 0 | 0 | 0 |
| 50104 | ENSMUSG00000112538 | 0 | NA | NA | NA | NA | NA | Gm47667 | predicted gene, 47667 [Source:MGI Symb | 0 | 0 | 0 | 0 | 0 | 0 |
| 50105 | ENSMUSG00000112539 | 63.67884692 | 0.282989044 | 0.291233744 | 0.971690438 | 0.331204575 | 0.774747658 | Gm40787 | predicted gene, 40787 [Source:MGI Symb | 68.264 | 46.256 | 57.356 | 64.587 | 66.528 | 79.083 |
| 50106 | ENSMUSG00000112540 | 0 | NA | NA | NA | NA | NA | Gm47461 | predicted gene, 47461 [Source:MGI Symb | 0 | 0 | 0 | 0 | 0 | 0 |
| 50107 | ENSMUSG00000112541 | 0 | NA | NA | NA | NA | NA | Gm9072 | predicted gene 9072 [Source:MGI Symbol | 0 | 0 | 0 | 0 | 0 | 0 |
| 50108 | ENSMUSG00000112542 | 0 | NA | NA | NA | NA | NA | Gm47840 | predicted gene, 47840 [Source:MGI Symb | 0 | 0 | 0 | 0 | 0 | 0 |
| 50109 | ENSMUSG00000112543 | 0 | NA | NA | NA | NA | NA | Gm48744 | predicted gene, 48744 [Source:MGI Symb | 0 | 0 | 0 | 0 | 0 | 0 |
| 50110 | ENSMUSG00000112544 | 0 | NA | NA | NA | NA | NA | Gm48089 | predicted gene, 48089 [Source:MGI Symb | 0 | 0 | 0 | 0 | 0 | 0 |
| 50111 | ENSMUSG00000112545 | 38.73327763 | 0.047207852 | 0.498336858 | 0.094730806 | 0.924528649 | 0.987546253 | 1300014J16F | RIKEN cDNA 1300014J16 gene [Source:MG | 35.336 | 39.061 | 40.25 | 68.442 | 38.172 | 11.138 |
| 50112 | ENSMUSG00000112546 | 0 | NA | NA | NA | NA | NA | Gm6992 | predicted gene 6992 [Source:MGI Symbol | 0 | 0 | 0 | 0 | 0 | 0 |
| 50113 | ENSMUSG00000112547 | 1.86681633 | -0.318764417 | 1.751994108 | -0.181943772 | 0.855626852 | NA | Gm47096 | predicted gene, 47096 [Source:MGI Symb | 3.2124 | 0 | 3.0187 | 3.8559 | 0 | 1.1138 |
| 50114 | ENSMUSG00000112548 | 0 | NA | NA | NA | NA | NA | Gm47338 | predicted gene, 47338 [Source:MGI Symb | 0 | 0 | 0 | 0 | 0 | 0 |
| 50115 | ENSMUSG00000112549 | 0 | NA | NA | NA | NA | NA | Gm35818 | predicted gene, 35818 [Source:MGI Symb | 0 | 0 | 0 | 0 | 0 | 0 |
| 50116 | ENSMUSG00000112550 | 1.711000422 | 0.145916439 | 1.5875776 | 0.091911375 | 0.926768454 | NA | Gm6627 | predicted gene 6627 [Source:MGI Symbol | 0.8031 | 3.0838 | 1.0062 | 0.964 | 2.1812 | 2.2277 |

| | | | | | | | | | | | | | | | |
|---|---|---|---|---|---|---|---|---|---|---|---|---|---|---|---|
| 50117 | ENSMUSG00000112551 | 0 | NA | NA | NA | NA | NA | Gm8219 | predicted gene 8219 [Source:MGI Symbol | 0 | 0 | 0 | 0 | 0 | 0 |
| 50118 | ENSMUSG00000112552 | 0 | NA | NA | NA | NA | NA | Gm36065 | predicted gene, 36065 [Source:MGI Symb | 0 | 0 | 0 | 0 | 0 | 0 |
| 50119 | ENSMUSG00000112553 | 0 | NA | NA | NA | NA | NA | Gm4556 | predicted gene 4556 [Source:MGI Symbol | 0 | 0 | 0 | 0 | 0 | 0 |
| 50120 | ENSMUSG00000112554 | 0 | NA | NA | NA | NA | NA | Gm47224 | predicted gene, 47224 [Source:MGI Symb | 0 | 0 | 0 | 0 | 0 | 0 |
| 50121 | ENSMUSG00000112555 | 0 | NA | NA | NA | NA | NA | Gm47027 | predicted gene, 47027 [Source:MGI Symb | 0 | 0 | 0 | 0 | 0 | 0 |
| 50122 | ENSMUSG00000112556 | 0 | NA | NA | NA | NA | NA | Gm47339 | predicted gene, 47339 [Source:MGI Symb | 0 | 0 | 0 | 0 | 0 | 0 |
| 50123 | ENSMUSG00000112557 | 0 | NA | NA | NA | NA | NA | Gm47626 | predicted gene, 47626 [Source:MGI Symb | 0 | 0 | 0 | 0 | 0 | 0 |
| 50124 | ENSMUSG00000112558 | 0 | NA | NA | NA | NA | NA | Gm32364 | predicted gene, 32364 [Source:MGI Symb | 0 | 0 | 0 | 0 | 0 | 0 |
| 50125 | ENSMUSG00000112559 | 0 | NA | NA | NA | NA | NA | Gm48599 | predicted gene, 48599 [Source:MGI Symb | 0 | 0 | 0 | 0 | 0 | 0 |
| 50126 | ENSMUSG00000112560 | 0 | NA | NA | NA | NA | NA | Gm19186 | predicted gene, 19186 [Source:MGI Symb | 0 | 0 | 0 | 0 | 0 | 0 |
| 50127 | ENSMUSG00000112561 | 0 | NA | NA | NA | NA | NA | Gm9035 | predicted gene 9035 [Source:MGI Symbol | 0 | 0 | 0 | 0 | 0 | 0 |
| 50128 | ENSMUSG00000112562 | 0 | NA | NA | NA | NA | NA | Gm48451 | predicted gene, 48451 [Source:MGI Symb | 0 | 0 | 0 | 0 | 0 | 0 |
| 50129 | ENSMUSG00000112563 | 0 | NA | NA | NA | NA | NA | Gm47608 | predicted gene, 47608 [Source:MGI Symb | 0 | 0 | 0 | 0 | 0 | 0 |
| 50130 | ENSMUSG00000112564 | 0 | NA | NA | NA | NA | NA | Gm47419 | predicted gene, 47419 [Source:MGI Symb | 0 | 0 | 0 | 0 | 0 | 0 |
| 50131 | ENSMUSG00000112565 | 0 | NA | NA | NA | NA | NA | Gm48473 | predicted gene, 48473 [Source:MGI Symb | 0 | 0 | 0 | 0 | 0 | 0 |
| 50132 | ENSMUSG00000112566 | 0 | NA | NA | NA | NA | NA | Gm32025 | predicted gene, 32025 [Source:MGI Symb | 0 | 0 | 0 | 0 | 0 | 0 |
| 50133 | ENSMUSG00000112567 | 0 | NA | NA | NA | NA | NA | Gm32899 | predicted gene, 32899 [Source:MGI Symb | 0 | 0 | 0 | 0 | 0 | 0 |
| 50134 | ENSMUSG00000112568 | 0 | NA | NA | NA | NA | NA | Gm47363 | predicted gene, 47363 [Source:MGI Symb | 0 | 0 | 0 | 0 | 0 | 0 |
| 50135 | ENSMUSG00000112569 | 0.160662974 | 1.126924175 | 4.080472857 | 0.276174898 | 0.782413736 | NA | Gm33111 | predicted gene, 33111 [Source:MGI Symb | 0 | 0 | 0 | 0.964 | 0 | 0 |
| 50136 | ENSMUSG00000112570 | 0 | NA | NA | NA | NA | NA | Gm47225 | predicted gene, 47225 [Source:MGI Symb | 0 | 0 | 0 | 0 | 0 | 0 |
| 50137 | ENSMUSG00000112571 | 0.335414862 | -1.779047242 | 4.030804179 | -0.441362855 | 0.658950329 | NA | Gm48207 | predicted gene, 48207 [Source:MGI Symb | 0 | 0 | 2.0125 | 0 | 0 | 0 |
| 50138 | ENSMUSG00000112572 | 0 | NA | NA | NA | NA | NA | Gm7530 | predicted gene 7530 [Source:MGI Symbol | 0 | 0 | 0 | 0 | 0 | 0 |
| 50139 | ENSMUSG00000112573 | 0 | NA | NA | NA | NA | NA | Gm48324 | predicted gene, 48324 [Source:MGI Symb | 0 | 0 | 0 | 0 | 0 | 0 |
| 50140 | ENSMUSG00000112574 | 0 | NA | NA | NA | NA | NA | Gm5750 | predicted gene 5750 [Source:MGI Symbol | 0 | 0 | 0 | 0 | 0 | 0 |
| 50141 | ENSMUSG00000112575 | 0 | NA | NA | NA | NA | NA | Gm31849 | predicted gene, 31849 [Source:MGI Symb | 0 | 0 | 0 | 0 | 0 | 0 |
| 50142 | ENSMUSG00000112576 | 0.267700746 | -1.497481649 | 4.057001436 | -0.369110456 | 0.712045396 | NA | Gm47621 | predicted gene, 47621 [Source:MGI Symb | 1.6062 | 0 | 0 | 0 | 0 | 0 |
| 50143 | ENSMUSG00000112577 | 0 | NA | NA | NA | NA | NA | Gm47170 | predicted gene, 47170 [Source:MGI Symb | 0 | 0 | 0 | 0 | 0 | 0 |
| 50144 | ENSMUSG00000112578 | 0 | NA | NA | NA | NA | NA | Gm33308 | predicted gene, 33308 [Source:MGI Symb | 0 | 0 | 0 | 0 | 0 | 0 |
| 50145 | ENSMUSG00000112579 | 0.17131964 | -0.796629176 | 4.080472857 | -0.195229623 | 0.84521319 | NA | Gm48450 | predicted gene, 48450 [Source:MGI Symb | 0 | 1.0279 | 0 | 0 | 0 | 0 |
| 50146 | ENSMUSG00000112580 | 0 | NA | NA | NA | NA | NA | Gm47673 | predicted gene, 47673 [Source:MGI Symb | 0 | 0 | 0 | 0 | 0 | 0 |
| 50147 | ENSMUSG00000112581 | 2.003866857 | -3.289735014 | 2.000576526 | -1.64439349 | 0.100094949 | NA | 1700022H01 | RIKEN cDNA 1700022H01 gene [Source:M | 4.0155 | 0 | 7.0437 | 0.964 | 0 | 0 |
| 50148 | ENSMUSG00000112582 | 0 | NA | NA | NA | NA | NA | Rpl19-ps2 | ribosomal protein L19, pseudogene 2 [Sou | 0 | 0 | 0 | 0 | 0 | 0 |
| 50149 | ENSMUSG00000112583 | 0 | NA | NA | NA | NA | NA | Gm4930 | predicted gene 4930 [Source:MGI Symbol | 0 | 0 | 0 | 0 | 0 | 0 |
| 50150 | ENSMUSG00000112584 | 0 | NA | NA | NA | NA | NA | Gm2398 | predicted gene 2398 [Source:MGI Symbol | 0 | 0 | 0 | 0 | 0 | 0 |
| 50151 | ENSMUSG00000112585 | 0 | NA | NA | NA | NA | NA | Gm47527 | predicted gene, 47527 [Source:MGI Symb | 0 | 0 | 0 | 0 | 0 | 0 |
| 50152 | ENSMUSG00000112586 | 0 | NA | NA | NA | NA | NA | Gm6729 | predicted gene 6729 [Source:MGI Symbol | 0 | 0 | 0 | 0 | 0 | 0 |
| 50153 | ENSMUSG00000112587 | 0 | NA | NA | NA | NA | NA | Gm47769 | predicted gene, 47769 [Source:MGI Symb | 0 | 0 | 0 | 0 | 0 | 0 |
| 50154 | ENSMUSG00000112588 | 0 | NA | NA | NA | NA | NA | Gm6859 | predicted gene 6859 [Source:MGI Symbol | 0 | 0 | 0 | 0 | 0 | 0 |
| 50155 | ENSMUSG00000112589 | 0 | NA | NA | NA | NA | NA | Gm48280 | predicted gene, 48280 [Source:MGI Symb | 0 | 0 | 0 | 0 | 0 | 0 |
| 50156 | ENSMUSG00000112590 | 0 | NA | NA | NA | NA | NA | Gm5186 | predicted gene 5186 [Source:MGI Symbol | 0 | 0 | 0 | 0 | 0 | 0 |
| 50157 | ENSMUSG00000112591 | 0 | NA | NA | NA | NA | NA | Gm4927 | predicted gene 4927 [Source:MGI Symbol | 0 | 0 | 0 | 0 | 0 | 0 |
| 50158 | ENSMUSG00000112592 | 0 | NA | NA | NA | NA | NA | Gm19972 | predicted gene, 19972 [Source:MGI Symb | 0 | 0 | 0 | 0 | 0 | 0 |
| 50159 | ENSMUSG00000112593 | 11.48053547 | -0.841123462 | 0.683377215 | -1.230833343 | 0.218385202 | 0.663735712 | Gm48882 | predicted gene, 48882 [Source:MGI Symb | 12.85 | 17.475 | 14.087 | 13.496 | 7.6343 | 3.3415 |
| 50160 | ENSMUSG00000112594 | 0 | NA | NA | NA | NA | NA | Gm48333 | predicted gene, 48333 [Source:MGI Symb | 0 | 0 | 0 | 0 | 0 | 0 |
| 50161 | ENSMUSG00000112595 | 0 | NA | NA | NA | NA | NA | Gm47340 | predicted gene, 47340 [Source:MGI Symb | 0 | 0 | 0 | 0 | 0 | 0 |
| 50162 | ENSMUSG00000112596 | 0 | NA | NA | NA | NA | NA | Gm48804 | predicted gene, 48804 [Source:MGI Symb | 0 | 0 | 0 | 0 | 0 | 0 |
| 50163 | ENSMUSG00000112597 | 0 | NA | NA | NA | NA | NA | Gm4777 | predicted gene 4777 [Source:MGI Symbol | 0 | 0 | 0 | 0 | 0 | 0 |
| 50164 | ENSMUSG00000112598 | 0 | NA | NA | NA | NA | NA | Gm40552 | predicted gene, 40552 [Source:MGI Symb | 0 | 0 | 0 | 0 | 0 | 0 |
| 50165 | ENSMUSG00000112599 | 16.35449307 | 0.473618333 | 0.541285164 | 0.874988572 | 0.381580124 | 0.8130053 | Gm47719 | predicted gene, 47719 [Source:MGI Symb | 13.653 | 10.279 | 17.106 | 16.388 | 23.994 | 16.708 |
| 50166 | ENSMUSG00000112600 | 0 | NA | NA | NA | NA | NA | Gm49918 | predicted gene, 49918 [Source:MGI Symb | 0 | 0 | 0 | 0 | 0 | 0 |
| 50167 | ENSMUSG00000112601 | 0 | NA | NA | NA | NA | NA | Gm47590 | predicted gene, 47590 [Source:MGI Symb | 0 | 0 | 0 | 0 | 0 | 0 |
| 50168 | ENSMUSG00000112602 | 0 | NA | NA | NA | NA | NA | Gm18557 | predicted gene, 18557 [Source:MGI Symb | 0 | 0 | 0 | 0 | 0 | 0 |
| 50169 | ENSMUSG00000112603 | 0 | NA | NA | NA | NA | NA | Gm18510 | predicted gene, 18510 [Source:MGI Symb | 0 | 0 | 0 | 0 | 0 | 0 |
| 50170 | ENSMUSG00000112604 | 0 | NA | NA | NA | NA | NA | Gm48518 | predicted gene, 48518 [Source:MGI Symb | 0 | 0 | 0 | 0 | 0 | 0 |
| 50171 | ENSMUSG00000112605 | 0.524667718 | -0.819976748 | 3.064824146 | -0.267544469 | 0.78904998 | NA | Gm7476 | predicted gene 7476 [Source:MGI Symbol | 0 | 1.0279 | 1.0062 | 0 | 0 | 1.1138 |
| 50172 | ENSMUSG00000112606 | 0 | NA | NA | NA | NA | NA | Gm46364 | predicted gene, 46364 [Source:MGI Symb | 0 | 0 | 0 | 0 | 0 | 0 |

| | | | | | | | | | | | | | | | |
|---|---|---|---|---|---|---|---|---|---|---|---|---|---|---|---|
| 50173 | ENSMUSG00000112607 | 0 | NA | NA | NA | NA | NA | Gm33979 | predicted gene, 33979 [Source:MGI Symb | 0 | 0 | 0 | 0 | 0 | 0 |
| 50174 | ENSMUSG00000112608 | 0 | NA | NA | NA | NA | NA | Gm46233 | predicted gene, 46233 [Source:MGI Symb | 0 | 0 | 0 | 0 | 0 | 0 |
| 50175 | ENSMUSG00000112609 | 0 | NA | NA | NA | NA | NA | Gm47480 | predicted gene, 47480 [Source:MGI Symb | 0 | 0 | 0 | 0 | 0 | 0 |
| 50176 | ENSMUSG00000112610 | 0 | NA | NA | NA | NA | NA | Gm30165 | predicted gene, 30165 [Source:MGI Symb | 0 | 0 | 0 | 0 | 0 | 0 |
| 50177 | ENSMUSG00000112611 | 0 | NA | NA | NA | NA | NA | Glipr1l3 | GLI pathogenesis-related 1 like 3 [Source: | 0 | 0 | 0 | 0 | 0 | 0 |
| 50178 | ENSMUSG00000112612 | 0 | NA | NA | NA | NA | NA | Gm7775 | predicted gene 7775 [Source:MGI Symbol | 0 | 0 | 0 | 0 | 0 | 0 |
| 50179 | ENSMUSG00000112613 | 0 | NA | NA | NA | NA | NA | Gm48175 | predicted gene, 48175 [Source:MGI Symb | 0 | 0 | 0 | 0 | 0 | 0 |
| 50180 | ENSMUSG00000112614 | 0.342639279 | -1.805976596 | 4.028547714 | -0.448294702 | 0.653940524 | NA | Gm9030 | predicted gene 9030 [Source:MGI Symbol | 0 | 2.0558 | 0 | 0 | 0 | 0 |
| 50181 | ENSMUSG00000112615 | 53.4899744 | 0.629185432 | 0.306134976 | 2.055254976 | 0.039854384 | 0.273460059 | Gm18337 | predicted gene, 18337 [Source:MGI Symb | 37.746 | 43.173 | 45.281 | 62.659 | 74.162 | 57.92 |
| 50182 | ENSMUSG00000112616 | 1.224468662 | 2.689446769 | 2.570056051 | 1.046454519 | 0.295351233 | NA | Gm47434 | predicted gene, 47434 [Source:MGI Symb | 0.8031 | 0 | 0 | 0 | 6.5437 | 0 |
| 50183 | ENSMUSG00000112617 | 0 | NA | NA | NA | NA | NA | Gm47937 | predicted gene, 47937 [Source:MGI Symb | 0 | 0 | 0 | 0 | 0 | 0 |
| 50184 | ENSMUSG00000112618 | 0 | NA | NA | NA | NA | NA | Gm48835 | predicted gene, 48835 [Source:MGI Symb | 0 | 0 | 0 | 0 | 0 | 0 |
| 50185 | ENSMUSG00000112619 | 0 | NA | NA | NA | NA | NA | 1700030E10 | RIKEN cDNA 1700030E10 gene [Source:M | 0 | 0 | 0 | 0 | 0 | 0 |
| 50186 | ENSMUSG00000112620 | 0 | NA | NA | NA | NA | NA | Gm18972 | predicted gene, 18972 [Source:MGI Symb | 0 | 0 | 0 | 0 | 0 | 0 |
| 50187 | ENSMUSG00000112621 | 0.995942328 | 0.3039145 | 2.42601415 | 0.125273177 | 0.900307287 | NA | Gm5173 | predicted gene 5173 [Source:MGI Symbol | 1.6062 | 1.0279 | 0 | 0 | 0 | 3.3415 |
| 50188 | ENSMUSG00000112622 | 0 | NA | NA | NA | NA | NA | Gm47164 | predicted gene, 47164 [Source:MGI Symb | 0 | 0 | 0 | 0 | 0 | 0 |
| 50189 | ENSMUSG00000112623 | 0.133850373 | -0.796629176 | 4.080472857 | -0.195229623 | 0.84521319 | NA | 4933436P19 | RIKEN cDNA 4933436P19 gene [Source:M | 0.8031 | 0 | 0 | 0 | 0 | 0 |
| 50190 | ENSMUSG00000112624 | 0 | NA | NA | NA | NA | NA | Gm19074 | predicted gene, 19074 [Source:MGI Symb | 0 | 0 | 0 | 0 | 0 | 0 |
| 50191 | ENSMUSG00000112625 | 0 | NA | NA | NA | NA | NA | Gm2628 | predicted gene 2628 [Source:MGI Symbol | 0 | 0 | 0 | 0 | 0 | 0 |
| 50192 | ENSMUSG00000112626 | 0 | NA | NA | NA | NA | NA | Gm18027 | predicted gene, 18027 [Source:MGI Symb | 0 | 0 | 0 | 0 | 0 | 0 |
| 50193 | ENSMUSG00000112627 | 9.267870868 | -0.206750377 | 0.816590291 | -0.253187405 | 0.800123399 | 0.962424862 | 4933412E12 | RIKEN cDNA 4933412E12 gene [Source:M | 2.4093 | 15.419 | 12.075 | 4.8199 | 13.087 | 7.7969 |
| 50194 | ENSMUSG00000112628 | 0 | NA | NA | NA | NA | NA | Gm47880 | predicted gene, 47880 [Source:MGI Symb | 0 | 0 | 0 | 0 | 0 | 0 |
| 50195 | ENSMUSG00000112629 | 0 | NA | NA | NA | NA | NA | Gm47771 | predicted gene, 47771 [Source:MGI Symb | 0 | 0 | 0 | 0 | 0 | 0 |
| 50196 | ENSMUSG00000112630 | 3.744256654 | -1.761616967 | 1.275103543 | -1.381548171 | 0.167110477 | NA | Gm36908 | predicted gene, 36908 [Source:MGI Symb | 11.243 | 3.0838 | 3.0187 | 2.8919 | 0 | 2.2277 |
| 50197 | ENSMUSG00000112631 | 0 | NA | NA | NA | NA | NA | Gm48555 | predicted gene, 48555 [Source:MGI Symb | 0 | 0 | 0 | 0 | 0 | 0 |
| 50198 | ENSMUSG00000112632 | 0 | NA | NA | NA | NA | NA | Gm4510 | predicted gene 4510 [Source:MGI Symbol | 0 | 0 | 0 | 0 | 0 | 0 |
| 50199 | ENSMUSG00000112633 | 0 | NA | NA | NA | NA | NA | Gm47627 | predicted gene, 47627 [Source:MGI Symb | 0 | 0 | 0 | 0 | 0 | 0 |
| 50200 | ENSMUSG00000112634 | 0.167707431 | -0.796629176 | 4.080472857 | -0.195229623 | 0.84521319 | NA | Gm47737 | predicted gene, 47737 [Source:MGI Symb | 0 | 0 | 1.0062 | 0 | 0 | 0 |
| 50201 | ENSMUSG00000112635 | 0 | NA | NA | NA | NA | NA | Gm47397 | predicted gene, 47397 [Source:MGI Symb | 0 | 0 | 0 | 0 | 0 | 0 |
| 50202 | ENSMUSG00000112636 | 0 | NA | NA | NA | NA | NA | Gm47223 | predicted gene, 47223 [Source:MGI Symb | 0 | 0 | 0 | 0 | 0 | 0 |
| 50203 | ENSMUSG00000112637 | 3.244386289 | 1.662649471 | 1.245000237 | 1.335461169 | 0.181725456 | NA | Gm48225 | predicted gene, 48225 [Source:MGI Symb | 1.6062 | 2.0558 | 1.0062 | 4.8199 | 2.1812 | 7.7969 |
| 50204 | ENSMUSG00000112638 | 0 | NA | NA | NA | NA | NA | Gm48463 | predicted gene, 48463 [Source:MGI Symb | 0 | 0 | 0 | 0 | 0 | 0 |
| 50205 | ENSMUSG00000112639 | 39.9179796 | -0.364508899 | 0.361625851 | -1.007972463 | 0.313467682 | 0.760152431 | A730063M1 | RIKEN cDNA A730063M14 gene [Source:N | 52.202 | 35.977 | 46.287 | 27.955 | 41.443 | 35.643 |
| 50206 | ENSMUSG00000112640 | 14.80034346 | 0.12065027 | 0.604868539 | 0.199465275 | 0.841898804 | 0.971854408 | Gm32687 | predicted gene, 32687 [Source:MGI Symb | 11.243 | 9.2513 | 22.137 | 15.424 | 20.722 | 10.025 |
| 50207 | ENSMUSG00000112641 | 2.467037139 | -0.430291143 | 1.410128884 | -0.305143131 | 0.760257143 | NA | Gm40709 | predicted gene, 40709 [Source:MGI Symb | 2.4093 | 3.0838 | 3.0187 | 1.928 | 4.3625 | 0 |
| 50208 | ENSMUSG00000112642 | 0.160662974 | 1.126924175 | 4.080472857 | 0.276174898 | 0.782413736 | NA | Gm17849 | predicted gene, 17849 [Source:MGI Symb | 0 | 0 | 0 | 0.964 | 0 | 0 |
| 50209 | ENSMUSG00000112643 | 0.331982614 | 0.16514309 | 3.986751476 | 0.041422971 | 0.966958701 | NA | Gm19688 | predicted gene, 19688 [Source:MGI Symb | 0 | 1.0279 | 0 | 0.964 | 0 | 0 |
| 50210 | ENSMUSG00000112644 | 0 | NA | NA | NA | NA | NA | Gm47220 | predicted gene, 47220 [Source:MGI Symb | 0 | 0 | 0 | 0 | 0 | 0 |
| 50211 | ENSMUSG00000112645 | 0 | NA | NA | NA | NA | NA | Gm48787 | predicted gene, 48787 [Source:MGI Symb | 0 | 0 | 0 | 0 | 0 | 0 |
| 50212 | ENSMUSG00000112646 | 0 | NA | NA | NA | NA | NA | Gm5654 | predicted gene 5654 [Source:MGI Symbol | 0 | 0 | 0 | 0 | 0 | 0 |
| 50213 | ENSMUSG00000112647 | 0 | NA | NA | NA | NA | NA | Gm5427 | predicted gene 5427 [Source:MGI Symbol | 0 | 0 | 0 | 0 | 0 | 0 |
| 50214 | ENSMUSG00000112648 | 0 | NA | NA | NA | NA | NA | Gm47793 | predicted gene, 47793 [Source:MGI Symb | 0 | 0 | 0 | 0 | 0 | 0 |
| 50215 | ENSMUSG00000112649 | 0 | NA | NA | NA | NA | NA | Gm47647 | predicted gene, 47647 [Source:MGI Symb | 0 | 0 | 0 | 0 | 0 | 0 |
| 50216 | ENSMUSG00000112650 | 0 | NA | NA | NA | NA | NA | Gm5184 | predicted gene 5184 [Source:MGI Symbol | 0 | 0 | 0 | 0 | 0 | 0 |
| 50217 | ENSMUSG00000112651 | 0 | NA | NA | NA | NA | NA | Gm18623 | predicted gene, 18623 [Source:MGI Symb | 0 | 0 | 0 | 0 | 0 | 0 |
| 50218 | ENSMUSG00000112652 | 0 | NA | NA | NA | NA | NA | 4921516A02 | RIKEN cDNA 4921516A02 gene [Source:M | 0 | 0 | 0 | 0 | 0 | 0 |
| 50219 | ENSMUSG00000112653 | 0 | NA | NA | NA | NA | NA | Gm36176 | predicted gene, 36176 [Source:MGI Symb | 0 | 0 | 0 | 0 | 0 | 0 |
| 50220 | ENSMUSG00000112654 | 2.901650798 | -1.246423469 | 1.406566135 | -0.886146366 | 0.375538661 | NA | 4930455C13 | RIKEN cDNA 4930455C13 gene [Source:M | 3.2124 | 1.0279 | 8.05 | 2.8919 | 0 | 2.2277 |
| 50221 | ENSMUSG00000112655 | 0 | NA | NA | NA | NA | NA | Gm48511 | predicted gene, 48511 [Source:MGI Symb | 0 | 0 | 0 | 0 | 0 | 0 |
| 50222 | ENSMUSG00000112656 | 0 | NA | NA | NA | NA | NA | Gm47929 | predicted gene, 47929 [Source:MGI Symb | 0 | 0 | 0 | 0 | 0 | 0 |
| 50223 | ENSMUSG00000112657 | 0.838537155 | -3.104405377 | 3.144633814 | -0.987207275 | 0.323541036 | NA | BC106175 | cDNA sequence BC106175 [Source:MGI S | 0 | 0 | 5.0312 | 0 | 0 | 0 |
| 50224 | ENSMUSG00000112658 | 0 | NA | NA | NA | NA | NA | Gm47395 | predicted gene, 47395 [Source:MGI Symb | 0 | 0 | 0 | 0 | 0 | 0 |
| 50225 | ENSMUSG00000112659 | 0 | NA | NA | NA | NA | NA | Gm40646 | predicted gene, 40646 [Source:MGI Symb | 0 | 0 | 0 | 0 | 0 | 0 |
| 50226 | ENSMUSG00000112660 | 0 | NA | NA | NA | NA | NA | Gm48761 | predicted gene, 48761 [Source:MGI Symb | 0 | 0 | 0 | 0 | 0 | 0 |
| 50227 | ENSMUSG00000112661 | 0 | NA | NA | NA | NA | NA | Gm6869 | predicted gene 6869 [Source:MGI Symbol | 0 | 0 | 0 | 0 | 0 | 0 |
| 50228 | ENSMUSG00000112662 | 0 | NA | NA | NA | NA | NA | Gm47922 | predicted gene, 47922 [Source:MGI Symb | 0 | 0 | 0 | 0 | 0 | 0 |

| | | | | | | | | | | | | | | | |
|---|---|---|---|---|---|---|---|---|---|---|---|---|---|---|---|
| 50229 | ENSMUSG00000112663 | 0 | NA | NA | NA | NA | NA | C630031E19 | RIKEN cDNA C630031E19 gene [Source:M | 0 | 0 | 0 | 0 | 0 | 0 |
| 50230 | ENSMUSG00000112664 | 0.331982614 | 0.16514309 | 3.986751476 | 0.041422971 | 0.966958701 | NA | Gm48076 | predicted gene, 48076 [Source:MGI Symb | 0 | 1.0279 | 0 | 0.964 | 0 | 0 |
| 50231 | ENSMUSG00000112665 | 0 | NA | NA | NA | NA | NA | Ryk-ps1 | receptor-like tyrosine kinase, pseudogene | 0 | 0 | 0 | 0 | 0 | 0 |
| 50232 | ENSMUSG00000112666 | 0 | NA | NA | NA | NA | NA | Gm48485 | predicted gene, 48485 [Source:MGI Symb | 0 | 0 | 0 | 0 | 0 | 0 |
| 50233 | ENSMUSG00000112667 | 0 | NA | NA | NA | NA | NA | Gm48270 | predicted gene, 48270 [Source:MGI Symb | 0 | 0 | 0 | 0 | 0 | 0 |
| 50234 | ENSMUSG00000112668 | 0 | NA | NA | NA | NA | NA | Gm48376 | predicted gene, 48376 [Source:MGI Symb | 0 | 0 | 0 | 0 | 0 | 0 |
| 50235 | ENSMUSG00000112669 | 0.956643647 | -0.817327529 | 2.26666353 | -0.360586174 | 0.718408827 | NA | Gm48689 | predicted gene, 48689 [Source:MGI Symb | 1.6062 | 2.0558 | 0 | 0.964 | 0 | 1.1138 |
| 50236 | ENSMUSG00000112670 | 1.468412105 | 1.945424723 | 2.004800999 | 0.970382958 | 0.33185564 | NA | Gm18042 | predicted gene, 18042 [Source:MGI Symb | 0.8031 | 0 | 1.0062 | 4.8199 | 2.1812 | 0 |
| 50237 | ENSMUSG00000112671 | 0 | NA | NA | NA | NA | NA | Gm18242 | predicted gene, 18242 [Source:MGI Symb | 0 | 0 | 0 | 0 | 0 | 0 |
| 50238 | ENSMUSG00000112672 | 0 | NA | NA | NA | NA | NA | Gm48118 | predicted gene, 48118 [Source:MGI Symb | 0 | 0 | 0 | 0 | 0 | 0 |
| 50239 | ENSMUSG00000112673 | 0 | NA | NA | NA | NA | NA | Gm7413 | predicted gene 7413 [Source:MGI Symbol | 0 | 0 | 0 | 0 | 0 | 0 |
| 50240 | ENSMUSG00000112674 | 0 | NA | NA | NA | NA | NA | Gm47459 | predicted gene, 47459 [Source:MGI Symb | 0 | 0 | 0 | 0 | 0 | 0 |
| 50241 | ENSMUSG00000112675 | 0 | NA | NA | NA | NA | NA | Gm48892 | predicted gene, 48892 [Source:MGI Symb | 0 | 0 | 0 | 0 | 0 | 0 |
| 50242 | ENSMUSG00000112676 | 0 | NA | NA | NA | NA | NA | Gm20108 | predicted gene, 20108 [Source:MGI Symb | 0 | 0 | 0 | 0 | 0 | 0 |
| 50243 | ENSMUSG00000112677 | 0 | NA | NA | NA | NA | NA | Gm48868 | predicted gene, 48868 [Source:MGI Symb | 0 | 0 | 0 | 0 | 0 | 0 |
| 50244 | ENSMUSG00000112678 | 0 | NA | NA | NA | NA | NA | Gm47492 | predicted gene, 47492 [Source:MGI Symb | 0 | 0 | 0 | 0 | 0 | 0 |
| 50245 | ENSMUSG00000112679 | 0 | NA | NA | NA | NA | NA | Gm40603 | predicted gene, 40603 [Source:MGI Symb | 0 | 0 | 0 | 0 | 0 | 0 |
| 50246 | ENSMUSG00000112680 | 0 | NA | NA | NA | NA | NA | Gm48760 | predicted gene, 48760 [Source:MGI Symb | 0 | 0 | 0 | 0 | 0 | 0 |
| 50247 | ENSMUSG00000112681 | 0 | NA | NA | NA | NA | NA | Gm47622 | predicted gene, 47622 [Source:MGI Symb | 0 | 0 | 0 | 0 | 0 | 0 |
| 50248 | ENSMUSG00000112682 | 0 | NA | NA | NA | NA | NA | Gm48874 | predicted gene, 48874 [Source:MGI Symb | 0 | 0 | 0 | 0 | 0 | 0 |
| 50249 | ENSMUSG00000112683 | 0 | NA | NA | NA | NA | NA | Gm18904 | predicted gene, 18904 [Source:MGI Symb | 0 | 0 | 0 | 0 | 0 | 0 |
| 50250 | ENSMUSG00000112684 | 0 | NA | NA | NA | NA | NA | Gm47958 | predicted gene, 47958 [Source:MGI Symb | 0 | 0 | 0 | 0 | 0 | 0 |
| 50251 | ENSMUSG00000112685 | 0 | NA | NA | NA | NA | NA | Gm40421 | predicted gene, 40421 [Source:MGI Symb | 0 | 0 | 0 | 0 | 0 | 0 |
| 50252 | ENSMUSG00000112686 | 0 | NA | NA | NA | NA | NA | Gm7034 | predicted gene 7034 [Source:MGI Symbol | 0 | 0 | 0 | 0 | 0 | 0 |
| 50253 | ENSMUSG00000112687 | 0 | NA | NA | NA | NA | NA | Gm18871 | predicted gene, 18871 [Source:MGI Symb | 0 | 0 | 0 | 0 | 0 | 0 |
| 50254 | ENSMUSG00000112688 | 0 | NA | NA | NA | NA | NA | Gm48514 | predicted gene, 48514 [Source:MGI Symb | 0 | 0 | 0 | 0 | 0 | 0 |
| 50255 | ENSMUSG00000112689 | 0 | NA | NA | NA | NA | NA | Gm47476 | predicted gene, 47476 [Source:MGI Symb | 0 | 0 | 0 | 0 | 0 | 0 |
| 50256 | ENSMUSG00000112690 | 0 | NA | NA | NA | NA | NA | Gm2446 | predicted gene 2446 [Source:MGI Symbol | 0 | 0 | 0 | 0 | 0 | 0 |
| 50257 | ENSMUSG00000112691 | 0.606727817 | -2.651277273 | 2.811524265 | -0.943003518 | 0.345679103 | NA | Gm8681 | predicted gene 8681 [Source:MGI Symbol | 1.6062 | 1.0279 | 1.0062 | 0 | 0 | 0 |
| 50258 | ENSMUSG00000112692 | 0 | NA | NA | NA | NA | NA | Gm19073 | predicted gene, 19073 [Source:MGI Symb | 0 | 0 | 0 | 0 | 0 | 0 |
| 50259 | ENSMUSG00000112693 | 2.205475424 | 0.868387188 | 1.406884894 | 0.617241106 | 0.537075706 | NA | Gm5512 | predicted gene 5512 [Source:MGI Symbol | 1.6062 | 2.0558 | 1.0062 | 1.928 | 2.1812 | 4.4554 |
| 50260 | ENSMUSG00000112694 | 0.339027071 | -1.79174149 | 3.919585585 | -0.457125237 | 0.647581032 | NA | Gm33037 | predicted gene, 33037 [Source:MGI Symb | 0 | 1.0279 | 1.0062 | 0 | 0 | 0 |
| 50261 | ENSMUSG00000112696 | 0.339027071 | -1.79174149 | 3.919585585 | -0.457125237 | 0.647581032 | NA | Gm48815 | predicted gene, 48815 [Source:MGI Symb | 0 | 1.0279 | 1.0062 | 0 | 0 | 0 |
| 50262 | ENSMUSG00000112697 | 0 | NA | NA | NA | NA | NA | Gm8137 | predicted gene 8137 [Source:MGI Symbol | 0 | 0 | 0 | 0 | 0 | 0 |
| 50263 | ENSMUSG00000112698 | 0.181769715 | 1.126924175 | 4.080472857 | 0.276174898 | 0.782413736 | NA | Gm7525 | predicted gene 7525 [Source:MGI Symbol | 0 | 0 | 0 | 0 | 1.0906 | 0 |
| 50264 | ENSMUSG00000112699 | 0 | NA | NA | NA | NA | NA | Gm48474 | predicted gene, 48474 [Source:MGI Symb | 0 | 0 | 0 | 0 | 0 | 0 |
| 50265 | ENSMUSG00000112700 | 0 | NA | NA | NA | NA | NA | Gm48757 | predicted gene, 48757 [Source:MGI Symb | 0 | 0 | 0 | 0 | 0 | 0 |
| 50266 | ENSMUSG00000112701 | 0 | NA | NA | NA | NA | NA | 2310002D06 | RIKEN cDNA 2310002D06 gene [Source:M | 0 | 0 | 0 | 0 | 0 | 0 |
| 50267 | ENSMUSG00000112702 | 0 | NA | NA | NA | NA | NA | Gm47571 | predicted gene, 47571 [Source:MGI Symb | 0 | 0 | 0 | 0 | 0 | 0 |
| 50268 | ENSMUSG00000112703 | 0 | NA | NA | NA | NA | NA | Gm18041 | predicted gene, 18041 [Source:MGI Symb | 0 | 0 | 0 | 0 | 0 | 0 |
| 50269 | ENSMUSG00000112704 | 0.133850373 | -0.796629176 | 4.080472857 | -0.195229623 | 0.84521319 | NA | Gm46329 | predicted gene, 46329 [Source:MGI Symb | 0.8031 | 0 | 0 | 0 | 0 | 0 |
| 50270 | ENSMUSG00000112705 | 0 | NA | NA | NA | NA | NA | Gm47552 | predicted gene, 47552 [Source:MGI Symb | 0 | 0 | 0 | 0 | 0 | 0 |
| 50271 | ENSMUSG00000112706 | 0 | NA | NA | NA | NA | NA | Gm46193 | predicted gene, 46193 [Source:MGI Symb | 0 | 0 | 0 | 0 | 0 | 0 |
| 50272 | ENSMUSG00000112707 | 0 | NA | NA | NA | NA | NA | D830005E20 | RIKEN cDNA D830005E20 gene [Source:M | 0 | 0 | 0 | 0 | 0 | 0 |
| 50273 | ENSMUSG00000112708 | 0 | NA | NA | NA | NA | NA | Gm8255 | predicted gene 8255 [Source:MGI Symbol | 0 | 0 | 0 | 0 | 0 | 0 |
| 50274 | ENSMUSG00000112709 | 10.24370875 | -0.10104059 | 0.830028364 | -0.12173149 | 0.903111673 | 0.983821605 | Gm19395 | predicted gene, 19395 [Source:MGI Symb | 10.44 | 10.279 | 11.069 | 6.7478 | 21.812 | 1.1138 |
| 50275 | ENSMUSG00000112710 | 0 | NA | NA | NA | NA | NA | Gm17829 | predicted gene, 17829 [Source:MGI Symb | 0 | 0 | 0 | 0 | 0 | 0 |
| 50276 | ENSMUSG00000112711 | 0 | NA | NA | NA | NA | NA | Gm32552 | predicted gene, 32552 [Source:MGI Symb | 0 | 0 | 0 | 0 | 0 | 0 |
| 50277 | ENSMUSG00000112712 | 0 | NA | NA | NA | NA | NA | Gm18755 | predicted gene, 18755 [Source:MGI Symb | 0 | 0 | 0 | 0 | 0 | 0 |
| 50278 | ENSMUSG00000112713 | 0 | NA | NA | NA | NA | NA | Gm48281 | predicted gene, 48281 [Source:MGI Symb | 0 | 0 | 0 | 0 | 0 | 0 |
| 50279 | ENSMUSG00000112714 | 0 | NA | NA | NA | NA | NA | Gm4798 | predicted pseudogene 4798 [Source:MGI | 0 | 0 | 0 | 0 | 0 | 0 |
| 50280 | ENSMUSG00000112715 | 0 | NA | NA | NA | NA | NA | Gm48476 | predicted gene, 48476 [Source:MGI Symb | 0 | 0 | 0 | 0 | 0 | 0 |
| 50281 | ENSMUSG00000112716 | 0 | NA | NA | NA | NA | NA | 4930471E19 | RIKEN cDNA 4930471E19 gene [Source:M | 0 | 0 | 0 | 0 | 0 | 0 |
| 50282 | ENSMUSG00000112717 | 0 | NA | NA | NA | NA | NA | Gm6362 | predicted gene 6362 [Source:MGI Symbol | 0 | 0 | 0 | 0 | 0 | 0 |
| 50283 | ENSMUSG00000112718 | 0 | NA | NA | NA | NA | NA | Gm48524 | predicted gene, 48524 [Source:MGI Symb | 0 | 0 | 0 | 0 | 0 | 0 |
| 50284 | ENSMUSG00000112719 | 0.367410362 | 2.078255025 | 3.800070157 | 0.54689912 | 0.58444804 | NA | Gm45925 | predicted gene, 45925 [Source:MGI Symb | 0 | 0 | 0 | 0 | 1.0906 | 1.1138 |

| | | | | | | | | | | | | | | | |
|---|---|---|---|---|---|---|---|---|---|---|---|---|---|---|---|
| 50285 | ENSMUSG00000112720 | 0 | NA | NA | NA | NA | NA | Gm19116 | predicted gene, 19116 [Source:MGI Symb | 0 | 0 | 0 | 0 | 0 | 0 |
| 50286 | ENSMUSG00000112721 | 0 | NA | NA | NA | NA | NA | Gm35608 | predicted gene, 35608 [Source:MGI Symb | 0 | 0 | 0 | 0 | 0 | 0 |
| 50287 | ENSMUSG00000112722 | 0 | NA | NA | NA | NA | NA | Gm3942 | predicted gene 3942 [Source:MGI Symbol | 0 | 0 | 0 | 0 | 0 | 0 |
| 50288 | ENSMUSG00000112723 | 0 | NA | NA | NA | NA | NA | Gm9110 | predicted gene 9110 [Source:MGI Symbol | 0 | 0 | 0 | 0 | 0 | 0 |
| 50289 | ENSMUSG00000112724 | 0 | NA | NA | NA | NA | NA | Gm47699 | predicted gene, 47699 [Source:MGI Symb | 0 | 0 | 0 | 0 | 0 | 0 |
| 50290 | ENSMUSG00000112725 | 0 | NA | NA | NA | NA | NA | Gm47600 | predicted gene, 47600 [Source:MGI Symb | 0 | 0 | 0 | 0 | 0 | 0 |
| 50291 | ENSMUSG00000112726 | 0 | NA | NA | NA | NA | NA | Gm47531 | predicted gene, 47531 [Source:MGI Symb | 0 | 0 | 0 | 0 | 0 | 0 |
| 50292 | ENSMUSG00000112727 | 0 | NA | NA | NA | NA | NA | Gm17794 | predicted gene, 17794 [Source:MGI Symb | 0 | 0 | 0 | 0 | 0 | 0 |
| 50293 | ENSMUSG00000112728 | 0.481988923 | 2.492401104 | 3.837077151 | 0.649557203 | 0.515978285 | NA | Gm48903 | predicted gene, 48903 [Source:MGI Symb | 0 | 0 | 0 | 2.8919 | 0 | 0 |
| 50294 | ENSMUSG00000112729 | 0.17131964 | -0.796629176 | 4.080472857 | -0.195229623 | 0.84521319 | NA | Gm46231 | predicted gene, 46231 [Source:MGI Symb | 0 | 1.0279 | 0 | 0 | 0 | 0 |
| 50295 | ENSMUSG00000112730 | 0 | NA | NA | NA | NA | NA | 4930567K20 | RIKEN cDNA 4930567K20 gene [Source:M | 0 | 0 | 0 | 0 | 0 | 0 |
| 50296 | ENSMUSG00000112731 | 0 | NA | NA | NA | NA | NA | Gm47739 | predicted gene, 47739 [Source:MGI Symb | 0 | 0 | 0 | 0 | 0 | 0 |
| 50297 | ENSMUSG00000112732 | 0 | NA | NA | NA | NA | NA | Gm47343 | predicted gene, 47343 [Source:MGI Symb | 0 | 0 | 0 | 0 | 0 | 0 |
| 50298 | ENSMUSG00000112733 | 0.160662974 | 1.126924175 | 4.080472857 | 0.276174898 | 0.782413736 | NA | Gm40650 | predicted gene, 40650 [Source:MGI Symb | 0 | 0 | 0 | 0.964 | 0 | 0 |
| 50299 | ENSMUSG00000112734 | 1.87377651 | -0.16401045 | 1.675160473 | -0.097907307 | 0.922005898 | NA | Gm47644 | predicted gene, 47644 [Source:MGI Symb | 4.8186 | 1.0279 | 0 | 0.964 | 1.0906 | 3.3415 |
| 50300 | ENSMUSG00000112735 | 0 | NA | NA | NA | NA | NA | Gm46191 | predicted gene, 46191 [Source:MGI Symb | 0 | 0 | 0 | 0 | 0 | 0 |
| 50301 | ENSMUSG00000112736 | 0 | NA | NA | NA | NA | NA | Gm48718 | predicted gene, 48718 [Source:MGI Symb | 0 | 0 | 0 | 0 | 0 | 0 |
| 50302 | ENSMUSG00000112737 | 0 | NA | NA | NA | NA | NA | Gm4898 | predicted gene 4898 [Source:MGI Symbol | 0 | 0 | 0 | 0 | 0 | 0 |
| 50303 | ENSMUSG00000112738 | 0.842329324 | -1.832359871 | 2.605893773 | -0.703159848 | 0.481956138 | NA | Gm48386 | predicted gene, 48386 [Source:MGI Symb | 0 | 3.0838 | 1.0062 | 0.964 | 0 | 0 |
| 50304 | ENSMUSG00000112739 | 0.133850373 | -0.796629176 | 4.080472857 | -0.195229623 | 0.84521319 | NA | Gm20597 | predicted gene, 20597 [Source:MGI Symb | 0.8031 | 0 | 0 | 0 | 0 | 0 |
| 50305 | ENSMUSG00000112740 | 0 | NA | NA | NA | NA | NA | Gm8264 | predicted gene 8264 [Source:MGI Symbol | 0 | 0 | 0 | 0 | 0 | 0 |
| 50306 | ENSMUSG00000112741 | 0 | NA | NA | NA | NA | NA | Gm47341 | predicted gene, 47341 [Source:MGI Symb | 0 | 0 | 0 | 0 | 0 | 0 |
| 50307 | ENSMUSG00000112742 | 0.367410362 | 2.078255025 | 3.800070157 | 0.54689912 | 0.58444804 | NA | Gm31793 | predicted gene, 31793 [Source:MGI Symb | 0 | 0 | 0 | 0 | 1.0906 | 1.1138 |
| 50308 | ENSMUSG00000112743 | 0 | NA | NA | NA | NA | NA | Gm49358 | predicted gene, 49358 [Source:MGI Symb | 0 | 0 | 0 | 0 | 0 | 0 |
| 50309 | ENSMUSG00000112744 | 1.006656641 | 1.034607824 | 2.256383849 | 0.458524743 | 0.64657549 | NA | Gm48880 | predicted gene, 48880 [Source:MGI Symb | 0 | 1.0279 | 1.0062 | 2.8919 | 0 | 1.1138 |
| 50310 | ENSMUSG00000112745 | 0.17131964 | -0.796629176 | 4.080472857 | -0.195229623 | 0.84521319 | NA | Gm48025 | predicted gene, 48025 [Source:MGI Symb | 0 | 1.0279 | 0 | 0 | 0 | 0 |
| 50311 | ENSMUSG00000112746 | 0 | NA | NA | NA | NA | NA | Gm19309 | predicted gene, 19309 [Source:MGI Symb | 0 | 0 | 0 | 0 | 0 | 0 |
| 50312 | ENSMUSG00000112747 | 0 | NA | NA | NA | NA | NA | Gm48721 | predicted gene, 48721 [Source:MGI Symb | 0 | 0 | 0 | 0 | 0 | 0 |
| 50313 | ENSMUSG00000112748 | 0 | NA | NA | NA | NA | NA | Gm6331 | predicted gene 6331 [Source:MGI Symbol | 0 | 0 | 0 | 0 | 0 | 0 |
| 50314 | ENSMUSG00000112749 | 0 | NA | NA | NA | NA | NA | Gm48428 | predicted gene, 48428 [Source:MGI Symb | 0 | 0 | 0 | 0 | 0 | 0 |
| 50315 | ENSMUSG00000112750 | 0 | NA | NA | NA | NA | NA | Gm47534 | predicted gene, 47534 [Source:MGI Symb | 0 | 0 | 0 | 0 | 0 | 0 |
| 50316 | ENSMUSG00000112751 | 0 | NA | NA | NA | NA | NA | Gm48084 | predicted gene, 48084 [Source:MGI Symb | 0 | 0 | 0 | 0 | 0 | 0 |
| 50317 | ENSMUSG00000112752 | 0 | NA | NA | NA | NA | NA | Gm48021 | predicted gene, 48021 [Source:MGI Symb | 0 | 0 | 0 | 0 | 0 | 0 |
| 50318 | ENSMUSG00000112753 | 0 | NA | NA | NA | NA | NA | 4930423D24 | RIKEN cDNA 4930423D24 gene [Source:M | 0 | 0 | 0 | 0 | 0 | 0 |
| 50319 | ENSMUSG00000112754 | 0 | NA | NA | NA | NA | NA | Gm9139 | predicted gene 9139 [Source:MGI Symbol | 0 | 0 | 0 | 0 | 0 | 0 |
| 50320 | ENSMUSG00000112755 | 0 | NA | NA | NA | NA | NA | Gm46190 | predicted gene, 46190 [Source:MGI Symb | 0 | 0 | 0 | 0 | 0 | 0 |
| 50321 | ENSMUSG00000112756 | 0 | NA | NA | NA | NA | NA | Gm48014 | predicted gene, 48014 [Source:MGI Symb | 0 | 0 | 0 | 0 | 0 | 0 |
| 50322 | ENSMUSG00000112757 | 0 | NA | NA | NA | NA | NA | Gm10743 | predicted gene 10743 [Source:MGI Symbo | 0 | 0 | 0 | 0 | 0 | 0 |
| 50323 | ENSMUSG00000112758 | 92.94890021 | -0.073167574 | 0.280811285 | -0.260557813 | 0.794433526 | 0.962046762 | Gm36172 | predicted gene, 36172 [Source:MGI Symb | 114.04 | 102.79 | 68.425 | 68.442 | 97.065 | 106.93 |
| 50324 | ENSMUSG00000112759 | 0.133850373 | -0.796629176 | 4.080472857 | -0.195229623 | 0.84521319 | NA | Gm47615 | predicted gene, 47615 [Source:MGI Symb | 0.8031 | 0 | 0 | 0 | 0 | 0 |
| 50325 | ENSMUSG00000112760 | 0.453341394 | -0.545391191 | 3.551113277 | -0.153583158 | 0.877938417 | NA | Gm47539 | predicted gene, 47539 [Source:MGI Symb | 1.6062 | 0 | 0 | 0 | 0 | 1.1138 |
| 50326 | ENSMUSG00000112761 | 0 | NA | NA | NA | NA | NA | Gm47972 | predicted gene, 47972 [Source:MGI Symb | 0 | 0 | 0 | 0 | 0 | 0 |
| 50327 | ENSMUSG00000112762 | 4.22127877 | 0.096379704 | 1.249545725 | 0.077131794 | 0.9385187 | NA | 4930459C07 | RIKEN cDNA 4930459C07 gene [Source:M | 8.031 | 1.0279 | 3.0187 | 0 | 5.4531 | 7.7969 |
| 50328 | ENSMUSG00000112763 | 0 | NA | NA | NA | NA | NA | Gm34304 | predicted gene, 34304 [Source:MGI Symb | 0 | 0 | 0 | 0 | 0 | 0 |
| 50329 | ENSMUSG00000112764 | 0 | NA | NA | NA | NA | NA | Gm48158 | predicted gene, 48158 [Source:MGI Symb | 0 | 0 | 0 | 0 | 0 | 0 |
| 50330 | ENSMUSG00000112765 | 0 | NA | NA | NA | NA | NA | 9430078K24 | RIKEN cDNA 9430078K24 gene [Source:M | 0 | 0 | 0 | 0 | 0 | 0 |
| 50331 | ENSMUSG00000112766 | 0 | NA | NA | NA | NA | NA | Gm40645 | predicted gene, 40645 [Source:MGI Symb | 0 | 0 | 0 | 0 | 0 | 0 |
| 50332 | ENSMUSG00000112767 | 0 | NA | NA | NA | NA | NA | Gm47532 | predicted gene, 47532 [Source:MGI Symb | 0 | 0 | 0 | 0 | 0 | 0 |
| 50333 | ENSMUSG00000112768 | 0 | NA | NA | NA | NA | NA | Gm48517 | predicted gene, 48517 [Source:MGI Symb | 0 | 0 | 0 | 0 | 0 | 0 |
| 50334 | ENSMUSG00000112769 | 0 | NA | NA | NA | NA | NA | Rpl17-ps2 | ribosomal protein L17, pseudogene 2 [Sou | 0 | 0 | 0 | 0 | 0 | 0 |
| 50335 | ENSMUSG00000112770 | 1.973079585 | -1.455016625 | 1.620074349 | -0.898117192 | 0.369123075 | NA | Gm48210 | predicted gene, 48210 [Source:MGI Symb | 1.6062 | 2.0558 | 5.0312 | 0.964 | 2.1812 | 0 |
| 50336 | ENSMUSG00000112771 | 13.98783781 | -0.334009371 | 0.610961186 | -0.54669491 | 0.584588352 | 0.908955096 | Gm47718 | predicted gene, 47718 [Source:MGI Symb | 19.274 | 8.2233 | 19.119 | 8.6758 | 15.269 | 13.366 |
| 50337 | ENSMUSG00000112772 | 0 | NA | NA | NA | NA | NA | Gm48400 | predicted gene, 48400 [Source:MGI Symb | 0 | 0 | 0 | 0 | 0 | 0 |
| 50338 | ENSMUSG00000112773 | 1.393881239 | -0.636148929 | 2.169171775 | -0.29326812 | 0.769317219 | NA | Gm3137 | predicted gene, 3137 [Source:MGI Symbo | 4.0155 | 0 | 1.0062 | 0 | 0 | 3.3415 |
| 50339 | ENSMUSG00000112774 | 78.31587152 | 0.296050768 | 0.291799981 | 1.014567468 | 0.310312053 | 0.759065371 | Gm36041 | predicted gene, 36041 [Source:MGI Symb | 64.248 | 78.122 | 68.425 | 63.623 | 78.525 | 116.95 |
| 50340 | ENSMUSG00000112775 | 0 | NA | NA | NA | NA | NA | Gm48722 | predicted gene, 48722 [Source:MGI Symb | 0 | 0 | 0 | 0 | 0 | 0 |

| | | | | | | | | | | | | | | | |
|---|---|---|---|---|---|---|---|---|---|---|---|---|---|---|---|
| 50341 | ENSMUSG00000112776 | 0 | NA | NA | NA | NA | NA | Gm6989 | predicted gene 6989 [Source:MGI Symbol | 0 | 0 | 0 | 0 | 0 | 0 |
| 50342 | ENSMUSG00000112777 | 0 | NA | NA | NA | NA | NA | Gm8723 | predicted gene 8723 [Source:MGI Symbol | 0 | 0 | 0 | 0 | 0 | 0 |
| 50343 | ENSMUSG00000112778 | 0 | NA | NA | NA | NA | NA | Gm33677 | predicted gene, 33677 [Source:MGI Symb | 0 | 0 | 0 | 0 | 0 | 0 |
| 50344 | ENSMUSG00000112779 | 0 | NA | NA | NA | NA | NA | Gm18029 | predicted gene, 18029 [Source:MGI Symb | 0 | 0 | 0 | 0 | 0 | 0 |
| 50345 | ENSMUSG00000112780 | 0.185640647 | 1.126924175 | 4.080472857 | 0.276174898 | 0.782413736 | NA | Gm48410 | predicted gene, 48410 [Source:MGI Symb | 0 | 0 | 0 | 0 | 0 | 1.1138 |
| 50346 | ENSMUSG00000112781 | 0 | NA | NA | NA | NA | NA | Gm9046 | predicted gene 9046 [Source:MGI Symbol | 0 | 0 | 0 | 0 | 0 | 0 |
| 50347 | ENSMUSG00000112782 | 0 | NA | NA | NA | NA | NA | Gm7940 | predicted gene 7940 [Source:MGI Symbol | 0 | 0 | 0 | 0 | 0 | 0 |
| 50348 | ENSMUSG00000112783 | 0 | NA | NA | NA | NA | NA | Gm48792 | predicted gene, 48792 [Source:MGI Symb | 0 | 0 | 0 | 0 | 0 | 0 |
| 50349 | ENSMUSG00000112784 | 1.630826856 | -1.883827287 | 1.788499039 | -1.053300698 | 0.292203199 | NA | Gm34868 | predicted gene, 34868 [Source:MGI Symb | 1.6062 | 4.1117 | 2.0125 | 0.964 | 1.0906 | 0 |
| 50350 | ENSMUSG00000112785 | 0 | NA | NA | NA | NA | NA | Gm47645 | predicted gene, 47645 [Source:MGI Symb | 0 | 0 | 0 | 0 | 0 | 0 |
| 50351 | ENSMUSG00000112786 | 0 | NA | NA | NA | NA | NA | Gm47696 | predicted gene, 47696 [Source:MGI Symb | 0 | 0 | 0 | 0 | 0 | 0 |
| 50352 | ENSMUSG00000112787 | 0 | NA | NA | NA | NA | NA | Gm48790 | predicted gene, 48790 [Source:MGI Symb | 0 | 0 | 0 | 0 | 0 | 0 |
| 50353 | ENSMUSG00000112788 | 0 | NA | NA | NA | NA | NA | Gm47821 | predicted gene, 47821 [Source:MGI Symb | 0 | 0 | 0 | 0 | 0 | 0 |
| 50354 | ENSMUSG00000112789 | 0 | NA | NA | NA | NA | NA | Gm19326 | predicted gene, 19326 [Source:MGI Symb | 0 | 0 | 0 | 0 | 0 | 0 |
| 50355 | ENSMUSG00000112790 | 4.04542702 | -0.398253763 | 1.046078994 | -0.380710984 | 0.703417717 | NA | Gm33378 | predicted gene, 33378 [Source:MGI Symb | 5.6217 | 5.1396 | 3.0187 | 3.8559 | 2.1812 | 4.4554 |
| 50356 | ENSMUSG00000112791 | 0 | NA | NA | NA | NA | NA | Gm47612 | predicted gene, 47612 [Source:MGI Symb | 0 | 0 | 0 | 0 | 0 | 0 |
| 50357 | ENSMUSG00000112792 | 0 | NA | NA | NA | NA | NA | Gm38407 | predicted gene, 38407 [Source:MGI Symb | 0 | 0 | 0 | 0 | 0 | 0 |
| 50358 | ENSMUSG00000112793 | 19.00233688 | 1.333135118 | 0.534907758 | 2.492271045 | 0.012692913 | 0.127461989 | Gm32872 | predicted gene, 32872 [Source:MGI Symb | 12.047 | 11.307 | 9.0562 | 38.559 | 18.541 | 24.505 |
| 50359 | ENSMUSG00000112794 | 8.919065917 | 0.542908005 | 0.724814564 | 0.74903021 | 0.453838997 | 0.857297539 | Gm48878 | predicted gene, 48878 [Source:MGI Symb | 5.6217 | 5.1396 | 11.069 | 9.6398 | 10.906 | 11.138 |
| 50360 | ENSMUSG00000112795 | 0 | NA | NA | NA | NA | NA | Gm2027 | predicted gene 2027 [Source:MGI Symbol | 0 | 0 | 0 | 0 | 0 | 0 |
| 50361 | ENSMUSG00000112796 | 95.28765638 | 0.139034207 | 0.254256943 | 0.546825608 | 0.584498548 | 0.908955096 | Gm40770 | predicted gene, 40770 [Source:MGI Symb | 104.4 | 69.898 | 97.606 | 114.71 | 94.884 | 90.221 |
| 50362 | ENSMUSG00000112797 | 0 | NA | NA | NA | NA | NA | Gm30034 | predicted gene, 30034 [Source:MGI Symb | 0 | 0 | 0 | 0 | 0 | 0 |
| 50363 | ENSMUSG00000112798 | 3.446058376 | 0.451548384 | 1.337461481 | 0.337615992 | 0.73565259 | NA | Gm40685 | predicted gene, 40685 [Source:MGI Symb | 5.6217 | 3.0838 | 0 | 8.6758 | 2.1812 | 1.1138 |
| 50364 | ENSMUSG00000112799 | 0 | NA | NA | NA | NA | NA | Gm48100 | predicted gene, 48100 [Source:MGI Symb | 0 | 0 | 0 | 0 | 0 | 0 |
| 50365 | ENSMUSG00000112800 | 2.803143896 | -0.256037238 | 1.402078149 | -0.182612673 | 0.85510194 | NA | Gm40604 | predicted gene, 40604 [Source:MGI Symb | 4.0155 | 1.0279 | 4.025 | 0 | 2.1812 | 5.5692 |
| 50366 | ENSMUSG00000112801 | 0 | NA | NA | NA | NA | NA | Gm48034 | predicted gene, 48034 [Source:MGI Symb | 0 | 0 | 0 | 0 | 0 | 0 |
| 50367 | ENSMUSG00000112802 | 0 | NA | NA | NA | NA | NA | Gm19005 | predicted gene, 19005 [Source:MGI Symb | 0 | 0 | 0 | 0 | 0 | 0 |
| 50368 | ENSMUSG00000112803 | 0.589026695 | 0.241472922 | 3.228838524 | 0.074786311 | 0.940384733 | NA | Gm36543 | predicted gene, 36543 [Source:MGI Symb | 1.6062 | 0 | 0 | 1.928 | 0 | 0 |
| 50369 | ENSMUSG00000112804 | 0 | NA | NA | NA | NA | NA | Gm30534 | predicted gene, 30534 [Source:MGI Symb | 0 | 0 | 0 | 0 | 0 | 0 |
| 50370 | ENSMUSG00000112805 | 1.923764 | -2.106737264 | 1.758021002 | -1.198357278 | 0.230777957 | NA | Gm34574 | predicted gene, 34574 [Source:MGI Symb | 3.2124 | 3.0838 | 3.0187 | 0 | 0 | 2.2277 |
| 50371 | ENSMUSG00000112806 | 2.977746313 | -2.227726531 | 1.424318406 | -1.564064974 | 0.117802311 | NA | Gm48146 | predicted gene, 48146 [Source:MGI Symb | 5.6217 | 2.0558 | 7.0437 | 0.964 | 2.1812 | 0 |
| 50372 | ENSMUSG00000112807 | 0 | NA | NA | NA | NA | NA | Gm47519 | predicted gene, 47519 [Source:MGI Symb | 0 | 0 | 0 | 0 | 0 | 0 |
| 50373 | ENSMUSG00000112808 | 0.301557804 | -1.64335916 | 4.042814367 | -0.406488899 | 0.684383412 | NA | Gm4739 | predicted gene 4739 [Source:MGI Symbol | 0.8031 | 0 | 1.0062 | 0 | 0 | 0 |
| 50374 | ENSMUSG00000112809 | 0 | NA | NA | NA | NA | NA | Gm47915 | predicted gene, 47915 [Source:MGI Symb | 0 | 0 | 0 | 0 | 0 | 0 |
| 50375 | ENSMUSG00000112810 | 0 | NA | NA | NA | NA | NA | Gm35101 | predicted gene, 35101 [Source:MGI Symb | 0 | 0 | 0 | 0 | 0 | 0 |
| 50376 | ENSMUSG00000112811 | 0 | NA | NA | NA | NA | NA | Gm7422 | predicted gene 7422 [Source:MGI Symbol | 0 | 0 | 0 | 0 | 0 | 0 |
| 50377 | ENSMUSG00000112812 | 0 | NA | NA | NA | NA | NA | Gm47415 | predicted gene, 47415 [Source:MGI Symb | 0 | 0 | 0 | 0 | 0 | 0 |
| 50378 | ENSMUSG00000112813 | 0 | NA | NA | NA | NA | NA | Gm34376 | predicted gene, 34376 [Source:MGI Symb | 0 | 0 | 0 | 0 | 0 | 0 |
| 50379 | ENSMUSG00000112814 | 0 | NA | NA | NA | NA | NA | Gm9040 | predicted gene 9040 [Source:MGI Symbol | 0 | 0 | 0 | 0 | 0 | 0 |
| 50380 | ENSMUSG00000112815 | 0 | NA | NA | NA | NA | NA | Gm48828 | predicted gene, 48828 [Source:MGI Symb | 0 | 0 | 0 | 0 | 0 | 0 |
| 50381 | ENSMUSG00000112816 | 0.531944269 | 2.618916564 | 3.285123409 | 0.797204926 | 0.425332026 | NA | Gm48798 | predicted gene, 48798 [Source:MGI Symb | 0 | 0 | 0 | 0.964 | 0 | 2.2277 |
| 50382 | ENSMUSG00000112817 | 0 | NA | NA | NA | NA | NA | Gm48122 | predicted gene, 48122 [Source:MGI Symb | 0 | 0 | 0 | 0 | 0 | 0 |
| 50383 | ENSMUSG00000112818 | 0 | NA | NA | NA | NA | NA | Gm35920 | predicted gene, 35920 [Source:MGI Symb | 0 | 0 | 0 | 0 | 0 | 0 |
| 50384 | ENSMUSG00000112819 | 0.167707431 | -0.796629176 | 4.080472857 | -0.195229623 | 0.84521319 | NA | Gm46332 | predicted gene, 46332 [Source:MGI Symb | 0 | 0 | 1.0062 | 0 | 0 | 0 |
| 50385 | ENSMUSG00000112820 | 0 | NA | NA | NA | NA | NA | Gm48298 | predicted gene, 48298 [Source:MGI Symb | 0 | 0 | 0 | 0 | 0 | 0 |
| 50386 | ENSMUSG00000112821 | 0 | NA | NA | NA | NA | NA | 1700104L18I | RIKEN cDNA 1700104L18 gene [Source:M( | 0 | 0 | 0 | 0 | 0 | 0 |
| 50387 | ENSMUSG00000112822 | 1.355901693 | -1.531138196 | 2.060135839 | -0.74322196 | 0.45734731 | NA | A130012E19 | RIKEN cDNA A130012E19 gene [Source:M | 0 | 2.0558 | 4.025 | 0.964 | 1.0906 | 0 |
| 50388 | ENSMUSG00000112823 | 0.17131964 | -0.796629176 | 4.080472857 | -0.195229623 | 0.84521319 | NA | Gm6125 | predicted gene 6125 [Source:MGI Symbol | 0 | 1.0279 | 0 | 0 | 0 | 0 |
| 50389 | ENSMUSG00000112824 | 1.813371089 | -0.779647222 | 1.791933429 | -0.435087157 | 0.663499165 | NA | Gm47917 | predicted gene, 47917 [Source:MGI Symb | 4.8186 | 2.0558 | 0 | 2.8919 | 0 | 1.1138 |
| 50390 | ENSMUSG00000112825 | 2.965098847 | -1.631462447 | 1.366305052 | -1.194068956 | 0.232450995 | NA | Gm9118 | predicted gene 9118 [Source:MGI Symbol | 3.2124 | 6.1675 | 4.025 | 0 | 3.2719 | 1.1138 |
| 50391 | ENSMUSG00000112826 | 0 | NA | NA | NA | NA | NA | Gm1559 | predicted gene 1559 [Source:MGI Symbol | 0 | 0 | 0 | 0 | 0 | 0 |
| 50392 | ENSMUSG00000112827 | 0 | NA | NA | NA | NA | NA | Gm48753 | predicted gene, 48753 [Source:MGI Symb | 0 | 0 | 0 | 0 | 0 | 0 |
| 50393 | ENSMUSG00000112828 | 0 | NA | NA | NA | NA | NA | Gm48495 | predicted gene, 48495 [Source:MGI Symb | 0 | 0 | 0 | 0 | 0 | 0 |
| 50394 | ENSMUSG00000112829 | 0 | NA | NA | NA | NA | NA | Gm29696 | predicted gene, 29696 [Source:MGI Symb | 0 | 0 | 0 | 0 | 0 | 0 |
| 50395 | ENSMUSG00000112830 | 0 | NA | NA | NA | NA | NA | Gm47765 | predicted gene, 47765 [Source:MGI Symb | 0 | 0 | 0 | 0 | 0 | 0 |
| 50396 | ENSMUSG00000112831 | 0.974138179 | -0.759835404 | 2.114820303 | -0.359290765 | 0.719377582 | NA | Gm35533 | predicted gene, 35533 [Source:MGI Symb | 1.6062 | 1.0279 | 1.0062 | 0 | 1.0906 | 1.1138 |

| | | | | | | | | | | | | | | | |
|---|---|---|---|---|---|---|---|---|---|---|---|---|---|---|---|
| 50397 | ENSMUSG00000112832 | 0 | NA | NA | NA | NA | NA | 4930484H19 | RIKEN cDNA 4930484H19 gene [Source:M | 0 | 0 | 0 | 0 | 0 | 0 |
| 50398 | ENSMUSG00000112833 | 0 | NA | NA | NA | NA | NA | Gm36595 | predicted gene, 36595 [Source:MGI Symb | 0 | 0 | 0 | 0 | 0 | 0 |
| 50399 | ENSMUSG00000112834 | 0 | NA | NA | NA | NA | NA | Gm48464 | predicted gene, 48464 [Source:MGI Symb | 0 | 0 | 0 | 0 | 0 | 0 |
| 50400 | ENSMUSG00000112835 | 0 | NA | NA | NA | NA | NA | Gm36173 | predicted gene, 36173 [Source:MGI Symb | 0 | 0 | 0 | 0 | 0 | 0 |
| 50401 | ENSMUSG00000112836 | 0.822613314 | -1.749415261 | 2.618121802 | -0.668194757 | 0.504009282 | NA | Gm47006 | predicted gene, 47006 [Source:MGI Symb | 0.8031 | 0 | 3.0187 | 0 | 0 | 1.1138 |
| 50402 | ENSMUSG00000112837 | 0 | NA | NA | NA | NA | NA | Gm48053 | predicted gene, 48053 [Source:MGI Symb | 0 | 0 | 0 | 0 | 0 | 0 |
| 50403 | ENSMUSG00000112838 | 0 | NA | NA | NA | NA | NA | Gm47098 | predicted gene, 47098 [Source:MGI Symb | 0 | 0 | 0 | 0 | 0 | 0 |
| 50404 | ENSMUSG00000112839 | 0 | NA | NA | NA | NA | NA | Gm47222 | predicted gene, 47222 [Source:MGI Symb | 0 | 0 | 0 | 0 | 0 | 0 |
| 50405 | ENSMUSG00000112840 | 0 | NA | NA | NA | NA | NA | Gm20049 | predicted gene, 20049 [Source:MGI Symb | 0 | 0 | 0 | 0 | 0 | 0 |
| 50406 | ENSMUSG00000112841 | 0 | NA | NA | NA | NA | NA | Gm48752 | predicted gene, 48752 [Source:MGI Symb | 0 | 0 | 0 | 0 | 0 | 0 |
| 50407 | ENSMUSG00000112842 | 0 | NA | NA | NA | NA | NA | Gm30232 | predicted gene, 30232 [Source:MGI Symb | 0 | 0 | 0 | 0 | 0 | 0 |
| 50408 | ENSMUSG00000112843 | 1.177744186 | -2.412679626 | 2.321486536 | -1.039282197 | 0.298673512 | NA | Gm46224 | predicted gene, 46224 [Source:MGI Symb | 0 | 3.0838 | 3.0187 | 0.964 | 0 | 0 |
| 50409 | ENSMUSG00000112844 | 0 | NA | NA | NA | NA | NA | Gm47036 | predicted gene, 47036 [Source:MGI Symb | 0 | 0 | 0 | 0 | 0 | 0 |
| 50410 | ENSMUSG00000112845 | 0 | NA | NA | NA | NA | NA | Gm47143 | predicted gene, 47143 [Source:MGI Symb | 0 | 0 | 0 | 0 | 0 | 0 |
| 50411 | ENSMUSG00000112846 | 0 | NA | NA | NA | NA | NA | 4930432O09 | RIKEN cDNA 4930432O09 gene [Source:M | 0 | 0 | 0 | 0 | 0 | 0 |
| 50412 | ENSMUSG00000112847 | 0.821015994 | 0.567017314 | 2.624106235 | 0.216080167 | 0.828925259 | NA | Gm47573 | predicted gene, 47573 [Source:MGI Symb | 0 | 1.0279 | 1.0062 | 2.8919 | 0 | 0 |
| 50413 | ENSMUSG00000112848 | 0 | NA | NA | NA | NA | NA | Gm32023 | predicted gene, 32023 [Source:MGI Symb | 0 | 0 | 0 | 0 | 0 | 0 |
| 50414 | ENSMUSG00000112849 | 0.133850373 | -0.796629176 | 4.080472857 | -0.195229623 | 0.84521319 | NA | Gm18040 | predicted gene, 18040 [Source:MGI Symb | 0.8031 | 0 | 0 | 0 | 0 | 0 |
| 50415 | ENSMUSG00000112850 | 0 | NA | NA | NA | NA | NA | Gm9176 | predicted gene 9176 [Source:MGI Symbol | 0 | 0 | 0 | 0 | 0 | 0 |
| 50416 | ENSMUSG00000112851 | 0 | NA | NA | NA | NA | NA | Gm7727 | predicted gene 7727 [Source:MGI Symbol | 0 | 0 | 0 | 0 | 0 | 0 |
| 50417 | ENSMUSG00000112852 | 0 | NA | NA | NA | NA | NA | Gm47767 | predicted gene, 47767 [Source:MGI Symb | 0 | 0 | 0 | 0 | 0 | 0 |
| 50418 | ENSMUSG00000112853 | 0 | NA | NA | NA | NA | NA | Gm47637 | predicted gene, 47637 [Source:MGI Symb | 0 | 0 | 0 | 0 | 0 | 0 |
| 50419 | ENSMUSG00000112854 | 0 | NA | NA | NA | NA | NA | Gm38403 | predicted gene, 38403 [Source:MGI Symb | 0 | 0 | 0 | 0 | 0 | 0 |
| 50420 | ENSMUSG00000112855 | 0 | NA | NA | NA | NA | NA | Gm47842 | predicted gene, 47842 [Source:MGI Symb | 0 | 0 | 0 | 0 | 0 | 0 |
| 50421 | ENSMUSG00000112856 | 0 | NA | NA | NA | NA | NA | Gm4305 | predicted gene 4305 [Source:MGI Symbol | 0 | 0 | 0 | 0 | 0 | 0 |
| 50422 | ENSMUSG00000112857 | 0 | NA | NA | NA | NA | NA | Gm35041 | predicted gene, 35041 [Source:MGI Symb | 0 | 0 | 0 | 0 | 0 | 0 |
| 50423 | ENSMUSG00000112858 | 0 | NA | NA | NA | NA | NA | Rnf212b | ring finger protein 212B [Source:MGI Sym | 0 | 0 | 0 | 0 | 0 | 0 |
| 50424 | ENSMUSG00000112859 | 0 | NA | NA | NA | NA | NA | Gm48522 | predicted gene, 48522 [Source:MGI Symb | 0 | 0 | 0 | 0 | 0 | 0 |
| 50425 | ENSMUSG00000112860 | 0.692375149 | -1.404276945 | 2.772265464 | -0.506544905 | 0.612474175 | NA | Gm47439 | predicted gene, 47439 [Source:MGI Symb | 0 | 1.0279 | 2.0125 | 0 | 0 | 1.1138 |
| 50426 | ENSMUSG00000112861 | 0 | NA | NA | NA | NA | NA | Gm47930 | predicted gene, 47930 [Source:MGI Symb | 0 | 0 | 0 | 0 | 0 | 0 |
| 50427 | ENSMUSG00000112862 | 0 | NA | NA | NA | NA | NA | Gm47094 | predicted gene, 47094 [Source:MGI Symb | 0 | 0 | 0 | 0 | 0 | 0 |
| 50428 | ENSMUSG00000112863 | 0.505131668 | 1.126696199 | 3.421159819 | 0.329331647 | 0.741905026 | NA | Gm19181 | predicted gene, 19181 [Source:MGI Symb | 0.8031 | 0 | 0 | 0 | 0 | 2.2277 |
| 50429 | ENSMUSG00000112864 | 0 | NA | NA | NA | NA | NA | Gm18596 | predicted gene, 18596 [Source:MGI Symb | 0 | 0 | 0 | 0 | 0 | 0 |
| 50430 | ENSMUSG00000112865 | 0 | NA | NA | NA | NA | NA | Gm40375 | predicted gene, 40375 [Source:MGI Symb | 0 | 0 | 0 | 0 | 0 | 0 |
| 50431 | ENSMUSG00000112866 | 0 | NA | NA | NA | NA | NA | Gm47704 | predicted gene, 47704 [Source:MGI Symb | 0 | 0 | 0 | 0 | 0 | 0 |
| 50432 | ENSMUSG00000112867 | 0 | NA | NA | NA | NA | NA | Gm48263 | predicted gene, 48263 [Source:MGI Symb | 0 | 0 | 0 | 0 | 0 | 0 |
| 50433 | ENSMUSG00000112868 | 0 | NA | NA | NA | NA | NA | 4930520K02 | RIKEN cDNA 4930520K02 gene [Source:M | 0 | 0 | 0 | 0 | 0 | 0 |
| 50434 | ENSMUSG00000112869 | 0 | NA | NA | NA | NA | NA | Gm48108 | predicted gene, 48108 [Source:MGI Symb | 0 | 0 | 0 | 0 | 0 | 0 |
| 50435 | ENSMUSG00000112870 | 0 | NA | NA | NA | NA | NA | Gm40689 | predicted gene, 40689 [Source:MGI Symb | 0 | 0 | 0 | 0 | 0 | 0 |
| 50436 | ENSMUSG00000112871 | 0 | NA | NA | NA | NA | NA | Gm35404 | predicted gene, 35404 [Source:MGI Symb | 0 | 0 | 0 | 0 | 0 | 0 |
| 50437 | ENSMUSG00000112872 | 0 | NA | NA | NA | NA | NA | Gm47058 | predicted gene, 47058 [Source:MGI Symb | 0 | 0 | 0 | 0 | 0 | 0 |
| 50438 | ENSMUSG00000112873 | 3.92075996 | -0.476403663 | 1.230737369 | -0.387087997 | 0.698691063 | NA | Gm48877 | predicted gene, 48877 [Source:MGI Symb | 2.4093 | 9.2513 | 2.0125 | 0.964 | 1.0906 | 7.7969 |
| 50439 | ENSMUSG00000112874 | 0 | NA | NA | NA | NA | NA | Gm34297 | predicted gene, 34297 [Source:MGI Symb | 0 | 0 | 0 | 0 | 0 | 0 |
| 50440 | ENSMUSG00000112875 | 0.294513347 | 0.165143496 | 4.080472857 | 0.040471657 | 0.967717103 | NA | Gm48074 | predicted gene, 48074 [Source:MGI Symb | 0.8031 | 0 | 0 | 0.964 | 0 | 0 |
| 50441 | ENSMUSG00000112876 | 0.684865378 | 2.990343487 | 2.966805827 | 1.00793367 | 0.313486306 | NA | Gm32443 | predicted gene, 32443 [Source:MGI Symb | 0 | 0 | 0 | 1.928 | 2.1812 | 0 |
| 50442 | ENSMUSG00000112877 | 0 | NA | NA | NA | NA | NA | Gm47537 | predicted gene, 47537 [Source:MGI Symb | 0 | 0 | 0 | 0 | 0 | 0 |
| 50443 | ENSMUSG00000112878 | 0 | NA | NA | NA | NA | NA | 1700042O05 | RIKEN cDNA 1700042O05 gene [Source:M | 0 | 0 | 0 | 0 | 0 | 0 |
| 50444 | ENSMUSG00000112879 | 0 | NA | NA | NA | NA | NA | Gm47726 | predicted gene, 47726 [Source:MGI Symb | 0 | 0 | 0 | 0 | 0 | 0 |
| 50445 | ENSMUSG00000112880 | 17.02463059 | 0.214197922 | 0.580916146 | 0.368724339 | 0.712333205 | 0.944239215 | Gm20337 | predicted gene, 20337 [Source:MGI Symb | 16.865 | 7.1954 | 23.144 | 17.352 | 13.087 | 24.505 |
| 50446 | ENSMUSG00000112881 | 1.727517405 | -3.017369124 | 2.365034573 | -1.275824531 | 0.202017554 | NA | Gm49344 | predicted gene, 49344 [Source:MGI Symb | 0 | 9.2513 | 0 | 0 | 0 | 1.1138 |
| 50447 | ENSMUSG00000112882 | 0 | NA | NA | NA | NA | NA | Gm47970 | predicted gene, 47970 [Source:MGI Symb | 0 | 0 | 0 | 0 | 0 | 0 |
| 50448 | ENSMUSG00000112883 | 0 | NA | NA | NA | NA | NA | Gm48023 | predicted gene, 48023 [Source:MGI Symb | 0 | 0 | 0 | 0 | 0 | 0 |
| 50449 | ENSMUSG00000112884 | 0 | NA | NA | NA | NA | NA | Gm47946 | predicted gene, 47946 [Source:MGI Symb | 0 | 0 | 0 | 0 | 0 | 0 |
| 50450 | ENSMUSG00000112885 | 0 | NA | NA | NA | NA | NA | Gm29807 | predicted gene, 29807 [Source:MGI Symb | 0 | 0 | 0 | 0 | 0 | 0 |
| 50451 | ENSMUSG00000112886 | 0 | NA | NA | NA | NA | NA | Gm18339 | predicted gene, 18339 [Source:MGI Symb | 0 | 0 | 0 | 0 | 0 | 0 |
| 50452 | ENSMUSG00000112887 | 0.160662974 | 1.126924175 | 4.080472857 | 0.276174898 | 0.782413736 | NA | Gm47961 | predicted gene, 47961 [Source:MGI Symb | 0 | 0 | 0 | 0.964 | 0 | 0 |

| | | | | | | | | | | | | | | | |
|---|---|---|---|---|---|---|---|---|---|---|---|---|---|---|---|
| 50453 | ENSMUSG00000112888 | 0 | NA | NA | NA | NA | NA | Gm47634 | predicted gene, 47634 [Source:MGI Symb | 0 | 0 | 0 | 0 | 0 | 0 |
| 50454 | ENSMUSG00000112889 | 1.608169416 | 0.832067133 | 1.844510406 | 0.451104602 | 0.651914161 | NA | Gm47073 | predicted gene, 47073 [Source:MGI Symb | 2.4093 | 0 | 1.0062 | 2.8919 | 0 | 3.3415 |
| 50455 | ENSMUSG00000112890 | 0 | NA | NA | NA | NA | NA | Gm18502 | predicted gene, 18502 [Source:MGI Symb | 0 | 0 | 0 | 0 | 0 | 0 |
| 50456 | ENSMUSG00000112891 | 1.981285638 | 1.792456683 | 1.756289902 | 1.020592717 | 0.307447442 | NA | Gm47391 | predicted gene, 47391 [Source:MGI Symb | 1.6062 | 0 | 1.0062 | 4.8199 | 0 | 4.4554 |
| 50457 | ENSMUSG00000112892 | 0 | NA | NA | NA | NA | NA | Gm8613 | predicted gene 8613 [Source:MGI Symbol | 0 | 0 | 0 | 0 | 0 | 0 |
| 50458 | ENSMUSG00000112893 | 0 | NA | NA | NA | NA | NA | Gm9583 | predicted gene 9583 [Source:MGI Symbol | 0 | 0 | 0 | 0 | 0 | 0 |
| 50459 | ENSMUSG00000112894 | 0 | NA | NA | NA | NA | NA | Gm47009 | predicted gene, 47009 [Source:MGI Symb | 0 | 0 | 0 | 0 | 0 | 0 |
| 50460 | ENSMUSG00000112895 | 0.990448572 | -3.34412486 | 2.6642679 | -1.2551759 | 0.209414912 | NA | Gm47567 | predicted gene, 47567 [Source:MGI Symb | 0.8031 | 5.1396 | 0 | 0 | 0 | 0 |
| 50461 | ENSMUSG00000112896 | 0 | NA | NA | NA | NA | NA | Gm8633 | predicted gene 8633 [Source:MGI Symbol | 0 | 0 | 0 | 0 | 0 | 0 |
| 50462 | ENSMUSG00000112897 | 0 | NA | NA | NA | NA | NA | Gm47831 | predicted gene, 47831 [Source:MGI Symb | 0 | 0 | 0 | 0 | 0 | 0 |
| 50463 | ENSMUSG00000112898 | 0 | NA | NA | NA | NA | NA | Gm47020 | predicted gene, 47020 [Source:MGI Symb | 0 | 0 | 0 | 0 | 0 | 0 |
| 50464 | ENSMUSG00000112899 | 0 | NA | NA | NA | NA | NA | 4930521O17 | RIKEN cDNA 4930521O17 gene [Source:M | 0 | 0 | 0 | 0 | 0 | 0 |
| 50465 | ENSMUSG00000112900 | 0.133850373 | -0.796629176 | 4.080472857 | -0.195229623 | 0.84521319 | NA | Gm48204 | predicted gene, 48204 [Source:MGI Symb | 0.8031 | 0 | 0 | 0 | 0 | 0 |
| 50466 | ENSMUSG00000112901 | 0 | NA | NA | NA | NA | NA | Gm18514 | predicted gene, 18514 [Source:MGI Symb | 0 | 0 | 0 | 0 | 0 | 0 |
| 50467 | ENSMUSG00000112902 | 4.912372442 | -0.868435087 | 0.962182792 | -0.902567676 | 0.366755388 | NA | Gm48884 | predicted gene, 48884 [Source:MGI Symb | 8.8341 | 5.1396 | 5.0312 | 3.8559 | 3.2719 | 3.3415 |
| 50468 | ENSMUSG00000112903 | 0 | NA | NA | NA | NA | NA | Gm5948 | predicted gene 5948 [Source:MGI Symbol | 0 | 0 | 0 | 0 | 0 | 0 |
| 50469 | ENSMUSG00000112904 | 0 | NA | NA | NA | NA | NA | Gm47030 | predicted gene, 47030 [Source:MGI Symb | 0 | 0 | 0 | 0 | 0 | 0 |
| 50470 | ENSMUSG00000112905 | 0 | NA | NA | NA | NA | NA | Gm47477 | predicted gene, 47477 [Source:MGI Symb | 0 | 0 | 0 | 0 | 0 | 0 |
| 50471 | ENSMUSG00000112906 | 0 | NA | NA | NA | NA | NA | Gm18189 | predicted gene, 18189 [Source:MGI Symb | 0 | 0 | 0 | 0 | 0 | 0 |
| 50472 | ENSMUSG00000112907 | 1.575476507 | -1.073397044 | 1.878444699 | -0.57142861 | 0.56770914 | NA | Gm47015 | predicted gene, 47015 [Source:MGI Symb | 2.4093 | 0 | 4.025 | 1.928 | 1.0906 | 0 |
| 50473 | ENSMUSG00000112908 | 0.133850373 | -0.796629176 | 4.080472857 | -0.195229623 | 0.84521319 | NA | Gm7392 | predicted gene 7392 [Source:MGI Symbol | 0.8031 | 0 | 0 | 0 | 0 | 0 |
| 50474 | ENSMUSG00000112909 | 0.672839099 | 0.267535205 | 2.816195809 | 0.094998794 | 0.924315785 | NA | Gm10120 | predicted gene 10120 [Source:MGI Symbo | 0.8031 | 0 | 1.0062 | 0 | 0 | 2.2277 |
| 50475 | ENSMUSG00000112910 | 0 | NA | NA | NA | NA | NA | Gm29685 | predicted gene, 29685 [Source:MGI Symb | 0 | 0 | 0 | 0 | 0 | 0 |
| 50476 | ENSMUSG00000112911 | 0.997777256 | 1.176137242 | 2.241393702 | 0.524734785 | 0.599767572 | NA | BC042761 | cDNA sequence BC042761 [Source:MGI Sy | 0.8031 | 1.0279 | 0 | 1.928 | 0 | 2.2277 |
| 50477 | ENSMUSG00000112912 | 0 | NA | NA | NA | NA | NA | Gm48160 | predicted gene, 48160 [Source:MGI Symb | 0 | 0 | 0 | 0 | 0 | 0 |
| 50478 | ENSMUSG00000112913 | 0 | NA | NA | NA | NA | NA | Gm30853 | predicted gene, 30853 [Source:MGI Symb | 0 | 0 | 0 | 0 | 0 | 0 |
| 50479 | ENSMUSG00000112914 | 0 | NA | NA | NA | NA | NA | Gm47702 | predicted gene, 47702 [Source:MGI Symb | 0 | 0 | 0 | 0 | 0 | 0 |
| 50480 | ENSMUSG00000112915 | 0 | NA | NA | NA | NA | NA | Gm18112 | predicted gene, 18112 [Source:MGI Symb | 0 | 0 | 0 | 0 | 0 | 0 |
| 50481 | ENSMUSG00000112916 | 0.346303622 | 2.004339653 | 3.884003423 | 0.516049919 | 0.605819544 | NA | Gm18622 | predicted gene, 18622 [Source:MGI Symb | 0 | 0 | 0 | 0.964 | 0 | 1.1138 |
| 50482 | ENSMUSG00000112917 | 0 | NA | NA | NA | NA | NA | Gm47331 | predicted gene, 47331 [Source:MGI Symb | 0 | 0 | 0 | 0 | 0 | 0 |
| 50483 | ENSMUSG00000112918 | 0 | NA | NA | NA | NA | NA | Gm18120 | predicted gene, 18120 [Source:MGI Symb | 0 | 0 | 0 | 0 | 0 | 0 |
| 50484 | ENSMUSG00000112919 | 0 | NA | NA | NA | NA | NA | Gm9049 | predicted gene 9049 [Source:MGI Symbol | 0 | 0 | 0 | 0 | 0 | 0 |
| 50485 | ENSMUSG00000112920 | 13.65290624 | 0.868261997 | 0.613281413 | 1.41576441 | 0.156844493 | 0.575786484 | Spata22 | spermatogenesis associated 22 [Source:M | 14.456 | 7.1954 | 7.0437 | 13.496 | 17.45 | 22.277 |
| 50486 | ENSMUSG00000112921 | 0 | NA | NA | NA | NA | NA | Gm48104 | predicted gene, 48104 [Source:MGI Symb | 0 | 0 | 0 | 0 | 0 | 0 |
| 50487 | ENSMUSG00000112922 | 3.159903107 | 0.308800049 | 1.159944456 | 0.266219686 | 0.790070023 | NA | Gm47572 | predicted gene, 47572 [Source:MGI Symb | 2.4093 | 2.0558 | 4.025 | 3.8559 | 3.2719 | 3.3415 |
| 50488 | ENSMUSG00000112923 | 0 | NA | NA | NA | NA | NA | Gm8768 | predicted gene 8768 [Source:MGI Symbol | 0 | 0 | 0 | 0 | 0 | 0 |
| 50489 | ENSMUSG00000112924 | 0 | NA | NA | NA | NA | NA | 4930579P08 | RIKEN cDNA 4930579P08 gene [Source:M | 0 | 0 | 0 | 0 | 0 | 0 |
| 50490 | ENSMUSG00000112925 | 2.901664159 | 1.970361695 | 1.423930865 | 1.383748147 | 0.166435585 | NA | Gm31763 | predicted gene, 31763 [Source:MGI Symb | 2.4093 | 0 | 1.0062 | 1.928 | 8.7249 | 3.3415 |
| 50491 | ENSMUSG00000112926 | 0 | NA | NA | NA | NA | NA | Gm7172 | predicted gene 7172 [Source:MGI Symbol | 0 | 0 | 0 | 0 | 0 | 0 |
| 50492 | ENSMUSG00000112927 | 0 | NA | NA | NA | NA | NA | Gm48332 | predicted gene, 48332 [Source:MGI Symb | 0 | 0 | 0 | 0 | 0 | 0 |
| 50493 | ENSMUSG00000112928 | 0.335414862 | -1.779047242 | 4.030804179 | -0.441362855 | 0.658950329 | NA | Gm47710 | predicted gene, 47710 [Source:MGI Symb | 0 | 0 | 2.0125 | 0 | 0 | 0 |
| 50494 | ENSMUSG00000112929 | 0 | NA | NA | NA | NA | NA | Gm47729 | predicted gene, 47729 [Source:MGI Symb | 0 | 0 | 0 | 0 | 0 | 0 |
| 50495 | ENSMUSG00000112930 | 0 | NA | NA | NA | NA | NA | Gm7086 | predicted pseudogene 7086 [Source:MGI | 0 | 0 | 0 | 0 | 0 | 0 |
| 50496 | ENSMUSG00000112931 | 0 | NA | NA | NA | NA | NA | Gm4303 | predicted gene 4303 [Source:MGI Symbol | 0 | 0 | 0 | 0 | 0 | 0 |
| 50497 | ENSMUSG00000112932 | 0.52105551 | -0.812028029 | 3.385364865 | -0.239864257 | 0.810435491 | NA | Gm48308 | predicted gene, 48308 [Source:MGI Symb | 0 | 0 | 2.0125 | 0 | 0 | 1.1138 |
| 50498 | ENSMUSG00000112933 | 0 | NA | NA | NA | NA | NA | Gm47491 | predicted gene, 47491 [Source:MGI Symb | 0 | 0 | 0 | 0 | 0 | 0 |
| 50499 | ENSMUSG00000112934 | 0 | NA | NA | NA | NA | NA | Gm32305 | predicted gene, 32305 [Source:MGI Symb | 0 | 0 | 0 | 0 | 0 | 0 |
| 50500 | ENSMUSG00000112935 | 0.596071152 | -1.224343926 | 2.897583865 | -0.4225396 | 0.672631203 | NA | Gm40983 | predicted gene, 40983 [Source:MGI Symb | 1.6062 | 0 | 1.0062 | 0.964 | 0 | 0 |
| 50501 | ENSMUSG00000112936 | 0 | NA | NA | NA | NA | NA | Gm33958 | predicted gene, 33958 [Source:MGI Symb | 0 | 0 | 0 | 0 | 0 | 0 |
| 50502 | ENSMUSG00000112937 | 0 | NA | NA | NA | NA | NA | Gm17746 | predicted gene, 17746 [Source:MGI Symb | 0 | 0 | 0 | 0 | 0 | 0 |
| 50503 | ENSMUSG00000112938 | 0 | NA | NA | NA | NA | NA | Gm48590 | predicted gene, 48590 [Source:MGI Symb | 0 | 0 | 0 | 0 | 0 | 0 |
| 50504 | ENSMUSG00000112939 | 0 | NA | NA | NA | NA | NA | Gm48251 | predicted gene, 48251 [Source:MGI Symb | 0 | 0 | 0 | 0 | 0 | 0 |
| 50505 | ENSMUSG00000112940 | 0 | NA | NA | NA | NA | NA | 4930511J24I | RIKEN cDNA 4930511J24 gene [Source:M( | 0 | 0 | 0 | 0 | 0 | 0 |
| 50506 | ENSMUSG00000112941 | 0.678286236 | 1.600482029 | 2.782216593 | 0.575254289 | 0.565119331 | NA | Gm48623 | predicted gene, 48623 [Source:MGI Symb | 0 | 1.0279 | 0 | 1.928 | 0 | 1.1138 |
| 50507 | ENSMUSG00000112942 | 0.181769715 | 1.126924175 | 4.080472857 | 0.276174898 | 0.782413736 | NA | Gm47597 | predicted gene, 47597 [Source:MGI Symb | 0 | 0 | 0 | 0 | 1.0906 | 0 |
| 50508 | ENSMUSG00000112943 | 0 | NA | NA | NA | NA | NA | Gm18983 | predicted gene, 18983 [Source:MGI Symb | 0 | 0 | 0 | 0 | 0 | 0 |

| | | | | | | | | | | | | | | | |
|---|---|---|---|---|---|---|---|---|---|---|---|---|---|---|---|
| 50509 | ENSMUSG00000112944 | 0 | NA | NA | NA | NA | NA | Gm48885 | predicted gene, 48885 [Source:MGI Symb | 0 | 0 | 0 | 0 | 0 | 0 |
| 50510 | ENSMUSG00000112945 | 0 | NA | NA | NA | NA | NA | Gm32151 | predicted gene, 32151 [Source:MGI Symb | 0 | 0 | 0 | 0 | 0 | 0 |
| 50511 | ENSMUSG00000112946 | 0 | NA | NA | NA | NA | NA | Gm9280 | predicted gene 9280 [Source:MGI Symbol | 0 | 0 | 0 | 0 | 0 | 0 |
| 50512 | ENSMUSG00000112947 | 0 | NA | NA | NA | NA | NA | Gm47493 | predicted gene, 47493 [Source:MGI Symb | 0 | 0 | 0 | 0 | 0 | 0 |
| 50513 | ENSMUSG00000112948 | 0 | NA | NA | NA | NA | NA | Gm48871 | predicted gene, 48871 [Source:MGI Symb | 0 | 0 | 0 | 0 | 0 | 0 |
| 50514 | ENSMUSG00000112949 | 0 | NA | NA | NA | NA | NA | Gm21939 | predicted gene, 21939 [Source:MGI Symb | 0 | 0 | 0 | 0 | 0 | 0 |
| 50515 | ENSMUSG00000112950 | 0 | NA | NA | NA | NA | NA | Gm32369 | predicted gene, 32369 [Source:MGI Symb | 0 | 0 | 0 | 0 | 0 | 0 |
| 50516 | ENSMUSG00000112951 | 0 | NA | NA | NA | NA | NA | Gm49929 | predicted gene, 49929 [Source:MGI Symb | 0 | 0 | 0 | 0 | 0 | 0 |
| 50517 | ENSMUSG00000112952 | 0 | NA | NA | NA | NA | NA | Gm48336 | predicted gene, 48336 [Source:MGI Symb | 0 | 0 | 0 | 0 | 0 | 0 |
| 50518 | ENSMUSG00000112953 | 0 | NA | NA | NA | NA | NA | Gm47984 | predicted gene, 47984 [Source:MGI Symb | 0 | 0 | 0 | 0 | 0 | 0 |
| 50519 | ENSMUSG00000112954 | 0 | NA | NA | NA | NA | NA | Gm49732 | predicted gene, 49732 [Source:MGI Symb | 0 | 0 | 0 | 0 | 0 | 0 |
| 50520 | ENSMUSG00000112955 | 0.492645588 | 0.9565034 | 3.447648281 | 0.277436479 | 0.781444974 | NA | Gm48889 | predicted gene, 48889 [Source:MGI Symb | 0 | 1.0279 | 0 | 1.928 | 0 | 0 |
| 50521 | ENSMUSG00000112956 | 0 | NA | NA | NA | NA | NA | Gm48645 | predicted gene, 48645 [Source:MGI Symb | 0 | 0 | 0 | 0 | 0 | 0 |
| 50522 | ENSMUSG00000112957 | 0 | NA | NA | NA | NA | NA | 4930547H16 | RIKEN cDNA 4930547H16 gene [Source:M | 0 | 0 | 0 | 0 | 0 | 0 |
| 50523 | ENSMUSG00000112958 | 0 | NA | NA | NA | NA | NA | Gm47443 | predicted gene, 47443 [Source:MGI Symb | 0 | 0 | 0 | 0 | 0 | 0 |
| 50524 | ENSMUSG00000112959 | 0.371281295 | 2.09253362 | 4.031852165 | 0.519000582 | 0.603760335 | NA | Gm32834 | predicted gene, 32834 [Source:MGI Symb | 0 | 0 | 0 | 0 | 0 | 2.2277 |
| 50525 | ENSMUSG00000112960 | 0.294513347 | 0.165143496 | 4.080472857 | 0.040471657 | 0.967717103 | NA | Gm47254 | predicted gene, 47254 [Source:MGI Symb | 0.8031 | 0 | 0 | 0.964 | 0 | 0 |
| 50526 | ENSMUSG00000112961 | 0 | NA | NA | NA | NA | NA | Gm48576 | predicted gene, 48576 [Source:MGI Symb | 0 | 0 | 0 | 0 | 0 | 0 |
| 50527 | ENSMUSG00000112962 | 0 | NA | NA | NA | NA | NA | Gm31711 | predicted gene, 31711 [Source:MGI Symb | 0 | 0 | 0 | 0 | 0 | 0 |
| 50528 | ENSMUSG00000112963 | 0 | NA | NA | NA | NA | NA | Gm6093 | predicted gene 6093 [Source:MGI Symbol | 0 | 0 | 0 | 0 | 0 | 0 |
| 50529 | ENSMUSG00000112964 | 64.12307746 | 0.170999289 | 0.289569452 | 0.590529449 | 0.554835749 | 0.897628185 | E430024I08F | RIKEN cDNA E430024I08 gene [Source:MG | 48.186 | 61.675 | 71.443 | 65.55 | 63.256 | 74.628 |
| 50530 | ENSMUSG00000112965 | 0 | NA | NA | NA | NA | NA | Gm34278 | predicted gene, 34278 [Source:MGI Symb | 0 | 0 | 0 | 0 | 0 | 0 |
| 50531 | ENSMUSG00000112966 | 0 | NA | NA | NA | NA | NA | Gm18499 | predicted gene, 18499 [Source:MGI Symb | 0 | 0 | 0 | 0 | 0 | 0 |
| 50532 | ENSMUSG00000112967 | 0 | NA | NA | NA | NA | NA | 4930428E07 | RIKEN cDNA 4930428E07 gene [Source:M | 0 | 0 | 0 | 0 | 0 | 0 |
| 50533 | ENSMUSG00000112968 | 0 | NA | NA | NA | NA | NA | Gm48530 | predicted gene, 48530 [Source:MGI Symb | 0 | 0 | 0 | 0 | 0 | 0 |
| 50534 | ENSMUSG00000112969 | 0 | NA | NA | NA | NA | NA | Gm48496 | predicted gene, 48496 [Source:MGI Symb | 0 | 0 | 0 | 0 | 0 | 0 |
| 50535 | ENSMUSG00000112970 | 0 | NA | NA | NA | NA | NA | Gm49370 | predicted gene, 49370 [Source:MGI Symb | 0 | 0 | 0 | 0 | 0 | 0 |
| 50536 | ENSMUSG00000112971 | 0.356960287 | 0.16514264 | 3.880370363 | 0.042558474 | 0.966053499 | NA | Gm33682 | predicted gene, 33682 [Source:MGI Symb | 0 | 1.0279 | 0 | 0 | 0 | 1.1138 |
| 50537 | ENSMUSG00000112972 | 0 | NA | NA | NA | NA | NA | Gm48417 | predicted gene, 48417 [Source:MGI Symb | 0 | 0 | 0 | 0 | 0 | 0 |
| 50538 | ENSMUSG00000112973 | 0 | NA | NA | NA | NA | NA | Gm48520 | predicted gene, 48520 [Source:MGI Symb | 0 | 0 | 0 | 0 | 0 | 0 |
| 50539 | ENSMUSG00000112974 | 0 | NA | NA | NA | NA | NA | Gm18026 | predicted gene, 18026 [Source:MGI Symb | 0 | 0 | 0 | 0 | 0 | 0 |
| 50540 | ENSMUSG00000112975 | 0 | NA | NA | NA | NA | NA | Gm21297 | predicted gene, 21297 [Source:MGI Symb | 0 | 0 | 0 | 0 | 0 | 0 |
| 50541 | ENSMUSG00000112976 | 0 | NA | NA | NA | NA | NA | Gm2654 | predicted gene 2654 [Source:MGI Symbol | 0 | 0 | 0 | 0 | 0 | 0 |
| 50542 | ENSMUSG00000112977 | 0 | NA | NA | NA | NA | NA | Gm34081 | predicted gene, 34081 [Source:MGI Symb | 0 | 0 | 0 | 0 | 0 | 0 |
| 50543 | ENSMUSG00000112978 | 0 | NA | NA | NA | NA | NA | Gm18601 | predicted gene, 18601 [Source:MGI Symb | 0 | 0 | 0 | 0 | 0 | 0 |
| 50544 | ENSMUSG00000112979 | 0 | NA | NA | NA | NA | NA | Gm8634 | predicted gene 8634 [Source:MGI Symbol | 0 | 0 | 0 | 0 | 0 | 0 |
| 50545 | ENSMUSG00000112980 | 1.029540662 | 0.20390886 | 2.204283627 | 0.092505727 | 0.926296241 | NA | D430020J02 | RIKEN cDNA D430020J02 gene [Source:M | 0.8031 | 2.0558 | 0 | 0 | 1.0906 | 2.2277 |
| 50546 | ENSMUSG00000112981 | 0 | NA | NA | NA | NA | NA | 5033424D13 | RIKEN cDNA 5033424D13 gene [Source:M | 0 | 0 | 0 | 0 | 0 | 0 |
| 50547 | ENSMUSG00000112982 | 0 | NA | NA | NA | NA | NA | Gm21007 | predicted gene, 21007 [Source:MGI Symb | 0 | 0 | 0 | 0 | 0 | 0 |
| 50548 | ENSMUSG00000112983 | 0 | NA | NA | NA | NA | NA | Gm32939 | predicted gene, 32939 [Source:MGI Symb | 0 | 0 | 0 | 0 | 0 | 0 |
| 50549 | ENSMUSG00000112984 | 0 | NA | NA | NA | NA | NA | Gm48419 | predicted gene, 48419 [Source:MGI Symb | 0 | 0 | 0 | 0 | 0 | 0 |
| 50550 | ENSMUSG00000112985 | 0 | NA | NA | NA | NA | NA | Gm48556 | predicted gene, 48556 [Source:MGI Symb | 0 | 0 | 0 | 0 | 0 | 0 |
| 50551 | ENSMUSG00000112986 | 0 | NA | NA | NA | NA | NA | Gm2123 | predicted gene 2123 [Source:MGI Symbol | 0 | 0 | 0 | 0 | 0 | 0 |
| 50552 | ENSMUSG00000112987 | 0 | NA | NA | NA | NA | NA | Gm5954 | predicted gene 5954 [Source:MGI Symbol | 0 | 0 | 0 | 0 | 0 | 0 |
| 50553 | ENSMUSG00000112988 | 0 | NA | NA | NA | NA | NA | Gm35298 | predicted gene, 35298 [Source:MGI Symb | 0 | 0 | 0 | 0 | 0 | 0 |
| 50554 | ENSMUSG00000112989 | 0 | NA | NA | NA | NA | NA | Gm48143 | predicted gene, 48143 [Source:MGI Symb | 0 | 0 | 0 | 0 | 0 | 0 |
| 50555 | ENSMUSG00000112990 | 2.048631249 | -0.292504249 | 1.743656719 | -0.167753346 | 0.866777327 | NA | Gm47372 | predicted gene, 47372 [Source:MGI Symb | 1.6062 | 5.1396 | 0 | 0 | 1.0906 | 4.4554 |
| 50556 | ENSMUSG00000112991 | 0 | NA | NA | NA | NA | NA | Gm36635 | predicted gene, 36635 [Source:MGI Symb | 0 | 0 | 0 | 0 | 0 | 0 |
| 50557 | ENSMUSG00000112992 | 0 | NA | NA | NA | NA | NA | Gm47869 | predicted gene, 47869 [Source:MGI Symb | 0 | 0 | 0 | 0 | 0 | 0 |
| 50558 | ENSMUSG00000112993 | 0 | NA | NA | NA | NA | NA | Gm48030 | predicted gene, 48030 [Source:MGI Symb | 0 | 0 | 0 | 0 | 0 | 0 |
| 50559 | ENSMUSG00000112994 | 0 | NA | NA | NA | NA | NA | Gm46312 | predicted gene, 46312 [Source:MGI Symb | 0 | 0 | 0 | 0 | 0 | 0 |
| 50560 | ENSMUSG00000112995 | 0 | NA | NA | NA | NA | NA | Gm47977 | predicted gene, 47977 [Source:MGI Symb | 0 | 0 | 0 | 0 | 0 | 0 |
| 50561 | ENSMUSG00000112996 | 0 | NA | NA | NA | NA | NA | Gm48064 | predicted gene, 48064 [Source:MGI Symb | 0 | 0 | 0 | 0 | 0 | 0 |
| 50562 | ENSMUSG00000112997 | 0 | NA | NA | NA | NA | NA | Gm49386 | predicted gene, 49386 [Source:MGI Symb | 0 | 0 | 0 | 0 | 0 | 0 |
| 50563 | ENSMUSG00000112998 | 0 | NA | NA | NA | NA | NA | Gm47656 | predicted gene, 47656 [Source:MGI Symb | 0 | 0 | 0 | 0 | 0 | 0 |
| 50564 | ENSMUSG00000112999 | 0 | NA | NA | NA | NA | NA | 9530050K03 | RIKEN cDNA 9530050K03 gene [Source:M | 0 | 0 | 0 | 0 | 0 | 0 |

| | | | | | | | | | | | | | | | |
|---|---|---|---|---|---|---|---|---|---|---|---|---|---|---|---|
| 50565 | ENSMUSG00000113000 | 0 | NA | NA | NA | NA | NA | Gm6049 | predicted gene 6049 [Source:MGI Symbol | 0 | 0 | 0 | 0 | 0 | 0 |
| 50566 | ENSMUSG00000113001 | 0 | NA | NA | NA | NA | NA | Gm40538 | predicted gene, 40538 [Source:MGI Symb | 0 | 0 | 0 | 0 | 0 | 0 |
| 50567 | ENSMUSG00000113002 | 0 | NA | NA | NA | NA | NA | Gm48620 | predicted gene, 48620 [Source:MGI Symb | 0 | 0 | 0 | 0 | 0 | 0 |
| 50568 | ENSMUSG00000113003 | 0.856470372 | -1.816219066 | 2.859255434 | -0.635207 | 0.525293418 | NA | Gm47138 | predicted gene, 47138 [Source:MGI Symb | 0 | 0 | 4.025 | 0 | 0 | 1.1138 |
| 50569 | ENSMUSG00000113004 | 0 | NA | NA | NA | NA | NA | Gm47951 | predicted gene, 47951 [Source:MGI Symb | 0 | 0 | 0 | 0 | 0 | 0 |
| 50570 | ENSMUSG00000113005 | 0.722877068 | -0.331317305 | 2.989821669 | -0.110815073 | 0.911762993 | NA | Gm34408 | predicted gene, 34408 [Source:MGI Symb | 2.4093 | 0 | 0 | 1.928 | 0 | 0 |
| 50571 | ENSMUSG00000113006 | 0 | NA | NA | NA | NA | NA | Gm48316 | predicted gene, 48316 [Source:MGI Symb | 0 | 0 | 0 | 0 | 0 | 0 |
| 50572 | ENSMUSG00000113007 | 0 | NA | NA | NA | NA | NA | Gm8700 | predicted gene 8700 [Source:MGI Symbol | 0 | 0 | 0 | 0 | 0 | 0 |
| 50573 | ENSMUSG00000113008 | 0 | NA | NA | NA | NA | NA | Gm48171 | predicted gene, 48171 [Source:MGI Symb | 0 | 0 | 0 | 0 | 0 | 0 |
| 50574 | ENSMUSG00000113009 | 0 | NA | NA | NA | NA | NA | 3732407C23 | RIKEN cDNA 3732407C23 gene [Source:M | 0 | 0 | 0 | 0 | 0 | 0 |
| 50575 | ENSMUSG00000113010 | 1.463326015 | -3.908076741 | 2.129442963 | -1.835257769 | 0.066467505 | NA | Gm34084 | predicted gene, 34084 [Source:MGI Symb | 1.6062 | 6.1675 | 1.0062 | 0 | 0 | 0 |
| 50576 | ENSMUSG00000113011 | 0 | NA | NA | NA | NA | NA | Gm1995 | predicted gene 1995 [Source:MGI Symbol | 0 | 0 | 0 | 0 | 0 | 0 |
| 50577 | ENSMUSG00000113012 | 0 | NA | NA | NA | NA | NA | Gm47196 | predicted gene, 47196 [Source:MGI Symb | 0 | 0 | 0 | 0 | 0 | 0 |
| 50578 | ENSMUSG00000113013 | 0 | NA | NA | NA | NA | NA | Gm47833 | predicted gene, 47833 [Source:MGI Symb | 0 | 0 | 0 | 0 | 0 | 0 |
| 50579 | ENSMUSG00000113014 | 0 | NA | NA | NA | NA | NA | Selenok-ps8 | selenoprotein K, pseudogene 8 [Source:M | 0 | 0 | 0 | 0 | 0 | 0 |
| 50580 | ENSMUSG00000113015 | 0.65103495 | -1.312578592 | 2.821398903 | -0.465222621 | 0.641772041 | NA | Gm6296 | predicted gene 6296 [Source:MGI Symbol | 0.8031 | 0 | 2.0125 | 0 | 1.0906 | 0 |
| 50581 | ENSMUSG00000113016 | 0 | NA | NA | NA | NA | NA | Gm47916 | predicted gene, 47916 [Source:MGI Symb | 0 | 0 | 0 | 0 | 0 | 0 |
| 50582 | ENSMUSG00000113017 | 0 | NA | NA | NA | NA | NA | Gm46401 | predicted gene, 46401 [Source:MGI Symb | 0 | 0 | 0 | 0 | 0 | 0 |
| 50583 | ENSMUSG00000113018 | 0 | NA | NA | NA | NA | NA | Gm9318 | predicted gene 9318 [Source:MGI Symbol | 0 | 0 | 0 | 0 | 0 | 0 |
| 50584 | ENSMUSG00000113019 | 29.27758891 | 0.614772509 | 0.434230256 | 1.415775386 | 0.156841278 | 0.575786484 | Gm47467 | predicted gene, 47467 [Source:MGI Symb | 21.684 | 16.447 | 31.194 | 27.955 | 32.719 | 45.668 |
| 50585 | ENSMUSG00000113020 | 0.167707431 | -0.796629176 | 4.080472857 | -0.195229623 | 0.84521319 | NA | Gm48063 | predicted gene, 48063 [Source:MGI Symb | 0 | 0 | 1.0062 | 0 | 0 | 0 |
| 50586 | ENSMUSG00000113021 | 0 | NA | NA | NA | NA | NA | Gm48440 | predicted gene, 48440 [Source:MGI Symb | 0 | 0 | 0 | 0 | 0 | 0 |
| 50587 | ENSMUSG00000113022 | 0 | NA | NA | NA | NA | NA | Gm9544 | predicted gene 9544 [Source:MGI Symbol | 0 | 0 | 0 | 0 | 0 | 0 |
| 50588 | ENSMUSG00000113023 | 0 | NA | NA | NA | NA | NA | Gm47686 | predicted gene, 47686 [Source:MGI Symb | 0 | 0 | 0 | 0 | 0 | 0 |
| 50589 | ENSMUSG00000113024 | 0.133850373 | -0.796629176 | 4.080472857 | -0.195229623 | 0.84521319 | NA | Gm48502 | predicted gene, 48502 [Source:MGI Symb | 0.8031 | 0 | 0 | 0 | 0 | 0 |
| 50590 | ENSMUSG00000113025 | 0 | NA | NA | NA | NA | NA | Gm49363 | predicted gene, 49363 [Source:MGI Symb | 0 | 0 | 0 | 0 | 0 | 0 |
| 50591 | ENSMUSG00000113026 | 0 | NA | NA | NA | NA | NA | Gm48669 | predicted gene, 48669 [Source:MGI Symb | 0 | 0 | 0 | 0 | 0 | 0 |
| 50592 | ENSMUSG00000113027 | 0 | NA | NA | NA | NA | NA | Gm7040 | predicted gene 7040 [Source:MGI Symbol | 0 | 0 | 0 | 0 | 0 | 0 |
| 50593 | ENSMUSG00000113028 | 0 | NA | NA | NA | NA | NA | Gm18065 | predicted gene, 18065 [Source:MGI Symb | 0 | 0 | 0 | 0 | 0 | 0 |
| 50594 | ENSMUSG00000113029 | 0 | NA | NA | NA | NA | NA | Gm40578 | predicted gene, 40578 [Source:MGI Symb | 0 | 0 | 0 | 0 | 0 | 0 |
| 50595 | ENSMUSG00000113030 | 0 | NA | NA | NA | NA | NA | Gm48778 | predicted gene, 48778 [Source:MGI Symb | 0 | 0 | 0 | 0 | 0 | 0 |
| 50596 | ENSMUSG00000113031 | 0 | NA | NA | NA | NA | NA | Gm34552 | predicted gene, 34552 [Source:MGI Symb | 0 | 0 | 0 | 0 | 0 | 0 |
| 50597 | ENSMUSG00000113032 | 0 | NA | NA | NA | NA | NA | Gm47578 | predicted gene, 47578 [Source:MGI Symb | 0 | 0 | 0 | 0 | 0 | 0 |
| 50598 | ENSMUSG00000113033 | 0 | NA | NA | NA | NA | NA | Gm47799 | predicted gene, 47799 [Source:MGI Symb | 0 | 0 | 0 | 0 | 0 | 0 |
| 50599 | ENSMUSG00000113034 | 0 | NA | NA | NA | NA | NA | Gm47812 | predicted gene, 47812 [Source:MGI Symb | 0 | 0 | 0 | 0 | 0 | 0 |
| 50600 | ENSMUSG00000113035 | 0.353089354 | 0.165142706 | 3.896102244 | 0.042386646 | 0.966190474 | NA | 5830428M24 | RIKEN cDNA 5830428M24 gene [Source:M | 0 | 1.0279 | 0 | 0 | 1.0906 | 0 |
| 50601 | ENSMUSG00000113036 | 0.342639279 | -1.805976596 | 4.028547714 | -0.448294702 | 0.653940524 | NA | Gm47021 | predicted gene, 47021 [Source:MGI Symb | 0 | 2.0558 | 0 | 0 | 0 | 0 |
| 50602 | ENSMUSG00000113037 | 0.160662974 | 1.126924175 | 4.080472857 | 0.276174898 | 0.782413736 | NA | Gm36550 | predicted gene, 36550 [Source:MGI Symb | 0 | 0 | 0 | 0.964 | 0 | 0 |
| 50603 | ENSMUSG00000113038 | 0 | NA | NA | NA | NA | NA | Gm48491 | predicted gene, 48491 [Source:MGI Symb | 0 | 0 | 0 | 0 | 0 | 0 |
| 50604 | ENSMUSG00000113039 | 0 | NA | NA | NA | NA | NA | Gm9265 | predicted gene 9265 [Source:MGI Symbol | 0 | 0 | 0 | 0 | 0 | 0 |
| 50605 | ENSMUSG00000113040 | 0 | NA | NA | NA | NA | NA | Gm9249 | predicted gene 9249 [Source:MGI Symbol | 0 | 0 | 0 | 0 | 0 | 0 |
| 50606 | ENSMUSG00000113041 | 0.844158739 | -0.371321695 | 2.388630557 | -0.155453799 | 0.876463574 | NA | Gm48302 | predicted gene, 48302 [Source:MGI Symb | 0.8031 | 1.0279 | 1.0062 | 0 | 0 | 2.2277 |
| 50607 | ENSMUSG00000113042 | 0 | NA | NA | NA | NA | NA | Gm47188 | predicted gene, 47188 [Source:MGI Symb | 0 | 0 | 0 | 0 | 0 | 0 |
| 50608 | ENSMUSG00000113043 | 0 | NA | NA | NA | NA | NA | Gm7753 | predicted gene 7753 [Source:MGI Symbol | 0 | 0 | 0 | 0 | 0 | 0 |
| 50609 | ENSMUSG00000113044 | 0 | NA | NA | NA | NA | NA | Gm47149 | predicted gene, 47149 [Source:MGI Symb | 0 | 0 | 0 | 0 | 0 | 0 |
| 50610 | ENSMUSG00000113045 | 1.477233843 | 0.510573446 | 1.660191398 | 0.307538906 | 0.758433214 | NA | Gm48621 | predicted gene, 48621 [Source:MGI Symb | 1.6062 | 1.0279 | 1.0062 | 1.928 | 2.1812 | 1.1138 |
| 50611 | ENSMUSG00000113046 | 0 | NA | NA | NA | NA | NA | Gm48579 | predicted gene, 48579 [Source:MGI Symb | 0 | 0 | 0 | 0 | 0 | 0 |
| 50612 | ENSMUSG00000113047 | 37.43589871 | -0.050865987 | 0.362655415 | -0.14025983 | 0.888454702 | 0.982382091 | Gm47469 | predicted gene, 47469 [Source:MGI Symb | 36.943 | 34.949 | 42.262 | 28.919 | 41.443 | 40.098 |
| 50613 | ENSMUSG00000113048 | 0 | NA | NA | NA | NA | NA | Gm48347 | predicted gene, 48347 [Source:MGI Symb | 0 | 0 | 0 | 0 | 0 | 0 |
| 50614 | ENSMUSG00000113049 | 0 | NA | NA | NA | NA | NA | Gm48544 | predicted gene, 48544 [Source:MGI Symb | 0 | 0 | 0 | 0 | 0 | 0 |
| 50615 | ENSMUSG00000113050 | 0 | NA | NA | NA | NA | NA | Gm8016 | predicted gene 8016 [Source:MGI Symbol | 0 | 0 | 0 | 0 | 0 | 0 |
| 50616 | ENSMUSG00000113051 | 0 | NA | NA | NA | NA | NA | Gm47672 | predicted gene, 47672 [Source:MGI Symb | 0 | 0 | 0 | 0 | 0 | 0 |
| 50617 | ENSMUSG00000113052 | 6.325769143 | 0.550385608 | 0.864812478 | 0.636421909 | 0.524501465 | 0.888334265 | Gm31513 | predicted gene, 31513 [Source:MGI Symb | 7.2279 | 4.1117 | 4.025 | 11.568 | 5.4531 | 5.5692 |
| 50618 | ENSMUSG00000113053 | 0 | NA | NA | NA | NA | NA | Gm18262 | predicted gene, 18262 [Source:MGI Symb | 0 | 0 | 0 | 0 | 0 | 0 |
| 50619 | ENSMUSG00000113054 | 0 | NA | NA | NA | NA | NA | Gm47514 | predicted gene, 47514 [Source:MGI Symb | 0 | 0 | 0 | 0 | 0 | 0 |
| 50620 | ENSMUSG00000113055 | 0 | NA | NA | NA | NA | NA | Gm48700 | predicted gene, 48700 [Source:MGI Symb | 0 | 0 | 0 | 0 | 0 | 0 |

| | | | | | | | | | | | | | | | |
|---|---|---|---|---|---|---|---|---|---|---|---|---|---|---|---|
| 50621 | ENSMUSG00000113056 | 0 | NA | NA | NA | NA | NA | 6030440G07 | RIKEN cDNA 6030440G07 gene [Source:M | 0 | 0 | 0 | 0 | 0 | 0 |
| 50622 | ENSMUSG00000113057 | 0 | NA | NA | NA | NA | NA | Gm48812 | predicted gene, 48812 [Source:MGI Symb | 0 | 0 | 0 | 0 | 0 | 0 |
| 50623 | ENSMUSG00000113058 | 0.160662974 | 1.126924175 | 4.080472857 | 0.276174898 | 0.782413736 | NA | Gm48311 | predicted gene, 48311 [Source:MGI Symb | 0 | 0 | 0 | 0.964 | 0 | 0 |
| 50624 | ENSMUSG00000113059 | 0 | NA | NA | NA | NA | NA | Gm46344 | predicted gene, 46344 [Source:MGI Symb | 0 | 0 | 0 | 0 | 0 | 0 |
| 50625 | ENSMUSG00000113060 | 0 | NA | NA | NA | NA | NA | Gm48041 | predicted gene, 48041 [Source:MGI Symb | 0 | 0 | 0 | 0 | 0 | 0 |
| 50626 | ENSMUSG00000113061 | 7.488665501 | -0.003253213 | 0.795400241 | -0.004090033 | 0.996736635 | 0.999154389 | Rps18-ps5 | ribosomal protein S18, pseudogene 5 [Sou | 7.2279 | 5.1396 | 10.062 | 4.8199 | 6.5437 | 11.138 |
| 50627 | ENSMUSG00000113062 | 0 | NA | NA | NA | NA | NA | Gm48895 | predicted gene, 48895 [Source:MGI Symb | 0 | 0 | 0 | 0 | 0 | 0 |
| 50628 | ENSMUSG00000113063 | 175.0591097 | -0.004281259 | 0.207530541 | -0.020629535 | 0.98354118 | 0.998580735 | Gm34667 | predicted gene, 34667 [Source:MGI Symb | 153.39 | 204.56 | 168.04 | 140.74 | 187.59 | 196.04 |
| 50629 | ENSMUSG00000113064 | 0 | NA | NA | NA | NA | NA | Gm40394 | predicted gene, 40394 [Source:MGI Symb | 0 | 0 | 0 | 0 | 0 | 0 |
| 50630 | ENSMUSG00000113065 | 0.614004368 | 0.315622926 | 2.898299649 | 0.108899342 | 0.913282328 | NA | Gm48870 | predicted gene, 48870 [Source:MGI Symb | 1.6062 | 0 | 0 | 0.964 | 0 | 1.1138 |
| 50631 | ENSMUSG00000113066 | 0 | NA | NA | NA | NA | NA | Gm47217 | predicted gene, 47217 [Source:MGI Symb | 0 | 0 | 0 | 0 | 0 | 0 |
| 50632 | ENSMUSG00000113067 | 0.301557804 | -1.64335916 | 4.042814367 | -0.406488899 | 0.684383412 | NA | Gm47166 | predicted gene, 47166 [Source:MGI Symb | 0.8031 | 0 | 1.0062 | 0 | 0 | 0 |
| 50633 | ENSMUSG00000113068 | 0.6996517 | 1.656076801 | 2.758183935 | 0.600422901 | 0.548224429 | NA | Gm48133 | predicted gene, 48133 [Source:MGI Symb | 0 | 0 | 1.0062 | 0.964 | 0 | 2.2277 |
| 50634 | ENSMUSG00000113069 | 0 | NA | NA | NA | NA | NA | Gm48541 | predicted gene, 48541 [Source:MGI Symb | 0 | 0 | 0 | 0 | 0 | 0 |
| 50635 | ENSMUSG00000113070 | 3.946571224 | -0.359452461 | 1.081777432 | -0.332279497 | 0.73967822 | NA | Gm48420 | predicted gene, 48420 [Source:MGI Symb | 3.2124 | 5.1396 | 5.0312 | 4.8199 | 4.3625 | 1.1138 |
| 50636 | ENSMUSG00000113071 | 0 | NA | NA | NA | NA | NA | Gm6353 | predicted gene 6353 [Source:MGI Symbol | 0 | 0 | 0 | 0 | 0 | 0 |
| 50637 | ENSMUSG00000113072 | 0.185640647 | 1.126924175 | 4.080472857 | 0.276174898 | 0.782413736 | NA | Gm47820 | predicted gene, 47820 [Source:MGI Symb | 0 | 0 | 0 | 0 | 0 | 1.1138 |
| 50638 | ENSMUSG00000113073 | 0 | NA | NA | NA | NA | NA | Gm48261 | predicted gene, 48261 [Source:MGI Symb | 0 | 0 | 0 | 0 | 0 | 0 |
| 50639 | ENSMUSG00000113074 | 0 | NA | NA | NA | NA | NA | Gm3772 | predicted gene 3772 [Source:MGI Symbol | 0 | 0 | 0 | 0 | 0 | 0 |
| 50640 | ENSMUSG00000113075 | 0 | NA | NA | NA | NA | NA | Rpl7a-ps1 | ribosomal protein L7A, pseudogene 1 [Sou | 0 | 0 | 0 | 0 | 0 | 0 |
| 50641 | ENSMUSG00000113076 | 0.321325949 | 1.911575114 | 4.048267893 | 0.472195804 | 0.636787034 | NA | Gm47088 | predicted gene, 47088 [Source:MGI Symb | 0 | 0 | 0 | 1.928 | 0 | 0 |
| 50642 | ENSMUSG00000113077 | 0 | NA | NA | NA | NA | NA | Gm49379 | predicted gene, 49379 [Source:MGI Symb | 0 | 0 | 0 | 0 | 0 | 0 |
| 50643 | ENSMUSG00000113078 | 0.305170013 | -1.657836477 | 4.041480335 | -0.410205257 | 0.681655385 | NA | Gm48734 | predicted gene, 48734 [Source:MGI Symb | 0.8031 | 1.0279 | 0 | 0 | 0 | 0 |
| 50644 | ENSMUSG00000113079 | 0 | NA | NA | NA | NA | NA | Gm48709 | predicted gene, 48709 [Source:MGI Symb | 0 | 0 | 0 | 0 | 0 | 0 |
| 50645 | ENSMUSG00000113080 | 0 | NA | NA | NA | NA | NA | Gm48928 | predicted gene, 48928 [Source:MGI Symb | 0 | 0 | 0 | 0 | 0 | 0 |
| 50646 | ENSMUSG00000113081 | 0 | NA | NA | NA | NA | NA | Gm3333 | predicted gene 3333 [Source:MGI Symbol | 0 | 0 | 0 | 0 | 0 | 0 |
| 50647 | ENSMUSG00000113082 | 0 | NA | NA | NA | NA | NA | D730050B12 | RIKEN cDNA D730050B12 gene [Source:M | 0 | 0 | 0 | 0 | 0 | 0 |
| 50648 | ENSMUSG00000113083 | 0 | NA | NA | NA | NA | NA | Gm48513 | predicted gene, 48513 [Source:MGI Symb | 0 | 0 | 0 | 0 | 0 | 0 |
| 50649 | ENSMUSG00000113084 | 0 | NA | NA | NA | NA | NA | A030003K21 | RIKEN cDNA A030003K21 gene [Source:M | 0 | 0 | 0 | 0 | 0 | 0 |
| 50650 | ENSMUSG00000113085 | 0 | NA | NA | NA | NA | NA | Gm47383 | predicted gene, 47383 [Source:MGI Symb | 0 | 0 | 0 | 0 | 0 | 0 |
| 50651 | ENSMUSG00000113086 | 0.462220778 | -0.685400455 | 3.191475091 | -0.21475977 | 0.82995462 | NA | Gm47376 | predicted gene, 47376 [Source:MGI Symb | 0.8031 | 0 | 1.0062 | 0.964 | 0 | 0 |
| 50652 | ENSMUSG00000113087 | 0.496077836 | -0.812642266 | 3.444120572 | -0.235950586 | 0.813471014 | NA | 4930473H19 | RIKEN cDNA 4930473H19 gene [Source:M | 0 | 0 | 2.0125 | 0.964 | 0 | 0 |
| 50653 | ENSMUSG00000113088 | 0.68303045 | 1.69421863 | 2.781340055 | 0.609137537 | 0.542433279 | NA | Gm36757 | predicted gene, 36757 [Source:MGI Symb | 0.8031 | 0 | 0 | 0 | 2.1812 | 1.1138 |
| 50654 | ENSMUSG00000113089 | 0 | NA | NA | NA | NA | NA | Gm46997 | predicted gene 46997 [Source:MGI Symbo | 0 | 0 | 0 | 0 | 0 | 0 |
| 50655 | ENSMUSG00000113090 | 0 | NA | NA | NA | NA | NA | Gm48190 | predicted gene, 48190 [Source:MGI Symb | 0 | 0 | 0 | 0 | 0 | 0 |
| 50656 | ENSMUSG00000113091 | 0 | NA | NA | NA | NA | NA | Gm9279 | predicted gene 9279 [Source:MGI Symbol | 0 | 0 | 0 | 0 | 0 | 0 |
| 50657 | ENSMUSG00000113092 | 0 | NA | NA | NA | NA | NA | Gm34662 | predicted gene, 34662 [Source:MGI Symb | 0 | 0 | 0 | 0 | 0 | 0 |
| 50658 | ENSMUSG00000113093 | 0 | NA | NA | NA | NA | NA | Gm36862 | predicted gene, 36862 [Source:MGI Symb | 0 | 0 | 0 | 0 | 0 | 0 |
| 50659 | ENSMUSG00000113094 | 1.131188945 | -0.206831842 | 1.968071741 | -0.10509365 | 0.916301498 | NA | Gm47447 | predicted gene, 47447 [Source:MGI Symb | 1.6062 | 0 | 2.0125 | 0.964 | 1.0906 | 1.1138 |
| 50660 | ENSMUSG00000113095 | 0 | NA | NA | NA | NA | NA | Gm7969 | predicted gene 7969 [Source:MGI Symbol | 0 | 0 | 0 | 0 | 0 | 0 |
| 50661 | ENSMUSG00000113096 | 0 | NA | NA | NA | NA | NA | Gm40884 | predicted gene, 40884 [Source:MGI Symb | 0 | 0 | 0 | 0 | 0 | 0 |
| 50662 | ENSMUSG00000113097 | 0 | NA | NA | NA | NA | NA | Gm47959 | predicted gene, 47959 [Source:MGI Symb | 0 | 0 | 0 | 0 | 0 | 0 |
| 50663 | ENSMUSG00000113098 | 0 | NA | NA | NA | NA | NA | Gm2912 | predicted gene 2912 [Source:MGI Symbol | 0 | 0 | 0 | 0 | 0 | 0 |
| 50664 | ENSMUSG00000113099 | 0 | NA | NA | NA | NA | NA | Gm34016 | predicted gene, 34016 [Source:MGI Symb | 0 | 0 | 0 | 0 | 0 | 0 |
| 50665 | ENSMUSG00000113100 | 0 | NA | NA | NA | NA | NA | Gm48754 | predicted gene, 48754 [Source:MGI Symb | 0 | 0 | 0 | 0 | 0 | 0 |
| 50666 | ENSMUSG00000113101 | 0.267700746 | -1.497481649 | 4.057001436 | -0.369110456 | 0.712045396 | NA | Gm33424 | predicted gene, 33424 [Source:MGI Symb | 1.6062 | 0 | 0 | 0 | 0 | 0 |
| 50667 | ENSMUSG00000113102 | 0.520796785 | -0.820103173 | 3.072141206 | -0.266948398 | 0.789508892 | NA | A930040O22 | RIKEN cDNA A930040O22 gene [Source:M | 0 | 1.0279 | 1.0062 | 0 | 1.0906 | 0 |
| 50668 | ENSMUSG00000113103 | 0 | NA | NA | NA | NA | NA | Gm48389 | predicted gene, 48389 [Source:MGI Symb | 0 | 0 | 0 | 0 | 0 | 0 |
| 50669 | ENSMUSG00000113104 | 3.014955729 | 0.688767407 | 1.398733409 | 0.492422218 | 0.622420896 | NA | Gm47806 | predicted gene, 47806 [Source:MGI Symb | 4.8186 | 1.0279 | 1.0062 | 5.7839 | 5.4531 | 0 |
| 50670 | ENSMUSG00000113105 | 0.160662974 | 1.126924175 | 4.080472857 | 0.276174898 | 0.782413736 | NA | Gm47371 | predicted gene, 47371 [Source:MGI Symb | 0 | 0 | 0 | 0.964 | 0 | 0 |
| 50671 | ENSMUSG00000113106 | 0.181769715 | 1.126924175 | 4.080472857 | 0.276174898 | 0.782413736 | NA | Gm47139 | predicted gene, 47139 [Source:MGI Symb | 0 | 0 | 0 | 0 | 1.0906 | 0 |
| 50672 | ENSMUSG00000113107 | 0 | NA | NA | NA | NA | NA | Gm3338 | predicted gene 3338 [Source:MGI Symbol | 0 | 0 | 0 | 0 | 0 | 0 |
| 50673 | ENSMUSG00000113108 | 0 | NA | NA | NA | NA | NA | Gm47832 | predicted gene, 47832 [Source:MGI Symb | 0 | 0 | 0 | 0 | 0 | 0 |
| 50674 | ENSMUSG00000113109 | 0 | NA | NA | NA | NA | NA | Gm7712 | predicted gene 7712 [Source:MGI Symbol | 0 | 0 | 0 | 0 | 0 | 0 |
| 50675 | ENSMUSG00000113110 | 0.267700746 | -1.497481649 | 4.057001436 | -0.369110456 | 0.712045396 | NA | Gm48528 | predicted gene, 48528 [Source:MGI Symb | 1.6062 | 0 | 0 | 0 | 0 | 0 |
| 50676 | ENSMUSG00000113111 | 0 | NA | NA | NA | NA | NA | Gm7754 | predicted gene 7754 [Source:MGI Symbol | 0 | 0 | 0 | 0 | 0 | 0 |

| | | | | | | | | | | | | | | | |
|---|---|---|---|---|---|---|---|---|---|---|---|---|---|---|---|
| 50677 | ENSMUSG00000113112 | 0 | NA | NA | NA | NA | NA | Gm8725 | predicted gene 8725 [Source:MGI Symbol | 0 | 0 | 0 | 0 | 0 | 0 |
| 50678 | ENSMUSG00000113113 | 3.219486852 | -1.028369825 | 1.275221188 | -0.806424669 | 0.419998027 | NA | Gm2614 | predicted gene 2614 [Source:MGI Symbol | 4.8186 | 3.0838 | 5.0312 | 1.928 | 0 | 4.4554 |
| 50679 | ENSMUSG00000113114 | 0 | NA | NA | NA | NA | NA | Gm47846 | predicted gene, 47846 [Source:MGI Symb | 0 | 0 | 0 | 0 | 0 | 0 |
| 50680 | ENSMUSG00000113115 | 0 | NA | NA | NA | NA | NA | 4930480M1 | RIKEN cDNA 4930480M12 gene [Source:M | 0 | 0 | 0 | 0 | 0 | 0 |
| 50681 | ENSMUSG00000113116 | 0 | NA | NA | NA | NA | NA | Gm47878 | predicted gene, 47878 [Source:MGI Symb | 0 | 0 | 0 | 0 | 0 | 0 |
| 50682 | ENSMUSG00000113117 | 0 | NA | NA | NA | NA | NA | Gm2544 | predicted gene 2544 [Source:MGI Symbol | 0 | 0 | 0 | 0 | 0 | 0 |
| 50683 | ENSMUSG00000113118 | 0 | NA | NA | NA | NA | NA | Gm3829 | predicted gene 3829 [Source:MGI Symbol | 0 | 0 | 0 | 0 | 0 | 0 |
| 50684 | ENSMUSG00000113119 | 0 | NA | NA | NA | NA | NA | Gm48883 | predicted gene, 48883 [Source:MGI Symb | 0 | 0 | 0 | 0 | 0 | 0 |
| 50685 | ENSMUSG00000113120 | 0 | NA | NA | NA | NA | NA | E330035G20 | RIKEN cDNA E330035G20 gene [Source:M | 0 | 0 | 0 | 0 | 0 | 0 |
| 50686 | ENSMUSG00000113121 | 0.371281295 | 2.09253362 | 4.031852165 | 0.519000582 | 0.603760335 | NA | Gm35161 | predicted gene, 35161 [Source:MGI Symb | 0 | 0 | 0 | 0 | 0 | 2.2277 |
| 50687 | ENSMUSG00000113122 | 0 | NA | NA | NA | NA | NA | Gm48644 | predicted gene, 48644 [Source:MGI Symb | 0 | 0 | 0 | 0 | 0 | 0 |
| 50688 | ENSMUSG00000113123 | 0 | NA | NA | NA | NA | NA | Gm40864 | predicted gene, 40864 [Source:MGI Symb | 0 | 0 | 0 | 0 | 0 | 0 |
| 50689 | ENSMUSG00000113124 | 0 | NA | NA | NA | NA | NA | Gm47193 | predicted gene, 47193 [Source:MGI Symb | 0 | 0 | 0 | 0 | 0 | 0 |
| 50690 | ENSMUSG00000113125 | 0 | NA | NA | NA | NA | NA | Gm47650 | predicted gene, 47650 [Source:MGI Symb | 0 | 0 | 0 | 0 | 0 | 0 |
| 50691 | ENSMUSG00000113126 | 0.952566124 | 1.285634123 | 2.448533191 | 0.525062975 | 0.599539413 | NA | Gm47588 | predicted gene, 47588 [Source:MGI Symb | 1.6062 | 0 | 0 | 1.928 | 2.1812 | 0 |
| 50692 | ENSMUSG00000113127 | 0 | NA | NA | NA | NA | NA | Gm48154 | predicted gene, 48154 [Source:MGI Symb | 0 | 0 | 0 | 0 | 0 | 0 |
| 50693 | ENSMUSG00000113128 | 0 | NA | NA | NA | NA | NA | Gm47813 | predicted gene, 47813 [Source:MGI Symb | 0 | 0 | 0 | 0 | 0 | 0 |
| 50694 | ENSMUSG00000113129 | 0 | NA | NA | NA | NA | NA | Gm21569 | predicted gene, 21569 [Source:MGI Symb | 0 | 0 | 0 | 0 | 0 | 0 |
| 50695 | ENSMUSG00000113130 | 0 | NA | NA | NA | NA | NA | Gm47721 | predicted gene, 47721 [Source:MGI Symb | 0 | 0 | 0 | 0 | 0 | 0 |
| 50696 | ENSMUSG00000113131 | 0 | NA | NA | NA | NA | NA | Gm48290 | predicted gene, 48290 [Source:MGI Symb | 0 | 0 | 0 | 0 | 0 | 0 |
| 50697 | ENSMUSG00000113132 | 0 | NA | NA | NA | NA | NA | Gm9231 | predicted gene 9231 [Source:MGI Symbol | 0 | 0 | 0 | 0 | 0 | 0 |
| 50698 | ENSMUSG00000113133 | 0 | NA | NA | NA | NA | NA | Gm21039 | predicted gene, 21039 [Source:MGI Symb | 0 | 0 | 0 | 0 | 0 | 0 |
| 50699 | ENSMUSG00000113134 | 0 | NA | NA | NA | NA | NA | Gm18795 | predicted gene, 18795 [Source:MGI Symb | 0 | 0 | 0 | 0 | 0 | 0 |
| 50700 | ENSMUSG00000113135 | 0 | NA | NA | NA | NA | NA | Gm48170 | predicted gene, 48170 [Source:MGI Symb | 0 | 0 | 0 | 0 | 0 | 0 |
| 50701 | ENSMUSG00000113136 | 1.663345318 | -0.615296061 | 2.022282288 | -0.304258246 | 0.760931153 | NA | Gm19951 | predicted gene, 19951 [Source:MGI Symb | 0 | 4.1117 | 2.0125 | 3.8559 | 0 | 0 |
| 50702 | ENSMUSG00000113137 | 0.321325949 | 1.911575114 | 4.048267893 | 0.472195804 | 0.636787034 | NA | Gm7511 | predicted gene 7511 [Source:MGI Symbol | 0 | 0 | 0 | 1.928 | 0 | 0 |
| 50703 | ENSMUSG00000113138 | 2.690921571 | 0.04996259 | 1.294373182 | 0.038599834 | 0.969209435 | NA | Gm47177 | predicted gene, 47177 [Source:MGI Symb | 4.8186 | 2.0558 | 1.0062 | 3.8559 | 2.1812 | 2.2277 |
| 50704 | ENSMUSG00000113139 | 0 | NA | NA | NA | NA | NA | Gm48011 | predicted gene, 48011 [Source:MGI Symb | 0 | 0 | 0 | 0 | 0 | 0 |
| 50705 | ENSMUSG00000113140 | 0.305170013 | -1.657836477 | 4.041480335 | -0.410205257 | 0.681655385 | NA | Gm48624 | predicted gene, 48624 [Source:MGI Symb | 0.8031 | 1.0279 | 0 | 0 | 0 | 0 |
| 50706 | ENSMUSG00000113141 | 0 | NA | NA | NA | NA | NA | Gm47748 | predicted gene, 47748 [Source:MGI Symb | 0 | 0 | 0 | 0 | 0 | 0 |
| 50707 | ENSMUSG00000113142 | 0 | NA | NA | NA | NA | NA | Gm18114 | predicted gene, 18114 [Source:MGI Symb | 0 | 0 | 0 | 0 | 0 | 0 |
| 50708 | ENSMUSG00000113143 | 0 | NA | NA | NA | NA | NA | Ly6e-ps1 | lymphocyte antigen 6 complex, locus E, ps | 0 | 0 | 0 | 0 | 0 | 0 |
| 50709 | ENSMUSG00000113144 | 0 | NA | NA | NA | NA | NA | Gm2754 | predicted gene 2754 [Source:MGI Symbol | 0 | 0 | 0 | 0 | 0 | 0 |
| 50710 | ENSMUSG00000113145 | 0 | NA | NA | NA | NA | NA | Vmn2r-ps95 | vomeronasal 2, receptor, pseudogene 95 | 0 | 0 | 0 | 0 | 0 | 0 |
| 50711 | ENSMUSG00000113146 | 0 | NA | NA | NA | NA | NA | Gm47406 | predicted gene, 47406 [Source:MGI Symb | 0 | 0 | 0 | 0 | 0 | 0 |
| 50712 | ENSMUSG00000113147 | 0 | NA | NA | NA | NA | NA | Gm47207 | predicted gene, 47207 [Source:MGI Symb | 0 | 0 | 0 | 0 | 0 | 0 |
| 50713 | ENSMUSG00000113148 | 0 | NA | NA | NA | NA | NA | Gm48456 | predicted gene, 48456 [Source:MGI Symb | 0 | 0 | 0 | 0 | 0 | 0 |
| 50714 | ENSMUSG00000113149 | 0 | NA | NA | NA | NA | NA | Gm49383 | predicted gene, 49383 [Source:MGI Symb | 0 | 0 | 0 | 0 | 0 | 0 |
| 50715 | ENSMUSG00000113150 | 0 | NA | NA | NA | NA | NA | Gm18263 | predicted gene, 18263 [Source:MGI Symb | 0 | 0 | 0 | 0 | 0 | 0 |
| 50716 | ENSMUSG00000113151 | 0 | NA | NA | NA | NA | NA | Gm48690 | predicted gene, 48690 [Source:MGI Symb | 0 | 0 | 0 | 0 | 0 | 0 |
| 50717 | ENSMUSG00000113152 | 0 | NA | NA | NA | NA | NA | Gm48422 | predicted gene, 48422 [Source:MGI Symb | 0 | 0 | 0 | 0 | 0 | 0 |
| 50718 | ENSMUSG00000113153 | 0 | NA | NA | NA | NA | NA | Gm48367 | predicted gene, 48367 [Source:MGI Symb | 0 | 0 | 0 | 0 | 0 | 0 |
| 50719 | ENSMUSG00000113154 | 0 | NA | NA | NA | NA | NA | Gm48713 | predicted gene, 48713 [Source:MGI Symb | 0 | 0 | 0 | 0 | 0 | 0 |
| 50720 | ENSMUSG00000113155 | 0 | NA | NA | NA | NA | NA | Gm48354 | predicted gene, 48354 [Source:MGI Symb | 0 | 0 | 0 | 0 | 0 | 0 |
| 50721 | ENSMUSG00000113156 | 0 | NA | NA | NA | NA | NA | Gm48223 | predicted gene, 48223 [Source:MGI Symb | 0 | 0 | 0 | 0 | 0 | 0 |
| 50722 | ENSMUSG00000113157 | 0.181769715 | 1.126924175 | 4.080472857 | 0.276174898 | 0.782413736 | NA | Gm48693 | predicted gene, 48693 [Source:MGI Symb | 0 | 0 | 0 | 0 | 1.0906 | 0 |
| 50723 | ENSMUSG00000113158 | 0 | NA | NA | NA | NA | NA | Gm31447 | predicted gene, 31447 [Source:MGI Symb | 0 | 0 | 0 | 0 | 0 | 0 |
| 50724 | ENSMUSG00000113159 | 2.217605442 | 0.243973147 | 1.500077021 | 0.162640413 | 0.870801565 | NA | Gm48771 | predicted gene, 48771 [Source:MGI Symb | 4.0155 | 0 | 2.0125 | 2.8919 | 3.2719 | 1.1138 |
| 50725 | ENSMUSG00000113160 | 0 | NA | NA | NA | NA | NA | Gm47440 | predicted gene, 47440 [Source:MGI Symb | 0 | 0 | 0 | 0 | 0 | 0 |
| 50726 | ENSMUSG00000113161 | 0 | NA | NA | NA | NA | NA | Gm47457 | predicted gene, 47457 [Source:MGI Symb | 0 | 0 | 0 | 0 | 0 | 0 |
| 50727 | ENSMUSG00000113162 | 0 | NA | NA | NA | NA | NA | Gm47554 | predicted gene, 47554 [Source:MGI Symb | 0 | 0 | 0 | 0 | 0 | 0 |
| 50728 | ENSMUSG00000113163 | 0 | NA | NA | NA | NA | NA | Olfr335-ps | olfactory receptor 335, pseudogene [Sour | 0 | 0 | 0 | 0 | 0 | 0 |
| 50729 | ENSMUSG00000113164 | 0 | NA | NA | NA | NA | NA | 4930559C10 | RIKEN cDNA 4930559C10 gene [Source:M | 0 | 0 | 0 | 0 | 0 | 0 |
| 50730 | ENSMUSG00000113165 | 0.935098222 | -2.037664976 | 2.285197225 | -0.891680138 | 0.372564403 | NA | Gm47863 | predicted gene, 47863 [Source:MGI Symb | 1.6062 | 1.0279 | 2.0125 | 0.964 | 0 | 0 |
| 50731 | ENSMUSG00000113166 | 0 | NA | NA | NA | NA | NA | Gm19047 | predicted gene, 19047 [Source:MGI Symb | 0 | 0 | 0 | 0 | 0 | 0 |
| 50732 | ENSMUSG00000113167 | 0 | NA | NA | NA | NA | NA | Gm48482 | predicted gene, 48482 [Source:MGI Symb | 0 | 0 | 0 | 0 | 0 | 0 |

| | | | | | | | | | | | | | | | |
|---|---|---|---|---|---|---|---|---|---|---|---|---|---|---|---|
| 50733 | ENSMUSG00000113168 | 0 | NA | NA | NA | NA | NA | Gm7614 | predicted gene 7614 [Source:MGI Symbol | 0 | 0 | 0 | 0 | 0 | 0 |
| 50734 | ENSMUSG00000113169 | 0 | NA | NA | NA | NA | NA | Gm6171 | predicted gene 6171 [Source:MGI Symbol | 0 | 0 | 0 | 0 | 0 | 0 |
| 50735 | ENSMUSG00000113170 | 6.828364388 | 0.091943187 | 0.952060846 | 0.096572806 | 0.923065654 | 0.987005368 | Gm36839 | predicted gene, 36839 [Source:MGI Symb | 9.6372 | 1.0279 | 9.0562 | 4.8199 | 13.087 | 3.3415 |
| 50736 | ENSMUSG00000113171 | 0 | NA | NA | NA | NA | NA | Gm48359 | predicted gene, 48359 [Source:MGI Symb | 0 | 0 | 0 | 0 | 0 | 0 |
| 50737 | ENSMUSG00000113172 | 0 | NA | NA | NA | NA | NA | Gm47836 | predicted gene, 47836 [Source:MGI Symb | 0 | 0 | 0 | 0 | 0 | 0 |
| 50738 | ENSMUSG00000113173 | 0 | NA | NA | NA | NA | NA | Gm48221 | predicted gene, 48221 [Source:MGI Symb | 0 | 0 | 0 | 0 | 0 | 0 |
| 50739 | ENSMUSG00000113174 | 0 | NA | NA | NA | NA | NA | Gm9292 | predicted gene 9292 [Source:MGI Symbol | 0 | 0 | 0 | 0 | 0 | 0 |
| 50740 | ENSMUSG00000113175 | 0.181769715 | 1.126924175 | 4.080472857 | 0.276174898 | 0.782413736 | NA | Gm9973 | predicted gene 9973 [Source:MGI Symbol | 0 | 0 | 0 | 0 | 1.0906 | 0 |
| 50741 | ENSMUSG00000113177 | 0 | NA | NA | NA | NA | NA | Gm7214 | predicted gene 7214 [Source:MGI Symbol | 0 | 0 | 0 | 0 | 0 | 0 |
| 50742 | ENSMUSG00000113178 | 0 | NA | NA | NA | NA | NA | Mylf-ps | myosin light chain, alkali, fast skeletal mus | 0 | 0 | 0 | 0 | 0 | 0 |
| 50743 | ENSMUSG00000113179 | 0 | NA | NA | NA | NA | NA | 1700030L22I | RIKEN cDNA 1700030L22 gene [Source:M( | 0 | 0 | 0 | 0 | 0 | 0 |
| 50744 | ENSMUSG00000113180 | 0 | NA | NA | NA | NA | NA | Gm46998 | predicted gene 46998 [Source:MGI Symbo | 0 | 0 | 0 | 0 | 0 | 0 |
| 50745 | ENSMUSG00000113181 | 0 | NA | NA | NA | NA | NA | Gm47954 | predicted gene, 47954 [Source:MGI Symb | 0 | 0 | 0 | 0 | 0 | 0 |
| 50746 | ENSMUSG00000113182 | 0 | NA | NA | NA | NA | NA | Gm20102 | predicted gene, 20102 [Source:MGI Symb | 0 | 0 | 0 | 0 | 0 | 0 |
| 50747 | ENSMUSG00000113183 | 0.133850373 | -0.796629176 | 4.080472857 | -0.195229623 | 0.84521319 | NA | Gm47664 | predicted gene, 47664 [Source:MGI Symb | 0.8031 | 0 | 0 | 0 | 0 | 0 |
| 50748 | ENSMUSG00000113184 | 0.898940922 | -0.606261109 | 2.77650979 | -0.218353672 | 0.827153559 | NA | Gm49654 | predicted gene, 49654 [Source:MGI Symb | 3.2124 | 0 | 0 | 0 | 2.1812 | 0 |
| 50749 | ENSMUSG00000113185 | 1.861265455 | 0.94739408 | 1.538763681 | 0.615685236 | 0.538102283 | NA | Gm47127 | predicted gene, 47127 [Source:MGI Symb | 0.8031 | 1.0279 | 2.0125 | 2.8919 | 1.0906 | 3.3415 |
| 50750 | ENSMUSG00000113186 | 0 | NA | NA | NA | NA | NA | A330076C08 | RIKEN cDNA A330076C08 gene [Source:M | 0 | 0 | 0 | 0 | 0 | 0 |
| 50751 | ENSMUSG00000113187 | 0 | NA | NA | NA | NA | NA | Gm5955 | predicted gene 5955 [Source:MGI Symbol | 0 | 0 | 0 | 0 | 0 | 0 |
| 50752 | ENSMUSG00000113188 | 1.509694657 | 0.5406839 | 1.764685395 | 0.3063911 | 0.759306884 | NA | Gm35638 | predicted gene, 35638 [Source:MGI Symb | 1.6062 | 2.0558 | 0 | 0.964 | 1.0906 | 3.3415 |
| 50753 | ENSMUSG00000113189 | 0 | NA | NA | NA | NA | NA | B020018J22I | Riken cDNA B020018J22 gene [Source:MG | 0 | 0 | 0 | 0 | 0 | 0 |
| 50754 | ENSMUSG00000113190 | 0 | NA | NA | NA | NA | NA | Gm47545 | predicted gene, 47545 [Source:MGI Symb | 0 | 0 | 0 | 0 | 0 | 0 |
| 50755 | ENSMUSG00000113191 | 0 | NA | NA | NA | NA | NA | Gm48343 | predicted gene, 48343 [Source:MGI Symb | 0 | 0 | 0 | 0 | 0 | 0 |
| 50756 | ENSMUSG00000113192 | 0 | NA | NA | NA | NA | NA | Gm18032 | predicted gene, 18032 [Source:MGI Symb | 0 | 0 | 0 | 0 | 0 | 0 |
| 50757 | ENSMUSG00000113193 | 1.522237856 | -3.977035884 | 2.067126455 | -1.923944167 | 0.054361582 | NA | Gm40576 | predicted gene, 40576 [Source:MGI Symb | 4.0155 | 4.1117 | 1.0062 | 0 | 0 | 0 |
| 50758 | ENSMUSG00000113194 | 0 | NA | NA | NA | NA | NA | Gm47998 | predicted gene, 47998 [Source:MGI Symb | 0 | 0 | 0 | 0 | 0 | 0 |
| 50759 | ENSMUSG00000113195 | 0 | NA | NA | NA | NA | NA | Gm48586 | predicted gene, 48586 [Source:MGI Symb | 0 | 0 | 0 | 0 | 0 | 0 |
| 50760 | ENSMUSG00000113196 | 0.9173971 | 0.153222854 | 2.48201849 | 0.061733164 | 0.950775329 | NA | D230049E03 | RIKEN cDNA D230049E03 gene [Source:M | 1.6062 | 0 | 1.0062 | 2.8919 | 0 | 0 |
| 50761 | ENSMUSG00000113197 | 0 | NA | NA | NA | NA | NA | Gm48318 | predicted gene, 48318 [Source:MGI Symb | 0 | 0 | 0 | 0 | 0 | 0 |
| 50762 | ENSMUSG00000113198 | 0.94782191 | 1.090542322 | 2.508664729 | 0.43471027 | 0.663772743 | NA | Gm6988 | predicted gene 6988 [Source:MGI Symbol | 0.8031 | 1.0279 | 0 | 3.8559 | 0 | 0 |
| 50763 | ENSMUSG00000113199 | 0 | NA | NA | NA | NA | NA | Gm9229 | predicted gene 9229 [Source:MGI Symbol | 0 | 0 | 0 | 0 | 0 | 0 |
| 50764 | ENSMUSG00000113200 | 0.784885323 | -1.683590003 | 2.624742731 | -0.641430485 | 0.521243032 | NA | Gm48632 | predicted gene, 48632 [Source:MGI Symb | 1.6062 | 0 | 2.0125 | 0 | 1.0906 | 0 |
| 50765 | ENSMUSG00000113201 | 0 | NA | NA | NA | NA | NA | Eif1ad13 | eukaryotic translation initiation factor 1A | 0 | 0 | 0 | 0 | 0 | 0 |
| 50766 | ENSMUSG00000113202 | 0 | NA | NA | NA | NA | NA | Gm48291 | predicted gene, 48291 [Source:MGI Symb | 0 | 0 | 0 | 0 | 0 | 0 |
| 50767 | ENSMUSG00000113203 | 0.160662974 | 1.126924175 | 4.080472857 | 0.276174898 | 0.782413736 | NA | Gm47587 | predicted gene, 47587 [Source:MGI Symb | 0 | 0 | 0 | 0.964 | 0 | 0 |
| 50768 | ENSMUSG00000113204 | 203.4940526 | 0.096466093 | 0.186607992 | 0.516945134 | 0.60519446 | 0.916376001 | Gm46430 | predicted gene, 46430 [Source:MGI Symb | 198.37 | 180.91 | 210.31 | 181.23 | 201.76 | 248.39 |
| 50769 | ENSMUSG00000113205 | 0 | NA | NA | NA | NA | NA | Gm48525 | predicted gene, 48525 [Source:MGI Symb | 0 | 0 | 0 | 0 | 0 | 0 |
| 50770 | ENSMUSG00000113206 | 0 | NA | NA | NA | NA | NA | Gm3742 | predicted gene 3742 [Source:MGI Symbol | 0 | 0 | 0 | 0 | 0 | 0 |
| 50771 | ENSMUSG00000113207 | 0 | NA | NA | NA | NA | NA | Cbx3-ps5 | chromobox 3, pseudogene 5 [Source:MGI | 0 | 0 | 0 | 0 | 0 | 0 |
| 50772 | ENSMUSG00000113208 | 0 | NA | NA | NA | NA | NA | Gm48421 | predicted gene, 48421 [Source:MGI Symb | 0 | 0 | 0 | 0 | 0 | 0 |
| 50773 | ENSMUSG00000113209 | 0.870506025 | 3.333445469 | 2.471995426 | 1.348483672 | 0.177502868 | NA | 1700081N11 | RIKEN cDNA 1700081N11 gene [Source:M | 0 | 0 | 0 | 1.928 | 2.1812 | 1.1138 |
| 50774 | ENSMUSG00000113210 | 0 | NA | NA | NA | NA | NA | 4930474N09 | RIKEN cDNA 4930474N09 gene [Source:M | 0 | 0 | 0 | 0 | 0 | 0 |
| 50775 | ENSMUSG00000113211 | 0 | NA | NA | NA | NA | NA | 4921525O09 | RIKEN cDNA 4921525O09 gene [Source:M | 0 | 0 | 0 | 0 | 0 | 0 |
| 50776 | ENSMUSG00000113212 | 2.262739465 | 0.318712649 | 1.515369644 | 0.210320069 | 0.833417872 | NA | Gm47435 | predicted gene, 47435 [Source:MGI Symb | 0 | 3.0838 | 3.0187 | 1.928 | 1.0906 | 4.4554 |
| 50777 | ENSMUSG00000113213 | 0 | NA | NA | NA | NA | NA | Gm19540 | predicted gene, 19540 [Source:MGI Symb | 0 | 0 | 0 | 0 | 0 | 0 |
| 50778 | ENSMUSG00000113214 | 4.727171008 | 0.70590445 | 0.998859229 | 0.706710645 | 0.479746313 | NA | Gm48010 | predicted gene, 48010 [Source:MGI Symb | 5.6217 | 2.0558 | 3.0187 | 7.7118 | 3.2719 | 6.6831 |
| 50779 | ENSMUSG00000113215 | 0 | NA | NA | NA | NA | NA | Gm18749 | predicted gene, 18749 [Source:MGI Symb | 0 | 0 | 0 | 0 | 0 | 0 |
| 50780 | ENSMUSG00000113216 | 0 | NA | NA | NA | NA | NA | Gm40841 | predicted gene, 40841 [Source:MGI Symb | 0 | 0 | 0 | 0 | 0 | 0 |
| 50781 | ENSMUSG00000113217 | 0 | NA | NA | NA | NA | NA | Gm47187 | predicted gene, 47187 [Source:MGI Symb | 0 | 0 | 0 | 0 | 0 | 0 |
| 50782 | ENSMUSG00000113218 | 0 | NA | NA | NA | NA | NA | 4933409F18 | RIKEN cDNA 4933409F18 gene [Source:M( | 0 | 0 | 0 | 0 | 0 | 0 |
| 50783 | ENSMUSG00000113219 | 0 | NA | NA | NA | NA | NA | Gm17890 | predicted gene, 17890 [Source:MGI Symb | 0 | 0 | 0 | 0 | 0 | 0 |
| 50784 | ENSMUSG00000113220 | 0 | NA | NA | NA | NA | NA | Gm35208 | predicted gene, 35208 [Source:MGI Symb | 0 | 0 | 0 | 0 | 0 | 0 |
| 50785 | ENSMUSG00000113221 | 0 | NA | NA | NA | NA | NA | Gm18880 | predicted gene, 18880 [Source:MGI Symb | 0 | 0 | 0 | 0 | 0 | 0 |
| 50786 | ENSMUSG00000113222 | 0.160662974 | 1.126924175 | 4.080472857 | 0.276174898 | 0.782413736 | NA | Gm48557 | predicted gene, 48557 [Source:MGI Symb | 0 | 0 | 0 | 0.964 | 0 | 0 |
| 50787 | ENSMUSG00000113223 | 0 | NA | NA | NA | NA | NA | Gm48147 | predicted gene, 48147 [Source:MGI Symb | 0 | 0 | 0 | 0 | 0 | 0 |
| 50788 | ENSMUSG00000113224 | 1.177510966 | 2.58959483 | 2.314810155 | 1.118707219 | 0.263265065 | NA | Gm48444 | predicted gene, 48444 [Source:MGI Symb | 0 | 1.0279 | 0 | 3.8559 | 2.1812 | 0 |

| | | | | | | | | | | | | | | | |
|---|---|---|---|---|---|---|---|---|---|---|---|---|---|---|---|
| 50789 | ENSMUSG00000113225 | 0 | NA | NA | NA | NA | NA | Gm36074 | predicted gene, 36074 [Source:MGI Symb | 0 | 0 | 0 | 0 | 0 | 0 |
| 50790 | ENSMUSG00000113226 | 0.181769715 | 1.126924175 | 4.080472857 | 0.276174898 | 0.782413736 | NA | Gm47808 | predicted gene, 47808 [Source:MGI Symb | 0 | 0 | 0 | 0 | 1.0906 | 0 |
| 50791 | ENSMUSG00000113227 | 0 | NA | NA | NA | NA | NA | 4933433N18 | RIKEN cDNA 4933433N18 gene [Source:M | 0 | 0 | 0 | 0 | 0 | 0 |
| 50792 | ENSMUSG00000113228 | 0 | NA | NA | NA | NA | NA | 4930471E15 | RIKEN cDNA 4930471E15 gene [Source:M | 0 | 0 | 0 | 0 | 0 | 0 |
| 50793 | ENSMUSG00000113229 | 0 | NA | NA | NA | NA | NA | Gm18442 | predicted gene, 18442 [Source:MGI Symb | 0 | 0 | 0 | 0 | 0 | 0 |
| 50794 | ENSMUSG00000113230 | 0 | NA | NA | NA | NA | NA | Gm48101 | predicted gene, 48101 [Source:MGI Symb | 0 | 0 | 0 | 0 | 0 | 0 |
| 50795 | ENSMUSG00000113231 | 0 | NA | NA | NA | NA | NA | Gm5189 | predicted gene 5189 [Source:MGI Symbol | 0 | 0 | 0 | 0 | 0 | 0 |
| 50796 | ENSMUSG00000113232 | 0 | NA | NA | NA | NA | NA | Gm48494 | predicted gene, 48494 [Source:MGI Symb | 0 | 0 | 0 | 0 | 0 | 0 |
| 50797 | ENSMUSG00000113233 | 0 | NA | NA | NA | NA | NA | Gm31063 | predicted gene, 31063 [Source:MGI Symb | 0 | 0 | 0 | 0 | 0 | 0 |
| 50798 | ENSMUSG00000113234 | 0.160662974 | 1.126924175 | 4.080472857 | 0.276174898 | 0.782413736 | NA | Gm49330 | predicted gene, 49330 [Source:MGI Symb | 0 | 0 | 0 | 0.964 | 0 | 0 |
| 50799 | ENSMUSG00000113235 | 0 | NA | NA | NA | NA | NA | Gm47872 | predicted gene, 47872 [Source:MGI Symb | 0 | 0 | 0 | 0 | 0 | 0 |
| 50800 | ENSMUSG00000113236 | 0 | NA | NA | NA | NA | NA | Gm40548 | predicted gene, 40548 [Source:MGI Symb | 0 | 0 | 0 | 0 | 0 | 0 |
| 50801 | ENSMUSG00000113237 | 1.412357418 | 0.098611615 | 2.395180442 | 0.04117085 | 0.967159692 | NA | Gm6109 | predicted gene 6109 [Source:MGI Symbol | 0 | 4.1117 | 0 | 0 | 4.3625 | 0 |
| 50802 | ENSMUSG00000113238 | 4.878781276 | -1.094803341 | 1.017037589 | -1.076463006 | 0.281720237 | NA | Gm47251 | predicted gene, 47251 [Source:MGI Symb | 4.8186 | 5.1396 | 10.062 | 4.8199 | 1.0906 | 3.3415 |
| 50803 | ENSMUSG00000113239 | 0.181769715 | 1.126924175 | 4.080472857 | 0.276174898 | 0.782413736 | NA | Gm18388 | predicted gene, 18388 [Source:MGI Symb | 0 | 0 | 0 | 0 | 1.0906 | 0 |
| 50804 | ENSMUSG00000113240 | 0 | NA | NA | NA | NA | NA | Gm48338 | predicted gene, 48338 [Source:MGI Symb | 0 | 0 | 0 | 0 | 0 | 0 |
| 50805 | ENSMUSG00000113241 | 0.435408177 | -2.179616019 | 3.531869654 | -0.617128103 | 0.537150234 | NA | Gm40663 | predicted gene, 40663 [Source:MGI Symb | 1.6062 | 0 | 1.0062 | 0 | 0 | 0 |
| 50806 | ENSMUSG00000113242 | 0 | NA | NA | NA | NA | NA | Gm48016 | predicted gene, 48016 [Source:MGI Symb | 0 | 0 | 0 | 0 | 0 | 0 |
| 50807 | ENSMUSG00000113243 | 0 | NA | NA | NA | NA | NA | Gm36525 | predicted gene, 36525 [Source:MGI Symb | 0 | 0 | 0 | 0 | 0 | 0 |
| 50808 | ENSMUSG00000113244 | 0 | NA | NA | NA | NA | NA | Gm31218 | predicted gene, 31218 [Source:MGI Symb | 0 | 0 | 0 | 0 | 0 | 0 |
| 50809 | ENSMUSG00000113245 | 0 | NA | NA | NA | NA | NA | Gm48574 | predicted gene, 48574 [Source:MGI Symb | 0 | 0 | 0 | 0 | 0 | 0 |
| 50810 | ENSMUSG00000113246 | 0 | NA | NA | NA | NA | NA | Gm36752 | predicted gene, 36752 [Source:MGI Symb | 0 | 0 | 0 | 0 | 0 | 0 |
| 50811 | ENSMUSG00000113247 | 0 | NA | NA | NA | NA | NA | Gm30155 | predicted gene, 30155 [Source:MGI Symb | 0 | 0 | 0 | 0 | 0 | 0 |
| 50812 | ENSMUSG00000113248 | 0 | NA | NA | NA | NA | NA | Gm18338 | predicted gene, 18338 [Source:MGI Symb | 0 | 0 | 0 | 0 | 0 | 0 |
| 50813 | ENSMUSG00000113249 | 0 | NA | NA | NA | NA | NA | Gm33312 | predicted gene, 33312 [Source:MGI Symb | 0 | 0 | 0 | 0 | 0 | 0 |
| 50814 | ENSMUSG00000113250 | 0.669251866 | -2.807208097 | 3.391018549 | -0.827836255 | 0.407763239 | NA | Gm48585 | predicted gene, 48585 [Source:MGI Symb | 4.0155 | 0 | 0 | 0 | 0 | 0 |
| 50815 | ENSMUSG00000113251 | 0 | NA | NA | NA | NA | NA | Gm18363 | predicted gene, 18363 [Source:MGI Symb | 0 | 0 | 0 | 0 | 0 | 0 |
| 50816 | ENSMUSG00000113252 | 0 | NA | NA | NA | NA | NA | Gm33489 | predicted gene, 33489 [Source:MGI Symb | 0 | 0 | 0 | 0 | 0 | 0 |
| 50817 | ENSMUSG00000113253 | 0 | NA | NA | NA | NA | NA | Gm48241 | predicted gene, 48241 [Source:MGI Symb | 0 | 0 | 0 | 0 | 0 | 0 |
| 50818 | ENSMUSG00000113254 | 0 | NA | NA | NA | NA | NA | Gm272 | predicted gene 272 [Source:MGI Symbol;/ | 0 | 0 | 0 | 0 | 0 | 0 |
| 50819 | ENSMUSG00000113255 | 2.606252026 | -0.039556485 | 1.437292157 | -0.027521534 | 0.978043765 | NA | Tes3-ps | testis derived transcript 3, pseudogene [S | 0.8031 | 4.1117 | 3.0187 | 0 | 4.3625 | 3.3415 |
| 50820 | ENSMUSG00000113256 | 0 | NA | NA | NA | NA | NA | Gm48527 | predicted gene, 48527 [Source:MGI Symb | 0 | 0 | 0 | 0 | 0 | 0 |
| 50821 | ENSMUSG00000113257 | 0.17131964 | -0.796629176 | 4.080472857 | -0.195229623 | 0.84521319 | NA | Gm30409 | predicted gene, 30409 [Source:MGI Symb | 0 | 1.0279 | 0 | 0 | 0 | 0 |
| 50822 | ENSMUSG00000113258 | 0 | NA | NA | NA | NA | NA | Gm7446 | predicted gene 7446 [Source:MGI Symbol | 0 | 0 | 0 | 0 | 0 | 0 |
| 50823 | ENSMUSG00000113259 | 0.465832987 | -0.698295221 | 3.18348759 | -0.219349126 | 0.826378096 | NA | Gm35190 | predicted gene, 35190 [Source:MGI Symb | 0.8031 | 1.0279 | 0 | 0.964 | 0 | 0 |
| 50824 | ENSMUSG00000113260 | 0 | NA | NA | NA | NA | NA | Gm53057 | predicted gene 53057 [Source:MGI Symbo | 0 | 0 | 0 | 0 | 0 | 0 |
| 50825 | ENSMUSG00000113261 | 0.160662974 | 1.126924175 | 4.080472857 | 0.276174898 | 0.782413736 | NA | Gm47404 | predicted gene, 47404 [Source:MGI Symb | 0 | 0 | 0 | 0.964 | 0 | 0 |
| 50826 | ENSMUSG00000113262 | 1.436617682 | -0.781446032 | 1.931604398 | -0.40455801 | 0.685802429 | NA | Gm48551 | predicted gene, 48551 [Source:MGI Symb | 2.4093 | 0 | 3.0187 | 0.964 | 0 | 2.2277 |
| 50827 | ENSMUSG00000113263 | 0.17131964 | -0.796629176 | 4.080472857 | -0.195229623 | 0.84521319 | NA | Gm4811 | predicted gene 4811 [Source:MGI Symbol | 0 | 1.0279 | 0 | 0 | 0 | 0 |
| 50828 | ENSMUSG00000113264 | 0 | NA | NA | NA | NA | NA | Gm19095 | predicted gene, 19095 [Source:MGI Symb | 0 | 0 | 0 | 0 | 0 | 0 |
| 50829 | ENSMUSG00000113265 | 0 | NA | NA | NA | NA | NA | Gm30198 | predicted gene, 30198 [Source:MGI Symb | 0 | 0 | 0 | 0 | 0 | 0 |
| 50830 | ENSMUSG00000113266 | 0 | NA | NA | NA | NA | NA | Gm19563 | predicted gene, 19563 [Source:MGI Symb | 0 | 0 | 0 | 0 | 0 | 0 |
| 50831 | ENSMUSG00000113267 | 0 | NA | NA | NA | NA | NA | Gm47969 | predicted gene, 47969 [Source:MGI Symb | 0 | 0 | 0 | 0 | 0 | 0 |
| 50832 | ENSMUSG00000113268 | 0 | NA | NA | NA | NA | NA | 4930477G07 | RIKEN cDNA 4930477G07 gene [Source:M | 0 | 0 | 0 | 0 | 0 | 0 |
| 50833 | ENSMUSG00000113269 | 0 | NA | NA | NA | NA | NA | Gm47494 | predicted gene, 47494 [Source:MGI Symb | 0 | 0 | 0 | 0 | 0 | 0 |
| 50834 | ENSMUSG00000113270 | 0 | NA | NA | NA | NA | NA | Gm48235 | predicted gene, 48235 [Source:MGI Symb | 0 | 0 | 0 | 0 | 0 | 0 |
| 50835 | ENSMUSG00000113271 | 0 | NA | NA | NA | NA | NA | Gm40653 | predicted gene, 40653 [Source:MGI Symb | 0 | 0 | 0 | 0 | 0 | 0 |
| 50836 | ENSMUSG00000113272 | 0 | NA | NA | NA | NA | NA | Gm34809 | predicted gene, 34809 [Source:MGI Symb | 0 | 0 | 0 | 0 | 0 | 0 |
| 50837 | ENSMUSG00000113273 | 0 | NA | NA | NA | NA | NA | Gm48423 | predicted gene, 48423 [Source:MGI Symb | 0 | 0 | 0 | 0 | 0 | 0 |
| 50838 | ENSMUSG00000113274 | 0.181769715 | 1.126924175 | 4.080472857 | 0.276174898 | 0.782413736 | NA | Gm41002 | predicted gene, 41002 [Source:MGI Symb | 0 | 0 | 0 | 0 | 1.0906 | 0 |
| 50839 | ENSMUSG00000113275 | 0 | NA | NA | NA | NA | NA | Tubb2a-ps2 | tubulin, beta 2a, pseudogene 2 [Source:M | 0 | 0 | 0 | 0 | 0 | 0 |
| 50840 | ENSMUSG00000113276 | 0 | NA | NA | NA | NA | NA | Gm33630 | predicted gene, 33630 [Source:MGI Symb | 0 | 0 | 0 | 0 | 0 | 0 |
| 50841 | ENSMUSG00000113277 | 0 | NA | NA | NA | NA | NA | Gm18116 | predicted gene, 18116 [Source:MGI Symb | 0 | 0 | 0 | 0 | 0 | 0 |
| 50842 | ENSMUSG00000113278 | 0.267700746 | -1.497481649 | 4.057001436 | -0.369110456 | 0.712045396 | NA | Gm33016 | predicted gene, 33016 [Source:MGI Symb | 1.6062 | 0 | 0 | 0 | 0 | 0 |
| 50843 | ENSMUSG00000113279 | 0 | NA | NA | NA | NA | NA | Gm47470 | predicted gene, 47470 [Source:MGI Symb | 0 | 0 | 0 | 0 | 0 | 0 |
| 50844 | ENSMUSG00000113280 | 0 | NA | NA | NA | NA | NA | Gm21559 | predicted gene, 21559 [Source:MGI Symb | 0 | 0 | 0 | 0 | 0 | 0 |

| | | | | | | | | | | | | | | | |
|---|---|---|---|---|---|---|---|---|---|---|---|---|---|---|---|
| 50845 | ENSMUSG00000113281 | 0 | NA | NA | NA | NA | NA | Gm11360 | predicted gene 11360 [Source:MGI Symbo | 0 | 0 | 0 | 0 | 0 | 0 |
| 50846 | ENSMUSG00000113282 | 0 | NA | NA | NA | NA | NA | Gm20368 | predicted gene, 20368 [Source:MGI Symb | 0 | 0 | 0 | 0 | 0 | 0 |
| 50847 | ENSMUSG00000113283 | 0 | NA | NA | NA | NA | NA | Gm35890 | predicted gene, 35890 [Source:MGI Symb | 0 | 0 | 0 | 0 | 0 | 0 |
| 50848 | ENSMUSG00000113284 | 0 | NA | NA | NA | NA | NA | Gm48213 | predicted gene, 48213 [Source:MGI Symb | 0 | 0 | 0 | 0 | 0 | 0 |
| 50849 | ENSMUSG00000113285 | 0 | NA | NA | NA | NA | NA | Gm36128 | predicted gene, 36128 [Source:MGI Symb | 0 | 0 | 0 | 0 | 0 | 0 |
| 50850 | ENSMUSG00000113286 | 0 | NA | NA | NA | NA | NA | Gm18627 | predicted gene, 18627 [Source:MGI Symb | 0 | 0 | 0 | 0 | 0 | 0 |
| 50851 | ENSMUSG00000113287 | 0 | NA | NA | NA | NA | NA | Gm40853 | predicted gene, 40853 [Source:MGI Symb | 0 | 0 | 0 | 0 | 0 | 0 |
| 50852 | ENSMUSG00000113288 | 0 | NA | NA | NA | NA | NA | Gm47982 | predicted gene, 47982 [Source:MGI Symb | 0 | 0 | 0 | 0 | 0 | 0 |
| 50853 | ENSMUSG00000113289 | 0 | NA | NA | NA | NA | NA | Gm48044 | predicted gene, 48044 [Source:MGI Symb | 0 | 0 | 0 | 0 | 0 | 0 |
| 50854 | ENSMUSG00000113290 | 0 | NA | NA | NA | NA | NA | A530058O07 | RIKEN cDNA A530058O07 gene [Source:M | 0 | 0 | 0 | 0 | 0 | 0 |
| 50855 | ENSMUSG00000113291 | 0 | NA | NA | NA | NA | NA | Gm19001 | predicted gene, 19001 [Source:MGI Symb | 0 | 0 | 0 | 0 | 0 | 0 |
| 50856 | ENSMUSG00000113292 | 0 | NA | NA | NA | NA | NA | Gm18413 | predicted gene, 18413 [Source:MGI Symb | 0 | 0 | 0 | 0 | 0 | 0 |
| 50857 | ENSMUSG00000113293 | 0 | NA | NA | NA | NA | NA | Gm48688 | predicted gene, 48688 [Source:MGI Symb | 0 | 0 | 0 | 0 | 0 | 0 |
| 50858 | ENSMUSG00000113294 | 23.92262566 | -0.471713244 | 0.560904426 | -0.840986847 | 0.400355303 | 0.827266295 | Gm49326 | predicted gene, 49326 [Source:MGI Symb | 20.881 | 16.447 | 46.287 | 28.919 | 8.7249 | 22.277 |
| 50859 | ENSMUSG00000113295 | 0 | NA | NA | NA | NA | NA | Gm49357 | predicted gene, 49357 [Source:MGI Symb | 0 | 0 | 0 | 0 | 0 | 0 |
| 50860 | ENSMUSG00000113296 | 0 | NA | NA | NA | NA | NA | Gm48032 | predicted gene, 48032 [Source:MGI Symb | 0 | 0 | 0 | 0 | 0 | 0 |
| 50861 | ENSMUSG00000113297 | 0 | NA | NA | NA | NA | NA | Rpl17-ps6 | ribosomal protein L17, pseudogene 6 [Sou | 0 | 0 | 0 | 0 | 0 | 0 |
| 50862 | ENSMUSG00000113298 | 0 | NA | NA | NA | NA | NA | Gm48559 | predicted gene, 48559 [Source:MGI Symb | 0 | 0 | 0 | 0 | 0 | 0 |
| 50863 | ENSMUSG00000113299 | 0 | NA | NA | NA | NA | NA | Gm35618 | predicted gene, 35618 [Source:MGI Symb | 0 | 0 | 0 | 0 | 0 | 0 |
| 50864 | ENSMUSG00000113300 | 0 | NA | NA | NA | NA | NA | Gm49371 | predicted gene, 49371 [Source:MGI Symb | 0 | 0 | 0 | 0 | 0 | 0 |
| 50865 | ENSMUSG00000113301 | 0 | NA | NA | NA | NA | NA | Gm48103 | predicted gene, 48103 [Source:MGI Symb | 0 | 0 | 0 | 0 | 0 | 0 |
| 50866 | ENSMUSG00000113302 | 0 | NA | NA | NA | NA | NA | Gm32219 | predicted gene, 32219 [Source:MGI Symb | 0 | 0 | 0 | 0 | 0 | 0 |
| 50867 | ENSMUSG00000113303 | 0 | NA | NA | NA | NA | NA | 4930423D22 | RIKEN cDNA 4930423D22 gene [Source:M | 0 | 0 | 0 | 0 | 0 | 0 |
| 50868 | ENSMUSG00000113304 | 0 | NA | NA | NA | NA | NA | Gm48618 | predicted gene, 48618 [Source:MGI Symb | 0 | 0 | 0 | 0 | 0 | 0 |
| 50869 | ENSMUSG00000113305 | 0 | NA | NA | NA | NA | NA | Gm47811 | predicted gene, 47811 [Source:MGI Symb | 0 | 0 | 0 | 0 | 0 | 0 |
| 50870 | ENSMUSG00000113306 | 0 | NA | NA | NA | NA | NA | Mif-ps7 | macrophage migration inhibitory factor, p | 0 | 0 | 0 | 0 | 0 | 0 |
| 50871 | ENSMUSG00000113307 | 0 | NA | NA | NA | NA | NA | Gm48516 | predicted gene, 48516 [Source:MGI Symb | 0 | 0 | 0 | 0 | 0 | 0 |
| 50872 | ENSMUSG00000113308 | 0 | NA | NA | NA | NA | NA | Gm48445 | predicted gene, 48445 [Source:MGI Symb | 0 | 0 | 0 | 0 | 0 | 0 |
| 50873 | ENSMUSG00000113309 | 0 | NA | NA | NA | NA | NA | Gm48408 | predicted gene, 48408 [Source:MGI Symb | 0 | 0 | 0 | 0 | 0 | 0 |
| 50874 | ENSMUSG00000113310 | 3.962074955 | -0.273512505 | 1.192737256 | -0.229314967 | 0.818624121 | NA | Gm47884 | predicted gene, 47884 [Source:MGI Symb | 0.8031 | 9.2513 | 3.0187 | 1.928 | 6.5437 | 2.2277 |
| 50875 | ENSMUSG00000113311 | 0 | NA | NA | NA | NA | NA | Gm47428 | predicted gene, 47428 [Source:MGI Symb | 0 | 0 | 0 | 0 | 0 | 0 |
| 50876 | ENSMUSG00000113312 | 0 | NA | NA | NA | NA | NA | Gm48052 | predicted gene, 48052 [Source:MGI Symb | 0 | 0 | 0 | 0 | 0 | 0 |
| 50877 | ENSMUSG00000113313 | 0 | NA | NA | NA | NA | NA | Gm32036 | predicted gene, 32036 [Source:MGI Symb | 0 | 0 | 0 | 0 | 0 | 0 |
| 50878 | ENSMUSG00000113314 | 0 | NA | NA | NA | NA | NA | Gm2721 | predicted gene 2721 [Source:MGI Symbol | 0 | 0 | 0 | 0 | 0 | 0 |
| 50879 | ENSMUSG00000113315 | 1.210023386 | -2.482020224 | 2.285496674 | -1.085987239 | 0.277484638 | NA | Gm8227 | predicted gene 8227 [Source:MGI Symbol | 3.2124 | 3.0838 | 0 | 0.964 | 0 | 0 |
| 50880 | ENSMUSG00000113316 | 0 | NA | NA | NA | NA | NA | Gm48253 | predicted gene, 48253 [Source:MGI Symb | 0 | 0 | 0 | 0 | 0 | 0 |
| 50881 | ENSMUSG00000113317 | 0 | NA | NA | NA | NA | NA | Gm48573 | predicted gene, 48573 [Source:MGI Symb | 0 | 0 | 0 | 0 | 0 | 0 |
| 50882 | ENSMUSG00000113318 | 6.105097927 | 0.190489649 | 0.915727287 | 0.208020065 | 0.835213301 | 0.971073158 | Gm18113 | predicted gene, 18113 [Source:MGI Symb | 4.8186 | 8.2233 | 4.025 | 1.928 | 8.7249 | 8.9108 |
| 50883 | ENSMUSG00000113319 | 0 | NA | NA | NA | NA | NA | Gm9628 | predicted gene 9628 [Source:MGI Symbol | 0 | 0 | 0 | 0 | 0 | 0 |
| 50884 | ENSMUSG00000113320 | 0 | NA | NA | NA | NA | NA | Gm47515 | predicted gene, 47515 [Source:MGI Symb | 0 | 0 | 0 | 0 | 0 | 0 |
| 50885 | ENSMUSG00000113321 | 0 | NA | NA | NA | NA | NA | Gm8075 | predicted gene 8075 [Source:MGI Symbol | 0 | 0 | 0 | 0 | 0 | 0 |
| 50886 | ENSMUSG00000113322 | 0.321325949 | 1.911575114 | 4.048267893 | 0.472195804 | 0.636787034 | NA | Gm48492 | predicted gene, 48492 [Source:MGI Symb | 0 | 0 | 0 | 1.928 | 0 | 0 |
| 50887 | ENSMUSG00000113323 | 0 | NA | NA | NA | NA | NA | Gm18625 | predicted gene, 18625 [Source:MGI Symb | 0 | 0 | 0 | 0 | 0 | 0 |
| 50888 | ENSMUSG00000113324 | 0 | NA | NA | NA | NA | NA | Gm48819 | predicted gene, 48819 [Source:MGI Symb | 0 | 0 | 0 | 0 | 0 | 0 |
| 50889 | ENSMUSG00000113325 | 0 | NA | NA | NA | NA | NA | Gm48080 | predicted gene, 48080 [Source:MGI Symb | 0 | 0 | 0 | 0 | 0 | 0 |
| 50890 | ENSMUSG00000113326 | 1.904382935 | 0.422883025 | 1.682042437 | 0.251410437 | 0.801496799 | NA | Gm47586 | predicted gene, 47586 [Source:MGI Symb | 0.8031 | 3.0838 | 1.0062 | 0.964 | 0 | 5.5692 |
| 50891 | ENSMUSG00000113327 | 0 | NA | NA | NA | NA | NA | Gm48615 | predicted gene, 48615 [Source:MGI Symb | 0 | 0 | 0 | 0 | 0 | 0 |
| 50892 | ENSMUSG00000113328 | 18.83770604 | -0.325654745 | 0.585043143 | -0.556633728 | 0.577777705 | 0.908070925 | Gm47260 | predicted gene, 47260 [Source:MGI Symb | 18.471 | 9.2513 | 35.219 | 20.244 | 10.906 | 18.935 |
| 50893 | ENSMUSG00000113329 | 0 | NA | NA | NA | NA | NA | 4930555G21 | RIKEN cDNA 4930555G21 gene [Source:M | 0 | 0 | 0 | 0 | 0 | 0 |
| 50894 | ENSMUSG00000113330 | 0 | NA | NA | NA | NA | NA | Gm34237 | predicted gene, 34237 [Source:MGI Symb | 0 | 0 | 0 | 0 | 0 | 0 |
| 50895 | ENSMUSG00000113331 | 0 | NA | NA | NA | NA | NA | 3110009F21 | RIKEN cDNA 3110009F21 gene [Source:M | 0 | 0 | 0 | 0 | 0 | 0 |
| 50896 | ENSMUSG00000113332 | 0 | NA | NA | NA | NA | NA | Gm9259 | predicted gene 9259 [Source:MGI Symbol | 0 | 0 | 0 | 0 | 0 | 0 |
| 50897 | ENSMUSG00000113333 | 0 | NA | NA | NA | NA | NA | Gm19011 | predicted gene, 19011 [Source:MGI Symb | 0 | 0 | 0 | 0 | 0 | 0 |
| 50898 | ENSMUSG00000113334 | 0.335414862 | -1.779047242 | 4.030804179 | -0.441362855 | 0.658950329 | NA | D030007L05 | RIKEN cDNA D030007L05 gene [Source:M | 0 | 0 | 2.0125 | 0 | 0 | 0 |
| 50899 | ENSMUSG00000113335 | 2.596195432 | 0.135680653 | 1.331840968 | 0.101874515 | 0.918856279 | NA | Gm48296 | predicted gene, 48296 [Source:MGI Symb | 2.4093 | 1.0279 | 4.025 | 4.8199 | 2.1812 | 1.1138 |
| 50900 | ENSMUSG00000113336 | 0 | NA | NA | NA | NA | NA | Gm16497 | predicted gene 16497 [Source:MGI Symbo | 0 | 0 | 0 | 0 | 0 | 0 |

| | | | | | | | | | | | | | | | |
|---|---|---|---|---|---|---|---|---|---|---|---|---|---|---|---|
| 50901 | ENSMUSG00000113337 | 0 | NA | NA | NA | NA | NA | Gm19220 | predicted gene, 19220 [Source:MGI Symb | 0 | 0 | 0 | 0 | 0 | 0 |
| 50902 | ENSMUSG00000113338 | 0 | NA | NA | NA | NA | NA | A530046M1 | RIKEN cDNA A530046M15 gene [Source:N | 0 | 0 | 0 | 0 | 0 | 0 |
| 50903 | ENSMUSG00000113339 | 0 | NA | NA | NA | NA | NA | Gm17915 | predicted gene, 17915 [Source:MGI Symb | 0 | 0 | 0 | 0 | 0 | 0 |
| 50904 | ENSMUSG00000113340 | 0 | NA | NA | NA | NA | NA | Gm7695 | predicted gene 7695 [Source:MGI Symbol | 0 | 0 | 0 | 0 | 0 | 0 |
| 50905 | ENSMUSG00000113341 | 0.699599566 | -1.425530384 | 3.053137229 | -0.466906751 | 0.640566595 | NA | 9430031K09 | RIKEN cDNA 9430031K09 gene [Source:M | 0 | 3.0838 | 0 | 0 | 0 | 1.1138 |
| 50906 | ENSMUSG00000113342 | 0 | NA | NA | NA | NA | NA | AA414992 | expressed sequence AA414992 [Source:M | 0 | 0 | 0 | 0 | 0 | 0 |
| 50907 | ENSMUSG00000113343 | 0 | NA | NA | NA | NA | NA | Gm34923 | predicted gene, 34923 [Source:MGI Symb | 0 | 0 | 0 | 0 | 0 | 0 |
| 50908 | ENSMUSG00000113344 | 0 | NA | NA | NA | NA | NA | Gm49251 | predicted gene, 49251 [Source:MGI Symb | 0 | 0 | 0 | 0 | 0 | 0 |
| 50909 | ENSMUSG00000113345 | 0 | NA | NA | NA | NA | NA | Gm48096 | predicted gene, 48096 [Source:MGI Symb | 0 | 0 | 0 | 0 | 0 | 0 |
| 50910 | ENSMUSG00000113346 | 1.309526411 | 0.956871372 | 1.855835071 | 0.515601514 | 0.606132753 | NA | Eprn | ephemeron, early developmental lncRNA | 1.6062 | 1.0279 | 0 | 1.928 | 2.1812 | 1.1138 |
| 50911 | ENSMUSG00000113347 | 0 | NA | NA | NA | NA | NA | Gm47323 | predicted gene, 47323 [Source:MGI Symb | 0 | 0 | 0 | 0 | 0 | 0 |
| 50912 | ENSMUSG00000113348 | 0 | NA | NA | NA | NA | NA | Gm8607 | predicted gene 8607 [Source:MGI Symbol | 0 | 0 | 0 | 0 | 0 | 0 |
| 50913 | ENSMUSG00000113349 | 0.181769715 | 1.126924175 | 4.080472857 | 0.276174898 | 0.782413736 | NA | Gm48369 | predicted gene, 48369 [Source:MGI Symb | 0 | 0 | 0 | 0 | 1.0906 | 0 |
| 50914 | ENSMUSG00000113350 | 0 | NA | NA | NA | NA | NA | Vmn2r-ps10 | vomeronasal 2, receptor, pseudogene 102 | 0 | 0 | 0 | 0 | 0 | 0 |
| 50915 | ENSMUSG00000113351 | 0 | NA | NA | NA | NA | NA | Gm48575 | predicted gene, 48575 [Source:MGI Symb | 0 | 0 | 0 | 0 | 0 | 0 |
| 50916 | ENSMUSG00000113352 | 0 | NA | NA | NA | NA | NA | Gm34639 | predicted gene, 34639 [Source:MGI Symb | 0 | 0 | 0 | 0 | 0 | 0 |
| 50917 | ENSMUSG00000113353 | 0 | NA | NA | NA | NA | NA | Gm18878 | predicted gene, 18878 [Source:MGI Symb | 0 | 0 | 0 | 0 | 0 | 0 |
| 50918 | ENSMUSG00000113354 | 0 | NA | NA | NA | NA | NA | Gm20789 | predicted gene, 20789 [Source:MGI Symb | 0 | 0 | 0 | 0 | 0 | 0 |
| 50919 | ENSMUSG00000113355 | 0.160662974 | 1.126924175 | 4.080472857 | 0.276174898 | 0.782413736 | NA | Gm48888 | predicted gene, 48888 [Source:MGI Symb | 0 | 0 | 0 | 0.964 | 0 | 0 |
| 50920 | ENSMUSG00000113356 | 0 | NA | NA | NA | NA | NA | Gm48012 | predicted gene, 48012 [Source:MGI Symb | 0 | 0 | 0 | 0 | 0 | 0 |
| 50921 | ENSMUSG00000113357 | 0 | NA | NA | NA | NA | NA | Gm19195 | predicted gene, 19195 [Source:MGI Symb | 0 | 0 | 0 | 0 | 0 | 0 |
| 50922 | ENSMUSG00000113358 | 0 | NA | NA | NA | NA | NA | 1700112M0 | RIKEN cDNA 1700112M02 gene [Source:N | 0 | 0 | 0 | 0 | 0 | 0 |
| 50923 | ENSMUSG00000113359 | 0 | NA | NA | NA | NA | NA | Gm48730 | predicted gene, 48730 [Source:MGI Symb | 0 | 0 | 0 | 0 | 0 | 0 |
| 50924 | ENSMUSG00000113360 | 0 | NA | NA | NA | NA | NA | Gm48355 | predicted gene, 48355 [Source:MGI Symb | 0 | 0 | 0 | 0 | 0 | 0 |
| 50925 | ENSMUSG00000113361 | 7.580820381 | 0.228683218 | 0.894429473 | 0.255674958 | 0.798201833 | 0.962424862 | B930059L03 | RIKEN cDNA B930059L03 gene [Source:M | 0.8031 | 8.2233 | 12.075 | 6.7478 | 8.7249 | 8.9108 |
| 50926 | ENSMUSG00000113362 | 0 | NA | NA | NA | NA | NA | Gm40557 | predicted gene, 40557 [Source:MGI Symb | 0 | 0 | 0 | 0 | 0 | 0 |
| 50927 | ENSMUSG00000113363 | 0 | NA | NA | NA | NA | NA | Gm48187 | predicted gene, 48187 [Source:MGI Symb | 0 | 0 | 0 | 0 | 0 | 0 |
| 50928 | ENSMUSG00000113364 | 0 | NA | NA | NA | NA | NA | Gm48059 | predicted gene, 48059 [Source:MGI Symb | 0 | 0 | 0 | 0 | 0 | 0 |
| 50929 | ENSMUSG00000113365 | 0 | NA | NA | NA | NA | NA | Gm48424 | predicted gene, 48424 [Source:MGI Symb | 0 | 0 | 0 | 0 | 0 | 0 |
| 50930 | ENSMUSG00000113366 | 0 | NA | NA | NA | NA | NA | Gm47259 | predicted gene, 47259 [Source:MGI Symb | 0 | 0 | 0 | 0 | 0 | 0 |
| 50931 | ENSMUSG00000113367 | 0 | NA | NA | NA | NA | NA | Gm47125 | predicted gene, 47125 [Source:MGI Symb | 0 | 0 | 0 | 0 | 0 | 0 |
| 50932 | ENSMUSG00000113368 | 0.185640647 | 1.126924175 | 4.080472857 | 0.276174898 | 0.782413736 | NA | Gm10054 | predicted gene 10054 [Source:MGI Symbo | 0 | 0 | 0 | 0 | 0 | 1.1138 |
| 50933 | ENSMUSG00000113369 | 0 | NA | NA | NA | NA | NA | Gm8504 | predicted gene 8504 [Source:MGI Symbol | 0 | 0 | 0 | 0 | 0 | 0 |
| 50934 | ENSMUSG00000113370 | 1.213558485 | -2.430141359 | 2.584716269 | -0.940196566 | 0.347116743 | NA | Gm46348 | predicted gene, 46348 [Source:MGI Symb | 0 | 6.1675 | 0 | 0 | 0 | 1.1138 |
| 50935 | ENSMUSG00000113371 | 0 | NA | NA | NA | NA | NA | Gm48588 | predicted gene, 48588 [Source:MGI Symb | 0 | 0 | 0 | 0 | 0 | 0 |
| 50936 | ENSMUSG00000113372 | 0 | NA | NA | NA | NA | NA | Gm18624 | predicted gene, 18624 [Source:MGI Symb | 0 | 0 | 0 | 0 | 0 | 0 |
| 50937 | ENSMUSG00000113373 | 0 | NA | NA | NA | NA | NA | | novel protein | 0 | 0 | 0 | 0 | 0 | 0 |
| 50938 | ENSMUSG00000113374 | 0 | NA | NA | NA | NA | NA | Gm33654 | predicted gene, 33654 [Source:MGI Symb | 0 | 0 | 0 | 0 | 0 | 0 |
| 50939 | ENSMUSG00000113375 | 0 | NA | NA | NA | NA | NA | Gm47838 | predicted gene, 47838 [Source:MGI Symb | 0 | 0 | 0 | 0 | 0 | 0 |
| 50940 | ENSMUSG00000113376 | 0 | NA | NA | NA | NA | NA | Gm48572 | predicted gene, 48572 [Source:MGI Symb | 0 | 0 | 0 | 0 | 0 | 0 |
| 50941 | ENSMUSG00000113377 | 0 | NA | NA | NA | NA | NA | Gm48446 | predicted gene, 48446 [Source:MGI Symb | 0 | 0 | 0 | 0 | 0 | 0 |
| 50942 | ENSMUSG00000113378 | 0 | NA | NA | NA | NA | NA | Gm47660 | predicted gene, 47660 [Source:MGI Symb | 0 | 0 | 0 | 0 | 0 | 0 |
| 50943 | ENSMUSG00000113379 | 0.342432689 | 1.990653086 | 3.900281263 | 0.51038706 | 0.609780321 | NA | Gm46436 | predicted gene, 46436 [Source:MGI Symb | 0 | 0 | 0 | 0.964 | 1.0906 | 0 |
| 50944 | ENSMUSG00000113380 | 0 | NA | NA | NA | NA | NA | Gm48682 | predicted gene, 48682 [Source:MGI Symb | 0 | 0 | 0 | 0 | 0 | 0 |
| 50945 | ENSMUSG00000113381 | 0 | NA | NA | NA | NA | NA | Vmn2r-ps97 | vomeronasal 2, receptor, pseudogene 97 | 0 | 0 | 0 | 0 | 0 | 0 |
| 50946 | ENSMUSG00000113382 | 0 | NA | NA | NA | NA | NA | Gm48429 | predicted gene, 48429 [Source:MGI Symb | 0 | 0 | 0 | 0 | 0 | 0 |
| 50947 | ENSMUSG00000113383 | 0 | NA | NA | NA | NA | NA | Gm47267 | predicted gene, 47267 [Source:MGI Symb | 0 | 0 | 0 | 0 | 0 | 0 |
| 50948 | ENSMUSG00000113384 | 0 | NA | NA | NA | NA | NA | Gm47859 | predicted gene, 47859 [Source:MGI Symb | 0 | 0 | 0 | 0 | 0 | 0 |
| 50949 | ENSMUSG00000113385 | 1.029540662 | 0.20390886 | 2.204283627 | 0.092505727 | 0.926296241 | NA | Gm48405 | predicted gene, 48405 [Source:MGI Symb | 0.8031 | 2.0558 | 0 | 0 | 1.0906 | 2.2277 |
| 50950 | ENSMUSG00000113386 | 0 | NA | NA | NA | NA | NA | Gm47357 | predicted gene, 47357 [Source:MGI Symb | 0 | 0 | 0 | 0 | 0 | 0 |
| 50951 | ENSMUSG00000113387 | 1.624093257 | 0.152393133 | 1.815115408 | 0.08395782 | 0.933089968 | NA | Gm31508 | predicted gene, 31508 [Source:MGI Symb | 1.6062 | 0 | 3.0187 | 2.8919 | 0 | 2.2277 |
| 50952 | ENSMUSG00000113388 | 0 | NA | NA | NA | NA | NA | Gm48111 | predicted gene, 48111 [Source:MGI Symb | 0 | 0 | 0 | 0 | 0 | 0 |
| 50953 | ENSMUSG00000113389 | 1.500382101 | -0.223605962 | 1.769303272 | -0.126380799 | 0.8994305 | NA | Gm9512 | predicted gene 9512 [Source:MGI Symbol | 0.8031 | 3.0838 | 1.0062 | 1.928 | 2.1812 | 0 |
| 50954 | ENSMUSG00000113390 | 0 | NA | NA | NA | NA | NA | Gm32635 | predicted gene, 32635 [Source:MGI Symb | 0 | 0 | 0 | 0 | 0 | 0 |
| 50955 | ENSMUSG00000113391 | 0 | NA | NA | NA | NA | NA | Gm48327 | predicted gene, 48327 [Source:MGI Symb | 0 | 0 | 0 | 0 | 0 | 0 |
| 50956 | ENSMUSG00000113392 | 0.133850373 | -0.796629176 | 4.080472857 | -0.195229623 | 0.84521319 | NA | Gm36262 | predicted gene, 36262 [Source:MGI Symb | 0.8031 | 0 | 0 | 0 | 0 | 0 |

| | | | | | | | | | | | | | | | |
|---|---|---|---|---|---|---|---|---|---|---|---|---|---|---|---|
| 50957 | ENSMUSG00000113393 | 0 | NA | NA | NA | NA | NA | Gm48415 | predicted gene, 48415 [Source:MGI Symb | 0 | 0 | 0 | 0 | 0 | 0 |
| 50958 | ENSMUSG00000113394 | 0.401551119 | -2.072982726 | 4.008251979 | -0.517178744 | 0.605031388 | NA | Gm48436 | predicted gene, 48436 [Source:MGI Symb | 2.4093 | 0 | 0 | 0 | 0 | 0 |
| 50959 | ENSMUSG00000113395 | 0 | NA | NA | NA | NA | NA | Gm48365 | predicted gene, 48365 [Source:MGI Symb | 0 | 0 | 0 | 0 | 0 | 0 |
| 50960 | ENSMUSG00000113396 | 0 | NA | NA | NA | NA | NA | Gm33467 | predicted gene, 33467 [Source:MGI Symb | 0 | 0 | 0 | 0 | 0 | 0 |
| 50961 | ENSMUSG00000113397 | 0 | NA | NA | NA | NA | NA | Gm9194 | predicted gene 9194 [Source:MGI Symbol | 0 | 0 | 0 | 0 | 0 | 0 |
| 50962 | ENSMUSG00000113398 | 0 | NA | NA | NA | NA | NA | Gm5194 | predicted gene 5194 [Source:MGI Symbol | 0 | 0 | 0 | 0 | 0 | 0 |
| 50963 | ENSMUSG00000113399 | 17.38546072 | -1.982136751 | 0.665676501 | -2.977627642 | 0.002904887 | 0.045719565 | Gm48199 | predicted gene, 48199 [Source:MGI Symb | 24.896 | 48.312 | 10.062 | 6.7478 | 8.7249 | 5.5692 |
| 50964 | ENSMUSG00000113400 | 0 | NA | NA | NA | NA | NA | Gm8605 | predicted gene 8605 [Source:MGI Symbol | 0 | 0 | 0 | 0 | 0 | 0 |
| 50965 | ENSMUSG00000113401 | 0 | NA | NA | NA | NA | NA | Gm18247 | predicted gene, 18247 [Source:MGI Symb | 0 | 0 | 0 | 0 | 0 | 0 |
| 50966 | ENSMUSG00000113402 | 0 | NA | NA | NA | NA | NA | Zfp996 | zinc finger protein 996 [Source:MGI Symb | 0 | 0 | 0 | 0 | 0 | 0 |
| 50967 | ENSMUSG00000113403 | 0 | NA | NA | NA | NA | NA | Gm9315 | predicted gene 9315 [Source:MGI Symbol | 0 | 0 | 0 | 0 | 0 | 0 |
| 50968 | ENSMUSG00000113404 | 0 | NA | NA | NA | NA | NA | Gm48622 | predicted gene, 48622 [Source:MGI Symb | 0 | 0 | 0 | 0 | 0 | 0 |
| 50969 | ENSMUSG00000113405 | 0 | NA | NA | NA | NA | NA | Gm4935 | predicted gene 4935 [Source:MGI Symbol | 0 | 0 | 0 | 0 | 0 | 0 |
| 50970 | ENSMUSG00000113406 | 0 | NA | NA | NA | NA | NA | Gm7762 | predicted gene 7762 [Source:MGI Symbol | 0 | 0 | 0 | 0 | 0 | 0 |
| 50971 | ENSMUSG00000113407 | 0 | NA | NA | NA | NA | NA | Gm47437 | predicted gene, 47437 [Source:MGI Symb | 0 | 0 | 0 | 0 | 0 | 0 |
| 50972 | ENSMUSG00000113408 | 0 | NA | NA | NA | NA | NA | Gm48540 | predicted gene, 48540 [Source:MGI Symb | 0 | 0 | 0 | 0 | 0 | 0 |
| 50973 | ENSMUSG00000113409 | 0.497389803 | 1.101799453 | 3.439268845 | 0.320358629 | 0.748696483 | NA | Gm8288 | predicted gene 8288 [Source:MGI Symbol | 0.8031 | 0 | 0 | 0 | 2.1812 | 0 |
| 50974 | ENSMUSG00000113410 | 0 | NA | NA | NA | NA | NA | Gm20043 | predicted gene, 20043 [Source:MGI Symb | 0 | 0 | 0 | 0 | 0 | 0 |
| 50975 | ENSMUSG00000113411 | 0 | NA | NA | NA | NA | NA | Gm48643 | predicted gene, 48643 [Source:MGI Symb | 0 | 0 | 0 | 0 | 0 | 0 |
| 50976 | ENSMUSG00000113412 | 0 | NA | NA | NA | NA | NA | Gm5191 | predicted gene 5191 [Source:MGI Symbol | 0 | 0 | 0 | 0 | 0 | 0 |
| 50977 | ENSMUSG00000113413 | 0 | NA | NA | NA | NA | NA | Gm21941 | predicted gene, 21941 [Source:MGI Symb | 0 | 0 | 0 | 0 | 0 | 0 |
| 50978 | ENSMUSG00000113414 | 0 | NA | NA | NA | NA | NA | Gm18504 | predicted gene, 18504 [Source:MGI Symb | 0 | 0 | 0 | 0 | 0 | 0 |
| 50979 | ENSMUSG00000113415 | 0 | NA | NA | NA | NA | NA | Gm3234 | predicted gene 3234 [Source:MGI Symbol | 0 | 0 | 0 | 0 | 0 | 0 |
| 50980 | ENSMUSG00000113416 | 1.10120282 | -0.134983118 | 2.137435647 | -0.063151898 | 0.949645548 | NA | Gm49328 | predicted gene, 49328 [Source:MGI Symb | 2.4093 | 0 | 1.0062 | 0.964 | 0 | 2.2277 |
| 50981 | ENSMUSG00000113417 | 0 | NA | NA | NA | NA | NA | Gm47478 | predicted gene, 47478 [Source:MGI Symb | 0 | 0 | 0 | 0 | 0 | 0 |
| 50982 | ENSMUSG00000113418 | 0 | NA | NA | NA | NA | NA | Gm47172 | predicted gene, 47172 [Source:MGI Symb | 0 | 0 | 0 | 0 | 0 | 0 |
| 50983 | ENSMUSG00000113419 | 0 | NA | NA | NA | NA | NA | Gm5789 | predicted gene 5789 [Source:MGI Symbol | 0 | 0 | 0 | 0 | 0 | 0 |
| 50984 | ENSMUSG00000113420 | 0 | NA | NA | NA | NA | NA | Gm7242 | predicted gene 7242 [Source:MGI Symbol | 0 | 0 | 0 | 0 | 0 | 0 |
| 50985 | ENSMUSG00000113421 | 0 | NA | NA | NA | NA | NA | Gm4166 | predicted gene 4166 [Source:MGI Symbol | 0 | 0 | 0 | 0 | 0 | 0 |
| 50986 | ENSMUSG00000113422 | 0 | NA | NA | NA | NA | NA | Gm47556 | predicted gene, 47556 [Source:MGI Symb | 0 | 0 | 0 | 0 | 0 | 0 |
| 50987 | ENSMUSG00000113423 | 0.321325949 | 1.911575114 | 4.048267893 | 0.472195804 | 0.636787034 | NA | Gm3791 | predicted gene 3791 [Source:MGI Symbol | 0 | 0 | 0 | 1.928 | 0 | 0 |
| 50988 | ENSMUSG00000113424 | 7.372778834 | -0.152665284 | 0.766417991 | -0.199193241 | 0.842111586 | 0.971854408 | Gm48161 | predicted gene, 48161 [Source:MGI Symb | 8.031 | 8.2233 | 7.0437 | 7.7118 | 6.5437 | 6.6831 |
| 50989 | ENSMUSG00000113425 | 3.448509517 | 0.53807749 | 1.261416364 | 0.426566124 | 0.669695376 | NA | Gm48653 | predicted gene, 48653 [Source:MGI Symb | 2.4093 | 2.0558 | 4.025 | 6.7478 | 5.4531 | 0 |
| 50990 | ENSMUSG00000113426 | 0 | NA | NA | NA | NA | NA | Gm6056 | predicted gene 6056 [Source:MGI Symbol | 0 | 0 | 0 | 0 | 0 | 0 |
| 50991 | ENSMUSG00000113427 | 0.528279927 | -0.837071592 | 3.369085264 | -0.248456636 | 0.803781118 | NA | Gm46378 | predicted gene, 46378 [Source:MGI Symb | 0 | 2.0558 | 0 | 0 | 0 | 1.1138 |
| 50992 | ENSMUSG00000113428 | 0.167707431 | -0.796629176 | 4.080472857 | -0.195229623 | 0.84521319 | NA | Gm7664 | predicted gene 7664 [Source:MGI Symbol | 0 | 0 | 1.0062 | 0 | 0 | 0 |
| 50993 | ENSMUSG00000113429 | 0 | NA | NA | NA | NA | NA | Gm20063 | predicted gene, 20063 [Source:MGI Symb | 0 | 0 | 0 | 0 | 0 | 0 |
| 50994 | ENSMUSG00000113430 | 0 | NA | NA | NA | NA | NA | Gm47086 | predicted gene, 47086 [Source:MGI Symb | 0 | 0 | 0 | 0 | 0 | 0 |
| 50995 | ENSMUSG00000113431 | 0 | NA | NA | NA | NA | NA | Gm6722 | predicted gene 6722 [Source:MGI Symbol | 0 | 0 | 0 | 0 | 0 | 0 |
| 50996 | ENSMUSG00000113432 | 4.278419969 | -1.908716912 | 1.190905998 | -1.602743555 | 0.108991283 | NA | 8430406P12 | RIKEN cDNA 8430406P12 gene [Source:M | 8.031 | 3.0838 | 9.0562 | 0 | 3.2719 | 2.2277 |
| 50997 | ENSMUSG00000113433 | 0 | NA | NA | NA | NA | NA | Gm47489 | predicted gene, 47489 [Source:MGI Symb | 0 | 0 | 0 | 0 | 0 | 0 |
| 50998 | ENSMUSG00000113434 | 1.423867364 | -0.698992261 | 1.944593905 | -0.359454105 | 0.719255406 | NA | 3200001D21 | RIKEN cDNA 3200001D21 gene [Source:M | 3.2124 | 0 | 2.0125 | 0 | 1.0906 | 2.2277 |
| 50999 | ENSMUSG00000113435 | 0 | NA | NA | NA | NA | NA | Gm19441 | predicted gene, 19441 [Source:MGI Symb | 0 | 0 | 0 | 0 | 0 | 0 |
| 51000 | ENSMUSG00000113436 | 0 | NA | NA | NA | NA | NA | Gm48392 | predicted gene, 48392 [Source:MGI Symb | 0 | 0 | 0 | 0 | 0 | 0 |
| 51001 | ENSMUSG00000113437 | 0 | NA | NA | NA | NA | NA | Gm47412 | predicted gene, 47412 [Source:MGI Symb | 0 | 0 | 0 | 0 | 0 | 0 |
| 51002 | ENSMUSG00000113438 | 0 | NA | NA | NA | NA | NA | Gm47369 | predicted gene, 47369 [Source:MGI Symb | 0 | 0 | 0 | 0 | 0 | 0 |
| 51003 | ENSMUSG00000113439 | 0 | NA | NA | NA | NA | NA | Gm49382 | predicted gene, 49382 [Source:MGI Symb | 0 | 0 | 0 | 0 | 0 | 0 |
| 51004 | ENSMUSG00000113440 | 20.40445312 | 0.72889545 | 0.517067026 | 1.409673047 | 0.158636247 | 0.579569253 | Gm46404 | predicted gene, 46404 [Source:MGI Symb | 17.668 | 11.307 | 17.106 | 35.667 | 25.084 | 15.594 |
| 51005 | ENSMUSG00000113441 | 68.19429746 | -0.178192776 | 0.341365933 | -0.521999294 | 0.601670822 | 0.915461118 | Gm40438 | predicted gene, 40438 [Source:MGI Symb | 53.808 | 74.01 | 89.556 | 50.127 | 94.884 | 46.781 |
| 51006 | ENSMUSG00000113442 | 0 | NA | NA | NA | NA | NA | Gm47973 | predicted gene, 47973 [Source:MGI Symb | 0 | 0 | 0 | 0 | 0 | 0 |
| 51007 | ENSMUSG00000113443 | 0 | NA | NA | NA | NA | NA | Gm48668 | predicted gene, 48668 [Source:MGI Symb | 0 | 0 | 0 | 0 | 0 | 0 |
| 51008 | ENSMUSG00000113444 | 0 | NA | NA | NA | NA | NA | Gm48672 | predicted gene, 48672 [Source:MGI Symb | 0 | 0 | 0 | 0 | 0 | 0 |
| 51009 | ENSMUSG00000113445 | 0 | NA | NA | NA | NA | NA | Gm48661 | predicted gene, 48661 [Source:MGI Symb | 0 | 0 | 0 | 0 | 0 | 0 |
| 51010 | ENSMUSG00000113446 | 0 | NA | NA | NA | NA | NA | Gm47455 | predicted gene, 47455 [Source:MGI Symb | 0 | 0 | 0 | 0 | 0 | 0 |
| 51011 | ENSMUSG00000113447 | 0 | NA | NA | NA | NA | NA | Gm48538 | predicted gene, 48538 [Source:MGI Symb | 0 | 0 | 0 | 0 | 0 | 0 |
| 51012 | ENSMUSG00000113448 | 0.185640647 | 1.126924175 | 4.080472857 | 0.276174898 | 0.782413736 | NA | Gm7042 | predicted gene 7042 [Source:MGI Symbol | 0 | 0 | 0 | 0 | 0 | 1.1138 |

| | | | | | | | | | | | | | | | |
|---|---|---|---|---|---|---|---|---|---|---|---|---|---|---|---|
| 51013 | ENSMUSG00000113449 | 0 | NA | NA | NA | NA | NA | Gm48073 | predicted gene, 48073 [Source:MGI Symb | 0 | 0 | 0 | 0 | 0 | 0 |
| 51014 | ENSMUSG00000113450 | 17.5318102 | 0.52701008 | 0.54276132 | 0.970979436 | 0.331558519 | 0.774747658 | Gm49359 | predicted gene, 49359 [Source:MGI Symb | 14.456 | 16.447 | 12.075 | 12.532 | 20.722 | 28.96 |
| 51015 | ENSMUSG00000113451 | 0 | NA | NA | NA | NA | NA | Gm47421 | predicted gene, 47421 [Source:MGI Symb | 0 | 0 | 0 | 0 | 0 | 0 |
| 51016 | ENSMUSG00000113452 | 0 | NA | NA | NA | NA | NA | Gm18616 | predicted gene, 18616 [Source:MGI Symb | 0 | 0 | 0 | 0 | 0 | 0 |
| 51017 | ENSMUSG00000113453 | 0 | NA | NA | NA | NA | NA | Gm30709 | predicted gene, 30709 [Source:MGI Symb | 0 | 0 | 0 | 0 | 0 | 0 |
| 51018 | ENSMUSG00000113454 | 0.674621893 | -1.426050892 | 3.091195043 | -0.461326727 | 0.644564212 | NA | Gm47373 | predicted gene, 47373 [Source:MGI Symb | 0 | 3.0838 | 0 | 0.964 | 0 | 0 |
| 51019 | ENSMUSG00000113455 | 0 | NA | NA | NA | NA | NA | Gm48825 | predicted gene, 48825 [Source:MGI Symb | 0 | 0 | 0 | 0 | 0 | 0 |
| 51020 | ENSMUSG00000113456 | 0 | NA | NA | NA | NA | NA | Gm30738 | predicted gene, 30738 [Source:MGI Symb | 0 | 0 | 0 | 0 | 0 | 0 |
| 51021 | ENSMUSG00000113457 | 0 | NA | NA | NA | NA | NA | Gm47152 | predicted gene, 47152 [Source:MGI Symb | 0 | 0 | 0 | 0 | 0 | 0 |
| 51022 | ENSMUSG00000113458 | 0 | NA | NA | NA | NA | NA | Gm33684 | predicted gene, 33684 [Source:MGI Symb | 0 | 0 | 0 | 0 | 0 | 0 |
| 51023 | ENSMUSG00000113459 | 0 | NA | NA | NA | NA | NA | Gm30655 | predicted gene, 30655 [Source:MGI Symb | 0 | 0 | 0 | 0 | 0 | 0 |
| 51024 | ENSMUSG00000113460 | 0 | NA | NA | NA | NA | NA | Gm48686 | predicted gene, 48686 [Source:MGI Symb | 0 | 0 | 0 | 0 | 0 | 0 |
| 51025 | ENSMUSG00000113461 | 0 | NA | NA | NA | NA | NA | Gm48186 | predicted gene, 48186 [Source:MGI Symb | 0 | 0 | 0 | 0 | 0 | 0 |
| 51026 | ENSMUSG00000113462 | 0 | NA | NA | NA | NA | NA | Gm48248 | predicted gene, 48248 [Source:MGI Symb | 0 | 0 | 0 | 0 | 0 | 0 |
| 51027 | ENSMUSG00000113463 | 0 | NA | NA | NA | NA | NA | 4930520P13 | RIKEN cDNA 4930520P13 gene [Source:M | 0 | 0 | 0 | 0 | 0 | 0 |
| 51028 | ENSMUSG00000113464 | 0 | NA | NA | NA | NA | NA | Gm47901 | predicted gene, 47901 [Source:MGI Symb | 0 | 0 | 0 | 0 | 0 | 0 |
| 51029 | ENSMUSG00000113465 | 0 | NA | NA | NA | NA | NA | Gm40271 | predicted gene, 40271 [Source:MGI Symb | 0 | 0 | 0 | 0 | 0 | 0 |
| 51030 | ENSMUSG00000113466 | 0 | NA | NA | NA | NA | NA | Gm48649 | predicted gene, 48649 [Source:MGI Symb | 0 | 0 | 0 | 0 | 0 | 0 |
| 51031 | ENSMUSG00000113467 | 0 | NA | NA | NA | NA | NA | Gm46350 | predicted gene, 46350 [Source:MGI Symb | 0 | 0 | 0 | 0 | 0 | 0 |
| 51032 | ENSMUSG00000113468 | 0 | NA | NA | NA | NA | NA | Gm48165 | predicted gene, 48165 [Source:MGI Symb | 0 | 0 | 0 | 0 | 0 | 0 |
| 51033 | ENSMUSG00000113469 | 0 | NA | NA | NA | NA | NA | Gm7065 | predicted gene 7065 [Source:MGI Symbol | 0 | 0 | 0 | 0 | 0 | 0 |
| 51034 | ENSMUSG00000113470 | 0 | NA | NA | NA | NA | NA | 4930448F12 | RIKEN cDNA 4930448F12 gene [Source:M | 0 | 0 | 0 | 0 | 0 | 0 |
| 51035 | ENSMUSG00000113471 | 0 | NA | NA | NA | NA | NA | Gm48116 | predicted gene, 48116 [Source:MGI Symb | 0 | 0 | 0 | 0 | 0 | 0 |
| 51036 | ENSMUSG00000113472 | 5.042048721 | -0.105450963 | 1.008261493 | -0.104586919 | 0.916703595 | NA | Gm47602 | predicted gene, 47602 [Source:MGI Symb | 1.6062 | 6.1675 | 8.05 | 6.7478 | 5.4531 | 2.2277 |
| 51037 | ENSMUSG00000113473 | 1.623570295 | 0.33491731 | 1.852249416 | 0.180816529 | 0.856511589 | NA | Gm7045 | predicted gene 7045 [Source:MGI Symbol | 3.2124 | 0 | 1.0062 | 0 | 2.1812 | 3.3415 |
| 51038 | ENSMUSG00000113474 | 0 | NA | NA | NA | NA | NA | Gm48110 | predicted gene, 48110 [Source:MGI Symb | 0 | 0 | 0 | 0 | 0 | 0 |
| 51039 | ENSMUSG00000113475 | 0 | NA | NA | NA | NA | NA | Gm49366 | predicted gene, 49366 [Source:MGI Symb | 0 | 0 | 0 | 0 | 0 | 0 |
| 51040 | ENSMUSG00000113476 | 0 | NA | NA | NA | NA | NA | Gm48309 | predicted gene, 48309 [Source:MGI Symb | 0 | 0 | 0 | 0 | 0 | 0 |
| 51041 | ENSMUSG00000113477 | 0 | NA | NA | NA | NA | NA | Gm45965 | predicted gene, 45965 [Source:MGI Symb | 0 | 0 | 0 | 0 | 0 | 0 |
| 51042 | ENSMUSG00000113478 | 0 | NA | NA | NA | NA | NA | Gm18405 | predicted gene, 18405 [Source:MGI Symb | 0 | 0 | 0 | 0 | 0 | 0 |
| 51043 | ENSMUSG00000113479 | 0 | NA | NA | NA | NA | NA | Gm47796 | predicted gene, 47796 [Source:MGI Symb | 0 | 0 | 0 | 0 | 0 | 0 |
| 51044 | ENSMUSG00000113480 | 0 | NA | NA | NA | NA | NA | Gm47413 | predicted gene, 47413 [Source:MGI Symb | 0 | 0 | 0 | 0 | 0 | 0 |
| 51045 | ENSMUSG00000113481 | 0 | NA | NA | NA | NA | NA | Gm48388 | predicted gene, 48388 [Source:MGI Symb | 0 | 0 | 0 | 0 | 0 | 0 |
| 51046 | ENSMUSG00000113482 | 0 | NA | NA | NA | NA | NA | Gm48167 | predicted gene, 48167 [Source:MGI Symb | 0 | 0 | 0 | 0 | 0 | 0 |
| 51047 | ENSMUSG00000113483 | 0 | NA | NA | NA | NA | NA | Gm47548 | predicted gene, 47548 [Source:MGI Symb | 0 | 0 | 0 | 0 | 0 | 0 |
| 51048 | ENSMUSG00000113484 | 0 | NA | NA | NA | NA | NA | Gm47157 | predicted gene, 47157 [Source:MGI Symb | 0 | 0 | 0 | 0 | 0 | 0 |
| 51049 | ENSMUSG00000113485 | 0 | NA | NA | NA | NA | NA | Gm48000 | predicted gene, 48000 [Source:MGI Symb | 0 | 0 | 0 | 0 | 0 | 0 |
| 51050 | ENSMUSG00000113486 | 0.294513347 | 0.165143496 | 4.080472857 | 0.040471657 | 0.967717103 | NA | Gm48676 | predicted gene, 48676 [Source:MGI Symb | 0.8031 | 0 | 0 | 0.964 | 0 | 0 |
| 51051 | ENSMUSG00000113487 | 0 | NA | NA | NA | NA | NA | Gm47422 | predicted gene, 47422 [Source:MGI Symb | 0 | 0 | 0 | 0 | 0 | 0 |
| 51052 | ENSMUSG00000113488 | 0 | NA | NA | NA | NA | NA | Gm47819 | predicted gene, 47819 [Source:MGI Symb | 0 | 0 | 0 | 0 | 0 | 0 |
| 51053 | ENSMUSG00000113489 | 0 | NA | NA | NA | NA | NA | Ubl5b | ubiquitin-like 5B [Source:MGI Symbol;Acc | 0 | 0 | 0 | 0 | 0 | 0 |
| 51054 | ENSMUSG00000113490 | 0 | NA | NA | NA | NA | NA | Gm18620 | predicted gene, 18620 [Source:MGI Symb | 0 | 0 | 0 | 0 | 0 | 0 |
| 51055 | ENSMUSG00000113491 | 17.73120522 | 0.154918734 | 0.518033831 | 0.299051384 | 0.764900839 | 0.954980327 | Gm19221 | predicted gene, 19221 [Source:MGI Symb | 16.865 | 18.503 | 15.094 | 25.063 | 15.269 | 15.594 |
| 51056 | ENSMUSG00000113492 | 0 | NA | NA | NA | NA | NA | Gm47792 | predicted gene, 47792 [Source:MGI Symb | 0 | 0 | 0 | 0 | 0 | 0 |
| 51057 | ENSMUSG00000113493 | 0 | NA | NA | NA | NA | NA | Gm7441 | predicted gene 7441 [Source:MGI Symbol | 0 | 0 | 0 | 0 | 0 | 0 |
| 51058 | ENSMUSG00000113494 | 0 | NA | NA | NA | NA | NA | Gm48102 | predicted gene, 48102 [Source:MGI Symb | 0 | 0 | 0 | 0 | 0 | 0 |
| 51059 | ENSMUSG00000113495 | 0 | NA | NA | NA | NA | NA | Gm19792 | predicted gene, 19792 [Source:MGI Symb | 0 | 0 | 0 | 0 | 0 | 0 |
| 51060 | ENSMUSG00000113496 | 0 | NA | NA | NA | NA | NA | Gm6575 | predicted gene 6575 [Source:MGI Symbol | 0 | 0 | 0 | 0 | 0 | 0 |
| 51061 | ENSMUSG00000113497 | 1.914476948 | 0.154752154 | 1.583546511 | 0.097725045 | 0.922150628 | NA | Gm18030 | predicted gene, 18030 [Source:MGI Symb | 2.4093 | 1.0279 | 2.0125 | 3.8559 | 2.1812 | 0 |
| 51062 | ENSMUSG00000113498 | 0.267700746 | -1.497481649 | 4.057001436 | -0.369110456 | 0.712045396 | NA | Gm4036 | predicted gene 4036 [Source:MGI Symbol | 1.6062 | 0 | 0 | 0 | 0 | 0 |
| 51063 | ENSMUSG00000113499 | 0 | NA | NA | NA | NA | NA | Gm18500 | predicted gene, 18500 [Source:MGI Symb | 0 | 0 | 0 | 0 | 0 | 0 |
| 51064 | ENSMUSG00000113500 | 0 | NA | NA | NA | NA | NA | Gm47249 | predicted gene, 47249 [Source:MGI Symb | 0 | 0 | 0 | 0 | 0 | 0 |
| 51065 | ENSMUSG00000113501 | 0 | NA | NA | NA | NA | NA | Gm18940 | predicted gene, 18940 [Source:MGI Symb | 0 | 0 | 0 | 0 | 0 | 0 |
| 51066 | ENSMUSG00000113502 | 0 | NA | NA | NA | NA | NA | Gm47411 | predicted gene, 47411 [Source:MGI Symb | 0 | 0 | 0 | 0 | 0 | 0 |
| 51067 | ENSMUSG00000113503 | 0 | NA | NA | NA | NA | NA | Gm47490 | predicted gene, 47490 [Source:MGI Symb | 0 | 0 | 0 | 0 | 0 | 0 |
| 51068 | ENSMUSG00000113504 | 0 | NA | NA | NA | NA | NA | Selenok-ps5 | selenoprotein K, pseudogene 5 [Source:M | 0 | 0 | 0 | 0 | 0 | 0 |

| | | | | | | | | | | | | | | | |
|---|---|---|---|---|---|---|---|---|---|---|---|---|---|---|---|
| 51069 | ENSMUSG00000113505 | 0 | NA | NA | NA | NA | NA | Gm48593 | predicted gene, 48593 [Source:MGI Symb | 0 | 0 | 0 | 0 | 0 | 0 |
| 51070 | ENSMUSG00000113506 | 0 | NA | NA | NA | NA | NA | Gm49377 | predicted gene, 49377 [Source:MGI Symb | 0 | 0 | 0 | 0 | 0 | 0 |
| 51071 | ENSMUSG00000113507 | 0 | NA | NA | NA | NA | NA | Gm48613 | predicted gene, 48613 [Source:MGI Symb | 0 | 0 | 0 | 0 | 0 | 0 |
| 51072 | ENSMUSG00000113508 | 0 | NA | NA | NA | NA | NA | Gm9742 | predicted gene 9742 [Source:MGI Symbol | 0 | 0 | 0 | 0 | 0 | 0 |
| 51073 | ENSMUSG00000113509 | 0 | NA | NA | NA | NA | NA | Gm19009 | predicted gene, 19009 [Source:MGI Symb | 0 | 0 | 0 | 0 | 0 | 0 |
| 51074 | ENSMUSG00000113510 | 0 | NA | NA | NA | NA | NA | Gm7239 | predicted gene 7239 [Source:MGI Symbol | 0 | 0 | 0 | 0 | 0 | 0 |
| 51075 | ENSMUSG00000113511 | 0.133850373 | -0.796629176 | 4.080472857 | -0.195229623 | 0.84521319 | NA | Gm19046 | predicted gene, 19046 [Source:MGI Symb | 0.8031 | 0 | 0 | 0 | 0 | 0 |
| 51076 | ENSMUSG00000113512 | 0 | NA | NA | NA | NA | NA | Gm9063 | predicted gene 9063 [Source:MGI Symbol | 0 | 0 | 0 | 0 | 0 | 0 |
| 51077 | ENSMUSG00000113513 | 0 | NA | NA | NA | NA | NA | Gm19118 | predicted gene, 19118 [Source:MGI Symb | 0 | 0 | 0 | 0 | 0 | 0 |
| 51078 | ENSMUSG00000113514 | 0 | NA | NA | NA | NA | NA | Gm18960 | predicted gene, 18960 [Source:MGI Symb | 0 | 0 | 0 | 0 | 0 | 0 |
| 51079 | ENSMUSG00000113515 | 0.342432689 | 1.990653086 | 3.900281263 | 0.51038706 | 0.609780321 | NA | Gm29675 | predicted gene, 29675 [Source:MGI Symb | 0 | 0 | 0 | 0.964 | 1.0906 | 0 |
| 51080 | ENSMUSG00000113516 | 0.860262541 | -0.538502919 | 2.596000693 | -0.207435584 | 0.835669695 | NA | Gm47368 | predicted gene, 47368 [Source:MGI Symb | 0 | 3.0838 | 0 | 0.964 | 0 | 1.1138 |
| 51081 | ENSMUSG00000113517 | 10.79818695 | 0.057404874 | 0.664412368 | 0.086399466 | 0.931148871 | 0.988420305 | Gm47905 | predicted gene, 47905 [Source:MGI Symb | 6.4248 | 12.335 | 13.081 | 8.6758 | 10.906 | 13.366 |
| 51082 | ENSMUSG00000113518 | 0 | NA | NA | NA | NA | NA | Gm48304 | predicted gene, 48304 [Source:MGI Symb | 0 | 0 | 0 | 0 | 0 | 0 |
| 51083 | ENSMUSG00000113519 | 0.321325949 | 1.911575114 | 4.048267893 | 0.472195804 | 0.636787034 | NA | Gm47541 | predicted gene, 47541 [Source:MGI Symb | 0 | 0 | 0 | 1.928 | 0 | 0 |
| 51084 | ENSMUSG00000113520 | 0 | NA | NA | NA | NA | NA | Gm4257 | predicted gene 4257 [Source:MGI Symbol | 0 | 0 | 0 | 0 | 0 | 0 |
| 51085 | ENSMUSG00000113521 | 0 | NA | NA | NA | NA | NA | Gm29787 | predicted gene, 29787 [Source:MGI Symb | 0 | 0 | 0 | 0 | 0 | 0 |
| 51086 | ENSMUSG00000113522 | 0 | NA | NA | NA | NA | NA | Gm48479 | predicted gene, 48479 [Source:MGI Symb | 0 | 0 | 0 | 0 | 0 | 0 |
| 51087 | ENSMUSG00000113523 | 0 | NA | NA | NA | NA | NA | Gm48366 | predicted gene, 48366 [Source:MGI Symb | 0 | 0 | 0 | 0 | 0 | 0 |
| 51088 | ENSMUSG00000113524 | 0 | NA | NA | NA | NA | NA | Gm47847 | predicted gene, 47847 [Source:MGI Symb | 0 | 0 | 0 | 0 | 0 | 0 |
| 51089 | ENSMUSG00000113526 | 0 | NA | NA | NA | NA | NA | Gm47453 | predicted gene, 47453 [Source:MGI Symb | 0 | 0 | 0 | 0 | 0 | 0 |
| 51090 | ENSMUSG00000113527 | 2.47160548 | -0.030941667 | 1.314839158 | -0.023532664 | 0.981225384 | NA | Gm48079 | predicted gene, 48079 [Source:MGI Symb | 2.4093 | 3.0838 | 2.0125 | 2.8919 | 1.0906 | 3.3415 |
| 51091 | ENSMUSG00000113528 | 4.393198481 | -0.134891438 | 1.165003135 | -0.11578633 | 0.907821884 | NA | Gm32699 | predicted gene, 32699 [Source:MGI Symb | 9.6372 | 0 | 4.025 | 3.8559 | 3.2719 | 5.5692 |
| 51092 | ENSMUSG00000113529 | 5.849492721 | -0.915821835 | 0.979056138 | -0.935412996 | 0.349575502 | 0.790905688 | Gm47484 | predicted gene, 47484 [Source:MGI Symb | 4.8186 | 9.2513 | 9.0562 | 8.6758 | 2.1812 | 1.1138 |
| 51093 | ENSMUSG00000113530 | 0 | NA | NA | NA | NA | NA | Gm8672 | predicted gene 8672 [Source:MGI Symbol | 0 | 0 | 0 | 0 | 0 | 0 |
| 51094 | ENSMUSG00000113531 | 0 | NA | NA | NA | NA | NA | Gm2915 | predicted gene 2915 [Source:MGI Symbol | 0 | 0 | 0 | 0 | 0 | 0 |
| 51095 | ENSMUSG00000113532 | 0 | NA | NA | NA | NA | NA | Gm47393 | predicted gene, 47393 [Source:MGI Symb | 0 | 0 | 0 | 0 | 0 | 0 |
| 51096 | ENSMUSG00000113533 | 0 | NA | NA | NA | NA | NA | Gm46992 | predicted gene 46992 [Source:MGI Symbc | 0 | 0 | 0 | 0 | 0 | 0 |
| 51097 | ENSMUSG00000113534 | 0 | NA | NA | NA | NA | NA | Gm19163 | predicted gene, 19163 [Source:MGI Symb | 0 | 0 | 0 | 0 | 0 | 0 |
| 51098 | ENSMUSG00000113535 | 0 | NA | NA | NA | NA | NA | AU015791 | expressed sequence AU015791 [Source:M | 0 | 0 | 0 | 0 | 0 | 0 |
| 51099 | ENSMUSG00000113536 | 16.05553277 | -0.022990534 | 0.605165053 | -0.037990518 | 0.969695242 | 0.994674947 | Gm49327 | predicted gene, 49327 [Source:MGI Symb | 15.259 | 8.2233 | 25.156 | 20.244 | 18.541 | 8.9108 |
| 51100 | ENSMUSG00000113537 | 0 | NA | NA | NA | NA | NA | Gm48807 | predicted gene, 48807 [Source:MGI Symb | 0 | 0 | 0 | 0 | 0 | 0 |
| 51101 | ENSMUSG00000113538 | 0 | NA | NA | NA | NA | NA | Gm48665 | predicted gene, 48665 [Source:MGI Symb | 0 | 0 | 0 | 0 | 0 | 0 |
| 51102 | ENSMUSG00000113539 | 0 | NA | NA | NA | NA | NA | Gm47684 | predicted gene, 47684 [Source:MGI Symb | 0 | 0 | 0 | 0 | 0 | 0 |
| 51103 | ENSMUSG00000113540 | 0 | NA | NA | NA | NA | NA | Gm47206 | predicted gene, 47206 [Source:MGI Symb | 0 | 0 | 0 | 0 | 0 | 0 |
| 51104 | ENSMUSG00000113541 | 0 | NA | NA | NA | NA | NA | Gm48409 | predicted gene, 48409 [Source:MGI Symb | 0 | 0 | 0 | 0 | 0 | 0 |
| 51105 | ENSMUSG00000113542 | 0 | NA | NA | NA | NA | NA | Gm19154 | predicted gene, 19154 [Source:MGI Symb | 0 | 0 | 0 | 0 | 0 | 0 |
| 51106 | ENSMUSG00000113543 | 73.5631478 | 0.750703018 | 0.369874912 | 2.029613235 | 0.042395869 | 0.283632674 | Gm36264 | predicted gene, 36264 [Source:MGI Symb | 27.305 | 46.256 | 91.568 | 86.758 | 98.156 | 91.335 |
| 51107 | ENSMUSG00000113544 | 0 | NA | NA | NA | NA | NA | 4833405L11I | RIKEN cDNA 4833405L11 gene [Source:M( | 0 | 0 | 0 | 0 | 0 | 0 |
| 51108 | ENSMUSG00000113545 | 0 | NA | NA | NA | NA | NA | Gm48238 | predicted gene, 48238 [Source:MGI Symb | 0 | 0 | 0 | 0 | 0 | 0 |
| 51109 | ENSMUSG00000113546 | 0 | NA | NA | NA | NA | NA | Gm18441 | predicted gene, 18441 [Source:MGI Symb | 0 | 0 | 0 | 0 | 0 | 0 |
| 51110 | ENSMUSG00000113547 | 0 | NA | NA | NA | NA | NA | Gm46389 | predicted gene, 46389 [Source:MGI Symb | 0 | 0 | 0 | 0 | 0 | 0 |
| 51111 | ENSMUSG00000113548 | 0.986449812 | -2.084851164 | 2.483799425 | -0.83937984 | 0.401256193 | NA | Gm34047 | predicted gene, 34047 [Source:MGI Symb | 0.8031 | 0 | 4.025 | 0 | 1.0906 | 0 |
| 51112 | ENSMUSG00000113549 | 0 | NA | NA | NA | NA | NA | Gm7739 | predicted gene 7739 [Source:MGI Symbol | 0 | 0 | 0 | 0 | 0 | 0 |
| 51113 | ENSMUSG00000113550 | 0 | NA | NA | NA | NA | NA | Gm29687 | predicted gene, 29687 [Source:MGI Symb | 0 | 0 | 0 | 0 | 0 | 0 |
| 51114 | ENSMUSG00000113551 | 0 | NA | NA | NA | NA | NA | Gm7768 | predicted gene 7768 [Source:MGI Symbol | 0 | 0 | 0 | 0 | 0 | 0 |
| 51115 | ENSMUSG00000113552 | 0 | NA | NA | NA | NA | NA | Gm48046 | predicted gene, 48046 [Source:MGI Symb | 0 | 0 | 0 | 0 | 0 | 0 |
| 51116 | ENSMUSG00000113553 | 0 | NA | NA | NA | NA | NA | Gm40932 | predicted gene, 40932 [Source:MGI Symb | 0 | 0 | 0 | 0 | 0 | 0 |
| 51117 | ENSMUSG00000113554 | 0.133850373 | -0.796629176 | 4.080472857 | -0.195229623 | 0.84521319 | NA | Gm36758 | predicted gene, 36758 [Source:MGI Symb | 0.8031 | 0 | 0 | 0 | 0 | 0 |
| 51118 | ENSMUSG00000113555 | 16.13840732 | -0.289116263 | 0.540524906 | -0.534880558 | 0.592732453 | 0.912925376 | Gm10095 | predicted gene 10095 [Source:MGI Symbc | 13.653 | 18.503 | 21.131 | 11.568 | 15.269 | 16.708 |
| 51119 | ENSMUSG00000113556 | 0 | NA | NA | NA | NA | NA | Gm48317 | predicted gene, 48317 [Source:MGI Symb | 0 | 0 | 0 | 0 | 0 | 0 |
| 51120 | ENSMUSG00000113557 | 0 | NA | NA | NA | NA | NA | Gm47657 | predicted gene, 47657 [Source:MGI Symb | 0 | 0 | 0 | 0 | 0 | 0 |
| 51121 | ENSMUSG00000113558 | 0.692607243 | 3.004183657 | 2.957557172 | 1.015765202 | 0.309741211 | NA | Tmdd1 | transmembrane and death domain 1 [Sou | 0 | 0 | 0 | 1.928 | 0 | 2.2277 |
| 51122 | ENSMUSG00000113559 | 0 | NA | NA | NA | NA | NA | Gm40987 | predicted gene, 40987 [Source:MGI Symb | 0 | 0 | 0 | 0 | 0 | 0 |
| 51123 | ENSMUSG00000113560 | 0 | NA | NA | NA | NA | NA | Gm48519 | predicted gene, 48519 [Source:MGI Symb | 0 | 0 | 0 | 0 | 0 | 0 |
| 51124 | ENSMUSG00000113561 | 0 | NA | NA | NA | NA | NA | Gm19176 | predicted gene, 19176 [Source:MGI Symb | 0 | 0 | 0 | 0 | 0 | 0 |

| | | | | | | | | | | | | | | | |
|---|---|---|---|---|---|---|---|---|---|---|---|---|---|---|---|
| 51125 | ENSMUSG00000113562 | 0 | NA | NA | NA | NA | NA | Gm47619 | predicted gene, 47619 [Source:MGI Symb | 0 | 0 | 0 | 0 | 0 | 0 |
| 51126 | ENSMUSG00000113563 | 0 | NA | NA | NA | NA | NA | Gm2492 | predicted gene 2492 [Source:MGI Symbol | 0 | 0 | 0 | 0 | 0 | 0 |
| 51127 | ENSMUSG00000113564 | 0 | NA | NA | NA | NA | NA | Gm48560 | predicted gene, 48560 [Source:MGI Symb | 0 | 0 | 0 | 0 | 0 | 0 |
| 51128 | ENSMUSG00000113565 | 0 | NA | NA | NA | NA | NA | Cbx3-ps2 | chromobox 3, pseudogene 2 [Source:MGI | 0 | 0 | 0 | 0 | 0 | 0 |
| 51129 | ENSMUSG00000113566 | 0 | NA | NA | NA | NA | NA | Gm40977 | predicted gene, 40977 [Source:MGI Symb | 0 | 0 | 0 | 0 | 0 | 0 |
| 51130 | ENSMUSG00000113567 | 0 | NA | NA | NA | NA | NA | Gm32300 | predicted gene, 32300 [Source:MGI Symb | 0 | 0 | 0 | 0 | 0 | 0 |
| 51131 | ENSMUSG00000113568 | 0.371281295 | 2.09253362 | 4.031852165 | 0.519000582 | 0.603760335 | NA | Gm46369 | predicted gene, 46369 [Source:MGI Symb | 0 | 0 | 0 | 0 | 0 | 2.2277 |
| 51132 | ENSMUSG00000113569 | 0 | NA | NA | NA | NA | NA | Gm48035 | predicted gene, 48035 [Source:MGI Symb | 0 | 0 | 0 | 0 | 0 | 0 |
| 51133 | ENSMUSG00000113570 | 0.371281295 | 2.09253362 | 4.031852165 | 0.519000582 | 0.603760335 | NA | Gm5953 | predicted gene 5953 [Source:MGI Symbol | 0 | 0 | 0 | 0 | 0 | 2.2277 |
| 51134 | ENSMUSG00000113571 | 3.299304355 | -0.521213311 | 1.229125751 | -0.424052063 | 0.671527844 | NA | Gm48137 | predicted gene, 48137 [Source:MGI Symb | 5.6217 | 1.0279 | 5.0312 | 4.8199 | 2.1812 | 1.1138 |
| 51135 | ENSMUSG00000113572 | 0 | NA | NA | NA | NA | NA | Gm7959 | predicted gene 7959 [Source:MGI Symbol | 0 | 0 | 0 | 0 | 0 | 0 |
| 51136 | ENSMUSG00000113573 | 0 | NA | NA | NA | NA | NA | AU017674 | expressed sequence AU017674 [Source:M | 0 | 0 | 0 | 0 | 0 | 0 |
| 51137 | ENSMUSG00000113574 | 0 | NA | NA | NA | NA | NA | Gm48037 | predicted gene, 48037 [Source:MGI Symb | 0 | 0 | 0 | 0 | 0 | 0 |
| 51138 | ENSMUSG00000113575 | 0 | NA | NA | NA | NA | NA | A930023M0 | RIKEN cDNA A930023M06 gene [Source:M | 0 | 0 | 0 | 0 | 0 | 0 |
| 51139 | ENSMUSG00000113576 | 0 | NA | NA | NA | NA | NA | Gm47658 | predicted gene, 47658 [Source:MGI Symb | 0 | 0 | 0 | 0 | 0 | 0 |
| 51140 | ENSMUSG00000113577 | 0 | NA | NA | NA | NA | NA | Gm19238 | predicted gene, 19238 [Source:MGI Symb | 0 | 0 | 0 | 0 | 0 | 0 |
| 51141 | ENSMUSG00000113578 | 0 | NA | NA | NA | NA | NA | Gm48642 | predicted gene, 48642 [Source:MGI Symb | 0 | 0 | 0 | 0 | 0 | 0 |
| 51142 | ENSMUSG00000113579 | 0 | NA | NA | NA | NA | NA | Gm47384 | predicted gene, 47384 [Source:MGI Symb | 0 | 0 | 0 | 0 | 0 | 0 |
| 51143 | ENSMUSG00000113580 | 0 | NA | NA | NA | NA | NA | Gm48237 | predicted gene, 48237 [Source:MGI Symb | 0 | 0 | 0 | 0 | 0 | 0 |
| 51144 | ENSMUSG00000113581 | 33.44305013 | -1.983522695 | 0.532664706 | -3.72377346 | 0.000196267 | 0.006211831 | Dio3os | deiodinase, iodothyronine type III, opposi | 22.487 | 56.535 | 81.506 | 20.244 | 7.6343 | 12.252 |
| 51145 | ENSMUSG00000113582 | 0 | NA | NA | NA | NA | NA | Gm33929 | predicted gene, 33929 [Source:MGI Symb | 0 | 0 | 0 | 0 | 0 | 0 |
| 51146 | ENSMUSG00000113583 | 0 | NA | NA | NA | NA | NA | Eif1ad9 | eukaryotic translation initiation factor 1A | 0 | 0 | 0 | 0 | 0 | 0 |
| 51147 | ENSMUSG00000113584 | 0 | NA | NA | NA | NA | NA | Gm46355 | predicted gene, 46355 [Source:MGI Symb | 0 | 0 | 0 | 0 | 0 | 0 |
| 51148 | ENSMUSG00000113585 | 20.36780788 | -0.100553324 | 0.522369109 | -0.192494775 | 0.84735466 | 0.972803041 | Gm19605 | predicted gene, 19605 [Source:MGI Symb | 19.274 | 27.754 | 16.1 | 10.604 | 25.084 | 23.391 |
| 51149 | ENSMUSG00000113586 | 0 | NA | NA | NA | NA | NA | Gm48606 | predicted gene, 48606 [Source:MGI Symb | 0 | 0 | 0 | 0 | 0 | 0 |
| 51150 | ENSMUSG00000113587 | 2.577092024 | -1.908291806 | 1.638943239 | -1.164342828 | 0.244285107 | NA | Gm36287 | predicted gene, 36287 [Source:MGI Symb | 8.031 | 3.0838 | 1.0062 | 0 | 0 | 3.3415 |
| 51151 | ENSMUSG00000113588 | 0 | NA | NA | NA | NA | NA | Gm48328 | predicted gene, 48328 [Source:MGI Symb | 0 | 0 | 0 | 0 | 0 | 0 |
| 51152 | ENSMUSG00000113589 | 0.843900015 | -0.389267667 | 2.389264755 | -0.162923622 | 0.870578572 | NA | Gm48374 | predicted gene, 48374 [Source:MGI Symb | 0.8031 | 2.0558 | 0 | 0 | 1.0906 | 1.1138 |
| 51153 | ENSMUSG00000113590 | 0 | NA | NA | NA | NA | NA | Gm48378 | predicted gene, 48378 [Source:MGI Symb | 0 | 0 | 0 | 0 | 0 | 0 |
| 51154 | ENSMUSG00000113591 | 0 | NA | NA | NA | NA | NA | 4930447K03 | RIKEN cDNA 4930447K03 gene [Source:M | 0 | 0 | 0 | 0 | 0 | 0 |
| 51155 | ENSMUSG00000113592 | 0 | NA | NA | NA | NA | NA | Gm7093 | predicted gene 7093 [Source:MGI Symbol | 0 | 0 | 0 | 0 | 0 | 0 |
| 51156 | ENSMUSG00000113593 | 0 | NA | NA | NA | NA | NA | Zp4-ps | zona pellucida glycoprotein 4, pseudogen | 0 | 0 | 0 | 0 | 0 | 0 |
| 51157 | ENSMUSG00000113594 | 0 | NA | NA | NA | NA | NA | Gm4052 | predicted gene 4052 [Source:MGI Symbol | 0 | 0 | 0 | 0 | 0 | 0 |
| 51158 | ENSMUSG00000113595 | 0 | NA | NA | NA | NA | NA | Gm35514 | predicted gene, 35514 [Source:MGI Symb | 0 | 0 | 0 | 0 | 0 | 0 |
| 51159 | ENSMUSG00000113596 | 0.342639279 | -1.805976596 | 4.028547714 | -0.448294702 | 0.653940524 | NA | Gm48452 | predicted gene, 48452 [Source:MGI Symb | 0 | 2.0558 | 0 | 0 | 0 | 0 |
| 51160 | ENSMUSG00000113597 | 0 | NA | NA | NA | NA | NA | Gm48631 | predicted gene, 48631 [Source:MGI Symb | 0 | 0 | 0 | 0 | 0 | 0 |
| 51161 | ENSMUSG00000113598 | 0 | NA | NA | NA | NA | NA | Gm48287 | predicted gene, 48287 [Source:MGI Symb | 0 | 0 | 0 | 0 | 0 | 0 |
| 51162 | ENSMUSG00000113599 | 0 | NA | NA | NA | NA | NA | 9530014B07 | RIKEN cDNA 9530014B07 gene [Source:M | 0 | 0 | 0 | 0 | 0 | 0 |
| 51163 | ENSMUSG00000113600 | 0 | NA | NA | NA | NA | NA | Gm7868 | predicted gene 7868 [Source:MGI Symbol | 0 | 0 | 0 | 0 | 0 | 0 |
| 51164 | ENSMUSG00000113601 | 0 | NA | NA | NA | NA | NA | Gm48735 | predicted gene, 48735 [Source:MGI Symb | 0 | 0 | 0 | 0 | 0 | 0 |
| 51165 | ENSMUSG00000113602 | 0 | NA | NA | NA | NA | NA | Gm5653 | predicted gene 5653 [Source:MGI Symbol | 0 | 0 | 0 | 0 | 0 | 0 |
| 51166 | ENSMUSG00000113603 | 2.319563167 | -0.297778695 | 1.369135694 | -0.217493924 | 0.827823442 | NA | Gm47004 | predicted gene, 47004 [Source:MGI Symb | 1.6062 | 4.1117 | 2.0125 | 2.8919 | 2.1812 | 1.1138 |
| 51167 | ENSMUSG00000113604 | 0.160662974 | 1.126924175 | 4.080472857 | 0.276174898 | 0.782413736 | NA | Gm19786 | predicted gene, 19786 [Source:MGI Symb | 0 | 0 | 0 | 0.964 | 0 | 0 |
| 51168 | ENSMUSG00000113605 | 0.59968336 | -1.233555632 | 2.892386901 | -0.426483618 | 0.669755482 | NA | 4930425K24 | RIKEN cDNA 4930425K24 gene [Source:M | 1.6062 | 1.0279 | 0 | 0.964 | 0 | 0 |
| 51169 | ENSMUSG00000113606 | 0.353348078 | 0.165142701 | 3.895042568 | 0.042398176 | 0.966181282 | NA | Gm47441 | predicted gene, 47441 [Source:MGI Symb | 0 | 0 | 1.0062 | 0 | 0 | 1.1138 |
| 51170 | ENSMUSG00000113607 | 0 | NA | NA | NA | NA | NA | Gm47444 | predicted gene, 47444 [Source:MGI Symb | 0 | 0 | 0 | 0 | 0 | 0 |
| 51171 | ENSMUSG00000113608 | 0 | NA | NA | NA | NA | NA | Gm48715 | predicted gene, 48715 [Source:MGI Symb | 0 | 0 | 0 | 0 | 0 | 0 |
| 51172 | ENSMUSG00000113609 | 0 | NA | NA | NA | NA | NA | Gm47190 | predicted gene, 47190 [Source:MGI Symb | 0 | 0 | 0 | 0 | 0 | 0 |
| 51173 | ENSMUSG00000113610 | 0 | NA | NA | NA | NA | NA | Gm47132 | predicted gene, 47132 [Source:MGI Symb | 0 | 0 | 0 | 0 | 0 | 0 |
| 51174 | ENSMUSG00000113611 | 0 | NA | NA | NA | NA | NA | Gm47750 | predicted gene, 47750 [Source:MGI Symb | 0 | 0 | 0 | 0 | 0 | 0 |
| 51175 | ENSMUSG00000113612 | 0 | NA | NA | NA | NA | NA | Gm48315 | predicted gene, 48315 [Source:MGI Symb | 0 | 0 | 0 | 0 | 0 | 0 |
| 51176 | ENSMUSG00000113613 | 8.519311899 | -0.970249356 | 0.905491661 | -1.071516612 | 0.283937205 | 0.736651636 | Gm40893 | predicted gene, 40893 [Source:MGI Symb | 18.471 | 3.0838 | 12.075 | 0.964 | 8.7249 | 7.7969 |
| 51177 | ENSMUSG00000113614 | 0 | NA | NA | NA | NA | NA | Gm49362 | predicted gene, 49362 [Source:MGI Symb | 0 | 0 | 0 | 0 | 0 | 0 |
| 51178 | ENSMUSG00000113615 | 0 | NA | NA | NA | NA | NA | Gm47150 | predicted gene, 47150 [Source:MGI Symb | 0 | 0 | 0 | 0 | 0 | 0 |
| 51179 | ENSMUSG00000113616 | 9.958809544 | -0.470961678 | 0.689263396 | -0.68328259 | 0.494428293 | 0.877473722 | Gm45941 | predicted gene, 45941 [Source:MGI Symb | 10.44 | 9.2513 | 15.094 | 10.604 | 5.4531 | 8.9108 |
| 51180 | ENSMUSG00000113617 | 0 | NA | NA | NA | NA | NA | Gm8655 | predicted gene 8655 [Source:MGI Symbol | 0 | 0 | 0 | 0 | 0 | 0 |

| | | | | | | | | | | | | | | | |
|---|---|---|---|---|---|---|---|---|---|---|---|---|---|---|---|
| 51181 | ENSMUSG00000113618 | 0 | NA | NA | NA | NA | NA | Gm48166 | predicted gene, 48166 [Source:MGI Symb | 0 | 0 | 0 | 0 | 0 | 0 |
| 51182 | ENSMUSG00000113619 | 0 | NA | NA | NA | NA | NA | Gm47118 | predicted gene, 47118 [Source:MGI Symb | 0 | 0 | 0 | 0 | 0 | 0 |
| 51183 | ENSMUSG00000113620 | 0 | NA | NA | NA | NA | NA | 7420701I03R | RIKEN cDNA 7420701I03 gene [Source:MG | 0 | 0 | 0 | 0 | 0 | 0 |
| 51184 | ENSMUSG00000113621 | 0 | NA | NA | NA | NA | NA | 2900060N12 | RIKEN cDNA 2900060N12 gene [Source:M | 0 | 0 | 0 | 0 | 0 | 0 |
| 51185 | ENSMUSG00000113622 | 0 | NA | NA | NA | NA | NA | Gm49749 | predicted gene, 49749 [Source:MGI Symb | 0 | 0 | 0 | 0 | 0 | 0 |
| 51186 | ENSMUSG00000113623 | 0 | NA | NA | NA | NA | NA | Gm48691 | predicted gene, 48691 [Source:MGI Symb | 0 | 0 | 0 | 0 | 0 | 0 |
| 51187 | ENSMUSG00000113624 | 0 | NA | NA | NA | NA | NA | Gm18879 | predicted gene, 18879 [Source:MGI Symb | 0 | 0 | 0 | 0 | 0 | 0 |
| 51188 | ENSMUSG00000113625 | 0.849425915 | -0.514303322 | 2.604813552 | -0.19744343 | 0.843480546 | NA | Gm3379 | predicted gene 3379 [Source:MGI Symbol | 0 | 0 | 3.0187 | 0.964 | 0 | 1.1138 |
| 51189 | ENSMUSG00000113626 | 2.26304814 | -1.784063009 | 1.698096229 | -1.050625388 | 0.293430676 | NA | Gm7240 | predicted gene 7240 [Source:MGI Symbol | 6.4248 | 4.1117 | 0 | 1.928 | 0 | 1.1138 |
| 51190 | ENSMUSG00000113627 | 0 | NA | NA | NA | NA | NA | Gm32526 | predicted gene, 32526 [Source:MGI Symb | 0 | 0 | 0 | 0 | 0 | 0 |
| 51191 | ENSMUSG00000113628 | 0 | NA | NA | NA | NA | NA | 1700020D12 | RIKEN cDNA 1700020D12 gene [Source:M | 0 | 0 | 0 | 0 | 0 | 0 |
| 51192 | ENSMUSG00000113629 | 3.311764931 | -2.36217311 | 1.364208256 | -1.731534096 | 0.083356549 | NA | Gm48558 | predicted gene, 48558 [Source:MGI Symb | 6.4248 | 3.0838 | 7.0437 | 0 | 1.0906 | 2.2277 |
| 51193 | ENSMUSG00000113630 | 48.51290964 | -0.513176577 | 0.341911955 | -1.500902703 | 0.133380729 | 0.535299351 | 4930404H11 | RIKEN cDNA 4930404H11 gene [Source:M | 56.217 | 70.926 | 44.275 | 49.163 | 37.081 | 33.415 |
| 51194 | ENSMUSG00000113631 | 0.267700746 | -1.497481649 | 4.057001436 | -0.369110456 | 0.712045396 | NA | Gm48745 | predicted gene, 48745 [Source:MGI Symb | 1.6062 | 0 | 0 | 0 | 0 | 0 |
| 51195 | ENSMUSG00000113632 | 0 | NA | NA | NA | NA | NA | Gm47446 | predicted gene, 47446 [Source:MGI Symb | 0 | 0 | 0 | 0 | 0 | 0 |
| 51196 | ENSMUSG00000113633 | 0 | NA | NA | NA | NA | NA | Gm29689 | predicted gene, 29689 [Source:MGI Symb | 0 | 0 | 0 | 0 | 0 | 0 |
| 51197 | ENSMUSG00000113634 | 0.133850373 | -0.796629176 | 4.080472857 | -0.195229623 | 0.84521319 | NA | Gm36756 | predicted gene, 36756 [Source:MGI Symb | 0.8031 | 0 | 0 | 0 | 0 | 0 |
| 51198 | ENSMUSG00000113635 | 0 | NA | NA | NA | NA | NA | Gm48816 | predicted gene, 48816 [Source:MGI Symb | 0 | 0 | 0 | 0 | 0 | 0 |
| 51199 | ENSMUSG00000113636 | 0 | NA | NA | NA | NA | NA | Gm8616 | predicted gene 8616 [Source:MGI Symbol | 0 | 0 | 0 | 0 | 0 | 0 |
| 51200 | ENSMUSG00000113637 | 1.150698365 | 1.470584334 | 2.140991259 | 0.686870779 | 0.492164162 | NA | Gm7049 | predicted gene 7049 [Source:MGI Symbol | 0.8031 | 1.0279 | 0 | 2.8919 | 2.1812 | 0 |
| 51201 | ENSMUSG00000113638 | 0 | NA | NA | NA | NA | NA | Gm9289 | predicted gene 9289 [Source:MGI Symbol | 0 | 0 | 0 | 0 | 0 | 0 |
| 51202 | ENSMUSG00000113639 | 0 | NA | NA | NA | NA | NA | Gm20288 | predicted gene, 20288 [Source:MGI Symb | 0 | 0 | 0 | 0 | 0 | 0 |
| 51203 | ENSMUSG00000113640 | 0 | NA | NA | NA | NA | NA | Adat3 | adenosine deaminase, tRNA-specific 3 [Sc | 0 | 0 | 0 | 0 | 0 | 0 |
| 51204 | ENSMUSG00000113641 | 0 | NA | NA | NA | NA | NA | Gm48164 | predicted gene, 48164 [Source:MGI Symb | 0 | 0 | 0 | 0 | 0 | 0 |
| 51205 | ENSMUSG00000113642 | 6.935878729 | -0.27390739 | 0.817404819 | -0.335093926 | 0.737554235 | 0.948137969 | Gm49329 | predicted gene, 49329 [Source:MGI Symb | 5.6217 | 6.1675 | 11.069 | 7.7118 | 4.3625 | 6.6831 |
| 51206 | ENSMUSG00000113643 | 0 | NA | NA | NA | NA | NA | Gm40598 | predicted gene, 40598 [Source:MGI Symb | 0 | 0 | 0 | 0 | 0 | 0 |
| 51207 | ENSMUSG00000113644 | 0 | NA | NA | NA | NA | NA | 4930448C13 | RIKEN cDNA 4930448C13 gene [Source:M | 0 | 0 | 0 | 0 | 0 | 0 |
| 51208 | ENSMUSG00000113645 | 0 | NA | NA | NA | NA | NA | Gm47076 | predicted gene, 47076 [Source:MGI Symb | 0 | 0 | 0 | 0 | 0 | 0 |
| 51209 | ENSMUSG00000113646 | 0 | NA | NA | NA | NA | NA | Gm46313 | predicted gene, 46313 [Source:MGI Symb | 0 | 0 | 0 | 0 | 0 | 0 |
| 51210 | ENSMUSG00000113647 | 0.587191767 | -1.12394062 | 3.226482889 | -0.348348545 | 0.72757844 | NA | Gm47210 | predicted gene, 47210 [Source:MGI Symb | 2.4093 | 0 | 0 | 0 | 0 | 1.1138 |
| 51211 | ENSMUSG00000113648 | 0 | NA | NA | NA | NA | NA | Gm48806 | predicted gene, 48806 [Source:MGI Symb | 0 | 0 | 0 | 0 | 0 | 0 |
| 51212 | ENSMUSG00000113649 | 0 | NA | NA | NA | NA | NA | Gm47665 | predicted gene, 47665 [Source:MGI Symb | 0 | 0 | 0 | 0 | 0 | 0 |
| 51213 | ENSMUSG00000113650 | 5.44760553 | -0.056138441 | 1.08796937 | -0.051599285 | 0.958847989 | 0.992027194 | Gm47826 | predicted gene, 47826 [Source:MGI Symb | 5.6217 | 3.0838 | 8.05 | 11.568 | 4.3625 | 0 |
| 51214 | ENSMUSG00000113651 | 0.167707431 | -0.796629176 | 4.080472857 | -0.195229623 | 0.84521319 | NA | Gm4900 | predicted gene 4900 [Source:MGI Symbol | 0 | 0 | 1.0062 | 0 | 0 | 0 |
| 51215 | ENSMUSG00000113652 | 0 | NA | NA | NA | NA | NA | 4930518P08 | RIKEN cDNA 4930518P08 gene [Source:M | 0 | 0 | 0 | 0 | 0 | 0 |
| 51216 | ENSMUSG00000113653 | 0 | NA | NA | NA | NA | NA | Gm48220 | predicted gene, 48220 [Source:MGI Symb | 0 | 0 | 0 | 0 | 0 | 0 |
| 51217 | ENSMUSG00000113654 | 0 | NA | NA | NA | NA | NA | Gm30893 | predicted gene, 30893 [Source:MGI Symb | 0 | 0 | 0 | 0 | 0 | 0 |
| 51218 | ENSMUSG00000113655 | 0 | NA | NA | NA | NA | NA | Gm47454 | predicted gene, 47454 [Source:MGI Symb | 0 | 0 | 0 | 0 | 0 | 0 |
| 51219 | ENSMUSG00000113656 | 1.588117572 | 1.132025104 | 1.923403777 | 0.588553021 | 0.556161156 | NA | Gm47370 | predicted gene, 47370 [Source:MGI Symb | 0 | 0 | 3.0187 | 0.964 | 1.0906 | 4.4554 |
| 51220 | ENSMUSG00000113657 | 0 | NA | NA | NA | NA | NA | Gm17821 | predicted gene, 17821 [Source:MGI Symb | 0 | 0 | 0 | 0 | 0 | 0 |
| 51221 | ENSMUSG00000113658 | 0 | NA | NA | NA | NA | NA | Gm46993 | predicted gene 46993 [Source:MGI Symbo | 0 | 0 | 0 | 0 | 0 | 0 |
| 51222 | ENSMUSG00000113659 | 0 | NA | NA | NA | NA | NA | Gm49248 | predicted gene, 49248 [Source:MGI Symb | 0 | 0 | 0 | 0 | 0 | 0 |
| 51223 | ENSMUSG00000113660 | 31.06012874 | 1.039624398 | 0.418163613 | 2.486166577 | 0.012912751 | 0.128577932 | Gm5626 | predicted gene 5626 [Source:MGI Symbol | 24.093 | 16.447 | 20.125 | 29.883 | 51.259 | 44.554 |
| 51224 | ENSMUSG00000113661 | 0 | NA | NA | NA | NA | NA | Gm48169 | predicted gene, 48169 [Source:MGI Symb | 0 | 0 | 0 | 0 | 0 | 0 |
| 51225 | ENSMUSG00000113662 | 15.90730255 | 0.049912601 | 0.612213933 | 0.081528039 | 0.935022027 | 0.988420305 | Gm35558 | predicted gene, 35558 [Source:MGI Symb | 14.456 | 11.307 | 21.131 | 16.388 | 6.5437 | 25.618 |
| 51226 | ENSMUSG00000113663 | 0 | NA | NA | NA | NA | NA | Gm48504 | predicted gene, 48504 [Source:MGI Symb | 0 | 0 | 0 | 0 | 0 | 0 |
| 51227 | ENSMUSG00000113664 | 0 | NA | NA | NA | NA | NA | Gm47184 | predicted gene, 47184 [Source:MGI Symb | 0 | 0 | 0 | 0 | 0 | 0 |
| 51228 | ENSMUSG00000113665 | 0.48903338 | 0.95637552 | 3.456690196 | 0.276673773 | 0.782030612 | NA | Gm47603 | predicted gene, 47603 [Source:MGI Symb | 0 | 0 | 1.0062 | 1.928 | 0 | 0 |
| 51229 | ENSMUSG00000113666 | 0 | NA | NA | NA | NA | NA | Gm35725 | predicted gene, 35725 [Source:MGI Symb | 0 | 0 | 0 | 0 | 0 | 0 |
| 51230 | ENSMUSG00000113667 | 0 | NA | NA | NA | NA | NA | Gm47853 | predicted gene, 47853 [Source:MGI Symb | 0 | 0 | 0 | 0 | 0 | 0 |
| 51231 | ENSMUSG00000113668 | 0.133850373 | -0.796629176 | 4.080472857 | -0.195229623 | 0.84521319 | NA | Gm47451 | predicted gene, 47451 [Source:MGI Symb | 0.8031 | 0 | 0 | 0 | 0 | 0 |
| 51232 | ENSMUSG00000113669 | 3.541402382 | 0.032203479 | 1.138495772 | 0.028285989 | 0.977434055 | NA | Gm36723 | predicted gene, 36723 [Source:MGI Symb | 2.4093 | 2.0558 | 6.0375 | 1.928 | 4.3625 | 4.4554 |
| 51233 | ENSMUSG00000113670 | 0 | NA | NA | NA | NA | NA | Gm49686 | predicted gene, 49686 [Source:MGI Symb | 0 | 0 | 0 | 0 | 0 | 0 |
| 51234 | ENSMUSG00000113671 | 0 | NA | NA | NA | NA | NA | Gm2233 | predicted gene 2233 [Source:MGI Symbol | 0 | 0 | 0 | 0 | 0 | 0 |
| 51235 | ENSMUSG00000113672 | 0 | NA | NA | NA | NA | NA | Gm34165 | predicted gene, 34165 [Source:MGI Symb | 0 | 0 | 0 | 0 | 0 | 0 |
| 51236 | ENSMUSG00000113673 | 0.321325949 | 1.911575114 | 4.048267893 | 0.472195804 | 0.636787034 | NA | Gm47486 | predicted gene, 47486 [Source:MGI Symb | 0 | 0 | 0 | 1.928 | 0 | 0 |

| | | | | | | | | | | | | | | | |
|---|---|---|---|---|---|---|---|---|---|---|---|---|---|---|---|
| 51237 | ENSMUSG00000113674 | 0 | NA | NA | NA | NA | NA | Hspe1-ps1 | heat shock protein 1 (chaperonin 10), pse | 0 | 0 | 0 | 0 | 0 | 0 |
| 51238 | ENSMUSG00000113675 | 0 | NA | NA | NA | NA | NA | Gm18259 | predicted gene, 18259 [Source:MGI Symb | 0 | 0 | 0 | 0 | 0 | 0 |
| 51239 | ENSMUSG00000113676 | 0 | NA | NA | NA | NA | NA | Gm18621 | predicted gene, 18621 [Source:MGI Symb | 0 | 0 | 0 | 0 | 0 | 0 |
| 51240 | ENSMUSG00000113677 | 0 | NA | NA | NA | NA | NA | Gm48487 | predicted gene, 48487 [Source:MGI Symb | 0 | 0 | 0 | 0 | 0 | 0 |
| 51241 | ENSMUSG00000113678 | 0 | NA | NA | NA | NA | NA | Gm6068 | predicted gene 6068 [Source:MGI Symbol | 0 | 0 | 0 | 0 | 0 | 0 |
| 51242 | ENSMUSG00000113679 | 1.034549114 | 0.072428415 | 2.202180142 | 0.032889414 | 0.973762774 | NA | Gm10432 | predicted gene 10432 [Source:MGI Symbo | 0 | 2.0558 | 1.0062 | 0.964 | 2.1812 | 0 |
| 51243 | ENSMUSG00000113680 | 0 | NA | NA | NA | NA | NA | Gm47405 | predicted gene, 47405 [Source:MGI Symb | 0 | 0 | 0 | 0 | 0 | 0 |
| 51244 | ENSMUSG00000113681 | 0.703108924 | -2.875459482 | 2.96384054 | -0.970180225 | 0.331956666 | NA | Gm45930 | predicted gene, 45930 [Source:MGI Symb | 3.2124 | 0 | 1.0062 | 0 | 0 | 0 |
| 51245 | ENSMUSG00000113682 | 0 | NA | NA | NA | NA | NA | Gm48630 | predicted gene, 48630 [Source:MGI Symb | 0 | 0 | 0 | 0 | 0 | 0 |
| 51246 | ENSMUSG00000113683 | 0 | NA | NA | NA | NA | NA | Gm47123 | predicted gene, 47123 [Source:MGI Symb | 0 | 0 | 0 | 0 | 0 | 0 |
| 51247 | ENSMUSG00000113684 | 0 | NA | NA | NA | NA | NA | Gm48418 | predicted gene, 48418 [Source:MGI Symb | 0 | 0 | 0 | 0 | 0 | 0 |
| 51248 | ENSMUSG00000113685 | 0 | NA | NA | NA | NA | NA | Gm21038 | predicted gene, 21038 [Source:MGI Symb | 0 | 0 | 0 | 0 | 0 | 0 |
| 51249 | ENSMUSG00000113686 | 0 | NA | NA | NA | NA | NA | Gm19189 | predicted gene, 19189 [Source:MGI Symb | 0 | 0 | 0 | 0 | 0 | 0 |
| 51250 | ENSMUSG00000113687 | 0 | NA | NA | NA | NA | NA | Gm49703 | predicted gene, 49703 [Source:MGI Symb | 0 | 0 | 0 | 0 | 0 | 0 |
| 51251 | ENSMUSG00000113688 | 0 | NA | NA | NA | NA | NA | A930027P06 | RIKEN cDNA A930027P06 gene [Source:M | 0 | 0 | 0 | 0 | 0 | 0 |
| 51252 | ENSMUSG00000113689 | 0 | NA | NA | NA | NA | NA | Gm29676 | predicted gene, 29676 [Source:MGI Symb | 0 | 0 | 0 | 0 | 0 | 0 |
| 51253 | ENSMUSG00000113690 | 0.321325949 | 1.911575114 | 4.048267893 | 0.472195804 | 0.636787034 | NA | Gm36501 | predicted gene, 36501 [Source:MGI Symb | 0 | 0 | 0 | 1.928 | 0 | 0 |
| 51254 | ENSMUSG00000113691 | 0 | NA | NA | NA | NA | NA | Gm38699 | predicted gene, 38699 [Source:MGI Symb | 0 | 0 | 0 | 0 | 0 | 0 |
| 51255 | ENSMUSG00000113692 | 0 | NA | NA | NA | NA | NA | Gm47482 | predicted gene, 47482 [Source:MGI Symb | 0 | 0 | 0 | 0 | 0 | 0 |
| 51256 | ENSMUSG00000113693 | 0 | NA | NA | NA | NA | NA | Gm48340 | predicted gene, 48340 [Source:MGI Symb | 0 | 0 | 0 | 0 | 0 | 0 |
| 51257 | ENSMUSG00000113694 | 0 | NA | NA | NA | NA | NA | Gm48747 | predicted gene, 48747 [Source:MGI Symb | 0 | 0 | 0 | 0 | 0 | 0 |
| 51258 | ENSMUSG00000113695 | 0 | NA | NA | NA | NA | NA | 9330159N22 | RIKEN cDNA 9330159N22 gene [Source:M | 0 | 0 | 0 | 0 | 0 | 0 |
| 51259 | ENSMUSG00000113696 | 0 | NA | NA | NA | NA | NA | 1700015C15 | RIKEN cDNA 1700015C15 gene [Source:M | 0 | 0 | 0 | 0 | 0 | 0 |
| 51260 | ENSMUSG00000113697 | 0 | NA | NA | NA | NA | NA | Gm47890 | predicted gene, 47890 [Source:MGI Symb | 0 | 0 | 0 | 0 | 0 | 0 |
| 51261 | ENSMUSG00000113698 | 0 | NA | NA | NA | NA | NA | Gm47987 | predicted gene, 47987 [Source:MGI Symb | 0 | 0 | 0 | 0 | 0 | 0 |
| 51262 | ENSMUSG00000113699 | 0 | NA | NA | NA | NA | NA | Gm48823 | predicted gene, 48823 [Source:MGI Symb | 0 | 0 | 0 | 0 | 0 | 0 |
| 51263 | ENSMUSG00000113700 | 0 | NA | NA | NA | NA | NA | Gm47387 | predicted gene, 47387 [Source:MGI Symb | 0 | 0 | 0 | 0 | 0 | 0 |
| 51264 | ENSMUSG00000113701 | 0.160662974 | 1.126924175 | 4.080472857 | 0.276174898 | 0.782413736 | NA | B230303A05 | RIKEN cDNA B230303A05 gene [Source:M | 0 | 0 | 0 | 0.964 | 0 | 0 |
| 51265 | ENSMUSG00000113702 | 1.018883997 | 1.207572996 | 2.096893663 | 0.575886616 | 0.564691823 | NA | Gm35558 | predicted gene, 35558 [Source:MGI Symb | 0.8031 | 1.0279 | 0 | 0.964 | 1.0906 | 2.2277 |
| 51266 | ENSMUSG00000113703 | 0 | NA | NA | NA | NA | NA | Gm47788 | predicted gene, 47788 [Source:MGI Symb | 0 | 0 | 0 | 0 | 0 | 0 |
| 51267 | ENSMUSG00000113704 | 0 | NA | NA | NA | NA | NA | Gm27528 | predicted gene, 27528 [Source:MGI Symb | 0 | 0 | 0 | 0 | 0 | 0 |
| 51268 | ENSMUSG00000113705 | 0 | NA | NA | NA | NA | NA | Gm48411 | predicted gene, 48411 [Source:MGI Symb | 0 | 0 | 0 | 0 | 0 | 0 |
| 51269 | ENSMUSG00000113706 | 0 | NA | NA | NA | NA | NA | Gm47201 | predicted gene, 47201 [Source:MGI Symb | 0 | 0 | 0 | 0 | 0 | 0 |
| 51270 | ENSMUSG00000113707 | 0 | NA | NA | NA | NA | NA | Gm10457 | predicted gene 10457 [Source:MGI Symbo | 0 | 0 | 0 | 0 | 0 | 0 |
| 51271 | ENSMUSG00000113708 | 0 | NA | NA | NA | NA | NA | Gm47584 | predicted gene, 47584 [Source:MGI Symb | 0 | 0 | 0 | 0 | 0 | 0 |
| 51272 | ENSMUSG00000113709 | 0 | NA | NA | NA | NA | NA | Hmgb1-ps11 | high mobility group box 1, pseudogene 11 | 0 | 0 | 0 | 0 | 0 | 0 |
| 51273 | ENSMUSG00000113710 | 0 | NA | NA | NA | NA | NA | Gm38577 | predicted gene, 38577 [Source:MGI Symb | 0 | 0 | 0 | 0 | 0 | 0 |
| 51274 | ENSMUSG00000113711 | 0 | NA | NA | NA | NA | NA | Gm9661 | predicted gene 9661 [Source:MGI Symbol | 0 | 0 | 0 | 0 | 0 | 0 |
| 51275 | ENSMUSG00000113712 | 0 | NA | NA | NA | NA | NA | Gm38428 | predicted gene, 38428 [Source:MGI Symb | 0 | 0 | 0 | 0 | 0 | 0 |
| 51276 | ENSMUSG00000113713 | 0 | NA | NA | NA | NA | NA | Gm48240 | predicted gene, 48240 [Source:MGI Symb | 0 | 0 | 0 | 0 | 0 | 0 |
| 51277 | ENSMUSG00000113714 | 0 | NA | NA | NA | NA | NA | Gm48635 | predicted gene, 48635 [Source:MGI Symb | 0 | 0 | 0 | 0 | 0 | 0 |
| 51278 | ENSMUSG00000113715 | 0 | NA | NA | NA | NA | NA | Gm6562 | predicted gene 6562 [Source:MGI Symbol | 0 | 0 | 0 | 0 | 0 | 0 |
| 51279 | ENSMUSG00000113716 | 0 | NA | NA | NA | NA | NA | Gm47518 | predicted gene, 47518 [Source:MGI Symb | 0 | 0 | 0 | 0 | 0 | 0 |
| 51280 | ENSMUSG00000113717 | 0 | NA | NA | NA | NA | NA | Gm40658 | predicted gene, 40658 [Source:MGI Symb | 0 | 0 | 0 | 0 | 0 | 0 |
| 51281 | ENSMUSG00000113718 | 0 | NA | NA | NA | NA | NA | Gm2645 | predicted gene 2645 [Source:MGI Symbol | 0 | 0 | 0 | 0 | 0 | 0 |
| 51282 | ENSMUSG00000113719 | 0 | NA | NA | NA | NA | NA | Gm9305 | predicted gene 9305 [Source:MGI Symbol | 0 | 0 | 0 | 0 | 0 | 0 |
| 51283 | ENSMUSG00000113720 | 0 | NA | NA | NA | NA | NA | Gm46995 | predicted gene 46995 [Source:MGI Symbo | 0 | 0 | 0 | 0 | 0 | 0 |
| 51284 | ENSMUSG00000113721 | 0 | NA | NA | NA | NA | NA | Gm21051 | predicted gene, 21051 [Source:MGI Symb | 0 | 0 | 0 | 0 | 0 | 0 |
| 51285 | ENSMUSG00000113722 | 6.762492068 | -0.785550758 | 0.845401528 | -0.929204327 | 0.352783205 | 0.793259753 | Snhg10 | small nucleolar RNA host gene 10 [Source | 6.4248 | 6.1675 | 13.081 | 3.8559 | 4.3625 | 6.6831 |
| 51286 | ENSMUSG00000113723 | 0 | NA | NA | NA | NA | NA | Vmn2r-ps10 | vomeronasal 2, receptor, pseudogene 10( | 0 | 0 | 0 | 0 | 0 | 0 |
| 51287 | ENSMUSG00000113724 | 1.017133346 | -0.93167857 | 2.468499519 | -0.377427082 | 0.705856244 | NA | Gm48398 | predicted gene, 48398 [Source:MGI Symb | 0 | 0 | 4.025 | 0.964 | 0 | 1.1138 |
| 51288 | ENSMUSG00000113725 | 0.17131964 | -0.796629176 | 4.080472857 | -0.195229623 | 0.84521319 | NA | Gm48358 | predicted gene, 48358 [Source:MGI Symb | 0 | 1.0279 | 0 | 0 | 0 | 0 |
| 51289 | ENSMUSG00000113726 | 0 | NA | NA | NA | NA | NA | Gm36099 | predicted gene, 36099 [Source:MGI Symb | 0 | 0 | 0 | 0 | 0 | 0 |
| 51290 | ENSMUSG00000113727 | 0 | NA | NA | NA | NA | NA | Gm47781 | predicted gene, 47781 [Source:MGI Symb | 0 | 0 | 0 | 0 | 0 | 0 |
| 51291 | ENSMUSG00000113728 | 0 | NA | NA | NA | NA | NA | Gm7622 | predicted pseudogene 7622 [Source:MGI | 0 | 0 | 0 | 0 | 0 | 0 |
| 51292 | ENSMUSG00000113729 | 0 | NA | NA | NA | NA | NA | Gm48589 | predicted gene, 48589 [Source:MGI Symb | 0 | 0 | 0 | 0 | 0 | 0 |

| | | | | | | | | | | | | | | | |
|---|---|---|---|---|---|---|---|---|---|---|---|---|---|---|---|
| 51293 | ENSMUSG00000113730 | 0 | NA | NA | NA | NA | NA | Gm47805 | predicted gene, 47805 [Source:MGI Symb | 0 | 0 | 0 | 0 | 0 | 0 |
| 51294 | ENSMUSG00000113731 | 0 | NA | NA | NA | NA | NA | Gm48650 | predicted gene, 48650 [Source:MGI Symb | 0 | 0 | 0 | 0 | 0 | 0 |
| 51295 | ENSMUSG00000113732 | 0 | NA | NA | NA | NA | NA | Gm18115 | predicted gene, 18115 [Source:MGI Symb | 0 | 0 | 0 | 0 | 0 | 0 |
| 51296 | ENSMUSG00000113733 | 0 | NA | NA | NA | NA | NA | Gm34785 | predicted gene, 34785 [Source:MGI Symb | 0 | 0 | 0 | 0 | 0 | 0 |
| 51297 | ENSMUSG00000113734 | 0 | NA | NA | NA | NA | NA | Gm48539 | predicted gene, 48539 [Source:MGI Symb | 0 | 0 | 0 | 0 | 0 | 0 |
| 51298 | ENSMUSG00000113735 | 0 | NA | NA | NA | NA | NA | Gm48607 | predicted gene, 48607 [Source:MGI Symb | 0 | 0 | 0 | 0 | 0 | 0 |
| 51299 | ENSMUSG00000113736 | 0 | NA | NA | NA | NA | NA | Gm40879 | predicted gene, 40879 [Source:MGI Symb | 0 | 0 | 0 | 0 | 0 | 0 |
| 51300 | ENSMUSG00000113737 | 7.800938216 | -2.369590043 | 0.930552068 | -2.546434665 | 0.010882958 | 0.115319492 | BB123696 | expressed sequence BB123696 [Source:M | 9.6372 | 12.335 | 17.106 | 0 | 3.2719 | 4.4554 |
| 51301 | ENSMUSG00000113738 | 0 | NA | NA | NA | NA | NA | Gm40910 | predicted gene, 40910 [Source:MGI Symb | 0 | 0 | 0 | 0 | 0 | 0 |
| 51302 | ENSMUSG00000113739 | 0 | NA | NA | NA | NA | NA | Gm48584 | predicted gene, 48584 [Source:MGI Symb | 0 | 0 | 0 | 0 | 0 | 0 |
| 51303 | ENSMUSG00000113740 | 0 | NA | NA | NA | NA | NA | Gm8598 | predicted gene 8598 [Source:MGI Symbol | 0 | 0 | 0 | 0 | 0 | 0 |
| 51304 | ENSMUSG00000113741 | 1.615109605 | -2.916473367 | 2.125979557 | -1.371825688 | 0.170117709 | NA | Gm48236 | predicted gene, 48236 [Source:MGI Symb | 2.4093 | 6.1675 | 0 | 0 | 0 | 1.1138 |
| 51305 | ENSMUSG00000113742 | 0 | NA | NA | NA | NA | NA | Gm48262 | predicted gene, 48262 [Source:MGI Symb | 0 | 0 | 0 | 0 | 0 | 0 |
| 51306 | ENSMUSG00000113743 | 0 | NA | NA | NA | NA | NA | Gm8712 | predicted gene 8712 [Source:MGI Symbol | 0 | 0 | 0 | 0 | 0 | 0 |
| 51307 | ENSMUSG00000113744 | 0.133850373 | -0.796629176 | 4.080472857 | -0.195229623 | 0.84521319 | NA | Gm47925 | predicted gene, 47925 [Source:MGI Symb | 0.8031 | 0 | 0 | 0 | 0 | 0 |
| 51308 | ENSMUSG00000113745 | 0 | NA | NA | NA | NA | NA | Gm19340 | predicted gene, 19340 [Source:MGI Symb | 0 | 0 | 0 | 0 | 0 | 0 |
| 51309 | ENSMUSG00000113746 | 0.339027071 | -1.79174149 | 3.919585585 | -0.457125237 | 0.647581032 | NA | Gm48242 | predicted gene, 48242 [Source:MGI Symb | 0 | 1.0279 | 1.0062 | 0 | 0 | 0 |
| 51310 | ENSMUSG00000113747 | 0 | NA | NA | NA | NA | NA | 1700105G05 | RIKEN cDNA 1700105G05 gene [Source:M | 0 | 0 | 0 | 0 | 0 | 0 |
| 51311 | ENSMUSG00000113748 | 0 | NA | NA | NA | NA | NA | Gm18628 | predicted gene, 18628 [Source:MGI Symb | 0 | 0 | 0 | 0 | 0 | 0 |
| 51312 | ENSMUSG00000113749 | 0.167707431 | -0.796629176 | 4.080472857 | -0.195229623 | 0.84521319 | NA | Mrto4-ps1 | mRNA turnover 4, pseudogene 1 [Source: | 0 | 0 | 1.0062 | 0 | 0 | 0 |
| 51313 | ENSMUSG00000113750 | 0.319491021 | 0.16514334 | 4.044653141 | 0.040830038 | 0.967431393 | NA | Gm5628 | predicted gene 5628 [Source:MGI Symbol | 0.8031 | 0 | 0 | 0 | 0 | 1.1138 |
| 51314 | ENSMUSG00000113751 | 0 | NA | NA | NA | NA | NA | Gm9293 | predicted gene 9293 [Source:MGI Symbol | 0 | 0 | 0 | 0 | 0 | 0 |
| 51315 | ENSMUSG00000113752 | 1.17805339 | 0.245060278 | 2.511372982 | 0.0975802 | 0.922265648 | NA | Gm47817 | predicted gene, 47817 [Source:MGI Symb | 3.2124 | 0 | 0 | 3.8559 | 0 | 0 |
| 51316 | ENSMUSG00000113753 | 0.556921942 | 2.679043159 | 3.638243063 | 0.736356289 | 0.461513899 | NA | Gm5791 | predicted gene 5791 [Source:MGI Symbol | 0 | 0 | 0 | 0 | 0 | 3.3415 |
| 51317 | ENSMUSG00000113754 | 0 | NA | NA | NA | NA | NA | Gm47401 | predicted gene, 47401 [Source:MGI Symb | 0 | 0 | 0 | 0 | 0 | 0 |
| 51318 | ENSMUSG00000113755 | 0 | NA | NA | NA | NA | NA | Gm46391 | predicted gene, 46391 [Source:MGI Symb | 0 | 0 | 0 | 0 | 0 | 0 |
| 51319 | ENSMUSG00000113756 | 0 | NA | NA | NA | NA | NA | Gm18025 | predicted gene, 18025 [Source:MGI Symb | 0 | 0 | 0 | 0 | 0 | 0 |
| 51320 | ENSMUSG00000113757 | 0 | NA | NA | NA | NA | NA | Gm47507 | predicted gene, 47507 [Source:MGI Symb | 0 | 0 | 0 | 0 | 0 | 0 |
| 51321 | ENSMUSG00000113758 | 0 | NA | NA | NA | NA | NA | Gm47407 | predicted gene, 47407 [Source:MGI Symb | 0 | 0 | 0 | 0 | 0 | 0 |
| 51322 | ENSMUSG00000113759 | 0 | NA | NA | NA | NA | NA | Gm47695 | predicted gene, 47695 [Source:MGI Symb | 0 | 0 | 0 | 0 | 0 | 0 |
| 51323 | ENSMUSG00000113760 | 0 | NA | NA | NA | NA | NA | Gm48222 | predicted gene, 48222 [Source:MGI Symb | 0 | 0 | 0 | 0 | 0 | 0 |
| 51324 | ENSMUSG00000113761 | 0 | NA | NA | NA | NA | NA | Gm4263 | predicted gene 4263 [Source:MGI Symbol | 0 | 0 | 0 | 0 | 0 | 0 |
| 51325 | ENSMUSG00000113762 | 0 | NA | NA | NA | NA | NA | Gm47386 | predicted gene, 47386 [Source:MGI Symb | 0 | 0 | 0 | 0 | 0 | 0 |
| 51326 | ENSMUSG00000113763 | 0 | NA | NA | NA | NA | NA | Gm49287 | predicted gene, 49287 [Source:MGI Symb | 0 | 0 | 0 | 0 | 0 | 0 |
| 51327 | ENSMUSG00000113764 | 0 | NA | NA | NA | NA | NA | Gm48617 | predicted gene, 48617 [Source:MGI Symb | 0 | 0 | 0 | 0 | 0 | 0 |
| 51328 | ENSMUSG00000113765 | 0 | NA | NA | NA | NA | NA | Gm47549 | predicted gene, 47549 [Source:MGI Symb | 0 | 0 | 0 | 0 | 0 | 0 |
| 51329 | ENSMUSG00000113766 | 0.321325949 | 1.911575114 | 4.048267893 | 0.472195804 | 0.636787034 | NA | Gm47374 | predicted gene, 47374 [Source:MGI Symb | 0 | 0 | 0 | 1.928 | 0 | 0 |
| 51330 | ENSMUSG00000113767 | 0.342432689 | 1.990653086 | 3.900281263 | 0.51038706 | 0.609780321 | NA | Gm48873 | predicted gene, 48873 [Source:MGI Symb | 0 | 0 | 0 | 0.964 | 1.0906 | 0 |
| 51331 | ENSMUSG00000113768 | 0 | NA | NA | NA | NA | NA | Gm9237 | predicted pseudogene 9237 [Source:MGI | 0 | 0 | 0 | 0 | 0 | 0 |
| 51332 | ENSMUSG00000113769 | 149.7310217 | 0.155375103 | 0.21429881 | 0.725039502 | 0.468427797 | 0.865461183 | 5033406O09 | RIKEN cDNA 5033406O09 gene [Source:M | 175.88 | 128.49 | 119.74 | 150.38 | 167.96 | 155.94 |
| 51333 | ENSMUSG00000113770 | 0 | NA | NA | NA | NA | NA | 1700019C18 | RIKEN cDNA 1700019C18 gene [Source:M | 0 | 0 | 0 | 0 | 0 | 0 |
| 51334 | ENSMUSG00000113771 | 0 | NA | NA | NA | NA | NA | Gm36529 | predicted gene, 36529 [Source:MGI Symb | 0 | 0 | 0 | 0 | 0 | 0 |
| 51335 | ENSMUSG00000113772 | 0 | NA | NA | NA | NA | NA | Gm47448 | predicted gene, 47448 [Source:MGI Symb | 0 | 0 | 0 | 0 | 0 | 0 |
| 51336 | ENSMUSG00000113773 | 0 | NA | NA | NA | NA | NA | Gm8278 | predicted gene 8278 [Source:MGI Symbol | 0 | 0 | 0 | 0 | 0 | 0 |
| 51337 | ENSMUSG00000113774 | 1.712571112 | 0.83037452 | 1.796818206 | 0.462136079 | 0.643983739 | NA | Gm36236 | predicted gene, 36236 [Source:MGI Symb | 1.6062 | 2.0558 | 0 | 0 | 3.2719 | 3.3415 |
| 51338 | ENSMUSG00000113775 | 3.003628513 | 0.178320171 | 1.491234223 | 0.119578916 | 0.904816723 | NA | Gm48027 | predicted gene, 48027 [Source:MGI Symb | 2.4093 | 0 | 6.0375 | 2.8919 | 0 | 6.6831 |
| 51339 | ENSMUSG00000113776 | 0.626495961 | 0.101295816 | 2.879413718 | 0.03517932 | 0.971936753 | NA | Gm19327 | predicted gene, 19327 [Source:MGI Symb | 0.8031 | 1.0279 | 0 | 1.928 | 0 | 0 |
| 51340 | ENSMUSG00000113777 | 0 | NA | NA | NA | NA | NA | Gm48060 | predicted gene, 48060 [Source:MGI Symb | 0 | 0 | 0 | 0 | 0 | 0 |
| 51341 | ENSMUSG00000113778 | 0 | NA | NA | NA | NA | NA | Gm40847 | predicted gene, 40847 [Source:MGI Symb | 0 | 0 | 0 | 0 | 0 | 0 |
| 51342 | ENSMUSG00000113779 | 0.160662974 | 1.126924175 | 4.080472857 | 0.276174898 | 0.782413736 | NA | Gm33785 | predicted gene, 33785 [Source:MGI Symb | 0 | 0 | 0 | 0.964 | 0 | 0 |
| 51343 | ENSMUSG00000113780 | 0.335414862 | -1.779047242 | 4.030804179 | -0.441362855 | 0.658950329 | NA | Gm33195 | predicted gene, 33195 [Source:MGI Symb | 0 | 0 | 2.0125 | 0 | 0 | 0 |
| 51344 | ENSMUSG00000113781 | 0.562214094 | -1.122836648 | 3.276281221 | -0.34271681 | 0.731811515 | NA | Gm35558 | predicted gene, 35558 [Source:MGI Symb | 2.4093 | 0 | 0 | 0.964 | 0 | 0 |
| 51345 | ENSMUSG00000113782 | 0 | NA | NA | NA | NA | NA | Gm48009 | predicted gene, 48009 [Source:MGI Symb | 0 | 0 | 0 | 0 | 0 | 0 |
| 51346 | ENSMUSG00000113783 | 0 | NA | NA | NA | NA | NA | Gm18617 | predicted gene, 18617 [Source:MGI Symb | 0 | 0 | 0 | 0 | 0 | 0 |
| 51347 | ENSMUSG00000113784 | 0 | NA | NA | NA | NA | NA | Gm46400 | predicted gene, 46400 [Source:MGI Symb | 0 | 0 | 0 | 0 | 0 | 0 |
| 51348 | ENSMUSG00000113785 | 0 | NA | NA | NA | NA | NA | Gm47975 | predicted gene, 47975 [Source:MGI Symb | 0 | 0 | 0 | 0 | 0 | 0 |

| | | | | | | | | | | | | | | | |
|---|---|---|---|---|---|---|---|---|---|---|---|---|---|---|---|
| 51349 | ENSMUSG00000113786 | 0 | NA | NA | NA | NA | NA | Gm49384 | predicted gene, 49384 [Source:MGI Symb | 0 | 0 | 0 | 0 | 0 | 0 |
| 51350 | ENSMUSG00000113787 | 0 | NA | NA | NA | NA | NA | Gm47807 | predicted gene, 47807 [Source:MGI Symb | 0 | 0 | 0 | 0 | 0 | 0 |
| 51351 | ENSMUSG00000113788 | 2.595155022 | 0.237648194 | 1.562504397 | 0.152094416 | 0.879112468 | NA | Gm47904 | predicted gene, 47904 [Source:MGI Symb | 4.0155 | 3.0838 | 0 | 1.928 | 6.5437 | 0 |
| 51352 | ENSMUSG00000113789 | 0.315620088 | 0.165143421 | 4.063300985 | 0.040642675 | 0.967580763 | NA | Gm47322 | predicted gene, 47322 [Source:MGI Symb | 0.8031 | 0 | 0 | 0 | 1.0906 | 0 |
| 51353 | ENSMUSG00000113790 | 1.472354105 | -3.941665183 | 2.621323271 | -1.503692897 | 0.132660459 | NA | Gm17872 | predicted gene, 17872 [Source:MGI Symb | 8.8341 | 0 | 0 | 0 | 0 | 0 |
| 51354 | ENSMUSG00000113791 | 0 | NA | NA | NA | NA | NA | Gm47216 | predicted gene, 47216 [Source:MGI Symb | 0 | 0 | 0 | 0 | 0 | 0 |
| 51355 | ENSMUSG00000113792 | 0.842381458 | 0.63517692 | 2.579321552 | 0.246257362 | 0.805483014 | NA | Gm10933 | predicted gene 10933 [Source:MGI Symbo | 0 | 0 | 2.0125 | 1.928 | 0 | 1.1138 |
| 51356 | ENSMUSG00000113793 | 0 | NA | NA | NA | NA | NA | Gm48553 | predicted gene, 48553 [Source:MGI Symb | 0 | 0 | 0 | 0 | 0 | 0 |
| 51357 | ENSMUSG00000113794 | 0 | NA | NA | NA | NA | NA | Gm36607 | predicted gene, 36607 [Source:MGI Symb | 0 | 0 | 0 | 0 | 0 | 0 |
| 51358 | ENSMUSG00000113795 | 0 | NA | NA | NA | NA | NA | Gm48119 | predicted gene, 48119 [Source:MGI Symb | 0 | 0 | 0 | 0 | 0 | 0 |
| 51359 | ENSMUSG00000113796 | 0 | NA | NA | NA | NA | NA | Gm36426 | predicted gene, 36426 [Source:MGI Symb | 0 | 0 | 0 | 0 | 0 | 0 |
| 51360 | ENSMUSG00000113797 | 0 | NA | NA | NA | NA | NA | Gm48043 | predicted gene, 48043 [Source:MGI Symb | 0 | 0 | 0 | 0 | 0 | 0 |
| 51361 | ENSMUSG00000113798 | 0 | NA | NA | NA | NA | NA | Gm47385 | predicted gene, 47385 [Source:MGI Symb | 0 | 0 | 0 | 0 | 0 | 0 |
| 51362 | ENSMUSG00000113799 | 0 | NA | NA | NA | NA | NA | Gm47487 | predicted gene, 47487 [Source:MGI Symb | 0 | 0 | 0 | 0 | 0 | 0 |
| 51363 | ENSMUSG00000113800 | 63.40847542 | 0.44590235 | 0.297853432 | 1.497052919 | 0.134379489 | 0.537569131 | 2210039B01 | RIKEN cDNA 2210039B01 gene [Source:M | 59.43 | 49.34 | 52.325 | 92.542 | 59.984 | 66.831 |
| 51364 | ENSMUSG00000113801 | 0 | NA | NA | NA | NA | NA | Gm18618 | predicted gene, 18618 [Source:MGI Symb | 0 | 0 | 0 | 0 | 0 | 0 |
| 51365 | ENSMUSG00000113802 | 0 | NA | NA | NA | NA | NA | 4933416O17 | RIKEN cDNA 4933416O17 gene [Source:M | 0 | 0 | 0 | 0 | 0 | 0 |
| 51366 | ENSMUSG00000113803 | 0 | NA | NA | NA | NA | NA | Gm18615 | predicted gene, 18615 [Source:MGI Symb | 0 | 0 | 0 | 0 | 0 | 0 |
| 51367 | ENSMUSG00000113804 | 0.339027071 | -1.79174149 | 3.919585585 | -0.457125237 | 0.647581032 | NA | Gm46424 | predicted gene, 46424 [Source:MGI Symb | 0 | 1.0279 | 1.0062 | 0 | 0 | 0 |
| 51368 | ENSMUSG00000113805 | 0 | NA | NA | NA | NA | NA | Eif1ad12 | eukaryotic translation initiation factor 1A | 0 | 0 | 0 | 0 | 0 | 0 |
| 51369 | ENSMUSG00000113806 | 0 | NA | NA | NA | NA | NA | Gm47688 | predicted gene, 47688 [Source:MGI Symb | 0 | 0 | 0 | 0 | 0 | 0 |
| 51370 | ENSMUSG00000113807 | 0 | NA | NA | NA | NA | NA | Gm6888 | predicted gene 6888 [Source:MGI Symbol | 0 | 0 | 0 | 0 | 0 | 0 |
| 51371 | ENSMUSG00000113808 | 0 | NA | NA | NA | NA | NA | Gm47988 | predicted gene, 47988 [Source:MGI Symb | 0 | 0 | 0 | 0 | 0 | 0 |
| 51372 | ENSMUSG00000113809 | 0 | NA | NA | NA | NA | NA | Gm48472 | predicted gene, 48472 [Source:MGI Symb | 0 | 0 | 0 | 0 | 0 | 0 |
| 51373 | ENSMUSG00000113810 | 0 | NA | NA | NA | NA | NA | Gm48259 | predicted gene, 48259 [Source:MGI Symb | 0 | 0 | 0 | 0 | 0 | 0 |
| 51374 | ENSMUSG00000113811 | 2.25705858 | -1.082420065 | 1.516326234 | -0.713843789 | 0.475323779 | NA | Gm47882 | predicted gene, 47882 [Source:MGI Symb | 4.0155 | 4.1117 | 1.0062 | 0 | 2.1812 | 2.2277 |
| 51375 | ENSMUSG00000113812 | 0 | NA | NA | NA | NA | NA | Gm48699 | predicted gene, 48699 [Source:MGI Symb | 0 | 0 | 0 | 0 | 0 | 0 |
| 51376 | ENSMUSG00000113813 | 0 | NA | NA | NA | NA | NA | Gm29968 | predicted gene, 29968 [Source:MGI Symb | 0 | 0 | 0 | 0 | 0 | 0 |
| 51377 | ENSMUSG00000113814 | 0 | NA | NA | NA | NA | NA | Gm9255 | predicted pseudogene 9255 [Source:MGI | 0 | 0 | 0 | 0 | 0 | 0 |
| 51378 | ENSMUSG00000113815 | 0 | NA | NA | NA | NA | NA | Gm48149 | predicted gene, 48149 [Source:MGI Symb | 0 | 0 | 0 | 0 | 0 | 0 |
| 51379 | ENSMUSG00000113816 | 10.28094782 | -0.36475443 | 0.748126228 | -0.487557335 | 0.625863422 | 0.919638611 | Gm47003 | predicted gene, 47003 [Source:MGI Symb | 19.274 | 11.307 | 4.025 | 10.604 | 10.906 | 5.5692 |
| 51380 | ENSMUSG00000113817 | 0 | NA | NA | NA | NA | NA | Gm21056 | predicted gene, 21056 [Source:MGI Symb | 0 | 0 | 0 | 0 | 0 | 0 |
| 51381 | ENSMUSG00000113818 | 0 | NA | NA | NA | NA | NA | Gm7119 | predicted gene 7119 [Source:MGI Symbol | 0 | 0 | 0 | 0 | 0 | 0 |
| 51382 | ENSMUSG00000113819 | 0 | NA | NA | NA | NA | NA | Gm48017 | predicted gene, 48017 [Source:MGI Symb | 0 | 0 | 0 | 0 | 0 | 0 |
| 51383 | ENSMUSG00000113820 | 0 | NA | NA | NA | NA | NA | Gm4012 | predicted gene 4012 [Source:MGI Symbol | 0 | 0 | 0 | 0 | 0 | 0 |
| 51384 | ENSMUSG00000113821 | 0 | NA | NA | NA | NA | NA | Gm48234 | predicted gene, 48234 [Source:MGI Symb | 0 | 0 | 0 | 0 | 0 | 0 |
| 51385 | ENSMUSG00000113822 | 0 | NA | NA | NA | NA | NA | Gm47151 | predicted gene, 47151 [Source:MGI Symb | 0 | 0 | 0 | 0 | 0 | 0 |
| 51386 | ENSMUSG00000113823 | 0 | NA | NA | NA | NA | NA | Gm270 | predicted gene 270 [Source:MGI Symbol;/ | 0 | 0 | 0 | 0 | 0 | 0 |
| 51387 | ENSMUSG00000113824 | 0 | NA | NA | NA | NA | NA | Gm47420 | predicted gene, 47420 [Source:MGI Symb | 0 | 0 | 0 | 0 | 0 | 0 |
| 51388 | ENSMUSG00000113825 | 0 | NA | NA | NA | NA | NA | 1700024I08F | RIKEN cDNA 1700024I08 gene [Source:MC | 0 | 0 | 0 | 0 | 0 | 0 |
| 51389 | ENSMUSG00000113826 | 0 | NA | NA | NA | NA | NA | Gm48577 | predicted gene, 48577 [Source:MGI Symb | 0 | 0 | 0 | 0 | 0 | 0 |
| 51390 | ENSMUSG00000113827 | 0 | NA | NA | NA | NA | NA | Gm47733 | predicted gene, 47733 [Source:MGI Symb | 0 | 0 | 0 | 0 | 0 | 0 |
| 51391 | ENSMUSG00000113828 | 0 | NA | NA | NA | NA | NA | Gm48252 | predicted gene, 48252 [Source:MGI Symb | 0 | 0 | 0 | 0 | 0 | 0 |
| 51392 | ENSMUSG00000113829 | 0 | NA | NA | NA | NA | NA | Gm18619 | predicted gene, 18619 [Source:MGI Symb | 0 | 0 | 0 | 0 | 0 | 0 |
| 51393 | ENSMUSG00000113830 | 0 | NA | NA | NA | NA | NA | 1700127F24 | RIKEN cDNA 1700127F24 gene [Source:M( | 0 | 0 | 0 | 0 | 0 | 0 |
| 51394 | ENSMUSG00000113831 | 28.2117968 | 1.035820094 | 0.468881594 | 2.209129355 | 0.027165645 | 0.21354341 | Gm49602 | predicted gene, 49602 [Source:MGI Symb | 20.881 | 11.307 | 23.144 | 37.595 | 23.994 | 52.351 |
| 51395 | ENSMUSG00000113832 | 0 | NA | NA | NA | NA | NA | Gm40392 | predicted gene, 40392 [Source:MGI Symb | 0 | 0 | 0 | 0 | 0 | 0 |
| 51396 | ENSMUSG00000113833 | 0 | NA | NA | NA | NA | NA | Gm48264 | predicted gene, 48264 [Source:MGI Symb | 0 | 0 | 0 | 0 | 0 | 0 |
| 51397 | ENSMUSG00000113834 | 0 | NA | NA | NA | NA | NA | Gm34022 | predicted gene, 34022 [Source:MGI Symb | 0 | 0 | 0 | 0 | 0 | 0 |
| 51398 | ENSMUSG00000113835 | 0 | NA | NA | NA | NA | NA | Gm19554 | predicted gene, 19554 [Source:MGI Symb | 0 | 0 | 0 | 0 | 0 | 0 |
| 51399 | ENSMUSG00000113836 | 5.698493151 | 0.530941497 | 0.888124312 | 0.597823401 | 0.549957775 | 0.895703112 | Gm3325 | predicted gene 3325 [Source:MGI Symbol | 4.8186 | 3.0838 | 6.0375 | 4.8199 | 7.6343 | 7.7969 |
| 51400 | ENSMUSG00000113837 | 0 | NA | NA | NA | NA | NA | Gm47902 | predicted gene, 47902 [Source:MGI Symb | 0 | 0 | 0 | 0 | 0 | 0 |
| 51401 | ENSMUSG00000113838 | 0 | NA | NA | NA | NA | NA | Gm46994 | predicted gene 46994 [Source:MGI Symbo | 0 | 0 | 0 | 0 | 0 | 0 |
| 51402 | ENSMUSG00000113839 | 0 | NA | NA | NA | NA | NA | NA | NA | 0 | 0 | 0 | 0 | 0 | 0 |
| 51403 | ENSMUSG00000113840 | 0 | NA | NA | NA | NA | NA | Gm18955 | predicted gene, 18955 [Source:MGI Symb | 0 | 0 | 0 | 0 | 0 | 0 |
| 51404 | ENSMUSG00000113841 | 4.312353249 | -0.755398375 | 1.0286115 | -0.734386476 | 0.462713227 | NA | Gm48701 | predicted gene, 48701 [Source:MGI Symb | 4.0155 | 7.1954 | 5.0312 | 1.928 | 4.3625 | 3.3415 |

| | | | | | | | | | | | | | | | |
|---|---|---|---|---|---|---|---|---|---|---|---|---|---|---|---|
| 51405 | ENSMUSG00000113842 | 0.729921525 | -1.61423457 | 2.706831497 | -0.596355766 | 0.550937584 | NA | Gm48138 | predicted gene, 48138 [Source:MGI Symb | 2.4093 | 0 | 1.0062 | 0.964 | 0 | 0 |
| 51406 | ENSMUSG00000113843 | 0 | NA | NA | NA | NA | NA | Gm6129 | predicted gene 6129 [Source:MGI Symbol | 0 | 0 | 0 | 0 | 0 | 0 |
| 51407 | ENSMUSG00000113844 | 0 | NA | NA | NA | NA | NA | Gm47943 | predicted gene, 47943 [Source:MGI Symb | 0 | 0 | 0 | 0 | 0 | 0 |
| 51408 | ENSMUSG00000113845 | 0 | NA | NA | NA | NA | NA | Gm39473 | predicted gene, 39473 [Source:MGI Symb | 0 | 0 | 0 | 0 | 0 | 0 |
| 51409 | ENSMUSG00000113846 | 0 | NA | NA | NA | NA | NA | Gm6217 | predicted gene 6217 [Source:MGI Symbol | 0 | 0 | 0 | 0 | 0 | 0 |
| 51410 | ENSMUSG00000113847 | 0 | NA | NA | NA | NA | NA | Gm48799 | predicted gene, 48799 [Source:MGI Symb | 0 | 0 | 0 | 0 | 0 | 0 |
| 51411 | ENSMUSG00000113848 | 0 | NA | NA | NA | NA | NA | Gm16419 | predicted gene 16419 [Source:MGI Symbo | 0 | 0 | 0 | 0 | 0 | 0 |
| 51412 | ENSMUSG00000113849 | 0 | NA | NA | NA | NA | NA | Gm47346 | predicted gene, 47346 [Source:MGI Symb | 0 | 0 | 0 | 0 | 0 | 0 |
| 51413 | ENSMUSG00000113850 | 0 | NA | NA | NA | NA | NA | Gm47191 | predicted gene, 47191 [Source:MGI Symb | 0 | 0 | 0 | 0 | 0 | 0 |
| 51414 | ENSMUSG00000113851 | 0 | NA | NA | NA | NA | NA | Gm48503 | predicted gene, 48503 [Source:MGI Symb | 0 | 0 | 0 | 0 | 0 | 0 |
| 51415 | ENSMUSG00000113852 | 9.983128684 | 0.287439701 | 0.836455366 | 0.343640214 | 0.731116878 | 0.94705581 | Gm48062 | predicted gene, 48062 [Source:MGI Symb | 5.6217 | 14.391 | 7.0437 | 7.7118 | 22.903 | 2.2277 |
| 51416 | ENSMUSG00000113853 | 1.513211182 | -0.959737254 | 2.154020921 | -0.445556143 | 0.655917896 | NA | Gm47583 | predicted gene, 47583 [Source:MGI Symb | 0 | 0 | 6.0375 | 1.928 | 0 | 1.1138 |
| 51417 | ENSMUSG00000113854 | 0 | NA | NA | NA | NA | NA | Gm2099 | predicted gene 2099 [Source:MGI Symbol | 0 | 0 | 0 | 0 | 0 | 0 |
| 51418 | ENSMUSG00000113855 | 0 | NA | NA | NA | NA | NA | Gm49374 | predicted gene, 49374 [Source:MGI Symb | 0 | 0 | 0 | 0 | 0 | 0 |
| 51419 | ENSMUSG00000113856 | 0.321325949 | 1.911575114 | 4.048267893 | 0.472195804 | 0.636787034 | NA | Gm10790 | predicted gene 10790 [Source:MGI Symbo | 0 | 0 | 0 | 1.928 | 0 | 0 |
| 51420 | ENSMUSG00000113857 | 0 | NA | NA | NA | NA | NA | Gm46338 | predicted gene, 46338 [Source:MGI Symb | 0 | 0 | 0 | 0 | 0 | 0 |
| 51421 | ENSMUSG00000113858 | 0 | NA | NA | NA | NA | NA | Gm4057 | predicted gene 4057 [Source:MGI Symbol | 0 | 0 | 0 | 0 | 0 | 0 |
| 51422 | ENSMUSG00000113859 | 0 | NA | NA | NA | NA | NA | Gm7979 | predicted gene 7979 [Source:MGI Symbol | 0 | 0 | 0 | 0 | 0 | 0 |
| 51423 | ENSMUSG00000113860 | 0 | NA | NA | NA | NA | NA | Gm45949 | predicted gene, 45949 [Source:MGI Symb | 0 | 0 | 0 | 0 | 0 | 0 |
| 51424 | ENSMUSG00000113861 | 0.181769715 | 1.126924175 | 4.080472857 | 0.276174898 | 0.782413736 | NA | Gm6190 | predicted gene 6190 [Source:MGI Symbol | 0 | 0 | 0 | 0 | 1.0906 | 0 |
| 51425 | ENSMUSG00000113862 | 0.371281295 | 2.09253362 | 4.031852165 | 0.519000582 | 0.603760335 | NA | Gm7537 | predicted gene 7537 [Source:MGI Symbol | 0 | 0 | 0 | 0 | 0 | 2.2277 |
| 51426 | ENSMUSG00000113863 | 0 | NA | NA | NA | NA | NA | Gm7126 | predicted gene 7126 [Source:MGI Symbol | 0 | 0 | 0 | 0 | 0 | 0 |
| 51427 | ENSMUSG00000113864 | 0.160662974 | 1.126924175 | 4.080472857 | 0.276174898 | 0.782413736 | NA | Gm48381 | predicted gene, 48381 [Source:MGI Symb | 0 | 0 | 0 | 0.964 | 0 | 0 |
| 51428 | ENSMUSG00000113865 | 0 | NA | NA | NA | NA | NA | Selenok-ps6 | selenoprotein K, pseudogene 6 [Source:M | 0 | 0 | 0 | 0 | 0 | 0 |
| 51429 | ENSMUSG00000113866 | 0 | NA | NA | NA | NA | NA | Gm47450 | predicted gene, 47450 [Source:MGI Symb | 0 | 0 | 0 | 0 | 0 | 0 |
| 51430 | ENSMUSG00000113867 | 0 | NA | NA | NA | NA | NA | Gm47456 | predicted gene, 47456 [Source:MGI Symb | 0 | 0 | 0 | 0 | 0 | 0 |
| 51431 | ENSMUSG00000113868 | 0 | NA | NA | NA | NA | NA | Gm34719 | predicted gene, 34719 [Source:MGI Symb | 0 | 0 | 0 | 0 | 0 | 0 |
| 51432 | ENSMUSG00000113869 | 0.572870759 | -2.577107685 | 3.182795569 | -0.809699407 | 0.418112959 | NA | Gm48869 | predicted gene, 48869 [Source:MGI Symb | 2.4093 | 1.0279 | 0 | 0 | 0 | 0 |
| 51433 | ENSMUSG00000113870 | 0 | NA | NA | NA | NA | NA | Gm48185 | predicted gene, 48185 [Source:MGI Symb | 0 | 0 | 0 | 0 | 0 | 0 |
| 51434 | ENSMUSG00000113871 | 0.853038123 | -0.516856556 | 2.386032976 | -0.216617524 | 0.828506428 | NA | Gm48349 | predicted gene, 48349 [Source:MGI Symb | 0 | 1.0279 | 2.0125 | 0.964 | 0 | 1.1138 |
| 51435 | ENSMUSG00000113872 | 0 | NA | NA | NA | NA | NA | Gm47474 | predicted gene, 47474 [Source:MGI Symb | 0 | 0 | 0 | 0 | 0 | 0 |
| 51436 | ENSMUSG00000113873 | 0 | NA | NA | NA | NA | NA | Gm46342 | predicted gene, 46342 [Source:MGI Symb | 0 | 0 | 0 | 0 | 0 | 0 |
| 51437 | ENSMUSG00000113874 | 0 | NA | NA | NA | NA | NA | Gm47868 | predicted gene, 47868 [Source:MGI Symb | 0 | 0 | 0 | 0 | 0 | 0 |
| 51438 | ENSMUSG00000113875 | 0 | NA | NA | NA | NA | NA | D230030E09 | Riken cDNA D230030E09 gene [Source:M( | 0 | 0 | 0 | 0 | 0 | 0 |
| 51439 | ENSMUSG00000113876 | 0 | NA | NA | NA | NA | NA | Gm48042 | predicted gene, 48042 [Source:MGI Symb | 0 | 0 | 0 | 0 | 0 | 0 |
| 51440 | ENSMUSG00000113877 | 0 | NA | NA | NA | NA | NA | Gm47208 | predicted gene, 47208 [Source:MGI Symb | 0 | 0 | 0 | 0 | 0 | 0 |
| 51441 | ENSMUSG00000113878 | 0.663785267 | -1.400021131 | 3.106754789 | -0.450637796 | 0.652250621 | NA | Gm19045 | predicted gene, 19045 [Source:MGI Symb | 0 | 0 | 3.0187 | 0.964 | 0 | 0 |
| 51442 | ENSMUSG00000113879 | 0 | NA | NA | NA | NA | NA | Gm48580 | predicted gene, 48580 [Source:MGI Symb | 0 | 0 | 0 | 0 | 0 | 0 |
| 51443 | ENSMUSG00000113880 | 1.661097209 | 1.356636461 | 1.723930125 | 0.786943996 | 0.431314651 | NA | A030005L19 | RIKEN cDNA A030005L19 gene [Source:M( | 0.8031 | 0 | 2.0125 | 3.8559 | 2.1812 | 1.1138 |
| 51444 | ENSMUSG00000113881 | 0 | NA | NA | NA | NA | NA | Gm18592 | predicted gene, 18592 [Source:MGI Symb | 0 | 0 | 0 | 0 | 0 | 0 |
| 51445 | ENSMUSG00000113882 | 0 | NA | NA | NA | NA | NA | Gm48550 | predicted gene, 48550 [Source:MGI Symb | 0 | 0 | 0 | 0 | 0 | 0 |
| 51446 | ENSMUSG00000113883 | 0 | NA | NA | NA | NA | NA | Gm36500 | predicted gene, 36500 [Source:MGI Symb | 0 | 0 | 0 | 0 | 0 | 0 |
| 51447 | ENSMUSG00000113884 | 0 | NA | NA | NA | NA | NA | Gm2270 | predicted gene 2270 [Source:MGI Symbol | 0 | 0 | 0 | 0 | 0 | 0 |
| 51448 | ENSMUSG00000113885 | 0.428363721 | -0.543730382 | 3.624677344 | -0.15000794 | 0.880758351 | NA | Gm47802 | predicted gene, 47802 [Source:MGI Symb | 1.6062 | 0 | 0 | 0.964 | 0 | 0 |
| 51449 | ENSMUSG00000113886 | 0.167707431 | -0.796629176 | 4.080472857 | -0.195229623 | 0.84521319 | NA | Gm48905 | predicted gene, 48905 [Source:MGI Symb | 0 | 0 | 1.0062 | 0 | 0 | 0 |
| 51450 | ENSMUSG00000113887 | 0 | NA | NA | NA | NA | NA | Gm35619 | predicted gene, 35619 [Source:MGI Symb | 0 | 0 | 0 | 0 | 0 | 0 |
| 51451 | ENSMUSG00000113888 | 0 | NA | NA | NA | NA | NA | LOC667541 | hypothetical LOC667541 [Source:MGI Syn | 0 | 0 | 0 | 0 | 0 | 0 |
| 51452 | ENSMUSG00000113889 | 0 | NA | NA | NA | NA | NA | Gm48501 | predicted gene, 48501 [Source:MGI Symb | 0 | 0 | 0 | 0 | 0 | 0 |
| 51453 | ENSMUSG00000113890 | 0.17131964 | -0.796629176 | 4.080472857 | -0.195229623 | 0.84521319 | NA | Gm48368 | predicted gene, 48368 [Source:MGI Symb | 0 | 1.0279 | 0 | 0 | 0 | 0 |
| 51454 | ENSMUSG00000113891 | 5.228789231 | 0.661915502 | 0.95169851 | 0.695509655 | 0.486735963 | 0.873681495 | Gm47683 | predicted gene, 47683 [Source:MGI Symb | 4.0155 | 2.0558 | 6.0375 | 3.8559 | 8.7249 | 6.6831 |
| 51455 | ENSMUSG00000113892 | 0 | NA | NA | NA | NA | NA | Gm48605 | predicted gene, 48605 [Source:MGI Symb | 0 | 0 | 0 | 0 | 0 | 0 |
| 51456 | ENSMUSG00000113893 | 0 | NA | NA | NA | NA | NA | Gm48800 | predicted gene, 48800 [Source:MGI Symb | 0 | 0 | 0 | 0 | 0 | 0 |
| 51457 | ENSMUSG00000113894 | 4.59628704 | -0.509727663 | 0.998655723 | -0.510413801 | 0.60976159 | NA | Gm48838 | predicted gene, 48838 [Source:MGI Symb | 8.031 | 4.1117 | 4.025 | 4.8199 | 4.3625 | 2.2277 |
| 51458 | ENSMUSG00000113895 | 0.321325949 | 1.911575114 | 4.048267893 | 0.472195804 | 0.636787034 | NA | Gm48383 | predicted gene, 48383 [Source:MGI Symb | 0 | 0 | 0 | 1.928 | 0 | 0 |
| 51459 | ENSMUSG00000113896 | 0 | NA | NA | NA | NA | NA | Gm18591 | predicted gene, 18591 [Source:MGI Symb | 0 | 0 | 0 | 0 | 0 | 0 |
| 51460 | ENSMUSG00000113897 | 0 | NA | NA | NA | NA | NA | Gm48227 | predicted gene, 48227 [Source:MGI Symb | 0 | 0 | 0 | 0 | 0 | 0 |

| | | | | | | | | | | | | | | | |
|---|---|---|---|---|---|---|---|---|---|---|---|---|---|---|---|
| 51461 | ENSMUSG00000113898 | 6.774495804 | 0.191083909 | 0.878624346 | 0.217480781 | 0.827833683 | 0.969794868 | Gm19144 | predicted gene, 19144 [Source:MGI Symb | 4.8186 | 9.2513 | 5.0312 | 11.568 | 2.1812 | 7.7969 |
| 51462 | ENSMUSG00000113899 | 0 | NA | NA | NA | NA | NA | Gm9595 | predicted gene 9595 [Source:MGI Symbol | 0 | 0 | 0 | 0 | 0 | 0 |
| 51463 | ENSMUSG00000113900 | 0 | NA | NA | NA | NA | NA | Gm40437 | predicted gene, 40437 [Source:MGI Symb | 0 | 0 | 0 | 0 | 0 | 0 |
| 51464 | ENSMUSG00000113901 | 0 | NA | NA | NA | NA | NA | Gm49381 | predicted gene, 49381 [Source:MGI Symb | 0 | 0 | 0 | 0 | 0 | 0 |
| 51465 | ENSMUSG00000113902 | 626.3911728 | -0.042034376 | 0.14471987 | -0.290453384 | 0.771469409 | 0.956965056 | Ndufb1 | NADH:ubiquinone oxidoreductase subuni | 697.09 | 650.67 | 557.46 | 532.12 | 628.2 | 692.81 |
| 51466 | ENSMUSG00000113903 | 0 | NA | NA | NA | NA | NA | Gm5655 | predicted gene 5655 [Source:MGI Symbol | 0 | 0 | 0 | 0 | 0 | 0 |
| 51467 | ENSMUSG00000113904 | 0 | NA | NA | NA | NA | NA | Gm32351 | predicted gene, 32351 [Source:MGI Symb | 0 | 0 | 0 | 0 | 0 | 0 |
| 51468 | ENSMUSG00000113905 | 0 | NA | NA | NA | NA | NA | Gm48872 | predicted gene, 48872 [Source:MGI Symb | 0 | 0 | 0 | 0 | 0 | 0 |
| 51469 | ENSMUSG00000113906 | 0 | NA | NA | NA | NA | NA | 4930529N2C | RIKEN cDNA 4930529N20 gene [Source:M | 0 | 0 | 0 | 0 | 0 | 0 |
| 51470 | ENSMUSG00000113907 | 0.349477146 | 0.165142768 | 3.911021834 | 0.042224967 | 0.966319359 | NA | Gm47167 | predicted gene, 47167 [Source:MGI Symb | 0 | 0 | 1.0062 | 0 | 1.0906 | 0 |
| 51471 | ENSMUSG00000113908 | 0 | NA | NA | NA | NA | NA | Gm10811 | predicted gene 10811 [Source:MGI Symbo | 0 | 0 | 0 | 0 | 0 | 0 |
| 51472 | ENSMUSG00000113909 | 0 | NA | NA | NA | NA | NA | Gm36377 | predicted gene, 36377 [Source:MGI Symb | 0 | 0 | 0 | 0 | 0 | 0 |
| 51473 | ENSMUSG00000113910 | 0.462220778 | -0.685400455 | 3.191475091 | -0.21475977 | 0.82995462 | NA | Gm48286 | predicted gene, 48286 [Source:MGI Symb | 0.8031 | 0 | 1.0062 | 0.964 | 0 | 0 |
| 51474 | ENSMUSG00000113911 | 0 | NA | NA | NA | NA | NA | Gm21930 | predicted gene, 21930 [Source:MGI Symb | 0 | 0 | 0 | 0 | 0 | 0 |
| 51475 | ENSMUSG00000113912 | 0 | NA | NA | NA | NA | NA | Gm21937 | predicted gene, 21937 [Source:MGI Symb | 0 | 0 | 0 | 0 | 0 | 0 |
| 51476 | ENSMUSG00000113913 | 0 | NA | NA | NA | NA | NA | Gm46323 | predicted gene, 46323 [Source:MGI Symb | 0 | 0 | 0 | 0 | 0 | 0 |
| 51477 | ENSMUSG00000113914 | 0 | NA | NA | NA | NA | NA | Gm48458 | predicted gene, 48458 [Source:MGI Symb | 0 | 0 | 0 | 0 | 0 | 0 |
| 51478 | ENSMUSG00000113915 | 0 | NA | NA | NA | NA | NA | Gm48019 | predicted gene, 48019 [Source:MGI Symb | 0 | 0 | 0 | 0 | 0 | 0 |
| 51479 | ENSMUSG00000113916 | 0 | NA | NA | NA | NA | NA | Gm18881 | predicted gene, 18881 [Source:MGI Symb | 0 | 0 | 0 | 0 | 0 | 0 |
| 51480 | ENSMUSG00000113917 | 1.332753434 | -0.599334234 | 1.909214978 | -0.313916578 | 0.753584399 | NA | Gm48382 | predicted gene, 48382 [Source:MGI Symb | 0.8031 | 0 | 4.025 | 0.964 | 1.0906 | 1.1138 |
| 51481 | ENSMUSG00000113918 | 23.7804096 | 0.148125861 | 0.455037011 | 0.325524864 | 0.744783868 | 0.949997329 | Gm6566 | predicted gene 6566 [Source:MGI Symbol | 28.912 | 18.503 | 20.125 | 28.919 | 26.175 | 20.049 |
| 51482 | ENSMUSG00000113919 | 0.133850373 | -0.796629176 | 4.080472857 | -0.195229623 | 0.84521319 | NA | Gm34466 | predicted gene, 34466 [Source:MGI Symb | 0.8031 | 0 | 0 | 0 | 0 | 0 |
| 51483 | ENSMUSG00000113920 | 0 | NA | NA | NA | NA | NA | Gm18783 | predicted gene, 18783 [Source:MGI Symb | 0 | 0 | 0 | 0 | 0 | 0 |
| 51484 | ENSMUSG00000113921 | 0 | NA | NA | NA | NA | NA | Gm48054 | predicted gene, 48054 [Source:MGI Symb | 0 | 0 | 0 | 0 | 0 | 0 |
| 51485 | ENSMUSG00000113922 | 1.22088717 | 1.402256566 | 2.259311288 | 0.620656646 | 0.534825561 | NA | Gm48887 | predicted gene, 48887 [Source:MGI Symb | 0 | 2.0558 | 0 | 1.928 | 0 | 3.3415 |
| 51486 | ENSMUSG00000113923 | 0 | NA | NA | NA | NA | NA | Gm10126 | predicted gene 10126 [Source:MGI Symbo | 0 | 0 | 0 | 0 | 0 | 0 |
| 51487 | ENSMUSG00000113924 | 0 | NA | NA | NA | NA | NA | Gm48493 | predicted gene, 48493 [Source:MGI Symb | 0 | 0 | 0 | 0 | 0 | 0 |
| 51488 | ENSMUSG00000113925 | 0 | NA | NA | NA | NA | NA | Gm7544 | predicted gene 7544 [Source:MGI Symbol | 0 | 0 | 0 | 0 | 0 | 0 |
| 51489 | ENSMUSG00000113926 | 0 | NA | NA | NA | NA | NA | Gm3565 | predicted gene 3565 [Source:MGI Symbol | 0 | 0 | 0 | 0 | 0 | 0 |
| 51490 | ENSMUSG00000113927 | 0.744190399 | -2.947713985 | 2.874421214 | -1.025498271 | 0.305128141 | NA | 5330409N07 | RIKEN cDNA 5330409N07 gene [Source:M | 2.4093 | 2.0558 | 0 | 0 | 0 | 0 |
| 51491 | ENSMUSG00000113928 | 0 | NA | NA | NA | NA | NA | Gm48209 | predicted gene, 48209 [Source:MGI Symb | 0 | 0 | 0 | 0 | 0 | 0 |
| 51492 | ENSMUSG00000113929 | 0 | NA | NA | NA | NA | NA | Gm3926 | predicted gene 3926 [Source:MGI Symbol | 0 | 0 | 0 | 0 | 0 | 0 |
| 51493 | ENSMUSG00000113930 | 0 | NA | NA | NA | NA | NA | Gm36235 | predicted gene, 36235 [Source:MGI Symb | 0 | 0 | 0 | 0 | 0 | 0 |
| 51494 | ENSMUSG00000113931 | 0.342432689 | 1.990653086 | 3.900281263 | 0.51038706 | 0.609780321 | NA | Gm5928 | predicted gene 5928 [Source:MGI Symbol | 0 | 0 | 0 | 0.964 | 1.0906 | 0 |
| 51495 | ENSMUSG00000113932 | 0 | NA | NA | NA | NA | NA | 2900024D18 | RIKEN cDNA 2900024D18 gene [Source:M | 0 | 0 | 0 | 0 | 0 | 0 |
| 51496 | ENSMUSG00000113933 | 0 | NA | NA | NA | NA | NA | Gm34611 | predicted gene, 34611 [Source:MGI Symb | 0 | 0 | 0 | 0 | 0 | 0 |
| 51497 | ENSMUSG00000113934 | 0 | NA | NA | NA | NA | NA | Gm48609 | predicted gene, 48609 [Source:MGI Symb | 0 | 0 | 0 | 0 | 0 | 0 |
| 51498 | ENSMUSG00000113935 | 0 | NA | NA | NA | NA | NA | Gm35732 | predicted gene, 35732 [Source:MGI Symb | 0 | 0 | 0 | 0 | 0 | 0 |
| 51499 | ENSMUSG00000113936 | 0.133850373 | -0.796629176 | 4.080472857 | -0.195229623 | 0.84521319 | NA | Gm8784 | predicted gene 8784 [Source:MGI Symbol | 0.8031 | 0 | 0 | 0 | 0 | 0 |
| 51500 | ENSMUSG00000113937 | 0 | NA | NA | NA | NA | NA | Rpl21-ps2 | ribosomal protein L21, pseudogene 2 [Sou | 0 | 0 | 0 | 0 | 0 | 0 |
| 51501 | ENSMUSG00000113938 | 0 | NA | NA | NA | NA | NA | Gm46996 | predicted gene 46996 [Source:MGI Symbo | 0 | 0 | 0 | 0 | 0 | 0 |
| 51502 | ENSMUSG00000113939 | 0 | NA | NA | NA | NA | NA | Gm47871 | predicted gene, 47871 [Source:MGI Symb | 0 | 0 | 0 | 0 | 0 | 0 |
| 51503 | ENSMUSG00000113940 | 0 | NA | NA | NA | NA | NA | Gm40477 | predicted gene, 40477 [Source:MGI Symb | 0 | 0 | 0 | 0 | 0 | 0 |
| 51504 | ENSMUSG00000113941 | 0 | NA | NA | NA | NA | NA | Gm35755 | predicted gene, 35755 [Source:MGI Symb | 0 | 0 | 0 | 0 | 0 | 0 |
| 51505 | ENSMUSG00000113942 | 0 | NA | NA | NA | NA | NA | Gm18632 | predicted gene, 18632 [Source:MGI Symb | 0 | 0 | 0 | 0 | 0 | 0 |
| 51506 | ENSMUSG00000113943 | 0 | NA | NA | NA | NA | NA | Gm47804 | predicted gene, 47804 [Source:MGI Symb | 0 | 0 | 0 | 0 | 0 | 0 |
| 51507 | ENSMUSG00000113944 | 0 | NA | NA | NA | NA | NA | Gm48847 | predicted gene, 48847 [Source:MGI Symb | 0 | 0 | 0 | 0 | 0 | 0 |
| 51508 | ENSMUSG00000113945 | 0 | NA | NA | NA | NA | NA | Vmn2r-ps99 | vomeronasal 2, receptor, pseudogene 99 | 0 | 0 | 0 | 0 | 0 | 0 |
| 51509 | ENSMUSG00000113946 | 0 | NA | NA | NA | NA | NA | Gm47250 | predicted gene, 47250 [Source:MGI Symb | 0 | 0 | 0 | 0 | 0 | 0 |
| 51510 | ENSMUSG00000113947 | 0 | NA | NA | NA | NA | NA | Gm10465 | predicted gene 10465 [Source:MGI Symbo | 0 | 0 | 0 | 0 | 0 | 0 |
| 51511 | ENSMUSG00000113948 | 6.985800806 | 1.401171282 | 0.890890432 | 1.572776215 | 0.115770645 | 0.500151317 | Rpl17-ps3 | ribosomal protein L17, pseudogene 3 [Sou | 7.2279 | 1.0279 | 3.0187 | 9.6398 | 7.6343 | 13.366 |
| 51512 | ENSMUSG00000113949 | 2.860148087 | 2.49826743 | 1.494657191 | 1.671465167 | 0.094629833 | NA | Scamp4 | secretory carrier membrane protein 4 [So | 2.4093 | 0 | 0 | 4.8199 | 4.3625 | 5.5692 |
| 51513 | ENSMUSG00000113950 | 0 | NA | NA | NA | NA | NA | Gm47080 | predicted gene, 47080 [Source:MGI Symb | 0 | 0 | 0 | 0 | 0 | 0 |
| 51514 | ENSMUSG00000113951 | 0 | NA | NA | NA | NA | NA | Ubl5c | ubiquitin-like 5C [Source:MGI Symbol;Acc | 0 | 0 | 0 | 0 | 0 | 0 |
| 51515 | ENSMUSG00000113952 | 0 | NA | NA | NA | NA | NA | Rpl31-ps2 | ribosomal protein L31, pseudogene 2 [Sou | 0 | 0 | 0 | 0 | 0 | 0 |
| 51516 | ENSMUSG00000113953 | 0 | NA | NA | NA | NA | NA | Gm48201 | predicted gene, 48201 [Source:MGI Symb | 0 | 0 | 0 | 0 | 0 | 0 |

| | | | | | | | | | | | | | | | |
|---|---|---|---|---|---|---|---|---|---|---|---|---|---|---|---|
| 51517 | ENSMUSG00000113954 | 0 | NA | NA | NA | NA | NA | Gm7822 | predicted gene 7822 [Source:MGI Symbol | 0 | 0 | 0 | 0 | 0 | 0 |
| 51518 | ENSMUSG00000113955 | 0 | NA | NA | NA | NA | NA | Gm48168 | predicted gene, 48168 [Source:MGI Symb | 0 | 0 | 0 | 0 | 0 | 0 |
| 51519 | ENSMUSG00000113956 | 0 | NA | NA | NA | NA | NA | Gm19368 | predicted gene, 19368 [Source:MGI Symb | 0 | 0 | 0 | 0 | 0 | 0 |
| 51520 | ENSMUSG00000113957 | 0 | NA | NA | NA | NA | NA | Gm9257 | predicted gene 9257 [Source:MGI Symbol | 0 | 0 | 0 | 0 | 0 | 0 |
| 51521 | ENSMUSG00000113958 | 0 | NA | NA | NA | NA | NA | Gm48020 | predicted gene, 48020 [Source:MGI Symb | 0 | 0 | 0 | 0 | 0 | 0 |
| 51522 | ENSMUSG00000113959 | 0.167707431 | -0.796629176 | 4.080472857 | -0.195229623 | 0.84521319 | NA | Gm46339 | predicted gene, 46339 [Source:MGI Symb | 0 | 0 | 1.0062 | 0 | 0 | 0 |
| 51523 | ENSMUSG00000113960 | 0 | NA | NA | NA | NA | NA | 4933412O06 | RIKEN cDNA 4933412O06 gene [Source:M | 0 | 0 | 0 | 0 | 0 | 0 |
| 51524 | ENSMUSG00000113961 | 0 | NA | NA | NA | NA | NA | Gm48498 | predicted gene, 48498 [Source:MGI Symb | 0 | 0 | 0 | 0 | 0 | 0 |
| 51525 | ENSMUSG00000113962 | 0 | NA | NA | NA | NA | NA | Gm47618 | predicted gene, 47618 [Source:MGI Symb | 0 | 0 | 0 | 0 | 0 | 0 |
| 51526 | ENSMUSG00000113963 | 0 | NA | NA | NA | NA | NA | Gm6554 | predicted gene 6554 [Source:MGI Symbol | 0 | 0 | 0 | 0 | 0 | 0 |
| 51527 | ENSMUSG00000113964 | 0 | NA | NA | NA | NA | NA | Gm48667 | predicted gene, 48667 [Source:MGI Symb | 0 | 0 | 0 | 0 | 0 | 0 |
| 51528 | ENSMUSG00000113965 | 2.806529524 | -1.560078386 | 1.322732683 | -1.17943588 | 0.238224655 | NA | A730091E23 | RIKEN cDNA A730091E23 gene [Source:M | 2.4093 | 5.1396 | 5.0312 | 0.964 | 2.1812 | 1.1138 |
| 51529 | ENSMUSG00000113966 | 0.829837731 | -1.7608533 | 2.399900965 | -0.733719152 | 0.463119924 | NA | Gm47980 | predicted gene, 47980 [Source:MGI Symb | 0.8031 | 2.0558 | 1.0062 | 0 | 0 | 1.1138 |
| 51530 | ENSMUSG00000113967 | 0.335414862 | -1.779047242 | 4.030804179 | -0.441362855 | 0.658950329 | NA | Gm31025 | predicted gene, 31025 [Source:MGI Symb | 0 | 0 | 2.0125 | 0 | 0 | 0 |
| 51531 | ENSMUSG00000113968 | 0 | NA | NA | NA | NA | NA | Gm48813 | predicted gene, 48813 [Source:MGI Symb | 0 | 0 | 0 | 0 | 0 | 0 |
| 51532 | ENSMUSG00000113969 | 0 | NA | NA | NA | NA | NA | Gm48731 | predicted gene, 48731 [Source:MGI Symb | 0 | 0 | 0 | 0 | 0 | 0 |
| 51533 | ENSMUSG00000113970 | 0 | NA | NA | NA | NA | NA | Gm7550 | predicted gene 7550 [Source:MGI Symbol | 0 | 0 | 0 | 0 | 0 | 0 |
| 51534 | ENSMUSG00000113971 | 0 | NA | NA | NA | NA | NA | Eif1ad4 | eukaryotic translation initiation factor 1A | 0 | 0 | 0 | 0 | 0 | 0 |
| 51535 | ENSMUSG00000113972 | 0 | NA | NA | NA | NA | NA | Gm48497 | predicted gene, 48497 [Source:MGI Symb | 0 | 0 | 0 | 0 | 0 | 0 |
| 51536 | ENSMUSG00000113973 | 0 | NA | NA | NA | NA | NA | A030014E15 | RIKEN cDNA A030014E15 gene [Source:M | 0 | 0 | 0 | 0 | 0 | 0 |
| 51537 | ENSMUSG00000113974 | 0 | NA | NA | NA | NA | NA | 4930442G10 | RIKEN cDNA 4930442G10 gene [Source:M | 0 | 0 | 0 | 0 | 0 | 0 |
| 51538 | ENSMUSG00000113975 | 0 | NA | NA | NA | NA | NA | Gm36372 | predicted gene, 36372 [Source:MGI Symb | 0 | 0 | 0 | 0 | 0 | 0 |
| 51539 | ENSMUSG00000113976 | 0 | NA | NA | NA | NA | NA | Gm18243 | predicted gene, 18243 [Source:MGI Symb | 0 | 0 | 0 | 0 | 0 | 0 |
| 51540 | ENSMUSG00000113977 | 0 | NA | NA | NA | NA | NA | Gm34276 | predicted gene, 34276 [Source:MGI Symb | 0 | 0 | 0 | 0 | 0 | 0 |
| 51541 | ENSMUSG00000113978 | 0 | NA | NA | NA | NA | NA | Gm48260 | predicted gene, 48260 [Source:MGI Symb | 0 | 0 | 0 | 0 | 0 | 0 |
| 51542 | ENSMUSG00000113979 | 0 | NA | NA | NA | NA | NA | Gm47012 | predicted gene, 47012 [Source:MGI Symb | 0 | 0 | 0 | 0 | 0 | 0 |
| 51543 | ENSMUSG00000113980 | 0 | NA | NA | NA | NA | NA | Gm47655 | predicted gene, 47655 [Source:MGI Symb | 0 | 0 | 0 | 0 | 0 | 0 |
| 51544 | ENSMUSG00000113981 | 0 | NA | NA | NA | NA | NA | Gm47011 | predicted gene, 47011 [Source:MGI Symb | 0 | 0 | 0 | 0 | 0 | 0 |
| 51545 | ENSMUSG00000113982 | 0 | NA | NA | NA | NA | NA | Gm34557 | predicted gene, 34557 [Source:MGI Symb | 0 | 0 | 0 | 0 | 0 | 0 |
| 51546 | ENSMUSG00000113983 | 0 | NA | NA | NA | NA | NA | Gm47742 | predicted gene, 47742 [Source:MGI Symb | 0 | 0 | 0 | 0 | 0 | 0 |
| 51547 | ENSMUSG00000113984 | 0.305170013 | -1.657836477 | 4.041480335 | -0.410205257 | 0.681655385 | NA | Gm48890 | predicted gene, 48890 [Source:MGI Symb | 0.8031 | 1.0279 | 0 | 0 | 0 | 0 |
| 51548 | ENSMUSG00000113985 | 0.294513347 | 0.165143496 | 4.080472857 | 0.040471657 | 0.967717103 | NA | 9130015A21 | RIKEN cDNA 9130015A21 gene [Source:M | 0.8031 | 0 | 0 | 0.964 | 0 | 0 |
| 51549 | ENSMUSG00000113986 | 43.56836447 | 1.400305013 | 0.408091094 | 3.43135401 | 0.000600576 | 0.014147778 | Gm31333 | predicted gene, 31333 [Source:MGI Symb | 34.533 | 24.67 | 12.075 | 59.767 | 50.168 | 80.197 |
| 51550 | ENSMUSG00000113987 | 0.160662974 | 1.126924175 | 4.080472857 | 0.276174898 | 0.782413736 | NA | 4930453C13 | RIKEN cDNA 4930453C13 gene [Source:M | 0 | 0 | 0 | 0.964 | 0 | 0 |
| 51551 | ENSMUSG00000113989 | 0 | NA | NA | NA | NA | NA | Gm47013 | predicted gene, 47013 [Source:MGI Symb | 0 | 0 | 0 | 0 | 0 | 0 |
| 51552 | ENSMUSG00000113990 | 0 | NA | NA | NA | NA | NA | Gm47263 | predicted gene, 47263 [Source:MGI Symb | 0 | 0 | 0 | 0 | 0 | 0 |
| 51553 | ENSMUSG00000113991 | 0 | NA | NA | NA | NA | NA | Gm48151 | predicted gene, 48151 [Source:MGI Symb | 0 | 0 | 0 | 0 | 0 | 0 |
| 51554 | ENSMUSG00000113992 | 0 | NA | NA | NA | NA | NA | Gm9158 | predicted gene 9158 [Source:MGI Symbol | 0 | 0 | 0 | 0 | 0 | 0 |
| 51555 | ENSMUSG00000113993 | 0 | NA | NA | NA | NA | NA | Gm47659 | predicted gene, 47659 [Source:MGI Symb | 0 | 0 | 0 | 0 | 0 | 0 |
| 51556 | ENSMUSG00000113994 | 0 | NA | NA | NA | NA | NA | Gm47375 | predicted gene, 47375 [Source:MGI Symb | 0 | 0 | 0 | 0 | 0 | 0 |
| 51557 | ENSMUSG00000113995 | 0 | NA | NA | NA | NA | NA | Gm47648 | predicted gene, 47648 [Source:MGI Symb | 0 | 0 | 0 | 0 | 0 | 0 |
| 51558 | ENSMUSG00000113996 | 0 | NA | NA | NA | NA | NA | Gm48455 | predicted gene, 48455 [Source:MGI Symb | 0 | 0 | 0 | 0 | 0 | 0 |
| 51559 | ENSMUSG00000113997 | 0 | NA | NA | NA | NA | NA | 1700040E09 | RIKEN cDNA 1700040E09 gene [Source:M | 0 | 0 | 0 | 0 | 0 | 0 |
| 51560 | ENSMUSG00000113998 | 0.349477146 | 0.165142768 | 3.911021834 | 0.042224967 | 0.966319359 | NA | Gm40909 | predicted gene, 40909 [Source:MGI Symb | 0 | 0 | 1.0062 | 0 | 1.0906 | 0 |
| 51561 | ENSMUSG00000113999 | 0 | NA | NA | NA | NA | NA | Gm47681 | predicted gene, 47681 [Source:MGI Symb | 0 | 0 | 0 | 0 | 0 | 0 |
| 51562 | ENSMUSG00000114000 | 0 | NA | NA | NA | NA | NA | Gm48762 | predicted gene, 48762 [Source:MGI Symb | 0 | 0 | 0 | 0 | 0 | 0 |
| 51563 | ENSMUSG00000114001 | 0 | NA | NA | NA | NA | NA | Gm48664 | predicted gene, 48664 [Source:MGI Symb | 0 | 0 | 0 | 0 | 0 | 0 |
| 51564 | ENSMUSG00000114002 | 0 | NA | NA | NA | NA | NA | Gm48637 | predicted gene, 48637 [Source:MGI Symb | 0 | 0 | 0 | 0 | 0 | 0 |
| 51565 | ENSMUSG00000114003 | 4.074539863 | -0.456840989 | 1.148085317 | -0.397915541 | 0.690692444 | NA | Gm9616 | predicted gene 9616 [Source:MGI Symbol | 4.0155 | 6.1675 | 4.025 | 5.7839 | 0 | 4.4554 |
| 51566 | ENSMUSG00000114004 | 0.990062021 | -2.088395155 | 2.279709817 | -0.91607938 | 0.359625262 | NA | Gm48552 | predicted gene, 48552 [Source:MGI Symb | 0.8031 | 1.0279 | 3.0187 | 0 | 1.0906 | 0 |
| 51567 | ENSMUSG00000114005 | 0 | NA | NA | NA | NA | NA | Gm24474 | predicted gene, 24474 [Source:MGI Symb | 0 | 0 | 0 | 0 | 0 | 0 |
| 51568 | ENSMUSG00000114006 | 0 | NA | NA | NA | NA | NA | Gm47997 | predicted gene, 47997 [Source:MGI Symb | 0 | 0 | 0 | 0 | 0 | 0 |
| 51569 | ENSMUSG00000114007 | 0 | NA | NA | NA | NA | NA | Gm48660 | predicted gene, 48660 [Source:MGI Symb | 0 | 0 | 0 | 0 | 0 | 0 |
| 51570 | ENSMUSG00000114008 | 0 | NA | NA | NA | NA | NA | Gm48033 | predicted gene, 48033 [Source:MGI Symb | 0 | 0 | 0 | 0 | 0 | 0 |
| 51571 | ENSMUSG00000114009 | 0 | NA | NA | NA | NA | NA | Gm47606 | predicted gene, 47606 [Source:MGI Symb | 0 | 0 | 0 | 0 | 0 | 0 |
| 51572 | ENSMUSG00000114010 | 0 | NA | NA | NA | NA | NA | Gm47432 | predicted gene, 47432 [Source:MGI Symb | 0 | 0 | 0 | 0 | 0 | 0 |

| | | | | | | | | | | | | | | | |
|---|---|---|---|---|---|---|---|---|---|---|---|---|---|---|---|
| 51573 | ENSMUSG00000114011 | 0 | NA | NA | NA | NA | NA | Gm47955 | predicted gene, 47955 [Source:MGI Symb | 0 | 0 | 0 | 0 | 0 | 0 |
| 51574 | ENSMUSG00000114012 | 0 | NA | NA | NA | NA | NA | Gm18974 | predicted gene, 18974 [Source:MGI Symb | 0 | 0 | 0 | 0 | 0 | 0 |
| 51575 | ENSMUSG00000114013 | 0 | NA | NA | NA | NA | NA | Gm48219 | predicted gene, 48219 [Source:MGI Symb | 0 | 0 | 0 | 0 | 0 | 0 |
| 51576 | ENSMUSG00000114014 | 4.500809932 | -0.678705006 | 1.04143188 | -0.651703697 | 0.514592337 | NA | Gm48350 | predicted gene, 48350 [Source:MGI Symb | 6.4248 | 7.1954 | 3.0187 | 4.8199 | 1.0906 | 4.4554 |
| 51577 | ENSMUSG00000114015 | 0 | NA | NA | NA | NA | NA | Gm47417 | predicted gene, 47417 [Source:MGI Symb | 0 | 0 | 0 | 0 | 0 | 0 |
| 51578 | ENSMUSG00000114016 | 3.175045262 | 0.051037538 | 1.21722431 | 0.041929443 | 0.966554945 | NA | Gm47258 | predicted gene, 47258 [Source:MGI Symb | 3.2124 | 3.0838 | 3.0187 | 0.964 | 6.5437 | 2.2277 |
| 51579 | ENSMUSG00000114017 | 0 | NA | NA | NA | NA | NA | Gm40662 | predicted gene, 40662 [Source:MGI Symb | 0 | 0 | 0 | 0 | 0 | 0 |
| 51580 | ENSMUSG00000114018 | 0 | NA | NA | NA | NA | NA | Gm36495 | predicted gene, 36495 [Source:MGI Symb | 0 | 0 | 0 | 0 | 0 | 0 |
| 51581 | ENSMUSG00000114019 | 20.97689493 | 1.065297192 | 0.502397732 | 2.120425959 | 0.033970139 | 0.24678257 | Gm47155 | predicted gene, 47155 [Source:MGI Symb | 15.259 | 7.1954 | 18.112 | 27.955 | 27.265 | 30.074 |
| 51582 | ENSMUSG00000114020 | 0 | NA | NA | NA | NA | NA | Gm48564 | predicted gene, 48564 [Source:MGI Symb | 0 | 0 | 0 | 0 | 0 | 0 |
| 51583 | ENSMUSG00000114021 | 0 | NA | NA | NA | NA | NA | Gm47109 | predicted gene, 47109 [Source:MGI Symb | 0 | 0 | 0 | 0 | 0 | 0 |
| 51584 | ENSMUSG00000114022 | 0 | NA | NA | NA | NA | NA | Gm17330 | predicted gene, 17330 [Source:MGI Symb | 0 | 0 | 0 | 0 | 0 | 0 |
| 51585 | ENSMUSG00000114023 | 0.898940922 | -0.606261109 | 2.77650979 | -0.218353672 | 0.827153559 | NA | Gm6474 | predicted gene 6474 [Source:MGI Symbol | 3.2124 | 0 | 0 | 0 | 2.1812 | 0 |
| 51586 | ENSMUSG00000114024 | 0.476489653 | -2.293616034 | 3.421506946 | -0.670352587 | 0.502633051 | NA | Gm46336 | predicted gene, 46336 [Source:MGI Symb | 0.8031 | 2.0558 | 0 | 0 | 0 | 0 |
| 51587 | ENSMUSG00000114025 | 65.90382873 | -0.264328058 | 0.279864992 | -0.944484184 | 0.34492228 | 0.786033695 | Gm49331 | predicted gene, 49331 [Source:MGI Symb | 63.445 | 69.898 | 82.512 | 53.983 | 65.437 | 60.148 |
| 51588 | ENSMUSG00000114026 | 0 | NA | NA | NA | NA | NA | Gm48480 | predicted gene, 48480 [Source:MGI Symb | 0 | 0 | 0 | 0 | 0 | 0 |
| 51589 | ENSMUSG00000114027 | 0 | NA | NA | NA | NA | NA | Vmn2r-ps10 | vomeronasal 2, receptor, pseudogene 10 | 0 | 0 | 0 | 0 | 0 | 0 |
| 51590 | ENSMUSG00000114028 | 0 | NA | NA | NA | NA | NA | 9630002D21 | RIKEN cDNA 9630002D21 gene [Source:M | 0 | 0 | 0 | 0 | 0 | 0 |
| 51591 | ENSMUSG00000114029 | 0 | NA | NA | NA | NA | NA | Gm4806 | predicted gene 4806 [Source:MGI Symbol | 0 | 0 | 0 | 0 | 0 | 0 |
| 51592 | ENSMUSG00000114030 | 0 | NA | NA | NA | NA | NA | Gm18921 | predicted gene, 18921 [Source:MGI Symb | 0 | 0 | 0 | 0 | 0 | 0 |
| 51593 | ENSMUSG00000114031 | 0 | NA | NA | NA | NA | NA | Gm48136 | predicted gene, 48136 [Source:MGI Symb | 0 | 0 | 0 | 0 | 0 | 0 |
| 51594 | ENSMUSG00000114032 | 0 | NA | NA | NA | NA | NA | Gm2139 | predicted gene 2139 [Source:MGI Symbol | 0 | 0 | 0 | 0 | 0 | 0 |
| 51595 | ENSMUSG00000114033 | 1.233017188 | 0.208113188 | 2.04700053 | 0.101667383 | 0.919020692 | NA | Gm47705 | predicted gene, 47705 [Source:MGI Symb | 2.4093 | 0 | 1.0062 | 2.8919 | 1.0906 | 0 |
| 51596 | ENSMUSG00000114034 | 0 | NA | NA | NA | NA | NA | Gm9267 | predicted gene 9267 [Source:MGI Symbol | 0 | 0 | 0 | 0 | 0 | 0 |
| 51597 | ENSMUSG00000114035 | 0 | NA | NA | NA | NA | NA | Gm31126 | predicted gene, 31126 [Source:MGI Symb | 0 | 0 | 0 | 0 | 0 | 0 |
| 51598 | ENSMUSG00000114036 | 0 | NA | NA | NA | NA | NA | Gm48228 | predicted gene, 48228 [Source:MGI Symb | 0 | 0 | 0 | 0 | 0 | 0 |
| 51599 | ENSMUSG00000114037 | 0 | NA | NA | NA | NA | NA | Gm48817 | predicted gene, 48817 [Source:MGI Symb | 0 | 0 | 0 | 0 | 0 | 0 |
| 51600 | ENSMUSG00000114038 | 0 | NA | NA | NA | NA | NA | Gm40999 | predicted gene, 40999 [Source:MGI Symb | 0 | 0 | 0 | 0 | 0 | 0 |
| 51601 | ENSMUSG00000114039 | 0 | NA | NA | NA | NA | NA | Gm48305 | predicted gene, 48305 [Source:MGI Symb | 0 | 0 | 0 | 0 | 0 | 0 |
| 51602 | ENSMUSG00000114040 | 0 | NA | NA | NA | NA | NA | Gm48319 | predicted gene, 48319 [Source:MGI Symb | 0 | 0 | 0 | 0 | 0 | 0 |
| 51603 | ENSMUSG00000114041 | 0 | NA | NA | NA | NA | NA | Gm9269 | predicted gene 9269 [Source:MGI Symbol | 0 | 0 | 0 | 0 | 0 | 0 |
| 51604 | ENSMUSG00000114042 | 0 | NA | NA | NA | NA | NA | Gm18473 | predicted gene, 18473 [Source:MGI Symb | 0 | 0 | 0 | 0 | 0 | 0 |
| 51605 | ENSMUSG00000114043 | 0 | NA | NA | NA | NA | NA | Gm48625 | predicted gene, 48625 [Source:MGI Symb | 0 | 0 | 0 | 0 | 0 | 0 |
| 51606 | ENSMUSG00000114044 | 0 | NA | NA | NA | NA | NA | Gm19134 | predicted gene, 19134 [Source:MGI Symb | 0 | 0 | 0 | 0 | 0 | 0 |
| 51607 | ENSMUSG00000114045 | 18.08070988 | -0.240662894 | 0.534582302 | -0.450188667 | 0.652574407 | 0.925756399 | Gm34220 | predicted gene, 34220 [Source:MGI Symb | 23.29 | 12.335 | 23.144 | 22.171 | 14.178 | 13.366 |
| 51608 | ENSMUSG00000114046 | 0 | NA | NA | NA | NA | NA | Snapc1l | small nuclear RNA activating complex, pol | 0 | 0 | 0 | 0 | 0 | 0 |
| 51609 | ENSMUSG00000114047 | 0 | NA | NA | NA | NA | NA | Gm7557 | predicted gene 7557 [Source:MGI Symbol | 0 | 0 | 0 | 0 | 0 | 0 |
| 51610 | ENSMUSG00000114048 | 0 | NA | NA | NA | NA | NA | 4930549C15 | RIKEN cDNA 4930549C15 gene [Source:M | 0 | 0 | 0 | 0 | 0 | 0 |
| 51611 | ENSMUSG00000114049 | 0 | NA | NA | NA | NA | NA | Gm48140 | predicted gene, 48140 [Source:MGI Symb | 0 | 0 | 0 | 0 | 0 | 0 |
| 51612 | ENSMUSG00000114050 | 0.85298599 | -3.123636412 | 2.795026974 | -1.117569326 | 0.263750975 | NA | Gm47666 | predicted gene, 47666 [Source:MGI Symb | 0 | 4.1117 | 1.0062 | 0 | 0 | 0 |
| 51613 | ENSMUSG00000114051 | 0 | NA | NA | NA | NA | NA | Gm48008 | predicted gene, 48008 [Source:MGI Symb | 0 | 0 | 0 | 0 | 0 | 0 |
| 51614 | ENSMUSG00000114052 | 0 | NA | NA | NA | NA | NA | Gm48616 | predicted gene, 48616 [Source:MGI Symb | 0 | 0 | 0 | 0 | 0 | 0 |
| 51615 | ENSMUSG00000114053 | 0.17131964 | -0.796629176 | 4.080472857 | -0.195229623 | 0.84521319 | NA | Gm35330 | predicted gene, 35330 [Source:MGI Symb | 0 | 1.0279 | 0 | 0 | 0 | 0 |
| 51616 | ENSMUSG00000114054 | 0 | NA | NA | NA | NA | NA | Gm47706 | predicted gene, 47706 [Source:MGI Symb | 0 | 0 | 0 | 0 | 0 | 0 |
| 51617 | ENSMUSG00000114055 | 0 | NA | NA | NA | NA | NA | Gm32089 | predicted gene, 32089 [Source:MGI Symb | 0 | 0 | 0 | 0 | 0 | 0 |
| 51618 | ENSMUSG00000114056 | 0 | NA | NA | NA | NA | NA | Gm5668 | predicted gene 5668 [Source:MGI Symbol | 0 | 0 | 0 | 0 | 0 | 0 |
| 51619 | ENSMUSG00000114057 | 0 | NA | NA | NA | NA | NA | Gm31450 | predicted gene, 31450 [Source:MGI Symb | 0 | 0 | 0 | 0 | 0 | 0 |
| 51620 | ENSMUSG00000114058 | 0 | NA | NA | NA | NA | NA | Gm46317 | predicted gene, 46317 [Source:MGI Symb | 0 | 0 | 0 | 0 | 0 | 0 |
| 51621 | ENSMUSG00000114059 | 0 | NA | NA | NA | NA | NA | Gm48153 | predicted gene, 48153 [Source:MGI Symb | 0 | 0 | 0 | 0 | 0 | 0 |
| 51622 | ENSMUSG00000114060 | 0 | NA | NA | NA | NA | NA | Gm18630 | predicted gene, 18630 [Source:MGI Symb | 0 | 0 | 0 | 0 | 0 | 0 |
| 51623 | ENSMUSG00000114061 | 0 | NA | NA | NA | NA | NA | Gm48626 | predicted gene, 48626 [Source:MGI Symb | 0 | 0 | 0 | 0 | 0 | 0 |
| 51624 | ENSMUSG00000114062 | 0 | NA | NA | NA | NA | NA | Gm48313 | predicted gene, 48313 [Source:MGI Symb | 0 | 0 | 0 | 0 | 0 | 0 |
| 51625 | ENSMUSG00000114063 | 0 | NA | NA | NA | NA | NA | Gm47555 | predicted gene, 47555 [Source:MGI Symb | 0 | 0 | 0 | 0 | 0 | 0 |
| 51626 | ENSMUSG00000114064 | 0 | NA | NA | NA | NA | NA | Gm9370 | predicted gene 9370 [Source:MGI Symbol | 0 | 0 | 0 | 0 | 0 | 0 |
| 51627 | ENSMUSG00000114065 | 0 | NA | NA | NA | NA | NA | Vmn2r-ps96 | vomeronasal 2, receptor, pseudogene 96 | 0 | 0 | 0 | 0 | 0 | 0 |
| 51628 | ENSMUSG00000114066 | 0 | NA | NA | NA | NA | NA | Gm9202 | predicted gene 9202 [Source:MGI Symbol | 0 | 0 | 0 | 0 | 0 | 0 |

| | | | | | | | | | | | | | | | |
|---|---|---|---|---|---|---|---|---|---|---|---|---|---|---|---|
| 51629 | ENSMUSG00000114067 | 0 | NA | NA | NA | NA | NA | Gm47065 | predicted gene, 47065 [Source:MGI Symb | 0 | 0 | 0 | 0 | 0 | 0 |
| 51630 | ENSMUSG00000114068 | 0.335414862 | -1.779047242 | 4.030804179 | -0.441362855 | 0.658950329 | NA | Gm48783 | predicted gene, 48783 [Source:MGI Symb | 0 | 0 | 2.0125 | 0 | 0 | 0 |
| 51631 | ENSMUSG00000114069 | 0 | NA | NA | NA | NA | NA | E130119H09 | RIKEN cDNA E130119H09 gene [Source:M | 0 | 0 | 0 | 0 | 0 | 0 |
| 51632 | ENSMUSG00000114070 | 0 | NA | NA | NA | NA | NA | Gm47727 | predicted gene, 47727 [Source:MGI Symb | 0 | 0 | 0 | 0 | 0 | 0 |
| 51633 | ENSMUSG00000114071 | 0 | NA | NA | NA | NA | NA | Gm47390 | predicted gene, 47390 [Source:MGI Symb | 0 | 0 | 0 | 0 | 0 | 0 |
| 51634 | ENSMUSG00000114072 | 0 | NA | NA | NA | NA | NA | Gm49230 | predicted gene, 49230 [Source:MGI Symb | 0 | 0 | 0 | 0 | 0 | 0 |
| 51635 | ENSMUSG00000114073 | 0 | NA | NA | NA | NA | NA | Gm30302 | predicted gene, 30302 [Source:MGI Symb | 0 | 0 | 0 | 0 | 0 | 0 |
| 51636 | ENSMUSG00000114074 | 0 | NA | NA | NA | NA | NA | Gm48150 | predicted gene, 48150 [Source:MGI Symb | 0 | 0 | 0 | 0 | 0 | 0 |
| 51637 | ENSMUSG00000114075 | 0 | NA | NA | NA | NA | NA | Eif1ad18 | eukaryotic translation initiation factor 1A | 0 | 0 | 0 | 0 | 0 | 0 |
| 51638 | ENSMUSG00000114076 | 0 | NA | NA | NA | NA | NA | Gm48896 | predicted gene, 48896 [Source:MGI Symb | 0 | 0 | 0 | 0 | 0 | 0 |
| 51639 | ENSMUSG00000114077 | 0 | NA | NA | NA | NA | NA | Gm48776 | predicted gene, 48776 [Source:MGI Symb | 0 | 0 | 0 | 0 | 0 | 0 |
| 51640 | ENSMUSG00000114078 | 0 | NA | NA | NA | NA | NA | Tspyl-ps | testis-specific protein, Y-encoded-like, pse | 0 | 0 | 0 | 0 | 0 | 0 |
| 51641 | ENSMUSG00000114079 | 0 | NA | NA | NA | NA | NA | Gm47855 | predicted gene, 47855 [Source:MGI Symb | 0 | 0 | 0 | 0 | 0 | 0 |
| 51642 | ENSMUSG00000114080 | 0 | NA | NA | NA | NA | NA | Gm47661 | predicted gene, 47661 [Source:MGI Symb | 0 | 0 | 0 | 0 | 0 | 0 |
| 51643 | ENSMUSG00000114081 | 0 | NA | NA | NA | NA | NA | Gm48904 | predicted gene, 48904 [Source:MGI Symb | 0 | 0 | 0 | 0 | 0 | 0 |
| 51644 | ENSMUSG00000114082 | 0 | NA | NA | NA | NA | NA | Gm40383 | predicted gene, 40383 [Source:MGI Symb | 0 | 0 | 0 | 0 | 0 | 0 |
| 51645 | ENSMUSG00000114083 | 0 | NA | NA | NA | NA | NA | Gm47860 | predicted gene, 47860 [Source:MGI Symb | 0 | 0 | 0 | 0 | 0 | 0 |
| 51646 | ENSMUSG00000114084 | 0 | NA | NA | NA | NA | NA | Gm48897 | predicted gene, 48897 [Source:MGI Symb | 0 | 0 | 0 | 0 | 0 | 0 |
| 51647 | ENSMUSG00000114085 | 0 | NA | NA | NA | NA | NA | Gm6841 | predicted gene 6841 [Source:MGI Symbol | 0 | 0 | 0 | 0 | 0 | 0 |
| 51648 | ENSMUSG00000114086 | 0 | NA | NA | NA | NA | NA | Gm18968 | predicted gene, 18968 [Source:MGI Symb | 0 | 0 | 0 | 0 | 0 | 0 |
| 51649 | ENSMUSG00000114087 | 0 | NA | NA | NA | NA | NA | Gm7046 | predicted gene 7046 [Source:MGI Symbol | 0 | 0 | 0 | 0 | 0 | 0 |
| 51650 | ENSMUSG00000114088 | 0 | NA | NA | NA | NA | NA | Gm47898 | predicted gene, 47898 [Source:MGI Symb | 0 | 0 | 0 | 0 | 0 | 0 |
| 51651 | ENSMUSG00000114089 | 0 | NA | NA | NA | NA | NA | Gm47116 | predicted gene, 47116 [Source:MGI Symb | 0 | 0 | 0 | 0 | 0 | 0 |
| 51652 | ENSMUSG00000114090 | 0 | NA | NA | NA | NA | NA | Gm9472 | predicted gene 9472 [Source:MGI Symbol | 0 | 0 | 0 | 0 | 0 | 0 |
| 51653 | ENSMUSG00000114091 | 0 | NA | NA | NA | NA | NA | Gm40655 | predicted gene, 40655 [Source:MGI Symb | 0 | 0 | 0 | 0 | 0 | 0 |
| 51654 | ENSMUSG00000114092 | 0 | NA | NA | NA | NA | NA | Gm47418 | predicted gene, 47418 [Source:MGI Symb | 0 | 0 | 0 | 0 | 0 | 0 |
| 51655 | ENSMUSG00000114093 | 0 | NA | NA | NA | NA | NA | Gm47979 | predicted gene, 47979 [Source:MGI Symb | 0 | 0 | 0 | 0 | 0 | 0 |
| 51656 | ENSMUSG00000114094 | 0.342432689 | 1.990653086 | 3.900281263 | 0.51038706 | 0.609780321 | NA | Gm48375 | predicted gene, 48375 [Source:MGI Symb | 0 | 0 | 0 | 0.964 | 1.0906 | 0 |
| 51657 | ENSMUSG00000114095 | 0.503095663 | 2.549494177 | 3.348715477 | 0.761334964 | 0.446457022 | NA | Gm47424 | predicted gene, 47424 [Source:MGI Symb | 0 | 0 | 0 | 1.928 | 1.0906 | 0 |
| 51658 | ENSMUSG00000114096 | 0 | NA | NA | NA | NA | NA | Gm48641 | predicted gene, 48641 [Source:MGI Symb | 0 | 0 | 0 | 0 | 0 | 0 |
| 51659 | ENSMUSG00000114097 | 0 | NA | NA | NA | NA | NA | Gm35615 | predicted gene, 35615 [Source:MGI Symb | 0 | 0 | 0 | 0 | 0 | 0 |
| 51660 | ENSMUSG00000114098 | 0 | NA | NA | NA | NA | NA | Gm40660 | predicted gene, 40660 [Source:MGI Symb | 0 | 0 | 0 | 0 | 0 | 0 |
| 51661 | ENSMUSG00000114099 | 0 | NA | NA | NA | NA | NA | Gm48695 | predicted gene, 48695 [Source:MGI Symb | 0 | 0 | 0 | 0 | 0 | 0 |
| 51662 | ENSMUSG00000114100 | 0 | NA | NA | NA | NA | NA | Gm48647 | predicted gene, 48647 [Source:MGI Symb | 0 | 0 | 0 | 0 | 0 | 0 |
| 51663 | ENSMUSG00000114101 | 0 | NA | NA | NA | NA | NA | Gm48152 | predicted gene, 48152 [Source:MGI Symb | 0 | 0 | 0 | 0 | 0 | 0 |
| 51664 | ENSMUSG00000114102 | 0.349477146 | 0.165142768 | 3.911021834 | 0.042224967 | 0.966319359 | NA | Gm48048 | predicted gene, 48048 [Source:MGI Symb | 0 | 0 | 1.0062 | 0 | 1.0906 | 0 |
| 51665 | ENSMUSG00000114103 | 0 | NA | NA | NA | NA | NA | Gm8557 | predicted gene 8557 [Source:MGI Symbol | 0 | 0 | 0 | 0 | 0 | 0 |
| 51666 | ENSMUSG00000114104 | 4.100855007 | 1.053148167 | 1.156807835 | 0.910391627 | 0.362616012 | NA | Gm47585 | predicted gene, 47585 [Source:MGI Symb | 4.8186 | 3.0838 | 0 | 6.7478 | 3.2719 | 6.6831 |
| 51667 | ENSMUSG00000114105 | 0 | NA | NA | NA | NA | NA | Gm47713 | predicted gene, 47713 [Source:MGI Symb | 0 | 0 | 0 | 0 | 0 | 0 |
| 51668 | ENSMUSG00000114106 | 0 | NA | NA | NA | NA | NA | Gm47042 | predicted gene, 47042 [Source:MGI Symb | 0 | 0 | 0 | 0 | 0 | 0 |
| 51669 | ENSMUSG00000114107 | 0 | NA | NA | NA | NA | NA | Gm48031 | predicted gene, 48031 [Source:MGI Symb | 0 | 0 | 0 | 0 | 0 | 0 |
| 51670 | ENSMUSG00000114108 | 0 | NA | NA | NA | NA | NA | Gm48459 | predicted gene, 48459 [Source:MGI Symb | 0 | 0 | 0 | 0 | 0 | 0 |
| 51671 | ENSMUSG00000114109 | 0 | NA | NA | NA | NA | NA | Gm18629 | predicted gene, 18629 [Source:MGI Symb | 0 | 0 | 0 | 0 | 0 | 0 |
| 51672 | ENSMUSG00000114110 | 0.335414862 | -1.779047242 | 4.030804179 | -0.441362855 | 0.658950329 | NA | Gm46379 | predicted gene, 46379 [Source:MGI Symb | 0 | 0 | 2.0125 | 0 | 0 | 0 |
| 51673 | ENSMUSG00000114111 | 0 | NA | NA | NA | NA | NA | Gm9321 | predicted gene 9321 [Source:MGI Symbol | 0 | 0 | 0 | 0 | 0 | 0 |
| 51674 | ENSMUSG00000114112 | 0 | NA | NA | NA | NA | NA | Gm48301 | predicted gene, 48301 [Source:MGI Symb | 0 | 0 | 0 | 0 | 0 | 0 |
| 51675 | ENSMUSG00000114113 | 2.326814214 | -1.756638759 | 1.481286972 | -1.185886862 | 0.235666982 | NA | Gm47639 | predicted gene, 47639 [Source:MGI Symb | 1.6062 | 6.1675 | 3.0187 | 0.964 | 1.0906 | 1.1138 |
| 51676 | ENSMUSG00000114114 | 0 | NA | NA | NA | NA | NA | Gm48499 | predicted gene, 48499 [Source:MGI Symb | 0 | 0 | 0 | 0 | 0 | 0 |
| 51677 | ENSMUSG00000114115 | 0 | NA | NA | NA | NA | NA | Gm30239 | predicted gene, 30239 [Source:MGI Symb | 0 | 0 | 0 | 0 | 0 | 0 |
| 51678 | ENSMUSG00000114117 | 0 | NA | NA | NA | NA | NA | Gm48795 | predicted gene, 48795 [Source:MGI Symb | 0 | 0 | 0 | 0 | 0 | 0 |
| 51679 | ENSMUSG00000114118 | 0 | NA | NA | NA | NA | NA | Gm48829 | predicted gene, 48829 [Source:MGI Symb | 0 | 0 | 0 | 0 | 0 | 0 |
| 51680 | ENSMUSG00000114119 | 0 | NA | NA | NA | NA | NA | Gm48039 | predicted gene, 48039 [Source:MGI Symb | 0 | 0 | 0 | 0 | 0 | 0 |
| 51681 | ENSMUSG00000114120 | 0 | NA | NA | NA | NA | NA | Gm4120 | predicted gene 4120 [Source:MGI Symbol | 0 | 0 | 0 | 0 | 0 | 0 |
| 51682 | ENSMUSG00000114121 | 0 | NA | NA | NA | NA | NA | Gm47728 | predicted gene, 47728 [Source:MGI Symb | 0 | 0 | 0 | 0 | 0 | 0 |
| 51683 | ENSMUSG00000114122 | 0 | NA | NA | NA | NA | NA | Gm48712 | predicted gene, 48712 [Source:MGI Symb | 0 | 0 | 0 | 0 | 0 | 0 |
| 51684 | ENSMUSG00000114123 | 0 | NA | NA | NA | NA | NA | Gm48416 | predicted gene, 48416 [Source:MGI Symb | 0 | 0 | 0 | 0 | 0 | 0 |

| | | | | | | | | | | | | | | | |
|---|---|---|---|---|---|---|---|---|---|---|---|---|---|---|---|
| 51685 | ENSMUSG00000114124 | 0 | NA | NA | NA | NA | NA | Gm19084 | predicted gene, 19084 [Source:MGI Symb | 0 | 0 | 0 | 0 | 0 | 0 |
| 51686 | ENSMUSG00000114125 | 0.622883753 | 0.113321311 | 2.884614835 | 0.039284729 | 0.968663382 | NA | 1700037F03 | RIKEN cDNA 1700037F03 gene [Source:M | 0.8031 | 0 | 1.0062 | 1.928 | 0 | 0 |
| 51687 | ENSMUSG00000114126 | 0 | NA | NA | NA | NA | NA | Gm48257 | predicted gene, 48257 [Source:MGI Symb | 0 | 0 | 0 | 0 | 0 | 0 |
| 51688 | ENSMUSG00000114127 | 0 | NA | NA | NA | NA | NA | Gm8231 | predicted gene 8231 [Source:MGI Symbol | 0 | 0 | 0 | 0 | 0 | 0 |
| 51689 | ENSMUSG00000114128 | 0 | NA | NA | NA | NA | NA | Gm47452 | predicted gene, 47452 [Source:MGI Symb | 0 | 0 | 0 | 0 | 0 | 0 |
| 51690 | ENSMUSG00000114129 | 0 | NA | NA | NA | NA | NA | Gm16476 | predicted pseudogene 16476 [Source:MG | 0 | 0 | 0 | 0 | 0 | 0 |
| 51691 | ENSMUSG00000114130 | 0 | NA | NA | NA | NA | NA | Gm18954 | predicted gene, 18954 [Source:MGI Symb | 0 | 0 | 0 | 0 | 0 | 0 |
| 51692 | ENSMUSG00000114131 | 0 | NA | NA | NA | NA | NA | Gm47460 | predicted gene, 47460 [Source:MGI Symb | 0 | 0 | 0 | 0 | 0 | 0 |
| 51693 | ENSMUSG00000114132 | 0 | NA | NA | NA | NA | NA | Gm4808 | predicted gene 4808 [Source:MGI Symbol | 0 | 0 | 0 | 0 | 0 | 0 |
| 51694 | ENSMUSG00000114133 | 3.134506211 | -0.606541295 | 1.365196753 | -0.444288557 | 0.656833974 | NA | Gm20075 | predicted gene, 20075 [Source:MGI Symb | 7.2279 | 0 | 4.025 | 0.964 | 4.3625 | 2.2277 |
| 51695 | ENSMUSG00000114134 | 0 | NA | NA | NA | NA | NA | Gm47408 | predicted gene, 47408 [Source:MGI Symb | 0 | 0 | 0 | 0 | 0 | 0 |
| 51696 | ENSMUSG00000114135 | 0.181769715 | 1.126924175 | 4.080472857 | 0.276174898 | 0.782413736 | NA | Gm7252 | predicted gene 7252 [Source:MGI Symbol | 0 | 0 | 0 | 0 | 1.0906 | 0 |
| 51697 | ENSMUSG00000114136 | 0 | NA | NA | NA | NA | NA | 4921511I17F | RIKEN cDNA 4921511I17 gene [Source:MC | 0 | 0 | 0 | 0 | 0 | 0 |
| 51698 | ENSMUSG00000114137 | 0 | NA | NA | NA | NA | NA | Gm48685 | predicted gene, 48685 [Source:MGI Symb | 0 | 0 | 0 | 0 | 0 | 0 |
| 51699 | ENSMUSG00000114138 | 0.863771898 | -1.920093103 | 2.569535405 | -0.747253025 | 0.454910842 | NA | Gm36423 | predicted gene, 36423 [Source:MGI Symb | 3.2124 | 0 | 1.0062 | 0.964 | 0 | 0 |
| 51700 | ENSMUSG00000114139 | 0 | NA | NA | NA | NA | NA | Gm20336 | predicted gene, 20336 [Source:MGI Symb | 0 | 0 | 0 | 0 | 0 | 0 |
| 51701 | ENSMUSG00000114140 | 0 | NA | NA | NA | NA | NA | Gm6943 | predicted gene 6943 [Source:MGI Symbol | 0 | 0 | 0 | 0 | 0 | 0 |
| 51702 | ENSMUSG00000114141 | 0 | NA | NA | NA | NA | NA | Gm34215 | predicted gene, 34215 [Source:MGI Symb | 0 | 0 | 0 | 0 | 0 | 0 |
| 51703 | ENSMUSG00000114142 | 0 | NA | NA | NA | NA | NA | Gm47264 | predicted gene, 47264 [Source:MGI Symb | 0 | 0 | 0 | 0 | 0 | 0 |
| 51704 | ENSMUSG00000114143 | 0 | NA | NA | NA | NA | NA | Gm48548 | predicted gene, 48548 [Source:MGI Symb | 0 | 0 | 0 | 0 | 0 | 0 |
| 51705 | ENSMUSG00000114144 | 0 | NA | NA | NA | NA | NA | Gm9309 | predicted gene 9309 [Source:MGI Symbol | 0 | 0 | 0 | 0 | 0 | 0 |
| 51706 | ENSMUSG00000114145 | 0 | NA | NA | NA | NA | NA | Gm48561 | predicted gene, 48561 [Source:MGI Symb | 0 | 0 | 0 | 0 | 0 | 0 |
| 51707 | ENSMUSG00000114147 | 0.797635641 | -1.749522449 | 2.643029821 | -0.661938218 | 0.508010818 | NA | Gm18939 | predicted gene, 18939 [Source:MGI Symb | 0.8031 | 0 | 3.0187 | 0.964 | 0 | 0 |
| 51708 | ENSMUSG00000114148 | 1.197506817 | -0.21630013 | 2.096469441 | -0.103173519 | 0.917825256 | NA | Gm47701 | predicted gene, 47701 [Source:MGI Symb | 0.8031 | 1.0279 | 2.0125 | 0 | 0 | 3.3415 |
| 51709 | ENSMUSG00000114149 | 0 | NA | NA | NA | NA | NA | Gm47646 | predicted gene, 47646 [Source:MGI Symb | 0 | 0 | 0 | 0 | 0 | 0 |
| 51710 | ENSMUSG00000114150 | 2.498418082 | 0.324089531 | 1.322615678 | 0.245036814 | 0.806427927 | NA | Gm46367 | predicted gene, 46367 [Source:MGI Symb | 1.6062 | 3.0838 | 2.0125 | 3.8559 | 1.0906 | 3.3415 |
| 51711 | ENSMUSG00000114151 | 0 | NA | NA | NA | NA | NA | Gm47442 | predicted gene, 47442 [Source:MGI Symb | 0 | 0 | 0 | 0 | 0 | 0 |
| 51712 | ENSMUSG00000114152 | 0 | NA | NA | NA | NA | NA | Gm5374 | predicted gene 5374 [Source:MGI Symbol | 0 | 0 | 0 | 0 | 0 | 0 |
| 51713 | ENSMUSG00000114153 | 0 | NA | NA | NA | NA | NA | Gm48692 | predicted gene, 48692 [Source:MGI Symb | 0 | 0 | 0 | 0 | 0 | 0 |
| 51714 | ENSMUSG00000114154 | 0 | NA | NA | NA | NA | NA | Gm48597 | predicted gene, 48597 [Source:MGI Symb | 0 | 0 | 0 | 0 | 0 | 0 |
| 51715 | ENSMUSG00000114155 | 0 | NA | NA | NA | NA | NA | Gm7986 | predicted gene 7986 [Source:MGI Symbol | 0 | 0 | 0 | 0 | 0 | 0 |
| 51716 | ENSMUSG00000114156 | 0 | NA | NA | NA | NA | NA | Gm48523 | predicted gene, 48523 [Source:MGI Symb | 0 | 0 | 0 | 0 | 0 | 0 |
| 51717 | ENSMUSG00000114157 | 1.79128324 | -0.140149002 | 1.601099042 | -0.087533 | 0.930247856 | NA | Gm47870 | predicted gene, 47870 [Source:MGI Symb | 1.6062 | 2.0558 | 2.0125 | 2.8919 | 2.1812 | 0 |
| 51718 | ENSMUSG00000114158 | 0 | NA | NA | NA | NA | NA | Gm49372 | predicted gene, 49372 [Source:MGI Symb | 0 | 0 | 0 | 0 | 0 | 0 |
| 51719 | ENSMUSG00000114159 | 0 | NA | NA | NA | NA | NA | Gm48866 | predicted gene, 48866 [Source:MGI Symb | 0 | 0 | 0 | 0 | 0 | 0 |
| 51720 | ENSMUSG00000114160 | 0 | NA | NA | NA | NA | NA | Gm48352 | predicted gene, 48352 [Source:MGI Symb | 0 | 0 | 0 | 0 | 0 | 0 |
| 51721 | ENSMUSG00000114161 | 0 | NA | NA | NA | NA | NA | Gm48662 | predicted gene, 48662 [Source:MGI Symb | 0 | 0 | 0 | 0 | 0 | 0 |
| 51722 | ENSMUSG00000114162 | 0 | NA | NA | NA | NA | NA | Gm3785 | predicted gene 3785 [Source:MGI Symbol | 0 | 0 | 0 | 0 | 0 | 0 |
| 51723 | ENSMUSG00000114163 | 0 | NA | NA | NA | NA | NA | Gm32296 | predicted gene, 32296 [Source:MGI Symb | 0 | 0 | 0 | 0 | 0 | 0 |
| 51724 | ENSMUSG00000114164 | 0 | NA | NA | NA | NA | NA | Gm48636 | predicted gene, 48636 [Source:MGI Symb | 0 | 0 | 0 | 0 | 0 | 0 |
| 51725 | ENSMUSG00000114165 | 0.963688104 | -2.04288252 | 2.26700855 | -0.901135781 | 0.367516131 | NA | Gm47513 | predicted gene, 47513 [Source:MGI Symb | 1.6062 | 2.0558 | 1.0062 | 0 | 0 | 1.1138 |
| 51726 | ENSMUSG00000114166 | 0 | NA | NA | NA | NA | NA | Gm18246 | predicted gene, 18246 [Source:MGI Symb | 0 | 0 | 0 | 0 | 0 | 0 |
| 51727 | ENSMUSG00000114167 | 0 | NA | NA | NA | NA | NA | Gm7002 | predicted gene 7002 [Source:MGI Symbol | 0 | 0 | 0 | 0 | 0 | 0 |
| 51728 | ENSMUSG00000114168 | 0 | NA | NA | NA | NA | NA | Gm47194 | predicted gene, 47194 [Source:MGI Symb | 0 | 0 | 0 | 0 | 0 | 0 |
| 51729 | ENSMUSG00000114169 | 0.321325949 | 1.911575114 | 4.048267893 | 0.472195804 | 0.636787034 | NA | Gm47075 | predicted gene, 47075 [Source:MGI Symb | 0 | 0 | 0 | 1.928 | 0 | 0 |
| 51730 | ENSMUSG00000114170 | 0 | NA | NA | NA | NA | NA | Gm47120 | predicted gene, 47120 [Source:MGI Symb | 0 | 0 | 0 | 0 | 0 | 0 |
| 51731 | ENSMUSG00000114171 | 0 | NA | NA | NA | NA | NA | Gm18313 | predicted gene, 18313 [Source:MGI Symb | 0 | 0 | 0 | 0 | 0 | 0 |
| 51732 | ENSMUSG00000114172 | 3.048093962 | -0.697652029 | 1.411100021 | -0.494402961 | 0.621021623 | NA | Gm36298 | predicted gene, 36298 [Source:MGI Symb | 3.2124 | 7.1954 | 1.0062 | 5.7839 | 1.0906 | 0 |
| 51733 | ENSMUSG00000114173 | 0 | NA | NA | NA | NA | NA | Gm48638 | predicted gene, 48638 [Source:MGI Symb | 0 | 0 | 0 | 0 | 0 | 0 |
| 51734 | ENSMUSG00000114174 | 0 | NA | NA | NA | NA | NA | Cycs-ps3 | cytochrome c, pseudogene 3 [Source:MGI | 0 | 0 | 0 | 0 | 0 | 0 |
| 51735 | ENSMUSG00000114175 | 0 | NA | NA | NA | NA | NA | Gm48051 | predicted gene, 48051 [Source:MGI Symb | 0 | 0 | 0 | 0 | 0 | 0 |
| 51736 | ENSMUSG00000114176 | 0 | NA | NA | NA | NA | NA | Gm48329 | predicted gene, 48329 [Source:MGI Symb | 0 | 0 | 0 | 0 | 0 | 0 |
| 51737 | ENSMUSG00000114177 | 0.167707431 | -0.796629176 | 4.080472857 | -0.195229623 | 0.84521319 | NA | Gm18362 | predicted gene, 18362 [Source:MGI Symb | 0 | 0 | 1.0062 | 0 | 0 | 0 |
| 51738 | ENSMUSG00000114178 | 0 | NA | NA | NA | NA | NA | Gm48656 | predicted gene, 48656 [Source:MGI Symb | 0 | 0 | 0 | 0 | 0 | 0 |
| 51739 | ENSMUSG00000114179 | 0 | NA | NA | NA | NA | NA | Eif1-ps2 | eukaryotic translation initiation factor 1, p | 0 | 0 | 0 | 0 | 0 | 0 |
| 51740 | ENSMUSG00000114180 | 0 | NA | NA | NA | NA | NA | Gm47989 | predicted gene, 47989 [Source:MGI Symb | 0 | 0 | 0 | 0 | 0 | 0 |

| | | | | | | | | | | | | | | | |
|---|---|---|---|---|---|---|---|---|---|---|---|---|---|---|---|
| 51741 | ENSMUSG00000114181 | 0 | NA | NA | NA | NA | NA | Gm46376 | predicted gene, 46376 [Source:MGI Symb | 0 | 0 | 0 | 0 | 0 | 0 |
| 51742 | ENSMUSG00000114182 | 0 | NA | NA | NA | NA | NA | Gm47019 | predicted gene, 47019 [Source:MGI Symb | 0 | 0 | 0 | 0 | 0 | 0 |
| 51743 | ENSMUSG00000114183 | 0 | NA | NA | NA | NA | NA | Gm47195 | predicted gene, 47195 [Source:MGI Symb | 0 | 0 | 0 | 0 | 0 | 0 |
| 51744 | ENSMUSG00000114184 | 2.226323441 | 0.878860199 | 1.515800712 | 0.579799305 | 0.562049967 | NA | Gm7143 | predicted gene 7143 [Source:MGI Symbol | 1.6062 | 3.0838 | 0 | 0.964 | 4.3625 | 3.3415 |
| 51745 | ENSMUSG00000114185 | 0 | NA | NA | NA | NA | NA | Gm47662 | predicted gene, 47662 [Source:MGI Symb | 0 | 0 | 0 | 0 | 0 | 0 |
| 51746 | ENSMUSG00000114186 | 0 | NA | NA | NA | NA | NA | Gm48681 | predicted gene, 48681 [Source:MGI Symb | 0 | 0 | 0 | 0 | 0 | 0 |
| 51747 | ENSMUSG00000114187 | 0 | NA | NA | NA | NA | NA | Gm18626 | predicted gene, 18626 [Source:MGI Symb | 0 | 0 | 0 | 0 | 0 | 0 |
| 51748 | ENSMUSG00000114188 | 0 | NA | NA | NA | NA | NA | Gm40849 | predicted gene, 40849 [Source:MGI Symb | 0 | 0 | 0 | 0 | 0 | 0 |
| 51749 | ENSMUSG00000114189 | 0 | NA | NA | NA | NA | NA | 9330199G10 | RIKEN cDNA 9330199G10 gene [Source:M | 0 | 0 | 0 | 0 | 0 | 0 |
| 51750 | ENSMUSG00000114190 | 3.513360438 | -0.694057777 | 1.26494835 | -0.548684677 | 0.583221863 | NA | 4930556H04 | RIKEN cDNA 4930556H04 gene [Source:M | 4.0155 | 2.0558 | 7.0437 | 5.7839 | 2.1812 | 0 |
| 51751 | ENSMUSG00000114191 | 0 | NA | NA | NA | NA | NA | Gm18631 | predicted gene, 18631 [Source:MGI Symb | 0 | 0 | 0 | 0 | 0 | 0 |
| 51752 | ENSMUSG00000114192 | 0 | NA | NA | NA | NA | NA | Gm33115 | predicted gene, 33115 [Source:MGI Symb | 0 | 0 | 0 | 0 | 0 | 0 |
| 51753 | ENSMUSG00000114193 | 0.562214094 | -1.122836648 | 3.276281221 | -0.34271681 | 0.731811515 | NA | Gm48139 | predicted gene, 48139 [Source:MGI Symb | 2.4093 | 0 | 0 | 0.964 | 0 | 0 |
| 51754 | ENSMUSG00000114194 | 0 | NA | NA | NA | NA | NA | Gm3800 | predicted gene 3800 [Source:MGI Symbol | 0 | 0 | 0 | 0 | 0 | 0 |
| 51755 | ENSMUSG00000114195 | 0 | NA | NA | NA | NA | NA | 2900042G08 | RIKEN cDNA 2900042G08 gene [Source:M | 0 | 0 | 0 | 0 | 0 | 0 |
| 51756 | ENSMUSG00000114196 | 0 | NA | NA | NA | NA | NA | Gm47547 | predicted gene, 47547 [Source:MGI Symb | 0 | 0 | 0 | 0 | 0 | 0 |
| 51757 | ENSMUSG00000114197 | 0 | NA | NA | NA | NA | NA | Gm47743 | predicted gene, 47743 [Source:MGI Symb | 0 | 0 | 0 | 0 | 0 | 0 |
| 51758 | ENSMUSG00000114198 | 0 | NA | NA | NA | NA | NA | Gm47039 | predicted gene, 47039 [Source:MGI Symb | 0 | 0 | 0 | 0 | 0 | 0 |
| 51759 | ENSMUSG00000114199 | 0 | NA | NA | NA | NA | NA | Gm49341 | predicted gene, 49341 [Source:MGI Symb | 0 | 0 | 0 | 0 | 0 | 0 |
| 51760 | ENSMUSG00000114200 | 0 | NA | NA | NA | NA | NA | Gm34721 | predicted gene, 34721 [Source:MGI Symb | 0 | 0 | 0 | 0 | 0 | 0 |
| 51761 | ENSMUSG00000114201 | 0 | NA | NA | NA | NA | NA | Gm18079 | predicted gene, 18079 [Source:MGI Symb | 0 | 0 | 0 | 0 | 0 | 0 |
| 51762 | ENSMUSG00000114202 | 0 | NA | NA | NA | NA | NA | Gm18078 | predicted gene, 18078 [Source:MGI Symb | 0 | 0 | 0 | 0 | 0 | 0 |
| 51763 | ENSMUSG00000114203 | 1.063656444 | 0.142299728 | 2.383976614 | 0.059690069 | 0.952402482 | NA | Gm35279 | predicted gene, 35279 [Source:MGI Symb | 0 | 1.0279 | 2.0125 | 0 | 0 | 3.3415 |
| 51764 | ENSMUSG00000114204 | 0 | NA | NA | NA | NA | NA | Gm47520 | predicted gene, 47520 [Source:MGI Symb | 0 | 0 | 0 | 0 | 0 | 0 |
| 51765 | ENSMUSG00000114205 | 0 | NA | NA | NA | NA | NA | Gm7480 | predicted gene 7480 [Source:MGI Symbol | 0 | 0 | 0 | 0 | 0 | 0 |
| 51766 | ENSMUSG00000114206 | 1.179805695 | 1.5225842 | 2.130518386 | 0.714654335 | 0.474822662 | NA | Gm30551 | predicted gene, 30551 [Source:MGI Symb | 0.8031 | 0 | 1.0062 | 1.928 | 0 | 3.3415 |
| 51767 | ENSMUSG00000114207 | 0 | NA | NA | NA | NA | NA | Gm41109 | predicted gene, 41109 [Source:MGI Symb | 0 | 0 | 0 | 0 | 0 | 0 |
| 51768 | ENSMUSG00000114208 | 0 | NA | NA | NA | NA | NA | Gm34307 | predicted gene, 34307 [Source:MGI Symb | 0 | 0 | 0 | 0 | 0 | 0 |
| 51769 | ENSMUSG00000114209 | 0.51014012 | 1.031855138 | 3.09002894 | 0.333930574 | 0.738431948 | NA | Gm48433 | predicted gene, 48433 [Source:MGI Symb | 0 | 0 | 1.0062 | 0.964 | 1.0906 | 0 |
| 51770 | ENSMUSG00000114210 | 0 | NA | NA | NA | NA | NA | A330084C13 | RIKEN cDNA A330084C13 gene [Source:M | 0 | 0 | 0 | 0 | 0 | 0 |
| 51771 | ENSMUSG00000114211 | 0 | NA | NA | NA | NA | NA | Gm47635 | predicted gene, 47635 [Source:MGI Symb | 0 | 0 | 0 | 0 | 0 | 0 |
| 51772 | ENSMUSG00000114212 | 0 | NA | NA | NA | NA | NA | Gm47985 | predicted gene, 47985 [Source:MGI Symb | 0 | 0 | 0 | 0 | 0 | 0 |
| 51773 | ENSMUSG00000114213 | 0.589026695 | 0.241472922 | 3.228838524 | 0.074786311 | 0.940384733 | NA | A430090L17 | RIKEN cDNA A430090L17 gene [Source:M | 1.6062 | 0 | 0 | 1.928 | 0 | 0 |
| 51774 | ENSMUSG00000114214 | 0 | NA | NA | NA | NA | NA | Gm41102 | predicted gene, 41102 [Source:MGI Symb | 0 | 0 | 0 | 0 | 0 | 0 |
| 51775 | ENSMUSG00000114215 | 0 | NA | NA | NA | NA | NA | Gm47038 | predicted gene, 47038 [Source:MGI Symb | 0 | 0 | 0 | 0 | 0 | 0 |
| 51776 | ENSMUSG00000114216 | 0 | NA | NA | NA | NA | NA | Gm48592 | predicted gene, 48592 [Source:MGI Symb | 0 | 0 | 0 | 0 | 0 | 0 |
| 51777 | ENSMUSG00000114217 | 0.17131964 | -0.796629176 | 4.080472857 | -0.195229623 | 0.84521319 | NA | Gm7054 | predicted gene 7054 [Source:MGI Symbol | 0 | 1.0279 | 0 | 0 | 0 | 0 |
| 51778 | ENSMUSG00000114218 | 0 | NA | NA | NA | NA | NA | Gm47423 | predicted gene, 47423 [Source:MGI Symb | 0 | 0 | 0 | 0 | 0 | 0 |
| 51779 | ENSMUSG00000114219 | 0 | NA | NA | NA | NA | NA | Gm31456 | predicted gene, 31456 [Source:MGI Symb | 0 | 0 | 0 | 0 | 0 | 0 |
| 51780 | ENSMUSG00000114220 | 1.532456365 | 0.492622301 | 1.75969763 | 0.27994713 | 0.779518068 | NA | Gm10353 | predicted gene 10353 [Source:MGI Symbo | 0.8031 | 0 | 3.0187 | 0.964 | 2.1812 | 2.2277 |
| 51781 | ENSMUSG00000114221 | 0 | NA | NA | NA | NA | NA | Gm47351 | predicted gene, 47351 [Source:MGI Symb | 0 | 0 | 0 | 0 | 0 | 0 |
| 51782 | ENSMUSG00000114222 | 0 | NA | NA | NA | NA | NA | Gm19320 | predicted gene, 19320 [Source:MGI Symb | 0 | 0 | 0 | 0 | 0 | 0 |
| 51783 | ENSMUSG00000114223 | 0.160662974 | 1.126924175 | 4.080472857 | 0.276174898 | 0.782413736 | NA | Gm19108 | predicted gene, 19108 [Source:MGI Symb | 0 | 0 | 0 | 0.964 | 0 | 0 |
| 51784 | ENSMUSG00000114224 | 0 | NA | NA | NA | NA | NA | NA | NA | 0 | 0 | 0 | 0 | 0 | 0 |
| 51785 | ENSMUSG00000114225 | 0 | NA | NA | NA | NA | NA | Gm47883 | predicted gene, 47883 [Source:MGI Symb | 0 | 0 | 0 | 0 | 0 | 0 |
| 51786 | ENSMUSG00000114226 | 0 | NA | NA | NA | NA | NA | Gm48412 | predicted gene, 48412 [Source:MGI Symb | 0 | 0 | 0 | 0 | 0 | 0 |
| 51787 | ENSMUSG00000114227 | 0 | NA | NA | NA | NA | NA | Gm5204 | predicted gene 5204 [Source:MGI Symbol | 0 | 0 | 0 | 0 | 0 | 0 |
| 51788 | ENSMUSG00000114228 | 0 | NA | NA | NA | NA | NA | Gm47605 | predicted gene, 47605 [Source:MGI Symb | 0 | 0 | 0 | 0 | 0 | 0 |
| 51789 | ENSMUSG00000114229 | 0 | NA | NA | NA | NA | NA | Gm47072 | predicted gene, 47072 [Source:MGI Symb | 0 | 0 | 0 | 0 | 0 | 0 |
| 51790 | ENSMUSG00000114230 | 0 | NA | NA | NA | NA | NA | Gm48239 | predicted gene, 48239 [Source:MGI Symb | 0 | 0 | 0 | 0 | 0 | 0 |
| 51791 | ENSMUSG00000114231 | 0 | NA | NA | NA | NA | NA | Gm40968 | predicted gene, 40968 [Source:MGI Symb | 0 | 0 | 0 | 0 | 0 | 0 |
| 51792 | ENSMUSG00000114232 | 0.874376958 | 3.338720136 | 2.469103386 | 1.352199407 | 0.176311533 | NA | Gm17747 | predicted gene, 17747 [Source:MGI Symb | 0 | 0 | 0 | 1.928 | 1.0906 | 2.2277 |
| 51793 | ENSMUSG00000114233 | 0 | NA | NA | NA | NA | NA | Gm48629 | predicted gene, 48629 [Source:MGI Symb | 0 | 0 | 0 | 0 | 0 | 0 |
| 51794 | ENSMUSG00000114234 | 0.181769715 | 1.126924175 | 4.080472857 | 0.276174898 | 0.782413736 | NA | Gm3131 | predicted gene 3131 [Source:MGI Symbol | 0 | 0 | 0 | 0 | 1.0906 | 0 |
| 51795 | ENSMUSG00000114235 | 2.237657286 | 0.098826104 | 1.846307917 | 0.05352634 | 0.957312545 | NA | F530104D19 | RIKEN cDNA F530104D19 gene [Source:M | 6.4248 | 0 | 0 | 4.8199 | 2.1812 | 0 |
| 51796 | ENSMUSG00000114236 | 0 | NA | NA | NA | NA | NA | Gm47876 | predicted gene, 47876 [Source:MGI Symb | 0 | 0 | 0 | 0 | 0 | 0 |

| | | | | | | | | | | | | | | | |
|---|---|---|---|---|---|---|---|---|---|---|---|---|---|---|---|
| 51797 | ENSMUSG00000114237 | 0 | NA | NA | NA | NA | NA | Gm48694 | predicted gene, 48694 [Source:MGI Symb | 0 | 0 | 0 | 0 | 0 | 0 |
| 51798 | ENSMUSG00000114238 | 0.492645588 | 0.9565034 | 3.447648281 | 0.277436479 | 0.781444974 | NA | Rps17-ps1 | ribosomal protein S17, pseudogene 1 [Sou | 0 | 1.0279 | 0 | 1.928 | 0 | 0 |
| 51799 | ENSMUSG00000114239 | 0 | NA | NA | NA | NA | NA | Gm48245 | predicted gene, 48245 [Source:MGI Symb | 0 | 0 | 0 | 0 | 0 | 0 |
| 51800 | ENSMUSG00000114240 | 0 | NA | NA | NA | NA | NA | Gm47360 | predicted gene, 47360 [Source:MGI Symb | 0 | 0 | 0 | 0 | 0 | 0 |
| 51801 | ENSMUSG00000114241 | 0.524667718 | -0.819976748 | 3.064824146 | -0.267544469 | 0.78904998 | NA | Gm30108 | predicted gene, 30108 [Source:MGI Symb | 0 | 1.0279 | 1.0062 | 0 | 0 | 1.1138 |
| 51802 | ENSMUSG00000114242 | 0.17131964 | -0.796629176 | 4.080472857 | -0.195229623 | 0.84521319 | NA | Gm31219 | predicted gene, 31219 [Source:MGI Symb | 0 | 1.0279 | 0 | 0 | 0 | 0 |
| 51803 | ENSMUSG00000114243 | 5.20992396 | 1.775402928 | 1.026771512 | 1.729111986 | 0.083789054 | 0.426583908 | Gm48163 | predicted gene, 48163 [Source:MGI Symb | 4.8186 | 1.0279 | 1.0062 | 6.7478 | 7.6343 | 10.025 |
| 51804 | ENSMUSG00000114244 | 2.6958474 | 0.202333605 | 1.282048038 | 0.157820611 | 0.874598157 | NA | Gm47119 | predicted gene, 47119 [Source:MGI Symb | 2.4093 | 2.0558 | 3.0187 | 0.964 | 3.2719 | 4.4554 |
| 51805 | ENSMUSG00000114245 | 0 | NA | NA | NA | NA | NA | Percc1 | proline and glutamate rich with coiled coi | 0 | 0 | 0 | 0 | 0 | 0 |
| 51806 | ENSMUSG00000114246 | 0 | NA | NA | NA | NA | NA | Gm48603 | predicted gene, 48603 [Source:MGI Symb | 0 | 0 | 0 | 0 | 0 | 0 |
| 51807 | ENSMUSG00000114247 | 160.4167864 | 0.266125588 | 0.319502631 | 0.832937079 | 0.404880216 | 0.829488307 | Gm32063 | predicted gene, 32063 [Source:MGI Symb | 236.11 | 84.289 | 115.72 | 197.62 | 200.67 | 128.09 |
| 51808 | ENSMUSG00000114248 | 0 | NA | NA | NA | NA | NA | Gm8352 | predicted gene 8352 [Source:MGI Symbol | 0 | 0 | 0 | 0 | 0 | 0 |
| 51809 | ENSMUSG00000114249 | 0 | NA | NA | NA | NA | NA | Gm47709 | predicted gene, 47709 [Source:MGI Symb | 0 | 0 | 0 | 0 | 0 | 0 |
| 51810 | ENSMUSG00000114250 | 0 | NA | NA | NA | NA | NA | Gm49142 | predicted gene, 49142 [Source:MGI Symb | 0 | 0 | 0 | 0 | 0 | 0 |
| 51811 | ENSMUSG00000114251 | 1.936049003 | -0.845455968 | 1.485215156 | -0.569248142 | 0.569187754 | NA | Tmed10-ps | transmembrane p24 trafficking protein 10 | 2.4093 | 2.0558 | 3.0187 | 1.928 | 1.0906 | 1.1138 |
| 51812 | ENSMUSG00000114252 | 0 | NA | NA | NA | NA | NA | Gm5630 | predicted pseudogene 5630 [Source:MGI | 0 | 0 | 0 | 0 | 0 | 0 |
| 51813 | ENSMUSG00000114253 | 31.56240584 | -0.230136565 | 0.450684429 | -0.510637932 | 0.609604609 | 0.916376001 | Gm47798 | predicted gene, 47798 [Source:MGI Symb | 35.336 | 37.005 | 30.187 | 47.235 | 22.903 | 16.708 |
| 51814 | ENSMUSG00000114254 | 0.535401493 | -2.48596965 | 3.684117979 | -0.674780141 | 0.499815459 | NA | Gm48067 | predicted gene, 48067 [Source:MGI Symb | 3.2124 | 0 | 0 | 0 | 0 | 0 |
| 51815 | ENSMUSG00000114255 | 60.59423396 | 0.119605695 | 0.294170695 | 0.406586031 | 0.684312058 | 0.935528319 | Gm10734 | predicted gene 10734 [Source:MGI Symbo | 68.264 | 53.452 | 52.325 | 68.442 | 67.618 | 53.465 |
| 51816 | ENSMUSG00000114256 | 0 | NA | NA | NA | NA | NA | Gm29844 | predicted gene, 29844 [Source:MGI Symb | 0 | 0 | 0 | 0 | 0 | 0 |
| 51817 | ENSMUSG00000114257 | 0 | NA | NA | NA | NA | NA | NA | NA | 0 | 0 | 0 | 0 | 0 | 0 |
| 51818 | ENSMUSG00000114258 | 0 | NA | NA | NA | NA | NA | Gm47074 | predicted gene, 47074 [Source:MGI Symb | 0 | 0 | 0 | 0 | 0 | 0 |
| 51819 | ENSMUSG00000114259 | 0.17131964 | -0.796629176 | 4.080472857 | -0.195229623 | 0.84521319 | NA | Gm49351 | predicted gene, 49351 [Source:MGI Symb | 0 | 1.0279 | 0 | 0 | 0 | 0 |
| 51820 | ENSMUSG00000114260 | 0 | NA | NA | NA | NA | NA | Gm19198 | predicted gene, 19198 [Source:MGI Symb | 0 | 0 | 0 | 0 | 0 | 0 |
| 51821 | ENSMUSG00000114261 | 0 | NA | NA | NA | NA | NA | Gm48243 | predicted gene, 48243 [Source:MGI Symb | 0 | 0 | 0 | 0 | 0 | 0 |
| 51822 | ENSMUSG00000114262 | 0 | NA | NA | NA | NA | NA | Gm49532 | predicted gene, 49532 [Source:MGI Symb | 0 | 0 | 0 | 0 | 0 | 0 |
| 51823 | ENSMUSG00000114263 | 0 | NA | NA | NA | NA | NA | Gm7201 | predicted gene 7201 [Source:MGI Symbol | 0 | 0 | 0 | 0 | 0 | 0 |
| 51824 | ENSMUSG00000114264 | 0 | NA | NA | NA | NA | NA | Gm48095 | predicted gene, 48095 [Source:MGI Symb | 0 | 0 | 0 | 0 | 0 | 0 |
| 51825 | ENSMUSG00000114265 | 0 | NA | NA | NA | NA | NA | Gm47521 | predicted gene, 47521 [Source:MGI Symb | 0 | 0 | 0 | 0 | 0 | 0 |
| 51826 | ENSMUSG00000114266 | 0 | NA | NA | NA | NA | NA | Gm10403 | predicted gene 10403 [Source:MGI Symbo | 0 | 0 | 0 | 0 | 0 | 0 |
| 51827 | ENSMUSG00000114267 | 0.449470461 | -0.545136533 | 3.562105917 | -0.15303771 | 0.878368537 | NA | Gm48600 | predicted gene, 48600 [Source:MGI Symb | 1.6062 | 0 | 0 | 0 | 1.0906 | 0 |
| 51828 | ENSMUSG00000114268 | 0 | NA | NA | NA | NA | NA | NA | NA | 0 | 0 | 0 | 0 | 0 | 0 |
| 51829 | ENSMUSG00000114270 | 0 | NA | NA | NA | NA | NA | Gm48746 | predicted gene, 48746 [Source:MGI Symb | 0 | 0 | 0 | 0 | 0 | 0 |
| 51830 | ENSMUSG00000114271 | 0 | NA | NA | NA | NA | NA | Gm32224 | predicted gene, 32224 [Source:MGI Symb | 0 | 0 | 0 | 0 | 0 | 0 |
| 51831 | ENSMUSG00000114272 | 0 | NA | NA | NA | NA | NA | Gm36704 | predicted gene, 36704 [Source:MGI Symb | 0 | 0 | 0 | 0 | 0 | 0 |
| 51832 | ENSMUSG00000114273 | 0 | NA | NA | NA | NA | NA | Gm34585 | predicted gene, 34585 [Source:MGI Symb | 0 | 0 | 0 | 0 | 0 | 0 |
| 51833 | ENSMUSG00000114274 | 0 | NA | NA | NA | NA | NA | Gm35333 | predicted gene, 35333 [Source:MGI Symb | 0 | 0 | 0 | 0 | 0 | 0 |
| 51834 | ENSMUSG00000114275 | 0 | NA | NA | NA | NA | NA | Gm48534 | predicted gene, 48534 [Source:MGI Symb | 0 | 0 | 0 | 0 | 0 | 0 |
| 51835 | ENSMUSG00000114276 | 0 | NA | NA | NA | NA | NA | Gm47849 | predicted gene, 47849 [Source:MGI Symb | 0 | 0 | 0 | 0 | 0 | 0 |
| 51836 | ENSMUSG00000114277 | 0.538988726 | 1.126003851 | 3.343747182 | 0.336749099 | 0.736306047 | NA | Gm48583 | predicted gene, 48583 [Source:MGI Symb | 0 | 0 | 1.0062 | 0 | 0 | 2.2277 |
| 51837 | ENSMUSG00000114278 | 1.497287341 | -0.202022933 | 1.918626684 | -0.105295592 | 0.91614126 | NA | Gm49027 | predicted gene, 49027 [Source:MGI Symb | 0.8031 | 0 | 4.025 | 1.928 | 0 | 2.2277 |
| 51838 | ENSMUSG00000114279 | 0 | NA | NA | NA | NA | NA | H2bc14 | H2B clustered histone 14 [Source:MGI Syr | 0 | 0 | 0 | 0 | 0 | 0 |
| 51839 | ENSMUSG00000114280 | 0 | NA | NA | NA | NA | NA | Gm2791 | predicted gene 2791 [Source:MGI Symbol | 0 | 0 | 0 | 0 | 0 | 0 |
| 51840 | ENSMUSG00000114281 | 0 | NA | NA | NA | NA | NA | Gm48320 | predicted gene, 48320 [Source:MGI Symb | 0 | 0 | 0 | 0 | 0 | 0 |
| 51841 | ENSMUSG00000114282 | 0 | NA | NA | NA | NA | NA | 5330431K02 | RIKEN cDNA 5330431K02 gene [Source:M | 0 | 0 | 0 | 0 | 0 | 0 |
| 51842 | ENSMUSG00000114283 | 0 | NA | NA | NA | NA | NA | Gm47057 | predicted gene, 47057 [Source:MGI Symb | 0 | 0 | 0 | 0 | 0 | 0 |
| 51843 | ENSMUSG00000114284 | 1.186850152 | 0.606173358 | 2.099620832 | 0.288706108 | 0.772806288 | NA | 9630015K15 | RIKEN cDNA 9630015K15 gene [Source:M | 0.8031 | 0 | 2.0125 | 0.964 | 0 | 3.3415 |
| 51844 | ENSMUSG00000114285 | 0.716887508 | 1.693828953 | 2.739549946 | 0.618287305 | 0.536385969 | NA | Gm47773 | predicted gene, 47773 [Source:MGI Symb | 0 | 0 | 1.0062 | 0 | 2.1812 | 1.1138 |
| 51845 | ENSMUSG00000114286 | 0 | NA | NA | NA | NA | NA | Gm47850 | predicted gene, 47850 [Source:MGI Symb | 0 | 0 | 0 | 0 | 0 | 0 |
| 51846 | ENSMUSG00000114287 | 0 | NA | NA | NA | NA | NA | Gm5197 | predicted gene 5197 [Source:MGI Symbol | 0 | 0 | 0 | 0 | 0 | 0 |
| 51847 | ENSMUSG00000114288 | 0 | NA | NA | NA | NA | NA | Gm32123 | predicted gene, 32123 [Source:MGI Symb | 0 | 0 | 0 | 0 | 0 | 0 |
| 51848 | ENSMUSG00000114289 | 0 | NA | NA | NA | NA | NA | Gm48453 | predicted gene, 48453 [Source:MGI Symb | 0 | 0 | 0 | 0 | 0 | 0 |
| 51849 | ENSMUSG00000114290 | 0 | NA | NA | NA | NA | NA | 4930435F18 | RIKEN cDNA 4930435F18 gene [Source:M | 0 | 0 | 0 | 0 | 0 | 0 |
| 51850 | ENSMUSG00000114291 | 0 | NA | NA | NA | NA | NA | Gm47471 | predicted gene, 47471 [Source:MGI Symb | 0 | 0 | 0 | 0 | 0 | 0 |
| 51851 | ENSMUSG00000114292 | 1.723595754 | -1.99756196 | 1.742046574 | -1.146675405 | 0.251515799 | NA | Gm6076 | predicted gene 6076 [Source:MGI Symbol | 3.2124 | 2.0558 | 3.0187 | 0.964 | 1.0906 | 0 |
| 51852 | ENSMUSG00000114293 | 0.181769715 | 1.126924175 | 4.080472857 | 0.276174898 | 0.782413736 | NA | Gm48078 | predicted gene, 48078 [Source:MGI Symb | 0 | 0 | 0 | 0 | 1.0906 | 0 |

| | | | | | | | | | | | | | | | |
|---|---|---|---|---|---|---|---|---|---|---|---|---|---|---|---|
| 51853 | ENSMUSG00000114294 | 0 | NA | NA | NA | NA | NA | Gm47577 | predicted gene, 47577 [Source:MGI Symb | 0 | 0 | 0 | 0 | 0 | 0 |
| 51854 | ENSMUSG00000114295 | 0 | NA | NA | NA | NA | NA | Gm2534 | predicted gene 2534 [Source:MGI Symbol | 0 | 0 | 0 | 0 | 0 | 0 |
| 51855 | ENSMUSG00000114296 | 0 | NA | NA | NA | NA | NA | Pou5f1-rs6 | POU domain, class 5, transcription factor : | 0 | 0 | 0 | 0 | 0 | 0 |
| 51856 | ENSMUSG00000114297 | 0 | NA | NA | NA | NA | NA | Gm30600 | predicted gene, 30600 [Source:MGI Symb | 0 | 0 | 0 | 0 | 0 | 0 |
| 51857 | ENSMUSG00000114298 | 0 | NA | NA | NA | NA | NA | Gm48371 | predicted gene, 48371 [Source:MGI Symb | 0 | 0 | 0 | 0 | 0 | 0 |
| 51858 | ENSMUSG00000114299 | 0 | NA | NA | NA | NA | NA | Gm47791 | predicted gene, 47791 [Source:MGI Symb | 0 | 0 | 0 | 0 | 0 | 0 |
| 51859 | ENSMUSG00000114300 | 0 | NA | NA | NA | NA | NA | Gm32243 | predicted gene, 32243 [Source:MGI Symb | 0 | 0 | 0 | 0 | 0 | 0 |
| 51860 | ENSMUSG00000114301 | 0 | NA | NA | NA | NA | NA | Gm48300 | predicted gene, 48300 [Source:MGI Symb | 0 | 0 | 0 | 0 | 0 | 0 |
| 51861 | ENSMUSG00000114302 | 0.709843051 | 3.034749291 | 2.67440342 | 1.134738786 | 0.256484781 | NA | Gm48683 | predicted gene, 48683 [Source:MGI Symb | 0 | 0 | 0 | 0.964 | 2.1812 | 1.1138 |
| 51862 | ENSMUSG00000114303 | 0 | NA | NA | NA | NA | NA | Gm4022 | predicted gene 4022 [Source:MGI Symbol | 0 | 0 | 0 | 0 | 0 | 0 |
| 51863 | ENSMUSG00000114304 | 0 | NA | NA | NA | NA | NA | Gm48099 | predicted gene, 48099 [Source:MGI Symb | 0 | 0 | 0 | 0 | 0 | 0 |
| 51864 | ENSMUSG00000114305 | 0 | NA | NA | NA | NA | NA | Gm48437 | predicted gene, 48437 [Source:MGI Symb | 0 | 0 | 0 | 0 | 0 | 0 |
| 51865 | ENSMUSG00000114306 | 0 | NA | NA | NA | NA | NA | Gm3227 | predicted gene 3227 [Source:MGI Symbol | 0 | 0 | 0 | 0 | 0 | 0 |
| 51866 | ENSMUSG00000114307 | 0 | NA | NA | NA | NA | NA | 4930526H09 | RIKEN cDNA 4930526H09 gene [Source:M | 0 | 0 | 0 | 0 | 0 | 0 |
| 51867 | ENSMUSG00000114308 | 0 | NA | NA | NA | NA | NA | Gm48390 | predicted gene, 48390 [Source:MGI Symb | 0 | 0 | 0 | 0 | 0 | 0 |
| 51868 | ENSMUSG00000114309 | 0.328370405 | 0.16514316 | 4.003149536 | 0.041253308 | 0.967093956 | NA | Gm48295 | predicted gene, 48295 [Source:MGI Symb | 0 | 0 | 1.0062 | 0.964 | 0 | 0 |
| 51869 | ENSMUSG00000114310 | 0.321325949 | 1.911575114 | 4.048267893 | 0.472195804 | 0.636787034 | NA | Gm48602 | predicted gene, 48602 [Source:MGI Symb | 0 | 0 | 0 | 1.928 | 0 | 0 |
| 51870 | ENSMUSG00000114311 | 0.160662974 | 1.126924175 | 4.080472857 | 0.276174898 | 0.782413736 | NA | Gm31452 | predicted gene, 31452 [Source:MGI Symb | 0 | 0 | 0 | 0.964 | 0 | 0 |
| 51871 | ENSMUSG00000114312 | 0 | NA | NA | NA | NA | NA | Gm18077 | predicted gene, 18077 [Source:MGI Symb | 0 | 0 | 0 | 0 | 0 | 0 |
| 51872 | ENSMUSG00000114313 | 0 | NA | NA | NA | NA | NA | Gm4066 | predicted pseudogene 4066 [Source:MGI | 0 | 0 | 0 | 0 | 0 | 0 |
| 51873 | ENSMUSG00000114314 | 0 | NA | NA | NA | NA | NA | Gm48413 | predicted gene, 48413 [Source:MGI Symb | 0 | 0 | 0 | 0 | 0 | 0 |
| 51874 | ENSMUSG00000114315 | 0 | NA | NA | NA | NA | NA | Gm48696 | predicted gene, 48696 [Source:MGI Symb | 0 | 0 | 0 | 0 | 0 | 0 |
| 51875 | ENSMUSG00000114316 | 0.160662974 | 1.126924175 | 4.080472857 | 0.276174898 | 0.782413736 | NA | Gm17740 | predicted gene, 17740 [Source:MGI Symb | 0 | 0 | 0 | 0.964 | 0 | 0 |
| 51876 | ENSMUSG00000114317 | 0 | NA | NA | NA | NA | NA | Gm49873 | predicted gene, 49873 [Source:MGI Symb | 0 | 0 | 0 | 0 | 0 | 0 |
| 51877 | ENSMUSG00000114318 | 0 | NA | NA | NA | NA | NA | Gm48781 | predicted gene, 48781 [Source:MGI Symb | 0 | 0 | 0 | 0 | 0 | 0 |
| 51878 | ENSMUSG00000114319 | 0 | NA | NA | NA | NA | NA | Gm47509 | predicted gene, 47509 [Source:MGI Symb | 0 | 0 | 0 | 0 | 0 | 0 |
| 51879 | ENSMUSG00000114320 | 0 | NA | NA | NA | NA | NA | Gm30839 | predicted gene, 30839 [Source:MGI Symb | 0 | 0 | 0 | 0 | 0 | 0 |
| 51880 | ENSMUSG00000114321 | 1.723285425 | 1.301727174 | 1.700594598 | 0.765454139 | 0.444001165 | NA | Gm8971 | predicted gene 8971 [Source:MGI Symbol | 0 | 2.0558 | 1.0062 | 2.8919 | 3.2719 | 1.1138 |
| 51881 | ENSMUSG00000114322 | 0 | NA | NA | NA | NA | NA | Gm47041 | predicted gene, 47041 [Source:MGI Symb | 0 | 0 | 0 | 0 | 0 | 0 |
| 51882 | ENSMUSG00000114323 | 0 | NA | NA | NA | NA | NA | Gm47758 | predicted gene, 47758 [Source:MGI Symb | 0 | 0 | 0 | 0 | 0 | 0 |
| 51883 | ENSMUSG00000114324 | 0 | NA | NA | NA | NA | NA | Gm18135 | predicted gene, 18135 [Source:MGI Symb | 0 | 0 | 0 | 0 | 0 | 0 |
| 51884 | ENSMUSG00000114325 | 1.819118595 | 3.284740217 | 2.05422546 | 1.599016408 | 0.109816957 | NA | Rpl13-ps2 | ribosomal protein L13, pseudogene 2 [Sou | 0 | 0 | 1.0062 | 0 | 5.4531 | 4.4554 |
| 51885 | ENSMUSG00000114326 | 0 | NA | NA | NA | NA | NA | Gm4938 | predicted pseudogene 4938 [Source:MGI | 0 | 0 | 0 | 0 | 0 | 0 |
| 51886 | ENSMUSG00000114327 | 0 | NA | NA | NA | NA | NA | Gm40988 | predicted gene, 40988 [Source:MGI Symb | 0 | 0 | 0 | 0 | 0 | 0 |
| 51887 | ENSMUSG00000114328 | 0 | NA | NA | NA | NA | NA | Gm48246 | predicted gene, 48246 [Source:MGI Symb | 0 | 0 | 0 | 0 | 0 | 0 |
| 51888 | ENSMUSG00000114329 | 0 | NA | NA | NA | NA | NA | Gm30489 | predicted gene, 30489 [Source:MGI Symb | 0 | 0 | 0 | 0 | 0 | 0 |
| 51889 | ENSMUSG00000114330 | 0 | NA | NA | NA | NA | NA | Gm47349 | predicted gene, 47349 [Source:MGI Symb | 0 | 0 | 0 | 0 | 0 | 0 |
| 51890 | ENSMUSG00000114331 | 0 | NA | NA | NA | NA | NA | Gm48056 | predicted gene, 48056 [Source:MGI Symb | 0 | 0 | 0 | 0 | 0 | 0 |
| 51891 | ENSMUSG00000114332 | 0.160662974 | 1.126924175 | 4.080472857 | 0.276174898 | 0.782413736 | NA | Gm8847 | predicted gene 8847 [Source:MGI Symbol | 0 | 0 | 0 | 0.964 | 0 | 0 |
| 51892 | ENSMUSG00000114333 | 0 | NA | NA | NA | NA | NA | 1700084D21 | RIKEN cDNA 1700084D21 gene [Source:M | 0 | 0 | 0 | 0 | 0 | 0 |
| 51893 | ENSMUSG00000114334 | 0 | NA | NA | NA | NA | NA | Gm31393 | predicted gene, 31393 [Source:MGI Symb | 0 | 0 | 0 | 0 | 0 | 0 |
| 51894 | ENSMUSG00000114335 | 0 | NA | NA | NA | NA | NA | Gm8456 | predicted gene 8456 [Source:MGI Symbol | 0 | 0 | 0 | 0 | 0 | 0 |
| 51895 | ENSMUSG00000114336 | 0 | NA | NA | NA | NA | NA | Gm18643 | predicted gene, 18643 [Source:MGI Symb | 0 | 0 | 0 | 0 | 0 | 0 |
| 51896 | ENSMUSG00000114337 | 0 | NA | NA | NA | NA | NA | Gm48704 | predicted gene, 48704 [Source:MGI Symb | 0 | 0 | 0 | 0 | 0 | 0 |
| 51897 | ENSMUSG00000114338 | 0 | NA | NA | NA | NA | NA | Gm4059 | predicted gene 4059 [Source:MGI Symbol | 0 | 0 | 0 | 0 | 0 | 0 |
| 51898 | ENSMUSG00000114339 | 0 | NA | NA | NA | NA | NA | Gm47974 | predicted gene, 47974 [Source:MGI Symb | 0 | 0 | 0 | 0 | 0 | 0 |
| 51899 | ENSMUSG00000114340 | 0 | NA | NA | NA | NA | NA | Gm18132 | predicted gene, 18132 [Source:MGI Symb | 0 | 0 | 0 | 0 | 0 | 0 |
| 51900 | ENSMUSG00000114341 | 0 | NA | NA | NA | NA | NA | Gm47104 | predicted gene, 47104 [Source:MGI Symb | 0 | 0 | 0 | 0 | 0 | 0 |
| 51901 | ENSMUSG00000114342 | 0 | NA | NA | NA | NA | NA | 4921520N01 | RIKEN cDNA 4921520N01 gene [Source:M | 0 | 0 | 0 | 0 | 0 | 0 |
| 51902 | ENSMUSG00000114343 | 0 | NA | NA | NA | NA | NA | Gm48322 | predicted gene, 48322 [Source:MGI Symb | 0 | 0 | 0 | 0 | 0 | 0 |
| 51903 | ENSMUSG00000114344 | 0 | NA | NA | NA | NA | NA | Gm17941 | predicted gene, 17941 [Source:MGI Symb | 0 | 0 | 0 | 0 | 0 | 0 |
| 51904 | ENSMUSG00000114345 | 0 | NA | NA | NA | NA | NA | Gm10873 | predicted gene 10873 [Source:MGI Symbo | 0 | 0 | 0 | 0 | 0 | 0 |
| 51905 | ENSMUSG00000114346 | 0 | NA | NA | NA | NA | NA | Gm5195 | predicted gene 5195 [Source:MGI Symbol | 0 | 0 | 0 | 0 | 0 | 0 |
| 51906 | ENSMUSG00000114347 | 0 | NA | NA | NA | NA | NA | Gm47913 | predicted gene, 47913 [Source:MGI Symb | 0 | 0 | 0 | 0 | 0 | 0 |
| 51907 | ENSMUSG00000114348 | 0 | NA | NA | NA | NA | NA | Gm41031 | predicted gene, 41031 [Source:MGI Symb | 0 | 0 | 0 | 0 | 0 | 0 |
| 51908 | ENSMUSG00000114349 | 0.349477146 | 0.165142768 | 3.911021834 | 0.042224967 | 0.966319359 | NA | Gm7591 | predicted gene 7591 [Source:MGI Symbol | 0 | 0 | 1.0062 | 0 | 1.0906 | 0 |

| | | | | | | | | | | | | | | | |
|---|---|---|---|---|---|---|---|---|---|---|---|---|---|---|---|
| 51909 | ENSMUSG00000114350 | 0 | NA | NA | NA | NA | NA | D14Ertd670e | DNA segment, Chr 14, ERATO Doi 670, exp | 0 | 0 | 0 | 0 | 0 | 0 |
| 51910 | ENSMUSG00000114352 | 0 | NA | NA | NA | NA | NA | 4931407E12 | RIKEN cDNA 4931407E12 gene [Source:MG | 0 | 0 | 0 | 0 | 0 | 0 |
| 51911 | ENSMUSG00000114353 | 0 | NA | NA | NA | NA | NA | Gm7896 | predicted gene 7869 [Source:MGI Symbol | 0 | 0 | 0 | 0 | 0 | 0 |
| 51912 | ENSMUSG00000114354 | 3.790915275 | 0.00713844 | 1.215825979 | 0.005871268 | 0.995315433 | NA | Gm34961 | predicted gene, 34961 [Source:MGI Symb | 3.2124 | 6.1675 | 2.0125 | 5.7839 | 0 | 5.5692 |
| 51913 | ENSMUSG00000114355 | 13.07467119 | 1.056381079 | 0.657242783 | 1.607292018 | 0.107990324 | 0.484761602 | Gm40916 | predicted gene, 40916 [Source:MGI Symb | 12.047 | 5.1396 | 8.05 | 14.46 | 10.906 | 27.846 |
| 51914 | ENSMUSG00000114356 | 0 | NA | NA | NA | NA | NA | Gm47265 | predicted gene, 47265 [Source:MGI Symb | 0 | 0 | 0 | 0 | 0 | 0 |
| 51915 | ENSMUSG00000114357 | 1.806916214 | 1.06222317 | 1.65810419 | 0.640625104 | 0.521766285 | NA | Gm48449 | predicted gene, 48449 [Source:MGI Symb | 2.4093 | 1.0279 | 0 | 1.928 | 4.3625 | 1.1138 |
| 51916 | ENSMUSG00000114358 | 0 | NA | NA | NA | NA | NA | Gm5200 | predicted gene 5200 [Source:MGI Symbol | 0 | 0 | 0 | 0 | 0 | 0 |
| 51917 | ENSMUSG00000114359 | 0 | NA | NA | NA | NA | NA | Gm47816 | predicted gene, 47816 [Source:MGI Symb | 0 | 0 | 0 | 0 | 0 | 0 |
| 51918 | ENSMUSG00000114360 | 0.798947607 | -0.296306415 | 2.620405875 | -0.113076534 | 0.909969879 | NA | Gm48596 | predicted gene, 48596 [Source:MGI Symb | 1.6062 | 0 | 1.0062 | 0 | 2.1812 | 0 |
| 51919 | ENSMUSG00000114361 | 0 | NA | NA | NA | NA | NA | Gm31104 | predicted gene, 31104 [Source:MGI Symb | 0 | 0 | 0 | 0 | 0 | 0 |
| 51920 | ENSMUSG00000114362 | 7.633467267 | -0.553142802 | 0.789863819 | -0.70030148 | 0.483739048 | 0.872271041 | 4933433G19 | RIKEN cDNA 4933433G19 gene [Source:M | 8.031 | 10.279 | 9.0562 | 9.6398 | 5.4531 | 3.3415 |
| 51921 | ENSMUSG00000114363 | 0 | NA | NA | NA | NA | NA | Gm48174 | predicted gene, 48174 [Source:MGI Symb | 0 | 0 | 0 | 0 | 0 | 0 |
| 51922 | ENSMUSG00000114364 | 0 | NA | NA | NA | NA | NA | Gm47426 | predicted gene, 47426 [Source:MGI Symb | 0 | 0 | 0 | 0 | 0 | 0 |
| 51923 | ENSMUSG00000114365 | 0 | NA | NA | NA | NA | NA | Gm9098 | predicted gene 9098 [Source:MGI Symbol | 0 | 0 | 0 | 0 | 0 | 0 |
| 51924 | ENSMUSG00000114366 | 0 | NA | NA | NA | NA | NA | Gm4165 | predicted gene 4165 [Source:MGI Symbol | 0 | 0 | 0 | 0 | 0 | 0 |
| 51925 | ENSMUSG00000114367 | 0 | NA | NA | NA | NA | NA | Gm48216 | predicted gene, 48216 [Source:MGI Symb | 0 | 0 | 0 | 0 | 0 | 0 |
| 51926 | ENSMUSG00000114368 | 0 | NA | NA | NA | NA | NA | Gm47953 | predicted gene, 47953 [Source:MGI Symb | 0 | 0 | 0 | 0 | 0 | 0 |
| 51927 | ENSMUSG00000114369 | 1.998057258 | -1.416831084 | 1.615124926 | -0.877226932 | 0.38036339 | NA | Gm41077 | predicted gene, 41077 [Source:MGI Symb | 1.6062 | 2.0558 | 5.0312 | 0 | 2.1812 | 1.1138 |
| 51928 | ENSMUSG00000114370 | 21.1773351 | 0.293726256 | 0.509995529 | 0.575938885 | 0.564656491 | 0.901542552 | Gm46411 | predicted gene, 46411 [Source:MGI Symb | 23.29 | 17.475 | 16.1 | 16.388 | 35.99 | 17.822 |
| 51929 | ENSMUSG00000114371 | 0.17131964 | -0.796629176 | 4.080472857 | -0.195229623 | 0.84521319 | NA | Gm48769 | predicted gene, 48769 [Source:MGI Symb | 0 | 1.0279 | 0 | 0 | 0 | 0 |
| 51930 | ENSMUSG00000114372 | 0 | NA | NA | NA | NA | NA | 2310067P03 | RIKEN cDNA 2310067P03 gene [Source:MG | 0 | 0 | 0 | 0 | 0 | 0 |
| 51931 | ENSMUSG00000114373 | 0 | NA | NA | NA | NA | NA | Gm48708 | predicted gene, 48708 [Source:MGI Symb | 0 | 0 | 0 | 0 | 0 | 0 |
| 51932 | ENSMUSG00000114374 | 0 | NA | NA | NA | NA | NA | Gm7911 | predicted gene 7911 [Source:MGI Symbol | 0 | 0 | 0 | 0 | 0 | 0 |
| 51933 | ENSMUSG00000114375 | 0 | NA | NA | NA | NA | NA | Gm31544 | predicted gene, 31544 [Source:MGI Symb | 0 | 0 | 0 | 0 | 0 | 0 |
| 51934 | ENSMUSG00000114376 | 0 | NA | NA | NA | NA | NA | Gm18807 | predicted gene, 18807 [Source:MGI Symb | 0 | 0 | 0 | 0 | 0 | 0 |
| 51935 | ENSMUSG00000114377 | 0 | NA | NA | NA | NA | NA | Gm48247 | predicted gene, 48247 [Source:MGI Symb | 0 | 0 | 0 | 0 | 0 | 0 |
| 51936 | ENSMUSG00000114378 | 0.17131964 | -0.796629176 | 4.080472857 | -0.195229623 | 0.84521319 | NA | Gm49355 | predicted gene, 49355 [Source:MGI Symb | 0 | 1.0279 | 0 | 0 | 0 | 0 |
| 51937 | ENSMUSG00000114379 | 0 | NA | NA | NA | NA | NA | Gm4241 | predicted gene 4241 [Source:MGI Symbol | 0 | 0 | 0 | 0 | 0 | 0 |
| 51938 | ENSMUSG00000114380 | 0 | NA | NA | NA | NA | NA | Gm47942 | predicted gene, 47942 [Source:MGI Symb | 0 | 0 | 0 | 0 | 0 | 0 |
| 51939 | ENSMUSG00000114381 | 0 | NA | NA | NA | NA | NA | Gm47361 | predicted gene, 47361 [Source:MGI Symb | 0 | 0 | 0 | 0 | 0 | 0 |
| 51940 | ENSMUSG00000114382 | 0 | NA | NA | NA | NA | NA | Gm4814 | predicted gene 4814 [Source:MGI Symbol | 0 | 0 | 0 | 0 | 0 | 0 |
| 51941 | ENSMUSG00000114383 | 0 | NA | NA | NA | NA | NA | Gm3963 | predicted gene 3963 [Source:MGI Symbol | 0 | 0 | 0 | 0 | 0 | 0 |
| 51942 | ENSMUSG00000114384 | 2.585222395 | -0.142737859 | 1.283034599 | -0.111250202 | 0.911417943 | NA | 4930455J16I | RIKEN cDNA 4930455J16 gene [Source:MG | 4.0155 | 2.0558 | 2.0125 | 1.928 | 3.2719 | 2.2277 |
| 51943 | ENSMUSG00000114385 | 0 | NA | NA | NA | NA | NA | Gm48344 | predicted gene, 48344 [Source:MGI Symb | 0 | 0 | 0 | 0 | 0 | 0 |
| 51944 | ENSMUSG00000114386 | 0 | NA | NA | NA | NA | NA | Gm47576 | predicted gene, 47576 [Source:MGI Symb | 0 | 0 | 0 | 0 | 0 | 0 |
| 51945 | ENSMUSG00000114387 | 1.111859485 | -1.041840268 | 2.15740327 | -0.482914012 | 0.629156792 | NA | Gm48767 | predicted gene, 48767 [Source:MGI Symb | 2.4093 | 1.0279 | 1.0062 | 0 | 0 | 2.2277 |
| 51946 | ENSMUSG00000114388 | 0.353089354 | 0.165142706 | 3.896102244 | 0.042386646 | 0.966190474 | NA | Gm48510 | predicted gene, 48510 [Source:MGI Symb | 0 | 1.0279 | 0 | 0 | 1.0906 | 0 |
| 51947 | ENSMUSG00000114389 | 0 | NA | NA | NA | NA | NA | Gm47734 | predicted gene, 47734 [Source:MGI Symb | 0 | 0 | 0 | 0 | 0 | 0 |
| 51948 | ENSMUSG00000114390 | 0 | NA | NA | NA | NA | NA | 1700062C10 | RIKEN cDNA 1700062C10 gene [Source:MG | 0 | 0 | 0 | 0 | 0 | 0 |
| 51949 | ENSMUSG00000114391 | 0 | NA | NA | NA | NA | NA | Gm48439 | predicted gene, 48439 [Source:MGI Symb | 0 | 0 | 0 | 0 | 0 | 0 |
| 51950 | ENSMUSG00000114392 | 0 | NA | NA | NA | NA | NA | Gm8961 | predicted gene 8961 [Source:MGI Symbol | 0 | 0 | 0 | 0 | 0 | 0 |
| 51951 | ENSMUSG00000114393 | 0 | NA | NA | NA | NA | NA | Gm47359 | predicted gene, 47359 [Source:MGI Symb | 0 | 0 | 0 | 0 | 0 | 0 |
| 51952 | ENSMUSG00000114394 | 0 | NA | NA | NA | NA | NA | Gm19153 | predicted gene, 19153 [Source:MGI Symb | 0 | 0 | 0 | 0 | 0 | 0 |
| 51953 | ENSMUSG00000114395 | 0.342639279 | -1.805976596 | 4.028547714 | -0.448294702 | 0.653940524 | NA | Gm47464 | predicted gene, 47464 [Source:MGI Symb | 0 | 2.0558 | 0 | 0 | 0 | 0 |
| 51954 | ENSMUSG00000114396 | 1.620608875 | -0.486635318 | 1.858099182 | -0.261899538 | 0.793398896 | NA | Gm31227 | predicted gene, 31227 [Source:MGI Symb | 1.6062 | 4.1117 | 0 | 2.8919 | 0 | 1.1138 |
| 51955 | ENSMUSG00000114397 | 0 | NA | NA | NA | NA | NA | Gm7621 | predicted gene 7621 [Source:MGI Symbol | 0 | 0 | 0 | 0 | 0 | 0 |
| 51956 | ENSMUSG00000114398 | 0 | NA | NA | NA | NA | NA | Gm20763 | predicted gene, 20763 [Source:MGI Symb | 0 | 0 | 0 | 0 | 0 | 0 |
| 51957 | ENSMUSG00000114399 | 1.572400321 | 0.521780432 | 1.939267349 | 0.269060598 | 0.787883051 | NA | Gm47994 | predicted gene, 47994 [Source:MGI Symb | 0.8031 | 2.0558 | 1.0062 | 0 | 0 | 5.5692 |
| 51958 | ENSMUSG00000114400 | 0 | NA | NA | NA | NA | NA | Gm47533 | predicted gene, 47533 [Source:MGI Symb | 0 | 0 | 0 | 0 | 0 | 0 |
| 51959 | ENSMUSG00000114401 | 1.238640426 | 1.454958615 | 2.28972858 | 0.635428421 | 0.525149037 | NA | Gm38604 | predicted gene, 38604 [Source:MGI Symb | 0 | 0 | 2.0125 | 0.964 | 0 | 4.4554 |
| 51960 | ENSMUSG00000114402 | 0 | NA | NA | NA | NA | NA | Gm49360 | predicted gene, 49360 [Source:MGI Symb | 0 | 0 | 0 | 0 | 0 | 0 |
| 51961 | ENSMUSG00000114403 | 0 | NA | NA | NA | NA | NA | Gm21021 | predicted gene, 21021 [Source:MGI Symb | 0 | 0 | 0 | 0 | 0 | 0 |
| 51962 | ENSMUSG00000114404 | 2.863896178 | -0.155207598 | 1.281735024 | -0.121091798 | 0.903618326 | NA | Gm40915 | predicted gene, 40915 [Source:MGI Symb | 4.0155 | 1.0279 | 4.025 | 4.8199 | 2.1812 | 1.1138 |
| 51963 | ENSMUSG00000114405 | 0 | NA | NA | NA | NA | NA | Gm47825 | predicted gene, 47825 [Source:MGI Symb | 0 | 0 | 0 | 0 | 0 | 0 |
| 51964 | ENSMUSG00000114406 | 0 | NA | NA | NA | NA | NA | Gm48323 | predicted gene, 48323 [Source:MGI Symb | 0 | 0 | 0 | 0 | 0 | 0 |

| | | | | | | | | | | | | | | | |
|---|---|---|---|---|---|---|---|---|---|---|---|---|---|---|---|
| 51965 | ENSMUSG00000114407 | 0 | NA | NA | NA | NA | NA | Gm48765 | predicted gene, 48765 [Source:MGI Symb | 0 | 0 | 0 | 0 | 0 | 0 |
| 51966 | ENSMUSG00000114408 | 0 | NA | NA | NA | NA | NA | Gm47810 | predicted gene, 47810 [Source:MGI Symb | 0 | 0 | 0 | 0 | 0 | 0 |
| 51967 | ENSMUSG00000114409 | 0.77079641 | 0.856159616 | 2.652840961 | 0.322733111 | 0.746897374 | NA | Gm9042 | predicted gene 9042 [Source:MGI Symbol | 1.6062 | 0 | 0 | 1.928 | 1.0906 | 0 |
| 51968 | ENSMUSG00000114410 | 0 | NA | NA | NA | NA | NA | Gm2048 | predicted gene 2048 [Source:MGI Symbol | 0 | 0 | 0 | 0 | 0 | 0 |
| 51969 | ENSMUSG00000114411 | 0 | NA | NA | NA | NA | NA | Gm48176 | predicted gene, 48176 [Source:MGI Symb | 0 | 0 | 0 | 0 | 0 | 0 |
| 51970 | ENSMUSG00000114412 | 0 | NA | NA | NA | NA | NA | 4930405A10 | RIKEN cDNA 4930405A10 gene [Source:M | 0 | 0 | 0 | 0 | 0 | 0 |
| 51971 | ENSMUSG00000114413 | 0 | NA | NA | NA | NA | NA | NA | NA | 0 | 0 | 0 | 0 | 0 | 0 |
| 51972 | ENSMUSG00000114414 | 0.167707431 | -0.796629176 | 4.080472857 | -0.195229623 | 0.84521319 | NA | A930014D07 | RIKEN cDNA A930014D07 gene [Source:M | 0 | 0 | 1.0062 | 0 | 0 | 0 |
| 51973 | ENSMUSG00000114415 | 0 | NA | NA | NA | NA | NA | Gm41030 | predicted gene, 41030 [Source:MGI Symb | 0 | 0 | 0 | 0 | 0 | 0 |
| 51974 | ENSMUSG00000114416 | 0 | NA | NA | NA | NA | NA | Gm47907 | predicted gene, 47907 [Source:MGI Symb | 0 | 0 | 0 | 0 | 0 | 0 |
| 51975 | ENSMUSG00000114417 | 0 | NA | NA | NA | NA | NA | Gm34586 | predicted gene, 34586 [Source:MGI Symb | 0 | 0 | 0 | 0 | 0 | 0 |
| 51976 | ENSMUSG00000114418 | 0 | NA | NA | NA | NA | NA | Gm18883 | predicted gene, 18883 [Source:MGI Symb | 0 | 0 | 0 | 0 | 0 | 0 |
| 51977 | ENSMUSG00000114419 | 3.238789681 | 0.181630572 | 1.156500981 | 0.15705181 | 0.875204016 | NA | 5430414B19 | RIKEN cDNA 5430414B19 gene [Source:M | 4.0155 | 2.0558 | 3.0187 | 4.8199 | 2.1812 | 3.3415 |
| 51978 | ENSMUSG00000114420 | 0 | NA | NA | NA | NA | NA | Gm48906 | predicted gene, 48906 [Source:MGI Symb | 0 | 0 | 0 | 0 | 0 | 0 |
| 51979 | ENSMUSG00000114421 | 0 | NA | NA | NA | NA | NA | Gm48526 | predicted gene, 48526 [Source:MGI Symb | 0 | 0 | 0 | 0 | 0 | 0 |
| 51980 | ENSMUSG00000114422 | 0 | NA | NA | NA | NA | NA | Gm30411 | predicted gene, 30411 [Source:MGI Symb | 0 | 0 | 0 | 0 | 0 | 0 |
| 51981 | ENSMUSG00000114423 | 0 | NA | NA | NA | NA | NA | Gm48404 | predicted gene, 48404 [Source:MGI Symb | 0 | 0 | 0 | 0 | 0 | 0 |
| 51982 | ENSMUSG00000114424 | 0.819128932 | -3.070895205 | 2.828495588 | -1.085699132 | 0.277612125 | NA | Gm47414 | predicted gene, 47414 [Source:MGI Symb | 0.8031 | 4.1117 | 0 | 0 | 0 | 0 |
| 51983 | ENSMUSG00000114425 | 8.340258578 | -1.128739376 | 0.813293861 | -1.387861671 | 0.165179171 | 0.590574884 | Gm33447 | predicted gene, 33447 [Source:MGI Symb | 12.85 | 17.475 | 4.025 | 5.7839 | 5.4531 | 4.4554 |
| 51984 | ENSMUSG00000114426 | 0 | NA | NA | NA | NA | NA | Gm10769 | predicted pseudogene 10769 [Source:MG | 0 | 0 | 0 | 0 | 0 | 0 |
| 51985 | ENSMUSG00000114427 | 0 | NA | NA | NA | NA | NA | F830002E08 | RIKEN cDNA F830002E08 gene [Source:M | 0 | 0 | 0 | 0 | 0 | 0 |
| 51986 | ENSMUSG00000114428 | 0 | NA | NA | NA | NA | NA | Gm47707 | predicted gene, 47707 [Source:MGI Symb | 0 | 0 | 0 | 0 | 0 | 0 |
| 51987 | ENSMUSG00000114429 | 0 | NA | NA | NA | NA | NA | Gm9025 | predicted gene 9025 [Source:MGI Symbol | 0 | 0 | 0 | 0 | 0 | 0 |
| 51988 | ENSMUSG00000114430 | 36.0517772 | 0.276567631 | 0.456848816 | 0.605381083 | 0.544925817 | 0.8954353 | Gm40264 | predicted gene, 40264 [Source:MGI Symb | 28.109 | 32.893 | 37.231 | 68.442 | 22.903 | 26.732 |
| 51989 | ENSMUSG00000114431 | 0 | NA | NA | NA | NA | NA | Gm48601 | predicted gene, 48601 [Source:MGI Symb | 0 | 0 | 0 | 0 | 0 | 0 |
| 51990 | ENSMUSG00000114432 | 0.167707431 | -0.796629176 | 4.080472857 | -0.195229623 | 0.84521319 | NA | Gm49391 | predicted gene, 49391 [Source:MGI Symb | 0 | 0 | 1.0062 | 0 | 0 | 0 |
| 51991 | ENSMUSG00000114433 | 9.948689789 | -0.560214119 | 0.758624503 | -0.738460354 | 0.460234751 | 0.859817816 | Gm48488 | predicted gene, 48488 [Source:MGI Symb | 15.259 | 5.1396 | 15.094 | 8.6758 | 3.2719 | 12.252 |
| 51992 | ENSMUSG00000114434 | 0 | NA | NA | NA | NA | NA | Gm48360 | predicted gene, 48360 [Source:MGI Symb | 0 | 0 | 0 | 0 | 0 | 0 |
| 51993 | ENSMUSG00000114435 | 0 | NA | NA | NA | NA | NA | NA | NA | 0 | 0 | 0 | 0 | 0 | 0 |
| 51994 | ENSMUSG00000114436 | 0 | NA | NA | NA | NA | NA | Gm48283 | predicted gene, 48283 [Source:MGI Symb | 0 | 0 | 0 | 0 | 0 | 0 |
| 51995 | ENSMUSG00000114437 | 0 | NA | NA | NA | NA | NA | 4930415C11 | RIKEN cDNA 4930415C11 gene [Source:M | 0 | 0 | 0 | 0 | 0 | 0 |
| 51996 | ENSMUSG00000114438 | 3.394766088 | 0.751451018 | 1.398221754 | 0.537433362 | 0.590968304 | NA | Gm47827 | predicted gene, 47827 [Source:MGI Symb | 1.6062 | 2.0558 | 4.025 | 11.568 | 0 | 1.1138 |
| 51997 | ENSMUSG00000114439 | 1.966293852 | -0.872872727 | 1.530085657 | -0.570473112 | 0.568356854 | NA | Gm48395 | predicted gene, 48395 [Source:MGI Symb | 1.6062 | 1.0279 | 5.0312 | 1.928 | 1.0906 | 1.1138 |
| 51998 | ENSMUSG00000114440 | 0 | NA | NA | NA | NA | NA | Gm47505 | predicted gene, 47505 [Source:MGI Symb | 0 | 0 | 0 | 0 | 0 | 0 |
| 51999 | ENSMUSG00000114441 | 0 | NA | NA | NA | NA | NA | Gm33472 | predicted gene, 33472 [Source:MGI Symb | 0 | 0 | 0 | 0 | 0 | 0 |
| 52000 | ENSMUSG00000114442 | 0 | NA | NA | NA | NA | NA | F630042J09 | RIKEN cDNA F630042J09 gene [Source:M | 0 | 0 | 0 | 0 | 0 | 0 |
| 52001 | ENSMUSG00000114443 | 6.11364124 | 0.784588329 | 0.896248703 | 0.875413628 | 0.381348887 | 0.8130053 | Gm19241 | predicted gene, 19241 [Source:MGI Symb | 7.2279 | 2.0558 | 4.025 | 5.7839 | 10.906 | 6.6831 |
| 52002 | ENSMUSG00000114444 | 0 | NA | NA | NA | NA | NA | Gm47008 | predicted gene, 47008 [Source:MGI Symb | 0 | 0 | 0 | 0 | 0 | 0 |
| 52003 | ENSMUSG00000114445 | 0 | NA | NA | NA | NA | NA | Gm7089 | predicted gene 7089 [Source:MGI Symbol | 0 | 0 | 0 | 0 | 0 | 0 |
| 52004 | ENSMUSG00000114446 | 0.167707431 | -0.796629176 | 4.080472857 | -0.195229623 | 0.84521319 | NA | Gm33524 | predicted gene, 33524 [Source:MGI Symb | 0 | 0 | 1.0062 | 0 | 0 | 0 |
| 52005 | ENSMUSG00000114447 | 0 | NA | NA | NA | NA | NA | Gm47636 | predicted gene, 47636 [Source:MGI Symb | 0 | 0 | 0 | 0 | 0 | 0 |
| 52006 | ENSMUSG00000114448 | 0 | NA | NA | NA | NA | NA | C630044B11 | RIKEN cDNA C630044B11 gene [Source:M | 0 | 0 | 0 | 0 | 0 | 0 |
| 52007 | ENSMUSG00000114449 | 0 | NA | NA | NA | NA | NA | Gm48325 | predicted gene, 48325 [Source:MGI Symb | 0 | 0 | 0 | 0 | 0 | 0 |
| 52008 | ENSMUSG00000114450 | 0 | NA | NA | NA | NA | NA | Gm18523 | predicted gene, 18523 [Source:MGI Symb | 0 | 0 | 0 | 0 | 0 | 0 |
| 52009 | ENSMUSG00000114451 | 0 | NA | NA | NA | NA | NA | Gm47912 | predicted gene, 47912 [Source:MGI Symb | 0 | 0 | 0 | 0 | 0 | 0 |
| 52010 | ENSMUSG00000114452 | 1.513184552 | 1.049050507 | 1.941143069 | 0.540429257 | 0.588901036 | NA | A530001N23 | RIKEN cDNA A530001N23 gene [Source:M | 0 | 0 | 3.0187 | 3.8559 | 1.0906 | 1.1138 |
| 52011 | ENSMUSG00000114453 | 0 | NA | NA | NA | NA | NA | Gm17805 | predicted gene, 17805 [Source:MGI Symb | 0 | 0 | 0 | 0 | 0 | 0 |
| 52012 | ENSMUSG00000114454 | 0 | NA | NA | NA | NA | NA | Gm48567 | predicted gene, 48567 [Source:MGI Symb | 0 | 0 | 0 | 0 | 0 | 0 |
| 52013 | ENSMUSG00000114455 | 0 | NA | NA | NA | NA | NA | NA | NA | 0 | 0 | 0 | 0 | 0 | 0 |
| 52014 | ENSMUSG00000114456 | 2.85932641 | 0.219086862 | 1.383288592 | 0.158381167 | 0.874156453 | NA | H2bc9 | H2B clustered histone 9 [Source:MGI Sym | 4.8186 | 2.0558 | 1.0062 | 4.8199 | 0 | 4.4554 |
| 52015 | ENSMUSG00000114457 | 0 | NA | NA | NA | NA | NA | Gm36346 | predicted gene, 36346 [Source:MGI Symb | 0 | 0 | 0 | 0 | 0 | 0 |
| 52016 | ENSMUSG00000114458 | 0 | NA | NA | NA | NA | NA | Gm47551 | predicted gene, 47551 [Source:MGI Symb | 0 | 0 | 0 | 0 | 0 | 0 |
| 52017 | ENSMUSG00000114459 | 0 | NA | NA | NA | NA | NA | Gm2736 | predicted gene 2736 [Source:MGI Symbol | 0 | 0 | 0 | 0 | 0 | 0 |
| 52018 | ENSMUSG00000114460 | 0 | NA | NA | NA | NA | NA | Gm10779 | predicted gene 10779 [Source:MGI Symbo | 0 | 0 | 0 | 0 | 0 | 0 |
| 52019 | ENSMUSG00000114461 | 0 | NA | NA | NA | NA | NA | Gm48155 | predicted gene, 48155 [Source:MGI Symb | 0 | 0 | 0 | 0 | 0 | 0 |
| 52020 | ENSMUSG00000114462 | 0 | NA | NA | NA | NA | NA | Gm8371 | predicted gene 8371 [Source:MGI Symbol | 0 | 0 | 0 | 0 | 0 | 0 |

| | | | | | | | | | | | | | | | |
|---|---|---|---|---|---|---|---|---|---|---|---|---|---|---|---|
| 52021 | ENSMUSG00000114463 | 0 | NA | NA | NA | NA | NA | Gm48162 | predicted gene, 48162 [Source:MGI Symb | 0 | 0 | 0 | 0 | 0 | 0 |
| 52022 | ENSMUSG00000114464 | 0 | NA | NA | NA | NA | NA | Gm18071 | predicted gene, 18071 [Source:MGI Symb | 0 | 0 | 0 | 0 | 0 | 0 |
| 52023 | ENSMUSG00000114465 | 0 | NA | NA | NA | NA | NA | Gm19139 | predicted gene, 19139 [Source:MGI Symb | 0 | 0 | 0 | 0 | 0 | 0 |
| 52024 | ENSMUSG00000114466 | 0 | NA | NA | NA | NA | NA | Gm19216 | predicted gene, 19216 [Source:MGI Symb | 0 | 0 | 0 | 0 | 0 | 0 |
| 52025 | ENSMUSG00000114467 | 0 | NA | NA | NA | NA | NA | Gm48430 | predicted gene, 48430 [Source:MGI Symb | 0 | 0 | 0 | 0 | 0 | 0 |
| 52026 | ENSMUSG00000114468 | 0 | NA | NA | NA | NA | NA | Gm48879 | predicted gene, 48879 [Source:MGI Symb | 0 | 0 | 0 | 0 | 0 | 0 |
| 52027 | ENSMUSG00000114469 | 33.51085451 | -0.124188209 | 0.499199199 | -0.248774857 | 0.803534941 | 0.963596811 | C730002L08 | RIKEN cDNA C730002L08 gene [Source:M | 55.414 | 34.949 | 14.087 | 24.099 | 46.897 | 25.618 |
| 52028 | ENSMUSG00000114470 | 38.39797878 | 0.108295364 | 0.401336653 | 0.269836716 | 0.787285875 | 0.960870081 | Gm49395 | predicted gene, 49395 [Source:MGI Symb | 55.414 | 29.81 | 25.156 | 38.559 | 45.806 | 35.643 |
| 52029 | ENSMUSG00000114471 | 0 | NA | NA | NA | NA | NA | Gm47891 | predicted gene, 47891 [Source:MGI Symb | 0 | 0 | 0 | 0 | 0 | 0 |
| 52030 | ENSMUSG00000114472 | 0 | NA | NA | NA | NA | NA | Gm38397 | predicted gene, 38397 [Source:MGI Symb | 0 | 0 | 0 | 0 | 0 | 0 |
| 52031 | ENSMUSG00000114473 | 0 | NA | NA | NA | NA | NA | Gm49398 | predicted gene, 49398 [Source:MGI Symb | 0 | 0 | 0 | 0 | 0 | 0 |
| 52032 | ENSMUSG00000114474 | 0 | NA | NA | NA | NA | NA | Gm34388 | predicted gene, 34388 [Source:MGI Symb | 0 | 0 | 0 | 0 | 0 | 0 |
| 52033 | ENSMUSG00000114475 | 0 | NA | NA | NA | NA | NA | Gm2726 | predicted gene 2726 [Source:MGI Symbol | 0 | 0 | 0 | 0 | 0 | 0 |
| 52034 | ENSMUSG00000114476 | 0 | NA | NA | NA | NA | NA | Gm21063 | predicted gene, 21063 [Source:MGI Symb | 0 | 0 | 0 | 0 | 0 | 0 |
| 52035 | ENSMUSG00000114477 | 0 | NA | NA | NA | NA | NA | Gm8795 | predicted gene 8795 [Source:MGI Symbol | 0 | 0 | 0 | 0 | 0 | 0 |
| 52036 | ENSMUSG00000114478 | 0 | NA | NA | NA | NA | NA | Gm34558 | predicted gene, 34558 [Source:MGI Symb | 0 | 0 | 0 | 0 | 0 | 0 |
| 52037 | ENSMUSG00000114479 | 0 | NA | NA | NA | NA | NA | Gm49375 | predicted gene, 49375 [Source:MGI Symb | 0 | 0 | 0 | 0 | 0 | 0 |
| 52038 | ENSMUSG00000114480 | 0 | NA | NA | NA | NA | NA | Gm3637 | predicted gene 3637 [Source:MGI Symbol | 0 | 0 | 0 | 0 | 0 | 0 |
| 52039 | ENSMUSG00000114481 | 1.080710637 | -0.022140248 | 2.593617264 | -0.008536436 | 0.993188992 | NA | Gm46415 | predicted gene, 46415 [Source:MGI Symb | 3.2124 | 0 | 0 | 0 | 3.2719 | 0 |
| 52040 | ENSMUSG00000114482 | 0 | NA | NA | NA | NA | NA | Gm7604 | predicted gene 7604 [Source:MGI Symbol | 0 | 0 | 0 | 0 | 0 | 0 |
| 52041 | ENSMUSG00000114483 | 0 | NA | NA | NA | NA | NA | Gm49144 | predicted gene, 49144 [Source:MGI Symb | 0 | 0 | 0 | 0 | 0 | 0 |
| 52042 | ENSMUSG00000114484 | 0 | NA | NA | NA | NA | NA | Gm8517 | predicted gene 8517 [Source:MGI Symbol | 0 | 0 | 0 | 0 | 0 | 0 |
| 52043 | ENSMUSG00000114485 | 0 | NA | NA | NA | NA | NA | Gm47330 | predicted gene, 47330 [Source:MGI Symb | 0 | 0 | 0 | 0 | 0 | 0 |
| 52044 | ENSMUSG00000114486 | 0 | NA | NA | NA | NA | NA | Gm48447 | predicted gene, 48447 [Source:MGI Symb | 0 | 0 | 0 | 0 | 0 | 0 |
| 52045 | ENSMUSG00000114487 | 0 | NA | NA | NA | NA | NA | Gm49393 | predicted gene, 49393 [Source:MGI Symb | 0 | 0 | 0 | 0 | 0 | 0 |
| 52046 | ENSMUSG00000114488 | 1.34908933 | 1.640931007 | 1.932083572 | 0.849306433 | 0.395710802 | NA | Gm5802 | predicted gene 5802 [Source:MGI Symbol | 0 | 1.0279 | 1.0062 | 3.8559 | 1.0906 | 1.1138 |
| 52047 | ENSMUSG00000114489 | 0 | NA | NA | NA | NA | NA | Vmn2r-ps10 | vomeronasal 2, receptor, pseudogene 10 | 0 | 0 | 0 | 0 | 0 | 0 |
| 52048 | ENSMUSG00000114490 | 0 | NA | NA | NA | NA | NA | Gm19866 | predicted gene, 19866 [Source:MGI Symb | 0 | 0 | 0 | 0 | 0 | 0 |
| 52049 | ENSMUSG00000114491 | 0 | NA | NA | NA | NA | NA | A230103O0 | RIKEN cDNA A230103O09 gene [Source:M | 0 | 0 | 0 | 0 | 0 | 0 |
| 52050 | ENSMUSG00000114492 | 0.17131964 | -0.796629176 | 4.080472857 | -0.195229623 | 0.84521319 | NA | Gm9984 | predicted gene 9984 [Source:MGI Symbol | 0 | 1.0279 | 0 | 0 | 0 | 0 |
| 52051 | ENSMUSG00000114493 | 10.99374497 | -0.189515501 | 0.692445068 | -0.273690304 | 0.78432263 | 0.960670212 | Gm47071 | predicted gene, 47071 [Source:MGI Symb | 16.865 | 6.1675 | 12.075 | 15.424 | 7.6343 | 7.7969 |
| 52052 | ENSMUSG00000114494 | 0 | NA | NA | NA | NA | NA | Gm48663 | predicted gene, 48663 [Source:MGI Symb | 0 | 0 | 0 | 0 | 0 | 0 |
| 52053 | ENSMUSG00000114495 | 0 | NA | NA | NA | NA | NA | Gm5664 | predicted gene 5664 [Source:MGI Symbol | 0 | 0 | 0 | 0 | 0 | 0 |
| 52054 | ENSMUSG00000114496 | 0.51034671 | -2.381743948 | 3.331081356 | -0.715006238 | 0.474605189 | NA | Gm47601 | predicted gene, 47601 [Source:MGI Symb | 0 | 2.0558 | 1.0062 | 0 | 0 | 0 |
| 52055 | ENSMUSG00000114497 | 0 | NA | NA | NA | NA | NA | NA | NA | 0 | 0 | 0 | 0 | 0 | 0 |
| 52056 | ENSMUSG00000114498 | 0 | NA | NA | NA | NA | NA | Gm48339 | predicted gene, 48339 [Source:MGI Symb | 0 | 0 | 0 | 0 | 0 | 0 |
| 52057 | ENSMUSG00000114499 | 0 | NA | NA | NA | NA | NA | Gm48489 | predicted gene, 48489 [Source:MGI Symb | 0 | 0 | 0 | 0 | 0 | 0 |
| 52058 | ENSMUSG00000114500 | 1.846944447 | 0.377174383 | 1.515561484 | 0.248867754 | 0.803463079 | NA | Gm8983 | predicted gene 8983 [Source:MGI Symbol | 0.8031 | 2.0558 | 2.0125 | 2.8919 | 1.0906 | 2.2277 |
| 52059 | ENSMUSG00000114501 | 1.764909323 | -0.101551649 | 1.652148297 | -0.061466425 | 0.950987753 | NA | Gm48582 | predicted gene, 48582 [Source:MGI Symb | 2.4093 | 3.0838 | 0 | 2.8919 | 1.0906 | 1.1138 |
| 52060 | ENSMUSG00000114502 | 0 | NA | NA | NA | NA | NA | Gm40975 | predicted gene, 40975 [Source:MGI Symb | 0 | 0 | 0 | 0 | 0 | 0 |
| 52061 | ENSMUSG00000114503 | 0 | NA | NA | NA | NA | NA | Gm20242 | predicted gene, 20242 [Source:MGI Symb | 0 | 0 | 0 | 0 | 0 | 0 |
| 52062 | ENSMUSG00000114504 | 1.195909497 | 1.367900466 | 2.270154055 | 0.602558431 | 0.546802481 | NA | Gm48321 | predicted gene, 48321 [Source:MGI Symb | 0 | 2.0558 | 0 | 2.8919 | 0 | 2.2277 |
| 52063 | ENSMUSG00000114505 | 0 | NA | NA | NA | NA | NA | Gm48471 | predicted gene, 48471 [Source:MGI Symb | 0 | 0 | 0 | 0 | 0 | 0 |
| 52064 | ENSMUSG00000114506 | 0 | NA | NA | NA | NA | NA | Gm5666 | predicted gene 5666 [Source:MGI Symbol | 0 | 0 | 0 | 0 | 0 | 0 |
| 52065 | ENSMUSG00000114507 | 0 | NA | NA | NA | NA | NA | Gm30747 | predicted gene, 30747 [Source:MGI Symb | 0 | 0 | 0 | 0 | 0 | 0 |
| 52066 | ENSMUSG00000114508 | 0 | NA | NA | NA | NA | NA | Gm47711 | predicted gene, 47711 [Source:MGI Symb | 0 | 0 | 0 | 0 | 0 | 0 |
| 52067 | ENSMUSG00000114509 | 0 | NA | NA | NA | NA | NA | 4930471G24 | RIKEN cDNA 4930471G24 gene [Source:M | 0 | 0 | 0 | 0 | 0 | 0 |
| 52068 | ENSMUSG00000114510 | 0 | NA | NA | NA | NA | NA | Gm48859 | predicted gene, 48859 [Source:MGI Symb | 0 | 0 | 0 | 0 | 0 | 0 |
| 52069 | ENSMUSG00000114511 | 0 | NA | NA | NA | NA | NA | 0710001A04 | RIKEN cDNA 0710001A04 gene [Source:M | 0 | 0 | 0 | 0 | 0 | 0 |
| 52070 | ENSMUSG00000114512 | 7.287466975 | 1.348859368 | 0.841989289 | 1.601991124 | 0.109157572 | 0.486100716 | Gm47348 | predicted gene, 47348 [Source:MGI Symb | 3.2124 | 7.1954 | 2.0125 | 12.532 | 7.6343 | 11.138 |
| 52071 | ENSMUSG00000114513 | 0 | NA | NA | NA | NA | NA | Gm47780 | predicted gene, 47780 [Source:MGI Symb | 0 | 0 | 0 | 0 | 0 | 0 |
| 52072 | ENSMUSG00000114514 | 0 | NA | NA | NA | NA | NA | Gm47010 | predicted gene, 47010 [Source:MGI Symb | 0 | 0 | 0 | 0 | 0 | 0 |
| 52073 | ENSMUSG00000114515 | 0.953031439 | -0.811520803 | 2.12857635 | -0.381250502 | 0.703017378 | NA | Aldoa | aldolase A, fructose-bisphosphate [Source | 1.6062 | 1.0279 | 1.0062 | 0.964 | 0 | 1.1138 |
| 52074 | ENSMUSG00000114516 | 3.208079518 | 0.817671582 | 1.293510892 | 0.632133511 | 0.527299639 | NA | Gm46440 | predicted gene, 46440 [Source:MGI Symb | 4.8186 | 2.0558 | 0 | 5.7839 | 4.3625 | 2.2277 |
| 52075 | ENSMUSG00000114517 | 0 | NA | NA | NA | NA | NA | Gm19290 | predicted gene, 19290 [Source:MGI Symb | 0 | 0 | 0 | 0 | 0 | 0 |
| 52076 | ENSMUSG00000114518 | 0 | NA | NA | NA | NA | NA | Gm48208 | predicted gene, 48208 [Source:MGI Symb | 0 | 0 | 0 | 0 | 0 | 0 |

| | | | | | | | | | | | | | | | |
|---|---|---|---|---|---|---|---|---|---|---|---|---|---|---|---|
| 52077 | ENSMUSG00000114519 | 0 | NA | NA | NA | NA | NA | Gm32376 | predicted gene, 32376 [Source:MGI Symb | 0 | 0 | 0 | 0 | 0 | 0 |
| 52078 | ENSMUSG00000114520 | 0 | NA | NA | NA | NA | NA | Gm41073 | predicted gene, 41073 [Source:MGI Symb | 0 | 0 | 0 | 0 | 0 | 0 |
| 52079 | ENSMUSG00000114521 | 0 | NA | NA | NA | NA | NA | Gm48050 | predicted gene, 48050 [Source:MGI Symb | 0 | 0 | 0 | 0 | 0 | 0 |
| 52080 | ENSMUSG00000114522 | 0 | NA | NA | NA | NA | NA | Gm32067 | predicted gene, 32067 [Source:MGI Symb | 0 | 0 | 0 | 0 | 0 | 0 |
| 52081 | ENSMUSG00000114523 | 0 | NA | NA | NA | NA | NA | Gm48394 | predicted gene, 48394 [Source:MGI Symb | 0 | 0 | 0 | 0 | 0 | 0 |
| 52082 | ENSMUSG00000114524 | 3.600707413 | -4.16927996 | 1.592401466 | -2.618234189 | 0.008838612 | NA | Gm48357 | predicted gene, 48357 [Source:MGI Symb | 3.2124 | 9.2513 | 8.05 | 0 | 1.0906 | 0 |
| 52083 | ENSMUSG00000114525 | 0.983224154 | -3.332395149 | 2.379709572 | -1.40033691 | 0.161412453 | NA | Gm47990 | predicted gene, 47990 [Source:MGI Symb | 0.8031 | 3.0838 | 2.0125 | 0 | 0 | 0 |
| 52084 | ENSMUSG00000114526 | 0 | NA | NA | NA | NA | NA | A930015P04 | RIKEN cDNA A930015P04 gene [Source:M | 0 | 0 | 0 | 0 | 0 | 0 |
| 52085 | ENSMUSG00000114527 | 0 | NA | NA | NA | NA | NA | Vmn1r-ps11 | vomeronasal 1 receptor, pseudogene 117 | 0 | 0 | 0 | 0 | 0 | 0 |
| 52086 | ENSMUSG00000114528 | 0 | NA | NA | NA | NA | NA | Gm46388 | predicted gene, 46388 [Source:MGI Symb | 0 | 0 | 0 | 0 | 0 | 0 |
| 52087 | ENSMUSG00000114529 | 0 | NA | NA | NA | NA | NA | Gm4939 | predicted gene 4939 [Source:MGI Symbol | 0 | 0 | 0 | 0 | 0 | 0 |
| 52088 | ENSMUSG00000114530 | 0 | NA | NA | NA | NA | NA | Gm48725 | predicted gene, 48725 [Source:MGI Symb | 0 | 0 | 0 | 0 | 0 | 0 |
| 52089 | ENSMUSG00000114531 | 0 | NA | NA | NA | NA | NA | A530065N2( | RIKEN cDNA A530046M15 gene [Source:N | 0 | 0 | 0 | 0 | 0 | 0 |
| 52090 | ENSMUSG00000114532 | 0 | NA | NA | NA | NA | NA | 2610318M1( | RIKEN cDNA 2610318M16 gene [Source:N | 0 | 0 | 0 | 0 | 0 | 0 |
| 52091 | ENSMUSG00000114533 | 0 | NA | NA | NA | NA | NA | Gm8416 | predicted gene 8416 [Source:MGI Symbol | 0 | 0 | 0 | 0 | 0 | 0 |
| 52092 | ENSMUSG00000114534 | 0.185640647 | 1.126924175 | 4.080472857 | 0.276174898 | 0.782413736 | NA | Gm32004 | predicted gene, 32004 [Source:MGI Symb | 0 | 0 | 0 | 0 | 0 | 1.1138 |
| 52093 | ENSMUSG00000114535 | 2.374140414 | 0.126621389 | 1.545395688 | 0.081934607 | 0.934698714 | NA | Gm47246 | predicted gene, 47246 [Source:MGI Symb | 0.8031 | 0 | 6.0375 | 1.928 | 4.3625 | 1.1138 |
| 52094 | ENSMUSG00000114536 | 0.803314871 | 3.228356836 | 3.175155518 | 1.0167555 | 0.309269756 | NA | Gm48837 | predicted gene, 48837 [Source:MGI Symb | 0 | 0 | 0 | 4.8199 | 0 | 0 |
| 52095 | ENSMUSG00000114537 | 0 | NA | NA | NA | NA | NA | Gm48569 | predicted gene, 48569 [Source:MGI Symb | 0 | 0 | 0 | 0 | 0 | 0 |
| 52096 | ENSMUSG00000114538 | 0 | NA | NA | NA | NA | NA | Gm31517 | predicted gene, 31517 [Source:MGI Symb | 0 | 0 | 0 | 0 | 0 | 0 |
| 52097 | ENSMUSG00000114539 | 0.538730002 | 1.11098254 | 3.037664833 | 0.365735722 | 0.714562288 | NA | Gm47921 | predicted gene, 47921 [Source:MGI Symb | 0 | 1.0279 | 0 | 0 | 1.0906 | 1.1138 |
| 52098 | ENSMUSG00000114540 | 5.36252085 | -0.299252206 | 0.929277451 | -0.32202676 | 0.747432422 | 0.950739825 | Gm6421 | predicted pseudogene 6421 [Source:MGI | 5.6217 | 6.1675 | 6.0375 | 7.7118 | 2.1812 | 4.4554 |
| 52099 | ENSMUSG00000114541 | 0 | NA | NA | NA | NA | NA | Gm29680 | predicted gene, 29680 [Source:MGI Symb | 0 | 0 | 0 | 0 | 0 | 0 |
| 52100 | ENSMUSG00000114542 | 0 | NA | NA | NA | NA | NA | Gm18142 | predicted gene, 18142 [Source:MGI Symb | 0 | 0 | 0 | 0 | 0 | 0 |
| 52101 | ENSMUSG00000114543 | 0 | NA | NA | NA | NA | NA | Gm48028 | predicted gene, 48028 [Source:MGI Symb | 0 | 0 | 0 | 0 | 0 | 0 |
| 52102 | ENSMUSG00000114544 | 0 | NA | NA | NA | NA | NA | Gm49146 | predicted gene, 49146 [Source:MGI Symb | 0 | 0 | 0 | 0 | 0 | 0 |
| 52103 | ENSMUSG00000114545 | 0 | NA | NA | NA | NA | NA | Gm32184 | predicted gene, 32184 [Source:MGI Symb | 0 | 0 | 0 | 0 | 0 | 0 |
| 52104 | ENSMUSG00000114546 | 0.167707431 | -0.796629176 | 4.080472857 | -0.195229623 | 0.84521319 | NA | Gm49350 | predicted gene, 49350 [Source:MGI Symb | 0 | 0 | 1.0062 | 0 | 0 | 0 |
| 52105 | ENSMUSG00000114547 | 0.85435009 | 0.823372753 | 2.383559862 | 0.345438252 | 0.729764924 | NA | Gm3226 | predicted gene 3226 [Source:MGI Symbol | 0.8031 | 1.0279 | 0 | 0 | 2.1812 | 1.1138 |
| 52106 | ENSMUSG00000114548 | 0 | NA | NA | NA | NA | NA | NA | NA | 0 | 0 | 0 | 0 | 0 | 0 |
| 52107 | ENSMUSG00000114549 | 0 | NA | NA | NA | NA | NA | Vmn2r-ps10 | vomeronasal 2, receptor, pseudogene 10( | 0 | 0 | 0 | 0 | 0 | 0 |
| 52108 | ENSMUSG00000114550 | 0 | NA | NA | NA | NA | NA | NA | NA | 0 | 0 | 0 | 0 | 0 | 0 |
| 52109 | ENSMUSG00000114551 | 0.181769715 | 1.126924175 | 4.080472857 | 0.276174898 | 0.782413736 | NA | Gm6035 | predicted gene 6035 [Source:MGI Symbol | 0 | 0 | 0 | 0 | 1.0906 | 0 |
| 52110 | ENSMUSG00000114552 | 0 | NA | NA | NA | NA | NA | Gm48570 | predicted gene, 48570 [Source:MGI Symb | 0 | 0 | 0 | 0 | 0 | 0 |
| 52111 | ENSMUSG00000114553 | 0 | NA | NA | NA | NA | NA | Gm7644 | predicted gene 7644 [Source:MGI Symbol | 0 | 0 | 0 | 0 | 0 | 0 |
| 52112 | ENSMUSG00000114554 | 0 | NA | NA | NA | NA | NA | 5430425K12 | RIKEN cDNA 5430425K12 gene [Source:M | 0 | 0 | 0 | 0 | 0 | 0 |
| 52113 | ENSMUSG00000114555 | 0.534859069 | 1.101323472 | 3.351640271 | 0.328592386 | 0.742463804 | NA | Gm48706 | predicted gene, 48706 [Source:MGI Symb | 0 | 1.0279 | 0 | 0 | 2.1812 | 0 |
| 52114 | ENSMUSG00000114556 | 0 | NA | NA | NA | NA | NA | Gm3895 | predicted gene 3895 [Source:MGI Symbol | 0 | 0 | 0 | 0 | 0 | 0 |
| 52115 | ENSMUSG00000114557 | 0 | NA | NA | NA | NA | NA | Gm8345 | predicted gene 8345 [Source:MGI Symbol | 0 | 0 | 0 | 0 | 0 | 0 |
| 52116 | ENSMUSG00000114558 | 0.642651897 | 2.906894826 | 3.440535777 | 0.844895974 | 0.398168923 | NA | Gm9570 | predicted gene 9570 [Source:MGI Symbol | 0 | 0 | 0 | 3.8559 | 0 | 0 |
| 52117 | ENSMUSG00000114559 | 0 | NA | NA | NA | NA | NA | Gm47429 | predicted gene, 47429 [Source:MGI Symb | 0 | 0 | 0 | 0 | 0 | 0 |
| 52118 | ENSMUSG00000114560 | 0 | NA | NA | NA | NA | NA | Gm35160 | predicted gene, 35160 [Source:MGI Symb | 0 | 0 | 0 | 0 | 0 | 0 |
| 52119 | ENSMUSG00000114561 | 0 | NA | NA | NA | NA | NA | Gm48515 | predicted gene, 48515 [Source:MGI Symb | 0 | 0 | 0 | 0 | 0 | 0 |
| 52120 | ENSMUSG00000114562 | 0 | NA | NA | NA | NA | NA | Gm18080 | predicted gene, 18080 [Source:MGI Symb | 0 | 0 | 0 | 0 | 0 | 0 |
| 52121 | ENSMUSG00000114563 | 0 | NA | NA | NA | NA | NA | 1700016K05 | RIKEN cDNA 1700016K05 gene [Source:M | 0 | 0 | 0 | 0 | 0 | 0 |
| 52122 | ENSMUSG00000114564 | 0 | NA | NA | NA | NA | NA | Gm10739 | predicted gene 10739 [Source:MGI Symbo | 0 | 0 | 0 | 0 | 0 | 0 |
| 52123 | ENSMUSG00000114565 | 0 | NA | NA | NA | NA | NA | Gm47651 | predicted gene, 47651 [Source:MGI Symb | 0 | 0 | 0 | 0 | 0 | 0 |
| 52124 | ENSMUSG00000114566 | 0.181769715 | 1.126924175 | 4.080472857 | 0.276174898 | 0.782413736 | NA | Gm48500 | predicted gene, 48500 [Source:MGI Symb | 0 | 0 | 0 | 0 | 1.0906 | 0 |
| 52125 | ENSMUSG00000114567 | 0 | NA | NA | NA | NA | NA | Gm47774 | predicted gene, 47774 [Source:MGI Symb | 0 | 0 | 0 | 0 | 0 | 0 |
| 52126 | ENSMUSG00000114568 | 3.760159908 | 0.670358593 | 1.243681856 | 0.539011315 | 0.589879047 | NA | Gm48732 | predicted gene, 48732 [Source:MGI Symb | 5.6217 | 1.0279 | 2.0125 | 10.604 | 2.1812 | 1.1138 |
| 52127 | ENSMUSG00000114569 | 0.181769715 | 1.126924175 | 4.080472857 | 0.276174898 | 0.782413736 | NA | Gm35215 | predicted gene, 35215 [Source:MGI Symb | 0 | 0 | 0 | 0 | 1.0906 | 0 |
| 52128 | ENSMUSG00000114570 | 0 | NA | NA | NA | NA | NA | Gm48058 | predicted gene, 48058 [Source:MGI Symb | 0 | 0 | 0 | 0 | 0 | 0 |
| 52129 | ENSMUSG00000114571 | 0 | NA | NA | NA | NA | NA | Gm35595 | predicted gene, 35595 [Source:MGI Symb | 0 | 0 | 0 | 0 | 0 | 0 |
| 52130 | ENSMUSG00000114572 | 0 | NA | NA | NA | NA | NA | Gm48457 | predicted gene, 48457 [Source:MGI Symb | 0 | 0 | 0 | 0 | 0 | 0 |
| 52131 | ENSMUSG00000114573 | 0 | NA | NA | NA | NA | NA | Gm47732 | predicted gene, 47732 [Source:MGI Symb | 0 | 0 | 0 | 0 | 0 | 0 |
| 52132 | ENSMUSG00000114574 | 0 | NA | NA | NA | NA | NA | NA | NA | 0 | 0 | 0 | 0 | 0 | 0 |

| | | | | | | | | | | | | | | | |
|---|---|---|---|---|---|---|---|---|---|---|---|---|---|---|---|
| 52133 | ENSMUSG00000114575 | 9.771297379 | 0.717797385 | 0.731050861 | 0.98187065 | 0.326163576 | 0.772018857 | Gm47814 | predicted gene, 47814 [Source:MGI Symb | 4.0155 | 6.1675 | 12.075 | 7.7118 | 14.178 | 14.48 |
| 52134 | ENSMUSG00000114576 | 1.797010217 | -1.181120951 | 1.966322222 | -0.600675178 | 0.548056355 | NA | Naip3-ps1 | NLR family, apoptosis inhibitory protein 3, | 2.4093 | 0 | 5.0312 | 0 | 0 | 3.3415 |
| 52135 | ENSMUSG00000114577 | 0 | NA | NA | NA | NA | NA | Gm35217 | predicted gene, 35217 [Source:MGI Symb | 0 | 0 | 0 | 0 | 0 | 0 |
| 52136 | ENSMUSG00000114578 | 0 | NA | NA | NA | NA | NA | Gm47158 | predicted gene, 47158 [Source:MGI Symb | 0 | 0 | 0 | 0 | 0 | 0 |
| 52137 | ENSMUSG00000114579 | 0.17131964 | -0.796629176 | 4.080472857 | -0.195229623 | 0.84521319 | NA | Gm4130 | predicted gene 4130 [Source:MGI Symbol | 0 | 1.0279 | 0 | 0 | 0 | 0 |
| 52138 | ENSMUSG00000114580 | 0 | NA | NA | NA | NA | NA | Gm48782 | predicted gene, 48782 [Source:MGI Symb | 0 | 0 | 0 | 0 | 0 | 0 |
| 52139 | ENSMUSG00000114581 | 0 | NA | NA | NA | NA | NA | Gm8277 | predicted gene 8277 [Source:MGI Symbol | 0 | 0 | 0 | 0 | 0 | 0 |
| 52140 | ENSMUSG00000114582 | 0 | NA | NA | NA | NA | NA | 3110040M0 | RIKEN cDNA 3110040M04 gene [Source:M | 0 | 0 | 0 | 0 | 0 | 0 |
| 52141 | ENSMUSG00000114583 | 1.933955351 | -1.364704768 | 1.603715128 | -0.850964579 | 0.39478903 | NA | Gm48881 | predicted gene, 48881 [Source:MGI Symb | 3.2124 | 3.0838 | 2.0125 | 0 | 2.1812 | 1.1138 |
| 52142 | ENSMUSG00000114584 | 6.510035202 | 0.595692539 | 0.863394324 | 0.689942617 | 0.490230274 | 0.875501998 | Gm47694 | predicted gene, 47694 [Source:MGI Symb | 2.4093 | 8.2233 | 5.0312 | 5.7839 | 9.8156 | 7.7969 |
| 52143 | ENSMUSG00000114585 | 0.328370405 | 0.16514316 | 4.003149536 | 0.041253308 | 0.967093956 | NA | Gm32401 | predicted gene, 32401 [Source:MGI Symb | 0 | 0 | 1.0062 | 0.964 | 0 | 0 |
| 52144 | ENSMUSG00000114586 | 0 | NA | NA | NA | NA | NA | Gm31804 | predicted gene, 31804 [Source:MGI Symb | 0 | 0 | 0 | 0 | 0 | 0 |
| 52145 | ENSMUSG00000114587 | 0 | NA | NA | NA | NA | NA | Gm48533 | predicted gene, 48533 [Source:MGI Symb | 0 | 0 | 0 | 0 | 0 | 0 |
| 52146 | ENSMUSG00000114588 | 0 | NA | NA | NA | NA | NA | Gm6411 | predicted gene 6411 [Source:MGI Symbol | 0 | 0 | 0 | 0 | 0 | 0 |
| 52147 | ENSMUSG00000114589 | 0 | NA | NA | NA | NA | NA | Gm18518 | predicted gene, 18518 [Source:MGI Symb | 0 | 0 | 0 | 0 | 0 | 0 |
| 52148 | ENSMUSG00000114590 | 0 | NA | NA | NA | NA | NA | 5930438M1 | RIKEN cDNA 5930438M14 gene [Source:M | 0 | 0 | 0 | 0 | 0 | 0 |
| 52149 | ENSMUSG00000114591 | 0 | NA | NA | NA | NA | NA | Gm47996 | predicted gene, 47996 [Source:MGI Symb | 0 | 0 | 0 | 0 | 0 | 0 |
| 52150 | ENSMUSG00000114592 | 0 | NA | NA | NA | NA | NA | Gm5042 | predicted gene 5042 [Source:MGI Symbol | 0 | 0 | 0 | 0 | 0 | 0 |
| 52151 | ENSMUSG00000114593 | 0 | NA | NA | NA | NA | NA | Gm19716 | predicted gene, 19716 [Source:MGI Symb | 0 | 0 | 0 | 0 | 0 | 0 |
| 52152 | ENSMUSG00000114594 | 0 | NA | NA | NA | NA | NA | Gm48402 | predicted gene, 48402 [Source:MGI Symb | 0 | 0 | 0 | 0 | 0 | 0 |
| 52153 | ENSMUSG00000114595 | 0 | NA | NA | NA | NA | NA | Gm48004 | predicted gene, 48004 [Source:MGI Symb | 0 | 0 | 0 | 0 | 0 | 0 |
| 52154 | ENSMUSG00000114596 | 0.535401493 | -2.48596965 | 3.684117979 | -0.674780141 | 0.499815459 | NA | Gm47523 | predicted gene, 47523 [Source:MGI Symb | 3.2124 | 0 | 0 | 0 | 0 | 0 |
| 52155 | ENSMUSG00000114597 | 0 | NA | NA | NA | NA | NA | Gm47067 | predicted gene, 47067 [Source:MGI Symb | 0 | 0 | 0 | 0 | 0 | 0 |
| 52156 | ENSMUSG00000114598 | 0.524667718 | -0.819976748 | 3.064824146 | -0.267544469 | 0.78904998 | NA | D130062J10 | RIKEN cDNA D130062J10 gene [Source:M | 0 | 1.0279 | 1.0062 | 0 | 0 | 1.1138 |
| 52157 | ENSMUSG00000114599 | 0 | NA | NA | NA | NA | NA | Gm48070 | predicted gene, 48070 [Source:MGI Symb | 0 | 0 | 0 | 0 | 0 | 0 |
| 52158 | ENSMUSG00000114600 | 0 | NA | NA | NA | NA | NA | Gm46387 | predicted gene, 46387 [Source:MGI Symb | 0 | 0 | 0 | 0 | 0 | 0 |
| 52159 | ENSMUSG00000114601 | 1.051247473 | -0.23417022 | 2.362313183 | -0.099127508 | 0.921037031 | NA | Gm48188 | predicted gene, 48188 [Source:MGI Symb | 2.4093 | 0 | 1.0062 | 2.8919 | 0 | 0 |
| 52160 | ENSMUSG00000114602 | 4.164853534 | -1.025156483 | 1.099459895 | -0.932418261 | 0.351120411 | NA | Gm49223 | predicted gene, 49223 [Source:MGI Symb | 9.6372 | 2.0558 | 5.0312 | 3.8559 | 2.1812 | 2.2277 |
| 52161 | ENSMUSG00000114603 | 1.974158331 | 1.890201242 | 1.911361095 | 0.988929432 | 0.32269767 | NA | Gm17938 | predicted gene, 17938 [Source:MGI Symb | 2.4093 | 0 | 0 | 2.8919 | 6.5437 | 0 |
| 52162 | ENSMUSG00000114604 | 0 | NA | NA | NA | NA | NA | 2010203P06 | RIKEN cDNA 2010203P06 gene [Source:M | 0 | 0 | 0 | 0 | 0 | 0 |
| 52163 | ENSMUSG00000114605 | 0 | NA | NA | NA | NA | NA | C130051F05 | RIKEN cDNA C130051F05 gene [Source:M | 0 | 0 | 0 | 0 | 0 | 0 |
| 52164 | ENSMUSG00000114606 | 0 | NA | NA | NA | NA | NA | Gm7633 | predicted gene 7633 [Source:MGI Symbol | 0 | 0 | 0 | 0 | 0 | 0 |
| 52165 | ENSMUSG00000114607 | 0.660353019 | -0.01500701 | 2.83315355 | -0.005296928 | 0.995773683 | NA | Gm8990 | predicted gene 8990 [Source:MGI Symbol | 0 | 1.0279 | 1.0062 | 1.928 | 0 | 0 |
| 52166 | ENSMUSG00000114608 | 0.622883753 | 0.113321311 | 2.884614835 | 0.039284729 | 0.968663382 | NA | Gm36161 | predicted gene, 36161 [Source:MGI Symb | 0.8031 | 0 | 1.0062 | 1.928 | 0 | 0 |
| 52167 | ENSMUSG00000114609 | 0 | NA | NA | NA | NA | NA | Gm29767 | predicted gene, 29767 [Source:MGI Symb | 0 | 0 | 0 | 0 | 0 | 0 |
| 52168 | ENSMUSG00000114610 | 1.001209504 | 0.17748899 | 2.219748584 | 0.079959051 | 0.936269824 | NA | Cyp2c52-ps | cytochrome P450, family 2, subfamily c, p | 0.8031 | 0 | 2.0125 | 0.964 | 0 | 2.2277 |
| 52169 | ENSMUSG00000114611 | 0 | NA | NA | NA | NA | NA | Gm47749 | predicted gene, 47749 [Source:MGI Symb | 0 | 0 | 0 | 0 | 0 | 0 |
| 52170 | ENSMUSG00000114612 | 6.078333999 | 0.153651387 | 0.962820375 | 0.159584686 | 0.873208243 | 0.979835844 | Gm47550 | predicted gene, 47550 [Source:MGI Symb | 3.2124 | 7.1954 | 7.0437 | 13.496 | 2.1812 | 3.3415 |
| 52171 | ENSMUSG00000114613 | 0 | NA | NA | NA | NA | NA | Gm10732 | predicted gene 10732 [Source:MGI Symbo | 0 | 0 | 0 | 0 | 0 | 0 |
| 52172 | ENSMUSG00000114614 | 0 | NA | NA | NA | NA | NA | Gm47745 | predicted gene, 47745 [Source:MGI Symb | 0 | 0 | 0 | 0 | 0 | 0 |
| 52173 | ENSMUSG00000114615 | 0 | NA | NA | NA | NA | NA | Gm48370 | predicted gene, 48370 [Source:MGI Symb | 0 | 0 | 0 | 0 | 0 | 0 |
| 52174 | ENSMUSG00000114616 | 0 | NA | NA | NA | NA | NA | Gm26754 | predicted gene, 26754 [Source:MGI Symb | 0 | 0 | 0 | 0 | 0 | 0 |
| 52175 | ENSMUSG00000114617 | 0.331982614 | 0.16514309 | 3.986751476 | 0.041422971 | 0.966958701 | NA | Gm48766 | predicted gene, 48766 [Source:MGI Symb | 0 | 1.0279 | 0 | 0.964 | 0 | 0 |
| 52176 | ENSMUSG00000114618 | 0 | NA | NA | NA | NA | NA | Gm41183 | predicted gene, 41183 [Source:MGI Symb | 0 | 0 | 0 | 0 | 0 | 0 |
| 52177 | ENSMUSG00000114619 | 0 | NA | NA | NA | NA | NA | Gm49141 | predicted gene, 49141 [Source:MGI Symb | 0 | 0 | 0 | 0 | 0 | 0 |
| 52178 | ENSMUSG00000114620 | 0 | NA | NA | NA | NA | NA | Vmn1r-ps96 | vomeronasal 1 receptor, pseudogene 96 [ | 0 | 0 | 0 | 0 | 0 | 0 |
| 52179 | ENSMUSG00000114621 | 0 | NA | NA | NA | NA | NA | Gm31107 | predicted gene, 31107 [Source:MGI Symb | 0 | 0 | 0 | 0 | 0 | 0 |
| 52180 | ENSMUSG00000114622 | 0.133850373 | -0.796629176 | 4.080472857 | -0.195229623 | 0.84521319 | NA | Gm18990 | predicted gene, 18990 [Source:MGI Symb | 0.8031 | 0 | 0 | 0 | 0 | 0 |
| 52181 | ENSMUSG00000114623 | 0 | NA | NA | NA | NA | NA | Gm34788 | predicted gene, 34788 [Source:MGI Symb | 0 | 0 | 0 | 0 | 0 | 0 |
| 52182 | ENSMUSG00000114624 | 0 | NA | NA | NA | NA | NA | Gm48267 | predicted gene, 48267 [Source:MGI Symb | 0 | 0 | 0 | 0 | 0 | 0 |
| 52183 | ENSMUSG00000114625 | 0 | NA | NA | NA | NA | NA | Gm48218 | predicted gene, 48218 [Source:MGI Symb | 0 | 0 | 0 | 0 | 0 | 0 |
| 52184 | ENSMUSG00000114626 | 0 | NA | NA | NA | NA | NA | Gm48266 | predicted gene, 48266 [Source:MGI Symb | 0 | 0 | 0 | 0 | 0 | 0 |
| 52185 | ENSMUSG00000114627 | 0.353089354 | 0.165142706 | 3.896102244 | 0.042386646 | 0.966190474 | NA | B230220B15 | RIKEN cDNA B230220B15 gene [Source:M | 0 | 1.0279 | 0 | 0 | 1.0906 | 0 |
| 52186 | ENSMUSG00000114628 | 0 | NA | NA | NA | NA | NA | Gm19018 | predicted gene, 19018 [Source:MGI Symb | 0 | 0 | 0 | 0 | 0 | 0 |
| 52187 | ENSMUSG00000114629 | 0 | NA | NA | NA | NA | NA | Gm48244 | predicted gene, 48244 [Source:MGI Symb | 0 | 0 | 0 | 0 | 0 | 0 |
| 52188 | ENSMUSG00000114630 | 0 | NA | NA | NA | NA | NA | Gm48105 | predicted gene, 48105 [Source:MGI Symb | 0 | 0 | 0 | 0 | 0 | 0 |

| | | | | | | | | | | | | | | | |
|---|---|---|---|---|---|---|---|---|---|---|---|---|---|---|---|
| 52189 | ENSMUSG00000114631 | 0 | NA | NA | NA | NA | NA | Gm5199 | predicted gene 5199 [Source:MGI Symbol | 0 | 0 | 0 | 0 | 0 | 0 |
| 52190 | ENSMUSG00000114632 | 0 | NA | NA | NA | NA | NA | Gm48399 | predicted gene, 48399 [Source:MGI Symb | 0 | 0 | 0 | 0 | 0 | 0 |
| 52191 | ENSMUSG00000114633 | 0 | NA | NA | NA | NA | NA | Gm46416 | predicted gene, 46416 [Source:MGI Symb | 0 | 0 | 0 | 0 | 0 | 0 |
| 52192 | ENSMUSG00000114634 | 0 | NA | NA | NA | NA | NA | F830210D05 | RIKEN cDNA F830210D05 gene [Source:M | 0 | 0 | 0 | 0 | 0 | 0 |
| 52193 | ENSMUSG00000114635 | 0 | NA | NA | NA | NA | NA | Gm49392 | predicted gene, 49392 [Source:MGI Symb | 0 | 0 | 0 | 0 | 0 | 0 |
| 52194 | ENSMUSG00000114636 | 0 | NA | NA | NA | NA | NA | Gm18275 | predicted gene, 18275 [Source:MGI Symb | 0 | 0 | 0 | 0 | 0 | 0 |
| 52195 | ENSMUSG00000114637 | 0 | NA | NA | NA | NA | NA | Gm47851 | predicted gene, 47851 [Source:MGI Symb | 0 | 0 | 0 | 0 | 0 | 0 |
| 52196 | ENSMUSG00000114638 | 6.207004933 | 0.116796658 | 0.868654004 | 0.134457053 | 0.893041169 | 0.982562372 | Gm31834 | predicted gene, 31834 [Source:MGI Symb | 5.6217 | 5.1396 | 7.0437 | 2.8919 | 7.6343 | 8.9108 |
| 52197 | ENSMUSG00000114639 | 0 | NA | NA | NA | NA | NA | Gm31946 | predicted gene, 31946 [Source:MGI Symb | 0 | 0 | 0 | 0 | 0 | 0 |
| 52198 | ENSMUSG00000114640 | 0 | NA | NA | NA | NA | NA | Gm48107 | predicted gene, 48107 [Source:MGI Symb | 0 | 0 | 0 | 0 | 0 | 0 |
| 52199 | ENSMUSG00000114641 | 0 | NA | NA | NA | NA | NA | Gm46476 | predicted gene, 46476 [Source:MGI Symb | 0 | 0 | 0 | 0 | 0 | 0 |
| 52200 | ENSMUSG00000114642 | 0 | NA | NA | NA | NA | NA | Gm5461 | predicted gene 5461 [Source:MGI Symbol | 0 | 0 | 0 | 0 | 0 | 0 |
| 52201 | ENSMUSG00000114643 | 0 | NA | NA | NA | NA | NA | Gm18064 | predicted gene, 18064 [Source:MGI Symb | 0 | 0 | 0 | 0 | 0 | 0 |
| 52202 | ENSMUSG00000114644 | 0 | NA | NA | NA | NA | NA | Gm47243 | predicted gene, 47243 [Source:MGI Symb | 0 | 0 | 0 | 0 | 0 | 0 |
| 52203 | ENSMUSG00000114645 | 0 | NA | NA | NA | NA | NA | Gm49399 | predicted gene 49399 [Source:MGI Symbo | 0 | 0 | 0 | 0 | 0 | 0 |
| 52204 | ENSMUSG00000114646 | 0 | NA | NA | NA | NA | NA | Gm47888 | predicted gene, 47888 [Source:MGI Symb | 0 | 0 | 0 | 0 | 0 | 0 |
| 52205 | ENSMUSG00000114647 | 0 | NA | NA | NA | NA | NA | Gm30127 | predicted gene, 30127 [Source:MGI Symb | 0 | 0 | 0 | 0 | 0 | 0 |
| 52206 | ENSMUSG00000114648 | 0 | NA | NA | NA | NA | NA | Gm19034 | predicted gene, 19034 [Source:MGI Symb | 0 | 0 | 0 | 0 | 0 | 0 |
| 52207 | ENSMUSG00000114649 | 0.160662974 | 1.126924175 | 4.080472857 | 0.276174898 | 0.782413736 | NA | 3110006O06 | RIKEN cDNA 3110006O06 gene [Source:M | 0 | 0 | 0 | 0.964 | 0 | 0 |
| 52208 | ENSMUSG00000114650 | 0 | NA | NA | NA | NA | NA | Gm47570 | predicted gene, 47570 [Source:MGI Symb | 0 | 0 | 0 | 0 | 0 | 0 |
| 52209 | ENSMUSG00000114651 | 0 | NA | NA | NA | NA | NA | Gm6200 | predicted gene 6200 [Source:MGI Symbol | 0 | 0 | 0 | 0 | 0 | 0 |
| 52210 | ENSMUSG00000114652 | 0 | NA | NA | NA | NA | NA | Gm41043 | predicted gene, 41043 [Source:MGI Symb | 0 | 0 | 0 | 0 | 0 | 0 |
| 52211 | ENSMUSG00000114653 | 0.346303622 | 2.004339653 | 3.884003423 | 0.516049919 | 0.605819544 | NA | Gm18398 | predicted gene, 18398 [Source:MGI Symb | 0 | 0 | 0 | 0.964 | 0 | 1.1138 |
| 52212 | ENSMUSG00000114654 | 3.283271261 | 0.858508728 | 1.229050087 | 0.698514029 | 0.484855787 | NA | Gm47918 | predicted gene, 47918 [Source:MGI Symb | 4.8186 | 1.0279 | 1.0062 | 2.8919 | 3.2719 | 6.6831 |
| 52213 | ENSMUSG00000114655 | 0 | NA | NA | NA | NA | NA | 4631422I05F | RIKEN cDNA 4631422I05 gene [Source:MG | 0 | 0 | 0 | 0 | 0 | 0 |
| 52214 | ENSMUSG00000114656 | 0 | NA | NA | NA | NA | NA | 2810403G07 | RIKEN cDNA 2810403G07 gene [Source:M | 0 | 0 | 0 | 0 | 0 | 0 |
| 52215 | ENSMUSG00000114657 | 0 | NA | NA | NA | NA | NA | Gm47764 | predicted gene, 47764 [Source:MGI Symb | 0 | 0 | 0 | 0 | 0 | 0 |
| 52216 | ENSMUSG00000114658 | 0 | NA | NA | NA | NA | NA | Gm30008 | predicted gene, 30008 [Source:MGI Symb | 0 | 0 | 0 | 0 | 0 | 0 |
| 52217 | ENSMUSG00000114659 | 0 | NA | NA | NA | NA | NA | Gm31191 | predicted gene, 31191 [Source:MGI Symb | 0 | 0 | 0 | 0 | 0 | 0 |
| 52218 | ENSMUSG00000114660 | 0 | NA | NA | NA | NA | NA | Gm47881 | predicted gene, 47881 [Source:MGI Symb | 0 | 0 | 0 | 0 | 0 | 0 |
| 52219 | ENSMUSG00000114661 | 0 | NA | NA | NA | NA | NA | Gm48707 | predicted gene, 48707 [Source:MGI Symb | 0 | 0 | 0 | 0 | 0 | 0 |
| 52220 | ENSMUSG00000114662 | 0 | NA | NA | NA | NA | NA | Gm31683 | predicted gene, 31683 [Source:MGI Symb | 0 | 0 | 0 | 0 | 0 | 0 |
| 52221 | ENSMUSG00000114663 | 0 | NA | NA | NA | NA | NA | Gm40976 | predicted gene, 40976 [Source:MGI Symb | 0 | 0 | 0 | 0 | 0 | 0 |
| 52222 | ENSMUSG00000114664 | 1.461775315 | -0.076374232 | 1.915478993 | -0.039872133 | 0.968195068 | NA | Gm48639 | predicted gene, 48639 [Source:MGI Symb | 2.4093 | 2.0558 | 0 | 0.964 | 0 | 3.3415 |
| 52223 | ENSMUSG00000114665 | 4.844777081 | 0.134023352 | 1.221958364 | 0.109679148 | 0.912663838 | NA | Lncbate6 | brown adipose tissue enriched long nonco | 9.6372 | 1.0279 | 3.0187 | 0 | 9.8156 | 5.5692 |
| 52224 | ENSMUSG00000114666 | 0 | NA | NA | NA | NA | NA | Vmn1r-ps10 | vomeronasal 1 receptor, pseudogene 101 | 0 | 0 | 0 | 0 | 0 | 0 |
| 52225 | ENSMUSG00000114667 | 0 | NA | NA | NA | NA | NA | Ctsm-ps | cathepsin M, pseudogene [Source:MGI Sy | 0 | 0 | 0 | 0 | 0 | 0 |
| 52226 | ENSMUSG00000114668 | 0 | NA | NA | NA | NA | NA | Gm47563 | predicted gene, 47563 [Source:MGI Symb | 0 | 0 | 0 | 0 | 0 | 0 |
| 52227 | ENSMUSG00000114669 | 0 | NA | NA | NA | NA | NA | Gm48384 | predicted gene, 48384 [Source:MGI Symb | 0 | 0 | 0 | 0 | 0 | 0 |
| 52228 | ENSMUSG00000114670 | 0 | NA | NA | NA | NA | NA | Gm18072 | predicted gene, 18072 [Source:MGI Symb | 0 | 0 | 0 | 0 | 0 | 0 |
| 52229 | ENSMUSG00000114671 | 0 | NA | NA | NA | NA | NA | Gm48211 | predicted gene, 48211 [Source:MGI Symb | 0 | 0 | 0 | 0 | 0 | 0 |
| 52230 | ENSMUSG00000114672 | 0.319491021 | 0.16514334 | 4.044653141 | 0.040830038 | 0.967431393 | NA | Gm48192 | predicted gene, 48192 [Source:MGI Symb | 0.8031 | 0 | 0 | 0 | 0 | 1.1138 |
| 52231 | ENSMUSG00000114673 | 0 | NA | NA | NA | NA | NA | 1700016H03 | RIKEN cDNA 1700016H03 gene [Source:M | 0 | 0 | 0 | 0 | 0 | 0 |
| 52232 | ENSMUSG00000114674 | 0 | NA | NA | NA | NA | NA | Gm19239 | predicted gene, 19239 [Source:MGI Symb | 0 | 0 | 0 | 0 | 0 | 0 |
| 52233 | ENSMUSG00000114675 | 0 | NA | NA | NA | NA | NA | Gm18834 | predicted gene, 18834 [Source:MGI Symb | 0 | 0 | 0 | 0 | 0 | 0 |
| 52234 | ENSMUSG00000114676 | 0 | NA | NA | NA | NA | NA | NA | NA | 0 | 0 | 0 | 0 | 0 | 0 |
| 52235 | ENSMUSG00000114677 | 0 | NA | NA | NA | NA | NA | Gm48733 | predicted gene, 48733 [Source:MGI Symb | 0 | 0 | 0 | 0 | 0 | 0 |
| 52236 | ENSMUSG00000114678 | 0 | NA | NA | NA | NA | NA | Gm48481 | predicted gene, 48481 [Source:MGI Symb | 0 | 0 | 0 | 0 | 0 | 0 |
| 52237 | ENSMUSG00000114679 | 0 | NA | NA | NA | NA | NA | Gm47256 | predicted gene, 47256 [Source:MGI Symb | 0 | 0 | 0 | 0 | 0 | 0 |
| 52238 | ENSMUSG00000114680 | 0 | NA | NA | NA | NA | NA | Gm47202 | predicted gene, 47202 [Source:MGI Symb | 0 | 0 | 0 | 0 | 0 | 0 |
| 52239 | ENSMUSG00000114681 | 2.904746087 | 1.337058977 | 1.492640995 | 0.895767288 | 0.370377064 | NA | 1700026N04 | RIKEN cDNA 1700026N04 gene [Source:M | 0 | 0 | 5.0312 | 5.7839 | 3.2719 | 3.3415 |
| 52240 | ENSMUSG00000114682 | 0.428363721 | -0.543730382 | 3.624677344 | -0.15000794 | 0.880758351 | NA | Gm48775 | predicted gene, 48775 [Source:MGI Symb | 1.6062 | 0 | 0 | 0.964 | 0 | 0 |
| 52241 | ENSMUSG00000114683 | 0 | NA | NA | NA | NA | NA | Gm47763 | predicted gene, 47763 [Source:MGI Symb | 0 | 0 | 0 | 0 | 0 | 0 |
| 52242 | ENSMUSG00000114684 | 0 | NA | NA | NA | NA | NA | Vmn2r-ps10 | vomeronasal 2, receptor, pseudogene 105 | 0 | 0 | 0 | 0 | 0 | 0 |
| 52243 | ENSMUSG00000114685 | 0 | NA | NA | NA | NA | NA | Gm48351 | predicted gene, 48351 [Source:MGI Symb | 0 | 0 | 0 | 0 | 0 | 0 |
| 52244 | ENSMUSG00000114686 | 0 | NA | NA | NA | NA | NA | Gm4573 | predicted gene 4573 [Source:MGI Symbol | 0 | 0 | 0 | 0 | 0 | 0 |

| | | | | | | | | | | | | | | | |
|---|---|---|---|---|---|---|---|---|---|---|---|---|---|---|---|
| 52245 | ENSMUSG00000114687 | 4.048581082 | -0.171953681 | 1.127541526 | -0.152503191 | 0.878790074 | NA | Gm5452 | predicted gene 5452 [Source:MGI Symbol | 0.8031 | 6.1675 | 6.0375 | 5.7839 | 3.2719 | 2.2277 |
| 52246 | ENSMUSG00000114688 | 0.133850373 | -0.796629176 | 4.080472857 | -0.195229623 | 0.84521319 | NA | Gm47381 | predicted gene, 47381 [Source:MGI Symb | 0.8031 | 0 | 0 | 0 | 0 | 0 |
| 52247 | ENSMUSG00000114689 | 0.695987358 | -1.412518496 | 2.768247659 | -0.510257271 | 0.609871234 | NA | Gm9465 | predicted gene 9465 [Source:MGI Symbol | 0 | 2.0558 | 1.0062 | 0 | 0 | 1.1138 |
| 52248 | ENSMUSG00000114690 | 0 | NA | NA | NA | NA | NA | Gm33321 | predicted gene, 33321 [Source:MGI Symb | 0 | 0 | 0 | 0 | 0 | 0 |
| 52249 | ENSMUSG00000114691 | 0 | NA | NA | NA | NA | NA | Gm48438 | predicted gene, 48438 [Source:MGI Symb | 0 | 0 | 0 | 0 | 0 | 0 |
| 52250 | ENSMUSG00000114692 | 0 | NA | NA | NA | NA | NA | Gm48157 | predicted gene, 48157 [Source:MGI Symb | 0 | 0 | 0 | 0 | 0 | 0 |
| 52251 | ENSMUSG00000114693 | 0 | NA | NA | NA | NA | NA | Gm41177 | predicted gene, 41177 [Source:MGI Symb | 0 | 0 | 0 | 0 | 0 | 0 |
| 52252 | ENSMUSG00000114694 | 0 | NA | NA | NA | NA | NA | Gm47995 | predicted gene, 47995 [Source:MGI Symb | 0 | 0 | 0 | 0 | 0 | 0 |
| 52253 | ENSMUSG00000114695 | 0 | NA | NA | NA | NA | NA | Gm48648 | predicted gene, 48648 [Source:MGI Symb | 0 | 0 | 0 | 0 | 0 | 0 |
| 52254 | ENSMUSG00000114696 | 0.505131668 | 1.126696199 | 3.421159819 | 0.329331647 | 0.741905026 | NA | Gm47479 | predicted gene, 47479 [Source:MGI Symb | 0.8031 | 0 | 0 | 0 | 0 | 2.2277 |
| 52255 | ENSMUSG00000114697 | 0 | NA | NA | NA | NA | NA | Gm30806 | predicted gene, 30806 [Source:MGI Symb | 0 | 0 | 0 | 0 | 0 | 0 |
| 52256 | ENSMUSG00000114698 | 0 | NA | NA | NA | NA | NA | Rpl31-ps3 | ribosomal protein L31, pseudogene 3 [Sou | 0 | 0 | 0 | 0 | 0 | 0 |
| 52257 | ENSMUSG00000114699 | 0 | NA | NA | NA | NA | NA | Vmn1r-ps13 | vomeronasal 1 receptor, pseudogene 131 | 0 | 0 | 0 | 0 | 0 | 0 |
| 52258 | ENSMUSG00000114700 | 0 | NA | NA | NA | NA | NA | Gm19010 | predicted gene, 19010 [Source:MGI Symb | 0 | 0 | 0 | 0 | 0 | 0 |
| 52259 | ENSMUSG00000114701 | 0 | NA | NA | NA | NA | NA | Gm47731 | predicted gene, 47731 [Source:MGI Symb | 0 | 0 | 0 | 0 | 0 | 0 |
| 52260 | ENSMUSG00000114702 | 0.160662974 | 1.126924175 | 4.080472857 | 0.276174898 | 0.782413736 | NA | Gm40828 | predicted gene, 40828 [Source:MGI Symb | 0 | 0 | 0 | 0.964 | 0 | 0 |
| 52261 | ENSMUSG00000114703 | 0 | NA | NA | NA | NA | NA | Gm47776 | predicted gene, 47776 [Source:MGI Symb | 0 | 0 | 0 | 0 | 0 | 0 |
| 52262 | ENSMUSG00000114704 | 0 | NA | NA | NA | NA | NA | Gm48003 | predicted gene, 48003 [Source:MGI Symb | 0 | 0 | 0 | 0 | 0 | 0 |
| 52263 | ENSMUSG00000114705 | 13.26477092 | -0.154594672 | 0.609786936 | -0.25352244 | 0.799864523 | 0.962424862 | Gm18760 | predicted gene, 18760 [Source:MGI Symb | 9.6372 | 12.335 | 20.125 | 15.424 | 9.8156 | 12.252 |
| 52264 | ENSMUSG00000114706 | 0 | NA | NA | NA | NA | NA | Gm47952 | predicted gene, 47952 [Source:MGI Symb | 0 | 0 | 0 | 0 | 0 | 0 |
| 52265 | ENSMUSG00000114707 | 0.818715751 | 2.030181432 | 2.59128962 | 0.78346373 | 0.433354844 | NA | Gm47558 | predicted gene, 47558 [Source:MGI Symb | 0.8031 | 0 | 0 | 1.928 | 2.1812 | 0 |
| 52266 | ENSMUSG00000114708 | 26.01865082 | -0.045451701 | 0.472578994 | -0.096177996 | 0.923379206 | 0.98705827 | Gm30177 | predicted gene, 30177 [Source:MGI Symb | 21.684 | 24.67 | 33.206 | 33.739 | 29.447 | 13.366 |
| 52267 | ENSMUSG00000114709 | 11.34764093 | 1.146294952 | 0.725266671 | 1.580515138 | 0.113988943 | 0.496457714 | Gm47920 | predicted gene, 47920 [Source:MGI Symb | 8.031 | 5.1396 | 8.05 | 26.027 | 15.269 | 5.5692 |
| 52268 | ENSMUSG00000114710 | 24.16509973 | -0.77444537 | 0.553672596 | -1.398742462 | 0.161890226 | 0.585248745 | Gm40923 | predicted gene, 40923 [Source:MGI Symb | 54.611 | 15.419 | 21.131 | 17.352 | 13.087 | 23.391 |
| 52269 | ENSMUSG00000114711 | 0 | NA | NA | NA | NA | NA | Gm47759 | predicted gene, 47759 [Source:MGI Symb | 0 | 0 | 0 | 0 | 0 | 0 |
| 52270 | ENSMUSG00000114712 | 0 | NA | NA | NA | NA | NA | Gm47569 | predicted gene, 47569 [Source:MGI Symb | 0 | 0 | 0 | 0 | 0 | 0 |
| 52271 | ENSMUSG00000114713 | 0 | NA | NA | NA | NA | NA | Gm9633 | predicted gene 9633 [Source:MGI Symbol | 0 | 0 | 0 | 0 | 0 | 0 |
| 52272 | ENSMUSG00000114714 | 0.342639279 | -1.805976596 | 4.028547714 | -0.448294702 | 0.653940524 | NA | Gm48763 | predicted gene, 48763 [Source:MGI Symb | 0 | 2.0558 | 0 | 0 | 0 | 0 |
| 52273 | ENSMUSG00000114715 | 3.977687929 | 0.184783648 | 1.058741363 | 0.174531434 | 0.861447829 | NA | Gm34471 | predicted gene, 34471 [Source:MGI Symb | 4.0155 | 4.1117 | 3.0187 | 3.8559 | 2.1812 | 6.6831 |
| 52274 | ENSMUSG00000114716 | 0 | NA | NA | NA | NA | NA | NA | NA | 0 | 0 | 0 | 0 | 0 | 0 |
| 52275 | ENSMUSG00000114717 | 0 | NA | NA | NA | NA | NA | Gm17832 | predicted gene, 17832 [Source:MGI Symb | 0 | 0 | 0 | 0 | 0 | 0 |
| 52276 | ENSMUSG00000114718 | 0 | NA | NA | NA | NA | NA | Gm47753 | predicted gene, 47753 [Source:MGI Symb | 0 | 0 | 0 | 0 | 0 | 0 |
| 52277 | ENSMUSG00000114719 | 0.353089354 | 0.165142706 | 3.896102244 | 0.042386646 | 0.966190474 | NA | Gm18134 | predicted gene, 18134 [Source:MGI Symb | 0 | 1.0279 | 0 | 0 | 1.0906 | 0 |
| 52278 | ENSMUSG00000114720 | 0 | NA | NA | NA | NA | NA | Gm41162 | predicted gene, 41162 [Source:MGI Symb | 0 | 0 | 0 | 0 | 0 | 0 |
| 52279 | ENSMUSG00000114721 | 0.17131964 | -0.796629176 | 4.080472857 | -0.195229623 | 0.84521319 | NA | Gm47938 | predicted gene, 47938 [Source:MGI Symb | 0 | 1.0279 | 0 | 0 | 0 | 0 |
| 52280 | ENSMUSG00000114722 | 0 | NA | NA | NA | NA | NA | Gm31392 | predicted gene, 31392 [Source:MGI Symb | 0 | 0 | 0 | 0 | 0 | 0 |
| 52281 | ENSMUSG00000114723 | 0.963977846 | 3.490969125 | 2.983271141 | 1.170181643 | 0.241927879 | NA | 5530402G07 | RIKEN cDNA 5530402G07 gene [Source:M | 0 | 0 | 0 | 5.7839 | 0 | 0 |
| 52282 | ENSMUSG00000114724 | 0 | NA | NA | NA | NA | NA | Gm47751 | predicted gene, 47751 [Source:MGI Symb | 0 | 0 | 0 | 0 | 0 | 0 |
| 52283 | ENSMUSG00000114725 | 0 | NA | NA | NA | NA | NA | Gm2854 | predicted gene 2854 [Source:MGI Symbol | 0 | 0 | 0 | 0 | 0 | 0 |
| 52284 | ENSMUSG00000114726 | 0 | NA | NA | NA | NA | NA | Gm48900 | predicted gene, 48900 [Source:MGI Symb | 0 | 0 | 0 | 0 | 0 | 0 |
| 52285 | ENSMUSG00000114727 | 0 | NA | NA | NA | NA | NA | Gm47269 | predicted gene, 47269 [Source:MGI Symb | 0 | 0 | 0 | 0 | 0 | 0 |
| 52286 | ENSMUSG00000114728 | 0 | NA | NA | NA | NA | NA | Gm6128 | predicted pseudogene 6128 [Source:MGI | 0 | 0 | 0 | 0 | 0 | 0 |
| 52287 | ENSMUSG00000114729 | 0 | NA | NA | NA | NA | NA | C330022B21 | RIKEN cDNA C330022B21 gene [Source:M | 0 | 0 | 0 | 0 | 0 | 0 |
| 52288 | ENSMUSG00000114730 | 0 | NA | NA | NA | NA | NA | Gm47014 | predicted gene, 47014 [Source:MGI Symb | 0 | 0 | 0 | 0 | 0 | 0 |
| 52289 | ENSMUSG00000114731 | 0 | NA | NA | NA | NA | NA | Gm47914 | predicted gene, 47914 [Source:MGI Symb | 0 | 0 | 0 | 0 | 0 | 0 |
| 52290 | ENSMUSG00000114732 | 1.045955322 | 2.382148677 | 2.401164967 | 0.99208039 | 0.321158312 | NA | Gm40918 | predicted gene, 40918 [Source:MGI Symb | 0 | 0 | 1.0062 | 1.928 | 0 | 3.3415 |
| 52291 | ENSMUSG00000114733 | 0.181769715 | 1.126924175 | 4.080472857 | 0.276174898 | 0.782413736 | NA | Gm4815 | predicted gene 4815 [Source:MGI Symbol | 0 | 0 | 0 | 0 | 1.0906 | 0 |
| 52292 | ENSMUSG00000114734 | 0 | NA | NA | NA | NA | NA | Gm48397 | predicted gene, 48397 [Source:MGI Symb | 0 | 0 | 0 | 0 | 0 | 0 |
| 52293 | ENSMUSG00000114735 | 0 | NA | NA | NA | NA | NA | Gm48156 | predicted gene, 48156 [Source:MGI Symb | 0 | 0 | 0 | 0 | 0 | 0 |
| 52294 | ENSMUSG00000114736 | 0 | NA | NA | NA | NA | NA | Gm47336 | predicted gene, 47336 [Source:MGI Symb | 0 | 0 | 0 | 0 | 0 | 0 |
| 52295 | ENSMUSG00000114737 | 0.133850373 | -0.796629176 | 4.080472857 | -0.195229623 | 0.84521319 | NA | Gm8483 | predicted gene 8483 [Source:MGI Symbol | 0.8031 | 0 | 0 | 0 | 0 | 0 |
| 52296 | ENSMUSG00000114738 | 0 | NA | NA | NA | NA | NA | Gm41041 | predicted gene, 41041 [Source:MGI Symb | 0 | 0 | 0 | 0 | 0 | 0 |
| 52297 | ENSMUSG00000114739 | 0.781711799 | -0.350673833 | 2.438876442 | -0.143784994 | 0.885670253 | NA | Gm47885 | predicted gene, 47885 [Source:MGI Symb | 1.6062 | 0 | 1.0062 | 0.964 | 0 | 1.1138 |
| 52298 | ENSMUSG00000114740 | 0 | NA | NA | NA | NA | NA | Gm47653 | predicted gene, 47653 [Source:MGI Symb | 0 | 0 | 0 | 0 | 0 | 0 |
| 52299 | ENSMUSG00000114741 | 0 | NA | NA | NA | NA | NA | Gm47971 | predicted gene, 47971 [Source:MGI Symb | 0 | 0 | 0 | 0 | 0 | 0 |
| 52300 | ENSMUSG00000114742 | 0 | NA | NA | NA | NA | NA | Gm47936 | predicted gene, 47936 [Source:MGI Symb | 0 | 0 | 0 | 0 | 0 | 0 |

| | | | | | | | | | | | | | | | |
|---|---|---|---|---|---|---|---|---|---|---|---|---|---|---|---|
| 52301 | ENSMUSG00000114743 | 0 | NA | NA | NA | NA | NA | Gm34672 | predicted gene, 34672 [Source:MGI Symb | 0 | 0 | 0 | 0 | 0 | 0 |
| 52302 | ENSMUSG00000114744 | 0 | NA | NA | NA | NA | NA | Gm38409 | predicted gene, 38409 [Source:MGI Symb | 0 | 0 | 0 | 0 | 0 | 0 |
| 52303 | ENSMUSG00000114745 | 0 | NA | NA | NA | NA | NA | Gm48802 | predicted gene, 48802 [Source:MGI Symb | 0 | 0 | 0 | 0 | 0 | 0 |
| 52304 | ENSMUSG00000114746 | 0 | NA | NA | NA | NA | NA | 4933436N17 | RIKEN cDNA 4933436N17 gene [Source:M | 0 | 0 | 0 | 0 | 0 | 0 |
| 52305 | ENSMUSG00000114747 | 0 | NA | NA | NA | NA | NA | Gm48547 | predicted gene, 48547 [Source:MGI Symb | 0 | 0 | 0 | 0 | 0 | 0 |
| 52306 | ENSMUSG00000114748 | 0 | NA | NA | NA | NA | NA | Gm6734 | predicted gene 6734 [Source:MGI Symbol | 0 | 0 | 0 | 0 | 0 | 0 |
| 52307 | ENSMUSG00000114749 | 0 | NA | NA | NA | NA | NA | Gm48442 | predicted gene, 48442 [Source:MGI Symb | 0 | 0 | 0 | 0 | 0 | 0 |
| 52308 | ENSMUSG00000114750 | 0 | NA | NA | NA | NA | NA | Gm18265 | predicted gene, 18265 [Source:MGI Symb | 0 | 0 | 0 | 0 | 0 | 0 |
| 52309 | ENSMUSG00000114751 | 0 | NA | NA | NA | NA | NA | 5430425E15 | RIKEN cDNA 5430425E15 gene [Source:M | 0 | 0 | 0 | 0 | 0 | 0 |
| 52310 | ENSMUSG00000114752 | 0 | NA | NA | NA | NA | NA | Gm35110 | predicted gene, 35110 [Source:MGI Symb | 0 | 0 | 0 | 0 | 0 | 0 |
| 52311 | ENSMUSG00000114753 | 0 | NA | NA | NA | NA | NA | Gm34059 | predicted gene, 34059 [Source:MGI Symb | 0 | 0 | 0 | 0 | 0 | 0 |
| 52312 | ENSMUSG00000114754 | 0 | NA | NA | NA | NA | NA | Gm8996 | predicted gene 8996 [Source:MGI Symbol | 0 | 0 | 0 | 0 | 0 | 0 |
| 52313 | ENSMUSG00000114755 | 0 | NA | NA | NA | NA | NA | Galr3 | galanin receptor 3 [Source:MGI Symbol;A | 0 | 0 | 0 | 0 | 0 | 0 |
| 52314 | ENSMUSG00000114756 | 0.160662974 | 1.126924175 | 4.080472857 | 0.276174898 | 0.782413736 | NA | Gm47273 | predicted gene, 47273 [Source:MGI Symb | 0 | 0 | 0 | 0.964 | 0 | 0 |
| 52315 | ENSMUSG00000114757 | 0 | NA | NA | NA | NA | NA | Gm48126 | predicted gene, 48126 [Source:MGI Symb | 0 | 0 | 0 | 0 | 0 | 0 |
| 52316 | ENSMUSG00000114758 | 0 | NA | NA | NA | NA | NA | Gm48005 | predicted gene, 48005 [Source:MGI Symb | 0 | 0 | 0 | 0 | 0 | 0 |
| 52317 | ENSMUSG00000114759 | 2.094514586 | 0.562471951 | 1.697832367 | 0.331288272 | 0.740426749 | NA | Gm10257 | predicted gene 10257 [Source:MGI Symbo | 0 | 5.1396 | 0 | 1.928 | 3.2719 | 2.2277 |
| 52318 | ENSMUSG00000114760 | 0 | NA | NA | NA | NA | NA | Gm48006 | predicted gene, 48006 [Source:MGI Symb | 0 | 0 | 0 | 0 | 0 | 0 |
| 52319 | ENSMUSG00000114761 | 0.528279927 | -0.837071592 | 3.369085264 | -0.248456636 | 0.803781118 | NA | Gm47242 | predicted gene, 47242 [Source:MGI Symb | 0 | 2.0558 | 0 | 0 | 0 | 1.1138 |
| 52320 | ENSMUSG00000114762 | 0.160662974 | 1.126924175 | 4.080472857 | 0.276174898 | 0.782413736 | NA | Gm47007 | predicted gene, 47007 [Source:MGI Symb | 0 | 0 | 0 | 0.964 | 0 | 0 |
| 52321 | ENSMUSG00000114763 | 0 | NA | NA | NA | NA | NA | Gm49354 | predicted gene, 49354 [Source:MGI Symb | 0 | 0 | 0 | 0 | 0 | 0 |
| 52322 | ENSMUSG00000114764 | 0 | NA | NA | NA | NA | NA | Gm20241 | predicted gene, 20241 [Source:MGI Symb | 0 | 0 | 0 | 0 | 0 | 0 |
| 52323 | ENSMUSG00000114765 | 0 | NA | NA | NA | NA | NA | Gm8514 | predicted gene 8514 [Source:MGI Symbol | 0 | 0 | 0 | 0 | 0 | 0 |
| 52324 | ENSMUSG00000114766 | 0 | NA | NA | NA | NA | NA | Gm5203 | predicted gene 5203 [Source:MGI Symbol | 0 | 0 | 0 | 0 | 0 | 0 |
| 52325 | ENSMUSG00000114767 | 0 | NA | NA | NA | NA | NA | Gm46392 | predicted gene, 46392 [Source:MGI Symb | 0 | 0 | 0 | 0 | 0 | 0 |
| 52326 | ENSMUSG00000114768 | 2.170151415 | -1.102059179 | 1.534968245 | -0.717968716 | 0.472776577 | NA | Gm48857 | predicted gene, 48857 [Source:MGI Symb | 4.8186 | 1.0279 | 3.0187 | 1.928 | 0 | 2.2277 |
| 52327 | ENSMUSG00000114769 | 0 | NA | NA | NA | NA | NA | Gm48312 | predicted gene, 48312 [Source:MGI Symb | 0 | 0 | 0 | 0 | 0 | 0 |
| 52328 | ENSMUSG00000114770 | 0 | NA | NA | NA | NA | NA | 4921522A10 | RIKEN cDNA 4921522A10 gene [Source:M | 0 | 0 | 0 | 0 | 0 | 0 |
| 52329 | ENSMUSG00000114771 | 1.129798215 | -0.382394472 | 2.319957523 | -0.164828221 | 0.86907919 | NA | Gm49064 | predicted gene, 49064 [Source:MGI Symb | 0.8031 | 3.0838 | 0 | 2.8919 | 0 | 0 |
| 52330 | ENSMUSG00000114772 | 0 | NA | NA | NA | NA | NA | 2210404E10 | RIKEN cDNA 2210404E10 gene [Source:M | 0 | 0 | 0 | 0 | 0 | 0 |
| 52331 | ENSMUSG00000114773 | 0.133850373 | -0.796629176 | 4.080472857 | -0.195229623 | 0.84521319 | NA | Gm48568 | predicted gene, 48568 [Source:MGI Symb | 0.8031 | 0 | 0 | 0 | 0 | 0 |
| 52332 | ENSMUSG00000114774 | 0 | NA | NA | NA | NA | NA | Gm35281 | predicted gene, 35281 [Source:MGI Symb | 0 | 0 | 0 | 0 | 0 | 0 |
| 52333 | ENSMUSG00000114775 | 0 | NA | NA | NA | NA | NA | Gm33172 | predicted gene, 33172 [Source:MGI Symb | 0 | 0 | 0 | 0 | 0 | 0 |
| 52334 | ENSMUSG00000114776 | 0 | NA | NA | NA | NA | NA | Gm18260 | predicted gene, 18260 [Source:MGI Symb | 0 | 0 | 0 | 0 | 0 | 0 |
| 52335 | ENSMUSG00000114777 | 0 | NA | NA | NA | NA | NA | Gm46427 | predicted gene, 46427 [Source:MGI Symb | 0 | 0 | 0 | 0 | 0 | 0 |
| 52336 | ENSMUSG00000114778 | 0 | NA | NA | NA | NA | NA | Gm9634 | predicted gene 9634 [Source:MGI Symbol | 0 | 0 | 0 | 0 | 0 | 0 |
| 52337 | ENSMUSG00000114779 | 0 | NA | NA | NA | NA | NA | Gm17940 | predicted gene, 17940 [Source:MGI Symb | 0 | 0 | 0 | 0 | 0 | 0 |
| 52338 | ENSMUSG00000114780 | 0.349477146 | 0.165142768 | 3.911021834 | 0.042224967 | 0.966319359 | NA | AI197445 | expressed sequence AI197445 [Source:M | 0 | 0 | 1.0062 | 0 | 1.0906 | 0 |
| 52339 | ENSMUSG00000114781 | 0 | NA | NA | NA | NA | NA | Gm47463 | predicted gene, 47463 [Source:MGI Symb | 0 | 0 | 0 | 0 | 0 | 0 |
| 52340 | ENSMUSG00000114782 | 0 | NA | NA | NA | NA | NA | Gm7920 | predicted gene 7920 [Source:MGI Symbol | 0 | 0 | 0 | 0 | 0 | 0 |
| 52341 | ENSMUSG00000114783 | 0 | NA | NA | NA | NA | NA | Gm48217 | predicted gene, 48217 [Source:MGI Symb | 0 | 0 | 0 | 0 | 0 | 0 |
| 52342 | ENSMUSG00000114784 | 0 | NA | NA | NA | NA | NA | Gm47754 | predicted gene, 47754 [Source:MGI Symb | 0 | 0 | 0 | 0 | 0 | 0 |
| 52343 | ENSMUSG00000114785 | 0.449470461 | -0.545136533 | 3.562105917 | -0.15303771 | 0.878368537 | NA | Gm41115 | predicted gene, 41115 [Source:MGI Symb | 1.6062 | 0 | 0 | 0 | 1.0906 | 0 |
| 52344 | ENSMUSG00000114786 | 0 | NA | NA | NA | NA | NA | Gm48594 | predicted gene, 48594 [Source:MGI Symb | 0 | 0 | 0 | 0 | 0 | 0 |
| 52345 | ENSMUSG00000114787 | 0 | NA | NA | NA | NA | NA | 4930455M05 | RIKEN cDNA 4930455M05 gene [Source:M | 0 | 0 | 0 | 0 | 0 | 0 |
| 52346 | ENSMUSG00000114788 | 0 | NA | NA | NA | NA | NA | Gm48899 | predicted gene, 48899 [Source:MGI Symb | 0 | 0 | 0 | 0 | 0 | 0 |
| 52347 | ENSMUSG00000114789 | 0 | NA | NA | NA | NA | NA | Gm31577 | predicted gene, 31577 [Source:MGI Symb | 0 | 0 | 0 | 0 | 0 | 0 |
| 52348 | ENSMUSG00000114790 | 0 | NA | NA | NA | NA | NA | 4921509O07 | RIKEN cDNA 4921509O07 gene [Source:M | 0 | 0 | 0 | 0 | 0 | 0 |
| 52349 | ENSMUSG00000114791 | 0 | NA | NA | NA | NA | NA | Gm47730 | predicted gene, 47730 [Source:MGI Symb | 0 | 0 | 0 | 0 | 0 | 0 |
| 52350 | ENSMUSG00000114792 | 0 | NA | NA | NA | NA | NA | Gm6416 | predicted gene 6416 [Source:MGI Symbol | 0 | 0 | 0 | 0 | 0 | 0 |
| 52351 | ENSMUSG00000114793 | 0 | NA | NA | NA | NA | NA | Gm48414 | predicted gene, 48414 [Source:MGI Symb | 0 | 0 | 0 | 0 | 0 | 0 |
| 52352 | ENSMUSG00000114794 | 0 | NA | NA | NA | NA | NA | Gm47782 | predicted gene, 47782 [Source:MGI Symb | 0 | 0 | 0 | 0 | 0 | 0 |
| 52353 | ENSMUSG00000114795 | 0 | NA | NA | NA | NA | NA | Gm48396 | predicted gene, 48396 [Source:MGI Symb | 0 | 0 | 0 | 0 | 0 | 0 |
| 52354 | ENSMUSG00000114796 | 3.590126731 | -0.198909176 | 1.252630344 | -0.158793196 | 0.873831809 | NA | A930028N01 | RIKEN cDNA A930028N01 gene [Source:M | 6.4248 | 3.0838 | 2.0125 | 6.7478 | 3.2719 | 0 |
| 52355 | ENSMUSG00000114797 | 239.5467138 | -0.008281589 | 0.157132825 | -0.052704385 | 0.957967446 | 0.991861398 | Gm49336 | predicted gene, 49336 [Source:MGI Symb | 238.52 | 241.56 | 240.49 | 229.43 | 236.66 | 250.61 |
| 52356 | ENSMUSG00000114798 | 0 | NA | NA | NA | NA | NA | Gm20769 | predicted gene, 20769 [Source:MGI Symb | 0 | 0 | 0 | 0 | 0 | 0 |

| | | | | | | | | | | | | | | | |
|---|---|---|---|---|---|---|---|---|---|---|---|---|---|---|---|
| 52357 | ENSMUSG00000114799 | 0 | NA | NA | NA | NA | NA | Gm47978 | predicted gene, 47978 [Source:MGI Symb | 0 | 0 | 0 | 0 | 0 | 0 |
| 52358 | ENSMUSG00000114800 | 38.49692421 | -0.543004097 | 0.435238324 | -1.247601756 | 0.212176935 | 0.656735971 | Gm31600 | predicted gene, 31600 [Source:MGI Symb | 63.445 | 42.145 | 31.194 | 32.775 | 44.715 | 16.708 |
| 52359 | ENSMUSG00000114801 | 0 | NA | NA | NA | NA | NA | Gm48549 | predicted gene, 48549 [Source:MGI Symb | 0 | 0 | 0 | 0 | 0 | 0 |
| 52360 | ENSMUSG00000114802 | 0 | NA | NA | NA | NA | NA | Cts7-ps | cathepsin 7, pseudogene [Source:MGI Syr | 0 | 0 | 0 | 0 | 0 | 0 |
| 52361 | ENSMUSG00000114803 | 0 | NA | NA | NA | NA | NA | Gm46443 | predicted gene, 46443 [Source:MGI Symb | 0 | 0 | 0 | 0 | 0 | 0 |
| 52362 | ENSMUSG00000114804 | 0 | NA | NA | NA | NA | NA | Gm18073 | predicted gene, 18073 [Source:MGI Symb | 0 | 0 | 0 | 0 | 0 | 0 |
| 52363 | ENSMUSG00000114805 | 0.133850373 | -0.796629176 | 4.080472857 | -0.195229623 | 0.84521319 | NA | Gm9626 | predicted gene 9626 [Source:MGI Symbol | 0.8031 | 0 | 0 | 0 | 0 | 0 |
| 52364 | ENSMUSG00000114806 | 0 | NA | NA | NA | NA | NA | Gm8963 | predicted gene 8963 [Source:MGI Symbol | 0 | 0 | 0 | 0 | 0 | 0 |
| 52365 | ENSMUSG00000114807 | 0 | NA | NA | NA | NA | NA | Gm32090 | predicted gene, 32090 [Source:MGI Symb | 0 | 0 | 0 | 0 | 0 | 0 |
| 52366 | ENSMUSG00000114808 | 0 | NA | NA | NA | NA | NA | Gm47947 | predicted gene, 47947 [Source:MGI Symb | 0 | 0 | 0 | 0 | 0 | 0 |
| 52367 | ENSMUSG00000114809 | 0 | NA | NA | NA | NA | NA | Atp5c1-ps | ATP synthase, H+ transporting, mitochonc | 0 | 0 | 0 | 0 | 0 | 0 |
| 52368 | ENSMUSG00000114810 | 0 | NA | NA | NA | NA | NA | Gm36101 | predicted gene, 36101 [Source:MGI Symb | 0 | 0 | 0 | 0 | 0 | 0 |
| 52369 | ENSMUSG00000114811 | 0 | NA | NA | NA | NA | NA | Vmn1r-ps13 | vomeronasal 1 receptor, pseudogene 133 | 0 | 0 | 0 | 0 | 0 | 0 |
| 52370 | ENSMUSG00000114812 | 0.167707431 | -0.796629176 | 4.080472857 | -0.195229623 | 0.84521319 | NA | Gm48254 | predicted gene, 48254 [Source:MGI Symb | 0 | 0 | 1.0062 | 0 | 0 | 0 |
| 52371 | ENSMUSG00000114813 | 0 | NA | NA | NA | NA | NA | Gm47510 | predicted gene, 47510 [Source:MGI Symb | 0 | 0 | 0 | 0 | 0 | 0 |
| 52372 | ENSMUSG00000114814 | 0 | NA | NA | NA | NA | NA | Gm48361 | predicted gene, 48361 [Source:MGI Symb | 0 | 0 | 0 | 0 | 0 | 0 |
| 52373 | ENSMUSG00000114815 | 0 | NA | NA | NA | NA | NA | Gm41118 | predicted gene, 41118 [Source:MGI Symb | 0 | 0 | 0 | 0 | 0 | 0 |
| 52374 | ENSMUSG00000114816 | 0 | NA | NA | NA | NA | NA | Gm10323 | predicted gene 10323 [Source:MGI Symbo | 0 | 0 | 0 | 0 | 0 | 0 |
| 52375 | ENSMUSG00000114817 | 0 | NA | NA | NA | NA | NA | Gm48356 | predicted gene, 48356 [Source:MGI Symb | 0 | 0 | 0 | 0 | 0 | 0 |
| 52376 | ENSMUSG00000114818 | 13.30671181 | 2.13058726 | 0.644389221 | 3.306367008 | 0.000945142 | 0.020103727 | Gm35164 | predicted gene, 35164 [Source:MGI Symb | 5.6217 | 5.1396 | 4.025 | 22.171 | 26.175 | 16.708 |
| 52377 | ENSMUSG00000114819 | 0 | NA | NA | NA | NA | NA | Gm48785 | predicted gene, 48785 [Source:MGI Symb | 0 | 0 | 0 | 0 | 0 | 0 |
| 52378 | ENSMUSG00000114820 | 0 | NA | NA | NA | NA | NA | Gm48200 | predicted gene, 48200 [Source:MGI Symb | 0 | 0 | 0 | 0 | 0 | 0 |
| 52379 | ENSMUSG00000114821 | 0 | NA | NA | NA | NA | NA | Gm3509 | predicted gene 3509 [Source:MGI Symbol | 0 | 0 | 0 | 0 | 0 | 0 |
| 52380 | ENSMUSG00000114822 | 0 | NA | NA | NA | NA | NA | Gm4813 | predicted gene 4813 [Source:MGI Symbol | 0 | 0 | 0 | 0 | 0 | 0 |
| 52381 | ENSMUSG00000114823 | 0 | NA | NA | NA | NA | NA | Gm31630 | predicted gene, 31630 [Source:MGI Symb | 0 | 0 | 0 | 0 | 0 | 0 |
| 52382 | ENSMUSG00000114824 | 0 | NA | NA | NA | NA | NA | 3110015C05 | RIKEN cDNA 3110015C05 gene [Source:M | 0 | 0 | 0 | 0 | 0 | 0 |
| 52383 | ENSMUSG00000114825 | 0 | NA | NA | NA | NA | NA | Gm19119 | predicted gene, 19119 [Source:MGI Symb | 0 | 0 | 0 | 0 | 0 | 0 |
| 52384 | ENSMUSG00000114826 | 0 | NA | NA | NA | NA | NA | Gm10000 | predicted gene 10000 [Source:MGI Symbo | 0 | 0 | 0 | 0 | 0 | 0 |
| 52385 | ENSMUSG00000114827 | 6.662393359 | -0.150242423 | 0.818024645 | -0.183664911 | 0.854276338 | 0.973825032 | Gm46432 | predicted gene, 46432 [Source:MGI Symb | 4.8186 | 9.2513 | 7.0437 | 6.7478 | 6.5437 | 5.5692 |
| 52386 | ENSMUSG00000114828 | 207.9336161 | 0.535363089 | 0.193426094 | 2.76779144 | 0.005643756 | 0.072368161 | AI463229 | expressed sequence AI463229 [Source:M | 156.6 | 167.55 | 185.15 | 205.33 | 235.57 | 297.4 |
| 52387 | ENSMUSG00000114829 | 0 | NA | NA | NA | NA | NA | Gm9130 | predicted gene 9130 [Source:MGI Symbol | 0 | 0 | 0 | 0 | 0 | 0 |
| 52388 | ENSMUSG00000114830 | 1.075966422 | -0.197496223 | 2.150415569 | -0.091840957 | 0.926824403 | NA | Gm47400 | predicted gene, 47400 [Source:MGI Symb | 2.4093 | 1.0279 | 0 | 1.928 | 1.0906 | 0 |
| 52389 | ENSMUSG00000114831 | 0 | NA | NA | NA | NA | NA | Gm18361 | predicted gene, 18361 [Source:MGI Symb | 0 | 0 | 0 | 0 | 0 | 0 |
| 52390 | ENSMUSG00000114832 | 0 | NA | NA | NA | NA | NA | Gm21388 | predicted gene, 21388 [Source:MGI Symb | 0 | 0 | 0 | 0 | 0 | 0 |
| 52391 | ENSMUSG00000114833 | 0 | NA | NA | NA | NA | NA | Gm41192 | predicted gene, 41192 [Source:MGI Symb | 0 | 0 | 0 | 0 | 0 | 0 |
| 52392 | ENSMUSG00000114834 | 0 | NA | NA | NA | NA | NA | Gm48705 | predicted gene, 48705 [Source:MGI Symb | 0 | 0 | 0 | 0 | 0 | 0 |
| 52393 | ENSMUSG00000114835 | 14.06797601 | -0.23526956 | 0.579605911 | -0.405912976 | 0.684806544 | 0.935737593 | Gm48194 | predicted gene, 48194 [Source:MGI Symb | 11.243 | 16.447 | 18.112 | 15.424 | 9.8156 | 13.366 |
| 52394 | ENSMUSG00000114836 | 0 | NA | NA | NA | NA | NA | Gm18517 | predicted gene, 18517 [Source:MGI Symb | 0 | 0 | 0 | 0 | 0 | 0 |
| 52395 | ENSMUSG00000114837 | 0 | NA | NA | NA | NA | NA | Gm48090 | predicted gene, 48090 [Source:MGI Symb | 0 | 0 | 0 | 0 | 0 | 0 |
| 52396 | ENSMUSG00000114838 | 0 | NA | NA | NA | NA | NA | Gm48703 | predicted gene, 48703 [Source:MGI Symb | 0 | 0 | 0 | 0 | 0 | 0 |
| 52397 | ENSMUSG00000114839 | 0 | NA | NA | NA | NA | NA | Gm40989 | predicted gene, 40989 [Source:MGI Symb | 0 | 0 | 0 | 0 | 0 | 0 |
| 52398 | ENSMUSG00000114840 | 0 | NA | NA | NA | NA | NA | Gm48212 | predicted gene, 48212 [Source:MGI Symb | 0 | 0 | 0 | 0 | 0 | 0 |
| 52399 | ENSMUSG00000114841 | 4.419803075 | -0.132298543 | 1.137748319 | -0.116281027 | 0.907429822 | NA | Gm47352 | predicted gene, 47352 [Source:MGI Symb | 5.6217 | 8.2233 | 0 | 3.8559 | 4.3625 | 4.4554 |
| 52400 | ENSMUSG00000114842 | 1.306617125 | 1.546876254 | 2.477123789 | 0.624464656 | 0.532322475 | NA | Gm47797 | predicted gene, 47797 [Source:MGI Symb | 0 | 2.0558 | 0 | 5.7839 | 0 | 0 |
| 52401 | ENSMUSG00000114843 | 0 | NA | NA | NA | NA | NA | Gm35733 | predicted gene, 35733 [Source:MGI Symb | 0 | 0 | 0 | 0 | 0 | 0 |
| 52402 | ENSMUSG00000114844 | 0 | NA | NA | NA | NA | NA | Gm19222 | predicted gene, 19222 [Source:MGI Symb | 0 | 0 | 0 | 0 | 0 | 0 |
| 52403 | ENSMUSG00000114845 | 0 | NA | NA | NA | NA | NA | Gm47511 | predicted gene, 47511 [Source:MGI Symb | 0 | 0 | 0 | 0 | 0 | 0 |
| 52404 | ENSMUSG00000114846 | 0 | NA | NA | NA | NA | NA | Gm29927 | predicted gene, 29927 [Source:MGI Symb | 0 | 0 | 0 | 0 | 0 | 0 |
| 52405 | ENSMUSG00000114847 | 0 | NA | NA | NA | NA | NA | Gm47892 | predicted gene, 47892 [Source:MGI Symb | 0 | 0 | 0 | 0 | 0 | 0 |
| 52406 | ENSMUSG00000114848 | 0 | NA | NA | NA | NA | NA | Gm35389 | predicted gene, 35389 [Source:MGI Symb | 0 | 0 | 0 | 0 | 0 | 0 |
| 52407 | ENSMUSG00000114849 | 0 | NA | NA | NA | NA | NA | Gm48595 | predicted gene, 48595 [Source:MGI Symb | 0 | 0 | 0 | 0 | 0 | 0 |
| 52408 | ENSMUSG00000114850 | 0 | NA | NA | NA | NA | NA | Gm8438 | predicted gene 8438 [Source:MGI Symbol | 0 | 0 | 0 | 0 | 0 | 0 |
| 52409 | ENSMUSG00000114851 | 0.995947842 | -0.043351441 | 2.675688395 | -0.016201977 | 0.987073258 | NA | Gm46409 | predicted gene, 46409 [Source:MGI Symb | 0 | 3.0838 | 0 | 2.8919 | 0 | 0 |
| 52410 | ENSMUSG00000114852 | 0 | NA | NA | NA | NA | NA | Gm47189 | predicted gene, 47189 [Source:MGI Symb | 0 | 0 | 0 | 0 | 0 | 0 |
| 52411 | ENSMUSG00000114853 | 0 | NA | NA | NA | NA | NA | D030051J21 | RIKEN cDNA D030051J21 gene [Source:M | 0 | 0 | 0 | 0 | 0 | 0 |
| 52412 | ENSMUSG00000114854 | 0 | NA | NA | NA | NA | NA | Gm34354 | predicted gene, 34354 [Source:MGI Symb | 0 | 0 | 0 | 0 | 0 | 0 |

| | | | | | | | | | | | | | | | |
|---|---|---|---|---|---|---|---|---|---|---|---|---|---|---|---|
| 52413 | ENSMUSG00000114855 | 0 | NA | NA | NA | NA | NA | Gm47909 | predicted gene, 47909 [Source:MGI Symb | 0 | 0 | 0 | 0 | 0 | 0 |
| 52414 | ENSMUSG00000114856 | 0 | NA | NA | NA | NA | NA | Gm47642 | predicted gene, 47642 [Source:MGI Symb | 0 | 0 | 0 | 0 | 0 | 0 |
| 52415 | ENSMUSG00000114857 | 0.267700746 | -1.497481649 | 4.057001436 | -0.369110456 | 0.712045396 | NA | Gm48770 | predicted gene, 48770 [Source:MGI Symb | 1.6062 | 0 | 0 | 0 | 0 | 0 |
| 52416 | ENSMUSG00000114858 | 0 | NA | NA | NA | NA | NA | Gm5790 | predicted gene 5790 [Source:MGI Symbol | 0 | 0 | 0 | 0 | 0 | 0 |
| 52417 | ENSMUSG00000114859 | 0.487198452 | -0.685759572 | 3.136894395 | -0.218610985 | 0.826953095 | NA | Gm47735 | predicted gene, 47735 [Source:MGI Symb | 0.8031 | 0 | 1.0062 | 0 | 0 | 1.1138 |
| 52418 | ENSMUSG00000114860 | 0.167707431 | -0.796629176 | 4.080472857 | -0.195229623 | 0.84521319 | NA | Gm49291 | predicted gene, 49291 [Source:MGI Symb | 0 | 0 | 1.0062 | 0 | 0 | 0 |
| 52419 | ENSMUSG00000114861 | 1.718587302 | -1.908667498 | 1.840722588 | -1.036912086 | 0.299776831 | NA | Gm47991 | predicted gene, 47991 [Source:MGI Symb | 4.0155 | 2.0558 | 2.0125 | 0 | 0 | 2.2277 |
| 52420 | ENSMUSG00000114862 | 0 | NA | NA | NA | NA | NA | Gm4817 | predicted gene 4817 [Source:MGI Symbol | 0 | 0 | 0 | 0 | 0 | 0 |
| 52421 | ENSMUSG00000114863 | 2.589300446 | 2.176055474 | 1.485335375 | 1.465026357 | 0.142913727 | NA | Gm48824 | predicted gene, 48824 [Source:MGI Symb | 0.8031 | 0 | 2.0125 | 3.8559 | 2.1812 | 6.6831 |
| 52422 | ENSMUSG00000114864 | 0 | NA | NA | NA | NA | NA | Gm41071 | predicted gene, 41071 [Source:MGI Symb | 0 | 0 | 0 | 0 | 0 | 0 |
| 52423 | ENSMUSG00000114865 | 0.133850373 | -0.796629176 | 4.080472857 | -0.195229623 | 0.84521319 | NA | Gm29776 | predicted gene, 29776 [Source:MGI Symb | 0.8031 | 0 | 0 | 0 | 0 | 0 |
| 52424 | ENSMUSG00000114866 | 0 | NA | NA | NA | NA | NA | Gm40922 | predicted gene, 40922 [Source:MGI Symb | 0 | 0 | 0 | 0 | 0 | 0 |
| 52425 | ENSMUSG00000114867 | 0 | NA | NA | NA | NA | NA | Cts8-ps | cathepsin 8, pseudogene [Source:MGI Syr | 0 | 0 | 0 | 0 | 0 | 0 |
| 52426 | ENSMUSG00000114868 | 0 | NA | NA | NA | NA | NA | Gm47335 | predicted gene, 47335 [Source:MGI Symb | 0 | 0 | 0 | 0 | 0 | 0 |
| 52427 | ENSMUSG00000114869 | 0.267700746 | -1.497481649 | 4.057001436 | -0.369110456 | 0.712045396 | NA | Gm48876 | predicted gene, 48876 [Source:MGI Symb | 1.6062 | 0 | 0 | 0 | 0 | 0 |
| 52428 | ENSMUSG00000114870 | 0 | NA | NA | NA | NA | NA | Gm49646 | predicted gene, 49646 [Source:MGI Symb | 0 | 0 | 0 | 0 | 0 | 0 |
| 52429 | ENSMUSG00000114871 | 0.160662974 | 1.126924175 | 4.080472857 | 0.276174898 | 0.782413736 | NA | Gm21370 | predicted gene, 21370 [Source:MGI Symb | 0 | 0 | 0 | 0.964 | 0 | 0 |
| 52430 | ENSMUSG00000114872 | 0 | NA | NA | NA | NA | NA | Gm18759 | predicted gene, 18759 [Source:MGI Symb | 0 | 0 | 0 | 0 | 0 | 0 |
| 52431 | ENSMUSG00000114873 | 0.301557804 | -1.64335916 | 4.042814367 | -0.406488899 | 0.684383412 | NA | Gm47052 | predicted gene, 47052 [Source:MGI Symb | 0.8031 | 0 | 1.0062 | 0 | 0 | 0 |
| 52432 | ENSMUSG00000114874 | 0 | NA | NA | NA | NA | NA | Gm48470 | predicted gene, 48470 [Source:MGI Symb | 0 | 0 | 0 | 0 | 0 | 0 |
| 52433 | ENSMUSG00000114875 | 3.663411951 | -0.407606399 | 1.103940658 | -0.369228541 | 0.711957384 | NA | Gm30363 | predicted gene, 30363 [Source:MGI Symb | 2.4093 | 6.1675 | 4.025 | 3.8559 | 2.1812 | 3.3415 |
| 52434 | ENSMUSG00000114876 | 0 | NA | NA | NA | NA | NA | Gm48026 | predicted gene, 48026 [Source:MGI Symb | 0 | 0 | 0 | 0 | 0 | 0 |
| 52435 | ENSMUSG00000114877 | 0 | NA | NA | NA | NA | NA | Gm33022 | predicted gene, 33022 [Source:MGI Symb | 0 | 0 | 0 | 0 | 0 | 0 |
| 52436 | ENSMUSG00000114878 | 0 | NA | NA | NA | NA | NA | Gm9086 | predicted gene 9086 [Source:MGI Symbol | 0 | 0 | 0 | 0 | 0 | 0 |
| 52437 | ENSMUSG00000114879 | 0 | NA | NA | NA | NA | NA | 4930467J12I | RIKEN cDNA 4930467J12 gene [Source:M( | 0 | 0 | 0 | 0 | 0 | 0 |
| 52438 | ENSMUSG00000114880 | 0 | NA | NA | NA | NA | NA | Gm49098 | predicted gene, 49098 [Source:MGI Symb | 0 | 0 | 0 | 0 | 0 | 0 |
| 52439 | ENSMUSG00000114881 | 0 | NA | NA | NA | NA | NA | Gm8588 | predicted gene 8588 [Source:MGI Symbol | 0 | 0 | 0 | 0 | 0 | 0 |
| 52440 | ENSMUSG00000114882 | 0 | NA | NA | NA | NA | NA | Gm48581 | predicted gene, 48581 [Source:MGI Symb | 0 | 0 | 0 | 0 | 0 | 0 |
| 52441 | ENSMUSG00000114883 | 0.587191767 | -1.12394062 | 3.226482889 | -0.348348545 | 0.72757844 | NA | Gm48860 | predicted gene, 48860 [Source:MGI Symb | 2.4093 | 0 | 0 | 0 | 0 | 1.1138 |
| 52442 | ENSMUSG00000114884 | 0 | NA | NA | NA | NA | NA | Gm47675 | predicted gene, 47675 [Source:MGI Symb | 0 | 0 | 0 | 0 | 0 | 0 |
| 52443 | ENSMUSG00000114885 | 0 | NA | NA | NA | NA | NA | Gm7371 | predicted gene 7371 [Source:MGI Symbol | 0 | 0 | 0 | 0 | 0 | 0 |
| 52444 | ENSMUSG00000114886 | 0.703108924 | -2.875459482 | 2.96384054 | -0.970180225 | 0.331956666 | NA | Gm48432 | predicted gene, 48432 [Source:MGI Symb | 3.2124 | 0 | 1.0062 | 0 | 0 | 0 |
| 52445 | ENSMUSG00000114887 | 0 | NA | NA | NA | NA | NA | Gm48894 | predicted gene, 48894 [Source:MGI Symb | 0 | 0 | 0 | 0 | 0 | 0 |
| 52446 | ENSMUSG00000114888 | 0.401551119 | -2.072982726 | 4.008251979 | -0.517178744 | 0.605031388 | NA | Gm49616 | predicted gene, 49616 [Source:MGI Symb | 2.4093 | 0 | 0 | 0 | 0 | 0 |
| 52447 | ENSMUSG00000114889 | 0 | NA | NA | NA | NA | NA | Gm29764 | predicted gene, 29764 [Source:MGI Symb | 0 | 0 | 0 | 0 | 0 | 0 |
| 52448 | ENSMUSG00000114890 | 0 | NA | NA | NA | NA | NA | Tpi-rs9 | triosephosphate isomerase related seque | 0 | 0 | 0 | 0 | 0 | 0 |
| 52449 | ENSMUSG00000114891 | 0.538988726 | 1.126003851 | 3.343747182 | 0.336749099 | 0.736306047 | NA | Gm47272 | predicted gene, 47272 [Source:MGI Symb | 0 | 0 | 1.0062 | 0 | 0 | 2.2277 |
| 52450 | ENSMUSG00000114892 | 0 | NA | NA | NA | NA | NA | Gm5463 | predicted gene 5463 [Source:MGI Symbol | 0 | 0 | 0 | 0 | 0 | 0 |
| 52451 | ENSMUSG00000114893 | 0.944152054 | -0.673133018 | 2.474469057 | -0.272031293 | 0.785597958 | NA | Gm48684 | predicted gene, 48684 [Source:MGI Symb | 2.4093 | 1.0279 | 0 | 0 | 0 | 2.2277 |
| 52452 | ENSMUSG00000114894 | 0 | NA | NA | NA | NA | NA | Gm7084 | predicted gene 7084 [Source:MGI Symbol | 0 | 0 | 0 | 0 | 0 | 0 |
| 52453 | ENSMUSG00000114895 | 0 | NA | NA | NA | NA | NA | 4930579J19I | RIKEN cDNA 4930579J19 gene [Source:M( | 0 | 0 | 0 | 0 | 0 | 0 |
| 52454 | ENSMUSG00000114896 | 0 | NA | NA | NA | NA | NA | Gm34245 | predicted gene, 34245 [Source:MGI Symb | 0 | 0 | 0 | 0 | 0 | 0 |
| 52455 | ENSMUSG00000114897 | 0 | NA | NA | NA | NA | NA | Gm5923 | predicted gene 5923 [Source:MGI Symbol | 0 | 0 | 0 | 0 | 0 | 0 |
| 52456 | ENSMUSG00000114898 | 0 | NA | NA | NA | NA | NA | Gm49390 | predicted gene, 49390 [Source:MGI Symb | 0 | 0 | 0 | 0 | 0 | 0 |
| 52457 | ENSMUSG00000114899 | 0 | NA | NA | NA | NA | NA | Gm48214 | predicted gene, 48214 [Source:MGI Symb | 0 | 0 | 0 | 0 | 0 | 0 |
| 52458 | ENSMUSG00000114900 | 0 | NA | NA | NA | NA | NA | Gm46396 | predicted gene, 46396 [Source:MGI Symb | 0 | 0 | 0 | 0 | 0 | 0 |
| 52459 | ENSMUSG00000114901 | 0 | NA | NA | NA | NA | NA | Gm47462 | predicted gene, 47462 [Source:MGI Symb | 0 | 0 | 0 | 0 | 0 | 0 |
| 52460 | ENSMUSG00000114902 | 3.440658399 | 1.721730972 | 1.2312036 | 1.398412879 | 0.161989118 | NA | Gm4118 | predicted gene 4118 [Source:MGI Symbol | 0.8031 | 2.0558 | 2.0125 | 4.8199 | 8.7249 | 2.2277 |
| 52461 | ENSMUSG00000114903 | 0 | NA | NA | NA | NA | NA | Gm46419 | predicted gene, 46419 [Source:MGI Symb | 0 | 0 | 0 | 0 | 0 | 0 |
| 52462 | ENSMUSG00000114904 | 0 | NA | NA | NA | NA | NA | Gm49352 | predicted gene, 49352 [Source:MGI Symb | 0 | 0 | 0 | 0 | 0 | 0 |
| 52463 | ENSMUSG00000114905 | 0.367410362 | 2.078255025 | 3.800070157 | 0.54689912 | 0.58444804 | NA | Gm48113 | predicted gene, 48113 [Source:MGI Symb | 0 | 0 | 0 | 0 | 1.0906 | 1.1138 |
| 52464 | ENSMUSG00000114906 | 0 | NA | NA | NA | NA | NA | Gm48098 | predicted gene, 48098 [Source:MGI Symb | 0 | 0 | 0 | 0 | 0 | 0 |
| 52465 | ENSMUSG00000114907 | 13.01904854 | -0.025251271 | 0.595275729 | -0.042419453 | 0.966164321 | 0.993855397 | Gm47794 | predicted gene, 47794 [Source:MGI Symb | 12.047 | 12.335 | 15.094 | 14.46 | 15.269 | 8.9108 |
| 52466 | ENSMUSG00000114908 | 0 | NA | NA | NA | NA | NA | Gm9547 | predicted gene 9547 [Source:MGI Symbol | 0 | 0 | 0 | 0 | 0 | 0 |
| 52467 | ENSMUSG00000114909 | 0 | NA | NA | NA | NA | NA | Csnk2a1-ps1 | casein kinase 2, alpha 1 polypeptide, pseu | 0 | 0 | 0 | 0 | 0 | 0 |
| 52468 | ENSMUSG00000114910 | 0 | NA | NA | NA | NA | NA | Gm48756 | predicted gene, 48756 [Source:MGI Symb | 0 | 0 | 0 | 0 | 0 | 0 |

| | | | | | | | | | | | | | | | |
|---|---|---|---|---|---|---|---|---|---|---|---|---|---|---|---|
| 52469 | ENSMUSG00000114911 | 0 | NA | NA | NA | NA | NA | Gm17878 | predicted gene, 17878 [Source:MGI Symb | 0 | 0 | 0 | 0 | 0 | 0 |
| 52470 | ENSMUSG00000114912 | 0 | NA | NA | NA | NA | NA | Gm48049 | predicted gene, 48049 [Source:MGI Symb | 0 | 0 | 0 | 0 | 0 | 0 |
| 52471 | ENSMUSG00000114913 | 0 | NA | NA | NA | NA | NA | Gm18276 | predicted gene, 18276 [Source:MGI Symb | 0 | 0 | 0 | 0 | 0 | 0 |
| 52472 | ENSMUSG00000114914 | 0 | NA | NA | NA | NA | NA | Gm48571 | predicted gene, 48571 [Source:MGI Symb | 0 | 0 | 0 | 0 | 0 | 0 |
| 52473 | ENSMUSG00000114915 | 0 | NA | NA | NA | NA | NA | Gm5667 | predicted gene 5667 [Source:MGI Symbol | 0 | 0 | 0 | 0 | 0 | 0 |
| 52474 | ENSMUSG00000114916 | 0 | NA | NA | NA | NA | NA | 4930572O13 | RIKEN cDNA 4930572O13 gene [Source:M | 0 | 0 | 0 | 0 | 0 | 0 |
| 52475 | ENSMUSG00000114917 | 0 | NA | NA | NA | NA | NA | Gm48250 | predicted gene, 48250 [Source:MGI Symb | 0 | 0 | 0 | 0 | 0 | 0 |
| 52476 | ENSMUSG00000114918 | 0 | NA | NA | NA | NA | NA | Gm9012 | predicted gene 9012 [Source:MGI Symbol | 0 | 0 | 0 | 0 | 0 | 0 |
| 52477 | ENSMUSG00000114919 | 0 | NA | NA | NA | NA | NA | Gm6122 | predicted gene 6122 [Source:MGI Symbol | 0 | 0 | 0 | 0 | 0 | 0 |
| 52478 | ENSMUSG00000114920 | 0 | NA | NA | NA | NA | NA | Gm19097 | predicted gene, 19097 [Source:MGI Symb | 0 | 0 | 0 | 0 | 0 | 0 |
| 52479 | ENSMUSG00000114921 | 0 | NA | NA | NA | NA | NA | Gm18578 | predicted gene, 18578 [Source:MGI Symb | 0 | 0 | 0 | 0 | 0 | 0 |
| 52480 | ENSMUSG00000114922 | 0 | NA | NA | NA | NA | NA | Ppifos | peptidylprolyl isomerase F, opposite stran | 0 | 0 | 0 | 0 | 0 | 0 |
| 52481 | ENSMUSG00000114923 | 0 | NA | NA | NA | NA | NA | Gm49345 | predicted gene, 49345 [Source:MGI Symb | 0 | 0 | 0 | 0 | 0 | 0 |
| 52482 | ENSMUSG00000114924 | 0.160662974 | 1.126924175 | 4.080472857 | 0.276174898 | 0.782413736 | NA | Gm38408 | predicted gene, 38408 [Source:MGI Symb | 0 | 0 | 0 | 0.964 | 0 | 0 |
| 52483 | ENSMUSG00000114925 | 0 | NA | NA | NA | NA | NA | Gm47212 | predicted gene, 47212 [Source:MGI Symb | 0 | 0 | 0 | 0 | 0 | 0 |
| 52484 | ENSMUSG00000114926 | 0 | NA | NA | NA | NA | NA | Gm2379 | predicted gene 2379 [Source:MGI Symbol | 0 | 0 | 0 | 0 | 0 | 0 |
| 52485 | ENSMUSG00000114927 | 0 | NA | NA | NA | NA | NA | Gm47114 | predicted gene, 47114 [Source:MGI Symb | 0 | 0 | 0 | 0 | 0 | 0 |
| 52486 | ENSMUSG00000114928 | 0 | NA | NA | NA | NA | NA | Gm47848 | predicted gene, 47848 [Source:MGI Symb | 0 | 0 | 0 | 0 | 0 | 0 |
| 52487 | ENSMUSG00000114929 | 1.010062259 | 3.555195121 | 2.385557701 | 1.490299362 | 0.13614554 | NA | Gm48441 | predicted gene, 48441 [Source:MGI Symb | 0 | 0 | 0 | 3.8559 | 1.0906 | 1.1138 |
| 52488 | ENSMUSG00000114930 | 0 | NA | NA | NA | NA | NA | Gm8447 | predicted gene 8447 [Source:MGI Symbol | 0 | 0 | 0 | 0 | 0 | 0 |
| 52489 | ENSMUSG00000114931 | 0.503302254 | -0.837877584 | 3.426698643 | -0.244514523 | 0.806832356 | NA | Gm32256 | predicted gene, 32256 [Source:MGI Symb | 0 | 2.0558 | 0 | 0.964 | 0 | 0 |
| 52490 | ENSMUSG00000114932 | 0 | NA | NA | NA | NA | NA | Gm4936 | predicted gene 4936 [Source:MGI Symbol | 0 | 0 | 0 | 0 | 0 | 0 |
| 52491 | ENSMUSG00000114933 | 0 | NA | NA | NA | NA | NA | Gm48040 | predicted gene, 48040 [Source:MGI Symb | 0 | 0 | 0 | 0 | 0 | 0 |
| 52492 | ENSMUSG00000114934 | 0.335414862 | -1.779047242 | 4.030804179 | -0.441362855 | 0.658950329 | NA | Gm48342 | predicted gene, 48342 [Source:MGI Symb | 0 | 0 | 2.0125 | 0 | 0 | 0 |
| 52493 | ENSMUSG00000114935 | 1.000898647 | -2.10615299 | 2.475207832 | -0.850899453 | 0.394825209 | NA | Gm47040 | predicted gene, 47040 [Source:MGI Symb | 0.8031 | 4.1117 | 0 | 0 | 1.0906 | 0 |
| 52494 | ENSMUSG00000114936 | 0.167707431 | -0.796629176 | 4.080472857 | -0.195229623 | 0.84521319 | NA | Gm47061 | predicted gene, 47061 [Source:MGI Symb | 0 | 0 | 1.0062 | 0 | 0 | 0 |
| 52495 | ENSMUSG00000114937 | 0.492645588 | 0.9565034 | 3.447648281 | 0.277436479 | 0.781444974 | NA | Gm30054 | predicted gene, 30054 [Source:MGI Symb | 0 | 1.0279 | 0 | 1.928 | 0 | 0 |
| 52496 | ENSMUSG00000114938 | 0 | NA | NA | NA | NA | NA | Gm48134 | predicted gene, 48134 [Source:MGI Symb | 0 | 0 | 0 | 0 | 0 | 0 |
| 52497 | ENSMUSG00000114939 | 1.010263336 | 1.395395317 | 2.689636066 | 0.518804508 | 0.603897074 | NA | Gm48566 | predicted gene, 48566 [Source:MGI Symb | 1.6062 | 0 | 0 | 0 | 0 | 4.4554 |
| 52498 | ENSMUSG00000114940 | 0 | NA | NA | NA | NA | NA | Gm21378 | predicted gene, 21378 [Source:MGI Symb | 0 | 0 | 0 | 0 | 0 | 0 |
| 52499 | ENSMUSG00000114941 | 0 | NA | NA | NA | NA | NA | Gm47779 | predicted gene, 47779 [Source:MGI Symb | 0 | 0 | 0 | 0 | 0 | 0 |
| 52500 | ENSMUSG00000114942 | 0 | NA | NA | NA | NA | NA | Gm49361 | predicted gene, 49361 [Source:MGI Symb | 0 | 0 | 0 | 0 | 0 | 0 |
| 52501 | ENSMUSG00000114943 | 0 | NA | NA | NA | NA | NA | Gm8947 | predicted gene 8947 [Source:MGI Symbol | 0 | 0 | 0 | 0 | 0 | 0 |
| 52502 | ENSMUSG00000114944 | 0 | NA | NA | NA | NA | NA | Gm48299 | predicted gene, 48299 [Source:MGI Symb | 0 | 0 | 0 | 0 | 0 | 0 |
| 52503 | ENSMUSG00000114945 | 0 | NA | NA | NA | NA | NA | Gm48265 | predicted gene, 48265 [Source:MGI Symb | 0 | 0 | 0 | 0 | 0 | 0 |
| 52504 | ENSMUSG00000114946 | 0 | NA | NA | NA | NA | NA | Gm47756 | predicted gene, 47756 [Source:MGI Symb | 0 | 0 | 0 | 0 | 0 | 0 |
| 52505 | ENSMUSG00000114947 | 0 | NA | NA | NA | NA | NA | Gm18577 | predicted gene, 18577 [Source:MGI Symb | 0 | 0 | 0 | 0 | 0 | 0 |
| 52506 | ENSMUSG00000114948 | 0 | NA | NA | NA | NA | NA | Gm6066 | predicted gene 6066 [Source:MGI Symbol | 0 | 0 | 0 | 0 | 0 | 0 |
| 52507 | ENSMUSG00000114949 | 0 | NA | NA | NA | NA | NA | Gm47530 | predicted gene, 47530 [Source:MGI Symb | 0 | 0 | 0 | 0 | 0 | 0 |
| 52508 | ENSMUSG00000114950 | 0 | NA | NA | NA | NA | NA | Gm47747 | predicted gene, 47747 [Source:MGI Symb | 0 | 0 | 0 | 0 | 0 | 0 |
| 52509 | ENSMUSG00000114951 | 0 | NA | NA | NA | NA | NA | Gm20784 | predicted gene, 20784 [Source:MGI Symb | 0 | 0 | 0 | 0 | 0 | 0 |
| 52510 | ENSMUSG00000114952 | 0.656740811 | -0.013525372 | 3.112905782 | -0.004344934 | 0.996533255 | NA | Gm29717 | predicted gene, 29717 [Source:MGI Symb | 0 | 0 | 2.0125 | 1.928 | 0 | 0 |
| 52511 | ENSMUSG00000114953 | 0 | NA | NA | NA | NA | NA | 4930519K11 | RIKEN cDNA 4930519K11 gene [Source:M | 0 | 0 | 0 | 0 | 0 | 0 |
| 52512 | ENSMUSG00000114954 | 0 | NA | NA | NA | NA | NA | Gm47746 | predicted gene, 47746 [Source:MGI Symb | 0 | 0 | 0 | 0 | 0 | 0 |
| 52513 | ENSMUSG00000114955 | 0 | NA | NA | NA | NA | NA | Gm48109 | predicted gene, 48109 [Source:MGI Symb | 0 | 0 | 0 | 0 | 0 | 0 |
| 52514 | ENSMUSG00000114956 | 0 | NA | NA | NA | NA | NA | Gm47893 | predicted gene, 47893 [Source:MGI Symb | 0 | 0 | 0 | 0 | 0 | 0 |
| 52515 | ENSMUSG00000114957 | 0 | NA | NA | NA | NA | NA | Gm47316 | predicted gene, 47316 [Source:MGI Symb | 0 | 0 | 0 | 0 | 0 | 0 |
| 52516 | ENSMUSG00000114958 | 0 | NA | NA | NA | NA | NA | Gm48612 | predicted gene, 48612 [Source:MGI Symb | 0 | 0 | 0 | 0 | 0 | 0 |
| 52517 | ENSMUSG00000114959 | 0 | NA | NA | NA | NA | NA | G630093K05 | RIKEN cDNA G630093K05 gene [Source:M | 0 | 0 | 0 | 0 | 0 | 0 |
| 52518 | ENSMUSG00000114960 | 0 | NA | NA | NA | NA | NA | Gm47649 | predicted gene, 47649 [Source:MGI Symb | 0 | 0 | 0 | 0 | 0 | 0 |
| 52519 | ENSMUSG00000114961 | 0.647861426 | 0.1905833 | 2.628676701 | 0.072501613 | 0.942202722 | NA | A930002C04 | RIKEN cDNA A930002C04 gene [Source:M | 0.8031 | 0 | 1.0062 | 0.964 | 0 | 1.1138 |
| 52520 | ENSMUSG00000114962 | 0 | NA | NA | NA | NA | NA | Gm6343 | predicted gene 6343 [Source:MGI Symbol | 0 | 0 | 0 | 0 | 0 | 0 |
| 52521 | ENSMUSG00000114963 | 0 | NA | NA | NA | NA | NA | Gm29010 | predicted gene 29010 [Source:MGI Symbo | 0 | 0 | 0 | 0 | 0 | 0 |
| 52522 | ENSMUSG00000114964 | 0 | NA | NA | NA | NA | NA | Gm18945 | predicted gene, 18945 [Source:MGI Symb | 0 | 0 | 0 | 0 | 0 | 0 |
| 52523 | ENSMUSG00000114965 | 0 | NA | NA | NA | NA | NA | Gm47906 | predicted gene, 47906 [Source:MGI Symb | 0 | 0 | 0 | 0 | 0 | 0 |
| 52524 | ENSMUSG00000114966 | 0 | NA | NA | NA | NA | NA | Gm18681 | predicted gene, 18681 [Source:MGI Symb | 0 | 0 | 0 | 0 | 0 | 0 |

| | | | | | | | | | | | | | | | |
|---|---|---|---|---|---|---|---|---|---|---|---|---|---|---|---|
| 52525 | ENSMUSG00000114967 | 0 | NA | NA | NA | NA | NA | Gm7473 | predicted gene 7473 [Source:MGI Symbol | 0 | 0 | 0 | 0 | 0 | 0 |
| 52526 | ENSMUSG00000114968 | 0 | NA | NA | NA | NA | NA | A630019I02I | RIKEN cDNA A630019I02 gene [Source:M( | 0 | 0 | 0 | 0 | 0 | 0 |
| 52527 | ENSMUSG00000114969 | 0 | NA | NA | NA | NA | NA | Hba-ps3 | hemoglobin alpha, pseudogene 3 [Source | 0 | 0 | 0 | 0 | 0 | 0 |
| 52528 | ENSMUSG00000114970 | 0 | NA | NA | NA | NA | NA | Gm49069 | predicted gene, 49069 [Source:MGI Symb | 0 | 0 | 0 | 0 | 0 | 0 |
| 52529 | ENSMUSG00000114971 | 0 | NA | NA | NA | NA | NA | Gm49162 | predicted gene, 49162 [Source:MGI Symb | 0 | 0 | 0 | 0 | 0 | 0 |
| 52530 | ENSMUSG00000114972 | 0 | NA | NA | NA | NA | NA | Gm49119 | predicted gene, 49119 [Source:MGI Symb | 0 | 0 | 0 | 0 | 0 | 0 |
| 52531 | ENSMUSG00000114973 | 0 | NA | NA | NA | NA | NA | Gm19156 | predicted gene, 19156 [Source:MGI Symb | 0 | 0 | 0 | 0 | 0 | 0 |
| 52532 | ENSMUSG00000114974 | 0 | NA | NA | NA | NA | NA | Gm21010 | predicted gene, 21010 [Source:MGI Symb | 0 | 0 | 0 | 0 | 0 | 0 |
| 52533 | ENSMUSG00000114975 | 0 | NA | NA | NA | NA | NA | Gm49090 | predicted gene, 49090 [Source:MGI Symb | 0 | 0 | 0 | 0 | 0 | 0 |
| 52534 | ENSMUSG00000114976 | 0 | NA | NA | NA | NA | NA | Gm2333 | predicted gene 2333 [Source:MGI Symbol | 0 | 0 | 0 | 0 | 0 | 0 |
| 52535 | ENSMUSG00000114977 | 22.24665689 | -0.244331612 | 0.46247041 | -0.528318369 | 0.597278385 | 0.914521894 | Gm9502 | predicted gene 9502 [Source:MGI Symbol | 26.502 | 23.642 | 22.137 | 18.316 | 26.175 | 16.708 |
| 52536 | ENSMUSG00000114978 | 0 | NA | NA | NA | NA | NA | Gm48925 | predicted gene, 48925 [Source:MGI Symb | 0 | 0 | 0 | 0 | 0 | 0 |
| 52537 | ENSMUSG00000114979 | 0 | NA | NA | NA | NA | NA | Gm31072 | predicted gene, 31072 [Source:MGI Symb | 0 | 0 | 0 | 0 | 0 | 0 |
| 52538 | ENSMUSG00000114980 | 0 | NA | NA | NA | NA | NA | 4933432I03F | RIKEN cDNA 4933432I03 gene [Source:M( | 0 | 0 | 0 | 0 | 0 | 0 |
| 52539 | ENSMUSG00000114981 | 1.066648353 | -0.993405617 | 2.381002554 | -0.417221567 | 0.676516352 | NA | Gm34093 | predicted gene, 34093 [Source:MGI Symb | 3.2124 | 0 | 1.0062 | 0 | 2.1812 | 0 |
| 52540 | ENSMUSG00000114982 | 0 | NA | NA | NA | NA | NA | Vmn1r14 | vomeronasal 1 receptor 14 [Source:MGI S | 0 | 0 | 0 | 0 | 0 | 0 |
| 52541 | ENSMUSG00000114983 | 0 | NA | NA | NA | NA | NA | Gm48997 | predicted gene, 48997 [Source:MGI Symb | 0 | 0 | 0 | 0 | 0 | 0 |
| 52542 | ENSMUSG00000114984 | 0 | NA | NA | NA | NA | NA | Gm48947 | predicted gene, 48947 [Source:MGI Symb | 0 | 0 | 0 | 0 | 0 | 0 |
| 52543 | ENSMUSG00000114985 | 0 | NA | NA | NA | NA | NA | 1810028F09 | RIKEN cDNA 1810028F09 gene [Source:M | 0 | 0 | 0 | 0 | 0 | 0 |
| 52544 | ENSMUSG00000114986 | 0 | NA | NA | NA | NA | NA | Gm21162 | predicted gene, 21162 [Source:MGI Symb | 0 | 0 | 0 | 0 | 0 | 0 |
| 52545 | ENSMUSG00000114987 | 0 | NA | NA | NA | NA | NA | Gm49022 | predicted gene, 49022 [Source:MGI Symb | 0 | 0 | 0 | 0 | 0 | 0 |
| 52546 | ENSMUSG00000114988 | 0 | NA | NA | NA | NA | NA | Gm49226 | predicted gene, 49226 [Source:MGI Symb | 0 | 0 | 0 | 0 | 0 | 0 |
| 52547 | ENSMUSG00000114989 | 0 | NA | NA | NA | NA | NA | Gm49303 | predicted gene, 49303 [Source:MGI Symb | 0 | 0 | 0 | 0 | 0 | 0 |
| 52548 | ENSMUSG00000114990 | 0 | NA | NA | NA | NA | NA | Gm49036 | predicted gene, 49036 [Source:MGI Symb | 0 | 0 | 0 | 0 | 0 | 0 |
| 52549 | ENSMUSG00000114991 | 0 | NA | NA | NA | NA | NA | 1700128I11F | RIKEN cDNA 1700128I11 gene [Source:M( | 0 | 0 | 0 | 0 | 0 | 0 |
| 52550 | ENSMUSG00000114992 | 0 | NA | NA | NA | NA | NA | Gm2150 | predicted gene 2150 [Source:MGI Symbol | 0 | 0 | 0 | 0 | 0 | 0 |
| 52551 | ENSMUSG00000114993 | 0.710308366 | 0.13927508 | 2.769785472 | 0.050283707 | 0.959896308 | NA | Gm6363 | predicted gene 6363 [Source:MGI Symbol | 0 | 1.0279 | 1.0062 | 0 | 0 | 2.2277 |
| 52552 | ENSMUSG00000114994 | 0 | NA | NA | NA | NA | NA | Gm49097 | predicted gene, 49097 [Source:MGI Symb | 0 | 0 | 0 | 0 | 0 | 0 |
| 52553 | ENSMUSG00000114995 | 73.07572286 | 0.030519087 | 0.260252038 | 0.117267429 | 0.906648135 | 0.984294725 | Gm49284 | predicted gene, 49284 [Source:MGI Symb | 66.657 | 76.066 | 74.462 | 79.046 | 68.709 | 73.514 |
| 52554 | ENSMUSG00000114996 | 0.167707431 | -0.796629176 | 4.080472857 | -0.195229623 | 0.84521319 | NA | Gm48958 | predicted gene, 48958 [Source:MGI Symb | 0 | 0 | 1.0062 | 0 | 0 | 0 |
| 52555 | ENSMUSG00000114997 | 0 | NA | NA | NA | NA | NA | Gm49038 | predicted gene, 49038 [Source:MGI Symb | 0 | 0 | 0 | 0 | 0 | 0 |
| 52556 | ENSMUSG00000114998 | 0 | NA | NA | NA | NA | NA | Gm2157 | predicted gene 2157 [Source:MGI Symbol | 0 | 0 | 0 | 0 | 0 | 0 |
| 52557 | ENSMUSG00000114999 | 1.032791295 | -0.925393667 | 2.656262528 | -0.348381855 | 0.727553428 | NA | Gm7962 | predicted gene 7962 [Source:MGI Symbol | 4.0155 | 0 | 0 | 0 | 2.1812 | 0 |
| 52558 | ENSMUSG00000115000 | 0 | NA | NA | NA | NA | NA | Gm49192 | predicted gene, 49192 [Source:MGI Symb | 0 | 0 | 0 | 0 | 0 | 0 |
| 52559 | ENSMUSG00000115001 | 0 | NA | NA | NA | NA | NA | 4930474H20 | RIKEN cDNA 4930474H20 gene [Source:M | 0 | 0 | 0 | 0 | 0 | 0 |
| 52560 | ENSMUSG00000115002 | 0.181769715 | 1.126924175 | 4.080472857 | 0.276174898 | 0.782413736 | NA | Gm10847 | predicted gene 10847 [Source:MGI Symbo | 0 | 0 | 0 | 0 | 1.0906 | 0 |
| 52561 | ENSMUSG00000115003 | 14.06957085 | 0.652176327 | 0.625371814 | 1.042861724 | 0.297012339 | 0.747778325 | Gm48933 | predicted gene, 48933 [Source:MGI Symb | 14.456 | 9.2513 | 9.0562 | 23.135 | 20.722 | 7.7969 |
| 52562 | ENSMUSG00000115004 | 0 | NA | NA | NA | NA | NA | Gm48920 | predicted gene, 48920 [Source:MGI Symb | 0 | 0 | 0 | 0 | 0 | 0 |
| 52563 | ENSMUSG00000115005 | 1.652451045 | -1.818234607 | 1.870471245 | -0.972073007 | 0.33101423 | NA | Gm49195 | predicted gene, 49195 [Source:MGI Symb | 1.6062 | 2.0558 | 4.025 | 0 | 0 | 2.2277 |
| 52564 | ENSMUSG00000115006 | 0 | NA | NA | NA | NA | NA | Gm49054 | predicted gene, 49054 [Source:MGI Symb | 0 | 0 | 0 | 0 | 0 | 0 |
| 52565 | ENSMUSG00000115007 | 46.11668766 | -0.364106459 | 0.383765672 | -0.948772875 | 0.342736141 | 0.784392005 | 5830448L01I | RIKEN cDNA 5830448L01 gene [Source:M( | 42.564 | 44.2 | 69.431 | 58.803 | 31.628 | 30.074 |
| 52566 | ENSMUSG00000115008 | 0 | NA | NA | NA | NA | NA | 4933429O19 | RIKEN cDNA 4933429O19 gene [Source:M | 0 | 0 | 0 | 0 | 0 | 0 |
| 52567 | ENSMUSG00000115009 | 0 | NA | NA | NA | NA | NA | G930009F23 | RIKEN cDNA G930009F23 gene [Source:M | 0 | 0 | 0 | 0 | 0 | 0 |
| 52568 | ENSMUSG00000115010 | 0 | NA | NA | NA | NA | NA | Gm20150 | predicted gene, 20150 [Source:MGI Symb | 0 | 0 | 0 | 0 | 0 | 0 |
| 52569 | ENSMUSG00000115011 | 0 | NA | NA | NA | NA | NA | Gm49013 | predicted gene, 49013 [Source:MGI Symb | 0 | 0 | 0 | 0 | 0 | 0 |
| 52570 | ENSMUSG00000115012 | 0 | NA | NA | NA | NA | NA | 9630009A06 | RIKEN cDNA 9630009A06 gene [Source:M | 0 | 0 | 0 | 0 | 0 | 0 |
| 52571 | ENSMUSG00000115013 | 0 | NA | NA | NA | NA | NA | Gm49231 | predicted gene, 49231 [Source:MGI Symb | 0 | 0 | 0 | 0 | 0 | 0 |
| 52572 | ENSMUSG00000115014 | 0 | NA | NA | NA | NA | NA | Gm49061 | predicted gene, 49061 [Source:MGI Symb | 0 | 0 | 0 | 0 | 0 | 0 |
| 52573 | ENSMUSG00000115015 | 0 | NA | NA | NA | NA | NA | D330046F09 | RIKEN cDNA D330046F09 gene [Source:M | 0 | 0 | 0 | 0 | 0 | 0 |
| 52574 | ENSMUSG00000115016 | 0 | NA | NA | NA | NA | NA | Gm33906 | predicted gene, 33906 [Source:MGI Symb | 0 | 0 | 0 | 0 | 0 | 0 |
| 52575 | ENSMUSG00000115016 | 0 | NA | NA | NA | NA | NA | AA536875 | predicted gene, 33906 [Source:MGI Symb | 0 | 0 | 0 | 0 | 0 | 0 |
| 52576 | ENSMUSG00000115017 | 1.122573798 | -0.362418364 | 2.131244112 | -0.170050142 | 0.864970703 | NA | Gm2573 | predicted gene 2573 [Source:MGI Symbol | 0.8031 | 1.0279 | 2.0125 | 2.8919 | 0 | 0 |
| 52577 | ENSMUSG00000115018 | 0 | NA | NA | NA | NA | NA | Ndor1 | NADPH dependent diflavin oxidoreductas | 0 | 0 | 0 | 0 | 0 | 0 |
| 52578 | ENSMUSG00000115019 | 0 | NA | NA | NA | NA | NA | Gm4278 | predicted gene 4278 [Source:MGI Symbol | 0 | 0 | 0 | 0 | 0 | 0 |
| 52579 | ENSMUSG00000115020 | 0 | NA | NA | NA | NA | NA | Vmn1r218 | vomeronasal 1 receptor 218 [Source:MGI | 0 | 0 | 0 | 0 | 0 | 0 |
| 52580 | ENSMUSG00000115021 | 0 | NA | NA | NA | NA | NA | Vmn1r173 | vomeronasal 1 receptor 173 [Source:MGI | 0 | 0 | 0 | 0 | 0 | 0 |

| | | | | | | | | | | | | | | | |
|---|---|---|---|---|---|---|---|---|---|---|---|---|---|---|---|
| 52581 | ENSMUSG00000115022 | 0 | NA | NA | NA | NA | NA | Gm49387 | predicted gene, 49387 [Source:MGI Symb | 0 | 0 | 0 | 0 | 0 | 0 |
| 52582 | ENSMUSG00000115023 | 0 | NA | NA | NA | NA | NA | Gm48967 | predicted gene, 48967 [Source:MGI Symb | 0 | 0 | 0 | 0 | 0 | 0 |
| 52583 | ENSMUSG00000115024 | 0 | NA | NA | NA | NA | NA | Gm49277 | predicted gene, 49277 [Source:MGI Symb | 0 | 0 | 0 | 0 | 0 | 0 |
| 52584 | ENSMUSG00000115025 | 23.03526443 | -0.072894961 | 0.445302157 | -0.16369775 | 0.869969089 | 0.978765244 | Gm4240 | predicted gene 4240 [Source:MGI Symbol | 24.093 | 20.558 | 26.162 | 22.171 | 20.722 | 24.505 |
| 52585 | ENSMUSG00000115026 | 4.364479356 | -0.564460563 | 1.002521571 | -0.563040816 | 0.573407091 | NA | Gm49041 | predicted gene, 49041 [Source:MGI Symb | 6.4248 | 4.1117 | 5.0312 | 2.8919 | 3.2719 | 4.4554 |
| 52586 | ENSMUSG00000115027 | 0 | NA | NA | NA | NA | NA | Vmn1r81 | vomeronasal 1 receptor 81 [Source:MGI S | 0 | 0 | 0 | 0 | 0 | 0 |
| 52587 | ENSMUSG00000115028 | 0 | NA | NA | NA | NA | NA | Gm9391 | predicted gene 9391 [Source:MGI Symbol | 0 | 0 | 0 | 0 | 0 | 0 |
| 52588 | ENSMUSG00000115029 | 0 | NA | NA | NA | NA | NA | Gm18885 | predicted gene, 18885 [Source:MGI Symb | 0 | 0 | 0 | 0 | 0 | 0 |
| 52589 | ENSMUSG00000115030 | 0 | NA | NA | NA | NA | NA | Gm49274 | predicted gene, 49274 [Source:MGI Symb | 0 | 0 | 0 | 0 | 0 | 0 |
| 52590 | ENSMUSG00000115031 | 0 | NA | NA | NA | NA | NA | Gm49051 | predicted gene, 49051 [Source:MGI Symb | 0 | 0 | 0 | 0 | 0 | 0 |
| 52591 | ENSMUSG00000115032 | 0 | NA | NA | NA | NA | NA | Gm49301 | predicted gene, 49301 [Source:MGI Symb | 0 | 0 | 0 | 0 | 0 | 0 |
| 52592 | ENSMUSG00000115033 | 0 | NA | NA | NA | NA | NA | Gm49206 | predicted gene, 49206 [Source:MGI Symb | 0 | 0 | 0 | 0 | 0 | 0 |
| 52593 | ENSMUSG00000115034 | 0 | NA | NA | NA | NA | NA | Gm19109 | predicted gene, 19109 [Source:MGI Symb | 0 | 0 | 0 | 0 | 0 | 0 |
| 52594 | ENSMUSG00000115035 | 0 | NA | NA | NA | NA | NA | Gm9614 | predicted gene 9614 [Source:MGI Symbol | 0 | 0 | 0 | 0 | 0 | 0 |
| 52595 | ENSMUSG00000115036 | 0 | NA | NA | NA | NA | NA | Gm49043 | predicted gene, 49043 [Source:MGI Symb | 0 | 0 | 0 | 0 | 0 | 0 |
| 52596 | ENSMUSG00000115037 | 0 | NA | NA | NA | NA | NA | Gm49262 | predicted gene, 49262 [Source:MGI Symb | 0 | 0 | 0 | 0 | 0 | 0 |
| 52597 | ENSMUSG00000115038 | 0 | NA | NA | NA | NA | NA | Gm48989 | predicted gene, 48989 [Source:MGI Symb | 0 | 0 | 0 | 0 | 0 | 0 |
| 52598 | ENSMUSG00000115039 | 0 | NA | NA | NA | NA | NA | Gm7043 | predicted gene 7043 [Source:MGI Symbol | 0 | 0 | 0 | 0 | 0 | 0 |
| 52599 | ENSMUSG00000115040 | 0.401551119 | -2.072982726 | 4.008251979 | -0.517178744 | 0.605031388 | NA | Gm18994 | predicted gene, 18994 [Source:MGI Symb | 2.4093 | 0 | 0 | 0 | 0 | 0 |
| 52600 | ENSMUSG00000115041 | 0 | NA | NA | NA | NA | NA | Gm48939 | predicted gene, 48939 [Source:MGI Symb | 0 | 0 | 0 | 0 | 0 | 0 |
| 52601 | ENSMUSG00000115042 | 0 | NA | NA | NA | NA | NA | Gm29742 | predicted gene, 29742 [Source:MGI Symb | 0 | 0 | 0 | 0 | 0 | 0 |
| 52602 | ENSMUSG00000115043 | 0 | NA | NA | NA | NA | NA | Gm32455 | predicted gene, 32455 [Source:MGI Symb | 0 | 0 | 0 | 0 | 0 | 0 |
| 52603 | ENSMUSG00000115044 | 1.200498726 | -1.245585098 | 2.300526566 | -0.541434781 | 0.588207939 | NA | Gm48978 | predicted gene, 48978 [Source:MGI Symb | 4.0155 | 0 | 1.0062 | 0 | 2.1812 | 0 |
| 52604 | ENSMUSG00000115045 | 0 | NA | NA | NA | NA | NA | Gm49028 | predicted gene, 49028 [Source:MGI Symb | 0 | 0 | 0 | 0 | 0 | 0 |
| 52605 | ENSMUSG00000115046 | 0 | NA | NA | NA | NA | NA | Gm49302 | predicted gene, 49302 [Source:MGI Symb | 0 | 0 | 0 | 0 | 0 | 0 |
| 52606 | ENSMUSG00000115047 | 0 | NA | NA | NA | NA | NA | 4930540I17F | RIKEN cDNA 4930540I17 gene [Source:MG | 0 | 0 | 0 | 0 | 0 | 0 |
| 52607 | ENSMUSG00000115048 | 0 | NA | NA | NA | NA | NA | Gm48921 | predicted gene, 48921 [Source:MGI Symb | 0 | 0 | 0 | 0 | 0 | 0 |
| 52608 | ENSMUSG00000115049 | 23.97105349 | -0.692317279 | 0.45141214 | -1.53367005 | 0.125110845 | 0.518510965 | Gm49172 | predicted gene, 49172 [Source:MGI Symb | 33.73 | 27.754 | 27.169 | 15.424 | 16.359 | 23.391 |
| 52609 | ENSMUSG00000115050 | 0 | NA | NA | NA | NA | NA | Gm21154 | predicted gene, 21154 [Source:MGI Symb | 0 | 0 | 0 | 0 | 0 | 0 |
| 52610 | ENSMUSG00000115051 | 0 | NA | NA | NA | NA | NA | Gm49049 | predicted gene, 49049 [Source:MGI Symb | 0 | 0 | 0 | 0 | 0 | 0 |
| 52611 | ENSMUSG00000115052 | 0 | NA | NA | NA | NA | NA | Gm9287 | predicted gene 9287 [Source:MGI Symbol | 0 | 0 | 0 | 0 | 0 | 0 |
| 52612 | ENSMUSG00000115053 | 0 | NA | NA | NA | NA | NA | Gm49275 | predicted gene, 49275 [Source:MGI Symb | 0 | 0 | 0 | 0 | 0 | 0 |
| 52613 | ENSMUSG00000115054 | 0 | NA | NA | NA | NA | NA | Gm49184 | predicted gene, 49184 [Source:MGI Symb | 0 | 0 | 0 | 0 | 0 | 0 |
| 52614 | ENSMUSG00000115056 | 0 | NA | NA | NA | NA | NA | Gm49103 | predicted gene, 49103 [Source:MGI Symb | 0 | 0 | 0 | 0 | 0 | 0 |
| 52615 | ENSMUSG00000115057 | 0.181769715 | 1.126924175 | 4.080472857 | 0.276174898 | 0.782413736 | NA | Gm8705 | predicted gene 8705 [Source:MGI Symbol | 0 | 0 | 0 | 0 | 1.0906 | 0 |
| 52616 | ENSMUSG00000115058 | 0 | NA | NA | NA | NA | NA | Gm19883 | predicted gene, 19883 [Source:MGI Symb | 0 | 0 | 0 | 0 | 0 | 0 |
| 52617 | ENSMUSG00000115059 | 0 | NA | NA | NA | NA | NA | Gm48983 | predicted gene, 48983 [Source:MGI Symb | 0 | 0 | 0 | 0 | 0 | 0 |
| 52618 | ENSMUSG00000115060 | 0 | NA | NA | NA | NA | NA | Gm49306 | predicted gene, 49306 [Source:MGI Symb | 0 | 0 | 0 | 0 | 0 | 0 |
| 52619 | ENSMUSG00000115061 | 0 | NA | NA | NA | NA | NA | Gm49290 | predicted gene, 49290 [Source:MGI Symb | 0 | 0 | 0 | 0 | 0 | 0 |
| 52620 | ENSMUSG00000115062 | 1.734639499 | 1.29570771 | 1.822379454 | 0.71099776 | 0.477085624 | NA | 4930544F09 | RIKEN cDNA 4930544F09 gene [Source:M | 0 | 3.0838 | 0 | 2.8919 | 1.0906 | 3.3415 |
| 52621 | ENSMUSG00000115063 | 0 | NA | NA | NA | NA | NA | 4930465K09 | RIKEN cDNA 4930465K09 gene [Source:M | 0 | 0 | 0 | 0 | 0 | 0 |
| 52622 | ENSMUSG00000115064 | 0 | NA | NA | NA | NA | NA | Gm49213 | predicted gene, 49213 [Source:MGI Symb | 0 | 0 | 0 | 0 | 0 | 0 |
| 52623 | ENSMUSG00000115065 | 0 | NA | NA | NA | NA | NA | Gm49109 | predicted gene, 49109 [Source:MGI Symb | 0 | 0 | 0 | 0 | 0 | 0 |
| 52624 | ENSMUSG00000115066 | 0 | NA | NA | NA | NA | NA | Gm49242 | predicted gene, 49242 [Source:MGI Symb | 0 | 0 | 0 | 0 | 0 | 0 |
| 52625 | ENSMUSG00000115067 | 0 | NA | NA | NA | NA | NA | Dpep2 | dipeptidase 2 [Source:MGI Symbol;Acc:M | 0 | 0 | 0 | 0 | 0 | 0 |
| 52626 | ENSMUSG00000115068 | 0 | NA | NA | NA | NA | NA | Gm48926 | predicted gene, 48926 [Source:MGI Symb | 0 | 0 | 0 | 0 | 0 | 0 |
| 52627 | ENSMUSG00000115069 | 0 | NA | NA | NA | NA | NA | Gm7382 | predicted gene 7382 [Source:MGI Symbol | 0 | 0 | 0 | 0 | 0 | 0 |
| 52628 | ENSMUSG00000115070 | 0 | NA | NA | NA | NA | NA | Gm4740 | predicted gene 4740 [Source:MGI Symbol | 0 | 0 | 0 | 0 | 0 | 0 |
| 52629 | ENSMUSG00000115071 | 0 | NA | NA | NA | NA | NA | Gm49026 | predicted gene, 49026 [Source:MGI Symb | 0 | 0 | 0 | 0 | 0 | 0 |
| 52630 | ENSMUSG00000115072 | 0 | NA | NA | NA | NA | NA | Vmn1r2 | vomeronasal 1 receptor 2 [Source:MGI Sy | 0 | 0 | 0 | 0 | 0 | 0 |
| 52631 | ENSMUSG00000115073 | 0 | NA | NA | NA | NA | NA | Gm49283 | predicted gene, 49283 [Source:MGI Symb | 0 | 0 | 0 | 0 | 0 | 0 |
| 52632 | ENSMUSG00000115074 | 0 | NA | NA | NA | NA | NA | Ndor1 | NADPH dependent diflavin oxidoreductas | 0 | 0 | 0 | 0 | 0 | 0 |
| 52633 | ENSMUSG00000115075 | 0 | NA | NA | NA | NA | NA | Gm49033 | predicted gene, 49033 [Source:MGI Symb | 0 | 0 | 0 | 0 | 0 | 0 |
| 52634 | ENSMUSG00000115076 | 0 | NA | NA | NA | NA | NA | Gm49106 | predicted gene, 49106 [Source:MGI Symb | 0 | 0 | 0 | 0 | 0 | 0 |
| 52635 | ENSMUSG00000115077 | 0.160662974 | 1.126924175 | 4.080472857 | 0.276174898 | 0.782413736 | NA | Gm46455 | predicted gene, 46455 [Source:MGI Symb | 0 | 0 | 0 | 0.964 | 0 | 0 |
| 52636 | ENSMUSG00000115079 | 0 | NA | NA | NA | NA | NA | Gm36147 | predicted gene, 36147 [Source:MGI Symb | 0 | 0 | 0 | 0 | 0 | 0 |

| | | | | | | | | | | | | | | | |
|---|---|---|---|---|---|---|---|---|---|---|---|---|---|---|---|
| 52637 | ENSMUSG00000115080 | 0 | NA | NA | NA | NA | NA | Gm5212 | predicted gene 5212 [Source:MGI Symbol | 0 | 0 | 0 | 0 | 0 | 0 |
| 52638 | ENSMUSG00000115081 | 0 | NA | NA | NA | NA | NA | Gm18714 | predicted gene, 18714 [Source:MGI Symb | 0 | 0 | 0 | 0 | 0 | 0 |
| 52639 | ENSMUSG00000115082 | 0.881705114 | -0.66605326 | 2.559736336 | -0.260203854 | 0.794706531 | NA | Gm49189 | predicted gene, 49189 [Source:MGI Symb | 3.2124 | 0 | 0 | 0.964 | 0 | 1.1138 |
| 52640 | ENSMUSG00000115083 | 0 | NA | NA | NA | NA | NA | Gm34756 | predicted gene, 34756 [Source:MGI Symb | 0 | 0 | 0 | 0 | 0 | 0 |
| 52641 | ENSMUSG00000115084 | 0.670829724 | -2.782124174 | 3.401449285 | -0.817923168 | 0.41340106 | NA | Gm49246 | predicted gene, 49246 [Source:MGI Symb | 0 | 0 | 4.025 | 0 | 0 | 0 |
| 52642 | ENSMUSG00000115085 | 0 | NA | NA | NA | NA | NA | Gm9198 | predicted gene 9198 [Source:MGI Symbol | 0 | 0 | 0 | 0 | 0 | 0 |
| 52643 | ENSMUSG00000115086 | 0.185640647 | 1.126924175 | 4.080472857 | 0.276174898 | 0.782413736 | NA | Gm49177 | predicted gene, 49177 [Source:MGI Symb | 0 | 0 | 0 | 0 | 0 | 1.1138 |
| 52644 | ENSMUSG00000115087 | 0 | NA | NA | NA | NA | NA | Gm48998 | predicted gene, 48998 [Source:MGI Symb | 0 | 0 | 0 | 0 | 0 | 0 |
| 52645 | ENSMUSG00000115088 | 0 | NA | NA | NA | NA | NA | Gm49165 | predicted gene, 49165 [Source:MGI Symb | 0 | 0 | 0 | 0 | 0 | 0 |
| 52646 | ENSMUSG00000115089 | 0.36353943 | 2.065989254 | 4.034137301 | 0.512126658 | 0.608562368 | NA | Gm4350 | predicted gene 4350 [Source:MGI Symbol | 0 | 0 | 0 | 0 | 2.1812 | 0 |
| 52647 | ENSMUSG00000115090 | 1.639785005 | -2.959199773 | 2.029700681 | -1.457948849 | 0.144854642 | NA | Gm49016 | predicted gene, 49016 [Source:MGI Symb | 0.8031 | 1.0279 | 7.0437 | 0.964 | 0 | 0 |
| 52648 | ENSMUSG00000115091 | 0 | NA | NA | NA | NA | NA | Vmn1r22 | vomeronasal 1 receptor 22 [Source:MGI S | 0 | 0 | 0 | 0 | 0 | 0 |
| 52649 | ENSMUSG00000115092 | 0 | NA | NA | NA | NA | NA | E130202H07 | RIKEN cDNA E130202H07 gene [Source:M | 0 | 0 | 0 | 0 | 0 | 0 |
| 52650 | ENSMUSG00000115093 | 0.556921942 | 2.679043159 | 3.638243063 | 0.736356289 | 0.461513899 | NA | Gm8047 | predicted gene 8047 [Source:MGI Symbol | 0 | 0 | 0 | 0 | 0 | 3.3415 |
| 52651 | ENSMUSG00000115094 | 0 | NA | NA | NA | NA | NA | Gm7106 | predicted gene 7106 [Source:MGI Symbol | 0 | 0 | 0 | 0 | 0 | 0 |
| 52652 | ENSMUSG00000115095 | 0 | NA | NA | NA | NA | NA | Gm49311 | predicted gene, 49311 [Source:MGI Symb | 0 | 0 | 0 | 0 | 0 | 0 |
| 52653 | ENSMUSG00000115096 | 0.133850373 | -0.796629176 | 4.080472857 | -0.195229623 | 0.84521319 | NA | Gm48940 | predicted gene, 48940 [Source:MGI Symb | 0.8031 | 0 | 0 | 0 | 0 | 0 |
| 52654 | ENSMUSG00000115097 | 0 | NA | NA | NA | NA | NA | Gm49150 | predicted gene, 49150 [Source:MGI Symb | 0 | 0 | 0 | 0 | 0 | 0 |
| 52655 | ENSMUSG00000115098 | 5.496416956 | 0.861578393 | 0.984719364 | 0.874948158 | 0.381602114 | 0.8130053 | Gm49134 | predicted gene, 49134 [Source:MGI Symb | 1.6062 | 7.1954 | 3.0187 | 6.7478 | 3.2719 | 11.138 |
| 52656 | ENSMUSG00000115099 | 29.51162674 | -0.099833468 | 0.427599393 | -0.233474297 | 0.815393126 | 0.967767553 | 1700087I21F | RIKEN cDNA 1700087I21 gene [Source:MC | 42.564 | 22.614 | 26.162 | 32.775 | 23.994 | 28.96 |
| 52657 | ENSMUSG00000115100 | 132.7731215 | 0.225350801 | 0.254749852 | 0.884596393 | 0.376374353 | 0.809739497 | Gm34934 | predicted gene, 34934 [Source:MGI Symb | 170.26 | 89.429 | 106.66 | 151.34 | 134.15 | 144.8 |
| 52658 | ENSMUSG00000115101 | 0 | NA | NA | NA | NA | NA | Gm7778 | predicted gene 7778 [Source:MGI Symbol | 0 | 0 | 0 | 0 | 0 | 0 |
| 52659 | ENSMUSG00000115102 | 0 | NA | NA | NA | NA | NA | Gm49270 | predicted gene, 49270 [Source:MGI Symb | 0 | 0 | 0 | 0 | 0 | 0 |
| 52660 | ENSMUSG00000115103 | 0 | NA | NA | NA | NA | NA | Gm49263 | predicted gene, 49263 [Source:MGI Symb | 0 | 0 | 0 | 0 | 0 | 0 |
| 52661 | ENSMUSG00000115104 | 0 | NA | NA | NA | NA | NA | Gm49029 | predicted gene, 49029 [Source:MGI Symb | 0 | 0 | 0 | 0 | 0 | 0 |
| 52662 | ENSMUSG00000115105 | 0 | NA | NA | NA | NA | NA | Gm49238 | predicted gene, 49238 [Source:MGI Symb | 0 | 0 | 0 | 0 | 0 | 0 |
| 52663 | ENSMUSG00000115106 | 0 | NA | NA | NA | NA | NA | Gm8043 | predicted gene 8043 [Source:MGI Symbol | 0 | 0 | 0 | 0 | 0 | 0 |
| 52664 | ENSMUSG00000115107 | 0 | NA | NA | NA | NA | NA | Gm18368 | predicted gene, 18368 [Source:MGI Symb | 0 | 0 | 0 | 0 | 0 | 0 |
| 52665 | ENSMUSG00000115108 | 0 | NA | NA | NA | NA | NA | Gm49228 | predicted gene, 49228 [Source:MGI Symb | 0 | 0 | 0 | 0 | 0 | 0 |
| 52666 | ENSMUSG00000115109 | 0 | NA | NA | NA | NA | NA | Gm49032 | predicted gene, 49032 [Source:MGI Symb | 0 | 0 | 0 | 0 | 0 | 0 |
| 52667 | ENSMUSG00000115110 | 0 | NA | NA | NA | NA | NA | Gm49269 | predicted gene, 49269 [Source:MGI Symb | 0 | 0 | 0 | 0 | 0 | 0 |
| 52668 | ENSMUSG00000115111 | 1.581182368 | -0.513551535 | 2.042803368 | -0.251395481 | 0.801508361 | NA | Gm4632 | predicted gene 4632 [Source:MGI Symbol | 1.6062 | 0 | 4.025 | 3.8559 | 0 | 0 |
| 52669 | ENSMUSG00000115112 | 0 | NA | NA | NA | NA | NA | Gm19207 | predicted gene, 19207 [Source:MGI Symb | 0 | 0 | 0 | 0 | 0 | 0 |
| 52670 | ENSMUSG00000115113 | 0 | NA | NA | NA | NA | NA | Smim40-ps | small integral membrane protein 40, pseu | 0 | 0 | 0 | 0 | 0 | 0 |
| 52671 | ENSMUSG00000115114 | 0 | NA | NA | NA | NA | NA | Gm48930 | predicted gene, 48930 [Source:MGI Symb | 0 | 0 | 0 | 0 | 0 | 0 |
| 52672 | ENSMUSG00000115115 | 0 | NA | NA | NA | NA | NA | | novel protein | 0 | 0 | 0 | 0 | 0 | 0 |
| 52673 | ENSMUSG00000115116 | 0 | NA | NA | NA | NA | NA | Gm46496 | predicted gene, 46496 [Source:MGI Symb | 0 | 0 | 0 | 0 | 0 | 0 |
| 52674 | ENSMUSG00000115117 | 0 | NA | NA | NA | NA | NA | Gm18094 | predicted gene, 18094 [Source:MGI Symb | 0 | 0 | 0 | 0 | 0 | 0 |
| 52675 | ENSMUSG00000115118 | 0 | NA | NA | NA | NA | NA | Gm36617 | predicted gene, 36617 [Source:MGI Symb | 0 | 0 | 0 | 0 | 0 | 0 |
| 52676 | ENSMUSG00000115119 | 0 | NA | NA | NA | NA | NA | Gm17756 | predicted gene, 17756 [Source:MGI Symb | 0 | 0 | 0 | 0 | 0 | 0 |
| 52677 | ENSMUSG00000115120 | 0 | NA | NA | NA | NA | NA | 4930425P05 | RIKEN cDNA 4930425P05 gene [Source:M | 0 | 0 | 0 | 0 | 0 | 0 |
| 52678 | ENSMUSG00000115121 | 0 | NA | NA | NA | NA | NA | Gm49236 | predicted gene, 49236 [Source:MGI Symb | 0 | 0 | 0 | 0 | 0 | 0 |
| 52679 | ENSMUSG00000115122 | 0 | NA | NA | NA | NA | NA | Gm49685 | predicted gene, 49685 [Source:MGI Symb | 0 | 0 | 0 | 0 | 0 | 0 |
| 52680 | ENSMUSG00000115123 | 0 | NA | NA | NA | NA | NA | Gm33299 | predicted gene, 33299 [Source:MGI Symb | 0 | 0 | 0 | 0 | 0 | 0 |
| 52681 | ENSMUSG00000115124 | 107.1526686 | -0.33895926 | 0.223489327 | -1.516668668 | 0.129350359 | 0.52627949 | Gm49201 | predicted gene, 49201 [Source:MGI Symb | 110.03 | 122.32 | 126.79 | 89.65 | 89.431 | 104.7 |
| 52682 | ENSMUSG00000115125 | 0 | NA | NA | NA | NA | NA | Gm6852 | predicted gene 6852 [Source:MGI Symbol | 0 | 0 | 0 | 0 | 0 | 0 |
| 52683 | ENSMUSG00000115126 | 0 | NA | NA | NA | NA | NA | Gm19223 | predicted gene, 19223 [Source:MGI Symb | 0 | 0 | 0 | 0 | 0 | 0 |
| 52684 | ENSMUSG00000115127 | 3.574244534 | 1.580260649 | 1.233416115 | 1.281206423 | 0.200121171 | NA | Gm46515 | predicted gene, 46515 [Source:MGI Symb | 3.2124 | 1.0279 | 1.0062 | 1.928 | 9.8156 | 4.4554 |
| 52685 | ENSMUSG00000115128 | 0 | NA | NA | NA | NA | NA | Gm49062 | predicted gene, 49062 [Source:MGI Symb | 0 | 0 | 0 | 0 | 0 | 0 |
| 52686 | ENSMUSG00000115129 | 0 | NA | NA | NA | NA | NA | Gm48916 | predicted gene, 48916 [Source:MGI Symb | 0 | 0 | 0 | 0 | 0 | 0 |
| 52687 | ENSMUSG00000115130 | 0 | NA | NA | NA | NA | NA | Gm49152 | predicted gene, 49152 [Source:MGI Symb | 0 | 0 | 0 | 0 | 0 | 0 |
| 52688 | ENSMUSG00000115131 | 0 | NA | NA | NA | NA | NA | Gm49268 | predicted gene, 49268 [Source:MGI Symb | 0 | 0 | 0 | 0 | 0 | 0 |
| 52689 | ENSMUSG00000115132 | 0 | NA | NA | NA | NA | NA | Gm18888 | predicted gene, 18888 [Source:MGI Symb | 0 | 0 | 0 | 0 | 0 | 0 |
| 52690 | ENSMUSG00000115133 | 0 | NA | NA | NA | NA | NA | Gm20167 | predicted gene, 20167 [Source:MGI Symb | 0 | 0 | 0 | 0 | 0 | 0 |
| 52691 | ENSMUSG00000115134 | 0 | NA | NA | NA | NA | NA | Gm49108 | predicted gene, 49108 [Source:MGI Symb | 0 | 0 | 0 | 0 | 0 | 0 |
| 52692 | ENSMUSG00000115135 | 0 | NA | NA | NA | NA | NA | Gm19157 | predicted gene, 19157 [Source:MGI Symb | 0 | 0 | 0 | 0 | 0 | 0 |

| | | | | | | | | | | | | | | | |
|---|---|---|---|---|---|---|---|---|---|---|---|---|---|---|---|
| 52693 | ENSMUSG00000115136 | 0 | NA | NA | NA | NA | NA | Ear-ps1 | eosinophil-associated, ribonuclease A fam | 0 | 0 | 0 | 0 | 0 | 0 |
| 52694 | ENSMUSG00000115137 | 0 | NA | NA | NA | NA | NA | Gm34794 | predicted gene, 34794 [Source:MGI Symb | 0 | 0 | 0 | 0 | 0 | 0 |
| 52695 | ENSMUSG00000115138 | 5.363886604 | 0.144797505 | 0.981997216 | 0.147452052 | 0.882775223 | 0.981769403 | Gm36899 | predicted gene, 36899 [Source:MGI Symb | 3.2124 | 2.0558 | 10.062 | 5.7839 | 3.2719 | 7.7969 |
| 52696 | ENSMUSG00000115139 | 0 | NA | NA | NA | NA | NA | Gm4251 | predicted gene 4251 [Source:MGI Symbol | 0 | 0 | 0 | 0 | 0 | 0 |
| 52697 | ENSMUSG00000115140 | 0 | NA | NA | NA | NA | NA | Gm32764 | predicted gene, 32764 [Source:MGI Symb | 0 | 0 | 0 | 0 | 0 | 0 |
| 52698 | ENSMUSG00000115141 | 0.77650227 | 1.950354251 | 2.938137078 | 0.663806418 | 0.506814217 | NA | Gm49256 | predicted gene, 49256 [Source:MGI Symb | 0.8031 | 0 | 0 | 3.8559 | 0 | 0 |
| 52699 | ENSMUSG00000115142 | 0 | NA | NA | NA | NA | NA | Gm33251 | predicted gene, 33251 [Source:MGI Symb | 0 | 0 | 0 | 0 | 0 | 0 |
| 52700 | ENSMUSG00000115143 | 0.133850373 | -0.796629176 | 4.080472857 | -0.195229623 | 0.84521319 | NA | Fkbp1a-ps1 | FK506 binding protein 1a, pseudogene 1 [ | 0.8031 | 0 | 0 | 0 | 0 | 0 |
| 52701 | ENSMUSG00000115144 | 0 | NA | NA | NA | NA | NA | Gm7968 | predicted gene 7968 [Source:MGI Symbol | 0 | 0 | 0 | 0 | 0 | 0 |
| 52702 | ENSMUSG00000115145 | 0 | NA | NA | NA | NA | NA | Gm35166 | predicted gene, 35166 [Source:MGI Symb | 0 | 0 | 0 | 0 | 0 | 0 |
| 52703 | ENSMUSG00000115146 | 0 | NA | NA | NA | NA | NA | Gm4681 | predicted gene 4681 [Source:MGI Symbol | 0 | 0 | 0 | 0 | 0 | 0 |
| 52704 | ENSMUSG00000115147 | 0 | NA | NA | NA | NA | NA | Gm5219 | predicted gene 5219 [Source:MGI Symbol | 0 | 0 | 0 | 0 | 0 | 0 |
| 52705 | ENSMUSG00000115148 | 6.351470627 | 0.03969972 | 1.013949446 | 0.03915355 | 0.968767967 | 0.994547305 | Gm49125 | predicted gene, 49125 [Source:MGI Symb | 9.6372 | 4.1117 | 5.0312 | 10.604 | 8.7249 | 0 |
| 52706 | ENSMUSG00000115149 | 0 | NA | NA | NA | NA | NA | 9330188P03 | RIKEN cDNA 9330188P03 gene [Source:M | 0 | 0 | 0 | 0 | 0 | 0 |
| 52707 | ENSMUSG00000115150 | 0.706178709 | 0.10198885 | 3.03884937 | 0.033561667 | 0.97322669 | NA | Gm10370 | predicted gene 10370 [Source:MGI Symbo | 0 | 2.0558 | 0 | 0 | 2.1812 | 0 |
| 52708 | ENSMUSG00000115151 | 0.771003 | -1.69715066 | 2.640552792 | -0.642725518 | 0.52040222 | NA | Gm19276 | predicted gene, 19276 [Source:MGI Symb | 1.6062 | 2.0558 | 0 | 0.964 | 0 | 0 |
| 52709 | ENSMUSG00000115152 | 0 | NA | NA | NA | NA | NA | Gm49253 | predicted gene, 49253 [Source:MGI Symb | 0 | 0 | 0 | 0 | 0 | 0 |
| 52710 | ENSMUSG00000115153 | 1.3993034 | 1.698869132 | 2.170302421 | 0.782779909 | 0.433756368 | NA | Gm49240 | predicted gene, 49240 [Source:MGI Symb | 0 | 0 | 2.0125 | 1.928 | 0 | 4.4554 |
| 52711 | ENSMUSG00000115154 | 0.476489653 | -2.293616034 | 3.421506946 | -0.670352587 | 0.502633051 | NA | Gm49067 | predicted gene, 49067 [Source:MGI Symb | 0.8031 | 2.0558 | 0 | 0 | 0 | 0 |
| 52712 | ENSMUSG00000115155 | 0 | NA | NA | NA | NA | NA | Gm49182 | predicted gene, 49182 [Source:MGI Symb | 0 | 0 | 0 | 0 | 0 | 0 |
| 52713 | ENSMUSG00000115156 | 0 | NA | NA | NA | NA | NA | Gm49279 | predicted gene, 49279 [Source:MGI Symb | 0 | 0 | 0 | 0 | 0 | 0 |
| 52714 | ENSMUSG00000115157 | 0.160662974 | 1.126924175 | 4.080472857 | 0.276174898 | 0.782413736 | NA | Gm49397 | predicted gene, 49397 [Source:MGI Symb | 0 | 0 | 0 | 0.964 | 0 | 0 |
| 52715 | ENSMUSG00000115158 | 0 | NA | NA | NA | NA | NA | Gm48912 | predicted gene, 48912 [Source:MGI Symb | 0 | 0 | 0 | 0 | 0 | 0 |
| 52716 | ENSMUSG00000115159 | 0 | NA | NA | NA | NA | NA | Gm17892 | predicted gene, 17892 [Source:MGI Symb | 0 | 0 | 0 | 0 | 0 | 0 |
| 52717 | ENSMUSG00000115160 | 3.469817334 | 0.070918412 | 1.132294272 | 0.062632492 | 0.950059155 | NA | Gm6532 | predicted pseudogene 6532 [Source:MGI | 4.0155 | 2.0558 | 4.025 | 1.928 | 5.4531 | 3.3415 |
| 52718 | ENSMUSG00000115161 | 0.181769715 | 1.126924175 | 4.080472857 | 0.276174898 | 0.782413736 | NA | Gm20555 | predicted gene, 20555 [Source:MGI Symb | 0 | 0 | 0 | 0 | 1.0906 | 0 |
| 52719 | ENSMUSG00000115162 | 0 | NA | NA | NA | NA | NA | Gm9501 | predicted gene 9501 [Source:MGI Symbol | 0 | 0 | 0 | 0 | 0 | 0 |
| 52720 | ENSMUSG00000115163 | 0 | NA | NA | NA | NA | NA | Gm18076 | predicted gene, 18076 [Source:MGI Symb | 0 | 0 | 0 | 0 | 0 | 0 |
| 52721 | ENSMUSG00000115164 | 0 | NA | NA | NA | NA | NA | Gm30159 | predicted gene, 30159 [Source:MGI Symb | 0 | 0 | 0 | 0 | 0 | 0 |
| 52722 | ENSMUSG00000115165 | 0 | NA | NA | NA | NA | NA | Gm48987 | predicted gene, 48987 [Source:MGI Symb | 0 | 0 | 0 | 0 | 0 | 0 |
| 52723 | ENSMUSG00000115166 | 1.448534708 | 4.05966489 | 2.110058612 | 1.923958352 | 0.054359804 | NA | Vmn2r-ps11 | vomeronasal 2, receptor, pseudogene 11( | 0 | 0 | 0 | 0.964 | 3.2719 | 4.4554 |
| 52724 | ENSMUSG00000115167 | 0 | NA | NA | NA | NA | NA | Gm49015 | predicted gene, 49015 [Source:MGI Symb | 0 | 0 | 0 | 0 | 0 | 0 |
| 52725 | ENSMUSG00000115168 | 0 | NA | NA | NA | NA | NA | Gm48910 | predicted gene, 48910 [Source:MGI Symb | 0 | 0 | 0 | 0 | 0 | 0 |
| 52726 | ENSMUSG00000115169 | 0 | NA | NA | NA | NA | NA | Gm30563 | predicted gene, 30563 [Source:MGI Symb | 0 | 0 | 0 | 0 | 0 | 0 |
| 52727 | ENSMUSG00000115170 | 0 | NA | NA | NA | NA | NA | Vmn1r38 | vomeronasal 1 receptor 38 [Source:MGI S | 0 | 0 | 0 | 0 | 0 | 0 |
| 52728 | ENSMUSG00000115171 | 0 | NA | NA | NA | NA | NA | Gm48943 | predicted gene, 48943 [Source:MGI Symb | 0 | 0 | 0 | 0 | 0 | 0 |
| 52729 | ENSMUSG00000115172 | 0 | NA | NA | NA | NA | NA | Gm2582 | predicted gene 2582 [Source:MGI Symbol | 0 | 0 | 0 | 0 | 0 | 0 |
| 52730 | ENSMUSG00000115173 | 0 | NA | NA | NA | NA | NA | Gm2638 | predicted gene 2638 [Source:MGI Symbol | 0 | 0 | 0 | 0 | 0 | 0 |
| 52731 | ENSMUSG00000115174 | 0 | NA | NA | NA | NA | NA | Gm35996 | predicted gene, 35996 [Source:MGI Symb | 0 | 0 | 0 | 0 | 0 | 0 |
| 52732 | ENSMUSG00000115175 | 0 | NA | NA | NA | NA | NA | Gm18525 | predicted gene, 18525 [Source:MGI Symb | 0 | 0 | 0 | 0 | 0 | 0 |
| 52733 | ENSMUSG00000115176 | 0 | NA | NA | NA | NA | NA | Gm49063 | predicted gene, 49063 [Source:MGI Symb | 0 | 0 | 0 | 0 | 0 | 0 |
| 52734 | ENSMUSG00000115177 | 0 | NA | NA | NA | NA | NA | 4930523O13 | RIKEN cDNA 4930523O13 gene [Source:M | 0 | 0 | 0 | 0 | 0 | 0 |
| 52735 | ENSMUSG00000115178 | 0 | NA | NA | NA | NA | NA | Gm49220 | predicted gene, 49220 [Source:MGI Symb | 0 | 0 | 0 | 0 | 0 | 0 |
| 52736 | ENSMUSG00000115179 | 0 | NA | NA | NA | NA | NA | Gm48927 | predicted gene, 48927 [Source:MGI Symb | 0 | 0 | 0 | 0 | 0 | 0 |
| 52737 | ENSMUSG00000115180 | 0 | NA | NA | NA | NA | NA | Gm30556 | predicted gene, 30556 [Source:MGI Symb | 0 | 0 | 0 | 0 | 0 | 0 |
| 52738 | ENSMUSG00000115181 | 0 | NA | NA | NA | NA | NA | Vmn1r10 | vomeronasal 1 receptor 10 [Source:MGI S | 0 | 0 | 0 | 0 | 0 | 0 |
| 52739 | ENSMUSG00000115182 | 0 | NA | NA | NA | NA | NA | Gm49128 | predicted gene, 49128 [Source:MGI Symb | 0 | 0 | 0 | 0 | 0 | 0 |
| 52740 | ENSMUSG00000115183 | 1.327769957 | -2.617282484 | 2.107202751 | -1.242064857 | 0.214212634 | NA | Gm49203 | predicted gene, 49203 [Source:MGI Symb | 4.8186 | 1.0279 | 1.0062 | 0 | 0 | 1.1138 |
| 52741 | ENSMUSG00000115184 | 0.167707431 | -0.796629176 | 4.080472857 | -0.195229623 | 0.84521319 | NA | Gm49197 | predicted gene, 49197 [Source:MGI Symb | 0 | 0 | 1.0062 | 0 | 0 | 0 |
| 52742 | ENSMUSG00000115185 | 0 | NA | NA | NA | NA | NA | Gm18715 | predicted gene, 18715 [Source:MGI Symb | 0 | 0 | 0 | 0 | 0 | 0 |
| 52743 | ENSMUSG00000115186 | 49.39184995 | 0.685159945 | 0.316416129 | 2.165376167 | 0.03035889 | 0.230640658 | Gm49417 | predicted gene, 49417 [Source:MGI Symb | 35.336 | 42.145 | 36.225 | 56.875 | 56.712 | 69.058 |
| 52744 | ENSMUSG00000115187 | 0 | NA | NA | NA | NA | NA | Gm49210 | predicted gene, 49210 [Source:MGI Symb | 0 | 0 | 0 | 0 | 0 | 0 |
| 52745 | ENSMUSG00000115188 | 0.185640647 | 1.126924175 | 4.080472857 | 0.276174898 | 0.782413736 | NA | Gm46522 | predicted gene, 46522 [Source:MGI Symb | 0 | 0 | 0 | 0 | 0 | 1.1138 |
| 52746 | ENSMUSG00000115190 | 0.167707431 | -0.796629176 | 4.080472857 | -0.195229623 | 0.84521319 | NA | Gm49542 | predicted gene, 49542 [Source:MGI Symb | 0 | 0 | 1.0062 | 0 | 0 | 0 |
| 52747 | ENSMUSG00000115191 | 0 | NA | NA | NA | NA | NA | Gm49071 | predicted gene, 49071 [Source:MGI Symb | 0 | 0 | 0 | 0 | 0 | 0 |
| 52748 | ENSMUSG00000115192 | 0 | NA | NA | NA | NA | NA | Gm18532 | predicted gene, 18532 [Source:MGI Symb | 0 | 0 | 0 | 0 | 0 | 0 |

| | | | | | | | | | | | | | | | |
|---|---|---|---|---|---|---|---|---|---|---|---|---|---|---|---|
| 52749 | ENSMUSG00000115193 | 0 | NA | NA | NA | NA | NA | Gm6533 | predicted gene 6533 [Source:MGI Symbol | 0 | 0 | 0 | 0 | 0 | 0 |
| 52750 | ENSMUSG00000115194 | 1.947733935 | 0.387128511 | 1.735393788 | 0.223078193 | 0.82347465 | NA | Gm48909 | predicted gene 48909 [Source:MGI Symbo | 0 | 3.0838 | 2.0125 | 0 | 4.3625 | 2.2277 |
| 52751 | ENSMUSG00000115195 | 0 | NA | NA | NA | NA | NA | Gm5671 | predicted gene 5671 [Source:MGI Symbol | 0 | 0 | 0 | 0 | 0 | 0 |
| 52752 | ENSMUSG00000115196 | 0 | NA | NA | NA | NA | NA | Gm6212 | predicted gene 6212 [Source:MGI Symbol | 0 | 0 | 0 | 0 | 0 | 0 |
| 52753 | ENSMUSG00000115197 | 0 | NA | NA | NA | NA | NA | Gm49075 | predicted gene, 49075 [Source:MGI Symb | 0 | 0 | 0 | 0 | 0 | 0 |
| 52754 | ENSMUSG00000115198 | 0 | NA | NA | NA | NA | NA | Gm49729 | predicted gene, 49729 [Source:MGI Symb | 0 | 0 | 0 | 0 | 0 | 0 |
| 52755 | ENSMUSG00000115199 | 0 | NA | NA | NA | NA | NA | Vmn1r15 | vomeronasal 1 receptor 15 [Source:MGI S | 0 | 0 | 0 | 0 | 0 | 0 |
| 52756 | ENSMUSG00000115200 | 20.65945725 | -0.225668089 | 0.552269103 | -0.408619797 | 0.6828187 | 0.935166147 | Gm46516 | predicted gene, 46516 [Source:MGI Symb | 36.943 | 13.363 | 16.1 | 14.46 | 16.359 | 26.732 |
| 52757 | ENSMUSG00000115201 | 0 | NA | NA | NA | NA | NA | Gm49276 | predicted gene, 49276 [Source:MGI Symb | 0 | 0 | 0 | 0 | 0 | 0 |
| 52758 | ENSMUSG00000115202 | 0 | NA | NA | NA | NA | NA | Gm9275 | predicted gene 9275 [Source:MGI Symbol | 0 | 0 | 0 | 0 | 0 | 0 |
| 52759 | ENSMUSG00000115203 | 0 | NA | NA | NA | NA | NA | Gm49178 | predicted gene, 49178 [Source:MGI Symb | 0 | 0 | 0 | 0 | 0 | 0 |
| 52760 | ENSMUSG00000115204 | 0.301557804 | -1.64335916 | 4.042814367 | -0.406488899 | 0.684383412 | NA | Gm36642 | predicted gene, 36642 [Source:MGI Symb | 0.8031 | 0 | 1.0062 | 0 | 0 | 0 |
| 52761 | ENSMUSG00000115205 | 1.041825665 | 2.372145788 | 2.219541612 | 1.068754817 | 0.285180164 | NA | Gm16374 | predicted pseudogene 16374 [Source:MG | 0 | 1.0279 | 0 | 1.928 | 2.1812 | 1.1138 |
| 52762 | ENSMUSG00000115206 | 0.371281295 | 2.09253362 | 4.031852165 | 0.519000582 | 0.603760335 | NA | Gm49286 | predicted gene, 49286 [Source:MGI Symb | 0 | 0 | 0 | 0 | 0 | 2.2277 |
| 52763 | ENSMUSG00000115207 | 0 | NA | NA | NA | NA | NA | Gm49132 | predicted gene, 49132 [Source:MGI Symb | 0 | 0 | 0 | 0 | 0 | 0 |
| 52764 | ENSMUSG00000115208 | 0 | NA | NA | NA | NA | NA | Gm49118 | predicted gene, 49118 [Source:MGI Symb | 0 | 0 | 0 | 0 | 0 | 0 |
| 52765 | ENSMUSG00000115209 | 0 | NA | NA | NA | NA | NA | Gm34250 | predicted gene, 34250 [Source:MGI Symb | 0 | 0 | 0 | 0 | 0 | 0 |
| 52766 | ENSMUSG00000115210 | 1.022057521 | 0.202542166 | 2.069425296 | 0.09787363 | 0.922032639 | NA | Gm49308 | predicted gene, 49308 [Source:MGI Symb | 0.8031 | 1.0279 | 1.0062 | 0 | 2.1812 | 1.1138 |
| 52767 | ENSMUSG00000115211 | 0.133850373 | -0.796629176 | 4.080472857 | -0.195229623 | 0.84521319 | NA | Gm18152 | predicted gene, 18152 [Source:MGI Symb | 0.8031 | 0 | 0 | 0 | 0 | 0 |
| 52768 | ENSMUSG00000115212 | 0 | NA | NA | NA | NA | NA | Gm8417 | predicted gene 8417 [Source:MGI Symbol | 0 | 0 | 0 | 0 | 0 | 0 |
| 52769 | ENSMUSG00000115213 | 0 | NA | NA | NA | NA | NA | Gm49315 | predicted gene, 49315 [Source:MGI Symb | 0 | 0 | 0 | 0 | 0 | 0 |
| 52770 | ENSMUSG00000115214 | 0 | NA | NA | NA | NA | NA | Gm49202 | predicted gene, 49202 [Source:MGI Symb | 0 | 0 | 0 | 0 | 0 | 0 |
| 52771 | ENSMUSG00000115215 | 0 | NA | NA | NA | NA | NA | Gm41148 | predicted gene, 41148 [Source:MGI Symb | 0 | 0 | 0 | 0 | 0 | 0 |
| 52772 | ENSMUSG00000115216 | 0 | NA | NA | NA | NA | NA | Gm49183 | predicted gene, 49183 [Source:MGI Symb | 0 | 0 | 0 | 0 | 0 | 0 |
| 52773 | ENSMUSG00000115217 | 0 | NA | NA | NA | NA | NA | Gm32729 | predicted gene, 32729 [Source:MGI Symb | 0 | 0 | 0 | 0 | 0 | 0 |
| 52774 | ENSMUSG00000115218 | 0 | NA | NA | NA | NA | NA | Gm46477 | predicted gene, 46477 [Source:MGI Symb | 0 | 0 | 0 | 0 | 0 | 0 |
| 52775 | ENSMUSG00000115219 | 77.70790828 | -0.112155013 | 0.279199733 | -0.401701721 | 0.687903558 | 0.936885029 | Eef1akmt4 | EEF1A lysine methyltransferase 4 [Source: | 87.538 | 70.926 | 83.518 | 70.37 | 95.974 | 57.92 |
| 52776 | ENSMUSG00000115220 | 6.443148336 | -1.372954652 | 0.907234265 | -1.513340825 | 0.130193108 | 0.528370682 | Gm49768 | predicted gene, 49768 [Source:MGI Symb | 10.44 | 10.279 | 7.0437 | 0.964 | 4.3625 | 5.5692 |
| 52777 | ENSMUSG00000115221 | 0 | NA | NA | NA | NA | NA | Gm49121 | predicted gene, 49121 [Source:MGI Symb | 0 | 0 | 0 | 0 | 0 | 0 |
| 52778 | ENSMUSG00000115222 | 19.4300249 | 0.263271235 | 0.490109485 | 0.53716821 | 0.591151428 | 0.912336134 | Gm49747 | predicted gene, 49747 [Source:MGI Symb | 18.471 | 16.447 | 18.112 | 26.027 | 16.359 | 21.163 |
| 52779 | ENSMUSG00000115223 | 0 | NA | NA | NA | NA | NA | Gm48985 | predicted gene, 48985 [Source:MGI Symb | 0 | 0 | 0 | 0 | 0 | 0 |
| 52780 | ENSMUSG00000115224 | 0.17131964 | -0.796629176 | 4.080472857 | -0.195229623 | 0.84521319 | NA | Gm49215 | predicted gene, 49215 [Source:MGI Symb | 0 | 1.0279 | 0 | 0 | 0 | 0 |
| 52781 | ENSMUSG00000115225 | 0 | NA | NA | NA | NA | NA | Gm7272 | predicted gene 7272 [Source:MGI Symbol | 0 | 0 | 0 | 0 | 0 | 0 |
| 52782 | ENSMUSG00000115226 | 0.401551119 | -2.072982726 | 4.008251979 | -0.517178744 | 0.605031388 | NA | Gm49229 | predicted gene, 49229 [Source:MGI Symb | 2.4093 | 0 | 0 | 0 | 0 | 0 |
| 52783 | ENSMUSG00000115227 | 0 | NA | NA | NA | NA | NA | Gm49057 | predicted gene, 49057 [Source:MGI Symb | 0 | 0 | 0 | 0 | 0 | 0 |
| 52784 | ENSMUSG00000115228 | 0 | NA | NA | NA | NA | NA | Gm48948 | predicted gene, 48948 [Source:MGI Symb | 0 | 0 | 0 | 0 | 0 | 0 |
| 52785 | ENSMUSG00000115229 | 0 | NA | NA | NA | NA | NA | Gm33525 | predicted gene, 33525 [Source:MGI Symb | 0 | 0 | 0 | 0 | 0 | 0 |
| 52786 | ENSMUSG00000115230 | 0.301557804 | -1.64335916 | 4.042814367 | -0.406488899 | 0.684383412 | NA | AU022793 | expressed sequence AU022793 [Source:M | 0.8031 | 0 | 1.0062 | 0 | 0 | 0 |
| 52787 | ENSMUSG00000115231 | 0 | NA | NA | NA | NA | NA | 4930402D18 | RIKEN cDNA 4930402D18 gene [Source:M | 0 | 0 | 0 | 0 | 0 | 0 |
| 52788 | ENSMUSG00000115232 | 0 | NA | NA | NA | NA | NA | Gm49378 | predicted gene, 49378 [Source:MGI Symb | 0 | 0 | 0 | 0 | 0 | 0 |
| 52789 | ENSMUSG00000115233 | 0 | NA | NA | NA | NA | NA | Polr2d-ps1 | polymerase (RNA) II (DNA directed) polyp | 0 | 0 | 0 | 0 | 0 | 0 |
| 52790 | ENSMUSG00000115234 | 0 | NA | NA | NA | NA | NA | Gm49130 | predicted gene, 49130 [Source:MGI Symb | 0 | 0 | 0 | 0 | 0 | 0 |
| 52791 | ENSMUSG00000115235 | 0 | NA | NA | NA | NA | NA | Gm49102 | predicted gene, 49102 [Source:MGI Symb | 0 | 0 | 0 | 0 | 0 | 0 |
| 52792 | ENSMUSG00000115236 | 0 | NA | NA | NA | NA | NA | Vmn1r11 | vomeronasal 1 receptor 11 [Source:MGI S | 0 | 0 | 0 | 0 | 0 | 0 |
| 52793 | ENSMUSG00000115237 | 0 | NA | NA | NA | NA | NA | Gm46474 | predicted gene, 46474 [Source:MGI Symb | 0 | 0 | 0 | 0 | 0 | 0 |
| 52794 | ENSMUSG00000115238 | 0 | NA | NA | NA | NA | NA | Gm19236 | predicted gene, 19236 [Source:MGI Symb | 0 | 0 | 0 | 0 | 0 | 0 |
| 52795 | ENSMUSG00000115239 | 0 | NA | NA | NA | NA | NA | Gm9264 | predicted gene 9264 [Source:MGI Symbol | 0 | 0 | 0 | 0 | 0 | 0 |
| 52796 | ENSMUSG00000115240 | 0 | NA | NA | NA | NA | NA | Gm5144 | predicted gene 5144 [Source:MGI Symbol | 0 | 0 | 0 | 0 | 0 | 0 |
| 52797 | ENSMUSG00000115241 | 0 | NA | NA | NA | NA | NA | Gm41322 | predicted gene, 41322 [Source:MGI Symb | 0 | 0 | 0 | 0 | 0 | 0 |
| 52798 | ENSMUSG00000115242 | 0 | NA | NA | NA | NA | NA | Gm49018 | predicted gene, 49018 [Source:MGI Symb | 0 | 0 | 0 | 0 | 0 | 0 |
| 52799 | ENSMUSG00000115243 | 0 | NA | NA | NA | NA | NA | Gm5207 | predicted gene 5207 [Source:MGI Symbol | 0 | 0 | 0 | 0 | 0 | 0 |
| 52800 | ENSMUSG00000115244 | 0 | NA | NA | NA | NA | NA | Gm49002 | predicted gene, 49002 [Source:MGI Symb | 0 | 0 | 0 | 0 | 0 | 0 |
| 52801 | ENSMUSG00000115245 | 0 | NA | NA | NA | NA | NA | 4930478E11 | RIKEN cDNA 4930478E11 gene [Source:M | 0 | 0 | 0 | 0 | 0 | 0 |
| 52802 | ENSMUSG00000115246 | 0 | NA | NA | NA | NA | NA | Gm48984 | predicted gene, 48984 [Source:MGI Symb | 0 | 0 | 0 | 0 | 0 | 0 |
| 52803 | ENSMUSG00000115247 | 0.678054141 | -2.792799308 | 2.990673428 | -0.933836266 | 0.350388359 | NA | Gm6201 | predicted gene 6201 [Source:MGI Symbol | 0 | 2.0558 | 2.0125 | 0 | 0 | 0 |
| 52804 | ENSMUSG00000115248 | 0 | NA | NA | NA | NA | NA | Gm49037 | predicted gene, 49037 [Source:MGI Symb | 0 | 0 | 0 | 0 | 0 | 0 |

| | | | | | | | | | | | | | | | |
|---|---|---|---|---|---|---|---|---|---|---|---|---|---|---|---|
| 52805 | ENSMUSG00000115249 | 5.954952634 | -1.146062542 | 0.967978048 | -1.183975757 | 0.23642266 | 0.685294058 | Gm49085 | predicted gene, 49085 [Source:MGI Symb | 10.44 | 9.2513 | 5.0312 | 7.7118 | 2.1812 | 1.1138 |
| 52806 | ENSMUSG00000115250 | 0 | NA | NA | NA | NA | NA | Gm49020 | predicted gene, 49020 [Source:MGI Symb | 0 | 0 | 0 | 0 | 0 | 0 |
| 52807 | ENSMUSG00000115251 | 0 | NA | NA | NA | NA | NA | Gm4774 | predicted gene 4774 [Source:MGI Symbol | 0 | 0 | 0 | 0 | 0 | 0 |
| 52808 | ENSMUSG00000115252 | 0 | NA | NA | NA | NA | NA | Gm48996 | predicted gene, 48996 [Source:MGI Symb | 0 | 0 | 0 | 0 | 0 | 0 |
| 52809 | ENSMUSG00000115253 | 0 | NA | NA | NA | NA | NA | Vmn1r20 | vomeronasal 1 receptor 20 [Source:MGI S | 0 | 0 | 0 | 0 | 0 | 0 |
| 52810 | ENSMUSG00000115254 | 0 | NA | NA | NA | NA | NA | Gm49140 | predicted gene, 49140 [Source:MGI Symb | 0 | 0 | 0 | 0 | 0 | 0 |
| 52811 | ENSMUSG00000115256 | 0 | NA | NA | NA | NA | NA | Gm49259 | predicted gene, 49259 [Source:MGI Symb | 0 | 0 | 0 | 0 | 0 | 0 |
| 52812 | ENSMUSG00000115257 | 0 | NA | NA | NA | NA | NA | Gm30467 | predicted gene, 30467 [Source:MGI Symb | 0 | 0 | 0 | 0 | 0 | 0 |
| 52813 | ENSMUSG00000115258 | 0 | NA | NA | NA | NA | NA | Gm32093 | predicted gene, 32093 [Source:MGI Symb | 0 | 0 | 0 | 0 | 0 | 0 |
| 52814 | ENSMUSG00000115259 | 0.181769715 | 1.126924175 | 4.080472857 | 0.276174898 | 0.782413736 | NA | Gm49073 | predicted gene, 49073 [Source:MGI Symb | 0 | 0 | 0 | 0 | 1.0906 | 0 |
| 52815 | ENSMUSG00000115260 | 0 | NA | NA | NA | NA | NA | Gm49005 | predicted gene, 49005 [Source:MGI Symb | 0 | 0 | 0 | 0 | 0 | 0 |
| 52816 | ENSMUSG00000115261 | 0 | NA | NA | NA | NA | NA | Gm35769 | predicted gene, 35769 [Source:MGI Symb | 0 | 0 | 0 | 0 | 0 | 0 |
| 52817 | ENSMUSG00000115262 | 0 | NA | NA | NA | NA | NA | Gm49135 | predicted gene, 49135 [Source:MGI Symb | 0 | 0 | 0 | 0 | 0 | 0 |
| 52818 | ENSMUSG00000115263 | 1.323847419 | 1.876094015 | 2.007969798 | 0.934323822 | 0.35013688 | NA | Gm5089 | predicted gene 5089 [Source:MGI Symbol | 1.6062 | 0 | 0 | 1.928 | 2.1812 | 2.2277 |
| 52819 | ENSMUSG00000115264 | 0.572870759 | -2.577107685 | 3.182795569 | -0.809699407 | 0.418112959 | NA | Gm49260 | predicted gene, 49260 [Source:MGI Symb | 2.4093 | 1.0279 | 0 | 0 | 0 | 0 |
| 52820 | ENSMUSG00000115265 | 0 | NA | NA | NA | NA | NA | Gm18153 | predicted gene, 18153 [Source:MGI Symb | 0 | 0 | 0 | 0 | 0 | 0 |
| 52821 | ENSMUSG00000115266 | 0 | NA | NA | NA | NA | NA | 4930557F08 | RIKEN cDNA 4930557F08 gene [Source:M | 0 | 0 | 0 | 0 | 0 | 0 |
| 52822 | ENSMUSG00000115267 | 0 | NA | NA | NA | NA | NA | Vmn1r76 | vomeronasal 1 receptor 76 [Source:MGI S | 0 | 0 | 0 | 0 | 0 | 0 |
| 52823 | ENSMUSG00000115268 | 0 | NA | NA | NA | NA | NA | Gm18524 | predicted gene, 18524 [Source:MGI Symb | 0 | 0 | 0 | 0 | 0 | 0 |
| 52824 | ENSMUSG00000115269 | 0 | NA | NA | NA | NA | NA | Gm5209 | predicted gene 5209 [Source:MGI Symbol | 0 | 0 | 0 | 0 | 0 | 0 |
| 52825 | ENSMUSG00000115270 | 0.185640647 | 1.126924175 | 4.080472857 | 0.276174898 | 0.782413736 | NA | 5430430K15 | RIKEN cDNA 5430430K15 gene [Source:M | 0 | 0 | 0 | 0 | 0 | 1.1138 |
| 52826 | ENSMUSG00000115271 | 0 | NA | NA | NA | NA | NA | 4930529K09 | RIKEN cDNA 4930529K09 gene [Source:M | 0 | 0 | 0 | 0 | 0 | 0 |
| 52827 | ENSMUSG00000115272 | 0 | NA | NA | NA | NA | NA | Gm49965 | predicted gene, 49965 [Source:MGI Symb | 0 | 0 | 0 | 0 | 0 | 0 |
| 52828 | ENSMUSG00000115273 | 0 | NA | NA | NA | NA | NA | Gm9335 | predicted gene 9335 [Source:MGI Symbol | 0 | 0 | 0 | 0 | 0 | 0 |
| 52829 | ENSMUSG00000115274 | 0 | NA | NA | NA | NA | NA | Gm49040 | predicted gene, 49040 [Source:MGI Symb | 0 | 0 | 0 | 0 | 0 | 0 |
| 52830 | ENSMUSG00000115275 | 2.056347911 | 0.129185567 | 1.642465361 | 0.07865345 | 0.937308272 | NA | 9630050E16 | RIKEN cDNA 9630050E16 gene [Source:M | 0.8031 | 4.1117 | 1.0062 | 0.964 | 5.4531 | 0 |
| 52831 | ENSMUSG00000115276 | 4.963548459 | 0.463984465 | 1.074543466 | 0.431796833 | 0.665889082 | NA | 9930017N22 | RIKEN cDNA 9930017N22 gene [Source:M | 6.4248 | 1.0279 | 5.0312 | 9.6398 | 6.5437 | 1.1138 |
| 52832 | ENSMUSG00000115277 | 0 | NA | NA | NA | NA | NA | Gm19016 | predicted gene, 19016 [Source:MGI Symb | 0 | 0 | 0 | 0 | 0 | 0 |
| 52833 | ENSMUSG00000115278 | 0.497389803 | 1.101799453 | 3.439268845 | 0.320358629 | 0.748696483 | NA | Gm34150 | predicted gene, 34150 [Source:MGI Symb | 0.8031 | 0 | 0 | 0 | 2.1812 | 0 |
| 52834 | ENSMUSG00000115279 | 0 | NA | NA | NA | NA | NA | Gm49273 | predicted gene, 49273 [Source:MGI Symb | 0 | 0 | 0 | 0 | 0 | 0 |
| 52835 | ENSMUSG00000115280 | 0 | NA | NA | NA | NA | NA | Gm7107 | predicted gene 7107 [Source:MGI Symbol | 0 | 0 | 0 | 0 | 0 | 0 |
| 52836 | ENSMUSG00000115281 | 0 | NA | NA | NA | NA | NA | Gm49743 | predicted gene, 49743 [Source:MGI Symb | 0 | 0 | 0 | 0 | 0 | 0 |
| 52837 | ENSMUSG00000115282 | 0 | NA | NA | NA | NA | NA | Gm49055 | predicted gene, 49055 [Source:MGI Symb | 0 | 0 | 0 | 0 | 0 | 0 |
| 52838 | ENSMUSG00000115283 | 0.17131964 | -0.796629176 | 4.080472857 | -0.195229623 | 0.84521319 | NA | Gm49252 | predicted gene, 49252 [Source:MGI Symb | 0 | 1.0279 | 0 | 0 | 0 | 0 |
| 52839 | ENSMUSG00000115284 | 1.045696598 | 2.377089708 | 2.217824212 | 1.071811596 | 0.283804664 | NA | Gm34678 | predicted gene, 34678 [Source:MGI Symb | 0 | 1.0279 | 0 | 1.928 | 1.0906 | 2.2277 |
| 52840 | ENSMUSG00000115285 | 0 | NA | NA | NA | NA | NA | Gm49068 | predicted gene, 49068 [Source:MGI Symb | 0 | 0 | 0 | 0 | 0 | 0 |
| 52841 | ENSMUSG00000115286 | 0 | NA | NA | NA | NA | NA | 4930429C20 | RIKEN cDNA 4930429C20 gene [Source:M | 0 | 0 | 0 | 0 | 0 | 0 |
| 52842 | ENSMUSG00000115287 | 0 | NA | NA | NA | NA | NA | Gm49111 | predicted gene, 49111 [Source:MGI Symb | 0 | 0 | 0 | 0 | 0 | 0 |
| 52843 | ENSMUSG00000115288 | 0 | NA | NA | NA | NA | NA | Gm9212 | predicted gene 9212 [Source:MGI Symbol | 0 | 0 | 0 | 0 | 0 | 0 |
| 52844 | ENSMUSG00000115289 | 0 | NA | NA | NA | NA | NA | Gm49164 | predicted gene, 49164 [Source:MGI Symb | 0 | 0 | 0 | 0 | 0 | 0 |
| 52845 | ENSMUSG00000115290 | 0 | NA | NA | NA | NA | NA | Gm19098 | predicted gene, 19098 [Source:MGI Symb | 0 | 0 | 0 | 0 | 0 | 0 |
| 52846 | ENSMUSG00000115291 | 0 | NA | NA | NA | NA | NA | Gm18605 | predicted gene, 18605 [Source:MGI Symb | 0 | 0 | 0 | 0 | 0 | 0 |
| 52847 | ENSMUSG00000115292 | 0 | NA | NA | NA | NA | NA | Gm33246 | predicted gene, 33246 [Source:MGI Symb | 0 | 0 | 0 | 0 | 0 | 0 |
| 52848 | ENSMUSG00000115293 | 0 | NA | NA | NA | NA | NA | Gm49333 | predicted gene, 49333 [Source:MGI Symb | 0 | 0 | 0 | 0 | 0 | 0 |
| 52849 | ENSMUSG00000115294 | 0 | NA | NA | NA | NA | NA | 1700047E10 | RIKEN cDNA 1700047E10 gene [Source:M | 0 | 0 | 0 | 0 | 0 | 0 |
| 52850 | ENSMUSG00000115295 | 0 | NA | NA | NA | NA | NA | Gm19129 | predicted gene, 19129 [Source:MGI Symb | 0 | 0 | 0 | 0 | 0 | 0 |
| 52851 | ENSMUSG00000115296 | 0 | NA | NA | NA | NA | NA | Gm48911 | predicted gene, 48911 [Source:MGI Symb | 0 | 0 | 0 | 0 | 0 | 0 |
| 52852 | ENSMUSG00000115297 | 0 | NA | NA | NA | NA | NA | Gm49045 | predicted gene, 49045 [Source:MGI Symb | 0 | 0 | 0 | 0 | 0 | 0 |
| 52853 | ENSMUSG00000115298 | 0 | NA | NA | NA | NA | NA | Gm18741 | predicted gene, 18741 [Source:MGI Symb | 0 | 0 | 0 | 0 | 0 | 0 |
| 52854 | ENSMUSG00000115299 | 0 | NA | NA | NA | NA | NA | Gm49168 | predicted gene, 49168 [Source:MGI Symb | 0 | 0 | 0 | 0 | 0 | 0 |
| 52855 | ENSMUSG00000115300 | 26.10306705 | 0.206030426 | 0.494078926 | 0.416999016 | 0.676679127 | 0.932891189 | Gm49188 | predicted gene, 49188 [Source:MGI Symb | 24.093 | 16.447 | 32.2 | 27.955 | 41.443 | 14.48 |
| 52856 | ENSMUSG00000115301 | 0 | NA | NA | NA | NA | NA | Gm6330 | predicted gene 6330 [Source:MGI Symbol | 0 | 0 | 0 | 0 | 0 | 0 |
| 52857 | ENSMUSG00000115302 | 0 | NA | NA | NA | NA | NA | Gm49394 | predicted gene, 49394 [Source:MGI Symb | 0 | 0 | 0 | 0 | 0 | 0 |
| 52858 | ENSMUSG00000115303 | 0 | NA | NA | NA | NA | NA | Gm4291 | predicted gene 4291 [Source:MGI Symbol | 0 | 0 | 0 | 0 | 0 | 0 |
| 52859 | ENSMUSG00000115304 | 0 | NA | NA | NA | NA | NA | 4930434J06I | RIKEN cDNA 4930434J06 gene [Source:M | 0 | 0 | 0 | 0 | 0 | 0 |
| 52860 | ENSMUSG00000115305 | 0 | NA | NA | NA | NA | NA | Gm30584 | predicted gene, 30584 [Source:MGI Symb | 0 | 0 | 0 | 0 | 0 | 0 |

| | | | | | | | | | | | | | | | |
|---|---|---|---|---|---|---|---|---|---|---|---|---|---|---|---|
| 52861 | ENSMUSG00000115306 | 0 | NA | NA | NA | NA | NA | Gm49159 | predicted gene, 49159 [Source:MGI Symb | 0 | 0 | 0 | 0 | 0 | 0 |
| 52862 | ENSMUSG00000115307 | 0 | NA | NA | NA | NA | NA | Gm49313 | predicted gene, 49313 [Source:MGI Symb | 0 | 0 | 0 | 0 | 0 | 0 |
| 52863 | ENSMUSG00000115308 | 0 | NA | NA | NA | NA | NA | Gm48944 | predicted gene, 48944 [Source:MGI Symb | 0 | 0 | 0 | 0 | 0 | 0 |
| 52864 | ENSMUSG00000115309 | 0 | NA | NA | NA | NA | NA | Gm49010 | predicted gene, 49010 [Source:MGI Symb | 0 | 0 | 0 | 0 | 0 | 0 |
| 52865 | ENSMUSG00000115310 | 0 | NA | NA | NA | NA | NA | Gm35360 | predicted gene, 35360 [Source:MGI Symb | 0 | 0 | 0 | 0 | 0 | 0 |
| 52866 | ENSMUSG00000115311 | 0 | NA | NA | NA | NA | NA | Gm35823 | predicted gene, 35823 [Source:MGI Symb | 0 | 0 | 0 | 0 | 0 | 0 |
| 52867 | ENSMUSG00000115312 | 1.757426182 | -0.101786716 | 1.629413579 | -0.062468312 | 0.950189896 | NA | Gm8518 | predicted gene 8518 [Source:MGI Symbol | 2.4093 | 2.0558 | 1.0062 | 2.8919 | 2.1812 | 0 |
| 52868 | ENSMUSG00000115313 | 0 | NA | NA | NA | NA | NA | Gm48941 | predicted gene, 48941 [Source:MGI Symb | 0 | 0 | 0 | 0 | 0 | 0 |
| 52869 | ENSMUSG00000115314 | 0.469265235 | -2.274672822 | 3.439794491 | -0.661281605 | 0.508431736 | NA | Gm48966 | predicted gene, 48966 [Source:MGI Symb | 0.8031 | 0 | 2.0125 | 0 | 0 | 0 |
| 52870 | ENSMUSG00000115315 | 0 | NA | NA | NA | NA | NA | Gnb1-ps2 | guanine nucleotide binding protein (G prc | 0 | 0 | 0 | 0 | 0 | 0 |
| 52871 | ENSMUSG00000115316 | 1.302947269 | 0.17976329 | 1.96941963 | 0.091277292 | 0.927272261 | NA | Gm49096 | predicted gene, 49096 [Source:MGI Symb | 1.6062 | 2.0558 | 0 | 1.928 | 0 | 2.2277 |
| 52872 | ENSMUSG00000115317 | 2.219099024 | 0.885511187 | 1.426418334 | 0.620793469 | 0.534735521 | NA | Gm32618 | predicted gene, 32618 [Source:MGI Symb | 1.6062 | 1.0279 | 2.0125 | 0.964 | 4.3625 | 3.3415 |
| 52873 | ENSMUSG00000115318 | 0 | NA | NA | NA | NA | NA | Gm49089 | predicted gene, 49089 [Source:MGI Symb | 0 | 0 | 0 | 0 | 0 | 0 |
| 52874 | ENSMUSG00000115319 | 0 | NA | NA | NA | NA | NA | Gm34417 | predicted gene, 34417 [Source:MGI Symb | 0 | 0 | 0 | 0 | 0 | 0 |
| 52875 | ENSMUSG00000115320 | 0 | NA | NA | NA | NA | NA | 4930524C18 | RIKEN cDNA 4930524C18 gene [Source:M | 0 | 0 | 0 | 0 | 0 | 0 |
| 52876 | ENSMUSG00000115321 | 0 | NA | NA | NA | NA | NA | Gm34060 | predicted gene, 34060 [Source:MGI Symb | 0 | 0 | 0 | 0 | 0 | 0 |
| 52877 | ENSMUSG00000115322 | 0 | NA | NA | NA | NA | NA | Gm49023 | predicted gene, 49023 [Source:MGI Symb | 0 | 0 | 0 | 0 | 0 | 0 |
| 52878 | ENSMUSG00000115323 | 0 | NA | NA | NA | NA | NA | Gm18762 | predicted gene, 18762 [Source:MGI Symb | 0 | 0 | 0 | 0 | 0 | 0 |
| 52879 | ENSMUSG00000115324 | 1.143707168 | -3.552083234 | 2.317588164 | -1.532663693 | 0.125358739 | NA | 4930527F14 | RIKEN cDNA 4930527F14 gene [Source:M | 0.8031 | 1.0279 | 5.0312 | 0 | 0 | 0 |
| 52880 | ENSMUSG00000115325 | 0 | NA | NA | NA | NA | NA | Gm48979 | predicted gene, 48979 [Source:MGI Symb | 0 | 0 | 0 | 0 | 0 | 0 |
| 52881 | ENSMUSG00000115326 | 0 | NA | NA | NA | NA | NA | Gm35019 | predicted gene, 35019 [Source:MGI Symb | 0 | 0 | 0 | 0 | 0 | 0 |
| 52882 | ENSMUSG00000115327 | 0 | NA | NA | NA | NA | NA | Gm7566 | predicted gene 7566 [Source:MGI Symbol | 0 | 0 | 0 | 0 | 0 | 0 |
| 52883 | ENSMUSG00000115328 | 0 | NA | NA | NA | NA | NA | Gm30691 | predicted gene, 30691 [Source:MGI Symb | 0 | 0 | 0 | 0 | 0 | 0 |
| 52884 | ENSMUSG00000115329 | 0 | NA | NA | NA | NA | NA | Gm49272 | predicted gene, 49272 [Source:MGI Symb | 0 | 0 | 0 | 0 | 0 | 0 |
| 52885 | ENSMUSG00000115330 | 0 | NA | NA | NA | NA | NA | Gm49221 | predicted gene, 49221 [Source:MGI Symb | 0 | 0 | 0 | 0 | 0 | 0 |
| 52886 | ENSMUSG00000115331 | 0 | NA | NA | NA | NA | NA | Gm9721 | predicted gene 9721 [Source:MGI Symbol | 0 | 0 | 0 | 0 | 0 | 0 |
| 52887 | ENSMUSG00000115332 | 0.685278559 | -2.809870862 | 3.376578027 | -0.832165239 | 0.405315682 | NA | Gm48952 | predicted gene, 48952 [Source:MGI Symb | 0 | 4.1117 | 0 | 0 | 0 | 0 |
| 52888 | ENSMUSG00000115333 | 3.135590699 | 1.797701434 | 1.454676831 | 1.235808116 | 0.216529898 | NA | Gm34590 | predicted gene, 34590 [Source:MGI Symb | 4.0155 | 0 | 0 | 4.8199 | 2.1812 | 7.7969 |
| 52889 | ENSMUSG00000115334 | 0 | NA | NA | NA | NA | NA | Gm17923 | predicted gene, 17923 [Source:MGI Symb | 0 | 0 | 0 | 0 | 0 | 0 |
| 52890 | ENSMUSG00000115335 | 0 | NA | NA | NA | NA | NA | Gm41279 | predicted gene, 41279 [Source:MGI Symb | 0 | 0 | 0 | 0 | 0 | 0 |
| 52891 | ENSMUSG00000115336 | 0 | NA | NA | NA | NA | NA | Gm48980 | predicted gene, 48980 [Source:MGI Symb | 0 | 0 | 0 | 0 | 0 | 0 |
| 52892 | ENSMUSG00000115337 | 0 | NA | NA | NA | NA | NA | 4930570B17 | RIKEN cDNA 4930570B17 gene [Source:M | 0 | 0 | 0 | 0 | 0 | 0 |
| 52893 | ENSMUSG00000115338 | 400.0629607 | -0.126311837 | 0.199443418 | -0.633321661 | 0.526523611 | 0.888334265 | Pnp | purine-nucleoside phosphorylase [Source | 361.4 | 326.88 | 564.5 | 336.43 | 404.62 | 406.55 |
| 52894 | ENSMUSG00000115339 | 0 | NA | NA | NA | NA | NA | Gm48913 | predicted gene, 48913 [Source:MGI Symb | 0 | 0 | 0 | 0 | 0 | 0 |
| 52895 | ENSMUSG00000115340 | 0.556921942 | 2.679043159 | 3.638243063 | 0.736356289 | 0.461513899 | NA | Gm48965 | predicted gene, 48965 [Source:MGI Symb | 0 | 0 | 0 | 0 | 0 | 3.3415 |
| 52896 | ENSMUSG00000115341 | 0 | NA | NA | NA | NA | NA | Gm48919 | predicted gene, 48919 [Source:MGI Symb | 0 | 0 | 0 | 0 | 0 | 0 |
| 52897 | ENSMUSG00000115342 | 0 | NA | NA | NA | NA | NA | Gm49095 | predicted gene, 49095 [Source:MGI Symb | 0 | 0 | 0 | 0 | 0 | 0 |
| 52898 | ENSMUSG00000115343 | 0 | NA | NA | NA | NA | NA | Vmn1r21 | vomeronasal 1 receptor 21 [Source:MGI S | 0 | 0 | 0 | 0 | 0 | 0 |
| 52899 | ENSMUSG00000115344 | 0 | NA | NA | NA | NA | NA | Gm49364 | predicted gene, 49364 [Source:MGI Symb | 0 | 0 | 0 | 0 | 0 | 0 |
| 52900 | ENSMUSG00000115345 | 0 | NA | NA | NA | NA | NA | Gm33497 | predicted gene, 33497 [Source:MGI Symb | 0 | 0 | 0 | 0 | 0 | 0 |
| 52901 | ENSMUSG00000115346 | 0 | NA | NA | NA | NA | NA | Gm19120 | predicted gene, 19120 [Source:MGI Symb | 0 | 0 | 0 | 0 | 0 | 0 |
| 52902 | ENSMUSG00000115347 | 0 | NA | NA | NA | NA | NA | Gm8238 | predicted gene 8238 [Source:MGI Symbol | 0 | 0 | 0 | 0 | 0 | 0 |
| 52903 | ENSMUSG00000115348 | 0 | NA | NA | NA | NA | NA | Gm49155 | predicted gene, 49155 [Source:MGI Symb | 0 | 0 | 0 | 0 | 0 | 0 |
| 52904 | ENSMUSG00000115349 | 0 | NA | NA | NA | NA | NA | Gm9348 | predicted gene 9348 [Source:MGI Symbol | 0 | 0 | 0 | 0 | 0 | 0 |
| 52905 | ENSMUSG00000115350 | 0 | NA | NA | NA | NA | NA | Gm49079 | predicted gene, 49079 [Source:MGI Symb | 0 | 0 | 0 | 0 | 0 | 0 |
| 52906 | ENSMUSG00000115351 | 0 | NA | NA | NA | NA | NA | Gm19077 | predicted gene, 19077 [Source:MGI Symb | 0 | 0 | 0 | 0 | 0 | 0 |
| 52907 | ENSMUSG00000115352 | 0 | NA | NA | NA | NA | NA | Gm49160 | predicted gene, 49160 [Source:MGI Symb | 0 | 0 | 0 | 0 | 0 | 0 |
| 52908 | ENSMUSG00000115353 | 1.707879031 | 2.196113525 | 1.788082773 | 1.228194554 | 0.219373943 | NA | Gm48968 | predicted gene, 48968 [Source:MGI Symb | 0.8031 | 0 | 1.0062 | 2.8919 | 1.0906 | 4.4554 |
| 52909 | ENSMUSG00000115354 | 0 | NA | NA | NA | NA | NA | Gm49083 | predicted gene, 49083 [Source:MGI Symb | 0 | 0 | 0 | 0 | 0 | 0 |
| 52910 | ENSMUSG00000115355 | 0 | NA | NA | NA | NA | NA | 4930445E18 | RIKEN cDNA 4930445E18 gene [Source:M | 0 | 0 | 0 | 0 | 0 | 0 |
| 52911 | ENSMUSG00000115356 | 0 | NA | NA | NA | NA | NA | Gm2581 | predicted gene 2581 [Source:MGI Symbol | 0 | 0 | 0 | 0 | 0 | 0 |
| 52912 | ENSMUSG00000115357 | 1.82908834 | -0.286008634 | 1.775665206 | -0.161071261 | 0.872037275 | NA | Gm49214 | predicted gene, 49214 [Source:MGI Symb | 4.0155 | 0 | 2.0125 | 3.8559 | 1.0906 | 0 |
| 52913 | ENSMUSG00000115358 | 0 | NA | NA | NA | NA | NA | Gm49138 | predicted gene, 49138 [Source:MGI Symb | 0 | 0 | 0 | 0 | 0 | 0 |
| 52914 | ENSMUSG00000115359 | 0 | NA | NA | NA | NA | NA | Gm49779 | predicted gene, 49779 [Source:MGI Symb | 0 | 0 | 0 | 0 | 0 | 0 |
| 52915 | ENSMUSG00000115360 | 0 | NA | NA | NA | NA | NA | Gm48932 | predicted gene, 48932 [Source:MGI Symb | 0 | 0 | 0 | 0 | 0 | 0 |
| 52916 | ENSMUSG00000115361 | 0 | NA | NA | NA | NA | NA | Gm49211 | predicted gene, 49211 [Source:MGI Symb | 0 | 0 | 0 | 0 | 0 | 0 |

| | | | | | | | | | | | | | | | |
|---|---|---|---|---|---|---|---|---|---|---|---|---|---|---|---|
| 52917 | ENSMUSG00000115362 | 0 | NA | NA | NA | NA | NA | Gm49088 | predicted gene, 49088 [Source:MGI Symb | 0 | 0 | 0 | 0 | 0 | 0 |
| 52918 | ENSMUSG00000115363 | 0 | NA | NA | NA | NA | NA | Gm49003 | predicted gene, 49003 [Source:MGI Symb | 0 | 0 | 0 | 0 | 0 | 0 |
| 52919 | ENSMUSG00000115365 | 0 | NA | NA | NA | NA | NA | Gm18582 | predicted gene, 18582 [Source:MGI Symb | 0 | 0 | 0 | 0 | 0 | 0 |
| 52920 | ENSMUSG00000115366 | 0 | NA | NA | NA | NA | NA | Gm9298 | predicted gene 9298 [Source:MGI Symbol | 0 | 0 | 0 | 0 | 0 | 0 |
| 52921 | ENSMUSG00000115367 | 0 | NA | NA | NA | NA | NA | Gm33301 | predicted gene, 33301 [Source:MGI Symb | 0 | 0 | 0 | 0 | 0 | 0 |
| 52922 | ENSMUSG00000115368 | 0.133850373 | -0.796629176 | 4.080472857 | -0.195229623 | 0.84521319 | NA | Gm48942 | predicted gene, 48942 [Source:MGI Symb | 0.8031 | 0 | 0 | 0 | 0 | 0 |
| 52923 | ENSMUSG00000115369 | 0 | NA | NA | NA | NA | NA | Gm49014 | predicted gene, 49014 [Source:MGI Symb | 0 | 0 | 0 | 0 | 0 | 0 |
| 52924 | ENSMUSG00000115370 | 0.79210974 | -1.697240781 | 2.618300099 | -0.648222403 | 0.516841114 | NA | Gm15667 | predicted gene 15667 [Source:MGI Symbo | 1.6062 | 2.0558 | 0 | 0 | 1.0906 | 0 |
| 52925 | ENSMUSG00000115371 | 0 | NA | NA | NA | NA | NA | Gm18146 | predicted gene, 18146 [Source:MGI Symb | 0 | 0 | 0 | 0 | 0 | 0 |
| 52926 | ENSMUSG00000115372 | 0 | NA | NA | NA | NA | NA | 1700100I10F | RIKEN cDNA 1700100I10 gene [Source:MC | 0 | 0 | 0 | 0 | 0 | 0 |
| 52927 | ENSMUSG00000115373 | 0 | NA | NA | NA | NA | NA | Gm49194 | predicted gene, 49194 [Source:MGI Symb | 0 | 0 | 0 | 0 | 0 | 0 |
| 52928 | ENSMUSG00000115374 | 0 | NA | NA | NA | NA | NA | Vmn2r-ps11 | vomeronasal 2, receptor, pseudogene 111 | 0 | 0 | 0 | 0 | 0 | 0 |
| 52929 | ENSMUSG00000115375 | 0 | NA | NA | NA | NA | NA | Gm49039 | predicted gene, 49039 [Source:MGI Symb | 0 | 0 | 0 | 0 | 0 | 0 |
| 52930 | ENSMUSG00000115376 | 0 | NA | NA | NA | NA | NA | Gm35766 | predicted gene, 35766 [Source:MGI Symb | 0 | 0 | 0 | 0 | 0 | 0 |
| 52931 | ENSMUSG00000115377 | 0 | NA | NA | NA | NA | NA | Gm4942 | predicted gene 4942 [Source:MGI Symbol | 0 | 0 | 0 | 0 | 0 | 0 |
| 52932 | ENSMUSG00000115378 | 0 | NA | NA | NA | NA | NA | Gm39653 | predicted gene, 39653 [Source:MGI Symb | 0 | 0 | 0 | 0 | 0 | 0 |
| 52933 | ENSMUSG00000115379 | 0 | NA | NA | NA | NA | NA | Gm48990 | predicted gene, 48990 [Source:MGI Symb | 0 | 0 | 0 | 0 | 0 | 0 |
| 52934 | ENSMUSG00000115380 | 0.335414862 | -1.779047242 | 4.030804179 | -0.441362855 | 0.658950329 | NA | Gm46453 | predicted gene, 46453 [Source:MGI Symb | 0 | 0 | 2.0125 | 0 | 0 | 0 |
| 52935 | ENSMUSG00000115381 | 0.849399285 | 3.303488339 | 2.515401219 | 1.313304738 | 0.189080283 | NA | Gm49266 | predicted gene, 49266 [Source:MGI Symb | 0 | 0 | 0 | 2.8919 | 1.0906 | 1.1138 |
| 52936 | ENSMUSG00000115382 | 0 | NA | NA | NA | NA | NA | Gm46475 | predicted gene, 46475 [Source:MGI Symb | 0 | 0 | 0 | 0 | 0 | 0 |
| 52937 | ENSMUSG00000115383 | 0.649696354 | 1.537929172 | 3.118369374 | 0.493183773 | 0.621882743 | NA | Gm49216 | predicted gene, 49216 [Source:MGI Symb | 0 | 0 | 1.0062 | 2.8919 | 0 | 0 |
| 52938 | ENSMUSG00000115384 | 0.321325949 | 1.911575114 | 4.048267893 | 0.472195804 | 0.636787034 | NA | Gm49205 | predicted gene, 49205 [Source:MGI Symb | 0 | 0 | 0 | 1.928 | 0 | 0 |
| 52939 | ENSMUSG00000115385 | 0 | NA | NA | NA | NA | NA | Gm19311 | predicted gene, 19311 [Source:MGI Symb | 0 | 0 | 0 | 0 | 0 | 0 |
| 52940 | ENSMUSG00000115386 | 0 | NA | NA | NA | NA | NA | Gm35909 | predicted gene, 35909 [Source:MGI Symb | 0 | 0 | 0 | 0 | 0 | 0 |
| 52941 | ENSMUSG00000115387 | 0.342432689 | 1.990653086 | 3.900281263 | 0.51038706 | 0.609780321 | NA | Gm49244 | predicted gene, 49244 [Source:MGI Symb | 0 | 0 | 0 | 0.964 | 1.0906 | 0 |
| 52942 | ENSMUSG00000115388 | 207.1495561 | -0.247465038 | 0.222700203 | -1.111202574 | 0.266481163 | 0.722551261 | NA | NA | 232.1 | 240.53 | 201.25 | 127.25 | 229.03 | 212.74 |
| 52943 | ENSMUSG00000115389 | 0.328370405 | 0.16514316 | 4.003149536 | 0.041253308 | 0.967093956 | NA | Gm48936 | predicted gene, 48936 [Source:MGI Symb | 0 | 0 | 1.0062 | 0.964 | 0 | 0 |
| 52944 | ENSMUSG00000115390 | 0 | NA | NA | NA | NA | NA | Gm49035 | predicted gene, 49035 [Source:MGI Symb | 0 | 0 | 0 | 0 | 0 | 0 |
| 52945 | ENSMUSG00000115391 | 1.144113709 | 0.823834277 | 2.553104362 | 0.322679437 | 0.746938027 | NA | Gm49123 | predicted gene, 49123 [Source:MGI Symb | 2.4093 | 0 | 0 | 0 | 0 | 4.4554 |
| 52946 | ENSMUSG00000115392 | 0 | NA | NA | NA | NA | NA | Gm17882 | predicted gene, 17882 [Source:MGI Symb | 0 | 0 | 0 | 0 | 0 | 0 |
| 52947 | ENSMUSG00000115393 | 0 | NA | NA | NA | NA | NA | Gm49052 | predicted gene, 49052 [Source:MGI Symb | 0 | 0 | 0 | 0 | 0 | 0 |
| 52948 | ENSMUSG00000115394 | 0 | NA | NA | NA | NA | NA | Gm49078 | predicted gene, 49078 [Source:MGI Symb | 0 | 0 | 0 | 0 | 0 | 0 |
| 52949 | ENSMUSG00000115395 | 0 | NA | NA | NA | NA | NA | Gm49224 | predicted gene, 49224 [Source:MGI Symb | 0 | 0 | 0 | 0 | 0 | 0 |
| 52950 | ENSMUSG00000115396 | 0 | NA | NA | NA | NA | NA | Gm48994 | predicted gene, 48994 [Source:MGI Symb | 0 | 0 | 0 | 0 | 0 | 0 |
| 52951 | ENSMUSG00000115397 | 0 | NA | NA | NA | NA | NA | Gm49232 | predicted gene, 49232 [Source:MGI Symb | 0 | 0 | 0 | 0 | 0 | 0 |
| 52952 | ENSMUSG00000115398 | 0 | NA | NA | NA | NA | NA | Gm534 | predicted gene 534 [Source:MGI Symbol;A | 0 | 0 | 0 | 0 | 0 | 0 |
| 52953 | ENSMUSG00000115399 | 0 | NA | NA | NA | NA | NA | 9430068D22 | RIKEN cDNA 9430068D22 gene [Source:M | 0 | 0 | 0 | 0 | 0 | 0 |
| 52954 | ENSMUSG00000115400 | 0.133850373 | -0.796629176 | 4.080472857 | -0.195229623 | 0.84521319 | NA | Gm49137 | predicted gene, 49137 [Source:MGI Symb | 0.8031 | 0 | 0 | 0 | 0 | 0 |
| 52955 | ENSMUSG00000115401 | 0 | NA | NA | NA | NA | NA | Gm49157 | predicted gene, 49157 [Source:MGI Symb | 0 | 0 | 0 | 0 | 0 | 0 |
| 52956 | ENSMUSG00000115402 | 0 | NA | NA | NA | NA | NA | Gm49243 | predicted gene, 49243 [Source:MGI Symb | 0 | 0 | 0 | 0 | 0 | 0 |
| 52957 | ENSMUSG00000115403 | 0 | NA | NA | NA | NA | NA | Gm49304 | predicted gene, 49304 [Source:MGI Symb | 0 | 0 | 0 | 0 | 0 | 0 |
| 52958 | ENSMUSG00000115404 | 0 | NA | NA | NA | NA | NA | Vmn1r31 | vomeronasal 1 receptor 31 [Source:MGI S | 0 | 0 | 0 | 0 | 0 | 0 |
| 52959 | ENSMUSG00000115405 | 0 | NA | NA | NA | NA | NA | Gm41307 | predicted gene, 41307 [Source:MGI Symb | 0 | 0 | 0 | 0 | 0 | 0 |
| 52960 | ENSMUSG00000115406 | 0.305170013 | -1.657836477 | 4.041480335 | -0.410205257 | 0.681655385 | NA | Gm49300 | predicted gene, 49300 [Source:MGI Symb | 0.8031 | 1.0279 | 0 | 0 | 0 | 0 |
| 52961 | ENSMUSG00000115407 | 0 | NA | NA | NA | NA | NA | Gm2140 | predicted gene 2140 [Source:MGI Symbol | 0 | 0 | 0 | 0 | 0 | 0 |
| 52962 | ENSMUSG00000115408 | 0 | NA | NA | NA | NA | NA | Gm4529 | predicted gene 4529 [Source:MGI Symbol | 0 | 0 | 0 | 0 | 0 | 0 |
| 52963 | ENSMUSG00000115409 | 0 | NA | NA | NA | NA | NA | Gm8139 | predicted gene 8139 [Source:MGI Symbol | 0 | 0 | 0 | 0 | 0 | 0 |
| 52964 | ENSMUSG00000115410 | 0 | NA | NA | NA | NA | NA | 2810457G06 | RIKEN cDNA 2810457G06 gene [Source:M | 0 | 0 | 0 | 0 | 0 | 0 |
| 52965 | ENSMUSG00000115411 | 0 | NA | NA | NA | NA | NA | Gm6986 | predicted gene 6986 [Source:MGI Symbol | 0 | 0 | 0 | 0 | 0 | 0 |
| 52966 | ENSMUSG00000115412 | 0.455176322 | 0.955156269 | 3.546674842 | 0.269310357 | 0.787690862 | NA | Gm49094 | predicted gene, 49094 [Source:MGI Symb | 0.8031 | 0 | 0 | 1.928 | 0 | 0 |
| 52967 | ENSMUSG00000115413 | 0 | NA | NA | NA | NA | NA | Gm49166 | predicted gene, 49166 [Source:MGI Symb | 0 | 0 | 0 | 0 | 0 | 0 |
| 52968 | ENSMUSG00000115414 | 0 | NA | NA | NA | NA | NA | Gm49257 | predicted gene, 49257 [Source:MGI Symb | 0 | 0 | 0 | 0 | 0 | 0 |
| 52969 | ENSMUSG00000115415 | 0 | NA | NA | NA | NA | NA | Gm49074 | predicted gene, 49074 [Source:MGI Symb | 0 | 0 | 0 | 0 | 0 | 0 |
| 52970 | ENSMUSG00000115416 | 0 | NA | NA | NA | NA | NA | Gm7313 | predicted gene 7313 [Source:MGI Symbol | 0 | 0 | 0 | 0 | 0 | 0 |
| 52971 | ENSMUSG00000115417 | 2.708861955 | 0.042578796 | 1.431613145 | 0.029741831 | 0.976272951 | NA | Gm48949 | predicted gene, 48949 [Source:MGI Symb | 0 | 3.0838 | 5.0312 | 4.8199 | 1.0906 | 2.2277 |
| 52972 | ENSMUSG00000115418 | 0.160662974 | 1.126924175 | 4.080472857 | 0.276174898 | 0.782413736 | NA | Gm49296 | predicted gene, 49296 [Source:MGI Symb | 0 | 0 | 0 | 0.964 | 0 | 0 |

| | | | | | | | | | | | | | | | |
|---|---|---|---|---|---|---|---|---|---|---|---|---|---|---|---|
| 52973 | ENSMUSG00000115419 | 0 | NA | NA | NA | NA | NA | Gm49113 | predicted gene, 49113 [Source:MGI Symb | 0 | 0 | 0 | 0 | 0 | 0 |
| 52974 | ENSMUSG00000115420 | 0 | NA | NA | NA | NA | NA | Rmrp | RNA component of mitochondrial RNAase | 0 | 0 | 0 | 0 | 0 | 0 |
| 52975 | ENSMUSG00000115421 | 0 | NA | NA | NA | NA | NA | Gm48917 | predicted gene, 48917 [Source:MGI Symb | 0 | 0 | 0 | 0 | 0 | 0 |
| 52976 | ENSMUSG00000115422 | 0 | NA | NA | NA | NA | NA | 4930452G13 | RIKEN cDNA 4930452G13 gene [Source:M | 0 | 0 | 0 | 0 | 0 | 0 |
| 52977 | ENSMUSG00000115423 | 0 | NA | NA | NA | NA | NA | | novel protein | 0 | 0 | 0 | 0 | 0 | 0 |
| 52978 | ENSMUSG00000115424 | 0 | NA | NA | NA | NA | NA | D130079A08 | RIKEN cDNA D130079A08 gene [Source:M | 0 | 0 | 0 | 0 | 0 | 0 |
| 52979 | ENSMUSG00000115425 | 0 | NA | NA | NA | NA | NA | Gm49305 | predicted gene, 49305 [Source:MGI Symb | 0 | 0 | 0 | 0 | 0 | 0 |
| 52980 | ENSMUSG00000115426 | 0.305170013 | -1.657836477 | 4.041480335 | -0.410205257 | 0.681655385 | NA | Gm19510 | predicted gene, 19510 [Source:MGI Symb | 0.8031 | 1.0279 | 0 | 0 | 0 | 0 |
| 52981 | ENSMUSG00000115427 | 0 | NA | NA | NA | NA | NA | Gm49174 | predicted gene, 49174 [Source:MGI Symb | 0 | 0 | 0 | 0 | 0 | 0 |
| 52982 | ENSMUSG00000115428 | 0 | NA | NA | NA | NA | NA | Gm41290 | predicted gene, 41290 [Source:MGI Symb | 0 | 0 | 0 | 0 | 0 | 0 |
| 52983 | ENSMUSG00000115429 | 0 | NA | NA | NA | NA | NA | Gm49153 | predicted gene, 49153 [Source:MGI Symb | 0 | 0 | 0 | 0 | 0 | 0 |
| 52984 | ENSMUSG00000115430 | 0 | NA | NA | NA | NA | NA | Gm41300 | predicted gene, 41300 [Source:MGI Symb | 0 | 0 | 0 | 0 | 0 | 0 |
| 52985 | ENSMUSG00000115431 | 98.06848309 | -0.173632654 | 0.231203778 | -0.750994018 | 0.452656256 | 0.856769845 | Gm3219 | predicted pseudogene 3219 [Source:MGI | 103.6 | 105.88 | 102.64 | 101.22 | 94.884 | 80.197 |
| 52986 | ENSMUSG00000115432 | 0 | NA | NA | NA | NA | NA | D130009I18I | RIKEN cDNA D130009I18 gene [Source:M( | 0 | 0 | 0 | 0 | 0 | 0 |
| 52987 | ENSMUSG00000115433 | 0.321325949 | 1.911575114 | 4.048267893 | 0.472195804 | 0.636787034 | NA | Gm49267 | predicted gene, 49267 [Source:MGI Symb | 0 | 0 | 0 | 1.928 | 0 | 0 |
| 52988 | ENSMUSG00000115434 | 0 | NA | NA | NA | NA | NA | Gm6479 | predicted gene 6479 [Source:MGI Symbol | 0 | 0 | 0 | 0 | 0 | 0 |
| 52989 | ENSMUSG00000115435 | 0 | NA | NA | NA | NA | NA | Gm18075 | predicted gene, 18075 [Source:MGI Symb | 0 | 0 | 0 | 0 | 0 | 0 |
| 52990 | ENSMUSG00000115436 | 0 | NA | NA | NA | NA | NA | Gm8353 | predicted gene 8353 [Source:MGI Symbol | 0 | 0 | 0 | 0 | 0 | 0 |
| 52991 | ENSMUSG00000115437 | 0.51034671 | -2.381743948 | 3.331081356 | -0.715006238 | 0.474605189 | NA | Gm49314 | predicted gene, 49314 [Source:MGI Symb | 0 | 2.0558 | 1.0062 | 0 | 0 | 0 |
| 52992 | ENSMUSG00000115438 | 0 | NA | NA | NA | NA | NA | Gm49193 | predicted gene, 49193 [Source:MGI Symb | 0 | 0 | 0 | 0 | 0 | 0 |
| 52993 | ENSMUSG00000115439 | 0 | NA | NA | NA | NA | NA | Gm36107 | predicted gene, 36107 [Source:MGI Symb | 0 | 0 | 0 | 0 | 0 | 0 |
| 52994 | ENSMUSG00000115440 | 0 | NA | NA | NA | NA | NA | Gm49091 | predicted gene, 49091 [Source:MGI Symb | 0 | 0 | 0 | 0 | 0 | 0 |
| 52995 | ENSMUSG00000115441 | 0 | NA | NA | NA | NA | NA | Gm34759 | predicted gene, 34759 [Source:MGI Symb | 0 | 0 | 0 | 0 | 0 | 0 |
| 52996 | ENSMUSG00000115443 | 0 | NA | NA | NA | NA | NA | Gm9395 | predicted gene 9395 [Source:MGI Symbol | 0 | 0 | 0 | 0 | 0 | 0 |
| 52997 | ENSMUSG00000115444 | 0 | NA | NA | NA | NA | NA | Gm48963 | predicted gene, 48963 [Source:MGI Symb | 0 | 0 | 0 | 0 | 0 | 0 |
| 52998 | ENSMUSG00000115445 | 0 | NA | NA | NA | NA | NA | Gm49791 | predicted gene, 49791 [Source:MGI Symb | 0 | 0 | 0 | 0 | 0 | 0 |
| 52999 | ENSMUSG00000115446 | 0 | NA | NA | NA | NA | NA | Gm49186 | predicted gene, 49186 [Source:MGI Symb | 0 | 0 | 0 | 0 | 0 | 0 |
| 53000 | ENSMUSG00000115447 | 0 | NA | NA | NA | NA | NA | Gm48964 | predicted gene, 48964 [Source:MGI Symb | 0 | 0 | 0 | 0 | 0 | 0 |
| 53001 | ENSMUSG00000115448 | 0 | NA | NA | NA | NA | NA | Gm21178 | predicted gene, 21178 [Source:MGI Symb | 0 | 0 | 0 | 0 | 0 | 0 |
| 53002 | ENSMUSG00000115449 | 0 | NA | NA | NA | NA | NA | Gm49175 | predicted gene, 49175 [Source:MGI Symb | 0 | 0 | 0 | 0 | 0 | 0 |
| 53003 | ENSMUSG00000115450 | 0 | NA | NA | NA | NA | NA | Gm4821 | predicted gene 4821 [Source:MGI Symbol | 0 | 0 | 0 | 0 | 0 | 0 |
| 53004 | ENSMUSG00000115451 | 0.946860264 | 0.386550567 | 2.714838108 | 0.142384389 | 0.886776392 | NA | Gm49025 | predicted gene, 49025 [Source:MGI Symb | 2.4093 | 0 | 0 | 0 | 3.2719 | 0 |
| 53005 | ENSMUSG00000115452 | 0 | NA | NA | NA | NA | NA | Gm18812 | predicted gene, 18812 [Source:MGI Symb | 0 | 0 | 0 | 0 | 0 | 0 |
| 53006 | ENSMUSG00000115453 | 0 | NA | NA | NA | NA | NA | BC048602 | cDNA sequence BC048602 [Source:MGI Sy | 0 | 0 | 0 | 0 | 0 | 0 |
| 53007 | ENSMUSG00000115454 | 0.481988923 | 2.492401104 | 3.837077151 | 0.649557203 | 0.515978285 | NA | Gm49086 | predicted gene, 49086 [Source:MGI Symb | 0 | 0 | 0 | 2.8919 | 0 | 0 |
| 53008 | ENSMUSG00000115455 | 0 | NA | NA | NA | NA | NA | Gm41219 | predicted gene, 41219 [Source:MGI Symb | 0 | 0 | 0 | 0 | 0 | 0 |
| 53009 | ENSMUSG00000115456 | 1.381343554 | 4.001184002 | 2.130088574 | 1.878412029 | 0.060324821 | NA | Gm35248 | predicted gene, 35248 [Source:MGI Symb | 0 | 0 | 0 | 3.8559 | 1.0906 | 3.3415 |
| 53010 | ENSMUSG00000115457 | 2.852980788 | 0.168888338 | 1.372865987 | 0.123018809 | 0.902092204 | NA | Gm2387 | predicted gene 2387 [Source:MGI Symbol | 4.0155 | 1.0279 | 3.0187 | 5.7839 | 3.2719 | 0 |
| 53011 | ENSMUSG00000115458 | 0.301557804 | -1.64335916 | 4.042814367 | -0.406488899 | 0.684383412 | NA | Gm48931 | predicted gene, 48931 [Source:MGI Symb | 0.8031 | 0 | 1.0062 | 0 | 0 | 0 |
| 53012 | ENSMUSG00000115459 | 0 | NA | NA | NA | NA | NA | Gm49280 | predicted gene, 49280 [Source:MGI Symb | 0 | 0 | 0 | 0 | 0 | 0 |
| 53013 | ENSMUSG00000115460 | 0 | NA | NA | NA | NA | NA | Gm49127 | predicted gene, 49127 [Source:MGI Symb | 0 | 0 | 0 | 0 | 0 | 0 |
| 53014 | ENSMUSG00000115461 | 0 | NA | NA | NA | NA | NA | Gm48988 | predicted gene, 48988 [Source:MGI Symb | 0 | 0 | 0 | 0 | 0 | 0 |
| 53015 | ENSMUSG00000115462 | 0 | NA | NA | NA | NA | NA | Gm49185 | predicted gene, 49185 [Source:MGI Symb | 0 | 0 | 0 | 0 | 0 | 0 |
| 53016 | ENSMUSG00000115463 | 0.160662974 | 1.126924175 | 4.080472857 | 0.276174898 | 0.782413736 | NA | Gm49356 | predicted gene, 49356 [Source:MGI Symb | 0 | 0 | 0 | 0.964 | 0 | 0 |
| 53017 | ENSMUSG00000115464 | 0 | NA | NA | NA | NA | NA | Gm49019 | predicted gene, 49019 [Source:MGI Symb | 0 | 0 | 0 | 0 | 0 | 0 |
| 53018 | ENSMUSG00000115465 | 0 | NA | NA | NA | NA | NA | Cyp2c73-ps | cytochrome P450, family 2, subfamily c, p | 0 | 0 | 0 | 0 | 0 | 0 |
| 53019 | ENSMUSG00000115466 | 0 | NA | NA | NA | NA | NA | Vmn1r3 | vomeronasal 1 receptor 3 [Source:MGI Sy | 0 | 0 | 0 | 0 | 0 | 0 |
| 53020 | ENSMUSG00000115467 | 0 | NA | NA | NA | NA | NA | Vmn1r37 | vomeronasal 1 receptor 37 [Source:MGI S | 0 | 0 | 0 | 0 | 0 | 0 |
| 53021 | ENSMUSG00000115468 | 0.503302254 | -0.837877584 | 3.426698643 | -0.244514523 | 0.806832356 | NA | Gm49217 | predicted gene, 49217 [Source:MGI Symb | 0 | 2.0558 | 0 | 0.964 | 0 | 0 |
| 53022 | ENSMUSG00000115469 | 0 | NA | NA | NA | NA | NA | Gm49227 | predicted gene, 49227 [Source:MGI Symb | 0 | 0 | 0 | 0 | 0 | 0 |
| 53023 | ENSMUSG00000115470 | 0 | NA | NA | NA | NA | NA | Gm18683 | predicted gene, 18683 [Source:MGI Symb | 0 | 0 | 0 | 0 | 0 | 0 |
| 53024 | ENSMUSG00000115471 | 0 | NA | NA | NA | NA | NA | C230086J09I | RIKEN cDNA C230086J09 gene [Source:M( | 0 | 0 | 0 | 0 | 0 | 0 |
| 53025 | ENSMUSG00000115472 | 0 | NA | NA | NA | NA | NA | Gm49070 | predicted gene, 49070 [Source:MGI Symb | 0 | 0 | 0 | 0 | 0 | 0 |
| 53026 | ENSMUSG00000115473 | 0 | NA | NA | NA | NA | NA | Gm49278 | predicted gene, 49278 [Source:MGI Symb | 0 | 0 | 0 | 0 | 0 | 0 |
| 53027 | ENSMUSG00000115474 | 0 | NA | NA | NA | NA | NA | 4930430F21 | RIKEN cDNA 4930430F21 gene [Source:M | 0 | 0 | 0 | 0 | 0 | 0 |
| 53028 | ENSMUSG00000115475 | 0 | NA | NA | NA | NA | NA | Gm49295 | predicted gene, 49295 [Source:MGI Symb | 0 | 0 | 0 | 0 | 0 | 0 |

| | | | | | | | | | | | | | | | |
|---|---|---|---|---|---|---|---|---|---|---|---|---|---|---|---|
| 53029 | ENSMUSG00000115476 | 0 | NA | NA | NA | NA | NA | Gm49047 | predicted gene, 49047 [Source:MGI Symb | 0 | 0 | 0 | 0 | 0 | 0 |
| 53030 | ENSMUSG00000115477 | 0 | NA | NA | NA | NA | NA | Gm20070 | predicted gene, 20070 [Source:MGI Symb | 0 | 0 | 0 | 0 | 0 | 0 |
| 53031 | ENSMUSG00000115478 | 0.167707431 | -0.796629176 | 4.080472857 | -0.195229623 | 0.84521319 | NA | Gm49198 | predicted gene, 49198 [Source:MGI Symb | 0 | 0 | 1.0062 | 0 | 0 | 0 |
| 53032 | ENSMUSG00000115479 | 0 | NA | NA | NA | NA | NA | Gm33203 | predicted gene, 33203 [Source:MGI Symb | 0 | 0 | 0 | 0 | 0 | 0 |
| 53033 | ENSMUSG00000115480 | 0 | NA | NA | NA | NA | NA | Gm49249 | predicted gene, 49249 [Source:MGI Symb | 0 | 0 | 0 | 0 | 0 | 0 |
| 53034 | ENSMUSG00000115481 | 0 | NA | NA | NA | NA | NA | Gm4940 | predicted gene 4940 [Source:MGI Symbol | 0 | 0 | 0 | 0 | 0 | 0 |
| 53035 | ENSMUSG00000115482 | 0 | NA | NA | NA | NA | NA | Vmn1r36 | vomeronasal 1 receptor 36 [Source:MGI S | 0 | 0 | 0 | 0 | 0 | 0 |
| 53036 | ENSMUSG00000115483 | 0 | NA | NA | NA | NA | NA | Gm9732 | predicted gene 9732 [Source:MGI Symbol | 0 | 0 | 0 | 0 | 0 | 0 |
| 53037 | ENSMUSG00000115484 | 0.181769715 | 1.126924175 | 4.080472857 | 0.276174898 | 0.782413736 | NA | Gm33121 | predicted gene, 33121 [Source:MGI Symb | 0 | 0 | 0 | 0 | 1.0906 | 0 |
| 53038 | ENSMUSG00000115485 | 0 | NA | NA | NA | NA | NA | Gm49126 | predicted gene, 49126 [Source:MGI Symb | 0 | 0 | 0 | 0 | 0 | 0 |
| 53039 | ENSMUSG00000115486 | 0 | NA | NA | NA | NA | NA | Gm49017 | predicted gene, 49017 [Source:MGI Symb | 0 | 0 | 0 | 0 | 0 | 0 |
| 53040 | ENSMUSG00000115487 | 0 | NA | NA | NA | NA | NA | Gm49148 | predicted gene, 49148 [Source:MGI Symb | 0 | 0 | 0 | 0 | 0 | 0 |
| 53041 | ENSMUSG00000115488 | 0.17131964 | -0.796629176 | 4.080472857 | -0.195229623 | 0.84521319 | NA | Gm49099 | predicted gene, 49099 [Source:MGI Symb | 0 | 1.0279 | 0 | 0 | 0 | 0 |
| 53042 | ENSMUSG00000115489 | 0 | NA | NA | NA | NA | NA | Gm49129 | predicted gene, 49129 [Source:MGI Symb | 0 | 0 | 0 | 0 | 0 | 0 |
| 53043 | ENSMUSG00000115490 | 0 | NA | NA | NA | NA | NA | 4930404O17 | RIKEN cDNA 4930404O17 gene [Source:M | 0 | 0 | 0 | 0 | 0 | 0 |
| 53044 | ENSMUSG00000115491 | 0.321325949 | 1.911575114 | 4.048267893 | 0.472195804 | 0.636787034 | NA | Siah1-ps2 | siah E3 ubiquitin protein ligase 1, pseudog | 0 | 0 | 0 | 1.928 | 0 | 0 |
| 53045 | ENSMUSG00000115492 | 0 | NA | NA | NA | NA | NA | Gm41277 | predicted gene, 41277 [Source:MGI Symb | 0 | 0 | 0 | 0 | 0 | 0 |
| 53046 | ENSMUSG00000115493 | 0 | NA | NA | NA | NA | NA | Gm5673 | predicted gene 5673 [Source:MGI Symbol | 0 | 0 | 0 | 0 | 0 | 0 |
| 53047 | ENSMUSG00000115494 | 0 | NA | NA | NA | NA | NA | Gm49293 | predicted gene, 49293 [Source:MGI Symb | 0 | 0 | 0 | 0 | 0 | 0 |
| 53048 | ENSMUSG00000115495 | 0 | NA | NA | NA | NA | NA | Gm38541 | predicted gene, 38541 [Source:MGI Symb | 0 | 0 | 0 | 0 | 0 | 0 |
| 53049 | ENSMUSG00000115497 | 0.822793274 | -0.447303882 | 2.407293551 | -0.185811939 | 0.852592242 | NA | Gm49207 | predicted gene, 49207 [Source:MGI Symb | 0.8031 | 2.0558 | 0 | 0.964 | 0 | 1.1138 |
| 53050 | ENSMUSG00000115498 | 0 | NA | NA | NA | NA | NA | Gm49104 | predicted gene, 49104 [Source:MGI Symb | 0 | 0 | 0 | 0 | 0 | 0 |
| 53051 | ENSMUSG00000115499 | 0 | NA | NA | NA | NA | NA | Gm48991 | predicted gene, 48991 [Source:MGI Symb | 0 | 0 | 0 | 0 | 0 | 0 |
| 53052 | ENSMUSG00000115500 | 0 | NA | NA | NA | NA | NA | Gm49147 | predicted gene, 49147 [Source:MGI Symb | 0 | 0 | 0 | 0 | 0 | 0 |
| 53053 | ENSMUSG00000115501 | 0 | NA | NA | NA | NA | NA | Gm48971 | predicted gene, 48971 [Source:MGI Symb | 0 | 0 | 0 | 0 | 0 | 0 |
| 53054 | ENSMUSG00000115502 | 0 | NA | NA | NA | NA | NA | Gm49222 | predicted gene, 49222 [Source:MGI Symb | 0 | 0 | 0 | 0 | 0 | 0 |
| 53055 | ENSMUSG00000115503 | 4.725905672 | -0.074734113 | 0.987104324 | -0.075710451 | 0.939649461 | NA | Gm6392 | predicted gene 6392 [Source:MGI Symbol | 2.4093 | 6.1675 | 6.0375 | 3.8559 | 6.5437 | 3.3415 |
| 53056 | ENSMUSG00000115504 | 0 | NA | NA | NA | NA | NA | Gm48946 | predicted gene, 48946 [Source:MGI Symb | 0 | 0 | 0 | 0 | 0 | 0 |
| 53057 | ENSMUSG00000115505 | 0.160662974 | 1.126924175 | 4.080472857 | 0.276174898 | 0.782413736 | NA | Gm9247 | predicted gene 9247 [Source:MGI Symbol | 0 | 0 | 0 | 0.964 | 0 | 0 |
| 53058 | ENSMUSG00000115506 | 0 | NA | NA | NA | NA | NA | Gm49237 | predicted gene, 49237 [Source:MGI Symb | 0 | 0 | 0 | 0 | 0 | 0 |
| 53059 | ENSMUSG00000115507 | 0 | NA | NA | NA | NA | NA | Vmn1r24 | vomeronasal 1 receptor 24 [Source:MGI S | 0 | 0 | 0 | 0 | 0 | 0 |
| 53060 | ENSMUSG00000115508 | 0.133850373 | -0.796629176 | 4.080472857 | -0.195229623 | 0.84521319 | NA | Gm49786 | predicted gene, 49786 [Source:MGI Symb | 0.8031 | 0 | 0 | 0 | 0 | 0 |
| 53061 | ENSMUSG00000115509 | 3.6438759 | -0.108883105 | 1.092287194 | -0.099683586 | 0.920595532 | NA | Gm49012 | predicted gene, 49012 [Source:MGI Symb | 3.2124 | 5.1396 | 3.0187 | 3.8559 | 2.1812 | 4.4554 |
| 53062 | ENSMUSG00000115510 | 0 | NA | NA | NA | NA | NA | Gm9334 | predicted gene 9334 [Source:MGI Symbol | 0 | 0 | 0 | 0 | 0 | 0 |
| 53063 | ENSMUSG00000115511 | 0 | NA | NA | NA | NA | NA | Gm49072 | predicted gene, 49072 [Source:MGI Symb | 0 | 0 | 0 | 0 | 0 | 0 |
| 53064 | ENSMUSG00000115512 | 0 | NA | NA | NA | NA | NA | Rpl17-ps7 | ribosomal protein L17, pseudogene 7 [Sou | 0 | 0 | 0 | 0 | 0 | 0 |
| 53065 | ENSMUSG00000115513 | 0 | NA | NA | NA | NA | NA | Gm2361 | predicted gene 2361 [Source:MGI Symbol | 0 | 0 | 0 | 0 | 0 | 0 |
| 53066 | ENSMUSG00000115514 | 0 | NA | NA | NA | NA | NA | Gm49009 | predicted gene, 49009 [Source:MGI Symb | 0 | 0 | 0 | 0 | 0 | 0 |
| 53067 | ENSMUSG00000115515 | 0 | NA | NA | NA | NA | NA | Gm5672 | predicted gene 5672 [Source:MGI Symbol | 0 | 0 | 0 | 0 | 0 | 0 |
| 53068 | ENSMUSG00000115516 | 0 | NA | NA | NA | NA | NA | Gm8341 | predicted gene 8341 [Source:MGI Symbol | 0 | 0 | 0 | 0 | 0 | 0 |
| 53069 | ENSMUSG00000115517 | 0 | NA | NA | NA | NA | NA | Gm48954 | predicted gene, 48954 [Source:MGI Symb | 0 | 0 | 0 | 0 | 0 | 0 |
| 53070 | ENSMUSG00000115518 | 1.391715992 | -1.483605466 | 2.084881866 | -0.711601693 | 0.476711458 | NA | Gm10791 | predicted gene 10791 [Source:MGI Symbo | 0 | 5.1396 | 1.0062 | 0 | 1.0906 | 1.1138 |
| 53071 | ENSMUSG00000115519 | 0 | NA | NA | NA | NA | NA | Gm48955 | predicted gene, 48955 [Source:MGI Symb | 0 | 0 | 0 | 0 | 0 | 0 |
| 53072 | ENSMUSG00000115520 | 0 | NA | NA | NA | NA | NA | Gm41335 | predicted gene, 41335 [Source:MGI Symb | 0 | 0 | 0 | 0 | 0 | 0 |
| 53073 | ENSMUSG00000115521 | 0 | NA | NA | NA | NA | NA | Gm34198 | predicted gene, 34198 [Source:MGI Symb | 0 | 0 | 0 | 0 | 0 | 0 |
| 53074 | ENSMUSG00000115522 | 0 | NA | NA | NA | NA | NA | Gm49173 | predicted gene, 49173 [Source:MGI Symb | 0 | 0 | 0 | 0 | 0 | 0 |
| 53075 | ENSMUSG00000115523 | 0 | NA | NA | NA | NA | NA | Gm6340 | predicted pseudogene 6340 [Source:MGI | 0 | 0 | 0 | 0 | 0 | 0 |
| 53076 | ENSMUSG00000115524 | 0 | NA | NA | NA | NA | NA | Gm49122 | predicted gene, 49122 [Source:MGI Symb | 0 | 0 | 0 | 0 | 0 | 0 |
| 53077 | ENSMUSG00000115525 | 0 | NA | NA | NA | NA | NA | Gm5213 | predicted gene 5213 [Source:MGI Symbol | 0 | 0 | 0 | 0 | 0 | 0 |
| 53078 | ENSMUSG00000115526 | 0 | NA | NA | NA | NA | NA | 4930413F20 | RIKEN cDNA 4930413F20 gene [Source:M | 0 | 0 | 0 | 0 | 0 | 0 |
| 53079 | ENSMUSG00000115527 | 0 | NA | NA | NA | NA | NA | Gm4355 | predicted gene 4355 [Source:MGI Symbol | 0 | 0 | 0 | 0 | 0 | 0 |
| 53080 | ENSMUSG00000115528 | 0 | NA | NA | NA | NA | NA | Gm8408 | predicted gene 8408 [Source:MGI Symbol | 0 | 0 | 0 | 0 | 0 | 0 |
| 53081 | ENSMUSG00000115529 | 0 | NA | NA | NA | NA | NA | 9630013A20 | RIKEN cDNA 9630013A20 gene [Source:M | 0 | 0 | 0 | 0 | 0 | 0 |
| 53082 | ENSMUSG00000115530 | 0 | NA | NA | NA | NA | NA | Gm4266 | predicted gene 4266 [Source:MGI Symbol | 0 | 0 | 0 | 0 | 0 | 0 |
| 53083 | ENSMUSG00000115531 | 0 | NA | NA | NA | NA | NA | Gm35419 | predicted gene, 35419 [Source:MGI Symb | 0 | 0 | 0 | 0 | 0 | 0 |
| 53084 | ENSMUSG00000115532 | 0 | NA | NA | NA | NA | NA | Gm41206 | predicted gene, 41206 [Source:MGI Symb | 0 | 0 | 0 | 0 | 0 | 0 |

| | | | | | | | | | | | | | | | |
|---|---|---|---|---|---|---|---|---|---|---|---|---|---|---|---|
| 53085 | ENSMUSG00000115533 | 0 | NA | NA | NA | NA | NA | Gm48929 | predicted gene, 48929 [Source:MGI Symb | 0 | 0 | 0 | 0 | 0 | 0 |
| 53086 | ENSMUSG00000115534 | 0 | NA | NA | NA | NA | NA | Gm7508 | predicted gene 7508 [Source:MGI Symbol | 0 | 0 | 0 | 0 | 0 | 0 |
| 53087 | ENSMUSG00000115535 | 0 | NA | NA | NA | NA | NA | Gm49006 | predicted gene, 49006 [Source:MGI Symb | 0 | 0 | 0 | 0 | 0 | 0 |
| 53088 | ENSMUSG00000115536 | 0 | NA | NA | NA | NA | NA | Gm32857 | predicted gene, 32857 [Source:MGI Symb | 0 | 0 | 0 | 0 | 0 | 0 |
| 53089 | ENSMUSG00000115537 | 0 | NA | NA | NA | NA | NA | Gm6704 | predicted gene 6704 [Source:MGI Symbol | 0 | 0 | 0 | 0 | 0 | 0 |
| 53090 | ENSMUSG00000115538 | 0 | NA | NA | NA | NA | NA | Hspd1-ps5 | heat shock protein 1 (chaperonin), pseud | 0 | 0 | 0 | 0 | 0 | 0 |
| 53091 | ENSMUSG00000115539 | 1.315438862 | 0.075736522 | 2.01805652 | 0.037529435 | 0.970062871 | NA | Gm6997 | predicted gene 6997 [Source:MGI Symbol | 0.8031 | 3.0838 | 0 | 2.8919 | 0 | 1.1138 |
| 53092 | ENSMUSG00000115540 | 0 | NA | NA | NA | NA | NA | 4930597G03 | RIKEN cDNA 4930597G03 gene [Source:M | 0 | 0 | 0 | 0 | 0 | 0 |
| 53093 | ENSMUSG00000115541 | 0 | NA | NA | NA | NA | NA | Gm49258 | predicted gene, 49258 [Source:MGI Symb | 0 | 0 | 0 | 0 | 0 | 0 |
| 53094 | ENSMUSG00000115542 | 0 | NA | NA | NA | NA | NA | Gm49046 | predicted gene, 49046 [Source:MGI Symb | 0 | 0 | 0 | 0 | 0 | 0 |
| 53095 | ENSMUSG00000115543 | 0.160662974 | 1.126924175 | 4.080472857 | 0.276174898 | 0.782413736 | NA | B230362B09 | RIKEN cDNA B230362B09 gene [Source:M | 0 | 0 | 0 | 0.964 | 0 | 0 |
| 53096 | ENSMUSG00000115544 | 0 | NA | NA | NA | NA | NA | Gm49281 | predicted gene, 49281 [Source:MGI Symb | 0 | 0 | 0 | 0 | 0 | 0 |
| 53097 | ENSMUSG00000115545 | 7.132644977 | -0.051402955 | 0.962310951 | -0.053416159 | 0.95740033 | 0.991861398 | Gm48908 | predicted gene 48908 [Source:MGI Symbo | 13.653 | 2.0558 | 6.0375 | 14.46 | 4.3625 | 2.2277 |
| 53098 | ENSMUSG00000115546 | 0.767390791 | -1.69152783 | 2.44773922 | -0.691057207 | 0.489529591 | NA | Gm49077 | predicted gene, 49077 [Source:MGI Symb | 1.6062 | 1.0279 | 1.0062 | 0.964 | 0 | 0 |
| 53099 | ENSMUSG00000115547 | 0.167707431 | -0.796629176 | 4.080472857 | -0.195229623 | 0.84521319 | NA | 1100001I12F | RIKEN cDNA 1100001I12 gene [Source:MG | 0 | 0 | 1.0062 | 0 | 0 | 0 |
| 53100 | ENSMUSG00000115548 | 0 | NA | NA | NA | NA | NA | 5430440P10 | RIKEN cDNA 5430440P10 gene [Source:M | 0 | 0 | 0 | 0 | 0 | 0 |
| 53101 | ENSMUSG00000115549 | 0 | NA | NA | NA | NA | NA | Gm2203 | predicted gene 2203 [Source:MGI Symbol | 0 | 0 | 0 | 0 | 0 | 0 |
| 53102 | ENSMUSG00000115550 | 0 | NA | NA | NA | NA | NA | Gm49225 | predicted gene, 49225 [Source:MGI Symb | 0 | 0 | 0 | 0 | 0 | 0 |
| 53103 | ENSMUSG00000115551 | 0 | NA | NA | NA | NA | NA | Gm49115 | predicted gene, 49115 [Source:MGI Symb | 0 | 0 | 0 | 0 | 0 | 0 |
| 53104 | ENSMUSG00000115552 | 0 | NA | NA | NA | NA | NA | Gm48959 | predicted gene, 48959 [Source:MGI Symb | 0 | 0 | 0 | 0 | 0 | 0 |
| 53105 | ENSMUSG00000115553 | 0 | NA | NA | NA | NA | NA | Gm49124 | predicted gene, 49124 [Source:MGI Symb | 0 | 0 | 0 | 0 | 0 | 0 |
| 53106 | ENSMUSG00000115554 | 0 | NA | NA | NA | NA | NA | Gm19111 | predicted gene, 19111 [Source:MGI Symb | 0 | 0 | 0 | 0 | 0 | 0 |
| 53107 | ENSMUSG00000115555 | 0 | NA | NA | NA | NA | NA | Gm49307 | predicted gene, 49307 [Source:MGI Symb | 0 | 0 | 0 | 0 | 0 | 0 |
| 53108 | ENSMUSG00000115556 | 0 | NA | NA | NA | NA | NA | Gm49133 | predicted gene, 49133 [Source:MGI Symb | 0 | 0 | 0 | 0 | 0 | 0 |
| 53109 | ENSMUSG00000115557 | 0 | NA | NA | NA | NA | NA | Gm40318 | predicted gene, 40318 [Source:MGI Symb | 0 | 0 | 0 | 0 | 0 | 0 |
| 53110 | ENSMUSG00000115558 | 0 | NA | NA | NA | NA | NA | Gm49000 | predicted gene, 49000 [Source:MGI Symb | 0 | 0 | 0 | 0 | 0 | 0 |
| 53111 | ENSMUSG00000115559 | 0 | NA | NA | NA | NA | NA | Gm49170 | predicted gene, 49170 [Source:MGI Symb | 0 | 0 | 0 | 0 | 0 | 0 |
| 53112 | ENSMUSG00000115560 | 0.17131964 | -0.796629176 | 4.080472857 | -0.195229623 | 0.84521319 | NA | Gm49191 | predicted gene, 49191 [Source:MGI Symb | 0 | 1.0279 | 0 | 0 | 0 | 0 |
| 53113 | ENSMUSG00000115561 | 0 | NA | NA | NA | NA | NA | Gm49285 | predicted gene, 49285 [Source:MGI Symb | 0 | 0 | 0 | 0 | 0 | 0 |
| 53114 | ENSMUSG00000115562 | 0 | NA | NA | NA | NA | NA | Gm48977 | predicted gene, 48977 [Source:MGI Symb | 0 | 0 | 0 | 0 | 0 | 0 |
| 53115 | ENSMUSG00000115563 | 0 | NA | NA | NA | NA | NA | Gm34562 | predicted gene, 34562 [Source:MGI Symb | 0 | 0 | 0 | 0 | 0 | 0 |
| 53116 | ENSMUSG00000115564 | 0 | NA | NA | NA | NA | NA | Gm49199 | predicted gene, 49199 [Source:MGI Symb | 0 | 0 | 0 | 0 | 0 | 0 |
| 53117 | ENSMUSG00000115565 | 0 | NA | NA | NA | NA | NA | Gm18082 | predicted gene, 18082 [Source:MGI Symb | 0 | 0 | 0 | 0 | 0 | 0 |
| 53118 | ENSMUSG00000115566 | 0 | NA | NA | NA | NA | NA | Gm49136 | predicted gene, 49136 [Source:MGI Symb | 0 | 0 | 0 | 0 | 0 | 0 |
| 53119 | ENSMUSG00000115567 | 0 | NA | NA | NA | NA | NA | Gm18148 | predicted gene, 18148 [Source:MGI Symb | 0 | 0 | 0 | 0 | 0 | 0 |
| 53120 | ENSMUSG00000115568 | 0.160662974 | 1.126924175 | 4.080472857 | 0.276174898 | 0.782413736 | NA | Tiaf2 | TGF-beta1-induced anti-apoptotic factor 2 | 0 | 0 | 0 | 0.964 | 0 | 0 |
| 53121 | ENSMUSG00000115569 | 0 | NA | NA | NA | NA | NA | Gm49169 | predicted gene, 49169 [Source:MGI Symb | 0 | 0 | 0 | 0 | 0 | 0 |
| 53122 | ENSMUSG00000115570 | 0 | NA | NA | NA | NA | NA | Gm48986 | predicted gene, 48986 [Source:MGI Symb | 0 | 0 | 0 | 0 | 0 | 0 |
| 53123 | ENSMUSG00000115571 | 0 | NA | NA | NA | NA | NA | Gm6616 | predicted gene 6616 [Source:MGI Symbol | 0 | 0 | 0 | 0 | 0 | 0 |
| 53124 | ENSMUSG00000115572 | 0 | NA | NA | NA | NA | NA | Gm18369 | predicted gene, 18369 [Source:MGI Symb | 0 | 0 | 0 | 0 | 0 | 0 |
| 53125 | ENSMUSG00000115573 | 0.633540418 | -1.320719359 | 2.623870149 | -0.503347835 | 0.61471974 | NA | Gm49131 | predicted gene, 49131 [Source:MGI Symb | 0.8031 | 1.0279 | 1.0062 | 0.964 | 0 | 0 |
| 53126 | ENSMUSG00000115574 | 0 | NA | NA | NA | NA | NA | Gm49245 | predicted gene, 49245 [Source:MGI Symb | 0 | 0 | 0 | 0 | 0 | 0 |
| 53127 | ENSMUSG00000115575 | 12.71008275 | -0.368099053 | 0.610556608 | -0.602890949 | 0.546581237 | 0.89549457 | Gm49024 | predicted gene, 49024 [Source:MGI Symb | 18.471 | 14.391 | 10.062 | 13.496 | 9.8156 | 10.025 |
| 53128 | ENSMUSG00000115576 | 0 | NA | NA | NA | NA | NA | Gm2675 | predicted gene 2675 [Source:MGI Symbol | 0 | 0 | 0 | 0 | 0 | 0 |
| 53129 | ENSMUSG00000115577 | 0 | NA | NA | NA | NA | NA | Gm49250 | predicted gene, 49250 [Source:MGI Symb | 0 | 0 | 0 | 0 | 0 | 0 |
| 53130 | ENSMUSG00000115578 | 0 | NA | NA | NA | NA | NA | Gm46506 | predicted gene, 46506 [Source:MGI Symb | 0 | 0 | 0 | 0 | 0 | 0 |
| 53131 | ENSMUSG00000115579 | 0 | NA | NA | NA | NA | NA | Gm49156 | predicted gene, 49156 [Source:MGI Symb | 0 | 0 | 0 | 0 | 0 | 0 |
| 53132 | ENSMUSG00000115580 | 0 | NA | NA | NA | NA | NA | Gm41230 | predicted gene, 41230 [Source:MGI Symb | 0 | 0 | 0 | 0 | 0 | 0 |
| 53133 | ENSMUSG00000115581 | 0 | NA | NA | NA | NA | NA | Gm20127 | predicted gene, 20127 [Source:MGI Symb | 0 | 0 | 0 | 0 | 0 | 0 |
| 53134 | ENSMUSG00000115582 | 0 | NA | NA | NA | NA | NA | Gm48953 | predicted gene, 48953 [Source:MGI Symb | 0 | 0 | 0 | 0 | 0 | 0 |
| 53135 | ENSMUSG00000115584 | 0.850737881 | 0.833803351 | 2.386603902 | 0.349368133 | 0.726812954 | NA | Gm5854 | predicted gene 5854 [Source:MGI Symbol | 0.8031 | 0 | 1.0062 | 0 | 2.1812 | 1.1138 |
| 53136 | ENSMUSG00000115585 | 0 | NA | NA | NA | NA | NA | Gm41271 | predicted gene, 41271 [Source:MGI Symb | 0 | 0 | 0 | 0 | 0 | 0 |
| 53137 | ENSMUSG00000115586 | 0 | NA | NA | NA | NA | NA | Gm48918 | predicted gene, 48918 [Source:MGI Symb | 0 | 0 | 0 | 0 | 0 | 0 |
| 53138 | ENSMUSG00000115587 | 1.038213457 | 2.372140906 | 2.221591093 | 1.067766662 | 0.28562578 | NA | Gm29697 | predicted gene, 29697 [Source:MGI Symb | 0 | 0 | 1.0062 | 1.928 | 2.1812 | 1.1138 |
| 53139 | ENSMUSG00000115588 | 0 | NA | NA | NA | NA | NA | Gm18533 | predicted gene, 18533 [Source:MGI Symb | 0 | 0 | 0 | 0 | 0 | 0 |
| 53140 | ENSMUSG00000115589 | 0 | NA | NA | NA | NA | NA | Gm18003 | predicted gene, 18003 [Source:MGI Symb | 0 | 0 | 0 | 0 | 0 | 0 |

| | | | | | | | | | | | | | | | |
|---|---|---|---|---|---|---|---|---|---|---|---|---|---|---|---|
| 53141 | ENSMUSG00000115590 | 0 | NA | NA | NA | NA | NA | Gm49219 | predicted gene, 49219 [Source:MGI Symb | 0 | 0 | 0 | 0 | 0 | 0 |
| 53142 | ENSMUSG00000115591 | 0 | NA | NA | NA | NA | NA | Gm31282 | predicted gene, 31282 [Source:MGI Symb | 0 | 0 | 0 | 0 | 0 | 0 |
| 53143 | ENSMUSG00000115592 | 0 | NA | NA | NA | NA | NA | Mcpt-ps1 | mast cell protease, pseudogene 1 [Source | 0 | 0 | 0 | 0 | 0 | 0 |
| 53144 | ENSMUSG00000115593 | 2.078778761 | -1.057551481 | 1.541866494 | -0.685890435 | 0.492782202 | NA | Gm4824 | predicted gene 4824 [Source:MGI Symbol | 2.4093 | 2.0558 | 4.025 | 2.8919 | 1.0906 | 0 |
| 53145 | ENSMUSG00000115594 | 0 | NA | NA | NA | NA | NA | Gm46497 | predicted gene, 46497 [Source:MGI Symb | 0 | 0 | 0 | 0 | 0 | 0 |
| 53146 | ENSMUSG00000115595 | 0 | NA | NA | NA | NA | NA | Rps12-ps2 | ribosomal protein S12, pseudogene 2 [Sou | 0 | 0 | 0 | 0 | 0 | 0 |
| 53147 | ENSMUSG00000115596 | 0.133850373 | -0.796629176 | 4.080472857 | -0.195229623 | 0.84521319 | NA | Gm48969 | predicted gene, 48969 [Source:MGI Symb | 0.8031 | 0 | 0 | 0 | 0 | 0 |
| 53148 | ENSMUSG00000115597 | 0 | NA | NA | NA | NA | NA | Gm49030 | predicted gene, 49030 [Source:MGI Symb | 0 | 0 | 0 | 0 | 0 | 0 |
| 53149 | ENSMUSG00000115598 | 0 | NA | NA | NA | NA | NA | Gm9290 | predicted gene 9290 [Source:MGI Symbol | 0 | 0 | 0 | 0 | 0 | 0 |
| 53150 | ENSMUSG00000115599 | 0 | NA | NA | NA | NA | NA | Gm49154 | predicted gene, 49154 [Source:MGI Symb | 0 | 0 | 0 | 0 | 0 | 0 |
| 53151 | ENSMUSG00000115600 | 0 | NA | NA | NA | NA | NA | Gm49299 | predicted gene, 49299 [Source:MGI Symb | 0 | 0 | 0 | 0 | 0 | 0 |
| 53152 | ENSMUSG00000115601 | 0 | NA | NA | NA | NA | NA | Gm49007 | predicted gene, 49007 [Source:MGI Symb | 0 | 0 | 0 | 0 | 0 | 0 |
| 53153 | ENSMUSG00000115602 | 1.788600534 | 2.413949509 | 2.016424207 | 1.197143687 | 0.231250555 | NA | Gm7908 | predicted gene 7908 [Source:MGI Symbol | 1.6062 | 0 | 0 | 5.7839 | 0 | 3.3415 |
| 53154 | ENSMUSG00000115603 | 0 | NA | NA | NA | NA | NA | Gm19099 | predicted gene, 19099 [Source:MGI Symb | 0 | 0 | 0 | 0 | 0 | 0 |
| 53155 | ENSMUSG00000115604 | 0 | NA | NA | NA | NA | NA | Gm7691 | predicted gene 7691 [Source:MGI Symbol | 0 | 0 | 0 | 0 | 0 | 0 |
| 53156 | ENSMUSG00000115605 | 0 | NA | NA | NA | NA | NA | Gm9643 | predicted gene 9643 [Source:MGI Symbol | 0 | 0 | 0 | 0 | 0 | 0 |
| 53157 | ENSMUSG00000115606 | 0 | NA | NA | NA | NA | NA | Gm17932 | predicted gene, 17932 [Source:MGI Symb | 0 | 0 | 0 | 0 | 0 | 0 |
| 53158 | ENSMUSG00000115607 | 0 | NA | NA | NA | NA | NA | Gm2803 | predicted gene 2803 [Source:MGI Symbol | 0 | 0 | 0 | 0 | 0 | 0 |
| 53159 | ENSMUSG00000115608 | 0 | NA | NA | NA | NA | NA | Gm48922 | predicted gene, 48922 [Source:MGI Symb | 0 | 0 | 0 | 0 | 0 | 0 |
| 53160 | ENSMUSG00000115609 | 0 | NA | NA | NA | NA | NA | 4930447J18I | RIKEN cDNA 4930447J18 gene [Source:M( | 0 | 0 | 0 | 0 | 0 | 0 |
| 53161 | ENSMUSG00000115610 | 0 | NA | NA | NA | NA | NA | Gm31251 | predicted gene, 31251 [Source:MGI Symb | 0 | 0 | 0 | 0 | 0 | 0 |
| 53162 | ENSMUSG00000115611 | 0 | NA | NA | NA | NA | NA | Gm4606 | predicted gene 4606 [Source:MGI Symbol | 0 | 0 | 0 | 0 | 0 | 0 |
| 53163 | ENSMUSG00000115612 | 0 | NA | NA | NA | NA | NA | Gm29800 | predicted gene, 29800 [Source:MGI Symb | 0 | 0 | 0 | 0 | 0 | 0 |
| 53164 | ENSMUSG00000115613 | 0 | NA | NA | NA | NA | NA | Gm48982 | predicted gene, 48982 [Source:MGI Symb | 0 | 0 | 0 | 0 | 0 | 0 |
| 53165 | ENSMUSG00000115614 | 0 | NA | NA | NA | NA | NA | Gm41276 | predicted gene, 41276 [Source:MGI Symb | 0 | 0 | 0 | 0 | 0 | 0 |
| 53166 | ENSMUSG00000115615 | 0.747854741 | -0.257859511 | 2.698313412 | -0.095563217 | 0.92386748 | NA | Gm49200 | predicted gene, 49200 [Source:MGI Symb | 2.4093 | 0 | 0 | 0.964 | 0 | 1.1138 |
| 53167 | ENSMUSG00000115616 | 0 | NA | NA | NA | NA | NA | Gm48923 | predicted gene, 48923 [Source:MGI Symb | 0 | 0 | 0 | 0 | 0 | 0 |
| 53168 | ENSMUSG00000115617 | 0 | NA | NA | NA | NA | NA | Gm48975 | predicted gene, 48975 [Source:MGI Symb | 0 | 0 | 0 | 0 | 0 | 0 |
| 53169 | ENSMUSG00000115618 | 0 | NA | NA | NA | NA | NA | Gm49298 | predicted gene, 49298 [Source:MGI Symb | 0 | 0 | 0 | 0 | 0 | 0 |
| 53170 | ENSMUSG00000115619 | 0 | NA | NA | NA | NA | NA | Gm49082 | predicted gene, 49082 [Source:MGI Symb | 0 | 0 | 0 | 0 | 0 | 0 |
| 53171 | ENSMUSG00000115620 | 0 | NA | NA | NA | NA | NA | Gm45924 | predicted gene, 45924 [Source:MGI Symb | 0 | 0 | 0 | 0 | 0 | 0 |
| 53172 | ENSMUSG00000115621 | 0 | NA | NA | NA | NA | NA | Gm8664 | predicted gene 8664 [Source:MGI Symbol | 0 | 0 | 0 | 0 | 0 | 0 |
| 53173 | ENSMUSG00000115622 | 0 | NA | NA | NA | NA | NA | Gm49117 | predicted gene, 49117 [Source:MGI Symb | 0 | 0 | 0 | 0 | 0 | 0 |
| 53174 | ENSMUSG00000115623 | 0 | NA | NA | NA | NA | NA | Gm41225 | predicted gene, 41225 [Source:MGI Symb | 0 | 0 | 0 | 0 | 0 | 0 |
| 53175 | ENSMUSG00000115624 | 15.29291793 | -0.028437413 | 0.608471725 | -0.046735801 | 0.962723796 | 0.992956886 | Gm49204 | predicted gene, 49204 [Source:MGI Symb | 12.85 | 23.642 | 10.062 | 23.135 | 9.8156 | 12.252 |
| 53176 | ENSMUSG00000115625 | 0 | NA | NA | NA | NA | NA | 2900040C04 | RIKEN cDNA 2900040C04 gene [Source:M | 0 | 0 | 0 | 0 | 0 | 0 |
| 53177 | ENSMUSG00000115626 | 0 | NA | NA | NA | NA | NA | Gm17772 | predicted gene, 17772 [Source:MGI Symb | 0 | 0 | 0 | 0 | 0 | 0 |
| 53178 | ENSMUSG00000115627 | 0 | NA | NA | NA | NA | NA | Gm41168 | predicted gene, 41168 [Source:MGI Symb | 0 | 0 | 0 | 0 | 0 | 0 |
| 53179 | ENSMUSG00000115628 | 0 | NA | NA | NA | NA | NA | Gm49084 | predicted gene, 49084 [Source:MGI Symb | 0 | 0 | 0 | 0 | 0 | 0 |
| 53180 | ENSMUSG00000115629 | 0 | NA | NA | NA | NA | NA | Gm49235 | predicted gene, 49235 [Source:MGI Symb | 0 | 0 | 0 | 0 | 0 | 0 |
| 53181 | ENSMUSG00000115630 | 0 | NA | NA | NA | NA | NA | Rpl19-ps5 | ribosomal protein L19, pseudogene 5 [Sou | 0 | 0 | 0 | 0 | 0 | 0 |
| 53182 | ENSMUSG00000115631 | 0 | NA | NA | NA | NA | NA | Gm19371 | predicted gene, 19371 [Source:MGI Symb | 0 | 0 | 0 | 0 | 0 | 0 |
| 53183 | ENSMUSG00000115632 | 0 | NA | NA | NA | NA | NA | Gm17922 | predicted gene, 17922 [Source:MGI Symb | 0 | 0 | 0 | 0 | 0 | 0 |
| 53184 | ENSMUSG00000115633 | 0 | NA | NA | NA | NA | NA | Gm29753 | predicted gene, 29753 [Source:MGI Symb | 0 | 0 | 0 | 0 | 0 | 0 |
| 53185 | ENSMUSG00000115635 | 0 | NA | NA | NA | NA | NA | Gm49271 | predicted gene, 49271 [Source:MGI Symb | 0 | 0 | 0 | 0 | 0 | 0 |
| 53186 | ENSMUSG00000115636 | 0 | NA | NA | NA | NA | NA | Gm49120 | predicted gene, 49120 [Source:MGI Symb | 0 | 0 | 0 | 0 | 0 | 0 |
| 53187 | ENSMUSG00000115637 | 0.17131964 | -0.796629176 | 4.080472857 | -0.195229623 | 0.84521319 | NA | Gm30970 | predicted gene, 30970 [Source:MGI Symb | 0 | 1.0279 | 0 | 0 | 0 | 0 |
| 53188 | ENSMUSG00000115638 | 0 | NA | NA | NA | NA | NA | Gm18074 | predicted gene, 18074 [Source:MGI Symb | 0 | 0 | 0 | 0 | 0 | 0 |
| 53189 | ENSMUSG00000115639 | 0 | NA | NA | NA | NA | NA | 4930404K13 | RIKEN cDNA 4930404K13 gene [Source:M | 0 | 0 | 0 | 0 | 0 | 0 |
| 53190 | ENSMUSG00000115640 | 0 | NA | NA | NA | NA | NA | Gm49247 | predicted gene, 49247 [Source:MGI Symb | 0 | 0 | 0 | 0 | 0 | 0 |
| 53191 | ENSMUSG00000115641 | 0.160662974 | 1.126924175 | 4.080472857 | 0.276174898 | 0.782413736 | NA | Gm9891 | predicted gene 9891 [Source:MGI Symbol | 0 | 0 | 0 | 0.964 | 0 | 0 |
| 53192 | ENSMUSG00000115642 | 0 | NA | NA | NA | NA | NA | Gm18977 | predicted gene, 18977 [Source:MGI Symb | 0 | 0 | 0 | 0 | 0 | 0 |
| 53193 | ENSMUSG00000115643 | 1.4790954 | -0.918267124 | 1.823178285 | -0.503662824 | 0.614498337 | NA | Gm49011 | predicted gene, 49011 [Source:MGI Symb | 0.8031 | 1.0279 | 4.025 | 1.928 | 1.0906 | 0 |
| 53194 | ENSMUSG00000115644 | 0 | NA | NA | NA | NA | NA | Vmn1r17 | vomeronasal 1 receptor 17 [Source:MGI S | 0 | 0 | 0 | 0 | 0 | 0 |
| 53195 | ENSMUSG00000115645 | 0.301557804 | -1.64335916 | 4.042814367 | -0.406488899 | 0.684383412 | NA | Gm49288 | predicted gene, 49288 [Source:MGI Symb | 0.8031 | 0 | 1.0062 | 0 | 0 | 0 |
| 53196 | ENSMUSG00000115646 | 0 | NA | NA | NA | NA | NA | Gm10926 | predicted gene 10926 [Source:MGI Symbo | 0 | 0 | 0 | 0 | 0 | 0 |

| | | | | | | | | | | | | | | | |
|---|---|---|---|---|---|---|---|---|---|---|---|---|---|---|---|
| 53197 | ENSMUSG00000115647 | 0.167707431 | -0.796629176 | 4.080472857 | -0.195229623 | 0.84521319 | NA | Gm49265 | predicted gene, 49265 [Source:MGI Symb | 0 | 0 | 1.0062 | 0 | 0 | 0 |
| 53198 | ENSMUSG00000115648 | 0 | NA | NA | NA | NA | NA | Gm49008 | predicted gene, 49008 [Source:MGI Symb | 0 | 0 | 0 | 0 | 0 | 0 |
| 53199 | ENSMUSG00000115649 | 2.71693415 | -2.00141214 | 1.614121038 | -1.239939318 | 0.21499784 | NA | Gm18813 | predicted gene, 18813 [Source:MGI Symb | 0.8031 | 7.1954 | 5.0312 | 0 | 3.2719 | 0 |
| 53200 | ENSMUSG00000115650 | 0 | NA | NA | NA | NA | NA | Gm48992 | predicted gene, 48992 [Source:MGI Symb | 0 | 0 | 0 | 0 | 0 | 0 |
| 53201 | ENSMUSG00000115651 | 0 | NA | NA | NA | NA | NA | Gm7595 | predicted gene 7595 [Source:MGI Symbol | 0 | 0 | 0 | 0 | 0 | 0 |
| 53202 | ENSMUSG00000115652 | 0 | NA | NA | NA | NA | NA | Gm18143 | predicted gene, 18143 [Source:MGI Symb | 0 | 0 | 0 | 0 | 0 | 0 |
| 53203 | ENSMUSG00000115653 | 0 | NA | NA | NA | NA | NA | Gm49059 | predicted gene, 49059 [Source:MGI Symb | 0 | 0 | 0 | 0 | 0 | 0 |
| 53204 | ENSMUSG00000115654 | 0 | NA | NA | NA | NA | NA | Gm19303 | predicted gene, 19303 [Source:MGI Symb | 0 | 0 | 0 | 0 | 0 | 0 |
| 53205 | ENSMUSG00000115656 | 0.160662974 | 1.126924175 | 4.080472857 | 0.276174898 | 0.782413736 | NA | 4930556J02I | RIKEN cDNA 4930556J02 gene [Source:M( | 0 | 0 | 0 | 0.964 | 0 | 0 |
| 53206 | ENSMUSG00000115657 | 0 | NA | NA | NA | NA | NA | Gm41325 | predicted gene, 41325 [Source:MGI Symb | 0 | 0 | 0 | 0 | 0 | 0 |
| 53207 | ENSMUSG00000115658 | 0 | NA | NA | NA | NA | NA | Gm19829 | predicted gene, 19829 [Source:MGI Symb | 0 | 0 | 0 | 0 | 0 | 0 |
| 53208 | ENSMUSG00000115659 | 0 | NA | NA | NA | NA | NA | Gm41138 | predicted gene, 41138 [Source:MGI Symb | 0 | 0 | 0 | 0 | 0 | 0 |
| 53209 | ENSMUSG00000115660 | 0 | NA | NA | NA | NA | NA | Gm48962 | predicted gene, 48962 [Source:MGI Symb | 0 | 0 | 0 | 0 | 0 | 0 |
| 53210 | ENSMUSG00000115661 | 0 | NA | NA | NA | NA | NA | Gm48976 | predicted gene, 48976 [Source:MGI Symb | 0 | 0 | 0 | 0 | 0 | 0 |
| 53211 | ENSMUSG00000115662 | 1.768368202 | -3.063643882 | 1.885230976 | -1.625076142 | 0.104146336 | NA | Gm7232 | predicted gene 7232 [Source:MGI Symbol | 2.4093 | 2.0558 | 5.0312 | 0 | 0 | 1.1138 |
| 53212 | ENSMUSG00000115663 | 0 | NA | NA | NA | NA | NA | Gm48999 | predicted gene, 48999 [Source:MGI Symb | 0 | 0 | 0 | 0 | 0 | 0 |
| 53213 | ENSMUSG00000115664 | 0.185640647 | 1.126924175 | 4.080472857 | 0.276174898 | 0.782413736 | NA | Gm32913 | predicted gene, 32913 [Source:MGI Symb | 0 | 0 | 0 | 0 | 0 | 1.1138 |
| 53214 | ENSMUSG00000115665 | 0 | NA | NA | NA | NA | NA | 4933425B07 | RIKEN cDNA 4933425B07 gene [Source:M | 0 | 0 | 0 | 0 | 0 | 0 |
| 53215 | ENSMUSG00000115666 | 0 | NA | NA | NA | NA | NA | Gm49031 | predicted gene, 49031 [Source:MGI Symb | 0 | 0 | 0 | 0 | 0 | 0 |
| 53216 | ENSMUSG00000115667 | 1.86681633 | -0.318764417 | 1.751994108 | -0.181943772 | 0.855626852 | NA | Gm10389 | predicted gene 10389 [Source:MGI Symbo | 3.2124 | 0 | 3.0187 | 3.8559 | 0 | 1.1138 |
| 53217 | ENSMUSG00000115668 | 0 | NA | NA | NA | NA | NA | Vmn1r25 | vomeronasal 1 receptor 25 [Source:MGI S | 0 | 0 | 0 | 0 | 0 | 0 |
| 53218 | ENSMUSG00000115669 | 0 | NA | NA | NA | NA | NA | Gm49151 | predicted gene, 49151 [Source:MGI Symb | 0 | 0 | 0 | 0 | 0 | 0 |
| 53219 | ENSMUSG00000115670 | 0 | NA | NA | NA | NA | NA | Gm2611 | predicted gene 2611 [Source:MGI Symbol | 0 | 0 | 0 | 0 | 0 | 0 |
| 53220 | ENSMUSG00000115671 | 0 | NA | NA | NA | NA | NA | Gm49289 | predicted gene, 49289 [Source:MGI Symb | 0 | 0 | 0 | 0 | 0 | 0 |
| 53221 | ENSMUSG00000115672 | 0 | NA | NA | NA | NA | NA | Gm7814 | predicted gene 7814 [Source:MGI Symbol | 0 | 0 | 0 | 0 | 0 | 0 |
| 53222 | ENSMUSG00000115673 | 0 | NA | NA | NA | NA | NA | Gm18884 | predicted gene, 18884 [Source:MGI Symb | 0 | 0 | 0 | 0 | 0 | 0 |
| 53223 | ENSMUSG00000115674 | 0 | NA | NA | NA | NA | NA | Gm49297 | predicted gene, 49297 [Source:MGI Symb | 0 | 0 | 0 | 0 | 0 | 0 |
| 53224 | ENSMUSG00000115675 | 0 | NA | NA | NA | NA | NA | Gm49080 | predicted gene, 49080 [Source:MGI Symb | 0 | 0 | 0 | 0 | 0 | 0 |
| 53225 | ENSMUSG00000115676 | 0 | NA | NA | NA | NA | NA | 4930449C09 | RIKEN cDNA 4930449C09 gene [Source:M | 0 | 0 | 0 | 0 | 0 | 0 |
| 53226 | ENSMUSG00000115677 | 0 | NA | NA | NA | NA | NA | Gm49241 | predicted gene, 49241 [Source:MGI Symb | 0 | 0 | 0 | 0 | 0 | 0 |
| 53227 | ENSMUSG00000115678 | 0 | NA | NA | NA | NA | NA | Gm49212 | predicted gene, 49212 [Source:MGI Symb | 0 | 0 | 0 | 0 | 0 | 0 |
| 53228 | ENSMUSG00000115679 | 0 | NA | NA | NA | NA | NA | Gm48995 | predicted gene, 48995 [Source:MGI Symb | 0 | 0 | 0 | 0 | 0 | 0 |
| 53229 | ENSMUSG00000115680 | 0 | NA | NA | NA | NA | NA | Gm35167 | predicted gene, 35167 [Source:MGI Symb | 0 | 0 | 0 | 0 | 0 | 0 |
| 53230 | ENSMUSG00000115681 | 0.603115608 | -2.642822926 | 3.106511347 | -0.850736608 | 0.394915683 | NA | Gm2682 | predicted gene 2682 [Source:MGI Symbol | 1.6062 | 0 | 2.0125 | 0 | 0 | 0 |
| 53231 | ENSMUSG00000115682 | 0 | NA | NA | NA | NA | NA | Gm46499 | predicted gene, 46499 [Source:MGI Symb | 0 | 0 | 0 | 0 | 0 | 0 |
| 53232 | ENSMUSG00000115683 | 0 | NA | NA | NA | NA | NA | Gm49021 | predicted gene, 49021 [Source:MGI Symb | 0 | 0 | 0 | 0 | 0 | 0 |
| 53233 | ENSMUSG00000115684 | 0 | NA | NA | NA | NA | NA | Gm6087 | predicted gene 6087 [Source:MGI Symbol | 0 | 0 | 0 | 0 | 0 | 0 |
| 53234 | ENSMUSG00000115685 | 1.327924942 | 0.217872788 | 1.987306221 | 0.109632217 | 0.912701059 | NA | D730044K07 | RIKEN cDNA D730044K07 gene [Source:M | 1.6062 | 2.0558 | 0 | 0.964 | 0 | 3.3415 |
| 53235 | ENSMUSG00000115686 | 0.513958919 | -2.394274821 | 3.756096013 | -0.637437066 | 0.523840191 | NA | Gm49310 | predicted gene, 49310 [Source:MGI Symb | 0 | 3.0838 | 0 | 0 | 0 | 0 |
| 53236 | ENSMUSG00000115687 | 0 | NA | NA | NA | NA | NA | Gm18962 | predicted gene, 18962 [Source:MGI Symb | 0 | 0 | 0 | 0 | 0 | 0 |
| 53237 | ENSMUSG00000115688 | 0.267700746 | -1.497481649 | 4.057001436 | -0.369110456 | 0.712045396 | NA | Gm49209 | predicted gene, 49209 [Source:MGI Symb | 1.6062 | 0 | 0 | 0 | 0 | 0 |
| 53238 | ENSMUSG00000115689 | 0 | NA | NA | NA | NA | NA | 4930435M08 | RIKEN cDNA 4930435M08 gene [Source:M | 0 | 0 | 0 | 0 | 0 | 0 |
| 53239 | ENSMUSG00000115690 | 0 | NA | NA | NA | NA | NA | Gm48924 | predicted gene, 48924 [Source:MGI Symb | 0 | 0 | 0 | 0 | 0 | 0 |
| 53240 | ENSMUSG00000115691 | 1.606107907 | -1.928626566 | 1.879766041 | -1.025992875 | 0.304894942 | NA | Gm48914 | predicted gene, 48914 [Source:MGI Symb | 1.6062 | 3.0838 | 3.0187 | 1.928 | 0 | 0 |
| 53241 | ENSMUSG00000115692 | 6.942580185 | -0.23589366 | 0.795652467 | -0.296478261 | 0.766864866 | 0.955295274 | Gm49163 | predicted gene, 49163 [Source:MGI Symb | 7.2279 | 8.2233 | 7.0437 | 4.8199 | 6.5437 | 7.7969 |
| 53242 | ENSMUSG00000115693 | 0 | NA | NA | NA | NA | NA | 1700085D07 | RIKEN cDNA 1700085D07 gene [Source:M | 0 | 0 | 0 | 0 | 0 | 0 |
| 53243 | ENSMUSG00000115694 | 0 | NA | NA | NA | NA | NA | Gm19235 | predicted gene, 19235 [Source:MGI Symb | 0 | 0 | 0 | 0 | 0 | 0 |
| 53244 | ENSMUSG00000115695 | 0 | NA | NA | NA | NA | NA | Gm48973 | predicted gene, 48973 [Source:MGI Symb | 0 | 0 | 0 | 0 | 0 | 0 |
| 53245 | ENSMUSG00000115696 | 0 | NA | NA | NA | NA | NA | Gm49292 | predicted gene, 49292 [Source:MGI Symb | 0 | 0 | 0 | 0 | 0 | 0 |
| 53246 | ENSMUSG00000115697 | 0 | NA | NA | NA | NA | NA | Vmn1r216 | vomeronasal 1 receptor 216 [Source:MGI | 0 | 0 | 0 | 0 | 0 | 0 |
| 53247 | ENSMUSG00000115698 | 1.748108113 | -0.579769056 | 1.670134211 | -0.347139201 | 0.728486744 | NA | Gm49076 | predicted gene, 49076 [Source:MGI Symb | 3.2124 | 1.0279 | 2.0125 | 0.964 | 3.2719 | 0 |
| 53248 | ENSMUSG00000115699 | 0 | NA | NA | NA | NA | NA | Gm6961 | predicted gene 6961 [Source:MGI Symbol | 0 | 0 | 0 | 0 | 0 | 0 |
| 53249 | ENSMUSG00000115700 | 0 | NA | NA | NA | NA | NA | Gm7517 | predicted gene 7517 [Source:MGI Symbol | 0 | 0 | 0 | 0 | 0 | 0 |
| 53250 | ENSMUSG00000115701 | 0.17131964 | -0.796629176 | 4.080472857 | -0.195229623 | 0.84521319 | NA | Vmn1r6 | vomeronasal 1 receptor 6 [Source:MGI Sy | 0 | 1.0279 | 0 | 0 | 0 | 0 |
| 53251 | ENSMUSG00000115702 | 0 | NA | NA | NA | NA | NA | Gm5206 | predicted pseudogene 5206 [Source:MGI | 0 | 0 | 0 | 0 | 0 | 0 |
| 53252 | ENSMUSG00000115703 | 0 | NA | NA | NA | NA | NA | Gm49112 | predicted gene, 49112 [Source:MGI Symb | 0 | 0 | 0 | 0 | 0 | 0 |

| | | | | | | | | | | | | | | | |
|---|---|---|---|---|---|---|---|---|---|---|---|---|---|---|---|
| 53253 | ENSMUSG00000115704 | 0 | NA | NA | NA | NA | NA | Gm49171 | predicted gene, 49171 [Source:MGI Symb | 0 | 0 | 0 | 0 | 0 | 0 |
| 53254 | ENSMUSG00000115705 | 0 | NA | NA | NA | NA | NA | Vmn1r28 | vomeronasal 1 receptor 28 [Source:MGI S | 0 | 0 | 0 | 0 | 0 | 0 |
| 53255 | ENSMUSG00000115706 | 0 | NA | NA | NA | NA | NA | Gm49143 | predicted gene, 49143 [Source:MGI Symb | 0 | 0 | 0 | 0 | 0 | 0 |
| 53256 | ENSMUSG00000115707 | 0 | NA | NA | NA | NA | NA | Gm2267 | predicted gene 2267 [Source:MGI Symbol | 0 | 0 | 0 | 0 | 0 | 0 |
| 53257 | ENSMUSG00000115708 | 0 | NA | NA | NA | NA | NA | Gm49187 | predicted gene, 49187 [Source:MGI Symb | 0 | 0 | 0 | 0 | 0 | 0 |
| 53258 | ENSMUSG00000115709 | 0 | NA | NA | NA | NA | NA | 4930431P22 | RIKEN cDNA 4930431P22 gene [Source:M | 0 | 0 | 0 | 0 | 0 | 0 |
| 53259 | ENSMUSG00000115710 | 0 | NA | NA | NA | NA | NA | Gm49208 | predicted gene, 49208 [Source:MGI Symb | 0 | 0 | 0 | 0 | 0 | 0 |
| 53260 | ENSMUSG00000115711 | 0 | NA | NA | NA | NA | NA | Gm41336 | predicted gene, 41336 [Source:MGI Symb | 0 | 0 | 0 | 0 | 0 | 0 |
| 53261 | ENSMUSG00000115712 | 0 | NA | NA | NA | NA | NA | Gm49255 | predicted gene, 49255 [Source:MGI Symb | 0 | 0 | 0 | 0 | 0 | 0 |
| 53262 | ENSMUSG00000115713 | 0 | NA | NA | NA | NA | NA | Gm18216 | predicted gene, 18216 [Source:MGI Symb | 0 | 0 | 0 | 0 | 0 | 0 |
| 53263 | ENSMUSG00000115714 | 0 | NA | NA | NA | NA | NA | Gm18926 | predicted gene, 18926 [Source:MGI Symb | 0 | 0 | 0 | 0 | 0 | 0 |
| 53264 | ENSMUSG00000115715 | 0 | NA | NA | NA | NA | NA | Gm33936 | predicted gene, 33936 [Source:MGI Symb | 0 | 0 | 0 | 0 | 0 | 0 |
| 53265 | ENSMUSG00000115716 | 0 | NA | NA | NA | NA | NA | Gm49044 | predicted gene, 49044 [Source:MGI Symb | 0 | 0 | 0 | 0 | 0 | 0 |
| 53266 | ENSMUSG00000115717 | 0 | NA | NA | NA | NA | NA | Gm49161 | predicted gene, 49161 [Source:MGI Symb | 0 | 0 | 0 | 0 | 0 | 0 |
| 53267 | ENSMUSG00000115718 | 0 | NA | NA | NA | NA | NA | Gm49179 | predicted gene, 49179 [Source:MGI Symb | 0 | 0 | 0 | 0 | 0 | 0 |
| 53268 | ENSMUSG00000115719 | 0 | NA | NA | NA | NA | NA | 4930538L07I | RIKEN cDNA 4930538L07 gene [Source:M | 0 | 0 | 0 | 0 | 0 | 0 |
| 53269 | ENSMUSG00000115720 | 0 | NA | NA | NA | NA | NA | 4930433E05 | RIKEN cDNA 4930433E05 gene [Source:M | 0 | 0 | 0 | 0 | 0 | 0 |
| 53270 | ENSMUSG00000115721 | 0 | NA | NA | NA | NA | NA | Gm49239 | predicted gene, 49239 [Source:MGI Symb | 0 | 0 | 0 | 0 | 0 | 0 |
| 53271 | ENSMUSG00000115722 | 0 | NA | NA | NA | NA | NA | Gm8472 | predicted gene 8472 [Source:MGI Symbol | 0 | 0 | 0 | 0 | 0 | 0 |
| 53272 | ENSMUSG00000115723 | 0.335414862 | -1.779047242 | 4.030804179 | -0.441362855 | 0.658950329 | NA | 1700087M2 | RIKEN cDNA 1700087M22 gene [Source:M | 0 | 0 | 2.0125 | 0 | 0 | 0 |
| 53273 | ENSMUSG00000115724 | 0 | NA | NA | NA | NA | NA | Gm49087 | predicted gene, 49087 [Source:MGI Symb | 0 | 0 | 0 | 0 | 0 | 0 |
| 53274 | ENSMUSG00000115725 | 0 | NA | NA | NA | NA | NA | 4930572G02 | RIKEN cDNA 4930572G02 gene [Source:M | 0 | 0 | 0 | 0 | 0 | 0 |
| 53275 | ENSMUSG00000115726 | 0 | NA | NA | NA | NA | NA | Gm30083 | predicted gene, 30083 [Source:MGI Symb | 0 | 0 | 0 | 0 | 0 | 0 |
| 53276 | ENSMUSG00000115727 | 0 | NA | NA | NA | NA | NA | 4930432J09I | RIKEN cDNA 4930432J09 gene [Source:M | 0 | 0 | 0 | 0 | 0 | 0 |
| 53277 | ENSMUSG00000115728 | 0 | NA | NA | NA | NA | NA | Gm49190 | predicted gene, 49190 [Source:MGI Symb | 0 | 0 | 0 | 0 | 0 | 0 |
| 53278 | ENSMUSG00000115729 | 1.254589243 | -1.392189065 | 1.938007023 | -0.718361207 | 0.472534599 | NA | Gm5468 | predicted gene 5468 [Source:MGI Symbol | 2.4093 | 1.0279 | 2.0125 | 0.964 | 0 | 1.1138 |
| 53279 | ENSMUSG00000115730 | 0 | NA | NA | NA | NA | NA | Gm7459 | predicted gene 7459 [Source:MGI Symbol | 0 | 0 | 0 | 0 | 0 | 0 |
| 53280 | ENSMUSG00000115731 | 0 | NA | NA | NA | NA | NA | Gm19050 | predicted gene, 19050 [Source:MGI Symb | 0 | 0 | 0 | 0 | 0 | 0 |
| 53281 | ENSMUSG00000115732 | 0 | NA | NA | NA | NA | NA | Gm9302 | predicted gene 9302 [Source:MGI Symbol | 0 | 0 | 0 | 0 | 0 | 0 |
| 53282 | ENSMUSG00000115733 | 0 | NA | NA | NA | NA | NA | Gm49001 | predicted gene, 49001 [Source:MGI Symb | 0 | 0 | 0 | 0 | 0 | 0 |
| 53283 | ENSMUSG00000115734 | 0 | NA | NA | NA | NA | NA | Gm7708 | predicted gene 7708 [Source:MGI Symbol | 0 | 0 | 0 | 0 | 0 | 0 |
| 53284 | ENSMUSG00000115735 | 0 | NA | NA | NA | NA | NA | Gm49180 | predicted gene, 49180 [Source:MGI Symb | 0 | 0 | 0 | 0 | 0 | 0 |
| 53285 | ENSMUSG00000115736 | 0 | NA | NA | NA | NA | NA | Gm8586 | predicted gene 8586 [Source:MGI Symbol | 0 | 0 | 0 | 0 | 0 | 0 |
| 53286 | ENSMUSG00000115737 | 0 | NA | NA | NA | NA | NA | Rpl19-ps3 | ribosomal protein L19, pseudogene 3 [Sou | 0 | 0 | 0 | 0 | 0 | 0 |
| 53287 | ENSMUSG00000115738 | 0 | NA | NA | NA | NA | NA | Gm7543 | predicted gene 7543 [Source:MGI Symbol | 0 | 0 | 0 | 0 | 0 | 0 |
| 53288 | ENSMUSG00000115739 | 0 | NA | NA | NA | NA | NA | Gm49254 | predicted gene, 49254 [Source:MGI Symb | 0 | 0 | 0 | 0 | 0 | 0 |
| 53289 | ENSMUSG00000115740 | 0 | NA | NA | NA | NA | NA | Gm49042 | predicted gene, 49042 [Source:MGI Symb | 0 | 0 | 0 | 0 | 0 | 0 |
| 53290 | ENSMUSG00000115741 | 0 | NA | NA | NA | NA | NA | Gm19146 | predicted gene, 19146 [Source:MGI Symb | 0 | 0 | 0 | 0 | 0 | 0 |
| 53291 | ENSMUSG00000115742 | 0 | NA | NA | NA | NA | NA | Gm48950 | predicted gene, 48950 [Source:MGI Symb | 0 | 0 | 0 | 0 | 0 | 0 |
| 53292 | ENSMUSG00000115743 | 0 | NA | NA | NA | NA | NA | Gm7004 | predicted gene 7004 [Source:MGI Symbol | 0 | 0 | 0 | 0 | 0 | 0 |
| 53293 | ENSMUSG00000115744 | 0 | NA | NA | NA | NA | NA | Vmn1r80 | vomeronasal 1 receptor 80 [Source:MGI S | 0 | 0 | 0 | 0 | 0 | 0 |
| 53294 | ENSMUSG00000115745 | 0.860314674 | 2.043280159 | 2.552981052 | 0.800350695 | 0.423507639 | NA | Gm46448 | predicted gene, 46448 [Source:MGI Symb | 0 | 0 | 1.0062 | 1.928 | 0 | 2.2277 |
| 53295 | ENSMUSG00000115746 | 0 | NA | NA | NA | NA | NA | Gm49218 | predicted gene, 49218 [Source:MGI Symb | 0 | 0 | 0 | 0 | 0 | 0 |
| 53296 | ENSMUSG00000115747 | 0 | NA | NA | NA | NA | NA | Gm49145 | predicted gene, 49145 [Source:MGI Symb | 0 | 0 | 0 | 0 | 0 | 0 |
| 53297 | ENSMUSG00000115748 | 0 | NA | NA | NA | NA | NA | Gm49093 | predicted gene, 49093 [Source:MGI Symb | 0 | 0 | 0 | 0 | 0 | 0 |
| 53298 | ENSMUSG00000115749 | 0 | NA | NA | NA | NA | NA | Gm41289 | predicted gene, 41289 [Source:MGI Symb | 0 | 0 | 0 | 0 | 0 | 0 |
| 53299 | ENSMUSG00000115750 | 5.265773193 | 0.010255318 | 1.043444652 | 0.009828329 | 0.992158254 | 0.99859176 | Gm49282 | predicted gene, 49282 [Source:MGI Symb | 1.6062 | 7.1954 | 7.0437 | 4.8199 | 9.8156 | 1.1138 |
| 53300 | ENSMUSG00000115751 | 0 | NA | NA | NA | NA | NA | 4930503F20 | RIKEN cDNA 4930503F20 gene [Source:M | 0 | 0 | 0 | 0 | 0 | 0 |
| 53301 | ENSMUSG00000115752 | 0.339027071 | -1.79174149 | 3.919585585 | -0.457125237 | 0.647581032 | NA | 1700010G06 | RIKEN cDNA 1700010G06 gene [Source:M | 0 | 1.0279 | 1.0062 | 0 | 0 | 0 |
| 53302 | ENSMUSG00000115754 | 0 | NA | NA | NA | NA | NA | Gm48972 | predicted gene, 48972 [Source:MGI Symb | 0 | 0 | 0 | 0 | 0 | 0 |
| 53303 | ENSMUSG00000115755 | 0 | NA | NA | NA | NA | NA | Gm48993 | predicted gene, 48993 [Source:MGI Symb | 0 | 0 | 0 | 0 | 0 | 0 |
| 53304 | ENSMUSG00000115756 | 2.480226141 | -0.084869757 | 1.33596723 | -0.063526825 | 0.949346999 | NA | Gm49519 | predicted gene, 49519 [Source:MGI Symb | 1.6062 | 4.1117 | 2.0125 | 3.8559 | 2.1812 | 1.1138 |
| 53305 | ENSMUSG00000115757 | 0 | NA | NA | NA | NA | NA | Gm48934 | predicted gene, 48934 [Source:MGI Symb | 0 | 0 | 0 | 0 | 0 | 0 |
| 53306 | ENSMUSG00000115758 | 0 | NA | NA | NA | NA | NA | Gm49294 | predicted gene, 49294 [Source:MGI Symb | 0 | 0 | 0 | 0 | 0 | 0 |
| 53307 | ENSMUSG00000115759 | 0 | NA | NA | NA | NA | NA | Gm18787 | predicted gene, 18787 [Source:MGI Symb | 0 | 0 | 0 | 0 | 0 | 0 |
| 53308 | ENSMUSG00000115760 | 0 | NA | NA | NA | NA | NA | Gm18909 | predicted gene, 18909 [Source:MGI Symb | 0 | 0 | 0 | 0 | 0 | 0 |

| | | | | | | | | | | | | | | | |
|---|---|---|---|---|---|---|---|---|---|---|---|---|---|---|---|
| 53309 | ENSMUSG00000115761 | 0 | NA | NA | NA | NA | NA | 1700015H07 | RIKEN cDNA 1700015H07 gene [Source:M | 0 | 0 | 0 | 0 | 0 | 0 |
| 53310 | ENSMUSG00000115762 | 1.245658253 | 3.845479106 | 2.213078452 | 1.737615358 | 0.082278612 | NA | Gm34907 | predicted gene, 34907 [Source:MGI Symb | 0 | 0 | 0 | 1.928 | 1.0906 | 4.4554 |
| 53311 | ENSMUSG00000115763 | 0.133850373 | -0.796629176 | 4.080472857 | -0.195229623 | 0.84521319 | NA | Gm48974 | predicted gene, 48974 [Source:MGI Symb | 0.8031 | 0 | 0 | 0 | 0 | 0 |
| 53312 | ENSMUSG00000115764 | 0.428363721 | -0.543730382 | 3.624677344 | -0.15000794 | 0.880758351 | NA | Gm18811 | predicted gene, 18811 [Source:MGI Symb | 1.6062 | 0 | 0 | 0.964 | 0 | 0 |
| 53313 | ENSMUSG00000115765 | 0 | NA | NA | NA | NA | NA | Gm19227 | predicted gene, 19227 [Source:MGI Symb | 0 | 0 | 0 | 0 | 0 | 0 |
| 53314 | ENSMUSG00000115766 | 0 | NA | NA | NA | NA | NA | Gm49139 | predicted gene, 49139 [Source:MGI Symb | 0 | 0 | 0 | 0 | 0 | 0 |
| 53315 | ENSMUSG00000115767 | 1.795592857 | 0.389474302 | 1.758275064 | 0.221509313 | 0.824695886 | NA | Gm10366 | predicted gene 10366 [Source:MGI Symbo | 1.6062 | 3.0838 | 0 | 3.8559 | 0 | 2.2277 |
| 53316 | ENSMUSG00000115768 | 0.167707431 | -0.796629176 | 4.080472857 | -0.195229623 | 0.84521319 | NA | Dpep2nb | Dpep2 neighbor [Source:MGI Symbol;Acc | 0 | 0 | 1.0062 | 0 | 0 | 0 |
| 53317 | ENSMUSG00000115769 | 0.36353943 | 2.065989254 | 4.034137301 | 0.512126658 | 0.608562368 | NA | Gm48938 | predicted gene, 48938 [Source:MGI Symb | 0 | 0 | 0 | 0 | 2.1812 | 0 |
| 53318 | ENSMUSG00000115770 | 0 | NA | NA | NA | NA | NA | Gm49056 | predicted gene, 49056 [Source:MGI Symb | 0 | 0 | 0 | 0 | 0 | 0 |
| 53319 | ENSMUSG00000115771 | 0.531944269 | 2.618916564 | 3.285123409 | 0.797204926 | 0.425332026 | NA | Gm49312 | predicted gene, 49312 [Source:MGI Symb | 0 | 0 | 0 | 0.964 | 0 | 2.2277 |
| 53320 | ENSMUSG00000115772 | 0 | NA | NA | NA | NA | NA | Gm48935 | predicted gene, 48935 [Source:MGI Symb | 0 | 0 | 0 | 0 | 0 | 0 |
| 53321 | ENSMUSG00000115773 | 0.328370405 | 0.16514316 | 4.003149536 | 0.041253308 | 0.967093956 | NA | Gm31458 | predicted gene, 31458 [Source:MGI Symb | 0 | 0 | 1.0062 | 0.964 | 0 | 0 |
| 53322 | ENSMUSG00000115774 | 0 | NA | NA | NA | NA | NA | 4933431J24I | RIKEN cDNA 4933431J24 gene [Source:MG | 0 | 0 | 0 | 0 | 0 | 0 |
| 53323 | ENSMUSG00000115775 | 0 | NA | NA | NA | NA | NA | Gm41318 | predicted gene, 41318 [Source:MGI Symb | 0 | 0 | 0 | 0 | 0 | 0 |
| 53324 | ENSMUSG00000115776 | 0 | NA | NA | NA | NA | NA | Gm49158 | predicted gene, 49158 [Source:MGI Symb | 0 | 0 | 0 | 0 | 0 | 0 |
| 53325 | ENSMUSG00000115777 | 0.877291758 | 2.066311913 | 2.565974136 | 0.805273866 | 0.420661659 | NA | Gm32815 | predicted gene, 32815 [Source:MGI Symb | 0 | 1.0279 | 0 | 0.964 | 3.2719 | 0 |
| 53326 | ENSMUSG00000115778 | 0 | NA | NA | NA | NA | NA | Gm10876 | predicted gene 10876 [Source:MGI Symbo | 0 | 0 | 0 | 0 | 0 | 0 |
| 53327 | ENSMUSG00000115779 | 0.465832987 | -0.698295221 | 3.18348759 | -0.219349126 | 0.826378096 | NA | 4930592A05 | RIKEN cDNA 4930592A05 gene [Source:M | 0.8031 | 1.0279 | 0 | 0.964 | 0 | 0 |
| 53328 | ENSMUSG00000115780 | 0 | NA | NA | NA | NA | NA | Gm49053 | predicted gene, 49053 [Source:MGI Symb | 0 | 0 | 0 | 0 | 0 | 0 |
| 53329 | ENSMUSG00000115781 | 0 | NA | NA | NA | NA | NA | Gm49319 | predicted gene, 49319 [Source:MGI Symb | 0 | 0 | 0 | 0 | 0 | 0 |
| 53330 | ENSMUSG00000115782 | 0 | NA | NA | NA | NA | NA | Gm49116 | predicted gene, 49116 [Source:MGI Symb | 0 | 0 | 0 | 0 | 0 | 0 |
| 53331 | ENSMUSG00000115783 | 0 | NA | NA | NA | NA | NA | Bc1 | brain cytoplasmic RNA 1 [Source:MGI Sym | 0 | 0 | 0 | 0 | 0 | 0 |
| 53332 | ENSMUSG00000115784 | 0 | NA | NA | NA | NA | NA | Gm30929 | predicted gene, 30929 [Source:MGI Symb | 0 | 0 | 0 | 0 | 0 | 0 |
| 53333 | ENSMUSG00000115785 | 0.349477146 | 0.165142768 | 3.911021834 | 0.042224967 | 0.966319359 | NA | Gm6740 | predicted gene 6740 [Source:MGI Symbol | 0 | 0 | 1.0062 | 0 | 1.0906 | 0 |
| 53334 | ENSMUSG00000115786 | 0 | NA | NA | NA | NA | NA | Gm4486 | predicted gene 4486 [Source:MGI Symbol | 0 | 0 | 0 | 0 | 0 | 0 |
| 53335 | ENSMUSG00000115787 | 0 | NA | NA | NA | NA | NA | Gm36146 | predicted gene, 36146 [Source:MGI Symb | 0 | 0 | 0 | 0 | 0 | 0 |
| 53336 | ENSMUSG00000115788 | 0 | NA | NA | NA | NA | NA | Gm48915 | predicted gene, 48915 [Source:MGI Symb | 0 | 0 | 0 | 0 | 0 | 0 |
| 53337 | ENSMUSG00000115789 | 4.970721499 | 2.056835676 | 1.148676667 | 1.790613263 | 0.073355377 | NA | Gm18147 | predicted gene, 18147 [Source:MGI Symb | 0.8031 | 2.0558 | 3.0187 | 10.604 | 1.0906 | 12.252 |
| 53338 | ENSMUSG00000115790 | 0 | NA | NA | NA | NA | NA | 9530086P17 | RIKEN cDNA 9530086P17 gene [Source:M | 0 | 0 | 0 | 0 | 0 | 0 |
| 53339 | ENSMUSG00000115791 | 0 | NA | NA | NA | NA | NA | Vmn1r217 | vomeronasal 1 receptor 217 [Source:MGI | 0 | 0 | 0 | 0 | 0 | 0 |
| 53340 | ENSMUSG00000115792 | 0 | NA | NA | NA | NA | NA | Vmn1r16 | vomeronasal 1 receptor 16 [Source:MGI S | 0 | 0 | 0 | 0 | 0 | 0 |
| 53341 | ENSMUSG00000115793 | 4.899907119 | 0.944737125 | 1.113907417 | 0.848128947 | 0.396366162 | NA | Gm48961 | predicted gene, 48961 [Source:MGI Symb | 0 | 6.1675 | 4.025 | 4.8199 | 4.3625 | 10.025 |
| 53342 | ENSMUSG00000115794 | 0.133850373 | -0.796629176 | 4.080472857 | -0.195229623 | 0.84521319 | NA | Gm49101 | predicted gene, 49101 [Source:MGI Symb | 0.8031 | 0 | 0 | 0 | 0 | 0 |
| 53343 | ENSMUSG00000115795 | 0 | NA | NA | NA | NA | NA | Gm8484 | predicted gene 8484 [Source:MGI Symbol | 0 | 0 | 0 | 0 | 0 | 0 |
| 53344 | ENSMUSG00000115796 | 0 | NA | NA | NA | NA | NA | Gm48970 | predicted gene, 48970 [Source:MGI Symb | 0 | 0 | 0 | 0 | 0 | 0 |
| 53345 | ENSMUSG00000115797 | 0 | NA | NA | NA | NA | NA | Gm48981 | predicted gene, 48981 [Source:MGI Symb | 0 | 0 | 0 | 0 | 0 | 0 |
| 53346 | ENSMUSG00000115798 | 0 | NA | NA | NA | NA | NA | | novel protein | 0 | 0 | 0 | 0 | 0 | 0 |
| 53347 | ENSMUSG00000115799 | 0 | NA | NA | NA | NA | NA | Vmn1r19 | vomeronasal 1 receptor 19 [Source:MGI S | 0 | 0 | 0 | 0 | 0 | 0 |
| 53348 | ENSMUSG00000115800 | 0 | NA | NA | NA | NA | NA | Frg2f4 | FSHD region gene 2 family member 4 [Sou | 0 | 0 | 0 | 0 | 0 | 0 |
| 53349 | ENSMUSG00000115801 | 3.822007903 | -1.348925248 | 1.140009383 | -1.183258022 | 0.236706904 | NA | | novel transcript | 3.2124 | 7.1954 | 6.0375 | 0.964 | 2.1812 | 3.3415 |
| 53350 | ENSMUSG00000115802 | 0 | NA | NA | NA | NA | NA | Vmn1r-ps13 | vomeronasal 1 receptor, pseudogene 136 | 0 | 0 | 0 | 0 | 0 | 0 |
| 53351 | ENSMUSG00000115803 | 0 | NA | NA | NA | NA | NA | Gm35496 | predicted gene, 35496 [Source:MGI Symb | 0 | 0 | 0 | 0 | 0 | 0 |
| 53352 | ENSMUSG00000115804 | 0 | NA | NA | NA | NA | NA | Gm49107 | predicted gene, 49107 [Source:MGI Symb | 0 | 0 | 0 | 0 | 0 | 0 |
| 53353 | ENSMUSG00000115805 | 0 | NA | NA | NA | NA | NA | Gm49110 | predicted gene, 49110 [Source:MGI Symb | 0 | 0 | 0 | 0 | 0 | 0 |
| 53354 | ENSMUSG00000115806 | 0 | NA | NA | NA | NA | NA | Gm49234 | predicted gene, 49234 [Source:MGI Symb | 0 | 0 | 0 | 0 | 0 | 0 |
| 53355 | ENSMUSG00000115807 | 0 | NA | NA | NA | NA | NA | Gm49100 | predicted gene, 49100 [Source:MGI Symb | 0 | 0 | 0 | 0 | 0 | 0 |
| 53356 | ENSMUSG00000115808 | 0.825760208 | 0.783249908 | 2.404539989 | 0.325737942 | 0.744622635 | NA | Gm18949 | predicted gene, 18949 [Source:MGI Symb | 0.8031 | 0 | 1.0062 | 0.964 | 2.1812 | 0 |
| 53357 | ENSMUSG00000115809 | 0 | NA | NA | NA | NA | NA | Gm48951 | predicted gene, 48951 [Source:MGI Symb | 0 | 0 | 0 | 0 | 0 | 0 |
| 53358 | ENSMUSG00000115810 | 6.279824072 | -0.258723655 | 0.949002791 | -0.272626864 | 0.785140059 | 0.960670212 | Gm49066 | predicted gene, 49066 [Source:MGI Symb | 11.243 | 2.0558 | 7.0437 | 1.928 | 8.7249 | 6.6831 |
| 53359 | ENSMUSG00000115811 | 0.185640647 | 1.126924175 | 4.080472857 | 0.276174898 | 0.782413736 | NA | Gm41293 | predicted gene, 41293 [Source:MGI Symb | 0 | 0 | 0 | 0 | 0 | 1.1138 |
| 53360 | ENSMUSG00000115812 | 0.531246861 | 1.101376188 | 3.36099124 | 0.327693858 | 0.743143147 | NA | Gm49309 | predicted gene, 49309 [Source:MGI Symb | 0 | 0 | 1.0062 | 0 | 2.1812 | 0 |
| 53361 | ENSMUSG00000115813 | 0.961910823 | -0.938554428 | 2.269016656 | -0.413639285 | 0.679138298 | NA | Gm46447 | predicted gene, 46447 [Source:MGI Symb | 0.8031 | 1.0279 | 2.0125 | 1.928 | 0 | 0 |
| 53362 | ENSMUSG00000115814 | 0 | NA | NA | NA | NA | NA | Gm49181 | predicted gene, 49181 [Source:MGI Symb | 0 | 0 | 0 | 0 | 0 | 0 |
| 53363 | ENSMUSG00000115815 | 0 | NA | NA | NA | NA | NA | Gm8328 | predicted gene 8328 [Source:MGI Symbol | 0 | 0 | 0 | 0 | 0 | 0 |
| 53364 | ENSMUSG00000115816 | 0.133850373 | -0.796629176 | 4.080472857 | -0.195229623 | 0.84521319 | NA | Gm34589 | predicted gene, 34589 [Source:MGI Symb | 0.8031 | 0 | 0 | 0 | 0 | 0 |

| | | | | | | | | | | | | | | | |
|---|---|---|---|---|---|---|---|---|---|---|---|---|---|---|---|
| 53365 | ENSMUSG00000115817 | 0 | NA | NA | NA | NA | NA | Gm30246 | predicted gene, 30246 [Source:MGI Symb | 0 | 0 | 0 | 0 | 0 | 0 |
| 53366 | ENSMUSG00000115818 | 0 | NA | NA | NA | NA | NA | Gm18580 | predicted gene, 18580 [Source:MGI Symb | 0 | 0 | 0 | 0 | 0 | 0 |
| 53367 | ENSMUSG00000115819 | 0.167707431 | -0.796629176 | 4.080472857 | -0.195229623 | 0.84521319 | NA | Gm9108 | predicted gene 9108 [Source:MGI Symbol | 0 | 0 | 1.0062 | 0 | 0 | 0 |
| 53368 | ENSMUSG00000115820 | 0 | NA | NA | NA | NA | NA | Gm41311 | predicted gene, 41311 [Source:MGI Symb | 0 | 0 | 0 | 0 | 0 | 0 |
| 53369 | ENSMUSG00000115821 | 0 | NA | NA | NA | NA | NA | 6330576A10 | RIKEN cDNA 6330576A10 gene [Source:M | 0 | 0 | 0 | 0 | 0 | 0 |
| 53370 | ENSMUSG00000115822 | 0 | NA | NA | NA | NA | NA | Gm33450 | predicted gene, 33450 [Source:MGI Symb | 0 | 0 | 0 | 0 | 0 | 0 |
| 53371 | ENSMUSG00000115823 | 1.266900876 | -2.548032382 | 2.280137557 | -1.117490642 | 0.263784599 | NA | Gm49004 | predicted gene, 49004 [Source:MGI Symb | 1.6062 | 0 | 5.0312 | 0.964 | 0 | 0 |
| 53372 | ENSMUSG00000115824 | 0 | NA | NA | NA | NA | NA | Gm49114 | predicted gene, 49114 [Source:MGI Symb | 0 | 0 | 0 | 0 | 0 | 0 |
| 53373 | ENSMUSG00000115825 | 0 | NA | NA | NA | NA | NA | Gm49048 | predicted gene, 49048 [Source:MGI Symb | 0 | 0 | 0 | 0 | 0 | 0 |
| 53374 | ENSMUSG00000115826 | 0 | NA | NA | NA | NA | NA | Gm49034 | predicted gene, 49034 [Source:MGI Symb | 0 | 0 | 0 | 0 | 0 | 0 |
| 53375 | ENSMUSG00000115827 | 0 | NA | NA | NA | NA | NA | Gm49261 | predicted gene, 49261 [Source:MGI Symb | 0 | 0 | 0 | 0 | 0 | 0 |
| 53376 | ENSMUSG00000115828 | 0 | NA | NA | NA | NA | NA | Gm18761 | predicted gene, 18761 [Source:MGI Symb | 0 | 0 | 0 | 0 | 0 | 0 |
| 53377 | ENSMUSG00000115829 | 0.455176322 | 0.955156269 | 3.546674842 | 0.269310357 | 0.787690862 | NA | 4921515G04 | RIKEN cDNA 4921515G04 gene [Source:M | 0.8031 | 0 | 0 | 1.928 | 0 | 0 |
| 53378 | ENSMUSG00000115830 | 0 | NA | NA | NA | NA | NA | Gm49176 | predicted gene, 49176 [Source:MGI Symb | 0 | 0 | 0 | 0 | 0 | 0 |
| 53379 | ENSMUSG00000115831 | 0.167707431 | -0.796629176 | 4.080472857 | -0.195229623 | 0.84521319 | NA | Gm48937 | predicted gene, 48937 [Source:MGI Symb | 0 | 0 | 1.0062 | 0 | 0 | 0 |
| 53380 | ENSMUSG00000115832 | 0 | NA | NA | NA | NA | NA | Gm19158 | predicted gene, 19158 [Source:MGI Symb | 0 | 0 | 0 | 0 | 0 | 0 |
| 53381 | ENSMUSG00000115833 | 0 | NA | NA | NA | NA | NA | Gm49196 | predicted gene, 49196 [Source:MGI Symb | 0 | 0 | 0 | 0 | 0 | 0 |
| 53382 | ENSMUSG00000115834 | 1.263441998 | 0.900942224 | 2.051072711 | 0.439254161 | 0.660477384 | NA | Gm10385 | predicted gene 10385 [Source:MGI Symbo | 1.6062 | 1.0279 | 0 | 3.8559 | 1.0906 | 0 |
| 53383 | ENSMUSG00000115835 | 0 | NA | NA | NA | NA | NA | Gm41253 | predicted gene, 41253 [Source:MGI Symb | 0 | 0 | 0 | 0 | 0 | 0 |
| 53384 | ENSMUSG00000115836 | 0 | NA | NA | NA | NA | NA | Gm49060 | predicted gene, 49060 [Source:MGI Symb | 0 | 0 | 0 | 0 | 0 | 0 |
| 53385 | ENSMUSG00000115837 | 2.796383678 | 0.088737621 | 1.621837268 | 0.054714257 | 0.956366111 | NA | 9130002K18 | RIKEN cDNA 9130002K18 gene [Source:M | 0 | 6.1675 | 2.0125 | 0.964 | 7.6343 | 0 |
| 53386 | ENSMUSG00000115838 | 0 | NA | NA | NA | NA | NA | Gm46507 | predicted gene, 46507 [Source:MGI Symb | 0 | 0 | 0 | 0 | 0 | 0 |
| 53387 | ENSMUSG00000115839 | 0 | NA | NA | NA | NA | NA | Gm49233 | predicted gene, 49233 [Source:MGI Symb | 0 | 0 | 0 | 0 | 0 | 0 |
| 53388 | ENSMUSG00000115840 | 0 | NA | NA | NA | NA | NA | 5830426I08F | RIKEN cDNA 5830426I08 gene [Source:MC | 0 | 0 | 0 | 0 | 0 | 0 |
| 53389 | ENSMUSG00000115841 | 0 | NA | NA | NA | NA | NA | Gm49105 | predicted gene, 49105 [Source:MGI Symb | 0 | 0 | 0 | 0 | 0 | 0 |
| 53390 | ENSMUSG00000115842 | 0.52105551 | -0.812028029 | 3.385364865 | -0.239864257 | 0.810435491 | NA | Gm49092 | predicted gene, 49092 [Source:MGI Symb | 0 | 0 | 2.0125 | 0 | 0 | 1.1138 |
| 53391 | ENSMUSG00000115843 | 0 | NA | NA | NA | NA | NA | Gm17873 | predicted gene, 17873 [Source:MGI Symb | 0 | 0 | 0 | 0 | 0 | 0 |
| 53392 | ENSMUSG00000115844 | 0 | NA | NA | NA | NA | NA | Gm49081 | predicted gene, 49081 [Source:MGI Symb | 0 | 0 | 0 | 0 | 0 | 0 |
| 53393 | ENSMUSG00000115845 | 0 | NA | NA | NA | NA | NA | 4933402J15I | RIKEN cDNA 4933402J15 gene [Source:MC | 0 | 0 | 0 | 0 | 0 | 0 |
| 53394 | ENSMUSG00000115846 | 0.781453075 | -0.369215669 | 2.439554878 | -0.151345507 | 0.879703174 | NA | Gm41144 | predicted gene, 41144 [Source:MGI Symb | 1.6062 | 1.0279 | 0 | 0.964 | 1.0906 | 0 |
| 53395 | ENSMUSG00000115847 | 0 | NA | NA | NA | NA | NA | Gm49050 | predicted gene, 49050 [Source:MGI Symb | 0 | 0 | 0 | 0 | 0 | 0 |
| 53396 | ENSMUSG00000115848 | 0 | NA | NA | NA | NA | NA | Gm9509 | predicted gene 9509 [Source:MGI Symbol | 0 | 0 | 0 | 0 | 0 | 0 |
| 53397 | ENSMUSG00000115849 | 0 | NA | NA | NA | NA | NA | Gm49149 | predicted gene, 49149 [Source:MGI Symb | 0 | 0 | 0 | 0 | 0 | 0 |
| 53398 | ENSMUSG00000115850 | 0.167707431 | -0.796629176 | 4.080472857 | -0.195229623 | 0.84521319 | NA | Gm48956 | predicted gene, 48956 [Source:MGI Symb | 0 | 0 | 1.0062 | 0 | 0 | 0 |
| 53399 | ENSMUSG00000115852 | 112.5525923 | 0.209389069 | 0.248196781 | 0.843641354 | 0.398869848 | 0.82594896 | Gm52969 | predicted gene, 52969 [Source:MGI Symb | 96.372 | 81.206 | 135.84 | 129.17 | 107.97 | 124.75 |
| 53400 | ENSMUSG00000115854 | 1.606287867 | -1.175097167 | 2.050416046 | -0.57310182 | 0.566575755 | NA | Gm36644 | predicted gene, 36644 [Source:MGI Symb | 1.6062 | 5.1396 | 0 | 2.8919 | 0 | 0 |
| 53401 | ENSMUSG00000115855 | 0 | NA | NA | NA | NA | NA | Gm34643 | predicted gene, 34643 [Source:MGI Symb | 0 | 0 | 0 | 0 | 0 | 0 |
| 53402 | ENSMUSG00000115856 | 13.62707407 | -0.381113131 | 0.613987129 | -0.620718437 | 0.534784896 | 0.889858671 | Gm18095 | predicted gene, 18095 [Source:MGI Symb | 8.8341 | 20.558 | 17.106 | 15.424 | 9.8156 | 10.025 |
| 53403 | ENSMUSG00000115858 | 0 | NA | NA | NA | NA | NA | 4930588J15I | RIKEN cDNA 4930588J15 gene [Source:MC | 0 | 0 | 0 | 0 | 0 | 0 |
| 53404 | ENSMUSG00000115860 | 0 | NA | NA | NA | NA | NA | Gm49426 | predicted gene, 49426 [Source:MGI Symb | 0 | 0 | 0 | 0 | 0 | 0 |
| 53405 | ENSMUSG00000115862 | 0 | NA | NA | NA | NA | NA | Gm41333 | predicted gene, 41333 [Source:MGI Symb | 0 | 0 | 0 | 0 | 0 | 0 |
| 53406 | ENSMUSG00000115863 | 0 | NA | NA | NA | NA | NA | Gm49485 | predicted gene, 49485 [Source:MGI Symb | 0 | 0 | 0 | 0 | 0 | 0 |
| 53407 | ENSMUSG00000115866 | 0 | NA | NA | NA | NA | NA | Gm49467 | predicted gene, 49467 [Source:MGI Symb | 0 | 0 | 0 | 0 | 0 | 0 |
| 53408 | ENSMUSG00000115867 | 255.7168555 | 0.259120866 | 0.243965058 | 1.062122861 | 0.288179913 | 0.73951397 | Gm17753 | predicted gene, 17753 [Source:MGI Symb | 242.54 | 205.58 | 249.55 | 163.88 | 314.1 | 358.66 |
| 53409 | ENSMUSG00000115868 | 1.01186617 | -0.80774997 | 2.233670792 | -0.361624449 | 0.717632691 | NA | Gm2999 | predicted gene 2999 [Source:MGI Symbol | 0.8031 | 1.0279 | 2.0125 | 0 | 0 | 2.2277 |
| 53410 | ENSMUSG00000115869 | 23.6864739 | 0.131112526 | 0.497658673 | 0.263458737 | 0.792197023 | 0.961434599 | Gm31814 | predicted gene, 31814 [Source:MGI Symb | 28.109 | 16.447 | 23.144 | 25.063 | 35.99 | 13.366 |
| 53411 | ENSMUSG00000115871 | 0 | NA | NA | NA | NA | NA | Gm49507 | predicted gene, 49507 [Source:MGI Symb | 0 | 0 | 0 | 0 | 0 | 0 |
| 53412 | ENSMUSG00000115875 | 0 | NA | NA | NA | NA | NA | Gm49408 | predicted gene, 49408 [Source:MGI Symb | 0 | 0 | 0 | 0 | 0 | 0 |
| 53413 | ENSMUSG00000115877 | 0 | NA | NA | NA | NA | NA | Gm49478 | predicted gene, 49478 [Source:MGI Symb | 0 | 0 | 0 | 0 | 0 | 0 |
| 53414 | ENSMUSG00000115880 | 0 | NA | NA | NA | NA | NA | 4930445N06 | RIKEN cDNA 4930445N06 gene [Source:M | 0 | 0 | 0 | 0 | 0 | 0 |
| 53415 | ENSMUSG00000115882 | 4.470074793 | 0.267007623 | 1.0090102 | 0.264623314 | 0.791299658 | NA | Gm5481 | predicted gene 5481 [Source:MGI Symbol | 4.0155 | 6.1675 | 2.0125 | 5.7839 | 3.2719 | 5.5692 |
| 53416 | ENSMUSG00000115885 | 0 | NA | NA | NA | NA | NA | Gm19811 | predicted gene, 19811 [Source:MGI Symb | 0 | 0 | 0 | 0 | 0 | 0 |
| 53417 | ENSMUSG00000115887 | 0.181769715 | 1.126924175 | 4.080472857 | 0.276174898 | 0.782413736 | NA | Gm49471 | predicted gene, 49471 [Source:MGI Symb | 0 | 0 | 0 | 0 | 1.0906 | 0 |
| 53418 | ENSMUSG00000115888 | 0 | NA | NA | NA | NA | NA | Gm31369 | predicted gene, 31369 [Source:MGI Symb | 0 | 0 | 0 | 0 | 0 | 0 |
| 53419 | ENSMUSG00000115894 | 0 | NA | NA | NA | NA | NA | Gm49453 | predicted gene, 49453 [Source:MGI Symb | 0 | 0 | 0 | 0 | 0 | 0 |
| 53420 | ENSMUSG00000115896 | 0 | NA | NA | NA | NA | NA | Gm36677 | predicted gene, 36677 [Source:MGI Symb | 0 | 0 | 0 | 0 | 0 | 0 |

| | | | | | | | | | | | | | | | |
|---|---|---|---|---|---|---|---|---|---|---|---|---|---|---|---|
| 53421 | ENSMUSG00000115898 | 0 | NA | NA | NA | NA | NA | Gm49524 | predicted gene, 49524 [Source:MGI Symb | 0 | 0 | 0 | 0 | 0 | 0 |
| 53422 | ENSMUSG00000115902 | 1.158156003 | -3.567567658 | 2.311061604 | -1.543692151 | 0.122662929 | NA | | RIKEN cDNA D730005E14 gene [Source:N | 0.8031 | 5.1396 | 1.0062 | 0 | 0 | 0 |
| 53423 | ENSMUSG00000115903 | 0 | NA | NA | NA | NA | NA | Gm17851 | predicted gene, 17851 [Source:MGI Symb | 0 | 0 | 0 | 0 | 0 | 0 |
| 53424 | ENSMUSG00000115907 | 0 | NA | NA | NA | NA | NA | Gm49432 | predicted gene, 49432 [Source:MGI Symb | 0 | 0 | 0 | 0 | 0 | 0 |
| 53425 | ENSMUSG00000115908 | 32.6081528 | -0.347309693 | 0.520679548 | -0.667031564 | 0.504751972 | 0.881592135 | Cyp2d35-ps | cytochrome P450, family 2, subfamily d, m | 41.761 | 24.67 | 43.269 | 44.343 | 33.809 | 7.7969 |
| 53426 | ENSMUSG00000115911 | 0 | NA | NA | NA | NA | NA | Gm3919 | predicted gene 3919 [Source:MGI Symbol | 0 | 0 | 0 | 0 | 0 | 0 |
| 53427 | ENSMUSG00000115912 | 0 | NA | NA | NA | NA | NA | Gm35569 | predicted gene, 35569 [Source:MGI Symb | 0 | 0 | 0 | 0 | 0 | 0 |
| 53428 | ENSMUSG00000115914 | 0 | NA | NA | NA | NA | NA | Cyp2d34-ps | cytochrome P450, family 2, subfamily d, p | 0 | 0 | 0 | 0 | 0 | 0 |
| 53429 | ENSMUSG00000115916 | 0 | NA | NA | NA | NA | NA | Gm8702 | predicted gene 8702 [Source:MGI Symbol | 0 | 0 | 0 | 0 | 0 | 0 |
| 53430 | ENSMUSG00000115917 | 0 | NA | NA | NA | NA | NA | Gm49406 | predicted gene, 49406 [Source:MGI Symb | 0 | 0 | 0 | 0 | 0 | 0 |
| 53431 | ENSMUSG00000115918 | 1.51708046 | -0.427560069 | 1.918708969 | -0.222837375 | 0.823662079 | NA | A930001M0 | RIKEN cDNA A930001M01 gene [Source:N | 3.2124 | 1.0279 | 1.0062 | 3.8559 | 0 | 0 |
| 53432 | ENSMUSG00000115919 | 1962.03281 | 0.574979377 | 0.194894407 | 2.950209739 | 0.003175583 | 0.048599236 | Gm31583 | predicted gene, 31583 [Source:MGI Symb | 1283.4 | 1335.3 | 2110.1 | 1760.2 | 2862.9 | 2420.4 |
| 53433 | ENSMUSG00000115920 | 0 | NA | NA | NA | NA | NA | Gm18283 | predicted gene, 18283 [Source:MGI Symb | 0 | 0 | 0 | 0 | 0 | 0 |
| 53434 | ENSMUSG00000115921 | 0 | NA | NA | NA | NA | NA | Gm18685 | predicted gene, 18685 [Source:MGI Symb | 0 | 0 | 0 | 0 | 0 | 0 |
| 53435 | ENSMUSG00000115923 | 0.819128932 | -3.070895205 | 2.828495588 | -1.085699132 | 0.277612125 | NA | Gm49521 | predicted gene, 49521 [Source:MGI Symb | 0.8031 | 4.1117 | 0 | 0 | 0 | 0 |
| 53436 | ENSMUSG00000115924 | 0 | NA | NA | NA | NA | NA | Gm19277 | predicted gene, 19277 [Source:MGI Symb | 0 | 0 | 0 | 0 | 0 | 0 |
| 53437 | ENSMUSG00000115925 | 0.160662974 | 1.126924175 | 4.080472857 | 0.276174898 | 0.782413736 | NA | Gm33432 | predicted gene, 33432 [Source:MGI Symb | 0 | 0 | 0 | 0.964 | 0 | 0 |
| 53438 | ENSMUSG00000115926 | 0 | NA | NA | NA | NA | NA | Gm8233 | predicted gene 8233 [Source:MGI Symbol | 0 | 0 | 0 | 0 | 0 | 0 |
| 53439 | ENSMUSG00000115928 | 0 | NA | NA | NA | NA | NA | Gm18930 | predicted gene, 18930 [Source:MGI Symb | 0 | 0 | 0 | 0 | 0 | 0 |
| 53440 | ENSMUSG00000115930 | 0 | NA | NA | NA | NA | NA | A930027H12 | RIKEN cDNA A930027H12 gene [Source:M | 0 | 0 | 0 | 0 | 0 | 0 |
| 53441 | ENSMUSG00000115931 | 0 | NA | NA | NA | NA | NA | Gm27825 | predicted gene, 27825 [Source:MGI Symb | 0 | 0 | 0 | 0 | 0 | 0 |
| 53442 | ENSMUSG00000115935 | 0 | NA | NA | NA | NA | NA | 4930573C08 | RIKEN cDNA 4930573C08 gene [Source:M | 0 | 0 | 0 | 0 | 0 | 0 |
| 53443 | ENSMUSG00000115936 | 0 | NA | NA | NA | NA | NA | 4833415N18 | RIKEN cDNA 4833415N18 gene [Source:M | 0 | 0 | 0 | 0 | 0 | 0 |
| 53444 | ENSMUSG00000115938 | 0.346303622 | 2.004339653 | 3.884003423 | 0.516049919 | 0.605819544 | NA | Gm17241 | predicted gene, 17241 [Source:MGI Symb | 0 | 0 | 0 | 0.964 | 0 | 1.1138 |
| 53445 | ENSMUSG00000115943 | 0 | NA | NA | NA | NA | NA | Gm49455 | predicted gene, 49455 [Source:MGI Symb | 0 | 0 | 0 | 0 | 0 | 0 |
| 53446 | ENSMUSG00000115944 | 0 | NA | NA | NA | NA | NA | AU021063 | expressed sequence AU021063 [Source:M | 0 | 0 | 0 | 0 | 0 | 0 |
| 53447 | ENSMUSG00000115946 | 12.86411065 | 0.132151209 | 0.740827311 | 0.178383285 | 0.858421975 | 0.975439379 | Mirt2 | myocardial infraction associated transcrip | 22.487 | 2.0558 | 12.075 | 16.388 | 15.269 | 8.9108 |
| 53448 | ENSMUSG00000115948 | 0.328370405 | 0.16514316 | 4.003149536 | 0.041253308 | 0.967093956 | NA | Gm49419 | predicted gene, 49419 [Source:MGI Symb | 0 | 0 | 1.0062 | 0.964 | 0 | 0 |
| 53449 | ENSMUSG00000115952 | 0 | NA | NA | NA | NA | NA | Gm18814 | predicted gene, 18814 [Source:MGI Symb | 0 | 0 | 0 | 0 | 0 | 0 |
| 53450 | ENSMUSG00000115955 | 0.845734943 | 0.605113486 | 2.575938408 | 0.23490992 | 0.81427865 | NA | 9530056E24 | RIKEN cDNA 9530056E24 gene [Source:M | 0 | 2.0558 | 0 | 1.928 | 1.0906 | 0 |
| 53451 | ENSMUSG00000115956 | 0.167707431 | -0.796629176 | 4.080472857 | -0.195229623 | 0.84521319 | NA | NA | NA | 0 | 0 | 1.0062 | 0 | 0 | 0 |
| 53452 | ENSMUSG00000115957 | 0 | NA | NA | NA | NA | NA | Gm49466 | predicted gene, 49466 [Source:MGI Symb | 0 | 0 | 0 | 0 | 0 | 0 |
| 53453 | ENSMUSG00000115958 | 1.909597977 | -0.719792747 | 1.789893897 | -0.40214269 | 0.687579017 | NA | Gm28040 | predicted gene, 28040 [Source:MGI Symb | 0 | 5.1396 | 2.0125 | 0.964 | 0 | 3.3415 |
| 53454 | ENSMUSG00000115959 | 0 | NA | NA | NA | NA | NA | Gm18494 | predicted gene, 18494 [Source:MGI Symb | 0 | 0 | 0 | 0 | 0 | 0 |
| 53455 | ENSMUSG00000115961 | 0 | NA | NA | NA | NA | NA | Gm41409 | predicted gene, 41409 [Source:MGI Symb | 0 | 0 | 0 | 0 | 0 | 0 |
| 53456 | ENSMUSG00000115962 | 0 | NA | NA | NA | NA | NA | 4930588D02 | RIKEN cDNA 4930588D02 gene [Source:M | 0 | 0 | 0 | 0 | 0 | 0 |
| 53457 | ENSMUSG00000115964 | 0 | NA | NA | NA | NA | NA | Gm6979 | predicted gene 6979 [Source:MGI Symbol | 0 | 0 | 0 | 0 | 0 | 0 |
| 53458 | ENSMUSG00000115965 | 0 | NA | NA | NA | NA | NA | Gm30564 | predicted gene, 30564 [Source:MGI Symb | 0 | 0 | 0 | 0 | 0 | 0 |
| 53459 | ENSMUSG00000115968 | 0 | NA | NA | NA | NA | NA | Gm38619 | predicted gene, 38619 [Source:MGI Symb | 0 | 0 | 0 | 0 | 0 | 0 |
| 53460 | ENSMUSG00000115969 | 0 | NA | NA | NA | NA | NA | Gm46519 | predicted gene, 46519 [Source:MGI Symb | 0 | 0 | 0 | 0 | 0 | 0 |
| 53461 | ENSMUSG00000115970 | 6.865166075 | 0.396118367 | 0.889585953 | 0.445283972 | 0.656114549 | 0.92651636 | 8430426J06 | RIKEN cDNA 8430426J06 gene [Source:M | 1.6062 | 11.307 | 5.0312 | 6.7478 | 9.8156 | 6.6831 |
| 53462 | ENSMUSG00000115971 | 0 | NA | NA | NA | NA | NA | Gm10856 | predicted gene 10856 [Source:MGI Symb | 0 | 0 | 0 | 0 | 0 | 0 |
| 53463 | ENSMUSG00000115973 | 0 | NA | NA | NA | NA | NA | Gm36002 | predicted gene, 36002 [Source:MGI Symb | 0 | 0 | 0 | 0 | 0 | 0 |
| 53464 | ENSMUSG00000115975 | 0 | NA | NA | NA | NA | NA | Gm49526 | predicted gene, 49526 [Source:MGI Symb | 0 | 0 | 0 | 0 | 0 | 0 |
| 53465 | ENSMUSG00000115976 | 0.51014012 | 1.031855138 | 3.09002894 | 0.333930574 | 0.738431948 | NA | Gm31462 | predicted gene, 31462 [Source:MGI Symb | 0 | 0 | 1.0062 | 0.964 | 1.0906 | 0 |
| 53466 | ENSMUSG00000115979 | 31.77070663 | 1.10461064 | 0.525229694 | 2.103100133 | 0.035457017 | 0.252635616 | Gm49410 | predicted gene, 49410 [Source:MGI Symb | 24.093 | 7.1954 | 29.181 | 67.478 | 38.172 | 24.505 |
| 53467 | ENSMUSG00000115980 | 61.61027414 | -0.024173827 | 0.29659803 | -0.081503667 | 0.935041409 | 0.988420305 | Gm49437 | predicted gene, 49437 [Source:MGI Symb | 60.233 | 57.563 | 68.425 | 52.055 | 54.531 | 76.855 |
| 53468 | ENSMUSG00000115983 | 0 | NA | NA | NA | NA | NA | Gm5476 | predicted gene 5476 [Source:MGI Symbol | 0 | 0 | 0 | 0 | 0 | 0 |
| 53469 | ENSMUSG00000115985 | 0 | NA | NA | NA | NA | NA | Gm18531 | predicted gene, 18531 [Source:MGI Symb | 0 | 0 | 0 | 0 | 0 | 0 |
| 53470 | ENSMUSG00000115986 | 0 | NA | NA | NA | NA | NA | Gm49439 | predicted gene, 49439 [Source:MGI Symb | 0 | 0 | 0 | 0 | 0 | 0 |
| 53471 | ENSMUSG00000115987 | 699.0338307 | -0.109052722 | 0.119091115 | -0.915708293 | 0.359819914 | 0.799452417 | Vps28 | vacuolar protein sorting 28 [Source:MGI S | 720.38 | 732.91 | 722.48 | 610.2 | 716.54 | 691.7 |
| 53472 | ENSMUSG00000115988 | 0 | NA | NA | NA | NA | NA | Gm49441 | predicted gene, 49441 [Source:MGI Symb | 0 | 0 | 0 | 0 | 0 | 0 |
| 53473 | ENSMUSG00000115989 | 0 | NA | NA | NA | NA | NA | Gm49533 | predicted gene, 49533 [Source:MGI Symb | 0 | 0 | 0 | 0 | 0 | 0 |
| 53474 | ENSMUSG00000115992 | 0 | NA | NA | NA | NA | NA | Gm10832 | predicted gene 10832 [Source:MGI Symb | 0 | 0 | 0 | 0 | 0 | 0 |
| 53475 | ENSMUSG00000115993 | 0 | NA | NA | NA | NA | NA | Zfp42-ps1 | zinc finger protein 48, pseudogene 1 [Sou | 0 | 0 | 0 | 0 | 0 | 0 |
| 53476 | ENSMUSG00000115996 | 0 | NA | NA | NA | NA | NA | Gm49462 | predicted gene, 49462 [Source:MGI Symb | 0 | 0 | 0 | 0 | 0 | 0 |

| | | | | | | | | | | | | | | | |
|---|---|---|---|---|---|---|---|---|---|---|---|---|---|---|---|
| 53477 | ENSMUSG00000115997 | 0 | NA | NA | NA | NA | NA | Gm49404 | predicted gene, 49404 [Source:MGI Symb | 0 | 0 | 0 | 0 | 0 | 0 |
| 53478 | ENSMUSG00000116000 | 0 | NA | NA | NA | NA | NA | Gm49497 | predicted gene, 49497 [Source:MGI Symb | 0 | 0 | 0 | 0 | 0 | 0 |
| 53479 | ENSMUSG00000116004 | 2.033199652 | 0.681633122 | 1.511303089 | 0.451023442 | 0.651972653 | NA | Gm49539 | predicted gene, 49539 [Source:MGI Symb | 1.6062 | 2.0558 | 1.0062 | 0.964 | 5.4531 | 1.1138 |
| 53480 | ENSMUSG00000116007 | 0 | NA | NA | NA | NA | NA | Gm4116 | predicted gene 4116 [Source:MGI Symbol | 0 | 0 | 0 | 0 | 0 | 0 |
| 53481 | ENSMUSG00000116008 | 0 | NA | NA | NA | NA | NA | Gm49440 | predicted gene, 49440 [Source:MGI Symb | 0 | 0 | 0 | 0 | 0 | 0 |
| 53482 | ENSMUSG00000116009 | 1.225890109 | 1.582632941 | 2.149231825 | 0.736371443 | 0.461504678 | NA | Gm49503 | predicted gene, 49503 [Source:MGI Symb | 0.8031 | 0 | 1.0062 | 0 | 1.0906 | 4.4554 |
| 53483 | ENSMUSG00000116010 | 0.401551119 | -2.072982726 | 4.008251979 | -0.517178744 | 0.605031388 | NA | Gm36026 | predicted gene, 36026 [Source:MGI Symb | 2.4093 | 0 | 0 | 0 | 0 | 0 |
| 53484 | ENSMUSG00000116011 | 0 | NA | NA | NA | NA | NA | Gm49501 | predicted gene, 49501 [Source:MGI Symb | 0 | 0 | 0 | 0 | 0 | 0 |
| 53485 | ENSMUSG00000116013 | 0 | NA | NA | NA | NA | NA | C87114 | expressed sequence C87114 [Source:MGI | 0 | 0 | 0 | 0 | 0 | 0 |
| 53486 | ENSMUSG00000116016 | 0 | NA | NA | NA | NA | NA | Gm49496 | predicted gene, 49496 [Source:MGI Symb | 0 | 0 | 0 | 0 | 0 | 0 |
| 53487 | ENSMUSG00000116017 | 0 | NA | NA | NA | NA | NA | Gm18154 | predicted gene, 18154 [Source:MGI Symb | 0 | 0 | 0 | 0 | 0 | 0 |
| 53488 | ENSMUSG00000116021 | 0 | NA | NA | NA | NA | NA | Gm49474 | predicted gene, 49474 [Source:MGI Symb | 0 | 0 | 0 | 0 | 0 | 0 |
| 53489 | ENSMUSG00000116024 | 0 | NA | NA | NA | NA | NA | Gm49527 | predicted gene, 49527 [Source:MGI Symb | 0 | 0 | 0 | 0 | 0 | 0 |
| 53490 | ENSMUSG00000116025 | 0 | NA | NA | NA | NA | NA | Gm49517 | predicted gene, 49517 [Source:MGI Symb | 0 | 0 | 0 | 0 | 0 | 0 |
| 53491 | ENSMUSG00000116029 | 0 | NA | NA | NA | NA | NA | Gm41414 | predicted gene, 41414 [Source:MGI Symb | 0 | 0 | 0 | 0 | 0 | 0 |
| 53492 | ENSMUSG00000116030 | 0 | NA | NA | NA | NA | NA | Gm49500 | predicted gene, 49500 [Source:MGI Symb | 0 | 0 | 0 | 0 | 0 | 0 |
| 53493 | ENSMUSG00000116031 | 0 | NA | NA | NA | NA | NA | D030024E09 | RIKEN cDNA D030024E09 gene [Source:M | 0 | 0 | 0 | 0 | 0 | 0 |
| 53494 | ENSMUSG00000116033 | 0.181769715 | 1.126924175 | 4.080472857 | 0.276174898 | 0.782413736 | NA | Gm49442 | predicted gene, 49442 [Source:MGI Symb | 0 | 0 | 0 | 0 | 1.0906 | 0 |
| 53495 | ENSMUSG00000116038 | 0 | NA | NA | NA | NA | NA | Gm46563 | predicted gene, 46563 [Source:MGI Symb | 0 | 0 | 0 | 0 | 0 | 0 |
| 53496 | ENSMUSG00000116039 | 0 | NA | NA | NA | NA | NA | Gm49483 | predicted gene, 49483 [Source:MGI Symb | 0 | 0 | 0 | 0 | 0 | 0 |
| 53497 | ENSMUSG00000116040 | 0 | NA | NA | NA | NA | NA | Gm49464 | predicted gene, 49464 [Source:MGI Symb | 0 | 0 | 0 | 0 | 0 | 0 |
| 53498 | ENSMUSG00000116041 | 0 | NA | NA | NA | NA | NA | Gm20369 | predicted gene, 20369 [Source:MGI Symb | 0 | 0 | 0 | 0 | 0 | 0 |
| 53499 | ENSMUSG00000116044 | 0 | NA | NA | NA | NA | NA | Gm49487 | predicted gene, 49487 [Source:MGI Symb | 0 | 0 | 0 | 0 | 0 | 0 |
| 53500 | ENSMUSG00000116045 | 0 | NA | NA | NA | NA | NA | NA | NA | 0 | 0 | 0 | 0 | 0 | 0 |
| 53501 | ENSMUSG00000116048 | 0 | NA | NA | NA | NA | NA | Septin2 | septin 2 [Source:MGI Symbol;Acc:MGI:97 | 0 | 0 | 0 | 0 | 0 | 0 |
| 53502 | ENSMUSG00000116056 | 0 | NA | NA | NA | NA | NA | Gm4544 | predicted gene 4544 [Source:MGI Symbol | 0 | 0 | 0 | 0 | 0 | 0 |
| 53503 | ENSMUSG00000116057 | 0 | NA | NA | NA | NA | NA | Vmn1r216 | vomeronasal 1 receptor 216 [Source:MGI | 0 | 0 | 0 | 0 | 0 | 0 |
| 53504 | ENSMUSG00000116058 | 0 | NA | NA | NA | NA | NA | Gm8843 | predicted gene 8843 [Source:MGI Symbol | 0 | 0 | 0 | 0 | 0 | 0 |
| 53505 | ENSMUSG00000116061 | 0 | NA | NA | NA | NA | NA | Gm49472 | predicted gene, 49472 [Source:MGI Symb | 0 | 0 | 0 | 0 | 0 | 0 |
| 53506 | ENSMUSG00000116066 | 0.730949792 | 3.072456834 | 2.934263219 | 1.04709653 | 0.29505506 | NA | Gm49477 | predicted gene, 49477 [Source:MGI Symb | 0 | 0 | 0 | 0 | 3.2719 | 1.1138 |
| 53507 | ENSMUSG00000116068 | 0 | NA | NA | NA | NA | NA | Gm49457 | predicted gene, 49457 [Source:MGI Symb | 0 | 0 | 0 | 0 | 0 | 0 |
| 53508 | ENSMUSG00000116069 | 0 | NA | NA | NA | NA | NA | Gm49510 | predicted gene, 49510 [Source:MGI Symb | 0 | 0 | 0 | 0 | 0 | 0 |
| 53509 | ENSMUSG00000116071 | 0.517184577 | -0.812115922 | 3.393570547 | -0.239310163 | 0.810865085 | NA | Gm49433 | predicted gene, 49433 [Source:MGI Symb | 0 | 0 | 2.0125 | 0 | 1.0906 | 0 |
| 53510 | ENSMUSG00000116072 | 0 | NA | NA | NA | NA | NA | Gm46526 | predicted gene, 46526 [Source:MGI Symb | 0 | 0 | 0 | 0 | 0 | 0 |
| 53511 | ENSMUSG00000116074 | 0 | NA | NA | NA | NA | NA | Gm49491 | predicted gene, 49491 [Source:MGI Symb | 0 | 0 | 0 | 0 | 0 | 0 |
| 53512 | ENSMUSG00000116075 | 0 | NA | NA | NA | NA | NA | Gm49476 | predicted gene, 49476 [Source:MGI Symb | 0 | 0 | 0 | 0 | 0 | 0 |
| 53513 | ENSMUSG00000116079 | 2.434905757 | 1.480975869 | 1.483054893 | 0.998598147 | 0.317989398 | NA | Gm36245 | predicted gene, 36245 [Source:MGI Symb | 0.8031 | 3.0838 | 0 | 1.928 | 5.4531 | 3.3415 |
| 53514 | ENSMUSG00000116081 | 0 | NA | NA | NA | NA | NA | Gm49593 | predicted gene, 49593 [Source:MGI Symb | 0 | 0 | 0 | 0 | 0 | 0 |
| 53515 | ENSMUSG00000116083 | 2.503807571 | -0.016359068 | 1.458764136 | -0.011214334 | 0.991052444 | NA | Gm5469 | predicted gene 5469 [Source:MGI Symbol | 2.4093 | 5.1396 | 0 | 1.928 | 1.0906 | 4.4554 |
| 53516 | ENSMUSG00000116088 | 0 | NA | NA | NA | NA | NA | Gm49460 | predicted gene, 49460 [Source:MGI Symb | 0 | 0 | 0 | 0 | 0 | 0 |
| 53517 | ENSMUSG00000116089 | 0.185640647 | 1.126924175 | 4.080472857 | 0.276174898 | 0.782413736 | NA | Gm7859 | predicted gene 7859 [Source:MGI Symbol | 0 | 0 | 0 | 0 | 0 | 1.1138 |
| 53518 | ENSMUSG00000116090 | 0.965523032 | -0.944628791 | 2.267086424 | -0.416670834 | 0.67691919 | NA | Rpl19-ps6 | ribosomal protein L19, pseudogene 6 [Sou | 0.8031 | 2.0558 | 1.0062 | 1.928 | 0 | 0 |
| 53519 | ENSMUSG00000116091 | 0 | NA | NA | NA | NA | NA | Gm49479 | predicted gene, 49479 [Source:MGI Symb | 0 | 0 | 0 | 0 | 0 | 0 |
| 53520 | ENSMUSG00000116092 | 0 | NA | NA | NA | NA | NA | Gm49422 | predicted gene, 49422 [Source:MGI Symb | 0 | 0 | 0 | 0 | 0 | 0 |
| 53521 | ENSMUSG00000116093 | 0 | NA | NA | NA | NA | NA | Gm3888 | predicted gene 3888 [Source:MGI Symbol | 0 | 0 | 0 | 0 | 0 | 0 |
| 53522 | ENSMUSG00000116095 | 0.133850373 | -0.796629176 | 4.080472857 | -0.195229623 | 0.84521319 | NA | Gm49549 | predicted gene, 49549 [Source:MGI Symb | 0.8031 | 0 | 0 | 0 | 0 | 0 |
| 53523 | ENSMUSG00000116096 | 0 | NA | NA | NA | NA | NA | 4933404G15 | RIKEN cDNA 4933404G15 gene [Source:M | 0 | 0 | 0 | 0 | 0 | 0 |
| 53524 | ENSMUSG00000116097 | 53.09966353 | 0.316311017 | 0.314551122 | 1.00559494 | 0.314610459 | 0.761014653 | Gm36738 | predicted gene, 36738 [Source:MGI Symb | 52.202 | 38.033 | 51.318 | 49.163 | 62.165 | 65.717 |
| 53525 | ENSMUSG00000116102 | 0 | NA | NA | NA | NA | NA | Gm20082 | predicted gene, 20082 [Source:MGI Symb | 0 | 0 | 0 | 0 | 0 | 0 |
| 53526 | ENSMUSG00000116104 | 0 | NA | NA | NA | NA | NA | Gm46524 | predicted gene, 46524 [Source:MGI Symb | 0 | 0 | 0 | 0 | 0 | 0 |
| 53527 | ENSMUSG00000116107 | 0 | NA | NA | NA | NA | NA | Gm49423 | predicted gene, 49423 [Source:MGI Symb | 0 | 0 | 0 | 0 | 0 | 0 |
| 53528 | ENSMUSG00000116109 | 0 | NA | NA | NA | NA | NA | Gm49498 | predicted gene, 49498 [Source:MGI Symb | 0 | 0 | 0 | 0 | 0 | 0 |
| 53529 | ENSMUSG00000116113 | 0.17131964 | -0.796629176 | 4.080472857 | -0.195229623 | 0.84521319 | NA | Gm49545 | predicted gene, 49545 [Source:MGI Symb | 0 | 1.0279 | 0 | 0 | 0 | 0 |
| 53530 | ENSMUSG00000116114 | 1.718380712 | -0.529318392 | 1.691001853 | -0.313020587 | 0.754265022 | NA | Gm35853 | predicted gene, 35853 [Source:MGI Symb | 4.0155 | 0 | 2.0125 | 0.964 | 1.0906 | 2.2277 |
| 53531 | ENSMUSG00000116116 | 0.528073336 | 2.60973587 | 2.979250867 | 0.875970499 | 0.381046072 | NA | Gm671 | predicted gene 671 [Source:MGI Symbol;A | 0 | 0 | 0 | 0.964 | 1.0906 | 1.1138 |
| 53532 | ENSMUSG00000116117 | 1.321838044 | 0.121187624 | 1.989776353 | 0.060905148 | 0.951434749 | NA | Cyp2d39-ps | cytochrome P450, family 2, subfamily d, p | 0.8031 | 0 | 3.0187 | 1.928 | 2.1812 | 0 |

| | | | | | | | | | | | | | | | |
|---|---|---|---|---|---|---|---|---|---|---|---|---|---|---|---|
| 53533 | ENSMUSG00000116118 | 0 | NA | NA | NA | NA | NA | Gm49511 | predicted gene, 49511 [Source:MGI Symb | 0 | 0 | 0 | 0 | 0 | 0 |
| 53534 | ENSMUSG00000116120 | 1.238723049 | 0.879930073 | 2.259902744 | 0.389366346 | 0.697005164 | NA | Eif2s3x-ps1 | eukaryotic translation initiation factor 2, s | 1.6062 | 0 | 1.0062 | 4.8199 | 0 | 0 |
| 53535 | ENSMUSG00000116121 | 0 | NA | NA | NA | NA | NA | Gm49486 | predicted gene, 49486 [Source:MGI Symb | 0 | 0 | 0 | 0 | 0 | 0 |
| 53536 | ENSMUSG00000116122 | 0 | NA | NA | NA | NA | NA | Gm49537 | predicted gene, 49537 [Source:MGI Symb | 0 | 0 | 0 | 0 | 0 | 0 |
| 53537 | ENSMUSG00000116123 | 0 | NA | NA | NA | NA | NA | Gm49451 | predicted gene, 49451 [Source:MGI Symb | 0 | 0 | 0 | 0 | 0 | 0 |
| 53538 | ENSMUSG00000116124 | 0 | NA | NA | NA | NA | NA | Gm49420 | predicted gene, 49420 [Source:MGI Symb | 0 | 0 | 0 | 0 | 0 | 0 |
| 53539 | ENSMUSG00000116125 | 0 | NA | NA | NA | NA | NA | Gm41361 | predicted gene, 41361 [Source:MGI Symb | 0 | 0 | 0 | 0 | 0 | 0 |
| 53540 | ENSMUSG00000116127 | 0 | NA | NA | NA | NA | NA | Gm8375 | predicted gene 8375 [Source:MGI Symbol | 0 | 0 | 0 | 0 | 0 | 0 |
| 53541 | ENSMUSG00000116129 | 0 | NA | NA | NA | NA | NA | Gm5477 | predicted gene 5477 [Source:MGI Symbol | 0 | 0 | 0 | 0 | 0 | 0 |
| 53542 | ENSMUSG00000116130 | 137.5632464 | 0.310090066 | 0.214240528 | 1.447392182 | 0.147787114 | 0.559179858 | Gm49431 | predicted gene, 49431 [Source:MGI Symb | 110.83 | 107.93 | 149.93 | 143.63 | 153.78 | 159.28 |
| 53543 | ENSMUSG00000116131 | 0 | NA | NA | NA | NA | NA | Gm49506 | predicted gene, 49506 [Source:MGI Symb | 0 | 0 | 0 | 0 | 0 | 0 |
| 53544 | ENSMUSG00000116132 | 0 | NA | NA | NA | NA | NA | Gm49428 | predicted gene, 49428 [Source:MGI Symb | 0 | 0 | 0 | 0 | 0 | 0 |
| 53545 | ENSMUSG00000116136 | 0 | NA | NA | NA | NA | NA | Gm3861 | predicted gene 3861 [Source:MGI Symbol | 0 | 0 | 0 | 0 | 0 | 0 |
| 53546 | ENSMUSG00000116138 | 825.2000794 | -0.028017924 | 0.111331022 | -0.251663223 | 0.801301386 | 0.96255706 | C030006K11 | RIKEN cDNA C030006K11 gene [Source:M | 870.56 | 824.39 | 803.99 | 794.32 | 863.77 | 794.17 |
| 53547 | ENSMUSG00000116140 | 0 | NA | NA | NA | NA | NA | Gm7748 | predicted gene 7748 [Source:MGI Symbol | 0 | 0 | 0 | 0 | 0 | 0 |
| 53548 | ENSMUSG00000116141 | 19.76927892 | -0.447404754 | 0.489939907 | -0.913182919 | 0.361146346 | 0.800074801 | Gm49495 | predicted gene, 49495 [Source:MGI Symb | 25.699 | 21.586 | 21.131 | 18.316 | 19.631 | 12.252 |
| 53549 | ENSMUSG00000116145 | 4.890166368 | -0.116158251 | 1.129133759 | -0.102873774 | 0.918063153 | NA | 5730521K06 | RIKEN cDNA 5730521K06 gene [Source:M | 3.2124 | 5.1396 | 7.0437 | 10.604 | 0 | 3.3415 |
| 53550 | ENSMUSG00000116149 | 0 | NA | NA | NA | NA | NA | Gm18476 | predicted gene, 18476 [Source:MGI Symb | 0 | 0 | 0 | 0 | 0 | 0 |
| 53551 | ENSMUSG00000116150 | 0 | NA | NA | NA | NA | NA | NA | NA | 0 | 0 | 0 | 0 | 0 | 0 |
| 53552 | ENSMUSG00000116151 | 0 | NA | NA | NA | NA | NA | Gm49535 | predicted gene, 49535 [Source:MGI Symb | 0 | 0 | 0 | 0 | 0 | 0 |
| 53553 | ENSMUSG00000116155 | 0 | NA | NA | NA | NA | NA | Gm34646 | predicted gene, 34646 [Source:MGI Symb | 0 | 0 | 0 | 0 | 0 | 0 |
| 53554 | ENSMUSG00000116157 | 0 | NA | NA | NA | NA | NA | Cyp2d67-ps | cytochrome P450, family 2, subfamily d, p | 0 | 0 | 0 | 0 | 0 | 0 |
| 53555 | ENSMUSG00000116158 | 0 | NA | NA | NA | NA | NA | Kiss1 | KiSS-1 metastasis-suppressor [Source:MG | 0 | 0 | 0 | 0 | 0 | 0 |
| 53556 | ENSMUSG00000116160 | 0.476489653 | -2.293616034 | 3.421506946 | -0.670352587 | 0.502633051 | NA | Gm41392 | predicted gene, 41392 [Source:MGI Symb | 0.8031 | 2.0558 | 0 | 0 | 0 | 0 |
| 53557 | ENSMUSG00000116162 | 0 | NA | NA | NA | NA | NA | Gm36329 | predicted gene, 36329 [Source:MGI Symb | 0 | 0 | 0 | 0 | 0 | 0 |
| 53558 | ENSMUSG00000116164 | 17.55086003 | -0.085620072 | 0.540299507 | -0.158467797 | 0.874088194 | 0.979986838 | 1110013H19 | RIKEN cDNA 1110013H19 gene [Source:M | 12.85 | 16.447 | 25.156 | 22.171 | 13.087 | 15.594 |
| 53559 | ENSMUSG00000116165 | 17.2908534 | 0.276435771 | 0.569812632 | 0.485134508 | 0.627580935 | 0.920328819 | Pdxp | pyridoxal (pyridoxine, vitamin B6) phosph | 9.6372 | 18.503 | 19.119 | 28.919 | 13.087 | 14.48 |
| 53560 | ENSMUSG00000116166 | 0 | NA | NA | NA | NA | NA | Gm49403 | predicted gene, 49403 [Source:MGI Symb | 0 | 0 | 0 | 0 | 0 | 0 |
| 53561 | ENSMUSG00000116167 | 0 | NA | NA | NA | NA | NA | 4930504C09 | RIKEN cDNA 4930504C09 gene [Source:M | 0 | 0 | 0 | 0 | 0 | 0 |
| 53562 | ENSMUSG00000116168 | 0 | NA | NA | NA | NA | NA | Gm49430 | predicted gene, 49430 [Source:MGI Symb | 0 | 0 | 0 | 0 | 0 | 0 |
| 53563 | ENSMUSG00000116170 | 0 | NA | NA | NA | NA | NA | Gm6332 | predicted gene 6332 [Source:MGI Symbol | 0 | 0 | 0 | 0 | 0 | 0 |
| 53564 | ENSMUSG00000116171 | 0 | NA | NA | NA | NA | NA | 4930543I03F | RIKEN cDNA 4930543I03 gene [Source:MC | 0 | 0 | 0 | 0 | 0 | 0 |
| 53565 | ENSMUSG00000116173 | 0.870712616 | 0.665767941 | 2.367602424 | 0.281199214 | 0.778557607 | NA | Gm32126 | predicted gene, 32126 [Source:MGI Symb | 0 | 2.0558 | 0 | 0.964 | 1.0906 | 1.1138 |
| 53566 | ENSMUSG00000116174 | 1.244960844 | 2.670927122 | 2.112382515 | 1.264414519 | 0.206081284 | NA | Gm10362 | predicted gene 10362 [Source:MGI Symbo | 0 | 0 | 1.0062 | 0.964 | 3.2719 | 2.2277 |
| 53567 | ENSMUSG00000116178 | 0 | NA | NA | NA | NA | NA | Gm49529 | predicted gene, 49529 [Source:MGI Symb | 0 | 0 | 0 | 0 | 0 | 0 |
| 53568 | ENSMUSG00000116179 | 0 | NA | NA | NA | NA | NA | Olfr290 | olfactory receptor 290 [Source:MGI Symb | 0 | 0 | 0 | 0 | 0 | 0 |
| 53569 | ENSMUSG00000116180 | 6.43708747 | 0.566650068 | 0.896008228 | 0.632416143 | 0.527114988 | 0.888334265 | Gm49492 | predicted gene, 49492 [Source:MGI Symb | 6.4248 | 1.0279 | 8.05 | 7.7118 | 8.7249 | 6.6831 |
| 53570 | ENSMUSG00000116182 | 0 | NA | NA | NA | NA | NA | Gm33666 | predicted gene, 33666 [Source:MGI Symb | 0 | 0 | 0 | 0 | 0 | 0 |
| 53571 | ENSMUSG00000116183 | 0 | NA | NA | NA | NA | NA | Gm49480 | predicted gene, 49480 [Source:MGI Symb | 0 | 0 | 0 | 0 | 0 | 0 |
| 53572 | ENSMUSG00000116184 | 0 | NA | NA | NA | NA | NA | 4632433K11 | RIKEN cDNA 4632433K11 gene [Source:M | 0 | 0 | 0 | 0 | 0 | 0 |
| 53573 | ENSMUSG00000116185 | 0 | NA | NA | NA | NA | NA | Gm49411 | predicted gene, 49411 [Source:MGI Symb | 0 | 0 | 0 | 0 | 0 | 0 |
| 53574 | ENSMUSG00000116187 | 0 | NA | NA | NA | NA | NA | Gm49456 | predicted gene, 49456 [Source:MGI Symb | 0 | 0 | 0 | 0 | 0 | 0 |
| 53575 | ENSMUSG00000116188 | 0 | NA | NA | NA | NA | NA | Gm49424 | predicted gene, 49424 [Source:MGI Symb | 0 | 0 | 0 | 0 | 0 | 0 |
| 53576 | ENSMUSG00000116190 | 0 | NA | NA | NA | NA | NA | Gm41386 | predicted gene, 41386 [Source:MGI Symb | 0 | 0 | 0 | 0 | 0 | 0 |
| 53577 | ENSMUSG00000116191 | 0 | NA | NA | NA | NA | NA | Gm19199 | predicted gene, 19199 [Source:MGI Symb | 0 | 0 | 0 | 0 | 0 | 0 |
| 53578 | ENSMUSG00000116192 | 0 | NA | NA | NA | NA | NA | Gm18869 | predicted gene, 18869 [Source:MGI Symb | 0 | 0 | 0 | 0 | 0 | 0 |
| 53579 | ENSMUSG00000116194 | 0 | NA | NA | NA | NA | NA | Gm49434 | predicted gene, 49434 [Source:MGI Symb | 0 | 0 | 0 | 0 | 0 | 0 |
| 53580 | ENSMUSG00000116197 | 0 | NA | NA | NA | NA | NA | Gm38619 | predicted gene, 38619 [Source:MGI Symb | 0 | 0 | 0 | 0 | 0 | 0 |
| 53581 | ENSMUSG00000116199 | 0 | NA | NA | NA | NA | NA | Gm49438 | predicted gene, 49438 [Source:MGI Symb | 0 | 0 | 0 | 0 | 0 | 0 |
| 53582 | ENSMUSG00000116202 | 0 | NA | NA | NA | NA | NA | Rpl10a-ps3 | ribosomal protein L10A, pseudogene 3 [Sc | 0 | 0 | 0 | 0 | 0 | 0 |
| 53583 | ENSMUSG00000116204 | 0.335414862 | -1.779047242 | 4.030804179 | -0.441362855 | 0.658950329 | NA | Gm36480 | predicted gene, 36480 [Source:MGI Symb | 0 | 0 | 2.0125 | 0 | 0 | 0 |
| 53584 | ENSMUSG00000116207 | 0 | NA | NA | NA | NA | NA | Nnt | nicotinamide nucleotide transhydrogenas | 0 | 0 | 0 | 0 | 0 | 0 |
| 53585 | ENSMUSG00000116210 | 0 | NA | NA | NA | NA | NA | Spn-ps | sialophorin, pseudogene [Source:MGI Syn | 0 | 0 | 0 | 0 | 0 | 0 |
| 53586 | ENSMUSG00000116212 | 0 | NA | NA | NA | NA | NA | Gm18979 | predicted gene, 18979 [Source:MGI Symb | 0 | 0 | 0 | 0 | 0 | 0 |
| 53587 | ENSMUSG00000116213 | 0 | NA | NA | NA | NA | NA | Gm5404 | predicted gene 5404 [Source:MGI Symbol | 0 | 0 | 0 | 0 | 0 | 0 |
| 53588 | ENSMUSG00000116215 | 0.185640647 | 1.126924175 | 4.080472857 | 0.276174898 | 0.782413736 | NA | Gm5470 | predicted gene 5470 [Source:MGI Symbol | 0 | 0 | 0 | 0 | 0 | 1.1138 |

| | | | | | | | | | | | | | | | |
|---|---|---|---|---|---|---|---|---|---|---|---|---|---|---|---|
| 53589 | ENSMUSG00000116217 | 1.751540361 | -1.167820134 | 1.970155728 | -0.592755241 | 0.553344966 | NA | Gm46495 | predicted gene, 46495 [Source:MGI Symb | 3.2124 | 0 | 4.025 | 0 | 3.2719 | 0 |
| 53590 | ENSMUSG00000116219 | 0.765090549 | -0.195763385 | 2.933488969 | -0.066733977 | 0.946793485 | NA | Gm28023 | predicted gene, 28023 [Source:MGI Symb | 2.4093 | 0 | 0 | 0 | 2.1812 | 0 |
| 53591 | ENSMUSG00000116220 | 1.678566237 | -1.865386216 | 1.90929337 | -0.977003453 | 0.328567443 | NA | A430088P11 | RIKEN cDNA A430088P11 gene [Source:M | 0.8031 | 2.0558 | 5.0312 | 0 | 2.1812 | 0 |
| 53592 | ENSMUSG00000116222 | 0 | NA | NA | NA | NA | NA | Gm34593 | predicted gene, 34593 [Source:MGI Symb | 0 | 0 | 0 | 0 | 0 | 0 |
| 53593 | ENSMUSG00000116223 | 0.133850373 | -0.796629176 | 4.080472857 | -0.195229623 | 0.84521319 | NA | Gm41396 | predicted gene, 41396 [Source:MGI Symb | 0.8031 | 0 | 0 | 0 | 0 | 0 |
| 53594 | ENSMUSG00000116226 | 5.113634657 | -0.999554065 | 1.194051405 | -0.837111418 | 0.402529943 | 0.828953151 | Gm49502 | predicted gene, 49502 [Source:MGI Symb | 6.4248 | 0 | 14.087 | 5.7839 | 3.2719 | 1.1138 |
| 53595 | ENSMUSG00000116227 | 0 | NA | NA | NA | NA | NA | Gm49513 | predicted gene, 49513 [Source:MGI Symb | 0 | 0 | 0 | 0 | 0 | 0 |
| 53596 | ENSMUSG00000116229 | 0 | NA | NA | NA | NA | NA | Gm49470 | predicted gene, 49470 [Source:MGI Symb | 0 | 0 | 0 | 0 | 0 | 0 |
| 53597 | ENSMUSG00000116231 | 0 | NA | NA | NA | NA | NA | Gm36330 | predicted gene, 36330 [Source:MGI Symb | 0 | 0 | 0 | 0 | 0 | 0 |
| 53598 | ENSMUSG00000116234 | 0 | NA | NA | NA | NA | NA | Gm49550 | predicted gene, 49550 [Source:MGI Symb | 0 | 0 | 0 | 0 | 0 | 0 |
| 53599 | ENSMUSG00000116237 | 0 | NA | NA | NA | NA | NA | Gm30339 | predicted gene, 30339 [Source:MGI Symb | 0 | 0 | 0 | 0 | 0 | 0 |
| 53600 | ENSMUSG00000116238 | 4.506624157 | -0.322817933 | 1.376752396 | -0.234477844 | 0.814614032 | NA | Gm49413 | predicted gene, 49413 [Source:MGI Symb | 0.8031 | 13.363 | 1.0062 | 9.6398 | 0 | 2.2277 |
| 53601 | ENSMUSG00000116239 | 1.606287867 | -1.175097167 | 2.050416046 | -0.57310182 | 0.566575755 | NA | Gm49520 | predicted gene, 49520 [Source:MGI Symb | 1.6062 | 5.1396 | 0 | 2.8919 | 0 | 0 |
| 53602 | ENSMUSG00000116241 | 0 | NA | NA | NA | NA | NA | Gm49458 | predicted gene, 49458 [Source:MGI Symb | 0 | 0 | 0 | 0 | 0 | 0 |
| 53603 | ENSMUSG00000116242 | 0 | NA | NA | NA | NA | NA | Gm18815 | predicted gene, 18815 [Source:MGI Symb | 0 | 0 | 0 | 0 | 0 | 0 |
| 53604 | ENSMUSG00000116243 | 0 | NA | NA | NA | NA | NA | Gm49552 | predicted gene, 49552 [Source:MGI Symb | 0 | 0 | 0 | 0 | 0 | 0 |
| 53605 | ENSMUSG00000116244 | 0 | NA | NA | NA | NA | NA | Gm49461 | predicted gene, 49461 [Source:MGI Symb | 0 | 0 | 0 | 0 | 0 | 0 |
| 53606 | ENSMUSG00000116245 | 0 | NA | NA | NA | NA | NA | Gm49452 | predicted gene, 49452 [Source:MGI Symb | 0 | 0 | 0 | 0 | 0 | 0 |
| 53607 | ENSMUSG00000116246 | 0 | NA | NA | NA | NA | NA | NA | NA | 0 | 0 | 0 | 0 | 0 | 0 |
| 53608 | ENSMUSG00000116247 | 0 | NA | NA | NA | NA | NA | NA | NA | 0 | 0 | 0 | 0 | 0 | 0 |
| 53609 | ENSMUSG00000116249 | 1.015555487 | -0.983278458 | 2.45507575 | -0.400508399 | 0.688782098 | NA | Gm49484 | predicted gene, 49484 [Source:MGI Symb | 4.0155 | 0 | 0 | 0.964 | 0 | 1.1138 |
| 53610 | ENSMUSG00000116250 | 0 | NA | NA | NA | NA | NA | Gm34939 | predicted gene, 34939 [Source:MGI Symb | 0 | 0 | 0 | 0 | 0 | 0 |
| 53611 | ENSMUSG00000116253 | 1.434761637 | 0.447626221 | 1.932695724 | 0.231607187 | 0.816843125 | NA | Gm41392 | predicted gene, 41392 [Source:MGI Symb | 1.6062 | 2.0558 | 0 | 3.8559 | 1.0906 | 0 |
| 53612 | ENSMUSG00000116255 | 0.428363721 | -0.543730382 | 3.624677344 | -0.15000794 | 0.880758351 | NA | Gm20564 | predicted gene, 20564 [Source:MGI Symb | 1.6062 | 0 | 0 | 0.964 | 0 | 0 |
| 53613 | ENSMUSG00000116256 | 0 | NA | NA | NA | NA | NA | Gm32405 | predicted gene, 32405 [Source:MGI Symb | 0 | 0 | 0 | 0 | 0 | 0 |
| 53614 | ENSMUSG00000116260 | 0 | NA | NA | NA | NA | NA | 1700041B01 | RIKEN cDNA 1700041B01 gene [Source:M | 0 | 0 | 0 | 0 | 0 | 0 |
| 53615 | ENSMUSG00000116262 | 225.5559706 | 0.054917356 | 0.212552847 | 0.258370363 | 0.796121087 | 0.962236316 | Gm49544 | predicted gene, 49544 [Source:MGI Symb | 173.47 | 195.3 | 295.84 | 245.81 | 227.94 | 214.97 |
| 53616 | ENSMUSG00000116263 | 0 | NA | NA | NA | NA | NA | Gm49473 | predicted gene, 49473 [Source:MGI Symb | 0 | 0 | 0 | 0 | 0 | 0 |
| 53617 | ENSMUSG00000116265 | 1.092148988 | -1.219782293 | 2.337692795 | -0.521788961 | 0.601817277 | NA | Cyp2d32-ps | cytochrome P450, family 2, subfamily d, p | 1.6062 | 0 | 3.0187 | 1.928 | 0 | 0 |
| 53618 | ENSMUSG00000116270 | 14.70234541 | 0.520362261 | 0.599715999 | 0.867681139 | 0.385568914 | 0.816437825 | Gm1600 | predicted gene 1600 [Source:MGI Symbol | 17.668 | 7.1954 | 11.069 | 12.532 | 16.359 | 23.391 |
| 53619 | ENSMUSG00000116272 | 0 | NA | NA | NA | NA | NA | Gm49540 | predicted gene, 49540 [Source:MGI Symb | 0 | 0 | 0 | 0 | 0 | 0 |
| 53620 | ENSMUSG00000116274 | 0 | NA | NA | NA | NA | NA | Gm18724 | predicted gene, 18724 [Source:MGI Symb | 0 | 0 | 0 | 0 | 0 | 0 |
| 53621 | ENSMUSG00000116275 | 0 | NA | NA | NA | NA | NA | Zc3h11a | zinc finger CCCH type containing 11A [Sou | 0 | 0 | 0 | 0 | 0 | 0 |
| 53622 | ENSMUSG00000116280 | 0 | NA | NA | NA | NA | NA | Gm49435 | predicted gene, 49435 [Source:MGI Symb | 0 | 0 | 0 | 0 | 0 | 0 |
| 53623 | ENSMUSG00000116283 | 0 | NA | NA | NA | NA | NA | Gm49412 | predicted gene, 49412 [Source:MGI Symb | 0 | 0 | 0 | 0 | 0 | 0 |
| 53624 | ENSMUSG00000116284 | 36.74673527 | 0.043374556 | 0.36742891 | 0.118048839 | 0.90602896 | 0.983991807 | Gm3787 | predicted gene 3787 [Source:MGI Symbol | 30.518 | 33.921 | 44.275 | 34.703 | 42.534 | 34.529 |
| 53625 | ENSMUSG00000116287 | 3.539023376 | 0.619736299 | 1.1743337 | 0.527734407 | 0.597683689 | NA | Gm3924 | predicted gene 3924 [Source:MGI Symbol | 3.2124 | 4.1117 | 1.0062 | 1.928 | 7.6343 | 3.3415 |
| 53626 | ENSMUSG00000116288 | 5.074839855 | -0.071233181 | 0.976058595 | -0.072980436 | 0.941821686 | NA | Gm41349 | predicted gene, 41349 [Source:MGI Symb | 6.4248 | 2.0558 | 7.0437 | 3.8559 | 3.2719 | 7.7969 |
| 53627 | ENSMUSG00000116289 | 0 | NA | NA | NA | NA | NA | Gm31342 | predicted gene, 31342 [Source:MGI Symb | 0 | 0 | 0 | 0 | 0 | 0 |
| 53628 | ENSMUSG00000116290 | 1.675856373 | -2.027189323 | 2.040621806 | -0.993417456 | 0.320506558 | NA | Gm49518 | predicted gene, 49518 [Source:MGI Symb | 4.0155 | 4.1117 | 0 | 1.928 | 0 | 0 |
| 53629 | ENSMUSG00000116291 | 0.181769715 | 1.126924175 | 4.080472857 | 0.276174898 | 0.782413736 | NA | Gm4383 | predicted gene 4383 [Source:MGI Symbol | 0 | 0 | 0 | 0 | 1.0906 | 0 |
| 53630 | ENSMUSG00000116292 | 0 | NA | NA | NA | NA | NA | Gm49551 | predicted gene, 49551 [Source:MGI Symb | 0 | 0 | 0 | 0 | 0 | 0 |
| 53631 | ENSMUSG00000116294 | 0 | NA | NA | NA | NA | NA | Gm49415 | predicted gene, 49415 [Source:MGI Symb | 0 | 0 | 0 | 0 | 0 | 0 |
| 53632 | ENSMUSG00000116295 | 2.61549222 | -1.02569364 | 1.421001135 | -0.721810571 | 0.470410949 | NA | Gm32885 | predicted gene, 32885 [Source:MGI Symb | 6.4248 | 2.0558 | 2.0125 | 1.928 | 3.2719 | 0 |
| 53633 | ENSMUSG00000116296 | 0 | NA | NA | NA | NA | NA | Gm49468 | predicted gene, 49468 [Source:MGI Symb | 0 | 0 | 0 | 0 | 0 | 0 |
| 53634 | ENSMUSG00000116297 | 0 | NA | NA | NA | NA | NA | Gm49546 | predicted gene, 49546 [Source:MGI Symb | 0 | 0 | 0 | 0 | 0 | 0 |
| 53635 | ENSMUSG00000116298 | 0 | NA | NA | NA | NA | NA | 4833412C15 | RIKEN cDNA 4833412C15 gene [Source:M | 0 | 0 | 0 | 0 | 0 | 0 |
| 53636 | ENSMUSG00000116299 | 0 | NA | NA | NA | NA | NA | Gm29904 | predicted gene, 29904 [Source:MGI Symb | 0 | 0 | 0 | 0 | 0 | 0 |
| 53637 | ENSMUSG00000116300 | 0 | NA | NA | NA | NA | NA | Gm49509 | predicted gene, 49509 [Source:MGI Symb | 0 | 0 | 0 | 0 | 0 | 0 |
| 53638 | ENSMUSG00000116301 | 0 | NA | NA | NA | NA | NA | Gm49547 | predicted gene, 49547 [Source:MGI Symb | 0 | 0 | 0 | 0 | 0 | 0 |
| 53639 | ENSMUSG00000116303 | 0 | NA | NA | NA | NA | NA | Gm49465 | predicted gene, 49465 [Source:MGI Symb | 0 | 0 | 0 | 0 | 0 | 0 |
| 53640 | ENSMUSG00000116304 | 0 | NA | NA | NA | NA | NA | Gm49488 | predicted gene, 49488 [Source:MGI Symb | 0 | 0 | 0 | 0 | 0 | 0 |
| 53641 | ENSMUSG00000116305 | 11.06338561 | -0.088676693 | 0.695290509 | -0.127539053 | 0.898513766 | 0.982798755 | Lncppara | long noncoding RNA near Ppara [Source:N | 16.865 | 6.1675 | 11.069 | 12.532 | 14.178 | 5.5692 |
| 53642 | ENSMUSG00000116307 | 0 | NA | NA | NA | NA | NA | Gm49490 | predicted gene, 49490 [Source:MGI Symb | 0 | 0 | 0 | 0 | 0 | 0 |
| 53643 | ENSMUSG00000116308 | 0 | NA | NA | NA | NA | NA | Gm49482 | predicted gene, 49482 [Source:MGI Symb | 0 | 0 | 0 | 0 | 0 | 0 |
| 53644 | ENSMUSG00000116309 | 0 | NA | NA | NA | NA | NA | Gm49418 | predicted gene, 49418 [Source:MGI Symb | 0 | 0 | 0 | 0 | 0 | 0 |

| | | | | | | | | | | | | | | | |
|---|---|---|---|---|---|---|---|---|---|---|---|---|---|---|---|
| 53645 | ENSMUSG00000116311 | 0 | NA | NA | NA | NA | NA | 4930478M13 | RIKEN cDNA 4930478M13 gene [Source:M | 0 | 0 | 0 | 0 | 0 | 0 |
| 53646 | ENSMUSG00000116314 | 0 | NA | NA | NA | NA | NA | Gm49459 | predicted gene, 49459 [Source:MGI Symb | 0 | 0 | 0 | 0 | 0 | 0 |
| 53647 | ENSMUSG00000116317 | 0 | NA | NA | NA | NA | NA | Gm49536 | predicted gene, 49536 [Source:MGI Symb | 0 | 0 | 0 | 0 | 0 | 0 |
| 53648 | ENSMUSG00000116319 | 0 | NA | NA | NA | NA | NA | Gm49429 | predicted gene, 49429 [Source:MGI Symb | 0 | 0 | 0 | 0 | 0 | 0 |
| 53649 | ENSMUSG00000116321 | 0 | NA | NA | NA | NA | NA | Gm49522 | predicted gene, 49522 [Source:MGI Symb | 0 | 0 | 0 | 0 | 0 | 0 |
| 53650 | ENSMUSG00000116323 | 0 | NA | NA | NA | NA | NA | Gm18092 | predicted gene, 18092 [Source:MGI Symb | 0 | 0 | 0 | 0 | 0 | 0 |
| 53651 | ENSMUSG00000116324 | 0 | NA | NA | NA | NA | NA | Gm33696 | predicted gene, 33696 [Source:MGI Symb | 0 | 0 | 0 | 0 | 0 | 0 |
| 53652 | ENSMUSG00000116325 | 0 | NA | NA | NA | NA | NA | Gm46501 | predicted gene, 46501 [Source:MGI Symb | 0 | 0 | 0 | 0 | 0 | 0 |
| 53653 | ENSMUSG00000116331 | 0 | NA | NA | NA | NA | NA | B230214G05 | RIKEN cDNA B230214G05 gene [Source:M | 0 | 0 | 0 | 0 | 0 | 0 |
| 53654 | ENSMUSG00000116335 | 0 | NA | NA | NA | NA | NA | Gm49516 | predicted gene, 49516 [Source:MGI Symb | 0 | 0 | 0 | 0 | 0 | 0 |
| 53655 | ENSMUSG00000116336 | 0 | NA | NA | NA | NA | NA | Gm49425 | predicted gene, 49425 [Source:MGI Symb | 0 | 0 | 0 | 0 | 0 | 0 |
| 53656 | ENSMUSG00000116338 | 0 | NA | NA | NA | NA | NA | 7530414M10 | RIKEN cDNA 7530414M10 gene [Source:M | 0 | 0 | 0 | 0 | 0 | 0 |
| 53657 | ENSMUSG00000116339 | 0 | NA | NA | NA | NA | NA | Gm35933 | predicted gene, 35933 [Source:MGI Symb | 0 | 0 | 0 | 0 | 0 | 0 |
| 53658 | ENSMUSG00000116341 | 244.9297355 | -0.119030634 | 0.529067614 | -0.224981894 | 0.82199336 | 0.969160361 | Cyp2d36-ps | cytochrome P450, family 2, subfamily d, p | 410.39 | 153.16 | 201.25 | 237.14 | 321.73 | 145.91 |
| 53659 | ENSMUSG00000116343 | 0 | NA | NA | NA | NA | NA | Gm6847 | predicted gene 6847 [Source:MGI Symbol | 0 | 0 | 0 | 0 | 0 | 0 |
| 53660 | ENSMUSG00000116345 | 0 | NA | NA | NA | NA | NA | Gm19141 | predicted gene, 19141 [Source:MGI Symb | 0 | 0 | 0 | 0 | 0 | 0 |
| 53661 | ENSMUSG00000116347 | 0 | NA | NA | NA | NA | NA | Gm49416 | predicted gene, 49416 [Source:MGI Symb | 0 | 0 | 0 | 0 | 0 | 0 |
| 53662 | ENSMUSG00000116348 | 0 | NA | NA | NA | NA | NA | Gm18722 | predicted gene, 18722 [Source:MGI Symb | 0 | 0 | 0 | 0 | 0 | 0 |
| 53663 | ENSMUSG00000116350 | 0 | NA | NA | NA | NA | NA | Gm35772 | predicted gene, 35772 [Source:MGI Symb | 0 | 0 | 0 | 0 | 0 | 0 |
| 53664 | ENSMUSG00000116352 | 0 | NA | NA | NA | NA | NA | Gm49447 | predicted gene, 49447 [Source:MGI Symb | 0 | 0 | 0 | 0 | 0 | 0 |
| 53665 | ENSMUSG00000116353 | 0 | NA | NA | NA | NA | NA | Gm49469 | predicted gene, 49469 [Source:MGI Symb | 0 | 0 | 0 | 0 | 0 | 0 |
| 53666 | ENSMUSG00000116354 | 769.6246749 | 0.116580914 | 0.208487039 | 0.559175832 | 0.576041725 | 0.907413147 | Cyp2d38-ps | cytochrome P450, family 2, subfamily d, m | 848.08 | 532.46 | 835.18 | 1044 | 772.16 | 585.88 |
| 53667 | ENSMUSG00000116355 | 0 | NA | NA | NA | NA | NA | Gm5217 | predicted gene 5217 [Source:MGI Symbol | 0 | 0 | 0 | 0 | 0 | 0 |
| 53668 | ENSMUSG00000116356 | 0 | NA | NA | NA | NA | NA | Gm49407 | predicted gene, 49407 [Source:MGI Symb | 0 | 0 | 0 | 0 | 0 | 0 |
| 53669 | ENSMUSG00000116358 | 0 | NA | NA | NA | NA | NA | Gm49450 | predicted gene, 49450 [Source:MGI Symb | 0 | 0 | 0 | 0 | 0 | 0 |
| 53670 | ENSMUSG00000116360 | 0 | NA | NA | NA | NA | NA | 9430014N10 | RIKEN cDNA 9430014N10 gene [Source:M | 0 | 0 | 0 | 0 | 0 | 0 |
| 53671 | ENSMUSG00000116365 | 0 | NA | NA | NA | NA | NA | Gm34531 | predicted gene, 34531 [Source:MGI Symb | 0 | 0 | 0 | 0 | 0 | 0 |
| 53672 | ENSMUSG00000116366 | 0 | NA | NA | NA | NA | NA | Gm49475 | predicted gene, 49475 [Source:MGI Symb | 0 | 0 | 0 | 0 | 0 | 0 |
| 53673 | ENSMUSG00000116367 | 0 | NA | NA | NA | NA | NA | Gm49414 | predicted gene, 49414 [Source:MGI Symb | 0 | 0 | 0 | 0 | 0 | 0 |
| 53674 | ENSMUSG00000116368 | 0.167707431 | -0.796629176 | 4.080472857 | -0.195229623 | 0.84521319 | NA | Cyp2d33-ps | cytochrome P450, family 2, subfamily d, p | 0 | 0 | 1.0062 | 0 | 0 | 0 |
| 53675 | ENSMUSG00000116371 | 0 | NA | NA | NA | NA | NA | Gm34284 | predicted gene, 34284 [Source:MGI Symb | 0 | 0 | 0 | 0 | 0 | 0 |
| 53676 | ENSMUSG00000116376 | 0.506966596 | 2.558866737 | 3.339834942 | 0.766165628 | 0.443577757 | NA | Tmem249 | transmembrane protein 249 [Source:MGI | 0 | 0 | 0 | 1.928 | 0 | 1.1138 |
| 53677 | ENSMUSG00000116378 | 0 | NA | NA | NA | NA | NA | Gcat | glycine C-acetyltransferase (2-amino-3-ke | 0 | 0 | 0 | 0 | 0 | 0 |
| 53678 | ENSMUSG00000116380 | 0.638982041 | 0.404325377 | 3.14176733 | 0.128693609 | 0.897600092 | NA | Gm39556 | predicted gene, 39556 [Source:MGI Symb | 1.6062 | 0 | 0 | 0 | 0 | 2.2277 |
| 53679 | ENSMUSG00000116383 | 0 | NA | NA | NA | NA | NA | Apol10c-ps | apolipoprotein L 10C, pseudogene [Sourc | 0 | 0 | 0 | 0 | 0 | 0 |
| 53680 | ENSMUSG00000116384 | 0 | NA | NA | NA | NA | NA | NA | NA | 0 | 0 | 0 | 0 | 0 | 0 |
| 53681 | ENSMUSG00000116385 | 0 | NA | NA | NA | NA | NA | Gm18334 | predicted gene, 18334 [Source:MGI Symb | 0 | 0 | 0 | 0 | 0 | 0 |
| 53682 | ENSMUSG00000116386 | 0.160662974 | 1.126924175 | 4.080472857 | 0.276174898 | 0.782413736 | NA | Gm49538 | predicted gene, 49538 [Source:MGI Symb | 0 | 0 | 0 | 0.964 | 0 | 0 |
| 53683 | ENSMUSG00000116388 | 0 | NA | NA | NA | NA | NA | Gm3749 | predicted gene 3749 [Source:MGI Symbol | 0 | 0 | 0 | 0 | 0 | 0 |
| 53684 | ENSMUSG00000116389 | 0 | NA | NA | NA | NA | NA | Gm49504 | predicted gene, 49504 [Source:MGI Symb | 0 | 0 | 0 | 0 | 0 | 0 |
| 53685 | ENSMUSG00000116395 | 0.17131964 | -0.796629176 | 4.080472857 | -0.195229623 | 0.84521319 | NA | Gm49499 | predicted gene, 49499 [Source:MGI Symb | 0 | 1.0279 | 0 | 0 | 0 | 0 |
| 53686 | ENSMUSG00000116396 | 0 | NA | NA | NA | NA | NA | Gm20556 | predicted gene, 20556 [Source:MGI Symb | 0 | 0 | 0 | 0 | 0 | 0 |
| 53687 | ENSMUSG00000116397 | 0 | NA | NA | NA | NA | NA | Gm49525 | predicted gene, 49525 [Source:MGI Symb | 0 | 0 | 0 | 0 | 0 | 0 |
| 53688 | ENSMUSG00000116398 | 0 | NA | NA | NA | NA | NA | Gm5216 | predicted gene 5216 [Source:MGI Symbol | 0 | 0 | 0 | 0 | 0 | 0 |
| 53689 | ENSMUSG00000116400 | 0 | NA | NA | NA | NA | NA | Gm49530 | predicted gene, 49530 [Source:MGI Symb | 0 | 0 | 0 | 0 | 0 | 0 |
| 53690 | ENSMUSG00000116402 | 0.328370405 | 0.16514316 | 4.003149536 | 0.041253308 | 0.967093956 | NA | Gm36246 | predicted gene, 36246 [Source:MGI Symb | 0 | 0 | 1.0062 | 0.964 | 0 | 0 |
| 53691 | ENSMUSG00000116403 | 0 | NA | NA | NA | NA | NA | Gm19571 | predicted gene, 19571 [Source:MGI Symb | 0 | 0 | 0 | 0 | 0 | 0 |
| 53692 | ENSMUSG00000116404 | 0 | NA | NA | NA | NA | NA | Gm49541 | predicted gene, 49541 [Source:MGI Symb | 0 | 0 | 0 | 0 | 0 | 0 |
| 53693 | ENSMUSG00000116406 | 0 | NA | NA | NA | NA | NA | Gm49489 | predicted gene, 49489 [Source:MGI Symb | 0 | 0 | 0 | 0 | 0 | 0 |
| 53694 | ENSMUSG00000116409 | 0 | NA | NA | NA | NA | NA | Gm34095 | predicted gene, 34095 [Source:MGI Symb | 0 | 0 | 0 | 0 | 0 | 0 |
| 53695 | ENSMUSG00000116412 | 0 | NA | NA | NA | NA | NA | Gm49505 | predicted gene, 49505 [Source:MGI Symb | 0 | 0 | 0 | 0 | 0 | 0 |
| 53696 | ENSMUSG00000116413 | 0.160662974 | 1.126924175 | 4.080472857 | 0.276174898 | 0.782413736 | NA | Gm49436 | predicted gene, 49436 [Source:MGI Symb | 0 | 0 | 0 | 0.964 | 0 | 0 |
| 53697 | ENSMUSG00000116414 | 1.013648964 | -2.157154189 | 2.470907524 | -0.873021013 | 0.382651625 | NA | Gm4432 | predicted gene 4432 [Source:MGI Symbol | 0 | 4.1117 | 1.0062 | 0.964 | 0 | 0 |
| 53698 | ENSMUSG00000116416 | 0 | NA | NA | NA | NA | NA | Gm32886 | predicted gene, 32886 [Source:MGI Symb | 0 | 0 | 0 | 0 | 0 | 0 |
| 53699 | ENSMUSG00000116419 | 0 | NA | NA | NA | NA | NA | Gm19267 | predicted gene, 19267 [Source:MGI Symb | 0 | 0 | 0 | 0 | 0 | 0 |
| 53700 | ENSMUSG00000116423 | 0.294513347 | 0.165143496 | 4.080472857 | 0.040471657 | 0.967717103 | NA | Gm7085 | predicted gene 7085 [Source:MGI Symbol | 0.8031 | 0 | 0 | 0.964 | 0 | 0 |

| | | | | | | | | | | | | | | | |
|---|---|---|---|---|---|---|---|---|---|---|---|---|---|---|---|
| 53701 | ENSMUSG00000116424 | 0 | NA | NA | NA | NA | NA | Gm7167 | predicted gene 7167 [Source:MGI Symbol | 0 | 0 | 0 | 0 | 0 | 0 |
| 53702 | ENSMUSG00000116426 | 0 | NA | NA | NA | NA | NA | Gm30567 | predicted gene, 30567 [Source:MGI Symb | 0 | 0 | 0 | 0 | 0 | 0 |
| 53703 | ENSMUSG00000116428 | 0 | NA | NA | NA | NA | NA | Gm36618 | predicted gene, 36618 [Source:MGI Symb | 0 | 0 | 0 | 0 | 0 | 0 |
| 53704 | ENSMUSG00000116429 | 0 | NA | NA | NA | NA | NA | Ddit3 | DNA-damage inducible transcript 3 [Sourc | 0 | 0 | 0 | 0 | 0 | 0 |
| 53705 | ENSMUSG00000116433 | 0 | NA | NA | NA | NA | NA | Gm49553 | predicted gene, 49553 [Source:MGI Symb | 0 | 0 | 0 | 0 | 0 | 0 |
| 53706 | ENSMUSG00000116435 | 0 | NA | NA | NA | NA | NA | Gm49548 | predicted gene, 49548 [Source:MGI Symb | 0 | 0 | 0 | 0 | 0 | 0 |
| 53707 | ENSMUSG00000116437 | 7.524272982 | 0.081613597 | 0.830156103 | 0.098311145 | 0.921685229 | 0.986685804 | Gm46545 | predicted gene, 46545 [Source:MGI Symb | 9.6372 | 6.1675 | 6.0375 | 5.7839 | 14.178 | 3.3415 |
| 53708 | ENSMUSG00000116438 | 0 | NA | NA | NA | NA | NA | Gm18911 | predicted gene, 18911 [Source:MGI Symb | 0 | 0 | 0 | 0 | 0 | 0 |
| 53709 | ENSMUSG00000116439 | 0 | NA | NA | NA | NA | NA | Gm49528 | predicted gene, 49528 [Source:MGI Symb | 0 | 0 | 0 | 0 | 0 | 0 |
| 53710 | ENSMUSG00000116441 | 0 | NA | NA | NA | NA | NA | NA | NA | 0 | 0 | 0 | 0 | 0 | 0 |
| 53711 | ENSMUSG00000116442 | 0 | NA | NA | NA | NA | NA | Gm5214 | predicted gene 5214 [Source:MGI Symbol | 0 | 0 | 0 | 0 | 0 | 0 |
| 53712 | ENSMUSG00000116445 | 0 | NA | NA | NA | NA | NA | Gm49481 | predicted gene, 49481 [Source:MGI Symb | 0 | 0 | 0 | 0 | 0 | 0 |
| 53713 | ENSMUSG00000116449 | 0 | NA | NA | NA | NA | NA | Gm18155 | predicted gene, 18155 [Source:MGI Symb | 0 | 0 | 0 | 0 | 0 | 0 |
| 53714 | ENSMUSG00000116450 | 0 | NA | NA | NA | NA | NA | Gm49534 | predicted gene, 49534 [Source:MGI Symb | 0 | 0 | 0 | 0 | 0 | 0 |
| 53715 | ENSMUSG00000116455 | 0 | NA | NA | NA | NA | NA | Gm35974 | predicted gene, 35974 [Source:MGI Symb | 0 | 0 | 0 | 0 | 0 | 0 |
| 53716 | ENSMUSG00000116461 | 0 | NA | NA | NA | NA | NA | Gm44502 | predicted readthrough transcript (NMD ca | 0 | 0 | 0 | 0 | 0 | 0 |
| 53717 | ENSMUSG00000116464 | 0 | NA | NA | NA | NA | NA | Gm49523 | predicted gene, 49523 [Source:MGI Symb | 0 | 0 | 0 | 0 | 0 | 0 |
| 53718 | ENSMUSG00000116467 | 0 | NA | NA | NA | NA | NA | Gm49515 | predicted gene, 49515 [Source:MGI Symb | 0 | 0 | 0 | 0 | 0 | 0 |
| 53719 | ENSMUSG00000116469 | 0 | NA | NA | NA | NA | NA | Gm49448 | predicted gene, 49448 [Source:MGI Symb | 0 | 0 | 0 | 0 | 0 | 0 |
| 53720 | ENSMUSG00000116472 | 1.648863812 | -4.095738768 | 1.984206651 | -2.064169458 | 0.039001652 | NA | Gm49421 | predicted gene, 49421 [Source:MGI Symb | 4.8186 | 2.0558 | 3.0187 | 0 | 0 | 0 |
| 53721 | ENSMUSG00000116475 | 0 | NA | NA | NA | NA | NA | Gm19945 | predicted gene, 19945 [Source:MGI Symb | 0 | 0 | 0 | 0 | 0 | 0 |
| 53722 | ENSMUSG00000116478 | 22.12378505 | 0.010363029 | 0.472916404 | 0.021913025 | 0.982517335 | 0.998580735 | Gm18890 | predicted gene, 18890 [Source:MGI Symb | 26.502 | 15.419 | 24.15 | 26.991 | 19.631 | 20.049 |
| 53723 | ENSMUSG00000116480 | 0.528073336 | 2.60973587 | 2.979250867 | 0.875970499 | 0.381046072 | NA | Gm49512 | predicted gene, 49512 [Source:MGI Symb | 0 | 0 | 0 | 0.964 | 1.0906 | 1.1138 |
| 53724 | ENSMUSG00000116481 | 0.17131964 | -0.796629176 | 4.080472857 | -0.195229623 | 0.84521319 | NA | Gm49514 | predicted gene, 49514 [Source:MGI Symb | 0 | 1.0279 | 0 | 0 | 0 | 0 |
| 53725 | ENSMUSG00000116482 | 0.133850373 | -0.796629176 | 4.080472857 | -0.195229623 | 0.84521319 | NA | Gm34152 | predicted gene, 34152 [Source:MGI Symb | 0.8031 | 0 | 0 | 0 | 0 | 0 |
| 53726 | ENSMUSG00000116483 | 0 | NA | NA | NA | NA | NA | Gm19782 | predicted gene, 19782 [Source:MGI Symb | 0 | 0 | 0 | 0 | 0 | 0 |
| 53727 | ENSMUSG00000116490 | 0 | NA | NA | NA | NA | NA | Gm46525 | predicted gene, 46525 [Source:MGI Symb | 0 | 0 | 0 | 0 | 0 | 0 |
| 53728 | ENSMUSG00000116491 | 0 | NA | NA | NA | NA | NA | 4930513L16I | RIKEN cDNA 4930513L16 gene [Source:M( | 0 | 0 | 0 | 0 | 0 | 0 |
| 53729 | ENSMUSG00000116493 | 0 | NA | NA | NA | NA | NA | Gm49445 | predicted gene, 49445 [Source:MGI Symb | 0 | 0 | 0 | 0 | 0 | 0 |
| 53730 | ENSMUSG00000116494 | 0 | NA | NA | NA | NA | NA | Gm49449 | predicted gene, 49449 [Source:MGI Symb | 0 | 0 | 0 | 0 | 0 | 0 |
| 53731 | ENSMUSG00000116498 | 0.133850373 | -0.796629176 | 4.080472857 | -0.195229623 | 0.84521319 | NA | Gm38563 | predicted gene, 38563 [Source:MGI Symb | 0.8031 | 0 | 0 | 0 | 0 | 0 |
| 53732 | ENSMUSG00000116499 | 0.267700746 | -1.497481649 | 4.057001436 | -0.369110456 | 0.712045396 | NA | Gm7318 | predicted gene 7318 [Source:MGI Symbol | 1.6062 | 0 | 0 | 0 | 0 | 0 |
| 53733 | ENSMUSG00000116500 | 0 | NA | NA | NA | NA | NA | Gm36560 | predicted gene, 36560 [Source:MGI Symb | 0 | 0 | 0 | 0 | 0 | 0 |
| 53734 | ENSMUSG00000116504 | 1.220680051 | -3.657308518 | 2.422931929 | -1.509455744 | 0.131182357 | NA | I730030J21R | RIKEN cDNA I730030J21 gene [Source:MG | 3.2124 | 4.1117 | 0 | 0 | 0 | 0 |
| 53735 | ENSMUSG00000116505 | 7.444125148 | 0.527207691 | 0.782915933 | 0.673389912 | 0.500699265 | 0.880928618 | Gm10865 | predicted gene 10865 [Source:MGI Symbo | 8.031 | 4.1117 | 6.0375 | 7.7118 | 7.6343 | 11.138 |
| 53736 | ENSMUSG00000116506 | 119.0664936 | 0.056197262 | 0.259509583 | 0.216551781 | 0.828557667 | 0.969955221 | 5730414N17 | RIKEN cDNA 5730414N17 gene [Source:M | 110.03 | 112.04 | 127.79 | 80.974 | 126.51 | 157.05 |
| 53737 | ENSMUSG00000116508 | 88.50483902 | 1.01574706 | 0.252729443 | 4.019108532 | 5.84E-05 | 0.002373508 | Gm49463 | predicted gene, 49463 [Source:MGI Symb | 57.823 | 58.591 | 59.368 | 117.61 | 138.51 | 99.132 |
| 53738 | ENSMUSG00000116514 | 0 | NA | NA | NA | NA | NA | Gm49443 | predicted gene, 49443 [Source:MGI Symb | 0 | 0 | 0 | 0 | 0 | 0 |
| 53739 | ENSMUSG00000116516 | 0 | NA | NA | NA | NA | NA | Gm49444 | predicted gene, 49444 [Source:MGI Symb | 0 | 0 | 0 | 0 | 0 | 0 |
| 53740 | ENSMUSG00000116518 | 18.88317968 | -0.230326886 | 0.559663133 | -0.411545576 | 0.680672529 | 0.934384556 | Gm49494 | predicted gene, 49494 [Source:MGI Symb | 30.518 | 12.335 | 18.112 | 18.316 | 23.994 | 10.025 |
| 53741 | ENSMUSG00000116519 | 0 | NA | NA | NA | NA | NA | Gm49409 | predicted gene, 49409 [Source:MGI Symb | 0 | 0 | 0 | 0 | 0 | 0 |
| 53742 | ENSMUSG00000116520 | 0 | NA | NA | NA | NA | NA | 4930414F18 | RIKEN cDNA 4930414F18 gene [Source:M( | 0 | 0 | 0 | 0 | 0 | 0 |
| 53743 | ENSMUSG00000116521 | 0 | NA | NA | NA | NA | NA | Gm4111 | predicted gene 4111 [Source:MGI Symbol | 0 | 0 | 0 | 0 | 0 | 0 |
| 53744 | ENSMUSG00000116525 | 0 | NA | NA | NA | NA | NA | Gm30371 | predicted gene, 30371 [Source:MGI Symb | 0 | 0 | 0 | 0 | 0 | 0 |
| 53745 | ENSMUSG00000116528 | 0 | NA | NA | NA | NA | NA | Gm49508 | predicted gene, 49508 [Source:MGI Symb | 0 | 0 | 0 | 0 | 0 | 0 |
| 53746 | ENSMUSG00000116529 | 0 | NA | NA | NA | NA | NA | Gm4600 | predicted gene 4600 [Source:MGI Symbol | 0 | 0 | 0 | 0 | 0 | 0 |
| 53747 | ENSMUSG00000116530 | 0 | NA | NA | NA | NA | NA | Gm32312 | predicted gene, 32312 [Source:MGI Symb | 0 | 0 | 0 | 0 | 0 | 0 |
| 53748 | ENSMUSG00000116531 | 0 | NA | NA | NA | NA | NA | Gm19082 | predicted gene, 19082 [Source:MGI Symb | 0 | 0 | 0 | 0 | 0 | 0 |
| 53749 | ENSMUSG00000116532 | 0 | NA | NA | NA | NA | NA | Gm5232 | predicted gene 5232 [Source:MGI Symbol | 0 | 0 | 0 | 0 | 0 | 0 |
| 53750 | ENSMUSG00000116533 | 0 | NA | NA | NA | NA | NA | Gm49561 | predicted gene, 49561 [Source:MGI Symb | 0 | 0 | 0 | 0 | 0 | 0 |
| 53751 | ENSMUSG00000116534 | 0.267700746 | -1.497481649 | 4.057001436 | -0.369110456 | 0.712045396 | NA | Gm49731 | predicted gene, 49731 [Source:MGI Symb | 1.6062 | 0 | 0 | 0 | 0 | 0 |
| 53752 | ENSMUSG00000116535 | 0.133850373 | -0.796629176 | 4.080472857 | -0.195229623 | 0.84521319 | NA | Gm5962 | predicted gene 5962 [Source:MGI Symbol | 0.8031 | 0 | 0 | 0 | 0 | 0 |
| 53753 | ENSMUSG00000116536 | 0 | NA | NA | NA | NA | NA | Gm2805 | predicted gene 2805 [Source:MGI Symbol | 0 | 0 | 0 | 0 | 0 | 0 |
| 53754 | ENSMUSG00000116537 | 0 | NA | NA | NA | NA | NA | Gm49698 | predicted gene, 49698 [Source:MGI Symb | 0 | 0 | 0 | 0 | 0 | 0 |
| 53755 | ENSMUSG00000116538 | 0 | NA | NA | NA | NA | NA | Gm49746 | predicted gene, 49746 [Source:MGI Symb | 0 | 0 | 0 | 0 | 0 | 0 |
| 53756 | ENSMUSG00000116539 | 0 | NA | NA | NA | NA | NA | Gm49603 | predicted gene, 49603 [Source:MGI Symb | 0 | 0 | 0 | 0 | 0 | 0 |

| | | | | | | | | | | | | | | | |
|---|---|---|---|---|---|---|---|---|---|---|---|---|---|---|---|
| 53757 | ENSMUSG00000116540 | 0 | NA | NA | NA | NA | NA | Gm49624 | predicted gene, 49624 [Source:MGI Symb | 0 | 0 | 0 | 0 | 0 | 0 |
| 53758 | ENSMUSG00000116542 | 0 | NA | NA | NA | NA | NA | Gm17783 | predicted gene, 17783 [Source:MGI Symb | 0 | 0 | 0 | 0 | 0 | 0 |
| 53759 | ENSMUSG00000116543 | 0 | NA | NA | NA | NA | NA | Gm36363 | predicted gene, 36363 [Source:MGI Symb | 0 | 0 | 0 | 0 | 0 | 0 |
| 53760 | ENSMUSG00000116544 | 0 | NA | NA | NA | NA | NA | Gm49626 | predicted gene, 49626 [Source:MGI Symb | 0 | 0 | 0 | 0 | 0 | 0 |
| 53761 | ENSMUSG00000116545 | 0 | NA | NA | NA | NA | NA | Eif3s6-ps3 | eukaryotic translation initiation factor 3, s | 0 | 0 | 0 | 0 | 0 | 0 |
| 53762 | ENSMUSG00000116546 | 0 | NA | NA | NA | NA | NA | Gm49755 | predicted gene, 49755 [Source:MGI Symb | 0 | 0 | 0 | 0 | 0 | 0 |
| 53763 | ENSMUSG00000116547 | 0 | NA | NA | NA | NA | NA | Gm7667 | predicted gene 7667 [Source:MGI Symbol | 0 | 0 | 0 | 0 | 0 | 0 |
| 53764 | ENSMUSG00000116548 | 0 | NA | NA | NA | NA | NA | Gm49773 | predicted gene, 49773 [Source:MGI Symb | 0 | 0 | 0 | 0 | 0 | 0 |
| 53765 | ENSMUSG00000116549 | 0.822793274 | -0.447303882 | 2.407293551 | -0.185811939 | 0.852592242 | NA | Gm49728 | predicted gene, 49728 [Source:MGI Symb | 0.8031 | 2.0558 | 0 | 0.964 | 0 | 1.1138 |
| 53766 | ENSMUSG00000116550 | 0 | NA | NA | NA | NA | NA | Gm3507 | predicted gene 3507 [Source:MGI Symbol | 0 | 0 | 0 | 0 | 0 | 0 |
| 53767 | ENSMUSG00000116551 | 3.864272391 | 1.40763962 | 1.107853231 | 1.270601178 | 0.203870568 | NA | Gm41517 | predicted gene, 41517 [Source:MGI Symb | 3.2124 | 1.0279 | 2.0125 | 4.8199 | 6.5437 | 5.5692 |
| 53768 | ENSMUSG00000116552 | 0 | NA | NA | NA | NA | NA | Gm34396 | predicted gene, 34396 [Source:MGI Symb | 0 | 0 | 0 | 0 | 0 | 0 |
| 53769 | ENSMUSG00000116553 | 0 | NA | NA | NA | NA | NA | Gm7307 | predicted gene 7307 [Source:MGI Symbol | 0 | 0 | 0 | 0 | 0 | 0 |
| 53770 | ENSMUSG00000116554 | 0 | NA | NA | NA | NA | NA | 4930563N14 | RIKEN cDNA 4930563N14 gene [Source:M | 0 | 0 | 0 | 0 | 0 | 0 |
| 53771 | ENSMUSG00000116555 | 0.301557804 | -1.64335916 | 4.042814367 | -0.406488899 | 0.684383412 | NA | Gm38477 | predicted gene, 38477 [Source:MGI Symb | 0.8031 | 0 | 1.0062 | 0 | 0 | 0 |
| 53772 | ENSMUSG00000116556 | 0 | NA | NA | NA | NA | NA | 1700007H22 | RIKEN cDNA 1700007H22 gene [Source:M | 0 | 0 | 0 | 0 | 0 | 0 |
| 53773 | ENSMUSG00000116557 | 0.167707431 | -0.796629176 | 4.080472857 | -0.195229623 | 0.84521319 | NA | 4933432I09F | RIKEN cDNA 4933432I09 gene [Source:MG | 0 | 0 | 1.0062 | 0 | 0 | 0 |
| 53774 | ENSMUSG00000116558 | 0 | NA | NA | NA | NA | NA | Gm49697 | predicted gene, 49697 [Source:MGI Symb | 0 | 0 | 0 | 0 | 0 | 0 |
| 53775 | ENSMUSG00000116559 | 0 | NA | NA | NA | NA | NA | Gm19867 | predicted gene, 19867 [Source:MGI Symb | 0 | 0 | 0 | 0 | 0 | 0 |
| 53776 | ENSMUSG00000116560 | 0 | NA | NA | NA | NA | NA | Gm2808 | predicted gene 2808 [Source:MGI Symbol | 0 | 0 | 0 | 0 | 0 | 0 |
| 53777 | ENSMUSG00000116561 | 0.160662974 | 1.126924175 | 4.080472857 | 0.276174898 | 0.782413736 | NA | Gm18335 | predicted gene, 18335 [Source:MGI Symb | 0 | 0 | 0 | 0.964 | 0 | 0 |
| 53778 | ENSMUSG00000116562 | 0 | NA | NA | NA | NA | NA | Gm46542 | predicted gene, 46542 [Source:MGI Symb | 0 | 0 | 0 | 0 | 0 | 0 |
| 53779 | ENSMUSG00000116563 | 0 | NA | NA | NA | NA | NA | | novel protein | 0 | 0 | 0 | 0 | 0 | 0 |
| 53780 | ENSMUSG00000116564 | 512.3168473 | 0.12608069 | 0.171145193 | 0.73668847 | 0.46131182 | 0.860071824 | Riok2 | RIO kinase 2 [Source:MGI Symbol;Acc:MG | 543.7 | 484.15 | 441.74 | 570.67 | 628.2 | 405.44 |
| 53781 | ENSMUSG00000116565 | 0 | NA | NA | NA | NA | NA | Gm49734 | predicted gene, 49734 [Source:MGI Symb | 0 | 0 | 0 | 0 | 0 | 0 |
| 53782 | ENSMUSG00000116566 | 0 | NA | NA | NA | NA | NA | Fkbp1a-ps3 | FK506 binding protein 1a, pseudogene 3 [ | 0 | 0 | 0 | 0 | 0 | 0 |
| 53783 | ENSMUSG00000116567 | 0 | NA | NA | NA | NA | NA | Gm49639 | predicted gene, 49639 [Source:MGI Symb | 0 | 0 | 0 | 0 | 0 | 0 |
| 53784 | ENSMUSG00000116568 | 0 | NA | NA | NA | NA | NA | Gm49790 | predicted gene, 49790 [Source:MGI Symb | 0 | 0 | 0 | 0 | 0 | 0 |
| 53785 | ENSMUSG00000116569 | 0 | NA | NA | NA | NA | NA | Gm49620 | predicted gene, 49620 [Source:MGI Symb | 0 | 0 | 0 | 0 | 0 | 0 |
| 53786 | ENSMUSG00000116570 | 0 | NA | NA | NA | NA | NA | Gm49754 | predicted gene, 49754 [Source:MGI Symb | 0 | 0 | 0 | 0 | 0 | 0 |
| 53787 | ENSMUSG00000116571 | 4.028036755 | 0.571023628 | 1.22558626 | 0.465918758 | 0.641273654 | NA | Gm49741 | predicted gene, 49741 [Source:MGI Symb | 5.6217 | 0 | 4.025 | 6.7478 | 1.0906 | 6.6831 |
| 53788 | ENSMUSG00000116572 | 0 | NA | NA | NA | NA | NA | Gm49695 | predicted gene, 49695 [Source:MGI Symb | 0 | 0 | 0 | 0 | 0 | 0 |
| 53789 | ENSMUSG00000116573 | 0 | NA | NA | NA | NA | NA | Gm36903 | predicted gene, 36903 [Source:MGI Symb | 0 | 0 | 0 | 0 | 0 | 0 |
| 53790 | ENSMUSG00000116574 | 0 | NA | NA | NA | NA | NA | Gm18870 | predicted gene, 18870 [Source:MGI Symb | 0 | 0 | 0 | 0 | 0 | 0 |
| 53791 | ENSMUSG00000116575 | 0 | NA | NA | NA | NA | NA | Gm49648 | predicted gene, 49648 [Source:MGI Symb | 0 | 0 | 0 | 0 | 0 | 0 |
| 53792 | ENSMUSG00000116576 | 0 | NA | NA | NA | NA | NA | Gm49725 | predicted gene, 49725 [Source:MGI Symb | 0 | 0 | 0 | 0 | 0 | 0 |
| 53793 | ENSMUSG00000116578 | 0 | NA | NA | NA | NA | NA | Gm49633 | predicted gene, 49633 [Source:MGI Symb | 0 | 0 | 0 | 0 | 0 | 0 |
| 53794 | ENSMUSG00000116579 | 0.973879455 | -0.768123796 | 2.429069234 | -0.31622145 | 0.751834428 | NA | Gm49701 | predicted gene, 49701 [Source:MGI Symb | 1.6062 | 2.0558 | 0 | 0 | 2.1812 | 0 |
| 53795 | ENSMUSG00000116580 | 0.160662974 | 1.126924175 | 4.080472857 | 0.276174898 | 0.782413736 | NA | Gm9525 | predicted gene 9525 [Source:MGI Symbol | 0 | 0 | 0 | 0.964 | 0 | 0 |
| 53796 | ENSMUSG00000116581 | 10.91391275 | -0.506844102 | 0.81055751 | -0.625303074 | 0.531772163 | 0.888991281 | Gm46565 | predicted gene, 46565 [Source:MGI Symb | 19.274 | 6.1675 | 13.081 | 19.28 | 5.4531 | 2.2277 |
| 53797 | ENSMUSG00000116582 | 0 | NA | NA | NA | NA | NA | Gm36169 | predicted gene, 36169 [Source:MGI Symb | 0 | 0 | 0 | 0 | 0 | 0 |
| 53798 | ENSMUSG00000116583 | 0 | NA | NA | NA | NA | NA | Gm2349 | predicted gene 2349 [Source:MGI Symbol | 0 | 0 | 0 | 0 | 0 | 0 |
| 53799 | ENSMUSG00000116585 | 0 | NA | NA | NA | NA | NA | Gm33086 | predicted gene, 33086 [Source:MGI Symb | 0 | 0 | 0 | 0 | 0 | 0 |
| 53800 | ENSMUSG00000116586 | 0 | NA | NA | NA | NA | NA | 4930556C24 | RIKEN cDNA 4930556C24 gene [Source:M | 0 | 0 | 0 | 0 | 0 | 0 |
| 53801 | ENSMUSG00000116587 | 0 | NA | NA | NA | NA | NA | Gm49753 | predicted gene, 49753 [Source:MGI Symb | 0 | 0 | 0 | 0 | 0 | 0 |
| 53802 | ENSMUSG00000116588 | 0 | NA | NA | NA | NA | NA | Gm49655 | predicted gene, 49655 [Source:MGI Symb | 0 | 0 | 0 | 0 | 0 | 0 |
| 53803 | ENSMUSG00000116589 | 0.167707431 | -0.796629176 | 4.080472857 | -0.195229623 | 0.84521319 | NA | Gm31323 | predicted gene, 31323 [Source:MGI Symb | 0 | 0 | 1.0062 | 0 | 0 | 0 |
| 53804 | ENSMUSG00000116590 | 4.898283528 | -0.4613886 | 0.978652018 | -0.471453174 | 0.637317151 | NA | Gm53028 | predicted gene, 53028 [Source:MGI Symb | 8.8341 | 5.1396 | 3.0187 | 5.7839 | 3.2719 | 3.3415 |
| 53805 | ENSMUSG00000116591 | 0 | NA | NA | NA | NA | NA | Gm49715 | predicted gene, 49715 [Source:MGI Symb | 0 | 0 | 0 | 0 | 0 | 0 |
| 53806 | ENSMUSG00000116592 | 0 | NA | NA | NA | NA | NA | Gm18770 | predicted gene, 18770 [Source:MGI Symb | 0 | 0 | 0 | 0 | 0 | 0 |
| 53807 | ENSMUSG00000116593 | 0 | NA | NA | NA | NA | NA | Gm49585 | predicted gene, 49585 [Source:MGI Symb | 0 | 0 | 0 | 0 | 0 | 0 |
| 53808 | ENSMUSG00000116594 | 0 | NA | NA | NA | NA | NA | Gm49601 | predicted gene, 49601 [Source:MGI Symb | 0 | 0 | 0 | 0 | 0 | 0 |
| 53809 | ENSMUSG00000116595 | 0 | NA | NA | NA | NA | NA | Gm30695 | predicted gene, 30695 [Source:MGI Symb | 0 | 0 | 0 | 0 | 0 | 0 |
| 53810 | ENSMUSG00000116596 | 0 | NA | NA | NA | NA | NA | Gm49798 | predicted gene, 49798 [Source:MGI Symb | 0 | 0 | 0 | 0 | 0 | 0 |
| 53811 | ENSMUSG00000116597 | 0.556921942 | 2.679043159 | 3.638243063 | 0.736356289 | 0.461513899 | NA | Gm536 | predicted gene 536 [Source:MGI Symbol;/ | 0 | 0 | 0 | 0 | 0 | 3.3415 |
| 53812 | ENSMUSG00000116598 | 0 | NA | NA | NA | NA | NA | Gm6438 | predicted gene 6438 [Source:MGI Symbol | 0 | 0 | 0 | 0 | 0 | 0 |

| | | | | | | | | | | | | | | | |
|---|---|---|---|---|---|---|---|---|---|---|---|---|---|---|---|
| 53813 | ENSMUSG00000116599 | 0 | NA | NA | NA | NA | NA | Gm2477 | predicted gene 2477 [Source:MGI Symbol | 0 | 0 | 0 | 0 | 0 | 0 |
| 53814 | ENSMUSG00000116600 | 0 | NA | NA | NA | NA | NA | Gm49707 | predicted gene, 49707 [Source:MGI Symb | 0 | 0 | 0 | 0 | 0 | 0 |
| 53815 | ENSMUSG00000116601 | 0 | NA | NA | NA | NA | NA | Gm49642 | predicted gene, 49642 [Source:MGI Symb | 0 | 0 | 0 | 0 | 0 | 0 |
| 53816 | ENSMUSG00000116603 | 0 | NA | NA | NA | NA | NA | Gm8118 | predicted gene 8118 [Source:MGI Symbol | 0 | 0 | 0 | 0 | 0 | 0 |
| 53817 | ENSMUSG00000116604 | 0.867048273 | -1.842698135 | 2.849929781 | -0.64657668 | 0.517905955 | NA | Gm49745 | predicted gene, 49745 [Source:MGI Symb | 0 | 4.1117 | 0 | 0 | 1.0906 | 0 |
| 53818 | ENSMUSG00000116605 | 0.695728634 | -1.425610641 | 3.05889997 | -0.46605337 | 0.641177299 | NA | Gm19142 | predicted gene, 19142 [Source:MGI Symb | 0 | 3.0838 | 0 | 0 | 1.0906 | 0 |
| 53819 | ENSMUSG00000116606 | 0.167707431 | -0.796629176 | 4.080472857 | -0.195229623 | 0.84521319 | NA | Gm10479 | predicted gene 10479 [Source:MGI Symbo | 0 | 0 | 1.0062 | 0 | 0 | 0 |
| 53820 | ENSMUSG00000116607 | 0.695728634 | -1.425610641 | 3.05889997 | -0.46605337 | 0.641177299 | NA | 4930534H18 | RIKEN cDNA 4930534H18 gene [Source:M | 0 | 3.0838 | 0 | 0 | 1.0906 | 0 |
| 53821 | ENSMUSG00000116608 | 0 | NA | NA | NA | NA | NA | Gm41495 | predicted gene, 41495 [Source:MGI Symb | 0 | 0 | 0 | 0 | 0 | 0 |
| 53822 | ENSMUSG00000116609 | 0 | NA | NA | NA | NA | NA | Gm5675 | predicted gene 5675 [Source:MGI Symbol | 0 | 0 | 0 | 0 | 0 | 0 |
| 53823 | ENSMUSG00000116610 | 0 | NA | NA | NA | NA | NA | Gm49678 | predicted gene, 49678 [Source:MGI Symb | 0 | 0 | 0 | 0 | 0 | 0 |
| 53824 | ENSMUSG00000116611 | 0 | NA | NA | NA | NA | NA | 4930404A05 | RIKEN cDNA 4930404A05 gene [Source:M | 0 | 0 | 0 | 0 | 0 | 0 |
| 53825 | ENSMUSG00000116612 | 0 | NA | NA | NA | NA | NA | Gm49621 | predicted gene, 49621 [Source:MGI Symb | 0 | 0 | 0 | 0 | 0 | 0 |
| 53826 | ENSMUSG00000116614 | 0 | NA | NA | NA | NA | NA | Gm46610 | predicted gene, 46610 [Source:MGI Symb | 0 | 0 | 0 | 0 | 0 | 0 |
| 53827 | ENSMUSG00000116615 | 0 | NA | NA | NA | NA | NA | 4930548J01I | RIKEN cDNA 4930548J01 gene [Source:M( | 0 | 0 | 0 | 0 | 0 | 0 |
| 53828 | ENSMUSG00000116616 | 0 | NA | NA | NA | NA | NA | Gm49688 | predicted gene, 49688 [Source:MGI Symb | 0 | 0 | 0 | 0 | 0 | 0 |
| 53829 | ENSMUSG00000116617 | 2.665582561 | -2.014010442 | 1.502224552 | -1.340685345 | 0.180022635 | NA | Gm49767 | predicted gene, 49767 [Source:MGI Symb | 1.6062 | 8.2233 | 3.0187 | 0.964 | 2.1812 | 0 |
| 53830 | ENSMUSG00000116618 | 0 | NA | NA | NA | NA | NA | Gm49719 | predicted gene, 49719 [Source:MGI Symb | 0 | 0 | 0 | 0 | 0 | 0 |
| 53831 | ENSMUSG00000116619 | 0 | NA | NA | NA | NA | NA | Gm49615 | predicted gene, 49615 [Source:MGI Symb | 0 | 0 | 0 | 0 | 0 | 0 |
| 53832 | ENSMUSG00000116620 | 2.185998148 | -1.629023848 | 1.608610941 | -1.012689773 | 0.311208365 | NA | Gm4828 | predicted gene 4828 [Source:MGI Symbol | 0.8031 | 3.0838 | 6.0375 | 0.964 | 0 | 2.2277 |
| 53833 | ENSMUSG00000116621 | 0 | NA | NA | NA | NA | NA | Gm7483 | predicted gene 7483 [Source:MGI Symbol | 0 | 0 | 0 | 0 | 0 | 0 |
| 53834 | ENSMUSG00000116622 | 0 | NA | NA | NA | NA | NA | Gm5221 | predicted gene 5221 [Source:MGI Symbol | 0 | 0 | 0 | 0 | 0 | 0 |
| 53835 | ENSMUSG00000116623 | 1.542388992 | 1.178831019 | 1.840581277 | 0.640466701 | 0.521869231 | NA | Gm49775 | predicted gene, 49775 [Source:MGI Symb | 0.8031 | 1.0279 | 1.0062 | 0.964 | 5.4531 | 0 |
| 53836 | ENSMUSG00000116624 | 0.640816969 | 1.600078776 | 2.832778663 | 0.564844263 | 0.572179697 | NA | Gm49706 | predicted gene, 49706 [Source:MGI Symb | 0.8031 | 0 | 0 | 1.928 | 0 | 1.1138 |
| 53837 | ENSMUSG00000116625 | 0.294513347 | 0.165143496 | 4.080472857 | 0.040471657 | 0.967717103 | NA | Gm8134 | predicted gene 8134 [Source:MGI Symbol | 0.8031 | 0 | 0 | 0.964 | 0 | 0 |
| 53838 | ENSMUSG00000116626 | 0 | NA | NA | NA | NA | NA | Gm49705 | predicted gene, 49705 [Source:MGI Symb | 0 | 0 | 0 | 0 | 0 | 0 |
| 53839 | ENSMUSG00000116627 | 3.901488147 | -0.273638222 | 1.271207547 | -0.215258494 | 0.829565788 | NA | Gm30124 | predicted gene, 30124 [Source:MGI Symb | 1.6062 | 9.2513 | 2.0125 | 3.8559 | 0 | 6.6831 |
| 53840 | ENSMUSG00000116628 | 1.090618289 | 3.652571688 | 2.872889547 | 1.27139301 | 0.203588868 | NA | 4930478L05I | RIKEN cDNA 4930478L05 gene [Source:M( | 0 | 0 | 0 | 0 | 6.5437 | 0 |
| 53841 | ENSMUSG00000116629 | 0 | NA | NA | NA | NA | NA | Gm49638 | predicted gene, 49638 [Source:MGI Symb | 0 | 0 | 0 | 0 | 0 | 0 |
| 53842 | ENSMUSG00000116630 | 0 | NA | NA | NA | NA | NA | Rpl7a-ps4 | ribosomal protein 7A, pseudogene 4 [Sou | 0 | 0 | 0 | 0 | 0 | 0 |
| 53843 | ENSMUSG00000116631 | 0 | NA | NA | NA | NA | NA | Gm7366 | predicted gene 7366 [Source:MGI Symbol | 0 | 0 | 0 | 0 | 0 | 0 |
| 53844 | ENSMUSG00000116632 | 17.70503689 | -0.207186945 | 0.557169068 | -0.37185651 | 0.709999687 | 0.944239215 | Magef1 | MAGE family member F1 [Source:MGI Syr | 14.456 | 16.447 | 26.162 | 17.352 | 22.903 | 8.9108 |
| 53845 | ENSMUSG00000116633 | 0 | NA | NA | NA | NA | NA | Gm18519 | predicted gene, 18519 [Source:MGI Symb | 0 | 0 | 0 | 0 | 0 | 0 |
| 53846 | ENSMUSG00000116634 | 0 | NA | NA | NA | NA | NA | Gm7531 | predicted gene 7531 [Source:MGI Symbol | 0 | 0 | 0 | 0 | 0 | 0 |
| 53847 | ENSMUSG00000116635 | 0.133850373 | -0.796629176 | 4.080472857 | -0.195229623 | 0.84521319 | NA | Gm49699 | predicted gene, 49699 [Source:MGI Symb | 0.8031 | 0 | 0 | 0 | 0 | 0 |
| 53848 | ENSMUSG00000116636 | 0 | NA | NA | NA | NA | NA | Gm20741 | predicted gene, 20741 [Source:MGI Symb | 0 | 0 | 0 | 0 | 0 | 0 |
| 53849 | ENSMUSG00000116637 | 0 | NA | NA | NA | NA | NA | Gm8130 | predicted gene 8130 [Source:MGI Symbol | 0 | 0 | 0 | 0 | 0 | 0 |
| 53850 | ENSMUSG00000116638 | 0 | NA | NA | NA | NA | NA | Gm49650 | predicted gene, 49650 [Source:MGI Symb | 0 | 0 | 0 | 0 | 0 | 0 |
| 53851 | ENSMUSG00000116639 | 0.603115608 | -2.642822926 | 3.106511347 | -0.850736608 | 0.394915683 | NA | Gm49730 | predicted gene, 49730 [Source:MGI Symb | 1.6062 | 0 | 2.0125 | 0 | 0 | 0 |
| 53852 | ENSMUSG00000116640 | 0.185640647 | 1.126924175 | 4.080472857 | 0.276174898 | 0.782413736 | NA | Gm8670 | predicted gene 8670 [Source:MGI Symbol | 0 | 0 | 0 | 0 | 0 | 1.1138 |
| 53853 | ENSMUSG00000116641 | 0 | NA | NA | NA | NA | NA | Gm41505 | predicted gene, 41505 [Source:MGI Symb | 0 | 0 | 0 | 0 | 0 | 0 |
| 53854 | ENSMUSG00000116642 | 0 | NA | NA | NA | NA | NA | Gm18168 | predicted gene, 18168 [Source:MGI Symb | 0 | 0 | 0 | 0 | 0 | 0 |
| 53855 | ENSMUSG00000116643 | 0 | NA | NA | NA | NA | NA | Gm30162 | predicted gene, 30162 [Source:MGI Symb | 0 | 0 | 0 | 0 | 0 | 0 |
| 53856 | ENSMUSG00000116644 | 0 | NA | NA | NA | NA | NA | 4933426B08 | RIKEN cDNA 4933426B08 gene [Source:M | 0 | 0 | 0 | 0 | 0 | 0 |
| 53857 | ENSMUSG00000116645 | 2.898139787 | -0.534663096 | 1.282538756 | -0.416878705 | 0.67676713 | NA | Gm49584 | predicted gene, 49584 [Source:MGI Symb | 3.2124 | 5.1396 | 2.0125 | 4.8199 | 1.0906 | 1.1138 |
| 53858 | ENSMUSG00000116646 | 0 | NA | NA | NA | NA | NA | Gm49763 | predicted gene, 49763 [Source:MGI Symb | 0 | 0 | 0 | 0 | 0 | 0 |
| 53859 | ENSMUSG00000116647 | 0 | NA | NA | NA | NA | NA | Gm7423 | predicted gene 7423 [Source:MGI Symbol | 0 | 0 | 0 | 0 | 0 | 0 |
| 53860 | ENSMUSG00000116648 | 0.983224154 | -3.332395149 | 2.379709572 | -1.40033691 | 0.161412453 | NA | Rpl31-ps12 | ribosomal protein L31, pseudogene 1 2 [S | 0.8031 | 3.0838 | 2.0125 | 0 | 0 | 0 |
| 53861 | ENSMUSG00000116649 | 0 | NA | NA | NA | NA | NA | Gm49780 | predicted gene, 49780 [Source:MGI Symb | 0 | 0 | 0 | 0 | 0 | 0 |
| 53862 | ENSMUSG00000116650 | 0 | NA | NA | NA | NA | NA | Gm49657 | predicted gene, 49657 [Source:MGI Symb | 0 | 0 | 0 | 0 | 0 | 0 |
| 53863 | ENSMUSG00000116651 | 0 | NA | NA | NA | NA | NA | Gm17844 | predicted gene, 17844 [Source:MGI Symb | 0 | 0 | 0 | 0 | 0 | 0 |
| 53864 | ENSMUSG00000116652 | 0 | NA | NA | NA | NA | NA | B830017H08 | RIKEN cDNA B830017H08 gene [Source:M | 0 | 0 | 0 | 0 | 0 | 0 |
| 53865 | ENSMUSG00000116653 | 0 | NA | NA | NA | NA | NA | Gm8900 | predicted gene 8900 [Source:MGI Symbol | 0 | 0 | 0 | 0 | 0 | 0 |
| 53866 | ENSMUSG00000116654 | 0 | NA | NA | NA | NA | NA | Gm10823 | predicted gene 10823 [Source:MGI Symbo | 0 | 0 | 0 | 0 | 0 | 0 |
| 53867 | ENSMUSG00000116655 | 0 | NA | NA | NA | NA | NA | Gm35455 | predicted gene, 35455 [Source:MGI Symb | 0 | 0 | 0 | 0 | 0 | 0 |
| 53868 | ENSMUSG00000116656 | 1.291619825 | -2.551117698 | 2.090478447 | -1.220351112 | 0.222331802 | NA | Gm49708 | predicted gene, 49708 [Source:MGI Symb | 1.6062 | 1.0279 | 4.025 | 0 | 1.0906 | 0 |

| | | | | | | | | | | | | | | | |
|---|---|---|---|---|---|---|---|---|---|---|---|---|---|---|---|
| 53869 | ENSMUSG00000116657 | 10.42826152 | -0.533418206 | 0.768870874 | -0.693768257 | 0.487827553 | 0.873910634 | Gm49774 | predicted gene, 49774 [Source:MGI Symb | 22.487 | 7.1954 | 7.0437 | 4.8199 | 6.5437 | 14.48 |
| 53870 | ENSMUSG00000116658 | 0 | NA | NA | NA | NA | NA | Gm49580 | predicted gene, 49580 [Source:MGI Symb | 0 | 0 | 0 | 0 | 0 | 0 |
| 53871 | ENSMUSG00000116659 | 0 | NA | NA | NA | NA | NA | Gm6551 | predicted gene 6551 [Source:MGI Symbol | 0 | 0 | 0 | 0 | 0 | 0 |
| 53872 | ENSMUSG00000116660 | 0 | NA | NA | NA | NA | NA | 9030404E10 | RIKEN cDNA 9030404E10 gene [Source:M | 0 | 0 | 0 | 0 | 0 | 0 |
| 53873 | ENSMUSG00000116661 | 0 | NA | NA | NA | NA | NA | Gm5966 | predicted gene 5966 [Source:MGI Symbol | 0 | 0 | 0 | 0 | 0 | 0 |
| 53874 | ENSMUSG00000116662 | 0 | NA | NA | NA | NA | NA | Gm46541 | predicted gene, 46541 [Source:MGI Symb | 0 | 0 | 0 | 0 | 0 | 0 |
| 53875 | ENSMUSG00000116663 | 0 | NA | NA | NA | NA | NA | Gm49671 | predicted gene, 49671 [Source:MGI Symb | 0 | 0 | 0 | 0 | 0 | 0 |
| 53876 | ENSMUSG00000116664 | 0 | NA | NA | NA | NA | NA | Gm18549 | predicted gene, 18549 [Source:MGI Symb | 0 | 0 | 0 | 0 | 0 | 0 |
| 53877 | ENSMUSG00000116665 | 1.052482331 | 1.09488462 | 2.207898318 | 0.495894494 | 0.619968851 | NA | E130310I04F | RIKEN cDNA E130310I04 gene [Source:M | 0 | 2.0558 | 0 | 0.964 | 2.1812 | 1.1138 |
| 53878 | ENSMUSG00000116666 | 0 | NA | NA | NA | NA | NA | Gm49649 | predicted gene, 49649 [Source:MGI Symb | 0 | 0 | 0 | 0 | 0 | 0 |
| 53879 | ENSMUSG00000116667 | 0 | NA | NA | NA | NA | NA | Gm49641 | predicted gene, 49641 [Source:MGI Symb | 0 | 0 | 0 | 0 | 0 | 0 |
| 53880 | ENSMUSG00000116668 | 0 | NA | NA | NA | NA | NA | Gm49568 | predicted gene, 49568 [Source:MGI Symb | 0 | 0 | 0 | 0 | 0 | 0 |
| 53881 | ENSMUSG00000116669 | 0 | NA | NA | NA | NA | NA | Gm49570 | predicted gene, 49570 [Source:MGI Symb | 0 | 0 | 0 | 0 | 0 | 0 |
| 53882 | ENSMUSG00000116670 | 0 | NA | NA | NA | NA | NA | Gm49752 | predicted gene, 49752 [Source:MGI Symb | 0 | 0 | 0 | 0 | 0 | 0 |
| 53883 | ENSMUSG00000116671 | 0 | NA | NA | NA | NA | NA | Gm49635 | predicted gene, 49635 [Source:MGI Symb | 0 | 0 | 0 | 0 | 0 | 0 |
| 53884 | ENSMUSG00000116672 | 0 | NA | NA | NA | NA | NA | Gm49630 | predicted gene, 49630 [Source:MGI Symb | 0 | 0 | 0 | 0 | 0 | 0 |
| 53885 | ENSMUSG00000116673 | 7.782357473 | 0.065167003 | 0.808464091 | 0.080605934 | 0.935755347 | 0.988420305 | A630089N07 | RIKEN cDNA A630089N07 gene [Source:M | 9.6372 | 3.0838 | 10.062 | 9.6398 | 9.8156 | 4.4554 |
| 53886 | ENSMUSG00000116674 | 0 | NA | NA | NA | NA | NA | 4930551I23F | RIKEN cDNA 4930551I23 gene [Source:M | 0 | 0 | 0 | 0 | 0 | 0 |
| 53887 | ENSMUSG00000116675 | 0 | NA | NA | NA | NA | NA | Gm49581 | predicted gene, 49581 [Source:MGI Symb | 0 | 0 | 0 | 0 | 0 | 0 |
| 53888 | ENSMUSG00000116676 | 0 | NA | NA | NA | NA | NA | Gm5768 | predicted gene 5768 [Source:MGI Symbol | 0 | 0 | 0 | 0 | 0 | 0 |
| 53889 | ENSMUSG00000116677 | 0 | NA | NA | NA | NA | NA | Gm49569 | predicted gene, 49569 [Source:MGI Symb | 0 | 0 | 0 | 0 | 0 | 0 |
| 53890 | ENSMUSG00000116678 | 0.717584917 | 3.048786435 | 2.951298099 | 1.033032358 | 0.301588756 | NA | Pgk1-ps1 | phosphoglycerate kinase 1, pseudogene 1 | 0 | 0 | 0 | 0.964 | 0 | 3.3415 |
| 53891 | ENSMUSG00000116679 | 0 | NA | NA | NA | NA | NA | Gm49739 | predicted gene, 49739 [Source:MGI Symb | 0 | 0 | 0 | 0 | 0 | 0 |
| 53892 | ENSMUSG00000116680 | 0 | NA | NA | NA | NA | NA | Gm49562 | predicted gene, 49562 [Source:MGI Symb | 0 | 0 | 0 | 0 | 0 | 0 |
| 53893 | ENSMUSG00000116681 | 0 | NA | NA | NA | NA | NA | Gm49618 | predicted gene, 49618 [Source:MGI Symb | 0 | 0 | 0 | 0 | 0 | 0 |
| 53894 | ENSMUSG00000116682 | 0 | NA | NA | NA | NA | NA | Gm49789 | predicted gene, 49789 [Source:MGI Symb | 0 | 0 | 0 | 0 | 0 | 0 |
| 53895 | ENSMUSG00000116684 | 0 | NA | NA | NA | NA | NA | Gm49735 | predicted gene, 49735 [Source:MGI Symb | 0 | 0 | 0 | 0 | 0 | 0 |
| 53896 | ENSMUSG00000116685 | 0 | NA | NA | NA | NA | NA | Gm49664 | predicted gene, 49664 [Source:MGI Symb | 0 | 0 | 0 | 0 | 0 | 0 |
| 53897 | ENSMUSG00000116686 | 0 | NA | NA | NA | NA | NA | Gm49681 | predicted gene, 49681 [Source:MGI Symb | 0 | 0 | 0 | 0 | 0 | 0 |
| 53898 | ENSMUSG00000116687 | 0 | NA | NA | NA | NA | NA | Gm49629 | predicted gene, 49629 [Source:MGI Symb | 0 | 0 | 0 | 0 | 0 | 0 |
| 53899 | ENSMUSG00000116688 | 0 | NA | NA | NA | NA | NA | Gm49720 | predicted gene, 49720 [Source:MGI Symb | 0 | 0 | 0 | 0 | 0 | 0 |
| 53900 | ENSMUSG00000116689 | 0 | NA | NA | NA | NA | NA | 4930405D01 | RIKEN cDNA 4930405D01 gene [Source:M | 0 | 0 | 0 | 0 | 0 | 0 |
| 53901 | ENSMUSG00000116690 | 1.097073163 | -0.155822922 | 2.139629041 | -0.072827074 | 0.941943726 | NA | Gm49736 | predicted gene, 49736 [Source:MGI Symb | 2.4093 | 1.0279 | 0 | 0.964 | 2.1812 | 0 |
| 53902 | ENSMUSG00000116691 | 0 | NA | NA | NA | NA | NA | Gm8363 | predicted gene 8363 [Source:MGI Symbol | 0 | 0 | 0 | 0 | 0 | 0 |
| 53903 | ENSMUSG00000116692 | 0 | NA | NA | NA | NA | NA | Gm49795 | predicted gene, 49795 [Source:MGI Symb | 0 | 0 | 0 | 0 | 0 | 0 |
| 53904 | ENSMUSG00000116693 | 0 | NA | NA | NA | NA | NA | Gm29805 | predicted gene, 29805 [Source:MGI Symb | 0 | 0 | 0 | 0 | 0 | 0 |
| 53905 | ENSMUSG00000116694 | 0 | NA | NA | NA | NA | NA | Gm6440 | predicted gene 6440 [Source:MGI Symbol | 0 | 0 | 0 | 0 | 0 | 0 |
| 53906 | ENSMUSG00000116695 | 0 | NA | NA | NA | NA | NA | 4930578N18 | RIKEN cDNA 4930578N18 gene [Source:M | 0 | 0 | 0 | 0 | 0 | 0 |
| 53907 | ENSMUSG00000116696 | 0.615839296 | 1.537131441 | 3.17677869 | 0.483864817 | 0.628481811 | NA | Gm49683 | predicted gene, 49683 [Source:MGI Symb | 0.8031 | 0 | 0 | 2.8919 | 0 | 0 |
| 53908 | ENSMUSG00000116697 | 2.423835382 | 0.454257838 | 1.609849114 | 0.28217417 | 0.777809963 | NA | Gm49694 | predicted gene, 49694 [Source:MGI Symb | 4.0155 | 2.0558 | 0 | 1.928 | 6.5437 | 0 |
| 53909 | ENSMUSG00000116698 | 0 | NA | NA | NA | NA | NA | Gm49676 | predicted gene, 49676 [Source:MGI Symb | 0 | 0 | 0 | 0 | 0 | 0 |
| 53910 | ENSMUSG00000116699 | 0 | NA | NA | NA | NA | NA | Gm49631 | predicted gene, 49631 [Source:MGI Symb | 0 | 0 | 0 | 0 | 0 | 0 |
| 53911 | ENSMUSG00000116700 | 0 | NA | NA | NA | NA | NA | Gm18694 | predicted gene, 18694 [Source:MGI Symb | 0 | 0 | 0 | 0 | 0 | 0 |
| 53912 | ENSMUSG00000116701 | 0 | NA | NA | NA | NA | NA | Gm30726 | predicted gene, 30726 [Source:MGI Symb | 0 | 0 | 0 | 0 | 0 | 0 |
| 53913 | ENSMUSG00000116702 | 0 | NA | NA | NA | NA | NA | Gm6553 | predicted gene 6553 [Source:MGI Symbol | 0 | 0 | 0 | 0 | 0 | 0 |
| 53914 | ENSMUSG00000116703 | 0 | NA | NA | NA | NA | NA | Gm46540 | predicted gene, 46540 [Source:MGI Symb | 0 | 0 | 0 | 0 | 0 | 0 |
| 53915 | ENSMUSG00000116704 | 0 | NA | NA | NA | NA | NA | Gm5491 | predicted gene 5491 [Source:MGI Symbol | 0 | 0 | 0 | 0 | 0 | 0 |
| 53916 | ENSMUSG00000116705 | 0 | NA | NA | NA | NA | NA | Gm29737 | predicted gene, 29737 [Source:MGI Symb | 0 | 0 | 0 | 0 | 0 | 0 |
| 53917 | ENSMUSG00000116706 | 0 | NA | NA | NA | NA | NA | Gm32431 | predicted gene, 32431 [Source:MGI Symb | 0 | 0 | 0 | 0 | 0 | 0 |
| 53918 | ENSMUSG00000116707 | 1.1400417 | 2.589352293 | 2.336715871 | 1.108116021 | 0.267811713 | NA | Gm49590 | predicted gene, 49590 [Source:MGI Symb | 0.8031 | 0 | 0 | 3.8559 | 2.1812 | 0 |
| 53919 | ENSMUSG00000116708 | 0 | NA | NA | NA | NA | NA | Gm49689 | predicted gene, 49689 [Source:MGI Symb | 0 | 0 | 0 | 0 | 0 | 0 |
| 53920 | ENSMUSG00000116709 | 0 | NA | NA | NA | NA | NA | Gm49722 | predicted gene, 49722 [Source:MGI Symb | 0 | 0 | 0 | 0 | 0 | 0 |
| 53921 | ENSMUSG00000116710 | 0 | NA | NA | NA | NA | NA | Gm6557 | predicted gene 6557 [Source:MGI Symbol | 0 | 0 | 0 | 0 | 0 | 0 |
| 53922 | ENSMUSG00000116711 | 0 | NA | NA | NA | NA | NA | Gm18495 | predicted gene, 18495 [Source:MGI Symb | 0 | 0 | 0 | 0 | 0 | 0 |
| 53923 | ENSMUSG00000116712 | 0 | NA | NA | NA | NA | NA | Gm49598 | predicted gene, 49598 [Source:MGI Symb | 0 | 0 | 0 | 0 | 0 | 0 |
| 53924 | ENSMUSG00000116713 | 0 | NA | NA | NA | NA | NA | Gm49613 | predicted gene, 49613 [Source:MGI Symb | 0 | 0 | 0 | 0 | 0 | 0 |

| | | | | | | | | | | | | | | | |
|---|---|---|---|---|---|---|---|---|---|---|---|---|---|---|---|
| 53925 | ENSMUSG00000116714 | 0.267700746 | -1.497481649 | 4.057001436 | -0.369110456 | 0.712045396 | NA | 2210020O09 | RIKEN cDNA 2210020O09 gene [Source:M | 1.6062 | 0 | 0 | 0 | 0 | 0 |
| 53926 | ENSMUSG00000116715 | 0 | NA | NA | NA | NA | NA | Gm40881 | predicted gene, 40881 [Source:MGI Symb | 0 | 0 | 0 | 0 | 0 | 0 |
| 53927 | ENSMUSG00000116716 | 0 | NA | NA | NA | NA | NA | Gm49643 | predicted gene, 49643 [Source:MGI Symb | 0 | 0 | 0 | 0 | 0 | 0 |
| 53928 | ENSMUSG00000116717 | 0 | NA | NA | NA | NA | NA | Gm49574 | predicted gene, 49574 [Source:MGI Symb | 0 | 0 | 0 | 0 | 0 | 0 |
| 53929 | ENSMUSG00000116718 | 69.64813627 | -0.018935209 | 0.271594522 | -0.069718671 | 0.944417581 | 0.990077979 | Gm49668 | predicted gene, 49668 [Source:MGI Symb | 77.901 | 68.87 | 63.393 | 74.226 | 59.984 | 73.514 |
| 53930 | ENSMUSG00000116719 | 0 | NA | NA | NA | NA | NA | Gm29719 | predicted gene, 29719 [Source:MGI Symb | 0 | 0 | 0 | 0 | 0 | 0 |
| 53931 | ENSMUSG00000116720 | 0.185640647 | 1.126924175 | 4.080472857 | 0.276174898 | 0.782413736 | NA | Gm31641 | predicted gene, 31641 [Source:MGI Symb | 0 | 0 | 0 | 0 | 0 | 1.1138 |
| 53932 | ENSMUSG00000116721 | 0 | NA | NA | NA | NA | NA | Gm9536 | predicted gene 9536 [Source:MGI Symbol | 0 | 0 | 0 | 0 | 0 | 0 |
| 53933 | ENSMUSG00000116722 | 0 | NA | NA | NA | NA | NA | Gm49627 | predicted gene, 49627 [Source:MGI Symb | 0 | 0 | 0 | 0 | 0 | 0 |
| 53934 | ENSMUSG00000116723 | 6.921608201 | -0.032601884 | 0.822960987 | -0.039615346 | 0.968399793 | 0.994245784 | Gm46559 | predicted gene, 46559 [Source:MGI Symb | 8.8341 | 5.1396 | 7.0437 | 10.604 | 5.4531 | 4.4554 |
| 53935 | ENSMUSG00000116725 | 0 | NA | NA | NA | NA | NA | Gm29686 | predicted gene, 29686 [Source:MGI Symb | 0 | 0 | 0 | 0 | 0 | 0 |
| 53936 | ENSMUSG00000116727 | 0 | NA | NA | NA | NA | NA | Gm8881 | predicted gene 8881 [Source:MGI Symbol | 0 | 0 | 0 | 0 | 0 | 0 |
| 53937 | ENSMUSG00000116728 | 0 | NA | NA | NA | NA | NA | Gm41467 | predicted gene, 41467 [Source:MGI Symb | 0 | 0 | 0 | 0 | 0 | 0 |
| 53938 | ENSMUSG00000116729 | 0 | NA | NA | NA | NA | NA | Gm49644 | predicted gene, 49644 [Source:MGI Symb | 0 | 0 | 0 | 0 | 0 | 0 |
| 53939 | ENSMUSG00000116730 | 0 | NA | NA | NA | NA | NA | 4930403O18 | RIKEN cDNA 4930403O18 gene [Source:M | 0 | 0 | 0 | 0 | 0 | 0 |
| 53940 | ENSMUSG00000116731 | 0 | NA | NA | NA | NA | NA | Gm18771 | predicted gene, 18771 [Source:MGI Symb | 0 | 0 | 0 | 0 | 0 | 0 |
| 53941 | ENSMUSG00000116732 | 0 | NA | NA | NA | NA | NA | Gm18769 | predicted gene, 18769 [Source:MGI Symb | 0 | 0 | 0 | 0 | 0 | 0 |
| 53942 | ENSMUSG00000116733 | 9.397591346 | 0.129868005 | 0.7368259 | 0.176253312 | 0.86009494 | 0.97570916 | 6330408M09 | RIKEN cDNA 6330408M09 gene [Source:M | 9.6372 | 7.1954 | 10.062 | 8.6758 | 16.359 | 4.4554 |
| 53943 | ENSMUSG00000116734 | 0 | NA | NA | NA | NA | NA | Gm8376 | predicted gene 8376 [Source:MGI Symbol | 0 | 0 | 0 | 0 | 0 | 0 |
| 53944 | ENSMUSG00000116735 | 0.17131964 | -0.796629176 | 4.080472857 | -0.195229623 | 0.84521319 | NA | Gm49555 | predicted gene, 49555 [Source:MGI Symb | 0 | 1.0279 | 0 | 0 | 0 | 0 |
| 53945 | ENSMUSG00000116736 | 0 | NA | NA | NA | NA | NA | Gm49692 | predicted gene, 49692 [Source:MGI Symb | 0 | 0 | 0 | 0 | 0 | 0 |
| 53946 | ENSMUSG00000116737 | 0 | NA | NA | NA | NA | NA | Gm49757 | predicted gene, 49757 [Source:MGI Symb | 0 | 0 | 0 | 0 | 0 | 0 |
| 53947 | ENSMUSG00000116738 | 0 | NA | NA | NA | NA | NA | Gm49565 | predicted gene, 49565 [Source:MGI Symb | 0 | 0 | 0 | 0 | 0 | 0 |
| 53948 | ENSMUSG00000116739 | 0.653308562 | 1.538013299 | 3.112409277 | 0.494155223 | 0.62119656 | NA | Gm49733 | predicted gene, 49733 [Source:MGI Symb | 0 | 1.0279 | 0 | 2.8919 | 0 | 0 |
| 53949 | ENSMUSG00000116740 | 0 | NA | NA | NA | NA | NA | Gm49659 | predicted gene, 49659 [Source:MGI Symb | 0 | 0 | 0 | 0 | 0 | 0 |
| 53950 | ENSMUSG00000116741 | 0 | NA | NA | NA | NA | NA | Gm49564 | predicted gene, 49564 [Source:MGI Symb | 0 | 0 | 0 | 0 | 0 | 0 |
| 53951 | ENSMUSG00000116742 | 0 | NA | NA | NA | NA | NA | Gm49665 | predicted gene, 49665 [Source:MGI Symb | 0 | 0 | 0 | 0 | 0 | 0 |
| 53952 | ENSMUSG00000116743 | 0 | NA | NA | NA | NA | NA | Gm49573 | predicted gene, 49573 [Source:MGI Symb | 0 | 0 | 0 | 0 | 0 | 0 |
| 53953 | ENSMUSG00000116744 | 0 | NA | NA | NA | NA | NA | Gm49764 | predicted gene, 49764 [Source:MGI Symb | 0 | 0 | 0 | 0 | 0 | 0 |
| 53954 | ENSMUSG00000116745 | 0 | NA | NA | NA | NA | NA | Gm41481 | predicted gene, 41481 [Source:MGI Symb | 0 | 0 | 0 | 0 | 0 | 0 |
| 53955 | ENSMUSG00000116746 | 0 | NA | NA | NA | NA | NA | Gm18825 | predicted gene, 18825 [Source:MGI Symb | 0 | 0 | 0 | 0 | 0 | 0 |
| 53956 | ENSMUSG00000116747 | 0 | NA | NA | NA | NA | NA | Gm41496 | predicted gene, 41496 [Source:MGI Symb | 0 | 0 | 0 | 0 | 0 | 0 |
| 53957 | ENSMUSG00000116748 | 0 | NA | NA | NA | NA | NA | Gm7295 | predicted gene 7295 [Source:MGI Symbol | 0 | 0 | 0 | 0 | 0 | 0 |
| 53958 | ENSMUSG00000116749 | 0 | NA | NA | NA | NA | NA | Gm49612 | predicted gene, 49612 [Source:MGI Symb | 0 | 0 | 0 | 0 | 0 | 0 |
| 53959 | ENSMUSG00000116750 | 0 | NA | NA | NA | NA | NA | 4931420L22 | RIKEN cDNA 4931420L22 gene [Source:M | 0 | 0 | 0 | 0 | 0 | 0 |
| 53960 | ENSMUSG00000116751 | 0 | NA | NA | NA | NA | NA | Gm49793 | predicted gene, 49793 [Source:MGI Symb | 0 | 0 | 0 | 0 | 0 | 0 |
| 53961 | ENSMUSG00000116752 | 0 | NA | NA | NA | NA | NA | Gm49622 | predicted gene, 49622 [Source:MGI Symb | 0 | 0 | 0 | 0 | 0 | 0 |
| 53962 | ENSMUSG00000116753 | 0 | NA | NA | NA | NA | NA | Gm36001 | predicted gene, 36001 [Source:MGI Symb | 0 | 0 | 0 | 0 | 0 | 0 |
| 53963 | ENSMUSG00000116754 | 0 | NA | NA | NA | NA | NA | Gm33255 | predicted gene, 33255 [Source:MGI Symb | 0 | 0 | 0 | 0 | 0 | 0 |
| 53964 | ENSMUSG00000116755 | 0 | NA | NA | NA | NA | NA | 6430553K19 | RIKEN cDNA 6430553K19 gene [Source:M | 0 | 0 | 0 | 0 | 0 | 0 |
| 53965 | ENSMUSG00000116756 | 0 | NA | NA | NA | NA | NA | Gm49770 | predicted gene, 49770 [Source:MGI Symb | 0 | 0 | 0 | 0 | 0 | 0 |
| 53966 | ENSMUSG00000116757 | 1.091781011 | 2.43899895 | 2.376659209 | 1.026229987 | 0.30478319 | NA | Gm4786 | predicted gene 4786 [Source:MGI Symbol | 0 | 1.0279 | 0 | 0 | 2.1812 | 3.3415 |
| 53967 | ENSMUSG00000116758 | 0 | NA | NA | NA | NA | NA | Gm19291 | predicted gene, 19291 [Source:MGI Symb | 0 | 0 | 0 | 0 | 0 | 0 |
| 53968 | ENSMUSG00000116759 | 0 | NA | NA | NA | NA | NA | Gm49666 | predicted gene, 49666 [Source:MGI Symb | 0 | 0 | 0 | 0 | 0 | 0 |
| 53969 | ENSMUSG00000116760 | 0 | NA | NA | NA | NA | NA | Gm1604a | predicted gene 1604A [Source:MGI Symb | 0 | 0 | 0 | 0 | 0 | 0 |
| 53970 | ENSMUSG00000116761 | 0 | NA | NA | NA | NA | NA | Gm36796 | predicted gene, 36796 [Source:MGI Symb | 0 | 0 | 0 | 0 | 0 | 0 |
| 53971 | ENSMUSG00000116762 | 0 | NA | NA | NA | NA | NA | Gm46560 | predicted gene, 46560 [Source:MGI Symb | 0 | 0 | 0 | 0 | 0 | 0 |
| 53972 | ENSMUSG00000116763 | 0 | NA | NA | NA | NA | NA | Gm41492 | predicted gene, 41492 [Source:MGI Symb | 0 | 0 | 0 | 0 | 0 | 0 |
| 53973 | ENSMUSG00000116764 | 0 | NA | NA | NA | NA | NA | Gm49748 | predicted gene, 49748 [Source:MGI Symb | 0 | 0 | 0 | 0 | 0 | 0 |
| 53974 | ENSMUSG00000116765 | 3.317701232 | -0.580528409 | 1.397042871 | -0.415540869 | 0.677746003 | NA | Gm49762 | predicted gene, 49762 [Source:MGI Symb | 8.8341 | 3.0838 | 0 | 5.7839 | 1.0906 | 1.1138 |
| 53975 | ENSMUSG00000116766 | 0.531944269 | 2.618916564 | 3.285123409 | 0.797204926 | 0.425332026 | NA | Gm49702 | predicted gene, 49702 [Source:MGI Symb | 0 | 0 | 0 | 0.964 | 0 | 2.2277 |
| 53976 | ENSMUSG00000116767 | 0 | NA | NA | NA | NA | NA | 4930517M08 | RIKEN cDNA 4930517M08 gene [Source:M | 0 | 0 | 0 | 0 | 0 | 0 |
| 53977 | ENSMUSG00000116768 | 0 | NA | NA | NA | NA | NA | Gm18269 | predicted gene, 18269 [Source:MGI Symb | 0 | 0 | 0 | 0 | 0 | 0 |
| 53978 | ENSMUSG00000116769 | 0 | NA | NA | NA | NA | NA | 8030451O07 | RIKEN cDNA 8030451O07 gene [Source:M | 0 | 0 | 0 | 0 | 0 | 0 |
| 53979 | ENSMUSG00000116770 | 0 | NA | NA | NA | NA | NA | Gm49576 | predicted gene, 49576 [Source:MGI Symb | 0 | 0 | 0 | 0 | 0 | 0 |
| 53980 | ENSMUSG00000116771 | 0.636946036 | 1.591416711 | 2.83781177 | 0.560790087 | 0.574940645 | NA | Gm49691 | predicted gene, 49691 [Source:MGI Symb | 0.8031 | 0 | 0 | 1.928 | 1.0906 | 0 |

| | | | | | | | | | | | | | | | |
|---|---|---|---|---|---|---|---|---|---|---|---|---|---|---|---|
| 53981 | ENSMUSG00000116772 | 0 | NA | NA | NA | NA | NA | 4930483P17 | RIKEN cDNA 4930483P17 gene [Source:M | 0 | 0 | 0 | 0 | 0 | 0 |
| 53982 | ENSMUSG00000116773 | 0 | NA | NA | NA | NA | NA | Gm49605 | predicted gene, 49605 [Source:MGI Symb | 0 | 0 | 0 | 0 | 0 | 0 |
| 53983 | ENSMUSG00000116774 | 0 | NA | NA | NA | NA | NA | Gm49690 | predicted gene, 49690 [Source:MGI Symb | 0 | 0 | 0 | 0 | 0 | 0 |
| 53984 | ENSMUSG00000116775 | 0.36353943 | 2.065989254 | 4.034137301 | 0.512126658 | 0.608562368 | NA | Gm49769 | predicted gene, 49769 [Source:MGI Symb | 0 | 0 | 0 | 0 | 2.1812 | 0 |
| 53985 | ENSMUSG00000116776 | 0 | NA | NA | NA | NA | NA | Gm46544 | predicted gene, 46544 [Source:MGI Symb | 0 | 0 | 0 | 0 | 0 | 0 |
| 53986 | ENSMUSG00000116777 | 0 | NA | NA | NA | NA | NA | Zfp520-ps | zinc finger protein 520, pseudogene [Sour | 0 | 0 | 0 | 0 | 0 | 0 |
| 53987 | ENSMUSG00000116778 | 0.181769715 | 1.126924175 | 4.080472857 | 0.276174898 | 0.782413736 | NA | Gm49713 | predicted gene, 49713 [Source:MGI Symb | 0 | 0 | 0 | 0 | 1.0906 | 0 |
| 53988 | ENSMUSG00000116779 | 0 | NA | NA | NA | NA | NA | Gm4829 | predicted gene 4829 [Source:MGI Symbol | 0 | 0 | 0 | 0 | 0 | 0 |
| 53989 | ENSMUSG00000116780 | 0 | NA | NA | NA | NA | NA | Dynlt2a3 | dynein light chain Tctex-type 2A3 [Source: | 0 | 0 | 0 | 0 | 0 | 0 |
| 53990 | ENSMUSG00000116781 | 0.328370405 | 0.16514316 | 4.003149536 | 0.041253308 | 0.967093956 | NA | Gm49712 | predicted gene, 49712 [Source:MGI Symb | 0 | 0 | 1.0062 | 0.964 | 0 | 0 |
| 53991 | ENSMUSG00000116782 | 0.185640647 | 1.126924175 | 4.080472857 | 0.276174898 | 0.782413736 | NA | Gm46562 | predicted gene, 46562 [Source:MGI Symb | 0 | 0 | 0 | 0 | 0 | 1.1138 |
| 53992 | ENSMUSG00000116783 | 0 | NA | NA | NA | NA | NA | Gm6048 | predicted gene 6048 [Source:MGI Symbol | 0 | 0 | 0 | 0 | 0 | 0 |
| 53993 | ENSMUSG00000116784 | 0 | NA | NA | NA | NA | NA | Gm49716 | predicted gene, 49716 [Source:MGI Symb | 0 | 0 | 0 | 0 | 0 | 0 |
| 53994 | ENSMUSG00000116785 | 0 | NA | NA | NA | NA | NA | Gm41434 | predicted gene, 41434 [Source:MGI Symb | 0 | 0 | 0 | 0 | 0 | 0 |
| 53995 | ENSMUSG00000116786 | 0 | NA | NA | NA | NA | NA | 4930570E03 | RIKEN cDNA 4930570E03 gene [Source:M | 0 | 0 | 0 | 0 | 0 | 0 |
| 53996 | ENSMUSG00000116788 | 0 | NA | NA | NA | NA | NA | Gm5674 | predicted gene 5674 [Source:MGI Symbol | 0 | 0 | 0 | 0 | 0 | 0 |
| 53997 | ENSMUSG00000116789 | 0 | NA | NA | NA | NA | NA | Gm49579 | predicted gene, 49579 [Source:MGI Symb | 0 | 0 | 0 | 0 | 0 | 0 |
| 53998 | ENSMUSG00000116790 | 0 | NA | NA | NA | NA | NA | Gm49596 | predicted gene, 49596 [Source:MGI Symb | 0 | 0 | 0 | 0 | 0 | 0 |
| 53999 | ENSMUSG00000116791 | 0 | NA | NA | NA | NA | NA | Gm18918 | predicted gene, 18918 [Source:MGI Symb | 0 | 0 | 0 | 0 | 0 | 0 |
| 54000 | ENSMUSG00000116792 | 0.335414862 | -1.779047242 | 4.030804179 | -0.441362855 | 0.658950329 | NA | D16Ertd519e | DNA segment, Chr 16, ERATO Doi 519, exp | 0 | 0 | 2.0125 | 0 | 0 | 0 |
| 54001 | ENSMUSG00000116793 | 0 | NA | NA | NA | NA | NA | Gm34256 | predicted gene, 34256 [Source:MGI Symb | 0 | 0 | 0 | 0 | 0 | 0 |
| 54002 | ENSMUSG00000116794 | 0 | NA | NA | NA | NA | NA | Gm49656 | predicted gene, 49656 [Source:MGI Symb | 0 | 0 | 0 | 0 | 0 | 0 |
| 54003 | ENSMUSG00000116795 | 0.626495961 | 0.101295816 | 2.879413718 | 0.03517932 | 0.971936753 | NA | Gm49600 | predicted gene, 49600 [Source:MGI Symb | 0.8031 | 1.0279 | 0 | 1.928 | 0 | 0 |
| 54004 | ENSMUSG00000116796 | 0 | NA | NA | NA | NA | NA | Gm49784 | predicted gene, 49784 [Source:MGI Symb | 0 | 0 | 0 | 0 | 0 | 0 |
| 54005 | ENSMUSG00000116797 | 0 | NA | NA | NA | NA | NA | 2600002D14 | RIKEN cDNA 2600002D14 gene [Source:M | 0 | 0 | 0 | 0 | 0 | 0 |
| 54006 | ENSMUSG00000116798 | 0 | NA | NA | NA | NA | NA | Gm49591 | predicted gene, 49591 [Source:MGI Symb | 0 | 0 | 0 | 0 | 0 | 0 |
| 54007 | ENSMUSG00000116799 | 0 | NA | NA | NA | NA | NA | Gm32357 | predicted gene, 32357 [Source:MGI Symb | 0 | 0 | 0 | 0 | 0 | 0 |
| 54008 | ENSMUSG00000116800 | 0 | NA | NA | NA | NA | NA | Gm29823 | predicted gene, 29823 [Source:MGI Symb | 0 | 0 | 0 | 0 | 0 | 0 |
| 54009 | ENSMUSG00000116801 | 0 | NA | NA | NA | NA | NA | Gm38551 | predicted gene, 38551 [Source:MGI Symb | 0 | 0 | 0 | 0 | 0 | 0 |
| 54010 | ENSMUSG00000116802 | 62.34513895 | 0.120284149 | 0.307718405 | 0.39089033 | 0.695878302 | 0.939992732 | Gm5165 | predicted gene 5165 [Source:MGI Symbol | 73.885 | 59.619 | 45.281 | 61.695 | 55.622 | 77.969 |
| 54011 | ENSMUSG00000116803 | 0 | NA | NA | NA | NA | NA | Gm49558 | predicted gene, 49558 [Source:MGI Symb | 0 | 0 | 0 | 0 | 0 | 0 |
| 54012 | ENSMUSG00000116804 | 0 | NA | NA | NA | NA | NA | Gm5223 | predicted gene 5223 [Source:MGI Symbol | 0 | 0 | 0 | 0 | 0 | 0 |
| 54013 | ENSMUSG00000116805 | 0 | NA | NA | NA | NA | NA | Gm49723 | predicted gene, 49723 [Source:MGI Symb | 0 | 0 | 0 | 0 | 0 | 0 |
| 54014 | ENSMUSG00000116806 | 0 | NA | NA | NA | NA | NA | Gm49680 | predicted gene, 49680 [Source:MGI Symb | 0 | 0 | 0 | 0 | 0 | 0 |
| 54015 | ENSMUSG00000116807 | 0 | NA | NA | NA | NA | NA | Gm7501 | predicted gene 7501 [Source:MGI Symbol | 0 | 0 | 0 | 0 | 0 | 0 |
| 54016 | ENSMUSG00000116808 | 0 | NA | NA | NA | NA | NA | Gm20164 | predicted gene, 20164 [Source:MGI Symb | 0 | 0 | 0 | 0 | 0 | 0 |
| 54017 | ENSMUSG00000116809 | 0 | NA | NA | NA | NA | NA | A630036G19 | RIKEN cDNA A630036G19 gene [Source:M | 0 | 0 | 0 | 0 | 0 | 0 |
| 54018 | ENSMUSG00000116810 | 34.9325841 | -0.202869754 | 0.375495466 | -0.540272181 | 0.58900934 | 0.911039598 | Gm32624 | predicted gene, 32624 [Source:MGI Symb | 33.73 | 45.228 | 33.206 | 27.955 | 32.719 | 36.757 |
| 54019 | ENSMUSG00000116811 | 0 | NA | NA | NA | NA | NA | Gm16385 | predicted gene 16385 [Source:MGI Symbo | 0 | 0 | 0 | 0 | 0 | 0 |
| 54020 | ENSMUSG00000116812 | 0.181769715 | 1.126924175 | 4.080472857 | 0.276174898 | 0.782413736 | NA | Gm2792 | predicted gene 2792 [Source:MGI Symbol | 0 | 0 | 0 | 0 | 1.0906 | 0 |
| 54021 | ENSMUSG00000116813 | 0.160662974 | 1.126924175 | 4.080472857 | 0.276174898 | 0.782413736 | NA | 4930579D07 | RIKEN cDNA 4930579D07 gene [Source:M | 0 | 0 | 0 | 0.964 | 0 | 0 |
| 54022 | ENSMUSG00000116814 | 0 | NA | NA | NA | NA | NA | Gm49636 | predicted gene, 49636 [Source:MGI Symb | 0 | 0 | 0 | 0 | 0 | 0 |
| 54023 | ENSMUSG00000116815 | 3.547521424 | -0.877468691 | 1.192158737 | -0.736033436 | 0.46171035 | NA | Gm49760 | predicted gene, 49760 [Source:MGI Symb | 1.6062 | 6.1675 | 6.0375 | 1.928 | 1.0906 | 4.4554 |
| 54024 | ENSMUSG00000116816 | 0 | NA | NA | NA | NA | NA | Cstdc3 | cystatin domain containing 3 [Source:MGI | 0 | 0 | 0 | 0 | 0 | 0 |
| 54025 | ENSMUSG00000116817 | 0.685278559 | -2.809870862 | 3.376578027 | -0.832165239 | 0.405315682 | NA | Gm5679 | predicted gene 5679 [Source:MGI Symbol | 0 | 4.1117 | 0 | 0 | 0 | 0 |
| 54026 | ENSMUSG00000116818 | 0 | NA | NA | NA | NA | NA | Gm7513 | predicted gene 7513 [Source:MGI Symbol | 0 | 0 | 0 | 0 | 0 | 0 |
| 54027 | ENSMUSG00000116819 | 1.699439758 | 1.391747046 | 1.860057236 | 0.748228075 | 0.454322599 | NA | Gm49586 | predicted gene, 49586 [Source:MGI Symb | 0.8031 | 0 | 2.0125 | 1.928 | 5.4531 | 0 |
| 54028 | ENSMUSG00000116820 | 0 | NA | NA | NA | NA | NA | Gm17928 | predicted gene, 17928 [Source:MGI Symb | 0 | 0 | 0 | 0 | 0 | 0 |
| 54029 | ENSMUSG00000116821 | 0 | NA | NA | NA | NA | NA | Gm8538 | predicted gene 8538 [Source:MGI Symbol | 0 | 0 | 0 | 0 | 0 | 0 |
| 54030 | ENSMUSG00000116822 | 0 | NA | NA | NA | NA | NA | Gm41504 | predicted gene, 41504 [Source:MGI Symb | 0 | 0 | 0 | 0 | 0 | 0 |
| 54031 | ENSMUSG00000116823 | 0.17131964 | -0.796629176 | 4.080472857 | -0.195229623 | 0.84521319 | NA | Gm49595 | predicted gene, 49595 [Source:MGI Symb | 0 | 1.0279 | 0 | 0 | 0 | 0 |
| 54032 | ENSMUSG00000116824 | 0 | NA | NA | NA | NA | NA | Gm18021 | predicted gene, 18021 [Source:MGI Symb | 0 | 0 | 0 | 0 | 0 | 0 |
| 54033 | ENSMUSG00000116825 | 0 | NA | NA | NA | NA | NA | Rps2-ps7 | ribosomal protein S2, pseudogene 7 [Sour | 0 | 0 | 0 | 0 | 0 | 0 |
| 54034 | ENSMUSG00000116826 | 0 | NA | NA | NA | NA | NA | Gm49682 | predicted gene, 49682 [Source:MGI Symb | 0 | 0 | 0 | 0 | 0 | 0 |
| 54035 | ENSMUSG00000116827 | 0.17131964 | -0.796629176 | 4.080472857 | -0.195229623 | 0.84521319 | NA | Gm5063 | predicted gene 5063 [Source:MGI Symbol | 0 | 1.0279 | 0 | 0 | 0 | 0 |
| 54036 | ENSMUSG00000116828 | 0 | NA | NA | NA | NA | NA | Gm18896 | predicted gene, 18896 [Source:MGI Symb | 0 | 0 | 0 | 0 | 0 | 0 |

| | | | | | | | | | | | | | | | |
|---|---|---|---|---|---|---|---|---|---|---|---|---|---|---|---|
| 54037 | ENSMUSG00000116829 | 0.535401493 | -2.48596965 | 3.684117979 | -0.674780141 | 0.499815459 | NA | Gm49704 | predicted gene, 49704 [Source:MGI Symb | 3.2124 | 0 | 0 | 0 | 0 | 0 |
| 54038 | ENSMUSG00000116830 | 0 | NA | NA | NA | NA | NA | Gm49714 | predicted gene, 49714 [Source:MGI Symb | 0 | 0 | 0 | 0 | 0 | 0 |
| 54039 | ENSMUSG00000116831 | 37.86458788 | -0.163602965 | 0.349585489 | -0.467991292 | 0.639790817 | 0.923932929 | Gm30505 | predicted gene, 30505 [Source:MGI Symb | 37.746 | 42.145 | 40.25 | 37.595 | 33.809 | 35.643 |
| 54040 | ENSMUSG00000116832 | 0 | NA | NA | NA | NA | NA | Gm49658 | predicted gene, 49658 [Source:MGI Symb | 0 | 0 | 0 | 0 | 0 | 0 |
| 54041 | ENSMUSG00000116833 | 0 | NA | NA | NA | NA | NA | Gm9584 | predicted gene 9584 [Source:MGI Symbol | 0 | 0 | 0 | 0 | 0 | 0 |
| 54042 | ENSMUSG00000116834 | 0 | NA | NA | NA | NA | NA | Gm18020 | predicted gene, 18020 [Source:MGI Symb | 0 | 0 | 0 | 0 | 0 | 0 |
| 54043 | ENSMUSG00000116835 | 0.767390791 | -1.69152783 | 2.44773922 | -0.691057207 | 0.489529591 | NA | Gm49594 | predicted gene, 49594 [Source:MGI Symb | 1.6062 | 1.0279 | 1.0062 | 0.964 | 0 | 0 |
| 54044 | ENSMUSG00000116836 | 4.194141363 | -0.228975697 | 1.105591675 | -0.207106929 | 0.83592635 | NA | Gm49727 | predicted gene, 49727 [Source:MGI Symb | 2.4093 | 8.2233 | 3.0187 | 3.8559 | 6.5437 | 1.1138 |
| 54045 | ENSMUSG00000116837 | 0 | NA | NA | NA | NA | NA | Vmn2r-ps11 | vomeronasal 2, receptor, pseudogene 11! | 0 | 0 | 0 | 0 | 0 | 0 |
| 54046 | ENSMUSG00000116838 | 0 | NA | NA | NA | NA | NA | Gm6475 | predicted gene 6475 [Source:MGI Symbol | 0 | 0 | 0 | 0 | 0 | 0 |
| 54047 | ENSMUSG00000116839 | 0 | NA | NA | NA | NA | NA | Gm49778 | predicted gene, 49778 [Source:MGI Symb | 0 | 0 | 0 | 0 | 0 | 0 |
| 54048 | ENSMUSG00000116840 | 5.51685215 | -0.513264624 | 0.95842342 | -0.535530136 | 0.592283324 | 0.912923013 | Gm21691 | predicted gene, 21691 [Source:MGI Symb | 11.243 | 2.0558 | 6.0375 | 3.8559 | 5.4531 | 4.4554 |
| 54049 | ENSMUSG00000116841 | 0 | NA | NA | NA | NA | NA | Gm49737 | predicted gene, 49737 [Source:MGI Symb | 0 | 0 | 0 | 0 | 0 | 0 |
| 54050 | ENSMUSG00000116842 | 0.838537155 | -3.104405377 | 3.144633814 | -0.987207275 | 0.323541036 | NA | Gm49588 | predicted gene, 49588 [Source:MGI Symb | 0 | 0 | 5.0312 | 0 | 0 | 0 |
| 54051 | ENSMUSG00000116843 | 0 | NA | NA | NA | NA | NA | Gm49750 | predicted gene, 49750 [Source:MGI Symb | 0 | 0 | 0 | 0 | 0 | 0 |
| 54052 | ENSMUSG00000116845 | 0 | NA | NA | NA | NA | NA | Gm46546 | predicted gene, 46546 [Source:MGI Symb | 0 | 0 | 0 | 0 | 0 | 0 |
| 54053 | ENSMUSG00000116846 | 48.6598618 | -1.311406524 | 0.358356352 | -3.659504061 | 0.000252704 | 0.007567512 | 1810044K17 | RIKEN cDNA 1810044K17 gene [Source:M | 76.295 | 63.731 | 68.425 | 40.487 | 19.631 | 23.391 |
| 54054 | ENSMUSG00000116847 | 0 | NA | NA | NA | NA | NA | Gm30251 | predicted gene, 30251 [Source:MGI Symb | 0 | 0 | 0 | 0 | 0 | 0 |
| 54055 | ENSMUSG00000116848 | 0.167707431 | -0.796629176 | 4.080472857 | -0.195229623 | 0.84521319 | NA | 4930542D17 | RIKEN cDNA 4930542D17 gene [Source:M | 0 | 0 | 1.0062 | 0 | 0 | 0 |
| 54056 | ENSMUSG00000116849 | 0 | NA | NA | NA | NA | NA | Gm49710 | predicted gene, 49710 [Source:MGI Symb | 0 | 0 | 0 | 0 | 0 | 0 |
| 54057 | ENSMUSG00000116850 | 0 | NA | NA | NA | NA | NA | Gm49742 | predicted gene, 49742 [Source:MGI Symb | 0 | 0 | 0 | 0 | 0 | 0 |
| 54058 | ENSMUSG00000116851 | 0 | NA | NA | NA | NA | NA | Gm35576 | predicted gene, 35576 [Source:MGI Symb | 0 | 0 | 0 | 0 | 0 | 0 |
| 54059 | ENSMUSG00000116852 | 2.615932559 | -0.946832249 | 1.336922884 | -0.708217549 | 0.47881017 | NA | 1700102H20 | RIKEN cDNA 1700102H20 gene [Source:M | 3.2124 | 2.0558 | 5.0312 | 0.964 | 1.0906 | 3.3415 |
| 54060 | ENSMUSG00000116853 | 0 | NA | NA | NA | NA | NA | Gm49609 | predicted gene, 49609 [Source:MGI Symb | 0 | 0 | 0 | 0 | 0 | 0 |
| 54061 | ENSMUSG00000116854 | 0 | NA | NA | NA | NA | NA | Gm49567 | predicted gene, 49567 [Source:MGI Symb | 0 | 0 | 0 | 0 | 0 | 0 |
| 54062 | ENSMUSG00000116855 | 0 | NA | NA | NA | NA | NA | Gm49608 | predicted gene, 49608 [Source:MGI Symb | 0 | 0 | 0 | 0 | 0 | 0 |
| 54063 | ENSMUSG00000116857 | 0 | NA | NA | NA | NA | NA | Gm49582 | predicted gene, 49582 [Source:MGI Symb | 0 | 0 | 0 | 0 | 0 | 0 |
| 54064 | ENSMUSG00000116858 | 37.6699438 | -0.099320929 | 0.414920276 | -0.239373525 | 0.810815958 | 0.966452061 | Gm49797 | predicted gene, 49797 [Source:MGI Symb | 51.399 | 40.089 | 25.156 | 34.703 | 50.168 | 24.505 |
| 54065 | ENSMUSG00000116859 | 0 | NA | NA | NA | NA | NA | Gm49651 | predicted gene, 49651 [Source:MGI Symb | 0 | 0 | 0 | 0 | 0 | 0 |
| 54066 | ENSMUSG00000116860 | 0 | NA | NA | NA | NA | NA | Gm46593 | predicted gene, 46593 [Source:MGI Symb | 0 | 0 | 0 | 0 | 0 | 0 |
| 54067 | ENSMUSG00000116861 | 0 | NA | NA | NA | NA | NA | Gm8253 | predicted gene 8253 [Source:MGI Symbol | 0 | 0 | 0 | 0 | 0 | 0 |
| 54068 | ENSMUSG00000116862 | 1.160043064 | -0.43591056 | 2.278443107 | -0.191319484 | 0.848275297 | NA | Gm49607 | predicted gene, 49607 [Source:MGI Symb | 0 | 2.0558 | 2.0125 | 2.8919 | 0 | 0 |
| 54069 | ENSMUSG00000116863 | 0.181769715 | 1.126924175 | 4.080472857 | 0.276174898 | 0.782413736 | NA | Gm49592 | predicted gene, 49592 [Source:MGI Symb | 0 | 0 | 0 | 0 | 1.0906 | 0 |
| 54070 | ENSMUSG00000116864 | 0 | NA | NA | NA | NA | NA | Gm3320 | predicted pseudogene 3320 [Source:MGI | 0 | 0 | 0 | 0 | 0 | 0 |
| 54071 | ENSMUSG00000116865 | 0 | NA | NA | NA | NA | NA | Gm49560 | predicted gene, 49560 [Source:MGI Symb | 0 | 0 | 0 | 0 | 0 | 0 |
| 54072 | ENSMUSG00000116866 | 0 | NA | NA | NA | NA | NA | Gm8938 | predicted gene 8938 [Source:MGI Symbol | 0 | 0 | 0 | 0 | 0 | 0 |
| 54073 | ENSMUSG00000116867 | 0 | NA | NA | NA | NA | NA | Gm49677 | predicted gene, 49677 [Source:MGI Symb | 0 | 0 | 0 | 0 | 0 | 0 |
| 54074 | ENSMUSG00000116868 | 0 | NA | NA | NA | NA | NA | Gm29721 | predicted gene, 29721 [Source:MGI Symb | 0 | 0 | 0 | 0 | 0 | 0 |
| 54075 | ENSMUSG00000116870 | 0 | NA | NA | NA | NA | NA | Gm49696 | predicted gene, 49696 [Source:MGI Symb | 0 | 0 | 0 | 0 | 0 | 0 |
| 54076 | ENSMUSG00000116871 | 0 | NA | NA | NA | NA | NA | Gm33912 | predicted gene, 33912 [Source:MGI Symb | 0 | 0 | 0 | 0 | 0 | 0 |
| 54077 | ENSMUSG00000116872 | 0 | NA | NA | NA | NA | NA | Gm6815 | predicted gene 6815 [Source:MGI Symbol | 0 | 0 | 0 | 0 | 0 | 0 |
| 54078 | ENSMUSG00000116873 | 0 | NA | NA | NA | NA | NA | Gm49577 | predicted gene, 49577 [Source:MGI Symb | 0 | 0 | 0 | 0 | 0 | 0 |
| 54079 | ENSMUSG00000116874 | 0.160662974 | 1.126924175 | 4.080472857 | 0.276174898 | 0.782413736 | NA | Gm34035 | predicted gene, 34035 [Source:MGI Symb | 0 | 0 | 0 | 0.964 | 0 | 0 |
| 54080 | ENSMUSG00000116875 | 7.051325058 | -0.147611331 | 0.828192928 | -0.178233025 | 0.858539975 | 0.975439379 | Morf4l1-ps1 | mortality factor 4 like 1, pseudogene 1 [Sc | 8.031 | 3.0838 | 11.069 | 5.7839 | 6.5437 | 7.7969 |
| 54081 | ENSMUSG00000116876 | 0 | NA | NA | NA | NA | NA | Gm49721 | predicted gene, 49721 [Source:MGI Symb | 0 | 0 | 0 | 0 | 0 | 0 |
| 54082 | ENSMUSG00000116877 | 0 | NA | NA | NA | NA | NA | 4921517D16 | RIKEN cDNA 4921517D16 gene [Source:M | 0 | 0 | 0 | 0 | 0 | 0 |
| 54083 | ENSMUSG00000116878 | 0 | NA | NA | NA | NA | NA | Gm29880 | predicted gene, 29880 [Source:MGI Symb | 0 | 0 | 0 | 0 | 0 | 0 |
| 54084 | ENSMUSG00000116879 | 0 | NA | NA | NA | NA | NA | Gm49794 | predicted gene, 49794 [Source:MGI Symb | 0 | 0 | 0 | 0 | 0 | 0 |
| 54085 | ENSMUSG00000116880 | 0 | NA | NA | NA | NA | NA | Gm49634 | predicted gene, 49634 [Source:MGI Symb | 0 | 0 | 0 | 0 | 0 | 0 |
| 54086 | ENSMUSG00000116881 | 0 | NA | NA | NA | NA | NA | Gm49674 | predicted gene, 49674 [Source:MGI Symb | 0 | 0 | 0 | 0 | 0 | 0 |
| 54087 | ENSMUSG00000116882 | 0.181769715 | 1.126924175 | 4.080472857 | 0.276174898 | 0.782413736 | NA | Gm49614 | predicted gene, 49614 [Source:MGI Symb | 0 | 0 | 0 | 0 | 1.0906 | 0 |
| 54088 | ENSMUSG00000116883 | 3.381933148 | 0.99204384 | 1.2393657 | 0.800444808 | 0.423453129 | NA | Gm49700 | predicted gene, 49700 [Source:MGI Symb | 0.8031 | 2.0558 | 4.025 | 6.7478 | 1.0906 | 5.5692 |
| 54089 | ENSMUSG00000116884 | 0 | NA | NA | NA | NA | NA | C130040N14 | RIKEN cDNA C130040N14 gene [Source:M | 0 | 0 | 0 | 0 | 0 | 0 |
| 54090 | ENSMUSG00000116885 | 0 | NA | NA | NA | NA | NA | 4930420G21 | RIKEN cDNA 4930420G21 gene [Source:M | 0 | 0 | 0 | 0 | 0 | 0 |
| 54091 | ENSMUSG00000116886 | 0 | NA | NA | NA | NA | NA | Gm49589 | predicted gene, 49589 [Source:MGI Symb | 0 | 0 | 0 | 0 | 0 | 0 |
| 54092 | ENSMUSG00000116887 | 0 | NA | NA | NA | NA | NA | Gm6029 | predicted gene 6029 [Source:MGI Symbol | 0 | 0 | 0 | 0 | 0 | 0 |

| | | | | | | | | | | | | | | | |
|---|---|---|---|---|---|---|---|---|---|---|---|---|---|---|---|
| 54093 | ENSMUSG00000116888 | 0 | NA | NA | NA | NA | NA | 7120432I05F | RIKEN cDNA 7120432I05 gene [Source:MG | 0 | 0 | 0 | 0 | 0 | 0 |
| 54094 | ENSMUSG00000116889 | 0 | NA | NA | NA | NA | NA | Gm49771 | predicted gene, 49771 [Source:MGI Symb | 0 | 0 | 0 | 0 | 0 | 0 |
| 54095 | ENSMUSG00000116890 | 0 | NA | NA | NA | NA | NA | Gm49617 | predicted gene, 49617 [Source:MGI Symb | 0 | 0 | 0 | 0 | 0 | 0 |
| 54096 | ENSMUSG00000116891 | 0 | NA | NA | NA | NA | NA | Gm32679 | predicted gene, 32679 [Source:MGI Symb | 0 | 0 | 0 | 0 | 0 | 0 |
| 54097 | ENSMUSG00000116892 | 0 | NA | NA | NA | NA | NA | Gm49787 | predicted gene, 49787 [Source:MGI Symb | 0 | 0 | 0 | 0 | 0 | 0 |
| 54098 | ENSMUSG00000116893 | 0 | NA | NA | NA | NA | NA | Gm49717 | predicted gene, 49717 [Source:MGI Symb | 0 | 0 | 0 | 0 | 0 | 0 |
| 54099 | ENSMUSG00000116894 | 0 | NA | NA | NA | NA | NA | 4930547E14 | RIKEN cDNA 4930547E14 gene [Source:M | 0 | 0 | 0 | 0 | 0 | 0 |
| 54100 | ENSMUSG00000116895 | 13.89166596 | -0.539265962 | 0.615227247 | -0.876531337 | 0.380741248 | 0.812628154 | Gm3435 | predicted gene 3435 [Source:MGI Symbol | 13.653 | 24.67 | 11.069 | 9.6398 | 8.7249 | 15.594 |
| 54101 | ENSMUSG00000116896 | 0 | NA | NA | NA | NA | NA | Gm49792 | predicted gene, 49792 [Source:MGI Symb | 0 | 0 | 0 | 0 | 0 | 0 |
| 54102 | ENSMUSG00000116897 | 0 | NA | NA | NA | NA | NA | Gm49647 | predicted gene, 49647 [Source:MGI Symb | 0 | 0 | 0 | 0 | 0 | 0 |
| 54103 | ENSMUSG00000116898 | 11.81918957 | 0.036242776 | 0.614204456 | 0.059007674 | 0.952945996 | 0.9915542 | Gm49785 | predicted gene, 49785 [Source:MGI Symb | 9.6372 | 13.363 | 12.075 | 11.568 | 10.906 | 13.366 |
| 54104 | ENSMUSG00000116899 | 0 | NA | NA | NA | NA | NA | Gm49777 | predicted gene, 49777 [Source:MGI Symb | 0 | 0 | 0 | 0 | 0 | 0 |
| 54105 | ENSMUSG00000116900 | 0 | NA | NA | NA | NA | NA | Gm8387 | predicted gene 8387 [Source:MGI Symbol | 0 | 0 | 0 | 0 | 0 | 0 |
| 54106 | ENSMUSG00000116901 | 0 | NA | NA | NA | NA | NA | Gm49667 | predicted gene, 49667 [Source:MGI Symb | 0 | 0 | 0 | 0 | 0 | 0 |
| 54107 | ENSMUSG00000116902 | 0 | NA | NA | NA | NA | NA | Gm49660 | predicted gene, 49660 [Source:MGI Symb | 0 | 0 | 0 | 0 | 0 | 0 |
| 54108 | ENSMUSG00000116903 | 79.47078531 | -1.249789467 | 0.350627269 | -3.564438871 | 0.000364635 | 0.010067457 | Gm19522 | predicted gene, 19522 [Source:MGI Symb | 82.72 | 133.63 | 119.74 | 53.019 | 65.437 | 22.277 |
| 54109 | ENSMUSG00000116904 | 0 | NA | NA | NA | NA | NA | Gm49640 | predicted gene, 49640 [Source:MGI Symb | 0 | 0 | 0 | 0 | 0 | 0 |
| 54110 | ENSMUSG00000116905 | 0 | NA | NA | NA | NA | NA | Gm49652 | predicted gene, 49652 [Source:MGI Symb | 0 | 0 | 0 | 0 | 0 | 0 |
| 54111 | ENSMUSG00000116906 | 0.160662974 | 1.126924175 | 4.080472857 | 0.276174898 | 0.782413736 | NA | Gm49572 | predicted gene, 49572 [Source:MGI Symb | 0 | 0 | 0 | 0.964 | 0 | 0 |
| 54112 | ENSMUSG00000116907 | 0 | NA | NA | NA | NA | NA | 4930553E22 | RIKEN cDNA 4930553E22 gene [Source:M | 0 | 0 | 0 | 0 | 0 | 0 |
| 54113 | ENSMUSG00000116908 | 42.534102 | 0.556134796 | 0.374531763 | 1.484880191 | 0.137575564 | 0.541762309 | Gm49599 | predicted gene, 49599 [Source:MGI Symb | 35.336 | 37.005 | 31.194 | 71.334 | 45.806 | 34.529 |
| 54114 | ENSMUSG00000116909 | 0 | NA | NA | NA | NA | NA | Gm49559 | predicted gene, 49559 [Source:MGI Symb | 0 | 0 | 0 | 0 | 0 | 0 |
| 54115 | ENSMUSG00000116910 | 0 | NA | NA | NA | NA | NA | Gm49670 | predicted gene, 49670 [Source:MGI Symb | 0 | 0 | 0 | 0 | 0 | 0 |
| 54116 | ENSMUSG00000116911 | 0 | NA | NA | NA | NA | NA | Gm5964 | predicted gene 5964 [Source:MGI Symbol | 0 | 0 | 0 | 0 | 0 | 0 |
| 54117 | ENSMUSG00000116912 | 0.181769715 | 1.126924175 | 4.080472857 | 0.276174898 | 0.782413736 | NA | Gm5487 | predicted gene 5487 [Source:MGI Symbol | 0 | 0 | 0 | 0 | 1.0906 | 0 |
| 54118 | ENSMUSG00000116913 | 0 | NA | NA | NA | NA | NA | Gm49566 | predicted gene, 49566 [Source:MGI Symb | 0 | 0 | 0 | 0 | 0 | 0 |
| 54119 | ENSMUSG00000116914 | 0 | NA | NA | NA | NA | NA | Gm20942 | predicted gene, 20942 [Source:MGI Symb | 0 | 0 | 0 | 0 | 0 | 0 |
| 54120 | ENSMUSG00000116915 | 0 | NA | NA | NA | NA | NA | Gm17900 | predicted gene, 17900 [Source:MGI Symb | 0 | 0 | 0 | 0 | 0 | 0 |
| 54121 | ENSMUSG00000116916 | 0 | NA | NA | NA | NA | NA | Gm34967 | predicted gene, 34967 [Source:MGI Symb | 0 | 0 | 0 | 0 | 0 | 0 |
| 54122 | ENSMUSG00000116917 | 0 | NA | NA | NA | NA | NA | Gm49653 | predicted gene, 49653 [Source:MGI Symb | 0 | 0 | 0 | 0 | 0 | 0 |
| 54123 | ENSMUSG00000116918 | 0 | NA | NA | NA | NA | NA | Gm49578 | predicted gene, 49578 [Source:MGI Symb | 0 | 0 | 0 | 0 | 0 | 0 |
| 54124 | ENSMUSG00000116919 | 0 | NA | NA | NA | NA | NA | Gm17809 | predicted gene, 17809 [Source:MGI Symb | 0 | 0 | 0 | 0 | 0 | 0 |
| 54125 | ENSMUSG00000116920 | 0 | NA | NA | NA | NA | NA | Gm49575 | predicted gene, 49575 [Source:MGI Symb | 0 | 0 | 0 | 0 | 0 | 0 |
| 54126 | ENSMUSG00000116921 | 0 | NA | NA | NA | NA | NA | Gm17879 | predicted gene, 17879 [Source:MGI Symb | 0 | 0 | 0 | 0 | 0 | 0 |
| 54127 | ENSMUSG00000116922 | 0 | NA | NA | NA | NA | NA | 4833419O12 | RIKEN cDNA 4833419O12 gene [Source:M | 0 | 0 | 0 | 0 | 0 | 0 |
| 54128 | ENSMUSG00000116923 | 0 | NA | NA | NA | NA | NA | Gm7559 | predicted gene 7559 [Source:MGI Symbol | 0 | 0 | 0 | 0 | 0 | 0 |
| 54129 | ENSMUSG00000116924 | 0 | NA | NA | NA | NA | NA | 4933401D09 | RIKEN cDNA 4933401D09 gene [Source:M | 0 | 0 | 0 | 0 | 0 | 0 |
| 54130 | ENSMUSG00000116925 | 0 | NA | NA | NA | NA | NA | Gm49776 | predicted gene, 49776 [Source:MGI Symb | 0 | 0 | 0 | 0 | 0 | 0 |
| 54131 | ENSMUSG00000116926 | 0 | NA | NA | NA | NA | NA | Gm18234 | predicted gene, 18234 [Source:MGI Symb | 0 | 0 | 0 | 0 | 0 | 0 |
| 54132 | ENSMUSG00000116927 | 5.996682994 | -0.576089711 | 0.979692906 | -0.58803091 | 0.556511546 | 0.898080805 | Gm30881 | predicted gene, 30881 [Source:MGI Symb | 2.4093 | 8.2233 | 11.069 | 7.7118 | 5.4531 | 1.1138 |
| 54133 | ENSMUSG00000116928 | 0 | NA | NA | NA | NA | NA | Gm18169 | predicted gene, 18169 [Source:MGI Symb | 0 | 0 | 0 | 0 | 0 | 0 |
| 54134 | ENSMUSG00000116929 | 0 | NA | NA | NA | NA | NA | 2210009P08 | RIKEN cDNA 2210009P08 gene [Source:M | 0 | 0 | 0 | 0 | 0 | 0 |
| 54135 | ENSMUSG00000116930 | 0 | NA | NA | NA | NA | NA | Gm49711 | predicted gene, 49711 [Source:MGI Symb | 0 | 0 | 0 | 0 | 0 | 0 |
| 54136 | ENSMUSG00000116931 | 0 | NA | NA | NA | NA | NA | Gm18496 | predicted gene, 18496 [Source:MGI Symb | 0 | 0 | 0 | 0 | 0 | 0 |
| 54137 | ENSMUSG00000116932 | 0 | NA | NA | NA | NA | NA | Fpr-rs5 | formyl peptide receptor, related sequence | 0 | 0 | 0 | 0 | 0 | 0 |
| 54138 | ENSMUSG00000116933 | 0 | NA | NA | NA | NA | NA | Atp5o | ATP synthase, H+ transporting, mitochond | 0 | 0 | 0 | 0 | 0 | 0 |
| 54139 | ENSMUSG00000116934 | 0 | NA | NA | NA | NA | NA | Gm3292 | predicted gene 3292 [Source:MGI Symbol | 0 | 0 | 0 | 0 | 0 | 0 |
| 54140 | ENSMUSG00000116935 | 0 | NA | NA | NA | NA | NA | Gpr31a | G protein-coupled receptor 31, D17Leh66 | 0 | 0 | 0 | 0 | 0 | 0 |
| 54141 | ENSMUSG00000116936 | 0 | NA | NA | NA | NA | NA | Bod1-ps | biorientation of chromosomes in cell divis | 0 | 0 | 0 | 0 | 0 | 0 |
| 54142 | ENSMUSG00000116937 | 0 | NA | NA | NA | NA | NA | Gm41480 | predicted gene, 41480 [Source:MGI Symb | 0 | 0 | 0 | 0 | 0 | 0 |
| 54143 | ENSMUSG00000116938 | 8.159499613 | 0.689755961 | 0.757780404 | 0.910231984 | 0.36270018 | 0.801664945 | Gm49645 | predicted gene, 49645 [Source:MGI Symb | 5.6217 | 4.1117 | 9.0562 | 12.532 | 8.7249 | 8.9108 |
| 54144 | ENSMUSG00000116939 | 0 | NA | NA | NA | NA | NA | Gm49619 | predicted gene, 49619 [Source:MGI Symb | 0 | 0 | 0 | 0 | 0 | 0 |
| 54145 | ENSMUSG00000116940 | 0 | NA | NA | NA | NA | NA | Gm7450 | predicted gene 7450 [Source:MGI Symbol | 0 | 0 | 0 | 0 | 0 | 0 |
| 54146 | ENSMUSG00000116941 | 0.856727441 | -0.738450537 | 2.821005776 | -0.26176853 | 0.793499902 | NA | Gm46578 | predicted gene, 46578 [Source:MGI Symb | 3.2124 | 0 | 0 | 1.928 | 0 | 0 |
| 54147 | ENSMUSG00000116942 | 0.572870759 | -2.577107685 | 3.182795569 | -0.809699407 | 0.418112959 | NA | Gm49744 | predicted gene, 49744 [Source:MGI Symb | 2.4093 | 1.0279 | 0 | 0 | 0 | 0 |
| 54148 | ENSMUSG00000116943 | 0 | NA | NA | NA | NA | NA | Gm18237 | predicted gene, 18237 [Source:MGI Symb | 0 | 0 | 0 | 0 | 0 | 0 |

| | | | | | | | | | | | | | | | |
|---|---|---|---|---|---|---|---|---|---|---|---|---|---|---|---|
| 54149 | ENSMUSG00000116944 | 0 | NA | NA | NA | NA | NA | Gm7477 | predicted gene 7477 [Source:MGI Symbol | 0 | 0 | 0 | 0 | 0 | 0 |
| 54150 | ENSMUSG00000116945 | 0 | NA | NA | NA | NA | NA | Gm49663 | predicted gene, 49663 [Source:MGI Symb | 0 | 0 | 0 | 0 | 0 | 0 |
| 54151 | ENSMUSG00000116946 | 5.85549587 | 0.560431129 | 0.962278497 | 0.582400137 | 0.560297189 | 0.899475387 | Gm41442 | predicted gene, 41442 [Source:MGI Symb | 4.0155 | 8.2233 | 2.0125 | 8.6758 | 2.1812 | 10.025 |
| 54152 | ENSMUSG00000116947 | 0 | NA | NA | NA | NA | NA | 2410003I16R | RIKEN cDNA 2410003I16 gene [Source:MG | 0 | 0 | 0 | 0 | 0 | 0 |
| 54153 | ENSMUSG00000116948 | 0 | NA | NA | NA | NA | NA | Gm4802 | predicted gene 4802 [Source:MGI Symbol | 0 | 0 | 0 | 0 | 0 | 0 |
| 54154 | ENSMUSG00000116949 | 0 | NA | NA | NA | NA | NA | Gm7177 | predicted gene 7177 [Source:MGI Symbol | 0 | 0 | 0 | 0 | 0 | 0 |
| 54155 | ENSMUSG00000116950 | 1.878190405 | -0.145453173 | 1.826145305 | -0.079650383 | 0.936515322 | NA | Gm18564 | predicted gene, 18564 [Source:MGI Symb | 0.8031 | 5.1396 | 0 | 0.964 | 4.3625 | 0 |
| 54156 | ENSMUSG00000116951 | 0.167707431 | -0.796629176 | 4.080472857 | -0.195229623 | 0.84521319 | NA | Gm49756 | predicted gene, 49756 [Source:MGI Symb | 0 | 0 | 1.0062 | 0 | 0 | 0 |
| 54157 | ENSMUSG00000116952 | 0 | NA | NA | NA | NA | NA | Gm6914 | predicted gene 6914 [Source:MGI Symbol | 0 | 0 | 0 | 0 | 0 | 0 |
| 54158 | ENSMUSG00000116953 | 25.06136621 | 0.147826546 | 0.469807866 | 0.314653195 | 0.753024986 | 0.952989051 | 9030025P20 | RIKEN cDNA 9030025P20 gene [Source:M | 16.865 | 22.614 | 32.2 | 35.667 | 19.631 | 23.391 |
| 54159 | ENSMUSG00000116954 | 0 | NA | NA | NA | NA | NA | Gm5676 | predicted gene 5676 [Source:MGI Symbol | 0 | 0 | 0 | 0 | 0 | 0 |
| 54160 | ENSMUSG00000116955 | 0 | NA | NA | NA | NA | NA | Gm49632 | predicted gene, 49632 [Source:MGI Symb | 0 | 0 | 0 | 0 | 0 | 0 |
| 54161 | ENSMUSG00000116956 | 0 | NA | NA | NA | NA | NA | Gm49610 | predicted gene, 49610 [Source:MGI Symb | 0 | 0 | 0 | 0 | 0 | 0 |
| 54162 | ENSMUSG00000116957 | 1.001209504 | 0.17748899 | 2.219748584 | 0.079959051 | 0.936269824 | NA | Gm34680 | predicted gene, 34680 [Source:MGI Symb | 0.8031 | 0 | 2.0125 | 0.964 | 0 | 2.2277 |
| 54163 | ENSMUSG00000116958 | 0 | NA | NA | NA | NA | NA | Gm6705 | predicted gene 6705 [Source:MGI Symbol | 0 | 0 | 0 | 0 | 0 | 0 |
| 54164 | ENSMUSG00000116959 | 0 | NA | NA | NA | NA | NA | Gm49604 | predicted gene, 49604 [Source:MGI Symb | 0 | 0 | 0 | 0 | 0 | 0 |
| 54165 | ENSMUSG00000116960 | 0 | NA | NA | NA | NA | NA | Gm49679 | predicted gene, 49679 [Source:MGI Symb | 0 | 0 | 0 | 0 | 0 | 0 |
| 54166 | ENSMUSG00000116961 | 0.48903338 | 0.95637552 | 3.456690196 | 0.276673773 | 0.782030612 | NA | Gm49662 | predicted gene, 49662 [Source:MGI Symb | 0 | 0 | 1.0062 | 1.928 | 0 | 0 |
| 54167 | ENSMUSG00000116962 | 0.160662974 | 1.126924175 | 4.080472857 | 0.276174898 | 0.782413736 | NA | Gm19000 | predicted gene, 19000 [Source:MGI Symb | 0 | 0 | 0 | 0.964 | 0 | 0 |
| 54168 | ENSMUSG00000116963 | 0 | NA | NA | NA | NA | NA | Gm5966 | predicted gene 5966 [Source:MGI Symbol | 0 | 0 | 0 | 0 | 0 | 0 |
| 54169 | ENSMUSG00000116964 | 0.481988923 | 2.492401104 | 3.837077151 | 0.649557203 | 0.515978285 | NA | Gm46555 | predicted gene, 46555 [Source:MGI Symb | 0 | 0 | 0 | 2.8919 | 0 | 0 |
| 54170 | ENSMUSG00000116965 | 0 | NA | NA | NA | NA | NA | Gm36742 | predicted gene, 36742 [Source:MGI Symb | 0 | 0 | 0 | 0 | 0 | 0 |
| 54171 | ENSMUSG00000116966 | 0 | NA | NA | NA | NA | NA | Gm18550 | predicted gene, 18550 [Source:MGI Symb | 0 | 0 | 0 | 0 | 0 | 0 |
| 54172 | ENSMUSG00000116967 | 0 | NA | NA | NA | NA | NA | Smim34 | small integral membrane protein 34 [Sour | 0 | 0 | 0 | 0 | 0 | 0 |
| 54173 | ENSMUSG00000116968 | 0 | NA | NA | NA | NA | NA | Gm49587 | predicted gene, 49587 [Source:MGI Symb | 0 | 0 | 0 | 0 | 0 | 0 |
| 54174 | ENSMUSG00000116969 | 0 | NA | NA | NA | NA | NA | Gm19771 | predicted gene, 19771 [Source:MGI Symb | 0 | 0 | 0 | 0 | 0 | 0 |
| 54175 | ENSMUSG00000116970 | 0 | NA | NA | NA | NA | NA | Gm49796 | predicted gene, 49796 [Source:MGI Symb | 0 | 0 | 0 | 0 | 0 | 0 |
| 54176 | ENSMUSG00000116971 | 0 | NA | NA | NA | NA | NA | Gm49766 | predicted gene, 49766 [Source:MGI Symb | 0 | 0 | 0 | 0 | 0 | 0 |
| 54177 | ENSMUSG00000116972 | 0 | NA | NA | NA | NA | NA | Gm6278 | predicted gene 6278 [Source:MGI Symbol | 0 | 0 | 0 | 0 | 0 | 0 |
| 54178 | ENSMUSG00000116973 | 0 | NA | NA | NA | NA | NA | AV205837 | expressed sequence AV205837 [Source:M | 0 | 0 | 0 | 0 | 0 | 0 |
| 54179 | ENSMUSG00000116974 | 2.023834424 | -2.29253971 | 1.782840093 | -1.285891943 | 0.198480776 | NA | Gm49761 | predicted gene, 49761 [Source:MGI Symb | 8.031 | 1.0279 | 1.0062 | 0.964 | 0 | 1.1138 |
| 54180 | ENSMUSG00000116975 | 0 | NA | NA | NA | NA | NA | 4930470H14 | RIKEN cDNA 4930470H14 gene [Source:M | 0 | 0 | 0 | 0 | 0 | 0 |
| 54181 | ENSMUSG00000116976 | 0 | NA | NA | NA | NA | NA | 4930500H12 | RIKEN cDNA 4930500H12 gene [Source:M | 0 | 0 | 0 | 0 | 0 | 0 |
| 54182 | ENSMUSG00000116977 | 0 | NA | NA | NA | NA | NA | Gm49637 | predicted gene, 49637 [Source:MGI Symb | 0 | 0 | 0 | 0 | 0 | 0 |
| 54183 | ENSMUSG00000116978 | 0 | NA | NA | NA | NA | NA | Gm49765 | predicted gene, 49765 [Source:MGI Symb | 0 | 0 | 0 | 0 | 0 | 0 |
| 54184 | ENSMUSG00000116979 | 0 | NA | NA | NA | NA | NA | Gm32865 | predicted gene, 32865 [Source:MGI Symb | 0 | 0 | 0 | 0 | 0 | 0 |
| 54185 | ENSMUSG00000116980 | 0 | NA | NA | NA | NA | NA | Gm49597 | predicted gene, 49597 [Source:MGI Symb | 0 | 0 | 0 | 0 | 0 | 0 |
| 54186 | ENSMUSG00000116981 | 0 | NA | NA | NA | NA | NA | Gm49611 | predicted gene, 49611 [Source:MGI Symb | 0 | 0 | 0 | 0 | 0 | 0 |
| 54187 | ENSMUSG00000116982 | 0 | NA | NA | NA | NA | NA | 4930567J20R | RIKEN cDNA 4930567J20 gene [Source:MG | 0 | 0 | 0 | 0 | 0 | 0 |
| 54188 | ENSMUSG00000116983 | 0 | NA | NA | NA | NA | NA | 4930553J12R | RIKEN cDNA 4930553J12 gene [Source:MG | 0 | 0 | 0 | 0 | 0 | 0 |
| 54189 | ENSMUSG00000116984 | 0 | NA | NA | NA | NA | NA | 4930549P19 | RIKEN cDNA 4930549P19 gene [Source:M | 0 | 0 | 0 | 0 | 0 | 0 |
| 54190 | ENSMUSG00000116985 | 0 | NA | NA | NA | NA | NA | Gm49669 | predicted gene, 49669 [Source:MGI Symb | 0 | 0 | 0 | 0 | 0 | 0 |
| 54191 | ENSMUSG00000116986 | 0 | NA | NA | NA | NA | NA | Gm17804 | predicted gene, 17804 [Source:MGI Symb | 0 | 0 | 0 | 0 | 0 | 0 |
| 54192 | ENSMUSG00000116987 | 2.230510745 | 1.82308047 | 1.545125885 | 1.179891223 | 0.238043481 | NA | Gm49625 | predicted gene, 49625 [Source:MGI Symb | 0 | 1.0279 | 2.0125 | 4.8199 | 2.1812 | 3.3415 |
| 54193 | ENSMUSG00000116988 | 0 | NA | NA | NA | NA | NA | Gm49673 | predicted gene, 49673 [Source:MGI Symb | 0 | 0 | 0 | 0 | 0 | 0 |
| 54194 | ENSMUSG00000116989 | 0 | NA | NA | NA | NA | NA | Gm49709 | predicted gene, 49709 [Source:MGI Symb | 0 | 0 | 0 | 0 | 0 | 0 |
| 54195 | ENSMUSG00000116990 | 0 | NA | NA | NA | NA | NA | Gm49738 | predicted gene, 49738 [Source:MGI Symb | 0 | 0 | 0 | 0 | 0 | 0 |
| 54196 | ENSMUSG00000116991 | 0 | NA | NA | NA | NA | NA | Gm6032 | predicted gene 6032 [Source:MGI Symbol | 0 | 0 | 0 | 0 | 0 | 0 |
| 54197 | ENSMUSG00000116992 | 0 | NA | NA | NA | NA | NA | Gm29719 | predicted gene, 29719 [Source:MGI Symb | 0 | 0 | 0 | 0 | 0 | 0 |
| 54198 | ENSMUSG00000116993 | 136.1178303 | 0.566901992 | 0.219116904 | 2.587212502 | 0.00967559 | 0.105940753 | Gm32461 | predicted gene, 32461 [Source:MGI Symb | 118.86 | 113.07 | 96.599 | 131.1 | 178.86 | 178.22 |
| 54199 | ENSMUSG00000116994 | 0.48903338 | 0.95637552 | 3.456690196 | 0.276673773 | 0.782030612 | NA | Gm49684 | predicted gene, 49684 [Source:MGI Symb | 0 | 0 | 1.0062 | 1.928 | 0 | 0 |
| 54200 | ENSMUSG00000116995 | 163.1551746 | 0.357570726 | 0.186237677 | 1.919969857 | 0.054861707 | 0.330220445 | Gm21926 | predicted gene, 21926 [Source:MGI Symb | 135.72 | 135.69 | 157.98 | 187.01 | 184.31 | 178.22 |
| 54201 | ENSMUSG00000116996 | 1.616939548 | 4.220104972 | 2.069621202 | 2.039071193 | 0.041442923 | NA | Gm8128 | predicted gene 8128 [Source:MGI Symbol | 0 | 0 | 0 | 1.928 | 1.0906 | 6.6831 |
| 54202 | ENSMUSG00000116997 | 0 | NA | NA | NA | NA | NA | Gm20040 | predicted gene, 20040 [Source:MGI Symb | 0 | 0 | 0 | 0 | 0 | 0 |
| 54203 | ENSMUSG00000116998 | 0 | NA | NA | NA | NA | NA | Gm21050 | predicted gene, 21050 [Source:MGI Symb | 0 | 0 | 0 | 0 | 0 | 0 |
| 54204 | ENSMUSG00000116999 | 0 | NA | NA | NA | NA | NA | Gm18723 | predicted gene, 18723 [Source:MGI Symb | 0 | 0 | 0 | 0 | 0 | 0 |

| | | | | | | | | | | | | | | | |
|---|---|---|---|---|---|---|---|---|---|---|---|---|---|---|---|
| 54205 | ENSMUSG00000117000 | 0 | NA | NA | NA | NA | NA | Gm32509 | predicted gene, 32509 [Source:MGI Symb | 0 | 0 | 0 | 0 | 0 | 0 |
| 54206 | ENSMUSG00000117001 | 0 | NA | NA | NA | NA | NA | Gm49858 | predicted gene, 49858 [Source:MGI Symb | 0 | 0 | 0 | 0 | 0 | 0 |
| 54207 | ENSMUSG00000117002 | 0 | NA | NA | NA | NA | NA | Gm32892 | predicted gene, 32892 [Source:MGI Symb | 0 | 0 | 0 | 0 | 0 | 0 |
| 54208 | ENSMUSG00000117003 | 0 | NA | NA | NA | NA | NA | Gm49937 | predicted gene, 49937 [Source:MGI Symb | 0 | 0 | 0 | 0 | 0 | 0 |
| 54209 | ENSMUSG00000117004 | 0 | NA | NA | NA | NA | NA | Gm38593 | predicted gene, 38593 [Source:MGI Symb | 0 | 0 | 0 | 0 | 0 | 0 |
| 54210 | ENSMUSG00000117005 | 0 | NA | NA | NA | NA | NA | Gm49825 | predicted gene, 49825 [Source:MGI Symb | 0 | 0 | 0 | 0 | 0 | 0 |
| 54211 | ENSMUSG00000117006 | 0 | NA | NA | NA | NA | NA | Gm9224 | predicted gene 9224 [Source:MGI Symbol | 0 | 0 | 0 | 0 | 0 | 0 |
| 54212 | ENSMUSG00000117007 | 0 | NA | NA | NA | NA | NA | Gm41556 | predicted gene, 41556 [Source:MGI Symb | 0 | 0 | 0 | 0 | 0 | 0 |
| 54213 | ENSMUSG00000117008 | 0 | NA | NA | NA | NA | NA | Gm49817 | predicted gene, 49817 [Source:MGI Symb | 0 | 0 | 0 | 0 | 0 | 0 |
| 54214 | ENSMUSG00000117009 | 0 | NA | NA | NA | NA | NA | Gm41597 | predicted gene, 41597 [Source:MGI Symb | 0 | 0 | 0 | 0 | 0 | 0 |
| 54215 | ENSMUSG00000117010 | 0 | NA | NA | NA | NA | NA | Gm49933 | predicted gene, 49933 [Source:MGI Symb | 0 | 0 | 0 | 0 | 0 | 0 |
| 54216 | ENSMUSG00000117011 | 0 | NA | NA | NA | NA | NA | E130008D07 | RIKEN cDNA E130008D07 gene [Source:M | 0 | 0 | 0 | 0 | 0 | 0 |
| 54217 | ENSMUSG00000117012 | 0 | NA | NA | NA | NA | NA | Gm18992 | predicted gene, 18992 [Source:MGI Symb | 0 | 0 | 0 | 0 | 0 | 0 |
| 54218 | ENSMUSG00000117013 | 3.333005541 | -0.199066431 | 1.198935149 | -0.166036029 | 0.868128597 | NA | Gm30531 | predicted gene, 30531 [Source:MGI Symb | 1.6062 | 6.1675 | 3.0187 | 4.8199 | 3.2719 | 1.1138 |
| 54219 | ENSMUSG00000117014 | 0 | NA | NA | NA | NA | NA | Gm9300 | predicted gene 9300 [Source:MGI Symbol | 0 | 0 | 0 | 0 | 0 | 0 |
| 54220 | ENSMUSG00000117015 | 0 | NA | NA | NA | NA | NA | Gm7947 | predicted gene 7947 [Source:MGI Symbol | 0 | 0 | 0 | 0 | 0 | 0 |
| 54221 | ENSMUSG00000117016 | 0 | NA | NA | NA | NA | NA | Gm6174 | predicted gene 6174 [Source:MGI Symbol | 0 | 0 | 0 | 0 | 0 | 0 |
| 54222 | ENSMUSG00000117017 | 0 | NA | NA | NA | NA | NA | Gm49844 | predicted gene, 49844 [Source:MGI Symb | 0 | 0 | 0 | 0 | 0 | 0 |
| 54223 | ENSMUSG00000117018 | 0 | NA | NA | NA | NA | NA | Gm7217 | predicted gene 7217 [Source:MGI Symbol | 0 | 0 | 0 | 0 | 0 | 0 |
| 54224 | ENSMUSG00000117019 | 0 | NA | NA | NA | NA | NA | Gm49896 | predicted gene, 49896 [Source:MGI Symb | 0 | 0 | 0 | 0 | 0 | 0 |
| 54225 | ENSMUSG00000117020 | 0 | NA | NA | NA | NA | NA | Gm49845 | predicted gene, 49845 [Source:MGI Symb | 0 | 0 | 0 | 0 | 0 | 0 |
| 54226 | ENSMUSG00000117021 | 0 | NA | NA | NA | NA | NA | Gm49826 | predicted gene, 49826 [Source:MGI Symb | 0 | 0 | 0 | 0 | 0 | 0 |
| 54227 | ENSMUSG00000117022 | 0 | NA | NA | NA | NA | NA | Gm17920 | predicted gene, 17920 [Source:MGI Symb | 0 | 0 | 0 | 0 | 0 | 0 |
| 54228 | ENSMUSG00000117023 | 0 | NA | NA | NA | NA | NA | Gm49876 | predicted gene, 49876 [Source:MGI Symb | 0 | 0 | 0 | 0 | 0 | 0 |
| 54229 | ENSMUSG00000117024 | 0 | NA | NA | NA | NA | NA | Gm49831 | predicted gene, 49831 [Source:MGI Symb | 0 | 0 | 0 | 0 | 0 | 0 |
| 54230 | ENSMUSG00000117025 | 0 | NA | NA | NA | NA | NA | Gm49950 | predicted gene, 49950 [Source:MGI Symb | 0 | 0 | 0 | 0 | 0 | 0 |
| 54231 | ENSMUSG00000117026 | 0 | NA | NA | NA | NA | NA | C87487 | expressed sequence C87487 [Source:MGI | 0 | 0 | 0 | 0 | 0 | 0 |
| 54232 | ENSMUSG00000117027 | 3.718757086 | 0.976594347 | 1.204642009 | 0.810692588 | 0.417542231 | NA | Gm49886 | predicted gene, 49886 [Source:MGI Symb | 2.4093 | 4.1117 | 1.0062 | 3.8559 | 9.8156 | 1.1138 |
| 54233 | ENSMUSG00000117028 | 0.435408177 | -2.179616019 | 3.531869654 | -0.617128103 | 0.537150234 | NA | Gm49874 | predicted gene, 49874 [Source:MGI Symb | 1.6062 | 0 | 1.0062 | 0 | 0 | 0 |
| 54234 | ENSMUSG00000117029 | 0 | NA | NA | NA | NA | NA | Gm36201 | predicted gene, 36201 [Source:MGI Symb | 0 | 0 | 0 | 0 | 0 | 0 |
| 54235 | ENSMUSG00000117030 | 0 | NA | NA | NA | NA | NA | Gm49959 | predicted gene, 49959 [Source:MGI Symb | 0 | 0 | 0 | 0 | 0 | 0 |
| 54236 | ENSMUSG00000117031 | 0 | NA | NA | NA | NA | NA | Gm49882 | predicted gene, 49882 [Source:MGI Symb | 0 | 0 | 0 | 0 | 0 | 0 |
| 54237 | ENSMUSG00000117033 | 0 | NA | NA | NA | NA | NA | Gm49912 | predicted gene, 49912 [Source:MGI Symb | 0 | 0 | 0 | 0 | 0 | 0 |
| 54238 | ENSMUSG00000117034 | 0 | NA | NA | NA | NA | NA | NA | NA | 0 | 0 | 0 | 0 | 0 | 0 |
| 54239 | ENSMUSG00000117035 | 0 | NA | NA | NA | NA | NA | Gm5094 | predicted gene 5094 [Source:MGI Symbol | 0 | 0 | 0 | 0 | 0 | 0 |
| 54240 | ENSMUSG00000117036 | 0 | NA | NA | NA | NA | NA | Gm33890 | predicted gene, 33890 [Source:MGI Symb | 0 | 0 | 0 | 0 | 0 | 0 |
| 54241 | ENSMUSG00000117037 | 11.28581669 | 0.077433579 | 0.677630317 | 0.114271126 | 0.909022871 | 0.985098284 | Gm50433 | predicted gene, 50433 [Source:MGI Symb | 14.456 | 12.335 | 6.0375 | 9.6398 | 17.45 | 7.7969 |
| 54242 | ENSMUSG00000117038 | 2.486981385 | 0.552866401 | 1.601191165 | 0.345284444 | 0.729880541 | NA | Gm49880 | predicted gene, 49880 [Source:MGI Symb | 0 | 2.0558 | 4.025 | 0 | 3.2719 | 5.5692 |
| 54243 | ENSMUSG00000117039 | 4.042455999 | -1.135047516 | 1.177651944 | -0.963822564 | 0.335134892 | NA | Gm32358 | predicted gene, 32358 [Source:MGI Symb | 6.4248 | 4.1117 | 6.0375 | 0 | 5.4531 | 2.2277 |
| 54244 | ENSMUSG00000117040 | 0 | NA | NA | NA | NA | NA | Gm49805 | predicted gene, 49805 [Source:MGI Symb | 0 | 0 | 0 | 0 | 0 | 0 |
| 54245 | ENSMUSG00000117041 | 0 | NA | NA | NA | NA | NA | Gm46574 | predicted gene, 46574 [Source:MGI Symb | 0 | 0 | 0 | 0 | 0 | 0 |
| 54246 | ENSMUSG00000117042 | 0.824421612 | 3.264491806 | 2.808832384 | 1.162223786 | 0.245144575 | NA | 2700054A10 | RIKEN cDNA 2700054A10 gene [Source:M | 0 | 0 | 0 | 3.8559 | 1.0906 | 0 |
| 54247 | ENSMUSG00000117043 | 0 | NA | NA | NA | NA | NA | Gm49822 | predicted gene, 49822 [Source:MGI Symb | 0 | 0 | 0 | 0 | 0 | 0 |
| 54248 | ENSMUSG00000117044 | 0 | NA | NA | NA | NA | NA | Gm35097 | predicted gene, 35097 [Source:MGI Symb | 0 | 0 | 0 | 0 | 0 | 0 |
| 54249 | ENSMUSG00000117045 | 0 | NA | NA | NA | NA | NA | Gm18365 | predicted gene, 18365 [Source:MGI Symb | 0 | 0 | 0 | 0 | 0 | 0 |
| 54250 | ENSMUSG00000117046 | 0 | NA | NA | NA | NA | NA | Gm19183 | predicted gene, 19183 [Source:MGI Symb | 0 | 0 | 0 | 0 | 0 | 0 |
| 54251 | ENSMUSG00000117047 | 0 | NA | NA | NA | NA | NA | Gm5230 | predicted gene 5230 [Source:MGI Symbol | 0 | 0 | 0 | 0 | 0 | 0 |
| 54252 | ENSMUSG00000117048 | 3.193940125 | 0.586135261 | 1.173847724 | 0.499328191 | 0.617548198 | NA | Gm41584 | predicted gene, 41584 [Source:MGI Symb | 1.6062 | 4.1117 | 2.0125 | 4.8199 | 3.2719 | 3.3415 |
| 54253 | ENSMUSG00000117049 | 0 | NA | NA | NA | NA | NA | Gm6540 | predicted gene 6540 [Source:MGI Symbol | 0 | 0 | 0 | 0 | 0 | 0 |
| 54254 | ENSMUSG00000117050 | 4.88910158 | -4.631522344 | 1.501196172 | -3.085221259 | 0.002034008 | NA | 1700023B13 | RIKEN cDNA 1700023B13 gene [Source:M | 4.8186 | 13.363 | 10.062 | 0 | 1.0906 | 0 |
| 54255 | ENSMUSG00000117051 | 0 | NA | NA | NA | NA | NA | Gm10512 | predicted gene 10512 [Source:MGI Symbo | 0 | 0 | 0 | 0 | 0 | 0 |
| 54256 | ENSMUSG00000117053 | 0 | NA | NA | NA | NA | NA | A730049N16 | RIKEN cDNA A730049N16 gene [Source:M | 0 | 0 | 0 | 0 | 0 | 0 |
| 54257 | ENSMUSG00000117054 | 0 | NA | NA | NA | NA | NA | Gm49869 | predicted gene, 49869 [Source:MGI Symb | 0 | 0 | 0 | 0 | 0 | 0 |
| 54258 | ENSMUSG00000117055 | 0 | NA | NA | NA | NA | NA | Gm9805 | predicted gene 9805 [Source:MGI Symbol | 0 | 0 | 0 | 0 | 0 | 0 |
| 54259 | ENSMUSG00000117056 | 1.202463136 | -2.416971838 | 2.318718553 | -1.042373959 | 0.297238335 | NA | Gm18270 | predicted gene, 18270 [Source:MGI Symb | 0 | 4.1117 | 2.0125 | 0 | 1.0906 | 0 |
| 54260 | ENSMUSG00000117057 | 0 | NA | NA | NA | NA | NA | Gm49846 | predicted gene, 49846 [Source:MGI Symb | 0 | 0 | 0 | 0 | 0 | 0 |

| | | | | | | | | | | | | | | | |
|---|---|---|---|---|---|---|---|---|---|---|---|---|---|---|---|
| 54261 | ENSMUSG00000117058 | 0 | NA | NA | NA | NA | NA | Gm49955 | predicted gene, 49955 [Source:MGI Symb | 0 | 0 | 0 | 0 | 0 | 0 |
| 54262 | ENSMUSG00000117059 | 0 | NA | NA | NA | NA | NA | Gm7912 | predicted gene 7912 [Source:MGI Symbol | 0 | 0 | 0 | 0 | 0 | 0 |
| 54263 | ENSMUSG00000117060 | 0 | NA | NA | NA | NA | NA | Gm7253 | predicted gene 7253 [Source:MGI Symbol | 0 | 0 | 0 | 0 | 0 | 0 |
| 54264 | ENSMUSG00000117061 | 0.401551119 | -2.072982726 | 4.008251979 | -0.517178744 | 0.605031388 | NA | Gm49958 | predicted gene, 49958 [Source:MGI Symb | 2.4093 | 0 | 0 | 0 | 0 | 0 |
| 54265 | ENSMUSG00000117062 | 0 | NA | NA | NA | NA | NA | Ldha-ps3 | lactate dehydrogenase A, pseudogene 3 [! | 0 | 0 | 0 | 0 | 0 | 0 |
| 54266 | ENSMUSG00000117063 | 0 | NA | NA | NA | NA | NA | Gm49936 | predicted gene, 49936 [Source:MGI Symb | 0 | 0 | 0 | 0 | 0 | 0 |
| 54267 | ENSMUSG00000117064 | 0 | NA | NA | NA | NA | NA | Gm49892 | predicted gene, 49892 [Source:MGI Symb | 0 | 0 | 0 | 0 | 0 | 0 |
| 54268 | ENSMUSG00000117065 | 0 | NA | NA | NA | NA | NA | Gm49941 | predicted gene, 49941 [Source:MGI Symb | 0 | 0 | 0 | 0 | 0 | 0 |
| 54269 | ENSMUSG00000117066 | 0 | NA | NA | NA | NA | NA | Gm49860 | predicted gene, 49860 [Source:MGI Symb | 0 | 0 | 0 | 0 | 0 | 0 |
| 54270 | ENSMUSG00000117067 | 0 | NA | NA | NA | NA | NA | Gm49819 | predicted gene, 49819 [Source:MGI Symb | 0 | 0 | 0 | 0 | 0 | 0 |
| 54271 | ENSMUSG00000117068 | 0 | NA | NA | NA | NA | NA | Vmn1r-ps14 | vomeronasal 1 receptor, pseudogene 140 | 0 | 0 | 0 | 0 | 0 | 0 |
| 54272 | ENSMUSG00000117069 | 6.866737881 | 0.130991156 | 0.892342769 | 0.146794663 | 0.883294097 | 0.981769403 | Gm49894 | predicted gene, 49894 [Source:MGI Symb | 6.4248 | 4.1117 | 9.0562 | 3.8559 | 3.2719 | 14.48 |
| 54273 | ENSMUSG00000117070 | 5.801354107 | -0.482398072 | 0.925672909 | -0.521132321 | 0.602274597 | 0.915492852 | Gm41541 | predicted gene, 41541 [Source:MGI Symb | 12.047 | 4.1117 | 4.025 | 5.7839 | 3.2719 | 5.5692 |
| 54274 | ENSMUSG00000117071 | 0 | NA | NA | NA | NA | NA | Gm9598 | predicted gene 9598 [Source:MGI Symbol | 0 | 0 | 0 | 0 | 0 | 0 |
| 54275 | ENSMUSG00000117072 | 0.783546727 | 0.691127367 | 2.662355566 | 0.259592436 | 0.795178172 | NA | Gm49926 | predicted gene, 49926 [Source:MGI Symb | 0.8031 | 0 | 1.0062 | 2.8919 | 0 | 0 |
| 54276 | ENSMUSG00000117073 | 0 | NA | NA | NA | NA | NA | Gm49843 | predicted gene, 49843 [Source:MGI Symb | 0 | 0 | 0 | 0 | 0 | 0 |
| 54277 | ENSMUSG00000117074 | 0 | NA | NA | NA | NA | NA | Gm49925 | predicted gene, 49925 [Source:MGI Symb | 0 | 0 | 0 | 0 | 0 | 0 |
| 54278 | ENSMUSG00000117075 | 0 | NA | NA | NA | NA | NA | Gm49922 | predicted gene, 49922 [Source:MGI Symb | 0 | 0 | 0 | 0 | 0 | 0 |
| 54279 | ENSMUSG00000117076 | 0 | NA | NA | NA | NA | NA | Gm41600 | predicted gene, 41600 [Source:MGI Symb | 0 | 0 | 0 | 0 | 0 | 0 |
| 54280 | ENSMUSG00000117077 | 0 | NA | NA | NA | NA | NA | Gm18735 | predicted gene, 18735 [Source:MGI Symb | 0 | 0 | 0 | 0 | 0 | 0 |
| 54281 | ENSMUSG00000117078 | 0 | NA | NA | NA | NA | NA | Gm4561 | predicted gene 4561 [Source:MGI Symbol | 0 | 0 | 0 | 0 | 0 | 0 |
| 54282 | ENSMUSG00000117079 | 2.710432117 | -2.764192485 | 1.537055857 | -1.798368271 | 0.072118667 | NA | Gm41611 | predicted gene, 41611 [Source:MGI Symb | 4.0155 | 6.1675 | 4.025 | 0.964 | 1.0906 | 0 |
| 54283 | ENSMUSG00000117080 | 0.462220778 | -0.685400455 | 3.191475091 | -0.21475977 | 0.82995462 | NA | 1700063J08I | RIKEN cDNA 1700063J08 gene [Source:M( | 0.8031 | 0 | 1.0062 | 0.964 | 0 | 0 |
| 54284 | ENSMUSG00000117081 | 0 | NA | NA | NA | NA | NA | Gm49948 | predicted gene, 49948 [Source:MGI Symb | 0 | 0 | 0 | 0 | 0 | 0 |
| 54285 | ENSMUSG00000117082 | 0 | NA | NA | NA | NA | NA | Gm18139 | predicted gene, 18139 [Source:MGI Symb | 0 | 0 | 0 | 0 | 0 | 0 |
| 54286 | ENSMUSG00000117083 | 0 | NA | NA | NA | NA | NA | Gm49800 | predicted gene, 49800 [Source:MGI Symb | 0 | 0 | 0 | 0 | 0 | 0 |
| 54287 | ENSMUSG00000117084 | 149.6067916 | -0.986610633 | 0.286736144 | -3.440831066 | 0.00057993 | 0.013773229 | Gm22146 | predicted gene, 22146 [Source:MGI Symb | 201.58 | 209.7 | 186.16 | 159.06 | 66.528 | 74.628 |
| 54288 | ENSMUSG00000117085 | 0.335414862 | -1.779047242 | 4.030804179 | -0.441362855 | 0.658950329 | NA | 4930542M0: | RIKEN cDNA 4930542M03 gene [Source:N | 0 | 0 | 2.0125 | 0 | 0 | 0 |
| 54289 | ENSMUSG00000117086 | 0 | NA | NA | NA | NA | NA | Gm31328 | predicted gene, 31328 [Source:MGI Symb | 0 | 0 | 0 | 0 | 0 | 0 |
| 54290 | ENSMUSG00000117088 | 0 | NA | NA | NA | NA | NA | Gm49875 | predicted gene, 49875 [Source:MGI Symb | 0 | 0 | 0 | 0 | 0 | 0 |
| 54291 | ENSMUSG00000117089 | 0 | NA | NA | NA | NA | NA | Gm49914 | predicted gene, 49914 [Source:MGI Symb | 0 | 0 | 0 | 0 | 0 | 0 |
| 54292 | ENSMUSG00000117090 | 0 | NA | NA | NA | NA | NA | Gm49921 | predicted gene, 49921 [Source:MGI Symb | 0 | 0 | 0 | 0 | 0 | 0 |
| 54293 | ENSMUSG00000117091 | 0 | NA | NA | NA | NA | NA | Gm5698 | predicted gene 5698 [Source:MGI Symbol | 0 | 0 | 0 | 0 | 0 | 0 |
| 54294 | ENSMUSG00000117092 | 0 | NA | NA | NA | NA | NA | Gm49934 | predicted gene, 49934 [Source:MGI Symb | 0 | 0 | 0 | 0 | 0 | 0 |
| 54295 | ENSMUSG00000117093 | 0 | NA | NA | NA | NA | NA | Gm49949 | predicted gene, 49949 [Source:MGI Symb | 0 | 0 | 0 | 0 | 0 | 0 |
| 54296 | ENSMUSG00000117094 | 0 | NA | NA | NA | NA | NA | Gm49956 | predicted gene, 49956 [Source:MGI Symb | 0 | 0 | 0 | 0 | 0 | 0 |
| 54297 | ENSMUSG00000117095 | 0 | NA | NA | NA | NA | NA | Gm49841 | predicted gene, 49841 [Source:MGI Symb | 0 | 0 | 0 | 0 | 0 | 0 |
| 54298 | ENSMUSG00000117096 | 0 | NA | NA | NA | NA | NA | Gm30420 | predicted gene, 30420 [Source:MGI Symb | 0 | 0 | 0 | 0 | 0 | 0 |
| 54299 | ENSMUSG00000117098 | 1.863042735 | 0.466864209 | 1.545291039 | 0.302120569 | 0.762560153 | NA | Gm49909 | predicted gene, 49909 [Source:MGI Symb | 1.6062 | 2.0558 | 1.0062 | 0.964 | 1.0906 | 4.4554 |
| 54300 | ENSMUSG00000117099 | 0 | NA | NA | NA | NA | NA | Gm3547 | predicted gene 3547 [Source:MGI Symbol | 0 | 0 | 0 | 0 | 0 | 0 |
| 54301 | ENSMUSG00000117100 | 0 | NA | NA | NA | NA | NA | A330072L02 | RIKEN cDNA A330072L02 gene [Source:M( | 0 | 0 | 0 | 0 | 0 | 0 |
| 54302 | ENSMUSG00000117101 | 0 | NA | NA | NA | NA | NA | Gm8950 | predicted gene 8950 [Source:MGI Symbol | 0 | 0 | 0 | 0 | 0 | 0 |
| 54303 | ENSMUSG00000117102 | 0.842149364 | -3.107408315 | 2.805524882 | -1.107603192 | 0.268033224 | NA | | Novel transcript | 0 | 1.0279 | 4.025 | 0 | 0 | 0 |
| 54304 | ENSMUSG00000117103 | 0 | NA | NA | NA | NA | NA | Gm49916 | predicted gene, 49916 [Source:MGI Symb | 0 | 0 | 0 | 0 | 0 | 0 |
| 54305 | ENSMUSG00000117104 | 0.167707431 | -0.796629176 | 4.080472857 | -0.195229623 | 0.84521319 | NA | Gm49893 | predicted gene, 49893 [Source:MGI Symb | 0 | 0 | 1.0062 | 0 | 0 | 0 |
| 54306 | ENSMUSG00000117105 | 24.29064882 | -0.074555763 | 0.493271097 | -0.151145615 | 0.87986085 | 0.98128671 | Dreh | down-regulated in hepatocellular carcinoı | 24.093 | 28.782 | 22.137 | 37.595 | 13.087 | 20.049 |
| 54307 | ENSMUSG00000117106 | 0 | NA | NA | NA | NA | NA | Gm49872 | predicted gene, 49872 [Source:MGI Symb | 0 | 0 | 0 | 0 | 0 | 0 |
| 54308 | ENSMUSG00000117107 | 0 | NA | NA | NA | NA | NA | Gm18738 | predicted gene, 18738 [Source:MGI Symb | 0 | 0 | 0 | 0 | 0 | 0 |
| 54309 | ENSMUSG00000117108 | 0 | NA | NA | NA | NA | NA | Gm49935 | predicted gene, 49935 [Source:MGI Symb | 0 | 0 | 0 | 0 | 0 | 0 |
| 54310 | ENSMUSG00000117109 | 0 | NA | NA | NA | NA | NA | Gm19969 | predicted gene, 19969 [Source:MGI Symb | 0 | 0 | 0 | 0 | 0 | 0 |
| 54311 | ENSMUSG00000117110 | 0 | NA | NA | NA | NA | NA | Gm49808 | predicted gene, 49808 [Source:MGI Symb | 0 | 0 | 0 | 0 | 0 | 0 |
| 54312 | ENSMUSG00000117111 | 0 | NA | NA | NA | NA | NA | Tpt1-ps7 | tumor protein, translationally-controlled, | 0 | 0 | 0 | 0 | 0 | 0 |
| 54313 | ENSMUSG00000117112 | 0 | NA | NA | NA | NA | NA | Gm4512 | predicted gene 4512 [Source:MGI Symbol | 0 | 0 | 0 | 0 | 0 | 0 |
| 54314 | ENSMUSG00000117113 | 11.00532065 | 0.330681325 | 0.643967096 | 0.513506554 | 0.607597027 | 0.916376001 | Gm49883 | predicted gene, 49883 [Source:MGI Symb | 12.85 | 7.1954 | 9.0562 | 11.568 | 11.997 | 13.366 |
| 54315 | ENSMUSG00000117114 | 0 | NA | NA | NA | NA | NA | Gm49867 | predicted gene, 49867 [Source:MGI Symb | 0 | 0 | 0 | 0 | 0 | 0 |
| 54316 | ENSMUSG00000117115 | 0 | NA | NA | NA | NA | NA | Gm9586 | predicted gene 9586 [Source:MGI Symbol | 0 | 0 | 0 | 0 | 0 | 0 |

| | | | | | | | | | | | | | | | |
|---|---|---|---|---|---|---|---|---|---|---|---|---|---|---|---|
| 54317 | ENSMUSG00000117116 | 0 | NA | NA | NA | NA | NA | Gm49868 | predicted gene, 49868 [Source:MGI Symb | 0 | 0 | 0 | 0 | 0 | 0 |
| 54318 | ENSMUSG00000117117 | 0 | NA | NA | NA | NA | NA | Gm49901 | predicted gene, 49901 [Source:MGI Symb | 0 | 0 | 0 | 0 | 0 | 0 |
| 54319 | ENSMUSG00000117118 | 0 | NA | NA | NA | NA | NA | Gm49878 | predicted gene, 49878 [Source:MGI Symb | 0 | 0 | 0 | 0 | 0 | 0 |
| 54320 | ENSMUSG00000117119 | 0 | NA | NA | NA | NA | NA | Gm49854 | predicted gene, 49854 [Source:MGI Symb | 0 | 0 | 0 | 0 | 0 | 0 |
| 54321 | ENSMUSG00000117120 | 0 | NA | NA | NA | NA | NA | Gm50238 | predicted gene, 50238 [Source:MGI Symb | 0 | 0 | 0 | 0 | 0 | 0 |
| 54322 | ENSMUSG00000117121 | 0.685278559 | -2.809870862 | 3.376578027 | -0.832165239 | 0.405315682 | NA | Gm49902 | predicted gene, 49902 [Source:MGI Symb | 0 | 4.1117 | 0 | 0 | 0 | 0 |
| 54323 | ENSMUSG00000117122 | 0 | NA | NA | NA | NA | NA | Gm9319 | predicted gene 9319 [Source:MGI Symbol | 0 | 0 | 0 | 0 | 0 | 0 |
| 54324 | ENSMUSG00000117123 | 11.2536201 | 0.476791848 | 0.71629735 | 0.665633968 | 0.505645089 | 0.881592135 | Gm49890 | predicted gene, 49890 [Source:MGI Symb | 16.865 | 3.0838 | 8.05 | 16.388 | 11.997 | 11.138 |
| 54325 | ENSMUSG00000117124 | 0 | NA | NA | NA | NA | NA | Gm49895 | predicted gene, 49895 [Source:MGI Symb | 0 | 0 | 0 | 0 | 0 | 0 |
| 54326 | ENSMUSG00000117125 | 0 | NA | NA | NA | NA | NA | Gm49836 | predicted gene, 49836 [Source:MGI Symb | 0 | 0 | 0 | 0 | 0 | 0 |
| 54327 | ENSMUSG00000117126 | 0 | NA | NA | NA | NA | NA | Gm49924 | predicted gene, 49924 [Source:MGI Symb | 0 | 0 | 0 | 0 | 0 | 0 |
| 54328 | ENSMUSG00000117127 | 0 | NA | NA | NA | NA | NA | Gm49812 | predicted gene, 49812 [Source:MGI Symb | 0 | 0 | 0 | 0 | 0 | 0 |
| 54329 | ENSMUSG00000117128 | 2.518207342 | 0.021038685 | 1.501942675 | 0.014007648 | 0.988823879 | NA | Gm49915 | predicted gene, 49915 [Source:MGI Symb | 2.4093 | 1.0279 | 4.025 | 0.964 | 0 | 6.6831 |
| 54330 | ENSMUSG00000117129 | 0.371281295 | 2.09253362 | 4.031852165 | 0.519000582 | 0.603760335 | NA | Gm49928 | predicted gene, 49928 [Source:MGI Symb | 0 | 0 | 0 | 0 | 0 | 2.2277 |
| 54331 | ENSMUSG00000117130 | 0.133850373 | -0.796629176 | 4.080472857 | -0.195229623 | 0.84521319 | NA | Gm49802 | predicted gene, 49802 [Source:MGI Symb | 0.8031 | 0 | 0 | 0 | 0 | 0 |
| 54332 | ENSMUSG00000117131 | 0.167707431 | -0.796629176 | 4.080472857 | -0.195229623 | 0.84521319 | NA | Gm41506 | predicted gene, 41506 [Source:MGI Symb | 0 | 0 | 1.0062 | 0 | 0 | 0 |
| 54333 | ENSMUSG00000117132 | 0 | NA | NA | NA | NA | NA | Gm32055 | predicted gene, 32055 [Source:MGI Symb | 0 | 0 | 0 | 0 | 0 | 0 |
| 54334 | ENSMUSG00000117133 | 0 | NA | NA | NA | NA | NA | Gm49806 | predicted gene, 49806 [Source:MGI Symb | 0 | 0 | 0 | 0 | 0 | 0 |
| 54335 | ENSMUSG00000117134 | 0 | NA | NA | NA | NA | NA | Gm18521 | predicted gene, 18521 [Source:MGI Symb | 0 | 0 | 0 | 0 | 0 | 0 |
| 54336 | ENSMUSG00000117135 | 0.17131964 | -0.796629176 | 4.080472857 | -0.195229623 | 0.84521319 | NA | Gm49811 | predicted gene, 49811 [Source:MGI Symb | 0 | 1.0279 | 0 | 0 | 0 | 0 |
| 54337 | ENSMUSG00000117136 | 0 | NA | NA | NA | NA | NA | Gm49835 | predicted gene, 49835 [Source:MGI Symb | 0 | 0 | 0 | 0 | 0 | 0 |
| 54338 | ENSMUSG00000117137 | 0 | NA | NA | NA | NA | NA | 4930471L23I | RIKEN cDNA 4930471L23 gene [Source:M( | 0 | 0 | 0 | 0 | 0 | 0 |
| 54339 | ENSMUSG00000117138 | 0 | NA | NA | NA | NA | NA | Gm9191 | predicted gene 9191 [Source:MGI Symbol | 0 | 0 | 0 | 0 | 0 | 0 |
| 54340 | ENSMUSG00000117139 | 0 | NA | NA | NA | NA | NA | Gm35692 | predicted gene, 35692 [Source:MGI Symb | 0 | 0 | 0 | 0 | 0 | 0 |
| 54341 | ENSMUSG00000117140 | 0 | NA | NA | NA | NA | NA | Gm18138 | predicted gene, 18138 [Source:MGI Symb | 0 | 0 | 0 | 0 | 0 | 0 |
| 54342 | ENSMUSG00000117141 | 0 | NA | NA | NA | NA | NA | Gm18214 | predicted gene, 18214 [Source:MGI Symb | 0 | 0 | 0 | 0 | 0 | 0 |
| 54343 | ENSMUSG00000117142 | 0 | NA | NA | NA | NA | NA | Gm46573 | predicted gene, 46573 [Source:MGI Symb | 0 | 0 | 0 | 0 | 0 | 0 |
| 54344 | ENSMUSG00000117143 | 0 | NA | NA | NA | NA | NA | Gm49807 | predicted gene, 49807 [Source:MGI Symb | 0 | 0 | 0 | 0 | 0 | 0 |
| 54345 | ENSMUSG00000117145 | 3.178126064 | 2.541472767 | 1.41124879 | 1.800867986 | 0.07172369 | NA | Gm4356 | predicted gene 4356 [Source:MGI Symbol | 0.8031 | 0 | 2.0125 | 9.6398 | 3.2719 | 3.3415 |
| 54346 | ENSMUSG00000117146 | 0 | NA | NA | NA | NA | NA | Fut4-ps1 | fucosyltransferase 4, pseudogene 1 [Sour | 0 | 0 | 0 | 0 | 0 | 0 |
| 54347 | ENSMUSG00000117147 | 0 | NA | NA | NA | NA | NA | Gm11166 | predicted gene 11166 [Source:MGI Symbo | 0 | 0 | 0 | 0 | 0 | 0 |
| 54348 | ENSMUSG00000117148 | 0 | NA | NA | NA | NA | NA | Vmn1r229 | vomeronasal 1 receptor 229 [Source:MGI | 0 | 0 | 0 | 0 | 0 | 0 |
| 54349 | ENSMUSG00000117149 | 0 | NA | NA | NA | NA | NA | Gm49919 | predicted gene, 49919 [Source:MGI Symb | 0 | 0 | 0 | 0 | 0 | 0 |
| 54350 | ENSMUSG00000117150 | 0 | NA | NA | NA | NA | NA | Gm9351 | predicted gene 9351 [Source:MGI Symbol | 0 | 0 | 0 | 0 | 0 | 0 |
| 54351 | ENSMUSG00000117151 | 0 | NA | NA | NA | NA | NA | Gm49820 | predicted gene, 49820 [Source:MGI Symb | 0 | 0 | 0 | 0 | 0 | 0 |
| 54352 | ENSMUSG00000117154 | 0.160662974 | 1.126924175 | 4.080472857 | 0.276174898 | 0.782413736 | NA | Gm7898 | predicted gene 7898 [Source:MGI Symbol | 0 | 0 | 0 | 0.964 | 0 | 0 |
| 54353 | ENSMUSG00000117156 | 0 | NA | NA | NA | NA | NA | Gm49904 | predicted gene, 49904 [Source:MGI Symb | 0 | 0 | 0 | 0 | 0 | 0 |
| 54354 | ENSMUSG00000117157 | 0 | NA | NA | NA | NA | NA | Gm49946 | predicted gene, 49946 [Source:MGI Symb | 0 | 0 | 0 | 0 | 0 | 0 |
| 54355 | ENSMUSG00000117158 | 0 | NA | NA | NA | NA | NA | Gm49866 | predicted gene, 49866 [Source:MGI Symb | 0 | 0 | 0 | 0 | 0 | 0 |
| 54356 | ENSMUSG00000117159 | 0 | NA | NA | NA | NA | NA | Gm49952 | predicted gene, 49952 [Source:MGI Symb | 0 | 0 | 0 | 0 | 0 | 0 |
| 54357 | ENSMUSG00000117160 | 0 | NA | NA | NA | NA | NA | Gm49859 | predicted gene, 49859 [Source:MGI Symb | 0 | 0 | 0 | 0 | 0 | 0 |
| 54358 | ENSMUSG00000117161 | 0.469265235 | -2.274672822 | 3.439794491 | -0.661281605 | 0.508431736 | NA | Gm49803 | predicted gene, 49803 [Source:MGI Symb | 0.8031 | 0 | 2.0125 | 0 | 0 | 0 |
| 54359 | ENSMUSG00000117162 | 0 | NA | NA | NA | NA | NA | Gm49840 | predicted gene, 49840 [Source:MGI Symb | 0 | 0 | 0 | 0 | 0 | 0 |
| 54360 | ENSMUSG00000117163 | 0 | NA | NA | NA | NA | NA | Gm5978 | predicted gene 5978 [Source:MGI Symbol | 0 | 0 | 0 | 0 | 0 | 0 |
| 54361 | ENSMUSG00000117164 | 0 | NA | NA | NA | NA | NA | Gm49951 | predicted gene, 49951 [Source:MGI Symb | 0 | 0 | 0 | 0 | 0 | 0 |
| 54362 | ENSMUSG00000117165 | 0 | NA | NA | NA | NA | NA | Gm49832 | predicted gene, 49832 [Source:MGI Symb | 0 | 0 | 0 | 0 | 0 | 0 |
| 54363 | ENSMUSG00000117166 | 0 | NA | NA | NA | NA | NA | Gm18961 | predicted gene, 18961 [Source:MGI Symb | 0 | 0 | 0 | 0 | 0 | 0 |
| 54364 | ENSMUSG00000117167 | 0 | NA | NA | NA | NA | NA | Gm6599 | predicted gene 6599 [Source:MGI Symbol | 0 | 0 | 0 | 0 | 0 | 0 |
| 54365 | ENSMUSG00000117168 | 0 | NA | NA | NA | NA | NA | Gm49908 | predicted gene, 49908 [Source:MGI Symb | 0 | 0 | 0 | 0 | 0 | 0 |
| 54366 | ENSMUSG00000117169 | 0 | NA | NA | NA | NA | NA | Vmn1r-ps13 | vomeronasal 1 receptor, pseudogene 139 | 0 | 0 | 0 | 0 | 0 | 0 |
| 54367 | ENSMUSG00000117170 | 0 | NA | NA | NA | NA | NA | Gm19197 | predicted gene, 19197 [Source:MGI Symb | 0 | 0 | 0 | 0 | 0 | 0 |
| 54368 | ENSMUSG00000117171 | 2.168063805 | 1.868054121 | 1.574610023 | 1.186359857 | 0.235480218 | NA | C230013L11 | RIKEN cDNA C230013L11 gene [Source:M( | 0.8031 | 0 | 2.0125 | 5.7839 | 2.1812 | 2.2277 |
| 54369 | ENSMUSG00000117172 | 0 | NA | NA | NA | NA | NA | A230051N06 | RIKEN cDNA A230051N06 gene [Source:M | 0 | 0 | 0 | 0 | 0 | 0 |
| 54370 | ENSMUSG00000117173 | 0 | NA | NA | NA | NA | NA | NA | NA | 0 | 0 | 0 | 0 | 0 | 0 |
| 54371 | ENSMUSG00000117174 | 0 | NA | NA | NA | NA | NA | Gm49939 | predicted gene, 49939 [Source:MGI Symb | 0 | 0 | 0 | 0 | 0 | 0 |
| 54372 | ENSMUSG00000117175 | 0 | NA | NA | NA | NA | NA | Gm20098 | predicted gene, 20098 [Source:MGI Symb | 0 | 0 | 0 | 0 | 0 | 0 |

| | | | | | | | | | | | | | | | |
|---|---|---|---|---|---|---|---|---|---|---|---|---|---|---|---|
| 54373 | ENSMUSG00000117176 | 0 | NA | NA | NA | NA | NA | Gm49879 | predicted gene, 49879 [Source:MGI Symb | 0 | 0 | 0 | 0 | 0 | 0 |
| 54374 | ENSMUSG00000117177 | 0 | NA | NA | NA | NA | NA | Gm49810 | predicted gene, 49810 [Source:MGI Symb | 0 | 0 | 0 | 0 | 0 | 0 |
| 54375 | ENSMUSG00000117178 | 0 | NA | NA | NA | NA | NA | Gm49852 | predicted gene, 49852 [Source:MGI Symb | 0 | 0 | 0 | 0 | 0 | 0 |
| 54376 | ENSMUSG00000117179 | 0 | NA | NA | NA | NA | NA | Gm49907 | predicted gene, 49907 [Source:MGI Symb | 0 | 0 | 0 | 0 | 0 | 0 |
| 54377 | ENSMUSG00000117180 | 0 | NA | NA | NA | NA | NA | Gm19399 | predicted gene, 19399 [Source:MGI Symb | 0 | 0 | 0 | 0 | 0 | 0 |
| 54378 | ENSMUSG00000117181 | 0 | NA | NA | NA | NA | NA | Gm31143 | predicted gene, 31143 [Source:MGI Symb | 0 | 0 | 0 | 0 | 0 | 0 |
| 54379 | ENSMUSG00000117182 | 0.455176322 | 0.955156269 | 3.546674842 | 0.269310357 | 0.787690862 | NA | Gm18068 | predicted gene, 18068 [Source:MGI Symb | 0.8031 | 0 | 0 | 1.928 | 0 | 0 |
| 54380 | ENSMUSG00000117183 | 8.226010487 | -0.774393109 | 0.865907907 | -0.894313475 | 0.371154192 | 0.807214774 | Gm20008 | predicted gene, 20008 [Source:MGI Symb | 8.031 | 8.2233 | 15.094 | 12.532 | 4.3625 | 1.1138 |
| 54381 | ENSMUSG00000117184 | 0 | NA | NA | NA | NA | NA | Gm49851 | predicted gene, 49851 [Source:MGI Symb | 0 | 0 | 0 | 0 | 0 | 0 |
| 54382 | ENSMUSG00000117185 | 0 | NA | NA | NA | NA | NA | Gm34510 | predicted gene, 34510 [Source:MGI Symb | 0 | 0 | 0 | 0 | 0 | 0 |
| 54383 | ENSMUSG00000117186 | 0 | NA | NA | NA | NA | NA | Gm46576 | predicted gene, 46576 [Source:MGI Symb | 0 | 0 | 0 | 0 | 0 | 0 |
| 54384 | ENSMUSG00000117187 | 0.954866367 | 0.067917755 | 2.273052348 | 0.029879539 | 0.976163124 | NA | Gm4708 | predicted gene 4708 [Source:MGI Symbol | 0.8031 | 1.0279 | 1.0062 | 2.8919 | 0 | 0 |
| 54385 | ENSMUSG00000117188 | 0 | NA | NA | NA | NA | NA | Gm49849 | predicted gene, 49849 [Source:MGI Symb | 0 | 0 | 0 | 0 | 0 | 0 |
| 54386 | ENSMUSG00000117189 | 0 | NA | NA | NA | NA | NA | Gm49897 | predicted gene, 49897 [Source:MGI Symb | 0 | 0 | 0 | 0 | 0 | 0 |
| 54387 | ENSMUSG00000117190 | 0 | NA | NA | NA | NA | NA | Esp5 | exocrine gland secreted peptide 5 [Source | 0 | 0 | 0 | 0 | 0 | 0 |
| 54388 | ENSMUSG00000117191 | 0 | NA | NA | NA | NA | NA | Gm49853 | predicted gene, 49853 [Source:MGI Symb | 0 | 0 | 0 | 0 | 0 | 0 |
| 54389 | ENSMUSG00000117192 | 6.708762 | 0.603177919 | 0.80700041 | 0.747431985 | 0.454802844 | 0.857337687 | Gm18649 | predicted gene, 18649 [Source:MGI Symb | 4.8186 | 5.1396 | 6.0375 | 7.7118 | 7.6343 | 8.9108 |
| 54390 | ENSMUSG00000117193 | 0 | NA | NA | NA | NA | NA | Gm36252 | predicted gene, 36252 [Source:MGI Symb | 0 | 0 | 0 | 0 | 0 | 0 |
| 54391 | ENSMUSG00000117194 | 0 | NA | NA | NA | NA | NA | Gm49913 | predicted gene, 49913 [Source:MGI Symb | 0 | 0 | 0 | 0 | 0 | 0 |
| 54392 | ENSMUSG00000117195 | 0 | NA | NA | NA | NA | NA | Gm49881 | predicted gene, 49881 [Source:MGI Symb | 0 | 0 | 0 | 0 | 0 | 0 |
| 54393 | ENSMUSG00000117196 | 0 | NA | NA | NA | NA | NA | Gm49823 | predicted gene, 49823 [Source:MGI Symb | 0 | 0 | 0 | 0 | 0 | 0 |
| 54394 | ENSMUSG00000117197 | 0.17131964 | -0.796629176 | 4.080472857 | -0.195229623 | 0.84521319 | NA | Gm6618 | predicted gene 6618 [Source:MGI Symbol | 0 | 1.0279 | 0 | 0 | 0 | 0 |
| 54395 | ENSMUSG00000117198 | 0 | NA | NA | NA | NA | NA | Gm18067 | predicted gene, 18067 [Source:MGI Symb | 0 | 0 | 0 | 0 | 0 | 0 |
| 54396 | ENSMUSG00000117199 | 0 | NA | NA | NA | NA | NA | Gm4566 | predicted gene 4566 [Source:MGI Symbol | 0 | 0 | 0 | 0 | 0 | 0 |
| 54397 | ENSMUSG00000117200 | 0 | NA | NA | NA | NA | NA | Gm49905 | predicted gene, 49905 [Source:MGI Symb | 0 | 0 | 0 | 0 | 0 | 0 |
| 54398 | ENSMUSG00000117201 | 0 | NA | NA | NA | NA | NA | Gm49834 | predicted gene, 49834 [Source:MGI Symb | 0 | 0 | 0 | 0 | 0 | 0 |
| 54399 | ENSMUSG00000117202 | 0 | NA | NA | NA | NA | NA | Gm9349 | predicted gene 9349 [Source:MGI Symbol | 0 | 0 | 0 | 0 | 0 | 0 |
| 54400 | ENSMUSG00000117203 | 0 | NA | NA | NA | NA | NA | Gm4701 | predicted gene 4701 [Source:MGI Symbol | 0 | 0 | 0 | 0 | 0 | 0 |
| 54401 | ENSMUSG00000117204 | 0 | NA | NA | NA | NA | NA | C330011F03 | RIKEN cDNA C330011F03 gene [Source:M | 0 | 0 | 0 | 0 | 0 | 0 |
| 54402 | ENSMUSG00000117205 | 0 | NA | NA | NA | NA | NA | Rps18-ps7 | ribosomal protein S18, pseudogene 7 [Sou | 0 | 0 | 0 | 0 | 0 | 0 |
| 54403 | ENSMUSG00000117206 | 0 | NA | NA | NA | NA | NA | Gm49809 | predicted gene, 49809 [Source:MGI Symb | 0 | 0 | 0 | 0 | 0 | 0 |
| 54404 | ENSMUSG00000117207 | 0 | NA | NA | NA | NA | NA | Gm18784 | predicted gene, 18784 [Source:MGI Symb | 0 | 0 | 0 | 0 | 0 | 0 |
| 54405 | ENSMUSG00000117208 | 0 | NA | NA | NA | NA | NA | Gm49945 | predicted gene, 49945 [Source:MGI Symb | 0 | 0 | 0 | 0 | 0 | 0 |
| 54406 | ENSMUSG00000117209 | 0 | NA | NA | NA | NA | NA | Gm49954 | predicted gene, 49954 [Source:MGI Symb | 0 | 0 | 0 | 0 | 0 | 0 |
| 54407 | ENSMUSG00000117210 | 0 | NA | NA | NA | NA | NA | Gm49857 | predicted gene, 49857 [Source:MGI Symb | 0 | 0 | 0 | 0 | 0 | 0 |
| 54408 | ENSMUSG00000117211 | 0 | NA | NA | NA | NA | NA | Gm49917 | predicted gene, 49917 [Source:MGI Symb | 0 | 0 | 0 | 0 | 0 | 0 |
| 54409 | ENSMUSG00000117213 | 0 | NA | NA | NA | NA | NA | Gm49814 | predicted gene, 49814 [Source:MGI Symb | 0 | 0 | 0 | 0 | 0 | 0 |
| 54410 | ENSMUSG00000117214 | 0 | NA | NA | NA | NA | NA | Gm49813 | predicted gene, 49813 [Source:MGI Symb | 0 | 0 | 0 | 0 | 0 | 0 |
| 54411 | ENSMUSG00000117215 | 0 | NA | NA | NA | NA | NA | Gm45926 | predicted gene, 45926 [Source:MGI Symb | 0 | 0 | 0 | 0 | 0 | 0 |
| 54412 | ENSMUSG00000117216 | 0 | NA | NA | NA | NA | NA | Gm49801 | predicted gene, 49801 [Source:MGI Symb | 0 | 0 | 0 | 0 | 0 | 0 |
| 54413 | ENSMUSG00000117217 | 0 | NA | NA | NA | NA | NA | Gm4766 | predicted pseudogene 4766 [Source:MGI | 0 | 0 | 0 | 0 | 0 | 0 |
| 54414 | ENSMUSG00000117218 | 0 | NA | NA | NA | NA | NA | Gm19283 | predicted gene, 19283 [Source:MGI Symb | 0 | 0 | 0 | 0 | 0 | 0 |
| 54415 | ENSMUSG00000117219 | 14.16665967 | -0.798798963 | 0.656662504 | -1.216452832 | 0.223812469 | 0.669614916 | Gm49911 | predicted gene, 49911 [Source:MGI Symb | 17.668 | 9.2513 | 27.169 | 14.46 | 11.997 | 4.4554 |
| 54416 | ENSMUSG00000117220 | 0 | NA | NA | NA | NA | NA | Gm49932 | predicted gene, 49932 [Source:MGI Symb | 0 | 0 | 0 | 0 | 0 | 0 |
| 54417 | ENSMUSG00000117221 | 0 | NA | NA | NA | NA | NA | Gm17830 | predicted gene, 17830 [Source:MGI Symb | 0 | 0 | 0 | 0 | 0 | 0 |
| 54418 | ENSMUSG00000117222 | 0 | NA | NA | NA | NA | NA | Gm49906 | predicted gene, 49906 [Source:MGI Symb | 0 | 0 | 0 | 0 | 0 | 0 |
| 54419 | ENSMUSG00000117223 | 0 | NA | NA | NA | NA | NA | Gm49842 | predicted gene, 49842 [Source:MGI Symb | 0 | 0 | 0 | 0 | 0 | 0 |
| 54420 | ENSMUSG00000117224 | 0 | NA | NA | NA | NA | NA | Gm49940 | predicted gene, 49940 [Source:MGI Symb | 0 | 0 | 0 | 0 | 0 | 0 |
| 54421 | ENSMUSG00000117225 | 0 | NA | NA | NA | NA | NA | Gm49862 | predicted gene, 49862 [Source:MGI Symb | 0 | 0 | 0 | 0 | 0 | 0 |
| 54422 | ENSMUSG00000117226 | 23.44257997 | 0.251200092 | 0.458897345 | 0.547399314 | 0.584104426 | 0.908955096 | Gm31084 | predicted gene, 31084 [Source:MGI Symb | 23.29 | 24.67 | 16.1 | 19.28 | 28.356 | 28.96 |
| 54423 | ENSMUSG00000117227 | 0 | NA | NA | NA | NA | NA | Gm34799 | predicted gene, 34799 [Source:MGI Symb | 0 | 0 | 0 | 0 | 0 | 0 |
| 54424 | ENSMUSG00000117228 | 0 | NA | NA | NA | NA | NA | Gm49877 | predicted gene, 49877 [Source:MGI Symb | 0 | 0 | 0 | 0 | 0 | 0 |
| 54425 | ENSMUSG00000117229 | 0 | NA | NA | NA | NA | NA | Gm35550 | predicted gene, 35550 [Source:MGI Symb | 0 | 0 | 0 | 0 | 0 | 0 |
| 54426 | ENSMUSG00000117230 | 0 | NA | NA | NA | NA | NA | Gm29748 | predicted gene, 29748 [Source:MGI Symb | 0 | 0 | 0 | 0 | 0 | 0 |
| 54427 | ENSMUSG00000117231 | 0.321325949 | 1.911575114 | 4.048267893 | 0.472195804 | 0.636787034 | NA | Gm41609 | predicted gene, 41609 [Source:MGI Symb | 0 | 0 | 0 | 1.928 | 0 | 0 |
| 54428 | ENSMUSG00000117232 | 0 | NA | NA | NA | NA | NA | 1600002D24 | RIKEN cDNA 1600002D24 gene [Source:M | 0 | 0 | 0 | 0 | 0 | 0 |

| | | | | | | | | | | | | | | | |
|---|---|---|---|---|---|---|---|---|---|---|---|---|---|---|---|
| 54429 | ENSMUSG00000117233 | 0 | NA | NA | NA | NA | NA | Gm36594 | predicted gene, 36594 [Source:MGI Symb | 0 | 0 | 0 | 0 | 0 | 0 |
| 54430 | ENSMUSG00000117234 | 1.020693421 | -3.383091396 | 2.599252854 | -1.301563021 | 0.193065808 | NA | Gm7818 | predicted gene 7818 [Source:MGI Symbol | 0 | 4.1117 | 2.0125 | 0 | 0 | 0 |
| 54431 | ENSMUSG00000117235 | 0 | NA | NA | NA | NA | NA | Gm49938 | predicted gene, 49938 [Source:MGI Symb | 0 | 0 | 0 | 0 | 0 | 0 |
| 54432 | ENSMUSG00000117236 | 0 | NA | NA | NA | NA | NA | Gm49829 | predicted gene, 49829 [Source:MGI Symb | 0 | 0 | 0 | 0 | 0 | 0 |
| 54433 | ENSMUSG00000117237 | 0.267700746 | -1.497481649 | 4.057001436 | -0.369110456 | 0.712045396 | NA | Gm18648 | predicted gene, 18648 [Source:MGI Symb | 1.6062 | 0 | 0 | 0 | 0 | 0 |
| 54434 | ENSMUSG00000117238 | 0 | NA | NA | NA | NA | NA | | novel transcript | 0 | 0 | 0 | 0 | 0 | 0 |
| 54435 | ENSMUSG00000117239 | 0.133850373 | -0.796629176 | 4.080472857 | -0.195229623 | 0.84521319 | NA | Gpr31c | G protein-coupled receptor 31, D17Leh66 | 0.8031 | 0 | 0 | 0 | 0 | 0 |
| 54436 | ENSMUSG00000117240 | 0 | NA | NA | NA | NA | NA | Gm49827 | predicted gene, 49827 [Source:MGI Symb | 0 | 0 | 0 | 0 | 0 | 0 |
| 54437 | ENSMUSG00000117241 | 0.167707431 | -0.796629176 | 4.080472857 | -0.195229623 | 0.84521319 | NA | Gm9214 | predicted gene 9214 [Source:MGI Symbol | 0 | 0 | 1.0062 | 0 | 0 | 0 |
| 54438 | ENSMUSG00000117243 | 0 | NA | NA | NA | NA | NA | Gm49944 | predicted gene, 49944 [Source:MGI Symb | 0 | 0 | 0 | 0 | 0 | 0 |
| 54439 | ENSMUSG00000117244 | 0 | NA | NA | NA | NA | NA | Gm9210 | predicted gene 9210 [Source:MGI Symbol | 0 | 0 | 0 | 0 | 0 | 0 |
| 54440 | ENSMUSG00000117245 | 0 | NA | NA | NA | NA | NA | Gm49887 | predicted gene, 49887 [Source:MGI Symb | 0 | 0 | 0 | 0 | 0 | 0 |
| 54441 | ENSMUSG00000117246 | 0 | NA | NA | NA | NA | NA | Gm17891 | predicted gene, 17891 [Source:MGI Symb | 0 | 0 | 0 | 0 | 0 | 0 |
| 54442 | ENSMUSG00000117247 | 0 | NA | NA | NA | NA | NA | Gm49923 | predicted gene, 49923 [Source:MGI Symb | 0 | 0 | 0 | 0 | 0 | 0 |
| 54443 | ENSMUSG00000117248 | 0.887948424 | 0.692526802 | 2.801019149 | 0.247241009 | 0.804721709 | NA | Gm29770 | predicted gene, 29770 [Source:MGI Symb | 0 | 2.0558 | 0 | 0 | 3.2719 | 0 |
| 54444 | ENSMUSG00000117249 | 0 | NA | NA | NA | NA | NA | Gm46577 | predicted gene, 46577 [Source:MGI Symb | 0 | 0 | 0 | 0 | 0 | 0 |
| 54445 | ENSMUSG00000117250 | 3.29553076 | -0.083411375 | 1.141987759 | -0.073040516 | 0.941773877 | NA | Gm49871 | predicted gene, 49871 [Source:MGI Symb | 4.0155 | 3.0838 | 3.0187 | 1.928 | 3.2719 | 4.4554 |
| 54446 | ENSMUSG00000117251 | 0 | NA | NA | NA | NA | NA | Gm49891 | predicted gene, 49891 [Source:MGI Symb | 0 | 0 | 0 | 0 | 0 | 0 |
| 54447 | ENSMUSG00000117252 | 0 | NA | NA | NA | NA | NA | Gm9514 | predicted gene 9514 [Source:MGI Symbol | 0 | 0 | 0 | 0 | 0 | 0 |
| 54448 | ENSMUSG00000117253 | 0 | NA | NA | NA | NA | NA | Gm49930 | predicted gene, 49930 [Source:MGI Symb | 0 | 0 | 0 | 0 | 0 | 0 |
| 54449 | ENSMUSG00000117254 | 0 | NA | NA | NA | NA | NA | Gm34567 | predicted gene, 34567 [Source:MGI Symb | 0 | 0 | 0 | 0 | 0 | 0 |
| 54450 | ENSMUSG00000117255 | 5.240568929 | 0.311160614 | 1.02309073 | 0.304137849 | 0.761022873 | 0.95402891 | Gm49864 | predicted gene, 49864 [Source:MGI Symb | 4.0155 | 1.0279 | 9.0562 | 9.6398 | 4.3625 | 3.3415 |
| 54451 | ENSMUSG00000117256 | 0 | NA | NA | NA | NA | NA | Gm5497 | predicted gene 5497 [Source:MGI Symbol | 0 | 0 | 0 | 0 | 0 | 0 |
| 54452 | ENSMUSG00000117257 | 1.006244586 | -3.367758763 | 2.959437494 | -1.137972595 | 0.255131926 | NA | Gm4948 | predicted gene 4948 [Source:MGI Symbol | 0 | 0 | 6.0375 | 0 | 0 | 0 |
| 54453 | ENSMUSG00000117258 | 0 | NA | NA | NA | NA | NA | Gm7773 | predicted gene 7773 [Source:MGI Symbol | 0 | 0 | 0 | 0 | 0 | 0 |
| 54454 | ENSMUSG00000117259 | 0 | NA | NA | NA | NA | NA | Gm19012 | predicted gene, 19012 [Source:MGI Symb | 0 | 0 | 0 | 0 | 0 | 0 |
| 54455 | ENSMUSG00000117260 | 0 | NA | NA | NA | NA | NA | Gm49818 | predicted gene, 49818 [Source:MGI Symb | 0 | 0 | 0 | 0 | 0 | 0 |
| 54456 | ENSMUSG00000117262 | 0.160662974 | 1.126924175 | 4.080472857 | 0.276174898 | 0.782413736 | NA | Gm9316 | predicted gene 9316 [Source:MGI Symbol | 0 | 0 | 0 | 0.964 | 0 | 0 |
| 54457 | ENSMUSG00000117263 | 4.838261327 | 0.476151945 | 1.033719942 | 0.460619869 | 0.645071354 | NA | Gm35290 | predicted gene, 35290 [Source:MGI Symb | 4.0155 | 3.0838 | 5.0312 | 5.7839 | 1.0906 | 10.025 |
| 54458 | ENSMUSG00000117264 | 0 | NA | NA | NA | NA | NA | 4930452A19 | RIKEN cDNA 4930452A19 gene [Source:M | 0 | 0 | 0 | 0 | 0 | 0 |
| 54459 | ENSMUSG00000117265 | 0 | NA | NA | NA | NA | NA | Gm49847 | predicted gene, 49847 [Source:MGI Symb | 0 | 0 | 0 | 0 | 0 | 0 |
| 54460 | ENSMUSG00000117266 | 0 | NA | NA | NA | NA | NA | Gm49947 | predicted gene, 49947 [Source:MGI Symb | 0 | 0 | 0 | 0 | 0 | 0 |
| 54461 | ENSMUSG00000117267 | 0 | NA | NA | NA | NA | NA | Gm49830 | predicted gene, 49830 [Source:MGI Symb | 0 | 0 | 0 | 0 | 0 | 0 |
| 54462 | ENSMUSG00000117268 | 0 | NA | NA | NA | NA | NA | Gm49884 | predicted gene, 49884 [Source:MGI Symb | 0 | 0 | 0 | 0 | 0 | 0 |
| 54463 | ENSMUSG00000117269 | 0.335414862 | -1.779047242 | 4.030804179 | -0.441362855 | 0.658950329 | NA | Gm9706 | predicted gene 9706 [Source:MGI Symbol | 0 | 0 | 2.0125 | 0 | 0 | 0 |
| 54464 | ENSMUSG00000117270 | 0 | NA | NA | NA | NA | NA | Gm49953 | predicted gene, 49953 [Source:MGI Symb | 0 | 0 | 0 | 0 | 0 | 0 |
| 54465 | ENSMUSG00000117271 | 0 | NA | NA | NA | NA | NA | Gm6149 | predicted gene 6149 [Source:MGI Symbol | 0 | 0 | 0 | 0 | 0 | 0 |
| 54466 | ENSMUSG00000117272 | 0 | NA | NA | NA | NA | NA | Gm49855 | predicted gene, 49855 [Source:MGI Symb | 0 | 0 | 0 | 0 | 0 | 0 |
| 54467 | ENSMUSG00000117273 | 0 | NA | NA | NA | NA | NA | Gm49815 | predicted gene, 49815 [Source:MGI Symb | 0 | 0 | 0 | 0 | 0 | 0 |
| 54468 | ENSMUSG00000117274 | 0 | NA | NA | NA | NA | NA | Gm17921 | predicted gene, 17921 [Source:MGI Symb | 0 | 0 | 0 | 0 | 0 | 0 |
| 54469 | ENSMUSG00000117275 | 1.389209159 | 0.935224046 | 2.008619442 | 0.465605393 | 0.641497982 | NA | Gm50047 | predicted gene, 50047 [Source:MGI Symb | 0.8031 | 2.0558 | 0 | 0 | 4.3625 | 1.1138 |
| 54470 | ENSMUSG00000117276 | 0 | NA | NA | NA | NA | NA | Gm49910 | predicted gene, 49910 [Source:MGI Symb | 0 | 0 | 0 | 0 | 0 | 0 |
| 54471 | ENSMUSG00000117277 | 0 | NA | NA | NA | NA | NA | Gm49828 | predicted gene, 49828 [Source:MGI Symb | 0 | 0 | 0 | 0 | 0 | 0 |
| 54472 | ENSMUSG00000117278 | 0 | NA | NA | NA | NA | NA | Gm36684 | predicted gene, 36684 [Source:MGI Symb | 0 | 0 | 0 | 0 | 0 | 0 |
| 54473 | ENSMUSG00000117279 | 0 | NA | NA | NA | NA | NA | Gm49850 | predicted gene, 49850 [Source:MGI Symb | 0 | 0 | 0 | 0 | 0 | 0 |
| 54474 | ENSMUSG00000117280 | 0 | NA | NA | NA | NA | NA | Gm49903 | predicted gene, 49903 [Source:MGI Symb | 0 | 0 | 0 | 0 | 0 | 0 |
| 54475 | ENSMUSG00000117281 | 0 | NA | NA | NA | NA | NA | Gm49848 | predicted gene, 49848 [Source:MGI Symb | 0 | 0 | 0 | 0 | 0 | 0 |
| 54476 | ENSMUSG00000117282 | 0 | NA | NA | NA | NA | NA | Gm49898 | predicted gene, 49898 [Source:MGI Symb | 0 | 0 | 0 | 0 | 0 | 0 |
| 54477 | ENSMUSG00000117283 | 0 | NA | NA | NA | NA | NA | Gm4471 | predicted gene 4471 [Source:MGI Symbol | 0 | 0 | 0 | 0 | 0 | 0 |
| 54478 | ENSMUSG00000117284 | 183.4940712 | 0.142711362 | 0.19023317 | 0.750191791 | 0.453139202 | 0.857073339 | Gm7072 | predicted gene 7072 [Source:MGI Symbol | 160.62 | 167.55 | 195.21 | 173.52 | 221.4 | 182.67 |
| 54479 | ENSMUSG00000117285 | 9.233708455 | 0.084830778 | 0.707730695 | 0.119863077 | 0.904591614 | 0.983865503 | Gm49863 | predicted gene, 49863 [Source:MGI Symb | 5.6217 | 12.335 | 9.0562 | 9.6398 | 8.7249 | 10.025 |
| 54480 | ENSMUSG00000117286 | 0 | NA | NA | NA | NA | NA | Gm1043 | predicted gene 1043 [Source:MGI Symbol | 0 | 0 | 0 | 0 | 0 | 0 |
| 54481 | ENSMUSG00000117287 | 0 | NA | NA | NA | NA | NA | Gm36200 | predicted gene, 36200 [Source:MGI Symb | 0 | 0 | 0 | 0 | 0 | 0 |
| 54482 | ENSMUSG00000117288 | 0 | NA | NA | NA | NA | NA | Gm49856 | predicted gene, 49856 [Source:MGI Symb | 0 | 0 | 0 | 0 | 0 | 0 |
| 54483 | ENSMUSG00000117289 | 0.956437057 | 1.282470111 | 2.271719443 | 0.56453719 | 0.572388597 | NA | 1700093J21I | RIKEN cDNA 1700093J21 gene [Source:M( | 1.6062 | 0 | 0 | 1.928 | 1.0906 | 1.1138 |
| 54484 | ENSMUSG00000117290 | 0 | NA | NA | NA | NA | NA | Gm34883 | predicted gene, 34883 [Source:MGI Symb | 0 | 0 | 0 | 0 | 0 | 0 |

| | | | | | | | | | | | | | | | |
|---|---|---|---|---|---|---|---|---|---|---|---|---|---|---|---|
| 54485 | ENSMUSG00000117291 | 0 | NA | NA | NA | NA | NA | Gm18271 | predicted gene, 18271 [Source:MGI Symb | 0 | 0 | 0 | 0 | 0 | 0 |
| 54486 | ENSMUSG00000117292 | 0 | NA | NA | NA | NA | NA | E330032C10 | RIKEN cDNA E330032C10 gene [Source:M | 0 | 0 | 0 | 0 | 0 | 0 |
| 54487 | ENSMUSG00000117293 | 0 | NA | NA | NA | NA | NA | Gm49821 | predicted gene, 49821 [Source:MGI Symb | 0 | 0 | 0 | 0 | 0 | 0 |
| 54488 | ENSMUSG00000117294 | 0.703108924 | -2.875459482 | 2.96384054 | -0.970180225 | 0.331956666 | NA | Gm49839 | predicted gene, 49839 [Source:MGI Symb | 3.2124 | 0 | 1.0062 | 0 | 0 | 0 |
| 54489 | ENSMUSG00000117295 | 0 | NA | NA | NA | NA | NA | Gm5492 | predicted gene 5492 [Source:MGI Symbol | 0 | 0 | 0 | 0 | 0 | 0 |
| 54490 | ENSMUSG00000117296 | 0 | NA | NA | NA | NA | NA | Gm20742 | predicted gene, 20742 [Source:MGI Symb | 0 | 0 | 0 | 0 | 0 | 0 |
| 54491 | ENSMUSG00000117297 | 0.678054141 | -2.792799308 | 2.990673428 | -0.933836266 | 0.350388359 | NA | Gm49888 | predicted gene, 49888 [Source:MGI Symb | 0 | 2.0558 | 2.0125 | 0 | 0 | 0 |
| 54492 | ENSMUSG00000117298 | 0 | NA | NA | NA | NA | NA | Gm9284 | predicted gene 9284 [Source:MGI Symbol | 0 | 0 | 0 | 0 | 0 | 0 |
| 54493 | ENSMUSG00000117299 | 0 | NA | NA | NA | NA | NA | Gm9288 | predicted gene 9288 [Source:MGI Symbol | 0 | 0 | 0 | 0 | 0 | 0 |
| 54494 | ENSMUSG00000117300 | 0.185640647 | 1.126924175 | 4.080472857 | 0.276174898 | 0.782413736 | NA | Gm18736 | predicted gene, 18736 [Source:MGI Symb | 0 | 0 | 0 | 0 | 0 | 1.1138 |
| 54495 | ENSMUSG00000117301 | 0.545309144 | 2.651974629 | 3.666219369 | 0.72335405 | 0.469462394 | NA | Gm35883 | predicted gene, 35883 [Source:MGI Symb | 0 | 0 | 0 | 0 | 3.2719 | 0 |
| 54496 | ENSMUSG00000117302 | 0 | NA | NA | NA | NA | NA | Gm34266 | predicted gene, 34266 [Source:MGI Symb | 0 | 0 | 0 | 0 | 0 | 0 |
| 54497 | ENSMUSG00000117303 | 1.17654932 | 1.687714554 | 2.567949793 | 0.657222566 | 0.511037817 | NA | Gm49889 | predicted gene, 49889 [Source:MGI Symb | 1.6062 | 0 | 0 | 0 | 5.4531 | 0 |
| 54498 | ENSMUSG00000117304 | 0.974421879 | -3.346724782 | 2.664166553 | -1.256199534 | 0.209043636 | NA | Gm49861 | predicted gene, 49861 [Source:MGI Symb | 4.8186 | 1.0279 | 0 | 0 | 0 | 0 |
| 54499 | ENSMUSG00000117305 | 0 | NA | NA | NA | NA | NA | Gm49963 | predicted gene, 49963 [Source:MGI Symb | 0 | 0 | 0 | 0 | 0 | 0 |
| 54500 | ENSMUSG00000117306 | 0 | NA | NA | NA | NA | NA | Gm8945 | predicted gene 8945 [Source:MGI Symbol | 0 | 0 | 0 | 0 | 0 | 0 |
| 54501 | ENSMUSG00000117307 | 0 | NA | NA | NA | NA | NA | Gm49837 | predicted gene, 49837 [Source:MGI Symb | 0 | 0 | 0 | 0 | 0 | 0 |
| 54502 | ENSMUSG00000117308 | 0 | NA | NA | NA | NA | NA | Gm20596 | predicted gene, 20596 [Source:MGI Symb | 0 | 0 | 0 | 0 | 0 | 0 |
| 54503 | ENSMUSG00000117309 | 0.167707431 | -0.796629176 | 4.080472857 | -0.195229623 | 0.84521319 | NA | Gm49960 | predicted gene, 49960 [Source:MGI Symb | 0 | 0 | 1.0062 | 0 | 0 | 0 |
| 54504 | ENSMUSG00000117310 | 33.93993526 | -1.036322814 | 0.429058134 | -2.415343588 | 0.015720373 | 0.146980912 | Ptp4a1 | protein tyrosine phosphatase 4a1 [Source | 53.005 | 54.48 | 29.181 | 16.388 | 30.537 | 20.049 |
| 54505 | ENSMUSG00000117311 | 0 | NA | NA | NA | NA | NA | Esp22 | exocrine gland secreted peptide 22 [Sourc | 0 | 0 | 0 | 0 | 0 | 0 |
| 54506 | ENSMUSG00000117312 | 0 | NA | NA | NA | NA | NA | Gm49931 | predicted gene, 49931 [Source:MGI Symb | 0 | 0 | 0 | 0 | 0 | 0 |
| 54507 | ENSMUSG00000117313 | 2.050052468 | -1.54357849 | 1.738878992 | -0.887685973 | 0.374709693 | NA | Gm49838 | predicted gene, 49838 [Source:MGI Symb | 4.0155 | 5.1396 | 0 | 0.964 | 2.1812 | 0 |
| 54508 | ENSMUSG00000117314 | 0 | NA | NA | NA | NA | NA | C030013G03 | RIKEN cDNA C030013G03 gene [Source:M | 0 | 0 | 0 | 0 | 0 | 0 |
| 54509 | ENSMUSG00000117315 | 0 | NA | NA | NA | NA | NA | 1600022D10 | RIKEN cDNA 1600022D10 gene [Source:M | 0 | 0 | 0 | 0 | 0 | 0 |
| 54510 | ENSMUSG00000117316 | 0 | NA | NA | NA | NA | NA | Gm4946 | predicted gene 4946 [Source:MGI Symbol | 0 | 0 | 0 | 0 | 0 | 0 |
| 54511 | ENSMUSG00000117317 | 5.320626972 | -0.01359691 | 1.067871414 | -0.012732722 | 0.989841032 | 0.99859176 | Gm49961 | predicted gene, 49961 [Source:MGI Symb | 8.8341 | 2.0558 | 5.0312 | 2.8919 | 11.997 | 1.1138 |
| 54512 | ENSMUSG00000117318 | 0 | NA | NA | NA | NA | NA | Gm18679 | predicted gene, 18679 [Source:MGI Symb | 0 | 0 | 0 | 0 | 0 | 0 |
| 54513 | ENSMUSG00000117319 | 0 | NA | NA | NA | NA | NA | Gm49833 | predicted gene, 49833 [Source:MGI Symb | 0 | 0 | 0 | 0 | 0 | 0 |
| 54514 | ENSMUSG00000117320 | 0 | NA | NA | NA | NA | NA | Gm18070 | predicted gene, 18070 [Source:MGI Symb | 0 | 0 | 0 | 0 | 0 | 0 |
| 54515 | ENSMUSG00000117321 | 0.653308562 | 1.538013299 | 3.112409277 | 0.494155223 | 0.62119656 | NA | Gm49927 | predicted gene, 49927 [Source:MGI Symb | 0 | 1.0279 | 0 | 2.8919 | 0 | 0 |
| 54516 | ENSMUSG00000117322 | 20.93430434 | 0.025428643 | 0.486201595 | 0.052300617 | 0.958289162 | 0.991891687 | 6330415G19 | RIKEN cDNA 6330415G19 gene [Source:M | 21.684 | 19.53 | 21.131 | 26.991 | 22.903 | 13.366 |
| 54517 | ENSMUSG00000117323 | 0 | NA | NA | NA | NA | NA | Gm36487 | predicted gene, 36487 [Source:MGI Symb | 0 | 0 | 0 | 0 | 0 | 0 |
| 54518 | ENSMUSG00000117324 | 0 | NA | NA | NA | NA | NA | Gm32432 | predicted gene, 32432 [Source:MGI Symb | 0 | 0 | 0 | 0 | 0 | 0 |
| 54519 | ENSMUSG00000117325 | 0 | NA | NA | NA | NA | NA | Gm49962 | predicted gene, 49962 [Source:MGI Symb | 0 | 0 | 0 | 0 | 0 | 0 |
| 54520 | ENSMUSG00000117326 | 0 | NA | NA | NA | NA | NA | 4930564C03 | RIKEN cDNA 4930564C03 gene [Source:M | 0 | 0 | 0 | 0 | 0 | 0 |
| 54521 | ENSMUSG00000117327 | 2.577124696 | -0.196262485 | 1.412177981 | -0.138978576 | 0.889467079 | NA | Gm6467 | predicted gene 6467 [Source:MGI Symbol | 3.2124 | 1.0279 | 4.025 | 3.8559 | 0 | 3.3415 |
| 54522 | ENSMUSG00000117328 | 0 | NA | NA | NA | NA | NA | 4930474M22 | RIKEN cDNA 4930474M22 gene [Source:M | 0 | 0 | 0 | 0 | 0 | 0 |
| 54523 | ENSMUSG00000117329 | 0 | NA | NA | NA | NA | NA | Gm49943 | predicted gene, 49943 [Source:MGI Symb | 0 | 0 | 0 | 0 | 0 | 0 |
| 54524 | ENSMUSG00000117330 | 0 | NA | NA | NA | NA | NA | Gm49824 | predicted gene, 49824 [Source:MGI Symb | 0 | 0 | 0 | 0 | 0 | 0 |
| 54525 | ENSMUSG00000117331 | 0 | NA | NA | NA | NA | NA | Gm49816 | predicted gene, 49816 [Source:MGI Symb | 0 | 0 | 0 | 0 | 0 | 0 |
| 54526 | ENSMUSG00000117332 | 0 | NA | NA | NA | NA | NA | Gm49942 | predicted gene, 49942 [Source:MGI Symb | 0 | 0 | 0 | 0 | 0 | 0 |
| 54527 | ENSMUSG00000117333 | 3.193733535 | 1.261677864 | 1.188270885 | 1.0617763 | 0.288337251 | NA | Gm16386 | predicted gene 16386 [Source:MGI Symbo | 1.6062 | 2.0558 | 2.0125 | 5.7839 | 4.3625 | 3.3415 |
| 54528 | ENSMUSG00000117334 | 0 | NA | NA | NA | NA | NA | A730006G06 | RIKEN cDNA A730006G06 gene [Source:M | 0 | 0 | 0 | 0 | 0 | 0 |
| 54529 | ENSMUSG00000117335 | 5.353708614 | 1.717473207 | 0.993233471 | 1.72917371 | 0.08377801 | 0.426583908 | Gm49870 | predicted gene, 49870 [Source:MGI Symb | 2.4093 | 1.0279 | 4.025 | 4.8199 | 9.8156 | 10.025 |
| 54530 | ENSMUSG00000117336 | 0 | NA | NA | NA | NA | NA | Gm49865 | predicted gene, 49865 [Source:MGI Symb | 0 | 0 | 0 | 0 | 0 | 0 |
| 54531 | ENSMUSG00000117337 | 0.17131964 | -0.796629176 | 4.080472857 | -0.195229623 | 0.84521319 | NA | Gm49964 | predicted gene, 49964 [Source:MGI Symb | 0 | 1.0279 | 0 | 0 | 0 | 0 |
| 54532 | ENSMUSG00000117338 | 0 | NA | NA | NA | NA | NA | Gm49804 | predicted gene, 49804 [Source:MGI Symb | 0 | 0 | 0 | 0 | 0 | 0 |
| 54533 | ENSMUSG00000117340 | 0 | NA | NA | NA | NA | NA | Gm49920 | predicted gene, 49920 [Source:MGI Symb | 0 | 0 | 0 | 0 | 0 | 0 |
| 54534 | ENSMUSG00000117341 | 0 | NA | NA | NA | NA | NA | Gm34684 | predicted gene, 34684 [Source:MGI Symb | 0 | 0 | 0 | 0 | 0 | 0 |

**Supplementary Table S1 |** Results of RNA-Seq for WT and KO livers from male DIO mice. DESeq2 results for all genes and the results of GSEA on the Hallmark gene sets are shown.

| NAME | SIZE | ES | NES | NOM p-val | FDR q-val | FWER p-val | RANK AT MAX | LEADING EDGE |
|---|---|---|---|---|---|---|---|---|
| HALLMARK_ALLOGRAFT_REJECTION | 196 | -0.66830415 | -1.5113256 | 0 | 0.103071444 | 0.045 | 2838 | tags=47%, list=16%, signal=56% |
| HALLMARK_TGF_BETA_SIGNALING | 54 | -0.48738965 | -1.4672534 | 0 | 0.14887583 | 0.151 | 1132 | tags=19%, list=6%, signal=20% |
| HALLMARK_INFLAMMATORY_RESPONSE | 198 | -0.56452847 | -1.4623072 | 0 | 0.13417637 | 0.201 | 3149 | tags=42%, list=17%, signal=51% |
| HALLMARK_KRAS_SIGNALING_UP | 199 | -0.6202797 | -1.458994 | 0 | 0.11488233 | 0.201 | 3027 | tags=46%, list=17%, signal=55% |
| HALLMARK_HYPOXIA | 198 | -0.55228394 | -1.4450331 | 0 | 0.103305764 | 0.201 | 3636 | tags=46%, list=20%, signal=58% |
| HALLMARK_ANGIOGENESIS | 36 | -0.565402 | -1.4265144 | 0 | 0.13318308 | 0.307 | 1574 | tags=33%, list=9%, signal=36% |
| HALLMARK_IL2_STAT5_SIGNALING | 199 | -0.49113852 | -1.4260441 | 0 | 0.12229977 | 0.307 | 3189 | tags=40%, list=18%, signal=48% |
| HALLMARK_APOPTOSIS | 161 | -0.52439725 | -1.4186269 | 0 | 0.12160982 | 0.307 | 2486 | tags=32%, list=14%, signal=37% |
| HALLMARK_UV_RESPONSE_DN | 143 | -0.47633144 | -1.4142033 | 0 | 0.11443098 | 0.307 | 2426 | tags=30%, list=13%, signal=34% |
| HALLMARK_EPITHELIAL_MESENCHYMAL_TRANSITION | 196 | -0.60128075 | -1.3967445 | 0 | 0.10868779 | 0.307 | 2745 | tags=42%, list=15%, signal=49% |
| HALLMARK_PANCREAS_BETA_CELLS | 40 | -0.4813735 | -1.3707412 | 0 | 0.12631413 | 0.353 | 3875 | tags=23%, list=22%, signal=29% |
| HALLMARK_APICAL_JUNCTION | 198 | -0.47533 | -1.368325 | 0 | 0.12053804 | 0.353 | 4048 | tags=41%, list=22%, signal=53% |
| HALLMARK_INTERFERON_GAMMA_RESPONSE | 195 | -0.4445092 | -1.3625617 | 0 | 0.120329805 | 0.411 | 2535 | tags=28%, list=14%, signal=32% |
| HALLMARK_MYOGENESIS | 200 | -0.49010175 | -1.3540645 | 0 | 0.12007616 | 0.411 | 4296 | tags=38%, list=24%, signal=49% |
| HALLMARK_COAGULATION | 138 | -0.41165316 | -1.3534292 | 0 | 0.11587118 | 0.411 | 3056 | tags=32%, list=17%, signal=38% |
| HALLMARK_COMPLEMENT | 193 | -0.43197158 | -1.3532228 | 0.0990099 | 0.11219171 | 0.411 | 3517 | tags=33%, list=20%, signal=41% |
| HALLMARK_IL6_JAK_STAT3_SIGNALING | 87 | -0.5543007 | -1.3511732 | 0.096525095 | 0.11446567 | 0.411 | 3066 | tags=45%, list=17%, signal=54% |
| HALLMARK_PEROXISOME | 102 | -0.4191325 | -1.3418498 | 0.11752577 | 0.12431385 | 0.452 | 4746 | tags=39%, list=26%, signal=53% |
| HALLMARK_GLYCOLYSIS | 200 | -0.37909767 | -1.3415395 | 0 | 0.12077097 | 0.452 | 5090 | tags=41%, list=28%, signal=57% |
| HALLMARK_FATTY_ACID_METABOLISM | 155 | -0.41911894 | -1.3296694 | 0.091649696 | 0.12057141 | 0.452 | 5046 | tags=45%, list=28%, signal=62% |
| HALLMARK_NOTCH_SIGNALING | 32 | -0.5379298 | -1.3271322 | 0.095703125 | 0.11754432 | 0.452 | 2273 | tags=31%, list=13%, signal=36% |
| HALLMARK_P53_PATHWAY | 200 | -0.40709466 | -1.3172717 | 0.11218569 | 0.14291735 | 0.548 | 2898 | tags=25%, list=16%, signal=29% |
| HALLMARK_UV_RESPONSE_UP | 156 | -0.38186383 | -1.3019066 | 0 | 0.15061219 | 0.548 | 2422 | tags=22%, list=13%, signal=25% |
| HALLMARK_TNFA_SIGNALING_VIA_NFKB | 198 | -0.49079376 | -1.296668 | 0 | 0.1492024 | 0.548 | 3055 | tags=38%, list=17%, signal=46% |
| HALLMARK_XENOBIOTIC_METABOLISM | 196 | -0.36678386 | -1.2785958 | 0 | 0.17687821 | 0.548 | 4547 | tags=36%, list=25%, signal=48% |
| HALLMARK_REACTIVE_OXYGEN_SPECIES_PATHWAY | 49 | -0.3914508 | -1.2618198 | 0.20656371 | 0.18302515 | 0.588 | 4614 | tags=45%, list=26%, signal=60% |
| HALLMARK_ADIPOGENESIS | 200 | -0.38541177 | -1.2602358 | 0.20689656 | 0.17835748 | 0.588 | 4881 | tags=38%, list=27%, signal=52% |
| HALLMARK_APICAL_SURFACE | 44 | -0.4897507 | -1.2494509 | 0.21230158 | 0.18377993 | 0.639 | 2414 | tags=32%, list=13%, signal=37% |
| HALLMARK_MITOTIC_SPINDLE | 199 | -0.36147326 | -1.2434647 | 0.11608961 | 0.18105571 | 0.639 | 5543 | tags=39%, list=31%, signal=55% |
| HALLMARK_HEDGEHOG_SIGNALING | 36 | -0.48961812 | -1.2231466 | 0.18490566 | 0.19763619 | 0.639 | 3087 | tags=36%, list=17%, signal=43% |
| HALLMARK_KRAS_SIGNALING_DN | 199 | -0.39965957 | -1.2220052 | 0 | 0.19506164 | 0.639 | 3831 | tags=28%, list=21%, signal=35% |
| HALLMARK_ANDROGEN_RESPONSE | 98 | -0.34832504 | -1.2089454 | 0.11264822 | 0.20822315 | 0.639 | 4723 | tags=40%, list=26%, signal=54% |
| HALLMARK_ESTROGEN_RESPONSE_LATE | 197 | -0.37003377 | -1.1538969 | 0.11306043 | 0.26121983 | 0.781 | 3555 | tags=32%, list=20%, signal=39% |
| HALLMARK_PI3K_AKT_MTOR_SIGNALING | 105 | -0.27428648 | -1.1125011 | 0.18737672 | 0.33430773 | 0.899 | 5244 | tags=30%, list=29%, signal=43% |
| HALLMARK_ESTROGEN_RESPONSE_EARLY | 197 | -0.34245452 | -1.0766778 | 0.09505703 | 0.3635614 | 0.951 | 3555 | tags=29%, list=20%, signal=36% |
| HALLMARK_OXIDATIVE_PHOSPHORYLATION | 198 | -0.25834903 | -1.0200375 | 0.5766129 | 0.43710467 | 1 | 6501 | tags=39%, list=36%, signal=61% |
| HALLMARK_G2M_CHECKPOINT | 196 | -0.2936401 | -0.9716138 | 0.3968254 | 0.5129486 | 1 | 5007 | tags=32%, list=28%, signal=44% |
| HALLMARK_MYC_TARGETS_V2 | 57 | -0.28414115 | -0.9668572 | 0.46341464 | 0.5039807 | 1 | 3965 | tags=25%, list=22%, signal=31% |
| HALLMARK_SPERMATOGENESIS | 134 | -0.28442773 | -0.95027345 | 0.5204918 | 0.52910936 | 1 | 3588 | tags=22%, list=20%, signal=28% |
| HALLMARK_WNT_BETA_CATENIN_SIGNALING | 41 | -0.3631463 | -0.94331646 | 0.51696604 | 0.52994657 | 1 | 4387 | tags=39%, list=24%, signal=51% |
| HALLMARK_PROTEIN_SECRETION | 95 | -0.1859828 | -0.93656486 | 0.6089494 | 0.522958 | 1 | 5755 | tags=31%, list=32%, signal=45% |
| HALLMARK_INTERFERON_ALPHA_RESPONSE | 95 | -0.2638043 | -0.76460844 | 0.69512194 | 0.8036836 | 1 | 3017 | tags=19%, list=17%, signal=23% |
| HALLMARK_E2F_TARGETS | 200 | -0.19604218 | -0.6652204 | 0.88667995 | 0.9429418 | 1 | 5007 | tags=29%, list=28%, signal=39% |

**Supplementary Table S1 |** Results of RNA-Seq for WT and KO livers from male DIO mice. DESeq2 results for all genes and the results of GSEA on the Hallmark gene sets

| NAME | SIZE | ES | NES | NOM p-val | FDR q-val | FWER p-val | RANK AT MAX | LEADING EDGE |
|---|---|---|---|---|---|---|---|---|
| HALLMARK_CHOLESTEROL_HOMEOSTASIS | 73 | 0.45226383 | 1.1578225 | 0.28024194 | 1 | 0.691 | 959 | tags=33%, list=5%, signal=35% |
| HALLMARK_HEME_METABOLISM | 191 | 0.28824356 | 1.1417108 | 0.11332008 | 0.8206479 | 0.892 | 2248 | tags=22%, list=12%, signal=25% |
| HALLMARK_UNFOLDED_PROTEIN_RESPONSE | 112 | 0.2420649 | 1.0237892 | 0.27 | 0.8225647 | 0.95 | 3410 | tags=28%, list=19%, signal=34% |
| HALLMARK_DNA_REPAIR | 149 | 0.23695563 | 0.93828595 | 0.48514852 | 0.8682633 | 1 | 3890 | tags=28%, list=22%, signal=35% |
| HALLMARK_BILE_ACID_METABOLISM | 112 | 0.2667276 | 0.9291706 | 0.5981132 | 0.719499 | 1 | 1547 | tags=17%, list=9%, signal=18% |
| HALLMARK_MTORC1_SIGNALING | 199 | 0.25237572 | 0.8980391 | 0.67061144 | 0.6607746 | 1 | 1606 | tags=16%, list=9%, signal=17% |
| HALLMARK_MYC_TARGETS_V1 | 200 | 0.208681 | 0.8459446 | 0.5962264 | 0.67479783 | 1 | 5642 | tags=41%, list=31%, signal=59% |
