## Supplementary Table 2 for "G protein-coupled receptor 151 regulates glucose metabolism and hepatic gluconeogenesis"

**Supplementary Table S2 |** Gene list used for custom GSEA on cAMP-regulated genes.

| | ensembl_gene_id | mgi_symbol | description |
|---|---|---|---|
| 1 | ENSMUSG00000000275 | Trim25 | tripartite motif-containing 25 [Source:MGI Symbol;Acc:MGI:102749] |
| 2 | ENSMUSG00000001435 | Col18a1 | collagen, type XVIII, alpha 1 [Source:MGI Symbol;Acc:MGI:88451] |
| 3 | ENSMUSG00000001542 | Ell2 | elongation factor for RNA polymerase II 2 [Source:MGI Symbol;Acc:MGI:2183438] |
| 4 | ENSMUSG00000001670 | Tat | tyrosine aminotransferase [Source:MGI Symbol;Acc:MGI:98487] |
| 5 | ENSMUSG00000003134 | Tbc1d8 | TBC1 domain family, member 8 [Source:MGI Symbol;Acc:MGI:1927225] |
| 6 | ENSMUSG00000005125 | Ndrg1 | N-myc downstream regulated gene 1 [Source:MGI Symbol;Acc:MGI:1341799] |
| 7 | ENSMUSG00000005514 | Por | P450 (cytochrome) oxidoreductase [Source:MGI Symbol;Acc:MGI:97744] |
| 8 | ENSMUSG00000013089 | Etv5 | ets variant 5 [Source:MGI Symbol;Acc:MGI:1096867] |
| 9 | ENSMUSG00000014867 | Surf4 | surfeit gene 4 [Source:MGI Symbol;Acc:MGI:98445] |
| 10 | ENSMUSG00000019232 | Etnppl | ethanolamine phosphate phospholyase [Source:MGI Symbol;Acc:MGI:1919010] |
| 11 | ENSMUSG00000020029 | Nudt4 | nudix (nucleoside diphosphate linked moiety X)-type motif 4 [Source:MGI Symbol;Acc:MGI:1918457] |
| 12 | ENSMUSG00000020034 | Tcp11l2 | t-complex 11 (mouse) like 2 [Source:MGI Symbol;Acc:MGI:2444679] |
| 13 | ENSMUSG00000020063 | Sirt1 | sirtuin 1 [Source:MGI Symbol;Acc:MGI:2135607] |
| 14 | ENSMUSG00000020085 | Aifm2 | apoptosis-inducing factor, mitochondrion-associated 2 [Source:MGI Symbol;Acc:MGI:1918611] |
| 15 | ENSMUSG00000020390 | Ube2b | ubiquitin-conjugating enzyme E2B [Source:MGI Symbol;Acc:MGI:102944] |
| 16 | ENSMUSG00000020415 | Pttg1 | pituitary tumor-transforming gene 1 [Source:MGI Symbol;Acc:MGI:1353578] |
| 17 | ENSMUSG00000020429 | Igfbp1 | insulin-like growth factor binding protein 1 [Source:MGI Symbol;Acc:MGI:96436] |
| 18 | ENSMUSG00000020644 | Id2 | inhibitor of DNA binding 2 [Source:MGI Symbol;Acc:MGI:96397] |
| 19 | ENSMUSG00000020718 | Polg2 | polymerase (DNA directed), gamma 2, accessory subunit [Source:MGI Symbol;Acc:MGI:1354947] |
| 20 | ENSMUSG00000020973 | Dnaaf2 | dynein, axonemal assembly factor 2 [Source:MGI Symbol;Acc:MGI:1923566] |
| 21 | ENSMUSG00000021453 | Gadd45g | growth arrest and DNA-damage-inducible 45 gamma [Source:MGI Symbol;Acc:MGI:1346325] |
| 22 | ENSMUSG00000021719 | Rgs7bp | regulator of G-protein signalling 7 binding protein [Source:MGI Symbol;Acc:MGI:106334] |
| 23 | ENSMUSG00000021779 | Thrb | thyroid hormone receptor beta [Source:MGI Symbol;Acc:MGI:98743] |
| 24 | ENSMUSG00000022106 | Rcbtb2 | regulator of chromosome condensation (RCC1) and BTB (POZ) domain containing protein 2 [Source:MGI Symbol;Acc:MGI:1917200] |
| 25 | ENSMUSG00000022124 | Fbxl3 | F-box and leucine-rich repeat protein 3 [Source:MGI Symbol;Acc:MGI:1354702] |
| 26 | ENSMUSG00000022270 | Retreg1 | reticulophagy regulator 1 [Source:MGI Symbol;Acc:MGI:1913520] |
| 27 | ENSMUSG00000022299 | Slc25a32 | solute carrier family 25, member 32 [Source:MGI Symbol;Acc:MGI:1917156] |
| 28 | ENSMUSG00000022462 | Slc38a2 | solute carrier family 38, member 2 [Source:MGI Symbol;Acc:MGI:1915010] |
| 29 | ENSMUSG00000022464 | Slc38a4 | solute carrier family 38, member 4 [Source:MGI Symbol;Acc:MGI:1916604] |
| 30 | ENSMUSG00000022684 | Bfar | bifunctional apoptosis regulator [Source:MGI Symbol;Acc:MGI:1914368] |
| 31 | ENSMUSG00000023150 | Ivns1abp | influenza virus NS1A binding protein [Source:MGI Symbol;Acc:MGI:2152389] |
| 32 | ENSMUSG00000024085 | Man2a1 | mannosidase 2, alpha 1 [Source:MGI Symbol;Acc:MGI:104669] |
| 33 | ENSMUSG00000024190 | Dusp1 | dual specificity phosphatase 1 [Source:MGI Symbol;Acc:MGI:105120] |
| 34 | ENSMUSG00000024294 | Mib1 | mindbomb E3 ubiquitin protein ligase 1 [Source:MGI Symbol;Acc:MGI:2443157] |
| 35 | ENSMUSG00000024785 | Rcl1 | RNA terminal phosphate cyclase-like 1 [Source:MGI Symbol;Acc:MGI:1913275] |
| 36 | ENSMUSG00000024900 | Cpt1a | carnitine palmitoyltransferase 1a, liver [Source:MGI Symbol;Acc:MGI:1098296] |
| 37 | ENSMUSG00000025076 | Casp7 | caspase 7 [Source:MGI Symbol;Acc:MGI:109383] |
| 38 | ENSMUSG00000025190 | Got1 | glutamic-oxaloacetic transaminase 1, soluble [Source:MGI Symbol;Acc:MGI:95791] |
| 39 | ENSMUSG00000025381 | Cnpy2 | canopy FGF signaling regulator 2 [Source:MGI Symbol;Acc:MGI:1928477] |

| | | | |
|---|---|---|---|
| 40 | ENSMUSG00000025764 | Jade1 | jade family PHD finger 1 [Source:MGI Symbol;Acc:MGI:1925835] |
| 41 | ENSMUSG00000025907 | Rb1cc1 | RB1-inducible coiled-coil 1 [Source:MGI Symbol;Acc:MGI:1341850] |
| 42 | ENSMUSG00000025981 | Coq10b | coenzyme Q10B [Source:MGI Symbol;Acc:MGI:1915126] |
| 43 | ENSMUSG00000026317 | Cln8 | CLN8 transmembrane ER and ERGIC protein [Source:MGI Symbol;Acc:MGI:1349447] |
| 44 | ENSMUSG00000026333 | Gin1 | gypsy retrotransposon integrase 1 [Source:MGI Symbol;Acc:MGI:2182036] |
| 45 | ENSMUSG00000026526 | Fh1 | fumarate hydratase 1 [Source:MGI Symbol;Acc:MGI:95530] |
| 46 | ENSMUSG00000026579 | F5 | coagulation factor V [Source:MGI Symbol;Acc:MGI:88382] |
| 47 | ENSMUSG00000026586 | Prrx1 | paired related homeobox 1 [Source:MGI Symbol;Acc:MGI:97712] |
| 48 | ENSMUSG00000026605 | Cenpf | centromere protein F [Source:MGI Symbol;Acc:MGI:1313302] |
| 49 | ENSMUSG00000026701 | Prdx6 | peroxiredoxin 6 [Source:MGI Symbol;Acc:MGI:894320] |
| 50 | ENSMUSG00000026781 | Acbd5 | acyl-Coenzyme A binding domain containing 5 [Source:MGI Symbol;Acc:MGI:1921409] |
| 51 | ENSMUSG00000027346 | Gpcpd1 | glycerophosphocholine phosphodiesterase 1 [Source:MGI Symbol;Acc:MGI:104898] |
| 52 | ENSMUSG00000027363 | Usp8 | ubiquitin specific peptidase 8 [Source:MGI Symbol;Acc:MGI:1934029] |
| 53 | ENSMUSG00000027513 | Pck1 | phosphoenolpyruvate carboxykinase 1, cytosolic [Source:MGI Symbol;Acc:MGI:97501] |
| 54 | ENSMUSG00000027792 | Bche | butyrylcholinesterase [Source:MGI Symbol;Acc:MGI:894278] |
| 55 | ENSMUSG00000027806 | Tsc22d2 | TSC22 domain family, member 2 [Source:MGI Symbol;Acc:MGI:1919283] |
| 56 | ENSMUSG00000027829 | Ccnl1 | cyclin L1 [Source:MGI Symbol;Acc:MGI:1922664] |
| 57 | ENSMUSG00000028211 | Trp53inp1 | transformation related protein 53 inducible nuclear protein 1 [Source:MGI Symbol;Acc:MGI:1926609] |
| 58 | ENSMUSG00000028266 | Lmo4 | LIM domain only 4 [Source:MGI Symbol;Acc:MGI:109360] |
| 59 | ENSMUSG00000028300 | C9orf72 | C9orf72, member of C9orf72-SMCR8 complex [Source:MGI Symbol;Acc:MGI:1920455] |
| 60 | ENSMUSG00000028426 | Rad23b | RAD23 homolog B, nucleotide excision repair protein [Source:MGI Symbol;Acc:MGI:105128] |
| 61 | ENSMUSG00000028542 | Slc6a9 | solute carrier family 6 (neurotransmitter transporter, glycine), member 9 [Source:MGI Symbol;Acc:MGI:95760] |
| 62 | ENSMUSG00000028645 | Slc2a1 | solute carrier family 2 (facilitated glucose transporter), member 1 [Source:MGI Symbol;Acc:MGI:95755] |
| 63 | ENSMUSG00000028655 | Mfsd2a | major facilitator superfamily domain containing 2A [Source:MGI Symbol;Acc:MGI:1923824] |
| 64 | ENSMUSG00000028967 | Errfi1 | ERBB receptor feedback inhibitor 1 [Source:MGI Symbol;Acc:MGI:1921405] |
| 65 | ENSMUSG00000029167 | Ppargc1a | peroxisome proliferative activated receptor, gamma, coactivator 1 alpha [Source:MGI Symbol;Acc:MGI:1342774] |
| 66 | ENSMUSG00000029267 | Mtf2 | metal response element binding transcription factor 2 [Source:MGI Symbol;Acc:MGI:105050] |
| 67 | ENSMUSG00000029695 | Aass | aminoadipate-semialdehyde synthase [Source:MGI Symbol;Acc:MGI:1353573] |
| 68 | ENSMUSG00000029817 | Tra2a | transformer 2 alpha [Source:MGI Symbol;Acc:MGI:1933972] |
| 69 | ENSMUSG00000030795 | Fus | fused in sarcoma [Source:MGI Symbol;Acc:MGI:1353633] |
| 70 | ENSMUSG00000031596 | Slc7a2 | solute carrier family 7 (cationic amino acid transporter, y+ system), member 2 [Source:MGI Symbol;Acc:MGI:99828] |
| 71 | ENSMUSG00000031700 | Gpt2 | glutamic pyruvate transaminase (alanine aminotransferase) 2 [Source:MGI Symbol;Acc:MGI:1915391] |
| 72 | ENSMUSG00000031889 | D230025D16Rik | RIKEN cDNA D230025D16 gene [Source:MGI Symbol;Acc:MGI:2443049] |
| 73 | ENSMUSG00000032078 | Zpr1 | ZPR1 zinc finger [Source:MGI Symbol;Acc:MGI:1330262] |
| 74 | ENSMUSG00000032079 | Apoa5 | apolipoprotein A-V [Source:MGI Symbol;Acc:MGI:1913363] |
| 75 | ENSMUSG00000032238 | Rora | RAR-related orphan receptor alpha [Source:MGI Symbol;Acc:MGI:104661] |
| 76 | ENSMUSG00000032633 | Flcn | folliculin [Source:MGI Symbol;Acc:MGI:2442184] |
| 77 | ENSMUSG00000032745 | Gpbp1 | GC-rich promoter binding protein 1 [Source:MGI Symbol;Acc:MGI:1920524] |
| 78 | ENSMUSG00000033107 | Rnf125 | ring finger protein 125 [Source:MGI Symbol;Acc:MGI:1914914] |
| 79 | ENSMUSG00000033760 | Rbm4b | RNA binding motif protein 4B [Source:MGI Symbol;Acc:MGI:1913954] |
| 80 | ENSMUSG00000033863 | Klf9 | Kruppel-like factor 9 [Source:MGI Symbol;Acc:MGI:1333856] |
| 81 | ENSMUSG00000034858 | Fam214a | family with sequence similarity 214, member A [Source:MGI Symbol;Acc:MGI:2387648] |

| | | | |
|---|---|---|---|
| 82 | ENSMUSG00000034903 | Cobll1 | Cobl-like 1 [Source:MGI Symbol;Acc:MGI:2442894] |
| 83 | ENSMUSG00000034936 | Arl4d | ADP-ribosylation factor-like 4D [Source:MGI Symbol;Acc:MGI:1933155] |
| 84 | ENSMUSG00000035372 | 1810055G02Rik | RIKEN cDNA 1810055G02 gene [Source:MGI Symbol;Acc:MGI:1919306] |
| 85 | ENSMUSG00000035530 | Eif1 | eukaryotic translation initiation factor 1 [Source:MGI Symbol;Acc:MGI:105125] |
| 86 | ENSMUSG00000035828 | Pim3 | proviral integration site 3 [Source:MGI Symbol;Acc:MGI:1355297] |
| 87 | ENSMUSG00000035992 | Fnip1 | folliculin interacting protein 1 [Source:MGI Symbol;Acc:MGI:2444668] |
| 88 | ENSMUSG00000036225 | Kctd1 | potassium channel tetramerisation domain containing 1 [Source:MGI Symbol;Acc:MGI:1918269] |
| 89 | ENSMUSG00000036353 | P2ry12 | purinergic receptor P2Y, G-protein coupled 12 [Source:MGI Symbol;Acc:MGI:1918089] |
| 90 | ENSMUSG00000036478 | Btg1 | BTG anti-proliferation factor 1 [Source:MGI Symbol;Acc:MGI:88215] |
| 91 | ENSMUSG00000037035 | Inhbb | inhibin beta-B [Source:MGI Symbol;Acc:MGI:96571] |
| 92 | ENSMUSG00000037112 | Sik2 | salt inducible kinase 2 [Source:MGI Symbol;Acc:MGI:2445031] |
| 93 | ENSMUSG00000037266 | Rsrp1 | arginine/serine rich protein 1 [Source:MGI Symbol;Acc:MGI:106498] |
| 94 | ENSMUSG00000037686 | Aspg | asparaginase [Source:MGI Symbol;Acc:MGI:2144822] |
| 95 | ENSMUSG00000037826 | Ppm1k | protein phosphatase 1K (PP2C domain containing) [Source:MGI Symbol;Acc:MGI:2442111] |
| 96 | ENSMUSG00000038500 | Prr3 | proline-rich polypeptide 3 [Source:MGI Symbol;Acc:MGI:1922460] |
| 97 | ENSMUSG00000038582 | Pptc7 | PTC7 protein phosphatase homolog [Source:MGI Symbol;Acc:MGI:2444593] |
| 98 | ENSMUSG00000039210 | Gpatch2 | G patch domain containing 2 [Source:MGI Symbol;Acc:MGI:1915019] |
| 99 | ENSMUSG00000039782 | Cpeb2 | cytoplasmic polyadenylation element binding protein 2 [Source:MGI Symbol;Acc:MGI:2442640] |
| 100 | ENSMUSG00000039989 | Cbx4 | chromobox 4 [Source:MGI Symbol;Acc:MGI:1195985] |
| 101 | ENSMUSG00000040093 | Bmf | BCL2 modifying factor [Source:MGI Symbol;Acc:MGI:2176433] |
| 102 | ENSMUSG00000042328 | Hps4 | HPS4, biogenesis of lysosomal organelles complex 3 subunit 2 [Source:MGI Symbol;Acc:MGI:2177742] |
| 103 | ENSMUSG00000042423 | Fbrs | fibrosin [Source:MGI Symbol;Acc:MGI:104648] |
| 104 | ENSMUSG00000042510 | AA986860 | expressed sequence AA986860 [Source:MGI Symbol;Acc:MGI:2138143] |
| 105 | ENSMUSG00000045636 | Mtus1 | mitochondrial tumor suppressor 1 [Source:MGI Symbol;Acc:MGI:2142572] |
| 106 | ENSMUSG00000045973 | Slc25a51 | solute carrier family 25, member 51 [Source:MGI Symbol;Acc:MGI:2684984] |
| 107 | ENSMUSG00000046434 | Hnrnpa1 | heterogeneous nuclear ribonucleoprotein A1 [Source:MGI Symbol;Acc:MGI:104820] |
| 108 | ENSMUSG00000047446 | Arl4a | ADP-ribosylation factor-like 4A [Source:MGI Symbol;Acc:MGI:99437] |
| 109 | ENSMUSG00000048271 | Rbm33 | RNA binding motif protein 33 [Source:MGI Symbol;Acc:MGI:1919670] |
| 110 | ENSMUSG00000049044 | Rapgef4 | Rap guanine nucleotide exchange factor (GEF) 4 [Source:MGI Symbol;Acc:MGI:1917723] |
| 111 | ENSMUSG00000051495 | Irf2bp2 | interferon regulatory factor 2 binding protein 2 [Source:MGI Symbol;Acc:MGI:2443921] |
| 112 | ENSMUSG00000052934 | Fbxo31 | F-box protein 31 [Source:MGI Symbol;Acc:MGI:1354708] |
| 113 | ENSMUSG00000054115 | Skp2 | S-phase kinase-associated protein 2 [Source:MGI Symbol;Acc:MGI:1351663] |
| 114 | ENSMUSG00000056429 | Tgoln1 | trans-golgi network protein [Source:MGI Symbol;Acc:MGI:105080] |
| 115 | ENSMUSG00000056501 | Cebpb | CCAAT/enhancer binding protein (C/EBP), beta [Source:MGI Symbol;Acc:MGI:88373] |
| 116 | ENSMUSG00000056673 | Kdm5d | lysine (K)-specific demethylase 5D [Source:MGI Symbol;Acc:MGI:99780] |
| 117 | ENSMUSG00000059811 | Atl2 | atlastin GTPase 2 [Source:MGI Symbol;Acc:MGI:1929492] |
| 118 | ENSMUSG00000060550 | H2-Q7 | histocompatibility 2, Q region locus 7 [Source:MGI Symbol;Acc:MGI:95936] |
| 119 | ENSMUSG00000062901 | Klhl24 | kelch-like 24 [Source:MGI Symbol;Acc:MGI:1923035] |
| 120 | ENSMUSG00000063406 | Tmed5 | transmembrane p24 trafficking protein 5 [Source:MGI Symbol;Acc:MGI:1921586] |
| 121 | ENSMUSG00000067279 | Ppp1r3c | protein phosphatase 1, regulatory subunit 3C [Source:MGI Symbol;Acc:MGI:1858229] |
| 122 | ENSMUSG00000069045 | Ddx3y | DEAD box helicase 3, Y-linked [Source:MGI Symbol;Acc:MGI:1349406] |
| 123 | ENSMUSG00000078650 | G6pc | glucose-6-phosphatase, catalytic [Source:MGI Symbol;Acc:MGI:95607] |

| | | | |
|---|---|---|---|
| 124 | ENSMUSG00000081766 | 2210409E12Rik | RIKEN cDNA 2210409E12 gene [Source:MGI Symbol;Acc:MGI:1919631] |
| 125 | ENSMUSG00000085054 | Gm15834 | predicted gene 15834 [Source:MGI Symbol;Acc:MGI:3802168] |
| 126 | ENSMUSG00000089809 | Rasgef1b | RasGEF domain family, member 1B [Source:MGI Symbol;Acc:MGI:2443755] |
| 127 | ENSMUSG00000095562 | | erythroid differentiation regulator 1 [Source:NCBI gene (formerly Entrezgene);Acc:170942] |
| 128 | ENSMUSG00000099474 | 1700097N02Rik | RIKEN cDNA 1700097N02 gene [Source:MGI Symbol;Acc:MGI:1914772] |
